# Independent origin of *MIRNA* genes controlling homologous target genes by partial inverted duplication of antisense-transcribed sequences

**DOI:** 10.1101/682591

**Authors:** Lydia Gramzow, Dajana Lobbes, Sophia Walter, Nathan Innard, Günter Theißen

## Abstract

Some microRNAs (miRNAs) are key regulators of developmental processes, mainly by controlling the accumulation of transcripts encoding transcription factors that are important for morphogenesis. MADS-box genes encode a family of transcription factors which control diverse developmental processes in flowering plants. Here we study the convergent evolution of two *MIRNA* (*MIR*) gene families, named *MIR444* and *MIR824,* targeting members of the same clade of MIKC^C^-group MADS-box genes. We show that these two *MIR* genes most likely originated independently in monocots (*MIR444*) and in Brassicales (eudicots, *MIR824*). We provide evidence that in both cases the future target gene was transcribed in antisense prior to the evolution of the *MIR* genes. Both *MIR* genes then likely originated by a partial inverted duplication of their target genes, resulting in natural antisense organization of the newly evolved *MIR* gene and its target gene at birth. We thus propose a new model for the origin of *MIR* genes, MEPIDAS (MicroRNA Evolution by Partial Inverted Duplication of Antisense-transcribed Sequences). MEPIDAS is a refinement of the inverted duplication hypothesis. According to MEPIDAS, a *MIR* gene evolves at a genomic locus at which the future target gene is also transcribed in the antisense direction. A partial inverted duplication at this locus causes the antisense transcript to fold into a stem-loop structure that is recognized by the miRNA biogenesis machinery to produce a miRNA that regulates the gene at this locus. Our analyses exemplify how to elucidate the origin of conserved miRNAs by comparative genomics and will guide future studies.

## Introduction

MicroRNAs (miRNAs) are short, non-coding RNAs which function as sequence-specific guides to control gene expression by post-transcriptional gene silencing. They are important factors for the normal development and physiology of plants and animals (He and Hannon 2004).

Genes encoding miRNAs (*MIR* genes) are transcribed by RNA polymerase II; the transcripts are capped at the 5’ end, polyadenylated at the 3’ end, and spliced (if they contain introns) (Xie, et al. 2005). The resulting primary miRNAs (pri-miRNAs) feature a stem-loop structure that is recognized and processed by the DICER-LIKE1 (DCL1) protein in plants (Park, et al. 2005). Generally, the 5’ and 3’ ends are first cut by DCL1 at the base of the stem leaving the precursor miRNA (pre-miRNA), which is then cut a second time by DCL1 near the loop to produce 21 nucleotide (nt) long miRNA/miR* duplexes. The miRNA/miR* duplexes are transported from the nucleus to the cytoplasm, possibly by HASTY in plants (Park, et al. 2005). One strand from the duplex, the mature miRNA, then associates with an ARGONAUTE (AGO) protein to form an RNA-induced silencing complex (RISC) (Rogers and Chen 2013). The target mRNA is recognized because it contains a target site which has a high degree of complementarity to the mature miRNA. The target mRNA is negatively regulated by the RISC, generally by cleavage or translational repression (Bartel 2004; Fabian and Sonenberg 2012; Huntzinger and Izaurralde 2011).

In plants and animals, some miRNAs are deeply conserved (Axtell and Bartel 2005; Berezikov 2011; Chavez Montes, et al. 2014; Pasquinelli, et al. 2000), whereas the majority of miRNAs are found only in a few lineages (Chavez Montes, et al. 2014; Fahlgren, et al. 2007; Hertel and Stadler 2015). Several hypotheses as to how new *MIR* genes may originate have been brought forward. Arguably the simplest hypothesis is that new *MIR* genes originate by duplication of existing *MIR* genes (Nozawa, et al. 2012). However, this mechanism predominantly gives rise to new *MIR* gene family members but not to new *MIR* gene families. Another possibility involves the evolution of new *MIR* genes from random sequences (Fahlgren, et al. 2010; Felippes, et al. 2008). According to this hypothesis, inverted repeat sequences in the genome may become transcribed. The resulting transcripts fold into stem-loop structures that may be processed by the miRNA biogenesis machinery to form mature miRNAs. New *MIR* genes may also evolve from transposable elements (Piriyapongsa and Jordan 2008; Piriyapongsa, et al. 2007). If a transposable element is inserted in both the sense and antisense direction in close proximity within the genome, a transcript of this genomic region will fold into a stem-loop structure. As described for the evolution of *MIR* genes from random sequences, this transcript structure may be recognized by the miRNA biogenesis machinery and processed to produce a mature miRNA. According to the inverted duplication hypothesis, a new *MIR* gene may evolve if a paralog of the gene it later comes to target is inversely duplicated (Allen, et al. 2004). If the genomic region of the paralog and its inverse duplication is transcribed, a stem-loop structure is formed. This stem-loop structure may first be processed into short interfering RNAs (siRNAs) due to an extended region of complementarity in the stem. Accumulation of mutations in the genomic locus of this inverted duplication may lead to a decrease in the complementarity in the stem of the resulting transcript and hence, a miRNA may be formed. The above hypotheses on how new *MIR* genes may originate are not mutually exclusive. Rather, examples of *MIR* genes which are thought to have originated according to each of these different hypotheses have been found (Allen, et al. 2004; Felippes, et al. 2008; Nozawa, et al. 2012; Piriyapongsa, et al. 2007).

In plants, the mature miRNA is usually almost completely complementary to its target mRNA over its whole length of about 21 nt. A number of *MIR* genes exhibit extended sequence complementarity to their target genes exceeding the complementarity between the mature miRNA and the target site (Allen, et al. 2004; Fahlgren, et al. 2010; Xia, et al. 2015). Hence, for a number of plant *MIR* genes, it is thought that they evolved according to the inverted duplication hypothesis. However, selection pressure mainly acts on the mature miRNA which is only 21 nt in length. Hence, over evolutionary time only the mature miRNA is conserved and extended complementarity to the target gene is lost. Consequently, especially for older *MIR* genes, it is hard to elucidate how they originated (Axtell and Bowman 2008; Fahlgren, et al. 2010).

The *MIR444* gene family was discovered in rice (*Oryza sativa*) (Lu, et al. 2008; Sunkar, et al. 2005; Sunkar, et al. 2008). The *MIR444* gene family is special because most of their transcripts derive from the antisense strand of their target genes. Hence, the miRNAs encoded by these *MIR* genes have been named natural antisense transcript miRNAs (nat-miRNAs (Lu, et al. 2008)). In fact, four of the six *MIR444* gene family members in *O. sativa* are natural antisense to one of their target genes. All known target genes are *AGL17*-like MADS-box genes. By regulating the expression of these genes, miR444 has important functions in the nitrate-signaling pathway for root and shoot development and in tillering (Guo, et al. 2013; Yan, et al. 2014). Furthermore, miR444 is involved in antiviral responses (Wang, et al. 2016). The *MIR444* family has also been found in other species of the monocot grass family Poaceae (order Poales), including wheat (*Triticum aestivum*), barley (*Hordeum vulgare*), maize (*Zea mays*), sorghum (*Sorghum bicolor*) and sugarcane (*Saccharum officinarum*) (Lu, et al. 2008; Sunkar, et al. 2005; Sunkar, et al. 2008).

The *MIR824* gene was initially described in *Arabidopsis thaliana* (Fahlgren, et al. 2007; Kutter, et al. 2007; Rajagopalan, et al. 2006). Interestingly, its target gene *AGL16* also belongs to the subclade of *AGL17*-like genes. The miR824-mediated regulation of its target gene has been shown to be important for stomatal complex development (Kutter, et al. 2007) and flowering time repression (Hu, et al. 2014) in *A. thaliana*. miR824 has also been identified in other Brassicaceae species, namely *Brassica rapa*, *Brassica napus* and *Brassica oleraceae* (Kutter, et al. 2007).

*AGL17*-like genes are one of 17 clades of MIKC^C^-group MADS-box genes that were present in the most recent common ancestor of flowering plants (Gramzow, et al. 2014). Only one other clade of MADS-box genes from flowering plants, the *DEF*-like (also known as *AP3*-like) genes, have been found to be regulated by miRNAs, namely by miR5179 (Aceto, et al. 2014). In order to gain a better understanding as to how miRNA regulation of *AGL17*-like genes by two different *MIR* gene families may have evolved, here we clarify the origin of *MIR444* and *MIR824* by systematic genome and transcriptome analyses and by thorough experimental verification in several plant species. We find that the *MIR444* gene originated early in the evolution of monocotyledonous plants about 120 million years ago (MYA), while the *MIR824* gene originated about 50 MYA in a common ancestor of the eudicot plant families Brassicaceae and Cleomaceae. Our studies reveal that in both cases the future target genes were likely transcribed in the antisense direction prior to the origin of the *MIR* genes. Subsequently, partial inverted duplications gave rise to the *MIR* genes and led to natural antisense organizations of the target and the *MIR* genes. Hence, we describe MEPIDAS (MicroRNA Evolution by Partial Inverted Duplication of Antisense-transcribed Sequences), a model describing the origin of *MIR* genes. Our model suggests that transcription of a newly evolved *MIR* gene is probably driven by a promotor that originated prior to the inverted duplication event which gave rise to the *MIR* gene. Our study also shows how thorough genome and transcriptome analyses can be used to elucidate the origin of old *MIR* genes and may hence guide the investigation of the origin of many more *MIR* genes.

## Results

### Phylogeny of *AGL17*-like genes

As the target genes of both miR444 and miR824 belong to the family of *AGL17*-like MADS-box genes (Kutter, et al. 2007; Lu, et al. 2008), we first reconstructed a phylogeny of *AGL17*-like genes from angiosperms. We find that *AGL17*-like genes of monocots and eudicots form distinct clades (Figure 1, Supplemental Figure 1), suggesting that the most recent common ancestor (MRCA) of both taxa contained just one *AGL17*-like gene. Monocot *AGL17*-like genes form a clade with one to five members from each of the monocot species studied. Nearly all monocot *AGL17*-like genes are putative targets of miR444 as based on the target genes previously identified (Lu, et al. 2008) and those identified in this study (see below). In eudicots, there are three major subclades of *AGL17*-like genes each with members from nearly all eudicot species studied (Figure 1, Supplemental Figure 1). We named these subclades *ANR1*-, *AGL16*- and *AGL21*-like genes after a representative from *A. thaliana*. The target genes of miR824 as described previously (Hu, et al. 2014; Kutter, et al. 2007) and in our study (see below) all belong to the subclade of *AGL16*-like genes.

**Figure 1.**
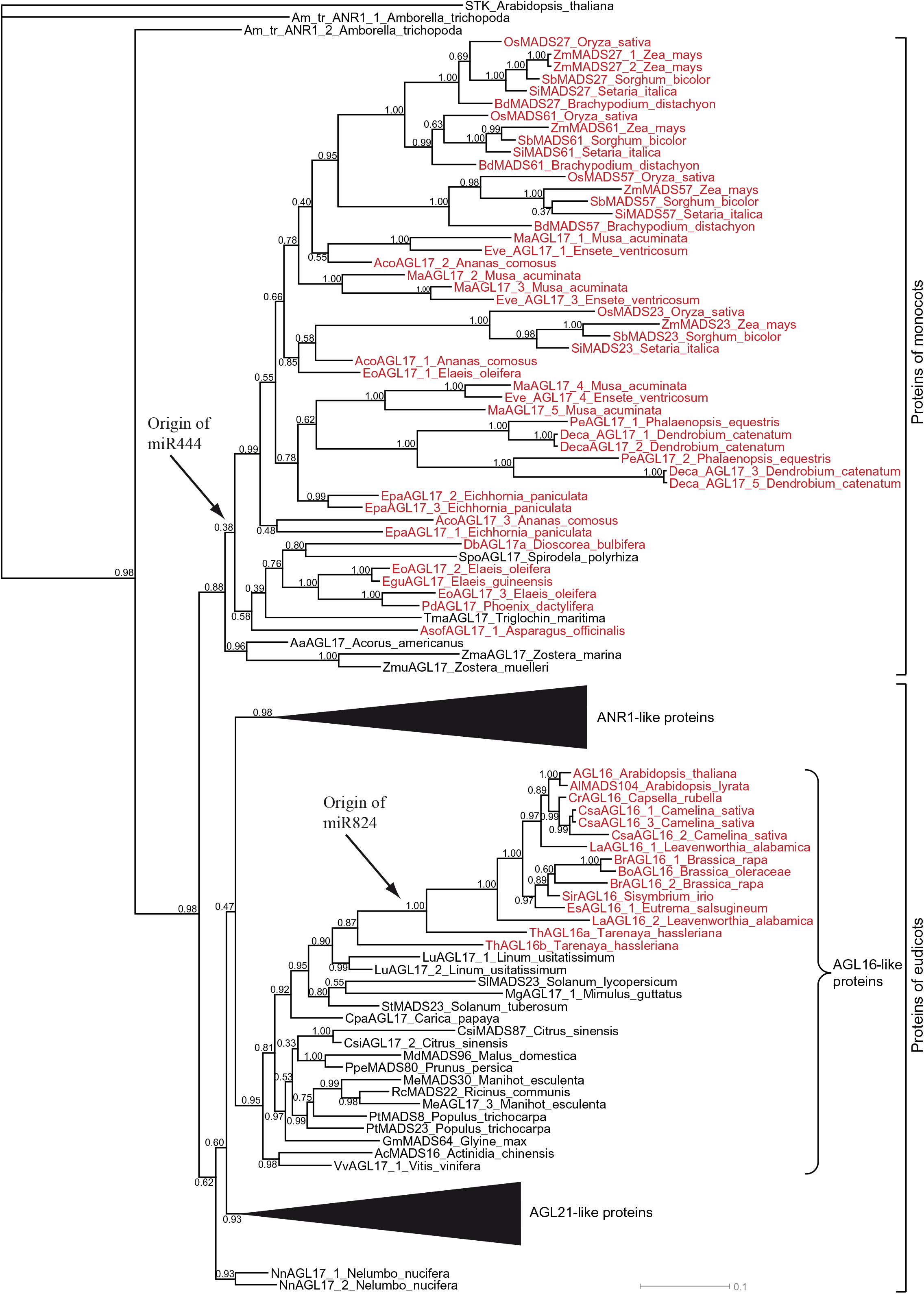
Phylogeny of AGL17-like proteins as reconstructed using MrBayes. AGL17-like proteins from monocotyledonous and eudicotyledonous plants form two separate clades (indicated on the right). AGL17-like proteins from eudicots are subdivided into three subclades which we named ANR1-, AGL16- and AGL21-like proteins after representative proteins from *Arabidopsis thaliana*. The subclades ANR1- and AGL21-like proteins have been collapsed and are represented by triangles. AGL17-like proteins of which the corresponding transcripts are targeted by miRNAs are highlighted in red. The putative origins of the miRNAs as judged by the species to which the AGL17-like proteins of the different clades belong, are indicated by arrows. Numbers at the different nodes represent posterior probabilities. For a complete version of the phylogeny see Supplemental Figure 1.

### Presence of the *MIR444* gene family in monocots

*MIR444* genes have previously been identified in a number of species belonging to the grass family Poaceae (Lu, et al. 2008; Sunkar, et al. 2005; Sunkar, et al. 2008). There are ten orders of monocots (Janssen and Bremer 2004). Apart from the order Poales, core monocots include the following orders (presented from most closely to most distantly related to Poales): Zingiberales, Commelinales, Arecales, Asparagales, Liliales, Pandanales and Dioscoreales. Alismatales and Acorales are early-branching orders of monocots. To get an initial idea on the distribution of miR444 we conducted Northern blot analyses. We obtained signals for RNA isolated from plants of the orders Poales, Commelinales, Zingiberales, Asparagales, Liliales and Dioscoreales (Supplemental Figure 2).

Whole genome sequences are available for species of all monocot orders except Liliales and Acorales (Al-Mssallem, et al. 2013; Cai, et al. 2015; Costa, et al. 2017; D’Hont, et al. 2012; Golicz, et al. 2015; Ming, et al. 2015; Singh, et al. 2013; Wang, et al. 2014; Zhang, et al. 2016). In all investigated genomes, apart from the genomes of the three Alismatales species *Spirodela polyrhiza*, *Zostera marina* and *Zostera muelleri*, we found at least one *MIR444* gene (Table 1, Supplemental Data, Supplemental Figure 3). In all species for which we identified *MIR444* genes, there is at least one *MIR444* gene that is orientated natural antisense to an *AGL17*-like gene (Table 1, Figure 2 illustrates one example for each order, Supplemental Data). We identified at least one *MIR444* gene per species in transcriptome datasets suggesting expression of these genes. Using RNA secondary structure prediction, we found evidence that all identified pri-MIR444 sequences are able to fold into a stem-loop structure from which the mature miR444 may be processed (Table 1, Figure 2, and Supplemental Data). By investigating small RNA data from the NCBI SRA database we found evidence of the generation of mature miR444 for all species for which this type of data is available (Table 1). Up to three mature miR444 species are generated from an individual pre-MIR444 (Table 1, Figure 2). The positions of the mature miR444 species were often shifted as compared to the positions of the mature miR444 species of osa-MIR444d (Table 1, Figure 2). As no whole genome sequence of a species of the orders Liliales or Acorales has been sequenced yet, we searched for evidence of pri-miR444 in these orders in the data of the 1000 plants (1KP) project. We were able to identify a putative pri-MIR444 for three species of the Liliales (Table 1, Supplemental Data). In contrast, we were unable to find a putative pri-MIR444 for any species of Acorales.

**Figure 2.**
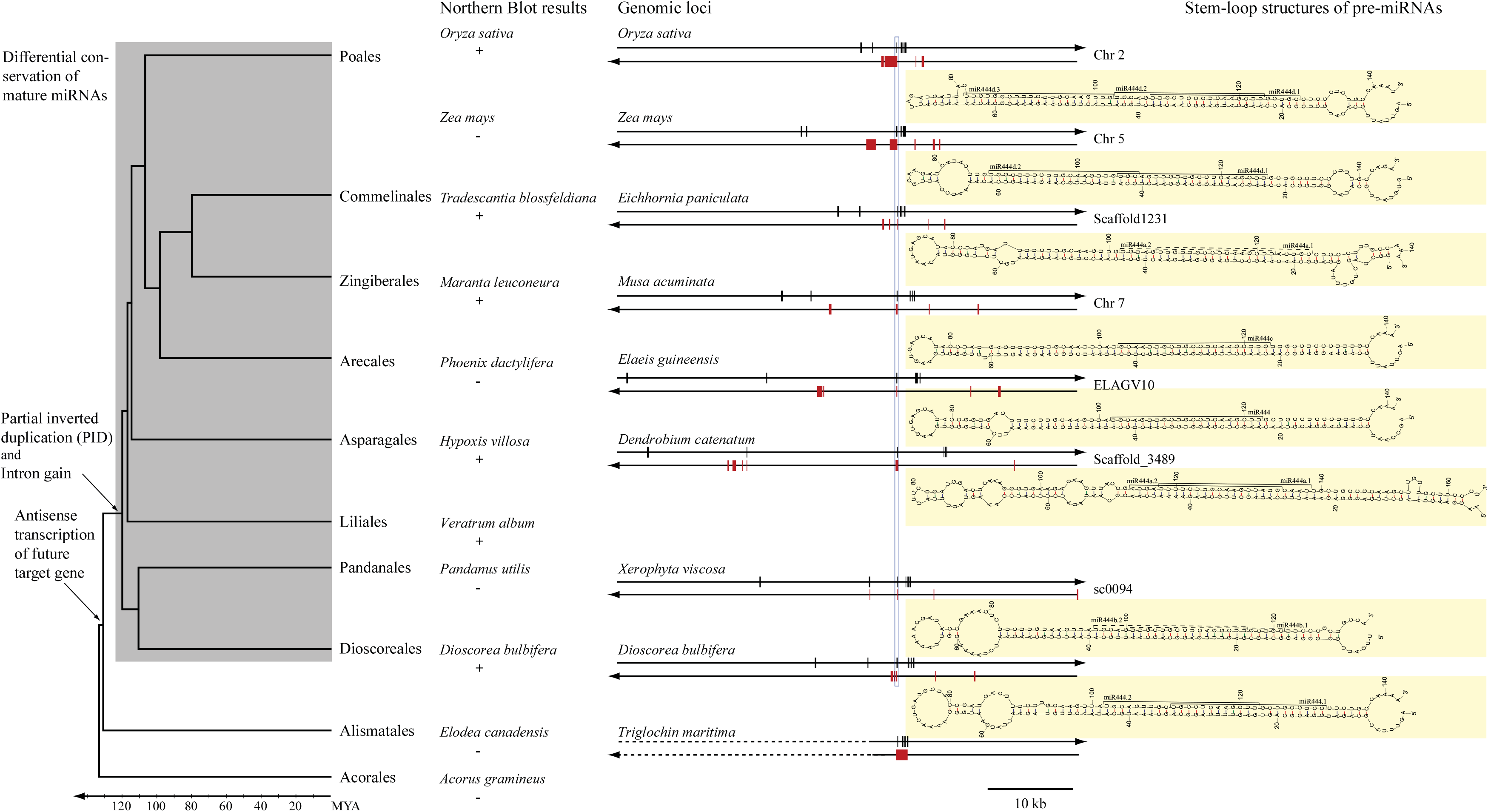
Evolution of the *MIR444* gene in monocotyledonous plants. On the left, a phylogeny of monocotyledonous plant orders is shown (Janssen and Bremer 2004). Approximate divergence times of the orders can be deduced from the time scale at the bottom. Presence (“+”) and absence (“-“) of a signal in our Northern blot analysis for the detection of mature miRNAs are indicated below the analyzed plant species names. The genomic loci of the *AGL17*-like genes and the *MIR444* genes or the antisense gene (in case of *Triglochin maritima*) are depicted by double arrows representing double-stranded DNA; the arrow heads indicate the 3’ ends. Exons are shown as boxes on the strand transcribed into RNA (i.e. coding strand), whereas introns are shown as lines. Exons of the *AGL17*-like genes are shown in black while exons of the *MIR* genes and the antisense transcript are colored red. The blue rectangle marks the position of the mature miRNA and the miRNA target site. In cases where the genomic loci were taken from public genome data, the corresponding identifier of the scaffold is shown next to the locus. The stem-loop structures of the pre-miRNAs are shown as predicted by mfold (Zuker 2003) and shaded yellow. The mature miRNAs are indicated on the stem-loop structures either based on sequence similarity to osa-miR444d.1 and osa-miR444d.2 (dashed lines) or based on small RNAs found in next-generation sequencing data of the SRA database at NCBI (Sayers, et al. 2012) (solid lines). If several *MIR444* genes were found in one species, one *MIR444* was arbitrarily chosen for the depiction of its genomic locus and stem-loop structure in this figure. The events in the evolution of *MIR444* by the partial inverted duplication mechanism are indicated on the phylogeny. Note that in the case of *Triglochin maritima* only part of the genomic locus of the *AGL17*-like gene was sequenced and we do not have sequence information for the rest of the locus (indicated by dashed lines).

**Table 1:**
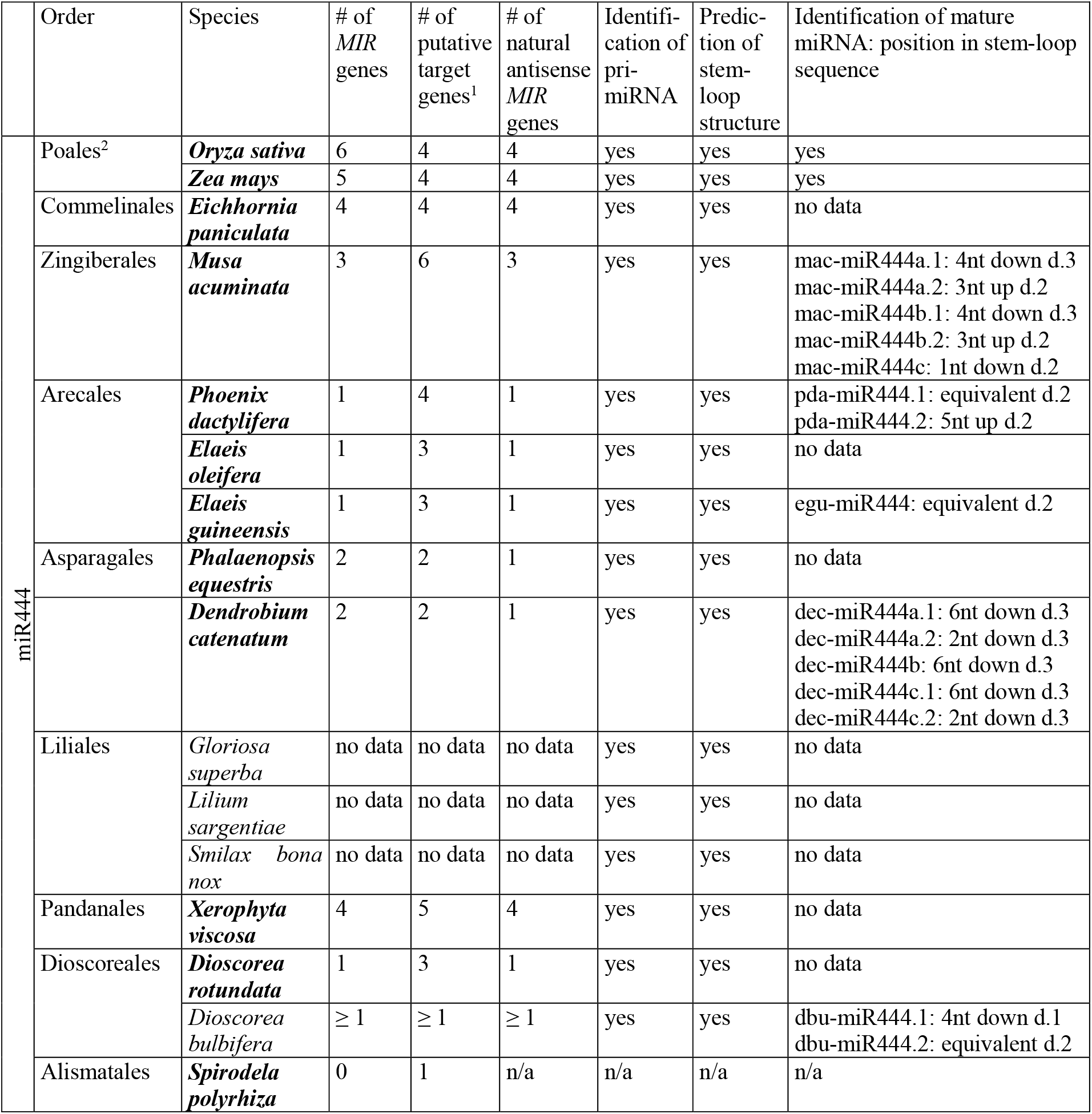
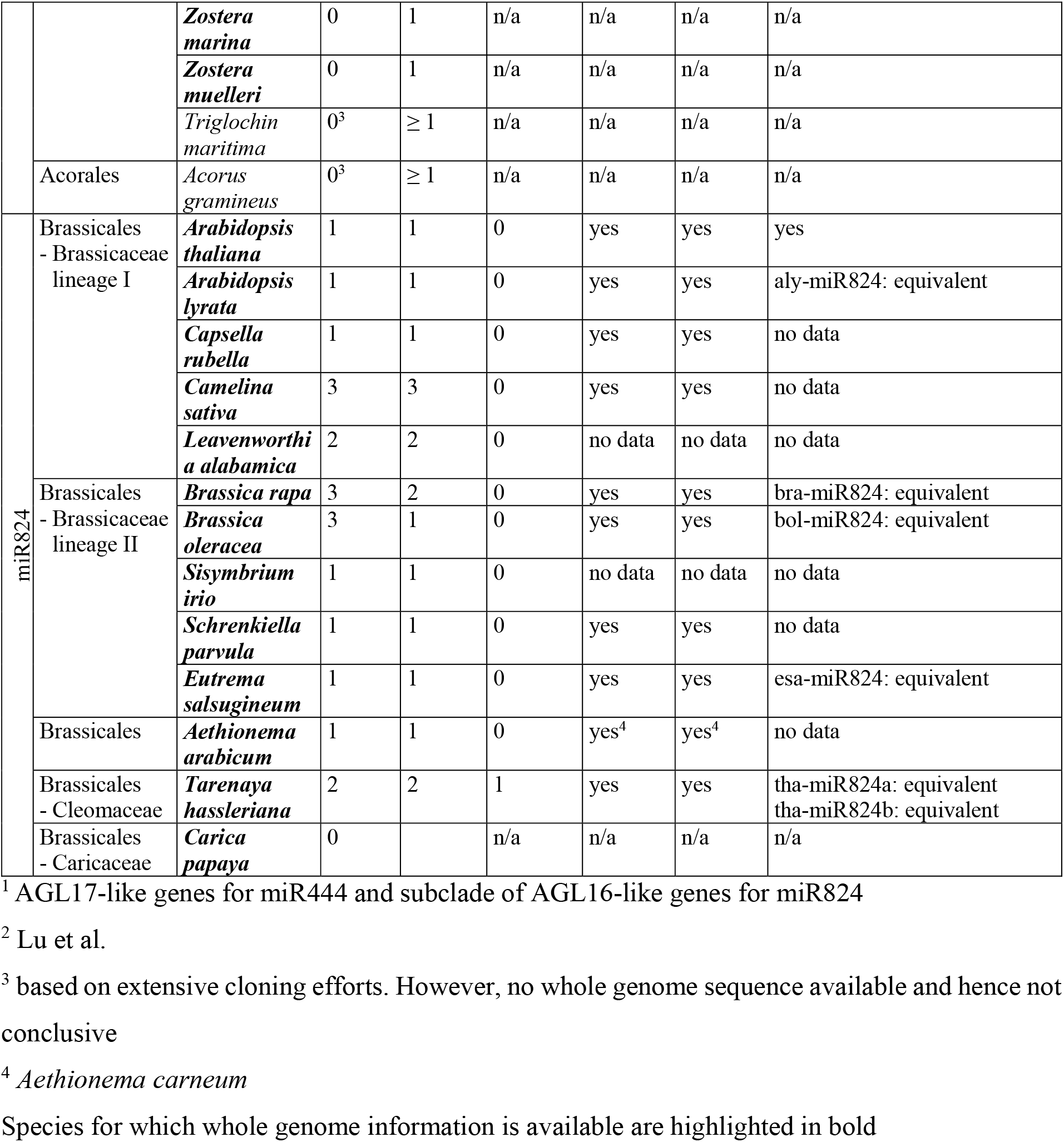
Presence of miR444 in monocots and miR824 in Brassicales.

### Regulation of an *AGL17*-like gene by miR444 in aerial yam

Dioscoreales are one of the earliest branching orders for which we found evidence for the presence of *MIR444* genes. Hence, we investigated *Dioscorea bulbifera* (aerial yam) in more detail. We first confirmed the expression of the *MIR444* gene and an *AGL17*-like gene (*DbuAGL17*). For both the *D. bulbifera* pri-MIR444 and *DbuAGL17* we found a high number of alternative splice variants, including some lacking the exon harboring the mature miRNA or the miR* (Supplemental Figure 4), and splice variants lacking the exon which contains the miRNA target site, respectively. Using genome walking we were also able to confirm that the *dbu-MIR444* gene is natural antisense to *DbuAGL17* (Figure 2). Using miRNA rapid amplification of cDNA ends (miR-RACE) (Song, et al. 2010a), we confirmed the generation of two mature miRNAs, dbu-miR444.1 and dbu-miR444.2. Sequencing revealed that the mature dbu-miR444.1 is shifted 4 nucleotides downstream in comparison to osa-miR444d.1 (Figure 2).

Regulation of the *DbuAGL17* by *dbu-MIR444* was confirmed using a transient expression assay in *Nicotiana benthamiana* leaves (Franco-Zorrilla, et al. 2007). The greater the concentration of *A. tumefaciens* containing the *dbu-MIR444* gene construct used, the more the signal of DbuAGL17 fused to GFP was reduced (Supplemental Figure 5). *N. benthamiana* leaves co-infiltrated with *A. tumefaciens* containing a construct with a miRNA not complementary to the *DbuAGL17* (*ath-MIR160a*), and leaves co-infiltrated with *A. tumefaciens* containing the empty vector, showed no reduction of the signal of DbuAGL17 fused to GFP. This observation indicates that the reduction of the signal is specific to the expression of the *dbu-MIR444* gene.

Using a modified 5’-RACE technique, we identified cleavage products of the *DbuAGL17* transcript in *D. bulbifera* leaves (Supplemental Figure 6). Surprisingly, we obtained numerous cleavage products but none with a cleavage site at the expected positions for dbu-miR444-dependent cleavage events. We also analyzed the cleavage products of the *DbuAGL17* transcript isolated from *N. benthamiana* leaves infiltrated with *A. tumefaciens* containing DbuAGL17 fused to GFP alone or together with *A. tumefaciens* containing the *dbu-MIR444* gene construct (Figure 3). One band on our agarose gel was found exclusively from *N. benthamiana* leaves which had been co-infiltrated with *A. tumefaciens* containing the *dbu-MIR444* gene construct. Analysis of this band revealed that most sequenced clones had a cleavage site at a position expected for dbu-miR444.1-mediated cleavage (Figure 3). We hypothesize that expression of *DbuAGL17* is regulated by dbu-miR444, either by translational repression or by transcript cleavage followed by rapid degradation of the cleavage products.

**Figure 3.**
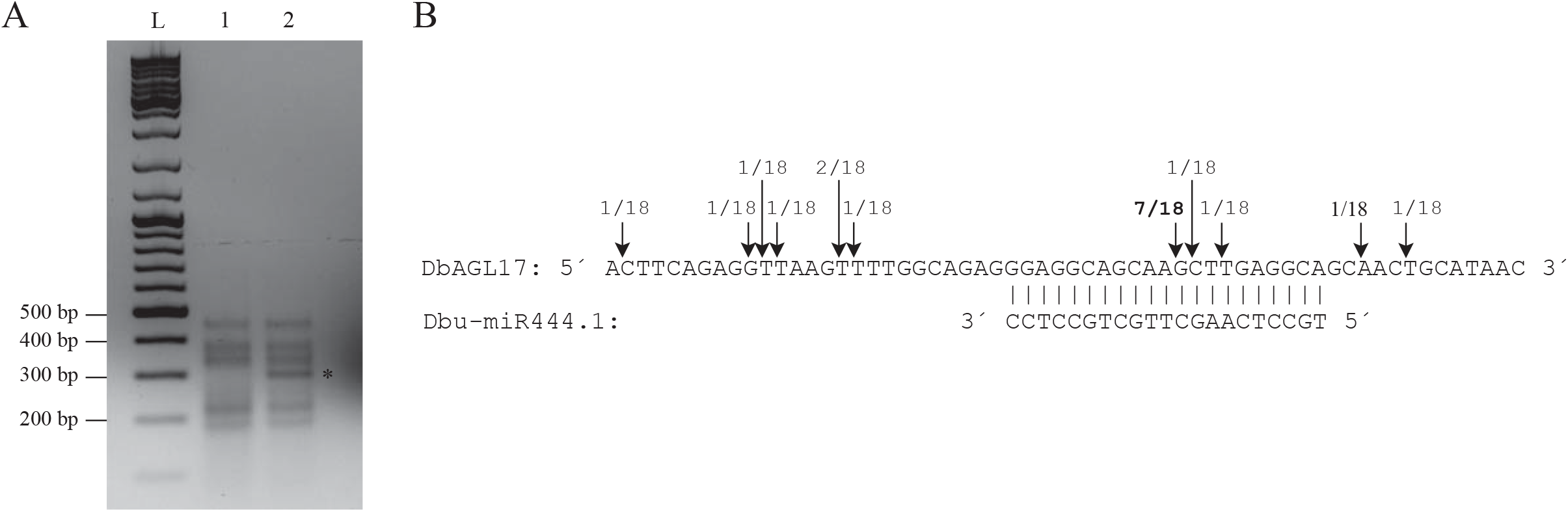
Cleavage of *DbAGL17a* mRNA by dbu-miR444. Modified 5’ RACE of *DbAGL17a* from *N. benthamiana* leaves co-infiltrated with *A. tumefaciens* containing the *DbAGL17a::*mGFP5ER and the *dbu-MIR444* construct was performed. (A) Agarose gel electrophoresis of the modified 5’ RACE products from *N. benthamiana* leaves infiltrated with *A. tumefaciens* containing only the *DbAGL17a::*mGFP5ER construct (1) or from *N. benthamiana* leaves co-infiltrated with *A. tumefaciens* containing the *DbAGL17a::*mGFP5ER and the *dbu-MIR444* construct (2). The 300 bp band which was only found when the *dbu-MIR444* is present is marked by an asterisk. The GeneRuler DNA Ladder Mix (L, Thermo Fisher Scientific) was used as a DNA size standard, lengths of some marker bands (in bp) are indicated on the left. (B) Identified cleavage products from *DbAGL17a* after analysis of the clones containing the 300 bp band. Shown is a partial sequence of *DbAGL17a* surrounding the miRNA target site and dbu-miR444.1. Perfect base pairing between dbu-miR444.1 and *DbAGL17a* is shown by vertical lines. The arrows indicate the cleavage sites identified by modified 5’ RACE, where the first number indicates the number of clones with this cleavage site and the second number represents the total number of sequenced RACE products. The site between the nucleotides that are complementary to positions 10 and 11 of the mature dbu-miR444.1 where cleavage would be expected is indicated by bold numbers.

### An antisense transcript of an *AGL17*-like gene is present in Alismatales

For the order Alismatales, we did not detect a signal for a mature miR444 in our Northern blot analysis (Supplemental Figure 2), and we did not identify a *MIR444* gene in the genomes of the species *Spirodela polyrhiza* (Wang, et al. 2014), *Zostera marina* (Olsen, et al. 2016) or *Zostera muelleri* (Golicz, et al. 2015). To further investigate the status of *MIR444* in Alismatales, we conducted detailed analyses in *Triglochin maritima*. We found several transcripts for potential *AGL17*-like genes as well as several antisense transcripts to the *AGL17*-like genes of *T. maritima*. The longest antisense transcript has a total length of around 850 bp and is antisense to exons 4 to 7 of an *AGL17*-like gene. However, none of these antisense transcripts contains a putative miR*, or would be predicted to fold into a stem-loop structure as typical for pri-miRNAs (Supplemental Figure 7). Additionally, genome walking of around 9.5 kb downstream of the exon containing the putative target site of an *AGL17*-like gene did not identify a sequence which could serve as a putative miR* (Supplemental Data).

### No *MIR444* gene in Acorales

Acorales is the earliest branching order of monocots. We did not detect a signal for a mature miR444 in Acorales in our Northern blot analysis (Supplemental Figure 2) and did not identify a putative pri-MIR444 in transcriptome data. We were able to amplify the transcript of an *AGL17*-like gene from *Acorus gramineus* (*AgAGL17*). *AgAGL17* contains a conserved potential miR444 target site (Supplemental Figure 8). However, we were unable to amplify a pri-MIR444 or a *MIR444* gene from cDNA or genomic DNA, respectively. Additionally, genome walking of around 9 kb 3’ of the last exon of the *AgAGL17* gene did not identify a sequence which could serve as a putative miR* (Supplemental Data).

### Presence of the *MIR824* gene family in Brassicales

*MIR824* has been identified in *A. thaliana* and other crucifer species (Kutter, et al. 2007). To elucidate as to how the *MIR824* gene originated, we searched whole genome sequences of the eudicot order Brassicales (including e.g. crucifers and *Carica papaya* (papaya)) (Cheng, et al. 2013; Haudry, et al. 2013; Hu, et al. 2011; Kagale, et al. 2014; Liu, et al. 2014; Slotte, et al. 2013; The Arabidopsis Genome Initiative 2000; Wang, et al. 2011).

We found evidence of *MIR824* genes in all species of the crucifer family (Brassicaceae) investigated and in the spider plant *Tarenaya hassleriana*, belonging to Cleomaceae, which is the sister family of Brassicaceae. However, we did not find *MIR824* genes in *C. papaya*, the most closely related species to Brassicaceae and Cleomaceae for which whole genome information is available (Table 1, Supplemental Data) or in genomes from other closely related plant species such as *Theobroma cacao*, *Gossypium raimondii* and *Citrus sinensis*. We found evidence of expression in transcriptome datasets for at least one *MIR824* gene for every species for which such data is available (Table 1, Supplemental Data). All identified pri-MIR824s potentially form a stem-loop structure which may represent the pre-MIR824. Information on mature miR824 was found in miRBase (Kozomara and Griffiths-Jones 2014) and in small RNA data for all species for which this kind of data is available. In all cases, the mature miR824 was identical in sequence and position to ath-miR824 (Table 1, Supplemental Data).

Three major lineages of Brassicaceae are distinguished, lineage I, extended lineage II and lineage III (Franzke, et al. 2011). However, no complete genome sequence for a species of lineage III is available. For all genomes studied of species belonging to lineage I (*A. thaliana*, *Arabidopsis lyrata*, *Capsella rubella*, *Camelina sativa* and *Leavenworthia alabamica*) the *MIR824* genes were found on different chromosomes to their corresponding *AGL16*-like genes (Figure 4). In contrast, the *MIR824* genes were located on the same chromosomes as their corresponding *AGL16*-like genes for all genomes investigated of species from extended lineage II (*B. rapa*, *B. oleracea*, *Sisymbrium irio*, *Schrenkiella parvula*, and *Eutrema salsugineum*). The distances between the *MIR824* genes and the *AGL16*-like genes range from 0.8 to 8.5 Mbp (Figure 4). In *Aethionema arabicum*, a species of the earliest branching lineage of the Brassicaceae, the *AGL16*-like and the *MIR824* gene were found on different, short scaffolds. Hence, it is not possible to determine from the current genome assembly of *A. arabicum* whether or not the *AGL16*-like and the *MIR824* gene are on the same chromosome. For *T. hassleriana*, we found two *AGL16*-like genes. These two genes are about 12 kbp apart. Interestingly, we found that one of the two *MIR824* genes in this species is natural antisense to one of the two *AGL16*-like genes (Figure 4). We thus hypothesize that the natural antisense organization of a *MIR824* and an *AGL16*-like gene may represent the ancestral genomic organization of the *MIR* and its target gene.

**Figure 4.**
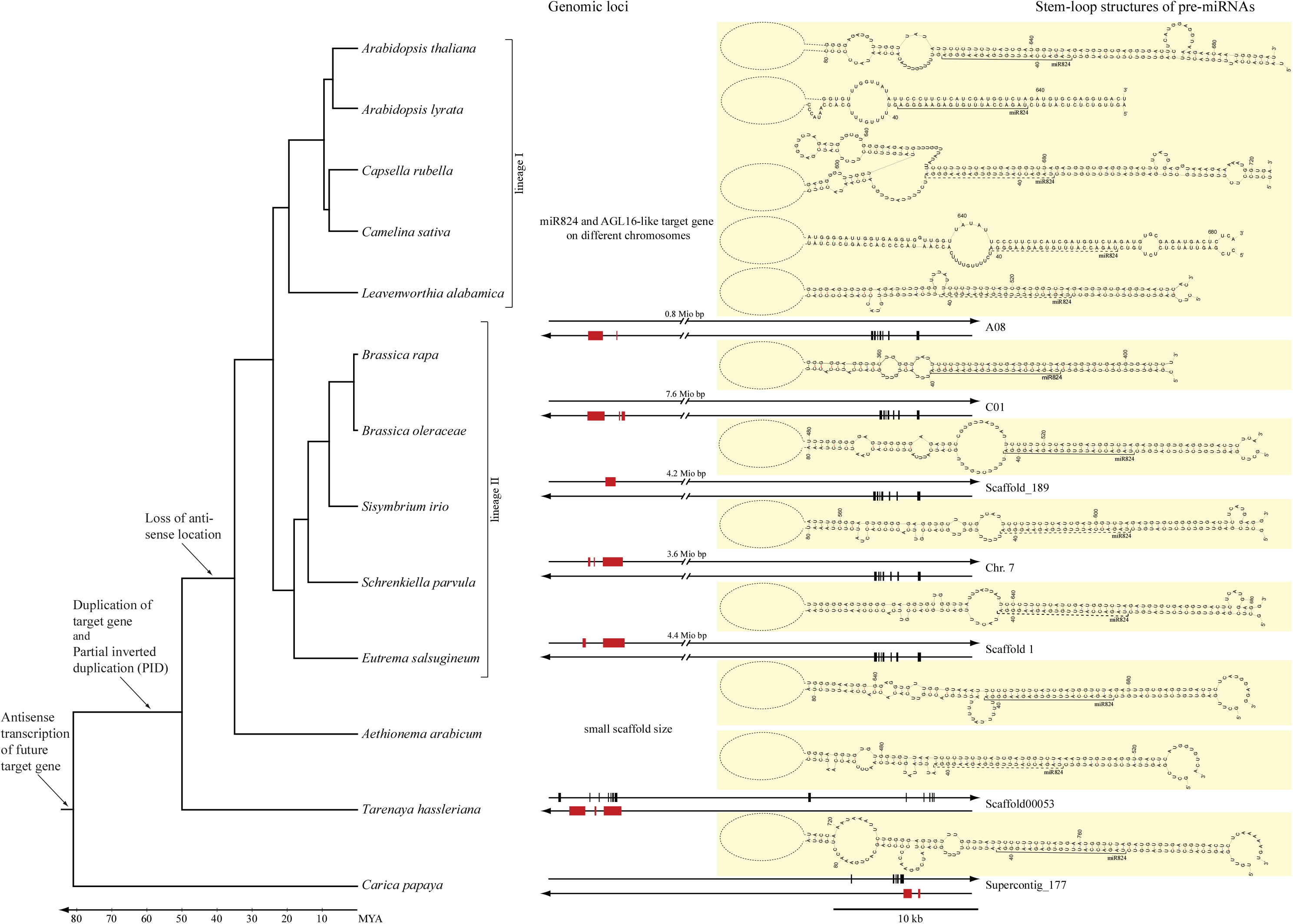
Evolution of the *MIR824* gene in Brassicales. Shown on the left is a partial phylogeny of Brassicales (Franzke, et al. 2011), with plant species from Brassicaceae, one species from Cleomaceae (*Tarenaya hassleriana*) and one species from Caricaceae (*Carica papaya*). The different lineages of Brassicaceae are marked on the right. Approximate divergence times of the species can be deduced from the time scale at the bottom (Franzke, et al. 2011). If occurring on the same chromosome/scaffold, the genomic loci of the *AGL17*-like genes and the *MIR824* genes or the antisense gene (in case of *Carica papaya*) are depicted by double arrows representing double-stranded DNA; the arrow heads indicate the 3’ ends. The corresponding identifier of the scaffold is shown next to the locus. Exons are shown as boxes on the strand transcribed into RNA (i.e. coding strand), whereas introns are shown as lines. Exons of the *AGL17*-like genes are shown in black while exons of the *MIR* genes and the antisense transcripts are colored red. The stem-loop structures of the pre-miRNAs are shown as predicted by mfold (Zuker 2003) and shaded in yellow. The mature miRNAs are indicated on the stem-loop structures either based on sequence similarity to ath-miR824 (dashed lines) or based on small RNAs found in next-generation sequencing data of the SRA database at NCBI (Sayers, et al. 2012) (solid lines). The events in the evolution of the *MIR824* by the partial inverted duplication mechanism are indicated on the phylogeny.

### Regulation of *AGL16*-like genes by miR824 in the spider plant

The spider plant *T. hassleriana* represents the earliest branching species in which we identified a *MIR824* gene. Hence, we investigated this species in more detail. Using 3’ RACE we confirmed expression of the *T. hassleriana* pri-miR824a. For the 3’ part of the *T. hassleriana* pri-MIR824a we identified four splice variants. We were also able to clone transcripts of the two *AGL16*-like genes. We named the *AGL16*-like gene which is natural antisense to the *MIR824* gene *ThAGL16a,* and the other gene *ThAGL16b*.

To confirm regulation of *ThAGL16a* by miR824, we used a modified 5’-RACE technique. We identified cleavage products of the *ThAGL16a* transcript in *T. hassleriana* leaves. For 12 out of the 16 clones sequenced, the cleavage site was found between the nucleotides in the target site that are complementary to positions 10 and 11 of the mature miR824, as would be expected for a miR824-mediated cleavage (Supplemental Figure 9). This suggests that miR824 regulates *ThAGL16a* by mRNA cleavage in *T. hassleriana*.

### An antisense transcript of an *AGL16*-like gene is present in papaya

We were unable to identify the *MIR824* gene in *Carica papaya*, the most closely related species to Brassicaceae and Cleomaceae for which whole genome information is available (Ming, et al. 2008). However, by searching short read data we were able to find evidence of an antisense transcript of the *AGL16*-like gene of *C. papaya* (Supplemental Data). The antisense transcript had a length of 1032 bp. It begins downstream of the *AGL16*-like gene and is antisense to the last exon of the *C. papaya AGL16*-like gene (Figure 4). The antisense transcript does not contain a putative miR* and is not predicted to fold into a stem-loop structure according to conventional *in silico* methods (Supplemental Figure 10).

## Discussion

### miR444 and miR824 regulate closely related target genes

Both miR444 and miR824 regulate *AGL17*-like MIKC^C^-group MADS-box genes (this study; (Kutter, et al. 2007; Lu, et al. 2008)). While nearly all monocot *AGL17*-like genes are targets of miR444, the target genes of miR824 belong to only one out of three subclades of eudicot *AGL17*-like genes (Figure 1). Seventeen clades of ancient and conserved MIKC^C^-group genes are distinguished in flowering plants (Gramzow, et al. 2014). miRNA-regulation has been shown for only one other clade of MIKC^C^-group genes in flowering plants: miR5179 has been described to regulate *DEF*-like genes in *Orchis italica* (Aceto, et al. 2014). The fact that 15 clades of MIKC^C^-group genes do not seem to be regulated by miRNAs, yet two miRNAs which regulate the clade of AGL17-like genes have evolved independently, may be due to chance. Another possibility is that the miRNA-dependent regulation of this clade of MIKC^C^-group genes is evolutionary advantageous, such that there is strong selection for the conservation of miRNAs regulating *AGL17*-like genes once these miRNAs evolved. Prominent phenotypes of *AGL17*-like genes are rare but this clade of MADS-box genes has been strongly maintained during angiosperm evolution (Gramzow and Theissen 2015). A number of *AGL17*-like genes, such as *ANR1* and *AGL21* from *A. thaliana*, *OsMADS25* from *O. sativa* and *GmNMHC5* from *Glycine max*, have been found to have functions in root development (Liu, et al. 2015; Yu, et al. 2015; Yu, et al. 2014; Zhang and Forde 1998). Other *AGL17*-like genes are involved in flowering time determination, e.g. *AGL17* and *AGL16* from *A. thaliana* (Han, et al. 2008; Hu, et al. 2014), in the regulation of stomata density on leaves such as *AGL16* (Kutter, et al. 2007) and in the determination of tiller number like *OsMADS57* from *O. sativa* (Guo, et al. 2013). These functions may benefit from an additional layer of regulation provided by miRNAs. Another explanation for the apparently preferential regulation of *AGL17*-like genes by miRNAs as compared to other MIKC^C^-group genes could be that *AGL17*-like genes are somehow prone to the evolution of *MIR* genes. However, the mechanism for such a potential predisposition to the evolution of *MIR* genes remains unknown.

### Independent origin of *MIR444* and *MIR824* genes

We found evidence of *MIR444* genes in all orders of core monocots but not in the monocot orders Alismatales and Acorales despite analyses of the genome of three Alismatales species *S. polyrhiza*, *Z. marina* and *Z. muelleri,* as well as extensive cloning efforts in the Acorales species *A. gramineus*. There are some reports of miR444 in eudicots, like in *Boechera* species (Amiteye, et al. 2011), *Citrus sinensis* (Xu, et al. 2013), *Gossypium hirsutum* (Ruan, et al. 2009), *Dimocarpus longan* (Lin and Lai 2013) and *Fragaria × ananassa* (Ge, et al. 2013). However, for the miR444 found in *Boechera* species, no stem-loop could be predicted (Amiteye, et al. 2011), making the presence of a genuine miRNA seem unlikely. Additionally, in the case of *C. sinensis* it has since been shown that the identified miRNA is a novel miRNA, rather than one belonging to the previously defined *MIR444* family (Taylor, et al. 2017). Likewise, the sequences for the miR444 of *D. longan* and *F. ananassa* provided by the authors exhibit at least five mismatches as compared to the most similar monocot miR444 (Supplemental Figure 11), rendering assignment of these sequences to the miR444 family questionable. In contrast, sequences of miR444 from *G. hirsutum* are identical to miR444 from *O. sativa* (Supplemental Figure 11) but their presence may have been due to contamination with *O. sativa* miRNAs. Hence, we are quite confident that *MIR444* genes evolved in a common ancestor of core monocots about 120 MYA (Magallon, et al. 2015) (Figure 2), and were not present in the much earlier stem group of monocots and eudicots.

We detected *MIR824* genes in different Brassicaceae species and in the Cleomaceae *T. hassleriana*, but not in other eudicot species. There are no reports of miR824 in species outside the order Brassicales, to which all the investigated species belong. Therefore, *MIR824* genes likely evolved in a common ancestor of Brassicaceae and Cleomaceae about 50 MYA (Guo, et al. 2017).

The miR444 target sites are found in regions encoding the K-domain while the miR824 target sites are present in sequences encoding the C-terminal domain of the corresponding MIKC^C^-group MADS-box genes (Kutter, et al. 2007; Lu, et al. 2008; Sunkar, et al. 2005). Additionally, pre-miR444 sequences are quite dissimilar to pre-miR824 sequences, e.g. an alignment of the precursor sequences of ath-miR824 with osa-miR444d reveals a sequence identity of only 0.1 (Supplemental Figure 12).

Taken together, our analyses strongly suggest that the *MIR444* and *MIR824* genes originated independently.

### Antisense transcription of future target genes

We found an antisense transcript of an *AGL17*-like gene in *T. maritima* (Alismatales). We hypothesize that antisense transcription of *AGL17*-like genes evolved in a common ancestor of monocots after Acorales branched off, but prior to the evolution of *MIR444* genes which originated in the remaining monocots after the Alismatales branched off. Similarly, there is an antisense transcript of the *AGL16*-like gene in *C. papaya* (Caricaceae). Hence, it is conceivable that *AGL16*-like genes were transcribed in antisense orientation in a common ancestor of Brassicaceae, Cleomaceae and Caricaceae before *MIR824* genes evolved in a common ancestor of Brassicaceae and Cleomaceae. Taken together, we suggest that antisense transcription of the future target genes of miR444 and miR824 evolved prior to the origin of the corresponding *MIR* genes. In general, acquisition of transcription from the antisense strand is not unlikely, as it has been shown that around 25% of plant genes exhibit antisense expression (Li, et al. 2006; Poole, et al. 2008; Yamada, et al. 2003). In fact, existence of antisense transcription would ensure immediate expression of a partly inversely duplicated genomic locus, potentially giving rise to a new miRNA. Furthermore, the antisense transcript may have already been involved in regulation of the sense gene (the future target gene) in a certain spatio-temporal pattern. Consequently, the newly evolved *MIR* gene, driven by the existing promoter, would gain the same transcription pattern as the antisense transcript had had previously, enabling it to regulate its target gene in the same useful spatio-temporal pattern. In this way, antisense transcription could truly facilitate *MIR* gene origin.

### Evolution of *MIR444* and *MIR824* genes by partial inverted duplication

In all species in which we analyzed the genomic location of the *MIR444* genes, they were found to be natural antisense to their target genes, and to have an intron separating the exons encoding the mature miRNA and the miR*, as has been shown for *O. sativa* previously (Lu, et al. 2008; Sunkar, et al. 2005; Sunkar, et al. 2008). It has been suggested that the natural antisense organization and the intron are best explained by an origin of the ancestral *MIR444* gene by a partial inverted duplication of its target gene (Lu, et al. 2008). In each investigated species the mature miR444 and the miR* are encoded by regions antisense to the third exon and to the downstream region of the *AGL17*-like target gene, respectively. The similarity between the target gene and its downstream region generally only comprises the parts encoding the stem of the pre-MIR444. We hypothesize that the ancestral *MIR444* gene likely evolved by a partial inverted duplication of a region including the third exon, where the copy inserted into a region downstream of an *AGL17*-like gene (Supplemental Figure 13).

The miR824 target sites are found in a region coding for the C-terminal domain in the last exon of their target genes. Sequence similarities between the *MIR824* gene and its target gene beyond the mature miR824 have previously been noted in *A. thaliana*: The region encoding the part of the stem-loop on which miR824 is found was shown to be complementary to a region from the last exon, while the region encoding the part of the stem-loop carrying the miR824* was found to be most similar to a region within the third intron (Fahlgren, et al. 2007). Our analyses of *T. hassleriana* reveal that one *MIR824* gene is natural antisense to an *AGL16*-like gene in this species. The region encoding the mature miR824a is natural antisense to the last exon and the region coding for the miR824a* is natural antisense to the third intron of the *AGL16*-like gene. The similarity between the genomic regions encoding the mature miR824a and the miR824a* only comprises the parts giving rise to the stem of pre-miR824a. Consequently, the ancestral *MIR824* gene presumably evolved by a partial inverted duplication of a region containing the last exon, where the copy is found in the third intron of the target gene (Supplemental Figure 13).

Evolution of *MIR* genes by inverted duplication of their target genes has been hypothesized for a number of young, i.e. species-, genus- or family-specific plant miRNAs (Allen, et al. 2004; Baldrich, et al. 2018; Fahlgren, et al. 2007).

For both the *MIR444* and the *MIR824* genes, the partial duplication of the target gene resulted in the evolution of *MIR* genes that were natural antisense to their target genes. Whereas the natural antisense organization was conserved for *MIR444* genes, it was only conserved for the *MIR824a* gene of *T. hassleriana* but lost for the *MIR824* genes of Brassicaceae. Beside the *MIR444* genes and their corresponding *AGL17*-like target genes in *O. sativa*, for which the natural antisense organization had previously been shown (Lu, et al. 2008), the *tha-MIR824* gene and its target gene *ThAGL16a* is only the second example of a *MIR*-target gene pair for which the natural antisense organization has been found so far.

### Position of miR444 but not of miR824 shifted during evolution

Relative to the position of the miR444 species in the stem-loop of pre-miR444d of *O. sativa*, the mature miR444 species are often shifted in other monocot species (Figure 2). The difference in the position of the mature miRNAs may be due to slight differences in the precursor structures of *MIR444* in the different species. These differences may lead to different processing of the *MIR444* by the miRNA machinery. Indeed, it has been shown that the precise recognition of the mature miRNA from the pri-miRNA depends on precursor structure-encoded processing signals (Bologna, et al. 2013; Mateos, et al. 2010; Song, et al. 2010b). In *A. thaliana,* members of the same miRNA family can be processed in different ways. For example, while *MIR171a* is processed in a base-to-loop direction, *MIR171b* is processed in a loop-to-base direction. Furthermore, the mature miR170/171a and miR171b are offset by three bases. It was hypothesized that this shift in their sequences may have originated as a consequence of diversification of the processing pathways of the family (Bologna, et al. 2013). This is quite similar to what we observe for *MIR444*.

In contrast, the mature miRNA of *MIR824* seems to be identical in all species as based on published data and small RNA data on *MIR824* (Kutter, et al. 2007). Hence, only one miRNA processing pathway seems to be possible for *MIR824* in all species studied.

### MiRNA Evolution by Partial Inverted Duplication of Antisense-transcribed Sequences

Based on our observations on the evolution of *MIR444* and *MIR824* genes we propose the scenario MEPIDAS (MiRNA Evolution by Partial Inverted Duplication of Antisense-transcribed Sequences) for the evolution of *MIR* genes. Our scenario represents a significant modification of the inverted duplication mechanism suggested by Allen, et al. (2004).

According to our hypothesis, a miRNA evolves at the locus of its future target gene (Figure 5A). First, antisense-transcription of the gene evolves (Figure 5B) by acquisition of a promoter. As discussed above, this is an important step, as for a miRNA to evolve the corresponding locus of the *MIR* gene must be transcribed. A partial inverted duplication (PID) at the genomic locus of the future target gene leads to an antisense transcript that folds into a stem-loop structure (Figure 5C). This stem-loop structure is recognized by the siRNA biogenesis pathway as proposed by Allen, et al. (2004) or it may even be recognized by the miRNA biogenesis pathway straight away if the PID was small enough. At the time of the PID, or subsequently, splicing of the antisense transcript may evolve (Figure 5D) as in the case of *MIR444*. It should be noted that if complete exons with surrounding intron sequences are inversely duplicated, at least the acceptor splice sites upstream and the donor splice sites downstream of these inverse exons would be present and hence the evolution of splicing might be facilitated. After the PID event, the antisense transcript folds into a stem-loop structure with an extended stem of perfect complementarity. Furthermore, the antisense transcript is perfectly complementary to its target gene over an extended region (the region of the antisense transcript which is overlapping with exons of the target gene). Hence, the transcript has the potential to give rise to several mature miRNAs as in the case of *MIR444*. Different mature miRNAs may then evolve in different lineages after speciation (Figure 5E). The natural antisense organization of the *MIR* to its target gene may be lost at any stage of the evolution of *MIR* genes by MEPIDAS (Figure 5F). The prerequisite for the loss of the natural antisense organization is the prior duplication of the target gene. Then, the target gene natural antisense to the *MIR* gene may degenerate (Figure 5F) and the miRNA will come to regulate the duplicate gene.

**Figure 5.**
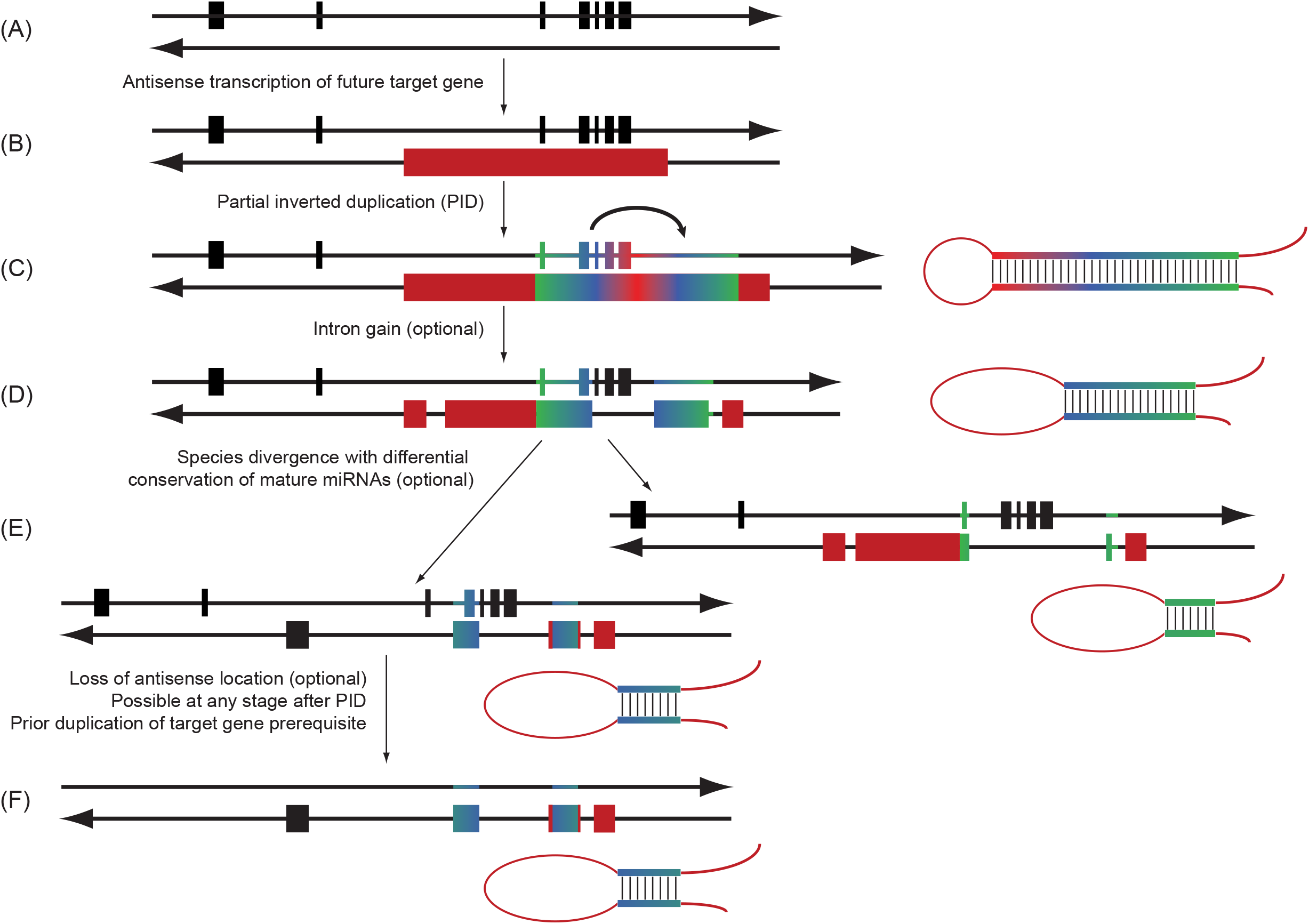
Schematic representation for the origin of miRNAs according to the scenario MEPIDAS (MicroRNA Evolution by Partial Inverted Duplication of Antisense-transcribed Sequences). As in Figures 2 and 4, the genomic loci of the (future) target genes and the *MIR* genes or antisense genes are depicted by double arrows representing double-stranded DNA; the arrow heads indicate the 3’ ends. Exons are shown as boxes on the strand transcribed into RNA (i.e. coding strand), whereas introns are shown as lines. Exons of the (future) target genes are shown in black while exons of the *MIR* genes and the antisense genes are colored red. Regions which are reverse complementary to each other due to a partial inverted duplication are indicated by a color gradient. Putative stem-loop structures of pre-miRNAs encoded by *MIR* genes are given on the right if applicable. For the stem of the stem-loop structure the same color gradient is used as for the genomic locus to indicate which genomic regions the stem is encoded by. (A) Future target gene with regular transcription of the coding strand. (B) Acquisition of a promoter (not shown) leads to antisense-transcription of the gene, indicated by an exon appearing on the lower DNA strand. (C) State of the locus after partial inverted duplication (PID) of the gene. After PID, the antisense transcript is able to fold into a stem-loop structure which is recognized by the siRNA or miRNA biogenesis pathway. (D) Structure of the locus after splicing of the antisense transcript evolved (optional) and partial sequence divergence of the inverted duplication locus. These events lead to shortening of the stem-loop structure. (E) Different genomic loci giving rise to diverse mature miRNAs in different plant lineages. This may evolve by conservation of different parts of the inverted duplication locus after speciation. (F) Genomic locus after the natural antisense organization of the *MIR* gene to its target gene is lost, which may happen at any stage of the evolution of *MIR* genes by PID. The prerequisite for the loss of the natural antisense organization is the prior duplication of the target gene.

MEPIDAS provides a detailed picture of the evolution of *MIR* genes. Our scenario suggests that the transcription of the future *MIR* gene is ensured by existing antisense transcription of the future target gene. It also shows how a partial inverted duplication may lead to a transcript that is immediately recognized by the miRNA biogenesis machinery. Furthermore, we show that a natural antisense organization of the *MIR* to its target gene may actually be a frequent outcome of a partial inverted duplication but may be rapidly lost in most cases. Our study also illustrates how comparative genomics helps to elucidate the origin of older *MIR* genes. It may hence serve as a blueprint for the clarification of the origin of many more *MIR* genes whose origin has so far remained enigmatic.

## Materials and Methods

### Phylogeny of *AGL17*-like sequences

Identification of *AGL17*-like genes is described in Supplemental Methods. Coding sequences were translated into protein sequences using Transeq (Rice, et al. 2000) and aligned using Probalign (Roshan and Livesay 2006). The alignment was trimmed with trimAL (Capella-Gutierrez, et al. 2009) with the parameters -gt 0.9 and -st 0.00001. The phylogeny was reconstructed using MrBayes (Ronquist and Huelsenbeck 2003). The protein substitution model WAG (Whelan and Goldman 2001) was chosen, 24 million generations were calculated sampling every 100th generation and excluding the first 25% of generations from the consensus tree reconstruction containing all compatible groups (contype=allcompat). The phylogenies were rooted using SEEDSTICK (STK) from *A. thaliana* as a representative of the outgroup.

To identify transcripts of *AGL17*-like genes of species of the orders Liliales, Pandanales, Alismatales and Acorales, we searched the 1KP data (www.onekp.org) using BLAST with the sequence of *OsMADS23* as query.

### Identification of *MIR444*

The monocot genomes listed in the Supplemental Methods were searched using BLAST (Altschul, et al. 1990) with the sequence of the pre-miRNA of *osa-MIR444d* as query sequence (Lu, et al. 2008). The best BLAST hits for all of those species were examined to determine whether the identified genomic loci may encode *MIR444*. To do so, the genomic sequences of the corresponding BLAST hits were extracted including 30 kbp of sequence up- and downstream of those BLAST hits. By comparison of those genomic sequences to the sequences of the pre-miRNAs of *O. sativa* and with attention to the canonical splice sites, putative exons encoding the precursors of *MIR444* in these species were determined. Subsequently, we attempted to obtain expression evidence for the putative miRNAs identified by querying the short read archive (SRA) at NCBI (Sayers, et al. 2012). We searched these expression datasets for each species independently, using BLAST with the sequences of the putative miRNAs as query. The identified expressed sequences were assembled using Sequencher 5.1 (Gene Codes, Ann Arbor, Michigan, USA) where the sequences of the putative miRNAs were included as well. If some of the identified expressed sequences assembled with the putative miRNAs with an identity of more than 98% covering (almost) the whole length of the putative miRNAs, we considered those miRNAs to have evidence of expression. For the miRNAs shown in Figure 2, we attempted to assemble whole transcripts. To do so, we iteratively searched the SRA database using the ends of the transcripts assembled in the previous step as query sequences. The RNA structures of the miRNA transcripts were determined using the RNA folding form of mfold (Zuker 2003). If the predicted RNA structure indicated a stem-loop structure that is typical for plant miRNAs, the corresponding genomic sequences were considered as *MIR444* loci. To find evidence of mature miRNAs, we searched small RNA data in the NCBI SRA database. Here, we used the sequence equivalent to the region ranging from the beginning of osa-miR444d.3 to the end of osa-miR444d.1 of the corresponding species as search sequence and chose the option “somewhat similar sequences (blastn)”.

As for the *AGL17*-like genes, we searched the 1KP data to identify transcripts of *MIR444*, using the precursor sequence of *osa-MIR444d* as query for BLAST.

Alignment methods used for visualizing conservation of the *MIR444* genes and their target sites are described in Supplemental Methods.

### Identification of *MIR824*

The Brassicales genomes listed in the Supplemental Methods were searched using BLAST (Altschul, et al. 1990). We took the sequence of the *pre-MIR824* from miRBase (Kozomara and Griffiths-Jones 2014) and the coding sequence of *AGL16* (Parenicova, et al. 2003) from *A. thaliana* as queries. The corresponding genomic sequences were downloaded and in the case of *AGL16*-like genes, the exon-intron structure was adopted as annotated or predicted using FGENESH (Salamov and Solovyev 2000). In the case of the putative *MIR824* sequences, the genomic sequence with similarity to the *MIR824* from *A. thaliana* was used to predict the RNA structure of the corresponding transcript using the program mfold (Zuker 2003). If the part with similarity to the mature miR824 sequence of *A. thaliana* was part of a stem, we assumed that the genomic sequence identified corresponds to the genomic locus of *MIR824* in the species investigated. We also searched and assembled transcript data and small RNA data from the short read archive (SRA) of NCBI (Sayers, et al. 2012) as described for *MIR444*. Structures of the transcripts were also predicted as described for *MIR444*.

To exclude the presence of *MIR824* genes in more distantly related species, we generated a Hidden Markov Model (HMM) from an alignment of the 5’-arm sequences of pre-miR824 as found in *A. thaliana*, *A. lyrata*, *C. rubella*, *E. salsugineum*, *B. rapa* and *T. hassleriana* using HMMER 3.1b2 (Eddy 1996). This HMM was used to search the genomes of *C. papaya*, *Gossypium raimondii*, *Theobroma cacao*, *Citrus sinensis* and *Citrus clementina* as downloaded from NCBI (Sayers, et al. 2012) via nhmmer (Eddy 1996). The presence of *MIR824* genes would have been identified as two results of the nhmmer search, one result in forward, one result in reverse direction, on the same genomic fragment where the two results have a distance of less than 3000 nt. These criteria were successfully tested in *T. hassleriana*.

### RNA isolation

*Dioscorea bulbifera* and *Acorus gramineus* plant material was kindly provided by the Botanical Garden of the Friedrich Schiller University in Jena. *T. hassleriana* was grown in a greenhouse under the following conditions: 16 h light with 19°C during the day and 17°C at night.

Young leaf material was harvested and ground under liquid nitrogen conditions either with a mortar and pestle or using the Mixer Mill MM 400 (Retsch) with 35 ml grinding jars and 20 mm grinding balls. Total RNA was isolated with QIAzol Lysis reagent (Qiagen) according to the manufacturer’s instructions.

15-20 µg of total RNA was digested with DNase I (recombinant, RNase-free, Roche Life Science) according to the manufacturer’s recommendations. Following digestion the RNA was purified using a Phenol/Chloroform extraction with subsequent precipitation. The RNA pellet was resuspended in an appropriate volume of RNase-free water.

### miRNA Northern blotting

Methods used for Northern blotting are described in the Supplemental Material.

### Cloning of transcripts

Methods used for 5’ RACE, RT-PCR and 3’ RACE are described in the Supplemental Material.

### Genome Walking

Genomic DNA from 1 g leaf material was isolated using the DNeasy Plant Maxi kit (Qiagen) according to the manufacturer’s instructions. Genomic DNA (2.5 µg) was digested separately with the following restriction enzymes: *Dra*I, *Eco*RV, *Pvu*II and *Stu*I overnight. The purification of the digested DNA and the ligation to the GenomeWalker adaptors were carried out according to the manufacturer’s instructions of the GenomeWalker Universal Kit (Clontech Laboratories). PCR reactions contained in a total volume of 50 µl 1x Long PCR buffer with MgCl_2_, 0.2 mM dNTPs, 0.25 µM Adaptor primer 1, 0.25 µM gene-specific primer, 1 µl of the DNA library and 0.4 µl Long PCR Enzyme Mix (ThermoFisher Scientific). The PCR program was as follows: 94°C for 2 min, 10 cycles of 94°C for 20 s, 68°C for 7 min, 25 cycles of 94°C for 20 s, 68°C for 7 min + 5 s/cycle, and a final step of 68°C for 10 min. The nested PCRs contained in a total volume of 50 µl 1x Long PCR buffer with MgCl_2_, 0.2 mM dNTPs, 0.25 µM Adaptor primer 2, 0.25 µM nested gene-specific primer, 1 µl of a 1:50 dilution of the first PCR product and 0.4 µl Long PCR Enzyme Mix (ThermoFisher Scientific). The PCR program was the same as for the first PCR. The PCR reactions were separated on a 1% agarose gel with ethidium bromide (EtBr) staining, all bands were excised from the gel and the DNA was extracted using the QIAquick Gel Extraction kit (Qiagen) according to the manufacturer’s instructions. The purified PCR products were cloned into the pJET1.2 blunt vector according to the manufacturer’s instructions of the CloneJET PCR Cloning kit (ThermoFisher Scientific) and sequenced (Macrogen). An overview of the primers used can be found in Supplemental Table 1.

### MiR-RACE

MiR-RACE was carried out as described before (Song, et al. 2010a) where 2 µg of DNAse I digested total RNA isolated from *D. bulbifera* leaf material was used as raw material and 5U poly (A) polymerase (NEB) was used for polyadenylation. HotStarTaq *Plus* DNA Polymerase (Qiagen) was used for amplification of cDNA where an annealing temperature of 63°C was chosen. miRNA-specific primers used for nested PCR reactions are listed in Supplemental Table 1. The nested PCR reactions were separated on a 1.5% agarose gel with ethidium bromide (EtBr) staining, bands were excised from the gel and the DNA was extracted using the QIAquick Gel Extraction kit (Qiagen) according to the manufacturer’s instructions. The purified PCR products were ligated with the pGEM^®^-T vector (Promega) as recommended by the manufacturer and subsequently sequenced (Macrogen). An overview of the primers used is provided in Supplemental Table 1.

### Reporter gene assay in *Nicotiana benthamiana*

First we created the plant transformation vectors pGREENII 35S and pGREENII 35S-mGFP5ER. To do so we digested pGREENII 0000 (Hellens, et al. 2000a; Hellens, et al. 2000b) using *KspA*I (*Hpa*I) to remove the multiple cloning site (MCS) of this vector. The digested vector was gel purified using the QIAquick Gel Extraction kit (Qiagen) and the 5’overhangs were filled in using Klenow fragment (ThermoFisher Scientific). The vector was religated using T4 DNA Ligase (ThermoFisher Scientific). The 35S promoter including MCS and octopine synthase gene terminator was released by a *Not*I digestion from the vector pMS37 and subsequently cloned into the *Not*I sites of the prepared pGREENII vector to create the vector pGREENII 35S. The mGFP5ER gene (Haseloff, et al. 1997; Siemering, et al. 1996) was amplified from the vector pBINmGFP5ER using primers which contained restriction enzyme recognition sites for *Bam*HI (forward primer) and *Bgl*II (reverse primer), respectively. The amplified and digested mGFP5ER gene was then cloned into the *Bam*HI sites of the pGREENII 35S vector to create the vector pGREENII 35S-mGFP5ER.

To test regulation of the *AGL17*-like gene of *D. bulbifera* by dbu-miR444, first the cDNA from exon 2 to 7 of *DbAGL17a* was fused to the 5’end of mGFP5ER. The chimeric construct was cloned into a binary vector under the control of a Cauliflower Mosaic Virus (CaMV) 35S promoter (pGREEN35S-*DbAGL17a::*mGFP5ER). A partial cDNA of the *dbu-MIR444* was cloned in a binary vector under the control of a CaMV 35S promoter (pGREEN35S-*dbu-MIR444*). As a negative control *MIR160a* from *A. thaliana* was also cloned in a binary vector under the control of a 35S promoter (pGREEN35S-*ath-MIR160a*). All constructs were transformed separately into *Agrobacterium tumefaciens* strain GV3101 by electroporation. *A. tumefaciens* cells containing a binary vector were incubated in YEB medium containing 50 µg/ml kanamycin, 20 µg/ml rifampicin, 2 µg/ml tetracycline and 25 µg/ml gentamycin at 28°C for 16 h. The *A. tumefaciens* cells were centrifuged and washed with 40 ml infiltration medium (10 mM MgCl_2_, 150 µM acetosyringone). After another centrifugation step the cells were resuspended in infiltration medium to adjust the absorbance of OD_600_ to 0.5. The *A. tumefaciens* cells were incubated for 1.5 h at 28°C. As positive controls *A. tumefaciens* carrying the empty vector pGREEN35SmGFP5ER and the vector expressing the DbAGL17a::mGFP5ER fusion protein (pGREEN35S-*DbAGL17a::*mGFP5ER), respectively, were used for infiltration of four-week-old *N. benthamiana* leaves. *A. tumefaciens* carrying pGREEN35S-*DbAGL17a*::mGFP5ER constructs were mixed with different amounts of *A. tumefaciens* carrying pGREEN35S-*dbu-MIR444* constructs and used for co-infiltration of *N. benthamiana* leaves. As controls *A. tumefaciens* containing pGREEN35S-*DbAGL17a::*mGFP5ER constructs were mixed with equal amounts of *A. tumefaciens* containing the empty vector pGREEN35S and pGREEN35S-*ath-MIR160a*, respectively, and used for co-infiltration of *N. benthamiana* leaves. Five days post-infiltration the GFP expression was monitored using an Olympus SZX16 microscope equipped with a fluorescent light source with a GFPA filter set (excitation 460 – 490 nm, absorption 510 – 550 nm).

### Detection of cleavage products by modified 5’ RACE

Modified 5’ RACE was carried out using the GeneRacer Kit (Invitrogen by life technologies) according to the manufacturer’s instructions without the enzymatic pretreatment of the RNA. The cDNA synthesis was carried out using a gene-specific primer. For the first PCR Platinum Taq DNA Polymerase High Fidelity (Invitrogen by life technologies) was used according to the manufacturer’s instructions. If necessary nested PCRs were carried out using HotStarTaq *Plus* DNA Polymerase (Qiagen). PCR products were separated on a 1% agarose gel with ethidium bromide (EtBr) staining, all bands were excised and the DNA was extracted either using the provided S.N.A.P. columns or the QIAquick Gel Extraction kit (Qiagen) according to the manufacturer’s instructions. The purified PCR products were either cloned into the provided pCR 4-TOPO vector (Invitrogen by life technologies) or into the pGEM-T or pGEM-T Easy vector (Promega) and subsequently sequenced (Macrogen).

## Supporting information

Sequence data

## Supplemental Material

**Supplemental Figure 1.** Fully resolved phylogeny of AGL17-like proteins.

**Supplemental Figure 2.** Northern blot hybridization analysis of the presence of miR444.

**Supplemental Figure 3.** Alignment of the stem-loop regions of *MIR444* transcripts.

**Supplemental Figure 4.** Alignment of the *MIR444* transcripts identified in *Dioscorea bulbifera*.

**Supplemental Figure 5.** GFP reporter gene assay to show the dbu-MIR444 mediated down regulation of the *DbAGL17* target gene.

**Supplemental Figure 6.** Identification of cleavage products of the *DbAGL17* transcript in *D. bulbifera* leaves.

**Supplemental Figure 7.** Secondary structure prediction of the antisense transcripts to the *AGL17*-like gene of *Triglochin maritima*.

**Supplemental Figure 8.** Alignment of the AGL17-like MADS-box transcripts identified in several orders of monocotyledonous plants.

**Supplemental Figure 9.** Validation of miR824 mediated cleavage of *ThAGL16a* in *Tarenaya hassleriana*.

**Supplemental Figure 10.** Secondary structure prediction of the antisense transcript to the *AGL16*-like gene of *Carica papaya*.

**Supplemental Figure 11.** Alignment of alleged miR444 from non-monocot species.

**Supplemental Figure 12.** Multiple sequence alignment of ath-pre-miR824 and osa-pre-miR444d.

**Supplemental Figure 13.** Scenarios for the origin of the *MIR444* and *MIR824* genes.

**Supplemental Table 1.** Primers used in this study.

**Supplemental Table 2.** Overview of the sequences submitted to GenBank including accession numbers and sequence descriptions.

**Supplemental Data.** Sequences used in this study.

**Supplemental Methods.** Detailed methods on access of genome data, identification of *AGL17*-like genes, alignments, miRNA Northern blotting, 5’ RACE, RT-PCR and 3’ RACE.

## Data Accessibility

An overview of the sequences submitted to GenBank including accession numbers can be found in Supplemental Table 2.

## Acknowledgements

We thank the members of the Theißen group for fruitful discussions and Ulrike Wrazidlo for excellent technical assistance. We are grateful to Anne Elisabeth Greßler and Tim Schmäche for experiments with *Triglochin*. We thank the Botanical Garden of our university for providing plant material, and especially Dr. Stefan Arndt for helping to find the different species in the Garden. The vector pBINmGFP5ER was kindly provided by Dr. Trevor Fenning, thanks for that. We are grateful to Dr. Rebecca Schwab for providing the pMS37 vector. We thank Dr. Phil Mullineaux/Dr. Roger Hellens, the John Innes Centre and the Biotechnology and Biological Sciences Research Council (BBSRC) for providing the pGREEN vector. We kindly thank Prof. Dr. Eric Schranz of the Wageningen University for the *T. hassleriana* seeds. We thank The 1000 Plants (1KP) initiative; Y. Zhang from BGI-China and E. Carpenter-US, who manage the 1KP database; and M. Barker, D. O. Burge, Tao Chen, M. Deyholos, P. Hollingsworth, A. Larsson, J. Leebens-Mack, C. Rothfels, M. Ruhsam, and S. Weststrand for providing transcriptomes of the 1KP project. The 1KP initiative led by G.K-S.W. and M. Deyholos is funded by the Alberta Ministry of Enterprise and Advanced Education, Alberta Innovates Technology Futures (AITF) Innovates Centre of Research Excellence (iCORE), Musea Ventures, and BGI-Shenzhen.

## Author Contributions

LG, DL and GT designed the research; LG, DL, SW and NI performed research; LG, DL and GT analyzed data and wrote the paper.

