## Supplementary material for "Independent origin of *MIRNA* genes controlling homologous target genes by partial inverted duplication of antisense-transcribed sequences": Sequence data

#### Marking of feature of AGL17-like genes and MIR444 genes

exons of AGL17-like genes<br>
exons of miR444<br>
mature miR444.1, mature miR444.2, mature miR444.3

#### miR444 in monocots

##### Poales

###### Oryza sativa

*OsMADS23 and osa-miR444a*

>Chr8

tccaaactactgaacgacgtgtttttgcaaaagtttttaatacgaaagttgctttaaaaatcatattaactcat<br>
ttttttaataatttaaactaatattatattagttatacggtaacgagtgtttcgtcgtatgtgctggacatcccg<br>
aacacagcctatgagAGTTGCTAGCTTGCTATACGCTCTCTGGTCTATGCTGCTCTGCCTGTCTTGCTCGTCCTC<br>
TGCCGCGGAGACacttgctccctctgttctgattttgacatcttgcttgtaatattttaccactcgtcttatttt<br>
aaataaatatataaatataaaaaacgaaaagttatgcttaaagtacttttaataataaagtaagtaaaaaataat<br>
tctaaatttttttaataagacgagtggtcaaacagtaaaatgtcaaaatccctaataACTTAtactttctctgtt<br>
ttatattataagcctttatagcattacccacgttcatatatatatgttaatgaatctaaacatatatgcgtacct<br>
agattcattaatatatatatatacatatacatatatatatacatatatatatacatatacatatatatatatata<br>
tacatatatatatatatatgtatatatatacatatgtatatatatatatatatatatatgtatatgtatatatat<br>
atgtatatatatatgtatatgtatatgtgagaatgtgaaggatgctgggaagttttataacctgaaacggaggta<br>
gtaTTAGTGACGGAGGGAGCATAATAGAT**AAACTCGTTGTCGTCGTCCCGTGCGGTGCGGCCCCCGCTACCCGCC**<br>
**GCGCGATGCCCCCTCTTCTCCGCCCGCCTGATCTAGGGTTTTACTGGAGCCGCCGCGCGCAGCGACCAGCCGACC**<br>
**ACGGACGAGGACGAGGGGATCAG**GTGAGCCCGCCGCCACTCTCTCCCCCACCCCTCCGAACCCGTTCAATTCAGC<br>
TCCCCGCGGATGTGATCTTGGTTAGGGGCTAGGGTTGGGTCGGTTAACGAAGATCGGTTCCGCGTCTCATCGTGG<br>
CGTGTGTCTTCGCGTGTGTGTGTTGGGTTGGGGGGAGTCCGCGCATGCGTTGTGTGGGTTTTTTTTTTTCAATGC<br>
TGGATGCGATTGAAATTGAGCAGAATGCCCAGTTTACTCATTCAGACCGAATGAGCTTCATTTACTCATTCATCT<br>
AGTGTTGGTGTTGGAAGTTTTGAAGACCTGAGTATGCGGTGGCTATATAATGTTTTTGTTGCAGATCAGTTAGCT<br>
TTGCTGGTTGGACAAATCGAGTCCTAGATATCTTGGATTTTTCTTTTTTCCATAGCAGAAGGTTGTGGGTAAAGG<br>
TGGTGCAAGACAACATGGAATATAAAAGGCCATTGTATTCATGCTGGCTGAATAAATACTAGTTTCTCGCACACA<br>
TTATCCAATGGACTCTGTTTTTTCATTGTGCATAGGCGAAGGGGGTTATGTTGTTTAAATAAAGCATTAGGGCAC<br>
CATTATATTTCCATATGCACGGACTATTTCTCATTATATATTCTATAATTGACACTGAACCAGTAGGGTCTGAAA<br>
AGGTCATATAAGCTTAGCTGTGAGTAGCTCAGATATTCTTTTCCTATTTATTTTGGCTGAACAAATTTAACAATT<br>
TTAGCTAATATTCGTTTCAATATATCTTAGATGCAAAAAAAAGATATGCCTTGATGCAAAATTTAGTCGCTTTTG<br>
CTTGCATCGTGTAGATTTATTGCTATTGTGCCCCTATGCTCTGGCCTCCAGGCTTACTCACGACAGGTGATGGAA<br>
CTACCAACATCTGTAGCTTTCCACCCATGCAGTCTTGTGCCAGGCTGGTTGCATGCTTTAGCTGTTTTGCATGTG<br>
TCGTCTGTCCTATTTTTGTTTTCTTGATGTCAGTTGACCAATGGAAAATCTTATGAGTATAGGATTAAACAATGG<br>
TCTAAAAGTCGGTGCTAGGTACTGGGTAGCCACTGTAatttggtgcttccctgccttgggggctctatattgcta<br>
tgtaatgttatattgtcttctttcatgtcatatgcttcccaactgcttgaaactccatagtcaagtttgttctag<br>
gctgtaggcaccgcctagttgacatctaggtgctaggtgccagccttgctgaccctctagCGCCTAGCGGCTTCT<br>
TGAACCAATAGTATTAAAGTGAATAAGTGTTTTAGCATTTGTTTTGGAAGATATAGCTTTACTACGGGGTATGAA<br>
TACCATTATCCCAAAGAGTTGATTGCATTTTTCAAATTTTTTTAGATGATTGATCCGTAATTTTCCTGGTTGATC<br>
ATGATATTCTTTACATTGCACATGCCATGATATAGATTTGAATGTTCCCAACCAATTAGTTAATTGCACTGCCAT<br>
CCTGTCCTATGAGAAGTTGAAAATACTGGCTGTATATGACTAATAAGCACTTGATTGAGTTCTTCCCTCTGTTAT<br>
TTCCTTTGGTATGTCTTAGAGGTAGATCTATAGGGTGCTTTTATGCTATTCTCATATCACCAAACCAAAAATTCT<br>
GCGCATTCTTACTGGATAACACTAAACAGATAACATGTCAACCACTTTTGCAGAGAAAATTATGTCTGTGACACG<br>
AGCAAGCATTAAAAGTATCACATGTACTTTTGCTGTAGTTGATATATGACGGAACTAACCTTTTAATTGTTCTGA<br>
TTAG**ATATGCAATTGGGGGCAGCAAGCTAGAGGTGGCAACTGCATAATTTGCAAGAAATT**GTAAGTTCTCTTGGA<br>
AAAGTCTGAAATGACATGTATTCATAATTTTATTGATCCTTTAGTTCTTTCTAGATGGAACCCACTTAGCGTGAT<br>
TATTGCTATAAAGAAGCCACCACATGAAGCGACATCGTCAAGTTTTCACACAGATGGTCCCTAAATGTGTAATTT<br>
CTTCCCGCACAAACCATTTTCTTGCTCTAGGATTAGGAACTTTTCTGTGAGACGACTGTCAATCCACATATCGCA<br>
ACCACAGTGCACATATCGATGCCCCACATCAATGCGCAAAGATTTATCTAGTTACAGCATCAGTACAAACCACCA<br>
TACACGCATACCAGTGAGGCAACCAAAATCAAATATATGTTTTAGGCATTAATCTTATTGGTTGGAACTATTTTT<br>
CTTGTTCATTCTTGTTGACGTACATGGTTCTAGTTGCTGTAGCTATATAATAACTTAGCTGTTGGTAAAATAATG<br>
TTCTTGTGACTTGTGAGAGGTGATTATTCAATGGCAGCCTTGATTCCAGGATCTGTTACGAACACCAAATTTTGA<br>
GCTAAGCGCATAAGCAACTTTTCTCGTATTTAAACTGCACTTGAAAGAGGAACCCTTTTAAGTTAAtactccccc

ggtcaaaagtatttgtcgtttaggacaagatttggtcaaaattttaaaattccaacaaccaattttatgttagaa
taaatgtataaaatctaataagtttatgatattacaatagtacttttcgaggcaaatttatgtatataatttctt
ttatttttaaactaagcatttaagaaaccaatgatggttggaattttcaaagtttgatcaaatcttggtctaaac
aacaaatatttgaccagaggtagtaTTAACGTTAAAAAATTAATATGAAGTAGCTTCATGGTACTTGGAAACAG
AATATAGAATGTAAGTTCTATATGTTTTCTATAgggactatctatatttatggcaccatatgaaaaaaatagtta
aatgttttagcatgtactatttacaccgtaaattccatatcttggtccatatatttacactgtaaattacaactt
acactgtaacttaaaacttacacagtaattttctCTAATACTTACATATATACAATAACTAGAAAGTACTAGTGC
AAACTTACATTAAAAGGTGATGGATCCAAAACAAGTAGCAAAAGGGCACCAATTTCAATCATCCAGATGCATGGT
GAACCTAAGCTGCATGTGAACCTAATCATATTGGCATGAACGTCCAGATGCATGGTGCCATAGGCTCAAGGCAAA
AACGAGCGGTAGAACTACGACTGAAAACAGACCAAAACAGAAATATAGCAAAATCGTTTTCTATATATTTATTCT
CCATATGTAAGATCATTGGGGGTGCCAAGTTACCCACTATCCCAATTCCGCATCCTTGGTCACGTGGGTGGCTCA
TTTGTGAACTTCTAATGTATCACTGGATATTATTCTGCTGTTTAGCATTTTAGATCATCGGCAATTTCCTTATAT
TCCAAGCAGCTTCATTGCATTTCAAAAGTAACCACATGCATCTACCAAGTTCAAATCAATACAAGATCAAGTGCC
TATGTTTGGCGCGATACAAATAAGTGCTGTGCACTCTCTGCCCTGCATTATTTTGAACCACAAGAACTACATAGT
CTTCAGAACAGCAGAATACATGACATAAACTCTGTTCTGGTTGAATGGGTTCCCAGCAACTTGAACAAAAAACTG
AACTAAACTGATAAATTGTTCATGCAGAAAATATTGCATGGGTAATGCTTTGAAGTTTAGCTTCGAGGTTTTATA
GTGACACCAAAGAAAATATGCCATTAGTCACTTAATCTTAGCGTTTGCCATCCTCACTTGGGCTGAAGTCTGCTG
CAATTTGAATAAACTTTTAACAATGTGTTGTAATGTGTGTCTTGGTATCTGAATTTAAAATAACAAGAACTCACC
CTGGATTTGGTGTTCCTGACTACTCCTTTTCTACATTCTCTGGCAGACTTAGCTCAAGATTAGGGCCAATTTCAT
TATCTGTAATCAGAAAGCTGTGCTGGATGGTAGAACCTGTTGGATGCCCATTACTCTTTGTCTTATATACCTGCT
AAAATAAATAAAAAAAAATGACATTTTGTAATTTGCATACACCAGAAAAGTGAATTGAGAAAATATCACTTGTAGA
ATCTGCATCATTGTTTGGCAAAATGTTTCTGCATGCCTCAACCTTTTTCTGTAAATTGATGTTTTCTTGATGGAC
AAGGCTGACTTTCTTGTGTAGTTCCATGTTTTCCTGATGTATGAGGCTTCCCTACAATTCAGTGAGAAGTTTGTG
CAGTGTCAAGCAAATTATCTCACTCCAATAAGATTTGGACCAAGTAGACTAGTGTACATAAGTAATGTTGAGTGA
ATGTGCTAAGTGGTAACAACCTTCCTGCTTAATTCTTGAATTTGATCCATCATGACATTGTCCTGTAAAGTGAAT
GTTAGTCAAAGTAGTTATTGATATTATGTCAAGGAAGAATTAGAAGTGGATGGACTGTTAACAAATTGTAATTTA
ACCTTCCTCAGACGGATATTTTTTAAGCTCATCTCCAGCTTACTTTCCAAATTTTGCAAGTCTTCTACATCCAGA
CCAGAAAGCTGTTGTCCCAACAACTGTCTGCAATATCACGTCATGCACAGTCATTATTTGAATGCTACTCCCTTC
ATCCTAAAATAAACCAACTTCTAGAGTTCAAACTTTGTCCCAAAATAAATCAGTTCCTACCTTCTTACCTACCAG
CGATTCTCAAATTGAGAAATTTTATCCCTTCAATACCCTTTACCTACCCAATGACCAACTCATTTACCTCCACAT
TGGGATGCTAGGAGCtttttttttttttttttttttttGGGGGGGGGGGGGGTGGGGGGGGAGCACAGGGAGTAA
ATGTTATCGCACTTGTTATAAACCTTTTGTTCAATTTTTAGTTTTTTACACGCACACTGTTTTTAATGTCGTTAG
ATGCAAATATTCATATACCTGAATTTAGACCTAGAAATTAAATTGCTATGACCAAAAACCCAAATGTCCTGGCAC
CTGCTACCATGATGCTGAGAAACGTATATATAGTTTATAAGATGCTTCTTAGTTTATCTTTGCAGAATCTGATTA
TCAAATCATACAGGTTAACTGAACTAACCATATTATCTAAAATTCTGTGTGCAAGGACACCACTTTAATCAAGTG
AAATGGACAATATCATTGCTTTTCACTCCAGTATTTTGTAAGGTTGTTTGTCCAAATATTGTATGCTAGCAACAC
CACTAATTTGGCCATCAAGGCATCAACTATCTATGGTGACAAGTTTAGAATTAAGCATGTACTAGGAGTATTACT
AAATGGGGAGAAGCATCAAACATATATAACTTATGTCAGTATCTCAAGTGATCTATTCAGCTCCTGATTATCAAA
TAATACAGGTTAACTGAACTAACACGTTATATTCAATTTCGTCTGTAAGGACACCCACTTTAATCAAGTAAGAAA
TGGGAACAGATAATTGCTTATTAGTGTGTTTGAAGGTTGTTTATCCAATACCATATGTTAACAGTGCCACTAATT
AACCATTTGCTATCTTGAGTGATAGGAAAGGCAAAGCATGTCCTAAGGGTGTTACTATCAGTGATTATGGAACAA
CTAGTGCAACTTGAGTTTCTTTGTTTTTATTGCTTGACTTGAGGACATTATATAAGTCTCGGTCTGTATACCTGA
AGACACGACCGAAACCTCCAAATTTCGTTTTTGGTAATATTGGTCTTATATGGTGCATAGCACAGTAGATGCCTA
GAGAGCCATAATGTATAGTTATAGTTCATGGTTACACTGATAGTTGTTTCATAGTGAAACCATGTACTCCCCTCT
TTCAAAATATAAGGCTTATTTTGAGTGGGCACAAGAATTGGGAAGAGTATAAAACCAAGAGAAAGTGGAGTTAAA
TATGAACATACACTCAAATATCTCAGGAATAGGGAGAACATGAGTAGTGAGATGGGAATAATACGGAGACAATTT
ATTTTTTTTAATGTTCTGTTTAATTTTTGGTACCTAAAACTTGTTTAATGGTTCTCAAGTGAATCATATGCACTA
GCATACTAGAAAACAATGAAAGACACAGTCCTCATTATTTATTTACTTTTATCAATTTATCAATTATTCAAATTC
ATAGAACTCATGCACATTTATGATTTATTGTAG**GTTGGCTGAAGATCATACCGATGATATTCTTGCAAGTTATGC**
**AGTTGCTGCCTCAAGCTTGCTGCCTCCTGTTGCCAAAGCTAGTAAAATGAAAACATTAAATTAGTATAGCTGGAC**
**GGTTTTTCTTGCAGCCTTATGTTCTTCTCTTCAGCTCATGAATTTCTATTTATCATAGGTAGAAAGTGGCAACAA**
**CATTAGTATGAGCTTTCCAGTC**AAAAAAAAAAAACACATTAGTATGAGCTAGCAACACTTTTGAGTTTACATGTT
CTGAAAATGCATAAGGATTTCAGATTAGTGAAATTAAGTATTCTCTAGTCAAATATATTGGACGTTGGGAGGACA
AAACTTAGCTCTCCCAACATCAATTATATTTGACTGGAGGGAGTATTTGGGTTTTTATTTTTTAAATTTCTGGTG
GTACATGTTTGAATCGTCAAACCGGTAGTGGTTCCTTTGACATCATTTGTCATCCTTCTTTAATTAACTAGAATA
TTCATCTTATGAACTACTCTGATAATATTCTTCTTAATTACATCAATTTTTTTTCTTTTCCATCTTTTGTGATGA
GAACCagggtttacgattccgacgaaatccggttggatttcgcgaaatttcgatattccgacgaggttcgaaagt
agaaaccgtctgagatttcgagaagtttcatccaaattcaaatttgaattgataaaaaaaagagaaaaacaaata
aaatcttataaatactatgatattcatagaacctattgggatataaatattcaaaaaaaaatgtattgtgtgtat
ttactgaaacttatatcaggaaagaaaaaaaaataaagaattaaaaaaaagaaaaacacttacaTGGGTCGGATTG
CATGTACTAGGCCTAGTTAGTAGTTGGTCCAAATTACAGATTACAGATCGAAATTTTGATGATTTCGATCGGAAT
TTTGCCAAAACGACTGGTTTTCAAGAACTTGTCAACAAACTTAGGGAATTTGTATTCAAATTTTGGTTTATGAAA
TTTGTTCGGAATATACCGGTTTTCCACCGGATTTCGCGAAATTCTGAATTCCGCCAGTGCCCGAAATGGAAGGGC

TTGTCGGAATCGTAAATCGTAAACCCTGATGAGAACTATTCTTTTCAAGCTTTCTTGTTGGGTGCTTGTATTCAC
TTGCTATTTTCAGCTATTTCACAAAAGCATACAAAAATACTCCCACTTCTCTATGGAATGAATTAAGATGATAGA
ACACGGAAATCTTTGCACAGTAAATTAAATAAAACATTGAGTGGGGGTTGAAGTAGCAGAAAACCCCCCGTGTTA
TGGCTTGTGAATTGTTGGCCTTGTGGCCATAATCTTCGATTTTATGGTTAAAGTTTTGACAAAACCCTAATCCAT
CCAAACCCTAAGTCTCATGGGGGCCAAGGTACCACTTTATGATAAATATTTTGCAGTGATCCAAACCATCCTTTT
GTTTTCCTTTGTTTTTTTTAATTGTACATTGTTTCTTAATAATAGTATAGAATCAAGTTTAAAATGTCCAATAGT
AACTTGTAATTGGCAGGGTTTACGATTACGACAGACCCTTTCGTTTCGGGCACTGGCGGAATTCAGAATTTCGCG
AAATCAGGTGGAAAACCGGTAAATTCCGAACAAATTTCATAAACCAAAATTTGAATACAAATTCCCTGAGTTTGT
TGACAAGTTCTCGAAAACCGGTCGGTTTTGGCAAAATTCCGATCGAAATCATCGAAATTTCGATCTGTAATCTGT
AATTTGGACCAACTACTAACTGGGCCTAGTACATGCAATCTGGCCCAtgtaagtgttttcctttttttaattctt
tattcttttctttcctgatgtaagtttaagtaaatacacacaatacatcatttttttttgaatatttatatccca
ataggttctatgaatatcatagtatttataggattttatttgtttttttctttttttatcaattcaaatttgaat
ttggatcaaagtcctcgaaatcttagacggtttctactttcgagcctcgtcggaatgtcgaaattttgtgaaatc
cgattttatttcgttggaatcgtaaACCCTGGTGATAGTAACAATCTTAGGTTGAGATCAACATATCCAATCCAA
CAATTGTAACCTACTTGGTACATATGCTCCATAACAGTAACTGCAATGAGTTAAGAAAAAAAGGTCACTGGAACG
TGTTGGTTCTCTACCCTACCAATTCAATGGTAGTGCCAAAATCTAGGTAAATTTTGACACTACCAATTATTTGGT
AGGGTAGAATTCATATAATGTGTTTGGTTTGGTACCATTTGCTACCAATATGCTTGTTATAGTGAAGAAAGTTCT
AGAATGCTACCAAATTAGCTATGGGTATtgccaatactttggctctaatccaaaccaatcaaactactatttgaa
ctgccaaacaattggtagggcacattttggcatcgatccaaaatggcccATtttggattgaagccaaaacatgac
cttaccaatctattggtatcttgaagatattgtatggttggattaaagccaaatcattggtaatgcccaTTAACA
ATTTGGCACTAATCTAGAACTTTCTTCTCtatatcatcaatatttggctttatattggtagcaattcaaatatgt
accatgcatctctaccctaccaaagaattggtagtgccaaaacatgccaaaaatttggcactagcaaagatttgg
tagggTAAAGAACCAAACAGCTTTTGTTTCAGTCAGAAAGTTAAGTGGGGCAAAAAGATTGCCCAAAATATTATG
AAATACCTCTCGATATACCTTTGATTTCACAATATGTTGTATTAATTTATGTTTTACAGGCATACAACAGATCCA
ACTTTCAATGACATGTTTATTGTTTAGTTTAGCTACAATAATTGTTGGGTAGTTGTAGAGCAGCACATGGTACTT
ATGCCACAAGCCCGCAAATAAAATAATTCGCATTTGCTCATCCTAATTCATGCTAGCATCTCTATTGCAGAACAA
TTTTGATACTATTTTTTTTTTAAAATTAGCCTTGAAAATGCCACTTCCCCTAAGGAATGACAAATGCATTTTTTT
TTACAAATGCAAAAACTTGCAGGTAGACACATATTTGGCAGGTGAATTCGTGTCATATGGACATATCTTTCATTG
GAAGATATGTCCATATGTTGGAGCTATTTCACCAGTTCTTTCATTGGAATAATGTTAGCATTTAATCAGTGATAT
AGGAGATGATTCACCTTCAACAAGCTCTCATTTATCATCATGACGAGTCAGAAGAATCCGATTGTCAAGTTACGC
CTTCTACCATATACTCCCCACCAAAAACAGCTACCACTGGTAAATAAATTGCTTTCTGTTCTAATTGGTGCCTGA
ATTTCTGTATGCTAGATCTGATTGTTTGTTGCTTTCCTGGTATTGTGTAGTTTTTAGTTATGATGTACATAATCC
TAAGTTAATCCAGAAATTAATTTTTGAGTTCTTTCTGGAGTTTATTCAGTAGTTTATAGGTTGTGACCACCAGTG
TCCTTTTATGGCTGAATACAACTCAAAGAATCTTCATGTTGGAGTTGGATTCACTTTGGCAGTTCATACAGTGGA
TGCTGATAGATGCGCTCGATTATTCCTGTCGATAGCATCCTAGTATTGTGTCTTTATATAGCTATATCTAGCATT
TTCATGTAAAAGCTATCTGTTCTGTGGATGGGTGCTCTTATTGGTTGCAGCAATCAGGATAATCAGCTAAAATAA
TGATATATTGCAGCCTTGTCAGCCTTTATTATATACTGGGTTCCCTAAATTGTCATCTAGTCATTAGTGCCAAAG
TGCATGTAGTCTTCTCTGATCGTTATTTATAACTTTATATGCCAATGGATGTTACAGACAACGTAGTGTATGTCA
ATCTATCTGGCGTTTTATTTATCAGAATACTAGAAGAGAATTTGCACCAGTGGTTCAAGTAAATCCCTGAACATT
GAAACTAGATAGTTTGGGCTTTTTGGCCCCTAATCTTGTTTGATTGATACAAACAAGTCAACGGGTCACAAGTGC
TTTTGACTGACAGACTACATCCGTTTCTCTCCTCTCATCTTCACTCTTCCCCATCGAATTCTCCTCTCATCTTCT
TCACTCTTCCCCATCTTTGACCAAATTAGCCGCATCCTATGCCTGGTGGAATATTCTGACACAACATAGCAGTGT
GGCTTGTTTCAGCAGAAAATTATTTGACAGGCGTTGATTGCAAAGCTGTCACAGCTGCACCCATCTGATCAGATA
TAGGTAAACTGAGATGTGGTTCTCAGCCAGTGGGAATGATGTACAACCACGTTGTCATTCAGGGGAACTAAAAAC
CATCTACTCCTGTTTTTAGCAACTCACATACAGTACAGATACCTAGAATCAACAGATATCCCTACTTTAACCACT
TTAAACAAGTCTGAGGACATATATTCCTAGTATCAATGTTCATACACTTGAACCAGTAATGCAAGGTTCAGGACA
TACAATGCAATTTACTCTTGTTTTCTTTTATAGTTCATTTGTCCTCATTTTGGTCCCTGATTTAAGTTGGACATT
CTTATCTCACTAGACGGTTGGGTAAGTAATGTTAGGGGTCAAAATATAGTGGAGTTATGCTATCCAATCTTGAAA
GCAATGTAGAGCTTTGATCTGTTACTATCCAGTACTACAGTGAACAGCTAGACTTGGAGCTAGTGATTTACTCCC
TTATTGGATACGGATATTGAAGTAGATATGGGCTGGATGCCTATACTATGTAAATTCACCCATTGACGGTTCATT
GGCTCAGTCAGATAGAGAACATGCCTGCTTAACTGAATTAGTGAAAGCAAGTCAAATAACTAAGTGGCAACTCTT
ATCAGCAAAACTAATTTGTGATTATGATCTGTGGCTTGTTTATTTCAACCATTTCTTTTATATGAATTGCGTGTA
ACTCAGAATTTCATGTGAATACTGAGTAACTAGCACATCTATTTTGATTTGGGCCCAATTTTTCAACTGGCAAGA
AAGAATTGGAGTACAAGATCAAAGGGTGGGAGTAACTCTTAGTCTTAGGCATCGGGATTGGGCAGGATTTGTTGT
GCACCTAATTTTCTTTCATTTCTCGTACTTGAACTTGTTTGTGCATATAAGTATGTGATCATCCAACTATTTCTA
AGAGGAAATTAGCCACAAGCAACAAATAGGAAGTTTTTATATTCATACCTTTGCCTCAGAACTTGCGTTCATGGT
TTGATGGGGATCTTCTTTCGTCTCATTGTATCTCTCAATTATGGATTTCATGCTGGTTGAAAAAAAAAATCAAGA
TCCCATTAGAACACATGTCAACACATGCCACAGTAGTGTTGAATGCTATCATATAGCTCCTGCAAGCTTATTTTG
TAGTACATGGGCATCAGTATGTAAGATGACAAATGCAAACTTTTAATGAAGGCCTAGTTCTTTGACAAGTAATCC
ACGTGGATAGATAACTGTAAGGTCAACACAAAGAATTAAGTGTGGAGTGTTTTGAGTTAATTGGTTCTTACTGCT
TAATTTCCTGTTATCTTAAAACTTCCTGTTATCTTAAAACTAGACACTGTTCTCCTAATTTGTCATCCTTTTCCT
ATTTAGTTTGTTGTTTTCTACGAATCTTGAGCTATTTGTATGGAGAAAAAGCAGTGACAATGGTAAAAGATGGTT
TTGAAGGCACTACAAGACATGCCTCAAGTTATAACCGTGCTGTTACGAGGATCCAAGCATCCAAAACACTTGGAA

GATTAAGCTTCAGGCGTTGTCAACATAATGCATAATTAGTTGCATACTTCAGTGGATGATTGAACGGGATGTTTG
GATATGCAAATATTTCTATGCATCCGGACTGAATACTTTCTGAACACAAAGCAAGCCAACCTGCCATGGCCATGA
CACGATGGTTCTATCCATATTTTCCTATTTGTCATGTACGGCTGGGCTGGGCCTAGCACCAAAACTATTCGTTGT
TGTTTATTATTATTAGGTAATTAGGTTATTCGTAGATCATGGTTGTTTATTTTATTATGAACATTTTGATAACAT
TTTTTTTAAAAAATTAAAGCCTTTATTTTTCCGATCATGTGGAGGACGCACACTGATATATACGTATATCACACG
TTATATAAGAAACGTGCTTGAACACACGGGCAAGAGATAGATACTATATAGGAGTAGCGATGAATCCATAACTTC
ACTAAATTAAACGTGCACCCATATGTGAGTAATCATGAGCATCAAGATTTAAATTCTATCAGAGTTGCTCGCCCA
CCCCCTTTCAAGAAAAATCCATTAATTAATCTTGTTACATAGGAGTATCGTTGTGAATAGCATGGCAAACAATAC
AGTTGTCTCCTGATATGTCCTAGCTCCTTGTATAGATTCGTTGGATAATGTGTCCCGGATAatgtgtcccatcca
gtttcaaattgctatattttgagacaaaaggagtAACTTTTTTTCCCCTTTGGATTTACTTTGAAACCTATTTTT
AAGTTTCAGAAAACAACGTATCTATAGATGATACATAATATCTTACTATATGTAGTTTAATAAGCCAGGATGAAC
AATGTAGAATTAATCATATTCTTTTGTACGATAACATAATAGACTTCGACAATTGTTGAGGGACATAAAGTAAAG
ATCTTGATAAAGCTGTTGTCCTAGCTACATGTATCTCTCTACCACCAGGTTGATTCATTGAGTAGCTAGTGAGGG
AGGAAGAAAAAAAAAACAAGCGTTCTTTCATTTCATTCTCATTCACCAAGAGATGGAAATTTTGAATATTGTACG
AATCCTTATACATTGTAGTTAACGACAAGTCCATTATCGCACTAAATTCAATTTGAAAGTACTGAGAATAGTGAT
AAATCTTTGATCATATTTCTATTAAAAACACACGTATATTACACATGTGTAATATTATTGGACATCAGAATTTAA
ATCCTATCATACAAAGACATGCACCTAGCTATGACTCAATCTTATCTATGATAGTTTCCATTGATTTTGAGGAAA
TTTTAGCACGTTCTGTTTGAGGATTCTCCTCATTCAAGGGTTGGAATGGTCTATTGATCGCGCAGAACCATCCAA
AAATTACTTAAATCAAACCAGTAAAGTCGATTTCAGAGTAAGAATCTAAGTGGTCCAATCACCGTACAGATCTCT
CCAAAGCAAACGCATCATGTACCAAGTTGGTACGAGGTATCTCTGCTACTTGCACATCATCACTGCCCTAAACTA
AAGTCAACCAAAGCGTTCTAACTTCCAAGACATGCTTAACCCCGTAATTAGCAGATCTAATTAGGAGCTTAGATC
CTAATTAAGCTAACTAATGTGGGAGTGAAGCAAAGAAAAAAAACAAATACCTGGAGCTGGCAAAGTCATAGAGAC
GGCTGGTGCTGGAGAAGACGAGGAGGCCGACCTCGGCATCGCAGAGGATGGAGAGCTCCCTCGCCTTCTTGAACA
GCCCGCTCCGCCGCTTCGAGAATGTCACCTGTCGGCTCGTCGCGTTGTCGAtcctctttatctctatcttccctc
tccccatcttccccttcgatctcctcctccttttccctctTACAGATCTACGCAGAAATAACAATTAGCAGCAGC
AACAACCTCGGCTATCTTTCAAGCCAAGCTCTCTATCTATCTCTAATCTTGTGCTAATCTCCTTAGAGTGCTCTT
TACAGGTAGCTCTATTTCTCTAATCAAGGCTGGTTAGCTAGCTAGGTAGCTAGCCCTGATGCACTACAACTTCTT
CCTTAAGAAAGTATTCCAAGAGAGGACATAAAATAAAAAAAATATATAGTTAAACAAGTACAGGGCGCGTTTGGA
GGCTGCACGATGCAGGCAGGGTCTTCCAGGATCAATCAATTAATCCCTTTTTTAGCTTTCGTCTACCTCGTCCTC
ATTTGTTAATGTGCCATGATAAACATATCAAAACCTTCATCGTATCGATGGCCGACGCGTGCATGAGAGGATGCC
CCATCATAATGCTGCAAATTAATTGATTAATTCAGCTGTGTTTGTCGTAAGCTATAGGTTAGCTTCTAATTTAAT
TAGGTTGCTAGCTAATGCGGTTTCCTTTATTGTTTTTCTGAGTTTCGTTTTTGGAAAGGAAGGTGTGATGACTGA
CGAGGTTCATGAAGATGTGGAATAATTAGCTAAGTAAAGCAAATCTTCATGATACTCTGTCACCCTAATTAGTCT

>OsMADS23

ATGGGGAGAGGGAAGATAGAGATAAAGAGGATCGACAACGCGACGAGCCGACAGGTGACATTCTCGAAGCGGCGG
AGCGGGCTGTTCAAGAAGGCGAGGGAGCTCTCCATCCTCTGCGATGCCGAGGTCGGCCTCCTCGTCTTCTCCAGC
ACCAGCCGTCTCTATGACTTTGCCAGCTCCAGCATGAAATCCATAATTGAGAGATACAATGAGACGAAAGAAGAT
CCCCATCAAACCATGAACGCAAGTTCTGAGGCAAAGCTTTGGCAACAGGAGGCAGCAAGCTTGAGGCAGCAACTG
CATAACTTGCAAGAATATCATCGACAGTTGTTGGGACAACAGCTTTCTGGTCTGGATGTAGAAGACTTGCAAAAT
TTGGAAAGTAAGCTGGAGATGAGCTTAAAAAATATCCGTCTGAGGAAGGACAATGTCATGATGGATCAAATTCAA
GAATTAAGCAGGAAGGGAAGCCTCATACATCAGGAAAACATGGAACTACACAAGAAAGTCAGCCTTGTCCATCAA
GAAAACATCAATTTACAGAAAAAGGTATATAAGACAAAGAGTAATGGGCATCCAACAGGTTCTACCATCCAGCAC
AGCTTTCTGATTACAGATAATGAAATTGGCCCTAATCTTGAGCTAAGTCTGCCAGAGAATGTAGAAAAGGAGTAG

>osa-miR444a_gi|297726460|

AAACTCGTTGTCGTCGTCCCGTGCGGTGCGGCCCCCGCTACCCGCCGCGCGATGCCCCCTCTTCTCCGCCCGCCT
GATCTAGGGTTTTACTGGAGCCGCCGCGCGCAGCGACCAGCCGACCACGGACGAGGACGAGGGGATCAGATATGC
AATTGGGGGCAGCAAGCTAGAGGTGGCAACTGCATAATTTGCAAGAAATTGTTGGCTGAAGATCATACCGATGAT
ATTCTTGCAAGTTATGCAGTTGCTGCCTCAAGCTTGCTGCCTCCTGTTGCCAAAGCTAGTAAAATGAAAACATTA
AATTAGTATAGCTGGACGGTTTTTCTTGCAGCCTTATGTTCTTCTCTTCAGCTCATGAATTTCTATTTATCATAG
GTAGAAAGTGGCAACAACATTAGTATGAGCTTTCCAGTC

#### Stem-loop structure

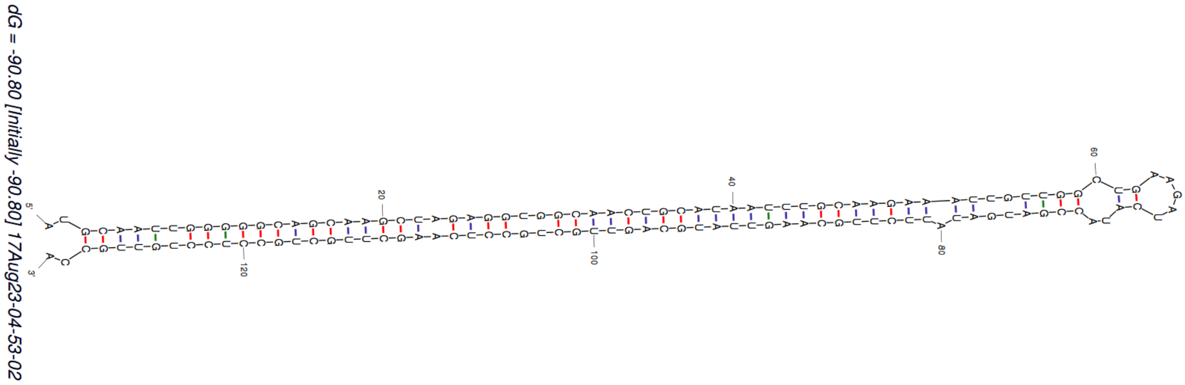


*OsMADS27 and osa-miR444b*

>Chr2

CATTCTCATATATACATCAAATCTTAATTAAATGAATGAATTTCTATATTGTCGATTTAAGTTTTTGATATCTGA
TTATGTGAATGCCAAAATGACATCTACTAAAATCTCAAACATTGGTGAGGATGAGACATCATCTAGTACAACGAA
TCTGGATAAATAGCTAGCTCATGTCTCATCCTCACTAAGGTTGTTTTTTTTGGGACAAATGGAGTATTTGTGAAC
GAAAGAAGGCCAAGCATGCACTTAATCACCACCTTTCGTGCGTGCAACGATTTTTCTCTTGGCTCTAGCTAACCT
AAGCCGGGACACTTTTAACAGGCCTTTTAAATGGCATCCAGCCATAAATTAAACTAATCGCCAATATCAGGCAGC
AGCCAATTGGGCGCTGTCTCCTCTTTCCTCTCTTCCCCCTCCCCCTTGCGTTTCATTGCTAAATAAAGCAAGTCG
CTGGCAATTTGAGGAAAGTCCACCCATACCCTGCAAGTTGCATCCACCACCAGAAAAAGCCTCTCCCGCAAACGC
CACACACCAAACCACAATTAATCCCATCAAATTCCCCTATAAGCTTTGGCCTCTTTGCAGCTTTCTTCTACCAGC
TCCCAGCTAGATCCCCTCTTGCCTTCTTTGGCACATTCGTTCTTGCTCTCTGCTGTGGATTATTTGCCTCCGGCT
CCGGCCAATCGGCGGCGTGGCGCGGGCAGATCGATCGATCTCATCGGCGGCGGGGATTGCCTGTTCTAGCTATAG
CTTGATAGCTTAGCTAGCTTGGTCATTTCTGGTGAGAGATCGATCGAGATAGCGGCGAAGAAGATGGGGAGGGGG
AAGATTGTGATCCGCCGGATCGACAACTCGACGAGCCGGCAGGTGACGTTCTCGAAGCGGAGGAACGGGATCTTC
AAGAAGGCCAAGGAGCTGGCCATCCTCTGCGACGCCGAGGTCGGCCTCATGATCTTCTCCAGCACCGGCCGCCTC
TACGAGTACTCCAGCACCAGGTCTGTTCTGTTCTTCATATCATCGATCATCCGTCCTCTCTCCTTGCTTGCTTGT
TGCTTGTTGATGATTAACTTTTGCTTGAAGATAATATTTGTTATTGATTCTTGTTCTTTTCCTGCGCTTAGCTTT
TTCTGCTTTAATTTCCTAGTATTTTGATATTTAGCCAGGGAGCCAGCATGGGCCATGGCCAGCTCTTCTCTGTGC
AGTCAGAAAGATGCATAAACTATGCAAAAACTGCCCTCTTATTGAGAAAAGAAATAAACTATGCAAAAACCGTAT
ATAGACATTCATGCACCTAACTAAAAAATGGTTATCACATATATATAAGAGCTGCACGGAGTAATGATTATATAT
GTTTTTGTCTTTATTTTTCTTTTGAAGGAGATCTGTATGTGTAGTTCTTCTATCTAATTTATCTTTTGTTTTGCA
TATGTATATTTTTTTTCTTCCTTGTTCTTGACTACTAATTAATTAATTAATTGCAAAACTAAACATTGTCTTTGG
TTTGTATTAATTTGTATATTACTTGGCCGTACGTTATCATGTGATCAACATATATGCATTGCAGAAACCATCAAA
TGAAAGAATATGTGGTGAATGGCAGCTAAAAAAATGATCTAGGGTTTCGTGATAATATGAAGTGTATAGTGTGTG
TCTCTAAATGAAAATTTGTATTACTTCGATCTCTCAGCTGTGAATAATTGCGTGGGTCTATACATCTCAACTACC
AATTAGGAGGTGTGTTAGATGGATGCTTATTTCTTGGTCAAAAAATAACACATAAAAATCACATGATTCACATAT
ATGATGTGTAATCTTAATTAACAGATTAAATTTTGTTACTTACCGTTATATATATGCAAATATGCAGCATGAAGT
CAGTTATAGATCGGTATGGCAAGTCCAAGGATGAGCAGCAAGCCGTCGCAAATCCCAACTCGGAGCTTAAGGTTT
GGTTCCCAAGATTGATTATTACAATTTATACAGTTTGATTATTCTCAGATTTAAGTGAGAAATATATCCACTCTA
AAAAAAATTAGATCATCCTGAAAGCAAATGCCACAGTGGAGTTTAGAAGTAATACTTAGCAAGCTAGCTTGCATG
CATCATAGCTCGAATAATCACACACTATATATTTCCATTTTGTCCGCCATATATAGGTTCCTAACTTCCAATTAC
ATATAGACACAGTAGAACACCATTTGTTTTAAATATAGTCCTTGCGCAGGAGTGTTGCCATTCGTATAAGTTTCA
TTAATTTCAAGGGAAATATGTGTCTCTTTTATGGTAAATTTTGGGGAGTGGGGCCATGGGGGGAGTATGTAGCAC
ACTGAATTTTGATGAAATAATAACAAGAATAAAATTATTTGATCTTTTAATAGATATAGGTGCACATATTATTTT
TGTGTGTGCTAGCTTTAGATATTTACACCATTGCAACATAAATATATATTGCATATGTCTCCTGGCAAACTAATT
AAATTGTCTAAACACAATGTTAATCTACTTATGTTCTATGTCTATCCAAAATGACCAACCTCTGATTGAGTATAT
ATATACATGGATAGTTCCCAAAAAATTGCAAGTCTGTTTGTACACATTTGTTATGGAAATAATAAACTAAGCAAC
TAAATAAACTTAATAATACATTATCAACTAAAGAAATATCAACCATATATACGTAAGACAATCTATATTATGCAA
ACAAATAAAGCAAGCAAATAATCAACTGCATAAGATGACCATATCTTTCTTGGCAATTAACAGCGTGGTTTTCAT
AATGAGTATATTTTAATCTTAGAGATTTTATGACTGATATGACTTGATTCATCTTAGCGTACTTCACACCTTATA
TAATCCCTTTTTCCATTGTAGTGTATTGTACAGTTTTGTTCTGTGTTATCCAAAATAATCACCTTGCCCTAGTGA
CGTATGAAAATTGCCTAACAGTTCAACTTTAGTTTTAGATTTGTGAAGAATGAAACTAAGTATCTAATTAAGATT
AAAACATAGATAAACCAAACATGGAGCATTGCATGAATAATTAAAACAACCTTCCTTATAAATAAAGTCATTGTG
GTTTGTCAGCATTTTGATTCTAATCTACCTATATTAATTAAGGAAGTCGATCTTACATGTAGTCTTTGATTAGCG
TGGAGCTATCAATCCTCGTGGTCAATTTTTAATGCATGGTTGATATCATCTTATATCTTTGATATCCTACTCAAT
GAAATCTGATAAGACGAAAAAAATCACCAACAAATTCACATGGGATAGGGGCCAAGACGATTAAACGTACCTTTC
GCATGAGAGAAAAACACATTGTTTATGATGTTTAtatagatcctgatgggcctgataaggtctggaaaaaaaaga

actgggagtaaaaatgatctttaagatcacttaagattaatggtaaacatagataaaccaaacatggatataaac
atcataacatgccgttgggcttgttaaccgatcatctttagtactggtttataattggtctgaaagatttatgat
ttgcctgaccgagtaaagatcggttctccagtagtAACTTAAGGACACGATCGCGAAGTCAGGGAAATTGAAGTA
ATAAAAAGAAGAATGATTCATCATGTTTTGTTTGAATATCGATTTTACAACCAGTCCTGATTCCTGAAAATTATA
ACTTCATTTTAAAATAAAATTGGATTTGAAAAAGTTGATTAACAAGGATTTATTTGGAAGGAGATAAGTTAAAAT
CCCAGTAAGAAGGTGAAAGTTTTATTGTAGTATTATGTGAGTCACAAGCTAGTAATCATGCTGGTTTTAATCTTC
ACAATCAGTACATTCTGAAATTCATGTTAATTTAAACATTTTCATACAATTTGTTCCAATCGAAAAGTAATATCT
ACTCAGAGCTTCAGTTGAGAATAATTTGAAACAAAATCAAGCATATGGTGTCAAAAGAGGTAAATTCGGGACTTa
ggccctgtttctttcagcttgggattattataatccagattattgggagtaagctgaaagaaacagacaacttat
tgaaatagcttattataatctgaagcccagcttattataatctgataagctcatttaggtgagcttttccaaat
tattgggtgaaaaattactaccatgctaccccactccctctatagacttgcgaacccaataatctagactctaat
aatttaggaaagaaacaactaacggcttattctactatagattataataatctagtttatagtaatctgacttaa
tcatttagattataataattttaaactaaaagaaacaggGCCTTAATTAAGGAAGAAGTGGGCAAATAAAGAAGC
TCTGACCAAAGCCCTACAGGGCCACATACCTAGTACGAGGTAGTTGGGTCAAACAAGTAACCAAATTATATCTTT
TAGATAAAGAAACAAGTAACCAAATTATAGTTGTTTAAGACCCATCAAACAAGATGTAAGTAAGTAGTATTTGCC
TTGGGCTGGGTTTTTAGGATGTCAATACAGACTTACCTTGGGCTGGGTTGAGTAGTCGCAAGTGCAACGGGCTGA
AATAAATTTAGGGCTTTCAGACTTTTGACCCACCTCAAATTGGCATGACTCTGTATCTATCAAAAATTGACTTGA
CAGGAAAAATAAAAGAACAGATCTCCACCAAAATGGAATCATGTACTCGATACTTACAAAGAGATGTGCATTACA
ATCACATGCAAACTTGGGTAGAAATTATGAAGACCAGAGTGGCAGCTCATGTTTTTCCACTTAAACAGAAATGAG
GAGTAAGTCCTCCTGTTTGCTCAACTATTCAGAGAAGATACAAGGAGAACAACACaaaaaaaaataaaaataaaa
atagaaatagaaaGTTCACAGTACATGCAAAAGAATAGCTAATCTTTTAACAAAGAACCCTGTTTTGCACCATCT
TAGCAGCGGTTTTTATGTGGTCAGTACGGCAGTCAATAGTCAATGAATTACACCCCTGCAATCTAAGAGTAACAA
ACATTCAAAGAAAATGTCATGTGCCTTTTTTGTCCGCAAAGGTGAATGCTGAACATGTGAGGATACAAAAATAGT
GAAATACTACATCCGTTCAAAAATGTTACTAACTACTACAGAAAGAAATATTTTATAGGACTATGAATCTAGACA
GGAAATGTCCAGGTTATATTTTGGGACATAGGTGGTTCAGTAGTAAACTACATACTCAAGAAAGCACCTAGGCAC
TGGTTTTCCTTTCTCAAATAGTAATCCCTACAAACCTTGCATTCTGCTGAATAGGTCAACCAAGGTTTAAGACAC
TAACTAGGATTCTAGGAATGCACTAGGTAGCATGTTAACAGCAGAAATTAACTTCATAAAAGCTGGAATTTTTCA
TTTGCTTGAGAACAGTCATACACATCTTTTTCCTTCCAAAAGTAAACATATATTATTCAGTCTCATTTGATTTAC
AAAAGTACAAAGCTGAGTAGCATGGGACAATGAACGTTCCGCTACCAAAGCACAGGTGACAGGAAAATTATAGTG
CTATTCTGATGAACAATCAACTCCAAAATTTACAGGATCACTCACTGGGCATTCCTAGAGAAATTCATGCAAACC
TACAAAACAAAGAAACCAAGGATCAAGCCATGAAACTAACATAATGCATTTAGAAATTCATAATTCACTACTAAG
AATTGTGACTGGCCTCATGATCAACTCAAAATTGCCCTTCCAAGCAAGCATGTCATGGGTAATAAGAGAAACATT
CAGCAGTGGAAGCCAATACACAATGTAGAGACGACGTAATCTGATCAGAGTAAACAACAATAGATGATCAGGATG
GACTAAACAGTTTACATGTGCAGCCTTCATTATAAGAACACTAGGAACAATTTTCATACATATTAGATGTTTGAC
TTCCTTACCAGACTTTACTCATCTAGCACCTTACCAGACTTTACTCACCTTGCTAGATGTTCATATGTTGATGAG
TTCATTTGCAATTGCCTGGTGGCACTAGCATCCATGGAAATAAATATATTTTTGTTAATCAAAAGATTTAGAAGA
ACCATTAAGATAATGTCAACCTTGTCAAAACATGTACCTGCCTCCATCTAGTTTTCATATTATAACTGTTGTAAG
CATCATGCTTCAATATTATCTCATGCCACAAGAGTTTTGTTATTAATCATTGTGTTTGTTGGTCAGACATGTTTA
TTACTTTAATTTTGTCTTGTGTGCCTGTAATTGAATAACTCTACC**CTGCTTTCTCACTTGATAGTTTTGGCAAAG**
**GGAGGCAGCAAGCTTGAGACAACAACTGCACAACTTGCAAGAAAATCATCGGTATGGTCTGAAAAACC**CTGCAAA
ATTTCCAAGCATATTATTCATTAGAAACTTGTTGGAAAGAGAAGCGCTAGCATGTACTATCTGAAACCAGCAAGA
TAGTAATAAAGACAAAAGCCAAGAATCAATGCCCTACAAGTAAGCATTATCATGTGCAAGATTACAAGCACCTAC
CAATCAACCCAAATACCTGAGACAACAAGGATACTATTAAACACAAATAAACCAAATCAGAACTACTCTGCTGTG
AACTCAGTTTCCAGTTCTAAGAAAAAAAAAATAGATCCTCCATATCAGTCTATAGATCAGTCCTATACTCCTATT
GGTCAGTATTGGTATCGCTCCCTATTGTATGGTGCATATTGATCCCATAAATGACTAGAAACTATTGAAAGGCCA
TTGATTTCACACGCAAGCATAGAGAACAAGATATTGCGAAGCCATAATTGCTACTAATAACAAGACAAGAAAAAG
GTATCACATATTGGTTTGCTATTATAGGCAGTTGATGGGCGAAGATCTATCTGGGCTGAATGTTAAGGAATTGCA
ATCTCTAGAGAATCAGCTGGAAATAAGTCTACGTAGTGTCCGTACAAAGAAGGTCAACCCACTTATCATATTTCT
CGTTATGCATTGATCATTTATAAAATAAACTTAACTGACATGGTGTTACTGTTACTGCTGCTGTTTTCAGGACCA
CGTCTTGATTGATGAAATTCATGAACTGAATCGGAAGGTCTCTATAGCATTTTAGTTTTTGTAATCCTCCATGAA
GGATGTTTATTATTATTATTATTACTTTAGTATACTGACTCTACTTGGTTGCAGGGAAGTCTAGTTCACCAAGAA
AACATGGAATTATACAAGAAGATCAGTTTAATTCGTCAAGAAAATGCTGAGTTATATAAGAAGGTAGCCAGCATG
TTAGATAAAGAAAACATAGTGTTGCCACCTAACCTGCTTAGCACATACTGACTTATATTGTAACAATGTTTTAGA
TCTACGAGACTGAAGGACCAAGTGAAGTCAATCGGGATTCACCAACTCCTTACAATTTTGCAGTAATTGAAAAAA
CAAATGTTCCTGTGCAACTTGGACTCAGCACACTACCACAACATAGTGACGCCGAACAATCAACTGCTCCTAAGC
TAGGGTAACACTTAGCTCTACGTATTTTCAGTCCATTTGCAAATCCAATCTTCATTGGAGTAACGAATGTTCAAA
CACATGCAGGTTACAGTTGAATCCATGACCAGAAAAACATTGTTTTGCTACACCCTTGTATGCTGAAGGGATACT
ATTTGCTGTCAAGAGTTGATACTCAGATCAAGAAAAAAAAAACTATTCATGCCATAGTGATTGAGAAATGGACTT
GTATACTGGAAATAAATGCCACAGTGGAATTGAGTTGTGGAATCCAGGCAATGAGTGTTTTATGATTCAATTTTT
CCATGTATTTTTCATATATTAACCAGTGCGGGCTTGTGTAAAAAATATAACCAGGCAAGGAAAATTAGCTGTATT
GTGCATATGGAGCAAATTTCAGGCGTTACTACCATTAAAATCATGACATGCTGTAAGCAAATATCTTGTAGACCT
TTATCTACATAGATTTTTTTCAGTAGGGGCCTCCCTACCGTTTCAAAAGAAAAAGAATGCAGATGATTGATATTT
CGAATGCCTTCAGACATGGATAATTCATAACTATTGCGATGTGGAACCTTCTTAATTTTCAATTTTTAACACGCT

ATCTACAAGTGGAATATCATAATCATTTGCTTGTCTTCATTAGTAAGACAAGTTTGAATGCACGTGCCAATGTGA
GACAACCGTACCATATAACGGCCATTAGTGCTAGATTGGCTTTGCACCATGAAAACATTGGTTTTACAACCTACT
GTGAGGACATTAAAAGCAGAGATACTCTAGCTGGCCTGTTTGGAGATAAACAGTCGAGAAATATTGTGAAGACAA
TCAAAGTGTTTATTCCTGGGATGCAGTCCAATCCCAAATGATGGATCCAGAACAAAGAAAATATACCTTCCATAT
GCCTATGTGGGGCTAGAAAGCCCAAGTGCCCAACAAAAGCAAGGGTCCAGTAATCCCTAATCCCAGAAGTATTGA
GGGCTTTTGAGTGAAATCCAGCACATGCGGGCCAATATCCTACCAGAATGGCTCAAGCAAGGGTTCTGGTCATAT
TGCAAGACAGCCCAGAAAGCTTTGAGGTTTGAGCTAAGAAAAAAAATATTTATACATGCAGGAATTGCCCTAGAG
GGATGGCCCAGAAAGCTTCGAGCTAAGAAAGAAAAAATATCTACCAAGAAAAACTGTCTAGAGTTATTATTCAAG
CTTCATACATGTACATACTGAAATCAGTGGCAAATCTAGGATAAAAATTGAGGTGCTTATGTAATTTGCATTGAT
AGGTCAAAATATGAAAAATTGATACATATTCCAATTAGCGCACCAAAATCTAAAATAATTCATAAAGCAGAGATG
TGACCACCTAATCAAATTATAAATTAATTCTATCCTTGGACGTAGAAAAAGAGTAACCATGGTAACAATTGCACC
ACAAGCCCATAACCAACAGGTACTAGTCTTCTAGAAGCTATGTAAACCCATCAATTACAGAAGCCATGATCTAAT
CCCATCACCCATTGGCCCTTGCCATTTTTGCTGTACTGTACGTAGGCACTAGGCAGTAGGCTACTAGCGCCCAGA
CTAGCCACAATCGTGGATTCACCCTGACTGAAATTTCAGTGGCACCCAACAAAATCCACAGTTGGGCATCAACTA
ACAAGGTTAACACATCTACAACAGCGTGGATATAAAAATACATGATCTTATAGATAAATTTCTATAATTTTCAAG
AATCGGTTATAAGATGAATCTACATCAATCCCAGTAACATTTCTAATTAAAACAAATCTTTACTGCTTCAAGAGT
TAGTGGTTGCAATGAGCCGTTTTGTACTCATCTCCTCTCTTCAAAGGTGTGAGAGTCATCTTTGCAAAAGCATGG
CTACATCCATTGCAATTCATGAAAACATCACTATCCATCGAAAAACACATGCACTTGGCTAGTTCAATACAAATT
AAAAACAAATATTTGCACATGCACTTGGCTAATTCAATAGAAATTTAAAACAAATATTTGCTCATTCTGAACCTT
ACAGGATTTTTTTAATGGAGAGTGAATTATGGAGGTAC**CTAAAGATTTTCTTGCAAGATGTGCAGTTGCTGCCAC**
**AAGCTTGTCACCATCCCCTTGCATGAATC**TGTCATAAAGCAGTAGATCGAATTTAGAGCAACATTTTGGGAGTGG
AGCTGAAGTAACTGGAACTAGTAGATGTAACAGATCAGCACAATCGCAAAGAATAACTAGATGCCATGTAGTATT
GGCTCACACCAAAAACTGTACTCTTATCAATTAAAAAAACTGCTGTTCTTTCGTGTTAATATAGCTGAGTTTGCC
ATGGAAGCAGTGAACCATGCATTCATACTGATTTTGCACATACAATAAAAGGACCTCCACAGAGAAGAAACATAT
GAATGCTTCCTCGCTCTATCTAATCATAGAGAGGAAAACTCTCAAAGTAAAGCCATACGGGTAATTTGGAAAACA
ATCATAGTTTCATCTTCCTCATTAAAACAGTAACATATGCATGCATCTGCCTGATAGCCTAGCGAAAATTGAACA
AGATCCAGCCCAATAATTGAATTTTCTAAAACATGCAGGCAAATTTCACCGTTTTCTAAAAGGAATTTATGATTT
AGATCTACTGCGTCCTACAGTTTAGCAAAAACACATCAAACTCCTAAGACCAATTCTTCAAATGCGCGCAGGTTC
ACAACTTGTTGGCACTATAATGTTAGTATCCAAGGGACAAAGCCATTAACAAATCTGTTACCAGAGAAACATCAC
ATGGAAGGATGATGTTAGAAGGGAAGCGGAGACTGGAGTTTGTGTTACATACTTCTTCTTATAGCACTGAAATGT

>OsMADS27

ATGGGGAGGGGGAAGATTGTGATCCGCCGGATCGACAACTCGACGAGCCGGCAGGTGACGTTCTCGAAGCGGAGG
AACGGGATCTTCAAGAAGGCCAAGGAGCTGGCCATCCTCTGCGACGCCGAGGTCGGCCTCATGATCTTCTCCAGC
ACCGGCCGCCTCTACGAGTACTCCAGCACCAGCATGAAGTCAGTTATAGATCGGTATGGCAAGTCCAAGGATGAG
CAGCAAGCCGTCGCAAATCCCAACTCGGAGCTTAAGTTTTGGCAAAGGGAGGCAGCAAGCTTGAGACAACAACTG
CACAACTTGCAAGAAAATCATCGGCAGTTGATGGGCGAAGATCTATCTGGGCTGAATGTTAAGGAATTGCAATCT
CTAGAGAATCAGCTGGAAATAAGTCTACGTAGTGTCCGTACAAAGAAGGACCACGTCTTGATTGATGAAATTCAT
GAACTGAATCGGAAGGGAAGTCTAGTTCACCAAGAAAACATGGAATTATACAAGAAGATCAGTTTAATTCGTCAA
GAAAATGCTGAGTTATATAAGAAGATCTACGAGACTGAAGGACCAAGTGAAGTCAATCGGGATTCACCAACTCCT
TACAATTTTGCAGTAATTGAAAAAACAAATGTTCCTGTGCAACTTGGACTCAGCACACTACCACAACATAGTGAC
GCCGAACAATCAACTGCTCCTAAGCTAGGGTTACAGTTGAATCCATGA

>osa-miR444b

GATTCATGCAAGGGGATGGTGACAAGCTTGTGGCAGCAACTGCACATCTTGCAAGAAAATCTTTAGGGTTTTTCA
GACCATACCGATGATTTTCTTGCAAGTTGTGCAGTTGTTGTCTCAAGCTTGCTGCCTCCCTTTGCCAAAACTATC
AAGTGAGAAAGCAG

Stem-loop structure

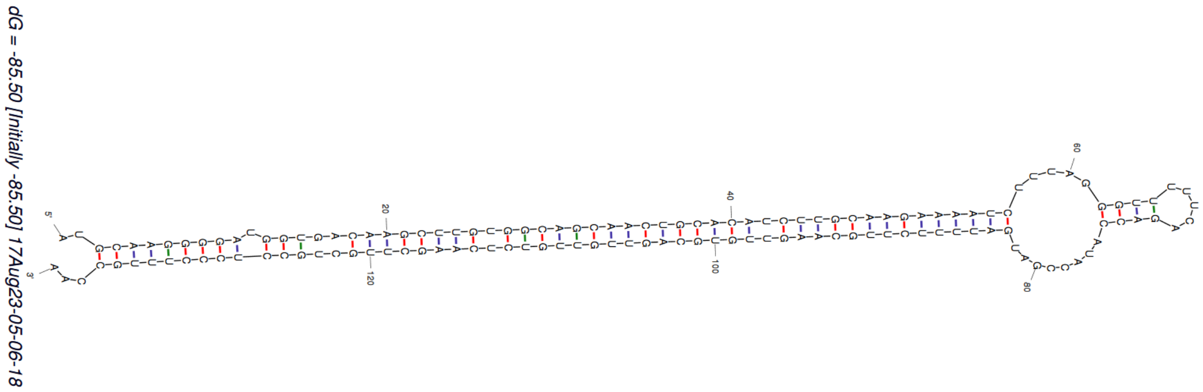


*OsMADS61 and osa-miR444c*

>Chr4

GTAATTTTAGCTGGCAATTTTTATGGTTGAGTAAGATTAAATTAAATTAGAGTATTCAGTTTGTATATAGTACAT
TTGTTCCAAAATATAGCAACTTTTGACATCATTATATTTTGGGACAAATGCAGTATTATATATATGCAAAAAAAA
AAGTAGAGGAAAAAAATTCAATGTGAAATTAATGAACTAGGCATGTGATTATTTTTTTCAGTATGTCTCTATATA
TGTCTAATATATAAATCATGGCAAATGATTGAAACTTTGAAGTAACGTTGGTGCAAATTAATACAAAATTCTAAA
AATTGCTAGAAAAATATAAAGATGAGTAATGGCATATATATTTTATAGAAAGTATATATATCAGTTCTTCATGAA
TAAAAAATATTAAGAGACCAAACATGGATTAATTAGTAGCACAACTATGGATGCACAACCTTAGCATCCCCTTTA
ACTTTCCTCAGTAGTGCGACCAATTCCAGGCACTGCACAATTTCTAGAAGCTAGCTACCTGTGCCTGGCCTTGCC
CTGGCCTCTTAATTGCAGCCATTTATTTAGCATTCACAGTTAAAATAACCCACCTTGTGATCCTAGCCAATGGGG
GGATATCTCTCCCCTCCTAACTAAATAAAGGAAGGAAGCAATTAAAGGAAAGCCAACTTGCAAAGGCACCAACAA
AAGAcccctccatttccattcctcctcttcctctcttctcccccccccctccAATAATTCAACCCCCTCTTCTGGG
AAACCTGGGAGCTTTCTCCTTCAAGAGCTCCCTGCTAGCTAGATCTCCTCctctctttctctctctctctctc
tctttctGTCCATTTCTATCTGTGCTCTCGCCTGAGCGGCGACAATCCAGCTGAGATCGATCGATCGATCGATGG
GGAGGGGCAAGATAGTGATCCGGCGGATCGACAACTCGACGAGCCGGCAGGTGACGTTCTCGAAGCGGCGCAACG
GGATCTTCAAGAAGGCCAAGGAGCTGGCCATCCTGTGCGACGCCGAGGTCGGCCTCGTCATCTTCTCCAGCACCG
GCCGCCTCTACGAGTATGCCAGCACCAGGTCAATGGCTTCTCCCTCCGGCCCTAGCTTATTAGCTTTTCTATAGA
TTCCGTTCTTGCTTTGGCTTATGATTATTAGCATGTTTTCCTCGATCGTCTTCAACTCTCCATGCGTAAACACTT
TATTTGGTTGGCATCTTTCTTCTTATTATTGTTCTCCGTTTCTACGGTTTTTTTTTCTCTCCATGAAAGATTAGA
ATCTTATGAATGAGTTTTACCTTTGACAGCTTCAGTAGTACAAAAGGTTATCGTGaaaattttatatatattttt
tttttgcATCTTTGCACCTATTATTCCCTGTCATATGTATACATATACATCGGCCATAAGTGTTTAAGAACTAAT
GCATGTGTCAGGTTCTTCAATTTCCTGTTAGTCTCAGATTAACTACTAGTTGTTCATCTTTGTTTTTAACCTTCT
TCTAGCTAGGGTCCCCCTATGTATGGCCATATGTCAGTTGTGTGTTTACTATTTTGTCATGCATATTTTTCGAAG
GAATAAATATGTCAACTATGCTTTGCAGTGAAGCAATAATTGATCTAGCACTCTTGGAATAAAATTGTGTGATAA
CTGTACACTTATACGTATGATTTCTGCTCACTTCATATAATTGTTAATTTACCCTTTTTGCCGTACTCTTACACT
CAAAAGCTAGCACACGCATGCATGTGCTTTCAGATTACAAAAATAATCTTAGTAAGAGTTGTGCCGGCCATTCGT
TTTGTGCAGCATGAAGTCAGTGATTGATCGATATGGGCGAGCTAAGGAGGAGCAGCAGCACGTCGCAAACCCCAA
CTCGGAGCTGAAGGTATGCCTTGATGACTGCAACTAAACATATTTCTTCTTCTGCCGATCTCTGTCATTTGTGTT
GTTTAGGAGTTTTGTAGCGTCTTCATATATATAACAGAGAACTAACCTTTCTAAACTCTAACCTGAGAGAAATTA
ACTCAACAGAGATTAATAAAAAAAGCATGCAGGAGCTAGCAAGTTGTTCTAATTAATGCAGTACTTAACTAAAAA
TAAGAACAGAGATCGAAATAATCCTTTGACATCATCTATTCATTTGGGTTAGTTCCAGTGTTTTGGTTGGACATA
TGCACATGCACTACACCCCCTGCCGGAATACTATTTATCTCTTTTCAAGCGTAATTGAACAAGATAAATTTGATC
ATCGCACAGATGAAAGCCTGATTAATTTGGTTTCCGGTGGAGATAAATTTGTCAGTGCTGTTTATTCAGAATTCA
CAAATTTCTGCGACAGTATGTCTGAAGGATAGTTTCCCGATATCTGACCGTCTATGATATGCATAATTAATTTTC
CCACTAATACTGTTTGATATTTCAAACATGCACAAACTTATAAATGTTCAATATTTTCATCGTGTTGACAACATC
CTTTATTTTCCTTCGTCTACATCATCTCAACTGTTACCAAGAAATTATTTTCGAGTAAGGACACATCAGAGTTTT
TCAGTTAAAATCTCCGTTGTTTTGAGTGAATTATGTAATAAGAGATAATTAGGCAAGTTTATATTTGTAGTTATA
CGTAGAGCGGGAGTGTTCACTTCAATTATCTTAAAAGAAAACACAATCAAGGTTCTCTAATTTTACTGTGCATCA
GTTAGGCAAGTTTATATTTGTAGTTATACGTAGCAAAACTAAGGCACAAAGTACAGGAAAATGGCTTTAATTAGC
TAGGCTTTTAATTTAAACAGGCCAGGAAAAAAAAAAACTACCTACTGCATGTGTACATTGCAGTAACTTTGTTATC
GGAGCTATTGCCTATTTAGCTAGCACTTAGCAGGATGCATTCCCATTTTGGTCCATTAGTGACACCTTTTCACCA
TTATTGCATGTAAGCACATTGCTTCATCCTCTACGTTATAATTCGCGTGATCTGCATGCATGTATGCATATACAA
AAACAATGGTCGTGTTTATTAGGTATACTAGAGGGGAACCCCCATGCGTGTTGCCGCGGAAAAAATGTATGAGAT
AAGTAAACATGAGATGCAAATTAGATTTGCTTTATTTTAAATGATTGGACATCATATAAGTAAATATCATTTATA
TATTATATATCATGCTAAAAATATTAAGATAATGTATCTTCATATGAGATCAAAGGGTACAAATAATTGAGATGG
TGTATTAAAAATGCTAGAAATAAATTAGATGTGTGTATACCTAGAGAACATAGACAATTTACATCGTAGATGAAT

GATCCAAAGGCTACAAATAACTAGGATGATGTGGCTTCAGGAGAGAAGAGAGAATTAACATTAACTATATATAGT
GATGATGAACCTTATAAGTACCGAATACTCTCACCAAAGACAATCTTTAGCTTGGTATCAACAAATGTAGATGGA
CATGGTAACCAAGCATCTCTATATTCATATAAGAGGAATTTCAAATAGTTCTTGATAGCCTATCCACGATGAGCT
TTTATGTTGTAGTGGTGTCTCCTCTGTCCTTACTCTTCAACCAACCTTCCAATAAGCGAAATGTTTCCTTAGTTG
ATAGACCTAACAATGCCTACCGTGCAAGCAAATTCTACCATTGGTTTTGCCCTATTGGGATATAAGGTAGCTTTG
GATGGTATATAAGGAGTTATCTTAAGAAGTAGTGCCAACTTAAGTATATCTAGCTCATCCCTTCATAGGTGCAAA
ATTATGACATAGGCATGTAATGGGTAGAGGCATGCTATATGAAGGTAGCATAGCTTGCAGAACATATTGCATTCT
CATCACCGTGACTCAATGATGTTGACACTAGTTAGTAAGCCAACCCTTTTTTTTTTCCAAAAAAAAAACTATAG
CTAGTAGCTCAACTCAGACTTAGTAATTTATTTGTGAGTTTTCTCGGGATAAACCCAAGGAATAACAAGGCTTAC
AACTTTTATTACTAACTACATTTGTGATGAAAAAAGTAAGCATTGTTTGTTGTAAGAGAAATTCTACGTTTTACG
ATTGACAAGCGACATGTCCTATAAATTGGCATCGACGAACGTGAGTTCTACGAAATACAAAATACCTTGGTACAA
GCATTTCTGTCCCTTTTTATCATCGCCACTAGGTTTCCATCCATAGACCTGCATTCTTCTAAAAGGACCATTCTG
CTCATAAAAAGATCTCCATTAAGTGTGCAAACATTTTTTTCATACAGTTTTTATCAAATTCTAGGGGCAAATTGG
TTTTCTAAAGACTTAATGGAGGTTTATGGATGGAAACCTAATGGCAGTGACAAAACGAGAAAAAAAACACTTGTA
CGATGATATTTTTCTAGAGCTCGAGCTTTCTAATGGCAACTTTTAAAATCGGTAGTTTGTCAATAACGAAACATA
TAATTTCTCTTGTTTTAAATGTTAGTGATGTGGTTTTTTGCATAGTAAATGTACATTGACAATGGTAAGGAATAA
ATATGTATTGGCCTTCTCAACTCTTAGTTGATTACACTTGAATCAAGTTTTTTTTAACAATAGAATAAAACCTAG
TCTCTACAGCAACAAAGATGGTAGCCAATAACTTCAAAATAAACAACCTCATCCATTTATATGCATCCACCTACA
CCCCAAAATCTTCTGAGAGCTCACCTTTTCACAATGGCGAAAGCCCTTGTGCCATTGAATTGGGTTTGCCAAATC
ACCACTTTCGCTATCTTTCATTTTGAGAGAAAATAGAAACATTTCTAGTCTTTACCAAGGTTACCAAAATCAGGT
GTTTGCTCCCCCTCTTTTGCTGTAGTGTGAGAAATAATCCCGCCTAAAACAAGTGAATCTATCTTACAACTAACA
AGTCACACACGTCATCTTTCTTAAAAAAAGGGAAATAAAGAGGACAACAAATCGGGTGAGGATCCTCTACCATAA
TATCACTATACCATTACGGTCACTTATATGTGAGTTCCGCTTTCTTAGTGTTACACATATCAATGTCTATAATGG
TAGTGTCATTATGGTAGAGGGTGTAAACCCCAACAAATCATTGTCGCGAACGCAGCTATTAGCAACTAGAATTAC
CCTATGTTGCCTTCCTCAAAGACTAGCATTGATTACCTTAATCGACTTTCTTGTCCCCGCATGGAGATTATATGC
TTACTGCAACTGTGCAATAAGGATCGAAGCAGTCTAGGGCAGATTCGCAACCCCAGATTTTGATTCGATGATCTA
GATTATGTATTAAAGAACATTTGTGTAAACCTAGATTTCTCATCATGCATTGCTAATGCAACTGCTAAGTTGTAG
TTATACTAGCAACCACACCAGATCTTGGTCCCCAGTCCCCAAGCCATAAAAACTACCACACATGTAAAGGAATAT
CTTACCTTCTAAATtactacctccatattttaatgtatgacgccattgacgttttgttcaacgtttaatcattcg
tcttatttaaaaaatttatgtaattatcatttattttattatgacctgattcgttatcaaatgttctttaagcat
gacataaatatttcatatttacataaaagttttgaataaaataaatggtcaaacattggtcgaaaagtcaacgg
tgtcgtttaaaatacggagggagtaCTTTTTATCATTTAAAATTTGAGGATTTGTCCAAACGTCGAAATTGGATA
ATAATGGGTGACAATCAACTTTGCCATTTTGTCTTTCCAAAAAATTTTCCGTTAAAGTACGGCTGAACcatttaa
gccctctcgctttggttcttttgaatatttttattgtttaagccgctgtttttttaaaccttaagtagtagactgt
tagcttcacatagaccagaaaagctactgtaacctagcttttagcttaacatagagtgacttatgaccttaaaaa
atacaaacgaaaaaatatctatttatttaggtttaaacATAAACAAAAGAGGGCCTTAATCAACAAACAGCCGGA
TGCTCATGCTCATGTGTCGTTGAGAGGATCCCACTCGATAATCAATTGGCTTAATAAACTCAGCTAGAGGGGTAA
AATATCGATCAGTAGATCACACAGATAATAATCCCTTTTATCAGAAGCTGATTTCCTATAAGAAAAATGTAGAGC
AGCCGCATCCACGATGCGCAAAACAACCAAAAGGGCAATATTCAATC**CACCGAACACCCTTCAGGATAATTCGTA**
**TACAAAACTTGCAGGTACACAACTAATTATGGTCCAATTCCTAAGCTGCGTAAACGGTGGAAATGTACATACAAA**
**TTGTCAATCGAACAGTCCTCATGCCATCGAAATGCTTCTGAATTTAGTTAAAATAAAATGCTTCTGAAGAGAACC**
**ATCCCATCACAGCTCGCATACAGAACACC**CTGCTTGAATTCAGAGGATAAAAACATCATTATAACAAGAAAACTA
ACAAAGAATCACAAGGGGCACGTAC**CTGTTAAATCTGTGTTAAATCCTCTTAAAAGGATCAGTACTGCATCTTTA**
**AGTTCACATTGAACCTTTTGCTTCGGTAACAGCTGACAGATCTGTACAGATTTCAGACTAACTGCTTATCTTTCA**
**CCGCAATAAAGCTTGTACCAGGATGATATAATCATCACTTACCTGACTTGTGCACTCTGCAGATACTTTGTAGTT**
**AATCTAGCTTGGTGGGACTTGGGAGCATTTTGTTTAGACAAACAAGTTTTTTATCTGACTTGTGCACTCTGCAGA**
**TACTTTGTAGTTAATCTAGCTTGGTGGGACTTGGAAGCATTTTGCTTAGACAAACAAGTTTTTTACCTGACTTGT**
**GCACTCTGCAGATACTTTGTAGTTAATCTAGCTTTGGTGGGACTTGGAAGCATTTTGCTTAGACAAACAAGCTTT**
**TTACAATAGGGGGTAACACCCCGCAAAACGAGGACTGATAATGGAGTACCAAATTCAATAGAGTTGGTAGCTCAT**
**CGTCTTATCTGGAAAAAAAACTGTCTTCAAACTGGAAAGATAATACTACTTCCCTTATTTGTTTACAACAAATTT**
**CTTTGTTTGCCTCTTGTTACTTCTTTTACCATACACAATAACTTCCTCAGTTCAAATAGTCTTCTTGTCTAATGT**
**AATTGAGTTTCCTTTTTTTTAAAAAAAAACTAGTGAGTTGATAACCCAGCATGTTTCTATGGGTATACACAACAT**
**ATTTATATTGCTGCTTTTGGCAAACCTGGATTCTGCATTTCTTGACTCTGTTTTCATATCCAATAGTTCTGGCAA**
**AGGGAGGCAGCAAGCTTGAGACAACAACTGCACAGCTTGCAAGAAAATCATCGGTATGGCCATAAAACC**CTGCAA
GTTTTTCAATGTTCCATCAGAAAATTCCTCAAGATAGAAAAACATAAACCATCATATATCTACTACTATGAAATA
CATGCATCATGAACCCTTGATGATGAACTTTTTTTTTTTGTGGGGACTTGCTCATGAATCCAGCAACTGCTAGCG
ACAATAATACTTCCAGCACCTGATTGTATAGTAAATATAAACTAGTACCAGATCATAAAAACAAGCACCTAGTGC
TTGGGTATATGATACTTAGAAAAAAGAAGAAAAAACAGAGTATGAAAGTGATATATGCTGATTCCAAATTTCCTA
TAGATAATGCATTGTTTAACCTTCACAAACAAAAGAAATGAAGAAGGCACGTGAATTTTCTATATCATTTAAAAG
TAAGAAATATATACCTTCAGCCATTCGATTTTTATGCATTTCCGGTTATCTCTAAAAGAAAAACAGAACTGAGCA
CCCATCAAAGGCGGACAGACCAAATTTACAAAGGAGAGTGAATATCACATTTTCTTCACCTGTGCTTTCTGCCTG
GTAAATTATTGTTCAAACACTCCATTTTGTCCTTCAGGCCTAATTTACAATACCTAAACTAGAAAGGTACAAATT
ATTTTTATTATCAATTAACAGCTGTCTGTTGATCAGCTAAAATTAAGTGTCAATATTCACTCTGTATATCATCAG

AACAACACTATAACATCCATTGGTTGATGGCAGGCAGTTGATGGGGCAAGATCTTTCTGGATTGGGTGTCAAGGA
ACTGCAAACTCTAGAAAATCAGCTAGAAATGAGCATACGCTGCATCCGGACAAAAAAGGTCAACCTACTAACCAT
ATTTTTTTGTTGGGCACCTGATTGATTATATAATGGATTAACTTAAGATCAATATTGTTGTTTTCAGGACCAGCT
CATGATTGATGAAATCCACGAACTGAATCGAAAGGTTCTACAACATCATGTACAATTGCTAAATCTTTGAATCAG
CTAGTAAGCTAACTCTATTGCTGGCTTATTTGCAGGGAAGTCTCATCCACCAAGAAAACATGGAACTGTACAGAA
AGGTCAACCTGATTCGCCAAGAAAATGCTGAGCTGTACAAGAAGGTACTTGATAAATGCAGATAGATGATTTCAC
AGGATATACGCTTCATTATATGATGACAGCTAATCTCTCTCCTCAGCTCTATGAGACAGGGGCAGAAAATGAAGC
GAATCGAGATTCAACAACTCCATACAACTTTGCGGTTATCGAGGAAGCCAACACTCCTGCTCGTCTTGAACTCAA
TCCCCCAAGCCAACAAAATGATGCTGAGCAAACCACACCTCCTAAACTAGGGTAACTCAGATTTTCTGCACATGA
TCCATCTATCCATTCTCAGTTCTTTAGTGAAACATCAATTATAACTTAATGAAATTATTGCAGGCTCCAATTGCA
TCCATGACAATTTATGGTGTGGAAGTGTTGCCTATATTTTCACTATTCTTAAGGGCTACAGTTCAGAGCTCTTAA
AATGTATTGGTCATAAAGAAGCTAACACATAACAAGACACAGTTTGCATGCTTGGATGATTAATATCTGATACTG
TAAAATCTATTCGAATAAAAATGTATTATCTGAAGAGCCCATGAAAATTATTTTCTTATGACTATTTGGCACTAT
CTGCAAGTTTGTTTACCCAAATGACAGCTACATGTGTGCAAACTTATTGGCAAAAACAAACAGTGCGCGAAGTGT
AGAGCATGCAAGATCAATGCTCTGTTTTATCTTAAAAAGCCATAAGCAGACAGAAGAGATGAAAAGGCATTATAA
TTTATCTGTGCAGATGTACCAGAACAAGAAAAGTATCAGCAAGAGCGACGCCAGGGGGCGGAGGGGCCTGGCCTC
TTCCCACCCCCTCCACCCTTCTATTCTGCTGATGGATTAGCTAGGCACTGTGCTTGCACGTATGTATGTTTAGCA
AGAGCGATAAGGTCTTTACTCTAGCCTTGTAGGCACACAACACCATCTATTTCCTTTTCCTTTTAGAATACCGAG
TAATATGTACTAATCATTTGTATTATTTACATCACTAGGTGGCTAACATTGAAATTTTAAATTTCTAAAATATTA
GTTAATACTTGTAACAGATGGATAGTTTGTTGCTCTTTTTGCCATAATCTAGTTCTTATTTTTTGTTTATTTTTT
TCCTGTTTACCTACACTTTTAGCTTGAATGGTACAATCACTGCTTATAATCTGTCTGCTCATAAGGCTAAATTAA
CCTGCCTGTGTTTTAAATTGTCGTGTGTTTTTTTATACTGATACGGTATATTACCTATGGGGAACATTTAGATTA
GCATAAACTTTTCTACGCTATTTTTTTTATAATCTTAATCACTTATAAAGCTGGGCCCCCTCATACTTAGATCCT
GGCTTCGCCACTGGGTATCAGCGTTCAATCTTTCTTAATAATTTTAGACACATGTGCTCATTAAAACTTTTCTTA
CTGTTCTAAGAAATGCTCGCAAATAGTACCTTTAGCAAAACTTTATACACATAAAATAATCAGAAGCACATCAAT
GTAATATGTCCAGAAAGTTTATAGCGAAAGAATCATTTTGTCCAGGACCCAAGAACTTGGACCTTACACCTCTAA
TTGTGCCTCTTAAATACCCATGCACTTTCATGTTACTAACTTATACCACCGGATAGAATGATTATTAGTTATGTA
TATCAGAAAAACTATTTTATACTAGGCACGAGACCTTGGGTAATAATTACATCATAACTTGTTTCAAATAATTCT
TGGAATTTGTAATGTGTACTTCACTAGGTTGGGAGCGCCTAATCCCCCGAGATATGAAAATGAAATAATGCATGC
AAAGACATGGAAAATTtcaaaaataaagtaacattcatgaacagttccaataataatattaacatactacctcca
tcccaaaatataacttttggctatgaatctggacacacagttgtccagattcatagctaaaaatgcttatatttt
gggacggagggagtaGAGTATGATGGCATCTACAAGGAGACAAGTGTCAAAGGGAGTGAATTAATCAAGTTTTAA
TAGTGTGTGGGGTAGAGAAAAGAATGTTAGATACTTAGAACATAATGGATGGTAATCCAGTTTGTTGAATTTTTA
AAGTTCTATGCAAACATAAATGCAGTATAGAAAGTAAAGACAAACGGTAAATACCAATTATGTACAAGACATAAT
GTAAATTAGATACTTAGAACATAGTAGTTTTATATAGAACGGTCCATGTGTTCATTCATATACCTTGATAGGTTG
AAGGTAGATCCTTCTGAAGATTACACCAAGGACTTGGTACAACACGTTCGTAATTATACAGGTACATGACTCTCA
CAAGGACTCTTACATAGACTCTCACACTTAGAATTTCATAAGTCTTTTATTTAGGTTATAACTATAACTATCTAC
TAACACTTAGATGGAGTGAAGTGTGATGGCTGGTGGAGAGGTGAAGAGGAATGGTTGAGTGAGATGTGTTGTGGT
TGTGTTCCTGAGCCCCTTATATAGGACTCGGTGAAGAATCTTTATGATATCTCCATTTGACTTTAAGGGGTTTTA
TGATTATCCCTTTAGACTTCACGGAGAGAATCTTTGGTGATGACGCTTTTGACCTTCAATAAAAGTATCATAATA
TCGCCTTTTGACTATAGGCATATCTTATCTTGAATAGTGTAGATTCCTTATCTTCAATAGTGTTTTTTGTCGAGG
ATAACAAGATCTTCATTCATTATCTTCACCGTATAAGTCTTCTACAGACATCTTTTACAACGTTCCTAAATCAGA
TTGAAGAATCTGATCTAGTGTTGCATAGATAGAATTGACAAGACCACGATCTATAATGGTTTGGAGAGGTTTGGA
AAGATATTTCATCTTGGTTGTtgtttacacccaaatttggcacctagggtcgaaataggaaaaattgcgaaagtt
tggggctctgccggtttcctccagaaagggccggtctgaccgccatatgtgggccggtcagaccgccgccggaga
gccggtcagaccggtggcatgtggccggtctgaccggtagagtctgagttagactccgttttcgtcaggtctcgg
gtggttggcgactccatatcaccgcaaggcgtttaggtgctgcgatcgtgcttgttgacttgtcacaaaaagttg
cTCTTGCCTTTGATACCAGCTCTTGCCCAAACACCATATATATTGCATAATCATCCCAATTCCCTCCTTCCACAA
TATATACTTCGGACCCAGCCAATAGCACCATGTAGACCGTGTGCCTACAAAGATTCTTACCAAAAGAAACTTAAA
GGCCAAAATCCCATGAGTACCGCATATGAATATCTCCAACCCAGATTGAGCTTCAAGACAGAGCAAATTTTATCT
TGCTTTGGGCAAGACATCCTCGACCTTGTTTGGCAAAAGTTTAAAGAGAGACAAATGAAAATATTTCAAGATTAT
TTCTTTCATCTCTTATAAGGCAGAGAAAGAAACCCAGCCATACAAGTAATCATATACATGCTTCCTATCTTAAGG
TTAGATGATCATCAGGTAATCTCTGACCCATCCTATAGATTCATGGTTCTCAAAGCCGAAATAAATCTCCAAAGA
TTCCGAAACTTACCCACATACAACCATGATGAAATATCTCTCCAAACCATTATAGATCATGGTCTTGTCAATTCT
ATCTATGCAACACTAGATCGGATTCTTCAATTTGATTTAGGAAGCGCTGTAAAAGATGTCTGTAGAAGACTTAGA
AATGGAAGATACAGGATTATCTTCTCCTCCATACCACCTAAATTCACACCACATGCTCGTCCAGCAATTCACtat
atatatatatatatatatatatatatatatatatatatatatatatatatatatatatatCATGAATAAACAATTTA
ATTTTCAAGAAGGCGGTCCTAGCAAATTATCAGATGAACAAGAAGAAATGTATACAATGATTGCAAATCATGAGA
ATTGGAGATTATTTTCTGAAGCTGCAGAAATTGAAGAAGCTATTACCATAGATTATGAATATCAGTTGGTTTTCC
AAAACAAGATCACAAGGATTTTTATTAGCAGATATTATAACCAAAGTTGTGAATATTTCAAGGGAGCAGGACGTA
TAATTAAACCAGATTTTACTACAGAAGCCTGGCTCAGTGGTATCAGAAAACAGAGCCATCAAAAATTGACATGAT
GCAATTGGCACCCATGATATACAGTGAAGATAATGAATGAAGATCTTGTCATCCCCGACAAAAAACACTATTGAA
GATAAGGAATCTACACTATTCAAGATAAGATATGCCTATAGTCAAAAGGCGATATTATGATACTTTTATTGAAGG

TCAAAAGCGTCATCACCAAAGATTCTCTCCGTGAAATCTAAAGGGACATCATAAAACCCCTTAAAGTCAAATGGA
TATATCATAAAGATTCTTCACCAAGTCCTATATAAGGGGCTCAGGAACACAACACAACCAAAACACATCTCACTC
AACCATTCCTCTCACCTCTCCACCAGCCATCACACTTCACTCCACTCCATCTAAGTGTTAGTAGTATAGTTATAA
CCTAAATAGAAGACTTATGAAAGTCAAGTGTGAGAGTCTATGTAAGAGTCCTTGTGAGAGTCCTGTACATGTATA
ATTATGAACGTGTTGTACCAAGTCCTTGGTGTAATCTTCAGAAGGATCTATCTTCAACCTATCAAGGTATATGAA
TGAACACATGGACCATTCTATATAAAACTACCATGTTATGGTATTTACAGTTTGTCTTTACTTTATGTACTGCAT
TTATGTTTGCATAATTATTTTTTAAGAAAATCAACAAACTGGATTACCATCCAGTTGCCACATACCAATCTCACA
TCCAACATTGCATCTAGATGCTGCATTCCTATCCTCATATCTGGTGACTAGACCCCTAGCACTGCCTGGGCAAGA
GCTGGTATCAGAGCCAAGTGCTCGTAACCTCCCCAAGCCATCTTCTCCGAAAgggggagggggagggggggag
agaggggTCGGGTAGCAAATAGTACTTTTAGCAAAACTTTATACACACAAAATAATCAGAAGCACATCAATGTAA
TATGTCCAGAAAGTTTATAGCAAAAGAATAATTTTGTCTAGGACCCAAGACCTTGGACCTTACACCTCTAATTGT
GCCTCTTAAATACCCATGCACTTTCATGTTACTAACTTGTACCACCGGATAGAATGATTATTAGTTATGTATATC
AGAAAAACTATTTTATACTAATCACGAGACCTTGGGTGATAATTACATCATAACTTGTTTCAAATAATTCTTGGA
ATATGTAATGTGTACTTCACTAGGTTGGGAGCGCCTAATCCCCCGAGATATGAAAATGAAATAATGCATGCAAAG
ACATGGAAAATTTCAAAAATAAAGTAAAATTCATGAACAGTTCCAATAATAATATTAACAtactacctctgtccc
aaaatataacttttggctatgaatctggacacacggttgtccaaattcatagctaaaaatgcttatattttggga
cggagtgagtaGAGTATGATGGCATCTACAAGGAGACAAGTGTCAAAGGGAGTGAATTAATCATGTTTTAATAGT
GTGTGGGGTAGAGAAGATAATGTTTTGGTGATTAACTACACTAGCAAAATAATTTAGCGATAGTAATAATAAGAC
TTAC**GATTTTCTTGCAAGATATGCAGTTGCTGTCTCTAGCTTGCCGCCTCCACTTGCAGCAA**CTAGCAGAGAAGG
CAAAGGAAGAATTAGTCAGCTGGATCAAAAGGGTCATATTTGAAATTAGTTGCTAGTGCTATTGGCAGTTAAAAA
AATTGCAAACATACAACAAAAAGTTCATTGGCATACTACACAAATATTTGCTTAACAAATTTAAATGTTGCAGTA
ATGCAGCCTTCCCCTTTCCCTTACTATGAACCGCGAGAGTTCGTGTTAAAAACAAGTTTATGTTTCAGTAAAAAA
CAAGTTGATGTTTGCTCAAAATGCAGAACTAGAAACCTTCAGTTGTAGCTCAAATGGCATGGAAAACTATTTCCA
TACAAATTGTTCTACAAACTAGCTTTTGCATTGCATGTGTTTCTAGTAGCGGTGCAGTGCTACTACAAAGTACAA
ACTGTCCTGAATAATAATAATCATCAGGAAAATAGCATGGAACTAGCAAAGGTTTTGTATGGGAAACTTGGTTAA
TACAATTGTATATTCAGTATTTCTTCACTGAATTAGAAACTTTAGTAAAGTGGTAAATGCTCAAAAGCAGAGGGC
ATACAATGTGTTCATCAAAGCATACAAAAGCTCATATCAGCGAAGAAAGAAAAACTAAAAATCAGAGATATGAAG
AATCAACTGCTGTTCTCAGCTAAATTCACCCCATAGTACCAGTAGGTCAACCCAcccccccccccaaaaaaaaa
aaaaaCTAATCTATTTGGCACTGGTAGGACATGCTCAAAACAACTATATTCATAATTTAGTGACTAATTTGACTG
GCCCATCAAACATTAAAAGACAATCATGATACAGGGCTGTAAAAATAGAAGGCCTGATCATGTTGCAAGACATGA
GTTCACAACCAAACAGACACTACAAAAACTAGTCAACTCAAATGCTCTGTTTCAAGAACTGCATAATGCAAGTGC
TGGTCCAAACTCCAAAACTGTGAAGATAACTTTGACATCAGATAAACAACACATTGGGTATGCAACTTAGTATCT
CTTTTGTTTTATTATCCTGCCAATAGCAAGGATGATATTTATATGATGAGACATGTATTAACACTAACCAGACTC
CATCACACTAGTGTCATGCTCATGTCAAAAAAGAATCCAATAATAGTATAGCGCTAATGGCAGAACAATCTGCCA
TGAAATTTTCATTAAGCTCAATTTGTGCAAGGAGAAAAGGTTGATATATAACACAGTCACAAAAATACAACCTTG
AGCACAAAATATTTTTCAACGTGTCTGTAAAAACTAAATGTTTCCTTGTGCTGACAGCAGTGGGGATAGGAACAT
GAAAGTCTTTTGGCAATGGGCTATTGAGAATGGAGTATGTTTGGGCCAAGGCCCATTTAGTGGTTGTTATAATCT
GAGGTGTTAAGATGAGGACCATACAAAAACTGAGCACTGCTTTGTCCAAAAGAGTAGATGAAATAAAATAGAGTT
ATTTCTGTATACTTTGCTTCTCTTCTATCTTCTCAACTCCATTCTGTATTCACATTGGTATTGTTGCATTGCGAT
TGGGCTCATCAGTCATCACCTCTTGGGACGTGACACAAAAATGCGGCAAAGAGATCTTTTAGACAATCTACTACT
AGTATCCCCACATCACACTCATACATCCAACCTAAAATTCACAAAGTTGTTGGTTATGTTTCAAATTGTTCACCA
TAATTACCATCCATCACATTCTTTTATTACTGGAAAGGTATTAGTGTTACGGAATCCTTTCCAAGTTCCAACAGT
GAAAACATTTTGGAAATAATAATGAAGCAACAAGTCTAAGCATGAAAACCATATGCTTTATATTTTTACTGAATT
AATTCATCAATTCTCAAATGGTCAGCCCACATGCCAATTTGACCTCCAGCATTGGCAGAATATTTTGCATTTTTA
ACTATTTATTCATGCACCTCTATGAAACTATTGCATGCCTATCAACAAAAGTAAACTACCATAAGATATACTGCA
TAAACTCTTGAGTTCTACAAAACCACCAAATAGACATTCTAACATGCAGACTGAAACTCAGTAAATGCATCTGAA
GAAATGTAACTACAGCCAGGAAGTTTCAGAATAAAACATTCTCTTGAATGAATTGTACTATATTATGTCATATAG
AAGGGAGTGTTAAACTTTAGCAGCATGCAAGAGTATAACTTTAGCATACTCATATGTCCGTTACGCATAATGAAC
ACATACATCCGACATATACACAGATCAGATAAACTGTGTATACATCCTCAGTTAGAAAACATATGTTACCACACT
TCACAAGCCAAAACAATTATAAATTAGTTCAAACTTAAGCTTTCATTATATCAGAGTGGATGTGCAGTACATGAC
TACATGTAGGGCGCTGACACTAAAGTCTAGAAAGAAGTACTAGAGGCAATCTCGTTAGAGACCAATACAAGAGTT
ATCCACTTATCCTAACCACGGTTAAAAGCAGCAAAATCCTCCCAAAACAAAACTAAGTAATGGCAATTCTAGTAT
CAGTGCAACTCTAAAAAAACAATTAATAATGCACTCCCTGATCAAGAAAGAAAAGAACTAAGCTAACATTAACAC
TGGCATTGACAACCGTACATGAAACAAAACGCATAAGCTGCTTGTAATTTCATGCTACAACGAAACTAATCAAAC
TTTTCACACAAATCACACCAAAACGACCGCACAGCTCGGGGCATTCAGTTAACTCAGTATCAGCAATTAACAACT
TAACTATGCAAAAAAAAATACATATATATGAGAGCCTGAGGAGAAGCTGTAAAATCGGTGGAACGTCTCAC**CTGA**
**GAAGCCGGCTCGAAGGTTGAACCCTAGATGAAGAGAGGCGGGTCGCTAGGCGGCACAGAATCAAAATCAACGCAA**
**GCATATCGGCTGGGGCTACAGCCTACAGGAGCGGCAATGGGAGAAGCTAATCGGGGAGCCAACGGCCACCACGAT**
**CCGAGGGTCGTGGTCATGCGACGGGCGGC**AGCAAGCTAGATAGCTACCTCCTACGGAAATCTAGGGGGGGGGGGG
GCAGATGCCATGGAGGACGAGACCGCGATTGGCGCGATCCTATCCAATCTAAGCGAGGGGGGGCTCCGAGAGAAC
GACGCGAAGACGACGACCGCCGCGAGGGGGGGGAGGCGATGATGGATTTTGCGCTGGATTTGGTCGGATGCGACG
GAGCGTGTGGTGGTGGTGGTGCTCTTTGTAGATGAAGAGAGAGAGAGATAACCCCCAAAGCAAAGCAGGCGCCCC
CACGCCAAGTGGGGTGGGGTTTCCGGGGGAGTAATAAAAAATGCGGCGGACGAGCGGCGCTCCGGCGGCCACGTC

GCCCCCGGCGCCATTTTCAGAGGGGGGGCGCTGCTGCGGGGCACCCCGACAGGTGTACCGTCCGCCCACCCGCCA
CCGCCGATCGCCGGCAAGTGGGCGCGTGGGTTGGGCCTGCGCCTCCTGCATGCTCTGCCTGTCGAGAAGGCTCGC

>OsMADS61

ATGGGGAGGGGCAAGATAGTGATCCGGCGGATCGACAACTCGACGAGCCGGCAGGTGACGTTCTCGAAGCGGCGC
AACGGGATCTTCAAGAAGGCCAAGGAGCTGGCCATCCTGTGCGACGCCGAGGTCGGCCTCGTCATCTTCTCCAGC
ACCGGCCGCCTCTACGAGTATGCCAGCACCAGCATGAAGTCAGTGATTGATCGATATGGGCGAGCTAAGGAGGAG
CAGCAGCACGTCGCAAACCCCAACTCGGAGCTGAAGTTCTGGCAAAGGGAGGCAGCAAGCTTGAGACAACAACTG
CACAGCTTGCAAGAAAATCATCGGCAGTTGATGGGGCAAGATCTTTCTGGATTGGGTGTCAAGGAACTGCAAACT
CTAGAAAATCAGCTAGAAATGAGCATACGCTGCATCCGGACAAAAAAGGACCAGCTCATGATTGATGAAATCCAC
GAACTGAATCGAAAGGGAAGTCTCATCCACCAAGAAAACATGGAACTGTACAGAAAGGTCAACCTGATTCGCCAA
GAAAATGCTGAGCTGTACAAGAAGCTCTATGAGACAGGGGCAGAAAATGAAGCGAATCGAGATTCAACAACTCCA
TACAACTTTGCGGTTATCGAGGAAGCCAACACTCCTGCTCGTCTTGAACTCAATCCCCCAAGCCAACAAAATGAT
GCTGAGCAAACCACACCTCCTAAACTAGGGCTCCAATTGCATCCATGA

>osa-MIR444c_gi|297602864|

GCCGCCCGTCGCATGACCACGACCCTCGGATCGTGGTGGCCGTTGGCTCCCCGATTAGCTTCTCCCATTGCCGCT
CCTGTAGGCTGTAGCCCCAGCCGATATGCTTGCGTTGATTTTGATTCTGTGCCGCCTAGCGACCCGCCTCTCTTC
ATCTAGGGTTCAACCTTCGAGCCGGCTTCTCAGTTGCTGCAAGTGGAGGCGGCAAGCTAGAGACAGCAACTGCAT
ATCTTGCAAGAAAATCGGTTTTATGGCCATACCGATGATTTTCTTGCAAGCTGTGCAGTTGTTGTCTCAAGCTTG
CTGCCTCCCTTTGCCAGAACTATTGGATATGAAAACAGAGTCAAGAAATGCAGAATCCAGGTTTGCCAAAAGCAG
CAATATAAATATGTTGTGTATACCCATAGAAACATGCTGGGTTATCAACTCACTAGTTTTTTTTTAAAAAAAAGG
AAACTCAATTACATTAGACAAGAAGACTATTTGAACTGAGGAAGTTATTGTGTATGGTAAAAGAAGTAACAAGAG
GCAAACAAAGAAATTTGTTGTAAACAAATAAGGGAAGTAGTATTATCTTTCCAGTTTGAAGACAGTTTTTTTTCC
AGATAAGACGATGAGCTACCAACTCTATTGAATTTGGTACTCCATTATCAGTCCTCGTTTTGCGGGGTGTTACCC
CCTATTGTAAAAAGCTTGTTTGTCTAAGCAAAATGCTTCCAAGTCCCACCAAAGCTAGATTAACTACAAAGTATC
TGCAGAGTGCACAAGTCAGGTAAAAAACTTGTTTGTCTAAGCAAAATGCTTCCAAGTCCCACCAAGCTAGATTAA
CTACAAAGTATCTGCAGAGTGCACAAGTCAGATAAAAAACTTGTTTGTCTAAACAAAATGCTCCCAAGTCCCACC
AAGCTAGATTAACTACAAAGTATCTGCAGAGTGCACAAGTCAGGTAAGTGATGATTATATCATCCTGGTACAAGC
TTTATTGCGGTGAAAGATAAGCAGTTAGTCTGAAATCTGTACAGATCTGTCAGCTGTTACCGAAGCAAAAGGTTC
AATGTGAACTTAAAGATGCAGTACTGATCCTTTTAAGAGGATTTAACACAGATTTAACAGGGTGTTCTGTATGCG
AGCTGTGATGGGATGGTTCTCTTCAGAAGCATTTTATTTTAACTAAATTCAGAAGCATTTCGATGGCATGAGGAC
TGTTCGATTGACAATTTGTATGTACATTTCCACCGTTTACGCAGCTTAGGAATTGGACCATAATTAGTTGTGTAC
CTGCAAGTTTTGTATACGAATTATCCTGAAGGGTGTTCGGTG

Stem-loop structure

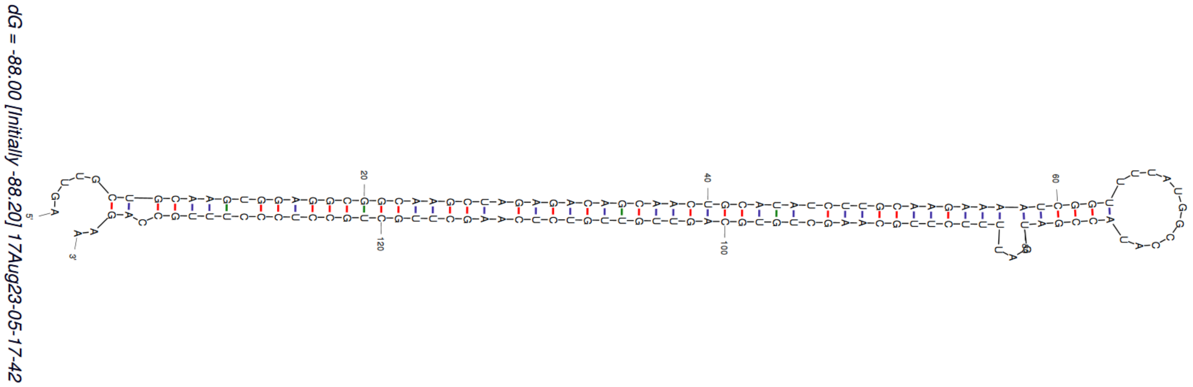


*OsMADS57 and osa-miR444d*

>Chr2

AAAATGAGATGTGTTGAGTCCTTTCCTATCACTTACATGACACTATACATATTAATTTTTAGCTTATATGCAATT
CTATAGCCAGTATTATGAATGGGGAGTGGTACCTATTTTGAGCAAACGATGAGGTGTACAATATCGTACTGGAAA
ATGTAAAATGTACTTATCATGAGCTGTAGCCGCACTCATAACCAATCAATAACTGGCACAAGGAAAGCCATGGAA
CATCTTTACATGGTCTTTCTTGCGGCATGGTTTTAAAGTAGCAAATTCGTGCTACCGTATATGAACATGTGACCG
TGACACAAAATCAAGAAATCCAGAATGAACATGGATCCTAAGCACATGACCTCACCGATCGATCACCACCAAAGC
CTTTTTCCATTTGTGTGGCAATAATAATTAAAGTTTTGTATAGTTTAGTACATGGCGCCGGCGGGTAATTAAGCT
GCAGCCACCTGCATCCGATCCATTCGTGCATATAACCTAACCACCCATTTCGCCTTCTCTCCACTCTCTCTCctc
tccctctctctctctctctgtctctctctctcGGGGTATTGCAAGATAGTAGCTAATTAACCAGCCTTTGGCCTT
CAGTTTCCCCATCGGTTCATCAGAGCACGCCTTCAATTTCCTAATCAATCTCCCATATAATTCCCCTTAGCTTGT

ATGCCTATCATTAGTACTTCTTGTGTTGTGTTGTCTTCTTGCCTTCTTTGAGCTAGAGCTAGCTAGGTTTGTGAC
CTGCTGTACGTGCGTGTGTTAATTAGATAGATGTCTTAGATCTGGGAGAGAGCGAGGAGATGGGGAGGGGGAAGA
TAGTGATAAGGAGGATAGACAACTCGACGAGCAGGCAGGTGACGTTCTCGAAGCGTCGGAACGGGCTTCTGAAGA
AGGCGAAGGAGCTATCCATCCTCTGCGATGCGGAGGTCGGCCTTGTCGTCTTCTCCAGCACCGGCAGGCTCTATG
AGTTCTCCAGCACCAAGTAAGCAATCCATCGGATTTATAAGTATATATGTATATTTCCCCTAATTTAATTGTAGT
TTCCTTAGTTGTTCTCGGATCTGCTTTGAAGTCTCCTTTTGTTCTTGATTTATTTTTGAAAGAACTCTCCTTGTT
CTTTCACTTGATAGGTATATATATCTGGTTCTGGTAATCTTTCATCCTTTTAAGTTTGAGACGTTTTTATCAACA
AAATGACCCAAGTATATGTCTGATCAGCTTTTCATCCTTTCAAGTTTTGAGGCATTTTTATCAACACAAAGACCC
AAGCTAAGCATATATGTATGCATGGCTATTAGTCATTCTTGTGATCTTTTAGTAAGCTATAGCTACATAGCAATC
AATATGTATGCATATAGTGTCAGTTTAATTGTGATATTTTCCCTCCGTTCTCGAAAACAATTGACACTTGGCACT
ACTAAGTTTTGATCACTCCTTTTCTTTCAAAAAATTTATAAACACTTAAAATATGTTATATTATTAAAGTATGTA
GCGTAGAAAATATAACCATACAACTTCCATTCATCATCATATAATTATTTAAAAGTAATTTGTGATCAAAGTAAC
AGCCGTAGTACTCAACCGTGTCAACCAAGTCAGTTATTCAAGAACTGAGGGAGTAAATATTGAATGCAAAATTAA
TATGAAAATTTTTTCTTCTGCTATATATCCTTTTTCAGCGTACGTATGTTCACTCTTCTGAATTGATTACAACTT
AGAAACTCAAGGTGGATATATAGCGAAGAAATATTAACATGTTAACGTTTAATAAAATTAGATGAATTTGTTCTC
TTTTTTTTTGTTTCAAGTGATGTTGATTTGTACCAAAATCAGATGGCATGTGTCACGGTCAAACTTATTAATTAA
AGAGATTTAGCATGCAATCTTGACAATCAAATCTTTACAGAGAATGGCATTTCTATATATGAAGCCTTACATGCA
TGTGCATAAATATTTTTTTGGAGCTAGAGTGTGGTAGCTTTACGTGCCACTGCATCAAAGACAGATAGATATTTC
CAGATTTTTAAATATCTATTTATCTGTGTGTTTTGTCTAAGTAAATTGAAGGTTTTTTAATATAAGCTGTATGTT
TTAGCTAAAAATGAAACAACTGACTCAACCATTTGGCTTTACTTGCATGCTAGCATGAAAACTGTGATAGACCGG
TATACCAACGCAAAGGAGGAGCTACTTGGCGGGAATGCAACTTCAGAAATTAAGGTATGGTTAAGAAGATAAGCT
ATCTTGTTAATTGTTACTATACGAGAACACGCATGTGATCTTGATCTGACGACCAATGACAGCTTTTAGTCTCTT
TAAGATTGATTTTATAAATAAGATGCTGATTGGCCAACTAGTTAAAGTGCTGAATGCACTGTGTTCTCATGCATG
CATGCAGCTGGGCATGGTGCACTGCTCACGCTGAAGAATATAGTGAGAAATTATTTCTGAACCCCTCTGGCTGAT
GCAGGCATGCAGCAGGGGTTGTTACTGAATTACATATACATTGTTTACTGTATATATAGACGCAAAcctttttat
aaaaaaaattcatgagaattttagatgttatcatttctaggagtttttttttctatatagtccttttaaatca
aaataatgaattatatgaaatcttacaaaattcctatggaatgtctttttctcgtataagttttggaggaattct
tgtgtttttccaatggcataatcaaacggtcatttatatatttttttcatgtgttttgtaatcatctgttttaca
cttagacttgcatttctgtgattcaaagcgccctcaacattttcctatcttctccTTCCAGATGCCATATATAAA
AAACTGATGCGTGTTGTGCCTAATGCTGATAGTACCAACTCCTATAACTGGAGAACTCCAGCAAACTCCCCTCAA
AGGAGCTGAATATGCGCACAATATATGCGCACAATCCGATAGTACAGAATCTTCTGAACTTGAGGTTGGCAAGAG
CTGCAAGCCACGCATGGGCCAAAGGGGGCAGATTGCTGCAGGTTCCTAGATTAGGACACTGGGCCCAACCATAAT
CTAACTAAGCCCCTGCAAACTCCTCAATTATCAAGCGTGTTCTGGAGCAGGAAGAAGCACTAGTCGATCTCCCTC
GGTTAATGCATCAACAGTAATGCATAGGACGAAAGTAACCGAGCAATTCG**TAATCTCAACATAAGCAAAAACAAG**
**CATACTTTAAACAAGATAAAACAGTGTACAATACAAGAGCTTGCAGAGTATTGTATAGGCAGGAGAAGCATTTGA**
**TGGTAGTGCAATATAATGCTGTACAAGTTGCTTGAAAATAACATGCTCAAGTCTTAATTTCTCAGTAATTAACTT**
**TCTTCATAGTGGCCACACTGGTACTCACTCCCGTTACTCCACTACTTAGCTTCTCACGTTTTCATGAACCACACA**
**CTTCATCAGCATCTGAGGGCATCAGGGCAGGCCATCCTCAAGTTTGCAAATGATGGAAAATCCATGCACATC**CTG
ACATCGAACGTTCAGCCAAAGAAGCGTATGAAGATCATAAGGCGATTCGCGAAAAAAAAAAGGGGCAGATTATAG
GTGCATACGTAC**CTTTTGGCACGATAAGGACTTCGCTGATTGCATTGGTCCAAGATACAGCGCAGTAATTTCGCT**
**GCATCTTATGACAACCACAAACTCAGAAAACAACACCCAACACATACTGAACCTGTCTGAGTATCTAACATCCAT**
**ATTTCAGAGGGGGGCAAACTTACTCGGTATATTTATGCATTATGTGGCAACTTGGAGGTTAAAAACAGTTGGAGA**
**CTCGAAAGGAAATCTAAAAACTGTTAGAAAATGGTTTTCTAAATAACAGATGCCAATAGCGTAAGCAGAATGGCT**
**GTAAAGCAACAAAGCCAGAACAACAAGGATACATAGCAAACCAAGTGATATTAACAGTTAAAAGTACCGTTGTTG**
**CTCAATTGACCAGTAAACTCGACAGACAATTGTAAACATAAAAAATATATACCTTAAGAAACAAACCGTCTACGC**
**ATAAAAATTTCACTGCTGCTGATGGTATGATATTCATGTTACTGAATCATGGGGGAATATGAAGTGATATTTTCA**
**GGAGTACTAGCTTATGTACTCCTTTATGAATTTCATTTCCATGTCATCATTTATTTAATGTTTATTCTTTCTATA**
**TGTTAAGGAAAAGGTCAAAACAAGCATCCACACCTACATGGTCTCAAAGAACTGTAGTACATAGCATGCCCCCAC**
**AGAGATCCTTCTCTTTCCCTCTATTTTGGGTTCAAAAATGCCATCGCTCATGAAAAATACAAAAAGATACCAAGC**
**ACCAAAAAGGGATAAATCTGATTTGCCCTGAGTTTACATTGGAGACTGTACTCATCATTACTACAGTATAACTGT**
**TATATCAACAAAATGGTTGAAAACAAAGTCCTGATTAAAGATACAATTATACATAAGTACCTCCTATGTCTATGC**
**GTAAGTTTTGTGCCAACAAAGTGATCAGCAGTTATGCATTATGCACTCCAATGGCCAGTCCATAATACAGATCAG**
**CAAGGTTATGTCTAGATTTTTTTTTTTCCTTAGAACACAGGGAAGTTCAGTGTGCAAATTTTTCAGCTAGGAGTTT**
**AGGACCAAAGGTTTAAAAGTCTAATGATGATGAAACAACCAAGCTACTGGAGCACTAGACAAAAACAAGTGTCAG**
**CTGAATTGACAAGTTGGTCCAAATTAATCTAGAATAATAAGTACTGAAGACTGAACAACTATAACACTCTGATGC**
**ACTTGTGCTACGCATATCCCACATCTTGAGATATTGGAACTATGTGAATCAGATAGCATTTTACCTCATCTCAAA**
**TGCATATGTGATTGAGTTGTGTGTGATTTTTGGCAACATACTGCGTTTCTAGATTAATCATATGTATATAGTTCA**
**AGGAAACTGACATGATTTTCTTTTTTTGCTGATAGATTTGGCAGAGGGAGGCAGCAAGCTTGAGGCAGCAACTGC**
**ACAACTTGCAAGAAAGCCACAAGTAATCATCTAATGATT**CTGCAATGGAAATTGATCATACATTAATATAAAATT
CATCAAGAAAGTAGTTTATCAACAATTTAAATGCCACATGGAATTGTATAAGAACCACCAAAAGATGCTTAAATA
CTTGAAGGAAAACTTTAGGAATATAATCAGAATATTCACTGTCAGATTTATCAAGAAAGTAGTTTATCAACAATT
TAAATGTCACATGGAATTGTATAAGAACCACCAAAAGATGCTTAAATACTTGAAGGAAAATTTTAGGAATATAAT
TAGAATATTCACTGTCAGATTTATTTTAATAGTATTCCTAAGTTTCTTTAACAAATAAGTTTGGCCTGTCTCAAA

CCATGTCGACAGAAGAGAGCTGTAGTGGCATATAGATTGGGACTTCTTGAACTGCTGAGATTACTCTTTTGCATG
GTTGTAGGCAACTGATGGGTGAGGAGCTTTCTGGCCTAGGTGTTAGAGACCTACAAGGTTTAGAGAATAGGCTTG
AAATAAGTCTACGTAATATCAGAATGAGAAAGGTTATATTCCTTGTGCTCTTCAATATCCGTATTTTTGCTTTTG
ATTTGCTATAGGCTAATGATGATACAGTAAACACCTATCAAACAGGACAATCTTTTGAAAAGTGAAATCGAGGAG
TTACATGTGAAGGTTTACCCCTTGCACTTAATGTCATTTCATTATTTCAATGCAAAAGAACATGCATGGAATATT
GTGAATTTTGTGTACTAAATATTAACTTTACTTCAGGGAAGCCTAATTCACCAGGAAAACATCGAACTTTCTAGA
AGCCTAAATGTCATGTCGCAACAAAAATTGGAACTGTATAACAAGGTTCGATGCAAAATTCTCTTAGAAACTTGT
GACTACATGTTAACACATTATCCCAAGCAACTAAAGGAACAACTTGATTTCAGCTTCAGGCCTGTGAACAGAGAG
GTGCCACAGATGCAAATGAAAGTTCCAGCACTCCATACAGCTTTCGTATCATACAAAATGCTAATATGCCTCCTA
GTCTTGAATTGAGCCAATCACAGCAAAGAGAAGGGGAGTGCAGCAAAACAGCTGCTCCAGAACTGGGGTAAGTAT
TTCTTATCTAGAATACATAAATTGCATTACTCAAAGAAGGGGAAGCTCAATCAAGCACGTATGTTTTCATATGTA
GCAGACTTCATCTGCCTTAAGACTATGCCGTACAAGCTGGACGATAAGTGATCAATTTCAAACAAGCACAAAGCC
TTTCAGATGCAAGTAGTATTTTTGTGATAGCTGAGACAATAAAAAAGTGTACAGACTACAGACTGGACAGATTAT
GTTGTCGGATGCTCATGTAATGACTGTGGTGTAATAAGAAATACGTAGTATAAGTTTGTAACCCTGCTTACTCTC
TATTGCTTTCTAGAAAAGCTCAACCAAAACTCTGAAAAGTCTTTGGACTGAAAAGGTTGTAACAATGTTTACCAG
GAAGGTGCTTTGGTGATGAAAAAAGAGTTGTCACTATGCCTAAGAAGGAAGGGGTGTGAGCTACTGGCGAAAACT
GTTATGTATTGCACATGGTGAATAATGGCTGTTGAACATGTCTACTCAGACTTAACAGTCAGCCCTTACTCATGG
CTGTTCAAGTGGTGTTACAGATGAAGGTGCATCCAACCCAATTCAAAGCAGCAACACTCATACTGGTGCTGCAAA
TACCGTGTATGCTAGAGAGAGGTGGTTGGATAGAAGGCCCATGTATCATGTCATGTCATTGATTCACAATTTCAC
GTGTCGCAACATTTATTATTAGTAGTATTCTACTGCAAGTCTGCAACTAAGAAATAAATCTTCTGTTAAACATGA
ACGGCTCCTGATGCAATACACTGGAGTTAACCACTAAAATCTCAGCAATTGAAAGTTCCACTTAGAAAAGCCCAT
ACCACATTAAAATGGTAATGCCTGTATTTAATTGATTTTGAATTAAAAGTGGTGCTATCAACTTTGAGTTCCAAA
ACTAATTATTTCCAAAGAAAAAGGTAATTCAATAATTACCATCACTTGGTATAGATTGGTCTTCTGTGACAACCA
CGGAAGAATTACACAATACATATAAGAATGAGCTACATGAGTTTACTGTCTTACTTGTGAGAGGTCTTAC**TTGTG**
**CCTTTCTTGCAAGTAATGCAGTTGTTGCCTCATGCTTGGTGCCACCATGTGCAATAA**CTAGTCAAGCAAAATATA
AAAAAATATATATATTTGAGAAATGCACATTGAGTGACCTACAAAGCAATAAGCACAGAATATAGCGTAGCTGCT
ACTTGAAAATTGAATCAGTCATAAACAAATAATTGCCTATGCTCGAATGGATTCTAGAATGTCGATAAACGGCAC
AATGCTAAATACTTTAATGGGTCATCAGCATGAGGCAAATTCAGCTAACAAAATGGAGCATGCAAACTGGCCAAT
CTCAGTTAGTTTATTTCTTTTCCCAGAATACAATACTGGATCAAATCAGATATTTGAGTACGAACTATGATTCCG
AAACAGATATCTGCAGTGCGGACTTACAGTTTGTAAGATCTGAAGGAATCTAGCAAATCATCTACAAATGTCGTT
AGAAACAACCCGAGGTTTCCAACCCCTTGAAGCCCGGAATCCCCCCTACTCCCCCATGATAATCCAAACGCGAAA
CCCTAGAAAACCAGACCCCCAGATCTCCCAACGAGCTCATGATCACCGTAACGCAAACCACGCCACACACGAAAT
CCAAACAAACGAAACCGAGATCTAGCTGCAGCAGAAGAAGCAGAATGGGGGGTCTAGAAACTTCTCTCCGTAC**CT**
**CGAATCCGCCGCGGGTTCCAGCCATGGTGCGGCAAGGGTGTGGGCGACTGCGACTGCGACCGCGAGCGATGAAGC**
**GACGGGTGGGAGGCCGGCGCACTTTAAAACGGGGGCGGAGAAGGGTCTAGAAGGATCGAATCGATGTTTCCAGAA**
**TTCCAAGTGGGGGGTAGAAAAGGCAGGAAAATTGGCGCAGTGGAGTTCGGCTTTTTTGGCGAAGGGGACGAGAGG**
**AGGGAGGGAACGGAGCGCGGCGAAAGGG**ATTAGGAGGGGAAGATTTGGGTTAAGCGGGGATTAGTTTAATTACGT
CGCGGGGAGAGGGTTGTGGCCGCGCAGGCGGCCACGTCGGCCCGGGCGGATTTTAAATGGAACAGGTGTCGGATG
CGTAGCGTGGAAAGAGAGATCTGTGCTCGTTAATCAATTCGGGTGTAACTTTTGTAATGTTGTCCTGGTGAAGGG
GACTCTGAACTGATGAGGGGAATAGAAAGTTCCAGAACATCTAAGAATGCGTGTAAAAAACAATGGATTCTACTC
TCGAAAGTTCGAGAACATATGGACATGAAATAATGTGGAATGACGAATCGTCGAATAAACTTGCCGCACTCGGGT
TTAGTCGAATCAGCTGCTCCAATAACAAGGCCCgtttaacttggtaaaatgaaaatttttggtgtcacatcgga
catttgatcggatgttggaagtgatttttggacacgaatgaaaaaactaatttcataactcgcttggaaactgtt
aaacaaatcttttgagcctaattaaaccgtcattagcacatgtgaattactgtagcactttatagctaatcatgg

>OsMADS57
ATGGGGAGGGGGAAGATAGTGATAAGGAGGATAGACAACTCGACGAGCAGGCAGGTGACGTTCTCGAAGCGTCGG
AACGGGCTTCTGAAGAAGGCGAAGGAGCTATCCATCCTCTGCGATGCGGAGGTCGGCCTTGTCGTCTTCTCCAGC
ACCGGCAGGCTCTATGAGTTCTCCAGCACCAACATGAAAACTGTGATAGACCGGTATACCAACGCAAAGGAGGAG
CTACTTGGCGGGAATGCAACTTCAGAAATTAAGATTTGGCAGAGGGAGGCAGCAAGCTTGAGGCAGCAACTGCAC
AACTTGCAAGAAAGCCACAAGCAACTGATGGGTGAGGAGCTTTCTGGCCTAGGTGTTAGAGACCTACAAGGTTTA
GAGAATAGGCTTGAAATAAGTCTACGTAATATCAGAATGAGAAAGGACAATCTTTTGAAAAGTGAAATCGAGGAG
TTACATGTGAAGGGAAGCCTAATTCACCAGGAAAACATCGAACTTTCTAGAAGCCTAAATGTCATGTCGCAACAA
AAATTGGAACTGTATAACAAGGCCTGTGAACAGAGAGGTGCCACAGATGCAAATGAAAGTTCCAGCACTCCATAC
AGCTTTCGTATCATACAAAATGCTAATATGCCTCCTAGTCTTGAATTGAGCCAATCACAGCAAAGAGAAGGGGAG
TGCAGCAAAACAGCTGCTCCAGAACTGGGGTAA

>osa-MIR444d_PUT-157a-Oryza_sativa-0160611
GAATCCCTTTCGCCGCGCTCCGTTCCCTCCCTCCTCTCGTCCCCTTCGCCAAAAAAGCCGAACTCCACTGCGCCA
ATTTTCCTGCCTTTTCTACCCCCCACTTGGAATTCTGGAAACATCGATTCGATCCTTCTAGACCCTTCTCCGCCC
CCGTTTTAAAGTGCGCCGGCCTCCCACCCGTCGCTTCATCGCTCGCGGTCGCAGTCGCAGTCGCCCACACCCTTG
CCGCACCATGGCTGGAACCCGCGGCGGATTCGAGTTATTGCACATGGTGGCACCAAGCATGAGGCAACAACTGCA

TTACTTGCAAGAAAGGCACAAAATCATTAGATGATTACTTGTGGCTTTCTTGCAAGTTGTGCAGTTGCTGCCTCA
AGCTTGCTGCCTCCCTCTGCCAAATCTATCAGCAAAAAAAGAAAATCATGTCAGTTTCCTTGAACTATATACATA
TGATTAATCTAGAAACGCAGTATGTTGCCAAAAATCACACACAACTCAATCACATATGCATTTGAGATGAGGTAA
AATGCTATCTGATTCACATAGTTCCAATATCTCAAGATGTGGGATATGCGTAGCACAAGTGCATCAGAGTGTTAT
AGTTGTTCAGTCTTCAGTACTTATTATTCTAGATTAATTTGGACCAACTTGTCAATTCAGCTGACACTTGTTTTT
GTCTAGTGCTCCAGTAGCTTGGTTGTTTCATCATCATTAGACTTTTAAACCTTTGGTCCTAAACTCCTAGCTGAA
AAATTTGCACACTGAACTTCCCTGTGTTCTAAGGAAAAAAAAAATCTAGACATAACCTTGCTGATCTGTATTATG
GACCGGCCATTGGAGTGCATAATGCATAACTGCTGATCACTTTGTTGGCACAAAACTTACGCATAGACATAGGAG
GTACTTATGTATAATTGTATCTTTAATCAGGACTTTGTTTTCAACCATATTGTTGATATAACAGTTATACTGTAG
TAATGATGAGTACAGTCTCCAATGTAAACTCAGGGCAAATCAGATTTATCCCTTTTTGGTGCTTGGTATCTTTTT
GTATTTTTCATGAGCGATGGCATTTTTGAACCCAAAATAGAGGGAAAGAGAAGGATCTCTGTGGGGGCATGCTAT
GTACTACAGTTCTTTGAGACCATGTAGGTGTGGATGCTTGTTTGGGACCTTTTCCTTAACATATAGAAAGAATAA
ACATTAAATAAATGATGACATGGAAATGAAATTCATAAAGGAGTACATAAGCTAGTACTCCTGAAAATATCACTT
CATATTCCCCATGATTCAGTAACATGAATATCATACCATCAGCAGCAGTGAAATTTTTATGCGTAGACGGTTTGT
TTCTTAAGGTATATATTTTTTATGTTTACAATTGTCTGTCGAGTTTACTGGTCAATTGAGCAACAACGGTACTTT
TAACTGTTAATATCACTTGGTTTGCTATGTATCCTTGTTGTTCTGGCTTTGTTGCTTTACAGCCATTCTGCTTAC
GCTATTGGCATCTGTTATTTAGAAAACCATTTTCTAACAGTTTTTAGATTTCCTTTCGAGTCTCCAACTGTTTTT
AACCTCCAAGTTGCCACATAATGCATAAATATACCGAGTAAGTTTCCCCCCTCTGAAATATGGATGTTAGATACT
CAGACAGGTTCAGTATGTGTTGGGTGTTGTTTTCTGAGTTTGTGGTTGTCATAAGATGCAGCGAAATTACTGCGC
TGTATCTTGGACCAATGCAATCAGCGAAGTCCTTATCGTGCCAAAAGGATGTGCATGGATTTTCCATCATTTGCA
AACTTGAGGATGGCCTGCCCTGATGCCCTCAGATGCTGATGAAGTGTGTGGTTCATGAAAACGTGAGAAGCTAAG
TAGTGGAGTAACGGGAGTGAGTACCAGTGTGGCCACTATGAAGAAAGTTAATTACTGAGAAATTAAGACTTGAGC
ATGTTATTTTCAAGCAACTTGTACAGCATTATATTGCACTACCATCAAATGCTTCTCCTGCCTATACAATACTCT
GCAAGCTCTTGTATTGTACACTGTTTTATCTTGTTTAAAGTATGCTTGTTTTTGCTTATGTGTGAGATTAA

Stem-loop structure

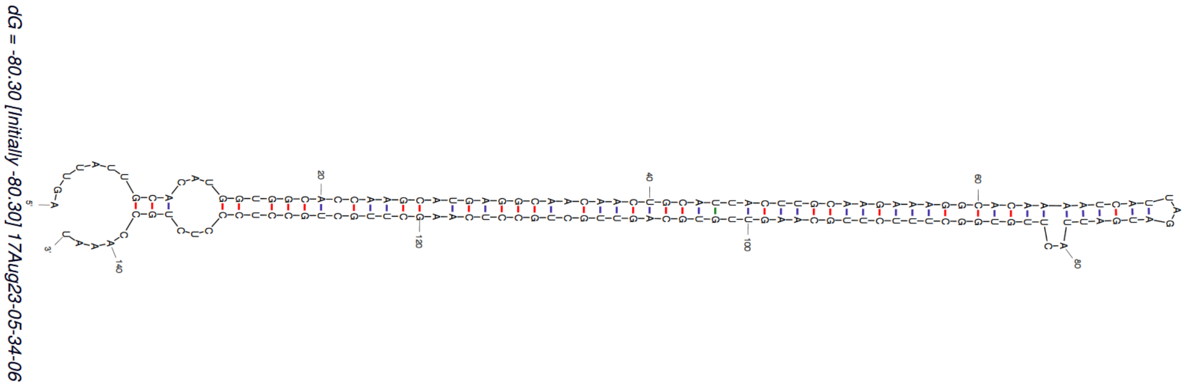


*osa-miR444e*

>Chr8

CCAAATGCTCGTGATGAAAAGCACAGCATGGGATGGTGCGCCCTGTTTAAGGGCAGCATGATGAGCAGAATCCAA
TAGGTGAAAAGATTCGGCACCATGGTGTCCCTCGCACGCAGCTTCAGCGAAGCTGCAGCCAGCCCAAATAGCAAG
AGTAAACAAATGCAGCCAAGTACACAGTATAGGTACATGTCTTATGCATGTGAATTTGTTATTATCAATTTACCA
TGTTCCAACACCTGTTACTACTATTAACCATGACTGGTGATTGTTGGCTTTATAATGATATTTATATGACCTCAC
CCTGTCCAGTTAGAAATATGTCTGATTTAGTCAGCCCCAATAAGCCACAGCCAAACAATATAGTTTGGTATTACC
CAGTATCACAGTATAAGTCTATCCTCCTGGCACACGATACAAGTCCATCGACAAAACCAAGAGGTTATTTAACCA
CTTGATTTTTCTTCAGGTTTTTGCACGCACAAGATCATACCTAACGTACAGGTACAAAAATATTTTTATAAGAGC
CATCTATCTGTCTGTCAGTACCTGTTTCTGCTAATTATTAGTTCTTGCTTATTTTTTTTCTCTGTACACAAAGCC
AGTGCAAATTAAAAAGATTGTACTAATTTTAAACTGGGCAGCATCCAAGCAGAGTGAAGCTACTCCACGCAAGTG
GGAGTTTTTCCTGTCAGCAAACTCAAACACATTGGAATTAATAATTGCATCCTATTGTTCCCTGTTCCCTTCCTT
GTTTTTCCCCCCGCTGCCTCCTCTCCTCATATCTGCCCATTCGGTTATAAAAGTAGACCTGGTCTCGTCCCCTCT
TT**GCACCAACACAGCATTCTTCCTGCTCCAGTTCTAGTCTTCTAGGGTTAGAAGTGATGAGAGGTCAAGATAAGC**
**TGTGGGTAAACAAGCTCAATCACTATGGTGTGGGTGTAGAGATCTGATCTATATATCCAG**GTAATTAATTAAACT
AATTACGTTAAGTTTACTTTCGAGCCATCTGTGATCTTGGCATGACTGCTGGTTTGCAGCTATGGCTTTCAAAAG
CTTGCTGTGTCAATCTCCTAGATCTAATGGTTGGCACAGCAAGTATATAACACATATTTGGTTTGTTCACCTAG
ATCTGCTTTCCTTTACTGTTCCATGTCAATATTGATCCTTGTGGAACATAGAATATAAGGGGCAGCACAGATCTT
GTTAAATTACATAGTGTGCTTGATCTGTGGTACCAGTTTGATTTTTTCTGATACATGGGAATCTCATACTAAATG
CCAACGTTACTTTTCCTTTCCAAAAATCTTGCTGGTTCAGGTATTATTTACATATCAGGTGCACTTAACATGCAT
GTACGCTTGATCTACCAAAGATGTCTTCAGCAAGTTTATTACCATCTATCATAAAAAATAAGTATATTACAAGAT

GCAACAAAGCAAAGGTGCCACTATACTGCATATGGAACCTTAAGTGAAGTCACCTCACTGGGGGTTTTAGAGTTG
GAGATGCCTCCATTGTGTGTGATGCATATTTTCCACTTCGGATGGCTAATAAAGTAATAATAACATAGCAAAACT
ATATTTGTTATTGACAATACTTGACACTTGATTGTGCTAACATTTACTAGTTCTTCTTGTAG**GTTTTGTAACCAG**
**GGGCAATAAGCTAGAGGCAGCAACTGCATAACTTGCGAGAAACTT**GTAAGATTTCTTTACAACCTATGGAAAAGA
TAGGTTTACTTGTCTTAATTTTTTGCTGTTCTTGTTAACTAGGAAACCAGAAGAATTATTAACAAGAGAGAATTT
ATAACAAACAAATTTGATGCCAATTCTATCCTTGACTCTTTTTTTTTCTATTTTTTTTATCTCTTGAGTCATCTA
GTCATTTCACTGTTTGAAAGTTGGGAACATCTGCATCACCTAAGATATGATCCTACCTATCCCTGTGGAATGTGT
TATCTGTGTGTTCAGTTTACAGGCTGTGTTACTAAAATAATGCATGTATTTGTGAGTAGTAGTGTTCTGATGTGA
TGACCTTCAAGCTAGACCTCCACATTTGCGAGAACTTTCACCATCAAGACAAAAAAACATGATGAAACAACCTAT
CCAATACATATAAAGCGCCATAGGATGAACTCACTCATACCAAACTGGCACTCTCATGGTATGGTATGACATGTG
GAAATGGGTATACGCATGAGATCTCCCGGAACTGAATTCACTTTGATATAATAACACCTTGGCATGTAATAGCAT
TAGTAAATGACGTCCAAGGAATATTAGAGTAATCATAGCGTTAATACTGTAAAAGGAACATATTACTTGTACTAG
TCCTTGATTGACATTCAGGGTCtgtttagttcccacaccacaccctgtcacatcgaacgtttgaacacctgcatg
aagtattaaatataggctaaaaaaataactaattgcacagattgcgactaatttgagatgaatcttttaagcct
aagtgctccatgatttgacaatgcgggattaattaggcttaataaattcgtctcgcggtttactaacggatttg
taattagtttttttattagtgcccaaacaccccatgcgacaccctatataatatccgatatgacacgctaaaact
ttacacctctagatctaaacaccccctcaatacgtactgaaacagcTACCGGAATCCAGAAGTAAAACTTTTTCA
CCTGAGTATCACATAAGCTCAGCCATTTCGCTATGGTACTCTAAGGTCTAAACATGGCAAATAGTCTAAGTCTGT
AACAATTATTATAGATATCAGTACAAACAACTTAATGGTGAGGAGACAGTTATATCATGATTTTCATGCTGATTA
TTATTCCTATATATGATGTGCTTCCTTCAG**GTTGGATGATGATCTTGCCAATGATTTTGCCGCAAGTTATGCAGT**
**TGCTGCCTCAAGCTTACTGCCTTTGTTTGCTAAACCATGAAACCAGCACAACAATAAGAGACTCCAATTGCAAGA**
**GACCTTCCAGGAAATCTTTAGTAGGCATCATATGTTCAATACAAATATACCTGAGGTTGTTAGAGCAAATTATCT**
**GCTCATGTACGATTACACTATCTCTATTATCAGAAAATCACAAGAACTTTTGTTGCTGGGTTTTGCATTAG**TACA
AGTATCTGGCTCCTGATTTTTCTTGACGTTTATCATTTTCCATTCTGAGCTATACGAGCAATAGGCTCCGGATTG
TAGTGATCTCCTGTGCAAGTTTTACCTCTTCGGCAAAAAATAATTGTCGGCCTACGGGGCAAACTATTCTGTCCC
CTCTAACTAATAATGTCTTTGAACATTTTGATCATATACCCTCCCATTTTATCAAAAATTGAGGATTTATTCTTC
TGTCACTGACATGTAGGGTTTATCTGAGGGCATTGACATGCGGACCATGAGGGTAAATCCTCAATTTTAATCTAA
TGGGAGGGCATATGATCAAGATCCACATTAAGGACCAGCTATTTAAAATACGGAAATTAAATTCTTAAAAAAATA
GTAAGTAGATCTGTATACTTGTAACATTTCAATTTATTAGGTTACTGCAACAAAATTGATGGAGAAAATACTCCA
AAGAGTCGTGTGTGAATTATGACCAGATATTTGTACCTTTCAATACGGTGGAACTTTCCTGCTTAATGACAGTAG
AAATCACAATGTAAATGAACTAGTCACTTGATTATAAAGCCTCATTTAGAGGATCACAACCCTTACATGGGTTTC
TTTGGAATTCAGATAGAATAGGCTCTTCCATAGGGATTTTGGATTTAAATTTTGTGAGGTCATATGTCATGTTTT
CATTTCTTTGTTTTTGTAATCTTATATTTCTAGACTTTGTTTCCTACAAATGCCTTGTAAAGTTTTCACGAGTTG
TTCATTCCTCTGGATTTTAAAGGATGTGACGCTATTATATTTTTTCTTTAAGCCACAAACAACTAAATTTCAACT

>osa-MIR444e_gi|115476535|

GCACCAACACAGCATTCTTCCTGCTCCAGTTCTAGTCTTCTAGGGTTAGAAGTGATGAGAGGTCAAGATAAGCTG
TGGGTAAACAAGCTCAATCACTATGGTGTGGGTGTAGAGATCTGATCTATATATCCAGGTTTTGTAACCAGGGGC
AATAAGCTAGAGGCAGCAACTGCATAACTTGCGAGAAACTTGTTGGATGATGATCTTGCCAATGATTTTGCCGCA
AGTTATGCAGTTGCTGCCTCAAGCTTACTGCCTTTGTTTGCTAAACCATGAAACCAGCACAACAATAAGAGACTC
CAATTGCAAGAGACCTTCCAGGAAATCTTTAGTAGGCATCATATGTTCAATACAAATATACCTGAGGTTGTTAGA
GCAAATTATCTGCTCATGTACGATTACACTATCTCTATTATCAGAAAATCACAAGAACTTTTGTTGCTGGGTTTT
GCATTAG

Stem-loop structure

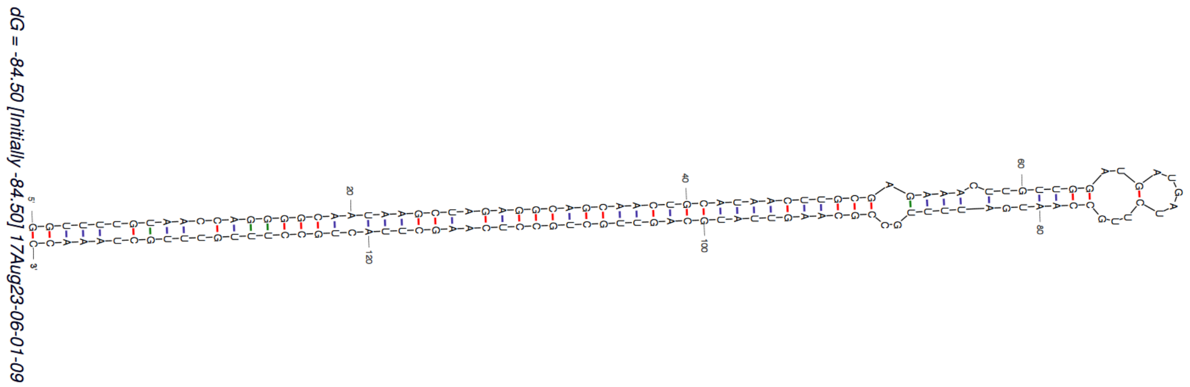

*osa-miR444f*

>Chr6

CGCTCTGGATGATGCTGATGTGTTGCCAATGACCGGGGCTGCTGCTCTCTTTCCTTTAGCCCGTGCTCGTGCAAC
ACCACTTCTGGTGATCAGTGGCACACCCAGTAACTCAATCACATGATCTCGTTGACTGTCACATCCAATCATCGA
ACACACACCCAGCTTCTCAATCACTAGATCTCGTTCTTTGTCACGGCCAAACACCTGAGACACACATATGACTGA
ACTTGTCTCTGGCATGTGTTGAACTGCCTCGATACTAACACCCACTGAATTAAGCACTCCTTCTAAATCAGCCAC
AGCCTTGTCGAGCTTCTCTCGGGCCTCCTTTATTTTTGTTTTGCTACCACGGATCAAAATTTTTGCAAGATTTAG
AATGGAAGAATAGCGTTTACCGAACCAGTTCCGGTCAGCATCCTCCATCTTTTGCCGCAGCGCCTGATCATCGGA
CTCGCGGAGAAGATCCTCGGCGTCGTAGGTGGTGTCCTTGAGCTGTGACAGAAGGAGCGCCATGTCCTTGTTCTT
GAACCTGCCCCACTCGCCGCGATTGATGAGGAAGCGAGCCTTGGGCAAGGTGGCCTGGAGGCGGGACAGTTCATT
GCTCAGCTGGGTCGCTTCATGGATGTTGATCCCGGTTGAGATGATCGTTTCCACGGCGTCCATCGCTGCTGCCGT
AGTCCTCGATTACTTCTCCGGATTTGATCTTGCCCTCGCTACTTCGATGAGATCGAACGAGCGTATCGATTGAGA
TCACAAGGAGATTATGGGGTGGAGAAGTAGGAGCAACTAGGTCAAGTCAATGGACCTCCTCGGTTATGGACTTCC
TGTAGAATCCACCAACTCCACAAGGTGACGTGGCCACCTTGACAGCCACGAGCTCTCCCTAACACCCCGAATTAA
TTAATCAATCAATTAATAAATATACTTAACACTAACACAGCTAACGACCCCACT**AATCTCTCCCTCTCCCTCTCG**
**TCGAAATTCCAATTCCACTCGCCTCGCCTCGCCTCCCCCACCTTCCGATTCGCTCGCGTATTATATTGCGCCGTC**
**TCGAGACTTCTAGATCCTTCCAGTGGAACCCGCCGCCGCTTCTGGATCCTTCTCCTGCCCCTCATTTTAAGCGCC**
**ACCTCCCCCGCCCCCCCACGCCACCGACCGAGCTGCCGGCGGAGGTGCGAAGCCATCGAG**GTGAGTGGAAGCTTC
TGGAACCTTACCCTCCTTTTTCCTTTCTATCTCGCTGCGGTTGGCTTAATTTGCCTCTGCGGGCGCGTCGCGGTT
TTGCTTCTAGAACCTTCTAGTTCTTGTGGTGGTATGAAGTTTCTCCGTGAGTGTCAAATGTGGATCCGCCATTAG
AGATGGATTTGAGTTCGCTGTGCTTTGTGACGATAGGCTTGTTTCGGTGGAGATTTCATGGATAGGTCGACGACG
AGTTCATAGAAGGAGGGGCTTTATATGCATTAGATGAACGGCTTCCAACAGTTTCCGTAGCAAGAAGCTTCTATG
GTTTAATCTAATTCTAGGAGGGCGGTAGGGGTGATGTTTGTGTGCCGAAGTCTTGCTAACCTGGATCAATGGAAT
GCGCTTGTTGTGCAATGCAGTTGGAAACTGTGGAGTAGCCCGTGTGACAGTGTGAGTTAATTACTTGGTTGCTTA
TTTGAGGATTAGGATACACACAGGAGGTCATGAGATGTTGTTTGGTTAAACTGAACATTTAATTTGTAGTCTATT
AGGCTGAAAGGAAAATAAAGAACCATTCAGCCTAGTTGCAAGACTAGTTTTTCTTTGTAACGTTGGTGAATTGTA
CTGTGTTTAAGCCAAACTAAGTTGTTAAAGGTTTAGACTGACGTCCCCTTATGTGATGTTTATCTTTGTTGTTGT
TGTCATATTTTGAAAATTGTGCATTCACCATCTCTAAAATAGCTCATAGGGCATTAACGAGTGATTTCAATCGAG
GTGCCTTGTTTACAACTATTGCTTTGCATATTCTGATCATTGCAATGTTTTCAACCAG**TTATTGCATATGGTGGC**
**ACCAAGCATGAGGTTACAACTGCATTACTTGCAAGGGAATCACAAGTCACAAGAAACACATGTTATTACAATGGA**
**T**GTAAGATTTTTTTATTCCCAGTTTATTTTCCCATGATTTTGCGACAGTGAATAGTTTTATTTTTTTTAAATAAA
ATGAACTTACATGATCATTGATGTTGTCTTTAACAACAACTCACCAATGAACAACCATACATTGTCATATGTGAA
CTCTGGTTAAAGAGTATCTTTTTGTAACTTAAGATAAATGGGCATCTGTCCTGCAAATGGAGATGGAAAGTTTGC
ATAGAAGTAACAGTGATATGCTATTCTGCTTATAATCACCTTTCCAGATATATGTTGTGCTGTACTAGCAGTCTA
ATACATACTATATGATGTCTGTTGTTAATATTCAGAAAATGACCCTGTTTTTTGCCACCATAAGTTTCCAACTTT
TCAGGCTATCATATTTCAGTATGCTGTGATAGATAAGTGAACCTCTTTGTGTGTAAATGTTTCATGGTCTGATGG
ATAAACAAGATAATTGTGTACAGAACCCAGCTTGCAGCATATCTGCATTAAGCAGATGTAGTTTTGCTGCAAGCA
AAGCATGGCTTATTGCTATACCATTGCCTTTGTGAGTGGAATGGTATATCTTCTAAATTAAGGAAAAGATGACTT
TCTCCTTTTCTGGAGCTATGTTGGGGTTGGCTCTGTTCAAGATCAGAGGGAGTATATTCCTCCTCTCAAGAGCTC
AAAACTTTTCTTGTGAATAAGTTTCCATGTTAAGGACTGCATAGAGCCCCTTTTCCATTGATAGGTACACCCAAG
GCAGCGGGCAGTCTCATTATATAAAATTATGTTGGTATGGTCAAAGCGCACCATCAGCAGGTTGAACCTTCCTAC
TTAGTTCTTCAATTCCTTGTGTCAAAGGTGGTCCTATGGCAGAAGATTGATTGCTTCAATCTAAATGGAATTAAA
ACAAAGTTGCAGATATTAATGAAGCACATGCTAAGGTGACCTTCATCTGAGATGTGACTATTGACATCACATACG
CTCATTTTGAGTTGGTTCTCTAAATTCCAGAGATCTCTTGCATTTAGGCCAGCAAGCTCCTGTCCATATTTATCA
ATAAACACAGGACGGAGTTGATTTAACATTCTGAGAAAAATTGTCACAATTTGATTTCTTATACATGTATTCTTT
CTCACTAACTTACTAATATTATGGAAGAGAAGCTGCTTCACTTTCCATTGCAGAATATTTCATTTACATATAAAA
CGATTCATTCCTGACCTGAACGTATTTATCTTTACACTTTTGTTTCATACAGTTCTTACATGATTTTGCTATTGC
CTGAAAAAATATAATTGACTTGAAAAAGCATTAGCTTCGTTATTGACCTGTGATTTAACTCTTTGCAG**AATCAAT**
**GGGTCATCATACTATGACTTTCTTGCAAGTTGTGCAGTTGTTGCCTCAAGCTTGCTGCCTCCGTCTGTCAGAACT**
**GCAACTAAGCTAGGAAGTTCAATCA**GTTTTTTTTTTTTTTATCTTGAACAGTAAATTCAATGTTGACAATTAATG
TAGTAATGATTTCTTAAGGAATGAATACTAATTTTGTGTTCATGGGGATCGATAGTCTTTTTTAAACATCATGTT
TAAACACGATCATTACTGGTGCTTTGAGTAGAGCCTAATACTGCATTACTATTTATTGCTTGCTGTTATAGTTTG
TTACGTACCATGTCCATTCCTTAAGGAAGCTAACAATAGATCTATCTTTTGTATTTTCCTCTTCGTGACACCTTT
TTCAATTGGATACGTGCAGTATAAAACTATCGAATTCACATTAAGAGTGCGATATATTTTGGGATCTTCATTCTC
CTTATTTGAATCATGATCTTGTTCACAGTATATATTCTATGTAATATGAGTATGACTACATGGAACACCTACTAT
GTCCTCGATAAATTATAAGTAGCTTTTGATCCTCCTAAGATTTTTATCTTATGAACACATGACTACCATTGACTG
TTTTCTTGGGAAAAAAATTAACATGACCCGCTTGTTTCAGGTCAGCTTGTGACCTGTATTGTACGTTAGTATTAT
TGTTTTATGGTGTAATTGCTAGTAGTTATCAACCCATGCACCTGTACGGGCTAGAAATTTAAAACATGTTTGCCT
AGTGCCGAATAATTTAGGATTTACTAAATACTACCACATCGTCTGGCAGATGTACTATTTCCACTACTATGCCAA
ATTTTCACAAGAATTCGAACTTTCTTATTCTCAAAGCCATGTGTGTGCTAACTGTCGAAACAGAACCAAGCGTAG
CATGCCTATGAACATTTAACTTAGGCATTTAAATATTATGCTAG**ATGTTCTGGCTGGACTGATCCCCCGATGATG**
**GTGTTCTGTGTTTTGTTTGATCATGGCTGTGTGCAACTGTGCATACATCCACAACTGGTTATGCACTTTTGATCG**

**TCAGCGTACGGTGGAAGATGATGAGACAGGACTGGCAAGTGAAGTGGTGACCAAATGATGTGAACGTCATTTGGC**
**ACCTCAGAGGTACATCAGAGGTACATGTATGAATCTTAGTGTTAATTGTTCAAACATAGGTTTTCTCTTCCTCCA**
**AGAGGAATTGGTTTATTTTGGGATGAAGGGTTTTTTTTGGCCGCCTCCTCTTCTTCTTCTTTTTTTTTTTTGAGA**
**TGTCTGTCATTTGGATCAGTAATGTTGAAACATGTGTTATGTGAAAACTCTGTAA**ATAGTTTTTTGTTGCACGCG
TCTATTACTTGGTACAGTAAATGGTAATGGATACATTCGGCCCAAATATGTGTACATTGGTCCAAAGGTGCGCAT
AAGCTGGCCCGATTCGGTCTAATTTCTCGAGTGGGCTTCGTGCGGTTCATTCAAAATTAGTTGTGGGCCAAATTT
CCTTGCAGCTACTATTGACTGTAGGCCAAAATTTGGCTCATATTTGACCATCTGTTGGCACGTGGACTAAAATAG
GAATGAGTCTATTTGCCTGACTAAAATTGGCATGATAGGAAGGGTGTTAGGTGCCTCCCTCATCTCTTAACTGCT
TATTGAATCGTCTCCCTCAATTTCAAAACCGCGTATAATGCCTTCTCCTCTTCAAAAACCAGAGCGGAGAGCAGT
TTCTGTGGACGTAGTAGGTTGATCACAGTTGACTTGTTAAGTCAGTAGGTGCCTAGGTGGGACATACATGCCAAT
GACTTTCTCCTTATATCATCCTTTTGCTCACCTCTCCCTGTTCTCTCTTTCCTCTCTTCCCTCTTCCTCTCTCTC
ACCTTCCATCAACAGTAGACGTCGGGAGGGTGGTGACAATGGTGAGCCAGACCTAAGGAAGGCGGTGGTAGAGGC
GAGCCGGAGCCGTGCGTTGTTAGGGGTTTGTATCGGGCTAGGCAACTGGCTGCGTCGCATGATAGGAAGGGTGTT
AGGTGGCTACTGGCAAAGGACTTGCAGTGAGTGTTGCTCGGTGGTCAAGCGTGAAGGTGAGCGGCTGTCCGCACC
CTCCTCACTTGTGCAGGAAGCCGAGGAGGCACGAGGGCATGCTGCAATTTGGTTTTCTGTAGACACGCCAAATGG

>osa-MIR444f_gi|150172133|

AATCTCTCCCTCTCCCTCTCGTCGAAATTCCAATTCCACTCGCCTCGCCTCGCCTCCCCCACCTTCCGATTCGCT
CGCGTATTATATTGCGCCGTCTCGAGACTTCTAGATCCTTCCAGTGGAACCCGCCGCCGCTTCTGGATCCTTCTC
CTGCCCCTCATTTTAAGCGCCACCTCCCCCGCCCCCCCACGCCACCGACCGAGCTGCCGGCGGAGGTGCGAAGCC
ATCGAGTTATTGCATATGGTGGCACCAAGCATGAGGTTACAACTGCATTACTTGCAAGGGAATCACAAGTCACAA
GAAACACATGTTATTACAATGGATAATCAATGGGTCATCATACTATGACTTTCTTGCAAGTTGTGCAGTTGTTGC
CTCAAGCTTGCTGCCTCCGTCTGTCAGAACTGCAACTAAGCTAGGAAGTTCAATCAGATGTTCTGGCTGGACTGA
TCCCCCGATGATGGTGTTCTGTGTTTTGTTTGATCATGGCTGTGTGCAACTGTGCATACATCCACAACTGGTTAT
GCACTTTTGATCGTCAGCGTACGGTGGAAGATGATGAGACAGGACTGGCAAGTGAAGTGGTGACCAAATGATGTG
AACGTCATTTGGCACCTCAGAGGTACATCAGAGGTACATGTATGAATCTTAGTGTTAATTGTTCAAACATAGGTT
TTCTCTTCCTCCAAGAGGAATTGGTTTATTTTGGGATGAAGGGTTTTTTTTGGCCGCCTCCTCTTCTTCTTCTTT
TTTTTTTTGAGATGTCTGTCATTTGGATCAGTAATGTTGAAACATGTGTTATGTGAAAACTCTGTAATT

Stem-loop structure

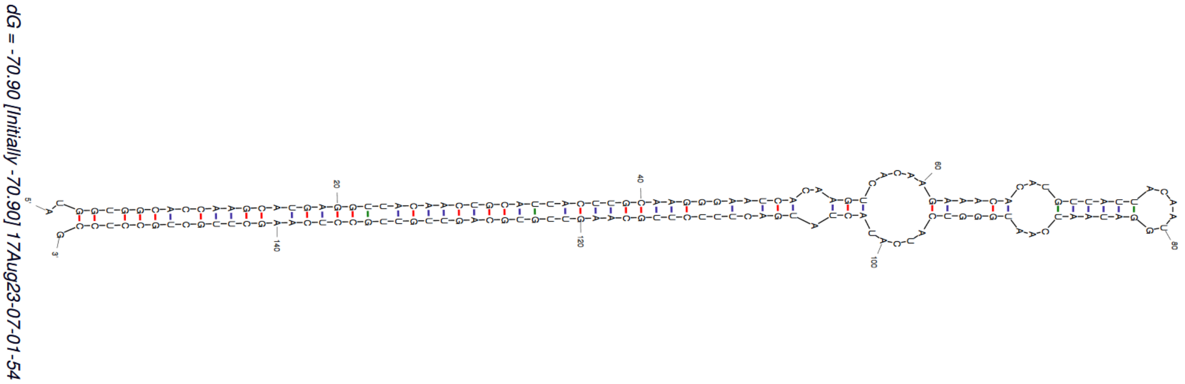


#### Zea mays

*ZmMADS23 and zma-miR444a*

>1

ATTGCCATTTTTCAATGAGCTAAACAACTATAGTGATATTATTACGTATCATTATATTAACTATCGAAAGTATTT
TTATAATAAATCTATTTAACAAACTAATTGTTCAAATAGAACTAAAACAAGGAGCAGCGTCAAATAGTCCACTTA
TATCACCTCAATTATATTATACTTGCTCTAGGTGCAAGTGCGCCTTCAAATCGCGCTACCCCAGCTGAAAGAACA
ATAATGGAGCaaaaaaaaaGCCGCTGCATTATATTAGCTGATGGAGTAGTGAGAAGGACAGCTACATTAAGGAAT
CCGCAGCGTGATGAGCATCGCCACGAACGCGCACGCGTCAGGCGCTGACTCGACAGAAGTAGTAAGCCGTGCAGA
TTTGTTTATTGCCAATGATGTAGTACTAATCAATAATTAGTGAACTAGGACGAGGTAGCTAAAGAAAGCAAAGAC
GTACGGTGAAGGGCCAGGGCCTTCCAAACGCTCTGTACTTGTATATATGGGTCTTTCCAGGTGGTCCTAGTATAC
TTTGTTGGGGAAGAAGCACAGGACAGGCATCTCAACCTACATAGGTAGCAACCTGGCCATTCTCAAAAGAGGATT
GAGCAGTCGTTGCAGGAGGATCATAATCTAGACTAccagagagagagagagaagtgagtgagtgagagagagagaga
gagagagagagagagagagagagagagagagagagagtagatagttaggtctcagctagctctttttctccctcTAA
GCTAGATCTGTGGTGTTGGAAAGGAGATGGGGAGAGGGAAGATAGAGATCAAGAGGATCGACAACGCGACGAGCC
GGCAGGTAACGTTCTCCAAGCGCCGGGGCGGGCTGTTCAAGAAGGCCAAGGAGCTCGCCATCCTTTGCGATGCCG
AGGTCGGCCTCGTCGTCTTCTCCAGCACCGGCCGCCTGTATCACTTCGCTAGCACCAGGTTTGATTCTTGCTCCT
CCTTTGCTAGCTTAATTGACGAATAGTTGTTAGATCTACTAACTAGTATTACGCACACGAATCAGGCATACAGCA

TATATATGCTCCAGAACCGATGAAGTATATATATTTCGGTCAGGAGAGAGAGATGCCAGTGGCATAGATTTTTGA
TACCTCGTACCTAGTACCATTTTTTATGGGGAGTATGTTAGTTTACCGGTGTGTGTATATATATACTCCTCTTGG
AAAGTAAAAAAGGGAGAGAGAGACTTGTGCAAATAAAGTTGTCATAGGTTGAGTACTTGCAATTTTGCAACGAAA
TTTGTATTAATTTGGGGATCAAATTAACATGAAAGACTAGAGAACGAAGGAATAATGATCTAGGTATTCGTTGTG
TGATGTGTCACATGCCTCACAATATTACAAAGATTTCAacccagctgctttgccgagtgcttcatatactcgaca
aagtctagaaaatactcgacgaagtctttgccgagtatgactttcagcaaactgtacatcgacaacggtttcttt
atcgaatacctttttgtcgaacactcagcaaaagctttgtcgagtgccacctggtactcagcaaagaaaagtcgtc
atcatggcgccaagtaacggtgacagagcctttgccgagtgtcctccgtgacactccgcaaaggctccatctttg
acaaatgtctgctggactagcgctcgacaaataagcccctataggcccatttgccagttcttttgccgaatgcac
taggaggcactcgttaaaatcttcgtctttgccaagtgcctgttggactaactctcggtaacgggaacactagtg
gcccctttgataatctctttgtcgagtgcattaggcaaaggttgtttctttatcgagtgcctgtttcggtaaaaa
gcatgtttccggtagtgGTTCCAGACATGAATTTATGGcttttattgtttttctttttattttacaactttttAG
TACGTTTATTCTTACCATGGAATGATGAGTTTGAACACTGGTCAATTTGTTTCACATGGAGTAAAATGAGTAAAT
GTATATTTCTTTGTTCTCGTGAATTACTATATATGTCGATTTCATGCCTATATATAAACTGAAACTGATTGAACG
ATTGCCACCAAGATAAGACTTTCACTTTGTCTCCCCCCAAAGTGCAATCCTTATCCTAGAGTCCTTACATGGTAT
AGTGTAATGTAAAAGTGGTTTTTATCGCACTACCTAGCTCAGCATGGCAAAATAGTTGTTTTCTAGATGGTTACT
TTTCACTCCGGAGATGATCCGATATCAATGCAATAACAATAAATGAACTTAATTTTTTGCTGATTTGGAAAGAAA
TAAACTGATAATGTATTTTGTCTATCTAGAGAACCATCTTTTGTGAACCCTTAGTTGTGCACAAGCCAGAAATAA
GCATTTTAATTTATCATGATGTTCGTGAACAGTAGGTGAAGCTTCACCAAGGTATAAATACTAGATCGCTCAATC
ATTTCAAGGTAATGAACGAACAATCTTCAGTAATTATACCTAGGTGAACCTTCCCCTAGGGTGATGAATAACACT
TATATATGCTATTTGTCACCTTAAATGATGGATCATACGGTTCATCACCCGTTTCAAATGTTACAATCAGATTAC
TAGATATATCAAAGCGACACAATTGTTTTCCTTCTGGATCCGCACAATCTAACTCTAATTTAAGTCATTTTAACT
AGTTTTAAGACGCCCTACTCTAATTTCTATTATTTTAACAAGTTTAATAAATTCCACTCTTTTCAGTTATTTTAA
ACATTTTAATACATTCAATCTTAATTCCAGTCATCTTTATTAGTTTAAGACACTCCAATCGAGTTTCAATCATTT
TAACCAGCTTAAGATACTCCAATCAAATTTCAGTTATTCTAGCTAGCTAGTTTATTATATTCAACTCTAAGTTTA
GTTCTTTTAACCAATTTTAAGATGCTTCGCTCTGATTTTAGTCATTTTAGCCAGTTTAAAACATCTCTCTCTTTT
TAGTCATTTTAAACAGGttaagacattcgacactaaatttggtccttttatttagtttaagacaacctactctaa
tttcagtcatttatcaagtttaagacattcAACCCTgatttttttagttttcaactaattttaagtcatttcatt
tatttatatttCATATTATTTAGTGTTTGTTTAATTGCCCTTTAGATTTCCAACTAATTTTAAGTCATTTTATAT
GTTTTCAAATATTTTCATCTTACTATATGCTTATTAATTGACACATATTTAGGCTACGACACTTTTAAGTAATTC
TATCTAATATTCAGCtattctaagtcatttcatattatttaagtcattatttattatATCATATTATTCAGCCCT
TAAATGTTATGGAAAACAAAGCAGGATGAGAACATTAAGTAATTTCAACTTTTCTATACCATCATGCCAATTCAG
AGTTGCATCTAAGTCATTCTTTTCAACTTACACTATTTAAGATATTTTATTTACATCTAAGTTAATCCTGCTATT
GCAAACTTGGATCTAAGTCATTCTTTCCAACTCACaagacatctttatattgcatctaagtcattcagatgttgc
ataattgaatctatgtcattattccttctaactcatagtatttaagacattatatattgcatctaagtcattcat
gtcattgtagagttgaatctaaaccattccttccaactcacactatttaagatattatatgttggagctaagtca
ttGCGGCCAATATAGTGTTTAATAACGAGTTGAATATCACAAGAGTAGATCGAACACATTTGGACAGGCGTGTGA
TGTGTTCGATCTTCGTTCAGACACGCGTGGCTAACTTGATTGGATACGTTCGGTCGGGTTTGATGAGTGGACTAT
TTGTTGTCACCAAGGTTTAAAATAGCGGGCTATGTAAAGTAGCGACATCATTTTTAGGCTACATGGCTATAGCGT
GGGGTGATCGTACTGACTAAATAACAGATTTAAAAAAACATAGAGTTCGAATAACAGAAGGTGTGCGTATTTGAT
TTGTATAGTAGGCGCTGAGTGGTGCGGCTGTAATAAACAGCATCAGCCCTACACAAGCACTTCCAATCACAGCCG
TAGAATTTGCACAGTAAAATATTTGTTACTGAGCCATTAGATCTTAGCAAATATGGGAGAGACAAATAATAGTTC
TCTCCAAATAGCGGAATTTACTCGATCTAGCGCAGAAAGTTACACAGAAAATAGACAATAATTTCGTATGGATAC
CAACGATTTCCAAAAGGGCCCTATTACCAAGTTGTACAAGGCCATGTGCATGCACTACCTTTTGctctctctctc
tctctctctctctTACATGATTATAATAAATTACATATAAGAACTTTAGCGAACTGCACTGCAGGTCCTCGATAA
TTAAAAAAGGTTAAAATGAGAAGAAAGAAACGTAACTTCCCTGCCTTATCTAGACATCTAGTACGGAGGTTTTCT
CAATATATATGAAAAAGTTTTTTAATATAACGTCCGGAACTGTTTTACCACCATGGGTCGAGTTTTGTTATTATT
GTCTTGACTTAATAGACAAGTAGATGCATTACTCTCAATCATGGTATACATTTTCGTCATATTTAAATTTTTCGT
ACGCTGCCCTGCCAAAAACGAAAAATGGAGCAAAATTTGTCGAAATTCGCCAATTATGATCGGAAAGGGTTTTTG
GATTaaaaaaaaGAAATTCGTAAATTTTGAAGTGAAATTTGTAAATTTCTCGATGGATTTTGTAGAATTTGGACC
AAAACGCTAATTGTGATGGAAAAATATTTTTTGTTAGGGTGTTGTTGGAATGTGTATAATTTATGCTACACAAGA
AATAATAATCACACATAATTTGTTCACTGGAGTCACAAAAACATATATAAAGTAAAAATGCAATCCGAAAGCAAC
ACAAAGAAAATTCTTCATATACATATATAACAAGTCTTTAGTTCGTGCTACATAATACAACTCCACAATGGTGAC
ATATACAAGTTTAAATAAAGTTTTAGTATAATATAAAGGTTCACACAGTTCAAAGGTCGAAGATAATTGAATACA
TATGACATAAAGAACTGAAATGTCCCCCTTTGCGCATGGTGACCATCGATGGTATGTTGAATCCTACATGTAGCA
CGTAAATAATACTAAAACTATAGCACGTAAATAATACAAAGCAACAAGGATAATACTAACACTACTATGTGAGGA
AAATAAAACTAAAACTATAGCACATGAATAATTATGCAAATGATACTACTATCTAATGTAAATACCTAGCCCTCC
ATGCTGTTAACTATCCCTAGTACTTGATAATATATACTAAGTACTAACAAATAAGCCTAATACTTGATAACATAC
ATAACAACTAACCCTAACATTTGAtaacatatagtatatgctatattgctatatataacaaatatgcattaacat
atatacatatatgcactatcataCGTATAACTATGCAGTAGCAGTAAATGAAATTATCAGTACACAATATAAATT
ACTTATTATATCATACATTCACTTGCGAAATCATTTAAAAATCATGAATACAACCATGCTTTCACCTTAGCCGCT
AGAAAAGGGGCGAAATCTGACCAAGGTATACCACGGATTTCGCTAAGTTTCTCGCTGCGGTTTGCTAtttttttt
atttttAACTTCTCAAATACGCTTAGATTTGACAGCACACTACAGAAGACTCAGAAAAACGTCCGattttttta

tcatattttctttggttttataatttttcattttAAAAAAATCATTTAACGAATTATACCGATTTTCCCAAAAA
CATTTCCGCAACATACCACATGCGAAAAATATGATCGGCATGTTGTTATGTCCTACGCACAGCAGGCTCGCAACT
GTGATCACACATAGTTCTTTATATGTTAATTACAGCACAAGTGATCGACGTGACTTTCTAACCGAAAAAACTGAT
TTTTATCGACCAGCATGGAATCTGTGATTGAAAGATACGAGGAAAGAGAGGGGCACCATCAGACTATGAGCGCAA
GTGCTGAGGCCAAGGTATGATAAGCACGTCGTTTGTATTCCTGCTAGTACTTCCAGTTTTTTTATTTGACTCTGG
ATAGTACAATTTTCATATAagatcatcttcaacaatgtctcaatttagtgcctcaaattgaaatatgggctctac
acaaaaactgccccaacaatgtctcatttcataaaatttgatcaaaaaattacactagtacatagaagctttata
gtagcggtcctagagctatttgccctggcgttttccagagacgtcagtgctaggggccagtagaaattagtcttt
tttacatgcggttgtttaccaaccgccagtgtaaataacattatcattggtggttttctttagcagcccgctagt
gtaaatctcattagcactgtcggtttgccgcatgtagaaatattattatcactggcggtgtcttaagcggcccgc
ctgtgaaaatgatatctccacatgcgctttcttaagaaaaccgtcagtgctaatgatattatcactgAGTAATGA
AGTAGTTAGTGATATATTGTTTATCTTAGATGTATTGATCACCTCTCTGGTATTGATGCAATGTATTGGGGTATC
GATCACTGAGTAATCACACAATGTATTGGGGTACTGAATCCTCTCTGGTACCGATTATTCatgcaatgtattgac
agtttcaataagtgaaccaccagtgtatatgatatttccactggcggtgtactttatacagccgccagtgtatat
acatcctttacactgacggtgtactttatacagccgccagtgtatatgatatttccactggcggtgtattttata
tagccgctagtgtatatacatcctttacactgacggtgtactttatacagccgccagtgtatatgatatttccac
tggcggtgtactttatacagccgtcagtgtatatacatcctttacactgacggtgcactttatacagccgccagt
gtatatgatatttccactggcggtttacttaataaagccgtcagtgtaaatacatcctttacactgatggttctg
ttaagtgaaccgccagtgtatatatatatttacactgtcggttcgttaaggcgggcccgcttgtttttttttactg
gcgtgctgtaactgaaaccgctagtataaatttctgcgtgccgccaccttagagctcttttctactagtgTTATA
CGGgtgcctcaaaaatactacaccgcagtggtctgccttataatataaatttgatgctttactgttggaccgaaa
tgttttgttggtaccttaaattctataaaatatactcattttaaattatatgacattttataggtcatgttgt
tggagcatctccaataacgtgacccataaaaatgccctataatttaaaaatgagtatatttttacagaatttaga
gcaacaataaaacatcttgctccaacagtaaagcctcaaatctaggttataagacagtacactacgatatagtat
atttgaggcagttgagagcgtgccttatatttttttgactaaattttatgaaatgaggcactgttagagtagttg
tttctgcatagagcctcatatttcaatttgagacatcaatttaatacatttttggagatgctctaaCGAGTACTA
TGTATAAAGTTGGATTATGGACATTCTAATATATGGTATATACCCCATCTGGCAGCCGATCTTCCGTTGAAAAAA
TATATATTCTTGGCCAATCCGtttttttcttcctttttttCCCCTTGAAACAGTTAATCCTAATTTCCTCTGTAA
AATGAGGCCGATGCCTGAATGCTAATTGTTTGCACTATGTGAAAAACTTGCAAGTAAAAAAGTCCGTGATCCAAG
TAAAATGCAAACCGACCTAAAATCACGGCCCACTGAACATTACCACAAAACCTTGATTTTCTAGAGGGACACCCT
TGCCGCCTTGCGTGGTGTTGGTTGTAATATTAAATTTCTATAATTATAATTAATAAAAAGTCCAGTCCGCTTTCT
TAAAAAAATACAGATTCATGATGTCCAAGCACTTCTACGTCAACCACTTGCAACACAATAAgcttgttcgtttta
gtgccaatccatatagattagattggattggattagtttaaattcataataagtcaaaattcttcttaaaatttt
ctaatcctttgtaatccatgtggaatagaaataaccgaacaagacctaagatggaAAAGTAGCTTGACGGCTATC
CAGATTCAACTTCAAGCTTTGCTAAGACCATAGTTGCATAGCTTGCAGAGAAAACATGTCACTTGGCTATCATCA
TTTCACAAATCACAAATAGGGTAAAAAATGATGTACATTATGATCTGAAGCTTCGAAAGCATGTTAATAGGCTGT
TAAATTTGCTTATAGTGTCAATGATTATGTGGGGCTAAGTTAAACGATAATGGCCTAGCATGTACCCACCTTATA
GTATCAATATTGAGttgacgaagcgatagatggcgttgacattgttgtagtacacaagcatacttcgggcgaggg
agttatgaagattttagaggagctgacgaactagcttgcatcagctaccccattagcaacaacacagtactcaac
ctcagcaatgaagcgggagacaatgggctaacgcttgaaggaccaaaccgtgtaggcagagatagaccgccaagt
gtccaggcacccgacctatttagcatcggtatagacaaccaactcatgcatggaggagtgccaaagagctcgtag
tcaacattaccttaaagatagtaaaggatctgcttgacCAGACTTATGCCAGTGCCCCTAGGTACGGGTACTTGA
GGCGTTACAACGAAACTAGGCATGTGCGGAGAGGGTAGGCTACGGGGCCCACCTTCAAGGTAGGCTAGGTCAACA
AGACCCCCTGAAGGGATGGAGTACATGATCGGGACCCTGGCTACGCATCAACCTAGGGGGTCCCTCACCACATAT
ACCTTTGCCATGCTTCCTTCAATAATAGCCCTCACTATCCTGTAAGCAGGGTCCGATTGCACCACGTGGCCCGAC
CGACGCCCTACCTGTGGTGGGGTTTGATACTACTACGTGGTTTCAGGTACCCGATGAGGGGCCCAGATCCCATAT
GGGCAGGCTCCAGACCCCCTAGAGGGTGtgaaggaaataagtgacgctaagagggggtgaattaggacttctaaa
actttcactaaactaggccacaaataaatccctagagcaaaacctatgcaaataatcaaactagaatgttcaaac
taggttttgtctaagtgttgctatctttaccgcaatgtctaagttccaatcctaaacaatataagtataaagaca
agattgaaacttaaatgcttaatataaatgtgaaagccaaagagcaaggtagagatgcaaactctcgtggatgac
gtcggtatttttattgaggtatccgaaaccacgcaaggtcccgactaatcctcgttggtgcccctacgcaaaggg
aagcccacgtgagggccaagcaccacggtctagtaactccgtagagagtcgcgggccttctccacgcgcaagtgg
tgctccgcttccagctcctctcggactctccccgccgtctcaactatcgagcttcaggccgaaaacgctgcgggc
ctcgttctctccagtacacggtgccggtcgtgacacaaacgcggttgtcacggtctcacaagactctctccccac
tcgatacaattacaacggctcgcgcaagagtcgaggggttgtgtggtttttctaaactcactcaactaactagga
ttcacctagagcaagcgctaaagcgatctaactaacctaagcacttcgcaaagcacctacgctaatcactgagtg
attctattaagcacttgggtgtatgagcacttggaaatgtctactatatgccttggtatgttgcttgggctccca
caccttgaaatggccggttggggtggtatttataggccctaacaccatttctagccattggacagaaagcagtag
ctttctgtcgatgggcacaccggacagtccggcgtgcaccggacagggcactgttcactgtctggtgcctgccac
attagccgaccattggggtctgcagcagtcgactgttggatccgaccattttctgactgtccggtgcacactgga
cagtctggtgctgcagcctgaggacgcctgttcgcgggcccctctgcgcagactgtccgattgtcccacccggca
gttactgttcactgtctggtgcgccaccaggcgctggcttactgcttttcttggatttcttcgccgattcttttg
ggcttcttttgttcttgagtcttggacttctatgcttctttttatgtcttcttttaaggtgttgcatcctcattg

ccttagtccaatcctcttagcatcatgtgaactacaaacataaacactagcaaatacattagtccacaggttgtg
ttaatcatcaaacaccaaaacctattgagccaaatggcccggggtccattttccttacaatctccccctttttgg
tgattgatgacaacacaaccaaagcaagcaaatataacaaacatttgaatgaaaatatgcaatctacttgctagg
atccatgagtgtccccataatgtgagattatggacttaagcatcccccctaactccataattcacttacttcctat
tttataccaaagaaccaaaaccacttgaaggatcaaaaatttaaattggtggtgtttggtgtttgataacatttt
attgctttggttcctacaaattctcctcttttggcatcaaccaccgaaaaggaagacattaaaagcacataggaa
gcaaataaaatttgccctcataaaggatttatcaattaataccaattgaagtactcaatattattacactcccta
aatgagtgttctccatttagaaagaattttctcctcctttagagacgaagaaatactcccccctaatagagatctt
ctacttaggaaaaagcatgtgagaaatagaggtgcaagacatgattttgattagactaacttatgcatttagagc
aacatgagaagtataatatatgaaatatagtctaaccaaatattggagaagacatgtaaatgatgtcaaatgtag
tctcaaagataccaattgaaaatcaatggcggaccatttgaagaccaatggcaggcttgtgagacataagaattc
ttgtagatttgcagtagaagcttgttgtctttctctggggacttcattttccttacaatgagactgctacatgaa
ataaacttgaaaatcggtattagtctcaaagacttagattatagagtaacctcccccctgaaagtgtgcataaaaa
ttttgaatacttgtaggatacatgcactttgattcgatatcaagaggggcagattatccataattcgagctttgg
cacatataagttaaatgataccaattgaagcataccaaatgaagtataattgaagtgttataccacttgaagcat
ttcactatatcaattgatgtatcatttttcttaggaatgttaagtaagctatgtgccttgcttatcattttaaacc
atgtagttttgcttaagagtacgaattgaaaatgagcaatcctaccatatgatttacctagtatgcatgcatttt
taatcaaaagtaaatacaaattgtattacaaggtatgaaaggtaaaactagatacatgaaaatacatacacatat
ctaaaaacaagaaatacagtcaatctagttaccttagtcatggacagggaaatttgggtccaaattatgtattaa
cccacttggcaataaacttgatccaatatggtgtatcccatgatatacatctcaattttgccaagtctccaaatt
ccctgtcgttctttggactcctcacttcccttttgggatctaaaccttcttggtgcttttcatgattgaaattat
ttcctttggcacccagatactttttggccccctttccctttgttgacttgcttgttgcccttgattgctatcatctt
tccattctttttcttcttgattaagtatagtgtagcagccttttttgtccatcttgttgatgtaggtgttggagat
tttgcatgttggctttttgcttgagcttttgcttatccccggtcttcaccttgcattggaaagccttgtggccttc
tttgtgacatagcctacaaatcacagtttcaccctccacaggcttgttcactcccgcagtggtgttatcctgtgg
aggttgaatttgtttggctttgtctttcttgtcatacaaatcctttccaagacgagccacctcttgttttaattg
ctcattctccattgcaatctcgtccgaacatgtttctataataacattcttaacaagaacttgattgcaggggtt
agaatcatcaattaaatcaaagcaggaagtagaagcatctttcttaataatttcaggtatttcaatcaaagatac
ttctgggatgataccaatgtttttatcttagtagctagctcatcattgtgtatgctaagaatttccatttttcct
agcaaattttcaatggcttcttaggagctagctaacttagccttaatttttttcattcttttttaataaggtcatca
tcggtatcagtggatagagcactagactctaattgctaaattttagttgatagctcacaatttaaatatataaat
gtttcaaattttttctagcaaacctttataatcattttgagagctaatcaacttattttttcaaagttttaagttga
gcctttttgcttagtgcaaacatcctaaaaaaattaacgacttctgcaagctcatctacagagggatttctctcacc
tttatcatcactactactatcatcactagaggatagaatactcatttacatcttgccataaggcacttgcatga
tgaccatgatgaggatcggtggctgcgtgcccttggaggttcatcttcgcttgaagagtcatcccaagttttgac
acttgtaagcatcttgcctttgctcctcccttttttgtgggttcgaccatagatggacagctatccctgaagtggcc
cttctccccgcatgcgaagcatcctctctttctttgtttttgcctgtcaatgttaaagagaagatcttctacctg
caggggcacacccattagattaatcttgcggatcatccccattacttctctgacacgtcgaattgtttttttcgtc
cttggaggatgatgttgatggttgattatcttcatcatcatcactttattcttcttcctcttcttcacttgagga
acttggagtgggagccttctttttatccttccttcatcacatgcaaaagcatatagccttgaggaagttggctcc
tctgcccgacacattcttcgcaacatctcaaaggccgcaattttcccaatcacaatgatcggggtcattgtactc
aagtcctccatgttgtgaagaatagtgatgatgctcgcatatctttgtcgtggtaatagggagataatcttcctt
acaatgtcaacatcacctagcttattgataccaatagaattaagctcattgatgattagattcaaacgagaatac
atatctctaacaagctcattatctttcatagcaaaagaattatactcatttaaaactaggcaatgtttttgttca
cggacattggatgtgctgccatggagctcatgcaattttaaccaaatctcattagcagtttttaatgtaaaaact
tgattaaaataatccatgctaagagattcatacaagcaatttttggctctagcatttaaatgaatttctttttcc
tcactcatggtgggtttagtgggattttttggtggtttcatcccatcacgagtgactctccaaacacctagatca
atagtctctagataacaagccattctagcactataatatgggaagttagtgtcgtcgaagtgtggtggcctatgg
gtatccatccccctttctctaaaaagagtcgactcttttagcagtgaagctaaagcatttcaaatgagccaaatca
ggttctgataccaattgaaggaaacaggtgactcctaagagtgggtgaattaggacttcaaaaactttcactaaa
ctaggccacaaataaattcctagagcaaaacatatgcaaataatcaaactagaatgttcaaactaggttttgtct
aagtgttgctatctctaccgcaatgtctaagttccaatcctaaataatataagtataaagacaagattgaaactt
aaatgcttaatataaatgcagaagctaaagagcaaggtagagatgcaaactctcgtggatgatgtcggtattttt
accgaggtatccagaaccacgcaaggccccgactaatcctcgttggtgcccctacgcaaagggaagcccacacga
gggccaagcaccacggtcgagtaactctgtagagagccgcgggccttctccacgtgcaactggtgcttcgcttcc
ggctcctctgggacgctccccaccgtctccactatcgagcttccggccgaaaacgccgcgggcctcgtttcctcc
ggtacacggtggaggccatgacacaaacgcggttgtcactgtctcgcgagactctcacccactcggtacaattac
aacgactcgcacaattggtatgttgcttgggctcccacactttgaaatggccggttcgggtggtaCACAgtgcct
tggtatgttgcttgggctcctacaccttgaaatggcaggtttggttggtatttattggtcccaaccccatttcta
gccattggacagaaattagcaactttctgtcaacgggcacatcgaaatgtctggtgcgcaccagacagggcattg
ttcactgtccgatgtctgccacgtcagtcgaccgttggggtctgcatcagtcgaccgttggatccgatagttgcc
tgactatccggtgcacaccggataatctggtgttgtagcttgagggcgcctatccgcgggcccctcaagtataag
tagataccatcccaactaaataagagatcacaattaataattcccacaatgcaatgcgtgacatattaaatttaa

ttccataagttactcatgtgagggtccaagccgctcttgaccgtgggcacgactgatatactagttttatactct
acagaggttgtgtatatttaccccacaagtcgtgatacccatctaccacgggtcatgaatcacatacacctctac
cgaggaggcaaggcggggtaacactatgagccctttacaaagttctaatagcttgagaaaactcgctagggtttc
gttgagaagtgggcaatatagcaatccctcgtccaaaaagtcatcgcaacaagaacggcccgagaacctccctat
accgacaactcctcatactgtggttgcccctttcgaataagctagtcctccactagctttcctaattagttagcc
aagtgcttctcataccaccctcgtggtagcactatttctcaggtggttctccatgttccaattaacataataat
cttatcatgaacaataattacgccaaaataatgatgaagcacgatcataattcacatttaatataatattgatct
caaaaccaggtagagcaatagcaagactactcaataattcatttgtatgcaaggtgtagggtaaacaaaacaggt
aaccaattaggtcccatcaaattaaccaattaggtctcattatcaagatcattattaaggtaaagaaagtgatca
aggacataacttttcttatacttgcgataaccttgttgtctatatgtgtcacaaccggttttggggcaaaccga
atgcataacatatgtgtgccaggatccatttttcacacatatgttgacgtcacaagtgtaatatatcaaaagaca
atgcaataaaggtgtaaagggactagagtaactttattacaccatccagacaatagtgtcttaaaaagtatcatc
atcaaagtacagtggaataaacatcgacactcttccacaggaaggtgactggcgcatcgttagactcaatactca
tcaacgtcatctacaaagtctacttcatcaccctctgatcaaaatattagcaaggggtgagctcacttatggtcg
gggcttagcaagtgagaggagtaatgcaagtttaacaaggtaggctatggataagcggtaagcattttagttggt
caatattttattaacaacgcctgacttactaattgtaagtgtatcccaaaatttcccaattaaacataagtacag
aattatatcaaaagcacttaaatgaaaccacttaagcaaaccatttatgaatcatcggatcacgatttattttcc
atctttaagtttaattatcatgtgagggtccaagttgctcttaatagtgagcacgactgacatatcagttttaca
ctctgcagatgaggtacaactttacccacaagtcgtgtatcccctctagtctaggttgatcggacccataaacac
ttcctaggtgaatggctagggatccactatgaggctttcacaaaatatccttagtacaaagccacccgctaaggt
ttccacattagtattgatggctctctggacgaggtaacttagccaagattaccaccccatgttaaatgagctgcc
acgaaaccactgccgcccccctcttgcccatctttcaggtagggttgctaagcactaagtctatagagctgactac
caaatccagagccattatagcacacgtggttgcactgttatcctgggtggtcaatccatgttccatttaatacaa
aatggtcttatttaaccatcgggttaacatcataatacaaattgcatttccaaaataatagtatcattaatgagt
gacattataattatcaaattgagcggtagcacaaaatgctaagcatagtatttatcccagggtaatcaaggatta
gggttggtaattctaagaaatcctatttaggcaaatcccatcaaattatgaaaatatttaaaaagtaaatattat
tatatgcattacttgggtacaaacaaaaatacagagggaaaatccacttgccttgctcgaacgtgtcatgtgggt
cgtcttcgaacgtgtttggttcgtctaccacacagtcagcttctaatatcgattcgcataacgtacatacaaaag
aaaacatacatcaaataaataaacatacaccattacatgggtaacattcaccattaaatgaacaatcacatacca
cgtaaactatacttgtaaataatgcctaaatcatagttattttattaatccactaaaatctccaatttgaattat
cactggtgaaagttatatgggcatgtgaccacaaaagtattttacctagcatagcttagatgaaatgacttaaca
cattctacaaaattaaattactaattaattattttctttatgactaataatattcttttggttttctcctttctt
ttctaaatagaaattatatgtatcattaaaattttatttataaatcacataacattaatattctaagaattaat
gtaaattactaattttagactttagtaacataatcatggttatttcgacataaatagatattctactaattctta
gtgaataagcaatatgcgtatgaacacatgtatcatgactaggaaaaatctatattttaacctgaccaagaagat
gtggcctagtacctctttcatttcttatatcagtaaaactatacttaaacaaatactaacatattattgtttta
tgcgttaaaatgagaagttaatcgcaaatgtttctagtattaaatcaccaagttaatgaatcgtgacataactta
tttattttcgaaatataaaaatacccatatccaaaatattgttgcagaggctagttcataactaaacatataCAT
CAGGGTGTAGCAAGAAATAAACAGGGATCTATTTTTTACAGTTTCTAGACTAGATAATATATGAACTAATTTTAC
ACAAATCCAGGGGCTAACACGTAAAATCATCATCTTCTACCTTGGGCTAGCTCATGTGCAGGGAGGTCTTGGCGA
GGCTCGGTCGGGACGGGGTGGCGCGACACGGGCGCGTGCGACGAGGCAGGGCTCGGCACGACGGACGGCGAGGCG
CGCAGGGGCTCCGACGAATGGAGGCGAGATCCGGTGAGTAGAGCGCGGGCGAGGAGAGAGGAGAGGGAGCCGGGC
GAGTTTGGTAGGGAGGGGTAGGGGCTCGGCTTGGCCTTTTATAGGGGAAGAGGGGGAAGGGAAGGGGCGCCGAGG
GAAGAGGAAGGGACGCCAACGTCCATCAAAGCCATTGATGGCAGATCCATCAATGGGGGCGAGAGTGGGGGAGAG
GAAGTGGGAGGGCTCAATAGTTTCCGGCTcagaggagatgaggggatgtgggccagggtcgggcacggccggagt
gcgcggtggctcggtcggggttgcgcgcgggtgcgtgcggactgtctgctggtgcaaggatgcggcgactggagg
taggagaagggcctgacgggtgggccccacctagcggtgagagagagagagaggaaggccggtagggctggtggc
tgggccggttcaaccggtcctgctaggatttctcttttttttttcttttatccatatttccaaaaatgtgaata
aatatattttaattatttctaaaagttcataataattatacccaaaaatattactattaaagagtttatttttg
gaccaaagtgctcatcttaattaattgggtttttaatataaaagaaattccacaaaaacaatgatcaatcaaatg
tgagcaaacagaaattattccaaaggtaaataagatcaaatatttgaaataaaaattcattactatatttgccac
taataatctaaatgcattattttagctgatggttttgagctgttgcaatatgctgccccAAAAGTTGTGCAtca
ggttgctcgtcagctcgcttctatgcgtagcaatacatatagacaaagaataggtaaaaataacattacacaaaa
tatgaaaacaaaaccatacaataatattctacgcattgctacaaaaccgcaggtttgagaatcactaaattcgga
gttaaaatgaattagatattgattttacaagttttctaggcattttggcattatttttaaaccttaaaatgatta
ttatacattttactattttcctgatttatcttatttgtttaatagagaaatgacagTGGGTTCTATTAATAAAG
ACTTTAGGGTTCAATTCATAAAGAAGGACCTAGATTGAatagttcactactaaccatggatcgcgggttgtttta
tagagaacacaggggggcttttgaaaatatgctcggccgaagggatatGGATGTTTTCTGGTCATCCATCCAGAGA
TCAGTATCCCGGATAAAACGACAAACTGGTGTGCATTGAAGGTCCTTCgatcccgatctaacggtcacagttcaa
ttagtcgaattgttttatgatcatctaatcctagcccttaccttggagatcaacgactctagttatctcttcttc
cctcaaatccaaacacggcggtgccaCCCATTCTGTGGCGGCAATATCGCCGGAGCATCACCCATTCGCAATCTG
GTGCACAAAATCTGGTATCCGACAACCGCAAAACATATCAGGGCTCATAACGAACATGGGGACGCGACTTACCGG
TGGTTGAGGCGGCGAAGGGGGCCTGCCCACATGCACAACAGCTCCACTACGATGCGCCCGCTTCAGCGAGCGATT

CCTGTTCCACCGTGATCTTCGGGCTCCTCTAGGCTCTGAACACAGGCGGTGAGGTCAAACAGATCTTCTCCACCG
CACAGCGCACgctcctcagcgtatgcctgaaagtcgcatagatggggtgaataggtgaaacctaaaaattacaaa
ctttgaacacaaacttcaccagaggttagcgttagaatgagtaataataaaatcggagtgcggaaTATTGttctt
gcaatgagttgctcaatctatgcggataacattaggagccaactcaaaatgattgcaagcaagagaacttagaga
gaggggagagaaagaatcaagtcgcaatacaagattaacacaaagatgcgggcgatttgtttcctgaggttcggt
tccaaagaacctacttcccgttgaggagaccacgtaggctaggtctctttcaaccctttcactctctcaaacggt
cacttagaccggttgagtttttctccttaatctcgagggtcacttagaccccgcaaggatcaccacacaattaga
tgtctcttgctagctttacaagtcacttgagagtttagaaggagatgaaggaagcacgatcaagaaacctagcaa
caaagagcaataagaaaaacacaagatcacattctctctagactttccaaggcacaaatcactaatgatcactag
tcacacttatggcacttggaggagtttcgtagctttgaatgtgtcttggaatgatgtgcttgctcctgtattgaa
tgttgatgaatggagtgcatggatgaagtgaatggaggtggttggggttgtatttatagccaccaaccattttct
agttgtttcccactttctgccaaccgcagacggtacgcgctgctggcccagacgatccccccctgcacatcaatga
ctaaaatcgcaacggtcagtagtaacggctacataaacggctataagtgcagtaaatgtgtcgtcagatgtcaga
taaagctgtcgtggacggttcgcaatgacactaaaaatgaattttaccgaaccccacaccttcgggtttttctgg
ttttcaacggaaggtccacccgcctgagagccggacggtccgcgcAtaggtttgttgccggtgagtagctcatat
gcagtcttcttgaggaggcgatgaagatagagacgattaatggcatggcaagccgtattcacggcttccgactaa
aactgttctggcgacttatattctcccagcatcgtccttgccatgtcaatgagcgtcatgttcttcctctccact
ataccattttgttgtggtgtgtagggagcgaagaactcatacttgatgcctttctccttaagatattcctccact
tgaagattcttaaattcagatccattatcactccttatatttactttagctcaaactcattttgagctcgtctt
aggaagcgcttgagagtttcttgggtttcagatttatcatgtaaaaagaatacccaagtgaagcgggaatagtca
tcaataattactagaccatacttacttccaccgatggtaaagtaaggaacaggcccgaagagatccatatgaaga
agttccaatggtcttgatgtcatcatcacattcttgcttggatgacttgtttccacttgtttacctgcttgacac
gttgcacaaggtctatctttctcaaaacatatatcggttagtcctaccacatgttctccctttagaagcttatta
agattcttcatctcaacatgtgctagacggtgatgccaaagctagttggtcttagcaagtaaacatgcatctaga
tcggcattttctttagtaaaatctactaaataaagcttgtcgtcagatacacttttaaatgctaatgaaccatca
cttcttctaaagacagttacatcaacatttgtaaatagacaattataccccatgtgacataactgactaacagaa
agtaaagtatatccaagcgattctaccaaaaacacattggaaattgaattctcggtagtaatcgctattttacct
atcccctttaccttgccttgattcccatctcaaagatgatcgtgtcatgagaatccttgttcttgacgtaggagg
tgaacatgttcttctccccgtcatgtagtttgtgcatccgctatcaatgatccagcttgatcccccggatgcata
aacctgcaagatatttaggcttgagatgtaggtacccaacttgcgtcgggtcctgtcaggttagttacataaaat
tttggaacccaaatacaagtcttacctttcttgcattttggtcccacacttgtagcaacaattttatcattctta
ctatgaattacataggatgcatcaaaagtatgaaatagaatggagggtccataagatgcattcatattcttaagc
gcatgagaaacaacatgagaagcatgacacctagttctactcctatcactagatgaagcaatcacagtgcgagga
gtaaaattatcattgcaaataacatgataagcattatgagaagtactcttgacatgagaatacaaataagcatgg
tttctcttctcatgaacagaatgcacactattagccataggcgacttccccttctcctttgtgaagttgaggtcc
ttttggctcttagtgttcttggttcccttttggaaaccaattccatccttaatggagggatgtctaccaatggta
aaagcgtccctagcaaatttaactttctctattgcacttttgcaacttttaagttgaacagttaaatgagcaatc
tcatcatttaattttgcaatagtagaagcatgattactacaaacattaaaacgtgatctttgcactttgcacaaa
ttgaaacatgctcaattgaagaacttgcaacatttactttctctattctagcatttaaatcatcatttaaactct
agaagctagtgatagaggaatgacaagaagataattcaaaagcaagcatttcattcttctttatttcaagcacaa
gagattttctacttcaactactttatcatgttcctcataaagtagatcttgtttttttagcaatctatcatttt
cattaagagcatcaattaattcattaattttacctaccttagatctatctaagcccttaaaaagattgttatagt
ctacatcatcatcatcacattcctcatcactagaagtatacttaggggtatcacaggtatgtactttcttctct
tttgccatgagtgaagtgtgatgttcgttgggtaagagataggatttgtcgaaggcagaggcagcaagtccctca
tcatcgaagacggatgtagagcagtctgagtcccattccttgccaatgtgtgcctcgccctttgtcctcctgtag
aatttcttgttctccttcttccctttcttatcttgtacctgatgattctcattattgggacattgagcaataaaa
tgaccagtcttaccatagaagcatgaatgctttccccttaatttgttcttgttgggctagtccttatgtcccttc
aatgtagtcttgaagcgctttattacaagcgacatttctttcggcagcctctacttgtgccaccttgtttgggag
cgtatccgtgttggttgttgccttgagtgcaacgggctatggctcataaagaggaagaagttcgttggtgatgtc
ttcaatgtaccttgcttctttcaccatcatgtgtccgcttacaaacttaccaaggatttcctcgggcgacatttt
ggtgtgtctagggttttcacgaataaggttaacaagatggggatcaattacagtgaatgaccttaacatgaggtg
cacgacgtcgtgatcaatccatcttgtgcttccttagctccgtattttgttcactagggttttgagcctattgta
tgttttagttggctcctctccccttttcatgtcgaatctccctagctcgccttccaccaattccatcttggtgat
catcgtggcatcattaccctcatgagagatcttgagggtgtcccatatttgtttggcgttgtccaaaccactaac
cttgttgcactcatccctgcatagagatgctagaacaacagtagtggcttgggcattttatgaatttgctcatt
aataaatatggcattatcactactatcaaaatgtattccattttctactacttcccaaatgctagaatgaagaga
gaataaatgaccgcacatcttatgactccaaaaagagtaatattccccatcaaaatggggaggcttgccaagagg
tatagatactaaatgagcattggaattgtacggaagaataatcaaaagaataatttggttcaccatctttttct
ttgtagaggaatcatcattgtcgacgtcttttggtgaagaagaggaggcatcactgccgtagtagatgatcttct
tgatgtgccttttcttcttcctgtcctttctcttgttgttcgagcctgagtcgatgggcttgtcaccttttggct
catcaatgttgagagtctccttgtcattgatcacaatccccttgcccttaggattcatctctttgggcggttagt
cccttgatgaagaaaactctgataccgattgagagcacctagaggggggtgaataggtgatcctgcaaaaatcaac
tctaaacaagaaaaacttggcttatgataagtgctattgaaaactaaaactgagttgttatgagtagaggttgag

aggagaagagaactcttcacttgattgttccttgtaaatgtctagttgataatcttgccaagaatggaagaggga
aagtgatagaattacatggacaatagcaggcttcaccatgggactaaatgtctcatcatagtaaactctaggaca
ctaagtgaagccctagagaacccatcgtgccttgtacctatctagagaaccatctgactttagaaccatccacct
tcctcttatgtgtccagactcacttgctagtttccacattactgtctgaggatgggacaccaagttacaggtgtt
gttggcaaggagggtctcatactcttcctccataacgggatactagtgaggatcaactagggcactgcgaacgga
agatggcataagggagaccgacgaagaggtcacaaagaggacaagataatcaaccagggaagaaacccagcgact
tgtcgtatcaccttgggttgacttggcgagggtcatggtgaaggggatgatacatcatcggcttggcctagcag
tagggcagtgcacccattcacactcttcgctaaataaacatgcataaagttagtgtagcagggagggatcaacgc
cagcgagggcattggagggggccctggtgatgagggtagggtcaaaaaagtcgaggttcaagacgaaccaactta
ggacgctagtggccaggcggaggcccagaggatgcacataatttgaccaggggcaatagggcagtccaaaggtgg
tagggaggcaccgaggccatatgcaagagtggcatgggtgaacctacaggcaaaaaacaagatgtggctaactaa
agagagctacatggtcaggaagaaagtccaactcgagaaggaagtcgatggtggagatggaagagggagaaagaa
aagtcatctcagcgaaaacaatgtgacaggagatcaagagacgatgagtgataagattaaggcaccattacccct
tgtgattagggaagtaaccaagaaagacacaacgggagaagtggggcaagattatgggagtaatggtagaaaggt
tgtggtagcatgcaaacccaaagatatgaagatggtcgtaggagagtggtgccaaacatagtgaagtagagtata
ggatcacgaaacgccttagtacaaagatgatcaaggatggaggtgatgtggaggatcttagccggtagccaacat
gaacgaaggcctgaaagaccaaggaacacatgatgttgttagtaatgcaaatcatacgatcatcctcgccattct
agaaagaagtgtaggggcagaacatccgaagttgaacaccataggagaagaatgactgagctgagttatcaaact
cacagtcgttatcacgctgaatgaccttaatggtgcgaacaaactaagtggaaaccaagcaaagaagtgcaggag
agtcacaaatgtatcgaacttcaaacacaagggaaaatccataaataatgagagtgatcatcaagaacgagtaga
taatagtataatacttatcactaaaaacagaagaggtccacaaatcatagtgtataagaacaaaattgtgcactg
tctaggaggagctagagaatggaagtctaacatgacaacctagctaacaagtgtggcatagatactcaaagaatc
ccctaatacaagagatgattgaactactacacaacttggataacatgctaggactaggatgaccaagctgacgat
gtcgaatgttggatatggtggaagtaatcaatgtgtgtggttgtagcaaggacaagggcgatgatggagcgaaaa
cggtaggtggagaagtaaatgaaggcagacgtaggatgtacatgggcctagagctatcacaacgggcgaacaggA
GGATGTTCATGTTGTCCACAATGTCAGAGGTGAACGTGAGGATGGTCATGTTGTCCATAACGACAGACGAAGGAT
GAGACATGAGATGGATTGGATGAGGATAGTATACCAGCATCCGAAGTGATATGGTAGAAGGCACCAAAATTAGCA
ACCTAGTCAATGACGATAAGAGTCAGGTCATCATTCTAAACGAGCTAGCAAGTGACTGTTGGTCCCACTCACTGT
GCCACATGTTCCAGGACAGCACCGAGTGGGGTGGGCGACTGGGGCCCGTCGGTCAGCGAGCACGTCCGGAGAAAG
ACACCGGCAATTGGGGGCATGTCGATAAGTTGGAGGAGAAAGGTagcgtctaccctagagaaacccgccctcgga
cgagcccaccgagcccaaggtcaagcgtggcgaagTCTAACAGGGGGTCGGCGTACCGAAGGGAAGCCACTCAA
GTGGATCCCTTGCCTGCCCTGAGACCAACCCGTCATAAATGATCGGGCACATTAACTGCACCGGGCATGGTCGCA
TTAACTGCGCCTGGCACCGAGCCACAGCGGAGGCACCGGTCCGCCTCCCACTTATAGCCTACGAGCCCCTTTGAC
CACAGAGCACTCATGACCTACGAACCCCTCAGACTACGGGGCATTTATGGTTCGCACACCGCGCGACATGCGGCG
CACTCAAGACCTACCGGGGCTAGGCTTGAGTGCGTAGACCTCCGGCAGGGTGCAGCACGACGAACCGTGTCAAGC
CCCAAGCTACGAAGGCCAGGGGTCGACCTAGCCCGAATGATAGCGCTCGAGCCTCATGATGCTCGCGTTATCCGC
CATAAAGAATACGAGATAGCCAGGCGACCCCGGGAACGCTTGGTTTACCGCCGCCTGGTGACTCCTTCCTAAGAA
AGCCACCACTAGCTGACCCTTCCTTGGCCTATAAAAGGAAGAGGTTGGTCCTTAGACCAAGAGGGACCTAGAAGG
ATGGACCCCAAGTGAGAAGACGCCCAAACGAAACGACGAACAGGAGGACGCACCGACGAGGATCCAGAGGCCAGA
GCCCTGCCGCCAAGCGAAGACTAGCTAGGACTCCACTTTGTAACCACCTCCACCACGAATaagagacatgggagc
gtcccctccctctctctcgccctcttataacccctactacgagctttaatcatcaatggtgtcggtagggtaagc
cttttatgactggaagtagggacgttctgcccgaaccagtataaatcttgcgtccaccgcgcacaccgtccggag
cccggaacgcgcacaataaatttacttgtcagagcaaggttcgaaacaccgaaagttggcgccaggtaggggcct
tgcgtgttcccgacgaagttcttcaagctccggatggcaaaccacgatgccaatcgatctcccggaacaacgatc
tatttcgagagcctcgacttctccctcaacgacgacgacgaAGCGGTCAGGGCCCCAGAAACCCGAcccccccc
ccccccGCTGCCGAGGAATCACGACATGACCGCAAGGGCCAGTAGCGGCTCCCATATCGACTACACCCCCGGAGC
GGCCATCCTCGGGAACCTTGATTTCGTCATCAAGCAAGAGAGCAAAATGGTCGGAATGGCACCTACCTGGCCTCC
TCTGCCCGAGAGCCCCGACACAGTTTTAGAGGTCGTTGGTGACCTCAGGATCGTCTCGCGCAAGCGACGTGAGAG
GCACAAGCCGACCCCGCCTTCAAAGGATGACAAAATAAGGCGATAGATCGCGACCTATATGGACCAAACCCCTC
ATATGACGATATATCAGCTTTCTTCGCGTCCTACACGGAGCTTTCCGGGTTATACAACGCAAAAACGATGTCCTA
CTACGGTGGCTCAAGCAACAGCCACGACGCCTTCCGTGTCGACTACGCCACGCCCTGGCGATGATCGCCGCCCTA
GTGGCCCATGGCCGTCCCCCACCTCTGATGACTTAGAGTTTGTAGGGATGGCCAACGGCAGTGGGGACCTCATCT
TCGACGCGCTCTGCATGGACTCGAACTCGGATCCGAGTTCCCCGCGAGCAGTGAGGAAGTAccccccccccACCG
ACGAGGAGAACGAAGACCGCCTGACCATTGCAACCCCATCGACGAGACAAGCAAAGTGGGGCGCTACGTGTGGTA
GGACACGGAGGGATCGTCGAACGTAATGCTTCCGTTGAGAGCCCATGGTAGATGCCCGACCGAAGTAGATCCGTG
CTCTACTTGTAAAACCCTCGAACGGTTATGAATAAAGCATGGTTTTCGAAATGCTCTCGACGTCAAACACGGTAA
ACAGGGGGAGAACCTCAGCTAAAGCCATGGtcctttctctcctttttttcccttttttttctctctccctttctcg
ttttccttAGGTGACTTTCAAGATAGGAAGCGTACCAGCAGACGAACGAGACACAATGAACAACGACGAACTTGA
CAAAGTAACGAGCTGGAACCCCCATCGCAAAACCAGCTGTCACTTTTTTCTCTTCTGAAATTGAGTGCTTCAAGT
TTTTCTTACCTTTCGTCGGAAGGGTCTTTGGCGAAACATGACACAGCATCGCTCTTTCTTTTTCTAGGGCACCCT
CGACGGGAAAGGGGCCTCGGGACAATGTAACCTAAGACCCCCGTGATCGAACCTTGAGGGAAACATTGAACGGGG
TGGCCATGTCTCCGTCTTCACTAGAAACGAATATTAGGGACAGCTAATAAGAACGAAAGGACCCACTGAGGAATT
GGGTCGTGACTGCGAACACCTACACCTCGAAGGCTACGATAATGGTCACCAGTGGAACCGGTGGACCCAAAAACT

CCCCAAGGTGGACATAACGCTCGTTGAGGCTGCGGAAAAGGTTGAAGACAAGCGTGTGATAaggccaactccagc
aatgtcgtgcatgcgagcgtgtataacactgttttacactatagatgcattctttatagagaggagtttgaaata
ggaagtgagataaggggtgggataaggagtctgctggagatagtctAAGAGacgaactccccaaggtcataaaca
gaggttgccatgcctttcatctagcaacagtatttgatgacagtgaggtcccccctggacaaaggtcctaaagacg
ggtcaagccggagagcattagccttgcaattgctaaggaactactcctcagtggcaatccagacctgaagatcag
tgccatcacactcgtggacgacgtccaccagctcaatagtgaccaatccaaggatctaagagattactacactat
ccatctaacactaggagaggagagtaagagacagtgtgccaatgaaaacatggacaacatagcgttgtagcgtga
ggacgaggctacgttagcgggtataagaggaccccggttcaaggacggagattagggagcaaatgttccaaatac
cctacacctacacgtgaaggttggtgatgACGACAGCCTCATATACTAAGGGGTGTCGACCATCGGGTATGCCGA
GGGAGTCAATGTCGAAACAGCCAAACAGATGGCGCTTAACCTCGACAAGTTGCCTAGCAAGGGTATCGACAACAA
CACATTCATACTCTTAGGCATTGCGACGACACGTTCATGCTCCTAAGCAACATCCACAATCGTCTTGGTAGCTCA
GTAGTGTGGGTTTCCTGAGGCAAAAGATGCCTAGGCCTATGGCAGCCAAACCTAAACCGACAATGTCGTGGTTGA
GCAAGCTAGGCCAACAGCAACCAGACCCATGCCCATGCCGGTGGCGCCGGCTAGGGCCAACCAGGACGACAATAC
CCAAGCCAAGGGTGACCATGCCAACGACGCTTAGGCCAGGGCCGGCTAGGAGCACCTAGCCTGGAGGAGGCTAGG
TCGCCACCTGCAATGGCAAGGACAAGAGGAGGCAAGCGATCAGGGAGCGCAACAAGGTTGGTGGAGGAAATCCTA
GCAGACTAAGGACCAACAAAAAGGGAGGAGAAAAGAACGGATGAGAGCCAACAAAGAGAAAGGAGGGGCACCAAC
CATGGTGGCGATAGCCTAGCGGCTAGCCAACAACTAGGAGGGGTAGAGGGAGAAAAAAATTGGTTGGATAGCATG
TTAGAAAGTGAAAACCCTAATCCATTGATTATAAGCTAAATATTACATACACTCGGCACTAGTTACATACACGGA
CTGGGCTCTAGGCTCATAACACTTAGACATATACAAACAATCTAACAAGCATGACCATAAACAATCAAATGCTCA
AGAAACTAAACTAAGAAATCAACCAGCAATGAAGTAATATTTTCCAATGTAGTGTGCATCTCAAGGCTGCTGAGT
GATCACTGTAGTAGATCGGAAAATGGATAGCAGCATCCAGAATTTTGATCACAATTCATTAGATTACATGCATTT
TCTGCTGGTATAAACAGATTGTTAACATGATTCAACCAAACTGAAACACGAACCATAACCAAATTCTCAATTGAA
TATACAGCTCCGTTTTGAGGCGGTAGGCAAGCAAAATACGAACACACGTTGCCAGTTCTTAACTATATGTTGTCT
TGATCAGAACATGCTCTAGCCTCCAATTTAGGCTGAGTTACAGAGCTCATCTGCACTGACCCTGATGGCAACAAG
TGTTATCTAACGATATTTGGAAAAAGAGAAGAATAGCTTCTGGATGTTTGTTAATGAACAGGTTGAAGCTGAAAT
TTGGCTTGAGAAAGAGCACCACGACATACCAATGAATAGCAAGAATAACATCCATTGCCTTAATCAATTCATAAA
CAataatctgaaatgattaaacacttgtcatggaccttcggtccatgaggatagatgtcttctccggcgaggtta
taccccctctgccgtaggcttggttctagggtagcagccctaggcgcttgagagtcattggtagagagagagattg
gtttgataatatatttcttattcttttgtctcctgcgtacatgcctatatataggccgccggttgaccaactagg
caaccactaattaggcaagtgactaattggaattatctcccctacaatttaggtctaccaatttagccaccaaat
cagtcactttcctttgccatgccttccttgcgcgcgctttcctttgccatgccttccttgcgcgcgctgctgcct
gcgcacgtgccttccttgcgcgcgctgctgccagcgcccccctgcggcctctcgcttttggcacgcataggtgcgg
ccccacatgagtggccactccctatgatgcccatgacatctcccccctcccttgaggaccagctcgtcctcgagct
gccatagacatacttgatacgacttaaggttaagccttttacatctcggcttacctttattatttaagacgaaaa
tacaaaataattgtggaactaaaatataaatttgaactgcacctatgtcctcctgtcctcctggatcccaaggaa
agagctgaactgcggccttcacgttggatgaaaggtggaggagaaatcagcccgttcttatgttgggtctgtcca
gcaccaaggcaggtgcccgaatgaaggagacgaccctctttggccgtgcaggggggctgcatacccagatctcga
cttctgcaggggggttcaaaaagtatagacgagacctggtggttgggcgcacaggctgcgagttggtgcagcgatg
agctgatcagcaagtgcagcttccgcaccgcccgcctaacaaggaagccgtgcactgcggcttgcaggcgcacca
cggcttgctcatgtgatgacagccatggggtggccctggaggccacagcaaggtgcttctccccacgaagggact
gtacctggcggcgtgccaggaaccctcgcgctgccgtctgaagccgcaccgctgcttggagttccttctctgtct
tctccttgtagcggtggaacatcacaacgaactgatcccactgctctgcgaagcggccaaaactctctttgagcc
gagtgaatgcatctggagttggagagggcgggtgagagggcgttgcggcggctgatggcgcggcggctggtggtg
gtgagggatgggcggctggtggtggtgaggataacctggagctgataccaggtgtcatggaccttcggtccatgg
ggatagttgtcttctccggcgaggttataccccctctgccgcaggcttggttctagggtagcagccctaggcgctt
gagagtcattggtagagagagagattggtttgataatatatttcttattcttttgtctcctgcgtacatgcctat
atataggccgccggttgaccaactaggcaaccactaattaggcaagtgactaattggaattatctcccctacaat
ttaggtctaccaatttagccaccaaatcagtcactttcctttgccatgccttccttgcgcgcgctgctgcctgcg
cacgtgccttccttgcgcgcgctgctgccagcgcccccctgcggcctctcgcttttggcgcgcataggtgcggccc
cacatgagtggccactccctatgatgcccatgacaACACTCTAATATCAGGACCAAACAACAAAAAAGGCTGTCA
TATATACAGAAGGATCCATAGCTAAGTGCCCTAGTAGTTGACTCCAGTACCTAGATTCAACTCATCATGGCAGCA
AATTTAAGGGTAGTTAAATATATACCCTCGTTGACCATGCCAGCAAATTtaagggctagtttggaaacttaaatc
ccctctgcaattggagaagaatgagagggaaactagttcattttcacctcaatcccctcaaatgccggagaggat
ttgagtttccaaactagccctaaAGGTAGTTAAAAATACACTCTCGCTAGCAACGCATAACggctagtttgggaa
ctctattttcccaaggatttctatttttccaagaatgaactaatttctattagaaaaatagaaatctcttagga
aaatggggttcctaaattagccTTAAAATGATAAAATAGCCACGGGGATTTTCCAGTCGGTTGGAAGACAGGTTT
CTTATTCTAAACCATACCACCGTGGGAGTGGTTTTCCCAATTATAGCCAAATTATTCTTCAAGGCTATCATATAT
CCTCATCGATGTGGCAAAGGATTCTGATTGCTGCAACTAAGTGCAGTAGAGAAATTATCTAGAAGACATTTAAGT
TGGTAAAGAAATATTTATGTAGTTATGAAACTGTCATTTGAAGACACAACTGTAGTCCCTCTCCTTAAAAGTCTA
AGAACCATCTAACTTCAGTTGCATTGAACAATACAGCAGAGGGAACATGCTGATTTGTTTTCTTGACCAAAAGAA
CTGCATCCAAACCATTTTTGGTGGAGGATATGGTTCGAGCTAAAATTGATAATGTAACAATACTACAAAGaaaaa
aaaaCTAGTAACACAGCTGGCAACAGAAAAATTACCAACAAAAGTTCAGATCTAACTAATAATATGAATCAAATA
TCTCAACTATTTTCTTTGCACAAAGTTTCATTTAGACAATACCTGCAAATGTTACATTTGAAAACTTTCATATGT

GTAATGCGTAAGCGCAAAAATGCACTCAGAATTAAACGTTGGGTGATTAATAAGTCAACAAGTTCAATGTTCAAT
GTCACTAATATATTTTGGTCGTAGCATGTATTACTGAATCAGAGTAAGAAAAATAGTTAGCGACCAAATTTACTG
TGGTTTAGCTAACATATGTGGTCTGTCTGCATTAGGTAGTACACTAGTACCTAAACACAGGTTTTCTAAAAGGCA
AATCTCTTGTAAATTGCCTTAAGAGTAGGGTTTGTAAGGAAAACTATTAAACTGCCATTTATAGTGTAAATGTAG
ACCAAGCCAAAAGCACCAGAAATTTGACTGGCATGTCCTGTATTTTGATTCTGATGTAGGGAGTTAAGCAAAACA
ACAGGAGCCATGAtgtcaaggaccttgaatccctggctcgccggacctcggctacgtagctcgtagccgtcggcc
gtcttctccggttggggttatgccccctaaccgcaggcttaaagaagctggaggtaggagaagagtaaattctt
tattgatttgaactgtgccgcatacatgcctatatatggccggcggcccattacaaatggagcaagtctccttga
ttgatagattgctactttcctaatcttagcctgattgcttctttcctaatcttagccacttgctatctgtggctg
atttgattgcttctttcctatctgtggctgatttgattgcttctttcctaatcctagccacttgctatctgtggc
tgattagcatgcgctgctgtggctgcgcgctcggctgctcgccggacatcacgcgcgcgcctgtgcctagtgaac
gttttaccgtacatgacatctctcccggcctcgaaagccagctcgtcctcgagctggaactgcggaaagctggcc
gtgaagtcgtcaagctcctcccaagttgctgccgtcgctggttcaccttgccactggaccaggacttgtttaatg
ccgcgagctaggcggccagccaagacacgtgcaggtgcagacactgcggcgccgttgagcgtggccggcagcacc
ggaggttcggaaggtggcgcccccacaaacttcctgaggacccccacgtggaagacgtcgtggagtctggcgccg
gccgggagctgtaggcggactgccacgtcgttgatgacttccttgacacggtagggtccgacatagcgcggcttg
agcttgcctgcggccgagcgtggcagtgacgcggacggccgctgccgtaagcgaagaagggcccagtcgcccacc
tggtagacgactctcctgtggtgctgatcgtaggcgcggcgctgggactcttgagcctggtggagacggaaacgg
acgtcgtcgaggaaggcagcgcgctcctccatctcttgcgccaccgcgggaacgcgcgcatcgcccggctcatag
gaccggatggtcggcgggtcgcggccgtacaccacccggaatggcgtgtcgcggagtgaggactggaaggcagtg
ttgtaggtgtactccgcccacggaagccacctgagccactgtcgagggcggtcgccggtgaagcagcgaaggtac
atggcgatgacgcggttggctgcctccgactggccgtctgtctgggggtggaaggcggaggacatgaagagcttg
gtgcccgtgagctccatcagttcggtccagaaggccgaagtgaacacggcgtcgcggtccgagacgatggactgc
ggaacgccgtgtagccgcaccacttcggcgaagaacgcctgcgccactgattccgccgtgtatgggtgcgccagc
ggaatgaaatggcaatacttgctgaagcggtcgacgacggtgaggatgaccgtcttgccttgggcgctgggcagc
gcctcgatgaaatcgatgccgatgtccgtccacaccgccgaggggattggcaatggctgtagaaggccggccggc
cgcaagtgatcagacttgtatttttggcacgtgccgcacgccttgacgaaatcctgcacaacacgacgcatgttt
ggaaaatggaagtcgcgctttagccggtgtagtgtgcgtaggacgccttcgtggccatcgtcgtgggtggcagcc
actatttcctgcagaagaggggacgacgccgggatgtagaggcgcccatgcgtggcaaccatcccgcccatgatg
gaccacggcgtcgtacgtgtacccgcgcgaatttcctcacagatggcgacgagggccgggtccgtagcttgtgct
tggcgcagccgtgcgatgaagtcgaagcggggtgcggatatggccatcagcgcgccctcctcggtgtcctttctg
gacagggtgtcggcgacggtgttcgtggcgcccgagcggtactccacctcgaagtcgaagcccaggagcttcccc
acccaatggtgttgagggatggtggccagccgctggtcgaggaggtacttgagactgaaatggtcagtcttgacg
gtgaagtgccggccccacaagtatggccgccaatggcgaacggcctggaccaagccgatgagttcgcgctcatag
gcggcgagggcgcgatgacgggggcgattggccggctgaagaacgcaaccgggtgtccctcctggacaagtacc
ccgccgaaccctgtggaggaggcgtcgcactccaccacgaacggcttgctgaagtccggcatggcgagcaccggg
gcggaggtcacggctgccttgagcgccgcgaacgctgccatagccgcgtcgtcccatacgaaaccatccttcttg
aggagggccgtcaacggggcagccacggtgccgtagttgtggacgaagcggcgatagtacccggcgaggccgagg
aagccgcgaactgcccggaccgagcgaggtggcggccagtcgcggatggcagccaccttggccggatccatcgcc
accccggctgccgagatgatgtggccgaggtaggagacggccggcgcgccgaaagagcacttcgagcgcttgacg
aagagctggtggcgctgcagctcgtcgagaaccgcacggagatggcggaggtgatcggcccaggtcttgctgtat
attagaatgtcgtcaaaaaataccaggacgaaacggcggaggaacggtcggagcacgtcgttcatcaacgcctgg
aatgtagcgggggcgttgcagagcccgaatgccatcacgaggaactcgtacaatccatcgtgcgtgcggaaggcg
gtcttgtgcacatcctccggccgcatccgcacttggtggtagccggaccgaaggtcgagcttggtgaagaatttg
gcgccgtggagctcgtcgagcagctcctctaccaccgggattggaaacgcgtctttgatggtaagtgcgttcagg
gctctgtagtcgacgcagaaacgccaggagccatcgggcttcttgaccaagaggaccggggacgaaaacggggag
tcgctgcggcgcacgatcccttgctcgatcatcgcggtgcactgacgctccagctcgtccttgtgggcagccggg
tagcggtacggacggactgccaccggcggcgcgtcgggcttgaggaggatacgatggtcgtgggcgcgcttggga
ggcaagcccgaaggggcggcgaagagcccccccaaacgagtccaggagagcctccatcagcgcatcccccccctgtc
atggtgcgcaccgccggttgcgctgcgcgtggggtgcccgtccaggagaccgagcgccctgggcgctggaacgtc
atgcggcgggtggcgaggtcccaaacgatcggccccagggcccctagccagcacgtgccgaggacgacgtcgtac
ccggccaacggcatgacgaagagatcggccgggtatggttcgcccgcgacgaggagcggcgcgttgcgcaggacg
ccggcacaggagatccgctcgccgttggccaccatggccgtgaggcgtggacggcgctggatggggaggcccgac
tggtgtgctgccgtcgcggagatgaagttgtgtgtgctgcccgagtcgagcagcgccaccagggaggccggaccg
agggtgaccgcgatctgcatggtttcagccggcgcaaccccggctaccgcttgtagggagaagagcggggcgttg
ggctcggccgcgcccgtgtccgcgtcgtccccggtgtcatcgatctccaccccatccacaaaaaagagacgtctg
caaaaacgattgtggccccgagaatacttctcgttgcagttaaaacataagccttgccgacgacgatccgccatc
tcctcgggtgacaggcggcgagggccctcgcctcgtgcttgtgctgcacctatcggcggcgctggcagggccaag
tgaggtagcggcgccggcagcgtcggagcggctggtcgtggagggggcgccggcagaagaccgcgcgctgccggc
cttgtcgccacctggatgcgagcgatctccagctgttccacctcccgcgccaggctcatcgcggcgctgagagtg
tcggggtggaggatccgcaccgagtggctgagcggcggcagcaaccctcccgtgtagagctggacgcgctgcgcc
tcctccatgggcccggcgtggggtaggagctcctggaaacggttggagtactcttcgaccgagcccgtgcggcgg
cactcggtgagctcgaagaaaggcgcggacctgctgttcggcccgaagcgcaggttgagctgctccttgaaccgg

cgccatgacggggtgccgtcatcttcttgcagctgaatgtaccagtgttgcgcgatgccctcgagatggtacgaa
gccatccacacttgctcctccggcgaagaacgatgttggcgaaagtaagactcacacttgttgatgaagaggaga
ggatcggtcttgccatcaaaccgagggaagtcccatttctggaacttaggcgggcagtcggtgtcgcgagggcgg
tcatgcagggcgccgccggaggacgacgacgcggacttctccttcaggccggagatgtcggcctgcatctcctgc
atcatcttcacgacgtcggcgagtgtgtactcagacatggcgctggaggcaggggggacgggtgggtgatggtggt
ggtgtgtggccggtggtggtttcggtggactgggacgtgctggacgaagatcgccggactcgtgataccaagctg
tcaaggaccttgaatccctggctcgccggacctcggctacgtagctcgtagccgtcggccgtcttctccggttgg
ggttatgccccccctaaccgcaggcttaaagaagctggaggtaggagaagagtaaattctttattgatttgaactg
tgccgcatacatgcctatatatggccggcggcccattacaaatggagcaagtctccttgattgatagattgctac
tttcctaatcttagcctgattgcttctttcctaatcttagccacttgctatctgtggctgatttgattgcttctt
tcctatctgtggctgatttgattgcttctttcctaatcctagccacttgctatctgtggctgattagcatgcgct
gctgtggctgcgcgctcggctgctcgccggacatcacgcgcgcgcctgtgcctagtgaacgttttaccgtacatg
acaCATGAACATCATGTTAATAATGCAAATGACTCCAACTCTGCATAAGTAGCCGCTATGAAGCATCACCTGCTA
TATTCATAGACTATGTTGACCCCAAAAATAGAAATCACAAAACTGAAACAATATTTCATATACTATGATACTGG
TTACCTTTCATCCTCAAGAACTTCTGTTGAACACACATTGTTGGCTCCAGGTTCTGCTCATCTCAAACACAACCA
GGATGAAGGTGGTTCTAAACTAGTCAGGTTTAAGTGAAACAAACATTAATCCACAAGCACCAAGCAATTTAACTT
TTCCATAATTTTTCCAAAATAGTACAAGATGGTGGAACTTTACAAAACTACCACGAAAATTATTTTGATAAGGTG
GAAAATATTTATAACTTAGAATGGTACATTTTTATCGTAGTGCCATATTGGCGCTCTATTTGTTTGAGCTTTTGG
CGACTTTTGGACTACTAGCTTTTGGACTTCCCAAAAGCCAAACAGCTCCTAGAAGCCAGAAGCCCACAAGCTCCC
AAACAAACAGAGCTGCATTTCAGACCAACTATTCTCAACTATGCTGAGAGCAGCTACTCAAGATGTGCAGCACAA
ACATAATGGGTGTACCAATTCCTATGCTATAGATAACGCTTCCACAGTTGTCTCAATTTATATGCTATAACATTT
TCAGGATGCAAGGCTTAATTTAAGTTCATGTTATATGGTGTCATTTTCTCATGTTTATGAATGTAGAAGGataaa
aaattgaacaaaactaaaataaaaaGGGTTCATACAACATATTTAAAGGAGCAATGAAAGTTTAGAACAAGTTAT
CTTGTGAGCAAGCATATTTTGATAAGTAAACAAGGATGCAAAATACAGTTTGTGTTATAGCATCTCCTCAGCAGG
GTGCTCCTCCCCACAAAGCCCCTCCCTGCCGCGGGCACCTCTCTTCATCACCGAGCCCCTCGATGCCACTGAACC
TCTCCCCACCAAGCCAATCCCTGAACATCGGTGGAAGATGGGAGAAAAAGAAAAGAAAACACACGCCACTGGTAG
AAGGGTCCCAATTGTCATTAAAAAAGTAGGGTCGCAAATTTGAGGCTTGTTGCTAGAGTTGGAGCAGAAATTTGA
GGGCTGAAAAAGTAAGGGTGACCCTCAAATTCTATGTAGGGGGCCAAGTCTGAGAACAAAAGAGTTCAAGAGATT
TTGGGAGCCAGTACTGCATAAACATTTAAACAGTTACACAATATCCAAAGACCTAATTGTTCTCGAACCATTTTT
TCTCTAGAATGTGG**AGAAAATTTCTCTTGAACCATTAATAGTTTCCAACCCAAAGAAGACTTCTTACAAGCTTAT**
**AGCTCACAAAACGATATTGTGTCTACATGTCCCTGTAGGATCTGACTGCTTGCTGCATTCTTATTTTCTTTAATT**
**ATTTGGCTCTTGTTGGGTACTTGGGTTTCAACTCTACTAAACCCCAGCTTCCTTTGAAGCAAAAGATTTTGTTCT**
**TTTCGTTGTAGTTGTAATCCCTATGCTAAAGTTGACCCATTTGGTTGTCCAGTCACTCTCATTTATGTTAATTGA**
**AAGCTTTCCCCATCTTACTAGCTTTGGCAAAGGGAGGCAGGAAGCTTGAGGCAGCAACTGCATAACTTGCAAGAG**
**CACCATCG**GTATGACCCTCAACCAACCTACGATAAATAGTTCAAATCATTGTGAGAGAAAAAATGTGCCTACTCA
AATTATTTTTTCCTAGTACTCTAACAGTACTATCATAGCAGAGACATTGTACTTACATCTGAGGAAACAAACACC
ACTGCAGTGAAAATAAGAACAAATTCTCAGCACATGATGACGAATTGCAAGATGCAATTGGTTCAATGTTCTATG
TGTATCTATTAATGCTAAGCACAGGAAGTTTGTCACCGAAAAAAGTTAAACCTTATACATTATGGCTCCAAAACT
TTCAGAAACAGCAATAAGCAACAATTTGCTTCATTATTCTATTTGATTAAGTGTGTGTCATTGCTCCTCCCACAA
AAGAAGTTATGTTATGCATTAGTTATTTCAACTGTACACATGATGGCTACAAAAATGGTATTTTATTTTAATGTC
TCTTAAACGTGAAATAGGATCGTACATTAGTACAATAACATGTTCATTTTCCGAAAAGCATAAGCTAATGTTTTA
TACAAATTTTCACTGCAGGAAGTTGTTGGGTCAGCAGCTCTCTGGCCTGGACGTGAGAGATTTGCAGAATTTAGA
GAATCAGCTGGAGACAAGCCTAAGAAATATTCGTCTAAAGATGGTGGCATTACGACTCCACATAACAACATAGAT
ACCAGTTGTTACTTTGTTTACTAACTTTCAATTTGCAGGACCAACTTATTTTTTATCAGATTCAAGAATTAAACA
GGAAGGTTTGTCCACACAACTAATTCAAAATTACTACTCCCTCCATCCCAGAAAACAAGGCATATAAAATTTTGA
ATTTGATTCCAAAAAATAAGGCGTGTTTGTGTTATTCTTGACGCATGTGCCCTTACAATATATTCGGGAGAGGGG
TTATATGCATGATTATATTCCTTCCTTCGGACCTTGGTTTTATCGCATGCATTCTTTCCTGTGCGCTATTCTAGG
TGAGGTGGAGCATGTCTGCCTAATTGTTCCTTGATTAGGAATGATGAAATCTTATCTTTCCTTGCATGCAGTGaa
aaaaaaGGCTTGCATGCAATTCAATTACGGATCCAGCAATTCCGCTTAGGGGCACCTGAGGGAAAAAGGCTATAA
TGATTAGCTGCTTTAATTTTCGTGAAAAAAACTATACGCCTACTTTTTTGGAATGGTGGGAGTATCATATTCTAG
GGTCTCGCTTAACTACACAAGTTTCATATTTGAATTACAGGGATACCTCATGCACCAGGAAAACATAGAACTACA
CAACAAAGTCAACCTTCTTCATCAAGAGAACATTAAATTACGTAGAAAGGTACATGCATCTGGAAATAGTTAACC
AAAAAAGGGTGCAATTTGTAGAACTCGTCTTTCGTTTTTATATGTCAAGTGTATGTGAATCACAAAGCATAATT
TGATTCAGGCGTATGGACAAGGAGTAAATGAGCATCCAACAAGTACTACAGTTAGACACAGTATTCTGAATACAG
AGAATGAAGATGTTCGGATCAATCTTGAGCTGAGTGTGCAAGGGGACAAATCAGAAACACCAAGTATAGGGTGAG
ATTTTTGTCTAAACTTCAGGGCAATTCAGAGTCAGTAAACCAGATGATTTAAGAAAGACCACTGAAAGTGCCAGA
ACGGTTCAAATGCAATGAGGATAACTAAAACTGGGAGACCAGAAATCTATATCATTGGTTGAATTGCAACACCTA
ACAGCTAAACTTTCAGCTGCTATATGCAGTAGTTTCTGTGTGAAAAATGTTGATTTTAATTTAGCTTTTCCGATA
TGTTTGCCAATTGCCATATATGCTCTTGTAAATTGTACTTTGAATTATAAGTCGTCTTGGCTTCTCTAGGTACAT
AGCATTTGCTATGCAGTATGCACctagatatgtgttatctcttgatccaaaacaacttataatttggaacggagg
gagtaACTTCAAAAGATGCAGAACTTAATATAAAACAATTTCTTTTGAACTCAAAATGAAAATGTGCACTATATG
ATTGAGAATAGTACAGAGGATTTTGGTAAAATGACAGTTTAAACTTCAACTACTGTCATATGACAAAAAAGATA
CATCCAAATCTACAGTAGGTCTCAAAACTGAGCCATTGCACAAATAAAAATATAAAGTGCCTGGTACTCTAAGCA

CAAGACCTGCTTACACATTCATTGTGTTTTGTAGTATTCTACAAAAATAAATAACTTAGAACCTAGATTGAAAAT
CAAGCCTAGTATTGTATAATTATCACATCATAATACATACAATAACAAGTTAGACCAACAAAAAAACATTGATCC
AATCAGTATCAATCTATCAATTGCTGTGGGAATAACAAAAGATAAATGATATTTCTCATAATACCAAAGAACACT
TAC**AAGCTCTTGCATGCTATGCAGTTGTTGTCTCGAGCTTGCTGCCCCCAAATACATAT**CTAATCAAAATAATTG
AAGGTTAGTAAAATCAATTATAGAAGGTATGAACATACTTATAGTGTAATTTTATTACAACTTGAGGATCATCAT
AATACTGTCAAATGCAATGTAAGAACAATATATATtgaaagaggatggctgggaaatgattgttggattgattat
tgagggagcccaaggctcagtacatataggcaaggatgtcccagaaataggaaagtgctaatctagagaatcagg
aaagtgctaatctaaggacaccttgccaatcagatccaatactaccaaaaaggggagaggggctgcagccgccac
tgctctagggttgcagccgaccaccctcctgggccaaccgaacaccctaaaggcccatgggcctatacagcatgc
taacacgcccccgcagtgtgaacatcggtgcttagccacggacattcagactggaccgaaattcagtgaagaccg
aagaaggtagccccttcgtgaagacgtcggcgaactgagatgtcgtgggaacatggagaacccgcacggcaccca
tggcgaccttttcacggacgaagtgaagatcgatctcgatgtgcttcgtccgttgatgttgtaccgggttggagg
acatgtagactgcgctgacattgtcacagtagacaatagtggcactgcgaagcgggatgtgtagctcgtggagga
gctgtcgtagccaagaagcctcagcaacaccattggcgacagcgcgatactcggcttctgcactggagcgagaaa
cagtattctgacgtttggaggaccaggagaccaagttgtcgccgaggaacacagcgtagcctgacgtggacttgc
gcgtatcgggacagccagcccagtcagcatcagagtaaaccaccaagtcatcttgagtcgacggtcggagatgaa
gacccaggtgaagagtgccccgaacgtaccgaagaatgcgcttgagagcagccaggtgaggctctcgcggatcat
gcatatgaaggcagacctgctgcacggcgtaagcaatgtcgggtcgtgtgaaggtcaaatactgtagggcaccgg
caagactgcggtaatctgacgggtcagtgactggagcgccagagtcggcagagatcttgggactggtgtcaacgg
gcgtggtacaaggtttgcattccgccatgccagcacggtccaagatgtccaacatatactgccgctgagagagaa
ccataccatcagaagtgcgctggacctgcatccccaagaagtgatgaagtgctcccaaatccttcatagcaaatt
cccgctggagagctaggatgaccttccgcagtagatgtgtcgaagaggcagtgagaacaatatcatcgacgtaaa
gtagtagataggcagtgtctgaaccttggtggaagacgaaacgaagtgtctgacttggcttctacgaagttcagg
gagaccaggtatgtggcgaagcggctgtaccaagcacgaggagcctgttttaacccatatagtgacttgttcaga
cggcagacaaagtccgggtgagcagggtcctcgaatccggagggttgagcagcgaacactgtctctgtcagggtg
ccgtgtaggaatgcattcttgacgtcgagctggtgaacaggccattgacgggagagcgccagagagagtactgtg
cgaacagttgcaggcttgacgacgggactgaaggtttcggcgaagtcaatgccagggcgctgggtgaagccccga
agtacccagcgtgccttgtagcgttccagggaaccgtcagcatgaaacttatgcttgaagatccacttcccggtg
acagtgttggcgtgactagggcgaggaaccaagtcccaagtcttgttctgaagaagcgccgtatgttcctcctcc
atagcggcccgccaatttggatcggccagggcgctgcggaaggtcttcggcaccggagataagggcacagcgtgg
tagaggaccggcacgcggaagccgtccttggcccgggtcaccatgcgatgatggttaaccgtgggaggaacggcg
acagcacctcgcggtagagggccgggttgtggagcagcctgcccggccggaactggttcagttggtgcagcggtg
gccggcacggggcgcctggagtagacgcgcagaactggaggctgcgcgcttggtggagacggtgccacgtgtggc
aacgcggcaatgtcctgctgtcccgcaggcgtggccgcgcgtggcgcggaggcggtcggaggcggcaccccggtc
ggtgaagagggagccgcgcagggcgcagttcccgcacgtggaggtacgtggaaccccgggggaactggaggcgca
ggaagaacaacgccacgggagcctggagatcctgcagaagacatggtcggcagtcctatagggggggactgtcagg
ttagggatttcatctaaaaaggtgaagtccgcagctgcgggcggtgagggctgcttggcgaaagggaaggtggac
tcgtcgaagataacatgacgagagatgatgattctattggtggtagggtcaagacatcggtatcccttatgatga
gcagagtagccgatgaagatgcaaagcgcagaacgaggtgcaagtttgtggggagcagtggccgagagattgggg
taacacttacaaccaaaaacgcggagatgatcgtaggacggcgcactaccaagaagggcagaatgcggagtcgag
aaatcgagcgtcttggttggtaagatgttgaggaggagagtcgcagtgtgcaacgcctcgtcccaatatggaggg
ggcatgctcgcctgaaacagaagagagcggatgacagcgttagtggtgcgaataatccgctcagctttaccattc
tgtggggacgtgtagggacacgacatgcgaaacaagacgccatgcgtgagtaaaaaggtgcgggtgctggagttg
tcaaattccttgtcgttgtcactttggacggccttaacgcggttgccgaactgggtagacacataggcaaagaag
ttagacagagtggaaaaagtgtcagatttcaggcgtaaagggaatgtccataagtaatgtgagcaatcgtcaaga
ataacaagataatatttgtaaccggagatactaggaatgggtgatgtccataggtcacaatgaataagatcaaag
ttatttgatgcccgagatactgaagtaccaaagggaagacgaacatgtcgtcctaactgacacgcatgacatata
ggatccaaacttttattacatgcaggaatggcagaagctagtttggagagggcctcgcgaccgggatggccgagg
cgacggtgccataggtgggatggtgcagcactggcgacaagggagtgagcgacgggcagctgcaaggggtagagc
ggaccggagctattgcacctgatgatcacgttccacgagagaagatccttcacagaacaaccatatgggtcaaat
tccacagagcaatggttatcgatagtaaattgtcgaacagaaataaggttcttgataagaggaggtgaaacaaga
acattatttaaaataagagggccaagttgcgccttgccggtggcagtgacgggcaggagcgaaccattaccgaca
acaattgatgtgggagaaggaaatcgaggagtggaggagtgagagagggtaccgtcatgagaggtcatgtgagag
gaagcaccggtgtcaaggtaccagtcgttgttcaccggtggagtgagcgtcatggtgctgaatgtagaggccagc
gagttttggtcccaaaccggagtgccaaccataggctggtattgcagcggcgcaagctgctgcggctggaaggcc
tgctggacggagggcgggtaagtctgctgagggagatgcatctgctgggccagcagagcttgctgcagagcctgc
tgttgagggccggcgcgtccctggggaagcgggacagtggaggggtctgggtgtcgggccacatttgaatggacc
cggtccacggattgtagacggagggccaggtcgcgccggacccagcaggagctccaggggcggaagagggagctg
agggagcgctggggccttgagcaggggtgctgccagccctgcggtccttcttcttcttgtgctttggcttggggc
ctccaccgccgatccgcggagagccactgttggtcgctgcaggctggcgagaagaaccagtggcgacaagagcag
tggagggggcaacagggtggtgtaccatgttgatttcctcgaggaggagttcgttgcggacggcggcgaaggagg
ggaacgggcgcccgcgccggagatggacgtcgatgtccttgtagcgctcgttaagcccacgaagcacgttgagga
ccaaggtgcggtcagtgacgtgctcgttgagctcagcgaggtcgtcggccatcttcttgaagcgcttgcaatact

cggtgatggagaggtctccttgagcaaaggcgcgaaattgagcgtcgaggtggagggcgcgagtctcctggttgc
caaggaagtggtcctcgatggcgagccaggcctcacgggcggatgcgcggtggtcgatcacggcgtcggcgaggt
cagcggagacagtgctgtagagccatgacttgacgacggcgtccatccgagtccagtcagcgagggctgagatgg
agtcgctgcggacgtggtccagaagcgagtacttggtgacggcgaggaggaattgctcgcgccaacgggggtagt
tgccagcggtgtcgagtaccgtcggcaccaggctgcgtgcgttctgcacggagactgcctgggcgtggagattga
ggacagcagcggcctcgtggagaagcatggcgtgggggacgctcatctcgccgtccgcgtgttgctggtcggcgg
gatctgcggcggcggcggcgtcggcccgggccttggccgcccgagcaagggcagcgcggacgcgctctgcggcgg
cgtcgcgctcgttggaggcgttggcggcgtcctgctcggcggtctgggcgacggctagggcggcctcgcggaggg
cgcggcgagcctcgacgtctggctgcgggggtggatcgtcgtcttggcggtggggtcggcaggatcgacgaggga
gagtggaggtgtcccaaccataccagcgcgcagggaggatgcgcagaggtgcagcggaagagtgagaggggaagg
actagtcccggtctcgtgataccatgaaagaggatggctgggaaatgattgttggattgattattgagggagccc
aaggctcagtacatataggcaaggatgtcccagaaataggaaagtgctaatctagagaatcaggaaagtgctaat
ctaaggacaccttgccaatcagatccaatactaccaaaaaggggagaggggctgcagccgccactgctctagggt
tgcagccgaccaccctcctgggccaaccgaacaccctaaaggcccatgggcctatacagcatgctaacaTATATC
ATTGCAGAGCCATGTACATATATTAGAACTTTTGTTAGGCAAATTACTTGCAGACAACAGAAAAAGTAGTTCACA
AGTCACAGGTGGCACAAAGCGTACATCCAGCATGGTGAATGGTGAAAATGCAGTGGAATCTAGTGGCACTAGCAG
CAACATGCAAAAATGGGGACTGAAATTGATGTTTATTGTCACACATGAAACACAGATGAACCCACACTAGTTATG
AGTATGCAACAGATCAACTTAACAAACCTGCCCACCCCACCGCCCACACTGCCAAGTGCCAACCCCATCCAACCC
CCACAAGGCCACAACCCCCAAACTTGTGACCATAAAGGCACTAAGCAAGAGTCTGCTATTTTTATATCAGATTGG
AACATCATCAAGGTACTCCATCAGTTCATGACTGAGAAGTCAAGATAAGAGTCCATATTCTCAAATTTCACATGA
TCTGATGCGTGGCCAATACTTCTTGACATTGAAATTATGCATGTATGTGGGTTATACAAATACCAAATCATATGA
AACAAATGTGCAAACTCTCTAATAATGAATAATTGAATATATAATGTATTTTACAATATAATGTAAATATTTACA
TTATGATATATACTTCAAAAGTGCACTCCTAACCTTAATAAATTTTTTAACTAAGTTAAAATATTTGGCAAGTCC
TTATAACATACTTTGACCAAAAAAAGAATGCAACGTATTGTGGTATGAACCATAGCCACAGGAGACGGGTATAGT
GGCAGAGACCCTCGACCTAAAAACGCCGTCCCCGACCCAGGTACACCTGGGTCATTTTCGTTCCCACCATGTAAA
AACCTCTCACAGGAAGTGTACCGCATTCCTCGACCCTTTTGATCTGGGGACTTTGTGACTATGGTCTAAACAATT
TATGAGACAATGAAGATTACTCCAACTTATGGGATATCATGATATTATTTTTTATCACACTATCATAAACTAAAC
TTCATTTAACTTTGTAAAAGTAAAACATGTTGGCCGTAAATGCATGTATCTTACTCCAATGCATAGCATAAAAAC
ATGCAATTGACCTTCATTATGTTAGGTACCaaaaaaaaGGGATGTATGGGCCAATATTCCTAGGTTCCAGAGTAA
TACGCTATAGCCTTTATATTCCATGTACTTGAGATATTTTGATTGGGAACaaaaaaaaTCCTACTTGGAGAAGAC
TATACCTAGAGATAGAGGCTCAATACAGAAGTATTGTAAGTGGAGCAAGCATATTATCCAGCTTGAAAAAGTTTC
CTCAGCTCATATTGCAACCTTCAGgatgcatacccggagtcccgaacacaaattctacaatctttggggttgttt
gacatggctacagagcaaaactccaaagagggactaaccataatcagccaaacatcctaacttcagagttgtaaa
ctttgtgaaattttaagagctacaatataaatgtttggagtaccttttaaattctctgaaactctttaaaaacaa
tctagacataaagtttgaaagtactttgtttggcatggctccagagcaaaactccaaggaggagctagccagagc
ctaccaaacactcttaactccagagttacaaacttcatggagtttttagagccaaacctcttttgctgcttggaa
atcactaaaaagcaacctagatacagagtttggagttatttgtctgccattgccactggttatgagaaagcaatt
taaactttggagcatagttgctccaccagagttttggagtagagctactttagagtggagcagagtcatgccaaa
taGCttacctaccattgtacttgttatgggaaagcaccatagactctggagcagaggtacttcaccTTCAAGTGG
TTGCGGGGCAGCTAGGACAGCCAGGGACAGCGACGTCACCCAGTGTCCCTCTCGTCCGCCTAATTCTGATGAACA
GCTAGGGACAGCACTGGGATGGTACTGTTCCAACCACTGACCCTAAACCAAAATGTCTTACGTAAGGGATCATCC
TATCCTGTcttgtctcatcccgtcattgcaaccaaactagggcaggaaacgagccgagccgagctcggctcggct
cggtgtctgaacgagctcggctcggctcggctcgtccatttacgagctggcgaaagaggctcggctcggctcag
ctcaactttggctcgcgagccggctcggctcggctcgcgagccatattctggtacttaccattgcattatttggt
gaattaacattatataaatttgaaatatagaatcatcattctacattatgagtaaattaaattataacttgtaat
atattcatcatcaaagactaaaaatgagctattatccataaaattatctaatattcattattattccataaatta
atcatttttgtgaggctcgcgagccggctcgcgagctggaacgagccggctcgactcggctcgctgcaaaaacga
gctcgaagaagaggctcggctcggctcgttcgaggctcgcgagccaccccgagccgagccgagccgctccgagct
cgagccagctcgcgagcctcgagctaattttccagccctaaaccaaaCGCACCCttagagaatagctgctctagg
atggagcagagtcatgccaaacacctccttagagaatatgttgctgatttttaaaaaaaatgttttattttataa
tgtctttgtcattagtaactttgagctctatgatatcaatttttttaggtataattttatatatttataagattat
atatttcttatattattggtttgttgaacggtttgataaataatattatcgagacaaactatacgacaaatatat
attatctgtccgagtaatatccgatatctttcttacccgcccgaattattaaccgatcccatcatgcattcgtcc
caaattttaattacttgtacccgattctgtatccgagaaaattatattagggtacgataccgaatgacctattat
ccgactgtattcgtcctattttcacccctacACCAAATGATGCAAGTTTCCTCGTGACAAAATCGATGAATCATA
TACACGCAGAGCAACATTTTCACTTCCCAAGTCCCCACCTACGAGTTCCCAAGTGCCCGCCTATACGAACACCAA
AACCGCACAACTACATCCTGTTTTACCCGATGGTCCGATCCATTGCGCCTGATGACATCAAGCAAACTGCAGAAG
TGAATCTATGGATCCAGTTCCAAGCTCCACACACTACGGATCCAATTCCAAACTCGGCGCACTACAAGCAGCACA
AAAACCAAAACCACATCGGGCACGAGGAGGACGGGCCGGTGACACCAACCACACAAGATCGTTCGAACTAAAGCG
GAATTCAACCTATCGGAGTCAGTTGAGCTTAC**CTGGATGGATCGGGCCCTCGCTTCGTCGCGAGGGCACTCCGCT**
**AAACCCTAGATCAGGTGGCGTCGGCAGTGTATTGAGTTCAT**CTGAGGTAATAGGCGAGTGCCCGTGCGTTGCAAC
GGAAACTGTTGAAGGAAGCGAGTAGAGGGGAGGGTTTATCGGCTAGACAGGGAAGAAGAGCACGGATCTGGGGAC
GACGACAACGGCGGTGGTGTGCAAGGGAGTGCGACGCGAGGGTCGGGGGGCAACGAACGTGTGCTACGATGTTAG

ACCGGTCagccaacaacccacgcatgcgagccaacaacacacgcatgcgagAGACAGAGAACAACATTATATGAA
TAATGATTAATATGATAAAATACATTAATATGCAAACTCCCTTTTTTGATCACTAACCAATCATGTAGTCCACAA
CAGCATGCAACATTTTATCACTCTCCATTCGTAGTCCGATAATACAGTCAAATTGCAAAAAGTTCATGACATACT
TTATTTCATAACATAAGAACCCTCAGTCCACCAAGCTTGCGTATAGCTCTCTCAGCGACACACCTCTTGTACCTC
TGGAACTTGAGTTGGGTGATACCCTGGTGCTTGGGCTTGCCATAGACAATACACTTGGGCAAAGGCCTCTTCCTA
CAATCCATCATATTTTTGTCACCCCTTGCAACAATAGAAAAGTTGTTTGATAAAATTATAGTGTGCAACGATTAC
ATAGAAATAAACAACATATGTATTCTCGACATCAATATATCACAAATCATTTGCACAAATGAATGCCACCACAAC
TATAGACAGACAGATCACACCCTATAATCCACCAGATTTTTCCCTTCCCTATGCGTCCAATGTAATACAACAATA
ATATAGGCTTGATATGGCAATACACCTATGCGACCTGTGCCATGGCCGATCTGGTCGATCAGATGGTCACCTTGG
TCCAAAGAGCTCCATGGCTTCTACACCAAGCGAGCATTCGTCCGTTGCAAAATCCATGTACTTTTGAATTATATG
TAGCTTCAAGTTAAAATAAACAGTATGGTAGCAACTAAAGAAAAATAGAATCAGCCAAAACTGAAATACTAACAT
CCAACATCTTGATCAGGCACAATATTCTCATATTCAATCCCAGTTAAGGAAGTAGTATCAGGCACAAACATAAGC
AGTAGCAGTTAAAACAACAATATCAAATTTGAGGCACCAAACTGGAGTTCTTGTTTCCAAACTTGGTTTGACGAT
CTAAAAGCCCCATGGAGATTATTTTCTATTTCAGACATGATGGCAGGAAGTTACAGAACATTAAACATGATGGTA
TTAATGTTGCATGTGAATCAATCCAAGTACTACTAACTAATAAAACAACCAATTTCTTATCGATACGTTAAATAC
TTAAATGTCATAGTACATAGCAAATCAGCTAATCAGGTACTTTAAATTGTAATTTGGCAACCTGTCAGCACAAAA
TGAGAGCTGCTAATGGGCTTAGATCCAAACCAAACCAACCTCAATCCATAAAAGCACAGCTACCTCACCGAATCA
GTCTAATCCACCATTAAAAAGGTTTGTTCCAAAATAAATAATATAGGGGTTCTTTAGTGAATACAACATGGAGAC
AACTTCAACAATGAGAAAGTAGCAAAGCAAAACACCTACCACCATCAGCTTGAAGAACTTGGACATGATATCAAT
CTGCAAAAAAGATATTATCCAACAGTAGAAAATTAAAAATGGATTTAGTCTTCCATTCTTAGTAAAATAAATAGA
GTAATCAATCCTACCGAGGCATTCCCACTCCGTTTTATATCTGTCTGTCGAAAATACGACCTAGATATGTGATGA
TGCCCCATTTGCCTTTGCACCGTTACCAGACTAAAAAATATTGTTTACATCAGATGAGAAAAAGACCCACAAAGG
AAGCCTATGCCACAAGCCCACAACAACGCCAGTCTTCCAATCTCAAGAGGCCAGTACAAGCaaaaaaaaTCTGAA
ACTCAAATCTAGAAATCAATCATCGTAATTTCCCTATCAAAATATCCACCAGTTCCAGTTACAAGATTAAAACAG
ATTCATTTTAGGAAAGTGGATCCCAGGAACAAGGAAATGTAAACAACCAGCGAGGTGAGTAGCTCTAGGGACCAA
GAAAATGTAAGCAACCAGCGAGCTGATGTACTCAATCTAAAATAATAACAAAATTGCTTAACTAAATACCCTCAA
CATGCCCGCAATGACCAGTCCCAAAAAAATCCTGGAGAATCATACCACCTGTGCCTGAATTGTAGATAACTATCA
AGGAATTCATGCAAAGAAGTGTTCTGTGCCACTGTGAAAAACAACAAAAGTTAGCATATGTCATTGACTAAGTTT
AGATGTAGCTGAATTTATGTACCAGTTGTGTAAGAACAACATAGTAAATTAATCATACAAACTAATCTATTTATT
GAACTATCGATTATTGAGGTAACTTCTGTTCGAGAATCTTGGGCACATGACATATCCTTATGCGGTTACATCTTA
ACATTTGTATTCCCAGCCAAAAAGTCTACAAACACCATCTTTACCCTGTTCCCTAACTGTCTATTCAGGAACGGG
CAAGAGTGCAAGACACTACATGAACAATATTTGATAATGTGAATAATAGGAAGTAACAGGTGCATACGATTATAA
TGAAAAATGCACAATTGCACACTCAATCTTGTGCCTAGGATCCAAGTGGGATCATCCCTATTATCCATCTTTTCA
TCTGACTAAATGCTAAATCTGATTGTCTTTAAGTCAGCCCAAGATAGGCGAGGGAATTTCAAACCGCACTTGGTC
CACAAAGATCAGATATGAGGCATGAGATTGAGTATGCTCAACTGCTCATAGGTTCACTAATATAATATTTTGCAC
CTTATATTTCCTTGCTAAATAAAATGTGATAGTCATTTGTCCGAAATACTGAATAAAGGACTCCAGATAGCATGA
CTCTTCTGTCACAATCACCATTATGTATATGCGAACACTTCAGGGACATTTCATGTGTAAAGAGAAATTGTGATT
GTATTTTCAAGATGTGAACAGATTAGCTGAGATTTCTCTACAATATGAGGAAAATTCAATATGAAATAACGAATA
GCTAGAGATATGTTTTTTTCTGGACCTGTCACTTGAATTGCCACAACAAACTGGAATAGTTTAAACAACCAGCAA
AACCAAATTTGTTAAGAGACTTGACCTGCAACTAAACATATAGGCTAGAATCTAAAACAATGCATGTAGTATTAA
GAAGGAAGAACCTAATTGTTAGGTGCACATCAAACTAATGTGGACTCATCATCTTACACTAATATCAACCAAAAA
ACACAAAGAGAAGGTCAGCTATTCCGAGCAAAACCATGGGATGACAATTATAACAAAATCTTAAAGCTATGAATG
GTTCTCAGATATAGAATACAAACCTAAATTGTCCTTTCACACAAGCCACAAACCAAATATTTTCCCCATGTATAC
GTACACAACGTTGCAGGCTCACTACCCATTGGTATATTTTTATACTGATATAAGTACTTGCTATTCTCTCTTGCt
tttcctttttttttCATTCACTGATGGAAGTCTGAACCAAAATCTACATGCATTGGTGGAATGGAAATGTGACCTC
CACTTTCGTAAGTCCGAGCGGGCTTACAATAGACCGCTAATCAGCTCATGTCATGACAATTTGTTAGAGGTTACT
TAACCGCACTTTGGTCCAGCCTAATATATATACAAGCAAAACACAAGAAGTGGGCTTTTGCAAGATAAGAGTAGG
GGGTGGATGAATGCAAGCAGCTAGTTGTCAGGCACCAACTTTCTATACTCTTGATAGAAATCTCTATCTAAACGA
GAGCAAGGGACAATCAAACACATGTAAATATAGAAACTAAAATTGATGAAACCACATGAAAACATTGCTTTGGCC
GCAATTTCAAACTGAACCTATTTTGTAAAGGAAATCAAACGCCTAATCTCCAACAAACTATCAACACCAAGAAAG
AAACCCAGTAGataatagctacagataaataatagctataggtaaaCCTAAGAACACCAAGAGAGGTTCACTAAG
ACTAACCATGGTATGAGCAGTTGCAGTACCCGCCGAAGAACTTGCGATATACCTAGGATGAAGACAGAACAATTA
TTTTTTAATCCTACCAAACAGATAGCACCCCACCTGACATGAGGTTACATGACATGTTCTCACCAATCACTGTTT
TCATTACCCAATAAATCTGCCAATGGATCTATCAGCGCCCAGATTTCCTACCAAATCGGATGCCTACCATCGGTG
AGAGACATGCGAGCCCAACAAAGCCATGTCCAGAAGGGAAAGCAACTGTTGGCTTGTCTTACTGTTGAGTTGATG
GTTCAGAATTATTCCTCGAAATAGTGGATTAGTGGTTGTGATTAGTGTGCCTTGGAGGCAGAATGGTGCCATCAT
TTACTACCAAAATGAGCAGGGAAAGTAAGAGGACTGCAACAGATGAATCTGGTAATTCTGAACTTGTGTGGGTGG
GGGCTCACCAGGCTCAGACTCGGACCTAAGGTAGATGCGGCAGCAAGCGACGNNNNNNNNNNNNNNNNNNNNN
NNNNNNNNNNNNNNNNNNNNNNNNNNNNNNNNNNNNNNNNNNNNNNNNNNNNNNNNNNNNNNNNNNNNNNNNNNN
NNgtcggcccacgggcgcgggagagagagaaaaaaaaagagagagaaagagagagagaaaagagaaagaaaagag
aaagaaaacattttcgaaatccgatctttctacatgaatgcatttgcactttcaaagcaatcaaaagaaatgcaa
ggttcggcatggtgcatcaaacaacataaagtatttagggtttttatacatacgggaattccaaaccgaatcccg
ctagaactttggaaaaggtcaaggtttagcgagggaaaaagaaaaagaaaggtaacgcccgaattttggcgagca

aagaaaagaaaaaaaaattcaactgcaaaattcggggcgttacaaacctatccccttaaaagaatctcgccctc
gagattcagggctggctagcaaagagctccgggtacttggccatcagatcatcttcacgctcccaggttgcttct
tcctcagagtggtgactccatctgactttgcacattctgacggtcttccttcgggtgactctatctgcaacctca
aggatctgagctggcttctcaacataggtcaagtcctcctggacttcaagaccttccactggcaactgctcttct
ggcatacgcaagcacttcttcaactgagacacatgaaagacatcatgcacagcagacaaattctctggcaaactg
agctgataagccacttctcctcgtcttgcaagaatctgatacggaccaatgtagcggggtgctagcttgcctttc
actccgaatcttctgactcctctgatcggtgacactttcagatagacaaagtctccgacttcgaaactcagctct
cttcttcttgtgtctgcatagcttcgctgccttgattgcgctatcttcagattctctcgaaccatcttgacgttc
tcttcggcttcgagcaaaatgtctggcccaaacacttgcttctctccaggctgatcccattgcaacggagttctg
caactccttccatacagcgcttgaaacggtgacatcttcaaactggcctgataactgttgttataggaaaactcc
gcataaggcaatcgcttgtcccatccggactgatcttgcaacgcacaggctctcaacatatcttcaaggatttga
ttggttctttcagtctgaccatctgtctgcgggtgataagctgaactgaaattcagatgcgtgcccaaggcttca
tgcaactgctgccagaaatgagaggtgaactgcgttcctctgtctgacactatcttctttggcacaccatgaaga
caaacgatccgagacatatacaattctgccaatactgcactgttgtagttggtcttgacaggtatgaagtgggct
gacttggtcaagcggtccactactacccagatggaatcgtagccggctcgagtgcgaggcaatccgactatgaaa
tccataccaatttcatcccatttccactgagggatctgcaacggttgcaacaatccagcaggtctctggtgttct
gccttaattcttcgacaactatcgcacatagccacatgctctgcgatttctcttttcattccgtaccaccagaat
ttcttcttcagatcctgatacatcttctcactgccagggtgaatcgaataagctgtctcatgagcttccttgaga
atcaactcccgaatggactggacattgggaacacacaagcggtctttgaaccataccacaccttctgcatcttct
cgaaaatctttgcctctgccatctagaatcaatcgccgaatctcactgatcttctcatcattcttctgcgcgtct
ttgatttcgcgctccaaggtaggttccaattcaactgtgactcctcgcgaattgttcagaaatccgagactcaac
ctgtcaaactccttggccaactcataaggcatcgggcgagcgaccatcagattgacttgactctttctgctcaaa
gcatctgccactacgtttgctttgcctggatggtaatgaatctccaactcatagtctttgatcaattctaaccat
cttcgttgcctcatgttcaattctgactgagtgaatatgtacttcagactcttgtggtctgtgtaaacatcgcat
ttctgtccatacagatagtgcctccatgtcttcagtgcgtgaaccactgctgccaactctagatcatggatgggg
tagttcttctcatgaaccttcagctgtcgggacgagtaagccacaactcttccctcttgcatcaacacacatccc
acacctgtgtaacaagcatcacaatacaccgagaagggcttgtgcacatcaggcaagactaggacaggcgctgta
gtcaacttctctttcagcgcttcaaaggcctcttggcatttctgggtccacttgaactcaaccttgttgcctagc
aacgctgtcattggcttcgcaatcttcgaaacccttcaatgaatcgccgataatatccggccattccaatgaag
ctcttgattcctcgagcatccgttggcgctttccagttcagaatgtctgccactttcttcggatccacagccaat
ccttctttgttgattatgtgacccaagaacaggacttcactgatccagaactcacacttgctcaactttgcatac
aactggtgctctcgcaatctctgcaataccatcctcaaatgatctgcatgctcttcttcactttgagaataaacc
agaatgtcatcaatgaataccaccacgaacttatcaagataatccatgaatacactgttcatcaagttcataaag
aatgctggcgcattggtcaaaccaaaagacatcactgtgaactcatataaaccatacttggtaatgaatgccgtc
ttcggaatgtccgaaggtcggatcctgagctgatgataacctgacctcagatcgatcttggagaacacactggct
cctctcaactggtcgaacagatcttctattctgggcaaaggatacttgttcttgatcgtgacttcattcaaagct
cgataatcgatacacatcctcttagaaccatctttcttttccacaaacaagacaggggcggcccaaggcgaggtg
cttggccggatgtaacctttctctgacagctcatcaatttgcttcttaagttcatccaactctggtccagatatt
ctgtaagctctcttgaagataggggcggttccaggaagaagctctatagcaaattcaactttccgctctggtggc
atacccggtaaatcctttggaaacacatctgggaattcagacacaaccttgatactctcatttgggtctgcttca
ctgctatcgacagccatctgataacaacttcctttctttggctcaggcgggactaactcggtcaccacttcctct
cctagtggggacactaacttgattgtccttttatcacaactgataactgcctgatacttatctagccaattcatc
cctaggatgacatctattccctgagtacccattactataaggttggcgggaaacgctatccccttattttcaca
cttatattcaaacaaatgctatcggctcgaattctaccaccggctgaatcaatttgaatggggttgacatggta
gttattggaagtttatgtgcttctacccatgatgcagtaatgaaagaatgtgttgctccagtatcaaataacact
tctgcaatatgggaatcgactgggaacatacctactgtcatgccggggtctcctgaactgcttcagcttccaag
tgattcaatcttccatgattatagcgctgctgagagcgattgcctgctccaggctgagacacattctgctttgct
ggggcattggggcctgactgctgctgggctgccttctttggacattgcatcacccaatggccttgctctccacag
tggaaacacgctctgtttccaacttgagctggtgctgcctgactgttctgctgatttgctggggcaggaagacga
ggtgcttgctgattctgcttctgaaactgacctcctgactgattgctctgacggttctggtactgattctgaggg
tactgcctttggaactgctgatactgctgacgctgctgatgctgctgaggtggacgctggttctgcctgaactgc
tgaggttgattgcctgaaaagcgagggcgactgctgcttccagactgcggtccactgatcttgcgcttacgatct
tccatctccttgcgcttcctttctgtcatgattgctctgtcaatcaggtgctggaatgtcgggaaggtgtgattc
atcagctgatactgcagagggtcaaccaagcctctcaggaaacggtactgtcgcttggcgtcggtgttgacatct
tcaggagcatagcgagacaattgcagaaacctgtctcgatactcactgacagacgatgacccttgcttaagggcc
aggaattcctccttcttcactgtcatcagacctgcaggaacatggtactgacgaaagctaccccctgaattcttcc
caggtgatggtgtcggggttggcatgggtggcgaggtaagactcccaccatgactgggctgctcctctcaacaga
cggggaccatacagaactttctccctgtcatcgcactgagcggtatgcaactcccgctccacagtgcgcagccaa
tcttcagcatccatggggtcagaagagtgagcgaacgttggtggatgacctctcatgaattcagcacgcttatct
ctaggcatctgaggcatctgaggctggggtggagcctgctgctgttgctgctgctgctgctgaatggcggcc
agagtctgaccgatggcttgaactgcctgagtctgcatcagaaacatctgctcgatggacatcgggggcgggggc
ggcaggtgctgctgctggggcacctcatcctgcggagcggctcgctcctgctgagcacgccttcctcctctacgc
ctgttctctgacatctgcagaacgcaaccacacatcagaactgatctggcaaagcttgcagcataagaaaagagt

atagaattctccaacagcactgaacagatgagcatcttcactgatctccaacacagaccacacagcttctcagat
aaagaggaaaggagaataatgggtttcccaactatataactaactttattgacatagtacgtaaaccaaaatgca
ggggatacccacactctggtgacaatcattacaaagatccaaaccaaacatagttcatcatgacaaacataaatg
cacaggatatagcaaactaccctgtctaactaagactaactaagagcgagactaacactaagaccgaagcttcta
tgtatatatttatttcgtattaattacaagatccaactctaacgatctatggttctaggttgatcttggtcttgc
aggcgggattgccataagactggtgtccacgctgaggtgagcggtacgggtgctgagtcccgacgggagcggggg
aaccaccatccagaaaacgacgttccgcgcgctcagcccgcagcagggcgatctcagcacgagccctactcagct
cgtctagtgcatggtccagctccgtgtttagcacggcggctaggttgactgtgctgctcaacctaggattgccct
caccgacaggtgagacaatcacgcctcctgtgctgccagatgaacggcgggggtaatacttcaggtcaagaccgt
cagctgccccaccgaaaaccgagcagtagtgcgaaagcgcacgccgtgcagcatcttgcatggctgcctcagctg
agtcccgctcagagatagaatagtgctctgaacaggcctctgcaccctggagactgtcctccgggcagcgcacca
agcaagtcgcctcccagcggtccgggtagaccccgcgactatgctggtagaccacacagcgatactcgacggacc
aagtatgccggtcaaatgcccgacgtagcagggtgtcgagcgcatcgtggaagtgacacccgcgagcagcgtcgc
gagtgatgggtcgagcaacccatccttccggctcagggttggcagagaagtcggtgtcgtggctcgagctgtcgt
cgccatctccgtcgtctgggtctcctccagcagctactccagaagctggggcgcccagtggtggtgcagggggcg
gctccagagaaagcacaggggagcagctcctcacagactctatctcctggtggagcgagagggaagagctccgct
gctcctgtcgtcgacgttcctgctcctcacgcaggcggtggtgtagtctctccaagtggctggactgtccggcca
cgggacggcgaagcggacgctcagcaagacgggagggtaagaaggggatgactgactttcgtgcagtgtgtctaa
gtcgagccatctacaaaagacatcgcaagcaaaagggtgagaacagaattaatatgaccagcaagtaataagtaa
atgaaggatcagaataaaacataattttcagcaaggtataatatgtagtagaacataggtttggtcggtaggacc
aacttttgaagagatatcaaagtcaaggaagagacagaggtctatagtccttagaacgaccattctactctaggt
tagcggtcctacagtcagcacggctttgataccacttatgtcacacccggttttaaaaaggcaaaccgaatgcga
accatgtacgtgccaggatcagttattcacgtacacagcagttacataatatggacatcatcacacagtgctcaa
aatagtattaataaggggaaataatcgattacatcatacgtctgagacgtccatatagttcttacaataaatcaa
agtgcggaaaagaaacgtagataacgcggccttcacaggcagccgactggggggttgccgctaacccacacctaga
actcgtcgtaatctcggaactcctggaagtctccttccacagcttcatcttcgcctgagcagtggttgcaatgct
gacaacctgggggggggtttggtgtgtagagcaagggtgagtacacatcaacatactcagcaagtatcctgttt
ggctgtagtggactagctttatgtggggataagtcgagcagtGCTAATCTCTAGTTCCGTANNNNNNNNNNNNNN
NNNNNNNNNNNNNNNNNNNNNNNNNNNNNNNNNNNNNNNNNNNNNNNNNNNNNNNNNNNNNNNNNNNNNNNNNNN
NNNNNNNNNNNNTGCAGGCGTGGCGGCGGCCTTGGGCGAACCACAACATGAGCTGCGACGGAGGCGATGAGAGGC
ATGAACCTGTCATGTCTGAGCAGTAAGTACTCTCTAGGGTtgggccgtgggccgggtttagtgtcctaatctggt
gttatgggtttatttggagtctgtaatggcctgttaagagaccggatatatactctgtacttggccagttggtaa
aactacgatcagaataatccagagaatcatatccgtaacaaactcacctgggttgtagcgactcgctgacgctcc
gacgaactcacgtcccggcggtggcgcgccggcggccacgggttgtctcagttggtatcagaggagtggtcctcg
gaggtttccgcgacggcgccgtccaccaaaccttccgtgcagacgcgtctgctgctggagacaatggaggtcgcg
gagcagaagagcgaggagcgctgggatcggatgatgaagatcctggagcgtttgggcgacaaggtggaggcgatg
gagattgggcagcaacgtcttcaacgtcaagcagaggtgatggaggcgggacagcaacgtcttcatcaccaagtt
gaggctgctgcaacatctgcgaggaaggcggaagaggaacgaaccatcctggcgcgtcaagtggatgaaaccggg
aaggcggtgtcgcgccttcgattggaatggatggcgaaggagatggaaggtatggaaactgagtctgagaaggaa
gggcgtaaccaaaggcggtctgagggtgatgatcgggagtatggagagcaggagtttcgtgtggatgatcagccg
gaaggaagtagagagcgtcgtgagcatcggggaggaaatatgaggaaccgggtgaatgacccacaaggagggaga
agagattatcggccttcagatttgccgaagatggcatttcccaagtttcagggtgctgaacctaaggtttggttc
gataagtgcttggagtattaccatatttaccaggtgcctgagttcttgtggccgtcctcagcttatctgcacatg
gaaggaaatgcagcacgctgggtgcgggaatacaagagaagtcaagggctgggaaattgggtccagtttactcag
gcagtacaggagaagtttggggcagaagaggattcccaagcaatgaggggtttgttgcagctgacgcagaggtcc
agtttagaggagtacattcatgcttttgaagaggccaagtatgctaataccttacacaaccctgggttgggagaa
tctttctttgtttctcagtttgtgaattgtttgacggggaattgcagggacatgtgttggcacattttacctcg
aacactggatcgggcgatacttctggcgcagattcagcaggaagtggtggagatgaacaaggccaaatttcagag
gaattatctaggtggcaaagtaatggtgaacacagtgaaaggagatcctagaggagtgactggccctgctgattt
ggctaaggagaggcagctgagagagtttcggaggctaaatggtttatgttttggttgtggagagaagtacgagcc
aggacatcaagctaaatgccttaaaagaggtggagcacaattacataatctgactgtagaggatatgggttggt
gttgtcagatgagacgttgcaacagctggatcaggaagataaggaagtggaggaagcatataagctgtctataaa
tgctatcagtggaactgaccgtgagaattgtatgagagttcgggcactgatccagaatcaagtgttggtgatgtt
ggtagactcgggcagttcctcaagctttgtcagccagaggatggtagatcagttggctttatcaactgaagttgt
ttcaccggtaagagtcaaggtggcaaatggagagttcatgttgagttcaaaaagggtggcagcgctggagtggtg
ggcagaagggcacacttatgccagtaacatgagggtactggatttagaagcctatgatgctatattggggtatga
ttggctaaagagccatagtcctatggagtgtgattgggacaacaaggtgctgacattctgggataaaggggtcag
agtgcagctgaaggggggatggcgtaggaagtacagaggtattgcaagtgtcagctgttcagctgagcaggtggat
gaagggaaatgaagtgtgggcttttgtgatgctcgaagcaatacaggaggaaggtgctgaaaaggaacaggaacc
gctgaggcagatacttaaggagtttgaagatgtgtttgctgtgcccactggactacctccggttagaatgtatga
ccatcatataccattgctcccagggtctattccggtgaattccaggccttataggtattcacctttccacaagac
agaaattgaaaggcaagtgacagagttattaactgcaggattgatagtacccagtgccagtccatttgcatctcc
cgtattactggttcagaagaaagatggcagctggaggttttgtattgattatagaaggttgaactccatgaccat

caagaaccgttttcccatgcccttggtggatgagattctggatgagttggcaggaactcagtttttcacaagtct
agacatgactgcggggtatcatcagattcggatggcagaggaggatgagggtaaaactgcgttcaaaactcatca
tggccattaccaattccgagttatgccttttggcttgacaaatgcaccagcaacattccagtgtgccatgaacac
agtgctagaaccatttctacggaaatttgttatggtattcttagatgatatattgatctacagtccgagtctgga
ggagcactacaaacatattaggagggtattgtccaagctcagagaacagaagtttttcttgaagcataagaagtg
tgtgtttgcgaagaaagagctgagatatcttggacatattatatccagtgaaggggtcgcgacggaccctgcaaa
gacagctgctatgctggcatggcctagaccacagaatatcactgagttgaggggattttggggcttacagggta
ctacaggaagtttgtgtgtgggtatggagttttggcacgacccctcactcagctgttgcaaaaacagcatggttt
caagtggaatgatgaagctcaagcagcatttgaggccctaaagaaagctatgacgactactcctgtcttggctct
accacgatttgatcttccttttgtggtagagacagatgcatgtgacaccggaatggggggctgtgctcatgcaaac
aaggaaagcccatagcttatatcagcaaggctttaggagaaaggaatagacacctatctatctatgataaagagt
ttttagccttaattcttgctgtggagaaatggaggccatatctgcagcggagtccttttgttatcaagactgatc
ataagtcacttacatttttaggagagcagcaacttcagtctgagctacaaaggaaagctatggcaagattaatgg
gccttcagtttcagatcgtttacaaaaaagggtagagaatcgagtagctgatgctttatccagggtgggggcta
tgatggagttggctgttctttctgaagtgcaaccagtgtggattcaagagattttgaattcctatgagacagatg
ctgaagcccaggagttaattactcagttgttggtgcatagtccaaatgaacaagggtttagcttgcagcaaggta
ttgtcaggaaaggtgaccaaatctggatagggtccaactctgcccttagaaccaaactgatagcagctttacatg
acagtgccatgggggggcattctggcatacatgccacttatcagagagtcaagaaacttttctggtggaagggac
ttaaggtggatgtgactcaatttgtacaacagtgcttgacctgtcaacaggcgaaaccagaaaggactcatccag
ccggactgttacagcctcttccgattccacatggcatttgggcagatgtcacaatggactttatcgaaggttac
caaagtctgagggttatgacaccatcatggtagtagtggacagattcaccaaatatggcatttcttccactca
ggcaccctttcactgcccagggagtggctcaaatatttttggatcaggtggtgaagttacatggacctcccaagt
ccattgtgtcagacagagacaagattttcattagctctttctggaagcacctgtttcaacttatggatgttaagt
tgctgttttccactgcctatcatccacagactgacggtcaaagtgaacgggtgaatcaatgtttggagacatatc
taagatgtgcagtgaatgctcagccggctaagtggaagttgtggttagcattggcagaattttggtacaatacta
cttaccatactagtctgggttgtacgcctttcaaagcactatatggatatgatgccccgatgttggcgttacctc
agtctggtgtcctggaggaggggtcagtaacagattggctggctgacagggttgctttctcaactcatctcaaag
aacaactcgctcgagctcaaaatcggatgaaacaaatggctgatcggggaaggacagcaagagagtttcaggttg
gggagatggtgctgctcaagcttcagccatatgctcaaaagactgtggtgaaccgtccttgcccaaagttggctt
tcaagtttttggtcctttcagagtgttggctaagattggtgcggcagcttacaaattggatcttccagccgatg
ctcaagttcatccggtgtttcacgtttcccagctgaaaccttattatccatcatatactccagtttttacttctt
tgcctccattggtggacctctcaagaactggcattgttcctgaagaaatcctggatcgacgtcttgtgcgcaaag
gccatcaagctgttccacaggttcttatccgctggtctggtgtgccagtggaatcagcaacttgggaagattact
atgtgttgaagaaccgttttccagacgccatcgcttggggacaagcggattctgaggcccggggaattgtcatgt
ctgagcagtaagtactctctagggttgggccgtgggccgggtttagtgtcctaatctggtgttatgggtttattt
ggagtctgtaatggcctgttaagagaccggatatatactctgtacttggccagttggcaaaactacgatcagaat
aatccagagaatcatatccgtaacaaactcacctgggttgtagcgactcgctgacgctccgacgaactcacgtcc
cggcggtggcgcgccggcggccacgggttgtctcaGAACCATGACATGAGCTGTGATGTGAGGTTCATCACGTAG
TTTCCTCTGGCCCCTTGGCGCTGCGTCTCATCCACAGAGGCAACAATCCACGTGGGTGGTCGAGGCTGTCTACGG
TTCCTAGATCCCATTGCGGTGCAACCTCCAGTGAGGGTGTCGAATCCAATGAGGGCTTCGATGAGGGTGCGAGAT
CTAGCGAGGGCTCAGACACCAAATCCGATGAGAGGGCCACACGAGCTCTAGCCACCAATTTATGAGAGGAATCAA
TTGCTGAGAGCTCTACTACAACATATGCATCAGATAAAATTATGATAGGAAATGACATCTCGTTGGTACCACTTC
TTCAATTAAGACAAACAGGAGTGTTACACCAATGACATGATCCTTTTCAACATTTTAGAAAACGAACGGCAAAGC
TGTATATGAGCACAGAAAATGCAACTCTTTCCTACTAAAAAATACCTTGTATTTTCTTTCTACAAAGCAATGTCA
TCCAATTGTAAGAAACTCTCACAACTGATCGAAATAAAATAAAATCTGCAGAACTAATTGTTTTTTACCCCTGGC
TTCTCTCCTAAAGAGCACCACCATGTCAATACAAAAAACAGGGTTCTTACTTCGATAAGCAATTAAGCATTAAGT
AAATATCATTAAGCATTGACTCCTAAATACTCAATAGCTAGTAGCTACCTAGTTGTTTCGGACCACTTACAAACT
CAATTTGCAGCGATGGAGATACGCTTGTCATGTAGAGGAACCACCGTGCGCCAAAAACACAACCCACTCCATACA
GTGTCCTCAGCGACATGACCAACAGGGTACCTATTCCATACAACAATGGCTTCCCGACCATTGTTTTTGCATGCT
TAGCAGCAAGGACAACTGCCCCCCTCATCTGCCAAAAAACAGAAACATGCCAAAATTTGTCAAGACCAAGAATTC
AAAGAGAACTCCGCTATTGTTAAAGTAATTCATAACATTAACCACTAATCTGGCTTCAGCTCGAGCGTAGGCCAC
AAGAGTTGCAGCATCTACTCCCTTAACTCTCTATGCATCTCCAGTTCGAGCTCCAATTGCTCTGCTTCCTCCCTT
AGGACCCGCTTTCTCCTTCAGGATCCAAAACCTGCAGGAACACAAACTTGTGTAACCAAAACTGAGCAATAACAA
CAATAAATATAAAACTGAAAGAATAGATTCTGGAGTTGTAGACTGAGGAGCACCACCATATTCCATCAGAGTGGA
GGCAACAAAAGATCTGGATTAGCCAAATCTCTTCGCCAAGCTCATTTGAGACATACGAAGTGATTTCTCATCATC
ACATCTAGCTTCTGCCATATAAAAAATGTCGTCATGAATACATGTTTTGGCCCCCCAAAATATACAAGAAAAACA
GCATGAGACAGAAAGATACACTTTTCAGGAACTGTTTTCCTGCCATATATTTCTCACCAATTTATACTGCTGGTC
TTCCCAATTGGTAACTGTTTTCCAAATTCTGCTTGATTAACTAAGCTTGCATTCCCACATCTCAAATCACTTAGC
ATAAGAAAGCAGAACAGAAAATCATTCCACAAACACTACTTGATTAACTAAGCTTGCATTCCCATATCTCAATTC
ACTTAGCATAAAGAAAGCAGAACAAAAAATCATTCCACAAACACTCACATCGATGCAGCCCTTATAAAGATGCTC
CCTCCAGAACCACCTCCACGGTCTCCTGGAGAGTGGAGACAACCTAACATAAATCCCCACAATAATaaaaaaaaC
AGGAAACTCCATTGAAATCTATAATTGAACCAATAACTTGAGTTCATCCATTAACTCATTTGTACGTGTAAGACA
TAACTTATGCCAATAAATAAGCATAACATAATTCTCCTAACACCTCACATTACAACAAAGATATCCATCCCATGA

CCCCAACCCAAACTTCTGTTGTCACCAGATTCTACaaaaaaaagaaaaCTTTGGATTATAGACCTAACAGGATCT
AGAACGAGCCAACAAAATAAACAAGTGGAACTGTACTCTAATTATCATGTGGAAGTCTTCACAGCCTTGAATGCT
GGATGAAGCAGCCCAAGCTCATCCTTCGTCTCAAGCGGGATGATAGGCATGACTAATTCCTAATTTGGGTGCTCT
AATCCTGTACTCCAGCTCACATGAACCATCCCTCTGAAAGCACATGATAGGCATGACTAATTCCACCAGTAGCAA
TAAACAACTATCACTTGTGAGTAAAACCGAAAAATCTCTTGTTTGTGTACCTTGTCGGCAAGCATCTGATGCAGC
ACCCTCACAACCTGCGCGCCAGCAGAGTGCCCCACAAAATGCACTGGGTTGTGCTCATAACAAACAGGATAGTGA
CCTGCACCAAATTAGAACAGCCATGTAAGAACTAATGGATAAATAAATTTCTGAAAGGCCATGGCCATGTATTGT
GTTCTTTGGAATCTCCTACCTGTGTGATATATTCTCCCAAACCTGTTGTGGCCGCAAGCCTTGCTGTGCTCCTCG
CCATAGTCCACCTGCTTACCCTTGAGGTAGTAGAAGAGTTCACGCGCCCTGCATCGCTAGGAACAATAAACTCTC
AGCGATCTATATTCAGGGGAACATAATCTTTCTTGTTCACAGACACGGAGAAGATACCAAATCATAATAAAAAAC
ATAATGGGCATGTGATTTCAGGGAAGCATTGGCACCAGTGATTTTGGTCTCCTTTGTTCTGTTGAGCACCTTCCT
CACCTTGTTTGCAAGCTCTATGATCGAATCATCTTCGTGGCCTGAAAAGGGGCAGCATAATCACAATTTGATGTT
TGAGTCTAGGATGTAAATACTGACTTTGCAGAAAGGACATACATGAACTACTCCCAGGTAATACCGTATTTTATG
TAACACATGGGGTATAAAAAACAGATATTCAGAGTGTTAACCACTTAAAAATATAAAAATAGGTTCTCTTACTAA
TGCTGGCTAATGCCTTCTCCATTGAGAACAGGTAATCCCTATTATAGCCAGTAGAGCCATTGGTTGTAGCAATTT
GCATGCCATTGCAAACAAGATTGTAGAACAAGTCACAATTTGATGCTTGCTCAGGTAAAATGCCCAAAAGAACAA
TAGAGGGGTGCAATGTCTCACCTTGCCATATTCTGCAAAGGAGCAGGGCCAAGATAGTACTTGTTGCCAATTGGA
TCTGGAGTGGAAACGAACCTGCATGAACACCAAGTTATAAGTTCCACAGCAACATCAGCGGATAGCATGGAATTG
GTTTTTTTGGTTTTGGAACTTCAGATCCTTACACTAAGACGCCCATCACAGCTGGTTTCAAGGGATCTCCTTCCT
GAACATGCATCAAGGAAACATGAGAGTTCTGAAAGAAAAGGCAGTTCAAAGCTAACTGTTCATACCTTGTAGAAA
TCAATGGATATCTTCTGGTCATACTCACACTCTCTTCTCTCCAAGTACTGAAATGAAGGTGAAATTTTAGTGTGA
TTCTAGTTTTTTTGGAACTGCAGAACACGGGAGAGAAAAAAAGTAGCGCACCTGCATTGCTTTTAGCTCTTTTTC
TGGACCACCCTTGACACAATATGCAATTCCCCACTACAAAAAAAGTTAACTGTAAATAAGACCAACATAACAAAA
TTATGGGAATCTTAAATGTGAGTAGCATCTGATGAAAAGAACTTGAAATGAGCATATCATCATTCATCAAAAACA
AAATCAGAACTAAAAACGAAACATTTTATACAAAAGAGTAACGAATGGAGTAGAAGACAAAAAGGCCAATCATGC
TCTCTAAATAGATAAGAGATCTTGAAAAGATGAGAAGCTGTGGGTGGATTCTAGATGTCAGCCACTGATCTTGCA
TGGGTTTCAGGTGCTATATAATGATGCATGTTGTGTCGTCACGCAAAGCTGATCCATGGGTCCAACATGCCAAAC
TGGTGTGCAAAGCTAACAACTATTGGAGTGGTTAAAGTAATGAGATTTTACACAGCTCACTATACAAGTAGAAAA
AGAAATTACTCCCTACCCTATTCTTCCACTCTTTTCTAATTCCATAGTCCAAAGTGAAGACCTCGATAAGATTTC
AAGGCCAATGTAGAGCTACATATTCATGCGAGAACATGGAAATTGTGACGCAAAACTGAATAACAATTCATTAAA
AACAATTTTATGGCTAATGGCAAGAAAACAAGTTGAACCAATATGTTGTCAGAAAGAAATTCATGACGACAGGCC
ATGCAAAGTACCTTGCCCATGTAACACTGTGTCACCGTGTAAAAAGGCGGGGCATGCAAAGATGCTACACAAACA
GAACATTATTTAGGTACATTTCAGTAAATGATTTTAGTCGTCCAAATCCATATCTAAATTGCACATGTAATACTG
TGTCACCACCTGCAGTAAAAGCTAGATAATTCATCTTCGCAACAAGAACACTGCAAACAGGGAAAATAAGGATAA
ACACGAGGTGGTTGGTTTCTCACGCATGGCCTCGTCGTCGGTTTCAAGCGTGCAGGTCCTCGCCGGATGCTCCGG
TGTGCCTCTGTGGTCAATGCAAGCTGCGGGGGTGGGAAACAGAACAAGGGAGGAAGTTATCTTAACTTTTGAGAT
TGAAATAACGACAACTAAAACAGGAAAAAAAATTTATAATATAGATGCTAGTTGGTTTGCTGTACGGGTTCTTCAT
TTCTATAATCATCAGGTATGTTTCGAGTTATCTCAACTTTTGAGATTGAAATAACTAATTCTGAAACCTTATTTT
CACTCTGAGCCTAAGAGAATAATTTATCCTCAGTTAATCAACACTTTATGACACTATAAGACCGAGAAGTTAGAT
AAACAGAACTGCACTTATAATAATACAATGTAGTCTTAAGTTATTGCATACACAATATAGCTGAAAATGCATCCT
TACACCTAAGTCGTCAAACTCAAGCACGCTAAACTCCATTATCAGAGGCCGATTGTTTGGAGCACCGTGGAGTAG
AGGGCTTATCTAGATTGGTAGTTTCTGTGAACTGGAGACATTTGTCCTCATCATTTTGGGGCCCTCATTTTGGAG
AGAACAGTAGGTAACTCAGTACTCGTACTTGCTTATCTCGTCACCGGACGAGCAAAAGTGTGGATCTCTTACCGG
GAGGGGGGCATCACAAATGGCTCCGACTGCTCCACAGCGATGCCTTGTGGTCGTGGTTGTGTTGTCCAACACGGC
TCTATGGGCTAGATAGTACCAGATCCGCGCAGAGTCgggggagcagggaggggaggggagggaagggcaggagga
ACAAGCCATGGGCTCACCTGTGCCTCCCGAAACCCCAAAAAATCCCAAGAAAGCAAAGTGTGGATCTCTTAACGA
GAGAAGTTAGATAAACAGAGCAGGGAGGGGAGGGAACGGAGAAGGATGCTTACCAGCTCCAAGCTGTTGACGTGG
TACTCCAAGGAAGCATAGTGCTTCTTCTATTTCTTCCCCTCGCCAGTGCTCTTGATCGGTTCCCCTACTACTACG
GGCACCACCTCCATGTTCTCCTCCTTGCAGGAATCCTCTGCATGCGGCGCACCTCCTCGAGCTCCTCATCCTCGC
TCCCTAGGTCGCGGCGGCGCTTGGGCCATCCCCGGTGGCGGCGGGTCTTCGTCTCAACCTCCTGCTCGCGTGTCG
CGCGCGAGTCACGTAAGTGGAGCGCGGGGTTGGGGGAGGGGGAGTCCGCTGACCCGGGGGAGGATTTCGTGGGGA
GAGCGGTTGCTCCGATAGTGGGCCTAGGTCTGGGCCACAAAAAGCAGATGGGACTCCCAGCACGCATGTACTTCG
TATTTGGAAGCTAAACTTAGTACCACATCGACTACTAAATTGCGCACACACTTGTTTATAAGTTCCAGTGTGAAA
CAACTTGTGTAGTCTTAGTTGGTCATGGCTGTCACCCCACACATCTAGCTTTTTTGGCGTTGATGCACAGGTGTG
GGAACGGAAAAGATGCGTGGGAATGGGAATGGAGAAGATGCCATGGTTGTGGGACCGGTTTTGGTGTCAGACAAC
AGTGTGTGGGTGGGAATGAGAAATGCCACGACCAGGACATCggaatggaaatggcagaaccaggaatggggcagc
ctacccccttagagccttaataagta**gtatagatttCCACAGACCGGCAGTGCACGT**GGAGTGCCTAACGAACGCG
ACCTGCAGAtatagatgtccaaatgggtgggccgtgccgatccggcccagattagctagcgggtcatgtcgtgcc
ggcccacgggctccaagcgaggcccaagcacgacctgctaggttaataacggttaataatcgtgtcgggccggtc
tacagcccgacagatgtaatgggctcataacaatatatgatgtttaaatgataaaaatatcctaaaagtaagcat
tttttatggtttaaaccatatttattaactctaaacatttatttacacccacataaccaacaaaaacacatgttt
cttgtgttttatactctatatttaagtataatatatgtatttatatgaaaaaatagaaaaaaaaataaaatcagg
ccgtgccgggcctagcccgtcgtgccacgcctttggcccaggcacggcccggtgaccgggtcaggccggcacggg

>ZmMADS23_XM_020546680.1
ATGGGGAGAGGGAAGATAGAGATCAAGAGGATCGACAACGCGACGAGCCGGCAGGTAACGTTCTCCAAGCGCCGG
GGCGGGCTGTTCAAGAAGGCCAAGGAGCTCGCCATCCTTTGCGATGCCGAGGTCGGCCTCGTCGTCTTCTCCAGC
ACCGGCCGCCTGTATCACTTCGCTAGCACCAGCATGGAATCTGTGATTGAAAGATACGAGGAAAGAGAGGGGCAC
CATCAGACTATGAGCGCAAGTGCTGAGGCCAAGCTTTGGCAAAGGGAGGCAGGAAGCTTGAGGCAGCAACTGCAT
AACTTGCAAGAGCACCATCGGAAGTTGTTGGGTCAGCAGCTCTCTGGCCTGGACGTGAGAGATTTGCAGAATTTA
GAGAATCAGCTGGAGACAAGCCTAAGAAATATTCGTCTAAAGATGGACCAACTTATTTTTTATCAGATTCAAGAA
TTAAACAGGAAGGGATACCTCATGCACCAGGAAAACATAGAACTACACAACAAAGTCAACCTTCTTCATCAAGAG
AACATTAAATTACGTAGAAAGGCGTATGGACAAGGAGTAAATGAGCATCCAACAAGTACTACAGTTAGACACAGT
ATTCTGAATACAGAGAATGAAGATGTTCGGATCAATCTTGAGCTGAGTGTGCAAGGGGACAAATCAGAAACACCA
AGTATAGGGTGA

>zma_miR444a_PUT-1-171a-Zea_mays-14134
ACGTGCCAACTGCCGGTCTGTGGAAGTTACCTCAGATGAACTCAATACACTGCCGACGCCACCTGATCTAGGGTT
TAGCGGAGTGCCCTCGCGACGAAGCGAGGGCCCGATCCATCCAGATATGTATTTGGGGGCAGCAAGCTCGAGACA
ACAACTGCATAGCATGCAAGAGCTTGTTGGTTGAGGGTCATACCGATGGTGCTCTTGCAAGTTATGCAGTTGCTG
CCTCAAGCTTCCTGCCTCCCTTTGCCAAAGCTAGTAAGATGGGGAAAGCTTTCAATTAACATAAATGAGAGTGGC
TGGACAACCAAATGGGTCAACTTTAGCATAGGGATTACAACTACAACGAAAAGAACAAAATCTTTTGCTTCAAAG
GAAGCTGGGGTTTAGTAGAGTTGAAACCCAAGTACCCAACAAGAGCCAAATAATTAAAGAAAATAAGAATGCAGC
AAGCAGTCAGATCCTACAGGGACATGTAGACACAATATCGTTTTGTGAGCTATAAGCTTGTAAGAAGTCTTCTTT
GGGTTGGAAACTATTAATGGTTCAAGAGAAATTTTCTGCAAAAAAAAAAAAAAAA

Stem-loop structure

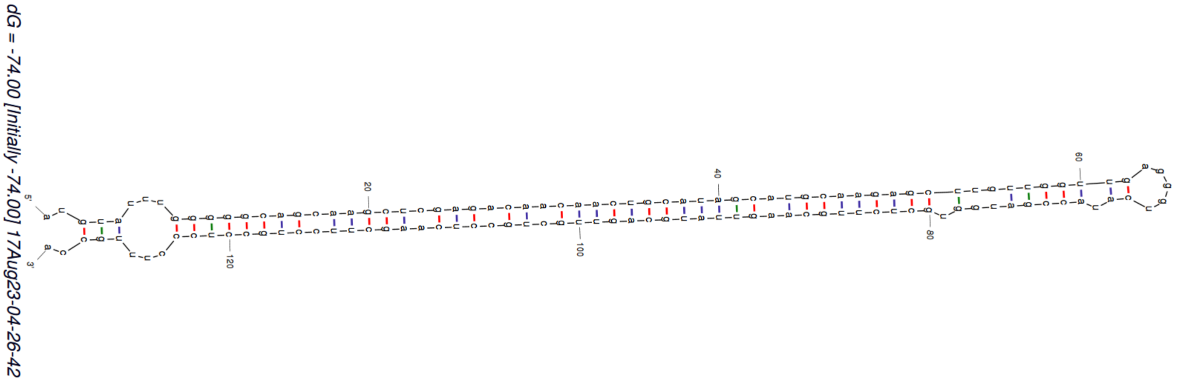


*ZmMADS27_1 and zma-miR444b*

>5
ACAATAGAAAAGAGCGCCCTCGTGCTCAGAGTGCTCGCCTTTAGTTTCGTGGCTACTATATACTAGCTGCGACGA
TTTTTATTTTCTTTCTCTTCTGGGCTATCATTAGCACGGCCATTTAGGGGTAATGAGCAGCGGCCAATAAACAAA
GCCCTGAGATGCATGGGGTAGCCAATCGGGTGTCGCCGTGCGCGCGCGCGTCTGGCCTCCTGCCCTCCCTTTC
CTTGCTAAATAAAGCAAGTAGCAGATAGGGGAAAGTCTGCTCGCAAGTTGCATCCGCCCTGCGCCAAGAAAAGCC
ATCGTTCTTCCCACAAACGCACACATAGAAGCATCATTCCCCTCTCGGCTAGCTTCTTCCTCTCTCTCCCTCCTC
CTCCTCTTCCTCTTCCTCCTCCCTTGGGAAACCTGCTGCCTTTGAGCTTTCTTCTTCGAGAGCTCCCACCAGATC
TCCTCCTCCTTACCTTCTTTGGCACGTTCGGCGGCGCGCGCGGAGAAAGATAGATCCCGCCATCGTCGTCGTCGG
TCCTTGCTTCCGATCGGAGGGCCACAACCACAACCTCTCGCTCCATAGCGTGCAAGCGCGAGCCAGGGTCAAGAA
GAGAGCTAGCTAGCTATAGGCCGGAGATCGATGGGGAGGGGAAAGATCGTGATCCGCAGGATCGATAACTCCACG
AGCCGGCAGGTGACCTTCTCCAAGCGCCGGAACGGGATCTTCAAGAAGGCCAAGGAGCTCGCCATCCTCTGCGAT
GCGGAGGTCGGCCTCGTCATCTTCTCCAGCACCGGCCGCCTCTACGAGTACTCTAGCACCAGGTCGTCGCCCCTTC
GATTATTATCCGTCCCTTCTTTAATTTAATACTTCAATAATGACAACTGTTTTCTTGCTCCTGTTGAAGCGATAT
ACTATATATAATAATAGAGTTGTTCTTGGCTTCTTGCTTTAATTAGTTTGGGTTATCGATTGCAATTTATGATTT
GAAACTACACACTACCTCCTGTCTCATATACATGGACTGGCGAGTGATTATGGACCTATACACCATGTATACACA
AGTCCTAAGGTAGACAAAAGATTTTAATTTACCTGTCGATTGTCGAATATTGACTGTACATAGCGCACAGCTAGC
AACACTCCTGGTTTCTTTGACGGGGCAAAGCAAGGTCTGTGTGTGCGGTCTAACAATAAAATACAATCTTTGGTA
CGGTTAGTATTTGTGTTTTTGCCCCTCTTCTTCCTTGTTGTTTGGTCTATTTGAACTATTATTCCTCTCCAAAAA
ATTCCCTATGTATCATCGGAATTTATACTCATCCCTTGCATTGTGTGCCATATATTGAGTGGCATATAGTAGGGA
ACATATGCAAAACTAAATTTAATTAGCTTTGTCATATTGTTTCATGTGATCAACTCGACACAGCATGTCATTGCA
GTAGGAAACGCATATGCTTTTGTAATGTAAGATATATGTATATATGTATGGGGTTTTATCAACAGGGGCTGGTAG
TTAGTGCCATCAGAAGAGTTTTTCCCCTAGAGATTGAGATCGAGATCGAGATCGAGAGAGAGAGACTAGATCTAA

TTCTGTAATTAAGATTTTAGCTTTTTTAGATAGCTATAAACTTAAATATTTTACAACGAATAGCATAGTAATAGT
TTTTTTTTGGATAAACTGACGTTTGTGCTGTGTGCAGCATGAAATCAGTTATAGATCGGTACGGCAAGGCCAAGG
AAGAGCAGCAAGTCGTCGCAAATCCCAACTCGGAGCTTAAGGTTTGCTCTAGATAATTTTTCCTCTCTATATTCT
GAAGTGTTTGCTTGTTTAACTAAAAGATTGCTATTTTTTTGCTTTCATTTTGATATATATATATATCCTTTAAGT
TTATTGGTTTTAAGGATGGGAAGAAATTACACAACTATATATGGTCACATCAACAAGCGTTCTACCTTTTTTATC
TTATCCAGCTGCTTGATCTTTTCTAGTTAACAAAAAATAATTTCAGTACTTGAGGATATCATCGGGTTGCTTGTT
TAGTTGGCCCATTATAAAAACGGTTCGACCAATCACGCTTTCATATTATATTATATTATATTATATTATATTATA
TTATATATGCTGCGTGGATATGTAATCCCTAAACAGCGCTAGATATTATTACAAAGTACAAACTAATGAAATAAA
GATTAATAATTAACCATGCATGGGATCGGACGCCCATCCTCAACGATGGTGTGAGTTATCTCACTTCTTGCTTCA
ACTCTTCTATTTGGTGTTATCTTATGCTCTCAAAAGTGGCACATATAGATGCAATTTAGTATTTCGGGCCCGTTT
GGCAAAGCTCCAGCTCCTGCTTCTTTAGCTCTAGATCCTGATCCTCGTGAGGAGCTGATCCTCCTGATTCTCCTA
GAGAATCAGAATCATTTTTAAAACTGTTTGGCAAATAAACTGATTTTTGTCTTCTGAAAGTTGGTGGTATGTGGC
GATTGTCTGTTTTACCCTTTACAGTTCAACTTTGTTACAAACTAATAAGTAGGATGTATATACGGAAAAAATTAT
GAAACAATAACATATAAAATATTTCCATCATAGTAGTGCATAAAATGGTCTCAAATGTTTAATCCATAAATTGGC
TCGTTATGTTTTTGTCCAATGGCAATTGCGGGTAATTTCGATCAAACTTTAAGGGGAGGTCCATATTTGGTTGGT
TGAGGAGCTGTTTTAAGGGAGGGTGGAGCAGATTTTTTTAGATCCCACCTTTCTTCACAAAAACGACTCCCGATC
TTTCGGCTCCACACGAGAATCATTAAAAAAAAGTACCCATTTGGCACGGCTCCTCCAGATCCTCGCTTAGAATCA
GGAGCTGGAGCTGTGCCAAACGGGCCCTTCATTTATGTTTTTCCAACACCTTTTAGTCACTAACAAGTTTGGATA
AGTTAGAGCCGAAACAGAACATGACCTACCAAAAAAGGAAAATAAAGATGAAAGGGAAATAAATAAATATATACA
TGACATCAATAATAGTCATATTATATTATTAGAAGATCTAAAGCTTGTTCATATTTCTCATGAGTTATTCCAGTT
AGAGCAGTACGATGAGAGGTCTCCCCTTGTCGGTCCACATTCAAATCCTTGTACAACCTTTTACTGCAATGCCTA
TAGGGCGCCCTATGATGGAGGTTTTTAAAGCCCGTTTATGGTTCAAACCATCTACTTTCTATACACTCCCATCCA
AGGACAATTTTTATAGGTATATTTCATACTCTCAAGTCCCTTTTTATTGCGACCATCCATGTCAAGTTTGATCGA
CCTTGGCCATCTTTAAATTGTTGTTGTATCTTATTGTTCTCTCGTGGCTCTAGTGGTCTCCACTGAACATATCCA
AACCATCTCAATCAATGTTTGACAAGTTTTCTTTAATGTGCTACCCTTCTCCTATCCCTTATATCATCATTTAG
GACTCGATGTGCTTTTGTATTATTTCAAATATAATGCAACATCTGCATTCACATGTGACACTTAAATGTTAAGCA
TGTGTCATATTATTAGTAGACCAACATTCTGCATGTTACAATATTAATATCATAAGTTTGATCAGGTTCTATAGA
ATTTACCTTTAGCTTCTATGGCACATACTTATCATGAAGAATGACATATATTTGGTGTCGTTTCATATGCTTCAC
TTTAAATATATTGCTAATATAATACCATCGATAATCCCATCTCTTTGTAACACTGATCCTAGATATTGAAAAGTG
TTCTCCTTGTGCACTAGCTTACCTTCCAAAGTAACATCTTCACTCGTGTATATAGTAGCTCCAAGGCATACCTCA
TGTACTCTATTCTAGTTTCAAGTCTAGTTTCAATTGCAGATTTTTCCACCACAAATCTAGTTTTCTATGTACTCT
CAACCATTTTTTCATTAATGGCTAGCACTACATCCATAAGCAAAAATAATATACCAAGGGATATCTCCTTGTATG
TTTTGTGGCCTCATCCATCACCAAGGCAAATAAAATAAGTTTTCAAAGATGGTCCCTAATGTAGTCATATTTTAA
TTGAGAACTCACATATGCCATTAATTGCTTGAATAATAGTCATAACATTATTGTATATGTCCTTCATGAGGCAAA
CATGATTTGTTTAGAATTTATGTTTGTATAAAACCCACCACATGGCATTTCTTAGCTACTTTTTATAGATCTTCT
ATAAATCTATAAAAATAACATTCCATAACTCTGTCTCAAACTTTATAGTATGTCTCGTCGATCACTTACTTTCTA
GATAATTAGCACACCTTTGATTGTGATCTAGCTAAATATCATTAGGATTAATTTTATTACATGACATGGACTTTG
TTAAGTTTTAATGATATTTTACTAAAGGATTACAAAAAATTTCAGGATCGACAAATTAAACCATAACAATCGCCC
CTAAATTTACCTCTACCCTTGAGGCATATTCATACTTTAATTAGGATACCATTAGGGCTCATCAACGTGCCTCGT
TTGCTAGACATTATATTCTTGAATCTTGTGCACCTAGAATCTATAATGTCATCAAAGGAGCCATCCAATTAAATA
GTTATGTTCTCATTCTCACCGTTGAATATTTTGTATAAACACTATTGTGATTTATGGTTGGCCTTATCCTCTTTC
ACCAACAATGAATCCATCATCATATTGATGCATTTAACTTGATTGGGGTCTCTCACATTCCTTTCATCAACCCTA
TCCATCATATAGGTATTTCTCTTCTTTGTTTGTACTTAGGTGTTGGTAAATATTCTCATAGGCATGACCCTCTCA
CTCACAACATGTTTTGCAGCCTTCTTTACCATCTTAAACTTCTCTACATAGTTCATATTATGTCATGGCACAACC
ACCTATATTGTTCTTTCTTCTTCATCCTAGTTCCTATAACAAGTATCTTTAGGTTTGCATCCTCTTGTTTTAGTC
ACTTCAAACACCTTTGAAGATACCTTCAAATATATGTTGCCATATTCACATGCATGTTATTTGCATCTTCTCCCA
AAATCCTTCTTTGATAACATTATTATTGAAGTCCTCTAAGGCCTCCTATTTGAGTTTCCACTACTTTGTCCTCGC
AACTGTGGTTTGTCCGTCCATATAAGCATGTAACTAAAAGCTCTCTAGACATCACCTTATTACAATATAGGCACG
CTCGTGTGTCATCTCTTTTTGGAAGGATGAAATCAACCTAACAATAGTATTTTACACTACTAAAGGTCATCAGAT
GGGCATATATTTGTTTAAAGTAAATGTTAGTTATCATCATATCATAAACTATTAAAAAGTTCAAGACTCTCCCTC
CGTCCTAATTTCTACCACCATACCCAAACACCCCATGTGCCACCTCAAACCCTATATTTGTAGCACCACATGACC
ATTAAGATCTACTCATATGAAAAGTTTCTTGCTAATAGGTACACCACTACTCATATCGTCATCTAATTCTTCGTA
GAAATGCCACTTTGCACTCTCATGTGTCCTAGATAAGGGGCATATTCCTTAATTGCGTTCAAGACTGAATCACCA
ACGCAGTAGGATTACTTATGTACTAACAATAAATACAATGGGGAGAATTATCTCAATTGAATGCTAACAGGATAG
TGTGTGAAAAATAAAAGGTCCCATTCTCGTGAGAAAGAAAATAACGAGTTTCTATCGATCTTACTTATGCTTTAA
TCGATTGAAAACACAATTAATTTCTTCTTTTCATTCATCATGTAATGTGGAGAATTACATCTATATCTAAAACTT
TTGGATATAGATAATTTCTATTTTTATTTTTTAAGACAAGACACTTCAATGCACACATTTTGCTAGAAAATTGGT
ACAAAGTACAAATTTTAGTGAAATAAAGGTTTTCCTTGCAAAACGCTATATGATGCCAAAAACCTATGAATAGGT
TTATGTATCCAATGCATGAAAATCATGATGCAAATAAAGTTTCAAGTTCTGAAAACAAAAGGTACAAGAATAATA
CCTAGTGAACAATGAACAAGAATTGATCATTAAGTATTTGCATTACATAGTATTCTAGATAGTGTTTATGGCGGT
GTAGAAACATGGTGAAGAAAAAATTGCAACAACTATATCTAGCTATAATGACAATTGAGAACTATTTGTTTTCAT
TTTAATATTAATAAATAGTATAAAATGACATATGTATAACAAATAAGTAACAACAAAAAATGTACGTTTAGGTTA
GGGCATTTACGTGCATTTGCATAGTTGCATGGCTGCCAAGATATTTCTTAGGAAACAAAAGATTGGTTCCGAATC

AGTGGCGGAACTCTCATTATGGTTCAGTATAGCATAAGCCATACCTAAAAAACATGTATGGTGTAGTGGTCATTG
CATATCACTTAAATTTAAGATAAGATTATGATAAGATAAGGTTTAAGATTTATTAAGAGGTCTCAAGTTCAAGTC
TTGCATGTGTAGTAATTTTCTTTTTTTTTTCTTTTTCAGCCATACCTCATTTATACGTTGTCCTCCGCCACTGTT
CGGAATACATTCACTTAACTAGCACCAGGAACACTGTCCATCCCTCTGCAGTACAGAACAGATCAACGTACCCCT
GGATGTGTTTTTGGGTTCTATAGTATGCTCTATATAAAGTTCATACAGTCATACATGTTTAGCTTAGATATTTAT
CATTATATGAGAACTGTGGCATTATTCTATCCCATTTGGTGATGTTTTCAAGTTAGCAAAGCCTTATTTAAATAT
ACTTGAGCTAATAGTTAGCTGCTAAAATTAATTGAAGGTATCCAAACAGACTAGCTAGTTCAGCTATTAGCTACT
TTTAGACAATTACCTAATAGATAGCTAATTCCACTAGCAATTCTTTTAGTCAACTAACTATTATGTCTAGTGCAT
TCAAATGGCCCTAAGTCAGCTAACTATTAGCTAGCTATAACTATTAGGTCTAGTGCATTTTATTAGCTAGCTAAT
TCTACTATTAATTTTTTAGTCAACTAACTATTAGGTCTAGTGCATTCAAAAGGCCCTAAAAGTAAGCTTCAAAAT
CTTGTTGATTTAGGTCTTACACAATCTGGACTCGAAACAAATGTTTTGGTTGGTCTTAAAATTCTTTAGACTCTG
TTTAAATGCACTAGATATAATAATTAGATGCTAAAATTAGTTAGAAGTATTCAAACAATCTAGCTTATAGTTTAG
TTATTACTCCCTCCGTTTCTTTTTAGTTGTCGCTGGATAGTTTAATTTTGCACTATCCAGCGACAACTAAAACGA
AACGGAGTGAGTACATATTTTTAGTAAATTAGCTAATAGCTAGCAAATTTTTTTAGTCAATAAACTATTAGCTCA
TGCATTACAATAATCCATGACTAATAATAACAGACAAATGTTCTGTCCCTTTTTGGTCTGGCCCACTCTATTCTA
CTAGATCAATTATTCTGTCGCTCGTTTCCTATGCACATGTTCGTGTCGACCTTCTCCTCACAACGCCAAAAAACG
TATCTTCGTTTTACCGGGTCGTCGGCGCTCCTCAGCGAATATATGCATGATTAATTGCTGTCTTGAGCTCGTTCT
TTGTGTACATGTGCTGCTGAAGTGCGAAGTCAAGCAACAAGCAAGTGGAATTGTTCCGAAATCAAATCAAGCATC
ATGATTTATGAATGAACCGTTTTGCAATAACCGTGAGACTTTAGCTGCAGGAAGTGAGGAACGCGGGAACATATA
GGCTTATTCGCAAGTGAAACGGAGGTGGTATTGGGAAGCTAAATTATTCTTGAGACAGCCTACGGATACTAAAAT
TTGCTATGCCTAAGTTTTTGCTGATAAAAACTATCAGTCTACTACAGATTTGTCAAGCCAAATGTAGAGGTTGGT
AAATTATGGTACCAAATTTGAGGCTTTCTTTAGAATGGGGGATTTTTAGAGGAAAAAACAAACAGGGTTCAGCAA
TCCAGGACCAGGACTATAACAGTGTAGATGTTCTTTTAGAGTTTCTTTGGAATGGAGAATTTTTACAGACTAGAA
CAGGATTCAGCAATCCAGGATTATGACAGTGTAGATGTTTTTTGTTTGTTAAAGAGGATCTCAGGAAAACTAAAG
GAATACTGTTGGAATGGAGGGTAGCATTTCTCTTTCCGTTTCTCATGAACATTTGATAGCTTTTTTCTCATGAAT
ATTTGCTAGTTTTTTTCCCGTTGCAACGCAGGCATGTTTGCTAGTCATTAGTAATAAAACATCAATGCATCTTAT
TTACATGTTCTCAAATCAAAATTTACATGTTGGTGACTAGGAATCCTTAACACACCAGTTAGCCTTTTCACAGAA
GTTAATTAGGAGATCCAACAGAGAACAGTGTGTCT**TACGAACTTGAATTCTACTGCTGTATGTTATCTGGAACTC**
**ATATTGCATCACGAAAACAATATGTCAGCAGATGCTTTATCCAGTAAATGTTAGGAATTCTTTTTGCCACTTCAG**
**TTGTGTCACATGACACTGGATTGTCAGAATTCTAGTGCTGTTGCAGAACATCCCATGAACTCCGAAGTACCAGGC**
**CGCAGGCCGTATAGTTTACATGCCAGAAGCTGCCCTCAAATATACCAGTCGTCTTCTACAACCTTGCAAAAAAAA**
**AAAGAACGTAGAATAAAACTGTGTATGCCAATCCAGCCGGTTGATTGTGCTCTAAAATATACGTAGCAGTGAAAT**
**TAAGTTGGAACAGGTCCAAGCTCTACTAACATCAAAGGTTGTCGTAGAATAGATACTGAACACTCTGAACACGTA**
**ACGGTAGAAGAACCAGCATTGACTAATTGACTAAAAGGCACTCGGCACCCTTCAAATCATAAAATCAGGAGATCA**
**TTTACTTATACTGACATGTTCTTGTAAGAATTAGCACTCGCCTTGTTATGACTTCATGAAGTCTGCTTGCAACAT**
**CCTAACACAACAGTACCCGCTGTAATATCGTACCGTACTGCTTGTTTATCCAGAAAGTAAACGAAGGCGCATTAA**
**AATAGCCTCCACGTTGCAAAATTGGACATGTCAGATTGCAATTTCTAGTTCTAGTAATAAATATTGTTCCCTCTT**
**AGTTTTGTACTCTTTCCATTCCAATTTATAAGATGTTTTGGCTTTTCTAGACATTAATTTTTGCCATGCACTTAG**
**ATATATACCATGTCTAGATGCGTAGTAAAATCAATGTATCTAGAAAACCCAAAGCACCTCATAATACCTAAAACA**
**TCTTATAATTTGGTATGAATGAAGTACCTAAGAGCTGCTGTGTGTAATTGTGTATCATGCTGCAAGATTCCAACT**
**AAACGTCATGTCTTACCAACATATACCAGTTAAAATCTGTGCCAATGTGTTCATTGATAACATAACCTGTTTCTT**
**CATAGTTTTGGCAAAGGGAGGCAGCAAGCTTGAGACAACAACTGCACAACTTGCAAGAAAATTATCGGTATGGTT**
**CGAAAACC**CTGCAAAAGTTTCAAGCAAGTTATTCATTAGAAACTTGAGTGTTCAGAGACAGAAGCACAATGCACA
AGCATGCAACATCCAAATGCAGGTACAGCTATTACACATTCATAGATAACAGAAACAATTAGCAAGAGTCAAATT
ATCATGTGAAAGGCTAGACATTAGACATCAAAATACTTCATACGAGATAAGTTGAACATGTGGAAATTATTCTAC
GGCCAACAATAAAATTAAAACAAAACAGGGATATCACAGTTAAACTACTTCCCCTGCTTCATTTTAAAGGTGCAT
TTGCATTTGCATTTGAACAAGTTTTTAAGGCTGTCTCTAACAGCTACCCTATCACATCCCCTATCTTACCTCCTA
TTTCAAACTTCATTCAGCAAACAGTGTAATCTAGAGTATAGTGCAAATCCCCTATTTTACACAATGTGCTGCGGA
TAGCCTAACATTGTACTTTGACCATGAATTCCTCTTATACAATAAATCAAACCCAGAAAATTCAATGTTGTTTCA
AAAGCTTTGTGAGATATGAATATTGATACAATTTTCATACACTAAGCCTGCATATAATTTGATTAACTGTTAGAC
AAATTTTGTAAAATTAAACAGCCTTTTTTCCATACAAAAGAAACAGAGGGTATATCAAAACATATCCTGATGATG
CTCCAAACATTTATAGAGAAAACACACACATATATTGGCAATTGGCAAACATAAGCTAGACTGTGTGTGTATAAT
TGTATAATACACAACATTAATTGCTGCTGCCTATTAATTTGTTACAGGCAGTTGACGGGAGATGATCTTTCTGGG
CTGAATGTCAAAGAACTGCAGTCCCTGGAGAATCAATTGGAAACAAGCCTGCGTGGTGTCCGCGCAAAGAAGGTT
TGATTCTTAATCAAATTTCTATAATCAATTGTTTAATAAATTACTAAAGATCGTTACTACTGTTGATGCTCTTAA
CAGGACCATCTCTTGATAGATGAGATTCACGATTTGAATCGAAAGGTCTGTAATATGGTGAATTTATAGTTCTGT
GTTATGTGTTTTATTTGATTTAGTAGCACTGACTAACCACTTTGTTGGTCACAGGCAAGTTTATTTCACCAAGAA
AATACAGACTTGTACAATAAGATCAACCTGATTCGCCAAGAAAATGATGAGTTACATAAAAAGGTAGCTGATGAA
TTGAACAAAAATACCACTAAGTAACTGCCTAACCACACATATGACATGATATTAATTGTTCTCAGATATATGAGA
CTGAAGGACCAAGTGGAGTTAATCGGGAGTCACCGACTCCATTCAACTTTGCAGTAGTAGAAACCAGAGATGTTC
CTGTGCAACTTGAACTCAGCACACTGCCACAGCAAAATAACATTGAGCCATCTACTGCTCCTAAGCTAGGGTATA
ACATTTATTCTCTCTATATATTTCAATTGGTTAACTAGTGTTGACCAGTGCTATAATGAGTTGCTAAACACATGC
AGATTGCAATTAATTCCATGAAGAAGAGTAAAACTGCCGTCTTATGATGCTGAAGGAAACTATTTATTGTGAAGA

GATGATACTCAGAGAAAGACATATTTGTGGCAGGGAGATTTGAGATATGAACTTATAAATGTAATGCAAATAATT
TTCAGACCGGAATGGGGTCGTGGAATTCAGAGGATGATTGCTTTCTAGTTGCATTTGATGTTTGATGAGACTTTT
CTCATGTGAAACGTTTATTAAAACTTCAAAATGGTGCAGATGTTGCTACTACACTTGTTAGGCTTGTATAAATAT
CTGACCATGTGTACTAGCATTACGAAAACAGAAAAGAAGACAGACAGCCCTGTCGAACATCTTGACGTGTTTCTA
AAAGGTTTGATAGGGGACAATGATGCGATCTAGGAGTACAAACAGAGTCTAGTTTCCACTCAGTTGAGTTTTATA
TTGCTCTCAAGCGTTATCAAGGGTTTAGTAGGATACAGATATAATATGAAGATGCAGTTAGATGTATTGGTCATT
TGTTGGGCCATATGGGAAAGCCAAAGCACAGCCTGATTCGACAGGAAAAATAATATTATACCCATGTAAAATACT
CTCAAGCCTGTGTTTATTTGAAGTATTAGGCAGGTTTATACAAATCAGGGCTAAAGGAAGCTACCGTCGAAAGGG
CTGGGCGTCATTCCACAGGCAGCCATCAGGATCGTGACATTGCAGGTGCAGGTGCATGAGACATTGAATCTGATC
AAGTGCTAGGAAGCAAGCTACTGTGTTCAGGTCCAGCACGGAAGTGGAATACAAAGTTCTAACCAATGCCATCGC
CAAGATCATGTGGGTACAGAAATTGTTTACTGAGTTGGGCATACAACATCCACAGATGGCTCATCCAGTAAGCGA
TAACCTGGGGGCTAAGTATTTGTTGTCAAATCCTATCTTCCATGATCAGCCTAAACATATTGAGAAAGAATTTCA
TTTCGTTCATGAAAGGGTGGCACAAAAGTTGTTGGATATGTCTTGATTAGTTCAGAGGATCAGATAGAGGATGCA
TTCACCAAAGAGTGTCTACTCGGCAGTTGGAAACTTCAGGCACAATCTCAACTTACCGGCATTGTGGAGTTGCTA
GGGTTGCGTGTGTGTTTGTTGTAAACTGAACCAGCCTATACTTTTGCATAAAGATGCAAGGTCTGGCAGTTCTAG
TCTAGATTGAGTTCGTTAGAGTTGTGTACACTTCGTGGTTAAGCCATAGAGATGATTGTAACTTGTAAGCCACAA
CAAGGACTCTTCCCATCTTATATAAGATGCAGCCGTCTGTATGTTTCGTGGAGCAGGTCTGGCTCTAGTTGCCTT
TTTTTTTGAACATAATGGCAGGAGCTCTGCCTCTCAATTAAGAAGAATAGAATTGATCCAGTTTTAAGGAAAACC
GGACCCAAAACCTACACAGGACGCACGGTAGAAAAACCGGCGAAGGCCCTAACAAACCACCTAGACAACGACAGC
CTCACACACACAAACACATCTAGAGTCATCGTTGCCAAGCTCAAACACAAGGGACGAGCAGCTCCGACTTCCCAA
GACGAGCCGCACACTGGCAAAGCAACTCGACGACGCCTAAATCCAGACACCGACGCAAGGAATAAGATCACGGCG
CTCCATCGTTGCCAAGCCTCGCACACTCCACACGAGGCAGCTCCGATTCTTCACCATGAGCAAAGATCCACAACG
CCGAACTCCGGGGGCAAGCAGCCGCCTAAAGCGACAATGGCCTCGACACCACCGAGCCAAGCGGAGTGACAAAGA
GCCTTCACAACGCCAACGAACACACAAACTAGAGGTCTTCTGGGCCGCGCCTTCAAGAAGGAGAAACGACGTCGA
AAAGTCGTCGCCGCCCGCCTGGTACCAACCAAGCAAAGTTTCCTCCGAAGACCTACTGTCGTCCGTCGACGCACC
AACCCAGCAGCCCCAACCCTCAGAGCCCCAACGTCGGACGAGCACCAGCGGGAGACCCCAAGAAGGCTGAGGCGA
GGGTTGCCACATGGTGGCGAGAGGTCCAAAGACGGCACCCTCAATAGGGTAGGGGTGGCGCCCGCGGGCGTCACC
GCTGTCCGCCCGCAAGCGGGCATGGCTTTCGCCAGAGGCCCTCCCACCACACGCCCCAACACACGAAGCAAGCCG
CCCCTGCAGAAGGCGCCGCTGACAGCTGCAGATCTGGCGACGGCCAACGCCCCCGACCTAGGCAAGCACAGGCAA
GCCACGCCGCCGCCACAGTGGGCCAGCCGCACGCGTGGGTGCCGACGCCGCTGGCCAGCCACGCGAGCGCGCGTG
CACCGCCACTGCCACCACGGCCGCAAGGCCACCGGCCCATGCCAGACCACACTGAAGCCCGAGCAGGCAGCAGCA
GCGCTGCGACCGCACCGCCAGAGCTCCACCGCAGTCACCACCTAGTGGTCTTCCTCGCCGCAGCCAGTGCACCGG
CTACGCTGAGTCGCCCATGTCGCCGTGCGCCGACCCCCGCATAGATCTGCCCGCTGACGCACACGTCGACAACCT
CCGCCTCGAAGAAGCACAGCTTGGCGTCGCCCTAGCCCCCACCGACAGTGCACGGCAGCGGCGACCGGAGCAGAT
CCTCAAGCGAGGCGCGGAGGCAGGCCACGACGTCCTCACAGCGACGCCCCGCCACACCAGGCCGCCTGGGCCTCA
GCCGCCGCCCGACGAGCTCCGCGGCGCGCGCCACGGCGCGCCCCAGCACGGCCACGACCACCCATAGGTACCGTG
CCGCGCGCGCTCCACCGCCGATGTGCTACGCTAACTCCCACCAGATCCGCGCGAAACACCCCGCCGTCGCCATCC
TAGGGGCCGAGCCGACCTCCGGCGGCGGCAAGGGAAGGGGAGAGAGGATTTGCGGGCGACGGCAGCAAGGGGAGC
GGGGCGCCGCCCGTGTCGCGCTCTAGTTGCCTAAGGCAGGCTAATTAGTGGTTGACTTGACCTGTCACAAGGTTG
CTTGTCTGGGATTTTTCTTATTTTCATCTTCCTGCTTCTGTTACTGTAAAAGATTAGTCCACCTATGTTGTCTTA
ACATAGCAGGTCTCACACCCTGTTAAAGGACAAGGAATGTGATAGGCCCAAGCCACGTTAAACCTAGCCATTCTA
ATAAAGATGAAACCCGAAAGAAAAGGTCACAAGGAACATATAAGATGATATCACATGGTATATGCTTTTTGCGGA
CGATGTAGAACTAATTGATGAAAGTTTGGCAAGAGTGAATAGGAAACTAGAGTTGTGACATAAGACTATTTTTTT
TCTCGAACGCGCAGGAGAGCTGCGCATCATTATATTAAGAGAGGAAAGGTCCAAAGTGGACCGAAATACAACACC
AAGACCAAACCCTCAAGCCTCCCCTACAAGGGGGGAGCCTGACGGGCCCTACTAGACCCTATTTAAAGTGACAAT
AATTCTCAAACTGTCTAAATGCTTCGCTCCCGCCTTACTCCAACTGATCAATTCATCTAGAAACAACCTCTTAGT
CAAACTAATGGAAGGGCTCACTCCATCAAAAACAGCCCTATTACGAGAGAGCCACAAGCACCAAGCTCCAAGGAT
AATGACACTATTGAAACCCCTCCTCTTGCACCTATGCACCCTTTTAGAGACCTTTTCCCACCACACATGAGACTA
TAGAGTTCAAAAGGTTTTAGACTTAATAGATTTAAAACCGAATATATGAGATGTGACTTCAGCACTCATACACAT
GAGAAAAGAGATTTTAGTTTTCAAGGTCAAGTAGTGCCTAAGAAGAATACCTTTCGTTATTTAGGATCAATGTTG
CAGAGATGAGGATATTGATGACGATGTTAGCCATAGAATCAAAGCAAGTTGGATGAAGTGGCGTCAAACATCTAG
AGTTTTATATGACAAGAGGGTACCACATAAGCAAAAAGAAAAGTTTATAGAATGGCGATTAGACCTTATAGGTTG
TACGTTGCAAAATGTTAGCCTACAAAAAGACAATATGTTCAACATATAAGTGTTGTAAAAATACGTATGTTGCGT
TGGATTTGCATCTATACAAGAAGGTATCGAGTCCGGAACAATGATATACGTAATAGATTAGGGGTAGCATCAGTT
GAAGAAAATTTGGCTAATATTGGTTGAGATGGTTTGGACATGTCCAACAGAGACCTCTAGAGGTAGTACGTAGTG
GGATTGTAAGGCACAATAGCAATGGAAAGAGAGGCAAGGGAAGACCGAAGTTGACATAAGAAGATACAGTAAAAG
GGTCTTGAAAGGATGGAATATACCTAAAAATTTAGTCTTAAGTAGGAGTGTATGAAAAACAACTATTTATGTGCT
TGAACCTTAACTTGTGGTTTTTGTTGAGAATCTGAAGAAGAATAGGATTGGTGTAATTGGATGGTTGTACTGCGT
TGGGCCCTTAGGCGATTTATATAGAGTACATGGCTTAGGAGATAAGGCAGCACATATATACATATGGCAGTCTAC
GAGGCACAATATCTAAACTACTATTTGTACTCTAACATCCCCCCATAGTTGTAGTAGCAGCACTGTAGATGGTCA
GACTGAAGAAGTTGATGAACGCATATCCCCCTGCAGTCGTGACGTTGTCGCACAGATGCAACAACTAGAAAAGAA
AACGATGAGTAGATCATGCATACGGTAGCCATTTGTGTCGATGTCGAGGTAGCCAAGTACGAGGACAAGCAGTCG
TTATTAATATGTCGTGCGAAGGATAGTCGTAGTTGTTGAGACCGCCGAAAAGTAGGAGCCACACAAGTCACGAGC

GCGCTAGGGCCGTCATGGTGATGGTGGCACACTGGAGGAAGCGCCATACATGAAGATGGTGACACGGTCAAGGCA
GACGATGAAGAAGGTCGATGCTGAAGTCGACATTGTCGAACCATCATGGCCAAGGCGTGCGCCAGGTTTGCCAAG
CCCGAGAAAACATCGGTGGTGAAGGCATGCAAAGGTGTTGCCAATAACGGACGTACTAGGTAGAGGGCAGCAACC
ATGTGTCAGTGTGTGAAGGGACTAGCAGAGAAGGCCTCCACGACGTTGAGATGCACAGAAGAACAATGAGGAAGG
TGTTGATGTAGTTCACAGTTATCAGAGTGGATTCTATTTGGTTTAACTATAGCTATCCCATGTACTCAAGTCATG
ACTTGTGGGTTCTATTTGGTTTAATTATATCCTACCCTAACTTGCTTGGGCTAAAAGGCTTTGTTGTTGTCGTTG
TACAATAGTCATATAAAACTCGGCCTGCAGGGGGGTAAGACAACACCCGAGTATTATATTAAGGAGACCTTCTCA
CGCAGGTCAAGAAAACCCCCGAACCCCTGCTCCACCCATACACAGCGGCACCGTAGCCCATGTGAGAAACGACCG
GGACCGAGCCTTATACATGTGCTTTGGCGTGGAACAAACGAGGGGATTTTTAACCACAACCTGAAATTCGCTCA
CATAGGAAGTCGAACCCAGGAATTGAGGAGTGTTACTCAGACCACCTAACCAACTCAGCTAGAGGCCCTTTCGCG
TACATTAGTCATATAACGCAGCTTCTCTACAATACTATTCACAAATTGTACTTCTTTAAAATAGTACTTAGAAGT
ATAGATCTTCTCTAAAATAATATTAAGGATCTTATTTTATTCATCTCTATTTTCTTTTTATATTTCCTAGACATT
TATATCCTTTTCTTCCTTTCTCTTGAAGACCGCTTCCTGGTACCTCCCTCCTAGCCACCGCCGCTACCAACAGTG
CACCGCGCACTCCCCGATGCTTTCCTCGGGTCCGCCCGCTCCCTCTAGATGCTTGCTATCTGCGGCAGTCTTGGG
CCAACTTCTAAGGCTACCATTCGTGAGCTTGGCAGTTGGGCTAGATATGTTCAGCAAACACTATAGTTCTAAGGC
TATCTCCAACAGCCTTCCTACCGCGCCCCTTATCTCACTCCTTATTTTAAACTCCATTATGTAAACAATGTGGTT
TCAGTGTTTTGCACGGCTGTATGCACGCGCGGTTTCAGTGTTTTGCATGGTTGTATGCACGCTCTGCTGAGCACG
GCCTAAGATAAAACACAATAATGAATCAAAAGCATTTTTGTAATACAGCCATTGTCTGAGACGTAGTGTGAGTAA
TTACGAAACAATTACTTAC**AATTTTCTTGCAAGATATGCAGTTGCTGCCTCTAGCTCGTCACCGCCCCTTGCATC**
**AAT**CTGCCGTAAAAACAAAATGCTGAGCTTAAAACACCAGTTTGAAACCAGATCTGAAGTAATTTGCTGCAGCTA
AATAAACAGCACTATAAAAAGACCAAATTCACACATGATTCCATAGCATATAATGCTGGTCACCAACATACATAT
TCATAGAAATATTCATAGAAAATTAACCGCACACCGACTAAATATCTGAACATGCCACAAGTAAAACAGAAAGAT
AGATGTTTTTTTCTTGCACGTTATTGTGCCTCACGTAAATGTACAGTGTAATGTTTCCAACCGAACTAAGAGATC
CCTACAAACAAAATCACAAAATTTATGACAACTAGTATTGCAAGTGACACTAAATCAAACAGTTAATAACTTGAA
AAAGGCCACGACAGCAATCTCTACTAACTATTAAGGGGGCAGTGTAGACTGCCCCCGCTCTCTACTCGCACGCGC
CAGCGAACGGCCTCCCGCACCTCACGCACCGCTCACCCGATGCAACTTCCCACGCCCACGTCACATCCTCATCAC
TCGCCCGCCTGCCTCCGTGTCCTCCGCATCCATGGCCAAGCCACCGCCACGGGACGACGGTGAGCGCCTGCTCGT
AGACGACGAGGAAGGCGAAGGCAAGGTTGGCGGCGACGCCGACGGAGACGACGAGCAGGCGGTCCGGGACGGGGC
AGTTGCGGAGGAGGTCGGGGTTGTCGGGCGCGAACTCGGACTTCGGGGCGTCGTCGGGGAAGTTGCGGCGAACAG
CCCGCACACATACGCACTCCCTTGCAGTTCAACGACGATTCAGGCGTTGTCCCCATCAATGTGGTGCTCGACCCC
GACGCGCCGCTCGACTCAAACGTCGACAGCAGCGGGTCACCGCTCCACCCGTTCCTCCCGAAGGGCGCGAGCTGT
TTCGCCGGGACCCGGATGAAATGGAGCACCCACCGGCGTTGCGCGCGTGCCTCAACTCGCCGCACCAGCTCCCCG
CTCCCGACGACCGACTGCTGAAGCCCGTCATCCACCGCCTCCTCCTGGAGCGCGCCTGCCAACCCGCCCAGGTTT
CGTTCAACACCGACGCAGAGGAGGAGGTTCTCACCACCACGCCCACCAAGTGTTCGAGAATGGGCCTCAACCGCG
GCACCTACTCTGTCCCCAACCGCCCCGTGGTCTGCATCCTGGGTGGGCCCAACTCCGACGACTACGGCTCCAACC
GCTCCATGCTCCTGGACATCCATGTTGTCGGCGGCGGCCTCTCCAACCCCGACCCGGACGACCCCAAGGTCATGG
GTCTCGAGGCCCGCGAACAAATGCGGTACATCGAGAAGCTGAAGCAGGCGCGCTATGACTCGCAACTGTCCTGTC
AGAGGGGAGACCTTGCCGCGACATGGCAGCCAGCACGGTCGCTGGCGTTCACGGTGGGCAACACGCGGCTTGCAG
TCCACGTCGCGTCGCAGTGCCAACAAGCGTCCTCGCGCGACACGCGGGCCATGGCCACCATGGCGAAGAAGAGCG
TCGGGGACCTCACGGAGGACACGATCATCGCGAGGCTCTGCACCTACACCACTTTCCAAGACCTCCTCCGAGTGC
ACGGCCTCGCTGTGCGACTATGCCTTCCCCAGAGCAGCTACGTTGCCACCCAAATCGTGCACACCTGCAACGGCT
ACGAACGCAAGGCAGACAATGATATTGGCCCTGAAATTGAGACGCTCGTGTTCACCTTGCCCTACAGCGCAAATC
GGCAGATGCTTAGTCAATATCCGGTGGCGACATATGTCAACTTTCCCCTTATGCGATGGAACTGCCAACTCGCCG
TGTTGGTCGGAGTGCTCTGGATCGAGGCTTGCTGCTGCACGCCGTGTTGGGAGGGAAGCTGTCGTCTCCTGCCGC
TGGGAGAAGCCGGGGACCCTCTGTTTGCTGAGGCCAAAATCTCCTTCCTTGCTCAGGCACTCGAGGACGATGCAA
TGGGTATCATACAGCGTTGTTTCTCGCCTTTATTCCATTGCAGCAAGGATGTAGTGTCGCGCCCCAATCGTGTCG
CGAGCATCACCGACATCCACATCGAGGAGGACAGCCCTGTGTGCAAGCTTGTTCCTGAGATCATGGGTTTCTCAA
TTTTTGCTGCACGGCTTGGATTACCAGTCAAGGACGAGGATGGGCCAGAGGGGATATTTACCAACAGAAAAGCTG
TCGAAGCCCATCAGTTCACAATATGGCCTTTCGGACAGACTGTATCTGGTCATTTCTTCCCAAATGGTGGCTACC
ATGCCGGGCAGAATATGGTGCTCATCTTGGATGTAGTCCGTTTCAAATATCCTCTGCATTGGATTCCATTGCAAC
TTATTTTGGAAGCCATGAATACAACTGATAACTCAACTGGAGGGTTCATGCTTAGATCAAGAAAGGTTGCAGCCC
CTTCATTGTTGCACACAGTGAGTTGCAGAGATGAAAACTGGAAAAGTGTGTCAAGGTTTTGTGTTGAGGATTTAC
CAAGTCTTCTGAAGGTAGGGATTGGTACTAGCGTGGGCTTGGTTACCTCTGGTTTGAGTGCTGTCAGAAGTCGCC
AAGGCAACTTTTCAATTTTTGTTTGGGATCCTGGTGATTGTACTGTCATATCGCCAGCTTCAAGTGTCGTTCCAG
TGTTGCAAGAGTTGTCGAATTTCAGATGGAGGGCTGCAGAGGAAACGTTATGCATTCAACAATGATGCAGCAGAA
GAAGCTGATCCTACACCATGAGCGACTTGGGGGCAAGCCACATTTTAAGGAGTGGGGAATGTTAGGTATACTATG
GGCTATGGTCATGTATACTAGATGGGCTATAGGTTATGTGGTATTAAGGAGTTATAAATACATGCTCAGACATCT
GTAAAAGGTATCAAAGAAGAAATATCAGAACTCTATCGAGAAGCCTCCGTGCTCCTCTGTCTCTATCTCTCTTCT
TCTCCAAGTCTCTATCCCAGACGATCCCTCTTCCGAAATCTCAATCTACAAGCTCTTCTTCCAGTTCTCTCGATC
CCCTTCCTAGGGGGCTGCTAATAGATGGCGCGGAGCGCGTACTCCACGGGGCCGAGGCGGAAGCGGGCGAGTGCC
GGGCCGAAGCCGATGGAGAACTGGGAGACATGGATGCCCCGGGACGCCGCGGCCAGGAAGTACCTAGACTCGTGG
ACGAGCACGATGGCGGCCAGGACGGAGGCCGTGGAGGCCACGGACTCGACGCTGGCGAGGAGGTCTGGCGCGGCC
GCGGCGGGCTTAGCGAAGTTGCGGTGGCGGCAGGGGAGCGGGAGGTGCACGGGAAGGTGGTGGTTGTGAGAGGTG

GAGTGGAGCGGGGACGGGAAGCGGTACTGCTTGAGGGATGGGGAGAGATGGGTGGCAATCATGGTGGGAGGGACG
GACGAAGAGGAGGAAGAGGCCGCAATGGATAATCTGACATTCTTGAAGGGAAGGAGTTTGAGGTGTGGGTTTGAC
GGTGAGTGAGCGAAGTGGACTCAACGGTTGTGTGGTGTGGACGAGGCGGAGGCGCCGAGCAATAGCTCGGCTAGG
TACTCCAACTAGGGAGATAATTGGGATTGTAACATGCTTGTCCGGTGGCTTCGTCACTCTATTATTCCGCCCCTT
CGCTCTTATAAGCCACTTGGCAGAAAATCTCTACTTCTCTACTAACTATTAAGACGATAGTGTAGACCACAGCCA
TGCCCTCGCCTCTCCTCCACAACCGCACCCGCACCCGCCCCTGTTTCCCTCGCCCGCACAAACCGCCACATCCGC
AAGCTACACATGGGCCATCTGCATGCCGCATCCGCCCCATCCTCTCACTGTCTATCCTCTCCGCACGACCACCAA
CCCAGTCCATCCCCTCCTCGCCCAGTGTCCGCTCCCTCCGCCCCTGCTCCCTCCGCATTCCCCCTCCCCCGCGCA
ATAGCCACGACGGTTGGGGAGGGGAGCAGCTGTCACAACATGTTTTGTTTTCCTGACATCTGTTTGTTCAATGGG
GAGCACATAGATAGAATGACGATTAATATTTCTGCATATTAATATATTTTATCATAACATATTGTCGCCGTAACA
ACGTACGGGCATACAGCTAGTTATTATTAAAGAAGATAAGCCACAAAGGTGGATGTGCAATGCAGTGCTATCTTA
GAACTGCCAAGACTTACCAAAACGACAATCTTATAGACAATTGTTAGATATATGGCAAATTGAAACTAGCATGTG
AAATATACAGTAACACAGTGCGAGCAGAGAAATAGAAAAATTATCCTCTCATATTCTCCCCTTCAGGCCTTGTTC
GGTTGGGGGTGGGGGGGGGGGGGCCTCCATCTACCCCTAACCGAACAGGGCCTCAATGTGTTGTATCACTGTCAA
ATTGCATGATATTATGATAACATCATTACATTGATTTTTTTATATACTAAGGGTGGTCTACTGTGCAACCATTTC
CAAAATGGTCATATCAGTCAAAACGACCATACAAACTACCTAAAAACACATATGACAGCATGGCATGAAGAGAAA
TAGAAAACAATTCCACGTGCACCATCAAACTTGTGGTAGTCTGATTTTGAACCTTGAACTATTAAAACAGGGTGA
CACACATACTCCAACTGTCGAAACCAGACTGATTTGGTCGAAGTGGATTTCTATTTCTTAATAATAATGAAAATC
TGGCTTGGTCTCTAGAAGTTCCATATGTAATTTATTTAAATCTTAAACGAAAATATGATACTAGTTCTAGTTTA
GGGTTTAGGGTTGATTTGGTTAGGATATTCCTTAGAAGTCAAATCAGTCATCTCTATAAAAGAGAGAAGCCAGTG
TCGATCGTAAGCAAGCAAGAAAAAAGAACCTTACTAGGATGAGGGCTCTCCTTCTGCCAATCCATATTAATTGGT
ATCAAAGCCCACTACCATGACATCGCCATCCATCACCGGTCTTGACACCGATCCTACGACCATCTCTAATAAGCT
CGACCTCATCTTGGCCCAGATCGTCACCCTCAACAAGCACCTTGACGCTCATGAACGACATATCGCAGGCGTGGA
ACATGTGCTCGGCGTCGTGTACGGCCTACTAGTATCTCCGACCCGACACCCCTACCCCAACAATTGTGACACCAC
CAGCAAAAGGCCATAGATGCAACAACTTCGGCAATCTCATCACCAACTTCGCCAACACTAGCATCACCAACACAG
GCGATCGCCCCTACACCAACATTGGAGGTCATGGTGCCTTTGTCATTGCCTCTACCAACCACATCACTGATCCTT
TTCATTTACCAACGGTGCAACTATCACGAGCAGGGTAGTGGCTATTTTTGCACGCGCGGATTCCCTACCGACTGA
GTGGCATGCCTTCAAAGCTCCAACGTCTGCTTGGAGTATGCCTCTAGCACAACCCCACTCTTTTGCACCCATAGA
GGGGTGGCTGCTCATGGAGGGCACTATCAACTCCATGTGCAGGCGCTCCTGAACGGATTAGATTTCGAGCAGATA
AGACGCAACAACCAATTCTCGGACCTCGAGGAGGAAGGAATGGATCTTGCACCATTGCAGAGGCAGATCCTATGG
ACCGAAGTCGATTTTGGGGAGCCTTATATCTTCCACGTCGACTACAACTACAACATAGGTCTCCTCCGGCGGGGC
TGCCATGGTGATCTTCTCTAGGCCACGTGTGCCCATCTATTCGAGCTCTTTACTTCTGATGGCACCCACTTGATC
AGGGATTTCGGTCCTTTCTTTCATGGTGGCTCGTCCGTGCGGCAGCGTCAGGGAAGCCACCACACCACCAGTTGT
GTGCATGCTTCATGAGCCACCTGACCCTACCTCGCTGCTACTTGCTATCTACCCCACATGGAGAGATATCCCAAC
AAACTCTACATGAGTGGACTACCAGGTCTTCCACTCCAGCAACCGCCACAGCTTTAGAACAAACTATCTTGGAGG
GGTGGAGTAGTGTTATGGCAAGTGGAGTCACACAAGGGCCCACACAACACTGTGGAGTGTGAACATTGTCAGATC
AGAGAGTCCAATTATGAGCAATATGTTGAATGTGTTCTCTAGCTCGATCAAGATCACCAAGTCTAGGCATATCCT
GACTAATTGCGCCCCGATCAAAGTACAACCCTATCCATGGTTGTTGCCTTCGGTCATAGACGCTATATCAATCAA
GGCAAGGGTCAGCACGGAAGCCAATGCTTCTTGCTCCGGTAACCCTAGACACCAAAATTCCTAAGAGGATCTGAT
CTCCTTTATCGTTTGAGTGGCCGAAAACGCCCAACTCTCCATGACGTCGGTAGACCCTAATGGTTCTACCCTACC
GGCCACCAACATGCCACCCCAAGTCCCTGATCCACAAGGAGGAGGAGAAACAAGTCTAGGCGATGGCGCTCCACA
CACGGACTCGACGTTGACGCCGACCGCCACAATCCACCCGGGCCCGAAGGTATAGACTGTATACTCCCCCACCTT
CTCTTCCACCTCACCATTGCACTAGTTCACGAGTCAATCAATCTATCGCTAAGAACCGGACGAGTCTAAGAGTGA
ATTCTGCTAGATCCTAAAGTCTATGACTCTATCACAATAAACCTAGTATGATACCTATATGTACATTGTAAAAAA
CTCGTCGGGGAGGGAAAGACAGCCCTCCTGGTATTATAATAAGAAGAGACCGAAACATAGTCTTGACCAAGAAAA
TCCCTGAACCATGTCCCCCAATCACTAGCGGGTCGTCACCGCGCACTCTGCAACGGCCCAGCCGGAGGGTACCGC
GCAGAACGTAACCCAGGAGCGGGCGGGGCATAGCGAGGGGATTTTTTAACCAAGCAAAAAATTCGCCCCCAAGGG
GGATCGAACCCGGGACCTGGAGGTGCTACTCGAAAGCCTTAACCATTACGCTAGAGGTCGTTTCTCCTATATGTA
CATTATAAACAAGTAAATAAACCAATTGGTACTAATTTCATGTTTTCTGATTTAAAACAAACTATTTTCAACCGT
AGATGCAATCATGGTTTGTTAAGGTTTTTTAATAAAGCTACACGAGGTGGTGGTGGAATTGAAGGGCTTAAGTGC
TAAATATGACATACGGTTGGATCTAGATCTAACGGGCCAATAAACTTGTTTGCACACTTGCACCTAATAGTTGCT
TGCACATTACCCGTTGCCTTTAAATAGTATATCAAATTTCATTTTTTTGTAAAAAAATACAAAACCACCTTTAAAA
CCAGCACAAAGACCAGATTTATCCTATTTCGATAATTGGAGAGTGTCACCCTTAATTATGTTTATCATCAAGTCA
AATGTATCATGCTTTGAGAGAACAACAGTGATTTAACATGCCTGCCCCCACCCTCCTCTCCAATATTCGAACTTC
CAAGACACACATATCATATACATTTAGAACATAAAAAAACAAATTTTTGCATCATGCACTATATAAACACTTTTT
CCCTTTCGAAGTTACATGCTGGATCTCTGATGTCATGGCACTATATAGAATCAAATTCTCATGTACCCATATTTA
TGATTATATAATTAAGAGAAATATTCAAATATATGAAATGTACTAGCGTTTATTCTTTGTATTAATGCTACATAA
ATAGCTAGCCTGTACCAACAAAAAATTCTGTGTGTTCTACATGACTACATGAGGTACACATAGTCAACAGGTACA
GCTAATTCTTTAACAAGTCCTATCAGAAGTAACGTTGGTTGAAAAATCAATATACCATCTTTGTTATTAGGAAGA
GCAATGAAGGTATACCCAGCAAGCATAATATAGCTGGACTCAAAAAACGTGGATCACGAATACATACAATACCAT
GCAATACACAACAACTTATTTGCCACATTCGCCTCAACGAAAGGGTCGCCTGCAAACACTACCAGGAATAATGCC
ACATCCACGGTGGCCGATGCTGACGATATAAACTCTCACCCGATTTAACACATCCCTAAGAATAATGCTATGCGG
CAGCGTGTGATATGCCCTAAAAGAGGGTTTCCATAAATCAACGAAAAAAATGAGAAAACGTTTCTAGACCCACCA

CATGTATATATATATCACCCATCTATCTAAAAGGCAAAAAGAAAAGTTGGTTTCCCGCACGCTCCTAATTCCTAAAT
CGTGAAAGCACATTACCATAAATCTGGTGCTGTGAAACTGAAATCCCGGTCTAAACATCAAGAAGGCAATGCAAG
TGACAATGGCACCAAATATAGCCCACCAACCAAAAAAAAACTAACTCGCGGAAAGGGACGGTGGGAGCTCTCAC**C**
**TGAGACGACGGCGCGCCGCTGCCGGTGATTGTGATGAAGGGGGGCTAGATACAAGAAGAAGGGAAGTGCGAGGAG**
**AAGGCACAAGGCAGTGCCGAGCAGGGATGCGGGTGAGCTAGAGAAGTTGAGGCAGGGTGCAGCGATTGCCGA**TTG
GCGGGGATCGCGCGGATCTGGCTCTATATAGACGCCGCGGCGCTGGGCCGCCGGGCGAGAGGGAGCTTGGATGCT
TGGCAGGACGCAGAGCCATGCGAGGCCAACAGGTGGCCTGGCCGTCGAAGCGAAGGAAAGGCAACGGGACACACG
TCGCCCGCGGGGGGCTCACCACGGCGGAGGGAGACAAGCGAAATCTGAGCTGGATTCCCATTCTCCCGGCTCGGC
TCGATGTGGTGTTGAACTGTTGGTGCGCCCGGGAGGTTGCGGGGCGGGCCTTCCCTATGTAGAGAGCAGGGACTG
CGGAAAGGCGTCGGCCTGGGGAGCCAAAAATATCTAGGGCTTTCCGGGACTAAAAAAACGCGGCGAGGAGTGTTC
CAGCTTGGGCCACGTCGCCCCCCTGCCCCCACGTCCGACAGCGGGCGGCGGCGGCAGCAGCAGCAGTGCAGCACG
CATCCGATCCGCCCACGGCCGGCTTGCCTACCGCCGCCGCACCTGCACGCCGTCGTGGCGCACACGCGATGCAGA
AGCCGGAAATCCGGATTTTCCTGTACGGCTTCTGTGTGCGTGGAGCGTGGTGTTCCCCGGTCCGGGCGGCGGGCG
CTATAGCGACGGGCGGCCCGAAGCCCCGAACACGAACACAAAGGCGATGATGGCACGGGAGAGGATGAGCGACGG
GGACGTGCGCTCGTCCCCGTCGGATTGAGGACGATGGCGAGGCGGATCTCTCAAGGCGCCGACAGGTGTCCCTCA

>ZmMADS27
ATGGGGAGGGGAAAGATCGTGATCCGCAGGATCGATAACTCCACGAGCCGGCAGGTGACCTTCTCCAAGCGCCGG
AACGGGATCTTCAAGAAGGCCAAGGAGCTCGCCATCCTCTGCGATGCGGAGGTCGGCCTCGTCATCTTCTCCAGC
ACCGGCCGCCTCTACGAGTACTCTAGCACCAGCATGAAATCAGTTATAGATCGGTACGGCAAGGCCAAGGAAGAG
CAGCAAGTCGTCGCAAATCCCAACTCGGAGCTTAAGTTTTGGCAAAGGGAGGCAGCAAGCTTGAGACAACAACTG
CACAACTTGCAAGAAAATTATCGGCAGTTGACGGGAGATGATCTTTCTGGGCTGAATGTCAAAGAACTGCAGTCC
CTGGAGAATCAATTGGAAACAAGCCTGCGTGGTGTCCGCGCAAAGAAGGACCATCTCTTGATAGATGAGATTCAC
GATTTGAATCGAAAGGCAAGTTTATTTCACCAAGAAAATACAGACTTGTACAATAAGATCAACCTGATTCGCCAA
GAAAATGATGAGTTACATAAAAAGATATATGAGACTGAAGGACCAAGTGGAGTTAATCGGGAGTCACCGACTCCA
TTCAACTTTGCAGTAGTAGAAACCAGAGATGTTCCTGTGCAACTTGAACTCAGCACACTGCCACAGCAAAATAAC
ATTGAGCCATCTACTGCTCCTAAGCTAGGATTGCAATTAATTCCATGA

>zma_miR444b_gi_195596206
TCGGCAATCGCTGCACCCTGCCTCAACTTCTCTAGCTCACCCGCATCCCTGCTCGGCACTGCCTTGTGCCTTCTC
CTCGCACTTCCCTTCTTCTTGTATCTAGCCCCCCTTCATCACAATCACCGGCAGCGGCGCGCCGTCGTCTCAGAT
TGATGCAAGGGGCGGTGACGAGCTAGAGGCAGCAACTGCATATCTTGCAAGAAAATTGGTTTTCGAACCATACCG
ATAATTTTCTTGCAAGTTGTGCAGTTGTTGTCTCAAGCTTGCTGCCTCCCTTTGCCAAAACTATGAAGAAACAGG
TTATGTTATCAATGAACACATTGGCACAGATTTTAACTGGTATATGTTGGTAAGACATGACGTTTAGTTGGAATC
TTGCAGCATGATACACAATTACACACAGCAGCTCTTAGGTACTTCATTCATACCAAATTATAAGATGTTTTAGGT
ATTATGAGGTGCTTTGGGTTTTCTAGATACATTGATTTTACTACGCATCTAGACATGGTATATATCTAAGTGCAT
GGCAAAAATTAATGTCTAGAAAAGCCAAAACATCTTATAAATTGGAATGGAAAGAGTACAAAACTAAGAGGGAAC
AATATTTATTACTAGAACTAGAAATTGCAATCTGACATGTCCAATTTTGCAACGTGGAGGCTATTTTAATGCGCC
TTCGTTTACTTTCTGGATAAACAAGCAGTACGGTACGATATTACAGCGGGTACTGTTGTGTTAGGATGTTGCAAG
CAGACTTCATGAAGTCATAACAAGGCGAGTGCTAATTCTTACAAGAACATGTCAGTATAAGTAAATGATCTCCTG
ATTTTATGATTTGAAGGGTGCCGAGTGCCTTTTAGTCAATTAGTCAATGCTGGTTCTTCTACCGTTACGTGTTCA
GAGTGTTCAGTATCTATTCTACGACAACCTTTGATGTTAGTAGAGCTTGGACCTGTTCCAACTTAATTTCACTGC
TACGTATATTTTAGAGCACAATCAACCGGCTGGATTGGCATACACAGTTTTATTCTACGTTCTTTTTTTTTTTGC
AAGGTTGTAGAAGACGACTGGTATATTTGAGGGCAGCTTCTGGCATGTAAACTATACGGCCTGCGGCCTGGTACT
TCGGAGTTCATGGGATGTTCTGCAACAGCACTAGAATTCTGACAATCCAGTGTCATGTGACACAACTGAAGTGGC
AAAAAGAATTCCTAACATTTACTGGATAAAGCATCTGCTGACATATTGTTTTCGTGATGCAATATGAGTTCCAGA
TAACATACAGCAGTAGAATTCAAGTTCGTA

Stem-loop structure

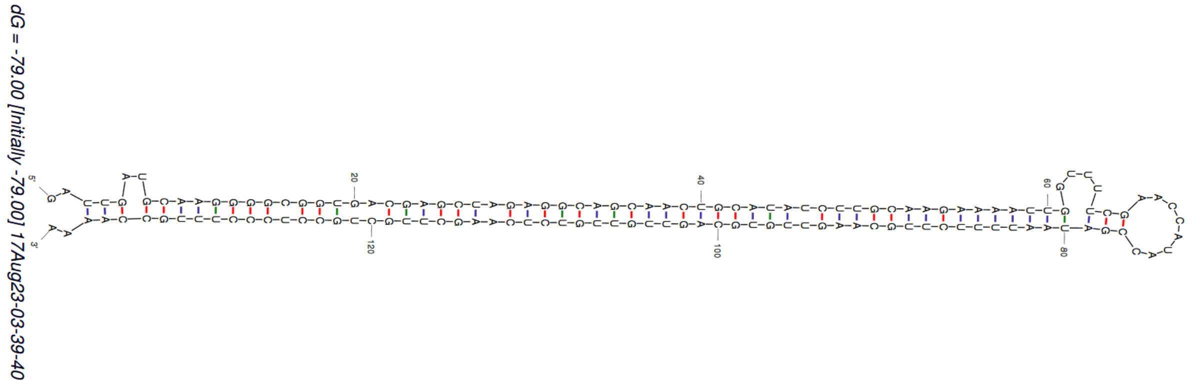

*ZmMADS61 and zma-miR444c*

>2

tccctacaaccagattctcagaaaagccggtaaaaaaaaactaaactaaataggcgcataggaaATACTTGATAG
CTAAGGTTCTAATTAATATGTTAGCTAGTAGTTTTAACAACTTCGTTAGCTGATTATCATATTCTCTCAAGCTAA
CATTCTAATAATAGCTCAGTTGAAAATCTTTACGAGCCCGGCAAATCATATCATAGCTGAGTAGAACACCACGTA
CTTCCTCCACTACCGTCGCGAATTGCGATCCAATTAACAACAAATCACTTCATCTCCCATCCGACGGTATCAGTT
TCACCAGTTCACCACTGACCCCAGTACAGGCAAGTGGTCAAGTGGACAAGAGAGCAGGCCGCACAGTCGCGCACG
CCGGCTCACATGCTCGGTTCCGGCGACACCCCGCAAAAGGCAGCCATGCCGCTTCGCCTCGGCTGTCCGCCTGCC
CCCGATCGGCGGAGGCGAAGGTACACCTGTCGGGGTGCGCGCAGCACAGCGGCAGCGCCCCCTTTGGAAATAGCG
CCGGGGCGACGTGGCCGGGCTGCGCACCGCTCCTCCGCACCATTTTTTACTC**CTCTTCTTCTCCCCCAAGGCCCC**
**AACCATCCCACTCTCTCTATACAACAGCCACCCCCGCGCCCTTCCGCCCTTGTCGAGTCCAAATCCAGGTTATAG**
**ATCCTCTCGCGCCGCCGCTCGCAGCCGCTCGCCGCCCGGTACATCTCCCCCGCGCCTCTCCCTGCTCCTGGATTT**
**CACCGATCTGGCATGCTTTACTGTCCCGTCATACGGTGGCAGCGGGAGCCGCCCCCTTCCTGCCTCCACAGCTCA**
**TGCCATGCATGGCCGATCCATTCCCTCAGATCGGGATTGCCGCATACATGCCGTTGCTTCCCCGTTAGCTTCTCC**
**CTTGTTACACCCCCTCGCTTCGTTCTGCGTAGTTTCGGGGTTTGACACCGCTTGTCTATCCCGTTCCTATAGCTT**
**TTAGGGTTTCGAGCTCAAACTCCGCGCCGGCGTCTCAG**GTGAGACATCTGTTTTGATCTTCTAATTTGCCTGCTA
TCATTCACCGTTAAGTGTTTGGTAGTATCGAGTAAGCTGTGGTCGTGTACAGTTGTGCCTAGGTTTTTGTCACCC
GTGTTTGAAGTTAATCCAGTTGTCTCACAGCATAAAATTACGAGGCTATTGCTTTATCCTGTCATGTGTGGATCT
TAGGTTGGAGGCTCGAAGTGAGCTTCAATAAGATCTATTTGTTGTTTTGTTAGTTTTCAGGAGTTATTGCCGGTC
AGTGTGCGTGAACATTGAACGTTTTGGGCTGCGATCTGATGTTAGGAGCTTGAGGTGTTATTTTGTTCTTGCATT
GATTGTGGATCTTTTCAAAACCAACATGGTAGATGATCTCTCTGTGTTTAATTTATGTAAATGCATTGTCTTTAC
GGTTTGCTTTCCTTTTGCAAGTGGTGGGTTATGTAACTTTGTAAGGCGACCAATTCTAAGGGTGAGGGCCTTGAT
AGTTGGTTTGGGTTACACCGGGCCTCAGTGTGGATAGTACTTGTACAAACAACGATGCGACATGGTGTTTGTTGC
TATATGCCACTAAAATTCAATTATCCCTTCTCTCCACAGCAATGTGTCAGTATGCAAATATGTTCTGTATTTGTT
TATGTGCGCATCTTGCATTTTACCTGTTTTTACAACAATCTGAACGGTCTGTGGTGGTACATGCCGCTGAAATTT
AACGGTTCCATCTCTATGCGGTGCATGCATGATTCAACACCGAATAAGTGCTATTGAAAGCTATACCTTACTGGA
CGGGTTACTCAGTGTCAATCAGAATATTGAGATGTCCTCATTGCCCCATAATTGAAAAAGATCAGATCCAATGCT
AGAGGCTCCCACATGAGTTGGAGGTATAGGGAAGGGACAAACTTATTTACTCGGTAAACTCATTTACTGCTCCGA
TGTTGCCCACTTGTTCTTATACCACATTTCTAAATTGCACGGGTGGGCTACCAATTCATATTTTGATGGATGACT
ACTGAGATGCCCGTGCATCGCTACAGTTCTATATATCATATGAGTAGACTTTGTTTTAATAAAACATAAAAATAT
GTGTTCTGTTGGTTAGGTGGTTCTAAATGTGGACTTGTGGAGCCAACAACAAGGTTCGAACCCCAATTTTACCTT
ATTTTTGCATAAAGAGAGAAGGGGAGACAACTGTAGAATCGTAGAACCATGGGAACTGGTGCTTTATATTTTGAT
GGATGAAGTGTTTCATATGTTGTCGTAAGTGGAACTGTTTTTTGGCTATTGAAGGCTCAAGTAAATATTGTCATA
TTACTTCATCCGTTCCAAATTGTAAGTCGTTCTTGCTTTTGTAGATACATAGCTTTTACTATGTATCTAGAAAAG
CCATAACAACTTACAATTTGGAGTGGAGGGAGTATATAAAAATTGATCGTATTTGTTCTACAAAAGAACTTAGTC
CTTTCTTTGCCATCTACTAACACATGTCACTAAAAATAGTAGGCAGTACACCAGTAGCAGAGAGAGTTATATCTT
ATCACACATTTCATTTTGTAATTGGCAATGATGCCACGTAAAAACACCAAATTTCGACCCACCCACCCATCCAAT
GTAAAGGTCTGTTTCTGCAGTGAGTTTGGACTAACCTGAATTGCTGATGACCACCTGTGCACAGCTGGATCCTTT
TTCCTTTTTAGGTAAAGATCTGCCTACGCTGTCAGTTTGGCCTTCCCTGGATTGCCgatggcactcagcaaacag
ctggatcctcttttctgatggccatcaacaaagatctggatcctttttctCCCTTCCTTGCAGTAGATATAAAT
TGATCCTTACGTAACATTCCTGTGATGTAGTGATAAATGACATGACCAAGTATATGTCATGACCTGTCAGAgccc
aaggggggctgtggccaggacctgtcagggcccaaaagggctacggcaagacgcaacaaggagcacaggtcaaag
gataagaagatcagttgagactatataggagattagttgagattatctaggaaggttatcagttagtagtttgtt
gagagttttaaggagataaggatctctagctagataggggcggccgaccggtctacttatgtaatcaatggatga
gaaataaagtaggcaagaattagaagggagaaaccctcctcttgctcggtcgtgggtagaggcccccggccggcg
ttcctgcccgttgacccccctcttcaatcccctttcattggcgggtactgtttaccgcgtcgcctccaccagcag
ctccaatccctcaccgatctagcaactataacatctggtatcggctagcttaggtttcgatttgatcctatggct
tctgctaccaatttcatggtgtcgtcctggcaaccgacgtccccatctccatgggttgcgctgcatgccaccttc
gtcacgtcgtcctggcaaccgacgtccccatctctatgggttgcgccgcatgccaccttcgtcacgccgtcctgg
caaccgacgttcccatctccatgggttgcatcgcatgccaccttcgtcacgtcatcctggcaatcgacgtcacca
tctccatggacaacaccgccttccaattttgtcatcacatcctggacgctggcgtgcccatctccagggggcagcg
ccaaatatcgtcacatcatcttcgttgacagcgtcttcgtccccatgggcgtcggggctgccatggctcaacaat
caccagctccatcaatggggggacttgtggccaacataagggctgatctcgcaaagatgcaatcctcgctcagat
ggttgagcaggggctgctgcaaatcaaggcaacaatgcaacatgagtagcagatggtgttgtccacacggctcta
gacgtcggcggctgtgggcctataggttgttgcttgtggtttccttgcatgacggagagtgcgggagatgctggc
agatgctcgaggcagccttggtgacagttgacctcggcacacgagggcgcgacctcaccctgtcggacgaccatc
agtagtgacgatgggccactatctccaagtgcgagcatggtgcttgtcccgcggacgacaagctgatgaacttca
actctacggcaacggcggtagggaaggtgctcccctcatcattggcaattgcactgcttagcgccaccacattcc
gctaccggtcgccacgagggcacttctgctgatcattgttgtgacctactctagatggccatccatgtgctcccc
tttcgtccagatggcgtccatgagatccaggtggctgcacacgtgcaagtccaatgcacggagggtgtccgcctt

atcttatggggtcaaacaaaattcaaatttgaaaaaaataagctgagatgtaaaaggcttgtcttctaggtgttt
ggttttaggtcaagtagagtcggtctttgggcactcctcatgctaaagctcgaggacgagctgtttgtccaggag
gggtgtagtgttaggacctgtcagggcccaaggggggctgtggccaggacctgtcagggcccaaaagagctacga
caagaagcagcagcaaggagcgcatgtcgaaggataagaagattagttgagagtatctaggagattagttgagat
tatctagtaaggttatcagttagtagtttgttgagagttttaaggagataaggatctctagctagaaaggggcgg
ccagccggcctatttatgtaatcaatggatgagaaataaagtagacaagaattagaagggagaaaccctcctctt
gctcgccgtgggcagaggcccccggctggcgttcctgcccgtcgatccccctctccaatcccctgtatgcatcta
cattgttcattggcaggtactgttcaccacgtcgcctccacccgcagctacaatccctcaccgatctagcaacta
taacaGTATACATAACATTAATTTTTTGTAATGTTATGTCTATAAAGTTCTCCCATACCTGGTATGTTGTAGTCA
TGATAGGGCTTTGTTTGTGCCGTGTTAACATTCGTGGTGGCTCTGAATCATGATTGTTCATTGTATCTTTGAGGC
ATAGCTGAGTTAGCTTCCAAATTGCACTTAGTGGGTTTctgaataaaaaaactcgactcgttgggagtaagattg
ctcccgtggtattatattaagaagaccttctcaagcaggtcgagaaaacccctaaacccATGTCTCACCTataca
cagtggcactgtagcccatatgagaacgatcgcagccgggaccaggccttagacctgtgctTTGGTGTGGGACAA
CCGAGAGGATTTTTTAACCCCAGcctgaacttcactcccacggggagtcgaacttaggacctgaggagtagtgct
acttaggccagctaaccaacttagctagatgtccttttgctAGTGGGTTTCCGAATAACCGTGAAATACTCTTAC
TGTCATAGAAGTTCTATGAATTCAGTTCTAAACAAAGGACCCTTGACGTTTCTTTCTATTATACTTGGTCTGTAG
TTAGTTTTTTTATTTAATGGAATTGAGAGTTTATGTTGACTTTGATGGGTTATATGTTTTCTGTTGTTTGTACAA
TGCACATCAGTTCACTTGTTTTAGTTGCTGTTAATTAACACCAGAGTACTTCTGAAACCTAGCAATATTCTAGGA
CTAAGTTCCTCTGTAATTAGGGACTACTGTCTTATCATGTTCAGTTTCAGGTTTCTGCAGCACCGTGGAATGCAA
CTAGTTTGTATTGCTTGTATGGTCTCGTAAATAACTGGTTTATATAAAAAAGTTTTGGTTGAATAGCAACTTTTA
GAAAACCTAAATAGATGAAGGGAAAACGACACTAATTTTAGTGCAAATTCTATGTGTGCTAAGTAAAAGATTGAT
ATCTCGTACTTTTAACTGCTCTGTTGGGTCATATTGCAATTTATTGCTGCTAATGGTACTAGGAGCTAATGCTAT
TAAATGGTCGGTTGGAATCCTTTAATTTGACGTTTTTGCCTCCCCTTTTCTGACAG**TTGCTGCAAGTGGCGGCAG**
**CAAGCTAGAGGCAGCAACTGCATACCTTGCAAGAAAAAATC**GTAAGTTACTTGATTGCTGTTCCTACCTGAAATG
TCTTCTGTTGTTTCGTATATGTATATGAATCGCTTTAGTCTTTACAAGTCCATCTTAGCTGAACTCAGGGTACCA
TGTTTTTCTCAAAGACCTCTGTTTACACATTTTTTGATCTGTACATGATTTGTGGATGTGCACTGAATCCATGAT
ATAGTAAAATAACTGAAGTCCCAAGTGTCTATTTAAGCCATTGATTTCTCTTTGTTGAGAGCTGACTTTTCTACA
TTCATGAATGCTATAATCATGTGGATTCAGTAGACAACGAGTTACTTGATTAGGATCCACTTGATTGTGATTGAG
CTGTCTAGTTTATAGATGGTTGGGCAGGATATGGAGTTCTTTTTATTGACCAGTGACAGCACATGCTTCCTCCAA
TTCATTTCTCTCTCCAGTGTGGTGTTTTGGACCAAGGTTGCAAAAGGTGCCTAGAGCCTAGTTGAGCTTAGGTAT
TTCCAAGGCGTTCTCTAGGTGGTGGCTAATACATGCATATATACTTAAAAGAATGAAAAAAAGGAGATCAGAGTG
GTCAATATCGGGGAGAAAgagaaggcccatttaacgtcacaagtctcctagcgtaacagcttatcctctcccctc
ccaaactcctttgttgatgcttgtcactcgcccacgcccttgacatcgcctgcttccccaccggccattgccgag
aaggcccatttaacgtcacaagtctcctagcataacatcttatcctctcccctcccaaacttctttgttgatgct
tgtcactcgcccatgcccttgacatcgcctgcttccccaccgtccattgccgCCCCCATCGATGTTACCTCCTCC
CCCACTGCCACATGACGTCTGTGCCGCCCCCATGCGTCCTTGCCTGGGCAGATGTGCCTGCCGCCAACGCCTAGT
GCAGCCTCTTCTCCCACTGGCCACCGCCGCTGCACGTTAGCATCCCTGCCTTGGCAGATCTGGCCACCTTTCACC
GGCAACAGTTGGGCGCTGCGTAGGTACCCACCTACCCCCTAGGCGAGGTGGTATGCCTCACCCAGCACATAAGTG
CACTTAGGCAAGCGCAGAATAGCGCCCTTTCGGACCTGTTTTGGTCATTGACATGGTCTTATGAACACTACTCTA
CCATGATTCCTTGAATGTGCGCCTTAATATATTTTAAATGCAATACAATTTTAGCAGAAAACTACGTACTGAGAC
GAATCTATACATATCATTTGCATGTCCAATACAAACATTATAGTCAAAGTTTTCGCCTGCGTGTTTGAGAGGTAT
TGCTTCATGCCTTCATGTCCAGGATAAAAATAGTTTGCTCCATACTCGTCTAATTATTGAATTCCAGTGTTTTCA
AGATCATAGGTTTAGCTTTCCAAGCTGTAGCAGTATACCAAATATTTCTTGGCTTCCTGTATAATCTTAAGCTAG
TGGAATTGTGTTCTCATATGGCAATTCGTTGTTTCTTTTTTATTTTCTCTGCATTTTGCTAAACTGTTATTATAC
TTCCATAAATGACGAAATGATTAGTGAACGACTATATGTTTATCTAGAATCCCTTTTATGGCACTTCTTTGTTCT
GGAATAAATGCGTAGATAAAAATGGGAAAAGATATCTGCTCTTCTGCCATACGTAACTTTCAAGACAAGCAGAAG
TTCAATTTTCCAGATACACACTTTATAGAATTTTTTCAATAAGTTCACTTGGAAATTTGTACATGTTACATACCA
TTTTGGTCAAACAAACTTGCATAACGTGTAAAAAACAAACCATTTGCTGGATTATTCAATATAACACAAGTTATT
TGAGATAGATTTGCACTACCATTGCCACTGTCTCTGGCATGCATACTTTATTTGTCTTGCTATGTTTTTAGGCCT
GTTACAACCAATCATTTTAGGCTGCTTTGAGTTGCAGACTACAAGGACAATGAGAACAAAGGCAATGTCCTGCAT
GTCTTCATTGATTTAATTGCAACCTGCAAGTTTTTAAATAATCTTAAATGAAATGCCACTAAAGAACTAAGAACA
GAGTCAATAGGTCCTATGCAGAAAAGGCAGGGTTACCCTAATTTAGGAGCCACAGGTTGCTCAACACCATTTTCT
TGCAGTGGAATATTAAGCTCAAGATGGATAGGAACGTTGGCACCCTCTGCAACTACAAAGTTGTACGGAGTTGTT
GAATCTCGGTTGACTTCACCTTCTGCTTCTTTCTCATAGAGCTGGGCATAGAAACAAATCATAGTCAACCATGTG
ATAGGGCTTATATCCTGAAAAATTGTATTTCTTTATAAAAAAACCTTCTTATATAATTCAGCATTTTCTTGACGA
ATTAGGTTGACCTTTCTGTGTAATCCCATGTTGTCTTGTTGGATGAGACTTCCCTACAAATGAGCCAGTACAGAA
TGAGCTTGCTGAAGTCCACTCAAAGACGTCATATTTCATAGAAATAGTGGAATGTTCCAGAACCTTTCGATTCAG
TTCGTGAATTTCGTCAACCAAGAGTTGGTCCTGAAATTGATAACCATGATCCTCAGTTTTACTTTTGGACAAAGA
AATCTTGTGAAGGGTAAAAAATGGTTAGCATGGTTACCTTTTTTGTCCGGATGCAACAAATGCTCATCTCTAGCT
GATTTTCTAGATTTTGAAGTCCCTTGACACCTAGTCCAGAAAGATCTTGTCCCATCAACTGCCTACAGCCAGTCA
AACAATCTCACTTAGTTTTTACAATTCGAGTTTGACTGTATTTTAATGTAATAGGGTCTTGTTGCTTGATCATAT
CGTGTTCCTTTTTAAGGATTCGGCTTCTGGCTATGTATGCGTTAGGGCACCTGGAGTGGTAAAGGCTCCCAATGG
CCGTAAAGTGAGCTCAGTGAGGCCTGTGTCAGATGAGGTTATTATCTGTTATGTTTGAGGCCAAGAGCTAACCCA

CATGGAAATATAAAAGTCACAGGCAACCTCTTCCTCTCTGCTGCTTTGTCTTCCTCGTCGCCAAGTGTCCTGCTC
ACAGCGCCACCTCCTTCCTCCATGCAAGGTCCTCGCCGCTAGCTGCAGTCTGTAGGAGGCCCTAGAGGCCCTTTC
CACTCCCTGCAGCGTGAGGCAGCTGCCTCCTCTCCTGTGGGAGGCCACAGTTGCCATTGTCTGCCACACACCTGT
GGAAGAGGAGTTTCTTGCCTTCGCTTGTGttttttttAATTTGCTGAGGCCAGGTATTAAGCTAATGGTCTGGTT
CTAAGTTTTTTGGGTTAAATACCGCTTTATGACCTTATCGGCTTTAAGGTCTTAGGACACCTTCATTTCCTACAC
TGCAGATGCCTTTAGTACTGTTACTTGGAATAGAGTGATTACAAATCAGAGCGCTCAAAAACTAGTAGGCAAAAA
ATTAAATACATTTAAATATGATGGTGTGTATCTGCTGCTTTGGGTTAATCATACAAAATTACAATAATGTTGTGT
ATCTGCTTCTTTGGACGTTAATAATAATGCATAAGATAACTAAATGTACAGTTTTCTATTGATAAATGGTTTGGC
ACATGAAGTTCACAAGCCTTTGCATCCTTTTACATTTCTCTTGCTTATGAAAACTGATGTCTGCATAAACCATGT
ATTATCCTATGGCATCATTACCACGTCCCTAAACTAGGTTCTGTTCTTATGCATTTTGTATGCTAAAATTATTCT
CACATATTGCTATACTAATTCATACACTATGCACAAGTTCGATGTTGGAGGTGTTCTAGTTACCTCGTGGTCCTT
ACTCCTTAAGGTTGATCCAGCAGCGCAGATTTGTTGTTGTTTTAAACCACTTTATATATTTTTCTTTAAAAAGCA
GAATGGATTTTCTTGTTTAAAAAATCCCTTGTGTTTCTGTACTCAGACGTTTTGTATTTGTGCTCATGCTGACAA
TTCTTTTATTGATCTTCCTTACATTTGGCAATTTTCTGAATGATGCATCTGTAAAACTTGCAG**GGTTTTGTTGAC**
**GTACCGATGATTTTCTTGCAAGTTGTGCAGTTGTTGTCTCAAGCTTGCTGCCTCCCTTTGCCAGAACTATTGAAT**
**GTGAAAGCACA**GTTAAGTAATGCCACGCCCAGAGAATCCAAAAGGTTGCTGATATAAAAAGTTGCTAATAAGTGG
TGGAACTTTTAGCGGAGTAAATTGTATGCCTTTATGAAATAACTGAAGTAGATATGTTGGCATGCTATTGTTTGA
ATCAAGAAATGATATTATAtactccctccgtcccaaaatattagtcgttttagcccttgaaattatgtctatatt
taaatgaatgacaatgaatctatatataatataaaacacatacattaagtattgtatgaatctagtaaaatgcta
aaacgactaatatttgggacggagggagtaCAAAATAACAAGACGAGAAATGACTGTATGAACTCAATTAAGGC
AAATACAAAACCTCTTCATCTTGTGTTGATATTTTATTTCAGACACATACTTAAATTAAGAAACAATTTTATCTA
ATGATATGAAGCTGGACGTGTATTGGCATGCTTTGATTATGTAATTTATTGCTGTTATCTACACTTCTCACATAC
AAGTGCTGACATCTCAATGGAGCTAAATAATCCCAG**AGCATGTGAAATCCTAATGACGCATGCGTCTTGAGGGAC**
**GAAACAGGACAAGTCAGAGATTGGTGATGAACTTGCTGGAACAAGCTTATGCATGAAACTGCTGATGGTTTTAAC**
**CTTTTTAGGATTTAGCTAAAGGCTATAATAATCTTCAAAGGTTCAATATAAATTCT**TGGACTATAGACTGAAGTA
TGGCTGCTGATCTTCCAAGATCTCTCCAGTTCAGAGGATGGGATTATGGGAGCTTAGACCAGGTACTGTGTTTCC
TGCTGACCACTGTGTCTTTTCGTATCTCCTATCACTTTTGTCTCAGGATGTTTGTATTTCAGGGTTGAAATTAGC
CTGAGGTTTTGGCTTCCTCCAGTaaaaaaaaaTTGCTTGGAGGGCCACTTTTGGCATATGAACCATCTGTGTTT
TGTGCTTAATGTTCAGAGTACTAGTTTGTATTAATTGCACATTTGCAGCCACTTTTTTTGAAACCGTAAATCCTA
CCATACATTGTTTTCTATTTTGCAAATGAAATTTGACTGTATTAAGGTGTATGCTGACTTTGCTAATGGGATAGT
TTATATAAAAACTTGACCATTTGGATCTTTAAAGAGGAAATAACCCACACAAACTAATTTAGTCCTGCATATTTG
TTTGGTATTTTATCTTCTCTATAAATAACTTATGTCAGGATGACTTCGATGTAGACCAGTTGTCTGCGTCCTGCG
GGCAGCACTAACCTTTGTTTTCCGATAATCTTCAGAGCTGTCTAAGCAATTCCAGAGTACAGAAGGGTATACTCT
GATCCTAGAAAAGATGGCATATTCTATCGGACCAGAGTCACCAGACCTTGGATGAAAGGGCTTCAGTTTAAGTCC
TACAAGAGATGGTACGATGTAGGCAGGAAAAAAATGGCATTCCTACCCTATCTTGGTTCTAAGCTGTAAACCTTG
GCCTAATTTGGTTAAATCTGTCTGAGCCTTTGTAGAACGATGGCAACCACCAAAAGTCCAACTACGTAGAACGAT
GGCAACCACCAAAAATTGCCCATTGTTCATCCGACGGGCAATTTGCCGTTTCTTCTCATGGTGAGACATGGGATG
CTGCAACATGACCCGAAAAACTGGATTTCAATTCAAGTGAACAGTTTCTCCATTTGTACGGTATTCCATTGCAAT
TTCTGCCTATGATGGTGCGCGGATACCTGGGCTAAGCCATGTGGAGACACAGACACCATATATGAAGTTGGTCTT
TTTGCTGCCAACCAATCATGAACACCAAATTCAGAGTTAGGAATGAACTATAACTATTCATAGTCTTGTTATTAC
CTTAAGCTCTTGATTTTGCAACGAACCTGTTGATGCGCAGTTGATCTAACACATGAGATGTGGTTTAATAGACAA
TACAAGTGTATTCTAGTTGTCTTTTTCATAGTGGGTCAGTGAAAAAAAGTGTTTTGAACCATTGTTCCTTGACCC
ATATGTTCGACGCACACAATTTGCCAAGAACTATCTGCACATATGGTTTTGTATCGGTTGCAAGGGCTGATGTTG
TACTAGTAAATGCTTTCTCCGTTCAAAGTTATTTTAACTATCTCAAAACATATTAAGTTGAACTAAAATTATAGA
GAATTATAAAGACTATAACATTTAACAATAAAATATAATTAATAAAAACTAATgatatttatttgatattataat
tatcgttattctataatatatatttaattaagtttcagatgttttgactttttttaaatgatttaagatagagaa
gttgggatgaatagaaataatttgggatatattacttataatttgagacggaggatttgcaaagcaggaagataa
tagagcgagtacttcattttagtgacaatatttatttgatattataattatcgttattctattatatatatttaa
ttaagtttgagatgttttgactttttttaaatgatttaagatagagaagttgggatgaatagaaataatttgggat
atattacttataatttgagacggaggatttgtaaagcaggaagataatagagcgagtacttcattttagtgccaa
tatttatttgatattataattatcgttattctataatatatatttaattaagtttcagatgttttgactttttt
aaatgatttaagatagagaagttgggatgaatagaaataatttgggatatattacttataatttgagacggagga
tttgcaaagcaggaagataatagagcgagtacttcattttagtgccaatattCTCAAGCAACCGATGTTTACAAG
GAACGTAGGGCTACTCATTCTCATATTTAATCATTTTCATATTCAAACTCTACCATACAAACATTGTAGTCTACC
ATGTAAAATAATGTTTTGGTGTGTATATTTAAACTTTATCATATGAACAACTTAGTTTACCGTGTAAAACAATAT
TTTGGATGATAAAAATAGTGTTTTAGATGATTATAGAGTGTATCGCTGAAAACGCACGTGTGGGCTGACTCTTGC
ATATGTCTGTGTCACATCAGCGTACatgagtccagatatgtttggattgtcgagaccaagaggatgagtccagat
ttgtttggattgtcgagaccaaaaggatctaaagaaggctgtttggTTATGAGACTATTTAATACTCGTAATTGA
TGTTCAAACATCTAATCTAAAACGTGAAACAGACTAAATTTTAGTCAGAAAAATCGAACGCCTCCAAACTAATGA
TTTACGATTTTAAGAATCAAAACGACATAATTGAGAATATATTTAAAAGCCATTGGTTGACCTTGGTTAAAAATA
CTTATAAATTTGAAAACTCTTAATACTGTATTCCAAATGAATTGACTAATACAATGACATGCACAAAATATGAGT
AAAGTAAAATGAGTAttttttttACTAGAGGGAGGATAGGTGAGAAGACATAATTTGTTTTGTACAACTGCATAT
ATATACATGCTTGAGATTAATGTCCATTCAAAGGTTAGTTTAATGATGACCCAACTTTTTAGGTGGAGCACAACA

ATTTTGGTAGCATAACATAGTTAACATGCATTATGCACAAAACCTATATGGTTGGACATAGCTACAATTAGTGTT
GTATATGTGAAAAGCTTGTTTAGATTTTAGCAATTAAGTGACAATATTTGTGAACTAATATATTTTGTGAGATGA
AATGTGATAATTGGTCCACAAAAGTAGAACATAAGGCCATGATGATGAAGATTGCGATGATCGTGATTGTGCAAA
AGATCGAGAAGTGAAAAGATAAAGACGAAGGATGACACATAAAGCAAAGGTATAATGATAAGGATTTGTCTTTCA
TCGTCCGTGTGGTGATCAGTGGAGAGGAAAAACTAGATTGATAGGAAACATAGTCATGCTATAAAGAGAGTGAAA
TTTGGTTGCTAATGTGATTGTCTAGTGCTAAAGTATTGGTCATCCCGTGAAAGATCTATTGGAGTTAGCCAATTT
AAAAGGTTCGAGAAGAAAACTTGATGGAGGATTTTTACCCACACTGGAGTCTCTGATGATAATTGGAGTTGGGTC
TTCGAATCCTAGAGCGCTGAATCGGCTCGACCCAATCATCCATCGAAATCCTCCGGTAACCACCAATGATCAGGT
CATCGGAGGTTAGAGCATTGACCTAACATAGTTGAAAACAATGGCATAGACCCTAACTTCTAGTGCCACTAGAGA
GCAGACATTGAAGCTTAGACTTTGATAGTTAGCAAAGTTTAGAAAGCATGTGGGAATTTCAAAAAGGCCACTATC
AAAGTTGTTTGTTGATCCCACCGAATTGCTACATCGAAGGAAGAGACGATGTTTGTGTGTGAGGGTGGGGTGGCA
GTGTATTCACTAGAACTATCTGGTGAGCACCGAATAGCAAGCACCAAAGGTTCGAGCATACAATGTGTAGTTCTA
ACATAGTGCCATCAACCATTGAAGAAGTTGTGGTGGTGGACTCTAGTGACCTACTAGATTCTCCTATAACACCAA
AAAAGTTGCCCCAAGCTTCCAATGACTAGTGTCATGAGAGGGCCTATGAAGTGTGTGGCCAAGAAAATTAGAACT
CTTCAACTTGTTGACACACACAAGAGTTCAAGAaacaccttcactcacatttgtttgctagtgattacatcctta
gtgaggtcagagtgatctagtgcattttctttgagagatagcatacaatagcactagaaaatcatatttgttgtg
agtttttatgttactcttggtggttgttgccacctatatagcttgTACTAGATATTTCTGTAGAAGAAATCACAA
TTTGAAATTGTGAGGTGATCCACAATTGATTGTGAGGAGCAACCACTGGAAACCACAATATGGATTTCAAAAACT
ACATTTGCTCATGATTTTAGGCATCACTCTTGCCCTCTATGGACAGTACAAGGAAAAGAAGAAAGTGCACATTGT
TTTGCACAATCAATGGACTTCAACCGATTATTCAGTTGGATAAGAAGAAGGCAACCAACAGGATGGGCAACTGAA
AATGGATAATGCCCCGACAGGCTCGACACTCATGCTTTCAGGTAAGCGTGGAGTTCATCCCACTATACTAAGTTC
GCGAGCCATGTCCAATTAGATGATAAATCTTCGTTTTTACATAGTTTCTTTCGAAACAGCCTACATTCACACTCT
TCCTTTTAGGAAAAAATAAACATGTGAAGAAAGCCCGTTAACCCACTCTTCCTTCACACAACCTAAAGAGGCAAC
AACCTCCCTTCTTTCTGCACAAGAAATGCAGCTACCAATGCTGCAATGTATTAGAAAGCCAATTCATACATATCC
CTTATTTTTTATTCAGCTCTATCGTCTCTAGCTTCATCTACTCTTGTGGAGGGCTTTTATTGAGATCACAATCTC
TCGCCGGCTCCGGCTCCTCATTCTGGTTCAACGCGAGGTTTGCCAAAACGTCTAGAGGAAAGTGTTTTTAGGGTA
ACCTTGTCGATTCCTTCGCGGGCCAGAAATTTGGCAACGACCCACCTTTCTTGAGGGTCATTAACGAACTTGAAA
ACGTCTTTTATTTTTCTGTCCATTTGCAGCTGGCTATGTACTTAAGGGCCAAATTTTGGACATCATAGTTTCATA
GCCTATCTAGCCAAGAGCAGGAGTCGGGTTTACCAACCACTCTAGTAGAATTGAGGCTTGTATACCCATCTTGTG
ATTGAGTTAACTCAAGATTAAGCACTCATGTCGAAGCCAACCTTGTGAGGTAGTTGAGcttgatagaggagcgat
gatccttaggctcgcctcaatatggatgtaggtgaccgataagttaccagaccacatagatatatcttatgtctt
cctgttgttgtattCGTTTTATCACTCTCTGTGCTTTACTTTGTGTTGTTCAATTCATATAGTGATATTAGGACT
TGTCATCCTAGAAGCAAACCTAGCTAGTTTATCATTTCTAGTTACATACTTAGCTTACATTCTTGATCACTAGAA
TTGTGTAGCTTGTCattaccctaggcttgcaagtcataacttggtgtagtgcttattaggcatcctttgtgtcta
acaagttttcctagtgtttgtggttgctagaattttgaattaggctatttacctcgcctcaggccatccaaagat
ccatacaagtggtattagagccaaatctctAATCAAATTAAGATTTAAGAAATATTTTGATTAAAAATGCTTAGT
AATATTCTTGGCAAGTCCTCATTCttcaagagcacaaattattattattattattattgttattgttattat
tattattattattattaGAAGAGAAGAATGTGTGCTAGTTTGAAAGGCTATGTGAAGGAATATTTAGGGAGTTGT
TGAAAAGGAGCTAGTACTACTTGATAGGAAGAATCCAATCCCAAGATAAGTTGAAAAATCTAGAACTTAATGATC
AAGATCTAAATGTTCGATATGAATTCTTAATCTCATGGTATTCGAGTGTATGAAAGACTTGGAAATGTCACACAA
GGCGTGGAAGACTTGGGAAAGCATATGAGGAAACCTTAACGATAAAGGATGGCAAGCTATATATCTTCAAGGACA
AATATGAAAAATTCAATATGCAAGGGGGGAGAGTGTGTCTTAGATATCTTCTAGAGGCTAAATGCCATTATGAAT
GAATTGAGGGCCTAGGACACAAGGTCGACGATTAAGACTTCTCCCACTATTTCTTGAGGTGTGTTCCCAAGAAAT
TCAAGATCATTATCATTGTGATCTTGAGAGGAGGATTGGAGCTTCTCACTCCTACACAAGTATTGGTAGAGATCA
TTACTCATGACACTTTTCATCATGAAAAATAGAGTTATGAGCAAGAGAAGGACAAGAAGAGCATAACATTCAAGG
TAAGATCTCCAATAACAATAAGAAGATTTAGGATGATGGTGATGATAAGGATGAGATGAAGTGATGATTAAGAGG
AGATGGCCTTCTTTATGTGTTGTTTTGGCAAAATACACAAAGGAGAGAGAGGCTTTGGTAGAATAAAAGGCCAAT
CATCATCATGGAGATCAAAGGAAGATCAAATATGTTATGATTTgggggggCTAGCCACATTGTAGCTGAATGCC
CCAAAAGAAGAAGGACAAGTATTCTAAGAAGGGAAACGATAAAAAAGATGATGACAAAAAGGATGTATTCTTCAA
TAAGAAGGGATACTCATATTTTCTGAGTTAGATTTCAATGCCTCAACATCAAGGGCAGATGATGATGAAGGTCAT
CCAAGTCTCTTATCAAGAACACCATCAATATTGATCCAACACTGTTAGACTAATTGGCAtgcctcatgtccaagg
gccctaaggtaaaatatgatgaaagtgatgatatctatagtgatattggagtttataaagaatattcttatgaca
accttgttgaagttctataagagtctcaatactttatgagctaggagatagaaaataggaactgaaaaggagatc
ccgagcctctagacaaaccctttgagaagctagaccaatcatgaggatcttttagaaactcatggtaggcttaag
gaagcttaaaacTCTATCTTTGAGCATAATGTTGAGTTCCATCCAAAAGTTGGTATGAGTGTTACATGTGTTTTA
CTTGAAAAATCCAACATGTACCCCTATTGTGACTAACCCCTTTTGTAGCACAAGCAAGACTTCTATCTAGGGTAA
TGAAATTTACATGTGATTCGTTTCTCACTATGAAGAACAACACACTCAAGAACGAGGTGGATGAGCTCACACATG
CACTAAGAAAGTGCTATGTGGGTGAGGCAAGGTTGCACCAATGCTTGGTAGCCTAAGGTTCTCCTTGAACAAGGA
AGGCCTAGGCTATGTTATAGAAAAGGAAAAAGTTTTTGCTCCTCACAAACAAACATTTGTGAAGAAAATGACAAT
TATTTTTACAACGAACAAGTaattacaaagccaaagggaggcattggacattggataatggttgcactcaacaca
taatcagagattgttaaattctcttataactcaaatgcaagtgaaggttatgacaacattgcactacacaacaat
tgctagggtaaatttgttggtttaggcaatattgcaataccaaatgatctctcaatatctaatgtgctattaatg
gagagtctcaacttcattttgctatttgtggcacatttgtgatttggagttcaaatatgtattactaactgaaga

tgctaagatcaaaggaatggattgtagtaatatcataggatttaggcatgtgaatcgatatcgagttgaatttg
gtgctagtgaggcaaggttatcaacatgattgttcaccaagtaatcaatggaatggttatgacatagaagacttc
catatgcaaggatgacgcaattggttatattaatcaagcatgatttagtcaatggtttgaagagtgtgacatttg
aggagggctggttgtgtagctcttatcaagagagaaaacaaattggaaacactcaccctagaaagatcatgatgc
aacttcaaagctatttgagttgtttcatatggatttatttggtccaaccatatatgttactattattggcaacaa
gtatgggttggtgatcattgatgacctctcaatattcacttgtgtattctttattagtgacaacagtaaggtttt
aacaaattcaaataattttgttaagagagtagagaaaaaattttaagcttaattttaagaaactgataagtgaca
atggaagtgagttccaaaatgctagatttttttatgacctatgtaatgatcatggcatcaagcatgagttctcggc
agtatacacctcAATAGACAACAAGGCTAGTGCTCTATGAGTCCAACTTTGTCAGACTGAGAACTAGATCCCCTT
TCCCTATATTATTGCCTAATGTTCATTGAGATGCAAAAATGAAGCTCCTACAAATCCCCTTTCTCTAAAGTTTTT
CCTCAATCCCACAAAATATCCCTCTCTCCCTCAAATAAAGGGAATCTAGTAGCACTTTTTCTCAATCACCTTTTC
TATCTTCTCCTTCATTGGCACATGGGCTCCATGTTACATTGTACCAAGCATGACCACCATGTTGCTCCCCACTGC
ACTAAGCTTGATGCCAGTGAGGTCTACATCGATGCGATTTGCATGCAAACCATCACAACCTTGCATGGTACATGC
CACCATTGCCAAGTGTTGAGCAGGTTGCCACCTTGATGCTCCGTGTGTTGCTCCTATATACCTCACCAACCACCA
TGCATTGCTCCTTAGAATCATGCCAACCACCGTTCGACAGAACAATCGCCCATTACACCATGCAATCGACATAGA
GACACTCCCATATCACACTTTGTgggaaattttaaagggaaacgaccttaaccatttctatatttgattttggt
gcttgacgaccatcacaaatcttatggactaactagtttgcctagttgattaattctcaggtgcataagtccatc
tctttcaattctaagtcgattgtccggaataccgtagattattccggacatgagaagctttttgaagaattcacc
tatgcgtggaccgttcgtgcccttccggcggaccgtccgcgacaacaaggtgtgcctcggataggaactatgcaa
aactcgcatttacactacggactgtctgaaggagaagagagcaccgcccgggactaagcttggaccgttcagcat
caggcacggactgtcctcccattgaaaaccaaagaaacccgaaggtgacaggttcggtaaaattcatttttagcg
tcctcgcggaccgtccagggtatacggttggaccgcccatgacttctttatctgacatctgacgacacatttaat
gcacttatagccgttatcgctaaacgttgcgatttcagtcattgatgtgcacggccggaccgtctggaccaggag
cgtggaccgtccggggttggcagaaagtgggcaatggctaggaagtggttggtggttataaatacaacccaacca
cctccattcacaagatccaagcatttcattcatcttcattcaatacaagagcaagcacttcatcccaagacgcat
tcaaagctcccaagctcatccaagtgtcatacttgtgactagtgatcattagtgattagtgccttggaaagtcta
gagagagtgtaatcttttgatttcttgttgctcttgttgcttggctttcttaatagtgctttcttcatctcctt
ataaattatcaagtaacttgtaaagctagcaagagacacctaactatgtagtgattcgtgcgaggtctaagtgac
cctcgagattaagaagaagcactcaaccggtgtaagtgaccatttgcgagagggaaagggttgaaagagacccga
cctacgtggcctcctcaactgggagtatgttctttggaaccgaacctcgggaaacaaatcaatgtgttcatttgt
gttgatctttatcttacgatttgtttcttcctctccgatttcattattactcattttaatcctaacctcgggtg
aagtttatgttcaaagtttatatttttaggtttcgcctattcaccccccctctaggagactttcaaaagttgtCA
CTTGGCCATGCCTCTCCTCTAAATGTCCCATACATCGTTTGTCTAGACCACCCACAACCTCATTATCCATCTACG
TTGGAGTCGGGAAGCCGTTGTCGCTCAAGATAAAATACAAAGAGTAAAGGGAGAGAGATTTTCTAACACTAACAT
GTGGAACTCATAAATAAAGGGGATTAGGTTTTTCTCAATCTGATGGAGCACATACCCTATCCTCCCCAATTCATT
TTTCTCAATCGCCTTTTCGGGGAAATGATCCCCGGTCCCTAGGACCCACCGGTCAGTGAGCGCGCTAAGGAAAGA
GCCCCTGTGGTCGAACCTTGTGGCACCGAGCCCAAGGCTGGGCGTGACGGAGCCTGGCAAAGGGAGACGGGTGTG
TCGGAAGGGAACCACTCGAGTGGATCCCGCGCCCGGACCCAAATTGACCCGCCATAAATGACTGGCCACATTAAC
CATGCCTAACATCAAGCCACAACGTGGGCACCGGTCCGCTTCCCACTCATGACCTACGGACCCTCTGCCTGCATA
GCACTCATGACCCACGAGCCCCCAGATTGCGGTGCATTAATGGCCCACGCGCCACTCGGTTTGCGGCGCACTCAT
GGCCTTTCGGACTAGGCCTGAGTGCGCAGACCTCCCGTAGGGCGCTACATGATAGGTCGTGCTAAGCCCTTGGCT
ACTGAAGCTAGGGGCCAGCGTAGCCAGAATGATGGCGCATGGGACCATCACCGCCCCCATGTTATCCACCACGAT
AAATACACGACAGCCAGGGAGCCCCAGGCCCCACGAGTACACATGGACTCACCCAGGGTAAAATGTTGGTTTTAC
CCCTGCCTTACGACTCCTTCCTAGAGAAGCCACTGTTGGTCGACCCCTCTCCTAGCCTATAAAAGGGACAGGGTC
GCCTCAGGGTAGGGAACGGAAAAAGACAGATCAAGATTGACCGATTGAGGAACAGAAGGATGGACTGAGGGACGA
CTGGATGCCTTCAGACAGAGATGAGACCACAAGGACGAACACGAGGAGGAGCGCCGACGAGGACTAGGAGGATGG
AGCCCCGCCAAGCCAATACTTAACTAGGACTCCACCTCTCAGCTTCTGAGCTCCACTTTCACCCCGATaagagat
atgggagcctctcctccctctctcggctatttgtaacccctactacgagtttgatcatcaatggtgccgatagtg
caagccaccgatgactggatgtagggacgttctgcccaaacaagtataaatcttgcgcccttcacgcacaccctt
cagagcacagaacgcgcacaataaatttacttgtcggagcgaggtacgaaacaccgacaCCCCTATACTCCATTT
TAAGAAAAAAGGGGAAGATGATCTAATGAAGTTCCTCTCAGTGATTCATTTTTGGTTGAGGCAATCAACATGACT
TATCATGCATAAAATAGTTGGGCACGACCATGATTTTCGAgcacttctcatcgtcaaaacaacacaattgttatt
tcatcggtttggatcctagcttaattccattaaggaatgctcgatgacttccctcgggatgcacatctcgtctga
gagtttccatgcaaatatgtttattccgcgtggaggaagtcgacaaccgagcttatttaggggcagaaaggcact
aatgtgcatccccttgccgctagagtcactagggtctagggatactTCCTTTTGTGTCCTTAATGTGTTGTGATG
GACCCTTCTGTGAAGTTGCATcattctacttcacactcataggcgtgttgaatggtcgtccctatggtgatggtc
cctgctgggctcgacatcatgagcttgaggtatatgtagttgggcatgaccatgatttcgtgtaacacgacctt
cccaatatggcgtggtaggacccacataatcctactaattcaaatgtgagggtttccatcctaaaatcgaatggg
gtgccaaaggtgatgagcatgtcgatctgccccaaaggcgtggccttgtagtggcatgatgccatagaaaggaac
tccagatggctgaagatgatcactgttgacgcacatggcttcgagggtagaagagtatagaatgttgatctcgct
gcctccaaccattagtaccttcgtgagtcgcaccatgccaatgatgggatcaatgataatgggtattgtctaggc
cgtgggatgtgctcggggtggtcagaatgaacgaatatgatggggtggtcggaccagtcgagaaaatatggggtg
gccagacgcacttggaacacctcccggcgctcgatcttgcgttgtcttttggtcgcgtagtagttgggcccaccg

aagatcatgaggcatcccatttcatcagaaaatagggtctttcctttccccgatgcatccatcgaatgcttagga
cgtttcccggatggttctctctagctcaatttttttgaggagcccacagtccttgtatgcatgtttaacagggta
gcggtggtttgggcaaggcgcctcgagcatcttctcgaagcgaccgttagctttttcaacgggggccttcttgcc
tttatcatcagtggcaacgaccatcatgcgctatgccgccatttgttcttatttatttttgtcaagatagtcag
tggtatcctcatgggcatccttgtcgcgttttgctttcctttgaggcgatcgaagacctcccaaccgcgtcttcc
ctagacatgtggttggtcgcaatgtcgagaagctccttcgtagtgcgggggattttgcgccctagtttgtgcact
agggtctcgctagtggtccgagagacaaaagagctgatgacgtccacatcggcgaggttagagagttcattgcat
tgccatgagaaaacttcggatgtaatcttggagagtctcccttgacttctatcagcaatttacgaggtcccaagg
gttcccggggtgtatgtacgtgctctagaaatctccaacaaaaatgttgacaaggtcggcctagctctggatgta
gctgggcctcatgtcttctagccaggctcgagctgagtcgaccaggtataaggggaggttatggatgataaatta
gtcatgatctggtcccctaggcaccctagcctagcacgctagcggtagtgttcgagccataggttggggtttgat
ttccctaagtacttgatgacattcgtaggtgcgcggtattgctgggggaccgatgcctggaggatatggtggtcg
aaggtgtgggtgcttggaggcgatgggcttgggcttcagtcttTATTTGGATCGTATTGCTCCTAGCATCGATTA
TACCCATGGCCTTCGATGGGATGCTCGTGCTCCTACCCACAACGCCAGGTGTCTAGGATATCTCGTGCATCCCGG
ATTGACCCAAGATATCGTCGAGCGGGAGCCAACTAATGTTCGTAAGATAGTTTTAAGTTGTTGAACTAGATAAGT
GTTCTCTCTTAGTCATGCTGTGGATGCACGCTGGCTGGCTTTAGCTTGCCAAGACATGGAGCTTTCGCCCTGTTG
TTGAGCAACAAGGATGAGAAGGCCTCATATTTTGCATTGGGTCTTGCGTTCATCCAGTGTTTTGGGTTTTGGAAG
TTTGTCAAGGAATGATGTCGCGACTGTGATGTTCTGACTATCCAGCCCCTTAGGGGCATCGACCACCTCAGGTCC
AGCTTTGACGGTCTTGCGGCTGTTTTTCATGTGTTTGACTGAGGCGCGGTAGGGTTGATGCTGCCGCAAGTACCG
ACAATTATATCGTAGTTTCTTTTGGGTCATCTATACTTGCAGTGTGGGCCTCGAGGGCACTCGTCATGAGGCATT
CCTGCATGGGGTTGTAGCTTGCCCCCACTTGATGTGGTGCCGAAGCTAGAGTCAGAGTCTGACAGGAGGATATCA
TAAACCGACTCCCTTGCGCAATCAGCCACCCATACAAACTCAGCATCTCCTAGGGTGGACCCTCGACCATGGACC
ACAGTCCGCAGGCACATCGATTATCTCCTCGACGTCACGAGTGTCGGGTAGGGACGATATAGTGGTGCCTATGCT
CAAAGCACCTAGGGGTCCGACCCCCTTTTACGAAGTGGACCACGGATCGTAGCCTTAACTCCCTAGGGACTAAGG
GGAGAGGCAGGCCATGTCCTAGGACGCGGTTTAGTGGGGTCCAAGTCGGAGATCCGTGAGCACCTCGATAACTGC
ACCAAGCTCATTCGCCTAACCCATGCaatgtcgctgctagccatccgatacctaatgagaggatgacaaaaacgc
gcaaggcccctacctagcgtgccaattgtcggtctttcgaacaagcgttgaggagtaaatttatgtatacgtgtt
ttaggatatgttgaacaagttttgcagcgaagatgcaaggaaaattagaacactagtgataacaaatgtacacct
atacttggatgaaagtagaaagctaatgagcaaaagctctatagctcaatgattggtagtttTGCTAAACTTGAT
gggaacatatcgaatgcaaaccacgttttggtgcaaactccgtgcaaacctactaaaaaagtttcaaaaaattct
gaaaaaacatgaatgtacatctcagcttacctcaactatatacaaaatttcatggtcaaattcatcttactctag
cagttacaaaaaagacaaaatttctgacaaacagtaatatttcaaatgcaccaaaatttgtctttttttgtaactg
ctagagtaagataaattttaccatgaaattttgtatatagataatttaagctgagatgtacattaatgtttttttc
cagaatttttttgaaactttttttagtaggtttgcatggggtttgcaccaaaacgtggtttgcattcaatatgttcc
cATACTTGATAGCATGAAAACTAGAtactccctccatcccataaaataatgcatatattttagcaaagtctttag
agcgaaactttgacaattaatttcttttaaaatatgacatcaacaaatacaaattgagcttcatattaaagtatt
tttgaatacaaattcaagtatataacatgtatacaataaaaatatatgtttttttttctgaattgttggacaaag
ttttaaaatgttgactttactaatacatgcatgccttactttatgggatggatggagtaatatgtttagtatgtg
tatgtgtgcaagatcccaaccatcaccctaagagtgtcattgtattatggtaaaataatgagatacttgaagttc
atccataacattaaactatggtatcccaaaggtgcacaatttgagttgattggttaatcatacttgagctatgcc
agttacttgaaaactagataaaaaGTTGGGGCATGtaaaaagttagtaggggtactgaaagggaaatgtgccctt
gggccatttctatattgttttggtgattacatgcacaacacattatgtttagactaacatgatgttgagcaaagt
tatataaagcaaattgaactcgaaagAGGACttagtgtaaatctatatggatcaagtaaaaaggtaagctctaga
cacttagtcaattgttttgtgtattaattatatttgtctaagtgtcagggactttatgaagtcaagtccaaaagg
agcaaaataaaacaaacaaagaaaaataaaaaggttgctggactgaaggcaccggacaatccagtgaccctcacc
agacagtctgatgcatggtccgtcgtactgctcgctcttgggaattttagccaggaccgtccgggcagagcatcg
gatagtccggtgtgccagcagccaacggccagtggccacgtcggccccgaaccaacgatcatgtgggacatcgga
cggtccggtgcccccagaagcagaaagtagcccattaggggattctgtgcccatgcactgttcactgtccggtgt
gcaccggactattcggtgcacccgcggacagaaggcaaccagggcctttcaaatggagctccaacggcttctagg
tcccttggggctataaaaggacctctaggcgtatggagctccacaccaagcacccattgaacattctacaatgc
cgagacttcgcaagcacacaattgtttcattgtgatagagatttgagcactctttgagttgtgttgtttcgtgtg
ctcttttcttgacttgtgtgcgtgtttttgctgtaactctagttcttctgtgtgtttctacccccttactcttgt
gtccatttgagatcatttgtgtaaggcgtgagaaactccaaattgtggagattcctcacaacggaatattgttga
gataaagagaactgtgatattcaagttgatctttggatcacttgaaaggggttgactgcaaccctcatccattgg
gtcaccacaacgtggagtaggtaagtattctatttgaccgaatcacaggataaaaatcatcgtgtcacttgtctt
tatcttactgcaatttctctctattcacttcacttacataattactcttagtttaatactcacttagtgaagag
aaattaagtgaagaagtttactcctctcatactcctcattcgaacttggctcttgctttcactaattcaatagta
gtccaagttcgttttgttgagctatttttataggatcacctattaaccccccctttaggtgctctcaGGTActag
tatcatagtcataaaaaaagcaaaactccatagtgctccatcaccgaagcagaatacatatgaacatgtagctat
tgtgctcaattgctacgtgtaaaggcaaccatgatggatttttttgtttgaattgaggaagcacctctcctatatg
acatggggaacatcatgatgattgtaagcaacctagtccaacactgaagaaccaagcacattgatacaagcgtaa
catcaATATTGAAAGTATTGGCACCAATGAGTAGTGAGACGACATCTTCTGATGTATTCTCTATGTCCTAATGTA
TAGGTTTGGAACCtaggttgcaaacctcatttgcgatagagatatcaacacttagtcaaacctttgttgcatgt

atgaacaatcagttatctacataggttttgattaagagaaaaatagaggtcatagtttaaagagaaattagattt
cctaattcaccctctcttacgcatcaacgttccctacTCTAGTGTTTGCCATTGATAGTCGTCCGAAGATCCATA
TAATATGCTACAGAATCTCTTAATTTAGCTTGAATCTAGCATGCCCATCAACCTCACAACACTAGGCACACTTGA
TCCAAACCAACCGATATGTTTAGGCCACCGTATGGGAAGACCTGAAGCATCCACAATTTTGCTCTAGAACCACAT
ACATAATATTTTTAAATCAATCACATAGATAGGCACTTTGAATAGAGAGTTTACATCAAGGTTAAACGAAAGAGT
ATAGAATACATCAGAAGAATAAAATAATATACTTGTTGAAGCAATTAAGTTCTTACATCAAGATCACGTAGTGGC
CAGGTACAAAAATACATAAGCATCTCTATAACTAATTCATAATTTATAGGACTTGAATGGCTAATAAGTGCACAT
TTGTGTCTAGTCTACATACAGAAAAGGAACACCAGCCCACCATTGTGCTCATCAATAATTCCCGAGTGACACATA
GGTTAGCCAAAGGAGTGGTTAAGTCTAAACACTAAGGATGACATTTGGATTATGTAGTTTCACTTGACAACTTTA
GAGTGTGCTATACTTGATGGGTTTTCTAGTCGATAGTATTAGAGCTATGCTACAAGCCTACAAATGAACTCAAGC
TTTCTCATTCTTAATTAACAACAATACAACTTACTTGGTGGCTCGAGGAATCTACTTGGTTGTACTCGGGCATGT
CATATGCGGGCTTAGACTTACTCAATTTTCAATTGTATACATTGGAGTTCTCTAAAGGATCTTATCATCTCAGTT
GTTTGCACACAATGCAATTTACTGATTATACATGTTTATCAGACATCTATATCTATACTATACTTAAAGCACCAG
TTTTAACGGTCGTCCCGCGTCATCTTTTTACAAATAACCTCTCACAGCTATTTCAAATTAATCTGTTGCACGCCT
ATAGATGGCCAAACGGTGGCCCGGCATGGGccagacgccagcgggccataactctggcccaggcatgtcatgccg
gcctgctgactgtgtcaggccagcccgttagaccgtcggcccatttgattaaatcgacgtaaaatattaaaaaaa
tagtgcagaaggtggggtttgaacccatgccctaatggaagaagggcggaagacactgagtgaagctgtctaacc
agtagaacatcatgctcaaatattttaatattaaatataaattgtatatatgtatataatttttttgtaaaatta
aaaaatataactgtgttgggtcaggacagcactataggtcgaggctacataccaagcacgacacGACGTTCTTGG
CTCTTGCAAGCATTATGTCGTTTCGGAGACCACATTGACGCAATGGACTCCATAGTGTTTGAGGTTGATGAATTG
GATGGAGCAACAATGATTGTCACACTAACAGTAAAATGAAAGGTTATTTGTTGGTTTTAAACGTTAGTAATTGCT
ACGAAGTAGCATAATTTATATGCAGCATATCCAGTTTTTATTGATGCCTGACTTTAGCAATCACTCCATATTTTG
ATCTATTTTTTTATAGGTTTGACTtcatgtgacttatttagaaaattgagctaacaaactttttcttatttggt
ctctatatggtgaaattatgtcattttataatatatgttcgttcagtcagtcgttgttaactctaatctaatcgc
tcactttattggtcgtgttgtaccaagacatattgcatggagtaaacaataacaacagttagccaaatcaaaaaa
tattatacagagagcggagacaattaataaaaaatcttgaatttttttgatagatagtttacgtgggtattattg
taagtcgtcgctatgcacgggcaaccgactagtgatacataatatataagctACAACTTTTAATTCTAGGTTTTC
TTATTCTGGGTAAGGCACTAGGGTCGATATTATCTTAGTTGACCTATTCGTTTCCTTAGCGCTGAGTAGGACAGG
GTGGAGTTTGACTAGGTTCTAAGTTCTATGGGTTAAGACTTTTCTTCACTCGCCGATATATATGTTCAAAGGCCT
TACACTACATTTATCAAGTCCCCCTAACTAGATTTTACTGTGGAATACTTAAAATATTGGGTAACTTATTAGTTT
GACTATATTCCAGTGGTTTCTATTGGGATCTATTTATACAATGCATACTTAGGTCTCCTAGATTGAACTCATGAC
AATTTGGATTAGGAATGGGGACATGAGTGTTGTGGTTGGCTATATTATTGCCTTTGACTATGATAGTCTAGGTCA
CTATGCCAATGAAGAATCATCCGTTCATTGGTAAGATTGTTTGGCATTCTAGTCGGTGAAATGCAAGATTCATAT
CAAGACTTAAAAGAAAAGTTGGCAAGGATGGAGAGGACTTATAATTTTTAATTCAACTTGGTTAACTAGATTTTA
GAGTATAATACTTAGTGAATTGCTTACCTTAAGGGTTCAACTACCTCATGTTGGTCAGACAAGATGTATATGATT
AAAAACTGATGGCAGTCTAGTTAGGGTGATCAGTATGACTCATATAGTAGGTTGCTCGATACTTCAGTTGACAAG
CAGAGAGGATAACACCTATTCTTCAATGTTTAATCTCGTTAAGAGGACAAAAGCTTATAATATCTATTTATAAGG
CATGGTAGTCGACAAGGCTGGTTGCAATGTACTTGATCTTTCAGTGGTTGCAGTAGTTCTCTAGTGAACGTGCAA
GCACATTAGTTCCACTAAGGATTCTTGGTGAAGCTAGATCTAAGGTGTCCGAAAAACGGGCAGGGTATGAATAGG
GTTGGTAGCATATGTACACAACAAAATGACACATAAGCATAAGTTGCTTACCTATGTCTTGGTTATGGTGAAGCT
AAATAGTAACTCCAAGATCCGATAGTCCTAGAATGGATCCTTCAACTTGTGTAGCTTGCGGTCATGTACTATTAT
AGAGAGAATGACCCTAAGACTCTAACAAAGGTCACAACCACCACCAAGATTATGCAAGTTTAAGCGACCAAGGAT
TATGATGGAACAAGCAAGATTGTAAAGGGAAAAGCATAGGACGACGGAGCTATACTAGATGGACTCAAGCACTAA
CACTAATGGTGGCCAACCATTCAAGGGTCCACTAGGGAAGGTCTTCACTCATGGAAAATTCTAATGTGGTTTGGT
TGGTTGTGAAACACAATGCTAGGTGAAACAATGGCTATGAGGGCTTGAACTACGATAGAGAAATTGGCATGCGTT
CTCTCTATGAATGATAGCAAGAAGAATGGAGGAAGAAGACAAGTTCTAGGAGGAAATAATGGTGTGGTACAATTA
CATAACTAAAGCAGTGGGGTGTCATACACGACGACAACACATGATATGAAGCATTTCCTTGTGCACATGGGCACA
ACATGGCAAAGAGGAAGGATATGTGACAACATGTGGCACTAAAGTTCCGAGTATGTCGAGCATCTAGTTGGGTGA
ACAACTAACCTAGTTAAATGACCAAATGATTGAATCATTTTTATTATGGTTCCAGACTTCACAACTTCTCTATTC
TTCTCATGATCAAGAATAAAATTTTATCGATATGTCTAAACGAAAGTTTTAAATTTGAACAACTCCAACTAAATT
AAGTTGGGATACCTTTGTATGCCATTTGTACCTAAAGCTCTATAAATCTTGGTTCAACTTCTTACTAAGTTTCCA
CATGGAATCAAAATGTGTTTGCTCATATATTTATGCATTTATATCTCCTGTCACAAAATTAGAGTTCCATACAAT
CTGAGTATACCACTCATATAGACTTTGGGATCACTTGGATTACCAACGAGATCTATGTCAAACTAAAGCTAGAGT
TTAGTGCTACAACTTTTATCTTTGAACTTTGGTAAGCGTTTAATTGAAATAAGCGGTTACTTGCCCTTAAAATGA
CATATTAATAACCATTCATGTTAGATATTTTTGCACTAACCTTATGTTGTTAGAGAATATGATATACTAGCTTCT
TAGTGAGGTTAACTGAAGGGTCTTTTAAGTCTGTTGGTGAAACAATATGCATTTGATGTGTGCAACTTGTTTTGG
AGCTTGTTCTTAATCATTTTTAAAATTTACTAGACTAGATATAGTGGGGATTAAAATGCATGAACATTCTCACTC
ATGCATGGTGATTGATGTTGTTCATGAAACACATACACTTAACATCGAGGTTGATATATTATACTAGCATACAAC
ACGTGCTAACGCTATGATTCCGGGGAGAGGAGGGAAGGGTTCGACGGTGGCGGCGGCACGAGTGAAGGGGGGTGG
CAGGGACGATGGGGAGGGACGAAGGATAGGGGACGATCTAAATAATATTGTTCATTGGATTTAAGTAAGGAAGAG
AGGGTACATCGATCTGAACTGTTGGttttgattgtctattaagtctgggtgtaatttgtttggtggatttagtgg
aggttatgcatgcgggggtgatttggttgggtaggttttgcaaagtttaaacgggtgggttatatagtggtataG
ATATCCGACTTGTGCCAAAATTTTATTTAAATGATTTGCAGTATATTAGTTATGATTTATCACTCTCTTTATTAT

CATCTTTATTTGCTAAATTGTTAATAGAAAATTCTAATTCTAAGGAACATAAGAAAAATCAAGAACATAGCGTTA
CTGCAAAACAACATGATGGAATTGATCAATATTTAGCTAAATTTTGTTGCTATCACTTTTAATGGCGTACCTTAA
GCTCTGTGTTGGGGTCTGCTACATGCTCCTCCTCCTTTGCTCGACCATATCGATCAATCACTGACTTTATGCTGC
ACAAAGACATGCTATTAATTATGCCAGACTCGTTTTTTGTGTAGAGTaaaaaaaactctaaaaaaTGTATGAGTT
ATACCGCTCGCACACACGTACATAGTACATGCGTGAGCTATATCCACATTTTCGATATCTAACATATTAAGTGGA
AGACCATGTGATGAATACACAAAAGTATGTTATACTTATTCATCTAGAAAAGTATGTAATCCttaggtcttgttc
gtttgtaccggattggtgggtcggaacgattcctaaccggattgcttctctaatttatataaattttgattagct
ggaacgatttcggatgcaatccgacagaaacgaacaaagccttaAATTTTCATAGCTTTAGATCTCCCTAGTCCC
TAGAGATACATGAATATAACCCCACTCCTTACCTTATTCCTCACTCTCACTCTAATGCTACCAAACCTCGATATA
GTTCATTCATACCACTATAATTGCTCACATGTAGCCAACTCCTCCAATGCTTCTGGGTACGATGGTATAATGCCC
CCTTTTCTCTATCCTATCGATCAAGTtagtacaaagaagctctatagtgacggttgtaaacctattttcattgat
gtttcttagaaccgtcggtgctatgggtcagtaaacatcattatttttacaggcggttaactaaggaccgccagt
gaaaatcgatttccaatggcggttgagttaagaaaaccgtcagtgaaaatcgatttccactagcggtcgatataa
taaatccgctagtaaaaattattttcagaaacattaaatggactttaaaaacagcaaaaaatatagaatacttca
tcctcatgcatacactccactactactctacctatggtataatttatgcctagtttgtagttatttctgcacata
ttacaaccatttgagtgtaaattgtttgtttgagaccttaaacgatttcaaataaaaaggttgtcaactacaaag
ttgaataacttttgaagttctacaactttcatttagacactttttcatctgaggtagtttgcaaaatttgaatt
ttaaatttgacatacttaaattcaattttgagaaccgaaatgagttcaaataaaaaaagttgtcaactacaaag
tttgataacttttagagatctacaattttatttgttagttttgtcatacgaggtcgtttgaaaaactcaaaaa
attaaggatagaaatgatttctagtggcgattccttaagaaaaccgccactagaaatcgatttctacaagcggtt
ttcttaaggaaccacctgcagaaatacgatttcaagtaacggtttttttaaggaaccgccactaaaaataacaca
ggcggttgataactgaatccgtctgtaaaaatatatcgtcccgcagtctttgagctttttctactagtgaCATC
ATCTGTCCTTCTATCTCCTATGCATTTTTGTGCGTCTAACTTTTAGTAGCTAGTAGATCTACTGCCCTTAGCCAC
TAACTATGGATCTGAATACATGACCTAAGTCACTAATTGATCTCCTTGCCCTTTTTATCTCTCATCGACAATCAC
CACAACTATAGATATCATCCCTAACCTAAGGCTAGGGTTCAAGCAAATTACAACCTTAATAATGTACACCACCTC
CATCGGATAGAGTGATGTCCATAGTAGAACACTAGTGGGTTTCAAAAGAACACCCAAAGGTGTGTAATTTAGAAT
TTTTTAGTACAAAAATATTTGGTGTGTATAGtagctaggggtggtgatatatcatgatccaaatgtttattcaca
aaatgttagagccctaaataaaaacttagtttaaaaataaatagaaataaggtccgatcctaacccaattcaatt
cttaaatcttatagtgtaaaatttagaacacattgtcagccctaacagtagcTAGGACTAGGTAAATATAGGCCA
GTGTCTAAGTACTGTCAAACTCTGGCCATGGATTAGCTAGCGGCTGCTGCTTAATTAGACACATATGTGATAACT
TGGGGGAAATCTATTGGATCCAATCCAAGAGATTCAACTTAGGTATGCACCAAGGTTTTCTAGAAAGAACAGTAA
CGAAACCTCACGAGCAAAAAGCACTCACATCTCCGGAGGGATCGAATATGAAGCATGCATGGAGACATGGAGTCA
ATTCTAACCTTGTGCTGGCGTATTCATAGAGACGACCGGTGCTGGAGAAGATGACCAACCCGACCTCTGCGTCGC
AGAGGATGGCGAGCTCCCTTGCCTTCTTGAAGATCCCGTTCCGCCGCTTGGAGAACGTCACCTGCCGGCTCGTGG
AGTTGTCGATCCGGCGGATCACTATCTTGCCACGCCCCATCGTTAAGCTGCGTCCGGTCTCACACGCACACACGG
CGCAAGCAGACTTCACGGATAGTAACCGAGAAGGAAAGAGGGAGATGGGAACTCCGGCCAGCAGGAGCCCGGCCT
GGAAGGAGAAGGGTTACGAATTATTGGAGGCGAGAGAAGAGGACGACGGGATAAGAGAGGAGTCTCTTGCTGGTG
GCTTTGCAAGTCGGCATTCGTTTAGCTGCTTTCCTTTCCTTTATTTAGCAAGGAAATCACGGAGACAGATTTGCC
TCCCCATTGGGCTAGGATCGCCAGATGGGCTAATTTATTCGAAATGCTAAACTAATGGTGGTAACCAAGCTAAGA
GGGTTTGGTAGGTTCTGGAAACAATGAGAGGTTGCCTGAAATTGGATGCACGCACAAGGCACAAAAGGTTATATA
TTAAAGCAACGCGCTTTGCCAAATGCGCCATATCTTATGCTGTGCCTAAACGTCACCACGAAATCTTaggtagtg
tttggttactagggactaattttagtccctttatttattccactttagtctataaattgtaaaatatggaaac
tataactctattttactttccatatttaacaatttagtgactaaagtggaataaattggagaaactaaaaattag

>ZmMADS61

ATGGGGCGTGGCAAGATAGTGATCCGCCGGATCGACAACTCCACGAGCCGGCAGGTGACGTTCTCCAAGCGGCGG
AACGGGATCTTCAAGAAGGCAAGGGAGCTCGCCATCCTCTGCGACGCAGAGGTCGGGTTGGTCATCTTCTCCAGC
ACCGGTCGTCTCTATGAATACGCCAGCACAAGCATAAAGTCAGTGATTGATCGATATGGTCGAGCAAAGGAGGAG
GAGCATGTAGCAGACCCCAACACAGAGCTTAAGTTCTGGCAAAGGGAGGCAGCAAGCTTGAGACAACAACTGCAC
AACTTGCAAGAAAATCATCGGCAGTTGATGGGACAAGATCTTTCTGGACTAGGTGTCAAGGGACTTCAAAATCTA
GAAAATCAGCTAGAGATGAGCATTTGTTGCATCCGGACAAAAAAGGACCAACTCTTGGTTGACGAAATTCACGAA
CTGAATCGAAAGGGAAGTCTCATCCAACAAGACAACATGGGATTACACAGAAAGGTCAACCTAATTCGTCAAGAA
AATGCTGAATTATATAAGAAGCTCTATGAGAAAGAAGCAGAAGGTGAAGTCAACCGAGATTCAACAACTCCGTAC
AACTTTGTAGTTGCAGAGGGTGCCAACGTTCCTATCCATCTTGAGCTTAATATTCCACTGCAAGAAAATGGTGTT
GAGCAACCTGTGGCTCCTAAATTAGGGTTGCAATTAAATCAATGA

>zma_miR444c_PUT-2-171a-Zea_mays-18627

CTCTTCTTCTCCCCCAAGGCCCCAACCATCCCACTCTCTCTATACAACAGCCACCCCCGCGCCCTTCCGCCCTTG
TCGAGTCCAAATCCAGGTTATAGATCCTCTCGCGCCGCCGCTCGCAGCCGCTCGCCGCCCGGTACATCTCCCCCG
CGCCTCTCCCTGCTCCTGGATTTCACCGATCTGGCATGCTTTACTGTCCCGTCATACGGTGGCAGCGGGAGCCGC
CCCCTTCCTGCCTCCACAGCTCATGCCATGCATGGCCGATCCATTCCCTCAGATCGGGATTGCCGCATACATGCC
GTTGCTTCCCCGTTAGCTTCTCCCTTGTTACACCCCCTCGCTTCGTTCTGCGTAGTTTCGGGGTTTGACACCGCT

TGTCTATCCCGTTCCTATAGCTTTTAGGGTTTCGAGCTCAAACTCCGCGCCGGCGTCTCAGTTGCTGCAAGTGGC
GGCAGCAAGCTAGAGGCAGCAACTGCATACCTTGCAAGAAAAAATCGGTTTTGTTGACGTACCGATGATTTTCTT
GCAAGTTGTGCAGTTGTTGTCTCAAGCTTGCTGCCTCCCTTTGCCAGAACTATTGAATGTGAAAGCACAAGCATG
TGAAATCCTAATGACGCATGCGTCTTGAGGGACGAAACAGGACAAGTCAGAGATTGGTGATGAACTTGCTGGAAC
AAGCTTATGCATGAAACTGCTGATGGTTTTAACCTTTTTAGGATTTAGCTAAAGGCTATAATAATCTTCAAAGGT
TCAATATAAATTCT

#### Stem-loop structure

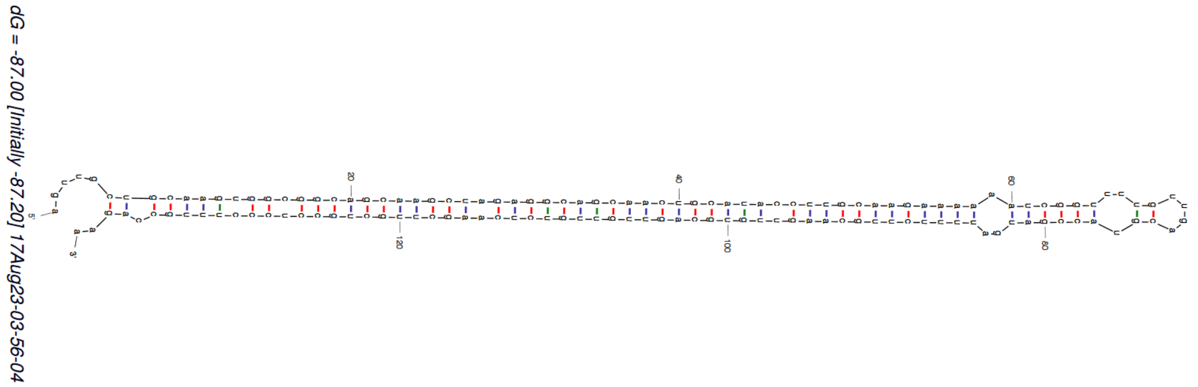


*ZmMADS57 and zma-miR444d*

>5

GTGAATGCCAAATTTATTGATTTCTGATTCACTCCATTTCATTAAGAAAACTATGGACCGTGGAATCTTTTGATT
GAGCAATCACATCGTGCTCGCTAACTATTTCAGCCCTTAGCTTAGTAGCCCCCTCTTGGATCAATCCTGGATCCG
CCACTGATTAAAAGTGTTGTGATGTAATCTCATTGTATTTGTCATTAATTGAGGAGAGCGAGGAAAAGACTAAAC
GGTGATTTTTTTGTTAACCTATACAACGTGGTGAAAAGGCTAAAGGATTGGTTGATTGTAGTGTAGTCCAAAGGA
ACGGTGATACACACGATCATTTATCCTTTCTCCACCAACCCCATTCCGGACTGTGCATACGGCGGCCACAAAACA
TCTTGAGAGAGAGGAAGGTAGGAATGCAAGTGACCCAACTAATATAATTATCTCTCCCGGCAGACACTTTGCGTA
TCATTCCACATGTCACTACCATACCACTAAGCGCAGAAAATTCGAAAATCAGGTACTACAAGTTCCAGAACCTTC
GAGGTTCCCAGGTCACTCGTAATATCGCCAGCTTAGTTTAGTCGCAACGAGGGGGGCCAGCATATAAAAAGCCGG
AGCCTGATTGGACCGAGCACTGCG**CTCCCACGCTCCACTCCCGACACCTGTTCCGTTTAAAATCTCCGGCGTCGA**
**CGTGGCCGC**GTGCGCAGCCACAAACCCTCCCTCCCGCCGCAATTAACAATCCGCTAATCAGCTCGGCGNNNNNNN
NNNNNNNNNNNNNNNNNNNNNNNNNNNNNNNNNNNNNNNNNNNNNNNNNNNNNNNNNNNNNNNNNNNNNNNNNNN
NNNNNNNNNNNNNNNNNNNNNNNNNNNNNNNNNNNNNNNNNNNNNNNNNNNNNNNNNNNNNNNNNNNNNNNNNNN
NNNNNNNNNNNNNNNNNNNNNNNNNNNNNNNNNNNNNNNNNNNNNNNNNNNNNNNNNNNNNNNNNNNNNNNNNNN
NNNNNNNNNNNNNNNNNNNNNNNNNNNNNNNNNNNNNNNNNNNNNNNNNNNNNNNNNNNNNNNNNNNNNNNNNNN
NNNNNNNNNNNNNNNNNNNNNNNNNNNNNNNNNNNNNNNNNNNNNNNNNNNNNNNNNNNNNNNNNNNNNNNNNNN
NNNNNNNNNNNNNNNNNNNNNNNNNNNNNNNNNNNNNNNNNNNNNNNNNNNNNNNNNNNNNNNNNNNNNNNNNNN
NNNNNNNNNNNNNNNNNNNNNNNNNNNNNNNNNNNNNNNNNNN**GCGTGCGCAGCCACAAACCCTCCCTCCCGCCG**
**CAATTAACAATCCGCTAATCAGCTCGGCGCAGCCCGAACCCTCCCTCTAATCGCCCGCTAAACCGCTCCCTCCCT**
**CCGCCAAAAGGCCAGTCCGGGGGCTTTTCCTATCTCCACTTGTTGGCGACTTGGTAGTAGACTCGCGGATACGAT**
**CCTTCTAGACACTTCGACGCGCCCGTTATAAAGGCCGGCGCATCCGGGCCATGACCACACAGCCCCTTGCTCATT**
**GGCTAGGGCGGGGAAGGAAATTGCGGGAGGATGCGAAG**GTACGAGAAGTTTCTAGACGCCACTGTTGTCTTCTCT
TCAATCTTCATCCATTCCTTGTGAGATCCGCTAGAGAGACTACGAGTTTTCGACGCCATTTTTCCTCTTCGGGTG
TGGTGTGATGAGCGGTGAGCGCGGCGGGGGTGCTGCAAGGGTGTTAGATGACTTAGATCCGAGGTTTGGCTCTGC
TATCCTGGTTCGAACGTGGCTAAATCTTCCTGTACGAAACCTAGCTGCTCAGCCCGCCGTCAGATAAGCTCCAGT
CCACAGTCGGATTTCAACTCAAATTTTGGCCGTCCGACAACATATATGTTTGCTAAATCCTTTAACGCCAGTCAG
ATTCTGACTCGAGTTTCGGCCGTTTGACAAGATATGTAACTGCGTACTTTGATGCCAATGAGCTTGCTCAGTCCT
AGTACCGTTTAAACCTGTGAAACCAATCACTATGATGTTGGTAATTTAATTATCGCATTTTGGTGATTTACCCCT
TGTTTGTTAGAAGTAGCACGCACTGGTTTACCTTCGTTGATTCATTGTCAGCATCAACGTTTAAATAGCTATAGC
CAGATTTAGCCTCTAAGAGGACTGGAGGCTAATCTAATCCTAGTGCTAGATGCTACTTGCCGCTATTTGCTAGAT
TGAGCTTGCTAAATCTGCTAAATAGCACTTAATTTAGCCCCCTAGTTAGGGGGGTATGAACTGATGTTTATGTCG
GGGACATATCCTGAGAAATGTTGGATATATATCCTGTGCAATCACTCATTTGGTGCAAGCCTAGAATCAAACCAT
TCAGAGTATCCCATCTAGATACAATGTTTTCATATGAAAGGGGTTAAATGTAAAATATGTACTAGATTTCAGGTG
GAAGGGATTAAATGAAAAACAACAATCAATGTTGGATTTTGATTGGATGCACTAGATCGCTTGATTGAATACAAA
TATGATTGATTGCACACGATATGTTTCCGTGTTTATCAAACTACCAAAAAATACTCGGCTAGTAGGGGGAAAGGC
TGTCCCCCACGATATTATATTAAGAAGAAGCTCAATCGGAGCTTCGACCGAAAAAGGTCATGAAAGCTAGCCCCC
TGTGCACCATGAGGGTCCGCCATCTATAGGCTAGCCTAGCCCCTACATGAGGATCCACCCCCTACAGGTCAGTTT
ATCTCGTGTGCACACAACCAGAGGAACCAAAAAGTGATCTTTTTTAATCTTAGGCTAAAATTTGCTTCCACTGGA

ATTCAAACTTAGGACCTAAGGAGTGCTACTTTGACCATCTAACCAACTTAGTTATCATTGGTTTGTTTGTGATAT
GTATCATGTAGGAATTGATGTGTTTGTCAAACTATCATTTGTTAGTTGCTATTTGTGATGTCCAATATATTTTAT
ATTTATGTGTAAAATTTCTTGATATTAGACATGTCTTTTGCTAAGCCCCTAAATGGGCTTAGCTGGCTATTTTGT
CTGTTATAGCATTCTTACAGCAACTCTAAGAGTGCATCTATATCCTTCCTTGATTTTGAACTCCAAATATTACCA
TTCTCTATTCTGAATTTCATCTCAACCAAAAATGGAGAGCGGCATTGACTTTTTAGAAGGAGTACAATATATTTT
GAATTTTTAGCTAACCAAAGATGCATCTCTGTAGAGGGAATACAAGATATAAAAAAACTGTTAGGGTAAAAAGAT
ATGAAACCGTTTTTTTATTTATAGAGTGAGTTTAAAAAAAGAAAGACTTCTGCTCTTGCCTCTTGGAAAGCTAGT
GTCTAAAAGTCATAGCCAGCTATTTAGAACATTCGTTAGCATAATGCGTTCGCTGCAGCTATTCAGCCCCAGCAG
CTGCTGCAGAGAAGGCCTACCGAAGCTATTCATTTTTTAAGCAGTTCAAAATTTATTTGCTGGCTGATCTAACAT
TTGTTTACTTTTTCATTCATTAG**CCATTGGTGTATTGCATATGGTGGCACCAAGCATGAGGCAGGAACTGCATTA**
**CTTGCAAGAAAGTCACAA**GTACACTGCATTACTTGCAAGAAAGTCACAAGTAAGAGACGAAAGCTTGTAGTTCAT
TTTAAGGTGTTCTATTGCTGCCTCTTTGGTTTCTCCAACCTCTTGTTCCACAAATAGCATTGTAGATTTTATATA
CTCCTGTACTGTGATCACAACCTTTATGTGCCTGGAGTTCACCATTATGGACCTTCCTCCTCCTGTGTATATCCT
GTACAGCTCAGGATTTGTAATTTCCCTGATGATAAAGCATGAGTTTTTTCCTAAACTATGTGGGACGAAAAAGC
AAACATGATTTCTGTTGAAGTGCGCTTCAACTAAAAAACTCGAACCCTTGCCCCACCCATGTGAGAACGACCGCA
ACCAGACAGACGAGGGGATTTTCTTTGACCACAGTCTGAAATTTACTCCCACGGGGATTCGAACTCAGGACCTCA
GGAGTGTTAATTAAACCACCTAACCAATTTAGCTAGAGGCCCTTTCGCAAGTGTGCTTCAACTAGAACTCATGAT
TCTTTCAAATAAAACTATTAAAAGAGCTGCTTAGGATTATGGACCAACTAATAGGATTTATCAAAAAAATAATGA
GTCCACTTTCGTGCTGGGCTGCTAGCTGTTGGGATGCTATCTGACCCACAGCCTGCCAACCTCCCTTACCCATTA
GAGTTCCAGTAGCANATTAGAGTTCCAGTAGCATTAGGAATAAATGTGCATCCAGTTTCAGTTCCAAACCATGAA
ACTTATACAATTCCTTATTGCAAAGTTCAATCATCAGTGAAGCAAATCCATATACATCATGCAAATGCCAAGTGA
AACCCAATACACTTTTGATGCAGAGGAGCCTCATGCCCTCATGTAACATTATTGGTCCTACACATTTTGTACACA
AACTATTTTGCTCATACAGCTGGTTATATATCATAAATATTCATGCAACCGGAGTGCTCTGCATTGTGTGCAATT
GGTGAAACACAATGGTATTTTCCACCTTGTACTGCATCTTCTTAGGGCAGTTGAAGTCTGTAACATATCAAAGCA
TAGATTTTTTTAGTATGCTTCTTCATGCTTGGAGTAACTCATTTTTTTATATTGGAATAAGAATAGTTACCCCAG
TTCAGGAGCCATAAATTCTTGGCACCGATTTCCCTCCTTTTGTTGCGAGTGCCTCTGTTCAAGATTAGCAGGGGC
ATCTGCTTGTTGTGTTGCAATACAAAATCTGTAGGGAGTGCTAGAGCTTGCACCAGCAGCACCTCTTGGTTCACT
GGCCTGGAGCTAAAATTAATTCATTTAGCTTCCTGTTGACAAAGCATCTTTATTTAGTTAAATTTTGTGCAAAGA
TCATTCGTCAAACCTTCCTATATAGTTCCTCTTTTTGCTGTGACATGAAATGTACTCTTCTACTGAGCTCCATGT
TTTCCTGGTGAATTAGGCTGCCCTGTTGCGATGTCCATGTCAGTAACTTCCATGCATGATCATCCTTTTGAAATA
AGAAAAAGGACATTGCTTTTAGGGATTAACCTTCCTCTGTAACACTTCAATTTCACTTTTCAAAGATGGTCCTG
TTTGACATTATTGCATATTAGATTGAAGCGAATAGAAAACTAAAACGTGAAAAATAAGTGGATAATGAATGACCT
TCCTCATTCTGATACTATGCAGGCTCATTCCAATCTGGTTCTCTAAACTTTGGAGGTCTCTTACACCTAGGCCTG
AAAGCTCCTCTCCCATTAGTTGCCTACAACATTGGAAAAGAGTTTCTCACATTATCTTAGGTAGGATACTAAAAC
TACCCAGCGTAAAATTACATGGGTACACAGGTGGAAGCAAAGGAACTTATTATTTTGTTAGGAGAAAGTCACTAT
CAGTTTCATGTCTTCAAATAGTGAATGTTGTTAATGTCTTCAATACTGGATATTTATTTCATCCTTCCCGGAAGA
GTTCCGTTTTGTGTATGTTTTGAATGTTCTTATGGAGTCCTTCAGGTGGCATAAAAAACTTTCTATTGATGAGTG
TTCCTTTTTTGTTGCAG**AATCCATTGCAAGATCATACTTGTGGCTTTCTTGCAAGTTGTGCAGTTGTTGCCTCAA**
**GCTTGCTGCCTCCCTTTGCCAGAGCTATCGTTAAAACAGGAAGTACTGTCAGTTTCCGAGCTACAATCAAGCTAT**
**AAGTTTAGAAAAATAATTTAGAAATGTACAACAAAGTCTA**GTAATACATTGCTAAAGATAAATCAATACTCTACA
AACAAGCTATAAGTTTAGAAAAATAATTTAGAAATGTACAACAAAG**TCTTTTAATACATTGCTAAAGATAAATCA**
**ATACTCAATAAGTTCCTTATGCTTTGGATTTTGGGATAACATACCTGTTTTATTTACATGCTATTTGGACCAACA**
**AAAAATACCTTAGGAACAGCCATTTGAGAAAAAAAACAAAGCCTTTTAGTCCTAAGCAAAAAAAACTTTTAGTCC**
**TAAGCAAGTTAGCATAGCCTAGAGTTGCAACTTCTACATGAGCACAATAAACAGACAAGAAAGAGAAAAGGAAAA**
**TAAATATATATGGCACCAAAAGCTATTACCATGTTATAATATAAATTTAATGACCGAAATTATAAACATTTTTAC**
**AATCTTAGCAAACCTCTCAACGTGTAGTCTGAATCGACAGAACATCAGTAGAGTAATTTTTGTTACATTTATAGT**
**CCCCAATGTCGTTTCAAGGCAAAGTTTGTTTGGTTTCTCTGTTTTCCTAAACATTGGCTTCTCAATGTGCATCTG**
**TACTAGTTTGACATCTGTGGCTATTGTGATTCTTCCTGGAATATCATCTGTCCATCACCTTTAAAGCATAACATT**
**ACCCTGATGGCCATGTGATCTGTAGGTCCCTGGAAAGGAAAATCGTGATTTGGAAAGGAAATCCATCAACAAACC**
**TGAGTCGGAAGGATGCTTAGGAAGCGG**GTACTCTCTATTTTCCAGATTATAGTTCATTTAGATTCAATGTAAGCT
TAACTTCTCTAACCTTGATAAGTTTATAGAAAAATGCACTAACGCATGCATCCTCGAATTAGTTTTACTAAATTT
CCATGAAAATATCTCGATGGTGCATTTAATTTAACTTATAGATGCTATTTTTTAAAACTTTGGTCAAAACTAGAG
AAGTTTGGAGCATGCTATACTGTCTGTCATGTGCGTGTAGTTTCCATGAAAAAATAGCATCTAAATTTCCATGAA
AAAAGCACTAATGCCAGCTTTGCTTGCCTGTCCCTACCAAGCTAGGCCAGCCTTGCCAAGTGCGAGCCGTGGGGA
AATCTTGAAATCACAATCAAAGACCAACGGCCAAAAAAAAACCAACCGTGTGCCAATTTTTTCACCTGGCTAGGC
AAGGAGAGGGAGCGGCAAAGGTTCCAAATGGCTCTTTAAAATCGACGTAAACTATAATTTGGAACAAAAATGAGT
GTATTGGTGGGCTTCGACTCCTGGAGAGGAGCCTTATCAAATGGTTCGGTAGAAATAGTTATTTACGGCCCTTTT
TTCATTAAAAAGGGAGGAGCTAGAGCCATTTGAAAAACTCTGAGGGCCTGTTTGGCAACAATGTTTCACACATGA
TAATACCGTAGTTTTGGCAGCTCCAAAAATAACATGGTTTTGAAAACCATTGTTTGGAGACAATATTATAAACCA
TGGTATTAGTCAAAAGCTAGTTATGCCATAAAAACTTTGGGTTGGAGTGGAATCTCCAATACTCTAAAAATACCA
TGGTATTTAGAAACCATGGTTTTAAAAACATAGTTTTCAAGAGTGCAACTAAACACCTCATGGCATAAAATACTA
TGGTATTGTCAAAACCATGGTATTGCATAGAAACTATAAAAAAAAATACTTTGCTCTCACACAGACCCTGAAACT
CTTCCAGCTCCTCCGAAGAAGCTTTGAGAAGCCCTACCAAACAAAGGCCTAATAAAGTTTCTCCAGGTTGAAATG

ACAATCTTAGGGCCTGTTTGGAAATACGGTTTTGAAATACTGTAATTTTGAGATACCACAGTTTATAATTGTACA
TGACACAAATACTACGGTATTACTTTACCACAGTAAAACCACAGTATTGCTCAAaaccgagatatgtttggttcc
attggaaaaacaaagtatatgtagagagaagagaagaaaactgaggtcccGAgtAGAGTTTCAAAAACTCCAAAA
ATACCACGGTTTTGGATAAACCATGGTATTTAAAACTGTGTTTTGCCTGTACAAACCAAACACCTTTTGAGTTTG
AATActatagtatcatcaaataccatagtattgtttCAAAACTGTAAAAATACCACACTTCCAAACAGGGCCTTA
TACTGTCTGTCATGTTGTACTATAAAGGCCATGGTTTTTCCATTGTTTGGTATTTTACTGTCCTGTCAACAGACA
ATTTCACTTTGAATTTTAAAAAGTTTTGACATAAAAATTTTAAATGGACACTTTGCCATAGCAGGTGTTATCACT
GTGCATACTCTAAAGCATAGCGGTTAAGATAGTTAACAAGCTTGGTGATATTCAATTAATCATTGCTGACATATT
TTGTTGTTAACAG**GATTCAATCAATTATTTCTGGGGTGGTCTACAGCCAATGCATAGTCTCTATCTGTGTGTGTG**
**TTTGTGTTTCCCGTACACTCTGTTCCTTTCGAGAGGCACTCGTGCCTGAAATATGGTTTAGCTTCAGGCAATGAA**
**GTGGACGTCGGATGGAGATCCGAGTGTCCACGATGATAGAGACCGGGTTTTGAATCTCTCGTCTCTCCATCTCTC**
**ATGCGTTCCCAACTTCCAACTCCAATGCCGTCTCCGCTGGCTGATCAGTGATCACCTCTGACCCTCTCTGGTGAT**
**GTGTACCCATTGTTAATCTCTCCAATGAAGCTACCCTGCCTGACCAGGCCCACAGGCTGTCTCCTGCATACGGTG**
**GCAACACACCAACACTGCTGGCCTGCCCTAAGACACATCTTCGCGTTGCGTCCACTGGATACCTGCTTTCTTAGT**
**CAGCAGCATGCCGAAACCCTCCATTTGTGAAACCTCAG**GTGCGCGTAGTTTAGGAGGTTAGATTTTACTTAATCG
ACCTTCGCATTGGATAATATGCCTAACATGGTGACTTTGGATCGCTTGTTTTCTAG**CGGTGATAGGCGGTGGTTG**
**TCGAACGCTGGCATGAAGGTGTGAGCTCCCAACTGCTGAATCGTGTTCCTCTTTTACATGCACGTGTCCAACAAG**
**AACCAAGGTGGAACTGTAAAGCCTTGTGTCTTTGAACTTTGCAGCCACGTATTCAGATTTTTTTTTGAGTTATTG**
**TAATTTATTTACATTCCCTCTAGAACCGAGATTAGCGCTACCTCAGTACACTGCCGGTGCAAGATAAACGAAGGA**
**AGCTGGGATTGCATGGCACTGCGGTCTGTTATGTAGCCGTGGAACAGCAAGACAAAAAGACATGGATATTCCTTG**
**TTTGGATTGGGGTGTTTGATCCAGAAACAGTCTCATTGCTAATTGTTTCATTATTACTACAAATCTTGGTCGAGG**
**CGTGAATATCCTTGCTCTTGATATAATTATAGTATTTTGAATCCCTATTGTGTTTTTTACTTAAGTGATAAATGT**
**TGCAAAGAAACCATTAGATAATTGTTGAATGATTAGTTATTGATTACATTGATTGATTTTGATTTAATGAATCTT**
**AAATAAACCAGAA**GATTCATGTGTATGAATATTTGTGTTGGGAATTTCTTGGTCACATGTTTTCCGCATAATAAT
ATCATAAAACTGTGGAATGAAACAGCTTTTGCATATTCCGCTATAACTTTTTATAGCACCCCTATCAAATTTATG
GGGTGTTTGGCATGCAAAACTCGAGTCAAATATTTCAACTCAAGAGTTACAAGTTTTGTGGAGTTTCTAGGAGCT
CTATTACAACTTTTTGGAGGGTCTTTTTGTTGCTGGTATGTAATCTTTTAAAAAGCAACATAGATATAGAAACTA
TCTAAACAATCATTGCCACTAGTTATAGACTCTAGAGCAAAGCTGTTGTACTAAAAGTTTATAGAGTAGAGCCGC
TCTAGGGTGGAGCAGAATCAATGCTAAACACGTCTTTATTGTTATTTTTACACTAGTTAGATTTATTTTGAAAGG
TTAGCTCTAGTCTCTTTCTATGCTAACAAATTTTAACAATATACCAACTTCTCCCAATAGAAAGGGAGCGTCTTA
GCAAACCGGGTCCACTTGGTGACTGCCCAGCCGGACCTGGACATCTCTAACCTCTTCCTTTATTTTCCGGCACGT
AGTTAGATGATATATTTGTGTGCGAAGTTAGCTTTTGTATGTGATTTCTCGTGAGAATTCCAAATTGTTTTGCAG
ATGTCCCCCCTAACGGGTAATGGTATTTGGTTTTTGGCTAAGAGCATCTCTAAGAGAGACTTTAAACATGGTCCT
ATTTTAAATATAGAGTTAAAATCATAAAACACTCTTCTAACCGTAGCCCTATTTTATAAAATTTCATCAAATAAT
TATAGGGCTCGGTCTCTCGGGTCCTTCTCCGAATCCTGGCCTCCCATCATTAACGGATAGGTGTCAACCATCCAC
ACAGGATATCGTAACCCGAGATCGAACCCAAGATTTGGAGGTGCTACTCAGAAGTCATTAATAGGGATGTAATAT
GGTTCGGAAAATATCCGTCCGGATCCGGATTCGAAGATGGTGAACACTggttcggatggaattttcggatatccg
gactttttttcggataacggatacgaatacggatattttgttcggatatcgaattcgaatacaatatgtctgata
tccgtcggatCGAATATCCGGATACTTTTTCGGATAGATCTTCTCGGATATCCGGATATCTTATCCGGATAgTTG
TGTGCATTTTGTTGATTTTTTGAAGAAAGAAATTAATAATACACAAATGGCCTAAAAAATTCATGAAAATTTATG
GAGGCATACCATATGTCCACATATAATCCTCCAAAATATttgtaccataaaatccacaatatgatagtgtttctt
tcatttcactttcacttgttgtgtgaattacacgtgaaatcactctaattatccaagttcactttatcattttca
tgtggagacttaatgttatattcttatagaatgtttattagttttggtaacttatttgcatttttgatgattttat
acaaaataaattaataacacgcacataggctaaaaattcatgatttttacaaaaaacgtgacatatgtacacata
taatactaaaaaatgtttgTACTCAAGGAGTGGATACATGATtacaaccatgatatgtgtggaatgatgtcttac
acgacatccgacaaaaaaattcgttaccgacaatatccgtgtccgactagttccgtattcgacaaatacctatccg
tatttgtatccgagaacatccgtattcgtattcatatccgaagctatccgtattcgaattcgaatccgaataaaa
atatgaaaacaaatatgatttcagtgatatccgtccgtattcgatccgattacatccctagTCATTAACCACTAA
GTTAGAGGACATTTCGCGATTGAATAATTAATCTTACACAAAAAAAATAAGTAAATTATGATAAGTTAGACAACA
TACTAAATAGACCTTATCTATTGTACTTCTTTTATATTTCTCACCACACGTCTGATTTTTTTTGTTATTCAAAAA
ATATATAATTTTTCCATTATCATTTTAATTGCACTAGTTGATCATCTAAAATTGTTATGTGAAATTTTGGACACC
TAAAATATAGCAACTTATGAAAGAGCCTCTAGGCTGGTTAGATGATATGAGCAGTACTTCTTAGGCTTGAGTGTA
ACTTTATGTAAGAGCAAATTTTAGATACAAGGTTGATTGAGTTTAAAAGCAAATTATAGGTTTTTAAACTCTATC
GTCTCCTTAGATTTGAGTTTAACTCTCATACTAGAAGGTTTTTAAATTCGATCAAAGTTTAAAAGGTTGACTTCA
ATCAACTCTCGGAATCTATTTATTCAAGGAATCTACTTATTCAAAGAGTGATAGAAAATGTATATACTATGCATT
CGATGTTTGGTAAAACGATTGGGTGTTTCTAGGAATACCAATGATCCATCCATGTCGCTACTAGATTATTTTTGG
TAGAGCTTTTTATTTGCAAGAAAATTAATTATTGCAAGAAATCAAGAGCAACTAGAATATACACATGGATCATTA
CCGGAATCCGGCTCTTTGCCGAGTGCTTTTTGTCGGGCACTCGGCAAACTCCTGCTCTCGGTAACGGCCGCATTT
GCCGAGAGCAGGACTCTCGGCACATAAATACACTCGACAAAGAAATCTTTGTCGAGTGCTAAACACTCGGCGAAT
GGCGGCGCTCGGCAAAGGGCCGTCAGCAGTCGTCTATAGCTGACGACCATTATCTTTGCCGAGCGCCAAGCTTTG
GCACTCAGCAAAGAAGCTTCTTTGTCGAGTGTCTTACGTCTGACACTCGGCAGAACATGCTTTACCGAGTGCCAT
GCTTGGGCACTCGACAAAGTATTTTTTATTTTTTTCTTTTCTCAACCAAACTTTTTGTGGTATGTTCCTACACTG
TATAGACCTACATGTATCATTTTGTCACAATTATAAAAGTGTTTTCTATAACTATTAGATTTAGTTCGTTTAATT

GAATTTCATTCTCAAATTCATATTTGAACTGCAAGTCACTCGAAACATGGAAAATTGTGCATACAAAAATTATAT
CCATGTTATATAACACAAGTTACGACTGATTTCAGGAACATACTGGAATCTTCGAGCACCATACTCACGAAACAT
GACTGTGAACTTGCCAGCAAGTTGTTTAAAAAATGTATAAAACACAAATAAAGTCAGAAAATCATGAAACTTGTC
CACATGTCATGATATCATATGTAGAGgTTGTGATAAAaTTTTGAAATGTTTCGAGAAAGTTGTGACGCACTATGT
GTAGAAACCTAAGTGACCTACACATGAAATCAtagagtttTATTGTAGACCAGCGAGGTTTCTACACATACTGCC
TCACAACTTTCTCAAAACATTTTCAAATTTTTATCACAACCTCTACATATGATATCATGACATGTGGACAAGTTT
CAGGATTTTCTGACTTTGTTTGTGTTTTATACATTTTTTAAACAACTTGATGGCAAGTTTACGGTCATGTTTCGT
GAGCATGATGCTCGAACATTCTGATATGCTCCTGAAATCGGTTGTAACTTGTGTTACCTAACATGGATATTATTT
TTTCATGCACAGTTTTCCGTGTTTCGAGCGACTTGTAGTTCAAATCTGAATTATCCTAAAAAATTCAATTAAACG
AACAAAATCTAATAGTTATAGCAAACACTTTTAGAATTATGTAAAAATAGAACATGTAGGTCTACATAATGTAGG
AACATACCACAAAAAGATTGATTGGGAAAAGAAAAAATATATATACTTTACCGAGTGTCCTAGGAGGACACTCGG
CAAAGAAGTCTTTGTCAAGTGCCTACGGAGGAACACTCGGCAAAGCTAAGTTTAACAAATAAATGAATCTTTGCT
GAGTGTCAGCCAGAGGACACTTCGGAAAGAAGGCTTTGCCGAGTGCCGGATCGTGGGCACTCGGCAAAGTATATT
TTAAAATTTTTAAAAAAATCTTTATCGAGTGTTAGATCGTGAGCACTCGGCAAAGAACACGTACATAGCACGTTG
CATCGTCCTTTTTCTTTCTAACTCTCTCACTCTCCTCTGAGCCCCACGCCGCCGCGCCCTCGGCCGCCGCCACCT
CCACCGCCGCCCCACGCTGCCACTCCCTCGGTCGCCAAGTCCGCCGCCACCTCACGCTGTCACTCCACACCGCCG
CCTCATCGTTGTCTTCAACCTCGGATCACACGGTGAGAAACTCCACACGAGTATCTATATTTTCTGTGGATGAAT
TAATGACTTGTTGTATATGTAGTTGATTTATATGTGGATTTGTAGTGCATAGATTGAAGTATATGTGGTTTAGGG
TTTAGGGTATATGTGGTTGGTTGCATGTTTTGGAAACCACCCCGTGCATTACATGTTTATATTAAACTATACTCG
GTTTAGGGTTTATATGAAATGATATGGTTTGATATTGAGTCAAATTTATATATGTTAGTTAGCTAGCTAGAGGCA
TCTCATTATAATATAAATGTAGTTTGCATGTGGCTATGTAGTTATGTAGTGTTGTTGTGCTATCTCTCATGGATT
GAATATATATAGATGTGATTGTCGGCCATCGTACCATTGAAATATGCGTGGATGTCAGCCATCGTGTTGATACTT
TTATTTGCAGGTTTTAGAAACCTCCCCGTGCAGGGGAGGTGCTGTCAAAATTTTCTATTGATAGGCTTCGGTTAT
ATATGCTAGAGGATGGAGGACCGTGAGTAGATGTACATGGGTCATGTAGGAAGGAATGATGTCACCCCTGAGTGG
ATTAGAAAGACCGATGATTTTGTGGAACGGGCATATGGCGAAGCTGCTAAAGGAGCGAGTctagtcccatgcccg
tACAACAAATGTGCCAAACAGAAAAGAAAATCAAAGAAGTCCATGGTATAACATATTTGGAAGTATGGATTTACG
CTGGGCTATACTAGGTGGATCTTTTATGGTGAAGCGCATCGTACGAGAGAGGAGGTGCTGAGACAACGCGTCGAG
CTTTATGATGCCGATGTAGGGGTAGTGGATATGTTGAACGACTATCATGAGGCACAGTACGCCGGAGGATGTACG
AATGATGAGCCAGAGCCGACTGCAAAGGCGTTCTACGACATATTTGACGCGACACAGAAGCCCCTTCACAACCAC
ACAAAGGTTTCTCAACTGGATGCCATTGGGCGTGTGCCCGATAAAAAGTACTCGGCAAAGAAGCCGTTGTCGATG
TACTGTTCGCCGAGATCTCTTTGTCGAGTGTCACACTTGGCAAAGACTTTGTCGAGTGTTTTTCAGTCACTCGGC
AAAGCGTTCGATTCCGGTAGTGGATGTACATACTATATACTCAATGTTTTGGTAAATTTATTGGATGTTCATAGG
AATACTAAGAATCTATTCATGTCGCTGCTTAGAGTCTGTTTGGTAAAGCTCTCCCTACTCTATATTCTCTGTAGA
GGAGGCTGTAGAGGAGGCTGATTATATAGGAAGCTGATTCTTTAAGAGAAATGATCCTCTCACTGTAATATGTTT
TTTTGTACATTTCTCTAGGATAAACTCTATGATAAGCTACTTTTTCCGGCTTCTGGTCTTCATTTCTAGAGAATC
ACTTCATAGAACTAGCATAATCACTTGCTAAGAAAAAAACCATTTGATAGAGCTCCCTCTAAATTCAACCTAGCT
AGAAGCTACTCTAGGAGCAATGACAAATAGGCCATTATTGTGCACAACAAATGTGCTACCATTCATTCTATCCCA
GCAAAAGTTTCATTAGGTAACATAATGGTTTTCAAATATAGTTTTGATTATTATTTTTATAATACATTAAAAAAA
TATAGCATAGATTTACTATTATATATAAACTTATACAAATATATTATGAAGTACCTTTTATTAAGACAAATTTAC
TAGTGTCAACTTTAAATATTGAAACTCAATACAAAAAAGTAAATTTATAATCAAAATTCTGAATAGCTCGAGTGT
TTGACCTATGGGTGGTAAGGCAGCTCTATACATTATTATATTAAAAATACATTCTCATGTAGATTGAGAAAACCC
TCGAATCTCTATCCTCATTATAGATGTCCAACGTGTCGTGCCGGGCCGAGTTGGCCCGCGGGCTGGCTGACGCGG
TCCAGGCCCAGCACGATTATTTTTACACGGGGCCGTGCCGGGTCGGGCCCGGACCCGTAAACAAAAAAACACTCC
AAAATTCAGAATTTAAAGGTAATATTTAAATTTATTTTAGATTCAATTAATTATATTTATAATtttagatacata
cttagaTAAAGAATTTCAGATACACATATAAACACATATCACAAATCAGAATGCTCTCGTATAATAACAGAATCA
TCAATATTATAGCTAAAAATAGACTGGCCGTTACAAAATTAGTCCCTCAAACGAGCTGTTAGTGTGCCTACCTTT
TAACCGTGCCGGGCCCGTGCATGCACCGCAGGCCATACAGTCGGCCCAAGCCTGGCACGACTATCGGGTCGGGCC
AGCCCGGACACGCTTCTAGTCGGACCGGGTCGTGCTGGGCCACGACCGTGCCGGGTCACGGGCCCACTGTCGGGC
CGACCCATTTGGACATGTATAATCCCCATGACTGAAATTTGCTTTCACGGGAAGTCGAACTCCGAGCACGAGGAG
TTCTACTTAGATCAACTAACCAACTCAGCTAGATGTACTTTCGCAAAATTTTTGAAAAGCTGATTAGCTAAACAC
AACAAATTGTCATTATTTGTGAGTAATGAAGGACTAAACTTCTGATCTTGGTCCGACCAACATACATCATATGGC
ACGTGTGTACTACGACCAATAACTGCTAGTTAGCTAGAAATAGGTATACATTTTCCTCACCGTACCATGAGTTCT
GAAGTTGCATCGTTCTTGCCAAGCTGCTCGTCTTTTGCCTTCCCATATCGGTCTATCACAGCTTTCATGCTAGAA
TAAAAAGTGGAACGAAACAGTGTCGTTAGTGCGTTAATCTGTATCTGAATTGTAGGCAATGTATGATCAGCTCTT
CCGGCCGGTTTATAATCGTGAATACTCAATAATCATATGTGTATATATAACCTGTTTTGATTTTACACATCGTAG
TTCTCGAGCAAACTCACTAGGATATATATAACTCTTCCCTGTTAAGATGAACATGGAGTGGAAATATTGGTCTCT
CGATCTAGTCTTAATTGTGTAAAGAAAACAAAAACTAATTAAGGAACTCAAGATCCGTCCTTGTTAACTGAACTT
AATTAGTTGAATAATAAAGGATCATGTACACACTAATATTTTTTGGCGAAAAAATAGTAGGCCCCTCAGATATTT
GTTTAATCTGCTCATACACAAGTTAATTATACGCAAAGCAAATTACAGATCTGTTAAAGTTGAACCAAGTCTGAT
CATCGGTAAGAAATTGCATACAGATATATGTGTAGCTAGCTAGGGAGAGTCACGACGTAGATGAATCGTACTTGG
TGTTGGAGAAATCGTAGAGCCTGCCGGTGCTGGAGAAGATGATGAGGCCAACCTCCGCGTCGCAGAGGATAGAGA
GCTCCTTCGCCTTCTTCAGCAGCCCGCTCCGCCTCTTGGAGAAGGTCACCTGCCTGCTCGTGGAGTTGTCTATCC
TCCTTATCACTATCTTCCCCCTCCCCATCCCTCCTTCTTCTGTCTCTCAGATCTAGTAGGCCGGACCGGATGGGA

AGGAATTGAAGGATCACACGGGAAGACGAAGAGCTCGCCGGCGGCCGGGGGGACGAGCGATGACTGGTCTGATGG
GGGAATTAAAGGCGGAAGCTATAGCCTAGCTACTAGAGGCCGGTTCCATTCCTAGCCACCTTCCAAAGAGCGGAA
CGGGAAGTGCCGATGATTTTTTTAGATAATGGGATAGAGTGTTCCCAGCCTACATCACGAGGATGACGGCCTGTC
ATGACaacaccacaccaacagatccaactcaaagagccacaagcggcgatacaaaccccaaacgcagttgggtgg
acagcaaaagtaaaaacaactctcacagagcgatcctattactaaactgccacccgtgaaaagcgaagatctgca

>ZmMADS57
ATGGGGAGGGGGAAGATAGTGATAAGGAGGATAGACAACTCCACGAGCAGGCAGGTGACCTTCTCCAAGAGGCGG
AGCGGGCTGCTGAAGAAGGCGAAGGAGCTCTCTATCCTCTGCGACGCGGAGGTTGGCCTCATCATCTTCTCCAGC
ACCGGCAGGCTCTACGATTTCTCCAACACCAACATGAAAGCTGTGATAGACCGATATGGGAAGGCAAAAGACGAG
CAGCTTGGCAAGAACGATGCAACTTCAGAACTCATGCTCTGGCAAAGGGAGGCAGCAAGCTTGAGGCAACAACTG
CACAACTTGCAAGAAAGCCACAAGCAACTAATGGGAGAGGAGCTTTCAGGCCTAGGTGTAAGAGACCTCCAAAGT
TTAGAGAACCAGATTGGAATGAGCCTGCATAGTATCAGAATGAGGAAGGACCATCTTTTGAAAAGTGAAATTGAA
GTGTTACAGAGGAAGGGCAGCCTAATTCACCAGGAAAACATGGAGCTCAGTAGAAGAGTACATTTCATGTCACAG
CAAAAAGAGGAACTATATAGGAAGGCCAGTGAACCAAGAGGTGCTGCTGGTGCAAGCTCTAGCACTCCCTACAGA
TTTTGTATTGCAACACAACAAGCAGATGCCCCTGCTAATCTTGAACAGAGGCACTCGCAACAAAAGGAGGGAAAT
CGGTGCCAAGAATTTATGGCTCCTGAACTGGGGTAA

>zma_miR444d_XR_002262532.1
CCTCCCACGCTCCACTCCCGACACCTGTTCCGTTTAAAATCTCCGGCGTCGACGTGGCCGCGTGCGCAGCCACAA
ACCCTCCCTCCCGCCGCAATTAACAATCCGCTAATCAGCTCGGCGCAGCCCGAACCCTCCCTCTAATCGCCCGCT
AAACCGCTCCCTCCCTCCGCCAAAAGGCCAGTCCGGGGGCTTTTCCTATCTCCACTTGTTGGCGACTTGGTAGTA
GACTCGCGGATACGATCCTTCTAGACACTTCGACGCGCCCGTTATAAAGGCCGGCGCATCCGGGCCATGACCACA
CAGCCCCTTGCTCATTGGCTAGGGCGGGGAAGGAAATTGCGGGAGGATGCGAAGCCATTGGTGTATTGCATATGG
TGGCACCAAGCATGAGGCAGGAACTGCATTACTTGCAAGAAAGTCACAAAATCCATTGCAAGATCATACTTGTGG
CTTTCTTGCAAGTTGTGCAGTTGTTGCCTCAAGCTTGCTGCCTCCCTTTGCCAGAGCTATCGGTAAAACAGGAAG
TACTGTCAGTTTCCGAGCTACAAACAAGCTATAAGTTTAGAAAAATAATTTAGAAATGTACAACAAAGTCTAGTA
ATACATTGTTAAAGATAAATCAATACTCAATAAGTTCCTCATGCTTTGGATTTGGGGATAACATACCTGTTTTAT
TTACATGCTAATATTTGGACCAACAAAAAATACCTTAGGAACAGCCATTTGAGAAAAAAACAAAGCCTTTTAGTC
CTAAGCAAGTTAGCATAGCTTAGAGTTGCAACTTCTACATGAGCACAATAAACAGAAAAGAAAGAGAAAAGGAAA
ATAAATATATATGGCACCAAAAGCTATTACCATGTTATAATATAAATTTAATGACCGAAATTATAAACATTTTTA
CAATCTTAGCAAACCTCTCAACGTGTAGTCTGAATCGACAGAACATCAGTAGAGTAATTTTTGTTACATTTATAG
TCCCCAATGTCGTTTCAAGGCAAAGTTTGTTTGGTTTCTCTGTTTTCCTAAACACTGGCTTCTCAATTGCATTCT
CAATGTGCATCTGTACTAGTTTGACATCTGTGGCTATTGTGATTCTTCCTGGAATATCATCTGTCCATCACCTTT
AAAGCATAACATTACCCTGATGGCCATGTGATCTGTAGGTCCCTAGAAAGGAAAATCGTGATTTGGAAAGGAAAT
TCATCAACAAACCTGAGTCGGAAGGATGCTTAGGAAGCGGGATTCAATCAATTATTTCTGGGGTGGTCTACAGCC
AATGCATAGTCTCTATCTGTGTGTGTGTTTTGTGTTTCCCGTGCACTCTATTCCTTTCGAGAGGCACTCTATGTC
AGGTGCCTGAAATATGGTTTAGCTTCAGGCAATGAAGTGGACGTCGGATGGAGATCCGAGTGTCCACGATGACAG
AGACCGGGTTTTGAATCTCTCGTCTCTCCATCTCTCATGCGTTCCCAACTTCCAACTCCAATGCCGTCTCCGCTG
GCTGATCAGTGATCACCTCTGACCCTCTCTGGTGATGTGTACCCATTGTTAATCTCTCCAATGAAGCTACCCTGC
CTGACCAGGCCCACAGGCTGTCTCCTGCATACGGTGGCAACACACCAACACTGCTGGCCTGCCCTAAGACACATC
TTCGCGTTGCGTCCACTGGATACCTGCTTTCTTAGTCAGCAGCATGCCGAAACCCTCCATTTGTGAAACCTCAGC
GGTGATAGGCGGTGGTTGTCGAACGCTGGCATGAAGGTGTGAGCTCCCAACTGCTGAATCGTGTTCCTCTTTTAC
ATGCACGTGTCCAACAAGAACCAAGGTGGAACTGTAAAGCCTTGTGTCTTTGAACTTTGCAGCCACGTATTCAGA
TTTTTTTTTGAGTTATTGTAATTTATTTACATTCCCTCTAGAACCGAGATTAGCGCTACCTCAGTACACTGCCGG
TGCAAGATAAACGAAGGAAGCTGGGATTGCATGGCACTGCGGTCTGTTATGTAGCCGTGGAACAGCAAGACAAAA
AGACATGGATATTCCTTGTTTGGATTGGGGTGTTTGATCCAGAAACAGTCTCATTGCTAATTGTTTCATTATTAC
TACAAATCTTGGTCGAGGCGTGAATATCCTTGCTCTTGATATAATTATAGTATTTTGAATCCCTATTGTGTTTTT
TACTTAAGTGATAAATGTTGCAAAGAAACCATTAGATAATTGTTGAATGATTAGTTATTGATTACATTGATTGAT
TTTGATTTAATGAATCTTAAATAAACCAGAA

#### Stem-loop structure

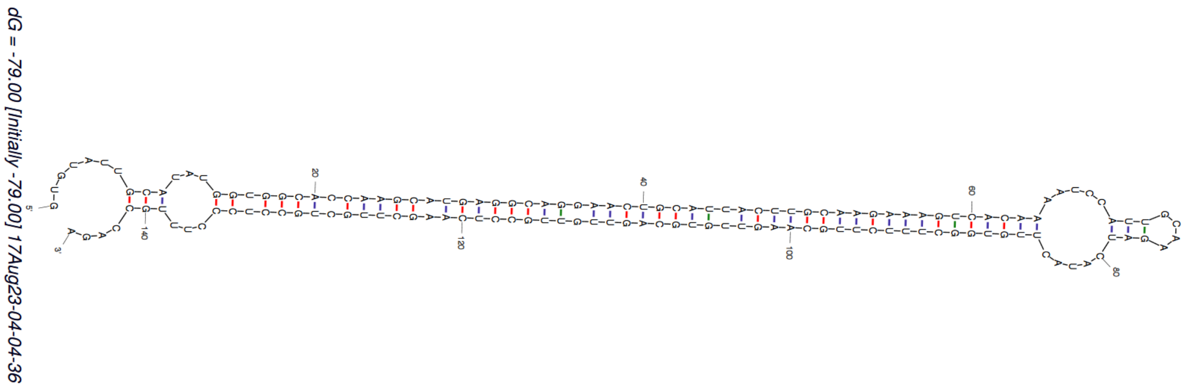


*zma-miR444e*

>1

aaattttgaaattacataaacaggtttgaaacacaataattttaactactaatctctttttcgtagctcagattg
tttgaataatacaccactgtatatatttactgaacatagtaatatatgctttgggtaaatctggtaatcactaa
atcataaagaataaaaaacaatatcctccactaccttctgcttgtatagagcacaaaactaatattacgtataga
aacctcagcaacattagtatttcattatttttcagcaatgggggaaaactcaaccatacattagcagctagctag
gtgagtaacaatctgatagattatattccttctgttttaatttataggtcgttttggctttttaaatacataac
atttattatatatctagacatatcttatatctagttgtatagtaaaagctatgttataaaaaagctaaaatggtc
tacaatttagaatagagggagaaaattgcattcttgcatcattgcttctgtttcatctcatggtaaatataggat
acatggctatggaaggtttctatacctatacggatgaacttcatcttattcctaca**ggtttggcaaagggaggca**
**gcaagcttgaggcagcaactgcataacttgcggtaaaatcatcagcacaccatggtccaac**ctgtagcaaataaa
tcatgctgctcatgaataaacacactcttttggatataccttaaaaacatcatcaattcaaaatacatttctcag
gcatagatttatactgcctgcaaggtttcgacttaacccatgggagttcatgagcatgtgttactcaggaagaag
aatggatatactagatcattaacatcacctacagttcgttacttcaatataaactttagagtaaggatgctgcaa
ctattcagatagtctgcttgaatcgtttactgataccatcatatatcagaggatacacatcatcacaagaatgtt
actagttttctttagtgcatatcatatcatatagaaattcatcctatggctcctttatgcattgtgtatatgagc
aaggttcgtgtgctctggtcagcaagagctccaatacatgtagagatattaaaaattgccataacattattatga
gaaccaaatctacggctacctttgacaatacattgtttgagtattggatagattgtttagttaagacttagtaa
tgtactccagtaatttatcagggctatcgtagagagccaatgccgatgctcaggaatttttgaaacttaattgttc
agaaaagagcactgcaataataatggaccaatagtcaaattgatgttttacttttccttaaaacctacctgcaaa
tgacagtttagtttctctgtgcgacattccctagctgataatgcaaacaagttttaaggggaataggccaaagaa
gagatcataaactatgtgtataattactctttaagttctgaaaattattggcaacatctattttcatagcttgtt
aagaaatcttac**aagtttatcgcaagttatgcagttgccgcctctagcttgctgcccccgcccaaatgcaaaac**c
tacaaaaagtaacacataaatatcagttgaatgaagtaccaacaattgctcatcagaaataattactgcttgcta
cgttatcactggcgaagtgttcatgttctagtggcatcttgtaaaaaaattatgttgggatcttcaacagacaca
ggttgggtgcatttggaagcaagaagactaaagacatacagatattcatcaagctagaatagctcatcaagcaca
cagcaatcaggacagaaaaaagacctataatgacttgtattttatgttccacaaggattaaaaaattgaaaccaa
acagaaaagataaaatgataaaaaaactaagagggctttatccatactgtccttgctgtgacatgggatctatga
gatcatcaacacaactacacatgcttcaaaaaaaaaaagctgaaagattcacaagtgaaggtgacaagatcatagt
actagctatatatgctcatgcataactaaatgagaaaaaaggttgaagatgaaacttatgttaaacaagtttagt
tagttacctggctagctagatcagatcttcaattccatgccatagcactaagatcactttttttacagcatagcc
cgaccccttataacgccagaaggcggattaggaaaacagagttgtgctcgtgcaaagaggggcgggaggccaagt
ctacttttatagccaaaggggcagagaccaggccacaagaagagttatgcagcagtgtgtgtgtgtgtgtgtgtg
tgtgtgtgtggagaggcgtgtgggggcacagacaaggcaacaagtcagaaaaacaattataccgtaccgtttggt
ttagtgacaatttggcaacctaacgttgattgtataaattagttaagtcacttcacatgtacagacaagaaagat
tccaacttgcgtggagcaacctaacgttgattggacgcctctctattagtaaaatgagtacattattttaattca
cactggccatgattgaacaaaaaaaatccctgagattgcctttcaaaattatcacaatcaagaactgacagacag
acggacacctctcatgtgtcatctaccatgagcgcatataataaaaaaaatccagtgctactgaatatcttctca
gttttgggatggactcctgattgttgcgtatggtataaagtagttgtctcaagacatctagctgccactgattct
taattggtaataacactttgaacatggttagcagacaataacaatttcatctatgtcagataggcagcttcgctg
aagctgcgtgcaagggacaccaggccatgagtgcttttgcatcttggattctgcctatccagcagcccttcaatt
gggctcaccatcctatgctgtggtttcaaacatgtgtgtcttgatcttggtccatggatcagcttcattagacaa
ggaaaagagtcattagagttttaagctgcattttgactgccaaaccgaacaactcagttgaacaatatgaaaatt
agtaattgcagctgtaacagcagattgtcaattattgagtagtcttgttccttgtgagaatgagtgtagagtttt
tattttctagtatatcagttcaaagcatcaatgctagaggttgcaacatcaaatccttgtgagaataatgttata
ctaccttcgttctcgattatttgtcgccggctagttcattttaactaaaccgcaacaaataaaaaagaacggag
ggagtaggttgcaagatcgaaacccaataaagctacaaattaaaaaggattcctgaaaacaaaggatatctaact

aaaactatggttttcataataggaaaccaggaaacttcttagttattgaaatactaaatattagcatggggaaat
cgaaatgcagatatgaaccaatatgcaaaaatatcatatgctaaatatgactatgattaaaatcctaaagtccag
gtagttgttagacgaatataattggtgggataatcggcatatactgcattgggcccTCATGGCATGATTATATAG
AGCACAAAAATACTAACACCCTATCGTCCTATTCTGATATTCAGGACCAAGAGGTCCTGGGATTCTCTTTTGGAC
AGGCCCAGAAAGGAGAAGAGAAGCTGAAATGACAAACAGACCtttggatttttggatatatttttatttgatt
tGGCATTATTCAACAAGGaaaaaaaattcgagactggataaaaaaaaGGAACTATGATTCATACTCACTATTTAG
ACCTCGCAACCAGACTTAAAAAAAATCAAGTTATTCTTAATAAAGAAAGAAATTTTCTTCCTTCCAACTTTGTTT
GCCCAAAAGAGAACTCTTTTCTCTTTCAAGATAGACATTGTGGTACTATTGTAATTTACTATAATATCCTCCTCA
TAGTGAGAGCAACACAGACGGCGAGACTGCAGAAGAAGCCAACAAGCTAATATCCCTTGGTCATAACGGTCGATG
CGTCTCTCATATGCTATGGCTGAAGAAGAAGCTGGCGAGTCCTCATGCAAGGCAGTAGCCCCATTGCACATGTTA
TGGCTAGACAAGAAGCTGATAAGGCCTCATGCAAGGCAGTAACCTTTAGGCCAATGTCAATTTAGCTGAGCGTGA
GAAGTCAAGCCATGGTCGAGGAGTCATGCTGGAGATAGTTGTCTCTGTTAGAAGTTGAGGTAAGTGATATCAAGT
TGACCACACATACGAATTCAAGCTTTTTTCATTACGGANNNNNNNNNNNNNNNNNNNNNNNNNNNNNNNN
NNNNNNNNNNNNNNNNNNNNNNNNNNNNNNNNNNNNNNNNNNNNNNNNNNNNNNNNNNTACGATCTAGAG
ATTCGTgcttacacctagtccctaattagttttggtggttgaattgcccaaaagaaaactttggactaactagtt
tgctctagtgtataagttatacaggtgctaaaggttcacacttagccagtaaaaagaccaagagttgggttcaac
aaaagagcaaaggggcagccgaaggcacctctggtctggcacaccggactgtccggtgtgccaccggacagtgtc
cggtgcaccaccggacatgtccggtgcaccagaggactccaactcgaactcgccaccttcgggaatttccagagg
cactcgcgctataattcaccggactgtccggtgtgcaccggacatgtccggtgctccaaggaagggcggcctcag
gaactcgccagcctcgggttttcattccagccgctccgctataattcaccggactgtccggtgtgcaccggactg
tccggtgcatctgcggagcaacggctacctgcagcgcatttattgcgcgccagcgcgcgcagaggacaggcacac
ccatgctggcgcaccggacactctacagtccatgtccggtgcgccaccggacatcaaggcgggcccagaagtcag
aactccaacggtcgatttccaacggcactggtgacgtggcgggggcaccggactgtccggtgtgcaccggactgt
ccggtgcgccatcgaacagacagcacagccaacggtcaagtttggtggttggggctataaataccccaaccaccc
caccattcattgcatccaagttttccacttcccaactactacaagagctctagcattcaattctagacacaccaa
agagatcaaatcttctccaaattccacacaacgccatagtgactagtgagagtgatttgcttgtgttctttcgag
ctcttgcgcttggattgctttcttctttcttgatcccttctttgcaatcaaactcacttgtaatttgaggcaaga
gacaccaaacttgtggtggtccttgtgggaactttgtgttccaagtgattgagaagagaaagctcactcgatccg
tggatcgtttgagagagggaagggttgaaagagacccggcctttgtggcctcctcaacggggagtaggtttgcaa
gaaccgaacctcggtaaaacaaatctccgtgtctcacttgcttattcgtttgggatttgttttgcgccctctctc
gcggactcgtttctttattactaacgctaacccggcttgtagttgtgtttatatttgtaaatttcagtttcgccc
tattcacccccccctctaggcgactttcaaCATGGTTTGCTACCAAGAAGTTGTGTGGAGGCAGACATGTCTACCC
AAGGTGATGATCTGCCTGACAGCAACTTGTTAGGATATATTCGTTGTCACCAACATATAATCAAAGGCACACAAA
CAAAGTGATCCAGCAAGCACTTTCTATGGATTTAACATTTTCCACCTTGAGAAAGTTGAATTATATGTAGTTTTG
TTTATCATCAACTTGTGGTTTAATTTCTATGCATCAAACAATGGCCAGCGTTCTGAGGACTGCTTTACTTTAACA
AGTAAACAACAAATTTTTTTGGCAGAAAAGGACACACACATATCAGTATATTACTGATAATCAGCTGAGCAACTG
AGATGGCAACACCAACAAGCTATTATAATTGTCTGAAGCCACGTCACATGCTTGATTCGAATGATGTGCAAGGCT
TCGCCAATATAGGGGCAACTCAAATTGCATGAATTTTTAATGGTGCATTATTGGAAGTGAGAGTGTGTTTGGTGC
TCCCAAGACATTCCTTGTCCTCACCTTTGTTTAAGCAAAGGAGAGTAGCTTTTTGACAATATGTTCAATCAAAGC
TTAACTTTAGGGCAGTACAGACTGAGTACTAGCAAGTGGTTGATTGATGACTACACATGGTTGAAAGCCTCTGTT
GATTTGGAAAATAATATGTGCTTGCTAAGCGCTTCAGAATTTTCAGAGCTAACTCGGGGGCAAAGGTGATAGCTC
TGAAAATAAACTGAGCTTGCTAAGCAGCCTAACTCGGGGAGGGGTTGCTGTAGAATGGCAACAGCTAATGCACCA
AGGATTCTATCCTTGTATCCGTGATCAACTGGTGACCTTCAACAGTTCAACTAAACAACTATTTCCCATCGTCAT
TCAACCTGCAAAAAGGGATAAAGCTACGGGGCCATCTAGCCGTCTTGATGCATCTTCGTGCGCCCTTCCTTGTCG
GCAACGCAATTCACGAGACTGGCAGAGAGTACCCCAGATCAAGTGGGCATCAAAAGGGCTTCAAAAAAAGTGAAC
AGAGGGTTGTTTTTTCCCTAAAAAACAGAGGGTAGTTACGAATACGAAC**CATACGTTGATCCGTCGGCACACAGA**
**CCACTCGTCCGTCCGTGTGTCATCTCCATGGCGGCATGGCAGTTGGCAGTCGACAGAGCCCGTGTCATCTCCGTG**
**CCCAGCGGGTGCAGCTCTCGGATCAGCACGAACCTCGCGGCGGCTGGCGGCTGACCCGTCGCCCTTGTAGACCGG**
**TCGAACACATATGCTGTCCGAGGGAGCGTTTCGTCGAACAGGTGATGTGCGCGGTGGTCGACGATTCTTGGGCAG**
**AGTTCTGTTAACACAGCCGAGGCGAAAATCAT**CCGAGTGGGTCATATTAGATTTGTACTGAGATAAAAGAACAAA
TGGAGAGAGAAAAATGGGTTATACAGTAGTACGCTCACGGACTTGTCACATTTTGTGACCACAAGAAATAGACCA
AATGTTAAATATATAACTTGTATAATGAAAGAGTTATTTGAACAAATTTAATAATATAATACTTTAGTTATTTTT
AACTAAAATAGAGAAGGAGTAGTTTATAGAAGACTTTATGATGTTATTATGTCACAAGTGTCGGACAACACACTA
CGAAATATATCATGTAGGACTTTTGGATGGTTGGCGGAGTCAACTATAAAAAGAGAGCTTTAGATCCAGAGAAAT

>zma_miR444e_FL182678.1

GAATGATTTTCGCCTCGGCTGTGTTAACAGAACTCTGACCAAGAATCGTCGACCACCGCGCACATCACCTGTTCG
ACGAAACGCTCCCTCGGACAGCAGATGTGTTCGACCGGTCGACAAGGGCGACGGGTCAGCCGCCAGCCGCCGCGA
GGTTCGTGCTGATCCGAGAGCTGCACCCGCTGGGCACGGAGATGACACGGGCTCTGTCGACTGCCAACTGCCATG
CCGCCATGGAGATGACACGGACGGACGAGTGGTCTGTGTGCCGACGGATCAACGTATGGTTTTGCATTTGGGCGG
GGGCAGCAAGCTAGAGGCCGGCAACTGCATAACTTGCGATAAACTTGTTGGACCATGGTGTGCTGATGATTTTAC
CGCAAGTTATGCAGTTGCTGCCTCAAGCTTGCTGCCTCCCTTTGCCAAACC

Stem-loop structure

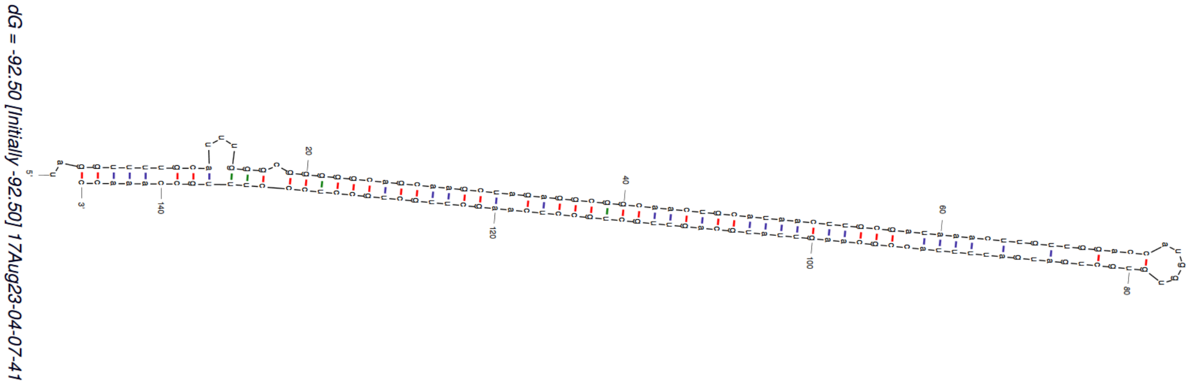


#### Commelinales

##### Eichhornia paniculata

*EpaAGL17_1 and epa-miR444a*

>LTAE01001227.1

AGAGGAAGCAAACAGCCTCCACCAAATAATAACCGCAAGAAGAGGAAAGCGATCAGGTGCATGCTCCACTGTGTC
CCATTTGTGGCCATCACCAATAAATAAATAGCTCAGGGTTTGTTTGCCTCTTCAGCCTTCACATATGTCATCTAT
ATCTCACCAGTATTCCCATTTCTATCAGTTTGTTCTTTGTCTTCTGCATCATTTTGGTTAATTCATCTGTATGTT
TAAACCTCGTTCCTTTAATTTATTGATCTGCAGTCGCTGAGGCTGCCTTTTACTTGTTTAATTACTTTCCGAGAT
CTGATTCGAATCTATGCACTGTTTCCTTGTTGATTTGGTGAAGCAGGAGAGAGAGAGAGATTGTTGTGCACATAT
AGATATTATCAGAAAGGAGGTAGAAGTATACAGAAACTGATCATGGGGAGGGGGAAGATCGTGATAAGAAGGATC
GACAACTCGACAAGCAGGCAGGTGACGTTCTCAAAGCGGCGGAATGGACTGCTGAAGAAGGCCAAGGAGCTGGCC
ATCCTCTGCGATGCCGAGGTCGGCATCATCGTCTTCTCCAGCACCGGCCGGCTCTACGACTACACCAGTAGCAGG
TCATCTCTTTCTCCCTCTTTATTGCCGGCCATGTATCCGTCTTCGACATGTTGTCTCCGTTTTGATTTTTCTCAT
GAGATCGATCCTTGAAAGTTGCATATTTCTCTGCACCTGAAATTTCCGAGATGTTTAAAATGGACATTTTGTAAT
TGTAGCTTATGTTCTTGTAGTAGCTTTTTAGGGTGCAAGATTCGCTCATAATTAACAAGACTTCTGTGTGTCAGA
AATAAGATAAGAAATTCATGTGATAGACCCCTCGATAATCTCATTCGTGGGGAGAAAAAAAAGATGGACTATCCA
ACTTTGGTACATCTTTGTTTTTTGTATCTATACATATATATATATGAATAGACTTATATGATATATCCGTATGTG
AATTACTATAAAGAGAATATACATGATTAAATGGTTAACTCTTTCATAAAAAAAATTAAGTCATTTCGGCATATC
TACAAGATTATTGTTTCATCACTGAGAGTTGATGCAATACCATATGACTACATGAATGATCTTATAACACTTGAT
ATAAATTCGTCAGTTAGATATATATATGACTTTGGACACTGTCTTGCATGTCATACGGGTTCTTCGAGGGCTGAA
GAAACGACCTTAATTAAACACCACTTAATTGAGGAGTTTCTAACCTTCAAATTTACGGGGCTGCTAGTCTTTCAA
ATACTTGTATTAATTAGATGAGCTATTGACCCATTTCTAAAAGTTTAAGCAATTAGGAATAACGTCATCTTTACT
AATACTAATCCACCGTATACTAATAGGTGAAAAATAACTTCTAATAAGAACTCCGTTGGAGTTTGAAATATGCAT
GATGAATACTAAAAAGAAATGTTCTCACCTTTTATTTGTTTAGCGTATGCATGTTTTATGTGCCTGGCTTAGCCC
TGGTAATGAATAACTTGTGTTTCTTAGCAGATTATTTAGAGTCTTAGAGAGATGCTTTTGCATTCATCAAAATGC
TTATTGTAGATTCATGTACGTACCAGAAGATTGTCATGAGGCTACTTACTGTATAACACATTTTGCTTGAATTCC
TTACTATAGAATAAGCGTACTTTATATATATACATCATAAATCGAAAATATATAGTTTGTTATGCTTGCTATAAA
TGATCAAAGTTATGCACCGTAAAATACAACGCAAAAAATGCGTTGTAACACTCGTATATTAATTTTTTTTGCAGC
TTGAGACGGTGATAACCCATGCTGTATATATCCTACTCGAGAGTGTTGCTCTATCACCTAATGGTTATCTCAAGA
GCATATGTGGTTCTTTCTGCAATATATATTGGTGCAGGACTTTCTTTTTCAACAATGTTTTACACTATACACCTT
CAGACATATATTAGAGGATAAACCTCCAATTAGCTTAAGTGATTAGTTCTTGTTGCTGAAAGAAGGAATTAGTAC
AACTGGCGGTTATGTTAACAGTTTATTTCATAGTTAAGTAGGAAGATTTCAAAGTTTACTTTTGTTTCAGATTAT
CATTATTAGCTAACAAAATGACATCACATTTCCATCATTTATGAGTTTTGTGTTATCATTTATGTTTCACTGTTA
TGGGACGGCTAAATGTGCAAGGAAACACTTACGTAGCATTGCCAGATTGAGAAATTTAATAAAAAAAGGAATAAA
ACATTATAATATCTTTATGTTCCTTGCCTACCTATGCTGTATTTAACTTGATTTCCCACTTACCCTCTTTTTATT
TCGGTATTGATGTGACAGCATCATGTGAATAGATGATGAATTTGAGGGAAAAGAAAACAAAATATATGGTTTTAC
CAAGAAAAAACCATAATAGTTCCAGGAAAAAAAGTTAATGAATTTTCAGAATTTTTTTCGTATTCCGTATTTAAT
AAGAGTCTCAGTTTGTTTTGACAGTGTTTTTTTGTAAAAATGATTTCTCCCATTATGTTATTTGATAATAGTCGA
TCGAAAAAATACTTTTTGTTGATAAGAAAAGTAGGCTAAAATTTTGAAAAATAAATTTCATTGTTTCAAATAGCA
AATCATCTTCTCCTCTCAAGTCTTCTATACTTTTAATTATAAGATAATTCCTTCATATTTGAAAATGATATTCCA
ACACATGAAAACAAATCACTTTTCAACAGCTCGTTTTCTTAAGAATTTATTTCCCTTATGCATTGGTATTTATCA
TAACAGATGGAATGCATTTCTTTTCTGCATTTAATTTATTCTCAACAAGGAAAGAAGTTTACTAACTAGTTAAAA
TATTTCTTTTTCCCCAGTCCTTATAACAAAATTTTCCAAAGTTTTTTTCTTCTTAACTGTTCTTCCTCTTCACAA
GTGCCACTGACCTGTGTGGGAGGAATTCTTGCAATTATATCATTAACTTTGTTGTATAAATCAAATTGTGCACGG
CATGCATCAACTAAATCTCTTACCCGAATTTTGCATTCAGTATGAACTCTGTGATAGAACGTTACAAAATGACGA

AGGAAGAGCCACAACTCGTTACTGATACAACTTCAGAAGTTAAGGTACGTGCTTCTTCCATCTCAAGTATTCATT
CAATTATCGATGTTAATTGAATTTCCATTTAGCCGATAATTGCTTATTGCATGATCCCTGCTATTTCTTTTGTTA
ATATTGTTGTTCATTCATGTCAAATTTCTTTTGATATTGTTCCATGTGAGTAGTGCTGAGTTCACCATATGTATC
AATCTCTAGATCTAGGTGCTCTCTTAAATTTGTTTAATTAATTAAAGCTTTGATTTGACTCCTATAAGATATCAT
TTTTAATTTCAGTGAACGGCAAAAGTTCAAAGATCCTTAATTTAATCCTACTTGTTCTTTCCTTTCATAAAAAAA
ATCTATTATATGTAAAATTGTTCTCTCTTTACTCATTTGATGTTGAACGTGGCAATGCCAAGTTGGTAAAATGTG
GAATGCCTCCTCATGGAAAAGAATTTTTTTTTTGACAAAAAATACAAATATTTGCCATTAACGAGATACTTTCTTT
TAAATAAAAGTAAACGTTGAATCTTTTGGTCGAACAAACTGATAGATATGAAATAAGATTAAAAAAAGGGTTTTT
CGACATATTTTCTGTTTGTTGGACTTCGAACTAGATATCATTGTGGACCAACTTGCTTTACTAGAGACAGAAGTA
CAAGACGATACTTCATATTTTTTTGCTTTGTTTATAGTTTATGATGCGTGTCCCTACACCTTCCTAAATATTCTT
TTAAACTTATCTTACTACTTGTAAGGCAAGAGATATTTTTTCCTACTTTTTAATATATTTTACTAATGTGAGGTG
AATTTCTCGAAGGAAGTTTTAGGAGGATTTTTCTTTATATTTTCTATAATTGATAAATTTTTTTGTAAGTGTACA
AAATCCTATTAAGTAATAAAAAGATCATATTTACATGGAGGGAGTTCAACTATTAAATATAGGATGAAATTGCTA
TTATCTAAATTAAAATTAATGATGAGGAAAGTAATTAAAAAAAGCAAGAATTAAAATCTAATAATATTACGACTA
AATAGAATGTCAACTAACATGCTTTTTCCAATGCAAGAAAAGATACCCTAGGGTTGAATTTATTTAGGAATATTC
GATTGTTTTATTACAACTCATATTATTATGTGATCTTAATTATGTCGTTTACTTAAGCTAATATGCATTTCTTTT
TCAAGATAACTCAACATACATGCATACATACATTCTCGTATCATGTGTTATCAATTTAATTCCATAAAGATCCTA
TGACAAAACCATATAAATGATGCTTATTTCTAGGGCAAACATTTATAGTTCACTTAGCAAAGCTACTAATCACAA
TATGGTTTTCACCAATTATTTATCTAGATCAATCCAAAGGATAAAGAATCACGATTGGATCTAAGAAATATGAAT
AGTCAAGAGAAAATCTTGAATCACATGAACATTTCAAAAATTCAATAAATCAAAATCCCTAAATTTTATCAAATC
CTAAGGCATAAAAGATATAGCCCCTCATCATGAAAAACACAATAATCGATTTTATCATGAAAAACAAACTGATCA
TATTTAAAAGTTAAGGAGTAGCAAGATGTCAAAGATTACAATGTAGGCCTCGAATCAACAAAAGTGTTGCGATGT
CTAGATCTATCCCTTGTCGTTCTCCGTTTTAGTCCCCCTCGACACAATGTTTCATCTCTTTTATATCTCCAAAC
CCCTAGGGCTTTACCCACTTAAAAATTATCAGAATATCCAAAAAAAATCAAAATACGCCATTATAGATCATCGAA
TGCGAGCGGCATCATGAGCAATACAATCATATCAAGTACATGTGCTTATTGGAATATTCAGGTTGATACGATCTT
TATCGACCAATACGGTCAGTATGAAATCTCTTCGGGAAGGATGATACGGCCGTATCGTCCGGTACGACCATGTTG
AAATCTTCTAAAAAAAACTTGTTTCAGTTGATCTTGATCAATACGACCATATCCTTAGCGTGGGTCCACCGTATT
GCTTCCTCCACATTATATTTTTGTTGTTTCTTCTGATGATCGTGTCCTGTCAAAACAAACACAACTATGTATAGA
TTTTCGAACAATAATCAATGTAGATATATAAAATCTTTGAATTTGACTCTCAAACGATGCCTAATTGTAATTTTT
ATGAACTTTGTGGAGAAAACCACTTAATATAAAAGGCCGAAACTTTGCGGAGAAAAGTTGACTAATTTAGACAA
ATATAATTAAGAATATTTTTTTAATTCTTTTAACATATAATAATTAAACATAAGAAGAATATTTCTATATCCTCC
TAATTATCCTTCTAAAATTCTTCTTAACCCCTCATGGTATCTTTTATAATTGATCAAGTTTTGAGAATAATTTA
ACTATGCGATTACTAAAATAATTCTTAAAAGAATAAGTTAGGAGAATTTTTTTATTTTTTCATATAAAAACGTAA
TAAATTTCATTGAAGTTGCTTCTTTACATTGCATTGTAACGATGTCCAAATAGGACAATGGTCTTGCGACTTTGA
AATGAACTAATGAAAACTTGATGCTTTGTCAAAGACATTTAGCATCTCCTAGTGAAGACCAACGGGA**GTTGTGGT**
**GGAGTGATAAGTGGCTCCTCCCCACTTAAGCAACCCCACTTAAGCAGGTCTCGAGTTCGAGTCCATATGAACGCA**
**GAATAATCCCCGCTGGGAGCCTCACCTGTCATAGTCAGACACACGACGCGGGACGACATGGTGGTGGGCCGAGAT**
**ACAACATGTGATCAGTGCACATAACCGTTAGTGTCGCGTGGGTTGTGGAATCAGCACCGCTTCTCGGACACTGGG**
**GTGACAACCGAAAAAAAGACATCTC**CTAGTGAAGAAAAAAGGGTTTGAAAAAAAAAAATCTAAAGAAAATAACTT
TTTAGGAAAATTAATTTAAAATAATTTATTTTATATGTTCAGTATATGACATGACTATATAAATAAAATTATTTT
GAAAATAGGAAAAAATATGACTCAGTGAGAGAAAAATAAATTTCCAATCTGGAAAGAGTAATTCACTTTCCCAAC
TATAACCTTATTTCTGATCAACAGAAAAATGATTTTCTATTGACCAATCGCTTTTAATCACTAAATATTGAAAA
ATAATGAAATTATTTTAACACTTCTTGAAGAATGAAAAGAGTTGCTAGATATAAACTTATAACTATCATCGTGTT
GATAGTTGATATCGCTCTTTAACACTTTGACCAATTAAATTTTTTTTTGTTGAGCCGATGATATGTACGAAATTA
GAACGTGCTTTAAACATTGAAAGAAATTGTCGCTAATAAACTCATATTTTGTTTGGCCTTTTGTTTTGTGCTCTT
ATAAATTTCATCAAAATAAATCGAACCTAAACATAAGCTATAGACACATAAACATAC**CATAGATAGGGCAATATT**
**AGCAGATATGTGCAAGCTTATTAAGATCCTCAATCAACCCTTAATTTCTTGGGAATTTAGAGTTTGCAACCGGAA**
**TAGGAGTCAACTCGCCTGGTAACC**CTGTGAGCATCAATATTTGTTAGCATGTTCTTCCTCCTCAAATAAAATAGA
TGCTTTATTAGTACGAAGTTAAATGTTCTTAATATCGAGCATTATATTTCTGCTTTGAGTACTAATTTTCAATAG
TGATGTATATTTTTTTTTTCCATAGTGATGTATATGATGGAACCATAAAGCTAATCATACAATTAAGTTGTCAAA
GGCGTATCTTATATACCTGTGCCCTTGTTCGAATTATACTATTAATGCAGTATTGTTTTGTATTTAAATTTGTTT
AGGAAAGAAAATCTATGAATTTCCTATCAAGTTTAGGTTGTATGACGTTAAACAAGTTTGCAAATAATATGGATA
AGTTGCATGCTCCATGCTCCCTTTTATATTGATCAGAAATTAGACCTTTGCAAGAAGCTTCAGGCATCTCATTAC
AATTTTTTGTTTGAGATATATGCAGAATATAATAGGGATCATTATCTTTGATTTAACCTACTTTTGTTCGAAGTA
TGAAACCTACTTTGAACCTTAATTTACTTAATTTTCATCTGCACTCTGAGGGTTCTTTTAGTAAAAAATAACTCT
TAAAAAATATTAAACATTAACCGGGAAACTGCCAACAGTATGAGATGTGAGATTTAAGAAAAGATATAGCAAAAA
TTTTACATATAAGAAGTGTAATGCAGTTTGCAAAGTTTCTGTTCAACATTCTCTGAAGACCCAGTGTCCTGTGGA
AACTTGAATAACCCAATACTACACTTTGAGGTGGATGCATAAATATAGAACTGCAATGCAAACTTAC**ATTCTTGT**
**AACTTTCCGCATGTCACTTAGTTTTGGCAAAGGGAGGCAGTAAGCTTGAGGCAACAACTACACAACTTGCAAGAA**
**AATCATAGGTATGCTCATTGTACC**CTACATAAGATCAATAAAAGGCAATGTTAATTCTATGGCATGCTTGAATAG
CTTTCATATTACTACTACTGATGAAATTCAATGATAGGCATTGATGATGGACAATATCCCAGTAATTATCAGCAA
AAGGAATATGAAACAAATATAAAGCAAAAGATAGGGGGTTATGTTTTTTTTTGTTTGGAGGATGATAAGGGTTAT
GCATTACCTAGATTTTTAAGATGCATATACCTAAAGTGGATCCATAAGAGTGAATAAACAGATTTGTCAACAATT

CAGGCAGTTGATGGGAGAAGAACTTTCAGGTTTAAGTGTCAAAGATCTACAAAATCTAGAGAACCAACTAGAAAT
GAGTCTTAAAGGCATCCGAATGAAAAAGGTTATTTTACTTCTCTCACTGAATATCCAATACTTTTCTTCTTATTG
ATCCTTAATATTCTTCATCTTCTCCATTTACAGGATGAAATTCTAATCAACGAAATCCAAGAACTGAATCGGAAG
GTCGATAACCACTTTGTCATCTTCATTTTCAATCAAATGATACATTCGAAGTTCCCTGATCTATGATAGGTTTCA
TTGCAGGGGAGCCTTGTTCAACAAGAAAATATGGAACTCTACAAGAAGGTAAATCTTATTCATCAGGAAAACATG
GAACTCTGTAGGAAGGTATAGAATTTAAAACTCTTCAAATCTGTGTACTACTAAAGAAGAATTACAACCAAAACC
CATCATGAATGGAATTACCTATGACAAACTATTCATTCATTAACGAATCTCAGGCTTATGGAACAAGGGAAGTCA
ATGGTGTTGAAAAAAACCAAATTTATCCACATGGTCTTAGCATTGCAGATGACATTAATGTTCCCACTCGCCTTG
AACTAAGCCACCCACATCAGCAAGATGGTTGTGCAACAAATTGCTCCTCCAAAACTAGGGTAACTTTCTTTCAGA
AAATAATACGATAGCTTCCTATAACAGTTTAAACATGCAGTGCTTAATGTAATGTGCAGCAGATTTCAACTACAT
TAAAAAGACAATGCACAGATCAGGATCATGTTCTCTAAAGGCCGCAGAATTCTCCAATCTGCGGTATGTTTCCAT
GTATAAGAGCAGCATGCTCAACAAATTATTTATAAGATGTATGAATCGGCAAGAATTTAGCTATTTGTGTAAAAC
TGTCAGAATAAAAGCAGTTGTGTAAGCTTGCATCTCAATGTTAACACAGTAATGTCCGTATGCATCTTAGTCTAA
AATGATGATTTCACAACTCAAGATGCATGATAAGGGATTATCAGATGCAAATGTTTCTAATGAATAAAGGTAAAA
AAATGCAATGTTAAAGAGGACAAATTCTCGAAAATAAAAAAATTCTCAAAGAACAAAATTCCCCACCAGATGCAA
AGTCAAAGAAGAAATTAATTGAATAAAATTTTCAACAAAAGTTTGTTAGACAGAACAAGTTCAAATGAAAAAGAA
TGAAAGAAATTTGCTGCCACTATAATATGTACAGGAGGCAACGGAAGGATCAGATTTCTGTAAGTTTGCTGCAAC
CACAAGAAAAATGCAATTGAAAATATTTTATTAAGATTTAGTCGTGAGTTAGGAGATAGGAAAGATCGACAATGG
CAGAACAAAACACCTTAGATCAAAGGGTACCAGGTAAGCAATTTTCCCATTAACAGCTGGCACATCCATTTTTAC
CGCAACGTGCATGCTTCATTAAATAAGAACCAATACTAAACATATTTCAAACAAATAGCTAAAAGAAACATAAAA
AAGCAATGAGGATTCTTCCATTATTACAGAGTAGGCTGATCCAAACAGAGCAAAAACATATTGATGGTGAACGAA
AGGCAAAATTCTGTTTCTTTCTCTTCTTATGATTCATTTGGTTTCCCTCACATGATCAGAAAAAAAATAAATAAG
AGGTTTGATAGAGTAAAGTGGCACTGTTGATCGAACTTCAGGCTAATATGAAACAGATGCTACAAGAAAGTGGAG
AAAATTTCGAGTAAGCTTTGTCCATGAATAAGGAAATTATAATGCACAGGGATATTACAAGAGAAAACACCAACA
GAGATCATCTAAGAAAAGATGGCATTTGAGTATTCAACCTTATTCCAACAAAAGTACTTGCCCTCGTGATCTGAG
GGAACATGGAAAAGGGTATTGAAGTAATAAAATAGAAAAGAAACACCCCCTTGTTGATCTAACTATTAGCTATGG
CAGCTCCATATTGGACCCACTAGCATTATATTGACTCCATAGTTTGATGTCAACAATACTGACGTGACTGTAGAC
ATTTTATTCCACAGCATAAAAGAAAATATACTTCAAAGTCGAGAACATTCAGGGAAGTATTTTCTTCGTCCTTAT
AAGTTGGAAGGTTTAAATGTAATGGAAGTTCAATACGTCTAGAACATGTGATCAAACAACTATCCCTAGTTCCAA
CAGTTTTTTTTAAAGAATTCCATCTCATTCTCGGAACTCCTAGTGCATGATTTTGAAGAAGACTCCTAGAACACG
TAAAAGATATATACTGAATTGCGAAGGACAATGGAATTACATATAACAAAAATGCAGAACTCTAAAATGAGTATA
CCCACAGAAAAATTACTTTTCAATGACAAAAAAGAATCATGTAAAAATAGTAAGCACATGTCCATTTGTTCACAA
CAACGACAAGAAAAAAATTAAAAAGAGCGGCAAACCTAGTAACCACCTACGATTGCACAACTTGTATAATGAGAAA
TGCACATACACATGAAATAAAAAATCACTATACAATCATATATGATAAAACATAATTGCTGTTATTTTTAGGATC
AATTAAAATGACCAATATGAATTAATACATTGCTAATGCTTTTGTTTTGGAAGAATCTTCACATATGTATCTGTT
TGTATATTTCGATGTGAAGAATTGCTTTTTCTAAGACAAATTTTTTAAGCCCATTACGAAGGCAATGGTGAGTGC
TCAGCAAGGAGTTCAAAGTACTAAGGTGGCAAGTTTATGGTAAGGCATTAACCAAAAATATGGCAGCTGATCTAA
AAGTTCTTTTTTGAGGGGTCAGTAAAAAAGTATAGCAACAATAAAACATTTATATCATTGATACAAACAATAATA
ACATAGTAACACATGTCACAGATGTAAGCATGGTAAATGATTATAAGCTAAGAGGAAAATCATTTGATCATTACA
ATGGATCTACATATATCCCAAATAAAGGAATGGAAATTAATAGTAATCACAGGAGATTGTATAGACATGCTACTA
TGCTAGATCTCTTGAAGAGTCTAATATGCAATTGGCGGAGAACTGTTTCTATCAGATTATTTCTTTTAACCATAA
CCTTCATCCTTCATATAATGGAGATGTTTTCCTGATCATGTCTGGCAGACGCAGACATCCACTCAAATATTGAAA
TACTACTGTAGTCAAGATTCAAGCAAACACTGGTTTAGATTTCAAATTCTCCATGCGCTTCTAAAAACTAGTAGT
AATTCTTGATACTTCCCAGTCACTTAATTCTGAGCAGTGAACAACCATATTATTCTCCATGATTTTGCAACAAAT
TAAAAGAAGCAACTTTTAAATGTTTTAATAGACTGTAAGTAGTAGTCTTGGGGAAAAAAAGACATAC**CAGAGCAT**
**TTCTTGCAGGTTGTGCAGTTCTTCCCTCAGGCTTGTTGCCTCCTAAACAGCTAGCCAAAAAATCACAATTT**CTGT
TCACAAAAAAAATGCAGAGAGCTTTAACAATATGGCTAGCTAGGACTGACACTTTTCAAATTATTGAATATTTAT
ACAAGAATTACTTATTATGCAATAACTTTATCGAAACCTCCGATGCTTATTGCATATTGCAATTATTGGTATAGA
GGAACAATATCATTAGAGTTCACACCCTCTGATCAATTTAAACAAGAGGGAAAAATCAACACGGGTAAAGGAGGT
TGCTGTCACGAATTTTCACTATGAGCAAAGAAATTGAAGTGGGCCCCAAGCTTGGGCCTACTGGCTTTTCTTATC
ACTTTCACAATTTTAGGTTTAAGGCAGTGGATGAGATAACAAAAATATTTAGACATGGTCGTTTCTTGCATAATC
TCCAAAAAATTCTAATGCACCTGGGTTTCACAATAGCAATGGCGTGTGAAATTTATCATAAATGTGTGGTAATCT
CACTTTTGAGGATTGGACTCCACTCATTTGAAAATGTTTCATAAGTGTAACCCTTGATTTGGTTCGAAGGTCCAA
ATGGAAGAAACTTAATTAACAGTAATATGTAAGGATAAAGCAAAGAATAGAAGCAAAAAAGATCCAAATGCATAC
AATCATATGGAGGTCCAAAAACGTTTTCATTCCCAGACTTTTTCTATGGATGCAGTGGGCCTGCTGGATTTGTGT
GCGTCCCTTATTTACCCCACGGTCCCCATCCACCCACCCCCCACCCCCAAAAACAAGATCCAAATACAGCTTTAT
TTTAGCATAACAAATTTGAGCTAAAATGGTATGAGCAATCATGTATACATATGCAGCAATTTTCTGTGGCATGAC
AAAACACGGTTATTTGGTTACCATATCAAAATTCATATTAGTATGTCCGAGTTTACCAACTTGGACCAAATTTAT
GTTATTACTGTTTCGTCGACATATCACAGGGTAAGATGATAATAAGGTTAAATGCTTTGAGTTTACCAAAATTTT
GATAAACAACCTACGTTTTTGGCTATGTTTTTCAGCATTTTTTCTTACGTATATTATTCTTTTTTGGCATTGCAT
ACACAGACATGGCTAGCAATTGCGAACATCCAAATGTCAACTTCATCGTGCATTGAGACCTAGAAGATCACAAGA
CAATGGTGATCTTACTCTAGTATGATTTCCAGAAAGCCTCAAAATGGGAAACCGACCTTTCAGAAACGCCAAAAA
CAGCCCCTCAACCTAAAATTATTTTCTAAAAATGTACATGACATTGTAGATGTATCATACCAGTAATTCACCAAA

CACAAGACCTTCATAGTCTCTGCAACTACTAGAGAAATTGTTGAACCGAAATAAACAAAAAAATGACAACCAAAA
TTCTCTCTTCAGAATCCTATCGTGATAGGATCAAAGGCAATAACAACAACAAAACCTTAATGACACAATCTTCTG
AAGAAGCATTCGAGGCTCCCTTTTGCTAACAACCAAATATGAAAGATTCCGATTAGAAACCTCAAATTTAGGTCC
CAAACACAGAAATAAACTGCTATGCATCAAAGACTGATAAAAATGCAAAAAGATCAAATGCAATATACAAAACAT
ACCAAATGCAGCGATCTTAAGCTACAACTACAAATATAAGGACAATTCTAACAGATTCAGAGGTAAAACAGAGGG
GACAAAGCAAATCCTTTCCCCAATCGAAGCAGATAAATCTCTGCAAAATGAATCAAAATCCATGGAAACAAAATA
CTCATATCAATCTATGAAAATTAC**CTCCGATTGGCTCTTCAAACGGAGGACGAGAGCGAGTGAAGTGAAGCATAA**
**GAGATCTTTTGGGGGACGCAGATAAATGTCTTCGGCAACCCGAGGGATTAAAAATAAACGTACGAACAAGTGGCT**
**CTCTCTGCT**CCACGTCGCTTCTCACATTTTTAAATGAACCCGAATCCATAATCTCGCTTTTTTATAATTACAATT
ATAATTCTTTTATTATTTTATTATTTTTCTGCACCTCCTAATTATATTTTTTTCTTCATCTAACGTAAATAATAA
ATTTATAAATTTGTATCCCAGTAATGTCATCCTCCCCTTAAATTTCATTTATGTTAACTTAATTTTTCGATTATC
ATCAATTATCATTTTCACTGATATTTTTATTAATATGATAAAAATATTTTTAAAAAAATAAAAAAATCAAACATA
AAATTAATATTTAATTATTTAAAATTATGCTCCATGTTATGATTTTTTAGTTCTAGAATACTTTTTGTATTAAAG
AAGTAAGGATAAAAATGATATCTTATAATTAGTAAAAATTGTTTTTATAGAATTTGACTGAAGGATCATTATAAG
TTTAAAAATTAAAAAAATGTCGATTTATAATTTAGAGTTAGATGCAAAAAAATATAATTAATTAAAGTGTTGAAA
AAAATAATTCTCCTTAAATATCATTAATATATTTTTTTGGGTGATAAATAATCTTTAATAAAATTTTGTTGAAAT
TCATCAAATATGAAACTCAGATTAGTTGAACACAATGTTCCTAAACACT

>EpaAGL17_1

ATGGGGAGGGGGAAGATCGTGATAAGAAGGATCGACAACTCGACAAGCAGGCAGGTGACGTTCTCAAAGCGGCGG
AATGGACTGCTGAAGAAGGCCAAGGAGCTGGCCATCCTCTGCGATGCCGAGGTCGGCATCATCGTCTTCTCCAGC
ACCGGCCGGCTCTACGACTACACCAGTAGCAGTATGAACTCTGTGATAGAACGTTACAAAATGACGAAGGAAGAG
CCACAACTCGTTACTGATACAACTTCAGAAGTTAAGTTTTGGCAAAGGGAGGCAGTAAGCTTGAGGCAACAACTA
CACAACTTGCAAGAAAATCATAGGCAGTTGATGGGAGAAGAACTTTCAGGTTTAAGTGTCAAAGATCTACAAAAT
CTAGAGAACCAACTAGAAATGAGTCTTAAAGGCATCCGAATGAAAAAGGATGAAATTCTAATCAACGAAATCCAA
GAACTGAATCGGAAGGGGAGCCTTGTTCAACAAGAAAATATGGAACTCTACAAGAAGGTAAATCTTATTCATCAG
GAAAACATGGAACTCTGTAGGAAGGCTTATGGAACAAGGGAAGTCAATGGTGTTGAAAAAAACCAAATTTATCCA
CATGGTCTTAGCATTGCAGATGACATTAATGTTCCCACTCGCCTTGAACTAAGCCACCCACATCAGCAAGATGGT
TGTGCAACAAATTGCTCCTCCAAAACTAGG

>epa_miR444a

AGCAGAGAGAGCCACTTGTTCGTACGTTTATTTTTAATCCCTCGGGTTGCCGAAGACATTTATCTGCGTCCCCCA
AAAGATCTCTTATGCTTCACTTCACTCGCTCTCGTCCTCCGTTTGAAGAGCCAATCGGAGAAATTGTGATTTTTT
GGCTAGCTGTTTAGGAGGCAACAAGCCTGAGGGAAGAACTGCACAACCTGCAAGAAATGCTCTGGGTACAATGAG
CATACCTATGATTTTCTTGCAAGTTGTGTAGTTGTTGCCTCAAGCTTACTGCCTCCCTTTGCCAAAACTAAGTGA
CATGCGGAAAGTTACAAGAATGGTTACCAGGCGAGTTGACTCCTATTCCGGTTGCAAACTCTAAATTCCCAAGAA
ATTAAGGGTTGATTGAGGATCTTAATAAGCTTGCACATATCTGCTAATATTGCCCTATCTATGGAGATGTCTTTT
TTTCGGTTGTCACCCCAGTGTCCGAGAAGCGGTGTCGATTCCACAACCCACGCGACACTAACGGTTATGTGCACT
GACCACATGTTGTATCTCGGCCCACCACCGAGCCTGACTAATCTGATGTCGTCCCGCGTCATGTGCCTGACTATG
ACAGGTGAGGCTCCCAGCGGGGATTATTCTGCGTTCATATGGACTCGAACTCGAGACCTGCTTAAGTGGGGTTGC
TTAAGTGGGGAGGAGCCACTTATCACTCCACCATAACAGATCGGAAGAGCGTCK

**Position of mature miRNAs based on comparison with osa-MIR444d and not based on experimental data!**

Stem-loop structure

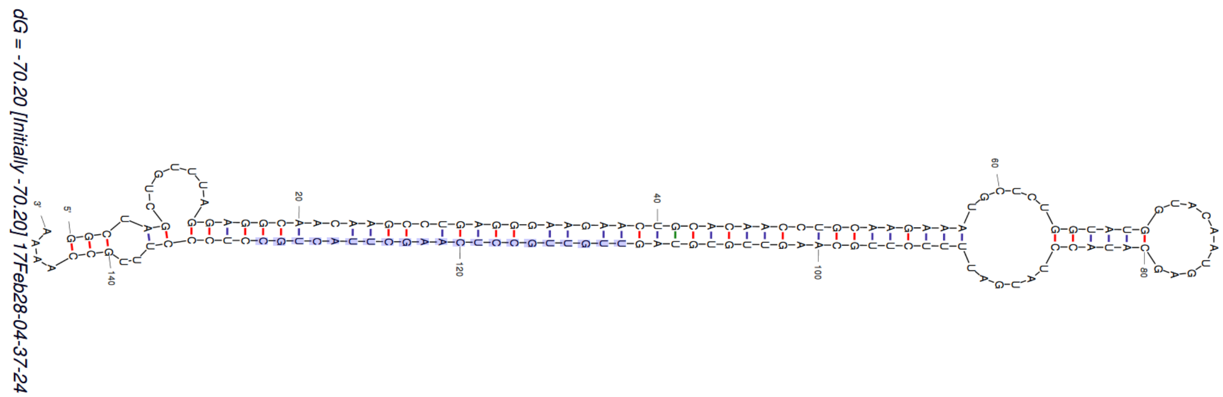

*EpaAGL17_2 and epa-miR444b*

>LTAE01016264.1 combined with LTAE01002067.1

ATCGTCTCCATCTTATAGCAAATTAAGGTTTCCTCTTTAATTTCCTTTCTTCTCTCTTCTTCTCCTCCTCGTCCT
CCTCCTCTTTCCTTCTTCGTCATTTTTTGCTCCCCCTTTTGTCCCCTCTCTGGTTGTCATGCCATTTATTGGTTT
GTTCGTTCTTTGCTTTTCCTAGCAAGTTTTGTTGTGAAGTAGCGACCATCGCCTGCCTTTTGGCTTTAAATCAAA
GTTGTTCTTCAACTGTTCATCTCTGATCTCTTCCGTATGTATTATTTTAATTTCTGATCGATTTTCATATGTAGA
CTTATTCTTTTCTACTTCCTTGGATTCGACCAAGATCGATCGCAAATTGAAGGTGTTTTGTTTTGTGTTTGCTGC
TTAGATCAGGAAGAAGTAGCTAGATAGAGAGATGGGGAGAGGAAAGATTGTGATAAGGAGGATCAATAACTCGAC
CAGCCGGCAGGTGACCTTCTCGAAGCGGAGGAACGGCTTGCTGAAGAAAGCCAAAGAGCTTGCCATACTTTGCGA
TGCTGAAGTCGGCCTCATGATCTTCTCCAGCACCGGCCGGCTCTATGATTTTTCTAGTACCAGGTTCATTTCCCT
CTACCTTAAGATGTATCTATATATGTATGAGATCTTGAGGGGTTTTTTTCCTCTTTTTGTTTCTTGCCCATATA
TTCTCTTTTTCCATTTGGCTGTTGATCAAATCTTCTCTTCAATTACAAGTTGCTTTGCTTAAAACGAACCCCTAA
AAGGAGAAATAAAAATTTTTAAGTGGGAGTAATTAGGAGAAGCAATGATGTATGAAATCTTAAGGGTCTTTCCTT
GTTCAAATCTTCTGGAATTTGTTTTTTTCTTATCAATAGAACTACTCGTCAAGAGAAAAAGAAAAGTAGGAAAGA
AATTAATTAAAAGGCTGTAATGGAGATTTATAGAATTATTCCTTGTAACTCTTCTAGAGTTAACTTTTATATGCT
TGCATCATAAGAATTAGAAATCAACTTGAATTCATGGTCGCGGAACACAGTAGATCGATCACATCATTTCTTTTA
TTGGTAGGGGGATCATTTTGTTTATCATATATGTTGTTTCTACTCCTTGTCATCTTTTACTTGAATGGCAGACTT
CATTCTCCAATTTATGTTCTAAATTCAATTTTTAAGGAATATTATACTATACTTAAATGAAGGACATGAACAAAA
AGCTTGTTTAACTCTAGAATTTAAGTAAAATTATTTTTGGCTCATAGAAGTGTGTCGTAAAATCATATTATAAAG
TTAGAGAATCTATGCACCAAACAAATATAACTAAAATAAGTAATATACTACCATCATCATAATCAAATAAATCAT
TTTATTGGACACACGAAATATATAAGGATCGCTATTTTGACTAATTGGTTCTTTTTGTTTTTGAACTTCATTTTT
AAAAACTAATCTAAATAAAAAATCTTAAAAAAATTATATAAATTTAAAAGTCATACAGGCGTGTATATTTTGTAT
AACAACAACAACAACAAAGTCTTCTCCCACTAGGTGAGGTCGGCTACATGGATCATACGATGTCATTGCGATCTA
TCAGTAACCAAATTTTTAGGTATATTATTCGCTAGTAGATCTCATTCTATTAACTCGTCAAGTGTTCATCTTGGT
CTCCATCTCCCTCTTTTAATCGGGCTCCAAATTGTTTGCTCTACTCTCCATATTGGCGCATCTGATGGTCTTCTC
TGAATATGACAAAATCACCTCAAACGATTCTCTACCATCTTCATGCTAACAGGGATCACTCCGACTCGTCCACGT
ATATAGTCATTCCTAATCTTATCCTTTCTAGTATGACCACACATCTATCTCAACATCTTCATTTCAGCTACCCCC
ACTTTATGTTCATATTGTGTCTTCATAGTCCAACATTCACTACCATATAACACTGCTGGTCTTATAGCAGTGCGA
AAGAATTTTTTTTTAACCTTATGGGGTACTTTTCTATCACAAAGGACTCAAGTAGCATTTCTCCACTTACACCA
ACCAGCTTGAATTCTATGATTAATATCATCATCAATCTCCTCATCTTGTCTTACAATTGATCTCAAATATTTAAA
CTTTGTCACTTGAGAAATTATATCATCTCCTAACTTAACTTTCAAATCGCATCTAAATTGTCTATTCGTGAAATT
ATAATGTATATACTCTGTCTTATTCCTACTTAAACAAAAACTCTTTGACTCCAACGTATTTCTCCATATTTCTAA
TTTTGCATTAATCTCATCTCTTGAGTCCGCTAAAAATACAATATCATCAGCAAAGAGCATGCACTTAAGTACCTC
CTATATGATACTCTCGGTTATAGCATCCACAACTAAGTTAAACAAATATGGATTTAAAGATGATCCTTGGTGTAG
GCTAATTATTGTAGGAAATGCTTTCGTCTCTCCTTCTAGTGTCCTAATCTTAGTCAATGCTCCATCATACATATT
TTGTATAAAAACATATAAATTTTATGACTAGAATTTTAGACTTTATGTTTTTCAAAAGGTTTTATACTGATTTGA
TTAGTTTTACTAATTTAAACTCATAAAAGTCGAGTTGACTTGAAAGTTTAACAAGCTTGAACAGGAATGTCAAGT
TACTTTCTTTTATTTTTTGAAGAGATATAGCACTTAAGGTTGCCTTGCCTCCACAGCCGGACACATTATACCATT
TTGGCTGTGCGATCACCAATATGATAAATGTTTCTTTTATTTTATCATTGATCACCAATATGGTTCATGTTTATG
GATTGTAGAACCTAAGGTTGCATTCGTTCTTCATCCAATGAGGATGGTTTTTCTTTTTTTTTTTTTTCTTCTTG
TGCTTTTCTGATTATCATCGCTATATATCAAACAAGAACTGTCCTCTTCCTGGCTCCGCAACTTAATTCTTGCAG
TGTATCTTAAGGAGGAGATGAGTGATGAATCGCATCGTGCTATGTTGGAACTATGACTTAATCACCTTTTTTTTA
TATTTTCAGCATGAAATCCGTGATAGAAAGATATCAGGAAACAAAAGAAGAACATCATCTAGTTGTGAGTACCAC
TTCAGAACTTAAGGTACATTCTCAGTTGAAGTACAAATAAAATTAATCAAGGTTAATTTCTATGTCTTCATGATA
AAATTTGTTTTTATATGGATGGATTTGATTTATTTGCTTGAGTTCTTTTGTAAGATTCCGCTTAAGATATTTCAC
CGTTAGAATATTCATGTGCTATATAATTCATCTGCTTTGATAGAGTTAAGAAGTGTGTAGCACACTTTTTGGTTT
AGTGCATGTAGAGCCCATTAAGTAATTAAGAATATCTACCGATCACATGAATAATAGATAAGGATACCTACTACG
TACATATTCATTTTTCTTATTTATGTTGCAGAACTAGATGATACTTTTTTGTACTTGATACCCTAACCTTTGTAC
CGAAATCATAATTCCATTAGTTTTTTATTCATGGAATAAAGATGGAAATCCACATTAAATATGATAGAATATCAA
GAAAATCAAGATAGCATCAACTTGAGTTTCACCTAGGACATCTCGTTGAACATATTACTAGAAGTTTCTACTTTC
ATAGTACATCTAAGCATATATTTTTCTCTGTGATTGATCATTTTAGCTCACGAAAAACATGGCGCAGTTTCTAAC
CATTTCCCGTACATGGCAACAACTTCACGACAGGCTAGTGGAAAAAAAAATGAGTGCATATAATTTTCAAGTAAA
TACTATTAAATTTCTTTTTTTTTCACATTTTGTTGAGTATTTCAATCTCAGAAAGATGCTTAGGTAAAGTTAGAA
GTCAGTCGACAAAAAAATTAATTCAATGAATAAAATAGAAAAAAGATTTGAAGTATTATTTGTACCAGGAACAAA
GAATAAGACGACGAAATAAAATAAAATAAAATGTGTTAACAATTAAAAAAAGCTCTCACAGACTAAATCTCAAGA
TTTAGACATTTCATGAAGTGATGACATAGAAGGATTGATTGATTTAGTGTGATTACTTAGTTAAAATGTTTAAAT
AATTGGAGCATAATATATGCTGTGGGATGGAGGCATACTACTCTTGAAACGATCTTGTAATCTTAAGATCGAAGT
AACAATTAGTCATGTGATTTGGAATGGAATGAAGCTTCGAAAATTGTGGAGGCAATCGAAAAAAGTATCAATTAA
TATCACAGAGATTAGGAGTGTTCACCGGGTCGGATCGATCCGATGAACCTGACGATCCGAATCGACCCAATCTCA
ATTAGGTCCGATCGGATCGTTGGTCGAGTCGAACCCGAACTGACTCGACCATAATCCGATCACTAACGAATCAGA
TACGGGTTTCAATTTTTCAAACCCGGTCAACCCGACCCAACCCGATGACCCGAATTTTTATATATTTTTTATTAA
ATTTGAACTAATTTATTCTTGATAAGAATATTATATTACTTTAGAATTTAAATATATTATAAATTTATACTTTAT

ATTCTTTAAATCATTATAAAATATTAAAAAATATGTTTCCTTATTTTTTTAAAATATAATTGATGTTAAGTATTA
TAAAAAAAAATTAAAAATATATGAAAAATTTAATCGGATAAATCGATGACCCGACC

NNNN

ATGAATTCGAATACTCAAAAAGACTTTATATCACTATAAATTTGCGATTATAAATATTAGAAAATCAACTGCAAA
TGCATATTTGCAGTAGACGTAATGACATGGTGGTAATAATGTCATTCGTGACGGAAAATTTCTTCAAAATTTAAG
ACTGACTCAAAATTCATCTTGCCCTACGTGGTGGTGCGTACACAGAATGAAAGAGATACGTAGAATGAAAGAGGT
CGCCTGTTGTGTTTGCGAGATATTTGTCATTACGAAGGATAATTTTGTGGGAGATCTGAAGGGAGGGAAGAAGAG
GGAGTGGCAATACATAAAGGGCTTTTGTGGAGAAATTTGCGGCAGAAAAATCCTTCACAAAAAGAAACAAAAGTA
CACTCTACTAATAAGTGTACCAAAAGATGTCACGGATGCCGCGAATTGCGGAGATTACGAATTGACCGCATTTGA
CGGTTCTTGACTACCGCATTTTGTACCGATATTAGAAATCACCGTAATTTATTGTCGTTAGTTAAACCGCCACAA
TTTGCATGAAAAAAAACCATTGGTTAGACCTCCACAAATTTTGGCGGGCATTCGATAGCTCAATTTTTCACACAT
TATAGTGAGGGAATTTTGTCCGTCTCACATTTAAAATGCGTCAATCTTATTTTCCGTTGAAAATCGTTGGCGAAC
GTAGTTGTTACGCCTCCAATTTTGATTCAAATTAAGGGGGATATTGAACTGCCCCAAAATTTCGCCGCAAGTTAG
TTATTTTCTTGTGGTGTACCTGTGCAAAGGTTTTGAATGTTCCATCGATTAGGGGTATAAAATGCAGAATAAATA
TTTATCCCCAACTCATACCAATGATAACTTTTATATTCAATTTTTTTCAAAGAACCATTATCATTAAGAAAATAT
AGCCAACTTGTGTGCGAAATTAGCCGAAAATTATTCCTGAACCCAAGCAAATATGGACATGTTTGGATATAAAGA
TTTTGACAAATCAAATATTTAATAACTAAAATTCAAACGCTCACAAAAAAAGAAATTAACAAAACAAATAAAACA
ACTCTTAAACTAAATATTACCTAAATATAAACAAAATTTCAAAATAGTTTATTTTACATAATAGTATTATTAAAA
CATAATTACTAAAATAATTTAGAAAAGGCATTTTCTTTTATATTTGATGAGATATTTATATTAAAAAATGGACAT
TAAAATGAGTTATAAGTATATAATATGTGATAATTATACGAGCTAAAATGAGTCTGCAAGGATATCTGTACTCAA
AAGAAAATGATGATACATTTTCCCATAAAATTGTCTCAAATTAGAGATACCAATCTTTTAAATCACAATTCTGCC
TTTTTTACACTAGACATCACTTTATCAACTTTTTAGAAATGTACAAAAGTGAGTTTGATTGAGCTGTAGTTTTTC
TTAAAATGTATCTAACAATAGAATTTGATCTCTGAAAATCATATATTTTTCTTTGAAATAAAATGTCATCAAGTG
AAAAGAATTTTACTCCTAAAATGTATGATGATCCTTTAAATTATCATAAATCACAAATTTAAATTTTATTTTTTT
AAATCTAAATTATGTGAACTCAATTTCTCTCCGCTCCAGTATTTATCTTTCCTCATCTATATAATGTTCAATTTC
TCTTTTCCCCTTTTTTTCATTCATCATTGCCTCCCTTATAATACTTTTTTTACACTCTATCACTGTCTCCCTTGT
AATACCTTTTTTTTCCCTCTTTTCTCATTTTTTCTCTACTATAACCTAATTTCATAACTTACTTTTCTTCACTTG
CTCATTTTTTTCTGCCATAAAGTTTCACTTTCTCTCTTTTCTAATATCTCATTTTTTTTTCTCACATCCAACTTAA
TCATTTTCCCCTCGTATGATTTTCTTTAATATCATAATTTTCACTCTCTTTAATATTTTTCATCTTCTCAATTCT
TTAATATTTCAATTTATCACTCCTTTAATATATTATTATATCTCCTCTAATATGTCATTTTTTTTTAGTTTTTA
AATTTTCTCATCTCTAGTCTTCTTACTTTATCACTTATAATTTTCTTTTATATCCTTATAATTCCCTTTTTTTCT
CTTTGTTCTAATATCATCGTTGTCTCTCTTCCAATTTTTTATTTTCCCTCTCCTGTTCGTCTCCTATATTCTTCT
AATATCTATCTTCATTTTGTCCATCTAATATACTTATTTATTATCTTCTCTATTCTCCTCTGATTATCTCTTTCA
CTCTCTCACATGATTATTATTTCTTCTCTAATCATCTTCCACCTATTGTCATCTAAGTTATAAAATAAAAATTAT
TATGAATAAATTTCGATAAAAAATCAAACATAATAATAATAATAATAATAATAATAATAATAATAATAATAATAA
TAATAATAATAATAATAATAATAATAACTGTAAAAATATAATTGCGTCAAATGTAATCTCTATTCAACACACCAC
TTTGACGATTAGGACCTCACAGGATGATAAACTCAATGCAGACTAGAGATAATGATATAGAACACATCAGCATTA
GATTTGTCAGTTCCTTTCTCGGAGGAACTGAATAGAGAACCCGTCCTTTTCATCCACTCCGTTTCAAAGCTGTGC
GTGTCAAATGTCCTACTAGCATCAAAATACATATCATGAACTGCATGGCAATTGAATTGTCAAGAAATCTCTTGG
AGGGAGATTATAGTTTAGCTCAAACTCATGCAATTTGTATTTTCTCTAGAGAACATATCTCAATCAGCCATTAAA
CCTGTCCTTGGCTCAAACTTTACTATGCTGTCCCAAAGAAGAGATCAAAAAGCTCGATCACATCTGAAAAGAGGT
TGGATCTTGCAAAAGAGAAGAAAATAGTCGTCGTGCCAATGTGCTTTAAGCAAGGCCTAAGTATCAACCGGAAAT
ACAATGCCCTTGGTAAACTTACTCCCGGATTCTGGTTGGCTCCTTGAAACACTCAATTGTTCAACTCAAGAATTT
GCAGGATTTACAGCCCAGTAGGGTGAAATCATGAATGGAGCCAATATCATTCGACAAAGTTTGCATAAAGCATCA
ATACCACTTTTAATCACTCAGTCTGGCACAACTTACAATGCCCTGGATCTTCTATATCATAGGAAGCACAAAATA
AGAGCTGATGATAAACCAAAAACATCCAGAACCTATAGTTAATTTTATTATTAATTTTCATAAAGGAATCAAGAC
ATAACTAAAAAGAAAGACAAATGTAGAAAGCTTTATATTTTGATGGATAAAAGGAGGAACGATTTGTGTCAAACT
TAAACTCTAGTTAACCTTAGTATCATATTAGCATAGTCCTACAAATTTCAAATGTTCATATCAATGCCTGTTACA
ATTGATCACAAAACATATGTTATATGTATATATCCATGTTGTCAGAACCAGACCGGACCGGCCTATTCGACCGGT
TCGACTGTTGATCGAAGTCTAGTCCGGTCCTAGTTTGATGCAAAAATCGACCAATGAGTTGGACTGGTCAAAACC
AATCAAAAATTGGAAAAACGAGAGTTGAACCGCTTGAACCGATCATTTTAATGAACCGGCCGGTTCAAGAAAAAA
AAGAAAAAAAAAATTGGGCTGGAAATGAGCTTTGGAAGTAAATGTGCCTGAAAAAAAATTATAGTTCATTGCCTTTT
AAATCAGTTTTTATTTCTTTTTCTTTTTAATTATAAAAGAATTTCTTTAATTGATATTTATATTTGAACAAATAA
TTATTTAAGATATATTTTTATTAATAATTTTTGTATCAAATAAAATATTAATTGATCATAACTTTTGAAAATATT
TTCATATTTAATCATTTATTTGTGATTCTTTATCTAAATTATAAATTATATTGTAGTTTAATATTTTTTAATAGT
ATCCGGTTCGACCGTCCGATTTAATCGGTTAAACCTGACCCGAAGGCTCTTCTGGGTCGATGTCCGATTCAATTA
TGACAATATTGGTATATATACATATACACACACTACTTGGGACTTTTGATGTGCTCCACAAAAACATCCCCAAAT
GCAGAAGAATATTTTAAAGTAGTTTCAAGTTCAACCACATGGCCAATAACTCTGTTTAGCTGATTATTATGAAGC
TTGATATAACCATTTTAATCTAATTGATGGAGAAAAATATATCATAATTAGGAAAAACATGTAATCAATTAAATA
GTCAATGGTACATATGCTGTTCATTGGAACCTGCAAGTATAGTAACATGTTGGCATTTTGATATTCACTGATTCT
GTTGCATTTCACATAAAGAATGAGCTTAATGCACAGAAAGAACGAAGATTTGGGTATCAAAAAAAAATCTTGATA

ACACCAGAATATAACATAGCGATGCTAAAAATAATAGGGTAAGTCAGATCATAGGACCTACGAAATGAATGCAAA
TTTGAGGCTTTATTTGTAAACCAACAGCGACAATAATAAATATAGATCTAATGTTGCACTAAGGAAATATCCAAT
CAGTTTTTGCTCGCCCCTTAACGGTCATAGTCCATTGCATATTCTGTTTTGGCTCCCATTCCAAGGAAATGAAAG
ACGCCTTGAGATATTCTTGAGCTAGCATTCTCAAAAATCAGCATACAGGAAATTATTCCAACTGGTATATAACAT
AGGTCTGACCATGCATGTGAAACTTTTTCATGGATAATTATGCTTATTCCCAAATCAGTTATCTTTTAACATTTT
ATTTCAGAATTCTCATTTCTATTTTCTTCTCTTTTGGAGATAGCATGTTATTTATTTTTCATTGTCATTCTTATC
CATGAAGGATATGCTGAAATGCAGGTCAGCATTCGCCAATTTTGGAACTCAAATGTTGGCAATGCATGATCATTA
AGGGTATACTTGTTCATCACTAACCTCATTATCCATATCTCATATGGCCCAGTTCATTATGCAAGATTTTGCAAC
TAGTTAATCATGCTTTGTTCGAACTGGATTTAGTTTATCACCGCTTACATGTTCCACAAGATGCCAGGCTTTCTT
TAGACATGTTAGATGCTAGTAGGTGCGGATAAACCAACTGGACATTGTGTAAGATTCTGTACTGATAATGCTAAG
ATCAAAAGTTATCATTCACATTGATGATAACATGCTCCTCTTACAAGCAGACATCAATTTCAAGCATGTTCCAAT
CAATCTGTGCAGATGTAGAAGGTAAACTTAGTTGATCATTCTCCTTGATACTTGGAATACAGAAAGAACAAACCA
CGCTTATAAATACAATACTGTACTCGTTCCAACTAAGCAGTAAGGGGCAAACTTCTAACTATATATGAACAAAAC
CAAGAACAATTATACATCTGTTAAAGAGGTTTAACATATAAAATAGACAATAGCAATCTAATAGACAGGATATCA
CAATTTTAAACTGTTCATTATTGCAATTATAATATCAACTAACCATGTGATATGATCACTTCCTGTTAACATGCT
CAAATTGAATTGCAGTCATATAACTAATACTGTCACTTACCTAATATAT**CATGTATTTTCTGCTTTCAATGCATT**
**AGTTCTGGCAAAGGGAGGCAGCAAGCTTGAGGCAACAACTGCATAACTTGCAAGAAAGTCATAGGTATAATCACT**
**GAACC**CTGTATTACATTTAAATGAAAGTAAAGTTACTTCCTCATTTGATTAATAGATATCCTGTAAAACCAAACT
GACAGATGTGCTTCAAACATTGCATAAATACAAACTTTTTAATTAACGACGTAACTCTAGGGGGGATATTGGGTC
TGACCCGACCTGCCAACCCGATCAACGCGACTAGATTTGAAAAGTTTTAGATCAAAATCCAACCCTACCCAAAAA
GTTCTGGGTCAGGTATGGATTGAAAAAATTATAACCCAATATCCAGACCCATTTGACCCAAAATTTTAATATTAA
TATTTATTTTTTTATAAAATTACTTAAGTGGCTAAATGATTTTAGTACTTATTAATCTTAATTACTTTAGTTACT
ATTATTTTATTTTATTTTAAACTGATTTTATTCTGAAATTTATTAAAATATTAAAAATTTCAGCCCATGTCCACT
TTTTGGACCTCAACCCAACCCGATCTAGTGTCTTGATCCGAGCTGAAAATTTTATTTTGAGATCAAGTTTGGGTT
TGAAACAAACCCGACCCAGTATCACCCTTTAGTAACTCTAGAGGCTACCATAAGTTTCCTGAGACCAAGGAGGGA
TGTGGGTAATGATGTTCACAACAAGAACTATGAATAACATATATAAGAAAAGGAATATTCACCAATGCAAAAGAT
ATCATTATTGCCTAGTGTACAGTATAAACTACTTTTGACACCACTTCTTATGTACATCTTACTTATAATGAGATA
TTAATCATCCTGACAATTGTTTCAGGCAGTTGATGGGAGAAGAACTTTCTGGCTTGGGCGTTAAAGATTTACAAA
ATCTAGAAAACCAATTAGAAAGTAGTCTACATGGTATTCGAAAGAAGAAGGTCAAAGATACTTCCTCACTTTATG
TCAGTATTTGTTCTTTTTTTTTATATTATATTCTTACTGCATGAGGCTACCTTCTTATAGGACCAACTTTTGGTC
GATGAAATTCAAGAATTGAATCAAAAGGTTGCAAATTATCTTCCTAGCTTTTTCATTTCATCCAAAGTGATAACA
TCAGAATTCCATGAACTTTGACTGATTCACTGCAGGGAAACCTTATTCACCAAGAAAATATGGAACTCTGTAGGA
AGGTAAACCTTATTCGTCAGGAAAACATGGAACTACATAAAAAGGTATCCAATTATAAATCTCCAGCATCCACAA
TAACTGTAGAAAAAGTTTGTCACAAACTCGTTTCGTCTCTGATAATGGGCAGAGTATCACCAACTAATCATTCAA
TCCAATCTATAACAGGTTTATGAAACAAGGGATGTAAATGATGCTAACAGAAGTCCAGTTGTTCCATTTGGTTTC
AGCATCATGGAAGAAGCAAATGTACCTATTCACCTTGAACTAAGCCAGCCAAAACAACAAGAAGATGAACTACAA
GCAGGACCAAAGCTAGGGTAACTTTCTTTTGAAAGTTCAGAACATTACAGCTTTGGGTTGAAACTTTAATCACAT
AATGATTTACTATGTGTGAAACAGACTTCAACTGAATTAAGGAGATGACTTGGAGGTACTTTTCATCGTCTTCTA
TGATAATAATTCAAGGCAAAAAGCGTATGGTATGAGCAGTATGGTTTGATGATAAGAAAGAGCTGCAAACTCAAA
TAGTGCGAACCCCACAATGTAAAACTGTCATGATAGAAGTATGAACACTCTGCCCAGTATTAGAATGTGATGTGT
GTTTATTAAGATGTCCTGTAGCCAGTGAGATGGGATGTGTATAGATGTAAGTTTGAATTGTAACAAATTCTTCAA
ACATCAAAAGTCTGGCAATAAAACCAACAATTTGATGTGTTTATTAAGATTTCCTGTTTCAGGATCAAATCCTAG
ATCTTCCACTGTTGAAAGAAAATTTAAACATTTACACTCCTGCCTCGAACTAAATTAATAAAAGGCTAGTAGCCT
CTTGGATACCATTGCACCGGAAAATGAAAAATAAAAAAATTTCTCCTTAAATGAATGAAACTTCTTAATTTTATA
TAGCATTCATCTAAATTTCAAGCTTTTAAATGAAGTAACAATTCATTCTTTTTATTTCAAGTCTTGACACACACA
TCGTTTTTATTTGTATGAATACATAACAAAATTGATCTTCAACCTTAAAGGCAGGTGAGATTCCATCCAGGACAT
TTCGCACCAGGACCATCTGTTTTAGATTAAGGCTAAGGTTATTCATTAACTGGTCTTTTTTCCCTCTAAAAAAGA
CTGGTATTGTTTTTTTTTAGGAAATAATTAAACGCCATTTGTGGGAACTGAACCCAGTTTTTAACAACAAAAAAC
TTGTTTCCATGACAATATAAGCAGAATATATTTACATAGACTTAATTAATTCTGCCCTAACAAATAGATGGTAGA
CAAGCCAGAAGAAAATAGGAAAGTACACACCACGATTTGCAGGTGAATCCTTTTTAAAAAATGCAATCAAGTAAT
TATCACAAAACCAAATTAACCTTCCCTTTCGGAACTGGACAATTCAGCATTGACCCAGATCAGACTGTCTCAACC
AGAATATTTTTTCTTCTGAAGGAAAGCCTTTCATCCAAAATATGATTTCCGACAACTAGGTCAAGGCAGAGCAGA
GTTTTTTCTTGTGTGTATTGGTAAGTGAATGTGATTGAAATGATATATGCAAATTTGTAGAGACAAAATGTAAAG
GAGTGCCATACAAGAAAGAACATATATTGATTTTGAACTTTCATCATGCAATGTATGAGATGGGGAAGAGGAAGA
AGTAGAGAAGAAACATTAAACATCAGGATTGAGAAAAAAAAAGCAACTTTAAAACCCAATGATAAATCGAGTAAA
TTACACAAATGGTCGACCAACTTTATTCGTTATACCAATATGGTCAATAAACTTCACTTAATAACACAATGGTCC
TCCAACTTATAAAATCTTTCACCCACGGTCATGTTGACTGACTATAGTCAGTTTTGCAGTTTACATTATGATGTG
GCTACCACACGTGCATTTCTCTCTCCTCCATGCACACATCTCTCTTCTACACATCAATATTCTCTCCTACGTGGC
AAAAATAAATTAGTAAAAATATCGATAGAGGAGAGAGGTTACCCCTTCTCTCTCCCTCATGGCCAGCGACGAGGG
GGAGAGAGAGGGTGGTCGTCACCCCCTCCTCTCTTTGCCCGACGACGAGTGGGAAAGAGAGGGGGGAGTCACCTC
CCTTCCTCTCCGTCGTCGTGCATGCAGAGAGAGGAGGGGGGCAAAGTCACTCCCCTCTTGTCTTTGCATGTGTCA
TGCGCAGAGAGGGGAGGGGGGCGAGCGACCCTCTCATCTCCCGTGCGCAGACAGAGGAGGGGGGCGCCCCCCTCC
TGTCTCTGCGAGCATCGTGCGCGCATAGAGGGAGGGGGAGCGCCCCCCCCCCCCCCCCCTTTCTCTCTTGTCT

CTGCTCGCGTAGTGCCCGCAGAGAGGAGGGGGAGCGGCATCGCCCCCCTCCTCCTATCTCTACATGCGTAGTGCA
CACAGAGAGGAGGGGGGCAAGAGGCCCCTCCCATCCGACAAGGGGCAGCAACAAGCTGCTGCCCCTTTTTTTAGA
ATTTATTGTTGTTTTAATACATTTTTTATGTGTATGAGGGATTCATGATGTGGCGAAGAGAGAAAACTATATATG
TTGCATCCACGTAAACATTTTAACCTAGAAACAGATAGTAACCGGTCAAAATGATCGTACCTGTAATTTTTTTGA
AGTTCGATAACCATGTGGTTACAAAGTGAAGTTCAGCGACCATTCTGAGAATGAATAAAAGTTGGATGGCCATTT
GTGTAATTCACTCGTATTATATCAAGAAACTGAGAAGATAAAAGAAGTTTCAAGTTCAAATAGGAAGTCACACAT
TAATGGGCGTCATTTCTATGACTAGACAATCGTGGAAAATGATGTAATCATTACATTTAGGATGCAAAATGTGTC
TCCATAATAAAAGATATTAAGAAGAAAGACATAATGAGGGAAAAATGATTCTTGAAAAGATAATGGTAAGAATAG
GTAATAGTGGTTGTTGATTACTCTTCATAACAAAAGACATCTAGGAGAAAGACATAATGAGGAAAAGGAATTCTG
AAAATAAATGATGGTAAGAATAGGAAACAGTGATGGTTGATTATGACATAAAAACTTGGAAGATTTGATTAGAAA
TATAAAAAAAGATTTGGAAGGCAAATAACCTGAAATATAATCAGATCTTGAAAAACATATATACATATGTTGTGG
TATAGTACCTTGGGTAAATAATGAACCACGTCAAAAGTAGAAGAAAATCAGAGGTACATGGACGATGAAACAGGA
AGATGAACTGTAAGATTCACCAATATCAGAGGTAATTTCAAGATGAAGATGGAAATGGTTCAAGAAATTATGGTA
TTAGTGAAGGACATTTTTTCACTCTCAATAATCAGGAAATAGAAACAATATGGGCAATTTACCTGGGAGATAAAC
ACCAACAATCAAGAAATGTTTGACAAATTATGACTCTGACATATGCAACAAAAATAGGGTTTAAAAGCATGAGAG
GTCTTCAAACCTTTACAAAATCAATTTTTAGGTAACTCAACTTTTGCTTTTAATATATAGATCACTCATCTTTTT
AAAAAATCAATATTTATGCCACTAGACTAATAAAATATTTAAGTTTAGCAATGCGTCAATGCAACATCACTAATC
ATATTAGCTAAAAAATATATTTGACAAATCATCATGTTTGCTTATTTTATTAGTGGAATGTCTTGAATATTAATT
TTTTTGAAAGATGTGTGATTCATAAGCTGAAAACAGATGTTGAGTGATCTAAACAATGATTTTGCAGAAGTTGTG
ACATAAAGCACTTTAAATCCTAAAAGATATGATCATATGATTTGCAAACTTACTTTTGCTTAGCATGGTATTTCT
AAGGGAGAAGTTGCCAAATATGAAATGATCACAACTACAAAGGGGAGTTTCTGTTAAGAAAGTGAGAAATATGCG
CCTGATCAAACAGTTTTTTACTTAAAACTATCATGGAAGTGCATAGCACTTATACTCACACAGGTGCAATTATCC
AAAGGAAAGAGTATACCTAGTTCTCATGCTAACAACATTCACTAGACCTCTCAAAAAAAGAAACATTACACTAGA
TGACACCATAGCACTATGTGGATATGCTACAAGTAGATGAAGTAGAATTTTAAAATGGAGATTCCAATTCTTGCA
TGTCTGTTATGGTTCACCTTTGAGCTATCTCCAAAAAGGTCAGGGACCCCATGCTGCAGCAACATATTCCAAGCA
AGGGACTTTTAATAACAAAAAAAGCTACTGCCTCTTGGTTTACAATAATCAAGGTTGCCCTTTCGGGCAAAAAAA
AACAATCAAGGTTGTGGCTCAAGCACAATCTTGCAAGAATACAAACTCAAAAAGGAGACCAGTTGTCTCTTAATA
ATCTTCTCAAAAAAAAGGTTGTTAATAATGCAACTGAACCACTTGATAACTATAGAATATGTCTACTATCATGTA
GATATTCCCCAATGTCAAAATGACATGGCAACAAAAAAAAACTACTTTGATATTTGTGTAGTGGGATCCTGAGTG
TTGGCAATTTCTTCTATATAAACTGATTTACATTCTCAGAACACCCTGAATAGTGACCTCTACTAATGCATTTTG
TAACTGAAACACAGGTCTTTGAGCATTATTTTTCAATAGGTAAATTAATTGATATTCATTTAATAAATAATTGTT
CCTTTTTCGATGTTCTCAAATACCAATGTATTATTTTCTGGTTAAAATTCTTCATGGTTCGTTGTTTAGTGCATC
TGAAAACACAATGTTCAACCATTTCTGAGTTAATTACTCAACAATCACTAATAATATGTTATTGTTATCTATTAA
AAGGCTCCAAGGAAACTAGGCATAC**CATAGATATTCTTGCAGGTTATGCAGTTGCTGCCTCAAACTTGGTGCCTC**
**CTACAACAC**CTAGTGAAACAATTGATATACAGTCTATAAGAAAAATGGAAAGTAAATAATGACTTGCTCAGGCAA
AATGTGTAAGCCAAAAGAAAGCACAAGTCTGCTTAGAGGCATATGCATGTCAAATATAAATATGCGTTCATATTT
GTATATCCAATATGCTTAATGTTATCTGTAGAACAACCCCATTTACTGCATGCCAGCATGTAATAATTCTCCTTT
GTAAGAATGTATAGTTATTGACAATATACCCATATTCTGAGAAGGATGACCACAAGCTTTTCGTTTGTGCAACAA
ACAAGACAGACGATAAAACATATTAAATCAAACGTGTTGTTTGCTCTTTTAATATTTTTTGTTCTCACAAGATAC
TTGGAGGAATCAGATTGAAGTTGTGTTAAAATTAGATAGAAGCTACAACAAATCTGTTAAGTACTAATGTATCCA
AATCCTCCTTTAGAAAAACCAAATGTACCAAAAACTAAGTGGAAAATATAGTTTGGAATATTTATAACTATACAA
AATATTGTTTCATTTTATTATAAGCATTATAATGATAACTTTCATTTGATAGAGTGTATAATCCTTCTCAAAAAA
GTATCCTGTTTATACTACATGCCCAGCATGGAACTCAATACCTCTGAAAAAACATAATCCTTAATGTGCAGTCTA
TAATTCATATTCTAGGATGTTAGAACATTTTGAACAAGCAATTATATTGCATGTTATTATATACTGGAAGTTTTAG
CATTATACAACAGTAGCAATTTGTTAGGTTATCATCTTTGAAGTTCATAGACAATGCCTTAAACGCAAATCTGTT
TGTCGATCGGTTGTGCTAGACTGATTCACTGGACTAATAGTGGTGATCACAGCTTTATATTTCCAGACAAAGAAA
GAAATCATAATGCAGAAGAAACTGGACACAAACTAAGAAAAGATGCCAAGAACACCACTTATGGTCTTCTGCAAC
ATGTACATTGCAAAACAGTGAAACACCGGCAATGTGTTCGTGTACTTTTATGTAGAATCTATTTACAATGGTAAA
GCATTTACCCATCATGACGCAATGTAGAATCTGGCAACAAATTAAGATCACTCCATTACACATCATGCAATGGAC
ACCAATTTTGGTTCGTGCAACATACCATTGCAAAACACATGCAACAATGTGTTCAAGTGTTCTTGTGCAGAATCT
ACAATGGGAAAACTGTTTAGAGTAAAGAAATCATTTTCAAGATTACAGCTTATTAATAGTAAATTAACAAATATC
AAAAGGACTCTGGATTTTCTTTCTTTAGTATTAATCTTATGTAGAGGCAGTCCAATCATCCTCAAACATATCATC
AAATATTTGATCATTTCTTCAATACCATAATGCCATTTTTTCAACTTTACATACAAATCAATGGGATATAATCCC
AAATCAAGTAGTACACCATGTACAAATTTGTTAAAAATGCACCACCTAGTGTTTAAGCTCGCTAGAAGCTTATCA
GTTAAGATAGAATGATTAAATGCAAATTTAGTTCTAACATTGAACACACTGAGACAAACATAGATCACATACATA
AGTGTTTGCTGATGCAAACTATAAACTAAACTAGAATGCTAAGCTACGCTGAGGATTTTTATCATTCAACTTGAT
GTCCATTACGACTGCAAAGTAGCCCTATTTATAAAGCTTTGGTGCTTCACAGAATACGACACCTAAGTTAAGTAA
GGAAACTAAATAAAGGTAATAGAAAATAAGTAAAGATTAAAAGGCAAATAAGGCAAGGAATCAATTTTGAGCTGC
CCAAATTGCAATTAATTTCAAATTTGAAAAATCAGCAGGAATAAAATGTTGTTTTTGAATTGAGGCTGATTTCAA
ATTAAAAAAATCAATGTTCGTCTTTTGTAAACTTCCTTAAATTAAGGCTCATTTTAGTCCATTCCCTCCAAGTGA
GAAGAACTCCGACGAGTTTTGAGACAAGTAGTGGTGCAAAGCAATGTTGTCAGAACCGGACCGGCCAGTTCGACC
ATCGACCGGGGCCTAGTCTAGTCCTAGTTCGACTGGGCAATGAGTTGGATTGGATGGAACCGGTCAAAACCGGAA
CAATAAGGGTTGAACTGCTTGAACCAGTCATTTTCAATGAACTGGTTGGTTCACTTGTAAAATAGAAAAAAAAAG

AAGGAAAAGTGGGCTTTGAGAGTAACTGGGCCATGAAAAATATTATAATTCAAGGCCTTTTAAACCAGTTTTTGT
TTTCTTTTTCTTTTTGTTTTTAATTATAAAAGAAATTCTTTAATTGATACTTGTGTTTGAATAAATAATTACTTA
AGATATATTTTTATTAATATTTTTTAGATCAAATAAAATGTTAATTGATCATAACTAATATTGTTTGAAAGTAAC
TTTAGTTCTTTTCATATTTAATCATATTATTTGTGATTCTTTATCTAAATTATAAATTATATTTTAATTTAATAA
TTTTTAATAATATTCAGTTCGACCATTCAGCTTAATCGGTTTAATCGATCCGATTGACCCGGAAGCTTCTCCGGG
TCGTCGTCCGGTTCGATTCTAACAACCTTGGTGCAAAGGGATTCATTAAGGCTGCAAAGTTCATCTCCTGTGATC
CAAGTAGCGTCGGTCGTAAGGCGACCAGTGCATTGAACAAGATAGCGGGTGATCGCACTAAAGTGGTTGAGGATA
CTTTCATTCTCCAAGATGGTGTCAATTTCATCATGTTGATGGCAGGGCACTTGTGGAGAATCAAAGAAGGTGTAG
ATGATGAAGATCCTTTAAATGATGAACTAGAAACAAATGCAGGATGCACAACAAGAAGCTCAAAGGTACCACGAC
AAGGACATAAATTCTCTACATTACATATTGAGCTTATATGCATGTCAAGTGGCAATTCAAGTACACAAGCATTGG
ATATTTGCCTAGAAATCACTCGAAACGAGCCACAAGCCCTACCAAGCAACTTTTTAAAAGAACCTCATGAAAACA
CTCAGGACGAACACAAGCCATTATCCAATCTCCAACATTAAATTCTAGATGTATGCATCGTAAATCAGCAATTGG
TTTGTAGTTTCAGTATGTAATGTTATTTTCTGCCGAATAATTTCATGTAACTCCTTAATGTGCTGGGCAAATGAC
AACTCTCTTAAGTGTGAGCATGTGTGGGAGGCGAAGTAGATCAATTGGAGTGCAAAACAATCTCAAAAAGGCTTG
TATTTGTGGATCTGTTGACAGAATTGTTATAAGCAAATTCAAACATGAGCAAAATTAGGTCCAAATTGCTAGTTT
ATCCCAGACTAATATTGAAGGAGGACACCCAAACTTTTGTAACTACTTCGATTTGTCATCAGTCAGAGAGTGGAA
TGATGAAGAATACTTAAGTTGTGTGCAAAAAGGTTTTTCAACCAGAAATAGCTTACAAATTTTACATCAGGATCA
GTGACTATCGACTGTGAGAAACCATTTAATTTAACAATTTCACTGAAGTTTGGTAATAAGGGACGCTCAAACTTG
AGTCAAAAAGCAAATAATCAAGAGAAAAGTCAGATGTTTTACTACAAAAAATGAAGTGTACATTTTAGAAAACCT
ATGAACGATCGCATGGAATCATGGAAGTGACGAGTACAAGGGAAAACAAGTACAAAGTCCATACTAAGGTCATGC
CACAAAGTGTGAGGTACAGGTAAAGGTGTATAAAAACTAGTATCTTGACATTTAACCTTGGCCATTTGACATGTG
TGACATTGTGTAACAATACCAATTATCTCTTTGTATAAAAAAGGCCGATCAGCAACTAAAGCAAACATCTTATCC
CTATTAAAGTGACCAGCGAGACCCCCAATGTAGTTCTCATACAAGAAAGTCTCGAAGAAATGTACGAGGAATGCA
CAATTGTGTGCCATGAAACACAAAATCATCATGAATGGCATAATCAGGGTGATCGATTCGTTCCTCATTTTAAAG
TGTCAATTTTTTTTCCTAAAATCAAGATAAGATGTTCATCACGTAAATGATCAAAACCACATATTTGTATGTGCA
TTGTGTGAAGGAGGATAATCAACGACTTAGGGCATCAATAGGTTTGTTATTGACACAAGAGCAATGTTTAATCAC
TATTTGATGTATTCAACCCACTTGATATGTCTAGGATTGAGCTTCGTTTATGTGTAGATGACAAAGAGCTTAATG
AGAAAAAAAAAACAAATTCATAAGGCACAAGATAATGACACTAATGTCGCAACTCGTAAAGTTTGAATAATGGCA
CAAAACTCCGAGTTGAGTATCTTTGTTTGGCATCATTAAAATTTTCACTAGTAGGTGAGAGGGTGACTTTCTTAA
CTAAGGACACAACTGATACCAACACCGAAGGTATCACAAGAATCCTCAAAAGGTGTAGAGAAGTTCAGAAGTCGA
AGAACAGGTGTCTTGGTCATTCTGATCTTAATTGTGATGAATGCACAATTAGCAAAAAGCATCCAGAGAAATTCT
TTACTTTTGGAATCCGTGAGGTGTCATGAATGTGCTAAGGTTCTTGTTAAATCGGCAATAGAAAAGTAGGTAAAC
CATGAAAACTTCTAACAACAGGCAAGGTTGTGTTTTGAGACCATTCGTGAATGGCTCGTACATTCTCAGGATCAA
CCTCAACCCCTTGAGAGGTAATTATGAAACCTAGGAAGGGGACATGAGGTTGTATAAAAGATTACTTCATAAGAT
TGACATATAATTTTCCTCGACGAAGTGTTTCAAAGATGGTGCGAAGATGACATATGCGGTCGTCTTGGTTGAGGT
TAAGGCAAGGAAGTTGTCAAAACAAACAACCAAAACTTGCCAATAAATGGTCGTTAGACATGTTCACAATCTACG
TGAAGGTGCTTGGGACATTAGAAGCCAAAGGGCATTATTATCCATTCATATAAACCATGTGGAGTTTTAAAGGCA
AGACGAATATTAATTCAATGATAATCACTAAAAAGGCCGATTTTGGAAAACCAAGCAGAGCCATGTAAAAGTAAA
TTATGTCATCAAGACGAGAGATTAGAAAGCAATATTTGATTGTGATTTTATTAATAGGTTGGCTATCAATACACG
TTCTCCAAGAACCATCCTTTTTAGGTGTTAAAAGAGAAGGTACCACACAAATGCTCATACTTGCTCTAGAAAGTC
TCGATTTGCTTATGAAGCTTAGTATTTTCATTAGGATTCACTCGATAGTGAGGACAATTAGGTAAAAATGATCCA
AGAATGAAATCAATGGCATGTTGGATATCTCTCATAGGCAAAAGTTCATCAAGTAACTCCTCTGGGGCTATATCA
TGGAATTCCTCAAGGAGAAGTATAAGAGATTTGTGTACATTTATGGCTGAGAATGATGGTATACAAACTTATTCA
AGACCTTTGCCAATATTAAGATCAAAAGTTTAGAAATTCTGGAAGTAATTGAAGAAGTACCTTGTTCGAGAGTTA
GGGCTTTATCAATTGGTTTGGCTGATGTGAGTTTAATAGCACAACAATTATGTGCAAATGTGAGGTTGTTACATC
ATCCTCAAATCCGCTGAGACAACAGCCCAAATCCATGTCTACCAAAATAATATTGCAACCACCATGGGAATGGCA
TGATACCAAATTTTAGCGATGTTCTAAACAACGCTCAGAGAAAAGTAGTGATTGTTTATCTACCTAAGCGACGCA
AAATAGGTGTGGATGAGGTTCAAATTTGAGTTTAAGACGAGCAACAACATGAGGTGAGATGATGTTAAGACAACT
ACCACTATCAATTATTACTTTGACGACTTTATCGTCGAGTCGAAGAAATGTGAAAGGTGGTGTCGTGCATCTCCA
ATCATTAGAGATTTGTGATGTGGACAACAATCGATGGACCAAATTAATGTTTGGCTAATGAGATTCTTCAAAAGA
GTCATCAAGATTAGATGGCATATCTTCGGAATCAATGATATAATTAATAGTATAATTAGGTAGATCCTCTTCATA
AATTAAGAGTGTGTTGAGTGGAAAACTTGTAGAGGTATGGCCAAATTGATGACAATTGAAACATTCCATATTTGA
GGAAAGTGGTTTGGAAAACAAAGGGTCATTAACTTTAGAAGGAGGCATGGGTACTAAATTCTTCGTGTGGTGCAT
AAGGATTGGTACAAAGGAAGTTATATTGTTGTTAGAAGGACAATTTGCCTTGGAAACCAACGGCACAACGAGACA
TTATAAGCTTTAAATATCGCCCAATTTCCTAAGCCTTGTGACAAGCCTCATCAAGGGTGGCAATATCGTGCAAGG
ATAGATAAAGGACGTATGTGGGGCGTAGCCCATTGGTGAAGTGAAAACTACTATCCAAGGGACCTCACACAAATC
AAAACTCGTTTTGAACTACTCAAATCGATATAAATATTCAACCACGATAATTGAACCTTGACGTAAATTCATACA
TTGTGTTAGCATGTATTGGCGATGATGTATTGAGACATAGAGAGATATGATGATATGTTTCATAGTGTGCCAATG
AGTGATAGAGTTTAAACCAAAATGTTCACGTTCTCCTTGCACACTTTGCCAGTAGCGTTTGACTAAGTCTATGAG
CTTGACTTTGGCAAATTGGGTTTGCTCAAGATCAATGATATGATATCATTCGAAGAAATCTTTGAGTTAAGCAAT
ATAATCAAGAAATATAGTACCATCAAGGCAACCATCAAATTCTTGAATATCAAACTTCACTAGACGAGTGATGTC
TTCATGAAGGTTGATGACGAGGAACATGTGTACCAGGACGAAATTGTGCAACTTGGCGAGGAGGGTGTAGAGGTT
CATTATGGAATCATTGGTAAGCAGAAAGTCGTGAAGCATTTGTGTAATCATAGTGATGATAGTGATGTTCATAAC

GCTTATACTCAGTATATTTTACGAAATTATTAGGAGATTGGGCAATATATGATGGTCGGTTGACATAATGGTGAT
GAGGATGATTGTTTGGCATCAAGGAATGCTTATCGCACAAGGTGTCAACTTCTTTAACCAAGCTGAAACACATTT
TGACAAGTCATGGAAGTCAATTATTCAATGATAACTTTTAATTGGTCAAACTTCATTAACCTTGGATGGAAGGGC
TTCCAAATCTTTTAGCTTCTTGAATATTCGGCAAATTATTCAATTAGGAGCAAATTAACACCAAAAGTACTGGAA
CATAATGCAAGAGACATAAGAAATTGTCCGAGTCCATGTGTAATATGGTAGTAGAAAAATAAAAGGCCGGTTATT
TATTGAATTTAAACAAAAAGGACCCATTGCTGGTGAACTAAAATAATCAAATGATAGCAAAAACACAGTAGTTAA
ACTAGAAAACTTATATCGAGTATCAAAACAATAAGAGCTTCATTGTTGCAGTATCAATATTGTTGGGGGTCCAAT
ATAGCGCAACAAGTTGAATATCTTGTGTTGGTGATGTAACCGATGGTGAAGATTTGAGGCCTAATAAAGAAAAAG
AGAAAGAATGACCAACCGCCCTTTCTTTTTTTCTTTTTTTCCTCTTTTTTGCAGATTGGTGGTCGCACACAAATA
AAACGGAGGAAAAGAGAGGAAAAAAAACTAAAGTAAAAGTAAACTAGCAAAGGAACAAAAGGAATATTTTTTTTT
TCTTTTTTTCTTCCTTTCTTGTGTTATTTAGAGACAAAAGAATAGTCTTTGATGCTCACTTGCAACTACATTCTA
GCCTTTGATTCCGAAGGTGAAGTCGAAAATTGACGGCGAACAAAGTTGATGAGTACTTGAACAAAGGGACGGTTG
ATGAGGACTAGAGGAGCAGGTAGTAGTTGTGTGAAGTGGATACTACAACAGCGGATTGTTTTAAGGAAGATTGG
AGGTATGGTGGAGAGGTGGTGATGCCTAAGAGAAAGAAGTAAGAGATCACCTATGTTGGATCTAGATTAGATGAC
TCTCATACTAAACTAATGCAAATTACAAACTAAACTAGAATGCTAAGTTATGATGAGTATTTTTACCATTCAATT
TGATGCCCATTACAACAACAAAGTAGCCCTATTTATAGAGCTTTGGTGCCTGAACGACATCTAAACGAAGTAAGG
AAACTAAGTAAAGGAAATATAAAATAAGGAAAAATTTGAAAATAAATTGGGCAAGGAACTAATAACAATACAATT
TTGAGTTTATTTCCTACCCAAATTGCAATTGATTTCAAATTTGAAAATTGAGTAGCAACAAATATGAGTTGGAAT
TGAGGCTTATTTCAAATTTGAAAAATCAACATTAATTTTGACCACGTATTTTGTAAACTTCCTTAAATTGAGGCT
TATTTCCATCATTTAGCTCATTGTAAGTCAAGTGCAAATCGACAAGAACTATGGTCCGGTGAGTGAACGAACTTT
CCTTGGCTAATATGGATGGTACGCAATCTCGGGTTCCATACTAAAGTTACCAAGCGAAGCGCTTGGTCCAGCTCT
AGTTCATATGGATGCTTCCAAATGTTTGTTAATCATTTCAGAACTTTTTATGATGATTTTCAGATATTTCTATAT
CAGTTAACATTTTATAGCATCTATAGTTAAATAAAATTTTGGGTAGTATTTTATACTAGGACTATTTTTCTTGAA
AACCTTTAGGATTAATTTTGTCCAATATCAATTTTCATTCACATAAAAAAGATTCGAGTATAGGAATCCAATCTT
TTACATCACAATTTCATAAATTCTTTTGTTTATTTTTTCAAGGGAATATAATATTTTGTTTCTTACTGGATAATT
TCATGTTGTTCGATCAAGAATAATTTCTTTTATGATGTACTTTATTTGAAATGGAGCTGTGTGAAGTTTATTTGC
TCACTTATTAATCACAAGATAAACTACTCCGCGTAAGTCATGCTTTTCCAAAATGATTCCTGCTGCATTTATAAT
ATAAATATTATAATACAACTCAATTAATATATGGTTTTAAATAAGAAGGGGAGGGTTGTAACTGTGTAACAACTT
GGATCCATGTTTGTCATCATACCCTGTCAGTCCACCATGGGGTTTATGTTCACAAAATTCATACAAACTCGTAGT
TTCCTACAACTCAGTCCAAATTTATTCTGAATCTAACCTCACCGTATTTTAGCCTCCACATGACTAGGAGCTTTG
GCAAAAGTATCTAAACAATCTCTCCGCAGTCTGTCCTTGATTTCTGTAGCACTTATCATCCAAACACCTATATTT
CTGAGTGAATTTGCAATGCGGCCACACATCCACTTCATGGTCAGTTGGTCACATCAGATACAATCAATTAGCAGT
ATAAATGAAAATGCTACAATCCTACCATACCAAGAGAATGCTTTGCAGTTGAAGTCACTATAATTCTCATTCTTA
ATAGTGCACCATAGTCATAACATTTTCTTCAGTATTATTATTTCTTATAACATCTTTCCGGTTACTGAAGTTAGA
AATTTGCTGTCAGCCAAAAATAGGGTATCTGAATTGTAGTCAAAGCCTCAAAGTCATGATTTCCTTCACTTTTTG
TTTGATTTATAGTTTATTCAAGTACATCCCCAATTTCATCTTTAATTTAAAAGTGAGACCATAGATGTCAAGGTA
ACTAAATTAGATATATATTCATCTTTTTCTTTTGTGATTCTTCTGCACCGCTTTGAATTGATATTCCAAATGGTT
GAGCATGCTAAGTAAAGCATGTGATAACGAGATCATCTAGACCAACCAGCATCCGAAACTTTCAATCTGTTATCC
TCCAAAGAAAGAAACTTGAGTTGTATGACCAATACTGCTATGATGCTAAATATAGTCCAGCAACGAGTGAATATA
GTTAGCCTGTCGCATTCAGTAATTTCAAAAGGAGAGTAATTTAACTCTAAGCTTCAAATTATTTAGCAGCAATCC
AGCATATTGTCATAAAATATTGTTGTTTTGTTAGAAAATGAGTCATTTGACATCCAAGTATAGTCATACTCTTAG
TCATACTAAGGTTCATGACACTTGCATTAGCAATATGAGGACACAAGATAGCTTTATAAGCATATCCAAAATAAA
CTGAGATTGCTGGATGTAGATCGACGATTGTAGTTATGGGAGTATCATCAAAATTTTAAGGTCTTTTGGAACTCC
AAAACCTGGACTCGAAACCCTTCTTTATTCAATTTGGAGTAGTAAATAACCAAGACATCAAACTCTATAATCAAG
ATATCTATGTTCTATTTTGAAAACGAAGTGGAATTGATGTTTCCAAAAAAAGATTGGAACTTCTCCTTCCATTTT
CAAGTTATCGAGGTCATGACAAGTTGACCAGTCAAAATCACATTTTACTAATCTTCAATGAACAAGGGAATGCAG
CACCGAATTCTTGAAGAACCCTGAATCCAAGATTCTCAGAATCAAGGCGACTTAAAACCCTAGAACAGAAAACCA
GTACACATCCAGGACAAAACCTGAGCTCTAAACAGCATAAGACACTAAGAACCGCACCAAATGAAAAAAAAAAAA
GATCATGAACAACCACACGAATCCTGCGTAAACGCAGGTTGATGACCATCAAAACCCCAAGCAAAATCAAACAAA
ATAAAACACGAAGCATTAATGGCAAAAGATTCAAGGTTCGTCACCATTAAAGCCAAACACAATGCCTAAAAAATG
TATAGAAAAAAGCATATAAATCTACTTCCACACACTACTTCTTTTTAC**CTGCTTCGATCG**AGAAACCAAGAATCC
GATCAGTCCACCGATCCAAATTGAGAAGAAAAAAAATTGCAGGAAGTAGAAGGAGAAGAGAGCGAGAGAACGATG
GAGATGGACGGGGGCGCTTATAAATATCTTGAATGGGGTATACCCGAGGGAGTAAAATAAGAAGCCCATGAGCCG
CTCGCGTCAGTCCACGTCGCCGCGCTCGTTTGTTTAATGCCTCGGTCTGACGCCCGTCGTTTATTTTGTTCCGCC
GGTACGCCAAACTGTCGCCCGGTTTAGTGGCACAAAACGAAGGGCTGCGCCGCAGGATCATTGTAAGGTCCGCCA
TTCCTCCGCTCGCGAGCCGTCCACGTCAGCAAGGGAAGCACGCAGGCTAAGGTTGGATCGGGGGTTCTGGCGGAG
CAACGTAAGTGGTAGTGGCCGGGGATGGGGGGACCGCTTCTTCGCTGCTCTCTCCTACGGCTACTCTTTGTTCAA
CTTTTTTCGGAAAAAAGATAGTGGAGGAAAAAACAATAATAAATTGAAGTGGAACACATTTC

>EpaAGL17_2

ATGGGGAGAGGAAAGATTGTGATAAGGAGGATCAATAACTCGACCAGCCGGCAGGTGACCTTCTCGAAGCGGAGG
AACGGCTTGCTGAAGAAAGCCAAAGAGCTTGCCATACTTTGCGATGCTGAAGTCGGCCTCATGATCTTCTCCAGC

ACCGGCCGGCTCTATGATTTTTCTAGTACCAGCATGAAATCCGTGATAGAAAGATATCAGGAAACAAAAGAAGAA
CATCATCTAGTTGTGAGTACCACTTCAGAACTTAAGTTCTGGCAAAGGGAGGCAGCAAGCTTGAGGCAACAACTG
CATAACTTGCAAGAAAGTCATAGGCAGTTGATGGGAGAAGAACTTTCTGGCTTGGGCGTTAAAGATTTACAAAAT
CTAGAAAACCAATTAGAAAGTAGTCTACATGGTATTCGAAAGAAGAAGGACCAACTTTTGGTCGATGAAATTCAA
GAATTGAATCAAAAGGGAAACCTTATTCACCAAGAAAATATGGAACTCTGTAGGAAGGTAAACCTTATTCGTCAG
GAAAACATGGAACTACATAAAAAGGTTTATGAAACAAGGGATGTAAATGATGCTAACAGAAGTCCAGTTGTTCCA
TTTGGTTTCAGCATCATGGAAGAAGCAAATGTACCTATTCACCTTGAACTAAGCCAGCCAAAACAACAAGAAGAT
GAACTACAAGCAGGACCAAAGCTAGGGTAA

>epa_miR444b

CGATCGAAGCAGGTGTTGTAGGAGGCACCAAGTTTGAGGCAGCAACTGCATAACCTGCAAGAATATCTATGGGTT
CAGTGATTATACCTATGACTTTCTTGCAAGTTATGCAGTTGTTGCCTCAAGCTTGCTGCCTCCCTTTGCCAGAAC
TAATGCATTGAAAGCAGAAAATACATG

**Position of mature miRNAs based on comparison with osa-MIR444d and not based on experimental data!**

Stem-loop structure

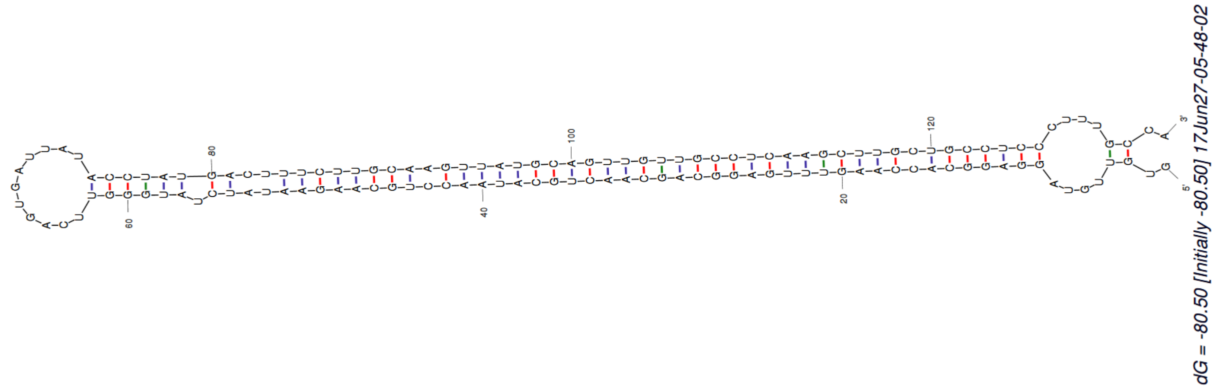


*EpaAGL17_3 and epa-miR444c*

>LTAE01003480.1 reverse

GTCTAAGAATATATAGAAAGGATTATAATAGCTAAAAGGAGAAGTAACCTCCACCAAATAGAGAGGCAAGAAAAG
GGAGTACCTCTAGTTGAGTCGTGACCATCCATCGACCAATAAATTGTTCATTGTCTCCATCTTACAGCAAAGGTT
TCCTCCTCTCCTTTGTTTCTTCTTTCTTCTTCTGCCTCTTGTTCTTCTTCTGGTAGTCATACCATTTCTGGATAT
GTTCTAATTGCTGTTTCTAAGCGAGTAATTTGTTGTGAAGTAGAGATCATGCATCTAATTGTTTTTTAGGCCTTA
AATCAAGGTTCTTTTTTCAGCTGCTGATCATCTCTAATTTCTTCTCGTCTGATATTGACTTATTTTTGACCTAGT
TTGTTGATTTGTTCTTTTTTTTTTTGCTTATGTTGAGGTTTTGAAGAAGTAGTAGACTAAGATGGGGAGAGGAAA
GATTGTGATCAGGAGGATCGACAACTCGACCAGCCGGCAGGTGACCTTCTCGAAGCGGAGGAACGGCCTGCTGAA
GAAGGCCAAGGAGCTCGCAATCCTTTGCGATGCCGAGGTTGGCCTCATGATCTTCTCCAGCACCGGCAGGCTCTA
CGATTTTTCTAGTACCAAGTTAGTTCTCCTCAACCTTGATTTATGTATGAGATCTTGAGGTTTTTTTACTTGCTC
AAAATTTTCAGGCCTTTTTTCTCATAAAAAAACCTGCTCAAGAGGAAAAAAAACTAAAAAGTAATTTAAGGAAGT
CGAGCTTGAAATAATGATGCGTCTTCTTCCATATTAAAGGAATTGAACGAAGGTAAAAACTAAAGAGAGGTCCTT
TTATTTGTTTTCTCAATTCTCAGACGCCTCTTCTTCTTTATTGTTCATTTTGATTGATGAGGGGATAAAGGTATG
CAAGAATACATAGTATATGTATTGGTGAATCCAAGATCTATATATACAGCAGGAGTGGATATATAGATGAAAAAA
TAATTAATTACAAGGAAGAATTCATAGAATAAACTACAAGCAATTAACAAGGAAGAGGGTGGAACAGCTTCTGTG
GATTAATACATAACTAACATCTGCTCTCATGCTGTTTAAATCATCGTGCTTTGTTGAGGTTAACCATTTCGTAAT
AAACTGGTGATCTTCTGAAGTTTTAACCTTTGGTAAATTTACCATCGAATTAATTCTTTCTGGATTCTGTAGATA
TCTGGAGGTACGAAACCTCTTGATGCCAGAAAGATAGTGCATAAAATCCTCTTGATTCCAGCTTTCTACAGTTTT
TATATGTTGAAAAAATGATTGACGACTATTTATCTGGAACATATATCAACTGGTTTTATGCTCTATGAACAGAGT
ATATGGATCGCATTGTTCTTTTTATAGGTAGTGGGAGCATATCGTTTTCCTATGTATTGTTTATACTTCCTGATA
TCTCTGCACTTGAATGAGAAATTTCATGGTGAGAACATTTTCCTTAATGTAACATGTATTCTATATTAATTATCT
TTTCATTCCTTTTTTTGGATAGAAGTTCCATGTTTATTAAATGACTGTAGAACATAAGTTTGCCTTTGTTGCTGC
AGGTCCAGTATTGAGGAATTCTCACTCTGATCAATTTTTTTCTTAATTTTCTGAGCACCATGATATAAGATAACA
AGAATAGGCCTCATGCTCAACTTAATTAAATTCTTGTGTTAAAAAAAATAATATATATCGTACATGTTAAGGAGG
ACATGAGTATGAACAAAACTGCTATATACGTTGGAACTAAAACTCGATTGATTTATTTTCTTTCAGCATGAAGTC
CGTAATAGAAAGATATCAGAAGACAAAAGAAGACAATCACCTAGTTATGAGTGCAACTTCAGAGCTTAAGGTAAG
TACATTAATCACAGCTGAAGTCCATAAATGAAACTATTCAAAAGCTGCTCTATATATTTTCATTGAGTGAAAATG

TACTTGATTTTGGTTTGAATCCCTTCATTAACTTTTGCCTAATATATAAATATATAAATAAATAAATAAATAAAT
AAATATATATATATATATATATATATATATCGTTGTATATTACATCACAATTCATCTACTTATATTTAAATATAAAT
ATCTTAGTACGAGTCTGTACTCATAGGCTCGTACTAACAAATTTGTATAAAATGTGAGAGAAAATGAGCTATTGT
AATTTAACTATAAGTAGCACACTTCTTGATATATACATATATATATATATATATATATATATATATATTACTTTT
AAAATATTTATGCAATATAATCGATCTGGTTGGCATGCTTATAAATTAATGAAAGATGGTTGGCATGATATCTTA
TGTTAATTGATTAATTATATCAATTTTAATTCTTTGAATGTAAGAGGGAAGTCAGCATTCAAATTATTAGAATCT
CAAGAAAATTATAGGAATAACTAGTTCAGCCTTAACTCGTCTGGATTAGCACAGTACATAAATCTATCTGCCTTA
ACATATTATTAGAAGTTTTCATCTTCTTGGGAGTTTCTCATTAGATATTATTTTCCTTCCATCCAAAAACTATAA
GTTCTATGTCATTAATACTTCTGACGACGAGTCCGGAAAAAAAAAATTGTTGAGAGGTGTCTGCATTGATAGAAAT
GTCATCAATAATTCTTTTATGTCTAAGAATTCCAATCACTTGATCGATTAATTTTTTTAGAGCATACAATTTCCA
TAATTAGACTTGTTACCACAAGCTTTACCAGATCAATATAGTATCAAAATCGGTGGAAGGCTTGATTGAATAATA
CTTGCATAGATTTGACTTGAAAAGAAGCAAATTTAATGAGTGTTTGGATTCAGAGAAAATGAGATTAACTAATAG
ACCAGATTAATTAGATTATAATAGTTTCGTGTACTAGTGAGTCTAATTGAAAAAAAATTAATTCCCTCCCTTTTG
TTAAATAAAAATTTTAATGATTGGGAAGGAAAAAAGTTAACCTAAGTTATTAATTTGTCAATATGACCATGATAT
ATGGTAGAGATAAATGCACATCAACAACGACCACGAGCAATAAAAACGAGATACTCTAAAAATCAATAGATTCGT
GATTGATTAATAATTAAATGTGTCAAAATACATAGAAACTAGCACTGCTTTTGTTTGTTATATGTGATCTTGTGT
CCCATTTGGTGAAAAAATAAAATTAAGATAGAGATCCTAATCCTCATATTTGAATCAAACAGTTTTTTTTCATCC
GGTGGAACCCATAGAGAGCAGATTATTCTCTCTATTCTTGACTGAACTTGTTTTCACTGGATTGAGGCAATGAGA
GTAACTGATGAATTAAAAAACTTAATCCATAGGCCTACAACCAAACCAGGCATTTGTCACACGCCACAAGATGTA
GTTCAATACTATAAGTAAATAGTTAATTAAAAGCCAATTGATGCATAAAAGGATCACAGTCCGAAGTACATATGA
TGCAACGTTATCCAGTGGACTTCATTGTCTTCAAGACATAAAAGTAAATTTGCTGCTGTTTTTCTTTTATAGCGT
CTCAGTTTTCATGCCTTCGTCACATGAGGACATATACTACAACCCTCCACTAACCACGAAACACAAGAAATAAGT
TAAAGTAGGTGCAATATTTTTTCTTTAAGCCACTGCCCAATTAGGGTGGTGTGTTTGGGTAGAGAAGAAGAGAGA
GAGGAGGGAAGGTATTGGGAGAAAGAGTAATTAAGAAAAAAAATTATTCTCCATTCGATTTCAATTGAAATTAGT
TAAAAGCAAAAGTAAGATTATTTTTTATTTCCTTCAAGTAAATATATACATATTTCAAATAAGATTATCGTTCAT
CTTATGTTAGTCAAATAAATTATTTTTTTTTCACTGAAAAAAATACTTTCTACAAGTGTAAGGACTCTGAAAGATT
CCTTGATTAGAGTGAATTTGCAATTGTAATTTTATTACAATTAATCTCGTTCCGAACAATTGATAGATGCATTCC
TCAACATGTTGACGAGACAGGTTGGACTGATGGGCATTATTTACTTTGCTTTATCGAAGAGATTGTAGAATATTG
GAGTATAATAATAGGACGACTGACGTGATCCTTAGAATATTAGTCAATAGCATCTGCTGCAACTAGAACTATTCT
TGAAGTAGTATGCAAGATACATCTATAACACAAGATAAATTTAAGTGTTAATAAAGCAATATAATACTATACAAC
ATGGTGATAATTTCAAGGATGCACTCTTACCATTCTAACGCTTGCTGTAAAGTAAAGATGTAGGTGGTTGTATTC
TTTCCTGGAAGACAGAACTGTTGCAGGTTTTAATTAACATCATATTGACCAAGTTTTAAATTTCAAAGTTTTAGT
TCTTTTTGTTTAAATATGCTTATTGTACAAGTTTTGACCACAAAAGAGTCTAGAGACATTACAAAGGGAGAAAGA
TGAAAAACAATTGAGGGGGAGGGTATGTGAAATGATCAGGTGCCAAATAAAAAAAACACACAACCCTCGCCAATG
TTGCTTTAGCAAAAGCCAAATTATCATAAGATTTAAAATGTTGGTGGTACACCAGATCTACTTCAGTTCATTTTG
CACGAGGTTCCATGAAAGCCTCCTGCTCCTATCAGAAATCTTATCCTCAATATGAACGGAGCCAATACCCCGACT
TCTAAATTTTCAGCAAAGTTTGCATTGATTCCCCTCTTATCACGGAGCAACTGTACATTCTTGCACAACTTACAC
TGCAAGGAGACTTATACCAAAACCGTAACGTCTGGTTTCATCCTTCATCTTCTGAGCCATAGGAAGCAACAAATT
TTATATGCTTAAACTGATATCATAGAAGACTTGTTAGTTTCCTTAGGCTAAGTTTGGATGTTTAGTGATTGAGGG
ATGATGATGAATAAACTGAAAATGATTAAGAAGATTGAAAAATAATGGGTTGAGAAAATTTGTTTTAATCTATGT
TCGGATAAAAAAACAAATGATAACTAATGGTATAAAATAATCAAACATTTTTTTATGGATAAAAATATAAAAGAT
GAATAGGAAAGGATAGTTTTATTAAAGAGACTCTTTTACCTTATAGAATTTGAGGAAGAGTATTTATGTATAAAT
AAATTGCAAATTAGAGAATCAATATTTTACATACTACTTACAATTACAAGAACTGTGTGCATTAAGAAGAACGAG
AAAAATAGTTCAAAATTGTAAATGAGTCGAAGGATAAATTTGAAAGAAAAAATTTAGTCCATCCCATTCTATGAA
ATTGATAGAATGAAATATGATACTTTTAAGAGTTGTCATAACCCTCCATAATCTTTTATGTCATCAAAAAATAAA
ATAGTCAAACTTGGATTATAATAACCATCATTAACCCATCTATCGTCATCCAAACAATTGCCTCACTGATTGGAA
GCATCAAATAATAGGCAGAAACATTTAGTACTTGGTTCATCTGCGGTTCACTTGATTTTGTTCAAAGGGAAACTA
CAATAGCAACTCCTAGATTTCATACAAAGAATGAATGATCACCTATAACAAGAACGAATAAGATTATAGCTCTTG
ATAAACCAAAATAAGGTGAGAAAGATTATAATTCCATGAAAGATTATAGTTCCATTGTAACTTTTCATAAAGAAA
AAAACTCCTTCATGAAAACATTTAAGATGGGATCAAGTCACCTCAATGGATCAAAAGAGAGTTTCTTCTTTTTTA
GAAAAAGAAGAAGGAAAGGATGACTAATGTTTAGTCATCCAGTATGTATTAGCATTATAAAAACATCTAGAATAA
AAAACAAATTGAGCAAACATTTTCTCGACCTTGCATTTTGCTAATTGGAAAGAGTTTTGAATCATTGTCCTACTT
AGGTTTTAATACATCACAACAAAGTAGAAAATTTCATGTCAGATCTAGAGCTCTAATTAAACTTTAGTATGCTAT
CAGCATGGTCCTACAAACTCAGATGTACCTACTGAAGTCTATCATAAATGAAGGATCTTAAGTTTTTTCTTGTAC
ATATAATTATCTACAGCAATAGCATTCATTGTAGCAAAATTTCTTAATAAAAAACAGTATCAAGTTCAACCTTAA
ATTATATCCAACATCATGTTTGACCAATATATTTTGAGGACTGATCGATCATTTCCATACAACAAATGGAAAAGA
AGATGCTTACTGAAAAATATTGGATCAATTGTTGGTCAATAGTAAATATGTTGTTCACTGGAGTATGCAAGTACA
GCAACATGTTCTGCTGCATTTCGCATGGAGATCTTAACATTAACTTTAGGTATCAAAGGGAACCTTGATGACATC
AGAATATGACAACATAGGGTAATTTTGTAGGACCTTATAATATGATTGCAAAGTTTATGTTTTATCGAATCAGCA
TAATTTGCAAATAAAGAATGCCAAAATAGTACATTGGCTTTGATCTTGCACTGGAATTTCTTTAATCATTTTTGT
TCACACAGTAGTCATAGAGCATATTGCAGATACCAATAGTTGGTCAGGTTCAATCGGTAGCAGATTAAGCTCCGG
CAAGAAGATGTCACAAAATTGAACCCCAAATATGCCTAACCTTGAATGGATTACTCAGGGTCTAGTAGGCCTCTA
GATACCAAGCTTAACATTGGAAAACAAGAATCTGTTGCATATCATGTTTTCATTCTCATCCCAAAAGAATGATTG

CACTTGCATTCTCCAGAATCACATTGAAGACATAGGATTCTTCATTTTCATTTTTTACCAAAAAACATGATGAAA
TGCAGGCAAGGAAGCTCTCTCTTTTTTCCTTGGGGGGGTGGGGGTTGGGTTTGGAGGGGAGGGAAGGGCATCTAT
GTTATAGAAATTGGATATGATTTTGTTGAAAAAAAAATGGATATGCATTTGCCAATTTTGGAGCTTAAAACTTAG
AATCGCATGATCACCAGGTCATTTTTTAGAGAGATCGGTGACCATCCATTTATTGCTGTGAAAGATGTCACATAC
TTAGGTTACAGTCAGAATAACAAAATTATAAGCACCTCCAATCAGCAATTAGCAGTTCTGCTACAGCTATGACAT
TCTAAATATAAACCCTCTCTCTGTAAGTATATCTATTTCACTGTACCCATATAGACTTAAAAAAGCAAATGGTGC
TGCCCGACGCTTCATGTGACATAAACCTGGACATTTTGGTCTTGACATTCACCATGACTTAAAGATTCAAAAAAA
TCTGTTGATGAGTTTAACTCGATCCATCTACACCGGATTGATGCTACACCTCCTCATGGGCCACATTATAATGGA
AAAATAAGTGATCAATATCTTCTTCAGATGCCTCGCATTTTAAAACCCAAAGATGCTCTTGAACCATAAATCAAC
CAGGCATTGTAAGATTAGTGGCAGAGAATTTTCACTTCGTATTGACAACGGGAAAAGTTATCATTCAGAGGTTTC
AATCTGTATTGGCAATGAAAAGGTCACATTGGTGAAAACATTACTCATCTTATAAGCAGGCATACCTTCAAGTAT
GTTCTGATCAATCTTTATGTAGGAGGAAGTTAGCCCACTTATATTTTAATACAGTATCTTCCAACTGTCCCCACT
AAATAGTTAAGGGAGAAACCACCACCTATCTAAGGACTTATCCATGCACAATTATACCTATGTTAAGAGGTTTAC
TATATAACATATCCATCAGCAATATAATGTAGATGAACTACTTCCTGCATTATATGATTTGAAATAGGATATTGC
AATTTCAAAGAATTATTCGCTACTACAGTTATAATATCAACTAACCTTGTAGTCTGAACAGAAGATATTCATATT
AAATTGCAGTCACACAAGAAATACTATCATTGCATACCAAGCATATCATGAACTTTCTGTTTTTTTCACACTAG**T**
**TCTGGCAAAGGGAGGCAGCAAGCTTGAGGCAACAACTGCATAACTTGCAAGAAAGTCATAGGTATGATCATTGTA**
**AC**CTGTATTGCATTCAAAAGAAAGTTACTACCCTGTTTGATTAGTAGATATCCTCGCAAATCAAAGTAACAAAAA
AGATATCCTCTCAAATCAAAGTAACAAAAAGGTGTCTGAAGGGTTAGAAACTAAAAATTTTTATGTTACAAGAGA
CAACTGTTCACATTGCAATTTTCAAACTCTAGTCTACTCCAAGTTTCTAGTTTTTTGACACTCATAATTAGCCTA
CTGGCTACTATAAGTTTGTTGCGACCAAAGAAGGATGCATGTAATGTAATGTCGACGACAAGAACTAAAAGTTAA
TATATGGAACAAGAAGAAAAATTGCAGATGAAAAGATATCATCTATGCCCTACCAGCAACGTATAAACTTAACAT
CCTAGTCTATAGTACATAGTATTATTCACATTCTGACAATAATTTCAGGCAGTTGATGGGAGAAGAACTTTCTGG
CCTAAACGTAAAAGATCTACAAAATCTAGAAAGTCAACTTGAAATGAGCCTGCATGGTATTCGAAAGAAGAAGGT
CAAAGATGCTCCCTCACTTAAACTCACTATTTATTATTTTTGATCTTCTATTCTTACTGCATGAAGCTATCTTCT
TATAGGACCAACTTTTAATCGATGAAATTCAAGAACTGAATAGAAAGGTTGTAAATTATCCTCCTATCTTTTTCA
CTTTAATCCAAAGTGATCACATCAGAACTTCCTAATGTATGAGTGGTTTCACTACAGGGGAGCCTTATCCACCAA
GAAAATATGGAACTCTATAGGAAGGTAAACATTATTCGCCAGGAAAACATGGAACTATATAAGAAGGTATAAAAT
AATAAAGCTTCAGCATCAACACAACCATAGAAAAAGTTTACGCCCAACTCATTTCATTTGTGATAATATAGGGGT
TGAGCCTAACCAACTAAGCATTTGATGTAATCTATAACAGGTTTACGAAACAAGGGAAGTAAATGGTGCTAACAG
AGGTCCAGTTGTCCCATATGGTTTCAGCATCACAGATGAAGCACATGTACCCATTCATCTTGAGCTAAGCCAGCC
ACAACATCAAGAAGATGGACAAGAAGTAGGGGCACCAAAGCTAGGGTAACTTTCTTTCAAACATTTAAAATATAT
TTTTTTTAAATCTTCAAACATTTAAAATGTTACACCTTTGGGTCAGTTTAATCAGACAATGTTTAATTCCATGTG
AAACAGACTTCAACTGCATTAAAGAGATGACTTGGTGAAGTTGTTGCCGTCCTCTATCACAGTATCTCAAGGCGA
GAAGAGCATGATATGAGCAATATGGTTTTATGTACAAGAAAGAACTGCATACTCAAAACAGCAATTAACACAACG
ATGGATTGTGTAAAGCTATGTTACAATAAAAGTATGAAGTCTCATCCTGGTATTAGAATGTAATAGCCTGGCTGA
AGCACGTAGCTATTAAGATGGGATGTTTACAGAAGTACAAGTTTTCTATGAAACAAATTCATCAAATACTAAAAA
AGCAAGCAAACAAATTTCAGTCTCTCCCTAGACATTCTTTATTTCATATACTAACTAATTCATCCAAATCTCAAG
CATCTAAAGAAAGAATTCATTCGTTCTCTTTTATTTCAAATGTTTTTTTTAGATGGAAAAGGATGTCTATTTTAT
TTCAAATGTTGGTACATGTGTTTTAAGTTGGTTGAAACATTCATCGACTCAAGTTGAACATAAGTGAGATTCCAT
CCAAGATATTCCACACATAGATGATACTGTCTCAAATATTATGGATCAACCGGTTTATGTGCTTTGTTTAGGAGT
ACGAAGTTGATTTACCATTGAAACAAAATAAAGACATTACTGGAGACCAAAGTAACTGAATGCTATGCAAGAAAT
AACATTATTCTAGAGAAGAAATAATTTTTTATTTCACTTTCCCTATGCAATCCATGCGATAGGGGAAGAGAGAAG
AATTAAACCAAGAGATCAAGGAAAAGCAAGTGGAAGCCAATAGAATTATCTCAGGGAACTGGGAAGATAGATATG
TCTGGAATGTAAATAGGTGATGTCACGTAAATGGATGGTTTCGTGCTGGACTTCTATGCAAGAATCATGTGACCA
TTACTTCTAGAATGCAAAAATGCATTGTTTTAATAACAAAGGGCATTAATAATGAGGAAAAGGAAAAAAATGACA
AAAAAGTAATTCACATTTCATTTTAAGCCGGAAGCTCTAGGGGGTACAAAAATATAATACGGGAAAAAAGGACTC
AGAAAAGAAAACAATGGTAAGAATAGGAAACTAAAGGAGTTAGATCATGACAACATTTACAGCACGTAGAAGATT
TTTTTCTTGAAATGTGAATAGAAATGGGAGACAAATCCTATAAAATTTATAAACAGGTTTAATATAAAAGAGTTC
AAATCTAATCACAAAATGATATCACATCTTGAATAAATGAAAGTGGTCGGGTGGAGGTATACAGGGTTAGATAAG
CCTTTATTTTCCATTTTATACCAAAGACACATTTTATGCCCCTTAAACCACTAGGGGAACTATCACTACAAAATT
ATTATTCCCATTTTGAGGACAAAAATTGGAGCAAGATTTCTATGGAGGGATGAAACTGTTTATGCCCATTTTTGT
CTCCTGGGTACAGGCTGTCCCTTGATCACAGTCTTGTTATGGGATAAAGCTCTTTGAACCAAACAGAGCCTAAAT
GTACAAGGAAATCAGAGGTACATGGACATGTAAACAGGATGTTGCACTCTAAGGTCCAAAAACATCAGAGACAAC
TTCAAGCTGAAAATGGAAATAGTAAGAAGAGTGGATAATACCTTGACAATCAGGAGACAGATAAAATGGGCAATG
TTCTAACGTCTAACGATTATTGATTTACGAGATCAAGACTGTTAATCAGGCAATTTTCAGCACCTTGCGAGTTGC
AGATATGCAATAAAAACTATGATCATGTGATTTCTTTTTGCTTTGCATGACATTTCAAAGGGAAAGCAGCTCAAC
AATTATAAAGAGAGGTTTTAGAAGTTTGTGCATGATCATGCATTTTGTTGATTTAAACAAATCATTGAAGTGCAA
GGCACTCAACTCACACAGGTGTCCAGTTATTTCTCAAAGGGAGGAATAACATAATTCTCATACTGACAAAATGAC
ACTAGATGCCACTACGACAGTATTGTGGATATACCACAAGTAGACAAAGTAGATTTTTAAAATGTAGATTCCATG
TCTGTTATCTCACCATTTAAGATATTTCCTTAAAATTTAATGATAGGTTTGCCACAGTAAAGTATTCACAAGCAA
GGGATTTTCCATTACAAAAAGAATTGTTGCTTCTTGATTGCCGACAATCGCTCAAGCACGACGTTTCAAGAATAG
AATCTCAGATAGGAGACCGATTTGCTGTTAAAAGGATGTCCTAGACGACCTTTCCATTCTAAAAAATGCTTTTAA

CAGGGATCAACAATCTAGTCTGTGAGTGGCCCACTAGGCAAGTACAAAACTGTAGCCTGGCATGGAAGATAAGAT
CAACTACCATAATATTAAAAAGGTAGATTAAGTAATATTATGGTAGTTGATCTTTATCTAGTTCGTAATGTAGAT
TGGTGCCTTTTTTTGGACGATTACCTGTTGATAAGTATCTCATTCTGCCGAAAAACATCTCATAATATTTTCTTG
TTAAATTTTCCATATTTTTTGCTCAACATATATATTAAACACAAATTAAAAATGTCCTCAGTTATTACTCGATCT
ACATAACATTTTGTATTAAAGGGCTCCAAGGAAACTACACATAC**CACAAAAGTTGTTGCAAGTTCCCTCTCAAAA**
**AACAGTTATTGCAGGTTATGCAGTTGCTGCCTCAAACTTGGTTCCTCCTAACACAC**CTAGTGAAACAATTCAAAT
ATACAGCTTCGTCAAAAAAAAATTTGGTTATACAGACAGTGAGAAAAAAATGAAAAAACAAATAATGGATGTGTT
GGTTAAAAAAAATACATATAACATAGATCAAAGAATAATGTCACCAAAATAAAATGACATCAAAAACATTCTAAC
GGATGTAAACTATAATACAAAAGAATTCTGGAATTCAAAACAGATAAGTTAAAACAATTTCCCTTTAGGAAAATT
CAGTTGGTAGAATGATTTTGTGTCAGGATTTTGAGAGGTTCAAGGTGTAAATTATTTAATTTCAAAATTCAGTAA
CTGATAACTTCCACTAATCTCTTTCCGACAAAGGACTTCTACTGGACATGGATGCTTTGTCTCTATAATGCTTGT
CTATCTTTGCAGAATAAATTCTGGTGATTTGCAGATACTTCTCTTATCAGCTAGTTTATGGAATTCAACATAGAA
GAACAGAAAACGTAAAAAAAATTTTCATCTAGGAAAATATAGTTCCGAGAACGACTTTCTTTCAGGGTTCCGAAA
AGTTCAAGTGTGACTTCTTAAAATCTAAATTGCAGATAATCGACAATTTCTAATCTACAGCCAACATAAGATCTT
CTGTTTTTTTTTTTGAGAGAAACATCAGATCTACTGCACATGGATGCTTCTAAATGCTCATTTATCTTTACAGGC
AAATTGTGGTATTTTTTAGATAATTCCTCGTATCAGCTAATTTTTCAACGCATATAAAGTTACCTAAATTTTGGA
TAGTATTTTTATACTAGGAATGTTCTTACCAGAATTTTTTAGAATGAATCTTGTTAATATAAATTCATATTCACA
TTTTTATCGTAAAAGGTGTCAGAATATATGACTTCTTTTACAACAAAATTTTAAATACTATTTTTTTTCTTTCTG
CATAATTTAGGTTTTCTAATCAAATGGCACATTGTGTCATGATGTAGAATTATCTTCTCATTGATTAATCACAAG
TGGTTCTTCCCGAAATAATTCCTGTAAAAATTTTTTTTTTGAGAGGGAAATTCCTGTCAACAAATATTTCGTTT
AAAATATAATAGAAGGATGCCTTAAATTAAAACTCAATTAAGCTTTGGCTCCAAACAAAAAGTGTGCAAGGACAT
GGATCCCTGTTTTCCATCATGTGTTCAATGTTCGTACAATTCATATGCACTGTTTTTAATAAGGGATATGACCTC
ATAATTTCTTACACCTCAGTCCAAATTATTCTAGATCTAACTTTACCCTCTTTCAGACTCCTCCATATGACTAGG
TGCGTCAGCCAAACTATCTAAAATAACCTCTCTCAGTCTGTCTTTAATGCCTGATTCCTAGCATACTAAGAGAAC
GCTAAATAATTGAAGCTATTACAATATGGACGATTAATAATACATCATATTCGCTACAATTTATTCAGAATATTT
AGTTCACCAATTTTTCCTGTATTCAAGTTACTGAAGTTCGATAATAACTTTAAGCCAAAAGCAGGATATGTGACC
TGTTTGTCAAAGACTTAAATCATGACTTTCCTCAGCAGAGTTTTTGGTCAATATTCAATTACAGACACAATTTAA
CATTTAATTTAAAAGTGATACCAGACATATGGTAACTTAACTACATGACATATTCTCCATCCAATTCTTCTGCAC
TTTTCTTCTTACATACTGAAGTCGACATTCCTAGTGACTCAACATCCCAAATGGAGCATGTGATAACAAAATTAT
CTAGACTTACCAGCCCCAAGAATTGTCAAAAAATGTAATTTTGCTAATAACTGCAATAATTTTAGTCTCCTTAAG
GATACATAATTATATGAGAGAAAGTGCTATGATGTTAAATTTAGTCCAAACAATAAAAGAGTGATGTAGCATACA
ATGATTTCAAAAAGGAGAGAAGTAATTTGACTACTCTTAAGCTTCAGCTTTGATTTTCCACCAAAATTTCATGCC
AACCTATCAAAATGATAGAACACAGGTGAAACAACTTTTGTCAACCAAAAATATCAGCTAGAGATGCATCCATGG
CAGTGATCCATCATGCTGCCAGATTTTTTTTCCCATATTATTTCTTTTAAGTCATGTTCATAGCCACAACTTTAA
GCCATACTAAGATTATTGACACTAGCATAGCAGAATCAGACCACAGGATAATTGGAGCAAGCATATTCAGAATCT
TCAATTCATCGCAGTTCAGGAAAAACCAGTGCATCAGTGGAGCTCAGTAACATGTACTAGTGATCTAAAGATCTA
TTTTTAATATCCAAGAATAAATTAAGATCCCTGGTTTGAGTTAGGCTAGAAAGTTCTCAACATAGTAATCCAACT
TCTAATGACAGAAAAAATGCATATTATGAACAGACCCGTCAAATGCTTTACAGTATATGACGAATATTTCCAGAC
AAATAATACTAATGAAACTCCAAAACCGGAGTTCAAAACCCTACTAGCTTCAATTTGGAGCAGTCAAACACTGCG
ACCCGATAAAATCAAACTCGAGGGTCAAGACATTCACATTCTCTTTGGAAGAGCAAACACGACGATGTTCCCAAA
CAAAGATAGAAAATTTCCCTCCATATTCCGATCATTGAGGTCGATAACATCGACAAGTCGACCCTTGAAGAGCAA
AATTTTGCTCAGCTTCCATGAACAAAGGGATGCACCGAATCTCGACCAACCCTAAATCTCGGTATCTAGGAGCGA
GAATTAACTAAATAAAACCCTAGAACATGTAACAGTACACTTCCAGCGCAAAACTTGAGCTTGAAACAACGCGCT
CAAGCACAAAGAAACACACGAAATGGAAGGAAAACGAACAATGAACCGCTGCATGAATCACTCATAATTGCAGAA
TGAAGAAATGTAAAACCTAACGAAAATCCAAGAAGAAAAGATGAAGCATTAGTCGCAAAAGATTCAAGGTTCATC
GCGACTGAAACCAACCTCATAATGCATCAAAAAAATTCAGAAAAAAGAAAGCATACAAAACTACGAAAGAACATTT
AC**CTGCGTCGATCAAACAAAGAACCGATCCATC**GACACCGGAATCTGAGAAAGAATTGGCAGGAGAAGCAAAACG
AAGAAGAAGAAGAGACCGAGAGAAGGAAGGCGACGGATGGGGGCCGCATAAATATCTTCGAGGGAAGGGTTGCGG
GGGAGGGGGAAGGTGATGCGGTGTACCCGAGGGAGTAAAATAAGAGCCCTCGTGAGCGGCTCACGTCCATCCACG
TCGACCCTTTCCTTTGTTTAATGCCTCGTCGGGACACAAGTGCTCCCCCATCCCTTATTCTATTCCGCCGGTACT
ACAAGTTGTCGCCCGGTTAGTGCCTTACGTTGGCACGGATCAAGAGGATTGCGCCGCAGGAGCAGCGGTAAGGGC
GCCGGTTCCCGCGAGCGCAAGCTGTCCACGTCGTCGGGAAAGTGTAAGAGGAAGGTGTGACCAAGGGGGGTTCCG
GCGGAGCAACGTAAGCGGTGGGAGGGGGATGAGGGGACCGCTTCCCCCTTCCTCCCCCACGTTGGACTCTACCCC
GCTCTCTTTTATTAATCACTTCTGTTCGCCCTTTTTTTTTTTTAAAATAATAAAAATGGAACTTTTCAAGTGGATG
ATATAAATATATATACACACGTGTCACGCTTCCATTGAACAAGTCTTTTTTATATAATAAATAAATAACATGATG
GATTTTTTTAACTCCTGTTTAATTTTAATAATAGATTCGACGTCATAATTTTCTTGATCAGAATGTTAAATATCA
AATATATAAACAGATTTTGGCATATTTATAGTTTTAGTGTTCATAAAAACAAAATTTCAGTTTTAAAGCTTCATA
AAATTTCGCGAATTAT

>EpaAGL17_3
ATGGGGAGAGGAAAGATTGTGATCAGGAGGATCGACAACTCGACCAGCCGGCAGGTGACCTTCTCGAAGCGGAGG
AACGGCCTGCTGAAGAAGGCCAAGGAGCTCGCAATCCTTTGCGATGCCGAGGTTGGCCTCATGATCTTCTCCAGC

ACCGGCAGGCTCTACGATTTTTCTAGTACCAACATGAAGTCCGTAATAGAAAGATATCAGAAGACAAAAGAAGAC
AATCACCTAGTTATGAGTGCAACTTCAGAGCTTAAGTTCTGGCAAAGGGAGGCAGCAAGCTTGAGGCAACAACTG
CATAACTTGCAAGAAAGTCATAGGCAGTTGATGGGAGAAGAACTTTCTGGCCTAAACGTAAAAGATCTACAAAAT
CTAGAAAGTCAACTTGAAATGAGCCTGCATGGTATTCGAAAGAAGAAGGACCAACTTTTAATCGATGAAATTCAA
GAACTGAATAGAAAGGGGAGCCTTATCCACCAAGAAAATATGGAACTCTATAGGAAGGTAAACATTATTCGCCAG
GAAAACATGGAACTATATAAGAAGGTTTACGAAACAAGGGAAGTAAATGGTGCTAACAGAGGTCCAGTTGTCCCA
TATGGTTTCAGCATCACAGATGAAGCACATGTACCCATTCATCTTGAGCTAAGCCAGCCACAACATCAAGAAGAT
GGACAAGAAGTAGGGGCACCAAAGCTAGGGTAA

>epa_miR444c
GTGTGTTAGGAGGAACCAAGTTTGAGGCAGCAACTGCATAACCTGCAATAACTGTTTTTGAGAGGGAACTTGCA
ACAACTTTTGTGGTTACAATGATCATACCTATGACTTTCTTGCAAGTTATGCAGTTGTTGCCTCAAGCTTGCTGC
CTCCCTTTGCCAGAA

**Position of mature miRNAs based on comparison with osa-MIR444d and not based on experimental data!**

Stem-loop structure

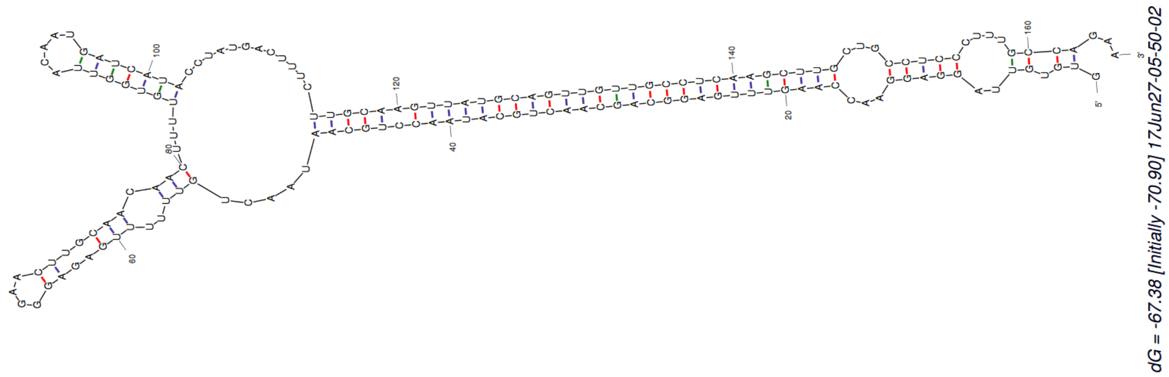


*EpaAGL17_4 and epa-miR444d*

>LTAE01004150.1
CGGCCACCACCTACAAGATAATATTCTTCAAATGTTCATTGCTCATATTATATGCACATATATATTTTAACAAAA
AGGAATCCATCAATCGAAGAAGAAGAAGGAAGGGGAGGACTTTTTCATGTAGCTTTATTTCCTCTACTTTGCTAT
TTGCTATCTATATATGAATGCCACAAACGGTAGAAATAGAGAGGGAAAGGCTTGATCTGAAGAAAAGGTTACCAC
CAGCCAGCTGCATGCTGTAGCCACCATTAGAGAAAGCTGGTCTGCCTGCACCTCTCCAAGAACAAAGGCAGGTTC
TTTTTTATGTAACCTTTCTCTTCTACTTCAGCTTCTCTTTATTTAAGGTTTCTTTCTTCAGGTAGAAGAAGATCA
TTTGTAGTGGTATCCACCATCTGTTGAACTTTACAATCTTTTGCGGTTTGCATATTTTGGTTGCAGATCAGGATT
CAGGAGGAAGCTGTGTAGATCGATATACATATAACAAAGAAAATGGGGAGAGGGAAGATTGTTATCCGCAGGATC
GACAACGCGACCAGCCGGCAGGTGACCTTTTCGAAGCGGAAGAATGGGCTGCTGAAGAAAGCGAGGGAGCTGGCC
ATCCTGTGTGATGCTGAGGTTGGTCTCGCCATCTTCTCCAGCACTGGTCGCCTTTATGACTTCGCCAGCACCAGG
TTTAATTTTTTTTTCTTTCGTTTTCTTCGCAATATCTGCTTCCAGCTCTCTATTACATATACGTAAGGCTACCT
AGCAAGATGATTTAGAAGGCGCTTGACGAAATCTGCCCTTTGGCTCCTGAAATGGAACTTTATCTAGCATATTTG
CTTGGCAGATATGTTTAGGATTTAGTTTTCATTTTAATTAGATCCACCAAAGATATAAAAAAAAGGAATTTCTGC
ATGCATGATGCATCTATGTATCTGGTGCCTGCTCTGCATGATTATACGTCAAGATCTCAATTAAGGAAAGGAGCA
TATAGCCTGAAATCTTGATCTCTCTCTCTCTCTCTCTCTCTCTCTCTCTCCACCAGTTTAGTTTATAGACGCA
GAAGAAAGGTAGGTAATCAACCTTCTAATTTTGCAAGTCATGTTTTTCAGCTTCCCTCATGGGGATGGTCCTAAG
TTATGCTCATCAATTTTTATACCATTCCCCGGGTCAATCTTATTGATGTATATCTGATTAAGGGTGAAGATGTTT
GTTATGTGAAAAAGTTGTAGATACACTTACAGAGACGGGAATTAGTGAAGTACTATTTTAGTTGTAGTAATCTTG
GTTTATTTCAACTATTCGGGCATGAATAGATGCATACATACATATCAATCTGATTTCTTAAAACAAAAAACAAAA
AAAATATAAGGAAGAAATAATCTTTGGTTTTCTTTTTGTCCTGATCACAATCTTTGGAATTTTGAGAATAATATA
CTTGTATAAAATACTTGCATTTTATTATAAAAAGTTCAATTTTGAGTGGATTTCCACATTCTAATGTAGCATGTC
TTTACTTATACTGTTTTACTTCGTGGTGCTCATATCTGATATATATGGTATATATAATACAAACAAGTATGTTAA
GGATATGCTTTTTCTCTAGTTCAAAAAGTTAATGGAAGATGCATTATAAATTATGCATTTTGAGTGTCATATCTA
TAATGACTTCACAAGTGGGTGTTATTATTTGTTCTAGGAGGCAAGATATATCTATAGATACGACCATTAGTTATC
TGCTTTTATTTTTTGCGGGTAATTATCTGCTTTCATTTTGTGTTACTCAAATATTTACTTAAACGTCAAAAGAGA
AATGAAACTATAAGCTACTGATCCTTTTGGAGCCAAAGTTATACATATCCATTTAAAGAAGATTTACCGTAGTAC
TTCATACATTGAAAGCTTTTCCTAACAAATTAACTTCAATATATGATCGTGTGTATTGTAACATACAATGTTTCT
TAACTATAGTTTAGAAGATCTTTGATAATGTCCAAGAAATGCTTGTCTTATAGTACATGTGCGATATATATACAT

ACATATTATGTATAAAGGTACAATCTTATGTTAGTTTTCTTTACTTCTTTCAGCATGAAGTCGGTGATTGAACGG
TACCAAAACACAAAAGAGGAGCCCAGCGCGGTTATGAACACAACTCACGAGGTATTTAAACTCAAGCAAATTTGT
TATATCTTTATTCCATTCATGCATTTCCAACTGTCTACAGTTTTTAATCTTTCTTCCGGAGAAGGAATTCTACAC
AATCAGCGGGAGATAAATAAAGAGTTTTATCATGGGGGGCTTAATTAGGGGTTAAAACATGAAGAATAAGTGAAA
CATTGTGTTACAACTAAAACCAGGCTAAGATATTAATTAAAAGTAAAATGAGATGTTCAGTATCATTGAACATCA
AACTCTCGATTAGATCTAAACAAAGAAGAAGAAAACATTTTATCCACTCATAAGATGTAATAATTAAACAAAGTA
GCCCAATCTTAATAGTCTTTGGAGATGAGAAGAGGTTATAATTAAGATTCCAACTTCCAACTTTTGAAACAATAT
AAGTGATGTTTGGAAGATTATAGGAAAATAGCATGCGAGTTCTTGTGGAATAAAATTAAGAAAGAGAGATCATAA
TCGTATGATGATCTGGGATAGCAGCTCAATGATTAAGGGAGGGATTTCTTTTTCTCCTGTTTGAATAGGAAGTTT
TAATGAAGGGGGATTAATTTAAAACTTTAACTTACGTTCCTGTCATATTATTTTTCTTTTTATCTTCAATGCTGA
ACTGAATGAAATTCCTTTCCATTATTTTTCCTCATATGTTAGCTTTTTCTCTTTCTCACTTCATTATATAACCAA
ATGAGATAGTAACATGGAATATAGCACCAGATCCTTCTCTGAGCTTGTTAGCATAGCAATTGAGGAAATGTAAGA
CCTAGATGGCACAAAGAGTTGGTTAATCCAACTCTCTAGCGGATTGGACAATAAAATAAGGGATCTCGGAATCAA
ATCTAGAAGTTGTCAATGTTGGGAGATGTTTGAATCTTTAGTCTTTACGCCATCAATTTTGAATTATAGACTAGC
TAGAAATTTGCTGATATAAGAATGTATATCAAAGAAAAACAATATGGAATATATATGGAAGGTAGATAAGATGAA
CAAAGCCTAGCCGAAGGCAATAAGGTAGAAAAGAGAAAACAAAGGTCAAGGGAGTAAAGCCATGACATGAGCCTT
TTGAACAGGCATTGCCAAGTTCATCGAAAGGGTGGATGGTTGTTCATCTTTAAGGTACTCACCTCGTGATTATGT
TAAGCTGATTGATATAAAAACAACTTGATTTTAGCTCACCTTGGAGGCTATGATGGCTAGTAAAGCTCTTGGACT
TGCGGACGGTAATGCCAAATAGAATCAAGCAATAAGAAATCTTATATGTATGCAACAAGAGCTAGTGAAGTTGCT
CAACATGGAGTGAAATACTCAACAAGAATCGATTTTCTTGCTAGGATTGAAAGGAAATGATAATGAAGGGAAAGG
AGGACCATCTTTTCCCTTCACATAGTAAAAGATAGTTTATCCTAATTTAAGATTAAAGAAAGGAAAAAGTTATCT
TGATTAATTTCCTCTTCCCTTTCTTTTCGTTTCCTTCTTCCATTAAGAATACCTTTAAAGGTTCAACAATCATTA
AAGGGAAGTGAAGAAAAATTAAAAAAAGAAAATGGAAGAGAACAAACATGAAAGAGGTTAATCTTGTTTAGATAA
AATAACTAAAAGAAAGTGAAGGGAATGATAGTTTTATTTGGACAAATATATTCCTTACATATGTGTTTATATATT
TAATTACGTAAAGTTAAAATTAAAAATATATTTTAGACATATTTTTTTCCTCCATCACACTGGAGTTGAAAGAAA
AAATGAACTATTGGAAGAAGAATGGAGAAGATATTTTCCTCTATTCCCCTATGTTAAGAATGCAATCAAACGAAC
TTAAATTAGTAATGTCTTTTTTTTCTTCCTCTCATTTACCCATCCTCAACCAAATAGACCCGTAAAGTCAAAATCA
ATTATATTTATTAGATAGACCTAAAAAAAGATAAGTTCATAGCTCAATTTTGTGATGAAAAGTTCTAACCACTTC
CACTACAAATTATATTATGAATGATATGATTATTGTATATCCTAGAGCCTAGTTTACTTAGTTAATAAAAATTAT
TTGATAAGCACCAATACTAAAAAGTATTTTAAGTGTTGAAACTTATTTAAAAAATAAGAAGTCACATACGCTTCT
ATAACAATATTGATAATAATAAATAATTATATTTAGGAAAAAATGTTGACTACGACACTCTTAAAATTAATTTTA
GCTTAAATGTATTAAAGAATTCAAATCAAATAAGCTTAATGCTTTAGTTTAATGTTACATTATAATTTATAGATT
TATTGAAAAAGACTTGTGAACTTTTAGAAACTTTAATTTATTTCATATCTTATTAAAAAAAATGAAAGGATGAGA
TATGAGATTGTCATATTTAAAAGAATAAGTGTAAAAAAGATAATATAATTGATTAAATAAAATAGAATTTAATGA
AAAGTACCACCTACTCTTCAATATTAGGGTATTAAATCTACAACGAATACAAAAAAAATTAGGAAAGAAAAAATT
TAATATCCACCACGATTCTAACATCTTGAAGTTGAGAAGATTAACATGATGTTACCTACTTAAATATAGCATCTT
CATTTTTTTCAATAAAATAAGTTGACAATTTTAAAAATGAGCTAAACTCTACTTTCTTTTTCAACTAACAACTTA
TAGATTCAATTTTTTATCAAGTGCGAAGATTTTCCAAAAATAACTCTTAATAATTTTGACCTACTCCCTATTGTG
CTTTTGGGAAGAGACATCATACTTTTTAATTGTATATGAATATATGATACGTTACTTGTGGAAATTATAATGCCT
GGAATAACACCTCATGTTACTAAGCATAACTAACCTTTTGTTCGTTAAATGTTTGGTGGAAAGCTTTGGAGGTTA
TTTTCCAAATTTTTTATAGATGACAACCTTTTGCATTATGCATAAAGTATGTGTTTCTCTGTCAATTTAAAGCAT
GTCAAACACAGGCTTACAATTGCAGACCATGATCCTGAAGCAAGAGCTTGATCATGAACCAAGTTGAAAGTAGCA
AGCAATATTTGTAGCAGGATTTTCAGATACCCCAAATAGAACACCTACTGAGCTACAGTAATCATATGAAGCCAA
TCTTAGCTATACAATATTAAGAGATTAAGGACCTATTTGAGAACTTTGTAAAAAAATTTGACGTTTTAAAAAATT
ATAGTAAGACACTTTTTTAAAAGTATCTTACAACAATTTTTTTTCAATTTAAAAATATTCTAAGTTTTTTAAATA
CGTTATAACTACAAAAGGATAAAAGAGAGAGGAGAAAAAAAATACGGAGATAGAAGTAAATAATTATTTAAAAGTG
CTTTCTCCACCAGGGAGTATATCCAACTTGCCTTGTCTCAAGTTGACATAAATATCGTCACTTGGAATCGACCAA
TATTGTAAACATTATTATAACAGAAAAGATTGTTGCATGCAACGGTATCAAATCTGATAATATACAGAACATAGC
ATGACTCAGGATCTTCGAGACTTCAATTTGCAAAATGCAGAGGTAAAATATTTATACAATAGAATATAAACATAT
CTTGTCACGTAGCTAAAAGTATAGTAACATCAGGAGCTCAAAGACTCCGAACCAAACAAAAAAAATTCAACACAA
GATAGTTTGCCTCTATTTCTCCTTCGGAAAGTAGTGTACCACATATATAACTCACATATAAGTAATTGAACGAAT
TCATCAGAACTATGATCCATATACACATTACACGTATAAAACGCTATATATGCATTATTACATCAGGACATTTAA
GAATCTTCTGCCACATCAGGTTACCTTCACCGATCAACTCTCATGTACAAGTTGAGAACACACATCAATGATTTT
TTTTTTACAGAATGAAATAGGGGATTGCCAAGTTTCATGCACTTAAACTTTCACTAGAAATATGCTCATTCTTCA
TTTCCACATTTAATTCCTCTGAACATTTTATCCAACTAAATGCCTTGTATAATAGGACAATGATTAGACACTAAA
GATTTTATTAAATTTCCACTAAATCGGCATTTTTTGCGGAATTGATCCTCAACCAAGCAGCATACAATTTCTTGT
TTAACTTGTGCATGAAGCTACAAGTGATCTAGCATTTTGGTGAGTGTTGATGTGACTAAACGGAACTTGATCACT
GAGTCATATTTACTCACAAATAGATAGAGGAAATACAATGTTTGCTTGAGGGTATGTTTGGATAGAGGAGGGAAA
GGGAAGGGAAGGGAGGGAAAAGTAATGTGAGGGAAGAGAATGGAGGGTAGAGAATTCAACCCCAAATTGGGAGTG
ATAATTTTTAATTAGTTAGGAGGGGAAAGGGAGTTAACTTTTAACTTCCCCTCCAAATAATTTTATATCCAAACA
AGGTCATGTTTACCCTCCCCTCCATTACCACCCCCTCCTTTCCCTTCAATCCAAAACACACCCTGAGTGAAAGCA
CTTGATTTGTGGTCGCATTTCAAAATTTCAACCTATGATTAAAGAGCAGACATGGGCCAACTTCTGAGGAAATGA
TGATAAAAAAAGTGCTCCTGCAGAAATAAGGCATATCTAGAATTTGTTTTCTTTAAAGGAGCATTATGGATTACA

TCACTTGAAGCAGTGCATTTGGATAAAATATACATTTACTGTTACTGCTGTAAATTTGATGCAGTAACACCAGAT
TATTGATGGGCTATCATAATGTCATGCTCGGACTTTAGTGGACAACAACAATCTCTTACCTAATATTTTCATATT
CTTCCAATAAAAACTCATCTTGTATTCTTGAGCATCAGGATGAAGATAATATAAAGGATTTGCCAGAGGTGGCAT
TTATGCCAATTTGATGGAATCAAATAAAATTATTAGAAATGTAGAAAGAAGAAATAACTAATTGGTCTAAAAGTA
ATTCAAATCTGTGGGTGTGGGTGGGTTGGGGTTAGGGTATGGGGAAACGAGGAATGGTAATTGCTTTAAGATGAT
ACTTTAAATTTGAAGCAATACTTCACATTTCATGCAAATCAGTTATCAAATCATGCTTATCTAACATGTCCAATT
GTTTGTTTGTACCTGTTAGTTCT**GGCAAAGGGAAGCAGCAAACTTGAGGCAGAAACTACATAACTTGCAAAAAGA**
**TCATAGGTAAGCCCTTGCTATTACC**CTGCATTATAATAAAATCATATTAAACATCCCTCACAAGAGAAGATTCTG
ATTAAATAACAAGATTTGCAATTTATCGTATGCAGAAACAAGTATGAATTTACTAGCATAGCCTTTAAAGGCACA
ACCTATACTGTATAACAAATGCATTATGCATGTGACATGTCACTCAGAACTGTATATCATTAACTATTCACTGAA
TTTCCCTATTGCTATGATTATCATAAAATCTCCCTCAATGATAAGAGGTAACATGTACTACTATCCATTGTGCAT
TGGCAAGGACAAACTAAAAAGTATATTGAAACCAAGAATGTTGCAGAAGCCATACAAAAATGTTAAGTAAGATGG
TTGAACAAAGCTCCACTATGACAAGTAACAAGTGCAACTATTAATTAAACAATATTCTATTTTATTATATGCATA
AATTGTACTCAATGGATCTCTGTAGATAATGGCAAAAACGAATCAAACATCTTTTGGAATAAAGAACGGAATATC
AAACAAATGACAATAAAAAAACGTCTTACAAATTTACCCTGATTAGCATGAGTGAAAGAATGAAGCCAAGAATTT
ATACAAATGTTATCAAACATACTTGCCAACAAATGTAAGAGAATACAACATCTTGTCAACAATTACAGGAAGTTG
ACAGGAGAACAGCTTTCTGGTTTAAGTGTCGAAGATCTACAAGCTCTAGAAAGCCAACTAGAAACGAGTCTACAT
AACATAAGAATGAAGAAGGTATTCCTAGCCTTGCAGCTAATTTTAGCGCATTGCATTTCCAGTTATTCTTATTGT
GATAATTTAATTTCACAGGATCAACTTTTGCTTAATGAAATTCAAGAGCTGAATCAAAAGGTTACAATCTCTACT
TCACACCCTTGTTTTGAGGTGAAGCGATGTTCTAAAACTGACTCAAGTTTGTGGGTACCACTGCAGGGAAGTCTT
ATGCATCAAGAAAATATGGAACTCTACAAGAAGGTAAACGTTATCCGAGAGGAAAACATCAAACTTTATGAAAAG
GTAACTAAACATCCGGCTTATAAACTTCACACAAAAAAGTAGAACTTTCCTAATCATCTCAAGTCATGGTAAATT
TGGAAGTTGTCAAGCACATGCTAACTATTCAAGCATCTGTAGGCTTATGGAACAAGGGAAGTAAATGTTGGAAGC
AGAGGTCCGGTTATCACATATGGTTTTACTATTACTGAGGCTAATGTACCAACCCATCTTGAACTAAGCCAGCCA
CAACATCACAAAGATTGGGCACAGACAGGAGGTCCAAAGCTAGGGTAACTTCTCTGTCCAGAAGCCATCAATAAA
TTGAGATCGTGCAGTGTTTAACTTATCTAAATTCAGCAGACTTCAACTGCATTAAGAAGGTTACAGGAGAAGTTG
TGGCTCTTCGCAGTTTGCAACATATTGTGTCAAGGTTTGGATACTTTATCGAGTCAACATTTCAAGGAGGGGAGA
TGGTTATCTGCGAAACCTTATTAAAATCGAAAAATAAAGTGAAGGCAATGACAATCAACAGCTCAACTTGTATTG
TTCCTGTTTAGACTTCTTTTCGACACTGCATTAATTATATGTTTAAATGAGATTGAAGTGATGCAGAGTAAAAGA
ATAATCTTTTCTGGAAACTTCTAGTCAGAACAAAAGTATAAAGTATGAAAAATCCAAGAACATCATCCACAACAG
TTATAAAAGACATAGCTCCACAACAGATGAGTCGAGAAACCCTGCTTTCGCAATGACCTCTAGACAGAACAGTGT
CAATATCAATTGCTACAAAGTAGGGGCACAATGCGAGAAGGGGATAATTAACAAAGAGAATTTGTTTAACTCAAC
AACGAAGAAACTATCTGCCAAAAAACAGAAAATCTAGAGCCAGGGCTCCAAGATGATGCATATCTCCACAAGACT
ATCTAACAAAAAAAATCTCCGCATGACCAATACTGTACTTTCATCTAGAGACAGTGCTGGGAGTTAGTAATAACA
TCCACAAAGATCTCATTAAACGAAGACATTACAGTGGATGAGGAGAAAAGAAAAACTATTATCTTCTTCACCAGA
TATACAAAAAAGAAGAAAAAATGGAGGCCCAAAATCTGCAGATGGAGAAAAGCCCCATACGATAAACAGGCCTAA
AAGTGGAAGGTTATTGAAAGGGATGGATGAAACCAGCAAACTAGGGAAAAAAGAGAAAAGAAGGCACAAATCTAA
AATTCCAAGCTCACTTTGATAGGACAAATGTCTTTAATCAATTTACTCAGATGTAGACATATTCACAAGTATCAA
GAAAGTAAAATTGTAAAATTAGGCCCCTCCTCAAGAATAATGCCATCTATCTCCCTTGTTCTCCAATAAAAAGGT
GATCAATGATATAGGATAAAAATTGACCAGTTAATTTTTTTTTTTTAAAAGTCGACTAGTTAAGTTCACAAACCA
ATTCAGACACAGGTCTTTATATGATCCTGAAGATGGGTTGTAGTTGCACTTTTAAGTGTTTATCGGATCCAAAAT
CGTAATGAGAGATTTCCACAAGCTTAGAAGATGACACCCGAGACACAGGGTTCAGTCTTACGAGAAACACGCAGA
TCGTATTTACAACAATTACTTTATCAGGGATTTAGTTCTATATACTGCTGTAAAGCCTCATTTATAGCACCAGAA
AATAGAAAAAGAAATAGAGAATTTATCCATATATATAATAATGATTGCTCAAAATAGAGTAATTGATTATGATAG
AGAAATAGATCCAAAGATTTAAGGTCAACTATAACAGAAAATATTAAAAGGTATACAACAATATAAAAGACAAAC
TTTGTTTAACTCAGTTTCCCGCCAACCATCCCTTCATAAGATAAAAAAAAAGGTTTCCTGTCAACCAATAGAAGT
GGACCCACTCAATCAATGAAGTTGCACACCATTCTTGTTCTCCAGTTTAGTCTTGTAGGTTCCAGTGAGGAGATG
AAATGAAAGAAAAAAATAGCATCTCAAGGTTGAAGGAACCATGGACATAATAAAAAATTGTCAAGTGCTTAGACT
AGACAACGAAAAAGCAGTATACAAACAAACAAGTTCACCCATAAGAATTCTGGAGAACAAGAATCTGAAGAAAAG
TAAGTCTTGCAAAAAGAGATGATAGTTCCCTCAGAAAAGTTGAGCAAGAGCATGAGGCTGCCATGAAAATTACTT
GTAGAGAAGACTTCAGAGTTCAATGATTACCAGCCCGCATTTCAAGATTGCTGCTTATTGTGGAAATGAATGGAA
AAAATTTATAGGTGTCCCTAGAGTCCATCAAAAGCATCTTCTCATGATTGACCATGACATACTGACTTAATTCTT
TAGTGATATAATGAATTATAAACAAATAAAAAAAATAGTTTAACTAATGAATTCATAGATTTTTTGGGGGGGGAT
GAAAATACGATGGTAGATTGGTAATCATGGTGCTAATGGAATTTTACTTTATAATTTAACGTCAAAAATCGGAAC
GTCTTTCCTCACTTGTACAGAAAGTTTATAACATAATGAAAGAAAAAATTAGCATTGGCTGGACTTGATTATTAA
ACAAATTTATAATATAATGAAAGGGTAAACAGTCTTGAAGCAAAAGGTAATTAAACACTCAGTAAAGCTTAACTT
GCCATTCTGAATCATTCTGCAAAATAAGTTTAGTAAATGAAAGGTTGCAAAAATGAAACCATCAGTCAACCCACT
CTCAGAAAAGAGGAAGAAAACATTGGCAAACTGGATTCGTTTGGGGGTTTATTTCAAGTTGTTGTATGTTGATGC
TTGAGTTTCCTTATGTGTGTTTCCTTCCTTTTAGCTGGCAGTTGCGAATGATGTTAGTTAATAGAAGGGAAGACA
GGGGTTTCCTCCCCCTTCCATTCATATCCTTTAGCTCAACGAAAATGTAGGGAAAATATAAAAGGTTTAGCAGCT
CCAACGGCAAAAGGAATTATGGTGAATACTATAATTAGCTTACTGGATACCAGGGTTTACACTGGAAAAGAAAT
TATGGTGAACAAAGTGGACTTACAAGAAATCACTTATTTCTTGAAAAAGTTGTGTTTCTTTTTCCAAGACAGTGG
ATCTACCCAACCACCACCTGAACTTCCTTGGTGTTTTTTTTTTTCATTTATAAAAGGAATAAAACAAAATAAGAAT

AATTCCATATTAGCTCAAATAGCTCCATATTTCAATGGTCATTCCAGATACCTTGACACTAACAGCATTCAAGTC
CTTAGTTATAGAAAATTGTTTATATACTAAGCTAAATTATATGCATATGTATACGCTTCCTCTTCTCATCGATTT
CTTTTTAGAAAGTAAACTATTTTACAATTGATATTGCAAAAAGGAATACCTAAAATGTGACTGTTCCCTAACAGA
TATGCGTTATAACTTCTCTAATATCTTTTAATTTATAGATAAAACAACAGTTAGTTAGATCACAATCTACTTAGA
CTTACATGTCTAGTTAAATAATGTTAAACATATAATAGCTTTCTAAAAAATTGTAGAAGAATTTAGGAACTAATG
AATAAATATATAACCTAATATTTGAAATAGTTTTTGAGAAACATTTGAAGTAGTTGCAAAGACATTAACTGATGT
ATATTTTGATTTTTTTCTTTAGACATTCTTCTTAATTTTTTCCCAGCACTCAGCTTCAGTACCAATTCATCTATT
GGTTAATTTCAATTCTTGATTGCCGATCAGGAAAAATTTATTTGTTATTTGATTCAGCAAACCGTTTGCATAGTT
CTGTGAGATATGTAAAAAGTGATATTCAAAGTTGATAACAATGGAACCAGACTTTTTTTTAGGGCTCACTTGAGA
AATGAAATAAACTCTTCTATAACAGTAACAGATTTTCTATAGCAATTACTCAAAAAACAAGAGGAAGACATGGAT
GTGAAACATCATAGTACATGTACAAGCACTTTTCAAATCTGCTCACATCCAATGAAGAAATTATGTGACAAAGGT
TAAAGCTCTTCATGTTAAAACTCAACAAAATCTCAATGATGGAAGCAATAACACCATTAATAAATGTCAATTGAC
AATTCGACGCTCCGATAATGTACATAATTCTACAAGGAAAATATATGTAC**CACAGGTCTTTATGCAAGTTATGTA**
**GCTCCTGCCTCAAGGTTGTTGCCTCCTCCTG**TGCCACAGAGAAGTGAAAATTACGAACAGATCATCTGAACCAAA
TCACCATATAAGGGAAGTTCATTCTAAATTATTCACAATGTACCAGCATTTGACGACATTCGATTGCCAATTTCT
TTCTCACCCTTAAACCTAATTGAAATTATATTACATTATGTATAATGGATTCATGTTGCACATAGTGCATTCCAT
CTTCTCAGTAAAAAATCCATCTACACTTTTAAAAAAAATGAATCCATCTACTATTCTTGCATATTTTTAGAACCA
ATCACTCTTTTAGGCTGAACGAACCAAGAAAAAACTGATCTATTACAATTCTGTGGAGCAATATCTTCATGAAAT
TGTTGAGGAATACACGAAGAAAGAGAATGCTGGGCAGATTATAACATTGTTTCACTATGCTAAGCTGCTCCAAAA
ATCCATTCAAAATACATTGTTTTCACCAATTTCTTAAGCATCTCCTACTGGTATACACATATAGCGATTCCCTTT
AATGTAGATGATATATTATTTGTAAAAAACAGAGTTTCAGAGAAAAAATGATTACTTTACTTGGAATGCCTCTGT
TAGATTATTGTGTCTATAACTTTCGATCAACATTTTCTCACAGTAATATGTGTTCTCCAACACAGTAAATTTGCA
TTTTGCTGATTAAACAATCAAGTTGTACATTCAAATTCGCATATGATTTCACTATGAATAGGTATGTGTAATATA
ATACTA

>EpaAGL17_4

ATGGGGAGAGGGAAGATTGTTATCCGCAGGATCGACAACGCGACCAGCCGGCAGGTGACCTTTTCGAAGCGGAAG
AATGGGCTGCTGAAGAAAGCGAGGGAGCTGGCCATCCTGTGTGATGCTGAGGTTGGTCTCGCCATCTTCTCCAGC
ACTGGTCGCCTTTATGACTTCGCCAGCACCAGCATGAAGTCGGTGATTGAACGGTACCAAAACACAAAAGAGGAG
CCCAGCGCGGTTATGAACACAACTCACGAGTTCTGGCAAAGGGAAGCAGCAAACTTGAGGCAGAAACTACATAAC
TTGCAAAAAGATCATAGGAAGTTGACAGGAGAACAGCTTTCTGGTTTAAGTGTCGAAGATCTACAAGCTCTAGAA
AGCCAACTAGAAACGAGTCTACATAACATAAGAATGAAGAAGGGAAGTCTTATGCATCAAGAAAATATGGAACTC
TACAAGAAGGTAAACGTTATCCGAGAGGAAAACATCAAACTTTATGAAAAGGCTTATGGAACAAGGGAAGTAAAT
GTTGGAAGCAGAGGTCCGGTTATCACATATGGTTTTACTATTACTGAGGCTAATGTACCAACCCATCTTGAACTA
AGCCAGCCACAACATCACAAAGATTGGGCACAGACAGGAGGTCCAAAGCTAGGGTAA

>epa_miR444d

ATTCGTCGCAGGAGGAGGCAACAACCTTGAGGCAGGAGCTACATAACTTGCATAAAGACCTGTGGGTAATAGCAA
GGGCTTACCTATGATCTTTTTGCAAGTTATGTAGTTTCTGCCTCAAGTTTGCTGCTTCCCTTTGCCAGAACTAAC
AGGTACAAACAAACAATTGGACG

**Position of mature miRNAs based on comparison with osa-MIR444d and not based on experimental data!**

Stem-loop structure

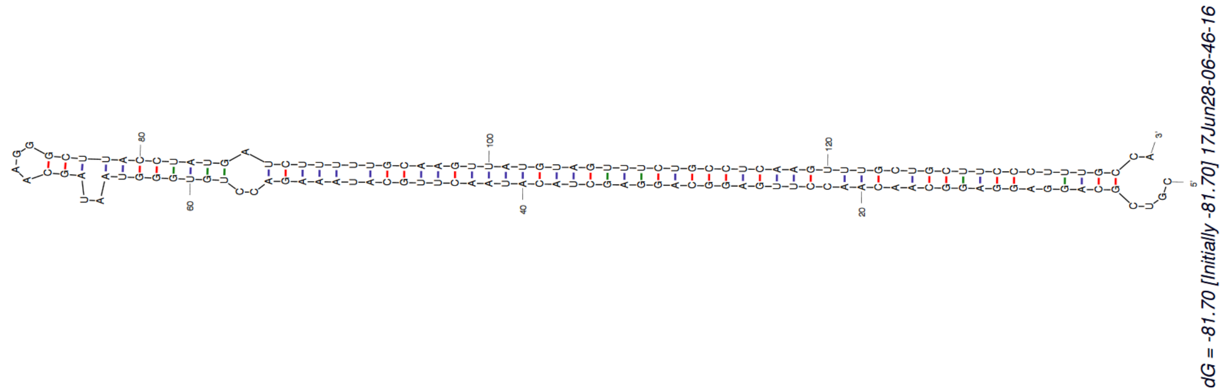

#### Zingiberales

##### Musa acuminata

*MaAGL17_6 and mac-miR444a*

>10

GATCCAGCTAGGAACTGCGCAGATCACAGCAGGATCAAAATTGTATATGCTTCTAATCCTATGTTTCCTTGTTGA
AATCAGCAACCTATTGGAATTTGTGTCGCATTCCTTGGATTAAGGGTTTTAACCTACTAATTTTGTTGAAACCGA
TGAAAATTCCCAGTTGAAGAGAGTGGGTTCTCAAGTAGCACACATGCACCGTATCGAGATATCTCTTCTTCCTCT
TCCTCTTCTTCCTTCTCGCGGATGAAAAATATTCTTAAACCAGATAATTAAATTAGTTTCTTTTTCGAAGAATCA
TCGGTCATACAGCACAAATAATTGAAGGTTTCTGCAGCTTAATTAAAAAAATTGATCTTAAATAACCCTAGAAAT
AATAAAGGAAAATAAATTTGATTCATAGTTGACCATTTCCTGAGAGAGTGAAATTGTTACATCAAACACTCGTAC
ATCTCTCTGACAAAATGAAATACAATTATTCCATGAGCTTTGAGCATGAACAGTAAGTACTCATCTGAAACATCT
TTATTACTCCAACCCATAGATTTGGAAAGAGGGGGACTTATTCATGATGGCATCACTATATGCCATTTCGTTGCC
ACCAAACTCCAATTTTAAGGTTGAAGCCGCGTCTCATGGAGAAACAACACTCTTTTGACCCTTAGGAAACGTCAC
ATTCATCAATGACAAATGGCCACAGGGTGTATGTACTGAGTCACCACGGACGACACACCAACCAAAAGCCAAACC
ATAGGAGGCTGACCAGGAAGATGAAACCCGGCACTTGCGTACAACCAGGTGTCAGTGATCTATACGGCAGGCGGT
GGGAGTTGTATCAAATTATAAGCAGCGAGGGAAACGTGCTTTGCGCCTCTCTGAACCACCAAAAGCTTCACACCA
TGGGAATCATCAAGCTGCAGAGATGTAATTAGTGGCTGCCTTTGTTTGACATCACTTCCGTTGTTGCTGTTAATG
AGGAGCAATGAGAAAAGGGTCATACTTAGGCATTTGCTGATAGCCTAACATCTTCCTTCCTTTCTATTCTGGTAT
AAAATAATAGGTGAAAGGGGGAAGACTGATACAAAGAGGAATCATCCTGAACCAAATAAAGGTAAGAGGTGTAGT
TGGGTCGTGGCACCAAGAAAAGGAAGCTTACCCTACTTGCAAAAGTTAAGGTTATATCTGCTGTAATCTCTCCTG
GCCGCATGTCTTTTCCCTTTCGTGGCAACCTCTTCTGAATTGGTTGTCTGCTAGCTTGTGTGGGCAGATCTGAGA
ACTTATTGCTTGGGAGACATGGTGAGAGGGAAGATCGTGATCCAGAGGATCGACAACTCCACCAGCCGGCAGGTG
ACCTTCTCCAAGCGGAGGAATGGTCTGATGAAGAAGGCCAAGGAGCTCGCCATCCTGTGCGACGCCGAAGTCGGC
CTCGTCGTCTTCTCCAGCACCGGCCGACTTTATGATTACGCCAGCACCAGGTTTTCCACTCCTTCCTGTTTCCAT
CTTTTAGTATCTCTTTGAGATGGGGAAGTTAATTAGCCCCAACATTTATTGAGTATCTACGTATTAGATTTCCTT
TTTCTTGTGTTTCCTCTGTGAGAGTGATTCTTGAATGATCCTCCCACAAGCTTTGTTATTCCTCTCTCTCTCTCT
CTCTCTCTCTCTCTCTCTCCTTTCTTCTGTATGCTCTCTTTTTCTCTCTCGAACTGAATGATGCATGCCAGAA
TTAGTCCATCCGATATACTTGCCTATATTTCTGGAGATATCTGCGCTTGTAATTGGATTTTCTTGACTGCCTCTT
TGACAAACCCTGGAAAGGAGCAAGGTGGTGTCCATTGTTCCCCAGCCAGCCAGTAGTCCCAAATTTAGGTTCTAC
TTTCCTTCCTATTGCATGAATCCTATGGATCGATATGGTGCCTCTCCTTGACAAGAAAGGGAAGCCAAGTAGGGG
GCTCCAAAAGTTGAAGGAAGATTTAGTTTTTATGTACAAAGCATTTGCATGCTCTCCAACCAATCAAAACACTTG
ATTAATAGATTTTTCCTTCTATTCATTGTGAACATTATGCTGGAGAAACATGACCCTGAAATGATCTAACTATAT
GTAGATATCATCAAACATGCGAGATCATGATGAAGGCATCATGCACAAGCATGCGAGAGCCCACCAGCTGGATTA
GACTATAATATTAAGCACTTAATGCAGCACTACTTACTCAGATAATCCATGAAGCGTACGGTACATCAGAAATTA
TTTTATGTGAATAAGGTCTGATCGAGGGAGCTGATTAAGCTTTGTCGGTACGATATGAGCACCACAGGGATGATT
TCAAATGGATACCTAGAGTTTGATTGATCTATAGAAGCCATCCATTGTAGTCTTTGGAACTTGAATTGTTCTTCT
AAAAGATATATCTTTTGTTTCACTGAGCCTCAAGTATTATTCTTGGACATAAAAATATAATTTTTTAGGAAAGGT
CGCTGCATTACTCATGATTTCCTTTTCATATCTTAGTATCGAGAGTGGAAAGAGAGTTCGAACTTCAAGTCAAAC
TTGGCTCCACTACATGATTTTGTTGCCCTATGTAACTCAATCGAATTGGATCTAGAAAGATGGATGGATTAGTTT
GCTAAGCTTGCCATTTTGGCTTTATGTTGATCGAGAAGCTTCGTTAATGATTTTTTACAAAATGTTCATTTTGTG
CTCATCATCTTGACACTTATGAGAGAGAGAGAGAGAGCTATATATATATACTCTATAGAATACCCAAAGCCATTT
TTGAGCATTATAGCTTAGATTCTGTTCTCAGCATGAAGTCTGTGATAGAGCGATACAAGAAGGCAAATGAGGAGC
ATCAACTTGTTCTCAATGCGACTTCAGAAGTCAAGGTATACCAACTTAAGCTCAGTCATGTATATGAGTTCATTT
CAAACACATGTTCATTCATTCCATGCTTTTTGATTTGCACCACACCTCTTCTCAAATTTTGCTTCGTTCACTACA
TGCATGCATTCTTGAACTACCAATTAATATCCTATTTCTAGATGGAAGGTGCATGAAAACAGATTAATTCTGATA
CAATACATAAATTAACATAACTTGCAACAATTTTCGGTAAGTAGTTTTTCCTCAGTGCTTTGTATTACGGTTCAT
ATGTCTACAATTTTTTAGCAAGATAATTAATGGTGAGCATAGTTATAATTTTCGACGAAGAATTATGTCCCAGTT
GATACTATTTGTTGCATGAAGCATGTCTGGTAATATGATTATACTATATATATATATATGTATGTATATTTTCTG
GATGATTTCTAATATTCTGAAACTTTCAGAAGCAAAAACATTTTGATTCTATAATTTTAGAAACACAAACAAAAG
AGAGCTTGTGACCTTTCTATTATTTAGGTTCCTGGCAACTTCTGAGAAAGAGAACTGTGTTCCAAAAGCAAAATC
ATGTTCAAATCAAGATCATTCTCGCCCACAAGAGTGGTGATTGTTTTAGAGCTCTAACTAAGTCTTCCCGAAGTG
ACTCATCATTAGTTACTTGATCTGACATGACTGTCGACGACTTTGCTAACTTAATCTTGTATTGGATACTAAAAT
AGATCTTACATCGTTTTCCAACTCCCTACTCTAATAAACATCAACTTTCGAGTTAGAAGATTAGGAAGCTTGTAT
ATGATAGTAGCTAAGATTTCTCGTAATAATAGCTGAGATTTTTTCAACTATATAAGAAAATATCATTACTCTGTC
GAGAAAAATCTTCCCTCATGTGTATAAATAGATAACACTATATCAAACACATATATTATCTTCGTTTCTTTCCTA
TTATTAATTTATGAAATTTGTGAAAATGGTTTAATATGGGCATCAAACTTCTTCACATCCATGCTAATTTGAGGC
TGACTTGCAATAGCAAACAGTCTAAACCTGTGGAGGAGTGCATATCTAATACCGTTGATATTGTACTATCATATC
AATATCACAATCTTATTGTAAATCTTTGCAAGAATAGAGCTAACATGCAAAAGGCATATAAACATTGTTCAAGAG
CTAGAATTAAATTTCATGACATTTTTATAGAACCTTTTTTTGTTTTCTTGATCAACTTCAATAATGAAAATGCTA
GCGAAGGTTTAACGAGGATTTTATTGCAATCATCCATCGAGAATCTTGGTATTTTGCAGAATTTGATTATGGTAA
GCATCTTCTTCGTATCATATCGTGTTCATGATATCCTTCCGTTATCAATTTCTTTGATTCAATTACTTCGAAACA

ATGCTTTACAAACTTACATTGGAGAAGCAGTAGTCTTTAGAGAAGGTGCTGCGTAACTATTGATTATGTTCTTGA
CAGGTCTTGACACATCGAATCAGTTGCTTCTTTGAGTTCATCATAGTTAGGGCCTTCTCTTTGTATCTATCTCCT
TTCAACATTATTAGCTCCAATCTTCTTTCTCTATAGGAAATTACAAATGAATGATGTGTTGTTCTTTCACATCTA
ACACAATTTAGAGAAGATATAACTCTTGATCTTGGTTTCCCTCCCCAAGAACATAGGTCTGAGCTTAGACCTTCT
CTTTTGCAGCATCATAGTTCTCTCCATTCTTGAAATTAGGCTGCACTATAAAAAACTAACACTTAAAGATAGTGA
TCAATTTGCAATTGAAGTCTAAAATTTATCAAATAAGAAAAAAAAATTTCATCTCGATCAATCTGGTTTTTTTTT
TTTTTTTTAAGGTTAGGTAGAGATGATCTAAGAAACTCATCACTATAAGTTTCAGGTGAGACTCAAATATCATCA
GAATCAATCAATTCCACTATGAGAATCTTGAGATTCATGTAGATACGTCCACCAGCTTGATCAGTCACCGAAGCT
TCAACTTGATGATGAAGACACACACATAAACTACAAACAAATTGGCAATGTATCTTAATTGCATCATATGCTCTT
TAAATTGTTCAAAAGAATAGATTTCATTGGCTTTTCACTTAAGAATGTATGATCATACTTGTACTTAGTCTTCGG
ACACAAGGCTTGATCCGTTCCCTCCTCATGAAGTTGATCTTGAGTCATGATCATGTCTTCGGTCTCAACATCTTC
GACCATGATTTCTTCATCATCAGACGTTTAATAGATGTTTCCATTTAGCTTTTCATATTTGGATTCTATCGAATT
CGGTGCAAGATTTCTCTTCGTGTTTAACACTTGATTGACTCTTTTTTGTTGCAAGCTACCTGAACATTCATGAAG
AAGCTCTCCATGTTGATCCTGCCATTAATTAATTAGGTTGTTGATGATACTGCTTTCCTCAACCATGCCATTACT
AATGAAAGATCTTTGAAATCATAGGCTTTAGATCTAACATCTTAAACCTTTTCTATTGAGACAGCATGAGAGTTT
CTTGTGTGTCATTAAATATAATGAATCAAAATCATATTTTAAAAAATAATACATCCTTAACCTCATATCTCGAGA
GTCTCGGATGATGACTACACATGCCAAGTGTATCATTCGATCCACTACTTGTCTTTTCGACTCATCATCAATTTA
TATGTTGGTCTTATCATATTCCATGGTATAACGATTTATTGAGGTTGATGTCATGAGCTCTCAATAGGCACCCAA
TCTAGGGTTGATACTTACAGCTATTGCATATCTAGGCTAGATTAGAATACCCATTAACTTAATTTGGGATAGTAT
TGGCAGAAAAAAATACACATTGAGTTGGGTTTGAGATGGTTTGGTATCCAAATAATTAGTTAGTCTTTTATTGGA
GTATAGTAAAATTTATGGATTCCTTACCTACATGACAACAAATTGACTTTTTTTTTATTTTTTTTGATACACTT
AAATATGGATTAAGATATTGGTTGTTCGAGGATGGGATGAATACATGTTAAAAAAATAAAGTGGTTAATTAATTA
AGTTAGAATAAAAATTATGGATGTCTTATTTATTATTATTTCTAATCAAAATTTTCATGCTAATGTAGTCCTTAG
GTCAATGCAAAAATATAATGAAGGGAGAGAAGGGGCAAAAGAAAACATGGGTAAAAAATTAAAATATACAGCTCC
TTATCTGAAAAATATGAAATATTTATCTTAAAATACTCATAGGGCTTAAATTTACAATCCTAATTTCGTAAGCAT
GGTTGTGAAACTTAAGGTACCGATAACGGGATAGTTATATCATTCTTACTCTAACCTTAATCCAGTGACTTAAAA
TGTTTAAATAAAAATAATAATAATACACCCATCAAGCTCTTATAAACGAACTTATAAACCTATGCTTTCGTTCCA
CATCAAAATTGTTGGAAGATTTCAACTAGCTTAAGCATATTGAATTAATAGATTAGACTTAATGAAGTTGATGGA
TTAATTATTTTATAGAGCCATCTTAATTGGGACCATATCTTGACGAGAATGTCATATAATGTTTTCATATAAAAT
TTCTTTATATGAAATTTTATATGAAAACATGGAAAATTTATAGGTTTTCATATAATGTTTCAAATCGAAAGCTTG
TGAGCATATTAAGGGTTATAATGGTAATCAACTTGTAGAACAATCATCTTTTATCGATAAGCAATATTTATAGGT
TTCTTAGATGTTGCAAATTGAAAGCCTTGTGAGTAAATTAAGGGATATAAAGGTAGAATCAACTTGTACAACAAT
TCGTGTTCTGAAATAAACATAGTGGCTTCACTGAAGATCTTCATCGCTAGCTTTTTCTACTGATCCATAGCACCA
GGCACATCAATATTTCGATTAATTAACTGGCTAATTTCTCTCTAAAAATTTTCACCCGGAGATATAATGAAATAC
ACGATTATTGGGTCTTTTTCTTTCCTTTTGTAATAACAGAAATAGCTACTGATCTCATTGTTTGAGCGAAAAGTA
TTTGAATAGCTACTGAAATAGCTACCTTTCTTGGTTGAATAGGCTACACTAGCATTCATCTCTCCGGTGGCCAAT
AAGCAAATCTCCCGGTCTCTTTAGTCAACAGCTAGCAGTTAAGACCACCAATCATGAAATTGACTTCTCAAGGAG
GGGTTAAAAAACACCTATAGATAATAAGGAGAACTAAATGTCAGTAATTAACAATGCTGAATTTTAAAGATACAA
TCAATTATGAAACACTGTCTAATCTGGAAAGGCAAGAGAAATAGACTTCGGCTGAGGAAGACAATGAGTGCCAAT
TGGTATCTTCAGCCTGCAATCCAATGTCTGGAGTTCTGCATAAATTTTCTTTTCATATGGGCATAAGGTAAAGCC
AGATAATAAATTTGTCAAGAAAACAAAAGTAAAATTTTAGATTAATTACATATAGGTTCCATCACATATTGTACT
TATCCATGCCTAATCACAATATATATATATATATATATATATATATAAACAACATCATCATTTTATCCAATATTTAT
TATATTTGAACCAGTATCATTTAATTTTGTTAGTTGTTTGACATATTCGACCAACATATACTCATGATTATTGAT
ACATGAAATGATATTATTATCTTTTAATTTAAACATAAATGTAATTAACAGAGATATACAACTTTTACTTGATTA
ACTTTCTTTATTTGTTCAGTTAATATTCCTTTCTATTTTAAATTAAAAATAAGAAAATAGTTCGTAAACATACTG
GAAAAGGCTAAAGAAATAATAAAAGCCCATCAGATTCTTTAAACATTCATTCTGTTGACATCTGAATAACATAAG
GTCTTAAATGTAATAAATTTAAATTAGTGGTGTCAGGTTCTTAAGAGAGGACTGAAAAGTGAATTAATCCAAATT
GTGGGTACCATATGTAAGTAACCCTGAAAATTTAGCTCAATAATTGTATCCCAGAAATAGGAGATGGTCCTTTCA
AGTCACCCAACAAATACCTTGTTAATCAGAGAGCCATCAGATTCTTCATTTTTAGTAGCTTGATCCTTTCAGCTT
TGTTATAAATATTGTTCAAGATCATTTATGGTGAGTCCTTAAGATTTGGAAAACCAATGTTAAAAAGAAAATTTA
GCATAACTAATAACTGCTGAGATCACAAACCCACCCCACCCATCATTACAATAATTAATTTAGAAAGTTATGGTC
ATTTAACTAAGAAGGAAATAAGTTATTTATTGGGTATGGAAAACAACTTATATACACAGTTATTTAACTGTAATC
TTGAAGAAATGCCCTGGGTCTGTATTCATGCAATCAAAGTTAGATTAAGCATGATAGTTGAAACACAGAACAATT
CGTAAGGTAGCAATGACTTAAAAAAGTATAAAACATGAGTTCAAGTAATAAATTTAGAAGAATTACTCAGAAAGA
ATACTGTCTAAAATCTAAACTATATATGTCGACAGTTCTAGAACAAGTACTTCAATCCTTTACTGTATGAGTAGT
TAAGAGTTATATGCAACATTTACCATCATATTTGAGATTCTAATTCACATCACTGTATATATTTGAAGTTTGCAA
ACTGTAGTCAAATAGATATAGCAAATCCAATCCACTTCTGACATGGATTGACTTTTGTGCTTTGTACTTACAAGT
T**TTGGCAAATGGAGGTAGCAAGCTTGAGGCAACAACTGCATAACTTGCAAGAAAATCATAGGTATGCTCATTTTG**
**CC**CTTCACACATCCAAATGAAAGCATTGTTAGCCAAAAATTCAATTACTTTAAACAAGAAGAAAAACTTCTCTAG
CAACATGAATGGTAATTCTGATCACATTTATTGGTCTCCTTTAGATCTGTAAAGATGAGGATATCATGACCAAAG
AAGTCTAATATCATCAGTATTTTCTTATGAAGTACATGATTCATGTATGTCCAGCTTTATAGGGTCAACACAAGT
GCACTAGCCCACCACATCCACTAATAGATCAATAAACCAACAAATAAACAGGAAACATAACAACAATCTTTTCAA
ATCATTTAAGCTTCAACACAAGATGGCAATAACATGTGTTTACCATTTCTCTGTCTCCCTAAGAACTTTCAAAT

TCACCAACTAAAGAAAAGCTTGGAAAACAGGATTTTAGCAACATCAATGGCTTGAATAAAAAACTAGAATTTTAA
ATTCTACATAAATTTCCCCTCTAAAGATCATCAGATTATTGTCAGATGCTAATTGGGCAACAGCAAAAAATAAAA
ATCAGACAAAAGGAAAGTTTAAGCTGAAGCAGCTGGTACCAAAACATTTACAACCAACATTTACCAAAGGAAAGA
AATCAACATACAATACTGAAATTCACACAATACAATACAGTAGAATGAGGTTCCACTTCACTAATGCATCCAAGT
CCTAGGAATAATCATACACGTTTTAAACAGTACAAAAATCATCAAATAGCATGAACAACTTTGATTCTTTAACAG
TTAAAGAAGGCAGCTTAGATCAAACCACTATCTCCAGATAGCAACTTTTCCAACTCACGACAAGAAATAAGTGAT
GCTCTAGCACCAAATATAGGGCAATAAGAAATAGTTGCATGCACAAGGATGTATGCAAAAGAGAATATATGAGAG
AAAATCATAAAAGCTATAGGGAGACATGCATTTAAGACTCTAGGAGAGCTGAAGAATGGAAAGTAATAGGCCAAA
GAATATACATAGCCGTAATTCACTTTTTACAGCATGAGTTGTGCAAAAATACGAACTATAGTCAAAAACTGTTTA
GAACTACACACAAAAAAGCAACTTTCCAGTATGCATTCCTGCAATCTTTATCACAAGTTAATAAACTTTTATCTT
TGTTATCATAAATTGTATTGAGATAGCAAGTTATTAGACATGGATACTTTGTCATGTAACGATATTTTTCTGCTT
GCCTGTTTTCCACAGTTGCCTTTAGACTAGTTGCAACATAGTTCTGATGGTTTTGCATTAGATCTATTTGCCAAT
CCTCTATTGTTTCTGGTCAAATTTTCAGTCATAATTTTCAAATGCATTTCATTACCTGGAGACCAAATAACTTTT
TTGTTGTCAAGCTCCTAACTGATATAATGTTCCTTTTGTTCACAAAATAGGTGACTTACATACTAAAGCCATCAG
ATAGTTAGCTGCAGTGTGTTTTTTCACCTTTTGATGGCACATCCACCTCATAGTAGCTTCTTGAACTATTACTGT
ATGTTTTCCACTATGAAATTTTTCTAGTTAACAGTTTTTATATGCTAGAATGAACTTGTTAAAAGTAATTAAGAG
GTTGATCTCTATGACTTATGCAAGTTCCTACTGAATTAATATGTTGATTGTAGTACTACATGTTACACAAAAAAA
GGGAACAATTATGTCACATAGCTTTTGCTTTGTGTTTTTCATAAACATGTCAACTAAAAGCTAAGGAATTTCATT
TGGTAACATTCAAACTTATTCCTAACAGACCATTGTCCTAAACACTAGAAGAAAGGATCAATAAGGTCATTGGAT
GTACCCAAGCAAAGTGGTTACTTTTGTTTCTCTAGAATTATTGACAGAACGAAAAGGAACCATTTCCATGCCATT
ACTTTTTAGAAAAAAAACTGAAAACAAAAACCCATGTTCCACTTTTCCTTCTGTAACTCTTCACACCTGCTAATA
GCTGCATATGCTACTTATTGCTTTCCTGTACTAATAGCAACATTGAAGGTGCCCTTGTTGCTTAGAGAATTGACA
AATGTCAAAATAAGGAGAAGAATTGTCCATGTAATACTCAGAAATCTGGAGAAATAGGAGCACTCTAGAAGAATT
CTCCTTTTATTAAACACTGCTCTCATGTAGTACTCAGAAATCTGCTCTCGTGTAGAACTCAAAAATCTACTCAGA
AAACTGATGAGCAGTGTTTACTAAAAGGAGGTTTTACACTAATGAGAATAGATCAACACTATAAGTAAATAAAGT
TATCAAGTCGCATGCTAAGAAACGACAACTACCAGAAGTTCTTTGAGGATATAAATGATTTTTCAAACTTGGTAA
AGACATCTGGTCAAAGTAAAAAATCATCCCTCTTAGTTCTAGTTGTTGATGATCTTAATGTCAATATCCAACTTA
AAATTAGACTTAGCTATCTGATTATCAGGAAATAACAATTGGATATATGGAATTACCCATATAAAGAAACATTCC
TCTTGACACTAAATATAAGCTTTAATTACTGGTTGATCTGTATCGGTCCACCCAATTTTGGTTTGTCCAACCAAG
CACCAGTAACAGGCCATACTATCTAGTAACATTCAGTACACCAGTCTTTTTTTTTTTCAGTGATTTTAAGCATCA
TAGATCAAATACATAACGATATATTAGTGCCACATTGGTTTGACAAACTTTCAATACACACCTTAGTGTAGACAT
CAAAGTCGTATTTTTAGGTATGTGATCTTCAAATCAAGGATTTTAATTTCGAATGATATCGCCCGTATCGGGTGG
TACGTACCGGTCCACCAGCAGACCGGCACGCGGACCCCCCGCTACCGGGCAATACCGTCGATTGGAGCTATTCCC
ACCCCGTTATCACTCGAAATCGGTCGGTGACGGTCGATTTTGACCGTCATCGCCCGTTACCGGGAGATATTAACT
GACGGAGAAGGAAGAAGAGGGAGAAAAAAAAGGGAGAACCTGGAGATCCGATGCTGCTCTCCCTCGATGATCCCG
ATCTGTCACCGCCCTCCCTCATCGGACGTCGAAGATGAGATGTCGCCTCCTCGTTTGAGGCGACGAGGCCTCGGC
GTCATCGGCGACTTCTCTTTAGCCACGCAGGGAGAGGAAGAGGCTTCACAGGGAGAAGAAAACGAGGTTTCTTCT
CCCCACGCGGGCCGAAGAAACGAGGCGATGTCGGCTCAACGATTTTGACCTTTTTTTAATTTTATTATTATTATT
ATTATTATATATATATATATATATATATATATATATATATATNNNNNNNNNNNNNNNNNNNNNCAAGGTTCGCAATACCG
TACCGTACCGGTATTTCGACCTGGGCTCGGTACCGGTACGGTACGGTGTACCGAGCGGTACACCCAGGTGTACCA
AGCGGTACACTCAGGTGCACCGAGCACTGTTGCACTGCTACAGTGCAACAGTGTACTGGTGCACTGTATTGCTAC
TGTAACAGTGCAGTGCATTGCTACAGTGCTGCATTTCACTGCTACAGTGCAGACCGATACTGGGTGGTCTGCGTA
CCGCTGGCCTGTCGGACCGGTACGTACCGCCCATACCGGGCGGTACGATTCGATATGGTAGACCTTGTATATATA
TATATATATATATATATATCGGTATGCCAGAGCATATCGCTCGGTACGCTCATACCGTATTGTATCGAGCTAAGC
TCGATACTTTGGTATGGTACGAAATTGCGAACCTTGCTTCAAATGACATCAATTAAAAAGTTGGAACTGCATCGA
CACAATCTTAAATTACATAAGCAGCAATTGGACCATAAAATCTTAAATTTTATAACAAAAAGCCAAACAATAAAC
CTCTATGCAAGAATGCCATTGAAGGGTGCCCTGAACTGAAGAAGAGTTCCAGATCAAGATGTGTTACGAATGTCA
TAATCTTAAGCTATCTAGAATTAAGTTGCCTAGGTAATCAAAATTCCTGTAGTAAAATTCTGTTAACGATAACAG
ACTGATGAGAGTGCAAATCAGCTCTTCGATTGCACAAAATATCCAAACACATAAATTTTATCACGAGAAGAAGAA
TTGTAAATGATATGATCCATTCATAAAATATATACAAGACATTCATGAAAGGAACAGTCATCAACCTAGGAAATG
AGGAGAATAAGAGTCAAGCCTATGGATCAAACAACCAAAATCAGATATGCAATGGAGTAGTGTCAAAATGCTACA
ATGAGTAGCCTGAATAGACTTTGTCAATGATCATTTATACATAATGGGAAAAAGTAACTCTAATGGAAGAGCACA
TATTACATCTAAACCAACTATTGCAGGCAGTTGATGGGAGAAGAGCTTTCTGGCCTGAGCATCAAAGATTTACAT
GATTTAGAGAACCAACTAGAAATGAGTCTACATGGTATCCGAACAAAGAAGGTCAGTAGTCTTTGCAACATTAAA
TGTCTCTTTTCTTTCGGCATTCCTAATGCTGTAAATTTCTGTTCTGCAGGATCAACTTTTGACGGACAAAATTCA
GGAGCTCAAATGGAAGGTTGTGCAGAAATTAAAACCCATCCTTCTCTGTTTAAGTTCAGATTGCAGCTAAGATGC
TTAACTATTTATTCAATGGAAACCTTACAGGGGAACCTCATCTACCAAGAGAATATGGAGCTCCACATGAAGGTA
AACCTCCTTGATCAAGAACATATGGAACTACATAAGAAGGTACATTAATTAGAAAGCTGGTAGACATACACACTC
AATCTAAAAACTCCATCAACAATCTTGTGGAATAACAAAAATTAATTTGTCTTGCACAAACTAAGCACTCAATCT
TCTACAGGTTTATGAAAAAAGGGTAGACAATGGTGACAATAGAGACTCAGCTATTTCATATGGTTTCGGCATTGC
GGAGAAAGCAAATGCATCCAATCATCTTGAACTAAATAGGCCACAACACCAAGCCAACAGGGAACAAACTGGAAC
ATCACGGCTATGGTAACCTTTCTTTCCAATACAACAACCTCACAGGTATCAAAACCTAATGTTAACTGTACAATG
TTCAACTGGATTTGTGGCAGTCTTGGATTGCATTAAGGAGATGGTTCAGAAATCAGGGTTGGGTGTGTGCACTTC

CATCATAATCATCTGAAGAATTCAACAATGTCATTTGATCAATCACATTTGGTGTTTAAGGAGGAGAAGCATCAA
TTGTAATTATGCATATCATATGACAGTAAGGTAAGCTTTGGTATTATGGTAAGAATGTTGATATACAACTTAGGT
AAGTAGGGTAAATCATAGGAACAAACTCTCTACTTGTAGGGGTAAGGTTGTGTACATTGGCTCTTTTTCAGACTT
CACAATGATAGTAGCCTCATGAACCAAGCTACTTTTTATATTATTTAATTAGAACTACCTATTTTTAGCAATTTA
ACAGAATAAAGTGGTGAAGACGAACCATGTAGTTAAGTTTAATAATTTTGGAAGTCGCTAGTTTGCTTTCTCTTT
TTCTCCTCCAATTAAACAAGACAAATACCACAAAATACCAGTTCAGATTGCCACAATAAAAGAACATTAGTTTTT
GAATGTTTCTAGAAATTGAACAAGTAAAAGCTTGGAACACAATTTCACGGACAACATTAGTCATCAAGCACCATG
AACACAAAAAGAAGAAAGTGAGAAAGTTTGTTTACAAAATAAAGCGCAGCAAGAAGTGTTTAATAATAGTTCAAT
CAGACAAAGAGAAGAAATTAACAGAAGGCTAGGTGGCAACCTTGCTGGATATAGGCAATCCAGTCAAGAAGACTC
TGAAAAGCAGTTGTCAGCTTTCACAAGAGAAATGCCAGGTCCTACCCACAAGAAGGAGACCTAACAGATTTTTAG
GCATGTTCCACTAATCCTTTAAAACAAATTAAAATTTAATTCACCCAACTACTTACAAAAGTTATTGAAATGTTG
GAGCTCAAAAAGGGGGAAAAAAAACCCAATTAAATTACTCGTCCAGTCAGCAGTCTTATCGGCCTCCCCGAGAAT
GTGCCCAATATTAAAATCTAGAAAGCTTTTTACTGAAATCATTTTAAATTTCAATGGTATCTAAGACAATCCAGG
GATGTTCAGAGTCTTTCACCCTGGAGATAGGAGATAAGTGACATGGATGCACCAACAACTGATTCTCTTGTGCCT
AGTTCAACAAGTAAAACTCTCTGAAAGGTAAAACTATTTCATTCATCCTCCTTGCTTTATATAGAACAGAAAAAC
ATTGTCCTGTATTGAGATCACCACTAAGAAAGGTATCTTTATTGTGTCATGTAAATCACTAGTCACAAATAGGAT
CCACGTATATGAAAAAAAACATGTGCCAGGATTCAGTAATTTCCATGCTTGAACAAAAAATTAAGGAATATTTCT
ATATTAAATGCATGAGGACCATTTGCATTATAAGCAGGATTATGAACAGAGCATCACAATTCACAATACCTGATA
GGGCATATTATGGATTCTGACAAGTGGACACGACCCCCACCACATCAGCCCAAGTGCGGAGCCTCGGGAGGCAAA
CGCAGCGCCACGTCGGATGCGTCACCAGAAATCCAGGATGGCGGTTGTTCACCTCCCGCGCCGTCTGGGGCCGCA
CCAACCAGCGTTGCCCCAACAAGGACCGCTTTGGGAAGCTCGGAACGGCACCACGTGACGCCATTCCACATATCC
TGGGCGACGAACGTTCGAAGGTACACTCGCCTCTCGGGGAGTCAGCCGCCACGCGAAGGCCGCATACGGCCGACC
CTCGGGCACTATGGTAAAAACCCCTTGCCAACATATGACAAGGGACACTTGGGACAATAGAATTCCACTAACTTG
ATCTTGAGGGGCCTAAGTCGAGATCCCCCTCCCCATCCCGACCTAAGCTAGTGTGCAGGAATCCGACGACGACAA
AGGCGGACGGGCTCGTTGAAGAGGCGCATCCCAACCAAATCTGAGCTCAACCCGATTAGGCGAAGACCTCGTCCC
CAAGATGGTCCTGGACATCGCCAATCGAACCAAGCCGTACTGACCCCACGGCCACGGCATAAGGTTCACCAACAA
TACCCTTGACATGACAATGCAGTACCAGCGCTTAGGATAGGTGCTTGCAGGAACCACGCCCTAGGTTAGTGCAAC
TGAATTCCCAGTTGCACTGTACAGTATCAGCCATTCCTTGGCAACACATGTAGTCCATAATGCTCATTATGTTTT
GGTCCACCCTCAAAGAAATTGGACTGCCTAATCTTTCTTCCAGAAGCTGACCCCTTCACAGCTTCCCCAATGTTT
TAAGAGAATACAAATATGGTAAGCTTCTCTTATAGGAGAAAATTATGCAGTATAGCAGGTCTTTTATGGCTTAGA
TATTTCCTGGTCTACCATGAGACTGAAAGAATAAGGGCTTATCTAGTGCACGAGGCTCCTGCCAATACGTGGTCT
GGGGAAGGTTAATATATGCAAGTAGTTTCCATGATTTGAGTCCTAGTCTCCTAGGTCAAAAAGGGGCAATTTCAC
CATTGTATCATAAGGAATCTACAGCTACATAAAAGTTATAATTTTCAGCTTCATACTACTAAAATTTACACATAT
TATCCAAGTTAAAAATTGATTGCTAACTACACCAAAAAAATTACAATATTTTATGAAAAAGTTCAATATTTCAGA
GGCTTCTAGCAAGCTCAGCAGCAAGCCTGATGGGGTCATCTAGAAACATATCCAATGTTAGCCAGGGCCTTTGCA
ATAAGAAATTATTAAAAAGGACTCCCAAAATATAATCTAAGTTTGTGTTTTCGTGTATTCTAACTATCAGAAGCT
TTAAAAAATATAATTCTTTAATAATGGTAGACACTTCATGTTTATGCATATCTGATGGGAAGAAATGTAACTGAA
TATTCATTAATGTAGTCAGAACCTTGGAGGTTGCACATTTTTCTGTTACTGTATTTCACGAAAAATATTGAAACT
TTGAGTTTATACTAAACTTGGAAGTACTTATTGAGAATAAAATAGCAAAGAACAGCTTCTAGGACTTTTATTATA
GAGAAAAACACTACAAGTCAATATTCATTGATGATTTTAAAATCATAAATCAAGAGACAATTAAACTTATAATTT
AAGAAGTTACAAATAATGATTTAAAATATTTCACATAAATTTCACAAATAGAAGTCAAGGAATATAGACATAC**CA**
**TAGAACTTCCTGCAAGTTATACAGTTGTTGCCTCAAGCTAGTTACATCCTCTTATCATAA**CTAGACAAGCAATCA
ACCAAATTTAGATACATGACATAGATAAAGTGCAAATGGCATCCTTATGAAAAAACACTGTTTCTAAGTTAAGCT
ATAAAATCTAAGTGCTCATGACAAGTCAAATGTAGACAATAATTTACTAAAATAAGTAAAGAAAATAGACATGCT
TACTAAGGTTTTCTATTAAGATAATTAGTCAAAAAACAAAGAGCAAATCTGACAGACTTAACGGGGATTTTCTTG
ACATTTCCTAGAGATATGTACAATTAAATTATAATTGTATTGAGAGGACTGTGGCCTCAAACCTGTTACTTCGAA
ACTATTCATGAAAGCTTCAAATGAAAGATTGACTCCATTGAGCATAATATACAGCAATCATGAGATGAAGAATCA
AACTTGTAAACAATATAACGTTTGCACCATGGACAAAAGAAGCAATTGTGATGGCATAAAATGTATTTTGATTTG
ATAAATAGTAAAATGTATCAATACTAGTTGAATATTGGCACACATTCATCGGTTAATTTATTTTCTCATTAACTA
AATTTTCTAGGTCAGATATTCATGAAAGAGTGCTGTTTACGCTGTTCCCCTTTTCATAGTTTGACAAAGACCCAA
AACATATCTAAATCATATGAATTTTAGTTCCTAAATATATTGCACGTTAGAAATCATGTTCAATGGGACTGAGAA
TCAGTTGGACCATAAGTTGCTACATAAAATACATAGCAAAAGGTATACTAAAAGATTTATTTTATCAGTTACATC
ATTCTGGACGTATATGTATGCCTACCCAGAACTTCAGTTTTTTCAGGGGATTGTAATACTTGATGCAAATTTGAT
AACAAGAAAAAAAAAAAGAGCCCTTACGCACAAAGGACTCCACAACATGCTACCCTCATTTACATAATTTTAGTGC
GTAACCTCATCCATCAAAGAACAATGATATCCACGCCATTGAGCAGCGATGGCGCAGATCCTACAATGCGAACAC
TAAAGTACTCCCCGAACTGTTATCGACCCTAAAATAACTCAAAAAGGAAAAGAAAATAAATAAATCAAAATAATG
GCTTTGTATCGCAACCCACAAGAGATCATTGAGGAAATGTACAAAACTTCAATCATCAGGAAGGGGATCGGGGGA
GAGGAGATCCGCTTCTCAC**CTCAGTTTCCACCAGCACGGCGTCCGCTCGTCTACGAGCAAACTCACAATTGAATC**
**AAGAGATCAAACTCGAGACGGGTGCAAGAAGCGAAAGAAGGG**TGGAGGAAGCAGATGGGTTTGGCTCTGGCAATT
GCAACGATAAATATCGTCTCCTTCTCGAGGAGGCGCATAATATAATAAGCAAAAGAAAGGCGAAACAAAGGGCTT
AGCATAGATCCACGTCTCCGCCTCCAATAATATATGTACTTCCGGTGCTTTTTATGTATCCATGCACTATTCGTT
AAACACACCAATTAGTCGATCGATCAACCAATCGGTCGGCACGTTACATCAAGGCATATGTTCCTTTGGATATAA
ACAAGGGAGACAACTGGGCTTAATCACAGCCGTCCACGTAGTTTGGTTGATTACTTAAATACTGGCCTCATGATA

TAAAGAAATATGTTCCATTAGATACAACACGGGGGAGATCTGGCTTTATCGCAGCCGTCCACGTCGACTTGATCA
CTTGGTTACTGGAGACGTCACACCAAGACAAGTTTATCCTCCCTCGAGTAGAAACAATGCGGGAGAATTGGACTT
CATCACTGCCGTCCACGTGGTGCGGTTGATTCAATCGATCTCTGGAATCGCGACAGCACGATGTGTTCGGAGAAA
AGGGTGACACGCCGTATCCTTTTTT

>MaAGL17_6
ATGGTGAGAGGGAAGATCGTGATCCAGAGGATCGACAACTCCACCAGCCGGCAGGTGACCTTCTCCAAGCGGAGG
AATGGTCTGATGAAGAAGGCCAAGGAGCTCGCCATCCTGTGCGACGCCGAAGTCGGCCTCGTCGTCTTCTCCAGC
ACCGGCCGACTTTATGATTACGCCAGCACCAGCATGAAGTCTGTGATAGAGCGATACAAGAAGGCAAATGAGGAG
CATCAACTTGTTCTCAATGCGACTTCAGAAGTCAAGTTTTGGCAAATGGAGGTAGCAAGCTTGAGGCAACAACTG
CATAACTTGCAAGAAAATCATAGGCAGTTGATGGGAGAAGAGCTTTCTGGCCTGAGCATCAAAGATTTACATGAT
TTAGAGAACCAACTAGAAATGAGTCTACATGGTATCCGAACAAAGAAGGATCAACTTTTGACGGACAAAATTCAG
GAGCTCAAATGGAAGGGGAACCTCATCTACCAAGAGAATATGGAGCTCCACATGAAGGTAAACCTCCTTGATCAA
GAACATATGGAACTACATAAGAAGGTTTATGAAAAAAGGGTAGACAATGGTGACAATAGAGACTCAGCTATTTCA
TATGGTTTCGGCATTGCGGAGAAAGCAAATGCATCCAATCATCTTGAACTAAATAGGCCACAACACCAAGCCAAC
AGGGAACAAACTGGAACATCACGGCTATGGTAA

>mac_miR444a
CCCTTCTTTCGCTTCTTGCACCCGTCTCGAGTTTGATCTCTTGATTCAATTGTGAGTTTGCTCGTAGACGAGCGG
ACGCCGTGCTGGTGGAAACTGAGTTATGATAAGAGGATGTAACTAGCTTGAGGCAACAACTGTATAACTTGCAGG
AAGTTCTATGGGCAAAATGAGCATACCTATGATTTTCTTGCAAGTTATGCAGTTGTTGCCTCAAGCTTGCTACCT
CCATTTGCCAA

Stem-loop structure

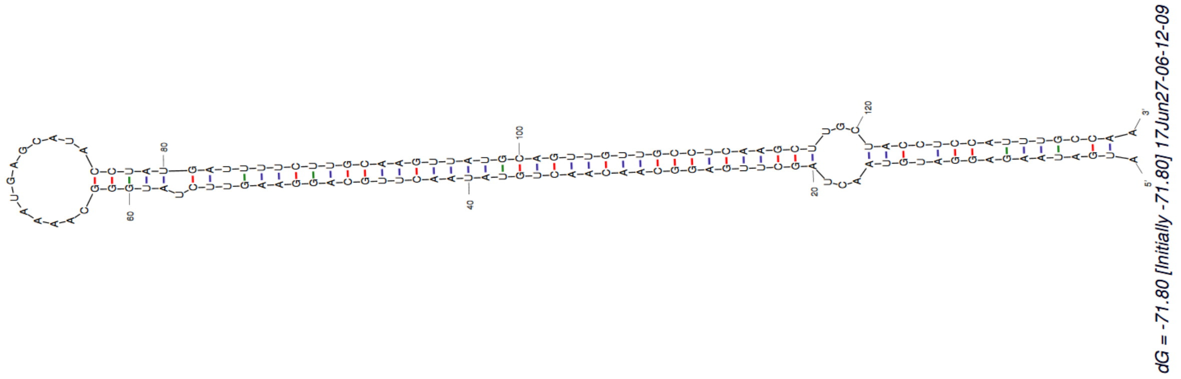


*MaAGL17_2 and mac-miR444b*

>3
CTTACATCCAACCTCTCCCCTCCCCTCCCCTCCCCTCCCCTCTCGGAGAGGTGGGGAAGAGGAGAGGAGAGAGTG
CGTGAAGGTATCTACCTGTCAGAATTCTATTAGCGTCGTCTCTCTCCTCCTCTCTCCCCCTTTCTATTTTTGTAT
TGGATAAATAGGTGAGGGAGGAAGGGGGGGAGATGTTGAAAGATAGCATCCGTCCTCACCAAAGAGAGGTGAGGA
GCAGCTCCATTTCGGCTTCTTGTCGTCGGGTAGAAGAAGGTTCATCCTCCTTCTTGTCTTCTGCTATGTGTAGAT
CTGTCGTCTGTGCTTCTTTCTTTCTTGTTCTTCTTGTTGATCCGTTGCATCTGATGGGGGTAGGTATAGTGGTGG
ATCATGGGGAGGGGGAAGATCGTGATCCGCAGGATCGACAACACCACCAGCCGGCAGGTGACCTTCTCGAAGCGG
CGGAACGGGCTGCTGAAGAAGGCCAAGGAGCTCGCCATCCTTTGTGACGCTGAGGTCGGCCTGGTCATCTTCTCC
AGCACCGGCCGCCTGTACGAGTTCTCCAGCTCCAGGTTCTCTGCCTCTCTCTCTCTTGCTCTATATCATCTTGAT
CTCCTGGATAGAGAGATCTATTAAACCCATGCATGAAAGCAGGAATACTTGGTCGGAGCCTCGATCTTATGGGTC
TTGGCTTCGGCTAATCGATCTATATCTTCTTCTCCTCCAAGCGGATCCTCTTTGAGGTCGCTGTAGTGATCCCTT
GAAATCCTCTAGGGTTTGGGTTGGAGGCTGTGCTTGCCTTGGAAGACGTTTTCATAACTCCATACGTCGTTGGAA
TCGAGTCTACAAGAGAAAGTCATTGGAAATGAGAATATAGCAGACGCTTTCTTTGCATTTCCCTCCGACCATTAT
TTTCTTCAGAGGTTTAGGTTTTAGAGATCTAGGAAGCTTTCTTGGGGGAGATGCGCACCTCCCCCAACAAACCTA
GGAAAACGACCGGTGAATCTCCAGTATTAGCATGCTTAGGTTCAGACTTAGTTGTCATTATTCTTCTCCATCAGA
AAAAATCCATGTAAGCAAAATATTTCCATTTTGAAACAAAAAAAAAGAGAAATACTGGCTCTCTCTTAGTTTGTG
CCCTCTGTTTAATCCAACTTCATTATTATCGATCCTAGAGACCTTAACAAATGCCTGATAGTAGAAACTCATTTC
ATTTGTATTGCAGCCTGCTGGTAATGACAGTCTACATTGGCAAAGTGACTGTGATTTAAGAAACTAAATAAAATT
ATTGGAAACATTCTACATCCGTTTCCAGATCTACTTTTATGAACAAATCGAAGTATGTATGTTACCTATGTTTTG
CATTGTTTTGTACAATTTTTTCTTGGAGATAACATTTGATGCTTCAAACTGTTATTTATTTTAGCAACCTAACCC
ACAACTGCTCTTTGCAGATGGAAGGTACTTTTGTAAAGTTGATATATATGTTTCCATATATGTTAAAATGACACG
AAATTTAGCTTTATAGTTTGTGACTCTTTTGGCATTGGCAAACAATTAAGTTCTTTTGTAACTATTTTTTGGTAT

CAAATATTTGTTTTGCAAAGATAATTGGTAATCATTGCTTGTTAATCGTCAAGAAAATGAACCATAAGCTTCTTT
TTTAGTAATCCTTAACGTATTGATTTAGGGGGAAAAGGAAGCTTGAGTTCAATCGTATGTACACGCGTTGGGTGG
ATAAGATGGAAGGATTTACTATGAGCTACCATGCATCTAGGTTTTTTTGTTCTATCATATGTTTTATTTTACAAG
AATTCAAACAATACATTTTGGAGTATATGGCATGATATCATCAGTTTTGTGAAATACATTATAAATGTTAATTTT
ACATTCTATTGATTAAATTCACTGCTATTGCCTAAATAACTTTCAGAAGAGAATTATTGAAAGATTATGTTATTA
GTTCACACTCTTAAATGTTGCAACGGATATAAATAAATGTTCTGAGTCAATCTCATGTTCTAAATAACAGCATGA
AGTCTGTCATAGAACGATATACCAAAGCTAAAGAGGAGCAACAGCAAATTGTTTCTGCAACTTCGGAACTTAAGG
TACGTACACTTCAAGCTTTAGTGTTTGTTATGTAGATAAGTTTATTAAGAACTGGATTATTTAACTCACTTGTTA
TTAACCAGTTGCATATGTCTTTCCAGTTTAGTTTGCTTAATTTATCATGTCATACTCTTACAATTTACTAGTAAA
TGTCATGAAACTTCATCTTCCAGTAGACTTCATGCTCAATTTTTTTGAGTTTGTCCAAGTGACAATGCTCTTCTC
TAAAATAATCCCAACACTTTGTATTTTTTGAAAGAGAGTCGTTCATTTCTCTAAAGTTTGATTGTAGGTTTTCCA
TGAACTAGTTTTAGGGAAATTCTTTGAGTGAAAATAAAAGTTAATTCTTTTCTCATGTTTGACCAACCACGAACT
GAAACATACGCTTGATTATTTGGTACTAAAAAGAGAATGTGAAATATTTGGTACTAAAAAGAGATGTGTAACATG
ATGAATATTCTCATTGATTTTCTCCATTTGCTTCATTTTTTTTCTTTCCAGAGGCATGGTAAGTTTATCAAAACC
TGCTTGAAATGGTTCATCTAAAATCATCCATGACCAAATTAATTTGACAAGAATTTGAGGCGAGAACCTCAAAAA
TCCTGATTTAAATTGATTTGAACTTGCTCAATCCCGTTAGAATTAGTGGCTTAATTAGCCTCATCAATGAGCAAC
TAGTTAGGAATTGATCAAACTGGGATAGAATTAACTTGAATCAATTGGTATTGAGTGTGATCTAGCTATGATGAT
CCATGGTCAAACTTAGTCAACCTAGGTAGATGAAACTCAAGCGATCAAATTTAGATGAATTAAACACAAGCGTTT
GTCAAGAAAGGAAGTCATGTAGATTTCACCATTCACATAAGCTAATATTGTTTGTGGAGACAATTGTGCAATTGA
TGTAGGTATAGTGCAAAGTTCAACGACACATCCATGGAGAGTTGGTTCATTTTTTTCACCATATCTAATGTAAATC
GATGGAAGTATAAGGTTGATAACGAAAAATTATTAAGAGCAAAATGTGAAGATAGATTGTACAAATTAGGAGCAA
ATTAGTAGCAAATTAGTAATTACTAATTACTAATTAGGAGCAAATTAGTAAGCTCCTAGTTGTACAAATTTGACT
TACGGCACGTCAAGGGCAATGATATATCAAACATATATGAGAACGAGGAAGGAACTTGTATTAAGTGTTGTAGGG
TTAATTGGCAACAAGTACAGTTCCCATGCATCATGAGTACGATAAAGGCATCACCATAGATAACTGATAGATGGA
GGACAAAAGTATAGTTGTAGGAAGGGAAGAAATAGACAATTGATAGCAAAATGAAAGGAAAATAATTTTCCATGA
AAAGGTGTCATGATCTTAAAAATATTTTTGATGATATAAATCTACCATGTTATGATGCAAATCTTGTTTAACCAT
ACCTATTATTTTAAAATCTTGAAACAAGGCGCTTATGCCAATAGATTATACGAGTACTATAAGTTTGTTTGTGAT
GATAATTTGATTTACATATTTGATCCAAATAAATTTTTTTATTTATTTATAGTTTATAAGATATTTCCCACATTA
CTTAAGTTAGCCCAGATGATTTTCTAGAAGTCGACCTGATAATCGAGCTAAGTTAGCCCTAATCCATAATAAACT
TGATATCACTTTAGGGGCCCATTAATAGGTTTCATTATCCAATCAATATCTTTTATTTATAATGGGATCTTCATT
ATCATCTTGCTTGAGATTGCTCTTCATGAATATAATAATTGGACTGCTGGATTGGCAACAAGATTCAATCATTGT
TGAAATATGGGATGATAACATTATTTTTATTATGTGCCTGATTTTAATTTATATTCAAATTAACTCAGAGATAAA
CCTAAACAATTGATAATATTTATATTCTAATTATGTTGAGGGTTTAACTTTTTTCATCATATTTTTTTGATACCT
AGACAATTAGTCCGATTGTCACGTGATCAACTATCACGTATCTTAGATTGAGAATAAAAGCTATCATCTTACAGA
TTTTGCACTTAAATAACAGATGATGGTGATATATCATATTCACGGACTCTAGGTTGAGGGTTTGAATCATTTAGA
ATACAAGCAATTATTCTGTGATGATATGCAATTTACTTACCTACCAAGGGTTTGAATCCTCTAGAATACAATCAA
TTAGTCTGTGGTGATATACAATTTACTTACCATAAGTTGGCTTAGGGTTTGAACCCTCTAGAGTATAAGCAATTA
GTCTGTGGTGACATACAAATTGCTTACTTTAAGTTGGGGATTTGAACCCACATCCTAAAAATTTATATATATATA
TATATAAATAATTAGTTTGATGATGGCATCCGATTCATTCACCTATAAATTATACACATAACAAATAGTTTTGTT
GAATAAATTTTATGCTCAAACAAAATCAAATAGTGATGTACTCGAATTATGTGATGTTATGTTTGACTCACTTTA
TCTTATCTTAGGCTGAGGATTCAAATCCTTATTATATAAATTGTAAATCTAAATAAGTAGTCTAGTTGTGATGTA
AGAGATATTATAATATAGGAATCTTAACAATTATCTAGCAATTCAGTCCTCGAGAGTCGTATAAAAAAGGGTACT
AAAAGCCTTGAGAAGTGCACCATACGTTAACCTACTGGTCACGATACTTTGGGAGACAATTAAATACCTAATTAT
AAGTGTTCGAGATAACATCTTTGTCCATTATTAATTCTTATAAAAAAAAGTTATTAGTACTTTACACTCATTTCG
ACCTTTAATTTATAATTTTCTTCTATAGTTTGGTAGCTTCTTTTATTTGTACTCCTAGCATGCCTTTAACTTGAT
TTTTAAGACCACAACAACAGATTCTGCCTATTCATGTAAAATATTAATATAAGGTCATTAACAGTCATAACACTA
AAACCCTTTTTGTAACCAACTAGTGGCAGAAGGTGATTTGATCAATTCAAAGTTGCAACATAGCAACATTGTATA
GCTTACAAATTTGGTTACACGTAAAATAGCATGGAAGGACCACGTCATACATTTACTTCACTGGATTTCAGATTT
CTCGAGGAACTTCTGCCATAAAGTTCAACAGAAAATAAATTTTCAGGCCTTGTATTGTTCAATAGTGGATAAAGA
TCACTTCTTGTTTGTTTGGGCATGACAACTTCAGATTCTGCTCAATTACACTAGGAATCATCATTAAGTAGCTCC
ACCGCATACTAATATGACCACAGATTCTCAAACTTCTGGCAAAGGAAACAGAATATCATCAAAGCAACAATATAG
AAAAAGTAAAAAAAAAAAAAGGTTTCTTCCAAAGACTCAGATATGAATGCTTATCAAATGTAAATGTGGGCAGTG
GTGGTAGAGGCTTAAAAGCAAGCCTAGATAGTGAAGCAAAAATTAGATGTGAAGTTGAGGATTAAAGAGCCCTAG
GACTGCATGCACATGATTATATATTGGAATTGTCATAAGCACTGTATGTTTGCAACAAGCACAATCTCAAAAGAC
AAAAAGAAAGAAAAAAAAAAGCACAACCGCAGAAGTTAATTCTATTTATGTTATACCACCTCAGGTTCTATTACTG
TCTGTATGATTAGCCTCTAGGTAAAATCAGTGCCTTCACCAAACTTCATCATATAACAATTTCATCAATTTGACT
ACAGGAACAAGAGAACATCTTGACAAATGACTTATGCCAATTAGCCCAATCATCATATGATGTTGAGTACCCTAT
GTCCCTAAACATATTTAGCAACGGAATTAGAGTTTTGCAAAAACTTTATCCTCTTAGAACAAACATTAGATGAGA
AACAAAATCCTGTACCATCAAAATCCTCCGGTGCAGAATGATGACCTTGATGCTCCGCTCACTGTCGTCTTCATA
TTAACAAAACAAATTATATGATGGCAGGGACATGGTAGTGATGATAAAACTTGGCATGTGGACCAGATGAACCAA
GATATTAAATGGAGCATTGCTTTATAACTTGCAAGGACAAGCTATTTTGATGTTTTGTTGGACAACTTGGCATCT
TCATTTTAGCAAGGCCAAGGATGAAAGGAGTGCACTCCAATGAGGATGAAAATCATACAATTATTATCTAATAGT
TTGGGCTGTTTAATATTTTTTAACCAATAGAATTTGAAGAAGATCCAAAATGCAAATAGTTCATTTCAGGTTAGA

GATGACCTATCCCTTATCCACAATGCATATGCTGGGGCACAGGGAACAGATCTACATTATTGAAGCTAAATATGC
AGTTGCCAATGTTCGAAGTCAAAGTTGGCAATTCATGTCCATCAAGAGTGTTAATATTTGACAAGGCCAGGGATG
AAAAGAGTGTCCTGCATTGAAGATGAAAATCATACAATTATTCTCTAATAGTTAGTCCTGCTTAATCTTTTTCAC
CAATAGAATTTGAAGAAGATCCAAAATGCAAATAGTTCATTTCTGATTAGAGATGACCTATCCCTTGTCCACAAT
GGATATATTGGGGTGCAGGGAACAGATCTTGAAGCTAAATGTGCAGTTGCCAATGTTAGAAATCAAAGTTGGCAA
TTCATGTCCATCAAGAGTATGTTAATATTTGACAAGTCAACTTCCATATGCTAAATTACTATCATCATGGAAGCT
TCTGTGACTTAGGAATCAATTGAAGGATTTCACTATAAGAACCAGATCTTATATCCTGTTTGGCTTTCACAATCA
GCTCAAAGTTACTTGTAGCTTCCAGTGTGCGCCTCCTAACCAACCATATTTTCAAAAATTTAGATTGTTGTTTAA
CTAATCTTTGGATCATTCAATTAGTTCCAGTAGAAAACTTTCGATCCTTTCATTGATATCATCAAAGTCAGATCT
TCTGCTCATTCTACGATATCACAATAATAACAGGTCACTTGAGTCAACAATTTGTCTGCCTCATTAAGAGGACTT
CTCCATCCACAAGGGTTGTAGGGCCAAGAGAATGGCGCATCCACATAGAGACTACAACATAGCTATATGGTAGTC
ATATCTTTGACAAAGTAGTAAAATACTAGAGATTTGCAAAAAAAGATGCAAATTCATGCTCGTAGCAAAACCAAA
TGCAAAGGAGCAGCATCTTTCCATACAACTAAGTTCCCTTCCACTGCAACCTTCATGAATTTTATCCTTTCTGAG
TTACAAGTTCTGGACTGCTGCTGTTTGTTCTATTGGTATATTTGTTGACATGTCAACTTTCTTCAGTTATAGAGT
AACCCAAAAAGAAAAAAGATAAAAATAAGGCTCTTGGAAACAATATGACAGTTATATTTCTGATATAGGATATAT
TCTTGCTACAGAAGTATCTCAGAAAACTACATACAATGGTCTATGATTGATCCCAATGTCATATCATTCATGGAT
GTTCTCCATAACGTCACTAGGTTGCCTAAGAAACTGCTCCTAAATCTGGCTAGGCTTGAAGATGAATAAGACATT
ATAGCAAACTAGAAAACTGCAAAGTTGCTTCTCCTTTCACAATGTACTGTAGATAGTAGTATATCCCCTTGTGCT
TCTTGTTTGAGGATCTTGTACGAGTATTGGTAATTGATTGTGAACAAAGAGCTTTCACATGTGTCTAACATGAAG
AATGGATCCTGGACATCTTGCAAAAATGTTGGATTAGCTCTATGAAATCTTCCGGTGCACAAAAGACAATTTTCC
AGGCCACTATAAGTAGATAATCATGATCTTAGATTACCATACTGGTCGATGCAGGTGGTATGTACTGGTCCAGTG
AATATTGGGATACATAATGTCCTGTACCTGTAAAATACTCTACTGAAAGTGGACCGTGGCACACCAACTGTTACA
GTCAGTGCAGGCGGTATGTACTGGTACAGTCAATATTGGAATACATAATGTCCTGTATTTGTAAAATACTCTGTA
CTGAAAGTGGACCATGGCATACCAAACTGTTACAGGTCGACATTCGTTCGATATAAGTTGTATACAACTAAGCAT
ATGTTCTTTTTTCAGACTCGACTACTTCTAAATCTAGCCCATAATTCAAGTGGGAATCCTCACTTTATCCACATA
AACCATCTAGCAGAGATCCAATATATCGACGTAACATGAAACACAATCTGACATAGATCCAACCTACTGACATAA
CAATAAACTAGCATTTGGAATACGTTCTTGTAGACCCCCCAACCCCATCTATCACAATGTGATTTTAAGCTTATA
AGATGGAGTTATATTGCTTTGGACACATAACATATTTTGTTTAAAATATTGGGTTATCATCATGTATTATTATAA
GTTCATATGTATTATTTTGTCTTCTTTCCATTTATTTACCATATAGTTGGGATATATGACTTGGTGTTGTTCTTG
TTTTTGTTGCAAATTATATTTCATTTAGATATTATGCACATGAGTATTATGATTCATATATTGTTGTTGTCGTCA
TTCTCATTCAATATCTTCAACAGATTTGAGATTATCTATTTGGAGCCTTAACGACCTTTTCTCACTCCTTTAGCC
TAGCCCTATCTGCAGTAGTGGGAATCAAAGAGAAACCTAAATACAAGTGCTTCTGAATCTCCATGATTCTGAAGC
TTCCTAGGAACCACAGTTCCTAGTATTTGGAATTATTATCCTCTAATATTTAGAATCAGGATTCAATGTATTCTA
AACCATATTCAGTATATTAACATATTTACACAATTCGAGTTCAAAATCTTTCTAATTCCTCCATAAGTTGTTGAA
TTCAAATAAAATGAAGGGTCAGATCAAGGATCACCATTTTGCCCAAACCAATACATACTGCCCCCTTTTGGGCAA
TCCTGGTCTGATTTATTTTTTCAATATGAGTGACAAATCTTAGGGCAGGGGGTCATGCAATTTGCTATTCATAGC
TCCCTGACCCTACAATTACCTCCAGCACCCTCTCCCCTACGCATGCTGCCACTACTATCATGTCCGCCTGTTGCC
TACCCGCCGATGGCCACAGCCAATCACTGCTGACCCATCTGATCTCCCTATACTTTCCTCATCATCTTCCTCATT
TGTCTCATCTTCCTCCTCCCCTATCCTTTGCCTCCTCTTCTCCTACTTCTCCCTTCCCCCTTCTTGGTACCATGG
TACACACCAACAAACTGTGTTATCATGGACTGGACCCATTTTGGTTTGGCCATAGATCTGCATGAGTTTGGTACA
TGAAACTCAAAAAGAAAAAAAAAAAACAAGAGATTACTTTGCCTTTTTAAAATTTCATCTTCTAAGACAAAGATG
AAGGATATTTTTAGGTGTTAGATCAGTTGATCTAACCTGGCTATAACCTGCATTGGCATGTTTAGACTAATCTTG
CATCATAACTACACAAAAAAAAATCAGGTTATCAAGAATGGTCAATTTTTAACTCTAGTACAACATTTCTAATAAG
GAAAGCTATAAGCCTATATCCTTTGTAAGAATCACAGATGATTATAATCACCCCTATTAATCTTCAGAACTTTGG
TAAATTTTCCTCTGACTTTACCATATCAGCTGTTAAAGCTAAAGAATTGACTAAAGAAGATGGCATAATATGCCA
TTTATTTAACAACTTCACTTGAACCTTCATGGCTGAAAATTTAAAATATGCACCTTTCTCAAAATCTCTCATTCA
CAAATCAACAAACAGACTAGAATGATGTCAGCTACGATATCAATTGAAGTCATTTTTACAGAAGAATAAGAAAAT
GGAGAACATTTTGTCTGGTTTGAATCATTTAGTTTAATTATATGCATTATCATTAATTATTCATTGTCAAAATTG
TACCAACTGTTGTCAATTTGTAGATTCCAAGTCCAACTGCAGCTTGAAAAGTTAAGG**CAGTAATTATTAAGATTA**
**TTTACTATACATACAAATATGCTTGTACAGTTATTGTCTCCAGGATGTAAAAGCAAATGCTCTGCGAAGATCTTC**
**ATGTCAACTACCGCACCAATCAGAGTTGTCATTT**CTAAAGCATAGAAAGAAAAAAAAAACCCACCAGAAGTATAT
GGGTAAATAAATAATGATGTAACTGCCAAAATGAAGAAAATGATAAATATAGCAAATTGCTATAGTTATATTCAT
GACCTTAACATTCATATATAAACTATTATCATTTAATAGTTTCTTACCTAGAACTTCTGCATGGACTTCTTTTAA
GATCTGCGGGTCTTTATTGTTACATTTGATCAAACAGCTCACCTCTAAAACATGTTTGCCTTTGGAGCTAGTTAC
TGAAGTTTTAAAAACTGCATATCAACATTGTAAGAGTATTGTGCTTTCTACTCAC**CAGTTCTGGCAAAGGGAGGC**
**AGCGAGCTTGAGGCAGCAACTGCATAACTTGCAAGAAAATCATAGGTATGCTCACTATACC**CTACAAATTCAAAT
AAAAGTATGGTCATTCAAAATCCATTTTTCTGTAGATAAGCACAAAAAATGTAGTAGCATTATCATGGATAGTTT
ACTTCTAATGTTCAGTTTTGCTTGAACTTTTGGACTTAGATATGTTCTCTGGTATATCAGTGTCTTGTATAACAT
GTATTATTCATGATATTGATGCCGCCGTCACAATTATGTGGAGTCAGCTTCATCAATGAATGCTCATGTTATTTA
ACAGTAAGAACTTAGTTACAGCACTAGAAAGACGTAGGAAACAATGTGCCACATCAAAACTCAAACCAATCAACG
AATTGATACACCTTAAACATTATAATCTAAGTGACCTCTTCTGAACTAAAATTACTAGGACTTCTGGGTGTCAGG
AAATTATTTCTACTATTCCAGTGGAGCTAATGAGCTCCATTCTTCATGTTATGACAGTATTATCAGTGTGTAAAT
CTTCTACAATGCATATGGTAAAGGGAATGCTAATCCAACTTAATAGAAATATAACAAGTGATATCATGTTTTAAG

GAAAACACAAAATTTATTTGTACACATTCATTAAGCAAGGAGAAAGTGAAATACAGCTAACTATAATGCAGCCCA
ATAGTTACCAAACACATGTACAGTAGCAAGCCAACCAGATAGACATGGGTAAACCAAACACATGGATTGTGCAGA
GACAAATAAAGGTATCATAAATCCGTATCTTTGGACTCAGCAAAAGTCCATGACTACCTCTGTACAATATATGCT
GGTACCAGTTCAGATAAGCAGTGCACAAACTGATGGAACAATGACAATTTGGATGACCAGTTACATGAAGCCAAG
TCACAGTGCACTACTGAGTCATTACATGCTACACAACAGAATTAGAACCCATAACTATTGCTATAAATTAGATAA
TGTCAACAAATGAAACTTAGATGTTTCTATAGTATAATTTAGTTATATAGTTTTAAGAGTACAAGACTATGCTTG
GAATATCCCAACCCACAAAATCAAACTTCTGTTTGATTGAATGCCAGAGCAAGTGGCACCCCAATGCCTTTCTGA
TGAAGGCTACAGAAATCCCAGACAAGAAGCACCGGACAAGAAGAAAGTCATAAAAGGAGTACCTGACAAGATGTT
TCAACAGGGAATTTAAACTCAATCTTCAAATACACCCGGTGAATACAAATTTCTTTCAATGTAATAGTTAATTAA
TGCATGCAAAAACTTACAAACAATTGTCTCTCCAGAATTCAAACTTGCATGACATGCCACTAAATTAGTTTTCTT
CCCCAAAAAAAGAGTAAAAACAGTATATATGCATCCATCAATCTGAGAATTCCAAGCAAAGGGTTCGCGAAACCAA
TTATCAGGCTGCACGTATCATCATTATGGATTTTATATTGGAAAATAATTTCATAAACACATAATGTCATAAATT
AGTGACAAGAATTACTGAGCCCATTGCTTCTGTGTCAACAACTGCAGGCAGTTGATGGGAGAAGAGCTTTCTGGC
CTAAATGTCAAAGATCTACAGAATCTAGAGAACCAACTAGAAATGAGCCTGCGTAATGTCAGAATGAAGAAGGTT
CTTGCTTCCTTTGCATCACTCAGTATTTCACACCTACTTTCTTCCAACAACTATTCTATAGCGTACAAATAATTC
TATTCCTGCAGGACCAACTTTTAATAGAAGAAATTCAAGAACTGGATCAAAAGGTTATGCCATAATTTTGCTACC
TCTCAACAAGTTTAGAATTGTACATCTAGTTCACCTATATTGATGCTGGTTTCATTGCAGGGAAGGTTTATACAC
CAAGAGAATATGGAACTGCACAAGAAGGTAAACCTTGTTCGTCAAGAAAACATGGAATTGCACAAGAAGGTATCA
GATACAAAAGCTATTTCATCATAGATCCATAAAAGTGTCCAATTTTATCCCTTGATGAGCTGTCAAATTATGTAT
AACATGAGTTAAGCTATCAATCATCATTGTCAGGTTTATGAAACAAGGGGAACAACTGGTAGTGATGGAGGTTCA
ATTATTCCATATGGTTTTAGTTTTACGGATGAAGCACATGTGCCCATTCATCTAGAGTTAAGCCAGCCACAACAG
CAAGCAGATGGAACCCAGACCCAACCTCCAAATCTAGGGTAACTTTTTTTTACAAACATTAGAAAATTCACACAT
TTAAAATTTCTAGTCTAACTGTACAACTTATTACTGAATTTGCACCAGACTTCAACTTCAGCTGACACGTGGAGA
GTGACTCTTCATGATCTTCCATCTATTATCTCACTGCTACAGTATCCCAAGGCTAGGAATCTTTAATTACGTAAA
AGTGCAGCAGTGACTCCTCAAAAACATTCTAGCCACCTATTGTATAAAGTTTATATCACCATGAAAATATGAATA
CTGTGCTAGGTATTAAGATCTGAGAAACTGAAACTGTATTTATTGCTTGCAATAAATGTATTGTGAATTTGCTAA
GTAGTCTTTTGTACCTAAATGAGGAATTCAAAGCTGAACTTGAACTTAAAGAACTACTCAGCAGGATCACATCAC
AACATTGCAAGCAGAACAAGCCAAGAGGAAGACTGGTTTATTATCCATCTGACAGAGAGAATGCAGTTAACAGAA
GGTAAGGTAGGCAATCTGTCAACAAATTGACTACTTATGACTGGAGCTTCGTCAGTCAACATGCACAACCTTTCA
AGAGTCATGCAGTAGGCATGTTCTGATAAAATGAAGTCCATTACTCACCCACTCAATTAAATCACCTCTAACACA
GACAAGTGGTAAAAAAAGAAAAGAAGAATAACGAATTGATCGTGTTGAAATGTGATGTGAATGGTGAAATATACC
ACACTAAATAAGAAACCAAAATAATAACTTCCTGCTGGCTCTCTTGGGTATATATTGCACAACAAATGATATCAA
TGCCATACAGGCTCTCATTCATGTATGACCAATTGCGGCATAGATCCCCCTTTTCTGGATATCCATAAATGGACC
GAGTGGAAATCCAAACAACAGCCACTTCTAACCTGGTAAGTGTCAATGTTCTGAAAAGTGGCCACAGTATGCAGG
CAGTTAAGGTCCCACATAGTGTATAAGCACTGGAAACATATGTGGCAGATACTGAAAAGTTTTGAAAAAATAACA
AAATAATAGCACGAAAAATATGGACAATATAAAGTTAATAGGATACATACAAAATGTAAGTGATCAAAATAGCAT
GATGATCTTATTTTATACTATTAAAATAATATATTAGCAGGAATAATAGAAAAAACATAAACCACATAGCGATCA
TATATTAAAGCTTGCACAGACCATATATGTGGCCACAATACATAGAGATTATGCATTGAGGTAATACGTCACTGC
ATACACCTATGCGACTGTATTTTTGAACAATGAGAATTCTTTCGGGTAGATTGCCACAAATGACCTATCCCATAT
CCTGTTGGAGCTCAAGTCATATACATCAGTTTGTTCTGACCAACCTCGAACTGATCGTAGGGTATTGGGAAAAAC
CACACCCGATTATAGGGCATCAATCATAGACCAATCCAGGTGTTCTAAATATTGTTCAAACCTAAAAGCCTGGCA
AGAAAATCTAGGTTTAGAAGAATTCAATTACACATGTTTATGTGCATGATGAGACCTGAGTCCTGACTACTCACA
GAATCCACGGAATCCAAACAAAATAGTCATGTAGGACAGGGAAAAAGTGGATTTCTAGAATAAATTAGACATCTG
ACAGATACAGAAGCCATAGCATACAGGGATTTCATATTTAGAGTTAGCACCTCAAGATGAAATTTCAACAAACAA
GAATATTCGGAGTCCTGTATTGATAAATCTAGTCCTGATTAAGGTTATAAAAAAAAATATGCAAATAAAGTCCAA
TAATGTTGGAGAAGCTATTCAGAAGCATAGGAATGATTTTAGAGTTCAATTATAATATAGCAAGAAAATTGTCAT
TGTGTGATTGGAAAAATTGAGTATGCAACGGACAAGGTCTGGCACATAACAGGACCATAAAAACTTATGAATACA
ATCAAGTTGAATCAGATCATAAAGAGTTTTTTGCAAGGAAATGTGCTGAGTCTAGAAATCCACTAAGTATGCACT
TAACCACAAGAACACAAAATTAAAGAAAGAAGATAGAAAACATTAAAGATGGTAAGAACTGGACTAATTCTAACT
CAACACATCTCATGCACAAGGGAACTGATGAACCATACGATATGAAAAGATGAAAGTGAAACACTTTTTTTCTTT
AGATTTTCATTAACTTCAATACAGCTATCTATTAAATTAAGAAATTAGCTTTTTGAAAGCAATCATTTAAATTTT
ATTCCGACCAAATGATCATAGAATGATAGAAACCAATACATTATGTGATAAGATAAGTAATTTTCCAAGATATAT
CTAAATCCAACCTTTTATTCATAGTTATAGATTTACAAGTCATGTGCAAACAAGATTTTTAAAATATAAAAGAGC
TCTAAGCTCATAAAAATAGGACATAAAGAAACATTCAGTTCTGATAAAATTTGATAGGAGTAAATATAAATAGAAT
CAGTGGTTTTGTTTGCATGGATTAATTGTGTCCAAAAATCTTTCCCTTTCCTAATTTTTCTGAAAATTTTGTCCA
AGTTGTTTGTTGTATGCCAGGAAGGGTAATTTAAATTGTTGGTCGATTTTAGTTGAAAAATAGTTGTTAAGTCAC
CATGGAGAATGTTTGCAGTTCTAAATTTGGATCGATTCCTTGTGTCTACAGGCTTAAATATCTAAATCATTTTTT
AAGACAACTTGCACGAATCTTGAGTTTATCAGTAAATTATGAAAAAAAATGACATTTCCTAAAACTCTGTAAGCT
TTGTTATCTTTCTTGCCTTTGACAAGTTTCCTTGAAGTTTCTTGACTATCTTGTTTCCAAAGCTTCTATTCCCTA
TCCTTTTAATTCCATCTTCATTCTTGGTTATCTTTAGTATTAGTTCTTTTGTTATGTCAGTTTGCTGTCCTATAT
ATTATTATAAAGCAAATCAGAATGTAGATTTGGATTACCAAACTACTACTGCCTTTTTTGAAGCATCTTAAGATC
TTTCTGACCAATATATATGGATCTCCCATATTTAGATAACCGATCAATAGGCACTAAAAATAATGATGAGAAAT
GAATTTAACAATACAAAAGAAAGATTTTTTCTCACTCGATCCAAAACCAATTTCTGCTAAATTCAATCCTAAACT

TGATTGGATAAACTGAAAATGCTATTGCACTTCTTATTAGTCAAGAGTCACTTCAACTGAGATCTTAGGTCATGC
TAATGGGTAGAAAGGTTCATGTGCAGTTAGAGTACACTAACAGAGTCCTTTAGGGCTACAGAAAATAAATAATGT
TCCATGTAGGTAGCCACCTCACTATAGAGGACCCAAGGATTAAAGGAGATTATTTGAGAAAACAAGCCACATGGG
CTCTTGAAGCATTCTCACTTAAATGCAGAAAGGCAAGGAAAAAAGAGAAAAGAAATTGAAGAGAAAATCACCAGG
AAGGACTCACCTTATTAGAGCAGAGGAAGGCTTTCAGTTTGAAAAAAAATGAGAGAAAAGAGGAAAAGAGGGTTG
CTAATTGGAAGAAAGGATCAGAGGGCTTAGAACACTAAGCAGTAGGCTAATTGCCCAAAAGACTACCAAGTAGTG
GTTATACAGTCAGGTGGTAAACCTGGTATTGACCCATTTTTACTGGTTCACTATTGAACCAGTAAACTGGTCGGC
AAATACCAGTCTGACTGGTTTTGAAAACCATGGTGATGACATTTACCAAAAATGACAAACTGCTATCAAGCTAAT
TTCAAAATAAAACAATGATAAAGTTTAACAAAGACGACTGAACTTGATAGAAGCAAAGTACCTTCTACTAGTTCT
AAATAAAAGACATCAAGACCTCGTTTTGTAATAACTAACTCCTGAACCAAGTTGTTCGCATCACGTTAGAGGAAA
TGAAATCTTTTTCCTATTATTTTCTTGTAAATAACTAACTCTGGAACCAAGTTGTTTGCAAGAATAATTCGAGCT
CTCTCCTTTTTCCTAAAAGATATTTGAGAAATCATCGGAAGCAACTCTTCATCAAAGAAAGAAAACCAAAAACAA
AAAGTTCACTTCTCCTGAATATATGAAACATAATATTTTTTTTTAAGTTTTCCCATCAAACACACTAGAAAGAAT
AGTTTTCTTGGAGAATAATTTTCAAGATTTTTTCCATGGGATCAAACAAATCCTAAGTGAAAAAAAAAAAGTAAT
TCAACAAAGACCAAGGGAAAGAACCTCCAAAATTTAAAGTATTCTTCAGCACAAGTAGAATAACTATCAACTTCA
TATCACAGTTACACAGAGCCAACACAGTGCAGCAAATACATTGAATCTGAAGTTTTCTGGTCCAAAATTTAAGTG
AAGCTAACACACAAGGAAAATGGAAAAAATATAAAATTGATAGACATTACAATGTTATTTAACATATTATCTGCC
TTAAGAAATGACAAAATCAAATCTAGTAACTTCAGGACCTTTATTTACAGCAAAAATTCAAATGATGGAGATTTA
AATTTTTGATTTCATAGAGTAAAAACCATTTCCAATTTGTGCTGTTGACTGTTGATGAGTGAACTTCTGACCTAC
GATGATGACATATGGCATTGTTTTGTTAAAAGCAGGTACAGGAAACAACTGAATCAGTTGCTAAAAATTGGCTGC
ATCCTTACAAGCAGAAATAAATAAAATGGTTATTTGACTGACGACTCCAGAATGGATAAAACTACCTGGCATGTT
GTCACATGACTCACACCAGATACGATAACACAAAATCTGATTGCTCACAAATAGAAGCATAAGTAGTAAGCAGAC
AGATCTTAAATAATGACTAGAAGTGCTCCTTTCATTATAAGTTTACATATAAACTTTGAACTTTTAATCATATAA
TTCTATGGGAACTGTTTTGCAAGTTTAAAGTACAACAAACAGGCGTCCAACAGCATTGCATTTCTACAATAAAAC
ATGCCAATAGCCTATGATTGTCACTTATCAAAACAGAAAAACAATCAGATTTCAATTGTAACAACTAATTTTATT
GTAAAGTTTTCATCTAAAGATAATAACATAC**CCAGATCTTCTTGCAAGTTATGCAGTTGCTGCCTCAGGCTAGTT**
**CCTCCCCTCTTGAAAGAT**CTAAAGAAAAAAGTCAAAATATATGCCATGAGTGATGTAGGTTTCAACCCAAATAAT
AGCTTATATACATACAAGATATTATATGTATGTATGTATGCATGTATGCATAAATGTATCAGTGTTGAATAAACT
TCAACATGCCTACAACATAAAATAATAAACTGAAGCATACATGTAAAAATTCTTTAAATGTGATGATATGAGACC
GTTTAAAGAAGGTGAAGAGACTTCATCACTCTGGTACTAGAAAATTATTCCAATGTCGGAGTGGACTATAACTTA
AAATAGTATATAATTCATGTCCAATAATAACCTTTTGTACTCTATTTGTGCATATTTTTCATATTATATTCGACT
CGAGCACTGATATGGTGTATTAATGTAATAATCATTTATAATTTGCGGATTATGTTAGAACCTATCATATATCCT
ACAAAATCTCCAGTCCATTGAGGATATGAACTTTGCAATTTGGACAATGGCAACAATGTGGAAGGAGGTAATATC
CCAAGGGTAACACATGCAAGGAGTAGATAAGAGAAGGGCAACTGTTGACTTCTCCAACATGAAATATAGCTTAGA
TCACATTGGTCAAAGTTTGTTGTGATCACTGCCATATTATCAATGGATTTCAAGGATATGAACGTGATGTCAAGT
TGTCAACATGTGTTCCAAATTCCCAAAGACTTTGAGGACTCTCCCAATAGCCTGACCAAACTTATCTTCTGAGCA
AATGCCTTCCTGTCATGGTTTTTATGTGGAAACCATGTCAAATGATTCTAGCACTTATGGATAGGTTTGATTACT
CATTCCTCCCCAGCCCCACAAAAACATTTCATATTGGCACTTGTCCCATCATGTGGAAGGCTAGGCTAATATTAT
TCAATAATATCAGGATGACAATGTCCAAGGCATGGCATAAGACATAGGACCAGTATAGTAGGTACATCTGCATCA
CATGTGGCCAAGTGTCAAGAAGCAAGTTCCCCCAACACAACATCCAAACATAAAAATTTTAGATTGATGGTACAT
GTTTTGTTGGTAGGTAAAAAACCATATGTCATTGTAGAGCTTGCTGATTTGACATGATTCCTTGTTCATCGATTG
TCATGTTCATGTTGTTGCATGCTTATAGTTAAGTTTGTGTGTCATGTAATTAAACTTTTTTTTAACTACATAATT
ATGACTTTGTTCCATTCCATGTTATCAACTTCACAAACTAATTCATTATATTTATGAATACTATCATGCAAATGG
ATTGACAATATTACTATGCACATTAAATTGTAGAACAAGAAAAACAAGTTTTTTAAATTAATCTCACACCACTAG
CCAAAGTAGCTGATGTACTAGCATAATGTATTGATGTAGAATAGAGTAGAGAAGTGTGAGATTAGAAATGAGAGT
AAGAGGCTAGAAGAGAGAAGATGCAATTCTTTTCATTCAAATCTAAAGTGTCTCATCCATCGACAAACTTCACTA
TTTATAAGGGTTTAGGAGCCTTAGTGCCTTAGGAGTTAGGATAATCAATGTAGACAATGATTCTCAAGAGTAACC
TCCTAGTAAACTTGTATGCCTTTTAGAAACCCCAAATGTGACTTTTAAGAGACCTAAACAGATTTAAAACAACTT
ATTACAGCTGATAAGAAAAATAGTGTATTTAATATAAGAGTACACTTAATACAAAAATAGAGTCCCAATGTGCTA
TAGAAAGACTTGAGTGCACTTAATACAAGAGTGCATTTAATACAAAAGTAAAGTCCCAATATTAGCACCTAGAAA
GACCAAAAGGCTCTTCAATTTTCCACACAACCGTTAGAATTTTTAGGCCAACTAATTGATGATTTTTTCATGAAT
TTCTTTGCCTAAAATAAGTTTTTATGAAATCCTTTATAAAACTAGATGGTTCTCCATCATGTATGTCAGTAGGTA
CGCCACCAACATCATAACTATATAGCCTATGAAAAGACACCAGTGTAAAATATGATCTAACTTTGGCTAGATGCA
AACATACCGTTGACAGGTGCATAACAATTACTAAACCCGTACCAGAGATTTTGAGCTCTAAATGAACACCATAAA
AGATATAATCCTTTGTTGCCACTGTGGTGGTTACGATCAGTGTTTTTAACTCAATTATTTTGCTTATTATGATAT
ATCTTTGTAGAATTTCAGAAATTGGCCAGTTGCTACATCAGTTTCAACATTAAAAAAAATCTAAATTTTAGTGAC
ACATCCCTCAATTGCCAACTCTATTCCATACTATCACTGAGGGCTAATCCTGGACAATGTATAATGCTTTCTATA
TAGACAAAACCGAGACCTTAACTTCTTCTTAATATGTTAATTTTGTTGTCCTCTTATGTGTGTTTCATAGAATGC
AGTTGAATTAAGGGGGGTCAAGCACATATGGTTTAGCCATGTTTTAACAAGGTTGAGATACCCCAGACATTACAT
GCAACCCTTATCCTTCAAAGTTTAAAATGGTCATCAAATAGTGTCTTTTTATATTATAACTTATTCTGTGATATA
TATTATATACAATGAAAGTTAGGCATGACAAGGTTTCTCATTTCTCTACAGTCATGTCTAGTCATACTATTATAA
ATGACCCATTACAATAGACTTTGGTAAGAACCACAGAATATATATACTTGTAGTATATATGGGCTTTCATTTTGA
AGTTTCGTGGAACTATCTATAAATGCTTAATGAAACCAATGGTCAGATCAAGTCTAATTTTCCTTTAAATGCTTT

AGAATTCGAGTCAGTCTTTTTCAAATTTCTTGTATTTTTGGAGTTATTTATACAAAATAGAAAGTGACCTTTTAG
ATTAGGGTTAACTTTAATTGTTTAAGAAGCATAAGTTTATTATGATTTATATTTTTTATAAGATAAAATTTTACT
TATTTATTTTTAAAAATTATTTTATATTTATTTATATGATTATATTTTTTATTTATTATATATTTGGGCCATACA
AAATTTTTTATAAGATTACTAACATTCATTTATTTGTTTTATTTATTTACTTATTAATTTAAATGAAAAATATGT
ATTCACTCATAAAAATATATTAAAAATATTAGAAGGATAAATTTGTCACGAAATATTCAACGAACTTTTATAATA
CAATTTTACCAAACAACACTTACGTCAATGTACATTTAAAATATAATTTTTATTTATTGTTTACCAAACATTCAA
AGCATTCTACAAGTACACATTTACTCAAATCTCTTTCTCAAATTGCACATTAAAAATATAAATGTGATCCCAAAT
GCAACAATAGAATTGTAGCATGAACGTTACCCCAAGAAAATTTGTCATGTACAAAATCAAATTTTGGAAGTCTGG
TAATGTGTTTGCAATTTTATCAGCCCAAGACTTCCACCAAGGTCTCTCGATGGAATGACACCAAGTTCACTTTTG
CAAACTTGAACGAAAAGCCACAATTTCAAACACTTAATGTAAAAGTGGGGTTCTGAACTATAGCTATCAAGGAAA
AATGGGACAAAAGATCATCCAATATCTTACTAGATATTGAGTGTTTGTAATTAAACTTTTTGCTCCAATAACAAT
AGAAGGGTCTGAATCAAAGTTGTTTCACGTCTCTTGGTATTGTGGAATGAATTATAGTTTGGGCTTGCTTAATGG
CCAAACAGGGTTCATATCTTCTCTCTATTTTTTCTCTAATCTTCTTTCTTGACTCCATTAAATCCTTATCATTCT
TCCAAAAACCACTGGGCTGCATGAGTAAAAATCGATGCTACCCATATGTATCTAGAAGCCATCTTTAAATGATAC
ATGCCTCTTACATACTACAGAACATTGAAATTCCATAAATAAGTTTAGTCATACTGCTATTTATTTTTTTTATTT
CATGTAAGAATGTGACAAATAACTACCTACATCAAAAAATATACAGGAAAATTATTTCCATGAAAAATATACTGT
GATATTTTCAGGATTAACAAAAGCATGTTGTAGTACTAGAGTACAATTTAGAATGCATCATATCTGATTTCTTAT
AACCATTCAGGACTTACATAACTTTATAGAGCTAACAGTTTAAGAGGAACTTCATTCTGAAATGCAAACAAAATA
GGGAATATTGTTCTACTTAGTTTTTAGTCCTATAATATTTTTAATTGAAATCATTGTATCCTTTTAATCGAACTA
TCTGAAATTATACATATAATAGGATACAACACAAATTACATCCATCAGAAATATAACTATCTGAATAGGTAATGT
CTTGTAAATCTTAATGTTCTACACATGGCATTTGCATGCTCGTGGTTGTCACATTACATGCCCAAATTGAGGTTA
TGTTGAACCAAACAAAGATAGTATATCATAAATATTCATAAATAATTACAGACATATTTAGAATACTTAGTATAT
ACATTCATACGTATACATATGTGCATATGCACATACATGTATTTATTCAAGGATTGTAATGAAGAAGCCTTTCAC
GCTAAAACAATGTTAACTTTTTAGGGAAAGAAAGTATTACAGACCAGTGATTTAGCACTTAAAACTAGCTTTCCT
GGAACTGACCTTAGACTTGTAGCCAAAGTTTAGCCAGGAAAACTGAAGGCAAGAAAAGTGCTACAGTGGACCAAA
AACAATCACTTAGCGGATCGTCTTTCATAAGTATGAACGTACGAGCTAAGAAAACTTGATAAGAAACAACAACAC
TTTTGATACCGGATAGGGGTTTCGACGTGACGACTTACAAGTTTTATCACTAAATATGTTTCAAAACATCTCGCA
GCGTATGACCACGAAACGTATCAAAGAACTCTTCCAAGATATGTCCATGAAATATATTTAATGCTCAACAACAAG
ACGCCCAAACTTTGAACAAAAACTTCTTACAGTCTAAAAAAATCAAGGAAAATACCTTATAATGTACAGCAATTC
CCTAGTAAGCCAGAGTTAAGTTAGACTGCAAGCCCATTCCAGTAAAGCAAGGTCCAAATGGCACAATCACGCACC
CGTCAGAAGAAGCAACAAAACTAGGTCGTACAAGCCCGTCGATAGCTATTTCATAGTCAAAGTAACACCCCAACC
TATCAAATCTGACCTAACCGATCAAAACCTTCAAGAAAACACTTCCGAATTCAAGGAATAATCGGCGATTACTCA
AAAGATCATGATAGGTAAAGAACTCGGTCTTTAC**CTCAGATTTCCGTCATTCGGCAACGAGGGCCAGCACCAATT**
**CTTCTCTCTCTCCTCCTTCCCTCCGAGGGTTTTCCGGCCATTGATCTGACAGCAAACCCAATTCA**AAACTAGAGC
TAATCAATCAAACACGAACTCCATTACCCGAAACAATGAACAACCGAGACCGGAAGAAATCAAGCAAGATCAGAT
CTATGAGATGTCAAGAGACAAAAAAGGATGCAGCAGTACCTGAAAGCAAGCCAACCACGAAGAGAGAAGGAGCGA
CGGTAGGGCGGTAGATATTTTGCAGGAGGAGCGTACGGTGCTCCCGAGGAGTAAAATAAGGGGTCTGAGGAACGG
TTCGCGGCCATCCACGTCGCCCGCCAAACTTATATATTTATCGAGGCGCGTGACAGAGAGGCGACGAGCAAAAGG
CGCATGATAACGAGAGGACAAACCATGAACAAAACGACTCGACAAATGCTCCCCGAAGAACCAACTTACCACGGA
CACGTGTCGAGACGGGATTGCTAAGGTGTAGTGTAACGACGAGGGTGGGGGTGCATGGGACCGCCTGGGTGGGTT
AAAACGGCTCGTGTGCCGTTTTGTTTATTGGCTCGCTGGACACGCGTCGGCGTCGGTAACGGCGTTTGGAGCCCA
CCTCATCCGTCCCACCGGCGTTCCCTACTCATCCTTTGATTAAGAAAACTAAATTCACATATATATATATATATA
TATATATATAATTGTTAGAGCAACAGGGATTAGACTGCTACATCAGCTATTTAGTTGACTCGTTGGCAGTCCATG
GCATTATTATTACTATCTATTGATCAAAATAT

>MaAGL17_2

ATGGGGAGGGGGAAGATCGTGATCCGCAGGATCGACAACACCACCAGCCGGCAGGTGACCTTCTCGAAGCGGCGG
AACGGGCTGCTGAAGAAGGCCAAGGAGCTCGCCATCCTTTGTGACGCTGAGGTCGGCCTGGTCATCTTCTCCAGC
ACCGGCCGCCTGTACGAGTTCTCCAGCTCCAGCATGAAGTCTGTCATAGAACGATATACCAAAGCTAAAGAGGAG
CAACAGCAAATTGTTTCTGCAACTTCGGAACTTAAGTTCTGGCAAAGGGAGGCAGCGAGCTTGAGGCAGCAACTG
CATAACTTGCAAGAAAATCATAGGCAGTTGATGGGAGAAGAGCTTTCTGGCCTAAATGTCAAAGATCTACAGAAT
CTAGAGAACCAACTAGAAATGAGCCTGCGTAATGTCAGAATGAAGAAGGACCAACTTTTAATAGAAGAAATTCAA
GAACTGGATCAAAAGGGAAGGTTTATACACCAAGAGAATATGGAACTGCACAAGAAGGTAAACCTTGTTCGTCAA
GAAAACATGGAATTGCACAAGAAGGTTTATGAAACAAGGGGAACAACTGGTAGTGATGGAGGTTCAATTATTCCA
TATGGTTTTAGTTTTACGGATGAAGCACATGTGCCCATTCATCTAGAGTTAAGCCAGCCACAACAGCAAGCAGAT
GGAACCCAGACCCAACCTCCAAATCTAGGACTTCAACTTCAGCTGACACGTGGAGAGTGA

>mac_miR444b

TGAATTGGGTTTGCTGTCAGATCAATGGCCGGAAAACCCTCGGAGGGAAGGAGGAGAGAGAGAAGAATTGGTGCT
GGCCCTCGTTGCCGAATGACGGAAATCTGAGATCTTTCAAGAGGGGAGGAACTAGCCTGAGGCAGCAACTGCATA
ACTTGCAAGAAGATCTGGGGTATAGTGAGCATACCTATGATTTTCTTGCAAGTTATGCAGTTGCTGCCTCAAGCT
CGCTGCCTCCCTTTGCCAGAACTGAAATGACAACTCTGATTGGTGCGGTAGTTGACATGAAGATCTTCGCAGAGC

ATTTGCTTTTACATCCTGGAGACAATAACTGTACAAGCATATTTGTATGTATAGTAAATAATCTTAATAATTACT
G

#### Stem-loop structure

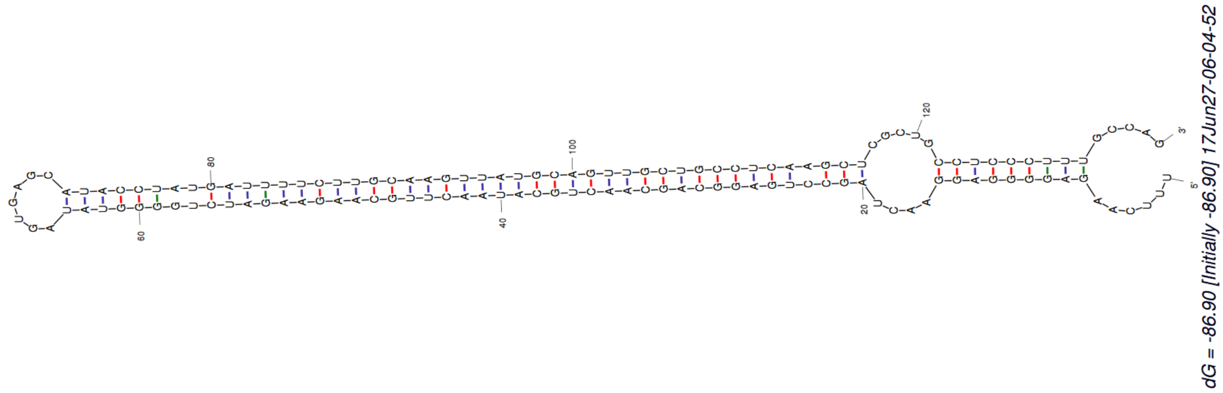


#### *MaAGL17_5 and mac-miR444c*

>7 reverse

AAGAAAAGATTAGGCGTCCGTCTCCGGGCCCGCGCGGGGCCCGCGAGCAATGCGGTCGCACCTTTTCTGCTTCCG
CCTCCGGTTTTCTGACAAGAGAAGGGGAGCCGAAACCGCAGGAAATCAAATGCCGAGGCAAATGTGCTTCGATTC
GACCCCCTGCGCACGATGGGATGCACGAACGAATCCGTCAAATCAGGTAACGGAGGGGTGGTTTGGTTTAAGCTT
TCTCCTTCTCTTAGGGCTGGCCTTTTTTGCATAAGATATACTGAGCAGAGAGACAGCGCTGACCTAAAAGGTGGC
ATCCCCACAAAATAAAGAGGGAGAGAGAGAGAGAGAGAGAGAGAGAGGGGTAATCAAGAGCTCCAGTTCAGTATT
TTCCTCATCCTCCTCTCCGGCTACAGGTTTGTACGCGTCTCTTCCTCTTGTTTATAATCATCTCTTCCATCTCTG
TTTTAACTGGTGATATGTTGTTGAGGTGGTCAGAGGGAGGAGGAATATGGGGAGAGGGAAGATCGTGATCCGCAG
GATTGATAACTCCGCAAGTCGGCAGGTGACCTTCTCGAAGAGGAGGAACGGGTTACTGAAGAAGGCCAAGGAGCT
TGCCATCCTTTGCGATGCTGAGGTCGGTGTGGTGGTGTTCTCCGGCACCGGCCGACTCTATGATTACGCCAGCAC
CAGGTTCTCTCTCTCTCTCTCTCTCTCTCTCTCTCTCTCTCTCTCTCTCTCTCCGCTTAAACTGATCTAAAGA
GGTTCTGCTTATACTTGTCTTGGTAATAGGGACGTTAGTGTTAGGGCTTGGTTGGGGAAAACGTGTGCTTGCTTG
TTATGCCACTACTAGACCCCTTTTAGTGGGTTTGGCGACGAGATCTGAAGCGAGTTGAAGTCGGGGAAAGAGAAC
AAATGTGGATCCGGTTCTCTCAACACTGGTCGTTCTTCAACTACGACACAACTCCATGTGTTAGAGATATTTATA
CTCGATGTAGACTTCTTAGGCACTGTCTCAGACTTCACCACTATGGAGACGACGAAATGATGTTACCTTCATCTC
TATAGTAACAACAACTAGGACAGTTCTTGTAACTTGGTGATCATTTGACGTAGATGATAGCGCAAGAATCCCCGG
GATGCATATGAATAAAGAAGAGCCATAGTTCACGTAAAAGGGTTTGATTAAGCTGTTTGTGTCTGAGTGATTATA
GCTCAGTTTTTTCCCTTCTCTCTTTGCCTTTATGCTGTCTAAATTGGTGCAAAAATCTGAAGGTTGTCAATTTTA
GTCTCAGCCTCTCATCTTTAGTACATATGTAACCAACTAAATCAGATAATTAAATATGCAAAATGGTATGATGAA
AGGCAATGATTGTGTTGACAAAACTTGGAGATTGGATTCTTGATATGATAACGAGGAAAAGTAGGTTAGCAGGCA
ATATTTCTTTGTGTTTGAATTGCTTTTTCCTACCTCTGAGGAAAGATAGACGTTTAAGTTGGATGATACAAGAAT
AGATCGATCATGCTGTAGTTTCATGATCTAGTAATTTGAACTAGTTAAACAAATAAGCTCTCACTCAAATTGGTT
GGTTACTTTGGCAAACATGTTGTACACCTTAAGATGTGTGTATACTACTGTGTTAGCTGAAACTTAATTTCTTTA
CCTCATTCAGATCTGCTAGTTATCAGCATGATATTTTGTGGACATAGAAGAACTGTTGGATCTTTTGGGCGAGTT
AATTAATCTACAGCAATGTTTGCTAGATCGATTGCTTCTGTCGAATCTTAAATTGGAGTAGTTGTCTTGTTATAT
CACACAACTATAAGCTGATGAGTTGTCTAGTTCTGTATCTCGAGTTTATTCCTATCTCGCTTGTTTTCAGTGTCA
TGAATGCTCCTTTACTTCAGTCACATAGTCAAAATACTAAAGACTAATGTAAACATTTAAAAGAAATATTTAATC
ACTTTGAACACTGTGTCACTTTGAAATCTCATGTTTCATTAGCATATATGTTTGCTCTTTGTTTTCCATTAATCT
CACCATTCTACAGTTTAATAGTCTGTCAAAAACAATTTAAACTTCATTGAATTAAAAGATGGAACACCTTAATGA
CAATCCACCCTGGAGTTTGTTGTCCTATCCAATATTCACATTCGGAATAAGCATGTACATAAACATAGTGGTTTT
AGGACATTCAGAATAAAATATTGGTTATCAACATATAGGAGAAGAAACCATTTTGGCTGTATTCCGAAAAGGAGA
TACATTTTCTCAAGTCATGAACACTTCCACAGATATTGATGATCACTTTTAAAGCATGACCTTCGTTTTCACGTC
TATCTTAATCCTAAATTTGTGGTATACTTTGCTTTCTTTATGGCAAAAGTACATGCAGGGAGGAATACGGTGTAA
TAACATATTCGTACATAGATGGACATTATATTTTTGTGTCGTGTGCTTGTAGATCTCTTCAATATCTCAAAAATA
AATTCTCTTATTATTACTATAGAAAAAAAAGGAATGTGTTTTTGTTTTCAAGTAGTTTTTGTTTTTTCACAATCT
TTGAGTGGTTGAATAAAACCATTTATTTTGCATTTGAGTGAAGGGAGGACCTCTTAATTATAAATTCGGACAATA
CAAAAACAAAATCGTAAAGCCCTAACTATTTGGTATCTACTACATAAAATGAATATGAAAACTTAAATAGTTTCA
CATACAAACTAAAAGGAAGAAAAAAAAATTTTTTCTTGTGTATGTATGTATGTATATGTGTACGTGTGTATATGT
GTTATACGTATGTATGTATGTATGCTTTTAGTTTTAAAATTATCAATTATATAGATAAAATCAGTGAGAGAAAAC
TTATATAGGTAAAATCTGTGAGAGGAAACTTGATATGGATGATTATAATCAACGCTGTTCAGTGAAAGGAAATGT

CTAATTGATGAACTTTATTAAGTTGTACGTTCCCTCCTTTAGTTTTAAAATTGTCAATTATATAGGTAAAATCTG
TGAGAGGAAACTTGATATGGATGATTATAACCAATGCTGTTCAGTGAATGGAAATGTCTAATTGATGAACTTTAT
TAAGCTGGGCATTCCCTAAAGTCAAATTTTGATGTCAATAAACAAATGCTTGCTACACCTGTCAATGGTCTTGCA
ATAAACCATTTCATTGGCCTGACCAATTTATAGACCTCAATTAAAGAGGTTTAGAACCCTAGATCGTTAAATGTT
TATGCATTTATGTCTGCGGTTGTGATAGGATTGTAAATTATCATGAAATCGAGATAAATTAGATTTTATTGTTTT
AGAATAGTGCTATGTCTACAATGTATTTAACTGCATTATTAATATTGTGGGAGTGGATTGGAAAGGTGAGGCTTC
CTTCATTTGCTTTCAATTCAAGTAGACACTGAGAAGCCAACTGCGACTCTACCAAATGCATGATTCCAGAAACAA
TATGGTGTGTATAATTAGCAATTAGAGGGGAGTGTGTAAGTGGGAGTTCCAGTTGTTAGAATCCTTCGTCTGTCA
TTGTATATCTAACTTCATTCAGTGAGTTATTCTAACCTGCCACTTCCCTAAGATGAAATATAGGAATTTGTAGAA
GTTTCAGTTTCTCCATATAGCCAGTTACTCTGACATCAGTTTTAGGACAATGGAACACAATATATAATAAATCAG
TTATGTATTCAGATCATAAAAAAAGTAAAATGTGAGCATAAGTTAGTAATCTTTATGTTATGTAATTTATTTATA
TTTTATGACAATAAGCTTCCTTTTTTTTTTCTCTATTTGGTGACAGAAAACTTCTTTACCTTAGTTTGATTGATC
CTCTGTCTTCTCAATATGCAAGCCAGTTACAACTAATCTTATTTTACACTCATCATATACAATGCATATGTTTCT
CATGTGTAACAATTGTATAAAATGTCAAAATTAAAGTTAATATAAGGTACTTGTGCAGCATGAAGTCTGTGATAG
AAAGGTATACTAGGGCAAAGGCAGAGAACCATCTGATTCTGGACACAGCTTCAGAAATAAAGGTATGTTTTATCC
TCAATCATAACAAGTATTTTTTTACTATAGATTTGTATGTTTGTAACCTCCTTGGTTAATTAACTCTGCTCATTT
CATCATCATGTTATCCTATTAGGGTACAACCAAAATAACTTTTCCATGAAAGGTAATACGCAAGCAGTCTACAAA
TATATTTTATAGTTCTATTAACCTTTTTCTTTTTTTCGAAAAAAAAATTTCTTTGCTTTTTGGGTCATATAAAGG
TTTGGTAACCCTTTAGTTAGATTATTTTAAGGAATATGTGTTTAAGCTATCTTAAGAAAGTCATTTTTTACAGAA
AAAGAAAGTATGCTTAACTAATGTGAAAGTACATCTAATTTTACTTCGAGTAATATCACATGAGAGGAAATAAAT
TTTCTTGAATAAACAAATCCCGTTACATGAGAGAGAAAAAAGCTGTATAAATACCTCTTCACGCCTTCTCTTAAG
GTTGTGGAGGTGATATTTATGCTTGGGATTCACCCCACTGTTAAAGGACTTCTCTAAATCATCCTTTCACCCTCT
TGTTAAGCAGCAACTTCTCTCTTTTACAACTAAATTGGTTGAAGATGTAGAATGGTTAAAAGTTCAAGGATTTCA
CAATTCAAAACAATAGTTAGATTCTTGAAACAACCACTTTTGACACTCTTAGATACCATGCTCAATATCCTGATG
AGGTAATAATGGAGGCCCTAGGTCTTCCATCCAGCTGAGAAATAGTGTACCAACAAGATTTTCGCCATGAGAGAC
GGTACAGTAGCTTTTTGGCAGTGCCACCATCAGATAGTTGTTGAAAATCAATGGTGCCACAACCTACAATAATGA
CTTCTGCCCTACTTAGGTGGTGCTACTAGCAACCTGGGCAATGAGTTTGGCAAGCAAGTAGGATAGTTTTAGGCG
GTGATACCATATCAACAATTCCATCGGTCTCCCATTATCCATCGCGTGGTTGAATTCATTGAACATCTTCCGATG
CCTCTTTATTCTTTAGAGTTCTTTTGGAATCAAACAGTTATTAAAGTAGAAAGGGTATCTAAGTTTTTACGATCA
TCATCCTCTTTGATTAAAAAAATAGCATATAAAATAACCATACGGTCAATTAGATTGAGGGTTAAAAATTAGAAA
CTATATGCACAAAAGAATGTTGAGATGGAGGCAAAGATTGATTGACAAAGGAGACCAATAGACTTGACTTCACGA
TGTAAGAAAGAGTGGAGGCAAAGATTGATTGACAAAAGATTTTACCAGAACCAATAAAAAATGATCTGAATGCTC
TTGATTTAATTAGTGATATCACATATTTTATAACTAATGTGAGAAAAGATCCATAAAACCAATTTTGATAACTTA
TTGTTATACGCTAAAAGGCAGTGATTGTAAATATATGAAACCTGCTAAACTCAACTTACAATGATTGTTGAACAT
TAGGACGCCTATAATTAGATGGTTATATCCTTACCAATAAGCTCCCTTAGTTATACCCTTTGTCTTTGATAAGCT
CCCTTAGTAATATCCTTCTCCAAGTCACACAAAGTTCAAGAAGAATATTAACAAATATGTAGCACAAACTAATGT
TACAGATGACGAATGTTATCTGCTACCACTAAATATGTATGTTTCACAAGATTATGCCTATTGTAATAGTTAGTT
ATAATCCCAGCGTATGAAATACAACCTGCCAGAATTAGAAGGCAAAGTCGGACCAAATATTTTGTTGATAAAAAA
CAATCTATCTTGGTACTCTGGTTCGAGCTTGATACAGAATGACAGTGAATATGCCAGACATGAACAAACAATAGA
ACCAAGCATGAGCAGTTTAGAAAACCAAAAACCAAAAAAAACAAACAAAGAAATGAATGTATAATACAAGTTGAT
ATCACAGTTATATGAGTACTGCTTATTCAGTAATTAATAGCAAGGTCCAGAGCTTCATGAATCTTTTATTACCAT
TGATCCCTGATTAGAGCAATATCTCTACTTCAATCAAGAGAATTCAAAATCTCTTTCTATTGTTTATGGTGAAGT
TACCGCTTATTTCGTAATAAAGAATCAAACCAGCAGGAGCTACCAAAGACAGCATATCTCATTCCATATTACATA
TAT**GTCACATCTCAATCACAGCGAACACAATGAAGTGCAGGGACTGTGGCTACATCATTCCAGTTGGGAAGCAGT**
**TGCCAATACTTAAATACAAAGTTGGCAATGCATAATAATCAAGGGTATGTTATAATAGGTGGTCAACATCTGTAG**
**TTCATTCTGGAAGCTCTTGCAAGGCCACTGAGGGTTAGGCAAAAGAAGGTATCAGCCTCCATTGTGCTATCATAA**
**TCCAT**CTTGCAAAGATACCAGAATAGCAGCTTCAGTCCGTCTCTCAACTCAGTCATCAAAGCCATTTCATTTAGG
TCAGTGCCCATTGACTCATTCAGTTCATTCTAAACAGACATTTAAATCTTCATGCCCATGAGATCGCATCTAATC
CCAGAGAATGCAAAGTAAAATGCTCATTTAGACCCTATAGCTAGCTTTGACAGAAATGCTTGCTGCTCCACTCAG
TAACAGATGTGTTGTCAAAAAGGCTAGGTGCTTTGCTGACAAACTATAAATTGCTCTTTACAATTTGAAAAAGCC
AAAACTTATACAATCCAAACCTGCTCATAAAATCCAGACTTGCTGGAAAAAGTAACAAATGAAACAAAAGAAAAT
ATTTTATGGCATTAAACTTTCAAAAGAAACAACACTACCATTTTCTTGCTGCTGATGTTGACTTCTATGATCCTT
AGAACTTACATATAAGCAATTGAATTGATCAATCTGTGTCTTAAATCACTGCCTATGTGGTTTCTTGTCATATTT
AGACAGATAAATATTGGCTTCTTGTTCTCACAAGACCTCAAAGGAGAGAACCAGCCACAACTACATTTTGCTTAC
AGTCATAGTCTCCTTTCAAGATACCGATACCGTGATGGCAGTTCAACAGACCAATATGGTTGAACGGTCCCACAT
GGCAAAGAAAATTTTGACAGAAACAGAGACGAAGACAAGAGGTAAATAGGGAAAGGATCATAAAGCTGTGCCAAC
AGAAATCTAGAGAAATCAACATATATTGATGGTGCTTTAGGAAATAAACAGCAAATAGTAGGAAATAGTTTTGTA
CGAACTAGATAATAGAATTTCAAAGATAATAATACAAGGCTTTTAAAATGTAGGATGAAGAAAGAAGACGTTCGG
AATCTGAATGGGAGTAACTTCACTAAGATTAAGAGTGTGGTAAAGGAGATTGTTGAAGAATCTAAAGGTAGAAAT
GATGATCCAGGAAAAGCGTTGGTGGGATTAAAAAAATTCAGAAGGCCATTAATACAAAACATGCATTAAAGATTG
GTAAAATCATAGAAACGAGAAGAATTTTAGAAGACATATAGTAGCTAAAAAGAGAAAAACTATATGGTTGGAAAA
ATAAAAAACATGAAAGTTTTTGCAGCAAGCTAAGTTCTAAGGATGGGGAAAAAGAAATTTATTAGATTGTAAAGC
TACAGATAGAAAAAAAGGAGTTCAGTAAAATTAAATGTATAAAGATATAGATGAAAAGGTGCTTGTTAATTGTAT

GAACAATAACAAAATAAAACATTGTATTCATAAATTATTTAGCAAGCAATAGACAACTTAAGCCCTTATTTAATC
CATAAAACATTATCATCCTTTCCTTAAAACCTTAATTCAATAGGCTTGAAAAACTGCATAATGTTTAATGATCTA
AAATAAGCTTATATTTATCCCTCAAATCAAAATTTCTGTCACAAGCAATAAATTCTTTAGAAAATTTGTACATAT
TTATTGTAGTTTTGTGTATTGACTTACCTTACGATCCTCCAATGTGTTAAGATATAATGTCATTTACCATTAAGT
GTATCTAGATTGCGTGAAAATATTTAACTGATTTTTTTCTTGTTTTGGATATTGAAAAGTTATGTATATTTTAAA
CAATAACTCATAGGTGTCTTTGCATATACACCAGCATACTGCATATGCAATATATGATGACATAACTACCTGCAC
AATGCTTTGCATTTCAAAAGCATGCTTTCATCAATGAAAATTAGAAGAAAACAAAATACATCATGAACGTTCAGC
CTGAAAATTATGAATGCAAAAATTGTAAAGACAAAAAACCCATAATTGAGTCATAATGTGAAATTACAGTGATCA
GACAATTGCTAACATTCCATTTGCTACTAATGCTAATAGCAATTCTCATACAATGTTAAAAACCAATCAAACTCG
TAATTTAGATTTGTATGGACAGGTGTAACCATGAACCAATATATGGCTCCATGGAAACCGATCAATACTGGTTCA
TGCCTGGCATTTCATTGATTTGAACCAGCATACTAGTTGTACCAAGTGGTAGGTTGTTCCCAAACTATGCCAATT
CTATTTCTTCAACTAGCAGCCCCCCTCGTTTTTATCAAACCAAAAGAACCCTTTTCCTTTCATTTATTCATTCTC
TTGGTGTATCATCAAAAGATTGGATTTCAAGCCAGAGTTGGCATTGTTCTTGAATCAAGGTAAATTTGAAGATAT
ATCTATAATTTCATTGCTTATTTTTCTTTTGAATCAATTTTTTACTTAAATTTGATTCATCTATACGTGATCTTA
CTATGTTTTGGTGTATCATACATAATTTTATTATCAAACCAAAGGATCCCTTTTCCTTTAATTTATTCATTCTCT
TGGTGTATCATCCAAAGATTGGATTTCAAGGCAGAGTTGGCATTGTTCTCAAATCAAGGTAAATTTGAAGATATT
CTATAATTTCATTCCTTATTTTTCTTTTGAATCGAATTTTTACTTAAATTTTTATTCATCTACACGTAATCTTAC
CATGTTTTGCTTGAAGATGCTTTCTTCATTCAATTGAAGCTTGATCTACACATAATTAACATTTTTCCAACAAAG
ATTACTTGAACATACATAGGATCTGGTCTACATCAGTGTCAGACTATGCTAGATTTTCATGTTTGACCTAAGTTC
ATGCAAATTCAATCTAGATACATACAGATCTAGGCTACTTCCATCAGGACCGACCTATATATATACAAATTGATG
ATAGACTTGTGTGGATCTGGTTTATCTCCTCGTAGATGACCAAAATTGAGGCAGATCTGGCCTGCACCCACAACA
ATCCAAGTGTATGCTAAATTTGTTGTTCATGTAGTATATAGGAATGTGTCAGAAAAGGTTTAGTAAAATACTGAT
CTAACATGGCAACACATTGATCACTTAGTGCACACCACAAGAATTAACATTTGGTATGAACTACTATGACTTTGT
TGCTCATGGTCGAGGAATAAGAAGGCTGAAGTAACATTTTCCTGATGGACATCTTGACATGGGCAAATGCACACC
AATCCCTTAAGAAATCAGAAAGACTTTCTAGGTAGAGTGGGAATAGAAAAAATCTCGAGCAGAACAAACATTTGC
AGAGGAGCTTGATAGGCAGGCATCTAAGCAGCATGCCTATGATCAGTATGAGTTAACGATGATGTTGTTAATGAC
CCTGAGATTGCTGTCAACATTTGTAATCTATGAAGCACTATTAAATGTATGAGCAGACGAAGAGGCACAGACATA
CTTTGCGACCATTCCAATGAGCATAACAGTGGCAATGGTAGCAACTACGTTATTGGACCTTAAATATGGTGCCCA
TCAGTTTATCAGGATAACCAATACGGGGAGTGGTGCTCCACATTCAAGAACAGTATATGATGAGGTCAATCACAT
TCCAATCCCTAGCAGGGGGTCCCTTTAAACAAGACATCAATACCATATATGGTTGATATTGACCCCATGGTATAA
TTCTCAGTGTCAAAACTACTAAGATCAATGATATACTCACGAGAACAAGAATGATAGACAGGCACTCTCATAGTG
ATTCAATTTTAATTCTCAACCCTCCTTTTGCATCTTCCAACACATTATATATCTATAATGTTCATCTAGAAATAC
AATTGATAAACAAAGCCTCCAAATCAATATTGCAGAAAGATATCTTGCTGATCAAGTTGGTGAATTCTGCATATG
GATAAAATCCTTCAAGATAGCCCTGACCCCGATTACAAAACAAGGAGTTTTGGTTTCCCAAAACCAAGAACCAAA
CCAAACTGGGCTGTTTTCTGGAATCAGAACTGGAACAGCTGGTTTCGGTTTGAACCGCTAATTTTTTTTAATATT
TTTAATATTTTTAATATTTTAATTACGAAATAACCAAAATACCCCTAGAAAATAACAATTATATGTCTATTGGAA
GGGACAAATTGGTCATTTGCCCCTCTATCCAAGCGGTGGACCATTGGACCGACCGTTTGAGCCCCAACAATCAAT
CAAGCCCAACATTGATCGGGCCTTTTTATTTTCTTTTTGAACCATCGGTTCCGGTTTCCCAACCCAAAATTCGAA
ACTGAACTACTCGGTACCGATTTCGACTCCATTTGGAGCCAGTGAACTGTTGGGCCAGTTCGGTTCTGGTTCCAG
ATCGATTCAAACCAAGCCGTGGTCTACTTCAAGATGCAACAAGATCTCTATGGTATGATTCTTACATGTGGTAGC
TAATCAAAACCCACTAGGGTAATAATCATCAACTTCCTAGCCATGATAGGAAGTCGTGTACCATAAGTCCATTGA
TTAATCACATCATTGACCCATCAGAGACTATTGTAGATGATATTGGAAATCACACTAAATACTCCAGGTGGCAAT
GGATAATGGCACCAATTAAGAAATCGATGAATTGATAAATACACATCTAAATCAATTTAGGCCATGTACCATATA
TCATATCGACCTTATGATCGACATCGATGACATCCCACGTGAAGAAAACAATGGACAAGGCACAAGAAATAATTT
GTTTTATCAACAACCATCAACGAGTGTATGCCTCAACGAAAATATTCATAGAAGGGAGATCCTACGAACGAGGGT
CACTAAATTTGATCAAGCAAAGAAAGATTGAATCGAATGTTATGTTTATTTCTAGTGATTGGACATGTTCTTGTT
ATGCTCTCTAAAATGAGCGAAAGTCAATTGAAAAGTGTATATAATCCTCCTCCATATATGTGGTTGAGATTATTA
TGACTATTGAGCCTATGTAACCGGAAGAGTTGAAATGAAAAAAGGATGAAGATTGGCTATGCTTCCCGCCAAATA
CACAATCTCACAAGTCACAACAATAAAACATTCAAGTGGCTTTTCCAACAAATGAAATAATCTTTACCTTCATAT
CATTTTTGGAGATGGGATAATCAAATTTCAAGCGATCTTAATTGTTTTAATCTCTTTATCCATAAAATTATTACA
TTTTCAATATCAAATGGTGGCTTATTGTCTCTTTTTTTGTTTTTTTTGGGTCAAAGTGGCCTATTGTCTGGCAAAA
GAAAAGTGCCTAATGTAGCTTACTACCTAAATTTAACATACCAAAATTGATATGACCCTGTCTTGGTGACTTGTT
CACTCTTTGCAGGGTCATCATCAGATAGAACCTTAGAACAAAATATAGAAAGTGCAACAGCTACTATTAGTGAGG
TGAAGTATCTGATCATGTTAATTTTTATCTCACAGAGATAGTATCAATTGACTTACCCTATTTTTTACCTTTAGG
GGCATCTATTTTGATTAGCCTACAACAATCTCAATAAGAATTCAACACAATTTCATATTGATATGACATGGATCC
CATAACAACATGACATTTCACATAATACTAAACAAAATGTGTTCCTATAATATGTTTTATAATTGTAATTGTGTA
GATGAGTAATGAATTCACGTGAAGGATCTGCACCTCAAAGAAAATTGTCCGATAATTCCTATCACAAATGACACA
AACTACGTCATCCAATGGTTGCAAGTATACTTAACCACTTTCTACGTCATACATATTGCAACCACCTTTTATATT
GACACTTGGAAAAGAAAGATTTTGGGCACTATTACATGCATAAATGGCTGCATCATATAACCAAGGATAGGACAA
CGACGAGGCCAAATAAAAACCATTTGACTCATAGTTTGTCACCAATGATTTATATCAAAGTTTCCAATATCCACC
CAATACAGTACGGTTCGATACAAGTGGTGAATACCAGTCTAATTGATTATTGACACAGGCAAATCGAGCATTATC
CCCATACCCAACTTGTACCCACACATACCATATGGTACAGGACCAGTACTTTTTTTTTACTATGCAGTCTTTTTT
AAGTGTTTTTTTATTTCTGAAACTCGGCACATACCGTATGCGAAATTGGAACCCTAGACTTATACCCTTTGATGC

ATTGAATATGTAGAGCTGAGAAACCAGATTATTATTTGATGAGATGGCAGATTGACCACACCCATCAAAGCTCAT
TTCTGAACAATTTATAGAGAGAACCAATAAGACGAGATGGCCCAATACGTGAACAATAATAGAATATCTGAACTC
AAATTGTAGGGCATGGTTAAGCCTCAAAAATCTCTTCCCTAGGTCATGCTATTGGAAGCTAGTTATAATCATCTA
TGTACTATTGGAAACTAGTTTTAATCATATACAAGGCGCAGCAAAGCTAGAGGCACATTGCAATCAGTTGACTCA
TCAGACATCAGTGATGACTAAGAGATCAGAAGGAAAGAAAAGGATGGTAGACTTCTCCCCAAGTTCTAATTGTGA
GATCCAAAATGACAATGGCGAACAGATGCAATAATGGTGACCAAAGATCCTATGTTTTTGCTGACAATAGATTGA
GCAATCATTTACTCATACCACCCAGGATGAAAAACATGATGATAGTGAATTGTGAGGACGGCAAGACTTCGCTCC
ATGGTTCCAACCAAGGATTGAAATATCGTACCGTACCGGAGTTTCGACATTCGCTCGGTACGATACTATATATTG
AGCGGTATACCGTTCGGTATAAATAAATAAATATATATATATATATATATTTATTTATTTATTTATTTATTTATAAA
AGGCGATGTCGCCTTTCATAAAAATATTTATACATTATATATATATATATATTCCGTCCGGTAATGGGCGATTCGTG
TACCGATCAACTGACCGACCGATACGTATCGCTCGGTACGGGCGGTATTATTCAAAACTGTATACCTTGATTCCA
ACTGTGAGGTCCAAATTGATGATAACAAACAACAGCAATAATGGTGACAAAAGATCCTATGTTTTTGTTGTCCAA
AGAGATTGACAATAGATTGAGCAATCATTTATTTGTATGATCCGGGACGAAAAACATGATGCCATATCTACCATT
ATTATATTAGATAGGCAAGGAAAGTCGAGAAGAAGACAGTTCGTATGGTTAATCACAAAAACACCAGAATCCTTG
CCAAGATCTCGTAAAGGAAAAATGAGCCAAGCTAAGTGGTTTGGAACTGTGAATCAATGAAGAGAACCTAAGGCT
TCTCCAAAGAATCACTAGGTTTTAGTAATGTATAGATTGCAACTTGAGCTCTTTTACTTGTTTGTAACTGCTACT
TCAGATCATATTTGTATTTGTTCCATGTATAGAAATTATTTATCTCTCAAAGTTTCTACCTTCTTCTGACTCATA
TGAATATCCAAGTTATACTACATGTCTACCACTCAAGCCATATAACATTTTCAAATGTCATGATCATACTCCTAA
ATCGAGATTACTCCACTTATTAGCTATCAGAAAAAATCCAAGTTTGCTGCATATCCAGATTGGTGGTTAATTTTA
CATGATAAACAAACCACCAAAAAGTGGGAACAATTGAATCACAATCAAAGGTCCAAAACATTGACATAAGACAAC
AAACTGATTAGAAAAACAAAATCATATTATTATAGGAATCTGAAATAACTTAACATGTTTTTAGGAGCTTTAACA
AATAGATTCTTCATTGTTTGATGCCATAC**CTCACAATTGAATTTCAGAATTCAGATTATAGTTCACCACTTCACC**
**TTATAGGATGATTATTTTCGAAGCCTAGTTAACTGTGAAAATATAAATCTAGCAATACCAAGAAATATCTAGTGT**
**GGCTAAACCTGGGCTTTGTACTTACAGTTTTGGCAAAGGGAGGCAACAAGCTTGAGGCAGCAATTGCATAAATTG**
**CAAGAAACTCATAGGTATGCTCACTTCACC**CTACATGACATGAAACATTAGCACTTAAAAGTCATCATTCTGTAA
CTTAGCCAATAACATGTAGCATCAAAAATATAATTGCAATCACATACAAAATAGTATCATTAGTAGACTTCAACA
AAAATCATCAGTAATATGAACCATGAACCACAGATGATTATATATCTGTTACATATATGTTCCACGTCAATCGCT
GCCCATAACTAGATAATATCATTAGGTACAAGCAACCAACATCTTTGCACAGTACAGGCTAATTAGTCAGTATTA
TGAACAAATAGAAGCCAAATGGCATCCTTTTTTAAGTATTAATAAAAAAAATATTAAATGTACTGAATGGAGTTCA
TCATTCATGTTCTTTTGATAAGGGGAAAAAGATCTGAAATCTCATGAAGATAGTTCAAGATAATATGAAAATACA
TAGGATGTATGAGTACAAGATAGACATTTTACAACATTTTTAACAATCAAACATGAAAAGTATAGTAAATACTAA
ATAGTAATCATGTTTTCAAAGTGATATATGAGATCATATCAACAAAAGCAACAGTCCATGACTGGCAGTACATAC
TTTCAAATTATCTGATGCCAAAATGTAAACGAAAGATAATTATCATATCATTTCATAATTTTATTCTCCCCGCTA
GGTAAGTACTCCTCTGCTTGCTTCCAAGTGTGCTAGAGACATAGATAGTATCTATGCAAAGTACAAACACACAGT
GAGCAAATGCAATCTAACATTATTACCATGATAATAGTAGTAATAGTCAAGGACTGCTGTACCACCTGAAACAGT
ATGAAACGATCACCTGGTCCAGTTCAGTAAAAATAATTAAAAAATCAGAAAATAGCAGTGGGGCCCATCCATCTA
GGCATTAGAAAAAAAATAAAATCCTATAATAAAAAAATTAGGGCTCGTAAACGCGAACCCTCTTCTCGCATTCAA
CGCTGCTGCCACTACCATTGCTTTGCAATCCTGCTGCCATCGTCGTTAAGTGACACCACTGCCAACATCATTGCC
GCCGCATTTCTTCTTCTTCCTAGATACGCACCTACCTCTTCTCCTCCTCTTCTTTCTTTGTCTTCCTTCTCCTCC
ACCATTTCCTCTTCTTCCTTCTGCCTTTGTCCTCCTTCCTATTCTATTCCTCCACCGCCTCTTCCTCTCTTCTTT
TCCTCTCTAAAACACCAGTACATACCTATGTACCAACACATGGTACACCAGTACTAACTGGTATGTACCAGTAAC
AGATCACCGAAATTGGTCTGATACCTGAAACAGCGATCTTTGGTAATAGTAACCATAAAAAGATATGGTCAATGG
CTTACCGGTATCCTAACTGCATGTGTGACGTCTATACTTGCAGAAGTGATACTGGGTGGCTATTCATATGTGTTA
TATCAGCCATATAAAAACTTGAGAACTTATATAAAAGAGCAACTTTTTCAACACTTGTAGGCAGTTGATGGGAGA
AGAACTTTCTGGTCTAAGTGTCAAGGATCTAGAAAATCTAGAGAACCAACTGGAGATGAGACTCCGTGGTGTTCG
AAAGACGAAGGTCATATTTACATTTGCCTTAATGCTGCTTAATGTTACTATTTATCTCATGTATACTTGACTGTG
TTGTTCATTTTACAGGAACAACTTCTAATTAATGAAATTCAAGAATTGAATCAGAAGGTATGGCAACCTACTGTC
TCTATGATACTCATTGTGTCTGAAGTAATGCAATTTCTACTGTTCTCATTGCAGGGGAGCCTTATTCACCAAGAA
AATATGGAGCTTTACCAGAAAATAAACATAATGCATCAAGAAAACAGAGAGTTACACAAGAAGGTATACTAATAT
CCGAAGAAAATTTGCAATCAAATTGTTTAAACTGGTGATAAACTATAGCACAAAACTAAGCATGTGCCTCCTCAG
GTTTATGCACCAACGGAGGAAAATGACCCTAATAGAAATTCAGTTATCCCACATAGTTTTCATATAACAAAGGAA
GTAAATGCACTTATTCATCTTGGACTACGCCAACAACATCAGCATGGAGAAGAGTTAAAACTTGGAGCTCCAAAA
CTAGGGTAATGTTTTTCTTCTAGTATGGATATTTGACATATATCAACTATTAGTTCAATAATACAATGTTTAAAA
TACCATGCAGCATATTTCAAGTGAAATAAGGAGAGAGAGAACAGATCAAGAATCAGCGTTCATGGTCTTCATCAT
ATTATCCCAAGGTTATAACAGTGTTGTAATTATCCTAAGCTATGGTGACAGCACCATACTATACAATGGCGACAG
CAGTACCTATTTATGGTCAACTTTGGCCATTTAATCCTAAATTTTCCTCTAATCATGACGAAATGCGACAAATCT
GTTCCCTTTAAACTCAATCAAACTGCACTTGGCTTGATCAGTCCTAGAGTCTAGAGATCCTTCTGTTAATCAAAA
CAGATCAAACCAAACTGATTTAACATATCAAAGGGATCAGTTGATCCAATTGAAACTCTCATAAGTTTCACCAAT
TCCAGTAGATGCTAGTGGGTTTCGAATATGATTTGGTGGCAAAACTGTAATAAGACTAGAATATGAAGTGAATCA
TGTCAGACTTTAATTTTACAAACTTTGCTCTTGATGTTTCAAGGCAGCACCTCTTTATCTTCCATTTATATTTCC
TCTTTTTTTTTGGCAAGGAATTCAATCTAGTAAGATCAACCACTCCATGGCCAAGGTAAATTCATAGTAAGTTTC
AAGGCACGCACAAGGACATAAATAGCAGATAAACTAGTGAATAACCTGGTAATCAAAGTAAAAGTCCATTTCTGT
AAATATAGTCCTGAGCCTTGGGATTTTATAAAATTTTATTATTACTTAATCTTCAAGATAGTATTGCTCATCACT

AGTTTTTTTTTAGTTATCTGATTCCATTTGCAGCTAGTTTGAAAACTGAAATAAACATTAAATTATTCACTGGTA
GCTATGTTCCATGAAAACACCTAGGTCGATGAAATTACTTCATGACAATAAGTATATCCAACAACTCTTCTCCAC
AAAAGGTGGTGGAACCATGTAAAGAACAAACAATAATGGCCATAACTTTATATCATATTTTTCTTGAGCCCATTC
TCTCTTCAAATTTAATCCCCTATTTCATTTTCTAGTTTGTCCTTCCTATCGTTCTAACTTGCTATTAAGGTTAAC
GAAATTGTCGTGCACAAATTACGAATTTGAAATCCAGTTAATACAAGAACATGATAGGGGCATGGATGTGAAACC
AAGATCAAGGTATTGTACCTTGAGCCAACGAGGCATTTAACAAAGACCATAATTAGGCTAGTGCCATGAATGTCT
CAACCATGCTCAATTTTTAAAGTCTTTTCTTTTGATTCATAATATTCTGTTTCTACTAGTCCTATTCGGTTCACA
ATTATCTGTACCTTTGTATAACAGAAAGCCTAACAAAAGCTACAATGATGTAGAATTCTTGTAGCTATGATAATT
GACCATACTCTACAGGCACAACCTGCTAACCGTATATAAACAGAACCATAGTGATTTTTGCATAAAACAGAGATA
CAGTCTTCCAGATTTTCAGGGTCACAAACCAAGATACTATTGGCACAACTTACAAATATTATCAACAAGTCTTAA
AAGAATATTAATATAC**CACAGAACTTCTTGCAGATTATGCAGCTACTACCTCAAGCTTGTTGCCTCCTTCTGCAA**
**TAA**CTAGTGAAACAAACCACAAATCAGATACATGTGAAATACAAATAAAACTTGATCGATATTTGACTTGGTTAA
GAGTTAAGGCAACAAAAGAATCAAAGCTACTTACTAATTGATGTGTGGCCTATGATTGTTGAGCACACTATAACA
ATACAACTAAACAAGAAAATATATTACAATCACAGGCATTTGTGACAACTGTCAGCAACAACAGCACGTCACACG
ACAAGACATAAAGCCCCAACAACAGCAAGCAACCCATATTGTACCAAAATTTGTGAGAATTAAGAAATACCAAAA
TCTTATTGCTATTTGGATCTCCAATGTGCCCTCAATCTATGCCTTCCCAAATTTTGCATACCGCTCGGTGTTTAC
CAGCCCAAGCAAGGATCAGTACCAAACCATATGATCAGGCACCGACCAGTACTACTTTTTAATATTTGCTTTAGC
TGATATTGATGGCACGAGTTGATATTCCGCCCAGTATACTGATATCTTTCTGGTCCGTTAGATTATTGAAATTGG
TATCAGAATAATTATTAAAATAACTATAAACCAAAATTCGATGCATATTATAAAAATATTTTTAATTTAGCAAAT
GAAAGGCTAAATATTTTTCTACCTGAAACTGTGAAGACAGTGATGAACCATATTATAACCCCAATGCTTCTCTAA
TCATCATAAGGGAAAAGTCCTAGGTAATGTGTTGAGAAGCAAAGAAAACCTTTATTTGAAATGAGTCCACAGAGA
TTGTGATTAATTATGAATCAACCCTATAGTCAAGGATTAAAAACAGATCTAACTAGATGGTACTGATTAATATTT
ACTAGTCCAACCACGTTTTGACACCGAGCCAAATTGCCTAGTCCAGTTCAAAACCAGGCTTGCAATCTGATTTTA
ATGATTAAAACCATGGCAGCCGGGTGGCAGCCAGGCAGCAGACTATATTTTTTTAATTAGTCCTCCTCTTTGTCA
TCTCAGTGTCGCCAAACAGCTACATCATCATCACCATCACTGGTTCCACCTTGGCCTCTGGTATTTATCATCCTT
CTCACCTCTTCCTCTACCTGTCCTCCTCTTCCCGTCTCCTCCTCTACCTCCCTTTTCTCCTCCATCTTTTCCTCC
CCCTCCATCAGTCCTTGTCTCTCTCCCCCCTCAACCTCCTCTGTACTCTCCACTTTTCTCTTGGTACGTTTAACA
CCGGCCCATATCGAGTTTCGATAAATCGATCCATACAGGTTCTGTCATCAACCAGTATCTTCAATCACCAATGAG
AGTTTAAAGATCTAGGATGATCAAAAGAGAAGTGATAAAATGAAAAGCTTTTCAAAAATATACATGAAATATATT
CCAATACATATTTATACATGTACACAAGCAAGACCATTCTATATTGCTCAAAATCCTAAAATGAATTTATTTTAA
AAATGTAGAATGTTTAGTGCAAAAGTCAAGATGTTTCCTAAGTTACCTTTCAAATTCAAATAGATTTTAAAGTTC
ACTTATTTTCTTTGTTTCATGGTAAATCTGAATTATCAACTCCAATAATTACTCACATTTACAAAAATAAAGAAA
TCTAATATGATTCATCAATCAATCATAAATATGGGAATTTAAAAGGGTCCTTCAGCATTTCATCAAAAGAGATCA
TTTGGTAACAAAAAAAAGAAAGCCCTATATGCAATTATATTTTCCCAAATTATGGTCTCCATTATAAGTCTTATC
TATTTATCAATTTCACTAAATTGCTTAATTTGATCAACTTATATTGATTATTTGGACATAAAGCAAAAAAATGAG
CCTTATGCTTAGGAGGTTAACTTATATTGGACAGGAACTAGATATGTTAAAACATGTTCTGTAATAATACTCTTT
TAGTACTATACATATAACATAGATTAAGTAAAGCTCAAGAAGAAAACTGTACCTTCTATATTCTACTATATAAGA
ACAATAGCAATAAGGTATGCACCTTTAAAAGTCTTTCACTTGTTATTGAAAATTATATTCCAAAATTCAAGACAT
ACTCTAAAAGATTATCGACAAGTAGAGCATGAAACCAAAAGTTTTTGTTTCAAAGAATTCAAATAATAAGTCCTA
CAACAGTTTGTTGAGTGGGGGGTCTGTCACATTTTTAGAGAAGGGAACCAAAGTGTGCACTGGCTTGCATCTTTT
ACCAAGCTCTCCAAATCTGATTTGATCTGGAGGGGAAACTGGCATCCTAATCTGATATCAATTTTATTCTGAGGA
GCAAGGCTTACATTGTACTAGAATTTTGAAGTAATAAATATCTATTTTCCTTTCGAAAAAGAAAAAACAAATCAT
CATTAATTTATAATGGCAAATATACAGATTAAGCAAAATCGACTGCCCATATTTATGGATCTTTTTTTTTTCCTC
CAAGAATGATGTACTATGGTTTCTCAACCTTCGGCCAGCATCCTCTACTGTCATGACCCAACTAAACCAGAATAG
CAACCCTACTTACGAGGATATCTGACCCATTTTTTGTGGAGTATAGCTCAAAAATGCTTTATCTCAAATTAACCA
GAGTTCATCCACCAAGCACTTATATATCAATTCAAGTGTTTATCCACTTTGGATATAGGACTAAATTCTGAGATG
TTATAGTCTCCCACTTGAGGTCTTTATGTCCTCTTTTAATGACCAACATGACCTATAATCAAGCCCTAACATAGT
GCACATGTGAACCAACCTAGTAGCTCCACGTTGTGACGTGCCCTCTAATTCCGTCCACGAGTAGCCCTTTGCTAT
TAGAAACTCTTATCATGCCCAATACTAGACTAGATCTAAGACCAATTGTCACGACCCATTAACCCATTTACTTTG
TTGAGTTTAAACATTTGGCCCAAAATGCTTTAACTCCACCCATTGACGGAAGACCCATCAAGCCCTTATAAAAAA
TTCAAGTTTTTAGCCACTACTAATGTGGGACTAAATTCTAGGGTATTACCTCTACTCTGGGAGTTAAGAAATACC
TTATAAGGATAATTAAAGTAAGAAAAGAGTTTTCTATACAAATAGAATAGTTAGAAGCTTGACAGACTCATAAAG
GCTTAATGATCTAGGCAATGATTTAAAAAGGCGCTCGGGCGCTCGCCTAGGCGCTTAGGCGAGGCGAGGCTCGAG
CGCCTCGTTTAAGGGCCAGGTGACGTGCTTCAAAGACGCTTCACCTGGAAGCTCGCCCAAGCCCAAGTGTCGAGC
GCTTCGGGCAAGCACTTAGTTCAATTAAGCGAACCGAACCAGACTTTTAAGTTTGGCTTGGTTAAATAACGTAGT
TTGTTACCTCAAAAGCCACCCTTCACACTAGCGCGACCCCAAGGAACCCTAGCACTACTTCTGCCTCCGCCGTCG
ATGCTTTCTATCGCTATCTCCGCTGCAAACGCAACTGACCGAGCCTTCCTCTGTCACCGACGGACGAGTTCGACG
ACCCCCATAGCTTCTTTCCGCTATCACTCCGTGTGCAGTTCCTTCCACTATCGAGGGAGTGGACCATAAGTTCCT
TTGCCACCGACATCGCCCTCTTTCACCACTGCCACTACTACTTTCCACAACTTTCGCCATTGTCGTCTGCAACTT
CCGTCGTTGTCGCTGCTGCTGTCCACAGCTTTTGCTGGTGTCACTATTGTTGTTCGCAGTTTCTGCCGCTACACT
GCTACTATTCGCAACTTCCGCCACTGTTGTTGCCATCGTTCGCAGCGTTGCCGCAACAATCTTAAACTAATCCTT
TGTTACTGTTAACATTTTTTCTCCCCTCTAACTACTTTAACAGTAAGTAATATATTGTTAACAGTATATTTTTAA
TTTAATATCATTTTTTATTTTAATAATTATATTTTTATTGTATTATATATATTTTATATTTTAATATTTCAGAGT

GCCTCGCTTCGCTCGGGCAAACGCCTAGCGCCTCAACCGTTTTGGGGCCTTGGCGCCTAGCGCTTTTAAAATCAC
TGGATCTAGGAAATTTAAGCTCGCAGCCTCGCAAAAGGTGAACTTTACAAGACAAGACTTGGCACCATAGTACGT
GGTAAGGTTGCTTCTTTACAAACTATGCGACAAAGGTGCAAGTGCTAGAAACAAGTTTCTTGTTTGCTAGGATAA
CCAAATCCCACATCAGTGCATGAACTCGAAAAAGGAGAAATTTGTGCACTGGGCCACTCTATTTTCGTAGTCAGC
AAATAACTTTCAAAGATTCTGTTTCAGCTGGAAAGTGTTCAATTAAGTGGCAAAGCAAGACGAGCTGCAAATAAC
ATTCCATGAGGAACATACGATCTCTAGTTATGATGTTCATTCCTCATCAACTAACAATTAGAAGCAATTCATAAC
ACAAAATTGTTGGTATAAAACGCATTAGGCATTCTATTATGAGATAACATATGAAGTAGTCATTTCCAACATAAA
ATTTTTGACTAGAGCATTAACCTTGTTCTGCCTACAACAGACTAACTTCTAACCAAGCAATAAAAGTGTATTTCT
AACATATAAACTAAAAACATAAGATCATCACAGACACTTCAACTACAAAAGGATGCATGCATGCGATACCATCCA
CATTTCAAGAACACCAATCTAATACCAATGAAAATCTAAGGTACACTATGTTTCCAAATCATCAAAGTTGAGATT
TTATTAATGACAAATACACTAACTCGGTAATAAAAGCTGATATCAGAAGCAACCTATAGTTCCGTTATATAATGC
ATAAAAAATAGCAGTAAGTCATTGATATGTGTCCACGAAAGGCCATCATGACCATTTGAAATCTGTGTCACACTA
ACAAAAGTGGCAGTTGGTACACCCCAAGTACAACAATTGCATGCTTTGAAGACGCAGACATATTCTACATCGAAA
TCAGTAATAATCGAACCCCATACTCCAAAGTTAAACTTAAAATAAATTATTGATTCAAATAGCCCGTACAAGTAT
GTAAAAACAGGTACTCCTAACTAACCAACACCAAACTACCTGTGTGCCCAAATCCTATGGTTCATTACCGAGAAT
GATGGTCCGCAGTTTCTTAAACTACCATTATCACCCTTGTGCTTGGGACAAAAGAACATCCTATGAACAAAAGTG
AAGCAAGAAAAAAGTCGTGTTAAAGCAGAAAAGCTAGTTGCTTGAACCAGCTTTGCAAAATTCAGTTGCGGCCAG
AAAGAGTGTGGAAAAGTGTCATAGGAAAACAAGTTGTGAAGAACTTAAGATGTGTTTTTTTTTTTTCCTCAACAA
ATTATGATCTATTACAAATATATTGGCTCAAAAGCCATCAAACAACATGTTATGTGTAACCATATTCAAAAGCTG
TTTGCAAGACAAACAAGTTCAAGTAATGCAGCACTCCTTACCACAGAAAACAAATACTTGACAAAGTGACAGACC
CCTCGACCAAAAACACGGAAACACCCAGATTATAACCCTCATTTTCACGAAAGCCAATCTCTGATGCCCACGGGA
ACCTAAACACTGGCACATTTGTGAATGACCAAGGTTGGGATCTCCGTCAACTCTAAATCTTTGCAGATACCGAAA
CAAAACTCAAAAATAAAAGAGGAAACCTTATCAAGATCGATCCGTGGGGAACCTTCGATTTAGCAAATCGAACCG
AACGAGAAAAAAGAATCAATGTTCAGATCGATCAGATCAAGCAAAACAGAACTTAC**GTTGGTGAATTACCTGATA**
**TCCTATCGTCGCTCCAAGTTAGGGCAACGCTCGCTCTCTCACTCTCGGGGGCGAAGTCAGGGAGCAGAATAAATA**
**TACTTGGAGATGTGCAATAAAGTTTCCCTAGAAATAAACAAAGGGGCCGTAGCTCAATAAATGGGCCATTGTCGC**
TTTTCATCGGTGGGGGATTCGATCCCCAGCCGCGTCGGTACATATATTCTATGTATGACAATAGCGGTTTGGTTT
GGTCCACGTCGCTGTCTAGGATCCCCACTTGCTGTTTAGAGTCTTAAAGCCTTTATTTATTGGCCACTGCGTACG
GTGGAGCACCAAAAGCCATGCTCGCCAAGCAACGTGAGCAGGCACTCGCGGTTTTCCGACGCCATCCGTCTCTGT
GGGCAGCAGCCAACGCACCTGCCTTCTGATTGGAGAGATCATCGTCGCACGACGGGGTCCCACGTGGTCTCCACC
ACCACGTCGATCGCGAGGCGTCGGCACCACTAATCATTCCGTTGCCGCTACATCTCCGTTGGCGCCCACCTTTCC
GCTGCGGGTTATTTGAACTTAGCAGCACAATGCGGAGGTGCGAATGACTCTGCTGTTTAGCTTGGATTACCAAGT
TTTGACTTGGAAGCGTTTCATTTTATATATTGGATTTTTATTTTATTTTATTATGATGCTTACTTCGTGATTCGA
AGGGCTGTTGGATTTTAAGTCCACTGTCGGAAACTCTCGTGTACTACTTCGGTAACACTACCCAATGCAGCGATA
GTGTATAGCATCGAGAGACGGTGCACCGATGGATACGTGATACTTCCAGTTCTTTATGCCTCGAATTCTGACACC
TTTAATGGAATAGACACTATAATATGTGCTCATTGCCAAATGACGTATCTCTTATTATGTATGAGAAACTCTGAT
CCACAACATCAATCTAGAGTGATCCAATGATAAAAATTCTTTATGCTTGCACATGGATCTTAAATTTTATATTAC
GTCGCCTGGTGCCTCATTCTGTTCTTGTATTGTTCCATAAAAGAACACGAAGGCAACTTACAGTTGTGAATCGGA
CTCGACATG

>MaAGL17_5

ATGGGGAGAGGGAAGATCGTGATCCGCAGGATTGATAACTCCGCAAGTCGGCAGGTGACCTTCTCGAAGAGGAGG
AACGGGTTACTGAAGAAGGCCAAGGAGCTTGCCATCCTTTGCGATGCTGAGGTCGGTGTGGTGGTGTTCTCCGGC
ACCGGCCGACTCTATGATTACGCCAGCACCAGCATGAAGTCTGTGATAGAAAGGTATACTAGGGCAAAGGCAGAG
AACCATCTGATTCTGGACACAGCTTCAGAAATAAAGTTTTGGCAAAGGGAGGCAACAAGCTTGAGGCAGCAATTG
CATAAATTGCAAGAAACTCATAGGCAGTTGATGGGAGAAGAACTTTCTGGTCTAAGTGTCAAGGATCTAGAAAAT
CTAGAGAACCAACTGGAGATGAGACTCCGTGGTGTTCGAAAGACGAAGGAACAACTTCTAATTAATGAAATTCAA
GAATTGAATCAGAAGGGGAGCCTTATTCACCAAGAAAATATGGAGCTTTACCAGAAAATAAACATAATGCATCAA
GAAAACAGAGAGTTACACAAGAAGGTTTATGCACCAACGGAGGAAAATGACCCTAATAGAAATTCAGTTATCCCA
CATAGTTTTCATATAACAAAGGAAGTAAATGCACTTATTCATCTTGGACTACGCCAACAACATCAGCATGGAGAA
GAGTTAAAACTTGGAGCTCCAAAACTAGGGTAA

>mac_miR444c

GCGACAATGGCCCATTTATTGAGCTACGGCCCCTTTGTTTATTTCTAGGGAAACTTTATTGCACATCTCCAAGTA
TATTTATTCTGCTCCCTGACTTCGCCCCCGAGAGTGCGAGAGCGAGCGTTGCCCTAACTTGGAGCGACGATAGGA
TATCAGGTAATTCACCAACTTATTGCAGAAGGAGGCAACAAGCTTGAGGTAGTAGCTGCATAATCTGCAAGAAGT
TCTGTGGGTGAAGTGAGCATACCTATGAGTTTCTTGCAATTTATGCAATTGCTGCCTCAAGCTTGTTGCCTCCCT
TTGCCAAAACTGTAAGTACAAAGCCCAGGTTTAGCCACACTAGATATTTCTTGGTATTGCTAGATTTATATTTTC
ACAGTTAACTAGGCTTCGAAAATAATCATCCTATAAGGTGAAGTGGTGAACTATAATCTGAATTCTGAAATTCAA
TTGTGAGATGGATTATGATAGCACAATGGAGGCCGATACCTTCTTTTGCCTAACCCTCAGTGGCCTTGCAAGAGC
TTCCTGAATGAGCTACAGATGTTGACCACCTATTATAACATACCCTTGATTATTATGCATTGCCAACTTTGTATT

TAAGTATTGGCAACTGCTTCCCAACTGGAATGATGTAGCCACAGTCCCTGCACTTCATTGTGTTCGCTGTGGTTG
AGATGTGACC

#### Stem-loop structure

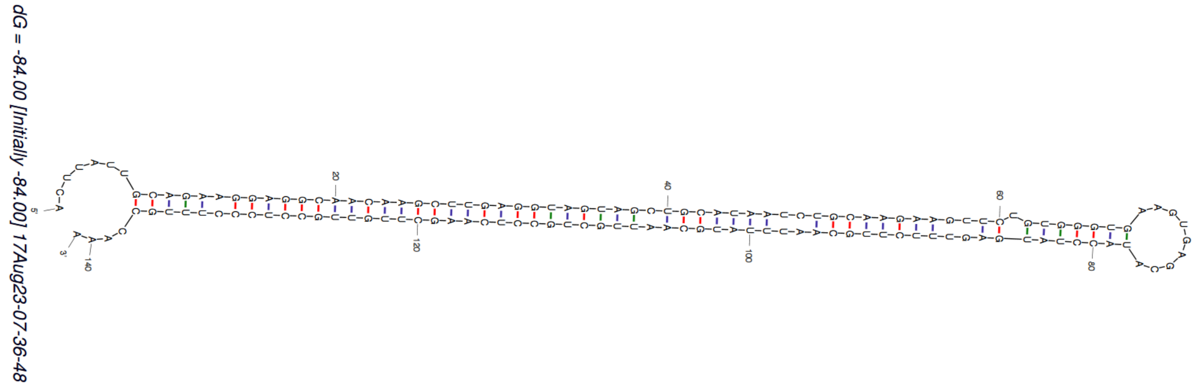


#### *MaAGL17_1*

##### >3_reverse

ACAGGTCCATTGACTGCGTACAGTGCTGCATATAGTTTCGAGCTGTTGCTTTAAGGTTTATGCGGCGTCACATCA
AGAGATAACCCTCCCTTCATCCCTAGGAAGGGTCACACTGTCAGGGGGACGGAGGGGGGGCTATCGAGTCAGTGA
CACACCAACCGACGGCAACAAAGACCATTGGCTACTCACGGAGAGCGGCACCAGCACGTAGGAGCAGGTGTCAGC
GATCCATGCGGCGGGCGGTGGGAGCTACCGAAGTATGCGTTTAGCCTGTTTTGCCATCGCTTCTGCAGTTGGTCT
CTCTGTGAGACCGCCACCAGCAGCGTTTTCTTTCGGTTTCTGTGCTCAAGAAGCAGTCATGAACTCATAATCATT
ATTATTATTCTCGCTCTATCTCTTTAGTCCTCTCATGTTAGAAAAATTCTACTAGACAAAGATATCTCGAGTAAG
ATTGATCGGAGCTAATATATATGAGATAGATATCGAGTCATCTAAATTTTGAATTGTAATAAAGAGAACTTTTGG
AAGCAAAAATTGGTGTGGATTTGATTGGAACAGTTCAAAAACAATAAACTATTTGTGGTTTTGTGTTTTACTTCT
ACTGACAAGAATTGAGCGAGCACAAGAGTAATGACAAAAGCACAAGCGGGTTTACCGTCTCCTCCCCCTCCTTTC
CGTTCCGGGCATAAAATAAGACATCAAAAGAGAGGGGACTGATATAAAGAAGATCCAACCTGCACCAGATAGACA
TAAGAAACTCTCGGCACCAAGAAAGAAAAGCTTACCTTCCTTGCTAAAGTTTAAGGTTTCTCCTCCACAGATCTC
GTGCTGCAGCCGCCTGTTTTTTCCCTTTCCTGGTAACCTGCTGCTCTTGGCTGGTTGTTTGGTAGTGCCTGCCGC
AAGATCTGCAGATCAGCTAGCTGAGATAGAGTCGTGAGGAGGAGGCATGGTGAGAGGGAAGATTGTGATCAGGAG
GATCGACAACTCCACCAGCCGGCAGGTCACCTTCTCGAAACGCAGAAACGGGCTGATGAAGAAAGCCAAGGAGCT
CGCCATCCTCTGTGATGCGGAGGTCGGCCTGGTTATCTTCTCCAGCACCGGCCGGCTCTACGATTTCGCCAGCAC
CAGGTTGTATGACTCTTGTTCCATCTTTGTCGGTGCTGGTTATGAGATTTGTGGGTTTCTTTCTTTCTTTCTTTT
CTTCTTTGTTTCTTGAATGGAAGATGAATATTGATGGGTTGAACGGATGCATAAATCTAAAGGGTAGAAAAAGTG
TAGGACATCAAACAGATCTCCTTTTTTGTATGCTTTTTCTGACTTTTATGATGTTGCAAGAATGATCTTGATTGA
TTCCCTCATTTGTTCCTCTTTCCTCCTCCCTCCTTCCATTCCATGCCTCTCTCTATCATTCTACCATCTTCTCGT
TCATCCATCTGTCTATCCACCTCCATATCTGTGCTTGTTGTTGAATTGTGGCTGCTATTTTCATATACCCTTGGA
ATGGATCGAGTTGTTCCCTTCTTTTGTTTCCTCGGAAGCACTACATTATGACAAGTACGAGGAACACGAAGAACA
ATCCCATTCGATGGTTGTGTTTGATGGCCCTACCAAATCTCATGAATGCTGTGAATCCCTACGGATGTTTCCTCT
CCAAGATCAGAAAGTGAAGCTGCACATGGAGTACTTTGATGTCCTCCGACCAATGAATGTGGTTCCTTAGATTTA
CTGTCAGCTGATCAGTGATTGCTGAACAAAACATGATCTAAAGTAGATCTTGACATCATGCGAGGCCATGATGAA
GGCATTCACATGCATGCATCAGATGTTTCCTCTTAGGTTTTTGCAGTAAACTGCTTGAATCATCTTTAATATCAG
CTGTGGAGATTGCTTGAGCAAAGACAAGGGATTTCTTTACCTTGGGTACCATAATTTCTAGTGTTAGAATTAGGA
AGCTTGCAAATGCCTTTGCCTATTTATATTTTCTACTAATTAGAAGGAAAAAAGGGGAATTAATTTCAATTTTTT
CCTCTTTGAGATAAACATATTATTTTAGCAGTAATTATTCTTGCCATACTCAATCATACTTGTATTTGAGCTAGA
AAAAATAGAACTTTTAGTTTGATAACTTGCTACACATAGCTAAGGCTGTCTTTTACAAGATCATAAATCCATAAT
AACTAAAAAATTCTTTTGCATAGTATGCTATATCGATCGAATATGTCCGTAGCATAGCTAATGTTTTAATTTTTT
TTTGCTCAACATCTCAATACAATTATAAAAACAGTTTAAGCATATTTAAATCGTTATAACTCATGTTCGATTTTC
AGCATGAAGTCTCTGATAGAACGATACAAAAAGGCCAAAGAGGACCATCATTTTTCTATGACTGCAACTTCGGAG
GTTAAGGTATATGCTAAACTTAGCGCGAGCTTATATATGAATTCATTTTTTATAAACACCTTCGATTAATGTTCG
CAACATTGGTTCCATACTATCTATCTATGTGGGAGATGCATAAAGAAGATTAGGTATCATATTTTTTCATAAATA
TTCATGAGATCTGCAAAAACTTAGATACTTAGAAGTTTTTTTCTTTTACCGATCTCACTGCTTAAGTATTAGGGG
GTATATGGTTGCAATTTTTCATAAGGTGAAATATTAGGAAGCATATTAATGAATCTCAAGGAAGAATTAATTCCC
AAATAACATTGTTTTAAGTACTTTGTTGTTAAAGAAAAAATAAACATAATGTTCATATTATGATTTTTTTGAGAA
AATTTATGTTCTTAAATCCTTAGAAAATTGTCTGCTTCTGATTTTTTTTCGGAACGTAAATGAGGTCACAAGCAC
ACAGTTTGTGAGCAAGAGTCAAGATCTAGGAAGAACTTAAAGAAAACTAGAACAAGAACATGTTCTTCATTTCAT
TTTTCTAAACCTCAAATGTGATACATTTAAGTGTTTCCTTTTTCACCAATAAGCCTTTTCCCATAAGAGTTTGCT
ATTTCTTAGTATCACATGCATGACATAAAGAGGGAAAAGAAAAAGATTGGCTATCCCTTGTAGATAAAGAGAGAG

GACAAAATAATAGAATGGAAATGCAAGGTATCAAAAATATATATCCAAAGAAATTAATGCAGAGGATATCTTAGC
ATTCATCCAACAAAAAGACCACTTTAGATTTCGATTTCAAAATGAAATTGCTCTTACATCCATAAATAGATCACT
GAACAAGAAATAACACATGAATATTCTCTAGAAAACAATTATGCTTAATCTCTCGTATCATTTGTATGTTGATAC
TTTATGTCGATAAGCTGGAGCATAATCCCTTAAACACTGTTTTAATCCTTGTAGAATTAGAAGTGTCAACCATTT
CAGAAATAAAATTCATGGAATTTTAAATAATCTCACCATACTTCAATGGAATGCTATTCTCCTACCATGAACTGG
ATAAAGGTAACTGTGGAAATTTTCTTGATGCTTCACTTGATAGGCATTTTGACATAAGTTTCTTGATTGTTTAAA
GACCTATAGACATGACTTAATGACATGGTGTGATCTATTTCATGCATCAAGCAGTTTATATTATTTTTGCATATA
ATTGTAGATTTTTAATAATTTGGTTCTTTTTTTCTAGTTGGAAGCATATATTAAAAAAATTAATTAGTTAAGAAA
GAGTTTTGTTAGGTAAGTATCTAACCTCGATCGAATTAATTGCTTTGTAATCGCAAGTTCTGGCAAAGGGAGGTA
GCAAGCTTGAGGCAGCAACTACGTAACTTACAAGATAATCACAGGTATCCTCGTTTTACTTTAATTTACGCACTC
AAACGAAACCATGATTGCTTGAAAGCCCTAGAACTTCTCTAATAACATTTCTTCAGTTTTCTTTAGATATTTCCT
TAATTTTGAGCAACACCATCAATATATATGGTTTATGTTAGTCCAATTAATACTATTATTACAAATATGCTTGTT
AATTGTTCCTTAACCAAGCTATAACAAATGAGTGAAAGTTAATTAACAATGTTAATTAGAAACATGTCTCCTTTT
AACAATATGTCATATGCAAAGTATTATAGAATTATAAAATCTATAGAACGAAGTATAAACAAGTAAACATGAACA
GCATCTACAAAAGAAAGAAATTAATGATATGAAAATACAATTTTAAAAGACACAAAATATGAAGTTTCTGCTAAT
TTGCTGTGTGTCCATCAAATAATATTTGCACACTGAGGTCTCTTTCATTAATGTGTGAGTTCTGGAAAAATTCAA
CCTCATTTTGAATGGAATAGACTCCAAAACCATCAGATAAAAATGCAGGATTCTTTGGAAAAAAATCATGTCTTC
ATCATCATGGATGATAGACATCAACACCTTCCTTGTTGTGTAAGATTAGCTTTGGTTCTCTTTGTCACAATTGAC
ATCTTAAGATTCGCTACAAAATAGTTATGTTGGTGTTGCATCAAATCTATATCATTTGTCAATAATTACTGGCCA
ACATCGAGCAGCAATTACTGTCCAATGGTAGACCATGTAGTGTAAATGTTTGTCATGTTTACAACCAAGAAATTG
TTCCTGTTGATCATTCAGTAATCGACTCACACACTATACCTCCTTGCTACAACATACGTAATATTTGGTTGATCA
GAGAATACACACGTCTCTAGTAGAAGAGCACATAGTACATGTTTGCCAACTTCTGCAGGCACTTGATGGGAGAAG
AGCTCTCTGGTCTTAGCATCAAGGACCTACAGGACTTGGAGAAACAACTAGAGATGAGTCTTCACGTTGTCCGAA
CGAAGAAGGTCTCAGTTACCTTTCAACAATGAGTTTCTCCTTTCTTGATCAGCATTCCTGGTGCAATGAATTGTT
ATTTTGCAGGACCAACTGTTGACAGATCAAATTCAGGAGCTTAAATGGAAGGTTCTGCCAATGGCAACCCTAATC
ATTCTGCTCTAAGCCTTACCTGCACCAAAAGTTGCTAATCGTTCTGTGGTCACACTGCAGGCAAACCTTATTTAC
CAAGAGAATGCGGAGCTTCATAAGAAGGTAACCCTCCTTAGTGAAGAAAACATGGAACTACACGAGAAGGTAGAA
CTTAGAAAGCTCATTTCGATCCGAAGTCTTTCCGGTGTATGTTGTCTCACACAAGCTAAGAATTCCAATCTTCTA
CATCAGGTGAAAGCAACAAGGGGTGAGATCGGTGACAGTGGAGACTCAGCTGTTCCATATGGTTGTCTTAGTATC
AGAGAGGAAGCAGATGCACCGAATCATCTCCAGTTAAGTAGGCCACTAGAGCAAGCAAATGGAGTACAAATTGGA
GCACCACAGCTACGGTAACTTCTCTTTCCTACAGTGGAACTTCATAGGTATCAAATGCTAGGTTTACTATAGAAT
GCTTAACCCTATTTATCACAGCCTTGGATTGCACTAAGAAGATCTCAAAGCTTCAACAGCATCATCTCCTGATGT
GGTGTACTGAGAGGAGCTTCAAGGAGAATTATATATATATATATGTATATCATGTGATTTAACTAAAACCAATTG
TTGTTCATTAGTTCTTCACTATGAAGTTATGAATACTATTGATCCAGTTGATAAGTTTAGCAATTTCTAAGATTG
TTTTTCGACGTGCCTAAACCGACGAATCGCAGTTTAGTTCGAGATAACGCAAGAAACTACAACAGACAAAGAGAT
GAAATTTAATGTATAACAGCTAGTCGAGGAAGTAAAATTCTCTCAAATCTCCAATACAATGAAAATAAAATTCAT
GTATCGTTCCCAGTCTTCATGTGAGAGTGGTTTGGTTTCTCCATGTAGATCGCATGAAGTTATGCATACCGACCT
AAGGAGTGTTGAGAGTTGACTTGTTGCGACAAGGAAAGAGAGATTCTACTGCTGTTGTCGAAGGAGAAAAGAGCT
GAGGAAATTATTTGTGTTTTGGGTCAAGAAAAATTCACGAGCGAAACATGTTCTGATCCAATCCAATCTGAGGCC
TCAGAAGCCAATATAATATCTCACCCTTTAATACAGGCCTTTAAAAGTCAAATTCATTGACTAGAATTGAGAGTC
ATGTATGATCCAATTAATCTTAAAGATGTTTGTTTGCTATGAAACATCATCCTATCAAATTTTATACCTAAAGCA
TTTAGTATGGATATGGTGATGATGCCTTAAATCTTGTTTGATGAAATCTTGAAGCTATCAATTAGTTTGACAGGT
AAAATTCTTATTTGATTCCTTTGAAAAATAAGATAAAATGGAATT

>MaAGL17_1

ATGGTGAGAGGGAAGATTGTGATCAGGAGGATCGACAACTCCACCAGCCGGCAGGTCACCTTCTCGAAACGCAGA
AACGGGCTGATGAAGAAAGCCAAGGAGCTCGCCATCCTCTGTGATGCGGAGGTCGGCCTGGTTATCTTCTCCAGC
ACCGGCCGGCTCTACGATTTCGCCAGCACCAGCATGAAGTCTCTGATAGAACGATACAAAAAGGCCAAAGAGGAC
CATCATTTTTCTATGACTGCAACTTCGGAGGTTAAGTTCTGGCAAAGGGAGGTAGCAAGCTTGAGGCAGCAACTA
CGTAACTTACAAGATAATCACAGGCACTTGATGGGAGAAGAGCTCTCTGGTCTTAGCATCAAGGACCTACAGGAC
TTGGAGAAACAACTAGAGATGAGTCTTCACGTTGTCCGAACGAAGAAGGACCAACTGTTGACAGATCAAATTCAG
GAGCTTAAATGGAAGGCAAACCTTATTTACCAAGAGAATGCGGAGCTTCATAAGAAGGTAACCCTCCTTAGTGAA
GAAAACATGGAACTACACGAGAAGGTGAAAGCAACAAGGGGTGAGATCGGTGACAGTGGAGACTCAGCTGTTCCA
TATGGTTGTCTTAGTATCAGAGAGGAAGCAGATGCACCGAATCATCTCCAGTTAAGTAGGCCACTAGAGCAAGCA
AATGGAGTACAAATTGGAGCACCACAGCTACGGTAA

*MaAGL17_3*

>9

TTTACAATAATGAAAAAAAATCTAACCATGGACAGCTCTCTGTTTAATTAGTTCAAATGTGAGGCTAAATTACTT
GATTTTGAATTTGAATTCAAGTTGATCTCATTAAGACAGTGAACCACTTCTTCAATCTATCTAAAATTATATGTA
GTAATGGCCTTAAATTATACTTACAAGCTGAGCAAGTCACAGGCTGATTTTTTGTTGGACTTTGAAGTTGGGACA
AGATCATCTTCCAAATATTGTCAAGTCTGAGAGGATCGAAATTGACATTCGTCCTATTTGGGAAGTTATTGTGTC

TCTTTCTGATAGCAGGTTGTGGATTTTGCCCTTTTTTATACTATAGGCTAAAAACTAAACAAATGTTGTTTAATT
CTGTACTAATTTGTATAGACCATTGATTCAAATGACATTAAAATATGGAAAACGAGAATGGTATTAGTATACGTA
GCAAAACATCTACAAAAGCTGGCTTGCTTTTATCTATTCTAGCAGCAAACAGGCTTTTCCCATCTTATTTATGAT
TATCACTAAATTTCATAAATTTCCAAGGTTCAGGTGATTAGGTGTTCTTGCTTTCATGAGCATCCAAGAACCATA
TAAAAAGTTCTTAAACATGACTGATGTTGCAGTCCTTCTTTAAGACATTACTGCTCAACTTGGAACTTGCTAGGT
TGGCCCATCATTTCACATGTGATTGTAAGTTCTGCTGCCTATGTTGACACTATATGATAGTCTTGGAAGATGATT
GATTGACAACATTTTGCCTTTTTTTGTTGTTAGAAAATTCATAAAGAAAGTCATTAGGAAGGATTTTTAGTGGAA
AATCAACATCATTCCAGTACTTGATGACTCTTGCATATACTATATTTTAGTGTCTTTCCTCGAAAATTAAAAAAG
ATATATTATAAAAAATCAGTCATTATATGACAAATATTAATCCAATAATTCACCAGATTTGGAGTATTTTTCCAC
ACCTTAAAGAAGTTACTAGTATGATTGGTTCCTTTATCTCACTGAAGCACTTTCAGCTTCTTCAGTGGGAGAATC
TTTTTTCTGGAAAACATAATCATGCATGGTTTCCTGCCTTACCTCTTTCCAGTCCATTCAAGAACAATTGCCTTG
TATCGTTCTGCAGGAAATCGCGGAAGATGGCCCCCCCTTATCGAGTGATCTTGCACAATGACAACTACAACAAGC
GCGAATATGTAGTCCAGGTGCTGATGAAGGTTATCCCAGGAATGACCCTTGATAATGCTGTCAATATCATGCAAG
AGGCACACCACAATGGCCTGGCAGTGGTAATTATATGTTCTCAAGTAGATGCCGAGGAACACTGTATGCAGCTCA
GGGGCAATGGTCTTCACAGTTCAATCGAGCCTGCAAGTGGTGGCTGCTGAAGCCAGTCCGACATATTATCTCTCA
ACTATTCTTGAATCCTTCTATTATGGTTGGTTGTAGTACTCACACGGGGAGTATGAAACAATGATCAAGGTGTAG
AAGGAGATGATTCAGGCTGTATATAAAGATTTCATCACAAGGTCTTTATACTATCACCAGTTGAACATATCAATA
ATATAATTGGTGTCTCGTACCTATCAAAATATTTCGTGATTTTGTGTAGGGTTCCAAGAGATTGAGATGATTAGT
ATTCACCATCTCCGCTCATCTTACTCCCCCCACAACTGCACACAAAAAAAGAAGTGTTTTTGACTTTTGTTCCTG
ATGATGATTGTTTGATGGACTCGTGCAGTTGCACGGACTCTAAATACATGCCAAAGTAGATTGCTGCTATTGGCA
ACTAAGAATACTTGTAAGATTGAACTAATGTCGACAAAGAGGAAACAAAATCATGGTTAATTATAGATTATTTTT
TATGATTAGTTAACTTTAATTTCTCGATCTCTATACTATAGTTATATCGAGTTATATTGAGATCATTATATTTAC
GAAAGTAAAATATTTAATAATCTTATATAAATATTTTATTTTTATAAATATAGGGATCCCAATATAATTTTTGAA
AATATAGATCTCAAAACAGCTGACTACATAAAGTAATATGTAATTAGTCCATAAAATCATCATGAAACACTTACG
ATGAACACAAAATTTTATTAGATAATAAAACTATGAGTCGAATTATATATATTATTATTTATTATATATACATAT
ATTAATTTAATAAATTTGAGATTTTGAATTAAAATGGTGTGTTAATCGGATGAATTCGTAGACATTATTATTTAT
TATTTCTTTGCTGGGTAGACGCCTCACCCTTAGTTTACTGAAATCGATAGTGACTCAAAATTTTCTTTGACTAAA
AATAAATTATCATGTCCAGTCCAAATATTAGAGAACAGGGAGCGGAATCTCGTGGCCCCGCCACGCGTCGGTGAT
CCATACGGCAGGCGGCGTCAGAGCTACCCAAAACAACATTGGGTCGTTCTCTCCGCTGTACCAGACGCTAAACGA
CACCGTCACAAAAGCTTGCAAAGGAATCTCACTCACTATCCTAGTATTCAAATCTCCAATCTCCTATCTTGAACG
TGGGGAAGGGGAAGAGTGTGTGTGTGAAGAAGACCTCTCTCTCTCTCTCTCTCTCTCTCTCCTTTCTATTTCCAT
ATTAGATATATAGATGATGGAGGAAGAAGGGCTGCCGCTGAAAGAGATTGCATCCACACCAGGTAACAGAAAGAT
CATCGTCGTCATTGTCCCCCCTATTCTCTTTGATTCTTTCCCCCCTTCCATCTTGAGCTTACCCTTGTCAGATCT
GATCGATGTACAGTTCCGGTGGGTAGAGGGAGAAATAAACTGGTGGATAATGGGGAGAGGAAAGATCGCGATCCG
CAGGATCGACGACACGACCAGCCGGCAGGTGACCTTCTCGAAGCGGAGGAACGGGCTGCTGAAGAAGGCCAAGGA
GCTCGCCATACTTTGTGACGCCGAGGTCGGCCTGGTGGTCTTCTCCAGCACCGGTCGCCTCTATGAGTTCGCCAG
CACCAGGTTCTGTTTCTCTATTTGTATCCCTTTCTTATCATCTTCTGATGGGTAGAAAGATCTTTCAAACCCATG
CATGCAAATAGGAACGCTTGGCTGAGGCCTCGATCTATGAGACAATTCTAGGTGCTGTCACTGATTCATCATCCG
ATGACCCTAGGTATTTCCCTTCTTCCAAGGTACAACCAAAAAAGTTTTGTGCTTGGCGTTCCTCCAACTATCTTC
TCTAAGAGTAACGTCTGTAAGCCGATACAAGGAGGACGACGCATTTTGCTTTCTAAGACCAAGCTTTGTGGTGGA
GGCTGTGCTTGCCTCGGACGATGGTTTGATACCTTGATTCGGATTGAGATCGCAAAAAAAGACCACATATAGCAC
AAGTGGAAGATATTTTCTCTGTCCCCCAACCTATCTGTAAAGTGACTGAAATATACAACATCTGTTTTCTTTGAG
GTTTAGAGTGCGGAAATTTTTGATTGTTCAAACCCCCCTTTAAACTGGAAAAATGACCAATCGATCCCAGCATGTA
ACATTCTTACATTCAGAATTAGCCACCATTCTTCTTCATTAGAACAATTTTATGTATCTCAGATTTTTCCAGTAT
ACATGAAGTTTGAGGAAGTCTTGCTTGCCTAAAGCTGAAATTTAGAGTTGTAGATGGATAAGTCTACCTAAGCAA
GTTGCTTTGGAATTTCTTCCACTGTTAGGCGCATTTGCTCGCTGAAAGCAGTGGATTCCATGATTGGAAGTTATA
TATTTTTGGTGATCTATTTGTTCAACATCATAAGGATTAGCATATATTTAGATCATCAAATCTGGTTCATTGGAA
CTACACATTTAATGCAATCTTCTTCTCGAAACCAAGATGGTTCATGAGCTGCTTTTGATCCATTATAGCAGAAGC
GTCATCGTCTCAGTATATTAGCTGAAAGTGTGACTCATTGATCAATCTAAGAGATTGACATTCCTTCATTTTATT
GGTTCAGTGACTATCTTTTTTGGGAGGTAAACTCACTTTGTTTTGACTGTGTTGGACAAAGATGGAAGGATCTTA
TGAACTACCGTCTGGGCTTTTATTACCTCGTATCTTTATTTTACAGGAAGTCAAACAATATCAAGCATTAGATTA
TTGATTTAGTGAAAATAAAATCTAAATGTTTCAATTTTTTAAAATTTTGTATTAAATTTTTTTTTTTTTCCATTT
TTTTGGTACTTCATTGATTAAATCTGCCTCTGTTGCCTAAGTGGACTTCAGAACAACAGTTACTGAAAATGTTTA
TTTTTCATGTTTGAATCAAGCTTACATTTTTTCTAAATATCAGCATGGAGTCTGTCATAGAACGATATAGCAAAG
CAAAAGAGGAGCGACAGCTAGTCATGTCTGCCGCTTCTGAGGTTAAGGTATGCGTATCTTGAGCTTTGGTTCATT
TAGAACTACACTATTAATTTCACTTGTCCGTATCCAGTTGCAAATTACTCCAATTGACTAATTATGGTATACTCT
TGAACAATTTAGTACTTTTTCTAAATATGATAAATTAAGTTTAGGATTGCTCAAATATGCTCTTCTCTAAGAAAA
CACGACATTTGGAAAATGTATGATTGTAGATTTTCTGTCAGTTAAGTTCAAAAGAAAAAAACTTTCTGCTTAAAA
AAAAAGATAAAAAAGCTAAATTTTTATTGGCAAACTAAAAATTGAAAAGTAGTATGATATTTATCCAAGAAAAGT
AACACTGTAGCCATATTCGTTTGTATTTGGTTTTTCCTCTTTTTTTTTTTTTTTTTTTTTCTGAGGCACATCAAC
ACCTTCATGAAATCAAATTTGACCAATAAGAAATCATGCATGACCAATCGAATTTGATCTAGAATTAGTGATGGT
CGTAACCTCAAAAATCCCAGTCCGGATCAGTTAGAACTTGTCTAGAAAATGATTGTGTTGGATCTAGTGGACTAA
TTAGCCTAGATAATGAGCAACAAGTTAAGAATTGATCAAACCTAGACATTACCGACTCGATTCAATTAGAGCTAA

GACTCATCTGGTTTTGATGATTCAAAAACATATATGTACAGAGGGTTAAAGTACACGAGATAATGAAGATGGTGA
TTAACAGGATAAAAGAAAGGGAAGGAATAATGTACTGTCTGATAGGAGAAAGGGAAGAAGCAGGTGATTAAGAAC
AAATATGGTAAGAATTACATAAGATAATGAAGATGTCGAGGGTTTGTAGGATTTCTACAAGAAAGCAACCATGAC
GGTAATACTCCTTGGAGTGTCTCGATGATATAAAACGAGATTTCAAGATGTCATTCATCGTCAAGTTCTAAAACA
AGGATTTAGACATTTCACTGAGATTTTGCAGTTGCAGAGTCAAAAACTTGACTCTACATGAGCTCGACAAGATGA
TCGAGGCGTTCTCATTGTTTTCCATCATATTCATAAATTAAATCTTGCTATAGCTTTGTAATTTTCTTCTGATAA
CATGCTCACTGGGCTTTACATTTTACCCCAAATTTGACATTCATTCTCTACAATCATTTGTGTATATGCTTGTAT
GTATATGGCCATGTGAGGACACAACCATAGTGCTAAGATACATATGACCATTTGGTAGGGATTGGAAGCATAATT
CTGGGCTAAACCGTACCTTATCTTGATAATCATCCTGTAAGCTAGCATAGCATTTGACTATCAATGTGAAGGGCC
CTATAATTGAATCTCACCTTCAACTAAATAAAATTACCTAAGGAAGGTTTCAAAAAAATTTCTGCCAATACTTCA
AATAACTTATCATTATTCAGATTATTTAATGAAACCAAATAACTTAGTTAGCAATGATCTTGAACTTGCTATATC
TCCATAAGAAGATCCACCTCCCTGGTTTAGTTGGTTATAATTTTTCTAAAAGAGTCTGGCCGAATGACCTTGATT
ATTAATAACAAGATCCTCGAACAAATTCTTAACCTATGACAATGTGGTTTTAAAAGAACCAATTTAGGTGATAAT
AATTTTCTAAAGATTGAAAAAACTATAATAAATGTAATAGTGCATTAAAAGTTGTTTTGAGGAGACTAATTCAGT
GATTTCCAAATATTATAATATATGTATATACATTATATGATATTTATATGTATATATATATATATATATATTTAA
AGAATAAAAAAGAAGCTAGTTAAAAAAAATAACTAGCTTCTTTTTTATTTTTATTTTTTCTATTTTATTTAATAA
TTCAGTGGGTCCATCTTTTGAGGTCTCAATTATTCATACATACATTATGACACCATAGTTAAAATAAATCCTATG
CTAAGTAGATGGAACAATCTGAGGGTAAAATTTTTACACTTTCTATAAGGAAAGAAACAAGGATGTTTGAGGAAT
AAAATTATGAAATTTATATTTATTAAAAATATTTATATGTGAGAGATGAATAAAGATAAATTAGAGGATAGTACG
GTATCACAAAGAATTGTAGAAGCCGGGTTATATATACTAGGTTTGATTCGTATTAAGTTTTTGGATTGAGCTGAT
ATCAAATTTGATACATATTAATCATTTAGATTTATTATAAAGTTTTTAATAATGCTTGGAGGAAAATAAAAACCT
TAATGTAAGTGATTGACAGTTTGTAATTTTGTACCAGATGGTTAGATTCAAAATCATGTGACCATAGTTTCTTGA
AAAAAGAAAGGCACAAATTAACCCACTATTTTTCTTTCTTTGACCTCTTTGTTTCAACCCTTCTGATTGCTTAAC
AATCAGATCAGAAAGTTGGACTGTTATAAGCATGCAAAGTCAAACAAATACTTATAGTTTAAAACAATTTCAGGC
TGTATACACAGGAAAATCGAAAAAACCTAAAGATTTGTCGTCTAGTTGGCAAAATAGATGTTTGATAAGTGATAT
TTTATTGGTCGACCAATCCCTAGCCAATGACTATGACCATGTTAATAGTATTTGCATCATTCTTTTTCCTTTCTT
AAGCATAGAACGAAGAAACCAGAGAAATAAACATGTTTCGTAACCATGTTATTGTCTTTTATTGTCGAGTAGCCA
ATGAAAATAAAATATTTCCATTTCCTTTTTCAAACCATATTCATTGGATGACCTTTTTCCCATTACCTCAAATTG
ATGGCTGTTTAGATCGAGATGATCGTTTGTCCCATCTGAGATATCTGCTCTCTTCAACTTGAGGCATCATAGGCA
CTTTAAGCCACCATTTATTGACCGGTAAAACAAATGGCATGTACATTCCTTCCAGACCAGATTCATGTGCAAAGT
ATGTGCATGGAATTAGTGGGCATATAAAGCGAACTAGGTTTCAAACAACCAAGAGTTGAACCTCCACAAGACTAC
TGCAAATAACAAGAGACTGACATCAGTTTTGTCCTCGGTTTGGTCCTCTGTTGTCAAAGTCAGGGGATAACCTGG
TGTGGAAATAGGGAAAAAGAAAGAAAGAGGAAAGAGAAAGAGAGCTGAATCCCTATGCTGTTGGAGGACCTTTCA
TTTATAACGAAGGATTCGTAATTTGCACTTAGAAAAATATTTACCATTGATCCGAAGAATATTTTTCGTTATCAT
TAGTAATACCATGATAAGACATAGGTCAGGTTTAACTATATGTCGTCATCCATTATATGTCGAAGTCTAAAAGTT
AAGTTTCTATGGAATGGATAATTTTGTTCTTATCAAGATCTTATCCTAATAATATAGGACATCAATCTACAAGGC
AAGAATTAATTTAATTATTATGCCTCAATTGGATTTTAGATTTGAGTTTCAAGTAATTGCGGAGAGTTTCAATTT
TGTTTATAATTTGCATGTTTATTTATTACCATCAGTCACCTAAATGCTCTCTAGTTTGATCTGCAATCAATTACC
TTAAGTAATATCTGGAGAGGACATTTTTCCACGGTTACCACTCAACTTGGAATCTCCAGCTAAGGACATATCCTT
CAAGATATGTTCAAGAATAGTAGTTAAATCTGGGTCACAAATGATTTTTAGAACGCCCTGTTGTTATAACTAAGT
GCACAAGGAAAGCTTTAGTTCCCAATCAAAAGATATCAAAGTATATGCTGTATTGAAAAAGCAGCACATTTATAT
CATATAATAATGAGTTCAATTAGACAAAAAGATCGTTCGGGTCCAACTTCAACTTACAAAGTTAATGTAGTAATG
ATTAAGATTCACAGAGTAATGATTAATGTAGTAATGATTAATAAAAATTAGCAACTCTTTAAATCTGCCTCTGCT
TGTACAGTCACTGCGTCTGCTTCAATATGTAAACGCAAAAAATGAGTTACCTACCCAGAATTAGGATCATCACCT
TTAAAAGCATTGGCACCACAGGAAACAAGGAGACAATGACCAACAATACGAATGCCATAATTAAAAAATGATAGG
AAATCAAATAAATTATAGCCAAGAATACTGATTGTAGCATTCATAAGTAATCTTTTAGCATTTACTAGTTTCTTA
ACTAATCTGTAACACTTCATTGTTATATTTCATCTCACAGTTCCATCTCAAAATAAAAATATTGTCTTCAGAGCT
TAGCTCCTGAACTGTGAATGATACATATCAACATTGTTTGAACCATGTGATTCTACTCATCAGTTTTGGCAAAGG
GAAGCAGCAAGCTTGAGGCAGCAATTGTATAACTTACAACAAAATTTGAGGTATGATCACCCTATAAACTCAAAT
AGATACATTGTCACTTAAAATCCATTTTTCTAGTAGAATAAACCGGGGTAATTTTACTGACAATTATATATATTT
TTCTTTCATGGCTCCAAAAATGATTATATATGATTATTCATGCATCCATCGTCCTGAGAATTACAACCAAAGGCT
ACTAAAAACCAATTTTAAAACTGCATGTTTCATTATTATGGACGTTATAATGAGGAGTTAATATCATGAACATCA
CATTTCAAATTTCAAATGAATTTTAACTGCAGGCAATTGACGGGGGAAGAAATTTCTGGTCTTAGCGTTAAAGAT
CTACAAAATCTAGAGAACCAACTCAGAATGAGCGTACGTAATGTCAGAATGAAGAAGGTCTTGACTGTCTCCGCT
CGCTGGGTATCTTATTTTTTGCTTCTTATAGCAGCTGTTCTATATGATGTACACAAGTTTTTTCTACAGGACCAA
CTCTTGGAAGAAGAAATACAAGAACTGGATCGAAAGGTTACAACAAAATTTTGCTCTCTCCAATATGTTTAGCAT
TCTTATTAATTTCACCAATGCTTATCCTGTTTTCACCGCAGGGAAGGCTTATTCACCAAGAAAATCTGGAACTAT
ACCAGAAGATAAACATGATTCGTCAAGAAAACATGGAATTATACAAAGAGGTACCAGACATAAAAGCAATTTCAT
CCCATGAAAAGCCAACTTACTGTCTAACACGTTACTCATTCAAGAATCAATGTCAGGTCTATGCAGCAAAGGGCA
AAAATGGTACTGACAGAGGCTCGATTGTACCATATGATTTCAGTATCACAGATGGGGCAGCATACAAGCCCATTC
ATCTTCGATTAAGCCAGCCACAGCCGAACGCAGACGGAATCAAAACAGAGCCGCCTCCAAATCTAAGGTAACTTT
CGTCACATAAAAGTAGGAGAGTGCTGCGTTACACATTTCTTATCTAACTATACAACTCACAACTGCAATTTTGAA
GCAGACTTCAGCTACACTACAGAGATGGTTAGTACGTCTATTATCTCAACGCTCATGTTTGAAGTCCCAGGCGAG

TGATCTTTAATTACGTGAAAGTGCAGCAGTGACATCCCAAAATAGTTTTTGCCACTTATTATGTGAAAAATGATG
TGTGCATGATTCTAACAATGATCGATTTAGTAGGAAGGGTGAGTGTTTGCTTATTAAGATTGTTTAATTTATGGA
CTCGTGGACACTGATATGATGTTGAATACCTTTCACTGTTGAACAAATTGAGAACCAAATCCATGATCACATACC
CCTACTTAGCTGTACTACGCAATGTAATCAAAATCTTGTCAGCCCCCGAGCAACGAAAAGAGTTCGTGTCCAGTA
AATCTCCCACCCCCCCCAGTCTTCTGATCCGAGGTGGCCGTGAGTGTCGTTGGGGGGGTCCACCTCGCGTAAGAA
ACTCCCTTGGTACGAAGCTTCAGTGCGCACTGCAGCTCTTCATGAGAACCTTTTGTCTCAACGCTGCAGCAGCTG
CTGCTGCTGCATGCTGCCCAACCTGCCCTTCTCAACCTCGCACAGTCTCTCTCTGTCGCCTGTCATGGCACGCTT
CACAGGGCGCCCAGATGTTCCCCGCCTCCCGAGGCGGGGCGGTCCCATCGGCAGAGCTCGGCAGCGGCCGTACCC
GCGTCGCGAGGGGGTCGCACACGTCCCGCGCTCGCGGCCACAGCCCGTCCGACGCACCCCACGGCACACACAAAA
ATACGCCCCAAAGGGGCCTCCGTACCGATACGCTGACACCCCCGAGCTCGAAAGCGATCCGCTTGCAGACACCTC
CAGCTTTGTCATTATTGTTCGCAGCTGCTGTTATTATTATCACTCCTTCAGAGCTGTGTTCAGACGATATCTCCA
TTATTTCCACCCACATTCCCATGCAAGCTCATTATATACACATATATCTACTGCACTGATCTCTATAAAAGCATA
ACTAAAAAGGTTGACGTGATAGGCACCCTGGAAATCGTATTTAAAACTGATAGATAAAGGATGGATGGATTGATG
GACTGATTGATGATCTCGCTGATGCGACACCGGGAGTCTCAGTTTAGTGAGAGAAAGGAAAACAAGGAACCTGCC
TTAATTTCTTGTTTGTGTGTATTCTCCGTTCCTACTTCTCTGTGCAGAGAATGCTGAGCTCCCATAGTCTCCTCA
TCCTCCGCTTCCCTTCGTCTCTTCCGCTGCTATGTACTCTTCTCATCTCTACCGGTACCGTACTTAATTCCATGT
GTATTGAATCAGCTGAAGTAGCACTGGTTTGAAGCAAATGACAGTGTTTTGCTTTTATGCTTTGCTTCCTTGTTT
CAGAGGAGAACAGGCCCCGTCTGGATTCAACATCGGCGATGCTGTTCTCGACCTCACCAGAAGTTTGTTGCTTCT
CTTATCCTGGTCCTGCAGCTTCATCCTGAATTACGACGGGGATTGATGCCTTTCTCCTTTTGTTTGTTCTGTTCC
ACTCTGAAGGTCCCATTCTTAGTCCCAAGAGCTCAAGCAATCTCACTGCTGTCAATTCCTCACCCCTACACCTGC
ACTTTGGGGCCATAAACCATGCGGTGTTTATCGTCACCTCTTTTTCTTTTAGTGGGCATTGTTAGAACGCGACGT
GTTTTGTGATGTAATTGTCTTGTAACAGAGTATGGCATCGCTGGACGTGGCTGCTCCGACGGCGGCGGCAGCGAC
ACATGTGGATGCTGGCAGGATGGCGTTCTCGAGAGGACAAATTGAGAACTGGGGCGACTCTGCGATGGTAGTGAG
CAGCCCAAGAACTGATACATCAACCGACGTCGAGATCGAGGATAGGACTAAGATGGTGGGCATTCTACCATCCTG
TTTTACTCTTCTAAGATTGACTTTGTACGTTTATGGAATTCTTTGTTGCTGTTCCACAGTTTGGTGGTGTTCAGC
AAGGTGCTATTATCTGTGCGGATTCTTCTGCAATGTCAAAGGGGAAAGCAGGGGAGCAAAAGGTGTCTTAAACGT
CAAGTTGAATTTATTTGTATTGCTTGGGTTATGTTAGAATGAATAGTGGATGATGCTTTCACTAATTTCTGATGC
AGACACTGCGTAGACTTGCTCAAAATCGTGAGGCTGCACGAAAAAGTCGCTTGCGGAAAAAGGTTTTTTTTTCT
TGCTCTAGGCAAACAAAAAATATTTTTTTTTGCACCTTAAAGAGTGTAATCATATGTGATACCTTATTAGTTTAT
ATCGCATGAAATTTAGTGTGTTAGCCTCTCAAAGATTGCTGTAGGCAAAGATGTTAGTTTTATGTTAGTTGGCAT
ATCCGTAGCAAGTTATTATTGTTTTAAGTTTCACTGATTGTTGATTGCTAGTTGCTACCGAGCATACTACGATAT
GAAAGTGAAGCTCAATAATCACTGTTTGTTGTATCCTAACATCAACTTCATGCATAGGTTTTCCTTTCCAAAGAG
TATGGCAATTGTACAGACCATGTTTAGTGTATTTTTAGTTAAGAACTACTACTTGTTTCTCTGTGTTTTGTCATT
CGAAATGGACACTATCAGCCTTTCATGCAGTGAAAACAGTCCTGTTTTAAATCAACGCCTTTTAGTTTATTGTAA
CTTTTCTTCTAGTTGTAGGTATATATTTCTGTTGTCTTCCATCTCTGTTTTTCTTTTTAAAAAAAGAATGTGATC
AAGCAGCCTGCCGAAAATTTTTAATAGAAAAGATTAGTTGAAAAAGCAGTAATCATTGCTACTTTGCTAAGCTCA
CATGTTCTTTACACCACGGAATAATGCACTGTCAACTGTGCTGTATAATTTGACATGCATTCAAGTTATCATCTA
TTCCCAAATAGCAAA

>MaAGL17_3
ATGGGGAGAGGAAAGATCGCGATCCGCAGGATCGACGACACGACCAGCCGGCAGGTGACCTTCTCGAAGCGGAGG
AACGGGCTGCTGAAGAAGGCCAAGGAGCTCGCCATACTTTGTGACGCCGAGGTCGGCCTGGTGGTCTTCTCCAGC
ACCGGTCGCCTCTATGAGTTCGCCAGCACCAGCATGGAGTCTGTCATAGAACGATATAGCAAAGCAAAAGAGGAG
CGACAGCTAGTCATGTCTGCCGCTTCTGAGGTTAAGTTTTGGCAAAGGGAAGCAGCAAGCTTGAGGCAGCAATTG
TATAACTTACAACAAAATTTGAGGCAATTGACGGGGGAAGAAATTTCTGGTCTTAGCGTTAAAGATCTACAAAAT
CTAGAGAACCAACTCAGAATGAGCGTACGTAATGTCAGAATGAAGAAGGACCAACTCTTGGAAGAAGAAATACAA
GAACTGGATCGAAAGGGAAGGCTTATTCACCAAGAAAATCTGGAACTATACCAGAAGATAAACATGATTCGTCAA
GAAAACATGGAATTATACAAAGAGGTCTATGCAGCAAAGGGCAAAAATGGTACTGACAGAGGCTCGATTGTACCA
TATGATTTCAGTATCACAGATGGGGCAGCATACAAGCCCATTCATCTTCGATTAAGCCAGCCACAGCCGAACGCA
GACGGAATCAAAACAGAGCCGCCTCCAAATCTAAGACTTCAGCTACACTACAGAGATGGTTAG

*MaAGL17_4*

>5
TCCTTTAAAGCATCTATAATCTAATGGTAACCTAATATCCAAAGGCATCTATAATCTATAATGTTCGAACTATCT
AAGATCACTTACATCAACCATGTACAAAACGACATCCTTAATTGAACTAAAAACATACAAATCTTTATATAAAGA
TTGGAGTTTAATTTTATTAGATTTTGTTGTATCTCTTCTCAAACCACTTTGAAACCTCCCAACAAATATATAAAT
TAGGTAGAGGTGACTTGATATTACCACCATCATAGTGAGCTTAAGAATTTTTTTTGTTTGAGTGCCTAACGTTTC
TTTTTCATGTGTATGATAATTATAATTTTAAAAAATATTTTTTTTTCAAAGGATAAAGGGTGAAGATAAATTTTT
ATCATTAGATCTTATGATGATCTAACAAAGTTTTACCAACCAGGTTAGTGCATCTTTAAAATTATTTCAAATAAG
GTTTAATATATAGTTGAAGAATGAAATGGTGGATATATAATTATTATGTTGCAATATATGATCGACTAGATGATC
CTTATATACCACATTAGCTATACATAACCAACACATGACTACCAAATTCATAAGAATATAATCACCACGGACGA
TATGTGATCGAACTATTATCCATTCGAACCTTCTGATTAGTATAATCAACCAATGTGTGCATCAGGTATAACAGA

TATGGTACCATCTTGATGCTTGAGACATTGCACATTACCTTCGTACTATATATGATTCTAAGTATAATCATCTCT
AGAGTTGATTATAAAAGGGGCTTCCAAGCACCTTATCATGAATACTTTATTTTAGTATTAGCCTACTAGTTTGAT
CGAATTGAACCAAAAACTTACTTGACTATCGAAGAGACATTTATCGAAAATACCCCCGACTTAGTTTTGAATGTT
TAGTACTATGAAAATAGTGAGTTGGATTGAGAATGACATAAAACTGACCTTAATTGGATCTGAATCACCTTCCCA
ACTATTGAAGTATTAGCCTACTATCGAAGATGCTGCACTAACATGCGCACTATTAAGAGACGATATGAAATTTTT
TAGATAAAGTTTTGAATCCTTAATCTTTAGTACTCCCAAATAATATTGTTGTATATTATTTCATGCTAGAATCTT
TTAACCTCAATTGATATATTTTTATATCTTAAACACTAGTATGATATACTAGATCAATTTATAAAAAGTGTAGTG
TTCAAAACCTCATTTAATATATTTTATATCTTAAACATTGTTCTAGTGGTATACCACTCTCTTATTAAGGATTAA
GAGTTCAAAATCTTAACTAATATATTTTTTATCAAACCATCTTGCCAAAGAGGGGTTAAGGGTTTGAAACCTCAA
TTAATATATGTTACACCTTAAACATTAGTCGAGTAGTGGTGTATAAGATCCTCTTATCGAAGATTAAGGATTCAA
AATCTCAACTGATATATTTTTATCATACTCTCTTACACCTTTTATACCCTTAAACATGAGTCTAGTAATGGTGTC
TCTAACCCTCATACAAGGTTAAGGTTAAGTGTTCAAAACCTAACACATTTTTTTNNNNNNNNNNNNNNNNNNNNN
NNNNNNNNNNNNNNNNNNNNNNNNNNNNNNNNNNNNNNNNNNNNNNNNNNNNNNNNNNNNNNNNNNNNNNNNNNN
NNNNATATATATATATACATATCTGGGCCCAAGTCAAAATCCATCAAGTAATGGGCCCAAATCAACCCAATTTCG
GACCTTGTGGACCAAATATGACTAGTATTTGGTGTATATATGTATCAACTTCTTTCTTTGACCTTGTACTCTGAT
GATGATGATATGTTTAATTAATTATCAATCATGATAATTATTTATCAAGAATGGGCGAAGGATAAGTGCAAGAAA
TCTTTAAAAGCATGATGGGATTTTGAGGGACTCACCAAGTCACATTGAGTCAATCATTCGCATTTTATATGGAAA
TAGTGCACTCGAAAGGATCTAAAACCTAATTATGGTGGGAATAGTGAACTATATCTCACAACTAAAATGGATGTG
ATAAGAACGAAAGATTCAATGCATCAACTATTTGCCTCCTCCACTCAAAGAGAATAATACACCCAAACAATCCAA
GCTGCAAAGGGTCATGTAAATCATCTTGCAACAACAGCTAATATGTCTCCTTTATACAAGTTCATGTCGCATGCA
TTATACTCAATCCGAAATCATATGATACATGAGAATAAGGTTCACTTAGGTCCTCCCTAAACCAAATTGTACCAA
CATATGCTTTAGTGGTAACACTCATTGGCTAAAGCATACAAGTCTTAGAGATGATATTGGCTATTCTTCTCATGA
ATATTTTTATTACAAATGGTAAGTAGGTCGAATACACCATCATCGTATTAGTCGAGTCAAGCATTATAATTTTTA
TGATAATAATTTAAACTCTCAATCTAAGGTAAGTTAATGGCATATCACTTTAGAGTAGTCGATTGGGTATATATA
TTTTTTTAAACGATAATTCGAACCCTTAATCTAAGGTAACTAAATCAAATATTATCACCAAACTAGTAGATCAAG
TATTTAAAATGTTTAGAAAAATGATTCAAACACTCAATATTTTAATCGGATATATCACTACTAAACTAATTAGTA
GAAGTATAATTTTTAAGATGATGATTTAGGCTCTCAGCTTAAGATAAGTAACACAAATAAACTATCACCAAACTA
GTTACTTAGTATATGTTATCATATGATAATCTATGTTACGCCATGCTACTTGTAACCCCGTCGAACTTACATTAG
AGGATCACTAGACCTTATATCACTTTTGATTATATATTATATTATTATCTCTATTGAGATAATCTGAATTAGTT
AATGTATTGTTGCACAAGTAATAGATCTAAGTTCATGTTCACCTACGAGCGTCACGTACACTATTTAATATAATA
ATTTTGACATTGTTCAAATTCTATATGTAGAAAAAAAATCATGAATATGAAAAAAAAAAAAAATAAGCTCTAGTTT
TGCAACTCTTGAGAAAAGAAAGTGAGCTTACATTTGCATATTCTTGTGAAAATAAAAGAGAAATCGATATAAACG
ATCTCATATCACTCCATCTTATTCTCATGACCTTTACTATAAATCAAACATAAGGGAAGAGAACACTTAATCTTT
AAGCTCGAAAACAATGATTGAAAGATTATATCCGAGTTTAGAAAAGAAAAACGATGAATTATCCCTATGCCTACA
AAAGAGAGTGAAAAAGATTTAATAGTAGTCCATGAAAAATAATAAGTTGAATATGATTTTGTTTTCATCAACTCT
CTCTCTCTCTCTCTCTCTCTCTTTATTTTTTTTTTATATTTCCTATATAAATAAGATATATATATTTTTTAAGGTGTAA
TTTGGGTGGATAAACTTTGACCATATTAATCGATATGAGTATCCTCGATATATAACATCGATGGCATTGAGTCAT
CAAAATTAAAACTTTGAGAAAAATAATAAATAAATAACAATAAGAAAATATTCACTCTCAATCATGTCGACACAT
GATTGTCATACAAATAGGTGAAACATCGGAATCATCACTCCAACCACGATAAAAAAAAAATATATTATCCAAATA
TTTTTTACTTACAAGATATTAAATCTTTACTCGAAGTGATTATGGATGCTCCCCATATAAAAAGGCACATCGATG
CCATCCATACTCGATGATAGATTATTTCTCTCATTACTATACTCTAATAAATCATATCGGGACTCTCTCTTATCA
TAGGTCTTCGTGCAAGGAGGGGATGTGGAGCTCACACCATCTCCAAAACTATGTTTCCAATATCTCTCTCTCTCT
CTCTCTCATTATCTTAGGCGAACCTATCACATCTAAAGTAATAATCAAACATTGGATCTTAGACTCCAACAACTA
ATCGATCGGGAGCATCCTCCAAACATGTATCTTCGTGCAAGAAACGAACACCGAGAGCTCGAACGTCTCCAATTC
GTCTTCGAAACACCCATCACATCTGAAACAATAATCGAACTTGGGGATCAAATCTCATCAAATAGAAACATCAGA
ATCAGGCATTTCACCTTTGTCACACAAGCTGAAACCTTATCTGCTGATTTCTTTAAGGCATCTCGCTCCAAGGAA
TCCATTAAAGAAGGAAAACGTACCGTCATGGGCGCACTGTCCGCCCCACATGGGGGGAGGGAGCTTCCCACCGCC
CCCACCCAAAACCGCCCTCGAATCCTTCCGACCTCTTCTTCCTCCTCCTCCTCCTCGTCACCGGCTGTGCCCCGA
CACCTGAAACCTTCGTCGCCCACAGAACCTCACGTCGCCTTATTATTATTATTATTATTATTCTTCCTTCACCAA
ACTCGTTCTTTACACTCTTGAGTTGTAAACTACTGCGAGTTCTCCACGAGGTTGTCTCTCCGTAGCAGCAGCACG
ACATGACGATGATTGACACTTGGCTTAGCACTCAAAGGTTATCTCGATGTGTTCCGAAAGCCACACATGGGAATC
TGCACCCCTTTGCTTACTTACAAAGCACTACGATGCAACGAAAAGAATAGTTACGATGATGTTGGCTTGTGACCA
TTCTGCTTGAAGTACGGATTCCTCGGCCTCAATCTTTATTGGCCTCACAGTACTGACACCCAGAACTCAGTGCAG
GGATGCCATCTGAGTGCCATCGATCGGAGATAGTGCTTTCAAAAGTGCATGCAAAGAGCTAGCTAGGGTTTTGAC
ACGTACCTTATGGTCTCGAGTGAACTTGGACTCGAATCCATTATGGTCTAAAGCTTGCTGCTTGTGTTTGGGACA
ATGTTCATGGTCAGGTGTTTCTTCATGTATAGCTCTTCCATCTTCCTGCTTATTCTGAGCACATATTTTGATGAA
TATTGATGAATATTTTCTTGAGCCTAAAAAGCTCTAATTTTTTTTTTTTGGGGATTCTCTATTTTCAGAATTTTC
GAATAAGTTTTCTAAATATCTAAAAAAACATAAATATATATTTTATGGACCGATCCATCCGTCATATTAATTTTT
TATACTATGTTATAGTGTTTTCAATAATATCGGAAATACATTGCGACACAATAAACAACAACATTATAACATACT
ATTTATTTTTTTTATAAATTTTGTAAAAATATTATGTTTATAGTGTTTTTATAAGAAAACATATTAATAAATATT
TTTAATAATATTAAAAACATATTATAACGTAATAAATATTTTTATATTATATTATGTATTTTTAGCATCATTAAA
AAAATACTATAATGTGATGTAAAAAATATTATAACTAATGTAGAAAAAAAAAAAAAGATAGATGAGTGGGGTGTT
TTAAGCATTAGGTAAAAAAAAAATATATTAAAGAGTTTTAAAAATGAGTGACACTTTTGTTAAAAAAAAAAGAAA

TTAAAACATGTAGTTATTAGTTCACATGAAACAATTGACTATTGTTGACATATATTCTTTCATTACTATTCATTT
TGTTTTCATGCGTTCATAATTGCCGTAAACGACAAGTGATTCAACAAAGAAAGCGGAGGTTAATATAAAGAAAGA
AGCAGGAAGACAAACACGAGGGCGACGTGCTGCCTCCCCTGCGCACCACGAGACATATATAATATGACGATCAAA
GGAATCAGTCAAGTTAAAGAACGGTTTAGTTTGCAGTACCAAAAGCTGCCTCGCCACCTCCCTCCCCTTCTACTC
TTTGCATCACATAAAGAATAGGAGATCCCGGAAGTGCCGATCTAAAAGGCAACAGTAGCAGCAGCAGCAGCAGCA
GCACCGCCACCCTCCACCATAAAGGTAAGATCAACGGGAAGCTCATCACTTGATCGATACAACCATCCATGTCTT
CTTCTTCTTCTTGTCGTAGCCATGTTGTTCTCTTTTCTTTGATGGTTTTCTGGGTTTGACCGCAAGATCTGGTAT
CTTGAGCTCAGGAGGAGTATGGGGAGGGGAAAGATCGCGATCCGCAGGATCGATAACTCCACTAGCCGGCAGGTG
ACCTTCTCGAAGAGGAGGAAGGGTTTACTCAAGAAGGCCAAAGAGCTCGCCATCCTTTGTGATGCGGAGGTCGGC
GTTGTTGTCTTCTCCGCCACCGGCCGAGTCTATGATTACGCCAGCACCAGGTTCTCTCTCTCTCTCNNNNNNNNN
NNNNNNNNNNNNNNNNNNNNNNNNNNNNNNNNNNNNNNNNNNNNNNNNNNNNNNNNNNNNNNNNNNNNNNNNNNN
NNNNNNNNNNNNNNNNNNNNNNNNNNNNNNNNNNNNNNNNNTCTCTCTCTCTCTCTCTCTCTTCTCTCGACGTTA
GTTCTGCTTGCTACTGTGTTCATGGTGGTGATGAGACTAGAAGCTAGAATAGTTTGAATGATGATGGGGAGTGCT
CTTCAACTGTGATCTATCTTCTTTCGCCTGTCATGAGGTTGAAGCTCCATTTGTCTTGACTACATCATCACTATG
GAGATGATGTTACCTTCATCCCCAAGCTTAACACCTGGGAACATCATCACTATGGAGATGATGTTACCTTCATCC
CCAAGCTTTAACACCTGGGAACATCATCACTATGGAGGTGATGTTACCTTCATCCCCAAGCTTTAACACCTGGGA
ACATCATCACTATGGAGATGATGTTACCTTCATCCCCCAAGCTTTAACACCTGGGAACTGGTACAAGGTTCATGT
GATCGAGTTGTACTTCGTTCCAAATATACATGCTTTTGGATTAGAGGTCATTTTGCCTAGTATGATCATTACTAT
AGACATGAGATCAGTGATCAAAGAAGAGCAGATCATAGGAAGGATTGGGCTGAGCTATTTGTTTTATTTAAGTTT
AAATTTTTATGTTAAATTGATATTTAATCTTATATATGTTAATCACATTTCAAAGAAGTATGGACATGAACATCT
TGATTCCAAGACAACTAATAGAGTGTCATGATGATAAGCATATGAAAGATTAAGCCCAAATGGATGCCACTAACA
TTTATGTCTACAACTTGGCAAATCCATGCAAAATATGTTATCCATGTTGTTCATTCTTTCATTCACAGATGATAG
TTGGTAATATTTTCATGTTCTTGCAATCTAATACACGCATTTCTCATTTCAATCTGAAATGTACATGTCAATCCT
AGATTTGTGGTTATCACTATTAGAACCGCAACATGCCTGTGCTAATGCTTCCATAAAAGAAATCATGTGTGTCAA
CATACCTCTACATAGATGAATAGCATATTTTGCATCTAATACATGCTTTTGTGCATCTTTCATACCTCAAAAAAG
AAACACCTATAGTGATATAAACAAAGAAAAATGTTTGGTTTTCAGAATGTTTATGGTTTATGAGAATATATAAAA
GTATGATGAGTTTAAAAGATAATACTTCCACAACATTTGTCTCTCCAAAATACAAGACCTTGTCACAATTAACAG
ATGCACTGTTAATAAACAGAATCAATAATCCATCAGATATGAAATCCGAGCTGACAATAATTTCTTCAGCTGCTT
GAAATAGAAACCTTATTGTTAGATTAAACCTTGTAAAAAAATCTTTGGCAAGATAAACTTGCCGAAACAAAAAAG
TTCATCAACCGTGTAAAAAACCTTGGGCAATGTAATTGTTCTAAAATCCTACATAAGCACATTCAGGCGTAAGCA
CAAGATAAACATTGTTGCAAATCTGTAAACATTTTTGCAATATAATTGTCATAGTAAATATTTTTGCAAATCTGC
AAATTGACTCTCAAATCGTACATAAGCACAGGATATTGCAGCAAACATTCAGGCATGAGATGTAACAAAAAAATC
ACTAAACAAGATACAATATAGAGAGAAGCAAATTTTATTCTGATTAGTTGTTAAAACTCTTCCCCTTTGTCCACT
ACTTGCATTCAGTAACCTGTTTTCTGATTGCCATTTCAGGACCAAATATAGGAATTGATGGATGTTGCAGTTTTG
TCTAAAGTTCTAATGTTTCAGGAGATATACACAACCAAATAAATGTTTCATTACTACTTGCAGTCAGTAACCTGT
TGTCTGATTGCCATCTCAGGATCAAAAATAGGAATTGATGGATGTTGCAGTTTTGTCTAAAGTTCTAATATTTTC
AGGAGATACACAACGAAATAAATGTTTCATAAGCTTGATCATCTTTCAAGATAATTTTATTCATGATATTGAAAT
TGTTTTTGTAGAAAATAAAAAAACCAAACCTAGTTTGATGTATCCCTTAAGCTTATCTAAAAGCAAAAACCATCT
TATTTCTTTTCCAGCATGGTTTATCATGATTTTGTAGCTCTATATTAGAATGTGTTTGATTCACTTATGTTAATC
ATGTTTTTCTACTAATAACAAACTGTTTTATCCACCAAACTTTTTCCACATCATATTTGATAAATTAATCATCTT
ACTGTTTACCTTTAATGTTTTGCAAGAACAAAAAACAAAGGTGATATCATGTACAACATTTGGTTCTGTAACTTC
TTTATTCCATTGAATAAAAAGGTACTAATGCATCCTTTTATTTGGTGAATAGAACAATTCTCTTACATCAGTTAA
ACCTCTTTCTTCTTAAGTACAGACGATCAATACCAATAGTTCTAAAAATGCACTTATCATTTGTGACACTAGTGA
TGATGCATATTCTTCTCATATGCAAAAAATATGTATAACGTTGAACTTAAAGCTTATACAAGGTGTTTATGCAGC
ATGAAGTCTGTGATGGAAAGATACACTAGAGCAAAGGGGGAGCATCATCTGGTTATGAACCCGAGTTCAGAAACT
ATGGTATATTTTATGCAAAGCATAACAACTAATTCATTTAATGTTAGATTAATCTACAATATAAGTTTTTAAGTT
AATGTTAGCTTTGTTAGTTTCATATCAATCTTAAACTATTAGGCATAACCAAAACCAATTTTAGGTGAAAAGTGA
TATGAAAAGAGCTATGAATAATTTTTGTTGCTTAATCAGCCTTCTCCTCTTTCACAAATTTAAATGCTTTTGCTT
TCTGGTTATAGGAAGGTAGACAACTTCTTTAGATCTACAATTTAATGAATCTTTTGATAAACTATTCTAGCTTAT
CCTTCTCCATAGGAGGAAAAGAAAGCAGCATGAGCCATTTATATGCAAGAAAGTGTCCAAGAAAAACTTCCTTTC
ATCATGCATGAGAGAGATTGGAAGTTCAATAAGAAGAGATGAGATGTGATGGTAGAATTCCATCACTCCAAAAAT
GCCAAATGGGGTGAAATAATAAATCACTAGTCTTGTCACCTAAGTGTGATGATTGTTAGTTCTCAATCTAATATT
GGTGGGACAATTATCATAGGTCACATGTAGCTGTAATAGTTTACATAATGCAAAACCTATTAAAGTTGACATTAT
ATATATATATATATATATATATATATATATATATATATATATATATAATATTCCTTTTCTATTGTAACGATAATCTGGA
ACTGAACTAGTTGCAAGATATCATGCACAGTGAAATTTAACAATTAATTCTGATGGTCTCCTGGTTGAACAAGTG
AATATATTTGAACCTTTATATATTTTAAAATGTTTTTGTTACCTATTTATGGACTTCCACACTCTTTTTTGCATG
TTGAATGAGATGACTATTGGCTAGTTTATTTGGTTAACAACAAGAAGACCCAGAATCAGCCTCATAATTGGCCTG
TCATTATGCTTAATTTTGGGTGTGGTAAGGAGGCAACTACAGAGAAATTTTATTTAGGCCTTCACCAAGTATAAG
CAGTTCGTCTTCCCAGACATCCAACTTTAAAAAATTAGAACTATTCTTCGTTTTAGTTTCACTTAACACTGAATT
GCTATTATGGTATAGATTCTCATTGGCACCTAACTAGAGAGGATGACCATAACTTGCAAAAAGAGTCACCTATGC
ACCACTATGAGGTGTGTATTTCTTTCCAATTAGAGTTATGTAACCCTTTCACTATGTGGATCTACTTGTTTCGAA
TATTCCAATGACAGAAAGTTCACTGCTGATAATAAAGAGATGAGGTCATCATAATAGATTTATTATTTCTATTGT
TGCCTATAGGAAAGCAAGAGGCCACAATGATGTAGGTCTACAAGTGCTCTGATGACGTGTCACATCTGTATGTTT

CTCTTGATATCAAAGAAAAGAAAGGGACAACAGTACTTCATTTGCTACCCATTACGTCTAATACAAATCCCAAAA
CTAGCAGAGTTTACCACCTATACCATCCATACAAGAAGCATAATCCTCAGCCAAAGGATAGGACTTAAAAGAGGA
ATTAATTTGGCATTCTCAACCCTTGGAAGCAAGATTAATAATGATTACATCAGCCAGAAATGCTTCCAGGGGTAT
CTTCACTGCACATATCATCTTTACTCAGTCTATTAATAACTGTACCTCGTTGCTACTAGGGATCACTCCAAAAAA
TGAGGCGCTCTTTCTTTTTCCTATAACCTATTACCATATCCATACTAGGTTCCCTAACTTCACGAGAATCTATCT
TGGTATACCAGGCTCCAGCCTTTCTTCTTACTCTTCACCCCCAAAGAAGGTTTCTTTGGAGTGCTAAATGTTGCT
CTTCCCTTGTTTTTTTTTATATGTTCCTATCATAGGATAATGTGTATGCCTTTCCTCTATTTGGTGAAGCTCAAT
GACATTCCAATTTCAGTCTGATGTGCCGCCAGTAAATTGGGCAATATATCAAATTTACCACCCAGTATCACCAGA
AAAAAGAAATTATAAAAATTAACAACTTACCAATACTAACCTATAAGGTCTTATAAAGTCTTTTGTTGGACCGAT
ACCCGCCAGTTTGACCAGTATTTGACTCCTTGATATCAAAATTTCTTATACTAAAGAGGATTAGGAATAAACTTT
GTAATAACTTCCCTTATATAAAGGCAATAGTTACTTGACTTCGTCCGCCTCCTCAAGGGTTGTATTCCATGCATC
GAGCTGGTTACTGACCTTCGCCTGCTCTTTGCTCACACTTCTGAAGCACTCGAAGTGTTTGCACTCCTTCCATTG
AGTTAGCTACTGTGATTCACCTTCTCAATGCCATCGAACTTCTGGAATGCAAGAAGTTTCACCCCAACTTGGAGT
AAGTCTCTAATAGTTTTGGTCGCCTCTAGGAATGTACCGTCTTCTCCATCAACGTTGTCGCCTACTCCACAGAGT
AGCAAAGGTACAGCACCGCGTACTGCCTGCTTCGTTCCTTGGTCGTGCACTCTTGCACGACCCGAAGTCCCTCAC
TTTCGGCTATCTTGATGAGAAGCTCGTTGACGCCGGTTTTACGAAGTTCCTTGGCCTCTGCCCTTCAGCCTTGTC
TCGGTACTTGAAGTTTGCCTCTACATGCTCCACCTCCTCGGCCCCTTTCACGACCAAGCACTCTCCCTCCATGAG
AGCAAGAGATCAATGACTTTCTCGGAAGTCCCGCCTCTGCGGTTCCATGGCGCTGCCATGCCCATGGCCCTACTA
TCCGTCGCCTCGCATCTGCATCCTTTTTCGTCACGATCAGTAGATATGTCTCTGTGACACTCCTCCGAGTCCACC
TCTATTCTAATTGATGCTTGATTTTGGATAGCTAAGTCCATCTGGACTCGTCGTCACTTTCTCACCCCTTTCGAC
CCCCTGCTTCAACACCTCTGTGTTCTCCAAGCAGCTCCCTTTGGTCGATGGAAAGACAGACTGTAACTCTCATGC
ATGGCCTCTACCATCATGTTATAGGGTTTGCACCGATTCTGTTCTCCTTAGCTTCCTTGGTAGCAACGTTCGCTT
ACTCGACCTTGTCCTCTGACTTGCCGGGCTCCCTTAAGCGAATATGGGCTCTGGAGCAGTCTAACTCTCCAGCTG
CTTCGATCATACCTCTATATGATCAAGTCCCTCCCATGGGACTCACAGGTACTTGCATTCGAACTTTTCTCTCGG
TGGTACACAGCCCCCATATGCTGATGACCAAGGCTTTCATCCGATGCAAAATTCGATGCATGCAGCAAGGCTCTC
ATCCTCTGCGGTACCACGACCTTCACTCCTTGAATCCATAGTCCTTCTTGCCGTTGTGTTGTTCACTGAAGTGGA
GCTCCTAGTAGCTCCCGATCATACCACTGTATGATATAGTCGCTCACGGGACTAGATTCGTGTGTATTGCATTGC
CACGAATTGTTCCACCACGATCCGCTGCACCATGTCGTCTCCTGGTAACATCTTCATTGCATTCTGATCCTTATG
GGATGAACTCGAATTGTAATACCTCAATATGTGGCCTCTGCCAACACATCGTAGGGTCTCTTCCACCTTCGATTT
TGTTCGTTCCTTTGGCAATCGACCTTCGTCCCCCCCGTTGTCGGGTCACACCTAGACAAAGCACTGCTCTAGGAC
AGCTCGTCGCCTACTAGCTCTCGAAGTCCACCGATTTCGTTGAAATTAGTGCACTATTGTTTGGATTCTAGGCCT
CTGCCCCTATCAGCACAATCTCCGCTATGCACCGCTTCCTTCATGGCAACACTGTAGCATATTCTTCAAGAGTAC
CCACCTCCGCTTTCTCTTACCCCGTGCCAAGGTCTTCTGAACTCAACTTCGCCTCTGCAAGTTGAGTCGCCTTAC
TTCCTCCATCAAATGCTTCTCCGAGATAAGATGCATGTGCCTAGAAGCTCCCTTCGTCTTTGGCACCATGCAAGA
TGAGTCTGCTCCATCCGAATGAATGACCCATATAACGACATGATCCCGCTCTTGCCTCTGTAAGAGTTCATGTCC
TTGACCCTTGTTCAAGGAAAGCTTTGTGCCTCCACTCCATGTTTCATCTTCTATGCTGGCTCCCTTCATGCGGCT
TGGGTACTTCTCTAAGTTACACCCAAGTTACTCCGCTCCTTGTTTTGCATTGAGTTGATGGTGGCTCTCGCGCCC
ACCATTCATCACGGGTCAGCCCTCCCTTGAGTCTGATCTCCATATCGACTCTAAGTATGCCTTCATTTGTATTGC
TTTGGGTCGCTCCCCCACTTGATCTCGCAATGCATCCACCAATGCATTCTCTCAAGCGAGATCATGCGACGATTT
CTCGCCGCTCACTCGGTTCATTGAGCTTTGTGGAGTTGTTGTTTTTTAGTTACTCCTCCTCAACATGTGCAGTCA
GTTACACATGATTCTCCCTCTAGAGAGCCGAGACTTATCCTTCCCAAGTATCTATCCTGTTGGAGCAACATCTCT
CTTCGTTTCCTTCCTTCTAAAAGCTTCGGAGACCACTATCCCCTTGGACTACTCCGACTTGTTGAACAAACTGTG
CATTGTTCTGCCTCTTGCAAACGCACTTGCTAGATTGCGACTCTACGTCAATACAGCCCCCGCTGCACTACTCAA
GGTCTAGCAACATACTGAACTCACTACACACTTCAACCTCCTATGGACGTATCCTTCTCGTGTCGACGAGAAAGT
TCCAATGCTTCATGGCACCGAGTTTCGGTCGCCTTGGCATGGCCACGAACATTCCATCGTCCGCATACAAGCCCT
TGCACGAGTACTGAATTCTTCGAGTTAGTAATTCCCCTTACTTCTGTGAGCTTTGCACAACTCTTTCGATCGTTG
AGCAACCCATTCCACATTGCATGGTCTCATCCTTTACCAAGCGCCTCGCTTGCCTTGAGCACCATCATGTATGGT
TGTCAACATTGAGCCGTAGCTCGAACTCAGCCATCCCAACATTTGTGCGCTACGCATTCTTTCAAGCTTGCTTGT
TCTCGTAGTGCCTCTTGCGCGAAGGGTTGGTCATTCCTCTAAATGTCAATCTCAGATGACCGCTCCTTCGAGCGA
CTCATTTTCCCTACATCTCCATGCTCGTTTACCCCCAAACAGTCGTGCGTGCTGACTGCCCTCAACGTAGTCCCG
TTGGGTCCCCCACATTTGCTTGCCAAGTGTTTCTATGAGTGCTTGTCCCGCTCTGATACCATATGTCACAGACTT
AGCTGATTTTGCCTAAGTCATGCGGCACTCTTGCGTGTCCGTCCGCAAAGGTCAGCCTCCCCGAAACCTCCCATG
GTCCCTTAGGACCTACAAAAGAGAAAACAAGTTAGAGAAAGCACCTCACTCGGGATCCACAAGTAAACATTTCAG
TAAACATTTCATAGACAATGCAAATTACAAACAGACTTTACAAGCTCTGAACGGTTGCACAACAAAGGGTCAAAA
TGGTCCATTACAGATCGAATATTTCTCACAAGTGTCCACATGACACAACCTTTATTTACAAGCCTAAAACGACCA
CCAAACCCAACTAAAATGAGGTTGTTAAACCTTCGACCGTCCCTCTACATGCTGTGCAAAGCATGAACAAATCAA
AATATACGGACATACATAAGTATTACATCAAACATCCTGCATGACTTGTATCAGAATTACTGTTTCTAGCACTTC
AATCCCAGTGAGTACAATGAAATGCAAGGAATTGAATGTGTTGTTCCAGTTGGATAAGCAGTTGCCAATGCTTAA
ACACAAAAGTTGGTAATGCATGACAATCAAGGGTATGTTATGATTGGTGGCAACATCTGTACCTCATTCAGGAAG
CTCTTGGAAGGCCATTGTCTTGAGCAGGTTTAGAAAAAGGCCTCAGCTTCCTCTGTGCTATCATAATCTACCTTG
CAAAAATGTCTGGACAGCAGCTTCCACTCTGCCTATCAAACTTGGCAATTGCTGTCATCCAGCATTAGGTCAGTG
CTCATCGATGCATAAATCTTATTATAATACAAATGCACTAAGTCGGTGCCCCAAGCTCATGCACTTCATCCTACT
TGACCTACATCCATAATATCTATATGCCAAATAGATCATATCTTCTTGAGAATGGAAAATGATGTGTCTTACTCG

TTATAATTAGACCCTCTAGCCAGTTTTGATATTCTCCTCAACTCATCAACAGATCGATTATCAAAAGGCCAGGTG
CTTCTTGAAACAATGACCACAAATTGCTGTCCACCATTATGAAAAACCATACCTCATAAATCAAAGTATGCTTTT
AAGGCCTATCATTGACCATATATAAAAACATGTGTAAAACTTCTAAAAGTGCAAGAAGTTTTCACATGATATATA
GATGATACATGCATTACCCATAGATCATGTACATGCAAGTCAAAAACTTATAAACGCATGAAGTTAATCATCAAA
TAAACCATACTTTATTGGTGTAGCTGTTACTACCACCAAAATGAATGTCATGTTGCCTAGAATGATAATCTCATG
TGGTTAGTGATCAAAATTATAATTTTGAATAACTCCTAACTGACATTTGTTCTTACGACATTTGATATTTTGATA
GACTGTGAGCCAATGTTGAATCAAACAAACAATAATTATGGAATCTAGGTCAGTAATTATTGGTAAATGCCAGCA
TTTGGTATAGTAGTGCACATGCACTTGCTATAACAACTAATATGGAAATATTTCTGAATGGTCTGACCATGATGA
CTGTGATCAGTAGAACTACTATGAATATATAGTTGTTGGTTGCCATATAAAGACTCTTATCAATGTATAGATGTT
GTTTTTGTTTCACATGATAGACAAACCACCAAAATGCAGCCGCAGTAGAATAGTAGTCAGAAAGTCTAAACTCTA
AAGCATTAATAAAATACTACAAACTGAGTAGAATAAGATTAAAAAAATGACTATAGTCAACAAACAATAGCAGAC
ATCATCAACAAGTAATCTAAAAGAGTTTTAACAAAGTTTCTAACTAGTGATTCAAGGAAAGTATTCTTCAGTGTT
TTATGTCATATGTTACTACTAGATGTCAGAACAGATAATAGTTCAGTTGACATGGTGCTTGCTTTCGAGTCTGGG
TAATTGTAGCAACATAAATCTAGCAGTACCAACATTTAATGGTTAAAATTGTTCTTTGTACTCACAGTTCTGGCA
AAGGGAGGCAACAAGCTTGAGGCAGCGACTGCATAAATTGCACGAAACTCATAGGTATGCTCAATTCACCTGCAT
GACATGAGCATAGTCGCTTTAAGTAATCATTTTGTAAGTTAGTAACTTGTAGCATCAAAAAGAGAATTGTGATAA
CATGCTTTGGTTTCATTCAAAGAATGATAGAGACAAAAACGAATCACAGATTGAGTTATCTCTAACATGAATGTT
CCATGCCAATTACCACTTACAACCAGATTGTCCTTAATTATGCATTAATGACCACATCCTTACACTGTTTAAACT
AGTCAAAGAACAATAGAAACTAAAATGCCTTAAAATCAAACACTTTGACTTGATTTATTTAGAAAGGCAATAGAT
ACAAATAAGTTACCCCAGAAGTTCCAACATTCACATTTATTCGTTGAGGACTTTGAAACCTGATGGAGATGGATT
TTGAGATAAGAAGAAACTATACAGGATTTAAGCATATAATATAGTTATTTACAACATTTTATGACAACCATACAT
GGTAAAAATATAGACAATAGTGATCTTGTCTTCAAAGTAATATATGACATCATATTAATAGAAAGCTTTGATCAA
TATTTTGGCAGTATTAACATCTAGACGTATATATATATATATATATATATATATATATATATATATATAATCTAATC
TTGTTTTCACATTGACATTCTGTGGTGAGAGAAAAATAGCCTTAATCATATGCTAATCTGTAACTAAAAGATTGC
ATCTGTGCCACTGAGAGAAGAAATGTGCAAGTCAAACTAGTTACTGCTGCCAAATAAATGAAACCAAATAAAGAG
GGGGCTATTATAAGAAATAGTAAAATCCTTCTTCAGAATCATTTGAAACTGAATGAAAGAATGAAACCAAATAAA
GAAGGAAACACTTCATATGCTACTGTTGCAAGTCACTTTACTATGGATGATAACCCAATGCTAGAATCCTATTAG
TAAAAATGAACCTAGGTCCTTATGTTTTTCTGTAGTAGTCAGTCAATGTAAAGAATGCATTGATTGGCTAAGGTG
TTTTGATTAAAACAATAACTATTCTTTATGTCAGCTGTCCCTGGTTTTCCCTTTCTTTCTTCTTTGTGTGATTTT
GTTCCACTTCTGATTTCTTTCTTCTGCAGTTTTCACACTGTATTCTTCTATTACCTTTGCCCACTAAAATTCTCA
AGTCAATTCCTACAACACATATTAAAGCCAATCATCAAACTATTATGATTGATGACAATGGCCTAACAATGCAAC
ATCTGGATTGTCTTCCCTACAAGACATTGGGTCCTCTGCATGCATCAAAGTATACGAGAGACTTATTAACAATAA
TATTTTTAACAAGAAACATCAAATAATAATTGTCTCATGACGACAGCAGTAACAGCAATAATAAAATAAATATGG
TCAATGGATTCCTAGTATCTTATAAGTATATTTGTTCTGAATATCCTTGTAGTTTCATATGGATTATACTAATTA
TATTTAATAAAAAGAACTTGAAAGCTTGAATAAAAACACACATCTCATATCTTTTTCAACACTTGCAGACAGTTG
ATGGGAGAACTTTCTGGTCTAGACATAAAAGATCTACAAAATATAGAGAACCAACTGGAGACGAATCTGGATGGT
ATCCGAAAGAGGAAGGTATTAGTTACATTAGCCCTAAAACTGCTCATACGGTATATCTGTGCATACTTATTTCGT
TGAGTTTTTCTCTCTACAGGAACAACTTCTACTCGAGGAAATTCAAGAATTACATCGGAAGGTTGGACAAACTAC
TGTCCCTATTATGCTCATTTTGAGTTTCAAATGTTGGACTTTGCACTAATGTCATTGCAGGGGAGCCTTGTTCAC
CAAGAAAATATGGAACTCTACAGGAATATAAACATCATGTGTCAAGAAAACACGGAATTACACAGGAAGGTACAG
AATTTTCCAAAGAAAACTGGCAAGCAAATTATTCTGACTGATGGTAATGCAGCACAAACTAACATACAATCATTC
ACTCAGGTTGATGCAACAAGGGAAGCAAATGACAAGAATTCAGTTATGCCAGACGGATTTAGTAAACCAGAGGAA
GCAAATGTACTCATTCATCTCCAACTGAGCCAACCACATCAGCAAGCCGAAGGATTACAACTGAAAGCTCCAAAG
CTAGGGTATTTTTTATTTCCAATATACAGATTTCACAGATCTAAACTACCAATTCAATAAGAAGAAGTCTAATAT
ACTTTCTGGCAGATTTCAACTGAACTGAGAAGACATAAAGATCAAGTACGGCTACTCAATGTCTTCGTCACATTA
TCACAAGACATATCAGCATTTTGTTAATTGGTGTATAAGGAGCAGCATTATTTCTCAAAATAGTAATCCATCTTG
TGAATTGACCAGAATAACCAATTACACAGTGAAGGTATGTGCAATTAGAATATTAAGATTATGTCAGGTGCTTTA
GACAATCTTTTTTGCTTAGTACACAATTAACAAGTATGCAAACTGATCATTCATGCTGAAAGGGTGTTCATTTGT
TCACATTGAAAGGTAGGAAACTCCTTGCAAAACGAAAGAAAAGAGGGAGATTTCCGACAACAACATAATTTGTCC
TTTCCAATTTGCTCCAGAATCGGCGCCCAAACCTGGCCCTACTGTCCAATCCGGTCTGGTTCTAATTTTTGTTCT
TCCAAAACACAGAAATATCTATCCAACCCAAAACATCGTGGTTACAAAAATCACCGTGTTTATCCAAGAGAAGCA
AAATGGAAAGTTCAGAAGGGGAGGTATGCTCCCCAAAATGCATGCCAATGATTCAACCTACCAAAATTTTTTGTC
GCCCGAAGGGCAACCTTTACCCAAAACACTTGGAACCCACCGGAGCATGGAGTGGAGCTCTCACTCGCGGCCACA
AATTCTACGCCTGCAAGAATAAAATAAGGGCACGAAGACGCACTTCTTAGTTCGTCGTTTTAGCAAATTACCTGC
GTATCCCTCATTTCGATGTAATCTTCTTATTTATCTATATAAATATAAATTTAGGCATACCAAACATAATTTGGT
ATTAAAATAGTTTACATTTTTCAATCAAATTTGATGTTGTTAAGAATGGTGAATCTGAAATTTATTGCTCGATAT
TAATTGGAAGAACATATTGCATGAAAAGTTATATGAACAAATATGCTATTTTCAATTTTCCAAACTATTTCAGCA
GATTTAATATCCTGTACCATTGGGTTCGGTTACTCATCCGAATACCTTGATGGTGATTGGTGAGTGAGGTGGCCC
CAGCCGTGGTCCACCGGAACCTTTCTCTCTGCCGCCGAATACAGACCAGAAGGCAAAACGTGTCCAACACAGCAT
CGTTTGAGTTCTTGCAGTGCATTGTGAATGCAAAAAGATAATGGAAGTTTGAATGAGGACGGAGGAGATTGAATC
GTCCAACTACTTATGAGGTCCCACCACTGAATCCGGAGAAAATTGACCAAATCTTACCATCTAGTGATTTTTTTA
TTATTTTTTTTGAAGGTTTCAAACATTTGACTTTCGGTAAGCCGCATACCAAATCTAATGTATCATAAGATTAAT
ATTTAAAATAAAAAAATTATTTTTTTAATATATGATTTAAAAATGATTTATGTTATCGAGGTGGTTCGATTCTTT

CTCTGTTCTTCTGCTCCTTTTCTCCATCATAGTCTCTTCAACTTGAATCTAAAGCAAATTCTATTAGGTACAACT
CGTTTACATGGATGCAAGGCAAAGTGAGCACGAAACTGAGCTGATCTCCAGCTATTGTGAATCGCTTGGTTCAAT
CATTAGTCCTATGCATCATAAACACCTCAAATCACGCATTTCCTTGAATTAAAGAATATGCCTACTTCAGTCAGG

>MaAGL17_4
ATGGGGAGGGGAAAGATCGCGATCCGCAGGATCGATAACTCCACTAGCCGGCAGGTGACCTTCTCGAAGAGGAGG
AAGGGTTTACTCAAGAAGGCCAAAGAGCTCGCCATCCTTTGTGATGCGGAGGTCGGCGTTGTTGTCTTCTCCGCC
ACCGGCCGAGTCTATGATTACGCCAGCACCAGCATGAAGTCTGTGATGGAAAGATACACTAGAGCAAAGGGGGAG
CATCATCTGGTTATGAACCCGAGTTCAGAAACTATGTTCTGGCAAAGGGAGGCAACAAGCTTGAGGCAGCGACTG
CATAAATTGCACGAAACTCATAGACAGTTGATGGGAGAACTTTCTGGTCTAGACATAAAAGATCTACAAAATATA
GAGAACCAACTGGAGACGAATCTGGATGGTATCCGAAAGAGGAAGGAACAACTTCTACTCGAGGAAATTCAAGAA
TTACATCGGAAGGGGAGCCTTGTTCACCAAGAAAATATGGAACTCTACAGGAATATAAACATCATGTGTCAAGAA
AACACGGAATTACACAGGAAGGTTGATGCAACAAGGGAAGCAAATGACAAGAATTCAGTTATGCCAGACGGATTT
AGTAAACCAGAGGAAGCAAATGTACTCATTCATCTCCAACTGAGCCAACCACATCAGCAAGCCGAAGGATTACAA
CTGAAAGCTCCAAAGCTAGGATTTCAACTGAACTGA

#### Arecales

##### Phoenix dactylifera

*PdaAGL17_4 and pda-miR444*

>NW_008246694.1
GTAAACCAAAAGCGGAATCGCCAAACCACGGGCCAGTTGGGACCGTAGGACGCACGTACTCGTCCAACCACGCGT
CGGCGATCTATACGGCAGGCGGCGGGTCTCAAAAGTCTCAATAAATCTAAAAGCAGTCGACGAGCGAAACGAGCT
CCGTCGTTCTGCGGACCACGAagaaaacataaacaaaaataagCATGATAAACAAGTGTATAAAGAAGGAAGCGA
GTGCAGAGCGGGGGCTCAGAATAGTTGGGTTTCatatggatttttttgtttttggtgaaCATATATGGATTCCT
TTCGCCATCCCAAACAATCGCCTCGTCCCTCCGTCAAAGGGAAGGAAAGCAGGGCCAGGGAGAGAGGGAACCAAG
CCCGCCTATTGGCCACCaccctttcctttcctttctatttttgtcTATAAAATAGTAGACTTGATAGGAAAGGC
ACCACCACCCACCACTTAGGCCAAAGCCAAATACAGAAAGAGGCTCAAACTCCAGTTGTCTTCCCGGAAATTAAG
GTTACAATCTCTCCACCAGATCTTCTCTCTgaccttctcttcctttccttttctctcccaCCTcgttgtctctct
ctctctctctctctctctctctcNNNNNNNNNNNNNNNNNNNNtctctctctctctctctctctctctctctc
tctctctctctacccctATCCTTCTGATTCCTTGTTGTTAGTTTTCCATAGCCAGATCTGATACATCTGATACTG
AGTTTTTGGTGGTTTGAgtgaaggagggaggagaagaaatatGGGGAGGGGAAAGATTGTGATCCGGAGGATTGA
CAACTCCACCAGCCGCCAGGTGACGTTCTCGAAGCGGAGGAACGGGCTATTGAAGAAGGCCAAGGAGCTCGCCAT
CCTTTGCGATGCCGAGGTCGGCCTCATGGTCTTCTCCAGCACCGGCAGGCTTTACGATTTTGCCAGCACCAGGTT
CGTGCTTGCTTCCTCTTCTACAAACTCTCTATGTCATCCTACATGATAGATATGGTTTCAATTCGATCAGTCTTT
CGTAAGGTCTTGTTGAGGTTATTGCATGGATGGACTGGGAGTAAGTCTAACCTATGGAAGAGTATTTTTACATTG
AAATGATAGATATGGATGGGACTTATCGATCATGTTAGAAAAGGTGGTCGGCTGCATAGTTTTGATTATCTTTGT
TGCATAGATGGTTTAGCTGATATGTTTTAGAGAAAGAACTAGAGAAAAGAGGAATGGTGTATCTCCCCCAccact
tctctcttattttatttatttcattttatttttttcgtagAAAATATCACTTCTCTTTTATAGCTTTcttattct
tagttttaatgTTAGCAAGAACTATTgtcatatgtttttttttattgtaacaTTGTCATGTGCTTTTATATGCTCA
TGTCGTATATCTCTCTTCCAATTTCTGTTCTCTCCGCATAGATTTTCATTAGAcaaactctctccctctctctct
ctcgctccccaCTCCATTCTACTTCATTTCTCGGATATTCAAGAGTGTATTTGGtggatttattttcttgtcttc
tATACTTACATGTTATAAGGCATGATATTGCTTCCTAGAGtactaaatcaaaaaaaattgcaagttGATCCAAGG
AATGTCAGTTGTAAGCCTGTTAGGTGCCTGAAGGTTTTTTCGAGTCTGTAAGAACACTGAAGTGggatagggaaa
aaaaaacaatataagaaTGTGGCAGTTTCTCATGCATCTGACACTTCTCATCTTTCCAAGTTTCTGATTTTCCCC
AATCAATCCAAAAATTATCAAATGATAAATGCTCCATGCACGTTCATCTTTAACCAAAGCCACTGATCATACCAT
CTTCTGCTTCAGAACACTGTTGGATGTGACGAAGGGGCACTGTTAAGGAAGATAAATAATCATCCATTCACGTGC
GCATTTGTGAAGTTAGGGTAGCGCTGAACATATATCGAGAATAACAAGGACGAAGTCCAGGGTAAAACTactaga
attaaatatttacttCTCACATTATTACTGTAGGCAACGAACATGAGGACGCATGTATTTTTTGAAGTGTGtga
aatgttattttaataaaacTGAACTTTTAGATTCAGAAGTGGAAGGGATGCTGAGGGTTAGAAGCTTGGATATAT
ATATGGCTATCCTCCAAAGGCATGCACCGCAATTGCTAAGCCGTAGCTGGATTATCATTGTTTTTGTGAATGCAT
CCTATTTATACAGATATGGTTTTGTCCAAGGAAGAGAGACAACCAATGATGGTACTGAACGTGGGAGTGGGGCAG
GTCTGACGTGCCTTGGACTATTTGATCATTTGATGCAATAACCGCTACAAGTTATGCCATCAGTATATGAAACTT
GGGAACAAGTTAATGGCTTGAGTCCAGTCCTGATCTAGCTTACTTTAAAGTGGGATGTGTAGCGGGAACCATATT
GGAAGATTAAATAGTCTAATGTATAGTCTAATGCATCATACTTCCCTTGCATCCATCTCACTATCAGATCCGATA
ACATTCAGAATATATGAAAGTGCGCAATAAAACCTCCTTCGAGAAGTACTGCCTACACAATTTTCTGTATGTGAT
GTTGATGTAACATTACATAAAACTAGGTAGTCATACATTATAACTTCACAAAGCAGTCTATAAGTAGCTGTTTAG
AGTTTCAAGATAAAACAAACGAAAAAGTGTCGTTAGGATCAATTGAGAGGAAACAGTATCATTTTAAGGTTAATT
ATTAGATTCTAATTGGAGAAATTTGTTAGTTTGTAAGGTATAGGTTaccatcattatttttgggtCTGAAAGGT
TGCCTTCATTATGACTGAAGAGTTAAACTAATTTATTTCGTCTTGTAGCATCATTCAGATGCATGCTCTTGATGG
AAAAAAATCGCCTACATATTGCGTGAACCTTGGAACTTCCTCTTGATATATACTTTTCTGTCTCTCCTTTTGATT
GTTGAAAGGAAAATACAATCTAAAACTACCTTCAAAAGACAAGAGAAGGAGAATAAACCCAAGCCATATTCAACT

TGGTATAGGAAAagtgtaataaaaaaattaccattTGAGCAACTTTGGTAGCATATATAGTTAACTAAGCAATCA
GTGGTTTAGTTACTTTCTTTGCAAATGTGAAAGTCTATATTTAGTTCAAACTACAAGATGAAAAACCTCCATCCG
TGATCTAGAAAAGATGATTGCTGAATTTGCTTCAATTTAAACATCCTAAGTATGTAAGACATTATCGATATATTT
ATAGCTTTCCTATTCTCAAAGTTTAACCAATGGGATTGAGCAGGTGAAGAGCTGTTTGGAATGTGACACAAGCAT
TCTTCCTACATGATCATGAATAATGATACAACATATATTGTTGAGATATGTCTAGAAGAGTTAGTGGACATGATA
ATAACTAGattctctttgtttttcatattctatgggttttgcatattttttttgatagaaaggaTAATCATTCTA
ATCAAGAAGGGTAGTTTGTTACTTGCTTCCATGTATCAGTAATAAAATCTGGTGAATCTTCAATTCAAGAATGAA
GATGCTTTGTTGGTTTATGCGCTGTTCAAATGCTTTATCTTCTAACATGACAAAATatgcatttttgaaaaaaaaa
tccttaaagctTTTGCATCCTCAAGAATGAAGGATGCAAGTAAGGAATCATATAGTGTCCACTGCATTATGAATT
CATGATGCTGATTAAGTCTCCTTCTAATATGACATCGTGAAGTCCCAATGCGACCCGGCCCATATAGAACATTCC
CTGGTTGTAAATGCTTTGATACATTCTGCTATTGTCCTAGCTCCAATAAGCTCCCCTTTGCTATCTCAAACTACA
CATCCAGTATCAGCTTTTCCATCTGCAAAAAAATACACCATCAGCATTGATTTATATACACCTTTTGGCAGGGTAG
ACCATCTTTCTTGTATTTGAACTTTTATACTAGTATTAATCTCTTGACTACTTTGGAATTCTTATAATAGCCAAC
AAGCTTTATTAACTATTTTCCATGGCTTGAGCTTTTATCCTCCAAAAATGAAGTCATTTATGCTTTTCCAAATTA
CCCAAGCAATAATAGCAAATGTAGTTAATCCTTCCATTTCCATTTCCAATTTGTTAAAGCTCTTATACCTAACCC
AAATGCTTTCAGCTTTGCTTCCCTTGGTTCTCATCTGGAGAGATGCAAGACTCCAAACTTGTAGTGCAAactaat
aatatcaaaaaattccACTCATGTATCATTCTGTTCGTAATATAGTTCACATAATTCATCCTTCAACTTGATTTT
CCTTTTACTCAAATTTGCTTTTATTGGAAGGATATTTAATTTTGCGAGcatatcacataaaaaaaattgatttttc
tgGATTATGTAGCTTCCAAAAATTACTATTCTCTACCAGAACTATGAAAATTAAAACTACTTGCCTTCTTTAAAT
CTTCTTGCTCCAGTATCAAGTGAAAAGCACTCTTAACAgtgaaaattctatttttttgtaTCATGTCATATCGATC
TATGATCTGTTAATCTATCACTAATTAGGATTTTAAGAATGTTCTGTGCAAAGAATACCTGATTAATAACTGTCT
CGTTATAGCACCCTCTATCCTTGCCCATCAAAATACCGACTGTAGCTTTAGGGCTCAAGATGCTAACAGGCCCCT
TGGAAGGCTGTTTTCACCGATAGCCATTTATACTTCCAAATTTTTACCCTATTCTCATGTTCAATTTGTGAGCAT
ACACCTGCTTCAGGCAACGGTTTTCTATATTTGAAACAACATAGAGTTCAACCGCCGTAGAAATCTCCATTCTTT
TTTTGCTAGTAGTGCTCTATTGAGAGCTTGTAATTCTCTGAAGCCacatctcctttctcttttaCTAAACAAAAT
TATTCCAGTTTATCATTCTAGTTTATCCAATGAACCTTCTCACTTTTCTTTTGCCCCTACCAAAACTTTGCTAGT
AATAAATTTAACTCCTTGCATGCTTGTTTGGGAACAAGAAAACAACTCACTACATATGTTGGAAGTGCTTGTGTG
ATCATcttcaataaataattttctctttgatttcccCATCATAGTTGGAAGCCCTAAATAATTTCACTCCTATTA
AATTGCTAATTTCTTCCTTTACATCTCGTGGTATCTTCTGCTGAATGTGATGGCTGACTTATCCAAGTTGATCAT
TTGCCCCAAAGCCTTCTCATATCTCTATAACAAGTTCAATATGTTTCTCGTTAGGATTAAGCCAAGAGAATGGGT
TGTGAAGAACTCTCATTATATCTTTTAGTATTGTATTAGGAGAGAATCGTATATACATGGTGGTACAAAAACATG
AGGTGAACTTACCACCAAGGTTACATGCATGTAATGTATACCCTATATACAATACCTAATACTCTAATTACACCC
CTCTCAAACTCATGGTGGGTTGTAGACCATGAGTTTGGAAAGAAGATAAGATGTTCGAGCTTTCGTCTGAGGCTT
AATTAGTAAGTAAATCAGCAGGCTGAGCCTCGGAAGATAATGAAGAGAGAGTGTCCCTGCAAGATGATGATAATG
GGTAAAGAAGGCATCAACACCAATTTGCTTGGTGAGCTCATGGTTGGTGGGGTTGGTAGCAATTTGAATGGCACT
GGTATTACCATAGTAGAGAGGAGTCGCCTCAGAAAGGATAGCACTCATATTTGCAAGAAGCTAATGAAGCCAAAT
GATCTCAGCAGTAGGAGTAGCCATAGCGCAGAGTACAGCTTTAGTAGATGATCATTTACTGAAGTGCCAAGAAAA
AATACATTGGAAAAAGCAGCCTTGAAAAATTGCATGGATCACTTGCCCAAGGAGCATTGGAAAAATTATAAAGCT
GAGGCTAGGAAGAATTGGATAAAAGCAAAGCTTGTGAAATGGTACATTGAAGAAAACACAACAGGTGTAAGAGAT
GGAAATAGAGAATTGTTGTGGAGGCTGAAACAAATTGGCTCAAAACATGCACAGCATGAACAATATATATCTAGC
CTGGAAATAATGAGATAAATGAGACTACCTATAAGATATTGATAGAGAGTACGATTAGAAAGAGGTGCACCAACT
CTAGGCTTTAAGTTATGATGGAGCACCAATGGAGTATTAGCAGTGTGGCAATCGACCAGCTCTAGAAAGAAGGTC
GGCATTGTATTTCTATTGGAATAAGAGGTAGCTCTTAGAGCTATATGCAACCTCAAGGTCCAAGAAATCCCAAAG
TGGTCATTGCAAGTTGCTGACTAAGATACTGCATAACAAATGCAATATGCTTTAGGTCATCCACggtaataatta
tattatcGACACAAAGCAGTAAAAAAAGTCTGACCACGTGAAGAAGAATTGACAAATGTGGTAGGATCATGATCA
AGGTAGGCCAAACTAGATCGAACTATTTAGTTCGGCCTATACCGAGCCAACCCGGTGAGAAATTGGGTCGGTTCG
TCTATAGAAATCAGTTTTGGCCCCGAACAGGCCGGAACATGTCTTGACTCGACCGAAAAAGACTCAAACCGGACT
GAATTGGCCGAAACTGATCTGAATCGGTCGGAACTGATTGGAACCAGCCAGTTCCGTTCGAACTTGGTGCGAACC
ATCCGCTTCGCACCGAGTCAATCCGGTTCGGAACTGGGCCCTgcccctctctttttttgttaaaaataattgTTT
CGGCCTTTCCAATAGTTTATTTTGAAAGTTCGTTCTAAACGACTCCCTTTTTTTACCAATCTATGAAATCACACT
GATTTAGTACGATTGAAGAAAGATTCAAGCCCAAATAAGATGTGCTTCTTCTTccctatctcatttctctattt
ttttgtttgaaaaaatagctcaaaaattataaaataagaaaaaatcttgccaaaatttttgatttggcattgatt
ttattaaatattgtagatctatagatgatctacacgatgacataaaaattttttaattttggattatatataatt
tttaaaaatttaaaatatactttggattaaaacataaaaaataaaaaaataaaaaatcagaagTATATTTAGGA
GCATGTAAGCTACTCTTCGGCTATATTTTGTGTCTATTTATTCAAGTTGATGCGATCTCGCGCATAGTAGCCgag
gcctaaatttgaaaaaaaatttaagaaataagtagcctttaaTGTATGCCTATTTGGCACTAGAGTGTATTTTTCA
GTGTGACTATTCAGGGCGTGCTCACGGCCTTATGTAGTGAGACGccattacaccaaaaaaaaaaaagcatgccta
catgcctagaaaaatatatgtagcatctatTATTGGATTCTACGTGGATCTGagctgaaattattttttttaaaat
atttacatttaaaattatttttaaatttaaaaaataattaaaaatatataattataccaaaacttatgaaaaatt
agtagtacattGGAAGTAAtttatgaagtagaaaacatatttttctgaatttatgatatttaaacatattttaaa
tttaaacattgttataaatatataaaagggGGTATTCATGTTTGGCATATGCCTGCCCTGACCGAAAGATTTTGG
gatgattaataaaaaatatgaaatattcatattatgtattagataaactACATGTATTTTCTGaggtaaaaaatt
atctttatcgtaacaaaattttggtatttttaaataattgaaaaaCTAACATACATGATAAACTCACAGAAATAG

AACCATTAAGAAAAAAGGAGCCAGAGTATGATATTGGTTGGAATCATGGGCAAATGTTCTTAGGTCGGCATCATT
GAaagtgcaactggtgcaacaTAGAGTTTAAAGGAGGAATAATCATgttgaagcagcacttagctggTGGTTACC
TCGACGTATCTATGTGCCGGAAATGCCGTCAGAAGGTTTGGTAGCGGATGAAGAAGCACTTTGCCGATTTCAGAG
CAGCGAAGGAGAGAGTTTCTAAGAAGAAGGCAGAGGTGGACTCCGAGCAGCAGAGCCACTTTTCTATTACTCTAG
GAAGTTAGAGAAGACATCCGCTCTAGATGACGAGGAGGCATAGATCTAAGCTGTCATGCAGGCGATCCTGGATGA
CAGTGACAGCAGGATGAGATAGCCAGGCAAAGGGCTTTATTAGGCCCCTCATACTACTAGTCAGGCTCTGGTTCT
ACGATCAGCAGGGCAAATCTAGAGTTCAAAAGGGCCTCCTCAGTTAAAGAGGCTGTCAGCAGAGGCGATGGCATA
TTGCATCTATGTTGGaagcttttggtagcaaaaacaAATCATCCAGAGAGATTCCATCAGGAGTATTAATCCATA
ATTTAGATCCATATGCCTTCGCCCGTAGAGATTCAAAGCAGTGAAGGATTGACAcaatgctgaagaaggataaCA
AGAaagatatgtggcgagctatttgATCATGATtctacttcagccacatccTAGCGAATGTGACAAACAATATGT
ACTATAGATCTGCTATTTCCTCCATATAGGCTGTCAGTCCCGATGTAGATCCTCCAGGCCCGAAGAACATCTACG
GTGAGCGTTTTGACAATAATAAAGAGGAGCTAGAGATATGGGTTGCCTCACACAAGAGTAAGTGACCCACATATG
GACTGACAAttatgtgtgatggttggaccggtcctacCAAATGGAGTATTATTAACTTTCCGACATATTGTGATA
CAAAGACTTTCTTCTACAAGTCAGTTCATACTTCTGATCAGGTGCACGACGCTACGTACATCCTTAAACTGATAG
AagaggtgattgatcaggtaggagagaagaatgtcgtgcaggtcatcactgataatgAGCCATAATATAACGCCG
CCGgaaagatcttgatggagcgacTGTCATATATTTTCTGGACGTCATGTGCTACACATTGTATCGACATCATGT
TGACAAATATTGTGAAGATTCATAGGGTGCAGCAGACAGTGGAGACAGCTCAAAGTATCACCAGATATATTTCTA
ACCATACGTGGGTTCTTTCACTAATGAGGAGGTATGCAGGGTGAGAGATCTTGAGACCAAGAGTCATatggtttg
ctaccaactacgttgcacttgatagctttcttgagaagaaagcGGTCCTACGTCAAATGTTTGTTCAATTTCGAG
TGgtagaaaagtagatatgctAGAGCATTATTAAGGACTTGGTGATGAGGCAGACATTCTGGCAACAAGCACTGC
AATAGTGAAGGTTATCAGACTATTGTATGAAGTGCTTCGGGTTCTAGACAACGAGAGATACCCCCAGATGGGCGA
TGGTGAgagcaaaaattcatatcatggaGGCAGATCCGATGTATGCTTAcgagtacatcgacatcattgagcggc
gATGGGTCTATTAAATGGATAGGGACTTGCATGTAACGAGTAAGTAggaacattgaaaattagactgctcctgta
cttgtacaattttactagACAATAACGAACTCTATGTGCAACATACTATTTGAACCCTCAGTTTTAGTATGCCAT
AGCTGGAATCGAAATGGATGATGAACTTCTGGCCGCTCTGCATAATGCGATTTATAAGATGGAGCATGATCCAAA
AGTCGCCGCCCTATGTCTAAAAGAGGTAAGTTAATTTAAGTGACATGTATAAGTTAACtagtatttttaattatt
aacatattacaatACAATTCTTTTTTATAGATCAAAATATTGAGAGGGAGCACTGATAGCTTTGGAGTTTCGACA
GCTGCCGTatgtaagaaaagtatgaatctaGATATATTACACATCGGGAGGATTTTTACCTGGCcatagatggtt
actaatataaCAATATCttatatacaatttttttttcaatatttttacagTCGAACGATGGATCCATTTTGGTTTG
TCCGCAAAAAATCTCAAGCAGTTCgctattcaaatatttttccaaacGGTCTCCTCGAGTAACTGTGAGCGCAAC
TAGTTAACCTTTGCCCTTATCCATAACAAGCAGAGAAACCGTTTGACATAGAAGTGTTTCAGTAATCTTGTATAT
ATTTCTATAATCTGAGGTTGACActgaagtgcattcaagagaaagtgGAGCTAAAGTATAGAAATTGAATCTATA
GTGCTTTTGTTGATGATgtggatgatcctatgatcgaaTGGCTTGTGGGCCAGTAGCAGGAGCTAGAGCTTGATA
AGCGAGGATTGCCTCCGCAACCAGCCAGCTTCATAGCCAGTGAGGCTAGGGTTGATGCAGGACAATGGGGAGAGA
GCAATATTTTATGCATCCTTCAGCCGATTAGCCACAACCATAGAGGAGTCGGTCACCATATGACTCTTTGTTCAA
GACAGCCTCTTAGAGATATAATCGAGAGATGTTTAGACGAGGCCACTATGCTACTCGATCGACTCGGTCAGGAGG
GCGACATACATCCTCCCAACATAGCCAGACAGACAGGATCACAAAGGATAAGCAAGCAGAGAGTgacaaatagaa
taaataggcAGCGAGTGACATAAAGGATAAGAAGATAGCTGCCCagcacaccaaaaaaagaaaataaaagacatc
tACCCAGCACACCAAATGGCCATAATGGATATTGAGTTTGAATTCTTGTAGCAaactttttataaatttggCATA
GTTGAGAGTGAGACTGCATAAAATAAACCCATGTAATGCGAGAATAATCAACAATTAATATGACATAATAGCAGT
TTCCATCTTTAGAAATAGAAGGGGAATAACTCTAGACAACTGGGTGAACTAAACCAAAAAGTGAAATAGCTATAG
AATTACTAGAGTGAAATGGAAGTCTCACATGCTTTCCAAGAACACAACCTTAGATATAGAGCCTATGACACCACT
ATGAATTAATGTGGTAAGATGGGAACCCTAAAGATGGTCAAGACGACAATGATAGATCTCGAAAGTAGAAGTGGA
AACATCTACCAAGGTAGAGGCAGGAGAATTGGGCACATGTGGGTGGTCCATCAGATACACTCCATTATGGCGATG
GCCAACACCAATCAGACGACCGATTTGTTGATCCATCACAAAGCAAGAGGATGCATCAAAGGAGACAAGGCAACC
ATGATTAGTGACACTACAGGAAAACTAACTTTACAAGAACATTTCAATGATCTTTATTATCTCCTTCCCTTTTGC
ATTTTCTGGAGCAATAAACGCATGATTTTTGCTCGCAAGACACACAAGCACCTAATAATTTGAGTAATGCATGAA
TAGTATCTTAGCAGCAGCATATAATGTAGTGAATCTGATCAAACCGAACGAAGTCAAGGGATCACCGCGAACaaa
aaaggggaagaggaaggagagaagatgGGAATcgaaaaaggggaagaacatAATACCTCTAGGGCTCTTCCGGTC
GGCgccgcagaggagggaggagaagagtgTTGAGGACCCCCGCAAACGCCGCCAACCAAGGAAGGCATCGAGCCA
CAAAGAAGNNNNNNNNNNNNNNNNNNNNNNNNNNNNNNNNNNNNNNNNNNNNNNNNNNNNNNNNNNNNNNNNNNNN
NNNNNNNNNNNNNNNNNNNNNNNNNNNNNNNNNNNNNNNNNNNNNNNNNNNNNNNNNNNNNNNNNNNNNNNNNNNNNNNN
NNNNNNNNNNNNNNNNNNNNNNNNNNNNNNNNNNNNNNNNNNNNNNNNNNNNNNNNNNNNNNNNNNNNNNNNNNNNNNNN
NNNNNNNNNNNNNNNNNNNNNNNNNNNNNNNNNNNNNNNNNNNNNNNNNNNNNNNNNNNNNNNNNNNNNNNNNNNNNNNN
NNNNNNNNNNNNNNNNNNNNNNNNNNNNNNNNNNNNNNNNNNNNNNNNNNNNNNNNNNNNNNNNNNNNNNNNNNNNNNNN
NNNNNNNNNNNNNNNNNNNNNNNNNNNNNNNNNNNNNNNNNNNNNNNNNNNNNNNNNNNNNNNNNNNNNNNNNNNNNNNN
NNNNNNNNNNNNNNNNNNNNNNNNNNNNNNNNNNNNNNNNNNNNNNNNNNNNNNNNNNNNNNNNNNNNNNNNNNNNNNNN
NNNNNNNNNNNNNNNNNNNNNNNNNNNNNNNNNNNNNNNNNNNNNNNNNNNNNNNNNNNNNNNNNNNNNNNNNNNNNNNN
NNNNNNNNNNNNNNNNNNNNNNNNNNNNNNNNNNNNNNNNNNNNNNNNNNNNNNNNNNNNNNNNNNNNNNNNNNNNNNNN
NNNNNNNNNNNNNNNNNNNNNNNNNNNNNNNNNNNNNNNNNNNNNNNNNNNNNNNNNNNNNNNNNNNNNNNNNNNNNNNN
NNNNNNNNNNNNNNNNNNNNNNNNNNNNNNNNNNNNNNNNNNNNNNNNNNNNNNNNNNNNNNNNNNNNNNNNNNNNNNNN
NNNNNNNNNNNNNNNNNNNNNNNNNNNNNNNNNNNNNNNNNNNNNNNNNNNNNNNNNNNNNNNNNNNNNNNNNNNNNNNN
NNNNNNNNNNNNNNNNNNNNNNNNNNNNNNNNNNNNNNNNNNNNNNNNNNNNNNNNNNNNNNNNNNNNNNNNNNNNNNNN
NNNNNNNNNNNNNNNNNNNNNNNNNNNNNNNNNNNNNNNNNNNNNNNNNNNNNNNNNNNNNNNNNNNNNNNNNNNNNNNN

NNNNNNNNNNNNNNNNNNNNNNNNNNNNNNNNNNNNNNNNNNNNNNNNNNNNNNNNNNNNNNNNNNNNNNNNNNN
NNNNNNNNNNNNNNNNNNNNNNNNNNNNNNNNNNNNNNNNNNNNNNNNNNNNNNNNNNNNNNNNNNNNNNNNNNN
NNNNNNNNNNNNNNNNNNNNNNNNNNNNNNNNNNNNNNNNNNNNNNNNNNNNNNNNNNNNNNNNNNNNNNNNNNN
NNNNNNNNNNNNNNNNNNNNNNNNNNNNNNNNNNNNNNNNNNNNNNNNNNNNNNNNNNNNNNNNNNNNNNNNNNN
NNNNNNNNNNNNNNNNNNNNNNNNNNNNNNNNNNNNNNNNNNNNNNNNNNNNNNNNNNNNNNNNNNNNNNNNNNN
NNNNNNNNNNNNNNNNNNNNNNNNNNNNNNNNNNNNNNNNNNNNNNNNNNNNNNNNNNNNNNNNNNNNNNNNNNN
NNNNNNNNNNNNNNNNNNNNNNNNNNNNNNNNNNNNNNNNNNNNNNNNNNNNNNNNNNNNNNNNNNNNNNNNNNN
NNNNNNNNNNNNNNNNNNNNNNNNNNNNNNNNNNNNNNNNNNNNNNNNNNNNNNNNNNNNNNNNNNNNNNNNNNN
NNNNNNNNNNNNNNNNNNNNNNNNNNNNNNNNNNNNNNNNNNNNNNNNNNNNNNNNNNNNNNNNNNNNNNNNNNN
NNNNNNNNNNNNNNNNNNNNNNNNNNNNNNNNNNNNNNNNNNNNNNNNNNNNNNNNNNNNNNNNNNNNNNNNNNN
NNNNNNNNNNNNNNNNNNNNNNNNNNNNNNNNNNNNNNNNNNNNNNNNNNNNNNNNNNNNNNNNNNNNNNNNNNN
NNNNNNNNNNNNNNNNNNNNNNNNNNNNNNNNNNNNNNNNNNNNNNNNNNNNNNNNNNNNNNNNNNNNNNNNNNN
NNNNNNNNNNNNNNNNNNNNNNNNNNNNNNNNNNNNNNNNNNNNNNNNNNNNNNNNNNNNNNNNNNNNNNNNNNN
NNNNNNNNNNNNNNNNNNNNNNNNNNNNNNNNNNNNNNNNNNNNNNNNNNNNNNNNNNNNNNNNNNNNNNNNNNN
NNNNNNNNNNNNNNNNNNNNNNNNNNNNNNNNNNNNNNNNNNNNNNNNNNNNNNNNNNNNNNNNNNNNNNNNNNN
NNNNNNNNNNNNNNNNNNNNNNNNNNNNNNNNNNNNNNNNNNNNNNNNNNNNNNNNNNNNNNNNNNNNNNNNNNN
NNNNNNNNNNNNNNNNNNNNNNNNNNNNNNNNNNNNNNNNNNNNNNNNNNNNNNNNNNNNNNNNNNNNNNNNNNN
NNNNNNNNNNNNNNNNNNNNNNNNNNNNNNNNNNNNNNNNNNNNNNNNNNNNNNNNNNNNNNNNNNNNNNNNNNN
NNNNNNNNNNNNNNNNNNNNNNNNNNNNNNNNNNNNNNNNNNNNNNNNNNNNNNNNNNNNNNNNNNNNNNNNNNN
NNNNNNNNNNNNNNNNNNNNNNNNNNNNNNNNNNNNNNNNNNNNNNNNNNNNNNNNNNNNNNNNNNNNNNNNNNN
NNNNNNNNNNNNNNNNNNNNNNNNNNNNNNNNNNNNNNNNNNNNNNNNNNNNNNNNNNNNNNNNNNNNNNNNNNN
NNNNNNNNNNNNNNNNNNNNNNNNNNNNNNNNNNNNNNNNNNNNNNNNNNNNNNNNNNNNNNNNNNNNNNNNNNN
NNNNNNNNNNNNNNNNNNNNNNNNNNNNNNNNNNNNNNNNNNNNNNNNNNNNNNNNNNNNNNNNNNNNNNNNNNN
NNNNNNNNNNNNNNNNNNNNNNNNNNNNNNNNNNNNNNNNNNNNNNNNNNNNNNNNNNNNNNNNNNNNNNNNNNN
NNNNNNNNNNNNNNNNNNNNNNNNNNNNNNNNNNNNNNNNNNNNNNNNNNNNNNNNNNNNNNNNNNNNNNNNNNN
NNNNNNNNNNNNNNNNNNNNNNNNNNNNNNNNNNNNNNNNNNNNNNNNNNNNNNNNNNNNNNNNNNNNNNNNNNN
NNNNNNNNNNNNNNNNNNNNNNNNNNNNNNNNNNNNNNNNNNNNNNNNNNNNNNNNNNNNNNNNNNNNNNNNNNN
NNNNNNNNNNNNNNNNNNNNNNNNNNNNNNNNNNNNNNNNNNNNNNNNNNNNNNNNNNNNNNNNNNNNNNNNNNN
NNNNNNNNNNNNNNNNNNNNNNNNNNNNNNNNNNNNNNNNNNNNNNNNNNNNNNNNNNNNNNNNNNNNNNNNNNN
NNNNNNNNNNNNNNNNNNNNNNNNNNNNNNNNNNNNNNNNNNNNNNNNNNNNNNNNNNNNNNNNNNNNNNNNNNN
NNNNNNNNNNNNNNNNNNNNNNNNNNNNNNNNNNNNNNNNNNNNNNNNNNNNNNNNNNNNNNNNNNNNNNNNNNN
NNNNNNNNNNNNNNNNNNNNNNNNNNNNNNNNNNNNNNNNNNNNNNNNNNNNNNNNNNNNNNNNNNNNNNNNNNN
NNNNNNNNNNNNNNNNNNNNNNNNNNNNNNNNNNNNNNNNNNNNNNNNNNNNNNNNNNNNNNNNNNNNNNNNNNN
NNNNNNNNNNNNNNNNNNNNNNNNNNNNNNNNNNNNNNNNNNNNNNNNNNNNNNNNNNNNNNNNNNNNNNNNNNN
NNNNNNNNNNNNNNNNNNNNNNNNNNNNNNNNNNNNNNNNNNNNNNNNNNNNNNNNNNNNNNNNNNNNNNNNNNN
NNNNNNNNNNNNNNNNNNNNNNNNNNNNNNNNNNNNNNNNNNNNNNNNNNNNNNNNNNNNNNNNNNNNNNNNNNN
NNNNNNNNNNNNNNNNNNNNNNNNNNNNNNNNNNNNNNNNNNNNNNNNNNNNNNNNNNNNNNNNNNNNNNNNNNN
NNNNNNNNNNNNNNNNNNNNNNNNNNNNNNNNNNNNNNNNNNNNNNNNNNNNNNNNNNNNNNNNNNNNNNNNNNN
NNNNNNNNNNNNNNNNNNNNNNNNNNNNNNNNNNNNNNNNNNNNNNNNNNNNNNNNNNNNNNNNNNNNNNNNNNN
NNNNNNNNNNNNNNNNNNNNNNNNNNNNNNNNNNNNNNNNNNNNNNNNNNNNNNNNNNNNNNNNNNNNNNNNNNN
NNNNNNNNNNNNNNNNNNNNNNNNNNNNNNNNNNNNNNNNNNNNNNNNNNNNNNNNNNNNNNNNNNNNNNNNNNN
NNNNNNNNNNNNNNNNNNNNNNNNNNNNNNNNNNNNNNNNNNNNNNNNNNNNNNNNNNNNNNNNNNNNNNNNNNN
NNNNNNNNNNNNNNNNNNNNNNNNNNNNNNNNNNNNNNNNNNNNNNNNNNNNNNNNNNNNNNNNNNNNNNNNNNN
NNNNNNNNNNNNNNNNNNNNNNNNNNNNNNNNNNNNNNNNNNNNNNNNNNNNNNNNNNNNNNNNNNNNNNNNNNN
NNNNNNNNNNNNNNNNNNNNNNNNNNNNNNNNNNNNNNNNNNNNNNNNNNNNNNNNNNNNNNNNNNNNNNNNNNN
NNNNNNNNNNNNNNNNNNNNNNNNNNNNNNNNNNNNNNNNNNNNNNNNNNNNNNNNNNNNNNNNNNNNNNNNNNN
NNNNNNNNNNNNNNNNNNNNNNNNNNNNNNNNNNNNNNNNNNNNNNNNNNNNNNNNNNNNNNNNNNNNNNNNNNN
NNNNNNNNNNNNNNNNNNNNNNNNNNNNNNNNNNNNNNNNNNNNNNNNNNNNNNNNNNNNNNNNNNNNNNNNNNN
NNNNNNNNNNNNNNNNNNNNNNNNNNNNNNNNNNNNNNNNNNNNNNNNNNNNNNNNNNNNNNNNNNNNNNNNNNN
NNNNNNNNNNNNNNNNNNNNNNNNNNNNNNNNNNNNNNNNNNNNNNNNNNNNNNNNNNNNNNNNNNNNNNNNNNN
NNNNNNNNNNNNNNNNNNNNNNNNNNNNNNNNNNNNNNNNNNNNNNNNNNNNNNNNNNNNNNNNNNNNNNNNNNN
NNNNNNNNNNNNNNNNNNNNNNNNNNNNNNNNNNNNNNNNNNNNNNNNNNNNNNNNNNNNNNNNNNNNNNNNNNN
NNNNNNNNNNNNNNNNNNNNNNNNNNNNNNNNNNNNNNNNNNNNNNNNNNNNNNNNNNNNNNNNNNNNNNNNNNN
NNNNNNNNNNNNNNNNNNNNNNNNNNNNNNNNNNNNNNNNNNNNNNNNNNNNNNNNNNNNNNNNNNNNNNNNNNN
NNNNNNNNNNNNNNNNNNNNNNNNNNNNNNNNNNNNNNNNNNNNNNNNNNNNNNNNNNNNNNNNNNNNNNNNNNN
NNNNNNNNNNNNNNNNNNNNNNNNNNNNNNNNNNNNNNNNNNNNNNNNNNNNNNNNNNNNNNNNNNNNNNNNNNN
NNNNNNNNNNNNNNNNNNNNNNNNNNNNNNNNNNNNNNNNNNNNNNNNNNNNNNNNNNNNNNNNNNNNNNNNNNN
NNNNNNNNNNNNNNNNNNNNNNNNNNNNNNNNNNNNNNNNNNNNNNNNNNNNNNNNNNNNNNNNNNNNNNNNNNN
NNNNNNNNNNNNNNNNNNNNNNNNNNNNNNNNNNNNNNNNNNNNNNNNNNNNNNNNNNNNNNNNNNNNNNNNNNN
NNNNNNNNNNNNNNNNNNNNNNNNNNNNNNNNNNNNNNNNNNNNNNNNNNNNNNNNNNNNNNNNNNNNNNNNNNN
NNNNNNNNNNNNNNNNNNNNNNNNNNNNNNNNNNNNNNNNNNNNNNNNNNNNNNNNNNNNNNNNNNNNNNNNNNN
NNNNNNNNNNNNNNNNNNNNNNNNNNNNNNNNNNNNNNNNNNNNNNNNNNNNNNNNNNNNNNNNNNNNNNNNNNN
NNNNNNNNNNNNNNNNNNNNNNNNNNNNNNNNNNNNNNNNNNNNNNNNNNNNNNNNNNNNNNNNNNNNNNNNNNN

NNNNNNNNNNNNNNNNNNNNNNNNNNNNNNNNNNNNNNNNNNNNNNNNNNNNNNNNNNNNNNNNNNNNNNNNNNNNNNNN
NNNNNNNNNNNNNNNNNNNNNNNNNNNNNNNNNNNNNNNNNNNNNNNNNNNNNNNNNNNNNNNNNNNNNNNNNNNNNNNN
NNNNNNNNNNNNNNNNNNNNNNNNNNNNNNNNNNNNNNNNNNNNNNNNNNNNNNNNNNNNNNNNNNNNNNNNNNNNNNNN
NNNNNNNNNNNNNNNNNNNNNNNNNNNNNNNNNNNNNNNNNNNNNNNNNNNNNNNNNNNNNNNNNNNNNNNNNNNNNNNN
NNNNNNNNNNNNNNNNNNNNNNNNNNNNNNNNNNNNNNNNNNNNNNNNNNNNNNNNNNNNNNNNNNNNNNNNNNNNNNNN
NNNNNNNNNNNNNNNNNNNNNNNNNNNNNNNNNNNNNNNNNNNNNNNNNNNNNNNNNNNNNNNNNNNNNNNNNNNNNNNN
NNNNNNNNNNNNNNNNNNNNNNNNNNNNNNNNNNNNNNNNNNNNNNNNNNNNNNNNNNNNNNNNNNNNNNNNNNNNNNNN
NNNNNNNNNNNNNNNNNNNNNNNNNNNNNNNNNNNNNNNNNNNNNNNNNNNNNNNNNNNNNNNNNNNNNNNNNNNNNNNN
NNNNNNNNNNNNNNNNNNNNNNNNNNNNNNNNNNNNNNNNNNNNNNNNNNNNNNNNNNNNNNNNNNNNNNNNNNNNNNNN
NNNNNNNNNNNNNNNNNNNNNNNNNNNNNNNNNNNNNNNNNNNNNNNNNNNNNNNNNNNNNNNNNNNNNNNNNNNNNNNN
NNNNNNNNNNNNNNNNNNNNNNNNNNNNNNNNNNNNNNNNNNNNNNNNNNNNNNNNNNNNNNNNNNNNNNNNNNNNNNNN
NNNNNNNNNNNNNNNNNNNNNNNNNNNNNNNNNNNNNNNNNNNNNNNNNNNNNNNNNNNNNNNNNNNNNNNNNNNNNNNN
NNNNNNNNNNNNNNNNNNNNNNNNNNNNNNNNNNNNNNNNNNNNNNNNNNNNNNNNNNNNNNNNNNNNNNNNNNNNNNNN
NNNNNNNNNNNNNNNNNNNNNNNNNNNNNNNNNNNNNNNNNNNNNNNNNNNNNNNNNNNNNNNNNNNNNNNNNNNNNNNN
NNNNNNNNNNNNNNNNNNNNNNNNNNNNNNNNNNNNNNNNNNNNNNNNNNNNNNNNNNNNNNNNNNNNNNNNNNNNNNNN
NNNNNNNNNNNNNNNNNNNNNNNNNNNNNNNNNNNNNNNNNNTGTTTTTCTTCAAGCCAAGGGTCGGGCCAAGCTGCAAC
CAGTAGACCCGCCCCTTGCCTTGATTACCACCATCACTTAGAATATCCGAACACCTTCCGGCGTTACCTGCATCG
GCCCAAATCATTCATCTTCGAGGTGCGCATAAGGAGCACGGACATAGAGTGGTGTGGAGGCTTCCGGGCCGAAA
GCCACCCTCCATCAAAGAGGAGGTTGCGGACGCGCGGTCGCTCGGagctggagaggagggaggaggagaggaact
TAGCGTCGGACGTTGCTCGGCGCTGGAGAGGACAAGGGTAGCTAGGGCATCGACCGAGAGTGGATGACTGAGGAA
AGCGAGCCAGGCGAGCactaaaaacaagaagaaaacaaagaggaggaggaggaggagacttACCTTAGAcgaggg
cgaggtcgacggcGACGGGGAAGCGGAATCTTCGATCGCCGATCAGGGCTCGGGATGGAGGTGAGAGTTTGATGT
CGGCAGCTAGGACATCGACCGAGAGAGGATGGGGAGAGGGGGCTGAACGAGCAGGTGGGTTTATGTTTAGTTTTT
TAACGACGTTTTTAAGCATCATCGTTTCTTaaacttttaacgacgcttttaagcctCGTCGTTTCTTAAGCTTTA
CCGACGCTACGGGGAAGCGTCGGCAAAGATTGGCGCTGAGCTAAACTTTCCCGATGCTTTCCTCTAGCATCGGAA
AAACTCAGATTTGCTGTAGTGTGAGTTGACAAACAGATAGTAAATTAACAGATAATTGAGGAATATTGAGGATGG
AAGGAACAGTAAAATGGATGGTCTTAAGATCACCCTGTTGAGATGGTGTAATTGGGTGCCATCACCTGTAACAAT
ATGAAGCTACATAGGAGGCGAACaggaaacaagaagagaagCATGTGGGGAAAGGATGCATCAGAGTCCAAGATC
CAAGTGAAACAGGATCTATCTGTAGAAAAGGCTGAAGATGCATGACCGGAGGGTAAAGCATCAGTGCTATGAAGT
TGTCAATGAACCTGTCATGGGAGACCTAGAGCTTGCATCAAATGAAGAATCCATGGCAGGCGCGGCAACAACTGG
TTCTTTAGGAGTAGCGCCTAGAGCTTGTTTGTAGCGTGGTGGTCGAATACTGCACTCCTCAATGATGTGTCCAGT
GCGTTTGCAGTAACGGCacaacttcttcctctttttcatagcatcagaagaaggaaggaaagtgGGACAAGAGCC
AATGCTAGTAGGCTGCATACTATGGGCAGCCAAAATAGAAGTAGGCTCGCCAATCTGAGATTGTAGGATACGAAG
ATGGGTCCCCTCAGCTATCAAGTCAGATAAAGCAGCATCCAAAGTAGAAAGTGGCGCACAATAGAGCAATTGACT
ATGGATGGTCTCAAACTCAAATCGGAGACCCATAAGAATCTGAAAAAAATGTTGCTGATCAAGAGCCTTGCGTTG
CTCATTAAAGCAAGTGCAGATTGACCAAGTAGAGCATAAATGATGCTGAACCGAGTGAAGTTGATGATGCCACAA
TTCCTGCATGTAATTGTAGAACTCCAATATAGAGTAGTCCTGTTGATGAGCAGTAGAGGGCTCTTGAAATATAAA
GTATGTACGAGCTTGATTAGCGAAGATAATCCCACAAGTCTTTGGTTGTAATATCCTGGCCTTCCATCAGATGTA
GAATTTGAAGCCTAATGGCATATGTGACACTTTAAGAAATTGCAGCAATGACACGACGATCAGCCATTGTCCATG
TAAATAAAAGATTAGGGTCCGAGCCAGCTGGCAGACAAGGCGTAGTCCCATCAATATGGCCCCATAGCTGATGAG
CAAATAGTTGAAGTTGAATAGATACTGCCCACAACTTTTAGGTGGCATAACTATCTAATTGAATGTCTAGAGTGA
TGGTGGAATTGGTTGAAGATCTCACAGCTGGAGGTGAGGAGGCTCCTAGAACTCCTGAAGACGAACTTGAAGGAG
TGCCCAAAGGAGCTAAAGAAGTTGGTTGTTGATTCATGATACCTTAGGTGGCTTTTGCCCTTGGAGTGACTGCTA
ACAAATGGCTTGCAGTGGCAGCTTCTAACAATGTCAAGCAAATGTATGATTGTAGGCTGTAGCAGTTAATTTGAG
GCAAATCATCAAACATGTAATCTGCAGTTGCTGTCACTGAAATGCAATGTTATAGCAGCTCATTCAGATATTCAT
AAACATGAGAGCTTGTCTATGGAAGACCATGGAAACCATGAAAAGAGCAAATGCGCAATTATTCTAGGTGTTGGC
TGGTCCAGGTGCCTTGGAGATATTTGGAACAAGGCATGGATCAGCTCCAATAGAGATCCAAGAATAATTCAGCAG
AAGTTACTTTAGTTCACATGCCAACATGGGCTGATGACTTTTAGATGGCCGCAGGCTGATATTATGCTTGGGTGG
TTTGGACCAATTATCGAGTAGCTGGCGGGTGCAGTATTGAGCATAACATAGGGATCTATAGATAAAGACTTGGAT
GTCCCAAAGCTTTAGCAAGATAGCGAACCATCGGAGATGAATGGTGGAACTGAAAATTCCAAAGAATTAGAGAGC
CCAAGATTGAGTCTGCGGCCAGCAAAGTTTGATAATGTCAAAATCTTCAAGCAGTATTACGAGGCTTTGTCTTGA
AGCTCATTTCCAAAAATGTGAGGAAAGCTTATTTTAGAGGTGCATGAAGCTAAGAGCATGGGCTATGAAGACCTC
TCATTCTATCTATTGGTATTGTATTAGAATTACCAAAAAAAGATGAGAATCATATAAAGATGATGGTTGAAATAA
CTATGAAGTGAGCTTACCATCAAGGTTACATGCATACAATATATACACTACATACAATACATAATACTCAACATT
TCTTCCCTCTTCATTATTGGCATTTGCAAATAGTAGAATGGTCTTTGCAAAGAGAAGATAAGATATTTTTAGCCC
TCTTTTGATAATAGAAATTCCAAGGACTTCTTTCCATTACTTTCATaggaccctttttttttttgggcttaaGCG
GGTTCTTCATACAacacgtgtacgaatacatccaattaagtcaaaaaACACCAACTCGTAGAGTGCCTGTAGCAG
CTCCTCTTTCCCTGACCAGAGGGTGTATCCAAAGTGGTAAGCCGCATAGGCAGCTGCCCAGTCGGCcaccccatt
ggcttctctgaatacatgtttggtCTAAAAGGCCACCCCATCCCTCATCATTATCCCTATGTCTCGGATCAGGGG
GTGGTCCGTGCTCTCACCCCTCGGTCTCCCTTAAATTCGCCCGACGATCGTGGCCGAATCGCCCACCAGAATGAT
CGAGCTGGCCTGCAGCACCTGCCGCACATGTCGAAGACCAGCCCAGGAGGCTCGCAGCTCTGCCTCCGGAACTGA
CGTGTCAAACAACTGGTAGCCACCCACTGCCACCATCCTGGAGTGCGGGTCCCATATAACAAACCCCGCACCACC
CCGCGTTCCTCCATCCAATACcgatccgtcgaagttgaccttgagaaaactcggggaTGGGAGCTCCcaagtgaa

aaacaccatctgagGAGCTGCCTGAGCAGAGAGGGAACCCTGAATGTCCCGAGCTGTTAAAGATCTATCTGAAGA
GAGAGTGGGTCTGATCTCCATCGTCTGTGCTCGAGCGAactccgcaacaaacctcggtgacattCCGCGCTCACC
AAAAGTGCGGGCGTTCTTCACCAGCTGGATCTGATGTGCTATGCAAGTCGCTTTGATGGCCTCCTAATATTTCTG
GGGGATGGCCAACCACTGCCGTATTACCTATAGAAACTGAAGCCTCTCACTCTAGGCCTCCTGCGGGATTCCTGT
CCACTGCCATGCCAACCTTGCCAATGTACACTGGAATAGTACATGATTCACCGACTCCTCAGCATCACATGTCTT
GCACTCCGCGGGAATCCCCCAGTTATGCCTATTCAGCACTATTTTCGTCGGAAGACGGTCCCATGTCACATTCCA
TAAGAAGAGTGCTGCCCTCAGATGAAGCCCCGACATCCAGATCTAAGTGTAGTCCTACCTCGGCTCATGCTCCTG
TTGGATGACACGGAAGAGGTCTCCCAATCTGACACTGGCCCTGCACGAGGTGCCCCAAACCCTGACATCTGGCCA
CGCATATCCGGGTATCGGAAGAgaccggatcctctcagctaggtgTACCCCGAACAACTGTCGTAGTTTGGCCTC
ATCCTACGCTGCCCCTCCTGAGGCtaggaggtcgcacacccgTAGCCCCTCCATTGCCTCAGTatcaaccatggt
cggccagcacCTCAGGAGAAGGATATCCACCCATGGGTCAGCAGTCACCTCGATGCTCCGCCCATCGCCTATCAA
CCACCTGGTGGTTGCCGACACAGTCGGCAAGTACCTCCTGATCtcgcgccacatgaaggagactcgaCGGCCACT
CCGTGCCACCTCTGAGCCCCCTCGACCATATCTGACTGCCATCATCTGATTCCAGAGCCCGTGTGGCTCTAACAG
AAACCGAACAGCATGCCGAGCAATGAAAGCCTCACGTCGCTCCAGGAGGGACTGCACCCCTAGACTGCCCTCACT
GGTGGGACGGCAGACCTGCTCCCAGACCACCAAATGCACCCTATGGCCTCCACCGTGTGACCCCCATAAGAAGCA
CTAAAGGAGTCGCTCGATCATCAGTAAAGTCGTCTTCGGTaccacagtgttggccatgagatatacCAGCATGGA
TTCGAACATTGAGCTGATTAGTGTCAATCTCCTCATCATGGATAGGGAGGACATCCTCCAACTTGCTCTGGACCC
GCTGCACCAGATCggagcactctgccacccgtaGTCTCCGGCCAGTGATGGGGACTCCCAAGTAGGTCAGTgtcc
cctcctgctctggtATCTGCCGATTCTTCCGGATTTCCTATCTGACTCTGCTCTCCGTACTCGGGCTGAAATGGA
TAACTGACTTCAGAAAGTTTACTCTCTGACCGGATGCCGCACAGTAGCCCGCCACCATTCTGTGGAGGATCCGTG
CATCAAGAACCCGCGCCTTGGCCAGAAGAAGATAATCATCAGCGAAAAGTAATGAGAATAGGCTCCCCCTTCTGA
GACcttttgatggaaaaaataatctGTTTGCTTTCCTTTTAATGAGAATAGAATGAGTCATGTAAGTAACGCATT
TCATAAGCAAGGCCAAATCTGTTTCGGTTTCAAACCTTGAGTTCCTGTTTTGTTATCTTAAGTTTCTCCGTAACT
CTTTCCATTCCTTGACTCATGTTCCGGCTCTCACGCAGCACTTCGTAAGTCTGGACCTATCAAATTTaaacattt
tccttttccttctttgatGACCATGGTGTTCAGCCTGggatattattaaaatttgggTATGATCTGAAAAATGTT
TTGCTTGTGCCTTTTGAGTACTTCATGCAccaatcaagagaagaaaggattctATCAAGTCTTTCCCTTGCAAGA
TAGTATGCCCATCATCTAAAGGTAATTCAGTTTATCATAGACTGAAATCAGGCCAAGGTGATAAGGTTAGGGCAA
GTGACAAGCTACCTCATTTCCAATAACATTAGCTATATTATTGGGCTTAAGGTGCCATAGAAGCAACAGGAAAGA
AAAGCATATAACTTAATGACTCTAGTTCATGTTTGATTGGAattattattagaaaaaaaacagCTGATATTTGTG
AATAAATACATAATGTTATGCATGCTTGGAAGGCCTAGATGCTTCTGAGCAAGTGAGTATGAGTCAGAAAAATTA
CTCTGAACCTTGTCAAGCATAAATTATCCaatcaaatatgaaagaaaTTGTTTCATTTCAATAAAGAACCTTGAT
TTCCCACACTATAGAATTAATGTGCAGAGAATGTCACAAAAATTGAACAAATACATTAATGCAATATCATTGATT
TGACTTGGTCCCTGTTAGAACAAACCAGTTATTCCATCCTACACATGTTCAGGAGACTAAGAAATTATGCTTAAG
CTGAATTGAGGACTTGGTTAGTAACTTGAAGAAGAACTTGGGAATTAGAGACACCCATTCTAAAAGGCAATAAAA
CGATCTTTAAAAATTCATCCCGGGTATTTGACTTGCATCTCATGAACTCGTAAACCTGCACAAGAAAATCATCAA
TTATATTTAGATGAAGCTTATACTTAATATGGAGGAATATATGAAGTATTATTTACAATTCTCAAGCAATAATCT
AGGCACTCGTAAGTTATAAGCATTCCTACAGTCAAAACATGGGCAACTAGCAACATGGTAACAATATTTTGTCAA
AGTATTATGAATTTTAAAACCACTAACTTTTTCTTTGACTGAAAGATACTATGCATTATGTCAAATCAAAAAGAA
CTATTAAATGTTGATAATTTATTCATGACTTATTCACCAAACATTCATGAATCAAAAGTAATGGCTTCAAGCTTT
ATCACTGGAAGTTATTTTGTTGCCATTGTTAAATGTCCTTGTCCACTAGATTTGCAGAGAAAATTGAGAAACCTG
GCCTGATTGGACAAGTCTAATATGCATGCTAATAGAATTTTAACTTTGATCATACCTATTACCATAAAGAGTTGA
AGTAACAAGTAGTCCAGGGCAAACCACTAGGCATGTACCATACCGGATTGACAAAACCATAACCATTTTTTTGAG
CTAGAAACTAATTCCAGCTAgataattttcttaattttatctttttttatatttcatcaaTTATCCAAACACTAAT
TCAAGAGTAGTCGGTAAAATTAAGGAAAACCCCCATAAGCTGAGATTAAACCTTCTCAGTTCACTTGCTCGATGC
CCAAAGTATGAATCCAAGatcaaaattaatgaattttgGGATTTTAGATTGATCCCCAAAACCCAAATTgtgtaa
attagaaaataacCCAAGCTATGCATTTGCTAAAGAAGATTTTCCGAAAATGCAGACAAGAAAGGTCCAATCTAG
AAAgctaccaaaatatttttttaggctATCTATTTAAAAACAGAATTTTCTAGAagttaatttatattaaaaatta
taatttctaGAGGAAAAATAAGTTGTATCAAAAGAAATACGGTAAATGCTCCAAAAGATGCATGTAATGTCTTTG
CACTTTAGGCAAGCTATACAAATTTTGAAAAGCTAAAAGTGATTTTGAATTGCAAGTTTCACTACTTCTAAATag
catttgaaatatatttttcttcgcCTAATTGCATTTCAATTGCAAATTTTAATACATATATCTTTTAAATACATT
TTAGATTACAACTTGTATGATCATAAATGATATTTGTATGTTTATCATACATAGAAATCCTCAAGTAGGGAATAG
CAATACTATATtagaatttcaatttttttcctttcttgtctttctttttttggtttgatattttttttatttttttt
ttcaaatgtttcTTTAGGCTTAGGTGAGTTTTTAAGCCTATGGTTGGGTTTATGATATTGTGCGAGTCCTAGTAC
AAATTTGTtatctagttttcttcttcttcctctttttctcccttcctcttcctccccatGATCCAATGCaataaa
agaggaaagaaaaaaaacaataaggAGTCACCTAACTACGTTCTGGTTAGTCTTGCATGAAATAAGTTTCCTATCC
CACTTGGTAGCCGATCAGCTCAGCACCAAGAACACTTATTTTTATGGCTCAGAGTCATGAACCCATCCTAGGACA
TGGTTGAGCACCAGTCCGATATAGGACACTAGGTATGAGCCAATATTGAGTGAGACCTAAATACTTGTATTTAGC
TTTATCTTTGTGCTCTAatcaaatttgataaaatataacattataCACCATCCTAGTGTTTTTTTACAATTTTCC
TTTTATGCCAGCATTTTTTACATGAAAAAGCATTATGTTGGGAAAAAAGTTACTTCTATGTGTTAAAAACATAAA
GGTCACAGAGATTTCCTTTCTGTAATATTTGTTTCATCTTTGTCATTTTGATATAGAAGGTAACCATTCATTTAT
CTTTTCAAAATGATAATGATATGATTATCTTCATTCATACTTATCTCGATTCTTGTGATTTTGAAGTAATAAACT
AACAAACCTTTTATTGTTAGCATGAAGTCAGTGATAGAACGCTACAACAAGACCCGAGAGGAGCATCATCTAGTT
ATGAATGCTAGTTCAGAGGCTAAGGTACATTTTAAGCACAAGTATATGTTACATCCCCTAAATAGTTTAAAGTTG

TAATCTTTTCTCTAACTCATGCTTTTACAATTAGCCAATACAACCACTTGACATTAGATTCTTGACTTTTATggt
tctcccactattttcaaGCAAAAATGCAACTAAATAGCTCGAACATGCACTGCTTAGGCTCaagtttctttttgt
tgctcaaATTTATATACTTCTCTTCTGAAAATATCACACCCTAAAATGTCCTCTTAATTTCATTCATAGCTCAAG
Caaataacatgaaaaaatg GGCTATGTttacaataattaattaagtaaTCATTCAATAAGGAAACCAAATacaga
tttttaaaaatacatattatctgactttttttttaagcataccaaggtatattttttttggatggtTGTAATAGA
CAGATCTTCACATATTAGAATAATCTTAACAATTTTTCTAGCTTTAAAGTATCAAACATAATTGTTTTAGATCAC
CACCTTGAAAAAGTCTATTTATGGAGTGGAGAATTAGTTCACTCTAAGTATCATTCTTGGCATATGAAAATTTAG
TTTATCCAGGTCCGACTATGAGTCATAGATTGGCATAGATGGAGTCTTCCTCCTTTACCCAAACAATTTTGGTCA
CTAATAAGAGCTTTCACAAGATCCCTGCTCATAGCTATGCTACATAATTGCAAGGTCAAGAGTAATGAGGCCTAA
GATTAGTGACAGACATCTTTCATGCTTAACTACAAGTCTACCTTCATGATTTCAAACCAAATAGTCTATGAATca
atttttaattgatcaaaatatttgaaataggaTCTTGATGTCAAAATAGAATAAACAAAGTGGAGAAAATGGAGA
ATTACCtagaattgaaggaaaaaatctATAGTTGTTGGTTATATTGGGTAAAATGTGAGTACTATGTGTAGTTCC
TAAATCAAAGTAGTCTAAAGCTAAGCTACATGGACTAAAATACAAAGCAGAGCACTAAACTAATTCACAACTAAG
TCTGATATCTAGACACTACGATCACAAGAAGCAAGAGATGAAGATAACATAATTTTAACAATGATTATCTACTTG
GAAGAGTTCTACTTATACATGTTTTATACCCTCTTGATCTCATTATAAATTTTACATCCCTAAgtatttttttgtt
actttcttaGCTCTGAAAGTAGCATGGTGGTTTGTGAACCAAACTATGTGCATGTTTGCATATGGTCATTCTATG
TCTTCTCAGATGTGATTTCTCACTTTTTGATGCTGCTATGGTGCTAGCTATTTCAAACTGATTCCTTTTTAATTC
ACTTTAATATTGGAagcttgatcattttttttttcaaccatatAGTAGCCATTACCATCAACCATTAAGTGGGCGG
TACTGCTAGTTGCTTGATTGTCACACTTAATTACTACAAGATTGCTACTTGTCGGTTGGTATAGTTAATTGCTGA
TTGCTAGTTATGATCATGGATAATTACCAATAGCCAGTAATCTCAATCATTACTAGTGTTGGCTAGCAGCTTTTC
AGTTGGATGATCACACCACTGCTTCTGCTTCTGTATTTCAGAGTCAAGTATCATACCATCCAATGGCTCCCACTT
GTTCACAAGTCGAGGGGAAGAATTCTAGATCATCCCCTAAATAGGGAATTTAGTTAGTAAAGTGAATAAGGACAT
TTTTGTGGTAAAATGTTCTCTCTTAGTATGTTTGATGGTGAGCTATTCATCATCCTTGATAAAAATGTTACTCTG
AAGTCATCAACAGTGGTCATCCAGAGTTATACGGCTAGAATCTACACCGATCTGTTGGCAATGATTTGGAGATGC
ATTCAGGTGTCCTAAATTATAAGTTGAAAAGTTATAGAAGCAAAACAAAGCAATCCTAACAACCACTGAGTAGTC
AAGATTGAAAAAAGGCAACTGGTGAAATAAGACCACATTAGGATTATTCCAGGAGACATAAGGCCACAGTGAAAA
ATCATGACCAGACATGTTTGCATGGGCAGTCAACCAGTTTTGCCTACTACTGGAGATTTGAACCATTGACTCACC
ATAAAAAATTAGTATACTATTAACTGCTATGTAGGTATTTGTTCGACCATCTTATTCAAAGGGTGGAAACTGACG
GGAAGACCGGGCTGCTTTCACTGAGTTGATCTTCCCAAGTCTTGTTAGGAGAGAGCAAAGGTTGAGGCCTCATCA
GGCTTTCACAAATTTGGTGCGTAATTGTTGGCCATACTAATTTGAAGTGAGTTGGTCTGTCTAATATCCAGGAAA
TGATGACCTTGAATTGGAAATGCTATGAAAGCATCTACCTCATTCCAATGCATTCCTATAACCCCAACCCACAGC
TTAAAAATCTATAAACAAGAGCTTGTGATGAAGAACCATGACAATTTTACAAGAGCTATTAACTACCAAGAATTG
GGACTTGGCAATTGACAGGACTTACAAGTGTAAAATCCATAATTGAGTTTTCCTTTGATAAATATTGAGTGAACA
TGGTGCAAAGTTGATGGTGGtgattttttgttttccaaaGGCCTAATAATCCATGGTTCAAAATCAGTTTTCAATT
TAGGTTAGGATTTTGCTCACTTTCCCTACAGAAAAGGATTGTACTACTTCTTCTCATGTAGAGTCAATTGTATAA
CACAATAGTTACTCAATATACATAACGTTTCCCTAGGCCTACGTATGGGCATTGGTCATTGATTGAGGACTACCT
GTCAAATTGGAGATAGCTTTGAGCCAAGGATGTATGACTTGCTGAAATGAGACAGCAGGGTTGGTACCATACCGT
TACTGGAAATCAGGACCCAGGACATCTGTGGGACACCGATCTATTTATGGGCTGAGACATCCTGCTCGTCTTGGT
TCAAACTAGTCAGGATCAGCCAGTACTAAGACCGATTCAGCCGGTACCAAGCATGAGTCGACTAGGAGGAAAGAT
CCACCTACCCTTGAGGTATTGGCACCGGTGCCAGCACTCGTTGATTCAAAACCATATCAATAATCATTCAAATAC
TCATTtaggacaaaaaataaaaattgaacaaaaaatTTGACATAATAGTATGGTACTGGGACACCAAACCATCCT
AAGAGTTAAGATGGTAGTGTCTTGATATCATCCCAACTCGGCCAAAATCGGTACAGGGCTTAGTACCGATATCTT
GAAGCATGCTTGGAGCTATGCGCGAAGATGCTAAATTACTAATTCAAGTCTTAATTTagttataatttattaatt
attttctatCATGTTGCACTAGTTCATCAATCTTAATAGACATTTGGATTGTTAATCTAAAGCCTTGGTCCAACT
TATAAGCTTAATAGCAATCATGGTATGATACAGGCTATAGCTAAAAGAATGCATGACAAacagaaattaataata
aactGGTGGAAGAGCTAAGAGATGTACAAGAGTTGCGTGACTTCAAGATGTCATACATTATTTAGAGGAAAATTG
GAAGCCTATAAAGTTCAACCTAGCTTCAATTGCTAAAAGTCACATTGCTCATGCTGGGGCGGATGACACGGCAAA
AGAGACAAGTAGACAACTTTAATGCTTGTAGATGCTCATGAGAAGTGACAAGAATTGCTCGTGAGATCATGAATG
ACTCGAATACATTAAAAGGCTAAGTGTAGAGATGCCCAAGTATCAGCTCTTGTCctttaaaatttatcataattt
acttgttattttattttatatcataaattcataacTAGTCATGACTCCTTGTGGTAGGTAGAAATGTAAGTTCAA
GCTGGTTTCAAACAAGCAATAACAAAGATACAACAAAGTGCACTAAGGTTtatcaaatagaaattttaatgGTGT
aagaaatgaataaaataataaatggagGGCCAATGGTCAAAAGTTCAGCATTGAGAAAAATTTGGCTATATCTGG
CAAATTCACCAAAATTAGGAAAGAGTGTGGATAACCAAAAATCTggaacatatataaaaaaataaaataaatgtg
agatttctaaaatatttaattttcattatgTAGAAACAAGGACTAGAAGGAGAGAATAATAAATGGTAGCAAACT
AACACCATCCCCAATGATAGATTGTATCTTTAGTCTGTAACACAGAATTTTAAAGTAATCAAGATTAACTCatca
ataaatcataaaatttatgtaaaattgAGAACAATTGATTAATTATGAACTTGTTTGTCTACAGGGAATAGTATG
GAAAACAGTTCAAATATGGAATAGTAAgagaaatttgaaaatttttcaatAGCTTGGAGAATATTGATACTATGT
GGAGGAGATATGGTTCTTTCACCAATTGGGGTTTAGTAATTCCATCAATATAGATCATCATTTTAACAAAATCTT
GAGttgttttcaataatttttaatgtCTTTTTAAAGAATTTCTCAGGTCTCAAAGAGTCCTAGGTCAGTGTGGAG
GTCTGAGATCTATCTTTCTAATCCCCAGTTCTTCAAATTACAAGTTAATGAAACTTATTCATTATACAATCAAAT
TTACCACTGTATATTATTTGTCAGCTTTTGCATGACAAATTCTTTGTCAACCCTGTGTTGGATCAAAAGAAGCTC
AAAGGAGGCTTTGGTGAATTTTTAAGGAGCTAACGAGCAAGTTTCTAAAGGCCACTTGATTTTCAAGAGCAAAAA
AGGCTTTGAAACACATGGATCTATATTGCTACTCTCCCCCTTCCTCATCACTTTGATTAGTTCTTTTCTGATCg

caaattatttattctcttATTATTTTGGCAACCCTTCTCAACTTGCATCCATATGTTCCTCTGGCCTCACGTGA
TTAATCAAATCTTGATGGTGCAGTTCATTGACATAGCTTAAAACTAATCGAACTTTTATCATCATTTAGTGTGAT
TCTCCAACCCTTAAATCACGGTTCGCTATCTCGGTACCAGACATCCCCTCGGTACCATTCTATTATAGTGTTGGT
ACTCTGGTATAGGAGCCAGTTTGGTATACCAAGTGCACCCTTTGTACAACGTATTGGTTcgatactagtatggta
cggtacggttgGCATGCACTAGTACCAACTGGTATGGCGAACCTTGCCTTAAATGGTAAGTTACCATTGATGGTA
AACTCATTTTCATCTTTAAAGAATTCTCTAATGCATGGAATTTGCaatcttgttttgctttttgccCCATGAGTT
TTATGTATCTCTTAGTCATTTAGACCTTAAAATGAATTCCAATATTAGTATTCAAACTGTGTTAAATCaacaaat
acaattaattagGTTAGTACATTCCTTGGATTGGCAATGAATATTCTTAGGAAAAGAACATCACATGGAGACCAC
AAGAAATGGCAATAAATCCAAATTTTGTAAATTAAATTGACAACATTTTGCAGTTGAATTTGCTAGACTTGAGTC
AGCTGCTTGACACCTCTCTCCTCCTTGATAGAAGTCCTGTGTTTGAAGCAGATGGTGTCATTGCCACCATgtttc
tcacaaaaaaaaagaaataaattaaaaaaaaactaagtcgTAGCTTATAAAGACCAATCGATTATATGGTTGATC
TGTTATTGTAGTTTTCATCCTCTTGTTATCTTTTTTACAAATTTTACTCATTTTATTTGCAAATGAGAAACTTGT
GGGAACTTTTAGGTTCCAAGCTCCCTTAGGAACCATATCATTAACAAGATGAGTAGCGGGGGAGGGCAATTTCCC
TATGATATGCTATTGCTTAAGTCTCATTTAGAACAACAAAATCAGGACATttgtttatgaaattttaataagaat
aaCCAAGGTAGCTTGATgagaaaatttcttatttacaAATAAATGGAGCCAGTATCTATAGCTCTTCAGAGTGA
ACTTTCATAGAAGATCTTCAGAAACTAAATCCAAGCTCTCGATCATCATATATGACAACCTATGATTTGGATAAT
TAAGATGTCATTGGAGCACTTCAATCTTGAATCTCCAATTTTCAATGATAAGTACAAGTGTCTCAAAATTAGACG
AAGTTATCGATTAACAATTTGCAAGCATAAAAAATTGTAATCATTATTCTCTTTTCTCTGAATATATGCAGTAAA
ACATAAAACCTCTTCAAAATATCAAAGTACCTCTGATGGCCTTAGAAGTCCCAATCATTGTCTTCGTCATCCATG
TGATCTTCCACCTCATGAAGGtcttatgaattatgatgatAGTCAATGGAATCCAAGCTATCTGACATCATAGAA
CATATTCAGAATTATCCAAAAACTCGTACTTTCCAACATTACAAcaccaaaagaaggaagaatcaAGAAGGCAGA
TTTTCAATACCTTTATTTTTCGATCATACCTGCTTCTTGTCCAAGGAAGAATGTAGTTACAGCATGTCAACATGA
AGGCATCGTGCCTCTCCCTCTTTCAATAAGTCATCCTACACTTCTCCATGTTGATTTGCTGGTTGCATTTTGTCC
ATACATCTGCCCTGAAAAAAGCATCTGTGGTATTTCTGTATATTATGGAAAAGCCATGCGGCATATGTTCCCTTT
GATCTATATTAATTAGAACCTGTGGTGTAAAATACCTTTGGCAAACTAATCTTTGCATATGGAGTACTTGATCCA
ATTAATGGTGCCAATGAGTAACTCCCACTATGGAATATGTTTCACGACATAGGAATGACTGATCAAAatgagaga
aataaaagaatgtAGATTTAGGATTGTGAGATAAGCGGAGGTAAGATTACAAAGCAAATGACGTGGGAAATGAAG
AAATGGTTATGGGAGGTTAGATGAAGGCATTCTCTCATTGTTTTTTGTCAAGGTTAACAAACTATGCATTGCAAC
TTCATGTCCATCTTGAGGTTTCTCTTGAGTTAGGTTCTCATATGGTTCTCTTAGTCTACAACAGGTTGCGGGTTA
AAGAAGAGTAAATGGAACAAAGGTCCCATTAGCCCATTTGTTCCATTTTAACTTGTCAAATGGTAAAAAAATACC
ATTTATTTGGAAATCAAAACaagtatttttaaaaagtttgtAAAGCCATTTTCCCACctaagcaaagaaaaatat
tgcacATTATAGTGTGgagtattttattttggaagaaCGAGGGTCAGGACCAACTAAACCAATTTCATCTAGGTT
GACTTATTCCACAAACCACAACCAAATGCAGCTTTCACGTGGAGTAGCAATTTCCTAGGAGGTGCAGGATGCTTC
CATCCAAGAGAAAGGAATTGCCTTCCCATTATAAATGGTCATGGCATTTCTAGTGAAGAAatttaatatgttttt
gatttAGATGCATAATTTGGAGGTAGccatgtaaaataaatattttattgctATTTCTCGTGCTTGCTTGTATCT
CTTTTGCTTGCTAATAACTTTTTGCATATCTTTAGGGATAGTTGTGTTTGAGCCaagttcgccgtctcggtaccg
gaccccgtatcGATGCCACACTAATATAgcgtcggtacggtatggtatacaGTTTATTTGgcataccgaatgtcg
gtacgccacTAGTACCGGTACGGGTACCGAATTGGTACGGTACGGTGCATCCGGTATCGTTTGGTTCAATACGGT
ACAACATACTATGGTTTGAGCCCATTAATAAAGAAAAGTATTACAGAACCTGCTGTCATGTGGTGCTGAGTACAA
GCTTAGGATCCACTTTTTGAAGAACTTGATCCTGAGACTTCTTTCTGTGATGTAATTAAGAAAATTTAGATTGTT
TAGAAAGATTAATGCAGTGCAATCAAAATGTTGAATCTATAATCAGAAATCATGTAATGTTTAAATATTCAACAA
ACGTAAAACCAGATACCGAAGCTTTTGAATGTTGCCTCTACATTCCATAATAATGATGCGGAAATTACACCATTT
ATTAGTCACGCAACATGAATAAATCATACAAATAATTTAAATGCAAATCTTatcctcctcccctcttcctgTCAA
CAGTCTCcttatcaaatatattattttcctttttttcttttgtgtgttgatttttttctttttcttttttttcttttt
ttgttggggaCTGGGTGGGTGGTTATGCAACAGATGTTAGGCAAAGCACCAAAAGTATCAGAATAGCTCATCAAC
TATTTAAACAGGAGCATTCAAAGATCACTACAGCATGCATGCATAACAATAAGGATTCAAAAAGTTATAGTTGAG
AAATTGTAGTTTACATCAGCAGTAAGTCATCTAGTAGAGGTTCAGAAGTATTTAACCTTGATGCTTTGAAACTGG
ATTTGTAATAGATTTGATGAAAAACTGGAGGAACAAACTTCAGATTATAAGTGATCTAACAGAATGTTTCTTTAT
TGAACCTGGTTACTGTAATACATAATCACTCCATAGCAGTAATATAGTATTTTAAAAGAATCTGTTGCATAGGCT
AAACTTTATCCACTGTACTCACTAG**TTCTGGCAAAGGGAGGCAGCAAGCTTGAGGCAGCAACTGCATAACTTGCA**
**AGAAAGTCATAGGTATGCTCACTTCACC**CTGcataaaataagatttttttaggCATTATTTCTCAGAAGGCGTTC
ATCGCAAAATAAGCATATTGTTACTTCTGCCAAGTGGTTTAAATTCAGGAAGTTGTGACATGACATATGGAGGCT
AAAAATTATTCATCATCTTGTAACatatataatactattttaATGTGACACTAGAAATGACATATTCCACTTCAG
AGGCACTAGCATATCACATGCCCTAAGAGTCAAGCTGATTAATAGTATCAGAAATGCCAATGGCAGACGGATAAA
ATGTCTCCAACAATAAATTACTTAAATATCTAACATACTAAGATCTAATACTTAAAAGAGAGTATTATTGAAGTC
TAGATAACAATTtcaagtttgtgggtaactttaGAATGAACTAGATAATTAGCAGTCTTCTCACCTAtcgtatca
aaaaaaaaagaaaaagaaaaagaaaagcttataTGAAGCAAATGTAGTAGAAGAAAATGTATCATTCCAGCACAA
GAAAAACGAAGGGGATTGGTTCACACATCACACGTACCGCTCCAAAAATCAGTTCTTAAGACCATAAGACTATGT
CTATGAAATATAGTAAAAGCATTTTCTCTATAATTGTAGAATTTCTATCTCACTTTATTATTCCGCAATTCAAAA
ACAATGAGGTATCTTAAATACAGCAGATGAATATAATTAGACATTTCTAGAAACCCATAGGACAAAAATAGAGAC
TGTAGAGCAACATCAACAGAAATGATAATGAAAGaagcacaaattcattgaaataACAGCAACAACATCTTAATC
AAAGTGCATGAACATTGCCAAGTTCTCAGTTTTGAtaacaatgaagaaaaagaaaatatttctaataaGGTTAGA
GCCAGTGTAACAAAACAAAGAGATGCTTCTTGGTCTAGGTGTCCACATATGCCTGGGAAACAACaactgattaaa

aaaaaagtatattagTGGAGGTGAAGTAGCATAATTGAGAATAAAGTGATTAAACTAGTTTAGATGGTAGGGATT
TGTGCAAAACCAAAATAACAGAACCCCATTTAGGAGGGGCTCATATCCCTTCTAATCTTCCATAGTGCATCTCTT
CATCTCTTTAtagtctctttcttttctttcttctgctgctgctttcttttttgtaacTTCTACTTTTGGTAAACA
CCATTATTTTGTAAATGCATATCACCTTTAGAAAATCACTTTTCTGAAAAGATCCAAATAATGTTCCACTCCATT
CTCTTCAATTAAGCACCTTGTACTTCtgcattctttatttttttaaaaaaaaactatagttTAATCTCATATGG
ATAACtgtattaatttttttgatcaaaaCTTGTATTTCTTTAAAATACAAAGAAATGGTTAGATTTATTTTGTAGT
ATCTCAAATTATTTTACAAAGATGCATCCTAGGAAGTTTGACATCCTAGATTTTCTTAGTTGTCTGTTTCCACTC
CCACACCAAAATACATCCCCAATGTTGTCCACCTAGAGCGACTAAGGTCTTCATCAGAGAAATTAGCTATTGACT
AGCTTTAGAAACACAAACAAAGGTCTCCAAAAAACCTTATCTGATGGCTAGGTCTATAGCAATAGTAAGTAGTGA
CTGATGAAATCGCAACAAAATAACCATGAGTCTTTAGTTCCCACTTGTATCCCAAGCGATACTCGGTGCACTCAA
TATACCAGACACTATGACAAGAAAAGAGTTCTAAAGACGATACTACATATACAATATCCATCTAGATACTAAACT
CTCATATTTACAATTGACAACATTTAATGTCCAACCATAAGAAGATTAAGTCCATCATGCACCAAAACCCAAACT
GAACTCATGGTTATGTCTCTGTGTAAGCAAAGCTCAAATAAGTGCTTAATATCAAATTGAGAGAAATCATTTGAA
ACATACCttaaatagaaatatataaCTGGCATAGCAAGGCATACACAACAACATTTCAATGATTGCAGGCAGTTG
TTGGGAGAAGAGCTTTCTGGCCTAAGTGTCAAAGATCTACAAAATCTAGAGAACCAACTTGAAATGAGCTTACGT
GGTATCCGAATGAAGAAGGTCTTGGTCATCTTTGCCTCATTCAATATTGATTTGCTGTCCTTTCTAACAATAGCC
ATTTTCAACGTATTAATTTTTTGCTTCTACAGGACCAACTTTTGATCGATGAAATTCAAGATCTAAACCGAAAGG
TTGGAAAACACGTGTTTACCACATCTCCTCTTCTAAGTTTTAGGTGATGAATATCAACTGCTTAACATTTAACTG
GTTCATAACAGGGGATCCTTATTCACGAAGAAAATATGGAACTATACAAGAAGGTAAACCTCATTCGTCAAGAAA
ACATGGAATTACAGAAGAAGGTACTGAATAAGAGAACGAATAGACTTGATACATCCATACAGAGACTTCTGTAAT
GAAATCCTTTCGACCATGGTAAATTATGTAAGTTGCAGGTATAGCACAAGCTAAGCATTAAATCTTGCATCTCAG
GTTTATGCAACAAGGGGCGTAAATGGCATTGATAAAGGTCCAATTACTCCATATGGTTTTAATGCCACAGAGGAT
GTAAATATACCCATTCATCTCGAACTAAGCCAGCCACAATTGCAAACAGATGGAGCGCAAGCAGCAGCACCAAAG
CCAGGgtaatttttctttcctaaacTCAAactgaataatatttttttaattactagTTTAATCATAGAATATTTCG
CTGAATGTTGCAGCCGACTTCAATTGCATAGACAAGACGATGCACAAATCATGGGTTGTTCCTTGTGATCTTTTG
TCAGTGTAGTTCTAAGGGTCACCACAGCATGTTATTAATGATAACCTGTGGTGTACAAGGAAGATAAGGATCTAC
TTCCGAAAAATGGTGTAAAATAATCAGAATAACTAATTACTCTGTAAATCTGTGTTACTATAAAAAGGTGATGAC
TGCGCCAGGAATTTCAGTTTAAGAGCTTCAAACAGCAATCATTTGCCTATTTAGAAGTGTATTACCATGTTTTAA
GTTAGTCATTAAGCCATATGATCAAGAAACTCCCAAGTTACATTGCAGGCATATGGGATTATCACTGCCTTAATG
ATGCTTACATTGAGAGGATTGATGCCTAGAGATGGACTGTATGCTGAAAACAAGTTTTTCACTTGTCACGTACCA
AAAATCATGAGAATTATGCAACCACGACAATCTGCACAAACTATTTTATTCACCATTAGTGAAAGAATAAGGTGG
AAAGGCATCCCTACTCTCCAGCCCAACAGAGTTATATACAAGCAAGCTAAATCATTTTGCTGAAATTATAAAACC
AACTCCATCTGAATTCAAGTTAGTAGGTTATACATAAGAAAATGTCGCGCCCTTCAAATGAACTTCCTTAAATGC
TTGATTTACTCTCCCTTCTAGAATATGCTGATAAGGAGCAACACGGTCATGCAGCCATACCAATAAATCTCTCTC
ATGAATATATAAAACGATAACAGAAGCAGCAGATTTCTTCCTGCAGAATTTGCCTTACTATAAAGCATAAAAGAA
GGGGCCTTCCtagaaaacaatttttttttatgtaaccACATGACCTGATTGAAATTCAAAAGAGAGGGGAGGCAAG
GGTTTGAAGCATCTGGAGGGGGAGGAGGTTATCATTTTGACGAGAACCCCTGAACCATGCAGTAACTGATCAGTC
CAATTCAATACGCCTCGAACCAGCTGGTTCGGCACGGTTCACTACACCTTGATGCAAAACATAGAAACGAACCAT
ATAAAATTGGAATCCTGAACATAAAACATGGTGGTACACATAAATTTCAAGGGCACATGACTACTGGTGCTTATT
TAGGAAGTGTGAAGCAAGCAACAAAAGACAATTACTATTACACAATAATGCCTCCTACCAGAAGGAAGATAAGAG
GTCTGCACTGATAACTGGATAGAATGTTCATGAAATCTTGGATTTCTTGTAATAATGTCGGGTTGTGTGCCCAAA
ACAAAAACCATTTTGGCAAAGGTTCACACAACCAGTAAAGCAACAAAAGAtaatgaataatttataatagcAGCA
AATACTGTTTGAGCAACATAAGAGGAGAATACTAGCAAAGCATGTAATGATATCTTGCATTTCTTTCCATCTCAA
TAACACTAGTCCTTGCAATAAATACTGGTAGATGTGCCAATAAAAGCCTACTTCAACAAGAAAGTACCAGCAGTA
GACATCTCTAAAGAGGATTCAACTGTAGTGCAACACTCTATCCATATCTCAAGCATCCACTCAAAGCTTGCTTCT
GTACTGTATAGAAGATAAAAGTCTCTAAGAAGATGCAGTAACACAATCAATCACATCCctaacacaatcaacata
tgACTAGGCATGGAGATGTAGTGAAGGATGAGACAAAAGGACTTCAACAATGGTGCTACTGAAAAACAGAAGATG
CAAACGACCATATAGAGTAATGATCATCCCTTTTTGTAACAAAAAACTAGAATGGAAAATGAACAACATAATTCT
AAATACCCAGCGCTTCAACAAGCTAGAAGGCACTCCTAATCATGCAGTCTTATAAACTTTCGTCTGTCAAGTGAG
CACAAGCAGTGATCTCAAAAGGATTACTGACAAGAAAGCAGATATCCCTAAGCAAATTCAACATGAACAGGAGAC
AGCTCTTCAAGGAAATTAAGGTGGAAAAGAATAAGAGAAGGAATGAGGACAGAAGAAaacaaacatggaaaagtt
cCAGTCCAGAACACTCAAAACTTCACTAAATACTTAACTCTCAGCAATCATCTCTACAACAAAAGGCCAATGTAA
GATTGTTTCTGCAGTCAACAAGGAactttataaactaaacaaagcatcAATTCTAACCATTGATAACAACCAATA
ATCCAATAATCcaaaatttcatttaattaccaaaaaaacacaaaatcCAACTTTGTCCTTAGGTCCTTTCTCGAT
ggcaaaaatcataattttaggCATCTCTCATAAAAAGGCCAACTGAGCCATCAGTCTGATTCTTAAGAATATTAA
GAAATTAGGATATAATTGATGCCAAATCTGACTATGTCTTGATGTATTCTCAACCTTAAATCTCTTGTAACTTTG
ATAGAGTTTTCATTTAAACCCCGAGTAGTTTGTATTGGTGATCTTCAACAGGACATTCTCATATTTATGACTTCC
ATCCACAAAATGTCCTTCAAATACAGGCCTACTCAAGGCCAAGCAAGAAAGGAAGTTGACCAAGATCATTCTCAT
CAAACTACCAATGAAGATATAATGCCTTCCAGTCTCACCTTAGGTAGAAAAGTGGATAAACATATCATTGTCATG
CTTACTATAGAATAGTCTTACCACAATAATCACCTAGATTGCCAAACAAGAACTGTAAACATGTTGAGGATGTAG
AAGACCATAAGGAGGAAGCATATGAAACTCGACTTGAACATTTATCATGAAGAGATTTATCACTTATATCTTGGT
AATAAAAGATCTGGTTGCAGAGCTGCAACAACTATCCAGAACCACAAAATAATCATCCAGGAAACAGGAGCAAAG
TCTCATTAATCTATAACATAGTTCTCAGACCAAATATGAATTACCAATTCAAATATAGTTTAGTAGACCTATTTT

GCAGGAGATAGGATTTCCCATGAGAAGGGAACAACATCACATGTCTCGGGGTGCAATAATGAACAAGTTTATTAG
CCATATCATGGTACTGATGAGTAATCCCAGCTTGTGGATgactacaaaaaaataaaaataaaaataggtgAAAGT
TGGTATATGAGTCTCGAATCACGAGATAGTCATGCCCAGTAGGAGTTACCAGAGAGCTGGGGTCCTCAGCTGTAG
AGAAAACAGAGAACCATGCAGTATTAATGGCATAAGTGTTCATAAACAGGTACATATGCAGTCTTATACAATGGA
AGATCAACAAGGAAACAGCACGTCAAAGCAATATAAGACTGATAACAATAGGAGAAAATAGGATCAGAAAGAGCA
TAATAAGATAGATCTACAAAGAAGGtattcccatgagaaatatgcAAACAACAAATAGCAGCATATAACATGTGT
ACTCTTTAAGCAGGCTATAATATCTGAAAAAGCACAAATGGCTGATCAAAATCCAAACGTTATCCACTAAACATG
AGGATGGACAAAATAGgtaaagcaaagaaaaaaagagaggacatGGGTAGGAATTAGAAAAGAGATTCTAAGAAA
CAGAAGACTAAAGTGACATTCAAGGTGCAAACATCacatcacaaaagaaaaactttgggTTCAAAAGTTTAGACA
GAAGCAGAggggaaggggaaaaaaaagaccaaATGACCATGAAACCTTTAGAGATTGATGGTTTTTATGCTTATT
GCTACCAAACATACAGCTTTATGAAAGTCATGGGGAACcagaaatatcaaatttttcatgcatcTAACACATTTG
AAATTCATTAATACAAGTGTATACAACTGATAAACAACATAAATACCACAATGTTTACTTCCGCATCTAGAAAGA
ATAGGTGGAGTCAGGACATTCATTTGAATGCATCAGATTAGCAGGTGGAGATACAAAACGATGCCACCAACATCA
CAAAGAAGGGTTGATTCATGAAGCAGTTTGCAGCCTAATAAAGATGccacaaaatattataataagaaaaaccat
cataagtcaaaaataagattcgccatctcggtactggagcccgtaccggtgccaccctagcacagtgtcggtatt
GTATGGTATAGGGCTTTTTTGGCGCACCAAGTGTTGGTACGCCCTGTACCATCCGGTTTGGACAGGTACGACGTA
CCTTGTCAACATATATCAGCTAACTTTTCCAAAGAATTATCTCCTAACAtcaaaaattcattttaaataagAAGG
CAAAAAaccgccaaaaaaaaaacaaaaataaaaaagaattcctCATACTGAAAAACTTAATTCACTTTTCCATGT
CAAGTTGTGTTTATCTCGCATACATCAACAATAATCTGCTTTATACAGTTCTCGCTCTATGCCATCTGCTCAAAG
CCATCCATAACTTTCTTAGTTGTACAAAGAGAGATTTGTGAACAACAATGTGCTTATcaatgttaatatttttat
gaaactttTTCATTAGctcacccaaaagaaaaaaataagcaatTTCCATTAAATGATTCCTGGCATAAGTATTTGG
CATCCCTCAACTTATTTTCGCCACGATAACATAAGAAACTAAagtaataatttaaataatagtgCATATAGCTGC
ATGTGGTCAACCATGCTAAAGTTAATGAATGGTTCattttcatggcatatattTGAAAATGTTGACCATGAAATC
AAATTTCCAGCTCACAATGTAGACACCAAACCAGTTGGTTTTTGAAAACATAATCATTGAGATAACAAAGTATTG
AATCTTCATTTAGTAATCATACATTCTCATACATTTCAAATTGATCATATTATTCTATCAATTAAAATTTCTGAA
TGGTTTTAGCAAGGTAACCTTGAACAATTTTCAACCATAGGCTTGAACCAAATTGAGAGGTACCATCGAGTGCCC
TAGACATACTCTTTTACCAAAAAGTTCAAAAGTTTTGACATTTGTAGGTCACGATAGAAATATCCTGATTCTAAT
TATTGTATTTAATGGACTAGCATGTTAAGGCCTTCAAGGAAAATAGATGTAC**CATTGAATTTCTAGCAAGTTATG**
**CAGTTGCCGCCTCGAGCTTGTTACGTCCACCTGCATTGA**CTGGTGAAAACAATCAATAGTTTAGTCCAACGAAAG
TATAGATTCAACTTGACAAATAATTACCATGCAAATAAAACTGTAAACTCCATTAAGTTCTTTATTGAATTTCTT
GCAAGTTATGCAGTTGCCACCTTGACCAATAACCACCACTAATTCATGTACagcttaattaatttttttgacca
tGTTTAGTTTCTGTGCATTAGACATGAGTGGTGGGAGTAACAATCTTCATAGAAGGTTGGCAAGTAGACAACAAT
CTATTTGAGAACTGCCTGACTCAAGAACTTGGCTTCCTAGCTATCTAAAGGAAAATCGCCTCCACCAACTGCATG
GCCTTGTCAGAATTTCCATCAGTGAGGATTGGGTTAAAGAGGGAAATTCCACCTTTATCTCTAGACACACTCCAT
GTCAGTAACGAGCTATGAAGCTCCCCTATATCTAAGATgttacatatttttataaggtCAACCACTCGATTTAAT
CCACTATGGTTGATGGTCCTTGTTTTGATTAGAGTTGTTCTCCAACTTGTTTCTATAATTGGGGGTAAAAATATA
TCTGAACCTCCTAACATTTTGATAGTGATGTTCTCAACATATCAATTGGCATAATGTTCTCACTGGATGAACCAC
ACCAAGGATCATGTCATCTTGTATCTAGATAAACAATCTTTTATTTCTACTAGCAATTTAGTAAAAGCATCAACA
CCTAGGTTAGAGACCTTAAATGGAAATGATTAAGCATATTTCCCTTCGTCACTAGGAAATCAAAGCTAGTCATCT
CAAATTTAGATTGAGGTTAGAATTAACTTCATTTGACTCCAAAACTATGTCAAACATTGGATATCAAGTGGTTGT
TTGATACTAGTGGCCGAGCTGATAGCTTGAAgctcttgaaaaattaaaatcaatttccAAAAGGATACCACTATT
TAGTGCATTTTTGCAAGCCGTATATTCAACATACTTCTCAAATTCACAATACACAATGCTCTGAGATCTTCAATA
GGGTCATTCATGGTTACATAAAATGGGGCAAAATAGAATAGTGAAAGCCTCGCATCTAAAGATTAAAAGTGTCCA
AAAGATGGACGTTAACTCAAATTACCGCTTCTAATTGCAGTGTCTTCTGTATATTGCACGGCTGACATGTCCAAT
GCATCAAATGTCTAGATATTAGGCATCAAATAGGGTTTCACTACAACCAATCCCCTCTATTAAGCATCAAATAGG
GTTTCACTACAACCAATCCCCTCTATTCCTTTATGATCAAATGCTATCGCCTTTCATCCAACAATTCCAATTTGA
CATTGTTTACAAAAGACATTTATGAAGGACTGgattacaaaatattagaaatttaaaatccaattcaactttTAAC
TCAAAGCCTGCTTTTACTCTTTGAGGATACAAGGACAAGATAACTGTGTTAGAGGAATCAAATGTTACTAGGTCA
CAATTACAAtttcctatttaaatatattatttagaaagGAATCTAATTTGAAGCAAGCAAAAATGCTTTGAAAAA
ACaatcaagcaaatttttttgtCATGCAACAAATGCAAAGGTCTGCTATTTCAGGCCAATGTTCATGCATGACAG
TTGCTAGCAAGTAAGACAAAGCCAAGATTTAACTATGCTTGGAATCCAAGAAGCCCTCGTCCTACCACACAAATA
GAGGTAGAAAGATTGGTTGCAATAGTGAGATAACAACCAAGAACAACTTCCTTTAATAGCAATCTCATGAAAGTT
CAGGTAGATTTCTCTTCCTATAAAGATTAAAGTAAAAACAATCACACAAGCAAAACAACAATAAAGTCAAGGATA
ATAATATATAGCAATTAAAGGTTAAGTTATACCAACAAGGATGAAGGTCTTGCACATCCTCTGTGCTATCAACAC
TCAACCTCACATTGTTAGTCTTCGTGGGACTTCAGTTAAAGACTTTGCCCCTCACCAAGTTTTCCTTCCACAGAC
TACCATTCTGAGTTGTCTCACAATGACACACCACACACAAGAATAGAAGAGAACTTGAATAAGATTGATTGTATG
AGAAAAGATGAGTAATTGGGGTTATTCATGGATGCATATAATGATTCCTAGTCCCTTAAAATTATATAGTTTGAC
TAATTCATGTTGAATTCAAATGACAACCAAATCAGattcaaaacaaataacaatTCACTACCATCTTCAATTTGT
AGATCTACGCATCATTCAATCTGAGCTATAAATTGAAATGGAACAAGCTTCAACATTCAAAAAATGGCTCTATTT
TAAGATTGGCATGCCAAGCATGAGAACATATTGTTTGATTGAACGTCTATAATCTATTATGGTATCTCTATCAAG
GTTTAATGAATATAATGAGAACATCCATTTTCAAAGCATAAGGCCAACACATCATATAACCTAACAATTGATATT
GTGAGTCATCCCTTTCTTAAATCCAAGACTTGCATGAGATATGATACGAGAAATTGCCCTTATATTAGACCAACA
TTGTCATATTAGATGAGTTTCTGAGGTTTTTATcatgattaattaattatgatttttttttcaaaagaaaagcta

tTGAACCATGTCTCATTTCCACACAGTTGCTTCTTAGATTCGTGTATCTTTCTTCAAAGTTTTATTACATTAAGT
GTTTCTGAATTCAGTTTATGCACcatttttaaattcatttctcATTCCACCAAGCAAGGATTCAAAATAATAGTA
GCTCTGGCTGGTCCAAGAGTAAGATGCCTAAGTTAACGATGAAGGTAGACTAGATTCTTCCCACTGTGTTGAGGC
ACCTTCTTGAGAAGAGTATGATGTTTCCTATTTTTGACATGTCAATTTCCGTCTacctcaaaaataatatatcag
aATTTGACACCATCAAAGCTATTCAGTTATATCATAGGCAATTAATGTGTCTCTTCTATCTCAGTATAACCAACA
AGACTTCTACAATATTTGGTGTTTCACAGTAGAACAATTTGAAATGAAATCTTTTAAACATATACCTTCTTGATT
AACCTTGCTTAGTTCTGTCGACTGACCATGATTATTAGACACAAAAGATTGATAAATCTGCCAGTTTTTCAAGTT
CAGACACAACAACATAtagaaatattgaaatatataacTAGTTTGTACCTAATTCACTTTGAGCCTTTACACATT
TTATGGATTAGCCAAaattagagctgatcatgggctggtcctcggatctaaactctattcattttttaatcaggCT
TCGGGCTAAgcttatgtaaaatttgatatttcttgTGTCCAAGCCCGgtccatgatcagctctagcCAAAACATG
TATAACATACTAATTTAGATCTAATTCACCATTGAACCTTGACTCATTTTATAGATTAGGCAAAACATACATAAT
GGAGGATTACTGACCATAACTTATCCTTTTGAATGTGTAGACAAGAAATTATCCTTATCCCTCAGGATGGAAGGC
ATTactggcctcctcctcctcctccatcacttACAAAAAACTGCATTACAAATTGATTTATCTTTTGCTCCCCAT
CTTGACAAAATATGTGTTTTAAAGTTGTCATTGCGCCGTACTATGCATGATGAATGTTAGATacaaatccaaatt
ttaggcCAACATGGTTAGTACGTTAAGATTTTGCTAGCTGTGGTCAACAAGATCCTAAATGTGCATCTCTTGCTG
TAATATTCTTGCTTCAATGTGAGTACATctataaaataatgaaacagaAATTTTATCCAAACAACAATTGCCAGG
AGTTACTCAGCAATCTAACAATAGCCATGCTACTACTAAGTGTTAGTAAAAGGGGTTACCCACCAATTTTAGTGT
GTGAACATGACTATTGAGTGTGACATTGACATGCAATGATGTCATGCATGTAACCTATGGACGGTTATGTAtgaa
atgaacaaataaatttagTGTTGCTTTAGAGAACATTACAAGAAGAGTGTCCGACAGTACATGTCCTAAGAAAAA
GGCAATATCTAATCTAAAAGGTTTGGCTTTCTAAATCAAGGTATGACCCATCAAAGTCCAAATATGTTTACAACT
AGCTCTCAAAGTGGCAATAGTTTCTATGAGGTCATCTTTAATTAATCCATACAGCCAAGTCTTAAAATCACTCC
ATAACAGTACTGGTAACTGATACTCACTAAAACTTGCATGCTTAAGTAAAATCAGGTTTCAAAGTAAATTTaatt
aaagcaaaaaataagCACATGGAACTACCCACTCAATGAAATTTATGCCAATACACTAGCTCACAAAAGCCAATC
TATGGCTAAAGAATTCTATCAAGAACACAGCAATCAGCCCATAATACATCTATTAATGTACTCTAGTTGGTTGCT
CAACATTGTGGTGCACAGATAACACTAATAGTATTGGATTAACAACAGATAaagcaaaagaatttttgtgcGTG
CGTgtaacacagagagagagagagagaaagtagtaATATAGGGGTTGAAGGAGATTAGCTTGTCACGGAATTGGA
AACTCTTCAAAGGGTAACATGACGGAATTGGTTGATCCTATATACATGCAAGAGggataaaagaagaagaaataa
agaaagcaaGTTCAGCCATTATTCCTTTCGAGAGGGGAAAAGGGGGTTGATGCATTAACTCAAGCCACACCAAAA
TCACAACCAAAAGAAGCGAACgagaataaatcaaatcaataacAACGAAACACATAACATATTCAAGAACCGAAT
TTTAATCTCGCACCAGCACATCTGAAGTCAACATATATATGAAATCCTCTTTAC**CTGAGATCCC**CTCCCGAGGCT
AAAGACTTCCACCCTCAGATCCTCTCCAtaactctccctctccctctctctctcttccatttctCAGACTCAA
ACTGATCGACCCTCCGGCCACAATCCGACAGCAAATTCACGCATAACCGCGACAACCGATCCAACAAGAAGCGAA
AGCCAGAGCAACGACGTATCAAACGTTCTACAAGAatgccgagagagagagagattggatccAAAAACCGGAAGA
GAATGGAGAGGGAAAGCCGAAGGGACGCCAAAATATCTCGGGGGGTCATGTGCTCCCAAGGGTATAAATAAATGG
AGGAGAGCAGCGGTCGATCTCCATCCACATCGCCGCCTCTATTTCTTTATAACCCTCGAAAGTTTTGAAACGGAA
ATGGTGCACACTTCAAGGTATTTTATCCACGGAATCGATAAAGTGTTACCCAGGAGCGATGCAGGGTGACAGCTG
GTGTACTGGATGCCTTCCACGTGGTCCATCTCCCGAGGGAGGGGATCGAGCTGTGGGAAAATATCTGGGCGATGC
CATGATGATGGAGGGTTCCTGCGGAGTATTATAATCGAAGTGGAGAAGCGGACCCGGTGCGCTCGTTTCGCTGCT
TTATTGAATACGTGTTGGCCGTTGAACAGGTAGTAACGCGTGCCCCCACTGCGCGCTTTCCACCTGGCAAGCGGG

>PdAGL17_4
ATGGGGAGGGGAAAGATTGTGATCCGGAGGATTGACAACTCCACCAGCCGCCAGGTGACGTTCTCGAAGCGGAGG
AACGGGCTATTGAAGAAGGCCAAGGAGCTCGCCATCCTTTGCGATGCCGAGGTCGGCCTCATGGTCTTCTCCAGC
ACCGGCAGGCTTTACGATTTTGCCAGCACCAGCATGAAGTCAGTGATAGAACGCTACAACAAGACCCGAGAGGAG
CATCATCTAGTTATGAATGCTAGTTCAGAGGCTAAGTTCTGGCAAAGGGAGGCAGCAAGCTTGAGGCAGCAACTG
CATAACTTGCAAGAAAGTCATAGGCAGTTGTTGGGAGAAGAGCTTTCTGGCCTAAGTGTCAAAGATCTACAAAAT
CTAGAGAACCAACTTGAAATGAGCTTACGTGGTATCCGAATGAAGAAGGACCAACTTTTGATCGATGAAATTCAA
GATCTAAACCGAAAGGGGATCCTTATTCACGAAGAAAATATGGAACTATACAAGAAGGTAAACCTCATTCGTCAA
GAAAACATGGAATTACAGAAGAAGGTTTATGCAACAAGGGGCGTAAATGGCATTGATAAAGGTCCAATTACTCCA
TATGGTTTTAATGCCACAGAGGATGTAAATATACCCATTCATCTCGAACTAAGCCAGCCACAATTGCAAACAGAT
GGAGCGCAAGCAGCAGCACCAAAGCCAGGGTAA

>pda-miR444
GGGATCTCAGTCAATGCAGGTGGACGTAACAAGCTCGAGGCGGCAACTGCATAACTTGCTAGAAATTCAATGGGT
GAAGTGAGCATACCTATGACTTTCTTGCAAGTTATGCAGTTGCTGCCTCAAGCTTGCTGCCTCCCTTTGCCAGAA

Stem-loop structure

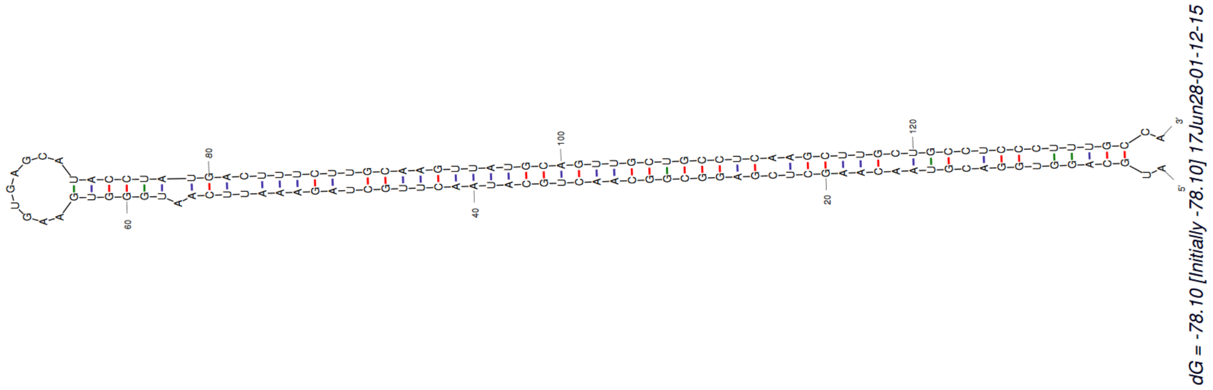


*PdaAGL17_1*

>NW_008246507.1

TGACCATCTATACGGCAGACGGCGCAATCAACAAGTCTCGGATGGGAAAAGTTATTATATGGACCAAGTCCGATG
GACCAATCAATCGACATACCGAAGCACATGTACGGTGGATAACAGGGACGTCGCATTTTATcttaaattgtttaa
atataAAAGGGGTAACGTGGTGAGTAGCTATTGGACGAGATCCGGTAGACATTCGAAGGGTAAACTAACGCCGTA
TGGTCCATTTGACTTGCCACAATTCTCGTTCCCCGCAGTAACGCAAACGAATTACATCAGAAACCGCTTCAGATT
CACCCAAGTTTACACCCTCTCACCATTATGCTTTATCCAGATGACGTAGATCTGCGTGCAGTGAAAGGCTGCAAC
GTTGGGTGTTTCTGGGGAGTGCACCCGAGTTCTTTGCAGAAAATGGAAGGCCTCCAGCCACTCCAAGAGTCCAAA
CGGGAAGTCTCCTCATGCCCTAAAGTGGCCGTCCTTGTCCTCTCTTATTGATGCATCCTAAAACCAAATGGCTGT
CCTTAAATAGGAGAGGACAGATATAGAGCAGCAGTCGATTCCTTCCATTATTTGCTCGGTAATCCACTCTTGTCC
CTTCCTTCCTCCCCAGATCTCTATCACCTCGGCTTCTTCATGGCGTTTCCTTGTTCTCCATCTGATGTTATTTCC
CTTCGTAAGACTTCTTTGCATTCTGTTATCAGATCTGAAGAGGATCGATCAAGTTATCGGCTGCTGCGAGCTAAG
TGGGGAAGGGAGAGATGGGCAGAGGGAAGATCGTGATCCGAAAGATTGAGAACTCGGCCAGCCGGCAGGTGACCT
TCTCCAAGCGGCGCAAGGGTCTGTTGAAGAAGGCCAAGGAGCTCGCCATCCTTTGCGATGCCGAGGTCGGCTTTA
TCATCTTCTCCAGCACTGGCAGGCTCTATGATTTTGCCAGCTCCAGgtcactcctctctctctctctctctct
cctccttcctNNNNNNNNNNNNNNNNNNNNTCCACGCTCCACGgtcactcctctctctctctctctctctcca
ccttcCTCACTATATCTGCATAAGTGCATTTTTTTTGTCTCATCCCATGCAAATTTTTCTCTTATTGCATGGTAT
GACATTTATGTGCAATCTTAGAGGCGTAGAAGTTTGGTAGATTTAATGGTAATTCTAGGTTTGGTAGAAGATTGA
TGCATCAAAAGGGTCGAAGAATAATGAGACCGAAGAAGCTGAATAAGTCCTTAGAGGAGAAGTAGTTCATTCGAA
AGAACAAAATGAGATATTTTTATCCCCTTCTGGCTTAATcaccttctttctcctctctcactTGGTCTCTTGGGT
TTTTCCTCCATCAAGTTCGGTGCTGAGGGATACAACTTCCCCTCACCAGGCACAGAAGtagtttttttaattgaaa
aggCTTGTGAAGCAATGACGAAATTAAAGGCTTGGAAGAGGtccctttttttgttttgttatgttttgttttgttt
gaaataGAAGAAGTGGAGGGTTCGTTATAAAGGTGTGGCAGAAGTTATGTTGAGAGTCTTCTATTGTAGATTTTG
ATTTTAACTTACGGTCTATTCAGTCAATATCAGCCGCCGAGGATAGAAAAACTAATTATGTGATTGATTTGTCTT
TTTCCCAAGAGGAAAGTAAAAGTTTTCGGTACGTCTTTGAAGGGCTTAGAGATTCTTCTGAGCCTAATTGTGGTA
GAGTAGTAGATGATGCTCTTGAGCCTAAAGATGATAACAGATTCTGAGAATACTGCATGAGGGATGCGATGAAGC
TTACCTTGGCTATTAGAATTAAATCATGATGAAGAGCTCCGGCACTTAATTCAGCAGCCATTGCGATGAAAACTT
TCGTGATGTTGACAAGGCTTGGTTGACTCGTATAAAACCTACGGGAACTTAAAAATGGTTTGTCTTCCATTGCTT
TGGACCGGCCGACGGGAATGCCATGATGCAGGGATGGTGATTTGACAagaaaccaaacaaaagaaCTGGAATTTT
CAGAAAGATAACAGAACTTAGTGTCCGtcaaattcatgaatcagaAAGAATGTGATGCGAGGATACATAtgccag
aaaacaaaaaagggacGTAGTTGGCACTGGCTGGAGCCTAGATTGTTTAACGCATATAAGCACTACTGGAGCCAA
GATTGTTCAGTAATCGATCCCATTTgcatttcaaatgcttcttcACACAATTATCTAATTGGATGTTGTAATAAG
GATAAGCAGTATCATCATAATTTAACTGATatgaaaatcaatttacaaGACAAAGGGGCATCTCCCACAGCACCA
TCGGAAGACACATTCATACCATTATAAATAGCTGTGAAATCTAGAAGTTACTGAGAATAGTATCTCGTGCGTCAA
GATCTTTTATCAACTCTGCAGTGAATtaactaactttttttttttgctgaataaTTGATGCCTTGACCAAGTTGC
TTATCTACTACATATATGGAGTTGAGCAAGCAGTGGATCCTTTCCGGTCTAGGTTGCAGGAACTTCCTGCAGTTC
GAGTACATGATAGTTTTTATATAGTTGGTTTAAGGAGAGGCAATTTGGCATTCTTTTCACCGTATTGCCTTTTTA
AACCACCCATCTGTCACAACAATTCACATACCTAAGATGTAAGAATTTCCTGCAGCCAAGATGTTGAGGATTCAA
ACTAGGACAACGAGTTTGAATTGATTGTGCATAAAATGGTAAAATTGCTCATAGTCTTTTCTGTTTTATTCTGGA
CACATCTCTATATTTTGATGTGATTATCCAAGTATAAAGGAAATCAAACTAAATTTTGAGTTTTGAATGATAACA
TGTGAATTACATTATATGCATCAGTGATATGCATATGCTACATTGAAGTATAGAGCTTTCCCACCCAAAAAGTGT
AGAGCTTATCTTGTTTCAAAATAGCTGTGATTATTTCAAAGTTTCTTCTTAAGAGGAAAAAACCAGGAAGTTCAA
TTATCTTATATCTTCATCATAACTAGTGAAAGAAAACAACCACAGAAGATGAGTTGCTTAGAGTCAAATGTTCCT
ATGGGTAACATATGACTTCTCTTTCCGATTGAGTATTTGACTTTTAGTCTTTAAGTTCTCTTGCTGGATTTTTTG
CAATGacaagatatttattttttttagattcaCAGTTTTTTTCATGTGAGAATCCTTATTGATAGTTCATGCATGC
CCAGTATGTTGTAGTTATAATTTCCAACCTTTTATGTGATTTATGTGAATTTTTTTCTCCAAGCTTTCGAAGGAAT

AGTATCTAAAGCATGCAAGAAGGAAAATCAATGATGGACAAGAACGACCATGCAATATTAGATAGGAAAAGGCTT
TTTTTATGTTATCAACTTTTCATGGAATAGTCCTATGGTGGAGTTTATACGCTCAAACAACTTCAGACTTCTGAT
TTGTATAGGATATCTCCGTCTGTATAGTCgtaattttaatttgaataggctgagttctgaaaaattcatTGTCCG
AAGACGatgaaaattagctatgaaggGGCAAAAAGCATAGATGGTATTTAAGGAGTTAATATGGTCATTGGAGTA
AGATGGAATCAGCAACTCATGGTCAAAGAGTACATCTCTGGAGGAAAAGCAAAGGGCAAATCCCAACCGATTTCT
TCAAAGTAATGAAGTACAAGACAAAAGTGAACAATTTGAGAACGGATCTTTCCTCTGGAAAACATGTAAAGTTAT
ATGTGCTCCCAAAGGATCTTCAGACTTGATATaaactactttggccactacgaataCGGTTGGACATAGTCTAGA
AGTAAGATCCGCCATATTGTGAAGCCAAATCATGTACTAAACTTCAagaggccataacttggtcatgCAACGccc
aattgagatgatcttttatttttgaaattgagGTTGAATCAGGCcaaaattctatcattttGTGTTCCAAAA
CGGGTTAGTCAAATCAAAACTTTCCTTTTcggaaaaaaaactttgaatAGTTGTTATCTCTTAATCTATGAAGAA
TAAGATGGAGATACAGAAGTCAAAGTTGATATAAAAATCAAGATCTAcctcttataaaattttatttttgaaat
tatttttattagttatatttatttttggaggtCTAGTTCTATTCGAATTAGGAGAAGAATCCAGtctaaattatt
aaagaaaggACCTCACCAATATTATAAATAAGAGCTATGTACCTCGATTTATGAGATGTGGAGTGTCAATAAAAG
CAAATTAGACTTAAGCCCTACTTTTGTAGCTCTtctatcttctatttttttaaaagaaatttgagTTGAGcgaa
ttgaaaaaaatatttctttttttcaatcgaatcaaattggtatcagagcttcgaTTCTTTACATCTATGAGAGT
TTTAGTTTATGGTCTAAACGCATAAGCAATACAAGCAATTAAAAGAAGAGCTATACGTGCTTCAAGCATATCGAA
GGAAGAAGGCGAGTATGGTTGTTAGTTCTTTCTCAATGGATGTTGAGATGAAGAAATGTCTCACATAACCAAAAG
AGAAGATTGATCAACTCAATGAGATGGTCTTTTGATTGGCGATACCTTCAGCATAAGTACTGGCAACTTTTGTTG
CGAAGCTATTTCTTTCATCTAAAAATGAAGTATACAACATCCAGTGAGAAGGTTGAGCAACCTACAAATATGCAA
AATCTTTTTTCAACTCTTTAAAGTTGAAGCTCGAATCGATATCTCTATATATGATAGAATTGTTAATATGGAGAT
GCTAGATAACTGGCTAGACCAGTTAGAAACCAACTTTACCATCTATGGATACTCCAATATCCAGAAGGTAGTTTT
TGCTCATCTCAAGCTTTCTAATCATGCTCTTATCTTGTGGAATTCATACATGAGAAACAACGATGCTTCTAATCT
GACATAGAATAAATTAAGTAGTTCCACCTAATCAACTACGAGAAGGATCGGTGGATTAAGTGGCAGTACCTGTGG
CAGAAGTATGATCAATCTGTCCATGACTACACCACCGAATTTCCCAAGCAAGCAATTTTACTTGAAATCTTCATC
AACGATCATGCAGTTTTTATGAAGTACGTTGGAGGTCTCCATGAAAGTACACAAAAAGAGCCGTAACTCTTTAAA
GTTGAAGATATTAGCAAAACCAACGTGAAGGCCATGAGGATTGAGGAGAAGAATTAGCTTAGGAAAGGTAAAAAG
GAcaacaagttaaaaaaaagacagTAGAAAATCAAACTAGAAGGAATGGGATAATAAGCAAGCCAATGtgactaa
aaaaaaagataatttagaaAAACATTATCATAACCTCGATGGACACATCAAGAAAAAGTATTGAAAACTTTATCC
AAAGTTACAGCCGAAGAGGAAGAAGCCAAAGGATGATAATTTGAACAAGAAAGGAAAGGTGATTCTCAATACTAG
AGGTGCAAGAAAAACTCTTCCACTTGAAAAATCTGGTGAAGCAGAGCATTATCAAGTTTATTATAGATCTTGAAA
GCTAGAAGAATCTCATCTCTGAAAATCTGGATCGAAAGCTGGGGTTGCAAACCAATCCTTATCCACATCCCTATC
TTCTTAGTTTGATTCATATCTACATTTTCTTATGAAAAACAAGTAAAATTTCTTCCTACCTCTAGACAAGTATGT
TAGACTGAATGCTTAATCTTAGTTTTCTAAATAATCTGTGAATTTCTAGGCACGTCGATATGTATTCCTAAAATA
AATTGTtcaactaattttcttttatgattctatccttaagggattttcttttaatctctAATCGTAAGTAATGAT
CTCTCTTGCTTATGGATTAGAGAATGTTACCATTCTATCATACAGATTATTTGAGCATGAGTTTAAATTGAAGGA
CTATTATATTAGAAGCGGTCATATGCAAAGAGGCTGGGCTATTGTTAGGAGCATGAATCAGGGCTTATCCAAGTA
GACATGTGTCAAATATTCTAGGTTCAAATCTTGGATGTTTCATCCCAAGTATCATGACCTAGGCAGTTATAGGTG
AGTCTTCCCTTTCAAAAACATGGTTGTTGTTAGAAGCAAATATTCTTAACGAAACTAGAAACTGttgataattat
aatatttttattataattaatctGCATCTTTATGTTTTGCTTATGGATTGTGCCAGCTGAGGAATTAACTTTCCA
AATGTGGATTCCTCTCCTTTAGGTTTTTGTTAATTAATCAAGTTTTTAATCCATCATGCATCTTCTATCTTCCTC
CCTTATGTTGATCCCTTTTCTGTTGTTAATTCGACTCTAATTGGACTCCTAAACATATTGGACTCTGTCTCTTAC
TCTATTGTCTAACCTCACTCTCTCCTCAAATGCGTGTTGCATGTGCCCAATTGCTAGTAAACTATAATGTTTGAA
TTAAACTAGCGCACTGTTTGTGTGGCAAAGGATTGGTTTTGAGTCGGATCAAATATGGATTAGGTCAAAGAATCT
TCAATGTGTACTTGATCGATTTACAACTATAAActgatttctatttcatttattttatatgcacACTTAGTTAAC
CTCCcaatcaaattataactttacTTATTAGTTATATatcttgataaataaaaattaaatgggTGTAGAAAGATA
AAACCTAGAGTTAGCTTCAATTTTTGAGAGATAAGAGGGAGACCCACCTATATTATGGTTAACTAGGCTCAAAAT
CTAAGAGATACACTATCCATACTTGAACCTAAAACCTTAAGTTGTTAGAAGCAAAGGAGCTATGCCTACTAGGAT
AATACCCAAATGGATTATTAAAAGTCTTTATACAAActcatttcttttaaaaagattCAAAAAGGTTGGAGAGCC
TAAGTTAAAAATTGCCATCTCTAATTTATTGAACTACCCAGAACATAGGCACATCTAATGCATATTTAGGTTCGT
CCAAGTCTCGAGCCTACGACCTTCCTATTATTATTTAATGGAAATAtaactaataatatatttttttcaataacat
ttttaaaaattgaagaaTGTTTTGGATAGTGAATTTGATAAATAGAATCTATTTGatatatttctcttttatttt
gcaataaggtatatatatatatataaggcaaCATAGAGGCAAAACATTCGGGTCAAGTCTCCTCCAATGGATGCAGG
ATGCCGTTAACTTGCGGGAATAGACCTCCCATCATCAACTTTACAATGCTTTTGCACTAGGATGAGTATATACAA
AggcaatggaaaagaaaaaaagaaaaccccaCTCTCATTTACAGTTCTAAAAGCAAAGAGCCATGGGTAAGTTGG
TCTCGAGCCATTGGCAGTGACTATAGTGGCAAGAACGACTGTTGAAAATGTTACTTTCACCCTCATCTGCCCTTT
AAGCAACAAAGAAGAGGGATCGCCCGAATGCAACTGTTCTAATGatgtaaattatttgaaattattGGAAGATAC
TATAGAAGCCACATGACGCAAATTAGCCGTGCAATTTCATCAGGGACCAAGTTGCCAAACCTGCTCTTAACCTGA
TCATCAATTCCTCGTTTCCCCTTTCAGTTCCTTTGAAAATTggagtttcaaatgaaaaaagaaaaagagtttggA
CGAGGAAAAATGCCTCAAAACAAGCATTTCCAAAGTTTGGCATGAAAGAATTGCTCCTAGAGCACCTTAGCACCA
TCACAAAATGAAAGAATTGTGTCATGCAACAGTAAATAGAGTTAGATGTATAATTTTACAAAAGTCACTATAATT
ATCTCTGAGTGAAGCAGTGAAATTTTTTTGTAACTCTGATTAATCTCCTTCAGTTCATTTGGATAATGAAGTTTT
AGTAAAAActattggaaagaaaaatatctccTAAAAGCACTTAAGTGTGTTTGAGTGTTAAGCATATGTCCATGT
AGACAAAAATGAGCAACAAAGCTTCATACTTAGCACCGAAAAATGAGTCCTCCGATGTTGGGGCATATGAAGTCT

TTGGTCACAAGCTATACATGGTTCCTCTCTTTTATGTGAAGTCTTTGatcatgagatttttttttaaacaagatG
GTGCAAATAGGCTCCTTGTTGAGAGATAGTCCCATGTTAGATAGGTTAAGAACTCCAATAGGTCTTATATATTCA
TGGGTCCTTCCACCTATCAGCTTAAGATTATATCAAAGTCAGCTTTCTTGCTTATTGATGATCATTGGCATCTCC
AAATCCATCATAACGTCTGTCCAAACGATCTATTGCAGATTGGCAAGGAACTCAGCCCACCaacaataagaagaa
aattaagaagaagaaagcttccATTAACTTGCATgttgaaaaaaattagtagacaacaaaataaaattgacTCGA
AATAAAGTATAAAATTTAGTCCAAACTTATCATTTATATGAATATAATTGTAAAATCGTAAATTCTTTGGTAAAA
TGGCTATATATAATTTGCTCAGTCTCTGAATCATGTACCCTTGATTTGTTAGATTGCTTGATTAGATATCCAATG
ACTTCGGTTTTGACACTTCAATTCACCATGAAATCTTTTGGTGATACATATCTTTTAGCTGTTGCATCCTTGACA
AGTCAAACTACTAAATTCACTAAGACTTGCAATTAAATCATATTAGATGGACTAATAATGACTAGTTGCAAATAG
GAAAGTAGTAACCAAGAAATTCCTAGTAAACTAACACAATTgaacaattattatatttaccaacaaaaaaaaaat
cttacgaGGAAGTCCAATTTCCACTCCGGTAGGAGTTTCCCGTAAAAGCAATTTAGGAAACCAATCTAATTCTTT
TAGGAAACTTCCTTGAGAGCCCAAATTTGGAAAGAAAGAGGACTTCTGAAGGTTGGGCTATCGAGCTATGAAGAG
GAGGATCAGTCAAGGAGGGTAGAGAGGGATGGAAGTGAAGGAGAAAAATCTAAAGTTAAAAATTCTCCCCGATGG
TCTTTTGATAAAAGCATTCTGATAAATCATTGGTTTGATTTGTTCAAGAGAGTAAATCTAGTGAAAGAGACATCT
TCATCAATATTTGTGAATTAATTGTTAGCTACTTATGAAATGGATTCCAACTTGATTAGTTAATTGAAATTCATA
TGTTTTCCTATGATTTTTATctagtttttgcatatgtatttaataatattttttctttatttatgacCATGTTTC
TTGTCTCTTGCATGAGAGTTGTTTTCTTCGCCATCATTCCACATTCCTTGCATTTGTATTTCTGTATTGGTTTCC
CTGATGACAACTGTGAATCGGCTCGGTTCCATAAGAAATATATGATCTGACAAGTCCTTTTTCCACGGGCAAAGG
ACAACAAAATTTGGCAAAATCACCTATATTTAGAGTATAGACAATAAAGATCGAGTCATCGACTATAGGGAATAC
TCTCCGAGGGAGAATGCTATGCCCATAAATAGAGGTTAGCAAAGAACTATTTATTAGCCCAAGCAAGCAATTGTA
AATAAGATTATTTAAAGCCACCCAAAAAGTGATGGAATCCTAGCACATAAAACAGGCCTGAATTGGGGTGTTGCC
TAGAAGGCTCTCAAAACAAGTGAGGTTTCatgttaagatttttttttcttttacaaaaaAGCTTTTATCCAGGGT
ACAAGGTAAATCATTGAGGGTTGCAATGACCTAGGTTTCGATCGCGCCTTTACCTGATTTTGACCGGGCTTCCTC
CCTCCTATTTCATACGCACAcaaagggagagaggagggaggggaagagagagagagagagagagatagatagataga
tagagagagagagagagagagagagagagagagagagagaggaNNNNNNNNNNNNNNNNNNNNgagagagagagaga
ctgggGGAGCGAGTACATGGCATGGTTAGTAATAAAAGGATTCCAAGGGATATGGGCAGAAGAGCTTATTGACTT
CTAAGATATTTTCTCATGTTATTGGAGTTACTCTTTTGCAAACAGGTAGCGTAAAGTTTGACACTAAATGCCTGC
CTTCCATGATGATTTAGAGGGAATCATTTATACGAAACAATCAGTAGGTTTCAAAGTCAACAACAAAGAGATCGT
TGTTATttctaccaagaaaaaaaaagaggagatcaCTCTTATGGCCTAAATCATATTAATAGGTAATGATGCAAAA
GGTTTTTCTTTATGGTTTGAGACCATGGATATCAGAGAACTACCTCAGATCATTGTTATCTGTCAACAAGTATCT
TGATGGTATACTTATGTCAATAATACATTGTTGGTCAAAATGCTAACAGAATTGAGAATAAAAGTTAGCAACTTC
TCTGTATAAAAGTTTAAAgccaagaaatcaaattttaagcataaatctTTCATAATACAAAGGCTAGGATGTCTC
AATTATCAGATCGATTCCCAAAATTATTTTGTGGGTCTTCCTCAATGTCCCCAATCACAGTACATTATAAGCAAA
TGATCAATCATATGCTTCCAAAAACATTAGAAAAATTCCTTAGGAATCCTGCAAATTCAACTTGTTCTTTCCCCC
TAACTGATGATTGGAACTCCACGCAGGCTTGAATCTTACTGAGAGGTTCACTGCAATTGAGGCTCCTTTAGTTAA
GTAAATCCTACCCAGACTGACATTGTGACGCTCCTTTAGTTAAGCATGcatttttttgaataacaaatcaTAGATC
ATGGTCAACGGACGATTCAGTTTCTCCAAAATGCATGGGGATTACTAGCAACAGAATTAACGGCAAAGCTGTAGA
AAAGCAGCAAAGTAATTGCTAACATTTTGAGAAACTGTTGTTTACTTCTTGCTCTCTGTTTACCGCATGGCATTT
TATAAACTTGTTGCTGGTAGTTAAATAAAGAGAGACCTTGATGTCCTTCAAGATTTGAATGAGTTACTAGCATCT
GTGCTATGAGATATGATTCATCCATCCACTGatcatttttcagaattttatttGGACTAATCTAGCTTAAAATAT
CTCTGAGCAAAGGAACTATGCTATTTTCTGAACCCCATCGTCTTCAGAACAAATAAGTAAAACACAAACTTTGGA
ATTAACTAATTGAAGATGCTTACCAATTTGGAACAAAGTCATGAGAGAATTTCAAGAAATTTGCACTAAGAAAAC
TAATAGACAAGGGATCATAAACTAACCCCAGATGGTTGGGAGAGACTGTTAATTCTAGTTCGTATTCCAGACATA
CAAGGGAAAAAGATTCAATAAATTAGATGTTGTAGAAGAGCAGTGGATGGACAACTAGTACATAACAAGACCACA
TTGAGACCATTTGTAGGGCTCGTTCGGTATTTAAGTCaccattttgaaaaaaaaactcaGTGGTTAGAGGATACAA
GGAACTCAAACTGTGTAATACTTATTAGAGAAAGCAGCActcttatatttaattatatattcatGTTCATTATCT
AATTCGATTATCATTTGCTACTCTGATAAGGCATAGTTTGTATCAACATTCAGTCTGATAGcataataaagaaaa
actaaCAGAAGATCCCTTATCATAGCAGTATGAAATCAATAATTGACCGCTACAGCAGGACAAAGGAGGAGAATC
ATCAGGTGCTAAATGCAACTTCAGAAGTTATGGTATGTCCAATTCTTGTGCAACACTATCAACAACTGATAACTG
AGCTTCTTCTTTGGTGGTGTACAAATGCACAATTTACTatgtttatattaattttgcCATCTCTTCTTGGATAAA
AGTAAAACTTTTTACTGCAAATGCCCTGGCATCTATTAAATTGCTGAGAACGAAAATAACACTCTTCCTTTTGCT
ATAGAAATTACTAAAATGCTGCATGATGAACTCTTATGTTACAGTTATGTACAAAAACATGCAAATTATTTGCAA
CACTTGTTTTGCATCATATATGTACTGCACCAGAAAAGTAGgggaaaaattagaaattacaCAAGTGTACATTAG
GTAATTATGTTTATCACAGTTCGATTGAAGAAGCTTAATTATGAAAACTTTATAATGTATAGTAGTAAATAGTTG
TTAAAAAGTAAATTACCAATATATGATGGAATACGGATATGACATAATGCAAAAACTATATAAAACTTATGATTA
TCCATGTCAGACATCTGTAAAAGCACAATAGTGAGGCTACTCAATAAATTACGTCCATGGAAATAGGATCTACAT
ATGGGTTATGTTGACTTGTAAAATGCATATGCTAAAGTGTTAAGattcatataaaataatgaatagCAAAGGCAG
TGGCACAATAATTTTTGATAAAGGTATTCATGATGGAGTTGAGAGGGGTGATAAGGGGCAGATGCTATAATGTAG
CAGGTCAACAATAAGGATGGTCCTAAAGTCTGTCATCACGAACAGGTGCTACAATTACTGATTATTGCTACTTAG
GTGAAGAATCAACTATGTTGCTCGTCTGTAAACAAAAACAAGGAGAAATTCTTACGGAACAAACTTggcattttt
gataaaaatatttatatggagtTGAGAGTGGGGACAAGGGGCAGATGCTATATTGTGGCAGGCCAACAATCAGGG
TGGTCCTAAAGCCTATCATCATAAACATATGCTATAAGTACTCTAGCAGATACTATACTTTGGCAGGCCCCTGAT
CAGGACCATCCTAAAGCCTGTCATCATAAACAGGTGCTAGAAGTACTGATTGTTGCTACTTAGGTGAAGAATCAA

CTCTCTAGCCCATTCATAACCAAAAACATTAAATTGTTGTGAGAAGATGAAAACAAGGAGAAATGCTATATGGGT
GTTATGAAAGAAACTTCTGCCAAATCATATTTATGAGCTTAGTTACTAGAAGCAGGTGCGGAAGCCAATTCAGGT
GTGTGTTTGGGGAAAGGATctcttaatctttttttttataataataaacgGGTGCAGAAACAAGTTCCTATGGAGA
TTGCAAAGCAGGTGTGGCACCCTATGCAGAAGATTCCTGCTACCCAGGTTATGAGTGTAAACAGCAGATCTAACG
AAAACAGATGAAGAATGCAACTGTTGTGGGCTGAATAATGAAAATTTATGCTTTGGTAATTGATGGCAAAAAGAT
GCTTCCAAAAGATTATTACAGTTCAGTTCTATTATGTTGAACAAACTCAAAGAAAGCAGTAGACATTCTCATCAA
GTTTTGAGCATGGAGAATGATGTAAAGATGTGTTTCTGGTGTGCTGCATGATTGTTACATATTATGGAGGAGACA
TAATACAAATTCTATCGAAAGGAAAGCCCATATGCTAGGTTGAGAAGTGGTAACTGAGACAGTCAGAAATAAGCA
TAACTACAAGAAGCTAAATACAACTGCAATCAGTAATTGGTATCTTGTTGAGACAGCAAGGCCATGTCATGACTA
AAGAGGGGCAATACACAATTAGAGGTACCTCAGACTTACATTAATGGGTTGAAGAGAGGATGGGAAGGCCTAGAG
AAACACAGATCGAAGTTGAaagaattaaattaatatattcgAGTAGAGGAAGCAGCCATGCATCTTTATAAGAAT
CAATGCTGAGTAATAATTCATAAATCTATTACAAGATAAGACTTGGAAACCATAGTTATGTTGCAATAGTTTCAT
ATTCAAGCAGGATTCTACAAGAGATAATGACCTTTCCGTGCTATTTTTCGTTTTCCTTGCTCAAAACAAATGCAA
TCACAGAGCCACTGAAGCAGTCAAAACAGTTTATACATTGGCCCAGAGGAAGCAAACTGTTCATGAACAGCTCAG
CTTGGCCAGCTAATCGGCTCTGTTGAGCTTGTTCAACTAAATGAGCCAAGCTTAAACATCAACTAAAGCTCAACT
CATAAAAGAGTGGAGCCTCAACACTTCAGGCTTGGAACCTTGAGCTTGGCTCAATAACAAGCTTGTCAGATTAAC
AAAAAAGGCAAGCTTGAACATAAATTATTTGGGCTCCTATAATCTTTCATACAATAAACTGCATCTGAACAGCTT
AGTAATTGGCTTGTTTGTATCCCTATTTATAAGAATTTATACCAAAGCATGACCAACAGGTTCTGTAGAATTGGT
AGTAATCATGTCATGTAGTGGGCTCATTGTGAATGGATTAACTGCAAAAACTGAGGAATTAATCATCAAACAATC
TCACCACTGAAAAGTCCTGAAAGCTGGCCCCAGCATATAGAACACAAAAAAGCTCCAACTGCTTGTTTGCATGAT
TCGTGTCATGCTAGGTTTGTATTGTTAGGAAAATTGGGGTTAATTGATAGCCAGTCCTTTGTTAGACTGCTATTT
TACAATCTATTTCTGTTGACTAAAATTTTGATTCCGGCCATTTAATTTCGATGGCCATTTACTGTTTAGTCCCAG
GTGTTGTCCATGCTTAATCATAAAGGTATGCAAAACAGATATAACTGCATACAGAAACTGGTGTAAGTCTGTCAA
AAATACAAGTGTCTATGCACAACATTGGCCCAACTTGGTGCAGGACTCAACACAAAATATGACATTAGGGCTAAA
CATAAAATGGTAGTCTGTCAAAAATACAAGTGTCCCTatgtcagatgtttgtcagtcccgAGTGGCAGGGAAGTA
GATATGCTCGGCACAGCAATGAAGGAAGCAGGGTCGAAGACTTGGTGACGAGGCAAACATTCTGGCAATGGGCCA
CTatgatagtgaaggctatcaaaccattatgtGAAGTGCTTCGGGCCATGAATAGCGAAAGATATCTCCAGATGG
GCtttctgtatcacatgatggagaggACAAAGGCTCAAATCAACGAGGCAGATCTAGCCCATGCCTAGAACTACA
TTAACATCATTGAGCGGCGGTGGGACTACCAAATGGTAGAGACTTGCATTTAGCaggtaagcaagaacattgaaa
atCAGACTGCTCCTGTACTAGTATCATTTTACTAAGACGATCTCCTCTATATGCAGCATACTACCTGAACCTCCG
GTTCAGTACAATGTAGctggaattggtatggatgacgAACTTTTGGCAGCTCTGCGTAATGTCATTTACAAGATT
GAGTTTGACCCAGAAGTTGCGACCCTATGTCTAGAAAAGGTAACTTAACTTAATTGGTATATGTAAGTTAACTGT
catcttctattattaacatagtacaaCACGGTTTTTTTTCACAgatcaaattatttagagagggcagcaaCAGTT
TTGGACAGCGGGCATCTGTCGTGAGTAAGATAACTATGAATCCAGGTATGTTACCCATTAGAAGGACATTCATCT
TgcattagatggttactaatatgatactatcttatatcaattttaaatcaacatttttgcagctgaatagTGGAA
TCATTTTGGGCTGtctgcaaaaaattttaaacgGCTAGCTATTCGGATCTTCTTCCAGACAGTCTtctcgagtgg
ctgtgagcgcgactggtccaccttcgccctcatccacaacAAGCAGAGGAACAGTTTGATACAGAAGCGCCTCAA
CAATCTTGTATATATCTACTACAATCTGAGATTGTGGCTAAAGTGCATTCACGAGGAAGTAGAGCTTAAGTATGC
AGATCTGATCAATAGTTCTTTCGTTGATGATGACAATGATCCTATGATTGGATGTCTTGTGGGCCAGCAGGAGGA
GCCAAAGCTTGACGAGCCAAGATTGCCTCCACGATCGATCAGCTTCGTAGCCACCAAGGCTAGGGTGGATGCAGG
GCAATGGGCTGAGTGCAATATTCCACACATTTCTACAGCTGATCAGCCACAGCCATAGGGGAGCCAGTCACCATA
TGACTCCTTGTCTGAGACATCCTCTCAAAGACATGATTGAGAGATGCTCGAACGAGGCCACTACACTATCCGATC
GACTCGGTTAGGAGGACGACATAGCCAGACAGATAGGGGCATAAAATGTAAGAAGGCAACTATCCAGCagaccaa
aaaataaaacaaaaggcacCTGCAGCCCCAATGAAGAGAGTCTTGAAGGAGTCTTGGAGCAGCAACGATGACGGT
tctggtggtcatggatctgctgGCACTGGAGGAAGTGAAGTCCCAGTTTATGAGACATAGCCGGAGGATGGTTGA
AATACTCTGGTGAGTCCTAGTTTACGTATGCTACACAAGATAGGAACTACGGAGCACGATCTTGTAGAGCCCGCG
ATGTTGAGATTTCATATAggagacgggctcctcgaggttGTTTAGGAGGACACGATATaactgcagatgacctcg
catgtGGAGTAGGATCTATAGACGTATCAGGTTCAGAATTGTatattgaattttattattcaCAGCTACAGacag
cctatgacccatacagatatggatatggatatggtgtgtctgaggcatcttccggtggctactatcctacaaAGC
CCGGAGCATCTCACGGGACAAATTTGCTGCCGATATATTCGGATAAGCCCTTCCACAGCCGTATTATCAgctaga
aaatacttctcaGCGCCAGAGCTTAATGAGAGAtgcgagagtgcttataacccggcACACATTTCTTATAgaatg
aacatacaagactacaacgctgCTTTAATAGAGGGATGgattgatgtgcctcctgattaTACTGATGATCCAAAT
ATCTACGAGAAACATCGCCACTTGATCCGATTTTAGAGATCATTATATATAATTGTTGGACTTcaaaagtatgta
attgattgtatctggatttgaagatatttttatgttattcattttatgatttgtatcattttaaagcaataggat
gcatcatctagcttaaaacgaaatcataaaaatatcgaaaaacataaaaaaatatcaaaaaattttaaattagac
ttaaattcataaaaaaggtttcaaaagagagagagctctccaaaaaaaaatttaggacaTGCCGGTGCAGGAACCA
TCATGCCCAaccggtttggttcggtatgAACCGGTACTGAACCGTACAAGGCGTCGACCCATTCAACACTCGATA
CCGGTTCAGCGATGCTTGCATCAAATAGTGAATACAATCCAATCCATTACTAGATTTATTTATAACCACACAGAT
TCTATCACTGACGAGGAGGTATACAGAGAGGAGATCTTGAGACCAGGTGTGATACAATTTGCTACTGACTATATT
GCACTCGGTAGCCTCCTTAAGAGGAAAGCAGGTTTGCAACAGATATTTGTTAGTGTCGAGTGGCAAGAGACAGAT
TTGCAAGGGCTGACACTGAGAGGAGCTATGTGGAGAACTTGGTGATGAGGCAGACGTCCTGACAACAGGATGAGa
agatagtgaaggctatcaagCAATTATACAAAGTGCTTTAGGCTGTTGGTAGCTAGAGGTACCCCAATAGGTttt
ctatattatataatgGAGAAGGCATAGAATCAGATCAAGAAGGTAGATCTGAAGCATGCCCAAgagtacatcaac

atcattgagcaGTAGTGGGACTACTAGATAGGTAGAGACTTGCATTTAGTATATAagcaagaattttaaaaattt
aactgCTCTtgtacttgtataattttactaataCAATTACGAACTCTATCTGCAGCTTGCTACCTAAATCCGAGG
TTCCAGTATATTGTTTTTGGGATCGATATGGACGGAAAACTTCTGGCGGACCTACATAATAAGATATATAAGATA
GAGCCAGATCTAGAAGTTACGCCCCATGTGTATAAAAGGTAAGTTAAGTGGCATGCGTAAGTTAACTTAGCTTGT
GTCTTCaatcattaacatattataACACATCTTTTTTATAGACAAATATTTAGGGAGGGCACTAGCAGCTTTGGA
GTCTCGGCATCTGTCATAAGTAAGAAAAGTACAAATCCAGATATATTAGATATTAACGATAGCTATTACTTAACG
TTAAatagttactaatatgatactatcttatatgtaattttctttaatatttttgcaaCTGCACTatggatttaa
ttttgatttgtcGGAGAGAAATCTTAACTAGCTAGCCATCCGAATCTTCTCCCGGACGGCTTCTTGAGTGGTACg
tgcaactggtccaccttcgccctcatccataACAAGTAGAGGAATCGTCTGACACAGAGGTGCTTCAACGACCTC
atatatgttcactacaatcaGAGATTAAGGTTGAAGtgcattcagaagaaagtgGAGCTGAAATATACGGATCCG
ATCCACAAAGTTTTTGTTCGGGATGAGCATGATCCTATAATCAGATGGTTTGTAGGCCAACAGTAAGAGCCAGAG
CTTGATGAGTCGGGATCGCCTCCACAACCGGCCATTTTCATAGCAAGCGAGGCTAGAGTGGATGTAGAGCAATAG
GCAGATCGGAACATTCCACAAATGCTTCTAGCCATGGTATGCTAAACCAGCCAGTATTGGCTGGTTTAGCTCATA
CTGGACTAGTCTAGTCTCTGACTGGTTCGGTACAGCCGTGGAAAACGGTTCCGGCCGGAACTGAACAGTACGGAG
CCCATACGGATCGGTACAAGAGCGAACCCGACGAAACTGGATCGATTTCATCCGATTCGGCCGATGCGAACCGGC
acccacagagtggcattaaagcCACTCTATGCCCCCGCGCCCATAGCGCGCACGAGGCAGTGCGAAAAAGCTTCC
TCCCGCGGAAAAAAGCGCCCAACGTGCGGTTACTCACGCCCGCGCGGGTGCGCTGGGCAGCTCATCTGCACGGGG
AGTGCGATGTCCCGTGCAAGATATTTAATGCCACATAGTGGCATTTAATGTCACTTTGGTATTTAACGTGGGGCG
CGCGCGTCCAGGCATGGCCCCCCGCACGCAGAGAAATTGCTTTTCGGTTCGGTACCATGTAAACCAATCTGAACC
GGTACCTTACCGGTCCGATAGGCGACCGGTGTGCCTACCAGTATCAGTACGGCATACCTTGCTTCCAGCTGACTC
ATTGTCCAAAACATCCTCCCAAAGATATGACCAAGAGTTGTTTAGACGAGGCCACCATGCTACCTAGTCGACTTA
GTCAGAAGGGTGACATCCATTCTTCCAGCATAATTAGGAAGAGAGGATCACGAAGGAAAAGCAGGCAGCTACTCA
GCACAgcctaaaagaaaaaggaaagacaaTTGTAGCCCTGATGGAGAGAGTAGAGGAAAAGTCAGTTCACTTAGA
TTTAGAGATCAAGAAGAGCAGTGATGATAGTTTTGATGATTATGGATCTGGTGGTCAGAAAGAGAGTGGAGTATA
TGAGACTATCATTTATAAGACATGTAAGGTGGTTTTCCATTTATTGGTGAGTCCCAGTTTACGCTTGCCATATAA
AATAGGGACCACGATGCACGAGCTGGTAAAACCCGcgaggatacgattgcatataagTCTTTTCAAGGTCATTCG
GAGGATACGATGCAGCTACAGATGACCTCGCTGTGAAAtaggatccatagacgtatTAGGTTCTAAGTCGTATGC
TAGATTCTATTACCTCCAGCCATAGGCAGCCTATGACTCATaaggatatggatatggtgcgtttAAGGCATCATC
TTCTAATGGCCCCTCCATAGCCATATCATCAACCAAAGAATACCTCTTAGAGCCAAAGCTTCAGGGAGAAATTCG
AGATCGATTATAACCCAGAGCGCATTTCTAATGGGATGAGTATACAAAACTACAATACCGTTTGGTTAGAGGAGT
GGTTTGATGTGcctgttgattatgttaaggaTCTGGATATCTACGAAAGACATTGCCACTTGACGAGATTCTAGA
TGTCGGTATGCATGTTGTACTTTAAAATACATGTAATTGAttatatcattttatgttttgtatctcactaattga
ttttaggatatttcatattACTTCTCCTAGCATGCAAAGCCAcgattgttgtttctttttacaTCTTAAATTTGG
CCTGATGTTAACACAAAAGGTTGGGGGTAGCAAATCCAGTGATAGATTTGTCAGATTTTGTCTCTCCAAGGTAGC
ATTATTTTAGAGCAATGCCATGATGACATAGAAAAATCACTCCAAACATCAAAAAGCATTATTTAGGGTGTACCG
GGAtgatagaaacataaaaaaaataatcaaattacacttaaaatctttcaaaaaaccaaaaaaaaattacttttc
attaaatcttggaaaaatttaaaaaaatattatgccgGTTTGGAACTGGCATGCCATAGTGTACCGTGTATCAGT
ACGGTACTGGTTCGAACCGGCACGCCACAGCGTACCGTGTATTGGTATGATACCGATTATTGTATTAAACCGGTT
GTCGAACAGTATGGGTTCTAGTACTAGTTCAGCTAACCTTGGCAgggtgctttttttttctttctttttttcatct
tgAGCAAAGGTGGGGATCCCAATATTTCATTGAAGAGATGAATTGATGAAACAAAAGGAAGCACCGGCAAAACAG
TGACTCAAGCcactaaaacataaaaaaaccaGCAGAAATACCCACCCAAGTGCATCACCATAATTAAGGTGTAAT
TGCAGGTTGCAATGTTTCATACATGAAATGTAGCTCACTATCTAAGCTTCCTAACTTTCAATTATTCATGGTACG
CGCTCAAAATTGGGACCCGCCCAAAAAGGTTCAGACCTAGTATGGTAAGAGATGTAAAGCATTGGCAGAACAAAT
AGGATTTAGAGCATTTGAGATAGATTCAGCTTATAATTAACATAGATATTAAGGCATATTTTACTTTAATGACTA
GGCCTTAGTTGTTCTAGCAATTCTAATTATTCAAGTACACCAAGGTACACCATACCATATCATACCGATATGTGG
TATGAGAGGCGTACTAACATGATACACCAAACCGGCCTCCGTACTAAATATATAGACGCAGTAATAGGGTGGGAC
CGATATAGGGCCCAATACCGAGATGGCTACCTTGAAgcaccccattttcatctaaccAAGATGAGCCTACTAACC
GATTCCAGTGGCTGACCGGACCCTCTCAAGCTTTTGACTTCTCTCTCTACATCCTTGTTTCAGTCTAATATACTT
CTTTGTTACTCATAAAAGACAAAAAGTTAGGATGATGGTCAACAATGTTCTGCTCTGGTGGTATTCCAATATCAG
CTGTACATTTtagcaacaaatattttaataaactactcaaaaaaaaagaatgtggtGCAAATGGCTTCCATTGTA
GgttctttcaaaatttccagatcaatattattattctattcCTGAAAAGATCCTCAGGATAATTCATGGACAGGA
AGAATCAGTCAGCTGGTCGAGATACTAGATCATGCCAGGCCTTTTATCTACAAAAGATTGGAACATACAGATGTA
AAAGTTAGTACTAACTGTAAACAGGTCACCAAATCCTAGATAATGATGCAAGGTAGCATAATGTAAAAGCAAGCA
TATGACTTCCCTTGAATACCTTGATCCTCCCAGTTACTTGTTCTGAACCTCTGAATGAGTCTCTGATTTTCATTC
AATCTCTAGGCTCTAATGATGCAGTTCCCAGAATTCAGCACAACTGCAGTGTCAAAAGTGCCTACAAGAGTGCTA
CCCAAACAAGCAGTAACAGCTAAATTTGACATGGGAATTGCATGGTAACAGAGAGGACATGACTGTATATAATGC
AAAAGGGTATTGGAAATGTCAGTTGACTTAGGGAAAAAAGTCAACAACTAGGATATACCTGGGAAATTGCTTAGC
TCAGATGCTTGATTGTCGTTTACTAAAGAATTTTAGGTCAGGGATCCATGAACTGCAGGAATTCTGTTCACATCT
AAGTTGGACACGCAATTGCTAAGCCTTGAGCTCAAATGCTCAAGGGCACGTTTTTAGCTGCTTGTCAACTTCCAT
GTTCACCAAGTAACACAGTTCTGCAACTTTCATTAACAAAATCATTGTGGACTCTCTTACACAGGCCTTCTCTAC
AAGCAGATGTCTACATTTCACTCGTTTGATTGAAGAGAGCCTTTGTAGCGGCTGGGTGTTCCATCAGCCGCTCCT
TATCTGCCCCTCTCAACTCAACCATCAATCCAGTTCCATCATTTCGGTTGGTGTCCTTTGCCTCTTTCTCATTCA
CAAAAGTATTTGTTTTACTCAATCAACTCATTCCAACAATAGTGGATCATCATAGCAATTGCATTTTAatagcaa

aaaacaaaataacaataaaagaaataatataaattagtGTGATACATGCATTACATCAGTCATAATATGAAAAGA
ATAGATAAGAAAGTGCAAAGATTTCCTCATAGATCATAAAACAATATCTTCTCCAAGACAGGAACTCCTGCCAGA
TAATATTAATTATGTCAACATATCATGCATGTTAACATATACAGGGAAGTTGTCAGAACTTGGGAGAAACATGTT
CTTCTGGTAATATAAAAGATGAGCAATAGATTCTTAGATCTTTTATAACCAATAGATCAGGAGTTCTGTGATCtc
aatttttagatttttctacAGCCTAGAGATAGGATACGTATGCATTATGCATTTTAGGCCTTTTAGTTACCTAG
AATAGATCCAAATTGCCAGCTTTTTGTACATGTAACAATAGCCACCCTtgattcaaaaaatttcaataaatgaGA
TAAGATAACAATGATATGTGAATGTTGAACTTTCTCAAGTCTCAACCTTCTGAATGTTTGAAAAGTATGTACTag
tgaaaaatcaaaaagcagcCTGTGATGAAAGCtataaaatataacatttagCTGCTATATGTTGAGGCAGGCAAG
AGAAGATTTAAGAGTCCTATTTGTTTAGACATTAGCATATAGAGACATCATAAGTACAATCTAGACCTAGCTAGC
ATTCAATATGAATATAGAAATTTGAAAGCACAGGTTTGAAGAACCTACTACCGAGAGCATTCAATATGAATATAG
AAATTTGAAAGCACCATCATCAGCCAGGGGCAAGAAGCCTTTGTGGGGGGGAGGAGCATTTCAGataatatttg
ctagtccAGGAGATGATGGTGGATTTACAGTGAGCCTCGAGGAGACGATGTTTGATGGCTGTgaagctggatatg
gagcgCGCTTATGATCGGATCCGGTGGGATTTTCTTCGGCGAGCGCTTGAGAGCTTCGGATTTCATCAACTTTGG
ATTGATTGGGTTCTAGGATGCGTGCAGGGGCCTAGCTTTTCCAATTTGGACAACGGCACGCCGTCGCCCTTCTTT
CGAGCTACTATGGGGCTTCGCCAGGGATGTCCTCTGTCTCCGTATATATTTATCATTTGCTCTGATGTACTTTTC
CGTGCTTTGCATGCAGCCTGTGGGAACCGGGAGTTGGCATCCTACTGTCCAGCCCCAGCGTCCGGCCCATTTCTC
ACTTGCTTTTCGCAGACGATTGCCTCCTACTTGCTAGGGCGAGAGCAACGGATGCGAGAGCCCTACAGGGGGTGT
TGACAAAGTATTGCGCGATGTCCGGGCAGAGAGTCAACCTCCAAAAGTCAGCCATATACTTCAGTCCGAGCATGG
AGCGGAGGGTGCGACGGGAGGTCAGGGCGATTCTGGGGATGCAAGAACAGGTGGGGCCCCTGGTCTATCTTGGGA
TTCCCCTCACAGGAAGGAGGTTGTGGGTGGAGTGTTCTACCCTAGTGGACATGGTGCGGAGCAGGCtagagggat
ggagggcatcaTCACTTTCCATGATGGGTAGATTGACGCTGATTAGGTCAGTTCTGTGCTCCATGCCAATCTATC
TGATGGCACACACTGTTGTGCCAAAGACGGTGTTGCTGGGGATTGAGCGGCTTATACGGAGCTTCCTTTAGGGCT
CCTTGGGGGGGCCATGGGGTACCTGGTGGCCTGAGAGCGTGTATGTCAGCCACTGCATGAGGGTGGTTTGAGGGT
GCCATCACTGGTGGAGAAGCGTGAGGCACTCCTGGAACGGCGGGCAGTGCAGTTCGTATTGGAGCCGCATGGATT
TTGGAGTCAGATCATGACGACTAGATATGGCCGAGCAGGGTTTGGGGGGCAGGCGAGGGGTGGCCGGAGGTTCTT
TATGTGGCGAGAGATTGGGGGATACTTACCTTTGGCCCTGGCGCACTCCATGTGGCTAATCAGCGACGGACAGAG
CATTGATGTGATGGACCCGTGGGTGGGCACTTTCATTGAGACAGTGGCCGACGATGGTCAGCGGTGAGGGGGAGA
TAGGGCTTCGGGTGTGTGACCTTCTCAGGCCAAGGGCGGCGGAGTGGGATGGCGATAGACTGGGCCAGTTGTTTG
GAGAGCATTTGGAGCACTACTTGTAGAGCTAGAGTCGGAATCGGCGATATTTCTCAGATTCTTCAGCAGGAGTCA
GAGCCGGGCTTAGACTGTCACTGGGTGTGGGGGTTTGGGCTACATCAGAGGATTGCTcctttcctctggaaggtg
gcctgagAGCGCGCCTGCCGACTTGCTCAGAGTTGAGCAGACGAGGAGTGAGCATCTCTCCCCTATGTCTGGATT
GTGGGGTGGACGAGTCGGTGGATCATATGTTATTCCAGTGTATGTGGGCGAGGGATGCTTAGAGCCTGCCAGGGA
TCCCGGAAGATGCTTGGAGTCGAAGGGATTGTTTTCTCGAGGAGGTTCGGCGATGGTCTGGTATCCCCTAGATGC
GTCCCGTGGCTATTAGGGCAACTTGTACAGCTTATCAAGTATGACTAGCCAGGAACATCCGGATATTTGGTAAGT
CCAGTCCGTCCTCGCATTTCGTCATAGAGCGAGCTCGTGCACAGGCAGCAGAGATCATCCATGTTGATCTAGACC
ATAGACTGTTGATAGCTCAGGATATCTGGAGCCCCCACATTGCTTCAGCAGTTCCTCATAGGGTATTCTTTACcc
gggagcccccacctccgagttttctcaaggtcaattttgacaGCTTAGTATCGGATGGAGGTAGGAGGAGAGGGG
TTGGCTTCGTCATCAGGAGCCCGAGCTCCAGAGTGGTAGCAACGGAGGGCTGTCAGATTTTTGATACCTCGGCTC
CTGGCGCAGAGCTGCAGGCGGCCTGGATAAGTATGTGTTACGTACGACGGGTGCTTCAGGCAACTTCAGTACTAC
TGAAGGGTGATCAGCTGGGTCCAACAGGGTCCAAGCAGTGACGGAGGAGTTCATCTCCTATTCCGCGATATTTGG
ATGAAGGTGAGAGTGTTGCATTTtaggctatgcatgtatattgtgaagctaatggggcggccgactggatGGCCA
CATTtgtagcgagccactcaggaggcacccTATGGACTGAGGGTGAGATGCCCAGAGTACTTCGGgaccttttat
tttctgattccattgggtgtattcgtacccgtgatgtatgaaatgcttgtttcagcaaaaaaaaaaaggtttgaa
tgGTAATAATGCATGAAAACttcaaacaaccaaaaaaaaagctGCATGCCACTATGATGCTCCTGGTGCCTATAAG
GAGACTCATAGAGCCAGTTGAGGTAGGATCCTAGGTAAACAAGGAAACCATCTCTAGGAAGGAACTGAAGTCTAC
CAATATACaagtcataaaataataaagaagacCTTATTGAAAAAGGATATTTTGCTAATTAAGGAAAACCACCTT
GTAAATAGAAGTTCAACAATAATCCATTTCATGCATATAACTTCCCTACAACTTACTCGCACATAAGGAGTAAAA
GCCAGCTACGGCAATTGGCATGGGCCAGCACACTTGGACTGATCTAatcaagaagagaaggaaacaaTTCCTTCA
AGTCATTTAATTTGGACCTCTCAAAGAAAATACTAATGAAATAATGGAGTGGCTAGACTAGGGACTACCAAACCA
TCCCGACCGAGCAATTCGAGGTATATCAAACCTACCAGCTGGTAACTGGGATGATTCGGTCGATCGATCCGGTCT
CAAATCAAAGGCTTCATTTAACTAAAAAGCGGATCACCTTGCGTCTCTCCTCAATCGCGATCTCTTCCGaattct
ctcccctctctctccctccaaatccatctcttccctctcccctctctccttctctctcaatctttctctcgtttc
ttccctctctcctgaTCCACCATACCGACTATGATCATGTTGATGTCGATGCGACGACAACTACGGCTAAGGTTA
GTTAGGGTTCCGCATTAGCGCAAGAACACTGCAGCCACCGCCTCCTCTTTTAGCTATGATGAAGATCCTCCGCTG
CCGCCTCCACCATCTTGTCTCCGCTGCCTTCGGCCCCTATCCTCCTCTATATTGGCACCTCCATAAGGCTGGGTg
aatcccctctctctcccctcctccctctctctaatTTCAATGTGTATCAGATCGGTATGGTATGCACCGATATCG
TACCGAACTATTCTCCGACCAGCACGTCCTCCAATACCACTTCAGTGAACCTTGGGCCAGACTCCTTTTGATTTC
ATTAATTGAATAATAAAACTGATTACAACTTGTCTTATTTATATTAGTGAGGTCCAAGTTCCCAAATTTGATTCA
AATTCCTTTTCTACTTAAAAGAGGatagaatttttctaaattagttaaggaaTTTGTGCTAAGTTGAAAGCCATC
CTAGAGATATCTCTAGATTCTTACATTAAATTTGCCGTTTGCTACATCGCCTAATTCCTCCAAGATTGGAAGATG
CACCATCAAAATCTTACCTGAAGAGGATGCTTTTACTTTGATCGGCCCACTTCAAGGCTTAAAACATCATAGCAC
CCCAACATTAGTAGGGTGCAATGAACCCTTAAGGTTTCCACATCCATGTAAAACTCAAAAAAGTtacccaatttc
aaaaaaagatcATTATAAAAAGGATGTCGTATGGGGAAGTTATGGCAGATCTTACTCCCAAACTTTATCCAAACC

TAGCCGAAGTGATCAAACTGGTCCACAATGGATCCAATGATCCTCCTAGGTCATGGACCAATATGTGGCACATCT
ATGTGCCAATGCATGGGGTTCAATGCATGCCAAGCATTTGAATAGCACAATATGTGCTGTGTTGGAACCAAATCA
ACACTATACCAATGCATGACACTTAAGAATTAAAACCAAACATCCATCGCTACTAGAATGGGGAACTCTTTGATA
GAATTTTTGGTTCCACACTAGAATCTTCACTTAAATGAAGTCACCTAGCCTCTTATGAAATGCTTATCAAAACTA
AACGTAGTTGGAGAAAAGTCTCTTACACCTTTAGAACTCCTTTCTAATTCATATAGTACTGAATTACAATAagga
aaataaatgaaacaatTAAAAACAACTTTTCTAAGCCTAATAGCTTGACTAAAGACCTTTCCTAATAGAGAAaac
tatgaaaaaataaaacaaaaagtttttaaaattcaTTTCTAATTCAAAGTAGCCAAGGAAAGGCAAAGTCATAAC
TACGTACAATCCTGTCGACAAATTTCAACAAAGTAGCCTTTCACTAAGATGACCATATGCCCCTCATTTAAGTTC
TGATTAACATGATTCAAGATTGCATTGGCAAGTTGTTTTGATGCTAAATACGATAGaagtattttgattttataa
ttcAGCTAGCTTTAGTAAGCATTCTTTTGGATTATGGTGTATGAAGTTGATCTTTGAAAGGCTAGCATTGGAAAG
TTGTTTTGATGCTTAATACGATAGAAGTACTTTGATTTTATAATTCAGCTAGCTTTAGTAAGCATTCTTTTGGAT
TATGGTGTATGAAGTTGATCTTTGAAAGGCAGCTAATCTATGGTATATTTTTCTCAACTAAAACAGCAATACGTC
AACATTGGTGAGCTAAGGGTCTAACAAGTGATGCCTGACAACTTTGGTAACTGATAGATTCAATGTGGTCTCAAA
GCCATTATGAAACCAATGTTGTTGTGGACCTCTTTTTGGACTCTAAAGCCCTCTTTGGTTCTAGCTTCATCTCTA
GCATGGATTAGTAGCATGTAGAGGACCCTTCACTCACATCAAACTCCCTGGCTTTAAAATTTGTCTTTTGATGAA
TTATTGTGGAATAAGATTTGAGAGCTAGCCTTAGCGCAATAAAAAGATTCCTCCATTATGACCTAGATGTCACAA
ATTCGAAAAATGGAAACAACCTCTCCCCACGCAACAGTAAGGTTGCGCAAATCAAACCATCCCCAGACCCTAGTG
TTGGGAATCTTGTGTACTAGGGAACCCTTTTAGGATATGATGGTCTTTCAACATGCTTGTGTCAAACACTTCACT
ATCATCAGCTTTCATGTGTTCCATCGCTTTCTTGTCCTTTTCCCATGATACTTTTGGATCTAAGCCTTCTCTATC
ACAAGCTTTCAGCATAATTTCATTAAtatatggattttgttttgattctgGGGTAGTTTTATACACTTTGATAGG
AAGAAGATTTTGATGTAAATCATCACTCAACTACGGAGTCATGCCATCCGCTCATATGGATTATTATTCACGTTG
CCTCCACTGCTACCCAAGGATTGCCGAACCGGTATCGGGTGCTGTACCGATCGGCGCCCGGTCCGGTTCGGATCG
GACCGAATCGATATCGAACTGGTCCGTCAAAAATGCTGCAGCGCGCGCTATGGCAAATCTAACTGGATTCGGTGC
AAACCAACTGGTTCGCACCGAGTTCTGGCTGAACTGGACGGAACCGGTCCAGTCCCGTCCGGTTCGCTTTTGTAC
GAGGCCAGAACCATTTTATATGTCCGAACCGACCCGGTTATAGACCGGGTCGACTCAGTTCGGCCCGGTTCGGCA
TTCCTTACTGTGATCTTTATTCTTTTTGGGATAGTTTTATACCCTTCGATAGAAAGATTTCGATGTAAATCATCA
CTCAACTACGGAGTCGGCTTCTTGCCATCCTCTCTTATGGATTGTTATCCATGTTGCCTCCACCCTCACCCTTAT
TCTTATGAATCGATACTATAGTGTTCTTCCTCCAATCATCAGGTATTTTCCTagtgtttaaaattttattaaaca
aatctCCATAGAGATTTCATTAGGCCCATCTACTTTtcctattttcatcttttccaTAGCCTTCTTTATCTCAAA
TactctaattttattatatgtatAGTTAATATTTCTAACCTCAATGGAACTAGTAAGGCATCATAAATCGTCTAC
TTGTCTTTCATTGAATAGCTTGTCAAAATAgtttctccatctttctttgatcTCATCTTTCTCAACTAAAACTTC
TTGATGTTCATCTTTGATGTATTTAACTTAAGTCAAGtctctagttttcttttctcctaGTCTTGCaatcatata
gatatttttttccctttccttagtTTCTCACTTGTGAAATAACTTTCCATACGTCTTGAATATAGTCAGTGAAACC
TGTTGCCGGAGCCGGAGATTAAGGAAAGAGAGTgcaagcgggggagggagggagtggGAGAGGGTAGAAGGCCGG
CGGAGGGTGGTGGACGCTGGAGGGCCGACGGAGGCCGCGAGAGCGAGGCTGTAGCCggtttcctctccctccctc
tatccccccttctctctctcttttcctctctcccatggtCTGCGGAGGGTGGTAGAACCGTTCGATTTGAGCTTCG
TACCGGTGTGAActggctcggttcgcaccggcaTGAGttcgaaccggtgcgaaccgagcatGGATCGCACCGGTT
CGTTGCTGAAAATTAGTGCTAGAAGTGCCACAGCGCTGGCCAGAGGGGGTCGAAACCAGGCAccggtacggtatc
ggttcggctcggttcggcaccggtaccgtaccgttaTCGAagccgaccggtacgtcggtacgatCGGTACCGCGA
ACCTTACTGTGACAACTTCATGAATGTTCTCAAGCTCTTGTGTTAGACATAGTACAGAATTTAAGTTTGCTAGCA
ACGTTCACTGTTTTGGTACCAAACTCCGTACCGGTATCATTCTATTACAGTATCGGTACATAGTACGGTACGAGA
CGATGAGGCATATTGAGTGTCAGAATGATACGAGATTGCGTACATAGTGCGGAGAAGTCGTGTTCGGGCCGGTTC
AGcgcataccgtaccggtaccaacCCCAACTGGTATGATTCGGCTGTGTACCAGTGCGGACTGGCCCGGTTCGCA
CCGGCCCAAACATGACTTCTTCGCACGGTCGTACGCAGGGCAGCGCGCCCtgcgtgggccatttaatgccactat
gtggcatttaacgcgtcCGTGCGTGGACGCGCGCGAACGCGTTGTTTTTTCGCGCTGCTTTATTTTTTTCGCATT
CGCGCGCAGACGCCCGAGCAGGAATCGCACGCCTAGCTCGCGCACACGCGCGATTCTTCGCGTAGGAAACCGCGC
TTGCAGCTGGTTTCCTAGCAAGGAATCATGTCCACGCGGGGCAGCGCACCCGCGCAGCCCGtatgggccatttaa
tgccacagagtggcatttaacgcgTCCGCACACGAACGCGAGTGGACGCATTTTTTTGCGTTCGCGCGTGAACGC
GTGTAGTTTTTTTCGCATCCGCACGCCGTGTCCGCGCGCGCGTCTCCAGTTCGATATTGGaccgataccgaaccg
gtaccataACAGTACTAGTGCctaccggtacgtcggtaccgTCGCTCCGGCGAACCTTGATACCAGTGCAGTATG
GTATGCGCGGTATGGCAAATCTTAATGTGGTACAGCAAATCTTGATTGCTAGCTTTTGTTGCTTGTAATACTTTG
ATCCTTatgcaaataaaaaagcaaaatctTACTTGTATACAAGGTAGTCACAAGgatctttcttgttttctaaaT
CATAAGCATGTTAAATGGATGAAGCTGCTAAAATGTAGcacctttttaaaaaatatagatcgTTGAATTTAAGAAG
TGGATGCATGTGTTTGTGAAGACAGTTCTTGATTGAAATCTACATTCCTGTCTGTAATGAAGCTGACCTTTTAGT
TATTTCTGAAAAGGATTGTAGATATCTAATTACTTGATTCAACACATCATGTTTACAGAGTCCAAGCTCGTGAAT
GGCTTCATTCATAAGATTGAACCTTTTTTcctaaatcaaataataatttttgtaaaatCATTGTTGGTGATTTGG
CATGCGGCTTTGAAATCAAACTTATTGTGCAATGCCCCAGAAAATACATAtgatagcaaaaaaataaacttagaA
AGTAGTAGCATACAACATCAAacctaaattatatttttcaaggtTTTTAGTCTTAGAGGTCAGGTCGAGATGTTG
CACCCCCATACCAGCACATGGTACACTACTGGAGGCAACATAACAGTTGCACATGTGTTGAGATACTATCGTATT
GAGTCCCTCTAATGTACCAATACATTATTGGGTCAGTATGATGCCCAGTACCAAGTGGgttcaaaaaaacttatt
tttttattaataaataattgttggttgcacatatatatgtaagGGCAAGAGAGACTTTCTATAGGTGTAGAATAC
AAAATTACAAATGATGCATGGTCATAGACATAAAGAATTTTCAGAAGAGAAAATTTGAGAAAGCATATTTCTTAT
AGAATATCAATACTCTAATTTATacttcaaaatattaatatattaaattgagattttattcaaatatttatgaAC

TGCTCATACTACACTTTGATGCTGGCTTTAAGGAATATTTCTTGTagaatttaactttttttcatttttatattg
ttaATTACTAGTACCTAAAGAACTAAATTATGTGCTCTGTTCTTTCTAGTTGTGGCAAAGGGAGGCAGAAAGCTT
AAGGGAGCAGCTGCATAACTTACAAGAAAATCATCGGTACCATCTCTTAACCTCGTTCACATTCAAAGATTGTTC
CTATGCCTGACTAGCCATTGGCACTATTCCCTTAGACATCATAccataaagaaaagatttaacTATATAACCATC
TATTTTGTTAGAAAAATAGTGTTATTCAACATGACGTAGACTTTCAAGCTTGTTTGAGTATAACTAATTCCCTca
tggaaaaatttaaaaattttcaccCAATTCCAGGGATCTAGAGCATCAAGATCATTAGGAAAATAATATGTAAAC
TAGTGAAAACatcaaattttgtaaatgacTAGAAATGAAGTATAAAAAGTATTTCTTAAGACTAATCACATGAAG
CAACATTTTCGTTTTTAAACACCACCTTGTCTTGTTGTTTACCATATTGTATGCATTGAATAAATCATTAATTGC
ATCATTAGTTATTGACAAAAGAGCAGGTTACTGAACTTCCAATAGTATGTCAAGATTTAGACAGCATAAAACAGA
TGGTACATCTAAGGACTTCATTTCTCACTTCAATACCCATGGACAAAGAATTGGTCCACAAGCCCACAGTTCACA
AAAACAGTGACAACTGCTGCTTCCACCATGGCAACTAACCTGTAACTATCAacttcatataaaattattatttgc
ttCAATTATCAAATTGAACCTCGGCATTTTTTATCCAAACATATAGCTTGGGCAGGAGCAAATGGTGGACAGTGG
CTTGAGTGCTTGTGAAAACAGTAAATGTGCCTAAAAACAACCTAGTAATAAAATTCTCTGACATTGACAATTTTC
CAATTATTCTCAATTAAGGTTTTCCCTAAATTTTTACTGACTTAATGAAATCCATGCTTGCCAAATATATAGTGC
CTTCTatgaataacttttttttttttgcaaatatttcatagaacatttcatttctttttctaaagttctccacttg
ttttatttttctttttattatcattCAGTCTTCATTTCTATTCATTCACTTTTCAATCCTCAACATATTCCAAAG
GTTTTCCAAAATGTATAGTagattataaatttttttccttGATAATATATTCTTGAATAACTATGCCTAAAAAGTA
TACAGACACTAAAACATATTAGAAGTCAAGATAAAACAAACGTTTCCTCCTACTAGTATACTGTGTAATGCAAGG
CTCGCCGTCTTAGTACCAGACCCATACCGGTGCCATGCTAGCACTATTGGTTCGGTGCGGTACGGTACGGGGTGT
TGTCGGCATGCCGAATGTCGGTATGCATCCCGTATCGTGTACCAATACTGAactggtatggtatgatatgcCCCG
TACTGCCCTGTTCGGCCAGTACGGCAAACCCATAATAATGTAATGTAGAATACTCAATAAGCTACACATACTATT
TGATTTCGCTTATGGATCCTTCTTTTTGAaacattctttttattttagtgATTGTTGTTATTGATATGTAATAAC
AGTACCAGAATTTAAAATCTATGACATTACATATTTATAGAATGGGATTATCACATTGATCTAGTAGCTTTATTA
CTATTCCTAAAGTTGTGCCTCACCCCCCAAGAAACCATGCttctatataaataaaaagaacacAGTTCTGACTGG
TCTTTTCTGTTATCTAATTTTTACGAGTTGAGAATCAGACTCTATGACATCATGAACTGCTCCAATTTCAATGTC
TAAAATTGCAAACCCCCAGAATGCTATAGAATACATGATCATGTGAAACACTGCTTGGATAGACCAAATTACCGT
GAAGATTTCTTAGAGTTTCACCAAGGGTGGACTACCAAGAGTGCAATCAGAAATCAATCTTATTTTGCCTATAGG
ATCTGAGAATAGCTGTTCTGGAAGAAAGTGAAAGCAAGACTCGGAGAACTGCTTCAGTACTTCAATTCCAGAGTC
TCAGCCTCCCAGCATATAATGCTAATAGCAGCAGCTGGAACAATCTTTTGGATAGGAGCACTGAGATCCTAAATA
CTTTTTTTCATCTCAGGCATCGCAATTTAGGATTTTGTCCCCATTTTACTTTGGTGGTAGCTTGGATATTAAAAG
AGTACAAGGCTTATACTCAGCCATACCTTGGTACAAGCTTTAATAGATAATTTCTGGTTGTATACTTCATGCAGA
GAATAAGGTTGTTAGAGAATTCTTTGATATGAATTTCTTACAAAGAAGTCTATGTGATGTTCTTGCCATGGAACA
TAAGGGAGCTGAGGTGTCAATTTTGGACGTCATAGCCCAATTACTTAAGAATTTACAAAATTGTGATTTAGAAGG
ATCACGAGTTCTATTGTGGACCAATTTAGCCACTTGGGGTAGGTCTAGACAAGGTTTGAGAGCAATATCCACCAC
ATCCAGAACTTGATCGTGAAGTGTAACTTCCGAAGGCCATAACTTTCTCGTATGATCTCCAGTTTCAACATGTCA
TGCGCAAAACAAAGCCCTTGACAAGTACTATGAGGATGCAACAAGGTTGTCACACCCCTATGGGTGTTCTACAAT
ATTGGGGCCTCAAAATGGGCCGATCAAAGTAAAACATTTCTTTTAAGATAAGACTTTGAACAAAAGCATCTTCTA
ATCTGTAAGAAATTAGGCGAAGGACAAAaggcaaagttgatgtagaagtcgagatctcaTTCCTAGATGAGTTTT
ATCTTAATATAATTGtcacaacccagaacctcacctaaaaagactAGCTGGAAGGTGTTATTTGGATTTCATGGC
CTTGTAACAAGCATCCACAATCTACCCAGCATTCTAGCTAGGTAAATCACCTCCAGCCCacccatatttttttaaa
aaatccaataaatacTTAATGTTAGGTTAAAAAAAACTTGGGGATTTATGATGCACCTAACAATATTTGATCACT
AGGTTTATGCTCTATGGCATAAATTTAGGACCAAAATCAAGGAAATTTGGGTTTCTGGCTTGAACCCCAAAACTC
actcaaaggaaaaaaaaaaggttttagaGGTATCTCTAATTCCAGATTATTCTGTCAAAAACATGCAACATAAAAA
CTTCAGCACACATCAAAAACCATGGCTAGAGAGTACTTAAATCAACATTGTTGCcatacttttcttaaaaaacat
aGCATTAAGACAacaaaacataagaaaaatacaaaatagaaTAGTTTCATAGCATAAATAGGCCATACCATCTGA
CCATGCAAAGCATATCAATCAATACCCTCTGCTAACATGATGTATCCCATACCAAACCTTCATCGATATGAGGTT
TGATATTCGATAATTTGCATCCtaataagagaaacaaaaatattaagggaaacaaaaagtaaataagtaaaatgg
GTAGAAAATGGTGAATGAAGTAGGATTGCTCAAGAAGGTTTCAGGGTACATTTGACAGAAATGAGAAAGAAGACA
ATGATATGGGAGTCAATAAGCAAGGTAAAAAGACCTACAACTGGCTGCTACACAAACAATGTCCTAATGCAAAAA
TGTAGAGGTGAAAAAAAGGAATAGGTGAATCTTAGTGTTCTATGATAAGATTAATAAAGTTTTGGTTCGATTAAT
GTTGGATTAAGTACATTACATGACTAGTTAGAAGAAGATATAAATAattatctcaaaagaaaaatggaaaaataa
gaagaatcaCATCTAAAGACAAGATGGCGGCACTGAAAACCTTTTTGGAGGTACTGAAAACCTTTTTGGAATGTG
ATATATCTTGTGTCCTTAAACTAGTTAATGATGGAAGTACTGAAAACCTCTTTGGAATGTGATACATCTTATATC
CTTAAACTAATGGTGGAATTTGGGTAGGGTATCACCATCCTAACATTCCTATTAGTTAGACTTGGATTAAGTACA
TTACATGACTAGTTAGAGAAGATATATATAattatctcaaaagaaaaatggaaaaatgagAAGAATCACATCTAA
AGACAAGATGGTGGTCCTTTTTGGAGGAATTGAAAACCTTTTTGGAATGTGATATATCCTTAAACTAGTTAATGA
TGGAAGTACCGAAAACCTTTTTGGAATGTGATACATCTTGTATCCTTAAACTAGGTAATGATGGAAGAACCAAAA
ACCTTTTTGGAATGTGATACATCATGTATCCTTAAACTAATGGTGGAATATGGGTAGGGTACCACCATCCTAACA
TTCCTATTAGTTAGACTTGGAATGTGATACATCTTGTATCCTTAAACTAATGTTGGAGTTTGGATAGGTACCACC
ATCCATAACATTCCTGTCAGTTGGAGCTTTTTTAACTGGTTACTCTTATCCAACACCTAGAATTCATGGCAGGAA
CATATATCCAAGTCGAACATGTGTTAGACACTTAAATACTTACCACCTTTTCTTAATTACAATTTTCCAAACTTG
GTAGGGAACTTGGAAGACTTCGATGAGTTCAAcgcatggaatgccgtaccggcccgtaccggccggtacgggccg
gtacgtaccggtccggccgtggaccggtaccggaacggataaaaaaccggtatttcccgatttttttatccgaacc

ggccggttcggggcccgtaccggccggttcggggcccgtaccggccggttcggggccgtaccggccggtaccggc
ctNNNNNNNNNNNNNNNNNNNNCTAATAATAAAAGTGAGATTTgttgtttttcaaaatattattagaatgccttt
gcaaaatattatttttcaaaatcctTAATATAATGGGAAGTTTGGGCAAGAAACATTCATTTCAAAAGATCttga
caaataataataatatctttcagtgctctatttcattacaaaatttacattttaacatcttttagttaaaaaaat
tgtttttcctacacatccttctcttttttttccatattgaTGAGTCGGACATTAGTAGACGAGTTCAAATCAGAT
TCTGGTGTCCCTTTCCATTCAACACCATAAGACACTAATTAATCTCGCATGTGAATTTGCATGGTTTaagtttta
gaaaatattttagatatagGATAAGTTTTAAAAAGATTCTAGGCTTTCTAGCATTGATTAAGTGCTAATATGCCT
TCCTCATAAAATTGTTTAAGATTTATTTCGTATGTTTTAAGAATTTTACACCATAAGACACTAATCAATGATaag
cttttaattttgttgtgACCAAAGTACTGCTCTGAATTTGGACTTTAGAGGATGTGAACGCTTATTACGTATAG
AATCATAGAAATCTAGGAGTTCACCCCTTGAACTCCTCCTTTGAGTCTATCAATGTTAGAATCCATTTTTATAAA
GTTTAGCATGGTTTAAGGGAGAAATCTAATTAGAGTTATGATGTAAAATTGGGACACCAGAATCTATAGGTCCTT
CAACTATATGGCTGCTTTTATCATACAATGTTAGTCTATAACATTAACCTTTAGGTCCTTCAACTATAAAAAAAA
CCTATAGGTCCTTCGAGCATACAGAACATGACATgctaacaatattcttagtgtTTTTCCACCATTGTTGATGCT
AACACCCTTTTTTTGTCATTAAATATAGGAAATGAAAAATCTTATATCATAGTTTATACTTGTGAGTTGTGACCA
CTTCTTAAGGTTAAGCATACATATCAAATAAATCACAATCATTGAAATGTCCAAATCAAGAATAATACAATCACT
AGTACGATAGTGCTTGACAATAAACAAATATTGTATGCATCTTCTATGAAAAAACTATCTTTTAAAGTTCTACTC
TACAACATGAGAATAATTAATAGTATAATTTAGTTAAAATCAAGGATTTAAATACCGTTGGAATGTTATGGGATC
AATATAGGTTTGGAACACTGGTTCtcacttgaatcggtatatacCAATGATACCAATTTGTACCAACAATCATAT
GTACTGGAAGTTGGTACATACTggttccattttatttttattgatacaAACCATTTCAATCCATACTGACCATAT
CGGTACCGTATCGCTCAGGTTCAAAACTGGTGTAAGGCTTGGTATTGGTCTTTTAAACCCCGGTACAGACCGGTC
CAACCTGGAACCGGTATCCGGACAACATAGATACCGGTTCTAGCTGTCTGGGTACCGGTCCAACTGTTTTCGGGC
TGGCACTGACCGGCATGCATAAGTACTGGCTGCATACCGGTCTCGTACCGGCCCGAGCCGGTGgccagttcgcac
cggtattgttttttttaaccaCAGTGCGCGCGTTGTGGTGTAGGAGCGCGTGGCCCTCCCTCGGGTCAGAGTGCC
CCGCGTGCGATTCTCCCGCGCGCGCTGTAGCCAGCTCTAGCGCGGACGCggatgcaaacaaaaaaaaaaaatttt
ctgcgCGCGTCcgtgaaccaaaaaaaaagtatataaacAAAGACCAGTtcgtaccggaccggttccgGTACGAATC
GGTCTCGAATCGGTCCGGTAGGCCAGACTGGTTCAGCGTACCTTGGTTAAAACTATCTCTTCCCACATCCCCTAT
ATATCTTTGCTACCTCTTTCCAAATCATACAATTAGGTATGCATCAAAATCTAATTGCAATATCCTTGTCCATAC
AAAACTTACAGCCAATGTTATTAATAGTTCTCACCGAGGATGCTACTTGATCCTCGAATATCATACTTGTTTCAA
ACTAATTTTTAGTACATCATAAGGCTTTCTCCGAGTCTATTATATGATCAAAAGTTTTTGTGGCAAACAAAGGGT
CTCAACATTATCCACGTGCTATAGACATAAGAGATAACAAAAGAGATATAtagtaattatatttataaatgattt
attgtaGCCATAGTTTGCCATACCGATCCGTATTGCCCGATACCAAGCATATGGAAACATTTGGTAAGGTATCAA
AACATGGTATCAAGATCCGATACAGCATAGAGGCTGAAAAGGCCTAGATGGAATGGTTGGTTCCACCCAGTTTCA
TGTTGTACCGATATATACTAGCCCAATTTTGGGTGGTGCCATGACTgggcataaaaaaaaactcggAACCTATGT
CAATACAAAGGGTGTAGCCGTACTATACCAGAACTTATCGTACCTTACTAGTAAGGTATCAATATGACAAACCTT
GGTTGTAGAACACTACTCAAAACTATAAGATATAATTGATAACAATGCATATAATGGCTCTCGTATGATAGTTTT
GTACAATGCATAAAACAtctttcattcacatcacattgGAACCGTAGTGTCATTActctaaaaaataatcatgga
AACCAAAGAAAGTTTCATACAGTAGTTTAAACCCCAACTCAAACACCTTGTTTTGGTAGCCAATTTACACTCACA
ATTCAACCACAAATCAATCTGCTGAATCAACCATTCTTTCTCAAAATCCTCAATTTTGTTCTTCCTTAGTGATCT
CAAAAAACCTTATATGAATATCTAAACCAAGAAAGGCGGAACTGTCCTAAATGGAATGGTTCTGCCTGTTCCGAG
CTGTACCGGTGGCTTACCAATACGGTTCCGGCTTTGAAAACAGCTCGTTGACGGCatcgaagaagaagagagaga
gagcgagcaggggaggaAAGAAGGATCGGCGGCGGACGCCAGCGGAGGGTCGCGGGGCGGCGGCGGACGCCGAGG
TCGCTACCACGTCCCTTGTTTCAAAAGTGGCCacggaaaaaatttagaaatttttaagtaaagtcggtaAATCGG
CCTTCGCAGGCCCCTCGCGGCCTCTGCCGACATCCGTCGCcgctcctccctctctcccccgctcgctctctcttt
ctcttctccttctctggcTCCGGCAGATTTTCTTTCTCGGTACGGGCCCATGACGCCAgacaaccggtacggtgc
ccggtaccgacAAAGTAAACCTTGACCCAAATGACAACAGCAGTGATGATTATGAAGGAGTTTGAACAAAGGTGT
GGAGTAATGAAGGTGATGGTAAGATTTTAGCCCAACCTAGTGGTGACAAGCCTGGAATCAAATAACCGTTTTAAG
TCTTACTGGACCTTGTACCAGTACCTTATTGGCATGAGTCAAATCAGTATAGTTTAGAGGACATGATATGCTATT
GGAGTTAGCATAGCTTGTACCATAACTCAGAATGGGCCTATGGGTGGCATATTGAACCTTCATGCCCTATTGGTA
CCGCATCAAGCTGATGTGATATGCCCTATACTAGCCAGCACTGGTCGGTATTTAAAACTGTTACAAATTAGGATG
GataaaaggaagggagaagaaaacaaggagatGGAGAGGATGATGACGAGATGGAATAAgagaagcatgatagaa
taaagAAGCTTAAGGCTTTTAACTCTTAATATTTTGTGAGGTCCATGCCTAACCACTGGACTAGATTAATTTTAG
CAATTCAGATGGTTGAATCTCCCAATTTAAAAACTCAGAATATCTTTTAGGAAATCCAAGTCCTGACATGATGTC
ATAGACTTAGACATAGACTCGTCGCAAGGGGTGCCGGCATGCGCATGGCACGATATTGCAAAAAGGGTTTGGCAT
GGGCCAAAAGTTGCAATGCTCCTCGATCATTTGTAACCCCCGAGGGTGGCCGTTGGGTGGGTGCCTAGTAGGGGT
ATCGAGGGGAGCAATGGTTAGCCAAGGCACGACAGAGCCATGGCATGATATCGGATGGCTCGGGAAGCATGATAT
TTGGCATGGTGGCGGACTACTAGGATATATATGGATACTCAAGAATTCCACTTGGTTGAGGGGTGCAAAACTTTG
TCCGTGTTCCCGACTCCCAAGCCCATAGGATATGGGTCCGCCATGGCACAATTGCAGGGCAAGGTAGTGCCATGC
CTGATGGTCTCGTGTGGAGGCATTTGGAGCAATGTTGGTTGGATGAGCTACACACGTTATGCGGAGGGGAGTTTC
GAGAGGTTGGTCCTTGAGGAATTTTCCGGACAACCCTGTGCGTTTTTTATGTCTATGATGGGGCGAAGAGGCGAT
AAAGGGCTGCTCGAGCAAGGGTGTGGCCTACCTTGACAAGCAAGTCTCATGAGCATTACAGGGCGATGTCTCATA
GGGATCCATGGAAGCTAGTCGACCGTGGAGTACAAATGTTGACTTATGGAATATCAATCGGATTATGTGTCTTAA
TGTTGGGCCAAGGATCAAGGACGGCCAAAAGGGTTGTGCCAAGGAGAGGACAACTAAATGCAATGATTGGTCGGT
CGAATATATTTACTTGATGGGCGGACTTGGGTGCTAATATCATGGTAGTGCACGATGTACGAGCAGGCCATGAAC

ATTGTCAGACAATATGGCCATGGATGGGGAAGGCATGCTGAGGATGCAAGACTAGGATTGTGGCATAAGAGCTTA
TTGGGGAGCTTGGATAGCAAAACCCTGAAGGGCTTGGACATGCACAATCAACCATAGCTTTGAGGGGTTGCAGGG
CATGTCAGCTTGGTGTTTTGACTGAGCTTGAGCCTTATGCATAGTAGTGGTGTTGAATGGTTGTGTCATGTGAAT
TTGGTTTGTGTATGCCCAAAGGGCAGCATGGTGGCTTACGGATCGGTGGCACAGGTTATGTTGTGGCACGAAAAC
ACTCTAAGAATCAAGACATCCTGGGTGAGCAGTACTTGGCTTGTGGGGCAAAAAGCCATAAAGCTAACATGGAGG
TGCCGAGTTGTAGAGCACTATTGGAAAGTGTCTATACTTTGGCATTGAGGTGCACAAGTGTTGCCATCGACCCGA
GGGGTCAGGTTGCATGCTTGGCTATGTAGCTAAACTGTTGCACATTTATATGTATGCCCGAGGGAAGGACTTGGG
AGGGCTAGCCTTGGTGTGTGGCATGTCAATGTGCCACATGGGTGTTTGGATCAGGAAAACACCCACCCCATGACA
ATGTGCCAACCTACATACATTATTGGGTGGCAAGATGAGCGCATCAAGTTGTATGGATCAAACATGTTGTGAAAC
AATAGTTTCATATACGATCTCTCCCACAAAACCATTAAACTCCACCTTAACAAGCAAAAGCAATGTACATACACG
GGAATGTACGTCTCTGGTGGAGGGTAACTCAGCCTACATTTGTTTAATGTCATCAAACTACCTCTTATGCTACTT
TATCATGAAATCCATTTGAAAGCCCACAAAAACCATGAACAAATCTTACAGGTGAATAGGTGTCACTAGTTAATA
TCACACTATGTGCAACAAGTTGATTATTACACTATTAAATAAGCTGATGATATAGCATATTCAAAATTTAGTCAA
TGCTTGCAGGAAATTGATGGGTGAACAGTTGTCTGGCTTAAGTGTCAAAGATCTACAAAATCTAGAAATCCAACT
TGAAATGAGTCTACGCAGTGTCAGAAAGAAGAAGGTTACCCTCATCAAATGTTGCTCCTCAATCCTTCTTATCCG
TATCTTCTCTACTTAATGTGTTTCTTTTCTCTGCTATAGGAACAAATTCTAACCGATCAAATTCAAGAATTGAAC
CGAAAGGTTTATCAATAGTTAGTTAAATCATCTTTAATGTTGAAAGTATTGCATTTTAAGTGCCTAAATTTCTAG
AATTTATTACAGGGGTACCTTATTCACAATGAAAACATTGAACTCTACAGGAAGGTAAACCTCATACGTCAAGAA
AACATGGAGTTACGTAAGAAGGTACTAAGTTTGAGACAGAGATATCTATTTTGCTCTTTGATCCAAACTCCTGAA
TGGAGCATTCATGAAGCTTGTCTTCATGTTTCTGCGgccacatgcattgcacatttCATGCTCTATGTAAAATAT
GTATGGATTTCATCTAAAACTAGATTAAATCAATCCCAAGGTTTTAAGTGCTGCTAATGCTGGTTTGGCCACCAT
ATTGGGCCAATATTAACTATAGCATTGGGTCAAATCAGGACTCATCAGAtctttttttgacattttttttgtttcg
acaaagtcaaaaaaaaatttgattttaagtcctttaatatttttggttgaaGCATACAAGTGCATTAGCACAG
CATAGCACGTATAGTGCTTGTCACTAGTTACTATGCCCAATGCCAACAATATTTAAAACCAAGATTCGCTGTCTC
GGTACCAAAATCCGTACCAATGCCATACTAGCATTGTGTCAGTACGGTACGGGGTTCTCttttggcataccaagt
gtcaATACACCATCCGTACTGGGTACCAGTACTAAACTGGCACGGTATACTCCATACTGCCCAGTTCGGGCCGAT
ATGGCGaacaatatttaaaaccttgttcAGTCCTTGAAAACATAACAGTTGTTTTTATAACATGCTAAGCGTGGA
ATTAGCATTGCAGATATACGCAACAAATGAGTTAAATGGAGGAACTAGTAAGGATACAAGTAATCCATATGGTTT
GTCTGTGATTGAGAATTCACATGTAGATGTACCTATTCATCTTGAGCTATGTCAACCTCAACAGCAAACAGATGG
ACCACGAGCAACAACACCGAAGCTAGGGTAATTATTTTGAAACAtcaaaccttttttatttttcaatatttatta
gtTTTACTTTAATATACTTAATTAGCCCTTGCTACAGACTTCAACTGCGTTAAGAAGATGACCGGAGATCAAAGT
GTATCGTCTATAGTGCGCATCAACCCAGTTGTCAAGGTTGCTATAGTTTGTTATCAGCTATAACATACAATGTAT
GAGCAAGAGCAGGAACAATACTCAGCAGTCATTTAAATGTGGTGTGgcatgatccaaaaaaaatcattactcTGT
AAAGCAATCTTCCAACTAGGACTATTAATAATATGACAGGTATTTCTGTTTAAGAACATTACTTGTACATGGAGA
CATGTCCTTCCATAGCATCCGAATGGTTGCTGATAACATCTACCAGAAAACTCACCTGCTAGATTGAAGGTACAA
GGGATTAGCACCACTCGAAGTGCAACAAGCTGCACAAAGGATGACTCACTATATACTGCTTATAAACTAGTAACA
ACAGGATATAAGACTCCAAAGGTTTATTGAAGTCCCACACAACTCCAATGTTGAATGAGGatataaaatcacaaa
aaaaaatgctaacaGCAGATTCAAATTGGTGTGATGACATCTTCAGAATGTGGCAGCCCAAGGAAGAAATATCAG
TGTAAAATGTTGAACAGTTGGGAACCTCATATTCCTAAAGGTGACAGATTCATAACTTGAAGAGTTGAAGGTGAA
AACCAAGTCCTCTACACATTAGTAAGGATGTTTGAACAGCAGACTGCTATGATAAAATCatgtcttcttttcttc
atcataAAGATTCATGATATGCAAAGTTGTTCCAATGATTGAAACACATAAGGGTAATTTAAGGTTGTTAAACAT

>PdAGL17_1

ATGGGCAGAGGGAAGATCGTGATCCGAAAGATTGAGAACTCGGCCAGCCGGCAGGTGACCTTCTCCAAGCGGCGC
AAGGGTCTGTTGAAGAAGGCCAAGGAGCTCGCCATCCTTTGCGATGCCGAGGTCGGCTTTATCATCTTCTCCAGC
ACTGGCAGGCTCTATGATTTTGCCAGCTCCAGTATGAAATCAATAATTGACCGCTACAGCAGGACAAAGGAGGAG
AATCATCAGGTGCTAAATGCAACTTCAGAAGTTATGTTGTGGCAAAGGGAGGCAGAAAGCTTAAGGGAGCAGCTG
CATAACTTACAAGAAAATCATCGGAAATTGATGGGTGAACAGTTGTCTGGCTTAAGTGTCAAAGATCTACAAAAT
CTAGAAATCCAACTTGAAATGAGTCTACGCAGTGTCAGAAAGAAGAAGGAACAAATTCTAACCGATCAAATTCAA
GAATTGAACCGAAAGGGGTACCTTATTCACAATGAAAACATTGAACTCTACAGGAAGGTAAACCTCATACGTCAA
GAAAACATGGAGTTACGTAAGAAGATATACGCAACAAATGAGTTAAATGGAGGAACTAGTAAGGATACAAGTAAT
CCATATGGTTTGTCTGTGATTGAGAATTCACATGTAGATGTACCTATTCATCTTGAGCTATGTCAACCTCAACAG
CAAACAGATGGACCACGAGCAACAACACCGAAGCTAGGGTAA

*PdaAGL17_2*

>NW_008246554.1

CATGGTTTGGAAGGTCCACTAATATCATTTCTGCAGCCATGAATCATGTAGACGAGAATGTAGTACTCTTCCATA
TCCATGAGATCGTCCTTTTAGAAGTGGCGCAGCTTAGTAGTGGTATTAGTGATGGTGGCAGCgagcctcctcctc
cattgGAACTAATAAATAATGTGGCAGACAAGAAGAGAAGGAGCATCCTTGGGTTGGGtttattttgaactttttt
ttatttttggtaaataCTATAGCTCTTTGTGGAAAATTTAGCTGAGCTAGTAGACCGGGTCTAGGATCTAATTTC
CAGAACCAAACTCTTCCTTGGccaccctcttcctttcttttatatttttgtgtaTAAAATAGTAGGATAGATAGG
AAAGACACCATCCACTTCCTAGGCCAAGGAAGATAGAGAGAGGCTCAAACTCCGGGCAGCCATTGAGAAAGCTCG

TCTTCCAGGAAAGTAAGGTTTCAATATCTCCACTAGATCTCCTCTCCGACCTTCTTGCTTTCCTTTTCTCTGCCa
ctcgtcttttttttttttctctctctctcttctcctttttaatTCCTTCTTGCTATTTTTCCATCGCCAGATCTAAT
ACACCAGATATTGGGTTTTAGGTGGAttgagaaagaaaggagaagaaatatgGGGAGAGGAAAGATTGTGATCCG
GAGGATTGACAACTCCACCAGCCGCCAGGTGACGTTCTCGAAGCGGAGGAACGGGCTGTTGAAGAAGGCCAAGGA
GCTCGCCATCCTCTGCGATGCCGAGGTTGGACTCATCATCTTCTCCAGCACTGGCAGGCTCTACGATTTCGCCAG
CACCAGGTTGTTTCTTAGTCCCCTTTCGACGCACTTTCTATCTCATCTTACATGATATACATGGTTACAAATTCG
ATCGGCCTTTCTTAAAAGATCTTGCTGCTAATGCATGGATGGATTCGGAGTAATTCTAACCTATGGAAGACTAGT
TTTATAAGGCAACGATAAAGATGATGGGACTTGTCCGACATGTTGGTAAAGGTGGTCGGGTTCGTAGTTTTGATC
ATTCTTTGCCTGGTTGGTTTTGGTGGTAGGGTTTAAAGCATGAACTAGAGGAAAACAGGAACGGTGCATCTCCCC
CcacttctttttcccttttttcctttgaGAAATCGCCATTTCCTTTTTGTAGCTTTCTTATTCATGGTTCCGACA
ATGTTACCAAGATTCATTTTCATAGGCTTTGATATGCTCATCTCATATATCTATCTTCGAAATAGATTTTCATTA
GACAACCCCCCcctccgctctctctctctctgtgacaACCACCCCTCATCTCATATATCTATCTTCCACTTAGAT
TTTCATTagacaccccccccccctctctctctctgactccttttcttttttttttactgtacttatttctcaaat
acttcgAGTCTGTATGATGTATTAATTTAACTTCGTCTTCTACATTTCCATGTTATGAGGCATTGTATTGCTTTT
CTAGAGTACCAAGTCAGACAAAATTGTGTTGGTCCAATGAATGTCAATTGTCAGCCTACTGCGAGGTTCTCCATG
GCTAAGTTTGTAAACAACAACAAAGTGCAAGTCATGGGCTTCTTAGGATTGGAAAAGAACATATAGCAAGTTGGC
AGTTCCTATAGACATTTGAGAGAGACTATTCATCTCTCCAACTTTCTCATTTCCCccaataaatctaaaaatatt
gcaTGATAAATGTTCCATCCATGTTTGTATTTAAACATAACCACTAATCATCTTGTTCTGTACACTATTGGATGT
TAAGGAAGATAAAgaataattcattagattaaatCATTCATTCACATGTGCATGATGAGATGTAATTACTGAATA
TGTGGGTAGTGCTGAACATATATAGAGAATAAAAGACGAAGTCCAGGGTAAAGCTATACTAAATATTTTACTTCT
CATATTCCTACTGCTGGCAACAAGCATCAAGACATAAGAATTTTCGCAGGATACagcaaattattttaataaaaa
ctgAACTTTTAGATCCAGAAGTAGGGGGATGGTGATGGTTAGATGCTTGTATATGGCTTGCTGTCTTCCGAAGTC
GCGCACAATAATTTCTAAGCTGTGGCTCACATTGTCGTCGTGCATTTTGTTCTTGCGGATACGTCTTCTTAAATA
GATGCAACTAAAGTACGGTACGAATTTAACATCAGGTTCACGTAGTGCCATTGTGGACTTGTCCATAATGCATTT
ATTGATATGGTTTTTTTCCACTCCCCATATTTTGTACGTTTTGAAGTGCGATGTGCAGCGGCCACCATGTCAGAA
TATCAAATAGTCCAAGGTTTATTCTTCCCTTGCGTCCACCTCACTATCAGATCTGACACATTGAGAaatttttttc
cccttaattttttttttttgagaagaggTGCCATTCATAAATTTATGTTCAGTTCTGCAATAAAACCTCCTTCGAG
TGGTATTGATGCACTACTGTATTTTGCATTTTCTCTGCCTACACAAATTTCTGGATGTGATATTTATATAACATT
ACATAGAACTACGTGTAACTTCATAAAAGCAGTATAGAAGTAATAGTTCAGAATTTCATGTTCATAAAGATAGCA
GAAAAGCTGTTATTGAGATCAAATGAGAGGAGATTGTGTCATTTCGAGGTTAATTATTAGTAATTATTGCTAAtg
gttttctctttttgatcttGCTAGTTTATGTGATGCTTACATACTTAGAATAGGGTTTATTAAAGTTTATTTGAT
TTAGATTCATTTGGATAAAAGCTGTgctttcaaaatttcttttacTCGAATTTAAATTGATCTCAATctgttttc
aaaaatttaatttgactTCAAGTAGGATTGTCACACCCCAGATCCAGAGCATGACTGGCCGTGCTACTGAGGGGT
ACAACCCACGATAACACAAAGTCAACATTacatttagctttcaaatcTAACTtgaatataatagaataaataaga
aTCCAATTTCATTATCCCTTACATGCAAGCAATACTAGTTCCGAGTATCTAAATCTAAACATAAATACTAAAccc
ataatcctcaatattcTGAAAATCATTGATTACAAAttcatagtcaatttaaaatactaaaatcctTCTACAAAA
TTGCCAAGCTTGCTAGGACGAATGCCAAGCCTGAACAAACCTTCATCCCTACTGATCTTATTCgtcttgagaaag
aaaaataaaaatagaagtatATGAGCGACATAGCTTAGTAAATAAACTTTTCACTATCTtactggatcaagcata
agtttttgtATGTTAATGTATAATTTAAAGAGAATAacatatattacaaaataaagttttcaTAGTACAATAT
ATCAAGTcataaattattcatatttcataaatatggttctaatgtatagcataaataaacTCATAAATTATTTAC
CATTTAACCATATCATTGATCAAACTGCTACTTTTAGACTAAATGCTAAGATCACTATTATACCTATGACAGGGC
CGTAATTGACAaataccaactactattacccatTGGTAGGATCGAATTTAAACTACTATTACCCACTAGCAGAGT
CATatcaactactattatccgctgGCAAAgtcatatgccaactactattatccactggcagggtgatggggacat
cagagcccttcattcaAATGATCATGATCCTCCGAATTGAGTCATacccggattgggtccatttgagttcgagtc
atttctttattgcattatttgcttttgccTTCGTCAAATCAATTTAGTCAGTTATTTTGGGATTGAGTCAGATC
CGGATTGGGTCTATTTGAGTttgaatcattttcttttattgcattatttgcttttatcttagttaagtcaatttg
gtcagttgttttgttattagactcgTCAATTAGTTTTATATAATTGAATCAATATGTAagatctatataaagacc
acccctctcatgtatgaggaattgatgattgaataagaaaaccctagcctctctcttattttttcttcttcctcct
cttctcttcttcttcctgtgtctctataacctcttctttcttctccctcccttgttctacctccttctcctcttc
ccttgcAGTGGTTctacatcaacttggtatcagagtcgTGTTTTGCCTCTATGCCAAAGCAACGGACCTTGGTGC
ATGGCAGAGCAGCCTTCGATCCTGCGTATCAGGTTGCCGCACGAGTTCTTCACCATAACCCCCACCCCGTGAACA
TCGGAATGTGTTGAAGGGCCACGCTCAACCCGCATTGCCAAAAGGAGAGAGAGCATCACCTCTCTTCTTGCATCA
CTGTCCTTGGGTCCCAAGGAGAATAGTCACCGGTGGTCTCAGGCCCACCGTAACAGCTGCCGGCTTCAACAGGGA
AGGAGTCCCTAGATCCACCGTTGCACCCACCTTCCACGGCCATGCACAACGCCAACAGCGCCACCAGCCCCTTCA
ATCGCGGCCGCCTTCACAATGACGCCATCGCTTCTTGGTTCACCGTCGGAGACCGTCGCGTCCCGAAGAAGAACC
AGCCGCCATCGTGACTGGCCGCCCCTCTCCGCCGCTGCTCCGTCGCTGGGGCCGCAACCACAGTTTGTCTCCGCA
GCCGCTGCCGAGGAGGTCGGCACGTCCTCCCGAATCCGTCgagaggttgaagaagaagTCAAGGTAACAGATAAG
TTTTAAACCTATTACCCACAACCCAGTAACCCggatccacaaaaaaaaaaaggtgcacGTGATCTGCACGTGCTGG
AGATAATGGGTATCGGAGCGGGTTGCGCCCGGTAATCCGAACTCgtaaagagtcaaaaaaaatagtagcaCGTGC
CttgcacacaaaaaaaacccTTACCTCTGTTCACCTGCTGTTGCCGCCGCCCGTGAAGCCCCGCTCCACCCTAGC
CGCCACGGCCGTCGCCCCTCCAGCCTCGCTGCCGTCTAGATCTGCCGCAACCCCACCTCCCATGCTGCACCGCCC
TTGCCGCCTTGCTATTTCTGCCTTTGCTCGCCGCCCACCGCTCCCTCCTCCATTAGCCACCGCCGAGTCCCACTG
CCCCTCCACCTTTTCCCCCCCTCGTCTTCTTCATTCCCTCCACTCCATCTAGCTCCCAAACaccctctctcctct

cactcgaGCCCCCTCGGCGCCCTTCCAGCACCACAAAACCCCCCCTCTTTCGGGATCCATTTCCCCCCTTTTCTC
AGTCACCGAGCGGGCCCAAATAGACCCAAGCAGACCTGCGGCCCACCCGCAACACCAAAACCGTAGGCCCTTTAG
GCGTTGTAGGCCCACGAATGCACCTCACTCCCCTCTCCACAGCTTCAACCTCCAGCCGTTCATCTCTCCTCGGCG
TCCACTTCAATCGTGCAACAGTCCGGATTCAACCGTCAACACCACTGAGTTTCTGCGCTGCGTCCGATGACGTCT
CCACTGCTTGAGAAGCCACGAGCGAACGACTCTGACCATTTACTTTCCAATAGGTGATAGGTTTTTACTTTTTct
agcttgttcatttaattatgttctagttctcttgcatgatggtcatattcttgaaacttacttttgctgtcaaaa
ttcagaacatagaACCCTtactttcaaacccatcctattcgcctttaaaatctagatattaaattaacACACCCT
AGCAACGAGACGTTTCTCAATATTTTTCGATCAGATTGATTTAGGATTAGAAtaactgtactacttgattgcaat
ctaatttcttaaattgaataatcttaatcctttttctgaataatcgaagtaaaaatcagcaacatttaaacttta
tGTTATTCTTATGAAAACTTACTGATCTctaaaatcataatagattgcattagtaggttgtcctgcatcaaattt
ggtacTGCATTATATTTATTAGGGTTccattagtgacactgcatttcacatcatcacctagTGTTGTCATTGCAT
GCCCACTCATAGTGATAGGAAGTACTATTTCAATTAACCTAAAACCCCTGTACCTACCATGAATTTCGACGTTTT
GAGATCTCTCATGGAACAGTTCGCTACCATGCGGGCTGGAAACgaggaattcaaacaagagatccaTGAGCTCAT
GCAAAATTTTAGGACCCCTAAACCGCCAACACCCGTTGCAGTACAACCGAAATTTGGTTTGGCTGCACCACATGT
AGCCCTTCCCCATCTTCCTAGGAACCAACCACCTCCCGATCACTAGCGCAATTTAGATGAGAGGCTGCTGCGTAC
CATAAGAGTAGAGGCCCCTACCTTTCCTAGTCAACCCAAGCCAAGCGTGTATCTCGATTGGAGAACCGCCATGGA
CAATTATCTTGAGTGGAACGAGATAACTGACGATAAAAAAGTGCGATTTggcaaaatgaagcttattggtcactg
gtactggcataatgttgaacatatATGTGAGACCCAAAGGCTTCCTCATAAAACTAcatgggaggatatgaaaga
aaagctaaatgagaagtatctgtCATTTTCATACTGACAATGCCTTATGGGTAGGTGGCAGAAGCTAACTCAAGA
GACATCGATGGTTGCTGATTTCATcgcacgatttgaggagtttcatatgAAGTGCGGTGCAATAGAAGAGGAGAC
TCTTACCCTCTCTAGGTTTCAGGCAGGACTCCgtgaggagatccagaaagaactaGTCCTTCGAGAGATTACTAC
TCTTAAccaagcataccagttggctcaagatatAGATAGTTTCTTAAGACTACCTGTGGTGAGGCCCATCGACCC
TTGACTATTGGCCCCAGGAGCTCGACCTAACCAAAATTATCTCCCACAACGTAGACCGCTTTTCAATCCTCCTAA
TCAGCCCGACCTAACCCAAGCATCGACTAGGACCTTCCCGATGCTTGCCCGAACCCTTCAACtgataagaaaaaa
ggaatattAGGTTCCAACCCTCCTTCTCgaagccaaacccaatgctTCTGGTGCCAAAATTCTGGCCACATCATG
CTCCAATGCCCTGGCAAGACTTATCTGATTACTGAATCAGAAGTTGGGACCCAAAAGatcgaatatgtcgaagaa
gtttATGAACTAGatatagaagactatgtagaagactttgaagacgaacccacAGGATTAGATTTTGTTCAAAAT
GATTTCCAAAGGAaaactattgatctaagtacaaccaagccacttcacatcactactgtggaagaggtagtgaaa
gatattgAGAGCTAGTCACCTGAGTTGGCACTTGTAGCTCAAGATACCAATGCAAAGTCCAACCTTGAACATTCT
TCTGTAATAGTtttccttctagaggagttttaTGATGTAGCTCCTGATGATCTTCCAGATGCACTTTCTCCGATA
CGTGATATCCAACACGTAATTGTTCTAGTTTTCGGAGTATCTCTGCCtaaccttccacatcataggtTCAACCCG
AATGCATATACTAGGACTTGAGATTTAATGAATCGATGGTTgagattacaaacctaggcaacccatagacctaTT
TTtatgtctcatcagtatagagtctctgagtttacacaatcatttgcatcacatccatattcattgtatcaaga
aatcagcaattgcattcactttaataacttaaaatataaatcccttgctgatttacacggacgttataaagagtt
aagtgaaagaagttacgtcatgataagaattaggatTGAACGACTTTTTTCGGAAACGTTCGAAAAATTGCAAGC
TTGTAGTACAGCACCGTTTGAATTTCTATGAAAAATTCGTTCGAATGTCtatgttgtggatctttctGATGACTA
TGGAATTAGTgcgatatttaatattgaagatttagtcctaTACAAAAAGATAACATTTATGCCGTTTTGGACCCC
TTTgttgatatacctgcccatgttgaTCACCCTGACCCATTATATGCTATTGAGCCACCACCTTCCAAATTTCAT
ACACACAGAGAACAactagaacatatattggatgagtaGATTATGTCAACCAGGAATGGTGGTTACCAatgttac
cttgttcggtggaaaggtccaTCAAAGTCTGACGTAATGTAAATTTTTAGAGcccagttgcagcgccttgacctc
gatcttctggagcagtacgacagccagCCAAATCTGTACTTGatggggtcgagtttttctcacccaggaggagtt
gatggggacatcagagccctttattcagatgatcctgagcTTACGGATTGAGTCACACCCtaattgggtccattt
gagttcgagtcattttcttttattacattatttgcttttatcttagtcaagtcaatttggtcagttattttggga
ttgagtcagacccggattggatccatttgagtccgagtcattttcttttattgcaatatttgcttttgttttagt
caagtcaatttagtcagttgttttgttattagactagtcaattgggtttatataattgggtcaatatgtaaggtc
tatattgaaagagtagatgccctataagccaatcatttgatgggtttctctttgtaatcctccaaatcatgtact
ttgacactcgctatatatcaataaaggcattgtgtttcatcatttgctgcttatctattttattattggatgatg
aaccccataaattaggacattgatTTTAGGGTNNNNNNNNNNNNNNNNNNNNNTTGGatccatttgagtccgagtc
attttcttttattgcaatatttgcttttgtcttagtcaagtcaatttagtcagttgttttgttattagactagtc
aattgggtttatataattgggtcaatatgtaaggtctatataaagaccacccctctcatgtatgagaaattgatg
attgaatgagaaaaccctagccttcctcttgttcttcttcttcgtcctcctcttctcttcttcttcctgcgtctc
tataacctcttcttccttcttcctcccttgttcttccttcttctcctctttcccttgCAGTGGTTCTACATCACA
GGGTCGTATGTCGATTtaccccaaaaaataaaaaaaaattaaaaataataaggagtGTAAAATCTACCTATCTCT
TTCATCTTAGACCGTCAAACTTCACTAGGTAATTTAGTTAAccctttttaaaaaatattaaaagtgaaattaaatt
ttgtttgatgattgaaataaaataatgaaaaataggaTGATTAACCAAATGATAGCGTTCGACGATTTTAGGTGG
GTCAGGCTTGAGCGATTCGATCAATAAATCATAGAGTATGGACTCGATTAGGATCAAGGTTAATCAACATGATTC
ATCAATGAAGGTTTCAAGGACATTCAACAGGATGGGGTGGTCGGTCGCGCCAGAGCGGTTTAGAATCTCTTTGTG
AGAGACAACTCAAGTCTATAGAACAAGTGGATGGGAACTCTGAGATCCATAGGTTCCGTTTACAAGGAGAAatag
agagatgagagagaagaaagagaaaataagagagagaaattagagagagatggctctctctctcctcctcttctt
cttctcttctttctttcttcttatttcttggCAAAACAAAGGATGAAGGAAGAGGGTGGACCAGTGGTTGTTGTG
GTCGTGGTCCAGCAAGGACGCAGCAGGCACCGATGAGGGGCAGTGAAGCGCAGCTGGCGACAGTAGCTCGTCGGC
TGAATTCAACCCCAAACAAAGCTGaacagaaatttttttggctcgGTTGTCCTCGGATCCCAGCCCACTCGCCGA
TAGTTATGGGATGGAAGAGAAGATTAGAGGAGTCTCTTATAGGCTAGAGGAGGTCGGAATCTGCCTCGACATTGG

ATTCCTCCGATGAAGTcggaaggaagaagactcccattaGGAGTCTTCTTCATTGCACGTGCGCCCaccccttcct
ttttttatttatttgaaccTGGCTGGCTAGCCGACTTTGGACTAGGACAGGCCCAGTTGGCTTGTTTCTGAAAAG
GATAACATGTActacttggagttggatttTAGAACCCAAGGGTTTATCTAGTTCACCAACTGGATTTTGTCTCAA
ATAGTTGATCGAGTTCCACCTTCTAACAAGGTCTATGACCGTTAAATTACATAGTACTAGATACTTCCAGGGTAC
CTGACGTGAGTTTTCTTCTTGTTAGAAGGGAGCTATTAGGCAATTCCTCCTTTGCTTGTTAGTTTTCTGGTTGAG
TGTTTGAGAACGATGAGCATCCTTAAACCCGTCAAGAGTGGAAGAGGCTGTGGCAATCCTACTAGGTTAACCTAG
TAGTGTGCATCCCGACTCCTCGACCACTTGCATGCATCCCCACATCACTCCTTGCTATAGTCTTGCATTAGCTGC
CTAACTTAGTCAATGGTAAGGCAATAGACACACCCTGACCCCCATGCTTACTACCAGTTCACTTTCTACTGCAAC
TAGGTTTTGCATGCACCTCAAGTATCCGATAGCCGCACTCTCACCTCTAGAACCCTATCGATGCATAATCTTCGC
ATTCGGAATGCCAACAGATGGTCAGCATCCACTATCAGATCAAAAGCAACTCCAGTCATCTGAGTCTTCGACAGC
ATTAACCAGCACATCATCAGAAGGTATCTGACTGCAGCAACTTCATCATGGATCCCCAATCGTAACCTCCTCACT
ATCGTATCCAATCAATAGATAGGAGCTTCGCCTACTGCAGGTATAGACTTTAGCCATGCATAAGACTATGCATGT
ACCTTTGCAATAGCTGCAAACTCTATCAAACAATATGCCAACAACAAAAGGCTTGAGAAAACATGGACCCAAACA
ACAGCATCAACTCCAAGTTTGAGTGTCTCTAGAAATAAATGGCTGTGAGCACCAAAACCATAAGGCAACTTTAGG
CCTCTTCACAAGATATTTTGGCCTGATCGATGACACTTCTAGCCTTGCAACTTGCAGTACCACAAGTCATACAAT
TCCGACAGGAGTTCAATGCTGATAACTGGATGCATAGATATTGCTGCTATAACAAACAGAACTAGCAACTTGATG
AGGATGAGCCCATTATGGATGATGAGGAGCCCACCCTCATTCTTCATCAAGTCTTATGACTTCCGAAGGAAGCAA
AGGTGGATTGGAGGTGCAACATCATCCACCATCAAATTGCTTGCCAAGAGcacatcatcttcctcatcaaCAGTG
GCAATAGTAGGAATATGGCCTTTAGGGAGCCTAAGCACAATCTTGGACTCATGACCATCTACAGCACTACGTATT
AGTGAGTTTTCCTAGTCAAAGACTAAACCATATTGGTCAATCACGGATGCCTAGTGTTCTTCAAATCAATAAGGG
CTCGAGGATGAAGTCTAGTTTGATGTCATCCCCATGAAAGTTAGCAATCTCCTTCATAGGTATCCATGGCTTTTt
gattacaatatcattgacCGAGATAATACCTATACCTTCACTCTCAATAGGAAGCCAaatctagagaagaagaaa
actgtTGAAACACAAGTTCTCTCCTTCTGATAATTCAAGGAGAAGAAAGCTAGTTTTACCTTGTCATTGGTAGCT
AAAATCTGAGATACCCTCTAAAGCTGACCAAGGGTATCAACTAGAGATCCAAGAGctcttaggccctatttggga
gagcttttggagggccaaaaagtactttttggccctccaaaaatacttttagatgaaagtgtttggtaaaaattt
cggaaagctgttttagctttttcaaaaaactaaaaacagctttctgaaagaagcttcattttggagctttccaaa
aaagctattttgatcTTTGTCAGAAAGCAATTTTTGTgaccaaaatatctataataaaatataacaattatatac
tttgtctctttataaacctaaaaatctgctggagccctaacaaagaactcCCTTCCGTGCAAAAAaagatattct
tattttctattattgtattatactataaatataatattatattacattatgatataatacattaatatgttatgt
taaataatttaatattatattaaattattatgctatagtatacaatattatattactatattataataatattat
attacattacaataatattatactatatcatatatttatgtattaatatatgttatcttttattatgctctgttg
tataatactatgcaatgttctttatggtaattttgccatacaaaagtactttctcagtttgttcaCCAAacacaa
tattttagaaatgtagttaccaaatagcaaacaacttttttataaaagctctacttcagaaagctttATTTTCAAC
AGCtttacttctaaaagctctactgccaatatCTCTCGCAAACAAGGCCTTAGACCCCGAGGATGTGGTCCTTAA
GGACTTATCAGGAGGTCTCCACCCAAGAGGGATATGGAACATACCATTGATTTAGTCCCATGAGCACTACTATTG
TAACATCTTCCTCACTATCACAGATTTCCCACTAAACATGAAAACCTGAAGCAACATGCCGAAGGATGGCTACAA
AGGGCTTCATTAAGGGAGCTTAAGCTCAAGTGGAGTACCAACTCTGCTTACATTCAAGAAAGATGCATCTTGGAG
GTCTTCATTAACAATCAAGCAATTAAAAAAGTAACTATCGAGTATTGGTTCCCTATAAGGAGACTTGGTGACATG
CTCTATTCCCCatacttctatttttttcttggtAGAGAAGGTTttcatttccttctttttggtTGACACCTAATTT
TTAAGTGCCCTTCATTGCCGATATTGTAATCGATTGATCTAGGTCACGGACCTCCAACGTCGCCCCATTGAAGCG
GCCAATATAGTCCATGAGGGACTCTTTCTCCTTCAACTTGATGTCGAGCAGTGCATTTGAGCTGCGATGTTGTCT
TCGGATGAAGGCGAAGTGTATCGCAAAGAGGTGgacgagctgctcgaaggagtgGATGGACTTGGGCCGAAGTCC
GGAGAACCAGAACCAAGCCTTCTTGTGCAGGGTTGCATGAAAGGCCTTGCAAAGCACGACGTCGATGGCTCCGTG
CAGCATCATAAGAGCCTTAAAATTCTCAAGATGATCTAGAGGATTAGAGATGCTGTCGTATGGCTCTACTTAGAG
CATCTTGAGCCGAGGTGGGATCGGTTTCTTTATGATCTGCCTGTTGAAGGGGAGTTCTAGGAAGAAGCTAAGGTC
GTTGTTCACCCTCAGATATTGTCCTCTAAGGGCTTGAATCTGCCGGCCTATCTCCTCAATTCTCCTATCAAGGTC
AAAGTTTCGGTGGGGCAGGGCCTCCACATAGTAAGGGGGCAANNNNNNNNNNNGAGGCTCGGAGCACGCTATTTTC
TATCCCGACCAAGACTGCTTTTCTGGGTGCAACGACTCTGGAGGATCATTCCGAGTCTCAGCATCGATAGTTGTG
GCTGACTAGGTGCATGCGGTGCGCCGGTGACTGAGAATGGTGGCTGGGCTGGCCAGACTCCGGCTAGACCTACTC
TGATCATGTTGTTGCCTACTGCAGGCTCTAGACTGCCGTGGTTAGCGCCTGCACTTGTTGAATaagcaggttgaa
ctgctcaTTGCCCATAGGTATAACAGATTGTTTAGGCAGAGCAGGTTCTAAGCTTTACAATGACTCTCGAGAGGT
CGGAATACGACATTTTGAAGCATGCGATGCACTGTGGGCTCCTCGTGGCCTCATGCTGGTGATCTCTCTCTTTGA
TTAATATTAATGTAGCTGGAGGGCAGGCCCTTTCTCTAGCGCCATTCTATTGCCACTGAATTTGGGACGAAGGTT
AGAGTGCAACTCGAATGACCTTGAGGTTGACTGAGGGTGAGCTAAGCAGGTTGATCGAGGTGTGGAAAAGTGGAT
CTCCTCCAAAGGTAGCATTCAAAAAGTTTGCTTGCTAGAGAGTTTTGGATAGGAGACCCTCCAATAcctaagtca
gaatagagaGGTAACAATGTGAAAGAGAGAAATTTCAGCAGAGTAGCGTTCTCTATGAATAGTAAATTACCTACC
TCTTAAGGTCCCCCGAGCTACCTCTTTATATAGGGGAGCCCGTTTTGTCATTATCTGGGTTGATGTGGCGTGCAC
TGATGGCGAAAAAATGACTGGTAGTACGACCGTGGTGTAGGAGCCAATCTATGTGGGCGACATGCAGCGTTGGTT
GGGCACAACCGTGTTGCTACCACGATTGCGGGCTGCCAAAGCTATCAGTTGTTAGAACTATCAGTCCTTGTCAAC
ATGTGCTGATTGTTCTTCTCGGCTGACTGGCTTCTCGGCCGGTAGCTCTTCGACTGATCGTCTCTTCGGCTGGGG
GCTCCTCAGTTGGTCAGCTCTTCGGTCGGGGTCGAAAtcattcttcttgatttatgTCTGAGGTCAAAGAGGTGG
TCTGGCACGAGATCGAGGTGGCCAATATTTTTCCCATCAGCTTTCATTTTAACTAGCATTAAGCTATCTTTTCAA
AGCTTCCTTAATCACCTCAACAAAAATTCAGATGAGAGAGATATGACAAAACTAACCACAATACATTAGTTTAGT

GAATCTCACAAGAAAGAGATGATATAGTACCAAATTTTCATCATCGTATACGACCTTGCCACACATTACTCATGA
CTTACCGACAATGTAGACCTAGTCCTAGTCAGAGTCATGCTATTAGATACAATCAAGAAcgttattatttatcat
tatatgcTTTAAATTTCATCTAGGAACCATCAAACACATAATTTGTTCCCATTAGTATTCTATGTCATATTACAT
AGTTGAGTTAAATAGTCAAATCTTAAAATACTCATGAGTCCTAATGGTCTAATTAGTCTAATGGACACTAAGACC
TATAAATGCCCAGGTTCACTAAAAGTATTGCTTTTCTATGAAATGCTAGTTATCCACCAGTTTTTCCTATCAAAG
CAAATAGTAAGGTTGATTTAGAAAAAGCTAACAAGCGGCGCTCCTGCAATCCATAGTACTGATTTTATATGTCAC
TATAGGTTTTAATAAAATCTTCAATTGCTCGTTATTTATCATGTTCATCATACTTGTTTTTAAAGATTGATCTTT
GATTTAAATCCTAAATACCTATTTAGGATAGAGACCAATTGACCATCATTATGTTTAGAACTTGAGCTTATTTTC
TATCTTCAAGCAATGTAACCCAAAATAACACACTAGCCACTACTGCTTCTTTGATGGCATCCGTAATTAGTTCTA
TCTCCATTCAGTTTAATCTCATGTTTGCCCTCACTCAGGTTCATGCAATATTATCCTTTTTTCAATTATGCATGA
CAATAAAGATCAACTAAAGATGACTTCTATCTAATGAAATGTGTTAACTCAGTTCATTAGTCAATCTCATCTTAG
GAGAGATCTAGTTAGGTCAGCATTGCCCAACATAACATCCCTTTCTTTTGTGCAACCACAACCCTTCTGCGCCTT
CTATGGGCTTGGCCATGATATTCATAGGTCTCAATTTATTAAATAGAGGACTTTGGTAAGAGTCATGTCCCATCT
TCTGGCTTTCATTGCCATGCTACATTAAAGTAGGTTGGGAGCATAACACTAGCTTCAGAGTGACGTAGTGGAAGG
CTAAATTATCTAAGCAATGATTTTAAACTTCCaagcaaaaattaaaataatactaGGAGCAAAATAActtcaaaa
actaaaggcttATCAAAGGAACCAAAGCACCTAAGAATAGCTAGAAGATCACCAAGGCCCCACCACCATGGACTC
CGCCAACTGACAGTGCTCTAGCATGCCTTCATAAAGGACTTCACCAACCAGCTTCGCCACTGCTTCACAAAATTC
AGTAATATTTTATAGCAAAATTCATAAAGGACTTCACCAACTAGCTTGCACAAAATCCAAGATGCAGAAATATTT
TATAGCAAGCATGAGCTCAAGTGCACTCAGGAAAAAgcaaacttcatttttttttttgctaaacaaaagcaaact
tcattgatcaatgaaaatatgaggTACACTGGAGGCTCTGGGACCCATACAAGGGATATGGATGACCAGGTATCC
TTGAAGTCCAAGTTAAACACAAAAATAAACTCGTATAATCACTACTAAGCAAAGTCATCAAAATAAAGCTTTTTG
AAGGAAAACCCTTCAGATGGTGACCTAATAATAGTTGGAACAATATGGGTAATAGTCATGTTGCCATTAGATGGT
TCTGATGATGAACATCATACCATTGGATCATGGTAAACTCCTTCTGCTTCCTTGAATGATGCTCTAGCAGAGGCC
CTCTTTTTTGAGCAATACCTTGGCTTCCATTGGATGAGGCAAATTTTGGGCTTGGACACAAACTTCTTGTGGTCT
TCTTAAAGACTTCGCTAGCGTTCTTTATTCTAATGGTGGTCTTCATGCTATCAGAATGGTTCTAATAATGCACTA
TATGACCTCCTCTAGGCCTCTTCTAGTTTCTGGACTGGGCCTAAACTGTATTAGCTTTCCTGCGTGAAGGAGTTC
GTTTCCTATGAACGAAGGGCTTGGAATTTCTCATAGAGGTCTAGGCTTATACATTACAATCTTCAGCTTACCTTC
CAAATTCAAGTGCCTCTTTCCACTTTACTAAAATTTTTAGTAGGTCCTCTTCCTTGTCAGCACAATTTCATGATC
AAGGATGGCCTtgatttctgattcttttgCCGTGTTGGAGCCATGAGTTGCAAGTCTCTAACGTTATCAACAAT
CGACCATCCTTCAAACTAGCATACGAGATTAGCAACATTAATGAAGGTGGGCTGAGTGTAGATCCTTGTCTCCCA
CTCAAAATGGTGCTAAAGCGTCTAAGATATTACCACGTGGCCCAAGTGGGTCTCCTTCAAGCCTATTATAGATGA
GGACTTTGTCGCCAGGCTTATATTGGATTCTTTTGATGTTCTAGTTGGCCAGtcttattaatatttttgcTATCT
TCTTTCAGACTTTCTCTTGCACTTACTAGATATGCCGCAAGAAAATTTGCCCCTTTGAGACAAGTCCTAATGGAT
TTCTGCATTGGGGCTAAAGCTATTGATTGCTAGGGTGCTAGTCCTATCAGGACTTCCAAGGGAGCGGTTGATGTT
ACTGCTGTAATCAATTAAACTCGACTTAGAAAGAAAGATGGTGTGttgtgtaacaacccaggacctcacccaaaa
tgactagccggaaggtattatttggattcctcgatcctgtataagtatccaagatctacccagcgaataatcaat
gtgggactaaacacacgcccgcacgggtccttacatactcccttcgttcaagccctgacgttctcgtcaggctaa
gtgttcaaatctattcaaatctaatcacaacacCATTATTGGCCTGTGGTCAACCCTAATGGATCCGTNNNNNNN
NNNNNNNNNNNNNtctacccagcgaataatcaatgtgggactaaacacacgcccgcacgggtccttacatactcc
cttcgttcaagccctgacgttctcgtcaggctaagtgttcaaatctattcaaatctaatcacaacacCATTATTG
GCCTGTGGTCAACCCTAATGGATCCGTACTGCAGTGtcccttagtccacataggttatgagccggGTCCGCTCGA
TACCATTTAtcacaacccaggaccttacccaaaatggctagccggaaggtattatttgggtttcttgatcctgta
taagtatccaagatttatccagcgaataacaaatgtggaactaaacacacgctcgcacgggTCCTTATATGTTGT
ATTTTCTGTGTGATAAGACTTTATAATAGGTTCTTCGAACTTCTATTTCAACTTCAATCTTCCCATGGTCAAGGA
TTGGAAGACACTAGGAAACTAGAATTCTCGGTTGACCCATTAAAAAATAACTCATGAAATTGATGTCATGGTTGA
ATACTACGTTATTCAGATCTTATGCAAACAAGCTAACTCTTTTAAGAAGAGTTTGGCAACATATGACATCAAAAG
CTTTCATATGGGTAATGACATGGGTCATCTTGGAGAGTCAATCCATCAACATATATATGGAATCATATCCTCCTC
TGTTCTTGGTAACCCTAAGATAAAGTTCATACCTGGATTTTCCCAAGGCCTTTCAGAGACTAGCAAGGGAATGTA
TGGTCTGGTTATCGATACTTTGAAGTTGGTTGATTGGTAGATCGGACACTTATTAGCTTTTCTTATATCCTTTTG
AATACTCAACCAATAGAAATGCACTTCACTAAAGTGACAGTTATATCTCAGCTGAAATGTCGAATTAGGCCTCTT
GTGTTTGATTTATGAATGATTTGATCATGCATAGATATCCCTAGAATGCAAAGACATGAATAAGTAGCCATCTTA
TAGAAGAAATCTGGATTTGCATATTTTACCAAAGCCACTTTCATAGAGCCCTATGACCTGATTGAAGCCAATTAA
TTTTACAGACATGTACCAAGCAGACAAGCCTATTGGCTTAGGGCACCGGCAACCTTGTTTCAACTCCTGCCTTGT
GGGCTAAATTCTTGCAGAAAGTTGCTCCACTTAGCATGCTTGAAGTTCAATATCCTTTAGGAGTTCAGATAGAGC
TCCATAGCTGAAAGCAAAGCATGAACTCCCTATAGATGGGATAATGTCTCTAGTGTTTTATAGTTTAGAACACAA
CATAGAATTCAATGTTATACATGGAGTAGCACCGCTTTACATCATTGAGTTTCTCGCAAAAGAGGACAATGGATG
TCCTTCTAGGCTAAGGACTCCTTCAATCCCAATGTGAGGTGTCAGATGAGACTTCGAAAGACTTTCAAAGTTAGG
TAGGCATAGTACCAGAGCCTCAACCACTTGATGCCTTAATGCTCGTTCACTAGCTTCGGTCCAGTTAAAGTATCC
CTTCTTCATGCATTTAGTGATCTCAACCAACACAATGTTGAAGTTGTGGATGAACCAATAGTAGAAAGTAGCCAA
CCTACGAAAGCTTTTGATTTTGTGGAAGTTTGTGGAGATTGGCCATTCAATAGTTGTTTGGACCTTTATGGGTCA
ACCTTGATGTCGTACATAGAGACGATGAACCTGAGTAAGGTTATCTTAGGTTGTATCCAAGTGCATTACTTCAGA
CtcatatacatttttttattaccaAAGGTTACTTAGTGATGAGAGCTAAAAGTTCTATTAGGACTAACAAAAGTG
TGTGCGCACGCTATGTATTAGAAGCTAGAAGTTCTATTAGGGCTAGTATTAGGTTGTACCCAAGTGCATTTCTTC

ATATTCATGTACATTTTCTCTCACCAGAGGATACTTAGTGATGGGAGCTAGAAGTTCTATTAGCGCTAGCAAAAG
TATGTGCGCCCTTTATAATAGGAGTTAGAAGTTCTATTAAGGCTCGCAAAAGTGTATGCGCATTTAGTGATGGGA
GCTAAAAGTTCAATTAGTGTGTGCGCACACGTGCTTAGTGATGGGAGCTAGAAGTTcatttgtgtgtgtgtaga
aGTTTAGTTAGTGTGTGTGAGCGCACTAAGTGATGGGAGCTAGAAGTTCTATTAGGGCTAGCAaaatgtgtttgt
gtgtgtgtatatgcaAGTTAGAAGTTCTATTAGGGCAAACAAAAGTGTGTGTTTCTATACATGTGCGCACGTGTG
CTTAGTGATGGAAGCTAGAAGTTCTATTAGGGCTAATAGAAGTGTTTGTGTATCTGTTATCTCTCATTTCagtta
aatacataaataacaTAGTGAGACCACCTAGTTGAAGGAACTatatagctaatttttttgaggaaaaccATATA
GTtataattgattttatttcttcactttgatataattaattgTATTAATTTGAGTTGAAAAGACTCTTTGCCC
ATGGTCAGGTTCTAGTTTCTACTAAAGTTTATGTTGCCAATCAGAAAGAGTCTGAAGGtgtcttttaaatttt
tttttttgcaagggAGTCTGAAGGTGTTATTGTTCTGTAAATTTTTGATTCCATTACAACTCCCCTAAAGGTGTT
AAAACATCACTCATTATCAGTTTCTTACTAATTtattattcataaaattatcaaaaaataacatttgctGATAC
GTTGAAGTTACctgttttaattatttcttaCTTATTCTTCATAAAATTGTGTTGAGACGCAGAATCTAATTTTAT
GCTTGGATCCGGCTCACCTTGAAACGGAAAAAGTCATCATTATCAAAAACTAGGTTCCATCACTCCGGTCAGAAC
TGGAACATATGAAATCTGATTGACTTGACTGTTGTAAACCCAACTTGATGTATAAAATGAACatagtctcatcaa
aaaaattCCAGTTGTATTGATGTTAGTGAGTATTGAATGTCAGCTTTCTATGGATACCACAAAGCCCAAGCTTAT
TGAACATTGCTAGGTGTACACTTAGTCTACCATCTAGTATTCACATGAATAATTTTTGTGTCAGTCTCTTGAGgc
attttaaaatgaattttaaTGTAAGATAAACTTTACACAGTACTAGGGACATTTATCAGAAAATATGGTGTTCCT
TGTTTTGCATTAGCTATACTTGGATTTAAGAATTTGATTTATGGCAATGTGAATATTGAAGGGCActaaaaatta
ttagataattTGTACCCAGGAAGGGTGCCAGTGTACCAGTTTGGGATTTCGACAATATGACACCAGAATTTCCAT
GTTCGATCCCACAGTGCACTATTGAAATGTAATGCAAAATTAGAACTCCAAAactttgttctttctttatctttg
tttttcatttgttttgtggttgtttttttggtttcttaCTTTCtacaatgatttttaaaatttggttAGGTTTTT
ATGCCTATGATTGGGTATATGCTACGTACCATACATGGGATCCAAggtttgttcttcttcctctctttttccctt
ccTCTTTTCATCCAACATTCCAATGCgagaaaagggaaagagagggaaaagggggaaaagaaaaaaagaaaaga
aaagtggtcAAATTCTTTTAGCTAAGTTCCAGTTAGTCTTGGATGAACTGAGTGGTTGCAATTGCCGTCCCGCTT
GGTCTCCAAAGCAGGACAGAAACAGGAAGTATGGTTCAGAGTTGTAGAGCAATCCTAGGACTTGGTCGAGCACCA
GCCTAACACTATGTATGAACCAGTAAAGATagaaagataaatccttgtatTTAGATTTATCTTTGTGCTCTAATT
GGATTTAGTagaatataacattatataccCTAGAGTGTGCAtgtttttctcattttcctttTTATGCCCCCATTT
TTGCACGAAaagttattatttgaaaaaaaaaagttccttgTATGTATTAGCAAATAAAACGTCATAGTGCTTTT
CCTTTCTTAATATTTGTTTCATCTGTATCATTTTGATGTAGAAGGTAAATGGATATGATTCTCTTTATTCTACCT
ATCTTGATTCTTGTGATTTTAAAGTAATAGACTTACAAATCTTTAACTGCCAGCGTGGAGTCGGTTATAGAACGC
TACAAGAAGACCAAAGAGGAGCATCATCTCGTTATGAATGCAAATTCAGAGGCTAAGGTACATTTGGAGCTCAAG
TACTTTTAAATACCATGCCCTAAATAGTTCGGTTATAATCTTTTCTAGTGCTCATGCTTTTCCAATTAGTCTTCA
CAACCATTTGACatgagtttctttttattattctccAACCATTTCGAAGCTAAAAGTGACTAAATGGCCAATTA
TCATAGAACATACATCTAATGCTTGAGTTTCCTCCTGCTGCTTGAATTGATATACTTAAGAAATTTCACAGCCTA
AAGCATCCTCTTCATTTGATGTATTAATAGCTTAAGAAAATACCATGAAAAAGGGgctattttatcataattaat
tAAGTAATCATTTATGGCTTTGACATTTCTTTTActaaataaaacataatatagaaaaatatatgctatTTGACT
ATACTTTTTCAAAGCATACCAAGGTAGAATTCTTGGATCATTATAATGGACTAATCTTTCTGTGATATTAgagta
atattaaaaaatatcatgGATGATGATCGTCATTCAGAACTTCAAGAAGGACCAATTGATGGAATGGAGAGTTAG
CTTATTGTAAGTATCATTCTTGTAAGTATCATATTCTTGGCACATGAGAACTCTCGTATACCTAGTTCTGCGTAC
AAGTCACAACTTGGCATGGACGGTCTGTTCCTCTTTTACCCAAACAACTTTGATCACTAATAAGAGCTTTCACAA
GATCTGACAACATGCAGACCTTTAAGATGTCAAACCAAACATTATGAATCAATTTTCAATTGATCGAGAGATTTG
AGATAATATCTTTGATGTGAAAATAGAAAACTACAAACATAGAGGAGAAAACAGCAAATTACCTAGAACTGAACG
AGAGATCTATCTCAAATGGTTATATTGGGTCAAACGTGAATACTACATCGATTCCATAGACAAAGCAGTCAAAAG
CTAAGTTACATGGCCTAAAATACGAAGCAGAATAGTAAACTAATTCACAACTAAGTCTAATATCTAGACACTAAG
AACATAAAGAGACAACGATAACATGATTTGACAATGATATATTACTTGCACGTAGTTCTAGTTCTCCCTACGGCG
CAGCCATTTGCCTACCGAATTTGACCTAACCCAACATATATCTGCCTTCTTGTTTGCATCAATTCTGGATCTGAA
AACCTTTTGTGAGTTTTACTTATACTCATTTTATAACCATTTCatcttattataaatttatattcctGTGTAAT
TTTTGGTGAGCTACTTTCTGTGTTCTGAAAGCAGCATGGTGGTTTGTGAACGAAACTAGCTATATGTTTGCATGT
AATCATTCTATCTTCTTAAATGCAGTTGTCTCACTTTTTGATTATCCGAACTGCTTCAAACTGAATCCTTTTTAA
TTCACTCAAATATGGGAAGCTTGATAATTGCTTCTTACGCCCACATGATCCTAATTATCATCACCCATTTAGTGG
TCAATACTGCTAATTGTCTTCCATACTAGAGTTAAACAATTATCTTTTGAAGCATTAACTCTTCAGGTGTTACTG
TGTTGTCATTAGGAAATAATATTACCATGTAGAGAGGCGGTCTAAGAATATTTCTTATGGGAGTTCGAGTGAAAT
TTGCATGGACAAATGGACAAGATTCTCTTGACAAGCTTCCAAGAAACTAGGCCGTTAATTCAAATGGTCCTAGGT
GGATTAATTGATTTAGTGATAGTGTTACTTACGAATGTAGCATCATTCAAAATCATCAAGGTGTTGAAACCTTAG
TTTACGAAATGGACTACTAAGAATTGAAATAGATCAAGAAAAGCTTTTGGTGTTTGTTGGACAGTTATCAAAGAT
TACCAAATTGTTTGAAGTAAGTTTTTTGGATATGGTATAATGCTTGGAACGGTACTCGCCAGGACCAtagtttta
agaaataaaacatCAGAAGAGGGAGATGAACCTGCTGAACCTAATTTAAACCTGCTTGCCATGGGCTAACTTGAC
AATTATTGAGTAGGctcacttaaaaatttttgAGCAATCAATTTAGGGCATCATACAACTAGAGAAGATTCATC
TTCTGGGTTCTGGATCAAGGATAAATTTGTGAAATCAAAGAAAGGTACTTCACttttaggaagaaaaagaagagg
gtaGAGGGGAAAACTTACAAGAGACATCTTTAAGGTTGCAgataagaggaggaagaattacAAAGCAAAGGATTT
AGAGAAATGAAcaaatggatatgaaaagtTAGCTGACAGTGTTCGAGGAAAGCTTGACAAACTATGTCATGCAAC
TTCACATACATCTGGTCATTTCTATTGAGTTAGGTTCTCATGTGGCTATCACATTTGGTTGTGCGTTAAAGAAGG
ATCAGATCAAATAAAGGTCCCTTTGGCCCATTTAACCCTTTTTACTTCCAACGACAAGAAAGACAACTAATATAT

TTCGAGCACAGAACGAGaattttcaaaagtttgaaAAAGCCACTTTTTAAAACTAACTTGGAGTAAATTATTCCA
CCCTATAGGGTGGAGTAATCTATTTCTGGGTTGGGAGCAAGTAAACCTATTTCATCTGAGTTGACTTACTCTGCA
ACCAAACATAGCCTATGGTTGGAGTAGCAATGTCTTATGAGGAGACTCCAGGAGATTTAATATGTTTCCGATTTA
GTTGCAAGTTGGGGGCAAAATGTCTAAATTCTTGAATATCTGATCACTTTTTCTCTTGCTCACTATATCTTTTTG
GTTTGCCATTAACTTCTTGCTTATGTCTAGGAACAGCTGTTTTTCTCTCCGTTAATAAAGNNNNNNNNNNNNNNN
NNNNNNNNNNNNNNNNNNNNNNNNNNNNNNNNNNNNNNNNNNNNNNNNNNNNNNNNNNNNNNNNNNNNNNNNNNN
NNNNNNNNNNNNNNNNNNNNNNNNNNNNNNNNNNNNNNNNNNNNNNNNNNNNNNNNNATGTAAGGTTTAAATAT
TCACCAGAAAAACAACCAGGTactgaaattttgaatattgacCACATACCATATTAATGAACATAGAACTATGC
AGAAATTGCACCCTTTATTAGTCATGCAGAATGAACAAACATCTCTACAAATAATGTATGAATATTTATAGGACT
TTAAGGCAGACTCCaccctcttcccctcttcctGTTGACAGTCCTCATATCCAATATTTTTGCCTGTGTTGATTT
TTTTCTGTGGGGTTCATTTGCTGGGGCTTGGGGGGTGGTGATACAACAGATAATTGACAGAGCACCGTAAGTGTG
AGAATAGCTCACCTACTATTTAAGAACCAAAACATTCAAAGATCATCGCAACATTCATCAACAAGATTAAGTAAT
CAAAAAGTTCGAGTCGAGAACTCCTAGTTTACATCAGCAGTAAGTGATCTAATATATGTTAGGAAGTATTTAACC
TTGATGCTTTGAATCTTGAGTTCCACTAAGTTTGAAGCAAAGTTGGTGCAGCAAACTTCGAGTATATGTGATCTG
ACTGAATGTTTCTTTCCTGAACCTGCTGAGTGTAATATTATTCACTCCAAAGCAATAATCCAGTACTTTAAATAA
TCTGTTGCATGGGCTAAACTTTGACCATTGTACTCACTAGTTCTGGAAAAAGGAGGCAGCAAGCTTGAGGCAGCA
ACTGCATAACTTGCAAGAAAGTTATCGGTATGCACACTTATCTTGCATAACATACAAGTTTTTAAGCATTATTCC
TCAAAAGGCATTCAtctcaaaaataagaatattgtcATAGGTGGTTTAGATTATTCAGCATCTTGTAACATGtat
ttattactttatttgaCGCTAAAATGACACTAGCACATCACATGTCCTAAAATTCAAACAGATTGGTAGTACTA
GAAATGGGAATGGCAGACGGGAAATATATCTTCGGCAATAAGTTACTTAAATTTCCAATGTACTAAGACCTAATC
ACCTAATACTTAAAAGATAGTACTAACAATGTCTAGATATCAATTCCAACTTGGTGTAATACCATAGAATGAACT
AGAAATTGGCATCCTTTACACTTGTCATCTCAAAGAAGATCTAGCAAGCTTATATCAAGCAACACTAGTAGAAAT
TGAAATACAAATGACTTGTATATGCAACCATGTGTATTTGTCTAGCCCATTGTAGGCCAAATCAGACCGAACCGA
CTTattcagcccgtaccgaaccaaCTGAAACCTGCCAGAAAATTGGGTCAGCTAGAATATTGGATTCAGTTCTGG
CCCCAAACCAACTGGAACCGGCCAGGAAATGCCTGAACCAGACCAAACTGGCTTGGAAATGGTCAGAACTATtcg
gaaccggccggttccatTCGAACTCGATGCGAACCATCCGGTTAGCACCGAGTCGACCAAGTTCAGAACCGGCCC
CCCTTCTCCTTTGCTTGTTTTGTTTAAAGAAGGTGTTGGACCAACACCTCCTTCTCTACTTCGTTTAGTCTTAAA
ACGCCTAAATGACTCCCATTCCTTCCAATTTACGAAATTACACTAATTTAACATGATTGAAGAAAAATCTAAGCT
TAAATAAGGTATTATTCTTCTCTCCTAtctcatttcctctattttattttttgagcttgaaaaaaagtccaaaa
tttgcaaaataaaggaagatcatgccaaaaatttttatttggacttaattttattatatattatagatctACAGA
TGATCTACATAAtgacatgaaaattttttaattttagattatatacaatttttaaaactaaaaatatatttttta
ttaaaaataaaaatataaaaaataattctttttaaaaaatataaaattagaagcGTATTTAGGGGCATACAAGA
TATTCTTTAGCAAAATTTAGGatctatttatttaagtttCAATGCGATCATGTGCCCAATGGCTttggtcaaaat
tttaaaaaaaattgagaaataagtagtcTTTGATGCGTACCCATgagcctagaaaaatatatgtagcatctatTA
TTAGGTTTTACATGAATCtaagctgaaattaatttttgaaatctttatatttaaaaattattttaaatttaaaa
ataactaaaaatatataaataatactaaaatttataaaaattagtagtatgtattacataatttagaagtaattt
ctaaagtagaaaacatatttttctaaatttatgatatttaaatatatttttaaaagtaaaaaattatataaataa
ataaataatataaaaattagtattatgtattagataaattagatgtatttctatggtaaaaaaaattatcttcat
cgtaacaaaatttactattttaaataattgataaactgacgtACTTGATAAATCTTCaagaatggagccatca
agaaaaaaagaaccaaaacgTGACATTAGTTGCTATCATGGGCAAATGCTCTTGGGTCGGCACCATTGGAAGTGC
AACTAATACAACATAGAGTTTAAGAGAGGAAGAGCAATCAGattgaagcagcacttagctaGTGGTTATCCTGAC
GTATCTATGTATCAGAAATGCCCACAGGAGGTTTGGTAGAGCTGATGAAAAATTACTTCGCTAATTTCAGAGCAG
CGAAGGAGAGGATTTCCAAGAAGAAGGCGAAGGTGGATGaccgagcggcagagccaccttctTATCACTTTAGAG
AGCCAAAGGAGGCGTTCGCTCTAGATGACGAGGCACGGATCCAAGCTGCCACGAAGgcaagcctggatgatcagc
GGCAGCAGGATCAAGTGGTCAGGCATAGAGCTCGATTTAAGCCCTCATACTATGAATCGAGCTCTGATTTAGCAA
GCAACAGGGAAGATCCAGAGTTCAGGAGGACCTCCTCAGTCAGAGAGGCCATCAGTAGAGGCGGGAGTCGTATTA
TAGCTATGCTGGAAATGTTTGGTAGCAAAAAGAAGTGATCAGAGAGATTCCACCAAGAGCAGTGATCCAAGATTT
AGATCGATATGCCTTCCCAAGCAAAGATTTGAAGCAGCAGAGGGTTGACAcaatgctgaagaaggataGATAGCA
AGCTATTGGATCTTgtttcacttcagccacattctAGCCAATATGGTAAACAATACGTATTGTAGCTCTACCATT
TCTGCCATGCAGGTTGTCGATGCTAGTGTAGATCATCTAGGCCCGAAggacatctacggtgagcttcttgacaat
aatAAGAAGCTAGAGAAATGAATTGCCTTATGCAAGAGTaagtggcccatatatggactGGCGGTGATTTGTGAC
GATTGGACTGGTCTTACTAGGTAGAGCATTATGAACTTTGACATATTGTAATATGAAGATTTCTTTCACAAGTTA
GTTGATGCTTCTGATCAGGTATACAATGCCACATACATTCTCAAattgatggaggaggtgattgacCAGGTAGAA
GAGGAGAATGACGTGCTTTATAATGAGCCACAATATAAGGTCACTGAGGAGATCTTGATAAAGCGGTGACCATAT
ATTTTCTAgaccccatgtgctgcacattgtatcaaCCTTATGTTGATGGATATTGTGAAAATTCATAGGGTACAA
TAGACAGTAAAGACAGCCCAAACCATcactagatatatttataattatatatggGTCTTTCACTGATGAGGAGAT
ATGCAAGAGAAGAAATTTTTAGATCAGAAGTCATACAATTTGCTACCAACTATGTTGCATTTGATAGCattcttg
agaagaaagcagCCCTACGTCAGATATTTGTCAATTCCGAATTATAGAAAAGTAGATATCCGAGGGTCGGCACTG
AAGGAACCATTGTCGAGGACTTGGTGACGAGGCAAATATTCTAGCAATGGGCCACTacgatagtgaaggctatca
gaCCATTGTATGAAGTGCTTCGGGCCATGGACAACAAGAGGTAACCCaagatgggcttcttgtatcacatgatga
agAGTAACGATTCAAACCAAGAAGGCAGATCTGACGTATGCCCAAAAATACATCAACATTATTGAGTGGTGGTGG
GACTATTAGATGGGTAGGGACTTACATTTAGCAAGTAAGCAAGAATATTGAAATAGACTGCTCTTGTATTTGTAC
AATATTACTAAGACAATAACAAACTCTATCTGTAGCATATTACTTGAACCCTCGGTTTCAGTACATCATAACTGG

AATCGATTTGGATGATGAACTTTTGGTCGTTCTatgtaatgtgatttacaagatggagcctgatTCAAAAGTTGC
GGTCCTATGTCTAGAACAGTTAAATTAACTTAAATGACATGTGTAAGTTAACTAATATCTTCcattattaacata
ttacaatACGGTTCTTTTttacaaactaaaatatttagagagggtACCGACAGTTTTGGAGTCTCGGCAGTTGTt
gtaagtaagaaaagtatgaatccaaatatttgcatatcagaaaaatatttatctGGCGTTAGATGGTTATTAAT
ATGACATTATCttatatacaatttttttaatatttttgtagctgaatggtggattcattttggtttgttcacaa
aaaaatttaagcgGCTAGCTATTCGAATCCTTTTCTAGACGGTCTTCTtgagtggctgtgagcgcaattagtcca
ccttcgccctcatccataACAAGTAGATGaaccgtttgacacaaaagTACCTCAATgatcttgtatatgttcact
acaatctgagaTTGAGACTGAAGTACATTCAGAAGAAAGTGGAGCTAAAGTACATAGATTCAATCCATAGTACTT
TCATTGATGAtaaggatgatcctatgatcggatggcttgtgggccagcatCAAGAGCAAAagcttgatgagcgag
gatCGTCTCCATGATCGGCTAATTTCATAGCTAGCGAGGCTGGGGTCGATGCAGGACAATGGGCAGAGAGCAATT
TTACACGCTTTCTTTTAGCTGATCAGCTACAACCACAGGAGAGCCGGTCACTACATAACTCCTTATTTGAGACAT
CTTTTTAGAGATATGATTAAAagatgcttagacgaggccacTACGCTAAACGGTCAACTTGATCAGGAAAACAAC
ATACATCCTGCCAACATAGCTAGGCAGATAGTGACATAAGGGTAAGCAGGCAGAGAGTAATACAAAAGATAAGCG
TGCAGAGAGTAGCACGAAGGATAAGTAGGTAGCGAGTAGCACGAAGGGTTAGAAGGCAGCTTGCTagcacaacca
aaaaaaaaacaaaaagcactTACAGGCCCAATGAAAAGTGTAGAGGAGtcagttcactcagattcagagaCTAGA
GTGGCAGTTCCGGTGGTCATAGATCTGCTGGCCAGAGAAAGATAGAGGACATGAGACCACTGTTTATAAGACAAT
CGCAGGAGTCTTTGATTCACCGATGAGCCTCAGTTTCGCATGCTACAAGGATAGGGACCATGGTGCACAATCTGG
TGGAGCCCGTAAGGATccaattgcatataggagacgggCTCTTCGAGGTCACTCAAAAGGACACGACGCAATTGC
AAATGATCTTATACGTAGAGTAAAATCTATAAGCgtatcaaattttgaatcatatactggatcctattatccaCA
GCCATAGGCAGTCTATGATCCATATGgatacgaatatggatatggtgcatctgaGGCATCTTTCGGTTACTACTA
TCCTACGTAGCCCAAAGCATCTTACGGTTCAGATTTTGCTGCTGGTTTATTCGAATGGGCTCCTTCACAGCTGTT
TTATCATCCAATTTATCAGGCAAAGGACACCTCTcaaagccagagcttcagcgagagatccaaAAGTGCTTATAA
CCCAAGCATGTTCCTTATGgaatgaacatacaagactacaacaccACTTGGTTGGAGGGATGGGCTGATGTGtct
cctgactatgctaatgattcGGATATCTACAAGAGACATCGTCActagataaaatttagatattgttATATATG
TCGTACTTTAAATACATataattgattttattattttatatttttttacctcactacttgatttaagaatatttta
tgttgcttcttctagcatgcaacatctcaataatcattttatgatttgtatcctTTTAAAACAATATGATACACT
ATTTAGGTTAAATCAGGATcgtagaaatatcaaaaaatattaaattagacttaaaattattgaaaatttttaaca
agattgattaccaatttaatcaacttgaaaattttaaaaaaataagcatgCTGGTACGAAATCGGCACGCCCAAG
CATACCATGTGTCGGTACTGTACCAAATTGGTACCGTACCAACCCGTCCACCGACCGGTATAGGTTTTGGTATTG
GTTCAGTGGACCTTGGTCTAGccaatatagaaaaaaaatgtagaggATTATTTCATATATCACATGTACCATTAA
AACACCAGTTTCAATAATTGCAGGCAGTTGATGGGCGAAGAGCTTTCTAACCTAGGTGTCAAAGATCTACAAAAT
CTAGAGAACCAACTAGAAATGAGCCTACGTGGTGTCCGAATGAAGAAGGTCTTGGTCACTTTTGCCTAATTCAGT
GTTAACTTGATGGCCTTTCTAACAATGGCAATTTAATGTATTTCTTTTAACTCTTACAGGACCAACTTTTGATCG
ATGAAATTCAAGAGTTAAACCGAAAGGTTGGAAAACATGTTTTTACCATCATCTCCTATCCTAAGCTTAAAGTGA
TGCCTATCAACTGCATGACATTTCACTGATTCATTGCAGGGGAAGCTTATTCACCAAGAAAATATGGAACTGTAC
AAGAAGGTAGAGCACATTCATCAAGAAAACATGGAATTACAGAAGAAGGTATTGAATAAGAGTACAAATAGACTT
GATACATCCACAGTGAAACTTCAGTAATGAAATCTTTTTTGGATTCTGACTAATGGTAAATCATGTAAGTTGTAG
TATAACAAAGCTAACCATCAAATGTTGCATCTCAGGTTTccacaatgaaggggacaaaagGAATTGATAAAGGTC
CAATTGTTCCCTATAGTTTTAATGTCTCAGAGGATGCAAATGTAGCCATTCATCTTGAACTAAGCCAACCACAAC
TGCAAAAAGAGGGGGGGCAAGCTGCAACACCAACGCTCGGgtaatttttctttccaaaagtcaaacaacatagat
ttttttaacTACCATTCTAATCACAGAATATTTCACTGATTGCTGCAGCAGGCTTCAATTGCATTGAGAAGATGA
TGCACAAATCAGAGGTGGTTCCTTGTGATCCTTCATTAGTGTAATTATAACGGTTACAACAGCATGCTACTAATG
ATAACCCATGATGTACAATGAAGAGAACGATATACTCACAACAGTCATTCATAAATGGTGTGAAACAATCAGACT
AGCCAATAATTATTGTAAAGCTATGTGATGGGAATTGCAACGGATTTCAAATCTACACAGGGCAAGCAAAGAAAC
ATGGTGCCATAACAAGAGCTTCATGAGCATAAGCATTAATGGCATAAGCATTTATAAATAGGTTCATATAAAATC
TTACGCAATCGAAAGTCAACAAAAGAGAAACAGGATGCCACTCAGAAAGGGGCACCAGAAAGATGGGGCCCTCAG
CTCTAAACGACGATGTATGCAGAACTAATGGCATAAGCATTCGCATATAAAGTCTTCGGTGATGGAAGATCAACG
AAGAGAGAAAATGGAACAGGTCCAGGTCACCAGAAATAGACTAAGTAAAATAAGACTGATACAATAGGAGAAAAC
ATAAGATCAGAAAGAGCATAATATGGTAGATCTACAAAGAAATTATTATCATGAGAAGTATGCAAATGCCACGTA
GCAGCATATAACATCTGTACTCTTTTTGCAGAACGTTCAATCCGAACAAACACAAATAGttgataaaaatcaaaa
cttaatcCACTGGACAGGAGGATGAACAACATGgatacagaaaagaaaaaaataaaatgtgagTATGGAGGTAA
TGAAGAAAGAGTCTCCTAGAGAGGGAAATCTATAGTAACATACAAGACGCAATTGAATGTCAGATTACAATAGAA
AAACTTtactccaaaaatttaaaatgcagGGGCGAATAGAAAGACCAAGATGGTTCTTATGTTTACTGCTACCAT
TTTATTGATGCCAAGATGATAGGTTTGGTTTAGTAGCTTGGAATAGTTACAGGAGTAAGACACCATtacaagggg
aaaaaagagatttacaaagaattgaaaaacagaaataacaatatttttatacattttGCACATTTTAGAATTTCA
TGGTTCTCAATATGTACGAGATCACCAAAATAGATACCATAATGTCTACTTCCATATTTAGATGGACATTTATTT
GAATGCAGCAGATCAGAAGGTGCAGAGACCAAAACAAGGCCACCGACATCAGAAAAAAGGCTAACTTGTCAGGGG
GCTGATAGCCTAAAAAAAGTTCATAGAATAGTATAATAAGCAAACCTAAGGGCCATGTAAGTCAAAAAGAATAAC
TTTTATATCACCTTACTTCTTCTCCAAAGAATTACCTCTTTCTAACATCAAACGTCCTTTTCACCGAGAAGGCAA
AATACCAAGAAAAACATCAACAAGAAATCCTCATATTGTGGCACTTATTTCACTTCCAATGCCAAATTTAATTAT
CTAGCACAAATCAACAATAATATCCTTTATAAGGTTCTTACTTTATGCTCGCTTCCTCAGGGCCATCCAAAACTT

>PdAGL17_2
ATGGGGAGAGGAAAGATTGTGATCCGGAGGATTGACAACTCCACCAGCCGCCAGGTGACGTTCTCGAAGCGGAGG
AACGGGCTGTTGAAGAAGGCCAAGGAGCTCGCCATCCTCTGCGATGCCGAGGTTGGACTCATCATCTTCTCCAGC
ACTGGCAGGCTCTACGATTTCGCCAGCACCAGCGTGGAGTCGGTTATAGAACGCTACAAGAAGACCAAAGAGGAG
CATCATCTCGTTATGAATGCAAATTCAGAGGCTAAGTTCTGGAAAAAGGAGGCAGCAAGCTTGAGGCAGCAACTG
CATAACTTGCAAGAAAGTTATCGGCAGTTGATGGGCGAAGAGCTTTCTAACCTAGGTGTCAAAGATCTACAAAAT
CTAGAGAACCAACTAGAAATGAGCCTACGTGGTGTCCGAATGAAGAAGGACCAACTTTTGATCGATGAAATTCAA
GAGTTAAACCGAAAGGGGAAGCTTATTCACCAAGAAAATATGGAACTGTACAAGAAGGTAGAGCACATTCATCAA
GAAAACATGGAATTACAGAAGAAGGTTTCCACAATGAAGGGGACAAAAGGAATTGATAAAGGTCCAATTGTTCCC
TATAGTTTTAATGTCTCAGAGGATGCAAATGTAGCCATTCATCTTGAACTAAGCCAACCACAACTGCAAAAAGAG
GGGGGGCAAGCTGCAACACCAACGCTCGGGTAA

*PdaAGL17_3*

>NW_008246539.1
CACAATTAATTGTTCTCTCCATTCTTCATCTTTATTccgtcaaaaattaaaaataaagacaaatcattgctataa
caaaatatatgatattactGCTAAAGTTGAAAAAAGAGATGTCTGCCAATGTTATCATATTTATAAACAATTCTT
ATGGATCATGCTCTTGTCTAGTGTGTGCCAGCTAAGAAGCGCACAATTTATGGGAGATCATCAAATTTCGTACAG
CACGCATTCGCACCATGATGTGCATCCATGTTTGCCATGGTTGATCCATCCATCACAaagccaaaagaaaatatt
gtcgctttttataaataattcttATGGATCATATTTTTGTTTAGCATGTGCCAGCTAAGAAATGCACATTTTAcg
aaaaatcatcaaatttttcttaaCATACATCCGTACCACGATGTGCATCCAAACCAGCAGGTGCCATGGTTGGTC
CATCCATTGCACACCCCAAATTTGAATGGGGCAGGTTGATGGAAAGGAGGGCAAAATGGTCAACTTGGCATAGGC
CGTCGGCAGCTCAATCAAATGATCAATCCCGGACAGGGGCCGACAGCGATATCTCGCCTCACGGAAGTCCGACGG
GGATCGTCGGATGTACGGAACAGACTCGTCGTACCACGTGTTGGCGATCCATACAACAGACGGCCGAAAGGGATT
TGATTTGAGTAAACCAACGCCGAGCGGTCTACTTGACTGCCCAATCCTCGTACCCCACAGTAATGCAACAAATTG
CGTCCggaccaaaaaaatataaatttgtagaGCCTACGCCACACCCTCTCATCTTTAGGCTTGGTCCAGACGGCA
TAATTCCGCGTGCACTGAAGGGTGCAAGATTGGGTGTTCTGTGGAATTAGGTGCATCTAGTTCTTTGCAGAAAAT
GGAAGGCCTCCAGCCATTGCAGACCGGAAACCTCCTGCCCTTATAGACGCGTCCTAAACAGACTGGTGTCCTTAG
CGGAGATCCATTTCCGACCACTGTTTGGTAGGTCATCCACTCTATTCCCTTCCTTCCTTCCCAGATCACTATCAT
CTTagcttcttcctcttgtttcctTGTTCTCCATCCGGTTTTATTTCCCTTCACAAGCCTTGTGTGCATTCTATT
ATCAGATCGGTAGAGGATCCACCGAATTATCGGCTGCTGCGAGGGGAAGGGAGAGATGGGGAGAGGGAAGATCGC
GATCCGAAGGATTGATAGCTCGACCAGCCGGCAGGTGACCTTCTGCAAGCGGCGAAGCGGTCTGTTAAAGAAGGC
CAAGGAGCTCTCCATCCTCTGCGATGCTGAGATCGCCCTTTTCATCTTCTCCAGCACCGGCAGGCTCTATGATTT
CGCCAGCTCCAGGTcactcccctctctctctctctctccacctccctctctctctttctctacaaGACGCTTTCT
CTTCGCATCctatatatgcaaaattttctattattgcaTGGTATGACATATNNNNNNNNNNNNNNNNNNNNNN
NNNNNNNNNNNNNNNNNNNNNNNNNNNNNNNNNNNNNNNNNNNNNNNNNNNNNNNNNNNCCTTTAAAAATTACTTA
AAGAAGAAATCCATTTTGAAGGCGAACGAAGGAAGTGGGATATTTTTATTGATCCCCTTTATCCCCCTTCTGGGT
TAATTACCATCATTCTCTTCCCTCGCTTGGTTTCTCggatttttcttctatcaagtGTGGTGATAAGGGAAAACA
TCCCCTCACAGGCGCAGAAGTAATTCTTTAATTGAAAAGTCTTGTGAAGCAATAACGACATTAAAGGTTTGGCGG
AGGCTCCCTTCGTTGTTTCATCTTGTTTGTTCTGAAGAAGGTGTGACAGAAGTCTCGTGAGAGACTTGTATTGTA
GATTTTAATTTTACGCCTAAggatggaaaagtaattttgtGATTGGTTAGtttgtgcttcttttttttttgtgt
tccaAAAGGGGAGAAAAATTAAGGTGGGCTTCTTATAATCTTTGGTAGGGCTTTTAGCGGGCTAAGGGATTATTC
TGAGCCTGAAGTTGATGACGGCTTCTGAGAGTAACGTATAAGGGATTCAATGAAGCTTATCTTGAATATCAGAAT
TAAATCATGATGAAGAGCTATAACACTTGGTTCATCAGTCATATGTCTGGTTTATCCTCCATTCCCTTGGACCAG
CCGATGGGAGTGCTATGATGTGGCGATGATGATTTTGACAAGAATCAATATCAGTCAAACTCATGAATCAGAAAG
AAAGGGATGCAAGAATATATACGCATGCctgaaaaaataaagcatatatAGTTCGCACTGGCTGAGTCAATAATT
GAAGATTTTAAGACCATTACGATCGACTCGTCGAACTTGATCAGATTTGGAGCATTTCAAAAGCTTCTTCACACA
ATTATCTAAATGGATATATGTTTTAGTAAGTGGTATCATCATAATTTCACTGAAATGAAAATTACAAGAGAAAGG
GCAGACACATGCTACCACTATTAATTGCTGTGACATCCAGGAGTTCAGTGAGAATATAATCTTGTGCATCAAGAT
TTTTTAAACTTCATCCCCTTTATTTTCCAACATGATGACCGTTGGTACGTTGTGATTTATATCTAGCCCAACCCC
GTAGTGAATCAACATGTTTTGTTGTTGAATAATTGATGATTTGAATCAACATTTATGGTGTAGAACTACCAATAG
ATCCTTTCTTAATTTGTAGGAATTTCCTCCAGCTTGAGTACATGATTGATAGTTCTAATTGGTTGAAGGAATGGC
ATTGGACGTTCTTTTCATTATATTTACCTTCTTAAACCATCACATACCTAGGAAATATTAGAATTTCCGACAACT
ACGTCGAATTTTAAattgcttatatatatatatatatattattctccatctttattcttttatgataaCAAAGAA
GTTTGGCTCTACACAGGAAAGGAGAGAGTTCTGCAAAAGTACATGTGGCAACTGTATATACCGCTGTCTTTGATG
TTACTTGCAGAAGTATAAAGGAAATCAAGCTAAATTTTGAGTTTTGAATGACAATATGTGAATTACATTTCATGC
ATTGATATGCATATATTACACTGAAGTGCATatattttctaacaaaaaaaaaatatatagagcttgaattgtttt
aaaataactGTGATTGTTTCCATATTTCTTCTTATGGAGACATGAAGTTCAATTATATTATATCTTCATCATCAT
TGGTGAAAGAAAACaaccataaaagatgtgttGTTTAGAATCAATTAATGTTGATATGTCCAACATATgtcttct
ctttcttatttgagTATTTGAGTCTTCGGTCTTTAAGTTGTCTTGCTAGCTTTTCATGTGATTATCCTTATATAA
CTTACACTTGCCCATTGCATTGTAGTTATAATTTCCTACATTCTACGTGATTCATGTGATTCttgtttccaaacc
ttgaaagTAATATCTAAAGCATACAAGGAGGATAATGAATGGATGGACAAGAAACATACAACATTTGGTTGGAAA
aggatttttttgaattactaATACACTTAAGGTCTATTTGGATATTTCCTGCAGGATTAGGCTGAAAATCCTATA

ATCTGGCCTAGCCAGTACATATCTTCTAAACTATGTGTATGAATTATGTTTAGTATTACAGAGAAATAGGCTTGA
CCCAACCCATATCAATTACATCTCTTCTAATCTATTTCTACAGGATGAAAAAAGGCCcgataatttcattttttt
ctctttatggaATTTGCGTTGGCTTACTTTgaaaaaaatcctttaaataTTGTGAATATGGATCTATACTGGCTT
GTGTTCTTGTTAATCCTATTGAATTAAACAAGACTTTATACTATATAACATCTCTAGACTTTGAGTCGTAGGACA
TCTTTGCTGGCGTGTACTCGTTACTTTTCTTTAAATAGGTTGGcttctaaaaaaattcatagtctagagacTTAC
TAAAAAGTTATAAGGAGGCAAAGAAGCATAGTTTGAGGTACTGAATGAGTAAATAGTCATTTGAGTAGGATGGAA
TCAACAATTCTTGGTGAAAGAGTACTTCTTTGGAGAAAAAGTGAAAGGTGAATTCCTATCTACTTCATTTGGATA
AGGAAGTACAAGATAAATACAAACAACTTAAGAATGAATGTTTCCCATTGATGACAATTATAACTATATGAGGGC
AATAGTTTGCACTTGATAAGATTCTTGCTTGGGTATCCATTGTTGTTTAACAAAATAAGTCAAATAATAAAGGAA
ATAATATTGAACACTTATttaagaatggttttgattATTTACTTAAATTTGCATTTCTACTTTAATGTAATGGCA
AAATTCTTATCTTCCAAGAAAATTGGGGATGCTTGGCAAAAGGTTCATAATCGGATGCATATTATATACACCTTA
AATTAATGATGTTGCATATATTGTTCATATAATTGGAATagatggcccactaatttgtttGCTATATCTTAGTTGC
TTTAGTGTCAAACTCTATAATCTGGAAATTTCATGATTTGATGTTTACAATATATTATtggtattttgtaaattt
aaaagCCAAAATTTAATGGGAAATAAATTTTCAATAGAAGTTTAGCTCTGCATAGGAAAGGAGAAAGTTCCGCAA
AAATACATGTGGCAACTGTATATATCGCTATCTTTGATGTTACTTGCAGAAGTATAAAGGAAATCAAGCTAAATT
TTGAGTATTGAATGACAACATGTGAATTATGTTTCATGCATTGATATGCATATATTACACCGAAGTGCAGatatt
ttctaacaaaaaatatatagagcttgaattattttaaaataactgtgatttttttctatgtttcttCTTATGCGGA
CATGAAGTTCAGTTATATTATATCTTCGTGAAAGAAAACAATCATAAAAGATGTGTTGTTTAGAATCAATGTTGA
TATGTCCAACATATgtcttctctttcttatttgGGTCTTCTGTCTTTAAGTTGTCTTGCTAGATTTTTACAAGAT
AAccagatttattttatttagattacATTTTTTCATGTGATTATCCTTATATAACTCACACTTGCCCATTGTGTT
GTAGTTATAATTTCCTACATTTTACGTGATTCATGTGATTCTTATTTCCAAGCCTTGAAAGTAGTATCTAAAGCA
TGCAAGGAGGATAATGAATGGATGGACAAGAAATATACAACTTTAGGTtggaaaaggtttttttttgaattacta
ATACACTTAAGGTCTATTTGGATATTCCTGCAGAATTAGGCTGAAAATCCTACAAGAATTAGACCTGGTCCGGCC
AGCACATATCTTCTAAACTATGTGCAAGTATTATGTTAGTATTATAGAATATGAGGCTTGATCCAACCCATATCA
ATTACATCTCTTCTAATCCATTTCTGCATGATGGAAAAATGGCCTggtaatttcattttttttctcctcatgGAAT
TTGGGTTGGCTTACTCTGAAAAAAATCCTTTGAATACTGTGAATATGGATCTATCCTTGCTTGTGTTCTTGTTAA
TCCTGTTGAATTaaacaaaattatccaaataagCCCTTAGGATGGAGTTTATACTATATAACATCTCTAGACTTT
GAGTCGTAGGACATCTTTGCTGGCGTGTACTGGTTACTTTTGTTTAAATAGGATGGCTTCTGAAAAAATTCATAGT
CCAAAGACTTACTAAAAAGTTTATAAGGAGGCAAAGAAGCATAGTTTGAGGTACTGAATAAGTAAATAGTCATTT
GGGCAGGATGGAATCAACAATTCTTGGTGAAAGAGTACTCTTTTTGGAGAAAGCGAAAGGTGAATTCCTATCTAC
TTCATTTGGATAAGGAAGTACAAGATAAATGCAAACAACTTAAGAATGAATGTTTTCCATGGATGACAATTATAA
CTATATGAGGGCAATAGTTGGCACTTGATAAGATTCTGGCTTGGGTATCCTTTGATGtttaacaaaataaatcaa
ataataaaggaAATAAAATTGAACACTTATTTAAGTACGGTTTTGATTATTTACTTAAATTTGCATTATTAATTT
AATGTGCTAAATAAGCAATAGGCATGGCAAAATTCTTATCTTCCAAGAAAATTGTGGATGTTTGACAAAAGCTTC
ACAATCAGATGCATACTACATACACCTTAAATTAATGATGTTGTATATATTGTTCATATCATTAGAATAGATGGC
CACTAAATTGTTTGCTATATCTTAGTTGCTTGAGTGTCAAACTCTATAGTCTGAAAATTTCATGATTTGATGTTT
ACAATAGATTATTggtattttctaaatttaaaagctaaaatttaatggaaaataaataagtaatgtTTCTTGCAC
TCTCTTCTTGGTCTTGTAAATGTTTTGCATACTATAGGATTTCATAAAGTACTTCTTGCATGTTCATATAATCAG
TTTTCTATATTCTTATATAAAGTTTCTCATTATGAGCTAGGATTAACTTTGCTCCATAAAATCTAAATTCAAGCT
ATATGACATCGataatttgaatattatgaCATTTTCATAAAgtaggaaaacaaaaattatgcaATTGGAAATGGG
CAGCGCTTGTATCGTACACCATCTAGATGTTGATGTAAATAGGACaaggatgtttttttttttttacactggCAG
TCACACATGTGTAATATATGCTtaaggaaaagaataaaaaaaaattcaaattatatttctataaattattaaaaa
tatgtaaataaaaatattattatatttaaaagaaatattaataGTAGAATATgatttgaataaatattttttctat
ttatccaATTACATTTTCAGAAGCAATTCAGAATTAATTTGTCATATTTTTGTTGGACCATTTTTCCAAGTTAGT
TTTAAGCTAGTTATACCCAAATCTGATTTGATTCGAAAGAATATCATTATTAGATTGCATTTCAAAagtaattca
aaatatttttatattttttaacaactAGAATTAATAAGAGGGAATaagttatatattatatatgaaaaatgatAA
ACGGAATCAAAATTCTTGATTTCAAAATTGTCATTTTATATAATATGGATAAGATTGATATATCTTTCATTATcc
tgtttgatttattttctttttgggaaaaaatactTGTAGTTGGATCAACTAGGACTGCCACTTTTCTAAGGTTT
TGTTTTGGATACCTAATATCCAAGCATCCTAGAATAGATTTAACCATGGGCATCTATCCCCTGACCAGCCAACTA
GTAAATAAGGTTATTGGGCGTTTGGTTGGTGTTCCCATGATTAGTGGATAAAGCTatagaaaagtgaaaaaatag
atAGAAAGCTATGGATAGTTAGTTCTATTTCACTCCGGTCCctgaataaaattatttaggttAAGTACCTTGAGT
CTTCGATTGGATGGAGACATTCACAACTTTTTTGGTGGTGTTTGGCTAGTTAAGCGAATAACTCTCCCTTACCCA
CCACTAAACCAGTATCTAAATAGCCTAATTGGTTTTGCTCCCCATCCCCACCCACAACCAACAAGCGGGGATCGG
GAATTCAAGTCTCGAGTGATGTTATACCTCAACTTTTCAATCCTAGTTAATTCCTTTTAGAGACTTTGTCCACAG
AAAATTTTGGGTTCATGTCCAAGACCTAGGTAACTTTCATGGATGGGACCTGGTCCATCTTAatctgaagaaaag
aaaggaaaaaaaaaacacgaaGCACCAAGTCTATCATTCAAGCTTTGCTTTAACTCCTACGCTACCTAGATTAGAG
AACTTTAGTCCAAAATTAGATTCGTAAAAGATTTTGGATCAAAGTCCACTAGTAGCAAAAACATATTTCACCATC
TTTGTATTTCTTGTCAgtgttaaaaaagaaatgtcTTCCACCATCTTGTCTTCGTCCCTTGCGCAGCCAACATAT
ATTAGCAtcagcttttcttttcctcGTTCCCTCTGCTCTGTCATAACATTTCTATCCAGTTTATGAGCTTGCACG
CTTCTCATGACCAGTCCATGGACTTAAACAAGTCTTCCACTTTCAAACCAGCCTCTACACTACCTAACCTGATTC
TTGGACAGGTAGAATTGGAGAAAATTTAGGCCTGATCCATAGCGTCTAGGCTGCATGGTTACTTGTCCTAACCTG
ATCCAATGGCACCCGTAGAAATGCTTAAGCAACTCAAATAAATCACTCAAAAAGTCACCAAGTTGGTCTGGTGTA
ATCTCATCGCTCGCTCTGAATTGGTACCATGAAGTGTGCCATTGAATGAAGCGAAGTGATTGGTGCAAGACACCT

GAGTGACTTTTTTTAAGTGATTTTTTGGAGTTACCGTAGCATTGCTCTTGTACAAGTCTTTCGATGTGAATATGA
TGGACATGGTCGGATATTGGAGCCAAACTCTTACCAGTAAGTTCAACTCTAGCTTGCTTTCCTCGTCTCAAGATT
TGAGTCCGCATAATGTACTGAAACAAGTCGCCAATAGCCTCCCTATTCCATGACCCGAGAATGCCTTTCATTTCa
gctttttcaaaaagatgAAACGAAAGAGGATGAGACAAGATTCACTCAAGGATTGCACGGACAGATCTACTACAC
ATTAAGGCAAACATGGTGATAATATTCCATGTTGGCATTTCGGTTTATTGAAATCTATCGCTTGAGGTATAGCCT
GTCCGACAATCTTTGGATCTCCCAGACATGAAAAAGATGGCATTCCTATAACACTTAGGCTAACAATTTAAGCAT
GATAATATCGAATCCATCAGTTCGCTGTTATTTTTTGCCTTCCTGATAAGTTCCCTCTCTTTTACCATTATCATC
TTCTggatgcttttgaaagaaaacaaCGTACAGTTGAATCTTCAGTCAACAAACCAAGCCTCCATGACATACTAT
CAAGCTTGTCACTGTTAACTCTTCCTCCATTGGACGGGCATGGCAAACAAACGATAGGCATATCCTTCATCAGAA
ACCTATTTTaggtttttccctttttctaattatttcaCTTTATATACTTGTCATACTTTTATCTATGCAATAAAT
GGGCTGGGTtataacaaaaaagaataagattAAATTCAGGTCTATATTACAATCACCCAAAAACCCTTACTAGGA
AGCTAATCCACACCTCAAACAATGTTTCTCTTAGAAACGGTCTTGAGCCCCTTAAGAATGCTTAAAACCTAGGAA
GGTAGTATGGAAACATATATAAGAATCAAGTTGATGTGTTACCTGAAGGCATTCAAAGTACTTGTGTGGGACCCG
TCTCACATTCGTTATTTGTTGAAAAGATTTTGAATAgttatacagaactaagaaatccaaaaatactttttcggct
agccattttgGTGAGGTagtgggttgttacaaatggtattagaacgaactcggcccatagcctatgtggactag
gcgATACTACAACATAGGTTCATGAATGCTGACCATGAGCTGATCGTGGtggttgtgattagatttaaatagatt
tgaatgaCATTAGGACTTGAACTGGAGGAGTATGTAATGACTCGTGcgaacgtgtgtttagtctcacatcggtta
ttcgccgaggagaACTTGGGTGCTTATACAGACTaaggaactcaaaaaatatctttcggctagctattttgagtg
aggtcctgggttgttacaatatgGTTGCATTACGGGTGTTGAAACCTTGACACTATATTTAAACTAAAATATTGT
AATACAATTAGTTTATGGGTGGCAATgtgttgggtttgggtttgggtcGAATCAAATATGAGTTGCTGGTCAAAG
GATCGTTGATGCAAACTTGATCCATTTTATAATAGGCCGATCAATAtctaattcatttattttacatGGACACAT
AATGAACTAAGAGTCCAATTAAAACTTCATTTTCtagttatatttattgataaataaaagcaaaataggGCCAAA
AAGAGAACCTATAGTTAGACCAGGGGAGAGAGACTGTGTGTGAGAGAAACAGTTGATGTTTTTGGCTCTCATAAG
TCTTTCTATCATCTAAACCAATTCTAAACTAGCTCATGGTTAATATACTATTGCCACCATTTTAAGGACTGGATT
CCATAATATATACCCAACTAGATTAttattgagaaaaagaaaagaagaactaCCCACATCATAATTACCCAGGAT
CGGTCAATTGGAGATGCACCGCAAGACTGGAACCTAGAAGCACTGACTATGGCCACTTGGATAATCCCTGACTGG
ACGGATTAAAATCTCTTATCCAAACTGATTTATTTTAGAATGAATAAAAAGGTTGAAGGGCCCGGGCTGAGAATT
GTCACATCTTTATTGAACTAGGGCATGCACGTATAGAAAACTTAGAACAATAGAGCATTAAAAgcaataattata
ttaaatagaatataaatataaaaaaagatattgatAATTTGATTGTGAAAAATCAAAGTTCGGGGAAGTTGAGAG
ACATGGACTGGATGGAGTGGTAGGAGAGCGTATTttcaagacattaaaatttgaaaattaatggAGAGGATGAGT
TAAGGTTGGAGATGGAGAACTCACTAACAGTTCTAGTCTGAAATTAATGGACGATCAGTCATCTTGAAGAGAAAA
TAATTAGTCTAGCTGACAAGCAGAACgataaaactaaaatatattacttcaagtatttaataaaatgacaTTAGT
AATATATTTTGGTTAGTAAATTTGATAGATTGGAactattcaatatttttttctttttcttttttttccaatgaaA
TATAGTGCTACCTTGATTAGGAACGCATGAAATTAGTAGAACTGGCATCACCTTTGACGGAAGCTGTGGATCCAG
CTTATGGTTGCCCTCTCTAAATTAATATATGATAGAATAATCATTCAAGACAGCTAGTGGACATTTAATCTTCCT
GGGATGCCCAAATTCCTAGCCTCCCTAGCCATTGATCCAGCTTATAGCTGTGATTTAATTGTAATATAGTATAGA
TGAGGACAAGAACTTTATTGTTTTAAGCTCTTGATAGAAGATAATTTGTGGGGATAAACTCCCCGTCATCAACAT
TAATGCAATGCTATTACACTGGGAGAAGCATAGACAAggacaaagaaacaaaagaaaaatcctctCATTTACTTG
GATCCTGCCCTACACAAGCTTGTGATGCTCCTTGCGTTGATAGGAGTGCATTTTTACATGAAAGAACATGGTCAA
GGATGATTCAGTTTTGACAAAATCTATGGGTTTCTAGCAAGAAACTAAGGGTAAAGCTGTAGAAAAGAAGCGAAG
TAATTGCTAACGTTTTGAGAAACTATTGCTAATTTCCTGCTCTATATTTACTGCATGGTATTctataaacctatt
gttgctAGCCTAAGAATGCAAGCTGAGCTAGAGAGAGTCCTTATGCCCTTGTACATTTGAATATCTAGTACCCTT
GCTATGAGATGCAATGAATCCACTAAGCATGTTCAACATTTTATTTAGGCTAATCCAGCTTAATATCTCTGAGTA
AAGAAAATACACTATTTTCTGAACCCCATCATCTTTCGAGCAAATAagtgaaacaaaaatattttggatggAAAC
AACTGACGATGCTAAACAATTTGGAACGAAGTCATGAGAGAATCTGAAGAAATTTACACTTAGAAACGAAACTAG
TAGAAAGAAATCATGAAGCAATGCCACACGGTTGGGAGAAGCATTGTTAATTCTAGTTCATATTCTAGGTAACAG
GGGGGAAACGTTTGATCAATTAGATGTTAGATGGTAGAATAGCAATGGATGGACAAATGCTACATAACAAGACCA
GATTGAGACCATGTGCGTGGCGGCTTAAATATTTAGGTCGCCACTTTGTATAAGAGGATACAAGAAACTCAAGCT
ATGTATTACTTATTAGAGAGGCAGCACTCATATTTATTCATGttcattatcttatttatttatcaattcCTGCTC
CTATAAGCCATATCATGTATCAACATCTACTCCGATAGTGCACTGAAGAAAAACTAACATAAAGTCCCTCCTCAT
ATCAGCATGAAATCAATAATTGAGCGCTACGACAGGACAAAGGAGGAGCATCATCAGGTTTTAAATGCAACTTCA
GTTAAGGTATGTTCATTTCTTATGCAGCATTACCATACAGATGTACAATTTACTAAATTTAAATAAGTTTTGCCA
TCTCTTCTTGGGTAAAAGTCAACCTTTTTAATTCAAATGTATGGTAACCACAAATTACTGAGTAGGAAAAGAACA
CCCTTCTTTTTGCTGAACATATGCAAATTACCAAGATGCTGCATACTCTTATTGTACCTTAACATACGAATAATT
TGTGACACTTCTTTTCCAGCATAGATTCCCTGTACCAGAAAAGtagggaaaaaaattagaaactgCACAGGAATA
GATCAGGCATATCCCAGTTCAATTTGAAGTAACATAACTATAAAAGCTTTAAAATGTATAGTAGTACATAGTTGG
TAAAAGGTGGATTACCAATAAATGATGGAATACATGTAGAATAAAAGGGGTAGACTCGTTACAAACATATGATTA
TCGGTGTTAGACATCCGGAAAAGCATAGCATCATAGAAGTGCTAGAGAAAAGTTGGCTGATTAAATTGTTTTGAT
GACGAGATTCGAAGGTGAATGCATGCCAAAGTGACATCAGTGAGGTTAACGAACAAAGTCAGCGGGAAATCATTT
ACATATGGTTTATGTTGGCTTACAAATACACACTGAAGGATTAAGATTCATGTGAAAGTGAATAGCGAAAGAAGA
TAATTTTTGACAAAGGTATTTACGATGGAATCGACAGTGGTGCTGAGGCGTAGATGCTATACTGTGGCAGGCCAG
CAATCAGGGGCATCTTAAAATCTGTCGATCAAAAACATTTGCTAGAAGGACTGCAGCAGATACTAGTTGTGGCAG
GGCCAATCATGGTCATCCTAGTCTGTCATCACTTGCAAACAAGTGCTAAGAAGTCCTGATTCTTTCTGCTTGGGT

GGAGAATCAACTATCTAGATCATCTGCAAATAAAAACATCATGACTGTCTGATGAAAACAAGGAGAAATGCTATA
TTGGCATTATGGAAGAAGCTTGAGTCGTGGTGACAAGCCTAGTTACTAGAAAGAGGAGTGGAGCAGATTCGGATG
CGGGTGTGGGGAAGCACTTCTCTTAAAActtttttgatgagaaaaaaaaacggGAGCAGAAACAAGTTTCCATGG
AGATTGCGAAGGAAGATTGCAAAGCGGATGTAGCACCCCAAGTAGAAGATTCCTGCTACACAGGTCATGCATGAA
CAAAAGATCTAACCAAAACAGACGACAGAACGCAGCTGCTGTCGGCTGAATAATGATGATGCTTTGGTGGTTGAT
GGCAAAAGGATACAAAAGATTACTTAAGTTCAGCTCTATTACGTCGAAcaaacttgaagaagaaaatgacaCAAG
GATGTCTTTCTGGGGTCTTGCACGATTGCTGTGTTATAGAGGAGACATGATATGAATTGTTTCAAACAGAAAGAA
AGCCCATAGCGCATGAGATGAATATTGCAAGAATAAGCATGCTAGGTTGGGACGTGGTAAAACAGAGAGAGTCAG
AAATGGACACAATCACAGGAAGCTAAATACAGCTGCAATTAGTAATTGTTGTTCTGTTGGCATGGCAAGGACATG
CCATGACTAAAAGGGGCAATATACAATTAGAAAGTGCCGGCTTTCACTAATGGTGGGTCCAAAAGAGGATGAGAA
GGCCTAAAAACACGGTTCAAAGTCAATAGAATGAATTTAATAAGTATTTGACTGGAGGAAGTAGCCACATATTTA
TAAGAAGGTTTTGCTActatagtttttatttattttagaaaagtcAGAATAGATTGTGGCCATTGGTTTGCAAG
GGGTTTTAAAGTGAAATAATAGGCAAATGGTTGGCATCCATTACCCCTTttcgttcctttttttaatccggAAA
CTCATCTTCGTCCCAAAAAACCTCTCGTCCTCTTAGGTTCATTAGTTGCCGGCTCCCATAACAGTCCATTCCGAT
CCCACCCCAACCaccaggggcggctcaatacattctggggcctaaggcgaattcaaTAAATGAggcttttttaa
taatattttttttaataataaaatacttaaaaaaatatatgaaaatagatagctatcaagtacactatttcaacc
acgaatctaacaaagatagacaagaagccttttcaatttaatataattcttgaatggaaacATATAAAAGTAATT
ATTCTAAATGAATGGTCATGAAAACTGNNNNNNNNNNNNNNNNNNNNNNNNNNNNNNNNNNNNNNNNNNNNNNNN
NNNNNNNNNNNNNNNNNNNNNNNNNNNNNNNNNNNNNNNNNNNNNNNNNNNNNNNNNNNNNNNNNNNNNNNNNNNN
NNNNNNNNNNNNNNNNNNNNNNNNNNNNNNNNNNNNNNNNNNNNNNNNNNNNNNNNNNNNNNNNNNNNNNNNNNNN
NNNNNNNNNNNNNNNNNNNNNNNNNNNNNNNNNNNNNNNNNNNNNNNNNNNNNNNNNNNNNNNNNNNNNNNNNNNN
NNNNNNNNNNNNNNNNNNNNNNNNNNNNNNNNNNNNNNNNNNNNNNNNNNNNNNNNNNNNNNNNNNNNNNNNNNNN
NNNNNNNNNNNNNNNNNNNNNNNNNNNNNNNNNNNNNNNNNNNNNNNNNNNNNNNNNNNNNNNNNNNNNNNNNNNN
NNNNNNNNNNNNNNNNNNNNNNNNNNNNNNNNNNNNNNNNNNNNNNNNNNNNNNNNNNNNNNNNNNNNNNNNNNNN
NNNNNNNNNNNNNNNNNNNNNNNNNNNNNNNNNNNNNNNNNNNNNNNNNNNNNNNNNNNNNNNNNNNNNNNNNNNN
NNNNNNNNNNNNNNNNNNNNNNNNNNNNNNNNNNNNNNNNNNNNNNNNNNNNNNNNNNNNNNNNNNNNNNNNNNNN
NNNNNNNNNNNNNNNNNNNNNNNNNNNNNNNNNNNNNNNNNNNNNNNNNNNNNNNNNNNNNNNNNNNNNNNNNNNN
NNNNNNNNNNNNNNNNNNNNNNNNNNNNNNNNNNNNNNNNNNNNNNNNNNNNNNNNNNNNNNNNNNNNNNNNNNNN
NNNNNNNNNNNNNNNNNNNNNNNNNNNNNNNNNNNNNNNNNNNNNNNNNNNNNNNNNNNNNNNNNNNNNNNNNNNN
NNNNNNNNNNNNNNNNNNNNNNNNNNNNNNNNNNNNNNNNNNNNNNNNNNNNNNNNNNNNNNNNNNNNNNNNNNNN
NNNNNNNNNNNNNNNNNNNNNNNNNNNNNNNNNNNNNNNNNNNNNNNNNNNNNNNNNNNNNNNNNNNNNNNNNNNN
NNNNNNNNNNNNNNNNNNNNNNNNNNNNNNNNNNNNNNNNNNNNNNNNNNNNNNNNNNNNNNNNNNNNNNNNNNNN
NNNNNNNNNNNNNNNNNNNNNNNNNNNNNNNNNNNNNNNNNNNNNNNNNNNNNNNNNNNNNNNNNNNNNNNNNNNN
NNNNNNNNNNNNNNNNNNNNNNNNNNNNNNNNNNNNNNNNNNNNNNNNNNNNNNNNNNNNNNNNNNNNNNNNNNNN
NNNNNNNNNNNNNNNNNNNNNNNNNNNNNNNNNNNNNNNNNNNNNNNNNNNNNNNNNNNNNNNNNNNNNNNNNNNN
NNNNNNNNNNNNNNNNNNNNNNNNNNNNNNNNNNNNNNNNNNNNNNNNNNNNNNNNNNNNNNNNNNNNNNNNNNNN
NNNNNNNNNNNNNNNNNNNNNNNNNNNNNNNNNNNNNNNNNNNNNNNNNNNNNNNNNNNNNNNNNNNNNNNNNNNN
NNNNNNNNNNNNNNNNNNNNNNNNNNNNNNNNNNNNNNNNNNNNNNNNNNNNNNNNNNNNNNNNNNNNNNNNNNNN
NNNNNNNNNNNNNNNNNNNNNNNNNNNNNNNNNNNNNNNNNNNNNNNNNNNNNNNNNNNNNNNNNNNNNNNNNNNN
NNNNNNNNNNNNNNNNNNNNNNNNNNNNNNNNNNNNNNNNNNNNNNNNNNNNNNNNNNNNNNNNNNNNNNNNNNNN
NNNNNNNNNNNNNNNNNNNNNNNNNNNNNNNNNNNNNNNNNNNNNNNNNNNNNNNNNNNNNNNNNNNNNNNNNNNN
NNNNNNNNNNNNNNNNNNNNNNNNNNNNNNNNNNNNNNNNNNNNNNNNNNNNNNNNNNNNNNNNNNNNNNNNNNNN
NNNNNNNNNNNNNNNNNNNNNNNNNNNNNNNNNNNNNNNNNNNNNNNNNNNNNNNNNNNNNNNNNNNNNNNNNNNN
NNNNNNNNNNNNNNNNNNNNNNNNNNNNNNNNNNNNNNNNNNNNNNNNNNNNNNNNNNNNNNNNNNNNNNNNNNNN
NNNNNNNNNNNNNNNNNNNNNNNNNNNNNNNNNNNNNNNNNNNNNNNNNNNNNNNNNNNNNNNNNNNNNNNNNNNN
NNNNNNNNNNNNNNNNNNNNNNNNNNNNNNNNNNNNNNNNNNNNNNNNNNNNNNNNNNNNNNNNNNNNNNNNNNNN
NNNNNNNNNNNNNNNNNNNNNNNNNNNNNNNNNNNNNNNNNNNNNNNNNNNNNNNNNNNNNNNNNNNNNNNNNNNN
NNNNNNNNNNNNNNNNNNNNNNNNNNNNNNNNNNNNNNNNNNNNNNNNNNNNNNNNNNNNNNNNNNNNNNNNNNNN
NNNNNNNNNNNNNNNNNNNNNNNNNNNNNNNNNNNNNNNNNNNNNNNNNNNNNNNNNNNNNNNNNNNNNNNNNNNN
NNNNNNNNNNNNNNNNNNNNNNNNNNNNNNNNNNNNNNNNNNNNNNNNNNNNNNNNNNNNNNNNNNNNNNNNNNNN
NNNNNNNNNNNNNNNNNNNNNNNNNNNNNNNNNNNNNNNNNNNNNNNNNNNNNNNNNNNNNNNNNNNNNNNNNNNN
NNNNNNNNNNNNNNNNNNNNNNNNNNNNNNNNNNNNNNNNNNNNNNNNNNNNNNNNNNNNNNNNNNNNNNNNNNNN
NNNNNNNNNNNNNNNNNNNNNNNNNNNNNNNNNNNNNNNNNNNNNNNNNNNNNNNNNNNNNNNNNNNNNNNNNNNN
NNNNNNNNNNNNNNNNNNNNNNNNNNNNNNNNNNNNNNNNNNNNNNNNNNNNNNNNNNNNNNNNNNNNNNNNNNNN
NNNNNNNNNNNNNNNNNNNNNNNNNNNNNNNNNNNNNNNNNNNNNNNNNNNNNNNNNNNNNNNNNNNNNNNNNNNN
NNNNNNNNNNNNNNNNNNNNNNNNNNNNNNNNNNNNNNNNNNNNNNNNNNNNNNNNNNNNNNNNNNNNNNNNNNNN
NNNNNNNNNNNNNNNNNNNNNNNNNNNNNNNNNNNNNNNNNNNNNNNNNNNNNNNNNNNNNNNNNNNNNNNNNNNN
NNNNNNNNNNNNNNNNNNNNNNNNNNNNNNNNNNNNNNNNNNNNNNNNNNNNNNNNNNNNNNNNNNNNNNNNNNNN
NNNNNNNNNNNNNNNNNNNNNNNNNNNNNNNNNNNNNNNNNNNNNNNNNNNNNNNNNNNNNNNNNNNNNNNNNNNN
NNNNNNNNNNNNNNNNNNNNNNNNNNNNNNNNNNNNNNNNNNNNNNNNNNNNNNNNNNNNNNNNNNNNNNNNNNNN
NNNNNNNNNNNNNNNNNNNNNNNNNNNNNNNNNNNNNNNGAGCTTAATTCATATTTGAGCTCGACTCGATTGGAC
TATTAACGAGCTTGAGTTGAGCTCGAGGAATTTGAATAGTTTTTCGAATTGAGCTCGAGTAGCAAAATACTCGAC
TCGAAGGATTGCAATCTTAatcgaatatatatattttcaataatattatattttatttataatatactcttaatt
taatattttaaatataatatttaatttttaatctgaTCTAACTCTTAgcttttaactatatttttttaattatga
ctaAAGCTCGAACTCGAACTCGAGTATAACTCGGTTGATATTCGAGTCAAGTTGAGTCGATCtcgagtattttt
tttaatcgaatTGAGTTCAGGCTAGGACATTAAAGCTTGATTAATCTCAAGTCGACTTTTGAGCTTGAGTATTTT

GAATCGAGTCGAGCCGGAGCCTCCAGCCACTCGACTCAGCTCGGCTCGATTGCACCCGTAAAAAATAACCAATGA
AGTTTAAGAATTTATGGTACTTACCATTCATACTTTGGGTCGGGAATTTATGGTACTTACAATTCACACTTTGAG
TTGGTGCAACCAGTGGTATCAGGTTAAAGATTGAGCTATCTTGGTAATTTACTTTAGACTGGATAGCTATGGTTG
CTCAAGTAAAACTAAGAGTATGGAATTTGAGAAGCACCGAAACAAAATCCTTGTAAGAATCAATAGCAAGTATGA
TTCATAAATCTATCAGAAATGCAGTGATAATTTTTTGCAATTATGGTTATTGTTGCAAAAATTTTCGTATTCAAG
TAGCAAACTGAATTCCAAAAGAGATAGTGTCCTTTAACtggtatttttttgttttctcttttcaaaacAAACGCAA
TCACAAAGATGGCCTCCAAAACCGTCAAAATGGTTTACAAGTAGACGCAGAGCAAGCAAACTATTCATGAACGAC
TTTGCTAGGCTAGCTAATAAGCTCTACGGATCTTGTTCAACTAAATGAGCTAAGCTTAAACATCATTTAAAACTT
AACTCATAAAAAAGCCGAGCCTCAACATTGGGCTTGGAAGTTCGAGCTTGGCTCAAGAACAAGCAAACAGGGCAA
GCTTGAGTACAGATTGTTCGAGCTTCAATAGCCTTTCATAAATTAGCTGCATCTGAGCAGCTTAGTAATTGGCTT
GTTTGTAGAACTGGGAGTAATCATCATGTGTGAATAGACTAATTGCAAAAACTGAGTTAGTACAGAATAGAAGTA
GGTTAAGTAAGTTTTAACCAGGCTAAATAGGTCTTAAAGGGGCTAGATGTGTTAAATAGATTACTTGATCTAACT
CGATTAGAATATTAAATGGGTCAGATGTCTGAAATCTGAACCCAATTTGAatgttaaacatgttaaacaaaTTGA
TCTATTTATGACCCAAACTTATTTAGCCTAAATCCAAATATATTTATGGCAGATCGAACACGGGTTGAGTTGACA
GGCTCGGTCATATTTTGCCAACCTAGAATCGAACCCTTGAACTCTTGTCTAAGAGGTGGGAGGTGGGAGGTGATA
CCATCCATGCTACCACCTGTTGGTGCATTAAATAAAGATCTTATGTAAATGAATCCTAAACCCAAAATAAATTAC
CCATTGATTGGTTCATTGAATAGTCCTACCAATGTCATAGGATGCACGCACCGGAAGGGGTCGTTACATTCTCTT
GTATTCATTCCTTAAAATTAGGTGCATACATCATACATGTCAATTGTCAAATGCTAATTCTTAATATGTTGCAAT
GCTGAATCAAATTTCATGTATTAAAGCATCATCTATTATGCTTGTGTTGCTGAGGGGTTATCATGATACCATGCT
TGTTGAGTACCAGCATGGAAAGGTACCATGTCCTTCAAAAGATCAATGAAAAAGATACTAAAGTGtgcttttta
attaaatgctTTTCACAGCTAATCATTTTCGAttgatttttgttattgttCAAATAAATGTAATAACTAATATAT
AATTATCCTTCATCATTGTGCTTTGTGTTTCGGGACACTTGTGTCTGTCTATAGAAAATTGTGTTGGAAACATAT
CTGAGTCAATACTTATCAGGCATTTGTATACTTATGAAGTTTTATGGTTGCAACTTTTAAGACTTGACAAAGAGT
TGAACTCTTCCAGTAAAGAATTTGACCCTCCATATTCGTGTTGGAAGTGACTATTCTTTCCTTTCAATGATTGGC
CAAGGATGTAGATATcgaaataattttataaaatgttatatttcttatattcttATAGGTTAATTGGGAGGTTCG
AATGACAAACATGTACTAAAGGatcttaacaaataatatattttttaatgtttatttttgatatttaaacaAAGAa
tattggatatatatatatttcagtcaaatatataatttatactaTGTATGTTTGcaccttctttttctctcttgc
taTTTTAGACACTTGGATGTACGCTCGAATCCAATTCTCATATTCGTTTCCATGCATCACTCGCTTTACGTTTTT
TTTCAAGTAGTTCATCATACAAGCTAATGTTCGAAGCTTAGAACTAAgtacatattaaaaattaagatcatAATT
CATGGTGATGGTTGTGGCTGgacataaatattttcatgcatTGATAATATGTTTCAAGTGTTGTATACGTACCAT
TGATATATAAGAAACTCATGTATTGATGGTGCAACTGCATGTCCATCATACGTACTCCATATATAATTGTCATGG
ATATGATGCACCATGGTGGACCCCTACTGTTACTAGCAACAAATTTGGTATTCGAGTCAATGTGCTTTTGATATT
CATTTTGGCTGGAACATCCCTATTCCTAGATCTTGCATCTGGATTAGTTAGAGGCCTTGATGGATCTAGTGCTTC
TGATTGAGGCTAGCAACCTAAATTGTCCATCACTTCCATGCTCCCTATACTCCATTCCACTTTCCTTTgacctaa
aacaaaaaaatgactTTTGGCTAGCATCATGATGATATCCTCCAAGATTTCCTGTAGTGATGCTGGGTGCCATAC
TAGTCAAAAGTCagatcggtatggtatggtatgatacGGATACTACATACAAAAGCATACTGAATTGGTGATATT
tcagagttttattttttttatcttcattatttttttgtttttagtgtTGATTAGAggtttgaaaacttggaattagg
gttttatttttttcttcatttttctatgtttttcaaaaaaaaattgttttatagTGGTCCTGGAGCACGATAAGTCAT
GTGCTTTGTGAAaagtttataaattaaaaaatagataatctaTAAAATTTAGCTAGAAAATGCATATTACTAGTC
CATTCTGGAGTATTTACAGCTTGCAAACATCGAGATCTCCTTCATAATTTGGGCTTGGACTTTGACTCCAACATG
GTTGCTGTGCCCCCACTTACGTCCGCTAATCGCTCTAACCTGGATAGCTACCCAAATACACATCCTCTGTTGACA
TTAGGCTGGCCATCCTACCTCAGGGAAGATCGAGGTAACAAACTCGCCAATGGCTGCAAATCCCCCCTCTCAATG
CTTTCCCAGACTCCCCACCTCTCCCATGTCCTCACTGTATTCAATCAAGAGGAGAAAGAGTCAGGAAGGCTCTTC
CTTGGCTCTCTCCTCTTTTGACTCAATTTTTGACCCAGGACCAGTATGTACTGGTTATAATTTAGCTGAACCAAG
CTGAACCAGGTAGCTCCATTTGGTTCGGACCAATTCTACTGCCTCAAATCAATAACAGGAACCATCTTGGTTCCG
CTCTGGTTCGATTCTATACGTCCCAAACCTTCTAGTTCGAGATGTTTCTACAGTCATTAAAGTAGTACACTACTG
AAAAATCCTAAAAGCAGGTCCTAACATTGAATGCAAAAAAGCTCCCATATGCTTGTTTGCATGATTCATGTCAAT
ATTGGTTTGTATTGATATGACAATTGGGGTACTTGATAAGCAGTCCTTTGTTAGGATGTCATCTTACAATCTGTT
CCTGTTGACTGAAAGTTTTATTTCTAGCCCTTGATTTTTCCATGGTCATTTAATGTTTAATCCATGGCCCTTTAA
ACATAATGGCGTGCAAAATAGTTATAGCCATGCACAGAAATTGGTGGATGCAAAAGAGGTGTTTGTGCATAACAT
TGTGCGCAAGACCAAACATATAACACTATGCAAGGGCTAAACAAAAAAATGGTAGTCAACCAAATGACCTAATATT
AAGTAGACTCATGAAAATTATCTTTATCTTCAATTATGACATTTGGGCCTCTTTTTGGACATTGGAGCATTCTCC
TAACTTCATTCTAGCATGGAATAGTAGCATGCAAAGACCGTTCACCCACATCTTACCCCCTAGCTTCAAAATTCA
TCTCTTAATGAATTACAGTGGAATAGGATATCACAGTCTTCAAATATGCTAACCTAACTtgacccaacctattta
atgAATGAAcgtttaaatttaaatttttgatttatttaataaataagccAGTTTCAAGATGATGATAAATTTTTT
ACCTGATCCACATCTAACACGAGCCATATCCAACCTGACCCACTTGCCACCCATCAAGCAGCACTTATAGTCATT
GTAGCCCCCATCTAAAAGTCCTAATTCTACGAGCACAATCTCGATGTGCCCAAAAGCCTAGTTTTCATCATAGGT
GAGAATAGGTTGGGATTAGGGATGAGGTGGCACTAAGCCTCAAAAGAGCTTTATTTAATGCAGCAAGAAACTTTA
TGCACAAGACTTGGGAGTATTTGATGGATGGTTACCATTCCATCAGCtaatgaaatattttagattAGCCATTCT
accatcaaattttatatttgtgaattgataaatatatttcacATTTATTGCGTCTCTTAGAAGTATTAGTCAAAA
AGAGGGATAGAATCAAGAGAGGTTATTTACAAAAAAGGGAGTGATTTATAGAGCATTGCAGACGTGTGACAATTT
CAGGCATAGCATTTATACATAGTAGATAAGAATAATTAGTAGCATTCATTAATTGAAGCAAGTTGATGCttatgc
aaataaataaagcaaaatCTTTCTTATGTATAAAGTAGTCACAAGCATTTTCCTCATTTTCTAATACATCAGCAT

GCTTAGTGGATCAAGCCAATAAAATTGCCAAACCTTTTTAACAGATATGGATGGTTGAATTTAAGAAATGAATGC
ATGTGTTTATTAAGACACTTATTGATTGGCATATATCATCCAATCTGTAAAGAACGTTCCTACCAGCAAGGACCA
TTCAGTCATTTCCAAAAAAGGATCATAAATGTCTCAATACTTGATGCAACACATCACATTTACAAAGTCCAAGCT
TGTGAACATATGGCTTGATTCATGAGATTGAACTTGTCTCTACCTAAATAATCATATATAGATGGACTTAAGTCA
TTGAAATCAAATGATAATACTTATGTCAATAAGTTGTAAAATcattattgatgattttaattTTGGCTCTAAATC
AATTTTATTATGCAATACTCCACATAATACATATGATAGCTAAAAATTAACAAAGCAAGTAGAAGAGCAGAAAAG
AACATCACATCtaaattaacttctccaagatTTTCAGCCTCGGCACCGGATAACATACTTAGTAGCTTGCCGTGG
TATATGTCGGCACATGATACATTGTTGGAGGCTAGATAGCAGTTGTACCATGTGTCAGGATACTAACATCTCGAG
CCCCATTATTGTAATGGTACCTTATCGGGTCAGCATAGTACACCCAATACCAAAATGTACTGGTCCATAGAAAAg
agctattaatttttttgttaacaaaTGATTATTAGATTCAAATATATAAGCATGCAAGGGCACGAGACTTTTGAc
cagtaaaaaaatataaatgatgcaTAATAATAGATATGTAGAATATTCAAAAGATTGAGAAAGCATCTCATCATA
AAATAGCAACTCTAATTTATATACTCCAAAACATTAATAAATTCATTGGAGATTTTCTTCAAAGAACTATGCATT
GTTCATACTACATTTGTAAGCTGGCTTGAAGGAATATTTCTCATAAATTTTAGCTTTTGCTCTTTTTCTACATTG
TTACTACTTACCATTATCTAAAAAGATATTGTATGAACTAAATTATGTTCTTTGTCCTTGCTAGTTTTGGCAATG
GGAGGCAGCAAGCTTGAGGCAACAATTGCAGAACTTACAAGAAAATCATCGGTACCATCTCTTAACTTGTTTACA
TCTAAAGATTGTTCCTACGCCCAACTAGCCATTGGAAATATTCCATTATATGTTATACCACAAAGAAGATTTAAC
TAAATAACCATATATTTTTGTCGTAAAAAAGGTTTGCGTTATTCAACATGTCTAAAAATTTAAAGCCCGTTCACA
TAAAACAAATCCCCTCATGGTAACATTTTACGAATATTCACTTATTTCCCAGGAATCTAGAGTGTCAAGATCACT
ATaaggaaaataatatataaactaGCAAAAACATCAAATTTCTACAATTGATTAGAAATGAACCGTAAAATCATT
AGTTCCCTAGCTGCCCATATCCTGAAAGTAAAATAGATTTCTTaagattaatcatataaacCAGGATTTTCTTGC
TGATACAGCACCTTGTTTTTGCAGTTTACCTTATTGCAATGCATTGACCAAATCATTAATGATGCCATTTGACAA
AAGAGCAGGTTACAGAACTTCTAATAGTCTGTCAATAATTAGATAGCATAAAACTAACGCCAGTGCATCTAAAGA
CTAGGTCTCTCACTTTAATACCTAGCCAGGCTTGGGAAgaaaaaatgtgaaattttaaTAGGCTTGGCCTAAAGC
CtaactaaaaatgaatatagtttaaGCTCAAGACCCGGCCCATGATCAACTCTAGTCACGGGTTAACCCGACCCA
ATTATAGCATTAATCACAAGCTTCGCATCACATTTTCAAATCTCAATACTGGACCccatatcaaaatattatcGG
TATGGTTAGATCAGTACGGTACATCCGTGCATCAACACATTGTATGATATTGGAGTTAGCACAAAATTGTACCAT
GTAACTACATAGGGGCATATCAAGCATTTGAACCATACCAATATTGTTAGGAAATTCATACTATACACAACTAGT
GTTAGTCAATACTCAAAATCGTTACGCCCCATGCCAACTAGTGTTAGTCAGTACTCAAAATCGTTGCAGAGTGGG
AAGGATGGGGGAAGCGGCAAGAAATcaaagtgagagagagaggggatgaTGAAAAGACTGCTAAGGGAATGCAAG
ATGGCCATAAAAGAGATGGGAGTAAgagaagcatgatagaataaagaagaaataagatatcaaaGCTTGAGGGtt
ttgcatattttaatatcatgTGGCCAGTGGCTAGCCATTGGACTAGAACAAATTTAGCAGTTTTGATGGTTTTTA
AACAACTAAAAATGAGAAATCCCAATCCTACCATGAAACTGACCTATATACATGTATGGGTAAAACAAAGATTTG
CTATACAAGTGCAAACTGCCCCATACTGGACCCCATACTGGATGTATCATATTGTATCCGTATCAGATTGACGCT
TAGTACTAGAAGTGTATCGAGTATCAGTACAGCGGATCAACACCCATACCGTGTGTACCAATATTGTACCAACAT
GGTACCGGTACATGCTCTGATACTAATATTGCAGACCTTAGTGTGAAACATGTTTTAGAACAATAGCATCATTTA
TTATCTTAGCCACGAAACCATTAAACTTCTAccttagctaaaaaaaaaacaaaataagtgCCTAGACTATGCATA
ATTATAGTTACTATAGACAAACTAACTAGTCCGGCGGTCATACAagtaaacaaaatttttataatttttcataat
agaTCTTTAGCCTACATTTNNNNNNNNNNNNNNNNNNNNNNNNNNNNNNNNNNNNNNNNNNNNNGAAagtccac
aaaaaaaaaagaataaatcatACTGGTGAACAAGGGCGCAACAATGTTGATTATTACACTATTAAATAAATTGAA
GATATAgcatattcaaaattttaatcaatGGTTGCAGGCACTTGATGGGAGAAGAGTTGTCTGGCTTAAGTGTCA
AAGATCTACAAAATCTAGAGAACCAGCTTGAAATGAGTCTACGTTCTGTccgaaagaaaaaggttttaagAACCT
TTCCCTCACCAAATGTTGTTCATTAATCCTTATCCACTCTTCCCTATTTGATGTGTTTCTTTTCTCTGTTGTAGG
AACAAATTCTGACCAATGAAATTCAAGAATTGAACCGAAAGGTTTATTAATGAATAATAGTTATTAACTCATTTT
TTAATGTTAACAAGTGCCTAAATTTCCAGAATTTGTTACAGGGTCACCTCATTCACAATGAAAACATTGAACTCT
ACAAGCAGGTAAACCTCACACATCAAGAAAACATGGAGTTACGTGAGAAGGTATTGAGTTTAAGACAAAAGATAT
CTACTTTGCTCTATAATCCAAACTCCAAAGTAGAGCACTCATGAAACTTATAATCTCCATGTTTCCACAACTTAT
AATCTCCATAAATCCACAGCCACATGCTTCGCATATGCCATGCtctatataaaatatgtatCGATTTGGTATAAA
ACTAGATTAATTCAATCACAAGATTTTTAGTGCTAGCCTATGTTGGTTTGGTCACCATACCGAGCCAATATCAAC
ACTAGCACTAGGTGAAAACAAGACCCATTGGATCCTTTTTCAATACTTTTGTTTTGACATGTTTAGAAAAGGagt
ttttgatttttaagtcctttcttaatattttttttgttctggcACATGACATGTGGATCAACAAAGCATGTAATA
TGCTAGCCACTAGCCAGCACGCCCCAGGCGAATGACATTTAAGATCCTGTTCAATccttgaaaaaataatagatt
attATTATAACATGCTATGCATGGAATTTGCATCGCAGGTGTGTGCAACAAATGAGTTAAATGGAGGAGCTAGTA
AGGATACAATTATTCCATATGGTCTTACTATGACTGAGAACTCAAATGCACCGATTCATCTTGAGTTATGCCAAC
CTCAACAGCAAACGGATGGGACGCAAGCAAGGGTTCTGAAGCTAGggtaattattttgaaatatcaaaCCTTTAC
ATATTTTCAATATTGATTACTTTGATTTGAACATAATCAATGCTTGCAACAGATTTCAACTGCATTAAGAAGATG
ACAAGTGATCAATATGCATCATCTTTAGAGACCATCAGCCTAGTTGCCAAGGCTTCTATGGCTTGTTATCAGCAA
TAACATCAATGTATgagcaagagagagaacaaTATTCAGCAGTCGCTTATGTGTGGTGTGACATGATCCAAAGAA
CAGTTACTTTGTAAAGCAAACTTCCAACTAGgactaataataatatgacaGGTATGTCTGTTTAAGAACATTACT
TGCATATGTAGAGACATGTCTCCTCATAGCATCCGAGTAGTTGCTGATGACAGCTACAGGAAACTCACCTACTAA
ATTAAAGTTACAAGGGGTTAACAGAAACTGAAGTGGAAAAAAACTGCACAAAGCATGACTCACTATCTTCTGAC
TACTGCTCAGTAACTAGCAgcaaaagatatataaaaatgTGACAGATTTGTTCCCCAAAAACCACTGTAGAATGG
TAATGACAGCATGTAGCTTAAGCTAGCTTACTAAACTAGAATGAACCAACTATTTTGAGTTTTACATAGGAACAC

TGCAACACTATAAATTGTGTGCTAGCACATAATTGGACAAGGGAAGTTTGCCTACAAAGAAATAACAAAGAGGCA
AGGCAGATCCCATTCTATTAACTATATAATGGATAAAAATACGGACCTGATCTTTCAAGCAACATGAAAGCGTCA

>PdAGL17_3
ATGGGGAGAGGGAAGATCGCGATCCGAAGGATTGATAGCTCGACCAGCCGGCAGGTGACCTTCTGCAAGCGGCGA
AGCGGTCTGTTAAAGAAGGCCAAGGAGCTCTCCATCCTCTGCGATGCTGAGATCGCCCTTTTCATCTTCTCCAGC
ACCGGCAGGCTCTATGATTTCGCCAGCTCCAGCATGAAATCAATAATTGAGCGCTACGACAGGACAAAGGAGGAG
CATCATCAGGTTTTAAATGCAACTTCAGTTAAGTTTTGGCAATGGGAGGCAGCAAGCTTGAGGCAACAATTGCAG
AACTTACAAGAAAATCATCGGCACTTGATGGGAGAAGAGTTGTCTGGCTTAAGTGTCAAAGATCTACAAAATCTA
GAGAACCAGCTTGAAATGAGTCTACGTTCTGTCCGAAAGAAAAAGGAACAAATTCTGACCAATGAAATTCAAGAA
TTGAACCGAAAGGGTCACCTCATTCACAATGAAAACATTGAACTCTACAAGCAGGTAAACCTCACACATCAAGAA
AACATGGAGTTACGTGAGAAGGTGTGTGCAACAAATGAGTTAAATGGAGGAGCTAGTAAGGATACAATTATTCCA
TATGGTCTTACTATGACTGAGAACTCAAATGCACCGATTCATCTTGAGTTATGCCAACCTCAACAGCAAACGGAT
GGGACGCAAGCAAGGGTTCTGAAGCTAGGGTAA

#### Elaeis oleifera

##### *EoAGL17_1 and eol-miR444*

##### >contig_KE504701

ccgcacgtactcgtcctacacgtgtcggcgatctatacggtaggcggcgcgttatcaaacgtcccaataaataga
agtaatcgacgagcgaaacgtgcccggctgttccccggaccacaaagaaatataaacaaccaataaacataataa
acaagtgtataaataaggaagcgagtgcagggcgagggctcagatatcaaaatggtatggttttatatgtattca
tttttggtaaagaaatccttcaccaccacctcattcctccgtcaaaggggaagaagaaaggcaagggagagtgg
gaaccaagcccacctattggccacctttccgttcctttctatttttgcctataaaatagtaagactagataggaa
aggcatcacccaccacctaggccaaagccaaatagagaaagaggctccaactccgtttgtggccattgataaagc
tcgtcttcccggaaattaaggttgcaatcgctccaccagatcgatctcctctccgtccttctctttctttccttc
tctctgccaccttggtctccctggcctccctcctttccttctgattccttgttgttagttttccatagcagatct
gatacatctgatactgggttttggtgggttgagaagaggagggagaagaagaaatatggggaggggaaagattgt
gatccggaggattgacaactccaccagccgccaggtgacgttctcgaagcggaggaacgggctattcaagaaggc
caaggagctcgccatcctctgcgacgccgaggtcggcctcgcgatcttctccagcaccggcaggctttacgattt
tgccagcaccaggttcttgcttctccctcttctacaaattactctctgtcatcctacatgatatatatggtttca
gttcgatcggtcttaatttcttaaggtcttgatggaggttattgcatggatggactgggagcaagtctaacatat
ggaagagtaattttacaccgaaatgatagaataatgttaggaaaggtggttggcttcttagttttgatatctttg
ttgcatagatggtttagctggtaggttttagagaatgaactagaaaaaaaaaaatgaggaatggcgtatctcccc
caccacttctcttttattttattttatttatttattttttaagaaaacatggcacttctcttttataggtttct
tattcctagtttaatgttaccaagagccattgtcatattctctctctctctnnnnnnnnnnnnnnnnnnnnnnnn
nnnnnnnnnnnnnnnnnnnnnnnnnnnnnnnnnnnnnnnnnnnnnnnnnnnnnnnnnnnnnnnnnnnnnnnnnnn
nnnnnnnnnnnnnnnnnnnnnnnnnnnnnnnnnnnnnnnnnnnnnnnnnnnnnnnnnnnnnnnnnnnnnnnnnnn
nnnnnnnnnnnnnnnnnnnnnnnnnnnnnnnnnnnnnnnnnnnnnnnnnnnnnnnnnnnnnnnnnnnnnnnnnnn
nnnnnnnnnnnnnnnnnnnnnnnnnnnnnnnnnnnnnnnnnnnnnnnnnnnnnnnnnnnnnnnnnnnnnnnnnnn
nnnnnnnnnnnnnnnnnnnnnnnnnnnnnnnnnnnnnnnnnnnnnnnnnnnnnnnnnnnnnnnnnnnnnnnnnnn
nnnnnnnnnnnnnnnnnnnnnnnnnnnnnnnnnnnnnnnnnnnnnnnnnnnnnnnnnnnnnnnnnnnnnnnnnnn
nnnnnnnnnnnnnnnnnnnnnnnnnnnnnnnnnnnnnnnnnnnnnnnnnnnnnnnnnnnnnnnnnnnnnnnnnnn
nnnnnnnnnnnnnnnnttttcttttgatgtaacattgtcatatgctttatatgcttagcactcaacatctcgatc
ttccatctgttctctccgcatagattttctattagacaacctctctctctctctcgctcgctcgctcgctcgc
tccccccctcctccatatattcctcagatacttcgagagtgcatttgatgtatttatcttcttgtcttctctatt
tccatggtataagacacgatattgcttcgtatacttaaataaaaaaactgcatgttgatcctaggagtgtcaatt
ataagccttgagctacttcatggtttttgaagtttgtaagaacactgaagtgcaggtcaagggattcttgggata
gggaaaaaacaatataagaatttggcagtttcttatgcatctgagacttttatctttccaaatttctcatttgc
cccaatcaatctaaaaatatttaaatgataaatgctccatgcatgtttatatttaaccatagccactaatcttat
catcttctgcttcagtacgctattggatgttagataatgaagggcactcttaaggaagataaagaatttcattaa
gtaaaatcattcattcacgtgtgtataatgtgatgcgtgggtagcgtacatcgaacatatatatatatatatata
tatagagagagagagagagagagagaannnnnnnnnnnnnnnnnnnnnnnnnnnnnnnnnnnnnnnnnnnnnnnnn
nnnnnnnnnnnnnnnnnnnnnnnnnnnnnnnnnnnnnnnnnnnnnnnnnnnnnnnnnnnnnnnnnnnnnnnnnnn
nnnnnnnnnnnnnnnnnnnnnnnnnnnnnnnnnnnnnnnnnnnnnnnnnnnnnnnnnnnnnnnnnnnnnnnnnnn
nnnnnnnnnnnnnnnnnnnnnnnnnnnnnnnnnnnnnnnnnnnnnnnnnnnnnnnnnnnnnnnnnnnnnnnnnnn
nnnnnnnnnnnnnnnnnnnnnnnnnnnnnnnnnnnnnnnnnnnnnnnnnnnnnnnnnnnnnnnnnnnnnnnnnnn
nnnnnnnnnnnnnnnnnnnnnnnnnnnnnnnnnnnnnnnnnnnnnnnnnnnnnnnnnnnnnnnnnnnnnnnnnnn
nnnnnnnnnnnnnnnnnnnnnnnnnnnnnnnnnnnnnnnnnnnnnnnnnnnnnnnnnnnnnnnnnnnnnnnnnnn
nnnnnnnnnnnnnnnnnnnnnnnnnnnnnnnnnnnnnnnnnnnnnnnnnnnnnnnnnnnnnnnnnnnnnnnnnnn
nnnnnnnnnnnnnnnnnnnnnnnnnnnnnnnnnnnnnnnnnnnnnnnnnnnnnnnnnnnnnnnnnnnnnnnnnnn
nnnnnnnnnnnnnnnnnnnnnnnnnnnnnnnnnnnnnnnnnnnnnnnnnnnnnnnnnnnnnnnnnnnnnnnnnnn

nnnnnnnnnnnnnnnnnnnnnnnnnnnnnnnnnnnnnnnnnnnnnnnnnnnnnnnnnnnnnnnnnnnnnnnnnnnn
nnnnnnnnnnnnnnnnnnnnnnnnnnnnnnnnnnnnnnnnnnnnnnnnnnnnnnnnnnnnnnnnnnnnnnnnnnnn
nnnnnnnnnnnnnnnnnnnnnnnnnnnnnnnnnnnnnnnnnnnnnnnnnnnnnnnnnnnnnnnnnnnnnnnnnnnn
nnnnnnnnnnnnnnnnnnnnnnnnnnnnnnaacatatatatatatatatatataagagagagagagagagagaga
gagagaataacaaggacgaagtcctcaaatatttacttctcacattattactgctggcaacgagctagcattag
ggtaagtcagcatttttacagataaaatattatttaataacattgaacttttagattcagaaagtgaaggggatg
ctgagggttagaagcatggatacggctatatattcttccaaagtcccatgcaccacaattctaagccatagcgca
ggatgatcattgttttctgaatcattgttcttgtgaatgtgtcctatttatatagatatggttgtcctaggaaga
gagacaactaattatggtactgaacatgggagtgggacagatctgacatgcctcttggactatttgatcatttga
tgcaataaccgccacaagttacgccatcattatatgaaacttgtgagcaagttaatggcttgagcccactcctaa
tctagtttattgtaaagtgggatgtgtagaggaaaccatgttggaagataaaatggtccaatgtataggatctaa
tgcatcatacttcccttgcatccacctcaccatcagatccgacaacattcagaatatatgaatgtgtgcaagaaa
accttcttcgagaagtacggcctgcacaatgttctatacgtgatgttgatgaaacattacataaaactagatggt
catacattataacttcacaaagcagtctacaagtacttgtttagagtttcaacgtaaaacaaataaaaaagtgtc
gttatgatcagaggagaggaaacagtatcatttaaggttaattatatattccatttggagcaatttgagtttgt
aaggtataggttaccttcattattttttttggtctggaaggttgccttcattatgatggaagagttaaactaaat
tgtcactttgtagcatcatttagatgcatgtgcttgatggaaaaaaaaaaatccgcctacatattgcatgcacct
gggaaaaaaatcctcttttgaagtggcttcacatcaagcacatgtcagaacttcatatcaagagggtcttagact
gttagcattgcatgccatcatatgtaggcttgggtaccatgtgaatggtagatcatatacttaacaagcctcgcc
attttcctagactcgccaagcttgttctcttttacctttcctttgcatcagaattgtaatgaaacaagtttttc
ctatcctgtctacattcaagctcaagtccccgttcatgccaacaactattctgagaggagtatttgttagacaaa
gccatctttacaagctgatggagaataagttgtggaaattgcattcaacatgatttatcaatttctatttattat
tacgaaattttccatctctagatttcaaatttttaatattataaagggctgggttgatttcaaaaagagaggtcc
acagtcttctatatttagtttaattactaaagtttcactgaagtcatttacatgatttgagttttaaagatcat
gccagacttgttatgttaaaattattatattaattgtagtttatatacactattgtctggataattgagtggaag
gtttatcattattttgaactcataacattcatttaactactctgtggtccctcttgcatgcatattcaagttcct
gaatttattctcactatattgtatctctcccaatcatttagcgatgaatctttatttctagttattggcaactct
tttctactccaagatggaactagaacatgcttctacttttcgattagaatgaagaaacgataaaaagaaacaatt
ggagtagacttcaacctaatggagaagcagctatggtggaaacgtcataagtcgatttgtgtattttttgctttt
tgtaggtaattaagcatgattaagagaatttgttatcacactacaactaaaggataggttttgttcgtaagaggt
aaagttgaaaccaaccaaccttagatcagttggtcaacatccttctccacttgtaagaggtctacaatttgaact
ctgctggcatgatcttatatattaatttgttggttttcaaaataaacagatataatgatgtctactaccatgtat
agttaaattgtatacttgcattcaatgatttctatatcttctttttttaagattttcaattttacaaaaaatgca
gtggataattgagttaatggtagagttcaagaataatatggcaaagctccaaggttgatgttaggttgctaccaa
attaatgcagcaaccacaagttggttgccattataaatgttactggtctagttttacttttgggtcagcacatct
agtgtacatagtgatttcaattttcggaatagtggttcgccttagtatgacaatattgtatgatttaggctacac
aaaccattgttaatgttgaaaggagagcttcaatatttgtaatcatttctaatatcttttggagcttttgcttga
ataattattttatgtaagccttaatcctttcataagttttatgtaattatttctcaggtcataatcagtctcct
taatttctttattcatccttataactaagtatagtattcttctaaattaaatacagacacatacatacatacata
cctacatatatacatatatattgtgactgagttaataacgtgccaaatgcaaaaaataatttttgagttcaaat
cttctttcatttgttaaaacaaatgaaattaatattactgattatttacttgatttttgatttctttaaacatt
gatttacaaaatatatgggtttattgcagtatatattaatcttaaatttaattttgaatcatcatgtatatata
aaattctcccaatatattttgtccaataaataataatataacatatttgccacatgtcaagaaaaatatttaatt
agatcttagctgaaattgtgatagtgcaaccattgaataatattttgattttaaaattaataataccattgaat
ggtaatttttcaatttaaaatttatgatgcgattatagttagcagatcatatcaagatcaaaatgatggcttca
agtttgtagatcaatttgtcatttcaatattatgtgattgataaagatgaaagcatataatttctatctttttct
ttggttgttgaaagaaaaaattacctaaaactaccttcaaaagacgagagaattagaacaaacccgacccatatt
tgacttggtataagagaagtgtattaacaaattgccatttgagtaactttggttgtataaatagttaactaagca
attagtggtttagttactctttttgcaaatgtaaaagtctgtatcatccaactgtaagatagaaagcctccatcc
acgatctagaaagatgattattgaattagcatcaattcaaacatccaaagcatttgatacattattgatatattt
atatcttttccatcaactcaaagtttaaccaatgagattaggtaggtgaagagccatttggaatgtgtcacaagc
attcttcatagatgatcatgaacaatgatatacaacatatatcgttgagatatgtctagaagagttactgataac
tagatttttttatttatatattctatgggttttgcattctctatataaaaaagataattattttaatcaagaagg
ggagttctttacttcaaactatcaatcaagtctgaaattcttcaacatgtattgcatgcttcgttgatttatgca
ttgtttcaaaatctgatccaaaatatgcattttggaaaaaaatttcttaagctttttcatccttaagaaggaagt
ttacaagtaaggaatcatagtattgattgctatttctcaaaaatggcaatccaagactggggaggtgaattgga
tttttttaaaaaaatgagaagcacttgtgccccccctaaaaaaatttgaaagtgtttgtgcagtgcttagcagta
actaaataagtactaaaaaataaatacaaaaccaatcattcaatcataaatataaatacaaaaatttataatggt
tcggtgctcaacttcacacctacgtccattccccagatcaacctattaaaattttataactataatcccataaga
ttataattcgattgtttttgtggattcacaatcaatctgattggtttttctaactcaccaactaaaatcttataa
ttgagtattttctaaggctcacaatcaatccagttgattttactgacttaccaatcaaaattttataatatcta
atttttggcaattgggccaatcctcaagaacccaggtttcttcccaaaaaaatttgtcatgatagataaagaaag
atgtataagaatttaaagcacaaagaataaaatctttaaactcttgaagaagtcgaagtggttgagcttttgaat

tacttcgatttcttacttgaatgcaaagaggggattgacagagaaataaatgcttttgctcggtgactgttgtct
caattaaatgcacatttcctctatattgagttaagtaaatatgaaaaggtgatgattgaaatacttttttgaatgt
ccttcttctctctctctcttttattttcacttaatagcttgaatattttcttcaataaatttaatagcttggttc
tcaattttgaattctctacttttatgacctcaaaaatgatcggagttaatacactagctgttggaatcaaaaata
gccgtttgacataaaatatgaccgtttgaaatattttgaaaactagttgttatgctatcagaattttatgagtcg
actcatccaaaatcttagttagctaagatgaagtgttgagttagctaagtaaagctctagattagctcaggaata
catactcatttttttatttctatctatctaaaattggatcagctgagatatctttttagtcggctaacttcatga
cttggtcagataatttattctttgggtcggctcaaaaaatttgtaagcttttactattttctatcttcttctaca
tctgagctcgaatcgactaacacattccttgagttggcttagttcctttttttagttggctatcttattttttatgg
tcggcttaggaggatatctgaattctttattcttctgtcttcaatttgagtctaaggctaaattagctatccatt
tatatgagtcaactatggcctgtcttgatcagctcagataaaacttgatcggctaagaaaaatatctgaatttca
cttctttctatcttttccttgaactgagatcggattggttaagtttctatcttggtcgactcaaaagcgtgtcag
ctatgccaaaaataatttttctattcctgagcagctttctttattcaaagatttttttaagctagatttaccttct
caaaattaatttttgagtagttaaacttgttgagaagtctttctgaactagatttttttttttttcaaattcattt
catatcgaacttagtttttttgctatcaaaatctgaaactttgattttagaaacatgaatttactcaaattttga
gaggacttcttttaaatgagataaagcattgagcattataagaaatcctacaattactttcacaagcattattag
taattataatttatactcaaattttttttgacatcatcaaaatcaattttaggactaacaactgcattataaattt
atgatgttgattaagtctcctaatgtagcatcatgaagtcccaatacgacccaaaagaaatatttttttggttata
aatacttcaatatattctgctattgttgtttggaccattttgtttgtcatagccgtattaagcctactttgcgat
ctcaaactacaaatgcaatactagcttttccatctccaaaaaatatactatcagcattgatttaaatactccatt
cttgtaattgaacttttatactagtactaatctcttgactactttggaattcttatgatagccaacaagctttat
taaaaattttctatagcttgaacttatatcctccaaaataaagtcatttatgctttttccaaactgcccaagcga
taatagcaaacattgttaatccttccattttcaatttgttcgaaacttcttgtatctaacccaaatgcttttagc
tttgcttcccttgattctcatttagagaagtgcaagactccatacttgtagtgcacactaatgacatttaaaaaa
atccattcatgtatcattttgttcatcatatagttcacataattcatcattcaagttcatttttcttttactcaa
attctttttttttagaaggatatttttgagagcatatcacgtaaaaaaatttttgatttttctaggttatgtacctt
ccaaaaattactcttccttatcaaaagtatgcaattaaaactacttgcttttgtcagatcttcttactttagtat
caagtgaaaaccaatagtcttaaccgtgaaaattctattttttttaccatgtcatattgatctataatttattgat
ctattcataattgggattttaagaatgttctatgaaaagaatatctgattaatacttgtctagtgtagcaccctc
tatccttgtccatcaaaacaccaactatagctttagggctcaagatgttaacaacagaggccatttggaggctat
tttcaccgatagtcatttatcctttaaatttttatcctattctcattttcaatttatgagtattcacctatttca
agtaatgattttatatacttgaaacaacatagaagtagactgttataggatctccattctttttttgctagtagt
gctcaattaagagcttgtaattctctaaagccacgtctcctttctctttacaaaattattctaatttataagtct
agtttatcccaaaaacctttttcactttttttttttgctccactttgctaatagtaaatttaactacttgcatgtatt
tttgaaataagaaaacaactcactagatatgttggaactgcttaagtaataatcttcaataaataattttacttt
gatttcctcatcatagttcgaagacctaaataatttcactcttattaaatttcactcctattaaattactattct
cttcctttacatctcatggtgtcttcttgttgaatgtgatggctggcttatccaagtcgatcattttctccaaaa
ccttctcatatttctccttgataggattaagccaagagaatgggttgtgaagaactcccattctatcttttggta
ttgtattaggagagaattatatatacatggtgattacaataaccatgagatgagcttacccctaagattatatgc
atataatatatattatatacaatagataatactctaacacctcccactcccctccaaaactcatggtgggtcata
gaccatgagtttggaaagaagatgagaatgttgagcttttatatgaggcttattaagtaaaatcagtaggttgaa
tctcagatggaagatagtgagagggcatccctacaagatgatgatgatgatgagtacagaaggcatcaacactaa
taagcttggtgagctcatgcttgataggtttggtggcaatatgaatagcattaatatgatcacaatagagtggag
tcagctcaaaaatgacagcacttatatatgtaagaagccaatgaagcaaaatgatctcatgagttggttggaaga
tggtagtctattctttattttcatgagaaaaggagtgccaagaaaatacagtagctaatgatagaatgataatga
acaaaatcacttgcccaaatagcattagagaaagaataaagctaaagttgggaagaactagataaaagcaaagct
catgaagtggtaccttgaagataatgcaacacatgtaagagatgggctaggtgaattcttgtgggattcgtaata
aattggcttaaaatatgacagcatgagcaatatctagtttgaacatagtgagataataagactacctacaagatg
tcgatagtgagtcggctcataaagagatacattaactccatgctttaagttatgatggagctccaatggagtatt
aacagtgcggcaattgatcaaactaactccagagagaaggtcagcaatgcatttctattggaataagtggtaatt
tttggagctatatgcaacctcaagctccaagaaatatcaaaataagacaacattaatcatcacacactgtgtact
aagatactacttaacaaaagtaatatgcttcgaatcatcctcaataataatcatatcatccttgctaataatcat
atcatcaatataaagcaataaaagaatccaactgcatgaagaagaatagacaaacatggtaagatcatgaccact
ctgagtgaacctaagagtaagaatggtttagctaaatcatttaaaccatgcatgaggagcttagcaaagagcata
aagagtctggcaaagatgacgcatatgtatggaaggatgaaaaattctagtggtgattttacatggacttcctca
tataaatctcaatgaagaaaagtattcttcacgtcaagctgataaaaagacattgatgaatagcaatcatagcta
taagagtccacacagtactcatatgggccaccaatgcaaaggtctccttgaagtcaatgtcattcttttgctcaa
attcatgagcaacaagacgagctttataatgtcaatgtagccatcaaagtgagtttcgatcttataaactaactt
ctagatgatgaaaattttgtcaagggaaagaggaacaagatcccaagtgcccgtatgcttaagagcatcaagctc
ctgagatattactacatgccactttgaaatatcaacaacctctttaaaagtcttgggctcttagtaagagtgaat
agtaataagaaagtatggaactctaaagaaaggcactgagaagtagcattgagtttagtagaagcagtgcgacag
tgtaaaggatagtgatgcttattaggtggagcaactgttgatggaggatcatgaatcgatagcttagatcagaca
ctcgtatatggaggaatatctatcgatgaggcagaggtgttatgaggagtaaaggagctagaaggcttaatgaga

aacataaaagacaagactagtggggcagaagtagaagtaataaaggaaagatcaacaaaaacaagttggagagca
gaaggtgaagggtctggtggatgaggagagagagtataaaaagaaagatgctcatgaaaaatgatatctcgagac
aaatacatgtatcatgaaacaaaatcatggcatctatacggctatgccctttatgctcaagactatatcctttga
aaacacaaagatccataagaaagtttggtatattcagagggaggtaaaatgacaaagcacaataaatacttgtag
attatcataggaaggggactaggaaaaagtctcttacgaggagtaagacctcaaaagatgaaagagggttgaag
attaataagataaacaacaatatgaatagcctcaacccaaaaatgaggggaacataggcaaagttaagaagtgag
ccagcaatctcaatgatgtatcaaatgagcattaggacaggagagttgagggagagtccccatgaggcaaggaag
tcaagagattcatgagatagatattcaccattagactaagaatgggaggccgtaagggatttagagaattgagtt
taaagcattgtggcagacttttataaatctggtatagttgagagcgagactacataaaaaatatatctatataa
agagagaataatcattaatgaatatgacataatagtagttgccatctttagaaacaaaagggaaggactctagat
atttgtgcaaactaaatcagaaggtgaaatagctacgggattactaaagtgaaatggaagtctcacatacattcc
aagggcataaccttcaaaaggcgacatatctgaattagatatagagcctaagacaccactacaaattaatataat
gagatgaagaccacaaaaaccaagataaatatagaaggaaaatatctacaaagatagaggcaaaagaaataggca
tatgcaagtggtccatcagatacactacattatggtgatggccaacactgattggacgatcagtttgttgattct
gcacaaagcaagaggatgaatcaaaggagacaaggcgatcatgatcaatgagttgactaacagatagtaaattaa
catataattgaggaatatagaggatggaaggaatagtttattaggtggtcttaagctcaccttgttgagatgtcc
gtaaataggtgccatcaattgtaacaatatgaggttgcatagaaggaggtgaacaggaaacaagaagagaagcat
gtaggatcatgtgaaaggatgcccgaagtctaagatccaagtgaaacatctgcggtgtatagttatttagttgtt
ggctgatcaaggtgccttggagatatttgggaaggagcggattaagctccattagagacccaagaataattcagc
agaagttagtctagttcacatgacaacatgggctgaagattttcgaatggatgctggctgatattatgcagattc
acaaacacttggtgcttggggtggtttggaccaattattcagcagttggtgggcgcagtcctaagcataacatag
agatccatggctgacatggataagattgtgaacaataggaaatgaatgatgaaactcaaatccaaagaaccagag
agtcccgaatagagtgcgcagctatcaaattttgataatctcaaaatctccaagcagtggtatgaggtcttgtct
tgaagctcatttcaaagatgagggaggcttatttagagatgcatgaaggaccagtgagttagctctgataccat
attaggattaaactaatagaatgggttgtgaaaagctcttattctatctattggtattgtattaggagagaatca
tacatatatacggtaattacaacaaccatgagggggcttaccaccaagattacattaatatatacactatatac
aatatatgatactctacatttcttctcccttattgacatttgtagataggaagagattagaaaacagagctcccg
cttcctaagacctcttgatgagacatatttactttctataacactcatttgattcatcggaagaattttcttca
ttggagtgttatttctgaaaagttgattcatgaaaatatgataccaagaaataattttgacatatttaattgatc
ataaaaaataattatttcaaattaatttatgtttagttgaatatttttttaaaaatatatatataatatctatta
tactcctaataaaggaagaaccttatcttattctatacattactattgtttgcaataaaaattatgatggttaag
agttttagtatagatcataatggaggagttgtgatggagagaatgtttatttcaattgtaaatttttgatggct
taaaagatattttgaaaaaaaagagcatctgattttcatcctatgaaaattcnnnnnnnnnnnnnnnnnn
nnnnnnnnnnnnnnnnnnnnnnnnnnnnnnnnnnnnnnnnnnnnnnnnnnnnnnnnnnnnnnnnnnnnnn
nnnnnnnnnnnnnnnnttactttctgttggcgaccttcaaacttcttttcttcttcttctttttcttctt
cttcttcttcttctttttgagaaagctcatctatcaagtcttctatttgctttagaacatggccaaacctgttt
ctgtttctgatcaagtgaggaagagtcagacaaataactttgaaccttatcaagcataaattatctgatcaaaga
tgaaagcaattgtttcatctcaataaagcaccttgatttcccacactacagaattaatgtgcagcaaatgtcaca
aaatcgaacaagtacatcaatgcaatatcattgactagactttgtccctgttagaaccaaccagttactcagtcc
tacacactttcaggaggctcatagatattattcttaagttgaacaagggacatggttagtagcttgaagaacaac
ttggaaattagagacacccattttaaaaggatataagaagattttaaaatttaacccgacatattatttgattc
ccatctcgtgaactcgtaaacctgcacaagaaaattaacaattatatttagatgaagtttatactgaatatgaag
gaacatatgaagtataattttcagttctcaagcaataatctagttattcgtaagttataagcattcccacaataa
gaacatgggcaactgccttagcatcatggtaacaatattttcaaattattacgaattttaaaactgctcacaact
ttttctttggttgaaagatactatgcattatactaatggtcaaataaaactaactattataattcattcatgact
tatttactaaacggtcatgaatcaaaataatggtttcaagcttaatcactgggagttatttttcgttattgttaa
atgtccttattagattagcagagaaattaagaaaagtggcctgattggacaagtctagtataggtgctaatagaa
tttaactttgatcatatctattactagaaagagttgaaataccaagtagtccagggcaaaacactgggaatgtac
cacagtggcttgacaaaaccacaaccatttttctgggctagaaactaatttcagctagataattttctgtgtttt
atcctttctttaaatttcatcaattatccaaacaccaattcaagagtagtcggtaaaattaaggaaagctcccat
aaaactaagattaaaccttctcaattcacttgatccacgcccaaattatgaatccaagatcaaaattaatgaatt
ttggaattttaggttgatccccaagacccaaattgtgtaaatttgaaaataacccaagcaatgcatttgcgcaag
tagatttcctgaaaatggagacaagaaaggccctacctagaaagccaccataatatttttttaggctatttattt
aaaaatggaaatttctagaagttaattttaaaataaaataataatttctggagggatatcatgtttacaccacta
atgtgtacatgctatcaaaattaggtatacaatcaatgaaacctagatgtgcttaggttccactagttttgaggc
cctcacaattttttaaatcaatccaataaaatgaccttcatttgagattgatttttttcacatgttgtgcacatggg
gccaatctatgatgccctaaacaaaacaatagcttctagtataattttaatttttaagttattttagaattaatt
ataagatatttaaaatgtaatgtgttgaaaatatagctataaacttagattattaaatactattttccatggtaa
gtagtgcattttcattaaaaatcattttattaattattattttaattatgtataaatgtttaaactttaaaacat
attttgaaatcataggaagaaaatataaattgtgtcaaaagaaatatggtaaatgctccaaaaagatgcatgtaa
tgtctttgcagtttcggtaagctaaacaaatttcaaaaggtgattttgaagtgcaagtttcactacattttaata
gtatttgaaattttgaatacatattttctttgcttactagcattcaattgctaattttaagacttatatcttta
aatacatcttagattacaagtcgtatgatcataaatgaaatttgtatgtttatcatacatagaaaacctcaagta

gggaatagcaatactatattggaatttcaaatttcttctttttttgtcttttttttggtttgatagttgtcaatc
tgtttatttttttatcttttatttttcaaatgtttctttaggcttaggtgagtttttaagcctatggttgggttt
atgatatcatacgggtcccactaccaagttggtatctggtttcttctgctttctctttttctcccttcctcatt
cttcccatgttccagtgcgacaaaaaaggaaggaaaaagcagttaggagtcttctaactatgtgctggttagtct
tggataaaataagtggttccaattcctctcccacttgtcagccaatcggttcagcaccaggaacacttactttg
tgacttgtcgctcagagtcacaaacccatcctaggacttggttgaccaccagcccgacctgggacactaggtatg
aaccaatatggagtgagacccaaagccttgtatttggcttataaattagtgctttaattggatttaataaatata
acattatacaccatcctaaagtgttttttttccccaaaaacagttttccatttatgcgaccatttttgcatga
aaagcattatgttgggaaaaagttacttctgtgtgttaaaacaaaagattatacagctttcttttgtagtatttt
ttcatctgtcattttgatgtagaaggtaatcaatcatttgtcttttcacaacgataacaatttgattctcttcat
tcatacttatcttgattcttgtgattttgaagtaataaactaacaaacctttcttgccagcatgaagtcagtga
tagaacgctacaacaagaccaaagaggagaatcatctagttatgaatgcaagttcagagactaaggtacatttga
agcaccagtatatcttgtaccatcccctaaatggtttaaagttgtcatcttttctctaactcatgctttgccaat
tagtctacacaaccagttgacattagattcttgactttatgattcctcaactattttcaagcaaatatacaacta
aatagctcatcatcatagaacatgcacagcttaggctcaagtttcttttagttgttcaaatttatgtacttctct
tatgaaaatatcataccctaaaatgtcctcttaatttcatttattcaatagcttaagcaaaaaaagaaagaaaga
aagaaaannnnnnnnnnnnnnnnnnnnnnnnnnnnnnnnnnnnnnnnnnnnnnnnnnnnnnnnnnnnnnnnnnnn
nnnnnnnnnnnnnnnnnnnnnnnnnnnnnnnnnnnnnnnnnnnnnnnnnnnnnnnnnnnnnnnnnnnnnnnnnnn
nnnnnnnnnnnnnnnnnnnnnnnnnnnnnnnnnnnnnnnnnnnnnnnnnnnnnnnnnnnnnnnnnnnnnnnnnnn
nnnnnnnnnnnnnnnnnnnnnnnnnnnnnnnnnnnnnnnnnnnnnnnnnnnnnnnnnnnnnnnnnnnnnnnnnnn
nnnnnnnnnnnnnnnnnnnnnnnnnnnnnnnnnnnnnnnnnnnnnnnnnnnnnnnnnnnnnnnnnnnnnnnnnnn
nnnnnnnnnnnnnnnnnnnnnnnnnnnnnnnnnnnnnnnnnnnnnnnnnnnnnnnnnnnnnnnnnnnnnnnnnnn
nnnnnnnnnnnnnnnnnnnnnnnnnnnnnnnnnnnnnnnnnnnnnnnnnnnnnnnnnnnnnnnnnnnnnnnnnnn
nnnnnnnnnnnnnnnnnnnnnnnnnnnnnnnnnnnnnnnnnnnnnnnnnnnnnnnnnnnnnnnnnnnnnnnnnnn
nnnnnnnnnnnnnnnnnnnnnnnnnnnnnnnnnnnnnnnnnnnnnnnnnnnnnnnnnnnnnnnnnnnnnnnnnnn
nnnnnnnnnnnnnnnnnnnnnnnnnnnnnnnnnnnnnnnnnnnnnnnnnnnnnnnnnnnnnnnnnnnnagacttaagc
aaaaaaaaaaaagaaagaaagaaaagaaaaaatgggctatgtttacactacttaataattgagtaatcattcaca
tggaaaccgaagacagatttaaaaataaatattatctgactttttttttcaaagcataccaaggtagattttttt
cgatggttgtaacagactgatcttcacatattagaataatcttaacaatttttctagctctaaggtatcatacat
aatgattccagctacatcaacacctgcaaaaagtctattgatggagtggagaattagtttactgtaggtatcatt
cttggcatatgagaattttgtttacccatgtccaactatgagtctttgctccacatagatggactcttcctcctt
tacccaatcaattttggtcactaatacaagcttttacaagatccctgttcatggctgcattacataattgcaagg
tcaagagtaatgaggaggcatctttcatgcttaacaacaagcggaccttcatgatttcaatccaaatattctaca
aattgattttcaattgatcaaaagatttgaaataggatctttgatgtcaaaatagaataaacaaacaagagaaaa
tggagaattgcctataactgagggaaaaaaaaactattgttgttggctatactgggtaaaatgtgagtactatgtg
tagttcctaaacaaagcagtctaagttacgcagcttaaaatacaatgcagaacactaaatgaatttacaactaag
tctgacatctagataccaagatcataagaagaaagagatgacgataacatgattttgacaatgatttttctactt
gcaagagttctacttatacacatttttatcctcttgatctcattataatttaatccctatgaaattttattac
tttctttgctctgaaagtagcatggtggttcgtgaatcaaactatgcacatgtttgcatatgatcattctgtctt
ctcaaatgtgatttctcacttgacactgctatggtgctacctgtttcaaactgattcctttaattcattttaat
atgggaagcttgatcattgcttctttttcttttcaaccatnnnnnnnnnnnnnnnnnnnnnnnnnnnnnnnnnnn
nnnnnnnnnnnnnnnnnnnnnnnnnnnnnnnnnnnnnnnnnnnnnnnnnnnnnnnnnnnnnnnnnnnnnnnnnnnnn
nnnnnnnnnnnnnnnnnnnnnnnnnnnnnnnnnnnnnnnnnnnnnnnnnnnnnnnnnnnnnnnnnnnnnnnnnnnn
nnnnnnnnnnnnnnnnnnnnnnnnnnnnnnnnnnnnnnnnnnnnnnnnnnnnnnnnnnnnnnnnnnnnnnnnnnnn
nnnnnnnnnnnnnnnnnnnnnnnnnnnnnnnnnnnnnnnnnnnnnnnnnnnnnnnnnnnnnnnnnnnnnnnnnnnn
nnnnnnnnnnnnnnnnnnnnnnnnnnnnnnnnnnnnnnnnnnnnnnnnnnnnnnnnnnnnnnnnnnnnnnnnnnnn
nnnnnnnnnnnnnnnnnnnnnnnnnnnnnnnnnnnnnnnnnnnnnnnnnnnnnnnnnnnnnnnnnnnnnnnnnnnn
nnnnnnnnnnnnnnnnnnnnnnnnnnnnnnnnnnnnnnnnnnnnnnnnnnnnnnnnnnnnnnnnnnnnnnnnnnnn
nnnnnnnnnnnnnnnnnnnnnnnnnnnnnnnnnnnnnnnnnnnnnnnnnnnnnnnnnnnnnnnnnnnnnnnnnnnn
nnnnnnnnnnnnnnnnnnnnnnnnnnnnnnnnnnnnnnnnnnnnnnnnnnnnnnnnnnnnnnnnnnnnnnnnnnnn
nnnnnnnnnnnnnnnnnnnnnnnnnnnnnnnnnnnnnnnnnnnnnnnnnnnnnnnnnnnnnnnnnnnnnnnnnnnn
nnnnnnnnnnnnnnnnnnnnnnnnnnnnnnnnnnnnnnnnnnnnnnnnnnnnnnnnnnnnnnnnnnnnnnnnnnnn
nnnnnnnnnnnnnnnnnnnnnnnnnnnnnnnnnnnnnnnnnnnnnnnnnnnnnnnnnnnnnnnnnnnnnnnnnnnn
nnnnnnnnnnnnnnnnnnnnnnnnnnnnnnnnnnnnnnnnnnnnnnnnnnnnnnnnnnnnnnnnnnnnnnnnnnnn
nnnnnnnnnnnnnnnnnnnnnnnnnnnnnnnnnnnnnnnnnnnnnnnnnnnnnnnnnnnnnnnnnnnnnnnnnnnn
nnnnnnnnnnnnnnnnnnnnnnnnnnnnnnnnnnnnnnnnnnnnnnnnnnnnnnnnnnnnnnnnnnnnnnnnnnnn
nnnnnnnnnnnnnnnnnnnnnnnnnnnnnnnnnnnnnnnnnnnnnnnnnnnnnnnnnnnnnnnnnnnnnnnnnnnn
nnnnnnnnnnnnnnnnnnnnnnnnnnnnnnnnnnnnnnnnnnnnnnnnnnnnnnnnnnnnnnnnnnnnnnnnnnnn
nnnnnnnnnnnnnnnnnnnnnnnnnnnnnnnnnnnnnnnnnnnnnnnnnnnnnnnnnnnnnnnnnnnnnnnnnnnn
nnnnnnnnnnnnnnnnnnnnnnnnnnnnnnnnnnnnnnnnnnnnnnnnnnnnnnnnnnnnnnnnnnnnnnnnnnnn
nnnnnnnnnnnnnnnnnnnnnnnnnnnnnnnnnnnnnnnnnnnnnnnnnnnnnnnnnnnnnnnnnnnnnnnnnnnn
nnnnnnnnnnnnnnnnnnnnnnnnnnnnnnnnnnnnnnnnnnnnnnnnnnnnnnnnnnnnnnnnnnnnnnnnnnnn
nnnnnnnnnnnnnnnnnnnnnnnnnnnnnnnnnnnnnnnnnnnnnnnnnnnnnnnnnnnnnnnnnnnnnnnnnnnn
nnnnnnnnnnnnnnnnnnnnnnnnnnnnnnnnnnnnnnnnnnnnnnnnnnnnnnnnnnnnnnnnnnnnnnnnnnnn
nnnnnnnnnnnnnnnnnnnnnnnnnnnnnnnnnnnnnnnnnnnnnnnnnnnnnnnnnnnnnnnnnnnnnnnnnnnn

nnnnnnnnnnnnnnnnnnnnnnnnnnnnnnnnnnnnnnnnnnnnnnnnnnnnnnnnnnnnnnnnnnnnnn
nnnnnnnnnnnnnnnnnnnnnnnnnnnnnnnnnnnnnnnnnnnnnnnnnnnnnnnnnnnnnnnnnnnnnn
nnnnnnnnnnnnnnnnnnnnnnnnnnnnnnnnnnnnnnnnnnnnnnnnnnnnnnnnnnnnnnnnnnnnnn
nnnnnnnnnnnnnnnnnnnnnnnnnnnnnnnnnnnnnnnnnnnnnnnnnnnnnnnnnnnnnnnnnnnnnn
nnnnnnnnnnnnnnnnnnnnnnnnnnnnnnnnnnnnnnnnnnnnnnnnnnnnnnnnnnnnnnnnnnnnnn
nnnnnnnnnnnnnnnnnnnnnnnnnnnnnnnnnnnnnnnnnnnnnnnnnnnnnnnnnnnnnnnnnnnnnn
nnnnnnnnnnnnnnnnnnnnnnnnnnnnnnnnnnnnnnnnnnnnnnnnnnnnnnnnnnnnnnnnnnnnnn
nnnnnnnnnnnnnnnnnnnntaccgataccggaccggtatgcccaggcgtaccgtgtgtcggtacggtatggt
accggtaccgtaccggtccgggaccggcacgccgtccggtaccggtacagcgaaccttatttcaaccatacagta
gtcattactcattaccatcaaccatctagtggtcggtactgctggttgcttgattgtcacacttaattaccgcaa
gactgctacttttagttggtatagttaattgcttattgctagttatgatcatgggtaattaccaataaccagtaa
tcttaatcactactagtgttggctagcagcttttcagtttgttgatcacaccactgcttctgttccagtgtttca
gagttaagtatcataccatccaatggctgctacttgttcacaagtcaagaggaagaattctagattgtcccctca
attgggaatttagttagtaaagtgaataaggacaactttgctgtcatatgttctctctcaggacgtttgatggtg
agccagtcatcatccttcttgataaaatttcactctgtagtcatcaacactgtagtcatccaaagtgatatgact
agaatctacaccgatctgttggcaatgatttggggatgcattcaggtgtcctaaattataagttgaaaagttata
gaagcaaagcaaagcaatcctaacaaccacttagtcaagattgaaagactactggtgaaataagaccacattagg
gtctttccaacagacataaggccacaccaaaaagtcatgaccagacatgtttgcatgggctaaagtgggggagcc
aaccggttttacctaccactggagatatgaaccaatgactcaccatataaaactgatatactctaactgctaagt
aggtgtttgtccaaccatcttattcaaagggtgaaactgactggcagaccaggctgctctcattatgagttaatc
ttacccgctctggtcaggaaaagaacaaagtttgaggccttatcaggcttacacaaatcttggtccataattgtt
ggccatactaaagatgccttcgaagtgagttggtctgtctaatattcaggaaacgtcgatcttgaattggaaatg
ctatgaaacatctacctcattccaatgcatccctataaccccattggatggcttaaaagtcaacaaacaagagct
tgcaaagaggaaccatgacaattttacaagagccattaactaccaagcatggtattctgtaccagcccaaaccgg
ccggtacaccccataccatactgtaccaacaaaaaatcagtccggttcaaaccagattttagaaaatgacctga
actggaccgaactgatcggttcggtatggtattggcccgaaccacccgatatcggatgatccggtccggttcggc
ccgattttttttaatgctgttgtgtgggatttttgatttttattatcggtatggtacggactggtatggtacg
gtacgataccgtatcggccggcaatcgacatgattttcgataccaagtctgcgaaccttgctaccaagcattgta
actttgcgattgaccagacatacaagtgtaaaatcaataattgagttttcctttgataaatattgagtgaacatt
gtgtgaagctgatggtggtgattttgttctccaaaggcctaataatccatggttcaaaatcagttttcaatttat
gttggggttttgcttacattccctacaaaaaaggcttgtattacttcttctcgtgtagagttaattgtatgacac
aatgattattcaatccacataatgttccctaggcctatgtatggtcattggtcattgatcaaagactacctgtca
aactggagatagctttcaagccaaggatttacatcttgcttaaatgagataacattgctggtactatactattcc
agtaggaagtcaggagctggagatctatgggataccgatctgtttatgggccgggacatccttcgtctcaactca
aactagttgggatcagccaataccaagcccgatttagcaggtaccgggcataagtcaactaggaggaaagatcca
cctacccttgcgggtgttggcatcggtgacatcaccatactcccgtcattgattcaaaagaatgttttgccgtac
tagtcggtactagtatgtacactgcaccgtatagtatcggtactaagtcggtatacagtacagggatgtttcaag
tattggtatgccgaaccaaccctcataccaagtttaccgacatagtaataggatggtaccaatatggggtctggt
accagatggcaaatcttgttcaaaaccatatcaatagtcattcaaataactatttaggattaaaaaaaaccaaaa
tttgatataatattatggtacagggacaccaaaccatcctaagagtcaagactgtagtgtcttgatatcatccta
acttggggcaaggctcaatatcaatatcttgaaccatacttggagctttgtgtgaagatgccaaatgactaattc
aagtccttatttagtcataattcataaattattctctatcattttgcattggttcatcaatcttaatagaagttt
ggcttgttaatctaaagtcttgatccaacttataagcttaatagcagtcatggtatgatacaagttgtagccaaa
agaatgcatgacaaacaaaattcataataaactggcggaagagctaagaactattaaattgcttgtatgataagc
ctatcaggttccgcctagcttcaactgccaaaagtcataattctgatggggcagatgaaaggggaaaggactcaa
atttaatgcttgtagatgctcatgagtgacaagaattactggtgaaatcatgaatgacttaaatgcattgaaagg
ctaagtggaaagatgccttagtgtcagcacttgtcctttgaaatttatcataatttacttgttattttatttata
tcataactagtcatgactttgtggtaggtagaaatataagttgaagctggtatagaacaagcaataacaaagata
caacaaagcacactaaggtttattgaatagaaattttaacggtacaagaaatgaataaaatgataaatggaggac
caatggtcaaaagtgcagcattgagaataaatggaggaccaatggtcaaaagttcagcattgagaaaattgggcc
atatctggcaaattcacaactttaggaaagagtgaggataatcaaaaatttggaacatatataacaaaacaaaat
aaatgcgagatttctaaaatagttacttttctttgtgtaaaagcaagtactagaaggagagaacaataaatggta
gcaaaataacaccatcatcaacaatatattgcatctctagtctgtaacacagaattatcatgtaatcatgattaa
ctcatcaataaatcatgaaatttatgtaaaattgagaacaattgattaattgtgaactaagtgatctacagggaa
tagtatggaaaacagttcaaatatcgaatagtacaagaaatttaagaatttttttcaatagttggagaatattgat
actatgtagaggagatatggttctttcaccaattaatgtttagtcattctatcaatatagatgataattttgacc
aaaccttgacttgttttcaagaatttttatgtctttttaaagaatttctaaggcctcaaagagtcctaggtcagt
gtggaagtcctagatctattgtcctaatcctcagttcttcaaattacaatgaaacttattcatgatacaatcaaa
tttgccatgttatattatttgtcagattttgcatgacaaattctttgtcaaccctgtgttggatcaaaaagaagc
tcaaaggaggctctggtgaattttgtaaggacctaacaagcaagtttctataagccacttgattttcaagagcaa
aaaagactttgaaacacatgaatttattattactagtcccccccttcctcgtcactttgatttagttcttttatt
tatcacaaattgtttaattctcttattattttgcaacccttctgaacttgtgtccatatgttcctctgtcaatc
aaatcttgttttgctctttgttacatgggttttctgtatctcttggtcatttagaccttaaaatgaattccaata

tcagcattcaaactgtgttaaatcagcacatacaattaattacattagtacatgccttggattggtaatgaatat
ttttaggaaagaacatcacctggagaccacaagaaatggcaataaatctgaagttttgcagattaaattgacagc
attttgcagttgaatgtgctatacttgagtcagctgcttgacacccctctcctccttgaggtcccacatttgaag
cagattgcataattcccaccatgtttctcacaaaaaaagagaaataaataaataaataaataaaataaaaagttg
gtagtttacaaagaccaatcaactatatgattgatctttattgtagttttagtcctctttttttacaagtttact
catttatttgcnnnnnnnnnnnnnnnnnnnnacaagttttactcattttatttgcaaatgaaaacttgtgggaac
ttttgggttccaagctctcatatcattaacaggatgaatagtggggagagaaatttccctatgatatgctatagc
ttaagtctcatttagaacaacaaaataaggacatttgtttatgcaattttaataagaataaccataggtagcttg
atgggaagtgtttctcatttacaaataaatggagccgatacctatagcacttcaaagtgatctttcatagaagg
tccttcagaaactaaaatccaagcttttgatcatcatatgtgacaagctatgatttggataattacaatgtcgtt
ggagcacttcaattttgaacattcaacttgcagtgataagtacaagtgtctcaaaattagaagaggttatcaatt
aatttaaattttgaaagtaagcattaaaaaattgtaaacactactctcttgtctctgaatttctgtagcaaaaca
taagaactcttcaaaagatcaaaatgaacgaagtacctctgatggccttaaaaaatcccaatcatagtcttcccc
atccatgtgatcttccaccttatgaaggtcttatgatgatgatcaacggaatccaagttatctgaccatttagaa
catattcaggattattagaaaaatcatggtttccaacagtacaacaccagaaggaggaagaatcaagaaggcaga
tttttaatacctttttttccctcttacgcctgcttcttgtccaaagaagaatgtagttacagcaatatgaacatg
aaggcatcatgcctctccctctttcagtaagtcatcctacacttctccatgttgattcactggttgcattttgtc
catccatcatcactgaaaaatgcatctgcagtacctctgtatattatggaaaagccatgtggcatatgttccctt
tgatctatattaattagaacctatggtgtaaaatgccttgggcaaactaatctttgtctaaaaagtacttgatcc
aattaatggtgtcaatgagtaactcccactatggacatgacataggaatgactgatcaaaatgagagagagagaa
ttacaatttacaattagataaatcgaacagctaggctaactgtcatttctactgctgcaaccttggaaccctact
tgctaaatcattacaatcagaatcattgattacaccactcatatcctttcatcagaaatctcatctaatcttaca
taacctcctcttacatgtgcctgtctaaattaaactacaaatttcacaatcatcaagtttaaatagtattttccc
tttcttgtcgtccataacatcatcatcatcatcaaatatactattaaaagcagtctgctccattttaatcttata
gcaagaccaccccacaaagatgatgctctagtcaaaatacctatcatagataggtgttccacaaggaatttcatc
tttcccaaccctcaatctcgcactgtagcaacagatatttacttattgtttcagtattttgcttctaacatcaca
cagatacaatcttccctcataacatactttttgtaattcatatggatcttagaggtctgttcatgtgttcctaaat
tagatagtcacactgttttgccaacaaaaattccacatattatgacagaagctctcctgctcaagctgtctttgc
tatgtctttctattcattatcaccaaaattaatacttaactataaatgcagacagatttaggactaaatcaacct
cacaaacatcacttctacagaaatagagaggcagatcacattcaggtagacagattgagagttcgactttccatg
ttctaactataccttaaatcaacatgactgcttctgagttttgaatttctgcaaatgctgtaaaatgcatcttgc
aaatttcacagtgatcgctagcatccatgtctgtgacaatcttcaatcctaacaatatgataccttctttcattg
aaaattatttgagacttatcagatggtccattcttcatagatttcaaccactcaatcaaacaacatccaacaatg
aactctgtagttggctgctccttttattaactagatatccttcacttagattttggtatttcattacaaggataa
gaaattcatgagcaaatgaaatcaatagactggtcaaacttttagattctgctacttaccatgaatatacaaaaa
tcaaaagcagtatacaggcaacaataattgcttggataatatactgctaatggcgagaaatgtggtcagtgttgg
aaatagatattcaacataaggttaagatgaaattcaactaaagaccacaatataatgcagaaaagctttaaaatt
aattataaaaatagcaggaggttttcttgcaatgaggcaatacaatctccagttctctcatttaccaatgtcag
aagaggcatatgacaaaatggtcataaatataaagatttaaaggtgacaagaaatgccatttggtttatcttact
agctccattgactatccactttgaatagcctataactgtcttctatgatggagttaaacaattctgtttagaagc
aagaatcctcatggcattgccgcattatcatgaggatattagataattgatagataggaggtctaataacaattc
ttatgtgagtttttagtgaaatttccatggacaaatagacgaatctctcaacaagctacaaagaaattaggcagt
gtatacaaaatagtcctaggtgtattaactgacactgtcacccatcgtctgcaatctcggctagagactgtaggt
caggtgccgctatggtatggtaccgagtcgaaccatattgacatgtggtatgtaggtgcataccaattcagtact
agtttgctattttaagagtgctagagtttttttttcatattgattctgagagactatatgattgattaatagata
caactgcaataattaaacataaacttgtaactcaatgcacagatgcatattaattaattccctgctctgtcatat
acatctttctttcaaccattaccttcttgaactcatgcttttgacttgtcatccaccctttactattgtttttg
ccttcatttcatagatttacataggccatattatcttgatatataggtggacatcatatcccaaggtctagttc
ttgtcttgctcctagtctcttatggttgcacctctgtatctctctctaattcccacttcccaatttagaatcagc
atccattagagtggcattagtctaaccaaatagtcttagggttttcttaagctttcattttgtttatgcaccatt
gttcccatcaagattaaaacattataggagaattctaaggttcctaaatgacaaaagtttatggttaaagtgcat
ttaattctatgacatactaatctgtgagaggttgccttctcgaatcattgttcatggaaatcatctgtatttgtt
aatggttcaattaagcaatacaaactcagatgatggtgagtttttcacctaagagcttggcataagttatgaaa
atgggtattaatggtgggcaagattttattcaaatattaaatattgtggttgtgtgttcaataatccctcttct
agatgaattgaagagaacctcttttagtgaggatccatataagatctatatgtcctaactttgtcattctctaca
gatttggaaactgattatgatcaggatcatctccccttcaatacctatcaattagtgcaaggattttgaactttt
attacaaacatattttacgatagaatatacaagttttccatcacaaaactcacccttcatgatgcaacaagagtg
cagtctgcggaaactataaccaactagaccatgccaagtttcatagaagccagcggttttagttgggtttctata
ttcctagttgaataattctagcatgtttccttattattggatcctatttccagcatgattatgtaccaggtttaa
tgcctgaatttggggtcatcctttatatgcttttagtacattgtatgaagggtgacttttgatatattaataaaa
aaattctcaaacagagctcccttcatcgaacaaagggttctgcaatccaatgtgccatcatcttgtctgttgatc
ttcaagtttaatcccgtcaatctaatcgctcttcactcctcatgctacatcactttccttcataattttgaaga
aaagataggagagtggcatcttgtggccttctctctttttcatcataggatggcccggaccaacccaaaccactc

taaaaccgagtgcaaccaagcaattccactcaattcagactgacttgggctggttcatgccagttcctgaaccaa
gaggttgaaccaacctagttccaactggtttgggttgatatgccccaaacagggagggcagttagagttggtttg
gcagaccatagtgtctatgcttgtcttcaaaaagtatcaccttcttacacttttacctgcataatgtagcatcgt
tcaaaatcataaagatgttgaaatcttagttaaaaatggactagtaagaaactaaatatatcgagacacaagctt
ttggtgtttgctcaaagacataccagagatgaccaatttgtttggagaagtattgtggagaatgctagaaatagt
tctcaacaagaccatagttttctgaaatagaacatcaggtaatggagatgtacctgctgggccttatttcaatct
acttgctgtggtccaactaagttataactattgagtgtgtttgcctgagaattctttaagaagccaatttaggac
atgataaactagacgagattcatcttagcaaccctctagagccaaacaaagctgtggaaacaaggacatatttct
gaaatcaaagaaaggtttttcaccttaagacaacaaaaagagaaaagaaaagaatatagatttacgattaagaga
taagtggatgtagaattacgaagcaatagaaaatgaagaaatggtttctagaatagcatgctaaatggagtagat
cgactgtgccagctcaacatccaaaacaaacaccaaacaaaaatatttgcaaagaatcatgagcaaaaaaaaaaa
ttatggaagaaaaacttgggagagaaacaaatttattcatcgaagaagaaaaatcaaaacctcacaaagacgcct
cacaaaagagtctccatatacacaaataatctctcctctctcctttttcccacacactcttctacataattttt
tcacggtcttccacacccaaaggagacatcaaaggtccctttaaatagagtaccacataggtgggagtttgactc
ctaccttgactcaaaatcaaagacccattagaacacaaactctaacaatctttagaaaagagtaacactagaaat
aaaatcttaaagtcattaaaactaatgatcaactaaaaatagaatcataattggattgagacttgacaatgaatc
tcaacactccctcttattgcaaagtccatactgacacaatgctgactttgtcgatgtcatgaaacgccatagtaa
gctagtagccatcgccatcaccgctgcagtaacagtaacatataacaacttattgaaccaacaattcaataatga
aactttttattgacttgaacattaattacttcgctggctcgtcctcttcgatgcatatatcaaaatcgacgatg
atgtcgtgcaatatttctctcagcattggttccatgatcatgatgtcgagatttcttcgagcatccagcttatc
ccaacgaacatcaaattatcggttacaataactgtgagcattgaccattacctgcacgcaagtaaaatcaaaagc
aaaactttgaccatatttatatacctccacttgagttgcttcaggtatctgtacacagttggtttctacatatgg
aattaaattttatgtatagaattttctaacagctcgaactttggagtagcttcaaaattaactgatgtatcaag
aatcttgttgatcaccaaagtacaataagaatcatctccattaagcatatttttctgttgttttattttcaaagc
taattcaaattagtcagtttttcatcctcgtagttgagtaaaccttcatatgaaatatccatatcttggatagct
tcaatatcaaattcttctccatgatttgactcctcaatattgatcttcataactgaatcggttgcagcatcatca
aactcttcttgagcaatttcagaatcaatagattcttcaatcatgaactgatttagacttgatgtggtagccatc
tgatcaagctttcttactccaaatttcttttgcacatgattgatccttaaattgatcgactttatcatattttga
gtcttccaaaattgatgtaatttaactgatcaagctccccatatttcaaattttcatctttaaactaattttgtt
tttttagctctttggaaaagcttgaaaccagaatttactcttcagataattttcgtttatgatttcttatattgc
aaaaattctagtagacaactgtaagaatatcccatactttctttggctcctttacatctcaaacatagaaaaaaa
tttctttatcaactgaaatttaaagaatgcatattaccttttagatttcatattccactttcttcttgcaatct
catctgttggcatcatttctgactcatccaaaatatcctacaaatcttggtctgtcaaataagacttaacaacag
ccttctaatatggataatagatattgttcaacttacgaaaaaaatagtagtaaaagcaatcttattgatattagc
acgatcagacatctcataactaaatctattgatagagatagtaatagcattgactatgatgatttacttcttcag
ttaccaaaaaataatctacacacttgcactaccgcacaacgaattactaaaaactctttaactacactgtgctat
gaatacggaaaccaccaataatcaagtcactactaaaaaaaaagcatcaaaactgcaaaccacttaaccactttc
caagcttatggctcaacatcatatagagccacaaacacacgagccaccaatctccgaaccttagctctgatacca
tgtagaatggcgtgctacataaagtagatcaatcgtgctgccccaagatctaaaacaaagaccaaacaaaaaaaa
atatttgtaaagaatcacaagaaaataattatcgaagaaaaattgggagagaaaaccaatttattcaaagaagaa
taagatcaaaacttcacaaagacacctcacgaaagagtctccacatgcacaaataatctctcttctctctttttt
tcattcttctacacaattttccatcatcttctacacctaaaggagacgtcaaaagtctctttaaatagagcacc
acgtaggtgggagtttgactcctatcttgactcaaaaacccaagacccactacaacacaaactctaacaaccttt
agaaggaaataattaaatagagtaacattaggaataaaatcctaaagtcattaaaactaacgttcaactaaaaga
atcctaaatggattaggacttggtattgaatcccaacatggttatgggaggttagatgaaggcattctcttggaa
aggttaacaaactatgtattgcaactacatgcccatcttgaggtttctcttgagttgggttctcatttggttcta
ttagtctacaactggctgcgggttaaagaagagtaaatggagtaaaggtcccatgagtcaatttattccatgtta
acttgacaaatgccaaaaatgaccagtttatttggagctcaaaacgagtaatttttaaagtttgtaaaagccatt
ttccccatctaatccaaagtaaatattggtcattatagggtaaagtatttatttggaggaaagagggttaggag
caactaaaccaattttatctgggttgacttgttccacaatcaaatgcagcctcaaggtggagcagcaattcctat
gaggtgaaggatgtttttgtccaagaggatggaattgtctttcaaatataaatggggatggtgtttctagtgaag
aatttaatatgtttttgatctggatgcataatatgtttttaatatggtgtttctatttctcttgcatgcttgtaa
ctacttttgcttcctaataactttttgcatatctttaggggtagttgtgcttgagcccattaataaagaaaagta
taacagaaacctcaccattaataaaaaaaaagtataacagaaacctcctgtcgtgtggtgatgagtacaagcaca
agttccatgttttgaagaacttgatctggaaactttttttgtgatgtaattaaggaaattcaaattgtttagga
agcttaatgaagcacaatcaaaaccaaagattcaagttccagcgggacgtcccacgggacaccggaacaagatat
catctcatgtgtcgggatagggcatctcttgagacatcccctatcccaagtttcaggatggggcaggacggtggt
gcatcccgttccatgagaaaatcaagagtcctgtcccacaggatttgaaatcttgatcaaaatattgaacctata
atcagaaatcatttaatgtttaaatattcaacacacataaaccagatactgaagcttttgaatgttgcctctaca
ttccatattaatgatgcagaaattacattgttcttagtcatgtaatatgaaaaaatcctacaaataatttaaatg
aagatcttatcctcttcccctttacctgtcgacagtctccatatccaatatattctttaattctttcctttttc
ttttgtgtgttgaattcttttctttttcaattgggaccgggcatgtggctatgcaacagatgttaggcaaagca
ccaaaagtgtgtgaataggtcatcaactatttaagggggaacattcaaagatcaatacagcatgcatcgataaga

ttaagtaatcaaaaagttatagtcgagaattcgtagtttacatcagcagtaagttatagtcgagaagtttttaaa
cctaatgcttttaaacttgatttgcaataaatttcatgaaaactggtgtatcaaacttcagattctaagtgacct
gacagaatgtttcttcctgaacctggttactgtaatccataatcactccaatagtaataatatagtatttaaaaa
tctgc**tgcatgggcttaaaactttgtccttgtactcactagttttggcaaagggaggcagcaagcttgaggcagc**
**aactgcataacttgcaagaaagtcaccggtatgctcacttcacc**ctgcataacatttttttttttaagcattatt
tctcaaaaggcattcatctcaaaaaggaaagcatactgtaacttctaccaagtggcttaaattcagaagattgtt
acttgttacataacacatggaggctaaaacttattcagcatcttgtaacatatataatactatttcaatttgaca
ctaataatggcatattacacttcaaaggcactagcatatcacatgccctgagagccagactgattaatagtatca
gaaatgcgaatggcagatgggtaacatgtctttgacaataaattacttaaatttcatgcatactaagacctaata
tgtaagttatagtactattaaagtctagataataatttcaagtttgtgtataactttagaatgaactacataatt
ggcagccttctcacttatcatctcaaaaaaaataaaataaatctaggaaagtttatatcaagcaaatgtagtagc
tactaaaatacaaacaagttgtaatgcaaccatgtatattagtcattccagcataagaaaatgaaggggactggt
aactggttcatatatcacatgtaacattccaaagatcagtagttaagaccatatgaccatgtttatgaaatatgg
taaaagcattctcactataattgtagaatttctatctcactttattattctggaattcaaaaacaatgaatataa
ttagacatttctagcaaccagtgggacaaagatagagactatgtggagcaacattaacagaaatgatcatgaaaa
gaagcacaaattcattgaaataacagtaacaacatcccagtcaaagtgaagagcactgccaagttatcagttttg
ataacaacgaagaaagagaaaatatttctaataaggttagagccagtgtaatgaaacaaatagatgcctcttggt
ctaggagtccacatatgcctgggaaactacaactgttttaaggaactagtatattagtggaggtgaagtagcaca
attaagaataaagtgattaagaaccagtaagatggtagggatttgtgaaaaaccaaaatatcagaaccccattta
ggaggtcctcatatcccttctaatcttccatagtagcatttcttcatctttttgtagtccctttcttcttttttt
tcttctgctgttgctttctttttgtaacttcttttggtaatcaccattatttttttcaatgcatatcacctttagc
aaatcacttttctgaaaagatccaaataatgttccacttcattctcttcaattgagcaccttgtacttctgtatt
ctttctttatttttatgttttaaaaacaactatagtttaatctcatatggataattgttattaattttgatca
aaatttgtatttctttaaaaaacaaagaaggtcagatttattttgtagtatcaaacattattttacaaagatgca
tcctaggaagtttgacatcctagatggtttgacaatctagattttcttagttgtctgtttccactcccacatcca
aatccattcccaatgttgcccatagattgataaggtcttcaacagtgaaattagccattgactagcttagaaata
gaaacacaaataatggatacagatgataaaaataccaaagctaacaaaaagtctcttcagacggctaggtataca
gcgatggtaagtagtaactcattgatgaaatcggaactaatttaaccataagtctctagttctcactggtatcac
aaaccatacccaggcactcaatataccagacactaatgacaacagaagagtttctaaagaagacactacatatac
aatatccatctagatactaaactcccatatttactattgacaacattaaatgcccaacaataagcagtaagtcca
tcattcaccaaatcccaaactgaactcactgttatgtatctgtgtaggcaaagctctaataagcgcttaatatca
aatttagagcaatcatttgaaacatatctaaaatataagtatataactgacatggcaaaacatacacaacatcgt
cttcaatgattgcaggcagttgttgggagaagagctttctggcctaagtgtgaaacatctacaaaatctagagaa
ccaactggagatgagcttacgtggtatccgaatgaagaaggtcttggtcatctttgcctcattcaatgttgactt
gctgtccttttaacaatagccatttttaatggattaatttttcactcccataggaccaacttttgattgatgaaa
ttcaagaactaaaccgaaaggttggaaaacatgtgtctaccacatcccctcctctaagttttaggtgatgaatat
caattgcttaacatttcactggttcatcgcaggggaaccttattcaccaagaaaatatcgaactatacaagaagg
taaaccttattcatcaagaaacatggaattgcagaagaaggtactgaataagagagcgaatagacttgaaaaatc
catacagaaacttctgtaatgaaatcctttcaaccaatggtaaattatgtaagttgcgggtatagcaaaggctaa
gcattaaatttgcatctcaggtttatgcaatgagggtgtaaatggcattgataaaggtccgattactccatat
ggttttaatgtcactgaggatgcaaacatatccattcatcttgaactcagccagccacaactgcaaacagatgca
acgtaagctgcagcaccaaaactagggtaattttctttccaaactaaaatggtatatatttttttaattactagtt
taatcatagaatatttcactgaatgttgcagctgacttcaattgcatggacaagatgatgcgcaaatcatgggtt
gctctttgtgtatcctttgccggtgtagttacaacagcatgctattaatgatcacctgtggtgtacaaggaagag
aaggatctattcccaaaaaatggtgtgaaataatcagaataaccaattactctgtcaagctgtgttactattaaa
aggtgatggctgtgctagggatttcagtttaagagcttcaaacagcaatcatttgcctatctagatgcgtattcc
catgttttaagttcatcattaaaacctataatcaagaaattcacaagttacattgcaggcatatgggattatcac
tacctaaatgatgcttacattgaaagagttgatacctagagatggactgtatcttgaaacaagttttcacttgtc
acgtacaaaaatcatgagaattatgcaactacgacaatctgcacaaaccattttatgcaccgttagtgaaagaat
aaggtggaaaggcatcccttctatccagcccaacagagttatcgacaagcaagctaaatcattttgcttgcgatt
ataaaaccaactccatttgaattcaagttagtaggttatacataagaaaatgtcatgccttcaaatgaacttcct
ttactgcttgattttactctcccttctagcagttttgtgcatatgctcatatagaagagcaacatagtcagccac
acaaataaatctctctcgtgagtatataaatataaaatgataacagacgtggcagatttctttctgcagaatttg
ccttactttaaagtgcaaaagaagggcctaagtagaaaccaatttatttatttagccacatgacctggctgaaa
tgcaaaagagagaggggaggtgagggttcgaagcattgggaggggaaggaccaaggtttgctatgccggtagacc
ggtcggccggcaatatggttcgttacccaccttgccgttccgtaccaagcccaaaccgacacaaggagagggtgg
agagagaatgggaggggagagacaaagagagggtggaaggagaatgggggggggagggaggggagagagagagg
agggcctttggaggacactagagagtagttgagcttgctgtgccattgccgagcttagtggaagctggagaaatc
tccaaaaaaattgaaaaaaattaaaaaaattaaccgacttcaacccaaaaatctacaaaaataatgaaacgaggg
ccaaagcccctatttcaatcaggaggagggtggaggccctcctttggccttcattgaccttggtggcggcatagc
cgcctcgccgacccttcggaggcctctgagagactcctctccctctccttccctccctccaccctcaccaagctc
gacagcggcgtagcaagcttggctgcctccaatggcctccggaggttatctctccttctccctctcttcctcccc
ctctctcttcttctcttccttctccctctccctctcgctcatttctcattccgaatgccggaaccatctcgattg

gccatggtacggttcggcatatcctgaactacgtgattcggggcggtttagcaaaccataaataggaggttatcg
tttcgactagagccagtttggttcaatacaccttgatccagctggttcgacatggttcaccagaccttgattcaa
aatatagaagagaaccatataaagaaaattggaatcccgggcataaaacatggtggtaaacataaatttctaggg
cacatgattattggtgcttatttaagaaatgtaaagcaagcaacaaaaggcaattaatattacacaataatgact
accactaccaaaaggaagataagagttctgcaccaataattggatagaatgttcatgaaatctaggattactcac
aatctgcatggcaactatttcttgcaataaagcgtggttggtgtgaccaaaaaaaaaaacattttggcaaaggtg
tgacaaccaataaggcaacaaagacaatgaatattttataaaagcaacaaatactatttaagatataagaggaaa
atactagcaaagcatgttaatgatatcttggatttcttgccatctgaataacagtagtccttgcaataatactgg
tagatttgccaataaaaacctatttcaacaagaagctaccaacagcagaccatctctaaagaggattcaaccatt
gcgcaacactctctccatatctcaagcatccacttaaatcttgctttcgtaccttataggagataaaagtctcta
agaagatgcagtaacacaatcaataacatcatcaacacaatcaacatatgaccaagcatagagttgtaggggaag
gatgagaaaatggacttcaacaatggtgctactgtgctactgaaaacaaagatgaaagagaccatatagaataat
gaacatcccctttcgtaacaacaaactagaatggaaaatgaacagcataatgctaaatactcagtgcttcaacaa
gctaaagggcgctcctgatcatgcagtcttataaactttctctgtcaggggagcacgagcagcgatctgaaaagg
actactgctgagaaagcagatatacctaagcaaattcaacatgaacaggagacagatcttcaagaaaatgaaagt
ggaaaagaataagcgaaggaatgaggacaaagaaacatggaaaaagcaccagtcctgactactcaaaacttcact
caatacttaactcccagtaatcatctctaccaccaaaagccaatgctagaattgtttctgtagccaacaaagaac
tttataaactaaacaaagcatcaattctaaccatcaataacaaccaataatccaataacccaaaatttcatttga
ttaccaaaaaacacaaaatccaacttagtgcttaggtcctttcttgataggaaaatcataatttaggcatctct
cataaaaaggccaactgatgatctgttccccaatctgtctgattattaagcaatagggatatcatttgatgccaa
atctgacaatgtcttgatttattctcgtaacttcaatggagctttcatttaaactcttagatggagtagcttgta
gtggtgatcttcaacaggacattctcatatttattactttcatccacagaatttccttcaaatacaggcttgctc
aaggccaaggaacaaggaagttgaccaaggtcattctcatcaacctaccaatgaagatataatgccttccggtct
caccataggtagaaaagtggataaacatatcattgtcatgctaactatagattagtcttaccaaataatcacct
aaattgccaaacaagaactgtaaacatgttgaggatgctggagaccttgaggaggaagcatatgaaacttgactt
gaacattatcatcaagagatttatcacttacatcccagtaagtgtaacaaaacaaaatctagtggcagagctgca
acagctattcagcaccacaggataatcatctgggaaacaggagcaaagtctcataatctatatcatagttctcgg
accaaatatggattaccaattaaatcatcatttagtagacctatattgcaggagataggatttcccatgagaagg
gaacaacattccatgtctcagagcacaagaaagaacaagtttattagccatctcattgtaccggtgagtaatacc
agccacttaatgtctatggtcatgatcattatagatgactaggaaaaaaggtgaaacatggtatatgagtctgca
atcacaagatagtcatgtccagtaggaggcatcagagaactggggtcctcaaccatagagaaaacagagaaccat
gcagtatcaatggcataagtattcataaacaggtacaaatgaagtcttagataatggaagatcaacaaggaacag
catgtcaaagtaatataaaactgataacaataggagaaaaaataggatcagaaagagcataataagatagaccta
caaagaagatatcaccgtgagaaatatgcaaacaacacatagcagcatataacgtatgtactctttaagaaggat
ataacatctgaaaaagcacaaatagttgatcaaaatccaaacttaatccactaaacctgaggatggacaaaacat
gtaaagccgagaaaagaaaaaaaagagagatacgagtggtaactagaaaagagattctaagaaacaggactatag
cgacattcaaggtgcaattaatacatcacattagaaaaggaaactttgggttcaaaagtttagactgaagcgcgg
gggggagcaggggggtgtgggtggaggttgggcgggcgtacctcaagaacgtgaagcctttagagattgatcgtt
ttatgctcatcgataccaaatacagctttatgaaggtcattgcgaactagaaatatcaaattttcatgcatctaa
cacatctgaaattcatgattacaagtctatacaactgattaccaacacagatacaacaaatgtctacttccacat
ctggatggaataggtggagtcaggacatttatttgaatgcagcagattagcaggtggagatacaaaataattcca
ccaaggcacaaacatcacaaagggctgactcatgaagcagttcatagcctaaaaaaggtgtcacaaaatactata
ataagcaaaaccataaataactcaaaatatatcatctaacttcttctccgagaattaccacctaacatcaaaatc
ctttagctaagaaggcaaaaaagccaaaaatcaaaaagagaccgtcaaactgaaacacttatttcactttccat
gccaagttgtgtttgtctagcacacattggcaataatctcctttataaagttctcgctttatgccatctgctcgg
acccatccataactttcttagttgcacaaagaaagctcatactccatgccttcagtcctatgaaaatagtaaaat
atgaaaatgacaaggaataccacatattgtacattggcatggatgaatccataatatcttccattcacaattacc
atgttcagaagctatgcccaattttgtcaaattagagaaccagaaggatgggggggggtgtggaggagattgag
ggagccaggagggaggaaggggggggggagtgagggagatttgtcgacaacaatatgcttatcaatgtgaatattt
tttagaaacttttcattagctcacccaaaagtaataagtgaagcaatttcactaaatgattccttgcataagcat
ttggcatcacatcctcacccaacctatttttgccatgataatataagaaactaaagcattgatttaaataatgga
acataagctgtatttggtcacccatgctaaagtaaataaatggttcattttcatggcatatattggaagtattg
actatgaaattgaatttccatcccacaatcaaaccaattggtttctgaaaacataattaccgaggtaacaaagta
tagaaccttgttttagtaatcatacattctcatacgtttcaaattgatcatagaattctatcaattaaaatttct
gaatggttttagcaaggtaaccttgaacaactttcaaccataggctagaatctaagttgagaggtaccatcgagt
ccctagacatcatctttaccaaaaagttcaaaagttttggcatttgtaggtcatgatagaaacatcttgattcta
attaaatgcattggcatgttaaggcttttaaggaaatagacgtac**catggaatttcttgcaagttatgcagttgc**
**cacctcaagcttgttacgtcctcctgcattgg**ctagtgaaaacaatgtatagtttagtccattgtaagtatagat
tcaacttgaccgataattaccatgatcataaaactgtaaaccccattaagttctttattgttgaatatattatgc
ataggtgttgcaaaaactccaaaaaataattcctcatgtaggttgatagtttatgagaaattgaggatgaagggc
tcttgtgctattttatttctaaatgtccatgagaatgcatctcaacaaataatcttccaaaagtcatattctct
tgcccatgcacagtcatcagcatctgcttatctttgtataaccacaattagtaaacgttttcagaaatataaag
ccatcatttatgttatctgaaattttgcatattatcatcgattcatgcacatgataaaacatagcaaaacaataa

agatgctctcaattgcttattatatatattttactagctccacttaactgactaaaaagggtggcctagtgcatg
aggctcccattactacggggtttggggacggtgagacatatgtagccttatccccacatgcagagaggctgtttc
tatgtttaaacccatgatacccaagtcacaaatgcacaatagtctatgtaactgacaattactatcatatttact
tggtacattatttcaaaagaaaacaacaatgaggcaattgattcaaatatagcaaaataacagtcactctaaatt
gttcattataattgttttgctggttatctatacttgattgctgcaaagaatttacatggtgcatcatttcaaaga
aaaactgatcctcaattgagaaaaattgatcctcatttgatgagattagaatgcaatactaccacatttccacaa
ctttccattgccacagaaaaccaagtgttggttatcacaggcagggcaggttagaatccaactatggaattagag
gcctcaccaactaggtggaaatataccctacgcaaagatgggaagagatggcacaatgtcatggccaagaaagta
tggaagcttaacatttttttatagagtgagaaaatttatatatcacaatcttgcacatgtttgaaagattacatt
gcattgaattcatgtacttcctagttgcagaacagttacatatctgctgatatgaatatttgtggtggctgacta
atgaatttagtatggcaaattgccccaaatctaaggaagattatagtttgcatcctctctcaatccatttatctc
gaaggtgcaagagaaactcgagcatgttctcattaatccaatggagagaaaagaatagctttgtacgttgttatc
ttgaaatgttactgtataaacattatagcatgaggttaaggctaaactacactaccaagcagcccaaggaagaag
tgtatattcattctctcaatgcaaatcccattcacaatgaggatgatctgaagatcaactagctcaagaatcatc
gaaaccaaagacaggccaatattagacaaaccaagtgatatgcgcaggcttccttactttttttcctcacgcatg
ccaataaattggaccttcatcaatgggctaaggagaagatgatggactcaaaggctgatcaagcacaaggagtaa
gagagacaaggctatgaaaatgggtagtctaagatggatccagatgatcaatcgaaagaactaaatgttgggctt
aggagactatcatggaatccaggatgaggaataaagtttgtctacttactaggggcagtcggtgcaccaaaacaa
gaagttagtcaatgctagtcaagggaagtactcaaataagtaaggctaaacatgatatcaaggctactcgatcag
tatggctagaaatttttttttcaaaaaaaaggtaaggatagcattaaaggaagacactgatagcttctttacaaa
caagaagttagcaaggctactcgatcagtatgcatgatgaatgttagacatactgcaagtccaaattttaggcca
acatggttcttacattaagatttgcttgctgtggtcaacatgatcctaaatgtgcatctcttgctgtaatattc
ttgcttcaatgtgagtacatctataaataatgaaatagaaatttttgtcccaacaataattgccaagagtcagtaa
ataacgccatctgacagtagccatgctactactacatgatagaaaaggagttactcagcaatttagtgtgtgaa
cttgacaactattgagtgtgatattgacatgcaatgatgttatgcatttaacctatagatgattatgtataaaat
gaacatacaaggaaacaaataaattaactgttgctttagagaatttctataacattacaagaagagtgtccaaca
gtacatgtcccaagcaaaaggcatcacctaatttataaggttttcttgcttgctaaatcaagatatgagccatca
aagtcccaatatgtttacaactagctctcaaaatggcaatagttttctatgaggtcatctttaattgatctagac
aactaagtcttaaaatcactccataccagtacagataactgataatcaccaaaacttgcaagcttaagtaaaaca
ggtttcaaataaatttaattaaagcaaaaaaaataagcacatagaactacccactcaatgaattttatgccaata
cgctaactcataaaagccaacctatggctaaagaattctatcaagaacaaagcaatcagcccataatagatctat
taaggtaccctagttgttgctcaacattgtggtacaaagacaacactcacagtattagattaagaacagataaac
caaaaactgtttttgtgcgtgtatatgtgacacacagagagagagagagagagaaagagcaaagtcgtgatataggg
gttgaaaactattagcctattacagatttggaaactcttaataaagcaagttcagccattctttcttcagagagg
ggggagatggacttaggcattaacttaagccgcaccaaaatcataaccagaagaagcaaaagaaataaatcaaat
caataataacgaaacacaatattttcaagaaccgaatttaaatctcgcgcagcacatctgaattcaagttataca
tgaaatcctctttac**ctgagatcccc**tccccaggccaaaaaaattccaccctcagatcctctccataactctccct
ctccctctctcttccatttctcaaactcaaactgacctaccctccggccacaatccgacagcaaataacgcataa
ccccaagaaccgatccaacaagaaggaaatgcgagagcaacgacgtagcaagagttctaaaggaaggccaagaga
gagattgaatccaaggagcggggaagaatcgagggggaaagacgatggaacaccaaaatatctcggccgtgtcat
gtgttcccaagggtttaaataaatgggggagagaagcggctgatctctatccacgtcgcccacccctatttcttat
atccctcgaaattttgaaaccgaaacgacacacacatcaaagtatttgtccacggaatcgataaagtgtaacct
aggagtggtcccgagtgacagctggtctattaggtctcttccacgtggttcatgtcgccagggaggggttggggc
tggaaaatatctgggcgacgccgtgatggtggacgattcctgcggagtattataatcaaagtggggagcggagcc
ggtgcactcatttcgcagctttccctttttttaagaatacgtgttggccgttgaacaggtagtagcgcgtgccc
cactgcccgctttccacgtggacagagaggcccgacgctggacccccgacgtgccaacttaggtcgtcgacacgt
gggtgttgtcggcggtgggaacgtgggttaaattaagtttgtaccattaggtacgttaatcccccttttatcct
tttaatcttgattacgatcatttttatcgtcttccgtttggatagccgtacgattggaggatggctgggttcttt
tgagggcgagcgtggtccccagccatgcctcaacgtgcttaacggagcaaggaatatatattaaattcccacttg
cacggaaaagcttcactttagaccaagtgcatcgtaccaagtaccggcgatttttttttttatatatatagaaata
ccaggtaccggagattgccagcaaaacggactaagttcattgtaccaagtgccggacattgcccgcaaaacaca
acggtcctcttcttttatgattcattcaaaagcttacacaaaaaaaggaaaaaatcttaattccatgccagttag
cttggctctatccatcctttccttcattaatcaaactaaattcgtattggttataaatagcgtagttcgtttta
gaggttgtttggttcgatgcagatgcctctaattagctcaagttatctgaaactttagttgctgataattgagtt
tcgtccagctgcaactaaaggttggtgctagatattttagagccatggtttttctattagaaaggttgcgtatag
gagatgtaagtaatattttagcgacaaagcaacctaatttttagtaaaataatccaaagtaaatttttaataaaa
gtaactaatttgtaagttaagaactccaactaagacacatattgaggattaatcttcggtctcaacatcactgtc
aatgtcaccaaacgcatgctgtatctaattgatgttgtaatcagacatgcacatcatgtgaaagcaattagttct
tataaaagaaaacatgaagtaaattaatccaacttggaagttaaaaactcagatcgatgcaagtgctgagaatgg
caatctttggcctgcctgtcattctcaattttgtttagactctagaagattttattaatgcttagtgctccact
tgttaattgttatgtcttgtttctaagaaaattaaattaaattagaaaaaaaatatagagatgttcctttaagtt
tggattatttatatttaaatcctaaaaatttacatcaaaatttcaaaattgttatgatatgtcccggccgtctat
ttttattaaaaaaaaagttagctaaatttttataaaatctcataaatgtactttccattaacttagcgag

>EoAGL17_1
ATGGGGAGGGGAAAGATTGTGATCCGGAGGATTGACAACTCCACCAGCCGCCAGGTGACGTTCTCGAAGCGGAGG
AACGGGCTATTCAAGAAGGCCAAGGAGCTCGCCATCCTCTGCGACGCCGAGGTCGGCCTCGCGATCTTCTCCAGC
ACCGGCAGGCTTTACGATTTTGCCAGCACCAGCATGAAGTCAGTGATAGAACGCTACAACAAGACCAAAGAGGAG
AATCATCTAGTTATGAATGCAAGTTCAGAGACTAAGTTTTGGCAAAGGGAGGCAGCAAGCTTGAGGCAGCAACTG
CATAACTTGCAAGAAAGTCACCGGCAGTTGTTGGGAGAAGAGCTTTCTGGCCTAAGTGTGAAACATCTACAAAAT
CTAGAGAACCAACTGGAGATGAGCTTACGTGGTATCCGAATGAAGAAGGACCAACTTTTGATTGATGAAATTCAA
GAACTAAACCGAAAGGGGAACCTTATTCACCAAGAAAATATCGAACTATACAAGAAGGTTTATGCAATGAGGGGT
GTAAATGGCATTGATAAAGGTCCGATTACTCCATATGGTTTTAATGTCACTGAGGATGCAAACATATCCATTCAT
CTTGAACTCAGCCAGCCACAACTGCAAACAGATGCAACGTAA

>eol-miR444
GGGGATCTCAGCCAATGCAGGAGGACGTAACAAGCTTGAGGTGGCAACTGCATAACTTGCAAGAAATTCCATGGG
TGAAGTGAGCATACCGGTGACTTTCTTGCAAGTTATGCAGTTGCTGCCTCAAGCTTGCTGCCTCCCTTTGCCAAA
ACTAGTGAGTACAAGGACAAAGTTTTAAGCCCATGCA

**Position of mature miRNAs based on comparison with osa-MIR444d and not based on experimental data!**

Stem-loop structure

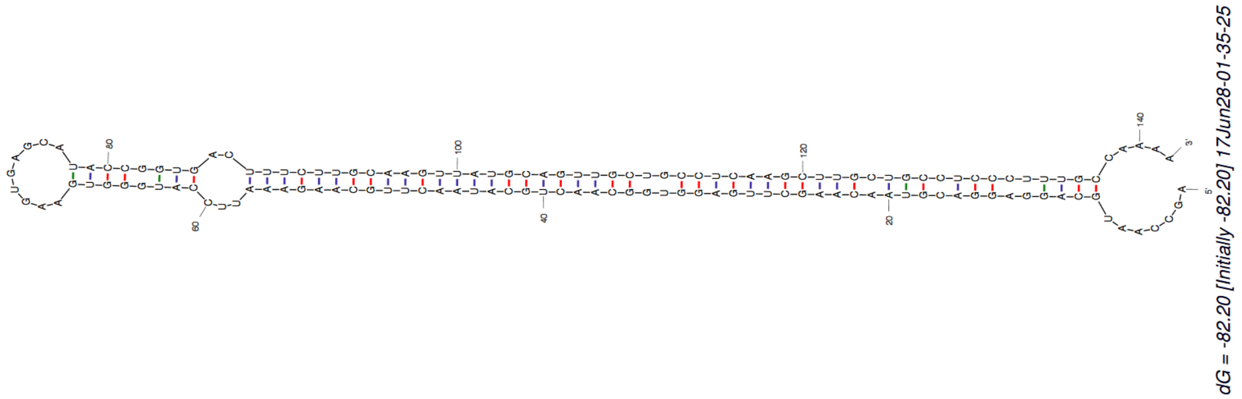


*EoAGL17_2*

>contig_KE504985
aaaatagttgcttcattctagtttagtgagctagcttaagatacatgttgtcattaccactctacagtagttttt
ggagaacaaatctgtcacattttatatacctcttgctggtagttactgagcagtagtcagtagatagtgagtga
tgcttttgtgcaaattttttttttctcactttagttggtgttattcctttgtaactttaatttagtaggtgggt
ttcctgtagctgttatcagcaactattcagatgctatgagaagacatgtctccatgtatgcaagtaattttctta
aacagagatacctgtcatattattagtagtcctagttggaagtctgcttgacagagtaaagtaactgttctttgg
atcatgtcacaccacatataaatgagtgctgagtattgttcttgctcctgatcatacattgatgttattgctgat
aacaaaccatagaagccttgacagctaggccgatggtctctaaagatgatgcacattgatcacttgtcatcttct
tactgcaattgaagtctgttgcaagaattgattatgttcatatcaaagtaataatattgaaatatataaaggttt
gatattcaaataattacctgagcttgggagctcttgcttgtgtgccatccgtttgctgttgaggttggcataact
cgagatgaataggtgcatgtgagttctgagtcatagtaagaccatatggaataatggtatccttactagctcctc
catttaactcatttgttgcacacacctgcaatgcaaattccgtgcatagcaatttataataataatctattattt
ttcagggattgaacaagatcttaaatgtcattggcatagggtggtgctaaagagtggttagcatattacagctgt
gttgacccatatgtcatgtgccaaaacaaaaaaaatattaagaaaggacttaaaaatcaaaaactttttttctaac
catgtctaaacgaaagtattaaaaaggattaaatgggttttgttttgaactagtgctagtgttgatattggctcg
gcagggtagccaaaccagcatagactagcacaaaaattcttgtgattgaattaatctagttttataccaaatcaa
tacatattttatatagagcatggcatgtgcgaagcatctggcggtggaaacatggagtttatagcttcatgagtg
ctagactctggagtttggattatagagcaaagtagatatcttgtatctcaaactcaatacttcttacgtaactc
catgttttcttgacgtatgaggtttacctgcttgtagagttcaatgttttcgttgtgaatgaggtacccctgtaa
caaattctagaaatttaggcacttgttaacatttaaaaatgatttaattgactattatatattgataaacctttc
ggttcaactcttgaatttcatcagttagaatttgctccctacaacagagaaaataaacacatcaaatagagaaaag
atggataaggattaataacaacatttggtgaggtcataacctttttctttcggacagaatggagactcatttcaag
ttggttctctagattttgtagatctttgatatttaagccagacaactcttctcccatcaattgcctacaaccatc

aattaaaatttgaatatgatatatcttaaatttatttaatagtgtaataatcaacatttttccacccttgttca
ccagtaggatttattcatttttttttgtggattttcaaatggctcttgtaatagcgtaaaagatagtgcatggtat
tctgtaccggcctgaaccgatcgagacatcccataccatatcataccgatggggaaccagttcggttcaggctag
ttttcagaaaatggtctgaatcaaatcgaatcggttagtttggtacagtatcagcccgaacccacccgatatcgag
cgattcggtacggtagtggttcggtccggttcggtttgatttttttctttttttctgaccattgggtgggatttt
gttgaattttttattatcggtacggtatggtatggcaccgtactaaccaggaacccatatgggcctcggtaccggt
tcctgaaatcttgaggtagtgtgatggcattgagcaaatgtaggctaaagatctagtatgattaattataaaaat
cttgttattggtatggctgcaacttgcaggactagttagtttatgtttataattatgtatagtctagacacttat
ttttgcctttttttgagctaaggtaggagttaatggttttgtggccgagatcataaatgaaactattgttctaaa
acatatttgacaccaaggttcacaatattagtatcaaagccgctcgtatcggtacgatgttggtatgcacagtat
ggggtttggtccgatgtactgatactcgatatgcttccagtatcgagcatcggttcaatactaatatggtatgat
acacccagtagttggtttgacccatacatgttgacccatatgtctagtcacccattagcatatacaggtcagttt
catgctaggactaggagtttctattttagttgcttaaaaaccatcaaaattgctaactttgatctagttcaatg
gctagccattggcatcaagaattaaagtatataaaaccctcaagctttgatttcttttgtcttctttattctatc
atgcttctcttactcccatcgcttctatgaccatcttgcactgccttagcagtcttttcatcatctctctctctc
tctctctccgatttcttatgccttccatcatccatcccactctgcaacaattttgagcattgactagtactagtc
gatatggggcgtatagtatgagcttcctaaccatattgatccgatacaaatgctcggttcaccccattccaatac
atggtataattttgtgctacctccaatatcatacaatgtgcctatgcgtggacataccatattgatctaaccata
ttggtaatatttcggtatgagttatagtatcgaggtttaaaaatttgatgctaggcttatgactaacattgtaaa
tgggtcaagtcaccccgtgattagagttgatcatggataggcctcgggcctaaactatattcatttttagctggt
tttaggccgggcctatttaaaatttgacctttttgtgcctaagcttggcccatgatcatctccacccatgatcca
attcaacctgacctggtgacgccagacctaacccattcttgaagacctgtggggccaagttgggttcaaatatta
gatttgttccacattttgagtcaggtctagatttcataaactttgacttgatcgactcgacctgatctatacact
ttatgagtttgttttggattttcatattatttaacaacccaacctgacttaactcgaattgaccggaaaaaagga
ttccaagtcatgtcttcttttcctctcttaagagagaagtgctcaaactaatctcactacattaagaaaagctag
ctattcgggttaaacttgttgaagcatattgcagttctggagaaacattaattcatcgacttttgtccatgttag
gaaacgattttatggcagtcaaaattcagtttgatttagaagtaagaacaaaatttttcttcatgaaatcttgtta
gtaacaaatgtagacttggacccaacttgaaccggacctagacttgaatatttgggttgatatgggtcatgcata
acctcaaccttatccaaactactcgtcaggtctaaaatatttccatgaacctgacccaacctgatcatctattgg
gttgggtttgagtccaatatcaagacccagacatgaaatcaagtcaaggtcaagcatgcctcaatccgacccatc
ccatttgcagccctacttgtgactgctaggttgggctagtgtctcaccatcaactcctttcctctacatcttggg
tcaaactcctctatgttcatcatagttgttgttgtcatcaagattttcataattggtgttgagatcacttaatca
tggtggaggagaatgaagagtgggttgagatggtggggtgaaaggagaggtgctgctcatgctaattttggggct
aatctgcttttgggtcaatgtctggagaggattggtaggagatagaggggggagggagattctaactaaagtgggc
ataaagattaaggaaggcttggagaaaggaggtttgagtcagtgggttgacttgttgaattgatgacttgcaaat
aggccaccaaaacaagtaggtgtttctgagctgaggtttaaactacttcatgaaactttcttttgtgttcatgat
catcattctaagtggcaaattggattgtggtattctagattgacaccttggcactactatgttccatgtcaatca
tgttatgtagactgctttccagaaaaattatgatcaaagaatattctaacagcttctctgcattaaatatataac
ccaaaattatctattaaagtttgtattcgtcatatctgagctatcatcaaactaaaatttggactaaatcctaaa
ttgcaatgctcgagatgaagaaaaaggtattcaggatttctaatgctcctatttaaaagattgttacgggtgctg
ctattggcaatatatgctatgagactgggaagttctggaatactgaagcagtcctctgagtcttgccttcacttt
ccatcagaacagctaggcagattttataggcaagataagatcgattctgattgcagtcctaatagtccaccttgg
tggattttgaagaaattttcacagtaatttggtctattcaaggggtgtttcacatattatgtattgtagggcatt
ctggtgatttgcaattttagatagtgaaattggagcagttcatgatgtcatagagtgtaattctcaacttgtaga
aactacataacagaaaagactggtcagaactgtggcttcttgttcattgatatagaatcatggtggcttggggtg
agacacaacttcagggatataatgaagcttctagatcagtgtaataatcccattccacaaatatgtaacttctga
gtataaaagctggtatataagactgtaactgttatcacatatcggtaacaatcattagtaaaataaaaaaatgtt
ttgaaaggattcatgaagtcaaatcacatagtgcatgcatcttgttgattattctaagtgcacatttcatgtgta
ccagcaggaagtaaggtctgttttatgatgatttctgatatttttgatatcatcgtatcaatgtattgccttct
agatacttttaaggaatagttacgcagaatctatcaccacggggagaaaatgtaagctaatatatgattcgagaa
actttggaatacgttgggattgaaagtgaatgaatctaaatgctgattgactgatactaaaaagaaaacaaaagc
tgagaaatttagaaaaggaagtgaaacatttttttgaaatatttggaaaaaaaatcataaaaggcactataaattc
aataaatgtgtgtttcattaagttgctataaatttgggtaaaaaccttaattgagaatgattggaatttttttga
atgttggagaattttgtacttggttgcttttagcgcatttactgtttgctaaatcactcaagccacagattgaca
cttgctcctgcccaagatttatgattagattatatgaaatattctgaggttcaattttgataattgacgtgtata
ataattttctatgaagttgataagtacaggttaattgcaatgatataggcagttgtcagtgtatttgtgaactgt
gggcctgtggaccaattctttatccagatattaaagtgagaaaccaattctttagaggtagtgtcactagtttta
tgttgtctcaatcttgacatactattagaggttcagtaacctgctcttttgtcaaatggcatcattaatgatttg
gtcaatgcattggaataagctacactgcagatacaaggtgccatatcagcaagaaaatgctggtttgtatgatta
atcttaagaaatacatttactttcaggatatggtcagctagagaactagtggttttatacttcatttctaatca
tttatgtaaatttgatgtttttgctagtttgtatattattttccttatagtgatcttgacactctagatcctggg
gaataagtgaataatcattaaattttaccatgagggaatttgttatatgtgaatggcttgaaataatgagacatg
ttgaataatgctaatttgttatagcaaaaatatatggttatttatttaaatcttctttatgctataaatataatg

gaatatttccagtggctagttgggcgtaggaacaatctttagatgtaaacaaggttaagagatggcaccgatgat
tttcttgtaagttctgcaattgttgcctcaagcttgctgcctccctttgccaaaactagtaagaacaaagaatat
aatttatttcttaaaatacctttttagataatggtaagtcacaatgtaaaaagaacacaaactaaaatctacaa
ggaatattccgtaaagccagcataaaaaatgtagtatgagcaatgcatagtttgttgaataaaagctccaatgaa
tttattaatgttttggggtataacttagagttgatattttatgaggagatgctttctcaatcttctaaatattct
gcatgtctatgactatgcatcatttatatttctttactggtcaaaagtctcttgtgttcttgcatgcttatatat
tgtatctaataattatttgttaacaataattaatagctttttctatagaccgagacctcttggtattgggtgtac
catactgacccagtaaggtactattattgggtgctattatggtacaagggcttgatatggtagtgtccatagtac
agctattatcttgcctccaacaatgtaccatgtgtcggtagggatggactgtaatgacctgacctatactggtaa
gctactggtacaggatctgatgctgagactgaaaatcttggaaaatataatttagatgtgatgttatttgctgtt
cttctactttcttacttttaaggtatcgtatgtattatgcggaacattgcttaataaaattgattcaaagtcat
atgctcaaatcatcaataatgattttacaacttattgacataagaattatcatttggtttcaatgacttaagttc
atctatatatgattatatagatcaaggtagtcaaaggtgaaaggcgtccttaaggcgcttggggggtctagcacc
ttgagatgagaggcgctagaaaagcactcatcagagtgaattaaggtgctaaaattgaaaaaaaataaaaatatg
atatataaataaataacaactgaagataactattgtaacatgaattaccatataattaacaatttaagcctaaaa
catatataaaacagtaaaatgctcgactagttcaagataaagtataagataacaagccatagacagctctaaatt
tatgaatgaaatagaaaatataataagtcctaaaatttatcaatggagtctcataaataaacagaaaacataaac
aaccttaaagtggaatcatataaatgaaacaaaaaagataaacagtcctaaagcctaaaattcatcaatggaatc
atcagatttagttctacttgcatagttgtcatcactttcatcttctcctccttcattggatctatatccttcaac
atctttttctgtctcatctgaattagcctctatttcttcaaaactaactttgccatactaattgaccattttacc
ctctgttagtaatagagctgagaagttaaatggatggcctttaggtttttatattttcaaccattaaaaaaataa
tcatagagttcactgaggcgccaaaggcaagcacctcgcctcactaagaaaggccctaaaaggtgccaaaggcat
gcgcctgagtgaaggtgcatcatctgaagcatcaaggggtgatgactcctagcgtcttgcttgagcgcctgaggt
gcgccttttggcgcctttgacttcactgatttagatcgaagcaagttcaatctcatgaatcaagccatatgttca
taagcttggactttgtaaatgtgatgtgttgcatcaagtattgagacctctacgatcctttcttagaataattga
atggtcctgggcctcctggctagtaggagcgttctttacagattggaccatatatgccagtcaataagtgtcttc
acagacacatgcattcatttcttatattcaacgatccatattttgttaaaaaggttcagcagtttatcagcttga
tccatttagcatgcttatgtattagaaaataaggaaatgagtgactactttatacataagaaagatttttgctttt
tatttgcataaggatcaacttggtcgaattaatgagtgcttctaattctacttctctactatatataatgctatg
cctgaaattatcacatgtctgtgatggtctatgaattcagtcccattttattaaataacctctcttgattttatc
cctctttttgattaatacttctaagggacgcaataaatgtgaactatatttatcaattcacaaatataaaaattg
atggtagaatgtctaatataaaatatttcattagatgatggaatggtaaccatctatcaaatactcccttagtct
tgcacctcaagcttcttgctgcattaaataaagatctctctcctctttttttttttttgctgtttgaggtaacag
aggcttagtgccattttatccctaatctcaacttatcccacttaagctggaacctaggcttctgagcatactgag
cctctgcttgcagaattagaacttttaaatggggggctacaatgagtgcagatgccactcagaatcaaacccctga
tctctttgtctaagaggtgagaggtgatccaagctaacacttgatagtccattaaataaaaatcttatgtaaatg
aatcctaaccccaaaataaattacagattgattggttcattgaatagccctacgcccctaccaatgtcgtacatt
gcatgtgaagggatcactacatcatcttgtattggtcacatgcaatagataaccaataaatctcataagcatctt
aaggtattcgttccttccaaattaggtgcatacatcatacatgtcaattgtcaaatgtaattgttaatatgtcac
attactgaatcaaatttcatgtattaaagcatcatctattgtgcttgtgtcattgaggggctatcatgttaccat
tcttgctgagtaccagcatggagaggtacaatatccataaaagatccatgaataacatactgaaatgtgcttttt
tatttaattcttttcgcagttaaccattttcaattgattgttgctactgttcaaacaagtgtaataatgaatata
taattatccttcataattgtgctaattgtacgctggatgcttgcatctgtctccaaaaaatcatgttggaaacat
atctaattcagatatgtgttaggcacttggatactaatgaagttttacggttgcaacttttttaagagttgaactc
ttccaacaatgagttaccatccatattcatgttagaagcgagtattcttcctttcattgattgcccaaggacata
tgtgttgaaataatttggaaatgttattattcttatattttaggttaattggacggtttgaatgacaaacatgt
tctaaagatcttaacaagcacaatcttcttaacatttatttttaatatttaaacaaagaatattgaagatatata
tttagttaaatttataatttatactaagtatatttgtacctccttcttctctcttgctgtgttagacacttggat
atgcattcaaatccaattctcatattcatgtccatgcaacactagcttaacatttttttcaagtagtttatcata
caagctaatgctagaagcttagaaccacgtgcacattaaaaataagatcacaattcatggtgatggttgtggctg
gacatatatatattttcatgcattaataatatgttgcaattgttgtttacctactgttgacactataagaaactc
atgtattgaaggtgcaattgcatgtccatcatacatagtccatataataatgttacagatatgatccatcatggt
gaatttccatcattactagcaacaaatttggtatttgagtcaatgtgcttttgatattgattttggctaggacgt
ccctattcctagatattgcttccggattagtgagaggccttgatggatctagtgcttatcatcgaggctagcttc
ctaaattgtctatcacttgcatgcaccctatactccatttcgctttcctttgaccaaaagggagaaaaaagact
tttggctagtatcatgataatatcctccaagatttcctatattggttattgggtgctttactagttgggaatcag
gttggtatggtatggtattgacatgacattcaaaagcacacttatttggtgctagtttgggttttaacccttta
tttttatatgatttatttatttggtttttagtgttgtttaaaggtttgaaaacttggatttagggtttattttta
tttgctatatttttcaaaaaatttgtcctgtagtgggaaaaaaagtagtggtcctggaacaccatacatcatgtg
cattatgcaaaatttataaattaaaaaaaacataatcaataaactttggcttgaaaatgtatattactggtccct
catggagtgtatgcagctcgcaaacattgagatcttcataatttggttgggactttgattccaacccagatgttg
tgcctacacttgcatctgccaatccctttaacacggatagccactcgaatacagatccttcgtctgatattaggc
tggccatcctacctcaaggaagttcaaggtagcaaactagccaatggctgcaaataccctctcttgaagctttcc

cagacttccacctctcccatcttttactgaattcaaagaagaggagaaagagttagaaaggctctttctcctctc
tttactcattctttgacccaaaactagtatgtgccagccatgattgagctgaaccaagcagaaccaggtagctct
gttggttcaaaccaattctacacttgctttaaatcaataggaggaacatcctggtaccattctggttcctttctg
tatgtcccaaacctcacagtttgagatgttttgacagttattggtcaaaaagcaggacttggagattgaagattc
tcctaggctttatagtaaatgtattagttggatggcactgtacaagtgagcaaacatttttttaataatttataa
cttataaaggtcaccttgaagaagtgtgatgaagaatgtggtgtgacatcgtgactgtcaaattcttccaggtta
gctgtggaaccaggaaagataggttacatataatgattaagataacacctattttgtattgctagatgattaaag
gtcactcttgattcttaatgggatatccaaccaaacataaacaagaatatgcaaaagagataacaagaagctgac
tcctcagttgagtgaagacttgcatcaaattttgccaattgaagggaataaaactgctccagaatctaaagaaat
ccatatatttataaaacttgggataaagaaggctcaaatccaaaagatcatgggagaaggacaagaaagcaatca
aatgcatgaaagctaatgatactcagtgattgacatgagcactttgaagctatcgtgtttgatgcaggtgaacag
tcgttacatgatactaatccatgctagaaatgaagctagatctagagggtttcaatgtccaaaaagaggcttaaa
tggcataattgaagataaagataattttcattaggaacttaatattaggtcatttggttgactaccattttctgt
ttagcccttaatagtattttatgttttgtcttgcacataatattaggtcatttggtttgacaccttgtttccaca
catgtgcaccaatgttttgtacacagttataactgttttgcacactgttatgtttaagtgtacagggccacggac
taaacattaaatgaccattgaaaaatcaagggttagaaataatggactggttatcaactacccaattttcatatc
aatataaaccaagcacgatatgaatcatgcaaacaagcagttgaagtttttcttcattctatgttggggcccact
tttaggatttgtactagtgatctggtttaatgattaactcagtttttgcaactagtctgttcatgatgagcacga
tacatgatgattactcccaattctatggaacatgttggtcatgctttggcgcatattcttataaataggtgtgca
aacaagccaattactgaacttctcagatgcagcttatttatgaaaggtttttggagcttgaacaatcagtattca
agtctgccctatttctcaattcagtattcaaatcttgttgttgagccaaggttgtacttccaagtccagtgttga
ggcttggctcttttatgagttaagcctttagatgatgtttcagcatagagtatttgcaaccgcaatcggaatcaa
ataggaatcgaaatggtcatatccttctatatgtttggttggtgactgaattggaatgggctttgaatactaagg
aagattagggttggatattaggggattagggcattcccatcctccctggaatcagaatgaaaatggaactctctc
taaccaattggctagaatgagagtgactcattcccattccagaattcaactccctccaaccgaacatgcttttgg
ttcatgtagttgataaagctccatggatcttgttagctaacctaatagaattgttcatgaatagtttgctttctc
tgaatctacttgtaaattgtttcaatcatggtctaccatactatcctgtaccacctggtacagagcgcaccatac
tgtacctgtctcctaccttactgaactgcgtaccttcgccataataagatggtattagtatggggtctcatactg
aaatggtggacctttgttttatgattttggaggccatctttgtgattgctttttttctttggaaaaaagaatgaa
aatagggcttcaaggatattgtctctgtggagtttatttctactttaatatgaaaatatttgcaacaataacca
tagttgcgaagtttacaaaatatttgtgacagatttatgaatcatgctcactattgattcttataaggatttgtt
acagtgcttctcaacttccatactcttagttttacttcagcaatactagctattaattacaaaagcaaagggcca
agatagctcaacggttaatctgacatcattggttataccaacccaaattatgaatggcgagataaattgtcaaac
ttcattggttatgttttctaatggtgcaatcgagccaagtccaatcaaacagatagaggctcgagcttaactcga
cttaaaacactcaagctcaaaactcaacttaagattggttgggtcttaatattcaagctcaaacttgactcgaac
gaaaaggctatgctcaagattgacttgacccgtctcgaatatcaaccgagcaatacttgagtttgaattcgaggt
caaatatgagcattggtcataacttaaaatatagttaaaagttagaagttggatcagattaaaagatgtaaatag
tatttttttaaatattaattaataatatatattataataattataatattatgaaaaatatatgttactcgatta
ggctcgtgagcctttaagattgagtattttattactcgagctcaacttaaaaaactgttcgagctacttgagctt
gatttgaactcgataatagttgagtggaacttagactcgagtcgaggctgagcaactcgtgagctgtttggctca
tttgcaccattattttttataaatggagtgatcgctatcatctttcttatgccatagtttgagattgtgacaat
acggacacaagcaaggatccactggttttgaatgccaccaccaatgccgatcgcaatggagaaggggtcatggtt
gacagtcacggacaactgcatcctcactaagtacctcaagagctaccccaaagtcctcaatctcaccctattaga
gttttgctttctaacaagttcccctatcaaggccctattagaggtggacctattggtgctccttgtcaacaagga
ggccatgatttgctgggtgttgttggcctatagagatcaatgcatttcggattggtgatggcagcgaggcagcaa
cgacaagggaagtccaagaggcgatcaaggtcgatgatatggtgtttggggtttttgacaaggagagggagatgg
ggaggtggttggggtggagggatgggattggagaggagttttattggagctagcagctagtgaggctaagaggat
tagaggggtttaggacaaaaatgagttttagggtctatttggatgattgagcccaaaatcttataggaaaaatg
gctttgggcctttacaaccacctaaataaagctgggttaggcctattttataaatacacaaagaaccactgtag
tgggccgggccgaatcccatagggaggaaaaaaaagacctattgagtctcttttccttcctatgggattcagac
cggcccactgcaatggatttctaggtatgtataaagacaggttgggctaggcctcatttaggtggttgtattggc
ccaaacctatgatttctatgggattttggccaattatctaaataggccctcaggatttttttttttaaagaaa
aaggtaatatatgccaagcagttgcctattatatcactttaaaccccttgcaaatcaatgaccataatctattc
tgacttttctaaaataaataaaaattatagtagcaaaccttcttataaatatgtggctacttcctccaatcaata
tattacttaaattctttctattgactttgatccatgtttttaggcattctcatctttgtgtcattcttggtgctc
aaaatttgatgagaatatctgtatatttcttgaaatttgttcaacatagtagaacgtttgtgtcttttggccatc
aactaccaaagtatcatcattactcagccaacaacagctgcattctgtcatctgttttggttagctctgttgttt
atgcatgacctgtgtagcaggaatcttctacttgaggtgctacacctgctttgcaatattggtggaaacttgttt
ctgctcctgctttcttattaaaaattttaagagaattgcttccccacacctgcatctgaatctgctccactccgg
tttcagtaactaagcttgtcactatgactcaattttcttccataacgccaatgtagcatttctccttgtttccat
aacacaaaaattgtgatgttttattttgcaaatgatctatatagttgattcccacctaagcagcaagaatcagta
ctttttagcacctgtttatgatgacagactaggatgaccatgattggccctgctacggtatagtatctgctatag
tccttgcacatgtttatgattgacagattttaagatgatcctgatgttggcctgccatagtatagcatctaccca

ctagcaccactgtcaatttccatcataaataccttgatcagaaattatctccctttgctattcatgttcatatga
atcttcatccttcagcgtgcacatttataagtcagcataaaccatatgtatatccttccgttgactttattcatt
agcctcactgttagtttggcccgcattcaccttagaacctcatcatcagccaatttaattagccaacctttctta
agcacttctatgatgctatgcttttacagatgtcagacaccgataatcgcaagtttgtaatgagtgtacccattt
tattctatttgtattccatcatttattggtaatccaccttttaccaactatttattactatacattttaaagctt
ttgtggttatgctacttcaaattgaactgtaatatacctgatctattcttgtgcagttctaattttccctactt
ttctggtacagggcatatatgctgcaaaacaagtgtcacaaattatttgtatgttaaagtacatgactataagag
tacatgatgcggcatcttgacaatttgcatatgttaagcaaaaagaaaggtgtactttcctacagtaattggtg
aatgccatacatttgcaaaaaggatgactttttacccaagaagaaatggcaaaacttatataaacatagtaatt
tgtacatctgtataccatcaaagaagttcatttgtcagtcggtgatgttgcataagaaggagcataccttgactg
aagttgcatttaaaacctgatgatgctcctcctttgtcctgttgtagcggtctattattgatttcatgctgatat
gatgagggactttatgttagtttttatcagagtagatgttgatacattttatggtttataggtgcttgaattaat
aatcaaatgagataatgaacatgaataaatatgagtgctgcctctcgaattagttatacacaggttgactttctt
ctgtcatccaaagtggtgacctcaatatttaactagcctcacaatggtctcattctggtttgtcatgtagcatct
gtccatccatagctattcgaccatctagctcctaattgattctgtatacctagaatacaaactagaatcatcaat
gcttctcccaactgattggcattgctttatgatttcccttctattactttcgttctctgtgcaagtttcttcaga
ttctctcatgactttgttccaaattgtttagcatcttcaattaactacaatatttttggttcacttatttatgat
ggggtttggaaaatagggtatttctttgctcatagatattttaagctagattaggctaaacaaattttaaacatg
cttggtggattcattgcatctcatagcaagggtactagaaattcaaaattccacgggcatcaagggctttctgta
gcttagcttgcattagccttatttaataataccatgcagtaaatatatagcaagaagttagctgtatggcaagaa
ttagtaatggtttcccaaaatgttagcaattactttcgctgcttttctacagctttacacttagtttcttggtag
aaacccatggattttgggcaaactgaatcatccgtggaccatgctctttcatgtaaaaatgcactcctatcaacg
gaagtagcatcacaagtttgtgtagagcatgatccaagtaaatgaggatttttgtttggtttttttgcccttcgtc
tgtgcttctcacagcatttcaatgttgatgatgggacgttcatctccacaaattatctgctatcccacatccgtt
gcagaacagctgaacacaaagtcttgccttcatgtttcttaaaaaacaataaaggaaagtcttgccctaatccac
gctatattgcaattaaatcacagatataagctggatcaatgtctagggagggtaagattttgagtatctcaggaa
gaataaatgtccacctgctgtctcgaatgattattctatcctataattaatatagaacgggcaaccacgttgtag
atttgcttgctagtatggcttgaaacttgtacgtagccataagctggatccacagctacccatccaaggtgatgc
aggagttactaattggatgcgttcctaatcagggtagcactttattccatcggaaaagaaaataataatattata
tagtttcaaactatgaaatttactaaccaaaatatatttaactctgagagtaatattgtttaaagaatataatgt
taatgacgttttaattaaatccttaaagtaacatattttagtttttatctttctgtttgttagctgtggtaatta
ttttctcttcaagatgattgattctccattaattacagatttaaactcttagaagttctctatctccaatcttaa
ctcatcctccagtaatttacaaattttaatattttaaaaatactttctcacaccgatccatcaagtccatgtatc
tctcaacttcccctgatttttgattttacacaattaagttatcaatattttatattatatttatattctatttaa
tataattattgcttttaatgatctatttgtttaattttgtatgcaaccatgcctgggttgctagtagtataataa
ataaagaggtgacaatttttaactcgggccatattaggaagtaattctttatgcactgcattcagtagagaaaaa
aatatactgcttcaccagattactcatagtgtaaccaataaggaaacaggcatttaatgtagcttagcatttta
ttaatttttttgatcaaattgcctttttctattgaatattttgacatctcaaataatgttacgacgtcctgttgc
tttttataggatacaacatgatatcatataatgcaacaggatatcataatattttgataattatttatgatgtc
ctgttaattttatgatatctcaataattatttataagatgcaacagaatgtcatagaatgtaacagaatattat
aatattatattataatatttattaattttataacattccaaaaaaatttatagcatgcaatagaatgttataat
attattatgatattttattggtttttatgatattctgataattatttacatgatgcaatataatatcataatatt
attatgacgtcctgttagttttatgacccccgataattatttatagaatgcgataggatgtcatagaatgtaa
catgatgtcataatattattatgatgtcatgttagttttatggcatccaaataattatttatgggatacaacaa
gatgttataggatgcaacagatgttatagatggaggatttttgtggagtgtccacatgttcaaaccacttcaaat
tgaatgccataatcagattatggcgtgagctcaattggattaaagtgggcacccaattggataggattggagaac
tcttgatcaacctggactctttggcgccaccttgcttgtgcttatccaatgttggcgccaacataggagagagca
tggggattcttnnnnnnnnnnnnnnnnnnnnnnnnnnnnnnnnnnnnnnnnnnnnnnnnnnnnnnnnnnn
nnnnnnnnnnnnnnnnnnnnnnnnnnnnnnnnnnnnnnnnnnnnnnnnnnnnnnnnnnnnnnnnnnnnnn
nnnnnnnnnnnnnnnnnnnnnnnnnnnnnnnnnnnnnnnnnnnnnnnnnnnnnnnnnnnnnnnnnnnnn
nnnnnnnnnnnnnnnnnnnnnnnnnnnnnnnnnnnnnnnnnnnnnnnnnnnnnnnnnnnnnnnnnnnnnn
nnnnnnnnnnnnnnnnnnnnnnnnnnnnnnnnnnnnnnnnnnnnnnnnnnnnnnnnnnnnnnnnnnnnnn
nnnnnnnnnnnnnnnnnnnnnnnnnnnnnnnnnnnnnnnnnnnnnnnnnnnnnnnnnnnnnnnnnnnnnn
nnnnnnnnnnnnnnnnnnnnnnnnnnnnnnnnnnnnnnnnnnnnnnnnnnnnnnnnnnnnnnnnnnnnnn
nnnnnnnnnnnnnnnnnnnnnnnnnnnnnnnnnnnnnnnnnnnnnnnnnnnnnnnnnnnnnnnnnnnnnn
nnnnnnnnnnnnnnnnnnnnnnnnnnnnnnnnnnnnnnnnnnnnnnnnnnnnnnnnnnnnnnnnnnnnnn
nnnnnnnnnnnnnnnnnnnnnnnnnnnnnnnnnnnnnnnnnnnnnnnnnnnnnnnnnnnnnnnnnnnnnn
nnnnnnnnnnnnnnnnnnnnnnnnnnnnnnnnnnnnnnnnnnnnnnnnnnnnnnnnnnnnnnnnnnnnnn
nnnnnnnnnnnnnnnnnnnnnnnnnnnnnnnnnnnnnnnnnnnnnnnnnnnnnnnnnnnnnnnnnnnnnn
nnnnnnnnnnnnnnnnnnnnnnnnnnnnnnnnnnnnnnnnnnnnnnnnnnnnnnnnnnnnnnnnnnnnnn
nnnnnnnnnnnnnnnnnnnnnnnnnnnnnnnnnnnnnnnnnnnnnnnnnnnnnnnnnnnnnnnnnnnnnn
nnnnnnnnnnnnnnnnnnnnnnnnnnnnnnnnnnnnnnnnnnnnnnnnnnnnnnnnnnnnnnnnnnnnnn
nnnnnnnnnnnnnnnnnnnnnnnnnnnnnnnnnnnnnnnnnnnnnnnnnnnnnnnnnnnnnnnnnnnnnn

nnnnnnnnnnnnnnnnnnnnnnnnnnnnnnnnnnnnnnnnnnnnnnnnnnnnnnnnnnnnnnnnnnnnnnnnnnn
nnnnnnnnnnnnnnnnnnnnnnnnnnnnnnnnnnnnnnnnnnnnnnnnnnnnnnnnnnnnnnnnnnnnnnnnnnn
nnnnnnnnnnnnnnnnnnnnnnnnnnnnnnnnnnnnnnnnnnnnnnnnnnnnnnnnnnnnnnnnnnnnnnnnnnn
nnnnnnnnnnnnnnnnnnnnnnnnnnnnnnnnnnnnnnnnnnnnnnnnnnnnnnnnnnnnnnnnnnnnnnnnnnn
nnnnnnnnnnnnnnnnnnnnnnnnnnnnnnnnnnnnnnnnnnnnnnnnnnnnnnnnnnnnnnnnnnnnnnnnnnn
nnnnnnnnnnnnnnnnnnnnnnnnnnnnnnnnnnnnnnnnnnnnnnnnnnnnnnnnnnnnnnnnnnnnnnnnnnn
nnnnnnnnnnnnnnnnnnnnnnnnnnnnnnnnnnnnnnnnnnnnnnnnnnnnnnnnnnnnnnnnnnnnnnnnnnn
nnnnnnnnnnnnnnnnnnnnnnnnnnnnnnnnnnnnnnnnnnnnnnnnnnnnnnnnnnnnnnnnnnnnnnnnnnn
nnnnnnnnnnnnnnnnnnnnnnnnnnnnnnnnnnnnnnnnnnnnnnnnnnnnnnnnnnnnnnnnnnnnnnnnnnn
nnnnnnnnnnnnnnnnnnnnnnnnnnnnnnnnnnnnnnnnnnnnnnnnnnnnnnnnnnnnnnnnnnnnnnnnnnn
nnnnnnnnnnnnnnnnnnnnnnnnnnnnnnnnnnnnnnnnnnnnnnnnnnnnnnnnnnnnnnnnnnnnnnnnnnn
nnnnnnnnnnnnnnnnnnnnnnnnnnnnnnnnnnnnnnnnnnnnnnnnnnnnnnnnnnnnnnnnnnnnnnnnnnn
nnnnnnnnnnnnnnnnnnnnnnnnnnnnnnnnnnnnnnnnnnnnnnnnnnnnnnnnnnnnnnnnnnnnnnnnnnn
nnnnnnnnnnnnnnnnnnnnnnnnnnnnnnnnnnnnnnnnnnnnnnnnnnnnnnnnnnnnnnnnnnnnnnnnnnn
nnnnnnnnnnnnnnnnnnnnnnnnnnnnnnnnnnnnnnnnnnnnnnnnnnnnnnnnnnnnnnnnnnnnnnnnnnn
nnnnnnnnnnnnnnnnnnnnnnnnnnnnnnnnnnnnnnnnnnnnnnnnnnnnnnnnnnnnnnnnnnnnnnnnnnn
nnnnnnnnnnnnnnnnnnnnnnnnnnnnnnnnnnnnnnnnnnnnnnnnnnnnnnnnnnnnnnnnnnnnnnnnnnn
nnnnnnnnnnnnnnnnnnnnnnnnnnnnnnnnnnnnnnnnnnnnnnnnnnnnnnnnnnnnnnnnnnnnnnnnnnn
nnnnnnnnnnnnnnnnnnnnnnnnnnnnnnnnnnnnnnnnnnnnnnnnnnnnnnnnnnnnnnnnnnnnnnnnnnn
nnnnnnnnnnnnnnnnnnnnnnnnnnnnnnnnnnnnnnnnnnnnnnnnnnnnnnnnnnnnnnnnnnnnnnnnnnn
nnnnnnnnnnnnnnnnnnnnnnnnnnnnnnnnnnnnnnnnnnnnnnnnnnnnnnnnnnnnnnnnnnnnnnnnnnn
nnnnnnnnnnnnnnnnnnnnnnnnnnnnnnnnnnnnnnnnnnnnnnnnnnnnnnnnnnnnnnnnnnnnnnnnnnn
nnnnnnnnnnnnnnnnnnnnnnnnnnnnnnnnnnnnnnnnnnnnnnnnnnnnnnnnnnnnnnnnnnnnnnnnnnn
nnnnnnnnnnnnnnnnnnnnnnnnnnnnnnnnnnnnnnnnnnnnnnnnnnnnnnnnnnnnnnnnnnnnnnnnnnn
nnnnnnnnnnnnnnnnnnnnnnnnnnnnnnnnnnnnnnnnnnnnnnnnnnnnnnnnnnnnnnnnnnnnnnnnnnn
nnnnnnnnnnnnnnnnnnnnnnnnnnnnnnnnnnnnnnnnnnnnnnnnnnnnnnnnnnnnnnnnnnnnnnnnnnn
nnnnnnnnnnnnnnnnnnnnnnnnnnnnnnnnnnnnnnnnnnnnnnnnnnnnnnnnnnnnnnnnnnnnnnnnnnn
nnnnnnnnnnnnnnnnnnnnnnnnnnnnnnnnnnnnnnnnnnnnnnnnnnnnnnnnnnnnnnnnnnnnnnnnnnn
nncgtaagataaaaatttaaatttattccgctccagatttggtgaaatctgaaagatcctacataggaaaaagg
cgatttttttttcaatagaaacaaagtagagtaaaaaaaaaaggcattttggtcaaaaaatttgaggaaaaaaaagtt
gagctagattaaatgtcaacctccttattaattgcactatgtggactaatccagtgaaacagtgcattttttttct
ttggtggtggtacaaaaaatttttcatttgcttttatttattaagcaatataattaataaatgaagattcaat
tagctcactattatgtgtccatgtaaaataaatgaactagacaccgactaacctattataaaatggatcaagtat
gcaccaacgatcctttgaccagcaactcataagtgatcggatccaaacccaaccaacacattgccacccataaac
tagttgcattgaaatattttaatttaaatgtagcgtcatggttccaacacacataatacagccaagtactttgaa
tacctgcaggtaactcatcaacttgattctaatatatgtttccataccctctcccaaggttttaagcatttttat
ggggctcaagaacagcttttctcagagaaacattgtttgagggtgtcggttagcttgctagtaaaagtttctgga
tgattgtgatagacctgaatttaaagcctatcattttgttttgagaaaaggaagggccaaaacccagctgtttta
ttaaatagataaaactatgacaagtatatgaagtgaaatagttagaaagagacttaaaactaaaatagatttctg
aggaaggatatgtctgttggtattgtctcgccatgcttgtccaatggaggaagagttcagtcacaagcttgacag
taagtcatggaggcttggtttgttgactgaagattcaactgttgttttctttcaaaagcatccaagaggtgataa
tggtaaaagagaggcaacttatcaggaaggcacgaaatagcagcgaaccgatggattcgatatcatcaggcttaa
attaattggaaccaaaatgttattgaaacgagcaccaaacttccatgacatgcatgtttgcaggtgccaagaggt
tgaatgtctcctgggtgagaatgcataggaatcccatccttctcatggctgggagacccatagattctgggacgg
gctgtaccttgttgtgcaaccgaaagattttaataaacggaaacgccaacatggaatcgccatgtttgccttaat
gtatggtagatcgagccatgcactcccacaattggcatcgtgccaactcaaatcttgaggcgaggaaagcaagca
agcgttgaactcagtggtaagaatttggctccatcccaccatgtccatcatgttcacattggaagacttgtataa
gagcaatgctaaatcactttaaaaagtcactaaggtgtcttacaccaatcacttagttcttcatgataccaattc
aaagggagaaatgagatggcatcagacgtcttacaccaatcacttagtttcttcatgatgccaattcaaagggag
aattgagatgacaccagaccaacttggtgaccttttgagcgatttacttgagttgcttgggcatttcttcttgtg
tgggtgctattggatcaggttgggacaagtaattaaccatgcatgcagcccagacgcatttgtcaggcctaaatt
ttcttcgagtctcccgtccaaaaatcatgttggtcagatagcgtgcaggcgggtatgaaattggaagacttgttt
aagtacatggactggttatgagaagcgtccaagctcggaaactggatagaaatcaaggttatgacagagctgggg
gaacgaggaaaaaagaaccgatatgctaatatatgttggctggatttcagctggcgatgtggaggtgtggaaggt
gacaaaaaggacgaaggcaagatggtgtgagacatttcctttcaacaaaccgacagaaaagtcaaagatgatgag
attttctttgcaacatgtggactttgatccaaaatcttttacgaatctaactttggactaatctaaacccccca
ccaccaccacgatcaagaaggaccaggtactatcggtaaaagttacctaggtctaggacatgaacccaaattttc
agtcgagaagatgtctaaaacgaattaactgggattgaaatcttgaggtatgacgtcactgaagacttgaattcc
caatcccccgcttgggatggggagctaaaccagttaggctgtttagatgctgcttcagtgtgtgggtaagggaga
gttatttgcttaaccagcccaacaccaccaaaaaggttgtgaatgtcttcgtccaaccaaagaatcaaggtgctc
aacctagataattttatgtcagcgactggaagtaaaataaagctaactatccatggctttctatctatttttcac
ttttcctatatatagctttatccactaaacgtggaaacaccaaccaaacactgaataatcttatttgctagatgg
ctggttagagaatagatgcacatggttaaatctattcagggatggttagatattcagtatccgaaacacaacctt

agataagtgacggatttagttgaatcaactacaaccttttttgttaaaaaaaggaaaataaatcaaacacaataa
tgaaagatataacaaatgatgatcatatcgatattttataaaatgacgatttcaaaatcaagaactttggttcta
tctatcatttttcataccaaaatgccctcttacctgttctatattatataacttattccctcttattaattctag
tcgttaaaaagatataaaaatattttgaattacttttgaaatccaatctaacaatgacattctttcaaatcaaat
catattttgggatgattagtttaaaactaatttggaaaatgatccgataaaatataccaattaattttgtattgc
atctgaaatatgatttgataaatagaaattttatttaaatcaaattctactattaattattatttaaatataaca
atattttattaaatattttataatttatagaaaatataatttgaaattttatagatggtgtaagatgcaagcact
actcatttccaattgcataattctagttttaatatttaagaaatatcgtaatattcaaattattgatgtgatatt
gcttgaatattggattttatggaacaaagataatcctatagctcataatgtgaaactttatgtaagaatatgaca
actacttatatgaatttgcaagaagtgctttatcaaatcctatagtacgtagaaccattgcatgatctagatgag
agttaaacattagttatttatttccctttaaaaaatttagcttttaaatttacaaaatttcaataatctatcgta
aacttcaagtcatgaaatttcaagactatagagtttgacactaaagcaactaacataaagcgaataatatagtgg
ctatctattctaattgtacaaataatatattcaaaatcattaatgtaatgcgtatatagcgtgcatctgattatg
aagcttctatcaagcatccacaattatcttggaagacaagaattttgccatacattttaattatttagtgcatta
aagtagtaatgcaaatttaaataaataatcaaaaccatacttaaataagtgttcgatattatttctttactgctt
gactttttttgtaaaacatcaagggatacgcatgccaaaatcttatcaactaccaattattgccctcatatagtt
ataattgtcatccatgggaaacattcattcttagattgtttacattgatcctgtacttcattattcaaatgaagc
atgtatgaattcacccttcactttttctcctttgaccatcgattgctgattcttactcaaatgactattaactta
tttggtatctaaaattgtgcttctttgcctccttataacttttaatatgtttctgtgctatgaatttttttagaa
gtcaacctatttaaactaaaataacgaatacacacaagtagagatgtcgtacaactcaacatctgaagatgttct
atagtatgaactccatcctaagggcttgtttggataatcttgtttaatccaataggatttataagaacataggcc
cggatagatctatattcatagtattcaaaggatttttccaaggtggaccggtcagaataccatattgagaaaaa
ataatattgccccatctttttccatcctaaggatggactggaagaggtctaattgacatagataggaccagacc
tcatattctatgatactagacataatatgtacacatggtttagaagatatgtattggataggccgagcctaattc
ctatagaattttcagcctcatcatgcagaaatatccaaataggccctaagcatattaataattcaaaagaaatcc
tttttcaatctaatgttgcatggatcttgtccatccattcattatcttccttgcatgctttagatactactttca
aggcttgaaaataagaatcacatgaatcatgtaaaatataggaaatcataactacactagagtggacaagtgtga
gttataaggataatcacatgaaaaaaatgtaatctaaataaaaataatatggttgtcttgaaaaaatcttgcaaa
caaacttacaccttggttgagagtttggagaagagaaaagaagaaaaaaaatattaataaggagagaaagaaaga
atgaaggtgtatttttttctcttggtggaaaatttaaagaatgaaaaaataattttcccttcatggttggaagc
aaagaagaaaggagaagaaaaaatattacaactttttttactattatatccttaaaggaacccaataacccaaga
acggaatcaagaaaagattttatctttaatttttctatctttcctttgttttcaccgatttttccgtccacat
tagtctcaatcaggtgggaggcttagatgggagacttaaaataacccttaaaattgaattgtaatattaatttgt
tcttcattattaaaagagtaagaataaaattataaaagattcactaactctcaatttctttctttccaactctac
ttgatttggaggagaaaaaactataaatttgagaacatttgtattcctctattttctctttttttcctttctttat
aaaaaattctaaccaagagaaagtattaacttttctcttattctttttcttctcttttattcttctctcccaacc
aaggcgctaaaagctaaagagtcaaatactcaaataacaaagagaagtcatatgtgttagacacatgaacatttg
attctaagtaatgcatcttttatgattgtttgctttcactaatgataatgatgaagacaaaatataatagaatgt
cctatcttcataagaagaaacatggaaacaatcacatctatttcaaaataagttaagctctatatatttttttgtt
aggaaatctctacactttagggtagcatatgcatatcaatgcatgaaatgtaattcacatgttgtcattcaatac
tcaaaatttagcttgatttccttttacttctgcaaatataacatcaaatacagaggcatatacagttgccacatg
tacttttgcataactttctcctttcttgtagagtcaactttcattgttatcataaaagaataaagacgaagattt
tttttaaaaaaaaaatagaagcaactctaaactcgttctcatagtttgaatccccacagttgtcggaaattctta
catttcttaggtatacatgtcaaggtttgtgacaggtggatggttaaagaaggtaaatatagtgtaaaaaacgtc
caatgccattccttccaccaattatagaaactatcatgtactcaagctgtcggaaattcctacaacctggactag
aaaggaagtgctaatagttctacagcataaatgttggttcaagccatcaattattcaacaataaaacatgttgat
tcaccacgaaggtgggcaggatttaaatcacaacatacgaacaaaaagaaggggaataaagttaaaaatcttgct
gcacaagataattattatactctcagtgaactggcctggatgtcacagcaattaatagtggtagcatgtgtgtcc
ccctttctcttgtatactgactttcatatcagtgaaattatgacgataccacttactaaagcatccatttagata
attgtgtgaagaagcttttgaaatgtgtgaaaactgatcaatttcaacaagtcgtcatggtcttaaagacttaaa
atattcaattatcgaataatctggacccagccggtgcgaactatatatgcttcatttttccaggcatgtgtgtcc
ttgcatcccttctttctgattcatgagtttgaccggcattaattcttgtcatctttctaaaactccagttcgggt
tcctatcaaaatcatcatagcgacatcatatatagcactcccatcggctggtccaaaggaatggaattcccttgg
tcttgaaacctatcaaacaagccatgccagcaccatcgaagctttcatcccaagttcatagcttgtagatactac
atgactgatgaactaagtgttatagctcttcatcatgatttaattctgatatccaagataagtttcactgaatcc
ctcatacagcacttcagaatctattatcaactttaggctccgtacaatctcgaggcccgccaaagccctaccaag
gaatatgactcaagaagcccagtttttttcctccctttgggaaaaaaaacacaaccaaccaatcacataattactt
ttctatcctcacgtgtgatatagttaaatctacaatacaagtctctcgcaagtggacttctgccacacctttgaa
agggaatctccattcttccattttcagaaaaaacgagatgaaacaataaaggggacctccgccaaacctttaatg
tcgctatggtttcacaagccttttcaattaaagaattacttctgtgcctggtgaggggatgctatcccttatcac
cgcacttgatagaagaaaaatcccagaaaccaagcgagagaagagaatgatggtaatttataagccagaagggggg
ataaaaatatctcacttccttccctttcaaatggatttctcctttaagtaattattgaaggacttgttcagcttc
ccttaatctttcattcttcttcatccctttaaattccatgcgtctatcttaaccgttaagcctatcccattaaaa

ctaccaagcttttgtgcctctaaagattgcacataaatgtcataccatgcaataatagaaaatcttgcgtatata
ggatgtggagagaaagcatctggctagagagagaaagagaggtggagcgagtgtgagagagagagaggagtgacc
tggagctggagaaatcatagagcctgccggtgctggagaagatgacaagggcgatctcagcatcgcagaggatgg
cgagctccttggccttcttcaacagaccgcttcgccgcttgcagaaggtcacctgccggctcgtcgagttatcaa
tccttcggatcacgatcttcccctgcccatccctctcttcccctcgcagcacccgataattcggtggatcctc
taccgatctgatgatagaatgcaaacaaacccatgaagggaaatctaaccaagatggagaacaagcaaacgagcg
gaagaagccaggatgcaagcgatcagggaaggaaggaagggaatagagtggattaccttccaaactatggtcgga
aaatgaacctcccctaaggacaccagtctgtttaggacgcgtctatgagaggggacgagggtagccacttaaggg
cagcaagattccggtctggaatggctggaggacttccattttctgcaaagaactagatgcacctgatttccgaca
cacccgaagttgcaccctttcagtgcacgcagaattatgtcatcaggaccaagcctaaaggtgagagggtgaggc
ctagggtgtcgagatatatattttatgctggacgtattttgttgcattactgtggggtacgaggattggcttgtc
caagagaccactcggcgttagtttactcaaatcaaatccctttcggccgtctgctgcatagatcgccaacacgtg
gtgcgacagagtggattccgtacgtccgacggtcccgatcgggcttccgtcgggcgacataccgctttcaagccc
tctggggggattgatcatttaattgagctgccgatggcctctgccaaattgaccgttttgccctcctttttcatcaa
tctgacctactcaaatttagggtttgtgaggggtgacattccttcctctgggctttgttatggatggaccaccag
agcacctgctaccgtagacgtacatcatggtgcatatgtgcaccgtctgagatctgatgatctctcgtaaattat
gcgcttcttagctggcatatgctggacaagaacgtgaaccataagaattgttataaagatgacaatatcaataaa
tatctatttttttcaatttaagcaataacgcatcatatatttgtttttctaagataagtaccatatatttttgtta
tagccatgattatttttgttttcaattttttgaaggaataaagacaaagaatggagagaaagtaaaatgtggcatta
gttccaccgacacacaataaacatatatatagcacatttcatactaactaataatctctagctgatgtatgggta
acatctagcattatatgccaattactatgacataaaagaccctaataggatataatttgcatttgaatgatttttg
atctagcttaatcattttttggatttaaaatttgctttacatcaatttcaacatatcccaccctgttgttttgtt
ccatacggtgttcttgttgctgcttgatacagtcatattaatgcctcaacttatcaaatctgactcctagaatta

>EoAGL17_2
ATGGGCAGGGGGAAGATCGTGATCCGAAGGATTGATAACTCGACGAGCCGGCAGGTGACCTTCTGCAAGCGGCGA
AGCGGTCTGTTGAAGAAGGCCAAGGAGCTCGCCATCCTCTGCGATGCTGAGATCGCCCTTGTCATCTTCTCCAGC
ACCGGCAGGCTCTATGATTTCTCCAGCTCCAGCATGAAATCAATAATAGACCGCTACAACAGGACAAAGGAGGAG
CATCATCAGGTTTTAAATGCAACTTCAGTCAAGTTTTGGCAAAGGGAGGCAGCAAGCTTGAGGCAACAATTGCAG
AACTTACAAGAAAATCATCGGCAATTGATGGGAGAAGAGTTGTCTGGCTTAAATATCAAAGATCTACAAAATCTA
GAGAACCAACTTGAAATGAGTCTCCATTCTGAGCAAATTCTAACTGATGAAATTCAAGAGTTGAACCGAAAGGGG
TACCTCATTCACAACGAAAACATTGAACTCTACAAGCAGGTAAACCTCATACGTCAAGAAAACATGGAGTTACGT
AAGAAGGTGTGTGCAACAAATGAGTTAAATGGAGGAGCTAGTAAGGATACCATTATTCCATATGGTCTTACTATG
ACTCAGAACTCACATGCACCTATTCATCTCGAGTTATGCCAACCTCAACAGCAAACGGATGGCACACAAGCAAGA
GCTCCCAAGCTCAGGTAA

*EoAGL17_3*

>contig_KE504736
tcgtcggcaagcatcctttttttgagttctacaaaggacccttcactctttttgtccgaaacatccagagaaaatc
atcaaattttgactcatatatccttttagtttcaccttaattgtttagatttttttataaatttattttaaaat
taattaattgtatgggaggagtaccttgcttcctccgcattccatcaaaaaagtatatatttatcttttttttt
caaaaaaaatgttgggatgacggaaaggagggcaaaatggtgaacttggtgggggctgaatcggaatacccgcc
tgacggaagccggatggagaccgttggacgtacggaagctctccgtccaaccacgtgttggcgatccatacggca
gacggcgctatcagccagtctcggaagggattttgatttgagtaaactaacgccgtgtggtttatttgaccttg
ccccaatcctcgtacgccacagatgcaaacaaattgcttccagaaccggtgacatgttgacccaatgttacaccc
gctcaccattacgctttatccggatgacgtagatctgcgtgcactgaaaggcagcaacgttgggtgtttctgagg
agtgcatccgagaactttgcaaaaaatggaaggcctccagcaactccaaacgggaagtctcctgccctaaagtggc
tgaccttgtcctctcttattgatgcatcctcaaacaaatgactgcccttaaataggggggagagaggaaaaggca
gcagcagcagcagtctattccttccatttttggtaggtaatccactcttatcccttccttcctccccagatctc
tatcacctcggcttccttgttctccatctgatgttattcccttcacaagtattctttgcattctgttatcagatc
taaagaggatcgatcaagttatcggcggctgcgagctaagggaggaagggagagatggcagagggaagatcgtg
atccgaaggattgagaactcgaccagccggcaggtgaccttctctaagcggcgcaagggtctgttgaagaaggcc
aaggagctcgccatcctttgcgatgccgaggtcggctttgtcatcttctccagcactggcaggctctacgatttt
gccagctccaggtcagtcctctccctctctctccctctccctccctctcccctcccccaccttactctctatat
ctacataagtggctttttttttttttttgttcataacccatgcaaagttttctattattgcatggtacatgtggc
atttatgtgcaatcttagaggcaagtttggtagttttaatggtagttctaggtttggtatttaagatagggttcg
aagaaaaatgagaccgaagaagctgtaatggtagttctaggtttggtatttaagatagggtcgaagaagaatga
gaccgaaaaagctgaagaagtccttagaggagaagttcattctaaagaacgaaatgagatatttttatccccttc
tggctcaatcaccttctttctcctctctcacttaattggtctcttggatttttcctccatcaagtgcggtgctaa
gtgatacatcttcccctcaccaggcacagaagtagttctttaattgagaagaattgtgaaacgatgacgaaatta
aaggtttggaagaggtccctttttatttgtttgtttggaatagaagaaatggagggttcttcataaagatgtggc
agtaagtgatgttgagagccttgtattgtagatttgattttaacttgtggtctattcatcgatatcatcccgcg

aggatagaaaactaatgacgtgattgatttgtactttttccaatagtaaaaatcttcagtaggtctttagtgggc
ttggagatttttctgagcctagtggtggctgagcactagatgcttttgagcctaaagatgatagcagattctaag
aacacagtacgagggatgcggtgaagcttatcttgtctattagaaacaaatcatgatgaggagctatgacactta
attcagcagtcaccatgatgagagctctggtaatgctgacaaggcttggttgactcgtataaagcctacgggaac
ttaaaaatggtttgtcttccatggctttggaccagccgatgggaatgtcatgatgcacggatggtgattttgaca
ataaaccaaacaaaagaactgaagtttctagaaggataacaagaactagtgtcagtcaaattcatgaatcagaaa
gaaagtgatgcgaggatacgcatgccagaaaacataaaaggcacgtagttagtactgcctggagacgacattggt
taaaatccatacgagcatcactggagccaacattgttcaataattgatccggtttggcatttcaaatgcttcttc
acagaactatcgaatggatgttttgatgaggataagtggtatcatcataatttaactgatatgaaattcatttta
caagacgaaggggcatctcgcacaataccatgggaagacacatactaccattataaatagctgtgacatccagaa
gtttactgagaatagtatctcgtgcaacaagatcttttctcaactctgcagtgaattaattagttctttgctgaa
ttattgatgccttgaccgagttacttatctgcactacatatatggagttgaacaagcagtggtccttctcagtgt
acgttgcgggaatttcctacagtttagagtacatgataacttttagttggttgaagtaaaggcaattggccgtag
ttaccttcttaaaccacctatctgtcacaaaccttcacatacgtaagattctacaactaagatgtggaggatcca
aactaggacaacgagtttgagttgattatgcatacaatggtaaaattgcttatagtctttctttttttgttccgta
atcacctctatatttggatgtgattagccaactacaggaaatcaaactaaattttgagttttgaatgataacatg
cgaactgcattttatgcattaatggaatgcatacgctacattgaagtatagagctttcccaccaaaagagtgtag
agcttatcttgtttcaatatagctgtgattatttcaatgtttcttaagaagaaaaatcgaggaagttcaattatc
tcatatcttcatcaacattggtgtaagaaacaatcacagaagatgcgttgcatagagtcaaatgttcctatgtgg
catatatatgactcctctttcctattggtgcatttgactctttagtctttaagttctcttgctcgtttttttacaa
cacaaccagattatttttttagattcacgtttctttttttcatgtgataatccttacttatagttcttgcatgccc
agtatgctgtagttaatttccaacctttttatgtgattcatttgattttttgttctccaagccttaaaataatagta
tctaaagcatgcacgaaggaaatgaatgatggacaagaacaaccgtgcaattaatattagatgggaaaaggatac
ttttaagtaatcaattatcatggaatagtcttatgatggagtttataccctcaaagaacttcagacttctgattt
gtagaatatctccatctttgtagtcgaaactttaatttgaataggctgacatatgaaaaatttgccgtctaaaga
caagataaaaagttatgaagaggtagcatagattgaggtattgaaggagtaagatggaatcagcaatttataatc
cctagctactgatttttcaaggtaatgaagcacacgataaaagtgaacaatttgagaatgaatcttcctatggat
aataagtatagttatatgaaggttggtgtggtcaaaaggtcggatatgactcagatggttgggttaactccgctc
ctacgtatgtgaggtattatccgctctaactcatgatcatttttggacttcagtagatcttggtcatttggagcc
tcacggttttgtcctttaaaaaaatatctcatgagggagagaaggtttcaatccatatgagctatacttcttttga
ctaacaaccgatgtgggactaaaaatgctctttctatgccacaacatagccccacagtcaagagggagtatgtgt
ggataggagatgttcctattgacggcaccaatattgcagtcaaagggttcgatggcttggatagttggattaact
ccgctcccgcatacgtgaagtattgttccctttggctcataattatctttggatttcaatgggccttggtcattt
agagcctcacattttatcctttaaaaagtacctcacagggaaaagaagattccaatctacataagctatactccc
tcttgactcataaccgatgtggaactaaagatatcctctctataccacaacattgccctccaattaagagagagt
atgtgtgggtaatatgttgcagcaagactccgaaagaggaatgatgaggttggtgttgaaccaatcgagtttgac
agttgctaagggtcaaaccaatgtatatcacgagtgaacctgtaaaagaagtcctaaaatcgataaagtaccctc
tggtttaggaccctccgattcttaattaagtcagctggggccttaaaaacagatagtgaaaatagagaaaatgaa
aggcaaagagagagatttgagtggaaggagtgagtgcttttgattccttcagtcggagaactgaaatgaattta
gcagagtccagactccatgagttctgatttatcttgcggagagtactccacctctatttatgtagagaggatttg
cttagatcgttcggagagtttattagaccattaagaatctgtcaggccgttagtttattataacagaatagctag
ctgtatgaaattgtggattgttcatccacttggtgattagctgtagtatggatggaaaacagttattgtagagct
agacgatagctgtaatgaaactacagagtaactgtttttgttctaggtcaaatcggctctagatcgatgtttctt
gtggcagatcggacaaaagctgagctaatcataatactcaattaaggattaaatcagatcaattcaggtcggtgt
ataactaaactgtgggttgaccagatcggcattccattgacctgaccaaaaattgactaatgggatttttaaatg
attctctttgattttgctacatggccagaggttggttatctataccaacaaaggcaataattggcactttaagat
tctagctttgggtattccttgatgtttcatataaaatcatctaaagaatatttttgaacacttatttgagtatag
ttttcattatctacattaaatttttcattattttaatgttttaaataagcaaaatattatggccaaattcttgtct
tccaagaaaattgtggatgcttgagaggaagtttcttggaagtgttctacgcaataattggaagcatattaagta
cactttaaattaatgattttgcagacaattgttgttagatattaattagaatagatggctcctaacttgtttgcc
atgtatattagttgctttaatgtcatacttgatggatatttcatgatttggtgtttagagtagattattgttata
tcatgaatttagaagctaaattttaaaaggaaagaaataaccaaatgtttctttcttctggattatgcaagtgtt
ctacatattgtatgacctcataaagcatttcttgcaaagtcatatatgcagttttacatatatattctcatacaa
ggtttcattaaactcattacaagtttcatgtgtttcatatataaaatccaatattcaaatcatatgtcattgata
attcaattatgacattcttaaatattaaaccaaacattatgcaattgagatgacacctatatttttacatgaactc
aattgttgaaatggttataattaccattataagcaaagtcatcgtatatcgatctctcatgtgtagttgtttta
aatcgattacaatatctgcttctagaattcaagatcaaacacctacttgatgtttgtcaatccatatttattttg
tattaatctcatcatcttcctttcaaatttgatgacaaataagaaacttatcagatcatatttggtgtttactac
atataattagaaatccattcacctaaatttcaaagtaactctgcatgtcactgaattatatagtctatttcagag
atttatatatacttgtatatacttcagaacctcatgaccaagctttttttcttcttttttttttttcttttgatggg
gggtgaagggtggggaatcctcttaacattgtgttgacatgattatgtgctaagtttcaagctatgcacaaggaa
acttggagtatcccttaacttgaatgctattcataaccatgagcgaggctgaacttgggcatcacaaaaccaaag
tctatataagcgcaacggaatggttatttctcaaaactatctttattattttttttttctcagaaaggaaaaacta

ggttccacttgcttcattaagaatataaagatcaaaagtacaaaggtacatagtgtaccaagaaaagttatttct
gagactatcaaaaaggtaccacgcgcacattgatttgcttctttcacagtgaatgccccaaaccgatgatcacaa
tccatcacaactaaaaacattctctccaaaagagatctgaaagaggacttgaatccgttcatttgttttgagggc
ctaacattaattgtcattctatttaagtcactttagggccatgctaactattgattttataatgcagtaactagc
ttggactgacttcaagtgaaagattcccaaagagaatcatacctaaactttctttcaaaataagtaaaatttctt
cctacatctggataagtatgttgttagactgaatgcttgatcttaattttctaaataatcagtgaatttctagga
atgtcaatatgtattcttgaaataaaatgactgactaattttcttttatgattctatccttgtgggattttcttt
tcatctctaatcctaagtagtgatctctcttgcttatggattggagaacattgcaattctatcatgaggattatt
tgagcattatattttaaattgaaggattattatattagaagcagtcatacacaagtactccgggctattgttaaa
agcatgaatataattgggtctacccacaaaacatgcaaggcctaaaaaaaaaaaaacttagtgccagtgcaagta
atctaggatacttaaaacttttaacttaataaacaaatccaacattattttcaaactcactccatgttcattaca
taatatttattttaaagaaaaaaaacttctaggttctaatccttccactcaaggcctattaatccaatctacact
ggatataggcatctgaaaaagggtaagaaaatgaaagagagctaaaagcccagtaagtcatccttattattattc
gatcaagcataatataacaaaactcaagattgtaatataagttttctatatttataatttaagacatattgagca
atataaaatataaaatgatcaacacatctgaataaaaaaagtgaaatttatgatcattgtatttaattgttcgca
cttttctcctagccttttgactatagccaggatctgcccgtggcaagccccaaaaaattttgattttaactataa
gatacacacatcagcatgcaccaatgatcatcctactagagatccaaaaaaactagctctgaaatcatacaaaaa
tatacccataatagtacgtacaagtgatcgtcctactggtgcggctcaaaagaaaactagctcctgattgcaaca
tgtcgttaaccagcatgtgctaatgatcacctaattggaggcctaaaaggagactaaatcctaaatcatatgatt
gcagttagactaaaaatgatacatcataatcaataaaatcaacaatcaaagtggtagaaaataatgtagttcata
ctcatgaaatttattttatgtttaatctagttgttcaaggtctacttatacagagcatacataatataattttaa
atatatataaatgttactcattgtaaattttaatacttaagttaacttctatatgtcaagttatgcgaagtgaac
caattcatatatataatcaaatactagatagatgtatgtatctttggcttgtaagatatagaaatatttccaagg
tactgcacatgacctaaataaatcaatgattaactttacatctaaggtaagcacatattgttattcaaaactcaa
gcatctaaagggattctagatcaatgaagcccactagaagtttcatctttcaaaaatgtctacatcaaggatgga
actccagatcagcttgaccaatagtcaggatcctagtacacatctcaaatctaacttagttcttctaatatcaca
tattcctaatctaggaatcatgaaaggtgcacgagtcataatcacctcagtagtgagatcggtccctggaaccta
tgcacgtcctagatctaatgctctgttcttatcttgatctataagataatacacctgattagacagaaaaagaga
aagatgaagaggcagagagagaaagctagagggatgtggacgaagataaagagaacgagaaaatggtgattggtc
ctatgcggataccccaagtgctaaaggacctgagagagagaaaaatgaagccagaaagagaggagaaagggaagg
gcggagagagagtgatgcagaggaaagtgatagagagaaattggaaagagtgaaggtggtctggtagagcttgac
aagcatggctaacaactcgactatggatggagacaaaggtggctactgaaggtggtccaaaacggagaagaagaa
tggggaaacgaatcatcttagagaagaacgagggtgatggagaggctcaacaatgttgtagctcaccaggcgtgc
aatgtgatggtgctcttatccatgatttgaggaaggagaagaagaaaagggaataaggtgatgagggagatgcga
agaagaggacaaagagctcgattttgaaagatatcatgggcatccatctcacccacccgtggaggaaaaggggag
aggagggcgcaatttctttctctcggttggcaacgataagcatggaaggaggctgtgacctttggtcatccttc
tttgtttgggctaggccaggaagaaagaggaggctgactttggaaggcagccaagcctcccaagataggcacgat
ccttcctcttgcagcctgccaaggatggcaaggcagtgggggaccatgtatcaaagattctagcttcaaatcta
taagtttcgtcccaaacatttgggacctgggtggttacaggtaagtcttccctttcaaaaacaatggttattgt
tagaagcaaatattcttaatgaatctagaaaatgttgataattataataatttcattataattagtccacatctt
tatgctttgcttaatgcttatggattgtgtcaactgcggaattaactttccagatgtagattcctcttcttcact
ttccagatgtagattcctcttcttcagccttttgttaattgatcaagtttttgattcatctatgcatcttctgtt
ttcctcccttatgttgatcccttttctgctgttaatttgactctagttggactcctaaacatattggactctatc
acctactctattgtctgacttctatctctcctcaaatgtgtgttgcatgtgcccagttgctaatgaagtataatg
tttgaatataactagtataccatctaggtggcaaaggattgagtttgagtcggatcaaatatgggttgggtcaaa
gaatcttcaacgaccactcaatcaatgtacaattataaacttatacctatttcctttatttcatatgcacaccta
gttaacctccaaatcaaattattacttattagtcataaatcttgataaataaaacaaaatggctggagaaagata
taacctggagttaggcacaatttttgatgtaccatccatatttgaacctaaaaccttaagttgttaagcaaaggc
ctataactaccaggataatacacaaccagattattaaaagcctttatacaaactcataccttttcggctgtgca
caatttgtttttgactggtttgggtcaattttcaaactaagctgatttgaatcagttttctaaaactaaaactaa
ttgaactaaccatataaaatatcagttcaaaccaaactgaaaaaattggtttggttaatcgaattgtaggctatt
agacaagttatgggatgggctatacaggctgctatccatttaattttttctaaccatacaaaatgggcaactt
atgcaatgagctaaatgaatagcttatgagatggactaaatgggctaaatatagaagcaacaaaccggtagggct
aaggcaaagaaaaaatctttatgcaccgcaggcagtgtagaaaatccgatatggagtgtactgctttatgtgatt
agtctacgtagctatcgctttttcacgcacatttaatgcttgcagtttcactttttcgtttgaaaattttgaatg
atgaaaatatccctgctctttgaaaaaaaattatagcatttatatttatattatgacatcctgtgtgcagcaagt
cataattttgacgcaagatgtcataatataccacaagatatcataatttttttttaaagagtaaggatattttgtc
attcaaaattttaaataaaaaagcgaaactgcagccattaaatacgtattgaaaagtgattggataaaattgtc
ccttctatattatgacatcctaaaccatattatgacatcctgagccatattatgacatcttgcatgtaaaaagtc
ataatttgacgtaggatattataatatgccataggatgtcataattttctggatcatattataacattctgcgtg
tagaaagtcataatatgtcataggatgtcataatttttctaaagagtagggatattttgtcatgcaaaatttta
acgaaaaataaactacagatattaaatgtgcattgaaaagcaatagttacgtggaccaatcatataaagcagtgc
gctccacgtcagattttctacaccacccatggtgcacaaagaatttctctaaggcaaatataagtaaatcggtg

gatggcagcccgtttagcccacaggtgaaggaaaaaatctgattcagatcaaattgccaggaaataattgaacag
atatattcttaaaattctcataaccattatgacttaccttgatgcgcagcggagatttgatcaagattgtttgaa
tcagttgcttcacgagatctggatgcgcaagttttctacttcacatgcacgtcaggcactgcagaaaaaatggat
gaacaccttgggcatcgctccttagaagggaagcaagaggaggccgtcaagatctgatctcacggctggaggct
agggagggattcttcacaccaagagagaaggatgccaagaactccacaccaatattagcgccctcccttttttg
tcacacaaaattactcttcctcatagtgatttcaaatctattcttcttttacgtgaagaaacacccctccttaaa
taattcatgctcaaaatcaaatcatagccctaagaaattaggtctctaattcaaatcaaatttgaattagataag
gctttatcttatccactaagagacccaacggagaattgacgccaacttgtgcatcttttatctttcatgcacgca
agggatgggatgcaattttcaatgccccctcttccttcttggtcagtcagataagatgagagagtcctagagaat
taaggctctgtggacctattccaatttggttcaatttgaatccatttcgaacctgattcgaatcaaatttgaaga
gactatttatcctaatctaattagaatttagatccaaatctaacttggataattaacccaattagtcttaatcaa
gttcaattaaatcagatctaatctaaaataattaagatttgaatccaatctctcaaaatttcttagatcacagat
tagatcaaatttatctctggaattaaacctacacctcatgtctagtgctaactaaatatagtcaatgtttaattc
atatgatctataatttaattcttaattaagtaaatctatttgttcaatcaagtcacatactttcaaattaaacac
ataaatgttaggtcaattcttttctacgttgcttgactttgtatgtgactctataggttcgaacactaagtcggt
agtgtagaaactatttttttatactaatcgaagtgaccatctagtaatgatatgatctgatgtcaggatggatcga
ataattacaaaataatattcaagaaccttgatatatggttaccacataattcatctctttaaccctcaatactca
ggatgacctcagagtaaactgtcaaatcttagatcagccatccatattgtatttcaaccttttaaatcctgtgat
aaatcacatggattaccctgatcaagatttactgaattaaaatacagtgaaacattatctcctgtaatccgaag
agatcaatcccatcttgactcatacacagacttcgcaagtatttgactgtgcccaataaccttctgtcactatat
taaaaatatagatagtccagcatcaaagtatagtgagttgcttgcaagttactatagtgatctcagatcgaaggg
acatttatgcccataccсttcatgagctgctcttgacagtaaagtgcttcacagttgaatcactattcagtaatg
atatacttctacatctcacctgcataccataccagtatctccacactctttgattacaagaacaaccaattcata
cggcatacaatgatctaaactcgataaacgttattgtcttattaataacgtattatttgatcgtgaacatagttt
aaaactattcgataagttctcattatctttatcgattgttttatagttctaaggacttcattatataattaaagt
tctataaggaatatgttaaaatatgatgaattgccaaaataatttttattattcatttataattcatatacaaag
tataattatttataattagataattgacttcaggacataattttcaacaacggatcattcggtttaaattaggtt
tcttaaaatcgaaaatgagatcgaaccaatttttttgattttgaaattttcaaactaaaatctaaatagaatct
ttaaaaaatcaatccaagccaaatcaaaattagtggttcaaattaattttacggtttaatcaatttcttgcgca
cccctatttcttttagaaagatttaaaaagttggatggcctatgctaaaaatttttatctctaattattgaacta
ctaacgacacaggcacatccgatgcatatttagaaaaacataaactagacatattacaaaaataactattacata
gtataaatattataattttaaagtgcaaaattgaaaattagggagtattgttaatgatttaaatctgaagttaat
ggaggaccaattattttgaagagaataaatagagatataaaagttaaatatatcatttagttatttgatgaaaa
tgaattaattaatgatccatgtaggtctcaaacttgcaacttttgcattattatttagcaaaaatattatcaata
atatattttaaaaatattttaaaaatgaagaatatttttttgatagtggatttattaatagaatctatttgatat
atttctcttttattttgcaatatggtatagataaagataagaccttttttcttaaggcaacatagaggcaagactt
tcggctcaagtctcctccaatggatggaggatgcttataacttgtggggataaacctcccatcatcaactttaca
atgcttttgcactaggataagtatatacaagggcaaaaaaaaaagaaaacctctacctctcatttgcagttgtaa
agaaagagtcatgggtacatcggtggtagttaaggctacaaatgggtcaagtgacccatgacccaacctgacttg
cttaaactcgaaatagatctgatttaaggacccatgggatgggttgtattgtggtatagaagggagggcttcatt
agtcctacattgattataaatcaaggaaaaatctggtatacattgattgtaagtcaagaaagaatccggtgtata
taagaagggaccatctttccatgtgagatatttttgaagggcaaaatcgtaaagcctatgaaccaaaatgggcca
tacgtcacaagtcgagccataagtcctagaccgcaacagtagtgcaccagaaagaggaccacccctacaattata
gagctcattctgtctatacaccaatctacaataaaataaggagggcatctctcatcctactcagattctctcttg
actgggaggctatattgtaatatagaagagagagagcttcattagccccatattggttgagtcaaggaggactctag
tttatatagaaaatgaccatccttcctatataagatgccttttgaagagcaaaactatgaagcccataggctaga
gcagacaatacctcaggtgtcgagccatgagcccttggttgcaactagttgctttccaaatttgaactcattcta
tattcaggtcggcctaagttattgagtttttatccaacctcgatcgattattgtatgggtctaatttgagtttc
taagatattacctaacttaacctgacccaaccaacccaatatacctattaacccaaggtgagtgggtggtattaa
gccagtttacagtggaccatctactcttctaaagtcctaattagaaattaatcctatataatgattgcgtccatc
accaatgaattttgaaataataatataattatcccaccaatccaagcttcttgatcagattttattcacatgaca
tagggatgtcgggaaaaactacatagttgagtgatattcttggattagaagtttgacttctagattatagattgc
tactctaactgatttagcattattcatgctctatttagtctattagatatctttaaacataataactcgtatta
tctatatggaatacataacttttagagctctttttaattatttttgagtcatattggaaaaattatcatgctaaa
tatatcgtagtatgcttccaaagagatatttatatcttatgatgaagctatgatttttctttgtctaaatttaac
tctttcactttaaaatccatatgtatgactaaatttttttagtttgaaaccagcttattcatagatccaacttgac
ctaaatgatgcgatcccaattcgaatttatcaggttgggaacgagttataaatggatctgactcaacctgaatga
cccattaagtcaaatattttagccatggatccgacccaacttgaactaatttgggttggatgttgatccaacatt
aggatccgaacaaggaattgggctaaatcgaggttacttgtgacatgatctaatcagacccttttgcacccctat
tggcaatgactatcatggcaacaaagaagagtgcttgcacagatgcatctgttccaatgatgtaaattatctgaa
attattggacgatgctatagaaaccactagaatttggcataagaggattgctcctacaacacctctgctcttaca
acaccttggcataatggtgaggtccacgtgccccaaattagctgtgcaatttcatgaggttccaagttgccaaac
ctactcttaacatgattatcaatgcatcatttcccattcggttcctaaaaaaaattggagttccaaatgaaaaaga

acaagagtttggatgaggaaaatgcctcaaacaaacatttcctaagatttggcatgaaagcattgctcctagaac
actatagcatcatcacaaaatgaaagaatgtgtcatgcaacattaaataatgttagatcgatgtataatgttaca
aaagtcaccattattatctctgggtggagcaaggaattttttttggacttaatgattaatctccttcaatttgg
ataatggagttttagtaaaacttgaaagaaaatatctcctaaaagcgcttaagtgtgtgcaaatgtaaagtatat
gtccatgcagacaaaaatgagcaatccaagcttcatactaagctgaaaaatgagtccgccaatgttgccggatat
gaagtctttggttataagcgatgcattgttcctcttttatgtgaagtctttgatcatagcattttttttccattt
ctttaaacaatatgatgctaatatgctacttgttgaggaatagccccatgttagataggttggggactccggtcg
gtcttgtatatccatgggtccttccacctatcagcttaagattttgagttgtctttcttccttattgatgatcat
tggcatctccaaatccatgatatcctctgtctacatcatttattgcagattgtcaaggaattcaatccaccaaca
ataagaagaaattaacgagaaaacttccattaacctgtacgtggaaaaataagtagacgactgttggtgcaaaaa
tccaccggcgtcggagaagctggagtcgagggagtcgcggtcgccgtcgggacctgcaaggaaagtctaaaccgg
agttggggttgctccggcaagaccctccgatgctcaagtcagttctctgcctcaacaagaatggagtgctcgaac
gaaattttagcagagttttgaaataagaattggagcttagaaaataacgtatctggagatccctatttatagga
gagggcgtaacagactgatagcgacgtctgtaaccgtctggtggtgggccgttcaggannnnnnnnnnnnnnnnnnnnnn
nnnnnnnnnnnnnnnnnnnnnnnnnnnnnnnnnnnnnnnnnnnnnnnnnnnnnnnnnnnnnnnnnnnnnnnnnnnnnnn
nnnnnnnnnnnnnnnnnnnnnnnnnnnnnnnnnnnnnnnnnnnnnnnnnnnnnnnnnnnnnnnnnnnnnnnnnnnnnnnn
nnnnnnnnnnnnnnnnnnnnnnnnnnnnnnnnnnnnnnnnnnnnnnnnnnnnnnnnnnnnnnnnnnnnnnnnnnnnnnn
nnnnnnnnnnnnnnnnnnnnnnnnnnnnnnnnnnnnnnnnnnnnnnnnnnnnnnnnnnnnnnnnnnnnnnnnnnnnnnnn
nnnnnnnnnnnnnnnnnnnnnnnnnnnnnnnnnnnnnnnnnnnnnnnnnnnnnnnnnnnnnnnnnnnnnnnnnnnnnn
nnnnnnnnnnnnnnnnnnnnnnnnnnnnnnnnnnnnnnnnnnnnnnnnnnnnnnnnnnnnnnnnnnnnnnnnnnnnnnnn
nnnnnnnnnnnnnnnnnnnnnnnnnnnnnnnnnnnnnnnnnnnnnnnnnnnnnnnnnnnnnnnnnnnnnnnnnnnnnn
nnnnnnnnnnnnnnnnnnnnnnnnnnnnnnnnnnnnnnnnnnnnnnnnnnnnnnnnnnnnnnnnnnnnnnnnnnnnnnnn
nnnnnnnnnnnnnnnnnnnnnnnnnnnnnnnnnnnnnnnnnnnnnnnnnnnnnnnnnnnnnnnnnnnnnnnnnnnnnnnn
nnnnnnnnnnnnnnnnnnnnnnnnnnnnnnnnnnnnnnnnnnnnnnnnnnnnnnnnnnnnnnnnnnnnnnnnnnnnnnnn
nnnnnnnnnnnnnnnnnnnnnnnnnnnnnnnnnnnnnnnnnnnnnnnnnnnnnnnnnnnnnnnnnnnnnnnnnnnnnnnn
nnnnnnnnnnnnnnnnnnnnnnnnnnnnnnnnnnnnnnnnnnnnnnnnnnnnnnnnnnnnnnnnnnnnnnnnnnnnnnnn
nnnnnnnnnnnnnnnnnnnnnnnnnnnnnnnnnnnnnnnnnnnnnnnnnnnnnnnnnnnnnnnnnnnnnnnnnnnnnnnn
nnnnnnnnnnnnnnnnnnnnnnnnnnnnnnnnnnnnnnnnnnnnnnngggagtccgggcctgtcggtcgaagctcgatag
gggcgtctggtggagcggggtagctctctgtctgagaggagctcggatatttcgaggttgctgacgagcccgct
gtcaacaggtgccttaggtgttggcgcggcaggctgaaatcattttgtcgtagaagctcgggcggagacttcccg
ttggtgaagtccggctggggttggcctgacgttgaacttattggaaagctgttcgccgggaggtctgatttcatg
aagagttcggctgaaggtcgaccgcttggctgaattcagaaggtgtcgtagaagcctgtccgcaggaggagcctg
tagaacactgttgaagttcggttggagccgtcgaaggtcggttgacgctgctgaaagtcggtggccggagaggta
tggaagacactagaggcagccgatcgccgtagaagttcatctgctggagctcgtccgtctgttgtagaagttcgc
ccggtacctatctgttacaggagctcggtcggagcctgacttttgtagaagtccggatgggaccgcttattgaag
aagtccgaccgaagtccgcggtagagattcgacgtggaccgcttgtagtagaagctcgacgtagatcgcttatag
tagaagctcggagtagatcgctcttcgtgggagctcggagtagaccgctcttggtgggagctcggagcaggccgc
tcttggtgggagctcggggtagaccgtttgtagtaggagcttggagtgttcaagctttgtcgaagtccggacgat
gtctacctgcaatagaaattcgaggaagaaatccggctcccgtaggagttcgggtgaagtctacctacagtagga
attcggggcagtggaccgtctcccgtaggagctcgggtggaatcttcctgcaatagaagttcgggacagtggacc
gtctcccgtaggagctcggagtagatcgtttgtagtagaagttcggagtaggtcgtttagctcccataggagctc
agggtaggtcgctcgtagtagaagttcggagtaggccgcctgtaggaggtcggacgaagtatacctgaaatgaga
attcgggatagaggtctggctcctgtaggagctcggatgaggtccggctcccgtaggagcttggacggagtccag
gaggcggtcgaccgccgtagagatttggagggagtccgaccgtcgtagagatctgctgttggggaagttcggcca
ccgtcgaagttttgaaagaagttcggattggagtcgatcgtggttggaaaagatctgaaagaggtccggagaacg
gtcgatcactgtagaaggtcggccggtagcggagcccggagagcacttggagcagcctgatagacgactaaagaa
gctcatcctggagaagcccggagggtgcggtaggagctcgtccgtcgaaggagttcggtcaacacttgtagtggg
caccgtagaagctcattcgtcagcgaggcttgggaacattgtggaagctcggcagctggaaaggtccgaaaagat
gtcgcccaagatcagaagccggaaggatcctggggaggccaatcctaatcacaaacttcnnnnnnnnnnnnnnnnnnnnn
nnnnnnnnnnnnnnnnnnnnnnnnnnnnnnnnnnnnnnnnnnnnnnnnnnnnnnnnnnnnnnnnnnnnnnnnnnnnnnnn
nnnnnnnnnnnnnnnnnnnnnnnnnnnnnnnnnnnnnnnnnnnnnnnnnnnnnnnnnnnnnnnnnnnnnnnnnnnnnnnn
nnnnnnnnnnnnnnnnnnnnnnnnnnnnnnnnnnnnnnnnnnnnnnnnnnnnnnnnnnnnnnnnnnnnnnnnnnnnnnnn
nnnnnnnnnnnnnnnnnnnnnnnnnnnnnnnnnnnnnnnnnnnnnnnnnnnnnnnnnnnnnnnnnnnnnnnnnnnnnnnn
nnnnnnnnnnnnnnnnnnnnnnnnnnnnnnnnnnnnnnnnnnnnnnnnnnnnnnnnnnnnnnnnnnnnnnnnnnnnnnnn
nnnnnnnnnnnnnnnnnnnnnnnnnnnnnnnnnnnnnnnnnnnnnnnnnnnnnnnnnnnnnnnnnnnnnnnnnnnnnnnn
nnnnnnnnnnnnnnnnnnnnnnnnnnnnnnnnnnnnnnnnnnnnnnnnnnnnnnnnnnnnnnnnnnnnnnnnnnnnnnnn
nnnnnnnnnnnnnnnnnnnnnnnnnnnnnnnnnnnnnnnnnnnnnnnnnnnnnnnnnnnnnnnnnnnnnnnnnnnnnnnn
nnnnnnnnnnnnnnnnnnnnnnnnnnnnnnnnnnnnnnnnnnnnnnnnnnnnnnnnnnnnnnnnnnnnnnnnnnnnnnnn
nnnnnnnnnnnnnnnnnnnnnnnnnnnnnnnnnnnnnnnnnnnnnnnnnnnnnnnnnnnnnnnnnnnnnnnnnnnnnnn
nnnnnnnnnnnnnnnnnnnnnnnnnnnnnnnnnnnnnnnnnnnnnnnnnnnnnnnnnnnnnnnnnnnnnnnnnnnnnnnn
nnnnnnnnnnnnnnnnnnnnnnnnnnnnnnnnnnnnnnnnnnnnnnnnnnnnnnnnnnnnnnnnnnnnnnnnnnnnnnnn
nnnnnnnnnnnnnnnnnnnnnnnnnnnnnnnnnnnnnnnnnnnnnnnnnnnnnnnnnnnnnnnnnnnnnnnnnnnnnnnn
nnnnnnnnnnnnnnnnnnnnnnnnnnnnnnnnnnnnnnnnnnnnnnnnnnnnnnnnnnnnnnnnnnnnnnnnnnnnnnnn

nnnnnnnnnnnnnnnnnnnnnnnnnnnnnnnnnnnnnnnnnnnnnnnnnnnnnnnnnnnnnnnnnnnnnnnnnnn
nnnnnnnnnnnnnnnnnnnnnnnnnnnnnnnnnnnnnnnnnnnnnnnnnnnnnnnnnnnnnnnnnnnnnnnnnnn
nnnnnnnnnnnnnnnnnnnnnnnnnnnnnnnnnnnnnnnnnnnnnnnnnnnnnnnnnnnnnnnnnnnnnnnnnnn
nnnnnnnnnnnnnnnnnnnnnnnnnnnnnnnnnnnnnnnnnnnnnnnnnnnnnnnnnnnnnnnnnnnnnnnnnnn
nnnnnnnnnnnnnnnnnnnnnnnnnnnnnnnnnnnnnnnnnnnnnnnnnnnnnnnnnnnnnnnnnnnnnnnnnnn
nnnnnnnnnnnnnnnnnnnnnnnnnnnnnnnnnnnnnnnnnnnnnnnnnnnnnnnnnnnnnnnnnnnnnnnnnnn
nnnnnnnnnnnnnnnnnnnnnnnnnnnnnnnnnnnnnnnnnnnnnnnnnnnnnnnnnnnnnnnnnnnnnnnnnnn
nnnnnnnnnnnnnnnnnnnnnnnnnnnnnnnnnnnnnnnnnnnnnnnnnnnnnnnnnnnnnnnnnnnnnnnnnnn
nnnnnnnnnnnnnnnnnnnnnnnnnnnnnnnnnnnnnnnnnnnnnnnnnnnnnnnnnnnnnnnnnnnnnnnnnnn
nnnnnnnnnnnnnnnnnnnnnnnnnnnnnnnnnnnnnnnnnnnnnnnnnnnnnnnnnnnnnnnnnnnnnnnnnnn
nnnnnnnnnnnnnnnnnnnnnnnnnnnnnnnnnnnnnnnnnnnnnnnnnnnnnnnnnnnnnnnnnnnnnnnnnnn
nnnnnnnnnnnnnnnnnnnnnnnnnnnnnnnnnnnnnnnnnnnnnnnnnnnnnnnnnnnnnnnnnnnnnnnnnnn
nnnnnnnnnnnnnnnnnnnnnnnnnnnnnnnnnnnnnnnnnnnnnnnnnnnnnnnnnnnnnnnnnnnnnnnnnnn
nnnnnnnnnnnnnnnnnnnnnnnnnnnnnnnnnnnnnnnnnnnnnnnnnnnnnnnnnnnnnnnnnnnnnnnnnnn
nnnnnnnnnnnnnnnnnnnnnnnnnnnnnnnnnnnnnnnnnnnnnnnnnnnnnnnnnnnnnnnnnnnnnnnnnnn
nnnnnnnnnnnnnnnnnnnnnnnnnnnnnnnnnnnnnnnnnnnnnnnnnnnnnnnnnnnnnnnnnnnnnnnnnnn
nnnnnnnnnnnnnnnnnnnnnnnnnnnnnnnnnnnnnnnnnnnnnnnnnnnnnnnnnnnnnnnnnnnnnnnnnnn
nnnnnnnnnnnnnnnnnnnnnnnnnnnnnnnnnnnnnnnnnnnnnnnnnnnnnnnnnnnnnnnnnnnnnnnnnnn
nnnnnnnnnnnnnnnnnnnnnnnnnnnnnnnnnnnnnnnnnnnnnnnnnnnnnnnnnnnnnnnnnnnnnnnnnnn
nnnnnnnnnnnnnnnnnnnnnnnnnnnnnnnnnnnnnnnnnnnnnnnnnnnnnnnnnnnnnnnnnnnnnnnnnnn
nnnnnnnnnnnnnnnnnnnnnnnnnnnnnnnnnnnnnnnnnnnnnnnnnnnnnnnnnnnnnnnnnnnnnnnnnnn
nnnnnnnnnnnnnnnnnnnnnnnnnnnnnnnnnnnnnnnnnnnnnnnnnnnnnnnnnnnnnnnnnnnnnnnnnnn
nnnnnnnnnnnnnnnnnnnnnnnnnnnnnnnnnnnnnnnnnnnnnnnnnnnnnnnnnnnnnngtcgcagccacca
cagaccacctcccagaggtgccgaattccccactgatccagaggtccgagacctctaaccttgtacttttccttc
attactgttttcttctttcttttgtcttcaaggaacattaagtcaataaattggtcttctccttatggggacttc
tcttcttgttcttcctttcttcttcttcttcttctttgtaatgagctgcatcttaacgcgttggtctcccgacg
gcagtatcctgccgatgctcttctcttgccacagaaaattttgctgaagtccacgatccgtcatctgaggaagaa
agttcgtcatttgacaaaaagatcgaaggagatagaggaagaacttcgcaggctgagggagagtcactcagaggc
caccgcggaggccactcacttccggaacctccacgtgaaaggaatcatggattatagtcggaagaaggcgaactt
tgagaaggagctcgaggaatatcggaaacgcaccagtgaccgatcctgggttcaagcctccaaaatcagctccct
tgaagcggagctggcgactgcacgggagaagatcggccagctggaggggagctcatcttggctcgcaactcaggc
cgaacgcgaacaagactggtcgaaaaaattctccgaccttcaacaacagctccaggatgccgagtcgaactacaa
tgcatatcgggcgggctggtgcaagtagatggatgactacagaaggaggctccaggcgtcgatcgacgaagttac
ccatcttcgaaggcggttatctgaaggagcccagctcgtctcagttcgggactccgacgaactccgatctctgag
gagaactatggaagagctatccgtcgccttgggagggaaggccgagctgcagcaattgaagatccaactgacgca
cgaacagcaggcggtcaaagatgcggaggcggagtccgaagtcctgaggaagagacttcgggagtcagagggcga
aagtcggcgattccatcggatgttccggaacatgctgctgaaaaagagagaactagaagaagaaattgaaaatct
gaagcagtccatgataaggagtgaagcagataactcgaggtcggaaggagctgagcttccctagagctcttttgc
cttccatttttttatatatatatttttcttttcagttcttgttattttgcttttctttctttagctattttgggc
tttgtagtatatacttcatgaatgaatgaaaaggagattttgtgatactttgtcttcactttgtattaaattgtc
gtccgccgaacttcttttgtcttttgacttgttcgacgtcgacgtcgtttgaagattcccagccattgttgtgtt
gatttactttcttatcatctttttttcgatcnnnnnnnnnnnnnnnnnnnnnnnnnnnnnnnnnnnnnnnnnnnn
nnnnnnnnnnnnnnnnnnnnnnnnnnnnnnnnnnnnnnnnnnnnnnnnnnnnnnnnnnnnnnnnnnnnnnnnnnn
nnnnnnnnnnnnnnnnnnnnnnnnnnnnnnnnnnnnnnnnnnnnnnnnnnnnnnnnnnnnnnnnnnnnnnnnnnn
nnnnnnnnnnnnnnnnnnnnnnnnnnnnnnnnnnnnnnnnnnnnnnnnnnnnnnnnnnnnnnnnnnnnnnnnnnn
nnnnnnnnnnnnnnnnnnnnnnnnnnnnnnnnnnnnnnnnnnnnnnnnnnnnnnnnnnnnnnnnnnnnnnnnnnn
nnnnnnnnnnnnnnnnnnnnnnnnnnnnnnnnnnnnnnnnnnnnnnnnnnnnnnnnnnnnnnnnnnnnnnnnnnn
nnnnnnnnnnnnnnnnnnnnnnnnnnnnnnnnnnnnnnnnnnnnnnnnnnnnnnnnnnnnnnnnnnnnnnnnnnn
nnnnnnnnnnnnnnnnnnnnnnnnnnnnnnnnnnnnnnnnnnnnnnnnnnnnnnnnnnnnnnnnnnnnnnnnnnn
nnnnnnnnnnnnnnnnnnnnnnnnnnnnnnnnnnnnnnnnnnnnnnnnnnnnnnnnnnnnnnnnnnnnnnnnnnn
nnnnnnnnnnnnnnnnnnnnnnnnnnnnnnnnnnnnnnnnnnnnnnnnnnnnnnnnnnnnnnnnnnnnnnnnnnn
nnnnnnnnnnnnnnnnnnnnnnnnnnnnnnnnnnnnnnnnnnnnnnnnnnnnnnnnnnnnnnnnnnnnnnnnnnn
nnnnnnnnnnnnnnnnnnnnnnnnnnnnnnnnnnnnnnnnnnnnnnnnnnnnnnnnnnnnnnnnnnnnnnnnnnn
nnnnnnnnnnnnnnnnnnnnnnnnnnnnnnnnnnnnnnnnnnnnnnnnnnnnnnnnnnnnnnnnnnnnnnnnnnn
nnnnnnnnnnnnnnnnnnnnnnnnnnnnnnnnnnnnnnnnnnnnnnnnnnnnnnnnnnnnnnnnnnnnnnnnnnn
nnnnnnnnnnnnnnnnnnnnnnnnnnnnnnnnnnnnnnnnnnnnnnnnnnnnnnnnnnnnnnnnnnnnnnnnnnn
nnnnnnnnnnnnnnnnnnnnnnnnnnnnnnnnnnnnnnnnnnnnnnnnnnnnnnnnnnnnnnnnnnnnnnnnnnn
nnnnnnnnnnnnnnnnnnnnnnnnnnnnnnnnnnnnnnnnnnnnnnnnnnnnnnnnnnnnnnnnnnnnnnnnnnn
nnnnnnnnnnnnnnnnnnnnnnnnnnnnnnnnnnnnnnnnnnnnnnnnnnnnnnnnnnnnnnnnnnnnnnnnnnn
nnnnnnnnnnnnnnnnnnnnnnnnnnnnnnnnnnnnnnnnnnnnnnnnnnnnnnnnnnnnnnnnnnnnnnnnnnn
nnnnnnnnnnnnnnnnnnnnnnnnnnnnnnnnnnnnnnnnnnnnnnnnnnnnnnnnnnnnnnnnnnnnnnnnnnn
nnnnnnnnnnnnnnnnnnnnnnnnnnnnnnnnnnnnnnnnnnnnnnnnnnnnnnnnnnnnnnnnnnnnnnnnnnn
nnnnnnnnnnnnnnnnnnnnnnnnnnnnnnnnnnnnnnnnnnnnnnnnnnnnnnnnnnnnnnnnnnnnnnnnnnn
nnnnnnnnnnnnnnnnnnnnnnnnnnnnnnnnnnnnnnnnnnnnnnnnnnnnnnnnnnnnnnnnnnnnnnnnnnn
nnnnnnnnnnnnnnnnnnnnnnnnnnnnnnnnnnnnnnnnnnnnnnnnnnnnnnnnnnnnnnnnnnnnnnnnnnn

nnnnnnnnnnnnnnnnnnnnnnnnnnnnnnnnnnnnnnnnnnnnnnnnnnnnnnnnnnnnnnnnnnnnnnnnnnnnnnnnnnnnnnn
nnnnnnnnnnnnnnnnnnnnnnnnnnnnnnnnnnnnnnnnnnnnnnnnnnnnnnnnnnnnnnnnnnnnnnnnnnnnnnnnnnnnnn
nnnnnnnnnnnnnnnnnnnnnnnnnnnnnnnnnnnnnnnnnnnnnnnnnnnnnnnnnnnnnnnnnnnnnnnnnnnnnnnnnnnnnn
nnnnnnnnnnnnnnnnnnnnnnnnnnnnnnnnnnnnnnnnnnnnnnnnnnnnnnnnnnnnnnnnnnnnnnnnnnnnnnnnnnnnnn
nnnnnnnnnnnnnnnnnnnnnnnnnnnnnnnnnnnnnnnnnnnnnnnnnnnnnnnnnnnnnnnnnnnnnnnnnnnnnnnnnnnnnn
nnnnnnnnnnnnnnnnnnnnnnnnnnnnnnnnnnnnnnnnnnnnnnnnnnnnnnnnnnnnnnnnnnnnnnnnnnnnnnnnnnnnnn
nnnnnnnnnnnnnnnnnnnnnnnnnnnnnnnnnnnnnnnnnnnnnnnnnnnnnnnnnnnnnnnnnnnnnnnnnnnnnnnnnnnnnn
nnnnnnnnnnnnnnnnnnnnnnnnnnnnnnnnnnnnnnnnnnnnnnnnnnnnnnnnnnnnnnnnnnnnnnnnnnnnnnnnnnnnnn
nnnnnnnnnnnnnnnnnnnnnnnnnnnnnnnnnnnnnnnnnnnnnnnnnnnnnnnnnnnnnnnnnnnnnnnnnnnnnnnnnnnnnn
nnnnnnnnnnnnnnnnnnnnnnnnnnnnnnnnnnnnnnnnnnnnnnnnnnnnnnnnnnnnnnnnnnnnnnnnnnnnnnnnnnnnnn
nnnnnnnnnnnnnnnnnnnnnnnnnnnnnnnnnnnnnnnnnnnnnnnnnnnnnnnnnnnnnnnnnnnnnnnnnnnnnnnnnnnnnn
nnnnnnnnnnnnnnnnnnnnnnnnnnnnnnnnnnnnnnnnnnnnnnnnnnnnnnnnnnnnnnnnnnnnnnnnnnnnnnnnnnnnnn
nnnnnnnnnnnnnnnnnnnnnnnnnnnnnnnnnnnnnnnnnnnnnnnnnnnnnnnnnnnnnnnnnnnnnnnnnnnnnnnnnnnnnn
nnnnnnnnnnnnnnnnnnnnnnnnnnnnnnnnnnnnnnnnnnnnnnnnnnnnnnnnnnnnnnnnnnnnnnnnnnnnnnnnnnnnnn
nnnnnnnnnnnnnnnnnnnnnnnnnnnnnnnnnnnnnnnnnnnnnnnnnnnnnnnnnnnnnnnnnnnnnnnnnnnnnnnnnnnnnn
nnnnnnnnnnnnnnnnnnnnnnnnnnnnnnnnnnnnnnnnnnnnnnnnnnnnnnnnnnnnnnnnnnnnnnnnnnnnnnnnnnnnnn
nnnnnnnnnnnnnnnnnnnnnnnnnnnnnnnnnnnnnnnnnnnnnnnnnnnnnnnnnnnnnnnnnnnnnnnnnnnnnnnnnnnnnn
nnnnnnnnnnnnnnnnnnnnnnnnnnnnnnnnnnnnnnnnnnnnnnnnnnnnnnnnnnnnnnnnnnnnnnnnnnnnnnnnnnnnnn
nnnnnnnnnnnnnnnnnnnnnnnnnnnnnnnnnnnnnnnnnnnnnnnnnnnnnnnnnnnnnnnnnnnnnnnnnnnnnnnnnnnnnn
nnnnnnnnnnnnnnnnnnnnnnnnnnnnnnnnnnnnnnnnnnnnnnnnnnnnnnnnnnnnnnnnnnnnnnnnnnnnnnnnnnnnnn
nnnnnnnnnnnnnnnnnnnnnnnnnnnnnnnnnnnnnnnnnnnnnnnnnnnnnnnnnnnnnnnnnnnnnnnnnnnnnnnnnnnnnn
nnnnnnnnnnnnnnnnnnnnnnnnnnnnnnnnnnnnnnnnnnnnnnnnnnnnnnnnnnnnnnnnnnnnnnnnnnnnnnnnnnnnnn
nnnnnnnnnnnnnnnnnnnnnnnnnnnnnnnnnnnnnnnnnnnnnnnnnnnnnnnnnnnnnnnnnnnnnnnnnnnnnnnnnnnnnn
nnnnnnnnnnnnnnnnnnnnnnnnnnnnnnnnnnnnnnnnnnnnnnnnnnnnnnnnnnnnnnnnnnnnnnnnnnnnnnnnnnnnnn
nnnnnnnnnnnnnnnnnnnnnnnnnnnnnnnnnnnnnnnnnnnnnnnnnnnnnnnnnnnnnnnnnnnnnnnnnnnnnnnnnnnnnn
nnnnnnnnnnnnnnnnnnnnnnnnnnnnnnnnnnnnnnnnnnnnnnnnnnnnnnnnnnnnnnnnnnnnnnnnnnnnnnnnnnnnnn
nnnnnnnnnnnnnnnnnnnnnnnnnnnnnnnnnnnnnnnnnnnnnnnnnnnnnnnnnnnnnnnnnnnnnnnnnnnnnnnnnnnnnn
nnnnnnnnnnnnnnnnnnnnnnnnnnnnnnnnnnnnnnnnnnnnnnnnnnnnnnnnnnnnnnnnnnnnnnnnnnnnnnnnnnnnnn
nnnnnnnnnnnnnnnnnnnnnnnnnnnnnnnnnnnnnnnnnnnnnnnnnnnnnnnnnnnnnnnnnnnnnnnnnnnnnnnnnnnnnn
nnnnnnnnnnnnnnnnnnnnnnnnnnnnnnnnnnnnnnnnnnnnnnnnnnnnnnnnnnnnnnnnnnnnnnnnnnnnnnnnnnnnnn
nnnnnnnnnnnnnnnnnnnnnnnnnnnnnnnnnnnnnnnnnnnnnnnnnnnnnnnnnnnnnnnnnnnnnnnnnnnnnnnnnnnnnn
nnnnnnnnnnnnnnnnnnnnnnnnnnnnnnnnnnnnnnnnnnnnnnnnnnnnnnnnnnnnnnnnnnnnnnnnnnnnnnnnnnnnnn
nnnnnnnnnnnnnnnnnnnnnnnnnnnnnnnnnnnnnnnnnnnnnnnnnnnnnnnnnnnnnnnnnnnnnnnnnnnnnnnnnnnnnn
nnnnnnnnnnnnnnnnnnnnnnnnnnnnnnnnnnnnnnnnnnnnnnnnnnnnnnnnnnnnnnnnnnnnnnnnnnnnnnnnnnnnnn
nnnnnnnnnnnnnnnnnnnnnnnnnnnnnnnnnnnnnnnnnnnnnnnnnnnnnnnnnnnnnnnnnnnnnnnnnnnnnnnnnnnnnn
nnnnnnnnnnnnnnnnnnnnnnnnnnnnnnnnnnnnnnnnnnnnnnnnnnnnnnnnnnnnnnnnnnnnnnnnnnnnnnnnnnnnnn
nnnnnnnnnnnnnnnnnnnnnnnnnnnnnnnnnnnnnnnnnnnnnnnnnnnnnnnnnnnnnnnnnnnnnnnnnnnnnnnnnnnnnn
nnnnnnnnnnnnnnnnnnnnnnnnnnnnnnnnnnnnnnnnnnnnnnnnnnnnnnnnnnnnnnnnnnnnnnnnnnnnnnnnnnnnnn
nnnnnnnnnnnnnnnnnnnnnnnnnnnnnnnnnnnnnnnnnnnnnnnnnnnnnnnnnnnnnnnnnnnnnnnnnnnnnnnnnnnnnn
nnnnnnnnnnnnnnnnnnnnnnnnnnnnnnnnnnnnnnnnnnnnnnnnnnnnnnnnnnnnnnnnnnnnnnnnnnnnnnnnnnnnnn
nnnnnnnnnnnnnnnnnnnnnnnnnnnnnnnnnnnnnnnnnnnnnnnnnnnnnnnnnnnnnnnnnnnnnnnnnnnnnnnnnnnnnn
nnnnnnnnnnnnnnnnnnnnnnnnnnnnnnnnnnnnnnnnnnnnnnnnnnnnnnnnnnnnnnnnnnnnnnnnnnnnnnnnnnnnnn
nnnnnnnnnnnnnnnnnnnnnnnnnnnnnnnnnnnnnnnnnnnnnnnnnnnnnnnnnnnnnnnnnnnnnnnnnnnnnnnnnnnnnn
nnnnnnnnnnnnnnnnnnnnnnnnnnnnnnnnnnnnnnnnnnnnnnnnnnnnnnnnnnnnnnnnnnnnnnnnnnnnnnnnnnnnnn
nnnnnnnnnnnnnnnnnnnnnnnnnnnnnnnnnnnnnnnnnnnnnnnnnnnnnnnnnnnnnnnnnnnnnnnnnnnnnnnnnnnnnn
nnnnnnnnnnnnnnnnnnnnnnnnnnnnnnnnnnnnnnnnnnnnnnnnnnnnnnnnnnnnnnnnnnnnnnnnnnnnnnnnnnnnnn
nnnnnnnnnnnnnnnnnnnnnnnnnnnnnnnnnnnnnnnnnnnnnnnnnnnnnnnnnnnnnnnnnnnnnnnnnnnnnnnnnnnnnn
nnnnnnnnnnnnnnnnnnnnnnnnnnnnnnnnnnnnnnnnnnnnnnnnnnnnnnnnnnnnnnnnnnnnnnnnnnnnnnnnnnnnnn
nnnnnnnnnnnnnnnnnnnnnnnnnnnnnnnnnnnnnnnnnnnnnnnnnnnnnnnnnnnnnnnnnnnnnnnnnnnnnnnnnnnnnn
nnnnnnnnnnnnnnnnnnnnnnnnnnnnnnnnnnnnnnnnnnnnnnnnnnnnnnnnnnnnnnnnnnnnnnnnnnnnnnnnnnnnnn
nnnnnnnnnnnnnnnnnnnnnnnnnnnnnnnnnnnnnnnnnnnnnnnnnnnnnnnnnnnnnnnnnnnnnnnnnnnnnnnnnnnnnn
nnnnnnnnnnnnnnnnnnnnnnnnnnnnnnnnnnnnnnnnnnnnnnnnnnnnnnnnnnnnnnnnnnnnnnnnnnnnnnnnnnnnnn
nnnnnnnnnnnnnnnnnnnnnnnnnnnnnnnnnnnnnnnnnnnnnnnnnnnnnnnnnnnnnnnnnnnnnnnnnnnnnnnnnnnnnn
nnnnnnnnnnnnnnnnnnnnnnnnnnnnnnnnnnnnnnnnnnnnnnnnnnnnnnnnnnnnnnnnnnnnnnnnnnnnnnnnnnnnnn
nnnnnnnnnnnnnnnnnnnnnnnnnnnnnnnnnnnnnnnnnnnnnnnnnnnnnnnnnnnnnnnnnnnnnnnnnnnnnnnnnnnnnn
nnnnnnnnnnnnnnnnnnnnnnnnnnnnnnnnnnnnnnnnnnnnnnnnnnnnnnnnnnnnnnnnnnnnnnnnnnnnnnnnnnnnnn
nnnnnnnnnnnnnnnnnnnnnnnnnnnnnnnnnnnnnnnnnnnnnnnnnnnnnnnnnnnnnnnnnnnnnnnnnnnnnnnnnnnnnn
nnnnnnnnnnnnnnnnnnnnnnnnnnnnnnnnnnnnnnnnnnnnnnnnnnnnnnnnnnnnnnnnnnnnnnnnnnnnnnnnnnnnnn
nnnnnnnnnnnnnnnnnnnnnnnnnnnnnnnnnnnnnnnnnnnnnnnnnnnnnnnnnnnnnnnnnnnnnnnnnnnnnnnnnnnnnn
nnnnnnnnnnnnnnnnnnnnnnnnnnnnnnnnnnnnnnnnnnnnnnnnnnnnnnnnnnnnnnnnnnnnnnnnnnnnnnnnnnnnnn
nnnnnnnnnnnnnnnnnnnnnnnnnnnnnnnnnnnnnnnnnnnnnnnnnnnnnnnnnnnnnnnnnnnnnnnnnnnnnnnnnnnnnn
nnnnnnnnnnnnnnnnnnnnnnnnnnnnnnnnnnnnnnnnnnnnnnnnnnnnnnnnnnnnnnnnnnnnnnnnnnnnnnnnnnnnnn
nnnnnnnnnnnnnnnnnnnnnnnnnnnnnnnnnnnnnnnnnnnnnnnnnnnnnnnnnnnnnnnnnnnnnnnnnnnnnnnnnnnnnn

nnnnnnnnnnnnnnnnnnnnnnnnnnnnnnnnnnnnnnnnnnnnnnnnnnnnnnnnnnnnnnnnnnnnnnnnnnnnnnnnnn
nnnnnnnnnnnnnnnnnnnnnnnnnnnnnnnnnnnnnnnnnnnnnnnnnnnnnnnnnnnnnnnnnnnnnnnnnnnnnnnnnn
nnnnnnnnnnnnnnnnnnnnnnnnnnnnnnnnnnnnnnnnnnnnnnnnnnnnnnnnnnnnnnnnnnnnnnnnnnnnnnnnnn
nnnnnnnnnnnnnnnnnnnnnnnnnnnnnnnnnnnnnnnnnnnnnnnnnnnnnnnnnnnnnnnnnnnnnnnnnnnnnnnnnn
nnnnnnnnnnnnnnnnnnnnnnnnnnnnnnnnnnnnnnnnnnnnnnnnnnnnnnnnnnnnnnnnnnnnnnnnnnnnnnnnnn
nnnnnnnnnnnnnnnnnnnnnnnnnnnnnnnnnnnnnnnnnnnnnnnnnnnnnnnnnnnnnnnnnnnnnnnnnnnnnnnnnn
nnnnnnnnnnnnnnnnnnnnnnnnnnnnnnnnnnnnnnnnnnnnnnnnnnnnnnnnnnnnnnnnnnnnnnnnnnnnnnnnnn
nnnnnnnnnnnnnnnnnnnnnnnnnnnnnnnnnnnnnnnnnnnnnnnnnnnnnnnnnnnnnnnnnnnnnnnnnnnnnnnnnn
nnnnnnnnnnnnnnnnnnnnnnnnnnnnnnnnnnnnnnnnnnnnnnnnnnnnnnnnnnnnnnnnnnnnnnnnnnnnnnnnnn
nnnnnnnnnnnnnnnnnnnnnnnnnnnnnnnnnnnnnnnnnnnnnnnnnnnnnnnnnnnnnnnnnnnnnnnnnnnnnnnnnn
nnnnnnnnnnnnnnnnnnnnnnnnnnnnnnnnnnnnnnnnnnnnnnnnnnnnnnnnnnnnnnnnnnnnnnnnnnnnnnnnnn
nnnnnnnnnnnnnnnnnnnnnnnnnnnnnnnnnnnnnnnnnnnnnnnnnnnnnnnnnnnnnnnnnnnnnnnnnnnnnnnnnn
nnnnnnnnnnnnnnnnnnnnnnnnnnnnnnnnnnnnnnnnnnnnnnnnnnnnnnnnnnnnnnnnnnnnnnnnnnnnnnnnnn
nnnnnnnnnnnnnnnnnnnnnnnnnnnnnnnnnnnnnnnnnnnnnnnnnnnnnnnnnnnnnnnnnnnnnnnnnnnnnnnnnn
nnnnnnnnnnnnnnnnnnnnnnnnnnnnnnnnnnnnnnnnnnnnnnnnnnnnnnnnnnnnnnnnnnnnnnnnnnnnnnnnnn
nnnnnnnnnnnnnnnnnnnnnnnnnnnnnnnnnnnnnnnnnnnnnnnnnnnnnnnnnnnnnnnnnnnnnnnnnnnnnnnnnn
nnnnnnnnnnnnnnnnnnnnnnnnnnnnnnnnnnnnnnnnnnnnnnnnnnnnnnnnnnnnnnnnnnnnnnnnnnnnnnnnnn
nnnnnnnnnnnnnnnnnnnnnnnnnnnnnnnnnnnnnnnnnnnnnnnnnnnnnnnnnnnnnnnnnnnnnnnnnnnnnnnnnn
nnnnnnnnnnnnnnnnnnnnnnnnnnnnnnnnnnnnnnnnnnnnnnnnnnnnnnnnnnnnnnnnnnnnnnnnnnnnnnnnnn
nnnnnnnnnnnnnnnnnnnnnnnnnnnnnnnnnnnnnnnnnnnnnnnnnnnnnnnnnnnnnnnnnnnnnnnnnnnnnnnnnn
nnnnnnnnnnnnnnnnnnnnnnnnnnnnnnnnnnnnnnnnnnnnnnnnnnnnnnnnnnnnnnnnnnnnnnnnnnnnnnnnnn
nnnnnnnnnnnnnnnnnnnnnnnnnnnnnnnnnnnnnnnnnnnnnnnnnnnnnnnnnnnnnnnnnnnnnnnnnnnnnnnnnn
nnnnnnnnnnnnnnnnnnnnnnnnnnnnnnnnnnnnnnnnnnnnnnnnnnnnnnnnnnnnnnnnnnnnnnnnnnnnnnnnnn
nnnnnnnnnnnnnnnnnnnnnnnnnnnnnnnnnnnnnnnnnnnnnnnnnnnnnnnnnnnnnnnnnnnnnnnnnnnnnnnnnn
nnnnnnnnnnnnnnnnnnnnnnnnnnnnnnnnnnnnnnnnnnnnnnnnnnnnnnnnnnnnnnnnnnnnnnnnnnnnnnnnnn
nnnnnnnnnnnnnnnnnnnnnnnnnnnnnnnnnnnnnnnnnnnnnnnnnnnnnnnnnnnnnnnnnnnnnnnnnnnnnnnnnn
nnnnnnnnnnnnnnnnnnnnnnnnnnnnnnnnnnnnnnnnnnnnnnnnnnnnnnnnnnnnnnnnnnnnnnnnnnnnnnnnnn
nnnnnnnnnnnnnnnnnnnnnnnnnnnnnnnnnnnnnnnnnnnnnnnnnnnnnnnnnnnnnnnnnnnnnnnnnnnnnnnnnn
nnnnnnnnnnnnnnnnnnnnnnnnnnnnnnnnnnnnnnnnnnnnnnnnnnnnnnnnnnnnnnnnnnnnnnnnnnnnnnnnnn
nnnnnnnnnnnnnnnnnnnnnnnnnnnnnnnnnnnnnnnnnnnnnnnnnnnnnnnnnnnnnnnnnnnnnnnnnnnnnnnnnn
nnnnnnnnnnnnnnnnnnnnnnnnnnnnnnnnnnnnnnnnnnnnnnnnnnnnnnnnnnnnnnnnnnnnnnnnnnnnnnnnnn
nnnnnnnnnnnnnnnnnnnnnnnnnnnnnnnnnnnnnnnnnnnnnnnnnnnnnnnnnnnnnnnnnnnnnnnnnnnnnnnnnn
nnnnnnnnnnnnnnnnnnnnnnnnnnnnnnnnnnnnnnnnnnnnnnnnnnnnnnnnnnnnnnnnnnnnnnnnnnnnnnnnnn
nnnnnnnnnnnnnnnnnnnnnnnnnnnnnnnnnnnnnnnnnnnnnnnnnnnnnnnnnnnnnnnnnnnnnnnnnnnnnnnnnn
nnnnnnnnnnnnnnnnnnnnnnnnnnnnnnnnnnnnnnnnnnnnnnnnnnnnnnnnnnnnnnnnnnnnnnnnnnnnnnnnnn
nnnnnnnnnnnnnnnnnnnnnnnnnnnnnnnnnnnnnnnnnnnnnnnnnnnnnnnnnnnnnnnnnnnnnnnnnnnnnnnnnn
nnnnnnnnnnnnnnnnnnnnnnnnnnnnnnnnnnnnnnnnnnnnnnnnnnnnnnnnnnnnnnnnnnnnnnnnnnnnnnnnnn
nnnnnnnnnnnnnnnnnnnnnnnnnnnnnnnnnnnnnnnnnnnnnnnnnnnnnnnnnnnnnnnnnnnnnnnnnnnnnnnnnn
nnnnnnnnnnnnnnnnnnnnnnnnnnnnnnnnnnnnnnnnnnnnnnnnnnnnnnnnnnnnnnnnnnnnnnnnnnnnnnnnnn
nnnnnnnnnnnnnnnnnnnnnnnnnnnnnnnnnnnnnnnnnnnnnnnnnnnnnnnnnnnnnnnnnnnnnnnnnnnnnnnnnn
nnnnnnnnnnnnnnnnnnnnnnnnnnnnnnnnnnnnnnnnnnnnnnnnnnnnnnnnnnnnnnnnnnnnnnnnnnnnnnnnnn
nnnnnnnnnnnnnnnnnnnnnnnnnnnnnnnnnnnnnnnnnnnnnnnnnnnnnnnnnnnnnnnnnnnnnnnnnnnnnnnnnn
nnnnnnnnnnnnnnnnnnnnnnnnnnnnnnnnnnnnnnnnnnnnnnnnnnnnnnnnnnnnnnnnnnnnnnnnnnnnnnnnnn
nnnnnnnnnnnnnnnnnnnnnnnnnnnnnnnnnnnnnnnnnnnnnnnnnnnnnnnnnnnnnnnnnnnnnnnnnnnnnnnnnn
nnnnnnnnnnnnnnnnnnnnnnnnnnnnnnnnnnnnnnnnnnnnnnnnnnnnnnnnnnnnnnnnnnnnnnnnnnnnnnnnnn
nnnnnnnnnnnnnnnnnnnnnnnnnnnnnnnnnnnnnnnnnnnnnnnnnnnnnnnnnnnnnnnnnnnnnnnnnnnnnnnnnn
nnnnnnnnnnnnnnnnnnnnnnnnnnnnnnnnnnnnnnnnnnnnnnnnnnnnnnnnnnnnnnnnnnnnnnnnnnnnnnnnnn
nnnnnnnnnnnnnnnnnnnnnnnnnnnnnnnnnnnnnnnnnnnnnnnnnnnnnnnnnnnnnnnnnnnnnnnnnnnnnnnnnn
nnnnnnnnnnnnnnnnnnnnnnnnnnnnnnnnnnnnnnnnnnnnnnnnnnnnnnnnnnnnnnnnnnnnnnnnnnnnnnnnnn
nnnnnnnnnnnnnnnnnnnnnnnnnnnnnnnnnnnnnnnnnnnnnnnnnnnnnnnnnnnnnnnnnnnnnnnnnnnnnnnnnn
nnnnnnnnnnnnnnnnnnnnnnnnnnnnnnnnnnnnnnnnnnnnnnnnnnnnnnnnnnnnnnnnnnnnnnnnnnnnnnnnnn
nnnnnnnnnnnnnnnnnnnnnnnnnnnnnnnnnnnnnnnnnnnnnnnnnnnnnnnnnnnnnnnnnnnnnnnnnnnnnnnnnn
nnnnnnnnnnnnnnnnnnnnnnnnnnnnnnnnnnnnnnnnnnnnnnnnnnnnnnnnnnnnnnnnnnnnnnnnnnnnnnnnnn
nnnnnnnnnnnnnnnnnnnnnnnnnnnnnnnnnnnnnnnnnnnnnnnnnnnnnnnnnnnnnnnnnnnnnnnnnnnnnnnnnn
nnnnnnnnnnnnnnnnnnnnnnnnnnnnnnnnnnnnnnnnnnnnnnnnnnnnnnnnnnnnnnnnnnnnnnnnnnnnnnnnnn
nnnnnnnnnnnnnnnnnnnnnnnnnnnnnnnnnnnnnnnnnnnnnnnnnnnnnnnnnnnnnnnnnnnnnnnnnnnnnnnnnn
nnnnnnnnnnnnnnnnnnnnnnnnnnnnnnnnnnnnnnnnnnnnnnnnnnnnnnnnnnnnnnnnnnnnnnnnnnnnnnnnnn
nnnnnnnnnnnnnnnnnnnnnnnnnnnnnnnnnnnnnnnnnnnnnnnnnnnnnnnnnnnnnnnnnnnnnnnnnnnnnnnnnn
nnnnnnnnnnnnnnnnnnnnnnnnnnnnnnnnnnnnnnnnnnnnnnnnnnnnnnnnnnnnnnnnnnnnnnnnnnnnnnnnnn
nnnnnnnnnnnnnnnnnnnnnnnnnnnnnnnnnnnnnnnnnnnnnnnnnnnnnnnnnnnnnnnnnnnnnnnnnnnnnnnnnn
nnnnnnnnnnnnnnnnnnnnnnnnnnnnnnnnnnnnnnnnnnnnnnnnnnnnnnnnnnnnnnnnnnnnnnnnnnnnnnnnnn
nnnnnnnnnnnnnnnnnnnnnnnnnnnnnnnnnnnnnnnnnnnnnnnnnnnnnnnnnnnnnnnnnnnnnnnnnnnnnnnnnn
nnnnnnnnnnnnnnnnnnnnnnnnnnnnnnnnnnnnnnnnnnnnnnnnnnnnnnnnnnnnnnnnnnnnnnnnnnnnnnnnnn
nnnnnnnnnnnnnnnnnnnnnnnnnnnnnnnnnnnnnnnnnnnnnnnnnnnnnnnnnnnnnnnnnnnnnnnnnnnnnnnnnn
nnnnnnnnnnnnnnnnnnnnnnnnnnnnnnnnnnnnnnnnnnnnnnnnnnnnnnnnnnnnnnnnnnnnnnnnnnnnnnnnnn

nnnnnnnnnnnnnnnnnnnnnnnnnnnnnnnnnnnnnnnnnnnnnnnnnnnnnnnnnnnnnnnnnnnnnnnnnnn
nnnnnnnnnnnnnnnnnnnnnnnnnnnnnnnnnnnnnnnnnnnnnnnnnnnnnnnnnnnnnnnnnnnnnnnnnnn
nnnnnnnnnnnnnnnnnnnnnnnnnnnnnnnnnnnnnnnnnnnnnnnnnnnnnnnnnnnnnnnnnnnnnnnnnnn
nnnnnnnnnnnnnnnnnnnnnnnnnnnnnnnnnnnnnnnnnnnnnnnnnnnnnnnnnnnnnnnnnnnnnnnnnnn
nnnnnnnnnnnnnnnnnnnnnnnnnnnnnnnnnnnnnnnnnnnnnnnnnnnnnnnnnnnnnnnnnnnnnnnnnnn
nnnnnnnnnnnnnnnnnnnnnnnnnnnnnnnnnnnnnnnnnnnnnnnnnnnnnnnnnnnnnnnnnnnnnnnnnnn
nnnnnnnnnnnnnnnnnnnnnnnnnnnnnnnnnnnnnnnnnnnnnnnnnnnnnnnnnnnnnnnnnnnnnnnnnnn
nnnnnnnnnnnnnnnnnnnnnnnnnnnnnnnnnnnnnnnnnnnnnnnnnnnnnnnnnnnngagataagaattaga
gcttagaaataatgtatctggggtccctatttatagggagagggcgtaacagactgatagcgacgcctgtaacc
gtctggtggtgggccgttcagggccatgagtagtttgttacggagagtagtgggatggagtcgtggctatcgccg
tggtctgtcacgtggagcctgttgtggagagtggagtggtgtccgttgtcgtgacttgccagagcatgatggagc
cgcatgggatctgttacagaaagtggagcagagtcgtggccattattgtggcctgttagggagtccgggcctgtc
ggtcgaagctcggtaggggcgtctggtggagcggggtagctctctgtctaagaggagctcggatgttttcgaggt
tgctaacgagcccgctgtcaacaagtgccttaggtgttggcgcggcaggctgaaatcattttgtcgtagaagctc
ggatggagacttcccgttggtgaagttcggctggggttggcctgacgtagaacttattggaaagctgttcgtcga
ggaggtctgattccatgaggagttcggctgaaggtcgaccgcttggctgaattcagaaggtgtcgtagaagcctg
tccgcaggaggagcctgtagaacactgttgaagttcggttggagccgtcgaaggtcggttgacgctgctgaaagt
cggtggccggagaggtacggaagacactggaggcagtcggtcgccatagaagttcgtctgctggagctcgtccgt
ctgttgtagaagttcgcccggtacctatctgttataggagctcgatcggagcctgacttttgtagaagtccggat
gggaccgcttattgaagaagcccggtcgaagtccgcggtagagattcgacgtggatcgcttgtagtagaagctcg
acgtagatcgcttatagtagaagctcggagtagaccgctcttcgtgggagctcggagtagaccgctcttggtggg
agctcgaagcaggccgctcttggtgggagctcagggtagaccgtttgtagtagaagcttggagtgttcaggcctt
gtcgaagtccggacgatgtctacctgcaataaaaattcggggaagaagtccggctcccgtaggagttcgagtgaa
gtctatctgcagtaggaattcggggcagtggaccgtctcccgtaggagctcgggtggaatcttcctgcaatagaa
gttcgaggcagtggaccatctcccgtaggagctcggagtagatcatttgtagtagaagttcggagttggtcgttt
agctcccataggagctcagggtaggccgttcgtagtagaagttcgggtaagccgcctgtaggaggtcggacgaag
tatacctgaaatagaaattcggggtagaggtccggctcctgtaggagctcggatgaggtccgactcccgtaggag
cttggacggagtccaggaggcggtcgaccgccgtagagatttggagggagtccggccgtcgtagagatctgctgt
tgggaagttcggccaccgtcgaagttctggaagaagttcggattggagtcgatcgtggctggaaaagatctgaaa
gaggtccggagaacgatcgatcactgtagaaggtcggccggtagcggagcccggagagcacttggagcagcctga
tagacgactgaagaagctcatcctagagaagcccggagggtgtggtaggagctcgcccgtcggaagagttcggtc
gacacttgtagtgggcatcgtagaagctcattcgtcagcgaggcttgggaacattgtggaagctcggcagctggg
aaggtccggaaagatgtcgcccaaggtcagaagccggaaggatcctggggaggtcaatcctgtcacgaacttcgg
ctgagggtatttatacccaacaacgaacaaaaaaaaaaaaaaaattaagaaatagaagtataaaattaggtccaa
attatcattgatagaacaacattgcaaaatcataaattttaatttctgcttttttttggtaaaatggctatatct
aatttgctcaatctctgaaccacgcattcttgatttgttagattgcctgattagatatccaatgactctggtttt
ggcagttcaattcatcatgaaatatgttggtgatacatatcttttagctgttgcatccttgacaagtcaaactac
taaatttgctaaagctttcaattaaatccacaaattagaattctgacattttttttgatagagagaattctaata
ttagaaggactaataatgactagttgcaaataggaaactagtaaccaagaaattccaagtgaactagcgaccggc
aattgaatgattcttgttttaccggtaaaagccaattcttatcaggaagtccaatttccactccagcaggagctt
cccgtaaaagcaatttaggaaaccagtctgattccttttggaaacttccttgggagcccaaatttagaaagaaag
aggacttctatgggttaggttatcgagctatgaagaggactggtcgaggagggtggagaggtttggaagtgaagg
agaaaaatgatcgcctgataaatcattggtttatttgttcaaggagagtaaatctggtgcaagagacatcttcat
cagtatttgtgaattaatttgtagctacttatgaaatggattccaactggagtagccgattgaaatttattgttt
cctataattttgtctattttacctacatatttaataatatttttctttatttatgatcattttttcttgtctcct
gcatgggagttttttcttcaccaacatttcatgtttcttgcatttgtatttcagtattggtttccctgatgacat
ctgtgaattggcgaatattcattcgtcatatatgatctgataagtagtcatttttccatgggcatgggacagcaa
aatttggtggaagcacctctatttagaaggtgttttgttcgcaattagaatcgaaagagaggaatcaaaatggcc
aaatcttctgaagcatttggttcatgactggaatcaaaatcagaatcaaaataaaaatttgaatctttaaaaaaa
atagagattgagttttatgtatattaagctaattccatttcatcctgaaattgaaattagaatgggattcctttc
aacaaaataattgaaatgaaagtcacccatttccattctcatttgaaaccccaactctccccacccagatatccc
ttagtataggcaacacagcatcgagccatcaactatataggaaatactctcctagggagaatgttttgcttataa
atggagtgtaccaaagagctactaactagatttaccaaaaaaggggagcaagcaagcaattggaaagaagattat
ttaaagccactcaaaaggggatggaatcctagcacataaagcatgcttggacactaaaatgcctgccttccatga
taacttagagggaatcattaggtttcagagtcaacaacaaagtgatcactgttatttcactcttatggcggcttg
accataaggctcccaatcccacccagaatattcaattcggaggataattttagaaagagaccctcccactatatg
ggcaaatccaatgagtgagtatgtcacttcagcaccatcgagatctggcaaactagaaactatttccaatacctg
cgctcaggccatctgtgatgacccggccaattgggcccagatcagccagccaaaccaacaaaaaaaaaaaaaaaa
agaaaacggagcagaggactcccgaagggagtcttcttcctcctctcgccgactcccaatcggagtcgggaagag
gcccttccagccctatttaaggaggagatccctcctccttcccttccaccagaaaattcgagctcaatcggcggc
aatcgnnnnnnnnnnnnnnnnnnnnnnnnnnnnnnnnnnnnnnnnnnnnnnnnnnnnnnnnnnnnnnnnnnnnnn
nnnnnnnnnnnnnnnnnnnnnnnnnnnnnnnnnnnnnnnnnnnnnnnnnnnnnnnnnnnnnnnnnnnnnnnnnnn
nnnnnnnnnnnnnnnnnnnnnnnnnnnnnnnnnnnnnnnnnnnnnnnnnnnnnnnnnnnnnnnnnnnnnnnnnnn

nnnnnnnnnnnnnnnnnnnnnnnnnnnnnnnnnnnnnnnnnnnnnnnnnnnnnnnnnnnnnnnnnnnnnnnnnnn
nnnnnnnnnnnnnnnnnnnnnnnnnnnnnnnnnnnnnnnnnnnnnnnnnnnnnnnnnnnnnnnnnnnnnnnnnnn
nnnnnnnnnnnnnnnnnnnnnnnnnnnnnnnnnnnnnnnnnnnnnnnnnnnnnnnnnnnnnnnnnnnnnnnnnnn
nnnnnnnnnnnnnnnnnnnnnnnnnnnnnnnnnnnnnnnnnnnnnnnnnnnnnnnnnnnnnnnnnnnnnnnnnnn
nnnnnnnnnnnnnnnnnnnnnnnnnnnnnnnnnnnnnnnnnnnnnnnnnnnnnnnnnnnnnnnnnnnnnnnnnnn
nnnnnnnnnnnnnnnnnnnnnnnnnnnnnnnnnnnnnnnnnnnnnnnnnnnnnnnnnnnnnnnnnnnnnnnnnnn
nnnnnnnnnnnnnnnnnnnnnnnnnnnnnnnnnnnnnnnnnnnnnnnnnnnnnnnnnnnnnnnnnnnnnnnnnnn
nnnnnnnnnnnnnnnnnnnnnnnnnnnnnnnnnnnnnnnnnnnnnnnnnnnnnnnnnnnnnnnnnnnnnnnnnnn
nnnnnnnnnnnnnnnnnnnnnnnnnnnnnnnnnnnnnnnnnnnnnnnnnnnnnnnnnnnnnnnnnnnnnnnnnnn
nnnnnnnnnnnnnnnnnnnnnnnnnnnnnnnnnnnnnnnnnnnnnnnnnnnnnnnnnnnnnnnnnnnnnnnnnnn
nnnnnnnnnnnnnnnnnnnnnnnnnnnnnnnnnnnnnnnnnnnnnnnnnnnnnnnnnnnnnnnnnnnnnnnnnnn
nnnnnnnnnnnnnnnnnnnnnnnnnnnnnnnnnnnnnnnnnnnnnnnnnnnnnnnnnnnnnnnnnnnnnnnnnnn
nnnnnnnnnnnnnnnnnnnnnnnnnnnnnnnnnnnnnnnnnnnnnnnnnnnnnnnnnnnnnnnnnnnnnnnnnnn
nnnnnnnnnnnnnnnnnnnnnnnnnnnnnnnnnnnnnnnnnnnnnnnnnnnnnnnnnnnnnnnnnnnnnnnnnnn
nnnnnnnnnnnnnnnnnnnnnnnnnnnnnnnnnnnnnnnnnnnnnnnnnnnnnnnnnnnnnnnnnnnnnnnnnnn
nnnnnnnnnnnnnnnnnnnnnnnnnnnnnnnnnnnnnnnnnnnnnnnnnnnnnnnnnnnnnnnnnnnnnnnnnnn
nnnnnnnnnnnnnnnnnnnnnnnnnnnnnnnnnnnnnnnnnnnnnnnnnnnnnnnnnnnnnnnnnnnnnnnnnnn
nnnnnnnnnnnnnnnnnnnnnnnnnnnnnnnnnnnnnnnnnnnnnnnnnnnnnnnnnnnnnnnnnnnnnnnnnnn
nnnnnnnnnnnnnnnnnnnnnnnnnnnnnnnnnnnnnnnnnnnnnnnnnnnnnnnnnnnnnnnnnnnnnnnnnnn
nnnnnnnnnnnnnnnnnnnnnnnnnnnnnnnnnnnnnnnnnnnnnnnnnnnnnnnnnnnnnnnnnnnnnnnnnnn
nnnnnnnnnnnnnnnnnnnnnnnnnnnnnnnnnnnnnnnnnnnnnnnnnnnnnnnnnnnnnnnnnnnnnnnnnnn
nnnnnnnnnnnnnnnnnnnnnnnnnnnnnnnnnnnnnnnnnnnnnnnnnnnnnnnnnnnnnnnnnnnnnnnnnnn
nnnnnnnnnnnnnnnnnnnnnnnnnnnnnnnnnnnnnnnnnnnnnnnnnnnnnnnnnnnnnnnnnnnnnnnnnnn
nnnnnnnnnnnnnnnnnnnnnnnnnnnnnnnnnnnnnnnnnnnnnnnnnnnnnnnnnnnnnnnnnnnnnnnnnnn
nnnnnnnnnnnnnnnnnnnnnnnnnnnnnnnnnnnnnnnnnnnnnnnnnnnnnnnnnnnnnnnnnnnnnnnnnnn
nnnnnnnnnnnnnnnnnnnnnnnnnnnnnnnnnnnnnnnnnnnnnnnnnnnnnnnnnnnnnnnnnnnnnnnnnnn
nnnnnnnnnnnnnnnnnnnnnnnnnnnnnnnnnnnnnnnnnnnnnnnnnnnnnnnnnnnnnnnnnnnnnnnnnnn
nnnnnnnnnnnnnnnnnnnnnnnnnnnnnnnnnnnnnnnnnnnnnnnnnnnnnnnnnnnnnnnnnnnnnnnnnnn
nnnnnnnnnnnnnnnnnnnnnnnnnnnnnnnnnnnaccatcagaatcaactctcttccgatcattcgcgtcctgg
catttaattgaagtcccgacatccatcgtgactcgaacttgagtcacttgcatggaagccaaggcccgtttcgcc
aggccactccctcgatggcaaaaaattttttctttatggtggaccatggatatcaaagatctacctcagaccatt
attatctgtcaacaagtatcttgatggcatactttgttgatttatgtcaataatacattgtcggtcaaaatgcta
acaaaattgagaataaaggttaacgacttctctatataaaagtttaaagataagaaatcaaattttaaacataaa
atctttcgtagcacaaagtctaagaggcctcaattatcaaatcgattcctaaaattatttttggatctttctca
atgttttgacttttttttttctgataaggcggataattatacaagcctattatgaatataaccaaaaaagtcaata
aataaaatatcacaaaaagataaagggggtgctctccccgcttgggcgctccatgacaaaaaaggtgacccagtca
gccatactgttcgccttccgataaacatgccagatgttcaggaaaaggcaccccttatcactctccagacatcac
gtaacaataggtgcacatgagctacccgacccgcctctgcaaccaggccacaataatggatgagtccccttcaat
atcgatatgatctacccctagggtccgcctcacacacatgccaacccaagccattcgaagctccgcatcagggac
agtgacatctactagatgactttcccctacagcaataaacattaagtccaggcctcaaattacaaacccaatttc
tcttctactgttatacatgctgccatcaaaattaaccctaataaagctcatatgtaggagctcgcaagtaatgaa
caccgaatgagtcactacaagaatagaaaaagagtcctaggtgtcctagattatcaagatcaaactagtagatag
ctgaagaatctctacagtctgagctagtgccccctctaagataaatctcgcatgctgcctccttaactcaaacac
tagactgctcttagccaaccaaatatggtacatgatataagctgcactaatgccaaaaaagcctggaagtctcgc
ttgaatactccttcctagtgcctcaagaaagaactaaatcgatatcgctgagtggacaaaagccggaaagaactg
gtcctccgacacacctgcacggtgctcggatattgaaaagaggacatggcccacaatctcatcatccatctcaca
accgaggtaggtgatgggggggtctatgcctctatctctcgaaccatcctgatggagagagggcctcgacccac
attccaaacaaaaggcacactcggtagtgagtgccccgccttcggatccaatctccatccataatgtcttggcca
ttcacagtaagtcataagcaaatgaacaatcatccactttccaaaaattgaaaatcattaggaatcctacaaatt
caacttgttcttttccctctaactgatgattggaacttcatccaggcatgaatcttactgaaaagttgaccacaa
ttgaggctctccctgcactaaacccatgaggatgatataggattcataagaacttttcctataggacacttacct
agacaatacttgaatcctacccagactgatgtcgtgacgctcctttagttaagtatgcatttttttgaatgacaga
tcatgattcagtttctccaaaatgtatgctattactagcaagagaattagcagtaaaagctgtagaaaagcagca
agctgttaacattctgagaaactgttgtttaccttttaatatgtttactgcatggcattttataaacctattgct
ggtagcagcattgcaagctgagctagagagaggccttgacatccttcgagatttgaatgagttactagtatcttt
actataagatatgctcatccactgatcattttggaattttatttagactaatctagcttaaaatatgtctgagc
aaaggtagtatgctaatttctgaaccccatcatcttcagaacaaataaatgaaacacaaactgtcgaattaacta
attgaagatgctaccaatttggaacaaagtcacgagagaatttcaagaaatttgcactaagaaagaaactaatgg
acaaggaatcacaaagtaaccccaaatagttgggagaaagattgttaattctagttcatattcaagatatacaat
ggaaaaattcaataaattagatgttgtagaggagcaatggatggacaactactgcataacacattgagaccatct
gtgtggctcgtttggtatttaggaaagaaactcgcagaggttagaagatacaaggaactcaagcggtgtaatact
tattagaaaaagcagcactcttatatttaattatttattcacgttcattatctcattcgattatcaattgctact
ctgatatggcatagaatgtatcagcatccagtctgatagcataataaagaaaactaacagaagatcccttatcat

ttcagcatgaaatcaataattgaccgctacagcaggacaaaggaggagaatcatcaggtgctaaatgcaacttca
gaagtgatgggtatgtccaattcttacgcaacattatcaacaactgataaatgagcttcttctttggcggtgtaca
aatgcgcaaattagtatgtttatattagttttgccatctcttcttggataaacttgatctttttactgcaagtaa
tctggtatctattaaatcaccgagtatgaaaatagcactcttcttttgctatagaaattgccaagacgctgcatg
atgtagtcttgtgttacggttatgtaccataacgtgcatattatttgcaacacttgtttcgcatcatatatgtcc
tgcaccagaaaagtaggggaaaatttagaaattacatgtgtacatcagctatatacgtttatcacagtttgattg
aagaagcttaattatgaaactttaaatgtatagttgaagatagttgttaaaaatgcattaccactacacaatgga
atataaatacgagacaaaatgcataaactatataaaacttatgattatccatgtcagacatctgcaaaaatatat
cattgaagttctggagaaaagttagtcaataaaataacgtccatgggaataggttctaaatatggtttatgttga
cattgaaaatgcttatgctaaagtactaagattcatataaataatgaatagcaaaggcacgatagttttgataaa
ggtattcatgatggagttgagagtggtaataagggaagatgccatatcgtggcaggccaacaatacgggtggtcc
taaagtctgtcatcataaataggtgctacaattactgattattgctacttaggtgaagaatcaaccatgtagctc
atctgtaaacaaaacattatcgttatggtgagataaaacaaggataaattcgtatggaagaacttggcattttga
taaaaatatttatacagagctgagagtggtgataaggggcagatgctatattgtggcagaccaacaatcagggtg
gtcctaaagtctgtcatcataaacatgtgctagtactctagcagatactatgctttggcaggcccccaataagga
tcatcctaaagcctgtcatcataaacaggtgctagaagtactgattgttgctacttagctgaagaatcaactctc
tagctcatttataaacaaaacaatataattgttgtgagaagatgaaaacaaggagaaatgctatatgggagttat
gaaagaatcctctgcaaagtcatatttataagcttagttactagaaacaggtgtggaagctaattcaggtgtggg
tttgggaaaggatctcttaattctttttttttaaataataaagggtgcagaaacaagttcccattggatattgc
aaagcaggtgtggcaccctatgcagaagtctcctgctactcaggtagtgagtgcaaacagcagatctaacgaaaa
cagatgacagaatgcaactgttgtggctgaataatgaaaatttatgctttggcaattgatggaaaaaaagatac
ttccaaaagattattacagttcagttctattatgttgaacaaacataaagaaagtattagacattttcattatat
ttcgagcatggagaactatgtaaagatgtgtttctgatgtgctgcatcattgtaatatattatggaagagacata
atataaattctatcaaaagaaagcccacacgctaggttgagaagtggtaaaactgagatggtcagaaatgagcat
aactacaggaagctaaataccactgcaatcagtaagatagcaaggccatgtcacaactaaagaggggcaatacac
aattagaggtactcagacttacattaatgggttgaagagaggatgggaaggcctagagaaacacagatcgaagtt
gaaagaattaaattaatatactcaagaagtggaagcagccattcatatttataagaatcaatgctgagtactaat
tcataaatctattacaagacaagacttggcaatcatagttatgttgcaaaaatttcatatccaagcagaatacag
gattccacaacagataatgacctttatgtgctatttttcattttccccgctcaagaaaaatgcaatcacagggat
agacactgaaacagtcaaaagttcaaaacaatttataaataggcccagaggaactagactgtccatgaactgctc
aacttggccagctaataggctctgttgagcttgttcaactaaactagctaagctttaacatcaactaaagcccaa
ctcataaaagagctgagcctcaacacttaaggccttgaatcttaagcctggctcaataacaagcttgtcaggtta
acaaataaggaatctagaacattaattgtttgggctcctgtaattttcatgcaataaactgcatctgagcaggt
cagcaattggcttgtttgtatccctatttataaaatttatgccaaagcatgaccatcaggttctgcagaataggt
agtaacatttcatgcagtgtgctcattgtgaatagattgactgcaaaaactgaggaattaagcatcaaatgagct
caccactgaaaagtcctgaaagctggccccagcatatggaagacaataaagccttaactgttcgtttgcatgatt
catgtcatgcttggtttgtattgctatgaaattggtgttgattgatggccagtcctttgttagactgccatttta
caatctcttcctgttgaccagaaattttactcgagccccttaacttcaatggccatttactgtttagtcccggtc
ctgtccatgcttaagcataaaggtgtgcaaaacagatgtaactatgtacagcacatagtgatgtctgccacaata
gatgaggacctagagcaagaaatcaaagagtaaacaatgagaaaatattcagaaacttataacgctaaaagtgac
aaaggtgcaggatgtttttttggttctttcttttattcattctgagcaattgtgggatcccagcaccatttcatt
gaataggtgaagtgatgaaacagagggagcatcggcaaaacagtgactcaagcctctaaaacataaaaatgagaa
ccagcagaaaatccacccaagtgcatcagttacaatgtttcgtacatgaaatataggtcactgcataagcttcct
cactttcaattattcatggtacgcactcaaaattgggaccttcccaaaaaaggttcagacctggtatggtaagag
gtaaagcataggcagcacaaataggatttaggtcctatttggctaagcttttagagtgtcagaaagtacttttttg
accattcaaaagtacttttggatggtacagtgatgtttggtaaaaaaatcaagaagctattttggctttgccaaa
aagctaaaaaggtttacttgggaaaagctccatttttgaagctttctcgagaagtattctctgactaatatcagaa
agctgtttaaaataaataaaatttnnnnnnnnnnnnnnnnnnnnatattataaatataatataatataataataa
ttaatattatgatatattataaacatataatataatttattatagtattattgtgttacaaatataatttgatat
aacaatatttaataataatatattatattatattatattacatcataattataataatattatattatagtataa
tataatattatattatattatatattattattattatataatccaatgatatttaatatataataatatataata
ttatattatattatattattattcactatgtaatacatgctatatcttttattatcattttatcataccgaaagt
acttttatttagttatcaaacatatattaaaattgtaaagtatttataaatgtggttaccaaacagcaaataa
cttttataatgactctacttccaaaagctctatttcttaaaagctctattttcaaaagctctactgctaacagca
atcctaaaagagccttagagcatttgagatagattcaacttatagtcaatatagatatttaaatccatggatgag
gttgtcctgatttttttatgaaataggatgcaccaccgttccacccatcttgacatttgggacaggagatatccca
aaacatcccgatcggaacgtcagaacgctagcaggatgaccatgtcttgacacatgagatggtaccccatcccgg
tatcccacaggatatcccgttgggacctgaatcatcataaggcattttacttaatgtctgagccttagttgttct
agcaattctaattattcaagcacaccattttcatataaccaagatgagcctactgatcggttccagtggctgacc
aaaccctgtcaagcatttgacttctctctctacatccttgtttcagtctaatataattctttgttccccataaaa
acacaagtttaggaggatggttaacaatgttcagctctggtggtgttccaatatcaactgtacattttagcaaca
aatattttaataaactactcaaaaagaaaagatggtgcaaatggcttccattctgggttctttcaaaatttccag
atcaatattattattcttttcctggaaagatcctcaggataattcatggacaggaagaggcagtcagctggtcac

atttctagaatcatgccaggccctttatgtataaaagattggaacatacagaagtaaaagttagtactaactgtc
aacaggtcactaaaatcctagataatgatgcaaggtaacataatgtaacagctagcatatgattttaccttgatc
ctcccaaatacttgttctgaatctctgagtacactgtctgatttccactcaatctcagggttctaatgatgtagg
tccctgaattcagcacaactagcagtgtcaaaagttcctacaagaatccgctacccaaacaagcaataacagtta
aatttgacatgagaattacacggtaacagagaggacatgactgtatataatgcagaagggtattggacaaggttt
gccataccaatcggtaccaatgcataccaatgtaccatagtgtaccgatactaaatcagtatgtggtatgaaggg
cgtgccaacacttggtatgccaaagcaagtaccataacggatgaaccaacactttaataggatgctatcgagaca
ggatttgatattgagatggtgaacctttattagaaatgtcaattgaattaggaaaaaaagtcgacaactatgatg
tacctgggaattatcgttgctttatcatcattgtttagctcaattgcttgagtgtcatttactaaagaattttag
gtcagggatccatgaactctgcaggaattttgtttacatctaagttggacgtgcaattgctaagacctgagctca
aatgctcaagggtatgtttttagctgcttgtcaacttccatatccaccaagcaatacagtttctgcaactttcat
ttacaaatcactgtggactctcttacgcaggccttctctacaagcagatgtctacgtctcactcgtttgattgaa
gagagcctttgtagtgtcagagtgtcccaactgcagcttcttgtctgcccctctcaactcaaccatcaatccagt
tccatcatttcggttggtgtcctttgcctctctatcatcatataagcatttgttttctcaatcaactcattccga
caatagcagaccatcatagcattgcgttttaatagcaaaaaacaaataacaattaaagaaataatatacataata
cgaaaagtatggataagacagtgcaaagatttccttacagatcataaacaatcttctttgttgagagtcttgaga
caggatctcctgcaagataattttagttatctcaacctatacagggaagttgtcagaacttgggagaaacatgtt
ctgctggtaatataaaagatgagttatagattcttagatcttgaatagtcaatagatcaggagttcattgatcac
aatttttagatttttctacagcctagcaataggatgcatatgcaatatgcatttagacttttatttatcta
gaatagattcaaattgccgggctccaagtatcaatcaatttgcagctttttgtacacataacaatagccaccctt
gcttcaaaaatttcgacaaatgagatcaagataaaacatatgtgaatgttgaactttatcaactttctgaatgt
ttgaaagtatgtagtagtgaaaataaacaagcagcctgtgatgaaagcaataaaatagaacatttagctgctata
tgtcaaggcaataagagaagattaaagagtcctacttgttcaaacattagcatatagagatatcacatacagtcg
agaccttgctagtattcatatgaatatagaaatttgaaaacacatgtttgaatggtactaatgaatgaaaaattt
caacaaactatgatgcttctggtgcccataaggagtactcaaggagagtgttgaggtgggatcctaggtcaacat
agaaaactatctctacgaaggaactaaagtctaccaacatacaagtcacaaaataataaagatgaccttattgaa
aaggatattttgctaaggaaaccaccttgtaaataaaagttcagtaataatccagttcatgcacataacttcctt
acaacttactcgcacataaggagtaaaagctagctatggcaacatgcacatttcatgggctagcacatgttgtgc
tatgcttggacttagggttcaacatctcggtaccaagctccacaactgtgctaacatgttactatgtcagtgcag
agccagtctaacatacggagtgtcagtacgcctccccatatcgcatgctggtaccaaaccaatatggtacgtcct
gtacctctagttagatacagtacaacaagccttgcttggactgatctaataaaaagcggaaacaattccttcaat
tagcttaactcggacctctcaaagaaaatactaataaaataatggagtggctatactagagaccacaaaaccatt
ccgaactgagcgattcgaggtataccaaacctatcggctagtaattgggatgattcggccgatcgatctagtctc
aaagcaaattgattccatttaactaaaagagaatcaccttgcccagtgtttgtcatactgccccatatcacttgg
tatgggacataacataccatatcgatatggcgtactatgttgaaccatgtattgatattgtaatagaattatgct
ggtatggggtccaatatcgagacggcaaaccttgacattacctctctcttatgtcatgatctctcctaaattctc
tccctctctctccctctctcccaatccatctcttcctcccccctctctctcgatcttcctctcctttcttccctgt
ctcctaatccaccaaaccagctatgatcacatcgatgccgatgtgacgatgactatgactaagggttttgcatta
gtgtaagaacaccacggccatcgccacctcttattgtgatgaagatcctcctttgccacctcaaccatctcatct
ccgctacctccggccccattccttgtctaaattggcacctccatcagactgggtgagggcccttctttcctccct
ctccctccatctagtttcaatatgtattggatccgtatggtatgcaccaataccataccgaaccattatctgact
agtacatccaccaataccagttcagtaaaccttaggctagactctttgtgctttcattaattgaaaacaaactga
ttacaatgcatcttatttatatcagtgacatttgagtttccaaatctgattcaaatttctttctacttaaaagag
gttagaacaagttttatgtcccgacgggataggggcatcccgtcctattcttattagaaatgggacttgacgggc
tcgggactttgaaacgcatccttgcagtgagagaaaagaagaagaaagaaggaaatgaaggaagataaagaaaag
ggaaaataaaaggaaagaagaagaaaatgaaaaaaaagaaaggaagggagaagaaaagaaaaagaaaaggaaaga
aaggaaagtagagaaaaagaaagaaaagaaggaagaaagagaaaagaaagaaaggaaagaagaaggagagaaaga
tggaggatatctactgggatacatgatcaggactgctatcagaatggaatcgaacgatggggcatcccgttccat
ggaaaatcggaatacctttgtcccatgggatttaaatcttggttagaaattttctaaggtagcaaggaatccatg
ctaagatgaaattcgtccaagagctctcttctcaattcctacatcaactttatcatttgttacatagcctaattc
ctcccagattggaagataagccatccatttcacccgaagaggatgcttttgcattggtcggcccatctcaaggct
caaacatcatagttccccaacattagtagggcacaatgaacccttgagcttgtcacatccatgtaaaactcaaaa
agttacccaatatccaaaaaaaggatcattataataggatgttttatgggtaagttatggtctctaaaagttgag
cttcatgatctagacttagatatggcggatcttgctcccaaaccctatccaaacttacctgaagtgatcaaattg
gtccacaatgggtccaatgatcctggaccaatatgcggtacatctatatgccaatgcatgggtctcaatgcatgc
caagcatcaaaatagcacaatatgtgccatgtcggtgccaaatcaacaacactgtaccaagtaccaacacatgac
acttaaaccaaacatccattgctgctagaatggggaaccctttgatagaattttgactccacactagaatcttca
cttaaatgaagtcacctagcctcttatggagtgcctatcaaaactaaacatagcctctcccctatagcctcacac
cataagaaggtccaggaaaagtctcttacacctagctccttgtctaattcatatagtactgaattacaataagga
aacaaatgaaataattaaaaacaacttttctaagccaaatagctttttctagcttgataaagacattcctgaata
gagaaaactatggaaaatagaacaaaaggatgttaaaattcatttctaattcaaaatagccaaggaaaggcaaag
tcataactatgtacaaccatgtcaacaaatttcaacaaagtagcctttcaccaagatgaccatatctctctcatt
caagctctaattaatatgattcaaaattgcattggaaagttgttttggtgctaaataagatagaagtactttgat

tttataattctgctagctttagtaagcattcttttggactatggtttacgaagttgatctttgaaaggcagctaa
tctatggtatgttttctcaactaaaataacaatactttgacattggtgagctaagggtctaacaagcaatgcctc
acaactatgtgatccaatcggagagaaggaagattttcttcaacccgcacaactcagatctctcaaaagaaaagg
tagaagaagaggagaattttgaaagcaagaatcaagtcgcttttctttctctttttttcattcacattgattaaa
aatgagatacatatcctctatttataataatgaggtccatccttacacaaattgagttggattcttctttagttc
taatgagactcttttacctaaatagatatgtcgagtagaaataatctgaaaaaagctaaaactcttcaaaaggta
gatctcgacttctacatcaactctaccttctatcatcctattggattcttctcggattagaagctacgctcgatc
aaagtctatccgaaaaaaatttttcaaattaatcaacccattttggagcccacacgaaggaattttgtcctcatt
gttggggcaccagcccctttaaaggatggcgtcacatcataaatctcaaaaaactatccaatttcaaataaaaaa
tatcattgaaatcagactcgtattaccaagtttggtttccagaagtttagcttgcaaattagcttcccaatgtgg
cgaatcttgctcttggaccctatccagccctatctatagtggccaaaatagtccacatcgagttcgtagatcctg
ttggatcactatggcaactgatagattcaatgtagtcttaaacctattatgaaacaaagtatgttgttttggacc
tctttttggactctggagccctcttcgggtcaagcttcatgtctagcatggatcagtagcatgtagagtaccctt
cactcacatcaaaccccccttggcttcaaaatttgtcttttgatgaattatagtggaataggatatgatggttttt
gagggcgagccttggtgatataataagattgctctattatgacctaggtgtcacaaattcgaaacatgaaaatag
ccgctccacatgcaataggaaagttaggtaaatctaacccttcgcaaactccattggcggaacctcatgcattag
ggcatccttttaggatatgatggtcttcaaacatgcttgtatcaaacacttcactatcataaactttcatgttcc
atcactatcttgtccttctcccatgatacttttggatctgtggcttctctatcacaagctttcaacataattta
ttaatatacggatttgtttagattctagggtagttctatacactttgataggatgattttgatgtaaatcatta
ctcaactatggagttgtgccgtcctctcttatggattgttattcatgttgcctcaaccctaaccttttcttaaat
tctaggatagtatataccctttgataggaatattttgatgtaaatcatcactcaactatagagttagcgtcttgc
catcctctcttatggattgttattcttgttgcctccaccctcacccttattcttatacattgatactatagttgt
aatacccaggccgaacaacttgggccttgtctagccataaaaaaatcaatccaaaaagaactgcacaattttaaa
gcaggaccaaataaaaattaaataagccagtacgtgcgttgcatgtgttgggagctgggaaaatacatgagaact
cccgaaggagttcttctccgattgaaatcctaaatccacttagaaattgagtggaaattaaactctagttctgtc
ccttaaatatctatttgaaggcctcaaccctcttgattctccctcaccaaagattggtcgtcatttgccacccctt
ttccccttcttttccttttgattgccagtggtttagtcttgatttggctgccaaaaattagatcaatactcatca
aaaatcaagtaagaactagcattatttaacttgccctcctctccttctgcagtccctaaatctcgaattctggcc
gatctattgttggatttcttcttgaaagtcgaaccatgctttttcctatttttttcattacttcctttggttttt
cgatgccttcagtcaccactaatcaccaacaagaggaccaccgactgcaccactgtgggcccaaccacgactagt
gatagccgtcgtcagtcttgagcctattgtctcgagaggtaaaggggaagtaaatcccctattcggttaaagaag
aaggaatgaagaagatggaaaaaaaaannnnnnnnnnnnnnnnnnnaagaaaagaaaaagagagagagaaagagc
ttatctctcttcctcttccatctctcttcccactttctctttccaccttctctctacttttttgtctttagaaaat
ttcactttctctctctataaatttctctctcaagaattatttctctctttataaaatttatctctctaagattga
tcgagtaaaggattttcttttgatgattttgatcgactttagtgaaggatcctattccatggtcatcaaatagcg
tgccagtacccatctcgattggatccggataccatcacatttgatctatttgatttttatggagccaattgtatc
ttaaatctcatcttgattaaatgaagttatttgattgaactgatctttgatcatcgagcactgtgttggcctcca
tcttgactttgttgcacactgttgatttgatcatctttgatctctattgaactacctagatcttaatatgatcc
tcgttagataaagttagatgactgactgactcagactgttggatgcaactatcgttcgtcatatctctcatgatt
atttgtatcctttctgattattggatttaattatgcataaggttatagttgtctcgatacaaaaatgtctagcaa
tgggatactgcgatgtgatctatgaattgaagattataaaaaaaatttatagaattatgtggatttgttggagta
cgtgtgagataagtaatgattcacttttttatagatttatcaataaattataaaatatttttctattgaattattt
atgatttataagttcaatgtatagtaattttggattgccaaatatgccttcttttatatcttaaatgatttatg
atcgattgtattaaaattgctatggattatatcgattgatttcgatatcgcctggtttatgacttttaaatttg
aatataaaatagggttttcaatcgagatggattttgactcatagcctgatcatgtattgaaccctactagtgggg
attaaatattggtacttcgataccctgccagtaagggttatgcacaggtattttgataccctacaggtggaggtt
atgcattgatattttgaaaccctgccagtggaggttgtgcactggtatttcgatatcctgctagtagagattgtg
cactagtatttcaataccctaccagtgagggtgtgcactaatatttgatacctcgtcattgggggttatgtagt
ggcattttgatttagtccatggctatcgaggtgaattggatgttttgaaagaattaatttatgaatgaaaatgaa
ttttgaaaagactaaatttgcatggatcttattttaattattgttatatattttgagttgatgcactttgcatgc
ttattttcagtatatcattattactgaatttatctagctaaagtgtacattatttattgagctatctagctcatt
actctatatttactattttcataaattcaaaggactagcttgatttggggattctatataggagcgtgaattag
aagcagaatttgataatctttattttagactaaggtttattgaaactgatggaaactatggaatattattttata
agacatttaatttagttatttgaataaagttgaacattaatttcaccgttgctttacgatatcatcataagatgc
cttgcatgcttgtggggaagttctccataagtacgtggcgattgccatgacctctggctcgtgatctcgggtcga
aggcatgacaatagtgctcctccaattgtcaagtattttcttagtgtttaaaatttttattaaacaaattagtcaa
cctctcaaatctccatagagatttcatcaggcccatcctcttttcctattttcatcttttccatagccttttac
cttagatactctagttctacgttgatagttaatatttctaaccttaatggacctcccaaattatctacatgtctt
tcattgaatagcatatgaaaatagtttctccatcttctttgatctcatctttctcaactaaaacttcttgatgtt
tatccttgatgcattaacctaagtctctagtttttttcctcctaatcttgcagtcttttagatatttttcacccct
agtttctcacctgtgatatctctccagatgccttgaatatagcctctcgtatcaccttcttaattccttcttgac
caccttatagttttcatacacttcctcattcctatatttaggtagtcttctatagcactatgtttgagccttaac
tgctcattgaactttcttattccaccaccaagtttcctcactagatggttttctttgattctcctcaaacctcta

tattcacattcttaatgacaatattatatggagtgtataagatggaacatgcggttgcacccaaagtattaagcc
tcaataagatggagcacatcccggctgtcccatcctgttctcatgagaaaataggattacaatggggttaagact
ctgaaacccatccatgcaaggagaaagaaaaaaggaaagatcaagaaaagggaaaatgaaaggaaagaagaagaa
gaagaagaaagaaaggaagggataaggaaagaaagaaagaaggaaagatggaagaaagaaaaagaaagaaggaaa
atagaaacaatggaagggataagaaagaaagaaaaacgaaaggaaagaaagaaagatagaaaaaagaaaggaagg
atagaaagaagnnnnnnnnnnnnnnnnnnnnaaaaagaaagaaaggaaaggaaaggaaagagaagggatgagaaa
tgggggatatctaccagatgcatgattaggactatcatcgagatggggtgggatagcgagacattcaattccatt
gagaaatcaggacaattctgtcccgcaagatttaaaaccttagttacaccatcgatatatgagtttctcatagta
tcaacaatacgaatacaacatgacacattgttaatgcatgaaaaagttgtgtcctaccataataatcaccatgaa
ttgtgatcgtatttgtgctttaatgtgcaaatatatctctaagcttctaacactagcttgcatggtaaattgact
gaaaaaatgtaaaccataagtatattcagcacaaatttatcgaatgattatataagtatcatgtagggctttcaa
aaccatgccgaaccacccagtacaccctgtatcgagccggacggataggaaactaagatggttccagcatccaaa
tcagttctcggagggatgaagaagaaagagaggaagagagagaaggggagagaaggagagggagaggaagaaag
ggcctccgacggccgtcgaaggtcgtctgacatcttgccaaggtgggtacacatggagggagggaagtagagaga
gtgggaggtccttaccagacctccggaggccgtcggaggcctttggacagacgacgcctccattgcgagatctcc
gatggccggtgagcccatgccgacaggcctgagggggtcgagctggaggagggcctccgtggcttcactccctt
cttcatttttgaaacaggggttcgcccctatttcagatttttttttttgattttaacatttgaaggttgcctactt
cataatttttttttaaaaaaaaataaagtggagttggcatatcatttgccaacttcgtccatttaaaagaacaaa
aatacggaatagggatgaatccctattccgcatgggagggcggaggccgtcctcgatgtgcttggcatcgaccgc
ccttgacactcgcttaccggcctatcacaggcttatccacaggcatgtcacaggggggtgacggcgtgggtgtggc
acggtgctgcgaaggatttggcatggccaaaatttgcaatgctactcgatcatttggaacccttgggtgagattt
ggggggatgccttgctggggtagtgagggagcaatggttcgccaaggagcggcatagccgtggcatgatatcggg
gggctcgggatgcatgatatttggaatgatggtggtccgccaagatatatgtggatacttaaggattcctcttgg
cttgaggggcgcggagctttggtcgcgtactcgactcctccagacccataagagacgggttcaccgtggcataat
tgtgtggacaatggtggtgccacacccgaatggtctcatatggaggtattcggggcaatgctggttggacgagct
ggccacactatgcagggaggggatatctcgaggttttatccttgaaggatttgtccagactactctatgcattct
tgatctttgtagtggagcgaagaggcgatagaggattagtcaggcaaaggcgtggcctaccaaacagacaagtct
cgtgagcatcgcaaggtgatgtcttgcggggatccacgggaaccgattgatcgtggagaacaagtaaggcttatg
gtatatcgatcgaattatgtgccttggagctgggccaaggatcatggatggccaagagggttatgccaatgagag
gacagcttgatgcatttatcaaccgatcgaagatatttacttgatggggcagacttggatgccgatatagtggca
gtgctcaatatgcgagtaagctgtggacctaggatgttaggtggtatggtcatgtatccccaggatcgtggtaat
gcacagtgatgggcgagcaggccatgaatattgtcaagcgatatagctatggatggaaaggtatatcgagaatgc
aagaccgggatcgtggcataagagcttgtggggaaagcttggatagtagaaccccagaggactcagatgtgtatg
atcaactgtagctttgagaagctgcagggcgtgttagcttggtagtttagctggacggcactagaggtgggtttg
cccaagcctcatgcatagtagtggcgttgcatggtcgtgctgtgtggatttagtatgtgtagtactgaaaggcag
cttggtgggttgcggattagtggcacagtggttatgtcatggcacgagggtatgtcaaggatcaagatatctggg
atgagcagtgcttggcttgaggggcagaatcttaaagtggacacagaggtgttgagctgcggagcactcttgaga
agagtatgttatggcacgaaagtgcgcatgagcattgcagttgatccgagggtcggattgcttacgggcacttaa
tgatgtcatctgggttcagtacaacagtgttgctaggcagaggcatagaggcgcatgcttggctatatggttgga
ctgctgcgtgcttattggtatgtccgaggaaggactaagagggctggccttggtgtgtggcatgctgaggcaccg
cacgggtgtttgagtaagaaacactcaccctgtgacaggccgtcagagaaatcgcatcgacgtcggcatccgatg
attcgatgagtctctccctctctctccttccctccctccacttgtaccgagctcgaaaagagcttggatggcctc
cgacgactgtcggaggccctctcttcctctccctctctctccctctcttcctctccttctccccctctcttcctc
tctttttttctccctctgggcccttctttgccgtgtcggtttgcaacattctaatccgatatggacccatatcga
ctcataccattggtcagccggcacggatcctggttctggctttgggaaccttggtatcatgtttagttgaacact
aatatattgaaattggtttagtgtaaaagaatttaaaaaatgctttagtatgttatttcggatctttcgatggg
tactatcaggggttcgctaaatcgggaaagcatgccaaaccatgccaatagcctacaggtatggttctgatgtaa
aattcggaacacgggcaagagggagaaggaaagagaggaaaagagagaggggaggaaaggagagggaaagagacc
ctctagaggctggctgaggctgctagagggccattatgcccactagagggctggcgaggaggggggagggagaaa
gaagagaagaaagagagagagagatggggtaggaaggagagcggagagggagataaagccttcgaaggcccgtcgag
ggctctggaggccgaagttggctaatcaaaacaatgtcttccgactttagttaactttttatttactagcttcg
ctgtgagttgacaaaaaatttaaaatcacgatcgaaaaagggcagtcgcccctatattcgtcggtagaaggcaag
gaggaggattcgagcctttgtgtccctccaatggccaccaccagcctctactccctctccttccctccctctccc
tccccctcccagtgatctctctctctcctccttcctctctcagttcagagtttcttactccttcgatcatctaag
ggccactacggctcttcaatggccaccactagcatctctgcagtctccctctccctccctcccctgcatctcttc
ctctttccagtttccctctcccttgctttctttgccatgttggttcaagcatactacagaatggtatgaaggcgt
actgaaccataccatcggccgattgacacggcaaaccttcgatactacaccttttcatgccggtttgcgacaagc
atggtatcatgatagcccctcgatgacccaagcataatacatgatgctttaatttatgaaatctaattaagcaat
gcaacatattaagaactagcatttgacaactgatataacagatatatgcacctaattcaaaaggagcgagtacct
taaaatgcttatgttacgagattcatcggtcatctattgcatatgactaattcaaaaggatgtagtgatgccttc
ccatataatgtacaacaccaattaagtggtaggcctatttgatgaaccaatcaagaggatgtagtataccttccc
atacaatgtacaacaccaattcaagtggtaggactatttgatgaaccaattgaatcagtaatttatgtgggttaa
gtttgtttacatagctttatttaatgcattagcttgtgttcgaaattccctaaaatatctcatctaattatagga

catttatatagttccctattttcctctctttccttgttcactgtaatcagatttttgggatttacacaggattt
gatcttatcccttatttcatgggatttatatgggatttgatcttatcccttatttcatgagatttataccagatt
tgatcttttaaaagatcatctttttatatatttatatgggattgattgcccacataaaaataaagaaattcacc
ctttttctcatggtatcagagccctttccatagggcctattttttttgccactgcttcttttctatctgtcttca
tcactaaagtccatagccttcatcgctgttggtcccattaccatcaacctttttgcttttattgcagaagtcgt
cattgttcgtagcttttctctgtcgaccgaatctgactgcactcccgttgtcagtcagtctttctattgttgctg
ttatttgatcccttctcctagtcaacatatttgaagcttctaaggccacggtgtccagccctttcgaacccgtct
ttgggataactcaatccattttaaatggtgaactgcgggccatgaacccttcctatcgtcttgatgggaagaatt
atcttcaatggtcacaataggtgaaaacatttctcaagagacgtggtaagctaaatcatttaattgctcttggtc
tgagtccctcagatccaaaatttctgcataggacaaagctgattccgtaatcatgtcatggctccgaaactcaa
tgcaacccgaagtcagccgcaattgcatgtttctctcatctgctaaagaaatataggacactgatcgacagacgt
attccaaagtccaagacgtagccgtcatttatgaactcaaaaataaaatctctgttgagaaaaatacctcgagat
tcaatccataataagcctagaataaagtctgcgatgaaaaatgaaatccaatgaacccaaggtcagcccaaatgg
agtccagatggaaaaactacgcgcgaaagaagattagaaggtggacaagtatggcatataggatggcagacgagc
ccacaggctgtgcgggcccgtaaacgcacgggctaggcccagcgggcatgcggcccagtgcgcaggcacatgcaa
gccagctgtggacgtgcgtgcgctaacacaggtcggcccagcgggtatgcgagcacgcggcccatggcgtatcgt
ggtccacgatggaccgaaggtccatgcgagagtgcatggaccaagagcccgtttccctaagtttcttgcggtcca
cgatggatcggagtcattttcgaggggggtttcttatggttcatgggatttttttgtggatcgggcgttgcagatgc
gcgatcagacggatgagagatgatccgagaggtgatccgacggctctgggaggcctcaatcttgatttgatggtt
ggaaagcgtgatcatgtgcaatcaagcgactgagagggcttgcagtgctggtcagatggtccaaacgcatctgga
gtctgaccaagagtctttctttgatgcagtttgggttttagggtccttaaacaggcctgtgggcgaacagagccg
aaagggagccgagagaggctctgaaagcagctgagaggagtttggcaagcgccaagcatgagcagccgacagagc
agaccaaagcaaagcatcaggcagagcagctcagaggcatgatgtaatgctccataggaagcgtccaggggcgtc
gaggatcctagtactggcaggcctgtgagaggatctccttaagagatttgtgagagcttttatgaggatagctt
ctagttgaaagagatctttgtacaagagttgagtggtgagtttctcttgtatttgtttgatttttttctctttca
tagtgaagcttgcatgcctcgtggaagtagcactaggactattccacgtacttgattgtgtgtttctgttttatt
ccttctttctgatgcaacggtggtactggatccacaacaagtggtatcaaagcaagtgatatcacaacaattggt
atcagagccggatcgataccagagtgcaggtgcggtggtagtgatcaagattgaagatggagaagacagatgcaa
tcaaagtggagatctatcacgaagggatgatcgatgctctcttatgcaaggagccgtctacatggaggatgccag
tgggagcccagtgtgaggtggagctctaggtggagatggattaggcacaagatgtgagactcgggaggacgatca
gcgacgtcgccaggaggctatcccgagcaggtctcaagtcatgcaggacggagatgggatcgatcggcctagctc
gactcccatgtttgcccatccatgagtaataggcgtttgccccaaggcatggggcgagaagatccaaaagctctc
aaagtcagatggaggccaaatatcgagtcgagatggagattgttaagaaaaatatctcgagattcaatccacaat
aaacctagaataagtctgtgataaaaaataaaatccaacgagcccaagatcagcccaaacaaatggagaaactac
gtgcaaagaaaatcagaggtggatgggtacggcacacaagatggcagacgagcccacaggccgtgcgggcttgca
gatgcataggcaggccgaccacgggtgcacgacctatggcataccacggtccacagtggatcgaaggcccatgcg
agagcgcatggaccaagagcccgtttctcctaggtttctcacaatccatggtggactgaagaaatttcccaaggg
ttttttggtggtttatgggatttttttatggatcgagtggtgcggatgcatgatcggatggatgagagatgatccg
agaggtgatccgatgactctgggaggctccgatcttgatctgatggttggaaagcttgatcaggtgtgatcaagc
ggttgagagggtttgcggtgctgatcagacggtccaaatgcatctggagtccgatcaggagtctttcttcgatgc
aatttgggttttaaggcctttaaacaggcttgtgggtgaacagaaccgagagggagctgagagaggctctggtag
caaccaagatgagttcggtgagtgccgagtgtgagtagccgacagagtagagcgtcagacagagcagctgcagag
gcatgatgtagcggtttgctccacaggaagcttctaagagcatcgaggaccttggtactggcaggcttatgagaa
ggtctccttggggggatttttgtgagagcttatatgaggatagcttttagttgagagagatctttatacaagggttg
agtggtgagtttctcttgtatttgtttgatttttttctctttcatagtgaagcttgcatgccccgtagtagcact
taggctattccacgtacttgattgtatgttgctattttattccttctttcttcctgccacaacagatgatatcgg
atctgcaacaagtggtattagagcaagtgatatcacaatatgttccacaccaaataaggttctctttctgtcact
gaatattataatcttatgaatggcttgtggcttgagttagatcactaccaagccatcaagatgtcttgccatgaa
gatactaatatccttctaaatcttgtggaatgagatcgtctcttcgagtttcttgcaggattgattgtcgaatat
gatcaagttcgtgtttagattctaggaagggagtgacttccataattaaaggaggtcttctctatcatccgggct
gaggaaagccttcgaagtgtaatgctctaacccctctcttcagaaggatccaccttggtatccatcaagaatctg
gatggtggtcccaagccaaatacttttggaacaaaatcagatgatcgtgcttcagacaatccttccaagcagtcc
aaccgtgatggtctttggtgtacttactgcaaaaagcctcaccacacaagggataattgtttaaagttcatggca
aagcacaaatcttgagtagaaccagaggattcaaagggcaatagtggaatcaagctaatttcaccgacaataagg
gacctatatctaaaaatttgaagggctgaattttgagaattcaacaaagaggaggttcaaaattacaaggtttct
aacctctctgtctactttctctggttcatgttcattgactcaatctggtaagtgccttgtttctaattctcttaa
tgccaatgcattgcctcgttcctcctcttggatgcttgactcaggagcaacagatcatatgacccatagctcatc
tcgatttgagacctacaaatcatgcccaaaatcaaacaattaccatggctgatgctttccctattaccatggctg
atggccaaggccaaatttccttagcccttcgtttatttaaaaatatgttacatgttccaaaattatccacttatc
taatctccattcatcagcttaccaaagatttaaattgttatatcacatttttgatacctattgcatctttcaag
acaaagtcatggggaggacgattggatctgctaaggaacggaatggcatttactaccttgatgctgccagtagtt
cttccactcctgcaaattggttcactctctcgtgcacatcgtccaattctgctcatcctttctcttccaatgaat
ctgaaatctgaatccaacatcttcgtcttggccatccacccttccttttgctgaaaatatatttctttattgtt

cgataatgttgatgtccaaagctttcattgtgatacttatgaatttgctaaacatcatcatgtgtcatttctttt
tgagtattaataaatcttccaaaccttttttcttatacatacagatatttgggtccatctacaattcctactcg
tactggggcaaagtggtttgtgtcattcattgatgactgtactaggaccacatggatatatcttatgaagaaaaa
tttgaggtcagccatataattccatcttttcaccaaatgataaagacacaattcgagatatccatacaaaagatt
cattctaataatgctcgagattattttaatttcacccttgcatcctttcacaaagaaggaatcattcatgaatcc
tcctgtgttgacacatcttaacaaataagatagcaaagagaaaaaccgacatctactaaatgtcactcgtgccct
tctttcctttctaacgccccaaaatttttttgggagaggctgtccttacagcaactttcctgattaataatccctt
cacgtgtccttggtaatcaatctcccattcacctattgtctatctttcattctgaacttcattacctgagcaatc
tccctcttcgcatctttgggtgtgtctcctatgttcatattcataaacaacgccagggaaagcttgatcctagag
ctctaaaatgcattttcatgggctattcgaatacacaaaagggtataagtgttatcatcctcctatccgaaaaat
gtttatctccatggatgtcacctttaacaagaatcaattttattttccaaaccccattcttctctttaggggggaa
aagcccaaagaagattcgaaatgggttgtccttgataatctcggaagtctacagcctatgcctcctccatatgaa
ccatcctcatatgagccaggatgagacactgaaaatcagcaacaaaagcagtctgaattacttcagtattcaagg
agaaaaaagatgattttcaaatcaatacagacccaagactccaatccaaaagaacaaaagtcaggtaaaccaatt
tctctcactaattctattcttaatgaatctgtttctgaagagcctacttctcaagacctagatctcccaattgcc
ttaagaaaggccaccagaaagtgcactcaacatctactctatctaatttcacagtttgtctcccttcatcgcctt
tcttcataacataaaagtttcctcactaccctaaactcaattgatatccctaaaaccatacaagaggcattaaaa
gacaaaaatttggtacaagccatggaagaagaaatgactgcactagagaagaataagacttaggaggttgtgact
aggccaaaagaaacaaccattggatgcaaatgggtttacactttaaaatataaagcagatgggtctcttgagaga
tacaaagcaagattagttggggggtatacacaaacctatgggatagactaccaagaaactttgcaccagtagcaaa
gatgaatactgtaagcattctattatctcttgctgctatgtttgattgggatttgtagcaatttgatgtgaagaa
tgcatttctacaaggagatttggaagaagagatctttgtggagccatcatccagatttaatgaagattttggagc
tgataaaatctaccgtttaaaaaaagctttgtacggcctaaaataatttctggagcatggtttgggaggtttaca
aaggcaatgctaagaatgggatacaaacaaagccagagagaccacacattattataaaacactcaaataaaggtg
aagttacagccttactagtgtatgtggatgatattattgtaacaaaaatgtaggtgaaagagaatctcttagaca
aagattagctcaagaatttgagatcaaagaactgaaaaaactgaagtacttcctaagcattgaagtggcttactc
aaaaaagggtatatttatctctcaacgaacatatgtactagaccttcttaaagaaactagaaatctttgcagcaa
aacagcaagtacgcctcttgattctaatcacaggataggaagaactaaagagaggatagctcggttgataagggg
agatatcagagacttgtaagaaaattaatttatctctcacatacaaggccagacatagcctactctgtgagtgca
gtcagctagtttatgcatgacccaaaagaagaacacttgcaagcaatttataaaatccttcactatcttaaagct
acatcaggtaagagaatcttgtttcagaaaagaggagagcttactcttgagtcctatactgacgctaattatgct
ggatcataagttgatagaaggtaacatctggacactgcacattcctaggaggtaatctggtgacttgaaggagta
aaaaaatagaatgtggtagcaagatctagtgctgaggctgagtttagagccatggctcaagggatttgtgaacta
ctgtgggtaaaataatcttagatgacttaaagatcaagagagagggaatgatgaagctatactgtgacaacaaat
ctaccattaatattgcgcataatccgattcagtatgatcggaccaagcacattgagatagacactttataaagga
gaagttggacaatggcttaattagtatcccatatatgccatcagagaatcaattagctgatgttctaataaagga
actacctacaacaaaattttaagaattattagcaagctgggaatggaagatatccactcaccaccttgaggggaa
gtgttgggaattttctaaaatctctaatctaattgtaggatatttatatagttctctattttctctctttccta
attcactgtaaccagatttttgggatttacacaggatttgatcttatcccttatttcatgggatttctacggga
tttgaacttttcccttatttcatgggatttacatgggatttgatcttatcccttatttcatgggatttataccag
atttgatctcttaagaagatcatccttttgtatatttatatgagattgcccacatgaaaaatagaagaaaatcac
tctttttctcagctcgaatttaccaccgttcacctcttagacaagtggtggtggttcaattctatgtagtgacta
tccttagtgactatccttattttggctcgcatttaaaaatcctagttctgcaagcacagtttcaattgcctagga
gcctaggttccaacctaggtggggtgccttagggttagggatgtaattaggagggagtatttgatggatggtggt
tgttccatcatctaataaatgttttaatttagacattccatgatcgccatctttatttatgaatgttgataaata
ttttccacaatttttatgtctcttgaaaatattagataaaaagagcgatagaatcaagagcttatttgctgaaaa
tggggtaaatttaagtggcattgtagtcgtgtgacaacttcatgaatgttctcaagctcttgtgtcagacagtac
agcagtaaagattgctggcttttgttgcttttaaaactttgctccttatgcgaataaattagcaatatcttattt
atacacaaagtagtcacaagcatctttcttcttttctaattcataaacatgctaaatggatgaagttgcgataat
gttgcaccttcttaacaaacatagatggccgaatttaagaagtggatgcatgtttgtgaagatagttcttgattg
agatctgcattcctgtccataatgaagctgatctttttgtgtcaaaaaacagaagctgatctttcagttatttct
gaaaaggattatagatgtctcattacttgattcaacacttcatgtttacagagtccaaacttgtgaacaaatggc
ttcattcatgagattgaacctttttcctaaataatcatgtatcaatggattattaagtcattgaaatcaaatga
tgattcttgtaaaataattattggtgatttcacatgtggctttgcaatcaactttattgttcaatgccccagaaa
tacatatgatagctaaaataacttagacagtagaagtagaatacagcatcaaacctaaattatattttcaaggt
ttttagtcttggagatcagcttgagattttgcacccctgtaccagcacatggtacactaccggaggcaagataac
agttgcaccatgtgtagggacactaccatattgaggccctctaatgtaccaatatgttattgggttagtatgatg
cccagtaccagtgggtgcaagacacttatttttatcaataaatagttatttgatgcacatatatatgtaaggaca
agagagactttctataggtaaagaatacaaatgatgcatagtcatagacatactgaatattcagaagaaaatttg
agaaagcatatttctcatagaatgtcaatactctaatttacacttcaaaatattaatatattgatatgagatttt
attcaactatttatgaactgctgatactacactttgatgcaggctttaaggaatatttcttgtagactttaactt
ttccatttttatattgttactaacttacttactagtaccttaagaactacatggggtttttgtttttctagtttt
ggcaaagagaggcagcaagcttaagggagcaactgcagaacttacaagaaatcatcggtaccatttcttaacctc

gtttacattcaaagattgttcctatgcctgactagtcattggcaatatgcctctagacatcgtaccataaagaaa
tattaaataaataaccatctattttgcttttaaaaaaatagcattactcaacatgactcagactttcaagcttgt
ttaagcataactagtttcctcaatgaaaaatttaacaagtattcatcaattaccaggatctagattctagagtgt
caagatcaccaggaaataatatataaactagtaaaacatcaaattttgtgaatgagtagagatgaagtatagaat
cactagttctctagttgcccattcatcctaagagtaaaatgtatttcttaagactaagcacatgaagcaacattt
tcatgctgatatgtcaccttgtcttcttgtttaccatattgcatgcattgactaaatcatcaattacaccattgg
ttatgacaaaacagcaggttactgaaattccaatagtatgtcaagatttagacaacataaaagtgatagtacatc
taaggacttcacttctcccttgaatacccacagacagaattggtccacaggcccacagttcacaaaacagtgaca
aactgacaactgtctgcaccatggctaataacctgtaactatcaacttcatataaattattatttgcttcaatta
tcaaattgaacctcagcatattttatccaatcatatagtttgggcacgagcaaatggactgtggcttgagtgctt
aagaaaacagtaaatgtgctaaaaacaacctagtaataaaatgctctgacattgaaaattttccaattattctca
attaaggtttccctaaattttactgacttaatgaaacctgcgcttaatgaatttataacgccttctttgaataac
ttttttgcaaatatttcataaacatttcattcatttttctaaagttttccatttttttttatgtttcttttattat
cattcaatcttcattttattcattcacttttcaatcctcaacatattccaaagttcttccaaaatgtaaaatag
cttacaatttttttccttgataacacattcttgaataaccatgcttaaaaagtatccagaaagcaatatgctgata
aggtgcatactaaaacatattagaaatcatgatcaaatgaacttttcctcctgctagtatactatgtaatgtaga
ctgaataatcaacaagctacatgtactatttgatttgacttatggatccttcttttcaaaacatcctctttattt
tagcgttggttgttattgatatgtgataacagttacagtcttctgatataccagattttaaaatctacgacatta
catattcatagaatgggattatcacactgatcttgtagctttattactattcctggagttatgtctcagcctcca
gacaccacacttatatataataaaaagaaggcagttctgactggtctttccgttatctaattttttacgagttgag
aattagactctatgacatcatgaactgctccaatttcaatgtctaaaattgcaaaccacagaatgctctagaata
gatgatcatgtgaaacactgcttggatagagcaaattactgtgaagatttcttagagttccaccaagggtagact
gccaagactgcaattagaaatcaatattatcttgcctataggatctcaatatctgttctggcggaaagtgaaggc
aacagtcagaggactgctccggtacttcacttccaatgtctcagcctcacagcatatattgctaatagcagcagc
tgtaacaatcttttagataggagcattgagatcctgaatacattttttttcatctcaggcatttcaatttaggat
tttgtcccaatttatttggtggtagcttagatattaaaagagtacaagctttaatagataatttctggttgtata
cttcatgcagagaataaagatgatatgttagagaattcttctatcatgagtttcctacaaagaagtctatgtgat
gtttttgccatggaacataggggtgctgtggtgtcaattttgaatgccatagcccaatttacttaagaatttaca
aaattgtgatctagaaggatcaccagttctattgcggactaatttggccacctgtggtaggtctagacagggtct
gagagcaaaattcaccatatctaggacctgatcatgaagagtaacttctgaaggccataatttctcatatattg
tccagtttcaacatgccatgcaccaaaatgaagcccttgacaaggactatgaggatgcaacaaggttgtcacacc
cgcatgaatattggggcctcaaaacaagctgatcacagtaaaacatttcttttagataatactttgaccaaacac
atcttctaatgaaggacacaaggcaaagttgatgttgaagttaagatttaactcctagacaagtgttatctttat
ataattatcacaacctagaacctcgccaaagaagactagctaaaagatctatttgggttcatcagccctgtataa
gcatccaagatctacccagcatcccagctaggtaaatcacctccacccacccatgttttggaaaaaaaatctaat
aataacttaacgttaggttgaaaaaaacttggggggtctatgaggggcctaataatcttcgatcaccaggttcat
gctctatggcatgaatgatcgaaaacaagaaaaattggatttttggcttgaaccccaaaactcaaaggataaaaa
aaggttatagaggtttctctaattctagattattctatgaacaacatgcaacgttaaaacttcaggatgcattaa
aaccatggctagagcacttaagtcaacattgttaccaatttaccatacttttgaaaagacatagcatcgagacaa
caaaacataagaaaaaacaaaaagaatagtttcagagcataaataggtcataccatctgaccatgcaaagtatat
cagtagcttgctagcaagtcacctttcggcatacatatcaaatagatcatagtaattgaaatgtccaaaatcaag
aataatacaatcactagtaagatagtgcttgacaataacaaacaccgtatgcatcttctatgaaaaagactatct
tttaaagttctactcttgttgccctatacccgtctatttattgtctaacaccattttactaaggtcttgcctact
gaagctggagcaacagttccatcagttcctctagggatctcactagggggcaagggtgtgcatctcctatcttg
ggaggtgctttgtcagccggttgacgagggcagtcttggcatttagtccttgatggtgaggcgagaggcccttttt
ggtgaggcatgcaaccaggctgctggtgtgtccggagagcccctggagtatgctgatggcggccaagtatggcct
atggacttctagtgctgacattcagacgaatagggccagctcctttatgtggaaggagatctcctcccatgccta
tgtggtggggactcagatcggatggatgctgggcgatggccacgcagcgagctttcatcatgactctcagctagc
tgaccttcccctaagtaggtggccgaccttcgttagtgttaaggtgggggaatctacgaggatctgttacctgtt
tcaacttggggtgggggctggagccctgaggcagtggctcagattttggggctgacttggtggataaagtgcta
ttgatcgctgtcccttccatcctagtccggatttgcgggtgtggagatctgcatgcaaaccgagggttgcaatga
gggatttatacaagatgtttcggagggagcctgtccaacatatagatagtgcttggatctagaggatagctattc
acctgagagtttgtttgttcatctggaaggtggtctggggccatctcctgtcgagagcggtcttgagggacaggg
gtctggctccttccgcttgcccttgttgtgatacgaaggaggagagacgatggagcatgtgttgttccagtgccc
gagagctgtccagatctggcagcagataggtccacttgcaactactattttgactgcaacatcgacccagtcctt
tatggaatttctccaacaaagttcggcttcagatttggttatgcttgatgggatccgtgtggcctatttcgcata
tcacatttggttatctaggaatagttaaatgttagagttgagacgatcatcagtggctgtagtggtggagagagc
tttagcccaagcaatggagtttgttcacttgttgctgcctgattttgttttgaggatggtggattgttgggactc
tcgctaggctcgtcgagcaacccatcgggtgttcacctcttaggagcccccaccccaacattcctcaagttcaat
ttcgatggcagtgtatttggctcgagaggaggtgcaggattcatggcccagactcgagactgctggcagcgggcg
agagttgcctagtggacactatggtcccgagggcagagctatgagcaatgtgggctggtattgcctatgcaaggc
tggctcttacagtagaccaactcatcattgagggtgactcagccactatgattagatggattcaaatgcgggata
cagaggatgctgctcacccacttctgagggatatcgcgatgctgctgaggggccaccatcatatgtgtccagca

tgtctacaggaaggccaacaccatagcaaactaggttgcctcctttatggcgaagcacccgagggatgtgatttg
gactgacatgtccactatgtcatgtggtttctgtgatgtttattttagatatttttgattgcgttcataccaaaa
tagtatgaatgtaccatcatatcaaaaaaaaagttctactctgcaacatgagaataattaatagtataatttagt
taaaatcaagtgtttaaatatgatcggaatgttatgggatcaataccaatttggaacatcggttcacatttgaat
tggtacataccaacgataccaatctgcatcaacaattcatatatattggaagttggtacatatcggttcccttta
ttttgattaatataaaccattcgatccataccaatcatagcggtatcatatagcccaagttcaaaactagtataa
gcaaagttcactgtgccaaaatcagatcctgtgccggtggtatgatttgttatgcccaaaccattctgtaccaag
ctcgaaccaacatggtgaagaagcggaggaaggaagaatcgaagaaagatacagagagagggagagagaggaaca
aagggaggagtggtcaaggatgccgatggtggctgccacctagagggcaggtagttgggtcgaaacaagggtgac
cacctctatttcgatcgcttaagtatttttatgctcatccttctctctctctgtctctctccattctcctctct
atggaggagggtggagctccgaggcctcgacacccctccgatggcctcgacgagcctccaatggcctctctaatc
ctccccccctctctcttttctctcttttttttctccctcattggtgtaccatttttagctgaaattgtaccgac
tgactgctggtatggttcagcacatgtcaaaccagtcggttcgcgatggtttggcaagtgctaggtataaggctt
ggtattggtcttttaaaccttggttaaaattctctcttcccatatcccttgtatattttattacctcttcctga
atcctacaattaggtatgcatcaaaatcagattcttatatccttgtcgacatagaatgtaccgctgatgtcatta
acactttccacgatggatgctactctcaaatattatccttgtttcaaactaattttaatacatcataagatgct
tcaagtctactacatgatccaaaagttagtgataaaaataggtctcaacattatccatgtactatagacattaga
gataacaaaagatatatatagtcattatatttataaatgatttgttgtagccatagtttgccataccgatctgta
cttgttgcccaaaaggttagcaacccaagagaggggtgaattaggttctttaaaaacttctatcaaatctaaatt
taaacgattatgtgcttcaaattaagctaatttgagtaattgtgaggtgtatgcaatgtgttacagcggaattag
gatataagaaacatgcaagaattaaaaataatgcaaagcaagtaagaaagcaaagcacaacacaaacacaataac
tttatagtagttcggtacaaccccacatctacatccactccccaagctttcttaggaattccactataatcagtt
tgattacgattcgattgttttatccgaacacacaatcagttgatttttttgggatcactaactaaaatcttaacac
cgattattttttgaattcacaatcaatctagttagttttgtgagattaccaaccacaatcttacaagtccaatttc
gggcttggatgagcctaacccaagtttctttcatcctaaagaaacttgaaagaaaatccaaaatacaagataaag
aaatttcagcacaaatgaaaatagaaaaatagaactcttctgtgtgtggagaagaagctgaaggattgctctttt
gaagcttgacttctcttctttgatgcttgagaggatactattgatgatggagaagatcactcttgaagactcact
gaattctatgtacaaatttccttctttttccctttttccaaagatattcaatgatgctcacacaaaataatgtgtt
ccactcttttaattctctttctcttatttttcaagtgattctttagccttttgtaagcttaaaagtggctggaga
gtgagttttgaccgttagaaggcttggactagctgttggagtagagaaatggccgttggagaagtacaaaaaatt
agttattacagcttcctcgtgtcaagagtcgactcgagctgtccacaagttgactcgaactcttcaagagtcgac
tcgacctcaggatccaaaattatgcattctatcattcctaagagagtcggctcgccatcttcatgagtcgacttg
acctgtccagggaccgactcgaccatttcatgggtcaacttgagcagggatcccaaaaatccagtcttctgtctt
tactaagagagtcggttcttgcttctcagtagttgattcatcgttcgtgtcagtgcgttccaatgtgccatagac
aactcgccactcccagggtcaactcctaaccttttttatctttattttctttgtttcaacctgttttgtttagggt
cggctccaatgcttttagggtcgactccactaagtcaaaaactaagaatttcatctttgaattattttgacttag
cttctttaagaggatttgtctcatggataaattcttcattttgagtacttagaaaatcctatatccatccttgaa
gcacatgattagtaacataattactcaaaattttgtgatcatcaaaaccaatcatacggtcaacaatctcttcct
ttttgatgatgacaaaactttgagtatatgcacaaataaacataatgtgtttcaattaatttgctaatatgaatc
atgaagtaaataactaagcacttgtaagcatatacttcatcaatatttcaagttatcttaaattcagatttcaaa
agtcaatgcacatttcaagagtattcaatatatacttattttatatgattgtctcaatgcaaacaaatgagcatt
tcaagcatctcaagtatgtgatcaactcaagcaattcaagcatacactcaacattacttcttccctttctgacag
tatcaaaaagtgcaaaaaaaaaagaaagatgtcatcaaaatcaactcaaatcaaattcaattttataacttctcc
ccctttttgaaagtttgtagcaaattcaaatggaatttcttccctttttggaattcaattttgaataatatagatt
caataactttttctcctcctttttaaataatatagatgaaacaataagttcttaaattcagttgattgatctttag
aaatccaggagagatagatctttccaaaaccaaggatattcacaagatatgtttagttttcaaaattcattcttt
ctttgcttgaaaagtatcaatcaattaagttcaaacaaggtatcattttcttagtttctctcaaatgatatcata
ttaaaaatgtatcaaagcaaaatgaacttgagaccaagcattctttcttttgacaagagataaagcaatggagta
taagacaaagataagctgtttgatacatgtgaaaattttcttaaagataatctacttgataccattgttttgaag
ttctaaccaaatagcaattttcttgaaattttttttgataccaagtagagatttaaaaagaacatttatcaaattc
atttctcgataataatgaatctaagtaaatatttcaaattcagtttgtaccaattaattatcaaaaatgataaca
agattaaacaatgcaaagtatgaataaacaaatcaaaaacacatgaatggtcaaatcaagataagcaagaataaa
tgtttatcaatgatctttcttattaagttccaaattcatttccccccctatttttgatagttaatgccaaaaaatt
tatttttcttaattctcccccttttttcttccaaattatgcacatttatttaatagtaattacaaaagcatatttt
tctcccccttaattttataagcaaagcaaatttcaataaatcaaacaaataatcacatcaaacacactgaatttt
catgaaagatcaaattttcattgattgccaaaatgtgatgacaaaaagtgattacaaagatagtaaatatcaaaa
catccaaaatgtgtgtaatttatatctatcaaaaggtagactgctagggcactaatgctttaagtaagaaatcct
agcattccatgagagagtaggtcagcatagccctatggagaaggtctaggctggctagatggacagggaggtcta
gtgtctccaaatgactgtccccgatcaaccctcactaatcacatgatccttataagtcacctctctcagcaatga
tatagcctatgaagtgtacctctgtatctcaaaatagactgcagtcacaatccgggtggaaaatctctcaataag
agcatcctcgttcaggataacaaatggtcgatctgagctctctagactagtctgtggaggtagtgctgtctctgt
ctactcgggctgtattgctagctcctcagatctgatgtctataataggcactattgcaggagcctcatcatcatt
agtgtcctcatctctatgcatctcctttgcgaagccaacgaccgtcaatctttgaaatcccatcctgcatagtga

accgtcattgtaagtatctgtgcgcatcaaaactctggtggtctccccgaaaatcattcccatgctcctggaaga
ccaaagtgagaaccatgccgtaaggtaaggatacctttgaccggttcactgcatcatgcatctgtccaatcatta
atgatggtagactagaaatctcctgaacaatgtgatacacaagggtgatgtcacactcagacaatcaaatctctc
catcttggggaagaaaatctggctgatcatatgatgcagtaatctcatctccacgctcagctagtttgcactgat
gtctctgataccccctaacatcatctcattccaatataactttcaatccaatgtctctctatataagatcatctga
aatgctcccatcagttggcatatgaagtgtgcgtccaaggctatccacatcaatctcaatttacatgccttttac
tttggagacaacacttcccacacccactttcatattactgtagaattctcttactagtcctgggtaagtgggcac
atccaaacaacaaagctactcccaacccatcctttttaatcttctgtccgctagaaaaccttcacgctctaaggaa
atcaatttcctaccaattatgacaatgcatgaagataagggtgccaaggatggaattggctgaggagctaatggt
ggttgtgaaggcactgaagctccttgagatggctgccctgggtcattcggctgtttggatcttcgttcacgcata
ggagaagggcttggtgaaactctagggtttgatgtttctgcacggatcctcttccttatttgggctagttgcttg
tggggtgccataagagaccaaaccaccaaaaaatcgggttaaaattggaggtggaagctttgatttaggagggaga
ggggtttgaatgaagttccaactcgaaagaaaagagaagaaatgagcagagccttgggttgccttgggttaaaaa
tcccatctccgacacaagggtcgactccagttagacacgagtcgactcctgataagaggtcgactctagtaaagc
tgggggtcgactcctcatagaactgcttttcttcttaaaaatataagtcaacttctcgagtctaagagtcgactc
aatacagaaataattcaagaagtttatatcaaaaaaggttggctcaaatagcgtgtggattaatggcaaatcaaa
tcaatgatcaaaaaaacgatatttgaatcagaataagtgaatcaatttgggggattaaaaaattaatcaatttaga
tatgatcaaatcaaggagaatctagatacaggctctaagggtgtagttcaactggaggattttcaaagccctaaa
gatcaatttaagatgcaatgatttatcatgcaaatcattgatatcttatacacaatcttttatgaataaatgtct
tatcttaaagttcttaggtcatgcaattttttgacaaatttgataattagatatcttatgaggatgtcaaatttt
agtttgtaagattttcacacaaataatttttgtatgcaatcaagcttattatgattcagctctcttatataattaa
taatcaaaataacatgatataaacaattaagagagcatgctgactttaaaataaacatatacttgttatgaacaa
tacatcaaaacattcaaagcaaagaataaagtcatcacaatatatattctagtatcatgcatgcatatgaagcat
caaaatcaatcaattaagcaaaataattgaagcaatcaatgatggtatgatgtcaaaggtttagcttttatcaaa
catcatcatccatccattttgagtccaaagcatccattcttcaagaatttactcctttcaaagcattggatgctt
ggcatacttgcaaaattatcatttttttagattttggtatctaaaatttcttggatccattgaagttagtgcacat
agtttcttttaaaattcaaatttgtttaaccaaagtatgaccaattcttttagtattgcattcaaaaattttgta
gcctattttattacatttaaagcaagtaatagaatgattttcatgtgaagctttagtaaacatatttttaacaaa
ttttatttcttgaagaattaaaaccaattccagctttattagaaataatgtttaattgtttaacattaatggaa
gcttttgagaactgaatgtgaacttttcaactataggctccaatatttcaatctctttcttaatttttatatttc
ttccactaaaatttttaaatttcaaaagatttcttttttcttcagccaaagaattcttttctttcaaaagttctt
tttctattaaaatctttcttttcttcagctaaaacaagtttgacttttaagctctttatttttcaagctttctt
tttgaaatcatccattagttcatgaaaggcttcaaatagttcttcaaaagtaaattcgaaagagttttcgatgtt
tacctcatcgttgtgagccacaaggcaaagattagctatacctccttcctctttcttttatccgaactagaaatg
tcactttctccccaaatagccatcatagtcttttttgcttcttttaggatttcttttgagtaggggcagtctgact
tctagtgttcctacttgttgtatttatagcatatgacatctctttcttcatcctccctatcttcctttcctttag
tcatttttcttgaggaattttttgaacttcctagtgagcagccctatgttctcatctccatcctcactttcttct
tcatttttatcttttgacccattgctttcttctgatttgatggttgattcaaatgcgatcttccttctttctttg
actcttcttcttcctcatttttggcacatgttcaactcatatgtcatgagtgaaccaattatcaattaattcttcc
atggatagcttgctaagatctcttgcttcctgtatcgctgttaatttagcttctcatgtctttggcaaacaccta
agaaccttgtaaataagctcattgttagtataggttttgccaagaccactcaaagcattaataatatcagtaaat
ctggaaaacataccagaaattgacttatgttgttccatttaaacaattcatacttgtgaataagcaaattaatt
ttagactctttaacttggcttgttccttctaagtaattctaatctgtcccaaatctcttttatggtagaacaagt
agatattctattaaattcattaatgtctaaggagcaatatagaatatttatggctttagcattcagctgagcata
tttttatcatgctcggctcaatctttttcgatttggaagtggacactccaactagcattgtgggtgtgtgtggtc
cgttagttatgactccccacatgtcatagtccaaagcttgcatgcaaattcttatgtgtgctttacaaatatgta
attcatgccattaaaaaagtgatggtttagaagtagagtggccttcagcaaatgaggaaccaactaagattgccaa
gatctttagcttaagaatgctaggtcaataacaaatataagagcacctactctgataccaattattgcccagaag
attagcaacccaagagggaggtgaattgggttctttaaaacttctaccaaatctaaatttagataattatgtgct
tcaaattaagctaacttgagtgattatgaggtgtatacaatgtgctgcaatggaattaagatacaagaaaaaaaa
agtaatgcaaagcaggtaaagcaagaaagcaaagcacaacataaatacaataaatttatagtggtttagtgcaat
cccgcacctacatccacttctcaagctttccttgaaaatttcattataatcagcccaattacagtatgattattt
tatccgagcgcacaatcaaacccatttgatttttttggaatcaccaataaaatatttacatcgattattttcggat
tcacaattaacctagttgattttatgggatcaccaaccataaccctacgagtccaatttcgggcttcaacaagcc
taacccaagttcttcactccaaagaaacttgaaaagaaaaatccaagaatacaagacaaagaaattttagtacta
attaaaatagaaaaaaagaactcttttgtgtgcggagaagctgaaggattgatcgtttgaaggttgacttctcttc
tttgatgcttgagaggatgttgttgatgatggagaagatcattcttgaaggctcactaaattctatgcataattt
cctctctttttctctttttcaaagatattcactgatgctcacataaaataatgtgttccactcttttaattccct
ttctcttgcttcctaagtgattctttagcctttgtaagcttgaaagtggctggggagtgagttttgaccattgaa
aggctaggaatagccgttggagtagagaaatggtcattggagaactacaaaaaactagctgtgatagcttcccca
tgtcaggagtcgattcgacctcttcaggagtcgactcaacctcaggatccaaaaattcttcattctatcattctt
gagagagtcgactcgatcatttcatgggtcaactcaagtaggggtgccaaaaatccagtcttctatcttcactga
gagagtcagctcttgcatctcaagagtcgacttgccattcatgccagtgtgttccaatgtgccagagttgacttg

ccgctcccagggttgactcctgaacttcctgatcttcatttttcttctattttcaacctatttctattcatgggt
cggctccaatgcttacgggatcgactccactaaatcaaaaacttggagtttcatctttgaattactttgcttagc
tcctttgagaggatttatcccatggataaattcttcattttgagtacctgaaaaatcctacatccatccttgaaa
cacatgattagtatcataattactcaacattttatgatcatcgaaatcaatcataggatcaacaatactgttaga
agcatattgaaatatttggtaaggtatcggcatggttatcaagattcaatatagcataggggccaaataggcctg
gttggagtggttgcttccacccagtttccaattttgggtggcaccatgcctgggcatgagaaaaaacacttccaa
ccctatgtcagtataaagggtgtagccataccatactaggaacacttctcagagctataagatataattgatgat
aatgcatatagtatcatgtgtagtccaaaccaaattcactccatctcttttaatacaattaatttgagaaccgaa
tccattggtttgttggcaccaaccagatggcacatataacattatctcttcaagtctctcgaagtcaattaattc
gggctacgtcagctggtggatcccaaggactcttaccaagcagaattcactccatctcttttaaggcaattaatt
tgagaattgagtccattggtttgttggaactgacgagatggcatgtatcatgctatctcttcaagtctcttgaag
tcaattaattggggttacatcagctggtggatctcgaggactcctaccaaaccagaatgcactccatcttttta
aggcaattaattctcataaatcccccaaagaggccctaattatgccacttttatttatcaacaaaccaattgatt
ctcaaaatctcaaccatggtttagggatgttaattagccaccacccgtttgatcttgtgaaatcaattttttacaa
ctcatggaattagagataaaatgcatcctatacattaaatggcaacccaaagaattcccgagtgcgacgcatttt
tgttcctcaaagggcaatctaaagggtttccaagtgcaatgcatgttcgttgctcatgccgattccaatatgtcc
ataaaagtgcataattaggtggaaattttcttagagatgctatcaaaaattgtaggcattcatcatcctatatag
ctccaacatgcttcatgtcttattgtttaaaagatgataaataaatttgaatatatataaataatcatatctttg
attaaaaaaattgtaaataaattttagtgtcattgcaccttgttttggtggccaattatactcatcaattcaaca
agtcaatccactgaaccaaacatcctttctcaattctccaaatcctccttaattttgttcttccttagattcata
atgcccactataatgatacactctatcctataattcctcatcaaaagttgacccaaaagcagcttaccccaaatt
tagcatgaggaacacccctctaaccccaccatctcaacccacttctcattctccttgccgtgattaatgaggga
acatgaaaatagatcataaacagcagagatttagtgcagatgatctcaaaaccttatatgaatatccaaacaaca
gcagcaatgatgatcatgaaggagtttgagcgaaggtgtggagtaatggaggtgatggtgagactatagtccaac
ctaatggtcacaacccttggaatcagataaatgctttaactcttactagaccctgtatcggtaccttattggcat
gagtcaaatcaatatggttcatccatgcaacaagacacaatatgctgtaggagttagcatagcaatcgtaccata
cctcagaatgggcctatgggtggcatattgagccttcatccatccatactaagctgatgtgatgtgccctatact
agccaatactgttggtacttaaaattgttacaaactaggacaaataaaggaagggagaagaaacaaagagatgga
gaggatgatgacaagactggtaaggtagcacaaaatagttgcaaaagagatggaataagagaagcatgatagaat
aaagaagcaatagtagaagtcatagcttaaggcttttgtctcctgatattgtgggttccatgcctaaccatcgga
ttagattaattttaacaactcagatggttgattctcctaagttaacaactcaaaatgtcatttaggaaatctagg
tcctatcaagatgtcacaaatttagccacaagctcgtcacaagaggtgctgacatgcgtatggcgcaatgctgca
aaggatttggcatgggccaaaagtctacaatgctcctcaatcatttagaacccacagggtggttgttgggtggat
gtctagttggagtattgaggggagcaatagttaaccaaggcatagcatagccatggtatgatatcgattgctcga
aatgcatgatatttggcatggcggcggaccaccaggatcatggtttgctataccgccttgtaccaccccatacag
gacatactgtaccatactaataccgcaccagtacagtgtctctaaccataccgacactcaatacgttgtcttgta
atgtaccgaactacatactgatattgtaataggatgatactagtatgggatctggtatagagacagtgaacgttg
gctaaaatataaatggatcctcaagaattcctcttggttgagtgtagaactttggtgatgttcccaactcctcca
agcctgtagaatataggtccacaattggcatgattgaggggaaaggcagtgccacaccagatggtctcacatgga
ggcatcatggtaatactggtcaagtgagctagccacattgtgtgaaggaaattttttgagagattggttctcaagg
aattgtccaagctaccctgtgcatttttttatctctatggagtaaagaggcaatagggacccgttgggcaaagggt
gtagcctaccgagataggcaagcctcatgagcataatagggtgatgtctcatggggatccactgaagctagttga
ttgtggagtacaaatgtggcttatggaatatcgattgtattatgtgccttggagctagggcttaaggacaaccaa
agagtcatgccaatgagaggatagctcgatgcaaggattggctagtcaaatatatttacttgataggcaaacttc
gatgccattatcataacagtgcacaatgtgcgagtgggctatgatcattgttgagtagtatggccatatggatga
gaagggatgttgtggatgcaagaccaggaccatggcagaagagcttacaggttagcttggatagcaaaaccctaa
agggctccaacatacacgatcaattgttgcttcgagggattatggggcatatgtcagtttggcagttcgactggg
caacacgagataggtttgccagagcctcatgcatagaagcggcattacatgggttgcttgcaagcacttgagggt
gtcgcatgtgttcagttcgacagagttgtcgagtaggggcatagttttgctacttggcaatatggctagactgct
gtgtttttacatatatgcccagaggaaggactcaggagggctggcattggtgtgtggtatgctagtgcattgcac
aagtatttgggtcaggaaacacttgacctatgataataagccaacccgcatataatatggtctaccataccacat
agtacagggcataccataccatatggggagcaaaccaatatgaaaccccacttcaagtcgttataacccatgtac
catccataccgagataagtaccgctccataccaaggcatgtaccatcccatattgaagcgtactgacctatacca
aggtgtaccaagataaaaattgaaaaaaataatttaggagctattttggcataggatgcataccataccataccg
actatatggaattgaaccgataaggtaccaatatgatatctgataccgagatagcagaccttactgcatatgtta
ataggtgataagatgaacatgtcaagttgtatggatcgaacatgttttaaaacaatattttcatatatcacctct
cccatgaaaccattaaactccaccttaggtcaatacaagcaaaagcgatgtacataattatagtcactttaaatc
ataaactagtcaatcccatagacatattaataaaacctttagcctacatttgtttaatgccatcgaactacctct
tatgctactcaatcatgaaatccatttgaaagcccacaaaaaccatgacaaatcttataggtgaataggggtcgc
tagttgatatcacactatgtgaaacaatgttgattattacattattgaataagtcgatgatatagcatattcaaa
attacgccgatgcttgcaggaaattgatgggcgaacagttgtctggcttaaatgtcaaagaactgcaaaatctag
agatccaacttgaaaggagtcttcattgtgtccgaaagaagaaggttttactcgcatcaaatcttgctcgttaat
cctgcttatccatatcttctctacttaatttgtttgtttgctctgttttaggaacaattctaaccgatcaaatt

caagaattggaccgaaaggtttatcaatgagtaatagttaaatcatttttaatgttgaaagtattgcattttaag
tgcctaaatttctagaatttgttacagggtaccttcttcacaatgaaaacattgaactctacaagaaggtaaac
cttatatgtcaagaaacatggagttgcgtaagaaggtattgagtttgagaaagagatatatattttgctctatga
cccaaacttcagagtggaccattcatgaagctctgatcttcatgtttccatgaccacatgcattgcacatcatgc
tctatgtaaaatatgtatggatttggtctaaatctaggttaaatcaaccccaaggttttaagtgctaccagtgct
gctttggctaccatgtcaacggatgtcaacactaacactaggtcaagtcaagactcatcaaatcctttttgaca
tttttttgtttcgacaaagtcaaaaaagggttttgattttaaatcctctaatattcttgtttagggcctttag
gcacacaagtgcatcaacatggcatggcacgaacagtgctagtcactagttaccatgcctcatgccaacaatatt
taaaaccttgttcaatccttgagaaataatagttattttataacatgctaagcatggaatttgcattgcagatgt
acgcaacaaatgagttaaatggaggagctagtaaggatacacttaatccatatgctttttctgtgattgagaatt
cacatgtagatgtacctattcatctcgagctatgccaacctcaacagcaaacacatggaacacaagcaacagctc
caaagctagggtaattattttgaaatatcaaacctttatatattttcaatatttattagttttacttgaatatac
ttaattagtccttgcaacagacttcaactgcattaagaagatgaccgtgatcaaagtgtatcatctatagtgcgc
atcagcccaactgtcaagacttctatagtttgttatcagctataacatgcaatgtatcggttatcaaatgcaatg
tatgaacaatgctcagcaatcatttctatgtggtgtgacatgatctcaaaaaaaatcattactctgtaaagcaat
cttccaactaggactattaataatatgacaagtatttctatttaaaacattacttgtatatgaggacatggccta
tcatagcatccgagtagttgttggtaatgctactggaaaactcacttgctagattgaaggtacaagggattagca
ccagttaaagtgcaacaagttttacacaaaggatgactcactatatactgcttagtaactagtaacaagaggcat

>EoAGL17_3
ATGGGCAGAGGGAAGATCGTGATCCGAAGGATTGAGAACTCGACCAGCCGGCAGGTGACCTTCTCTAAGCGGCGC
AAGGGTCTGTTGAAGAAGGCCAAGGAGCTCGCCATCCTTTGCGATGCCGAGGTCGGCTTTGTCATCTTCTCCAGC
ACTGGCAGGCTCTACGATTTTGCCAGCTCCAGCATGAAATCAATAATTGACCGCTACAGCAGGACAAAGGAGGAG
AATCATCAGGTGCTAAATGCAACTTCAGAAGTGATGTTTTGGCAAAGAGAGGCAGCAAGCTTAAGGGAGCAACTG
CAGAACTTACAAGAAATCATCGCCCACAAAAACCATGACAAATCTTATAGGAAATTGATGGGCGAACAGTTGTCT
GGCTTAAATGTCAAAGAACTGCAAAATCTAGAGATCCAACTTGAAAGGAGTCTTCATTGTGTCCGAAAGAAGAAG
GAACAAATTCTAACCGATCAAATTCAAGAATTGGACCGAAAGGGGTACCTTCTTCACAATGAAAACATTGAACTC
TACAAGAAGGTAAACCTTATATGTCAAGAAACATGGAGTTGCATGTACGCAACAAATGAGTTAAATGGAGGAGCT
AGTAAGGATACACTTAATCCATATGCTTTTTCTGTGATTGAGAATTCACATGTAGATGTACCTATTCATCTCGAG
CTATGCCAACCTCAACAGCAAACACATGGAACACAAGCAACAGCTCCAAAGCTAGGGTAA

#### Elaeis guineensis

##### *EguAGL17_1 and egu-miR444*

>REF_ELAGV10
AAAAGGATGGATAGAGCCAAAGCTAACTAGCATGGAATTAAGATTTTTTTCTTTTTTTGTGTAAGCTTGAATGAA
TCATAAAAGAAGAGGACCGTTGTGTTTTGCTGGCAATGTCCGGCACTTGGTCCAATGAACTGAGTCCCGTTTTGC
TGGCAATCTCCGGTACCTGGTATTTCTATATATATATAAAAAAAAAAAATCGCCGGTACGTGGTACGATGCACCT
GGTCTAAAGTGAAGCTTTTCCGTGCAAGTGGGAATTTAATATATATTCCTTGCTCCGTTAAGCACGTTGAGGCAT
GGCAGGGGACCACGCTCGCCCTCAAAAGAACCCAACCATCCTCCAATCGTACGGCTATCCAAACGGAAGACGATA
AAAATGATCGTAATCAAGATTAAAAGGATAAAAAGGGCGATTAACGTACCTAATGGTACAAACTTAATTTAACCC
ACGTTCCCACCGCCGACAACACCCACGTGTCGACGACCTAAGTTGGCACGTCGGGGGTCCAGCGTCGGGCCTCTC
TGTCCACGTGGAAAGGGAGCAGTGGGGGCACCCGCTACTACCTGTTCAATGGCCAACACGTATTCTTAAAAAAAA
AAAAAAAAGGGAAAAGCTGCGAAATGAGTGCACCGGCTCCGTTCCCCCCCCCCCCCCACTTTGATTATAATACTC
CGCAGGAATCGTCCCCCATCATGGCGTCGCCCAGATATTTTCCAGCCCCCAACCCCCTCCCTGGCGACATGAACC
ACGTGGAAGAGACCTAATAGACCAGCTGTCACTCGGGACCACTCCTAGGTTACACTTTATCGATTCCGTGGACAA
AATACTTTGAAGTGTGTTTCGTTTCGGTTTCAAAAATTTCGAGGGATATAAAGAAATAGGGTGGGCGACGTGGAT
AGAGATCAGCCGCTTCTCTCCCCCATTTATTTAAACCCTTGGGAACACATGACACGGCCGAGAT**ATTTTGGTGTT
CCTTCGTCTTTCCCTCTCGATTCTTCCCCGCTCCTTGGATTCAATCTCTCTCTTGGCCTTCCTTTAGAACTCTTG
CTACGTCGTTGCTCTCGCATTTCCTTATTGTTGGATCGGTTCTTGGGGTTATGCGTGATTTGCTGTCGGATTGTG
GCCGGAGGGTAGGTCAGTTTGAGTTTGAGAAATGGAAGAGAGAGGGAGAGGGAGAGTTATGGAGAGGATCTGAGG
GTGGAATTTTTTGGCCTGGGGAGGGGATCTCAG**GTAAAGAGGATTTCATGTATAACTTGAATTCAGATGTGCTGC
GCGAGATTTAAATTCGGTTCTTGAAAATATTGTGTTTCGTTATTATTGATTTGATTTATTTTCTTTTGCTTCTTC
TGGTTATGATTTTGGTGCGGCTTAAGTTAATGCCTAAATCCATTTCCCCCCTCTCTGAAGAAAGAATGGCTGAAC
TTGCTTTATTAAGAGTTTCCAAATATGTAATAGGCTAATATTTTTCAACCCCTATATCACGACTTTGCTCTCTCT
CTCTCTCTCTCTCTGTCACATATACACGCACAAAAACAGTTTTTTGGTTTATCTGTTGTTAATCTAATACTGTGA
GTGTTGTCTTTGTACCACAATGTTGAGCAACAACTAGGGTACCTTAATAGATCTATTATGGGCTGATTGCTTTGT
TCTTGATAGAATTCTTTAGCCATAGGTTGGCTTTTATGAGTTAGCGTATTGGCATAAAATTCATTGAGTGGGTAG
TTCTATGTGCTTATTTTTTTTGCTTTAATTAAATTTATTTGAAACCTGTTTTACTTAAGCTTGCAAGTTTTGGTG
ATTATCAGTTATCTGTACTGGTATGGAGTGATTTTAAGACTTAGTTGTCTAGATCAATTAAAGATTACCTCATAG
AAAACTATTGCCATTTTGAGAGCTAGTTGTAAACATATTGGGACTTTGATGGCTCATATCTTGATTTAGCAAGCT
AGAAAACCTTATAAATTAGGTGATACCTTTTGCTTGGGACATGTACTGTTGGACACTCTTCTTGTAATGTTATAG

AAATTCTCTAAAGCAACAGTTAATTTATTTGTTTCCTTGTATGTTCATTTTATACATAATCATCTATAGGTTAAA
TGCATAACATCATTGCATGTCAATATCACACTCAATAGTTGTCAAGTTCACACACTAAAATTGCTGAGTAACTCC
TTTTTCTATCATGTAGTAGTAGCATGGCTACTGTCAGATGGCGTTATTTACTGACTCTTGGCAATTATTGTTGGG
ACAAAATTTCTATTTCATTATTTTATAGATGTACTCACATTGAAGCAAGAATATTACAGCAAGAGATGCACATTT
AGGATCATGTTGACCACAGCAAGCAAAATCTTAATGTAAGAACCATGCGGGCCTAAAATTTGGACTTGCAGTATG
TCTAACATTCATCATGCATACTGATCGAGTAGCCTTGCTAACTTCTTGTTTGTAAAGAAGCTATCAGTGTCTTCC
TTTAATGCTATCCTTACCCTTTTTTTTTGGAAAAAAAAAATTTTCTAGCCATACTGATTGAGTAGCCTTGCTATC
ATGTTTAGCCTTACTTATTTGAGTACTACCCTTGACTAGCATTGACTAACTTCTTGTTTTGGTGCACCGACTGCC
CCTAGTAAGTAGACAAACTTTATTTCCTCATCCTGGATTCCATGATAGTCTCCTAAGCCCAACATTTAGTTCTTT
CGATTGATCATCTGGATCGATCTTAGACTACCCATCTTCATAGCCTTGTCTCTCTTACTCCTTGTGCTTGATCAG
CCTTTGAGTCCATCATCTTCTCCTTAGCCCATTGATGAAGGTCCTATTTATTGGCATGCGTGAGGAAAAAAAAGT
AAGGAAGCCTGCGCATATCACTTGGTTTGTCTAATATTGGCCTGTCTTTGGTTTCGATGATTCTTGAGCTAGTTG
ATCTTCGGATCATCCTCATTGTGAATGGGGTTTGCATTGAGAGAATGAATATACACTTCTTCCTTGGGCTGCTTG
GTAGTGTAGTTTAGCCTTAACCTCATGCTATAATGTTTATACAGTAACATTTCAAGATAACAACGTACAAAGCTA
TTCTTTTCTCTCCATTGGATTAATGAGAACATGCTCGAGTTTCTCTTGCACCTTCGAGATAAATGGATTGAGAGA
GGATGCAAACTATAATCTTCCTTAGATTTGGGGCAATTTGCCATACTAAATTCATTAGTCAGCCACCACAAATAT
TCATATCAGCAGATATGTAACTGTTCTGCAACTAGGAAGTACCTGAATTCAATGCAATGTAATCTTTCAAACATG
TGCAAGATTGTGATATATAAATTTTCTCACTCTATAAAAAAAATGTTGAGCTTCCGTACTTTCTTGGCCATGACA
TTTTGCCATCTCTTCCCATCTTTTGCGTAGGGTATATTTCCACCTAGTTGGTGAGGCCTCTAATTCCATAGTTGG
ATTCTAACCTGCCCCGCCTGTGATAACCAACACTTGGTTTCTGTGGCAATGGAAAGTTGTGGAAATGTGGTAGT
ATTGCATTCTAATCTCATCAAATGAGGATCAATTTTTCTCAAATGAGGATCAGTTTTTTCTTTTGAAATGATGCA
CCATGTAAATTTCTTTGCAGTAATCAATTATAGATAACCAGCAAAACAATTATAATGAACAATTTAGAGTGACTG
TTATTTTGCTATATTTGAATCAATTGCCTCATTGTTGTTTTTCTTTTGAAATGATGTACCAAGTAAATATGTTAG
TAATTGTCAGTTACATAGACTATTGTGCATTTGTGACTTGGGTATCATGGGTTTAAACATGGAAACAGCCTCTCT
GCATGTGGGGATAAGGCTACATATGTCTCACCGTCCCCAAACCCGGTAGTAATGGGAGCCTCATGCACCAGGCCA
CCCTTTTTAGTCAGTTAAGTGGAGCTAGTAAAATATATATAATAAGCAATTGTGAGCATCTTTATTGTTTTGCTA
TGTTTTATCATGTGCATGAATCGATGATAATATGCAAAATTTCAGATAACATAAATGATGGCTTTATGTTTCTGA
AAACGTTTACTAATTGTGGTTATACAAAAGATAAGCAGATGCTGATGACTGTGCATGGGCAAGAGAATATGACTT
TTGGAAGATTATTTGTTGAGATGCATTCTCATGGACATTTAGAAAATAAAAATAGCACAAGAGCCCTTCATCCTT
AATTTCTCATAAACTATCAACCGACATGAGGAATTATTTTTTGGAGTTTTTGCAACACCTATGCATAATATATTC
AACAATAAAGAACTTAATGGGGTTTACAGTTTTATGATCATGGTAATTATCGGTCAAGTTGAATCTATACTTACA
ATGGACTAAACTATACATTGTTTTCACTAG**CCAATGCAAGAGGACGTAACAAGCTCGAGGTGGCAACTGCATAAC**
**TTGCAAGAAATTCCATG**GTACGTCTATTTTCCTTAAAAGCCTTAACATGCCAATGCATTTAATTAGAATCAAGAT
GTTTCTATCATGACCTACAAATGCCAAAACTTTTGAACTTTTGGTAAAGATGATGTCTAGGGACTCGATGGTAC
CTCTCAACTTAGATTCTAGCCTATGGTTGAAAGTTGTTCAAGGTTACCTTGCTAAAACCATTCAGAAATTTTAAT
TGATAGAATTCTATGATCAATTTGAAACGTATGAGAATGGATGATTACTAAAACAAGGTTCTATACTTTGTTACC
TCGGTAATTATGTTTTCAGAAACCAATTGGTTTGATTGTGGGATGGAAATTCAATTTCATAGTCAATACTTCCAA
ATATATGCCATGAAAATGAACCATTTATTTACTTTAGCATGGGTGACCAAATACAGCTTATGTTCCATTATTTAA
ATCAATGCTTTAGTTTCTTATATTATCATGGCAAAAATAGGTTGGGTGAGGATGTGATGCCAAATGCTTATGCAA
GGAATCATTTAGTGAAATTGCTTCACTTATTACTTTTGGGTGAGCTAATGAAAAGTTTCTAAAAAAATATTCACA
CTGACAAGCATATTGTTGTCGACAAATCTCCCTCCCTCTCCCCCCCCCCACTCAATCTCCTCCACGCCCCCCCC
CCATCCTTCTGGTTCTCTAATTTGACAAAATTGGGGCTTAGCTTCTGAACATGGTAATTGTGAATGGAAGATATT
ATGGATTCATCCATGCCAATGTACAATATGTGGTATTCCTTGTCATTTTCATATTTTACTATTTTCATAGGACTG
AAGGCATGGAGTATGAGCTTTCTTTGTGCAACTAAGAAAGTTATGGATGGGTCCGAGCAGATGGCATAAAGCGAG
AACTTTATAAAGGAGATTATTGCCAATGTGTGCTAGACAAACACAACTTGGCATGGAAAAGTGAAATAAATGTTT
CAGTTTGACGGTCTCTTTTTGATTTTTGGCTTTTTTGCCTTCTTAGCTAAAAAGGATTTTTGATGTTAGGTGGTA
ATTCTCGGAGAAGAAGTTAGATGATATATTTTGAGTTATTTATGGTTTTGCTTATTATAGTATTTTGTGACACCT
TTTTTAGGCTATGAACTGCTTCATGAGTCAGCCCTTTGTGATGTTTGTGCCTTGGTGGAATTATTTTGTATCTCC
ACCTGCTAATCTGCTGCATTCAAATAAATGTCCTGACTCCATCTATTCCATCCAGATGTGGAAGTAGACATTTGT
TGTATCTGTGTTGGTAATCAGTTGTGTAGACTTGTAATCATGAATTTCAAATGTGTTAGATGCATGAAAATTTGA
TATTTCTAGTTCGCAACGACCTTCATAAAGCTGTATTTTGGTATCGATGAGCATAAAAACCGTCAATCTCTAAAG
GCTTCACGTTCTTGTGGTACGCCCGCCCAACCTCCACCCACACCCCCCCGCTCCCCCCTGTGCTTCAGTCTAAAC
TCTTGAACCCAAAGTTTTCCTTTTCTAATGTGATGTATTAATTGAACCTTGAATGTCACTATAGTCCTGTTTCTT
AGAATCTCTTTTCTAGTTACCACTCGTATCTCACTTTTTTTTTCTTTTCTCGGCTTTACATGTTTTGTCCATCCT
CAGGTTTAGTGGATTAAGTTTGGATTTTGATCAACTATTTGTGCTTTTTCAGATGTTATATCCTTCTTAAAGAGT
ACATATGTTATATGCTGCTATGTGTTGTTTGCATATTTCTCACGGTGATATCTTCTTTGTAGATCTATCTTATTA
TGCTCTTTCTGATCCTATTTTTTTCTCCTATTGTTATCAGTTTTATATTACTTTGACATGCTGTTCCTTGTTGAT
CTTCCATTATCTAAGACTTCATTTGTACCTGTTTATGAATACTTATGCCATTGATACTGCATGGTTCTCTGTTTT
CTCTACGGTTGAGGACCCCAGTTCTCTGATGCCTCCTACTGGACATGACTATCTTGTGATTGCAGACTCATATAC
CATGTTTCACCCTTTTTTTCCTAGTCATCTATAATGATCATGACCATAGACATTAAGTGGCTGGTATTACTCACC
GGTACAATGACATGGCTAATAAACTTGTTCTTTCTTGTGCTCTGAGACATGGAATGTTGTTCCCTTCTCATGGGA
AATCCTATCTCCTGCAATATAGGTCTACTAAATGATGATTTAATTGGTAATCCATATTTGGTCCGAGAACTATGA
TATAGATTATGAGACTTTGCTCCTGTTTCCCAGATGATTATCCTGTGGTGCTGAATAGCTGTTGCAGCTCTGCCA

CTAGATTTTGTTTTGTTGCACTTACTGGGATGTAAGTGATAAATCTCTTGATGATAAATATTCAAGTCAAGTTTC
ATATGCTTCCTCAAGGTCTCCTGCATCCTCAACATGTTTACAGTTCTTGTTTGGCAATTTAGGTGATTATTTTGG
TAAGACTAATCTATAGTTAGCATGACAATGATATGTTTATCCACTTTTCTACCTATGGTGAACCCGGAAGGCATT
ATATCTTCATTGGTAGGTTGATGAGAATGACCTTGGTCAACTTCCTTGTTTCCTTGGCCTTGAGCAAGCCTGTAT
TTGAAGGAAATTCTGTGGATGAAAGTTATAAATATGAGAATGTCCTGTTGAAGATCACCACTACAAGCTACTCCA
TCTACTCAGGGTTTAAATGAAAGCTCCATTGAAGTTATGAGAATTTTAAGTTTGAGAATACATCAAGACATTGTC
AGATTTGGCATCAAATGATATCCCTATTGCTTAATAATCAGACAGATTGGTGAACAGATCATCAGTTGGCCTTTT
TTATGAGAGATGCCTAAAATTATGATTTTTCCTATCAAGAAAGGACCTAGGCACTAAGTTGGATTTTGTGTTTTT
TTGGTAATCAAATGAAATTTTGGGTTATTGGATTATTGGTTGTTGTTGATGGTTAGAATTGATGCTTTGTTTAGT
TTATAAAGTTCTTTGTTGGCTACAGAAACAATTCTAGCATTGGCTTTTGGTGGTAGAGATGATTACTGGGAGTTA
AGTATTGAGTGAAGTTTTGAGTGGTCAGGACTGGTGCTTTTTCCATGTTTCTTTGTCCTCATTCCTTCTCTTATT
CTTTTCCACTTTCATTTTCTTGAAGATCTGTCTCCTGTTCATGTTGAATTTGCTTAGGTATATCTGCTTTCTCAG
CAGTAGTCCTTTTCAGATCGCTGCTCGTGCTCACCTGACAGAGAAAGTGTATAAGACTGCATGATCAGGAGCGCC
CTTTAGCTTGTTGAAGCACTGATTATTTAGCATTATGCTGTTCATTTTCCATTCTAGTTTGTTGTTACGAAAGGG
GATGTTCATTATTCTATATGGTCTCTTTCATCTTTTGTTTTTCAGTAGCACAGTAGCACCATTGTTGAAGTCCAT
TTTTCTCATCCTTCCCCTACAACTCTATGCTTGGTCATATGTTGATTGTGTTGATGATGTTATTGATTGTGTTAC
TGCATCTTCTTAGAGACTTTTATCTCCTATAAGGTACGAAAGCAAGATTTAAGTGGATGCTTGAGATATAGAGAG
AGTGTTGCGCAATGGTTGAATCCTCTTTAGAGATGGTCTGCTGTTGGTAGCTTCTTGTTGAAATAGGTTTTTATT
GGCAAATCTACCAGTATTATTGCAAGGACTACTGTTATTCAGATGGCAAGAAATCCAAGATATCATTAACATGCT
TTGCTAGTATTTTCCTCTTATATCTTAAATAGTATTTGTTGCTTTTATAAAATATTCATTGTCTTTTGTTGCCTT
ATTGGTTGTCACACCTTTGCCAAAATGTTTTTTTTTTTGGCACACCAACCACACTTTATTGCAAGAAATAGTTGC
CATGCAGATTGTGAGTAATCCTAGATTTCATGAACATTCTATCCAATTATTGGTGCAGAACTCTTATCTTCCTTT
TGGTAGTGGTAGTCATTATTGTGTAATATTAATTGCCTTTTGTTGCTTGCTTTACATTTCTTAAATAAGCACCAA
TAATCATGTGCCCTAGAAATTTATGTTTAGCACTACGTTTTATGCCCGAGATTCCAATTTTCTTTATATGGTTCT
CTTCTATATTTTGAATCAAGGTCTGGTGAACCGTGTCGAACCAGCTGGATCAAGGTGTATTGAACCAAACTGGTT
TAGGGGCTCTCGTCGAAACGATAACCTCCTATTTATGGTTTGCTAAACCGCCCCGAATCACGTAGTTCAGGGTAT
GTCGAACCGTACCACGGCCAATCGAGATGGTTCCGGCATTCGGAATGAGAAATGAGTAAGAGGGAGAGGGAGCAG
GAAAGAGAAGAAGAGAGAGGGGGGAGGAAAAGAGAGGGAGCAGGAGAGATGACCTCCAGAGGCCACTGGAGGGCA
GCCAAGCTTGCTACGCCACTGTTGAGCTTGGTGAGGGTGGAGGGAGGGAAGGAGAGGGAGAGGAGGCTCTCAGAG
GCCTCCGAAGGGTCAGCGAGGCGGCTATGCCGCCACCAAGGTCAATGAAGGCCAAAGGAGGGGCTCCACCCACCT
CCTGATTGAAATAGGGGCTTTAGCCCTTGTTTCATATTTTTTGTGGATTTTTGGGTTGAAGTCGGCAATTTTTTT
TGGAGATTTCTCCAGCTTCCACTAAGCTCGGCAATGGCACAGCAAGCTCAGCTACTCTCTAGTGGCCTCCAAAGG
CCCTCTCTCTCTCTCCCCTCCCTCCCCCCCCCGGTTCTCCCTCCACCCTCTCTTTGTCTCTCCCCCTCCCATTCT
CTCTCCACCCTCTCTTTGTGTCGGTTTGGGCTTGGTACGGAATGGCAAGGCGGGTAACGAACCATATTGCCAGCC
GACCGGTCTACCGGCATAGCAAACCTTGGTCCTTCCCCTCCCAATGCTTCGAACCCTCACCTCCCCTCTCTCTTT
TGCATTTCAGCCAGGTCATGTGGCTAAATAAAATAAATTGGTTTCTACTTAGGCCCCTTCTTTTGCACTTTAAAG
TAAGGCAAATTCTGCAGAAAGAAATCTGCCACTTCTGTTATCATTTTATATTTATATACTCATGAGAGAGATTTA
TTTGTGTGGCTGACTATGTTGCTCTTCTATATGAGCATATGCACAAAACTGCTAGAAGGGAGAGTAAAATCAAGC
AGTAAAGGAAGTTCATTTGAAGGGCATGACATTTTCTTATGTATAACCTACTAACTTGAATTCAAATGGAGTTGG
TTTTATAATCACAAGCAAAATGATTTAGCTTGCTTGTCGATAACTCTGTTGGGCTGGATAGAAGGGATGCCTTTC
CACCTTATTCTTTCACTAACGATGCATAAAATGGTTTGTGCAGATTGTCGTAGTTGCATAATTCTCATGATTTTT
TGTACGTGACAAGTGAAAAACTTGTTTTCAAGATACAGTCCATCTCTAGGTATCAACTCTTTCAATGTAAGCATC
ATTTAGGTAGTGATAATCCCATATGCCTGCAATGTAACTTGTGAATTTCTTGATTATAGGTTTTAATGATGAACT
TAAAACATGGGAATACGCATCTAGATAGGCAAAAGATTGCTGTTTGAAGCTCTTAAACTGAAATCCCTAGCACAG
CCATCACCTTTTAATAGTAACACAGCTTGACAGAGTAATTGGTTATTCTGATTATTTCACACCATTTTTTGGGAA
TAGATCCTTCTCTTCCTTGTACACCACAGGTGATCATTAATAGCATGCTGTTGTAACTACACTGGCAAAGGATAC
ACAAGGAGCAACCCATGATTTGCGCATCATCTTGTCAATGCAATTGAAGTCAGCTGCAACATTCAGTGAAATATT
CTATGATTAAACTAGTAATTAAAAAAATATATACCATTTGAGTTTTGGAAAGAAAAATTACCCTAGTTTTGGTGC
TGCAGCTTACGTTGCATCTGTTTGCAGTTGTGGCTGGCTGAGTTCAAGATGAATGGATATGTTTGCATCCTCGGT
GACATTAAAACCATATGGAGTAATCGGACCTTTATCAATGCCATTTACACCCCTCATTGCATAAACCTGAGATGC
AAAATTTAATGCTTAGCCTTTGCTATACCCACAACTTACATAATTTACCATTGGTTGAAAGGATTTCATTACAGA
AGTTTCTGTATGGATTTTTCAAGTCTATTCGCTCTCTTATTCAATACCTTCTTCTGCAATTCCATGTTTTCTTGA
CGAAAAAGGTTTACCTTCTTGTATAGTTCGATATTTTCTTGGTGAATAAGGTTCCCCTGCGATGAACCAGTGAAA
TGTTAAGCAATTGATATTCATCACCTAAAACTTAGAGGAGGGGATGTGGTAGACACATGTTTTCCAACCTTTCGG
TTTAGTTCTTGAATTTCATCGATCAAAAGTTGGTCCTATAGGGGTGAAAAATTAATCCATTAAAAATGGCTATTG
TTAAAAAGGACAGCAAGTCAACATTGAATAAGGCAAAGATGACCAAGACCTTCTTCATTCGGATACCACGTAAGC
TCATCTCAAGTTGGTTCTCTAGATTTTGTAGATCTTTCACACTTAGGCCAGAAAGCTCTTCTCCCAACAACTGCC
TGCAATCATTGAAGACGATGTTGTGTATGCTTTGCCATGTCAGTTATATACTTATATTTAGATATGTTTCAAAT
GATTGCTCTAAATTTGATATTAAGCGCTTATTAGAGCTTTGCCTACACAGATACATAACAGTGAGTTCAGTTTGG
GATTTGGTGAATGATGGACTTACTGCTTATTGTTGGGCATTTAATGTTGTCAATAGTAAATATGGGAGTTTAGTA
TCTAGATGGATATTGTATATGTAGTGTCTTCTTTAGAAACTCTTCTGTTGTCATTAGTGTCTGGTATATTGAGTG
CCCTGAGTATGGTTTGTGATACCAGTGAGAACTAGAGACTTATGGTTAAAATTAGTTCCGATTTCATCAATGAGT
TACTACTTACCATCGCTGTATACCTAGCCATCTGAAGAGACTTTTTGTTAGCTTTGGTATTTTTATCATCTGTAT

CCATTATTTGTGTTTCTATTTTCTAAGCTAGTCAATGGCTAATTTGACTGTTGAAGACCTTATCAATCTATGGGC
AACATTGGGAATGGATTTGGATGTGGGAGTGGAAACAGACAACTAAGAAAATCTAGATTGTCAAACCATCTAGGA
TGTCAAACTTCCTAGGATGCATCTTTGTAAAATAATGTTTGATACTACAAAATAAATCTGACCTTCTTTGTTTTT
TAAAGAAATACAAATTTTGATCAAAAAATTAATAACAATTATCCATATGAGATTAAACTATAGTTGTTTTTAAAA
CATAAAAAATAAAAGAAAGAATACAGAAGTACAAGGTGCTCAATTGAAGAGAATGAAGTGGAACATTATTTGGAT
CTTTTCAGAAAAGTGATTTGCTAAAGGTGATATGCATTGAAAAAATAATGGTGATTACCAAAAGAAGTTACAAAA
AGAAAGCAACAGCAGAAGAAAAAAAAGAAGAAAGGGACTACAAAAAGATGAAGAAATGCTACTATGGAAGATTAG
AAGGGATATGAGGACCTCCTAAATAGGGTTCTGATATTTTGGTTTTTCACAAATCCCTACCATCTTACTGGTTCT
TAATCACTTTATTCTTAATTGTGCTACTTCACCTCCACTAATATACTAGTTCCTTAAAACAGTTGTAGTTTCCCA
GGCATATGTGGACTCCTAGACCAAGAGGCATCTATTTGTTTCATTACACTGGCTCTAACCTTATTAGAAATATTT
TCTCTTTCTTCATTGTTATCAAAACTGATAACTTGGCAGTGCTCTTCACTTTGACTGGGATGTTGTTACTGTTAT
TTCAATGAATTTGTGCTTCTTTTCATGATCATTTCTGTTAATGTTGCTCCACATAGTCTCTATCTTTGTCCCACT
GGTTGCTAGAAATGTCTAATTATATTCGTTGTTTTTGAATTCCAGAATAATAAAGTGAGATAGAAATTCTACAAT
TATAGTGAGAATGCTTTTACCATATTTCATAAACATGGTCATATGGTCTTAACTACTGATCTTTGGAATGTTACA
TGTGATATATGAACCAGTTACCAGTCCCCTTCATTTTTCTTATGCTGGAATGACTAATATACATGGTTGCATTAC
AACTTGTTTGTATTTTAGTAGCTACTACATTTGCTTGATATAAACTTTCCTAGATTTATTTTTATTTTTTTCTGA
TGATAAGTGAGAAGGCTGCCAATTATGTAGTTCATTCTAAAGTTATACACAAACTTGAAATTATTATCTAGACTT
TAATAATACTATCTCTTACGTATTAGGTCTTAGTATGCATGAAATTTAAGTAATTTATTGTCAAAGACATGTTAC
CCATCTGCCATTCGCATTTCTGATACTATTAATCAGTCTGGCTCTCAGGGTATGTGATATGCTAGTGCCTTTGAA
GTGTAATATGCCATTATTAGTGTCAAATTGAAATAGTATTATACTATTATATATGTTACAAGATGCTGAATAAGT
TTTAGCCTCCATGTGTTATGTAACAAGTAACAATCTTCTGAATTTAAGCCACTTGGTAGAAGTTACAGTATGCTT
ACCTTTTTGAGATGAATGCCTTTTGAGAAATAATGCTTAAAAAAAAAATATGTTATGCAG**GGTGAAGTGAGCATA**
**CCGGTGACTTTCTTGCAAGTTATGCAGTTGCTGCCTCAAGCTTGCTGCCTCCCTTTGCCAAAACTA**GTGAGTACA
AGGACAAAGTTTTAAGCCCATGCAGCAGATTTTTTAAAATACTATGTTATTACTATTGGAGTGATTATGGATTAC
AGTAACCAGGTTCAGGAAAGAAACATTCTGTCAGGTCACTTAGAATCTGAAGTTTGATACACCAGTTTTTCATGA
AATTTATTGCAAATCAAGTTTAAAAGCATTAGGTTTAAAAACTTCTGAACATATACTAGACCACTTACTGCTGAT
GTAAACTATGAATTCTCGACTATGACTTTTTGATTACTTAATCTTATCGATGCATGCTGTATTGATCTTTGAATG
TTCCCCCTTAAATAGTTGATGACCTATTCACACACTTTTGGTGCTTTGCCTAACATCTGTTGCATAGCCACATGC
CCAGTCCCCAATTGAAAAAAAAAGAACAAGAATTCAACACACAAAAGATGAAAAGGAAAAGAATTAAAGAATATAT
TGGATATGGAGACTGTCGACAGGTAGAGGGGAAGAGGATAAGATCTTCATTTAAATTATTTGTAGGATTTTTTTC
ATATTACATGACTAAGAAACAATGTAATTTCTGCATCATTAATATGGAATGTAGAGGCAACATTCAAAAGCTTCA
TGATTTCTGATTATAGGTTCAATATTTTGATCAAGATTTCAAATCCCGTGGGACAAGACTCTTGATTTTCTCATG
GAACGGGATGCGCCACCGTCCTGCCCCATCCTGAAATTGGGATAGGGGATGTCTCAAGAGACGTCCTGTCCCGAC
ACATGGGATGATATCCTGTTCTGGTGTCCCGTGGGACGTTCCGCTGGAACCTGAATCCTTGGTTTTGATTGTGCT
TCATTAAGCTTCCTAAACAATTTGAATTTCCTTAATTACATCACAAAAAAAAAGTTTCCGGATCAAGTTCTTCAAA
ACATGGAACGTGTGCTTGTACTCATCACCACACGATAGGAGGTTTCTGTTATACTTTTTTTTTTTATTAATGGTGA
GGTTTCTGTTATACTTTTCTTTATTAATGGGCTCAAGCACAACTACCCCTAAAGATATGCAAAAAGTTATTAGGA
AGCAAAAGTAGTTACAAGCATGCAAGAGAAATAGAAACACCATATTCAAAACATATTATGCATCCAGATCAAAAA
CATATTAAATTCTTCACTAGAAACACCATCCCCATTTATATTTGAAGGACAATTCCATCCTCTTGGACAAAAACA
TCCTTCACCTCATAGGAAATTGCTGCTCCACCTTGAGGCTGCATTTGATTGTGGAACAAGTCAACCCTGATAAAA
TTGGTTTAGTTGCTCCTTACCCTCTTTCCTCCAAAATAAAATACTTTACCCTATAATGTCCAATATTTTACTTTG
GATTAGATGGGGAAAATGGCTTTTACAAACTTTAAAAAATTACTCATTTTGAGCTCCAAATAAACTGGTCATGTT
TTGGCATTTGTCAAGTTAACATGGAATAAATTGGCTCATGGGACCTTTACTCCATTTACTCTTCTTTAACCCGCA
GCCAGTTGTAGACTAATAGAACCAAATGAGAACCCAACTCAAGAGAAACCTCAAGATGGGCATGTAGTTGCAATA
CATAGTTTGTTAACCTTTCCAAGAGAATGCCTTCATCTAACCTCCCATAACCATGTTCGGATTCAATACCAAGTC
CTAATCCATTTAGGATTCTTTTTAGTTGATCGTTAGTTTTAATGACTTTAGGATTTTATTCCTAATGTTACTCTA
TTTAATTATTTCCTTCTAAAGGTTATTAGAGTTTGTGTTGTAGTGGGTCTTGGGTTTTTGAGTCAAGATAGGAGT
CAAACTCCCACCTACGTGGTGCTCTATTTAAAGAGACTTTTGACGTCTCCTTTAGGTGTAGAAGATGATGGAAAA
ATTGTGTAGAAGAATGAAAAAAAAGAGAAGAGAGATTATTTGTGCATGTGGAGACTCTTTCGTGAGGTGTCTTTG
TGAAGTTTTGGTCTTATTCTTCTTTGAATAAATTGGTTTTTCTCTCCCAATTTTTTCTTCGATAATTATTTTCTT
GTGATTCTTTACAAATATTTTTTTTGTTTGGTCTTTGTTTTAGATCTTGGGGCGGCACGGTTGATCTACTTTATG
TAGCATGCCATTCTACATGGTATCAGAGCTAAGGTTCGGAGATTGGTGGCTCGTGTGTTTGTGGCTCTATATGAT
GTTGAGCCATAGGCTTGGAAAGTGGTTAAGTGGTTTGCAGTTTTGATGCTTTTTTTTTTTAGCAATGACTTGATT
ATTGGTGGTTTCCGTATTTATAGCTCGGCGCAGTTAAAGAGTTTTTAGTAATTCGTTGTGCGGTAGTGCAAGTGT
GTAGATTATTTTTTTGGTAACTGAAGAAGTAAATCATCATAGTCAATGCTATTACTATCTCTATCAATAGATTTA
GTTATGAGATGTCTGATCGTGCTAATATCAATAAGATTGCTTTTACTACTATTTTTCTTCGTAAGTTGAACAATA
TCTATTATCCATATTAGAAGGCTGTTGTTAAGTCTTATTTGACAGACCAAGATTTGTAGGATATTTTGGATGAGT
CAGAAATGATGCCAACAGATGAGATTGCAAGAAGAAAGTGGAATATGAAATCTAAAAAGGTAATATGCATTCTTT
AAATTTCAGTTGATAAAGAAAATTTTTTTTATGTTTGAGACGTAAAGGAGTCAAAGAAAGCATGGGATATTCTTA
CAGCTGTCTACTAGAATTCTTGCAATATGAGAAATCATAAACGAAAATTATCTGAAGAGTAAATTCTGATTTCAG
GCTTTTCCAAAGATCTAAAAAAACAAAATTAGTTTAAAGACGAAAATTTGAAATATGGGGAGCTTGATCAGTTAA
ATTGCATCAATTTCGGAAGACTCAAAATATGGTAAAGTCGATCAGTTTAAGGATCAATCATGTGCAGAAGAAAAT
TTGGAGTAAGAAGAGCTTGATCAGATGGCTACCACATCAAGTCTAAATCAGTTCATGATGGAAGAATCTATTGAT

TCTGAAATTGCTTAAGAAGAGTTTGATGATGCTACAACCGATTCAGTTATGAAGATTAATATTGAGGAGTCAAAT
CATGCAGAAGAATTTGATATTGAAGCTATCCAAGATATGGATATTTCATATGAAGGTTTACTCAACTACGAGGAT
GAAAAACTGACTAATTTTGAATTAGCTTTGAAAATAAAGCAACGAGAAAATATGCTCAATGGAGATGATTCTTAT
TGTACTTTGGCGATCAACAAGATTCTTGATACATCAGTTAATTTTGAAGCTACTCCAAAGTTCGAGCTGTTTGAA
AATTCTATACATGAAAATTTAATTCCATATGTAGAAACCAACTGTGTACAGATACCTGAAGCAACTCAAGGGGAG
GTATATAAATATGGTCAAAGTTTTGCTTTTGATTTTACTTGCGTGCAGGTAATGGTCAATGCTCACAGTTATTAT
AACCAACAATTTGATGTTCGTTGGGATAAGCTGGATGCTCGAAGAAATCTCGACATCATGATCGTGGAACCAATG
CTGAGAGAAAATATTGCACGACATTATTGTCGATTTTGATATATGCATCGAAGAGGACGAGCCAGCGAAGTAATT
AATGTTCGAGTCAATAAAGAAGTTTCATTATTGAATTGTAGGTTCAATAAGTTGTTATATGTTACTGTTACTGTA
GCGGTGATGGCGATGGCTACTAGCTTACTATGGCATTTCATGACATCGACAAAGTCAGCATTGTGTCAGTATGGA
CTTTGCAATAAGAGGGAGTGTTGAGATTCATTGTCAAGTCCCAATCCAATTATGATTCTATTTTTAGTTGATCAT
TAGTTTTAATGACTTTAAGATTTTATTTCTAGTGTTACTCTTTTTAGTTATTTTCTTCTAAAGATTGTTAGAGTT
TGTGTTCTAATGGGTCTTTGATTTTGAGTCAAGGTAGGAGTCAAACTCCCACCTATGTGGTACTCTATTTAAAGG
GACCTTTGATGTCTCCTTTGGGTGTGGAAGACCGTGAAAAAATTATGTAGAAGAGTGTGTGGGAAAAAGGAGAGA
GGAGAGATTATTTGTGTATATGGAGACTCTTTTGTGAGGCGTCTTTGTGAGGTTTTGATTTTTCTTCTTCGATGA
ATAAATTTGTTTTTCTCCCAAGTTTTTCTTCCATAATTTTTTTTTTTGCTCATGATTCTTTGCAAATATTTTTGT
TTGGTGTTTGTTTTGGATGTTGAGCTGGCACAGTCGATCTACTCCATTTAGCATGCTATTCTAGAAACCATTTCT
TCATTTTCTATTGCTTCGTAATTCTACATCCACTTATCTCTTAATCGTAAATCTATATTCTTTTTCTTTTCTCTT
TTTGTTGTCTTAAGGTGAAAAACCTTTCTTTGATTTCAGAAATATGTCCTTGTTTCCACAGCTTTGTTTGGCTCT
AGAGGGTTGCTAAGATGAATCTCGTCTAGTTTATCATGTCCTAAATTGGCTTCTTAAAGAATTCTCAGGCAAACA
CACTCAATAGTTATAACTTAGTTGGACCACAGCAAGTAGATTGAAATAAGACCCAGCAGGTACATCTCCATTACC
TGATGTTCTATTTCAGAAAACTATGGTCTTGTTGAGAACTATTTCCAGCATTCTCCACAATACTTCTCCAAACAA
CTTGGTCATCTCTGGTATGTCTTTGAGCAAACACCAAAAGCTTGTGTCTCGATATATTTAGTTTCTTACTAGTCT
ATTTTTAACTAAGATTTCAACATCTTTATGATTTTGAACGATGCTACATTATGCAGGTAAAAGTGTAAGAAGCTG
ATACTTTTTGAAGACAAGCATAGACACTATGGTCTGCCAAACCAACTCGAACTGCCCTCCCTGTTTGGGGCATAT
CAACCCAAACCAGTTGGAACTAGGTTGGTTCAACCTCTTGGTTCAAGAACTGGCATGAACCAGCCCAAGTCAGTC
TGAATTGAGTGGAATTGCTTGGTTGCACTCGGTTTTAGAGTGGTTTGGGTTGGTCCGGGCTGTCCTATGATGAAA
AAGAGAGAAGGCCACAAGATGCCACTCTCCTATCTTTTCTTCAAAAATTATGAAGGAAAGTGATGTAGCATGAGT
GAAGAGCGATTAGATTGACGGGATTAAACTTTGAAGATCAACAGACAAGATGATGGCACATTGGATTGCAAAACC
CTTTGTTCGATGAAGGGAGCTCTGTTTGAGAAATTTTTTTATTAATATATCAAAAGCCACCCTTCATACAATGTA
CTAAAAGCATATAAAGGATGACCCCAAATTTAGGCATTAAACCTGGTACATAATCATGCTGGAAATAGGATCCAA
TAATAAGGAAACATGCTAGAATTATTCAACTAGGAATATAGAAACCCAGCTAAAACCGCTGGCTTCTATGAAACT
TGGCATGGTCTAGTTGGTTATAGTTTTCGCAGACTGCACTCTTGTTGCATCATGAAGGGTGAGTTTTGTGATGGA
AAACTTGTATATTCTATCGTAAAATATGTTTGTAATAAAAGTTCAAAATCCTTGCACTAATTGATAGGTATTGAA
GGGGAGATGATCCTGATCATAATCAGTTTCCAAATCTGTAGAGAATGACAAAGTTAGGACATATAGATCTTATAT
GGATCCTCACTTGAAGAGGTTCTCTTCAATTCATCTAGAAGAGGGATTACTGAACACACAACCACAATATTTAAT
ATTTGAATAAAAATCTTGTCCACCATTAATACCCATTTTCATAACTTATGCCAAGCTCTTAGGTGAAAAAACTCA
CCATCATCTGAGCTTTGTGTTGCTTAATTGAACCATTAACAAATACAGATGATTTCCATGAACAATGATTCGAGA
AGGCAACCTCTCACAGATTAGTATGTCATAGAATTAAATGCACTTTAACCATAAACTTTTGTCATTTAGGAACCT
TAGAATTCTCCTATAATGTTTTAATCTTGATGGGAACAATGGTGCATAAACAAAATGAAAGCTTAAGAAAACCCT
AAGACTATTTGGTTAGACTAATGCCACTCTAATGGATGCTGATTCTAAATTGGGAAGTGGGAATTAGAGAGAGAT
ACAGAGAGGTGCAACCATAAGAGACTAGGAGCAAGACAAGAACTAGACCTTGGGATATGATGTCCACCTATATAT
CAAGATAATATGGCCTATGTAAAATCTATGAAATGAAGGCAAAAAACAATAGTAAAGGGTGGATGACAAGTCAAA
AGCATGAGTTCAAGAAGGTAATGGTTGAAAGAAAGATGTATATGACAGAGCAGGGCATTAATTAATATGCATCTG
TGCATTGAGTTACAAGTTTATGTTTAATTATTGCAGTTGTATCTATTAATCAATCATATAGTCTCTCAGAATCAA
TATGAAAAAAAAAACTCTAGCACTCTTAAAATAGCAAACTAGTACCGAATTGGTATGCACCTACATACCGCATGT
CAATATGGTTCGACTCGGTACCATACCATAGTGGCACCTGACCTACAGTCTCTAGCTGAGATTGCAGACCATGGG
TGACAGTGTCAGTTAATACACCTAGGACTATTTTGTATACACTGCCTAATTTCTTTGTAGCTTGTTGAGAGATTC
GTCTATTTGTCCATGGAAATTTCACTAAAAACTCACATAAGAATTGTTATTAGACCTCCTATCTATCAATTATCT
AATATCCTCATGATAACGCGGCAATGCCATGAGGATCTTGCTTCTAAACAGAATTGTTTAACTCCATCATAGAAG
ACAGTTATAGGCTATTCAAAGTGGATAGTCAATGGAGCTAGTAAGATAAACCAAATGGCATTTCTTGTCACCTTT
AAATCTTTATATTTATGACCATTTTGTCATATGCCTCTTCTGACATTGGTAAAATGAGAGAACTGGAGATTGTAT
TGCCTCATTGCAAGAAAACCTCCTGCTATTTTTATAATTAATTTTAAAGCTTTTCTGCATTATATTGTGGTCTTT
AGTTGAATTTCATCTTAACCTTATGTTGAATATCTATTTCCAACACTGACCACATTTCTCGCCATTAGCAGTATA
TTATCCAAGCAATTATTGTTGCCTGTATACTGCTTTTGATTTTTGTATATTCATGGTAAGTAGCAGAATCTAAAA
GTTTGACCAGTCTATTGATTTCATTTGCTCATGAATTTTCTTATCCTTGTAATGAAATACCAAAATCTAAGTGAA
GGATATCTAGTTAATATAAAGGAGCAGCCAACTACAGAGTTCATTGTTGGATGTTGTTTGATTGAGTGGTTGAAA
TCTATGAAGAATGGACCATCTGATAAGTCTCAAATAATTTTCAATGAAAGAACGTATTATATTGTTAGGATTGAA
GATTGTCACAGACATGGATGCTAGCGATCACTGTGAAATTTGCAAGATGCATTTTAGAGCATTTGCAGAAATTCA
AAACTCAGAAGCAGTCCTGTTGATTTAAGGTATGGTTAGAACATGGAAAGTCAAACTCTCAATCTGTCTACCTGA
ATGTGATCTGCCTCTCTATTTCTGTAGAAGTGATGTTTGTGAGGTTGATTCCGTCCTAAATCTGTCTGCATTTAT
AGTTAAGTATTAATTTTTGGTGATAATGAATAGAAAAGACATAGCAAAGACAGCTTGAGCAGGAGAGCTTCTGTC
ATAATATGTGGAATTTTTGTTGGCAAAACAGTGTGACTATCTAATTTAGGAACACATGAACAGACCTCTAAGATC

CATATGAATTACAAAAGTATGTTATGAGGGGAAAGATTGTATCTGTGCGATGTTAGAAGCAAAATACTGAAACAA
TAAGTAAATATCTGTTGCTACAGTACGAGATTGAGGGTTGGGAAAGATGAAATTCCTTATGGAACACCTATCTAT
GATAGGTATTTTGACTAGAGCATCATCTTTGTGGGGGTGGTCTTGCTATAAGATTAAAATGGAGCAGACTGCTTT
TAATAGTATATTTGATGATGATGATGATGTTATGGACGACAAGAAAGGGAAAGTACTATTTAAACTTGATGATTG
TGAAATTTGTAGTTTAATTTAGACAGGCACATGTAAGAGGAGGTTATGTAAGATTAGATGAGATTTCTGATGAAA
GGATATGAGTGGTGCAATCAATGATTCTGATTGTGATGATTTAGCAAGTAGGGTTCCAAGGTTGCAGCAGTAGAA
ATGACAGTTAGCCTAGCTGTTCGATTTATCTAATTGTAAATTGTAATTCTCTCTCTCTCATTTTGATCAGTCATT
CCTATGTCATGTCCATAGTGGGAGTTACTCATTGACACCATTAATTGGATCAAGTACTCTTTAGACAAAGATTAG
TTTCCCAAGGCATTTTACACCATAGGTTCTAATTAATATAGATCAAAAGGAACATATGCCACATGGCTTTTCCAT
AATATACAGAGGTACTGCAG**ATGCATTTTTCAGTGATGATGGATGGACAAAATGCAACCAGCGAATCAACATGGA**
**GAAGTGTAGGATGACTTACTGAAAGAGAGAGAGGCATGATGCCTTCATGTTCATATTGCTGTAACTACATTCTTC**
**TTTGGACAAGAAGCAGGCGTAAGAGGGAAAAAAAAG**GTATTAAAAATCTGCCTTCTTGATTCTTCCTCCTTCTGG
TGTTGTACTGTTGGAAACCATGAGTTTTCTAATAATCCTGAATATGTTCTAAATGTCAG**ATAACTTGGATTCCGT**
**TGATCATCATCATAAGACCTTCATAAGGTGGAAGATCACATGGATGGGGAAGACTATGATTGGGATTTTTGAAGG**
**CCATCAGAGGTACTTCGTTCATTTTGATCTTTTGAAGAGTTCTTATGTTTTGCTACAGAAATTCAGAGAAAAGAG**
**AGTAGTGTTTACAATTTTTTTAATGCTTACTTTCAAAATTTAAATTAATTGATAACCTCTTCTAATTTTGAGACA**
**CTTGTACTTATCACTGCAAGTTGAATGTTCAAAATTGAAGTGCTCCAACGACATTGTAATTATCCAAATCATAGC**
**TTGTCACATATGATGATCAAAAGCTTGGATTTTAGTTTCTGAAGGACCTTCTATGAAAGATCACTTTGAAGTGCT**
**ATAGGTATCGGCTCCATTTATTTGTAAATGAGAAAACACTTCCCATCAAGCTACCTATGGTTATTCTTATTAAAA**
**TTGCATAAACAAATGTCCTTATTTTGTTGTTCTAAATGAGACTTAAGCTATAGCATATCATAGGGAAATTTCTCT**
**CCCCACTATTCATCCTGTTAATGATGAGAGCTTGGAACCCAAAAGTTCCCACAAGTTTTTCATTTGCAAATAAAA**
**TGAGTAAAACTTGTAAAAAAAAAAAAAAAGAGGAC**TAAAACTACAATAAAAGATCAATCATATAGTTGATTGGTCTT
TGTAAACTACCAACTTTTTTATTTTATTTATTTATTTATTTATTTCTCTTTTTTTGTGAGAAACATGGTGGCAAT
TATGCAATCTGCTTCAAATGCGGGACCTCAAGGAGGAGAGGGGTGTCAAGCAGCTGACTCAAGTATAGCACATTC
AACTGCAAAATGCTGTCAATTTAATCTGCAAAACTTCAGATTTATTGCCATTTCTTGTGGTCTCCAGGTGATGTT
CTTTCCTAAAAATATTCATTACCAATCCAAGGCATGTACTAACGTAATTAATTGTATGTGCTGATTTAACACAGT
TTGAATGCTGATATTGGAATTCATTTTGAGGTCTAAATGACCAAGAGATACAGAAAACCCATGTAACAAAGAGCA
AAACAAGATTTGATTGACAGAGGAACATATGGACACAAGTTCAGAAGGGTTGCAAAAATAATAAGAGAATTAAAC
AATTTGTGATAAATAAAAGAACTAAATCAAAGTGACGAGGAAGGGGGGGACTAGTAATAATAAATTCATGTGTTT
CAAAGTCTTTTTTGCTCTCGAAAATCAAGTGGCTTTTAGAAACTTGCTTGTTAGGTCCTTACAAAATTCACCAGA
GCCTCCTTTGAGCTTCTTTTTGATCCAACACAGGGTTGACAAAGAATTTGTCATGCAAAATCTGACAAATAATAT
AACATGGCAAATTTGATTGTATCATGAATAAGTTTCATTCTAATTTGAAGAACTGAGGATTAGGATGATAGATCT
AGGACTTCCACACTGACCTAGGACTCTTTGAGGCCTTAGAAATTCTTTAAAAAGACATAAAAATTCTTGAAAACA
AGTCAAGGTTTGGTCAAAATTATCATCTATATTGATAGAATGACTCAACATTAATTGGTGAAAGAACCATATCTC
CTTTACATAGTATCAATATTCTCCAACTATTGAAAAATTTCTTAAATTTCTTGTACTATTCGATATTTGAACTGT
TTTCCATACTATTCCCTGTAGGTCACTAAGTTCACAATTAATCAATTGTTCTCAATTTTACATAAATTTCATGAT
TTATTGATGAGCTAATCATGATTACATGATAATTCTGTGTTACAGACTAGAGATGCAATATATTGTTGATGATGG
TGTTATTTTGCTACCATTTATTGTTCTCTCCTTCTAGTACTTGCTTTTACACAAAGAAAAGTAACTATTTTAGAA
ATCTCGCATTTATTTTGTTTTGTTATATATGTTCCAAATTTTTGATTATCCTCACTCTTTCCTTAAGTTGTGAAT
TTGCCAGATATGGCCCAATTTTTCTCAATGCTGAACTTTTGACCATTGGTCCTCCATTTATTCTCAATGCTGAAC
TTTTGACCATTGGTCCTCCATTTATCATTTTATTCATTTCTTGTACCGTTAAAATTTCTATTCAATAAACCTTAG
TGTGCTTTGTTGTATCTTTGTTATTGCTTTTTCTGTACCAGCTTCAACTTATATTTCTACCTACCACAAAGTCAT
GACTAGTTATGATATAAAATAAAATAACAAGTAAATTATGATAAATTTCAAAGGACAAGTGCTGACACTAAGGCA
TCTTTCCACTTAGCCTTTCAATGCATTTGAGTCATTCATGATTTCACCAGTAATTCTTGTCGCTCATGAGCATCT
ACAAGCATTAAATTTGAGTCCTTTCCCCTTTCATCTGCCCCATCAGAATTATGACTTTTGGCAGTTGAAGCTAGG
CGGAACCTGATAGGCTTATCATACAAGCAATCTAATAGTTCTTAGCTCTTCCGCCAGTTTATTATGAATTTTTGT
TTGTCATGCATTCTTTTGGCTACAACTTGTATCATACCATGACTGCTATTAAGCTTATAAGTTGGATCAAGACTT
TAGATTAACAAGCCAAACTTCTATTAAGATTGATGAACCAATGCAAAATGATAGAGAATAATTTATAAATTATGA
CTAAATAAGGACTTGAATTAGTCATTTGGCATCTTCACACAAAGCTCCAAGCATGGTTCAAGATATTGATACTGA
GCCTTGCCCCAAGTTAGGATGATATCAAGACACTACAGTCTTGACTCTTAGGATGGTTTGGTGTCCCTGTACCAT
AATATTATATCAAATTTTTGATTTTTTTTTAATCCTAAATAGGTATTTGAATGACTATTGATATGGTTTTGAACA
AGATTTGCCATCTGGTACCAGACCCCATATTGGTACCATCCTATTACTATGTCGGTAAACTTGGTATGAGGGCTG
GTTCGGCATACCAATACTTGAAACATCCCTGTACTGTATACCGACTTAGTACCGATACTATACGGTGCAGTGTAC
ATACTAGTACCAACTAGTACGGCAAAATATTCTTTTGAATCAATGATGGGTGTATGGTGATGTCACCGATGCCAA
CACCCGCAAGGGTAGGTGGATCTTTCCTCCTAGTCGACTTATGCCCGGTACCTGCTAAATCGGGCTTGGTATTGG
CTGATCCCAACTAGTTTGAGTTGAGACGAGGGATGTCCCGGCCCATAAACAGATCGGTGTCCCATAGATCTCCTA
GCTCCTGACTTCCTATTGGAATAGTATAGTACCAGCAATGTTATCTCATTTAAGCAAGATGTAAATCCTTGGCTT
GAAAGCTATCTCCAGTTTGATGGATAGTCTTTGATCAATGACCAATGACCATACATAGGCCTAGGGAAACATTAT
GTGGATTGAATAATCATTGTGTCATACAATTAACTCTACACGAGAAGAAGTAATACAAGCCTTTTTTGTAGGGAA
TGTGAACAAAACCCCAACATAAATTGAAAACTGATTTTGAACCATGGATTATTAGGCCTTTGGAGAACAAAATCA
CCACCATCAGCTTCACACAATGTTCACTCAATATTTATCAAAGGAAAACTCAATTATTGATTTTACACTTGTATG
TCTGGTCAATTGCAAAGTTACAATGCTTGGTAGCAAGGTTCGCGGACTCGGTATTGAAAGTCATGCCGATTGCCG
GCCGATATGGTATCGTACCGTACCATACCAGTCCATACCATACCGTACCATACCGATAATAAAAAAATCAAAAAA

ATTTCACACAACAGCATTAAAAAAAATCAGGCCAAACCGGACCGGATCATCCGGTATCGGGTGGTTCGGGCCAAT
ACCATACCGAACCGATTGGTTTGGTCCAGTTCAGACCATTTTCTAAAAATCTGGCCTGAACCGGACTGATTTTTT
GTCGGTACGGTATGGTATGGGGTGTACCGGCCGGTTTGGGTTGGTACAGAATACCATACTTGGTAGTTAATGGCT
CTTGTAAAATTGTCATGGTTCCTCTTTGCAAGCTCTTGTTTGTTGACTTTTAAGCCATCCAATGGGGTTATAGGG
ATGCATTGGAATGATGTAGATGTTTTCATAGCATTTCCAATTCAAGATCGACGTTTCCTGAATATTAGACAGACT
AACTCACTTCAAAGGCATCTTTAGTATGTCCAACAATTATGGACCAAGATTTGTGTAAGCCTAATAAGGCCTCAA
ACTTTGTTCTTTTCCTGACCAGAGTGGGTAAGATTAACTCATAATGAGAGCAGCCTGGTCTGCCGGTCAGTTTCA
CCCTTTGAATAAGATGGTTGGACAAACACCTACTTAGCAGTTAAGAGTATATCAGTTTTATATGGTGAGTCATTG
GTTCATATCTCCAGTGGTAGGTAAAACCGGTTGGCTCCCCCACTTTAGCCCATGCAAACATGTCTGGTCATGACT
TTTTGGTGTGGCCTTATGTCTGTTGGAAAGACCCTAATATGGTCTTATTTCACCAGTAGTTTTTCAATCTTGACT
AAGTGGTTGTTAGGATTGCTTTGCTTTGCTTCTATAACTTTTCAACTTATAATTTAGGACACCTGAATGCATCCC
CAAATCATTGCCAACAGATCAGTGTAGATTCTAGTCGTATCACTTTGGATGACTACAGTGTTGATGACTACAGAT
CGAAATTTTATCAAGAAGGATGATGACTGGCTCACCATCAAACGTCCTGAGAGAGAACATATGACAGCAAAGTTG
TCCTTATTCACTTTACTAACTAAATTCCCAATTGAGGGGACAATCTAGAATTCTTCCTCTTGACTTGTGAACAAG
TAGGAGCCATTGGATGGTATGATACTTAACTCTGAAACACTGGAACAGAAGCAGTGGTGTGATCAACAAACTGAA
AAGCTGCTAGCCAACACTAGTAGTGATTAAGATTACTGGTTATTGGTAATTACCCATGATCATAACTAGCAATAA
GCAATTAACTATACCAACTAAAAAGTAGCAGTCTTGCGGTAATTAAGTGTGACAATCAAGCAATCAGCAGTACCG
ACCACTAGATGGTTGATGGTAATGAGTAATGACTACTGTATGGTTGAAAAGAAAAAAGAAGCAATGATCAAGCTT
CCCATATTAAAATGAATTAAAAAGGAATCAGTTTGAAACAGGTAGCACCATAGCAGTGTCAAGTGAGAAATCACA
TTTGAGAAGACAGAATGATCATATGCAAACATGTGCATAGTTTGATTCACGAACCACCATGCTACTTTCAGAGAA
AAGAAAGTAATAAAAATTACATAGGGATTTAAAATTTATAATGAGATCAAGAGGATAAAAAATGTGTATAAGTAG
AACTCTTGCAAGTAGAAAAATCATTGTCAAAATCATGTTATCGTCATCTCTTTCTTCTTATGATCTTGGTATCTA
GATGTCAGACTTAGTTGTAAATTCATTTATTGTTCTGCATTGTATTTTAAGCTGTGTAACTTAGACTGCTTTGTT
TAGGAACTACACATAGTACTCACATTTTACCCAGTATAGCCCACAACAATAGATTTTTTTCCCTCAGTTATAGGC
AATTCTCCATTTTCTCTTGTTGTTTATTCTATTTTGACATCAAAGATCCTATTTCAAATCTTTTGATCAATTGAA
AATCGATTTGTAGAATATTTGGATTGAAATCATGAAGGTCCGCTTGTTGTTTAGCATGAAAGATGCCTCCTCATT
ACTCTTGACCTTGCAATTATGTAATGCAGCCATGAACAGGGATCTTGTAAAAGCTTGTATTAGTGACCAAAATTG
TTTGGGTAAAGGAGGAAGAGTCCATCTATGTGGAGCAAAGACTCATAGTTGGACATGGGTAAACAAAATTCTCAT
ATGCCAAGCATGATACCTACAGTAAACTAATTCTCCACTCCATCAATAGACTTTTTGCAGGTGTTGATGTAGCTG
GAATCATTATGTATGATACCTTAGAGCTAGAAAAATTGTTAAGATTATTCTAATATGTGAAGATCAGTCTGTTAC
AACCATCCAAAAAAAATCTACCTTGGTATGCTTTGAAAAAAAAAGTCAGATAATATTTATTTTTTAAATCTGTCT
TCGGTTTCCATGTGAATGATTACTCAATTATTAAGTAGTGTAAACATAGCCCATTTTTTCTTTTCTTTCTTTTCT
TTTTTTTTTGCTTAAGCTAAGAATAAATGAAATTAAGAGGACATTTTAGGGTATGATATTTTCATAAGAGAAGT
ACACAAATTTGAACAACTAAAAGAAACTTGAGCCTAAGCTGTGCATGTTCTATGATGATGAGCTATTTAGTTGTA
TCTTGGCTTGAAAATAGTTGAGGAATCATAAAGTCAAGAATCTAATGTCAACTGGTTATGTAGACTAATTAGCAA
AGCATGAGTTAGAGAAAAGATGACAACTTTAAACCATTTAGGGGACGGTACAAGATATACTTGTGCTTCAAATGT
ACCTTAGTCTCTGAACTTGCATTCATAACTAGATGATTCTCCTCTTTGGTCTTGTTGTAGCGTTCTATCACTGAC
TTCATGCTGGCAAGAAAAGGTTTGTTAGTTTATTACTTCAAAATCACAAGAATCAAGATAAGTATGAATGAAGAG
AATCAAATTGTTATCGTTGTGAAAAGACAAATGATTGATTACCTTCTACATCAAAATGACAGATGAAACAAATAC
TACAAAAAGAAAAGCTGTATAATCTTTTTGTTTTTAACACATAGAAGTAACTTTTTTCCCAACATAATGCTTTTT
CATGCAAAAAATGGTCGCATAAAAGGAAAACTGTTTTTTTGGGGGGGGAAAAACACTTTAGGATGGTGTATAATG
TTATATTTTATTAAATCCGATTAAAGCACTAATTTTATAAGCCAAATACAAGGCTTTGGGTCTCACTCCATATTG
GTTCATACCTAGTGTCCCAGGTCGGGCTGGTGGTCAACCAAGTCCTAGGATGGGTTTGTGACTCTGAGCGACAAG
TCACAAAAGTAAGTGTTCCTGGTGCTGAACCGATTGGCTGACAAGTGGGAGAGGAATTGGAACCACTTATTTTAT
CCAAGACTAACCAGCACATAGTTAGAAGACTCCTAATTGCTTTTTCCTTTCCTTTTTTGTCGCACTGGAACATGG
GAAGAATGAGGAAGGGAGAAAAAGAGAAAGCAGAAGAAAACCAGATACCAAATTGGTACTGGGACCCGTATGATA
TCATAAACCCAACCATAGGCTTAAAAACTCACCTAAGCCTAAAGAAACATTTGAAAAATAAAAGATAAAAAAAATA
AAACAGATTGACAACTATCAAACCAAAAAAAAAGACAAAAAAAGAAGAAATTTTAAATTCCAATATAGTATTGCT
ATTCCCTACTTGAGGTTTTCTATGTATGATAAACATACAAAATTCATTTATGATCATACGACTTGTAATCTAAAA
TGTATTTAAAAGATATATGTCTTAAAATTAGCAATTGAATGCTAGTAAGCAAAGAAAATATGTATTCAAAATTTC
AAATACTATTAAGATGTAGTGAAACTTGCACTTCAAAATCACCTTTTGAAAATTTGTTTAGCTTACCTAGACTGC
AAAGACATTACATGCATCTTTTTGGAGCATTTACCATATTTCTTTTGACACAATTTATATTTCTTCCTATGATTT
CAAAATATGTTTTAAACATTTATACATAATTAAAAATAAATAATTAATAAAATGATTTTTTAATGAAAATGCACT
ACTTACCATGGAAAATAGTATTTAGTAATCTAAGTTTATAGCTATATTTTTCAAAACATTACATTTTAAATATCT
TATAATTAATTCTAAAATAACTTAAAAAATTTAAAATTATACTAGAAGCTATTGTTTTATTTAGGGCATCATAGA
TTGGCCCCATGTGCACAACATGTGAAAAAATCAATCTCAAATGAAGGTCATTTTATTGGATTGATTTAAAAATTG
TGAGGGCCTCAAAACTAGTGGAACCTAAGCACATCTAGGTTTCATTGATTGTATACCTAATTTTGATGGTGTACA
CATTAGTGGTGTAAACATGATATCCCTCCAGAAATTATTATTTTATTTTAAATTAACTTCTAGAAATTTCCATTT
TTAAATAAATAGCCTAAAAAAATATTATGGTGGCTTTCTAGGTAGGGCCTTTCTTGTCTCCATTTTCAGGAAATC
TACTTGCGCAAATGCATTGCTTGGGTTATTTTCAAATTTACACAATTTGGGTCTTGGGATCAACCTAAAATTCCA
AAATTCATTAATTTTGATCTTGGATTCATAATTTGGGCGTGGATCAAGTGAATTGAGAAGGTTTAATCTTAGTTT
TATGGGAGCTTTCCTTAATTTTACCGACTACTCTTGAATTGGTGTTTGGATAATTGATGAAATTTAAAGAAAGGA
TAAAACACAGAAAATTATCTAGCTGAAATTAGTTTCTAGCCCAGAAAAATGGTTGTGGTTTTGTCAAGCCACTGT

GGTACATTCCCAGTGTTTTGCCCTGGACTACTTGGTATTTCAACTCTTTCTAGTAATAGATATGATCAAAGTTAA
ATTCTATAAGCACCTATACTAGACTTGTCCAATCAGGCCACTTTTCTTAATTTTCTCTGCTAATCTAATAAGGAC
ATTTAACAATAACGAAAAAATAACTCCCAGTGATTAAGCTTGAAACCATTATTTTTGATTCATGACCGTTTAGTA
AATAAGTCATGAATGAATTATAATAGTTCTTGTTTATTTGACCATTAGTATAATGCATAGTATCTTTCAACCAAA
GAAAAAGTTGTGAGCAGTTTTAAAATTCGTAATAATTTGAAAAATATTGTTACCATGATGCTAAGGCAGTTGCCC
ATGTTCTTATTGTGGGAATGCTTATAACTTACGAATAACTAGATTATTGCTTGAGAACTGAAAATTATACTTCAT
ATGTTCCTTCATATTCAGTATAAACTTCATCTAAATATAATTGTTAATTTTCTTGTGCAGGTTTAAGTTCACGAG
ATGGGAATCAAATAATATGTCGGGTTAAATTTTTAAAAATCTTCTTATATCCTTTTAAAATGGGTGTCTCTAATT
TCCAAGTTGTTCTTCAAGCTACTAACCATGTCCCTTGTTCAACTTAAGAATAATATCTATGAGCCTCCTGAAAGT
GTGTAGGACTGAGTAACTGGTTGGTTCTAACAGGGACAAAGTCTAGTCAATGATATTGCATTGATGTACTTGTTC
GATTTTTGTGACATTTGCTGCACATTAATTCTGTAGTGTGGGAAATCAAGGTGCTTTATTGAGATGAAACAATTG
CTTTCATCTTTGATCAGATAATTTATGCTTGATAAGGTTCAAAGTTATTTGTCTGACTCTTCCTCACTTGATCAG
AAACAGAAACAGGTTTGGCCATGTTCTAAAGCAAATAGAAGACTTGATAGATGAGCTTTCTCAAAGAAGAAGAAG
AAGAAGAAGAAGAAGAAAAGAAGTTTGAAGGTCGCCAATAGAAAGTAAACATAAGTGAAGACTGGTTACCTGGGA
GACAATTAAATAAGGCTTTGGCACATGGGGAAGCAATGTGGAAGCAAAGGCCTACAAATTTGTGTTTGGGGAGCA
TAGAGACGCATCATTTCTTTCATGCAAAGGCATCTCAAAGACACAGGATGAATGAGATTTCAAGGCTCAAAGATG
ATGGATGAGAACAAAGTTGAGGATGTTGCAGTTTCAGGAATCTGAATGAAAGAAGAAAAAAAAAAAGCTTGAATC
CACACAGTTCTATTTCATGAGGTGGTAGAAAGATAAGTCTCAGGTGAAATGAATGGAAACAAACCTCAAAACTTC
ATGACCAACGAGTAACATCTTGTCTTGTTCCAACTCTACCTTTAAAAGCACCTGGGCCTGATGGCTTATTTGCGC
CTTATTTCCAAAAACATTGGAATAATGTTGGTCCTAATGTTATCAAAGCTTCCCTTGATTTTCTAAATTATGGTC
ATTTCTTGGATGAAATAAATTTTACCTATGGTGCTCTCATTCCTAAAAGCAAGAATTCTGATTCAATGACTCAAT
TTCGACCTGCAGGTCTTTGCAATGTGATCTACAATGTGATTTCAAAAACCATTAGCTTGCTAAGATAATTTCAGA
AAGCCAATGTGCTTTTGATTCTGGTCATTTAATCACGAACAACATTCATAAGGCTTTTGAAGGAATCCATTATTT
GAAGAATAAGTTCAGTGCTTCATTTAACTCTGAAACTAGACATTAGCAAAACTTATGATAGTGGTATGGCTTTTT
GGGGAGCAAATAATGCTCACAATGGATTGGTGGAAAATGGGTGGCCTTGGCTGTGAAATGTGTTACTTTTGTGAC
TTATTTTATTCTTGTTAAGGATTTGTTTGGCTCACGAAAAGAGGGGAAGAAATATGGTCAATGGAAAGTAAAAAA
AAAATCACCCTTGTTCGGTTAGAAATTTCAAAGTTGAGAGATTGAGAAAAGGTATTTTTATGGGAATATAATTTT
TATATTTTATAAGAAACAAAAATCATGTAGAACATGAATTTTTTGAATTTTCATAGGATGAAAATCAGTTGCTTT
ATTTTTTTTCAAAAATATTTTTTTAAGCCATCAAAAAATTTATAGTTAAAATAAACATTCTCTCCATCACAACTC
CTCCATTATGATCTATGCCAAAAACTCTTAACCATCATAATTTTTTATTTGCAAACAATAGTAATGTATAGAATA
AGATAAGATTCTTCCTTTTATTAGGGATATAATAGATATTTTACATATGTTTTCAAAAAAAAAAGAAAGCTCAAC
TAAACATAAACTAATATGAAATAAACTACTTTTTTATGATCAATTAAATATGTCAAAATTATTTTCTCGGTATCA
TATTTTCATGAATCAACTTTCCAGAAGTAACACTTCAGTGAAGAAAAAATTCTTCCGATGAATCAAATGAGTGTCA
TAGAAAAGCAAATATGTCTCATCCAGAGGTCTTAGAAAGTGGGAGCTCTGTTTTCTAATTTCTTCCTATCTACAA
GTGTCAATAAGGAGGAAGAAATGTAGAGTATCATATATTGTATATACTGCATATATTAATGTAACCTTGGTGGTA
AGCCCCCCTCATGGTTGTTGTAATTACCGTATATATGTATGATTCTCTCCTAATACAATACCAATAGATAGAATG
AGAGTTCTTCATAACCCATTCTATTAGCTTGATCCTAATATGGTATCAGAGCCAACTCACCGGTCCTTCATGCAT
CTCTAAAATAAGCCTCCCTCGTCTTTGAAATGAGCTTCAAGACAAGGCCTCATACTACTGCTTGGAGATTTTGAG
ATCATCAAAATTTGATAGCTGCGCACTCTATCCGGGACTCTCTGGTTCTTTGGATTTGAGTTTCATCATTCATTT
CCCATTATTCACAATCTTATCCATGTCAGCCATGGATCTCTATGTTATGCTTAGGACTGCGCTTACCAGCTGCTG
GATAATTGGTCCAAACCACCCCAAGCACCAAGTGTTTGTGAATCTGCATAATATCAGCCAGCATCCATTTGAAAA
TCTTCAGCCCTTGTTGTCATGTGAACTAGACTAACTTCTACTGAATTATTCTTGGGTCTCTAATGGAGCTTAATC
TGCTCCTTCCCAAATATCTCCAAGGCACCTTGATCAACCAACAACTAGAATAACTATACACTGCAGATGTTTCAC
TTGGATCTTAGACTTCGGGGCATCCTTTCACATGATCCTACATGCTTCTCTTCTTGTTTCCTGTTCACCTCCTTC
TATGCAACCTCATATTGTTACAATTGATGGCACCTATTTACGGACATCTCAACAAGGTGAGCTTAAGACCACCTA
ATAAACTATTCCTTCCATCCTCTATATTCCTCAATTATATGTTAAGTTACTATCTGTTAGTCAACTCATTGATCA
TGATCGCCTTGTCTCCTTTGATTCATCCTCTTGTTTTGTGCAGAATCAATGAACAATTGTCTAATCAGTGTTGGC
CATCACCATAATATAGTGTATCTGATGGACCACTTGCATGTGCCTATTTCTTTTGCCTCTATCTTTGTAGATGTT
TTTCCTTCTACATTTATCTTGGTTATCTTTGTGGTCCCCATCTCATTATATTAATTTGTAGTGGTGTCTTAGGCT
CTATATCTAATTCAAATATGTCGCCTTATGAAGGTTGTGCCCTTGGAATGTATGTGAGACTTCCATTTCACTTTA
GTAATCCCGTAGCTATTTCACCTTCTAATTTAGTTTGCACAAATGTCTGGAGTCCTTCCCCTTTTGTTTCTAAAG
ATGGCAACTACTATTATGTCATATTCATTAATGATTATTCTCTCTTTATATAGATTTATTTTTTATGTAGTCTCG
CTCTCAACTGTACCAGATTTATAAAAAATCTGCCACAATGCTTTAAACTCAATTCGCTAGATCCCTTACGCCCCC
CATTCTGAGTCTAATGGTGAATATCTATCTCATGAATCTCTCGACTTCCTTGCCTCATGGGGACTCTCTCTCAAC
TCTCCTGTCCTAATGCTCATTTGATACATCATTGAGATTGCTAGCTCAGTTCTTAACTTTGCCTATGTTCCCCCT
CATTTTTGGGTTGAGGCTATTCATATTGTTATTTATCTTATTAATCTTCAACCCTTTTTCATTTTTTGAGGTCTT
ACTCCTCGTAAGAGACTTTTTTCTAGCCCCCCTTCCTATGATAATCTACAAATATTTATTATGCTTTATCATGTT
ACCTCCCTCTGAATATACCAAACTTTCTTATCGATCTTTGTGTTTTCAAAGGATATAGTCTTGAGCATAAAAGGC
ATAGCCTCATAGATGCCATGATTTTATTTCATGACACATGTATTTGTCTCGAGATGTCATTTTTCATGAGCATCT
TTCTTTTTATGCTCTCTCTCCTCATCCACTAGACCCTTCACCTTCTGCTCTCCAACTTGTTTTTGTTGATCTTTC
CTTTATTACTTCTACTTCTGCCCCACTAGTCTTGTCTTTTATGTTTTTCATTAAGCCTTCTAGCTCCTTTACTCC
TCATAACACCTCTGCCTCATCCATAGATATTCCTCCATGTGCGAGTGTCTGATCTAAGCTATCGATTCATGATCC
ACCATCAACAGTTGCTCCACCTAATAAGCATCACCATCCCTTACACTGTCGCACTACTTCTACTAAACTCAATGC

TACTCCTCAGTGCCTTTCTTTAGAGTTCCATACTTTTCTTATTACTATTCACTCTTACTAGGAGCCCAAGACTTT
TAAAGAGGTTGCTGATATTTCAAAGTGGCATGCAGTAATATCTCAGGAGCTTGATGCTCTTAAGCATACGGGCAC
TTGGGATCTTGTTCCTCTTTCCCTTGACAAAATTTTCATCATCTAGAAGTTGGTTTACAAGATCGAAACTCACTT
TGATGGCTACATTGATGTTACAAAGCTCGTCTTGTTGCTTGTGAATTTGAGCAAAAGAATGGCATTGACTTCAAG
GAGACCTTTGCATTGGTGGCCCATATGAGTACTGTGTGGACTCTTATAGCTATGATTGCTATTCATCAATGACTT
TTTTATCAGCTTGACGTGAAGAATGCTTTTCTTCATTGAGATTTATATGAGGAAGTCCATGTAAAATTACCACTA
GAACTTTCTCATCCTTCCATACATGTGCGTCATCTTTGCCAGGCTCTTTATAATCTTTGCTAAGCTCCTCATGCA
TGGTTTAAATGATTTGGCTAAACCATTCTTACTCTTAGGTTCACCCAGAGTGGTCATGATCTTACCATGTTTGTC
TATTCTTCTTCATGCAGTTGGATTCTTTTATTGCTTTATATCGATGATATGATTATTAGCGAGGATGATATGATT
ATTACTGAGGATGATTCGAAGTATATTACTTTTGTTAAGTAGTATCTTAGTACACAGTGTGTGATGATTGATGTT
GGCTTATTTTGATATTTCTTGGAGCTTGAGGTTGCATATAGCTCCAAGAATTACCACTTATTCCAATAGAAATAC
ATTGCTGACCTTCTCTCTAGAGTTAGTTTGATCAATTGCCGCACTGTTAATACTCCATTGGAGCTCCATCATAAC
TTAAAGCATGGAGTTAATGTATCTCTTTATGAGCTGACTCACTATCGATATCTTATAGGTAGTCTTATTTATCTC
ACTATGTTCAAACTAGATATTGCTCATGCTGTCATATTTTAAGCCAATTTATTTCGAATCCCACAAGAATTCACC
TAGCCCATCTCTTACATGTGTTGCATTATCTTCAAGGTACTACTTCATGAGCTTTGCTTTTATCTAGTTCTTCCC
AACTTTAGCTTTATTCTTTCTCTAACGCTATTTGGGCAAGTGATTTTATTAATTATCATTCTATCATTAGCTACT
ATATTTTTCTTGGCACTCCTTTTTCTCATGGAAATAAAAAAAATAGACTACCATCTTCCAACCAACTCATGAGAT
TATTTTGTTTCATTGGCTTCTTACATATATAAGTGTTGTCATTTTTTAGCTGACTCCACTCTATTGTGATCATAT
TAATGCTATTCATATTGCCACCAAACGCATCAAGCATAAGCTCACCAAGCTTATTAGTGTTGATGCCTTCTGTAC
TCATCATCATCATCATCTTGTAGGGATGCCCTCTCACTATCTTCCATCTGAGATTCAACCTACTGATTTTACTTA
ATAAGCCTCACATGAAAGCTCAACATTCTCATCTTCTTTCTAAACTCATGGTCTATGACCCACCATGAGTTTTGG
AGGGGAGTGGGAGGTGTTAGAGTATTATCTATTGTATATAATATATATTATATGCATATAATCTTAGTGGTAAGC
TCATCTCATGGTTATTGTAATCACCATGTATATATAATTCTCTCCTAATACAATATCAAAACATAGAATGAGAGT
TCTTCACAACCCATTCTCTTGGCTTAATCATATCAAGGAGAAATATGAGAAGGTTTTGGAGAAAATGATCGACTT
GGATAAGTCAGCCATCACATTCAACAAGAAGGCACAATGAGATGTAAAGGAAAAGAATAGTAATTTAATAGGAGT
GAAATTTAATAGGAGTGAAATTATTTAGGTCTTCAAACTATGATGAGAAAATTAAAGTAAAAATTATTTATTGAA
GATTATTACTTAAGCAGTTCCAACATATCTAGTGAGTTATTTTCTTATTTCAAAAAATACATGCAAGTAGTTAAA
TTTACTATTAGCAAAGTGGGGCAAAAAAAAAGTGAAAAGATTTTTGGGATAAACTAGACTTATAAATTAGAATAA
TTTTGTAAAGAGAAAGGAGATGTGGCTTTAGAGAATTACAAGCTCTTAATTGAGCACTACTAGCAAAAAAGGAAT
GGAGATCCTACAACAGTCTAATTCTATGTTGTTTTAAGTATATAAAATCGTTACTTGAAGCAGGTGAATACTCAT
AAATTGAAAATGAGAATAGGATAAAAATTTAAAAGGATAAATGACTATCGGTGAAAATAGCCTCCAAAGTGGCCT
CTGTTGTTAACATCTTGAGCCCTAAAGCTATAGTTGGTGTTTTGATGGACAAGGATAGAGAGTGCTACACTAGAC
AAGTATTAATCAGATATTCTTTTCATAGAACATTCTTAAAATCCCAAGTATGAATAGATCAATAAATTATAGATC
AATATGACATGGTAAAAAAATAGAATTTTCACAGTTAAGACTATTGGTTTTCACTTGATACTAAAGTAAGAAGAT
CTGACAAAAGCAAGTAGTTTTAATTTGCATACTTTTGATAAGGAAGAGTAATTTTTGGAAGGTACATAACCTAGA
AAAATCAAAAATTTTTTATGTGATATGCTCTCAAAAATATCCTTCTAAAAAAAAGAAATTTGAGTAAAAGAAAAA
TAAAGTTGAATGATGAATTATGTGAACCATATGATGAACAAAATGATACATGAATGGATTTTTTTAAATGTCATT
AATGTGCACTACAAGTATGGAGTCTTGCACTTCTCTAAATGAGAATCAAGGGAAGCAAAGCTAAAAGCATTTGGG
TCAGATACAAGTTCCGAACAAATTAAAAATGGAAGGATTAACAATGTTTGTTATTATCGCTTGGGCAGTTTGGAA
AAAGTATAAATGACTTTATTTTTGGAGGATATAAGTTCAAGCTATAGAAAATTTTTAATAAAGCTTGTTGGCTAT
CATAAGAATTCCAAAGTAGTCAAGAGATTAGTACTAGTATAAAAGTTCAATTACAAGAAAGGTGTATATAAATCA
ATGCTGATGGTGTATTTTTGGAGATAGAAAAGCTAGTATTGCATTTGTAGTTTGAGATCGTAAAGTAGGCTTAAT
ACGGCTATGACAAACAAAATGGTCCAAACAGCAATAGCAAAATATATTGAAGCATTTATAACCAAAAAATATTTT
TTTTGGGTCGTATTGGGACTTCATGATGCTACATTAGGAGACTTAATCAACATCATAAATTTATAATGCAGTTGT
TAGTCCTAAAATTGATTTTGATGATGTCAAAAGAGTTTGAGTATAAATTATAATTACTAATAATGCTTGTGAAAG
TAATTATAGGATATCTTATAATGCTTAATGCTTTATCTCATTTAAAAAGAAGTCCTCTCAAAATTTGAGTAAATT
CATGTTTCTAAAATCAAAGTTTCAGATTTTGATAGCAAAAAAATTAAGTTCGATATGAAATGAATTTGAAAAAAA
AATCTAGTCTAGAAAGACTTCTCAACAAGTTTAACTACTCAAAAATTAATTTTGAGAAGGTAAATCTAGCTTAGA
AAATCTTTGAATAAAGAAAGCTGCTCAGGAATAGAAAAAATTATTTTTGGCATAGCTGACACGCTTTTGAGTCGA
CCAAGATAGAAACTTAGCCAATCCGATCTCAGTTCAAGGAAAAAGATAGAAAGAAGTAAAATTCAGATATTTTTT
TTTAGTCGATCAAGTTTTTATCTGAGCTGATCAAGACAGATCATAGTTGACTCATATAAATGGATAGCTAATTTA
GCCTTAGACTCAAATTGAAGGCAGAAGAATAAAGAATTCAGATATCCTCCTAAGCCGACCATAAAAATAAGATAG
CCAACTAAAAAAGAAACTAAGCCAACTCAAGGAATGTGTTAGTCGATTCGAGCTCAGATGTAAAAGAAGATAGAA
AATAGTAAAAGCTTATAAATTTTTTGAGCCGACCTAAAGAATAAATTATCTAACCAAGTCATAAAGTTAGCCGAC
TAAAAAGATATCTCAGCTGATCCAATTTTAGATAGATAGAAAGTAGAAAAATGAGTATGTATACCTGAGCCAATC
TAGAGCTTTACTTAGCTAACTCAGCACTTCATCTTAGCTAACTAAGATTTTGGATGAGTCGACTCATAAAATTCT
GATAGCATAGCAACTAGTTTTTCAAAATATTTCAAACGGTCATATTTTATGTCAAACAGCTATTTTTGATTCCAA
CAGCTAGTGTATTAACTCCGATCATTTTTGAGGTCATAAAAATAGAAAATTCAAAATTGAGAACCAAGCTATTAA
ATTTATTGAAGAAAATATTCAAGCTATTAAGTGAAAATAAAAGAGAGAGAGAGAGAGAGAGAAGAAAGACATTCA
AAAAGCATTTCAATCATCACCTTTTCATATTTACTTAACTCAATATAGAGGAAATGTGCATTTAATTGAGACAGC
AGTCACCGAGCAAAAGCATTTATTTCTCTGTCAATCCCCTCTTTGCATTCAAGTAAGAAATCGAAGTAATTCAAA
AGCTCAACCACTTCGACTTCTTCAAGAGTTTAAAGATTTATTCTTTATGCTTTAAATTCTTATATATCTTTCTT
TATCTATCATGACAAGTTTTTTTGGGAAGAAACCTGGGTTCTTGAGGATTGGCCCAATTGCCAAAAATTAGATAT

TATAATATTTTGATTGGTAAGTCAGTAAAAATCAACTGGATTGATTGTGAGCCCTAAAAAATAATCAGTTATAAG
ATTTTAGTTGGTGAGTTAGAAAAAACCAACCAGATTGATTGTGAATCCACAAAAACAATCGAATTATAATCTTAT
GGGATTATAGTTATAAAAATTTTCAAATAGGTTGATCTGAGGAATGGATGTAGGTGTGAAGTTGAGCACCGAACC
ATTATAAATTTTTTGTATTTATATTTATGATTGAATGATTGATTTTGTATTTATTTTTTTAGTGCTTATTTAGTT
ACTGCTAAGCACTGCACAAACACTTTCAAATTTTTTTAGAGGGGCACAAGTACTTCTCATTTTTTAAAAAAAAAT
CCAATTCACCTCCCCCAATATTGGATTGCCATTTCTGAGAAATAGCAATCAATACTATGATTCCTTACTTGTAAA
CTTCCTTCTTAAGGATGAAAAAGCTTAAGAAATTTTTTTCCAAAATGCATATTTTGGATCAGGTATTGAAACAAC
GCATAAATCAATGAAGCATGCAATACATGTTGAAGAATTTTAGACTTGATTGATAGTTTGAAGTAAAGAACTCCC
CTTCTTGATTAAAATAATTATCTTTTTTATATAGAGAATGTAAAACCCATAGAATATATAAACTAAAAAAAAATC
TAGTTATCAGTAACTCTTCTAGACATATCTCAACGATATATGTTGTATATCATTGTTCATGGTCATCTATGAAGA
ATGCTTGTGACACATTCCAAATGACTCTTCACCCACTTAATCTCATTGATTAAACTTTGAGTTGATGGAAAAGAT
ATAAATATATCAATAATGTATTAAATGCTTTGGATGTTTGAATTGATGCTAATTCAATAATCATTTTTTTCTAGAT
CGTGGATGGAGGCTTTCTATCTTATAGTTGGATGATACGGACTTTTACATTTGCAAAAACAGTAACTAAACCACT
AATTGCTTAGTTAACTATTTATACAACCAAAGTTACTCAAAGGGCAATTTTGTTAACACACTTCTCTTATACCAA
GTCAAATATGGCTCAGGTTTGTTCTAATTCTCTCATCTTTTGAAGGTAGTTTTAGGTAATTTTTTCTTTCAACAA
CTAAAGAAAAAGATAGAAAATTATATGCTTTCATTTTTTATCCATCACATAATATTGAAAATGACAAATTGATCT
ACAAACTTGAAGCCATCATTTTGATCTTGATATGATCTGCTAACTATAATCGCATCATAAATTTTAAATTTGAAA
AAAATTATCATTCAATGGTATTATTAATTTTTAAAATCAAAAAATATTATTCAATGGTTGCACTATCACAATTTC
AACTAAGATCTAATTAAATATTTTTCTTGACATGTGGCAAATATGTTATATTATTATTTATTGGAAAAAATATAT
TGGGAGAATTTTATATATACATGATGATTCAAAAATTAAAATTTAAGATTAACATATACTGCAATAAAACCCATA
TATTTTGTAAATCAATGTTTAAAGAAATCAAAAAGTCAAGTAAAATAATCAGTAATATTAATTTCATTTGTTTTA
ACAAATGAAAGAAGCTTTGAACTCAAAAAATTATTTTTTGCATTTGGCACGTTATTAACTCATTCACAATATATA
TGTATATATGTAGGTATGTAGGTATGTATGTGTCTGTATTTAATTTAGAAGAATACTATACTTAGTTTATAAGGA
TAAATAAAGAAATTAAGGAGATTGATTATGACCTGAGAAATAATTTACATAAAACTTATGAAAGGATTAAGGCTT
ACATAAAAATTAATTTATTTCAAGCAAAAGCTCCAAAAGATATTAGAAAATGATTACAAATATTGAAGCTCTCCT
TTCAACATTAACAATGGTTTGTATAGCCTAAATCATACAATATTGTCATACTAAGGCAAACCACTATTCCGAAAA
TTGAAATCACTATGTACACTAGATGTGATGACCCAAAAGTAAAACTAGACCAGTAACATTTATAATGGCAACCAA
CTTGTGGTTGCTGCATTAATTTGGTAGTAACCTAAACATCAACCTTGGAGCTTTTCCATACTATTCTTGAACTCT
ACCATTAACTCAATTATCCACTGCATTTTTTGTAAAATTGAAAATCTTAAAAAAAGAAGATATAGAAATCATTGG
ATGCAAGTATACAATTTAACTATACATGGTAGTAGACATCATTATATCTGTTTATTTTGAAAACCAACAAATATA
TAAGATCATGCCAGCAGAGTTCAAATTGTAGACCTCTTACAAGTGGAGAAGGATGTTGACCAACTGATCTAAGGT
TGGTTGGTTTCAACTTTACCTCTTACGAACAAAAACCTATCCTTTAGTTGTAGTGTGATAACAAATTCTCTTAAT
CATGCTTAATTACCTACAAAAAGCAAAAATACACAAAATCGACTTATGACGTTTCCACCATAGCTGCTTCTCCAT
TAGGTTGAAGTCTACTCCAATTGTTTCGTTTTATCGTTTCTTCATTCTAATCGAAAAGTAGAAGCATGTTCTAGT
TCCATCTTGGAGTAGAAAAGAGTTGCCAATAACTAGAAAATAAAAGATTCATCGCTAAATGATAGGGAGAGATAC
AATATAGTGAGAATAAATTCAGGAACTTGAATATGCATGCAAGAGGGAACACAGAGTAGTAAAATGAATGTTATG
AGTTCAAAATAATGATAAACCTTCCACTCAATTATCCAGACAATAGTGTATATAGACTACAATTAATATAATAAT
TTTAACATAACAAGTCTGGCATGATCTTTAAAACTCAAATCATGTAAATGACTTCAGTGAAACTTTAGTAATTAA
AACTAAAATATAGAAGACTGTGGACCTCTCTTTTTGAAATCAACCCAGCCCTTTTATAATATTAAAAAATTTGAA
ATCTAGAGATGGAAAATTTCGTAATAATAAATAGAAATTGATAAAATCATGTTGAATGCAATTTCCACAACTTAT
TCTCCATCAGCTTGTAAAGATGGCTTTGTCTAACAAATACTCCTCTCAGAATAGTTGTTGGCATGAACGGGGACT
TGAGCTTGAATGGAGACAGGATAGGAAAAACTTGTTTCATTACAATTCTGATGCAAAAGGAAAAAGGTAAAAAGA
GAACAAGCTTGGCGAGTCTAGGAAAATGGCGAGGATTGTTAAGTGTATGATCTACCATTCACATGGTACCCAAGC
CTACATATGATGGCATGCAATGCTAACAGTCTAGGACCCTCTTGATATGAAGTTCTGACTTGTGCTTGATGTGAA
GCCACTTCAAAAGGGGATTTTTTTCCCAGGTGCATGCAATATGTAGGCGGTTTTTTTTTTTCCATCAAGCACATG
CATCTAAATGATGCTACAAAGTGACAATTTAGTTTAACTCTTCCATCATAATGAAGGCAACCTTCCAGACCAAAA
AAAATAATGAAGGTAACCTATACCTTACAAACTCAAATTGCTCCAAATGGAATATAATAATTAACCTTAAAATGA
TACTGTTTCCTCTCCTCTGATCATAACGACACTTTTGTATTTATTTTACGTTGAAACTCTAAACAAGTACTTGTA
GACTGCTTTGTGAAGTTATAATGTATGACCATCTAGTTTTATGTAATGTTTCATCAACATCACGTATAGAAAATT
GTGCAGGCCGTACTTCTCCAAGAAGGTTTTCTTGCACACATCCATATATTCTGAATGTTGTCGGATCTGATGGTG
AGGTGGATGCAAGGGAAGTATGATGCATTAGATCCTATACATTGGACCATTTTATCTTCCAACATGGTTTCCTCT
ACACATCCCACTTTACAATAAACTAGATTAGGAGTGGGCTCAAGCCATTAACTTGCTCACAAGTTTCATATAATG
ATGGCGTAACTTGTGGCGGTTATTGCATCAAATGATCAAATAGTCCAAGAGGCATGTCAGATCTGTCCCACTCCC
ATGTTCAGTACCATAATTGGTTGTCTCTCTTCCTAGGACAACCATATCTATATAAATAGGACACATTCACAAGAA
CAATGATTCAGAAAAACAATGATCATCCTGCGCTATGGCTTAGAATTGTGGTGCATGGACTTTGGAAGAATATAT
AGCCGTATCCATGCTTCTAACCCTCAGCATCCCCTTCACTTTCTGAATCTAAAAGTTCAATGTTATTAAAATAAT
ATTTTATCTGTAAAAATGCTGACTTACCCTAATGCTAGCTCGTTGCTAACAGTAATAATGTGAGAAGTAAAATAT
TTGAGGACTTCGTCCTTGTTACTCTCTCTCTCTCTTTCTCTCTCTCTCTCTCTCTCTCTCTCTCTCTCTC
NNNNNNNNNNNNNNNNNNNNGAGAGAGGTTGTCTAATGAAAATCTATGCGGAGAGAACAGATGGAAGATCGAGAT
GTTTGAGTGCTAAGCATATAAAAGCATATGACAATGTTACATCAAAAGAAAAAAAAAAAGGAGAGAGAGAGAGAGA
GAATATGACAATGGCTCTTGGTAACATTAAACTAGGAATAAGAAACCTATAAAAGAGAAGTGCCATGTTTTCTTA
AAAAAATAAAATAAATAAAATAAAATAAAAGAGAAGTGGTGGGGGAGATACGCCATTCCTCTATTTTTTTTTTTT
TTCTAGTTCATTCTCTAAAACCTACCAGCTAAACCATCTATGCAACAAAGATATCAAAACTAAGAAGCCAACCAC

CTTTCCTAACATTATTCTATCATTTCGGTGTAAAATTACTCTTCCATATGTTAGACTTGCTCCCAGTCCATCCAT
GCAATAACCTCCATCAAGACCTTAAGAAATTAAGACTGATCGAACTGAAACCGTATATATCATGTAGGATGACAT
AGAGAGTAATTTGTAGAAGAGGGAGAAGCAAGAACCTGGTGCTGGCAAAATCGTAAAGCCTGCCGGTGCTGGAGA
AGATCGCGAGGCCGACCTCGGCGTCGCAGAGGATGGCGAGCTCCTTGGCCTTCTTGAATAGCCCGTTCCTCCGCT
TCGAGAACGTCACCTGGCGGCTGGTGGAGTTGTCAATCCTCCGGATCACAATCTTTCCCCTCCCCATATTTCTTC
TTCTCCCTCCTCTTCTCAACCCACCAAAAACCCAGTATCAGATGTATCAGATCTGCTATGGAAAACTAACAACAA
GGAATCAGAAAGAAAGGAGGGAGACCAAGGTGGCAGAGAGAAGGAAAAGAAAGAGAAGGACAGAGAGGAGATCGA
TCTGGTGGAGAGATTGAAACCTTAATTTCCGGGAAGACGAGCTTTATCAATGGCCACAAACGGAGTTGGAGCCTC
TTTCTCTATTTGGCTTTGGCCTAGGTGGTGGGTGATGCCTTTCCTATCTAGTCTTACTATTTTATAGGCAAAAAT
AGAAAGGAACGGAAAAGGGTGGCCAATAGGTGGGCTTGGTTCCCACTCTCCCTTGCCTTTCTTCTTCCCCTTTGA
CGGAGGAATGAGGTGGTGGTGAAAGGATTTCTTTACCAAAAAAATGAATACATATAAAACCATACCATTTTGATA
TCTGAGCCCTCCCTGCACTCGCTTCCTTATTTATCCACTTGTTTATTATGTTTATTGGTTGTTTATATTTTCTTT
GTGGTCCGGGGAACAACCGAGCACGTTTCGCTCGTCGATTACTTCTATTTATTGGGACGTTTGATAACGCGCCGC
CTACCTTATAGATCGCCGACACGTGGTAGGACGAATACGTGCGGGCCACGGTCCCATCTGGCCCGGCGATTCTAC
CATTGGTTTGCGGTGTTTTGTTGTCCATCTCTTTGAAATGGCCGATTTTTTCATCCCAGTGTTGGAATAATTAGA
AACATCATAGCCGCATTTATACAAGAGATTATATTTGGAAATATATATAGTTGATCACTAAAACTAACTTTCTAG
AAAGAATTATTAGTATAATAAAGAGTTATAGATAAAATCCTAAACTCTAACCTTTAAACCACCGTCAAAGTAAGT

>EguAGL17_1

ATGGGGAGGGGAAAGATTGTGATCCGGAGGATTGACAACTCCACCAGCCGCCAGGTGACGTTCTCGAAGCGGAGG
AACGGGCTATTCAAGAAGGCCAAGGAGCTCGCCATCCTCTGCGACGCCGAGGTCGGCCTCGCGATCTTCTCCAGC
ACCGGCAGGCTTTACGATTTTGCCAGCACCAGCATGAAGTCAGTGATAGAACGCTACAACAAGACCAAAGAGGAG
AATCATCTAGTTATGAATGCAAGTTCAGAGACTAAGTTTTGGCAAAGGGAGGCAGCAAGCTTGAGGCAGCAACTG
CATAACTTGCAAGAAAGTCACCGGCAGTTGTTGGGAGAAGAGCTTTCTGGCCTAAGTGTGAAAGATCTACAAAAT
CTAGAGAACCAACTTGAGATGAGCTTACGTGGTATCCGAATGAAGAAGGACCAACTTTTGATCGATGAAATTCAA
GAACTAAACCGAAAGGGGAACCTTATTCACCAAGAAAATATCGAACTATACAAGAAGGTAAACCTTTTTCGTCAA
GAAAACATGGAATTGCAGAAGAAGGTTTATGCAATGAGGGGTGTAAATGGCATTGATAAAGGTCCGATTACTCCA
TATGGTTTTAATGTCACCGAGGATGCAAACATATCCATTCATCTTGAACTCAGCCAGCCACAACTGCAAACAGAT
GCAACGTAA

>egu-miR444

ATTTTGGTGTTCCTTCGTCTTTCCCTCTCGATTCTTCCCCGCTCCTTGGATTCAATCTCTCTCTTGGCCTTCCTT
TAGAACTCTTGCTACGTCGTTGCTCTCGCATTTCCTTATTGTTGGATCGGTTCTTGGGGTTATGCGTGATTTGCT
GTCGGATTGTGGCCGGAGGGTAGGTCAGTTTGAGTTTGAGAAATGGAAGAGAGAGGGAGAGGGAGAGTTATGGAG
AGGATCTGAGGGTGGAATTTTTTGGCCTGGGGAGGGGATCTCAGCCAATGCAAGAGGACGTAACAAGCTCGAGGT
GGCAACTGCATAACTTGCAAGAAATTCCATGGGTGAAGTGAGCATACCGGTGACTTTCTTGCAAGTTATGCAGTT
GCTGCCTCAAGCTTGCTGCCTCCCTTTGCCAAAACTAATGCATTTTTCAGTGATGATGGATGGACAAAATGCAAC
CAGCGAATCAACATGGAGAAGTGTAGGATGACTTACTGAAAGAGAGAGAGGCATGATGCCTTCATGTTCATATTG
CTGTAACTACATTCTTCTTTGGACAAGAAGCAGGCGTAAGAGGGAAAAAAAAGATAACTTGGATTCCGTTGATCA
TCATCATAAGACCTTCATAAGGTGGAAGATCACATGGATGGGGAAGACTATGATTGGGATTTTTGAAGGCCATCA
GAGGTACTTCGTTCATTTTGATCTTTTGAAGAGTTCTTATGTTTTGCTACAGAAATTCAGAGAAAAGAGAGTAGT
GTTTACAATTTTTTTAATGCTTACTTTCAAAATTTAAATTAATTGATAACCTCTTCTAATTTTGAGACACTTGTA
CTTATCACTGCAAGTTGAATGTTCAAAATTGAAGTGCTCCAACGACATTGTAATTATCCAAATCATAGCTTGTCA
CATATGATGATCAAAAGCTTGGATTTTAGTTTCTGAAGGACCTTCTATGAAAGATCACTTTGAAGTGCTATAGGT
ATCGGCTCCATTTATTTGTAAATGAGAAAACACTTCCCATCAAGCTACCTATGGTTATTCTTATTAAAATTGCAT
AAACAAATGTCCTTATTTTGTTGTTCTAAATGAGACTTAAGCTATAGCATATCATAGGGAAATTTCTCTCCCCAC
TATTCATCCTGTTAATGATGAGAGCTTGGAACCCAAAAGTTCCCACAAGTTTTTCATTTGCAAATAAAATGAGTA
AAACTTGTAAAAAAAAAAAAAAAAGAGGACA

Stem-loop structure

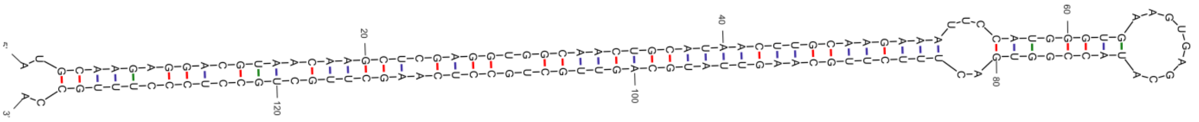

*EoAGL17_2*

>gi_527177216_gb_ASJS01001197_1_p5_sc00001_2392

GAGAGAGAGAGAGTTCCTTCTTGTAGTTCTAGGAGTCAGATTTGACAAGTTGAGGCATTATTACGACTGTATCAA
GCAGCAACAAGAACACCGTATGGAACAAAACAACAGGGTGGGATATGTTGAAAATTGACGTAAAGCAAATTTTAA
ATCCAAAAATGATTAAGCTAGATCAAAATCATTCAAATGCAAATTATGCCCTATATATTAGGGTCTTTTATACCA
TAGTAGTTGGCATATAATGCTAGATGTCACCCATACATCATCTAGAGATTATTAGTTAGTATGAAATGCGCTATA
TATATGTTTATTGTTTGTCGGTGGAACTAATGCCACATTTTACTTTTCTCTCTATTTTTTGTTTTTATTCCTTCA
AAAGTAAAAAACAAAAATAATTATAACTATAACAAAATATATGATACTTATCTTAAAAAAAATAAAATATATGGT
GCGTTATTGCTTAAATTGAAAAAAAGTAGTTGTTTACTGATGTTGTCATCTTTATAAACAATTCTTATGGTTCAC
GTTCTTGTCTAGCATATGCCAGCTAAGAAGCACATAATTTACGATAGATCATCAGATCTCCACGGTGCACATATG
CACCATGATGTACGTCTACGGTGGCAGGTGCTCTGGGTGGTCCATCCATAACAAAGCCAAGAGGAAGGAATGTCA
CTCCTCACAAACCCTAAATTTGAGTAGGTCAGATTGATGAGAAGGAGGGCAAAACGGTCAATTTGGCAGAGGCCA
TCGGCAGCACAATTAAATGATCAATCCCCCAAAGGGGCTTGAAAGCGGCATCTCGCCCGACGGAAGCCCGATCGG
GACCGTCGGACGTACGGAATCCACTCTGTCGCACCACGTGTTGGTGATCCATACAGCAGACGGCCGAAAGGGATT
TGATTTGAGTAAACTAATGCCGAGTGGTCTCTTGGACAAGCCAATCCTCGTACCCCACAGTAATGCAACAAAATA
CGTCCAGCATAAAATATATATCTCAACACCCTACGCCTCACCCTCTCACCTTTAGGCTTGGTCCTGATGACATAA
TTCTGCGTGCACTGAAAGGGTGCAACCTCGGGTGTGATGGAAATCAGGTGCATCTAGTTCTTTGCAGAAAATGGA
AGGCCTCCAGCCATTCCAGACCGGAATCTTGCTGCCCTTAAGTGGCTACCCTCGTCCCCTCTCAGAGACGCGTCC
TAAACAGACTGGTGTCCTTAGGGGGAGGTTAATTTTCCGACCATAGTTTGGTAGGTAATCCACTCTATTCCCTTC
CTTCCTTCCCTGATCGCTTGCATCCCGGCTTCTTCCGCTCGTTTGCTTGTTCTCCATCTTGGTTAGATTTCCCTT
CATGGGTTTGTTTGCATTCTATCATCAGATCGGTAGAGGATCCACCGAACTATCGGGTGCTGCGAGGGGGAAGAG
AGGGATGGGCAGGGGGAAGATCGTGATCCGAAGGATTGATAACTCGACGAGCCGGCAGGTGACCTTCTGCAAGCG
GCGAAGCGGTCTGTTGAAGAAGGCCAAGGAGCTCGCCATCCTCTGCGATGCTGAGATCGCCCTTGTCATCTTCTC
CAGCACCGGCAGGCTCTATGATTTCTCCAGCTCCAGGTCACTCCTCTCTCTCTCTCTCTCTCTCACTCGCTCCAC
CCCTCTTTCTCTCTCTAGCCAGATGCTTTCTCTCCACATCCTATATACGCAAGATTTTCTATTATTGCATGGTAT
GACATTTATGTGCAATCTTTAGAGGCACAAAAGCTTGGTAGTTTTAATGGGATAGGCTTGACGGTTAAGATAGAC
GCATGGAATTTAAAGGGATGAAGAAGAATGAAAGATTAAGGGAAGCTGAACAAGTCCTTCAATAATTACTTAAAG
GAGAAATCCATTTGAAAGGGAAGGAAGTGAGATATTTTTATCCCCCTTCTGGCTTATAAACTACCATCATTCTCT
TCTCTCGCTTGGTTTCTGGGATTTTTCTTCTATCAAGTGCGGTGATAAGGGATAGCATCCCCTCACCAGGCACAG
AAGTAATTCTTTAATAGAAAAGGCTTGTGAAACCATAACGACATTAAAGGTTTGGCGGAGGTCCCCTTTATTGTT
TCATCTCGTTTTTTCTGAAAATGGAAGAATGGAGATTCCCTTTCAAAGGTGTGGCAGAAGTCCACTTGCGAGAGA
CTTGTATTGTAGATTTTAATTATATCACACGTGAGGATAGAAAAGTAATTATGTGATTGGTTGGTTTGTGTTTTT
TTTTTCCCCAAAAGGGGAGGAAAAAAACTGGGCTTCTTGAGTCATAATCCTTGGTAGGGCTTTGGCGGGCCTCGA
GATTGTACGGAGCCTAAAGTTGATAATAGATTCTAAAGTGAAGTGCTGTATGAGGGATTCAGTGAAACTTATCTT
GGATATCAGAATTAAATCATGATGAAGAGCTATAACACTTGGTTCATCAGTCATGTAGTATCTACAAGCTATGAA
CTTGGGATGAAAGCTTCGATGGTGCTGGCATGGCTTGTTTGATAGGTTTAAAGACCCAGGGAATTCCATTCCTTT
GGACCAGCCGATGGGAGTGCTATATATGATGTCGCTATGATGATTTTGATAAGAACCCGAACTGGAGTTTTAGAA
AGATGACAAGAATTAATGCCGGTCAAACTCATGAATTAGAAAGAAAGGGATGCAAGGATACACATGCCTGGAAAA
ATGAAGCATATATAGTTCGCACCGGCTGGGTCAAGATTATTCGATAATTGAATATTTTAAGTCTTTAAGACCATG
ACGACTTGTTGAAATTGATCAGTTTTCGCACATTTCAAAAGCTTCTTCACACAATTATCTAAATGGATGCTTTAG
TAAGTGGTTTCGTCATAATTTCACTGATATGAAAGTCAGTATACAAGAGAAAAGGGGACACACATGCTACCACTA
TTAATTGCTGTGACATCCAGGCCAGTTCACTGAGAGTATAATAATTATCTTGTGCAGCAAGATTTTTTAACTTTA
TTCCCCTTCTTTTTGTTCGTATGTTGTGATTTAAATCCTGCCCACCTTCGTGGTGAATCAACATGTTTTATTGTT
GAATAATTGATGGCTTGAACCAACATTTATGCTGTAGAACTATTAGCACTTCCTTTCTAGTCCAGGTTGTAGGAA
TTTCCGACAGCTTGAGTACATGATAGTTTCTATAATTGGTGGAAGGAATGGCATTGGACGTTTTTTACACTATAT
TTACCTTCTTTAACCATCCACCTGTCACAAACCTTGACATGTATACCTAAGAAATGTAAGAATTTCCGACAACTG
TGGGGGATTCAAACTATGAGAACGAGTTTAGAGTTGCTTCTATTTTTTTTTTTTTTAAAAAAAATCTTCATCTTTA
TTCTTTTATGATAACAATGAAAGTTGACTCTACAAGAGAGCAGAAAGTTATGCAAAAGTACGTGTGGCAACTGTA
TATGCCTCTGTATTTTGATGTTATATTTGCAGAAGTAAAAAGGAAATCAAGCTAAATTTGAGTATTGAATGACAA
CATGTGAATTACATTTCATGCATTGATATGCATATGCTACCCTAAAGTGTAGAGATTTCCTAACAAAAAATATAT
AGAGCTTAACTTATTTTGAAATAGATGTGATTGTTTCCATGTTTCTTCTTATGAAGATAGGACATTCAATTATAT
TATGTCTTCATCATTATCATTAGTGAAAGCAAACAATCATAAAAGATGCATTACTTAGAATCAAATGTTCATGTG
TCTAACACATATGACTTCTCTTTGTTATTTGAGTATTTGACTCTTTAGCTTTTAGCGCCTTGGTTGGGAGAGAAG
CATAAAAGAGAAGAAAAAAGAATAAGAGAAAAAGTAAATACTTTCTCTTGGTTAGAAATTTTTTTATAAAAGAAA
AGAAGAAAAAAGAGAAAAATAGAGGGAATACAAATGTCTTCAAATTTATAGTTTTTTTTTCCTCCAAATCAAGTA
GAGTTGGAGAGAAAGAAATTGAGAGTTAGTGAATCTTTTATAATTTTTATTCTTACTCTTTTAATAATGAAGAAC
AAAATCAATATTACAATTCAATTTTAGGGATTATTTTAAGTCTCTCATCTAAGCCTCCCACCCGATTGAGACTAA
TGTGGACGGAAAAAATCGATGAAAACAAAAGGAAAGATAGAAAAATTAAAAGATAAAATCTTTTCTTGATTCCGT
TCTTGGGTTTATTCGGTTCCTTTTAAGGATATAATAGTAAAAAAAGATTGTAATATTTTTTTCTTCTCCTTTCTT
CTTTGCTTCCAACCATGAAGGGAAGAAATTATTTTTTTTTCATTCTTTAAATTTTCCACCAAGAGAAAAGAAATA
CACCTTCATTCTTTTCTTTTCTCTCCTTACTAATATTTTTTTTCTTGTTTTCTCTTCTCCAAACTCTTAACCAAG
GTGTAAGTTTGTTTGCAAGATTTTTTCAAGACAACCATATTTATTTTTATTTAGATTACATTTTTTTCATGTGAT

TATCCTTATAACTCACACTTGTCCACTCTAGTGTAGTTTTGGAAAAGGATTTCTTTTGAATTATCAATATGCTTA
GGGCCTATTTGGATATTTCTGCATGATGAGGCTGAAAATTCTATAGGAATTAGGCTCGGCATGTCCAATACATAT
CTTCTAAACCATGTGTACATATTATGTCTAGTATCATAGAATATGAGGTCTGGTCCTATCTATGTCAATTAGACC
TCTTCCAGTCAATCCTTAGGATGGAAAAAAGATGGGGCAATATTATTTTTTTCTCAATATGGTATTCTGACCGGT
CCACCTTGAAAAAATCCTTTGAATACTATGAATATAGATCTATCTGAGCCTATGTTCTTATAAATCCTATTGGAT
TAAACAAGATTATCCAAACAAGCCCCTAGGATGGATTTCATACTATAGAACATCTTCAGATATTGAGTTGTACGA
TATCTCTACTTGTGTGTATTAGTTATTTTAGTTTAAATAGGTTGACTTCTAAAAAATTCATAGCACAGAAACATA
TTAAAAAGTTATAAGGAGGAAAAGAAGCACAATTTTAGATAGCAAATAAGTTAATAGTCATTTGAGTAAGAATCA
GCAATCGATGGTCAAAGGAGAAAAAGTGAAGGGTGAATTCATACATGCTTCATTTGAATAATGAAGTACAGGATC
AATGTAAACAACCTAAGAATGAATATTTCCCATGGATGACAATTATAACTATATGAGGGCAATAATTGGTAGTTG
ATAAAATTTTGGCATGCGTATCCCTTGATGTTTTACAAAAAAGGTCAAGTTGTAAAGAAAATAATATCGAACACT
TATTTAAGTATGGTTTTGATTATTTATTTAAATTTGCATTACTACTTTAATGCACTAAAAAATTAAAATGTATGG
CAAAATTCTTATCTTCCAAGATAATTGTGGATGCTTGATAGAAGCTTCATAATCAGATGCACGCTATATACGCAT
TACATTAATGATTTTGAATATATTATTTGTACAATTAGAATAGATAGCCACTATATTATTCACTTTATGTTAGTT
GCTTTAGTGTCAAACTCTATCGTCTCGAAATTTCATGACTTGAAGTTTACGATAGATTATTGAAATTTTGTAAAT
TTAAAAGCTAAATTTTTTTAAAGGGAAATAAATAACTAATGTTTAACTCTCATCTAGATCATGCAATGGTTCTAC
GTACTATAGGATTTGATAAAGCACTTCTTGCAAATTCATATAAGTAGTTGTCATATTCTTACATAAAGTTTCACA
TTATGAGCTATAGGATTATCTTTGTTCCATAAAATCCAATATTCAAGCAATATCACATCAATAATTTGAATATTA
CGATATTTCTTAAATATTAAAACTAGAATTATGCAATTGGAAATGAGTAGTGCTTGCATCTTACACCATCTATAA
AATTTCAAATTATATTTTCTATAAATTATAAAAATATTTTAAATAAAAATATTGTTATATTTTAAATAATAATTA
ATAGTAGAATTTGATTTAAATAAATATTTCTATTTATCAAATCATATTTTCAGATGCAATACAAAATTAATTTGG
TATATTTTTATCGGATCATTTTTCCAAATTAGTTTTAAACTAATCATCCCAAAATATGATTTGATTTGAAAGAAT
GTCATTGTTAGATTGGATTTCAAAAGTAATTCAAAATATTTTTATATCTTTTTAACGACTAGAATTAATAAGAGG
GAATAAGTTAAATAATATAGAACAGGTAAGAGGACATTTTTGATATGAAAAATGATAGATAGAACCAAAGTTCTT
GATTTTGAAATCGTCATTTTATAAAATATAGATATGATCATCATTTGTTATATCTTTCATTATTGTGTTTGATTT
ATTTTCCTTTTTTTAACAAAAAAAGGTTTTGTAGTTGATTCAACTAAATCCGTCACTTATCTAAGGTTGTGTTTT
GGATACTGAATATCTAACCATCCCTGAATAGATTTAACCATGTGCATCTATTCTCTAACCAGCCATCTAGCAGAT
AAGATTATTCAGTGTTTGGTTGGTGTTTCCACGTTTAGTGGATAAAGCTATAGGAAAAGTGAAAAAATAGATAGA
AAGCCATGGATAGTTAGCTTTATTTTTACTTCCAGTCGCTGACATAAAATTATCTAGGTTGAGCACCTTGATTCT
TTGGTTGGACGAAGACATTCACAACCTTTTTGGTGGTGTTTGGCTGGTTAAGCAAATAACTCTCCCTTACCCACA
CACTGAAGCAGCATCTAAACAGCCTAACTGGTTTAGCTCCCCATCCCAAGCGGGGGATTGGGAATTCAAGTCTTC
AGTGACGTTATACCTCAAGATTTCAATCCCAGTTAATTCCTTTTAGACATCCTCTCGACTGAAAAATTTGGGTTC
ATGTCCTAGACCTAGGTAACTTTAACCGATAGTACCTGGTCCATCTTAATCGTGGTGGTGGTGGGGGGGTTTAGA
TTAGTCCAAAGTTAGATTCGTCAAAGATTTTGGATCAAAGTCCACATGTTGCAAAAGAAAATCTCATCATCTTTG
ACTTTTCTGCCGGTTTGTTGAAAAGGAAATGTCTCACACTATCTTGCCTTCGTCCTTTTTGTCACCTTCCACACC
TCCACATCGCCAGCTGAAATCCAGCCAACATATATTAGCATCGGTTCTTTTTTTCCTCGTTCCCCCAGCTCTGTC
ATAACCTTGATTTCTATCCAGTTTATGAGCTTGGACGCTTCTCATAACCAGTCCATGTACTTAAACAAGTCTTCC
AATTTCATACCCGCCTGCACGCTATCCGACCAACATGATTTTTGGACGGGTAGAATCGGAGAAAATTTAGGCCTG
ACAATGGCGTCTGGGCTGCATGCATGGTTAATTACTTGTCCCAACCTGATCCAATAGCACCCACACAAGAAGAAA
TGCCCAAGCAACTCAAGTAAATCGCTCAAAAGGTCACCAAGTTGGTCTGGTGTCATCTCAATTCTCCCTTTGAAT
TGGCATCATGAAGAAACTAAGTGATTGGTGTAAGACGTCTGATGCCATCTCATTTCTCCCTTTGAATTGGTATCA
TGAAGAAACTAAGTGATTGGTGTAAGACACCCTAGTCATGACTTTTTAAAGTGATTTAGCATTGCTCTTATACAA
GTCTTTCAATGTGAACATGATGGACATGGTGGGATGGAGCCAAACTCTTACCACTGAGTTCAACGCTTGCTTGCT
TTCCTCGCCTCAAGATTTGAGTTGGCACGATGCCAATTGTGGGAGTGCATGGCTCGATCCACCATACATTAAGGC
AAACATGGTGATTCCATGTTGGCGTTTCCGTTTATTAAAATCTTTCGGTTGCAGAACAAGGTACAGCCCGTCCCA
GAATCTATGGGTCTCCCAGCCATGAGAAGGATGGGATTCCTATGCATTCTCACCCAGGAGACATTCAACCTCTTG
GCACCTGCAAACATGCATGTCATGGAAGTTTGGTGCTCGTTTCAATAACATTTTGGTTTACAATTAATTTAAGCC
TGATGATATCGAATCCATCGGTACGTTCGCTGCTATTTCGTGCCTTCCTGATAAGTTGCCTCTCTTTTACCATTA
TCATCTCTTGGATGCTTTTGAAAGAAAACAACAGTTGAATCTTCAGTCAACAAACCAAGCCTCCATAACATACTG
TCAAGCTTGTGACTGAACTCTTCCTCCATTAGACAAGCATGGCGAGACAATAGCAACAGACATATCCTTCCTCAG
AAATCTATTTTAGTTTTTAAGTCTCTTTCTAACTATTTCGCTTCATATACTTGTCATAGTTTTATCTATTTAATA
AAAACAGCTGGGTTTTGGCCCTTCCTTTTCTCAAAAACAAAAATGATAGGCTTTAAATTCAGGTCTATCACAATC
ATCCAGAAACTTTTACTGGCAAGCTAACCGACACCCTCAAACAATGTTTCTCTAAGAAAAGCTGATCTTGAGCCC
CATAAAAATGCTTAAAACCTTGGGAGAGGGTATGGAAACATATATTAGAATCTAGTTGATGAGTTACCTGCAGGT
ATTCAAAGTACTTGGCTGTATTATGTGTGTTGGAACCATGACGCTACATTTAAATTAAAATATTTCAATACAACT
AGTTTATGGGTGGCAATGTGTTGGGTTTGGGTTTGGATCCGATCACCTATGAGTTGCTGGTCAAAGGATCGTTGG
TGCATACTTGATCCATTTTATAATAGGTTAGTCGGTGTCTAGTTCATTTATTTTACATGGACACATAATAGTGAA
CTAAGAATCTAATTGAATCTTCATTTATTAATTATATTGCTTAATAAATAAAAGCAAATGAAAAATTCTTTGTAC
ACCACCACCAGTGAAGAAAAAATGTACCATTTCATTGGATTAGTCCACGTAGCGCAATTAATAAGGAGGCTGATA
TTTAATCTAGCTAAATTTTTTTTCCTCAAATTTTTTTAACCAAAATGCCCTTTTCTTTTTTTTTACTCTACTTTG
TTTCTATAACATCTATTACATCCTATAACATCTTGTTGTATCCCATAAATAATTATCTGGATGCCATAAAAACTA
ACATGACATTATAATAATATTATGACATCATGTTACATCCTATGACATCCTGTCGCATTCTATAAATAATTATCG
GGGGTCATAGAAACTAACAGGACGTCATAATAATATTATGATATCATATTGCATCATGTAAATAATTATCAGAAT

ATCATAAAAACCAACAAAATATCATAATAATATTATAACATTCTGTTGCATGCTTTAAATATTTTATGAAATATT
ATAAAAATTAATAGAATATGATAATATAATATTATAATATTCTGTTACATCCTATGACATTCTATTGTATCTTAT
AAATAATTATTGGGATATCATAAAAATTAATAGGATATCATAAATAATTATCGAAATATTATGATATCCTGTTGC
ATTATATGACATCATGTTGTATCCTATAAAAAAAGCAACAGGGCGTCATAACATTATTTGAGATGTCAAAATATT
CAATAGAAAAAGGCAATTTGATCAAAAAAATTAATAAAAAAATTGAGCTACATTAAATGCCTGTTTCCTTATTGG
TTACACTACGAGTAATCTGGTGAAGCAGTATATTTTTTTCTCTACCGAATGTGGTGCACAAAAAAATTACTTCCTA
ATATGGCTCGAGTTAAAAATTGTCACCTCTTTATTTATTGTACTGCTAGCAACCCAGGCATGGTTGCATACAAAA
ATTAAAACAAATAGATCATTAAAAGCAATAATTATATTAAATAGAATATAAATATAAAATAAAATATTGATAACT
TAATTGTGTAAAATCAAAAATCAGGGGAAGTTGAGAGATACATGGACTTGATGGATCGGTGTGAGAAAGTATTTT
TAAAATATTAAAATTTGTAAATTACTGGAGGATGAGTTAAGATTGGAGATAGAGAACTGCTAAGAGTTTAAATCT
GTAATTAATGGAGGATCAATCATCTTGAAAAGAAAATAATTAGCACAGCTAACAAACTGAAAGATAAAACTAAAA
TATATTACTTTAAGGATTTAATTGAAATGTCATTAACATTATATTCTTTAAACAATATTACTCTCAGAGTCAAAT
ATATTTTGGTTAGTAAATTTCTTAGATTGAAACTATATAATATTATTATTTTTCTTTTTCCAATGGAATAAAGTG
CTACCCTGATTAGGAACGCATCCAATTAAGAACTCCTGCATCACCTTGGATGGGTAGCTGTGGATCCAGCTTATG
GCTACGTACAAGTTTCAAGCCATACTAGCAAGCAAATCTACAACGTGGTTGCCCGTTCTATATTAATTATAGGAT
AGAATAATCATTCAAGACAGCAGGTGGACATTTATTCTTCCTGAGATACTCAAAATCTTACCCTCCCTAGACATT
GATCCAGCTTATGTCTGTGATTTAATTGCAATATAGCGTGGATTAGGGCAAGACTTTCCTTTATTGTTTTTTTAA
GAAACATGAAGGCAAGACTTTGTGTTCAGCTGTTCTGCAACGGATGTGGGATAGCAGATAATTTGTGGAGATGAA
CATCCCATCATCAACATTGAAATGCTGTGAGAAGCACAGACGAAGGGCAAAAAACCAAAACAAAAATCCTCATTT
ACTTGGATCATGCTCTACACAAATGATGCTACTTCCGTTGATAGGAGTGCATTTTTACATGAAAGAGCATGGTCC
ACGGATGATTCAGTTTGCCCAAAATCCATGGGTTTCTACCAAGAAACTAAGTGTAAAGCTGTTGAAAAGCAGCGA
AAGTAATTGCTAACATTTTGGGAAACCATTGCTAACTTCTTGCCATACAGCTAACTTCTTGCTATATATTTACTG
CATGGTATTATTAAATAAGGCTAATGCAAGCTAAGCTAGAGAAAGCCCTTGATGCCCGTGGAATTTTGAATTTCT
AGTACCCTTGCTATGAGATGCAATGAATCCACCAAGCATGTTTAAAATTTTGTTTAGCCTAATCTAGCTTAAAAT
ATCTATGAGCAAAGAAAATACCCTATTTTCCAAACCCCATCATAAATAACTGAACCAAAAATATTGTAGTTAATT
GAAGATGCTAAACAATTTGGAACAAAGTCATGAGAGAATCTGAAGAAACTGGCACAGAGAACGAAAGTAATAGAA
GGGAAATCATAAAGCAATGCCAATCAGTTGGGAGAAGCATTGATGATTCTAGTTTGTATTCTAGGTATACAGAAT
CAATTAGGAGCTAGATGGTCGAATAGCTATGGATGGACAGATGCTACATGACAAACCAGAATGAGACCAATTGTG
AGGCTAGTTAAATATTGAGGTCACCACTTTGGATGACAGAAGAAAGTCAACCTGTGTATAACTAATTCGAGAGGC
AGCACTCATATTTATTCATGTTCATTATCTCATTTGATTATTAATTCAAGCACCTATAAACCATAAAATGTATCA
ACATCTACTCTGATAAAAACTAACATAAAGTCCCTCATCATATCAGCATGAAATCAATAATAGACCGCTACAACA
GGACAAAGGAGGAGCATCATCAGGTTTTAAATGCAACTTCAGTCAAGGTATGCTCCTTTCTTATGCAACATCACC
GACTGACAAATGAACTTCTTTGATGGTATACAGATGTACAAATTACTATGTTTATATAAGTCTTGCCATTTCTTC
TTGGGTAAAAAAGTCATCCTTTTTGCAAATGTATGGCATTCACCAATTACTGTAGGAAAAGTACACCTTTCTTTT
TGCTTAACATATGCAAATTGTCAAGATGCTGCATCATGTACTCTTATAGTCATGTACTTTAACATACAAATAATT
TGTGACACTTGTTTTGCAGCATATATGCCCTGTACCAGAAAAGTAGGGAAAAATTAGAACTGCACAAGAATAGAT
CAGGTATATTACAGTTCAATTTGAAGTAGCATAACCACAAAAGCTTTAAAATGTATAGTAATAAATAGCTGGTAA
AAGGTGGATTACCAATAAATGATGGAATACAAATAGAATAAAATGGGTACACTCATTACAAACTTATGTCTGACA
TCTGTAAAAGCATAGCATCATAGAAGTGCTTAAGAAAGGTTGGCTAATTAAATTGGCTGATGATGAGGTTCTAAG
GTGAATGCGGGCCAAACTAACAGTGAGGCTAATGAATAAAGTCAACGGAAAGGATATACATATGGTTTATGCTGA
CTTATAAATGTGCACGCTGAAGGATGAATATTCATATGAACATGAATAGCAAAGGGAGATAATTTCTGATCAAGG
TATTTATGATGGAAATTGACAGTGGTGCTATGGGGTAGATGCTATACTATGGCAGGCCAACATCAGGATCATCTT
AAAATCTGTCAATCATAAACATGTGCAAGAAGGACTATAGCAGATACTATACCGTAGCAGGGCCAATCATGGTCA
TCCTAGTCTGTCATCATAAACAGGTGCTAAAAAGTACTGATTCTTGCTGCTTAGGTGGAGAATCAACTATATAGA
TCATTTGCAAAATAAAAACATCACAATTTTTGTGCCATGGAAACAAGGAGAAATGCTACATTGGCGTTATGGAAG
AAAATTGAGTCATAGTGACAAGCTTAGTTACTGAAAACCGGAGTGGAGCAGATTCAGATGCAGGTGTGGGGAAGC
AATTCTCTTAAAATTTTTTAATAAGAAAGCAGGAGCAGAAACAAGTTTCCACCAATATTGCAAAGCGGGTGTAGC
ACCTCAAGTAGAAGATTCCTGCTACACAGGTCATGCATAAACAACAGAGCTAACCAAAACAGATGACAGAATGCA
GCTGTTGTTGGCTGAATAATGATGATACTTTGGTAGTTGATGGCCAAAAGACACAAAAGTTCTACTATGTTGAAC
AAATTTAAAGAAATATATAGATATTCTCATCAAATTTTGAGCACCAAGAATGACACAAAGATGAGAATGCCTAAA
AACATGGATCAAAGTCAATAGAAAGGATTTAAGTAATATATTGATCGGAGGAAGTAGCCACATATTTATAAGAAG
GTTTTGCTACTATAATTTTTTATTTATTTTAGAAAAGTCAGAATAGATTATGGTCATTGATTTGCAAGGGGTTTT
AAAGTGATATAATAGGCAACTGCTTGGCATATATTACCTTTTTCTTTTTAAAAAAAAAAAAAATCCTGAGGGCCTA
TTTAGATAATTGGCCCAAAATCCCATAGAAATTATAGGTTTGGGCCAACACAACCACCTAAATGAGGCCTAGCCC
AACCTATCTTTATACATACCTAGAAATCCATTGCAGTGGGCCGGTCTGAATCCCATAGGAAGGAAAAAGAGATTC
AATAGGTCTTTTTTTTCCTCCCTATGGGATTCGGCCCGGCCCACCACAGTGGTTCTTTGTGTATTTATAAAAATA
GGCCTAACCCAGCTTTATTTAGGTGGTTGTAAAGGCCCAAAGCCATTTTTTCTATAAGATTTTGGGCTCAATCAT
CCAAATAGGTCCTAAAATTCATTTTTGTCCTAAAACCCCTCTAATCCTCTCAGCCTCATTAGCTGCTAGCTCCAA
TAAAACTTCTCTCCAATCCCATCCCTCCACCCCAACCACCTCCCCATCTCCCTCTCCTTGTCAAAAACCCCAAAC
ACCATATCATCAACCTTGATCGCCTCTTGGACTTCCCTTGTCGTTGCTGCCTCGCTGCCATCACCAATCCGAAAT
GCATTGATCTCTATAGGCCAACAACACCCAGCAAATCATGGCCTCCTTGTTGACAAGGAGCACCAATAGGTCCAC
CTCTAGTAGGGCCTTGATAGGGGAACTTGTTAGAAAGCAAGACTCTGATAGGGTGAGATTGAGGAATTTGGGGTA
GCTCTTGAGGTACTTATGAGGATGCAGTTGTCCATGACTGTCAACCATGACCCCTTCTCCATTGCGATCGGCATT

GGTGGTGGCATTCAAAACCAGTGGATCCTTGCTTGTGTCCGTATTGTCACAACCCCAAACTATGGCATAAGAAAG
ATGATAGCAATCACTCCATTTATAAAAAAGATAATGGTGCAAATGAGCTAAACAGCTCACGAGTTGCTCAACCTC
GACTCGAGTCTAAGTTCCACTCAACTATTATCGAGTTCAAATCAAACTCAAGTAGCTCGAACAGTTTTTCAAATC
GAGCTCGAGTAACAAAATACTCAATCTTAAAGGTTCACGAGCCTAATCGAGTAACATATATTTTTTCATAATATT
ATAATTTATTTATAATATATATTATTAATTTAATATTTTTAAAAAAATACTATTTACATCTTTTTAATCTGATCC
AACTTCTAACTTTTAACTATATTTTTAAGTTATGACCAACGCTCATATTTGACCTTGAATTCAAACTCAAGTATT
GCTCGGTTTATATTCGAGACGAGTCAAGTCAATCTTGAGCATAGCCTTTTCGTTCGAGTCAAGTTCGAGCTTGAA
TATTAAGACCCAACCAATCTTAAGTTGAGTTTTGAGCTTGAGTGTTTTAAGTCGTGTTAAGCTCGAGCCTCTATC
TGTTTGATTGGACTTGGCTCGATTGCACCATTAGAAAACATAACCAATGAAGTTTGACAATTTATCTCGCCATTC
ATAATTTGGGTTGGTATAACCAATGATGTCAGATTAACCATTGAGCTATCTTGGCCCTTTGCTTTTGTAATTAAT
AGCTAGTATTGCTGATATAAAACTAAGAGTATGGAAGTTGAGAAGCACCATAACAAATCCTTATAAGAATCAGTA
GTGAGCATAATTCATAAATCTGTCACAAATATTTTGTAAACTTCGCAACTATGGTTATTGTTGCAAATATTTTTC
ATATTAAAGTAGAAAAATAAACTCCACAGAGACAATATCCTTTGAGCCCTATTTTTCATTCTTTTTTCCAAAAAA
AAAAGCAATCACAAAGATGGCCTCCAAAATCATAAAAACAAAGGTCCACCATTTCAGTATGAGACCCCATACTAA
TACCATCCTATTATGGCGTCGGTACGTGGTTCAGTAAGGTAGGAGACTGGTACAGTATGGTACGCTCTGTATCAG
GTGGTACAGGATAGTATGGTAGACCATGATTGAAACAATTTACAAGTAGATTCAGAGAAAGCAAACTATTCATGA
ACAATTCTATTAGGTTAGCTAACAAGATCCATGGAACTTTATCAACTACATGAACCAAAAGCATGTTCGGTTGGA
GGGAGTTAAATTCTGGAATGGGAATGAGTCACTCTCATTCTAGCCAATTGGTTAGAGAGAGTTCCATTTTCATTC
TGATTCCAGGGAGAATGGAAATGCCCTAATCCCCTAATATCCAACCCTAATCTTCCTTAGTATTCAAAGCCCATT
CCAATTCAGTCACCAACCAAACATATAGGAGGATATGACCATTTCGATTCCTATTTGATTCCGATTCCGGTTGCA
AATACTCTAAGCTGAAACATCATCTAAAGGCTTAACTCATAAAAGAGCCAAGCCTCAACATTGGACTTGGAAGTA
CAACCTTGGCTCAACAACAAGATTTGAATACTGAATTGAGAAATAGGGCAGACTTGAATATTGATTGTTCAAGCT
CCAAAAACCTTTCATAAATAAGCTGCATTTGAGAAGTTCAGTAATTGGCTTGTTTGCACCCCTATTTATAAGGAT
ATGCGCCAAAGCATGACCAACATGTTCCATAGAATTGGGAGTAATCATCATGTATCGTGCTCATCATGAACAGAC
TAGTTGCAAAAACTGAGTTAATCATTAAACCAGATAACTAGTACAAATCCTAAAAGTGGGCCCCAACATAGAATG
AAGAAAAACTTCAACTGCTTGTTTGCATGATTCATATCGTGCTTGGTTTATATTGATATGAAAATTGGGGTACTT
GATAACCAGTCCATTATTTCTAACCCTTGATTTTTCAATGGTCATTTAATGTTTAGTCCGTGGCCCTGTACACTT
AAACATAACAGTGTGCAAAACAGTTATAACTGTGTACAAAACATTGGTGCACATGTGTGCAAACAAGGTGTCAAA
CCACATGACCTAATATTATGTGCAAGACAAAACATAAAATACTATTAAGGGCTAAACAAAAAATGGTAGTCAACC
AAATGACCTAATATTAAGTTCCTAATGAAAATTATCTTTATCTTCAATTATGCCATTTAAGCCTCTTTTTGGACA
TTGAAACCCTCTAGATCTAGCTTCATTTCTAGCATGGATTAGTATCATGTAATGACTGTTCACCTGCATCAAACA
CGATAGCTTCAAAGTGCTCATGTCAATCACTGAGTATCATTAGCTTTCATGCATTTGATTGCTTTCTTGTCCTTC
TCCCATGATCTTTTGGATTTGAGCCTTCTTTATCCCAAGTTTTATAAATATATGGATTTTCTTTAGATTCTGGAG
CAGTTTTATTCCCTTCAATTGGCAAAATTTGATGCAAGTCTTCACTCAAGTGAGGAGTCAGCTTCTTGTCATCTC
TTTTGCATATTCTTGTTTATGTTTGGTTGGATATCCCATTAAGAATCAAGAGTGACCTTTAATCATCTAGCAATA
CAAAAATAGGTGTTATCTTAATCATTATATGTAACCTATCTTTCCCGGTTCCACAGCTAACCTGGAAGAATTTGA
CAGTCACGATGTCACACCACATTCTTCATCACACTTCTTCAAGGTGACCTTTATAAGTTATAAATTATTAAAAAA
ATGTTTGCTCACTTGAATTACAGTGCCATCCAACTACTGCATTTACTATAAAGCCTAGGAGAATCTTCAATCTCC
AAGTCCTGCTTTTTGACCAATAACTGTCAAAACATCTCAAACTGTGAGGTTTGGGACATACAGAAATGAACCAGA
ATGGTACCAGGATGTTCCTCCTATTGATTTAAAGCAAGTGTAGAATTGGTTTGAACCAACAGAGCTACCTGGTTC
TGCTTGGTTCAGCTCAATCATGGCTGGCACATACTAGTTTTGGGTCAAAGAATGAGTAAAGAGAGGAGAAAGAGC
ATTTCTAACCCTTTCTCCTCTTCTTTGAATTCAGTAAAAAGATGGAAGAGGTGGGAAGTCTGGGAAAGCTTCAAG
AGAGGGTATTTGCAGCCATTGGCTAGTTTGCTACCTTGATCTTCCTTGAGGTAGGATGGCCAGCCTAATATCAGA
CGAAGGATCTGTATTCGAGTGGCTATCCGTGTTAAAGGGATTGGCAGATGCAAGTGTAGGCACAACATCTGGGTT
GGAATCAAAGTCCCAACCAAATTATGAAGATCTCAATGTTTGCGAGCTGCATACACTCCATGATGGACCAGTAAT
ATACATTTTCAAGCCAAAGTTTATTGATTATGTTTTTTTTAATTTATAAATTTTGCATAATGCACATGATGTATG
GTGTTCCAGGACCACTACCTTTTTTTTTCCCACTACAAGATAGAATTTTTTGAAAAATATAGCAAATAAAAAATA
AAACCCTAAATCTAAGTTTTCAAACCTTTAAACAACACTAAAAACCAAATAAATAAATCATATAAAAAAATAAAA
AGGGTTAAAACCCCAAACTAGCACCAAATAAGTGTGCTTTTGCATGTCATGTCAATACCATACCGTACCAACCTA
TTCCCAACTAGTAAAGCACCCAATAACCGATATAGGAAATCTTGGAGGATATTATCATGATACTAGCCAAAAGTC
TTTTTTTCTCCCTTTTGGTCAAAGGAAAGTGAAAAAGGAGTATAGGGTGCATGCAAGTCATAGACAATTTAGGTG
GCTAGCCTCGATGATAAGCACTAGATCCATCAAGGCCTCTAACTAATCCGGAAGCAATATCTAGGAATAGGGACG
TCCTAGCCAAAATCAATATCAAAAGCACATTGACTCAAATACCAAATTTGTTGCTAGTAATGATGGAAATCCACC
ATGATGGATCATATCTGTAACATTATTATGTGGACTATGTATGATGGACATGCAATTGCACCTTCAATACATGAG
TTTCTTATAGTGTCAACAGTAGGTAAACAACAATTGCAACATATTATTAATGCATGAAAATATATATATGTCCAG
CCACAACCATCACCATGAATTGTGATCTTATTTTTTAATGTGCACGTGGTTCTAAGCTTCTAGCATTAGCTTGTA
TGATAAACTACTTGAAAAAAATGTTAAGTTAGTGTTGCATGGACATGAATATGAGAATTGGATTTGAATGCATAT
CCAAGTGTCTAACACAGCAAGAGAGAAGGAGGTACAAATATACTTAGTATAAATTATAAATTTAACTAAAATATA
TATCTTCAATATTCTTTGTTTAAATATTAAAAAATAAATGTTAAGAAGATTGTGCTTGTTAAGATCTTTAGAACA
TGTTTGTCATTCAAACCGTCCAATTAACCTAAAAATATAAGAATAATAACATTTTCCAAAATTATTTCAACACAT
ACGTCCTTGGGCAATCAATGAAAGGAAAGAATACTTGCTTCTAACATGAATATGGATGGTAACTCATTGTTGGAA
GAGCTCAACTCTTAAAAAGTTGCAACCGTAAAACTTCATTAGTATCCAAGTGCCTAACACATATCTGAATTAGAT
ATGTTTCCAACATGATTTTTTGGAGACAGATGCAAGCATCCAGCATACAATTAGCACAATTATGAAGGATAATTA

TATATTCATTATTACACTTGTTTGAACAGTAGCAACAATCAATTGAAAATGGTTAACTGCGAAAAGAATTAAATA
AAAAAGCACATTTCAGTATGTTATTCATGGATCTTTTGATGGATATTGTACCTCTCCATGCTGGTACTCAGCGAG
AATGGTAACATGATAGCCCCTCAATGACACAAGCACAATAGATGATGCTTTAATACATGAAATTTGATTCAGTAA
TGCGACATATTAACAATTACATTTGACAATCGACATGTATGATGTATGCACCTAATTTGGATGGAATGAATACCT
TAAGATGCTTATGAGATTTATTGGTTATCTATTGCATGTGACCAATACAAGATGATGTAGTGATCCCTTCACATG
CAATGTACGACATTGGTAGGGGCGTAGGGCTATTCAATGAACCAATCAATCTGTAATTTATTTTGGGGTTAGGAT
TCATTTACATAAGATTTTTATTTAATGGACTATCAAGTGTTCGCTTGGATCACCTCTCACCTCTTAGACAAAGAG
ATTAGGGGTTTGATTCTGAGTGGCATCTGCACTCATTGTAGCCCCCATTTAAAAGTTCTAATTCTGCAAGCAGAG
GCTCAGTATGCTCAGAAGCCTAGGTTCCAGCTTAAGTGGGATAAGTTGAGATTAGGGATAAAATGGCACTAAGCC
TCTGTTACCTCAAACAGCAAAAAAAAAAAAAAGGAGAGAGATCTTTATTTAATGCAGCAAGAAGCTTGAGGTGCAA
GACTAAGGGAGTATTTGATAGATGGTTACCATTCCATCATCTAATGAAATATTTTATATTAGACATTCTACCATC
AATTTTTATATTTGCGAATTGATAAATATAGTTCACATTTATTGCGTCCCTTAGAAGTATTAGTCAAAAAGAGGG
ATAAAATCAAGAGAGGTTATTTAATAAAATGGGACTGAATTCATAGACCATCACAGACATGTGATAATTTCAGGC
ATAGCATTATATATAGTAGAGAAGTAGAATTAGAAGCATTCATTAATTCGACCAAGTTGATCCTTATGCAAATAA
AAAAGCAAAATCTTTCTTATGTAATAAGTAGTCACTCATTTTCCTTATTTTCTAATACATAAGCATGCTAAATGG
ATCAAGCTGATAAACTGCTGAACCTTTTTAACAAAATATGGATGGTTGAATCTAAGAAATGAATGCATGTGACTG
TGAAGACACTTATTGACTGGCATATATGGTCCAATCTATAAAGAATGCTCCTACTAGCCAGGACCATTCAATTAT
TTCTAAGAAAGGATTGTAGAGGTCTCAATACTTGATGCAACACATCACATTTACAAAGTCCAAGCTTATGAACAT
ATGGCTTGATTCATGAGATTGAACTTGCTTCGATCTAAATCAGTGAAGTCAAAGGCGCCAAAAGGCGCATCTCAG
GCGCTCAGGTGAGACGCGAGGAGTCATCGCCCCTTGATGCTTCAGATGATGCACCTTCACTCAGGCGCACGCCTT
TGGCACCTTTTAGGGTCTTTTTTTAGTGAGGCGAGGTGCTTGCCTTTGGCGCCTCAATGAATTCTATGATTATTT
TTTTAATGGTTGAAAAATATAAAAAACCTAAAAGGCCATCCATTTAACTTCTCAGCTCTATTACTAACAGAGGGT
AAAGTGGTCAATTAGTATGGTAAAGTTAGTTTTGAAGAAATAGAGGTTAATTCAGATGAGAAAGAAGAAGATGTT
GAAGGATATAGATCCAATGAAGGAGGAGAAGATGAAAGTGATGACAACTATGCAAGTAGAACTAAATCTAATGAT
TCCATTGATGAACTTTAGGCTTTAAGACTGTTTATCTTTTTTTGTTTCATTTATATGATTCCACTTTAAGGTTGT
TTATGTTTTCTGTTTTATTTATGAGACTCCATTGATAAATTTTAGGACTTATTATATTTTCTATTTCATTCATGA
ATTTAGAGCTGCCTATGGCTTGTTATCTTATACTTTATCTTGAACTAGTCGACCATTTTACTATTTTATATATGT
TTTAGGCTTAAATTGTTAATTATATGGTAATTCATATTACAATAGTTTTCTCTAGTTGTTATTTATTTATATATC
ATATTTTTTTTCAATTTTAGCACCTTAATTCACTCGGATGAGCGCTTTTCTAGCGCCTCTCATCTCAAGGTGCT
AGACCCCCCAAGCGCCTTAAGGACGCCTTTCACCTTTGACTACCTTGATCTATATAATCATATATAGATGAACTT
AAGTCATTGAAACCAAATGATAATTCTTATGTCAATAAGTTGTAAAATCATTATTGATGATTTGAGCATATGACT
TTGAATCAATTTTATTAAGCAATGTTCCACATAATACATATGATACCTTAAAAAAGTAAGAAAGTAGAAGAACAG
CAAATGAAATCACATCTAAATTATATTTTCCAAGATTTTTAGTCTCAGCATCAAATCCTGTACCAATAGCTTACC
AGTATAGGTCAGGTCATTACAGTACATCCCTACCGACATATGGTACATTGTTGGAGGCAAGATAACAGCTGTACT
ATGGACACTACCATATCAAGCCCTTGTACCATAATAGCACCCAATAATAGTACCTTACTGGGTCAGCATGGTACA
CCCAATACCAAGAGGTCTCGGTTTGTAGAAAAAGCTATTAATTATTTGTTAACAAATAATTATTAGATACAAATA
TATAAGCATGCAAGAACACAAGAGACTTTTGACCGGTAAAGAAATATAAATGATGCATAGTCATAGACATGCAGA
ATATTTAGAAGATTGAGAAAGCATCTCCTCATAAAATATCAACTCTAAGTTATACACCAAAACATTAATAAATTC
ATTGGAGCTTTTATTCAACGAACTATGCATTGCTCATACTACATTTTTTATGCTGGCTTTACGGAATATTCCTTG
TAGATTTTAGTTTGTGTTCTTTTTTACATTGTGACTTACCATTATCTAAAAAGGTATTTTAAGAAATAAATTATG
TTCTTTGTTCTTACTAGTTTTGGCAGAGGGAGGCAGCAAGCTTGAGGCAACAATTGCAGAACTTACAAGAAAATC
ATCGGTGCCATCTCTTAACCTTGTTTACATCTAAAGATTGTTCCTACGCCCAACTAGCCACTGGAAATATTCCAT
TATATTTATAGCATAAAGAAGATTTAAATAAATAACCATATTTTTTTGCTATAAACAAATTAGCATTATTCAACA
TGTCTCATTATTTCAAGCCATTCACATATAACAAATTCCCTCATGGTAAAATTTAACGATTATTCACTTATTCCC
CAGGAATCTAGAGTGTCAAGATCACTATAAGGAAAATAATATACAAACTAGCAAAAACATCAAATTTATATAAAT
GATTAGAAATGAAGTATAAAACCACTAGTTCTCTAGCTGACCATATCCTGAAAGTAAAATGTATTTCTTAAGATT
AATCATACAAACCAGCATTTTCTTGCTGATATGGCACCTTGTATTTGCAGTGTAGCTTATTCCAATGCATTGACC
AAATCATTAATGATGCCATTTGACAAAAGAGCAGGTTACTGAACCTCTAATAGTATGTCAAGATTGAGACAACAT
AAAACTAGTGACACTACCTCTAAAGAATTGGTTTCTCACTTTAATATCTGGATAAAGAATTGGTCCACAGACCCA
CAGTTCACAAATACACTGACAACTGCCTATATCATTGCAATTAACCTGTACTTATCAACTTCATAGAAAATTATT
ATACACGTCAATTATCAAAATTGAACCTCAGAATATTTCATATAATCTAATCATAAATCTTGGGTAGGAGCAAGT
GTCAATCTGTGGCTTGAGTGATTTAGCAAACAGTAAATGAGCTAAAAGCAACCAAGTACAAAATTCTCCAACATT
CAAAAAAATTCCAATTATTCTCAATTAAGGTTTTTTACCCAAATTTATAGCAACTTAATGAAACACACATTTATT
GAATTTATAGTGCCTTTTATGATTTTTTTTTCCAAATATTTCAAAAAAATGTTTCACTTCCTTTTTCTAAATTTC
TCAGCTTTTTGTTTTCCTTTTTAGTATCGGTCAATCAGCATTTAGATTCATTCACTTTTCAATCCCCAACATATT
CCAAAGTTTCTCGAATCATATATTAGCTTACATTTTCTCCCCGTGGTGATAGATTGTGCGTAACTATTCCTAAAA
AGTATCTAGAAGGCAATACATTGACACGATGATATCAAAAAAATATCAGAAATCATCATAAAACAGACCTTACTTC
CTGCTGGTACACATGAAATGTGCACATAGAATAATCAACAAGATGCATGCACTATGTGATTTGACTTCATGAATC
CTTTTCAAAACATTTTTTTTATTTTACTAATGATTGTTACCGATATGTGATAACAGTTACAGTCTTATATACCAG
CTTTTATACTCAGAAGTTACATATTTGTGGAATGGGATTATTACACTGATCTAGTAGCTTCATTATATCCCTGAA
GTTGTGTCTCACCCCAAGCCACCATGATTCTATATCAATGAACAAGAAGCCACAGTTCTGACCAGTCTTTTCTGT
TATGTAGTTTCTACAAGTTGAGAATTACACTCTATGACATCATGAACTGCTCCAATTTCACTATCTAAAATTGCA
AATCACCAGAATGCCCTACAATACATAATATGTGAAACACCCCTTGAATAGACCAAATTACTGTGAAAATTTCTT

CAAAATCCACCAAGGGTGGACTATTAGGACTGCAATCAGAATCGATCTTATCTTGCCTATAAAATCTGCCTAGCT
GTTCTGATGGAAAGTGAAGGCAAGACTCAGAGGACTGCTTCAGTATTCCAGAACTTCCCAGCCTCATAGCATATA
TTGCCAATAGCAGCACCCGTAACAATCTTTTAAATAGGAGCATTAGAAATCCTGAATACCTTTTTCTTCATCTCG
AGCATTGCAATTTAGGATTTAGTCCAAATTTTAGTTTGATGATAGCTCAGATATGACAAGAATACAAACTTTAAT
AGATAATTTTGGGTTATATATTTAATGCAGAGAAGCTGTTAGAATATTCTTTGATCATAATTTTTCTGGAAAGCA
GTCTACATAACATGATTGACATGGAACATAGTAGTGCCAAGGTGTCAATCTAGAATACCACAATCCAATTTGCCA
CTTAGAATGATGATCATGAACACAAAAGAAAGTTTCATGAAGTAGTTTAAACCTCAGCTCAGAAACACCTACTTG
TTTCGGTGGCCTATTTGCAAGTCATCAATTCAACAAGTCAACCCACTGACTCAAACCTCCTTTCTCCAAGCCTTC
CTTAATCTTTATGCCCACTTTAGTTAGAATCTCCCTCCCCCTCTATCTCCTACCAATCCTCTCCAGACATTGACC
CAAAAGCAGATTAGCCCCAAAATTAGCATGAGCAGCGCCTCTCCTTTCACCCCACCATCTCAACCCACTCTTCAT
TCTCCTCCACCATGATTAGGTGATCTCAACACCAATTATGAAAATCTTGATGACAACAACAACTATGATGAACAT
GCAAGAGTTTGACCCAAGATGCAGAGGAAAGGAGTTGATGGTGAGACAATAGCCCAACCTAGCAGTCACAAGTAG
GGCTGCAAATGGGATGGGTCGGATTGAGGCATGCTTGACCTTGACTTGATTTCATATCTGGGTCTTGATATTGGA
CTCAAACCCAACCCAATAGATGATCAGGTTGGGTCAGGTTCATGGAAAATATTTTAGACCCGACGAGTAGTTTGG
ATAAGGTTGAGGTTATGCATGACCCATATCAACCCAAAATATTCAAGTCTAGGTCCGGTTCAAGTTGGGTCCAAG
TCTACATTTGTTTACTAACAAGATTTTAAGAAGAAAATTTTGTTCTTACTTCTAAATCAAACTGAAAGAAAATTT
TGTTCTTACTTCTAAATCAAACTGAATTTTGACTGCCATAAAATTGTTTCCTAACATGGACAAAAGTCGATGAAT
TAATGTTTCTCCAGAACTGCAATATGCTTCAACAAGTTTAACCCAAATAGCTAGCTTTTCTTAACGTAGTGAGAT
TAGTTTGAGCACTTCTCTCTTAAGAGAGGAAAAGAAAACATGACTTGGAATACTTTTTTTCGGTCAATTCGAGTT
AAGTCAGGTTGGGTTGTTAAATAATATGAAAATCCAAAACAAACTCATAAAGTGTATAGATCAGATCGAGTCGAT
CAAGTCAAAGTTTATGAAATCTAGACCCGACTCAAAATGTGGAACAAATCTAATATTTGAACCCAACTTGGCCCC
ACAGGTCTTCAAGAATGGGTTAGGTCTGGCATCACCAGGTCAAGTTGAATTGGATCATGGGTGGAGATGATCATG
GGCCAAGCTTAGGCACAAAAAAAATCAAATTTTAAATAGGCCCGGCCTAAAACCAGCTAAAAATGAATATAGTTT
AGGCCCGAGGCTTAGCCATGATCAACTCTAATCATGGGGTGACTTGACCCATTTACAACGTTAGTCGTAAGCCTA
GCATCAAATTTTTAAACCTCGATACTACAACTCATATCGAAATATTGCCAAATGGTTAGATCAATATGGTATGTC
CATGCATAGGCACATTGTATGATATTGGAGGTAGCACAAAATTATACCATGTATTGGAATGGGGGTGAACCAAGC
ATTTGTATCGTATCAATATGGTTAGGAAGCTCATACTATACGCCCCATATCGACTAGCACTACTCAATGCTCAAA
ATTGTTGCAGAGTGGGATGGATGATGGAAGGCATAAGAAATCGGAGAGAGAGAGAGAGAGAGAGAGAGATGATGA
AAAGACTGCTAAGGCAGTGCAAGATGGTCATAGAAGCGATGGGAGTAAGAGAAGCATGATAGAATAAAGAAGACA
AAAGAAATCAAAGCTTAAGGGTTTTATATACTTTAATTTCTCGATGCCAATGGCTAGCCATTGAACTAGATCAAA
GTTAGCAATTTTGATGGTTTTTAAGCAACTAAAAATAGAAACTCCTAGTCCTAGCATGAAACTGATCTGTATATG
CTAATGGGTGACTAGACATATGGGTCAACATGTATGGGTCAAACCAACTACTGGGTGTATCACACCATATTAGTA
TTGAACCGATGCTCGATACTGGAAGCATATCGAGTATCAGTACATCGGACCAAACCCCATACTGTGCATACCAAC
ATCGTACCGGTACGAGCGGCTTTGATACTAATATTGTGAACCTTGGTGTCAAATATGTTTTAGAACAATAGTTTC
ATTTATGATCTCGGCCACAAAACCATTAACTCCTACCTTAGCTCAAAAAAAAGCAAAAATAAGTGTCTAGACTAT
ACATAATTATAAACATAAACTAACTAGTCCTGCAAGTTGCAGCCATACCAATAAACAAGATTTTTATAATTAATC
ATAGTAGATCTTTAGCCTACATTTGCTCAATGCCATCACACTACCTCAAGATTTCAGGAACCGGTACCGAGGCCC
ATATGGGTTCCTGGTTAGTACAGTGCCATACCATACCGTACCAACAATAAAAAATTCAACAAAAATCCCACCCAA
TGGTCAGAAAAAAAAAATCAAACCGAACCGGACCGAACCACTACCATACCGAATCGCTCGATATCGGGTGGTTCG
GGCCGATACTGTACCGAACCAACCGATTCGATCTGATTCAGGCCATTTTCTGAAAAACTGGCCTGAACCGAACTG
GTTCCCCACCGGTATGGTATGGTATGGGATGTCTCGATCGATTCAGATCGGTACAGAATACCATGCACTATCTTT
TATGCTATTACAAGAGCTATTTGAAAATCCATAAAAAAATGAATAAATCCTATTGGCGAACAAGGGTGGAAAAAT
GTTGATTATTACACTATTAAATAAATTTAAGATATATCATATTCAAAATTTTAATCGATGGTTGTAGGCAATTGA
TGGGAGAAGAGTTGTCTGGCTTAAATGTCAAAGGTCTACAAAATCTAGAGAACCAACTTGAAATGAGTCTCCGTT
CTGTCCGAAAGAAAAAGGTTATGACCTCACCAAATGTTGTTATTAATCCTTATCCATCTTTTCTCTATTTGATGT
GTTTATTTTCTCTGTTGTAGGAGCAAATTCTAACTGATGAAATTCAAGAGTTGAACCGAAAGGTTTATCAATATA
TAATAGTCAATTAAATCATTTTTAAATGTTAACAAGTGCCTAAATTTCTAGAATTTGTTACAGGGGTACCTCATT
CACAACGAAAACATGGAACTCTACAAGCAGGTAAACCTCATACGTCAAGAAAACATGGAGTTACGTAAGAAGGTA
TTGAGTTTGAGATACAAGATATCTACTTTGCTCTATAATCCAAACTCCAGAGTCTAGCACTCATGAAGCTATAAT
CTCCATGTTTCCACCGCCAGATGCTTCGCACATGCCATGCTCTATATAAAATATGTATTGATTTGGTATAAAACT
AGATTAATTCAATCACAAGAATTTTTGTACTAGTCTATGCTGGTTTGGCTACCCTGCCGAGCCAATATCAACACT
AGCACTAGTTCAAAACAAGACCCATTTAATCCTTTTTTAATACTTTCGTTTAGACATGGTTAGAAAAAAAGTTTT
TGATTTTTAAATCCTTTCTTAATATTTTTTTTTTGTTTTGGCACATGACATACGGGTCAACACAGCTGTAATATG
CTAGCCACTCTTTAGCACCACCCCATGCCAATGACATTTAAGATCTTGTTCAATCCCTGAAAAAATAATAGATTA
TTATTATAAATTACTATGCACGAAATTTGCATTGCAGGTGTGTGCAACAAATGAGTTAAATGGAGGAGCTAGTAA
GGATACCATTATTCCATATGGTCTTACTATGACTCAGAACTCACATGCACCTATTCATCTCGAGTTATGCCAACC
TCAACAGCAAACGGATGGCACACAAGCAAGAGCTCCCAAGCTCAGGTAATTATTTTGAAATATCAAACCTTTATA
TATTTTCAATATTTATTACTTTGATATGAACATAATCAATTCTTGCAACAGACTTCAATTGCAGTAAGAAGATGA
CAAGTGATCAATGTGCATCATCTTTAGAGACCATCGGCCTAGCTGTCAAGGCTTCTATGGTTTGTTATCCGCAAT
AACATCAATGTATGATCAGGAGCAAGAACAATACTCAGCACTCATTTATATGTGGTGTGACATGATCCAAAGAAC
AGTTACTTTACTCTGTCAAGCAGACTTCCAACTAGGACTACTAATAATATGACAGGTATCTCTGCTTAAGAAAAT
TACTTGCATACATGGAGACATGTCTTCTCATAGCATCTGAATAGTTGCTGATAACAGCTACAGGAAACCCACCTA
CTAAATTAAAGTTACAAGGGAATAACACCAACTAAAGTGAGAAAAAAAAAAATTTGCACAAAAGCATCACTCACTA

TCTACTGACTACTGCTCAGTAACTACCAGCAAGAGGTATATAAAAATGTGACAGATTTGTTCTCCAAAAACTACT
GTAGAGTGGTAATGACAACATGTATCTTAAGCTAGCTCACTAAACTAGAATGAAGCAACTATTTTGAGTTTTGCA
GAGAAGCACATGCAACACTATAAATTGTGCCAAGCATATGATTGGACAAGGGAAGTTTGCCTACAAAGAAATAAC
AAAGAGTTGAGGCTGATCCTGTTCTATTAACTATATAATGGAAAAATACAGACCTGATCTTTCAAGCAATGTGAA
AGCATCAGTTAACATGTGCAATAGAAATTCAACTTTACAAAGATAAACTTAAGAACTGTCAGAATAATTGATTCA
TGAGCAACGATAGGACATACTTTAAACTGAAAACTACTTCTTGAAGGTATTTTTTCTTAAAAATATATGGTCATT
CATATACACATGGTTCACATAGATGCAAGAGACACAATCGAAAAGCAGCAAACAGATAAATTCTCTTTACATGTA
AACAGTTACTGTAGCCTTTAAAAGTAAACAGATTACTGTAAACAGATTATAGTGAAGACAAAAGCTATAACATAT

>EguAGL17_2
ATGGGCAGGGGGAAGATCGTGATCCGAAGGATTGATAACTCGACGAGCCGGCAGGTGACCTTCTGCAAGCGGCGA
AGCGGTCTGTTGAAGAAGGCCAAGGAGCTCGCCATCCTCTGCGATGCTGAGATCGCCCTTGTCATCTTCTCCAGC
ACCGGCAGGCTCTATGATTTCTCCAGCTCCAGCATGAAATCAATAATAGACCGCTACAACAGGACAAAGGAGGAG
CATCATCAGGTTTTAAATGCAACTTCAGTCAAGTTTTGGCAGAGGGAGGCAGCAAGCTTGAGGCAACAATTGCAG
AACTTACAAGAAAATCATCGGCAATTGATGGGAGAAGAGTTGTCTGGCTTAAATGTCAAAGGTCTACAAAATCTA
GAGAACCAACTTGAAATGAGTCTCCGTTCTGTCCGAAAGAAAAAGGAGCAAATTCTAACTGATGAAATTCAAGAG
TTGAACCGAAAGGGGTACCTCATTCACAACGAAAACATGGAACTCTACAAGCAGGTAAACCTCATACGTCAAGAA
AACATGGAGTTACGTAAGAAGGTGTGTGCAACAAATGAGTTAAATGGAGGAGCTAGTAAGGATACCATTATTCCA
TATGGTCTTACTATGACTCAGAACTCACATGCACCTATTCATCTCGAGTTATGCCAACCTCAACAGCAAACGGAT
GGCACACAAGCAAGAGCTCCCAAGCTC

*EoAGL17_3*

>REF_ELAGV07
TTTGTCCGAAACATCCAGATAAAATCATCAAATTTTGACTCATATATCCTTTTTAGTTTTAGATTTTTTTTATAA
ATTTTATTTTTTAAAAATTAATTAATTGTGTGGGAGGAGTACCTTGCTTCCTCCACATTCCATCAAAAAAGTATA
TATTTATCTTTTTTTTTCAAAAAAAAATGTTGGGACGACGGAAAGGAGGGCAAAATGGTGAACTTGGTAGGGGCT
GAATCGGCATACCCCGCCTGACGGAAGCCGGATGGAGACCGTTGGACGTACGGAATCTCTCCGTCCAACCACGTG
TTGGCGATCCATACGGCAGACGGCGCTATCAGCCAGTCTCGGAAGGGATTTTTGATTTGAGTAAACTAACGCCGT
GTGGTTTATTTGACCTTGCCCCAATCCTCGTACGCCACAGACGCCAACAAATTGCTTCCAGAACCGGTAACATGT
TGACCCAATGTTACACCCGCTCACCATTACGCTTTATCCGGATGACGTAGATCTGCGTGCACTGAAAGGCTGCAA
CGTTGGGTGTTTCTGAGGAGTGCATCAGAGAACTTTGCAGAAAATGGAAGGCCTCCAGCAACTCCAAACGGGAAG
TCTCCTGCCCTAAAGTGGCTGACCTTGTCCTCTCTTATTGATGCATCCTCAAACAAAATGACTGTCCTTAAATAG
GGGGGGAGAGAGGAAAAGGCAGCAGCAGCAGTCTATTTCCTTCCATTTTTTGGTAGGTAATCCACTCTTATCCCT
TCCTTCCTCCCCAGATCTCTATCACCTCGGCTTCCTTGTTCTCCATCTGATGTTATTTCCCTTCACAAGTATTCT
TTGCATTCTGTTATCAGATCTAAAGAGGATCGATCAAGTTATCGGCGGCTGCGAGCTAAGGGAGGAAGGGAGAGA
TGGGCAGAGGGAAGATCGTGATCCGAAGGATTGAGAACTCGACCAGCCGGCAGGTGACCTTCTCTAAGCGGCGCA
AGGGTCTGTTGAAGAAGGCCAAGGAGCTCGCCATCCTTTGCGATGCCGAGGTCGGCTTTGTCATCTTCTCCAGCA
CTGGCAGGCTCTACGATTTTGCCAGCTCCAGGTCAGTCCTCTCTCTCTCTCCTCCCCCCTCCCCCACCTTACTCT
CTATATCCACATAAGTGGCTTTTTTTTTTTTGTCATAACCCATGCAAAGTTTTCTATTATTGCATGGTACATGTG
GCATTTATGTGCAATCTTAGAGGCAAGTTTGGTAGTTTTAATGGTAGTCCTAGGTTTGGTATTTAAGATAGGGTT
CGAAGAAGAATGAGACCGAAGAAGCTGTAATGGTAGTTCTAGGTTTGGTATTTAAGATAGGGGTCGAAGAAGAAT
GAGACCGAAAAAGCTGAAGAAGTCCTTAGAGGAGAAGTTCATTCTAAAGAACGAAATGAGATATTTTCATCCCCT
TCTGGCTCAATCACCTTCTTTCTCCTCTCTCACTTAATTGGTCTCTTGGATTTTTCCTCCATCAAGTGCGGTGCT
AAGTGATACAACTTCCCCTCACCAGGCACAGAAGTAGTTCTTTAATTGAGAAGAATTTTGAAACGATGACGAAAT
TAAAGGTTTGGAAGAGGTCCCCTTTTTTAGTTTGTTTTGTTTGGAATAGAAGAAATGGAGGGTTCTTCATAAAGA
TGTGGCGGTAAGTGATGTTGAGGGTCTTGTATTGTAGATTTTGATTTTAACTTGTGGTCTATTCATCGATATCAT
CCCCGCGAGGATAGAAAAACTAATGACGTGATTGATTTGTACTTTTCCAATAGTAAAAATCTTCAGTAGGTCTTT
AGTGGGCTTGCAGATTTTTCTGAGCCTAGTGGTGGCTGAGCACTAGATGCTTTTGAGCCTAAAGATGATAGCAGA
TTCTGAGAACACAGTACGAGGGATGCGGTGAAGCTTATCTTGTCCATTAGAATCAAATCATGATGAGGAGCTATG
ACACTTAATTCAGCAGTCACCGTGATGAGAGCTCTGGTAATGCTGACAAGGCTTGGTTGACTCGTATAAAGCCTA
CGGGAACTTAAAAACGGTTTGTCTTCCATGAATTTGGACCAGCCGATGGGAATGTCATGATGTAGGGATGGTGAT
TTTGACAATAAACCAAACAAAAGAACCGAAGTTTCTAGAAGGATAACAAGAACTAGTGTCAATCAAATTCATGAA
TCAGAAAGAAAGTGATGCGAGGATACGCATGCCAGAAAACATAAAAGGCACGTAGTTAGTACTGCCTGGAGACGA
CATTGGTTAAAATCCATACGAGCATCACTGGAGCCAACATTGTTCAATAATTGATCCGGTTTGGCATTTCAAATG
CTTCTGCACAGAACTACCGAATGGATGTTTTGATAAGGAGAAGTGGTATCATCATAATTTAACTGATATGAAATT
CATTTTACAAGACGTAGGGGCATCTCGCACAGTACCATGGGAAGACACATACTACCATTATAAATAGCTGTGACA
TCCAGAAGTTTACTGAGAATAGTATCTCGTGCAACAAGATCTTTTCTCAACTCTGCAGTGAATTAATTAGTTTCT
TTTGCTGAATTATTGATGCCTTGACCGAGTTACTTATCTACACTACATATATGGAGTTGAACAAGCAGTGGCCCT
TCTCAGTGTACGTTGCAGGAATTTCCTACGGTTTAGAGTACATGATAACTTTTTAGTTGGTTGAAGTAAAGGCAA
TTGGCCGTAGTTAACTTCTTAAACCACCTATCTGTCACAAACCTTCACATACCTAAGATTCTACAACTAAGATGT
GGAGGATCCAAACTAGGACAACGAGTTTGAGTTGATTATGCATATAATGGTAAAATTGCTTATAGTCTTTTCTTT
TTTTGTTCCGTAATCACCTCTAAATTTGGATGTGATTAGCCAACTACAGGAAATCAAACTAAATTTTGAGTTTTG

AATGATAGCATGCGAACTGCATTTTATGCATTAATGGAATTTATACGCTACATTGAAGTATAGAGCTTTCCCACC
AAAAGAGTGTAAAGCTTATCTTGTTTCAATATAGCTGTGATTATTTCAATGTTTCTTAAGAGGAAAAATCGAGGA
AGTTCAATTATCTCATATCTTCATCAACATTGGTGTAAGAAAACAATCACAGAAGATGCGTTGCATAGAGTCAAA
TGTTCCTATGTGGCATATATATGACTTCTCTTTCCTATTGGTGCATTTGACTCTTTAGTCTTTAAGTTCTCTTGC
TCGTTTTTTACAACACAACCAGATTTATTATTTTAGATTCACGTTTCTTTTTTCATGTGATAATCCTTACTTATA
GTTCTTGCATGCCCAGTATGCTGTAGTTAATTTCCAACCTTTTATGTGATTCATTTGATTTTTGTTCTCCAAGCC
TTAAAAAGAATAGTATCTAAAGCATGCAAGAAGGAAAATGAATGATGGACAAGAACAACCGTGCAATTAATATTA
GGTGGGAAAAGGATACTTTTAAGTAATCAAATTATCATGGAATAGTCTTATGATGGAGTTTATACCCTCAAAGAA
CTTCAGACTCCTGATTTGTAGAATATCTCCATCTTTGCAGTCGAAACTTTAATTTGAATAGGCTGACATATGAAA
AATTTGCCGTCTAAAGACAAGATAAAAAGTTATGAAGAGGTAGCATAGATTGAGGTATTGAAGGAGTAAGATGGA
ATCAGCAATTTATAATCCCTAGCTACTGATTTCTTCAAGGTAATGAAGCACAAGATAAAAGTGAACAATTTGAGA
ATGAATCTTTCCTATGGATAATAAGCATAGTTATACGAAGGCTGGTGTGCTTAAAGGGTCGGATGGCTCAGATGA
TTGGGTTAACTCCGCTCCTGTGTATGTGAGGTATTGTCTGCTCTAGCTCATGATCATCTTTGAGCTTCAGTGGAC
CTTGGTCATTTGGAGCCTCACGGTTTTATCCTTTAAAAAATACCTCATGAGGAAGAAAATGTTCCAATCTATATG
AGCTATACTTTCTTTTGACTAACAACTACTGATGTGGGACTAAAGATGCTCTTTCTATACTACAACATAGCCTCC
CAGTCAAAAGAAAGTATGTGTGGATAGGAAATGTTCCCACTGACAGTGCTAATATTGCAGTCAAAGGGTTGGATG
GTTTGGATGGTTGGATTAACTCCGCTCCCGCATACGTGAGATATTGTTCCCTTTGGCTCATAACCATTTTTGGAC
TTCAGTGGGCCTTGGTCATTTAGAGTCTCATATTTTGCCCTTTAAAAGATACCTCACTAGGAAAAGAAGGTTTCA
ATCTATATAAGCTATACTCCCTCTTGACTCACAACCGATGTAGAACTAAATAGCCCTCCAATTAAGAGGGAGTAT
GTGTGGGTAATACATTGCAGCAGGACTTCAAAAGAGGAATGATGAGGTTGGTGTTGAACCGATCGAGTTGGAGAG
TTGCTAAGGGTCAAATCGATGTAGATTATGAGTGAATCTGTAGAAGAAGTCCTAAAATCGATAAGGTACTCTCTG
GTTTAGGACCCTCCGGTCTTTAATTAAGTCAGCTGGGGCCTTAAAAATAGATAGTGAAAATAGAGAAAATAGAAG
GCAAAGGGAGAGATTTTGAGTGGAAAGGTGAGTATTTTGAATTTCCTTCAGTCAGAGAACTGAAATGAATTCAGT
AGAGTTCAGGCTCCATGAGTTCTGACTTATCTTGCGGAGAGTACTCTGCCTCTCTTTATATAGAGAGGATTTGCT
TAGGCCATTAGGAGAGTTTATTAGACCATTAAGAATCTGTCAGGTCGTTAGTTTGTTATAACAAAATAGCTAGCT
GTATGAAAATCATGGGTTATTCACCCACTTAGTGATTAGCTGTAGTATGGATGGAAAACAGTTATTGTAGAGCTA
GACGACAGCTATAATGAAACTGTAAAGTAACTGTCTTTGTTTTAGGTCAAATTGGCTCTAGATCGATGTCTCATG
TAGCAGGTCGAACAAAAGCTGAGCTGATCATAATACTCAATTAAGGATTAAATCGGATCAATTCAGGTCGGTGTA
TAACTAAACTGTGGGTTGACCAGATCGGCATTCCATTGACCTGACTAAAAAATTGACAATGGGATTTTCAAATGA
TTCTCTCTGATTTTGCTATATGGCCAGAGGTCGATTATCTGTACCAACAAAGGCAATAATTGGCACTTTAAGATT
CTAGCTTTGGGTAATCCTTGATGTTTCATATAAAATCATCTAAAGAAATATTTTTGAACACTTATTTGAGTATGG
TTTTCATTATCTACATTAAATTTTCATTATTTTAATGTTTTAAATAAGCAAAATATTATGGCCAAATTCTTATCT
TCCAAGAAAATTGTGGATGCTTGATAGAAAGTTTCTTGGAAGTGTTCTACGTAATAATTGGAAGCATATTAAGTA
CACTTTAAATTAATGATTTTGTAGACAATTGTTGTTAGATATTAATTAGAATAGATGGCTCCTAACTTGTTTGCC
ATGTATATTAGTTGCTTTAATGTCATACTTGATGGATATTTCATGATTTGGTGTTTAGAGTAGATTATTGTTATA
TCATGAATTTAGAAGCTAAATTTTAAAAGGAAAGAAATAACCAAATGTTTCTTTCTTCTGGATTATGCAAGTGTT
CTACATATTGTATGACCTCATAAAGCATTTCTTGCAAAGTCATATATGCAGTTTTACATATATATTCTTATACAA
GGTTTCATTAAACTCATTACAAGTTTCATATGTGTTTCATATATAAAATCCAATATTCAAATCATATGTCATTGA
TAATTCAATTATGACATTCTTAAATATTAAACCAAACATTATGCAATTGAAATGACACCTATATTTTACATGAAC
TACAATTGTTGAAATGGTTATAATTACCATTATAAGCAAAGTCATCGTATATCGATCTCTCATGTGTAGTTGTTT
TTTAAATTGATTACAATATCTGCTTCTGGAATTCAAGATCAAACACCTACTTGATGTTTGTCAATCCATATTTTA
TTTTGTATTAATCTCATCATCGTCCTTTCAAATTTGATGACAAATAAGAAACTTATCAGATCATATTTGGCGTTT
ACTACATATAATTAGAAATCCATTCACCTAAATTTCAAATTAACTCTACATGTCGCTGAATTATATAGTCTATTT
CAGACATTTATATATACTTGTATATACTTCAGAACCTCATGACCAAGCTTTTTTTCTTCTTTTTTTTTTTTCTTTT
GATGAGGGGTGAAGGGTGGGGAATCCTCTTAACATTGTGTTGACATGATTATGTGCTAAGTTTCAAGCTATGCAC
AAGGAAACCTGGAGTATCCCTTAACTTGAATACTATTCATAACTATGAGCGAGGCTGAACTTGGGCATCACAAAA
CCAAAGTCTATATAAGCGCAACGGAATGGTTATTTCTCAAAGCTATCTTTATTTTTTTTTTTTTCTCAGAAAAGGA
AAAACTAGGTTCCACTTGCTTCATTGAGAATATAAAGATCAAAAGTACAAAGGTACATAGAGTACCAAGAAAAGT
TATTTCTGCGACTATCAAAAAGGTACCACGCGCACATTGATTTGCTTTTTTCACAGTGAATGCCCCAAGCCGATG
ATCACAATCCATCACAACTAAAAACATTCTCTCCAAAAGAGATCTGAAAGAGGACTTGAATCCGTTCATTTGTTT
TGAGGGCCTAACATTAATTGTCATTCTATTTAAGTAACTTTGGGGCCATGCTAACTATTGATTTTATAATGCAGT
AACTAGCTTGGACTTACTTCAAGGGTAAGATTCCCAAAGAGAATCATACCTAAACTTTTCTTTTCAAAAATAAGT
AAAATTTCTTCCTACATCTGAATAAGTAAAATTTCTTCCTGCATCTGGATAAGTATGTTGTTAGACTGAATGCTT
GATCTTAATTTTCTAAATAATCCGTGAATTTCTAGGAATGTCAATATGTATTCTTGAAATAAAATGACTGACTAA
TTTTCTTTTATGATTCTATCCTTGTGGGATTTTCTTTTCATCTCTAATCCTAATTAATGATCTCTCTTGCTTATG
GATTGAAGAACATTGCAATTCTATCATAAGGTTTATTTGAGCATTATATTTTAAATTGAAGGATTATTATATTAG
AAGCAGTCATAAACAAGTACTCCGGGCTATTGTTAAAAGCATGAATATAATTGGGTCTACCCACAAAACATGCAA
GGCCTAAAAAAAAAAAACCCTTAGTGCCAGTGCAAGTAATCTAGGATACTTAAAACTTTTAACTTAATAAACAAAT
CCAACATTATTTTCAAACTCAATCCATGTTCATTACATAATATTTATTTTTAAAGAAAAAAAAACTTCTAGGTTC
TAATCCTTCCGCTCAAGGCCTATTAATCCAATCTACACTGGATATAGGCATCTGAAAAAGGGTAAGAAAATGAAA
GAGAGCTAAAAGCCCAGTAAGTCATCCTTATTATTATTCGATCAAGCATAATATAACAAAACTCAAGATTGTAAT
ACAAGTTTTCTATATTTATAATTTAAGACATGTTGAGCAATATAAAATATAAAATGATCAACACATCTGAATAAA
AAAAGTGAAATTTATGATCATTGTATTTAATTGTTCACACTTTTTCTCCTAGCCTTTTGACTATAGCCAGGGCCT

GCCCATGGCAAGCCCCAAAAAATTTTTGATTTTAACTATAAGATACACACATCAGCATGCACCAATAATCATCCT
ACTAGAGATCCAAAAAAAACTAGCTCTGAAATCATACAAAAAAATATACCCATAATAGTATGTACAAGTGATCGT
CCTACTGGTGCCGCTCAAAAGAAAACTAGATCCTGATTTCAATATGTCGTTAACCAGCATGTGCTAATGATCACC
TAACTGGAGGCCTAAAAAAAGACTAAATCCTAAATCATATGATTGCAGTTAGACTAAAAATGATACATCACAATC
AATAAAATCAACAATCAAAGTGGTAGAAAATAATGTAGTTCATACTCATGAAATTTATTTTATGGTTAATCTAGT
TGTTCAAGGTCTACTTATACAGAGCATACATAATATAATTTTAAATATATATAAATATTACTCATTGTAAAATTT
CAATACTTAAGTTAACTTCTATATGTCAAGTTATGCGAAGTGAACCAATTCATATATATAATCGAATACTAGATA
GATGTATCTTTGGCTTGTAAGATATAGAAATATTTCCAAGGTACTGCACATGACCTAAATAAATCAATGATTAAC
ATTACATCTAAGGTAAGCACATATTGTTTTTCAAAACTCAAGCATCTAAAGGGATTCTAGATCAATGAAGCCCAC
TAGAAGTTTCATCTTTCAAAAAATGTCTACATCAAGGATGGAACTCCAGATCAGCTTGACCAATAGTCCGAATCC
CAGTACATATCTCAAACCTAACTTAGTTCTTCTAATATCACATATTCCTAATCTAGGAATCATGAAAGGTGCAGG
AGTCATAATCACCTCAGTAGTGGGATCGGTCCCTAGAACCTATGTACGTTTTAGATCTAGTGCTCTGGGCTTATC
TTGATCTATAAGATAATAAACCTGATTAGACCGAAAAAGAGAGGTGAAGAGGCAGAGAGAGAAAGCTAGAGGGAC
GTGGACGAAGATAAAGAGAACAAGAAAATGGTGATTGGCCCTATGCGGATACCCCAAGTGCTAAAGGACCTGAGA
GAGAGAAAAATGTAGCCAGAAAGAGGAGAAAGGGAAGGGGCGGAGAGAGAGTGATGCAGAGGAAAGTGATAGAGA
GAAAATTGGAAAAGAGTGAAGGTGGTCTGGTGGAGCTTGACAAGCATGGCTAACAACTCGACTATGGATGGAGAC
AAAGGTGGCTACTGAAGGTGGTCCAAAACGGAGAAGAAGAATGGGGAAACGAATCATCTTAGAGAAGAAACGAGG
GTGATGGAGAGGCTCAACAATGTTGTAGCTCACCAGGCGTGCGCGTGTAATGTGATGGTGCTCTTATCCATGATT
TGAGGAAGGAGAAGAAGAAAAAGGGAATAAGGTGATGAGGGAGATGCGAAGAAGAGGACAAAGAGCTCGATTTTT
AAAGATATCATGGGCATCCATCTCACCCACCCGTGGAGGAAAAAGGGGAGAGGAGGGCGCAATTTTCTTTCTCTC
GGTTGGCAACGATAAGCATGGAAGGGAGGCAGTGACCTTGGGTCATCCTTCTTTGTCTGGGCTAGGCCAGGAAAG
AAAGAGGAGGCTGACTTTGGGAAGGCAGCCAAGCCTCCCAAGATAGGCACGATCCTTTCTCTTGCTGCCTGCCAA
GGATGGCAAGGCAGTGGGGGGACCCTGCATCATAGATTCTAGCTTCAAATCTATAAGTTTCATCCCAAACATTTG
GGACCTGGATGGTTACAGGTAAGTCTTCCCTTTCAAAAAAAAAATGGTTATTGTTAGAAGCAAATATTCTTAATG
AATCTAGAAAATGTTGATAATTATAATAATTTCATTATAATTAGTCCACATCTTTATGCTTTGCTTAATGCTTAT
GGATTGTGTCAACTGCTGAATTAACTTTCCAGGTGTAGATTCCTCTTCTTCAGCCTTTTGTTAATTGATCAAGTT
TTTGATTCATCTATGCATCTTCTGTTTTCCTCCCTTATGTTGATCCCTTTTCTGCTGTTAATTTGACTCTAGTTG
GACTCCTAAACATATTGGACTCTATCACCTACTCTATTGTCTGACTTCTATCTCTCCTCAAATGTGTGTTGCATG
TGCCCAGTTGCTAATGAAGTATAATGTTTGAATATAACTAGTATACCATCTAGGTGGCAAAGGATTGAGTTTGAG
TCGGATCAAATATGGGTTGGGTCAAAGAATCTTCAACGACCACTCCATCAATGTACAATTATAAACTTATACCTA
TTTCCTTTATTTCATATGCACACCTAGTTAACCTCCAAATCAAATTATTACTTATTAGTCATAAATCTTGATAAA
TAAAAACAAAATGGCTGGAGAAAGATATAACCTGGAGTTGGGCACAATTTTTGATGCACCATCCATATTTGAACC
TAAAACCTTAAGTTGTTAAGCAAAGGTCTATAACTACCAGGATAATACACGACCAGATTATTAAAAGCCTTTATA
CAAACTCATACCTTTTCGGGGCTGTGCACAATTTGTTTTTGACTGGTTTGGGGTCAATTTTCAAACTAAGCTGAT
TTGAACCAGTTTTCTAAAACTAAAACTAATTGAACTAACCATATAAAATATCAGTTCAAACCAAACTAAAAAAAT
TGGTTTGGTTTGGTTTGGTTAATCGAATTGTAGGCTATTAGACAAGGTATGGGATGGGCTAAACAGGCTGCTATC
CATTTTAATTTTTTTCTAACCATACAAAATGGGCAGCTTATGCAATGAGCTAAATGAATAGCTTATGAGATGGAC
TAAATGGGCTAAATATAGAAGCAACAAACCGATAGGGCTAAGGCAAAAGAAAAAATCTTTATGCACCGCAGGCGG
TGTAGAAAATCCAATATGGAGTGCACTGTTTTATGTGATTAGTCTACGTAGCTATCGCTTTTTCACGCATATTTA
ATACCTGCAGTTTCACTTTTTTATTTAAAAATTTTGAATGATGAAAATATCCCTGCTCTTTGAAAAAAATTATAG
TATTTATGTTTATATTATGACATTCTGTATGCAGCAAGTCATAATTTTGACGCAGGATGTCATAATATACCACAA
GATATCATAATTTTTTTCAAAAAGTAAAGATATTTTTATCATTCAAAATTTTTAAATAAAAAAGTGAAACTGCAG
CCATTAAATACGTATTGAAAAGTGATTGAATAAAATTATCCCTACTATATTATGATATCCTAAACCATATTATGA
CATCCTGAGCCATATTATGATATCTTGCATGTAAGAAGTCATAATTTGACATAGGATATTATAATATGCCATAAG
ATGTCATAATTTTTTGAATCATATTATAACATTATGCATGTAGAAAGTCATAATATATCACAGGATGTGATAATT
TTTTCTAAAGAATAGAGATATTTTTATCATGCAAAATTTTTAAACGAAAAAAAAAACTACGGATATTAAATGTGC
ATTGAAAAGCAATAGCTACGTAGACCAATCATATAAAACAGTGCGCTATACGTCAGATTTTTTCTACACCACCCA
TGGTGCACAAAGAATTTCTCTAAGGCAAATATAAGTAAATCGGTGGATGGCAGCCCGTTTAGCCCACAAGTGAAA
GAAAAAATCTAATTCAGATCAAATTGCTAGGAAACAATTGAATAGATATATTCTTAAAATTCTCATAACCATTAT
GACTTACCTTGATGCGCAGCGGAGATTTGATCAAGATTGTTTGAATCAGTTGCTTCACGAGATCTGGATGCGCAA
GTTCTCTACTTCACATGCACGTCAGGCATTGCAGGAAAAATGGATGAACACCTTGGGCATCGCTCCTTTAGAAGG
GAAGCAAGAGGAGGCCGTCAAGATCTAATCTCATGGCTGGAGGCTAGGGAGGGATTCTTCACGCCAAGAGAGAAG
GATGCCAAAAACTCCACACCAATATTAGCGCCCTCCCTTTTTTTTGTCACACAAAATTACTCTTCTTCATGGTGA
TTTCAAATCTATTCTTCTTTTTACGTGAAGAAACACCCCTCCTTAAATAATTCATGCTCAAAATCAAATCATAGC
CCTAAGAAATTAGGTCTTTGATTCAAATCAAATTTGAATTAGATAAGGCTTTATCTTATCCACTAAGAGACCCAA
GGGAGAATTGACGCCAACATGTGCATCTTTTATCTTTCACGCACGCAAGGGATGGGATGCAATTTTCAATGCCCC
CTCTTCCTTCTTGGTCAGTCAGATAAGATGAGAGAGCCCTAGATAATTAAGACTCTGTGGACCTATTCCAAGTTG
GTTCAATTTGAATTCATTTCGAACCTGATTCGAATCAATTTTGAAGAGACTGTTTATCCTAATCTAATTAGAATT
TAGATCCAAATCTAACTTAGATAATTAACCCAATTAGCCTTAATCAAGTTCAATTAAATCAGATCTAATCTAAAA
TAATTAAGATTTGAATCCAATCTCTCAAAATTTCTTAGATCACAGATCATATCAAATTCATCTCTGAAATTAAAC
CTGCACCTCATGTCCAGTGCTAACTAAACATAGTCAATGTTTAATTTCGTATAACCTATAATTTAATTCTTAATT
AAGTAAATCTATTTGTTCAATCAAGTCACATACTTTCAAATTAAACACATAAATGTTAGGTCAATTATTTTCTAT
GTTGCTTGACTTTGTACGTGACTCTATAGGTTCGAACACTAAGTCGATAGTGTAGAAACTATTTTTTATACTAAT

CGAAGTGATCATCTAGTACTGATATGATCTGACGTCAGGATGGGTTAAATAATTACAAAATAATATTCAAGAACC
TTGATATATTGTTACCACATAATTCATCTCTTTAACCCTCAATGCTCAGGATGACCTTAGAGTAAACTGTCAAAT
CCTAGATCAGCCATCCATATTGTATTTCAACATTTTAAATCCTGTGACAAATCATATGGATTACCCTGGTCAAAA
TTTTATTGAATTGAAATACAGTGAAACATTATCTCCTGTAATCCGAAGAGATCAATCCCATCTTGACTCATACAC
AGACTTCGCAAGTATTTGACTGTGACCAATAGTCTTCTATCACTATATTAAAAATATAGATAATCCAGCATCAAA
GTACAGTGAGTTGCTTGCAAGTCACTATAGTGATCTCAGATCGAAGGGATATTTATTCCCATACCCTTCGTGAGC
TGCTCTTGATAGTAGAGTGCTTCACAGCTGAGTCACTATTCAGTAATGATCTACTTCTACATCTCACCTGCATGC
CATACCAGTATCTCCACACTCTTTGATTACAAGGACAACCAATTCATATGGCATACAATGATCTAAACTCGATAA
ACATTATTATCTTATTAATAATGTATTATTTGATCACGAACATAATTTAAAAACTATTCGATAAGTTTTCATCAT
TTTTATCGATTGTTTTATAGTTCTAAGGACTTCATTATATAATTAAGAGTTCTATAAGGAAGATGTTAAAATATG
ATGAATTGCCAAAATAATTTTTATTTATTCATTTATAATTCATATACAAGAGCATAATTATTTACAATTAGACAA
TTGACTTTAGGACACAATTTTCAATAATGGATCATTCGGTTTAAATTAGGTTTCTTAAAATCGAAAATGAGATCG
AACCAATTTTTTTGATTTTTGAAATTTTCAAACTAAAATCTAAATAGAATCTTAAAAAAATCAATCCAAACCCAA
TCAAAATTAGTGGTTCAAATTAATTTTATGGTTTAATCAATTTTCTTGCGCACCCGTATTTCTTTTAGAAAGATT
TAAAAAGTTGGATGGCCTGTGCTAAAAATTTTTATCTCTAATTATTGAACTACTAACGACACAGGCACATCCAAT
GCATATTTAGAAAAACATAAACTAGACATATTACAAAAAATAACTATTACATAGTATAAATATTATAATTTTAAA
GTGCAAAATTGAAAATTAGGGAGTATTGTTAATGATTTAAATCTGAAGTTAATGGAGGACCAATTATTTTGAAGA
GAATAAATAGAGATATAAAAAGTTAAAATATATCATTTTAGTTATTTGATGAAAATGAATTAATTAATGATCCAT
GCAGGTCTCAAACTTGCAACTTTTGCATTATTATTTAGCAAAAATATTATCAATAATATATTTTTAAAAATATTT
TTAAAAATGAAGAATATTTTTTGATAGTGGATTTTATTAATAGAATCTATTTGATATATTTCTCTTTTATTTTGC
AATATGGTATAGATAAAGATAAGACCTTTTTCTTAAGGCAACATAGAGGCAAGACTTTCGGCTCAAGTCTCCTCC
AATGGATGGAGGATGCTGATAACTTGTGGGGATAAACCTCCCATCATCAACTTTACAATGCTTTTGCACTAGGAT
AAGTATATACAAAGGTAAAAAAAAAAAGAAAACTTCTAGCTCTCATTTGCCGTTGTAAAGAAAGAGTCATGGGTA
CATCGGTGGTAGTTAAGGCTACAAATGGGTCAAATGACCCATGACCCAACCTGACTTGCTTAAACTCGAAATAGA
TCTGATTTAAGGACCCATGGGATGGGTTGTATTGTGGTGTAGAAGGGAGGGTTTCATTAGTCCTACATTGATTGT
AAGTCAAGGAAGAATCTGATATACATTGATTGTAAGTCAAGAAAGAATCCGGTGTATATAAGAAGGGACCATCTT
CCCATATGAGATATTTTTTGAAGGGCAAAATCGTAAAGCCTATGAACCAAAATGGGCCATACGTCACAAGTCGAG
CCATAAATCCTTGGCCGCAACAGTAGTGCACCAGAAAGAGGACCACCCCTACAATTATAGAGCTCATTCTGTCTA
TACACCAGTCTACAATAAAAATAAGGAGGGCACCTCTCATTCTACTCAAATTCTCTCTTGACTGGGAGGCTATAT
TGTAATATAGAAGAGAGGGCTTCATTAGCCCCATATTGGTTGAGTGAAGGAGGACTCTAGTTTATATAGAAAAGG
ACCATCCTTCCTATATAAGATGTCTTTTGAAGAGCAAAACTATGAAACCCATAGGCTAGAGCAGACAATACCTCA
GGTGTCGAGCCATGAGCCATTGGTTGCAACTAGTTGCTTTCCAAATTTGAACTCATTCTATATTTCAGGTCGGGC
CTAAGTTTATTGAGTTTTTATCCAACCTCGACCGATTATTGTATGGGTCTAATTTGAGTTTTCTAAGATATTACC
TAACTTAACCTAACCCAACCCAAACCCAATATACCTATTAACCCAAGGTGAGTGGGGTGGTATTAAGCCGGTTTA
CTGTGGACCATCTACTCTTCTAAAGTCCTAATTAGAAATTAATCCTATATAATGATTGCGTCCATCACCAATGAA
TTTTTGAAATAAATAATATAATTATCCCACCAATCCAAGCTTCTTGATCAGATTTTATTCACATGACATAGGGAT
GTCGAGAAAAACTACATAGTTGAGTGATATTCTTGGATTAGAAGTTTGACTTCTAGATTATAGATTGCTACTCTA
ACTGATTTAGCATTATTCATACTCTATTTTAGTCTATTAGATATCTTTAAACATAATAACTCATATTATCTATAT
GGAATACATAACTTTTAGAGCTCTTTTTAATTATTTTTTGAGTCATATTGGAAAAAATTATCATGCTAAATATAT
CGTAGTATGCTTCCAAAGAGATATTTATATCTTATGATGAAGCTATGATTTTTTCTTTTGTCTAAATTTAACTCT
TTCACTTTAAAAATCCATATGTATGACTAAATTTTTTTAGTTTGAGACCAGCTTATTCATAGATCCAACTTGACC
TAAATGATGCGATCCAAATTCGAATTTATCAGGTTGGGAATGAGTTATAAAATGATCTGACTCAACCTAAATGAC
CCATTAAGTCAAATATTTTAGCCATGGATCCGACCCAACTTGAACTAATTTGGGTTGGATGTAGATCCAATATTA
ATTAGGATCCGAACAAGGAATTGGGCTAAATCGAGGTTACTCGTGACCTGGTCTAATCAGACCCTTTTGCACCCC
TATTGGCAATGACTATCATGGCAACAAAGAAGAGTGCTTGCACAGATGCATCTGTTCCAATGATGTAAATTATCT
GAAATTATTGGACGATGCTATAGAAACCACTAGAATTTGGCATAAGAGGATTGCTCCTACAACACCTCTGCTCTT
ACAACACCTTGGCATAATGGTGAGGTCCACGTGCCCCAAATTAGCTGTGCCATTTCATGAGGTTCAAAGTTGCCA
AACCTACTCTTAACATGATTATCAATGCATCATTTCCCATTCGGTTCCTAAAAAAAAAATTGGAGTTCCAAATGAA
AAAAGAACAAGAGTTTGGATGAGGAAAAATGCCTCAAACAAACATTTCCTAAGATTTGGCATGAAAGCATTGCTC
CTAGAACACTATAGCATCATCACAAAATGAAAGAATGTCTCATGCAACATTAAATAATGTTAGATCGATGTATAA
TGTTACAAAAGTCACCATTATTATCTCTAGGTGGAGCAAGGAATTTTTTTTTTTTGGACTTAATGATTAATCTCC
TTCAATTTGGATAATGGAGTTTTAGTAAAAACTTGAAAGAAAAATATCTCCTAAAAGCACTTAAGTGTGTGCAAA
TGTAAAGTATATGTCCATGTAGACAAAAATGAGCAATCCAAGCTTCATACGAACCTGAAAAATGAGTCCGCCAAT
GTTGCCGCATATGAAGTCTTTGGTTATAAGCTATGCATTGTTCCTCTTTTATGTGAAGTCTTTGATCATAACATT
TCTTTTTCCATTTCTTTTAGACAATATGATGCTAATATGCTACTTGTTGAGGAATAGCCCCATGTTAGATAGGTT
AGGGACTCCGGTCGGTCTTGTATATCCATGGGTCCTTCCACCTATCAGCTTAAGATTTTGAGTTGTCTTTCTTCC
TTATTGATGATCATTGGCATCTCCAAATCCATGATAACCTCTGTCTACATCATTTATTGCAGATTGTCAAGGAAT
TCAATCCACCAACAATAAGAAGAAAATTAACGAGAAAACTTCCATTAACCTGTATGTGGAAAAAATAAGTAGACG
ACAAAAAAAAAAAAAGGACTAGAAATGAAGTATAAAATTTGGTCCAAATTTATCATTGATAGAACAACATTGCAA
AATGATAAATTTTTAATTTTCTGCTTTTTTTTGGTAAAATGGCTATATCTAATTTGCTCAATCTCGGAACCACGT
ATTCTTGATTTGTTAGATTGCCTGATTAGATATCCAATGACTCTGGTTTTGGCAGTTCAATTCATCATGAAATAT
GTTGGTGATACATATCTTTTAGCTGTTGCATCCTTGACAAGTCAAACTACTAAATTCGCTAAAGCTTTCAATTAA
ATCCACAAATTAGAATTCTGACATTTTTTTTGGTAGAGAGAATTCTAATATTAGAAGGACTAATAATGACTAGT

CGCAAATAGGAAACTAGTAACCAAGAAATTCCATGTGAACTAGCGACCGGCAATTGAATGTGTTGTGGGTAGGCT
GTGGGAGGACCTCTCTTAATAGTCCCACATCGGACAAACTCTGAAGTGTCATGTGCTTAAAAGGGATACGAGCAC
ACCTTCCCTCGTGAGGCGTCTTTTGCCCATAAGACGAAGGACAAAACTGTGAGGCCGTCTGCATGACGCTGGGCC
GGGGAGGTACAACCCCCGGTGGAGCCCGATCAGGGGCTCCAGGGACAACCCCCGGTGGGGACTACGGGGCACACT
CCTTGTCTGGACCTAGCCAAAGCGGGCAATACTTCGTGGGGCGGAGCGTGGTTGACTCCGACGACCTGCTTGACA
CATGGACACATCAGTGGCGCGTCAGGTAAGGGGCACCCGCCCCCTGTCCGCACATGACCCCTACAGAGTGGGGGG
CATGTTGTGGGTAGGCTGTGGGGGGACCTCCCTTAATAGTCCCACATCGGGCAAACTCTGAAGTGTCATGTGCTT
AAGAGGGATACGAGCACACCTTCCCTCACGAAGCGCCTTTTGCCCATAGAGCGAAGGACAAAACCGTGAGGCCGT
CTGCGTGACGCTGGACCGGGGAGGTACAACCCCCGGTGGGACCCGATCAGGGACTCCAGGGACAACCTCCGGTGG
GAACCACGAGAGCTGATGTGTCCATGTGTCAAGCAGGTCGTTGGAGTCAACCACGCTCCGCCCCACGAAGTATTG
CCCGCTTTGGCTAGGTTCAGACAGGGAGCGTGCCCCGTGGTTCCCACCGGGGGTTGTCCCTGGAGCTCCTGATCG
GACCCCACCGGGGGTTGTACCTCTCCGGCCCAGCATTACGCAGACGGCCTCACGGTTTTGTCCTTCGCCCTATGG
GCAAAAGGCGCCTCATGAGGGAAGGTGTGCTCGTATCCCTCTTAAGCACATGACACTTCAGAGTTTGTCCGATGT
GGGACTATTAAGGGAGGTCCCCCCACAGCCTACCCACAACATGCCCCCCACTCTGGAGGGGTCATGTGCGGACAG
GGGGCAGGTGCCCCCTACCTGGCGCGCCACTGATGTGTCCATGTGTCAAGCAGGTCGTCGGAGTCAACCACGCCC
CGCCCCACGAAGTATTGTCTGCTTTGGCTAGGTCCAGACAGGGAGCGTGTCTCGTGGTCCCCACCGGGGGTTGTC
CCTGAAGCCCCTGATCGGGCCCCACCGGGGATTGTACCTCTCCGGCCCAGCGTCACGCAGACGGCCTCATGATTT
TGTCCTTCGCCCTATGGGCAAAAGGCGCCTCGTGAAGGAAGGTGTGCTCGTATCCCTCTTAAGCACATGATACTT
CAGAGTTTGTCCGATGTGGAACTATTAAGGGAGGTCCCCCCACAGCCTACCCACAACAGAATGATTCTTGTATTT
ACCGGTAAAAGCCAATTCTTATCAGGAAGTCCAATTTCCACTCCAGCAGGAGCTTCCCGTAAAAGCAATTTAGGA
AACCAGTCTGATTCCTTTTGGAAACTTCCTTGGGAGCCCAAATTTAGAAAGAAAGAGGACTTCTATGGGTTAGGT
TATCGAGCTATGAAGAGGACTGGTCGAGGAGGGTGGAGAGGTTTGGAAGTGAAGGAGAAAAATCATCGCCTGATA
AATCATTGGTTTTATTTGTTCAAGGAGAGTAAATCTGGTGCAAGAGACATCTTCATCAGTATTTGTGAATTAATT
TGTAGCTACTTATGAAATGGATTCCAACCGGAGTAACCGATTGAAATTTATTTGTTTTCCTATAATTTTTGTCTA
TTTTTACCTACATATTTAATAATATTTTTTCTTTATTTATGATCATTTTTCTTGTCTCCCGCATGGGAGTTTTTT
CTTCGCCCACATTTCACGTTTCTTGCATTTGTATTTCAGTATTGGTTTCCCTGATGACAACCGTGCATTGGCAAA
TATTCATGAGTCATATATGATCTGACAAGTAGTCATTTTTCCATGGGCATGGGACAGCAAAATTTGGCGGAAGCA
CCTCTATTTAGAAGGTGTTTTGCTCACAATTAGAATCGAAAGAGATAGGAATCAGAATGGTCAAATCTTCTGAAG
CATTTGGTTCATGATTGGAATCGAAATCAGAATCGAAATAAAAATTTAAATCTTTAAAAGAAAGTAGAGATTGAG
TTTTATGTATATTAAACTAATTTCATTTCATCCCGAAATTAAAATTAGAATGGGATTCCTTTCAACCAAACAGTT
GAAATGAAAGTTACCCATTTTTATTTCCATTTGAAACCCCAACTCTCCCCACCCAGATCCCCTTAGTATAGATAA
CACAGCATCGAGCCATCAACTATATAGGAAATACTCTCCTAGGGAGAATGTTTTGCTTATAAATGGAGTGTACCA
AAGAGCTATTAACTAGATTTACCAAAAAAAAAGGGGAGCTATTAATTAGCACAAGCAAGCAATTGGAAAGAAGAT
TATTTAAAGCCACTCAAAAGGGGATGGAATCCTAGCACATAAAGCATGCTTGGACACTAAAATGCCTGCCTTCCA
TGATAACTTAGAGGGAATCATTAGGTTTCAAAGTCAACAACAAAGTGATCACTGTTATTTCACTCTTATGGCGGC
TTGACCATAAGGCTCCCAATCCCACTACAGAATGTTCAATTCGAAGGATAATTTTAGAAAGAGACCCTCCCCACT
ATATGGGCAAATCCAATAAGTGAGTACGTCATCCCAGCACCTTCGAGACCTGGCAAACTAGAAACTATTTTTAAT
ACCTGCACCCAGGCCGTCAGGATCAACTCTCTTCCGATCATTCGCATCCTACTATTTAATTGAAGTCCCGATGTC
CATCGTGACTCGAATCTGGATCACTTGCATGGAAGCCAAGGCCCGTTTCGCCAGGCCACTCCCTCGATGGCAAAA
AGTTTTTCTCTTTATGGTGGACCATGGATATCAAAGATCTACCTCAGACCATTATTATCTGTCAGCAAGTATCAT
GATGGCATACTTCGTTGATTTATGTCAATAATACATTGTCGGTCAAAATGCTAACAAAATTGAGAATCAAGGTTA
ACGACTTCTCTATATAAAAGTTTAAAGATAAGAAATCAAATTTAAATATAAAATCTTTCATAGCACAAAGTCTAA
GAGGCCTCAATTATCAAATCGATTCCTAAAATTATTTTTTGGATCTTTCTCAATGTTCCGACTTTTTTTTTTTCT
GATAAGGTGGGGTAATCATACAAGTCTATTATGAATATAACCAAAAAAGTCAATAAATAAAACATCACAAAAAGA
TAAAGGGGTGCTCTCCCCACTTGGGTGCTCCATGACAAAAAAGGTGACCCAGTCAGCCATACTGTTCACTTCCGA
TAAACATGCCGGATGTCCAGGAAAGAGCACCCCTTATCACTCTCCAGACATCACATAACAATTGGTGCACATGAG
CTACCTCGACCCGCCCTTGCAACCAGCCCACAACAATGGATGAGTCCCCTTCAATATCGATATGATCTACCCCTA
GGGTCCGCCTCACGCACATGCCAACCCAAGCCGTGCGAAGCTCCGCATAAGGGACAGTGACATCTACTAGATGAC
TTTCCCCTACAGCAATGAACATTAAGTCCAGGCCTCAAATTACAAACCCAATTTCTCTTCTACTGTTATACATGC
TGCCATCAAAATTAACCCTAATAAAGCTCATATGTAGGAGCTCGCAAGTAATGAACACCGAATGAGTCACTACAA
GAATAGAAAAAGAGTCCCAGGTGTCCCAGATTATCAAGATCAAGCTAGTAGATAGCTGAAGAATCTCTGCAGTCT
AAGCTAGTGCCCTCTCTAACATAAATCTCGCATGCTGCCTTCTTAACTCAAACAATAGACTATTCTTAGCCAACC
AAATATGGTACATGATATAAACTGCACTAATGCCAAAAAAGCCTGGAAGTCTCGCTTGAATATTCCTTCCTAATG
CCTCAAGAAAGAACTGAACCGATGTCGCTGAGTGGACAAAAGTCGGAAAAAAAACTGATCCTCCGACACACCTGCA
CGGTGCTCGGATATTGAAAAGAGGACATGGCCCACAGTCTCACCGTCCATCTCACAACTGAGGTAGGTGATGGGG
GGTCTATGCCTCTGTCTCTCGAAACCATCCTGATGGAGAGAGGGCCTCGAGCCATATTCCAAACAAAAAGGCACA
CTTGGGAGTGAGTGCCCCGCCTTCGGATCCAATCTCCATCCATAATGTCTTGGCCATTCACAGTAAGTCACAAGC
AAACGGACAATCATTTACTTTCCAAAAAATTGAAAATCATTACGAATCCTACAAATTCAACTTGTTCTTTTCCCT
CTAACTGATGATTGGAACTTCATCTAGGCATGAATCTTACTGAAAAGTTGACCACAATTGACGCTCTCCCTGCAC
TAAACCCATGAGCATGATATAGGATTCATAACAACTTTTCCTATAGGACACTTACCTAGACAATACTTAAATCCT
ACCCAGACTGATGTCGTGACGCTCCTTTAGTGTGCATTTTTTGAATGACAGATCATGATTCAGTTTCTCCAAAAT
GTATGCTATTACTAGCAAGAGAATTAGCAGTAAAAGTTGTAGAAAAGCAGCAAGCTGTTAACATTCTGAGAAACT
GTTGTTTTACCTTTTAATATGTTTACTGCATGGCATTTTATAAACCTATTGCTGGTAGCAGCATTGCAAGCTGAG

CTAGAGAGAGGCCTTGACATCCTTCGAGATTTGAATGAGTTACTAGTATCTTTACTATAAGATATGCTCATCCAC
TGATCATTTTTTGGAATTTTATTTAGACTAATCTAGCTTAAAATATGTCTGAGCAAAGGTAGTATGCTAATTTCT
GAACCCCATCATCTTCGGAACAAATAAATGAAACACAAACTGTCAAATTAACTAATTGAAGATGCTACCAATTTG
GAACAAAGTGACGAGAGAATTTCAAGAAATTTGCACTAAGAAAGAAAACTAATGGACAAGGAATCACAAAGTAAC
CCCAAATAGTTGGGAGAAAGATGGTTAATTCTAGTTCATATTCTAGATATACAATGGAAAAAATTCAATAAATTA
GATGTTGTAGAAGAGCAATGGATGGACAACTACTACATAACACATGGAGACCATCTGTGTGGCTCGTTTGGTATT
TAGGAAAGAAACTCACAGAGGTTAGAAGATACAAGGAACTCAAGCGGTGTAATACTTATTAGAAAAAGCAGCACT
CTTATATTTAATTATTTATTCACGTTCATTATCTCATTCGATTATCAATTGCTACTCTGATATGGCATAGAATGT
ATCAGCATCCAGTCTGATAGCATAATAAAGAAAAACTAACAGAAGATCCCTTATCATTTCAGCATGAAATCAATA
ATTGACCGCTACAGCAGGACAAAGGAGGAGAATCATCAAGTGCTAAATGCAACTTCAGAAGTGATGGTATGTCCA
ATTCTTACGCAACATTATCAACAACTGATAAATGAGCTTCTTCTTTGGCAGTGTACAAATGCGCAAATTAGTATG
TTTATATTAGTTTTGCCATCTCTTCTTGGATAAAGTTGATCTTTTTACTGCAAGTAATCTGGTATCTATTAAATC
ACCGAGTATGAAAAATAGCACTCTTCTTTTTGCTATAGAAATTGCCAAGACGCTGCATGATGTAGTCTGGTGTTA
CGGTTATGTACCATCACATGCATATTATTTGCAACACTTGTTTCGCATCATATATGTCCTGCACCAGAAAAGTAG
GGGAAAAATTTAGAAATTACATGTGTACATCAGCCATATACGTTTATCATAGTTCGATTGAAGAAGCTTAATTAT
GAAAACTTTAAAATGTATAGTTGAAGATAGTTGTTAAAAATGCATTACCACTGCACAATGGAATATAAATACGAG
ACAAAATGCATAAACTATATAAAACTTATGATTATCCATGTCAGACATCTGCAAAAATATATCATTGAAGTTCTG
GAGAAAAGTTAGTCAATAAAATAACGTCCATGGGAATAGGTTCTAAATATGGTTTATGTTGACATTGAAAATGCT
TATGCTAAAGTACTAAAAGTCATATAAAACAATGAATAGCAAAGGCACGATAGTTTTTGATAAAGGTATTCATGA
TGGAGTTGAGAGTGGTAATAAGGGGAAGATGCCATATCGTGGCAGGCCAACAATACGGGTGGTCCTAAAGTCTGT
CATCATAAATAGGTGCTACAATTACTGATTATTGCCACTTAGGAGAAGAATCAACCATGTAGTTCATCTGTAAAC
AAAAACATTATCGTTATGGTGAGATAAAAACAAGGATAAATTCGTATGGAAGAAACTTGGCATTTTGATAAAAAT
ATTTATATAGAGCTGAGAGTGGTGATAAGGGGCAGATGCTATATTGTGGCAGACCAACAATCAGGGTGGTCCTAA
AGTCTGTCATCATAAACATGTGCTAGCACTCTAGCAGATACTATGCTTTGGCAGGCCCCCGATAAGGATCATCCT
AAAGCCTGTCATCATAAACAGGTGCTAGAAGTACTGATTGTTGCTACTTAGCTGAAGAATCAACTCTCTAGGTCA
TTTGTAAACAAAAACAATATAATTGTTGTGAGAAGATGAAAACAAGGAGAAATGCTATATGGGAGTTATGAAAGA
ATCCTCTACAAAGTCATATTTATAAGCTTAGTTACTAGAAACAGGTGTGAAAGCTAATTCAGGTGTGGGTTTGGG
GAAAGGGATCTCTTAATTCTTTTTTTTTTTTAAATAATAAAGGGGTGCAGAAACAAGTTCCCATTGGATATTGCA
AAGCAGGTGTGGCACCCTATGCAGAAGTCTCCTGCTACTCAGGTAGTGAGTGCAAACAGCAGATCTAACGAAAAC
AGATGACACAATGCAACTGTTGTGGGCTGAATAATGAAAATTTATGCTTTGGCAATTGATGGAAAAAAAGATACT
TCCAAAAGATTATTACAGTTCAGTTCTATTATGTTGAACAAACATAAAGAAAGTATTAGACATTTTCATTATATT
TCAAGCATGGAGAACTATGTAAAGATGTGTTTCTGATGTGCTGCATCATTGTAATATATTATGGAAGAGACATAA
TATAAATTCTATCAAAAGAAAAGCCCACACGCTAGGCTGAGAAGTGGTAAAACTGAGATGGTCAGAATTGAGCAT
AACTACAGGAAGCTAAATACCACTGCAATCAGTAAGATAGCAAGGCCATGTCACAACTAAAGAGGGGCAATACAC
AATTAGAAGTACTCAGACTTACATTAATGGGTTGAAGAGAGGATGGGAAGGCCTAGAGAAACACAGATCGAAGTT
GAAAGAATTAAATTAATATACTCAAGAAGTGGAAGCAGCCATTCATATTTATAAGAATCAATGCTGAGTACTAAT
TCATAAATCTATTACAAGACAAGACTTGGCAATCATAGTTATGTTGCAAAAATTCATATCCAAGCAGAATACAGG
ATTCCACAACAGATAATGACCTTTATGTGCTATTTTTCATTTTCCCCGCTCAAGAAAAATGCAATCACAGGGATA
GACACTGAAACAGTCAAAAGTTCAAAACAATTTATAAATAGGCCCAGAGGAAGTAGACTGTCCATGAACTGCTCA
ACTTGGCCAGCTAATAGGCTCTGTTGAGCTTGTTCAACTAAACTAGCTAAGCTTTAACATCAAGTAAACCCCAAC
TCAATAAAAGAGCTGAGCCTCAACACTTCAGGCCTTGAATCTTAAGCCTGGCTCAATAACAAGCTTGTCAGGTTA
ACAAATAAGGAATCTAGAACATTAATTGTTTGGGCTCCTGTATTTTTTCATGCAATAAACTGCATCTGAGCAGGT
CAGCAATTGGCTTGTTTGTATCCCTATTTATAAAAATTTATGCCAAAGCATGACCATCAGGTTCTGCAGAATAGG
TAGTAAACATGTCATGCAGTGTGCTCATTGTGAATAGATTGACTGCAAAAACTGAGGAATTAAGCATCAAATGAG
CTCACCACTGAAAAGTCCTGAAAGCTGGCCCCAGCATATGGAAGACAATAAAGCCTTAACTGTTCGTTTGCATGA
TTCATGTCATGCTTGGTTTGTATTGCTATGAAAATTGGTGTTGATTGATGGCCAGTTCTTTGTTAGACTGCCATT
TTACAATCTCTTCCTGTTGACCAGCAATTTTACTCTACCCCTTAACTTCAATGGCCATTTACTGTTTAGTCCCGG
TCCTGTCCATGCTTAAGCATAAAGGTGTGCAAAACAGATGTAACTATGTGCAGCACATAGTGATGTCTGCCACAA
TAGATGAGGACCTAGAGCAAGAAATCAAAAAGAGTAAACAATGAGAAAATATTCAGAAACTTATAATGCTAAAAG
TGAGAAAGGTGCAGGATGTTTTTTTTGGTTCTTTCTTTTTATTCATTCTGAGCAATTGTGGGGATCCCAGCACCA
TTTCATTGAATAGGTGAAGTGATGAAACAGAGGGGAGCATCGACAAAACAGTGACTCAAGCCGCTAAAACATAAA
AATGAGAACCAGCAGAAAAATCCACCCAAGTGCATCAGTTACAATGTTTCGTACATGAAATATAGGTCACTGCAT
AAGCTTCCTCACTTTCAATTATTCATGGTACGCACTCAAAATTGGGACCTTCCCAAAAAAGGTTCAGACCTGGTA
TGGTAAGAGGTAAAGCATAGGCAGCACAAATAGGATTTAGGTCCTATTTGGCTAAGCTTTTAGAGTGTCAAAAAG
TACTTTTTCATCATTCAAAAGTACTTTTGGATGGTACAATGATGTTTGGTAAAAAAATCAAGAAGCTATTTTGGC
TTTGCCAGAAAGCTAAAAAGGTTTACTTGGAAAAAGCTCTATTTTGAAGCTTTCTCCAAAAGTATTCTCTGACTA
ATATCAGAAAGCTGTTTAAAGTAAATAAAATTTTTTTAATGATCAAAATGCCCACTAACAAATCTTATAATTACA
TTTTATATCATATCATATCATATTATTATATTATGAATATAATATAATATAATAATAAATTAATATTATGATATA
TTATAAACATATAACATAATCTATTATAGTATTATTGTGTTACAAATATAATTTGATATAACAATATTTAATAAT
AATATAATATATTATATTATATTACATCATAATTATAATAATATTATATTATAGTATAATATAATATTATATTAC
ATTATATAATAATAATATTATATAATCCAATGATATTTTAATATTTAATAATATATAATATTATATTATATTATT
ATTCACTATGTAATACATGCTATATCTTTTATTATCATTTTATCATACCGAAAGTACTTTTTTATTTTAGTTACC
AAACATATATTAAGATTTCAAAGTATTTTATAAATGTGGTTACCAAACAGTAAATAACTTTTTATAATGACTCTA

CTTCCAAAAGCTCTATTTCTTAAAAGCTCTATTTCCAAAAGCTCTACTGCTAACAGCAATCCTAAAAAGAGCCTT
AGAGCATTTGAGATAGATTCAACTTATAGTCAATATAGATATTTTAAATCCCATGGGATGAGGCTGTCCCGATTT
TTTTATAAAATGGGATGCACCACTGTTCCACCCCATCCTGACATTTGGGACAGGGGATATCCCAAAACGTCCCGA
TCGAGATGCCAGAATGCTAGCAGGATGACCATATCTTGACACATGGGATGGTACCCCATCCCGGTATCCCACGGG
ATATCCTGCTGGGACCTGAATCATCATAAGGCATTTTACTTTAATGTCTGAGGCTCTGAGCCTTAGTTGTTCTAG
CAATTCTAATTATTCAAGCACACCATTTTCATATAACCAAGATGAGCCTACTAATCGGTTCCAGTGGCCGACCAA
ACCCTATCAAGCATTTGACTTCTCTCTCTACATCCTTGTTAGGAGGATGGTTAACAATGTTTGTTCCCCATAAAA
AACACAAGTTTAGGAGGATGGTTAACAATGTTCAGCTCTAGTGGTGTTCCAATATCAACTGTACATTTTAGCAAC
AAATATTTTAATAAACTACTCAAAAAAGAAAAAGATGGTGCAAATGGCTTCCATTCTGGGTTCTTTCAAAATTTC
CAAATCAATATTATTATTCTTTTCCTGGAAAGATCCTCAGGATAATTCATGGACACGAAGAGGCAGTCAGCTGGT
CAAATTTCTAGAATCATGCCAGGCCCTTTATGTATAAAAGATTGGAACATACAGAAGTAAAAGTTAGTACTAACT
GTCAACAGGTCACTAAAATCCTAGATAATGATGCAAGGTAACATAATGTAACAGCAAGCATATGATTTTTACCTT
GATCCTCCCAATTACTTGTTCTGAATCTCTGAGTACACTGTCTGATTTCCACTCAATCTCAGGGTTCTAATGATG
TAGGTCCCTGAATTCAGCACAACTAGCAGTGTCAAAAGTGCCTACAAGAATCTGCTACCCAAACAAGCAATAACA
GTTAAATTTGACATGAGAATTACACGGTTACAGAGAGGACATGACTGTATATAATGCAGAAGGGTATTGGACAAG
ATTTGCCATACCAATTGGTACCAATGCATACCAATATACCATAGTGTACCGATACTAAATCAGTATGTGGTATGA
AGGGCGTACCAACACTTGGTATGCCAAAGCAAGTACCATAACGGATGAACCAACACTATAATAGGATGGTATCGA
GACAGGATTTGATATTGAGATGGTGAACCTTTATTAGAAATGTCAATTGAATTAGGAAGAAAAGTCGACAACTAT
GATGTACCTGGGAATTATCGTTGCTTTATCATCATTGTTTAGCTCAAATGCTTGAGTGTCATTTACTAAAGAATT
TTAGGTCAGGGATCCATGAACTCTGCAGGAATTTTGTTTACATCTAAGTTGGACGTGCAATTGCTAAGACTTGAG
CTCAAATGCTCAAGGGTATGTTTTTAGCTGCTTGTCAACTTCCATATCCACCAAGCAAAACAGTTTCTGCAACTT
TCATTTACAAAATCACTGTGGACTCTCTTACGCAGGCCTTCTCTACAAGCAGATGTCTACGTCTCACTCGTTTGA
TTGAAGAGAGCCTTTGTAGTGTCAGAGTGTCCCAACTGCAGCTTCCTGTCTGCCCCTCTCAACTCAACCATCAAT
CCAGTTCCATCATTTCGGTTGGTGTCCTTTGCCTCTTTATCATCATATAAGCATTTGTTTTTCTCAATCAACTCA
TTCCAACAATAGCAGACCATCATAGCATGGCATTTTAATAGCAAAAAACAAAATAACAATTAAAGAAATAATATA
CATAATACGAAAAGTATGGATAAGACAGTGCAAAGATTTCCTTACAGATCATACAACAATCTTCTTTGTTGAGAG
TCTTGAGACAGGATCTCCTGCAAGATAATTTTAGTCATCTCAACCTATACAGGGAAGTTGTCAGAACTTGGGAGA
AACATGTTCTGCTGGTAATATAAAAGATGAGTTCTAGATTCTTAGATCTTGAATAATCAATAGATCAGGAGTTCA
TTGATCACAATTTTTTAGAATTTTTCTACAGCCTAGCAATAGGATGCATATGCAATATGCATTTTAGACTTTTTA
TTTATCTAGAATAGATTCAAATTGCCGGGCTCCAAGTATCAATCAATTTGCAGCTTTTTGTACACATAACAATAG
CCACCCTTGATTCAAAAAATTTTCGACAAATGAGATCAAGATAAAAACATATGTGAATGTTGAACTTTATCAACT
TTCTGAATGTTTGAAAAGTATGTAGTAGTGAAAAATAAACAAGCAGCCTGTGATGAAAGCAATAAAATAGAACAT
TTAGCTGCTATATGTCAAGTTAATAAGAGAAGAGAAGATTAAAGAGTCCTACTTGTTCAAACATTAGCATATAGA
GATATCACATACAGTCGAGACCTTGCTAGTATTCATATGAATATAGAAATTTGAAAACACATGTTTGAATGGTGC
TAATGAATGAAAAATTTCAACAAACTATGATGCTTCTGGTGCCCATAAGCAAGGAGAGAGTTGAGGGGGGATCCT
AGGTCAACATAGAAAACTATCTCTACGAAGGAACTAAAGTCTACCAACATACAAGTCACAAAATAATAAAGATGA
CCTTATTGAAAAAGGATATTTTGCTAAGGAAAACCACCTTGTAAATAGAAGTTCAGTAATAATCCATTTCATGCA
CATAACTTCCTTACAACTTACTCGCACATAAGGAGTAAAAGCTAGCTATGGCAACATGCACATTTCATGGGCTAG
CACATGTTGTGCTATGCTTGGACTTAGGGTTCGACATCTCGGTACCAAGCTCCACACCTGTGCTAACATGTTACT
ATGTCAGTGCAGAGCCAGTCTAACATACGGAGTGTCAGTACGCCCCCCCATATCGCATGCTAGTACCAAACCAAT
ATGGTACGTCCTGTACCTCTAGTTAGATATAGTACAACAAGCCTTGCTTGGACTGATCTAATAAAAAAGCGGAAA
CAATTCCTTCAATTAGCTTAACTCGGACCTCTCAAAGAAAATACTAATAAAATAATGGAGTAGCTATACTAAAGA
CCACAAAAACCATTCCGAACTGAGCGATTCGAGGTATACCAAACCTATCAGCTAGTAATCAGGGATGATTTGGCC
GATCAATCTAGTCTCAAAACAAATTGATTCCATTTAACTAAAAAGAGAATCACCTTGCCTAGTGTTTGTCATACC
GCCCCATATCACTTGGTATGGGACATAACATACCATATCGATATGGCGTACTATGTTGAACCATGTATTGATATT
GTAATAGAATTATGCTGGCATGGGGTCCCATATCGAGACGGCAAACCTTGACATTGCCTCTTTCTTATGTCATGA
TCTCTCCTAAATTCTCTCCCCTCTCTCTCTCCCTCTCTCCCAATCCATCTCTTCCTCTCCCCTCTCTCTCGATCT
TCCTCTCCTTTCTTCCTTCTCTCCTAATCCACCAAACCAGCTATGATCACATCGATGTCGATGTGACGATGACTA
CGACTAAGGGTTTTGCATTAGTGTAAGAACACCATGGCCATCGCCACCTCCTCTTATTGTGATGAAGATCCTCCT
TCGCCACCTCAACCATCTCATCTCCGCTACCTCCGGCCCCATTCCTTGTCTAGATTGGCACCTCCATCAGACTGG
GTGAGGGCCCCTCTTTCCTCCCTCTCCCTCCCTCTAGTTTCAATATGTATTGGATCCGTATGGTATGCACCAATA
CCATACCGAACCATTCTCTGACTAGTACATCCACCAATACCAGTTCAGTAAACCTTAGGCTAGACTCTTTGTGCT
TTCATTAATTGAAAAATAAAACTGATTACAATGCATCTTATTTATATCAGTGACATCTGAGTTTCCAAATCTGAT
TCAAATTTCTTTTCTACTTAAAAGAGGTTAGAAACAAGGTTTTATGTCCCGACGGACAGGGGCGTCCCATCCTA
TTCTTATGAGAAAATGGGACCTGGGACTTCGAAACCCATCCTTGCAGTGAGAGAAAAAAAAGAGGAAAAAAGGAA
ATGAAGGAAAGATAAAGAAAAGGGAAAAATAAAAGGAAAGAAGAAGAAAAATGAAAAAAAAAGAAAGGAAGGGAG
AAGAAAAAGAAAAAAAAGGAAAGAAAAGTAGAGAAAAAGAAAGAAAAGAAAGAAAGGAAAGAAGAAGGAGAGAAAG
ATGGAGGATATCTACTGAAATACATGATCGGGACTGCTATCAGAATGGAATCGAACGATGGGGCATACCGTTCCA
TGGAAAAACCGGAATACCTTTGTCCCATGGGATTTAAAATCTTGGTTAGAAATTTTCTAAGGTAGCAAGGAATCC
ATGCTAAGATGAAATTCGTCCAAGAGCTCTCTCTCAATTCCTGCATCAACTTTATCATCTGTTACATAGCCTAAT
TCCTCCCAGATTGGAAGATAAGCCATCCATTTCACCCGAAGAGGATGCTTTTGCATTGGTCGGCCCATCTCAAGG
CTCAAACATCATAGTTCCCCAACATTAGTAGGGCACAATGAACCCTTGAGCTTGTCACATCCGTGTAAAACTCAA
AAAGTTACCCAATATCCAAAAAAAAGGATCATTATAATAGGATGTTTTATGGGTAAGTTATGGTCTCTAAAAGTT

GAGCTTCATGATCTAGACTTAGATATGGCGGATCTTGCTCCCAAACCCTATCCAAACTTACCTGAAGTGATCAAA
TTGGTCCACAATGGGTCCAATGATCCTCTTGGATCATGGACCAATATGCGGTACATCTATATGCCAATGCATGGG
TCTCAATGCATGCCAAGCATCAAAATAGCACAATATGTGCCATGTCGGTGCCAAATCAACACTGTACCAACACAT
GACACTTAAACCAAACATCCATTGCTGCTAGAATGGGGAACTCTTTGATAGAATTTTTGACTCCACACTAGAATC
TTCACTTAAATGAAGTCACCTAGCCTCTTATGGAGTGCCTATCAAAACTAAACATAGCCTCTCCCCTATAGCCTC
ACACCATAAGAAGGTCCAGGAAAAGTCTCTTACACCTAGCTCCTTGTCTAATTCATATAGTACTGAATTACAATA
AGGAAAACAAATGAAATAATTAAAAACAACTTTTCTAAGCCAAATAGCTTTTTCTAGCTTGATAAAGACATTCCT
GAATAGAGAAAACTATGGAAAATAGAACAAAAGGTTGTTAAAATTCATTTCTAATTCAAAATAGCCAAGGAAAGG
CAAAGTCATAACTATGTACAACCATGTCAACAAATTTCAACAAAGTAGCCTTTCACCAAGATGACCATATCTCTC
TCATTCAAGCTCTAATTAATATGATTCAAAATTGCATTGGAAAGTTGTTTTGGTGCTAAATAAGATAGAAGTACT
TTGATTTTATAATTCTGCTAGCTTTAGTAAGCATTCTTTTGGACTATGGTTTACGAAGTTGATCTTTGAAAGGCA
GCTAATCTATGGTATGTTTTTCTCAACTAAAATAACAATACTTTGACATTGGTGAGCTAAGGGTCTAACAAGCAA
TGCCTCACAACTATGTGATCCAATCGGAGAGAAGGAAGATTTCCTTCAACCCGCTCAACTCATATCTCTCAAAAG
AAAAAGGTAGAAGAAGAGGAGAATTTTTGAAAGCAAGAATCAAGTCGCTTTTCTTTCTCTTTTTTTCATTCGCAT
TGATTAAAAATAAGATACATATCCTCTATTTATAATAACGAGGTCCATCCTTACACAAATTGAGTTGGATTCTTC
TTTAGTTCTAATGAGACTCTTTTACCTAAAATAGATATGTTGAGTAGAAATAATCTGAAAAAAGCTAAAACTCTT
CAAAAGGTAGATCTCGACTTCTACATCAACTCTACCTTCTATCATCCTATTGGATTCTTCTCGGATTGGAAGCTA
CGCTCGATCAAAGTCTATCTGAAAAAAAATTCCAAATTAATCAACCCATTTTGGAGCCCACACGAAGGAATTTTG
TCCTCATTGTTGGGGGCACCGGCCCCTTTAGAGGATGGCATCACATCATAAATCTCAAAAAACTATCCAATTTCA
AAAAAAAATATCATTGAAATCGGACTCGTATTACCAAGTTTGGTTTTCATAAGTTTAGCTTGCAAATTAGCTTCC
TAATGTGGCGAATCTTGCTCTTGGACCCCATCCAGCCCGATCTGTAGTGGCCAAAATAGTCCGCATCGAGTTCAT
AGATCCTGTTGGATCACTATGGCAACTGATAGATTCAATGTAGTCTCAAACCCATTATGAAACAAAGTATGTTGT
TTTGGACCTCTTTTTGGACTCTGGAGCCCTCTTCGGGTCAAGCTTCATGTCTAGCATGGATCAGTAGCATGTAGA
GGACCCTTCACTCACATCAAACCCCCTTGGCTTCAAAATTTGTCTTTTGATGAATTATAGTGGAATAGGATATGA
TGGTTTTTGAGGGCGAGCCTTGGTGATATAATAAGATTGCTCTATTATGACCTAGGTGTCACAAATTCGAAACAT
GGAAATAGCCGCTCCACATGCAATAGGAAAGTTGGGTAAATCTAACCCTTCGCAAACTCCATTGGCGGGAACCTC
ATGCATTAGGGCATCCTTTTTGGATATGATGGTCTTCAAACATGCTTGTATCAAACACTTCACTATCATAAACTT
TCATGTTCCATCACTATCTTGTCCTTCTCCCATGATACTTTTGGATCTGTGGCTTCTCTATCACAAGCTTTCAAC
ATAATTTTATTAATATATGGATTTTGTTTAGATTCTAGGGTAGTTCTATACACTTTGATAGGATGATTTTGATGT
AAATCATTACTCAACTATGGAGTTGTGCTGTCCTCTCTTATGGATTGTTATTCATGTTGCCTCAACCCTAACCTT
TTTCTTTAAATTCTAGGATAGTATATACCCTTTGATAGGAATATTTTGATGTAAATCATCACTCAACTATAGAGT
TAGCTTCTTGCCATCCTCTCTTATGGATTGTTATTCTTGTTGCCTCCACCCTCACCCTTATTCTTATACATTGAT
ACTATAGTTGTAATACCCAGGCCGAACAACTTGGGCCTTTTCTAGCCATAAAAAAATCAATCCAAAAAGAACTGC
ACAATTTTTAAAACAGGACCAAAATAAAAAAATTAAAATAAGCCAGTACGTGCGTTGCATGTGTTGGGAGCTGGG
AAAATACATGAGAACTCCCGAAGGAGTTCTTCTCCGATTGAAATCCTAAATCCACTTAGAAATTGAGTGGAAATT
AAACTCTAGTTCCATCCCTTTGATATCTATTTGAAGGCCTCAACCCTCTTGATTCTCCCTCACCAAAGATTGGTC
GTCGTTTGCCACCCTTTTCCCCTCCTTTTCCTTTTGATTGCCAGCGGTTTAGTCTTGATTTGGCTGCCAGAAATT
AGATCAATACTAATCAAAAATCAAGTAAGAACTAGCATTATTTAACTTGCCCTCCTCTCCTTTTGCAGTCCCTAA
ATCCCAGATTCTGGCCGATCTATTGTTGGATTTCTTCTTGAAAGTCGAACCATGCTTTTCCCTATTTTTCCATTA
CTTTCCTTTGATTTTTCGATGCCTTTAGTCACCACTAATCACCAACAAGAGGACCACCGACTGCACCACCATGGG
CCCAACCACGGCTAGTGACAGCCATCGTCAGTCTTGAGCCTATTGTCTCGAGAGGTAAAGGGGAAGTAAATCCCC
TATTCGGTTAAAGAAGAAGGAATGAAGAAGATGGAAAAAAAAGGAAAAGGAAAAGGAAAAAGAAAATAAGAAAGA
AAAGAAAAAGAGAGAGAGAGAAAGAGCTTATCTCTCTTCCCTCTTCCATCTCTCTTCCTACTTTCTCTTTCCACCTT
CTCTCTACTTTCTGTCTTTAGAAAATTTTACTTTCTCTCTCTATGAATTTCTCTCTCAAGAATTATTTCTCTCTT
TATAAAATTTATCTCTCTAAGATTGATCAAGTAAAAGGCTTTCTTTTGATGATTTTGATCGACTTTAGTGAAGGA
TCCTAATCCGTGGTCATCAGACAGCATGCCGGCACCCATCTTGATTGGATCCGGATACCATCACATTTGATCTAT
TTGATTTTTATGGAGCCAATTGTATCTTAAATCTCATCTTGATTAAATGAAGTTATTTGATTAAACTGATCTTTG
ATCATCGAGCACTGTGTTGGCCTCCATCTTGACTTTTGTTGCACACTATTGATTTGATCATCCTTGATCTCTGTT
GAACTATCTAGATCTTAGTATGATCCTCATTAGATAAAGTTAGATGACTGACTGACTCAGACTGTTGGATGCGAC
TATCGTTCATCATATCTCTCATGATTATTTGTATCCTTTCTGATTATTGGATTTAATTATGCATAAGGTTATAGT
TGTCTGGATAAAAAGATGTCTAGCAATGGGATACTACGATGTGATCTATGAATTGAAGATTATAAAAAAAAATTA
TAGAATTATGTGGATTTGTTGGAGTACGTGTGAGATAAGTAATGATTCACTTTTTATAGATTTATCAATAAATTA
TAAAATATTTTTGTATTGAATTATTTATGATTTATAAGTTCAATGTATAGTAATTTTGGATTGCCAAATATGCCT
TCTTTTTATATCTTGAATGATTTATGATCGATTGTATTAAAATTGCTATGGATTATATCGATTGATTTCGATATC
GTCTGGTTTATGACTTTTTAAATTTGAATATAAAATAGTATTTTCAATCGAGATGGATTTTGACTCATAGCCTGA
TCATGTATGGAATCCTACTCGTGGGGATTAAAAATTGGTACTTCGATATCCTGCCAGTAGGGGTTATGCACAGGT
ATTTTGATACCCTACAGGTGGGAGTTATGCATTGATATTTTGAAACCCTGCCAGTAGAGGTTGTGTGCTGGTATT
TCGATATCCTGCTAGTGGAGATTGTGCGCTAGTATTTCAATACCCTGCCAGTGAGGGTGTGCACTAGTATTTCGA
TACGTCGTCATTGGGGGTTATGTAGTGGCATTTTGATTTAGTGCATGGCTATCGAGGTGAATTGGATGTTTTGAA
AGAAATTAATTTATGAATGAAAATAAATTTTGAAAAGACTAAATTTGCATGGTTCTTATTTTAATTATTGTTATA
TATTTTGAGTTGATGCACTTTGCATGCTTATTTTCAGTACATCATTATTACTAAATTTATCTAGCTAAAGTGTAC
ATTATTTATTGAGCTGTCTAGCTCATTACTATATATTTTACTATTTTTATAGATTCAAAGGACTAGCTTGATTTA
GGAATTCTATATAGGAGTGTGAATTAGAAGCAAAATTTGATAATCTTTATTTAGACTAAGATTTATTGAAGTTG

ATGGAAAACTATGGAATATTATTTTATAAGACATTTAATTTAGTTATTTGAATAAAGTTGAACATTAATTTCACC
GTTGCTTTACGATATCATCATAAGATGCCTTGCATGCTTGTGGGAAAGTTCTCCATAAGAACGTGGCGATTGCCA
TGACCTATGGCTCATGATCTAGGATCGAAGGCATGACAATAGTGCTCCTCCAATTGTCAAGTATTTTCTTAGTGT
TTAAAATTTTATTAAACAAATTAGTCAACCTCTCAAATCTCCATAGAGATTTCATCAGGCCCATCCTCTTTTCCT
ATTTGCATCTTTTCCGTAGCCTTTTTTACCTTAGATACTCTAGTTCTACATTGATAGTTAATATTTCTAACCTCA
ATGGACCTCCCAAATTATCTACATGTCTTTCATTGAATAGCATATGAAAATAGTTTCTCCATCTTTCTTTGGTCT
CATCTTTCTCAACTAAAACTTCTTGATGTTTATCCTTGATGCATTAACCTAAGTCTCTAGTTTTTTTCCTCCTAA
TCTTGCAGTCTTTTAGATATTTTTCACCCCATCCTTAGTTTCTCACATGTGATATCTCTCCAGGTGCCTTGAATA
TAGCCTCTCGTATCACCATCTTAATTCCTTCTTGACCACCTTATAGTTTTCATACACTTCCTCATTCCTATATTT
AGGTAGTCTCCTAATAGCACTATGTTTGAGCCTTAACTGCTCATTGAACTTTCTTATTCCACCACCAAGTGTCCT
CACTAGATGGTTTTTCTTTGATTCTCCTCAAACCTCTATATTCACATTCTTAATGACAATATTATATGGAGTGTA
TAAGATGGAACATGCGGTTGCACCCAAAGTATTAAGCCTCAATAAGATGGAGCATATCTCGGCTGTCCCATCCTA
TTCTCATGAGAAAATAGGATTACAATGGGGTTAAGACTCTGAAACCCATCCATGCAAGGAGATAAGAAAAAAAGG
AAAGATCAAGAAAAGGGAAAATGAAAGGAAAGAAGAAGAAGAAAATAAAGGAAGGGATAAGGAAAAGAAAGAAAG
AAGGAAAGATGGAAGAAAGAAAAAGAAAGAAGGAAAATAGAAACAATGGAAGGGATAAGAAAAGAAAGAAAAACG
AAAGGAAAGAAAGAAAGATAGAAAAAAGAAAGGATGGAAAGAAGAACAAGAAAGAAAGGAAAGGAAAGGAAGAGA
AGGGATGAGAAATGGGGGATATCTACCAGATGCATGATTAGGACTATCATCGAGATGGGGTGGGATAGCGGGACA
TTCAATTCCATAGAGAAATCAGGACAATTCTGTCCCGCAAGATTTAAAACCTTAGTTACACCATCAATATATGAG
TTTCTTATAGTATCAACAATATGAATACAACATGACACATTGTTAATGCATGAAAAAGTTGTCCTACCATAATAA
TCACCATGAATTGTGATCGTATTTGTGCTTTAATGTGCAAATATATCTTTAAGCTTCTAATACTAGCTTGCATGA
TAAATTGACTGGAAAAATGTAAACCATAAGTATATTCAGCACAAATTTATCGAATGATTATATAAGTATCATGTA
GGGCTTTCAAAACCACGTCGAACCACCCAGTACACCCTGTATCGAGCCGGACAGATAGGAAACTGAGATGGTTCC
AGCATCCAAATCGGTTCTCGGAGGGACAAAGAAGAAAGAGAGGAAGAGAGAGAGAAAGGGGGAGAGAAGGAGAGG
GAGAGGAAGAAAGAGCCTCCGACGGTCGTCGAAGGTCGTCCGACATCTTGCCAAGGTGGGTACACACAGAGGGAG
GGAAGGAGAGAGAGTGGGAGGTCCTTACCAGACCTCCGGAGGCCGTCGAGGCCTTTGGACAGATGATGCCTCCGT
TGCGAGATCTTCGATGGATAGTGAGCCCATGCTGACGGGCCTGAAGGCGGTCGAGCCGGAGGAGGGCCTCCGTGG
CTTCACCCCCTTCTTCATTTTTGAAACAGGGGTTCGCCCCTATTTCAGATTTTTTTTTTGATTTTAACATTTGAA
GGTTGCCTACTTCACAATTTTTTTTTAAAAAAAAATAAGGAGTCGGCATATCATTTGCCAACTTCGTCCGTTTAA
AAGAACAAAAATACTGAATAGGGATGAATCCCTATTCCGCATGGGAGGGCGGAGGCCGTCCTCGATGTGCTTGGC
ATCGACCGCCCTTGACACTCGCTCGCCGGCCTGTCACAGGCTTATCCACAGGCATGTCACAGGGGGTGCCGACGT
GGGTGTGGCACGGTGTTGCGAAGGATTTGGCATGGCCAAAATTTGCAATGCTACCTGATCATTTGGAACCCCCTG
GGTGAGATTTAGGGGGATGCCTTGCTGGGGTAGTGAGGGAGCAATGGTTCGCCAAGGAGCGGCATAGCCGTGGCA
TGATATCGGGGGGCTCGGGATGCGTGATATTTGGAATGATGGTGGTCCGCCAAGATATATGTGGATGCTCAAAGA
TTCCTCTTGGTTTGAGGAGGGGCGCAGAGCTTTAGTCGCGTACCCGACTCCTCCAGACCCGTAAGAGATGGGTCC
GTCATGGCACAGTTGTGTGGGCAATGGTGGTGCCACACCCGAATGGTCTCATATGGAGGTATTCGGGGCAATGCT
GGTTGGACGAGCTAGCCACACTGTGCAGGGAGGGGATATCTCAAGATTTTATCCTTGAAGGGTTTGTCCAGACTA
CTCTATGCGTTCTTGATCTCTGTAGTAGAGCGAAGAAATGATAGAGGACTAATCGGGCAAGGGTGCGGCCTACCC
AAACAGGCAAGAGTCTAGTGAGCATCACAAGGTGATGTCTTGTGAGGATCCACGGGAATCGATCGGTCGTGGAGA
ACAAGGCTTATGGTATATCGATCAGATNNNNNNNNNNNNNNNNNNNNCATTTATCAACCAGTCGAAGATATGTAC
TTGATGAGGCAGACTTGGATGCCGATATAGTGGCAGTGCTCAATATACGAGTAGGCTGTGGACCTAGGATGTTGG
GTGGTATGATCATGTATCCCCAAAATCGTGGTAATGCACAGTGATGGGCGAGCAGGCCATGAATATTGTCAAGCG
ATATGGCCATGGATGGAAAGGCATATCAAGGATGCAAGACCGAGATCGTGGCACAAGAGCTTGTGGGGAAAGCTT
GGATAGTAGAACCTCGGAGGACTCGGATGTGTACAATCAACTGTAGCTTTGAGAAGCTATAGGGCATGTCAACTT
GGTAGTTTAGCTGGGCGGCACTAGAGGTGGGTTTGCCCAAGCCTCATGCATAGTAATGGCATTGCATGGTCGTGT
TGTGTAGATTTGGTATGTGTAGTATTGAAGGGCAGCTTGGTGGGTTGCGGATCAGTGGCCCAGTGGTTATGTCAT
GGCATGAGGGTATGTCGAGGATCAAGATATTTGGGATGAGCAGTGCTTGGCTTGAGGGGCAAAACCTTAGAGTGG
ACATAGAGGTGTTGAGCTGCGGAGCACTCTTGGAAAGAGTATGTTATGACATGGAGGTGCGCATGAGCATTGCAG
TTGATCCGAGGGGTCGAATTGCTTACGAGCACTTGATGGTGTCATCTGGGTTCAGTACAACAGTGTTGCTAGGCA
GAGGCATAGAGGCGCATGCTTGGCTATATGGTTGGACTGCTGCGTGCTTATTGGTATGTCCAAGGAAAGGACTCA
AGAGGGCTGGCCTTGATGTGTAGCATGCTGAGGCACCGCACAGGTGTTTGAGTAAGAAAACACTCACCCCGTGAC
AGACCGTCGGAGAAATCGCATCGATGTCGGCGTCCGATGATTCGATGAGTGTCTCCCTCTCTCTTCCTCTCCTTC
TCCCCCCTCCTCTCTCTCTCTTTTTTTCTCCCTCTGGGCCCTTCTCTGTCGTGTTGGTTTGCACCATTCCGATCG
GTACAGACCCATATCGACTCATACCGTTGGTCGGCCGGCACGAATCCTGGTTTTGGCTCTGGGAACCTTGGTATC
ATGTTTAGTTGAACACTAATATGTTGAAGTTGGTTTAGTGTAAAAGAATTTTAAAAAATGCTTTAGTATGTTATT
TCGGATCTTTCGATGGGTACTATCAAGGGTTCGCTAAATCGAGAAAGCATGCCAAACCATGCCAATAGCCTACAG
GTATGGTTCTAATGTAAAATTCGGAACACGGGCAAGAGGGAGAAGGAAAGAGAGGAAAAGAGAGAGAGGGAGGAA
AGGAGAGGGAAAGAGACCCTCTAGAGGCTGGCTGAGGCTGCTAGAGAGCCATTATGCCCATTAGAGGGCTGGCGA
GGAGGGGGGAGGGAGAAAGAAGAGAAGAAAGAGAGAGAGAGAGATGGGGTAGGAAGGAGAGCGGAGAGGGAGATA
AAGCCTTCGAAGGCCCGTCGAGGCTCTGGAGGCCGAAGTTGGCTAATCAAAACAATGTCTTCTAACTTTAGTTAA
CTTTTTTTATTTTACTAGCTTCACTGTGAGTTGACAAAAAATTTAAAATCATGATCAAAAAAGGGCAATCGCCCC
TATATTCGTCGGTATAAGGCAATGAGGAGGATTCAAGCCTTTGTGTCCCTCCAATGGCCACCACCAGCCTCTGCT
CCCTCTCCTCCTTCCATCCCTCTCCCTTCCAGTGATCTCTCTCTCTCCTCCCTTTCCTCTCTCAGTTCAGAGTTC
CTTACTCCGTCGATCATCTAAGGGCCGCTACGGCTCTTCAATGGCCACCACTGGTGTCTCCGCAGTCTCCCTCTC

CCTCCCTCCCCCGCATCTCTTCCTTTTTTCGGTTTCCCTCTCCCCTGCTTTCTTTGCCATGTTGGTTCAAGCATA
GTACGGAATGGTATGGTGGTGTACCGAACCATACCATCGGCCGATCGACACGGCAAACCTTCGATACTACACCTT
TTCATGCCGATTTGAGACAAGCGTGGTATCATGATAGCCCCTCGATGACCCAAGCATAATACATGATGCTTTAAT
ATATGAAATCTAATTAAGCAATGCAACATATTAAGAATTAGCATTTGACAACTGATATAATAGATATATGCACCT
AATTCAAAAGGAGCGAGTACCTTAAAATGCTTATGTTACGAGATTCATCGGTCATCTATTGCATATGACTAATTC
AAAAGGATGTAGTGATGCCTTCCCATATAATGTACAACACCAATTAAGTGGTAGGACTATTTGATGAACCAATCA
AGAGGATGTAGTATACCTTCCCATATAATGTACAATAATAATTCAAGTGGTAGGACTATTTGATGAACCAATTGA
ATCAGTAATTTATGTGAGGTTAAGTTTTGTTTACATAGCTTTATTTAATGCATTAGCTCGTGTTGGAAATTCCCT
AAAATATCTCATCTAATTATAGGACATTTATATAGTTCCCTATTTTTCCTCTCTTTTCTTATTCATTGTAATCAG
ATTTTTTGGGATTTACACAGGATTTGATCTTATCCCTTATTTCATGGGATTTATATGGGATTTGATCTTATCCCT
TATTTCATGGGATTTATACCAGATTTGATCTCTTAAGAAGATCATCTTTTTGTATATTTATATGGGATTGATTGC
CAACATGAAAAATAAAAGAAATTCACCCTTTTTCTCATGGTATCAGAGCCCTTTCCATAGGGCCTATTTTTTTTG
CCATTGCTTCTTTTCTATCTGTCTTCATCACTAAAGTCCACAGCCTTCATCGCTGTTGGTCCCATTACCATCAAC
CTTTTTTGCTTTTATTGCAGAAGTCGTCATTGTTCATAACTTTTCTCTGTCAACCGAATCTGACTGCACTCCCAT
TGTCAATCAGTCTTTCTATTGTTGCTGTTATTTGATCCCTTCTCCTAGTCAACATGTTTGAAGCTTCTAAGGCCA
CAGTGTCCAGCCCTTCCAGACCCGTCTTTAGGATAACTCAATCCATTTTAAATGGTGAACTGCGGGCCATGAACC
CTTCCTATCATCTTGATAGGAAGAATTATCTTCAATGGTCACAATAGGTGAAAACATTTCTCAAGAGACGTGGTA
AGCTAAATCATTTAATTGCTCTTGGTCTGAGTCCCTCAGATCCAAAATTTTCTGCATGGGACAAAGCTGATTCCG
TGATCATGTCATGGCTCCGGAACTCAATGCAACCCGAAGTCAACCGCAATTGCATGTTTCTCTCATCTGCTAAAG
AAATATAGGACACTAATTGACAGACGTATTCCAAAGTCCAAGACGTAGCCGTCATTTATGAACTCAAAAACAAAA
TCTCTGTTGAGAAAAATACCTCGAGATTCGATCTATAATAAGCCTAGAATAAAGTCCGCGATGAAAAATGAAATC
CAATGAGCCCAAGATCAGCCCAAATGGAGTCCAGACGGAGAAACTATGCGCGAAAGAAGATTAGGAGGACGAGTA
TGGCATATAGGATGGCGACAAGCCCACAGGCTGTGCGGGCCCGTAGACGCACGGGCTAGGCCCAGCGAGCATGCG
GCCCAGTGCGCGGGTGCGTGCAAGCCAGCTGCGGACGTGCGTTAGCGCACACAGGTCGGCCCAGTGGGTGTGCGA
GCACGCAGCCCATGGCGTATCACGGTCTACGGTGGACCGAAGGTCCATGCGAGAATGCATGGAGCCATGGACCAA
GAGCCCATTTCCCCTGAGTTTCTTGCAGTCCACGATGGATCGGAGTCATTTTCCAGGGGATTTCTTATGGTTCAT
GGGATTTTTTGTGGATCGGGCGATGCAGATGCGCAATCAGACAGACGAGAGATGATCCGAGAGGTGATCCGACGG
CTCTAGGAGGCCTCAATCTTGATTTGATGGTTGGGAAGCTTGATCATGCGCAATCAAGTGACTGAGAGGGCTTGC
AGTGCTGGTCAGATGGTCCAGACGCAGCTGGAGTCTGACCAAGAGTCTTTCTTTGATGCAGGTTGGGTTTTAGGG
CCCTTAAACAGGCCTGTGGGTGAACAAAGCCGAAAGGGAGCCGAGAGAGGCTCTGGAAGCAGCTGAGAAAAGTTT
AGCAAGCACCGAGCACGAGCAGCTGGCAGAGCAAACCAAAGCAAAGCGTCAGGCAGAGCAGCTCAAAGGCATGAT
GTAGTGGTTTGTTCCATAGGAAACGTCCAGGGGCGTCGAGGATCCTAATACTAGCAGGCCTGTAAGAGGGTCTCC
TTGAGAGATTTTGTGAGAGCTTTTGTGAGGATAGCTTCTAGTTGAGAGAGATCTTTGTACAAGGGTTGAGTGGTG
AGTTTCTCTTGTATTTGTTTGATTTTTTTCTCTTTCATAATGAAGCTTGCATGCCCCATGGAAGTAGCACTGGGA
TTATTCCATGTACTTGATTGTGTGTTTCTGTTTTTATTCCTTTCTGATGCAACGGATGGTACCGGATCCACAACA
AGTGGTATCAAAGCAAGTGATATCACAACAATTGGTATCAGAGCTGGGTCGATATCAGAGTGCAGGTGCGGTGGT
AGTGATCAAGATTGAAGATGGAGAAGACAGGTGCAATCAAGGTGGAGATCTATCGCGAAGGGATGATCGATGCTC
TCTTATACAAGGAGCCGTCTACATCGAGGATGCCAGTGGGAGCCCAGTGCGAGGTGGAGCTCTAGGTGGAGATGG
ATTAGGCACAAGATGGGAGACTCGGGAGGACGATCAGTGACGTCAACAGGAGGCTATCCCGAGTAGATCTCAGGT
CATGCAAGATGGAGATGGGATCGAGCAGTCTGGCTCGACTCCCATGTTTGCCCATCTATGAGTAGTAGGCGTTTG
CCCCAGGGCATGGGGGCGAGAAGATCCAAAAGCTCTCAAAGTCAGATGGAGGCCGAATATTGAGTCGAGGTGGAG
ATTGTTGAAAAAAATATCTCGAGATTCGATCCACAATAAACCTAGAAAAAAGTCTGTGATAAAAAATAAAATCCA
ACGAGCCCAAGATCAGTCCAAACGGAGTTCAGACGGAGAAACTACATGCAAAAGAAAATCAGAAGGTGGATGGGT
ACGGCACGCAAGATAGCAGACGAGCCCACAAGCTGTGCGGGCTTGCAGACGCATGGGCAGGCCGACTGCGGGTGC
ACGACCTATGGCATACTATGGTCCATAGTAGATCGGAGGTCCGTGCGAGAGTGCATGGACCAAGAGCCCGTTTCT
CCTGGGTTTCTCACGGTCCAAGGTGGATCGAAGTCATTTCCCAAGGGTTTTTTGGTGGTTTATGGGATTTTTTAT
GGATCGAGTGGTGCGGATGCATGATCGGATGGATGAGAGATGATCCGAGAGGTGATCCGATGGCTTTAGGAGGCT
CCAATCTTGATCTGTTGGCTGGAAAGCTTGATCAGGTGTGATCAAACGGTTGAGAGGGTTTACGGTGCTAATCAG
ACGGTCCAGATGCATCTGGAGTCCGATCAAGAGTCTTTCTTCGATGCAATTTGGGTTTTAAGGCCTTTAAACAGG
CTTATGGGTGAACAGAATCGAGAGGGAGCCGAGAGAGGCTCTGGTAGCAATCAAGAGGAGTTCAGTGAGTGCCGA
GTGTGAGTAGCCGACAGAGCAGAGCGTCAGACAGAGCAGCTGCAAAGGCATGATGTAGCGGTTTGCTCCACAGGA
AGCTTCTAAGAGCATCGAGGACCTTGGTACTGGCAGACCTATGAGAAGATCTCCTTGGGGGATTTTGTAAGAGCT
TATATGAGGATAGCTTTTAGTTGAGAGAGATCTTTATACAAGGGTTGAGTGGTGAGTTCTTCTTGTATTTGTTTG
ATTTTTTTCTCTTTCATAGTGAAGCTCGCATGCCCCGTAGTAGCACTTGGGCTATTCCACGTACTTGATTGTATG
TTGCTATTTTTATTCCTTCTTTCTTCCTACCGCAACAGATGGTACCGGATCTGTAACGAGTGGTATTAGAGCAAG
TGATATCACAACATGTTCCACACCAAATAAGGTTCTCTTTCTATCACTGAATATTATAATCTTATGAATGGCTTG
TGGCTTAAGTTAGATCACTACCAAGCCATCAAGATGTCTTGCCATGAAGATACTAATATCCTTCTAAGTCTTGTG
GAATGAGATCGTCTCTTCGAGTTTCTTGCAGGATTGAATGTCGAATATGATCAAGTTCATGTTTAGATTCTACGA
AGGGAGTGACTTCCACAATTAAAGGAGGTCTTCTCTATCATCCGAGCTGAGGAAAGCCTTCGAAGTGTAATGCTC
TAACCCCTCTCTTCAAAAGGATCCACCTTGGTATCCATCAAGGATCTGGATGGTGGTCCCAAGCCAAATACTTTT
GGAACAAAATCAGATGATCGTGCTTCAGATAATCCTTCCAAGCAATCCAACCGTGATGGTCTTTGGTGTACTTAC
TGCAAAAAGCCTCGCCACACAAGGGATAATTGTTTTAAAGTTCATGGCAAAGCACAAATCTTGAGTAGAACCAGA
GGATTCAAAGGGCAATAGCGGAATCAAGCAAATTTCACCAACAATAAGGGACCTATATCTAAAAAATTTCAAGGG

CTGAATCTTGAGAATTCAACAAAGAGGAGGTTCAAAAATTATGAGGTTTTCTAACCTCTCTGTCTACTTTCTCTG
GTTCATGTTCATTGACTCAATCTGGTAAGTGCCTTGTTTCTAATTCTCTTAATGCCGATACATTGCCTCGTTCCT
CCTCTTGGATGCTTGACTCAGGAGCAACAGATCATATGACCCACAGCTCATCTCGATTTGAGATCTACAAACCAT
GCCCCAAAAATCAAACAATTACCATGGCTGATGCTTTCCCTATTACCATAGCTGGCAAAGGCCAAATTTCCTTAG
CCCCTTCCTTTTATTTAAAAAATATGTTACATGTTCCAAAATTATCCACTAATCTAATCTCCATTCATCAGCTTA
CCAAAGATTTAAATTGTTATATCACATTTTTTGATACCTATTGCATCTTTCAAGACAAAGTCATGGGGAGGACGA
TTGGATCTGCTAAGGAACGGAATGGCATTTACTACCTTGATGCTGCCAGTAGTTCTTCCACTCCTGCAAATCAGT
TCACTCTCTCGTGCACATCGTCCAATTCTGCTCATCCTTTCTCTTCCAATGAATCTGAAATCTAGATCCAACATC
TTCGTCTTGGCCATCCACCCTTCCTTTTGCTGAAAAATGTTTTCTTTATTGTTCGATAATGTTGATGTCCAAAGC
TTTCATTGTGATACTTATGAATTTGCTAAACATCATCATGTGTCATTTCCTTTTGAGTATTAATAAATCTTCCAA
ACCTTTTTTTCTTATACATATAGATATTTGGGGTCCATCTACAATTCCTACTCGTACTGGGGCAAAATGGATTGT
GTCATTCATTGATGACTGTACTAGGACCACATGGATATATCTTATGAAGAAAAAATCTGAGGTCAGCCATATAAT
TCCATCTTTCCACCAAATGATAAAGACACAATTCGAGATATCCATACAAAGGATTCATTCTAATAATGCTCGAGA
TTATTTTAATTTCACCCTTGCTTCCTTTCATAAAGAAGGAATCATTCATGAATCCTCCTGTGTTGACACACCTTA
ACAAAATAAGATAGAAAAGAGAAAAAACCGACATCTACTAAATGTCACTCGTGGCCTTCTTTTCCTTTCTAACGC
CCCAAAATTTTTTTGGGGAGAGGCTGTCCTTACAGCAACTTTCCTGATTAATAGTCCCTTCACGTGTCCTTGGTA
ATCAATCTCCCATTCACCTATTGTCTATCTTTCATTTTGAGCTTCATTACCTGAGTAATCTCCCTCTTCGCATCT
TTGGGTGTGTCCCCTATGTTCATATTCATAAACAATGCCAAGGAAAGCTTGATCGTAGAGCTCTAAAATGCATTT
TCATGGGCTATTCGAATACACAAAAAGGGTATAAGTGTTATCATCCTCCTATCCGAAAAATATTTATCTCCATGA
ATGTCACCTTTAATAAGAATCAATTTTATTTTTCCAAACCCCATTCTTCTCTTTAGGGGGAAAAGCCCGAAGAAG
ATTCGAAATGGGTTGTCCTTGATAATCTCTAAAGTCTACAGCCCATGCCTCCTCCATATGAACCATCCTCATATG
ACCCAAGATGAGACACTGAAAATCAGCAACAAAAGCAGTCTGAATTACTGCAATATTCAAGGAGAAAAAAGATGA
TTTTCAAATCAATACAGACCCAAGACTCCAATCCAAAAGAACAAAAGTCAAGTAAACCAATTTCTCTCACTAATT
CTATTCTTAATGAATCTGTTTCTGAAGAGCCTACTTCTCAAGACCTAGATCTCCCAATTGCCTTAAGAAAGGCCA
CCAGAAAGTGCACTCAACATCCACTCTATCCAATTTCACAGTTTGTCTCCCTTCATCACCTTTCTTCATAACATA
AAAATTTCCTCACTACCCTAAACTCAATTGATATCCCTAAAACCATACAAGAGGCATTAAAAGACAAAAATTTGG
TACAAGCCATGGAAGAAGAAATGACTGCACTAGAGAAGAATAAGACTTGGGAGGTTGTGACTAGGCCAAAAAGGA
AACAACCATTCGATGCAAATGGGTTTACACTTTAAAATATAAAGCAGATGGGTCTCTTGAGAGATACAAGGCAAG
ACTAGTTGCAAAGGGGTATACACAAACCTATGGGATAGACTATCAAAAAACTTTTGCACCAGTAGCAAAGATGAA
TACTGTAAGCATTCTATTATCTCTTGCTACTATGTTTGATTGGGATTTGTAGCAATTTAATGTGAAGAATGCATT
TCTATATGGAGATTTGGAAGAAGAGATCTTTGTGGAGCCATCATCCGGATTTGATGAAGATTTTGGAGCTGATAA
AATCTACCGTTTAAAAAAAGCTTTATACGGCCTAAAATAATTTCCTGGAGCATGGTTTGGGAGGTTTACAAAGGC
AATGCTAAGAATAGGATACAAACAAAGCCAGGGAGACCACACATTATTTATAAAACACTCAAATATAGGTGAAGT
TACAGCCTTACTAGTGTATGTGGATGATATTATTATAACAAAAAATGATGTAGGTGAAAGAGAATCTCTTAGACA
AAGATTAGCTCAAGAATTTGAGATCAAAGAACTCGAAAAACTGAAGTATTTCCTAGGCATTGAAGTGGCTTACTC
AAAAAAGGGTATATTTATCTCTCAACGAACATATGAACTAGACCTTCTTAAAGAAACTAGAAATCTTTGCAGCAA
AACAGCAAGTACGCCTATTGATTCTAATCACAGGATAGGAAGAACTAAAGAGAGGATAGCTCGGTTGATATGGGG
AGATATCAGAGACTTGTAAGAAAATTAATTTATCTCTCGCATACAAGGCCAGACATAGCCTACTTTGTGAGTGTA
GTCAGCTAGTTTATGCATGACCCAAAAGAAGAACACTTGCAAGCAAGCAATTTATAAAATCCTCCACTATCTTAA
AGCTACATTAGGTAAGAGAATCTTGTTTCAGAAAAGAGCAGAGCTTACTCTTGAGTCCTATACTGACGCTAATTA
TGCTGGATCATAAGTTGATAGAAGGTAACATCTAGACACTGCACATTCCTAAAAGGTAATCTGGTGACTTGAAGG
AGTAAAAAAATAGAATGTGGTAGCAAGATCTAGTGCTGAGGCTGAGTTTAGAGCCATAGCTCAAGGGATTTGTGA
ACTACTATGGGTAAAAATAATCTTAGATGACTTAAAGATCAAGAGAAAGGGAATGATGAAGCTATACTGTGACAA
CAAATCTACCATTAATATTGCGCATAATCCGATTTAGTATGATCAAACCAAGCACATTGAGATAGACACTTTATA
AAGGAGAAGTTGGACAGTGGCTTAATTAGTATCCCATATATGCCATCAGAGAATCAATTAGCTGATGTTCTAATA
AAGGACCTACCTACAACAAAATTTTAAGAAATTATTAGCAAGCTGGGAATGGAAGATATCCACTCACCACCTTGA
GGAGTGTTGGAAATTTTCTAAAATCTCTAATCTAATTGTAGGATATTTATATAGTTCCCTATTTTTTCTCTCTTT
CCTAATTCACTGTAACCAGATTTTTTGGGATTTACACAGGATTTGATCTTATCCCTTATTTCATGGGATTTATAC
GGGATTTGATCTTTTCCCTTATTTCATGGGATTTACATGGGATTTGATCTTATCCCTTATTTCATGGGATTTATA
CCAGATTTGATCTCTTAAGAAGATTATCCTTTTGTATATTTATATGAGATTGCCCATATGAAAAATAGAAGAAAA
TCACTCTTTTTCTCAGCTCGAATTTACCACCGTTCACCTCTTAGACAAGTGGTGGTGGTTCAATTCTATGCAGTG
GCTATCCTTAGTGACTATCCTTATTTTGGCTCGCATTTAAAAATCCTAGTTCTGCAAGCACAATTTCAATTGCCT
AGGAGCCTAAGTTCCAACCTAGGTGGGGTGCCTTGGGTTAGGGATGTAATTAGGAGGGAGTATTTGATGGATGGT
GGTTGTTCCATCATCTAATAAATGTTTTAATTTAGACATTCCATGATCACCATCTTTTATTTGTGAATGTTGATA
AATATTTTCCACAATTTTTATGTCTCTTGAAAGCATTAGATAAAAAGAGAGATAGAATCAAGAGCTTATTTGCTG
AAAATGGGGTAAATTTAAGTGGCATTGTAGTCGTGTGACAACTTCATGAATGTTCTCAAGCTCTTGTGTCAGACA
GTACAGCAGTAAAGATTGCTGGCTTTTGTTGCTTTTAAAACTTTGCTCCTTATGCGAATAAAATTAGCAATATCT
TATTTATACACAAAGTAGTCACAAGCATCTTTCTTCTTTTCTAATTCATAAACATGCTAAATGGATGAAGTTGCG
ATAATGTTGCACCTTCTTAACAAACATAGATGGCCGAATTTAAGAAGTGGATGCATGTTTGTGAAGATAGTTCTT
AATTGAGATCTGCATTCCTGTCCATAATGAAGCTGATCTTTTTGTGTCAAAAAACAGAAGCTGATCTTTCAGTTA
TTTCTGAAAAGGATTATAGATGTCTCATTACTTGATTCAACACTTCATGTTTACAGAGTCCAAACTCATGAACAA
ATGGCTTCATTCATGAGATTAAACCTTTTTTCCTAAATAATCATGTATCAATGGATTATGCAGTCATTGAAATCA
AATGATGATTCTTGTAAAATAATTATTGGTGATTTCACATGTGGCTTTGCAATCAACTTTATTGTTCAATGCCCC

AGAAAATACATATGATAGCTAAAAATAAACTTAGACAGTAGAAGTAGAATACAGCATCAAACCTAAATTATATTT
TTCAAGGTTTTTAGTCTTGGAGATCAGCTTGAGATTTTGCACCCCTGTACCAGCACATGGTACACTACCGGAGGC
AAGATAACAGTTGCACCATGTGTAGGGACACTACCATATTGAGGCCCTCTAATGTACCAATATGTTATTGGGTTA
GTATGATGCCCAGTACCAGTGGGTGCAATACACTTATTTTTTATCAATAAATAGTTATTTGATGCACATATATAT
GTAAAGACAAGAGAGACTTTCTATAGGTAAAGAATACGAATGATGCATGGTCATAGACATACTGAATATTCAGAA
GAAAATTTGAGAAAGCATATTTCTCATAGAATGTCAATACTCTAATATATTGATATGAGATTTTATTCAACTATT
TATGAACTGCTGATACTACACTTTGATGCAGGCTTTAAGGAATATTTCTTGTAGACTTTAACTTTTTCCATTTTT
ATATTGTTACTAACTTACTTACTAGTACCTAAAGAACTACATGGGGGTTTTGTTTTTTCTAGTTTTGGCAAAGGG
AGGCAGCAAGCTTAAGGGAGCAACTGCAGAACTTACAAGAAAATCATCGGTACCATTTCTTAACCTTGTTTACAT
TCAAAGATTGTTCCTATGCCTGACTAGTCATTGGCAATATTCCTCTAGACATCGTACCATAAGGAAAATATTTAA
ATAAATAACCATCTATTTTGCTTTTAAAAAAATAGAATTACTCAACATGACTCAGACTTTCAAGCTTGTTTAAGC
ATAACTAGTTTCCTCAATGAAAAATTTAACAAGTATTCATCAATTACCAGGATCTAGATTCTAGAGTGTCAAGAT
CATCAGGAAAATAATATATAAACTAGTAAAAACATCAAATTTTGTGAATGAGTAGAGATGAAGTATAGAATCACT
AGTTCTCTAGTTGCCCATTCATCCTAAGAGTAAAATGTATTTCTTAAGACTAATCGCATGAAGCAACATTTTCAT
GCTGATATGTCACCTTGTCTTCTTGTTTACCATATTGCATGCATTGACTAAATCATCAATTACACCATTGGTTAT
GACAAAACAGCAGGTTACTGAAATTCCAATAGTATGTCAAGATTTAGACAACATAAAACTGATGGTACATCTAAG
GACTTCACTTCTCCCTTGAATACCCATAGACAGAATTGGTCCACAGGCCCACAGTTCACAAAAACAGTGACAAAC
TGACAACTGTCTGCACCATGGCTAATAACCTGTAACTATCAACTTCATATAAAATTATTATTTGCTTCAATTATC
AAATTGAACCTCAGCATATTTTATCCAATCATATAGTTTGGACACGAGCAAATGGCCTGTGGCTTGAGTGCTTGA
GAAAACAGTAAATGTGCTAAAAACAACCTAGTAATAAAATGCTCTGACATTGAAAATTTTCCAATTATTCTCAAT
TAAGGTTTTCCCTAAATTTTTACTGACTTAATGAAACCTGCGTTTAATGAATTTATAACGCCTTCTTTGAATAAC
TTTTTTGCAAATATTTCATAAAACATTTCATTCATTTTTCTAAAGTTCTCCATTTTTTTTATGTTTCTTTTTATT
ATCATTCAATCTTCATTTTTATTCATTCACTTTTCAATCCTCAACATATTCCAAAGTTCTTCCAAAATGTATAAT
AGCTTACAATTTTTTTCCTTGATAACACATTCTTGAATAACCATGCTTAAAAAGTATCCAGAAAGCAACATGCTG
ATAAGGTACATACTAAAACATATTAGAAATCATGATCAAATGAACTTTTCCTCCTGCTAGTATACTATGTAATGT
AGACTGAATAATCAACAAGCTACATGTACTATTTGATTTGACTTATGGATCCTTCTTTTCAAAACATCCTCTTTA
TTTTAGCGTTGGTTGTTATTGATATGTGATAACAGTTACAGTCTTCTGATATACCAGATTTTAAAATCTACGACA
TTACATATTCATAGAATGGGATTATCACACTGATCTTGTAGCTTTATTACTATTCCTGGAGTTATGTCTCAGCCT
CCAGACACCACACTAATATATAAATAAAAAAGAAGGCAGTTCTGACTGGTCTTTTCTGTTATCTAATTTTTACGA
GTTGAGAATTAGACTCTATGACATCATGAACTGCTCCAATTTCAATGTCTAAAATTGCAAACCACAGAATGCTCT
AGAATAGATGATCATGTGAAACACTGCTTGGATAGAGCAAATTACTGTGAAGATTTCTTAGAGTTCCACCAAGGG
TAGACTGCCAAGACTGCAATTAGAAATCAATATTATCTTGCCTATAGGATCTCAATATCTGTTCTGGCGGAAAGT
GAAGGCAACAGTCAGAGGACTGCTCCAGTACTTCACTTCCAGTGTCTCAGCCTCACAGCATATATTGCTAATAGC
AGCAGCTGTAACAATCTTTTAGATAGGAGCATTGAGATCCTGAATACATTTTTTTTTCATCTCAGGCATTTCAAT
TTAGGATTTTGTCCCAATTTACTTTGGTGGTAGCTTAGATATTAAAAGAGTACAAGCTTTAATAGATAATTTCTG
GTTGTATACTTCATGCAGAGAATAAAGATGATATGTTAGAGAATTCTTCTATCATGAGTTTCCTACAAAGAAGTC
TATGTGATGTTTTTGCCATGGAACATAGGGGTGCTGTGGTGTCAATTTTGAATGCCATAGCCCAATTTACTTAAG
AATCTAGAAGGATCACCAGTTCTATTGTGGACTAATTTGGCCACCTGTGGTAGGTCTAGACAGGGTCTGAGAGCA
AAATCCACCGTATCTAGGACCTGATCATGAAGAGTAACTTCTGAAGGCCATAATTTTCTCATATATTGTCCAGTT
TCAACATGCCATGCACCAAAATGAAGCCCTTGACAAGGACTATGAGGATGCAACAAGGTTGTCACACCCGCATGA
ATATTGGGGTCTCAAAACAAGCTGATCACAGTAAAACATTTCTTTTAGATAATACTTTGACCAAACACATCTTCT
AATGAAGGACACAAGGCAAAGTTGATGTTGAAGTTAAGATTTAACTCCTAGACAAGTTTTATCTTTATATAATTA
TCACAACCTTGAACCTCACCAAAGAAAACTAGCTAAAAGATATTATTTGGGTTCATCAGCCCTGTATAAGCATCC
AAGATCTACCCAGCATCCCAGCTAGGTAAATCACCTCCACCCCACCCATGTTTTGGAAAAAAAATCTAATAATAA
CTTAACGTTAGGTTGAAAAAAAACTTGGGGGTCTATGAGGGGCCTAATAATCTTTGATCACCAGGTTCATGCTCT
ATGGCATGAATGATCGAAAACAAGAAAAATTGGATTTTTGGCTTGAACCCCAAAACTCAAAGGATAAAAAAAAAG
GTTATAGAGGTTTCTCTAATTCTAGATTATTCTATGAACAACATGCAACGTTAAAACTTCAGGATGCATTAAAAA
CCATGGCTAGAGGACTTAAGTCAACATTGTTACCAATTTACCATACTTTTTGAAAAAGACATAGCATTGAGACAA
CAAAACATAAGAAAAACACAAAAAAGAATAGTTTCGGAGCATAAATAGGTCATACCATCTGACCATGCAAAGTAT
ATTAGTGGCTTGCTAGCAAGTCAGCTTTCGGCATACATATCAAATAGATCATAGTAATTGAAATGTCCAAAATCA
AGAATAATACAATCACTAGTAAGATAGTGCTTGACAATAAACAAACACCGTATGCATCTTCTATAAAAAAAGACT
ATCTTTTAAAGTTCTACTCTTGTTGCCCTATACCCGTCTATTTATTGTCTAACACCATTTTACTAAGGTCTTGCC
TACTGAAGCTAGAGCAATAGTTCCATCAGTTCCTCTAGGGATCTCACTAGGGGGGCAAGGGTGTGCATCTCCTAT
CTTGGGAGGTGCTTTGTCAGCCGGTTGACGAGGGCAGTCTTGGCATTTAGTCCTTGATGGTGAGGCGAGAGGCCC
TTTTGGTGAGGCATGCAACCAGGCTGCTGGTGTGTCTGGAGAGCCCCTGGAGTATGCTGACGGCGGCCAAGTATG
GCCCATGCACTTCTAGTGCTGGCATTCAGACGAATAGGGCCAACTCCTTTATGTGGAAGGAGATCTCCTCCCATG
CCTATGTGGTGGGGACTCAGATCGGATGGATGCTGGGCGATGGCCACGCAGCGAGCTTTCATCATGACTCTCCGC
TAGCTGACCTTCCCCTAAGTAGGTGGCCGACCTTCGTTAGTGTTGAGGTGGGGAAATCTACGAGGATCTGTTACC
TATTTCAACTTGGGGGTGGGGGCTGGAGCCCTGGGGCAGTGGCTCAGATTTTTGGGGCTGACTTGGTGGATAAAG
TGCTATTGATCGCTGTCCCTTCCATCCTAGTCCAGATTTGCGGGTGTGGAGATCTGCATGTAGACCGAGGGTTGC
AATAAGGGATTTATACAAGATGTTTTGAAGGGAGCCTGTCCAACATATAGATAGTGCTTGGATCTAGAGGATAGC
TATTCACCTGAGAGTTTGTTTGTTCATCTGGAAGGTGGTCTAGGGCCGTCTCCTATTGAGAGCGGTCTTGAGGGA
CAGGGGTCTGGCTCCTTCCGCTTGCCCCTATTGTGATGCGGAGGAGGAGAGACGATGGAGCATGTGTTGTTCCAG

TGCCCGAGAGCTGTCCAGATCTGGCGACAGATAGGTCCACTTGCAACTACTATTTTGACTGCAACATTGACCCAG
TCCTTTGTGGAATTTCTCCAACAAAGTTCAGCTTCAGATTTGGTTATGCTTGATGGGATCCGTGTGGCCTATTTC
GCATATCACATTTGGATATCTAGGAATAGTTAGATGTTAGAGTTGAGACGATCATCAGTGGCTGTAGTGGTGGAG
AGAGCTTTAGCCCAAGCAATGGAGTTTGTTCACTTGTTGCTGCCTGATTTTGTTTGGAGGATGGTGGACTATTGG
GACTCTCGCTAGGCTCGTCGAGCAACCCATCGGGTGTTCACCTCTTAGGAACCCCCACCCCCAACATTCCTCAAG
TTCAATTTCGATGGCAGTGTATTTGGCTCGAGAGGAGGTGCAGGATTCGTGGCCCAGACTCGAGACTGCTGGCAG
CGGGTGAGAGTTGCCTAGTGGACACTATGGTCCCGAGGGCAGAGCTGTGGGCAGTGTGGGCTGGTATTGCCTATG
CGAGGCTGGCTCTTACAGTAGACCGACTCATCATTGAGGGTGACTCAGGCACTATGGTTAAATGGATTCAAATGC
GGGATACAGAGGATGCTGCTCACCCACTTCTGAGGGATATCGCGATGCTGCTGAGGGGGGCCACCATCGTATGTG
TCCAGCATGTCTACAGGAAGGCCAACACCAAAGCAAACTAGGTTGCCTCCTTTATGGCGAAGCACCCGAGGGATG
TGATTTGGACTGACATGTCCACTATGTCATGTGGTTTCTGTGATGTTTTATTTTTAGATATTTTTGGTTGCGTTC
GTACCAAAATAGTATGAATATACCGTCACATCAAAAAAAAAGTTCTACTCTGCAGCATGAGAATAATTAATAGTA
TAATTTAGTTAAAATCAAGTGTTTAAATATGATCGGAATGTTATGGGATCAATACCAATTTGGAACATCGGTTCA
CATTTGAATTGGTACATACCAACGATACCAATCTGCATCAACAATTCATATATACTGGAAGTTGGTACATATCGG
TTCCTTTTTATTTTGATTAATATAAACCATTCGATCCATACCAATCAGAGCGGTATCACATAGCCCAAGTTCAAA
ACTGGAATAAGCAAAGTTCGCTGTGCCAAAACCAGATCCTGTGCCGGTGGTATGATTTGGCATGCCCCAAACCAT
TCTGTACCAAGCTCGAACCAACATGGTGAAGAAGCAGAGGAAGGAAGAATCGAAGAAAGATACAGAGAGAGGGAG
AGAGAGGAACAAAGAGAGTGGTCAAGGATGCCGATGGTGGCTGCCACCTAGAGGGCAGGTAGTTGGGTCGAAACA
AGGGTAACCACCTCTATTTCGATCGCTTAAGTATTTTTTATGCTCATCCTTCTCTCTCTCTGTCTCTCTCCATTC
TCCTCTCTATGGTGGAGGGTGGAGCTCCGAGGCCTCGGCACCCCTCTGGTGGCCTCGACGAGCCTCCGATGGCCT
CCGACCCTTCCCTCCCCCCTCCCTCTTTTCCTCTCTTTTTTTTCTCCCTCATTGGTGTACCATTTTTTAGCTGAAA
TTATGCCGACTGACTGCTGGTATGGTTCAGCACATGTCGAACCAGTCGGTTTGCGATGGTTCGGCAAGTGCTAGG
TATAAGGCTTGGTATTGGTCTTTTAAACCTTGGTTAAAATTCTCTCTTCCCATATCCCTTATATATTTTTATTAC
CTCTTCCTGAATCCTACAATTAAATATGCATCAAAATCAGATTCTTATGTCCTTGTCCACATAGAATGTACCACT
GATGTCATTAACACTTTCCACGATGGATGCTACTCTCAAATATTATCCTTGTTTCAAACTAATTTTTAATACATC
ATAAGATGCTTCAAGTCTACTACATGATCCAAAAGTTTGTGATAAAAATAGGTCTCAACATTATCCATGTACTAT
AGACATTAGATATAACAAAAGATATATATAGTCATTATATTTATAAATGATTTGTTGTAGCCATAGTTTGCCATA
CCGATCTATACTTGTTGCCCAGAAGGTTAGCAACCCAAGAGGAGGGTGAATTAGGTTCTTTAAAAACTTCTATCA
AATCTAAATTTAAACAATTATGTGCTTCAAATTAAGCTAATTTGAGTAATTGTGAGGTGTATGCAATGTGCTACA
GTGGAATTAGGATATAAGAAACATGCAAGAATTAAAAATAATGCAAAGCAAGTAAGAAAGCAAAGCACAACACAA
ACACAATAACTTTATAGTAGTTCGGTACAACCCCACATCTACATCCACTCCCCAAGCTTTTCTTAGGAATTCCAC
TATAATCAGTTTGATTACAATTCGATTGTTTTATCCGAACACACAATCAGTTGATTTTTTGGGATCACTAACTAA
AATCTTGACACCGATTATTTTTTGAATTCACAATCAATCTACTTAGTTTTGTGAGATTACCAACCACAATCTTAC
AAGTCCAATTTCGGGCTTGGATGAGCCTAACCCAAGTTTCTTTCATCCTAAAGAAACTTAAAAAGAAAAATCCAA
AATACAAGATAAAGAAATTTCAGCACAAATGAAAATAGAAAAATAGAACTTCTGTATGTGGAGAAGAAGCTGAAG
GTTGCTCTTTTGAAGCTTGACTTCTCTTCTTTGATGCTTGAGAGGATGCTATTGATGATGGAGAAGATCACTCTT
GAAGACTCACTGAATTCTATGTACAAATTTCCTTCCTTTTTCCCTTTTCCAAAGATATTCAATGATGCTCACACA
AAATAATGTGTTCCACTCTTTTAATTCTCTTTCTCTTACTTTTCAAGTGATTCTTTAGCCTTTTGTAAGCTTGAC
AGTGGCTGGAGAGTGAGTTTTGACCGTTAGAAGGCTTGGACTAGCTGTTGGAGTAGAGAAATGGCCGTTGGAGAA
GTACAAAAAAATTAGTTATTACAGCTTCCTCATGTCAAGAGTCGACTCGAGCTGTCCACGAGTTGACTCGAACTC
TTCAAGAGTCGACTCGTCCTCAAGATCCAAAAATTCTACATTCTATCATTCCTAAGAGAGTCGGCTCGCCATCTT
CATGAGTCGACTTGACCTATCCAGAGACCGACTCGACCATTTCATGGGTCAACTTGAGCAGGGATCCCAAAAATC
CAGTCTTTTATCTTCACTAAGAGAGTCGGTTCTTGCTTCTCAGTAGTTGACTCACCGTTCGTGTCAGTGCATTCC
AATGTGCCAGAGTCAACTCGCCACTCCCAAGGGTCAACTCCTAACCTTTTTTATCTTTATTTTTCTTTTGTTTTC
AACCTGTTTTTGTTTAAGGGTCGGCTCCAATGCTTCTAGGGTCGACTCCACTAAGTCAAAAACTGAGAATTTCAT
CTTTGAATTATTTTGACTTAGCTTCTTTGAGGGGATTTGTCTCATGGATAAATTCTTCATTTTAAGAACTTAGAA
AATCCTATATCCATCCTTGAAGCACATGATTAGTAACATAATTACTCAAAATTTTATGATCATCAAAACCAATCA
TACGGTCAACAATCTCTTCCTTTTTTATGATGACAAAACTTTGAGTATATGCACAAATAAACATAATGTGTTTCA
ATTAATTTGCTAATATGAATCATAAAGTAATTAACTAAGCACTTGTAAGCATATACTTCATCAATGTTTCAAGTT
ATCTTAAATTCAGATTTCAAAAGTCAATGCACATTTCAAGAGTATTCAATATATACTTATTTTATATGATTGTCT
CAATGCAAACAAATGAGCATTTCAAGCATCTTAAGTATGTGATCAACTCAAGCAATTCAAGCATACACTCAACAT
TACTTCTTTCCCTTTTTGACAGTATCAAAAAGTGCAAAAAAAAAAAAGATGCCATCAAAATCAACTCAAATCAAA
TTCCATTTTTATAACTTCTCCCCCTTTTTGAAAGTTTGTAGCAAATTCAAATGGAATTTCTTCCCTTTTGGAATT
CAATTTTTGAATAATATAGATTCAATAACTTTTCTCCTCCTTTTTAAATAATATAGATGAAACAAATAAGTTCTT
AAATTCAGTTGATTGATCTTTAGAAATCCAGGAGAGATAGATCTTTCCAAAACCAAGGATATTCACAAGATATGT
TTAGTTTTTCAAAATTCATTCTTTCTTTTCTTGAAAAGTATCGATCAATTAAGTTCAAAAAAGGTATCATTTTTC
TTAGTTTCTCTCAAATGATATCATATTAAAAATGTATCAAAGCAAAATGAACTTGAGACCAAGCATTCTTTTCTT
TTTGACAAGAGATAAAGCAATGGAGTATAACACAAAGATAAGCTGTTTGATACATGTGAAAATTTTCTTAAAGAT
AATCTACTTGATACCATTGTTCTGAAGTTCTAACCAAATAGCAATTTTCTTGAAATTTTTTTTGATACCAAGTAG
AGATTTAAAAAGAACATTTATCAAATTCATTTCTCGATAATAATGAATCTAATAAAATATTTTCAAATTCAGTTT
AGTACCAATTAATTATCAAAAATGATAACAAGATCAAACAATGCAAAGTATGAATAAACAAATCAAAAACACATG
AATGGTCAAACCAAGATAAACAAGAATAAATGTTTATCAATGATCTTTCTTATTAAGTTCCAAATTCATTTCCCC
CCCTATTTTTGATAGTTAATGCCAAAAAATTTATTTTTCTTAATTCTCCCCCTTTTTCTTCCAAATTATGCACAT

TTATTTAATAGTAATTACAAAAGCATATTTTTCTCTCCCTTAATTTTATAAGCAAAGCAAATTTCAATAAATCAA
ACAAATAATCACATCAAACACACTGAATTTTCATGAAAGATCAAATTTTCATTGACCGCCAAAATGTGATGACAA
AAAGTGATGACAAAGATAGTAAATATCAAAACATCCAAAATGTGTAATTTATATCTATCAAAAGGTAGACTGCTA
GGGGCACTAATGCTTTAAGTAAGAAATCCTAGCATTCCATGAGAAAGCAGGTCAGCATAGCCCTATGGAGAAGGT
CTAGGCTGGCTAGATGGACAGGGAGGTCTAGTGTCTCCAAATGACTGTCCCCGATCAACCCTCACTAATCACATA
ATCCTTATAGTCACCTCTCTCAGCAATGATATAGCCTATGAAGTGTACCTCTGTATCTCAAAATAGACTGCAGTC
ACAATCCGGATGGAAAATCTCTCAATAAGAGCATCCTCGTTCAGTCTAACAAATGGTCGATCTGAGCTCTCTGGA
CTAGTCTGTGGAGGTAGTGCTGTCTCTGTCTACTTGGGCTATATTGCTAGCTTCTCAGATCTGATGTCTATAATA
GGCACTACTGCAGGAGCCTCATCATCATTAGTGTCCTTATCTTTATGCATCTCCTTTGCGAAGCCAACGACCGTC
AATCTTTTGAAATCCCATCTTGCATAGTGAACCGTCATTGTAAGTATCTGTGTGCATCAAAACTCTGGTGGTCTC
CCCCGAAAAATCATTCCCATGCTCCTGGAAGACCAAAGTGAGAACCATGCCGTAAGGTAAGGATACCTTTGACCG
GTTCACTGCATCATGCATTTGTCCAATCATAAATGATGGTAGACTAGAAATCTCCTGAACAATGTGATACACAAG
GGTGATGTCACACTCAGACAATCAAATCTCCCCATCTTGAGGAAGAAAATCCGGCTGATCATATGATGCAGTAAT
CTCATCTCCACGCTCAGCTGGTTTGCACTGATGTCTCTGATACCCCTAACATCATCTCATTCCAATATAACTTTC
AATCCAATGTCTCTCTATATAAGATCATCTGAAATGCTCCCATCAGTTGGCATATGAAGTGTGCATCCAAGGCTA
TCCACATCAATCTCAATTTACATGCCTTTTACTTTGGAGACAACACTTCCCACACCCACTTTCATATTACTGTAG
AATTCCCTTACTAGTCCCGGGTAAGTGGGCACATCCAAACAACAAAGCTACTCCCAACCCCTCCTTTTAATCTTC
TGTCTGATAGAAAAACCTTCACGCTCTAAGGAAATCAATTTTCCTACCAATTATGACAATGCATGAAGATAAGGG
TGCCAAAGACGGAATTGGCTGAGGAGCTAATGGTGGTTGTGAAGGCACTAAAGCTCCTTGAGATGGCTGCCCTGG
GTCATTCGGCTGTTTGGATCTTCATTCACACATAGGAGAAGGGCTTGGTGAAACTCTAGGGTTTGATGTTTCTGC
ACGGATCCTCTTCCTTATTTGGGCTAGTTGCTTGTGTAAGAGACCAAACCACCAAAAATCGGGTTAAAATTGGAG
GTGGAAGCTTTGATTTAGGAGGGAGAGGGGTTTGAATGAAGTTCCAACTCGAAAGAAAAGAGAAGAAATGAGCAG
AGCCTTGGGTTGCCTTGGGTTAAAAATCCCATCTCTGACACAAGGGTCGACTCCAGTTAGACACGAGTCGACTCC
TGATAAGAGGTCGACTCTAGTAAAGCTGGGGGTCGACTCCTCACAGAACTACTTTTCTTCTTAAAAATATAAGTC
AACTTCTTGAGTCTAAGAGTCGACTCCTCAATACAGAAAATAATTCAAGAAGTTTATATCAAAAAAAAGTTGGCT
CAAATAGCGTGTGGATTAATGGCAAATCAAATCAATGATCAAAAAACGATATTTGAATCAGAATAAGTGAATCAA
TTTGGGGGATCAAAAAATTAACCAATTTAGATATGATCAAATCAAGGAGAATCCAGATACAGGCTCTAAGGGTGT
GGTTCAACTAAAGAATTTTCAAAGCCCTAAAGATCAATTTAAGATGCAATGATTTATCATGCAAATCATTGATAT
CTTGTACACAATCTTTTACGAATAAATGTCTTATCTTCAAGTTCTTAGGTCATGCAATTTTTGACAAATTTTGAT
AATTAGATATCTTATGAGGATGTCAAAATTTAGTTTGTAAGATTTTCACACAAATAATTTTGTATGCAATCAAGC
TTATTATGATTCAGCTTTCTTATATAACTAATAATCAAAATAACATGATATAAACAATTAAGAGAGCATGCTGAC
TTTAAAATAAAACATATACTTGTTATGAACAATACATCAAAACATTCAAAGCAAAGAATAAAGTCATCACAATAT
ATATTCTAGTATCATGCATGCATATGAAGCATCAAAATCAATCAATTAAGCAAAATAATTGCAGCAATCAATGAT
GGTATGATGTCAAAGGTTTAGCTTTTATCAAACATCATCATCCATCCATTTTGAGTCCAAAGCATCCATTCTTCA
AGAATTTACTCCTTTCAAAGCATTGGATGCTTGGCATACTTACAAAATTATCATTTTTTAGATTTTGGTATCTAA
AATTTCTTGGATCCATTGAAGTTAGTGCACATAGTTTCTTTTAAAATTCAAATTTGTTTAACCAAAGTATGACCA
ATTCTTTTAGTATTGCATTCAAAAATTTTGTAGCCTATTTTATTACATTTAAAGCAAGTAATAGAATGATTTTCA
TGTGAAGCTTTAGTAAACATATTTTTAACAAATTTTTATTTTCTTGAAGAATTAAAACCAATTCCAGCTTTATTA
GAAACAATGTTTTGATTGTTTAACATTAATGGAAGCTTTTGAGAACTGAATGTGAACTTTTCAACTATAGGCTCC
AATATTTCAATCTCTTTTCTTAATTTTATATTTTCTTCCACTAGAAATTTTTTAAATTTTAAAAGATTTTCTCTT
TCTTCAGCCAAAGAATTCTTTTCTTTCAAAAGTTCTTTTTCTATTAAAATCTTTTTTTTTTCTTCAGCTAAAAAC
AAGTTTGACTTTTTAAGCTCTTTATTTTTCAAGCTTTTCTTTTTGAAATCATCCATTAGTTCATGAAAGGCTTCA
AATAGTTCTTCAAAAGTAAATTCGAAAGAGTTTTCGATGTTTACCTCATCGTTGTGAGCCATGAGGCAAAGATTA
GCTATATCTCCTTCCTCTTTTTCTTTATCCGAACTAAAAATGTCACTTTCTCCCCAAATAGCCATCATAGTCTTT
TTGCTTCTTTTAGGATTTCTTTTGAGTAGGGGCAGTCCGACTTCTAGTGTTCCTACTTGTTGTATTTATAGCATA
TGACATCTCTTTCTTCATCCTCCCTATCTTCCTTTTCCTTTAGTCATTTTTTCTTGAGGAACTTTTTGAACTTCC
TAGTGAGCAGCCCTATGTTCTCATCTCCATCTTCACTTTCTTCTTCATTTTTATCTTTCGACCCATTGCTTTCTT
CTAATTTGATGGTTGATTTGAATGCGATCTTCTTTCTTTTCTTTGACTCTTCTTCTTCCTCATTTTGGCACATGT
TCAACTCATATGTCATGAGTGAACCAATTATCAATTAATTCTTCCATGGATAGCTTGCTAAGATCTCTTGCTTCC
TGTATTGCTGTTAATTTAGCTTCTCATGTCTTTGGCAAACACCTAAGAACCTTGTAAATAAGCTCATTGTTAGTA
TAGGTTTTGCCAAGACCACTCAAAGCATTAATAATATTAGTAAATCTGGAAAACATACCAGAAATTGACTTATGT
TGTTCCATTTTAAACAATTCATACTTGTGAATAAGCAAATTAATTTTGGACTCTTTAACTTGGCTTGTTCCTTCT
AAGTAATTCTAATCTGTGCCAAATCTCTTTTGTGGTAGAACAAGTAGATATTCTATTAAATTCATTAATGTCTAA
GGAGCAATATAGAATATTTATGGCTTTAGCATTCAGCTGAGCATATTTTTATCATGCTCGGCTCAATCTTTTTTC
GATTTGGAAGAGGACACTCTAACTAGCATTGTGGGTGTTTGTGGTCCGTTAGTTATGACTCCCCACATGTCATAG
TCCAAAGCTTGCATGAAAATTTTTATGCGTGCTTTACAAAATATGTAATTCATGCCATTAAAAAGTGATGGTCTA
GAAGTAGAGTGGCCTTCAGCAAATGAGGAACCAACTAGGATTGCCATGATCTTTAGCTTAAGAATGCTAGGTCAA
TAACAAATATAAAAGCACCTACTCTGATACCAATTATTGTCCAGAAGATTAGCAACCCAAGAGGGGAGGGTGAAT
TGGGTTCTTTAAAACTTCTACCAAATCTAAATTTAAATAATTATGTGCTTCAAATTAAGCTAACTTGAGTGATTA
TGAGGTGTATACAATGTGCTGCAATGGAATTAAGATACAAAAAAAAAAAGTAATACAAAGCAGGTAAAGCAAGAA
AGCAAAGCACAACATAAATACAATAAATTTATTGTGGTTTAGTGCAATGCCGCACCTACATCCACTTCTCAAGCT
TCTTTGAAAATTTCATTATAATCAGCCCAATTACAGTATGATTATTTTATCCAAGCGCACAATCAAACCCATTTG
ATTTTTTGGAATCACCAATAAAAATAAAAAATATTTACATCAATTATTTTTCGGATTCACAATTAACCTAGTTGA

TTTTATGGGATCACCAACCATAACCCTACAAGTCCAATTTCAGGCTTCAACAAGCCTAACCCAAGTTTCTTTCAC
TCCAAAGAAACTTGAAAAGAAAAATCCAAGAATACAAGATAAAGAAATTTTAGTACTAATTAAAATAGAAAAAAG
AACTCTTTTGTGTGCGGAGAAGCTGAAGGATTGCTCGTTTGAAGCTTGACTTCTCTTCTTTGATGCTTGAGAGGA
TGTTGTTGATGATGGAGAAGATCACTCTTGAAGGCTCACTAAATTCTATGCATAAATTTCCTCTCTTTTTCTCTT
TTTCAAAGATATTCACTGATGCTCACATAAAATAATGTGTTCCACTCTTTTAATTCCCTTCCTCTTGCTTCCTAA
GTGATTCTCTAGCCTTTTGTAAGCTTTCCACTCTTTTAATTCCCTTCCTCTTGCTTCCTAAGTGATTCTCTAGCC
TTTTGTAAGCTTGAAAGTGGCTGGAGAGTGAGTTTTGACCATTGGAAGGCTAGGAATAGCTGTTGGAGTAGAGAA
ATGGTCATTGGAGAAGTACAAAAAAACTAACTATGACAACTTCCCCATGTCAGGAGTCGATTCGACCTCTTTAGG
AGTCGACTCGGCCTCAGGATCCAAAAATTCTGTGTTCTATCATTCTTGAGAGAGTCGACTCGATCATTTCATGGG
TCAACTCAAGTAGGGGTGCCAAAAATTCAGTCTTCTATCTTCACTGAGAGAGTCAGCTCTTGCATCTCAAGAGTC
GACTCGCCATTCGTGCCAGTGTGTTCCAATGTGCCAGAGTTGACTTGCCGCTCCCAGGGATCGACTCCTGAACTT
CCTATTCTTCATTTTTCTTCTATTTTCAACCTATTTCTATTCATGGGTCGGCTCTAATGCTTCGGGGATCGACTC
CACTAAATCAAAAACTTGGAGTTTCATCTTTGAATTACTTTGCTTAGCTCCTTTAAGAGGATTTATCCCATGGAT
AAATTCTTCATTTTGAGTACCTGAAAAATCCTACATCCATCCTTGAAACATATGATTAGTATCATAATTACTCAA
TATTTTATGATCATCAAAATCAATCATAGGATCAACAATACTGTTAGAAGCATATTGAAATATTTGGTAAGGTAT
CGGCATGGTTATCAAGATTCAATATAGCATAGGGGCCAAATAGGCCTGGTTGGAGTGGTTGCTTCCACCAGTTTC
CAATTTTGGGTGGCACCATTCTTGGGCATGAGAAAAAACACTTCCAACCCTATGTCAGTATAAAGGGTGTAGCCA
TACCATACTGGGAACACTTCTCAGAGCTATAAGATATAATTGATGATAATGCATATAGTATCATGTGTAGTCCAA
ACCAGAATTTACTCCATCTCTTTTAAGACAATTAATTTGAGAACCGAATCCATTGGTTTGTTGGCACCAACCAGA
TGGCACATATAACATTATCTCCTCAAGTCTCTCGAAGTCAATTAATTGGGGCTACGTCAGCTGGTGGATCCCAAG
GACTCTTACCAAAGCAAAATTCACTCCATCTCTTTTAAGGCAATTAATTTGAGAATTGAGTCCATTGGTTTGTTG
GAACTGACGAGATGGCATGTATCATGCTATCTCTTCAAGTCTCTTGAAGTCAATTAATTGGGGTTACATCAGCTG
GTGGATCTCGAGGACTCCTACCAAACCAGAATTCACTCCATCTCTTTTAAGGCAATTAATTCTCATAAATCCCCC
AAAGAGGCCCTAATTATGCCACTTTTATTTATCAACAAACCAATTGATTCTCAAAATCTCAACCATGGTTTAGGG
ATGTTAATTAGCCACCACCCGTTTGATCTTGTGAAATCAATTTTTACAACTCATGGAATTAGAAATAAAATGCAT
CCTATACATTAAATGGCAACCCAAAGAGTTCCCGAATGCGATGCATTTTTGTTCCTCAAAGGGCAATCTAGAGGG
TTTCCAAGTGCAATGCATGTTCGTTGCTCATGCCGATTCCAATATGTCCATAAAAGTGCATAATTAGGTGGAAAT
TTTCTCAGAGATGCTATCAAAAATTGTAGGCATTCATCATCCTATATAGCTCCAACATGCTTCATGTTTTATTGT
TTAAAAGATGATAAATAAATTTGAATATATATAAATAATCATATCTTTGATTAAAAAAATTGTAGATAAATTTTA
GTGTCATTGCACCTTGTTTTGGTGGCCAATTATACTCATCAATTCAACAAGTCAATCCACTGAACCAAACATCCT
TTCTCAATTCTCCAAATCCTCCTTAATTTTGTTCTTCCTTAGATTCATAATGCCCACTATAATGATACACTCTAT
CCTATAATTCCTCATCAAAAGTTGACCCAAAAGCAGCTTACCCACAAATTTAGCATGAGGAACACCCTTCTAACCC
CACCATCTCAACCCACTTCTCATTCTCCTTGCCATGATTAATGAGGGGAACATGAAAAATAGATCATAAAACAGC
AGAGATTTAGTGCAGGTGATCTCAAAACCTTATATGAATGTCCAAACAACAACAGCAATGATGATCATGAAGGAG
TTTAAGCGAAGGTGTGGAGTAATGGAGGTGATGGTGAGACTATAGTCCAACCTAATGGTCACAACCCTTGGAATC
AGATAAATGCTTTAACTCTTACTAGACCCTGTATCGGTACCTTATTGGCATGAGTCAAATCAATATGGTTCATCC
ATGCAACAAGACACAATATGCTGTAGGAGTTAGCATAGCAATCGTACCATATCTCAGAATGGGCCTATGGGTGGC
ATATTGAGCCTTCATCCATCCATACTAAGCTGATGTGATGTGCCCTATACTAGCCAATACTGTTGGTACTTAAAA
TTGTTACAAACTAGGACAAATAAAAGGAAGGGAGAAGAAAACAAAGAGATGGAGAGGATGATGACAAGACTGGTA
AGGTAGCACAAAATAGTTACAAAAGAGATGGAATAAGAGAAGCATGATAGAAAGAAGCAATAGTTGAAGTCATAG
CTTAAGGCTTTTGTCTCCTGATATTGTGGGTTTCATGCCTAACCATCGGATTAGATTAATTTTAACAACTCAGAT
GGTTGATTCTCCTAAGTTAACAACTCAAAATGTCATTTAGGAAATCCAGGTCCTATCAAGATGTCACAAATTTAG
CCACAAGCTCGTCACAAGAGTTGCTGACATGCGTATGGTGCAATGCTGCAAAGGATTTGGCATGGGCCAAAAGTC
TACAATGCTCCTCAATCATTTAGAACCCACAGGGTGGTTGTTGGGTGGATGTCTAGTTGGGGTATTGAGGGGAGC
AATAGTTAACCAAGGCATAGCATAGCCATGGTATGATATTGAGGGCTCGGAATGCATGATATTTGGCATGGCGAC
GGACCACCAGGATCATGGTTTGCTATACCGCCTTGTACCACTCCATACAAGACATACTGTACCATACTAATACCA
CACCAGTACAGTCTCTAACCATACCGACACTCAGTACGTTGTCTTGTAATGTACTGAACTACATACTGCTATTGT
AATAGGATGATACTAGTATGAGATCCGGTATAGAGATAGTGAACGTTGGCTAAAATATAAATGGATCCTCAAGAA
TTCCTCTTGGTTGAGTGTAGAACTTTGGTGATGTTCCCAACTCCTCTAGGCCTGTAGAATATAGGTCCAAAATTG
GCATGATTGAGGGGAAAGGCAGTGCCACACCAGATGGTCTCACATGGAGGCATCATGGTAATACTGGTCAAGTGA
GCTAGCCACATTGTGTGAAGGAAAGTTTTGAGAGATTGGTTCTCAAGGAATTGTCCAACCTACCCTATGCATTTT
TTATCTCTATGGAGTAAAGAGGCAATAGGGACCCGTTGGGCAAAGGGTGTAGCCTACCGAGATAGGCAAGCCTTA
TGAGCATAATAGGGTGATGTCTCATGGGGATCCACTGAAGCTAGTTGATTGTGGAGTACAAATGTGGCTTATGGA
ATATCAATTGTATTATGTGCCTTGGAGCTAGGCCTTAAGGACAACCAAAAGGGTCATGCCAAGGAGAGGATAGCT
CGATGCAAGGATTGGCTAGTTAAATATATTTACTTGATAGGCAAACTTAGATGCCATTATCGTAGCAGTGCACAA
TGTGCAAGTGGGCTATGAACATTGTTGAGTAGTATGGCCATATGGATGAGAAGGGATGTTGTGGATGCAAGACCA
GGACCATGGCAGAAGAGCTTACAGGTTAGCTTGGATAGCAAAACCCTGAAGGGCTCCAACATACACGATCAATTG
TTGCTTCGAGGGATTATGGGGCATATGTCAGTTTGGCAGTTCGACTGGGCAACACGAGATAGGTTTGCCAGAGCC
TCATGCATAGAAGCGGCATTACATGGGTTGCTTGCAAGCACTTGAGGGTGTCGCATGTGTTCAGTTCGACAGAGT
TATCGAGTAGGGGCATAGTGTTGCTACTTGGCAATATGGCTAGACTACTGTGTTTTTACATATATGCCCAAAGGA
AGGACTCAGGAGGGCTGGCATTGGTGTGTGGTATGCTAGTGCATTGCACAAGTATTTGGGTCAGGAAAACACTTG
ACCTATGATAATAAGCCAACCCACATATATCATGGTCTACCATACCACCTAGTACAGGGCATACCATACCATATG
GGGAGCAAACCAATATGAAACCCCACTTCAAGTCGTTATAACCCATGTACCACCCATACCGAGATAAGTACCGCT

CCATACCAAGGCATGTACCATCCCATATTGAAGCGTACTGACCTATACCAAGGTGTACCAAGATAAAAATTGAAA
AAAATGATTTAGGAGCTATTTTGGCATATGATGCATACCATACCATACCGACTATATGGAACCGAACCGATAAGG
TACCAATATGATATCTGATACCAAGATAGCAGACCTTACTGCATATGTTAATAGGTGATAAGATGAACATGTCAA
GTTGCATGGATCGAACATGTTTTAAAACAATATTTTCATATATCACCTCTCCCATGAAACCATTAAACTCCACCT
TAGGTCAATACAAGCAAAAGCGATGTACATAATTATAGTCACTTTAAATCATAAACTAGTCAATCCCATAGACAT
ATTAATAAAACCTTTAGCCTACATTTGTTTCATGCCATCGAACTACCTCTTATGCTACTCAATCATGAAATCCAT
TTGAAAGCCCACAAAAACCATGACAAATCTTATAGGTGAATAGGGGTCGCTAGTTGATATCACATTATGTGAAAC
AATGTTGATTATTACATTATTGAATAAGTCGATGATATAGCATATTCAAAATTACGCCGATGCTTGCAGGAAATT
GATGGGCGAACAGTTGTCTGGCTTAAATGTCAAAGAACTGCAAAATCTAGAGATCCAACTTGAAAGGAGTCTTCA
TTGTGTCCAAAAGAAGAAGGTTTACCCGCATCAAATCTTGCTCATTAATCTTTCTTATCCATATCTTCTCTACT
TAATTTGTTTGTTTGCTCTGTTTTAGGAACAAATTCTAACCGATCAAATTCAAGAATTGGACCGAAAGGTTTATC
AATGAGTAATAGTTAAATCATTTTTAATGTTGAAAGTATTGCATTTTAAGTGCCTAAATTTCTAGAATTTGTTAC
AGGGGTACCTTCTTCACAATGAAAATATTGAACTCTACAAGAAGGTAAACCTTATACGTCAAGAAAACATGGAGT
TGCGTAAGAAGGTATTGAGTTTGAGAAAGAGATATATATTTTGCTCTATGATCCAAACTTCAAAGTGGACCATTC
ATGAAGCTCTGGTCTTCATGTTTCCATGACCACATGCAGCACATCATGCTCTATGTAAAATATGTATGGATTTGG
TCTAAATCTAGGTTCAATCAACCCCAAGGTTTTAAGTGCTACCAATGCTGCTTTGGCTACCATGTCAACGGATGT
CAACACTAACACTAGGTCAAGTCAAGACTCATCAAATCCTTTTTTTGACATTTTTTTTGTTTCGACAAAGTCAAA
AAAAGGTTTTTGATTTTTAAATCCTCTAATATTTTTTTGTTTAGGGCCTTTGGGCACACAAGTGCATCAACATGG
CATGGCACGTACAGTGCTAGTCACTAGTTACCATGCCTCATGCCAACAATATTTAAAACCTTGTTCAATCCTTGA
GAAAATAATAGTTATTTTTGTAACATGCTAAGCATGGAATTTGCATTGCAGATGTACGCAACAAATGAGTTAAAT
GGAGGAGCTAGTAAGGATACACTTAATCCATATGCTTTTTCTGTGATTGAGAATTCACATGTAGATGTACCTATT
CATCTCGAGCTATGCCAACCTCAACAGCAAACACATGGAACACAAGCAACAGCTCCAAAGCTAGGGTAATTATTT
TTGAAATATCAAACCTTTATATATTTTCAATATTTATTAGTTTTACTTGAATATACTTAATTAGTCCTTGCAGCA
GACTTCAACTGCATTAAGAAGATGACCAGTGATCAAAGTGTATCATCTATGGTGCGCATCAGCCCAACTGTCAAG
ACTTCTATAGTTTGTTATCAGCTATAACATGCAATGTATCGGTTATCAAATGCAATGTATGAACTATGCTCAGCA
ATCATTTATATGTGGTGTGACATGATCTCAAAAAAATATCATTACTCTGTAAAGCAATCTTCCAACTAGGACTAT
TAATAATATGACAAGTATTTCTATTTAAAAACATTACTTGTATATGAGGACATGGCCTATCATAGCATCCGAGTG
GTTGTTGGTAATGCTACTGGAAAACTCACTTGCTAGATTGAAGGTACAGGGGATTAGCACCAGTTAAAGTGCAAC
ATGTTTTACACAAAGGATGACTCACTATATTCTGCTTAGTAACTAGTAACAAGAGGCATATGAAATGTGATAGAT
TTATTCTCCACAATAGTCACCACTAAATGGTGATGACAGCATTGTATATTAAGCTAGCTTGCTAAACAAAAACAA
AACAACTAGTACTGGAACCGGCACTGATGATGGTAGGCCAGCAGTACAATATAGTCCGATACTGTGCCATTCCGG
CACGTACCGATGGAAAACCAGCAAGGGAGGGGAATAGGGAGAAAGAAAGAGAGGAAAAGAGAGAAAGAGAGAAAG
TGAGGAAGAGAGGGAGAGTGGAAGAGAGGGAGAGAGCAAGAGGGCGGCAAAGATAGAGGGAGGTTGAGGGAGAGG
GAGAAGAGGAAAAGGGAGAAAGAAACAAAGGCTCCGCCCGTGTGTGGATTCTCTCCCCCTCCCTCAGGCTCTCTC
TCTCCGCTGCCCTCTCCCTCTCCCTCCCTCCCTCCCTCTTTCGCTCTTTTCTTCTCTCTCCTCCCTCTCCCTCTC

>EguAGL17_3

ATGGGCAGAGGGAAGATCGTGATCCGAAGGATTGAGAACTCGACCAGCCGGCAGGTGACCTTCTCTAAGCGGCGC
AAGGGTCTGTTGAAGAAGGCCAAGGAGCTCGCCATCCTTTGCGATGCCGAGGTCGGCTTTGTCATCTTCTCCAGC
ACTGGCAGGCTCTACGATTTTGCCAGCTCCAGCATGAAATCAATAATTGACCGCTACAGCAGGACAAAGGAGGAG
AATCATCAAGTGCTAAATGCAACTTCAGAAGTGATGTTTTGGCAAAGGGAGGCAGCAAGCTTAAGGGAGCAACTG
CAGAACTTACAAGAAAATCATCGGAAATTGATGGGCGAACAGTTGTCTGGCTTAAATGTCAAAGAACTGCAAAAT
CTAGAGATCCAACTTGAAAGGAGTCTTCATTGTGTCCAAAAGAAGAAGGAACAAATTCTAACCGATCAAATTCAA
GAATTGGACCGAAAGGGGTACCTTCTTCACAATGAAAATATTGAACTCTACAAGAAGGTAAACCTTATACGTCAA
GAAAACATGGAGTTGCGTAAGAAGATGTACGCAACAAATGAGTTAAATGGAGGAGCTAGTAAGGATACACTTAAT
CCATATGCTTTTTCTGTGATTGAGAATTCACATGTAGATGTACCTATTCATCTCGAGCTATGCCAACCTCAACAG
CAAACACATGGAACACAAGCAACAGCTCCAAAGCTAGGGTAA

#### Asparagales

##### Phalaenopsis equestris

*PeAGL17_2 and peq-miR444a*

>Scaffold000428

TCTAGAAAGAATGCAAATACATGCAAATTGAATTAGCAATGTATTTTAAATTTCATATCGAGTTTATTTTTATAA
ATGTAATTTGAATATACCAAATAAATTTTACTTCTAAACTAGACAAAAATAATTAATGTCCTGAATATATTTTAT
TTAGTGTATTTCTTTTACCATCCATGTGTATTTCAAAGAAACAAACATACCCAAAGTTAACTTATATGCAAATCA
ACATCATAGAGATTAAATAGTCCTCTCATTGAAAAAATCATCAAATAATTAATGCATGCCTCATATCATTGGCCT
CCTTATAACTTCCTAATTATAGACAAAATTGCATACTTGCAGTGCACTTTGATTAATTAATTAACAACCAATCTT
GGCAAGACAAACACTCTCCCCTCACCAATCAAGCTCTAAATATAGGGGTGTCAATTCTGGTCAGCAGGTCGGGTT
CGAGTTCGACTTGAAAATGATCTGACCTGAATAGGGTCAAGACGAACCCGACCTAAATAAAAAATGGGTCAAAAA
TTAAGACCCGAACCCAAGCTTGACCCGGGAACCCGCCAGACTTGAACTTTCGTGCTTATTCGAGTCATGTTTGAG
TTTTCAAGTTCAACCCGAACCCGACTTGAATAATAAACGGGTCAAGCTCTCTTACTCGAATCCAACTCAAAAAGT

GTTTGGGTCAACTCTGATCCGGATTTTTCAAGTCAAGTTCGACACCACTGCCCAAAAGAGGGTTTAAATGAAATT
AAAATAAAAGTCCCTCTTCTAACTCCCCCCTAGTATGGATCTTCCATTAGCTCCCCCTTCCTATAAATAAAGCCT
CAACAATTTCTTATCCTCCAGAAAGTTGAGAGAGAGACGGGGGGAGACAGTGTTTGGAGATGGAAGAGGGAAGA
TTGTGATGAGCAGGATCGACAACACGACGACAAGACAGGTGACCTTTTCGAAGAGGAGGAATGGGTTGCTGAAGA
AGGCGAGAGAGCTCGCCATCCTCTGCGATGCAGAGGTCGGACTTATCGTCTTCTCCAGCACCGATAGGCTCTACG
ATTTCGCCAGCACCAGGTTCTTCTTTCTCTCTTTAATTTCTCTTCCTTTTCATCTGTTCACTGCTGATGATTTAG
GGTACAATAGGCTGATCGGCTCCGTAGGGCTAATCCCTGAATCGATTTTCTTGCTTGATTAGACTCGTAATGGAA
AATGGGGGGAGAAATATAGATGGATTAAAGGATAGGGAATTTTTTTTTAAAAAAAAGGAGGAAGTCATCGGATT
CTCCGGACAGATCTGAAGGTTAGGCGGCTTAGAAGGATGAACATTAGAGTTCAGGAGTATTGAGTTTTTTGATGG
TTTGTTGTTTTGAGGACAATATGCAATCACGAGATACTCTTAATACTCTCGCTGTTAGGAAGTAGATCGATATAT
AAAATATCTGTAAATTTATCAAAGAACAGGTGCAGCGAGTATTCCGAGCGGAATCTTCATGAGAATATCCTCAAT
GATTCATATATATATATATATATATATATATATATATGAAAGAAAATAATTTCGTTTGACACGCATATCAGAAATTT
TTTTGAATAAGAAATATATAAATTTTAATTAAATATCATCTAGAACTTCGATTTCACTAAACCGAGCTCCACTAT
ACAACCACAATGACACCAAAATTATTCCAATCCAAATAAAATCAAATATGGATCGAAATCATTAAACTGCTTTTG
AAAGCTTTTGGTACTAATTTGAAAGATCAAACAAATTCCAAATCTGATAAACTTATAATCGAATCGAGTATCTCG
ATGAGACGAGTACAACAAGTCTAAGATTACTCAACCATTATGACACTGATGGAGGGAATCTCTTTGGGCAGTCCG
ATGGAATAGTTCAGATTCATTTCTTATTTGCAGAGAGACTTTTCCCAAATTTTATAAATAAAATCTTACCCACTT
CATGAAAAAGAACCTATGTCTTTAACCATGCAAATAGCTAGACAATGAGCGTGTTTGCGAATAAGGTGAATTTTA
GTGTCATTCTGGACCGAAGATCCGACGAATGCTTCCGGCTCTTGCTGCTGCCAACCACTGCTTGCTAACTGTCGT
CGATCGACTCGTGCTCATTGCATTGAAACCTTGTGCAGATTCCAAGACGTATGTTGCAAATTAGTGAAACATTGC
AAAGTAACCACTAACGCCACGAAGTAACAAAATATCTTACAAAACAACACATTATCCTGAGCACGGCGCAACAAC
ACTTTCAAAGTCGGTATACCGAGTTTTTATAAACTACTCCTAAATGCAACCTCCGGAGGAACTGTATGACCCTTT
CCTTCATGTGGGACAAAAGTCTGGTAATGTAAACACTAATTCATAATAGATTATTCCGTCTGGACAAAGTCTCCT
AGTTTTCTCTTTTGAGTTTTTCATGTAAAATCAATGATATCTCTACTCGATGCTTCTATTATCCAAGTGTTTGTT
AAAATTTTGTATAACGCTAAGATACTAAATAATCACTCTACTATCGTAAATCTTCCGACAAACGTAAAGTTAACT
TTAGGGTGAAAAATTCTCTCCTAAAATTTACTATTTCTTACTATTTCTCCATATAAACAATAAAATTTATTTTTT
TCTTATATTCTGCCGTTCCCACACACCCACCCACCNNNNNNNNNNNNNNNNNNNNNNNNNNNNNNNNNNNNNNNN
NNNNNNNNNNNNNNNNNNNNNNNNNNNNNNNNNNNNNNNNNNNNNNNNNNNNNNNNNNNNNNCACACACACACACACACA
CACACACACACACACACACACACATATAGATATATATATAATATTTTATAAATGGAGGAAGAGTACACACACACACA
CACACACTTATAAATGGAGGAAGAGTATAATTTAAAAAACAGAGGAAACGTAGTTAATTTAAAAAACAGAGGAAA
CGTAGTATTAACAATGCATTAAAAAGGTAAATATAACTATTTTGGTCCTTAAACATATAAACCCATTGGGTACAT
AATAAGGATGATTTTATTAATTTATTTTTCTGCTTTGCTGAGTCCTACCTGCCCGGGAAAGCAATGAATAATATC
CCTGATATTCGAAAGGAATGAATTTCTGTCATATTTAATAAATGATGAAATTTAAAGTTCCCAATGTTCTTTCTC
AAATGAACCATCGGACTCCAATATATATCTGCCGGCAACAGTTGGGAAATTTACGTGTATAGCACACAGTTCTCA
TGTCTAACACAGTTCGATATTTAAAGCTGAGGAAATTTACAGAAGTTGCACTCAATGCGGATTTGCCTATCTTAA
ACTGAAACTTTCAAATTTACGGGCTCAACATTTTACTTATGTATGAAAGAGTGAATTTGATCATTATAAAATGTA
ATTCGTGCTATTTTTTTATGTTATATGAAAGAGTTGGGTGCACTTCATGTAAAAATAAAAGTTATAGTGAGACGTG
GAAGTTTATGAGAATTGAGTGCTATGTCCATAAATGCCCCCCTTTTCCCCCTTTAAACTTATTACACTTTTTGGC
TTATAAATCCATACTTGAAAATGTGTATTATATTTTTAAGTTAATAATTAGGGTTGTAAACGATTCAAACACGAA
CGGACAGAGCTCATGTTTGTATTCCGTTCGTTTGAAATTAAGTGTGTTTGCAATAGGCTCGTCTTTAAAATTGTT
TGTGAAACTACTGTTCGGATCCGGTTCGATTTGTTTGAAATATGTTTGAAATGTTTGAATTATTCGAATTGTTTA
AATTGTTCAGAATCTTATAAATTTGCTTATATTGTTTGAATTGTTTTAAATATTTCAAAAACTTCAACATTTATA
AATCACATTACAAATGCATAATAACTGAAACACAACTCACAACAAAACCATGATATCAGATCCCATCCATAAAAA
TTCAGATCTAGAGGCAATATAAGAGAGTCCACTCCCAAAAACTTCAGAAACTAGGTTTCACTAAAAAAAACTAAA
TAAATTTAAAATAAATACAACAAAATTTAAGCAAAGACGTGAACTTGAAGATCAGATTATAATTTAAAGGATTTA
AATTTTTAAAATACAATAAATTTTGAGTATATAATTTTTTTTAAAATATGATAAATTATATATATATATAAACAT
TCCAAACATGATCGAACAAATGACTGTTCCTGTTTGGTTCATTTATAAACTCAACGAAAAAAAATTATTTGTGAT
CGTCTCATTTACTTATCAAACAAATCAAATGCTATCTCCTTGACGAACCAAACTCGAACTGTTCGTTAACAATGA
GCGTTTTTGCAGCCTTTTTAATAATAATGATAATTTTGATGTGATTTTAAATAGTACGATAAAAAAAACTTGTTC
GCAAATAATTCATTTCAGCTCGTTAAGTAGATTATGTTTGAATCATTCATTAACTGAGCTAAGCACAAGTAAATT
TTTTTATACTCAAATTATAAATGAACCAACTTTTTGTTCGACCGTGTTTGGCTCGTTTTAGTTCGAAATAAGAAA
GTTAGATATATTATTTATATAATAGAATAGTATATATATTTATTTCAAGCTTAAATCACTATAGTTTGTTAGTGA
TTGTGTTTGTTAATGCACAAGAATAGTGCATGTTTGAATTCAACCTTTCATATATTTTTTTTAATTAATTTAAAT
GAGCCAATTCGAGTCGAACATGAGTTGATAACAAACTCGAAAGTTTTTTTGAGTTGAAAATGAATAAAAGCCATT
TTGAACGTGAGTCAATTTTAAACAAAAAAAAAAAAAAAAAAAAAAAAAAAAAAAAAAAAAAAAAAAAAAAAAAAAAA
AAAAAAAAAACTTGAATACTAACACATTTGCTCAAGCTCGGCTCATTTAAACCCCTAATGTTATAATTTTATGTT
ATAAATGTGAATATGAAAATTTAGGTGTTAGATATGTAAATTCATAAGAATGGTGTGGTATGTATGTAAATACCC
CTATTTAATTATTTATTGAAGTGAATATATATTTTGTGTATAATTCATTATGGTTCAAAGTGATTTACTTTTATA
TTATCGGGCTTCAGAATAGTTTGGTGATTTATAGAAATATTAAGGGGTATGCATTTTAATTTAGAATTTACCTTA
ATCAAGCAATGCTAACTAAGATAGTTGGTTAGATTACAGCATAGGCACATAGCTTTACCTGTCAATTTTTATATA
TCTTCATATGCTCTATATATTAACCTTATTTACTTAAATTATTAATATCGTTTGTTAAATCTGAGCTTGTGTTGT
GAATCACACCCAATGATATAACGTAGAACAGGAGGAATAGAGAAAGTGGGAGAAGAGTGTTCTACTTAACAATAA
AATGAGAGAATTGCATAATATTTTATTCATGAAATAATTCGAGCTCAAAAAGTATGTTCGTGTTCAACACATTTG

CTCAACGAGCCAAACACGAGTTGTTCGCAAACAAAGAATTGATTTGACAACCCTAATTACATAACCATAAAATGT
TAAGGTTTGTGACAACTTAAAAATTATTATTTACCTTAAAATATCATTTCAATCTAATTAATAATTTTTATTCTA
TTTTCTTAATTAATATTTTTAATTGATAATAATTGTTTGTAAGCTTGTAGTCAATTAATTTTTTTAAAAATACTA
TTCATCAGTTATATATATATATATATATATATATAATTGTTGAGTAATATTTAACTTAAGAGCCAAAACGAACCGAA
CAAATGCCGAAACTAAATCTCGTTATAACCGGGAAAGGGCCTTATGCTTTAAATTTTCGTCAAAAATGAATATCG
GGTCGGGTCGGAATACATGAAAAAGCTTATTTTTAAGCTGTAAAGCCACGTGAAATCGCATTCCACGTGGCGAAT
TTTTATTCGGTACCGCATGTAAGGCTTCATGCGGTGCACGTACCGAATAAAAATTCCTCTACCAGGGCGCGCACA
TTGTTTACACCCATATAAACCAAAATCAATGTGCGATCATACAATTAATTAGCGGTTTAGCCTAAATTATATAGT
TAGCAATAATTTCACAGGCTCCCGCATAACAATTATCAAACATGCAATTATAAAGAGCTAGAAACCACTTTTGAT
CCCCTCCTGCTTAAATTATCACTTTTTTGTCAACGAGTCCCACATAACATGACCTAATTTAAACTTCATTCCAGT
ATCAATGTCATAAATTATTTAAAAACTATGAAAAACATTATTAAAAACGTATCAACAATAATCTGCATTAATATG
CAGGCAAAACATAAGCAAAGCACATATGAAGAGACTTGAACAATATAAAAATATGCAATTATCCTAAACAAAAAA
CATGAAAGACGATGGCTTTTATGAACATATTAAACATGATTAACTATTATATTAGCGAGTTAACAAGCATATGAC
CTAAAATACTTAGCTCCCCGTAATATAAGTTTTTTTTTGGAAAGTCATACTTGAAGCCGAATATAGGGAGCGTCA
GTGAATTTTTTATTATACTATCGAATGATTCTAATGGGAGCATTTGATGTGTTGATACCTAAGTGGACAACAGGT
TTTAACAAATTAGAGGGAGTTGAGATTCAATAATAGAAAGATCAACATCAATGCTATTGTTTGTAAAAGTAGGGA
CCCTTTCTATGGGCTAGCAGGTGCAAAGGACATTAAGATCAAACTTCAAGGTGGTGTCTAACATTAGTTCCAAAT
TATAACAGCAAAATTGAACTCAAGGAGGACCATCAAAACCTAAAAATATGATTCCCCTATTTCAACAAGCCTTAA
AACATTCATAGGAAATGGTATATAAATATTTTGAACTACTAAATATTTGTATATGAAAATTCAGCTACTTGAAGA
AGTAAAAAATAGCCGATTATCGCTAATGTAGCACAAAAGTGAAACTCATTGTCCTGGCCCATGAATTTTGTTAAG
GAGCTGTAATAAATCTTTACTGGCACTTCTTGGGCACTCATGGTCATAAATTAGGTGCTATAATAATGGAATCCA
GGTTGACTTAGGTTTAGCTCCACCACCCAATCCTAGGATGAAAAATTCTAGCACATGCTCCTAATTCCTATCTCT
ATTCGATAAGAATACCAGCATTATGTAATTATTATGCTTATATTTTTTCTTATTTTTTATCTTATCTCATGTTTT
TCCCTACATTTTATTCCTGCTTAGACAAGCAGTTGTTTAGTCTCATTGGAATAACAAAAAAGAATTGTTCTCTTT
CCTTTCAAGACATTAGACCTCCCACCGTGAGTAATATTTATTTCATAAAGCTAACGTCTGCCCAAAATAGCATAA
GCAATGTCCAGAAGCAATACATCACACTAAATCTATAAATTAACTAATCAAAGCGAAAATAAAAATCTCTAAGAT
AAAAAAATATCAATTTTATTCAACCATATATCTTTTACATAATTGGGGTATGATGTGATTCATGTTCCAAATTCA
TACAATTAGGCATGAAATTCATGCAACCCTCTTTATCAATTATGAATTCGCAAAACTAATCATTTTAGAATAAAC
TAAATAAATTAAAGTATTCTTCCAACAAGACTCAACATTGTAAGCATCCATGGTGCATTCTATGAAATTCAACAA
AATTATCAACTAAATCACAACCCACATGCATATGCAAGTTCCATTGCTCAAATCTAGTTGGACCAAAATATAGAT
GCAGTTAATAGATTACTAGATGTAACAACTAAATTGATTGCCATGATGAAATGATGCATTTCTTGAATAAACAGT
GAGCCACAGAAGCAAAATAATATTTTCGCTATCAGTTTAAAATTGTGCTCCATTGATGGAATATGATGAGAGATG
TAAAATCTCTTAATAGTTTTGGTCTCATACAAAACTTAAATTTGCTCATTTATTCAACTATAGTTTGTCATCTTT
TAAAAGATTAAATTCATCAAGCAATTAATCCCTTATAGGATGAAAAATCATTTTCATTTAAAAACTCTTCAATTG
AGCTCATATAATAGTATATTCCTTTCATTTTATACCTCCACGCTATACAATAATTTTAACTTGTCCAATGAATTT
AATTTTAGCAAATCCTATTCATTGTTTATTAGATATGATTTGTCATTCATAAAAGACATCTAAATCTAGCATCTA
CTTATTAAGAAATTTTAAGACCAATTTCATCATATTATTAGTATGCTATAAGTTTTTCATCATGTACATCAAAAT
TGACTTTTATAATTTTATCAGCATCATTATTAAACAACCTAAGTTATCTAAGCATCATATTTTCTCTCTTTTCAG
AAATACAATTCAAAATAATGTTTCCTTTATATCGAGATTGGAATCTACTTTTTCACATGCATAAATTTTGATTAT
TTTCTACGTGTAAAACTATGTTTTAAGCTCATTAACTTCATCAAAGAATTGATTTCATCTATTAAAAAATTCAAG
CGCTTTATTTTACAAGATTTCAACCGGTTACTTTTCAGGACAACCTCGCTAGTTTGAGTGGTATCAAGTAGTAAG
ATTAATTCCAAAAATTCAGAATGCCCTTGAAAAGCGCTCCTTCACTAATTCTATCCAAACAAATCAAAGTTAATG
GACAGATATATTTGCTTAAAATTTGGTCAATTGATGGAATAAATTAAACTAGTACAACAAAGTCAGCAATTTAAA
ACAAATCTAGGGACTACGTTGGACATTTACTAATTAACAAGATATTAAGTTTCATATATTTAGGATAGCAATATA
CTTTTGAGTGAAAAACAGTTGAAAATGTGATGATATGACAAAGATCACAAACACAAGAGACTAGTTTTGCATGGC
TATAAGATCCACCGCCTACATAAAACCCACAGAGATCAAATATTTCAGAGAGCTTCAAGATTGGTGCATAAAGGG
ATTATACAATACAAAGTAGAAGATATAGATAGAGATATCATTTTGCCAAATTCAATGGCTGATAAGA**AGCACAAA**
**CACATGCTGCAAAGATGATGGGTAAACATAAGGAAATGCTTTAATGTATGTACTCCCTTACCCTTTTGTTGTAGG**
**AGTTTAAACGAGCACTGTCTTTCTATTGTTGATAGCTCCTCCTCTTGCTAGAACAGAAAAAGAGGAAACAACTCC**
**TTTCTCTCTTGAAGCAACCGGACACTGCTCTTATAAGCACTAAAAAACTTTGCTTTCTTGGATGGCTGCCCTCTC**
**CTTTATCTCATCCGCTCTTCTCTTTGTTTGTTTCATGGTTGTGAGAAACTAAACCCCTCTTAAATAGTT**CTATTT
GTAGTGGATGCGAGAGTAGGATATTAATTAAAAACTTTATAAGGACAATTAAAATTAAAAAAATTAAATCACCCA
AAAAAATGATATTAACATAATGCAATATTCACTTCCTGCACAAGTGAAAATAGAGAATAGAGACAACAAGGGAGT
TTGTTTGTTATGAAATTCCAATGTTGCAACACGTAGCAAGTAATAAAAAATAAGAGAAAATGCAATATTGTTATA
AATTCATTCACTTCCTGCACAACCACAAAATTGAAGCTGCATATGCCAATTTCAAACAACCTCTAGAAGTGACAT
TTTAAGGCTTGCAATTCCAAAGTGGCAACAACCCCCCGAAGTTTGTAGAAACAAGGAAGCCTTCTAATTGTAAAT
AACAATACACGATCTGTTGGAACTGTATGAACTGCCAAGAATATCATAATTTACTTTACGTCACCAATCAATAGT
TTCAAGTAGCCAATCAATAGTTTCAAGTAGCAGAATTGATAATCAAACATGATTTCGGAAGAAGCATTTGTCTTT
CTTTTGCATTAAATGGATTAAAAGCACTTTTATGCATTTTTTTGGTTTTCTTGAACCAATGAACATTGTTCAGGT
CTATTCAAGAAAGGATCGCCCGAAAAAAAAAAAAAAGAAAAAGAAAATAGAGCCAACAAGGTAGTTTGTTGACGGA
AGAGAGAGAAAAGAGAAACATTTAGAATAGAAAGGACAACAACATACATTGTTTTTCCCTAGTTAATAAACATTA
AGCATGAATGAAGAACGGGGAATTGTAGGTCTGACCCCGAAAACTTTAAAATAGTGCTCATGATGATAATTATGA
TAATGATAATGAAAGCTTAACACAAGTAATTAAGGAAATACTTTTAAAAAGTTCTAGCAGGGTAGAGGCTAAAAA

GAAAAGAAAGAAAATCTATGGATGTCAGTACATCTAATATTAACACAAGTACATGCGAATTCATCTAAGCATGTT
TATTACTAAAAATAGTATTCTAACCAATACAACCATGAAAATTTTAAAAAACTAAAAATCATGATGTTCTCTATT
TCAGCATGAAGTCAGTGATTGAACGCTACAGCAAGGTGAAAGAGGATCATCAGCTTGTGATGCATTCAACTTCTG
AGATTAAGGTACATAAACATGAGATTTGCATTTAATCGGTTGTCATTTTGACTGTGAAGATGATAGAGCCTTTAT
CCTCTTTCTGAGTTCAATAAATTATAAATTAGGCATAAAGAGCTCTCACAATTTTTTCTATGACAGAGCATAAGA
TTTTAGAAAAGATATGAAAAGTATTTAAATTCCTGCGTAGTCATCCCTGCTGAAGAAAATCGAAAAAATTAACTT
TTTATGAATAAAAACCTCATTAAACTTATACGCAACATTATAATTTTCATGTATTATTTTTCAGTTTTCACCACA
GCAATTTGCATAATCCTTGAGCAATTGCTCTAAACTCACATATTGTACTTGATTTTGAGACTAAAGGTTAATTTT
AGGCAATTGCTTTTAACTCAAATTTCGCACTCAATCATGAGAAATCAATATTTTTTTACTCCACGTCATGAGATT
TTCTTTGAACAACACAATATCCTGAAGTTGATCTATGGAATAATATAATCCTATGGTCAGCCATTAGTATGGTTT
ATTAATTTTAGAGTGTGTGTTTTCTGTAATTTTTTCAGGATTAAATTTTAAATACCTTAGTATAGTTTCTTGGAG
GAAAAAAATAATTATTCCTCTTCAATTTCTTAAGATTCAGTTTCAAGAAAAATAGCAGGGGGAGGCCACCACACC
AGCAAAGCCACAAAATCCTCATGCTAGATGGATAGAAAGACACAATCGATGGCCACGGTTGGCTGTTGAATGGCT
GAAAGGTCGGTGGCAGAGATGGTTGCAGCAAGTGTATGGTGAGGTGGCTGGCAGGGTGTTGGAAGGATCGAGGAG
GTTCACGAAAGCAACGAAGGCGGCTGGAAGGGGGTCGTTGCAAGGTGGGAGGTGGTGAAAGGGATGACAGCAAGA
TAGGAGGGGGTTGTGTGGTAAGGAGCTATTTTTCGGGTAGATGCAACAAACACAGTGACGAAAACTGTAAAGGGA
GGGTCGTGGGGATGCAGGAAAAGGTAAATAAACATCACCTCATGACGAGTTATAGTCGGAGCACCTTCTCACCAT
TGAATATATAACCATAGGAAAGAGAAATAGAGACATCAACGGCTACCCTGACAAAAAATAAATCAAAAACTGACC
TAAACACAACAAATAACATATATATAAGATTAGGGTGACTAAGGGGGGACCCATTCCTATTGGGTGACAAACCCT
ATGCCCAAAGGAGGGTCATCCAGTCGACCCTATTCCCGCTCTACTAAATTCATCCAGGTTTTTGGTTCTCCTGCT
TCTAGCCCAGGGAAGGAGCATATAAGTTAGTCTCACATAATTGGGGAAGCTTCGACCTCATTTATTGAAGTTTTT
ATTACGTCATAGTATGAAATCAGATCAAGCATGACTGTAACCTATTGGGATTGAAGGGTTCCAACTACATCGATA
TGCGCATCTTGATAACATATGAGAAGTCTTCCCACAACATATGAGAAGTCTTCCCCATGCGAGCAAACATCCATA
TTGGTCCAAGTATATGGGGATACACCGCGAAGCTAAAGAAGGATTTCATGCCAAAGTGTATTTTGTAGAATCCTC
CAAAAGTAACCCAAGCTTGGTTCAACCCGATTTAAAGTGATGGAAACTTGAATCTCCCAGATCAGTGGTGGTCGT
GGCGATAAGCTATCAAATATCATGTCCCAAAACACCCTATTTAATGATTGTCATGTCTGAAACTAGCTCTTCTAT
CATGTTCTCCTCTGTCATCACCAAAATTGATCCGGCTAGCACTTCTTTGATGGCAAGGACACATGCACTTTCTTA
CTCTCCAAAAAGTATGACTATGAGTTTACCATCTTGCGTAAAGGAGCGGCAAAACTATTAGGAATCACACAACAA
ACAATTTCCAACTTTACCATGGCTTAAGGTCAATTAGAAACTAAACCACACATCGAAATAACGGTATTGCACCAA
TTTGATAAAAATTTCATCAAAAAATTAAAACATGAGCACAATCAAAACAACCAAAAAAGCACACGAGAATTTATG
TGAAAAACCCCAGAACGGAAAAAAAATCACAGAGTCACACCACCAGAAATTTCCAATATGAAGCTCAAAAGGTTTT
ACAAAGAACTCACAAGTTCTCTCAACACACAAAGAAAACTTACAGAACTCATTCTACAATCCTTCCCTCTGCGAT
CCAACATAAAAACCACGCTGAATCACCAAAATCCATCTCTTAGACACTCCTAACCTTCACTGGGATTCTCATCGA
AGTACACCGAAAGAAGACAACAACAAGATCTCCGATCCATGCTAAAAAATTGGCTTAAAAACCACTTTTTTGTGT
GGTAACAACACTGGACGCTTAGGCACCCAAACATGGCCCCTAAGCGCCAAACACACTATATTGATGCCTCATAAT
AGGTGCCTAGGCGCCAATAGACCAACCCCTATGCGCTGGTCTCTCGGGACTAAGAGCAGCGCCCAAAGTTGGTGT
TTAGGCACTGGCTCTTTTGTTCGAGCAGCACCCAAGTGGGCGCTTAGGCCCACAACCCGATGCATACGTGTTGGG
GTTTTACCTCGTTCAATAAAATCAGTCTCCACTCATAACATAAACTTTATTGCATACTCGACCGACAAGTCCTGA
TGTAAGTATTGGAGCCTCATGAAGAGATCTACCAACAGGGTTAGGTAGGCACTATCGCTGCAACTCTTTCTTCAT
CCTTTCCAAGATAGTAATCTAACTCCTCTCTCTATTTCTCTCCCTATTCAAATTTACCCACCACAACGGGCGCCC
TAGCAGTTAGTTTCACTTTTTGGTCAAAGGCCCTCTGAACGGTATGCAACTATTCTATGAAATCATCCAGATTCA
TTCTCCCCTTGAAATCCAGTATGTCTACTTTCACTCTTGAGCCTCTGTATCCAACAAACACTTCATCTTTGTATC
CTTTAACACGCACTTCATTTCTATATCCACTAACGGGCATTTCATCTATGCATCCTCTAACGAGTACTTCATCTC
TTGACCTCCTCTGATAATGTATCCCAAAGCATCGCGGAATGTCTTCTTCTCCATTCGAATCATCCCTATGGAAGG
GATTCAGGTGTCCTTGCCTCTCCTCTTCGGATCTTCCTACCATTCTCATTGGTAACGTCTTCATGGGTCATTTCC
TCAATGTCATCGAGTAGTTGTAACACTTGATGGAGCTACCGCTGTAACTTTTTCATGGTAAACTCCTGATCAGGT
TCTTGCAATTGCCTTTGTTGCTAGGGCTTTAAATGAACAAATGCACTCGATAAACAACCCGAACTCAACTTGAGT
AAGGCTCGATCATGTTCGTTCAAATTAGTAAATAAACATGCTTAAACAATTTTTAAAACCTCAAAGCTTAAATGA
ACTAAATATGAATGTAGTCGAATTTGTTCATTTATGCTCACAAATAAACTTCTTTGTTATTCAACAAGTAACTTG
TTAAATACTCGATATTAAGTTCAATGAAGTTTCATATTTAATATATATATATATATAAAAAATAAAATTTAATTA
ACTAGAATTTATTTAATTTAATATTTAATAGTAGTATATTATTATATAAATAATTTTTTTAGATTAATTATTAAT
TATATATTTATTTATTAAACTAGGTGTTAAATGTGACATTACTTGAGACTAATTTGTTGTATTAAATGCCCCTTA
AAATTAATTTTAAACCCTCACATACCTCACTTTCCTTCTAACCTCGTGTAAGTTCACCTTTTCCCTTTTTGTCTG
AAACATATGAAACCCTGAGTCGCTAGGGCATGCAAGTAACAAAAGCATGATGAGATTGCAGTGTGCAAGGATCTT
TGAAACGCTGGCTCGGTATATCTCCCCTTTCCACTCTGCTTTTAAGTCCTACCTGATTAACCCTCATTTCTCATC
CTTCCTCGCACCACACTTCACACGCTTCAACTAATTTAAATCTTATAAATCCTTCTTCAAATTTCATTTACTATA
TCCGAGATGTAACACAAAAAAGAGGAGAAATGGAATTTAAGATCCTGTATGGTTTAGATTTGTCATTTTTGTTGT
TTAGACCTATACGGGATTCAAGGAAAAATGGAGGATAAATCAGATTGAAGATCTGGTAGGGCTTTATTTGTCATT
TTTGTTGATTGTTAGGATTTAGTTTAAATTATTTATTAAATTTCTTATATGAAAATCTAGATATTCATTTGAACA
TTTAGCTCTAAATACTCATTCACTTTTCATAAACAAATTAAGTACAAATCAAGTTCAAACTTTGAAATATATAAA
AAATTCAAGCTGAACAATATATGAAAGCTCGACATAAACTCAAGTCAAACATTAACAACTTATAGTTACACTAAG
CCAAGTCCGAATAAGGATATACTTGGCTAGATTACGACCCTATTTGTGGCGGATGATTAGGGTGGGGAACTGAGT
CGGTGGAAGGAGATGTAGAGATTATGTAAAGTAAGAAAGAAAGATTTTGTGTCGGGACAACACTATAATGGTGGA

TGCTAGGAACCCTAGGATAAGACTAGGGTTAGTCTAGCTCTAATAATAAATGATATCCAAGTTAACATAGTATGA
AAATATAGATGTAACTTCTATTATTGAAATCAAATAGGGATTACAACAACGCTATAATGGTGGATGCTAAGAACC
CTGGGATAAAACTTAGGTTAGTCTAGCTCTAATAATAAATGATATCCAAGTTAACATATATGAAAACATAGATGT
AACTTCTATTATTGAAATCAAATAGGGATTGCAACCAAAGATATTCAAAACTAGGAAATAACAAGAGACATAACA
CATACCCAACTAGGGCTTAGGTTGAGGGTGGTAATGCTGCAACTTGCAAAGAAATAGTAAGTGTGGAGGGTTAGA
TGGTCAGAAAACAATTATAGAAGATCATGATTGGCTGTCAGATGGTTGGCGAGGAGACTAGCAGGGTGTTGGAAG
GGTCGGGGAGGTTCACGGAAACAAGGAATGCGGCTGGAAGCAGGTGGGAGTTGGTTGGAAGGTAAGGCAGCAAGA
TGGGAGGCGGTTGGGTGGTAGGCGACATTAGAAAAACAACAAAGGAGTTGTTTTCCGGGTGGCCACAGCAAAAAC
AGTGACAAAGGAAGTGGAGGCAAGGCTGTGGGATGCAGAAAACGATCGATAAACACTATCCCAGAATGAGTTATA
GCCAAAACAACTTCTAGCCTTTGATACAGCCCTAGAAGGAAGAATTGAAGACGCTGGCAGCTGTCTCAGCAAAAA
ACCACATCAAAAACTGACCTAAACACAACAAATAACATATATATAATACTGAGGTAACCAAAAGGGGTGCCGCCC
TTGGCCCAAAGAAGGGTCGGCCGACCCTATTCCTGCCCTGCTGAATCAGTGTAGGTTCATAGCAGTCCCACTTCT
GGAAGGAACACTAGCATCATAAAGCTATTCATAAATTTTATGGGTAATTAATGCATAAAAAACACTCATTTAAAT
GGATATAACATTACAGTTCTACCTAAATAAATAAGCAGAAGGAAAAAAATCTTTCATAAGAAACAAATGACAAAT
GCGACATGGAATGCCTTAAATACTACCATTAGGGAAAAAAAAAGATTTTTTACATGGATTAAACGTCATTAAAAA
GCTATTTAACCTCTGCATCATAATTTTATATGTAAACACCCAAGTTATAAATATTCAAAAAACAATATCACCTGT
ACAATAATCACAATAGCATGACTAATGGTGATTTCCAATGAAGAATGTTAGTAGAAAGGGCCTCGAATGACACAA
AATCAAGCCAACCTTAAACTATGCCAAGTAAAAGAAAGGTTCTATGACAGCAAAACCTTTAGCAAGCTTAAAAAC
ATGTTTTAGCATGCAAACAGGTTTTTAAATATACATTCAGCTAAAAATATTACAAAGTTTACACAATATCTATTA
TTTAATTTGACCAAATCGGTAATATAGCAAAATCTTTCTTGATACTATGCACTAGTTCTTTGGAACATCCTGGCA
GATAATTTTCACAAATAATATAGTTTTTATTTGATTTACCTGCACCAAATCTGCATTAGAATTATCAAACGATTT
TGTTTTGTGCCCACTTGAATTGGCAGAAGGAAGCGGCGAGCTTGAGACAGCAACTGCATAACTTGCAAGAAAATT
ATCGGTAATTTCTCTTCTCTTAATATCTGAACAGCACGTTGAATGCATGGCATGTTAGAGGATAAAAAAAATCAA
AAGCATAAATTTACCTGTGTGAATTTTATAGTTAATGTTTAATTCAACAAGAAAAAAAAAGATGAAAAAAGTCTC
ATTTAATCAGAAAACAGTTTACAAAATGATCGGTCGAAATTAAAATAAAATAGTAGAATAAATTTTGCTCTTGTA
AGATTGAAAAATTAAAAATGCAACAATTCAAAGTGCTGCAACTATACTGGCGGATATAATTTAAACTGCGCTTCC
TTGGAATAAAAAATAAATAAATAAACAAGCCTCCTTAAATAGAACATGGCAAACATTTATGCCACAATTTACATG
CATTAATCACCGGGGCCACTCATGACAAGAATGATAAAAATTAGCAAATAGAACAGGGAATATCATGATAGAGCC
ATCATGAATCACTTTGTTATCCGAGAAGTAAGGATTCCTTTCACAATGAATCAATAATAACAGCCTTGAGACTAT
TGGAGTAATCCAATTAAGTCAACCTCTTCAACCCTGCTTGTCCTAAAAATAACATTTGACACACATGCTAGCCTT
TATTACTATTAAGGGTAATAACAATATAAAAGAACTATTCACAGCAATTCTAACTTTAATTGTGCTACAAAATAT
AAGCTTGAATAAAGAAAAACGAAAAGAAAAAAAGAATAATATCTGTATGATTAAAAACAGCTGGTAAACTAGAAG
TCTAGAATTACATACAGTTTTTATTAAACTTAATTTATATCACCTACAATTTTCTTCTAAAGTCTACATGTGCAC
TCAATAAAAACAAAAAATATAAATACAAATATATCCTATTTAAGGAAAATAACCAATCAGCTTCTATGAAAACAG
GCATTCAACAGAATGTAATTCTTGAAATATTTAATATTCAGTGTTTCTAAAATGCATGAACCGCTCAACCATTCA
TGAAAATGTTTGAATTACATGTTAATCAAAATTTAGTCCTCCATATAACTTCTTCTACAGATCATGACTCTGTTT
TATAAAGCCTTCGAAAAATTACCATGACTTTCGATCAAAATAATCCAGTTTATGACGCCGTTTTAAAAGTATATT
TTAACTACTAAACAATAGAGAACATTTTCTTTCCCATTTGACTAAATTCGGACTTCAAAATCAAGTATCTTCGCC
AATCTCCAATTTAATAAATTAAATCTCCATTTTCAGAACCAGCAACCTTAATTAATAAAGGACAAAATAAATTAA
GTAGAAAGATTTATCAAAGATCCTTTTGCCAAAAAACAATATTTTCCTTCTGTAGTTGGGAGAGAGATTAGTGAA
GAGCAGAGGTAGATACTCTCATAATGTCCAATGAATTTTAAGCCCTCCACATATACTCTGCTAGTAGTGCACTCT
CAACCGAGACTGTAGGAACACAAGCTGATGAACCATCCTTTTCAAAGAAAATATCAAATATCAGAAGCCTCCTTC
CCCTGACCATGTCAAAGTCAGAGTAACAAGGACAAAAAAGTATTTATGAAAATTTAAGCACATAACAAAGCATCA
TAAGTGTGTGTAAATCTAAGAGGCCTATTGACAAGTAAAATGAGAATATATTTAAATTCTTCAAGCTATTTTTGT
CAATATTTTCTCTACGCCATTAAGTTTGCAAGTGTAAACCAAAATTTTTTTTATTTGAGAATTTGAGTCATGAAT
AACTGCATGATCACATCAACTGCAGATTATTCTGACAAATGAAACCAATCATAGAATGAGATCTTGAAAAATTTT
TACCAACAGTTAACTTGATAATTACAGGCAGCTGATGGGAGATGTTAATGGGCTCGGTCTCAAAGAGCTACAAAC
ATTAGAGAGCCAGATGGAAACAAGCTTAAGAGTGATCCGACAGAAAAAGGTATTTCGATCAAATACTTTAAGCGC
AATGTCTAAATACAGCCATCATTTTCATTATTTCTATATAATCATATATTAATCTCTCTGGCAGGATCAAGTTTT
AATCGATGAAGCTCAAGCACTACATAGAAAGGTAGCCCCTCTCATCTCATCTGAGATCAGATGTTTACAACATCA
ACATCTCAACTCAATATATCCATTGCAGGTCGATGATATGGAAGAAGAAAACATGAGGCTCTACAAAAAATTAAA
ACTTTCTCAGCAAGAAAACATGGAATTGCACCTAAAGGTAATTATAAGTACATTAAAAGGCTTGATGTTTGTTTT
CAATCATATTCGGTCCAATAATTAGCCAAATAGCACTTGAGCTTGCCAAAATTTGCCTCATATACAGCGCTAATA
TTCATCAAAGTGTCCAGCATCATTAACACTAAAACTTAAGCCAAGTATTTCCAGATAGAAGTTCTATGTACTGCC
TGAACATGAAAATCCATTTTTAGGTCGCCGCAATTGCAAGATTTATGGGGAGATCTACTAGTAGTGGCTCTCTTT
GTCCGACATCAACTGACCTTGAATTAAGCCCGCCACAGAAACAAACAAGTGAAATGCAAGAAGACGTACTAAAGC
TAGGGTAAACTCTCATAACTTCATTCATTGAACAAAATTTAATGTTGACAGACATTGTTCTTTATTTAATCATAA
TAAATTTCCTTGTATCCTGCTCCAGACTTCAATTGCTTTAAGAAGCTCCGATACATCAAGAATATCAATTAAGAA
GTGTGAGATTACCTCTAAAGGATCACTTCATATGTGGGGTCCTGCTGTGCCAACAGAATGAATTTTCTTTGCAGT
TTCAAAGTCTCGGTTTAGTTATTTTTGTTAGCGGAAAACATGTTCTCACCGATTAATAGTGGCTTATTTTACAAA
ATGGTGGTGTAGAAGAGGAATTGTCTTTGTCAAATTTCAAGATGTCAATTCAATTATGTTTCAAGGAATGCTAAA
CATAATTTCATCACCACTAAATACTAGAAGGCATTAATCTTCTCGCATTCCCCTGAATCTAATGCTGGTCATTCA
CATGTGCTGCATTTACATACATACCACCAAATTCAAACTACTAACATATCACACCTAAATATTATCCTAAAACTT

ATATAAAACCAAAAGACCAATTTGGATCAAAATATTACAATAACAACAAGCTATAAATTTAAAATACGACAATAA
TGCAAAAAGAATGAGAAACAACCAAATATAATAAAATCTGATGAAATTTCTCAAAATCTCCTAAAACCAAAAAAG
TCTGAAAATCATAAATTCTATCAGATAAAATATCAAAAAAAAAAACTTTATGAGTAATAAAAATGAAATATACAT
TTAATTCAAGGGAAAAAACGCCAAAAATATATAGTTTTTTTAGAAAAATCAAAAGCCTCACAAGTCGCTTAGTTA
AGAGCACTCGCGCGCAAAGAAAAGAGAGAAATAAGTGTTTCAACACGTCGTTCGTTATACGAAACTTGACCAATG
CCTTCTAGCCTAAGCCATCAATTTTAAGTAAATTAAGAACCGACAATCACAAACTGTCGGTTTCTCGTTGGCCCT
TGTGTTTGTAGATTTTCATTGCTTTAGTCATCGAAAATAACCGACGATCATGAGTCTACCGGTTATTCATCAATT
TTCTCAGACTTTCCTATAGTGCAAGTTGAAGTCGACAAGGTCGGAATAATACATCATTCAAGAACGAGTCGGATC
AAAAACATATAAGCTAAATACTCATAATGGCAATGGCAATGAAGTGCCAAAAATATGGCCAAGAGATAACTTCCA
AAAGATTGTATCCTTAAAACTCTTCTAATCTTTATATGCTTTGCACTTATAATACTTGTCCAGGCAAATTCACTC
TTAAAGAATAAATATTTGTTATTAAAATACATAATATCTTAGTCAACTTTAGAGAACTTCTTCACTTGCAATTTA
GTTAAAATAGAACAAAGATGCATTGGCTTAAGGATAAATTGAAAATTCGGATGACAACTTACAGATGGAGAACAA
ACACAGATTAGAGAAATCATTTAGTCTTTTTTCTTTCAAGATTGTTATATATATATATATACTTTTATCACCTTA
GATATTTTTTTATTTAGCATGGATGTCAGATCAAGACGTATCAACTGTAATTCTTCTAATATAGACTCTAGGGGT
GTAAATGAGTCCGAGCCGAGCTGGAGAAACTCTCAACTCGAGCCTAGTTTGAGCTAGATTAATTTTCCAAATTTG
ACCTGGAACTGCTTATTTATTATAATAGAGAAAATAATATTAATTTAAACTAGTGTTAAAAAAAATGAAAAAAAA
ATGTAAAAATTATGAGATAAAGAATAAGAAATCATTTTGATATAAAATATAAAGAATTAATTTTTTGAATAGTTT
AAATAATTTAAATATAAAGTCTTCAAATTTAATAAAGTTTGCATATATTTTATATTAATATTTTTTTTAAATAAC
ACCCTTGAACGGCTCATTTGATTTTAAAATTCGATTATGCATCAATTAATTATATAATTGGTTTTTACTTATTAA
ATTCACTATTGATCAGCTATTATTCATTACAAATTATAATTGTTCTCAAGCATCTAAAATCTCAAGCACTTAATA
TACTCTTAGACCACAAATCAAGCCAAGAAGATAGAGCTCCCTCTTCTCGCGACAAGATCAGTAATGAGTAGCGTG
CAAGTCTCTTCTACAACAATCTTAAACAATTGTCTCCATGAAATTATAACAAGATTCACATTAAAAAAAAATAAA
AAAATTAAAAGATGTTGCTAATTAAACGGAATAATTTCACCAAAAAAGTATATCGAGTGTTCATTAATTCGAAAA
ATTTATATATAAATAATTCAAAATTTGTCAATCCTATTATTAAACTCGATAAATTCACAATTAAATTCATTTATT
TTCAATTCATGTGGTTAATTCAAATACATAGTGCTCATCAAATCATCATTCATTTATACATACATATCACTTCAA
AAAAAAAATCAATGTTAAAAAAAATATTGATTTCATTTCCAAAAGATTACAATATTCACAATTCAACCATACAAAG
AAATAAAAAGGATAAAGAACTGAGGCGTTTGTCAGGCTTTCTTCTGAACTAGTCTACATCATATTTTTAAAGGGC
TTTTTTTATCTTTCCATACAGAAGTGAGTATATTGTTTCTTACTAAATTCTCTGAACATAACTATTGGGACTTCA
AGTTTTAACTATTGTACAAAGGAATACCTCCTTAATCTCTCCAAAGTCTACAACTCACCAAACATTTCATAATTG
TTAACCTTTTTCCAGTGGCGAGTTTCCCTAATCTTTAGCCAGAAAACGTTACACAGCACATTGCATTACAAATTT
GTGTTGTATAGTGCTCCTGAGTCAACCCAAATCTATACATAAAGAAGACCACCCAAGGGGGAAATGATCCTTTTC
TTTTTCTTCCAAGACTTTACTGTGCCTCTTTCTTATTTAAAATTCTCTATCATGCTTTATACTCGAGAGGGAGCA
CTTTTTTTTTTTTGGCATTCATACTATTCCTCTTACAAGCCTTACCCAAGTGATGTAAGACCACTAACAAATATA
TATATTGATACTTTTTTAAGTTTTTTTTTTTATTATTATTTTTCCATGCAGGAGCAAGTATATCCTTTCTTACTAA
ATTCTCAAAACACAAATCTTGGGACTTCTCTTTAAGAAGAATCTGATATACTAAAGTTTTAACGACAGTCCTACG
TAATTTCTCCTTAATTTTCATTATGGGCATATTCCTTTCAAACTACAACTCACCATACGTTTCATAATTTTTAAC
TTCTAATCTCCTTTTCAAGTGGCGAGTTTCCTTTAAAACCTATAGCCAGACAATTCTACACAACACATTGCATTG
CATATTTGTGTTTACGGTGCTCTTGAGATAAAGAATTACACCCAAGGGTAGGGGACTAGCAAGGCCTTACCCAAG
TGATGTGAGAACACTAATATATATATATATATGTGTGTGTGTGTAGGCATAACATGAATACATGTTTTACTCGATAG
CATTATCGGCTAAAAAAGTGAGCCAATTCTTTTCCAACTGATTAAATGGTTAGATTATAATTTTAATTGTAGCGG
TTTCTAAACCAACGGATTGATCAGCCGGTCAAGCAAACCGACTGAAAAATCAGCTGTCATTAACAAGCCAGTTCG
GATATAACTTAAAAATAGCTAGGTTACACTCTGAATTCAGTTGGCAATCCGCTAGAATGATCTAATATAACGACG
ACACGCACCAAAGCTCACTTCCTCTCCCTTTATCTTGTTCCAATTTCTCCTCTCGAGCAATGACACAAAACCACA
ATTCTTGGACGGCAAATATTCGGCGGTGACAAGGCCTGAGGCTTCGACTCAATAAACTTCACTTTTCTCCATCTT
CTTTCTCCAGTTCAACACAATAAACTTTACATGTGTTTTGGCTCAAACATGTGGTTTTGCTTATAGATTGCTTGC
TTGGATGGCAATCAGAGCAAGCCACCCAATTCCTATGTATATATATGTCTAAGACCTAATAGAAAATTAGGTTAT
CAATGTTTTTGTGGTTTCACTCGGATTGCATGCTCAATCAGCAAGAAATCTCAAGCATTCCACCCAATTCTTGTG
TATTTAATGTCAAAAACCTAATTTTGCTATAATTCATGGTTTTGCTCAGATTCCTTGTGTTGGAATACATGCTTA
GGACTGTTTATTGAGTTATAGGTTTAATTCAGCCAAGTTTGTTCCTTAATCTTAAAAGTTAGTTGCTATTTTTCT
TGACTTTGTTTCTTGTTATTTTCTTGGCTGATTTTTAGATCTTTTTCTTGGAAGTTTGTTATTAGCTCTAAGTTT
AAATATTGTAAGGCTACACAAAAGATGCAGTTCTAATTTTCCCTTCAATCTACAACAATTCTCTTAAGTTTTCTC
TTCTTTCTCATTTCAAGAAACCAATAATTTGGTATTAGAGCTCTCTTATTAAGGGACATGTGCGTGAGTTTGAGC
AACAATGGAGATCGAAGCTAGTTTTTTCGCAGTTGCTCCAATGTTTGACGGAGACAACTACCGGATCTGGGCAGT
TCTCATGGAGACATACCTGGAAGTTTTGGATCTCTGGGAAGTAGTGGAAGAAGACTGTGACATTCCTCTTCTCCT
TGAAAATCCCACCGTAGCACAAATTAAAGCACAAAAGGAGAAGAAGACAAAGAAGTCCGAGGCAAAAGTTTACCT
ATTTGTTGCGGTTTCCTCCACGATCTTCACTCGTATCATGTCGTTGAAAACGACAAAGGGGATTTGGGATTATCT
CAAATCCGAAGAGATGAAAAAAATTCGAGGGATGCAGGTGTTAAACTTGGTTCGTCGTGACTTTGAATTGCAAAAG
ATGAATGAGATAGAAACCATATATGGGTACTCTGACAGACTACTCAACATAGCAAATCGTGTCAAATTATTAGGT
TCTTCTCTGGAAAATTCAAGGATTGCTCAAAAAAATCCTTGTAACAGTGCCTGAAATATTTGAGACTACCATCACC
ACCTTAGAAAACACGAAGGATCTGTCAAAGATTATCACTCTTAAATTTCTATACAAGCAAAAGAGCAGAGGGCGT
AATGAGGGAAGGAGCCGTCGAAGGAGCACTACCAGTCAAGCATCAAGATGATGAAAAGAATAGGTGCCCAAGAAT
AAGAAGTTTCAAGCTGCAAGTAGACAAAGCTCTGCACCTGAGAACAGAAGCAAAATTGAAAATTTGAAAATAAAA
GATTAAAAGATTGAGAGTAAAAAAAAGATTACCCTCCATGTCAACATTGTAGCAAGAAAAGTCATCCTCCTTTCA

AATGTTGGAGAAGACCAGATGCAAAGTGCAACAAGTGTGGAGATGAAGCTGTTATTTGCAAGAATAAAAATCAAC
AGTAAGTAGATACAAGTGGCAGATCAAGAAGAAGACGACCAACTTTTTGTTGCAATTTGTTTCACAAGTTATGAA
ACAAGTGAAAACTGGCTCATTGATAGTGGTTGCACTGATCATATGACACATGACAAGAAGTTGTTCAAGGAATTG
AAGAGCACCAAGTACAAGAAGGTCAGAATTGATAATTGTGAATATATTGCAGTCAAAGAAAAAGGGAACCGTAGC
CATTGCAAGATGTTCAGGAACCAAGTTGATCACTGATGTTCTCTATGTACCAGATATTGATCAAAACTTACTCAG
TGTTGGTCAATTAATAGAAAAGGGATTCAAAGGGATTCAAGATTCAAAGTGATGTTCATGGAGAAGGTCCGTGTG
ATAGAGGATGCTACGGGCGAAATGATGTTTGAAGTTAAGATGGCAGGGGGGAGTTTTTCACTCAATCCAATGCAT
GAGGAACAAATTGAAAGCACTATGAAGACTCATCAGCAATTGAGATGGTTGAGCATGTGAAAGTTGATGCAGGGC
TAAGGAAATTAGAACTGGAGTCAACCTTACAGAAAACAACTGCAGAACTTTTAGAGTTGAAAGCTCACCTTTTGA
GTATGGAAACAGAACTCCCAACCATGTCGCAAATGAATGACGAATTGAAAGATAAAATGGATAAGGCAATGGCGA
GCCAATTTGATTCAGAGATGAGGCTGATAAAGGCTACTAAAGAAGTCACAGAACTGAAAGCAAACCTCTTGGATA
AGTAGACACAGTTGCAAAGCATTATGGATGAGAATGAACAACTAAAATCGGAGCTTAAGAGAGACAGGGAGCATT
AGGAGAATTATGTTGCAGCTATATTTTTGGATGCTGAGAGAGCAAAAGCTCTAGAATAATAAACATTGATGAGAC
TAGGTGTTGTCAATGGGGATGCTGAAAAGAATAACAATAAGGCAACAAAGGTTTTGGAGCAGCTTGAGGCTACTC
AAGCAGTGCAATTAGAGATGGAGGCTGAGCTGAGGAGGCCATCTGAAACAGCAGCTGAAACAGTTCTCACACCAG
GGAACAATGGCATAATAGTTGAGAAGACGAGCTCTCTTAAATCATACGACAATTCAGGAAGCTGCTAAGCTCCCC
TTTCTCCAACGACTAGGATGATGATTCTCCAAAGAGGAAGAATGACAATATGCTAGGGAAAATTTTAATGGATCC
CAATCTGGAGCAGAAAGAAAGCACTGAGAATTTTGTTGACAACCAAGCTGTCATTTTCATTTTCCAAAATTCTGT
GTTTCATGGGAAAACTAAGCATTTCAACATCAAGTTATTTTTTCTAAGAGAAGTGCAGAAAGAAGAATTGGCGAC
TGGTTTACTACAAATCTGAAGACCAGTTGGCAGATTTGTTTACAACGTCACTTTCAATCAGCAAGTTTGAGTTTC
TAAGGCAGAAATTTGGTGCACAGCTCCTAAAGTAAAGAGGAGTGTTGGAATAATGCTTAGGACTGCATATTGAGT
CTAGGTTTAATTGAGTCAAGTTTGTTCCTTAATCTATGGAATTAGTTGCTATTTTTCTTGATTTTGTTTCTTGTT
ATTTTCTTGACTGATTTTTAGGTCTTTTTCTTGAAAGTTTGTTATTAAAAGTTTGTTATTAGCTGTTATTAGCTC
CAAGTTTAAATATTGTAAGACAACATAAAAGGTGTAGTTCTAATTTTCCCTTCAATCTACAACAATTTTCTCAAG
TTTTCTCTTCTTTCTCATTTTAACATCGGTCAATAAGAGATTCGAGCAAGCCACCAAATTTTTATGTATTTACAT
GTCTGACACATACTTTATCACCTAAAATTTCATATTTACAAACATAATATTTGGGTGCTTTAAACATATGCCTAA
AAAAGTGCATTTCACTTTTAAAATGATTTAGAAACAAATGAAATGCTTCATAAATTTTTAAGCAAATGATTGTCA
TTTTGTTGAGTGGTTACCCAAAAAGTTATGTGTGAATGTGGGCTTAGTTTGGAGAATTAATCCACGTTTATCTCA
TTAAATTTTAATTTGAAGAAATGGGCTTCGGTGCAACCAGAGCTCAATTTGTTAAGCGAGTAACTGAGAATGAGC
CAACGAATAGAGAAACAAAAAATTCTATGAAGTATTTTATGTTGCACTACAACTATTGTGAAAAGATTGCACCATA
TATTTTGAACTTTTAGTGACTCCTAAAATATTTGGTACTAAATCAAATTATAATGTGTCCAAAGAATATTTTAAG
CAGGATGATTAAAAGAGAAACTGAAAAAAAAAAATTGTATCAAAAGACTTGTAAAGAATGAAAAAATTAATGTTG
AAGTTAAGATTGACATCAAAAAAGATATATTTCTATGTTAATGGTCGAATGTTTTATTTGTCGACGGTATCACAC
TATACAGTCAGCCAACTTCCACATACTCATGTTAGCCAGTGATAGTTATGTTGCATAACCTATCACATGAAATGT
GCCTGACTGAATATTTCATATTCTTCACATTGATCATATCCGATCTACATAAGCCAAAGAAAAGATTTGATGTAT
ATTTACAATATTTAATAGATGAGCTTCAAATATTATAAGAAAGAAATGCGATCACTTATGACATTACAACAAAAT
AGAAATTTCAAATGAAAGTCATTTTGAAAATGACAATAAGCAGTTTTCTTGCAAACAAATTGCCATCGGAATGGA
GCACTGCACAAAAAATTAGAATGTCCATATAATATAGATCTATCAAAAGCGTTTATTTTAAAATAAGATAGGAAAA
ATTCGTGATTCAACTATTCTCGAGTATTCTTACCTATTAATCAACCATTCACAATAAATAACATTGAAAAATTAA
GATCATTGCTAATAATATCTGAGGATGTTTTGTGGCAAATGGTTAGTGCAATTTTAAATATTACTAAAATTATCT
TCACATCTTTCTTATTTCGGTGTTACTCATAACTAGCCAAAGCAAAATATTTTTGAGATGTGTTATACTAGGGGA
ACAATTTAATTATACATATTTTTGATGTCGTGCACATTTAGAAAAATAAGTTCGATAATATTTTCAATATTGTGA
TGAATATTAAAGACAAAACCGATTATAACATTAAAATTAGATTAAACTTAAAAAATTGTTGCAAGAAAAAAACAC
TTGAAACTTCAAGAGACAACTATGAGTAGACTGATGAGATATAAAACTAAGTTTCTTACTTTAGATCCGCAAAA
AATAATTTCTTAATGATTGCAATCTTTGAATTTTTCTAGTACGTATGCATCTAACTTAAATTATTCTGTCGCCTA
AAGAATAAGGAATATATTTGACATGAAAAGTTATGACAGTCATATTTTCTTACAATGTTTGATATATGTTGCATT
TAATACTTTTTCAAGTCATACATTAATTAAATTCAATGTATTTTTCAAATATTTATATGCAATTTATTTGAAAAT
AAATGCATTGAGATAACTTGAAGTGAGCATTTCAGTGACTATACAAGCTTGAGTGAATACTTTATTATGACTTCT
TTAATTTGATGGAGCTACATCTAACTTATGATTAAAAGCTCGGTACAGTACCTATGAATGTATCCGTTTGAAAGA
TGCGTCATTTAAATGTCTGATTATATAATTTACAAAGAATTATTACAATACGGTACTAAATAATTAGCACTATCT
CATACTTTTGAATAAAATAAAAAAAGGAAAAATAAACAATGAAAAAAAATATTAAAATGTGGATGTGAGATAAAT
TTGCTACATAAGAAATACTTTTTAGTATTACTTCAAGCACCAACGATCAATAGAATAGGCTGTAGTGTGGGGTCA
TTGCAATGTGTGGTTTATGTATTTCAAAAATCACGTGAAGTATTAAGGTGATGCAATTCATGTTATCTCAACGAC
AACAGTGAACTTTGACCAACGACTGTCTATTCAATTTTAATTGTGATGAATTTTAATTATTTCTTAAATAAAACA
TATAATATCTAAATAATATTTAAACTAATATTGAAAGTGTTACATTCACTTGATAATCAAAGTTTTAATTACATG
TTCAAGTTATTTGTCAACTCTTTAAATATTGAGGATGAGAGTATTAGCAAAAAGAAGATTGAAGAAGGGTTCACA
ATATGGTACAAATTATATGGGATTTCTACTAACTCAAAATTTCACTAGACATAAATTGAATGCTCAAATTACATA
TCTTTTGATATTTTTAAAAGATTCATAATTAGGATGCATATAGTAGATTTTTACTATGCGGACAATAGTTGATGC
GAAAAGATACAAATTTCACTTGAGATTCTGCTCCTTCGAAAAAAAAAAAAATTTGATATTTGTGCCAAAGGAAAAA
TTCATTCCGATAATAAGATATACGAATTTTTAAAAAATTGTGCAAATGCAATATTCAGGTTCTATTGAGACCAAT
GTATTTCTATTCAACTATGATGAGTGTGATCATGTTCTCAAACAAAAAACAAACATATAAATCATATGATATCAC
TAAAGTTAAAATAAATAAAAGATAAGTCAAATTTGATACATTCGTTTTAGCAACTCAAGTCAATTAGATATATTA
ACTTTAATACCCTAATAGACAAAGTTGATTGGGCAGTAACAATAACGACTAAAACTAAAGCACCTATTAAAATCC

AAAGTAAAGTTAAACCAACTCATCAGAAACAAAAAGAAAAAAGTTTGATTTTTGTATTCTCTATGCTTTTATTGA
GTAAAGAATTACCTTAAAAGATGCTATTAATGCATGTGAAGTCAATCGTGAGTGGATGAATCTGATGATGATACA
AGCGAGAAAATCTTAAATTTTAGCATTTACTCACTCCTCTTACGAAACAGAATAAGATGGGTAAATAATTTAGTT
TTCAATTTCTCATGTTTTTATTGCTTGCATTTTTTTTTTCACTATTTGTTACAGTGTTGACTGCATTTATGTTGT
TTTATGCATTTATTTCTATGTTTTTTTTTTACTACATTATTTTTTGCTTATAATTATTCTAATCAATTTATTTCT
TATTACAAATGACTGCCAACGACAACTTCCACAAAGGTTCTTGCAAGATAGCAATTTTTCATAGTTTATATCAAA
GGAAGAAGCAAACAATTTTGCGCAACACCCACAGATGAGCAGGACTGTGACTGTCATTTCCACGACTGCCACCAA
GTTACTCATAAAGCATGTTGCTCCACCTTCAATGAAACATGCGATCCACACCCCTTAAGCTCCCACCACATTGCC
ACCTCTACCCTCCCCTTCAAGGTAAGCCCCTCCACTCTCATCAAACTTGGTTTGAAAGGTACCTCAAGTCAAGAA
AGATACAAATCCGAAAGTAATTGAAATTATAACAGCTGAAATTATTTTTTATCATATTCTATATCTAAAATCCTT
TCTATATGTTTCATCTCGCCAAGTAGATATTTCATAACATCACGAGTAGTCATGTGATTCAAAGATTTTGAGGTA
AATCTAAATAACATTGAATTACAATTACAATCTTAAATTATGCTTTCATAATAACAAATATTATTTATATTTATT
TTGCAGAAGAACTTCTGTTGATTGCCTTAGCATTATCGACAAATTTGGAGAAACTTTGAGATCCGAAGACAACCT
GCTTATGACATATGTTTATTGTTATACATAAAAGATAGGGTAGGATGTTTGGGCCATGTTGTACAAAGCCAAAAC
TTCTCCTGAGTATCTCAAAAAGAGGCAGACGGTGCAATAGAAACGAGCCTCAGAATGTGGGGGCATGAGTGCCTC
TGTACACATCAGAGCCTTGGTTCCCCCACAATGTATAGAAGGCGGATGATAAGATTTATTTATTGAATTAAATTG
ATATATATTATTATTTAATTTCCTGAATATTTAAAGTGTTAACAGAAGCAAACGGTTGGACACAAGTCTATCATT
GCTGAGTTGCATGCTCGCTCACATCAACATCCTGAGTATCCCAACATGTCCCCATAAATTAAATTATTTTTTATG
TATAAATTTTATTTGCAATCTTATTTCAATGCTAAAGAGCATTTATAGACTTTTGTAGGAGTATGTGCACCTGAA
GAATGTCTGAAGTAGGAGTCTACATGAGTTTGTAGCTCTAGTGGTCAGTCTAACAACTCAAATATCGAGATCTAA
AGGGAGTGAAGCTGCTTGTGGGAGAAAGAAAGGAAATGTGTACAGTATCAATGCACAGGGATATGCATACGAGAC
AACAATCCGGCTCTGGTGAGACTTCATCATCATTTTATGATGTAGTGAAGCGTGTGGTCATAATCATCGAAGAAT
TTAGAGCCAACTTTGCTATTGATCTACGATGGCTTGAAGAGACAGTGCAAGCTATAAATGCCAAGGTTGAGAGGC
TTGAGGCCAGCATTGAGCCGTTGTAGCACAAGTCACATATGAATGACAAGCGTCCTATGATCGTGGAGGAGCACG
AGGAGCAACTAGCCCTTTATTTTCCCAATGTAAAACTTGATTTTTAGTATTTTCATTTATTTTGTATATATTAAT
ATATTGGATGATTTGTTGTGTTATAGTGTTATATTTGTAATCTACGATTCTATATTTATTTTTGTTTATTTATTA
TGAGTTATTATTATTTGCATCATTTTGTCATTGTAAGTATGAAAATTTTAATCAACTTCAACACGCTTTTATTTT
TGCACAAGTTTCTAGCATATCACCAACGTGTTTTTCAGTCGCTAGACAGATAACAGTTTTTCAAGCCACATATTA
ACATTTGGACTGTCAGCTGCTAATCCATCAATCGAATTAACAATAGAAAAATTCAACATGTATTTAAATATCAAA
TGAAAAGTTAGCAAGTTCTTCAGTCCACACATTTATTGTTGAAATTTTGATGCTAATCCGCCAATAAGCCAATAA
TGGCGAAAACTACTCAAGCCAATAAAACATTTCAGACAACCACCTTTCTAGCTGGCAAAAAATACAACGTAAATA
AGTTATTTAACTAAATTACAAACATACATATACCTTAGCACTCTCTAATTTATCTGTCGGCAATCGTGGAATTTT
CTATAGTAATACAATACGTGTGCGCGTGCGCACGCACACGCACACGCACATATATGATTCTATGTATATTACATC
ACCTAATTAAAAAAGAAGTATGCGTGCTTAAAAAATTATTAGGGTGCAAATAATCCAACAATATTTGCATAAAAA
TGTAATCAATTATAATTAATACTCAAAAGACAAAACAATTCCATTATAACCATGCATTAAACCAAGTAAGAAATA
ATGATTGGCATAAAGACAATGGGTCTATTTCCAATTAGAAACAATTTTTTTCTTTGCAGATGATCCAATACTTGT
AAGTTTTTTATATCCTCAACATAATGAGATTTTGATCGATAAAAAAGGGTACTTCTAGCACAAATTTAATAAGAA
TTTTAAATAATTTGAAGAATGATTGATTCAATCGACAAGAAGCAAAACATAATAACAAGCCATGGAAAATATAAT
TTGAATTAGATTGTTGAACTAATGGATCAACCTGCCAATGCAAAACAAGAAAATAAAGCTAAAAAGCTTTAAACA
AATGATCATTGCATTTATATGTAC**CAAAAATGATTTAACATTGGAAGAAGAGAATTTTCTCTTTTTCTTGCAAGT**
**TATGCAGCTGCTTTCTCAAGTCTAATCACCTTGACAAAC**CTGAATACAAATAGAAAAAATAACAGCAATAAATAA
ATTCAGCCAAATCATTCATAAAAACAAATAGAAGGTCCTGCAAATGCAAGGAAGAAAAACTTAATAAATGGTTGG
TGAAAACAAGAGTAATTATACATTGGATAAATGATTTTTAAATTACTAACCAAAACAAATATTGAAACATGATGA
TGTTTTGATAATAGAAAACATGCCTTGGAAATACTAATGATTTTCTAGAGCATAGATTGGAAATTCAAATCAATA
TGATCCAAACAGCTCAACTTGAAGTGCTTGAAAGATCAAGCATCACCAAAAATATTTTAACAAGGTTTTTTATTA
TTATTATTATTTTGCTTAGACTACTTTCTAAAAACTTTATAAAAGAATATGCTAACCCACAAACACATTAAATAA
ATGTCCTACCAATACACATTAGAAAACATAAATTATTTTCAAAAAACATCTGATGACTAATGAATACGATCAAAA
TTATGCATATGATGCTCCAATGGCTCTTTTGCTATAAGGGATAGGGATATGACCATTATAAGTGTTTATTACCAA
TCATAGCCAGTTTGCTAAAAGAAGGTTTACCAACAATTAAATCAAACATTCAATATAGCCATCATTCAAATGATA
CGTTAAAAGCTCAAAAGAAATTAAACATGTTATACATAAGACCCATGATTGTACGTATGATCATATAGAACTTAG
AATTTTGTGGAATAAATTAGATGTAATGGCAACCTTGACTATACATTGGCAATTGTCAACAATCATATGACATTA
TTTAAGCTATGATTTAAAATCCTAGCAACTATAAATACCCATTGGAACCTTTAAAAGCCATCCTAAGGCTTACGC
GTATATTTCAAACTTCTCAAAGCTATAGAATTTCACTTGAAATCTCTTTGGTGAGACTAGCTAGTGAGCTATGTA
AATTTAGGATTGGAATACCAACTTCTTTCCACTAATTCATCACAACCTATCAACGTATACTATAAGTGAAATGCT
TCTCCTAGAAGATAAAACTCAAACTTCTCTACCCAATTAAGGTTTATGCCCTGCGATTCTCTAAGAAATGGATTA
ATGAAACTCTCAAAGGAAGAAAAATCATGACATCAATGTAGATTGATTGAGACTTAATTATCATACATCGAATTG
CATTTTCTACCTTCTAATTTTGCATTTTAATTTCTTAACTTTATTTACTAGTCTTTCTAGCTATAATATATAGTT
ATTCACATTGCTAGCTAGATAGTATACACATTCATTACGAGGTATTTTCAAAGTAGCACAAAATCTTGGAAAAAC
TAGGCCTGGCAATCATAAGAAAAGATCATTCCTATCACACACTTTGGTAACAATAATTCTTTTAATATCTACTAA
ATTCATAATGAAAACTTTTCTAATTCATCAATAAGTAGTACTTAGGCTAAGATAAAACATCCTTCCTAGGGTGGA
AGATGAAATTTCAATTGAGAAGCAGTCATTGAAAAGACTCAAAGTGAGAAAAGCAAACTATCAAATGCAAGAGCA
CTTATTTTGTTTATTTTGTGCACCCAAATTCAATGATTATACAATGGCTTCATGTAAAAAATTGATAATGAAATA
TTGCATAAGCTTATGTCATGAATGAAGAAACTAAGTTCACCAATTTTGAATTCATGCAAGTTCATAAATACCAAT

TATTTCCATGAAACCAGATGAATCAATCTCAAATATGTACACTTATTTTAATAAGATCATCATTAGTTTACATTC
TCATAGAAATGAATCTTATTTAAATCAACAGATAACAAACAAGATCCATTTCCATTCTAAGAAATGGGATGCCAA
GGTCTCAGTCCTCAAGTAAGCGAACATTACCTATATACTTATTTTATTTATAGGACCATGAAATAGAGATGATTC
AACAAAGAAAATGTTGCATTTAGCTGAATTATGGATAGATAGATAGAAATTTATATTCCTACAAATACAGGGCCA
TTAGAATGCCATTCTTGAGGCCATCAAATATATTCCTCCAAAAATCTACTTTTATATCCAAATTGTCATCGTCTC
TCACCTAAAAGCTAGACACCATAAACCCTACCTTACAAAAATGCAGCCTAGCAACAATCAACCGATTCAACCAAG
CATTTCCCTCAACAGCAAAAGTAAATCAATCCACCGATGCCAAAGATATCCTTCAGAGACACGACGATGGAGAGT
TGAGGCATTTAGACAGAAACTCTATAAATAATTCCTTGCAAAAAATCTTACAGAAGAATTCTAACCTCTAGCAAC
CCTACATTAAGATTTGTGTACTTCTCTAAATATAAAAAATCGTTATTATCACTAACATAGTTAAAAACGCAAATC
TTCCCAGTAATTCCTACTCAGCATTCACAAACAATAGCGAGACATCAAAACAACAAACAACCTACAATAATCAAA
CCAAAAGAAAGGCCAAATCAAATAATCAACCCTAAATCAGGGATCAATTCTTAAAATTTGCAATAACTCCAAATT
CTTTCCCTTAC**CTACTGTCTTCAACGACGAACACATTTCTTTCCATCGTTCGTTCAGAAAGAAAACACAAGCAGA**
**ATCAAGGGAATCGAGTCTCTATATAAAGCATGCGCATCTCACCTGCATCGAAAAAGCACCGGTAGATATTATCTT**
**ACCCGGCCTCACCGCGACCGTAGCGGCAGTTCGTGAGCC**ACGTCTCCTCCGCTGCCCAGACTACATTAACTAGCA
GGGGATGGAGTAAACTCCTGACCGGCAATGGTGGAAACAAAGGACCACGTTGCGGGCTGTGATTCAAACAATGGA
CTTCTTTAAGGAAGGGAATGGAGATGATGTCAAGTGTCAGTACGGGGGCGGTGATCGGCGCACACGTCAGGCTGG
TTCTGCTTCCAAGGAACCTGCATTGGTTCGAAGATCAAAGGGAGGCTACGTGGATCGTAGCAGAAGGTCTCTAGC
GAATATTATATATGGAATCCTTACATCTTTATTGAAGTAAGAGCACATGGAAATGTTGTTAAAATTAACTTAAAA
AGTTGGTTTCTTAATTAATCAAATATCATGAATTTCAAATTCTTGGTATATTTGGTTGAATAAAATGTTTAATCT
AACTAGAACTTAAATAGTAAATTAGTTAATATGAGTTATTCGGGTCAAATTGAACTTGATTGAAATTATTCTGAA
TTTATAAGTTTTGACTCAAATCTGACTCAAATAGCAATCGGCTAAATCTTAATCTAAATTTTTTTAGAGTAAATT
TCATTATTTTTTTAATCATATTTGGATGGAATTAATACAATCAATACAATCAACACCAAAATATTTTTATTCTA
AGTTTAGTATTATTAATTCAAAAGCTTTTATGCTTTATTTTTTTAATAAATTTATCAAGGATATGACTATATTTT
TTCAAAATTAGCGTGAATTCCAAGTCATAGTTTCAGGAAGGTTGATTGGTCGCTCGACAATTTAATTTGGTTTGA
TTGGCTACCAAGGTCTGATCAATATGAAATATGAAATTAATTCTTATAAAATACATGATATAGGATTTTAAGCTA
ATCTCATGATTTAACTAAAACCTCTTTGTCGTTTAAGCCACTAAAATCTCATTATGTGCTTGGAACTTTGAGGCA
ATTTCTTTAATTGTAAGAGAATAGGAATCAAAATCATAATGGATACAACTAGTTCTAAGACTTGATTAATCTTTT

>PeAGL17_2
ATGGGAAGAGGGAAGATTGTGATGAGCAGGATCGACAACACGACGACAAGACAGGTGACCTTTTCGAAGAGGAGG
AATGGGTTGCTGAAGAAGGCGAGAGAGCTCGCCATCCTCTGCGATGCAGAGGTCGGACTTATCGTCTTCTCCAGC
ACCGATAGGCTCTACGATTTCGCCAGCACCAGCATGAAGTCAGTGATTGAACGCTACAGCAAGGTGAAAGAGGAT
CATCAGCTTGTGATGCATTCAACTTCTGAGATTAAGAAGGAAGCGGCGAGCTTGAGACAGCAACTGCATAACTTG
CAAGAAAATTATCGGCAGCTGATGGGAGATGTTAATGGGCTCGGTCTCAAAGAGCTACAAACATTAGAGAGCCAG
ATGGAAACAAGCTTAAGAGTGATCCGACAGAAAAAGGATCAAGTTTTAATCGATGAAGCTCAAGCACTACATAGA
AAGGTCGATGATATGGAAGAAGAAAACATGAGGCTCTACAAAAAATTAAAACTTTCTCAGCAAGAAAACATGGAA
TTGCACCTAAAGGTCGCCGCAATTGCAAGATTTATGGGGAGATCTACTAGTAGTGGCTCTCTTTGTCCGACATCA
ACTGACCTTGAATTAAGCCCGCCACAGAAACAAACAAGTGAAATGCAAGAAGACGTACTA
AAGCTAGGGTAA

>peq_miR444a
GGCTCACGAACTGCCGCTACGGTCGCGGTGAGGCCGGGTAAGATAATATCTACCGGTGCTTTTTCGATGCAGGTG
AGATGCGCATGCTTTATATAGAGACTCGATTCCCTTGATTCTGCTTGTGTTTTCTTTCTGAACGAACGATGGAAA
GAAATGTGTTCGTCGTTGAAGACAGTAGGTTTGTCAAGGTGATTAGACTTGAGAAAGCAGCTGCATAACTTGCAA
GAAAAAGAGAAAATTCTCTTCTTCCAATGTTAAATCATTTTTGATATTAAGAGAAGAGAAATTACCGATAATTTT
CTTGCAAGTTATGCAGTTGCTGTCTCAAGCT**C**GCCGCTTCCTTCTGCCAATTCAAGTGGGCACAAAACAAAATCG
TTTGATAATTCTAATGCAGATTTGGTGCAGAACTATTTAAGAGGGGTTTAGTTTCTCACAACCATGAAACAAACA
AAGAGAAGAGCGGATGAGATAAAGGAGAGGGCAGCCATCCAAGAAAGCAAAGTTTTTAGTGCTTATAAGAGCAG
TGTCCGGTTGCTTCAAGAGAGAAAGGAGTTGTTTCCTCTTTTTCTGTTCTAGCAAGAGGAGGAGCTATCAACAAT
AGAAAGACAGTGCTCGTTTAAACTCCTACAACAAAAGGGTAAGGGAGTACATACATTAAAGCATTTCCTTATGTT
TACCCATCATCTTTGCAGCATGTGTTTGTGCT

Position of mature miRNAs based on comparison with osa-MIR444d and not based on experimental data!

Stem-loop structure

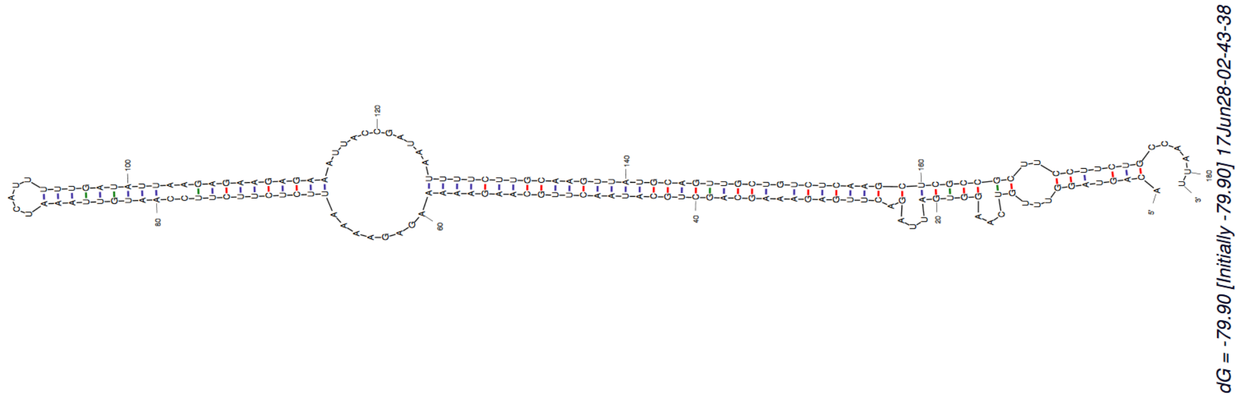


*peq-miR444b*

>Scaffold000212

TTTACAACATTATCTTTTAAATTCTAATCTATAATTACACCTACATTTATATTGTGGTTTTTACCTGAATAACAA
AGTTTAAAACGTAAGCTATCAATTTTTTTAGTTTTTTTTCTCTAACCAATTTAGTTTTTTGAATATATAAGATAT
TTATCTTTCTATAAATTATAATATTAACTAATTTCATTAATTTACTTGTGATGGTACCTATATTTCAAGATTCTA
TCATAATATTAGTTTCATGAACTAATTTCTTTACTCGCATTCACCCGTAATGTAGGAACTATTGTTTATTTGACA
CTACACCCGGACGCCGGTGTAGCGGCCCTTGCACACTTTTCACTGTATTATGGTCATACAATGCAATGCTTTCAT
GGGACTCCATAACATTTAAACAATGATTAAAATGATTCATGTCATGATTTTTCAATAAATTTTTCGCTGACTATC
AAAATTATTTTTAAGATTTAAATTTGATTAGAATGGAATGTGATTCAGAACTATTTGTAATCCTTGGTCCCATAC
AATGTAACTAAGAATGGGCTACAAAAATTGGGTTGATTGATAAGGAAAATAATTATAACAAGAGACCACGTCAAC
ATAGCTTCCGCGTATTACGAGTGAAGTATAGATAGGCATAATTTTGTTATATTTAACAATAAGAGATAATGCAAT
TACAAAACTATCCTTTCATCCTATCTCAATGAAATTCAAACAAAAAAAGAAAAGCAAATATTGTAATTATTTTTA
TATATAGAAAAAAAACCTTAGTGCCACCCTACCTGAAATGAGTGCATTATGTAGTTCTAGATATTGCATTAGACA
TAACCTTTAGTTTATCATTTATTCACCAAAACCTATGCCTATCTTACTCTGTTATATTTTCAATTAATTTTTATT
TTGAATGTTCATCTCTCATTTATTCGCATTGTAATGGAAGAAAGCATGCTCCTTCTTATCAATCTGCTGTTCAAT
TTTTTTCTCCGCAAAATACATATGAGAAATGAATGGACTGTACAACTATTATGTTCCCTTAATATGATTACCGAT
ATCCATATTTTTATATTTTGGCAATTTGATAATTAATACTAATTTATTGCTTCACGTGAGGTTTTTTTTTTTTTT
TTTTCCTCAACAGGTTCCATTTATGAATTTGTAATTTTGCTAGCTAACTCATTTTTTTATAAAACCACTATAAGCA
ACATGTGAAATGAAATTGGACACCCTGCACAATATGATGTATCAAATTGGAAGTTTTAATTGAAAGAAGTTTCAA
GGCATTCCAAATGTAGCTATTCCAATGTGCTGAATACCAACAACTCAAAGAATAATTCTGTAGTTATTTATTATA
ATGTGGTGCTTTTTAAAGTGTTGTCTTTAAGAAATTACGAAACATTTATCATATTAGGCAGCGAGATTCATGCAT
CTTTTTGAGATTGTTTAATTAAAGATCATAAATCAAATTGTTTATTTCTATAGATGCTTTCTAGATT**TGGCATAT**
**GGGAGTAGGCTTGAGGGAGTTGCTGCATAGCTTGTAAGAAGCCATCGAAACATCTCTTCATCTAATG**GTAAACCA
TTTTCTACTCTTTTTAATTCAGTATTTAATTCTTTTAAGGAAGTTTGAGCGTGGTTATCATTCTTATTTGAACTT
TGCATGCATGGGAAACCCATCAAACCTAAATCCTATTCTGACTAGCCCAATAATTTTTTCATAGTTTTATCAGAC
TTGTTGGGCATGATACGAAGAAATTTTCAGGCAATTTTTTTTATTCAAAGTGTTGGGCTGGATTAAGCTTAAGCA
AAGGTTTTTAACGGGCCAATTTATTTTAATATGGCTTGAATTTGACACCCATTTTTCTAGTCTAATCGTCATCAT
TATTTGTCTTGTCTCAATGCTTAAGTGTCAAATATATGGATATATCTTCTCCATTCAATTTTGTTGGGTGTCATA
TATGAATTTTCTAACTGTGTTCTATAAATTTAATTGCTTAGTGCAGCATAAATTTTCCTCCAATTTTTTATACTT
TTCTTTCAGCCATTTTTTTCGCGCGTTATTAATGTCATGGAAGTTAGTTTCATGCCTGTTTGAATTGTTCGTTGT
GTGGTCTTGTTTGTGTACAACAACAAACCTTTATCACATTAAGTGGGGTCGGCTACGTGAATCTTTTTTCTCCAT
TGTTTGTGTTCCAATAGATTTATTTTATATTTTATTATGTTTTAACCTTTTTCTATTGGCCTTTAACTATGACTT
TCTAAGCGAGTCAGCTGTTATATGAGATTTGAGTTATGACTTGGACTAGTATTTGAATTTGGAGTAGACATTGCG
GGATTATCTGTAAGACCAACGTTGATAGGGTTGTTGTTATGCATGTGGTAAGAGAGATGTTTACAACAACCATTC
TAACGTGGTGCTGTAAGTTTAGTTTCTTTGTTATTCTGCAAAACTTTCATATTATCTGAGTGCTAATGATCTTA
AAATCATATTAGTTGGCCCTTCTTAACTATAGAAGCAATTTATGATGTTGTGCTTTACTCATCAACCTAGAAAAG
ACCGCGCAACCTAAAATGTTAATTATAATTATGTGTATTGGTCATCTTTGCATCTATAACACCGAAAAGTATAAT
AATATACATGACCAAAAAACAAATTAATTAATTTTGAATATTGAAGCTTTGATCTTGATTTCATTTAAACTAACT
TGTACTAATCAAACCATCAACTTCTTCTGGCTCTCGAACGTACTTTAGAATTTTTCTCTAGATGTACCCTTTCAA
TCTCCACTTGAAGTACCTTGTCTCATGCCTTCACAAACATAGCTTATATTAAATTAAATTTGTGATAATCAAAAT
ATTAAATATATGTTATAAATTATGGGGTTGACATATTCTACATTTGTCACACTTATGTGATACAGGCATTAATTT
TAGTTTTACCACAACTTTTAAACATGTTATCTTTTGTTATACATTTGTTTTATTTTACATTATTTTTATTAACTA
GCTTGTTGTTGATGATTTATGACACATCAAAAATGGGGATGCCAGGGATTTGTATTTTAAAATTTTTATTGTGAT
TAGGGGAAGTAAATGTAGAAGTGAGAAGTATAATAATTATAATTATGTGTTTTGGTCATCTTTGCATCTATAACA
CCGAAAAGTATAATAATACACCGAAAAGTATAATAATATACATGACCAAAAAACAAATTAGTTAATTTTGAATAT
TGAAGCTTTGATCTTGATTTCATTTAAACTAACTTGTACTAATCAAACTATCAACTTCTTCTGGCTCTCGAACGT

ACTTTAGAATTTTGCTCTAGATATACCCTTTCAATCTCCACTTGAAGTACATTGTCTCATGCCTTCACAAACGCT
AATCAATAATATTAGCTTATATTAAATTAAAATCTTGATAATCAAAATATTAAATATGTGTTTAAAATTATGGGG
TTGACATATTCTACATTTGTCACACTTAAGTGATACAGGCATTAATTTTAGTTGTACTACAACTTTTAAACATGT
TATCTTTTGTTATCCATTTATTTTATTTTACATTATTTTTATTAACTAGCTTGTTGTTGATGATTTATGACACAT
CAAAAGTGGGGATGCCAGGGATTTGTATTTTAAAACTTTTATTGTGATTAGGGAAAGTAAATGTAGAAGTGAAAA
GCTTGCAAGTGTGTAATTAAACATCAAATGAAAAGAATAAGGAGGCGGTCAAAATATACACAATATTTATGTGGT
TCAACCCAATTGGCCCATGTCTACTAAGCCCGCAAAGAAACCATCACACCTTACAAACTCGCAATTTGGGCTAGT
GCAATCCAATTATAAGATTCCTAGATGAACCTCATTAAGCCTTTATAAATATTTAATAACTGAAAGGGAAACTAA
AACTACATAGTTTACTTTTTGATCGAACAAAAATAGTAAAACAAGAATGAAAATAAAAAATAAAGCGATTTTACA
AAGTCTTTGATAAATGTGCAATTTCATAAATAGCTTACAATTGATTCATCTTTGGTGCTCTTAAGCTTTGAAAGA
TGTTTTAGCTCTCAAAATTGAATTTGATAGAGTTTATCATGAGAGAACGTTAGAGAAATCTAGATAATGAAGGAA
TTTAGCTATAGTTTCATTACTCTATTTATTATTTAACTTTCTTTCTATTTATAACTCCCAATCTTCTTCTAAAAT
AATTTAAAATTTATTAAAACCCTCCCTAATTTTTTTTCAAGGAGCAGCAATTGTTTAATGACTATAAATCTAGTT
ATTAGTAGTGTTAGAACATTGATAGATAGAACCTTGTCACACAGAGCTTGCCTCGGTGTGCTTTGTGAGAGCAAC
CTTTTGTTCAGTATTAGTAGCACTGTTTTAATTTTTTTTAAGGCTTGTAATTTTCTTTTAATTAATATTGTATTA
AAAAATACTTATATGATATTATATAATTTTGCATGTGAAAAATATGATGGATTTAGTCATTTACAAAACTAGAAG
CTAATTTTTGAAAATTTAAATTAATTGTATATTTAAATGTTAAATGATACTAATTAGTTAATGACAAAAATTGGA
GAGATTGAGTTTTTCATTCATGGATATGTACAGTATTTATAATATCTTAGATCTTCTGATGAGAAGCATGTAAAA
TCTTGGAAAAAAAAAATTTTATAGTCCACGCCTCCACTTTTGATGATGTGACACAAATAATCTCATAGATTATTGA
GTTGACATAAAGTAAATAAGAAGGGTGGACTGTTGGATCGTTTAAACATTAATAGAGGCAGAATTGATGGAGATT
AATTTTTCCACTACTAACAAGGGAATAATGATCAAAATGACTTGAAAGTTTTTTGCTAAAGTTTATTTATCCAAC
TTTCAATTTCATAATATTAGAAATACTTTTGAGCAATTTATAAATTTAAGTTAAATGAGTTCTTGTTGAGAGATT
GGGCTAAAGCTGTAATGATATTAAAATTACTTCCAATGATATCCCTGAAGATTCTAAGTAGACCTTGAAATCTTG
GAGTTGACTTGGTCATGACATTGGTTCACTTGATCATGCATTGGAAAATGGGGTTGAAGCTTGTATTAAGATCTA
TGATACTTCCCACTCTAATGACTCTTAGATAAATTTTTAGTACTTTTAATATATGCTTGTAGTTGCCAAAATGAG
CTCAAAATACTATCAAGCCTACGGAGTCGAGAATCTTTCTTTTTGATGATGACAGACATATTACATTGATTGTGC
TGCAAAGTGACATCCCCCCTTACATGAAGGCATAGGTAAATAGAGATCATTTGGAATAAAAAATTTGCAATATGT
GGGGACTAGCTCTGTCTCTAAGTACATGCAGCTGAAGTTAGTAAACTTTTCAACCAATTTAATAACAATTTTGCT
GACATATGTTACTTCCTATTTTACCAACATCAAAAGAGAGATACGCCTAAAGGAAAACTTTATTTTGACAATGAG
AAAAGAGTATGCAGGCTCATAATTAAACAAGTGAACACAAGAATAAAGTTTCATATGTCAAACGAGTTCATACAT
CATTTTATCCAACCAATAATTGAAACATGGGAACTTCAACATATTACATTCTCTTTAAGATCAATGAATTTTGCC
ATGATGTAGAATATAAATGTCTTTTGAAAAGGGTGGTCTATGAATATGTCAATGTTTTCACGTGGAATATCATAT
TTTACTTCAAAAGGATAACAATTTTGATGATATGACTTAGAGAGAAAGATTTGTTGGTCGTTCCATTTCAGAACT
TGTAACTCAGAAATATTTCCAATGTTAAAAATTCAGAGATTAATGTAGTTGTTTTGTTTTTTCTCTTTAATCAGA
TAGGAGAGAGAGATTAATGTTTTTGTAAGTTCATTTTGATGTGAGTTAAGCAAATTGTAATTAGGTAGAATAAGGGT
AGCAGTGATAGAGTTAGAATTACTATCTTTGACCAAGACAATTTTTTGTAGGAAATATTGGATGGTGTCTTGGTC
ATCTTTGACCAAGACAGTTTTTTGTAGGAAATATTGGATGGTGCAATCCTCGTTAGATAGTCTAATCATATAATG
GAGATGTGATTTTGTAGATGTGGAATGCACTACTCTTTTCATGACAGTGCAATTAAATTCTAGCCGATTTTCATT
GATAGACAACATGTCAGATGCTCCATTTGGAATTAAGAAAAAAAAAAAAAAACTTACCTGATCTTCACAAGTAATG
ATATCAAGATGACCATTTTTGAAAGAAATAACTTTATTCGTTCAGAGGTATAAACTAATTTGTTGTAGAAGATAT
TTGTAAAGGATAACCAAAATTGAGTTCTACACTAAAGAGTGTTTTCAAATCTGGTTCCATGGAATAGGCTAAATA
CATTTGCATTGAATGGAAGAAGTGGCCATGGTACTTTAAGCCTTACATTTTTTATAAGAGGATAAGCTTGGCTTT
GAAAACGATTATGAACCATTGAGGAAGATTCATCATGTGATTGTATTGAGCCCTTAGGTAGTCAAAGGAATATGA
ATATCTATAGGGTGTAAAGATTTAAACACATTTTAATGAGCAGAAACTAATTTTTGAACGGATTTTTTATGTTTT
AAACACTAATTTTGCAATGAAGGATAAAGGAATAAAAAAGTAAATGGAAGATTAAATTATATAAAAAAAATCTTAA
CAAGCAAATTTATTAGATATAGATCTTGGTATAAACACTAATGGTCTAACAAAAAATGAGGATATTTTGTTATCT
TGACATTAAGAAATAGGAAACTTAAGTTGAACGGCTGGGGTTCAGCAACGATCAAAACAAAAAGGATAGGAAGAA
ACAAAATACATATTTGACCAAGGATCTTAAAAAATGTCGAGTTCAATGTTGCTTTGCAATCAATCTTTCATCTTG
CAAAGTCATCATTTGCCTGAAAGTGAGCATAAAAACATACAACAAATATATTAGTAGTGCTACTAGCAGTAAGTT
ATTATAATTACCCCATCAAATCCTTATAAAGCTTAGGGTCTGTTTGTGGCAACCGCAGCGGTTTGTTGAAGCAGC
TGAAGAAGAAGCTATTGTTCAAAAAGCTGCGGTGGCAATTATAATTTTTTTCTTTGAAAAGGTCAGATCGCAGCT
TTTTAACAGTCACAGGAGATAACCGGGTTTGGTGGGAAAACGTAGCTTTTAGCTGAAGCTTTTTGGCAAAACGTA
GTTTTGCCAAATAGGAAAAAATTGCGGTGTGGCTCCGCACCGCCTCCTTAAACACACCCTTAGTCTTTCTAGGTTT
TCCATCTTCATCTAAATCTACTGTCAGAGCCTATAGAGAACCTTTGGTTGAAGTTTTTGGTGGTCTGTCATTGTC
TCCTATTGCTTTGGTGGCCTCTCTATTAGTGCTTCACTTATGGATCCTCCTCAAAAATTAAATCAATGTTTTTTT
ATGCTTCTGTTGGGTGTGCTTTTGTTTTTTCACCGAAGTTGTTGGGTGGGGGGAGAATGTATTACGTTTTATGTC
TCCATCATCACTTTTTTTGTATATTTTTGTAACAAGGCTCAAAATTATCTCAACATAAGTTGCCCTCCACTGTCC
AAATTAAATCACTCTCATTTATTTATGAAGAGGTTTTTTGAGTTGGTGCCTCTATTTACCATTTTTATATTTGAT
AATTGTCATCCCAAGCTTAAGCTTATTTGCAGATTCATAGCTGACCTATGTTTGATTGTTGATTCTTTGGGCAAC
TTGACCTTTGCCATGCCCTTATTTATTAATTCTAAGAGCTTATTATTCGAGGCTATGCTAACATAGAGCCTTCAC
TTAGACCATTTTTAGGTATGCCCATGTTTTGATTTTTTTGAGGAACCAACTATTAACTTCAATTTGCATAGCTTT
TTCTCAAATTCAAGTCCACTACTTTCACCAATACACTCTAAACTCCCTTGGTTTTATTTTTGGTAAACATTTACC
AGTGAATCATATTATTGTTAGGCTTTAGAGGTCGTTTGTAGTTAACATCTCAATAGAAATGGACATTACTTCTTC

TTATATATATATAAAATTTATTATCAAATTTTAATTTAAATCATGTTTATTAAATTTTAAGACATTTAAAGGAAT
TATAGATAATTATTTTGTACTTAATGCTATTAGTATCATTTGTAATTATTTTGAATTACTTTATCACATGAAAAC
ATAAATTTTTAAGTTATGTTGGGAACTAGAATCTGATAAGTTATATGCTTATGATCAAATTCTCTTAGGCATTGA
AAAACAAGAGGATAGAAGATTTCAAATTATTTTTATTTGGAGATGTAAGAATAAGAAAAATGTATTTTTTTCATT
CATTAAAGTTTATTCAAAATGGAGCAAGACCCAATAATTTGTCTCATGACTTTGATAGTCTTGACTTGTGAGGAG
TGAAGCTATCCTTTGTTTATTGAGTATTGGTTAATACTGTCTTTTGGCTGTTCTTCGTAATTGCTTAGTGTTGTT
AAAAAAAAGTATTGAACATTGGGGACTTTATGTTTGTCACAAACATTTATAAGGTTAGTGAACGTGCTTTAAATG
TCTGTAGTAAGCATTCAATTCAAGTATTGTATTGCTCACAGGAGTAAGATCTCAATCTTTATCATTCTATTTTCA
GTATAGTCATTATTTGGATCATTTCCTTTTCTTTTATTTTAAAAGCTTTCTTGATGGACGATATTGCATTTTAGG
CTTGTAATTGAAGAATAAGATTTTGTGCTTACCTAAAGTTATAGGTTCTGATAAAAAGATAGACCTTGTTTGAGA
ACAGCTCTATTTGCTTTTTTGAATAACTTGAGTTCTACTGATGTAGTTCTTGAATTTTATCGCTCAAAATTTGAT
CTGCAAGTTGAATCAATAGAAGATTAGAAACAAACAGAAGGATACAAATAACTGGGGATGCCTAAATCCTTTGAT
TAATAACATGCTTTTGTCGTATAGCTTGCAAGCTTGTTTCTAGTTGGTTCTCTGTGGGCTTGTTTGGAATCACTT
CAATTTATTATTTTTTATTTTATTTTTTCTTTTATTTAAAAACATATTTGTTTAGTGTATTTTTTGCATCTTTTT
ATTTTTTTATTTTGTTTTTAAAAGCATTTTAACTTGTTTTAAAAACTCTCAAAACATCTAAAAGCTGTTTTGAGT
GTCTTTAACCATCAGCTGCTACGAACTAATTGTCTTAAAGGAACAATCACTTAGGTTTTGGTTGATTACATGTGA
TTGCTCACCTTTCCCTTGGTTGGATGTCATAGATGCTGTCTCTTGTCAATTTTCCAAAATGAGTTACTATGTTTT
TTTTTCTTGGTTGAGGTTTTATTTCTATATTATGTCACATTTTCTTTGTTTTTTCTGCTAATATGCTTCATATTT
TTTTTTTTAAATTCAAG**GTTGAAGAATTGTTACCGATGATTTTCTTGCAAGTTATGCAGCTGCTGCCTTAAGCTT**
**GTTACTTCACCCTGCAA**GAACTTGTGTTTTCAACAGACAAAGTTAGTTGGTACTGTACATTCTCATATTTGAGTG
AAAATAATAAATAAACAAGCATCATTAGTTAAAATTATCAAATAATTAAGCTTATTTGAGGAATAATTATTTTTT
CTGATGTTGATGAAGTAGTTTTTTTTTTTCTTTTTATCACCAGAAAACATGGAAATTGTTAGCTAGCACTTTGTG
CCCCACTGAATGTAAGATGATATTTGAGCTTTTAGTGTAATTTCTATTTATATTTTCGTTTCCTTTTTTATTTGC
CTGGAAAAATATGGGAACTAAAACTCTCATACTTAAATGTTTTTTTTTTAATTTTTTCTATTATTTATATATTTT
AGAAGCTGATTGATATTGTCTTTAACAGAGCTAGTCATAATTTGACAATTGCTTTGTTTATTTCCATTGTCCAGC
ATTCATTGTTTGACATCTTTCGGCCATCATTCAGCTTTAATGAAGCATACCTGGAAGGTTCAATTAAATATCCTA
CCGTGATAGAGTTTAAATTTAGGCAATTTTGTTTTCCTTTAATTAATTTTTAAGTATGTTTTCTTCTAATAAAAT
CTAACTTACAGCTATTGAGTATCATCATTTGGTCTCAAGGAAAATCCTAATAATGTTTTCAAGCTGCTTTGTTCC
TAAATCCAGGTCAGTAGTTACTCATATATCTATATTAATTTTTTTATTTGAATATTTTTTTAATATATATTTTG
TTAATCTTCATTTAAACTTTTTTGTAGCTTTCTTTGATTTAAATAATCATTTTAGACAGCTGGTATATATTTTAG

>peq_miR444b

GAGATGCTTTCTAGATTTGGCATATGGGAGTAGGCTTGAGGGAGTTGCTGCATAGCTTGTAAGAAGCCATCGAAA
CATCTCTTCATCTAATGGTTGAAGAATTGTTACCGATGATTTTCTTGCAAGTTA**TGCAG****CTGCTGCCTTAAGCTT**
**GTTAC**TTCACCCTGCAAGAACTTGTGT

**Position of mature miRNAs based on comparison with osa-MIR444d and not based on experimental data!**

Stem-loop structure

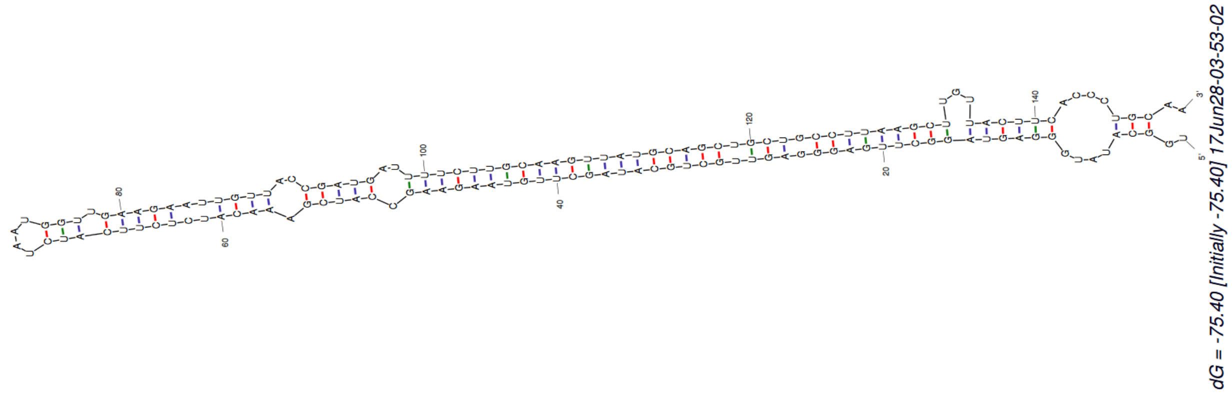


*PeAGL17_1*

>Scaffold000039_reversed

AGGCAAAAAGTTCTACCAACTTCCTACTCCCACCTAATTTACCCTTTACCTACAATTGAATTAAAACCATGGAAA
TAAACTTCTTTTACCAAAATTAAACTCATCAATAAATTAAGCTCTTGAATAAAAAAGATAATTTTACACATATTA
CACAATTCTTACATATATTACATATCCAACTCCCAAAGCTTACACTTATAACACTTTGGTGTCATAAATCTATAC
ATAAAAAAATTATATTTCATTTTTTTTTTTGTAAATAATGATAGTGATACTTTTCTTACACATTAAAAATTCAAAT

ATAAGTATGAAGCTTATGTGCTAGATACATAAATGCATAGAAATTACACCCACCCCGCCCAAACTTTAATATTTA
CACTTATACCGCCTCGATATCAAAAACATCTGCATGAAAATACACATTTCATGAAAATTTATATTTTAATTTTTA
AAATATAAAATAAAAATGAAAAATATTACACTGAAATAAAAATTTTATTGAGGCCCAATAATGAGACAAAATGAA
CTGGCTTCAATTAGTAAATTACTTAATTGTCATAAGGTACCGAACTTCCATATCTGTATTATAAATATACTTTTT
GCACTCCACATTGATTAATGAACAATTATATGGTAATGACCTCGTCGTATGGGGAAAGCAGCACCCCTCCTCCCT
TTATCAGGCTAAAACCAGTAGAGAGATAAGAGGAGGAAGAAGAAGAGATAGAGAGAGAAAGGGGGAAGAGAGAGA
AGATGGGGAGGGGGAAGATAGAGATAAGGAGGATTGACAACATAACAAATAGGCAGGTGACGTTCTCCAAGAGGA
GGAATGGGCTGCTGAAGAAGGCAAGAGAGCTGGCTGTGCTTTGTGATGCAGAGATTGGGCTAATCATCTTTTCCA
GCACAGGAAAGCTTTATGATTTCACCAGCTCCAGGTTTCTCTCTCTATCTCTTTCTCTCTCTCTCTACTTCAC
CGAAATAGAGGTTAATTATCTTCTTAATCATAATTAATCTCTTTTTAATCATGATTTGAAAGCTTAAGTACTGCT
ATAATGCTTTGATTAATATTAATTGTTGGTTTTAACTAAGTAGTTTTGTGTTTATAAGTTGTTTTGTATGTCAGA
TCTGAAGTGGGATTAAGCTAATTAGGAGGTAATTAAGGGATGATTACGTTCTTAAGATTGATAAAAAGGGGGAAG
AATTAACATTATCTAGCCGGTTTTTTAGGTTAAATGATTACATGATAATGAATGACATATAGTTAGCCAACAAAG
ATTAGTGGTCTTATCCAATTTGATGTTGGATTTATTCATTTCTATTTTTCATTTATGTTTTTAAATGTCACTCAA
TTCCTATGTATTTACATGTCTAATACATATATATATATATATATATATATATATATTAATTTCTATCTATGTCAT
TTTGTTTTGCTTCTTCAGCTTCGCAATCATTTAATCAAAATTTTTTATATTCATTTTATTGATGCATTTTGTATA
GATTATTATAAATGCATAAATCATACACACAAACTTATCTTTTGTGTATATGTTTTCTTCTGATTTCTGAACAAT
TAGAAGCTAACAACAAAAATGAGAAGAGATTATGAAGAATAGGAAGAGTAAGTAGTCCAAGGGAAAATGAAGACT
CTCTGACACTAAAAAAAATGATATGTTGCATTATAAATTAATGATGAAAATATGCCTAATTATGGATGAATATTA
CTATGCATGATCTACACAATTTTAAAATATTCATAATTGAAATATTTATATAAACTCTAGCTATTAAAACCATTC
AACAAATTGCCTTTTATATTTAGCTTTTGGAATTTATCCAAACACGACATTTTTGAAAATAATAGAGTGATTAAT
TACTTTCAACATCAATAGATTTCTAATATATTCATCTTTTAATCTACTGTGTTTAAACCCTGACAAGTACGACAT
TCTTTGAGTTAAGTGAGTTCTTTCATAAACCAAGTTAGTGCCTTTTTTGAGAAAGGTGAGTGATCTCTTGGGTGT
GTGCTCTTTGGAGTGGTCGCTTTGAGTTGGGTGAGTTATTTCCTGGCTAGGCAAATTATTCCTTTAAGCCAAGTG
AGTTCCTTTTTAGGTTTAACATGTGGTTTTTTCAATTGGGCAAGTTATTTTGTGGGCCAATTAGGTGTTCATTTG
AATTGGTCAAGTTCCTTTCTGTCATAATCCAAAAATAATACTCCCATTTAATACTTCAAAATCAATATTAATGTG
ACTTCTCAGTATGCTAGTGATCATGAATATCTATAAACATAGATTGGTTCTTCACCTCATAGTAGCTTCTTTCTT
CATGCCTTGTTTGTTTGCAAGATATTTTATTGCTCGTGAGCAATAGTTGAGTCACCATAACTGAAGATTATGAAT
ACCACCAATTATTAGTTGCACTGAAGTGCTATGCACAAACACCTCCATGCATATATAAGATGTCAAGTAGTGGTC
AACTCACTGTATTAAAAAAAAAAAGGACAAGGAGATATGTTAAAGGTGACCTGAGTAGGGCAGTGACAAATCTTG
AAGTTGATTTTAATAAGTACCACTTGGAATAATATTCCTTGATAACCCTTTATTGATTGCTCCAACGATAATTTG
GGATGTTTGTCTCTAATGGCCACGACTTAGAAAACCTTATTTTACCAAAAATAGATAAAAGAAGATTATGCGAGC
AAATTAAAAAATTTTTTGTAAGAAATTAACTTCATTTATTTACATATTTATTTTTTTGGATGAAATGATTTGATA
CCATAGACAATTTGATGGGGAGACTAGTAGCGATGATTTAACATAGAAAGTAAGACTATTCAATTTCTGTTACAT
AAAAGGGTTACTTTGACAAGTCCTATAATGATCATTTTAAGTAGGCATATGTTGCATCTAAATTTAATTCAAGTT
TTAATTCAACTACCAAAATTAAAGATTACAAATTTCATAGGTTTTGTCTTTAACTTTGCCTTGGGATATATGGCA
ACAACTTGTTGAGGTTTCCTTTGATAAAATTTTGGTTAGAGAATTTTATTTTCCTTTTATTTCAATTAAAATAAT
AGGGATGGATTTAGTGAGGTAGCCTAAGGGAAATGAAGGCTATCAACTTGTTAAAATTATTGTGGAGACCATCTT
AATGTTTGGGCTAAATGCATGTGTGGAAGCCGAGTTACTTTAACCTACTCAACCTCAAAAGTAAGTTTAGAAAAC
TCATGATGCTTCTTCTTGAATGATACCTAGATCTCATGGATCTTTGCCACCTTTTTTTTCTAGAGAGTTTGACTA
TCAAATTTGTTAAGCTAAAGGCATCATTGGTCTTTATGATAAAATACACACACATTCCTATGGAATGATCAGTGA
CGCTAGTGTAAGCATTAAAGAAGGAAAAGAGAAACATCCCCTCCTACAAGAATAGATGGGATCCTCCGCCTCTGA
GTTCATCGCCTCTATTTTTTGAAAAGTCTAAGATTTAAAGGGAAAATCCATGGCCACTAGCAAGTTTACGAAGTC
TTCAAAATCTAGTCAATTTATTGTTGGAGGAGCATTCATTTTGCCACCTTTTGTCAAAATAGAGACGTCAACATT
GATGTGTATTATATCAAATTATTAAGTTTCTTCTTTACATTGATCACTATAGGTGGATTCACCTTTATGTAAATG
TAATCTTGATTTATAATTCTATCATCCCACTTCTTATATGAATATTTTATTTCAGACTAGCTATATTAACTGTCA
TGTAATTTGTATAGTTATTGACTCAATTTTTTTAATTATTCTCCTTTTGTTCACACTCATTAATTTTTCCATTCA
AATAATATCAGTTAAAATAAATCATTTATGGTGGATGTGATATTTTCTTGAAAAACTATATAATCTTTCTCTATA
AATATCCTCCTCAATATTTAATGATTTATTATTCTTTTCATTTTCACCTCATGACTTTTTTCACTTGCACTTATA
ACACACTTTGAGTCATGGAAGCTTCCTAGAAGAAACGTTTCAAAGCATTCTAAAAGGGCAAGTAGACATAAACTA
AGATTTTCATATGGTTGTTAGTAGAATGAGTCCACTATCAACGCCAAAAATTTTAAGGTGTGGTACCCAAAAAGG
CTCAAGTTCACAAAGTCCTCCGAAAATAAGTTGAGCTTTCACTTGGAAATTCAAATTTGACCTATAACTACCATT
ATGCTAAGAAGTTGCCAACGAATTGAACTTATGGACTTTTCGCTTCTCAAATCATGCCAAATTTATTTAGAATAT
TCCATGATTTCTTGTATGAGGAATTAAGAATATCATTTTGAACATCAATTTCTTCAAACATCTCCTAATGATGAG
GGCCTTCAAATTCAATTGCATTCAATTCTTCGTGTTTTATATTTGAATTTTAATATTAGAGCACAAGGATGGTGA
GTGGAATGGGAAGTATGTGCTTACAATATCAATGAATGGTTGGATAGTTCTAATAATTCCTATCCTTTCACTACA
AAACTTATGCTCTTTTTGGTAGCCTATAGTTGTAAGAAATATTGATTTAGCCATAAGTTATTAAATTTTATGATA
GCTTTCTGATGGGCACTTAGTTGTTGCATTTCTCTTGGCAGAAAGACAATTTATGATAGTTTTCTAATGGGCAAT
TTTTTTGACAATTTTTTTACTGGCATCTACCCATTGATATATAGTCTAACGGCGATGTACCTGTCAGTTCATACC
TATCAATAAACTAATGGAGAGTTATTGGTTGGCCATCAATTTAAGACTCAACGAGCTATTGCTCATTGGTATAAG
GATATTGATAGTCCGCTGTAAGGCCATCAATTGGCTATTAAACAATTAGCTAAAAAAAACTTCTTTATTTTATTA
GAATTTATGCACGTACAATTGTATCCCAAAAACACATGAAATTTACATCACCCAAATCTAATTTACTTATCCATA
CAAGCAATCACAAATTAAAAATAAATGCAATCATAATGTATTAAATAGCTAAAATTGAAATAATTAGTACATAAC

TAATACATAACAAGTACATAAATGTCTTAGAAAAATAAATGAAATGCTAACTTAAAAAATTCATCATCAATCTCC
TCATCTTCCTCCTCCTCCTCATCCAAATCCAATGGCTACTACTCTCTCTAACGTGCCTCACGTCACACCTCCTCC
GATTCCATCTTCATTCTAACCTCGATCTGAGTGAATTGAGTTGAAATTCGAGCCTCCATATCTTTAGTGAAAGAT
GAGACCTATCTGCAAAGTTCATTGCATTGCATAGATGTAGAGCTATGCGCAAAAATTGAATGGTGTAAATCATTA
GCATGTCCTTGTGAGCTCAAGCCATAAACCCTTCCTTCTTTCTTGCCCTCATTGAATTCTTTCCAAAATTCATTA
TTCAACAAATCAAAATCATTACCAGGGACACGAACTCCTTCTAAGCCTCCTTCAGTCCTTTGTACTCCTACAAAA
ATATAACAAAGGTAAAGGTAAAATTAAATTTCAAAATATTTGTAATGATATAAATGTTACTTTTTCACTACATGT
GTTATATGTGATCTATACCCATCGGTGGTCTGACTGATGTTGATGTGCACAAGCTATAGCACGATGGTGATGGGC
TCTCTTCCAAGCTACAATGTCTATTAAGAAGAGAAATTTTTTGTTAAATAATTAATATCCACTTAAATTCAATAC
ATAAAATAACTCTTATCATTCTCCTTCTATACTGAGTATGTGGGTTTTCTCCTCTAATGTGTTTGGAAGTGACCA
TACCCCCAAAATTCAAATGTCAGTTTTATTTTGCCTTTTGGCTTTTCTTGATATAACCACGTTGTTTGTAGATGT
CCTTCTATAGAAGGCACAAAATAGTGTTAACACAACTTGTGTATCTCTATGAAAATTTCACACAAACAGGAGACC
TCTAGACTTAAGGCTACTTCTGATATGTGCATGATGCTTTGGACCATTAAGAGAACTGCATATGCATGAAAAATA
TTCATATGTTAGATTGATAAAGTTATATAACTTAAGATTAACATTTAAGGTTTATCTCACTATTGAAATCACATA
TCTCATACATCAATCGTTTTCTCCTTTCATGAAAAGTCCCAGAACATCAAACTTTATCTATAAGATGAAAAAAAT
AAATTATTAGGTTTATAACAAAAAATTGTAATAAAATTGTATTTTCCTGAAATCTATATTCTTTATGTATACCAG
CGAGCTCTAGATTGAATATGCATTTTGAGTTCCCAACCTAAATCACCTAGATTTGTACAAGTACAGGGACCCACA
CATGCATGGAATGTAGGCATTAGTGGTGGTATGGTAAGAGGTAAAGGCCTGTTTGGAGGGAGTAGAGATCATGGT
AGTCACAGATATTGTAGTTTTCCTCTGTCCGGTGATATGACATAGAGCTACACTCTTCTTCTTCCCAAATCTTTG
GATTAAATCTCGGCGAGGAGGGAGATCATCATCACCACCGCCCAAAGCCATCCACATCAAGAAAAAAATGAATTGA
TGATTGCACATAACAAAAAAATATATAAAAATTAATAATAGTAAACAAAAATATGATAAAGATAAAGTGAACTTT
ACAAACTACCACTACTAAAATTAGAAGTAAGAATAATCAATAGAAGTGTGATCAAATTATAAAGTGCAATCTACA
AACTATAACTACGAAAAATTGAAGTGAGAGTAATAAATAGAAGTATGATCAAATTATAAAGTAATCTAAAAACTG
TCACTATTGAAAAAAGAAGTAAGAATAGTAAACAAAAGTCTAAATATTAAACATAAGGAAAAGTAATAATCAAAA
GATAAAATAATTAATAAAAGAAGTTAAAAAACATACACATCTTATTCTACATCATCAAATGCGTGACTAGAGGAC
AAACCTATAACTTTTTCATCTTAACCATCATCGTAGTCATACTTATTTTGAGTAATTTCTTCATGTGTGCCACTA
ATATTCTCTAAAGTTATCTATTCCTGGATAGAATTAATTAGGATGCAAAATATGATATTTCTATTTCTTGATGAG
CTAGTTCAATACAATTTTTATTCTCAATAGTATCTCTTGGTTTACACTTGATAACTATTGCCCAATCTAATTTAT
ATATTGCTCTATATGCATGTCTAACATAGTATACCTCCATCGCTTGTTTTGCTATAATGAAAGAATCAAATTTAA
CATATTTTATGTTTATTTTGACTTTAGCGATAATATATAACTTATTTACTCTTATTCCACGACTAAGAAAATGAT
TAATGAATCTTGGATGAACATGATGTCACTACTCTCAATAAAGTTATTGTATTATAGTTGAACTACTTCAAGAAT
CTTGTAATAATTTATTTCATCATATCAATTATTATTTTATTTAACACAAAACACAAATATATCAGTTCTTTTTTT
TTTCCTTCGAACTAGGATCCTATATGAATATATAGTCATTCACATCATAAATTAGCCACACTTTCGCAATGTGCA
TATGACCTAAATTAAGTTCTATCAAGCCTAGATTTGTAACTCTACTATTCCAATTGTGAACCTATGAAGTTCGTA
CAGAATCATGTTTTTATTATTTACCTCAATATACATAACAATAAATATGTGAGATGGGTATAATTTTTATGCACT
TTAAACCATTTGAGAAAGTCTTCTTCAATCTTATCACATATTTTACTCAGCTCATTTAAATTCAATGTATTTACA
AATAATTCTCACACACTTTGAAAAACTTATAAATCAAGTAAATATTGCATTCAAATTACAGAGTTGAAAAAATAC
TAATAATAACATGTCTTACTTAAGATTAGGTTAAACTTCATCATAATTGACAAGAACATAAAGAATCATTGCTCG
ATGCTCCTAGTCATTGAGATGTGCATCTCTCTGATGCTTTATTAAACTGCTGAAATTAAAAATTATAGAGGAAGG
CAACACAACTACTACAACATCGGCGGGTTATCGTTGATCAAAGCAATTAGAGCAAAAAATAGTATTCTTTTGTAA
TACATATGTTTCATATATTAAGACTTCTTGTCAGGCTTTATTCTTCCCAATTCTTTACTTAAAATTTTATAATAA
AATAATAATTTAGCATAATTTATCAATACTTGTAACTAAATAGTTTTCTAATATATGTTATTACCTTTCAAATAC
ATACATAGAGTGATATTATATTAAAGCACCAACTATAGCCTCATATGATATATGCATGGGCAAATGCTCCATCAA
GTCAAAAAAATAAAGTGAAAATATTTTTTTGAGCTTACACAATATTATTGGAATGCCTTCTTCTAGCTGCTTTAA
TGTATCAAGTTTTAAATGAGTTACAAAATAATTCTTGAGGAGTGCACTAATTTTATTTAATGAATTAAATATTTT
ATATGAAAAATAACTAAATGCAGTTGGCAACAAACATTGCAATAAAATATGGCATTTGTTACTTTTTCAATCCAA
ATATCTTTCCTACTCCCAGGTCAACACAATGAATCAAATTTTAAACATAACCACTCAGAAGTTATAATGATTATA
ACCACCCACAAATTATTCTTCGTTGAACCAAATTAAGATAATATTTTATTTTAAGCTCAATAATTTTACCGTTAG
TAGTTTAAGAATAGAACTCCTTTCTTCCATAAAACTCCTTCAAGTCTAGTTTAGCATTAACATTATCCTTTGTTA
CCCTTAAAATCTATTGTAATGTTGAAAATATTTCAGAAAAACAATTCTCCATGTGCATTATATCAAGATAAGCCA
ATTAATTTTTTCTTCTAGTAAGACACATCCAAGAAAGTAGCCTATTTTATCCAATTATGATTAACTCCAAAATTA
AGGAGATATTAATAGATAACTCATGAAAATCAATAATTTTTGGAATTGTACTACTAATATAGAATAATTCATCCT
TAAGTAATCTTGGTGGTGACTATGATTTCATAAATCTCTTTTGAATGAAGGTATTTCTATTTCACCTAAAAGGAT
GATCAATAGGTAAAAATATATGATAAAAGTGAAACCATGAATTTTTTTACTATATTGAAGAGTAAATGCTTTTGA
TGAATCAATGCAGTATTAGCATGCTAATTTTTACATAGGGCCGCATCTCGACAACACTCCATACATTAGAAAATC
ACATATTGCTTACATTAAAGCTACATTCATTTGAAAGTTTTGTATCATAGAAACATCATAAGTGACAACACATGT
TTCCTAGGGATATTTTATTTCATCTTTTAGTGGCTACAAATATACATCAATTCCTCATTTTTGACTATTAAGAGT
TGAAATTATTAATATCAGAACATGAAGAGAGTAGTCATGCACATCTAAAGTGAAAGATTATGTTAAGTGACAATA
ACAATCTAATATGAATAGGGTGAGGTGGTTTGATTATATGGTGTGGACCCATCAGCACGAAGCCACACCTAACAT
TATGTGACTTCGTAAGTAAATTCAGGATAAGTATGATCAAAATATTTTCATGTTTCTCCAGAAAATATATGAGTC
AATGAAATGATTTCTACTAGACTGTTATAATATCATCTCATATACGGAGCATTATACATAGAACAAATAGTTTTT
AAAGAAGCGGAATGATTAGCCAAAAAAATGGTGTTTTGATAGAACTTTTTTTTATCTATCACTTTTCTATTTCGAG
GTTTGAATCTATGAGCCTTACAAAATCTACACTGAGTATCTATTTTATCAACCATATAGAATAAGATACTTCATT

TGACACAACAATCCATCTTCCTTGATTTTAAATTTAACTTTGAGATCAGCTTTTTATTCTGTATAATCTTGTGGC
ATTAAATTTTCTTATGAATTTATCTCTTTAAGAAATCTTGCAATCCTATTAAAGTTTTCCCAAGGTATGTTAAAC
TCTATTTTATTTTTTATATTCTAAGAGCCATATAGTAATGGGAAATGCATCATTCCTACATTGGTTTTTAGGTCA
TATTTAGTAGTTTATAAATATGTTGGTTTCTCTATTTGGTGGCTTTTCCTCTATTTTACTCAAATAACCAGTTCT
TGTTCTTTTAAAAATCACTTCACAAATCAGCTTAAGTCATACTCAATTGGAGGATCATAGGTCATTAAAGGAGCA
AAGCTAATTGAGAGGCCAACTTCAGTATAAATGTGATGTTCAACCACATCATGTGAAGTCCAAAATTAATAATTT
AGTTAAAAACCTCTACAGTATAAATCATACATGATTTTATCTAGCATTGAGTATGCTTGATTCTTGTTTAGAACA
TGGGCATCAAAATTTACTCTCAGATACATATGTCAGAAGCTAAAATATCGATTGCACAAATGACTCAACACCATT
CAACAACCCCTCAGTTAGTCCTTTTTTGCTCGACTGTTGTCTATTTTACATTTAGGATCTTACAAATCAAATATC
TATAGGTAACCTATATTCTTAATAAAATAATTTATTAAGTATAGTACACATCTTATGCAATTATTTCTTAGAGGT
ATTAATTGATTCATATATTATTTGATAATGAAGCACAAAAACTCATAAATATGAAAATTTGCATATATATCAAAA
AAAAGAACTTCATTGGTTTGTCTTGTGTACAAGAAGGCATTAGATTTCTTACCACTAAATTCTTATATAAGTTTT
TTACATAAACACATGAACCTGCATATTTTCATATAAACCTAACAATTAAGTGGCTTACATACATACCACAAAATT
CTTAGTTATTAATATATCTAACTGCAGCACTCACATATTTAATCCTAAACAGGGATTTAAACAATCCAATTTCAA
TTTAAATTGGGTTCAGCTTGATATCTGCGAGAATGGAAATGGCCTATCTCAAATTGAGCAACTTAAATTCATTCT
AGCTCAAAAATCACAATGACCTAGCTCAATTCAATTAATTGAGCTTAAAATAACTGATATATAACTAAATTTCAC
AAAAATCACAACACTTATACATTATAAAATTTTATGAAAAGTAAAAAAATGATTGAGGTTGGAGAAGATTTAATT
GAGCGCTATAGCATCGGTCGGCTCAAAGCTTTCGCACACAAGAAAAAGCATAATTGAGAGCCTTTCATTGTCGGT
TTGATATATAAATAATAACTGACACATTTAATGAGGTCAACCTATGAGTTTTTAATAAATTAGTAATTGACGGCT
TGCCTATAGGTTACCCATCGATTATCTGTCAATGTTAGTCCCTGATAAATTTCATCAATAATACTATGGAGAAAA
ACAAATGCCCATATGACCATCATTTTTCCAATTTTTTATCAGATATTTATATAGTATTTGATCCCTTACCAAAAG
ATAAGGTGGGCATTCCTACTTTAGATTTTATGAGCACAACACCTATCATCAACTTATGGGTTAGGCATTACATGT
TAACTTGTACATCTCCCATGATTTCATTGTGAAAAACTTATATAGTAAATTACATACCTTAAGGTTTTCTTGTGA
ATTTATATTTTTAGGATTACTTTGTAATTGTTAACAATTTTTCTAATTGTAGGTGAGCCAAATATCCATGATCTC
TCTGACAATCTGAAGGTTAACTTTCTTAGGTTGGTGGTACCAAAGGTTTAAGAAAATAAGGTGAAGAATAAAGAT
CCATCATAGATAAAGTTCAATGATGTAGCTTTATCCAATGTATTTTTATTTTTTTGCAACATTGTTTTCATACCC
ATGATCCCCATAATGGCATGAAATAGAATTTCTTAATTGTGAATCAACTCACTCCATGAGCTCATGATGATGTTT
TAGAGGTAGCCACTTTCTCCCTAAACACAATAATTATGAAAAATTGCTCTCTTGGCAGAGCAGAGTGTGACCCAA
CCCGCAATTACGCGGATATTTAGTCTCTTGGCCAAAAACTTGAACCGATTTCTTTAGGCAACTATTGTGCCAAAT
GGATATGATATAGAATTTAGAATTGATTTATATTTCACACATCAGGGGTTCTACGCAAGTATATGTATCATTTTA
AGTAATATAGAACTTTCTTTAAAAACACTAGTATTGATCCCACAGGGAGGTAAGGTGAGAAAGCCCTAAGCTATT
TCTAGTATACTACTATTAGGCCTCAGTTTGTGATTTTTGAACAGTTTGTTTTTCCTAAATTCACTGATTTACTCT
TAAATTAAAGTGAATTAAATTAAATAAAAACCAAATTGAATGAGACTGGGTAAATTTTAGAAAGTTCTTATATAA
TTAAAGTTTCTTTTCCCTTACGAAGAGATTATAATCAGCGTTGGAATCAAATGTTAAAGGTCAAAATGCACTTTT
GTGACAAATAGGAGCATACATATTCTTTACCCTCTCCCAATTTTATAAAGAAAATTGGATAATATAAAGTTAACT
TCCTAATTATATGGTTGTGCTGAGTAGATTCTAGCTTCTCACTTCTAGATTACTCACTGTGTCCTCTTTATAGGA
CCATCCAAGGACTCAGGTTCTGAACTAAGACCAATTTGTAAAGGTTCTAACTTGTCACTCTAGAATCCTCTATTA
TACGCTCATCAAAATTTATCTCCCTATTTATTTGCACTCCCACTTGGACGCGGCATCCCCTCCATAAAAGGTTTA
GTGTGGTACTCAGTAATTGAGTGAAATCATTGCCCTCCCGGGTCATTATGTTTTGCAACAGATTTTAACATTATA
AGTGATTAGTTTGTAAGTAACTTAAGAACTGCTGAAAATACCATCGCAATTTTGGAATAAAATAAATCTTCAAAC
CATTATAATCCCACATATAATTAAATCTCATCACAAAGAGAAAGAGTAAGGAAAACTACTCACTAAGAATAAAGA
AAATACAACATAAAACTTCATTTAAATCAAAGAAAATTCAGTAGAAAAATTTTTACAACAATTATAAGATAATGA
AGAACACAATACTTAACTTTTTCCTCTGAAAGAAAGATGAAAATCCCTCCAACAATGTCTCTTGCTGATTCACAT
AAAGTTCATACAATTCCTTTCAAATATCTCTCAAAATCCCTCACCAATCTCTACATCTCCTATCATATCTATGAC
CTCTGGACCTTCTTGGAAGTTCCAAGAGTTTGAAATGCCAAAATGTGCCTTCTCAAGTGTCAAAAGGTGCTTAAA
AATTCCTTGGGTTGGCTAGAACCTGTCCATATTGTTTCTTATGCCTACCATTTTCTGGGCAAAACTAGCACCTTG
GCGTCGTTTTTGGTGCCTAGAAGCCACCTTGGTGGCAACACACCCGTTTCACTTTGTTTCGCTACATTTTGCCCC
GTTTGAGCTCCAAACTGCTCCAAGTATTATACCTATAAGATAAACATGAAAATAACATTTTTTTAAAGTATAATT
TTATTTAAAAATATGTAATAAATTTAAAGGAGAATGAGCATAGAAAGACATGAAAATACGTCAGGTCGGAATTTT
ATTCTTTGCTCAATTTGAATGCAAATTGCACCATTTTTTCTCTATTTTTTCCAAAACTCCAACTCTTGTATCTGG
AACCAAAATCATGCAAAATATCTTTCTTTAAAGTACAATATCATAAAAAATGAGCATAAAATGTATATAAAAAAG
TAGGTAATTTTTAGGTGTGTCGGGGTACTTGTCGCAATCCTCCATTTATAGATGGCTCCATCTCACCCAATTAAT
CTTGCCAAGTTGGTGTGCATGTCATCTTACGGCATCTTCATCAATGAAGCAATAGAGCAATAATGGATTGGTAAG
AGGTTATGACATTAAGCAAATAAAATCAGTAGGGAGGGATAGGCTAATAAGAATAAGATTTAATTATATTATTTA
TCTTATAATAACTCACCTACTTTGACCATGAATCTGTATAAATCTATTATTGAAGTTGATATTTCTTATAAATAA
AGGAGTAAAATAAGTTAATCCATCAAACTTGTTGAAGACTCTAATTTATTTCTTATTGCTTTGGCTTTTTCTATT
TTAAGCCCTAACAATCTTAGGCAGATAAGATCAAGATTCTTAACCTCCGAGAGTCATTAATGAGCTGATTCAATT
CCCCTTTGATTTCCTTTATTTTGAAGTATTGAATTTGGGCATTTTTATTATCACTACGAATATATATATATATAT
ATATATTGAAAATCAAACAATGAAAAGTTTTGTCTTGATATTATTTACTAAGAAATATTAAATAAGATTTTCTTC
TTAAAAATATTAAATAAGAGTTGATTTTAATAAAAAATAGAAGCAAAAATATTCTTTTTTTTTACTAACACTTTT
TGTAATCACGCATGAATGATAAATAATAAATTAACTCCAATACTAAAAACGCAACTAATATTTTTAGATAATATT
TAATTTTTTTTAATATATCCAAACTAAAGCACCATACTTTGATCATGTGCAAGGGCAAAGCAATCTGGCTGGTTG
TGCCAGCCCAAGCTAAATCATTTTAATTCAGTGTTCAGTTATTTTATNNNNNNNNNNNNNNNNNTATATATATGTA

TATATATACATACATACATTATATATATATATATATACATACATACATATATACATATATATTGTATATATATAT
ATATATGGCTCCTTACATGTTAATCGATAAAAATTGTTACAAGAAAATAGGGAAAGGAAGCCCCGCTGCCAAAAA
CCTAAAAAATATTACAATACTCGCTCAAAATATGTGGGTTCCTCATCCACAAGACAACTCGAAGCAAAAAAAAAT
CCCAAAAATAACACACTAATACCAGCTTCAAAAGTATACTCAACACAAGAGAAATATACCCAAAAGTCTGTAAGT
GTAGCGAAAAAGAAACCAACAACCAAGAGGATAGGATCTTCGACTGTAGAGCTCCCCGCAGCTATGGGGAATAGT
ATATGGAAAATCTCCGGAAGATACCTACACAAAGAAGGAGAAACTCCAAGCCAAAAGAGCATCAAAGTGATAAAT
CGATGAGCCATAAAAAGGTAGAGCTTCAGGCCGAGGTAAATGATAAAAAAGCATATATATATATATATATATATA
TATTGGGAAGCCGTAGGAAAATTTATTGCTCAAGGGTAAAAGTTACAGTGTTAGGTGAAAGGATCCCACTACGAA
AGCAAAAAAAAAAAAAAATAGAACTCACACCTAAGGTTCTATTCACCAGGTATACAACACACCCAAAAGAAAGTCT
CCAAAATACTAATACATGTATACCTAAAATAATTCTCCAAAAATGGTGGAAAAAACCAAAGTATGTTAGATAAAG
AGAGAAAAAGTAGCATGTATATATCTATCTTATGTAAGGTATTTCCAACTTGTCTTTATGTACGACACTTGTAGA
TCTAGAGAGGGTCTTCTACTTACAAATACCAATAAAAAAAGAGACTTTGAAAGCCCATATTTAATTTATAAATTT
AATATCATAAATCCCTTACAAATTCTCTCAAATTCCCCTCTCAAATCTAACTTGAACAAGAATTCCTACTCCTAC
CTACAATTCTTGGAGAATTCCAGGACTTACTCCTAAGATTGTTAAAACTCCCAGTCTGAATCATCAACACTCATA
AAATAACTCCTGGAGTCACGTAAAAAGTAACTGTCTGAACGAACCCAGTTTGTGATGCTTAAGTGTCGTGCTCTC
TGTCTATTCAAGAAGCTTGGATGCCTAGATGCCCAAAACCTAGTGCCATGGTCTTCATCTGCAGCACTTTGGCGC
CCAAAAGTGGGTGCCTAGATATTAGACCACAATCAGCACCATTGCTTCATTTGGTTCTATTTCGCTCTGATTCAG
TCCAAATTTGTTCAAATGCTCCTACCTGTTATACCTACGTCATAAATATAAAAATACCTTTCATTAAATTACAGT
TTCATTCAAAAAATATGTGTAAAATAAAGAAGAAATTAAGTATATCCAAAGACATATCAATTACAAAATATAATA
CGATTAAGAGAGTGTTGATAGTGATCAAGTGAAAAAAGGAAGAGAATTTGTATATCATAATATTTTGATTAAAAT
GAAACTAAAAAATGGGGCCCAACCTAAATTATGTAACTATATAAAAAAAAATTTTAGTTTTATAAATATCCTCGTT
ATGGAAAAGATAGTAATTCATTAAACTCTATTAATAAATCTCTAGAAAAAGTAGTTTTAACTCCTATCTTTTACA
TAATAGAATGATAAATGTTTTTAATTAAAACACCCTTATATTTATAAAAAGGCCAAAACTAGGAGTATGATCAAG
TGATGAGATTTGATGAAGCATAGTTTTCACCAGAATTCAATTAATTGGTTGAAATATTTTTTTTGATGGCCCAAA
TGCACAATAAGCAAAGAGTTAACCAAGATCATTAAAGTTTATCATGAGTTCAAAATCATTTTAATATGACATAAC
TCAGAGATCATAGTTTTCATCCACAATCATGTTTTTATAGAAATATGTAATATATAACCAAAAAAAAAAAATAGAT
GTGTGTTACTGTAAAGTGAACAATCTTGATCTCTCAAACAAATGAAGGATTTCAAAATTAAAATTGGTAAGAGCT
CAATGTATCAGAAAGATTAATTATCGCACAATTTTGGTTTCGATAGGAGGCAGTAGGAATGAATTGGGTGGTGTA
TGCCCAACCATAATTTATATGACTTAATCATTTGGAATTGATCGATGTTTGATCCAAATAAGAGAGAGGGAGAGA
GTAGGATAAACAATAGGCAATAATACACAAAATAAGAAAATTTAAGTAGTGATTATAGAGATAAATATTACATAA
AATAATCTTAAAAGATGAATGTAATACTAATAATATACTTAAAAATACAGCAAACTTTTTTTTTTTTTTGAGGCT
ATGATAGCTAAATTAAAAAAATTGAAATGGTTGAGAAATGAAAATGTATATGATGTAGAACATAAGGTGATTAAG
ATAGAAGGGAAGCCTTCAAAAAAAAAGATAATGGAGAAGAAGAAATGCATTAATAAAGAGGCTCCTGAAACTAAT
ATGATGCTTTTATATAAATATGGATTGATGACAACAACTTGTTAGTATAGAAGGCTAGAAGGATAGAGTTCAATT
TGCATTTGACTTAGAGGATACATGAATTTTATAAGAGAAAAAATGTTGAAGGATTTAAGGTTTACAAATTTAACA
ATCTAAGTAATTCTTTTTATCAAGTAATTTAAACATGAGAAAGGTAAGATGTAGTGGTAGTGAAAATCAAAGTGA
TATTAAGAAAATATGAGATATGTAAAAATTTTCATAATAGCAATTTCTTTCAACAACTAAGATTTATCTTGATGT
CATCAATTATGACACATAACAAAAAAATTATACTACCCTTGGACAAATGTCTTTTGTGGAACCCAAAGTTAGTTT
TGTGTGATAGATTGAAAATAATAACATCAATAATGTTCAGTTCAAATCAAAAGAGCATGACATAATACTTAGAAT
AATGTACATCTACTACCAAAACAAGAAGCTAAAGATTCACCTGAGGTCATTGTGAGTTAGGTTTTTTAGGCACTA
GAGTACTGTGTCTTTTCATGTCTTAACATCACAAGTTAAATGTTAGAATTTTATTTCATGTCTTTTCTATCGTTG
TAGCTAATATATGCCATCCAAATAGCCTTAGACAATTTTTATTTGATATGAAAACATTCAATTCTTTCATTTATA
CAGATTTAGTTAAATTGATGAGTTCTATGTGTTTATTATTTTATCAATCATTATAGATGTTATCAATATATGAAA
AGAAATTTAAATTTTTGTGAGAGAAATTTTGAAACAAATGATTGTATTGTCCCTGCCAAATTTTGATTCACCATT
ATGTATGGATTGATTGTTTAACCATTTCATAACAACATATTAGTTTAAATTTCTAATAAAAATTTAGCATAGCCT
TTACGAAATTTAATTTTTCTCAAAATTAGTTAAATTAAGTTTCTCAAAGAAAAAGTTTTAATTTCAATTTATATA
ACTTTATTAAAGGCCTCCATTAACTTAATTGACATTCACAAATATTATCAATGCTTGAACATTTAAAAGTATTAA
AATTTCATTTTAAATATTAGAAGTCATCCTCAAAGTTTAATATACCTTTACTATCCCTTTTAATGGTCACTATTA
AATGTTAATAAATCCTTATTAACCCAATTTATTAGTCGAACACTATACACAGTTAGTTATCCAGTGATTAATACA
ATTTATTTTCTTTAGTTAACACTTGCAATTTGATTTTAAAACATCATAATAAACTCTATATAAAACCTATTTTAT
TAGTTAATATTACTTATTTTTTTAATCCAATATCACTTTTAATTCTATCTTATAAGTTCTTTCCATTTTAGGAGA
GAAAAAAGGCACACTTACTTGGTGATACTTTTTATAATTATTGCAGAGTTAGGTTAGATCATTGAAGTAATCCAA
TTGATGGATGAGAGGACTACTAAGATGCTTGAGAAACCATAAGAGGGACAATAAGCCAATTTTTTAATTATTGGG
GCTAAGTATATAGTGAAAGTGGTATTTTGAAAGGAGAATTTTGAATTTATTTCAGGAATATATAATTCACACTTA
TTTATTTAGAAACTTTGTCTATAAAAATCCAAATTTGATATATTTATGCTTAAATTATAGTAAATGGAGTTGAAT
ATGGATCTTGAAAGTTTTACTAAGTCTGGATAAGGTTTCTAGTGGAGTTTACTTCTAGAAGAGATAGAGGTCGAT
AATGAAAAAATAAATCATGTTTAACATATTTATTAGAATTTTTAATTGGTTGAGGGAAATCCAAGTTTTCACTTT
GAAAAGTTTATAGACCCTAATTTTATGAAATGAAGCTTTTGGAATTTTTTTAACTTATGTTATTATTTTTCAAGG
AACTAAATGATTGAAATGATGGAAAAAATAAATTACAAGCTTCTAACATTCAACTTTTAGGGTTTTAGCTCTCTA
GAATTTTGAATACAATAATCCTAAATAAATTAGAGATATAGGAGATGGATATGACATAAAATGATCTGAAAATTT
TTGATGTGCTTAGCCTTGAATAGTTGGGTCAATTGAATGATATTCTAAACTTAATATAAAATTTAGTCACTTGA
TTAACTTTTCAGTTAGGCTTTAGAATAGGAATTAAGAAAGAATTAATTTTTATTTACAAATATAGTAAGAACATG
GATTTTTAAAATTAACATAAAATTTTGAAAGGAAGTTGCAAGAATGTAAAAATCTTATATGACAAATTAATGAGT

TCAATGGTTCACCCAAACTCTTATTAAAGCTAGCCACCAAGCTGCCTTCAATTACCTTAGAATAACACCAATTTT
AAACTTTTTTTTTAATTTAAAAATTAAGAGTCATGTACAAATTATTTCTTCTACATAAGGCATATTTCTTGACTT
TACAAAATCAAAGACTAAAAGGAAGGAGAAGTTAGAGTGGTTAGAGTGGTTAGAGTGCTCCTTTGACATGATCAA
CATAATTCCACTCTTAAATCCTTAATTTTACCTCCACCTAAGCATGAAACATGACACATAAGAAAGAAAGTTAAC
TTGTGGTCAATACTTATATGAGATAAATTCATAAAATCATGAATATTTAAATATAAATATTGCAACATGAAATAA
ACACAAATGCTAAAAGCACAATAAGTAAATAGTTTCAAGTTAATAATCTTCATGCTTATCATTTGGTTTGTCTCT
TGATATTTCATCTTTCTAGTTGTGAAGCAGTTATCCTGGACTCTATTTTTTAGTAGGTCTCAGTGCTTCAAAGCC
CAATATTGGATCAACCTCTTATGATTAGAGGACCCTCCTTGAATCAATCGTGATGTATTGTGCTATATCACTTCC
ACCATATTTAAAAAAAACAATTTTTTTAAAGATTTAACGGCAAGGTGGCATTTTAATATAACCGACGCTAATTGG
TGTAGCTTCTCCCAAGTTACCCAAACATCCTCACTAGATAGATTGTCCTTTCATCAAGTCAAAAACCTTTGAGTA
GTACCTTCATCTAAAAGTACCTCAACTATAATTGTGATATCTAATTTAAAGAATTTTGGTTAGATGGGTATCTGC
AAATGATTTTTATTTCTTCATACAATTGTATTATACAAAAAATTTGGAGGTTGGGTAATTATTTTAAAAGAAATT
AAGTATTTCACGTTAAAAATTAGACTAATTAGCATATGAGGGGAAATTCATGAGCAAAAGCATTAGTACCAAATT
TTTTAATAAATTTAAATGGCCCACATGCTCTCATTTGTAATTTTTTATATTTGGTGAAAGGGAATCTCTTTGGAC
AAATACTATATACCATTATATAATCACCAAGCGCATAGCCCATAAGTTTTTAATTTTTTTCTTTTGTTATGCATA
CTTAGCATTTATGCTTATAATACTTTGTATATATTCATGAAGGCTATGAAAATCACACGCAAATATGAATTTTGA
ATAAAAAAATGTAAAACTGTGTTATTTATATCAATTAGTTTCGTAATATTATATTATTATATTACTTTAAATGAT
ATTTTATTGTTAGTTCTAGAAATGGTACTATTAAGACAAACTAAGTTATGAGGACATATAACTTCCAGTTGGCTA
AAATTTTACCAACTAAACTACTTATAAAGTTTCATGAGGTTTTATTGGTAACTTTAATTTGGCCATCAATTTAAT
TGTGAAAGGTAAGGAGAAGTTCAATAACTTTTTTTCAAGAAAATGTTATAAATTTAAAATTCATATAAAAAGCTA
TGGTAGATGGAATCTCATGTAATGTCACTATTTCTACATAGAAGAATTTAGCAATATGAGATGCATCATTTGTTT
TGGAGAAAGGTAAAAATTGTAGCAACTTATCGAACCTATCAATAACCATAAGGATATTGTCATGACCTCAAAGTT
TCATGGGTAACTTTAAAGTAAAATCCATAGATAAATCTTCCCAAAGCGTAGAAGGAATAGGCAAAGTTGTGTGTA
AGCACCAATATTTTTTTCTTTGCAACTTTAACTTGAAAGTAAGTATAACTTTGACTTATAATTTTCTCGACATTT
TTATGTGCAGATGGCCAATAAAGTTTTTGTTCTACAAAATTCTCAGTATTATTTTACTAAATTTCAACCAAGTCT
CATAGAATGAATTTTTATGATAATAAATTATCCTATGGAGGTTCTAAGAATACAATCTTTTACTTTCATAAAGAT
AACCATCTTAAATGCGAAAGAGCTATCAAATGCTAATGATTTAAACTCATGCTCTAAAGAGATAAGATGATATTC
AAGGAGAATTTTATTCTTCTTTCATGTTATTAAATCCAAGAATTTGAAACCTTAATTATGTGACTATTATAGATG
CCCCAGATAAAGCATATGCTCGCTTATTCTCCACTTCAATTTTATGTTTTGTTGAATTTATAATCTTGTAGGTGT
TTCACTCCTTTGATACGATAATGATTTAACTTTTCTAGGTGTTCATAAGTTCAAGAGCTTAGTGGTCCATGTTAA
GATCATTTTTTAATGCAAAAGATTGTACATTCGGTGTTTAAATATTTGAGTGATTGCAAAAAATTCTCTATTATG
TGTAGAATATCTATGTTAGAATTATTCAATTTCCCACTAAAAATGGCAACTAGATGGTTATCTTGGCTTAAACTC
TCCCTATCCATATGCTTGTAGTGTCATAGGTTATCAAAATATTTTTTTTAAAACCAAGAGGTTTAAAACATGTGA
TTTGATCATTAAAATTTTTACAACGTCTAAAGTCCAGAAAAATTTATTTGTCTTTAGTATTTATGTGATTGGAGT
TACCAAAATGTTGAAATTTTTTATAAATATTTGATAAAAAGTGTCTAAGCCATGAAAATTTTTATTCGTGAATGA
TTTTGGAATTTTCAAGTCTATAATAGATTATATTTTTTTATGTATCCACTTTTAGATCATCATATAATACAACAAA
AAGTATGAATATACTTCAAGCTAGTGAAAGAAGCACTTTTTTAGATTAATGGAAATTTTTTTCCAATTTTTTTTC
AAAACTAAGCACAAGTGATCTAGATGTGAAAGCTCATCTGGGCTATAAATAACAGTGCCATTATATTATACTATG
AAAAATATCTACTACAGGATTCTAGAATAGTGTTCATCAATCTCATGAATGTACTGGTAGCATTAAATAGACCAA
ATGAAATAACTATACCTTCATAAAGACCATCTAGGGTTTTCACTACAGTCTTCCACTCATCCCTTGGTTTAGTCC
TAATATGATGGTACCTACTTTTCAAATCTATTTTGGAAAAAATAGCTGCATTTGAAACAAATTCTAGCATGTTAT
CTATTTTAGGTAGAGAACATATACTTGATAATAATCTTATTTATAGCTCTTTTTTCTACATGCATTCTCTAAGTT
TCATCATTATTTGGGTCAGAAGTGTAACTACATTAGAAGTACTTAAACTCTCTTCAATATAACATTCTTTTTGGG
GTTCATCATCTTATTTCTTTAACTCAGCATTCTTAGAGAGATTCATTCTATGAAGACGAAAAATTGGAATTTAAG
AACCAATAATTAAATCTATACCATGTTGAACTTCCCTTAATGTTAGTAACTCATCTAGAAAATCATTAGGGACAA
TAACACTAAAATCAATATGAAGTTTGAGACCACAAGTATTTGGTGCTAACTCTAATTAGATGTAACCTCTCTAAA
TTTCATAACAAACAAAGACGTTCGGATAGAAAATTTTTTGGCTAATTTTTTAAACCCTAATGATTCAACTGTAAT
CTCTATGTTAGCTGGAATCTTAGAATTTAGTTAGAAATTTCATCCTTAGTATTAAGGGTTGATAATACTATTAAT
AAGACTTTGGCATTTTTCTTCTATGTTTTAGTTACATTTCAAAGTTCGGTTTCTGAAGTGGTGTGAGAAAAGTAT
TATGAGTAGGTGGGATTGGCCTAATTTCTATAAACTTTATAACAAATAATTTTATCAGCTTATTATTATACTTTA
TCACATAAGTATATTCTCAACCATAATCAAAACAGTGAAGTCCTAACCATATCTAATAAAGTTCTATTTTTGTGT
CCTACCTTTGGCATGGTGTTCCACCAAATTTATTTGGTATGGATCTTACTGGCTAAACAATTATTTTTAATGCAA
TGTCATAAAAAAATCTCATCCATAAATTCACCTTCATCAACACTTCAAATATTTTGGCTTATTAATATTTTTTTC
TCTTTGTGATAAAAAAATAAACTTATAAAACTCTAACAAAGCCTTAATCTTTTCAAGTATGATCACGATCCAAGA
TACCTAAAAAATTCTGCAAATAAAAAATTTAATTCTTACCTCAAGATTAATGGGTCCAAATAATCCATGTGCAAG
AATTCCTTGGTCTAGTGGTTGAAAGTTGTTTAATGATTTGAAATAATTTCCTTATTTTTTTTCTAGTTAGGAAAC
ATCACATAGAATATATATAGGGTATTGAAGCTTAGACATGCCAGATACAAGTTTATTCCTTAAAAAATGTATTTA
TATGCATAGAAGCATGACAATCTTTTATCTCAAAATAATGAGTCATTAAAATTAAAATTTAAATGTGTTATGGTA
TTATGATATAAATAATCAAAATAAATTAGAAAACATATAATTATGCATGTGTTCAAGAAGCATAAGATAGCCATT
CAATTTAATCAGTCAATGTGAAGATAAAAAGAAAATATATTTATACTTTTATCATATAGTTGGTTGTTATTTATA
CATTACCAATGTAACTTATATTAAGATCAACATGTTTTTTATTCATCATGTTATTTATGAATTTTCAAATAAGTT
GCATAAATTTCATGCAAGATGACATAAACAACATATAGGTAGCCCTCCTTTGACCTCAAATAAAAGTTACACATG
CATAGGAGGTTTATTTAAGATTTTGTTAGTGTAACTCAAACTAAATTAAGAGCAAATAGAAAATTTCATCCATGC

GTGGAAGAAAGAGAAGGGATGCAAGGAGGAGTGACAAATGTAAGAAATGAATATATACTTTTTTATTAAAACATG
ATATGATGAAGAAATGAAGAAACAAATATTTATAACTCGTATATTTAATTTTATTTAATCTTAAGTTTTATTATA
AATCAATATCTCTTTGTATTCTTATGCAATTCAATTCTTCATAACTTTAAGTCCATACACGAAATTCTAGATCTT
AAAGCACTATAGTTTAAAAGGCATTTGAGATCAAACTTCTTTCCCAAATTTTAGCAAAGAGAAAATTTTACTTTT
TCTTCGTAGAGATTGAGAGATAAACTTTCTTGGTTAGAGAATGGTGGTACTCCCATTATTCTTCAATTACTTTAT
TGTTGCTTAAGTTCTTCCATCATGCTTTAGCTTAACAATCCACTTATCAATTGATATAAGAGCTTTGGTTGGTGA
TTATGCAAGTAAAGCACAAGTGGTACAAACATTCCAAAACATATTTTTTTTTCGAAAAAATTATGTATTTTTAGT
GATTTCCTTCACAATCATGTTGGAAAGAATAGTGAGAAAATACTTATGTATCTTTCGGGCATTATTCTTAATCGT
GCGTTAGAACTCATCCAAACATTTATTTCCTAGTTTATTGTCCATTATAGATAACTCAATATTAAAAGAAGCTGA
AAAAAGACAACTTTAATGATACTATGTTACAACTAATTGAAACTTAATTATAACAATATCAAATTATGTTTTTAG
TCTTCAAAGTAGAAAAAATGTTTCTACTAAAGTAATTTATTCATTATTTTAAGAAAATTGTAAAAGCACTTCTCAT
TTCATCCATACTCTCACTTTAAACTTTTGAATTTCTTGTCATCTCCCTTCAATATTTTAATGCTAGTGTTCATAG
TATAATCCTTAATATATATAAAAATTTTCCAACTAATCTAATTAAATAATAAGATTATTTTTTTAACTTATTAAT
CTCTACTTAAATTTATGCATCTAACAAACTTTCAACTTTTCAAAAGAAGTTCATTTTCATGTGGGGGGAGAAAGG
ATGCAACGTAAGTATAACAAAAAAATGAAGATTGAAAGGCATTAATAACTTAAAATGGATGAAATTTACCATAAC
TATTACATAAAATTATGTTTCTTCTGTCAAGTAATAATATAAACACCTTATATTTATAACCATATCATTTTATTT
TATTTTTATTTTTAGGAAAAAAAATAAGTTAGCACATTAAAATTTACTTCACTAACTTCTCAAGAAATTTTCATAT
AAGTTGCATAAATTCTATGCATGATGGCATGAATGATATTTCATTAACCCCACCTTTGATATTAGATCTCATTTG
GAACTTATAAATACATGAGAGTTGTATTTAAGCTCTTCTTAGTTCAACTCTCTAAATTAAAGTGAGAAGTAATTT
AATTCATGTATAATGTAAAATAAAGTGAGAAAATTTACCCTTTTTTATAACATGATATGTTGAATATATAAAAGA
ATAAATACTCCCAAGTCATATATCTATGTAATTGTGTTTAATTTTGGCTCTTCAAGCGAATTTCTCTTTATGTTT
GTACAATATTTGTGATATCAAACCTATATATACCTATGGGAAATTTAGTTGTATGACAACATATTTAATAATAAT
TGACTCTCTTTTCCTAACATTTTAGATTGTTGGAAAGTTTTCTATATTTTACTTTGATTAAGTTAAGAGTGATAT
ATGAGGGTATGATAGTTATCCATTTACTCTTTAGTTATTTGCATAGCTTCTGTCATTCTTCCAAAATTATTCAAA
GATTTATCAAAACATTTGTATAATGAGAAATATTTCAAAATTAAATTCTAAATACACATCATATTCCAAATGCAT
CATCATTATAGCCTTACAAAAGTCAAAGAAGATGTGATCAAATAAAAGAGGTCCAGTGTTCAACAAACCTTATTG
TGCTTTATCAATGGGGTTTTGTGTCATGTCATGAGTGTAAATGAAAAAAAAAAATCTAAGATTAAGTGATACATTA
TTAGTAAGAAATTAAAGCTAAAGCTCTAATACTATTTGCTCACACACAACCCAAGATAGAGGAAGAAGGAAATTA
AAATTTTAAATGATAAACTAATTATTTTATGAAGATTTATATGGCTCTTTGTAAGCTCGTGCAATATTTTTATGA
ATCTAAAGAAAAAGCAAAAATAAAGTTCATGAAATAAAAAAGATGCTAATTTTTAGATGGTTTAGCTCTCTAAAC
CTTATTTCCACATCTAAGTATAACTCACACATATAATTCTCATACTCAAAGAACATACTTCTTATGACTTAGAAG
TGAAAAAAATAATTTTACCATCTCTTATTTTAATGGGAAGAGTACTTTGGCCTTATTTAGGGCAAGGGAGCTACC
TCTAAGTCTTCTCTAGAATACTTAGTTACATATATATAAGATTTCTGATATAGCACACCATGGTAGATTAAATCT
AGGACCCATCAACATAAAAGGTAAGCCAAGATGGGGCATAAAGCATCAAAGTGGGATCTGCTACAAAGGTTAAGT
ACAACAGAACTACTTCACATTAAAGAGATCCAATTACTAGAAGCTTACGTCATTAGTGATTGTATTTATTCCTTA
AAATGTGTTTATTTAAGTTTTTTTTGTAGTTTAGGACTTTATTATAAGTCAACATTTTTTACTTAGGTGTCAAGT
TCTATGCTTAGGTGGAGGTCAAAGTCAAGCAAGGGATTAAGAAGTAAGAATGTAAGAGTGGCATTATGCTGACCA
CTATAACTATTCAACTTCTCTCTCCTTCTAGTATTTGATTTATAAACTCAAGACATATTTCGTTTGGAAAATAAA
TGCTTTATATAAGATCCTTAATTTCTAAAGTAAAAGGGGAGCTTATTTTTTGGTGAAAAAATGGAGTTATTCTAA
GGTGATTGAAGGAGGTTTAATGGCCAACCTTAACAAGAGTGTTGGTGAACCCTTAAAATCCTTTCTTTGGAAAAT
AAGGTTCTTTATATTATTGCAACTTTTGATCAAAGTTTTGTCTTATTATTGTGAATCCATGTTACTAATAAGTTT
CTAAGAAAAAATCTAAAGTTTTTTCTTTCTTAATACCCATCTTAAAACCTAATTAAAAATTTATTAAAGTGACTA
AAATTGTACACTAGGTTCATAGTCTCTTTCAATTGACTGAATAATTTGTTGGTGTTTTTTGTTCTTTTTTCGATT
GTTTCCACTCTAAATCTCAAAAGATTTCTATCTAAAAAATTAGATACATCATTTGAAGAATAATATACTACAGTA
AGGGAAGAAAATATTAGCTTAAAATAGATTTAATCATTTAGATGATGTTCCTTAAAATTTATTATTGGTGTGTTT
ATGCTCAGAGGTAGCATAACCACACCAAAGGGTGGTGTGGTTACGCCAACCATGATTAGGGGTTCAAGTTCTTTT
CCACTCGTTTTGAGCTTGGTAACCTCACCAACATTTAATGTCCTTCCCATGCGAGGACAGGTGAGGTTTTGGTCA
AGAAATGGTTGACCAAAATGAAGTTAACAAATGAACCACGAATATATGTGGCGAACTGTTCAACGTATGATGCAC
ATATTGACTGTGGCGATCAATCAAGCCAGAAATAAAGGTAGTGATATTCATTATTTTAATTTTGAAATTTGAAAT
ATGATATGAGGATTGAATTTTTTATTTGAGTGATTAAAAATTTAAACTAGAACATTTGTGAGTTTTGTGTTTCCT
ATCAATAGGATCCATTGAAGATTCAAGCTGAGGATTTATGCAAATGATTGTCAAAACATTTTGAATTGATTTTTT
TTGTTCAATTTTATTTTATATTGTATCACTGTATGCCATTCCATATCATTTTGTATCACATCCTATTAGATAACA
TCATATCACATCACGTCTGTCATGTCATGTCATATCATATCATACAACCGAACACATTATGAAGGAAAACACAGG
AACTCAACTTCCACAAAAGAAGCCAAGGTTGCTTTGTAAAGACGTGTATTTGGGGCCTTGACCAAATAATGAAGA
ACTTAGTTACTTTGTGTCGGGGACATATCAATGGACTTATCCAAAAAGCAAAGAACCATGTCAGCATGTTTTCTT
GTAGTCCATACTGGAAAAGCATAATTCTAACCCTAAGGTGTGCAATTAATTTACTAAGACCAATCCTAGATTTAC
TTAGCTAAGATGCGAGTTTTATTTGAGTTTGGTATCAAAAATCAACAAAAATGACAAACCTGAGAATGGAACATA
ACATATCTTTGTTGTACAATGAGCTAAACTAAGAATAGGAATCATTGAGAGATTTGAGTTAACTCAAATTTATTA
ACAAAAGCCTTAACCTGCATTAGCAATCACCATTTTTGGGAGCTAAATGAACATGATCACATCGTGATGGTCTCA
TAGAGAGTGTACTCCAAATAAATCATGAGCAAAACCTGTAGAAAGGAATGGTCAAGGTCATACCATGTTTAACAA
CCTAATCCTGGGGTGTTCAATGGTTGCATTAAAGAATATTTATACTGATCATATGCTAAATTTACTAAGGAAGCC
TTTGAAATGAAAGAGAAAATAATGGATGGGCTACAAATGCAAACCATCCACAAAGGAATTACATCTTATCTAGGT
TAAGGGAGCCTTTAACATCCCTGTCATTCTAAACAACCACTTGAGCATATAAATCTATACTTGGCCAGCCAGTTT

GGAAAAGTAAGAGTACTTCTCCTCATTACATGTAAATATTTATGTAGAGGCCTCATTAATATAACTTTGGATATA
TCCATGCATGTGATGCCATTTTAAACATGGCCTAAAGGCCTTACAAGAAATGTAAACCATAAATTAAGTTGTTGA
TTCAAAAAGTGATACAATTATAATCATAAATTTTGAATAAGATAAAATTTTTATTATTAAAATGATGTAGAAATG
ATTTTATCCTATAGAGCCTAGGGCCTAAATTCAAAGAGCCCAGAAAATCCACAATCAAAAGAAAGGGAGAATTCA
ACACTCTATGTAAGGAGATAAATAAAACATGTAAAAAAAATTAACAACAGTTCAGATGTTGTTGAGTCGACAACA
TATCCCAAAAAGATTAATTGGGATCAGGCTTAACATATGCATTATTCTAAAAAACAATATGAATCCAGAATTAAG
ATACAAGTCACATACATACTATTGTTAATCCTCACCACATGGTTCCACATATCGATCACATTGAAAACCCATGCT
TGTACATTATTTCAGATATAAACTCACGCTTATGGATTGTCAAAAAGTACGGATAAAAATTAAGAAAAAAACCGA
CCAACCATTGTAAGGTTAGCATGAGTTATTTCATGTAACTTATATTGAACCATGCTGACAGGAGGAGAATTGAGC
AATGTCTAAAAATACGTGTTAGTTTAAATTATAATGCTCTTCCCTATTGAACATCACTCCTTGCAAAAAGATCAC
CTTTAATACAAAAGTGAGTATGTCTCACACCCAGTAAATATGTTAAAGAAAACATCCAAAAATGTTTGGATAGGC
AGAAACCATAGACAAATAGTCAAACAAATGAAAAACCTAACTACACCCTACCCTATTATAAACAAGCATCTAAAA
TAAGGTGCTCTAATTACATACATCTACGTGTGTGCAAAAAATGTGATTTGAGGCTGATGTATAATTTTGAAGGGG
TCATCAACACTGGCCCCATCCTAAAGACCCAAGAAATTTAAAGTGTATGACTAGAAGATGATAATTTTAAGCAAA
ATATATTTCAAATATTAACCTTATGTATAGTCTAAATAATGTATACTTTTTGTTTTAAAATAGATTTTAAAATAT
GATGGATAGGATTAATATAAAAATACCTTTTTTTTATATTTTGAAATAAGACCTCATGTACAGGTGTAAATAACC
TATGTGGCACATTTTAGTTTTTTGGATCCTAAGCAAAGTGATTCACGCGTTGATAACAATTCAATAATTTGCATT
GATCACTTAGATATACTCATAAGAAAAGCTCCTTTAACCATTGCAACTAGCAACGGGTCACACATCTAAAAGGGT
GACTTGCCATAAATAACATGCTAGGAGAAGAAGATGCAAAATTCAAGGTGGGTCAATATGATATGATTGCTCAAG
CAAGTTGTAAGAAATAAAATTATGATATGCTAAGTGGTTTTACATGATACAAAAGCTCGATCTAGGTATGAGCAA
CCATAACATGAATGAAATTACATATTAAACATGTGTACTAAGTTAAGTATGAGTAGCCACTTCAAATGACATGTA
AATATAGATATAATTTGCCTTCTCAACATGATCATATGAGGTAAACGATCACACATGAAAGCAAGCAAAATTTAT
GCAATATAAAATCACAATAGGTGAAGTGATCACGTAAAGAAAGACATGATATATCATGTCTATGATGATTTTAGG
TGATGTCACTGATGCTAGGTAACTCGACCTGCAACTACGCAGATATTTATGTTCCTAATTATGGCCAAAATCTTG
ACCCAATTTATATTTAACGGCTAGTGAAAAGAAATGTAATATTATAAAATTTAGATTATTAGTTCTCTCACTATG
GGTACTACACAAGTATACGTATCGTTTCAAGTATTATAGAGTTTTCTTTAAAAACCCCAGTATCGATCCCACGGA
GAAATTAAAGGAAAGAAGTAAATTTAGGAATCTTAACTATGAAAAAGTTTTACATAAGACTGCAAAAAAATATTTA
AAGTAGTATATGCAATGAAATAAAGTTTACTAATTCTTACAAATGCAAGTTAAAAGTAAATGTAGAGTTAAAAAA
TTCAAGATATATATATATATATATATATAAAGAGACATTCTCCTTACAAATAATTCATAATCAGTGTATGATTTCAA
ATGTTTAGAGGTTTAAATGTGATATTGTGACAAATAAGAGCATACCTATTCTCTTCCGATCGTAAAGAACATTGG
CTAACCTAAAGTTGGTGTTGACATGTATAGAGTGTTGGGTACGTTCTAGTATCTCACTCCAAGTTTATCAACTGT
GCCCTCTCTAAAAGAACCTTCCAGTGACCCTAGAAACATCAATTCTTTTCCCATCAAATCTTCTTCCTATCTATC
TGCATTACCCCTCGGACGGTCACCATGGCTCTTTCTAAAGTCAAATGTGACACTCACTAGTTATAGAAATCTTAG
CCCTTTCAGTTATTTAAGTTTCTCAACAAATTTTAACAATATAAGTGGTCAAATTGTAAAATAAATGAAGGACCG
GTGAAAATATCATCACAATTTTTTGAAATAGAAATTATAATCAAGTCAAAACCACCACTTGATTACTTTATATTG
CAAGGAGAAAAACAAACTACTTAGAATGCATGGAGATAACAAAACAAAACATAATTATAAAACTCAAATCCAAAG
TACAATACTTAAAGAAAAGAAACAAATACTTAACTTAAATATTCCCCCAAGCACTCCATTGAAGTTTTTTGATTT
CTCTGACTCCAATAGACTCTCACAAGACTAAAAAAACTCCACTAATCACAAAGACTCCCAAAACCTTACTAAGGT
AGCCACCCAGTCTATATGTGTGCCTCCCCTTTACAAAGCGACTCTAGAACTCCCCCTAATCGTGTAGAATTAATC
TATATATATAGAGCTTTTAAAATCACCAAAAGTTACTTTAGACATCTCTTCAAAATTCGGTCTTATTCAATGAAT
CTTCTAGAAACCGACTCTGAAGCCTAGGCGCCAGCCGCGATCTTCATCTGACGTATAGGTGCCAAATTCTGGCGC
CTAAGTGCCAGTTTGGCTATTTTGACAATTTTTCTTAGTTTGACCCATTTTGCTCCTTTTTAGTTTATTTTTGCT
CCTTTTTTCTTTTAAGCTCTTTTTCTTGCACTTGTAACCAAAAACACACAATACATTTCATTAAAACATAATATC
CTCATAAATGGGTATAAAACGTGCATAAAAAATAGGTAATTTTCAAGTGGATCAATTTCCCCAAACTTAAATGTT
ACTTGTCCTCAAGTGACTCATTTATTAAATAAAATATTATGCCTCATAAATTTCACAAAGGCCTCATCATATTAT
GCCTCATAATCACTCTTCCTCCATCTGCCATATGATCAAATTTCTCTTCCATATCTTGCATGTGACATTGCATTA
TAAACATATAGGACTGGTGCTCCAAATATCTCTCACTCAAAACATCCCAAGAACTAGATGGTGGCTCATTAAAAG
TTAAAGTAGAAGTTCTAGAAGTAGAAGGCCTAGAAGTAGTAGTAGAAGGTGCATCAGAAGCTCTACCTTTAGTTC
GTTCATAAACATCTGCTTTAGCCTTCCCAATAGGCTTAAGTGGTTTTAACTGTTCTTCCCAAGGATCAAACTTTA
CTCCAGAATCTTTGTAAATAGTAAAAATCAAAGAAGGAAAGCCTAGACCTGAAGTACTACTAGCTTGGGCAGAAA
TATTAATAGCCTCAAAGATAAGACAGCCAAAAGCAATAGGTTTGATTTCTTGGATGCACTAGTTCAAGATGGCTC
TTTCTCGTGTCACTTCACTCAAATGATTTGAAGGAAGGAAGCAAGCTGAAATAATGTCAGGCCTTTGAGGTGGCA
TTAAGCATGGAAGAAGAAAAAATTTACCTTATGATCATTTTGGATCTTGCAGGATCCACATCCATCAGTCAGATAA
TTCTACTTCAGATGATGTCTACTTAGAATAAAAACAATCTATAGCAGGTTGCAAACCATATACCACATTAATCCG
CTCTGAAGTGAATTCAATTCCTCTTCCTCGAACAAATATCTCATCATCAGACGCTTCAGGTAAGTTGGCACAAAC
CTTTCAAACTACAGCAATGACAACTATAGTGGTGAAATAACAAAAGCTCTCCTAGTGTTGAGCCACAATAACATT
TTTTATGGCACCACGGGGTTTGTAAAACTTTCACTCTGTAATGAAGGTCCTCAAAAATGCAAACTTGTTGTAGAA
AATTTTAGCTTTAAAATTTATGAATTTTGTATTGTCATAAGCTTTCTTCCCCTTTTCTCGCTTCATCTTAGGAGC
CATCTAAAAAAATTCAAGAAAAAATAAATTCCAACAAATTTTAAAATTAAGATAAATGGCTTTCTACAAGCTTTT
CAAGCAAACGAGGATTAAATCCATGGATTTTTCAACACCCAATCCAAAAAAATTTGCAATTTTTGAACTACACTA
AATGTATGGAAAATCCATAATGAAAATAACTATGTCAATCATAGCGCGTCTGGTCGCAAGGAAAGGTCTCCCCAT
ACTAATAGGAATTTTTTTATCTTCTTCCATGTCTAGCACAATGAAATCCACAAGAAAGATAAATTTATCCATCTT
AATGAGTATATTTTCTATAACCCCTTTAAGATAAGTGAGATCGATCAGCTAACTACAGGGTTACAGTAGTGGGAG

AAATCTCTCCAAAGTCTAACCTTCTAAAAATCGATAAAGGCATGAGATTAATTGCTAGCTCCTAAATCGCATAAT
GCTTTTCCATAAAATTATTCTTCAATATCACAGGGTATAGTAAAACTACCAGGGTTTTTAAGCTTTTGAGGATGC
TTATTTTGAATTATTGCACTACATTCTTCACTAGTGCAACAGTTTCATAAGCTTCAAACTTCCTCTTTTTGTTTA
ATATGTCTTTCATACATTTGGCATAACTCGGCATTTGTTTCAAAGCTTCTGCAAAAGAAATGTTGATTGTGCACT
TCTCAAAAATCTCGAAAAACTTGGAAAATTTTATATCAAGCTTTTGTTGTCTAAGTCTTTGAGGAAAAGAAAGTG
GTGGACTTTCTGAATCAATTTTAGTTGGAGAGACATGTTGTTCTTTATTCATAGAAATAATTTCTTTTGGTTTTT
CAATTTCTTCAGTTTCTCTCTCAATTTCAATCTCTTTTTGAATTCCAATTTTTAATTCATTTTCACTCCTAAGTT
GAATGACATTAATTTGTTCTTGTTTTTATTGTTCTCTTTCAGATTTATCTCAACATTTCTAGGAAGTGTTATCTA
GGGTCTATTGTTAAGTACGTTAGCAATATGTCCCAATTGATTCTCAATAATTCTTATTGCAACACCATGACTTTA
TAATTTATTCCCCATTTGTAAAACTACTTTTCTATAAACATCTTGTTTTCCTTAAATTGCTAGTTATTGAAGCCA
GGAGGATAAGAAGGTCGACAATTAGAACTACCTTAGTTGTTCCAATTTCCTATTGGGTTCTCTTGAGTATTCCCC
CATTGCTGATTTTCACCACCAGAATTATTATTATTCCAGTTAAAGTTGGGGTGATTTTTCCAACCTAGGTTATAA
AAGTTAGAATTGGGTTATTGGGTTTATTAGTAACTGTATGTAAAGTCTCTAATGATTGAGAATCATTCAATTGTC
CTAATAAATTTTCCTCAGTCACTTGAGGTGTTTGAGAAAAGTTTAGTGCAAAATTATCTAATTTCCTAGATATCA
CAGCAACTTGAGATGCTGAATAAATAAAGCATCATTATCATGAATACTAGCTACTCTCTTAAAATTTACCCTTTC
TAAAGGCCATTGAAAATTATTAGTAATTATCTCATCAATTAAAGCTTAAGCTTCATCAAATGGTTTTGACATTAA
AGTCCCTCGAGCAGCAGCATCAATTATAAATCTAAGAGATCCTATTAACCCATTATAAAATATTTGAATGATTAA
CCAGTCAGATAGAGCACAATGTGGACACTTTCTAAGGAGTTCTTTAAATCATTCTCGAGCTTCATAAATTTACTC
TCCTCCATACTAAGTAAAAGAAGTAATATCATTTCTTAGCTTAGCAGTTTTTTAAGGAGAAAAGTATTTAACCAG
GAATTTTTGTATCAAATTCCCCAAGTTCTAATAGATCCAGGTGCTAACGAATTTAACCACATTTTAGTTTTATCT
CTTCAGGAAAAAGGAAAATTATAACAGCATCATTTGATACACCATCATATTTGAATGTAGCACAGAGTTTTAGAA
AATTTGCAAGATGTAAATTTGGATCCTCAGATGGTGTTCCCCCAAACTGATTACTTTGCACCATTTGTATTGTTG
TCGGTTTAAGTTCAAATTTGTTAGTTGCTACAGTAGTAGTTTGAAAACTTAAGTGAATTTCATCTACTATAAGGC
TTGCATAATCCCTTAAAGGTCTTTGTATATTATGATTTCATTTTCAGTAATTAAAATCTTAGTTGTCATTTTGTG
ATGCGTTTGTCATTTCTTAAGAATTGTGAAGTTTTTTTCATTTTCGAATTCTTCTTATAGTTTGTTCTAATTCAG
AATCATAGGGCACTAAATCTTCAGATCCCAAAATTTGCATGCAAAAGATAAAATTCAAAAAACAAAATACTCAAA
ATATCAAAGTAAAAATTTTTATATTTCTCGGTTTTTGCAGATTTTGTTGTGTTTTTGTAATTTTTGAAAATTGAA
AAAACTCTAAAATTATTTTATAGTTATTCTACAATTCAAATAAGGTTTCTAAGTCAAAAAGTTTTGAAATAAAAT
TTTTATTTAAGTTCTCCAGCAATTGCTCCAAAAACTTGACTCAACCCACAATTACGCAGATATTGATGTCCCGAA
TATGGCCAAAATCTTGACCCAATTTATATTTAGTGGCTAGTGAAAACAAATGTAATATTATAAAATTTAGATTAA
TAGTTTTCTCACTAGGGGGTACTACGCAAGTATATGTATATTTTCAAGTAACATAGAGTTTCTTTAAAAACCCCA
GTATTGATCATACAGGGAAATTAAAGGAAAGAAGTAAATTTAGGAATCTTAACAATGACCCTTAAGACTGTAAAA
AAAAATTTTAAGTTGTTTATGTTATGAAATAAAGTTTACTAATTCCTACAAATCTAAGTTAAAAGTAAATGCAAA
GCTTTAGAAAATTCAAGTTATATATATAAAGAGACTTTCTCCTTGCAAATAATTCATAATCAGTGTATGATTTCA
GATGTTTAGAGGTTTAATTGTGGTATTGTGACAAATAGAAGCATACTTATTATTTACCTCTTCTGAGCATAAAGA
ACATTGGTTAATATAAAGTTGGTGTTGGCATGTAGTAGTAATGAGTACGTTCTAGTCTCTCATTCCTAGATTACC
CACTGGCCTCCTATAGGACCTACCAGTGACCCTTAGTTCCAATCTCTAGATCGAATTGATAAGAAAATATAGCCT
ATCAGCCCAGAAACATCAACTCTACACCTATCAAATCTTATCTCTCTATTTATTTGCACTCCCCCTCGGACGATT
GCCACAGCATTTTTCCAAGTGAAATGTGACACTCACTAGTAAGAGAGATCTTTGCCCTCTCGGGTCTAAGTTTCT
CAACAATTTTTAACAATACAAGTGGTCAAATTATAAAGTAAATGAAGGACCGGTGAAAATATCATCACAAATTTT
TAAAATAAAAGATGTTCTAATCAAGTCAAAACCATCACTTGATTACTTCATATTGCAAGGAGAAAGAAGAAAACT
ATTCAGAACACATGAGTATAACAAAACAAAAGATATTCCAAAGTACAATACCTAAAGACAAGATATAAATACTTA
ACTGAAATGTTTCCTCAAGCACTCCTTTGAAGTTTCCTAACTTCGATAACTCCAATAGACTCTCACAAAACTCCA
GAAATCACAAAGACTCACAAAACCTTACTAAGGTGGTCACCCACTCTATATGGATGTCTTCCCTTCTCAAAATGA
CACTAGAACCCCCATATATATATATAGCTTTGAAAATCACCAAGAGTTTCAAACATCTCTTCAATATCCGGCTTC
ATTCAATGAATCTTTTAGAAACCGACTCTGGTGTCTAGGCGCCAAAACTATCATGCCTAGGCTCAAGCCGCGATC
TTCATATGGCACCTAGGTGCCAGACTCTGCCACCTAGGCGCCATTTCGGGTGATTTTGACATTTTTTTCTTGGTT
TGGTCTATGTTGATCTTTTTTTATGTATTTTTGCTCATTTTTGCTTTTAAACTTTTTTTCTTGCACCCGTAACCA
AAAACACACAAAAAACCTTTCATTAAAGCATAATATTCTCAAAAATGGGTAGAAAATGTGCATAAAAAATAAGTA
ATTTTAAAGTGGATCACTAGGTAGCATTATTCATTCTAGGAAGAGTTCTGCATAAGGTGTTCTTTGTAGGCTAGC
CACCCCCATATGATACATGGATATTAGACATGAGCATATGGGGACAGATGCTCATGTGAGAACATCAAATGTGTG
AATTCAATTGCATAACTTAATGTGTGCAATGACACACAATGATATATATGTACATGCAAATGCTACAATCTGAAA
TAACTGTATGGAAAAAAATACTTATGTAGGGATACCAAATATGTAAGTGCATATGCACAATGTGTCACACACATA
TGATCCATAATATGATGCTACCATTGCAAGCAATGTGAAAATATATATTTTTTACATATGCATGATGCTAAATCA
CAAGCATAACATAAAATGTAATTTTCACATATGCACCATGCTAAATGAGAAGCGGAACATAAAATACTTTTTTCA
CGTATACACCATGTTAAATGAACAGAATAATACAAAATTCATTTTCACATATGTTATCATAAATGCAATACAAAA
TTGATTTTTTACATATGTCATCACGAGCACACTACAAAATTCATTTTTAAGTAGTATTTAAAAAATCACACATAC
AGATGATGATACATACATCAGTTTGATATGCAATCCTAAATATACTTAATGGAAAATCATCCTACCTAAGTGACC
TCTCATGGGTTTGAGTTTCAGTGATGGTAATGTGTACCAACACATCATTATGCAAGATAATCATGTGCTCAACTC
CATTTGGGTGATTTAAGAATTAGAGGTTTTCATGAAGAAAACTCTCGTGACCTAAAGAACAGAAGATCAAAGGAG
TTCTTAGATTCCTTGAGCTCAAAGGCTCAATCATACCATCAAAAAGATGGAGCCATTACAATAAATGCAAGATTA
CTTCAATCTTGAATATATAGGTCAAAGGAGAGGGGTTTTCGGAAAAACCCTAAAGGATCTTTGGGAAAAGTAAGA
GAACGATTTCTATCTTATAGGGTTTTTGATAACAACAACTGAACCTATAACTCTGAAATCATCCTAGAAAAGGCT

TTTCTAAGAAAAATAAAAATAAAGATGCAAGAAGATCTCATTATCATAGTGTTTGTCAAAGCGTTCTCTTTACTT
GTTTGACCTTCCTTATGTAAGTAAGATGATGCCTCTCTCATCCAAAGAAACTCCAAAAACTTCTCTCTTTTATCT
CAATCTCAGAAAGAGATGAAGTGGAATAAGAAGATGAGAAAGAAGAATAAAAAGAAAATCAAGGGGGAAGAAGAA
GGCGGAAGAAGATGACACTTCTTTTCTCTCCAAGTCTCCCTAGAATCCCACTTTTCAACTCCCAAGTGATTTAAA
AAGCGATTAAAATCAAAGTATAATGAAGCCAAAATGTTTTTTGCAAGTTCAGCCCTTCTTTGACCATGAAAATCC
TCTAAGTGGAAATATTGCAATAGGTTTTAATCTGCATGACATGATTGCAATTAGATACCTTTTGAGGTGAGGGGA
CTACTGAGCTTTCTATGGTATCTTTCATGATCAATTTTGGCCTCATGTGGCTTGATTAGCCAAAAAGTATAATAA
AAAATAGCCAAAATTGAAGCAACTCAGCGTAATCATGCCAAAACTAATGTAATTATGTGTGTTGCTCTGTTTGGC
ATAGTTATGCCAGTTTTGGGTCTAGTTACACCCAGACAGGCACACCAGTTCTGGGCGTAGCTACACCAATTTAAA
CTCTAGTTTCATGCCCAAAATGCAAGTCGAAGGGATTTTATGGATGTCTTAGACTATGTGAAGCTATAAATGAGT
ATAATATAGTCCCATAACATCTAGTAGACATCAATCATGTGTAAAATTAAGAATTGATGCTAATGAATATGTACT
CTCTAACACGAGTGTGTCGGTTCGGACTCTTATTTCATGCCCTGAACGTGCACCTGAGAATTGTTATGGATGTTT
TGAACTATGAGAAGTTATGTATGAATGAAATTAAGAATTGATGGATGTTTTAAACTATGCAAAGCTATGTATAAA
TGAGTATAATATAGTCCCATAACATCTATTTAGACATCAATCATGTGTAAAATTAAGAATTGATGCTAACGAATA
CGTACTCTCTAACACAAGTGTGTCGGTTCAGACTCTTATTTCATGCTCTGAACGTGCACCTGACAGTTGTTATGG
ATGTTTTGAACTATGTGAAGCTATGTATGAATGAAATATACTCTCATGTCATTCAAATGGACATCCCACTTGTGC
AACATAAAAAGTTGATGCCAGCATTGGGAAAAAGGAGAGGGGAAATGAGACGATTAAGAAGAAGAATAGAAGTAG
CATGAAAGTGGTGTTCTAATTAGGATAATGGAGAGAAGAGAGGAAGAAGACAGTAATGATGGTTTGAGTAGGGGA
AAAGGGTGGGGATATGATTGTGAAAGTGGTGTAGGGATTTAGCTAAGAGAAAAGTGAGAGGAAGAATTAAAGCTA
AAAGAAGAGAAGGAGAAGTAGAAGGGAAGAAGAACAAGAAAGGAAGAAGAGAGTAAGTGAAAGTGTAAGAGAAGA
TGAATGAGGTGAAGAGGAAAAAGTGTAGAAAGGAGAAGTGGTGAAGAAAAAGAAGTGTGGAGAAGTAAGTAGTGT
GGAAGAAGGAGAAGTGCGAGGAAGCATGTAGGAGATGAGAAGAAAGTAAAAAAAGTAGGAGAAAAGATCGAAGAA
GAAGCATGCAGGAAGAAGTGGAAGAGGAATTAAGTAGATAGAAGAGAGAATAAAGGAGAAGAAGTAAAGGACCTA
CAATAAAAGGCCAAAGAACATCGGAGGTGGAGTAGACTAAATGAACAAGATCACCAGAGTGAAGAAATCACAAGG
GATGGAGGAGAGGCTGAGTTATTTTGTATAAACATGAGGGGTCTGATATAGAGAATATAGGCAGTGGTGGCTGAA
GGAGGGAGCATTTGGGGCTTTGAAATTGAGTGAGAGAGGTTGGGATCGGCGATGGCGTGGTCAATCCACCCCAGT
GACATAATTATGTTACTCTCTTGGGTTGGATTTTGTGGTTATGCCACCTTGAGGTGTAACCATGCCACAAAAGGG
GTCCTAACATAATGTGGTTATGCCCAGGGGTAGCATAGCTATGCCCAAGGGTGGTGTGGTTACACCATCAAGGGC
CAGTGTTTTAAGAATTTTTTTCTTGCTTCAAGCTTGGTAATCTCATCATCATGGAATGTCATTCTACGGAAGAAT
AAGTAAAGTTTTGGTCAACAATTGGTTGACCAAATGAGGTCAACACAATTCAAGGCTAATTACATGAAACGTTTT
CAAATCATTTTGCCTAACATTAATCTGCAATATCATTAATTTATGTTGGCCTATTCTAGTCAAAATTTTAGAGAA
TTAAAATAAAAAATTATAATCCCCAATTATCAATAATTTCTATTTTCTATATTTTTAGTCATTTATTTCCTATGA
AAAAAAATATCAGAAGTAATACCAAAAAATCTTATTTAAAAAATTAAGGTATTAATGACCTTTTTCAAGTAAAAT
CTCTAAAGTTCATTCAACCAATTGAATGCTCTAATCAAAATCCTAAACATGGTCTCTTATAATTAAAACTGGCAT
TTATCCCTTCTAAGTTAAGCCCACTATCAAACCTAATACCAACTTAATCAAACTTTCTAGATCTATATGAGACTC
AATTTCATATAAATAATGCCTAAGATTATCAAGCTTTGACCTCTATAAACAAAGCTCTTAATAAACAAGCTAAAA
TTAAAAATTCTAGATAAAAATTCAGAATCATTCTTATAAATCATCACTTATTTATATACTTAGCCACACAAATGA
ATTATATTTACCTTCTACGCATCTTAGGAGGCCCCCTCATCCACCATATTGGTTACTTAAATGGTCTAAGACAAA
TTCTATAAATTTTTTTTTTTGTTAGAGACAAGTTGTAGTAGACATGGGAAGGTTGAAGCTAGTTTGAAACTACAA
AGCTTGGAAGTTTGGATAAGAAATTTGTAAGTGATATAAATTGAATACTAGCAAGGTTAGTCCCTATATTGTGCA
TTAGGGTAATTTACTTATTCAATTTAAGTATTTTCATTGGCATAACTCATGCTAGAATTCATGTGGTAAGTGATA
TTATGATTGTTTGCTATTGTTACCAAATAATTAACTCCATATGATAAAGTGAACATCACCCCACTAAAATCACTA
TGAAACTATGTCATTTGAGTGAAAATTTTGGCAAGGGAGAGTTCCCTTATCAAGCCTAACATTAATTATGATCCA
TACATAAAAATGTAAATTATATTTTTTTTGGATTTTACACTAGTCTTTCGAAAAACTTACTTGTTGCTTTGTGCA
ACAACTTGTTAGTTTATATTTGTTTAAAGGATAAATTTATTTGAAAATTTTCACATCACAAATCAATTCAATTTC
AAAAAATTAACCTTCATATTTTAAAAAAAATTGTTGGAGAATTGAATCATTTTACATTGCATATCAAACATATCA
AACATGCCACTAAAATCAGCTAGCTTTGATAAAAAAAAATTCTAAAAGGGTAGCGAAAATTAATCTACAACGTAA
GGAAAAATTACTAGATTAGAAAAGCTCATATGGATTTAGCTACCAAACTCCAAAGCATGAATACTAGGATTATGA
ATACACCTACAAATTGAAACCACGTGTGAAGGCTTTGTAGAACATCTTAATGATGATTATCTTTTTAGGAGACAT
GTGAGGAACCCCCACATAAATGAAAACTATAGGTCTACTAAACATGAACCTACTGGAAAAGTTACAATATAAGCC
CCAATATATGATGATATCTATGTCCACAATAATTTTTTTATGTAGCCAACAAGTTTTTAGATTGGCTTACCCATA
TGTTTGAATTTTTTAATAATATTCTATATCAACTATCAAATACTACTTTTTCTAAGATGAAATTTATAGGCCATG
CAAAGATTTATTGAAATAAAATTTTAACTGAGAATGATTTGGTAGAAACATTATGACATGGGTAGTGATGAAAGA
GATTTTGTGATCCAACCATATAAGTTTTTGAAGACATGAATACAATTGAGAATTTAATAGCTAGAAAACTAAGAT
CAGAAATGATAGTTAGCAAAAATATTTGTAAATGATCTTAGAACTAAAATTAGTAATGAAATTGAGCATCATGAT
TCCCAAACATTAGAAGATGACTTTTAATATGCTTTACTCTTAGAAAAGCACAAAAAAACAAGTGCCCACATCTAA
AACAAGAGGATAAACTTCATAGTAAAACATGCATTTCAGAATTAAAAACCTTTGAGTAGACCGAAGCAATCTACA
CCTAATACTTGAAGTCAAAAGACTCATAATAAATCAACAACTACAAATGCTAGGAACATAGTCTATTTCAAATGA
CATAAGAAGGAAACATGTAAAAATAAAATAAGGTTAAGATTATGTTTCTAGTCAAAGAAGAAGAGTTAAGATGGT
GAAATTGAAGAAATGATGATGATACATCGCTTTCTGCTGAAGAGAGAGAACATGTAGAAGATACTGAGCGTCATT
AAATGCATGACTGCATCCAGGAAAGAAGAGGATTGGATGAGGATAAATACTTTCACCATAATCATTTTTGTAGAG
AGAAGAAATAAAAGCTACTAATTGATGAAAGTAGTTCCATGAATATTATATTTGTTAATGTCATTACTAAAGGAA
ACCTCAAGGCTAAACCTCACCATAAATCATTTCACATAATTTGGGTGAACTATGTATCTTTGTTTATTACATCTC

ATTATCATGTTCCTACCAAGATGCAGTTTTGTTGACAATCCTTAGTGTGATATCCTCTCTATTGAGATTGTATAT
ATCTTGTTATGAAGGTGTTGCTTTATAAGATTTAACTATAATTCATCATGATTGGGAAAATACTTTTGTATTCAA
ATATAATAATAAGTCTAAATTTTTTTCCTATAAACCCAATAGAAAAGAGGCCAATCACACCTCATAAGATTTCTT
GTAACACAGCCTCACAATTTGAATTTTGAAATTCAACTAAAAACCTCGTTTGAATAAAAATTCCAACTAACATAA
AGATTACGATTGAATAATTATGGTTTAGAAAATTATCCCAACATTTTTTGTATCCAACTTCTTTATTTGTTCTAC
AGCTTAGAGCAGATACATGCAACTATAATGGGCACCACATATGTATCATTTTCTCAAAATGTCTTATTAAGTTTA
ATGATCTTATCCTTAATGATTTTGTGGATGAGTTACCGCCATAAAGTAATAACTATATGTATAAGGGTCTGTTTG
GGGCAGCTGTAGCTTATCCAGCTTATCCAGATAAGCAGCTTATCCTGATAATCTGCTGATCAATAACTTATCCTA
ATAAGCTGGCTAGTTGTTTAGTTTGAGTGTTTCGGAAAAAGCTGATTCTGGATAAATGATCATTCTGCCCTTTTT
CTAAGCACAATAAAAAATCAAAACACTTAATAACTTGCGGACGAGGTCCAAATCTTCATCTCCGCAAAATCATCA
AGTCCTTCTAATCTAAGCACAATAAAAAATTCTTACTATTCTTTTATATCCTCGCCAACTCAGAAACAAAAATAA
ATGAATAAATCTTTTCACTAAAACCCAAAGAAAACAGAATTTGACAATGAGAAAACAGAATTCATCAATGAATGA
GAATTTGATTCTACTAAACCAAAATTTGATTTTACTATGCAGTAACAACGATTAATCAATGCCCCTGATTCTCTA
AATCCCCTAATTTCGTAGATGCAGAATGTATTCTCCATACAAATAATGTATAATAAAACAGTGAAATAAATGATC
GATGGACTAATAAAAATCATCAAAACTAGTTCCAATGAGAAGACTGATGAAGAATCGCAAGGGTTTCACTTCCAG
TCTTTAATTGGGAAATCAGGAAGAATAGGTACAGCTTCGTACCTGAAATAAGGGCAGAGAAGAGGCGATATCCGA
TTAGGAAGAAGCCCACCATTTTTCTCACTCTACTTTTTAATCCCCTTCTTTGATTTTCCTGCCTGCAGACCTCAA
GCGGGACTCTCGATTTTTCGGCTCCGGGTACTCTTCCTTCGGCCTGAGGAAATCGTCGTCTGTGCTCCGCCCGTT
CTCCGAGCGTGCGAGAGGAGGAGGATAAGAGATGGCGAGGGGGAGAAATCCCTTCTCTGATTTATAAACTTGAAA
GGAAACTATATGATTTTTCCATACTTTCAACTTTTCTATTTAGAGAACAACTGCGAAAAAATAGCATTTGTAGTT
CTTTTCTCATGATAAATGGTTATTTTCACCAAGGTAAGAGACTATGTATCCCTATAACCTCCATAAGAGTTTTTT
ATCATAGAATTTCAATCTGCAGGAATTACTTGATATTTTGAAAATTTTAAATCCTATAAATTGGTAGAATAATGA
ATTTATTAGCCATCTATACATAGGGATGTTGAGAGAATTATAACCCAATGTTCTACTTGTCTTCAAGCAAAAGCT
AGAAAGAAAAAAATAGGTCGTTATACACCTTTGCCTAGTCCTTCTAGGCCTTCTAGGCCTTACAAAGATACATCT
ATGAAATTTCTTCTAGGTTTACCTAGGATACTTTGAGGTCATGAAATTATGTTTGTGGTTATTGATAGATTATCA
AAGATGCCACATTTCCTAACTTACTTCAAACTAAATGATGCATCTCACAATGCTAAAATCTTCTTTAGAAAAATT
GTGAGATTACATGAGATTCCATCTACCATAATTTCTGATAAGGATGTTAAATTTATGTAAATTTTTTTAAACATC
TTATAGGAACTAATAAGGACACAATTGAAATTCTTCTCAGCATTTCATCCTTAAAATAATGGCCAAACTAAAGTT
ACCAATATAACACCAAAAAACTTATGAGGAGTTACACCTCCTCACAGTTGAGTTCTTCTATAGTAGTACCATTTC
TAGGTAATTAGGATTTACGCCTCCTCGCGCTTGAGTTCTTCTATAATAGTATCATTTCTAGAACTACCAATAGAA
GATAATTTGAGTAATATATGAATATAATCCTATGAATTCAATTAATCTAATTAACGCACCTACTCATGTATTTTT
AATTTAAAATTTACATTTACATGTTATTTTTATAGCCTTCATGAAGATATGAAAAATAATTAGAAGTACAAAATG
CTAAATATGTTAATAAAGAAACAAAGTAAAATTTTTAAGTAATATGAGCTAGGAGATTTTGTATTCATACACATT
CACTAAGAGAGATTCCCTTCCACTAAATATAATAAGTTCTAAAAGCATGTGGGCCATTTAAAATTATTATGAAAG
TCTCTACTAATTCTTATATCCTTTAATTGTACCCACGCACATTTATTAGTCCTCCTTTAAATTTCTTTTAAACGA
GATGCCCAATCTATAAATTTATAGGTCAACACTACTGTAAGAGGAGATATACAAAATTTGCATGTATCCATCTAA
TTGAATTGGAAAATATAGTTATAGCTAAGAGACCTTTAGATGGAGATCCTACTTAAAGGTTCTTTATTAGATGGA
AGGATAAGCCATCAAGAGAGGATGTTTGGCTAACCTGAGAGAAGCAAACTAGTTAGCACCATAGCAGAATCACCT
CTAACTCTTCAGAGAAAAGTTCTTTGGTGTTCAGTGGAATTTGTAATGCATATCATGGTAGATTCAAGTTGGGGC
TCATTTACGCAGAAGGTGAACCCCAGATAGGGCCTAGAAGCAAAAGTGTGGGATCTACTATAAGAGCCTAACCAA
TGAAAACTACTTCACATGCACGAGGTCCAATTATTGGAAGTCAAACTATATGAAAAGCATGAAGATTAACTTAAA
GATGTTTTTCTATTTTATTATTAGTATTTGTATTTATTCCATATTGTAATGTGTATGTTTAAGTATTTGTTTACG
GTATTGCGACTTTATTTCATAGAAGTGTTTGCTATAAATCAACATTCATTCATAGGTGTCATATTCCATGCTTGG
GTGGATTTCAAAGTAAAGCAATGAAGAGTGGTATTATGTTGGACAAGTCAAAGAGATGCTCTAACCACTCTAATG
TCTCCTTCCTTTTAGTATTTGATATTTGAAATGAAGAAGTGTATCTTTCACGAAAGGAATAATTTGTGCAAGACA
CTTAATTTCTGAAGTAAAGGGAGACTTAATTTTCTATGTGAAATTAGAATTATTCTAATATGATTTAAGGAGGCT
TGGTGGCCAGCCTTAATAAGGGTTTAGGTGAACCATTTAACATTCATATTATCTGCATCAACCCTGATATATGAT
TCTATCGGACTTAAATAAGGGATCTAAGGAAGAACAAGTGCTAAGAATTATTTGTAACAAGTAAAACTTTTGTAT
TTTTGTATTTAATACAATCAATATATTGTGGGGATAAATACTAACCAAAAGATATGAGCCAATTCAAAAAGCACT
TGATCATGATCTAACCTCGTATAAACTTAAACTTTTGAGATCCATATTAGACTTAATTTTCTATATATTAAGCCT
AAGTTTATCAAATTTAGACTTCATTAGATAAAGCTCTTACGAAAATAAGAGAGAGTTGTAGGTTATGGACATAAA
TCCAAAATCCTTTTTATAAATCACCATTACACATATATACTTAGCACAACTAATTAGTTAGCTTTAATATACCTC
TTACACTTTCTTAAGCAACTTAAGAAGACCACCTCATCCACCAATTGGTTCACTTTAATGATCTAATCTAACTCT
GCATCAATTTCAAAGAAGTATCTCTAAGTAGGCATACCTTTCTTCTCTCTTAAAATGAAAAGAACTTGCAAGATA
GAATTCAAAGAGATCTTGGATTCACAAAAATAAGTAACATTAGCCAAGGGAATAAATTTTGTATAGAGTTCATTA
TGATGTTCAAGAATCAAATTGCAACTATTAACCCAAAAAAAAATGGATTAACCACTAGAAAAATAACTTTGTATA
TTGTTCAGTTTACAAATTAGGTCAGTAGATATTTATTAATATTTAATAGATAACCTTAAAGGGTTAGTAAAGGCA
TACTAAACTTATAGTAAGAGTAATAATATTAAAAAGGAAAGGTCAAACTTTCTAAATATTCAAGCAACACTAAGT
TTTATGAATGCGGATTAAGCTAAATCAGGTCTATAACAATGTCATATAAATTGACATTACAAGTACTTTATTTTG
TTTAGGAAACTATATTTGACTAATTTTGAAAAAAAAATTAAATTTCATAAATGGATAATCTATATTTTTATCATA
CATTTAATTAATATATAATTATTGAATGGTTAAAAAACAAATCCATACATACAAGTGAATCAAAATTGGGCAATG
ACAATACAATCAACTGTCTCAAAATTTCTCAAGTAAATTTACAATTTACTTCACATTTATTGCTAACATCCATAA
TGATTCATAAAATACTAAACATATAAAAGCATAAATTTAACTAATCTTTAAAAATGAAAGAATGAAGTGCTCCCC

CATCAAATAAAATTTGTACATCACCCTTACTTATATTGAGTATAGAACTTTGGATGACCTTATATTTTTTGACAT
AACCATTTGAATAACTTAGTTCAACATGTCCACATCCTTAGTCACCTCACTAATACTCTTACCAGATATCTACTT
AGAAGTCTACTTAGCGGAAATTAAATAATGGCAAACCTTTGTCTCAATAATCGCATATGAGGTAAAAAAAGGATT
GATTTTTCATCTCTTATCTTCATTGGGCTTGGGACTCTAAACTAATAATAAAACTAAATTTCCTCAAAATCTATG
TCTCCATTATAAAATTTTATTAATTCATCATATTTTATGGTTTATACTTATAGGCCAAATTTATTTTAACAATAC
CAAAAGTAATCACTTATAACACAAGCCTTATCGTTTCGGTCTACTTTTGCATTGACAAAAAATTTCTTAACTACT
TGGACAATGTTTGCATCAGGTTCACTGGTGAAATACTCTCATCCATATTTTACAATCATACTTTTAAACCTTTTC
CAATGACTTCTAAACCCTCACTCAGGGATCAAGTTCTCATGGAATATGAACTGCTCAAATCTTCCCTTAGCATCT
ACACTAATAAATTTTCTAAAATCACAAAATGTAAATATCCTCTTCTTCCATATAATCGTCTTTGCCAATGACATG
TTTGCTAAAAGAAGAAAAAAGTAAGATGCTAAACTAGATGTCTAAAGTTTTGCTATTCTGAGGATTCTCTCTCAA
TCTTATGCCCAACTTAAATGAAATGAATATGAACCACATGGTTATTTATAATCTAGCATAGCATTGTGAGTCTTT
AAGGAGAGCACTGCGGTGCTACTCCCTTAAGCATTATAGAGCTGCAATGCCAAATCTTAGTGGTTTGGTGCTCTG
ACCCGATAATCCATAGCTCTAAAAAGGAGTCAGCAATAATTCAATACATTATAAATTCTCATTAAGCAATTATTC
AATTTCACAGAAACCCTTATCAACTCATTATGAATTTTGTTATAGCTTTGTGGCACTATGCCTAAGCATTGTGGA
TCTATTGCTTGGGTTTCCAAAATTTACATTAGTTAACCATCAAACCATTGACTATACTAGCATTGTGACACTAAT
CAAAAGCATCAATGTGATGTTATAACAAAAAAGTCAAAAGTCAATAGCAATATTACAAAATTAACCAATCAACAA
GATACAAGATCTCAAACTGCTTTTTGGTGCTACACAACCACACTGCAATGCTACACAACATTTCAAGGTAGTTGA
AATACTTACCACTTCTGATACTACCATAATTCAGTGCTACATCATTAATTAATCAGCACTTAGGAATTGTACCTG
CCACACAAAATTTAGTTAATTTTTGAAAAATAACAATAATACGCAAAGAATATGATACCACAAAAATCTTGAATT
GCATCCCAAGAATCTATTAGATTATAGCCCTTAGCCTAATTGTCCTATGTTATCTAACAAATGTGAACCACATTA
ATTGAAATCCATTAACAATCTATAACCCATCTTAAATGCGGTGTTAATCTCTATTTATTAACTTTAAATTTTGGG
CATTTTTTCTATTCTCAATATTAATATGTTATAGTAGCAGACTTTCACTACAATGAAAAGACCTGATCACCTGGA
CTTCAACTTTATTGGTAATATAATATAGAATTAATTAGCAAAAATTCTTATTCATCTTCAAAATGCTTCTTAATA
AATTTTCAATGTCATATCATTTTGTAGTTTGCTCTTTATAAATCTTAACATTCTTATAAGCCTCCAGTCTAATTT
TTTTTTAACACATTCAAATATAGTAATATTTTCTCTCCAAACTATTTTCAATTATAATTCAAATTCTTAATTACC
TAAAAAACCTTATATTCTAATTTTATGCTTAACTGAGGCTTTTTCAAATATCATTTATAAGGGGAGATTCCTATT
AGAGTTTTATAAGTTGTTACTATAAGTTCATAAAGCGTCGACCAATCTCGATGAATAATCTTTTCTAAAAAGATC
GACTATTTTCTCTAATATCTACTTTGTTTCCCCATTCGATATCTCAATTTATCCACTAATTTAAGGGTTAGTAGC
TATTATGTATTTAATTACCATTTTAGCTACTTAAGTCTCAAATTTTTCTTTGACAAAAATGACTTCCTCCATCAT
TGATTATAACCCTAGGTGTTCCAAATCTTACAAAAATATTCTACTTCAAAATTTTAATTACTACCTTAGAGTCAC
CGCTAAATACAACCAATCACATTGGGACATAAAATTTTCCTCCTATTGTCACTAGATGATTGAATTGAACTCAGA
TAAGCTATAACTGGAACCCACTTGAACTAAACTCGAGCCAACCCTAATTTAATCTAAAACTAACATCTAAATGAA
TAGAAATAAAAATTAAATGAACTAAAAATGGAATTTTAAAAATTTAAATTTTCATGTTATAATAAAATTGGGCAT
TTTCTCTCTTTTAGCTATACAATTTATCATTTGACATTAAGCATGCCTCTTAGCAGATTCATTAATAGCCTTAAA
TAGGGTATGCCAATCAAAAATTTGATAGTGAAATCTTACGACTCATTTTATAACCTATAAAATGGCCATCATATT
TAGATTCATGACATTTATTGACTATACTTCAAACTTATTCTAAGGATATACATCTTCGAATGATATAATCCATAT
ATTTATTAAGCAAATAAGGTTTATCCTAAAAATGAATTTAATATCATGGAAAATTTTGTTATTACAAGTCAATTC
ATAAGTTGTAATGTTACTAATGAGGAAATGATTGTATCTGCATACAAAGACACATTCAAAGCAAACATACTCAAC
AAGAAAAAATATCAAATCATTAACTTATTTTCAGCAATGTTCCATTTAGTCAAGAAATGTGATCAACCATTTTTT
TATAGCTTTTATCACAGATTCTTAAGTCAAACTCTTATTATAATAAAATTCATCTAATAAATTTAGGTTTCATTA
TCATTTTAGGTGCAAGATATTTAATTATAATATAATTTGTATACACCATAACCTTAATACCAATCAATGTAATTA
AAATTTGTCAATAATAAAGACTGATTAATAATACTTTTTTAGAGCCTGCTTGGGGCAACTGCAACTTATTAAAAA
GTTATTGCGTAGTAGTTGTGTCAAGAAGTTATTGCAAGTAAACTGCATGTCAATTTGTTTGAACAATTGTACAAC
AGCTGTGCTGCAGCTGTTTATGATAAAAAGACTTTTTTACCCCTGTATTATTATGTTATTTATAGCAGAATCAGT
CTCGGTTGCACATTTCATAAATTAAAACGGATATTCAATTTCAATATTATATAATAAAACTAACATATCACAAAA
CAGTGAGTAAAAGAACAATCAAAATAAAAATGAAATCATAAAATAATAAATAATATTATCATAATGTTCAATTTA
AAATGTCGATGATTAGTTTCACCAGATAATCTCTGCTCCACGAAACTCTTCATGTTTGTACACTATCGGAAGTGG
AATCATCGAACTGATCCACCCACCTGAAAAAGTCACAGTTCTCTTATATATTCCTTGGACCCCTATAAAACTACC
TCCTATAGTTATACTCATGTCGCAAAGTAAGCAGAGAACACTACAGTCAATTGTGGTTGTGCAATATTTTAGAGA
CGTCGGATGGCCTCGACCTCATCTTTTCTAAAGGTTTAAAACCATAAAGCACAGACCCTTAGATGCAACACTAGT
CAAAAAAGAATAATAGAAAATATAAAAATAAAGTTTTATAAATTTGTGAAGTATAAACCAACGAATAATCAATCA
ATGCAAGAAGACGGTATACTTTAAAAAAGTTTCTTGGTTGTATTTAACTTATGTAAAACCGAGCTCATTTGTGAA
AATAGGGAATCTTAGTTGTATTTATACGTTGTAAAACCGAGCTACATATGTGTCATAAAGTCATATCATCACAAA
GGCCATCCAAAAAAAATCCCAACAATCATATGACTGAAATAGAAGAGGATCTTGGTTGCCTTTAAACATAATATCG
AGCTCGTATGTGTAACAAAGTCATATCACCACAAAAGACATCCAGAAGAACCTCTAAGGTCATTTGCGTAATCAT
AAAACTCAAAACCTAAAAAAATGCAATATATATANNNNNNNNNNNNNNNNTATATATATATATTAACTCACTATTG
CCTAAGGTCATTCATGTGGGAAAAAAATACATGCAGAAAGAATGAAAAAGTAATTTGCACCGAATCAAATTTCCAC
ATGAACTTCATTGCATATAACATCATTACAATAAGTAATAGTTGTCATTGCAACACATTACATAGATAAAGAAAA
TGTAGCCACAAAATCCTTTCACCAATAAACTGTTGACATGCAATTTGCTCTCAACTCCATAATCTTTCTAGAACC
ATCACCCATCTTGTGTACAAGACTCATGTAATCAGATTCAGTTTTTGGTGTTGAATGACCTAGGGGATATCAGAA
TGAATCGACAGGGACATGTGATTGGTCCTCATGCATTATTGAGCTCAATGTCATTACATGCGGGATGTCGACGAA
AATAGTTGTGGAGAGTCATCATAGTGACCACAGTTTTCACTTATTTCTTAAACGGATAGGTGGACATATCATGAA
AAATATATCATTTTTTTTCCAAACACAAAATGTCCTTTCAAGAACACCATGAAAAGATGAGTTCCACTTATTATA

AAATCTCCTTTCTACCCGAGCTAAATTGTTCGCTTTTTTAAATCAAAAAAATGATATTTTGTATCAGAAAATGGC
TTGACATATCCTATTTCTAATGGATATCCAGCATCAGTCCAATAATATTTCCCTGTTATAAGTTTGTTCATTCAA
ATGTATTGCAACAAATATGTTACTTATTCATATGTTAGCAAATATGGAGGCGGCTTCGAAAAGTTGTATTTGAGT
TCACGTGGCATATTCAAATATACCACTATCATGGGTGCTTCCTTCTCATCTAACGATCACAAAAGTTAAGAACAC
ATTGAAATCACATACATTCATAATGTTTTGTGTTTGTGACCCACACGTCCAATATATGCAATCTTTTCACTGTTG
AACACACGAGCATCCATGTGTGTTTCATCAATTTCCTCAATACAATCCTCGATCAATTACGTAAGGTATTAAGGT
CAACCACGCTCAAGTAAATTTTGAATAGATAAACACAAAAATCTACCTTGAAATATGACATTTAGTGTTGATCAT
GTCAAAGCTCTACAGAGGTGTTTTGTAATGTAGCGTTTTCAGGTCTTATATTTTTTTTAAGGTAAGTTAGTGAGA
GCTTCCAAACCATCCAAAAAAATATCTGCTAATTTTTTCTGATAAGTGTTGGAATCGCTTCATGGCTCCTCACATA
GACTCATTTTGTAAGAAAAAAATAGAGGGTTATAGCTAACACCTCTTTTGAAGTCATCCAAGATGAGCCATGGAGT
CTATAATATCTGTGTAGTTCTTGTAATAAATCAACAAATATTGGTTGTCATATAAAACATATTGATAAATCTCGT
AGGCTAACCCAGAAGGAACTCTTCAACCCACTTGTTGCCTATGTATGTGGATGTCATGCAATGAGAGCAAGCAAT
ATATGAGTTGTAAAACTCAAAAATTAAGCATATTACAGTTGCAATTTTTCTTTTCCAACTTTCTCTATCCAAATT
ATTCATTTTTTAAGCATCTAACTCATATGTTGGTTTTCTTTTTATTTTTAAATGAAAATAATTTAATTTAAACTA
TATATTTTTACTCTAAAAATGTGAATTTTTTATTTTGATAAAAGAGACCGAACATATCTCATATATTCAATGGTT
TAAATAATCGAGAACGCTATTCCGAGTAATGCTTCATCTCAAATAGATAAGAATATCACTCGATAATAGTTTCAA
TTATTATAAATACATAGGAATTGAATGGTATAAAAAAAGAATTGAAAGAAATAAGTAATATTAACTATAGCCAAA
ATCCAATTGATGCATGTGAATTCAGACTGGTATGTATGTAAATCTTCCCAAAAAGAATTGAAAGAAATACTAACT
TGATCTCTATTTCAGCATGAAAACAGTGATTAACCGCTACAACAAAGCAAAACAGGATCATCTGCAAACGATGCA
TGCGAATTCAGAGCTTAAGGTATGCACATATATCTGTTGATTATCTATTCACATTTCATTCTATGTACTTACTTA
TCCGATACGGATGAGATTTTGTATAGCTTTGAAAAAGAAAATCTGACCCGGGGATGAAATATTGGCATAATTCTG
TATAAGAATAAACTTGTGTAGCACCTTAGACAAGATATAAACCATCTATTTAAAGAATGCAAGTAGCAGTGATAA
AAAGCAAGCAATTGTACCAATTCAATGTAGCTAATGTTAAACCAAATCATGAAATTTGAACCAAAATTTAGCTCA
GAGCAAGAGCTGAACACTAGCACTACAATGCAATGCGGTAATGTGGTAAAACATCAAGCTTTATTGCAAACTTGC
ACTTCAAAGGGATGTGACGCACAAAACATGTACAACACTTTGCTTTGTTTTTCCTTGATCCTAAAGCGCACTTCT
AACAAGAAAAGATTTGTTGATACACTTCTATTGTCCAAGATTTATCAATGGGCACGATCCAAAGGTTAAAAAAGA
TGGTAGCAATCGTTGCCCTTAGCATTGTTATGAAACTTCGCTACGATTGCTAGTTGACTTCTTAAAAGCCATGGG
TTCCTCTATGCAAAGACAGTCTCATAAATTGAGGGCGGAAAACATGCTCGGCTCATCATCAGAAACAATGCATTA
TTTCCTAACATCTTTCAAGCAGACTGACAACGTCTTCACAGTTCTTGGAAATGATTGGGCCATTCAACATGCCTT
CAAGTGTCATAGATGGAGTAGTCTCTGTAGGAATTGTTTATTGACTCCACTAGAGATCGTCATTCATGAAAAAGG
ATTTGTTTATGCTGAGAATAAAAGGATCACTGTTTAAGGGAGAATAAGGTTTATCATGTTTGGGGACAGCTTACT
TGGCATTAATGCAAGTGCTCCATTTCATTGTGCACATCCCAACAATATAAATTTATAATTTTTACTACCTATAAG
TTGTTAACTTAATGGAACATCACACAAGATGATATCATTTATGCTAATTCCAAATTGTTTTGATCATGGTGAAAG
AAAAATCAAGCCATCCAGGACCATAAGAATAATTGGGTGAGCACAATTTTATCACGACCAACTGTAAACTTTCGG
GTTTGAAAAGGCCCTGTATCAAATAACTCGCAAAAGTCAAAGGGTAATACTTCGGGATTTTAGGCTATTTTCCTG
ACAATCAGAAAGATAAGCAACCATAAAAATTTTGCTTTAGACCTGCAGGAATTCATCAAGAGATGACAAAAACAA
ACCACTAGACAGCTTTCTGGGTACTTTTTTTTCTTTCTTCCAAGGACCACAATAAACCCAACGACAGCTGCCCAC
AGAGTTCTTGCAGCAGGNNNNNNNGGAATTGACATTTCAATCACTAGGGGCAGCAGGAACAACCTTTGTTTTCAC
TTCCTGCCTCTAAAATTCTGTAGGTGACTTCTATGACAGCCAGTGGGGATGGATTCTGAGTGTCGGAATCAATTT
TAGCATAATTTTGAACTTCATGGTGACCACTGATCCTAATAAATGAATTTTGCTGCAGCTCTGTTTGCATTTTTT
GGTGGAAGATATCTGCTCCTTTGAAATCACGATGGTCATAGAAGAGGAGGAAATGTGAAGAGGAAATGTGAAGCT
TAAGAGATCGAATATGGTGCAATCGTAGACTCCGTTACATTTCAACCCTTTTTAGGCTCTCCAAATTTAGGGAAT
GTCACAACCCGTCGTGCGACTTAGAATTAATTAAGGCAGGAACAGACATTCGGACGGTGAAATTTAGTATTCTAT
TATTCTAATTTTATCACTAACCATTTAATATCAAAAGAATCATGAAAGAATCCCTTAAAAAAAAGAAGTTTAATT
TATAAACTTGTCCAAAAAAGAGAGATTGGCTCCAAAAATAACTTCAAATACAAGCCCATCGTGCCTTCTCCAATA
GAACTGGCCAAAGTCCATAAGCTCTAAATCATCTGATTCATGGCCGTCATCATCTTCGCAAAAGAAAATGTAGTC
AAAGTAATCCGTGGAGTTGTATCAATGCTGCCATATTATCTCTAACTTTCTCCCCTTGATTGTCCATATTTTTGA
AATAACATAAGATTATAATTAGACATAATTTAACCCTTCATTAATATTATTTATGAATTAATTTAATCAATCTAA
TGACCTATTTTAAATTGATCCAACCTAAAATCTTTGATACCACTTTTGTCACGACACGACCTGATTGATTTGAAT
TAACCAAATCCAGAAATGCATTCAGATGGTATCAAATCTTAGAGAAATATAAAAATTTCTATAAACTTCCAATAT
ATATAAAAAGTCGCTAAGAAATTATCACAACACGGCCCGAGGGTTAGAATTAACCAAGGAGGGATAGACAATCAG
ATGGTAGGAATTAAGTTTTTTTATATTTATTTATCATTTAAAAAATAAAAGTGCCAATAACCAAATTAAATTAAT
ACAATATATTTAAATTTACCTGCAAGTGTCTTCTATAAACATTTTTCTTTCTATCGCGTATCCTTTAATCTTAAT
CAAACATAAAATATTAATTTAGAACGACTATAAGTTTAAACAAAACTAGTGAATACTCAAACTCCTAAAATATAG
TAATACATGAGGAAAAAAAGTAGTATGCGTGACAATTAAGTTTTTACTAAATTAATATGAACATTTGATGGTTTT
TAAGGAATCATGCTATATCAATAGTATATATGTGTATATGAATCATTTTATTTTGTACAAAAGTGATAGAAGCAC
CGTATAAAAATAATATCACTTAAAGTACTTCAATATCAATATAATTTTCTATTTACATGCATGAAAATCTTAGTG
TTTCTCCACTAGTATCATAAAAGGAGTAACTTCATCATAGTCAATCATGCACATACAAGCAGTCAGTATAGACAC
AAACACAGACACAGACACTGAGCTATATAATGCAANNNNNNNNNNNNNNNNNNNNNNNNNNNNNNNNNNNNNNNN
NNNNNNNNNNNNNNNNNNNNNNNNNNNNNNNNNNNNNATATATATATATATGATGTCCCGCCGCTCTGCTAGGGTTTC
GCCCCATGGCGAACCCTAGCGTTTCGATTGTCGTTGCTGCCCCGAACTCCTCTCCCTCACGCGTCCATGCGCGTC
GTCGATTGCGTCGGAGAAGACGCGACTCTGCTGCGTCGTCTTCGTAGCTTTGTCGACGACTCCACTGATGCTCGC
TAGCCCGCGCTCGTCTCCTCTGCGATTGTCGTTCCTTCGCGCCCAGGAACGCAAGGCTTTTCCTCCTTCGCGCTG

CTTCGTGCCCTGCGACGAAACTGCGGCTTGTCTTCCAATCGACGTGCGCTGCTTCTACACCTGCATCTGTGTCGA
TCGACATTTCTTACTGTAGAAATGGCATCAGAGCGTTCCTACGCCGCCGCCCTCGCGAGTGATAACCCCGTCTTC
ACTCTTTTTTTCGAATATGTTTCTGCAACGATAATTTTGCTCTTAGCACTGAAACCAGATTTCTCAAAAAAGCTT
CCCGTAAAAATGGCAAACCTGTAATCTTTCTCACTGAGGCAGAGATCTCCACCCTAGCTGCGCCCTGCCGACTGT
TTTTAGTTGGTAAGTTTCTTTATGGAAACCCTAAACTTGAGCTACTTTGAAAGAACTTTGAACAAATGAAGCTGA
AAGGGGGTTGGAAGATCGGTAAATTGGATGGTCGCCACATTCTTATTCAATTATCAAATGATGAAGACTACACCC
ATTTGTTTGCGAGAAGACTTCTTCGAATAGAAAACCAGGTGATGAAGGATTTAAAATGGACGGTGGATTTTGACC
CTTGTATTGAACCTCCTACCATTCCTGTTTGGCCGCGCTTTCCAGGTTTGAAGATCCACCTTTTTGATCAAGAGC
TTCTCTTTGCTTTAGCAATGGTGTACGGTAAACCCCTCAAAACTGACACACCTACAGTGAATCTAACTCGGCCTT
CAGAAGCCAGAATTTTGATAGAAAGGGATATTACTTTACCTTTTCCGGGAGAAATATGGATTGAAACATAACTGC
AGGGATTTTGACAGCAAGCAGAACTGGAACAGATACCTCATGAACAGATACCACTTTATTGTACAAACTGCCGTA
TGTTTGGACATTTCTCAATACAATTTTATCAGTTGCATCCAAATTTAAAACCTTCAAGTAAAAACGCTTCTGGAT
CTGTCCCACCTCATACATGTCCCTCCTCTGCTCCGGAGTGGATTCCACAGACTATTGAAACACCGGTTGACACGA
TGTCACCTCCCCCGCCCACGACCGGAGACAATACTATCTCTGTTCTTCCGACTGATATATTGGCAGATGGTTTTC
CCCAGGATGCATCAGGTGTTGAAGAGACATGTTTACCATCTCTTCAGAATCAGAATGTTGTTACTACTAGTCAGG
AGCTAGGGGCTCATTCTCTCCTACATGCCCAATAATCTGTTGACCACACCGATCAGATGAGTCAGTCTTCGGATG
AGGAATGACAGGTTTACTTATCTAGAAAGAAAAAGAAGAATGAAGACACACTCCAGCTTGGGAACTTAATGGATG
ATACTCTAGTAACTTCGGGTAATGGAATGACACTCAGCAATTTGTTAGCTGTCTCCGCTCCTAGAACCAGAGGCA
AAGGCAAGACAAAGCTGGCGCGTGCCAAACAGCTTATTTCATGCAAGTTTTGTTTTTTTTTTTGTGTTTTATCT
ATGTTTCATCCTTTTACGATGCTAGTGTTTGATGTATCATATGGGTAGGACACCTTTTTTGGGTCCTCTCATACA
ACATCTATGTATCTTTTCAATTTCAATAAAATTAGCCCTTAGGCGTTTATAAAAATATATATATATGTGTGCGTG
TAGCAGTAATCAATGCATAATAGTAAGTAGCTATAAATAGATAACATTCATAACAATGTGACAAATAAATAAAAT
ATATCAATCAAAATAAAACTAATGCAATTATCATTAACTGAAAAATGAATGAAGAAAAAAATATTACTATATATT
AAAAGTATGAGTTCTCAACTCGATGCGATTAATTTGTTATTTGATTTGTTTGAAAACCTAGCATAAACATGGCAT
TCATATTACTAAAATTTAATTTATTTATTATCTTATTACATATATATACAAATTTAATAATTTATATTATTTTAA
TTCGTTATTTACTTTACATTATTTTTAAGAATAAAATTTTTATTATTTTACTTTATGTGATTCTGCTTAAATTTT
CTATTTTATACATACATACATATATACATACATACATACATACATACATACATATATACATATATATATATATAT
ATATCTATATATATATATATAATCAATTAACTTTTAACTTTCTTAAAACTCACCTCGTCGAGCCCTAATCCACAT
GTAGATGTGATTCTGCCTGCTCTTCTCCTCTTTTCTGCTCCCATTATATGGCAACAACGAACGTGAACCCCTCCT
CTGTGCATGGGAGAGCTCCAATGGCTTCGGCTCCCTGTCACATGAGCTCGGCCTTTCTTTTCCATTCCTATCTGA
ACACAAGATCAAAATTGCACAGGGACTATAGCTTTGGCTCCCTTTCTACAAAGCCACAACGTTTACCTTCCTCGA
CTTCTTCAGTCACCTCCGTCTTCGCCTCCTCCTTCTTCCTCCTCTTTTTGCTTAGTTGAGGTTCGGGAATTCTCT
CAATAAAATGTAGAGGCGTTCAATCAATCCTTGCGAGCCATACACACCTCCAACACACGAGACAAATGGATGTAC
AAGGAGAGTCGCACACTTACTTTCCCCCTCTCCTTTCCTCTTCTTCTTCTAGACTTGCCTCAACTTCCTCATATG
TATTGTTTTATAGAACATATGTAATATCTGAGTTCTCGATTGTTCGATTCCTCTCCCTCCTTTGACTGGACCCAA
CTCCTCACTCTCTCCTTTGGTCTCCTTGTTTCCTCTTCCCCGGGTAACTGAAATGGGATGTTACAGAAACCACAA
ATTATACCATACATAAATGAAAATATCCAAACATAACTACAAAATATGTAGACTACATCTCACAGCAACACATAA
CAACGCATCACTACCCTAAAAAAAGTATTGCAAAAAAAAAACTACAATCACATCATAACATTATACCTATTAGTT
GTAACTCTCTTCTTATAATTGTTAAATCGTGAAAACTTTTTAGATTTAATAGTTTTATGTATGAATGGCCAAAAG
AAATGTAATTCAATATTTTATACAACTGTAAAATTTCAGTATAAATTTATTATTTCTCGTTTCAACTCAAGTATT
TACTCTTTTAAATCATTAAGAGTCATATCAGGATTATCGTCCTAATCAAATGTATTCTCAAATTATACCCATAGG
TATCTTCAAAAAGTTGCACCTAAAAGTACACATTTATCATTTCTTTTATCGTGCACGATTACATCACATCCCGTC
GGTAAAATCTGAGGGTAATGTTGTATGACTTAAAACATTATATTTTATTTCTTTTAATTATTTCCATTTAAGATC
AATGAAACTATCATTTTTCCTATTCAATTTTCATTTCATTCATTTCAACAAATTTTTCATTCACAATATATCACA
GATTATAAAATCGAATATAAGAAATCATAATGCAATTATTTGTTAATAAAACATTAATTTAAGAAATACAAGAAA
ATAATTACTCAACTGAAGCCTGAATGATTTTTTCCTTCCCTTGCTTTTCTTTTACTTCAATTAACACCCATTGTC
TTTTAAATCAACAAACCTAATTAAAACTCATATATTTTCTAAACAATTAATAATCAACGAAAATTTAACTACGTA
AGAATTTTATGATTAAAAACACTCACCTTTTTTAACCTTAAGATTCTATCTTCTTCCTCAAATCTCAAAAAGTTA
TCAGCCTTCTATCGCACTTCCTAATTGAAAAAAAAAATTAATAGCATTAGTTTAAGGAATTATATCACTTATATA
ACAATATAATTATCAACAATTATCTTATAAGTTTCTCGAATCAATCTTCCCACTCACTCCCATGCTTCAACTTCT
CCTTCCAATCTGAAAAATTTATATATGCTTCTTCTCCTTCCAATTCGAAATTTATATATTTGAGTCCTGAATGA
GTACAACTTTGCATTAAGGCTGTGAACCAAGCCAAATATTGCTTTGTAACCTTATTTAATTTTAACATGTAGGGA
GTCTCTGTCACTCCTTTATATATATATATTTATATATATATAATATTTTATTAATTTACTTAATAAAACTTTTCA
TTAAACATTTTTTGTTACTTACCTTTGTAACTGGTCGGCTGCCCCACCAGCACACACGAAGGTCCTTTTCTTCCT
CCAGCCTCCGCTCAACCAAGCACGGCGCGGTCTTCTCCTTCCCTCTTTTCATCTGAAAGGGAATCGAGTTCCTCT
TCCTTCATGCAGCTAAACACGGAATCTCGACTTCAACACTGCAAGTACACACACGACGCCGCTTTCTCCTACCGT
GAGTACAACGTTGTTGCCTCCTACAGCTAGGGTTTCATGTCATATCTCAGGTACAAGAACACACGTGACTCCTCT
TATTCCTCCTCTTCTTCTCCTCTGCTTCACTCACAGACAAGCCGCAAACTAGGGTTGGAAATGACCCCTCTTCCC
TCCTTCCTTCTCCTCTGTGACTTTGTTTCTTCTAATAACCAAAACACTTGAAGATTATGCTTCGATTTGATTATT
ATTATTAATATCATTATTTTTAGTTATTAAAAAATACGCATTAAAAGAATAAAAATTTCAACCTTTTCTAGTCGC
TTCTTCTCCACATAATATCAGAGTGATTAAGATCCTATTTTTAACCATTGTTCTTATCTTTCATTCTTCTAGATG
CCAAGCACAATTTCTTTGATTTTCCTTCCAAGAAGTCAATCAGCATGAATACTAAAAATAAGATAAAACAAAAGC
TCCATTGACTACATGTTTGTCACCAACAGGCCCATAGACTTGGAATTAATAATGAGAGAACAAAACATCAAATGA

TGAAATTAATCAAAATTTTAGTTTCAACCACTAACAAATAAAATGATCATCCTAATGAATTGAAAAGCTACTAAC
AAAAGTAGCTTAAATTAACATAAAACAACACGACTCTACTCTACTTATTCCAGGTACAAGAAGGCGCGACTCCTC
TTATCCCTCCTCTTCTTCTCCTCTGCTTAACTCACATACAAGCCGCAAACAAGAACTAGAAATGACCCCTCCTCC
CTCCTTTCTTATCCTCTGTGGCTTTGTTTCTTCAAATAACCAAACCACTTGATGATAACGCATCGGTTTGATTAT
TATTATTATTATTATTATCATTAGTTTTAGTTATTACAAAATACCCATTAAAAGAATAAAAAAATTCAACCTTTC
CTAGTCGCTTGCTCTCCACATAGTATCATAGTGCGATTAAGATCCTATAACCATTGTTCTTATCTTTTATTCTTC
TTTATTTCATGCACAATTTCTTTGATGTTCCTTCCAAGAAGTCAATCAGCATGAATAGTAAAAATAAGATAAAAC
AAAAGCTCCATGGACCAGGCCCATGGACTTGGAATTAATAATAAAAGAACAAACTAATAAAAATTTTAGTTTCAA
CCACTAATAAATAAAATGATCATCCTAATGAATTGAAAAGCTACTCACAAAAGTAGCTTAAATTAACATAAAACA
ATACAATAATGATATTACTTTTTTTCTATGTTAGTATTGTCCTCTATACATATTGTCACTTCTTTCACAAATTTT
TTAATCTCAATCTTAAACCAAAATATTAGTAGTTTAAAATATGATGCGCAATCAACCTGCCAGTAAATACTTATA
GTTTTAAGTTGCAATTTTCAATCAGGAAAGATGGAAATCAGACAGTTAAATTAGGAGTTTTCTATAATTAGGAGT
TTCTTTATTTAAATTATTTCCAATATCTTTATGATGATTTATTTTCCTTTTAGCTGAATATTTAAATTAGGAATA
ATATTTAATGAATATTTAAATTAGGAATATGTATTATTAAGTTACCAAAAATAGATTATACAGAATATTTTCTAC
CTAAAATAGATTGTAAGATTATTATTTAATGGAATGAAAACAATGGGAAAAGGCATCGAGCATTTTCATTATTCA
GCCTATTTTTCTCAACTTTGACATGATATCAGAGTCCAATTTTTGTCCTAACCACCGATAATTTTTGAAATACCG
TCAAGGCTCTCTTACCGTCGGACTTCTTGTAGAACGGGAAGGGAGGAAATATTTCTTTGCCTTGATTTTCACTCG
TCTGTTCTTTGTCTCATTATTTCTTCAAATCTCCTTCGCTAGCCATCGCTACTTCTGTGGCAACTTCGACCGCTT
CTTCCACTTCGGTTTCATCCGCTTTGACTTCTGCTTCTTTGTCGCACTCAAGGATTGAAGGCTACGACACTCCTT
CATAGACCGACGACTTTGGCAAACCCTAGATTTTCGGATTCGGCAATCTCGATGTCGGCACTCACGGCATCGACA
TCTCAGCTTCAACAGTTTTGGCATGAAGACCAGGAAACAAATCGTCCTCGTCTTCGCAAGAAGAATCAATAGCAT
TTGGATTGTTTGGCAGGGCGTGGATCCTAATTGTGGGCTATGATCAGTAGCATTTGGATTAGGATGCGGATCCTA
CTTGTAGGCTATAACCGATAGCATCTAGATCATGTAACTGGATCCTATTTGTGGACTATGATCGGTAGCATTTGG
ATCGGGATGCTGATCCTACTTGTAACCTATGATCGATAGCATTTGGATTGTGTAACTGGATCCTATTTGTGGGCT
ATGATAGGTAACATTTGGATCAGGATGTGGATCCTACTTGTAGGCTATGATAGATAGCATCTGGATCGTTGAACT
GGATCTTAATTGTGGGCTATGATCGGTAGCATTAGGATCGTGTAACTGGAAGCGAATCCTTCTTATGGCTATGAT
TCGAATTTCTGGGCGATAATGATAAAGGATTTCTTGCTTTTCTTTGCTCAAATCATGTTATGGTTCAGAAGTTGT
CTCTAAGGCTTCAACATTTTTGGCTTCGACACTCACGGTTTGAGGATTTTTGCTCCCGACAGTGTTATTAGTTTC
TTGTTGGCAAATTGATGTTTCCTGATCAGTCGACATCAATTTGAGGAGGGATGTCAATCAGGAAAAACGAAAATC
AGGCAGCTAAATTAGGAGTTTTCTATAATTAGGAGCTTCCTTAATTTTTTTTCAGTATCTTTATTGAGATTTATA
TTCTATTTAGCTGAATATTTAGATTAGGAATATGTATTATTAAGTTACTTAAAATATATTATACATAATAATAGA
TTATAAGATTATTATTTAATGGAATGAAAGCAAAGGGAAAAGGCATCGAACCTAGTCAGTATTCAGTCTATTTTT
CTCAAAACTTCGACAACAATAATGTAACTGAAAATAAAATATGTATGCTAATTAAAAGCAATGAATCAAAATAAA
AAATAAAACATTTTATTATTCAATTATGATCATTCGGAGAAGTTCAGAGCATTTTATCATATAATATAATATCTT
ATTTTATCTATATATTATTTACATTCATCGCACATGGCAAGCATACTACTAACATATATCAATATCAATATCTAT
TTCATTTTACGTCCAAAGATACCTTGGCATCTCCAACTATAGTAACATAAAGAGATGATATTCATCAATCTATCA
AACACAAGCAAGCATACATTTTATCTGTCCTTTCATAACAGTAAATGTACTTGCATAGTAATAAATTTCTACAAT
AAATAATTTAAACAATGCTAAGTGTATACATCCATTCATTTAATATGATAAAACATTTAGTAAAATTAGGTAAAT
CATAAATTTGCAGCGATAATCAATCTAACAAGTTCATTAATATTAATTTTGTTAACTTTTGAAGAAGGAATAATA
TTATAGCTTAGAAACATAAATACTCACTATAGTTGAAAGATTTGCATTCAACATTTCATTCTTTGGTCAATTGTA
TTTTAAAAATTTTAAATTTTACAAATCGACACCATGTTAAATTATTTCTTCTAAATCATGTACATTTATCTCCTC
TCATGATTTATTGATCTTTTTTACTTTGTGATATTGAAAATTTTCATTTCTATCTTTATACAAATTCTTATGTGA
ATTATCTAATATAACTGTTTTATTTAATTAGTATTTTTCATAGTAATATTCTCAAAAATTTATTTTACTCAAAAG
ATTATAGTATCCAATGTCCAATTAATTTCTGCTTTTATTTAAATCCTTACAGATTTTAATTCAATATTCTAAATA
TATATTCAATTCTCACTCAAATTTATTCATAACATAATTCTATCTTTAAGTAATAAATTAATCATCATAATTAGA
TTATTATCATTTTTAATTCCATTACAATTATCTAAAATCATTGATTTTAAAAAAATATAAATCCACAAAATCTAA
TCATTGTGATTACTATAAACAAATGATAATAAATTTAACAGAAAAATCTCATTATTAGTATTTTTCACATAACTT
ATATCCTACCATAAACCTCCTAAGAAATAATGACTGACCAGGAAAATAAAGTTCAAAGCTAGGACACGGTTTAAT
AATTTCAAAGATTCTTCAATTGCTATAAACGAAAATGTTTTAACAATTTACAAAATATTTTGCCGAGCGCCGATA
TAAGTAATGAAATTTACATCATAACCAGAATGTATCTAGTTCAGCAAGTACCTAGAAGTAATTTAAACAAGAAAC
TGAAGACCTTCCTTCACGTTAGGCCACATGCTAAGTATGTCCAAATCAATGTAGATGCTTTACCTCTGAATAATT
ATGAGTTCATTTGCTTTTCGTACAGTTGAGCATTCCCTGACTTAATTAATTTATTTGGTGTCTGCTCCAGATCTG
GCAAAGGGAAGCAACAAGCTTACGGCAGCAATTGCATAACTTGCAAGAAAAGCATCGGTAACATCTCTTCACCCT
TTAGAACATCAAGAATAAAATACAAAATGTCATAGAAATAGAACATAAAAATTACTAGCATGCATTTTAAAATTC
GTTACTCTTATTAAAGATGTAACTAGCAAACTATAGTTTATTTAATGGAATAAAAGACTGTAGTCCTCCAACTAC
TTATATGTGGTCAATTCAGTCTCAAGGTGTCACTAATACTAGAGTTAACCTAAATCACACATGAGAAAATAAATA
ATACTACTAAAACAAATAAACAATTACAACAAACACACTTGAAAGGGGAAATAAAACACACGCGACATGCAAACG
GCAGCTCCCGGGTGATGATAGTGATTGTTAAGAGGTTGGATGCGGTCGCGGCGAGGTTAAGAAGGGTGTTTCCAG
GCTATGATAGAGGAGTAACAGCAATTATGGGTGGCTGGAATGGCCAAACAGCAAGGGACAGCAGTCCAAACCGCA
GCTATTTGCTGTTTAGGGCAGCTGGAAGGCGGGAAGAAGCAAGGGGAGGCTGACCGGGTTGCTGGAACAGCAAAA
AACAGGCCGGAATGGCTGTTTCCGAGCGGCTGCAGTGAGCAGGATTGGCGGCGGCGGCAACGGTTAAGGCATGGG
ACTGCGGGACTGCCAAAAAAGGACCGTCCAGAGGCTGCGATAATGAAAAGAATGGCAGGACAGCAAGGAGCTGCA
AGGAGATAGCAACTGGACTCTTGTCGGGGCGGCCTGCAGCGATGGGGAGAGATGCAGATAAGTTAGGGAAANNNN

NNNNNNAAGATAAACACCACCTGAAGACTAAGATACAGCCGAAGCACCGCCTGGAAATCAAGATACAGTTTTCGA
AGGGAAGAAAAGATGCAGATGGTTGTCTCGGTACAAAGCACACAAAAAATTATTAATCCAAACCGTAATACACGT
ATACAGTGGACTAAAATGGAACGAAGGAAAATAAGAAAGAAAGAATAAGAAAAGAAACCGTGATTGTTTCGGCAG
CCCGACTCCGTCTCTTGTGGTACTATACCAGCTAAAATACCAGGAGAGAACTCTGCGGGAAAAAAGCTCCCGCAT
CAATCACATTTCTCCAATTCAATCCATCAACCATAAAAAATGGTCCTCCGGTTGAAATGGATTAAAACTAGGATA
CAAAATTAATCTCACTTTAATCACTCAACTACATAAAAGTGATCCATTTTAATCCCTAAAAATCTATTTTGGATC
AGATTTTTATAATCAAAGAACTAAATTGGAGAAATGTGATACTTTAAGTTTAAATTGGTCACATATGAATAATCA
AGATGACTATTTTGGGCTTTTCCAATATGTAATGTATTGGTTCTGTTTTGTGCCAAAAAGATGAAGTGCTAGAAT
ATAATAAGCCAACAGCTTAGAGAACACTATATTTAGTTAATGACTCATAAAATAAACAACGAGACAATGTTCTGG
CAAAACTACCACACATTAGAATGCAACAACATGGAATATCAAAATTATACTTCAGATTATGTCTCAGCAAAATCA
AACTACAAAAGTGTTCATAAATTCAGCTATTTAACCAACAATTATAAAGAAACTTAAAAGGAAATAAAAAATTCT
GTTGACTTTGATAATGAGTGGTGAAGATAAGCAAGGAAGAACAAAATTTATAAGTCAATAATTACAGCTGCTAAC
TAAATTTATAACTAAGATTAATCTTTTTAGCTATGTTTTCTATAATAACTACAACAATGTAAACTTTAACAAATA
TAATTATTGATGGATAATTAATTTACCAGTTCTATCTTCAAAAGCTTTTAATAGACAAAAAAATACTGCAGAAAC
TCATTAATTTCTATAATAAAATGAATAGAGAAGCCTTAATGACTAGTAGTTTAATAACAGAAGTGCTCAACAACA
TGTATAAAATATTTATTTTCAGGTAACTGAAAGCTTGCCACTCCATAAAAAGAAAACAAAAGGAACTCTTCATAT
AAGAATAACCTTTCGAAAAAATATACATTTTTCAATTGCACAGTTCGACTGATTAGAATCCTGCTAAATCTTCAT
TAATGACTGAATTACAACTTCAATATTCAACTTGCTTTTATCTTTTTTGTTGCGTCAACAAGATAAGTGCTTTTA
TAAGTGCCTCTAACCGAAGAATAGAACACCTGTGAAATATATTTACTTCTTTTATCATGTTTGTTGAATCTACAA
AATCACTATTTTGTGCCTCTAGCCGAAAAATAAAACACCTGTGCATTATATTTACTTAATTTTTTCCTTTTCTAT
CTATCCAACTATTTGGATTTATGACTGGCAGGTCTACTACGGAAGCTATACATTTATTGAGAGGTTTGATGGAGA
AGTTCAGAGATAATAAGGAGGATCTATATTTTATTTTTATTCATTTAGAAAAGCTTACGATAATGTTCCAAGAGA
TATCCTTTGGAGAGTTCTTGAGAAGGAAGGTGTTGATATGAAGTCATTAAAGATATGTATACTAGAGTTGTTGTG
TTCGAAATGAAGGAGGATTGACCAATTATTTTTCTATTTATGTTGGATTTCATCAGGGATCGACTTTAAGCCCAT
ATCTTTTTGCTTTAGTTATGAATGGTTTTACTAGACATTTACAGGAGGATGTACCTTGGTGTATGCTTTTTGCAG
ATGAAATTCTTCTCGTAGGTAAGACGAGAGAAAAAGCCCAAGAAAAACTTGAATTATGAAGGTCTACGTTGGAGC
CTAAAGGCCTTCATTTAGTAACTAAAACTGAATACATGGAATGTAATTTTAGTAGTAGTAGCTCTAGTGACAGAA
GTGTAAATTTGAGTGAGCAAATTATTAAAATGTGTGAACAATTTAAATATCTCGGGTATGTTGTTCAAAGCGATG
GTGATATAGCTAAGGATGTTAATAGCAAAATTCAAGTTGGTTGGTTGAAGTGGAGGAATGCTTCAGTGGTGAGAC
CTGCAATGCTTTATGGCGTTGAATGTTGGCCTTTAAGAGAGAAGCATACTACTAAGTTAAATGTTGCCGAGATAG
AATGATTAGGTTGATGAGTGGCTTTACACATAGAGTCAAAATGCGAAATGATTATCAAATTTTTGTTTTTCCCAT
CCGCGATAGGGTTGGTGTAGCTCCAGTGATGGACAAAATTAGAGAAAGTTATCTTATATGGTTTGGTCACATTAA
ATGTCGACCATCTGATGATCTGATTAGGAGTGTATATATTTTAGATTCGACTTATGTTAAGAAAGGCAGAGGTAG
ACCTAGGAAGACTTGGGCATATTGTATAAAAAATGATTTTAATTTATTAGAACTAAATGAAAATATGACGCACTA
AATAGAACACAATGGAGAAAAAGGATTCATGTAGTCGAGCCCACGTAGTGGGATGAAGACTTTGTTGTTGTAGTT
GCATTTATCCAACTAATTGTAATCATTTTAGATCCAACTTATGTTAAGAAAGGCAGAGGTAGACCTAGGAAGACT
TGGGCATATTGTATAAAAAATGATTTTAATTTATTAGAACTAAATGAAAATATGACACACTAAATAGAATGCAAT
GGAGAAAAAGGATTCATGTAGTCGAGCCCACGTAGTGGGATTAAGACTTTGTTGTTGTAGTTGTATTTATCCAAC
TAATTGTAATCATTTATTGAATCCTGAAAAGTTTGTTTAAAATTTTCCTTCTATTTTATACTTAATATCATAAAT
CATTTGTTTTAATTTAATACATAGGACCACAATTAACTTTCCTTTATGAAATCATTTTGTAACATAATATTATAA
CTTATAAACAAATTTCTAATTTTTACCTTTCAAATTTTTTTATAATGCAGTATTATAATTTCCTTTCAAAATTTA
AATTATTTTTTTCTTGCAATCTATAATTTAACTCATCTTAAATAATATTGTTATTTTAAATTTATGTTCATCTCG
ATTTCGAGTACACTACATCCACACAACTGCTCTATGCACTAAACCTTACTGCTCATTTTAATCGTTCGCCTCTAT
TTGACCTACCTGCAAAATGTGTTTGCGCACAAGCTCCCACCTTTTACCAACTCATCTTAGTCAAGATTCACTCAC
ACGATCACTCGCTCGGTCCGGCGTGCCACTCCAACAACTTCTTGTTCCCCCCTATGTTCTACTTCATCTCTTTCT
CTCTGTTATGATCTCGTCATGCCCGCGCCCGTCACCATGCACAGACATATTCTCACTACTCATGCATGTAGTTCC
CCTTCACCTTCTTTGCCTCACCTTCTTCTCTTATACCATTTGATTTTGTGCTCTCATTTCTCACCAGATAAATTC
AACCTTCTCTATTTGTTCTCGCACAAGCTCCCACCTTTCACCAACTCATCTTAGTCAAGATTCACTCACACGATC
ACTCGCTCGGTCCGACGAGCCACTCCAACAACTTCTTGTTTGCCCCCTATGTTCTACTTAATCTCTTTCTCTCTG
TTATGATCTTGTCATGCCCACGCCCATCACCATGTACAAACATATTCTCACTACTCATGCATGTAGTTCCCCTTC
ACCTTCTTTGCCTCACCTTCTTCTCTTACACCATTTGGTTTTGTGCTCTCATTTCTCACCCGATAAATTCAACAT
TTTTTATTTGTTTCGCCTCTCCTTTATTATAATAAAATTGCTTGTTGTCTCCCTCATCTTATTGTTTCTTCTTCA
CATCCCTTTCTTCCCCTACGTCTTCATCATCCTTGTTGAATCAAACTATCATTTTTTTTTATTGTTATAGGTCGT
TATAATCTTCATCTTATAAAAATTTCATTCTCAAAATTTATATTTCCTAAGACATTTATAGTGTGAATTTTTCCT
AACCCAAAATCACCATGAAATAGTTTAAATTTTGTTACTCATGTTATAGGAAATATTTAACATTAAAATTAGTAG
CAATTGTAAATCTAACTTAGTAAAAAAAACACCTACTCACAAGATTAGATCGCCATCTTTCCTCTAATACAAAAC
ATCATAACTAAAATAACACCTGCTATATGAACCAGGATAATATAATTAGAACAACACCCAATATTTAAAATAGGA
TAGCATAACTAGAACAACACTCAACACATGATAACTAAAAGAACACCCAAATGACGTGAAATAACATAAAACTCA
TTAAATTTTATCAGCATCTGCATCAATAGTAATAGAATGTTTGGGCCCTTGATGGGTTAAGGCATAAATCCATTG
TCGAATGTTCTATCGTGCTTGCTCCGTATGAGTTACTACGCCAAAGTTTAATCTTTTGTGGACCATTTCTGATAA
TAGATCAATCTTGAAAACTCCTCTGAGTATGATGCCTTCCTGAGTTAGCAATTTGTGTGGATCTTTGAGTAAAAT
TGGGACAGTTTCTCATATGATGTCCACGCAATCAACATAAATAACAAACCACAGAGCCAAATAAGCACTCACCAG
TTGTAGCTTCTTACACTTAGCATAATCGTTTGTAGTGCTACTGCCAATCTATCAAATCCATTGGCATGTAATTTT

GTTTACTCCTATTAAATCCTCCACAATGAAAGTTTGATGCCACCTCTTCCGAGTTCTTTCTCTTGTGCCTCTCTC
TATATTTTTGAATGAATTGAAAATTAATCACCACTGACCAAACCCTTATCAAGAACACTAAATAATCCTCAATCG
AAATCTAATTGAATCATTCAAACCAAACAAGAAATTCTATCATTTCTCCTATGTAGATGATACCAGCTATGGTGC
ATTTTTAGATAATGTTGTAAATTCTATCTCATATTACATGACTAATATCTTCTCTTGCACCAATCTAATAAATTT
ATTCTGCACCTGCCTCCGTCAATAATGGGGCACGTCAATCTCTAATATCTCCACAAAATCTCAATTAACCAACCC
ATTTGGTATATTTCTCAACCTGTCTTGACACTGCTCTGGTTACCGTCTTTTAGCTTCGCTATCCAATACAAATCC
AACTACCTATCACCCGAATAAATATATTATTTCCTTCACGAATTAAATATATGATTCAAAATATTTTCTATTCTC
AATAGTCAGTTTTCATCTTATATTGATCCTTTAATACCCTTGAATAGTGTTGGTTGCAAATTCTAAAACATCTTC
AAAAAGTCTCAACTATTTATCATTAGTAGATACTGATAGTTGTACAAGTTTTATTTTGATTTATATGTTATTTTT
TTAACGGTCGCATTTTTATAAATGTTTGTAGCTGTTTGATATCATGCATTTATGTATGCAATAAATAATATATTA
AAATGAATGCGATGTGCACATTAATTATATATTTATAGTGCTTCTATATGCATAACTGACTATGATTATATGTTT
CTATTAACTGTTTCGATATACATGACAGACTATGATGAAGTCGTCTATTTATGGCACTAGCGAGGGGAAAGCTAA
GGGACCCGTGCACCTAAATGGAAAAAAACAATGATATTGAAGTGCCTTAATTGAAATAATTTTTACTTGGTGCTT
TTCTGACAGTAGTACAGATAAAAAATAATATAAAAATATTATTGATATAATAAAACACATTGATAACACCTAAGT
GATCATATGTTTTATTCTGATTTACTATAAACTTAACTAGCATGTGCATTGTATCTTTTTTATTTATATTATTGT
ACTATAAGAGGATGAGCATTCACTAGCTTTGTTTAGCTTATTGTTGTTATAAACTAATATTACAGGTTTGCTTAA
GTGAATAAAATATGATTGATAGAAATAAAAATATTTATAGAAGACGCGGACAAGTTACAAGATAAATTTAATTAT
GTTGTCATAATTTATTTTTTTTATTGCCTCTTTCATCTGTAAAATATTATTTATAGAAAATACATATATAAACTT
AGTTCTCACCGGGTCGTGACAAAAGTAGTATTAAAGCCAGTATTTAGGATTAATTAATGAGATCTTGACGAAAGA
ACCAATAGGATAATTAAAAAAATTTAATAATAAAAAAATTAAAAATTTAAGTTAACTCAAATTAATAATTATAAC
CATTTAGGCATAGAAGTGCGAATGTCCTGGCCCAGATGATGGCTTTGATGACTCAACTGATTTTAAAATCGAGAT
CACAGCCTAACATAAGTGATACAGTTGTTATAGATAAGTACTTGAAGATATTCAAAGATTTGAGTCCACCATTGT
TTAAGGGAACAAAAGATTTGATTGAAACAGAAAATTGATTGATGCAAATTGTGAAGATAATATATATTATGATGT
GCTCAGCTAAAATAATTCATTCACTACGTCTATACTAGATGGAAAAGCTGAAAAGTGGTGGCATGGGTACTGTCA
GGATAAGCTACGGGACAAACCCAATATAATGGTCAGTTGGGAGGAGTTGGGTTTGTGCCATATCAGCTCAAAGAC
AGATGAAAAAGGTTCAACAACTTGGTGTAGAGAGAGAAGACAAGCATGCAGTATAAAGTAGAATTCATTGTATGC
GTCACATTTGATCGTTGCTGTAGAAGAAATTGAGATACGTCCCATAGGCAACTTAACGATCTTTGCAAGGATTGA
TAAGTTTATAAGCACTCACGGAACATATTTTCTAACATAATAGCGTCAATTCTCTAAATATAACTGTCTCAGTTC
CCCAAGTATAATTGCATCAGTTCCCCAAGTGTAACTCTGTCAGTTTCTCGAGTATAAAAACAATCAGAGGAAGTA
CTATTATCAATATTTAATATGTGAAATCTCAAATTTTTAGATGTAATGAATATAAAAAAAACACTTTTTTATATT
GTGTTTTTCAAAGATAAGAAGGGTTATACAAAACAATTATGACAGTATTTCTTCTACAAGTAGGAATAACTTTAA
GGAAGGAATTACTTTAGCTCGTCACCACTCCAAGTTCGAGGGATGTCGTTCCCTGCAAATCTCTTCAGTCTGTAT
GCTTCAGTTACAACTTTTTAGTGGATGATGTAGGGGTCATTCCAGGTTTGATCCAGTTTCCCGATAGCTTGCCTC
CTAGCGTCGAGGACAGACTTCTTCAGCACCAGGTAGCCTAGGGCAAACTTCTCTTCGTCTACGTGTTTGTTAAAA
TATTTGGCTATCCTTTGCTAAACCTCTTCTTGTTGGAGTGCTGTGTTGGCTTAATTCTCCTTGGCAAGATCTAGA
GTCAGTCTGACTCCTCCGCCATTTGCGCCACACTCCTAGATCTTCCTCGTCGATAGCTGAGGCGAAGTGTACCAT
CTGGTTATGATTGTTGTATTTGACCGGTAACATGGTTTCCATTCCATGGGACAACATAAAGAGGGTCTTTCCGGT
GGATGCAGACGGTGTTGTTTGGATTGCCCAAAGTACCGAAGAGATTAGGTCTGCCCAATTTCCCTTTGCTCATTA
TACATTTTTCCTTAGCAAATTTATTATGATTTGTTTGTAGTCTTCGTTTGTCTGTTAGTTCGAGGGCTTTATACA
GAAGCAAATTTGAGGTTGTTCCCTTAACTTTTACAGAAATCTTAAACTCTTTTACCGGCGAACTGGGTCACATTA
TCTGTAACTATGACGAGGGTAATTCGTATCAGCAAATCAGGTTAGCCAGGATGAAATTTTGGACTACCTCCGTCG
TTATGTGTGCCAGAGGCTTTTACTTCTACTCACTTGGAGAAATAACGTTTCATCTCTCCTTTATATTAAACAAAT
TGCTTGTTGACTCAAGCGCCTTCTTGTTTAAACTTTCCTTTCATTTCTTGATGCGGAAGCGCCAATGCTCGCATG
GGCAAGGCAATTGCTGCATGCAGATCGGCTCAGTCTAAGCGGGAAACGAACCTAGGGTTCTTCCTTGCGCAAGTT
TCTCTCTTATTTGGAGCGGGGCCCACTCGAATTTACTCTAGGACTACGCGTATGTTATTTATTATGTTTTTAAGT
CAATATTGTTTTGCTTCAAATTTTCTATCATTGGTGTATTTTTCCCTTGTTGGAGTTGTATACAATGATAGGTGG
TGCTCCGACTGTATCTTGCTCCAAGGTGGTGATTATCTACTCGTCTTCTTCCCTCTCTCACCTCTCCTCTTGTTG
TCGCAACCTCCAACGGTCCGTTGCTTGTTGTAGCCCTTGCAAATCCACTTACCATTTGTTGTGGCACCCTTTAAC
ATCCTAACTTGTTGTTGTCGCCCCTAACAACCCATTTATTGCTTGATAACGCCCCTACCAATCCTCGCCTTCTTG
CTATCCAACCGCAAGACAAATCTTCCCCTCGTGCCGCTAGACCTCTACCACCTACAACTTTCAATTTCCACTATC
TCAACACATTGCCTACCACCTTTGAACCTCCCCCTAAGGGGTTTGGTGTTTAGGTTGTTGTTTTCCTGTTCTCAA
GTGTCTTTGTTTCTAACACCCCTTTTAATTCCAACAGTAATATTATCATTCCGTTCGGTATCATTTGGCATCAAA
GCTAGGCTCACCATAGTCATATCCTAGAGGTTTCAAGCATCCGTCATAACTCATACCTATCTCCACACTCCCAAA
AAAAAAAAAAATCGTTCTCATTCCCATACCCTCTCTGACTCTCACATATCCAAATCCTAACCATGCCACCAAGGA
GACACCTGTCGAAACTTTCCCGGGAATAGAAACAACAAGAAAAGTGTTGACTTTAGCAGAGATTTGTCGAGATTG
ATGATAGGACCCAGGGGAAATTACCCTCGAGCATTATATAGAATAGGGTTGTAAACTAGTCGAGTCAAGTCGAGT
TCGACCCACCCTTAACTTGAAATTGAGCTGGACTAATTTTATTGAGTTCGAGATCAAGCTTAAGTCTACGAGTTC
GTGATCGACTCACAAGCCGACTTTTTTATTATAAAAAAGGGAAAATAATATTAATTTAAGATAGTGTTACAATAA
AATGACAAATATAATTAGCAATTAGTAAAGAAATAATAAGAAATTACTCAATATAAAATATATAAAATATTTTTT
AAATATATTTGAAATCTTTAAATTTAATATATTTTGAGTATAATTTATATTTATATTTTTTAAACGAGCCAGCTC
ACGAGCTAATAAGTTGAATTGGGGCTAGCTTGAGCTGTGTTGTTAGCTTATTGAGCTCAATTCACGAGCCACTTA
TCAAGCCAGCTCACGAGTCGGCTTGACTCACACCTCTAATATAGAGTCGAAGAGGAGAAAGCGATGAGAATGAGA
ATATTTATTAAAATCTTTTATTATCGGGCATTACACACCTAAAATTATCAAAGGAGCGCATAATCATTAAGGGCA

TAACATGCCCTAATAGTTAAAGGTCATTATGCTCTGCCTACCTATACATGATGAATTTTTAAAACCACTCAAGTC
AATAATGTGGTCCAATTAGGCTAAAACTCTGTTGATTATATGTTCTGACGTCAAATTCTTGAACTTTCCAACCCA
AAATATGTGTTAGAAGCAGCACAACCGTTATCAATTCACGAGCCACTTATCAAGCCAGCTCATGAGTCGGCTTGA
CTCACACCTCTAATATAGGGTCGAAGAGGAGAAAGCGATGAGAATGAGAATATTTATTAAAATCTTTTATTATCG
GGCATTACACACATAAAAACATCAAAGGAGAGCATATACATTAAGGGCATAACATGCCCTAATAGTTAAAGGTCA
TTAGGCTCTGCCTACCTATACATGATGAATTTTGTAAAACCACAAGTCAATAATGTGCTCCAATTAGGCTAAAAC
TCTGTTGATTATATGTTCTGACGTCAAATTCTTGAACTGTCCAACCCAAAATATATGTGTTAGAAGCAGCACAAC
CGTTATCACACAGCTTAACATTATAACCAACAGAAAAACATACTTTGTAGGAAGATATGAGGCACTGCAAAAAGC
AATACTAAAAAGCTCACCGTATGGATCAAATCAATTGCAGTTAATCAAAAAGAATGTCAAATAAACAGCTTATGC
TATGAGATGTGGTGACATCCTTGTCAATCATAACAGGCAGCTGATGGGAGAAGATATTAACGGGCTGAGTATCAA
AGAACTACAAAACTTAGAGAACCAGCTAGAAACAAGTCTCCATGCGATCCAGCAAAAGAAGGTCTTTAGATCCAA
CATGTCAAATTATGCCCAGTTTATCCCTGATTTCAATTATTTTTATAATGTTTTGATCTCTTTTGTAGGAACAAA
TTTTAATCAATGAAGTTCAAGAACTAAATCAGAAGGTAGTTCGCACAAATCTGATTAAACATGTCAGATTATTCT
GCAATCTGCAATACTTTTTCATTTTTTCTTTTTACATTTTTTTTATTTTTTTTATTTTTTTGCAGATTGCCCTCAT
GCATCGAGAAAAAATGGAACTCTGTAAAATGCTAAACCTTTGTCGCCAAGAAAATATGGAATTAAACAAAAAGGT
ATTCATCAAAGATCAAACCAACCACTAAATAGCAATAAATTTCAAAGCCAGTAAAATGTAAACGATGTTGATAGC
ATTGAAGCTCATATTTTATTTGTTGCATGACATAAGAATTGTTTGTCAGGTTAATGCAACTAAAGGATGCAGAGG
AGGATCTACTTCTGCCAATTCTGGTAGCCCAATATCAATCTGTCTTGAGTTAAGCCCTCCACAACAGAAAATTAA
TGAACTGCATGAGGACTGCCCGAAATTAGGGTGAATATCTTTTTCCTTCCTTTAAGAGTACAGCATATTGACGTC
ATACAAGATTTAATTGAGAAAAACAATTTGAATTTGCAGCAGACTCCAATTGTATTAAGATGCCCCAAGAAGAGT
AGATGATTCTCCATGGTCAAGATTATAAACAACTGGTATATTACATTCATTAGTGAATGTAAAAGCAACACTCAG
TTCAACTGAAATAAAAATAAGAGATTATCATAAAAGAACGAATTCATATGTAATGCAATATAGCACTTAGTAAAG
ACAATTTCTTTTGACATTTGAGTATGACGTTATGACACTTAATGGTAAATGAACGCAAGTTAATTCTCAGAAAAA
AGACACTTCCAAAATGGCAGTTCAATAACAATTCATTTCCACCAATTAAATATGAGCCGCATTAACAAGTATGCT
AGTTTGCAGACAGTATGGCATGCATTCAAAGCGAGGCCTTTTAACTCCATCTGAATAAATTGGACCTAATAATGA
TGAATACAGAACTGCAAGATGGTAGCTTTTTTACTTCATATACATTAACTAAACTTCACCAAAACGAATAAAGAA
CTACAAGATGACGACACCAACTCGTTTCAAATGTACAAACTTTTATTACTGGATATTTGATTGGAGGGATTTTAA

>PeAGL17_1
ATGGGGAGGGGGAAGATAGAGATAAGGAGGATTGACAACATAACAAATAGGCAGGTGACGTTCTCCAAGAGGAGG
AATGGGCTGCTGAAGAAGGCAAGAGAGCTGGCTGTGCTTTGTGATGCAGAGATTGGGCTAATCATCTTTTCCAGC
ACAGGAAAGCTTTATGATTTCACCAGCTCCAGCATGAAAACAGTGATTAACCGCTACAACAAAGCAAAACAGGAT
CATCTGCAAACGATGCATGCGAATTCAGAGCTTAAGATCTGGCAAAGGGAAGCAACAAGCTTACGGCAGCAATTG
CATAACTTGCAAGAAAAGCATCGGCAGCTGATGGGAGAAGATATTAACGGGCTGAGTATCAAAGAACTACAAAAC
TTAGAGAACCAGCTAGAAACAAGTCTCCATGCGATCCAGCAAAAGAAGGAACAAATTTTAATCAATGAAGTTCAA
GAACTAAATCAGAAGATTGCCCTCATGCATCGAGAAAAAATGGAACTCTGTAAAATGCTAAACCTTTGTCGCCAA
GAAAATATGGAATTAAACAAAAAGGTTAATGCAACTAAAGGATGCAGAGGAGGATCTACTTCTGCCAATTCTGGT
AGCCCAATATCAATCTGTCTTGAGTTAAGCCCTCCACAACAGAAAATTAATGAACTGCATGAGGACTGCCCGAAA
TTAGGGTGA

#### Dendrobium catenatum

##### *DecaAGL17_1 and dec-miR444a*

>JSDN01S052128.1 reverse
CATATGTTTAATTATGTATATAACGATAATTTATTTAAAATAGTGAACAACATTTCAAATAAGTATCCAAAAACA
CAACTCAACCGAACCCATGTTATTCAAGGCAAACTTTAACTAAGTAGCTCGTGATCACTTAATCCGAATTAAACA
TTAAACACACTTCCGAAAAAATTTAACGGTCCTATTTCCTCAATTACAAAGCATATAAATCAAACTTCCTTCTTA
TTTATTTTTCTCAAATTAGAGGTTAAGATGAAATAAACTCCAAAAACTTAGGTATACTTATGATTTACAATGCCA
AGTCAGTAATATATTAATTTCTAATTTCCCATTTACTCATTTAAGATAAAATCAAAAGCTATGCCTCACTTGAAA
TAAAGATCAATAATAGAAATTAGCTTCATATATATATATATATAATAATAATAATAATAACTGAAATGAAAATCT
AAAACTGAAAAAATAAAGTTAGAATATGACTTTTATTGAGGGACAAAACCAATTGTCTCTTGGTGATCAATAATG
GAAAAAATGAACTGGCTTCAATTTGTAAATTTCTTAATGTCCAAAAGGTACCAAACTTCCATATTTCTATTATAT
AAATATATTGCACTCCACATTAATTAATGAACAATTATCTCTTAAGAACCCCGTCGTATGGGGAAGGCAGTAGCC
CTCCCTTTACCAAGCTGAAACCAGAAGAAGAAGAAGAAGAAGAGAGAAAGGGGGGTTAGAGGAGAGAGGAGATGG
GGAGGGGGAAGATAGAGATAAGGAGGATTGAGAACATAACAAATAGGCAGGTGACCTTCTCCAAGAGAAGAAATG
GCTTGCTTAAGAAGGCGAGAGAGCTCGCTGTCCTCTGTGATGCAGAGATTGGGCTGATCATCTTCTCGAGCACAG
GAAGGCTTTATGATTTCACCAGCTCCAGGTTCTCTCTCTCTCTCTCTCTCTCTCTCTCTCTCTCTCTCTCTCTAC
TGTACAGATTCATGATCCATGCATATTTATATCTCCTCCTAATCTTAATTAATCGGCTCCCAGTCTTAGCTTAGC
CGCTTAATTACTGCTATAAAACTTTGATTAAGGTTAGTTGTTAGTGTTAATAAAGTAACAATCAGATCTGAATAG
GGATTAAGCTAATTAGAAGATGATTATCGGAGAGAATATTCTTTCTCTCACTAAGTTTCTTAGGTAATTAAAAGA
TGCAGAATAAAATATGCCATCTAATTACTCGTTCTTATCAAATTTGAGGTTTGATTTATGAATCTTCATTCTTCA
TTTGTTTCTTTAAATATTCTACAAGTTGTTAAAAATAAATTTTGTATATTATTTTTATTTATGCTGTATTTGATC

TGCCATTTTTTCTTCATGATCATTTAATAAAAATTTTATCTATTTATTGTACTGATGCATCTTCTATTCATTATT
ATTATTATATATGCTCAAATTGTTTAAATATTTTTTAACAGATGTTTAATTTCTCCTAATTATTGAATAATTAGG
AAGCGACCAACAGTTAAGAAACTTAAAAAAAAAGTTATTATCTACAAATTTATCACCTTGGGATCAAACTGTACA
GAGGTCTAATAGTTGTAAATATCTGCTTACAAAAATGGGTGAGAAGAAAAAGAAGAAATACAATCAGAATAGTTC
AGAAGTTAGATCCTCTGGAAGAAATCTTCGTCTAGTTCACAAGTGAGAATGGAAGCAAGATATAAGGCTTTAATT
GTTTGGGACGACATTCTTACTGCTTTTTAAAGCATCGTTCTAGTACAAAAATTTTAATTTTTATACTAACAAATT
GCTTTAAGAAGCAATAAAAAAGCCGTCCCGAACGAAACCTAAGAAGAGAACATATTATAAGATACCTCATAGAAA
CTTAACCAAGTCATTAGCTAGAAATGAGGCAAACTTGATGGGATATGGTGGTAGGTTTAAATAAAGCTTGGGTGA
TGCATGGTATGAATATCCTATGAAAGTCAAATCAATTTGTGAGGAATGGGTGGTTAAGATTTGATCATAGTTTGG
GAAGGTTGAGAGGTCAGAGAAGAAGATTATGTTATACTGGTTGTCAATGAGAGAGGACAAAAATAAAAGTAGAGG
TTGTAAAACTATGGAAGGAGAGAGTGAGGCTAAATTTGATGAAAATGGTTTGAATTAGAGGTTCAAGTTCATTTG
GGATAGCTTTCTCGCTACTTTCTGAAGCAGTTTTTTAGTATAAAAGATTTTTAAATTAAAAATCTTTTATACTAA
AAAACTATTTTAGAAAGCAGCAAGAAAATTGTCTGAAACAAAGCCCCACTATCAGATATAAAAAACTATGATGAG
CCCCAAAAGACTTACCTGGGTTAGTTTCCAAGCCAAGTCAAGAATGCATGCAAGAGAGAAAATGATAATTGGCAA
GCAAGAAAAAATAATAATTGCATTATGTTTAGCTTACCAAGGAAATTATTTGTAACTTGCCAAAATGTGATCAACT
TTGATGATGCTTTTGTTGATCTTGAAGCCCCACAATTTAAATTTGATTTCTCGAAATAACCAGCCCTAGAATAAT
ATAACATGATATGATGAACAAATGAAGAGATAAATATTTGCAACTCATATGTTTATCTAATTTAGTTAATTTTGA
CTATATAATTTGATGTCTCATTTTAATCTTATAAAACTTTTTTTTTTTCATGTTGAAAATCTTATATATAGCCAT
AATGTAAAAATATTTTTAAAAATACAAGTGAGATTACTCTATTTTTCTCCAAACTTAGTTTGGTTTAGAGTCATA
GAAATCTTTTAGTGAGGGTGTGGCAGTTATCCAATTAAGCTTTAGTTATTATCTAAGTTGCTTGGATTCTTCTTC
CATAATCCATCAAAGCTAAAATTAAATAGCACTTAATCAATTGATATTTGAGCTTTGAATGATAATTTTGTGCAA
CTCTCTAATCATTGTTGTTAATAGTGGATCAGATCTTATCTTTTTTTTTTTCCATAAATCTTCCTTCCTTCCATG
ATAAAATAATATGGTATAAGAGAGGACAGGAAAAACAACTTTCAAAGAATAATATAAAACTTTAATAAAATCAAA
TTGATTGTAAAAATTAGATTATAATAATTTCAAATTGCATTTTTAATATTTTAAAAAGTAAAAAATGAGGAAGAA
AACTATCCATCATTTAAAAAAAAAAAAAAACTAAAAAGACACATCATCTTCTATTATAAATCCTCAAGGCCTTAGT
GCTACTATTTATAACACAATAATTATTAATATTTGAAAGGATTTTCAAGTAGATCTGTTTGTAAGAGAAGCATCC
TACAAAGCTTTCAAGTTTTCAGAAAAGAGTTATTTTAAGTTGTGGGATGCTGATGCAACACCAAAAGACTTACCT
GGGTTAGTTTCCAAGCCAAGTCAAGAATGCATGCAAGAGAGAAAATGATAATTGGCAAGCAAGAAAAAATAATAAT
TGCATTATGTTTAGCTTACCAAGGAAATTATTTGTAACTTGCCAAAATGTGATCAACTTTGATGATGCTTTTGTT
GATCTTGAAACTCCACAGTTTAAATTTTATTTCTTGAAATAACCAGCCATACAATAATGTAACATGATAAGATGA
ACAAATGAAGAGATAAGTAGTGGGATGCTGATGCAACACAAGAATGCTAAAAAAAATGTTGTAATGATGGCCTCCT
CTATAAATATTTCATAAAAATATTATAAGTATAGCTTGATATGAATTTTTTTTTTAAAGAAGCTTATTTTCATAA
GCTTATCATTATAATTTATTTGATACATCCATGAAAAAAAAAAGTTGGCCATTTTAAACTTTTCATGATTCCTTC
ACTGCCTTTTACATGAATTTCCATATAAGAAATGTGAATTATAAATGCATGAATTGCTTCATAACTTCCTTTAGT
GTAACTCCCCAAATTAATTAATAAAGAAGAGAAAATTTCATTCTAGCATATGGAAGTAGAGAAAGTGGGCTTAGT
ACATAGCAAAGAGGAAGTGTAAAAACTATACCACTAGTTATAACATGATTTGCAGAAAAAAAAATGAAAATATTTA
TGTAATTTTAACTTTTAATATAATTTATTGTCTCTTTGCATTTCGATAACTTTTTTAATTTATGATGTAAAACCT
AAATGTAGCCATAAAATTTTAGGATTTAAGATACTACATTTTAAAAAGCAATTGAAATTACTCTTTAAAGAAAAA
AAAAACTTAGATTGATAATTATTATTAGCAATGTGCATTAGACTTGATCTAAATATTTTTTCCCTAAATTTTCCT
TCATTCCATGAAAAAGCATTAAGAGATGAAGTGAGAAAATAAACTTGCAAAGAATAATACAAACTTTAAATCAAA
TAAAAAAAAATTATCTTTAGTCCCAATAAATTCAAACTTGATTATTAGAAATTCAAATTGTATTTCCAGTATTCG
ATGAAGCAAAATATTTTTAACGAAGAAAATTATCCATTATTCGAAAAAACTTGTCCAAAATATTTCTTTCTTAAA
ATTTACTTCCATGATAAATAACACGCTTTAGGAGATTAAATTAGAAAATTAACTTTTAAAGAATAATACTAAATT
TATTCAAATAAAAAAAAATTAATGAAAACAAACCACAATCAAATGAAACTTGATTATAATATAACTTGTATAAAT
TTCATGCATGATGATATGAATGACATATCATTAACCATTTCTCGACAATTTGATCACATTTGGAAATTGCTAGTG
CGTGAATGTTATAGATGCATGGGCTCGAAGAAAAATTATTCTGTGCGGTCACCGAACGACATCCGGATACTAACC
AAAACGTACAACATCATTTTTTGACTCGGTATGCATCAAGGTATAGAGACGAATGCATCAAGGTATGGATTAGAG
GTATTGCTTATGCAAAATCACATGTTCATAGCTTACTTCAACTAGGTGATGACAGCTAGAGATACACTGAAGTCA
GTTTATGAGAGGGAGTTGTTATTTAAATGAATGTAAATATTAAAACAAAAATAAATTGCAAGAAACAAAAGAGAA
ACCTTGATTTTTGTGTGGATTGTTCTCCAAAGCTTTAGGTTAATTCAGACCTAAGTACAACTCACACTTAGAATT
CCTGTCTCGAGTAACCTACTCCCATATAACTCGTGGGTAGAGAAACCTTTACATTCTCTAAAGGGGAAGAGTGGC
AAAATGAACCTACATATTGTATTTAAAAGTGCATGGAATGATAACATGAATCAAATTCAAATAACTACCACCAAG
CAACTCATTCTTCATCTCATTCAAAACAAATAGATTTCATAAGATAAGGGGGAGACCCTCCTCGGTACAACCGCT
TATCCAGCAGTTCAGTGAGATTTTTCAGCTATTACAAGGATTGCCACCATAGAGAGAGCAGGAGCGCGCTATCAC
CTTGAAAGAGGGGGTGTCAACAATCAGCGTCAGGCCATATCGATACTCTCATATCCAAAATGATGAGATTGAGAA
GCTGATCCAGTAGATGTTGGAGGCAGGAATAGTTAGGCTGAGCATGAGTCATTTTGTAGCCCAGTCTGACTTGTC
AAGAAGAATGATGGGAGTTGGATATTTTATGTGGATTATCTGACATTGAACAAGGAGACTATGCCAAACAAGTTT
TTCATTCTGGTGATAGATGAGTTGTTGGAAGAGCTTTGTGGAGCGGCATTGTTCACAAAGATGGATCTGAAGTCT
GGGTACCATCGGATCAGGATGAGTGAGGATGTTCCAAAGACAACATTCGGGACACACGAATGACACTATAAGTTC
TTGGTGATGCCGTTTGGCTTCACCAACACCCATACTACATCCCAATCTCTCGTGAACAGGATATTCCGACCAAAT
CTCAGGAAATTTTTGCTGGTTTTCTTTGATGATATCTTGGTATATAACAAGTTGGGTGATGAGCATTTGGATCAT
TTAAGAGTGGTGTTAGAGATGTTGTAGAAATATCAGCTGATAACCAATCTGAAGAAGTGTTGTTTTGCACAGGCT
AGTGTGGAATGCTTAAGCCATGTGGTTTCTCAGGAGAGGGCGACTACTAATCAAGCCAAAATTGAGGTAATGATG

AAATGACCTCTACCTAGGGATTTGACGGAGCTGAGGGATATCTTAGAGCTGATTGTATACTATAAGAGATTTTTG
AGGGGATACGACATCGTAGCCTGGCTACTAACAGAGCAGCTGAACAATGATAGCTTATTGTGAGGAGAGGCAGCT
ACTCAGGCCTTTAAGAAATTGAAGCAGGCCATGACCATAACACCAATCTTGGCCCCACCAGATTTCAATCAAACT
TTTGATATAGAGACGAATGCATCAAGGTATGGATTAGAGGTATTGCTTATGCAAAATCACAGGTTCATAGCTTAC
TTCAACTAGGTGATGACAGCTAGAGATACACTGAAGTCAGTTTATGAGAGGGAGTTGTTGGCCATTGTGTTGGCA
ATTCAAAAGTGGCGGCCTTATTGGGTTGACACTTCATCGTTAGGATAGACTAGAGGAGCTTGAACCAGGGGCGAA
TCAAAGCCTTATTTTTTAGGGTGGGCAGACTTATTATATATAATTCTAAACAAAATCTTATATTTCATTATAATT
TATGAATTGAAAAATAATTTAAAATTTGCATCATATACATTCATTATATATTTTATATTATTTGGAAATGAATCT
ATATAAAATTGATGATTTTTTTCTCTATCAACCGGCCTAAATCATCATGGTCAATGAAAAGTTAGACACTTTTGA
TGGATCACAAACAACTCGAGATGGGTCATATTAGGGCAAACAAAAATTTTTGAGAGGGGGGTGGGGGGAGCAAAA
TTCCAAAATTACCCATCTTCTACATAATTTTTTTTTTTAGGGTGGTGGGCATTTGCCCACCGTGAATAGAGTGGG
GATCCACCAATGGCTTGAAGTACATGTTAGAGCAAAGGATGGCGACCGAGGAGTATTAGACGTGGCTATCAAAAC
TGTTGGGCTATGACTTTGAGATTTAGTATAGGCCAGGGAGAGGGCGGGCTATTAAAGAGCAGAGTATGCGGGATG
AGGAGCTGAGAAGGATTAGAGGGGGATCTTCTTAAGGATGAGAGTAGCCACCATGTCTACTCCCTAGAGGGGGAG
CGACTACTGTATCAGATGAGATTTGTACTTCTACAAACTTCGGTCCATATTCCACATATGTTGCAATAGTTCTAT
AAGAGCACCAAGAGGGGTCATTCAGGTGTGCATAAAACATATAGGAGATTAGCAGAGAGTTGCTTTAGAAGGGGA
TGCACAAGGATGTGGAGGCGATGGTAGCTAGGTGTGAGATATGTCAGATGAACAAATACATGATCATGGCTCTAG
AGGGATTACTTCAACCCCTAACTCTTCCATATAGAATATGGGAGGATGTCACTATATATTTTATAGATGGGCTAC
CCCGATTTAAGAGGGTTAACGATGATTCTAGTGGTGGTTGATAGATTGAGCAAGTATGCTCATTCATTCCCCTTC
GACATCCTAACATGACATTGACAGTGACAGTGGCAGCAACGTCCATAAGAAAGGTGGTCAGGTTACATGGAACAC
TAGAAAAGATAATATCCGACCGAGATAATGTGTTCCTCAAACATTTTTTGAGGGAGCTGTATAGGTTGCAGGGAA
CAGTGTTGAAGAGGAGCACCACTTACCACCCCCAGACAGACGAACAGACGGAGAATCAGAGATGGTAAATAGGAG
CCTGATGTTCTGCTTCAGAGACATCGTTGTTAGGCCCAAGTGGTTAGCATTGGAGGAGTATTGGTACAACACTTT
GTATCATTCGGCTTCCCATATGACACATTTTAAAGTGTTGTATGGAAGAGACCCACCTCACTTGATGCATTTTAG
TCAGAGCAGTGCGTCGATGTATTCAGTGGAGTAGTATTTAGAGGAGTGTGGTAGGTGTTGCATGAATTGAAAAGA
CACTTTTGAGGGATTATAAAAAAAAACATATGGATGGGAACATGAAGGGCATCCAGTTTGAAATTGGGGATAGGG
TGTTCTTAAAGCTTCGACCCTACAGATAAAGGACAGTAGCTAACATAATAAATGAGAAGCTAGCTCCAAGATACT
TTGGCTTTTCTGATATATCAGAGAGGATTGGAGTAGTGGCTTATAGTTTAAGGTTATCCGTGATCGCTACCATAC
ATCCAATGTTCCACGTGTCACAGTTGAGGAAAGTTAGTGGGGATCATGCAGCTAGTTTAGAGTTATGGGTGACAT
TGGCTAAGGATTTAGAGGTGGTGCTAGAACCATTGGAGCTAAAAGGGTACGATAGAGAAGAGGGATCAAAGAAGT
GCTGATCAAATAAAAAAACCTACCTAACTATGACTTTAGTTGGAAGTTGTATGATAGGGTGAAGCATCAGTTTCT
GTCTTTTCATCTTAAGGACAATGATCGATCCAAGAGAGATCGGTCTAGCCAATCCAGATGGACTAACTCTCTATT
ATAGGCAGAAGTGTTCCACCTCCAGAGCAGTCCACCACCTTTGGGGATGGGGTCCACGAATTAATGCAAAACTTA
AAGGGGAGACTAGGAGGAAAATTCAATTTAAGAGGGTTGTTTTGGGTAATTTGCAGATTTTGTATGCTCTAAGGA
GGATTGTAAACCTAGGGTGGGGTACCTTAATTTTGGTTGAAATTTCGAGAGTTATTAGATTTTGTTTGGGCTCTA
GCCACCGGCTATATATTGGGAGAGTTAGATCCTCCTCTCAAATTTGGGATCCATTGTATCTTGTTCATTTCTTCC
CTACTTCATCTATTACAAAGTTTTTATCTTCCTATTCCTCTTGTTCTATTTGTGATTGTGTGTTGGAGATTGCTG
AAATTAATTTCAGTCCTGTTTTATCTTGAGTTTTACAAGCACTACCTAACAATCATATAGTCCTCCACATTAATA
TGATTATTAGCAAATTCATGCAATTTACCCAACTTTCACAGATTAAATTAACTTCTAAATAGGCATTCTAATAAT
TTGTCAACACATCTTAGACTATTTGCGGAACCTAAGTCTTTGACTTAAAGAATTCTGATGGAAATAAATATGACA
TGGCAAGAAATAGTATCTCCATAGAAGTTTCTTCTATAAAATAAGTTTCATCGTTTACTTGAGGCAAGTAGCCTT
ACTATGGCTAAATGTTGATGAACATAGCTAATGTTGGTAAATAAGTAGTCCCTTCTCCTGAGGAGATTCAACATG
TTGATCAATTCTCAAAAATTGCTCTAGTTGCCAAACGATCTTCCATTTGATTAAAATTATCTAATAATTACAAAA
CCTTATTGTTAAACTTCTAAAGATCTTTTCATTTCAATAGGATGCTCCATGACATAATTTCTAATCCTCAATATC
ATCAATGCTCATCGCACTTTATATGATATTAAATTAAAAGTATCCAAGAATAATGTGAGGACCTTACTGACAAG
CATGACTCACCCCCAAATAAAGAAAAAAATAAGAGAATTTATAGAAGAAAAAAAATTAAGAATTTTATTGAATTT
TAGCAAAAAAAAGAATATAACCAATATAACACAAATCCGTCAGAATCTGCTAAGCTTGCCCTTTATGTATTTATA
AGATTTTGGTCGAGGTCTGATCCAACCTTAATTTTAAGTGGGGCAAACTTATAAGTAAAAATGTAAATTTTTTTT
TTCCATAAATATCCTTACAACTAAAATTTGTTCCGCATGTACTGCCTCTCGACCCCTATTTGTTCATTGTTGTAT
GCTTCACATATATGTCAGGGATCCAAGTGGGGCGTGCACCGGGGGGGTGGGGGGGGGGAATTGCCGAAAAACCTC
TGTAATAAAGACTTAAAAATGGATGGAGAGGAACAAATTACCTAAGGAATCATATAATCCAAATACTTGATAACA
TCTTTAGAATCTCAAAGTACAATCAAGAGTTCAATGGAATAAATTCTCTTAGCCAATCCTCTTGAACTAACAAGC
ATGTTACTTCAACAAGCTATTCTGGGGGA**CTTAGCTGAATTGTAAAATAGTATTGGTTTGTTGATGATTTCAATT**
**GACATTCCATAGGCAAATATAGCCTTTTATTGGTAACTTGAGCAAAAATAAATTCAAATGACTTGGAGGCTTTGG**
**AGTTTGGGGAGCTACAATACCACTTTAAGCAGCCTTTAAC**CTATATGGATATCAACTTTTTCATTATAATTCATA
GTTATATCACTACTTGATAGACATGACCCAACTGACCATAAGTTTAATATCAATAACACAATTGACATTTTATTA
ATATTCTACATTAAAGATATCTATATATATAATTTGAATCTGCTAAGTAAGCTTAATACATTAGCATTCTTCTTG
TTTTTTTTACAAATTAAACTAGCTTAATAAGTTAGCATTCTTCATTTTTTTTCTCACAAATTAAATATAGGCTTTA
GTTTGAAAGATTTGAGAGTAAACTAAGGTTTTGTTTGGGACAGTTTTTTTACTGCTTCATAAAACAACTTTCTAG
TACAAAAATTTAAATTTTTGTAATAAAAAGATGCTTAAGAAAGAAGCAAAAAAATCATCCCAAACAAACCGTAAG
AGCGTACATAC**CTATTCTGCTCAAAGCGCTTTCTTAAGGGCCATGATTTTCCCAGTCTTCGATATCGTAGAAATA**
**ATCATAGTTCCCTCAGAAAGCAAGTAAAAGTATACCACATCCCAATCCTTGTATCTTTGCCTGCTCCAAAAACAA**
**AATACCAAACTAAGACTTAGTTTGAGACGGCTTTTTTACTGCTTTTTAAAACAGCTTTCTAGTTCAAAAATTAAA**

**TTTTTTTTACAAGAAATCCGCTTTAAGTAGTAGTTAAAAAACCGTCCCAAACAAAGCCTAAGGTACTTTGTAGTT**
**TGAGTTTATCGTGGCTTGATTGGAATTTTGCATTGTATAGATGGCTTGAATGCTTTGGCCCATGTCCGAACCAGA**
**GCAGGGCCTAAAAGATGGACAAGAAC**CTGGAAGATGAAATTTAGTTGCCATTAAAAACCAGCTCAAAACTCAATA
CCTTGAGAAAGTGAACATTAGTTTCCTTTACATAACTGATAACAAGTAGAAACTTGGGATCTCTTAAATAAAGAA
AAGAATATCAAAATAGTTGATGGACTATTTACAGGGAGCTGTTTTAATTCTAGACTATTTTATGTATCCAATTTA
AATATCTGAAATAGTCCTTTGACTCTTCAAATTTGATCTAAAATAGTTTTTTGACACTCCAAATCTGATCCAAAA
TAATCAACAAAGGAACCGTTTTGGACCATATTTGAGAAATCCAAAGATCATTTGAACTTTTTCTTCAAAATAGTC
TGGGATTAAAATAATCCCATATAAATAATCTATGGATTATTTGACCTTCTTATCCCCCTAAAAATTACAGCCTCA
ACAGGGAAGTGAAGTGGAAAGACTTTTTTTTTTTTGGACATTTAGGCCATTTAAGGCAAACTCTCACACACAAC
GCATACACAAAGACTCGAACCCAAGACCTCTTTATTGAAAGCCTAAGCTCTCAACCAACAGGCCAAGGTGAACTG
TGTGTGGAAAGACTTTTTACTGTTTGATTTTTTCAATGAAACTTAGAAATCACTTCAGTTTATGAGGAAGTTAAA
GACTGCAGAGCAATTTCGTTGAATTCACCCGGAAATGGATATTATTATCATCAAAATCATAGACTTCTACGCGGC
AAAAACATTGAAATGGGAGGAAAGAGTTAGCAC**CTGACTGATAGACGACCAGAAAACCACCGAAATGTGCTTCTT**
**CAGCTCCCTTTCTGAAGTAGGAGATGCAAGTTCCAGTTTTTGAAACCAAACAAATGGG**CTATGATTATATAAAAT
GATGGGTGATTTTCCAGTTTGAAATGTATTAAGTATCAATAATAGCATATCATCTATCAACAGAAAAGAAACTGA
AACCCAAATTTAGAACAAAAGCATTACTCAGTACTTCCAGCTTTTTTTTCAGAAAAATTGCATCAACATAATATA
AAATAAAGGGACTAGAGGAAGAGGACGAGAGAGGACCAGGGTAGAGTGAAAACTTTCATAAAATATTAGAGAACA
CTAAAATATCCATTGTTTTTATTACACTCCATTCAGCGAATGTCGTTATTAAAATGTAGATTAGTAAAGTTAACC
ATAGTTTTAAATTGCAAATCATTGTCAATTAACCATGCATACGTGAAAAAAAAAAAAATAGAACTGAGAGAAATAC
TTAC**TTGATCTCTATTACAGTATGAAATCAGTGATTGGCCG**CTACAACAAAGCAAAGCAGGATCATCAGCAAGTG
ATGCATGCAAAGTCAGAGCATAAGGTATAAAACTATCCCATGGTTATCAATTTACGTGTCATTTATGTACTTAGT
AATCTGTTACAGATGAAATCTTAACATAATTTTGAATAAGAAAAACTGATCATATGAAAGATTGTCGTAATTTTT
AATAAAAAATAAACATGGGTAGTTCCATGGACATGATATAAGCCATCTAATTATAAAAAAGTTTAGGCTCCAAAA
ATTCGTGCCCGGCCTCTCTGCGGACCTCTTTCTTTTTCGGGTCATCGAGGAGAGTTATAACGCTCAAGTTCCAAG
GTAAATTTTCAAAAGGAAGATTGGGTTTAAAATAAGACACTGCCCATAGATTTCTCCCTCATTTTACGAGGACTT
CAGATCTGAGTTCAAGTTTCCTATATGCGCAAGCCCTTATAGTTAATTACCCCTCTCAATAACGAGAGAGCACTT
TCAATTAAGACTTGACATCAAATAGATACGTAGGGATACGTGGTATCTGGTGTATAGTAGCAACATAGGTTTAGC
TTTTTTAAAGATTTAATATTTAAAATTAAAAATTTCACCATCTTGCATTTAAAGCAGATGGTGGGGGTAGTGATG
ATGTACCCGATCAAGATCCCATGGTTCCGTCGTCCCAGGTCATTATGGTCATTGATCTCCTTAGTCAGAGGAGGT
AGTCGATCCAATCTAATAGAAGGATCCTTGTTGGCAGCCACGAATTGTGTCATCCAAGGAAACCTTGTAGACTTG
GGTTTAAATATTTGTCTAAGGGGACAAATTTGAATCCACCTTAGCTAACATAGGGCTTATGTCCTATCGATTATA
TTATATTGATTCATATCTCAAAGATTTGAAGTCATTAGGGAACATATAAACTATGACCATAACCATAATACAGTG
GAATCATTGGACCAAGCCACTAGGTCAGAGGGATGCGATCATGATAGTTCCCACATGGAGCCGTTTGACAAAACC
ACATGTTATTTGTGGATTCATATGGCTAAACCACTGATTAGATAATCAATGTCATGTCCTTACCGATAATAGACA
GATTGAGAATGAGTCCTAATGATATTATCGACCAGACTGGCTCTTTGTGAGCTAAAGTAATTTTTCAATTTGTCC
ACCATGGCAAATTGTTATGGAGGGTGGATTGAGATTTGGCTTTAAGTATTTTAGTGATTTATTTCAATTTTAGAA
ATTTTAGACTTTTAATGACTTTTAAGCCTAAATAAAGTGACAAGTGTGCAAGGGCCGCTGCACCGGCGCCTGGGT
GCAGTGTTAAATAGGCAAGGGTTCTCACATCAAGGACGGATGTGGGCAAAGAAGCTAGTCCATGAAACTAAAATT
AGGATAGGAACTTGGAATATAGGTTCCCTTACAGGAAAACTAATGGAAATAATAGATATTATGATTCATAGGAAG
TTAAATATTTTGTGCATTCAGGAAACTAAGTGGGTTGGAGAGAAGGCTATGAAAATTGATAGTTTAGGATTTAAA
TTTTGGTATTCGGTTAGGAGTCGTATTAGAAATGGTATAACACCTATAGATAAGAGTTTAAAGGATAATATTATA
AAAATTAATAGGATTGGTGATAGAATTATAACTTTAAAGATGATTGTTGATGATAATATTGTTAATATTATAAGT
GCTTATGCTCCTCAAGTAGGTTTGGATATAAATTCTAAATTAAATTTTTGGAATGACTTAGAAGAATTAATACAA
AAGATACCTCTTGAAGAAAGAGTTTTTATTGGAGGAGATTTAAATGGACATGTTGGTAGAGAGGTAGATGATTTT
AAAAGCATTCATGATGGTGATCATGGATTTGGAACGACGAATGTAGGTGTCAATATCATATTAGATTTTTGTTTA
GCTTATAATTTTTTAATTGCTAATACCTTATTTAAGAAAAGAAATGAGCATCTCATCACTTATAAAAATGGATAT
AATACAACCCAGATTGACTATTTTCTTATTAGAAAATTAAACCGTTTAGATTGCAAAGATTGTAAAGTTATTCCG
AGTGAAAGTTTATGTACCCAACATCGATTATTAGTTTTGGACTTTAAGATAAAGGGGCATAAAATAAAAAATAAA
GTAAAACGTTTTTCGAGGATTAGATGGGGGCATTTAAAGAATATTAACCAAAATATTTTTGTGAGTAAGATTTTA
GATCTTAAGATTTTGTACGTGCATGAGGATGCTAATATTATGTGGGAAAAAGTGAATAGTGATATTAAGAATATT
GCAAAAGATGTCATTGGAGTATTAAAGGGAAAAGCTCCTAAAAGTAAAAATAGTTGGTGGTGGAATCCTAATGTG
CAAGAAAAAATTAAAAATAAGAAGCTTTGTTATAAAAATTGGTTTAATTTTAAAAATTAAGATAATTGGAAAATA
TATAAAGTGGCTCATAAGGATGCTAAGAAAGCAATATGCGATGCAAAGGCATTAGCTTTTAGTGATTTATATAAT
AGATTAGATACTAAAGAGGGAGAAAGATATATGTATAAGTTAGCCAAACAAAGAGATAGGAGTAGTAAGGATTTA
GTTCAAGTGAAGTGTATTAAAGATGAAGATCAAAAAATACTTTTCTTAGAAGATGATATTCAACAAAGATGGATG
AGATATTTTGCTAAACTTTTTAATGAAGATTGTGAAAATAAGATTGACTATAGTGACCTTAGCACCCTAAATGAA
AATAAAGATTTCTATTTTTATCGGCGTATTAGTAAAACTGAAGTTAGAGAGGCTTTAAAGAAAATGAAGGGTGGA
AAAGCTGTTGGACCTGATGATATCCCAATAGAGGTGTGGAAATGCTTAGGAGATGAAGGAATTTCATGGCTTACT
AAACTTTTTAATACTATTTTGAAGACTAAAAAGATGCCAGATGAGTGGAGAATGAGTATGTTAATTCCTATTTTT
AAGAATAAAGGTGATGTACAAGATTGTGCGAATTATAGAGGTATTAAGTTAATGTGTCATACTTTAAAACTATGG
GAGCGAGTTATGGAGCATAGGTTGCGGCGTGATACTAATGTTTCTCAAAATCAATTTGGATTCATGCCGGGTAGG
TCGACTATGGAAGCCATACATTTATTAAGAGGTTTGATGGAGAAATATAGAGAAAATAAGGAGGATCTACATATG
ATATTTATTGATTTAGAAAAAGCATATGATAAAGTTCCAAGAGAAGTGCTATGGAGAGTTCTTAAAAAGAAAGGA

GTCAATAATGCTTATATTCAAATCATTAAAGATATGTATGCAGGAGCTATTACTTGTGTTCAAACACAAGGTGGA
TTGACCAAGTATTTCCCTATTTCGGTTGGGTTACATCAAGGATCAGCCTTAAGCCCATATCTTTTTACTTTAGTC
ATGGATGTTCTCACTAGACATTTGCAAGAGGATATACCTTGGTGTATGCTTTTTACTGATGATATTCTTCTTGTT
GATAAGACTAGAGAGGGAGTTGAAGGTAAACTTGAATTATGGAGGTCAACCTTAGAGTCTAAAGGTTTTCGCTTG
AGTAGATCTAAAACAGAATATATGGAGTGTAATTTTAGTAGTAATAGACCTACTGAAGGGATTGTGACTTTGGGC
GATCAAGTCATTAATAAGAGTACACGTTTTAGATATCTTGGATCTATTGTTCAGAGTGATGGTGAAATAGATGGG
GATATTATCAACAGAATTCAAGTTGGATGATTGAAGTGGAGGAATGCATCGGGCTTGTTATGTGATAGAAATGTA
CCCCTCAAACTTAAGGGAAAATTCTATAAGATGATAGTGAGGCCTGCAATGCTTTATGGTGCCGAATGTTGGCCA
TTAAAAGAGAAGCATAATACTAAGTTAAGTGTGGCAGAGATGAGAATGCTCAGGTGGATGAGTGGCTTTACCCTT
AGAGATAAGATACGAAATGAACATATACATGAAAAGGTTGGAGTAGCTCCAGTAGAAGATAAAATTAGAGAGAGT
CGCCTTAGATGGTTTGGCCACATAAAACGACGGCCTTTCGATGACCCGGTTAGGAGAGTAGAGGTATTAGATTTG
ACTTATATTAAGAAAGGTAGGGGTAGACCTAAAAAGACTTGGTTAGAAAATATTAGGAATAACCTTTCTTTATTA
GATTTAAACNNNNNNNNNNNNNNNNNNNNNNNNNNNNNNNNNNNNNNNNNNNNNNNNNNNNNNNNNNNNNNNNNNN
NNNNNNNNNNNNNNNNNNNNNNNNNNNNNNNNNNNNNNNNNNNNNNNNNNNNNNNNNNNNNNNNNNNNNNNNNNNNNNNN
NNNNNNNNNNNNNNNNNNNNNNNNNNNNNNNNNNNNNNNNNNNNNNNNNNNNNNNNNNNNNNNNNNNNNNNNNNNNNNNN
NNNNNNNNNNNNNNNNNNNNNNNNNNNNNNNNNNNNNNNNNNNNNNNNNNNNNNNNNNNNNNNNNNNNNNNNNNNNNNNN
NNNNNNNNNNNNNNNNNNNNNNNNNNNNNNNNNNNNNNNNNNNNNNNNNNNNNNNNNNNNNNNNNNNNNNNNNNNNNNNN
NNNNNNNNNNNNNNNNNNNNNNNNNNNNNNNNNNNNNNNNNNNNNNNNNNNNNNNNNNNNNNNNNNNNNNNNNNNNNNNN
NNNNNNNNNNNNNNNNNNNNNNNNNNNNNNNNNNNNNNNNNNNNNNNNNNNNNNNNNNNNNNNNNNNNNNNNNNNNNNNN
NNNNNNNNNNNNNNNNNNNNNNNNNNNNNNNNNNNNNNNNNNNNNNNNNNNNNNNNNNNNNNNNNNNNNNNNNNNNNNNN
NNNNNNNNNNNNNNNNNNNNNNNNNNNNNNNNNNNNNNNNNNNNNNNNNNNNNNNNNNNNNNNNNNNNNNNNNNNNNNNN
NNNNNNNNNNNNNNNNNNNNNNNNNNNNNNNNNNNNNNNNNNNNNNNNNNNNNNNNNNNNNNNNNNNNNNNNNNNNNNNN
NNNNNNNNNNNNNNNNNNNNNNNNNNNNNNNNNNNNNNNNNNNNNNNNNNNNNNNNNNNNNNNNNNNNNNNNNNNNNNNN
NNNNNNNNNNNNNNNNNNNNNNNNNNNNNNNNNNNNNNNNNNNNNNNNNNNNNNNNNNNNNNNNNNNNNNNNNNNNNNNN
NNNNNNNNNNNNNNNNNNNNNNNNNNNNNNNNNNNNNNNNNNNNNNNNNNNNNNNNNNNNNNNNNNNNNNNNNNNNNNNN
NNNNNNNNNNNNNNNNNNNNNNNNNNNNNNNNNNNNNNNNNNNNNNNNNNNNNNNNNNNNNNNNNNNNNNNNNNNNNNNN
NNNNNNNNNNNNNNNNNNNNNNNNNNNNNNNNNTATATATATATATATATATATATCACATATTTATATACGTTCATTCA
TAAAATTTCAGTCATTTTCAATTATAACAATTTCTATATCAATCTATGCATTTGAATTTATATTACCTTACTCAT
AATAGTCTTCAAAAAGAGTAAAATAATTATATAAAAAACTTCTTTCATATTTGAAATATTAAAATTTACATAACT
ATAATTACTCACTTTATTTGCCTTTTCAAATATTCGTCTTGAATTGAGAGCGTTTAACTGTTCACAATAACATAA
TGATTTAATTAATATTAAATTCTAATTTGAATGTCTCAACAAACATTTGATTTCTTCCATATAAAAGTACCTAAT
AAAATTTTTCCCTTCCTTGCAAATTTTCTTCTTTCATTGTCTGATTTAAATTCGCTTTAATTCAATTTTTCCTAC
AACATAATGTCTTGAATTAAGAGAATTCTAAGATTTTCTTTCTCCATTGCTTAAATAGAAATTCTTATGAAATAA
TTTCATAAGAATAAAAGAATCACAATCAAATCCTTCTTATTCACCTTTTTCTTCCTTTGTCGTTCCCCTTTGGCA
CGTACGTGATTTCCTCCTACAGGAAGAAGCGAACCGAAGACATGTCGCGTGAGGAGTCGTTAGTTTTCTCCCTCG
AACGAAATTACTTAAAATAGTGAAGTATACATCCAGAAGGCACTCGCACGAATTCGGCCTGGCTCAAATCATAGT
TCCGTCACTAGTTTTCCCGTCTGATTACGGGATTAATATTAGGGCTGGCCGCTTCAATTGAACCTGCCAATTAAA
CCGATTATTACAACCTCCATTTCTTATAAAAATTTCATCCTCAAAATTTAAAATCAAGTATAAGTAAATGTACCT
TAATTGAATATATGTGGATATTGAGCTTTCATTTATGCTTCTTTTTCCTAGGTGGCATCTCTTTTTGATCAATGT
TTCCATAAAATCTTTACAAAAAAATTTTCTCGACTCCTCAGTTGTTTCACATCATATTTCAGTATTCTTTCAGGT
TCTTTAACATATTTTAATCAAGCTGCAAATCAATTAATTTTGACTGGTACATACTTCTGAAGCATTGATATGTGA
AACATATTATAAATTCCTTGCAAATGCACTACCAGATCCAATCTGTAGGTCATATTTCTAATCCTCTCAATGATT
TCAAACGAACATATGAATCGTGAGCTTAATTTTCATGTCTTCCCAAATCATTGTACACCTTTCATTGGACTGACT
TTAAAGAAAACAAATTCTCATACTTCAAACTTAACTGGCCTATGCTTTGTATCATAATACTTTCGTTGTCTGTTT
TGTATTGCCTTTAGTCTATCTCTGATCAATATGATTTTTATGGTAGCATCATCCACCAGTTCAGTATCAATTATT
TTTCTTTCACATGTTTCTGCCCAACAAAAGGGGGTTTCGCATTTTCTTCCATAAAAAGCTTCGTATTAAGCCATT
CCAATGCTGGATTGAAAACTGTTATTATATGCAAATTCAATTAATGGCATACAGGTCTCCCAATTTTCTCCAAAA
TCTAATACACATAATCTAAGTAAATCCTCTAAAGTTTAAATTGTTCTTTCTAATTGCACATCAGTCTGAAGATGA
AATGTTATACTGAAAATCAATATTGTACTCATAACTTCTTGAATCTTCTTCCACAATCTAGAAGTGAATCGGGCA
TTTCTATCAAAAACTATTGATTTTGAAATTCCATACAATCTCACAATTTCATCTAATATATATCTTTGTCAAATT
CTCCATTGAATCTATTTGTTTGAATGGTAAGAAATAAGCTAACTTGGTTAATCTAACTATAACAATCCACATAGC
ATCCTTACTCTACCTTGACCTTGGTAATACAATTACAAAATCCATAACAATTTCTTCCCATTTTCATTCAAATAT
CGGTAATGGTTATAATTGACCACTTGGTTTTTGATGTTGAGCTTTAATCATCTGACAGAAAATATATTTTGCCAC
AAATTCGGCATCTCCTTTTTAATCCTGACCACCAAAATAACTCTCTCAATTCTTGATACATCTTTCTACTGCTTG
GATGAATTTCATGGCCAATGAGGTGAGCTTTTTGTAATATATCTTCTCCTAAATTGCCAACATTTCAACACATAT
TCTACCTTTATACTTTAGCAATCCTTCATCATCAACTACAAATTCTCTAACTATTTTGCTTGCAATTTCTTTATG
ATGTCTTCAAATTTTACATCTTGAGATAATGAATTGAATACTCTATCTTTCAAAGTGGATTGTATCACCAGCTGA
TTTAATAACATATCTTTAATATCCTTCTGATTGACCACTAATATCTCTAGTCTTTTCATGTCTCTGATTATCTGC
TCCTGCTGGGCGATTATCAAATTTACACATCCAAGCATTTTTCTGCTTAGTGTATCTGCATCTATATTTGCTGAT
GAGGACGTCCGATTGAGAAAAGTTTATCAGAGAAGGGGCCGAAGATCTATGGCTATGATTGGACCATTTTATTTT
TGAGCTTTCATGTATATTTGATTTATTTATGTATCTTTTTCGTTTTTACCATCACTTGAGGGAGTAGGCTTATGT
GAACCCAGGGGGCACTTTTGTCATTAGACATGTTATTGTATTTTGGCTATTTATAAACTCATCCCCCTCTTTGGA
GAGGATGATGATTATTATGAATGAAAATTTTCTGCACCTTGTATCTTGGAGAGTTTTGAGAGTTTGAAGTCCCTA

AGTGAGGGCAAGGCATCACCCGTCAACCACAGGAAGCGTCCGAGCTCACGGTCAATCTCCTTCACAAACTTCCCT
CTCCTCCTTTCCTCCAAGCCGCCGGCCTCCCTCTCCCCCTACTCTTGGTCGACTCCCTAGCAAACCGCCGGCTCA
ACCTCCCTTCCTAGCCGCCGGCCACCACCCCTTCCCTTGACTCCCAATATTTCAAACCTAGCATTCAATACTTTC
CTAACTTGACTCATTATAAATTTGGAAACTCAATAATTTCATATTCAAGTAAATTTGAATTTGAATTTTCTTTCC
AAAATGAGAAGATTTATGTTAGGAAGGTTTTCCATAATCATGCTAACATTAATTTGAATAGAGTTTACCCTAAAT
TTTCTTCTCCATTTAGTGTCCTAAATTTAGCAACCTCTTTAAACCCTCTCATTAATTCTCTCTCCCTAGCTTTGC
CGCCCCACCCCTCTCTCCTCTAATTATTACTTTCCCATTAATGCTCCCCAATCAATTCCCCTCCCATTAGAATGT
GCTGCCCCCTCCCTACCCCTAATTTAGGGACATTCAAATTCAAATATCCCCCAATCTCTCCACTTAATGCAACCA
TTAATTCCTATTAAACCCCAATTGATTCTTGTCTTAATTTGCATGCCACCTAGACAAGAATTCAATTCCAGAATT
TTGTTAATTGCTGGAATTAAATTCAAAGTTTCCTATCATTCCTTACTCATCACCCTAAGTAAGGGATTAAACACC
TCTCTAGGACCGACCATAAGCCCCCCCCCCCACCCCCCCAACCCNNNNNNNNNNAACCCCTCCCCTCCCCTCCC
CCAACCTCCCTACACTTCTCCTTCGCTTGATGTTTTCACCAAAGAAGAAGAAATAAATCCTCTTCCCTTGGCTGG
CCACATCCCAACCATTCCGGCACTTTAATCCCTTGTTGACTACAAGGTGTGTTTGCTTCAAAGCGCATCCACGAA
CAATCGACAGCCGGATCATGGAGAATGTTGGAGCCAAGTCCAAAAGCTTGAGATGATAAACTCTTCTCCTAAACC
TTTTATCTTTAACTCATCGATTGTAGCATCGTTAAAACTATGAAGGCTTCTCCGCTTATCCCATATTATGAGCAT
GTATGAGTGGATGCTTATAGTGACTATTATAATTATAAACTGAGGGATTTTCTTTGAAAGTCCCACAACAGTAGG
GTTTGATTTAATTTGATTGTTTTGTGCTCCTAATATTATCATATAATTGATTGTCTGAGCTTAGACACTATCTTA
TTGGCTTCAATCTTTTGTTTAATACGTGAGAAGTTGATTAATTGTTATTGTTCTTTTCATGAGCGTGAAATTTCA
AATTTAATAACATCGCATGTGCATGTTGTCCTACATCATTTGCCCTTCCAGGATGATATTGTATATCTAATTCAT
AATCTTTGATAAGCTCTAACCATTTTCTCTGCCTCAAATCAAGCTCTTTTTTTATTGAAAATATATTTGAGGCTG
TTATGATCTGTAAACACTTCACATCTGACTCCATATAAATAATGTCTCTATATTTTGAGTGCAAAAATCACAGTT
GTATGGGACTAAATTTTGTTGTTTAAAAGCAAAACCTATATGCCAACTACTGGACAAGGATGCCGGTGCGGCGTG
CAGCGGACTAAGGACTACACATAACCGGGTAGAGCAAGGTGGGTCGATCAGACCACAGGTGACCGGACAGGGCAA
GCTATGCAGGTGGGTGAGTCGGGGTAGCGCGGGTGAGCTGCAAGGTAGCACAGGTCAGCGGGCATGGGCAAAAAG
AATTCTAATTAAAAAATTAAATTTGCTGATTTTATTTTAATTTTGTGGCGGCAATGGCCGCCAGAAAAATGAGTT
TTTTCACATATATAATAGTTTTTGTATGCGCGTTTTACCGGCGAAGAAAATTTTCCGACGGAAATCCTCTAGAAA
TACTACTTAACGATGGATTTTGTTCTTTTTCTGACGAAAATGACAGTCGAAAAATGTTGTTTTTTTTTTGTAATG
ATTCCTCGAGAAAAAACTAAAAATGGAGGGACCCCTTAAGGTGACTAGGATAACTACTTAATTGATTAAACATAA
AAAAACCCTCTTGGTTGGATTTGCTAAACCAGTTGGGTGAAACAACCGGTCTTGGGACATCTCCGACATCACACC
ACTTACAGGTTTCAACAATCATCACCATATCACCGTAAGCTTCTCTTTGCTCACAAACTAAACATAAAAAACTAC
AAATTGGCATGTCATCTTCTTTATAATTAATTTCAACAACACTTAATACAGCTTAATTATTAAACAATCACTGAG
AATTCTTCTTAACCAAATTCCTTAAAATACTCTAAGAGTCTACCCTAAACTCAACTTCAACACTGCTTAGGACAA
CCAAGGGTTGCCTTTTAAAGCAACACCCTATCACCACAGATGAGCCTGTCTAATTAACATCAGTGAAAACCTCAA
TCTTCCTATCTGAGTTCCTTTTGAAGAATAATCTTTTTCATCGATTCCTTTTCAATATCGCTAGATTCTGAAAAC
AATATCCATATTTAAGTTGGGATATTCATAAATCAATTTACCACACCTCTTGGCAATTCAATATTAAGTCGGGTG
TGAAACAGATAAATGAGCCTTAACTATAAGTCTTCGATAAAATTTTTTATCTACTGGAGCACAATCATCTTTGGT
TCCCAGTTTGCCATTCAAATCCATAGGTTTAATTGCTAGTTTACATCCCAGCATTCATGTTTATTTCAAGAGATT
TGAAATATATTTGCATAGTGAGACCTAAATAACCCTTATGAGATCGAGCCACTGCATTCATGGGAAATATTTTAG
ATTTCCTATGTTTGATTTGAGTTTGCTTAATTCTTGTACAAATTCACCTCTTCGAATGATATCATCAATATACAT
AATCAGAATAGCAATTTTACATTCACGTAATCATTTAATAAAGATAATATAATCAGCTTGGCCACTTTGGTGAAT
CTATAAAAAGAAATATGTGGGAATTAAGACTAAAACCAAATACAAAGAACTTGATTACACCAGAAAAAGAGAAAA
TGATGTCAGTCCTGTTAGATATCCAAGAATGGAGTTCTTGCTACTCCCATGTTTCCAGCTTACACTCCTATATAT
TTTGGTCCTCGTAGAATCTTGTTTGGCGTTAGAAGGAAACGTTGAGGCTTGTCATCATCCTTGCTAGTTCAATAA
TTCCCTTCCAAAGGTCAAGAGATGCATACAAGATCATAAATACCCGGTTTTGATGCTGCCCGTGACCGAATTGTT
GGTGTCGGAATCTAAACAGTGACCTTCCATTGGGTTTTCAAAAAGGTAAATTAAGGGACAACGCGAAGAGAAGGA
TTGTCAAGTGTTGGAATCAATTTTGGCAGCGCTCTTCTATAATAATTTTTCATGTTGATATTTCTGGATTTTCTG
CAGCAAGAATATGGCAGACGGTTACTTGAATTGCAACGATAGTACCAAGTTAGATGAGAAAGAATAGAAGCTTTG
GTGATCAAGCGTGACGAACTCTGTGATCTCCTCTTCCCACCAAAATAGGAAAAATATTCCCCAACTTCAAGAACT
AAAACAAACTTACCACCAAAGAGTATCTTATAGACCTAGGGTGAATATTTGTTTAAGGGGATAAATCTAAATCTA
GCTTAGCTAACAGAGGGCCTTGTGCCCTATCAATTATATCCTATTGATTCATATCTCAAGAATTTGAAGTCCTTG
GGAAGCATACAAGCTCGGAGCATGACCGTAATTTACTGGTGACATTTGGTGGCAAAATCACCCGATTCAGAGGGT
CTCGGTCATAACAGTTTACCTATGATGCCATTAACGGCGAGCCATCGGTGTGGTGCCATTTGGCAGCGAAACCCT
TGGTTAGAGGACATGTGTCCCGTCTTTACTGACAAGAGACAATTTGAGAATGAATCCTAATGATATTCTCGATCA
AACATTCGCTCTTTGTGAGCTGAAATGATTTTCACATTTGTCCGTTTCGGCAAATTGTTGTAAAGGGTGGATGGG
TATTTTGTTTTGAGTGTTTTGGTGTTTTATTTTATTTTCAAATATCTTTATTTTGGTGAATTTTGGGTTGAGTAT
TCCTGGGAATTTGGGAGGTATATATAAATACTTGGGGATCCCCACTGGTTGAGATTTTCCATCAACACGGCCATC
ATCGGGATACTTATGAGCAATTTCATTTTATTGGTTGAACCTTGTCGACTCTGCTTACTTGCCAACAGAGGATCA
GATTCATTGTCAGCTTTATGCATTTATTATTGAATTTTCTTTGTTATAGACCTTGAAGCCCTAAAATGAACTGTA
TAAAATAAATAACTCAGTCGATGCTGAGACCTTATAACTATTTGATAGCTCGAGCTGTGTAGACTCTTATCCATC
TACATAGAACGATCGAGCTCGAACTTAAGCTTCGTTTAGGACAGCTTTCTCACTGCTTCTTAAAGCAACTTTTTA
GTATAAAAATTTTAATTTTTGTACTAGAAAGCTGCTTTAAGAAGCAGTAAAAAAGCCGTCCCAAACGAAGCCTTA
GATTCATTTATTCACTCGCACAGCATCCTCAGACGTCACGTGTGTACCTATAACATGCTCGTCGGGGACAATGGG
GGTATATGCTTTTACATGTACTTTGCATGATGTGCTTGTATAGTGGTTGTATGTATGCAGATATCTATGATCAAC

ATGCATACTCGCACACACACCTATACCTTCATCTGCATATTAATATACTTGTATGTGCATTTATGTATCTTTATG
TATGAATGTATGTTGGCGTGTGCATTTGCATTTTGTATGATATGCATGATATGCTTGATATGCATTATTGTGGTT
TAGCGCGTTGACATGGCATATAATGCGTTTCTACAATGTATGTATAATGTGTCCATGCATCATCAATTTTATTTG
CACATCGATTCAGTACTTTTATACAATGCATAAAGCATTTGTATAATGATGTAGCGTGTTGTCACTACGCAATGC
ATTTTGCATGTTTAATGGCATGTATGCATGATGTGTTTTGTATTTTGCAACATGTGTATGCATGATTTGGATGCA
TTAGTGCACATTTCAATGTCTGGGGTGTTTTTGCAATATTCAAATATGACTAATTGCAAAGTTACATTTAACACC
TACAAATGCCTTCGATTAACTTGTATGAGTGTTGCGGATCCAGTGGTGTAGCATTGCACTATGATTATTTGCTGC
TTCTTGCAGATGAGTTGTTAAGAAATGCACAGGGCTCCACTTGAAAATTTATAAATGTCTTATGTCCACTTGTAT
GCATTTGTGAATTCGGTGTTGTTTAGTAGTGCACTAGGTCTCATTTCATCTTCTGCATATTACTATCTTCTTCAG
CATTTAGCCTGCGAGATTATTTTTGCCTGCAATGTGTTGCATAACGCATCAATGCCTTCCATCTATCTTTTATAG
ATAAACTGATTTATCTAGCATGATAACCTAGTGTCGTAATTTTACGACCTCAATTTCTTAGCATATATCCCATAG
TTTTTACCACTGATTCCCTACTTTGCTTACAGGTGAATCACAGCGTTGTGATATAGGTCACCCGTTGCTTCTTAC
AAGCGAGTTATTGTGCATTGCAGGCTTTGTTGAATTTGTTCGTCAGTAGTGCGCCATCAGCAGGCGTTAAATAAT
CGTCAACGGTTGCATGCAAGTGGACTACATTCAACATATATTTTGCTCGTAGGTGGGTCATAGAGCTGTGTTGTG
GATTATCAGCTGCTTTAAACAAATAGACTGTCGTGTAGTAATGAATCTCCTACAGTTGCTTGTAGATATTTGTAC
CCATCACAAGCTTCTGTGTTTGCTTGTAGATAGCTTGTTGTGCATATGACGAGTTTTCTACAACAGCTCGTAGAT
GACATTTAGTATATATGGCGGGCCTCCAACAACGGCTCGTAGGCATTTGCATTTGTTTCCAATCGTTCACCCTTT
CCACGTCCTTGCTGGCAAGAGGCTGTTTGAGAATGAATCCTAAAAAATATTCTCGACTAGACATTAGTTCTTTGT
GAGCTGAAGTGATATTTGAATTCCTCCACCTCGACAAACTATTGCAGAGGGTAGATGGGTCTTGTTTTAGGTGTT
TGATAGTTGAATTCCAATTTTTAAATATTTTGAATAGTTTTGAAGTTTCAAATTTTGTGGGCAATTAGGAGGCAT
TTTGAATACTTGGAATTAATTCCTCCAGTTGAGATCTACGTTCAACAAAACCATCATCAGTATATTATGGGCATT
TGTATTTGGTTGATTGAACACTGTCAACTCTATTTATTGTCGACAGAGGATCGGGTTCAAGTCGGCTGTATGCTT
TTATTTAAAAAAAAAATCTGGTTATAGGACTTGAAGGTCTAATATGAATGTATTAAAATAAATATCTCAGTCGAC
GCCATGATTGTAGATTTGTTTGTTAGTTCAAGTCATGTCAACTCCTATTTACTAGCATATAAGGGTCGAGCTTGA
ACCTTTGTTTTTCCAACGCAACATCCTCGGGCGTCACCTGAGTACCAACAACACGCATGTCGGGGCCCATGCAAG
TATACGATTGCACACGTATTTTCTTGTATTATGCTTGCATACTCACATATACATTTACACTTTCATTTGTGTATT
AAGATGCTTGCACGTGCACTTCTGTATTTATGTGTATACATGTTGTAATGTACGCGTGGGAGCATTTTAAAATGT
TTAACTCTGATCAGTTTTGCGAGCAACTCTTGTTTGAGTGTGCATAGCATTTAATTTGAGAGCATATAGACGTCT
TATGTTCGATTATATGAGTATTGCAGATCCGGACGTGGCATCACGTTGCGTTATTAACCTATTGCTTTATACAGT
TGGATAGTTGTATTCATCGTCAGCTTCTGTATTTTTGCCTACAAGAAATATATTGCGCTAGCTGCAAATCTTTTT
CATTCGTGTTTGCAGGATATTCATTATATTTGCACCACATTAGTTGTAGGTGACTTGATGTATTCATTTACAATT
AAAAGGTGTTGATAGCAGGGTGTTGTTGCGGTCTCCTTAAATGTGTTCGTAGGATAGCTGACATATAAATGACAC
ATCTAGCATTATCACTTCCTGATAAGCTAGGAGGATTGATGTGTTCTCTTGAAGTTGAATTGTTGTCAATTGCAA
TTAGGAGACGTTGATCATATGTAGGTTGTATCAATCTTCATTATTCATAGATGTTGTATTGCATTCATCCCACTT
TTTTTTTACACTTTGCTTGCACTTAGATTATATAGTTCCCTCCATTCGCTCGATTGTCATATGTTGATATTCAAT
TGCAAAGCCGATACGAACATAGCACTTCAATCATCAATACCCTTGAATATTTTGCAGTGCTTGTTGGAGTATCTT
TCTGCATTCGTTTGCAACTAGTTCACATGTTATTTTTAATTATTGTTGATCATTGCAAGCAAGTCAAGTTTGGGA
TTTCATCCAGCATTTCTTATTCATTTGGATGATAGGGATCATTTGAATTTTCATTAAGATATATATATATATATA
CCTCCGCGCTAGAGAAAAGCATGTACAAGGCCTACCTGCCTCATCGTAAGCATTGCATTGGATTCTCTTCCCACT
ACGCTAGAAAAAAGCGTGTATGTGCAATTTTTTTGTGCGTCGCAAATGCTGAAAGCGGCAATTTTGAGTTAAGAG
CATTATACCTGTGTGACGAGAGGTTGACGATGGTGATAGTGTCGATGGAAATGACAAAAAAAATCCCTCTCTTTT
TCTTCTCTCTTTCTTTTTATTTTCTTACTTTTTTTTATTTTTCATTTTTTTTGCATATGCGTATATATATATATA
TATATATATATACACACATACATACATGAATGAAATTTACAAATATGTTTATACAACTATTGTGTTATAATTATA
ATGATTGTACTAAAAAAATCTTTTCAATCATGCTTCTATGTGATTTTAGTTATGTTAGTGCTGTTGTCAATAACA
CTTAGCATGTCACTATAAAATGTCTGATAAGAATCTTATTCATCCCTCTTTAAACAATCAATCAAACCTTTTAAT
AATCATGTCAAATTATCTTTGATAAAAGGACGAACATATGTTTTATTAGGTGTTCAAATAAGTTTCAACTATTTT
ATAAATTTTTTCAATAATGTGTTTTAATCACCTTATTCAAATATGTTCAAACTTAGTTGATTCTATGTGCGACCG
TTTGATCGATTACATCAGATTATCCTCTGATGGAAGGGCCGACCATATGTTCAATTTAAGTGTTCGAATAAATTT
CAGCTATTTTATAAAATGTTCTAATAATTTATTTTAAGAACCTTATTTCAACATGTTCAAATACTTAACTGAATT
TATGTTCAACCCTTTTATCAATCAATTTCAAATTAACTTCAATAAAATGGTCAAATATAGATTCAATTAAATGTT
CGAATAAGCTTCAACTATTTTAAAACTTTTAAGAGCAAAGTAACAATGATATTCAAATTTTGTTCAATATTTATT
CGACACTTAATTGAATCAATATTAGCCCGCAACAAAAAAGAGGGTATTTTAGTTTGAAGGGAAATTTTGGGCATA
TCTTTCAAAGAAATAAAATAATAAACACATTTCTCCAATTTTAAGCATTGTGAGGCATAGAAACTCAATTTTTCC
AATCTATTTATCTCTCTATATATAATCCTAAAATAGTTTATATCACCTTCTCTCCATCTGTAACGTGTGTCAACC
TCCTAAGAAAGGAATATATTTTAAATTCAATACAGAATCTGGACAAAAAATCAGCATTGGGACTTAATAGAAAAG
GAAAATGGAAAATGGAAAATCGCTAGAAATACATGTACATGACTAATGAAATAATTAACATTTATATATGTGGAC
AATTGAATAACCTCAAAGGGTCTAAAGTCGCTTGGGTATTCAGTTTGCTGCCCTAATAATGGACATTTGAAAATT
AAGAGAAATCATGCATGGAACAAAGATCCTAGAGAATAATGATAGGCCAAGAAAATAAAGTTCAAAACTATAACA
CGGTTTTCTTGGTGTTAATGACTCTGCAATTGCTATAGAAGAAAACTTTTGCACAATTTACAATATATTAGCATG
CACGAATATGAGTAATGAAAACTACATCCTAACCAGAATGTTTTAAGTTCAGTCAGTAC**CCATAAAAATTTTAAA**
**CATGATCTTGTAGACCTTCATTCTAGGCCATATGCTAGTTATGTTGAAATCCATCTAGATAAGTTGTGTCAGTAA**
**TTCTTATTTTATATGCTTTTTGCGCATAATTGAGCATTCCACTTTACTAATTGATTTGTTTGGTGTCCACACTAG**
**ATCTGGCAAAGGGAAGCAACAAGCTTGCGGCAGCAATTGCATAACTTGCAAGAAAATCATCGGTAACTTCTCTTC**

**ACCCTTTGAGTCCATCAGAAATAAAATA**CTGCATGTCAGAGAAAAAATAGCATAAATGAAAATTCATCTCTTAAA
TTACAAGCTTATTTATCTTCTATGGAAAGATGCAAAATAACCCTATTTCATGTTTTGATTCTGTTTTCAGCCAAA
AATAGCAAACGAATATAGCAAGTCAACAGCTTATATCATTATATTTGGTTAATGGATCATCAAATAAACCAAAAA
AACGATGCCCTGGCAAAACTACAATATATTAAATTGCATCAATACGGTATATAACAATTATATCAGTTGCAAAAT
CGTTAGACTAGGTTTCTGCAGAATCAAATATTAAAAGTATTCATTAATTCAGTTACTTCACCAAAAACTACAGAA
AATTTTAAGAAAGCAAGAGAAAGTTCTGTTGGCTAAAATTTTAAATAATGAGTGGTGAAGATAAACAAGAAACCA
GAAGATAATGAGTTAATAATTACAGCTGCAAACTAATTTACAACTACGGTTAATCTTTTTAGCTATGTTTTCTAT
AATAACTATAATGAATGTAGATTTTAACGAGAATAAATATTTATTGAAAATTAATTTACCCAATCTATCTTAAAG
CATAAGAAAGGAGTTAATAACTCTTATATTCAAATCATTAAAGATATATATGTAGGATTTATTACTTGTGTTCAA
ACACAAGGTGGTTTGACCGAGTATTTCCCTATTTCGGTTGGGTTACATCAAGGATCAGCCTTAAGCCCATATCTT
TTTGCTTTAGTCATGGATGTTCTCACTAGACATTTGCAAGAGGATGTACCTTGGTGTATGCTTTTTGCTGATGAT
ATTCTTCTTGTTGATAAGACTAGAGGAGTTGAAGGTAAACTTGAATTATGGAGGTCAACCTTAGAGTCTAAAGGT
TTTCGCTTGAGTAGATCTAAAATGGAATATATAAAGTGTAATTTTAGTAGTAATAGATCAAGTGAGGGGATTGTG
GCTTTGGGCGATCAAGTCATTAATAAGAATACACGTTTTAGATATTTTGGATCTATCGTTCAGAGAGATGGTGAA
ATAGATGGGGATTTTATCAATAGAATTCAAGTTGGATGGTTGAAGTGGAGGAATGCATCGGGCTTGTTGTGTGAT
AGAAAAGTACCCCTCAAACTTGAGGGAAAATTCTATAAGATGGCGGTGAGGCCTGCAATGCTTTATGGTGCCAAA
TGTTGGCCATTAAAAGAGAAGCATAATACTAAGTTAAGTGTGGCAAAGATGAGGATGCTTAGGTGGATGAGTGGC
TTTACCCTTAGAGATAGGATACGAAATGAGCATATACGTGAAAAGGTTGGAGTAGCTCCAATAGAAAATAAGATT
AGAGAGAGTCGTCTTAGATAGTTTGGTCACATATAACGGCAACACGGCAATACATTTTTTTATATAGTGTATATA
TATTAATGTAATATCATATTAAAAATGACAAACAATAATAAATGAATGACATGATAACTTGTCTAGCTTATAATA
TAAAATATCTAGTACAACCACAAAAATATAAGCAAATAATCACTTACTACAACATCAATCATTCAAGCTTCAACT
ATTAAGATTAACTTTAGAGGTTTGTTTTTGGCCTTCAATATCATCATCATTGAATATCACTACTTTCAACTCTGG
TTCAACAAGAGAAAGATTGTACTTGTGCAAAAAGAGAGGAGATGAAAATACCTTCACTGACAAAGAGAAATAAAA
ATGAGAATATTTGATAAATTGGGGTATTAAATGCAGAAGTTGATTTGGTGTTGTGGATATGAAATAAGAAAGTTA
GTTTAAATCTCGTCAAATATAAGAAGTTATATTTAAAAAAAATATATAAAATATTAATATACGTGTGTCGGACAT
GCGTGTCCAATTTGTTTTTAATATTTTTTTAATCGGACACGTTTTCGGCGTGTCCGACACGTGTCGGAGGAGTGT
CGGACGAGTGTCGTGTCGGACATGGTGTCGGACACTGCACGCCTCTTTGAGGAGAAGTGTCAGTGCATCATAGCT
ACTAGTTTAACATTCAACATTCTGCTTCCAGCCTCTAGTTATGGCTTCCTAACGCATCCCGTGCCAACACCCTAA
CTACATGTTGCCTTACACTATTATGATGATTTCTCCAACCACATGACCTGGACTTTGTCTGAGCAGCCAAGCCTC
CTCGTGAAAATCCTCCAGCTGCTGTGTTATGTTCGCATACCAGAATCACTCTCTATATCACATGCTACAAGAGGA
ACTAAGTTATTGAACCAATCAAGCCCCGCAACTTACCTAACTTTCTGAAACTGCCTTATCCCAACCCTTCATGGT
GTAAGGTTATGGGCATTGTTATTCTCATACCATTTGAGTGTTAGCAAAGCTTAAGGATAAGGAGCTCGAAAATTG
GATTGAGGCCAATAATAAACTGACGATATATGATGATCTAATGCAATGCCGCTCTTTTACCTTTCAACTAAGGCA
TTAATTATTTGGGAAATTATCAGAAATCAGATTTGGATGTTAAAGTTTTTATTGCCTTTATGTCATTGAAATCAA
AAGAATAATTCTTGAAGCTCTATATGATGATGATCTAATGGATGAGAATATGAGATTATTGGACCGATTGCTTGA
GCTGCTAAGAATTTGAAAGAGATGTTAAGCCATTCATTTTCTAAGAGAGTTTCAATATGTGAAACAATTGTCAAG
ATTTGACCTTGTAACTAAATGCTAGATCTTTAGTGTAATGATTTGATTTTGGAAGAGTTGCATCACTTTTTCAAG
ACCATAAGATCAAACCGTTCAGATTATTTTTTCTTTACAATGGAAATTCTCAAGACACTCGTTGAGAGGGAAAAT
ATTTCAGCAAGGCTATTTCATGTCTCTTGAATAATCTTAAAATTGATGAAAGATATGTATGACATGAATATTATG
CCATCAACAAAGAAGCTTGAAAAAATGTTGGCTGATTAAGCCTTGAACCTGAATCATTATTAGGCATAGTAATTC
AAAGGCACATGTGCACGTTGAAGTGCTCATAGCAAAATAGTTACTATTGTATATTAAGAGAACGTTAAAACAAAA
TGCAAAATGATATGAATAGCTTTAGGGGCTATCAGTTGAGTGTATTTAAACTCAAGTGTATTTTAAATGTAGGGT
AAAGTAAAATCTAATAGAGTTATTTGGTTGATTAAATAAATTCAAGAACCCATTTTTCTAGGCTAAAAGAAAATT
AAATTATAAATGCTTGAATGCAAAGAGTTAGAGCACAATGATGTTGTCTTTCCTGATGAGGTTGACAATGTTAGT
TGTGGTTACGACAATGGGTGGTCTATGATGAGCATTGGCGGTGGGACGTTGCCATTGGAGCTGAAATGAGCAGTA
TATGCAAGCGTTAAACATAGGAGGAGCATGCACAGAGTGGGGACTAGCTCTCCACAAATGCCAGGACACACGGGG
ACAAGCATATTGTGGGGAGGAGCACTTCGACAAACATGGGAAGTTGAAATTCGCTACATTCAAGGGACTTTCAAA
TTCCACAAAAACGAATTAAAGGATTAGTTCAAGTTCCATAAAAAAATTACGCTGAATTTATTTTTCACTCAACCA
AAAAGTCAAACGTGTAAAATCTCCAAATTTGAATGCAATACACTCCTAAAACTCTCAACCAAATAGCACCTTCCT
GAAACTTAGGTAGATGACATCAAATTATCTGAATAAGGCTCCTTGCGATCAACTGTCTTAGAGATCTGCAAAATA
AGAACACTAATTGACCTTCGTTAAAAATGTATCATAATGTATAACAGTCATGCATCCAATTCAAGGATGCCTGAT
GAGCAATGGAAATTTACGAATGGGGAAAAGGGATTGCAGGGCTTCAGAATAAAAAGCAGAATTTTTTCAAGCCTA
GATATGAGAAAGAAGCACTTATTAGGTCTCGTGATGAAATATTTAATACTAAATTTGATTGGATCTGGATCATAA
AGGAAATAGAAGTCTCTAGTTTCTTTGTCTTGTTGCTCCATTCACAACACTCTCCCAGATTGTGTGAACCCTTGG
CTAATTTTTTCCTTAACCATATGCTAGAGGATGGGTCTTAGCCAAGCTTTTGCACAGGTGATCTTATTAGAATTT
AGAAATAAAAATAAACGAACTTGAGTTTGAGGAGCCTCCGGCCACTATTGTTGCTCGACAACCTATATAGATGAA
AATGCATGTTGTTCCCCAATGATTATTGCTCACTTCATCACTTGTTGACTGTCTTTCCAATTTTTTTCACCATCA
GATTGTCATATTGTTCATTAAAAAATGGCATAAGTATGGAGATATCAAAATTCACAACCAACAATTGTGATTGTT
GATAATGCTAATAAATTCCTTATATTATTAATGAATTCTTCATTTGTAATGGTTAACTCTACATTCACACCACCA
TCGCAATCAAATTGTCATGGGATTTTAGCCATCTAATTCGGGATGTACGTCATCGTTCAAAGCTAAAGAATAATA
ATCACAGGATATCAGTTTCTTATTACTCTACAATTCACAATACCCATGTTCTAGCTCCACCTCCACCTCCACCTC
CGCTGCCGCCACCACCACAGTCACCCCCAACAACACTAGAAAATTTTCTTTTGAGAAGATTGAACTAATATGAAT
CTCAATCATAAGATTCTTCTTCAATGATCTTCACTCAGTCCATTTAGTTTTTGCACATGTTGTAACTATGTTTCA

AATATGTAATAAGTACAAATTATTCAATTTATTTCTTCCTTGATGCATTTTCGCAGCATTTAATAAGCCCTGTTT
TAGGAATGTTCCATGATCCCTAATTCATAAAAGGAGCTCTAGTCTTATTATATAGGATCTAATGGCTTGATAAAA
TGTTAATTGAACATTACGCTTATTTTAATTTTTTTAATTTCATCTAAGTCATTAGCGGGTATTAAAGTAATCGCA
TAAAAAGCATCAAAGCTTCACCTATCTCTCCATGATGGATTTATTAATACCTCCTAAGTGAATCATTTAGTATGC
TTAGGCTACAACCCTAATTTGTTCAATACTCTGACCCCATGTCCTTTGAACTTTCCAACCAAACAATTTATCTGT
GTTACAACTGTTGTCACCCAGCTTAACATTACAATCAACAGCAAAACATAACTTGTAAAAATATATGAGGCACAG
TAAGAATCAATGCTCAGCACGTACCGCAAGGATCACATTAATCGCAGATTATTTTAACTATATATCAAGAGGAAT
TTGAAATAAACTGTTTATGCTATAAGATGCTGTGACATCCTTGTTGATGCTAACAGGCAGCTGATGGGAGAAGAT
CTTAATGGGCTAAGTATCAAAGAACTACAAAACTTGGAGAATCAGCTAGAAAAAAGCCTCCAGGTGATACAGCAA
AAAAAGGTCTGTAGAGCAACCACGTCAAACCATGCCCAGTTATATCTATCATTTCAACTATTTTTAATAATTTTT
TGACCTCCTTTGTAGGACCAAATTTTAATCAATGAAACTCAAGAACTGAATCAGAAGGTAGTTCACACAAATCTG
ATTTTATAATTAATCAGCTGCTCAGACTAAATAATAAAATCTTTTTTTTCAGATTGCCCACATGCAACGAGAAAA
CATGGAACTCTACAGAATGATAAACCTTTGTCGCCAAGAAAATATGGACTTAAACAAAAAGGTGCTCGTCAAAGA
AGAAATCAACTCTAAATAGCAATAAATTTTAAATCCAGCAAAATGTAAATGTTATAAGTAATTAAGCTGATTTTC
ATATGCTGCCTGACATAAGAATTGGTTCTCAGGTTAATGCAACTAAAGGATTCAAAGGAGGATCTACTTCTTCAA
ACTCTGGCAGTCCGGTAACAATTTATCTTGAATTGAGCCCTCCGCAACAGAAAATTAGTGGAATGCACGAGGACT
TCCCAACATTAGGGTGAATTTCTTTTTTATTCCTTTAAGGGTACTGCACATAGTCATCATCCTAGATTTAATTGA
GGAAAACTCCTTGAATTTGCAGCAGACTCCAACTGCTTTAAGATGTTCCAGGAAAAGTGGATGATTCTCCGTGAT
CAAGATTCTGTACAATAGGAATATTACATTCCTTGGAGAATGTAAGCTCAACACTCACAATTCAATTAAAGTCAA
AGTTTGAAATTACCACAAAAGAACGAGTACATATGTAATGCTATATAATGCTTAGTGGATACAGTATTTTCTTTG
ACAAACTTGAGCTGTCAACTTCAGTATGACTTTAACAAACTTTTCTTGAGTATGACTTTAACAAACTTTTCCTAA
TGTATGGTAAATGAGACCTACGCAACTTCAATCTCAGAGGAAAAAAACATCCCATGGTGGCAATTCAGTAATTAT
TCTTTTCCACCAAATATGAATCACATTAAAATGTATTCTAGTAAGTAGATTGTCCGACATACATTAGACTTCCCA
ACATGTGATTTTTTAACACCATCTGAATTAACTAGACCTAACCAGCAGTACAGATCTCCAAGATGGTACCTTCAT
AACTTCTTTCCAAACAAATTTGACCTAACCAGGATGAATAAAGATCTGCAACATGACCACACCACCTCGTTTCAT
GTTTACAAATTTTTCTAAGGGGCTTTTTGGATGTAGAAATTTTGAACGCACTGCATTTCAAATGTAGGAGTTTTA
AGTTTTTGTGCGTTCGGTTCAAGGTAAGCTAAATCTAAAGTATTGAGAAAAGGATCAAAATGATCCTTCAACTCT
TTGAGAAGGACCAAAATGGTCTCCAACTCTTTTGAAGGACTAGGTCCCTCAACTCCTTTAAAAGGAGCATCTTTT
GTTCTAACATCAAAATTTTGGGATTAATGGTGTTAAATCAATGAGCATTTTAGTCATTTTTAATTAAATTATTTA
TTAAATATAATTTTCAAGTAATAGTATTTCAAAAAAAATTATTAAACATTTCTTTAATTAAAAAATTATTAATAA
TTCTAGTCTATAATGATCCTAAAATAAAACTAGATCCCCCCTCCCCTCTTTTAAAACAAGAGCACAAAAACATTG
CCTCCAATGGGGAACCAATAAATGTGCAACACATACCACCTCATATAGATTGAAGCAAAAGGTCCATTTTGAACA
AACCAGCACCCCACTCTTCTCACAATGCACGTTAGTTAAGAGTGTTTCTACTAAATAATCAAGAAACCTAAACAA
ATTCTCTTGAATTCTTCAAATTATAATTCAACATAACATTATACAGATTACATTGAATATTTCTTGTAGGATAAA
ATCATCAATGATCATGAACAAAAAACTGGAAAATGGCGCTAAACTATGTCCTTCCCTTAATTATCCTTCAAAGCT
ATTATTACATCTAATAAGTAGATGTCAATTCTAAGCTGCCATTCTCAAGACATCATCAATCATTTTTAAATAAAA
CTTTAAAACAAGATAATGAGATAAAAATAACATGAAACAATACAAGAGATTGTAAAATGTAAGTGTTAAAATCAA
AAACAAAACTTAAGGTGACATGATGATATCAGAAAGAAGTATCTATTTGAAACAATGATGAGCTTCTCAAGAAAA
AACCCGCATTTGTAGAGTCTTATTTCTAGCCTAGTGAATTTTAGAGTAGCAATAATTTTTTTTTCTAATTCCTAC
CCCTCATGTCAAATAGGACATGAAATCTCGCATATACTTCGGCTTCTATGCTCATAATCTAACATGAAAATGATA
CAATTGCATAAAATTTAGGAATCAAAATCATCAAAAATGGGAGGAGGATGAGGGAATGACTAGTCTGGATGAACT
AGACCAATGTCCTTGGACGCAATATGAGATGTTGATCACATCTGTGGGGTTAAGGAAAACAAGAGAAGACTCTGA
AAGCTTTTATACCATGATAGGGATCAAGAGAAGATTCCAAATTTTATGTGAAAGCTCAGCATCCAACAAACATGA
AGATAGAACTCGGTCATGTGTATACAAAAATTTGATCACATCTCCTTATTCCAGAATGTAAGGATGAATAATAGA
AGGTTAGTTTCGGATGAGTTTTGAAATATTTGGAGTGGCTATGTCCAAAAAATGTTACATAAACAATAGGATGAAT
GCCTTCAAAGTTCAAATTTGAAATTTTGTCAATAACCACAAGAAGAACCAATCTAAACGCCCTTATCCAAAAGAA
TAGTCATGTCCATTTGAATTACAAGCCTAGCCATAGATGATCTAGAAAGACAAGCAGTAACATCATTCATTTGCA
AGGGTTTACCAAAAACAGAACCCAGAGAATATAAAATAAACGAAGCTGAGAAAAAATAGCCCAAACATAAATGAT
AGGTGAATTCCACATATGAGTTAAAAGAGAGATAACTTTAACACATGCATTGGATAGCCACTCACATAATATTAT
TGATGTGTGATAAATCTAGAATAATCTGGTTACCAAACCCAACAGAGTATTCTCCAATCAAGCAAAAAGTTTAAA
TAAATAAATGTATAATCTCAAGGACGGGATAGCCCAATAAAAACTTGCCCATAATACAGAATTGCTACCTCAAAA
ATCTCAACTTGAGTGAGAATAAGAGAATGCTTATTTGATTCCAAATAACCAACAAATTCACATTTAAATTTAATC
CAAGATAAAACAAGAATTTAGGCCGACAAATGAATCTTTCCCTATGAATAGAAGCTTGTAACTAGAAGCTAGCAC
ACATGCAAAGATTAAAAATTGCAAGAAGATAAAACCTCAAATTATCTATTTATGCATTTCTCATCCCTAGGCTGA
TAGATATGATTGGCACAAGATCAACTCTATCTGATTCCAAAGAACCTTCAAATTAACATTTAAATTGTATCCAAG
AAAAAAAAAGAATTTAAAAAGTCACATGAGTACTTTCCTTTCTAAATAGAAGCCCAAAAATAAACTCTTGAATAA
AAAATCTCGGATAAATCTTCTTTAGTCCTAGAAGTAAATAAGATCGATATATACAATGATTGTTACAATCTTGGA
AATAACGAGAGGACCTACAATTATGCATCAATCTACCTAGATTGAAAACATTCAACTCAAATGAATTGAACCATT
ACAATCTTGGTGATAAAGAGAGGCCCTACAATTATGCATCAATCTACCAAGATTGAAATCATTCGAATCAAATGA
ATTGAACCATTAATGTTACATAACAACTCAATAAGAACTGTAGGTAACCTTCAATTGAACATAAGGGTTGATCTT
TAAGGTATCCAATAGCTAAGACAACCTATTCTCAAAACAATCATATGGGCATAATGATCATTAAGAAATAAATAC
AATTTCATAAGGCCAACATTATAAGCATGCTCTAATTGCAATTAAAGAAGTAGCAAACTAAAATAAAATAAACCA
ACTAAATGAAATCAAATAAACTTTCTCAAATAATTGCTTATTGCTTCTAATGAGCATAAACATCAATAAGATTAT

ACATATGAAAATTTGATTATCTCAAAATACACGTAATTGATCAAATATTCCCACTATTTTTAGTTGGAGAAATAT
ACATGAAAGTTCATTTCGAATGAACATACCCTCAATAAGAGATAACACCTTGCATATGTGAACTCATTAAACTCC
AAAGTCATTAGAAACAAGTTTAAACTAGTTTAAATTTGAAATTAGTTTGAACTACAAGGAAGACATGGCAGGGAC
AGAGAAGAGTTCCCAAGGAAAAAAAAAGTTTTTGGAAAAGGCATCAAAGAATCGGCAGCATGTGAAACGAGGTTC
GCATGATAATGCATCATAGGACTAGTGGCACATGCGAAGAGTTAGCTAGAAGGCATCACAAGACCAGCACTTGCA
AAGAAGAAAGGAGTTCAAAAGGAAGGCAACGCGAGACCCACGACATGCAAAGCAAAAGAGGAATTCAAGAGGAAG
GCATCGCAAGGCCAACAATGTCTGTGATGGAGTTCATTAAGGCATGGCAAGTGCAACAATGTGCGTGAAGAGTTT
GTGGAGAAAGCATTGTGGGACCAACATCGTGCAAAGGAGAAGAGGAGAACGAGGGAAAGGCATCACCAGACCAGC
GGCATACAAAGGAGAAGCATAGTTCATGTAAAAAGGTAGCGCAATACAATTGACTGGTGAAATAAATCTTTCATA
GCAAGCGTTCGAGAACGAAGAGAATGAGGTTCAAACCACATTTGAATGACATTCGAAACAATGTTTGAACTAAGA
TCTAATGAAATTCGGTATCATGTTTGAATTAAGGTTGGAATTTGTACTAGGTTGAGCCAATTTTGTTTGCAACTG
ACTTATATATACAATTTGCTAATACAAAAGCGATTAATTTCATCCGAAATAAATTTTATGGAAATCATAAATTTA
GCGAAATTAGCAATATATTCAAGACATTTAAAGTTGATTTCTCATGAAACTACATCCTTCTTAGGCCAATGTCAT
CGAATTCAATTAACTCAACATTTAAATAATATATCATGGTCAAAATCATTCATTTTACCTGCCCAATTCACCTAA
ATCTAATTTACACAATAAAAAAGTGCACTTAATTATTAAGCCTAGCAACAAATTGCAACACATGCCTCAAAACTC
TTTTTTCATTCAATTCAACAATGTCAACCATCATGGTGCAATCTGTATGATATGCCAAAAAACTTAAAGGGCTCA
AAACTTAACAAGAAAAAAGGGATACACCAAAAATAATGGAGATAGAAGTTGATTACAAGAATGATATATATTGAT
GTAGGCAAGGAATGGGTCAAAACAAGTTGAGTTGAAAATTTTGTAAGTTCTACTTGAATCAGCTCATTAATGAGC
CTCTTCTTATGTTTCAAGCTCATAAATGCTTTCCTCTATTAGATATCAGCTCATTAATGTCTAGTAGAGAGCCTT
TTTAGCTCATAAATTGAGGTTCATTCCCCCATTTGTAAATTGAGGTTCATTCCCCCATTTGTTGTCATTTTGGTT
CCCGTGGATGGCTCTTTGGTTCCCGTGGATGGTTTCTTGGTTTTCCATGCCTTGGCTCCTTGGTTTCCCCGGTTG
GCTGCTGGTTTTCCGTGTTGGGTGGTCTTTCCCGTGCCTTGGTGATTCGGTTCCATGCGGGCCTCCTTAGTTCTT
GCGGCTGCCTCTTCGGGTTCCGCCTGGGCTGTTTTCCCGTGACTTTTTGGGGGCTTTTCCGTGGCTATGGTGCTT
CGTTTCCGGGAAGTGACTGTTGGTTTTTCATATTCCCTATTTAAATAAGCTTAAATTAATGTTATGTGACTGTTT
GTATTCCCCCTTTCATACCTATAAATGAGGGGTTATGTGATGTTTTGGATATGTTATGTGAATAATTTTTGGAAA
TGTAAACCTGCGGGTTTCATGGAAAAGCTTGCGAGCTTTGTCCACTTCTTCTTCTCATCCCCTCCGGAAATCTTG
CGAGCTCTTTCCAACCTTCGTCTTGGCTTCTAACCTCTTTCGTCTCTCCGCAAGTGCCACTCCACCTGTATCCCT
CTTCTCTAAGTGAAGCAATTTAGGGTGTCTTTAAAATTAGACTAATAGTGAAATGGATATGCTAAGTAGAACAAA
CTCAATCTACATGGATGAAAAGAGCTAGATAACCGTTCCTCAAGCAATAATGATGATGCACCACAGAGGAAATTT
TTTTATAGTAATCCAACTTCAGGATAGCAATCATTGATCTTGAATTCATAAATGCTAACTTTGTTATCAAACTTT
ATGACAGATGAAGTATAAATGGGTGATGGGTAGCCATCATCCAAACAAAACCATTTAATTAAATGCACGCATGCA
GATTTTATTTGATTAAGTGCTTTTAATTAAATAAAAAACAGTTCACAAACAAAGATTAGCGTGCTAAGAAGTTCA
TAAAACGTGAGACTTAAGTTTCTAAATAAAAAAAAAAAATTGTAGGATAGCACTATATGGATCATTCAATAAAAAT
ATCTTCAACAAACTTCATAGAGACCAAACCTAATTTGCTGAGGAAAAAACACAAAAAATAAATCAACTTCAGCAA
GTTGGGTGGATCCCAGTCATTGGCCTACTGTCATTGGCCCTGCAAAGTTGGGTGGATCCCTTGCCCTTTCGGTCT
AGCTCATAGATTTCCAATTTTGTCAACGTTTCCATACATGATAATGATACCTTGTTTCCCTTTGCTTCTTTAACA
TCTTCCATACATGATCATGTTACCTTGTTGCCCTTACCTTTTTGGTCTGGCTCATAGATGCGCTCATGTTCATAC
TGTCTTTGGCCCTGAAGACTAGGATGGCTCCCTTGCCCTTTCAGTCTGGTTCATAGATTTCCAATTTTTTTTAAC
ATCTTCCATAGAAGATCATGATACCTCGTTTTCCTTGGCTGTACAGCAATGCCATCTTCTCTGTTAACTGCACCT
AAAGATTCATCACCACCTTCAATTCTCACGCCCTTTGTTTCTCCTAAGGTTCCTTCTCATATATTATCCAAGCAC
CTATGATTGTTATTTCTACTGCTGGCAATGCCCCCCAATCATTTGCTTAATTTTATTGGAGAGTTGATGCCTTCA
AGACCTCAATAAATTTTGTTTATTCAATTCCCATTATTTTTTCCAATAAAGATTAAATCCCACCACTTTTCAAGG
TAAACATGTTGTTCTTTTTACAGAATCTGAGATTTCTATGTTAGCAACTCAATTCGGTTTCTTTATAGCGGCAAA
TTCTCAATAGGGTGCCAAATCTTGAGATTATAAGTATATTTCATATTTGAATTTGACTAGATAATATTTTATTAG
GCCTATACATCCTCACCAAGAATGAATCCAAGATTAATGATTACCTTCAACCCAAGTAAGAAAATTAGCTAGATC
TACAAAAATCAAAACAAACAAATTAAGACAGAAGACCTAAAGCAATAATTAATATGCCCCCATTGAGACCTTAAG
GTCGTTGGTTCCAACAATAAAAATAGTCCCAAATGTAGTGTAAGACCCCAAACACTAATGGGAGCCTCAGGCACT
TCATATGCTTCCTTAAAACAAATTAATTAACCCTAAAAACAGTTTTGATGCATGGCTAGAAAATTTTTTCAGAAC
TAATGTTTTATGAATCAAGTTTTCAATCATGATAATGAATCTGTGCAAGCTTGCAGAAGAATACCAAATTCCTTC
CCAAACTATTGGTTCAAGACATCTTAAGCTAAATATGGTAAACAATAATAATTATAATAATATTATTATTATTAT
TATTATTATTATTATTATTATTATTATCAGTATTAATGAACTTGCATTACTAAAGATGCAACTGGAGATAGCATA
GGAAACAGATTAAGGAACACCAATTTCTATACATCCTCTATCTGATGGAGTCATCCTCTACTAGCCAGAATTATA
GCAGAATTTTCTAATACTCCGTGAGTAATGAATCTAAGCAACAAGATAAGTGTATTTAAAACACCACATAAGTAG
GATTTCATGAATTTTAAAGCTAATACAAACACATGCAACACATTAGAATAAATTAGATAAATTTGAAAAGCTATT
GACGATAATATAATTGCACTTATTATTCAACAACATCTAAAACAAGAATAGCAAACTAAGTCTTTCAACAAAATG
ATCAAATTTAAAGGTCCAAAAAAGTGATCTAC**CTTTGGATGAAGATATATTGCGTTGATTTTCTTGCAAGCTATG**
**CAATTTCTCCCTCAAGCTTACTTCCTTTTCCCAAACCTA**CTGACCAGTTATAAGATATAAAAGTGAGTTGTGACT
TTCAAGTGAACCGATTCAAAGAAGAAAAAAAAATTAATGTTATTTCATGGGGTACAACTTCTAACAACTTTAAAA
AAATAAAAAAATAAACTAATTTCACAACAAAAAGGGTATACAAGTATTACATAATTTAACATTCTCAACAAATGA
ATTTAAATCACTTTAAGCTTGTTAAACGTGCAAGTTGCTCTTTACGTTAAATAAAATATATCTACCTTTTAAGAC
ATACATTTTAAGATCCAAAATTTATGAATGAAGGTATTCCTCATAAGCAACCCATTTCTACCGATTGTGGTGAGA
AGTTGGCCGAGTTTTGCTAGAACACTATGGATATTGTCTCATCTAGTCATGTTCAGCATCAAACAAAAGGTTAAT
TGTTGATTTTTGCTTTCTTAGTAGTTTGATGTATTGTGACTTATATTTGATTTTTTTTTTAATTTCTTTTGAGTT

TATTTGGTAATTGATGATGAATTTTATTATTTATGGAGATTTCTTTGTAAAAATCCCTAGAGGTCTTTAATTCTA
TGAAAACGACCGAGGGGATTAGCATGAGAGCATATATTATGATAACAAGAGTTTGATTACAATTGCCTCTTCATA
TCTCTTGGTGACCACCTCATATCCCTCTATTCTCCAATATTACCCCTTTCGTGCATCTACTCTTCTTTTCCCATC
TCTTCTTCCTTTTCCCCCTTGCGACCAACTGCATAACACATAAGAAACCAAATAACTCTTTGATTAAAATCCCCA
TACTACATCAGAGGAAAATAATTCATGATGCATTTTTTTTCATTGCAAAAAGAATGTTCCTAGAAAATCCTGTAC
TATCATGTTGTCCAGAACTTGGACAACCAAAGCAACAATTTCGGGTTGTCGCCTACCTCCCCAAATCCTCATTCC
CCTCAGGTTGCCACCTACCACCCCAAACTATCATGGATTGATTCTCAGTGAAAAGTGAACATAAACAACTCCTTG
TTATCCATAAAATACAAACAATGGCATCCTTCCAAACAAGTAATGAACTATGATGTACCGACACACCTTCATAAG
AGGGTATGCAGTGTTGGACACGAACACGCGCCCAACACTGCTGGGACATGTGTTGGACATGGAAGAGCCATGTTG
AATTAAAATAATATTAAAATTAAAACATGCATGCATGTCGAACACGCGTTTCCTAATATTTGATGATTAAATTTA
TTCAAATCACATAATTTTAACATTTTAATATTCTTTTAATAAAATGAAATTGCGTCTGACATTCAGTATATCCTG
TATTCAAGCGTTAGTGGCGTTGTATGCTCTAGGGTTTCACTTTTCACGATCTCGTCTCCATCCTCTTCTTCCTTT
ACGCGACCAAGCAAGTTAGGCAAACTAACTAAGCGAGCGACTGTTCGATTGAAGCCAGATACATCCACATTTCCT
TTTCGTCTTCTCCAATTCATCTTCTTCATTCATTTGTTTTCGACTCTTCATCCATTCTGCAACGAGGTGAGGCCA
ACTAAACAAGCAACTGCTCAATTTTAAGTCTGGTACGTTCGATTTCATCCATTCTCTCATACTTCTCTTCATTTC
CGCACAAGCATATGACTATTCGTCTGCGCCATTTCCGCAAGCACCAAGGGCATTGAAAGTTTCAATTCTTGTGCC
TATTTTGCCGGCGGTAGCGGAGTGATGCGAGCAAAAATTTGGGAAGGGCCCCAAATTTTTGCAAGCAAAGACTAA
ACTTTCCTAGGTAATAAAAATTACAAAGATTGCCCCAAAATAGACAAATAACAACAAAAACAAACTTTAATATTA
TAAAAGAAGCTTACAAGACGCTTACAAACTTTGACGACATAAAGACAACTTAAACACCTAAAGTTGCTTTGCTTT
GAGAAGAAGGATACAGGTGGAGTCGAACTTGCGGATAGAAAAAAGAGATAAGAAGTCCAAGACGAAGACAGGACA
AGCTCACAAGCTTTCCTAGATAGAGAGGAGAAGAGGTGGAACAAGCTCGCAAGCCTTTCCAAGAAAACCCGTAGG
TTTTATCCAAAAATCACTCACATATCATCTCTTCTTCATACAAAACATCTCCTATTTATAGGCAAAAGGAAGAAG
CTAAACAGCCACAAAACCCCCAACTAAATCCAATCAGATTTAGGGAATATGGAAGACCAAAAGACAGCAAACAGC
TGCAGGAAAACTGCACAGAAAATAGGGAAAATTCCATGGAATTCCGCCTCAAACAGCCACAGGAATCCAGCCAGC
AAGGGACCCGAAAAATCAGTACAGGAGCCAAAGCAACAATCCACGGGAAACCTGCCAGCAAGGGACCCGAAAAGC
CAGCAAAAGAGACGAAGAGGCAGGGCACGGGAAACAAACAAGCAAGGGGAAACCGAGAAGACAAAGGGAGGAGAA
GAGGCACAAGTCAAAGTAGGGGAAACCAAAGAGACTAAAGTAAAGGAGAATGAACCAAATAAAGATAATTTATGA
GCTGAATTAAGTAGTGGGAAAGTATTTATGAGCTGAAACATAAGTAGAAAAAGCTCATTAATGAGCTAATTTTAG
AAGCTCACCAAAATTTAAACTCAACTTGTTTTGACCTCACCCTTGCCTACATCAATTCTCCTCCACTTAAAAAAG
CTCATCTACTGATGAGGAAGCTTCAAAATTTTGGAGATCCGGCATTTCAGATTCTTGGATTAGAAAATCATACTT
TCGTTTGATGGCTTCCCTTTCTCGTTCAAATTCTGCTAGAAGTAAACGTTTCTTGGACTCGAATCTCTTGTCACA
TAATTGCAGTTGTTCTTGAATCCTGAGTAGTTCGATCAGTATCGGGTTAGAAGAACTTTTTGTGGCGAGGTCATT
GGTTTTGAGTCAAAGCTTTTTCAAAATCTTTTGGCTTACTTTCATGATTGCTTTCCACTTGATATTTATCAATCT
TAACTTAAAAAGACACAAGACTTAAGAAGACACAAACCTAGGCTCACGCACAACAATCATGACACACCTAGACTC
ACTAGACAAAAACTTGATTAAATGACCTAGGATATACCAACTTGCTCTGATACATAATTGATGAAAACAAAAATT
TGGGAATGGCCCCAAATTTTTGCAAGCAAAGACTAAACTTTCTTAGGTAATAAAAACTACAAAGATTGCCCCAAA
ATAGAAAAATTAGACAAATTGAAACAAAAACAAGCATTAATATTATAAAAGACGCTTACAAACTTTGACGACATA
AAGACAACTTAAACACCTAAAGTTGCTTTGCTTTGAGAAGAATGATACAGGTGGAGTCGAACTTGCGGATAGAAG
AAAGAGATAAGAAGTCCAAGACGAAAAGAGGACAAGCTCGCAAGCTTTCCTAGATAGAGAGGAGAAGAGGTGGAA
CAAGCTCGCAAGCCTTTCCAAGGAAACCCGCAGGTTTTATCCAAAAATCACTCACATATCATCTCTTCTTCATAC
ATAACATCTCCTATTTACAGGCAAAAGGAAAAAGCCAAACAGAACCCCCAACCAAATCCAATCAAATTTAGGGAA
TATGGAAGGTCAAAAGACAGCAAACAGCTGCAGGAAAACTGCACAAAAAGCAGGGAAAATTCCATGAAATTCCCC
CCCCCCCCAAACAGCCACAGGANNNNNNNNNNCCCCCCCCAAACAGCCACAGGAATCCAGCCAACAAGGGACCCG
AAAAATCAGCACAGGAGCCAAAGCAACAGTCCACGGGAAACCAGCCAGCAAGGGACCTGAAAAGCCAGCACAGGA
GACAAAGAGGCAAGGCACGGGAAACCAACCAGCAATGGGAAACCAAGAAGACAAAGGGAGGAGAAGAGGCACAAG
TCAAAGGAGGGGAAACCAAATAGACAAAAGTAAAGGGGAATGAACCAAATAAAGATAATTTATGAGCTGAATTAA
GTAGTGGGAAAGTATTTATAAGCTGAAACATAAGTATAAGAAGCTCATTAATGAGCTAATTTTAGAAGCTCACCA
AAATTTAAAATCAACTTGTTTTGACACCACCCTTGCCTACATCACGGAGAGACCTGTTAGTATTTTTTTATTGTG
AATATTTTTTTTTCTTAATGCTAATTTCAAGCACATAAAAATTTTTCCTTACTATTTTCTTAATGCTAATTTCAT
CCAAGGTTCCGACTATGTTTAAGTGGAGAGATCTGTTCTACGCTATTCCTATGTTTCTTGTTTTTTTTTTTCTTT
GCTCTTGCAAAGTTCTTTGGATATTTAGAGGATTTTTGTGATCAAGAATCTTTGGTTGATAGATGGAGGATGGAT
CCAAAGATGCGGTGGTTCATTCATAGAGCTTCAGCATCGCTTTTACAAACAATCGTAATAAAACTTCTTGGTCAA
GCTTCATCCTCCTCTTATTATGAGAAATTGAATTGCATACTCATTCACTCAATCGATGAGGAACAAATTACTCCA
ACAGGCTAAAGATTTATTTTATGTGCATACAAATATTCGTCTTCTATCAAGAAAGACCACTTTATATATAAAGTG
ATAACAAGATGTGCGATATTGCTTGAGATGGGTTCGAGTGCCTTGATGATATGGGAATATTGGAGATTGTAAATC
TTTCTCTTGAAGAACCAGAGTAGGAAGCAATGATATTCAATGATGATGATATTGAAGACCAAAAACAAACCTCTG
AAGCTAATCATATTATTCCTTTTAATGAGCAATTGAAGTTTGATGTTATACTAATTGATTTTTTGCTTATATTTT
TGTTGTTGTAACAAACATTTTGTACTAAACATTTTATATTATAAACTGGACAAGTGTTTGCTCAAGAAAAATGAC
GCATCGGCATGAATCCCGCTTTGCAGACATAGGATTTCATTCTCCTTAAGTCAGATTGTAATTAACCAGATTTTT
CCTGAGGGTAGTATCAGGGTTTAGTTAGATCTAAACCATTATTTCCCATGAGCGCAAGCACTCATAGTTAATTAC
CCCTCTTAATGCGTAAAAGAGAACTCTCGATTAAAGATAAATAATTTTAGGTGTTTTTAATCTTAATTGTACGAA
AAAATAGAGGTAAACCCGATCAAGATCTCATGGTTCCATCATCCTTTATATACTGTAGGTTTCAATCTACTAACA
CTAGTGATGGTTAATCATAATGTCCAGGGAATCTGATTAGCAGTCATGAGTTACCTCCTCTTAGGAATCCTGAGT

ACACTGAGGTGATTCAATGTAAGTTATAATTTTAAAAAGGTGGTTTGGGTTTACATCTATTAGAATACACAGTAA
TCGATTTAGTTTGGGTTTAATTACTTCATTTAGAAGTTACCCGTTTAATTGAATTAGTTCGGGACTTACGGCTTC
CGTTCGAAGTAACTCGATTATTTTAGAAGTGGGTCTTACGATTATCATTGAAAGCAGCCCGAATAACATGTCTTA
CAACTTCCTTTGGAAGAGTGTCTATTTTGGATAGGTCTTACGGCTTATGGATTTAGAAGTAACTTGAAGGAAAGT
CTTACAACCTAGCCCTATAGGCAAGACGATCCATTTTGTTTGTTTGGGTAGGTCTTACAGCTTCCCTTTCGAAGT
TGTCTGTATGCACATTTAGATAGATCTTACAACATTCATTGCAAGTTACCTGTAGTTTATTTGTTTGGGTAGATA
GGTCATACGACTCTCTTTAGAAGTTACCTATAATTTTGTAGGACTTATTTCTTACCTTTGAAAGTAGCATGTTTT
TTTTAGTTTAGGGAAGATTAGACGGCAAGTATTTTTAGTTTGCAGTTTGTTTGATAACTTGATGCAAGACCTTAG
TTTCGCAATTATTTACTAGTAACTCTAGCTTGACACTAATGTTTCTGCTACAGTACCCCGCTTTTTTATTTCAGT
ACATGCTTTTCTTTCAATACTGTAGTTTATCTTAGTACTCGTTCTTTTCTTGCTTTAAAAGTTTAGTTTTCAATA
CTATGCATTTTCTTCTTTTAGCAGTTTAGTTCCAATACTTATGACTTTTGTTTGCTCATTTACAGGAGTCGTGCT
ACTTCAGAGCTTACCCAGGAGCACTAGGGAGCTTGCACAAGTATTTTGAGTGTTGTAGAGTTTTTTTTATTTGCG
AGTTTTGTATTTCAATGTTGGTGTAATATTTTAGTTTTAGGAGTTTGGTTTGTATTAGGATGATTAGTGTGAATA
TTATTTTCATTTTGCATTTTATTTGATTTATATTATTTATTCAATCAATTAAACGAATTTTAAATTAGTTAATTA
ATTTAATTAAATGAAATGAATTTTAGAATGGTTAAATGAATTAAATGAAGTTACAAATTTTGACATTAATTAAAT
AATTATATTTTAAATTTATTTTTGAATTAAATGTATTATTAAATAAATCAATGTAAGGAAAATCCAAAGTTACTT
TTAAATTAATTAATAAATTTTAATTACATATTTTTGATTAGTTCATTCTTCAATTGAATAAATAAAATGTCATTT
TAAAGTTATTTATTCAATAAATTTAATTTAATTTTAATATCATTTCTAATTCAATATTAGAAATTATCTTTCTTT
GTTAAATTTAAACGAGAATTTGATTTTTAACAGGACATAATGAAGAAAAAGGATTCGTGTAACTGACCCCACATA
GTGGGACAAAAGCTTGATGTTGTTGTTGTAATGTCATTTTTGAATTAAATAAATTTTAATCTTATTTTTATTTAA
TTTTTATTTGAATTTAATCAATTACTTAAATGAATTAATTTTAATTTAATTTTAAAATTAATTAGTTCATTATTT
TGAATCAAATAAAAATTAATTATTCTTAATTAATTTTTAAACGACTTAATTGAATTAATTTTTTAAAAATAAATT
CAAATGCCCAAAATTACTCTCCAATGGCTTTTGAAAATTTGATAGCCTTCAGGGACAATTTTGGCAACCTCTCAA
TGCCCCTTTCCCGCAAGGGAAAGTCGGGATGACACCTGTCGTTGATTTCTACCTACTAGAAACAAAGATCGGCAG
CCCCAGGTCTTCAAATTGGAAATCGTCCTACTCGGATCAAAGATCAATGGCGTATGGCCTTCAATTTCAAAATTT
CACCATCTGGATCAAAGATCAACAGCACGGGTCTTCAAATTTAAATTCATCCCACCCAGATCAAACATCAACGGC
GTAGAGCCTTGAATTTTAATTTTCATCTCACCAAAATCAAATATCAATGGCGCGAGTCTTCAAATTAGAAATCAT
CTGCCCAGATCAAAGATCGATGTCACGGGCTTCAAATTAAAAATAATCTGCCTAGATCAAAGATTGTTGGCGCGA
TCTTCAAATCAGAAAATACTTACTGATTTACAATTGCCAAGATTTTATATTCAATTACCTGAAGATTTATTATAA
TAATCTTCTGTGAACAGAGAAAATGACAGGTATTATTACTGTCTTAATTATAATAATTCCAAATTTAGGCGAACT
GTTATTCTTGACTAAAGTAAATGTTGGCTCAAGAAAAATGGCATGTTGACATGAATCCCAATTTCAGTTGTAGGA
TTCTATTCTCGTTATGTCAGATTATAATTAACCAGATTTCCCCTGAGGGTAATGTCAGTGTTTAATTAGATCTGA
GTCCTTACTTCCCATGAGCGCAAACATTAATAGTCAATTACCCCTCTTAACATGTAAGAAAGAACTTTTAATTAA
GGGTGAATAAACTTTAGGTGATTTTAATCTCAATTGTGGGAAAAATGGAGGCAAACCCAATCAAGATCCCATGGT
TCTGTTGTCCCGGGTATATTGTAGGTTTCAATCTACCAACACTAACAATCATTGACCGTGATATCCAAGAAATTT
GATTAGCAGTCATGGGTTGCCTCTTATTAGGGATTACGAGTACATTGAGTTTTAGTTTTATTGAATTTTGGGGTT
TACATCTATTAGAATACCTAATAGCTAGTTTGGTTTGGGATTTATGACTTCATTTAAAAGTTACCCTGAAATTTG
AATTGAAGCGGGTCTTACGGCTTCTCTTGAAAGTAGACCAGGTGTTTGAATTCGGGTCGGGACTTATGACATTCA
TTAACAGTAGCCTGTAAGGAGAGTCTTACAGCTCCTATTAGATGTAGCCCATTAGTGGATCTTACAACCTAGTCC
TATAGGTTAAGTTGCCCGTTATTTTTAATTTGGTTGACAGTCTTACGACTCGCTTAAAAGTTGCCTGTAAGTTTA
ATTTAGAGTGAGACTTACGGCTCTACTCATGAGTTATCAAAAGATGGCTCTAACAACTTAGCCCTACAGCAAGAC
TGCCTGTGGATTTAATTATTTGGATAAATCTTACAGCTTCTCTTTGGAAGTTAACTAAAAGCGGATCTTATAATC
TAATCCTTTAGATTAAGATGCATGTTAATTTTTATTTGAGGGTTTGATATTTCATTTAAAAATTTTCTAAATAAT
AGATCTCACAACTCCCTTAGAAGTTGTCTAGTTATTAGGTAGGATGGGTCTTACTGCTTCCCTTTAGAAGTAACC
CAATGGTGGGTCTTACAACCTAGCTCTATAGGCTAAGATGCCTAAAGATTTAATTGTTTGTCCGTTTGAGTGAAG
TGGACAGTAGTTGTTTCTTAGCTTGACACTAGACTGTAATTTTCAATTGTTTATTAGTTCGACACATATTAGCCA
GCAGTTGTTATTGTGTGACACTAGTATTTCTTGCTATAGTACTTTGCCCTTTTTTCTTTTCAGTACCATAGATTG
TACTAGTAGTTACTTTACCAGTGTCTGTTCTAAGACTCGGCATGGTGAACACATAAAGTTCTTAGCAAAGTGGAT
AAATCGCCCTTTGATGGACTGTACTTGGATGAATGCAAAAGAGCTTGAAAAGTGTCATCCGGAGCTTAGGGGGGA
ATTGATGCATTCAGTTATTTTGATTTATTTATGTTTTATTGTATTTTGAACAATATTAGTTGTGTTTTATTTTTG
TTTATGCTAATCCTATGATGGTTGGGCTTCCAATGATCATGGTAGGAAGGTGTTTGGCATGGCAAACTTTGGGTT
GCTTGCTTTTGGGTCCACTTCGTCCTTGGGCTGCTTGCTGATTTTTGTTTTGGTGTGGTGACTGCTCTCAGGTGG
GCCTTGGATGCTTGGTGTTGAGCTTAAAAAAATGTTGTATTCTACTTGGTGCTAGCTCATAAATGCTTGAGCTGG
AGTTTAGCTCAATGATTGGGGGATTTATTGCACTTCCTAGCTCATTAATTCATAAAGGTATGTTATTATGTCTTG
GATTCATTTTGGAGTGGACTTTGCCCCTTCTCCTCCCTTTGTCTCTTCGGTTTCCTCTTGGCTTGTGTTGTTCCC
GTGGCCCCTGGCTGTTTTCCCGTGCTGTTGTTCTGTTCTTTGTGCTGACTCCTCGGACCCCTTACTGGCTGGGTT
TCCCGTGGCTGCCTGCTTGGTTTCTCGTGGCCTTCTTTTCTCTGTTTACTGTACCGGGTTGGGGTTGCTGTGTTG
TGGTTGTTGGCCTTTCATTTCCATAAATCTAATTGGTTTGGTTGGAGGTGTTATGGCTGTTTGGTTCCTCTCTTT
TGCCTATAAAATGGAGATGTTTTGGCATGGAAAAGGTTGTTGAAATTGTGAGTAAAAATTCTACAAAAACCTGAG
GGTTTCTTTTTGGAAAAGCTTGAGAGCTTGTTCCATCCCCTTCTTCTCTTCTCTATTTCGTCTTGGCTTCTAACC
CCTTCTTCTCTACGCAAGTTCCACTCCACTTGTATACTTCTTCTTGAGCTTTGGAAGTGTATGTTGGCTGAAATT
TGGAGACATTTGCTGGAGGAGTTTGGAAAGGACGTTGGACTCTTTTGCAATGGTGGACCCTTGGTTTGCAAGCTT
AAGCAGAATTTTCTTAAATTCCTACTTTCGTTGCATGTGAGGAGACTGTTACACGGTTATACTGAGGATAGTGTT

TTATATTTTTATTTTGTTTTCGTTCGATTTGGCATTTGTTTTTCCTAAGTTTGTTTGGATGTTTGCAAGTTACAA
TTGAATTGAGTTTGGTTTAAGTCTTGTCCTGATGTTCTCCAATTTGTCTCTTTTCTCTAGTCTTTGCTTGCATCA
TGTATTATGGTAAGAGTCGTACTAGAAATTGCGTAGGTATGATTATAGATAAGAGTTTAAAGTATAATGTTATAA
AAATTAATATAATTAGAGATAAAATTATAGTTTTAAATATTATTATTGATACTAAGATTATTAATATTACAAGTG
CTTATGTTCCTCAAATAGATTTGGATATAGGTCCAAAATTATAATTTTGAAATGACTTAGAAAAATTAATGCAAA
AGATAACTCTTAAAGAAAAAAAAATTATTAGAGGAGATTAAATGGACATGTTGGTCGAGAGGTAGATTATTTTAA
AAGCATTCATGGTGGTTATGGGTTTAAAGAAACTAATGAAAGTGGTAAGACCATATTAGATTTTTCGTTAGCTTA
TTATTTTGTAATTGTTAATATCTTATTTAAGAAAAAAAATGAGCATCTCATCACGTATAAGAATGGATATAGTAA
AACTCAGATTAATTAATTTCTCATAAGAAAAATAGGTTATTTAGATTATAAAGATTGTAAAGTCATTCCAAGTGA
AAGTTTATGTACCCAATATCACTTATTAGTTTTAGAATTTAAGATGATGAGGTATAAAATAAAAAATAAAATAAA
ACTTTTTTTAGGGTTAGATGGGGACATTTAAAAAATCAATTTGGATTCATGCTGATTTGGTCTACTATGGAAGCT
ATACGTTAATTGAAAGGTTTGGTGGAAAAGTATAGAGAGAATAAGGAGGATCTACATATGATATTTATTGATTTA
GAGAAAGCATATGATAAAGTTCAAAGAGAAGTGCTATGGAGAGTTCTATAAAAGAAATGGGTTAATAATTTTTAT
ATTCAAATCATTAAGGATATGTATATATGAGCTATTACTTGTGTTAAAACAAAAGGTGGTTTGACCAAGTATTTT
CTTATTTTTGCTATATTACATCAAGGATCAGCCTTAAGCCCATATCTTTTTGTTTTAGTCATAGATGTACTCACT
GGACATTTGCAAGAGTTTGTACCCTGGTGTATGCTTTTTGTTAATGATATTATTCTTGTTCATAAGACTAGAGAG
GGAGTTGAAGGTAAATTTGAATTATGGAGGTCAACCTTGAGTCTAAAGGCTTTCGTTTGATTAGTTCTAAAATTG
AATATATGGAGTGTACTTTTAGTAATAATAGTTTTAGTGTGGAAATTGTGACTTTCGGTGATTAAGTCATTAATA
AGAGTACACGTTTTAGATATCATGGGTCTATTGTTCAGAGTTATAGTGAAATAGATGGGGATATTATTAGTAAAA
TTCAAGTTGGATGGTTGAAGTAGAGGAATGCATTGGATTTGTTGTATAATAGAAAAGCCCCCTCAAACTTAAGGG
AAAATATTATATGATGGTGGTTAGACCTACAATCCTATATGGTGCAGAATGTTGGCCATTAAAAGAGTAGCATAA
TATTAAGTTAAGTGTTGCAGAGATGAGGGTGCTTAGGTGGATGTGTGGTTTTACTCTTAGAGACAGGATACGAAA
TGAGCAAATACGTGAAAAGGTTGGAGTAGCTCAAGTGGAAGATAAGATTAGAGAGAGTCGCCTTGGATGGTTTGG
CCACATAAAATGGCGGCCTTCCGATGACCCGGTTAGAAAAGTAGATATATTAGATTTAACTTATGTTAAGAAAGG
TTGAGGTAGACTTAAAAAGACTTGGTTAGAAAATATTAAAAATAACCTTTTTTATTTGATTTAAAAGAGAATTTA
ATTTTTAATAATGCACAATGGAGAAAAAGGATGCTTTTAGATGACCCCACATAATGAGATAAAGACTTGATGATG
TTGTTGTTGTTATTGTAATATTTTTATAATTTAATAATTTAATCTCAACAGATCATTTGAAAGTCAAACTAAATT
TTAAAAATCAGGAAATTTCCTGATAAATTTTTCTTCTTTATTTCCTTATTTCTTCTCTACATTTTATTCCTTCCT
CCTTTTCTTACTTTTATTTTTCTCTTATTTGAACAACTCCAACCACTTATCCAGCAAGCCACCGCAGCCCTCGAA
AATCTCTCACAATAGTCCTCACAACTCAAAAGTAGAAAGAAAAAATAAATAAATAAAGTCACAAAGTAGAAAGAA
ATTGATAAGAATTTAATTGGATTTTTACTTATATATAACTAAAAATTTAAATTTCATAGCCAACTCCTTCCTTCA
ATGTCAAAACTTTGTCAACTGAATCACCGTCTCTGCTATTACCGCTCTAGGGAGAAAGATTAGGATCATTCGCTT
GCTACAGTACTGCTACATTAATTCTACTCCAATAATTATTTTGTTATAAATCCCCCCTCCTTTAGGAATTTCATC
AATGAAATTCCTTTCATCTCCATTTACAAAATATAAATAGGAAAATCTTTGTTTTATTTCATGCTTCTTTTCCAA
TTTGTTTCTTTTCTACTATATTTCTACATTTTATCTTAGACATTCGAAATTTGTTTGTTTATCAATTTTTTAATA
TTATATTTCAATATTTTTCATGTTCTTTAGCATATGTAAGATCTTTTTGCACTTCAATTTCTTCATGTGAGAATC
TCTGACTTTCAATGGAAAACTATTTTCTCAGCAATGAGATACATACAAAGACATTATGTAAACCTGATAAATTAG
CGGGTAATGCTAACATGTAAGCAACTTTGCAAATTTATTCAATAATCTCAAATGATCCTATGAATATTGGATTTA
GCTAACGTCATATACAATCTTATAACAAAAAATATTGTAGCAGTACTTTGGCGGTTGACAAAACTTTGGCAATAA
AGGGAGGGGTTGGCGGTAGAATCCAAAAGGAAAATTGAAAGAATATATATATATATATATATATATATATATATA
TATATATATATGTGTGTGTGTGTGTGTTAAAACCCAACCAAATTCTTTTCAATTTTCTTCCTACTTGTGAATTGT
AAGGGCTAACTGGCTAGAGAAGTGGCTGCAATTGGTAAAACCAAAGAAAAAAAATGACAAAGAAGAAGGAATTAA
AGAAAAAGAAGCGAAGGAAAAGAAGAAGAAATTAGACAAAATTTTGTCAGAAATTTTTCTTATTCTTAAGATTCG
GGTTGATTTTCAAATTGTAGTAAATTAAGGTTAAAATACTATATCCTACTCAGATTAAATTATTGAATTTGAAAA
ATATTACAGGTTGAAGAAATTATTATTTTGGGTTATTAGGAAAAGATGAAAGCTTTGGAGATGAAAAAATAAGAA
CATCCACTGAGGAGAAGACCCAAATTTTTAGGTGAGGTACTTAACTTCTAATATAATAATCGAATATATCATAAT
ATTTAAGTTTACACATTTATCATTCTTATAAATAATGAGAATGAAATTTAAGAATCCTATGAAAAGTCACATCAT
GTGAATTAAGATGGTTGACTTTGATGACCTAGGCAAGAAGAGGTGATTCCTAATAATTGACAGTTGGTTAGCCTA
ACATTTATCAGAAAAAGATGAATGAAAAGGAATTCTTAATTCTCGATGCTGAAGATTTGGTTAACCGAGCATTTA
TCAAGACAATAAAAAAGAAATTCCAATCTCTTGATAATGGAGAGTTGTTTAACACAACTTTTGTCAGGGAAAGTT
AACGCAAAGGGGCTAAAACCTATAATAAGCATGAAATGGCATTAAAGATAAATATGTTATGACAATCATAATGAA
TTTGATATGATATTAAAAATGAATATAGTATGATATTAAAAAATGAATATGATATGGTATGTCAAAAGTATTAAT
AATCAAGAATAAGTAATAAAATGAAAATGAGTAATTCATTAGTAATTTCATGATTATGTAACATTCTTGCTTGCT
ATTCCTGATTTGCTCATTTGTACTAATGATTCGTAACTAATATAATAAACAGACATCTACTTCACGAAACTAAAT
ATAATTGATTTTATGATTTTATTTATGAACGCTTCCAATTAAATACTTCTAATGGTTACTTAAATGATGTTAAGT
GCTATGTAAAACATTTAGACGATAACAAAAGACTATTTTCTCTCATCTGATCCTGATGAGTTAAATCAGAATAAA
TGTAGTGAGAATTTTATCTGCCCATATTGTCTGCTTTTGTTATACCTCATGATTGCATAGAAGGGGTGAAAATGG
GGGATGGCACATCCTTCCTTAAACATAAATAGAGTGTCAATATTAGACCCGACACAAAACTAACTTTGAATCCAA
CAAAAAAATTCGGGTTTGGGTCAACAAGGATAATATTCGAGTCGAGTTGGGTTAACATAAATGACTCAAAAAAAT
ATGGTGGGTTCAAGACATGTCAATTCAGGTTTGGGTTTGCGTAAATAACTCATAAATAATGATGGATTCAAGTTG
TATTGGGCCGAGGACAGGTCAACCCACAAAAAAAATTACTTAGCTATTGAATTCAATTTTTAATGAAAACCCCAT
ATAAATTGTATCTTTTATTATTTTAAGCATTTTCTTGTCCAAATAGAGTTAGCATGTTCAATTTGGGTTGAGTTG
ACATGACACTAAACCATCACAAATTAAATGATTTTCGTGGTGTGTCAAGTCAAGTTCAGGTTGGGAACCAGTTA

ATAAATGAGTCAGTTCATATTAGATATATTTGTGTCATTTTTTAGATATGCTTGAATATGATCCATGAACCCAAA
TTAACACCCACCCATAGCTAGGGGATTTAACTATATATTCTTTTCCTCCAAATTATCTCAAGTGCAAGTTTTCTT
CCCCCAATCACGATTATAAAGCTCCAATCCTAACTTCTTCCTTACCTCCATGAGTCGCATAATTATTATGAGTTC
TTCCTTCCATCCCACATGAGAGATTCCCTAGTTTTAGATGTTGATTGGTCATTGCACGATCATTACATTTATAAC
ATTTCTGTAGAGTTATATTGATGCATACTTTGACCGCAAGGACGTACACATCTATAGCTTTTTTGTCCTTCAATG
TAAGCAAATAATAATTGCTCGAACTACATATAATTTGATCTCTTTGGGAGATGGTATGTAGGCAATGGGTATAGG
GTCTCAAGTATTTGCAATTTCCTCTTAAATCTTTTGTTATGAAATTATATAATAAACAAATTATATTCCTTCCAT
CCCCATTTGTTGCCCGAGAGTATGCTAACCTTCTCTTCTTCCCAGGTGTTCTATTCCTTCCCACACTGCCTCCAA
CATGTAGTCATTGCTACCCTCTTTCTATCAATACATTGTTGTATTTGCCCACATAAATTTCAAGGTTGAACATCA
CATTGGTTAAATCTTCCTTTTGAACTTCTACAACCTACCATACTCTTTTTTTCCCCTCAATTAGTTTTCATTGAT
TTACTTTATATCAATCTAGTAGTGGGAATGCTATCTTTACTAATACTATAACCAACAACTTGTTAATCTATTGTA
GGGATTAATTTAGGTTATTTGGTAAGTCAGGATTGAATATGTTTAGAGTTTAGCCTTTTGAGATTATGTTTATTT
GGCATTACTAGTTTAAACTAGTTCAACTTGTGTTTATACATGTGTAATTAATTTAAAATTCTATTTAAATTTAGG
TTTTTTTGGATCGTATTAGTCTGGTATTATGTTTTTTTGGAGCTAGAGGAGCATTGAACTCGCTTCTATATATTA
AGGTAAGTAATTTGTATTTATTAAAGTGATAAGCCTACCATTATTTCTAAGGAGAAATATCCTTGATGCTTGTTG
CAGGTTTCTCGAGTATTTAATAGAATGTTAGTCATGCCTATCATATTCAAACTTGTCTTACACATTCATTGAAGC
TTTATGGTCATGTTGTTAAGTCATGAGATGTGCATCTAGCTTTACATTGATAATTTGTTGCATAATAATTACAAT
CTTGAAATGAAGCATGTGATGTAATGACTCATTCTAATATTAGAATCTACTGAAATTAAAAAAAAAAAGCTTTTA
GTCGTATTTTGTGTGTGATGGATGCAGTACCCTCGTGTTGATATTATTGGCTAACCTAGTTAATGGATTCAGAGT
AACAACTCTCTTAGGGTCACTAAACTCCATTAAGTAGGTAAAAATGTTCCTCTAGATCATGTGTTAAAGCATGCC
GACTAGGTAAAAAAGACATTATATGATAAATGATTAATAGGTCATTTCTGCCTGTATAAATATGGTATGCATTGA
GTGATCTAACAATTGAATCCAATATTTGTCAACTTGCATGCTTCTTGTGAAATTGCTTGATAGATATAGCTACCA
TATTTGATAATATTTTTTATGAAAATTTAAAATAATTTGATGTCCTAATCAATGTTTAAAATTGTGTATTCGGGT
ATTAAAGCGTTTTTCGTTCGTGAGACGAAGCGTAAGCCTCAAGGCGATTTGAGGCGTAAGCCTCAAAATCAACAA
ATTGCGTAAACATAAAATAATAAATTTTGTATATACTAATGAGTTAATAATATATAAAACATAAAAATGCTACTT
ACAAAATCTTTGAGAATAAATAAGGGAAACATATCACAAAAGACCTCACAAGTCATCAGCACTGTCATCACTCAT
CAGTCATCAGCACTGTCATCACTCATCAGTTATCACAAAAGGGGTCTTCATTTGAATCATCATTATGATGAACTG
CTCTCTCCTCTTCCCCCTCTCCTATAGAGTCCAATATCTCCCAATCAGCTTCTTCTACATTATTAGTTATCTTCA
TTTTCCCTAAATATTTTTAAACAATATCATAATGAATAATAGTGTAATACATTAATAAAAGTCTAAGAAAAAAAC
TATATTATTTATATACCTTTTTTGATTTGTTCTCGAGTTATGGTGGATGTGGATGTTTTAAGACTATCAAAGAAA
TCAACATTTAAATCTTCACCTTGTAATTCAGACGGCATCACTTCAACTTCATCATCTACAAATCACTCATCATCA
GATGCCACATCATTGCAAAGCAAAGGATCTTCATCATCCATTAATCCTTTTTTCTTCAAATTTCTATCTCTGAGT
CTCCTATTGTACATAATGTACACCAGTGAGTTCATCCTCAATGTCGACAAGCGATTTCTCCTCTTTGTATGCACT
TGGTTGTAGGTACTCCAATTCCTTTCACAGCCAGATGAAGAGCATGTCAAGCCCAAAACTTTCACAGCAAAACGC
TGTAATTCTGGAGTCTCATCACCAAATTGAATCCACCACTCAACTGGAGATCGTTTGGCAATTGTATCTTTTGCT
TATTGTAATCCAAAAAAACCTTCTTTGTTCCTAAAAGGTACCAATTGCAAGTCAGCATTTGATCTTTCAGTTGCA
TCACTTATTAGACGATCCATGCAATGATATAATCCCAACTTTATTTCAGGGTTTGTAGACTTGTTCTCTGCATAT
TGATACTGAGGGTTTAGGTAGTAGCCAGCGGCATGCAAGTGTCGGTGAAGCTGTAGTTCCCACCTATCATCAATA
ATATTCCAAATTTCCCTAAAGCTTCCCTCACTGCCTCCTAAGTTGGAGGCAATCATCTCTTTGGCTTCATCCATA
GCATTGTAGATGAAACCCATTGCAGGCTTCTTCTCAGCATCCACCATCCTTAAAACATGAACCAAAGGTCTAGTG
GTTGTAATGGCATATAAGACACTTTGCCAAAATCTGTCATTTAAAACAATCTTTCTAATCTCTTTTCCCTCAGCC
TTCTTCGCCCAAGCACAACTCAACCACTGTGTAGATGTGAACATAGCCTCTAAAGGTTGTCTGGCATCCAGCATG
CTCTCTAAGGTCAAGTATGCGGTGGCAAATCTTGTTGCGGCTGGCCTTAGAATTTCTTTCTTCGAAAACTTTCTC
ATAAGAGCTAGAATCCAAGAGTGGTTGTAAATAAACTTGGTTATTTTTTTGGCTTTAGTGAGTGCATTTTTATGT
TGAGGTAGATCCCCAAGCTTCTCAAGAATCAGATCTATGCAATGTGCAGCACATGGAGTCCAATACAAATGTTTC
CTCTTCTCCATAAGTTTATTACCTGCAGCCTTATAAGAACTTGCATTATCTGTCACAATTTGTACAACCAAATGT
TCTCCAATCTCTTCAACAATGCTATCTAATAATCTAAATATGAACTCAGCATCTTTGACCTGGTCAGATGCTTCA
ATTGACCTTAAAAACACAGTACCATAAGGATTATTAATGAGGATGTTGATCAGACTTCTATGTTTGATATCAGAC
CACCCATCTGACATAATTGTTACTCCTGTTTGGGCCCAAGTCTTCTTAATTTCTTCAATGCTTTTATCAGTGCTT
TCCAACTCCTTCAAAATCCATGTTCTTAAATCATACATGGTTGGAGGTTTGAACCCTCTCCCAAATGCTCCAACT
GAACGAATCATGTTGTAATAAGCAGGTGATGTTGCAACACTAAATGAAATTGCATTTTCGTAGAAGAATCTGCCT
ATATCCTTACACACTCTCTCCGCGCCTCTCTTGCATTTTCTGGGGCCATCTTCTGTGTAGTAGAAGGCCTATCCT
CTGATGTGTCAACCATATAATGATCCATAGGCCCTCTAGCACCCCTAGATGAAATGGGCTTCATAGAAGAAGATG
AATCCCTGACGCTGCGCTCACTCCCGTAGTAAGATCCAGCATCAACTCTATCCTCAAATTGCTTCTGTGCAAGAT
GTTTTGATGTGATAGACTTATTCATGTATGACTTTATCTCTTCCCTTACAGCATCAGGAACTTTGGAGCATGGAG
CAACATTCTTGTGAGAGCCTGAAAGGTGTTCCTTCATCCTAGTCACACCACCTTTCACAACTTTATCACAGTAAT
TGCATTTCAGATAGACATATGTTTTATCACTTCCTCCTACATCCACCTGAGTGCTATATTTCCATGCTGGATCTT
TTCTATGACCTGATGCAGCCTTATCCTTATTTTTGACCATCCTGATTGAGAGAAATTTTAGGAAGCAAACGAGTT
CTTACTAATCTTTTATATGACCTGATGCAGGTCGGAGGAGACAGCAGCCTCGGAGGCTCGGACAGAAGCGTACGC
TGGTCGGACGGAGAAGACAGCAACCCAACTCGCAGTCTCGCAGTAGAAGAAACCTCCAAGCAAATAAGTTTAGGG
TTTAATATGCGGGCTGGATGTGTTCAGTATTATATAATGTCGAGCCGTCGATAAAGCAGAACACTGCGTACTCTA
TGCGTATTTATGCGTATTTTGTGCGTACACCCTACGTGCACATTGCGTGTTTTGGAGCAGATGTAATTTAATGTA
TGATTTTGTATTAGTAGTTAATAAAACATGATTTTCTAGGTATTGTTTGAGTACCAGGAGTGTTAATTATTTTAT

TGTGCGTACATTATGCGTATCTTATGCGTTATATATTGGTAAACTATGCGTTCACCATACGTTTTTTGTGCGTGG
TCTCTGCGTCAACATTGCGTATTTTTAATGAAAGATAAAGTCAAAGAGAAAAAAGAACATTTAATGTACATCTTT
AAATACAAAAAAAAGAACATTTAATGACTATGTACAATATAAAAAATATAAGAACAAATTAAAAGAACATTTATA
ACAAATACAATTTTACAAGAGACCCTAATGTACATCTTCAACAAGACAACAACATTTTATACATAGTCACCTTGA
CAACAACATTTATAACTTGCATTACCTTCTGTTGTCCAATATAAATAATCTTTGACAACCAAGCCTTCCATTCAT
TACTGCGATACTTATCCTCCTCCCATGTTCCTGATGCCAATAACTTGTTAACAGAAACCTGGTCAGGTTAGAGAG
GTTTAAGTTTACAAGTTATTGGCATCAATTTCATGGGAGGAGGATAAATATCGTAGTAATGAATGGAAAGCTTGG
TTGTCAAAGATTATTTATATTGGACAACAGAAGGTAAGATTTTTTACAAGAGAAGCTTGCTTTGACACTTTGCCA
TAAGGCAAACGTTCCTGCATTATATAGTGAAAAATTTAATGTTGAGCAATAGTAAGTAATAAATTGTAAATAAAG
GCTCATAAAAATTCAACCTACCCTATTATCCTCTCAACCATATTTGCCACCCGTCAGTCCACAGCTAACGAAAAT
TATTATTCATGTCACGGCCCCTCAGTCCATACTCCATACTAAATGTAAGTAATCAAAAATATTTAATAGAATGTG
TGCGTGTTTTATGCGTACTAAATGCGTAATTTATGCGTTTGTTGTGCGTGCCTTAAGCGTATGTTATGCGTATTT
TATGCGTTCGTTGTGCGTACTATGTAAAAAACATTTCAAAATCATTTATAAAATTATTATTTTTTGCTTTAAAAC
AAATAAGAAATTCAAACACTAATCTTTGAACGAAATTGCCAAACACAAACTTCCAAACAAACATCATATTCCTTA
TTCCTCTTCCCTAAATACAATCCAACAAGAGCCATAAACCATTTGTACTTGTTCAAGTATCATGTTCAAGATTAG
AAAATGAAATCATACAAATCTAACAAGAGCCATAAACATCATATATTCATATTCCTTATTTCTTCATCATTTTAA
TGTAAGTAACTTACAAACAAAGATATATATTAATTAAAATATTTAATAGAATTTTATGAATAAAATTTACTGCTT
ATGTGTGCGTGTATCATGCTGCGTTATGCGTACGATATGCGTTCGTAATGCGTATTTTGCGAGTTCATTATGCGT
ATTTTATGCGTTCGTTGTGCGTTCTATGTTTAATATGTAAATTTTTCGTTGTGCGTTCTAGTAAGTAATAAATGC
TCATAAAAAATCAATCTACCCTATTATCCTCCCAACCATNNNNNNNNNNNNNNNNNNNNNNNNNNNNNNN
NNNNNNNNNNNNNNNNNNNNNNNNNNNNNNNNNNNNNNNNNNNNNNNNNNNNNNNNNNNNNNNNNNNNNN
NNNNNNNNNNNNNNNNNNNNNNNNNNNNNNNNNNNNNNNNNNNNNNNNNNNNNNNNNNNNNNNNNNNNNN
NNNNNNNNNNNNNNNNNNNNNNNNNNNNNNNNNNNNNNNNNNNNNNNNNNNNNNNNNNNNNNNNNNNNNN
NNNNNNNNNNNNNNNNNNNNNNNNNNNNNNNNNNNNNNNNNNNNNNNNNNNNNNNNNNNNNNNNNNNNNN
NNNNNNNNNNNNNNNNNNNNNNNNNNNNNNNNNNNNNNNNNNNNNNNNNNNNNNNNNNNNNNNNNNNNNN
NNNNNNNNNNNNNNNNNNNNNNNNNNNNNNNNNNNNNNNNNNNNNNNNNNNNNNNNNNNNNNNNNNNNNN
NNNNNNNNNNNNNNNNNNNNNNNNNNNNNNNNNNNNNNNNNNNNNNNNNNNNNNNNNNNNNNNNNNNNNN
NNNNNNNNNNNNNNNNNNNNNNNNNNNNNNNNNNNNNNNNNNNNNNNNNNNNNNNNNNNNNNNNNNNNNN
NNNNNNNNNNNNNNNNNNNNNNNNNNNNNNNNNNNNNNNNNNNNNNNNNNNNNNNNNNNNNNNNNNNNNN
NNNNNNNNNNNNNNNNNNNNNNNNNNNNNNNNNNNNNNNNNNNNNNNNNNNNNNNNNNNNNNNNNNNNNN
NNNNNNNNNNNNNNNNNNNNNNNNNNNNNNNNNNNNNNNNNNNNNNNNNNNNNNNNNNNNNNNNNNNNNN
NNNNNNNNNNNNNNNNNNNNNNNNNNNNNNNNNNNNNNNNNNNNNNNNNNNNNNNNNNNNNNNNNNNNNN
NNNNNNNNNNNNNNNNNNNNNNNNNNNNNNNNNNNNNNNNNNNNNNNNNNNNNNNNNNNNNNNNNNNNNN
NNNNNNNNNNNNNNNNNNNNNNNNNNNNNNNNNNNNNNNNNNNNNNNNNNNNNNNNNNNNNNNNAGAAGAAG
TTGATTGGGAGATATTGGACTCTATAGGAGAGGGGGAAGATGAGAGAGCAGCTCATCATAATGATGATTCAAATG
AAGACCCCCTTAGTGATGACTGATGAGTGATGATAATGCTAATGACTAGTGAGTTCCTAACTTCTTATGTGATAT
GTTAATGTCATTGCTATTTATCTTGTGTATTTTTATGTTTTATATATTATTTACTCATTAGTATATACAAAATTT
ATTATTTTATGTTTACGCAATTTTTTTATTTTGAGGCTTACGCCTCAAATCGCCTTGAGGCTTACGCCTCGCCTC
ACGAACGAAAAATGCCTCAATACCCGAATACGCAATTTTAAACCTTGGTCTCAAAAATGCCCATCTTCAAATCCC
AAATTATCGATTCAAGAACTATAAATCCTATGGCAAACTCCTCTCATAAATTAATTCCATATACCCATTGCGCTA
GCCACAAGAAAATAACCATGGTCAATAGATCATTGATCACCATGGCAGGTCGAGATATTTTTATAATAAATCCCT
CACATTTTAAAATATCCCCCATGTACTTAAATTATAAACTAACATATTATCCATTCAAATGCTAACTAAAGATTT
CAAATGCAATGTGTTATATCCTTTATATTCCATCTTTTGAGAGCTGTCACGAGGAGGGTGATTGATCTTGATAAA
GTAATAGATGGTCACTACCATACAGATAATCCAATAATCGAAAAACTTCAATACGACCTTTAATATTATCTCTTA
TCTCAATAATTTTTAAGCCCGATAAAAAATGACTTCAACACCCATAGCCAGGCTACCCATCATTTATTAGTCTCA
AAACCATGATTCATTCAATGCTTTAAGAAATAAATAAATAGAGAATTTTATTGTGATTTTTGTTAATTAGCGAAA
TAAAAATATATTAATTTTCTAAAGAGTGACACAAAAAACCCTTATTCAGTTATTCCTTTTTCTCACATTCATAAC
AACATTTGGGAACCACCATCTACACCTAACATCACTCTCGATGGTTTGGGTTTTTAATCGATGATCATACCCATG
TGTTATGGATATTTTTACTAAAAATCAGAAGTGATGTAAGTATAATATTTTCCAACTTATTTAACCTAAACAAAA
ATCAATTTGGAGTTGTAATTAAAAGAGACATCCCATAATTCCAAAAGACTACTTTAATCATCCCTCCTCATTCTT
CCAAAAAAAAAGGAATAATATGAGTCCTTTTGTGCACACACTTCAACAAAATAGAGTGGTTGAATGAAAGAACGG
TCATCTATTACCAAATAGCTCTATTTATTCAAAAAAATGTCCCACAGCAAGTCTTAACAGCCAATCACATCATAA
AAAAATTAGCAACAAAGACCTTTTAGTTCAAGAGTCCAACACAAGCCTGAAATAAATTATTTTACAATTTCATCG
CCTAAAACAATATTGTTCCAAAAATATTTAAAAGTGTTGCGTTTGCTCATAAAAATTTCTAAAACAAACAAGACA
TGAACCTCAAGCAATAAAATGTATTTTATTAGCTATTACAAATTCTACCATTTTATCGTGAGAAAGGTGGTTGGT
TCACTGATGTCACTTTCAATAAAAAAATATCGTTATTTTGTTATTGCATATCATAACAAAGAATTATCATTAACAA
ATAAATTTTCCTCCTAGATTTCTCTTGTTGTTTATCCAAACAAATAACTCTATTCCAAAACCAACCACTCATACC
CAAGTTGAGTTCAAACCAAAACAAATTGTCGAAACATTCACAGCGGTAGAGAGATGTAGAGTTTAACGTCCAAGC
TTTGTACGAAGATCCATCACAAAAATAGATAAGAGTGAGGGGTGCAACAATCATAAGAGACACTAGATTTTTTAT
GTGAAAAACTCAAGAAAGAGAAAAATAACGGGTCTGGCCTAGTCAAAAAGATTTCACTAGTGTTTGTTTTCGATT
ATAAATGATGTCCTTTATCTGGTGAAGGTGTTAGCACATTACAGCAAATATACATTGTATAAAGCAAATTATATG
ATTGGAGTTGCAGAATCAATTGACATGAGAGCTTTTCTCAAAAAAATGTGGGCTTGTTTGTTTGGTAGATAAAAC

CATAGCCGAGCAACTGAGGTCAGACGGATCATAGTCGGTGAGCGGTGAAGGCCAGAAGAATCGTAATCGACAGTC
AACGAAGATCAGAAGGACTATTGTCATACGACGGCGGTCAAAGGGATAGTAGACGGAATCAAGCGATCTTGGGCT
CATTGGATTTGTCTTATTGAGCCACTCGATTTGACTATAAGCATGTCATATTTAGATATCATTGGTCACCCAATT
AATATTCTAAAGCTTTCAAATATAAATTTGAGCTTAAATTGGATCGATTTTGGTGTTGTTGAGTTCGCCTCATCG
AGACGTCCAGTTTGATTATTAGTTTGCTAAGATTGGAGTTAAGATGATCCTGAAATTAATTTTGAAAGCTCCCAA
TTCAAATATGCCTCAAGATTCACGTCCAATCAAGCCATTTTATTATGATGTGTCAAGCATGCATACTATTTTTAT
TCCCAAAATAATCTCTGACTTAAAATAATGAGAATTGAAATAATATTCTGAGAGAAAAATGGAATCCTTTAAAAA
AAATTAAAATAAAAAATCTGAATTGGTAAAAACACTAAAAAGAAAAAATATTAATAGGTTACGGATGGGTCTTTA
TTAGAAAATATTAATCTAACAGTTCACCACAAAAGGTATAAAGCCCAATTAGTAGCTAAAGGCTACACTCAAACA
TATGAGATTTATTATCTTGAAATCTTTAGTCCAAAATGAACACGGCTAGAAACCTCTCAACTTTAGTAGCTAACA
AGGACTAGAAATAACATTTTCTATGTTCAACTTACATAGATACATGGTGATCTACAGAAGAACGTATATAGAGAA
GTCACATCGAGATTTAGCTAAAATACAGAATAAATTGTATGTATACTAAAAAAGATATTGTATGGATTAAAGCAA
TATCCAAGGGTGTGTTTCAAAATATTCACCAAGCTAAAGCTCAATATATAATACAAACAAAGTTAAAAGGACCAA
AATGTTATTTCCGAAACACATCTTCAAGAGAAGTCACCGCACTGATAGTTTATGATAATTTCATTCTCATGACAA
CTAATTGCTTCGAAAGAAAAGACAATTTAAAGAAAAGTTAAATCTAAGAATTTGTTAATATGAAGTCTCCACGTC
ACATGAAAAATACATAACAAACATTTTAAAAGAAAAGGGTAAGTGAAGATGCAAAAATGTCACGAGCCCCATGGA
GTTTTAATTATAAATTTGGAGATGCCCCAAAAGATAGTAGTAAAAATTCTCCTATCAAAAACTAGCGAGAAGACT
TATCATATAATAGAGTTGACTGTAGCGACTCAATTCATGCATAATTCTAGAAAAATAAATCTTAGGACTAAATTC
AAAATTCTATAGTATTTAAATGGAACTTCAGGCAAAAGAATTTAATTCCAAAAAGGGGAAGATTTGATGCTGGAA
GCTAGGAATTATAACATACAACATCATGACTTTGAGAAATCAAAAATAAAATGTAGTAACAAAATCTAAAGCAAA
AGCTAAATTAAATCAATGGCACTAGAAAACTGCAAATTGATAGTGTTAAAAGTTGGATTTAAAGGTTGAATGAAA
ACATCTTACGAGGCTCTATCATTATAATAAGTCACTCATCAATATTGTATATAATCTCGTGTAGTATGACCGTAC
AAAGCATATTAAAATAAATACGGACATTTAATAAAAAGGAATTTTGACCATGATCGAATATGTGCTCAACAGGCT
AGCATTTTAACCAAATGATTATAACAGGTAGTCTCAAAAATTATCAAGAAAACTAGCAATGAATGATATCCAATC
ATTTTCGTAAAGAGTGTGAAACATTCTCATATTTTTTGTCAAATATGATCCAATACCAAGATTTCCAACCCCCTT
TTCTTCATTTTACATGGCACCTTAGTGATTCGTATGATCCTTCTTGCCTATCTATGAGCTATTATAATTGTTTTA
TAATTATTATCCTGCCAACTTATCAATATATATGCTATTTTAATTATTCTATAATTATTATTCTACCAACATAAC
CACCTATAGCCAATTGTAACTATTTTGCACTTAGTATTCTTTGATTATGTATTTTTCCTATTAGCTCTCCTAGGA
TTATGTAAATACTTCACAAATGTTAGAATGAAATACTTAACCAATTCTTGCCTTTCTTTAGCTCAATATTTATTT
TTTTAGAAGTATTTTAATTTTTAGGTGAGTTGAAAACTAGAAGAGTTGCATAAAAATAGTTGATCCCTTTCTTTT
TAATTGTTTGGTGTAATTGATCATTTTCATGCTATTAATTTGATGTTTTAATTCTCCTTAATGATAAGTCTATCT
AAATTTACATTGAATTAATAAAGATGTATTTTTCATTTTATATATATATATCATATCTCTTGAGATTTATCTCAT
AAGTTAATATTAAAACTCTTCATCATGTATCTTGCTTTTGGGTTTATCATTTTCTACATTCGAGGCATGAGCAAC
AACATACCAACAATGCAACAATAACTAAAACACACATTAAAGGAACATGCATACACACACTAATCAAAAATATAT
CTTTTATCTATATGTTAAAAAGGTAAATTTTTAACAAATCTAAATGTAAAGATGAATAGATATTCAAATATTTTT
TATGTTAAAATCTAAATGTGAATACACTGTGGTGAGTGATAAAATACTATTTACTTAGGGGAAACAAACCAAACT
ATTAATGATTTTAACCTCAAAACATCCTAAATGCAACATGAGAATATAGAGAAAATAATGCATATGGGTTATTTA
CAACTAGTTCTTTCAATTAAATTTAGAAAATTAGGAGGATATCACAAGATTGGAAAAGAAATATTTTAATATCAC
TATTGAAGAACATGAAAAAATTTGGGTTTGTGATATTTATAGGAAATCAAACTTATGATTCATAGAATTCATTAT
GTGAGAGGGAGTTATGGAAAAAATTTGGAAGAATTATTTTAATAACAAATAATCAATTTGATTTTATGTGTGAAA
ACTTCTACTTTTTTGCTTTTATTGTTAAAAGCTACTTTTCCCATCCTAGTTTCTATTATGCTTTACCAAAAACAA
TTTTTCAAAAGCAATGTTTGAAAAAGTTGTGAGCTAAAGTGTGTTTGCGTAAACTTTTTTAGCTATTTTCTTCTC
TGCTTCGCTTTTATACCATTGTCACGTTGTTGTTTGACTACAATTTTCTAGTTATTTTTAGAAATTATTGTAGTG
AGAAGTAATGTTTAACAACCATATCTGCATGTGAGGTAGTTATAAGAAAATAATTATCTAACTTATTCAAAATAT
ATATATATATAAAACCCATAACTAGTTTCAAAAAAATCAACTTATACCACAACAAATTTACACACACTATTAAAA
ATAGTTTTAGCTGCCCCAAATGGACCCTAAATGAACACATTATTCTTGATTTTTCTTTATAACCTTCCTATGGTG
AAGGTTGTTAAAAAAAATAAAAAATTTCTTTTTACCACAAAAAATAGTTAGAAAGTTGTCTCAAACAAAACTTTA
ATTAAAAAGAGAACAAAGATGTGAATAGATCATAAATTATCAAAAATTAGCATAAAAGAGATACTTACTGAAGAT
ACTCAAAGCAAGGTATATTGGTATGCAATTATTACAATGATATAAGTTTTATTGGAGAGGCCTTTGTTGAAATAA
GTAGAAATTAAGATATGGATAAAAAATTTGTGGGACAAAAGCCTAAGTTATTCCAAAGTAAAGAATGAGTATTCA
TTATGCAAATTTAATGGGTGTTACTTAGAATTATCAGGCAACATTAATATAAAATAAATAAAGACTTAAGTGTGA
TTAATTTAATAAATTGGATCTCTTGTATGGAAGAATGTGACATTTTTTATGATAAAGATTATAGGGTTAAATTTG
AGTGGTTAAAGGAGTGATAACTAGGGTGTACAACATAATATGAGAATGCATCGAGACAATGGTATTTTTAAAATT
GTAATTAGGTTATTTTGTTGTATTGGTTCAAGTTTTGGAAGGAAAAGAAGTCACTAATAGCCAAATTGAGTATGA
AAAAAATGAGAAACATTAAGATAGATCTGTAAGAACATTACACAGGATATATTAAAAAATCTTTTATGTTGAATA
TAACAAAATTAAAAATAGAAACATGAAAAGTTATTTTACACCTTGTGAAGCATTCAGCATAAACAAAATGACAAG
AGTGATTCATAAAGTGAGGCCTAAATGGTTGAGATAAAGCGAAAGATAATGATTTTGATGGTGATTTGATCTAAC
AATATGTACACTCTTGTTACTTTAGATTTCTTCCTCTTTCTCCAGCATGTGTTGCTTTCCTCCTTTTTTTTATTT
TTGTCGTAAATTGTTCTCAAAATCTTAAATGCTATGATCAAACTTCATATCTACCTGCAAGATTATATTAGTGAG
CATTTACACGTGAATGAAACAATTAAATTTTTCAAATTAAGGAAGTCGTGTACAAAGTGCCTCAAAATTTTTTTA
TTTAAATTTTTAAATTGTATAAAGTTGAAATAAGTTAAGCTCTAGAAGACAAAAACAAAACAATCGATTAGCTAA
TCTTTCAAATAAGCTTCACAATAAGAATCTAACATATTGTAAGCACTACCACTGACCACTCAATTATATGTAATT
TAATAACATGGAAGAAGTAAAATCTGTTACCACATTTAAGGGAGCCTTCATGGATGTCAGATACCTTAGCAATAT

TTGATAATCTGAATAGTCCATGGTATTAAATCAGACATTCCATTGGGCATACTCCATTTTTTTTAATTACGTTTT
TAACATTTACAATGTACTAGAATTTCAGAATATTTGAGGTATGAAAGGGATATAGAGCAACAAAAATTAACAAAG
CCATTAAATTACAAAGGGCAGACACAAACGCCATCTTCAAAAAGTTCCGTATATACATTCCCGATACGAACATCC
TAGGCCTCTAACGCGATGATAATTTATTCAAAAAATAAGACATTTCATCAAAATATCCCGTTCCGTTTCGCCCTA
AACCCTAAAACATCAAACACTAAAAGATAACCCAAAATCCCATGTAAAAGGCATCCAAACAACAAGCAAACAGTA
CAAAACTGAGTTCGCCCAAGAAAGCAAACATCAAACAAACAACACCATCACGAATTTTTTAAGCTGAAGTACCAA
TGGCGAAGAAACTAGTCCCGGAGAAATCAAAATCAAAAGCAGTCAACAACTATCTAAGGAAGCCGAGTGCCATGC
TTCACCCCAGTAAGAAACGAACAAGACGAAATACCCAAAGAGAGTCCAAAGAGAGAGACAGCGAGAGAGAAAAAA
AAATCTTATTGCGCAACCTAAAGCATGCAAAAGTACTTCCACCGCATAAATTCATTGAGAAGCTCTTCGAAC**CTC**
**TTTCAATTCTCTGGCCAAAGAACCCATCCCTCTCCAGCCAGCGCAAAGCTTCTCTAGAAGGAGAAATGTGGAAAA**
**CAGGAGGAGAAGACGAGCGCGAAAGGTACGGAGCGCACAGGGGAAGGGCATATATATAATCATACGGACCTCGCC**
**AGCCCCGCCCGGCGGTGGATCCTAGCCTTTGATATTAGGTATTATACACACG**CGGCGCATGCGAAAACCCTTTGG
AGGCAAGGGACTTTTAAGCTTACTAAACGAACTTTTAAAAAATACTGATCGCGCAATTTTTAATCGCGAAGGCGC
GCGCCGCGCGGCACTTGACCTGCTGTGGTAGGGCCAGATGGACACGTATCGGGCAGGTATTCAGCGTCCAAATCT
TTAACGAGTTTAATGCCGAATTTTATAAGTTAATAAGTCAAATTTTAGCTAGCTCTGATGGATAAGTTTAAATAA
TATTAATATAAAAAATATATAATGTATATTAGAATTTAAAATTATAAATAGGAAGTTTTAAAAAATATTTAAAAA
TCATCTCATTTTTTATATTAAGTATAATTTTATTATTTGTTTATAATTCCACATTCTTTTTCTCATTTTATTTAA
ATAAGTAATTAATCATAATTATTTTCACTCTTTCATAGTAAATGAGCAGCTCATGAGCCGCCTCTTGAGTCGGCT
ATAACTCATAGAATAAAGCTTGAGCTCGAACTCAATTAATTAGTCAAATTCAAGCTCAAGCTAAGGGTGAATTAA
ACTCAGTTGGACCTGTTTACATTACACCTCGAAAATGGACTACTGAGAAGAGTTTGAGCCCAATTTGAAATTGAC
TTGATCTAAATTAAGTCAATTTGAATCAGATCGATTTTCAGTCAATCTGAATTCAACCGGTCAAAAAAATAAATA
AATCCGAACTAAACATGCTGGCAACCTAAATTATCAAAGC

>DecaAGL17_1

ATGGGGAGGGGGAAGATAGAGATAAGGAGGATTGAGAACATAACAAATAGGCAGGTGACCTTCTCCAAGAGAAGA
AATGGCTTGCTTAAGAAGGCGAGAGAGCTCGCTGTCCTCTGTGATGCAGAGATTGGGCTGATCATCTTCTCGAGC
ACAGGAAGGCTTTATGATTTCACCAGCTCCAGTATGAAATCAGTGATTGGCCGCTACAACAAAGCAAAGCAGGAT
CATCAGCAAGTGATGCATGCAAAGTCAGAGCATAAGATCTGGCAAAGGGAAGCAACAAGCTTGCGGCAGCAATTG
CATAACTTGCAAGAAAATCATCGGCAGCTGATGGGAGAAGATCTTAATGGGCTAAGTATCAAAGAACTACAAAAC
TTGGAGAATCAGCTAGAAAAAAGCCTCCAGGTGATACAGCAAAAAAAGGACCAAATTTTAATCAATGAAACTCAA
GAACTGAATCAGAAGATTGCCCACATGCAACGAGAAAACATGGAACTCTACAGAATGATAAACCTTTGTCGCCAA
GAAAATATGGACTTAAACAAAAAGGTTAATGCAACTAAAGGATTCAAAGGAGGATCTACTTCTTCAAACTCTGGC
AGTCCGGTAACAATTTATCTTGAATTGAGCCCTCCGCAACAGAAAATTAGTGGAATGCACGAGGACTTCCCAACA
TTAGGGTGA

>dec_miR444a_SRA

CGTGTGTATAATACCTAATATCAAAGGCTAGGATCCACCGCCGGCGGGGCTGGCGAGGTCCGTATGATTATATAT
ATGCCCTTCCCCTGTGCGCTCCGTACCTTTCGCGCTCGTCTTCTCCTCCTGTTTTCCACATTTCTCCTTCTAAAG
AAGCTTTGCGCTGGCTGGAGAGGGATGGGTTCTTTGGCCAGAGAATTGAAAGAGTAGGTTTGGGAAAAGGAAGTA
AGCTTGAGGGAGAAATTGCATAGCTTGCAAGAAAATCAACGCAATATATCTTCATCCAAAGTATTTTATTTCTGA
TGGACTCAAAGGGTGAAGAGAAGTTACCGATGATTTTCTTGCAAGTTATGCAATTGCTGCCGCAAGCTTGTTGCT
TCCCTTTGCCAGATCTAGTGTGGACACCAAACAAATCAATTAGTATAGTGGAATGCTCAATTATGCGCAAAAAGC
ATATAAAATAAGAATTACTGACACAACTTATCTAGATGGATTTTAACATAACTAGCATATGGCCTAGAATGAAGG
TCTACAAGATCATGTTTAAAATTTTTATGGCGGCCAATCACTGATTTCATACTGTAATAGAGATCAACCCATTTG
TTTGGTTTCAAAAACTGGAACTTGCATCTCCTACTTCAGAAAGGGAGCTGAAGAAGCACATTTCGGTGGTTTTCT
GGTCGTCTATCAGTCAGGTTCTTGTCCATCTTTTAGGCCCTGCTCTGGTTCGGACATGGGCCAAAGCATTCAAGC
CATCTATACAATGCAAAATTCCAATCAAGCCACGATAAACTCAAACTACAAAGTACCTTAGGCTTTGTTTGGGAC
GGTTTTTTAACTACTACTTAAAGCGGATTTCTTGTAAAAAAAATTTAATTTTTGAACTAGAAAGCTGTTTTAAAA
AGCAGTAAAAAAGCCGTCTCAAACTAAGTCTTAGTTTGGTATTTTGTTTTTGGAGCAGGCAAAGATACAAGGATT
GGGATGTGGTATACTTTTACTTGCTTTCTGAGGGAACTATGATTATTTCTACGATATCGAAGACTGGGAAAATCA
TGGCCCTTAAGAAAGCGCTTTGAGCAGAATAGGTTAAAGGCTGCTCAAAGTGGTATTGTAGCTCCCCAAACTCCA
AAGCCTCCAAGTCATTTGAATTTATTTTTGCTCAAGTTACCAATAAAAGGCTATATTTGCCTATGGAATGTCAAT
TGAAATCATCAACAAACCAATACTATTTTACAATTCAGCTAAG

Stem-loop structure

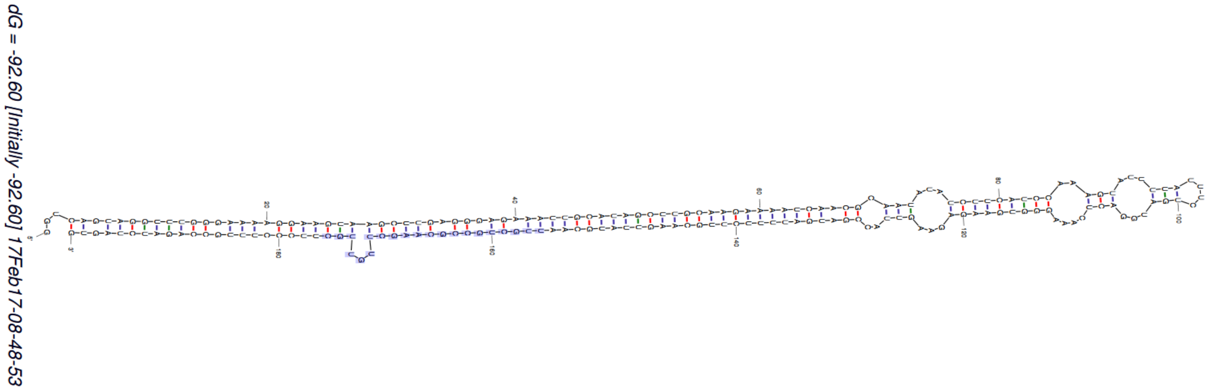


*dec-miR444b*

>JSDN01S015150.1

CCAGATAGTTTTCAATTATTCGTAAAATATATAACTACCGATATAGGATGAGTATTAAATTCAAAAGAGCC
ATAAGGATGTTTAGGAGCGGATTGAAAAAATATATACAGATCAATACAAATATTCAAAAAAAGAATGAAAA
TGATAAAAGATATCATTTTCTGTATTTAAAACCAAATACAACTATTGACCTGGACTTAGAAACTATGCAAC
CTGAGAACATTGCTGATACTCAATACCTGCAAGTTAGATTTTATTAGAAGGAAACAAACAAATATGTAGGA
ATAAATTAAAGTAAAACAAAATTGGCTTAATATCATCTTACATTCAGTAGGGCTCAATGTTCTGGTTAATT
CTTTCCATGCTAGTAGTTGATATCAAGGGGGAAAACTACAACACCATTTTTTTTTTTTTTTTTAGAAAAA
ACTACAGCACCATTAACAACAAAAAAATATTTTTTTCTCAAGCAATCTTAATTAGTTGATATTTTAACAAA
TGGTGCTTTTTTCTTCATTATTTTCACAGAAATCATGAGCATGTCAGTACCAACTTTGTTTAATGAATAAT
CTAGTTCTGGCAGGTTGAAGTAAAA**AGCTTGCGGCAGCAGCTGCATAACTTGCAAGAAAATCATCTGTAAC**
**ACCTCTTC**AACCTTGAGTTTGCAAAAAGAAAAAAATTAAACATATTAGCCGCAAAAAAAATAAAAAAAAG
CGGCATGATATATAAATAATCTCAATCAACAAAAAAAAAACATAGATACCCATTTTGGATAATTGAAGCAA
GAAATCTTTATTTTGGATAATCAAAAGCGTTCATGAGTGAGACGGAAGTAATATTTGGATAAGCAGAAATA
ATTTTTAAGTTTATTTTACTTATATTTATTTTTTTAAAATATATCTCAGGTCTAAATTACACACATCTCAA
CAAAACTAAAGTGAGGACTCATCTAAGGTGTAAGCTTCACATTGGATGCAAATTTTCCAGAAAAGCGTTGA
GGCACACCTCGTATTACAAATTTTAAACCATGCTTGAGTAAAATACAAAATCTGCTAAATCAATCATCTTT
AGTATTGTTCTTCTATTACAGGCCCAAAAATGCAATATCATCCTTCAAAAAAGCTTCTGTAATGAAAAAGT
ATGGGAACTAGTCCAAAATAATTATCATACTTGAAAATGGAGTAAAAAAGACTCAAATCTTACTTCTATGA
GCAATATCATAGCTGAATCGTATGCTTACTATAGGAAACTAAAGTTCATGCACTTGACCTTATAAATATTT
GTACAATGAACATAAAGGCGCCAATATTTGAATATTATTTTATAAAAACAGATTTTATAGCAAATCAATGA
AAATATTACACACAAAAAAACGAAAAGATTACAATATAATTACAACGAGCAACAAAAAGACAGCATCAACC
AAAATACAAGGAACAAAAGAAAACATGATAGCAGGGAAGAAAATTTTCATCAAAATTATGTGAAGAAACCC
CATTAACTCCTCGCAAGTCAAGAATAAAAAAATAGTTAGACAAACTATAGTCTCTTGTTTGAAATCTAGTA
AACCTTGATAAAGGAAGAAATATATATTTTTGTAATTATTGTGTCTCCTAGTAAAAAGAATTTGAAATCTT
TTATCCATTTGTTCCTCAAGACTTATAGATTTGAGATTTCTATCAAAAGCATATGAATAATCAAATCTTGG
GCCACCTACTTCCTTTACAACAAAAAGATCATGAGGAGTTTCACTAACACAACTGAAAAAGTCTAAATCAA
TGCTGACATTTAAAGAATAAGTTCCTCCTACAGCTATGATACTTAAAGAAGGAAGCATCAACAATAAGGGA
GGCTCTGATTCCAAGGCAAGAGGTATAGTAGGTGTAATATCCAAATTTTTCAAAAGATGATATGAAACTTT
CTCATAAAACTTGGACCCCATTTTATGCATGTCCTGAAAATCATCAAAACTATTTTTATTAAATTTTATAT
TTCATTTTGATTATTTTGATATATAAAAGTACATATATAAATATTCAACTTTTCATTTAAATTTATTTGAC
CTAAATTAGTAAACAATATGTGCCGAATGTTGGCCATTAAAAGAGAAGCATAATACTAAGTTAAGTGTGGC
AGAGATGAGGATGCTCAAGTGGATGAGTGGCTTACCCTTAGAGATAGGATACGAAATGAGCATATACGTGA
AAAGGTTGGAGTAGCTCCANNNNNNNNNNNNNNNNNNNNNNNNNNNNNNNNNNNNNNNNNNNNNNNNNNNN
NNNNNNNNNNNNNNNNNNNNNNNNNNNNNNNNNNNNNNNNNNNNNNNNNNNNNNNNNNNNNNNNNNNNNNN
NNNNNNNNNNNNNNNNNNNNNNNNNNNNNNNNNNNNNNNNNNNNNNNNNNNNNNNNNNNNNNNNNNNNNNN
NNNNNNNNNNNNNNNNNNNNNNNNNNNNNNNNNNNNNNNNNNNNNNNNNNNNNNNNNNNNNNNNNNNNNNN
NNNNNNNNNNNNNNNNNNNNNNNNNNNNNNNNNNNNNNNNNNNNNNNNNNNNNNNNNNNNNNNNNNNNNNN
NNNNNNNNNNNNNNNNNNNNNNNNNNNNNNNNNNNNNNNNNNNNNNNNNNNNNNNNNNNNNNNNNNNNNNN
NNNNNNNNNNNNNNNNNNNNNNNNNNNNNNNNNNNNNNNNNNNNNNNNNNNNNNNNNNNNNNNNNNNNNNN
NNNNNNNNNNNNNNNNNNNNNNNNNNNNNNNNNNNNNNNNNNNNNNNNNNNNNNNNNNNNNNNNNNNNNNN
NNNNNNNNNNNNNNNNNNNNNNNNNNNNNNNNNNNNNNNNNNNNNNNNNNNNNNNNNNNNNNNNNNNNNNN
NNNNNNNNNNNNNNNNNNNNNNNNNNNNNNNNNNNNNNNNNNNNNNNNNNNNNNNNNNNNNNNNNNNNNNN

NNNNNNNNNNNNNNNNNNNNNNNNNNNNNNNNNNNNNNNNNNNNNNNNNNNNNNNNNNNNNNNNNNNNNNN
NNNNNNNNNNNNNNNNNNNNNNNNNNNNNNNNNNNNNNNNNNNNNNNNNNNNNNNNNNNNNNNNNNNNNNN
NNNNNNNNNNNNNNNNNNNNNNNNNNNNNNNNNNNNNNNNNNNNNNNNNNNNNNNNNNNNNNNNNNNNNNN
NNNNNNNNNNNNNNNNNNNNNNNNNNNNNNNNNNNNNNNNNNNNNNNNNNNNNNNNNNNNNNNNNNNNNNN
NNNNNNNNNNNNNNNNNNNNNNNNNNNNNNNNNNNNNNNNNNNNNNNNNNNNNNNNNNNNNNNNNNNNNNN
NNNNNNNNNNNNNNNNNNNNNNNNNNNNNNNNNNNNNNNNNNNNNNNNNNNNNNNNNNNNNNNNNNNNNNN
NNNNNNNNNNNNNNNNNNNNNNNNNNNNNNNNNNNNNNNNNNNNNNNNNNNNNNNNNNNNNNNNNNNNNNN
NNNNNNNNNNNNNNNNNNNNNNNNNNNNNNNNNNNNNNNNNNNNNNNNNNNNNNNNNNNNNNNNNNNNNNN
NNNNNNNNNNNNNNNNNNNNNNNNNNNNNNNNNNNNNNNNNNNNNNNNNNNNNNNNNNNNNNNNNNNNNNN
NNNNNNNNNNNNNNNNNNNNNNNNNNNNNNNNNNNNNNNNNNNNNNNNNNNNNNNNNNNNNNNNNNNNNNN
NNNNNNNNNNNNNNNNNNNNNNNNNNNNNNNNNNNNNNNNNNNNNNNNNNNNNNNNNNNNNNNNNNNNNNN
NNNNNNNNNNNNNNNNNNNNNNNNNNNNNNNNNNNNNNNNNNNNNNNNNNNNNNNNNNNNNNNNNNNNNNN
NNNNNNNNNNNNNNNNNNNNNNNNNNNNNNNNNNNNNNNNNNNNNNNNNNNNNNNNNNNNNNNNNNNNNNN
NNNNNNNNNNNNNNNNNNNNNNNNNNNNNNNNNNNNNNNNNNNNNNNNNNNNNNNNNNNNNNNNNNNNNNN
NNNNNNNNNNNNNNNNNNNNNNNNNNNNNNNNNNNNNNNNNNNNNNNNNNNNNNNNNNNNNNNNNNNNNNN
NNNNNNNNNNNNNNNNNNNNNNNNNNNNNNNNNNNNNNNNNNNNNNNNNNNNNNNNNNNNNNNNNNNNNNN
NNNNNNNNNNNNNNNNNNNNNNNNNNNNNNNNNNNNNNNNNNNNNNNNNNNNNNNNNNNNNNNNNNNNNNN
NNNNNNNNNNNNNNNNNNNNNNNNNNNNNNNNNNNNNNNNNNNNNNNNNNNNNNNNNNNNNNNNNNNNNNN
NNNNNNNNNNNNNNNNNNNNNNNNNNNNNNNNNNNNNNNNNNNNNNNNNNNNNNNNNNNNNNNNNNNNNNN
NNNNNNNNNNNNNNNNNNNNNNNNNNNNNNNNNNNNNNNNNNNNNNNNNNNNNNNNNNNNNNNNNNNNNNN
NNNNNNNNNNNNNNNNNNNNNNNNNNNNNNNNNNNNNNNNNNNNNNNNNNNNNNNNNNNNNNNNNNNNNNN
NNNNNNNNNNNNNNNNNNNNNNNNNNNNNNNNNNNNNNNNNNNNNNNNNNNNNNNNNNNNNNNNNNNNNNN
NNNNNNNNNNNNNNNNNNNNNNNNNNNNNNNNNNNNNNNNNNNNNNNNNNNNNNNNNNNNNNNNNNNNNNN
NNNNNNNNNNNNNNNNNNNNNNNNNNNNNNNNNNNNNNNNNNNNNNNNNNNNNNNNNTTCAGCTCACAAAA
AGCAAATGTCTGGTCGAGAATATCACGTGTATTTATTCTCAAACAATCTCTTACTAGCAAGGATGTGGCAC
ATGTCCTTTAACCGGTGGTTTCGCCACCAAATGACACCACAGGGTAAATGGTTATGACCGGGACCCTCTAA
ATCGGGTGATTTGCCGCTAAATGGTACCACAAAATTACGGTCATATTCAGAGCTCACATGTTCCCCAATGA
TTCCAAATTCTTGAGATATAATCGATAGGGCACACATCCCTCTGTTAGCTAAGGTGGATTTAGATTTATCA
CCTTATTCACTCCAAGTCTACAAACTACCCATAGATGAAGCAACCCATAGTTGTCGATGAGAATTCTTCTA
TCAAGTTTGGATCGATTACCTCCTATGACTAAGTAGATCAGAGACTACGATGGCCCGAGAAGACGGAACTA
TGGAATCTTGATCGGGTCTATCATCACTTCCCCCACCATCCGCTTTAAGTGCAAGATGGCAAAATTTTTAA
TTTTAAATATTAAATCTCAAAAACTAAACCCATGCTGCTACTATATACCAGATATGTATATGTATCTGTTT
AATGACTAGCCTTAATCGAAAGTTCTCTCATGTTATCGGGAGGGGTAATTAATTATAGGGCTTGCGCCTAT
ATGAAACCTGAACTCAGGTCTGAAAGTTCTCATAAAATGAGGGAGAAATCTCTAGGTGATGTCTTATTCAT
AATCCAATATGCCTTCTGAAAAATTTTCCCAGGAACTTGAGCGTTAAAACTCTCCTCGACGACCCAAAGAA
GAAAGAAAGGGTTTGCGAAGAGGCCGGGGTGCGAATTTACGGAGCCCACATTTAAGTTTTGCATAAAGGTT
AAAGAAAGATAACTTGATTAAAAAATAAATTGTAGTAATAAAGAAACTCAATTAAACTGCCAAATCAAACT
ATCACAACAATAAATAATACTTGAGCAAAATATAAAACCTAAGAGCTTAGCCTTACATAACTATTCCAGAG
AATACTTATGCTAAAAATAAAAGCAATAGCTTTTTATAGATTCCATATAGGGGTGTCAATATTTGACACGA
CCCAAAAACCCGACTCGAACCCGACCCCAAAATTTCGGGTTAGGGTTAACCCAAACCCGAAACGGGTCGAG
TTCGGGTTCAACATTTTGACCCGTTTCAAAATTCGGGTCGGGTTCGGGTTAACCCTTTGACCCGAAAACCC
GATTTTGACCCGGCAACCCGAGATTGACCCGTTTTTACCCGAAAAAGAGGAGATAGACCCGTTTTTTACCC
GAAAAGATGAGATAGACCCGTTTTTTACCCAAAAACTAGCAATTTTTTGTTGGAAATGGGTCATTCGGGTC
GTGTCGGGTTGGCCCGGACCCGAAGCGGGTCGGATTCGGGTCAGGAAAACTGACCCGTTTTCAAAAAGGGT
TGGATTCGGGTTGACCCGAATAGTGCCGGATCAGTTTCGGGTTCGACCCGAACCCGACCCGAAATTAACAC
CCCTAATTCCATATGCAAATGCACAAACAAAACTAGTCATCATGAGATTTAAACTTAAACTACACATAGTC
CATGTTATTTTGATAGCATTCACCTTGCATGTATGATAGTGCTTGAAGTACAGTCGCTCCACCATTGAGTT
CTACTGCCTACTATAATACTCATGGTAATCCCATATAACTCTGATACGGGGGAGAGGGCAGCCAGCCCAGC
AGGGGTCGGTCTAGCAAAATCAGGTGGACAACCTTAGGGGTGGAACTCGCAGAAATTTCGAGCAGGTTATG
AAGGATACACTTGTGTAAGAATCGAGACCTTCAGGCCTTGATGGTTTGTGAAGGGAAGGAGGAGAGTGGGA
CAAGCTGGAGGAGGTTATGAAGGAGATAGAGCAGTTACATTTAGATGTAGCAGAGGAATCTCTGAACTCAA
TGGTGGGATTTATCCCAAATCACACCATGAGGTAAGGGGACAAATTGGAGAGCAAGCATTGGTGGTTTTGG
TAGATGGTGGAGCCACTCATAACTTCATCTCCCACCGGGTCGTGGACCAGTTAGGATTGAAGCTAGTGGAT
ACAAGGAGTTCTGGAGTCATTAGGGGGGCATGAAAGGTAGAGAGGAATAAGGGTATTTGCAGAGGGTTGGT
ATTAGCCCTCCAAGGCATATAAATAGTGGAGGAATTTTCTTCTATTGGAATTGGGAAGCACTGATGTCATC
ATGGGGATGAACTTGTTACAGACGATTGGAGAGACTAAGGTGGACTTGAAGTCTTTTTCTATGGAGGTACA
GTTGGAAGTGAAGGTGGTGCTACGAGGGGATGCCAGCTTATGCAGGACCATAGTTTCACTCACAACCATGA
TCAAGGATATATAGGAAGCAGGACAAGGCTATTTGGTAAAACTGCAGAGTTTGGAGGGGATCCGAGAGTCA
TAGGCAGACCGGATCCCTGACTGTTTATATCAGTTGCTACAACAATTTCAGAGGTCTTTTAGCAACCCAAC
AGATTAGCACCACCTTGAGAGCATGAGTATGCAATTTCATTGACAGGAGTTTTCCTATCAGTGTTCAACCT

TACCGGTACCTGCAGATCTAAAAAGATAAGATTGAGTGGCTCATCAAAGAGAAGCTAGAGGCATAGATTAT
TAGATCCAGTGTGAGCCCTTTTTCCAGTCCTATTTTACGGGTGAAAAAGAAGCACAAAAGTTGGAGGTTCT
ACGTGGATTATAAAACTTAAAACAAAGAAACTATGTTGAACAAATTTTCTATTCCTTTGATTGATGAGTTA
TTGGATGAACCATTTGGGGTGATTATCTTCTCAAAGATTGACCTGAAATTCGGATACCACGAGATCAGAAT
GAAACAAAGGGATGTGCAAAATACAACTTTCAGAACCCATGAGAGTCACTACGAATTTTTAGTTAAGTCTT
TCGGCCTCACCAATGCTCCAGGCACTTTTAAGTCACTCATAAATCGAGTTATTCAACCCTATCTTAGAAAG
TTTGTCATGGTAATCTTCAATGACATCCTAATCTACAGTTGAATGGAAGCATAACATATGGGACACTTGAG
GACAGTGTTGGAGACATTGAGGGAGCACCAGCTGTATGCCAACCTGAAGAAATGTTCATTTGCACAACCAT
AGGTAGAGTACTTGGGACATGTGGTGATCAAGACAAGGGTAGCAATGGACCAGTCCAAGATTGAGGCTATG
CTAAATTGGCCTTTGCCCAAGAGCATTAGGGAGTTAAGAGGATTCTTAGGCCTCACTTGGTACTACTAGAA
GTTCGTGCAAGATATGGCACAATAGTAGGGTCATTGACTGAGTAGCTGAAGAAGGAAAGTTTCAAGTGGGG
GGAGACAACTACACATGCTTTCGAGTTGCTAAAGAAAGTAATGACTAGCAGTGGCGGATCCACGGTTGGCT
GAGGGTGGGCAAATGCCCACCCTAAAAAAAAAATTATTATAGAGAAGGGGCATTTTAGACCTTCTGTCCCC
CCTAAAAAAAAATTCACTAAATCTTCCGATCGTGAGCCATTTTTCCCCATTCAGTTCGCGATTTCCTCACG
CCGTCACCCGTGCTCCAGTCGAGTTCGCTGCCTCGCAAAGTCACAGTCGCAGGACGAGCGGACCGGCGGCG
GCGGATCCCTCTTGTTCACAGCCTGCAGCCTTCATCTTCGTGTCTACGTGAAGAGCCGAAGAGGTTGCCGG
CTCGCCGCTGCCCTCTCTTTCCAGTTCCCGATTTGCAGCCTTCTTCACTGTTTTGAAGGTAAGTTTTGATT
GTTTGGCTTCTCATTACTACTCCATAATCTCCACTTGCTCTTTTTTTTGCTTCCATTTTTCCTTATCTTGT
TATCTGCATCTTGAAAAAATTTGGATATTTTACTGTTTATCTTCTTCACTGCATTATCTTCACTTGATTAT
CTTCTTCACTGCATTTTTCCTTATCAGGTTGCTATTCATCTTCAAATTTTATATTTTTACTGTTTCTAACC
GAGAATTCCCAGTTGGAAGAAATGTGGAAATGTAGTGTTTCTAAGTTCTAACTACTCTTTATATTACTCAA
ATGAATTAACTCAATTAATTAGGTTTAACTCAACTAAATGAATTAATTCAGTACCAATAAGGTAATTGGAT
GAGACAATTAGATTTTATATATATGCTTTGCTCACACAGATTTTGAGTTATCTAAACCTTTGTTTTTGTCA
TTTGTATTGAGACATAGTCCTATTGTAGCTTTTCAAATTAAAGATTAATTTTTTTTTTTTTTTTTTTTTT
TTTTTTTTAATGNNNNNNNNNNNNNNNNNNNNNNNNNNNNNNNNNNNNNNNNNNNNNNNNNNNNNNNNN
NNNNNNNNNNNNNNNNNNNNNNNNNNNNNNNNNNNNNNNNNNNNNNNNNNNNNNNNNNNNNNNNNNNNNN
NNNNNNNNNNNNNNNNNNNNNNNNNNNNNNNNNNNNNNNNNNNNNNNNNNNNNNNNNNNNNNNNNNNNNN
NNNNNNNNNNNNNNNNNNNNNNNNNNNNNNNNNNNNNNNNNNNNNNNNNNNNNNNNNNNNNNNNNNNNNN
NNNNNNNNNNNNNNNNNNNNNNNNNNNNNNNNNNNNNNNNNNNNNNNNNNNNNNNNNNNNNNNNNNNNNN
NNNNNNNNNNNNNNNNNNNNNNNNNNNNNNNNNNNNNNNNNNNNNNNNNNNNNNNNNNNNNNNNNNNNNN
NNNNNNNNNNNNNNNNNNNNNNNNNNNNNNNNNNNNNNNNNNNNNNNNNNNNNNNNNNNNNNNNNNNNNN
NNNNNNNNNNNNNNNNNNNNNNCCCCAAAAAAAAAAAAAGGCTGGATCCGCCCCTGATGACTAGTGTATCG
GTATAACCACTTCCTGACTTCAGCAGGCTGTTTGTGGTAGAGACAGATGCGTCAGGGATATGAACTAGGGA
TAGTACTCATACAGAATAAGAGACAGATTGCCCATTTTAGTCAGGTGCTCACAACCAGGGCGATACTCAAG
TCAGTGTACGAAAGAAAGTGGCGACCCTATTCATTGGGGAGACACTTCCTAGTTAGAAAAAATCAGAGGAT
CTTGAAGTATTTGCTGGAATAACGGATGGTCACCGAAGAGCATCCATGATGGCTATCCAAGATACTGAGAT
ATAATTTTGAGATCCAGTACTGACCGGGCCTAGAAAACAAAGAAGCTAATTCCTTATCCAGGAGAATCGAG
GAGCCTCAGTTAACAACTTTGTCAGTACATCTAATCCTAGATTGGGAAGTTCTAAAATAGAAGAGTGAACA
GGATGATAGACTAATGAAGATCATAGACGAGTTGTTGTAGAACCCGAATAATCATCCTAGTTATATGTTGA
GAGGAGTCCAATTGCTACATAAGGGGAGATTTGTTCTACCAAAAGGATTGGGATACATCCCCAAGCTGCTA
CAGAAGTTTCATGGCAGTGTAGTGGGAGGGCATTCAGGACTGCAGAAAACCTACAATCGATTATCAACAGA
ATTATACTAGGTGGAGATGCATCGTGATGTAAAGGATTGGATAGCCAAATGTGATGTTGAACAAAGGCAAA
AGTACCTATCCATGGCGCCGACAGTGTTACTACATCCCCTAGCCCTACCGAAGAGGGTGTGGGAGGAGGTG
ATGATGGATTTCATCAAGAGATTGCCATGATCAGAGGGGATTTCAGTGATACTAGTAGTAGTGGATTGTCT
GAGCAAAAATGCACACTTCATTCCTTTGAAGCGCCCTTATACCATCGTTATTGTGGCTACTGCCTTCATAA
AGGTAGTAGTGCGCCTACATGGAATTCCATAAGCCATTGTGTCAGACAAGGACAAGGTGTTCCTTAGTCAC
TTATGGAGAGTTGTTTCAATTACAGCGGACAATTCTAAAGAGAAATGTAGTCTACCACCCTCAGACATATG
GCCAGATAAAAGTGACCAACATAAGTATAGAGACCTATTTGAGATGCTTTGCAGCAAACACGCCAAAGAGC
TGGGTAAGGTGGATTGCTTGGGCGGAATATTGATACAATACTTCATATCATACAGCAGCGAAGACCACTCC
TTTCAGGATTCTCTATGGGAAAGATCCACCCCATCTGATTCATTATGGGCTGAGCATTATGCCTGTGTCTA
GTGTAGAGCAGTACTTCGAGGAGAGGAATAGGGTATTATCTGAGCTCAAGGTACATTTGTTGAGAGCCCAA
CAGATCATGAAGAGGAAGGCTGATGAGCACAGGAGGGAAGTAAACTTTAAGCCAGGAGATATGGTTTACAT
AAAATTGAGGCCCTACTGATAGAAAACACTAGCCAAAAGGAGTAATAAGAAGTTATCTGCATGGTAATTTG
GGCCTTATGAGATCTTGGAGCGTATTGGAACAGTAGCTTATCGATTGACGCTACCACCCTCATCAGCTTTG
CACCCAGTCTTTCATGTCTCACAGCTAAGGAAGGCGATAGGATTATTGTATCCTGCTTTGATGTTGAAGTA
TTAAAACAAAAATCTGATGAGGACATCATTGACACGTGTACCACCGGAGATCAACGGCCACGGTCATGAGT
TTATATTTTGAGTCATGTTATGTAGTAGTTTATTTTATTTTTCAGTTTTACTGTTTTACCCTTATGGATGG
TCATTCGATATTGTAGTCCCGATAGCATTTATGTCTTTTAGTTTTAGTTTTATTTTGGCCTATTTATAGGA
GGCCTCATGCATTGTATGAGGATGATTATTGATGAATGAAAATTTCTTTTTCTTGTTGATCTCTTTGTGAG
TTCATGAAATTTCCCAAGATCTCTTTGAGAGTTCTTGAGAAGATAGTATTGTGGATCTCTTTGTGAGTTCT

TGTGGAAGTGGTGTTCAAGGTTTGTAGTTTGATAACTTTCTTTTCCCTACCCAATCCTTTTTTCCCCTACT
TAACCGAAAATTCCCCTCCTCCCCAAAATTTCCCACCCCACCCGCTCCACCGCCGGCCTCCCTTGGCTGAT
CCAGCCACCGGCAGACCTCCCAGCTCCCTGTCGCTAGCCGCCGGGTCCAAGCCCATTATTGAATTAATTTG
AAATTTCAAATTAATCTCCCTCCAAAACCAATTGAAGCTGAAATTATTTTCAGTTTTCTAAATCCTTCCCT
CTCCCTCTAGCTCACGGCCAACCCCTCTCTTCCCCAAACTGCCACTTCCACCTCCCGTCCCCCTCCCTCTT
CTCTTCTCTATTTATTACGAACATGAATGGCGGATCCGGAATTTGATCACGGATTTGCGTATGACTTTCAA
GGAGATATTGACGTCGTTCGTTCGCCATTCTTTGACGTTGGGTTTGGCTTCAACCACACCGTTGAGGATTA
CTTGAACTGTATCCTTCCCACTCTCGTGGATGTCATCGACGAACAACTTTCGGCCGACGATTAGATCATCA
ACGGGCGTCCACCATCTTCCCCAAATCCGGCTTCTTCCCCGGCGACTAGTCTTCTCAGCGTCACTTGTGTG
ATAGTGGCATCGTTGAGTTTCTGCGCCATCCTCTTCCGCTAACCGAGTGAATCCAAGAGTGAAGTGAGGGA
CTTCAAGGTGAAGGATCTATATCTTAAGCCGTGGATTTTTGTTAAGAGCCATTTGCTTTATTACTTATCCA
TTGGATTAATTTTGTTGGTTTCATGTCGCTTCTAAGATAATTGATTGACGTTGCTTAGTTCTTCCATGGAA
ATTTCTTACGATCTATCTTGGAACACATTTGTTTTAAATTAGCAATGCAAGTTCACACGTTATTAATTTTC
TAACAAGCAATTCTTGTCTATCGCATTTAGTTTCACATCAAAATATTCTATCATTTCTCAATTTCTTCTTT
TTATTATCTTTTCTCGCTGATTTCATCCCCAAGGAGTATCAAACTCCTATAGTGTCCAGATAGTTTTCGAT
TATTCGTAAAATATATAACTACCGATATAGGATGAGTATTAAGTTCAAAAGAGCCTTAAGGATGTTTAGGA
GCGGATTGAAAAAATATATACAGATCAATACAAATATTCAAAGAAAGAAAGAAAATGATAAAATATATCAT
TTTCTGTATTTAAAACCAAATACAACTATTGACCTGGACTTAGAAACTATGCAACCTGAGAACATTATTAG
GCTTTTACTTAAGACCATTTGCTGATACTCAATACCTGCAAGTTAGATTTTATTAGAAGGAAACAAACAAA
TATGTAAGAATAAATTAAAGTAAAACAAAATTGGCTTAATAGCATCTTACATTCAGTAGGGCTCAATGTTC
TGGTTAATTCTTTCCATGCTAGTAGTTGATATCAAGGGAAAAAACTACAACATCATTAACAAAAAAAAAA
CATTTTTTTCTCAAGCAATCTTAATTAGTTGATATTTTAACAAATGGTGCTTTTTTCTTCATTATTTTCAC
AGAAATCATGAGCATGTCAGTACCAACTTTGTTTAATGAATAATCTAG**TTCTGGCAGGTTGAAGTAAAAAG**
**CTTGCGGCAGCAGCTGCATAACTTGCAAGAAAATCATCTGTAACACCTCTTCAAC**CTTGAGTTTGCAAAAA
GAAAAAAATTAAACATATTAGCCGCAAAAAAAATAAAAAAAAAAGCGGCATGATATATAAATAATCTCAAT
CAACAAAAAAAAAACATAGATACCCATTTTGGATAATTGAAGCAAGAAATCTTTATTTTGGATAATCAAAA
GCGTTCATGAGTGAGACGGAAGTAATATTTGGATAAGCAGAAATAATTTTTAAGTTTATTTTACTTATATT
TATTTTTTAAAAATATATCTCAGGTCTAAATTACACACATCTCAAAAAAACTAAAGTGAGGACTCATCTAA
GGTGTAAGCTTCACAGTGGATGCAAATTTTCCAGAAAAGCGTTGAGGCGCACCTCGTATTACAGATTTTAA
ACCATGCTCGAGTAAAATACAAAATCTGCTAAATCAATCATGTTTAGTATTGTTCTTCTATTATAGGCCCT
AAAATGCAATATCATCCTTCAAAAAAGCTTCTGTAATGAAAAAGTATGGGAACTAATCCAAAATAATTATC
ATACTTGAAAATGGAGTGAAAAAGACTCAAATCTTACTTCTATGAGCAATATCATAGCTGAATCGTATGCT
TACTATAGGAAACTAAAGTTCATGCACTTGACCTTATAAATATTTGTACAATGAACATAAAGGCGCCAATA
TTTGAATATTATTTTATAAAAACAGATTTTATAGCAAATCAATGAAAATATTACACACAAAAAAACGAAAA
GATTACAATATAATTACAACGAGCAACAAAAAGACAGCATCAACCAAAATACAAGGAACAAAAGAAAACAT
GATAGCAGGGAAGAAAATTTTCATCAAAATTATGTGAAGAAACCCCATTAACTCCTCGCAAGTCAAGAATA
AAAAAATAGTTAGACAAACTATAGTCTCTTGTTTGAAATCTAGTAAACCTTGATAAAGGAAGAAATATATA
TTTTTGTAATTATTGTGTCTCCTAGTAAAAAGAATTTGAAATCTTTTATCCATTTGTTCCTCAAGACTTAT
AGATTTGAGATTTCTATCAAAAGCATATGAATAATCAAATCTTGGGCCACCTACTTCCTTTACAACAAAAA
GATCATGAGGAGTTTCACTAACACAACTGAAAAAGTCTAAATCAATGCTGACATTTAAAGAATAAGTTCCT
CCTACAGCTATGATACTTAAAGAAGGAAGCATCAACAATAAGGGAGGCTCTGATTCCAAGGCAAGAGGTAT
AGTAGGTGTAATATCCAAATTTTTCAAAAGATGATATGAAACTTTCTCATAAAACTTGGACCCCATTTTAT
GCATGTCCTGAAAATCATCAAAACTATTTTTATTAAATTTTATATTTCATTTTGATTATTTTGATATATAA
AAGTACATATATATAAATATTCAACTTTTCATTTAAATTTATTTGACCTAAATTAGTAAACAATATGTGCCGA
ATGTTGGCCATTAAAAGAGAAGCATAATACTAAGTTAAGTGTGGCAGAGATGAGGATGCTCAAGTGGATGA
GTGGCTTACCCTTAGAGATAGGATACGAAATGAGCATATACGTGAAAAGGTTGGAGTAGCTCCAGTAGAAG
ATAAGATTAGAGAGAGTCGCCTTAGATGGTTTGGTCACATAAAACGGCGGCTTTCCGACGACTCGGTTAGG
AGAGTAAAGGTATTAGATTTGACTTATGTTAAGAAAGGTAGGGGTAGACCTAAAAAGACTTGGTTAGAAAA
TATTAGGAATGACCTTTCTTTATTAGAGAATTTGACTTTTAATAGGACACAATGGAGAAAAAGGATTCATG
TAGCCGACCCCACGTAGTGAGATAAAAGCTTGATGTTGTTGTTGTTGTAGTAAACAATATCACCCTATTAA
TCTACCATATCAAAATCCTATTTTTCCAAAATTTCATGCCAAACAAGGGTAAATCGCTAAATTGAGGGTGG
GGATAGTTTATAGGATAGTTTAACTAAGAGTGAAGCCACTCATCCACTTGAGCACCCTCATCTCTGCAACA
CTTTACTTTTATGCTTCTCTTTTAATGGCCAAAATTCTGCACCATAAAGCACTGCAGATCTCACCACTATG
TTATAGAATTTTCCCTTAAGTTTGAGTGGTTGATGAGGACATCGAAACAAGCCCCATGCATTCCTCCACTT
CAACCATCCAACTTAAATTCTGCTGATAATATCCCCATCTATTTCACCATCACTCTGAACAATAGATCCAA
GATATCTAAAACATGTATCGTTCTACATCAGTGGTACTTTTCTATCACATAATAGACCTATTGCATTCCTC
CAATTTAATCACCCTACTTGAATTCTACTAATAATGTCGCCATCTATTTGCAAATGCACCTTTGCACATAT
CAATTTCTAGGATCACCACCCTAAAACTATGCAAAAAATGATTGGGCTAGGTCAGACTCAGACCATATCAT
AATTCAATGTTGAAATGCCACTAGTAATGAAAATTAGGTTTGATGGGTTTCCCATCCATATCAAAGTTAAA
TAATATTGATGATCAAGATCAAAATTACTTCAAAAAATAAATAC**TAAATTTCAATGTGCAGAAAATGATTT**

**ACCATTAGATGAAGAGATGTTTCGATGGCTTCTTACAAGCTATGCAGCAACTCCCTCAAGCCAAATCCCGT**
**TTGCCAAAT**CTAAATAGCATCTATGCAATGTAAACAATTTGTTTCATGAGCATTAAATAAATCAATCTTGA
AAAGATATATATTTCTTGCTACATAATATGATAGGTGGATAGTAATTGCTATCAAAATCAAAAACTTTACA
AGCATGACAATATAACTAGGATACAATATTATACCTACTAAGCAATAAAAAAAATAGAGATATAATATTAT
ACCTCCTTAACCCATTATAATCATTTGATGTGCTAATGTGCACACTTTCGATAGTTACTTTTCTTATCCAT
CAAGCCAAGTTTTTATTGCCGATTCTTAGTTACATTGAGTCTAAGGATGACAATTAGTTCCAAATCAAATT
CCATTATTATTCAATTTCAATCTTATAAAACAATTACAACTAATTCAGTGTGTGCCAGCAAGGGCCAAACC
AATCTGACCCATCCACACCCTTTTTCTAAAGGACGGCAAATAATTATGCATTGTACAAGGTTATTCTTCTC
AATATTACATGCCATGAAAAAAAAAATTAAAAATACAATCACATATTCACATGATAACTTAAAAAATAAAA
CAAATACCATTAAGACATGCACATGTGAGAATAAGAAGACACATAGAAGAACATTTAAAATTTACAAATGT
AGTAAGCATAATAAACACAAAGGATCAAATCATCATTCAAAATAAAAAGTTAAAAACCCAAGAATATTACT
TAGCAAATTAAGTAATTAGCTTTTCATAAAAGACCTAGCAGCTTTCAAAGATAACCAAAAAAAAAAAACAA

>deca_miR444b

ATTTGGCAAACGGGATTTGGCTTGAGGGAGTTGCTGCATAGCTTGTAAGAAGCCATCGAAACATCTCTTCA
TCTAATGGTTGAAGAGGTGTTACAGATGATTTTCTTGCAAGTTATGCAGCTGCTGCCGCAAGCTTTTTACT
TCAACCTGCCAGAA

Stem-loop structure

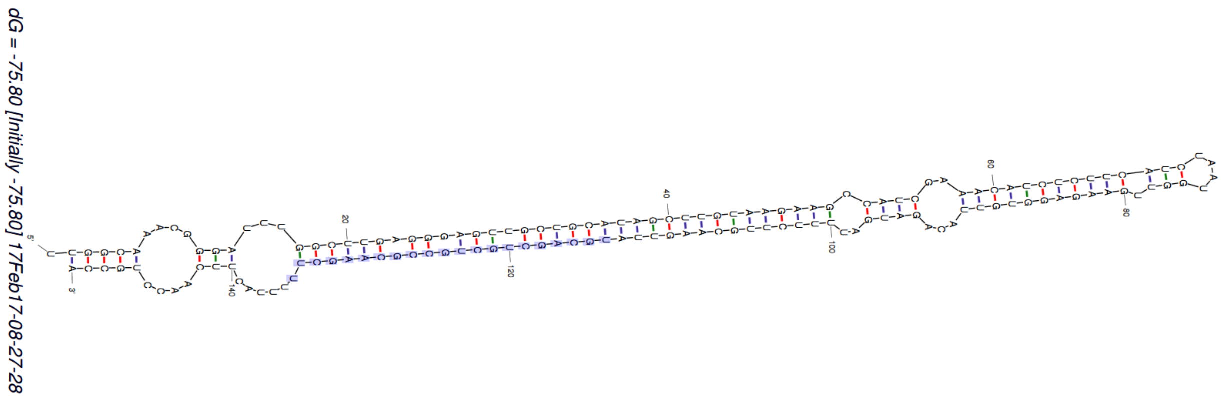


*DecaAGL17_3*

>JSDN01S008597.1

TTTACAAAGCAGTAAGAAAGCTATCTCACATGAAGCATCGGCAACTTTCTGCTTCGCTTTTTAGATCATAA
TCATAATTTCTATTTAACAACTACTTTTTTATTATTATTTTTTTAAAATATGTAATAAAAAATTGATTTCT
ACAACAACTTTAAAATTGTACTCTCTACAACAGTTTTACAAAAGAGTAAGAAAACTACCCCAATTATTAAA
ATCTTGGCTTCATTCAACCATGAATATAACTTGAAGATTTAATGCTTCATAGGATTTATATTATAAAATAG
TTAATTTAAGCTCATATATTTGCAGAAGTAATTAATAGATTTAGGCTTCATTTGAGACAGCTTTCTTATTA
CTTTTTAAAGTAACTTTTTAATAAAAAAAACTAAATAATAGCAACTTTTTAATACAAAATTTTTAATTTTT
GTATTAAAATTTTGTTTTAAGAAGCAATTAGAAAGCCGCCCCAAACGAAATCTTAGAAACATAGTTGATAT
GTAGGGTTTTATACATATTATTTATATTTCAAGTTGATTACATACTTTTTTTTTAAAAATAGTTACATAGA
TGATAGCCAAAAGAGAATTAATACCAACAAATTTAACCTAATCAAGCCCAACAAATCTCCCATGAATCACT
AATCACCATCATAGAAACTAGGCACCTCTCTTCAAATTATCATCAAATAAAACAACATCGCTAGCCCATAG
GGCGCCCTCCTTGTGCGGACGTCAGACACAACTCTACGTCCGGAGACGAATCAGATTCAACCACATCATCG
GTATCGAGCCACGGTACCAAGCAATACCGAGATGTGGTACCGAGCAAATGAATGACGTAGCACATTCTGGC
TGGTCTCCGAACAGCATTCAACATCTCCACAAAATAATTTTTTCCAAACCTGACCAAAACAAAGAGAGAAA
ACAAGAAAAAAATAAAATAAAATAAAATAATCTCTCCCTTCAACTTGAAAGAGTCCTTCCTTCATTAGCTC
CTCTTCTTATAAATAAAGCCTAACTTAGAATCTTTCATCTCCCCGCCTCCATTGTTTCCAGTAGAGAGAGA
AAGGTGAAAGGAAAGACAGAGAGAAAGAGAGAGGGGAGATGGAAGAGGGAAGATTGCGATAAGGAGGATC
GATAACACGACGACGAGGCAGGTGACCTTTTCGAAGAGAAGGAATGGGTTGCTGAAGAAGGCGAGAGAGCT
TGCCATTCTCTGTGATGCAGAGGTCGGCCTTATCGTCTTCTCCAACACAGATAGGCTCTATGATTTCGCCA
GCACCAGGTTCTTTTTTCTTCGCTTATGTTACTGTCATCTCTTCATAGGTAGACTATCCATAATCTATATA
AAGGGGATTAAGTAAACAATATGATTTTCTTGCCTGATTAGGCCCTCAAATTAATGTATTAGAAGGAGAAC
GAATGTTCATTTTTTATCTAGGTTTAATTTATTAATTTTTCTTAGTATGGAGAAAGAAGTCTGGACCTCTG
GGAAAATTATTAGGTCCGGTACCGGACGTAGAGATTGGTCCAGACACCAATAAAATATTGGCATGTGGAAT
GTACACATGTCTGCTGAATTTTAAGTCAAGCGCGTCAAAAGCACTGGTAAAGCGCAGTCAAAAAGCACAGT

CAAAGCGCAGTCAAAGCGCAATCCACGTGACAAGTTTTAATTGAATGTTCGGACCTATTTTTAGGTCCGGT
GACCGGACCTAATAATTTTCCGGACCTTTGGCCAGATCTGAAGGAGGATGAAAATCAGAATTAAGGAGCAC
AGGCTCTTCTTTTTTTAATTTAATGTTCTTTTGAGTTTATTGATGATGCTCACTTTGTTAAGGAAAAAAAA
ATCAAACTACTAAAATTATATATATATATATATATATATATATATATATATAGTTTAAAGGTAGGGGTAGACC
TAAAAAGACTTGGTTAGAAAATATTAGGAATGACCTTTATCCAATTTTGAGGGGGCAATTTTAATATAAAA
ATAATTAAATAAAAAATTAAATAATTAAATAATTAAATATTAAAAATATTATATATAATATTATGAAAAGT
TGTTTTAATTTTTTTTAGCTTTATATTTGATCTAAAAAAAAAATTTTAGCTTCCAAATTCAAACTAAATGT
CAATCAAAATCATCAACATATTTTCAGCCACAAAAAAAAGTAAAAAAAAAAAAATATTGCAAGAACATGAA
ACAAATATACATAATTGATATTCTTAATCAAATCCTAGAGATTTTTTTGTGCACTAATTAACTCACTTTCA
AAAGGTATTTCAAAACTAAAATTATTGCATAGAAATATGAGCCTTCATTACTACCTATTGTCATAGCAATT
AAAAAATAGAGAATATAGATACATTGGGAGAAGATAAAGCAAAAAATGCAATGTATGGTTGAGATGAATCA
AGCAATGACAAGTAGGGGCAAAAAAAAGCTTTTTAGGGGGGCAAAATTGATTTTTTAGGAGGGGCAAAATG
GATTTTTTACACTCCTATAAGTTAAAAAATTTTTTTTGAGGGGAGGGGGGGGGGCAATTGCCCCCCCTAA
GCATAGGGTTCCGCCCTTGGGGCGAATGATAATAATGATGATGAAGATGAAGGATGCTTAGCACAAATGAA
TAAAGAAAAGTTCTCAAAGTTTTCTAGCAGAGTAAAGTCTAAAAGGAAAAGAAATAAACTCTTCTAATTTG
ATTATATTCAGGTAGTGTACATGCAAGTTCATCTTACCATGCTTCTGATAGAAAGCGTCCATAAACTAAAC
AATGCAAAAACGAAAATTTAACAAAACTTAAAATCCTTATCGTGTCTATTTCAGCATGAAATCGGTGATTG
AACGATACAGCAAGATAAAAGAGGATCGTCAGCTAGTGATGCATTCAAATTCAGAAATTCAGGTAAATTAA
CCTATTGAGATTTGCATTTAATCCCCATTTTGATATCTTTGTAATTCATATAATTGCAAAAAGATGTTAGA
ATTATTTATTATTGTTGTGATTGGTCCAAGTTTAGGTTTGTAGTTCATTAGGTACATTTTATCTCTTTAGT
TCTATTGTTCTATAAATAGAGATGATTGTAATAAGTTTGGTATCCAATCCAAGTAATAAAGATTTCTCTAA
TTCCGTGTTGTCTCTCTCTCTCCTCCATTTTCTCTCTCTTTCATCCAAGAAAGTTTCCTTCCTCACCAAGA
TCCTACAACTGGTATCAAAGCAAGTTAGAGTTGATCAAGGGGAAGGAGAACTTGTAATCAAGAAAAAGAGA
GGGAAGGGAAATTCCTATTGTGCTTGGATCAAAGATGAGCGCCTAACTCTCGTGGGTGAGGTGATCAGGGC
ACGCAAAAGTGGCACAAGCAGGCACTGCGCCTGGCGCGAGCATAGCCTAGTCGAGACCGACAGGCAGACAC
TCGCACGGGAGCTCGCATGTCTAGGTTCGCGCATGAAGGCGTGGAGTGCGGGTCATGCGGATGTCGCAAGC
GGCGACGGGTGACATATGCATGGTGCGGGAGCGCGAAAGTGTTGCATGGGTGCGTGGTTGGATCAGGCGTT
GTGCTCGCGCGAAAGCATGAGTGGAGTTGGGCCGAGTGTGCGGAGGCATCATCTCGGGTGTTGTTCACGTA
GGTGATCACATAAGTAGGCCAAGACCGCAAGTTGGAACGGGATTCTCTATTTGCAATAATTAGGAGCTAAA
CCCGATCCTCTATTTGCAAGCAAATATAGTTGACAATGTTCGACTAACGAAATGATAATACACATAAACCC
TCGATGATAGCCGTATTGAATGCAAATCTCAACCCGGAGGGGTCTACGTAAATTTCAAAGAAATGCCCCAA
AAATTCAAAGTTAATTATTTGGGGCTCAAATTTAAATAAGAATTTTAAATCTATCGTTCTTAAAAAATTCA
ATCTATATAATCAAAATAAATCACCTAAACACTTAAATAAAAAATTTATCCACCTTTCACAACAATTTGTC
GAGGTAAATAAATTTAAATCTATTGCTCTTCAAAAATTCAATATATATAATCAAAATAAATCACCTAAACA
CTTAAAGTAAAAAATCCATCCATATTTCGCAACGGTTTGTCGAGATAAATAAATTTAAAAATCACTTCAGA
TCATAAAGTGCGAATGTTTAGTCGAGAATATTATTAGGGTTCATTATCAAATTATCTCATATCAGTAAGGA
TCTGACATGTGTCCTCTAACCGGTGATTTTGCCACCAAATAGCACTATATATAAAAGGTTTTGATCGGGGC
TCTCTGACCTGGTGGTTATATAGTCAAATGGTACCATTAAATTAAGGTCACGACTAGAGCCCACATGTTCG
CCAATGATTCTAAATCCTTAAGTAATGAATCAACATAATATAATCGATAGGGCACAAAGTCTTTTATTAGC
TAAGATGAACTCAAATATGTCCCCTTAGACAAACATTTACCCCAAGTCTACAATGTACCCTTATGCGAAGC
AACCCATTGCTATCAAAAGGGATGTATAAAACCAAAATATTTTTTAATGTAGGATTGCAAAGCCCTTTAAT
GAATATATTCTCCAGTTGACTATTTGTTGAAGCATAAGAAGAGAAAATCAATTCACTTTTCACTTTTTTTA
CGCGCCTATGTACCACCATGAGTTTAGTTTGAAAATGTTGTACCGGATTATGAACAATGCTAATAGCACAC
ATTGCTATCACTATAAAAGTCGTTTCCATTTAATCTTTAGATCTTCAAGTATAATTTTTTTAACAATAATA
GTTTACATATTCGATGGGCAATTACTCTAAAATTTAATTCAGCACTTTATCTTGAGATCACATGTTATTTC
CTACTTCTCCATGCTACAAAATTTCTCAAACAATGCAATATCCTGAAGTTGATTTATCCACTTATGATCTC
AAATAGTCAGTGAGGATTTTTAGCATTTTTTCTCTTCATTATCATCTGCTATAATAATTATTTTATGAACA
TGTACAATATGAACTGTCATTCTCTGTTAAATAGAGTGTTTTATAAAGACTGTATGATCTAATTGACTTCG
TCTATATCCTTTTTTTTTTATTAAGCTACCTGTGAAAACAATTAATTACATGACAACAACAAAAAAGTACAA
TACATATAAAGATTGACCATGTTTCTATATTTGTAGCTATGAAAATAAGAAATATATTCTAGAACTTGCAT
ATATTTACTTCCTAAATATACAAGGATTAATCATTTTTCTATATTTGTGGCTATGAAAAAAAAAAAAGGAAG
ATATTCAACAAATTTGACACGTTTATTTTCCTGGAAAAAACTGGAGATTTTTTCCCTTGTAAAGGCCCATG
AATTGCCACCTTGGTTTCTTTAAGGTTCAAAAAGATCAATTTTTATATTGAATCTAAAATCAATTTAACAA
GATCTATAAGAGCCCTACGCTACAAGAGTTCTTGCAATTGTTACTGAAAAAAAGAAAGCCTCGCATTTTCT
GAAGATTTGCACAAGATAGTTCAACTTCAAGAACCCATCATTACTTTGATCATATAAAATGCAACGAAAGC
ACTCATTTAAAAAATAAGATCCAAATCTAAAAAATTTATACCAATGACATCTTAATGCTGAGCCTACAACA
TAAGAGAAAAACAAATAGAAGTTAAAGAAAATCAAAACTAATAATTAAGTCATAGCTACATTACATGTACC
CTAACTCGATAAAAAAAATAAAAATAAAAAGAATAGTCCCTAAAAAGTAGATTTATTTTCTACATGAAGTT
GCCAGCCATCATTGAGGGTTAGTGCTCACATATAAATCACTACTAGATTCTAGTTTTGCATGAGTAAATAA
CACATAAAGAATACATGAGAAATATAACATGGAATGTTCTCAAAATTTTAGTAGCAGGTTAAAGTTTGTTT

TTAACAAGTAAACGTCCATAAAATGATATTGAACTTCTGCATCGTAAATTCATATTCAAACAAGACAAAGA
TCTAATGATTCAAAAAACATTATCATTTACATAATAATCAAAATAGTATAGATTATTTGCGATTTCCAATT
AAGAATGTTAGCATACAGGGCCTCAAGTCTTAATTAAAGAAGAATGATGAGTTAACCGAGAACAAGAAAGC
TTCTATGGCAGAATTAGCCTTTTGCAAGTTCGAAAAGATGCAAACAGTCTTTAGATATACATTATAGCCAG
AATATTACAAAGTTCAACACAATACCTGTTATGTAATTTGATCAAATAGCTGATACAGCAGCATCTTGTTT
TATAAAGTCATCCACTAGTACTGTAAAAATTCCCAGTTGATATTATTTGCAAACAATGTTGTTTTGATCTG
ATTGCCTCACCGTATCTGCATTAGACGTACTAACTGATTTTGTTTTTTGTCCACTAGTTCTGGCAGAGGGA
AGCAACATGCTTGAGGCAGCAACTGTATAACTTGCAAGAAAATTATCGGTAACTTCTCTTATCCCTAGAGT
ACCAAAAAATGTTAAGACACACGGCAGAATTGTAAGATTTTAAAAAATGCAACAATGCAGGTTGTCACAAT
TATACCGTCTGAAAATACTTTAAACTAAGCTTTTGATAGAATCAAAGTTGAAAAATATTTATGTCACATGG
TTAATAGTCATCAGGTACCTCATGGCATGAATAATAAAAATTATCAAATAGAAATGGGAAAATTATGAAAG
AACCATCCTCTTTGTATGGAGAAATTATTGTGCGGAATCCGAATGCATTCGGAGACGAATCTGGATGCGCC
ATGTCACCGGATACACTGGACCCCGGTGACGTGGCATGCCCGGATTCACCTCCGAACACGTTCGGATTCCG
CATACAATAATTTCTCCTTTGTATGAGGTACTCATTAATCACTTCATTACCGGAGAACTAAAATAGATTCC
TTTCACAAAGAGTCAACTATAACAGCTTTGAGACTACTGGATTAATGCAATTCATGCTTCGTTTGGGACGG
CATTCTTACAGCTTCATAAAGAAGCTTTTTAGTACAAAAAATTAAAAATTTTGTACTAAAAGCTTCTTTAT
GAAGCGGTAAGAAAGCCGCCCCAAACGAAGCCTCAGTCAAACTCCTGAACTTCGATTGTCCTATACATATT
GGACCCACGTGCTAGATAGTTACCAATATTATTCTCATCATCAAGGTTAAAAAAAACTCAGCATGTTATTG
TGGCTGTTACTTTACAGCAACTTCTATTGACAAAAGGGTAAGGGGGTAAAAATGTTCTGAGTTTAATTGTC
GATATAGATGCTTGACAAATGAAAAGAAAATGAGAAATCTCAATATAACTTAAAAATGACTACTAAAAAAT
GGAATTACATCCTATTTTTACATAATTTTTTTCTGATTAACTACAAATTTCTTCTTAACATCAGAAGAAAA
TATAAATTTTTACAAAAAACAAAATCAAAATCAGTAAGTTTTTACAAATATATCCAATTTAAGAAGATTAC
CAATCTGTTTCTATCAAAACAAGCATTTGACAGATTGTAATGCCTGAGATAGTTAGGATTGAGTCTTTCCA
AAATGCATGAAGCGCTCAATCATTCATATGATTGTTTAGATTACATGCTAATCAAATTTTAATCCTCCAAA
TTAACTTCTTCCATAGCTTGTGAATCCACTTTATGAAAAGCTTCAAAAAAACAAGAATGACATTCTACCAA
AATGAGCCAGTTTCAAAAGTATAGTTTTAACAGTTAAATAATAAACAAACTATTCTTTCCGTTTTATTGAA
TTAGGACTTCAAAATGAAATCTCTTCCCCGGTCTCTAACCTAATAAGCATTAAGCCTCAATTTTCCAAAAA
CATCAAGTCCGATTATTAAAGCAAAAATAAATTAAGGGTATTTGCAATCTTATGTATTCGCACGACTAACA
CCAAAATTTTGTATTTACTCATATACCACTGTCTAAATCTGATGCAGGACTCAGTATGATTTAGATGTATA
CGATAAGAACTTGCACGATTTAGATAAGAGCCAATAACAATTTAGACTGTTAGACACTAAATAATAACAGA
AAAATGTAAGAAGGGTAGCTACACTTAGGCCAAATCACATGCTAAGAACTTGAATCATCAAAGCATTTTAG
AATTTCAATAAAATACAAATTAAGTAGCCATCCAATAAAATCCTCACGTTAACAATGAGATTAGAAACATA
GAAAGTAAAACTCAATTCCTATGATTACTAAGTGGAGGCAGAGGGTGAGGCACGACTCCTTGCAATTGGAT
CACCACCTACAAGGTTAGCGGAATAAGAAGATAAATGAATAGGTGAAGAAGTTCACCCTTGTTGATGACCC
GAAACAATTTGCCACTGGACGTTGACAAGTTGTTCTTCGACGGCTTTGGAAAGTGTGAAGATGATGCGGTT
CAAGTATTCTTCAACGAAGTTGTCGGTTTCCGGGTTGAAGTTGAAGAAGGTTGACCGGAGAATGTCGACGT
CTCCTTTCTCGTTGTAAATGAATCCGAAATCAAGATCGGGATCGGCCATTCATTGGTTTCAAAGAGTTGTA
ATGTTTGAAGACGATCGTCGTTGCTTCCCTTGAAGACTCCCATTGAGAGGGAGGAGTTTATCTTGAAATTC
AAGAGGGAAAGAGAGAGGGATGTGACCAGCCAAGGGTGAGGATGGGGAGAGTGAACTAGGTTCAGATTTTA
GGAGAAAAGAGGTGTAGGGAAATCAAGCAATTGTTTCTCTAATTAATCCCTTTTGTTTGGAAAGGATTTGG
GTGTATTTAGGGAAAGGTGGGGGCGGCTAGAAGAGAGGGAAAGAGGATTTGAAATCTTCCCAAAGGAGGTG
CTAGCAATCAGGAAAGAAATCAAAGGGGTAATGGGGGGGTGGGGCTAGCGGCTTCCATAGGAGGGAGAGAG
ATTTAGGGAAAGCATTTAAAGTTACTTGATTTAGATTTAAATATTTGGAAAGGTGTATTTAAATCCAATTG
AATTTTAAATTCAAATCGAATTTGGAAAGGTATATTTAAATTAGAAAACTTTCCAATAATAGGTTTTGAAA
TTTAGGAAACAATTCAAATTCAAATTAAATTTGAATATGAATTAAATAAGTTTCTATTCTAGAAAGACCA
AATTGGAAAGGTTTGAACACAAGCCAAGAGGGGAAATCCCCAAGGGGGGCCGGCAGTTAGGGCTAGGGTTT
TGGTGAGAGGGGTGGCGGCTAGGTGGGGAGAAAAGGATTGATTGGTGGTGGTGTGGGATGAGACCGACCGG
GGGGGGGGGCGGGGGGGGGGAAGGCTTGGGAATTCGGTGGTGGGGCTTGGCGAACTGTGGATTAGGGGAAG
AACAAGGGAGAGAAAAATTGAAACTCAAAATGATAAACTAACCATGAATCCGAACTCTTCCAAGGTTGGTG
GGTAATCTGGTTCCCTCTCAAGGGAATATCTCTCCAATCTCTCTCAAGCTCTCACAAGAACTCAAACACGG
GGAAAATGTTATCATTCATCCATTAACCATGGAAAGAGAACATAAGGGCTATATATAGCCTCATACAATGG
ACTCAAAAATAAAAGACGACAATGCCCTTTGGGCTTACACATGACCACCCTTAAGTAAGGGGTAAAAGGGA
AAATAACTAAAAGCAAATACAAACATAAATAAATAAAAACAAATACATATGACGACTTAAAAATATATTGG
CGCGATCGTGGCCGTAGATCGGCGGCCGCGCCTCTGATAGACTTTCCAAGCTCCGATGTCCCCATCAAAAT
CACATCAAAGCAAATGCATAAAAAAGGATGGTTTTGCTTTAAATAAGTTAATATTAATGATATTTTTTTAG
CTGATTTAGAGATTTGAATGAAAAATAAGTAAATATAAAAATTTAGGAGTTATATATAAACACAAACAAAT
TTTGTGTTATGTATGTAAATCCCCTAATAAATTAAGTAGAAAGATCTATCAAACAAATGCCTTTCCTTCTG
TCACTTCAATTTATTTTCTTTTGTAGTAGGGAGAGAGATTTGTGAAGAGTAGAAGGTAGATACTTTCAATA
ATGTCAAATAAATATTTAAAGCACTAAGACAAACAGTTTCTCTAAAAAGATCTTGAATTTTAAGGCCATCA
GGTTCACGTATGATTATTCAATATTCACAAATGCAAATGAAAGATGTTGCATAGAGCACTCCTAGTAGTGC

GCTCTCAGCAAAAAATGATGTAGGCGCACAAAAGCTGATAAATTCAACTATCCCTGTCAAAAAATATATCA
GAAGTCTCTTTCCTCTGACAGTGTCAGAAACCAATATTAAGGTCATGAAATACTCATGAAAATCTACGCAC
ATAACAAACATGGTAAGTGGGTGTATAGATGTAAGGACACTGAAACACAAGCAAAATGATAGAATATAGTT
AAATTCATTCGGCTTTGTCTAGCTATACTTTTATCAATATCTTCTTTAAGCACTAAGAACACAAGCGTAAA
CTGGAATTCTGTTTATTTGAGAATTTGATTTCATGAAGATGATGCCACAAAACTGAGAAATTATTTTGTGC
GAATGCCGTCTAGAGACCAACCAAAATGCGCCGCATCACTCCTTCGTTCGGTACCGCGTCTCGGTGCCGCT
GATGTGGCGCTGGTTGGTCTCCGGAAGCGTTCTGCGTTTGGTCTCCACATAGAATAATTTAAACCAGAAGG
TTTCATAACCATTTTATCCGAGTTTGGGAACATAAGAGACAGAATCACATGGAAAAAATCATGCCTAATAG
AAAGAATATAATGCATGTATGGAACAGTAAGAAACTATTCTCAAATCTCAATGGATAACTACATGATCACA
TCAAAAGAAGATTATTTTAACAAATGAGAATGTTACTCAAATGAATCTATTCATATAACATGAGATCTTGA
AACATCTTTTCCCCCTCTGAACAACTTAATGATGATTACAGGCAGCTGATGGGTGATGTTAACGGGCTAGG
TCTCAAAGATCTACAGACCTTAGAGAATCAGCTAAAAGCAAGTTTACAAGCAATACACAAAAAGAAGGTCT
TTAGATCTGATACTTTAAGTCATGTCTAGTTACAGTCATCATTTTCATTGATTCTACTCATCTCTTAATCT
CTGGCAGGATCAAATTTTAATTGATGAAGGTCAAGAGCTAATTCAGAAGGTATTTTTTCATCGTAAATCTG
ATATACAAGATTCGCTCAACTGCTCCACTGAATATATTCATTACAGGTCAGGATTATGGACAGAGAGTACA
TGGATATCTACAAAAAGATAAACCCCTATGTTCAAGAAAATATGGAATTACACCAACAGGTATTCATTAAG
GATGTTAGAGAAGTTAGTTGTTTGTATTTGGTTCAATATTTAGCTCTATAATAAGTCACAAACACAAATAA
TAAAATTTTACTTAATTGCTTAACAAGTTGAGCCGAACGACACTTGAGCTTGCACAAGCTTCAGCTCATTT
ATGCCCCTTAGTGTTCATCAAAGAAATAAAAATCATTAACAAAAAAACTGCAAGGCCAGTATATCCATAAT
AGTGAAGATAGAAAGTTTTATATATTGCCTGAAGATGACAATCTATTGTTAGGTTGATGCAAGTGCAAGAT
TTAATGGAGGATTTACCACTTGCGGTTCTCTTAGTCCCACATCAACTGATCTTGAATTAAGCCCACCACAG
CAGCAAACAAGTGGAACGCAAGATGACGTGCTGAAGCTTGGGTAAACTCTTATAACTTCATAGAAAAAACT
GAATGTTGATAGACATTTTTCTAGATTTAATAAAGAAACTTCTTGAAATCTGCACCAGGCTTCAATTGCTT
TAAGATGTTCCAAGCAATGTTCAGGATACTCCACAGCTTCAAAAATAACGACGGTACGAATATTATAAGAT
TTCATGAATGTAAGAGCACTCAAAATTCAATTTAAGTGTGAGATTATCTTTAAAGAATCACTTCAGATACA
CGAAGAGGGATTTTCCTTGTCAATTAACATAGATGATAGCCAAAAGGGAATTAATACCAACAAATTTAACC
TAATCAAGCCCAACAAATCTCCCATGAATCACTAATCACCATCATAGAAACTAGGCACCTCTCTTCAAATT
ATCATCAAATAAAACAACATCGCTAGCCCATAGGGCGCCCTCCTTGTGCGGACGTCAGACACAACTCTACG
TCCGGAGACGAATCAGATTCAACCACATCATCGGTATCGAGCCACGGTACCGAGCAATACCGAGATGTGGC
ACCGAGCAAATGAATGACGTAGCACATTCTGGTTGGTCTCCGAACAGCATTCAACATCTCCACAAAATAAT
TTTTTCCAAACCTGACCAAAACAAAGAGAGAAAACAAGAAAAAAATAAAATAAAATAAAATAATCTCTCCC
TTCAACTTGAAAGAGTCCTTCCTTCATTAGCTCTTCTTATAAATAAAGTCTAACTTAGAATCTTTCATCTC

>Deca_AGL17_3

ATGGGAAGAGGGAAGATCGCGATAAGGAGGATTGATAACACGACGACGAGGCAGGTGACCTTTACGAAGAGAAGG
AATGGGTTGCTGAAGAAGGCGAGAGAGCTTGCCGTCCTTTGTGATGCAGAGATCGGCCTTATCGTCTTCTCCAAC
ACAGATAGGCTCTATGATTTCGCCAGCACCAGCATGAAATCGGTGATTGAACGATACAGCAAGATAAAAGAGGAT
CGTCACCTAGTGATGCATTCAAATTCAGAGATTCAGTTCTGGCGGAGGGAAGCAACATGCTTGAGGCAGCAACTG
TATAACTTGCAAGAAAATTATCGGCAGCTGATGGGTGATGTTAACGGGCTAGGTCTCAATGATCTACAGACCTTA
GAGAATCAGCTAAAAGCAAGTTTACATGCAATACGAAAAAAGAAGGACCAAGTTTTAATTGATGAAGCTCAAGAG
CTAATTCAGAAGGTCAGGATTATGGACAGAAAGTACATGGATATCTACAAAAAGATAAACCCCTATGTTCAAGAA
AATATGGAATTACACCAACAGGTTGATGCAAGTGCAAGATTTAATGGAGGATTTACCACTTGCGGTTCTCTTAGT
CCCACATCAACTGATCTTGAATTAAGCCCACCACAGCAGCAAACAAGTGGAACGCAAGATGACGTGCTGAAGCTT
GGGTAA

#### Liliales

##### Gloriosa superba

>gsu_miR444_scaffold-GDKK-2013839-Gloriosa_superba

GGGACCACTCGCACCCATCCTCACCTCTCGACACGTAACCACGCCTCCGCACGCGGCCCCACGACGCGCGC
TGCGCTCGAATACATTAACACCCCCCTTGCGCCTTATTTTAATCCCAAAACCCCTACAGCGGATCCATGCG
TCGCCTAGATATTTTCCCACCCGCCCCATGCTCCATCTTCATCCTCGCTCTCGCCACGTCTCTCCGTCCGA
ATTTCAGTTGGAGGCAGGAAGCTTGAGGTGGCAACTGCATAATAGCAAAAAAATCAACGGTAAAAGATGTT
ACCGAAGATTTTCTTGTAAGTTTTGCAGTTGCTGCCTCAAACTTTCTGCCTCCCTCTGCCAGAACTAGTGA
GTTAAATCCAAAATTTTTAGTCTGCCGAGTATGTTTA

Position of mature miRNAs based on comparison with osa-MIR444d and not based on experimental data!

Stem-loop structure

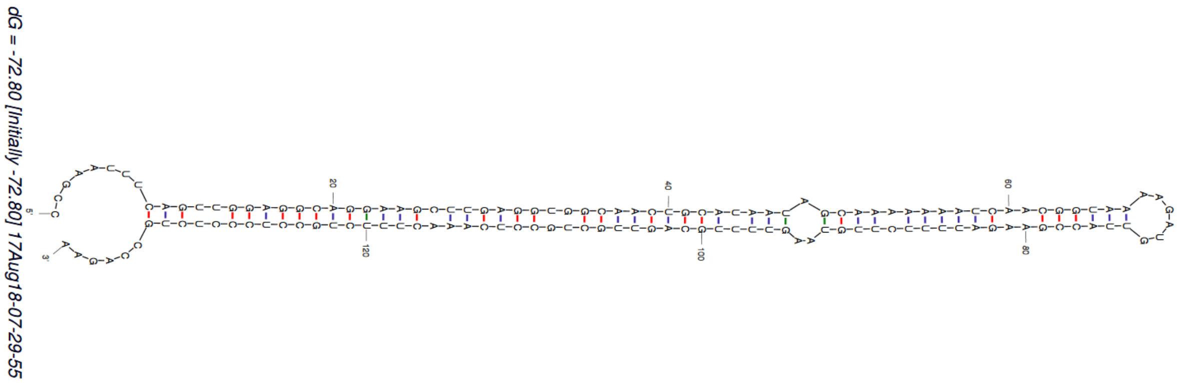


#### Lilium sargentiae

>lsa_miR444_scaffold-THEW-2000710-Lilium_sargentiae

AGAATAATATATCAATTATTACTTGTTTATTCGACTCACTAGTTCCAGCAGCGAGGGGCAACAAGTTTGAG
GCAACAACCGCATAACTTACGAGAAAATCATAGGTAGCATCCATTCACCAGTGATTTTCAAGCAAGTTATG
CAGTTGCTACCTCATCCTTGCTGCCTCTTTCTGCGAACGGGACCAGAGAGAAAGAGAGATAGAGAGAAT

`Position of mature miRNAs based on comparison with osa-MIR444d and not based on experimental data!`

Stem-loop structure

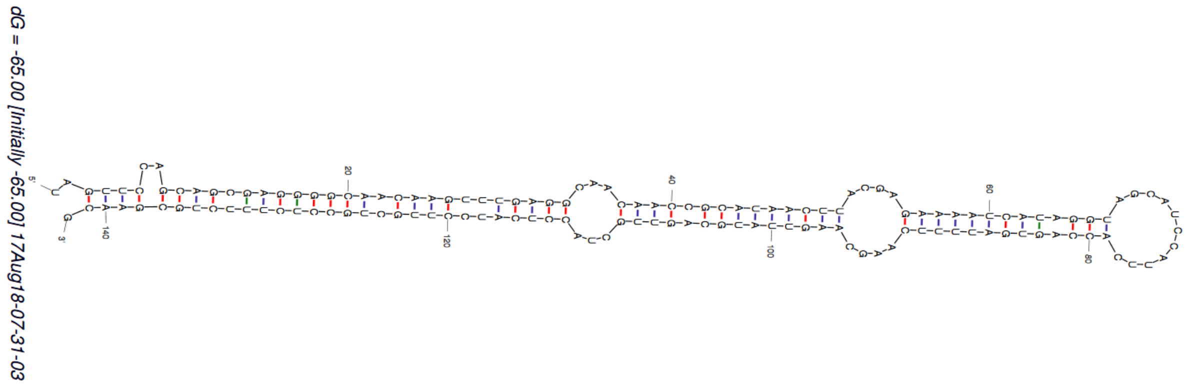


#### Smilax bona nox

>sbn_miR444_scaffold-MWYQ-2052889-Smilax_bona_nox

CGGGGATTATGACTTCTGCATTTTACTCACTAGTTCTGGTAGAGGGAGGCAACAAGTTTGAGGCAACACCT
ACATAGCTTACATGAAAATCATCGGTAACATCTTTTTACCAGTGATTTTCTTATTAGATATGCAGTTGCTA
CCTCAAGCTTGCTGCCTCCGTCTGCCAAAATAATGACTGCAAACCGGACGCTCCCGAACC

`Position of mature miRNAs based on comparison with osa-MIR444d and not based on experimental data!`

#### Stem-loop structure

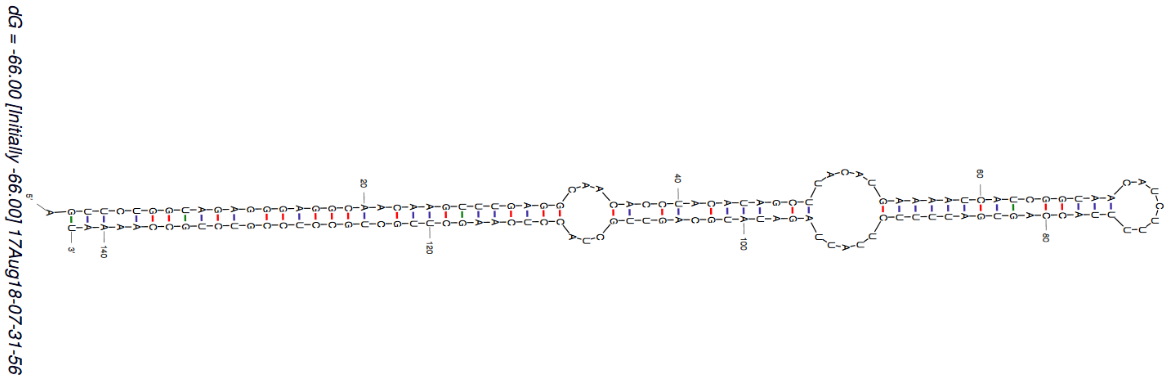


### Pandanales

#### Xerophyta viscosa

*XviAGL17_1, XviAGL17_2, xvi-miR444a and xvi-miR444b*

>MJHO01000094.1

TCTTGATTTGCTCCCTTGTAGAATTTGATCAAGGCTTTGAATCTTTGTTGAATTCTTGAATCTTCCGTTTG
CATCGTTGATTTCGCCTCTAACAACATCGATttcatttcttataatttgttGCACTTcaagttgagagaga
gagagagagagagagagagagagagagagagagagagagagagagagagagagtgagtcgCGCGGAATTTCTA
GGTGAGCATTTGATTACTGTGACAGATGTAATAAATTAGTAGGTTATAGTACATTCTGGTTTGATTGATCA
GATACTTGACAGTAAATTTGGACATGCTGATAACCTTGTTTCGATGTTACTTGATGTTGGAcatgttttag
tttttattgttttaatggCAGTCTGAATGTTCAAtatcaattctgagttccgcatTGGTAAACATTTCCTT
TTGACAATACCTTAGCCATAAGGTGTTATACTGTGTACACTTTGTTTATTAGATTAGGGATTTCTTGGAAC
CCTTTCCAGAGAGTGCTAGGGATTGGTGATAATTGAAGAAAGTCTATAGTGTTTCTCAGGAAAGATGTTAG
TACCGTGGTGCAAGTATCTGAAAACTCGTAGAAGTAATGTGCGTATATGTGCTTTTCgttatttattaaac
taaGCATACAATGAAGTTAACGTAAACAAAGTTTATGTTGCTACATGAAGCACATTCCAAGTTCATTCCAA
gttcacataaaaaaagaacacagtTTTTCTGTGACTAGTTTTGAGTACATTCTTGAAAGGTGTTCATGAGT
TTTCACAACGATgagctttttgttttatttctttcaagtCGATTAGAACGTTTATGTTTTGGAACTTGGAG
ACCTTGTTTGAGCTCGTGTTCGTCGAAAGAATAGAGTTACGACTTCTGTTATTCCTCTTACGTGCTGTTTG
GATTTTTCCTCCTGGATTGGCAAATTTCATAGTAGCTCGATCATTGTTCATAAATCGCTTTGGGTTATAGT
ATTGCTAGTTTGTTTGGCTTTTCACTGTAGATTAAGCAGTCCCTAATAACTGTTCACTGTAACTGCAG**TTC**
**TGGTAGAAGGAGGCAGTGAGTTTGAGGCACCAAGTGCATAACTTGCGAAGCTATTGGACG**GTAAGTTTTAA
TCAAggttcttatattttattgcacTGCATTGTTAAAGGAGTTAGCTTTTGAACTTGAGCTCAATACTAAC
TTTTCCAagcaattaaaattgaaagtcCAGAGAAATTGAACAAGCtgttataacatattaataaacttgac
ataagaaagagaaagttcattaacttttaaaacatttcAATTTCTTGAATTCAGGTGCTAGCTAGCACTCG
ATCGTGTTCTCTCTTGAGGATTAGCTAGGTTCTTTGGGATGGTGTGTCCTTGTCATCGTTGCATGTCCACA
AGATCGTGTTTCTTGAGACATTCTAAGTTGCTTGATTGATTTGAACTGCTTTTTGATGCTTTTGcgagttt
tttttaatcattcatCTTCTTAATGTTATCAAAACTGTACATGCTGGCTTTTCTTGGGTCGCTTTGTCTTA
GTAATAAGAATTCAATGAGTGTGTATACTTGACATTGGcataataccaaaaatatcCAAGATAGGTGTAAG
TTAATGCACCAAAAAACGTCTGGATGATTAATGacatatttgcaaattttatcattcaaaataaataatac
attgtctatttatagagaACATCCCTTTAGATTCCTACCAtatttttctcctaaatattctcctaataATG
CTCTCATAATTAACtctttcctaatacaaatattcctttatttttctacttttctacTACTACAATAGTTC
CTATTACCTAAAATTTTCCTACTAATACAATATTCTTTTCTGCTTCTCATTGCTCACTCgcttttttgtaat
atatctttaaataccCCTTCCAAGTTTGTCAGTtatgttttcaaattaaaaaaaaaaagagcatcaTAAAC
AAGATCTGGCATATGAGCTGGCGGCTGCGGATGGTGGTGTGGTTGCTGTGTATGTGGTATCGGTGAAGAAT
CTATAACTATGTGGTGGTTGGGATGTATAGTAACTCTCTTGGGGTGGCTACTAAGCGTACGATAACGAGCA
AGGAGATCAATATGTTAGTTGTGGTGGTTGCTCGTAATAAGGCTTCAAGTATTGCGGTGGTGACTGCAGGC
GCGGTTGGTAGGAGGCGTGGGATGTAGGAGGTGGTGAGTAGTTGCGAAACAGCGGCGGAACAAAAGGCTCT
GGTTATGGTGTGTTCCAATCGGATGCCagtgatacttttttttttctttttttctttttttttgatgaatttgg
tGGGAGTGGTGGATGTTGTGAGGATAAGATTAGTACAGTGCAACGGCGCGAGGGTTCGCGATCGCTGGctc
tgatatttttgttttagaatatttccTATACGTGCATATTTGTTACCTTATTAATCTGATTagtttaaaaa
caaagaaaagaaactccCACATATGATCTAGCAAAATCGCTTCTAGACCAAGGACGACTGTTAGCCATAGC
AATCTTGAGATCTGAAGTTTTGGATTGAAAGGAGAAGTCGCTAACAAGCAACTTAGATGACTTGGTTAGCG
ATTTGTGAGTGATTGGGTATTAGTTGCTATAATGGATCTAGGGTAGAATGCTTAAGGATGGTTGGGTGACT
GGGAAAGGATTTAGTgctagtataaatatatactaatatgttTATTGTACATGACTTTTGTTACGACATTG

TGTCTAGACATCGGACTTGAACTTGAAGGTAGTACCAAACCCTCCCTACCAATTAGGACTAGTTAGATTCA
CCGTTTTATTTGCCAGATGACACGCTCGATGCGGATTAGACGCGGCAACGGCCATTTTCCGGCCATGCGGA
ACATCTTCCTCTCCcatactctatttttttaatgattaatgAGGAAAATGAATATTCACGCACCGTGCGA
CCCTAAAAGTCGCGCGCTGccaaatctttttattaatttttttttttaaaaaaatatgaatgtgttttat
ttgttctATCTAATATGGTGATCAATAATGTTGAGCACCAATCTGggacattattatttaaaaaatgatta
atcGGACGCCATTTCCGGTGTTGCAAACTTTgaggaaaagaagaatagCGCCCGACTGGCGTTGCTTCTGT
TCCCACCATTGTTCAatactgttatttttatttataaattataaaaaataattgattatcGACATGATA
GGCATGGTATGATCGGCACATTGGGCATGTATCGAATTTGGAAGTTACCAGAATGGCACGGTATGTAAAAT
CTTGATTAGGTATGATCCAATAGGGTTGGGTAAGGCATACGGGTTGGGTTTTCAACACCCTAGTCTTTCAG
TTACATGCCATTATCTTGAGCTATAACCTATATAGTTGAGTTTGTGTTATAAAGAAGCTGGCAAAAGAGCC
TATACTCGACGGAAACTGAATTCACAATGTAGCAACTAGAGATGATTCGGTCAATTAGCTTCAAGAAGAAC
AACAACCAATGCCAGTGTCAACAGTTGGAGGTAATCAATCAGTTAACAGATGATTTGGCTAGTCACATGAA
TTGGAATCCGTAACAGCTGTATGATCATCCATAATCTATaatgttgttttatgtttggtctaaataatttt
cgtttttttaaaaaaaattgttttaataacCGGGCGTCATATTGATCCTTGATAAATTATTAGCACAGTTG
GCTGAGTTTTTATGCTTCACTATTCATCGTGGCTTCATTTTGTATCTTTTGATGAGTTGATAGGTTTGATG
TACCCGTTTGCACAATTCACAACCCAACAATCAATAACTCCTTGCATGTCCAccataaacaaattttctgc
CAACCATTGTCCTTGCTTGCTTCTAGTGCTTTGAGCTTTTGCAACTATATGATATCTCCCATGTAGTTAAG
ATGTGGAGGTTGGCTAGTCATTGGAGTGTTCGATTGACCGtcaagttaaataaaatccacCAAGATTATAG
CAGCTTATTGTTgcactttttttaattagttagaTTAAACGATTGGAGTCATACACTAAGTGGTCTAATATG
AGCATTCGTTTAATTACTTAGATTAAAACGATTGGAGTCATGCACTAGTGGCCAAGTATCAGCATTCTAAC
CACCCTGACACCAAATGGTTCCTTTAGAATGCGGGAATGTGCAGGGTTCTTCCCAAAAGGGGGTGCAAAAG
GCGCACAAACCCCAAGAAGGGGGTACGGGATACGAAATGGTCAATGTTATCTGTCTATATCTAGTGGTTTG
CCAAGTGACTTGAAGAGGAGTTTGTTGCTATAACTGGCGTTCAAAGTATTACATATAAGCACCATAGAAAT
TGGTTAGTGATTATCGTGTTGTGACGATCCATATGAGTTGTTATTAGTCTAATGTCAGCCTTTGGTTTTTG
TTTGTCTATTTGtatgtttaataatatttgtataagaaTTAGGTGCTTGTTGAAGCGATAGTAAAGTCAGG
TGTTCTTGAGTTGCCTCTGTTTGAATCACGGAATGTAGATGATATCTGGTGCTTATCTTTGATTTGTTATA
GGTATAGTAATAATTAAGGTTAGTATAGCCCCAAGACACTGGTTTCTGCTGTACGTCACTGTGTCGGCTTG
CCAACAAAATTTTCTCGCAGCATTTCTTTTTGGattgaaattttagaaagtgaaaaagtgtATAGACATGT
GATGTGTATAAACCAAAACTCAAGGTAGCTCATGTAATTGTAAATTAGAAGTTAGATGCTTTGACTAAATC
TAGTAACTACTAGAAAGTTAATTAGGTGAGCAATGATAAGGCAATGCATCTAAACCATGTAGCAAAGTACC
TTAATAAGTTAATGATAGTTTTAAGACAATAGGTTAATCTAACAATCAAAATTTCCAAGGTGAAATTGCTT
AATAAGTTGGGGTTAAGTGCTTTGTGTTGTTGCGATTGGTCTTGGACTATGAATAAGGCCCTTATCggact
aaaaaattaacttaaaccATTGGTCAACCAGCTATTTATCTTTTGGTGGGCTCATTTAGCTTGGCCATTAT
GTCTTGACCTATCCCCATTTTAGCGGTTGGGACAAGAGTGAACACTTGGGATTTCAACTTTCCAACAAAAA
GAGCGTTGTAGCGGTGGCGTGATTAATATTCAACTCTTCAAGACTGGGTTTTATCGATGTTTGGAGTAATA
GCTAATTCAAACGAGACGTGACTGGAGTAGCTttcttttgacaattttttttctcattaattTGAGTTGAC
TTGATTGGTTGAGGGAAATTTCGCATCCGTCTTTGTCTAAAGGCATGCTTGTTTAGTGGCTATGTACTTAA
CCGTTAAGATATAGCCTGTCTTACTAAGACTATGGTAGTCGTTGCTTTTATAAGCTTGTGAAGAgattaat
ctaatatttttttgccATCATTTTACTGTTAGTTTGTACTGATTTTGCGGGAACACTAGCTTCTTTCTGACT
TCTCTTTTAGGAATTTTGGAGGGTTTGTTCATTCACAGCTAGTTTTAGCTTCTTGATatctgaattttaaa
agtttagagATCCCAACCCTATATCGTTATTCCTTAGGTCCACATTTTGTCCTACCTGAATTTACTTCCAC
AAATTATCTTGAATACAACAAGAAGAATGGAAATCTACTAATAAATCCCGCCCGATCGGGTCTGGGAAGGT
TAGGATGTAACAGTCATACCTTTCGGAGGAGATTATTTTCTTGGGACTCTCGGCTTGGGTAATCAACACTT
GAACACAAATCCACGCCAGCATACAATCATACAAGTCCTTGGAGCATGCATTCAGACTACTTACACCCAAT
CAGGCAAATAGACCAAGTACACAACAAGTAAAGcattaaataaactataacCTAGGACAGAGCAAGGAGGC
AGCAGGAGGAGAAAATAAGTACGAAAAGCCAAAATGCTTAATCCACTAAGCAATGGTATCCGTTAACACGG
CTTCTTAGACCCACTCAGCTTGCTCGATAGCGAGGTCCTCATGGATGTTTAAATAGGTTAGATCTTGAATG
AATAGCCTTAAACACATTTCTCAAACGTCTAATAATTTGGTAACCAAGTCCGGAAATTGCTTTGTGGAGTT
GGTTAGCATTAAGTTTCCAATCTTATCTAAAATCATGTCATGAGCAAAGTTAAAAAGAGATTGTATGTGAG
ACAACGTTTGTTTTTTCTCAACACATTATAAACTTCTCCTATCCTGACAAATGATCGATGGTTTAGGATCT
TAAAGAGAATTATCATGATATTACGTGCTTACAATGCCGACATTGGCTTTGTCTATCGGTAGGTTGTTGAG
GTCTTGTGATGACCTTTCCATTGTCTTAATGAAGTTGTAACCTGTTGAAACACCAGGTAAATGCTCATGAT
TATAATCTAATAAAAGACTGATGCaaactattttagaaaagtaattaCCCTAGCTTAGGTGCTCCTGCTGG
GTCTGGATCAGCTTGATGGGGCAGGTTGTTTAACTCAAGCTGCGGTTGCAAATGTAAATTTCCCATGATTC
TCACGCCTAATGGATTGACTGGTAGATGTGAAGCTTCTCCTTGCAGCCCTGTTGATGCATAAACCTACAAA
AATACCCTGTTAGtgtgaaatttgaatatcagAGGTTATACAATGTTCTTCTCATTATGTCACTTAAAGAC
ATGAGGAAGGCTTCTTATCACTTTTACTTTAGGGATGTGTCACAGGATCTTTTACCTTCTTGTAGAGTTGC
ACATTTTCTTCGTGAATTATCTTTCCCTGCAATTGGAAGACTTAATCTGATAAGATGCATCACATGAATTC
CAAATCCAGCAGATGGATGATACCTTTTGTTTTAGTTCTTCAATTTCATCTGTTAAGATTCGGTCCTGCAA
AAAGTGTAAGAATTGTTTATCAATATTGTTGTCTATGACAGATTCAATGATTGTTCTGATTTGATAGATAA

GGATAGAAGGGAGCAATCATTATTAGTGTGGTTGGTAAATAGTTTGCAGACCTTCTTCATTCTGACAGTTC
GAAGAGTTATTTCGAGTTGGTTCTCTAAATCCTGCAGATCTTTAACGGAAAGATCAGAAAGATTCTCTCCT
GTCAATACCCTGTAAacattaatttaaactttgaaTATCCTGTGCTATCATTGTTGAGGATACAAAGTAGA
TTTTACGTATAATTACTGTTCTACAACATACAATCATGCAAATAGTTCAACGTATAGAATTTTGCAGGAGC
GAATATATCAATCCTTAGAAGAGTTATGAAAAGAATAGgatgctttttttattataattcatttattGCCA
GCAACACTGCAACATAATCAGCTTTTAAGAACCCACGTCAAATTCGTGTAGTTATTAGTGTTGAGTAGAAT
CCCATATTTATCATAACGGGTACATCATCTCATTCTCTAAATCTGTCTAAAACATGTATTAGACGAAATTT
CTGGCGAAGTTGTAACAAacgaaaattttcatgttgctTGGTGGATGTCATATCTAAATTGATTTGACTGT
GGTGAATAGTCTAATGCATGATTCTTAATTTTGTGCAACAG**GTAGAAAAAAATACCGGTGATCTTCTTGCA**
**AGTTATGCAATTGTTGCCTCAGACTTGCTGCTTCCCTTTGCCAAAACTAGTGAGTACAACATTCAAGAATT**
**AGCTTATTATACTAGTAAATACTTTCACATGTAACTTAAAAATGATTACTAAACAACTAAAACCAAATTAC**
**CAACGTCTTTTTTCAtgtatcaaattatcaatGCATGGTTTATCCTTGTCACAAGCTTTTTGACAGAAAAT**
**TGGATTCAGAATCGGATACTCTGGAGTTGACTCTGCTTCTCACTAAATAACATTTAGCAATTGCCCCTTGA**
**TTGTATGGTAACTATTAATTTCCTATATTCAAAGTAGCTGTTGGACATATTCTGCCATTACAGTGTAATGA**
**ATTCACACATGTTTAACTATGTGCTTACATCAGAACATGCTTATTGGGGAA**TTATGCTTTGTATTTTCTGG
AATTACTATTTCTTACTATATATCTCTTAGATGGACATCTTTTGCAACATGTGTGACACTGCTGATCTATA
GCATTATCTCATCTCTAGTGTTAGCTACAAATagcatctatttttttgtttctctattttcttttttacc
cTAAAACTTGAGAAATATCTATACACCATTAACAATGAAACAATTATAGGAGAAATCTGTAGATCATTGTT
GGGGGAATCCTTTGAAGATCAAATGGAAAAATAGCtcatgaattttatataaatctaaagTAGATCCAGAA
CAAGTCACCTTTTTCAAGTAAATCATTCTGTTATGTTGATTCTGCTAGAGCAAATTTTGGAACGAAATTCT
CGACCAAAGAGTTCATTTTATAGAAAACAGAACTATCAACAAGCATGACCTACCTTTTTCTGAATGTATGT
TATGGAAGGCCACTCATCTATccatataaaaatgatgttgTAGAAGTAGAAAGGGCAATAACAAGAAACAG
ATGGTGACAAAGGCCTTACAAAGTCAATAAAGGAATTACTCGCAGAAAAGAACTGGTTGGAGGAAAATTTC
TAGAAGTTTAATTTCATTATAACAAGAAATGGAGTAGGATTCCAAGAAAATTCCATAATGGTATAAGAAAA
TGTTCAAAGCACCATTATTAGGTCCTATTTTGGATGAACTTGAAACGCTGAAACTCGATGAGCAACTTGAA
TGGATGGATAATGGAATTGTATTTAGGTTCTTGCATACATAGGGGGTATTTTGTAGAGAGTCGGCACCTTA
CTTCTTATGCTATTTGGTAGAGAGTTGGCACCTCCTTAGTCCATTTCGTAGATGAGTTCGGGTAAGGGTGT
TCATTGGGTCGATTATGATTTGATCTAACAACTAGACTGGGGATCGGATCGAATAAGTCTCATAAGGTTAA
TTGGCCAAACCAGACTATACCCCGTTTGATCGAATTAGcattatctttaatatttaatttatgaagcTATT
TTCAATACAtgtgaatttaaatattttacaattattgaaaataataattcaaatattattttaaatttact
aattgtATAGTAACTTAAAGTTAAATACAGGTCTTCTGTCAAGTCCATCAATTTAACTGGTTCGGATTGAA
GACCGGTATGCGTTCGGTCTGAACCAGACTGAACCGTACATGTCGATTCGATTTTCATACCGAACCAAACA
TGTCAGTTCGATTTTCCAGTCCGCATTGTCATTTGAGGCAGATTATCGCGTGGGTGACGGTTGAAATATTG
ACCCTGTGACAAATGACATTGAAGCTCAATATTGAGGGGATGACATCATATATCGAGAGCTAGATGATCGT
GGGGGACAACAGAAGGGTGAGAATCATCACACAATGCCACTGCAATCTTGAACATTAAGCATGGAAAGTTA
GTGGATAAGATGACGAAATTCAAGCACCACGATCGAAAAGATTGACTTGGATGATGGCTTGGAGGCTGGCA
TAGTGGCCACACTCAACTAAGTTGTGGGCGACTAACAGCAGTGGTTGGCTGCATTGGAGGTACAATCCGAG
GCACTTTGCTAGGAGAATTAGGAGTTGCGTGGTGATCTGTGTAGCAACTTGCCTTGGATAGGGTTAGGTAC
AAAATTGCTGGAAGGCTCGAGTTGTGCCAGGCAGCAGCTACAAATTATGGCGATATGTGCAACTTCCTTTA
CCATATGGAGAGGTACtttgatatgtatttttacttaatatgCTTCAAAGtatgtacctttttttttttcc
ttagcCGGTGTCGCGACACTATGATAACGACGAGTGTATGACAATGTGGCAAGAGACTGGTGGCAATCATC
ACATGGCAAGTATTCCAAGTAGAATTCAAGAGGCAGTTATACCGGGAGAATGCCTTAGACAAAGTTAGGCT
TAGGTTGAAGCGACTGAAGCAAACCGACATAATTTTCTGGGTATGTGAAGGTGTTTGCAGAACTCAATTTT
CGAAATATCGAGCATGTCGGACCAGGACGTGTAACTTGCGTTTGTGAACTATCTAAATAAGTGGGTTGGAG
CTTAAATGGCAAGGCATGCAGAATCTTATGGGTGCCATCCATTTGATTGAGTGCTAGGAATTCAGGGGAGT
TATGATGAGGCTGAAAAGAGGACTATGATGACAATGGTGGTGAGAGAAACACCACtcaggaaaaaagaaat
tcgcAAACGAGGCTAAGGCCTGGAAGAGGCAAAGACTCAAAGCCAATAGTGTGATTCTTGTGTGATGGATC
GCACAACCCACGGAACTGTCCAAGTTGTACGAAGCTTGTCGCATTGTTGGTGCAGAAGGCACAAGAACAAA
AGCACAAGCGCAACACAACTTAGTGCGACACTTGGCGGAGGTTCAGCTGGAGAGCGAGCCAATGAAGGTGC
TTCTTGAGAAACGGGCTCATGCAACATTCCTATGACGGATCAAGTTGCTCTCAAAATGGGTTCATAGTGTA
GCGAGCTGAGGGATCCATGAATGCTTTAACTCCTTCCCAAGCCCATCGTTGGCATGGCACGGCAGGTCAAC
AGCATGTTAGACGACTACACACGTGTTAGTTGTTTATTGCCATAGTTGTTGCCACAATTCCGGGCATAGTG
ACAGGATTAAGAAGTTACCTTATGTGCAGCTAAATTATGTAAGTTGTTTATTTCCATAATTGTTGCggttt
tttgtaaattttatgcaCTGATAGTATTTCAATACTAGTTTTTacttagattttatcaaagaccatcataa
ttaattttggactAGAAAAGCTGGTTCTACTGTGAACCAGCGGCCAGCCCGGTCTGGAACAACACAAAAAC
CGGAATCGTTGTTGAATcagaaaaaaacatttaaaccACTCGCCCAGATGAACCAGTCGGCCAGGACCAGA
CCTGGTTGAACCATTTTCTAACCTAGCCTTTTTATGCTCGGGCCTTAAACCCTTTCttctctccaaaaatt
tttgtctAAGGTTTCTCCAACGGGCGATCCAATCAACGGTTACAATGTGCACCACTCCAGTAAAATCGAT
GACAATGATACTGACCCTTGTTTTTTACAGTGTCCTTTGCAATCACCTATTCTGCTTAGGTCTGCTACTAA
CTTTCTGGATTGTTTTCCACCATTCTATTATTGTTGTTTCAGATTTTTGTTGCTTCTTGTTCTTTATGCTT

```
TGTGATTTTTAAGatctttttgatgttttgttcttcttgttttcttgttCTCGTATTTGTTGAACTGttag
ttaactttttttttttttgaatttgacaaGCTTAACAATTAATACTATGACTTCAAAGGAAACTAGTGATG
ATTGATCATGCTATGGTGATTCTCTGTTCTCACTCCTAGAAATTTTTAGATTGAATTCATTGGTGTACTGT
ACcatattaacttttcttttgttgtctCTGACATACAGCTTTTGTAAATGAGTTAAACTGTTAATTGAAAA
TGTTCCAGTTTAATAAACATAGATTTAATGAAAgacattcataatatataagttgtcAAAGTGGGGACTGC
CGGTTCAACCATTTGGTTCAACAAGTTGAATCAGGAACCACTGGACTGGCTGGTTGTCGACCTGTCCAGTT
TTCAAAACATTGTAAAAGACATAATCTAAAGGCATCAAACCTTCAGCTAAAGCTAAAGTTTAATGTTCGAA
AACGCTTAAACATTCAACAATATGATGCTATTAATCAGAACTTGTATTCATTAAGCTGATGACAAAGCAGT
TTTACTTTCTAGCTGTCAGTTTATACAAACCCACGGTCtgtactttaattattatttatcttctgTAGACT
AGAGAAAGGGTGTTGAATTTGCAGTTTGTGTTCTACTATGTATTCTgcaacatttttaatttctcagTGTT
TTTTACTTATCTTTGTCACAAACATAGTTAATATTCATATACTCTATAAGTCCATATTACAAATGATTCAA
TTTGCCAATCTGAGTCTGTTGAATGTAATATTTTCATGCTGTACTTTTAAGGTCATTTAGACAAAATTGAC
TGTATTCTAATGTCTTGGATGTCTCAGATATATACCTAAGAAATTACTCTGACATGTCATTTAAAGATCAT
CTCACACACTGTAGTAGCTgacttcaatatttctcattATTATGTTGCCCATGGTGATATTTTGTCCACT
AGCACTTCTAAAAGAAGATAGAACCACAAACAAAGTGAATTATAACATCCTATCTTCTTTCAGAAGATAGA
ACCACCCTACATTTTGGCAAAATTAAAATGTGTTAAACATGATGCTACTTGATGTTCAGTAGCTATCATAT
TTGTGTAATATCGTCAACTTATCTCGAGTTGCTTATAGTTCCTGAAATCTTTCAACCAAAAACTTtttcag
taatttttctttgattcaGTTTGAATTAGTTCATAGATAGACCaaatttaatactttaatttCAGTGAAGA
atgcaaatatttgttttctagTTTCTGGACTTTACACTATTCAATACAATCAGCAAGCAACTTGGCATCTC
TCTTATGCCTTAAAAAAGCAacttggaaaagaaaaggtaaaaaagaatttgaaaacttttaagtGCTCACC
TAGAGATGTTTCTGAAATGAAGTCCATTAATTGTATAACCTTTGCCTTGGTTATATCTTTATAACAGGTTA
TGGATATTTTAAGCACCAGATTTCGCCGCTTTTATCTGTAGATATGTGCGTTATTCAGGGCTGCAGAATGG
GTGTAAGGAGATAGAGGGTAAAGATCGAGGCGTGAGAAGTGGAGGGAAAGATTTGCTCAGTCtcttttata
atcttttttaagtttattttcaatttctttctttctttcattccTAGGTTTTTGTGTAACTTCTATGTCAT
TCAGTTTTCTCATAGAAATTCAAAGTTAACTAACAACTTATCTTCTGTCATTGTTTTTTGAATTGCAGATT
TTTCAAAGAGATTATTCCAACACTTTTATGGTGATACATGACTTCTCTTTCTATGTAAGACCGCGTGTGAC
TGCATTATTTCTACTGTCTGTAGGCTATCTGTCACTCACATAACACTATCATTACTCTTCCTGCAGCAACT
CATAGCAAAGCaggatcctttttttttattaagatgaACATCGTTGTTACCTTTCTCCAATACGAACTTGG
TAAGTTCTCTCTCAGCTAAGATAAACATATTCTATAGATTACTTCTATATTCATTCATGGGCTAAGCGCAG
CGGCGGTGGTAAAGGTTATAGCTCACCTAGCCAAAGGCAATGATGGAAACATCCTCCAGTAGTAGCAGACA
TAGGTGCAGTGCGTATGCCGTCCCTCCCCTTCCCCCCTTTCGAAAAAATCCTCGTGAGTTAAGTGGGCTAG
GGAGTAGCATTGTGCCCTTGGGTTTACATTTATACTCATTCATATCCCCCTTCCAAAAAATTGCTCGCGAG
TTAGGCGGTGGCTTTGTGCCCTTGGGTTCGCCTTATATTCTTTCATATTGCCACTATAATTTTAGTCTTca
ctagtgttgcaaaccgtgcttGTCCGAGCCGGCTCGTATTGTTTGATGTGGGCCGCTCGTGGCGGCGggct
gcgggccggcccgcacggctaCTACGGCGGGGTGGGTCGAGCCGGCCCGTCCTACTAAAAGCGCCAGTTCG
GCCGGCAATCCGCCGAGCCGGCcgcaattaaaaaatgaataaataaataaaaaacatattataggCGGGCC
GGCCCGTCCCGCCCCGCGCGAGCGAGCTAGTTTTCCCAGCCTGGCTTGCGCGGCCGGTAAGGCGGGCTGGC
CTGCAGCTCGCACTGTAGGTAAGGTCGGCTAGCCCGCGGTCCGGCGAGCCGGACTATGGTACGGGCCACGG
GCTAAATTAGCATCTCTAGTCCTCACCCTCTAAAAATGGAAAGATTTGAACATAAAGGTGGCATGAAAAGC
TAATTGTATTTTGTCATTGGCAACTGCTGTCGATGGAAAAAAGGCTTTTCTAAGTCAAGACATGGATGCAT
TCATGATGATGATTGATGAAGGGAGACCTACTGATTTTGCTTGAATTCGAAGAAAAGCATGCTATGTGATC
AGTAATTCTGTAAAGCAGTTGGTAAGATACCTTAGCTTCTGAATCGGCATTTGGCGCTTGGCAGTGCTCAT
CTTTCGCTTTGTTGTACCGTTCAATCGTAGATTTCATACTGAGCAACAGAAGTCATATTAGAAATTCGAAT
TCAGAAGCAAATTAAACTATAACTTGTAAAaaatgaaggagaagaaatctaAAGAGCCGCACAATGTTGTA
GATTAAATCAAAAGTGTTAGAGATTTGACTATAACCAAAGTTGAAATAACCTGAATGTCGTGTGGAAAGTT
TTAtcttgttattaaaaaaaaaagggattccAATGACAAACCATTAATGAGTGCAATTAAATGCGTCATAG
TTGAATAGAAAGTGTGGTTCATTAGATCGGAATGAAGTATACTAGAATCAGAACGGGAATCGATATTTTTC
CACATATTATAATGGAGGTCtcaggaataaaaaaattcataaccATCGGAATTACATTTCAATTTGAATGG
AAATAATTTCCAATCATAACGGTTATAACgtaatcaaatattatcaacGGTTAATTGGAATGATTTCTATT
TTCGgctgttatttattttaccaaataccatttaaatgttttacaaCAATTATTAACAAATGTATCGGTta
tggacataatttttacgataattaagtttttttttttttttttttttttttttttttttgagataactagG
CGAAAAAATTTCGGTGCACTAACTCCTAATAGTGTTAAAGATATCGTTTTTAATCACGATCCATCagttca
aatataatactGATTTTAGCTGTCACGTGTTAGCAAAAGAGCTGGCCACCAGATAAGGAGTTCAAGAATCT
CATATGACTGAGGTGCATACGTTGTCAAGGTGTCCATCCGCATACTTTGGGATCGTCATGTCCAGACTGAT
CTCGGCCCACAGCTCGACCTGCTCGACTCGCACTCCGCATGCTGAGACCGACATTGGCCCACATTCGCATA
CCTCTGGAGACGCATTAACATGCCGACCTCAGCTCGACCGTAGACCCAAACTCTGAATCGTCCCCAGCCTC
AAGATCACCAGGCCGAGGCCGACCTCTGGTTGATGATGACTTGCAGACACCTCGGATCGAATCGAGATCTT
GGCAAACTACACGAAGTTCTGTATCCAGCATCGGCCAATCGAGACTAACTTCAGATATAATCTTAATCAAG
GTATTAAAAATCTGACCAAACTCATCCTACCGATGTGAGGAAAGATGAAGTCAGCCATATAACCACTGTGC
GGACGTGGATGAGTCAATTAAGCACTAAGCTGTCCACCAACAACGTAGGAGCTTATTTGGCACAATGAGAA
```

CCAGTCAATCCACGATTCTTGTCAGATTCACTATTTCCGCAACCGTCAAGATCTAGGCATGCGCCCAGCAA
CACCTATAAAAGGCTGTCCGGGGCACTGTTTATAGGGGCTCCAACTTCACGCACTCGAGAACACCTAAAGA
GGATTGAGATTTTCAATTCCACGCATCGAGAGCACTTAGAGAGGACTGAGATTTCTAGCGTTGTGCTAAGT
CTAGTCGTGTGCTTAATTGTATTGTAATATGAGGTGTTTTAGACCTCGACAGAGAGTTTGATGTGTCTAAG
ACCTTGacaaagagagaaaaatgagAGATCTTGTATTCTTCATAATCTAGTAATATCTATACGTTGGCTTA
CGCGATCGCCCGTTGTTTTTCCTTGTATTCCttaagggtttccacgtaaaaactTTCGTGTCTATTCCCTA
ttccttttaatatatttctgctTTCGAATACTATTGGTCGGCTGACGTCAACTCGATCAATTGGCATCGT
CTATAGGGAAATCAACACGAAAGCAATTGCATATAAGAGAAAGGCAGGCAGAGCGGCTCGCACCACGGCGA
GCGAGGAGCATAGATCCATGGAGGAGGCTTGAGGCTGCGAGCAAGGCTTTCtgtaccgagccgagccgagt
ttcggtaaactcagatatcggcacgtgccgaccatgccgatggtgtcgtgccaaccgtaccgatatatatc
tatttgttatttttttaaaaaaaaaaattaatttaaaattcaaatataagatttttttgacacgtgtcatG
TCGGTTCGTGCCAACAACTGTCGGAAAAAAACACGTGCCATGCGGCActgggctgtgccgaccgatACAGA
AAATCATGGCTGCGAGCGCTCCCTATAACATATTGACTGAAGCCTCGCAAGTGCTCGCAGGTAGCAGTGCT
GCACTTGCGACCAACAAATGTCACTAGCAATCTGGACGACACTCCCGCCAATTTGAGGGCAGTCCGTGGCC
TTCTACGCGAAGAATTGCCCAACCTGTTGCAGCCATCAGCATTGGCTGCACCAGAACAAGTACCGGTATGA
AAACGACACTTGACTTGCAGAATTCAAGACCGCGGCTCGACATTCAGCAGACTTGGACCATCTAGGATCTA
AGAACGTCTGGGAACTATGGTGGTCGACTGGCGAGGGAGAGGGACGAAAAACCTACATAGAGCTACCAGAA
TTCCTCGCCCAAACAAATCCGAGGACTGCGCAGAGACAAATGGCCGAAGCCCCACGGGACAAAGAACCTCG
ACATCGAAAAGTGCGGACCGAGCCCACATACTTCATAGCTCGAGGTCCCAGCACGGAGCAGCAGACGAAGA
AGGTGGAACTCCCGATCGACACGCTTGACGTCAAGGAGGAAGAGCCAACAACGATGTAGGGACGTAATCCA
ATCATTTCAAGAAGACCGACGCCACCAATCTGGGTGACTCCTTGCACTCCTTATTGTCACGGGTCTTAGAG
TAGGAGGTGCGGGCCAGAAAATTACTTAAGTGTCGAGGGACGTCCTTTGCATCCCGAGCCGATGCGAGCTT
CGAGATGAAGGGCGTAAGTCAATGACCTGGTTGACGACCTCTGCACTTTTTATCATCACGAGTCTCAGAGT
GAGGGACGTGGGTTAGAAAACTTACTCGTAAGTCCTGGGAGACATCTATTACACTCCTTACCGTTACAGGT
CTCGAAGCGGAAGTGTAGGCTAGAAAATGACTTAAGTGCAAGGAACGTCCTTTGCATCCTGAGTCTATGGG
CGCCTCGTGATGAAGGGCGTGAGCCAATAAACTTCTTCTTAGGTCCTGGGAGACATGCCTTGCACCCATTA
CCATCATAGGTCTCGGAGTGGAAGATGTAGGTTAAAGAATGACTTAAGTGCCAAGATTGTCTTTTGCATCC
TAAGCCCATGGGCGCCTTGAGATGAAGGGTGTGAGCCAATAAATCTTTCTCTTAAGTCTTGGGAGACACCG
CTTGCACTCCTTACCATCACGGGTCTCGGAGTGGGAGGTGTGAGCCAGAAAATGACAAGTGTTGAGGGATG
TCCTTTGCATCGAGAGCCCATGCGCGCCTCGGGATGAAGGGCGTGAGCCAACAAATCTTCCTTTCAAGTCC
TGTGGGATAATTCTTACACTTCTTACCATCACTGGGCTTAGAGTGGGAGGTGTGGGCCAGAAGATGACTTA
AGCGCCGAATGACATCCTTTTGCATCCCTTGTGGGCTAGAAAATGACTTAAGTGTCAAGAGACATCATTTA
CATCCCGAACCCATGCGTGCCTCAGGATAAAGAACGTGAGCCAACAAATTTTCCTCTTAAGTCATGGGGAA
ATACCTTTTTACTCTTTACCTTCACGGGTCTCGGAGTGGAAGGTGTTGGCCAGAAAGCGACTTAAGTGTCT
GAAgacgtcctttttttttatcccgAGCCCATACACATCTCAGGATGAAGGGCGTGAGTCAACTAGGTAACT
CCGCTGTACTTCCATTGCAAATCTCGGAGTCGGGCGTGAGCCAAAAGATTTTCTCCTGTCAATACGGAGAA
GACCTCGGAGTTGTGATGCTTGAGCTATTGAGGGGACATCCAGTAACGTCATTGGATCAAGGCAACATGTA
AGGTTCCAGCGGACGCACCGACATATGTCGCCACCCCCGTCGAGGGACAtctaatatagtatatatatttg
aactgCCACTTGTTAGCAGAAGAGCTGGCCACCAGATGAGGAGTCCAAGAATCTCATATGATCGAGGTGCA
TATGTTGTTAAGGTGTCCATCCGCTTACCTTGGGATCATCATGTTCAAAATCAATCTCAGCCCACAGCTCG
ACCTGCTCGACTCACACTCCGCATGCTGAGACCGACCTCGGCCCACATCTGCATACCTCTAAAGACGCATT
AACCTGCTAACCTTAGCTTGACCGTAGACCCACACTCCGTATTGACTCGAGGCCCCAGCCTCGAGATCACA
AGGCTGAGGCCGACCTCTGGTCGACGTCTTACATACACCTAGGATCGAACCGAGGTCCTAGCAAACTACAC
AAAGTTTTGTATCTAGCATTAGCTAACCGAAATTGACGTCAGATAGAAGCTTAATCAAGACCTTGGCAGCC
TGACCAAACTCATCCACTAAGATGAGGAGATAACAAGTCATCCGCATAACCACTGTGCGGACGTGGAAGAG
TTAGTTACGTATTATGGCCTCTACCAATAGACGGTGGGAATTTACTTGGCGTAAGGAGTACCACTCAACCC
ATGATTCTTGTCAGATCCACCATTTTCGCAATCGTCAAGATCCAGACGTGCGCCCAGCAACGCCTGTAAAT
GGCCGCGTAGGGAACTGTCTACATGGGTCTCAACTCATGTACTCGAAGGCACCTAGAGATGACTGAGGTTT
TCGGTGTTGTGCTAAGCCTAGTTGTGTGCTTGAGTGTATTGTAATATGAGGTGTCTTAGGCCTCGGCAAAG
AGAGTCCGGGGTATCTAAGAGCTCGACACAGAGAGAAAAATGAGAGTTTGTATTCCTCATAACTCTAGTAA
CATCCCTGCAATAGTTACCGTAATCGTTGGTAATTTTTCCTTGCATTTCTCAAAGCTTTCTACGTAAAAAT
TTTCGTGTCTATATCTCTTCCTCTTAGCCGTTTTCTGCTTTCAAACACTTATTGATCGACTGAGGTCAACT
CGATCAAGTTCGTATTGGATCAGAATCGCGGttggaaaataaaacttaacaCTTAATTAAGGAAAACTCAA
TGCttccatttttctaaaacaaatcaaacgCAACTAGACTgatctcccaaaaaaatttcgtatttGCAGAT
TTGACTTAATCTCTCACACCAATCGTTGTTATCCATTGGGAAGATGTGCTCATCCCCTACTTTTTCATTGG
TATAATCATCTCTCACACCAATCGCTGTTATCCTTTTCATAATGCCTCGGCTTTTGACTATAGGTAGTGTA
ACAGCGACTTCCCTAAAACAATTTAACGCATTTAATGAAGTTGGAGGTTTGTCAGGGAATCCATGGAGTGG
TAGATCTGTTATTTCACACTTAAATTTAGTTATGACTAGTTatacctgaaaaaaaaaaaaaaaaaaaaaa
aaaaaaaaaacgtactagtccagttggttagcaaGATTTGATGCTGAAAGACGACTGTGACCTTTTACTCG
TGAAGTATAGATCGGAAGTTTGAATCTTAACAAGCGCCTAGTAGGAGGTGCTAGCGTTTCTTTTAGATTCG

GATCGAGATATAGAATGGAAATCGAACTCAACGTTcgtaaattcaaaaaaataaaataaaataaaataaaa
taaataaaagaaaaaggttataACAAGTTCTGTCACTGATCAGGATGGAAGGTTGTGTTCTTCAATCGTGA
ATTTGACGTTAGTAGCTTCGTAGTAAATTTCGTTCTTCCATCCTAATAATCATATCGTATATGGACTCCTG
ACGAAATGGGTGCTTAGGCATAGACAAGCTCTAGATTTGGTTACATATGAATCCAGTTGTGTATATAACGA
GGTAAAAGTTGATTGAATCAGACACTAGGCCTTCGGGACTCAAATCCGATGTCCTACTCTCTATCCTGTTA
TCTATTCCAATCccttaaaatacatacttataaatatttttttgaaaaaattttcaacataaaagttgtaaa
tcttatcaagatctacaatttttatgttttaagtttttctaaaaaaatacttccaatatgtatttttagac
ctctaagtgttaacggatttcaaaatagttttaaaatctgttaaaatgttaagattttgaaatctaaaaaa
catatatttagaaaaaaaatttggaaaaatttttaacataaaagttatagatttgatgagatctacaactt
tcatattgaattttttttcaaaaaatacttgtaaacatatattttgagggaTTGGAACAGAGGACATGATAG
AGAGTGGGGGCAGAAGATTTGCGGCCAGGCCTTCGAATGTCCTAGGTAACTGATTTCTTAATTGCACGAAC
ACGGAGGAAAAAGTGATCGTGTAATATCATCTCAAAATGGTGAATCTCTAGGTGTACGCCGTTGAACGTGC
TATTCTTACGGTGCCTTGTCGGCCCATGTTGTTCATTGCGATCACCTCCCACGATCTGCTAGGGTGACTTA
GCGCCTTCATGTATGACTTGGCTTAAGCTAGCTTCTTCTCCACGGTCCACCTGATACGTTTGTACCATGGT
GAAGTGACCACGTAATCTCGATATCTTTCGACATGCTCTTGTTTTGGAAGGTTGACGTGCTTGATATCGTT
CGATCACGGCTTCTCAATGTCTTCGATCATCGTCCCAAGCTGCCGAACTTGAATATTCACAGCTCTTTTGA
TGTTAAGAACTGCTAATCATCAGGGGCCCCAAAGCTCCTGATGCGATCTGAACCGTTCAGAGCACCTCGAG
ATGCATTTGCGTGCATCCCGAGGTACTCTGGACGGCCTTGATTTCATCAGAGGGCGTCAGGGttccctgat
gaatggcatttTCCTTTCATGTTAGGGGTTATAGGTCCCTACACATAGTGGGAGCTCGAATAGCATCTCGT
ATGTGTGCTAGTGGTTGGAGCACTACACTTCTACACTTGACAAGGGAAAATCAAGGAATCCTCATTTGTCG
TAGGTGAGAAGCGAAACGAGGGTTGGAGAAGAATGTAAGAAGCAACccaaataactattttactcatgatt
tcaaaagttttacgTGGAAATTCTCAAACTTGTCAAGGTAAAACAAAGTGCAATTGTTGTCGGCATGAGAT
GTTTTCGACGTCCCATACCTAATATCACGAGCTTTTCTATCTAATGCATGCTACAATCGCCATTATGTACT
TTAGATAGTGGGTAGAGCAATTACTCATACCAAATAAtctggattttattttattctaggACCTCTGATAA
CATGATGTGAACgcaattttgatgagatattactatttattattataaaaataatacttgttCCTCTACAA
GAGGGTAGCAACGTTTTAGATTGGTCCACGTGTCCCAGGGCCTGCTGGTTCTAAACCTGATAAAAACCGTA
AGTTTGTAGCGGTTTCTTTGGAAGGTGATCTCATTGCTTTGGAGTCGGACAGAACTTAGGGCAAGCCATgg
tttaattagtttattttaagagatatatttgaacccatatatacatatatatatatatatatatatatata
tatatagttttctgCTTCACATGAATCTCATATATACATCATGCAATGCAGAATCTTTACGGAGAAATACA
ATAGAGGTTTGTAAATGTTTCCAAAaccatatatatcaaataaatttatgaaaactatagttaaaatttat
ttcgtgaactacaatttttttacgACTAATTTGGCacttagtttttaaatttttaattacaaaattaagaat
attttaaaatatatttaatatttaaaacttttctaaagatatctaaaaatacatactttCGCGATTACTGC
ATAACGGACCTGTACTATATGGAGCCCACTCCTTTGCGCACCTATTATCCACCGCATTGATTTGCTGATCA
TCATCTATCATTTGCTAATTGTTATCAACCAGTTCGATCTCGCACTACTCGCCGATGTGGTAATTTGCTGG
CGCTCTTTCGTTGGCCACAATTATATTTGGGTAAGGTGGTGAGATTAAGCAAGCAAGAGGGTTGTAGAGGT
GGAGGAGATGGTTACCTGTGAATGATCACAGCTGCAACGGAGGATAGTGACTTGAGATGGTATATAGgggc
agaaaaaaaaaagaatataatagatGGTTGCACAGATGAGGATCCTCAGCACCGGCTTTCTTGCACTGTCC
CACACACcgacatctcaaaaatatagcgacatatatttttttatatagcgacatatattcaatataagttat
agattttgataaaatctacaattttcatattgaacgtttttctaaaataggtGTTCTGCGACTTCAAACTc
ctaacagattttaaaagagttttaaaactcgttaagACGTTAAGACTTTTGAGCTtaacaaaaatctattt
agaatcaattttaaaaaatcgttcaatatgaaaattgttgtcatcaaaattgatcatatatatatatata
tattatatatatatatatatatatatatgtttttaaaatcgaTGCACGGAACGCTGTGTAAAAGTCGGTAC
AGAGGATTTTCATCCTGATTGCATACTAGTTGAGCCATCCACACTTAAGCAAAATAAGAGTCGCGCATATA
GCTTTAGTAGTGCACCTCGTCCCATAAGATTTATCATTTCTACGGTGTCGGAGTCGCAAGCCCGTGCAGCC
AAGACAGCCTCTAATGTGCGGGGAAGAAGTAATTTAAGTTTCTCAAGatatccaaatattattttagtccc
ttaatcaaaattttcatttgtatgTCAAATCTACGTGATATGCGAAAAAGCTTACCTCGTCGCCAACTCAT
GTATAGCTATAAATGAAATCCACGTGCATGTGAGTAATTAATGCATAGCTATAAATGAAATCTACGTGCAT
GTGAGTAATTAACTGATAGTATGATTAGTCATCCTTTCTTGAGGCGCAAAATGTTCAATGATATTCATTCA
AGcgactaaaaataattctgtacttagattgctttttttttttaatacgttAAAGGCTAAAATCGCATTTG
GGCTGTAACTTAATGGActataagtatataattaatctataaaataaaaatatatcaaccAAGAGTTGATT
AGCGGAATTGAatgctagaaaaaaaataggacatggCTCTCATGAATAAGTCGAaggtttgaattttaaca
aacgTGTGTTATAAGGGATTAACTGTACCACTCCGTTTCGAAAAGTAAGGTAGATTAAACTCGGCGgtcaa
atgtcaaaaataaaaaaaaataaaagataaaaaaggtGATGAAAACGAAAAGACTACTGCTTCAGTACCTCA
CAAGTTAATATACATAACTTAGTTCAGAAATTCAACATTCTTCtcataattcctttttttctcccCATATT
GAACTTTGAGTTCAATTGTTGCACTAGCTTCGgactaaaaattagaaaagaaaacaccTACAAAGTAACTC
TAATTACGATAATCGtggtttgattgatttttttttttaaatgtaaaaaaaaaggataaaaaaagacggggg
aaaaaaataaaaaaaaaaaaccaatgcaTGCGAGGCTGAAAGTACAGCCTTTCAGCTCGTGCGTTCACAGTT
GACATGATAATGTAGGATGCCTGTATAAACGTGAGAGAAGTAGAATCAATTATTCTGTGTCTCTTGTTGAT
TAggttatttaatataaaatacaaacccTAAAATACCTTGTCGCTACCTGGATCTGCTAATATTTCCTTAA
TTAGCAGTTATCAAATGATTTATCATGCTAATCGccgtaataataaaatttagcaCCGTGAACAAATTAGG

GTGTCGTCAAATACAATCATtagagaaggaagaagatgaagcaaATTAGGTCTATAAGAAATCCGATATTT
TCAGCCTCGCTTAAGCAAGAGTCGAcggtaaaatttattatctaataatcTTATTAGTAAATCAGTAAATA
AGGACTTAAGAGAAGCAATTAAGGGCTTGGTAGACCTACCTTGAGCTGGCGAATTCGTAGAGCCTGCCGGT
GCTGGAGAAGACGACGAGGCCGACCTCGGCGTCGCAGAGGATCGCGAGCTCCTTGGCCTTCTTGAACAGCC
CGTTTCTGCGTTTCGAGAATGTCACCTGCCTGCTCGTCGAGTTATCGATCCTCTTGATCACAATCTTGCCC
CTCCCCATCCCTTCCCAATCAAGAAATCTGCTAGATCTAGAACAACCAAgtacaaaaatagtattatatga
atttggaaaaatagtCAACAAAATCAAGAGGATCGATCGAGTAGgacaaaataacatacaaaatCAAAGGA
TTGAAAGATTCCCTTGCGGCGAGGAGAGATCTGGAGAGCGAGATGGGGAAGGAATAGGTGGTGAGGTGTTT
TGGTTGTGCTCGGTGGTGGGAGGAATGATGGGATTGCCTGATTTGAGAATGACTTTCTACTTTCCGAGGTT
GCAACTGCAAGTGGAAGCAAAGTGGGGGTGGTCCATTatggaaattaaattttttgttattataacgAATTT
TCGAAGTTGAAAATCTGTAAAATAGCCAAGTTTAgataatatttggaaaatttactaattaacccactata
taaatatttcttactaaTGAGCTTtcctcaaaatattcatacaatTCAACCCCCTattactattttgaatt
tttcacggacgaaaatattctcagAATATTTAATaccctaaaataaatatttgaaaaaactttcaacatg
aaaattgtaaatattatcaaatcctacacttttcattttgaaagtttttcaaaaatattcttctaaataat
tattttaggtgttaaaaatatcatttaaaattaatttttaagcgttatttttaacaccacgaataaatatt
tagaagtatttttagaaaatttttaacctaaaagttgtaaatcttataaagatctataactttatattgat
tttttttttcaaaaaatacttttaaataattatattgatgtgttgaaaataatacTGAAAATGAGATTAAA
ATCGACATTGTATGAAACGTGAGGGGGACTTTTTTGTTGGATAAGTTTGTCGGGGACTGATCGGCAACAAC
GAAAGTTTGGTTGCTGATTAGCAAGAACACAAGTTTGATGAAGCTgaatagcaatttttttaaatatatttt
ggattaattacatataagtttttttaaagatttttacgTATCGATCTCTGAAAATAGCAAAGTTGAAAAA
GGTGGATGGCTGATATGTATCaactccaaatatttttttatgtttttgaaacCATTTAGCTAcgacttaatt
atttattaatttttttaaatataaaaaaaggctgtcaaaattatagtttcaaatataaatttttgtatttata
tatcctTATACTCCACTAATATGCTGGGGCTGGAATTTGATTCTGGATGTAGGCCCCCTTCTTGCCTGCCT
GCCCCATCTACCATCCCAGAggttctcttttttattttctactttattttgcattttaggaagtatacaaga
taaataaaaaaatattatttctaagaaatttctaattatctttatatatatataatccaaatcacatattat
atcAGAAAATACATGATCGAGTTACACCTCTGACCGATGTAATTAATAAacgaatatataaaatggcttga
aagagaaaaagcaaatatacaaacacacaAGAATTTTACAAGAACAGGCTCGatctaaaaataagtaaaac
taCGGACGTTTAAACATCACCGATAAAAAATTCACTAGGTGAAAATAAGTTATAACTCTCCCACACTCTCT
CTATGAGCGGAGGTCAAAGAAATACCTCGCCTTGATTATAAGAATTATCTCTAAGTactattttattcttt
ctaaGTCTCGAATGTTGGATCTCGCATGCCACTAGTTTGCTTTGCTATTTATAAGCGTTCCCTGCGATTCG
TTTTGGTACCGCAGTCATGGCGAGAAAAAAAAGGATCTAGCAGAGATCGTGGGTTGAGTGGTCAACACCATC
CTCTGCTCATCACATGACCCCACGTTCTCAAGCAGTTGGTTTTCTGATGAGCGTCGTCCCACGACCTCACG
GCGGTGGTAGCTGCTAACTTATTCTCCAAAAGTTATGCATGCGGTCACGGTTTAGTCTCATAGCTTCGATC
AGAGCAAGGTGGATCTTGATCCATCTTTTGATAAATCTACATTTCATCAAGAGATCTTAGTAAAGAGCAGT
GCTTTAAATAAGAAGAATGCGTGAGAACACGAATCTGCGCACAGATTTATGCACGAAAATGGtagaaacgt
attttcgtggctgtaaaatatatttttataatcacgaaaaatatatatttttttataccattTTCGTATGCAA
TTCGTGAGCTCCTCCACATTCTCAAGCATGGtcctttaaataaattcaaggTGAAGATGGAGGGTTTTTGT
ATTTGTGGGTATCTCGGCAGTGCCGAGGTTTCTCcccttccaaaaaaataaaaaataattacatatcagcc
ctccaaaatTTCACTTATACATTTCAGCTCCCGAAAATGCGAAAACACGCGACGTGGCGGCATGTTTCACT
CatacatttcagcccctcgaaaatatattttcgctTTGACATGGTGCGCTCATGCGCCACCACGTCGCGTT
TTCAGGGGTTGAAAtacacaattattatttttagagtggctgatatgcaattatcctaaaaaaaaaaaaaa
gaaataagaaagagagaggcaGCCACCCACCGCGTCGGTTGTCGACGCGCGGCCGAGATGGGGACAAAACG
CACGAACTACACCTCGCCAGGTTGGTGGAAAACGTGACATGAGATTCCATTTCTGGCCCGGCAACTCCTAG
TAACCTGAGCACTATCGGtacaacccaaaaaaagaccGAACCAAAActaatgatcttttttttttttcggctC
GGTCTGGTTTAAGATTTTAGCTCGGAAGATTTACATGTCAGCCCATTattcaaaagttaatttacgaataa
aattttcatttacattCGTtcctaaaagatatatttaaaaataattttataaaaaaatttaaatataaaaa
ttataaattacagtaaaatgtgttttattttttgtatttaattttttttttttaaaaatatttttaaatataag
tttgagacttcaaagtcttaggagattagtagattaatttttaaatacactaatttttttgaatctcaaaat
atatatctagaaataatttttaaaaatttttaaatacaaaaattataaattatattaatttttttttattcat
actcaaagttttttttcaaaataatattatataataattaaaatgtccTCGTTAAACATCgagaaggatttt
tttttttggttgggaCATGTAAAGACTCAAGTTATaccgatatttttttaaacattaggTGGCTTAactat
taacaaataatataatccgATTacgcaaaaataaaaaagacgaATTAACCAAATCTCAAAAAGTGGGACAA
TATAGATGTAGAAAATTTGTGGAAGCACATATGCGCCTGTGCAGGTATCATTTTCAACGATTTAGAGCTTG
CACGAGCTTCCGTCGTCTCAGAATGACGGTGTTCAAATCTAAGTCATTCAGAATTAACGTCGTCTGCCGCC
AGCAGGCGGTGTGGTATGAAATTCTTCCCTTGATTTGTGGGATTGTGGGTTCGAACCCACGTGGGTGAAGG
GGGGTAAATTAAGTTGATTGGACTAACCGCCAACTTCActggacccaaaaaaaaaaaaaaaaaaaaaaaa
aaaaaaaaaaaaaaaaaaaaaaaaaaggaaaagaaagaaaaaattaacgtCACTGATCTAAAAGCATCAAAG
TATATCAGTAGATATCTGTAGATGATGAGTTTGccttttttttggaagaaaaaaaaaaagtaataaaaaagaa
aaagctactATTGGCTCATGTATTTTTgctgaaataattatttggaatGACATATCTGAAAGTTCTGACCT
GTTCCTCTGCTACAATTTTTTGGGGAGTTAcataaccttttttttattatgccaATAAACCTCttattttat

cctttaattttaatattaaatatttatttaaaaaatatataacacaaaagttgtaagttttaataagaata
atttaatataattttaaaatatattttaaatagctattttgaggctccaaaattcagcatattaattga
ttgattttgaaaccaACTAATACtttgaaaactcaaaatatccctttagaatttttttttttttgaaaaa
ataatgtacatgaaagttataaatctcataaaatttataatttttatattgaaaatttttcaaaaaatac
ttatataatagagaaaataGCTCTTGCACGAATATcgaaaaataagttttttttaaagcttaagtagtaac
taaaaaatttaggagtgATTTATATAAGCACGGATAATCTAGtagttgatttgtaattatcgcATTATTGG
GTTTCTAATctttcttgtctttttttcttcaaaattattgttaaaaaattctatctaaagtcgatttattt
attatataacagTCGTAAGAGATAAACCCACTATTAATATCAACCTATATTTTACCgaccgaaaaattta
atatttcgagtctaaatttttttttggattatcgGGAGAAatcctgcccgtaaccggcacTCATTcgaacg
accgtcgtaacaggtaaatccaCGGGTGATATCGATTCctattccaccaaccgaggGGTTCCATCCTTATG
GATAGGGCAGATTCTTTGAGCCAATTTCACTCCCCACAGAATACAGGAGGGTCTTCAGCTCTCACATTGGG
TCTTCAACCCTCACATTTTTACGGGACGTCTGCTGGAGTCGAACCTGGGACGCTTActtcgtgaggcgagc
gtCCAGGCCACCCACGCTGTGGTCGCCCGGGCCACCATTTCGAGTCTAAATTTTACATTCCGTTTCAGTGC
GAAACGTGGCGTCACCAAACCAGCAACCCGTTAATAGCATGGGATTGTCAACACCATGTTACTTTAGCCCT
CCAAAACCATACTAAGTGTTCCATACATGCTAGTGTTACATCAACATCCATATCACAAcaatctattaaaa
cttttcacTTAAAACAGTGTTATACTATTCATGAAAATTATCTAAACTAGTGTacagttttgtcaaaaaaa
aaatgtttggatAATCTCGAATCAAGCCTTACTTTTCAAAAGCCGTTCGAATCAGTCCctaactttctatt
tttttcaaatcaattcCTAATCAGATATTTGAAAGTGTTCCATACGGTAAAAAAGTATTGCAAGCATTACA
TTTAGCAATTAGACTTGAACGTTTTGgttgagctaaaaaaatatttgtaccaGCTAAAAAAGGGTGtgat
ataatttatgaaagttttgctaaattataatttgtgttaCCGCAcgtaaagttttttttttttgagattat
cgtggtggaactcgtaatacgagcgcacgcccccgtaaGAACCTCAAACTTGCGAAAAATTCCTACGTCCA
ATTCCGACCGACATTTTGAAATCCAATTATagaccttccatgaacggaaACTCCTATTATAgggtacataa
aattttcttcaaaaaattctacgaGAATTTCATCCATACGGCCTCGAATTCGGATGGTGAAGCTTTCTGA
GGCACTTTCCAACCAGTCAAACTAGCCGCGCTCAGGTGTTACGAAtggctaaaaaaatgttgacaACTGAG
GGGTCGATTTGAGAGAGGTTTGCAGATCGGGggcttattttgaaaatattccaaatttcttAACCGAAAAg
gaaataatcttttttttccaCTTGTAGAACGATTGTGGCAACGAGATAATGTGGTTCAACGGGCGAGGTGCT
GACGAAACTTGCAAAGAAAGGCAAGGAAGCAGAATCAGTTCTGACTCCCACCACGCGTCGGTGATCGTCGA
AAAGGGCGGCTCCTTTCCAATTTTATAATCTCCCACGCCGCTATCTGATTGGTCGTCAAACGCTAATCTGG
AAATAGTAATCACCATCGCCTGCTTCGTCATCCTAAAAAACCAAACAACGTATGTCAAGTCTACGGAATCA
GATCCCATGCACCCAATTTTCTGAACCCACGTCCGAAATAtataccaatttatatttttttttaaaaaaaaa
ttaaataaaaattataaattttaatattatatataattttttatgttgaaatttttttttttagtggttTAGGA
CTCCaatattaacagattttaaaataattttttaaaatccgttatgAACCCATTAActttagagttctaaaa
tattaatttataagtatttttgaaaaaattttaatataaaaaatgatatataacatcaaaatctataatttt
tatgtttaaaatttttttaaaaaaaatataagttggtaTGCGTTTAGATATGGGAGCAGAGAGTTGGGTGTT
GGGGATTTCAACTCCAAGTCTACGTATACAACCAATCTACGTACAAACTGTTCAATTAACAAATTGTTTtt
gcaattaataaaataaattagtattagttttaatatctaaaatttttatatttttatttgggatGATTTCGGT
TCAGTTCCGATTGCTTTCTATCTATTTCAGATACTTTTGGGTTAGTAAACCAGATCAATCCGAATTGATCC
AAATTAACCCAAATTGTAATTTGTTCAGCTTAGATTGCaattaaattcttatcaattcaacgtcaatttga
tttttgttaacTTTTTACAGTTCAGATCAAGACGgtcagttttaaaattttatattatttattatattgat
cATATAAAGTAGACCAATTTCGGACAGACTTTCACCCCAGGGTAAGAGAAATGGCAAAGTACTGGATAGTT
CTATAGATATTTGTTGGCTTTTATCCCGTTGGGGCGGCTACATGTGGAGCGAGCTCTTGTTGGCGGCgcat
taaatagatatatagaaTATGCATTCGGTTATTCATTTTTAGCAGTCACCCACCAACATAACCCACCATCC
TCTCCAGATCTCCTACGATCGCCGCTGTTCATCTTCTATTAATTTCTCCGAAACTTTCCTCTTATAAGTAA
GCAAAAATCTCCTGCAGATTTGGATTCTTCTGTATACCTTCCAGATTTGGATTTATGTACGCtaattattc
ttataattAATGGCAGTAATTTGCTGATAAAAGTGATGGGGAGAGGAAAGATTGTGATCAAGAGGATCGAT
AACTCGACGAGCAGGCAGGTGACATTCTCGAAACGCAGAAACGGGCTGTTCAAGAAGGCCAAGGAGCTCGC
GATCCTCTGCGACGCCGAGGTCGGCCTCGTCGTCTTCTCCAGCACCGGCAGGCTCTACGAATTCGCCAGCT
CAAGGTCTGTGTTCTCCAATATCACCACTGAGCTTTCGTTATCCGTCGTCTTTCCGATTAAGACTATGCTG
AAATTCTTTCGTGCGATTAATCGAAACTTGGTTTTATACCAGAGCTAATAACAACTGTGTGGTAGAAAATC
ATAGCAGATCTAGACAGTGTCTATCGAGAATTTTTCTTATGATCATCAAGAAATGAGAACCCTGATCAGGA
AGTTCATAAATAATCAGACAGAGATAGACAAATAGAGCCCATACTGCTTGCTTACACCTAGGAATTTGTTA
GTGCTCTAAACTAGATTTGGTGCCAAAACCCAAGCATTAGAGGTTGAAAATGGTTTCAATGCATCAAGCTC
ACTGCCAagtttcctttctttttctttttcttttctttttcttttttttgataaaaacgAAGATGAGCTTCA
TGGAGCTCATCACAGGCTGCGAAGCTTAGATCGCAGAAAATCTTTTACTGCTGTATTAAGCTTTCATCTGA
GACTTTGAGCTCCTGAAGCTTGATCTAAGGAGGCTAAAAAGGTGTTGAGATATCGTGAAAAGGGATCTGAA
TAAAGCTCTCCATGTAGATGGACGGATTCGCTTCAGCTCAAAGCTTGAacacagaaaacaaaaagccTCCA
TTGTTGGAGGTGAATAAAGCTATTTGTTGTAGCTGCTCTGAGAGGAGTTTCGGGAGCTCGATTTTGGTATC
GTAGTAGATAGTAATACAGTTATAGAATAATGGTAACTGGAGCTTAGGAGGTAAGGTCCGCCATTAATGGA
AAGAAGAGAGCTCGTTTAGCTCTAACTAAGGTAAGGGTTGAGACAAAAATCCCATGAGATAACACTCGTGT
AAGAGTTGGTTCGAAATTTAGTTCTATATATACAACTAGGATGGATGAGCACTGTTAGAAATAAGGTTGGc

cttgaaaaattaattaatgtcaaaaatgatttttgcatatttatattttatagaatatatttcatatac
tttaatcttttaataattaaatttctaaaagtttctGAAAAAGACCGCACCGAATTTGAACCAATGgttt
tttcctttctatttGGTGGTATAATTGCAATTCAGCCACACTGCTTTTCAAAAGTCGTAAAATCAaccttt
aacttttaaaatttcttcaaattagCGCATCGACAAAAACTCGAAAGTGTTCTAGAcgcctaaaaaaatat
tgcaagtgattgtaaaatatttatagcaCAATATCTAGTATTTGGACATAATTATTTGGGGCAAGCTAAAA
AGTTATtgtgaaataaattataaaagtgttGTGGCATTACCTTTTGTGTTATTGCAAGCAAAATTAGTATG
GTAAAAGGGTAAAAAAATATGGCGGAGTGTGCAGtgatctttttataatacaaagtaCCGCCCGAACAAA
CTTTATTGTATCCAATGACACTCCCAATACATTTTTAAGCTATTTCTATGAAGACCAAGATTGTCGTCATC
TTGAACAAGCTTTATTGTATCAGTTTCTActatctttaaattaaatttctaaaactatGTCAAAAGCATG
TTTATGGATCCacccaaaattatataagtaaaaactCATTTACATTTGAATTACAAGTTTCCGGGCCCATT
AGGATGTGTCATGTGTAATACGCTGCCAAAGTTGTATCAGAtgctaaaagtaattttacattttgaagcGC
GGCTATTCACACCTATTAACTTActattttgtacatttttttacCATAAAGTTGTGAACTACTAATTGTAAA
ACACTAGCTGCTTTATTGTCGCCTTAGGCTACCAACTTCAGTGTATTTTGCCATCATAAGTGcaacaatta
tataattcaatcGACCACATAAGTATGCCTAGTAGGCAATATTAGAAGTAACTTAATAGGTAGTATTATGT
ATGTAATTCATTgtttctattataatattatatgaatgtGTATTAATACGATATAATGTAGAAAACATAGG
TTCCATCTCTCTTCTATATTATTCAGCATATAATCTTCTATTCTCCCACATATCTCTTTCGCTCTTATCTA
AGCGTCTGGCTAAGCTTAGGTCGATTGAGGTCTGAATGCCGCACGCATGGATCAAAACTCTAATATCATGA
CAAGTCGTATCAGAGCTTAAGATCGTGGCAATCATGTTAGATAACGATAACAACGTTGCTACAGACAATCA
TAGAAGATGGCGAGGAAACTATCATGCGATGTGATGGCAGCCATGGACATCAAGTTTGAGAACATGGTGGA
CAAGGCGGCAAAAGTCGAGTTTGCGGTTGAGGGGACCACGAACTTGGATAATAGCCTAGAGGCTCACATGG
TGGCAACAGTAAGCAAAATCGTAAGCGACCAGCAGTAGCTGTTGGCGTAAGAACCTGCCTTAATCAAGCTA
AAGTAGAGCTTTGTAGTAAGACTCGAACTATGCCGCGTAGCGGTTGCTATTGATGGATGACTGAAAAAACT
GACATACATGTAGAAATTCTCAAGCCAAAGGTATTCAATTACTTATGAAGTACACGTGACATCGTTAActt
cttataatatatgaagaGGTACTTTAGTGTGGTTTTTTTGCTTGATTCGTCCAAACTTCGTGCGATCATTC
TCTTGCTAGGCGATATCCTGACATTGGTGGTGACGAGTATTCGACGATATGGCAATGAGAATCGGGAAGCC
ATCACCCCGATACAGTTCATGAATATATGGAGGTGTTCATAGAACTCATATTTGAGATTCTGGACATATTA
GATTAGGATGCATTGTTCATGTTCGTGAGCAGATGGGTCCAACTAGAGCTCAAGCGACAGGGCGCGCAGAC
TCTCGTGGCTGCCATCCCTGAGATAGAGTCTATGATCGAGTTCAAGTTAGGTCAGAGTGAAGAGGAGCCAA
AGGTAAGAGTACGATGATAATGGTAGGGGAGGAGAACCACCACTGGCGAGGAACCTACAAAGAGGCTAAGG
GCGAGAGAGCTAAAATCTAGAAAGTCAATATCATGCTTCTTATGCGACAGATGCACATAGCATAGGAGTGC
CCAAATCGTAAAAAACTTGCCACATTGCTATTACAGGAGGAATTGAAGCAAGAATGACAATGCAACACAAC
CTATTATAACCGCTTGGTACTTGAAAAACCAAGTCACGAAAGAGTCATGACGGACAAATGTTCACGAAAGT
CTAGCCGGAGAGCCAGCCAATCATGCGCTCATTGACATATGTAAGAAACAACCTTATGGCTGAACGATCTC
AAATTTGGGCTCCTAGTGTGAAGCGGTGAGGAATCCATGAAGATAATGAAATCCGCCCCCAAGCTAATTGT
TGGTATCACACGAAAAGTCAGCACCAAAGTGGGCAATTGAATTAAGTGCATCTGCATTCTGGTAATTCAGA
TTACGACTACGACATGGTGCTTAGATTAGAGTTGCTGGAGGAAGAGTTGGATACAATATTGTTCCAAAAAG
TAACAATATTCTCATCTAAGTTGGGAGACCATGCACTATTCAATCTTACACGTAGGATCAGTTATAGGTAT
AATGTTATCTATAATGTGTAACGCccctttatttgaattaaaaaagacCTTAAAGCCTTTAGAGAGCGTAT
GGATCCAAGGCCCATATACGAGAAGTGGTGATTGTTGGATTTAATCCCACATTGATTAAAAGATGAATAAA
TTAAGGGATTTAAATGCTAGAGGCTTATAGCGACAATTCATAATCCAATCATTTTCAGAGAATAACTCCCG
GTGGGGCAAAACTTACTAGACCAATTATAGATCAGGTAGGTGCGGGGCATTAcatatggtatcagagctga
CCACGCTCACATATGTACGTTTGGATTATAAGAAGGTGTGCCCTTTGGGTTCTAGATCGCCTTGACAAtgt
gtaaatttatatttctaaattgccTATCTCTAATCTCACACCCGATTAGGGATAAGTTTGGTATTATTCTA
ACGTATGATAAGGACGCATTTTAAAAATGGAAGAGTGTGATTCCCCTTGATTTGGATTAAAAAGAccttaa
aagatttttgtaaaCCTTATGTATACTAAACTCATATAGAAAAATAGTGATGGTCGTATTTAGTCCTACATT
CGTAATGGATGAATAAATTTAGGCGTTTATATGCGAAGGGCTTATAGCTGTTAGTAACTCcagatcaaatt
attttcggAGAATTTCGGCCATTGGATTGAGAGTTATTGGGCATCCTATTGATTGAGTAAGTGCGGGACAT
TACACAATACAAATATCAAAGGAGATTTAGAGGATCGATGAACATCACAAAATCGAGGGTGCCAATAAGTA
CTTGACTCTTTGGAAGGGACGTGAGACACAATAGGGCAACACTATTCGTAGGACACAAAGATGTAGCTGCT
ATTTGTGTTAGTGGGGTTAGGGGAAGAAATGTCACAGAGTAAGATTCATGACTAAGTCGTTTGCTTATGCT
CCCACCAACGAGTCGGAATAAGGTAAGAGTCGTAGAGTGGGTTATGTGGATTTGATCATGATAAGGGTAAA
AGTTACAAGAGTTGCATACTTTGATGAACAATATTTAGAAGAACAACATGATGCGTAACGTTATGTATGTA
ATTTGTTGTTTGCTGGTTACAATTCCTTAagaatcaatataaatatgacCAATTTGTTGAGAATTTGGCAT
CTTtcactttatctttttttcaatattagaacaatattatattttacacatCTCTCTCATTCTTGTTTAA
GAATCTGCCcgagttaatataattaaggTCAATATTAAGTTGTCGGATTAAAGTTCTAGCACTATAACAAT
GGTAGGCCTCgctagattataaaaaaaaagaatataagtacgattaataaaaatcaagagaaaaccaaaaa
gataaaaaataaaaataaatcatgatTCAAAAATTGAGAAGAAAGGGGGTCAACACGAAGTAGAGAATCCA
TAAGTCATCCTTTGGTACAAATACAAGAGAAACCTCTCCTAACCGATATCATTACAATAGCCGACCTTCTT
CCCAACAATCTCATTTTCTTGGTTCTAAAATAAGTGGTataattttgaccaaaaaaaaagtggtatAAATA
CAAGAGAAACCTCCTTAATCAGTGTCATTACAAGAGCAGACCTTCCCGATAGTCTCATTTTCTTGGTTGTA

AATTAAGTCTCGTCCATTTAACGTCTTAAAGAAGTTTTCTTCGAGATTTGCTCTTCATAATTTCACGCATT
TGGAGATCCCTTAGGGACtatttacttcgcgcggaactcgGAGAAGTGAGAACTGATGAACTATATAGTTA
GAAAGTGGTAGAGAAGTGGTATTTTCAATTTCTCTATTTACTTAGAATTGTGAAGTGGAAGCTAGAAGATA
ATTTCGTTATTTACTTAATATTGGAATTGAAGgtagaaaatagttttattgtttattttgatgatgaagaa
ttagaataaaaatgatacatattttaatttactactgttaataatatgtaatttttagttatatatatata
tatatatatataatatatatatatatatatatatatattatatatatatattattaatatatatatata
ttatatatatatatatatatatattgtattcttTTACTATTGCATtgagatgaaaataaaaaatcacaata
atatttttaaatagaatataatttatagatgattataaattacaatgtATGTACAAcacatttgtttataa
atatacactaACTACAAACAAACCCTTAAGATCTtatgcaattttaaattactactcaaggataattttaa
aacatttacatGGTTAGACGAAGAAGCTACTTTTAaggacaattttaaaaatactacatTCAGATCTTAAG
GGAGAAGCTTGGGCTCTTGCTTGCTGGCGGCAACAAAGGGTTGGACGAACGAGAAAGGTGAAAGCCAAATC
TGTTATTGATCAGTCACCATGACAACAAATTCGGTCCGCTCGTCAAAAAATCTCAATCGTTGCCGTCGTTA
ACCTAGCAGGCGACGATCATCGATTGGGGTCGAGAGATGTTAGATCAATCGGTGATCGCGAGAGTGGCAGT
GAGAAACATGAACTACAATGATCTCGTAAAAGGGCAGAATTACAAGAGCTCGTGGGCGACGAGCTCACCCA
GTCTGCGAAGTGGAGGAAGTCGTTGATATATCTATACTAGCAGAACTCAAATCCGAGTTTTGCGAGTTTGT
GGTGGCCGGGTAAACAACACAACTCATATCTGACTATCCAGTCCTGCGAGTTCATCCAGATCCGAACGAAG
TAGACAACCCCTTAAAGCCGAAAAagcataatttcatatcaatcAAACCTTCCAAGCACTACACGATTTTG
ATACCTGAATATTTGcttgatattttatgtaatactATAGCTACATGAGTCTTCCTACTAGTTGAAAACTT
GGTCGAGCTCCAATATTACAAAGCGCAAGACTAAGTTCGAGCAAAGATTTAAACACGATTAGTTAAACCAG
TCAAGCCTGAGCGTCGAGTTACTCGACTCACTAGCTTGCGAGTCTAAACAAAGTTTCCTACTTTACTCGAG
CATCATTTGCTACAAACTTTTGTCAGCAAGTTTGATCTCGAGATCGAGCATGAGGTTTTCGAAATCGAGTC
AAGTTCATCCACTCAACTCGAACTGGGTTATTTTCACCCAAGTTAGCCATCGGAAAGCACCAATTTTCTTG
CCATTTGACATTTAGTACAGAGCCTTTCTAGATTGCCACTCATTCAGTTATGGAGAGATCCTACTAAGACC
AATGCTATAACTTTGGCCCGCGGCTCGTGGGGCTAACTTTACGACCCACTAACAGCTTGCAAGGTCGAGCC
GCGTGAGTCAGTCCACAGATTTTGGCCCGCATGCCAGCCTGCACAACCTATTGATCGAGCGTAGGATGCAC
CTCTAGGGCTATCGCAGCGCCTTGTCGGTCTGGAACTCTGAGCTCTAGGTGTTGATGTCACGACCTGCAAG
CCGAGGGCCGCGAAGGAGGgaaaacatttttttcgaaaataaatgACAATCGAAATATAtgctttgcggaag
cttaatcgatcaaggagttcattttaaataccaaatcttgaaatcaaaatctcaaaagtcCAAATcaagga
agagccgcaacctaAACATTCGGTTCaggacgagaacccgcgggagAGGCGGTTCGGTCCCAACCCGAACT
CAGTTTGTGAAGGATACTAATGCATCAACCTCCCCAGATGATACAAGAGGATCCTAGGAAATCTATGGTCC
aaactctttaaaatttatgaaaacatacttaaaaaccatttttttacgaaataccggttcgacctgtattt
ttccaaaagattcGGAAATGCCAAACCTCTAGTAAACATTTATAAGATCTTCCTAAAGcattgaaatcaaa
tttgaaaaccGAATGCGCAAAATAAAACGTCATATTCCagagctattttcggtcgaccgaaatatcggact
ggtcgaccgaaagtcgacTAGAAACTTAAAACAGAACGACtataggtttcggtcgaccgatccaaatttc
ggtcgacagATACCTGCAGGATCTGCAAAAACATAATTGTAGCAAACAAGAAactcaacaccccattttt
caaatatttttccaaaacatccAAACATTATAtgaatacatatacatattccgaatacacatgcaggtatc
gatttgCACCCTTAAAACACATCCAACCTCATTAACcgaacatttttccaaaaacccTTCAAACACTTAAC
CAAAAGAGCTCATATCATCAATTTTGGTTACAACCACATATCCATTCATTCTTAAGCACCAAAGGCATATA
AAAGCCTAAATACACATACGACGAAGCTCGAATCTCAAATTGAAAATgcttttatacatatatatcataat
atgatcatttacaacttaaaatccaaaatacacaacttttagagtttaaactttcttaactaaggtttaca
accctaaaaacttacaaatcatcccaaaacataaaatgtaagttagtacTTCCGAATATTAAGTCTTTATA
TGTATGAGCTATATTACcaaaagttatatacaataaaagcCCCAccaaaatatctttccaaTTCCTTTCCA
AGCTAAACGACGAGAACTTAGCACCAATTTGCCGGTGCCTTTCCACGATCTAatggatctgaaaaagatta
aaagcaATGGAGTGAGTCAAATATGACTCAATGAGCGGAGCGAATAACgatcgcggcacctctcaccacaa
cacaaccaaaCCATAACACATGTCAATAAGCATTAAGTAAAgcatcatcataatcatcaaaGAACATGTAC
TAGAACGAATTCCTATAACCAAACCAATTATGTTTTACTTTCTTAAGTTAGCGTTTGCTCGATTAATCACC
CATTAGTCCTTTCACATTTgtaacgcaacacaccctttaggtaccgcagcaccaaggatctggatcaaaca
cggtgcacgtcacgtcccccagtaccctacagtggggacgatccctttggccttaaggaaGTTATTTGAT
ATACCTGCCTTTTCGAAATTttcgattcaagcctcgcctggtctcgtagctgCCACCCTgacctcgggctg
gccttaactacgcccacctaTCTCTTTCCAAGACAACAAATCTGTGACTCTGGCCCACATCGTCAAATTTC
GATtccggtcgatatccactcgactcacACAGAAGAGCTAATCTAGGCGATGCaatccaacgaaagagcct
cACCACAATATATTCActtatcaaatctcattagcactcacgttgtcgatgtctcttatcacaacaatact
catatTGGCAACACTAATcacataagtactagtgatagacatacaacggttttcacgcattcacaatttt
tttccaatacaTATCCAATATATATCATGAAATAAGAAAGTGTTCCCATAATAAGTCCAAGTCACATAATC
ACCACCAATATGATCATAAGTACCAAAACATACTAGTAAGTTGCAATAACACATCACCATCATActagcat
cacaacatgctcataaatcacatatcacaaaaatcTAGGATTTCTACTCGAAAAGTCTAAATATCACTTAA
TAATACTtgaatcacataaatcatctcaattaCATTCATCACAACCATAATCACACCAAGCAAGAAGTGCA
AACAGTTCATCTCGAGGGCTCGGACGAAATAACACAAAAACGCAAGCTCACCACAAACCGCACGCACCCGA
GAGAGGGGTTAGGTCAAGAGAAGCTAggagagtgagggagagagagatgggatgaTCAAGAATCCAAGAAT
CAAAGAGAGATGAGGGGAGGAACTCGCCTAGGTCAGTTGGGTCCAGGTCAGgggaagcagaagaagaagtg

aAGGGATAAGCAAATAAACTCAAGTCCCGCACTTAAACACGGGCTCCACGCAactatcagtcgaccgaaaa
tacaaatcggtcgaccgaaactgttcTGCACAGTACAAATAGCCCAAAGTCAAGCAACCCAACTCCATCtc
gcacacaaccaagcactctCACGTGCcgacggggtcaccaaccacgtgacacacccgcggtacgcccgaac
acgcatgcacgggTAACGAAAACTCATCCAAAGACATCTCACAAACTCAAATCATAAATCACAAAACAACG
CAAATAAAATAACGAACCTCACAGTtgaagctcttttttttttaattttaatgtatggtGGCCGAGCCAA
GCCAGCCCGAACACATAGACTCCGATTTGGCAGATCACTGAGCCGAGCCAAGCTAGGATTCCTTCCTATGC
AGGCCGAGCCGGCTCGGCCCATAGAGGAAGGgctagctaaaaaaaaaatttctagataaCCACACAGAAAC
ATGGCCCAACAAGCCCTTGTGTCCAGATGGGTGGCCCAACCAATCAGTTGTGTAACATTTGCAAACTTCTT
GCTTAGTCCTTGCTGATGACCATATGCTAGAGTTGGGTTGTTGGGCATGCTTATTTAGCAGGAGTTGAAAT
GTGCGATTTGTACTATCTAACTGTATTACAATGATCTAAAGGTGGTTATATTTGGCTCGGGAccaatatgc
atataagtgaTGCTGGGTAGGTTTAACTTGGAGTTACATTACGAATTTGTAAGGTAATGGATTTACTTaag
atgttttatttaatatgtaaagTTTTCATACAAAGTTTGTTTGATGATATCGGCTTCAAGATGCCGAGTAC
TGATCCTGTAGAGTCGACTACACCATATCacggtgcagtcgaccctATATACGcgtaaaaatacatatttt
tttaaaaacttccaatataaaaattatagattttgatgaaacaaataattttatgttaaaattttttttt
taaaaaaaatacttctaaataggtgttttacaGCTCGGAAGCTTAACGGTTTTACAAGTTCATTTCAGAAC
ACATTAAGTTTCGGAAATctaaacacttatttagaaatatttttttgaaaaactttcaatataaaaatttt
atatttcatcaagatctacaactttcatattaaaattatttatagaaaaaaaaacacaaataagtgtttta
ggagGTCGACAGCACCATAGGAACGGTTTTGCAAGATGCCATACATCACAAATAATTAGAGTGATCTAGAT
AGTTCGAGGCTTTTGGGGGCAGGCAGCCTCTAGGTATACGTGATATTCTAGAACGGCACATCACCGAGGAA
ACATTTATACGTCATAAAGAAAAAGCCACCTCCAGAACAATTGTAAATGTTCATGTAGCTTATGTTCACGT
TACGGTTTTACTTTGGTTCGTTAGTAACTAGTATTTATATCTCCATTGGATTACCTTATCTCACAATTGCT
AGGTAGTTGTATAAACACCCATTTGGGACTCATTCATAGCAAATAAATTAGGTAAAATTCGGCGCTCTCCT
GAAAAAATCCTCgcaagttagcggagaccAGAAATTGCCTTGTGCTTGGCTTCAGTCTGCCTTTTTGGCAG
TCGGAATCCCAACTCAATTGAAATATTAACATAGTGGATGTTGCTAATATGACTACTTAGATGCTCTTATG
TTGTTGATTACAATAACGATAGTTGAGGGGTGAAAAAGTTCCAACTCGAGCAATCTTGAAATTCAGGAAAG
TTGCATCTCCAACtgaattagataaataaaaaaaaaaaaaagttcgaaTTAACTTAATGTATATGTCTTATT
ACAATGGCAAGAGAGTTGAGGGGGGAAAAAAGTTACTTCCAGCTCAAGCAATCTTACAATTCATGAAAATT
GCATTATCCTAATGGATTTGCATAAGATTTGGGATTTACTTAATGATTATGTctcagtaaaatatttatca
agtCATAGAACATTAGAAAACAAGCGGagaagggaaaagaaaaggtatcACTACTGTTCATGAACATACAA
GTCAGATTTCAATGAAAACCCTAGTCAGAGTACAAATCTTTCGAGATTACAACTACAATAGTCAAGTTTCT
GAAGAAATAGATCCTAAATAATGAACGAAGGACACCACGTGACGAGCTTTTAAACAACCATCTAAGATGTC
AAACATGACCTGAAAATGCAAGTAGACTACTTCAAAAGCACAGTTATATTCTATGATAAAGACCTAAATAA
TGATCCAACAGCCGAAAGTTTTGTGAAGAAATATAGCCAAAATAACCAACAAGACCATGACCTTTAAACAA
GAACCACATGCTTatgtaacaaaaatatattcaaagatGTGCTGCGATGATTGATGCACCAGGAGATGACA
CTAAGATGCAACCAGACTACTTGAAAAAACCTTCTACATCATGATACCCTACCTAACAAGAAGAACTACCA
CATGTTAATGTGTGTCTTGAGTTTTGCAAGTTGCCAATCAAATTTAAGCTGAACGAAAGTTTTTGAGGCAC
TGACATACACAAGGAAGAGAAAAGTGGTCAGTAggtcattattttaaaacataaaacaaaatacggatcgg
cacagtcggcacacTCGGTATGTCTCAAGTTTTGAATTTACCGAAGCGCGGCTTGAggcggcaccttaaac
ctcgCATAAACCAAGTGGTATATAATCAGAAATCATGTCAGATTCATCCACGCTAAAGACGTGAACATCga
caatttaaaaagaaaaagcaaatgaataaataacCATGATTTCTATAGCTAAGTTTGCAGTAAATCAAAAC
ATAGGGTATGACTTTTACTATAGTAATCCTAAATTACATTGCATCCAAGCATCATATGTATGACTTGTAGT
ATCTACAGAGAACAAtttctatgataattttgttgtataatAATCCTCCCAGAACATTTAATCACCATTAT
GTCTGCTTTTGTCTTTCAATATATGAAAACTAGTTTCCGACATTGTTATTGTTGGTGTAGGTGTGCTTATT
GTTACTTCtacacattattttttttattcaccaTTCTGATGCTTGAATTAGGAAGCAGTAATTACCTTGAC
TATGATGTTAAGCATGTCGAATTTAACATAATGTTTTCTGCCAGCATGCAATCAGTTATTGACCGATACAG
CAAGAACAAAGGAGAACATTAC**CAAGGGCAAAATCCACACACAGAAGTAGAGGTATGTTTTATGCCACTCG**
**TAATGAAGCATCTGGCCTAGGAGTTCGTTTGCATGCAACGCA**CTGCATGATACCAGATTGCATAAGTCACA
TTAAACATACACAAAACAAAGAGAATAAGTTCACTGACTCCAACAAGTATGGCAATAAGGCTGTCTTGTGA
ATGTAACTTTCGCCTCCCATAAGTTGAGGAAAGCATGAGGAAGTGTTTTCTATATTctgtactattatttt
aaaaaaaaagggatactAAAAGCAGGTGTGTTCTACATACCAAAATTACTTGGAAAATGTGATAATGCTCT
GCTCCTTTAGGCTTGTACTGCTGAAGATACTCATGAAATGTTACAGTAGTAATAATACACATGATGACCAG
CTGGCCAGTTTTTACCAATGCTTAAACGTCAGTTAAGGTCTACCAACTAGCACACAGCCAGAAGCCACAAT
GGATTTAGGGATACCTATAAAACTAAGATAAACTTAAAAGGGaaagatagatagatagaatACTTGATAAT
AACTATGTGAATTCAGGTTTGGTAGGCTTCCTTGATCGTAAATTGACCTACTACTAGATATAAATAGAACT
CAAAACTTAGCAGTACTAAGATCAAGACCTATTACTTCAAAGGAAGAGCCATCTTGAACTTGccctttcaa
aaataattttaaatatcatatatgaaaattatccatttttcTCCTCAACCTTTACTGTTTGTTGAACGGCA
TCGTCTAGTGGCCAACCTAACAGCACTAGGCATTTAGTTGAATGGCATTCACGTTTTGCAtcatactaaat
aaaattttaaaaagtctgAGCagcaaaataaacattttaaagcaCAGGGATTGATCTACAACACACATGGG
TAATAAGAACTCGCACGACAATACAGAAAAAAATGTTAGCCAGTTAATGGAGGAGTGTTCAAGTTTGAGCT
ATATAGCAagtatctttcttctttcttagcTCGGATTTGGCGGTgcagagaagaaagaaggaagaaggaTT

GGATATCTAAGTATAACACAACAGGATTCAAGGAGAAGGAAGGAGGATTGGATCGCTAAGTAAAATACAAG
AGGATTCAATGACAAACATGCAGCTGGATTGGTGAAGAGGAGTTCGACTATAGTGGGGGACCAATAGTCCA
CAAATCCCCTCAAAGGGCTAGTTAGTCAAACCATCTACTAAGTTTCAGTCCACACTTCTTCCGGTTCTGAG
CACAATGACATGGCTATCCACACTATGGACTATGATCGCATAATGTATCATGGCTTGCTGGCAAGTCCATC
AAAAAGAAGATTCGATTTCCCATATGCCAGAtgcatgaaagaaaaaatcacataagaAAGCCAAGTGTCGC
TAACAcccttaaatttaaaaaaaaaaaaaaaaaaagtaaactgagagaggagagaggatAAGGTGCCAAAG
GCGATGACAACAAGATCGGAGGCGCAATAGAGAGATCCCATGGTACCCAACACTTCCATTATCTTTCTATT
CCCCTTTGTCTTTGAAAACCTCAATCCATATGGCTAGCACGACCAATAACCTTCTGCATTGTCACATATCT
AGCGGCACCACAGGTGTAGCACGGTCAAAGCTTGGTTGGACCACATCAAGTTGATTAGTTATAAAGAGAT
TCAAAGTCTATCACAGGGTTCAACCAACAACAGGAAACAATTGTCATACGCTCTTATCACGCGCACTGTGC
TTTCGTTGCATAGTTGCTAGTAAAACATGCCCATACTTGCTAAGCTTGGTATTTGTGAGGTGATCTAACGT
CGACTCATGGAGTTCCAAGGCAAAAAGCCGAAACGATAGCATGATGATTTGAGACTTAGATGTGAGGTAGA
GATGAACCTGAGTTTGTTTCCAAGATAGATACGCGAGAGGGAGGTTGCAGTTTGATATGAGAAAGAAGATG
ATGgaccgagagagagagagagagagagagagaggagagagagagagagagaggtaggAAGGCTAATCAAA
ATGTATGTCCGAGAATTGTATTGATTTTGCCTTATCAAATATAGTTATGATCAAACCATCATATATGTTTA
CAATACTGAGGAAAGCATTACACCTAAGTAAGGAGatcaagaaaaagataagattGCAAAATGAAGCAAAT
TGGCTTATATTCCTAAGCTTAGCTTTCTATGTGTTCTCCTAAATAGATCCGCAGAAGGTCAAAACCGACAT
TGTCAGTACTAATGCTACAGAATTGATCTACATCTATTCACATCTGCTTTAAGGGTGGAGATGTGTCAATC
ACAAAAACATACGTATAGGACAAGGACTTATCTATGTTTGGTTCCCATACACTGTTAACCTTGCGGATAAG
AATGCTTCCGAAGAgcaatagaaatttataaacaagAGTGGAGAGGACAGAGCATATTTGGGTCAGTTATA
GtttgatagaaatttagtAGAAAAATGTATTTGATACGAGTCTGATTTGTCAAAGTCCACATCATAAGTTT
ATAGATATCCATGGTTGTAATCGTTGACTTATGCATGAATAGATATGTATCTTAGTATGAATAGATACTCT
ATTCATGACTTATGCATGAATAGAGTATATTGTTTTTAGAGGgtatagattaatttaaaatcgCCAAAGAA
CCGCCGGTGTTTGTGCTGATGCTATCAAGGGCCAACACAATTTGGTTCATACATGAAACTTGATCAATGAT
TGGTGATGTCGCTATATTACAATTTGATTCTTTAAGAATCTTCCACGCACATATCCATCCTAGTTCAAAAA
GCAAGCAAACAAGACAATTTGTAGTATGATGATTTGGCTTATAAATCCTATTTATGGACAAATATTATACT
GTCCATGATCTTATCCGAATCTAATAGTAGAATTAATATCTACTAGATTATGTACCTACAAATTGTTTCCT
CATGCATGCTTGCATTTGATGGTTGTGACAAAATATAGCATGATATAACCCACATGAGTATGTTTTGATAG
CACCTAAACTTGCACTATTGAGACTTTAGCAGAACAACAAACAACCAAAATATGACATATAAACATGTTCA
ATTGTTGGTTAACTAATTCTACATATATTGTCTAGCATCATTTTAC**CTCATATGATCTGAACCTAGCATGT**
**TGTATGTACCAGCTTTGGCAGCGGGAAGCAGCAAGCCTGAGGCAGCAACTGCATAACTTACAAGAAGAGTT**
**TCGGTATCGTTTTTAAC**CTGTCATTCATAGTTTCAAAGTCATAACATAATCACTATTGCAAACAAACTGAT
AAGAGAACTGCAATTGCTAAtttgtgattaaaaaaaaacatgtttggATCAATTGGCATACGGACAAACAC
TTTGACAAATGTAACAATTgttctttgaaaaaatggtTCCTTCGTAAACGAAAGagcatcttatatttttc
atgacaTACTAGCCACATGACTCATCCAATCACTCAGCTAGCTTCATCAATTGCATACAGAAATAGTTTTA
CAATCTTTTCCTAAATATTAACTTACCCACTAGTAATTTGCATTAACTAATATTGAATGCTCCTTTATGGA
CACATGCAAACTTAGGCAAAACACTGAAAGATCCCACTTAAGCTTTATGAAGCAGTTAACATGGACCATTT
GGCAAATAACCCAATGTGGTCCCTAGTGCATTACAAAACATAGTAGACTAATAGCTATTAATtgcaaatat
agaaaaatatagttcATATTTGGATATACATTTTTTGGTCTAATTGTCATGGGATATAGTGCAGATGAGTT
TCTAAGCCCATAGCTTGATCAATTGCCCCGGAAAAAGCATACggaaaattatacatcaagAACAGACTTCA
AAAATAGGAATCCATTCATCAAGTTAACTATGTATAGTTGTTActctctaatttaaaaagaagataagagT
GAAGGCTagcaaaaccaacaaaaaggCGCCATCATGAAAGAATATTGGACAAGTAAAGATGctgaaatatt
tgatttctCGTGGGCACTGTTTCTTTGAACAAGCTGTAGCAATTGACAGCATGGAGCATATAAGCATCTGT
CAAATTGTTCCGGTTAATGTCTTTTGCCATGATTTAATCATGACAGTAACACACAAGATTTAGTagtaata
tctataattacaGGAAAATCACAGGAGAGAATTTATCGGGACTAGCTATTCCAGATCTGCAGAATCTGGAA
CACCAACTAGAAATGAGTCTTAGATCAATCCGTATGAAGAAGGTATTCAAACTATTGATCAAAGCCATTGC
TGAATgcatcaaattttatcatgaaatCATCGTAtttcatttattcattttttgcAGGAGCAAATCTTAATT
GACGAAATTGAGGAATTAAAAAGCAAGGTATTCGTAACAAATTTGTCTTTCAGATGGTGCGCATCATACTT
GacttttatgttttgtgtGTAGGGAAAGGCAATTCTTGAGGAGAATTTGAGCCTCTACAACAAGGTTAGCC
ATTTGACAATGTTGATATAAAATCTGATACTGCCATCAGATCTTCaagaaatttccaaataaatttacaag
caTCAAGCATCATCTCAGAAATCATCTAATCGAAGTTAACTTTCAGGTTTATGCCACAAGTGGATCGACCG
TAGAAACTACCAGTGCTCCACCTCTACCGTTAGGATTGAGAATCACTGAAAACTTAAATGTGCAGCCGCAG
CTGGAGTTGGGTCATTTGCAGCAACTAGAACAGGCTCAAGAAGAACCAAAGCTACGGTAACcaactttcct
tttttttttttgggaaaactaTTACATATGGATCTTATCTATAAgactataataatttaaatacattgATG
TTGCAGCAGACTTCAACTTCATTAAGAAGATTGCTGAGTTGGAAATGCCACTACTTAGTCTCATCCATTTG
TGTAATATAGAGTGGTGGAACAAACAACattcatgttaaaaatgcAAACATATGACATGTAAGATTTCAAG
GCATGCAAGATTTCAAGGCATGCATAtgggaaaaaagaagtttgagtAAAAAGATCCCCAAGAAGCAGAGA
ATGCAACGATGGCAGCTGTTATTTACCAAGAATCTCATCTAGTAACTTCATGTTTCTGGAATACTACAGCA
TGTAATAGATTCCCATGAAAGGAGCTGTTTTggttttattcaaattatcaaaatatcatgCAAAGAAAGCC
ATACCAAGCCCAACACATATGTGTTGGAAACATTACATTCTTCAAATCCATTATTCTGAACTCACAAAATT

TGTTTGCTAGGGTTCAAGAGAAAATGAGTTCCTAGAGAAGACCAAGACGGCAAAATCAAatgcaaaaaaca
aagaagatatGTCAATTCCAAAAGCATTGTCTACGTCAGGTCTATAAGTCGCAATGTTCTAACAAGTTTAA
AGATTGTAATTCCTGCCCTATCATATCTAAGGCAAGGCATATTATTTCATGTGTCCTCTATTGGagattcc
tatcaattaaaaaaaaaaaaaaaagttagccTCTACATTATTCCACGCAACAtcaattcaaaatagtttGA
CCTCTTAATTTATGTTGCTACTGTCATCTATTTTCAAAACCTTTTTCAGCACTTTATTAAATAGGCCTCTT
TTTTGTCAATAATGTCCTCTTTCTTGTCAATAATGCACATTTGGCAAACTAGAAAATGAACGAGATTTAGA
ACATAGGCAAGCCTTGATAATTTCATAACACATTAACTTATCTGTGTAAGTCAATTAAATACTTCAATATC
AGATTAGTACAAAAGGTGGCATCTTTTTGATAGTTTCATAACACAGTAACTTATCTGTGTAAGTCAATCAA
ATACTTCAATATCAGATTGGTAACAAAGGTGGcatctttttaaacttaaatcAGCTCAAGCATTGAGCCT
GCTTCAGGCAACTACACTAGAGGTGGGTTGGATGTAAATTCAAACCTATGTATTGATGCTAACGTTGATTA
TAATGATAACGAAAAATTAGCTATGTGATCTAAGATATTGAGTCAAGTCTGAAAGAACTAGTTTGATTTGA
ACTCTCGGGTTTAGGGTGCCTCGCAACGGAAAAGGCAAGGAAGGTGCAATTTTTTGCTATAGAGAAATTAT
AACAATCTTAGGGAGTCGATAGAGGCAGGGGGATTATTATTCCAATGCACCCCTTTTACTTGCGTATGTAA
TTATCCCCTCATCTTTATACACACAAACACATAGTCAGCTATCAACATTTTCTCCAGAAGACTAGTTGTAg
tcgaaatttaaaataccctttttaaaaactgtatTGACTTGAAAATCTAATGGGAAGTTGATTAAATTAAA
GAGCTCAAAATCCTAAATTAAAAAGCATGAAAAGTACTCGAATATTTACATTGTTGTGTCTTCTCTATTCT
GCTACTCCTTGATCTTTTCATACTTgcaatagaaaattaattggGTAGAACACCAGTTTACTTTACTTGTT
TAGCTTGCAATCACAACTAATCTCATGAACTTAGAATGGAGACTTTAGAGTTGCATGACGTGTCAACAATC
TAACTTGAAAGAGTTATCTAATTGAGTAGGCCCATTAATGGGTGTCCATTAGCTTTATTGCAAACTTTAAT
CCCAATCATAATTGACTAATTACTTAAGGTATGACGACTTTCAAGCACAAACATTGAGTATCTCAGATCAT
CTaaccaaaatattcatttcaacCATCTCATACATCATTCATCTTCCAGAAAGAATGTAACACTGTGTGAA
TTAAGATCCCAAAATAAGTGACATCACCTGGAATTCTATGCTAAGTCTAACATCTTACAAAAATTAAGTGC
AATGATGCTACTGGCACCTTCACAATATTTAGAACACAGTTAGCTAGCTACAATGTCTTTAGTATTTCCCT
CTGGCAAGTCTTAGTACTAAGATATTTGGAGCAGTTCATGATTCACATTGCAATTAGAAAAAAGGGGGGAG
AAATTTCAACAAGCTAAACAAAAATCCATTAAAGTCATTTTAGACTTTTGAAAACATATGCTAAAGAGAAC
ATAATAGAACCAAATAATTAGCATGTAAGACACTATTTCCAAGCATTAGAACCAAACAAAGTTTTAATAAC
TAACAAGTATTCCCCAAATAAATGCTACGAATCTCCAAGCTGAACTGAGATGATAGcatacacaaaaaatt
taaaaaaaaaaaagaggatacATGTGAACAGCATGTTAgattggggaaaaaaaaacaactagaAGCACTAG
CCTGACTAGGTAGAGTAAATGCAACCTAGAGAAGCTTACTAGCAAAATACAATGACAAAAGCAACTATAAC
CATTGCCTGACTTGGTAGATTGAACTAACAGTTGCACCTTTTGCACAAACATTAACAGTTCAGCTGAATGT
ATAC**CTTTTAGATTCTTGCAAGTTATGCAATTGCTGCCTCAAACTCGCTGCCTCCCCCAATCAAAA**CTGCA
CAATATAGTCAAAAATGATTAGGCAGTTCATGACGCAATAGCAAGTAAGTCGATACTACCATAATCAATTA
ACTAATTATCAGAGTAAGAACCCAACAAGACATGCAGCTCCAAATTCAGATCTTAAATCTTGCCAATCATG
CACCATATAAAGGAAATCAAAGGATCAAACATCATTAAATCTGAAACTATCACCCTGCATGGAAGATTTAA
TCTAGACCTTCAATAATCCAGTAAAAAGGAAGTTAATCAGTGTTCAGAGAATATTACTGTCCTTTTCTCTA
TTGGAGGGACAATTAAACAGAGGTAGATCTCTAGCTAAGTAAGTATTAGTTTAGCAGAGATAATATCAGCA
CAATCACAAGACAAACACAAGATTGAAACACTtgtcaaaatagaaatttattatttgtctttTCAAATGCA
TAATTGGTAGAGGTTACAAATACATCAGAATCTACTTGCTATCACAATTTCAAGTTGCAGAAAAACATTAG
CATTCATAAAACAAGCATTGAAGGCAATACTACCCAAATTAGTTACACCATGTAATAGTGAGATATCATGA
CCATAAGCTTCATGTGgttcaaaaacatttttctaggtttttgcaaaatcaacaCAAACAGGGTAACTCA
GTAAATCCTCACTTGGCTTATAATTAGTTACCTGTCTCAAATTAAATCTTTTACATAAATACATGTTACTG
CTGGAGCATGAATACATATGGGCAAAGGTAAAGTAAGTGACAGGGACTGATTAGAAGATGGCTTAAACTAA
AACCGAGCATTAGGGAATAGAAATGAGTAGGAATAGGGTGTAGAAATTTTGAAGACGTAGATAAATATCCC
AACAATTCCTGGGACAAGTGTAGGAGTAGGGTGTATCCAAACCCTTCTTCCAACCAAGTGCCTTAAGTTTG
GGCTACGAAAGCCATTGGATTATGTACCTGGAATTGTGGAGACATAGATAAGAATCTTATGAAGAACAGCA
GCTAAACGGAGCAAACACACATAAGAATCTTATGAAGGGAAGCCAAACTTATAGGGAAAGAATAAACCATA
AATTTCTCATGAAATACGAGGTTCGCTTAAGGTACATTATATGTTGAAGTCTTTAACTAAGAAGGACTTCC
GTGCCAGGCGGCACGCACGTGCTGGTGCCGGCCAGCTTCGGCACCGGCACGGGGCCGGCCAGCTTTGACAC
CGGGCCGGCACGGTGCCAAACCGGCCATGCCGGTGGGTGCCGACCATGTCGAGcagcaagctttttttttaa
gacaacccGTGACGGTCAAATCACAggcggattttttttttattacaacaGACCGCCCATGCAAGGCGATCA
AAATTGACCACCCTTCACGGGAGTCATCATCCCATGTAGTTGGGATGCATGGGTGTACTACGTCCACATGA
CATATAGGGTAGATTTAAATAGTTTGGGTTTTACAGATCTCTTATATCAGCCACATCAACACTCTATGTGA
CAATGAATTTGTACTATCTGTTTTGGagtcattaaattttatcttagtatttttattttattaatattttt
tattaccatttaaataatttaatatttaaatataataaattgtatggATCGCtacagtcggcacgtgccg
agtttcggattCACCAatactcggcacggcacagaaAACTTTGTTAAATAAACACCTAACAAGCATCACTA
CATTCAGTGAGTTTGTACAGTAGCAATCAAATGTGCAGAAGTGGTTCAGAAGAATTTCAACACAGCCACCT
TGACATCTAGTGGTCCTCTAATATTTCACACCCCTCGTTTAAAGCAAGAAAACCAACTGTGTTTTACAGCA
ATGATACTTAGTGATCTTTATGTATTACAGGCAGAATGCCTGAAAATACTGCTTACTTCGCCCAAATCAAA
ATTAGGTATAGCATACCAAATGATAACCTTCTACTAAAGGACTTGGCTTCaatcacaaaaattttattcta
ttctgCACAAGATGCAGTTAAAGATCACtgcataaatattacaaattgcGATACAAGAACAAAGAACTGAA

TACTCATGGTGATGGTGTATAAGCAAAGCCTTCCAAAACAACGACAGAATAGCATGTATAATTACGTAAAT
TGCTTAAGCAAAAAAAGGACAGGGATATGAGCCATTATCTTGTATGAATTTCTTGTCCATTCacttgttta
atttatgttcgttagatttccaaaaaaggTCAGCTTGATGAAATCCTTAGAACACGTCATGGATATCTT
CAAATGATAGAATTACGTCGAATAATTAATCTTTGGCAAGTTTTGAGTAGTTTGACAGGCTTCTACtgttt
ttatcataatttctaaaccGATAACACATATCCACACAACGGAACGCTTGTTGGGAAGCTGTTGAGAAAA
ATCACATTAGTCCAAATTGTCATGGTGAAATTCTATAGTTGGTCAAGggagaaaagaagatattgttacag
atttttttttcctttttttccttaaagCTGTTTTAGTGGAAACTTGgtacattttaaatttcccCCCCAA
CATTGCTCCTATCTTGTGAAATAAGTGGACAAGTCACCCTACACTCACAATCAGTTGTTTTATTACTAGTA
AGTTATTTATTCTTCCCTTCTGTTGaccctaatgtgattttgatgataacaaaacactTAGCAAAAGTGTT
ACTaacaaatttgttcaaattgcTAAGTATAGGTGTTGTCTTGGTGAGCTAAATATGTCAAACAAGCAACA
AAACACTTGGAACAAACTTGGATGAACTTAGTAAGCCTAAAGAACATGCGCGAAGCAAAGAAATCACTTTG
GAGACGCTTAAAGCTTGAAGTCAGATTTTTTCGAAAACGACGTTTCTCCAACATTCACTGTTCAAGagcta
tttataattttatgtgtaaTATGTATTCTCTAGTACTAGAAATAAAGAGTTAAAGTTAAACTCAAAGTTTA
GGATGATTTAGAgaccaaaatcaaaagttatatattctCTAAATCATTGCTGAAAAGAGAggatttccggt
cgaccagtCATTCTGACGGTGATTCGAAAACGCATAAACTGATCGGTCAACCGAATATTTgaaccggtcga
ccggaaattcagaaaaatagcCGTTAATAGCCGttggaaaagaaagttttaaataggGGAAGAATTGTTTT
GGCCTTTCATGATTTTTAATAGTGCAAAAACATTTCATATCCCTTAGAGAAGTTCCAAAGCCCCCTTGGAA
CATAGAGATTGAACTTAAACTTGCAAACCATTAAATTTAGCCCTAGACCTCTTGCCTACCATTAAAATATC
TTGAGAGAGGCTGTAAATtggattttggtttgtttttctCCTTTGTGAGAGATTGTATAGGTTGCTTGGTA
AGCCACAAAACCAAAAACTACAGGTTGCTAGGTTAGCCACTGAAACCTAGGGAGATTATGGGTTATCGCTA
AGCTTGTAAAAGCGAGGTTGTAAACAACTTGATCAACGCCCACAAGTTGATTGTCTAGTGAAGTCATCGGT
GGAGGCAATGGAAGTGGACATAGGCGGGGTTTCGctgaaccactataaatcaCGGTGTCGTATCTCCTTCT
CCCttgctttttttatattcttactAAACGCATGTGTAGGAATTAATTGATATATGTATACACGCATTTGTG
CTTGAATTGTTATAGAGCTAATATTGCTTTAGAGTGCTAAACTTGTTTAATTAGCtcttaattttacaatt
acccaattcacccccccccctccctcttgggacccaacaccttccaatttttgttttgtgcttCTTCTCCC
AACGAACATGATATGTGTGAAATGGTTGTACATTTGCAGCTACTTTCTTACATAGtttgacgtttcggtcg
gcacaaggcgtgccgaccgaaacgtctctgtgccgacagatatcggcacggcacggtcggcacgaccgaat
ttttttttttagtaaggaaaaattgttcatctttcaCGAATGAAGATAACGGTGCTCTTTTTTTGCGTTCC
ACGCGACACGACTGGTATTTCCAGCTGCATGGATGCTTTTTTTCGTATTATGCCGATGGGCGCGAATCAAA
GGTCAcacgctcttaattttttttttaagtttttcaattttaaaattgaaaatcttttaaaatattaaaata
tataataaaatataaaaaattaatattttatcatcggcacagtcgaaacgacacgatcggcacgtctcgag
tttggagttcactgaaactcgaaacggcacggcacggtaaactttGCTTTCTTATATAGTATATGAGATCT
AGATTCAGTTGTGGTTCTTTGATATTCTTTTAGAAGGTGGAGAACACACTATATTTGGTTCTACCCTATTA
GTAAAAGgcacttttttaagaaaatgaatacaaaatagaacaaataaattaactttcCGTTGGCAATATAGA
AGAGAcatttaatttacaaaaatatattaactaaacAAGCCAAAAAGAGTTTTTGTCAGTAACAACTTATA
GGCTCTATTCAATCAATTAAAGTCAACATGTACTTTCCCATTTTCATTCTCCTAGTGGTATCATGTTCATT
ATATCCTTATGtatagagatgtcaaaaaaagcTCGCGGCTTGCTACTCCGCCCGTTGCGCGCATAAATTGT
CAAGCTAAGCCCGGCCCACTCGGAAGTGGTCTGTATTGGCAATCCGAGCTAGGCCGCGACTTCTAGCCCGC
TCAAACAGAGCAACTAGGCCAGCCCACACTACTTATCCATTTGTCCATTCCTCTATACGCGATCCAAGATC
TTAAATTATTCGCTTATTTCTTTATCTCGGAGGCTCAACACTTCATTGTTATCATTTTCTTCCAAAGCTAA
ACATACATCACATATACTTGACCTTAGATCTGCTCACATTCATGATATCCTCCCCTATTTGTCCATTCATC
AATGATCAAACGAGTTAAATACATAGAAAATTTCTTGTTCTAGTGCCAAGTAATGATGGGGGCTTGCCTAC
ATTGTGGCTTCCAGGACTTGACTATTTAGTAATGTGCGTGCATAGACAACTAGAAAAAGTAGATAGAGAAA
GAACAATAGATCAGTTCTTGAGTCAGcttgtatttcttttacaCAATCAAACCTAAATCCTACTTCATAGT
CAATTGCTAGTGAAACAACCCAAATAATGTGACTATCCGAGGCAAAAGTTCAGATGTTGTTCCTAGCTACA
AAAATCGCCTCATGTAAATGACAACGTACTACAATCTTCTAAATTTATGCACAGCGGAAGCTAAGTGGCAC
CatccacaaaaagaaaaaatgaaacaaacaTAAAGTTCAGTTGTGGCACCTTCCACAGAAGTGAAATGAAA
CAAACATGAAGTTTAGTTGTGGCAAAACATAAGCAATAGTTACTCGCCACCTGGCACCTATTACGTAAGAT
GGTTAGCATTTCAACTTGCtggcataaaataaaaaccccCTTTAGTTCTTTACttagaattaaaaagaacT
TCCTCATGTACCTGCTTTAAGATGCTTAGCATCCGTCGGCTTCCTGgcattaaatcaaaatataccgTCAT
GTTTGCTACTTGGTATAAAAGAGCctcttctttaaaatttcttatattactAGTAATAACGAAAACTTCTT
GGTTGATAagtttgcaaaaattttctttcgaCCATATTATGCAGAATCAAACGACCATACAATTTGACGTA
CAGAAGATACAGGTCGATTGTTAACTAGTGGAGCTCCAAACCGTTAGATAAAACACATTGCCTTATTTCTC
CTCAAGTCATGCATTATCAAAATTCACTTACTTACATGCGTTCATTAAAGAGAACAACCTCAGATTTCCTT
GCTTCTACAATTTGATATTGTTGGATTATGTGTTTTCACGATGACCTAAGTTAGAGGAATTTCTCAACCAC
ATCTgctataattgtaaaatttgatacctttgaattatgtgttttgatgataacaaaatttgaaggGTGAT
AGCATAATTGACACAAGATATCGATATGACTAATCAAAGATGATATTCGGCATACAATTGGTTTACTTTCA
GGTAAATTTTGTTGTGGATCAACATAGAGTATCTAATAGCTGCCATAAAACAAGTAAGATAGAAATTCTAG
CTTGCTAAGAAAGCGactaaatctatatattattttacaatctACTCAACAAACAATTTGAAGGAAATCAT
AGCTTTGGAgagaactaaaattttaagaggCGTATCAAATCATCTTGTACAACATTTTTACACAGAAAAAA

TTCGCTCACAAGTCAAGCTTTACAAAGCTTCATCTACATGAACCGAGGATTGAAAGCAAGAGATCTCTACG
GTCTAATACTTCACCTATCGtaagatgtatttttttatataggtaGGCTGAAAAACTAAATAGATCAAGAT
CTCTCACAATAGCATTGAAAATACATCTAAACGCCAGTTGGCACAGCCAATACCGAgaggcacgtgcctgt
tccagcagttgttggcacggcacgtgccgaaaagtcttaatatttgaatgctttaattaattttacatttt
atttttatttttcaaaaaaatgtatatcggcACCACATCATCAAAACAATCGGCACATGatcagtgttgt
taaaggcgcgccaaggcgcgccttgccatgcgcctaggctcgccacacgaatctgcgcccgccaagaacaa
ggcaaggctcccgttcactaggcgcagcctTATCGCGCCTTGGCGCGCCtaggtgcgccttgcgcctaggc
gcgccttgctgcgcctagtgaacaagccccatttataaactatgtgagTCCAATTAAACCATCAGAACTTA
CTATTAAGTCCATATGAGAGCTCCTTACTCACTACAGTGggctaaatagaaaaaaaaaaaattcctaatttt
tctcaCAAGCAACAAGCCTGGTGGATTGGCAGAGGACGGCGAAGGCTGGAGACGagctaatttttcataat
gaacatgttcatcatgtgaacatctccaaacaagctagttattcatcatttttcacactttgagaCTTTGA
GTCAACATGTTCATGTAATGTGACTATGTGCTATTCCAATATGGCTTGTCCAGAATCCATCAATAAGCTAA
CTTCCGCAAACTTTAGAGGTAATTAGCAacttatacatatgttaagaactaagagtcttaagaatgatgta
agaaatttacctgtgtattttgatgtgtgtaAGATCTTTGCTAATAGGCTTGTTCTGATCTTTTCATgaca
atttgatttaaatttatatttttttatggagatatttatatctaaggaaGCAGCGGGCCTTGCGTTAATAA
GGCCCGCGCCGCGCCGCGCtttgcgccttgcgcctaggctctgGGACCACCCTTGCGccttggtgcgcctt
gcgcctttgacaacattgcaCATGATAGGATCTGAGTTTGCCGAAATTCTCGGCTCGAACAAAAAACCTTC
ATtctctttaatttgtttaaggGTGGATCTTGCGGATTCAAGGTCGTGTACAAGTTGTGCATTCAAACTTG
GTGAAGGTTGAACTTGTTGACTCAATTTCGGCTTAGTTTTTTGTGCATtgctatttagaaaaaaaaaaaaat
ctagctCGAGCGGTTATTCGAGAGTTCGGATGGTGATCATATGTCGAGTCTATTAGAATCCCACTTGATCG
AGATTTGGTTTGTGGATGTAGGTGGGCTTCAAATCATGTTATATCCGGCATTTATAACCATTTTTTTGTAA
GCTTGGTTTGTTCTTGTTAATATTGTAGGTATGTTACAATTCACCATCCATCAGCCTGCTAAACCTATCCA
ATTAtttcttatgatttttttggtgaCCAACTAAATTGATTAAGTACATGTTTTGTGACTATCGCGTGCGA
GCTTTGCCTAAGGAATTCGAAAAGCCCCTTTCATAGGTGTTTTCCCTTAGGAAGTCTAAAAGATGCATAAA
CAACATCTCGCTAAAGCAATTCAAGTGTCACGATTTTGCTAATAAACTACATAGATCAAGATCTCTAACAA
TAGCATTAAAAGTACAAACTTCTCTTCATCGTTTTAAGCGCCGATTGGCACAGTCCGTACCTGAGGGCCCG
TGTCCATTTCAGTAATTGTCGGCACCGACGTGCCGAAAaggattaatatttaaatgttttaattaattta
aattttattttatttccaaaaaaaatgtatatcggTAAGGGAGGTACGAAATCATCGGCACGATCGACAC
GTGCCAATATCAGCTTCGTTTGTCGACACTCCTGGCTCGATAGAAAACTTTCattctctttaattttatta
taggtGGATCTTGCAGATTCAAGGTCGGGTAGAAGTTGTACATTCAAACTTGGTGATGGTTCAATTTCAGC
TTTGTTTTTAGAGCATTGcgatttaggagaaattttttaaaataaataataataataacaatctaGCTCG
AGACCTTGCTCGAGAGTTGGGATGGTGATCATATGTCAAGTCCATTAGGACCCCACTTGATCGAGATTTGG
GTTGTGGACGTAAGTTGGCACCAAATCATGTTATATCTGGTGTTTATAACCATTTTTTTCGTAAGCTTGGT
TTGTTCTTGCTAATATTGTAGGTCTGTACAATTCAACCATGAAGGTTGAACTTGCGGACTCAATTTCAGCT
TCGTTTTTTGAGCATTGCGATttaagagaaaatttaaagaaacaatCTAGCTCGAGCGATTGCTCGAGTTG
GGATGGTGATCATACGTCGAGTCCATTAGAACCCAACTTGATCGAGATTTGGGTTGTGGACGTAAGTTGGC
ACCAAATCATGTTATATCTGGtgtttataacattttttttttggtaagctTGGTTTGTTCTTGCTAGTATTG
TAGGTCTGTGTTATGTACAGGTTGGAAGCGGATCGAGCTGCAAGAGGACGTGGATCGTGAGCAAAGTTTGg
aatgcatcaaaatattattaggaatatatttagtaggaaataattgtaatagaaagaatttatatttggag
atttttattagtataataggtagaaaaaaaatatagaggaatagtttctataattatatattttgtatcc
tTTATTGTTAAGtatggaagaaatatagtaattttaattacaatagtttgtagaAGTAGAGCCGCCATCGG
ACCGCCGGTGTTAtcctcgccgcgtccgggccgccgacgtgagccgatccaagtcggttcataacagtCTG
TTACAATTCAACCATCCAATCAGCCTACTAAactatctaaatatttttttgtaatttttttggtgactAAAT
GAGTTGATTAAGTACATGTTTTGTGACTATCACATGTGATTTGCCTAAGGAATTCGAAAAGTCCCTTTCAC
TGGTGTTTTCCCTTATGAATTCTAAAAGATGCATAAACAACATCTCGCAAAGCAATTCAAGTGTCACGATA
TTTCTATAAAACctcaacttttgtatttctttccCAAGAGACCAAGGTCAAAACATCTACTCACACCAGAC
GAGGAGATGTGCACTAAGCCACCAACTTTCACATCAAGCTAAAGTGTCGTACCTATCAAAAGGCGGTTCGC
GCAGATATGTCAAGGAGAAGAATCGCTAAATGTAGTTATCCCACTCAAAGTGCTTGTTCAAATTGTTGGCT
ACTCGGCTATCAATGTGCGAGCTTTGCCTAAAGAAGCCAAAGGATGTATACACCCAATTCGCCTCTTCCAT
GAGTGTTTTTGCATATTTAGTCTAATAAATGATTAAGCAATATCACCTTTAAGCATTTAGAGTGCCACGCT
ATCGTTATAAAATCTCAACTGCAGCTCCGTTCGTTATATCTTTTCCCGAGAGACAGGTCAAAGATATTGAC
TCGCATACCACAAGGAATGTATAATAAGTACCCAAAAGTGTTGTACTACCTATCAAAGGGTGtctttttttc
gaaaatctaTTATAAGAATGTATGGACCTATCAAGGTTGGTTTGCAGAGATATGTTAAGGAGAAGAATCGT
CAAATTTTGTTAACAAGTTCAAAGCATATGTTCTAGTTCTTGACTGTTAATGTgcacttttccttttttttt
ttgggtcaagaTGTGCAACTTTTGCAAAAGAAGTCTGTATACACCCAAGTCGCCCCTTTCATGGGTTTTTT
CGCTTATGAAGTTTAAAAAGATGCTTAAGCATCATCTTGCACTTCGTTCTCGAGAGACCGAgaccaaaaat
atcaactcGCATCCCACAAGGAATTGTGTACTACAAAGTCCCGAACTTCTAGATGCGCTAAAGTCATACCC
ATCAAAGGATGGTTCGCTCAGATCTGTTAAGGGTAAAGAAGAATCGCCAAATGTAGTTAACCCGTCCAACC
AGCTAATTCTTAACTAAAACCCGCATGGAACACCTTTTTTCGCtcaataattttagttattacATCCAAGA
CCAAGTTTCAAAATGGAACCCAATCAAATCCTATACAATACATATCTTGAGGTATTTATCAGCAATCTATA

TTTCAGACTTCATTTCATAGTATTGGCAATCTAATGCAAACAAATCTAACACAACAGATAACTACTATATC
TATAGTTCAGgcttcattaaattttatgtttatcaGCAATCTATCGGAGAAGCTAATTGCATAACTGCCAG
GCAGGACACTATAAGttactttgaacataaaatataatgatcGACAAATACGTTGAAACGAAATATAACAC
CAAATGTTCAATGTGCCGATGGAATAAGTTCTATCATCCAAAAGGTAATACGATAAGATAATCAACGTAGA
TAAGATACAACTGAATGAAATCATCAAcatcaaagaaagaaacatcTAAACCACAAGCAAAAGCTATCCAT
ATAATGTTGTACGCTCAATTCATCTAACCAAAACGCAGTTACAGACTACATATTGTATGCATTGTATGCAA
TAAGAGAAACCTCGCAAGAGATAGTCCAAAGGCAGATCGTCCTATCTACAACAGAAAGTAGCTTTAAATCG
TTTTCCAAacgaaagaaaaacaaaagctcCAAAGTTGACGTATATGAAATTTCCTGACGCCTTGGAAGTCT
CAGATGGAACCGTAAATGGTTAAGGCCACTCATGATGATGAAGAGTAAGAAAGCGATAGAACTGCGGCCAT
ACAGCTTAAGAAAAGCCAAGCCAGGAATATCAtcataattatgatataaaaatgagttcgatgaaaaaaat
ttcgtgttCTTCATGAAATGTATTAACATGTTAACAAACAATCACCGTAGTCACTCTctgaaacttttaac
aattagTCTCTCGACAAATCCCGAGCTTTATTGaattgaaaaaggaaaagaacaaTGAAAAAAAGCCTCAA
GACTTATCAATTTAAACGCTAAAATCCCTCTACAAATTacagaaaacaagaaaatcaaaCCTAAAAGTTTA
AATGCTAACCTGAAAATGCCGCGACAAACAATATAACGACCTCCATCCACGAAGAACTCACTCACTCAATT
TTTTCGAATATAAGAAGGATTATCGACCCACTCCTGATCTTTAATAACGTCAAAAGGAGAGTTCGCGACGG
TTCTCGAAGAGAGAAATACAGTAGCTTGGACTAAAGGGATTTCGGAGCTTTCTAGAAGCAGAAGGATCtag
aaaaagcaaaatcaagTACCACCAATTCTCCATTAATCTATTGGAGAACTAACAAATGAAAATGCAAAGCT
CAGCGATGAAGAAAGACACAATGGTCAACGGAGAGATCACCAAGGCCCGATGGATCATACTCATCGATCAG
GAGAAACCAAGAATCAAGAACAAGACAAGATGTAGAAGAGCAACAGTCGCAGCAAACATTCATATGcgtat
tattattacaataaattaCTGTAGGAAGGCTGTTATATCTCTGGTCACGTTCATCCCTCTCGTTCATTTAA
AAATCTCCATCGTCCAACCGAAGCGAAGCATCTGTGGCTACAACCAGGGTCCCAAACTCATGTGTAAACAC
AGAGCAAAGATGTCAGGATTTCCATTGGTCCAAATAGCAAGCCTGATTAAGAAATATTGGTGGGTCCCAGT
AAACCAGctaataaaacaacaaattttcatgtttcgTGTTCTGTTCTTTTCAGCTACATCCACCCTCCGCT
CCCACTACTTAAACGTAGTAGTTAATTAAacgttttaaagaaaaaaagctagGCTCTGCGGCTATCTTGCG
GACACGTCGCCcagtattttctttaaagtcgttctttttttttatttttaaacaaaaaattatccctTGGC
ACAAAtgaagttttcaaaattttaaccttACTTTAGATTTCGACATAATGTGCACGACGAATAGGATCCCG
GCCGCAACCGAGTTGATAAAATAATCAGAAATCCAGGCAGGAACATGGGACACGTAGTCACGCCACAAGAC
TGGTCCGTTGCGGTGGGACACGTCTGCACCGCGGAGACTGGAGGCCTAGTGGTCCCCCACTACGAAACAAT
cagttttacaatttaaaactaCCAACTTGGCTGCGCctttacaatttaaaacatCTACAGCGAGCTGGTGC
AACGTACCCACCTGGACACTTGGACAGCTTCGTGGTTATATTAATAGACAACTTGTGAGTTTAAtatcaag
tatatatattgatattaactGCGAAAACTCTTTCAAATAGCAGCATTGTTATTCTTCATAAAGAATGTACT
AGTAGTGgcagatttaaaattttacgcaaaaatattaaaatacaatataaaaaaaaaatcggttaTATTAAT
AGACAACTTGTGAGTTTAATATCAAGGTATTGATATTAACTGTGAAAGCTCTTTCGAATAGCagaattgtt
atttttcataaatagtgTATTTACGGTAGcggatctaaaatttaaaacaggaatatcaaaattgaatgtaa
aaaaaaaaagtcggttatatataataatcacaactttaacaaaataaaatcgattATACATAACAAACGAT
CACAactaacacaaaaataaaatcgataatattataattgacCTCGTCAAGATATTGATATTAACTGTTAA
AGCTCTTTCTAATAGCGGCAATGTTATTCTTCATAAAGGATGTGCAAGACACACATCCTGAGATACTGTAT
CAGGCCCATCATGACTCTGAAGAGCTTTAGCTAGTCTTCTATGACTTAAATGTCCATGGGACTCTGAAGAG
CTTTAGCTAGTCTTCTATGACTTGATATAGACACAAATAGAAAccttgatattttattttctgaacaaact
actcacctccaacaaactacataacaaaccacacacaaccaatcttcACTACACACGTATCTCGTaatccg
cacaattttctcgttATCCgtaacaatatcaaaattacaaatacacccatcgtgatgtgtgatcGTAATTG
TGCTACGCtagtatgtgtgattttggaagtacatagcaaagccctttatTTTCAGCACTATGCGTTGGAT
ACTCTCACTACGAAGGATTTCGaactaaaagaaattaaagaaatgcGGGTTGTGTTCCTTCCTAGCTTAGT
TGTATAAACCAAGGTAAATATCaccttcttaaaaaaattgggattTCAACGTAGCCTAGTTAGTTTTAGct
ataaagtaaatataaattatctttccaagCGGTTGTAGAATGTAGAAGTTCtaaattaggaaaaattatga
agagtggtgaaaaattatgaagagTGGTGAAAAATTGTAGCCAGCAAATCTTATAACGAgaccatataaat
tatttaactttaatgaattattaatttttttttaaaaaaaaatatcgtaaaaaaattctgcatgTAATCTG
CCcactttttatttcttaaagaaaaaaaaacttcaagaaatataaaatatatagagcaGAGAGTGGTTGGTTG
AAGTTACTCCTCATAATTTACTTCTCACACTTACCCTTCAAATGAATTGagaagatggaaaaaaaatttat
actccAATAACTCCTAAACTTCTAAGTTATAGCCGTCGATGATACAAATGTTAAagtaattattcttaaaa
ttaacttAAGAGTTATTGAATCCAATGAAGGTCTTAATGTCTAGCTTTTCAGTTCCcctaaattatagttt
ttatttatatttatagcatttttaggagaaattaTTTACCTTTGTAATTATGTGCTAATATATTAACGGA
GAAGGTTATTAGAATCAACCTATAAATATACAGTCAATGCAACGTTTATATTAGGTTGATTTTGGTTATGA
TAATTTacatttgtttcttattttcctGAGTGCTGCAAAGTGTCTATCAAAATCCTAAGAGTGGCCCTGAA
AtggagaatatataatttcagcGAGTTTTCAACGAAAATTCTAATATGGTGGCCGGACGGGACACTAATAC
TGCACGCCTGGCTCAGCGTGCATATCAAGCTGGGATTTTACTTCTTAACCGCGGTTCTGGTCCAATATGAA
CTGATGGTTCGTGATTAGAAACGGTATCATCAGTAACATCGGGGTTTGTGCATTGGAAATGTTTCATCtcg
ttatctccaaaaaaaaagatagaatttAAAGATTATCTATCCTtccactttatatatatatatatatatat
aatatatatattatatatattatatattatatatatatatatatatatatatataacaaaattccaatt
tcCGGATGGTGAGAAAAAGCTACTCCCTATAGtggtaataataaaataatacaaatatcctATGTTGTCAT

TATAATAAATCCAGACGCATCTCACatataaaatgagaaaaatccAGGCCTATTAGAATGAAAAAAAGttc
cccccaaaaaaaaggaatatattcttctatatgatagaaatttagtACTTCGAATATTCCAACAAATGGTA
GTTACTGAAGGATAATAAACTTATGATCACTAATTAAGCTCTAAGCATTCAACTCAAGCTCTAATAAACTT
ATACTCATTTAAAAACTATCGAGCCTAGTTAATCCAAACGAGTCAATGACTCAAGCTCGTTATAAGTCGAG
CTTTGGATCTTCTTTCTCGTTTACTAACAAATTAACCTGACCTTGAGCTGAATCTAtaccgagctactcgt
gagctgTTTGGCTCATTTACATCCCTTAAAGAGGATAGTCCGTTGAAAATAGCCTATAATGCAGGAGTCCA
TAGAAATATTTCGCACCGGTGAATTTTCACAGAGTTTTtctttgtcaaattttatcaaagtaTAGTTCGCG
ACGTACTCATTGCCAGTCCTAAAACCAAGAACTTGTAGCAAGAAAATGGGGCCAATGCAAGTCAAAATTTG
AGTTCAATAAGAACAAACAACGAACAAATAATTTCAAGTTTGCCTAGTTTTGGTATCGCTCTCGTGACACG
CATCAAATAGTGCTTTAGCCCTAGATGTTCAGTCAACTGCGCTACACCTAAATGCATTTTACTCATGTTGG
CATGTGACTCGAGTCGAGTCTGAATCAAACAAGCATCCAATTGGATTCGGTAACCGCCAATTGGGTTCGTA
AcagaccacatatatatacataataatagatCTATGGGACGCCATAAGTTGCATTTACAATATCTAAAGGA
ACTTGGATAGTGCATTAGTGAGTGTCTCTGGTAGTCACCCACGTCGCCGTAGAGTTTGCCGGAGCATTGCC
GGAGTTTGGGAGTAATCACGACTAGATCTTCGTTTCCACTCCAACACGAGGTATTTCGTGATATTTCATTC
TCTAGATTCAAAAATAGTAGGATTGTCTTGTATCTTCAACTCCTAACTAGCGCGCAGCAGGAATTGTATCG
AAACTAAGTGCAAAATAGGATAAATTGGCTACATTAAGATTATATCTGCTTAGCTCTGGGTGGATAATCTT
GGGGTGAACCAAAAGATACTAGGTAGAGCCGGAGCCGGGTCTAACTGTTGACTAGATAAGCGGCTTGCAAC
AAGAGTAGTAATTATGCACCctgtaaatagtttttataggTTCGTCGACGACGGAAACGAGCTTCTGGAAG
AATGATTTTTTGAGCTCATGCAAATTAACCTAGTTCTCCACTGAGAAGCTACAATAACATACGGAACTTTT
ATAAGAGCTCAACATAATTGGTTCCATCTCGATGCTTCTACACTCCTATAGAAGAGGAAGCGTTGACACAT
GTTTTGATTAAGCGTTGACATCGAGAAGTGACCTCGTTTCAAGGTCActgaatttaataaataataggaTT
CATATTTAAGGCATGCATGTCccgattctctctctctctctctctctctcttctctctcaaaaaaaaaa
aaaaaaaaaaaaaaaagaagttatgtCATTCTGGTTAGCGTCCATGATCGAAGATGCTTCTGGAAATGTTT
AGACAGTTAAACAAGATATTCtcaatattgaaaaaataatcaacatatatatacttcaacTGGAAAAGAGC
TAATGCAGATGAACTGAAGGTTATATATGCGCTCTAACGATGACACACATTGCAAATGATGACGACCAAAG
TAgataaaaaacaagaagaatatAATTCTTAC**CTCGAAAAGACGTCGCTGAAGCAGGAGGAGGTCTACGAT**
**CATCTTCACCAGGACGTTTGCAAGATGCAGagagagatgaagaagatgaacgaAGATGGGAATGGAGACCT**
**ccatactctctctctctctctctctctctctctctctctctctctctc**tctcgctctctctctctca
agcgAACGCAACGGCGCCTTCGAACGAATATGAAAACCAGAATCAAAATCTGCATAAGGATTTGgtggctt
ttaaaattatatatacaaggaTCACTGAGAGGTCTACGATCATCTTCACCAGAACGTTTGCA

>XviAGL17_1
ATGGGGAGGGGCAAGATTGTGATCAAGAGGATCGATAACTCGACGAGCAGGCAGGTGACATTCTCGAAACGCAGA
AACGGGCTGTTCAAGAAGGCCAAGGAGCTCGCGATCCTCTGCGACGCCGAGGTCGGCCTCGTCGTCTTCTCCAGC
ACCGGCAGGCTCTACGAATTCGCCAGCTCAAGTATGAAATCTACGATTGAACGGTACAACAAAGCGAAAGATGAG
CACTGCCAAGCGCCAAATGCCGATTCAGAAGCTAAGTTTTGGCAAAGGGAAGCAGCAAGTCTGAGGCAACAATTG
CATAACTTGCAAGAAGATCACCGGGTATTGACAGGAGAGAATCTTTCTGATCTTTCCGTTAAAGATCTGCAGGAT
TTAGAGAACCAACTCGAAATAACTCTTCGAACTGTCAGAATGAAGAAGGACCGAATCTTAACAGATGAAATTGAA
GAACTAAAACAAAAGGGAAAGATAATTCACGAAGAAAATGTGCAACTCTACAAGAAGGTTTATGCATCAACAGGG
CTGCAAGGAGAAGCTTCACATCTACCAGTCAATCCATTAGGCGTGAGAATCATGGGAAATTTACATTTGCAACCG
CAGCTTGAGTTAAACAACCTGCCCCATCAAGCTGATCCAGACCCAGCAGGAGCACCTAAGCTAGGGTAA

>XviAGL17_2
ATGGGGAGAGGAAAGATTGTGATCAAGAGGATCGATAACTCGACGAGCAGGCAGGTGACATTCTCGAAACGCAGA
AACGGGCTGTTCAAGAAGGCCAAGGAGCTCGCGATCCTCTGCGACGCCGAGGTCGGCCTCGTCGTCTTCTCCAGC
ACCGGCAGGCTCTACGAATTCGCCAGCTCAAGCATGCAATCAGTTATTGACCGATACAGCAAGAACAAAGGAGAA
CATTACCAAGGGCAAAATCCACACACAGAAGTAGAGCTTTGGCAGCGGGAAGCAGCAAGCCTGAGGCAGCAACTG
CATAACTTACAAGAAGAGTTTCGGAAAATCACAGGAGAGAATTTATCGGGACTAGCTATTCCAGATCTGCAGAAT
CTGGAACACCAACTAGAAATGAGTCTTAGATCAATCCGTATGAAGAAGGAGCAAATCTTAATTGACGAAATTGAG
GAATTAAAAAGCAAGGGAAAGGCAATTCTTGAGGAGAATTTGAGCCTCTACAACAAGGTTTATGCCACAAGTGGA
TCGACCGTAGAAACTACCAGTGCTCCACCTCTACCGTTAGGATTGAGAATCACTGAAAACTTAAATGTGCAGCCG
CAGCTGGAGTTGGGTCATTTGCAGCAACTAGAACAGGCTCAAGAAGAACCAAAGCTACGGTAA

>xvi_miR444a
TTCTGGTAGAAGGAGGCAGTGAGTTTGAGGCACCAAGTGCATAACTTGCGAAGCTATTGGACGGTAGAAAAAAAT
ACCGGTGATCTTCTTGCAAGTTATGCAATTGTTGCCTCAGACTTGCTGCTTCCCTTTGCCAAAACTAGTGAGTAC
AACATTCAAGAATTAGCTTATTATACTAGTAAATACTTTCACATGTAACTTAAAAATGATTACTAAACAACTAAA
ACCAAATTACCAACGTCTTTTTTCAtgtatcaaattatcaatGCATGGTTTATCCTTGTCACAAGCTTTTTGACA
GAAAATTGGATTCAGAATCGGATACTCTGGAGTTGACTCTGCTTCTCACTAAATAACATTTAGCAATTGCCCCTT

GATTGTATGGTAACTATTAATTTCCTATATTCAAAGTAGCTGTTGGACATATTCTGCCATTACAGTGTAATGAAT
TCACACATGTTTAACTATGTGCTTACATCAGAACATGCTTATTGGGGAA

Position of mature miRNAs based on comparison with osa-MIR444d and not based on experimental data!

Stem-loop structure

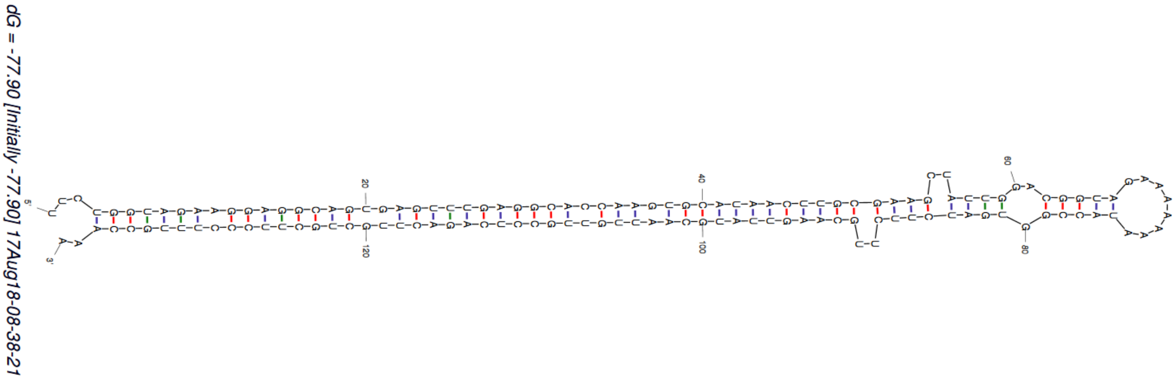


>xvi_miR444b

GGCGCCGTTGCGTTCGCTTGAGAGAGAGAGAGAGAGAGAGAGAGAGAGAGAGAGAGAGAGAGAGAGAGAGTATGG
AGGTCTCCATTCCCATCTTCGTTCATCTTCTTCATCTCTCTCTGCATCTTGCAAACGTCCTGGTGAAGATG
ATCGTAGACCTCCTCCTGCTTCAGCGACGTCTTTTCGAGTTTTGATTGGGGGAGGCAGCGAGTTTGAGGCA
GCAATTGCATAATTTGCAAGAATCTAAAAGGTTAAAAACGATACCGAAACTCTTCTTGTAAGTTATGCAGT
TGCTGCCTCAGGCTTGCTGCTTCCCGCTGCCAAAGCTGGTACATACAACATGCTAGGTTCAGATCATATGA
GTGCGTTGCATGCAAACGAACTCCTAGGCCAGATGCTTCATTACGAGTGGCATAAAACATACCTCTACTTC
TGTGTGTGGATTTTGCCCTTGG

Position of mature miRNAs based on comparison with osa-MIR444d and not based on experimental data!

Stem-loop structure

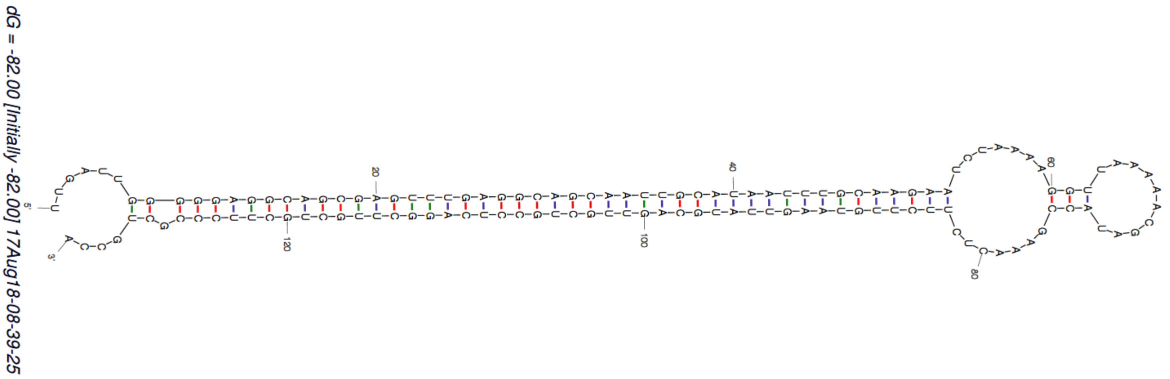


*XviAGL17_3 and xvi-miR444c*

>MJHO01000017.1

taattttaatttattaaattaatttttttggcgCACGTATTTATGTATTGTCCTAGTGGAAATAATTACCT
AAacaaatgaatatattattcaagaCAAAAGCCTTGCGGGCTTTCATCAGTTCGACCCTGCAATGCTTTCT
TTGTTCTGGAAGTTTCCCACAAAGGTTTCCAGACGTCGATGGTTAAAAGGACCTCTGTGGTCCCCACTTCC
GTCACCTGAAACCCAAAAGCGGAATCTCATACGGACGTGTCAGGCCCTCGCACGCGTCGGCCATCGCGACG
GCGAGGACGAAAAGATTCGTTTTTAAGTCCAGAACCAGCGACCCTGCGAAACCTAAGCACGATAGTAATAG
AGTAATATTGGAAGTTTGAAGAAGTAAAATGaaagtggagaagaagatgtcgctaaaaatataaaacaaaa
gaaaaagataaggtAGATGAGGTTGATCTGTCGCTCCCCTAGGAAGTAGGAGAAAGGAAACCGACAAGGTT
TTCCTATTGGCTGCTCATTGATGATATATTGATCAGCTAGCTGTCGCAGATCTGAAAAAAGACAGCCGCCG

ACGAATTTCAGGTTCTcggatttatttttcttttattctgtGTTTTTGTGATCCTCTTCTGATCTTTGTTA
GCTTGTACTGATTTGTTTAGTTTAGATTTATTGATGTTTGGAGTATAATCAGTCAGGCGGAGAGATGGGAA
GGGGGAAGACTGTGATCAAGAGGATTGACAATTCGACGAGTCGCCAGGTGACATTCTCGAAAAGAAGGAGC
GGGTTGCTGAAGAAGGCACGGGAGCTCGGCATACTCTGCGATGCGGAGGTGGGTGTTATCGTCTTTTCCAG
CACCGGCAGGCTCTATGAATTCGCCAGCACCAGGTTTGTCATCAATACttccttatatatattctactcgt
ttcttctaattattttattgcatatcatctctctctctctctctctctctctctctctctctctctctctc
tctctctcttaagaacgaagaaaatttaagaaattgttttctaaaattagaaacagttcgtttgaaagaaa
attcctTTGACTTCCTCTCTTCCAAAGACGGAAACTCATAAAATACTCCTCGTAAAAGATTAAACTCGAGA
CATAGACCCCAAGTTAGAACAGTAAAGCTCATGTGGTGAATCTTCAATAGTTGTTTCTAGCGCTTATCTTC
TCCTCCACATGTTTCTTCGATCTATGAGTTATACATTCAACCAATCGGAATGATATGAATGTAAGTTCAGC
GGATAAGTCATGAGTTAAAGGAAGATCTGATATTTTATGCGCAGAACAAAGTCTCAAATATCACGAAACAC
TTCTCAGCATGTACTTGAATCATATTTAGACTTCTTTTTGAGGGGGAAAGGAAAGCAAATGTGTTTAGATT
TCTAAAGTAACTTGTTACTGCTTTGTTGCAGTCATCTATTAGGATTTCTCCTACCTAGCTTCGATACTATC
TACAGTTTAGATAGATGATCATCAAAGTGATGAAGTAGAACTCCTAGTAGCACCTTTGGCCAAGTATTCAC
GACGTAAACAATCCCGTTAGTGGTCGACATAGCACTGGGCCCGAAAGAGTTTAGATGAATGTATAAGTCTA
TAAAGATGGAAAGAAAACATTAATGCTCAATTCTATAGAGATGGAAAGAAAACAATCTAGGTTTGGTCCAT
GCAAACATAGGAAACCACAACTTCCAAGTTGTTGCAATAGAAGGTTTTAAAGACTATTTTGTAAGAGGACG
ATTATTGTAGGAATTTTCGTTGGAAGAGCTAAAGTAGTATGCAAAAGAAATAGGAAGGTTTGAAGTGAACT
TGTATGCAACTCTTATTACTTGGCTCGTAAATGTAAGAGTACAAGTGTATTAATAGGCACCTAAGAACGTT
TTACCTActagaaaaatatctaaaatagtttACATGAATAAAGAAAGAGATAGTGTCTATCCTAAGTGGCA
TGATACTAGAATTCTTCCAATATATCTAACTTTAGAATTCTCTCAAAAGTATCTCCATATACAGATTCTTC
GCAAGCTCTATGGCGGACTTGTTATCAACATGTATATTTGTTGCTATATCCTATGGGACTTTAAGCTCATT
GAGAATGTTGCGTAGCTAAATGACATATGCATAATGTAGCCGCTATATATTCCGCTTCATATGTATAGAGT
GCGAAAATCGGTTGCTTTTTGGATGACTATGTACAAGCATTCCTCCAAAGTAGAAGATAAAACTTGATGTG
CACTTCTGGTCATCATAATCTCCGGCCCAATCGCTATAACTACAACCATCAAGATTCAAATTGTTAGAGAT
GagtaaaataaactatatgttGCATACCTTTGATGTAGCGGAGAATGCGATGGTAGTATTCATATGCAACA
TTTGTGGAGTCCCCACATATCAGCTAATTAGTCTAACTCCATATAAAATATCAGGTCTAGTGCATTTAAAA
TATCGAAGGCTTCCAACTAAGCTCTTGAAATATGTGGGACTTATATGCTGACCTTCACCTTCCTTCCTCAA
TTTGGTGCCATAATCAACCGGTGTGTCGATTAGATGACAATCTTCCATACAAAATTTCTTAAGAATTTTct
ctgtataattttattgagagATGAAAAtgccttttttccttttgatgTACTTTATTTACCAAGAAATAAGAC
ATAAGACCAAGATCAGACAAATCAAATGATTTGATCTTGAATCCTTGATTAATCACCGATGAATCATTGCT
TGTAAAGATGAAGATACACTTTTCACCCTTGTTGTCAATATGCATGTGTTCtcatttcttccaaaattttg
caatAAGCAATGctacaaaaaattcttaaataatcCCTTTTGGGTTTGTCAAGACAAGTTTGTTCCGGTGT
AATGTTATAGAGtctttttttatagaaatctatttaataaatagacgACATTCCTATGGCTTTGCTCTAGAA
TTCTTTAAGAACAAATTTTTGCTTAACATACATCTCACTGTATTAAGAGTACTTCTATTTCTCCTTTCATA
AACACCATTTTGAGATGGTGAATAAGGCAAGATAAGTTGATGGTTAGTACTATTTTTTccgcaaaaatttt
caaaatcatgtGATTTGTATTATCctctatatgattttggGACAATAGTTTACATTCACTTTCATTttcc
accaaaatttgaaggctTTGAATTTCTCCAgcaattcatcatttttttgcaaaaaaaaatatatttatgtc
ttACCCCTAAAGTCATcggaaagaaaggaaaatacttGTTCCCTTCAATTGATTTTAGCTCAATGGATCCA
CAAATGTCTGATATACCAATTGTTGTTTCCTTGTAGCTCTACTTACTCCGTTCACTTTCAAGCTCCTCTTA
TGTAGCTTTCGgagaatacatttttttatttcttattttctttcactaATCCATAAAAGACCTGTcacca
ttttttttgacaCAAAAACTTTAGCCCTTCATAATTAGAATGACCATATATAAGATGCCATAGAGTATCAA
CATCCTCAATTATAATGTTAAGGTATCTTCTTTGATCAACTTCAATAGTTAGATTGAACATTCGATTTTTA
GTCATTTTGACATGGGCGATTAGTCTTTTATTTCTATCTACATAGATAATCGAAGATCTTTCATGTATATT
TCATAACATTTTTGTAGTAATTGGCccaaactcaaaatattactTCAAATGTCTGGCACATAATAAACATT
ggaaatatattgatatgatCCATTTCTCAGTTTGATAAGAATCTTATATTTGCCTTTCATAGGTCTTTGAG
taaattatccaaatataatattactttcaaatttttcattgAGCTCcacaaatatattgttatcaCCACAC
ATATAGTTGCTTCCACCAGTATCGAGATACCATGTCGTACTTAAGAGTGGGCCCATCTTCTTTACCTTAAG
TAaggtttcttcttcattttatttttatacaaaatacactTCTTAACCATTGTGGTTAGTATCTCTGTTCC
AACAATTAGAAGAGGTGTGTCCGTATTTCTTGCAAATATGACATTGAGTGGTAGAATTGCCATATCTTGGt
cttttatatagttttaaaaaattagctcTTTTTCTGTCTCTTTGATAAGATGGTCTTGTAATTACTTCTTG
CAATTATTGCCTTGATGTTTCATAgaatcttcttcctcttcgtcCATGACCATGTCCTTGAAAGCCCTATC
CTCTTGGttaattgtttgaatttgtttttttgtttctctctctGTCAATAATCTTTTGTTCACAGGCCCATG
AGTTGCTCTACCATTATTTTCTCCATGATCTTGGGTTATTCGATggctacaaaaataaatcaaacttgGGG
GGTCCAAAGACCTTAAGATTTGTCAACTTGCACATGGGTTAGTTTTTGCCTATATCTTCTAAGTTGGTTAA
CGATAAGTCACTTATGGATTCACCattcttcattttcaaaagtttatattgAAATCTTAGAGCTTGGAAGC
GAATTTTCTTTACCTTCTCGACTCCTCCAGTAGACTTTTGTAGGATATCCCATACTTCTTTAGAAGACTTG
GTGCTGGCTGGCTTTAAGTTCAAATGTAGGTTCATCAAGTTCTTAGTAAATGATGAAAAAGAACttcttgt
cctttttttttttgtaagtaagAACTTGTattaacacaaaataaaatctttacaaACAATCAGCTTTGAAC
CGAAATAATCGGCGCTCAAAAGCTGCAAACAGACTcggaaaattaaaagatagaaAACTCGTTCTAACAAA

CAGAacaaagacaacaaaataaaatatccttCAGTGACTCTTTACTTGAGAAGCTCTTCTTTAAGCTCAAC
TCCAATCAGCCCAGTAGGAAAAAACACACTATCCGAGACTCTATACTttgaaaaacactttttttaagcCC
AAATCCAATCAGCAGAGAAGGAACATCACACCtgaaaaaacacaaaaaactaCAAGTATCAAGATTATaac
caaacacaaaatttgcataaaacTCCGGAACAAAATCAGACTCTCCAATTgtaaagaacataaaataaaga
caaaatcAATTAATGTGGATCAGATTATAAAACATCAAGAAAGTAAAGTATAAAAATCTGACAAATAAAGA
GTATCCCAAAACACTACAAGTAGTAATGAAATGCCTAGATCTAAGATATGGCACGACCTTTGCCAGTCTAC
ATTAATCAACAACACTCAAAGTTAAAATGTCATCTCTAACTCATAAATGCCTCTGACTATAAATGGAAAAG
CTATGACAGAGAGACAAATGGGGTAAAGTAAGTCTTGAAAAATAACTGTAGACTAGAGACAACCAAACTCC
ATATTGGAGAAAACGACCACCAGATATTATAGGCGtccaatactcgggtcttatcCAATTTTTTCCCAGGC
TATATCACATGAATTCTTTATGCCTGAGGCCCACATAGttaccaaaatcctttttggaaGAACACTACCAG
ATTTTAAAGACATCCAATACCGGGACTTAACCAAATTCTTTACCCAGGCTATACAACAATActtcttctaa
aaatagattgaGGTTCACATAGacaccaaaatccttttttggtaGAAGTCTCCCCGAAAAGGTGAGGTCGAA
AAATGGGACTTTAACCCAAACAATTAAGAGAAACAAACATACCCGAATCCCAAGATATAGCAAACCAGAAT
ATGATCAAAGAGCAGTAGTACAAGAAAGAGACACTGCACTTTTACTTGAGAGAGTTTGAAAGTGAAACTAA
GCTAAGGTCATCTATAAATAAGTTAAGGAGGATTCATAGCTAACTAGCATGTGTTTAGGAACAcctttaaa
aacatattcatATACAATACTCATCATACTATGTTGACTGATTATCATATCCAGACACAAGCATCATCATA
TcaactcataaaaaaattagcattataCCCACTAGCTATGAAAAACCAGTTCCAAAAccttttttgatatta
aattgatAGATGAAAAAAGGAAGCAGATACAACTGGAGCAACATTTCTTTGAGCAGAAATAAGAATCATCT
ACTCTAGAACATTGAGAGAGAGCTTAACCCCTTTAATCAGATTCATTACAGAACCTGGAGGAGCAGAAAAA
ACGCAGCCGTCCTTCTTGTCCTTTTTTTCGGTTTCTCTTTAAGAGTATTCTTCTCGAAGCTACTCATAGTT
TCTTCTACTTTGGTGGATTTCGGCTTCTCATAGCTTTCTCCAAGAAGCTCCTACATATCAAGAGAtcctaa
taataatttcatcagAATTAGTTATCTTTGGTTAATTTTGGGACTTGTGGCTGGATAAGATTTGTGGTGGC
TATTATTAGAACTcatctattattaaaatagatgAGTTTTGCTAGCATGCTTAAGTCCATGACTTGAATAG
CTGTGCTCataccaacttttttttctgtctACTGCTCGTTGCTGCTTGCACTCGTGTGCTTTGTCTGTCAG
CACGTACCCTTGCACCAGAGAACAACACATTCGCGCATGCCTTTGCACCTGCTTATTGCTTGCTTGCTCCA
CGTCTTTTCGTTATGCCGTGCCAACTCATTGTTGCCCCCCCAGCCGCCTGCATGTCTTCGCATACTTGAAT
TGCCGCTCGTTGTCCATGCCAATGTAGGCTAGGCTAATGCTTGGGAAGCCTTTAATCTGCCTAGACCACTC
TGCAAAACACGCGATCTTGATCTTCGGGAAGTAGACAAACATTTTACCATCCACGTAACAAGGGAAGGATT
TGGGTTATCAAAGGGGTGCAAAGCAGCGGGAGAGGAGAGAGGTCAGACctaaaaaaatcctaaaagaAGCA
ACACTCCAAATCAAGGACTAGAAGCCATTGTCGCCTTGTCCATTACTCCTCTCAACTACCCACTATTACTT
CATTCTATGTAATAACTTCTAGAGTTTATAAATGGAGTTAGATGATTTTCTCCCCTAGATGTAGGTTAATT
GACTGAACTACGTAAATCGTTGTCTATctctcaatttatttttctttttgctccTTTCCATGTAATTTGTC
ATCAATGCATCCTAACCAGTCTAGGGTTTAAGAGTTTTGAGAAAACCAAAGGAAAAAAGTTGTGCGAGAGA
TGAAAACCAAAACTCAACAAGAGAAACGAAGAAATTGTTGACAAGAAGATCCAGGTAATAAAAGTCAGTGG
CACATGCAGCTGAGCAGCAGTAGAAGGAAGCCACacaaagaggaagaagaagggttCCTCTTCCTGCACAA
GCCTATTTAAAGGGATAGGATGGCGAATAGGGATGAAAACCGAGGTGATGTAATTTGGCATGGGAATCGAG
TGGATATAGCATTAGATGTGTGGTTCCTTGTCAGTTGCACACTTGCACCAACCTCGATGCGCCTTAGGAAC
AAGCCGCATGCCTTTACTAGGTACCACTGCTAACACACCGACACTTGCCCTTCACATCGTCCCAAGCATTT
TGAATTGAAGGCTCCCACATCGGGAAATTCCGGCGAACAACCACAAGGTACACTAACATGGAGCCCAAAAT
TTCAAAGCGGATACCATGGCACATCATAATGCAATACGTCGTCTTGTCTTAAATGGGCGAAGATCTAGACC
CCTATGTAAGAGATATTTCGCACTTTATGTTGTGTTATAATGTATCAAGGTACATATCAAATTTGACCACT
TTCTTATGTGGGTAGCAATGATTCATCATCAAAGAATAATAGAATTCTATGGATGGTTCATTCGATGACCa
cttaaaattacaatatgaCAGCCCGCAACCCGGGCTCGTTAACTCACTAATGCCAAATCATACACGCAAGT
AGAAGacttatatatgtgattttagAAACTTGAGAGACTTTAAAATACTTTGATTCGCTACGGCACATGAC
AAGCATCTATGAGGAAGATGGTTCTCGCCACTCAACATGAACGTAACTTGCTATAGATTTAAATGCATCAT
TTTTTCTCATTGATACTTAAgaatgtataatattatatggtcCAAAACCACTTCCTGTTTCAAAccgtttt
ataaaatttcggctcaatatatttcaaacattcGGAAATGCCATAATTGATATATACCTTTTGTAatcata
aaaacttaaaaccaaaatttaaaaatacttaaatcagttttaaaaatcttacgCAAGAGGACTTTTTGGTC
GGCTGGCCAAAATTATTGGTCAACCGATAGTTCTAAAATCAGATTACCGCAACTATACCGGCAAGATCAGT
CGACCACAACAACACTTTCGGTCTATCGAAAGTTCAATATTATGCATATCGGCATTCAAATTTGGGAACTA
CCCAAGTATCCAAAATGACCATAAAACATTCAATAAGGATATCGGACATGTATACACCTGTATCATGGTCA
AATACACTTGCCCTAACTTGAGtgattttaaacataaaatcacCATTTATGCATCAAATgtattttgccaa
aaatacaaaaaaaataattaaaaaaaat**gccaactttTTGCATTTCCATCTAAAGCATGCGTTTCAAACAAG**
**CATAACCACTTAGATAAGGGCtcatatacatgcatacaacAAAATATGAGTTAGTGCTTCCTTTATATTCC**
**TTTACACTACATGAAGTTGGCCATATAGAACTAGGTATACATAAAGGGTTACATGTTACACTTAACCCCAG**
**CATTATACCTCAAAAGGAAATATAGTCTAGTCTTTCCTCTTTTGCTATACGTATAGGACTCTCGACTGGTG**
**CCTTCCCACAATCTC**CTGGgtttggaaaaaagaaaataaaagaatgtgAGTTAAACAGACTCGGTGAGCGG
CACAAAGCATTCCCCAACCAATGCTTATCGTAGCACCACCTGCAGCATATTCACATATACAATACATAACA
TGATTTGCACGATAACATAACTTGTCAACCAAGCCATAAAGCAAATATGCTAGTCAACTAGGTAATGATCA
Tccatccaaatcaattatgttg TTGCTTATTTGGAATCTTAGAGTAAAATCATTGGGtactaacccaaacc

agttagtaGTCCTTTCGTGTTGAATTAGATAACAGAATGACCTAGATCAGGGTAACCCGATCCAGTACCAA
GATCAGGTACCTAAACCACAAGTGTACATTTTGTGATGACAGACCTTACAGCAGGAGCGATCCCACAATAT
CCAGCGGGAACTACTTGGCAGCTATGCAACCTTCGGACGGTGTGACtctgagcctcgcactcccTAGTTAT
ACACTTGGGCCCCAAGCGTTGTCTATTATGCTCATTGGTCCCATAAAAACATTCATATATTAAGATTAAGC
ATTTTAGTTGTAATCCGTTGGTCATACCATCCCTTTTCACAAACCCGCCGATATCTTGGGACTAGAACTTA
CAACACACTTGTGCTAATCTCTTTAGCCTTTATAATCAGTCTACCATGATTATCGCTAGAGCAAACATGCT
CACGTACTCACATAAGTAGAACCATCTTCATAGGCACACCATTACATATGTCAAATAATGTTTATACAAAG
TACTACTAGAGCAATCCGCTCACATACTTATTCAGGTAGACAACAtcataaacatattaaaatatataaat
acaattaattCTACCACAAGTATTGATAGAGAATACCGTTCATATTCATCGCTAGGTAGAACAACATCATA
AGCATATTATAAGAACACAACATTCTACTAAGAGAATTTCCCAAGCAAGCTTGTTCTAATACTCATCCAAA
TAGAACACCATATCAATAACAAGTTAATCAAACAACATATTCATTTGTTATAGCATAAATCAACAACATGG
GTAGCATATAAACTATGGTAAACATACCAACATGTTTAACACCCAAACAACATGCTAGAATCCTATAACAA
GCATAATACAACTTTGTTAAgtcaacaaaacaaatcatGCAACATGTGCAAACCGCTCAACTTAGGTTTTT
TTCAGCCCTAGCCCCATGCTTCAAACCTCGACGAAAACCAAAGAATCTAGTATCCACGCGTATACAATGAT
CTAGTGAACCcaaatttaagtttaattgGATCTCCAAGCTTCTACTAAGAAGATATCAAAGTTTAATTGGA
TCTCCAATGATCTAGTGTCCATATTGGAGTTTGAATGTTAGATCCTGCAAGGAGGATTGGCTCTCAGCACT
AACTGAGTACTTACATCCCTCATCCATTGCATATATGCTTCCTGTAaatgaaactatatatatatatatat
atatatatatataatatatattaatatattatatattaatatatatatataataatataataatatatatt
atatatattattatatattataagagagagagagagagagagagagaggctattcttcAGCGTCGACCTGA
ATACCTTGGTAAGCAAACATCTGCAGCcgtgatctccttgatctaacgctgatatttaattttacacttaa
cggaataacgattcacgccgttacgTATTTAATGGtgccgttaagtgtaaaattaaatctcagccgttacg
TTAGATCAatgagatcaacggctgagatattagcttaccaAAGGTTATTCTcaagtcgagctgagaatagc
ctatatatataagcacaCTTATAAAACTAATACTAACATGTCTTTCCAGATCTTTTAACCACCCAAGCATT
GTCATTCTCCTTTCAATAGACCAACCTGAGCTACCCATCTACTGTTGGCTCTCTACCcagttcttttttaa
ttaatttaacaaacGCAAAAGTCGTCATGTCAATAAACGATTACTAAAACGTCAATGAAAAGATAGTACTA
TTCTCCTCTTGTCACCAGTTGTTTCGATCGAGCCATTCTAATGAAGTAATTCACCAAAATCTGAAGCTTGG
TTTATCTTAGCATGTCCACTTTAAGCCAAATGCctacaacatattaaatatctatgtaTCCATCTCCATTA
GGCTCCAACAATTACATAACATCTATCTGTGCATCCTCAACCGCCTTGCTAGCGGTAGATTTCAATATAGG
GTGCATGAAGGGTGGCTGTGTAGAGGGACCACGAGTGTCGCGAATCGGATAGGCTGAAACCTATAGAGAAA
GGTTCTAACAATATCCAATTAGATGGACGATAATGGTGTTGAACTGAAAGGATGAAGGAGAAAGGTGGTTG
TGGTTGCGGTGGTATTTGCATAGAGCAGCAAGGTGGTGGAATGACAAAGGTAGGCGAGGGCAAGTGGAGGA
AGACAGTTGACTTTTCTTTCCCACATATCCCGAAAGGCTATTCTAACTCTACCATGACTTAAAAATGACAT
CGATAACAGATTGTTGAAGAACATTAACATTTGCCAAAAGCAATAAGcctgaaaatttaaagaaaataaga
gataATAAATCTCATTGGTAGGTTTACATAC**ACATACGAACGAACTTTGacttccaaatttttgactttc**
**tgACCGCCCGTCCATTGCATGAACCATTGTGCCCATACGTCACGTTGCAC**CTGAATATAATCACTGTTATG
AGATACAACTTGCACAAGACAATGTTATCACAATGAGAATTCAGGACGCATTTTAAAATGCATAACATTTA
TATCATCTCCAAAATCTGCAAGAGAAGGTAAAAGACAATTCTGATAATCAAGCAAGTTTCAATTATATGTA
CAACCACCATATAGTGCATTAGCATATTGAAGAGATAAATTTGAAGCAGTAGGTAATCTACTACATTACTA
TTAGATGTTCATGGACCAGGGTTTATATGCTAACCTCTTGTATGGAGGTTCATAACTCTTtaagtagaaaa
aataaaaagataatttaacaatattgATACCTAatacacacaaaaaatattttaacatcaacCATGATAGG
ATAGAGAAGGTGATATATTGTGATAGTAGGAGAACCCCTATTGAAGATATgacaagaagaggaagaagtaA
CTATGCGACGCACCCCAATAGGAGATTCAAGGATGGAAGTGAGAGAAACAACCATGATGGTGATGCTGCTT
GTTAGACTAGGAAGAGGCAAAGTTGATAGGCTCTGGCAATTATTTAGTTTGGCTAGCATGTAAAACTAAGG
GGCTTGCTCATGAGCAAGTGCATTGCAAGTAAACACAGAACTTACTGCCCTGACACCATATTTAGAATAGC
CAAcacttatatcaaaattaagcatattgaaaattagatACTTAAAGgtcttttatacattttaaccTCTC
ATATTGAGGTTAATTAATCGTAAGTAGGAAGCAACTAAATAggaataatagtaaaagtaatatttgcaaaa
tgttactttactatttttatatgtttgcaTAAGTTAAGTTTCCCAATAAGCTCCATTCCACATGTTTTGA
GGGTCTTCCACACGTTTGGCTTGCAGGTGAGTGATTCAACAAATGATTTTCATTCATCATACCTTATAACG
TATCATTGCCATTAGGGTCTCTACAGAACTCTAACCAATGTAGATTTCAAGGTGGGTCATTACAAGTTACA
ACCAATACGACAAGCCACCTCATCCACCAATGAGTTTCTTTACATGCTTCGGTCATAGTTATGTCCTTATC
GTTCGATCACTGATATTTAACACAGTTatgtttggaaaaagaaatgatttaGACCAAGAAACTATCAAACT
GGTCCGAAATTGTTCACCTCAAACATTTCAGGACCCCTTTAAGAACAAACAATAATCACATCATCCACTTA
AAGCAGATAAGAATAAAACGTCTATGTTTTATAATAGAGTAAGTTGAATTGTGAGAACTCGACAGCGAATG
AAACCTGAAAGAATTGAACTTGTGAAGTTCACGAAACTGCTTAAGAGCATATACAGAGAACATTGAACCAC
AAACTTTATTTGGTTTAATGGAATACCTGGAGGGGACTCTGTATAAATTTCATCCACTAAGATCACCATCA
AGGAATGTGCTTTGGGCATTCATCTACCTAATTTCACCTTTACATGCAATCCTGAAAAGTTAAAGAGAAAT
ACAAACAAGTGTTTCCTAATATTCAATACCATATTCCTAACTGTAAACTTTGTCAACTAGTTGAGTTTATT
GTATTCAAGTCATCCAATAGCCGTAATCTTATTGATCCTTGCACATTCCACAGCCAACAATCTGGAGTTAC
TTTAACCTAATCCAAGTCCGATTATTCTCCAAGGCGTACAATCTACTTCCCTTGCTTGTCCAAAACAGGAT
CTGTTGCTTCtttataaacaagaaaacataaaCGGAACGAAGTAGCTTggaattaaatttgttaaactatc

aagaaaagatttttacgTAAGCGCAAATAAGATATCATGATGTAGAAGGGTTTAAGACTCAAC**TAAAGAAG**
**TTGCTGAGGGAAGATTCATTGTGCGTCACTTGGTGTGAAGGTATGATGAGAATCTGAATCAGGATTCTTGA**
**ATGCAACCAGAATAAGAAAATGATACACTTAATCTCCAACCATGCACCAATCTGACTGGGCAGTTACAATT**
**CACATGTC**CTGACAATGCCATACAAGTGTAAGCTAATGGTAGCTGTGGAAAATAGGGCAAGATGGCAGAGT
AGTAGGAGCGTAAAACTTGTGAAATTGTTAGCTACGAAACGACTGCGGTAAGAATATCAAACTCGGGATGT
AATAGCACTAGTGCATAAACATTATCATGTCTATTGGAAGAATAGTAGTACAACaatacaagaaataaaaa
gatggaCATAACGTCAACAACCATCTTGGTTCACGCCACATTCTAAAACATGTAAAAATAGCAACAATTTG
TAGATCACTTTCCCGTGGAGAAGAAAAGACAGAGAAAGTCAGCCACAACCACAACCAATAAATCAATTGTA
TGCAAGATTCATAAAAATGGTGTCGATCGGTATCAGATTGTAAACAATAGAGATCTGATATAGGGTATCTG
TTTATCGGTATTGccaaaattgtttgtttttaaaaaaaattaaaatattttaaattgtaaaaaattatta
taaacatatcaAAACTATTGTTTGGTAATATTGGTATTAGCGTATGGACCAGCATAGGTGCTTATTGAGCG
ATACTGGCGAAAGAGCCGATACGGACTTGTAAAAGTTTATCCGGCGGGAGTTGGTTGGGGCAAACgacaat
atttgaaatcttgcTAAAGTTTGGGCCAAAAACACCGACttcgaattatttttccaagtCTTGAGGGGT
TTAGAAGTTGATTTTAAACTCGCAAAAACTTTAGGGCCCACATCACTTGCTTCGAGTCaatgctaaaaaaa
actttggttacaaaaagatatagcatactaaattctttttttttattcatttttttttccaaacgaTCATC
TATCTTAGGTGTTTGGGGATCTGTACCGAATTACGGCACGGTTGAACCACCGAATATCACTCATCAGCGAA
TCTTGTCCTAGTTGATGAAGATATTTTAAGAGTTTGTCTAGGGTTTTTATAGCATATCGATGTTTGAGAGA
AAGGACATTTAGATTTAAGAAAATCTCCAAACTCATACACACCTGAATCCTACCAGATAGTTTGAACAGAC
TTGAAGAATTCTGCAATACAAGCATACTATAAAAGCAAAAGATAAATCCAGATATACAGAAACTAATCACT
AGGAGTTAACATAATATTCCATCCCACCATAATTAGATTACAAGATGGActcgaaataaaaaatgcaaaat
atcaAGACGTGTGAAAGGAATAGacaatattattgaaaagGAAATTTTACAAGTTTTCTGAATTTCCAAAC
TACGACAATCAAAATATAGGAAAGCAGAATCCAATAGTGACCACTtgagaaaaacaaagatacaTGCTAAA
CTAAGATGCCAAAGCAGTGGAGAAGATAACTTCAAGTATGTGGTTGTGGTAACTAGCAAATGACGCATGCT
AATTCTACATTTGATTAGAAACATGCCTGATCCTACTTACATTTTTCTTGTGTTAAGTCTAGAATTGACAA
TGAAAACTACAAAAGATACATGTTCAAATTCAGCAAACTCTCCAACGTAGAATTGTCCATATGTGAATGGT
AGCGAGAGAATAACCgcaaacaatattaatattaaaatccaGGAAAAACCAACAAGGAGAGGTGAAATGGG
GTAGGGAAGGCCTAAGAAGAATTGTTATAAAATCTAAACATTGACCTCATTTTGCTGATTTGAATGAAGAC
ATGTCTCAAAGTAGAGAACAATGCCATACAACTCAC**CTAGTGGGAAAGTCTTTGTGTTGTTTCTTAGTGTT**
**GCTGTAGAATGAATGACATGATGCATTTGATGATGGCTTTCAATC**CTAAAAAATGGAGGTAAACAAACCAA
ATAACCCTCTGTTATCAGCATGAAGTCGTCGATTGATCGGTACTACAAGACTAAGGAATGCAACCACCTAA
CCAATGCCAATGCGGAGCTCAAGGTATGTAGTATGTCAGTTTGTAATAAATCATTGGTCAATGTACTTTGT
TAATGCCATTTAAACTGTAAATCACTAGATAAAGCATAATTCTTGTGACTGCTACGTTACACTCACTTTGA
CTGATTATGAGTTGAATGAACTTGGATCAACATAGTCATGGATCTTTGTGTAAGAGCtttgtaaattaaaa
tgaataagaaacaagaaaaaggaCCAAAATGGAATAAAACAACATAGTAGGAATAGATAGAGAAAAAATGA
AGGTAGCAATTATATGTTAATCAAGCAAATGGTTGCcgtaaaattgtaattttgctATATTACTGTAGCAG
CTTGCTCAAATAATTTGAATCCATTCCCTAGCATTACTGTCCCCACATATTCTATATTCGAATCACATTAG
TTGAGTCGTCTATGAACATTTGGGAAGGGCTAAAACCCAAATAAAGCTAAAGGGGCGGCAAAGCAGGAGAT
TCGCAACTATCAtagaatataaagataaagCTACTACTAAAGCACATCAtctaaaagtttcatattactct
cctaataaATTTAACTGTTCAGCATTGAAAGTTCCATATGTTTGAATCACGTTGGAACGAATCTAGAAGTT
TATGCTTGTTAAGGACCACAGTTGcagttgaaacttgaaaattactaaaatagaaTGGATCGGAGAACTTA
TGGTGCAGAAATTATTGAACCAGCTAGCTGAAATGGTGGCTTGATATCTTAGACATATGCCATGTTACTAT
AGAACATCTCAAACACCCTGCTTTACTGATTAATGTAGAATATCGAAAAGAATGGAATGCCTGTAGTGATG
TGAACATTGTTGCTGTCTGGTTTAACATTGAAATAAAATCTGGTTGAGGCAGCACTTGGACATCAGACACG
TTTGGAAGACGCCAAGATTGGTTGAAAGTTTAAACCCACTGACCTATTTGTCACAAGACATCATGGCAAAT
TGTTTACTACAATTAGACATGGGacctaaattaaaattttgccaCTAGATAGCTGAACACTTccagagaga
gaaaaaaggagCAAGATTGTAAAAACGCAGCTTGGCAAATTATTCATTGATACTTTGCCTATGCTTGCACA
TGCTTCTTACCAGACAATTATACATGGAATGAAAAAGATGGCTATTATTGCACAgaatgaaacaaaaagaa
gaagcattTTTGCCATAATTCAAATAGCAAGTTACTCGTTGGATTTTACATACGCTTGCATATAAAGCCGG
TATCATACTTTATAAGCAAGAAGTAATGAAACTCTCAC**ATTATTCCAACCGGAAAGAGAACTAGCAGAGAA**
**TTCAGGCATATCTTCACAAGGAGGGGCATATCTGCATTTGAAATTCTGCTTAGAGACACAAGCTATCAAAT**
**TTCTGCACCATAACTAACATGAttagaagaaaacaacaataCTGAAATTGAACAATCAAAAAGATGCTATG**
**AACTTCCCCTCAGTGGCATGTTGTTAGATGTCCTGCATGGACATAGAATTTCACATTAATGTGCGGCTGCA**
**TC**CTGCATGGACACAGAATGTGTCACATCAATGGAGTATTCTTTGTAAAAGAGCTTACTAAAGGTAGAAAA
ATGTTGCTTAC**AATACTTTGCTTCCTGAGCTTAGTTAACTGCTTGATTAGCTCGCACATACTTATCTATTA**
**AACCTTGATGGTTTTGCAATCCAGTTTTGGCAAGGTGAGGCAGCAAGCTTGAGGCAACAACTGCATAACTT**
**ACAAGAAAGTCATTGGTATTGCACTTTGACC**CTgcaatagattttattaactatCAAATTTTGAGTATATA
TCGAACTTTTTACAAAGAAACAGGAGAAAAGCATGTATAAGTGCCTGCTATGGAGTcaatactttctaatt
attggTTCAGTATAGGAAATATGTTTGGACTTGTAACTATAGTCATGATGCGCAACACTTGAAACTCAAAG
CTAAAAGTGCTTCGCATCACAAAATGTTATGTTTGTCAAATACGCGAAGATGCATCTAGAATGATTTACAA
GTTACtttcagaattatttattagtacttatattttaataacttttcttGACCTTATAGCTAAATCAGTAA

TCACAATTTCACCATCTGAGGAAAAAGATCAAGTACAATATAGTCTAATTCCTGTTACTATGATAATAGTC
AGATATaacatcaattaaaaaaaaaaaaaattaacagattatgcttgaattacaatatatatataaagaga
gagagagagagagagagagagagagagagggaagcGGTACCAATCCTCGTTTAGAATGTCATAAGTCAAAG
TTGAATTGCCTAATCCACTCCAGGATGAGATTGATCATATCTTAacctcccccctctctctctctctctc
tctctctctctctctctctctctctctcttctctctttctctgtattatatatatatatatatatattata
tatatatatatatatatatatattatatataatatatatatatatattatatatcttatatatataatg
tgtgtgtgtgtgcacgCATGTGTGCGCACacatcattcaaattatttggaTTCCAATTCCTACTAAATTTT
GTACTTAGAAGAGTAGATGGATTCAACAGAGCTTTGGGAGTCATTTTAGGAACTTTAACCAAAATACTTTT
ACTGGTAATTCTACCTAACTAGGATAAGTACACTGatattcaacatcaaaaatCATTTTGGCATGTGATTC
ATTATGAACCATCTATCTGCTGCGATGTCAAAGGGAGTGCTCTTGTAAGAACTAAGATTTAATGAAACGAT
TCATAGGTAGATTGATCATTGTTTTAGACTTGCTTTAGTTGGTGGAACCATAATTTTGCTAATAGTGGTAC
ACTATTACGTACTAGCTATAGTAGCTTTAATGAAATGAATGTCTAAAGATAATGTCGGGTTGGAGAAATTA
TCTGTTGTatagattttttgttgttgattaATACAATCCCACTGGAAGTGGCTATATGAAAGGAGTTTCAAA
AAAGTTCGGGATATGTTGCCGTCTTACTCCTAGTTACCCATTAgcatataaatagttttaaaattagaagt
aaAAAGTTCCAATTGTCTAAAAGGATCAGTTAGCAGCAAAAAGACTAGCCATGTGCAAATCCATATAATAG
CAacaatgtgaaaaaaaaaaaatgtaactagATAAAGAAAGCCAAAGAAGACGCAATCGCCAAAACCAGAAG
TTTTGAAGGAGTTGAAAGGACTCATTCACACTGATGATTAGTTTTGCCTCAAGTTTACTCTTAAATAGGGA
CTTGGCCATAGAACTAATGAATCCATCGGATATCCTATCTAGAGCATGGTTATATATGCCGGGCAGCACGT
GCAGTTATCGGCAGTTATCGGCAAAGCACGAACCAGCACAACATGCTAaagaatcttatatttgaatttttt
aaattaattttgcatttCGTATTTTCgtttttggaaaacaaaaaataaatgttggtGCGTTTAGCAAGGCA
CCATTTGCACCGTCGGCATGCACTAGGATCTGAGTTACTGAAACTCAGCTCAGCTCggcattaaaaatttg
atctaaAGTGATTTAAACTAATCTCTGAAGAAGTATCTAGCTTCACTAAGTATTTGCAGCCACCTGCTGGG
AGAGGAGCTAGCTTCATTAAGTATCAATGATCTTCACAACATTGAGAGACAGCTAGAGACAAGCCTACGTG
GTGTCCGAGCAAAGAAGGTACTAAAAGTTTTGATCAATCTCAAGTTAAACATGTTATCATCGTTTAGTATG
TAATATCATAAATGCACATCTGAAACTTGTAGGAACAACTTTTGATGGGAGAAATTGAAGAACTTAGAAAG
CAGGTCTTTCAGATAATCTCATCTTCTTTTCACCAAAAACTATTGAAACTAATCATTACTCACCTAAACTT
CATTTGTCATATTACAGGGTAATATTGTGCAGCAACAAAATGCAGAACTGTACAGCAAGGTAAAGTCTGAT
TCAGAGGTAATTTATACGAAAAGTCTAAAATCGCGAGTTAGAATTACTCAAATATACTTTGAACGAGGCAG
ATATAAAGTCGTAAACATGATGCGCATTAGCACAATATTCACGAGCAACTAACAAATCATATCGTGCTTGC
AGGTTTATGCAACAAAGGGGTTAAATGAAATCAATCAAGGATTGGCTATTCCATTTAACTTCTGTTTGTCA
GAGAACTTAAATACTCCTATTCTTATCGACATTAGTCAGCCACAAAAGCAACTTGAGCAAACTCCAGATGG
GGCTCTAAATCTAGGGTAAAGTTTGTCTGGATTCAAGTCACAAATTCCATCACTGCTAGTTATAAATGTGT
CCCTTTACGTTGCAGCAGACTCCAACTGCACTGATCGATGGCTGTTACATTGCAACAGACTTCAACTGCAC
TGATCGATGGGTAACAACAAGTGCCATCATGTATCTTCTCGAAAGCGAGCATCAATGCTAGATATGCTTGC
TAGTATTTGCAACTAGTTGTTTATATGCATGACCAGGATGATTAGAAAAACTGGCAGCATCCATAAAATTT
GATCACAATAACCATAAAGTAGATTTATCTTGTATTATAGAATAAAACAGCGGTGGTCATGAGCCCTATAA
CTACCGTTGTATgtatattatagaataaaacAGTGGTGGTCATGAGCCCTATAACTACCGTTGTATGTATC
TGCATGGAGCATGATGCTTTCCTCGTGTAACCAAAAAATTGAGACCGTTTTTCGTCTTTTACGAAATTATG
GAAGTATAGGAGCTCGTAACAAAAGGAACATGATCTGCGTTCAGTTTAAGGACCTGTAAATGTCATAATAG
CCTATCAAATAATGTTTGATCTGTTTCGAATATTAGCTTGAGTTGGTGAAGTCACTGCTATAGAGAATTCA
ATTGGAATGAACAAAAGGCGCTTTGGGGTGAATGCGTAGTAGCCTTCAAATTGCTACAAATTCTTTTCCAA
CATCGAGATCAACTGGAAATTGGACAAGGAATCCTAAACAcctttctgttttttatttttgcaaatatacACA
GCTTTGCATCTAGTTTCATTCAGCCCCTTCTTCACGCCTTACTAGAAGCATTCCTTTTAACTGTAGTTTCA
CAGAAATCTTTAGATCACATAGATTACCCTACTTCGGTGCCAACATTACTAGAAATTGTTTCATAGCCTAC
AATCCATGTGGCggatttcttttgttttagagTTTGTAGTTAAGCCCACCCAATGATAAATATCGACAATT
CAATTtgcttcaaaaacaaaaatgaaaccCAAGTATTGCACGATATTTTAGTTGTCTTCTATTTTTGGAAA
CATAGCATGCATCTATAATTGGCAAGTCCATAATCTTACTTGCATAAATATAATGCAAAATTACAGTATTG
GATTACTGTTAATTGTTATGATGTATAAGTTCTGAAGATTTTCTGGGTTTATGATATTTAGTGAAGGATTA
TTGCTTAGACATGTTGACTTTGTACAAGTacatgatattaaaaaaaaaaaaaaaagaaccttaTCATTTAGG
GTCAAAGAAAGGCAAAGTGTCAACTGTATGTTTTGGTTTCCTATTCAATTAGTACTAACTTTACCCATTTC
CACAGCAGCTCGTGAAATATCTTTTCCGGCAGCAAACTAGTAAAGAtgtcattcaaattatttatagaaat
caTTTGGTTTTTAGCCCACCCACTTCACCAAAGATTCTTGCTTTAAgccaatatcaaaatatgaataataa
tcaTATCACGGTATCTAAGGATCTTCTTGGCCTCCCTACTGTCCGTCATATAGCTTGCCCCACTACAGTTG
CAACTCCATTGATGTTCCATCTAACTACGAATCAACCGCTTCCGGACTTAAATTAGTTCCATCTTTTCGCA
AGCCCAACCATGATGGATaggttttaacaaaataagacAGTTAGCACTTTTCCAAGTATATGTAGGTAAAA
TCCTATTCCACTACCCTCATAAATACCAAACATGGTATATGCCTCTATTGTTCAATTCATTATAATTGTTG
TCCAAATTACCGTTTTATTTTACTGAACTTTCTTAGCTACGcccattcaaatttttgaccCTTGatcattc
aaaaatataaaactcaaTATGTGTCAAGGGTCTATTTGGATAAGGTTTAACGAAGGGCATTTTATAAGGAG
AATTTGGAAGTATCCGTTCCAGTTTTGCTGTTTGGATGTCCATATATTCCACTCCTTCAAAATACTTCTTC
TAAATAACATCTACTCGGGTGTTATTTGTAaaatcttctttaaaaaatcctCTCCCAAAATTCTCATCTCT

CAATCACCAACTTTCCTTTATGTGTTGTTCAGGGCAGCACGGAGGCTCGGTGCCGCCCAGCACATGCCAGG
TCTGTAGGGATTCAGCACTGAGCTGGTGCTGTTCTGGCCAGCACGGAGGCTAGGTGCTGCTTGGTACTGTT
TCGGCTGGCACGGAGACTCGGTGCCATCCGAGACAAGCAGCTCGGCAGCACCAAGCCtgataagagagaga
gagagagagagagagaagaggcactgccgccgccgctgccgagATCTTAGcccatcctctctctctcct
ttctttCTCAATCCTTTCATGTCTTTTTTCGCTCCAGGATAGAGCTGGGGACAAAAGCTGCCAACAACTAC
CCTCCACCTTAgggctaaaattttttttagttttaaaagtttttagttttttttaataaataaattta
aaataaaaagtacagATCGGCACGCTCAGCACATCTCGAAGAAGTCACCGAAACTCGGCAAGCACGGCACA
AAAAAACATGTGTTGAATGGAAAAATCTCAGCTCCTAACACCAGCCGAGTTCTTTGTATGTGCTTGTTGTC
GAATCTCATAGGACATTGTCGCAGACGGTCTCACCGatcatcatcattatcatCATTTATTCTTGCCAGAC
ATCATCATCCATTGCAATTCTCACCAAACGGCGTCGTCATTATATGTAATGTTAATCGTGTCgaaccccccc
cccccactttgatctttttcattttaatatttttatctcataGATAAGTTCTATCAATATAAGTCAATGTAA
AGAAGATTTGATTGTTTAGAcctaaatatttctttcatctaagaattttttttaagaatcatTTTCAGCTC
TTCTGTGTGCTTTGATGGTAAATGGATATACTGGTCACATTCTTTGTAATGCTATTGTgtcaattaatttt
tgttacatCCACTGTatacttggtttttttttgggtattggATCGAatgtttttgttcaatttcCCATTAG
AATAGTAACACTAGTAGTATATTTTTGTCTAatcatttatgtttgttttggttctgatcccttttttttgt
catctAGTTTAAAATTGTCTCTATTGTTACAACattcaaatatagaaattCAAACACTTCATCGAATTACG
AAGTGAAAATGtgtaattaaaaagaagaggaTTTGGAAATCAAGCATTTGCAAGTCAATGCATTTATGAAT
ACTAAGTTTGAAAGTTCTAAAGTACCCTGAGCAAATGATTTTAGTTAGAATTTGTTAAAGATAAAACGTTT
CCCTCATGGGAATTGACAAGCAATGATAACAAGTACAAAACCTTATCACTACTACTATACGCAAGCTCATG
CATTTTTTGAACAAGGATGGAACTATAGCTCAATTGTATGATATCCCTTCGATAATTATAGTGCAGAGAAC
ACTGCCTGACAATAAATCCTACCcttctgttttaatttatcaaatactaaaccaaggttttccgtgccagA
TCCGTGAAGCTCAGACGCCggcacggtgccgagccgcCGTGCCGCCCGTGCCAACTGTGCGACCGGCATGG
TCAAGTTGTGCCAAgccgtgccggtcggcacataatttttttatttttttaatgttttgctTTGGGTTTTAC
ATATCCCTTCTATAAGCTATATTGAtactctatataataataaactatatactatatatttttactatgta
ATTACGACTTTtgaatatagttttataaattttttttattattgcttaaataatttagtatttcaaaaaaa
aaattatacagatcGGCgatgtcggcacggcatggtCGGCATGTCTTGAGTTTCTGATTTACCGAAACGTG
GCtaggcacagaaaaccttgtactaaaataaaaatagtgtaCTCTATTATGTACTACAAGATAATGAACAC
TCCCTCCCGATTGTAGCTCCTaatcttctatttttatttttttttatttttttggtcaaataaCGAACAACCA
TGGGGTAATTCATTCTATTCCAAAAACTGgagtttttttcgaaaatatataaaaaaccttTCTTTGCTGGCCT
TTGAGGTTGTTCGTTCAAAACTGCAATAGCGTTAGGTTATATCCaacttaatatttttggtatttttttaaa
atatttttgatcaaTGAAGAACTAAGAAAATGGTTTATCtggttctacaacttttttaattcaacttgttt
taagaaaataaaaaaggtatcCAATTGTCTTTCGAAATGGTGGAGCTGCTTCCATATCTTACgtcaaataa
actaaatatggtgtattatatgttattccACAGATTTTTAGTTTAACGGTTTTTGATAAACAAATATTGTA
GAATGTGAAGTTCGTAGGGTGAACTGTTAATAACAGAAATAATGTTTTGTTACTTTATCACTAATGTTTTA
ATACTTCTTTCAATTCTGATTAAATGATGAACTAAAATGTTGACAGTTTGTTTTAACCACAACATTATGCA
AAGAGGCCTTCATAATCCTAGAGTTCCAAAAGATCCTATGACAACCCTCAACTTCTAAAAAGGACCTCCGC
TAGCCTTGTAGTCagatttttttgcttttacaCAGTTGGAACTATTGATGTAGCTATAACGTGGCAATCATT
TGGTaaagagcaaaataattaaaaagtaatacatTTTTACCTAATTATTGTCACGTCAATAGTTGAAAAGG
CATTAAGGCAAAATGGTTGTAATGTAACTGAAggttcttttcaaaatttgaggTTTATCATGAGaactttt
aaaacttcaagATATAAGGGTCCATTTATATAAAGTTCAGGGGTCAAAACATACTTGTacctatatttatt
atctaattgTTCAATGTTTTTAGTAAAGAATAATTAGAAGTTACAGATTTGTTAGTTATGTTCTCTAGGA
GTAGATTGGTTGTTCTAGCCGCCATTGGGCCATTGGTATTAATCTTGCCGCTAAAAGAAGCCAATAATACT
TGTTTCAtgaaaattggtatttttttgagCAGTTTGTCAATGCCCTCGCACTGTTCACCATCTCAACGCCAA
GGATCATTGCTCGGTGAATAGTTACATACCAAAAGAAACACTATATACCTTTTTGTGAAATGCTCTTTCTA
CATACCAAAAGAAACACTTCTGGTTCGGAGAGGGAGAACAATGCTTTGCTTATGGAGACAAGCAATGGAGG
GTATCTTTGCCCAAGGATGAGACTCAGATTAATCGTAGGGAGAATATGGACACAAGAATAGTAAAACACAC
GAATTTACGTGGTTTGGTCAAGTGCCTACGTCAATGGGCAGTCACAAAGAAGTATTCGCACTAAGTAATGA
AGAGTAAAGAGAGTTGATTgcaaggttttacgtgccgggcggcacacgcggtgccgagccggcacaggccc
tccctctctctttctctccctccctccctttctctctcttgatctcgatttctctctctctctctctctct
ctctctctctctctctctctcttctctctctcgtctctctctctctctctctctatctcaaTCTCTTCGgg
gtttttgtaaaatttgggtgtcttcctcctcgcatTCCGAGGAGGACGCAGGTCGCCTGCggcgacctttt
tttttttaaagcgcCGTGTGTTCGGCGTGAccccatttctttttttaaaaaaaaaaattaaatacctttaa
aaaattaaaaaaatcttatataatatataataaataatataaataataatatttataatcggcacgtcttg
aatttaaatttaccgaATATCCGACACGAAATACATCCGTACTATCTgttttagatatgtaattaaagttt
ttttattttttttttattttttttatttcaatccataattggtttattattatttacataatttaatatttt
aaatataataaatttatacggatcggcacggtcggcacgtgccgagttttggatttaccaaTACTCGACTc
agcacggaaaaccttggttgatTGTGCCCCTAGCTGCTCAATTACATTCTTTTATAATCCTCTCATCTCCC
CTAACTAACCATCTTCGTTTTGTAGAGAATTATTAGAAAACTACTAGATAATAGTGACTATGTAATGGGAT
TGACTTGCACGTTAGTGTGCTCCTCTTTGGCTTCAAATCAGCTGACCAAACTTTTTAGGATGTGCAACACA
TGAGCCTCATATCATGCTCGGGCTCGTCTGTGCACATGGTCCCGCACGGATGGACGTAGTTGAATGTGACG

CTTGCTCTTCCAAGGTTAGGTGCTTCACATTGCTCCTTTAAGGGGTCCGGAGTTGGATATTGCAGCATCCA
AGGTCCACATAAATGGTTCAGGACTAGAGGTGGACGTTGCTTCAGCCGGAGTCGGTATGGAAAGTTCTCAG
TCGAACACTTAACATTGTTCTGTCTGCGGTTGGTATGCACAGTTCCTGTTCAGGCACTTGATATTGCTCCG
ACCGGAGTCAACATGGGTAGTGCTGGGTTGGGCGTTGGTCGGTATAGGTAGTCTGAGCGGGGAGACATTCC
AATCAAGGTTAGGGTCACGCACCAGTTTCTCCGGACGAAGTACGTTCACATAGCCACCTAGACCAGACATA
TATCCCACGTACCATCACACTAGCAACTGCCAGTAGTAGCCTTTCGGCCAATAGACGACTACGAGCACCAT
AGCCAAAGATGGTACAAAGCCCAGTAGGACAATAGTAGGAACCAGGATTATTGGTTGAAAATAAACTGAGA
CTTAATTCCATGTGGTGGGTTGcatgttgaaaatataattttgaatactgGTTAGATGGATGAATCATTTC
CAAACATGCAGCCAGTCCTATttcatttaacaaaaaatgtgATTTCACCAATATTTGAAGTTCTAATTGGT
AGTTCGTGTAttagatcacaacaaaatagggataatatgtagattatttattttgatttcaatATTCTTAT
TCTcataattcaatttaaaaatgccaatgaataatttaacaaatcgcatattctttttttataaatcagcag
AAATATATGGCATAAGTTCAACTATTACGTGATAATCAAAGAAGAGAACAATACATCAAATAAAGGATTAG
TAC**CATTGACTCTCTTGCAAGTTATGCAATTGCTGCCTCGACCTGGTGCATCCATCTACAT**ATGAAGAAGT
AACAAGGAATAAAACACTACCAATTTACAATGAATCgaaagattataaaatttccaGAATATAAAGTACAC
GTTAGGACCTTTAGACTTCACGGTGCAATAATAACAAAGTCTTCTCTCACAAAGTTTGTTTGGCTATTTGG
ACACTTGTTCGCCACTATGCTACCAAGTCTTCAAAAtgccttattttaaaaatatttaaagaatggCAAGG
TccaaaatgccacatggcatccACCTAAGCAGATCAGTCCACTATAGAAACTTGACTCTAGCAAATGACAT
ACTACAATGAAGGCCAAATATGGACCTGTTAGCGTGGTATCTTCGCATTTAGTTTAGAGATGGTGCACATA
TTGAAATTATGAGAAAGGTTTAAGGAGAAATGTCAACTAAAGAAAAGAGACAAGATATAGGTGGAAAAATC
CTAAAAAGGTTGGAGAAAAAAACCACAGATTCAAGGAGGCACcttgtaaaaaatttcactatATAATGTTG
TAGTGCTACAATATTTTCTTGTCCTTtccctacattttttttctaaaaagtgaGAGTTAATGAGAATTTTG
TCCTTTTGTACAGCAAATGATTGTCCTATTTATAGGAGTTTAGCAAGATATAGTAGAATTTAGTAATACAT
AAATTGACTAAATACTAAATTCCTCTATAAAGGGTGCGAGTGGACCTTGATGTGAACTATGAATAGTTGTA
TCGGAACGGATTGCTTTGACGTGGATATGTACGAGGCAAGGCTCGTAAAGAGGGGTCAATGAAGGGCGGCC
GATTAAAGCCAGCTCTAGGGCTGGCATTAATCATGACTAAAGCCAGCCCTAGGGCTGGCTTTGCATGCATA
GtcgaattagtttttttaaaaatttttaataaaaattaaaaaaacttttaaaagccTATAAATAATGATGAGT
TTCTAAAAGTTTCTCATTCAAATCACTTCATACTCAAAGTCtcaaattagttcaaaagttCAACTATTTAA
CTCGATGGCACAACCCAAAAGTCAATCAATCAATAATATTAGTTGGCAACATAGTACCCTTGTAGAAGGCA
ATatgttttattgaaatttcCACAAAAAAATGGAGGTGGAGTTTCTCGACGAAAACAATATTTAGCCGGAGG
TTTTCCAGATGTTAGATCTTGCGATAAATGCGGTGTTAAGATTCGTGACACTGTGAAAGCGCACCTATGTT
CGGGGAAGACCGCAAGAGAGCATAAGCAAAAGCAATGATAGTACATTGATTGTATGGTTGCGAAGGAGGAT
TATATTGCTGACGAGGGCCGTCCTACAGAAGAATAATACGATGATTTTGACTTACGAGATCCAAGAACACC
GCCAAGAGAAATAGAGGAACTAAAATAAGCAAGGGCTGAAATTTTAAGAGAAACGGAGATTAAGGAGCAAA
GAGACGGCTTCGCACAAAGCTCGTCTTACGCTCCTAGTAGTTCTTCAGGTTCTTCGAGCCAATGAGGGAGA
AGTAAAAGTGGTTGGACGTCATTCTTCGGTAGATCGGTATCCATGAGGACATTCGATATTAGACAGGTTAA
CCCACATGCATCTAAGCCAGTGGACTTCCAACAACCCATAATTAATGAATTGGAAGAGGGGGAAGGAAGTC
CAAATTGGCCAAGTCGATTGGCAAATTGTTCCATTTCAATCACAAAGCTCCTAACGTTGCGTCAAGTCCAT
CTTATTGCAATATGATCTCAATGGCAGAGGAGGTAGGAGAGGGTATTCAACCTCTCAAGCCATGGGAGATT
GTTGGATCCCTCTTCgatgaaaatgagaaagagCTACGCATCTACATCAATAGTTTGAAGAATCAATGGAA
AGAGTATGGTGTTACGATTACGTCTCATGGATAGACAAGTCCAACAAGATTGAAAATtatcaacttttaat
aaactaCGACAGAAAGACAATGTTTCACGACTCTATAGATGCTACCCTGTACGTTTAAGACCATGTCTACA
TAAAGTTgaagatagaaaaaattattgctGATGTTGGCATGAAGAATGTGGTGCAAATTGTAACTAACAAT
GGTTCCAATTATAGACGTACATGAATATCGATGATGGAGGCCTACCCACATTTATATTGGACAACTTGTGC
AGTTTAGTCTATCGACTTGATGATGGCCAATAGTGGTAAACTCAAACATGTGAGCAAGTTGGTTCGAAAGG
GGCAGACTATCaccaaatttgtatataaccACGGACCACTTCTGGCACAGATGAGAGCAAATCGAAGGCAA
TCGACTGAGACCGGGCATCACCCGATTTGCTACCAATTTTATGTCTCTAAATAGTATCAAGGAGAAAAAAA
GTGGTACTAAAAGCTCTATTTACAAGAGAGTAGTCGTCTGAATCTAAATATCGATTGACGGACCACGGAAG
AAAGGTGGATAGCATCATACTCGACAGGaggttttggaaaaatataattgtattcaTTGACATTATCGCCC
CTCTTTATGATGTACTTAGAAAAGTTGACACCGAAGATCTTCCACATATGGGATACATCTACCACTCGATA
AATAATGCAAAGAAGCAGATCCAAGAAAATAACCAAAAGGGATATAAtgactattataatataattgacAG
GTGGAATCACCAGATAGGCCACGACATTCATCTTGTAGGTTATATCTCAATTCTCATTTCCAGTACCAGCA
TAATCTGAGAACGAAGAATGACCTTATTATGGTGCTACGGAATGTTATTATCCGATTAGAGCCAAATAAAG
CCGTACAAGCACATGCTATCGAGAAGGTATAACATTGTATAGCTATTAACTTAATTAGCATTAATCTTTAA
GcataactattaaattaattagcatTGGGCtttaatgctaattttttttctaaataattattacatgGCAA
GCTATTTAGATATGTTAAAGGctcatttgaaaaatatacagCGAAGATGTCAAGATTAAATATGGTGCCTT
GTAAGATTGTTGtcatacttaaaaaaaaaaaactatgacaACATAGGACCGATCAATTTTTGAGGACCGAT
CAATGTTTGAGGATCCTAGTGatacacatataatataaaaaaaaaaaaaaaaaaaaaaaaaaaaaaaaaaa
aaagtaatgaagGTATGATGGATAGCAAACATCACGAGCAATAGATCATAACCACACAGACcttcaatttt
gaaagaagCTACAAAAAAGTGAAACATGTAGGCCCGCTTCAAATCATTGTTGAATCACATTTGTATTTCCT
AAGTAAGATTATCTACTTTCAAGTTCGAAGTCTTTTTATGCTAATTGCACAGGAGCTAATGATCTTATAGA

CCTCCTTTCGTTCACATGAAGCGAAAGCATCATCACTACTAGTTCCTTGAAACCTAAATATTGTTCTAGGT
TCCTTTATATGCCCCCGATCCAAATATTCGGATCAAATAATCAAGTAAAAGAAACATACTATCAGATAACC
AAAACTTGCAAAGCAATGTTGTCGAGCATTCTACAGAACAACGAATAAACATAAGCAAAGTATATCCAACg
agaaacaattttaaaattgaaggaTATCAATGTGAtaagaaaacgaaaaaaaaaaaaaaagaagattgcaA
TCAC**CTAAAATCTGGTTTCAGAACTCTTCCCCTGGTTGTTTATTCTCCAGAGAAAGGCCGCTTACAAATAC**
**TTGCAGATGTAGTAGATGAAGACGGAATCTTgactaaaattataaagaaacgGGGACAAGACGACGACAAT**
**CTGGGAACTTTCCTCAATTCAATGGATCAAATCAGAGGCAAGAACAGAGAGAGATTGAAAACATAGC**CAGA
TTAAAATACCGCCGCAATGcggcatatttttatttacattcaaATACGAGCGTACGCACGTGTCGAGTGCG
TGCTCAACGCGGCCAACAAAGATCAGGCAGCCGGTGGGGCCCAGAAAGGATCCCGTTAAAATGGAAAGGTC
CGAGCGTAGCCAAGTTGGTCCGGGTGCAGCCGCTGCTGCAGCTAGAGACTAGATGAGGTTCAGGTGCAATG
TATTGACCCTGCTTCCGGGAATAAAGAACCCCCCACTTGGGAAGAGTTGGTAATGGCCGCTTCAGAGGAGC
AACGTATACCTGTCCTTTCGTAACATTAGagctttcaaaatttcttcacaAGTTCACACAAAACAACCAAG
TTTATTTCCATATGGTGCAACTAGAACGCCTTCATGATCGaacaattttaatgaaatgaactgtattaatg
aaataattaattaattttattttaactacaaatatatatatatatatatatatatatatatgaattttttct
gaaataatcatgtttagtaaaaattatttacagaAATATGTGTGGTATTTACAAGCGTCATGGAAAGATAT
GTAGTAAATAACTGAGCTAGAGAAATAATCGTGAAGGTGTGAATCGTCAAGAAAATAGCATATACAATTAG
GgagtcaaaagaaaatatttgggtCAATACTGAACTGTAGATCAATTTTTGTGATGGTGTTATTATGTGTG
GATTATGTGTATAGCCATAGGAATATTTACAGTAAAGATATGAGAGAAGTTTAAGTTTGCCTGgaggcaat
ttttttttttttctggtatccgaCCCCTTTCTAagtggcagtccagaccgagCGATATAAATGCTAATTTA
GGTATTGTGACACCTCCTTCCAAAGAATTGTGTTGGggctcaaactcgagacctTTCCCTGTCTAGCCGC
ACTGATTTGCCTAGAGGCAATAGCTTATttacatatctatataatatacatgtatataatcTATAGTtgtc
gtgtgtgtgtgtgtgtgtgtgtgtgtgtgtgtgtagtaCAAGTAACAAGGAGACACCTAGATGGCTAGATGTG
TGATCCTTATAAGTTAAGGGCCTATGAGCATATTTCATTCACATATCGATTTATCGAGGATCAAATCATTC
CACCGAGTATGAGTGGCAAATTCTATCTTTGGGATACCTTATAGTTTcctacttattattttattatgatt
ggATCTGTATTGGTAGGAATAATCATGCTTTCAGTATcgctaaaaaatttatgctttcagtataaatttt

>XviAGL17_3
ATGGGAAGGGGGAAGACTGTGATCAAGAGGATTGACAATTCGACGAGTCGCCAGGTGACATTCTCGAAAAGAAGG
AGCGGGTTGCTGAAGAAGGCACGGGAGCTCGGCATACTCTGCGATGCGGAGGTGGGTGTTATCGTCTTTTCCAGC
ACCGGCAGGCTCTATGAATTCGCCAGCACCAGCATGAAGTCGTCGATTGATCGGTACTACAAGACTAAGGAATGC
AACCACCTAACCAATGCCAATGCGGAGCTCAAGTTTTGGCAAGGTGAGGCAGCAAGCTTGAGGCAACAACTGCAT
AACTTACAAGAAAGTCATTGCCACCTGCTGGGAGAGGAGCTAGCTTCATTAAGTATCAATGATCTTCACAACATT
GAGAGACAGCTAGAGACAAGCCTACGTGGTGTCCGAGCAAAGAAGGAACAACTTTTGATGGGAGAAATTGAAGAA
CTTAGAAAGCAGGGTAATATTGTGCAGCAACAAAATGCAGAACTGTACAGCAAGGTAAAGTCTGATTCAGAGGTT
TATGCAACAAAGGGGTTAAATGAAATCAATCAAGGATTGGCTATTCCATTTAACTTCTGTTTGTCAGAGAACTTA
AATACTCCTATTCTTATCGACATTAGTCAGCCACAAAAGCAACTTGAGCAAACTCCAGATGGGGCTCTAAATCTA
GGGTAA

>xvi_miR444c
GCTATGTTTTCAATCTCTCTCTGTTCTTGCCTCTGATTTGATCCATTGAATTGAGGAAAGTTCCCAGATTG
TCGTCGTCTTGTCCCCGTTTCTTTATAATTTTAGTCAAGATTCCGTCTTCATCTACTACATCTGCAAGTAT
TTGTAAGCGGTCCTTTCTCTGGAGAATAAACAACCAGGGGAAGAGTTCTGAAACCGGATTTTAGATGGATG
CACCAGGTCGAGGCAGCAATTGCATAACTTGCAAGAGAGTCAAWGGGTCAAAGTGCAATACCAATGACTTT
CTTGTAAGTTATGCAGTTGTTGCCTCAAGCTTGCTGCCTCACCTTGCCAAAACTGGATTGCAAAACCATCA
AGGTTTAATAGATAAGTATGTGCGAGCTAATCAAGCAGTTAAGTAAGCTCAGGAAGCAAAGTATTGATGCA
GCCGCACATTAATGTGAAATTCTATGTCCATGCAGGACATCTAACAACATTCCACTGAGGGGAAGTTCATA
GCATCTTTTTGATTGTTCAATCTCAGTATTGTTGTTTTCTTCTAATCATGTTAGTTATGGTGCAGAAATTT
GAATAGCTTGTGTCTCTAAGCAGAATTTCAAATGCACATATGCCCCTCCTTGTGAAGATATGCCTGAATTC
TCTGCTAGTTCTCTTTCTGGTTGGCATAATGATTGAAAGCCATCATCAAATGCATCATGTCATTCATTCTA
CAGCAACACTAAGAAACAACACAAAGACTTTCCCACTAGGACATGTGAATTGTAACTGCCCAGTCAGATTG
GTGCATGGTTGGAGATTAAGTGTATCATTTTCTTATTCTGGTTGCATTCAAGAATCCTGATTCAGATTCTC
ATCATACCTTCACACCAAGTGACGCACAATGAATCTTGCCTCAGCAACTTCTTTAGTGCAACGTGACGTAT
GGGCACAATGGTTCATGCAATGGACGGGCGGTCAGAAAGTCAAAAAATTTGGAGGTCAAAGTGCGTTCGTA
TGTGAGATTGTGGGAAGGCACCGGTCGAGAGTGCCTATACGTATAGCAAGAGAGGAAAGACTATATTTCCT
TTTGAGGTATAATGCTGGGGTTAAGTGTAACTTGTAACCCTTTATGTATACCTAGTTCTATATGGCCAACT
TCATGTAGTGTAAAGGAATATAAAGGAAGCACTAACTCATATTTTGTTGTATGCATGTATATGAGCCCTTA
TCTAAGTGGTTATGCTTGTTTGAAACGCATGCTTTAGATGGAAATGCAAAAGTTGGC

Position of mature miRNAs based on comparison with osa-MIR444d and not based on experimental data!

Stem-loop structure

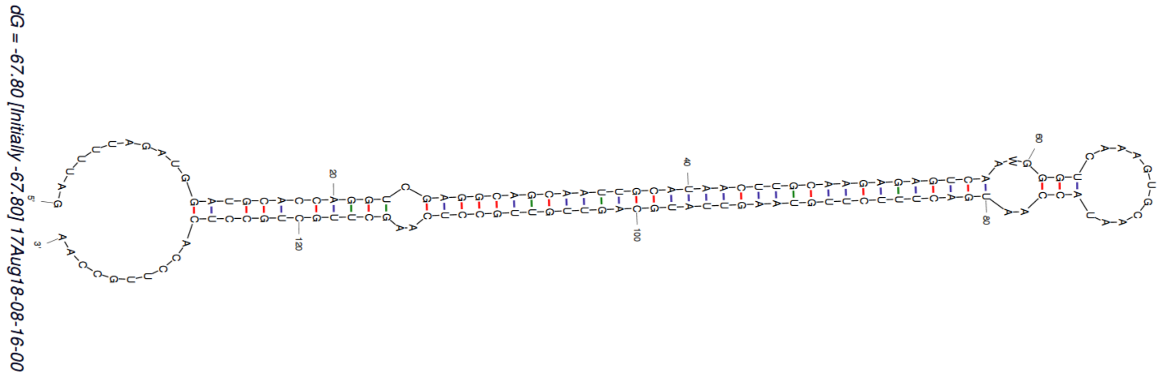


*XviAGL17_4 and xvi-miR444d*

>MJHO01000134.1

AAGCTCAACTGACCATAAATAGAGATTTGACCAAATTAGAAAAGGCCTAAAAAGagagattttaaaagtta
tagataaaaaattatttagaatacatgataaataacgcagagaaacaaaaaaaaatatatttagaatat
atgataaaaacactaattttaatttattaaattaatttttttggcacaCGTATTGATATATTGTCCTAAGT
GGAAATAATTACCTAAAcaaatgaatataattaaattaagacAAAAGCCTTGCGGGCTTTCATCAGTTTTT
CGACCCTGCAATGCTTTCTTTGTTCTGGAAGTTTCCCCACAAAGGTTTCCCAGACGTCGATGGTAAAAGGA
CCTCTGTGGTCCCCACTTCCGTCACCTGAAACCCAAAAGCGGAATCTCATACGGACGTGTCAGGCCCTCGC
ACGCGTCGGCCATCGCGACGGCGAGGACGAAAAGATTCGTTTTTAAGTCCAGAACCAGCGACCCTGCGAAA
CCTAAGCACGATAGTAATAGAGTAATATTGGAAGTTTGAAGAAGTAAAATGaaagtggagaagaagatgtc
gctaaaaatataaaacaaaagaaaaagataaggtAGATGAGGTTGATCTGTCGCTCCCCTAGGAAGTAGGA
GAAAGGAAACCGACAAGGTTTTCCTATTGGCTGCTCATTGATGATATATTGATCAGCTAGCTGTCGCAGAT
CTGAAAAAAGACAGCCGCCGACGAATTTCAGGTTCTcggatttatttttcttttattctgtGTTTTTGTGA
TCCTCTTCTGATCTTTGTTAGCTTGTACTGATTTGTTTAGTTTAGATTTATTGATGTTTGGAGTATAATCA
GTCAGGCGGAGAGATGGGAAGGGGGAAGACTGTGATCAAGAGGATTGACAATTCGACGAGTCGCCAGGTGA
CATTCTCGAAAAGAAGGAGCGGGTTGCTGAAGAAGGCACGGGAGCTCGGCATACTCTGCGATGCGGAGGTG
GGTGTTATCGTCTTTTCCAGCACCGGCAGGCTCTATGAATTCGCCAGCACCAGGTTTGTCATCAATACTtc
cttatatatattctactcgtttcttctaattattttattgcatatcatctctctctctctctctctctt
aagaacgaagaaaatttaagaaattgttttctaaaattagaaacagttcgtttgaaagaaaattcctTTGA
CTTCCTCTCTTCCAAAGACGGAAACTCATAAAATACTCCTCGTAAAAGATTAAACTCGAGACATAGACCCC
AAAGTTAGAACAGTAAAGCTCATGTGGTGAATCTTCAATAGTTGTTTCTAGCGCTTATCTTCTCCTCCACA
TATTTCTTCGATCTATGAGTTATACATTCAACCAATCGGAATGATATGAGTGTAAGTTCAGCGGATAAGTC
ATGAGTTAAAGGAAGATCTGATATTTTATGCGCAGAACAAAGTCTCAAATATCACGAAACACTTCTCAGCA
TGTACTTGAATCATATTTAGACTTCTTTTTGAGGGGGAAAGGAAAGCAAATGTGTTTAGATTTCTAAAGTA
ACTTGTTACTGCTTTGTTGCAGTCATCTATTAGGATTTCTCCTACCTAGCTTCGATACTATCTACAGTTTA
GATAGATGATCATCAAAGTGATGAAGTAGAACTCCTAGTAGCACCTTTGGCCAAGTATTCACGACGTAAAC
AATCCCGTTAGTGGTCGACATAGCACTGGGCCCGAAAGAGTTTAGATGAATGTATAAGTCTATAAAGATGG
AAAGAAAACATTAATGCTCAATTCTATAGAGATGGAAAGAAAACAATCTAGGTTTGGTCCATGCAAACATA
GGAAACCACAACTTCCAAGTTGTTGCAATAGAAGGTTTTAAAGACTATTTTGTAAGAGGACGATTATTGTA
GGAATTTTCGTTGGAAGAGCTAAAGTAGTATGCAAAAGAAATAGGAAGGTTTGAAGTGAACTTGTATGCAA
CTCTTATTACTTGGCTCGTAAATGTAAGAGTACAAGTGTATTAATAGGCACCTAAGAACGTTTTACCTAct
agaaaaatatctaaaatagtttACATGAATAAAGAAAGAGATAGTGTCTATCCTAAGTGGCATGATACTAG
AATTCTTCCAATATATCTAACTTTAGAATTCTCTCAAAAGTATCTCCATATACAGATTCTTCGCAAGCTCT
ATGGCGGACTTGTTATCAACATGTATATTTGTTGCTATATCCTATGGGACTTTAAGCTCATTGAGAATGTT
GCGTAGCTAAATGACATATGCATAATGTAGCCGCTATATATTCCGCTTCATATGTATAGAGTGCGAAAATC
GGTTGCTTTTTGGATGACTATGTACAAGCATTCCTCCAAAGTAGAAGATAAAACTTGATGTGCACTTCTGG
TCATCATAATCTCCGGCCCAATCGCTATAACTACAACCATCAAGATTCAAATTGTTAGAGATGagtaaaat
aaactatatgttGCATACCTTTGATGTAGCGGAGAATGCGATGGTAGTATTCATATGCAACATTTGTGGAG
TCCCCACATATCAGCTAATTAGTCTAACTCCATATAAAATATCAGGTCTAGTGCATTTAAAATATCGAAGG
CTTCCAACTAAGCTCTTGAAATATGTGGGACTTATATGCTGACCTTCACCTTCCTTCCTCAATTTGGTGCC

ATAATCAACCGGTGTGTCGATTAGATGACAATCTTCCATACAAAATTTCTTAAGAATTTTctctgtataat
tttattgagagATGAAAAtgccttttttcctttgatgTACTTTATTTACCAAGAAATAAGACATAAGACCA
AGATCAGACAAATCAAATGATTTGATCTTGAATCCTTGATTAATCACCGATGAATCATTGCTTGTAAAGAT
GAAGATACACTTTTCACCCTTGTTGTCAATATGCATGTGTTCtcatttcttccaaaattttgcaataagca
atgcaaaaaattcttaaataatcCCTTTTGGGTTTGTCAAGACAAGTTTGTTCCGGTGTAATGTTATAGAG
tctttttatagaaatctatttaataaatagacgACATTCCTATGGCTTTGCTCTAGAATTCTTTAAGAAC
AAATTTTTGCTTAACATACATCTCACTGTATTAAGAGTACTTCTATTTCTCCTTTCATAAACACCATTTTG
AGATGGTGAATAAGGCAAGATAAGTTGATGGTTAGTACTATTTTTTccgcaaaaattttcaaaatcatgtG
ATTTGTATTATCctctatatgatttttggGACAATAGTTTACATTCACTTTCATTttccaccaaaattttg
aaggctTTGAATTTCTCCAgcaattcatcatttttttgcaaaaaaaaatatatttatgtcttACCCCTAAAG
TCAAcggaaagaaaggaaaatacttGTTCCCTTCAATTGATTTTAGCTCAATGGATCCACAAATGTCTGAT
ATACCAATTGTTGTTTCCTTGTAGCTCTACTTACTCCGTTCCTTTCAAGCTCCTCTTAAATGTAGCTTTCG
gagaatacatttttttatttcttattttctttcactaATCCAAAAGACCTGTcaccattttttttgacaC
AAAAACTTTAGCCCTTCATAATTAGAATGACCATATATAAGATGCCATAGAGTATCAACATCCTCAATTAT
AATGTTAAGGTATCTTCTTTGATCAACTTCAATAGTTAGATTGAACATTCGATTTTTAGTCATTTTGACAT
GGGCGATTAGTCTTTTATTTCTATCTACATAGATAATCGAAGATCTTTCATGTATATTTCATAACATTTTT
GTAGCAATTGGCccaaactcaaaatattactTCAAATGTCTGGCACATAATAAACATTggaaatatattga
tatgatCCATTTCTCAGTTTGATAAGAATCTTATATTTGCCTTTCATAGGTCTTTGAGtaaattatccaaa
tataatattactttcaaatttttcattgAGCTCcacaaatatattgttatcaCCACACATATAGTTGCTTC
CACCAGTATCGAGATACCATGTCGTACTTAAGAGTGGGCCCATCTTCTTTACCTTAAGTAaggtttcttct
tcattttattttatacaaaatacactTCTTAACCATTGTGGTTAGTATCTCTGTTCCAACAATTAGAAGA
GGTGTGTCCGTATTTCTTGCAAATATGACATTGAGTGGTAGAATTGCCATATCTTGGtcttttatatagtt
ttaaaaaattagctcTTTTTCTGTCTCTTTGATAAGATGGTCTTGTAATTACTTCTTGCAATTATTGCCTT
GATGTTTCATAgaatcttcttcctcttcgtcCATGACCATGTCCTTGAAAGCCCTATCCTCTTGGttaatt
gtttgaatttttttttgtttctctctctGTCAATAATCTTTTGTTCACAGGCCCATGAGTTGCTCTACCATT
ATTTTCTCCATGATCTTGGGTTATTCGATggctacaaaaataaatcaaacttgGGGGGTCCAAAGACCTTA
AGATTTGTCAACTTGCACATGGGTTAGTTTTTGCCTATATCTTCTAAGTTGGTTAACGATAAGTCACTTAT
GGATTCACCattcttcattttcaaaagtttatattgAAATCTTAGAGCTTGGAAGCGAATTTTCTTTACCT
TCTCGACTCCTCCAGTAGACTTTTGTAGGATATCCCATACTTCTTTAGAAGACTTGGTGCTGGCTGGCTTT
AAGTTCAAATGTAGGTTCATCAAGTTCTTAGTAAATGATGAAAAAGAACttcttgtacttttttttttttgt
aagtaaaaCTTGTattaacacaaaataaaatctttacaaACAATCAGCTTTGAACCGAAATAATCGGCGCT
CAAAAGCTGCAAACAGACTcggaaaattaaaagatagaaAACTCGTTCTAACAAACAGAacaaagacaaca
aaataaaatatccttCAGTGACTCTTTACTTGAGAAGCTCTTCTTTAAGCTCAACTCCAATCAGCCCAGTA
GGAAAAAACACACTATCCGAGACTCTATACTttgaaaaacactttttttaagcCCAAATCCAATCAGCAGA
GAAGGAACATCACACCtgaaaaaacacaaaaaactaCAAGTATCAAGATTATaaccaaacacaaaatttgc
ataaaacTCCGGAACAAAATCAGACTCTCCAATTgtaaagaacataaaataaagacaaaatcAATTAATGT
GGATCAGATTATAAAACATCAAGAAAGTAAAGTATAAAAATCTGACAAATAAAGAGTATCCCAAAACACTA
CAAGTAGTAATGAAATGCCTAGATCTAAGATATGGCACGACCTTTGCCAGTCTACATTAATCAACAACACT
CAAAGTTAAAATGTCATCTCTAACTCATAAATGCCTCTGACTATAAATGGAAAAGCTATGACAGAGAGACA
AATGGGTAAAGTAAGTCTTGAAAAATAACTGTAGACTAGAGACAACCAAACTCCATATTGGAGAAAACGAC
CACCAGATATTATAGGCGtccaatactcgggtcttatcCAATTTTTTCCCAGGCTATATCACATGAATTCT
TTATGCCTGAGGCCCACATAGttaccaaaatccttttttggaaGAACACTACCAGATTTTAAAGACATCCAA
TACCGGGACTTAACCAAATTCTTTACCCAGGCTATACAACAATActtcttctaaaaatagattgaGGTTCA
CATAGacaccaaaatccttttttggtaGAAGTCTCCCCGAAAAGGTGAGGTCGAAAAATGGGACTTTAACCC
AAACAATTAAGAGAAACAAACATACCCGAATCCCAAGATATAGCAAACCAGAATATGATCAAAGAGCAGTA
GTACAAGAAAGAGACACTGCACTTTTACTTGAGAGAGTTTGAAAGTGAAACTAAGCTAAGGTCATCTATAA
ATAAGTTAAGGAGGATTCATAGCTAACTAGCATGTGTTTAGGAACAcctttaaaaacatattcatATACAA
TACTCATCATACTATGTTGACTGATTATCATATCCAGACACAAGCATCATCATATCcactcataaaaaaat
tagcattataCCCACTAGCTATGAAAAACCAGTTCCAAAAcctttttgatattaaattgatAGATGAAAAA
AGGAGAGATACAACCTGGAGCAACATTTCTTTGAGCAGAAATAAGAATCATCTACTCTAGAACATTGAGAG
AGAGCTTAACCCCTTTAATCAGATTCATTACAGAACCTGGAGGAGCAGAAAAAACGCAGCCGTCCTTCTTG
TCCTTTTTTTCGGTTTCTCTTTAAGAGTATTCTTCTCGAAGCTACTCATAGTTTCTTCTACTTTGGTGGAT
TTCGGCTTCTCATAGCTTTCTCCAAGAAGCTCCTACATATCAAGAGAtcctaataataatttcatcagAAT
TAGTTATCTTTGGTTAATTTTGGGACTTGTGGCTGGATAAGATTTGTGGTGGCTATTATTAGAACTcatct
attattaaaatagatgAGTTTTGCTAGCATGCTTAAGTCCATGACTTGAATAGCTGTGCTCataccaactt
tttttctgtctACTGCTCGTTGCTGCTTGCACTCGTGTGCTTTGTCTGTCAGCACGTACCCTTGCACCAG
AGAACAACACATTCGCGCATGCCTTTGCACCTGCTTATTGCTTGCTTGCTCCACGTCTTTTCGTTATGCCG
TGCCAACTCATTGTTGCCCCCCCAGCCGCCTGCATGTCTTCGCATACTTGAATTGCCGCTCGTTGTCCATG
CCAATGTAGGCTAGGCTAATGCTTGGGAAGCCTTTAATCTGCCTAGACCACTCTGCAAAACACGCGATCTT
GATCTTCGGGAAGTAGACAAACATTTTACCATCCACGTAACAAGGGAAGGATTTGGGTTATCAAAGGGGTG

CAAAGCAGCGGGGAGGAGAGAGGTCAGACctaaaaaaatcctaaaagaAGCAACACTCCAAATCAAGGACT
AGAAGCCATTGTCGCCTTGTCCATTACTCCTCTCAAACCCACTATTACTTCATTCTATGTAATAACTTCTA
GAGTTTATAAATGGAGTTAGATGATTTTCTCCCCTAGATGTAGGTTAATTGACTGAACTACGTAAATCGTT
GTCTATctctcaatttatttttctttttgctccTTTCCATGTAATTTGTCATCAATGCATCCTAACCAGTC
TAGGGTTTAAGAGTTTTGAGAAAACCAAAGGAAAAAAGTTGTGCGAGAGATGAAAACCAAAACTCAACAAA
GAAACGAAGAAATTGTTGACAAGAAGATCCAGGTAATAAAAGTCAGTGGCACATGCAGCTGAGCAGCAGTA
GAAGGAAGCCACacaaagaggaagaagaagggttCCTCTTCCTGCACAAGCCTATTTAAAGGGATAGGATG
GCGAATAGGGATAAAAACCGAGGTGATGTAATTTGGCATGGGAATCGAGTGGATATAGCATTAGATGTGTG
GTTCCTTGTCAGTTGCACACTTGCACCAACCTCGATGCGCCTTAGGAACAAGCCGCATGCCTTTACTAGGT
ATCACTGCTAACACACCGACACTTGCCCTTCACATCGTCCCAAGCATTTTGAATTGAAGGCTCCCACATCG
GGAAATTCCGGCGAACAACCACAAGGTACACTAACATGGAGCCCAAAATTTCAAAGCGGATACCATGGCAC
ATCATAATGCAATACGTCGTCTTGTCTTAAATGGGCGAAGATCTAGACCCCTATGTAAGAGATATTTCGCA
CTTTATGTTGTGTTATAATGTATCAAGGTACATATCAAATTTGACCACTTTCTTATGTGGGTAGCAATGAT
TCATCATCAAAGAATAATAGAATTCTATGGATGGTTCATTCGATGACCacttaaaattacaatatgaCAGC
CCGCAACCCGGGCTCGTTAACTCACTAATGCCAAATCATACACGCAAGTAGAAGacttatatatgtgattt
tagAAACTTGAGAGACTTTAAAATACTTTGATTCGCTACGGCACATGACAAGCATCTATGAGGAAGATGGT
TCTCGCCACTCAACATGAACGTAACTTGCTATAGATTTAAATGCATCATTTTTTCTCATTGATACTTAAga
atgtataatattatatggtcCAAAACCACTTCCTGTTTCAAAccgttttataaaatttcggctcaatatat
ttcaaacattcGGAAATGCCATAATTGATATATACCTTTTGTAatcataaaaacttaaaaccaaaatttaa
aaatacttaaatcagttttaaaaatcttacgCAAGAGGACTTTTTGGTCGGCTGGCCAAAATTATTGGTCA
ACCGATAGTTCTAAAATCAGATTACCGCAACTATACCGGCAAGATCAGTCGACCACAACAACACTTTCGGT
CTATCGAAAGTTCAATATTATGCATATCGGCATTCAAATTTGGGAACTACCCAAGTATCCAAAATGACCAT
AAAACATTCAATAAGGATATCGGACATGTATACACCTGTATCATGGTCAAATACACTTGCCCTAACTTGAG
tgattttaaacataaaatcacCATTTATGCATCAAATgtattttgccaaaaatacaaaaaaaaataattaaa
aaaaatgccaactttTTGCATTTCCATCTAAAGCATGCGTTTCAAACAAGCATAACCACTTAGATAAGGGCt
catatacatgcatacaacAAAATATGAGTTAGTGCTTCCTTTATATTCCTTTACACTACATGAAGTTGGCC
ATATAGAACTAGGTATACATAAAGGGTTACATGTTACACTTAACCCCAGCATTATACCTCAAAAGGAAATA
TAGTCTAGTCTTTCCTCTTTTGCTATACGTATAGGACTCTCGACTGGTGCCTTCCCACAATCTCCTGGgtt
tggaaaaaagaaaataaaagaatgtgAGTTAAACAGACTCGGTGAGCGGCACAAAGCATTCCCCAACCAAT
GCTTATCGTAGCACCACCTGCAGCATATTCACATATACAATACATAACATGATTTGCACGATAACATAACT
TGTCAACCAAGCCATAAAGCAAATATGCTAGTCAACTAGGTAATGATCATccatccaaatcaattatgttg
TTGCTTATTTGGAATCTTAGAGTAAAATCATTGGGtactaacccaaaccagttagtaGTCCTCTCGTGTTG
AATTAGATAACAGAATGACCTAGATCAGGGTAACCCGATCCAGTACCAAGATCAGGTACCTAAACCACAAG
TGTACATTTTGTGATGACAGACCTTACAGCAGGAGCGATCCCACAATATCCAGCGGGAACTACTTGGCAGC
TATGCAACCTTCGGACGGTGTGACtctgagcctcgcactcccTAGTTATACACTTGGGCCCCAAGCGTTGT
CTATTATGCTCATTGGTCCCATAAAAACATTCATATATTAAGATTAAGCATTTTAGTTGTAATCCGTTGGA
CATACCATCCCTTTTCACAAACCCGCCGATATCTTGGGACTAGAACTTACAACACACTTGTGCTAATCTCT
TTAGCCTTTATAATCAGTCTACCATGATTATCGCTAGAGCAAACATGCTCACGTACTCACATAAGTAGAAC
CATCTTCATAGGCACACCATTACATATGTCAAATAATGTTTATACAAAGTACTACTAGAGCAATCCGCTCA
CATACTTATTCAGGTAGACAACAtcataaacatattaaaatatataaatacaattaattCTACCACAAGTA
TTGATAGAGAATACCGTTCATATTCATCGCTAGGTAGAACAACATCATAAGCATATTATAAGAACACAACA
TTCTACTAAGAGAATTTCCCAAGCAAGCTTGTTCTAATACTCATCCAAATAGAACACCATATCAATAACAA
GTTAATCAAACAACATATTCATTTGTTATAGCATAAATCAACAACATGGGTAGCATATAAACTATGGTAAA
CATACCAACATGTTTAACACCCAAACAACATGCTAGAATCCTATAACAAGCATAATACAACTTTGTTAAgt
caacaaaacaaatcatGCAACATGTGCAAACCGCTCAACTTAGGTTTTTTTCAGCCCTAGCCCCATGCTTC
AAACCTCGACGAAAACCAAAGAATCTAGTATCCACGCGTATACAATGATCTAGTGAACCcaaatttaagtt
taattgGATCTCCAAGCTTCTACTAAGAAGATATCAAAGTTTAATTGGATCTCCAATGATCTAGTGTCCAT
ATTGGAGTTTGAATGTTAGATCCTGCAAGGAGGATTGGCTCTCAGCACTAACTGAGTACTTACATCCCTCA
TCCATTGCATATATGCTTCCTGTAaatgaaactatatatatatatatatatatatatatatatatatatat
atatatatatatatatatatatatatatatatatatatatatatatatatatatatagagagagagagaga
gagagagagagaggctattctcagctcgacctgaatagccttggtaagcaaacatctcagccgttgatctc
cttgatctaacggctgatatttaattttacacttaacggaataacgattcacgccgttacgTATTTAATGG
tgccgttaagtgtaaaattaaatctcagccgttacgTTAGATCAatgagatcaacggctgagatattagct
taccaAAGGTTATTCTcaagtcgagctgagaatagcctatatatataagcacaCTTATAAAACTAATACTA
ACATGTCTTTCCAGATCTTTTAACCACCCAAGCATTGTCATTCTCCTTTCAATAGACCAACCTGAGCTACC
CATCTACTGTTGGCTCTCTACCcagttctttttaattaatttaacaaacGCAAAAGTCGTCATGTCAATA
AACGATTACTAAAACGTCAATGAAAAGATAGTACTATTCTCCTCTTGTCACCAGTTGTTTCGATCGAGCCA
TTCTAATGAAGTAATTCACCAAAATCTGAAGCTTGGTTTATCTTAGCATGTCCACTTTAAGCCAAATGCct
acaacatattaaatatctatgtaTCCATCTCCATTAGGCTCCAACAATTACATAACATCTATCTGTGCATC
CTCAACCGCCTTGCTAGCGGTAGATTTCAATATAGGGTGCATGAAGGGTGGCTGTGTAGAGGGACCACGAG

TGTCGCGAATCGGATAGGCTGAAACCTATAGAGAAAGGTTCTAACAATATCCAATTAGATGGACGATAATG
GTGTTGAACTGAAAGGATGAAGGAGAAAGGTGGTTGTGGTTGCGGTGGTATTTGCATAGAGCAGCAAGGTG
GTGGAATGACAAAGGTAGGCGAGGGCAAGTGGAGGAAGACAGTTGACTTTTCTTTCCCACATATCCCGAAA
GGCTATTCTAACTCTACCATGACTTAAAAATGACATCGATAACAGATTGTTGAAGAACATTAACATTTGCC
AAAAGCAATAAGcctgaaaatttaaagaaaataagagataATAAATCTCATTGGTAGGTTTACATACACAT
ACGAACGAACTTTGACCTCcaaatttttttgactttctgACCGCCCGTCCATTGCATGAACCATTGTGCCCA
TACGTCACGTTGCACCTGAATATAATCACTGTTATGAGATACAACTTGCACAAGACAATGTTATCACAATG
AGAATTCAGGACGCATTTTAAAATGCATAACATTTATATCATCTCCAAAATCTGCAAGAGAAGGTAAAAGA
CAATTCTGATAATCAAGCAAGTTTCAATTATATGTACAACCACCATATAGTGCATTAGCATATTGAAGAGA
TAAATTTGAAGCAGTAGGTAATCTACTACATTACTATTAGATGTTCATGGACCAGGGTTTATATGCTAACC
TCTTGTATGGAGGTTCATAACTCTTtaagtagaaaaaataaaaagataatttaacaatattgATACCTAat
acacacaaaaaatattttaacatcaacCATGATAGGATAGAGAAGGTGATATATTGTGATAGTAGGAGAAC
CCCTATTGAAGATATgacaagaagaggaagaagtaACTATGCGACGCACCCCAATAGGAGATTCAAGGATG
GAAGTGAGAGAAACAACCATGATGGTGATGCTGCTTGTTAGACTAGGAAGAGGCAAAGTTGATAGGCTCTG
GCAATTATTTAGTTTGGCTAGCATGTAAAACTAAGGGGCTTGCTCATGAGCAAGTGCATTGCAAGTAAACA
CAGAACTTACTGCCCTGACACCATATTTAGAATAGCCAAcacttatatcaaaattaagcatattgaaaatt
agatACTTAAAGgtcttttatacattttaaccTCTCATATTGAGGTTAATTAATCGTAAGTAGGAAGCAAC
TAAATAggaataatagtaaaagtaatatttgcaaaatgttactttactatttttttatatgtttgcaTAAGT
TAAGTTTCCCAATAAGCTCCATTCCACATGTTTTGAGGGTCTTCCACACGTTTGGCTTGCAGGTGAGTGAT
TCAACAAATGATTTTCATTCATCATACCTTATAACGTATCATTGCCATTAGGGTCTCTACAGAACTCTAAC
CAATGTAGATTTCAAGGTGGGTCATTACAAGTTACAACCAATACGACAAGCCACCTCATCCACCAATGAGT
TTCTTTACATGCTTCGGTCATAGTTATGTCCTTATCGTTCGATCACTGATATTTAACACAGTTatgtttgg
aaaaagaaatgatttaGACCAAGAAACTATCAAACTGGTCCGAAATTGTTCACCTCAAACATTTCAGGACC
CCTTTAAGAACAAACAATAATCACATCATCCACTTAAAGCAGATAAGAATAAAACGTCTATGTTTTATAAT
AGAGTAAGTTGAATTGTGAGAACTCGACAGCGAATGAAACCTGAAAGAATTGAACTTGTGAAGTTCACGAA
ACTGCTTAAGAGCATATACAGAGAACATTGAACCACAAACTTTATTTGGTTTAATGGAATACCTGGAGGGG
ACTCTGTATAAATTTCATCCACTAAGATCACCATCAAGGAATGTGCTTTGGGCATTCATCTACCTAATTTC
ACCTTTACATGCAATCCTGAAAAGTTAAAGAGAAATACAAACAAGTGTTTCCTAATATTCAATACCATATT
CCTAACTGTAAACTTTGTCAACTAGTTGAGTTTATTGTATTCAAGTCATCCAATAGCCGTAATCTTATTGA
TCCTTGCACATTCCACAGCCAACAATCTGGAGTTACTTTAACCTAATCCAAGTCCGATTATTCTCCAAGGC
GTACAATCTACTTCCCTTGCTTGTCCAAAACAGGATCTGTTGCTTCtttataaacaagaaaacataaaCGG
AACGAAGTAGCTTggaattaaatttgttaaactatcaagaaaagatttttacgTAAGCGCAAATAAGATAT
CATGATGTAGAAGGGTTTAAGACTCAACTAAAGAAGTTGCTGAGGGAAGATTCATTGTGCGTCACTTGGTG
TGAAGGTATGATGAGAATCTGAATCAGGATTCTTGAATGCAACCAGAATAAGAAAATGATACACTTAATCT
CCAACCATGCACCAATCTGACTGGGCAGTTACAATTCACATGTCCTGACAATGCCATACAAGTGTAAGCTA
ATGGTAGCTGTGGAAAATAGGGCAAGATGGCAGAGTAGTAGGAGCGTAAAACTTGTGAAATTGTTAGCTAC
GAAACGACTGCGGTAAGAATATCAAACTCGGGATGTAATAGCACTAGTGCATAAACATTATCTTGTCTATT
GGAAGAATAGTAGTACAACaatacaagaaataaaaagatggaCATAACGTCAACAACCATCTTGGTTCACG
CCACATTCTAAAACATGTAAAAATAGCAACAATTTGTAGATCACTTTCCCGTGGAGAAGAAAAGACAGAGA
AAGTCAGCCACAACCACAACCAATAAATCAATTGTATGCAAGATTCATAAAAATGGTGTCGATCGGTATCA
GATTGTAAACAATAGAGATCTGATATAGGGTATCTGTTTATCGGTATTGccaaaattgtttgtttttaaa
aaaaattaaaatattttaaattgtaaaaaattattataaacatatcaAAACTATTGTTTGGTAATATTGGT
ATTAGCGTATGGACCAGCATAGGTGCTTATTGAGCGATACTGGCGAAAGAGCCGATACGGACTTGTAAAAG
TTTATCCGGCGGGAGTTGGTTGGGGCAAACgacaatattttgaaatcttgcTAAAGTTTGGGCCAAAAACA
CTGACttcgaattatttttttccaagtCTTGAGGGGTTTAGAAGTTGATTTTAAACTCGCAAAAACTTTAGG
GCCCACATCACTTGCTTCGAGTCaatgctaaaaaaaactttggttacaaaaagatatagcatactaaattc
tttttttttattcatttttttttccaaacgaTCATCTATCTTAGGTGTTTGGGGATCTGTACCGAATTACG
GCACGGTTGAACCACCGAATATCACTCATCAGCGAATCTTGTCCTAGTTGATGAAGATATTTTAAGAGTTT
GTCTAGGGTTTTTATAGCATATCGATGTTTGAGAGAAAGGACATTTAGATTTAAGAAAATCTCCAAACTCA
TACACAC**CTGAATCCTACCAGATAGTTTGAACAGACTTGAAGAATT**CTGCAATACAAGCATACTATAAAAG
CAAAAGATAAATCCAGATATACAGAAACTAATCACTAGGAGTTAACATAATATTCCATCCCACCATAATTA
GATTACAAGATGGActcgaaataaaaaatgcaaaatatcaAGACGTGTGAAAGGAATAGacaatattattg
aaaagGAAATTTTACAAGTTTTCTGAATTTCCAAACTACGACAATCAAAATATAGGAAAGCAGAATCCAAT
AGTGACCACTtgagaaaaacaaagatacaTGCTAAACTAAGATGCCAAAGCAGTGGAGAAGATAACTTCAA
GTATGTGGTTGTGGTAACTAGCAAATGACGCATGCTAATTCTACATTTGATTAGAAACATGCCTGATCCTA
CTTACATTTTTCTTGTGTTAAGTCTAGAATTGACAATGAAAACTACAAAAGATACATGTTCAAATTCAGCA
AACTCTCCAACGAAGAATTGTCCATATGTGGAATGGTAGCGAGAGAATAACCgcaaacaatattaatatta
aaatccaGGAAAAACCAACAAGGAGAGGTGAAATGGGGTAGGGAAGGCCTAAGAAGAATTGTTATAAAATC
TAAACATTGACCTCATTTTGCTGATTTGAATGAAGACATGTCTCAAAGTAGAGAACAATGCCATACAACTC
AC**CTAGTGGGAAAGTCTTTGTGTTGTTTCTTAGTGTTGCTGTAGAATGAATGACATGATGCATTTGATGAT**

**GGCTTTCAATC**CTAAAAAATGGAGGTAAACAAACCAAATAACCCTCTGTTATCAGCATGAAGTCGTCGATT
GATCGGTACTACAAGACTAAGGAATGCAACCACCTAACCAATGCCAATGCGGAGCTCAAGGTATGTAGTAT
GTCAGTTTGTAATAAATCATTGGTCAATGTACTTTGTTAATGCCATTTAAACTGTAAATCACTAGATAAAG
CATAATTCTTGTGACTGCTACGTTACACTCACTTTGACTGATTATGAGTTGAATGAACTTGGATCAACATA
GTCATGGATCTTTGTGTAAGAGCtttgtaaattaaaatgaataagaaacaagaaaaaggaCCAAAATGGAA
TAAAACAACATAGTAGGAATAGATAGAGAAAAAATGAAGGTAGCAATTATATGTTAATCAAGCAAATGGTT
GCcgtaaaattgtaattttgctATATTACAGTAGCAGCTTGCTCAAATAATTTGAATCCATTCCCTAGCAT
TACTGTCCCCACATATTCTATATTCGAATCACATTAGTTGAGTCGTCTATGAACATTTGGGAAGGGCTAAA
ACCCAAATAAAGCTAAAGGGGCGGCAAAGCAGGAGATTCGCAACTATCAtagaatataaagataaagCTAC
TACTAAAGCACATCAtctaaaagtttcatattactctcctaataaATTTAACTGTTCAGCATTGAAAGTTC
CATATGTTTGAATCACGTTGGAACGAATCTAGAAGTTTATGCTTGTTAAGGACCACAGTTGcagttgaaac
ttgaaaattactaaaatagaaTGGATCGGAGAACTTATGGTGCAGAAATTATTGAACCAGCTAGCTGAAAT
GGTGGCTTGATATCTTAGACATATGCCATGTTACTATAGAACATCTCAAACACCCTGCTTTACTGATTAAT
GTAGAATATCGAAAAGAATGGAATGCCTGTAGTGATGTGAACAATTGTTGCTGTCTGGTTTAACATTGAAA
TAAAATCTGGTTGAGGCAGCACTTGGACATCAGACACGTTTGGAAGACGCCAAGATTGGTTGAAAGTTTAA
ACCCACTGACCTATTTGTCACAAGACATCATGGCAAATTGTTTACTACAATTAGACATGGGacctaaatta
aaattttgccaCTAGATAGCTGAACACttcaagagagagaaaaaaggagCAAGATTGTAAAAACGCAGCTT
GGCAAATTATTCATTGATACTTTGCCTATGCTTGCACATGCTTCTTACCAGACAATTATACATGGAATGAA
AAAGATGGCTATTATTGCACAgaatgaaacaaaaagaagaagcattTTTGCCATAATTCAAATAGCAAGTT
ACTCGTTGGATTTTACATACGCTTGCATATAAAGCCGGTATCATACTTTATAAGCAAGAAGTAATGAAACT
CTCAC**ATTATTCCAACCGGAAAGAGAACTAGCAGAGAATTCAGGCATATCTTCACAAGGAGGGGCATATCT**
**GCATTTGAAATTCTGCTTAGAGACACAAGCTATCAAATTTCTGCACCATAACTAACATGAttagaagaaaa**
**caacaataCTGAAATTGAACAATCAAAAAGATGCTATGAACTTCCCCTCAGTGGCATGTTGTTAGATGTCC**
**TGCATGGACATAGAATTTCACATTAATGTGCGGCTGCATC**CTGCATGGACACAGAATGTGTCACATCAATG
GAGTATTCTTTGTAAAAGAGCTTACTAAAGGTAGAAAAATGTTGCTTAC**AATACTTTGCTTCCTGAGCTTA**
**GTTAACTGCTTGATTAGCTCGCACATACTTATCTATTAAACCTTGATGGTTTTGCAATCCAGTTTTGGCAA**
**GGTGAGGCAGCAAGCTTGAGGCAACAACTGCATAACTTACAAGAAAGTCATTGGTATTGCACTTTGACC**CT
gcaatagatttattaactatCAAATTTTGAGTATATATCGAACTTTTTACAAAGAAACAGGAGAAAAGCA
TGTATAAGTGCCTGCTATGGAGTcaatactttctaattattggTTCAGTATAGGAAATATGTTTGGACTTG
TAACTATAGTCATGATGCGCAACACTTGAAACTCAAAGCTAAAAGTGCTTCGCATCACAAAATGTTATGTT
TGTCAAATACGCGAAGATGCATCTAGAATGATTTACAAGTTACtttcagaattatttattagtacttatat
tttaataacttttcttGACCTTATAGCTAAATCAGTAATCACAATTTCACCATCTGAGGAAAAAGATCAAG
TACAATATAGTCTAATTCCTGTTACTATGATAATAGTCAGATATaacatcaattaaaaaaaaaaaaattaa
cagattatgcttgaattacaatatatatataaagagagagagagagagagagagagagagagagagagaga
gagggaagcGGTACCAATCCTCGTTTAGAATGTCATAAGTCAAAGTTGAATTGCCTAATCCACTCCAGGAT
GAGATTGATCATATCTTAacctcccccctctctctctctctctctctctctctctctctctctctctctctc
tctctctctctatatatctatctctctctctctctctctctctatatatatatatatatatatatatat
atatatatatatatatataatgtgtgtgtgtgtgcacgCATGTGTGCGCACacatcattcaaattatttgg
aTTCCAATTCCTACTAAATTTTGTACTTAGAAGAGTAGATGGATTCAACAGAGCTTTGGGAGTCATTTTAG
GAACTTTAACCAAAATACTTTTACTGGTAATTCTACCTAACTAGGATAAGTACACTGatattcaacatcaa
aaatCATTTTGGCATGTGATTCATTATGAACCATCTATCTGCTGCGATGTCAAAGGGAGTGCTCTTGTAAG
AACTAAGATTTAATGAAACGATTCATAGGTAGATTGATCATTGTTTTAGACTTGCTTTAGTTGGTGGAACC
ATAATTTTGCTAATAGTGGTACACTATTACGTACTAGCTATAGTAGCTTTAATGAAATGAATGTCTAAAGA
TAATGTCGGGTTGGAGAAATTATCTGTTGTatagattttgttgttgattaATACAATCCCACTGGAAGTG
GCTATATGAAAGGAGTTTCAAAAAAGTTCGGGATATGTTGCCGTCTTACTCCTAGTTACCCATTAgcatat
aaatagttttaaaattagaagtaaAAAGTTCCAATTGTCTAAAAGGATCAGTTAGCAGCAAAAAGACTAGC
CATGTGCAAATCCATATAATAGCAacaatgtgaaaaaaaaaatgtaactagATAAAGAAAGCCAAAGAAGA
CGCAATCGCCAAAACCAGAAGTTTTGAAGGAGTTGAAAGGACTCATTCACACTGATGATTAGTTTTGCCTC
AAGTTTACTCTTAAATAGGGACTTGGCCATAGAACTAATGAATCCATCGGATATCCTATCTAGAGCATGGT
TATATATGCCGGGCAGCACGTGcatgttccggcagttatcggcaAAGCACGAACCAGCACAACATGCTAaa
gaatcttatatttgaattttaaattaattttgcatttcgtattttcgtttttttggaaaacaaaaaataaat
gttggtGCGTTTAGCAAGGCACCATTTGCACCGTCGGCATGCACTAGGATCTGAGTTACTGAAACTCAGCT
CAGCTCggcattaaaaatttgatctaaAGTGATTTAAACTAATCTCTGAAGAAGTATCTAGCTTCACTAAG
TATTTGCAGCCACCTGCTGGGAGAGGAGCTAGCTTCATTAAGTATCAATGATCTTCACAACATTGAGAGAC
AGCTAGAGACAAGCCTACGTGGTGTCCGAGCAAAGAAGGTACTAAAAGTTTTGATCAATCTCAAGTTAAAC
ATGTTATCATCGTTTAGTATGTAATATCATAAATGCACATCTGAAACTTGTAGGAACAACTTTTGATGGGA
GAAATTGAAGAACTTAGAAAGCAGGTCTTTCAGATAATCTCATCTTCTTTTCACCAAAAACTATTGAAACT
AATCATTACTCACCTAAACTTCATTTGTCATATTACAGGGTAATATTGTGCAGCAACAAAATGCAGAACTG
TACAGCAAGGTAAAGTCTGATTCAGAGGTAATTTATACGAAAAGTCTAAAATCGCGAGTTAGAATTACTCA
AATATACTTTGAACGAGGCAGATATAAAGTCGTAAACATGATGCGCATTAGCACAATTCACGAGCAACTAA

CAAATCATATCGTGCTTGCAGGTTTATGCAACAAAGGGGTTAAATGAAATCAATCAAGGATTGGCTATTCC
ATTTAACTTCTGTTTGTCAGAGAACTTAAATACTCCTATTCTTATCGACATTAGTCAGCCACAAAAGCAAC
TTGAGCAAACTCCAGATGGGGCTCTAAATCTAGGGTAAAGTTTGTCTGGATTCAAGTCACAAATTCCATCA
CTGCTAGTTATAAATGTGTCCCTTTACGTTGCAGCAGACTCCAACTGCACTGATCGATGGCTGTTACATTG
CAACAGACTTCAACTGCACTGATCGATGGGTAACAACAAGTGCCATCATGTATCTTCTCGAAAGCGAGCAT
CAATGCTAGATATGCTTGCTAGTATTTGCAACTAGTTGTTTATATGCATGACCAGGATGATTAGAAAAACT
GGCAGCATCCATAAAATTTGATCACAATAACCATAAAGTAGATTTATCTTGTATTATAGAATAAAACAGCG
GTGGTCATGAGCCCTATAACTACCGTTGTATgtatattatagaataaaacAGTGGTGGTCATGAGCCCTAT
AACTACCGTTGTATGTATCTGCATGGAGCATGATGCTTTCCTCGTGTAACCAAAAAATTGAGACCGTTTTT
CGTCTTTTACGAAATTATGGAAGTATAGGAGCTCGTAACAAAAGTAAGGAACATGATCTGCGTTCAGTTTA
AGGACCTGTAAATGTCATAATAGCCTATCAAATAATGCTTGATCTGTTTCGAATATTAGCTTGAGTTGGTG
AAGTCACTGCTATAGAGAATTCAATTGGAATGAACAAAAGGCGCTTTGGGGTGAATGCGTAGTAGCCTTTA
AATTGCTACAAATTCTTTTCCAGCATCGAGATCAACTGGAAATTGGACAAGGAATCTTAAACAcctttctg
tttttatttgcaaatatacACAGCTTTGCATCTAGTTTCATTAGCCCCTTCTTCACGCCTTACTAGAAGC
ATTTCTTTTAACGGTAGTTTCACAGAAATCTTTAGATCACATAGATTACCCTACTTCGGTGCCAACATTAT
AAGAAATTGTTTCATAGCCTACAATCCATGCGGCggatttcttttgttttagagTTTGTAGTTAAGCCCAC
CCAATGATAAATATCGACCATTCAATTtacttcaaaaacaaaaatgaaaccCAAGTGTTGCACGATATTTT
AGTTGTCTTGTCTTTTTGGAAACATAGCACACATCTATAATTGAGTCCATAATCttacttgtataaatata
atgcaAAATTACAGTATTGGATTATTGTTAGTTGTTATGATGTATAAGTTCTGAAGATTTTCTGGGTTTAT
GATCTTTAGTGAAGGATTACAAGTACatgatattgttaaaaaaaaaaaaaagaaccttaTCATTTAGGGTC
AAAGAAAGGCAAAGTGTCAACTGTATGTTTTGGTTTCCTATTCAATTAGTACTGACTTTACCCATTTCCAC
AGCAGCTCGTGAAATATCTTTTCTGGCAGCAAACTAGTAAAGAtgtcattcaaattatttatagaaatcaT
TTGGTTTTTAGCCCACCCACTTCACCAAAGCTTCCTGCTTTAGcccaatatcaaaatatgaataataatca
TATCATGGTAGCTAAGGATCTTCTTGGCCTCCCTACTGCCCGTCATGTAGCTTGCCACACTGCAGTTGCAA
CTCCATTGATGTTCCATCTAACTACGAATCAACCTCTTCCGGACTTAAACTAGTTCCATCTGCTCTCAAGC
CCAACCATGATGGATAGGTTTTAACAAAGTAAGACAGTTAGCATTTTTCCAAGTATATGTAGGTAAAATCC
TATTCTATTACCCTTATAAATACCAAACATGGTAATATGCCTCTATTGTTCAATTCATTACTTGTTGtcca
aattatgttttatgttaCTGAACTCTCTTAGCTACGcccattcaaattttgaccCTTGATccttaaaaac
tataaaactcAATATGTGTCGAGGGTCTATTTGGATAAGGTTTAACGAAGGGCATTTTATAAGGAGAATTT
GGAAGTATCCGTTCCAGTTTTGCTGTTTGGATGTCCATATATTCCACTCCTTCAAAATACTTCTTCTAAAG
AACATCTACTTGGGTGTTATTTGTAAAATCTTCTTTAAAGAATCCTCTCCCAAAATTCTCATCTCTCAATC
ACCAACTTTCCTTTATATGTTGTTCCAGGCGGCACGGAGGCTCGGTGCCGCCCAGCACATGCTGAGTCAGT
AGGGATTCAGCACTGAGCTGGTGCTGTTCTGGCCAGCACGGAGGCTAGGTGCTGCTTGGTACTGTTTCGGC
TGGCACGGAGACTCGGTGCCATCCGAGACAAGCAGCTCGGCAGCACCAAGCCtgataagagagagagagag
agagagagaagaggcactgccgccgccgctgccgagATCTTAGcccatcctctctctctctctctctctct
ctctcctttctttCTCAATCCTTTCATGTCTTTTTTTCGCTCCAGGATAGAGCTGGGGACAAAAGCTGCCA
ACAGCTACCCTCTACCTTAgggctaaatttttttagttttaaagtttttagtttttttttataaataaa
tttaaaataaaaagtacagATCGGCATGCTCAGCACATCTCGAGTTTAGAATTCACCAAAACTCGGCACAG
CACGGCACAAAAAAACATGTGTTAAATGGAAAAATCTCAGCTCCTAACACCAGCCGAGTTCTTCCTATGTG
TTGTTGTCGAATCTCATAGGACATTGTCGCAGACGGTCTCACcgaacatcatcatcatttattCTTGCCAG
ACATCATCATCCATTGCAATTCTCACCAAACGGCGTCGTCATTATATGTAATGTTAATCGTGTCGAACGcc
agccccccccccccccaactttgatctttttcattttaatattttatctcataGATAAGTTCTATCAATATA
AGTCAATGTAAAGAAGATTTGATTGTTTAGAcctaaatatttctttcatctaagaatttttttaagaatc
atTTTCAGCTCTTCTGTGTGCTTTGATGGTAAATGGATATACTGGTCACATTCTTTGTAATGCTATTGTgt
caattaattttgttacatCCACTGTGTACTtggttttttttgggtattggATCGAatgtttttgttcaat
ttcCCATTAGAATAGTAACACTAGTAGTATATTTTGTCTAatcatttatgtttgttttggttctgatccct
ttttttgtcatctTGAGTTTAAAATTGTCTCTATTGTTACAACattcaaatatagaaattCAAACACTTC
ATCAAATTACGAAGTGAAAATGggtaattaaaaagaagaggaTTTGGAAATCAAGCATTTGCAAGTCAATG
CATTTATGAATACTAAGTTTGAAAGTTCTAAAGTACCCTGAGCAAATGATTTTAGTTAGAATTTGTTAAAG
ATAAAACGTTTCCCTCATGGGAATTGACAAGCAATGATAACAAGTACAAAACCTTATCACTACTACTATAC
GCAAGCTCATGCATTTTTTGAACAAGGATGGAACTATAGCTCAATTGTATGATATCCCTTCGATAATTATA
GTGCAGAGAACACTGCCTGACAATAAATCCTACCcttctgttttaatttatcaaatactaaacCAAGATTT
TTCGTGCCGGATCTATGAGGCTCCGACGCCGGCACAGTGCCGAGCCGCCGTGCCAACTGTGTGACCGGCAT
GGTCAAGTTGTGCCAAgccgtgccggtcggcacataatttttttatttttaatgttttgctTTGGGTTTT
ACATATCCCTTCTATAAGCTATATTGAtactctatataataataaactatatactatatatttttactatg
taATTACGACTTTtgaatatagttttataaatttttttattattgcttaaataattttagtatttcaaaaa
aaaaattatacagatcGGCgatgtcggcacggcatggtCGGCATGTCTTGAGTTTCTGATTTACCGAAACG
TGGCtaggcacagaaaaccttgtactaaaataaaaatagtgtaCTCTATTATGTACTACAAGATAATGAAC
ACTCCCTCCCGATTGTAGCTCctatcttctatttatttttttttttttttggtcaaataaCGAACAACCA
TGGGGTAATTCATTCTATTCCAAAAACTgagtttttcgaaaatatataaaaaacctTGTTTGTTGACCTT

TAAGGTTGTTCGTTCAAAACTGCAATAGCGTTAGGTTATATCCaacttaatatttggtatttttttaaaa
tatttttgatcaaTGAAGAACTAAGAAAATGGTTTATCtggttctacaacttttaattcaacttgtttta
agaaaataaaaaaggtatcCAATTGTCTTTCGAAATGGTGGAGCTGCTTCCATATCTTACgtcaaataaac
taaatatggtgtattatatgttattccACAGATTTTTAGTTTAACGGTTTTTGATAAACAAATATTGTAGA
ATGTGAAGTTCGTAGGGTGAACTGTTAATAACAGAAATAATGTTTTGTTACTTTATCACTAATGTTTTAAT
ACTTCTTTCAATTCTGATTAAATGATGAACTAAAATGTTGACAGTTTGTTTTAACCACAACATTATGCAAA
GAGGCCTTCATAATCCTAGAGTTCCAAAAGATCCTATGACAACCCTCAACTTCTAAAAAGGACCTCCGCTA
GCCTTGTAGTCagatttttttgcttttacaCAGTTGGAACTATTGATGTAGCTATAACGTGGCAATCATTTG
GTaaagagcaaaataattaaaaagtaatacatTTTTACCTAATTATTGTCACGTCAATAGTTGAAAAGGCA
TTAAGGCAAAATGGTTGTAATGTAACTGAAggttcttttcaaaatttgaggTTTATCATGAGaacttttaa
aacttcaagATATAAGGGTCCATTTATATAAAGTTCAGGGGTCAAAACATACTTGTacctatatttattat
ctaattgTTCAATGTTTTTAGTAAAGAATAATTAGAAGTTACAGATTTTGTTAGTTATGTTCTCTAGGAGT
AGATTGGTTGTTCTAGCCGCCATTGGGCCATTGGTATTAATCTTGCCGCTAAAAGAAGCCAATAATACTTG
TTTCAtgaaaattggtatttttttgagCAGTTTGTCAATGCCCTCGCACTGTTCACCATCTCAACGCCAAGG
ATCATTGCTCGGTGAATAGTTACATACCAAAAGAAACACTATATACCTTTTTGTGAAATGCTCTTTCTACA
TACCAAAAGAAACACTTCTGGTTCGGAGAGGAGAACAATGCTTGCTTATGGAGACAAGCAATGGACGGTAT
TTTGCCAAGGATGAGACTCAGATTAATCGTAGGGAGAATATGGACACAAGAATAGTAAAACACACGAATTT
ACGTGGTTTGGTCAAGTGCCTACGTCAATGGGCAGTCACAAAGAATATTCGCACTAAGTAATGAAGAGTAA
AGAGAGTTGATTgcaaggttttacgtgccgggcggcacgcgcggtgccgagccggcacaggccctccctct
ctctttctctccctccctccctttctctctcttgatctcgatttctctctctctctctctctctctctc
tctctctctctctctctctctctctctctctatctcaaTCTCTTCggggtttttttttaaatttgggtgt
cttcctcctcgcgttcCGAGGAGGACGCAGGTCGCCTGCggcgaccttttttttttttaaagcgcCGTGTG
TTCGGCGTGACcccatttctttttaaaaaaaaaaaaattaaatacctttaaaaaaattaaaaaaatcttata
taatatataataaataatataaataataatatttataatcggcacgtcttgaatttaaatttaccgaATAT
CCGACACGAAATACATCCGTACTATCTgttttagatatgtaattaaagtttttttatttttttttttatttt
ttttatttcaatccataattggtttattattatttacataatttaatattttaaatataataaatttatac
ggatcggcacggtcggcacgtgccgagttttggatttaccaaTACTCGACTCAAGCACGGAAACCTTGGTT
GATTGTGCCCCTAGCTGCTCAATTACATTCTTTTATAATCCTCTCATCTCCCCTAACTAACCATCTTCGTT
TTGTAGAGAATTATTAGAAAACTACTAGATAATAGTGACTATGTAATGGGATTGACTTGCACGTTAGTGTG
CTCCTCTTTGGCTTCAAATCAGCTGACCAAACTTTTTAGGATGTGCAACACATGAGCCTCATATCATGCTC
GGGCTCGTCTGTGCACATGGTCCCGCACGGATGGACGTAGTTGAATGTGACGCTTGCTCTTCCAAGGTTAG
GTGCTTCACATTGCTCCTTTAAGGGGTCCGGAGTTGGATATTGCAGCATCCAAGGTCCACATAAATGGTTC
AGGACTAGAGGTGGACGTTGCTTCAGCCGGAGTCGGTATGGAAAGTTCTCAGTCGAACACTTAACATTGTT
CTGTCTGCGGTTGGTATGCACAGTTCCTGTTCAGGCACTTGATATTGCTCCGACCGGAGTCAACATGGGTA
GTGCTGGGTTGGGCGTTGGTCGGTATAGGTAGTCTGAGCGGGGAGACATTCCAATCAAGGTTAGGGTCACG
CACCAGTTTCTCCGGACGAAGTACGTTCACATAGCCACCTAGACCAGACATATATCCCACGTACCATCACA
CTAGCAACTGCCAGTAGTAGCCTTTCGGCCAATAGACGACTACGAGCACCATAGCCAAAGATGGTACAAAG
CCCAGTAGGACAATAGTAGGAACCAGGATTATTGGTTGAAAATAAACTGAGACTTAATTCCATGTGGTGGG
TTGcatgttgaaaatataattttgaatactgGTTAGATGGATGAATCATTTCCAAACATGCAGCCAGTCCT
ATttcatttaacaaaaaatgtgATTTCACCAATATTTGAAGTTCTAATTGGTAGTTCGTGTAttagatcac
aacaaaatagggataatatgtagattatttattttgatttcaatATTCTTATTCTcataattcaatttaaa
aatgccaatgaataatttaaaaaatcgcatattctttttttataaatcagcagAAATATATGGCATAAGTTC
AACTATTACGTGATAATCAAAGAAGAGAACAATACATCAAATAAAGGATTAGTAC**CATTGACTCTCTTGCA**
**AGTTATGCAATTGCTGCCTCGACCTGGTGCATCCATCTACAT**ATGAAGAAGTAACAAGGAATAAAACACTA
CCAATTTACAATGAATCgaaagattataaaatttccaGAATATAAAGTACACGTTAGGACCTTTAGACTTC
ACGGTGCAATAATAACAAAGTCTTCTCTCACAAAGTTTGTTTGGCTATTTGGACACTTGTTCGCCACTATG
CTACCAAGTCTTCAAAAtgccttattttaaaaatatttaaagaatggCAAGGTccaaaatgccacatggca
tccACCTAAGCAGATCAGTCCACTATAGAAACTTGACTCTAGCAAATGACATACTACAATGAAGGCCAAAT
ATGGACCTGTTAGCGTGGTATCTTCGCATTTAGTTTAGAGATGGTGCACATATTGAAATTATGAGAAAGGT
TTAAGGAGAAATGTCAACTAAAGAAAAGAGACAAGATATAGGTGGAAAAAATCCTAAAAAGGTTGGAGAAAA
AAACCACAGATTCAAGGAGGCACcttgtaaaaaatttcactatATAATGTTGTAGTGCTACAATATTTTCT
TGTCCTTTccctacatttttttttctaaaaagtgaGAGTTAATGAGAATTTTGTCCTTTTGTACAGCAAATG
ATTGTCCTATTTATAGGAGTTTAGCAAGATATAGTAGAATTTAGTAATACATAAATTGACTAAATACTAAA
TTCCTCTATAAAGGGTGCGAGTGGACCTTGATGTGAACTATGAATAGTTGTATCGGAACGGATTGCTTTGA
CGTGGATATGTACGAGGCAAGGCTCGTAAAGAGGGGTCAATGAAGGGCGGCCGATTAAAGCCAGCTCTAGG
GCTGGCATTAATCATGACTAAAGCCAGCCCTAGGGCTGGCTTTGCATGCATAGtcgaattagtttttttaaa
aattttaataaaaattaaaaaaactttttaaaagccTATAAATAATGATGAGTTTCTAAAAGTTTCTCATTC
AAATCACTTCATACTCAAAGTCtcaaattagttcaaaagttCAACTATTTAACTCGATGGCACAACCCAAA
AGTCAATCAATCAATAATATTAGTTGGCAACATAGTACCCTTGTAGAAGGCAATatgttttattgaaattt
cCACAAAAAATGGAGGTGGAGTTTCTCGACGAAAACAATATTTAGCCGGAGGTTTTCCAGATGTTAGATCT

TGCGATAAATGCGGTGTTAAGATTCGTGACACTGTGAAAGCGCACCTATGTTCGGGGAAGACCGCAAGAGA
GCTAAGCAAAAGCAATGATAGTACATTGATTGTATGGTTGCGAAGGAGGATTATATTGCTGACGAGGGCCG
TCCTACAGAAGAATAATACGATGATTTTGACTTACGAGATCCAAGAACACCGCCAAGAGAAATAGAGGAAC
TAAAATAAGCAAGGGCTGAAATTTTAAGAGAAACGGAGATTAAGGAGCAAAGAGACGGCTTCGCACAAAGC
TCGTCTTACGCTCCTAGTAGTTCTTCAGGTTCTTCGAGCCAATGAGGGAGAAGTAAAAGTGGTTGGACGTC
ATTCTTCGGTAGATCGGTATCCATGAGGACATTCGATATTAGACAGGTTAACCCACATGCATCTAAGCCAG
TGGACTTCCAACAACCCATAATTAATGAATTGGAAGAGGGGGAAGGAAGTCCAAATTGGCCAAGTCGATTG
GCAAATTGTTCCATTTCAATCACAAAGCTCCTAACGTTGCGTCAAGTCCATCTTATTGCAATATGATCTCA
ATGGCAGAGGAGGTAGGAGAGGGTATTCAACCTCTCAAGCCATGGGAGATTGTTGGATCCCTCTTCgatga
aaatgagaaagagCTACGCATCTACATCAATAGTTTGAAGAATCAATGGAAAGAGTATGGTGTTACGATTA
CGTCTCATGGATAGACAAGTCCAACAAGATTGAAAATtatcaacttttaataaactaCGACAGAAAGACAA
TGTTTCACGACTCTATAGATGCTACCCTGTACGTTTAAGACCATGTCTACATAAAGTTgaagatagaaaaa
attattgctGATGTTGGCATGAAGAATGTGGTGCAAATTGTAACTAACAATGGTTCCAATTATAGACGTAC
ATGAATATCGATGATGGAGGCCTACCCACATTTATATTGGACAACTTGTGCAGTTTAGTCTATCGACTTGA
TGATGGCCAATAGTGGTAAACTCAAACATGTGAGCAAGTTGGTTCGAAAGGGGCAGACTATCaccaaattt
gtatataaccACGGACCACTTCTGGCACAGATGAGAGCAAATCGAAGGCAATCGACTGAGACCGGGCATCA
CCCGATTTGCTACCAATTTTATGTCTCTAAATAGTATCAAGGAGAAAAAAAGTGGTACTAAAAGCTCTATT
TACAAGAGAGTAGTCGTCTGAATCTAAATATCGATTGACGGACTACGGAAGAAAGGTGGATAGCATCATAC
TCGACAGGaggttttggaaaaatataattgtattcaTTGACATTATCGCCCCTCTTTATGATGTACTTAGA
AAAGTTGACACCGAAGATCTTCCACATATGGGATACATCTACCACTCGATAAATAATGCAAAGAAGCAGAT
CCAAGAAAATAACCAAAAGGGATATAAtgactattataatataattgacAGGTGGAATCACCAGATAGGCC
ACGACATTCATCTTGTAGGTTATATCTCAATTCTCATTTCCAGTACCAGCATAATCTGAGAACGAAGAATG
ACCTTATTATGGTGCTACGGAATGTTATTATCCGATTAGAGCCAAATAAAGCCGTACAAGCACATGCTATC
GAGAAGGTATAACATTGTATAGCTATTAACTTAATTAGCATTAATCTTTAAGcataactattaaattaatt
agcatTGGGCtttaatgctaattttttttctaaataattattacatgGCAAGCTATTTAGATATGTTAAAG
GctcatttgaaaaatatacagCGAAGATGTCAAGATTAAATATGGTGCCTTGTAAGATTGTTGtcatactt
aaaaaaaaaaaactatgacaACATAGGACCGATCAATTTTTGAGGACCGATCAATGTTTGAGGATCCTAGT
GATACacctataataataataaaaaaaaaaaaaaaaaaaaaaaaagtaatgaagGTATGATGGATAGCAAA
CATCACGAGCAATAGATCATAACCACACAGACcttcaattttgaaagaagCTACAAAAAAGTGAAACATGT
AGGCCCGCTTCAAATCATTGTTGAATCACATTTGTATTTCCTAAGTAAGATTATCTACTTTCAAGTTCGAA
GTCTTTTTATGCTAATTGCACAGGAGCTAATGATCTTATAGACCTCCTTTCGTTCACATGAAGCGAAAGCA
TCATCACTACTAGTTCCTTGAAACCTAAATATTGTTCTAGGTTCCTTTATATGCCCCCGATCCAAATATTC
GGATCAAATAATCAAGTAAAAGAAACATACTATCAGATAACCAAAACTTGCAAAGCAATGTTGTCGAGCAT
TCTACAGAACAACGAATAAACATAAGCAAAGTATATCCAACgagaaacaattttaaaattgaaggaTATCA
ATGTGAtaagaaaacgaaaaaaaaaaaaaaaagaagattgcaATCAC**CTAAAATCTGGTTTCAGAACTCTTC**
**CCCTGGTTGTTTATTCTCCAGAGAAAGGCCGCTTACAAATACTTGCAGATGTAGTAGATGAAGACGGAATC**
**TtgactaaaattataaagaaacgGGGACAAGACGACGACAATCTGGGAACTTTCCTCAATTCAATGGATCA**
**AATCAGAGGCAAGAACAGAGAGAGATTGAAAACATAGC**CAGATTAAAATACCGCCGCAATGcggcatattt
ttatttacattcaaATACGAGCGTACGCACGTGTCGAGTGCGTGCTCAACGCGGCCAACAAAGATCAGGCA
GCCGGTGGGCCCAGAAAGGATCCCGTTAAAATGGAAAGGTCCGAGCGTAGCCAAGTTGGTCCGGGTGCAGC
CGCTGCTGCAGCTAGAGACTAGATGAGGTTCAGGTGCAATGTATTGACCCTGCTTCCGGGAATAAAGAACC
CCCCACTTGGGAAGAGTTGGTAATGGCCGCTTCAGAGGAGCAACGTATACCTGTCCTTTCGTAACATTAGa
gctttcaaaatttcttcacaAGTTCACACAAAACAACCAAGTTTATTTCCATATGGTGCAACTAGAACGCC
TTCATGATCGaacaattttaatgaaatgaactgtattaatgaaataattaattaattttattttaactaca
aatatatatatatatatatatatatatatatatatatatatatatatatatatatatatatgaatttttct
gaaataatcatgtttagtaaaaattatttacagaAATATGTGTGGTATTTACAAGCGTCATGGAAAGATAT

>XviAGL17_4

ATGGGAAGGGGGAAGACTGTGATCAAGAGGATTGACAATTCGACGAGTCGCCAGGTGACATTCTCGAAAAGAAGG
AGCGGGTTGCTGAAGAAGGCACGGGAGCTCGGCATACTCTGCGATGCGGAGGTGGGTGTTATCGTCTTTTCCAGC
ACCGGCAGGCTCTATGAATTCGCCAGCACCAGCATGAAGTCGTCGATTGATCGGTACTACAAGACTAAGGAATGC
AACCACCTAACCAATGCCAATGCGGAGCTCAAGTTTTGGCAAGGTGAGGCAGCAAGCTTGAGGCAACAACTGCAT
AACTTACAAGAAAGTCATTGCCACCTGCTGGGAGAGGAGCTAGCTTCATTAAGTATCAATGATCTTCACAACATT
GAGAGACAGCTAGAGACAAGCCTACGTGGTGTCCGAGCAAAGAAGGAACAACTTTTGATGGGAGAAATTGAAGAA
CTTAGAAAGCAGGGTAATATTGTGCAGCAACAAAATGCAGAACTGTACAGCAAGGTTTATGCAACAAAGGGGTTA
AATGAAATCAATCAAGGATTGGCTATTCCATTTAACTTCTGTTTGTCAGAGAACTTAAATACTCCTATTCTTATC
GACATTAGTCAGCCACAAAAGCAACTTGAGCAAACTCCAGATGGGGCTCTAAATCTAGGGTAA

>xvi_miR444d
GCTATGTTTTCAATCTCTCTCTGTTCTTGCCTCTGATTTGATCCATTGAATTGAGGAAAGTTCCCAGATTG
TCGTCGTCTTGTCCCCGTTTCTTTATAATTTTAGTCAAGATTCCGTCTTCATCTACTACATCTGCAAGTAT
TTGTAAGCGGTCCTTTCTCTGGAGAATAAACAACCAGGGGAAGAGTTCTGAAACCGGATTTTAGATGGATG
CACCAGGTCGAGGCAGCAATTGCATAACTTGCAAGAAAGTCAATGGGTCAAAGTGCAATACCAATGACTTT
CTTGTAAGTTATGCAGTTGTTGCCTCAAGCTTGCTGCCTCACCTTGCCAAAACTGGATTGCAAAACCATCA
AGGTTTAATAGATAAGTATGTGCGAGCTAATCAAGCAGTTAAGTAAGCTCAGGAAGCAAAGTATTGATGCA
GCCGCACATTAATGTGAAATTCTATGTCCATGCAGGACATCTAACAACATKCCACTGAGGGGAAGTTCATA
GCATCTTTTTGATTGTTCAATCTCAGTATTGTTGTTTTCTTCTAATCATGTTAGTTATGGTGCAGAAATTT
GAATAGCTTGTGTCTCTAAGCAGAATTTCAAATGCACATATGCCCCTCCTTGTGAAGATATGCCTGAATTC
TCTGCTAGTTCTCTTTCTGGTTGGCATAATGATTGAAAGCCATCATCAAATGCATCATGTCATTCATTCTA
CAGCAACACTAAGAAACAACACAAAGACTTTCCGACTAGAATTCTTCAAGTCTGTTCAAACTATCTGGTAG
GATTCAGGAGAAGATTCGCTGATGAG

Position of mature miRNAs based on comparison with osa-MIR444d and not based on experimental data!

Stem-loop structure

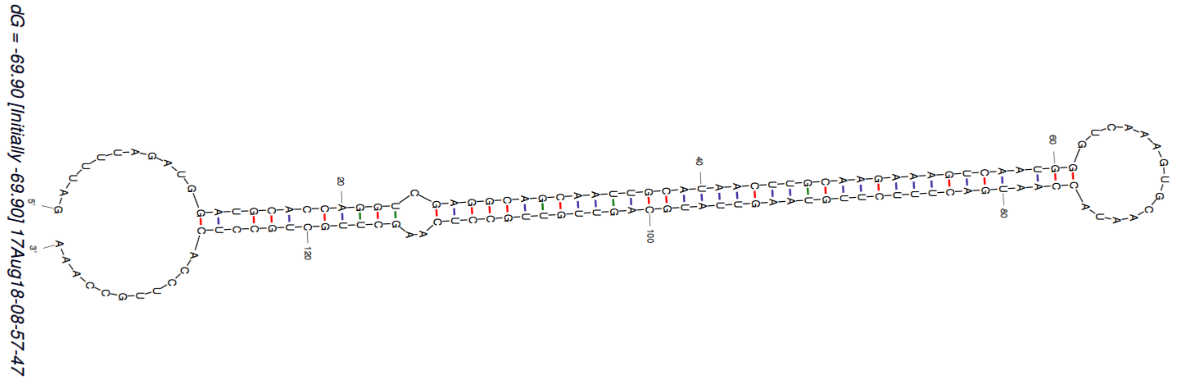


*XviAGL17_5*

>MJHO01000243.1
CAGTTAAAAATAGTAAGTACCTTAATTGAAGTTCATAATTACCTCTTAGCTATGACACTGGCCTGCGGGCT
TCAttctctccctttttttaaatcctgcaatatttttttcatctgGAAGTTTCCCACAAAGGGTCTCCAGA
CCAGACGTCAATAACTGAAAGAACCACTGTGGTCCCCACGTGCCTCACCTGAAACCCAAAAGCGGAATCTG
AATTTCCTACGGACGTGTTGGTACGCTTCACGTGTCGGCGATCGCGATGCCGAGCGGGAAAAAAGAATTAC
TTTTAAGTCAAGAACCAGCTAGTCTACGAAGCCCTAAGCACATCAGTAATAGAGTAATAAGTAAAACGaaa
gtggagaagaagatatcgctaaaaaaataagaaaaagatataaagaaagaagaaagtagGAAAATCATAAG
GTAGACGAGGTTGATCTAATCGCTACGCCTCTATCTGTCGCTCCCCTGGGAAGTAGGAGAAAGGAAACCGA
CAAGGTTTGCTTATTGGCTGCTCACAGTTGATATATTCATCAGCTAGCTGTCGCAGATCTGAAAAAAGGCC
TCAGCCGCCTACGCAAATTAGTTCCGGTTTCTTGGATTTATTTCTCCTTGAttcttatattatcttttcat
tCCGTGATCTCTTCTGATCTTGattagtttttcttttgcaCTGAATTTTTCTGGATTTAGATTATTGATTGA
GTAAAATCGATTAAACCACTGAGGAGCGATGGGAGGGGCAAGATTGTGATTAAGAGAATTGACAATTCGA
CGAGTCGGCATGTGACCTTCTCGAAACGAAGGAGCGGGTTGCTGAAGAAGGCACGGGAGCTCGCCATACTC
TGCGATGCAGAGATCGGTGTTATCGTCTTCTCCAGCACCGGCAGGCTCTATGAATTTGCCAGCACCAGGTT
TGTCATAAAAACCTTTCTTGATATGCCTTCTAAACGTTTCgtcagattttatttattgcataTCACCTAAA
TATCACCCTCTCTCActccctttctctctttcgctctctttaaaacaaagaaaatacaaaaaggaaCTTTC
TAAAACTAGAAAAGTTTCgtttaaaagaaattgttcCTTTCTCTTATTAGAAGAACTACGGTTTAAaccag
tgttgttaaaggcgcgcctaggcgcgcctcgccccgcgcctaggcgcgccccGGGGGTCTTGCGCCCGCCT
CGGTTAGAGGCGGGGGCGCACCTGTAATAGGCGCGCGCCCGAGGCGCGCCCGGGCGCAAGGCGCCAGGGCG
CGCCTCGGGCGCGCCTATTGCTGGCCGGGCTAGGTATTGTCAAAAAACATAAGATTATGGTCCGTGTGACT
CAATCCCCTCCCCCTCTCTCACCCACGCCCTCTCCTCTCCTTCCCAAAACAAAACTCTAGCACTCTCCTCA
TCCCCAAAACAAAATCCTAGCAAAAAATGAAgcgagaaaagaaagagaagaggaagaagaaaaagaggaga
agaagatgttgaagaagaaagcaagTTTCTTTTTCCCAGCTATGCTGTTGAGTGCTGAGCTACTTTgctgt
agcttttttttttgctgtttgcTGCTGTACTTTGCTGTATTTGATGTTGCTGCGGCTGTGTTTTGGctgtt
tttccttttccctttttgttttggtcaGCTAGCTCCACTGCCTACTTACTGTTGTTATTTTGTCCAAGCta
gcttctttcttcttttgctaTTTGATGtgctgtttttgttttattttgattcaACAGCTGGCTCACTGCCC

CTACTGCTCTTACAGCTGGCTATGCCCACTGCTATACTttgatgctgctgctgctatGTTTTGAACTGATT
ATtgctatttgatatatataactttaaatgtttggtttggtttaaatttatctatgttatgcatatattta
gataCTAGAAAATAATGCGCCTCGCTTGCAAAAGGCCCGCGCCTCGccttgcgccttgcgcctaggctacA
GAACCACCGTTGCGCCTCaatgcgccttgcgcctttgacaacattggtTTAAACTGCCCTTAGTTATTTGT
TTCCAACTTGATTCTTTGTTGTGTTCAAACTTCATTAGGTAGCTATACCATCTATATATCTGCTTTATTAT
AAAGTTCCTAATTAATAACATCTAAACGTCGCACTTTGACTTGCTATTTTCCAACTTCTAAGGAtggaaaa
ttattaattttactcGAACAATTAATGAGTAACGTAAAAGGCATAGACACGTAGTTATAGGCGGTAAGTCC
ATGTGGTCTTCAATAATCATTTCTCTCGTGGTTCATGTAGGTTCTTCATACATGTGACTGTAAATCTAATC
AGTCACAATTTTGTACGTGTAGGTTCACCCGATAAGTCATGATTTAAAAGAAGATGTGATATTTTATGTGC
AGTACGAAGTCTTAAATGTCACGAAGTTCTCCTCAGCATGAACttgacttattttagatttgtgAAATAG
TTTTTGCTTATGCTTTTGTTGCATTCATCCATTAGGATATCTCCTACATGGCTTTCGTACAATCAAAGTGG
taggttttattttcattgtgCTACTCTCTAGGATAGATGATCGTCAAAGCAAAGAAGTAGCAATGATTTC
GTGAGGGAACTGCAATCAACTACTAAAAGATGGAAAACATCGAGGTTCTACTTTTACCAACTCCTAGTAGA
ACCTTTGGCCAAGTATCTAGTGCTGGCCTGAAAAGGAGTTCTAACACGCTCATGATCCGCGAAGTAGCTAG
GAAACCACAGCTTCCAATTTATattaggatttatttttgttgtcagTATGTTTCAATTGATAAGTTAATG
GAGTTTGCTTTACAGatgcaaaaaagaaagaaatccaAGTGCAGTTTTAGACATTAGTGATTCTTCACCAT
TTAGTACATGTTATCTATAGCGGGGTAGAAGATTACTATCAAGTCCAAGGATGCATGTAGGGTGCTTACTA
TTGCATTTTggattaagaatttttaagcaACTTTGAAAATGATAACTAGATTAAAGACCTTTATAGGAAA
ATCTGTTATtgttttaagtatttgaaagatgttagaaatatttcttgcACATACAAGGTGCTGTATACCTC
TTTTAAGTGACTCCTTTGGTCTTCTTGTGCTACATAGTGGAGAGGTAATTCGCTTGGTTGAGGATtgagta
atttataaatggtCACCAAAACACCAAAATATGCAAGATggagaataaattttatctttcctTCGTCTTTA
ACTTCCCATGATATAATCCTTGACGCTTTGGCGACGGGATTTCTAAACAATTTAGTTGCCTTCTTATGCCA
TCTTCTTCATGATGTTCTCCATttcccatcttttttttttctttcttcgaTGTCTTTCCTTATGTCTCTTA
TTAGCTCTCTTTTGCACGTGGTTTGTAATCTCCAAGTTAGATAGAGTGAACTCACATGTGGGTCATAGTTG
GGAATTTGCTGGATGATGACTACCACCTTCTCGAAGATGAGATAGACAAATCTCTCATACTTACTAACAAA
CGATGCTGTTATATTCATTCAATCGAGGCTGACATCTATGTCCAACTTCATTATGATGTGAACTAGCTTAG
ATCCAAATCTCTCAAACTCGCTAACACACGATGTGTTCTTTTCCACCCGATCGAGGCCGGCATCGGTGTCC
AACATCATTATAATGTGACCCAGCTTATATCTGTTCCTTTCGGCATCCTGAAAGCTTTTAGTTCAAagtat
atctttatttttggacacATGATATCTATAGACTTATGACTCGTAATTGATGACCAATAAAAACGcctatc
aaatataatttctagttgATAGAACTCCTAAAAACATCTACTCTAGGTTGGGCTCTATATAATTCCTCCA
ATCCATTCTATAACATGATTACTATCAAAAGCAAACTTATTCACAAGATTGTCAATAGATACGAACCGACA
CAGATTGGCCCTTGATTGTGATTGAAACAAAGAATCCTTGAAGCGGATGTATATTATCGTAGactatttct
caaataatagaAATGAGACATATCAGTTCCAGCTTATGCAAATCGGTCTTAATTTGTTAGGACAGATGCAG
ATTCACCTTGAAACAGATTTGTATTGATGGAGATcgattatgatatatttttacagaCTCAAAATCAATTG
TAACTGATACAGAACGATCTCCAACTTTTTATTGATCAACGTAGATTGATATCAAAGATCCATCTAGACTA
AAGCATATTGGTCACTGAGCTAAGGATTGGtcagatctttttttttttttttccagcagTGTTCTTTAG
ATTGACGTAGATCGATCCTAGATATACtaccaaatacataaaaaagaataatactaATTTGGACTCTAGAT
TACTAGTTTGGCTGATTTAGTATGCCCTGGTCCCTTCTAGCCTAGTTCCTTaccatttactttttttaagat
GTTTATTGAACTTGACTTTGTCGCTAACTCATCTTTACCCCAATACCTGGAAATCGCTTTTAAAACCGTTG
TTTTCAAAGCTGGGGTTACTTGCTTTTATTGCATGTTTTTAGTGCTTTTGTATGACTAGGATTCGTAGCAT
GACACCTGCTATTATTTCCGATCTCCTAATGGAATACCCAAAACTTAGAACTTATTCTATACGATTTGGAG
ACAAGTTCACTTCCATACATAGACTCGGAGCTTCGTTCCGTTTATTCCCACAGCTCTTGGATGATTCCTTT
CTCCTAATATCTAGAGATCAGTGCAAACCTTCGAAGGATAGGGCTGTCACCATGTCAGTCAGAAAGATAGC
ATGTATCCAAGTCATTGCCCCAAGCTAATCATTGTCAAAGTTATTctatctgctgctgctgctataATGAA
CTTTTTAGGATGAAGGTGCCGAAAATAAGATGGTAGTCGTGATGTAGAAGGCTCACACTATCGATCGTACA
AATATCAATTGGCACATCCAAGGCACTTGAGCATGGCCAGGCGACGGCGGCATCAAGGTGACTAGGACTTG
CTCCGCACCCATAATACTCTATAGCCGCATGCTATTTCTGTCGAAGGCATCTCTGACACATACTTGAGAAT
GAAAAAGAATGCATCTTCCTCCCCACTTTTCTTTATCTCCTCTCTACCACTTCGTCACCGTTGCTAAAAcc
aagctcaagctcgaacgCTAGTGGTGAATAGCCATTCCTAACATCCTTTGTCAATCCAAACCTTCAATTAG
CGTCGTATTCTTTTGGCCAACATGTGCATATCCTTTCATATAACTCTACAAATCCTCTAAAATGCATCTCT
AGCTCAACTAACCTAGCCGAGCTTACTACATTTCCTAGGAGTTGGAGTTGCATCTTGATCTGATTTTTAAG
CCAGCTagaataaatagataatgtaGTCTGTTTTCGAGTAATGCTGGAGGATCAATCTCTAAAGAGCTTCC
ACCAGCCTTTTGGAGCAATGCTAGAGGAGAAGCATTTGAGGAACTGCTTCCAATCTTTTGTAAAGCTGGAG
GAGCATTACAAACAAGGAGGAATCCTTGAGGATCTGGTAGCTCCATTTTTGTGAGCATATTGGGATTGGCC
TATATGGACAAGCATCCATCAAAGCCTATTAAagggtgtatatatatatatatatatatatatatatat
tattatatttatttattttatttttagattatcgtggtggaacccataatgcgagcgcacgcccctgtag
ggacctcaaactcacgGGAAACCTCcacgcccagccccgaccAACGCTCTGGGATACGAGTATAGGTCTTT
TATGAACGGGGCTCCTGTCACgccttcaaaaaattctactgGGGATTTCACTCCACGCGGTCTCGAACtcg
gatggtgacgcttcctgagccacctcccaaccggtcgagctagccggGCTCGGCTAAAggttaacatatat
aaatacatattgataGTCAAACCCTATGTAAGCCACATGATAACCATGGATCTAGAGAGTATGCAAGAAGC

TGGCTCTTGTAGTCGTCCTTGCCATCGACGAGATCGTAGGAGTTCGCTGGAGATTGCTGGCGCATCGCCAA
TGGATGTCAGCATGAGCGCAAGCAAAGATTCGCCTCCACAAACGCATATCTAGGGTATTCTAATATCTCCA
CCAGTGGATGCTTCACCATCACATGGTCAACTTGGTGCTGCCTTTGGAATCCGCAATTTTCGGCCACTATT
GCAATCAATAGCTATGAATGCAACTTAGGTTGCTAACCTTTATTAAACACGATCTCCTCCACCTCTCTTGA
ACGACATTGGTTTAAGGGTATACCACGATCAAGCGGTTGAAGAGAACTTTATGATCCTCCTCATGCACATG
GTAGGTGTGGCCTATCGACATGAATCCAAGTTGACAAAACATTGTTCCATCTGATTCAAGATGATGAAACA
GAAGAGTTTCAGAGCATATCCATCAAGTTGAACTAGGCAATGAAGCAAGCTCAAGTAGGTCTCGTGGACAA
AGTCGATGATCAAGAGGTAACTGGAATTCCTCCACCTACATTATGGTGCAGGTACCTCAAGTAGAGAGTTC
TTCTCCTTTCTAGAAGAAATTGATTCGTCTCCTTATCTAGTTAGACATACAGCACCTTAGACAATCGCTAT
TATAGGCTGACTGACCCTTCTCATAATATTCTTCACTTTGTTCATACTATGAAAGCTTATCCACCAGGTGA
AATGCTCTACTGAAAATGTTCTTTCAATGTTGTTTGATGGGTATCTTTTGATATGGTTTAACAAACTCCCA
CCATAAAGCATAACTACTTAGGAGTAGTTCGTCGAAGTGATTGTGTCAATATTCCAGTCGAGCACTTCCAA
CCGGAAGAATTCTAACATAATAATGAATGTCAagcaagaaaaggaaaaaaaagtcattaatGCGTTACTAC
AAAAGCTTCACAACTGAGCAAGTTAACTTGGATGCCACAAGTGGTGACCTCATTTGCCTAAAACTTTGCAT
TTCTAAGTCACAGCTTGTATATGAGAGTTCCTACGTAATATCCCCAAATCAGATAGAGATGCTCCAAGCTG
TCTTAAAGATTCAACAAAAGCTGAGGAGGTCTTTGTCAATTGTGGAATCATCAAAGGCAAGCACTTTGTTA
ATATGAAGTCCACGCTCAGCTCATTTGATATGGGATAAATTTCTTCTAGAGGGAAGTAATGAGGTCGAAGT
GACCAAGATTGAAACAGGAAAAGAAAGATCATACTTCTTGAAATCGTGATTGGATTGAAGATTAGATACTA
TAGGCTTGTCCCGCTAGAAGGGAATGCTCTAATCTTAGATGCACTTAAACAACACATAAGCACATTATCCT
TTTAGCCACAAGCAGAGTTGGGATAATCTAGAGATTGTAGTGTTAACTGCACAAGTCACGTTGTACGCGAC
CCAATCTACATGCCGGCCCATGCTATGTAATTTTAGGCCCTCCTGCCTCCATAATTGACACCACATGACTG
TCGTTAGTGTGGCACAACCCACCAAGTGTTGACACCACATGAAGGATAGAATTGACTTGTCCTACCAGAAA
ATGGAATGCTCTGATTTCAAAGACGCTTAATTCacatataagtaaaatatCCTTCCAGCCACCTGCTGATG
TGTGACAATCTAGAGGCTGAAGTTTGATCTGCACAAGTTAGGGTCCAAATGGCCCAACCTTAGGTTCATAT
TGGCCCACACAATGCAATTTTTCAGCCAGCTTGTCTCCTCCTAGATTCCGTCAGTTCCGCCTCAGCAATAG
GATCGCCCACGACCTACTAACCCTATATCTGTGAGTTCTTATAACGTCTTCCAAATCAAGCTGCTCTTCGT
CATAAAGATAACTTATAACGTTCTTATAATATCTTTGATGGCATCAAGATAACTTATACTAATGGTCTCGT
CATAAAAATGATCTTCAATTGTTTGAACATGCTTCGTGCCTTAGTAGACAATAAGGTTGACATTCTCTTCT
ACAATTGCTtaataaaaatggaaattaCAGAAGAACTTGTTTTGGTATACTCAGATTTGGTGGGGTTCAAT
GGCTCTACAATTACCCCACTTGGAGCGATTGATTGGAGGGTAGTAATATGCAAACCTCCAAGGTTAAGGAT
GACTAAATTTAACCTCCTTTGTTGTAAAAATTCCCTATCTATACCTTAGAAGTTCTTGGAAGTTGTATCCA
CTTCTCACCTCCAAATGAAATTCATACCCAAAGCTGATGTCTACACCTTGTCCATGAAGCAAGAAAGTTAT
ATTGAGACTCTGTGACAAGACAAACCTTTCCAATTGAAGGCCCATTTCAAGTCAAAGACCCAACCGAGAAA
AGGCTAAGAGCATCACCAGCCAAATACCATTTAAAAATCCAAGAACTCCTGAATTAGATACCAAGCTCTTG
AACCTTGGCTCAACATTAATAGCCCTAGAACACACCGCCTTGAAGTCTtgcttacaaataaatactgaCAT
TTTGACCTTTTAGGATTCGACACGTGTCTTACATGTCAAAGATGACACGCAAAATGAGCAATCATGCTTGT
GAGGCCCATGTGCATTATTGTACAATGAAAGTTTCTAGGAGTGCTTTCGAAAAGTATGTCTGTTCGGGAGA
CATTTAATACTTCCTTGATTTCTACTGAGATAGCAGTTCAAATATATCATGAGGTCCAGGTGCTATACCCT
CACAAGGCCTAAATGTTTCTGTAGAAGACTGAAAGTTATGAGATGGCCCACTAAGCTAACCTCAATACTCC
TCATACAAGCCATTTCATGCATGCTTTTCACGTCGCTTGTGAAAACGGACAACTACTAAAACATTTCCTCA
TAATATAGTAAAACCTTTATGAGGTGTCTCCTTGTTCCCATAATTTTTCCCatctttttttaagctttCCG
TGAAATTTCTTTGtctcttcttttatttgacgTTTATTAGGCATCATACTGCATGACTAGATCTGAGGCCG
AGTCATGCAGGACGGATGCAGTGGCTTTATATAGATTAAGCACCTCTATTTCATTTAGGTGGAACAAGCC
AGCCCTAAAACATTCCTCCTCTCTTCTCTCAACTCCGACACAAGTATAATAACTAACCTTAGCGTGGGTCA
TcctaaaattacttaaaagttTTCTGTTCGGCTCTATGGCACCCTAAGAGGAGGGAGGGGTGAATTAGGTA
AATGTAAATTAAGagctattaaataattttgcatgTATAATTAGTTCTTGGCAAGATAAGAATACTTAGCA
AGATATGTGATCAAATTGTCAATCTCAAGCTAAGTAATATACTAGCATGCAAAGATAATCAATTTAATACT
AAGCATGCTTCTAGTGAATCTAGTAAGTAAAGACAAGAGGTAGAAAGTTGACACCATATTTATGGTGGTTT
AGTGAAGCCCGCCTACATCCACTCTCTCGCCTCCTCGAGGTTTTGTTCTAATCCACTAAATGATCAACTTT
CGGGTGTTGATCAAATTGTTTACACTTCACTTTTTTCGGGCTAAGTGATAGCCGTCTTCCCTAGCTCTTAG
CGATGAGCTGACTACCTAATTCCCTAGCTTTTCTGTTAATTAGCTAGAAACCCATTTTTACAATTCTCTCA
AATAAACCATAAACCCACAAACAATCTTTCTAAGATTCTCACTTGGAGGGCAAGATATTCTAGggctaaaa
ataaagatttacaAGTTTATGCTCAAATCTAGGTTCTTGGGAAGTAAGAACTTTCTCACTAGAGTTTAGAA
GTGTTTAGGCAAGTTATGAACTATTAGAGAAGCCAAAATACCAAAACtcgtatttataaatttctctcACA
ATGGCTAGTTTTTCTGAACTTTTGGAACCGGAATGACTTTCCGATCGACAGGACTTGCCAATTTAACAGCT
TTAGAAATGCTGTTGAGAGTTTGGTCAACTGGACTTCATCCTGCATTACTGCAAATCCTCTATGTTCAGCC
atgatttatataaactatatatataattatatatatatataatatatatatatatatatataatttatata
tatcatactttATTTGGTCCGTGAAATCATCCCAAACTTTGAgtttaactttaaaatctatatttctaGC
ACTAGATAATACcaatttttctcataaaatataaagaggtCCCGAACAATGAGTGTTGGAAAGACGTCATC
TTCGTAAAATTCGGCCTTCAATCTATAAGCGTATCCAATGTGATTTCCTTGTTCCGCGCATGTTCTCTAGA

CTTACTATGCTTATCTAAATTTGTTCAATTGAGTGGTGAGgttttgtttaaattgttttgcTCTTCTTGGC
TCATTAAGCAACACCTATACTTagcaaaacttgaacaaacttattagttatatttatgctaattatttg
ttatcattaaaATCGCAATAGGGCCAACAATCTCTCCCTTTTTTATAatgacaaaattttagtttggtTAG
GCACAAAACAATGCAAAAGTGTggtttagtaaaaaaaagatatacaaatCAAGCGCATTCTCCCCTTTTTT
GTCATATAAAAACGATTTATCAACTAGAACCATGATAGGCATTTATTCTCCTCTTTAAGTATGCAActtta
caaagttatataaagTGAATTCATGTCATACATACTTGGTTATCATGTGAAGTTCAAGTAAGACTGTAGAC
AAATTTGATCTAAGAACTCACATGCTAATattctaatacaataatataaatatatctaagcATGCTTTAGT
TGCTATTTAAGAAATGAGGTTTGGAATCATATAGGACAAGTTAGAGCAAAACTATCTGAATATAAAGCAAA
TAATCAGTGCAGCAATTTTTCGATTAACTGCCCAGATTTTTTGGTCGATCAAAAATTAATCGAgtcaaatt
ttcataaatgaatttaaaaataggccaataaatccaaataaatttcagaaaaatgtaaaaacgCTTCATGT
TTATTACCCAAAAAAGCTTGTGTCTCCTCCTTTCTTCGGTAAAATTagcatgatttttaaaatttttgacc
aAAGTCAAGATAGGAGCTAAAACAAAGGCTCAAGGGGTCTAGACTTAAtctaaacatcaaaatcacacttt
aAGCATATGAAATGGTCCATAAGGGATAAATacacaaatcaaccccttatCAAAGTctatttatgcatatt
tttcgAGATTTAGCGTGTTAAGTCGgatatatgcatattatcATTTAAGCATTAGCTTTATTATGCAAGAA
TTTCTccaaacatattattttatttcatattctaaCAATATCTAACATAGTCAGAAGAAGTAtagagtta
aaaatatttcacacCTCACCCCTTTGGCCTTGGGTCCACATGTTTATTGTCTAATCTCCACTCCAAAGGCA
CctatatctaatatttaggTTTTTAGTTTGGTACCCAAGTTTGCTTGGGTCCGTTGAGATTAGCCCTTAAA
CATTTTTGAGCCATGtagttttgacattttgtttcCTATACAGGCATCCATATGCTATATGTTATAGTCCC
TCACAATAATAGTAAATGATTTGAGGCTTACTACAATTATGaactatattagtttttttgaaCATACCAAAT
ACCATGCTTTTGAGTTGGTCTAGATGGAGCAAACATAGTCTTACTATGTTTCGTTTTTAATGAAGATTCAG
CTAGTGAGGTGCTCAAGtcgatatttatttctctaagaCTACGCGCTCACTTTATTATAGCTCATTGCCT
TTGGCCATTTGATCATGTTTGACTTGCATAGATAAaagttcattttcaaatttgcaagCAATGTCAGTTTG
ATCTTGGAAGCCTCTAACTCAATTCATAAGCACTCTTTTTTCACTAAGCAAGTCAACAACCATAGCGTTTA
ATTCATCATATAAGGATGATAAGGCTGACATgcttttcttatatttcttatagaGTACTTAAGCTTCTCTT
AAGCATCAAGCAATTCATCAAATCTTTTGCAATTGTCGATACGTAGGAGCATCGCTAGAGCTAGAGCTTAC
CTCATCATCACTACTTCTATGGGCCATGAAAGCAAAGAATCCACTTGACTTATCCTTTCTCGTTTGAGTTT
CATGAGGTAGGATCTTCTTCACTCCAAGTTGCCACCATTGCCTTCTACTTTTTTggctttgtatttttata
ttttggaaaatccGGTTGGATGTGTCCGGACTTTTTGCACCTATAGTAAATTATTTCATCCACTTCGCTTG
AGgtgtatttcttcttcttcttcttcgtctccaTCCTTTTCCTAAGCGTCTTCTCGAATTTCCGGGAAAGC
ATTGCAATATCttcctcattttcttctttacttGAAGAGCATCTAGTTTTAGCTTTAAGTGCTATGCTCTT
AGCTTTTAGAATCCTACTCTCATCTCCTTTATTTCCATCTCCTTTTTCATGAGCTTGCTAAGTAAGATATC
AAATGTTAGCTTGTCAAAGTCCCTTCCTTCCTCTATAGCATCGATCTCTAAGGTGTAGAGAGGAATGACTG
ACTTCATGAACTTCTTTATGAGGTCAATTGAGGTGAAAGTTTTGCCCAAGGTCTTTAAGAAATGAATAATC
TCCAAGAACCTCTTAGGCATTTCATCTATAGCTTCTCCTATCATCATCTTAAACTTTCTCGAAGCTATCCA
TGAGAATGCTAACCCTTGAATCATTTAGCTCTGGAGTCCCTTCATAAGTTATTTCCAATctattccaaatt
ttcttcGCGCTTTCATAGATTAACACACGGTTAAATTCCTGGGATTATTCAGCGagcatttaaatcaaata
atctaGCTTCATTATGGCTAAATATTCTTTtcacttcaaaataattttatgatcaaTATCCTTAGTAGGAT
AATTGAATCCATTTACTTGTTCTACATAGTAATCTATCTTAGgattagataattattttcattttaatttt
tcatatccataatttttgcCATTAAAATAAGGAGGTCTACCAAATGATCTATCTTCAACTACTCTATTACC
CTTATAATTtgccatatatatagaagaacctaatatcttaatttttatatgattaactATATGAATCAAGA
GCTCAaactctgataccaattgttgaTATTTAGAGACCCTAAGAGTGGGGAGGGGGTGAATTaggtaatta
taaattaaaagctaattaaacaattttgcATGCAAAGCAAATTAGCTCTTGGCAAGATAAGAATACTTAGC
AAGATATGTGATCAAATTGTCAATCTCAAGCTAAGTAACATACTATCATGCAAAGATAATCAAGTTAATCC
TAAGCATGCTTCTAGTGAATCTATTAAATAAAGACAAGAGATAGAAAGTTTACATCgaatttatagtggtt
cggtaAAGCCTGTCTTcgtccactctcttgcctcTTCGACGTCTTACTCTAATCCATTGGAGAATCAACTT
ATAGGTGTTGATCAAACTGTTTACACTTCACTTTTCTAGGCTACGTGATAAACGTTTTCCCTAACTCTTAC
TAGGGAGCTAGCTATCTGATttcctagtttttttttttttttttggttaccTAGAAACCGACTTTTACAA
TTCTCTCAAATAAACCAAAAGTCCTAAAGTAATCTCTCTAAGATGTCTCACTTGGAGGGCAAGAGATTCTA
GGGTTAAATATGACAGCCCACAACTCTGGGTCGCCACGAAATGCATTCACAAACCAACCATACATGCATGC
GGAAGCTTTACAAGATAAATGCATggtttcaaattcaaaatacaataagtaGTTCTTCTTAACTCAATATT
TTCAAGCTAAGATTCTAGACTTCTGGGCACATGGCAAGCATCTGCGAGGGAGGTGGTTCACATCACATGAC
CCAACATAGCTTGCTAAAGATTCAAGGTATCAATCTCTTTCATTGATACTTAAAAATCTTAAGTAATCTAA
GGTTTACGACCAAATTCCCTGTCTAAAaccatttccaaaatataagcaaaattaatatttttctcaaact
ttGTAAGTGTTTTATATCTTCGATATGTCCCTTTTAGCTtcccaaaagcttaaaatcCACTTTAAAgcca
tttaaataaaatttgaaaaaaaatcttattctaGAGGACTTTTCGGTCGACTAACCAAAATTATTGGTTGA
CAATAGTTCATACAAAGGCATAATGTAGGTTATCCATCATTTTTGGTCGACCGCTTAACAATTTTCGTCGA
CCTAAAATTCagaaacatacatatatagatttgaattttgggACTACCCAAGTAACCAAATTCCACATTTT
CAtcaattatatcttttaaacatGCACACATCTGTAACATACATAAGTACATATGCACAAGCTTTCTAAAG
TGATTTTGAACCCCAAATCACTACTTATgcaataaaaagtatttttaccaaaaaatacagaaaatacaaCA
TTATAAgcccttttttttcatttagaacttcaagttcaatttctaaaacaaacatattCCTATACGCAAGGG

TTTATACATgtttacaaatacaaaatatgagTTAGTCCTTCTATTTGTATCCTTTACACTATATGTTTATT
ATCTATACAAGGTATACAAGACTAGGCTACAAGTGTTACAACTACAGTTAGTCCTCAAGATCTATACctca
aaagtaaatataaacccaGCTCACAAGCTAGCCACCTAAACACCTACCGCCGGTGCCCTCCCACGATCAGC
GGGATTCGGGATAAGTTCAAACaaagagcgtgagtcaaacaAATTCAGAGCGGGTTAAACACAAACCTATG
CCCATGCTCACCAAACAACATgcgtatcaaaatatatatgtgaataCAAAGCATTACTTAGCATAACAACA
TAACATGTCATCTAAGACATAAAACAAACTTTCATACTATAACAACTTGGTAATTGATCACCCATCCCAAT
CAATTATGCTATTTATTATCTGGGACTATGTTGGCATGCAACCAATAAGCACTAACCCAAACCCAATTAGT
CCTTTCatatttagtttaggtactAGAACACTCTAGACTAAGGTAATCCTTTCTAGTACCAAGATCAAGAT
CTGCCATAAGTGCACATACCAGACCTTACAATATGGGCGATCCCACAGCGTCTTCTGGAAACTACTCGGCG
GCCATGCAATCTATAGATCATGTGTGATGCCCCGTACCTCGGGGGTCACGAAAAATAGAACagggtgattt
ttttttttttttttttttttttttttttttttttttttttttttttttgtaaactaaaagacaagacgtt
atgagctagcacttccaattctcaatagtacaatagatttacaaattaattaaattacacagcgcaagata
aataaatttacatattttaacccAAACCCATCTTTTGAACCTCatgctcgctctacgatcgtcgatAGACC
CTGCAAGGTCTAAGGGGAAGAaggggggtgagtatataaaatatactcagtgagtgacccaccctggcaaaG
GCAACCAACAGGactgcggccacacaccaaacataTGCAAGCACAGAAGGCATACATGCAGATgcgaatat
gcatataaatagaagtcctgcatattaaacttctatatgtCAATACAATCTGCACGCGTAACATATGTCCA
GGCACACAagttgtattctaatcatcttctagcaacagatgaacatacaacaagtcatcacaacatgctgg
agtaaacaactcactacaagcatgatctcacaaatccataaattagcttgcaaacaaaataatacagCATG
CTTTCACATCATAGCATAAGTCTGTCATGCACGtcacaacaaacaagcattgccaccaatagtggacttac
acaaaaccGACCACATAACCCATACACATAACCATACTCAATATGTTGCCAAAATGGACAAACTATAAACC
ATATATCCATATGCTATCACAAATAGTGAAGCATCAACTCAAATAAAGGCTTACAAGACAATACATCACAA
TAAAGATAATCACCACCACAGCCATAGGTTATGctatttatctttcttatgtagtgtgatctaacccatag
ttagtcctctcatattccgttctggtagagcggacccgcACCAGAGACTGTCTCCCAAGGCCGAGGAAACC
TATAACCCCTCCTCGCTaccccgagatagtcctctcaatacGAGCGCTCTCCGCCACGCTACCAGGTCCTG
GTAACGCGCCGGCCGGGCATTCCGTTGCGACTCAAAACCCACGCGGAAGGCGATCCCAACatgcgcaaaat
gggggctactcgtgAACTCTTACGCCTCATGGCAGCCGAGCTCGCACCCGTAAGCCCCTCCCTGGGCCAGT
GAGGCGAATCCAGACTACGTCCACCGACATCTTCTAGAGgccatcatattttattacgTTTGCTTGTGTTT
CTAAATTGGTCCAAATCGCTTTCCCCCACGAGCTTTTGCCCTCCGGGTTGTATTTCAAACCGTTCAGACTA
AACGTTAACTCGAGGATCCCGCGACCCAATTTTTCGACCACAATGGCCCCGTAAGGCAGCAAAACCacata
cgagtgtcacaaatgtattttaaaaaccaCCCACTCAATTCAAACGCTCCATTAGCAattaaggcacaacc
aatCACGATTGTGTCCCTAGAGTATTTTACTACTACACTCGCACtatgctcaataacatcaatatcatcaa
aatgGATAGGCATAGTTAAGAGCAACCCAAGCAAAGGTTCATAATTATAAACCATCACATTGTACATATCA
TAGGAAGTGTACATCCACATGCATGCATTTTAGCAATCATATGAGCAATCACAAAAGCCATATAGATGAAC
CTCATAACAACTACATCAAAAACTTAGGGTTCATAACCCGGTAAATAcacatgaaatatttataacatccA
TTAATACACCAtaattcaaacataaataaGCTTGCTACTATATAGCAATACGAAATCCTGCAGTGCTTTAA
CCGCTCACAACACAttaacaagataaaaaataccatTCAAGTgtttaacatataacacaaatcttAACACC
TACCAATTTATATACATCAAACGATACAACTcaaatgcatcgtaaggtaggttcaaacacttataaataaa
CGTAACATTCATAACAAAGCACTCTCCCATTAACAATATAGTTTAAGGTCGCACACGACTAGGAGTCGTTT
CCCACGGCAGGCAACACAAGGTGGACTATGTTGGTCTAATTGCAACtaacttacataatatttctaacgTT
CCTAATAACATGCAAACCACAGAACTCTTGTTCACAAGCACgactaggcaacgtcaaaTACATACACAATT
ATATTGGCATAATATAGCCGAGCACAATTAGCCCTAATCCGCCCTAATCGAGTCGACAAGCAATGCCACAC
GAAACATGTTCATACAACAGGGAGCATATATGAGGATGTCCAAACATGATTACAAGCACACCCAATCACCA
TGCATGACCATATACCTCTAATCGACTAACATTTATATGTTCgatgcacatacatctacatTGCCACTTCT
GCTATCCGTCCGTACACTATCCGGCTGGTATTTACCGTCTAGGAAAACGTGAACCACTACCGAGCGCCCAA
TATCGGACAATATCAATGTCATGATATCGAACAAAGTCACACAAGACGATCTGTATAGCTAGCATGAATAT
TACTACTATCTAGCATGAAAGTCATTAATGTCGAATACATGCTATCACAACCTAGGAGTCATATGTTATTC
TTAACAtaattcctagcatgctttTCTAACGGATAACATCTTCTAATACATGCTATCGCAACCAACAAATC
ATATCATGCCGACATATCAAGACAGGCTACTATATCATAGCATACAGTACTAACATAAtttctaacaatat
ttttaacaaccaaGTGAGACCGTAgggggcactcaccttaatccgaacgtaaaccttctagggtttaccga
tcGGCGCGTAAACGTCGAAGCATTTCCGGAAACGAACCCACGAACGAATCCACGATGTCGAGACGATGTGG
CAGAGCGGTCGAGTTCGCCAGAGACTTtgccggaggcggccggacggcagctacagtggcgacggcggcgg
cctTGGAGCTcaggaggcggcggagacgtcGGTGACGACCGGAGAGGGGGggcagccggcggcggcggtgg
ccgAACAGATGAGCCGAGATTAGACGGTGGAGATCCAGGCGATCGGCGCTTCAATCGGCGGCGATGAGGTT
CGGTGGGGCTCAGTTGACGTCGGGAGGCAGCGGTGGTCGACGGCGAGCGGTGGAGACACCGGAATCACGGC
GGCGAGGATTAGGGTTCGCAACCGCGCGAGCGTCCGCGCGCGGGGGAGAGGGTGATAGCAAGCCAGGGGGA
GAGAGGGGTTGCTGGAGGAGGTGGCGCGCACGCCGGCGATGGGACGAGCGGCGGCCGGGCGGCGCGGCGAA
CGGGCGAGCAGATCGTGAGAGAGACGCGGGCGGGAGAGAAGGAGGGGAGCACgcgggcgagagagagagga
aggagACCGGGCGACTCGGGCGAGTCAACCTGGCTCGCCCGGGTCAGTTGGGCTCCCGGCCCTGGGGCCGA
GTCCAACTCAAAtagggcccaaattggcccaaaatgcaaccaaatcaatccaaatgcaaatcaagcatGCA
AATAATCAACTAACTCTCGATCGAAGGTAATTAATCCGAAAATAAATTCgtttctaactaatcgaacacAG

GGACCTCACATCATATGTCTTCGAGCATCGCACTATGTAGTAATACAGCTAGCCAAAGATGGTCTAAACTA
CGCCCACTGATCTCTATATAGAGACATCACATCTTTAACCACTCTACTAAAATTATCGCTAGAGCAATTGG
CTCATATACCCATCCATAGTAGAACGAATCTCAAAGGCACACCACACTTATGTCACACTAATATCCCCACC
AAGATTAAGCATAATGGCATATATATCATGTCATaatcctaccacatgtatcaCTAGAGCTACCGGCTCAC
ATACATAGCCAGGTAAAACACAACCACAAGCACGTTATAAAGCAAACATATTCTACTGAGGATCTTACCCG
AGCACATCCACTCTAATACTCCTCTAAGTAAATTTCCATAACAATAATATGCTAGCATACAACATGCTTCG
CTTGTTAGCCATGGTAAAATACACCAACATATTCAATCACCCAAAATCAACTAACATGCTAGGATCACATA
ATAAGTATAACAAATCTTTTTCACACAAGCAAATCAAACACACCACACATGTAATACATCGGCTGGACAGC
TTTCGATCGACTAGACTTGGAGTTTTGGTCCACCGAAAATCTGCAGGACCCACGGATTGGGCAGGAAATGG
ACTTGAGCGCGTGAGTTTAGTCCTTTACCACTTGTGCGGGGCAATTTGGTTAATATACCGGGAGACCCTCA
GAGTGTTACCTCTCAAATGGATCCATCATTTTCGGGCTAGAAAAAATGGACCCGACCTTGTGGTGCAGCCGG
GACTTGGGCGGTGCTTGGGCCCAGGGCTGGTCGGGGTGTGACAGAAATGGTATTAGAGTGACCCTGTAGCC
TATACGCTAGGTTATGCGGAGGCCAGACGGGAGGAGATGCTGACTGGGATTTGTAGGAGTGCCTCGCTGTG
TTAGACTAAGTGCAATAGTGGTACACCAACAAGGGAGAGTTTGGGCTAGAGCTTCTGCCAGCGACTCGTGC
TACACATTGAGGATGTTGTATCCTAAGGATGGGGAAATGTAATTCCCTGGTTGGACAGCTTTCGTTCGATC
GGACCTAGAGTTTCGGTTGACTGAAAATCTGCATGATGCATGGAAGGGTTGGAATTGGGCCTGTGCGCGCG
AGTTTAGTCCCACACCGCTTGTGCGGGAGGGCAATTTGGGTTAATATGCCTGGAGGCCTTTAGAGTGTTAC
CTCTCAAGTGGATCCACCATTTTCGGGCTAGCAAAAAGGAACCCAGTCGTGTGGTGCGGCTTAGGTGGTGC
GGCAAGGTCTTATGTGGTTCGGGGTCTGGGATCGGTCGGGGCATGTCAACACAAGTGcaaaaccgctcacc
tcaaacGCAGCCCTAATCGTTCCACGCGTCAAACTTTGATGAAAATCGAAGAAGTTGGTGTCCACGGGTAG
ACAACAATCTGAAGagttcaaatttatgtttgatttAGCTCGAAGGCTCCCCACACAAGTAGAATAGCTTA
GGAACTCCAAGCAAGCACTTAGATAGGTGAGGTGCAGGTGGAAATGAAATATTCTCTCACTAGAGTTTAGA
AGTGTTTAGGAAAGTTATGAACTTTTAGAGAAGCCAAAACACCAAACttgtatttatagatttctCTCGCA
ACGGCTATACGACTAGTTTGTTCGAGCTTTCGGGTCAATCAAAACGACTTTTCCGTAGTCAGGACGTGCCA
ATTAGTGATGAGCGGCTATACGactagttcttttttttttttttttttttttttttgagattatcatggtg
gaCCTGCCAATTTAATAGCTTCAAAAAATGCAGTTGAGAGTTTGGTTAATCGGACCTCATTCTGCAGTAGA
TAAAAAATTCTCTCAGTTCAGCCATgattttgagaatatataatatttgattttggtgtCTAAAATCATCC
TAAACTTTGACCTTAACTTTAAATCTATATTCCTAGCACTATAGAATACAAATTTCACtcactaatatata
aaaaattcttgaacGGTGAGTGTTGGAAAGATGCCATCTTCGTAAACATCCGACTTTAAGCTATAAGCGTC
TCCAATGTGATTTCCTTGTCCTGTGCAGGTTCTCTAGACTTGCTAAGCTTATCTAGTTTTGTTCCATTGAG
TGGTGAGGCTTTGTTTGAATTCTTTTGATCTTCTTGGCTCATTAAGCAATACCTATACTTagcaaaacttg
aacaaacttATTCGTAAtacttttactaattatttattatcatcaaaatcacaataGGATCAACATTCTTA
TTTGGACTTACTTTCcatcaaatttttagttgaGGACTTCTAATCCTATTTTACAAGTAGGAAGGCTTTTT
TGATTTAACAATCccaacaaataaaatctttcaagGGTTTTAATTGAAGCTATTTTCACATTTATGGCTAA
GAACAATTATAGTTGTAGTTGTACCATATGTTTCATGAAATCATGAAACCAATTCATCTAAATACAGATAC
ATATAAAGCATATGAATCAATATGTAAATtaaattcatacaaaaattCACCATTGTTACTTAGTTtgcaaa
tttttgtaaaaattaatttattcgggCTAATTCGCAAAAcaattcaaatcaatttaaaaatttactcaTTT
TCATCCTCATCATTAGTGGAAGGCTGGCTGGCTTTACCATTTCCTCAAGAACAGAGGTGGCCTATGTTCCT
CAAACAAAAGCGTTGGAAGAAACGTGTTTTTGCTCCATTTCGTCTCCTTGCCAATAAGTTCTACCTTGTGT
TCGGCCTGTTTGTCCTTTCACTCTTCTAATTCAGTAGGTTACAAGTATGCATCCTTATCTGTTCATGTTCT
TGAGGGCTCACAACTGGAGGCAATGCAAATGAGCCAAACCAAAATGGCATGTAAGAACAGTGGACTCCGTT
TCCGTTTATAAAGCCACTCCTCCTTCCTGAAGCCCCTCTGTCTATATTTGGAGCTTGGATGTTAGATCCTG
CTAGTTTTCATCACCACTTGAGTCTTAATGACTCTCCTCCCTCATCTGCTGCATATATGCTTCTTGTTAAT
GAAACattaataattgtaaataagtGTGCAACAATATATCAAAGCAGACTATaagtaatgaaaattattgG
TTCAAACTTCAACAAAACTATTGCTAACATATGTTTTTGGGGATATTCGTCCACCGAAGCATTGACATGAT
AACATtacattatttaaaacatcaaCGGAAATAGTACCATTCTCCTATTATGCTCAGTTGTGTTGGAGTCA
TCCTACTGAAGTGAGCCACCCGAGTGtccacgtttttttttgccaaatgcCTCGGAATTCCAAATTTCAC
ACTAGTCATTGTACATTCTCCTTTAGCAATCCCACTTTGCCAATTGGCTCAGATCACAAGATAATTGGTAG
GTATATACGTATCATCTCCCTTAAGCTTCAAGGATCACATTCCATCTGCATGTGCTTCCTCGACCATCTGC
ATGTGCTTCCTCTAACATGCTGCATCCTACTGAAGTGATACAGTAAATGCTGTAGTTAGGATTGATACACA
TGCATGCTTACACATAGAGTAAAACCTCCAAGTTTCTAACTTTCTAACCGCGCATCCTATTCGATTTGCA
CCAGCACATCAAATCTGCAGAgatgcgccagtgccgcgtgCATGTGAACATAAAGTTCATTGTTATGAGAT
ACAAAATGCACATGAAACTATTATTAAACGAGAAATATAGACACATcattaaatatgcataacaTTTACAT
AATCTGAAATATTCAAGATCGTGTAAAGGAGTTCCACGTAAGTCCAATAATAAAGCAAGTTTCAATAATTT
GTACAACCTCAATCGTAATGCTTTAACATTATTCGAAGGGATAAAAGAGAGCAACATAAGACTGATTATGT
ACATGAGCTAACCTCTCATATAGAGGTTCATAACTCTTGGGTCACAAACAACTAAATAGAGAATGACAATA
ATAACGATACCTAatacatatctaaatatattctaacatattCCAAGCTCAAGAACTAAATCTTAAGTTTG
GAAGTGTTAATTATCGATCAAACACATGTCGATGCAGTTTTAATAGGACTGGCAAGGTGGCACATTGTGAT
AGTTGGAGCATTGAAATATATGGGCTTGTTCACGAGCAGAAGATTCAAAGAACTTCTGCTTGATACCATAT
TtgggaaaattaaaaaaatagagattgtTATTATCACAGATGCTAACCTCTTATATAGagactaataaata

ttaagtagGAAACGATTAAATAGGACCTATAACAATTCTAAAATTGCAACAATATTTTCTATGACtccatt
tttatgttttctaacATAGCATTCGTGCTGTAAGTTTCTTGGATTCCTCCACTACGTTAGCTTGCTAGTGA
ATGAATCAAATGTTCAACAACATCAACATGAATCAAATGTTCCATTGTAACCTACAATTTAAATGAGGATT
ACAAAACTAACAAACTCTTTATTATCAGCATGAAGTCGGTGGTCGATCGATACTACAAGACTAAGGAAAGC
AACCACCTAATCAATGCCAATGCAGAGCTTAaggtatatattatttccCTTTGTGATAAATGTATTGGTCT
ATGCATTTCATTAATGCCATTTCTGCTGTAAATCCATACATAATGTATACCTAGCACTCTGACTACGATAT
TCCACCCACTTTGAATGATTGTGAATTGACAAAAACTTGGATCGACATAGTTATGCAACaataaattatga
actttataaaatttaaaaaaaatgaaaataattgtaGGATTACAAAGTGAAAAGTTTAAGCTAGCATATGG
TTGCGATCTTATAGGACCAACttgcttaaatatatatggagaCAGTAATGCTTGATATTATGGACCATGTG
ATTCTATGTCACAATACAACTTGGCAAATTGTTCACTGTTTGTATAgaataagaaagaaagcatTATTGCC
ACAATACAACTAGCAATCGTTATAATTTTACGTACACTTGCGCATACAATCCCATATCAGTTATTCagata
aaaatcaacattatACCAAATGGAAACAGATCTAGACAGAGAATTCTGACATACCATTACGAAGAGGGAAC
ATATCTGCATTTGAAATTCTTCTTAGAGAGACAAGATACTCAGATTTCTCCACTATACCACTAACATGTTT
AGAAGAAGACAGCAAACAATACTGAACAATTAACAAGATGAAAATTAATCACTGAACGTTTCCTAGGTGGG
TGTGGTTGCATATCCTGCATGGACATAGCATGTGTCAGATTAAATATGTGGTTGCATGTCCTGCTTGGACA
TAGGAAATGTCAGATCAATggagtaattttgtaaaatagctCACTAAAGAGGTAGCAAAAACAATATtgct
aattatatttgtttcctGAGCTTACTTAGCTGCTTGATTTACCGAGCACATCTAGCAATTATAATTTGAAC
CTTGATGGTTTTGCTCCAGTTTTGGCAATGTGAGGCAGCAAGCTTGAGGCAACAACTGCATAACTTACAAG
AAAGCAATCGGTATAGTACTTTGACCCTGCAATCAACTTTATTTCACTAGCAAATTTGAGTATGTACTGAA
CTTTCTATAAGAACCAGGAGAAAGCATGCAGATGCTATATAGTGCTGAACGGAACCAATCTTTCTAGCAAT
TGCTTTTTGTATCAGAATATATGTTGGGACCTTAACTGTAGCCATGATGCACAACACTTGAAACTCAAGGC
TAAAAGGCATAGCATCACAAAATGTCACATTGGTCAAATACTGCGAAGATGCAGTCCAGGAACAACGTACA
AATcactttcaaaattacttGGTAATACTTGCTTTTAAGGAAACAAACAATTCTCGACCTTATAACTAAAT
CTTAAATAAAGACatttttttgaaggaaaaattccaagtaaaataaaatatagttccTGTTACCAAGATAAC
AGTCACCAgttcagtatatatatatatatatatatatatatatataatagttggGTGTGATTGCTGCAAAA
TTGTCTACTTAAAAGATCTTGCAGATGGATTAAATAGATTAGGGAGTGCCTCAGTGGTTACCTTAGCAAGT
TGAATCACAGTACCATTGCTGGTTGTACTACCTACTACCTATCTAGGATACTAAGAAAAGAGTGCATTTTC
gaccacaaaagaaaaagaagatgagaGTGCAGTGatttcaacatcaacatCAGCATCATGTAGGGTTGTG
CATTCAATATAAAACCATCTATCTTCCACAACGTCGAGGGGAGTGCTCTTGTAAGAAAACTAAGATGTAGT
CttcaatgatttttttaaacagaataAACATTGTTTCATACTTGCTTCAGTTGGTGGGACCCTAATTTGTTT
AGATTCTATTACAAACTGATGCGATTCGGGCGCGTAAGAGATCCTTCGCCTTGTGTCCAAGTGTTGCGATC
CCGACTTGCTACGATTCACTTGTTGCATCAACTCCCAAGTCTACTATAATCAGGCTTCACATCttctaaaa
ctatttctaaggATAAATTTCCTGTGTGGTTGGTTTCATGACCCGTTGTCTATAGGAGGCAAAAACCTTGA
AATGTGTCAGGCTCCTCCAtccgttgcctatagagagaggcaagaacctcaAAATATACCTTTGGATGCCA
CAATCGACTACTGGGGTCGATTGATAAGTCGGTGGTGGATGCcacggatctctccgataagttCTTAAGTT
CTCTAAGAACCTAAGAGGTTTATAAAACCTCACAATATTGAACACAGAGTTGTCTTCATTGATCCCTTAAA
GGGGTATAGAATACATAagaggaggggggggggggggagaagAGATCTGCCAGCCCTAGTGGCCTTGAC
ATAGCCAAAATCTCTAAGCAAGAatcaaggaaaataaaatgtagAGAATCTATCAAATATCCCTAATGTAG
TTCGGTCTTTAGTGAATATTAGCTGATTTAGCAACTTTCGGCCCTAATACATACTTAGCCGAATTAGATGA
TTTGGCCTTAAATTTTAGCCGAATTACGTGGATGCACATGCTTGGCCGGTTTGAAGACTTAGCCAAATAAC
TCTTGGAAGGTGGCCGAAATATACTTGGATGTGTAGAGTCATATCTTAGCCAAGTTGGATAACTTCTTGGC
CGAGATTTGCAATGCAAGGCTGGCATATTCGTTGATATCTTTCCAAGATCTTTGTGGTCAACTTGTTGGTG
TTGTAGCAGACTTGTAGGATCATGATTCTTGCCGAACTTGAATGAGTAGCCAAATTGTGTAACATGTCTTT
GCTCTGCCATCCTGGAACAACTCGTGATGtagccaaaataaattcttgCTCAACATCTTCAAGGATCTTCA
TGGACTGGGTTGCTTGAGGCCAAAAGCTCCAACTTGGCTCATTTCATGTCTTCATGAATCTTTTAAGTCGA
ACCCTCGTAATTGGCCCAATCGAATTTTGATCACGCTCCACATCATCATTGGTTGTGCCAATGGTGCTCGC
ATCATATACTAGCTGCAGCCATAGTCAAATGGATGTCTAACGATAATGACGGATTGGAGATATAATACGTT
GGAAAGATACATACTTTCATTTAAAGATAGAAGCTGGAATAAAGTTTGGACAGACAACCTTTTGCTAATAT
AAATGCAGTGGAAGTGGGTTTGGAAAGGGTTAGGAGAAGTTTGAGATTAAGGTCCCCTCTAATTCCTAGTT
GGATATTGAAGCTTTTGacctaaaattttgaaaaaaaaaagggaagaaaatcagaaaatgtAAAGTTCCAA
TGATCTAAAAAGATAAGCTAACAGATCAAAGATTAGCCAAGTGCAAAATCCATGGAATAGCAACAACGTGG
AAAAAGACTAGGAAACGAGCAAAATGACAAGTTCAGGCAAATATGAGATGACAAAAGAGGATAGGATTGTC
AAATCTAGATAGGGAATTACTAAAGACTCAGTTATACAGATGATTAGTTTGACTTAAGTATCTACTTAAAG
AGGGTGCATGGCCAACTGACGTCATGAATCCGTCGGATATTCTATCTGGAGTATTTAGATTAAGATCTTGA
GAAGCATCTTACTAATCCAACCttcctctattttctaattgcaGCCACCTGCTGGGAGAGGAGCTAGCCTC
ACTAAGTATCAGTGATCTTCACAACCTTGAAAGACAGCTAGAAACGAGCCTACGTGGTGTCCGAGCAAAGA
AGGTCCCAAAAGTTTTGATCGATCtcaacttaaataatttatactagtTCAGTATGTAATATCGTAAATGC
ACATCTGAAACGTGTAGGACCAACTTTTGATGGGAGAAATTGAAGAACTTAGGAAAAAGGTCTTTCAACGA
TTTTGTCTTCTTCTCTAATACTATTGAAAGTGATGATTTAGTTGTCTAATTTCATTTCTTCATACAGGGGA
ATATCGTGCAACAACAACATGCGGAACTATACAGTAAGGTAAGTCAGATTCATAGGCATTATAAtgtgaat

ttttaaatcaagcGTTATGAGCtactgaaatataatttgaagaaGGCAGACATAAATATGACATAGTAGCA
CAGTATTCACGAGCAACTAACAAGTGCTATTGTGCTTGCTAGGTTTATGCAACAAGGGGGTTAAATGAAAT
CAATCAAGAAATGGCTATTCCGTTTAACTTCTGTTTGTCAAAGAACTTAAACACTCCTATCCTTATCGAGC
TTAGCCAGCCACAAAAGCAACGAGAGCAAACTCCGGATGGGGCTCCAAATCTAGGGTAAGGTTGTGTAGAC
TCAAGTTTCAAAACTTTCATAGTTTCTAGTCATAAATATGTTCATTTACCTTGCAGCAGACTCCAACTGCA
CTAAGTATGATCAATGGTTAACAATGAGCTTCAGCATGTATCTTCATCGGAAGCTGCCATCTGCTTGCTAT
GAGATTCAAATACTTGTTTATATGCATGACCAGGATAATACAAATTGGCAGTACATATCAATTGGTCACAA
AAAATGTAATGTAGATCTATCCTGTATTATAGAATAAACCTTGGTTATCATGAGCCATACAACTATCCTTG
TATGTATATGCAAGAAGTATGATGTTTTTCTCTTctacctaaaaaaattaagacccTTATCATTTACATC

>XviAGL17_5
ATGGGGAGGGGCAAGATTGTGATTAAGAGAATTGACAATTCGACGAGTCGGCATGTGACCTTCTCGAAACGAAGG
AGCGGGTTGCTGAAGAAGGCACGGGAGCTCGCCATACTCTGCGATGCAGAGATCGGTGTTATCGTCTTCTCCAGC
ACCGGCAGGCTCTATGAATTTGCCAGCACCAGCATGAAGTCGGTGGTCGATCGATACTACAAGACTAAGGAAAGC
AACCACCTAATCAATGCCAATGCAGAGCTTAAGTTTTGGCAATGTGAGGCAGCAAGCTTGAGGCAACAACTGCAT
AACTTACAAGAAAGCAATCGCCACCTGCTGGGAGAGGAGCTAGCCTCACTAAGTATCAGTGATCTTCACAACCTT
GAAAGACAGCTAGAAACGAGCCTACGTGGTGTCCGAGCAAAGAAGGACCAACTTTTGATGGGAGAAATTGAAGAA
CTTAGGAAAAAGGGGAATATCGTGCAACAACAACATGCGGAACTATACAGTAAGGTTTATGCAACAAGGGGGTTA
AATGAAATCAATCAAGAAATGGCTATTCCGTTTAACTTCTGTTTGTCAAAGAACTTAAACACTCCTATCCTTATC
GAGCTTAGCCAGCCACAAAAGCAACGAGAGCAAACTCCGGATGGGGCTCCAAATCTAGGGTAA

#### Dioscoreales

##### Dioscorea rotundata

*DroAGL17_1 and dro-miR444*

>BDMI01000006.1
ggtTCGTAATGCATAGGAGAAATGTAGGCTGTAACTTTTTGACTTGCTTGGATTCTAGTCAAAAGGGTTGGGGAT
AAAAGGTTAAAATAGAGTGGGGATAAAGTTATCCGTATTTTATCACATTTGGTTTAACATTAGATTGTATTAAGG
TTATCATCATGTGGGTAGTTGGGCATGTATCTTCGGATATCCCAGTCTAAAATCGGATTCAAACATCTTTATGtt
tgtgttatatatatttttcatgttaCTTTTTGCTTGGCATTTtgaaccaaatttttttttttggcttgccAT
TGTttaaatgaaagataaaaatgCCAATGCTGCGTAGTATGGTTAAGCTTTGTACTTAGCAGTCCTATGAATGCA
CTAGGTTAGATTGTTGGGTGATATAAGTTGATGCAgagaaccaaaaaaataaaaataaaaataaataaaaaaata
aaagactagGATGAATGGAGAGTGTTAATCTTTCATGCCAAGTTATTATCAACTCAGACTTcattaatgattata
ttaagctgattaattaaatttgtaaaatgggAGGATTCATGTAGCCGACCCTAAATAGTTGGGACTAATGGCTT
GTTGTGGTTGTAGGAAATGTCATWCTTTTTCCAAATTATTCAAATCATTTTGTTTGGCAGGAAGTCATGGGAATA
ATTTGTTCCCTGCCTTATAATGTACGATCTAACTTAAGCAATGTTGAATATGGCTCTGGTGAAATTGTAACATTG
CTGTTTGTTTGAAGCtggtatctttttttttgtatctttGTGCTCTATTTAGAGGTATGTAGGTGTCCTTCATGT
CTCTTAGTGGTGTTTATCGTGCTCTGTTAATTCACCCATCTTTTAACTTATTCAATGATCCCTTATTTGCCTTCT
TAATGCGTCACATAATAAATTAGATACCTGGTTATGATTGTCTTGTCAATATGcttaaatgaattattttttcagT
GCTTTGCATGAACTaagctttatttttcttattaac**cAGTTTACGCAGATGGAGGCAGTGAGTTTGAGGCAGCAA**
**ATGCATAACTTCAAGAATTATAAATG**GTATGATTTCGATCAGCTTGAGttcattttaaatattgaattcaGTTCA
TAMGTagtcaataaaaatattaaattgggGTGCTYAGAAATAGTTAAAGTCAAAAATGGGGATTGATTGGCATTT
TTTGAgtgttttctttatgtttttRWTATATTGTGTGGAAATGATATTTTAGAACACCAGTTTAAGATATGCCAA
GGCAACATATTTTGGYATTATCTCTTAAAAATCGTAGTTTAAATACATTTTCACTTCAGTGCCTYCAAATCTTTT
TGATGATTAATGCATCATCGGCATCTGGTTatagtataatttttaatatccTGAGATGTAGAGGTGATTCCAGGA
TAATGGCAGAACTATGTTCTTTTAATCAATTAAGTTTATTGTTGATGTGGGTACTGTTATGGAGTTTATGTATGG
ACTTCTATTGGGCTAGAAGATGATGGCATACAATGTTGgaggaaattataaaattaatacataaatgaAGTGTAA
TTGTTGATTTTCGCTATGYATAGGTTAATTATCAAATGATCTACATTTTCTGAAGAAACATTAAAGTCCGAAAAT
GGTGGCTTTACAGCTTGAAGCATGCTCTGCTGTTTTCTAACTCTTCAGCTCCACATGTGACTGTTAGTATTGGAA
TAGATCTCACAATCATGATTATACAAAAGCATTCACTATTTGCAGTTTACCTGTTTTTGTCGCAAAACTTACTGA
TGGTTCTTAAATTGCCTTAGTGCTTAATCGAAGAGGCCCCCATAATCAACTAACATACCCTTGTAAGAATAGTTT
GTCCTTGTTTTTTTGGTTCTgttgtttttttccttgtttggGGTGTTTCTGTTGGTTTGTTGTGTTCCTGCTTGT
TCTGTCTTTAGTTTTTTCTAATAaagtagtggtttatccacttttattaaaaaaaagtttaactTTGCTTAAAAA
AGAATGGTATTGATATTCTGACATTTGGCATCTTCTAGAATATTCTTACAAATTACAGAKTCACAAAGAACTTTT
GAGCAAGATAGGTTTATATGATTGGGACAAAGTTTAWTTCATTGGTCATGTGGAGGATACCAAGTTGTGGAGACA
CCACTTAATGTATAATTCGTTGTATTTGTTAAATGATTGCAATCAGAAGGAAATAAGATATTTGTGCRTTACCCT
TAATATTTGGAAGAGCATATCTTGGCATGTGAACTGCAAAATATAAGTCAACCATTTGCTATTTGTTTTATGCTT
ATTCATCATGTTGCTTTGCGGATGTCATTTCTGTGCTTTGAGAGGCGGTTGAAGGACTAGAGGTTAGAGATCTTG
GGTAGGCTTTTCTAATAATTCCAAATGAAGCTGTTGTATCACACCTTGGGTTGTTGACcttgaaattttccttaa
aaggaTTGTGAACCTAAATACTGTATGTTAATTTTATGGCTTAAATATTGTGTTAGTGTGTGATGACtttaatta
gatttttatgattatctatctacttatttattttttgaaaactgGGGCAAGCATGCTRTTGAGAGGTACCGGGGG

TAATTATTGATGGAATTTCattcttgtatatatttagaaatgcCGGGCCAACTTAGTGTGATTATTATTKGGCT
GGTATACTTATAGTTTGGTAGACAAAATGGGGCAATTTTGTGATGCTAGAAGAGTGATAGTTATTGCTGTGGTGG
TATTTATTGTAGTTATTTTGAGATAATGGGATAACATGGAGTTATGGTTGCTGGGTGGCTTCTATGTTGTTTGAA
AAGTTGGAATCRCTCTAGAATCACAGAACTAGAACAGGATTttcatataaacatatattgtAGTATTTCTAGATA
GTTATTGTTGAAGGCCTGTTKCAgtctttttgaaatttcaaggACAGTCTTTGGATTTTGGCATTAACAAGTTG
GTAATTTCACTGggaagtaaataaatattttgggaAACCTGTGGCCATTTACTGTCTCATTGTGCATCTTGTGGT
GTTTTCCTTTATAGCAATACTTYRTAAGAAATTGTCGGTAGATTCAAAGAAACTCATTGCTATTTCCTTTGGAAT
GCCTCTTTTCATCCATATGATTTTGTARTCTKYTGTCCTTCTGCAACATGAGACTTGTTAAGATGCATTTTCATC
TATGGCCATGACCATTTTTATGATCCAGCTCTCTTTGACACCTCATGTGCGTCAGAATAATCATGACTCAGTATT
AGTAATCTGAAGATACAATTATAGTGAAGCAAATAATCAGTTTTCACATCATTGCTGCAGTTGAACAATACATAC
TTCTTGGTTATTGTGCTATATTGTTGCCACAAATAAGTGATTGCACTCGGTTGTCATTGCTTCTCTTGTTTACAA
ACCAAATCATACTTAAATAGTGTACGAGTTCATTGTGAAGTAATAAAACATAACCTTGTTTTTAGTTCTTGATCA
CATAGTTCAATCCTGCTTTAAGCTTCTTCAAGTACTAGGTTTTGAATTCCATTTAAGATCACCYCAATTTTGAAG
ACCTTGGTTGCACGCGATCCTTTTGATGTTGTGGTTGGCTTAGTTCAAGGTGAACTCGGACATCTSTGTTCTCCA
TAAAACTGAACCCATGTGGAATAACTGGGGCATTGTCAACTCCTGTGCGTCTTACGTYATAAATCTGATGCAAAA
TTCACATATAGCAAGTAAGACAATAGAACCTAatgttatttatcttttatttatcatcCTTTAAGATTTAAAGGT
TCCCGGTATTTCCTTTCAATTTAAGTGTCTTGCTATAGCTTTATAcccatttatttaataaatataatatgtttt
GTATTATCTCCTTAATAAAGGGTTACCTTCTTGTAGAGTTCCACGTTTTCTAGATGGAGTAGATTTCCCTGCGAG
GWAATCATAGAAAGTTAGGCTGCTAATACTCATCACCCTAAGTTTAAAGACACAATATAACATCTTTAAATGACC
TTCTGCTTTaattcttcaatttcttcaatcaGAAGTTGCtcctacaacaacaacaacaacaaaagccaTTAGTCC
CAACTATTTGGGGTTGGATAGAAGTTGCtcctagaaaaacaaaaatcaaaagaaaaagaagcaactATTTAGGGT
TGGCTAGAAGTTGCTcctagaaaaaacaaaaatctaaagaaaaagaagcaattaTAGTGGcagtttttttttgtg
aagagATGAATCTTGAGAGCAGAGCTATATTTTATTACCTTTTTCATTCTCACACTGTGTAAACTCATATCGAGT
TGGTTCTCGAGATTTTGTAGATCCTTAATGTTTAGGCCAGAAAGCTCTTCTCCCAGTAATTGCCTGCAGTTAGCA
TTAGAGATATGCTGGCATGATTTGCTGATGTTTGTCGAAGTTTATAAGATTGCATTCGGGTAGGATTTTTGGGGA
TGATTATGCAGACTTCATACATAGTTGCTTTGCCTGAGCGGAATTTTAGTGATAGAGAATTCCTATTTGCGATAA
TTTGTGGGATTGGTGGAAGGTGTTACACTGAATTTTATGGAAACATGGTTAAATATACCTTACCATTAGCAATAG
TATGACAAtaagtattagggttttgattgaaggaTGCAAAATGTACTATGCATACATTTGATGATGTTTAACACT
TTGTATCACATGTTTGAAAACACTTTTTAGTGATAGAAAATAAGCAAATTACTAAGTGTGGATTTTGAATGTCTT
TGCTTGACATGTTTTCATAAATTTGCAATAAATGCAGTCAAGCTTGCTTAAAAACAATTGCACTAGGTTTTTAGA
CAATTATCATTTTATGTAGTGAAAGCATGCAATATCATTACGATATGTTTTTAAGGCTTGTCTGAGCCCAAGGTT
TTAGTTTGAGAATCTTGCTAACATCAAATCCCTTCAATAAGCTTACCGATAATAGTAATGGTAAAACCAAAGATT
GAGAAGTTAAGCAAAACGGTCAAATtgatcttttctcttttacacAATTTGTTTAATATTCAAACTTATACtata
atttagatttatttttgtgatagTATTACTTCTAGATGTGACTTTAAATGACACATTTTAATTACTTGTGTTTTA
AGTGCGTCTAACATGCTAAATCAAATTGGTTCACAGTAAACTTGTTTCGATTGATAAAGAAATTGTGTGTCAAGC
TAACTTTATCACTTCCTACTTCTCTAAGATTAGTATGTAAAAATGAAACATCATTACAATTATgagttttagtat
tttttattagatataGCTCAATTAGATTACTCCATTTGATTGGAAGGACCTATAGTGTAGCTTTAACTGACAATG
TCTGGTTAACAAAGTTTATAATTAGGATTGATAaatggggatttttttttactagttcGTTATATGTTTTAAAGC
AAAGTGTGATATCCTATGTTCTTRATGGGAACTAGTTTTCGGtatgaaattttataacatGAGATGTGGCATTTG
CTCTATTACTTCTGGTTATAGATTGGCTGAACACTATTTGTTGTGAAATGATTGCATACGATCAGGAAAGATTTC
TAATGTATAAAGTTGAATRTATTCCAG**GGCAAAGTGATGGTACCGATGACTTTCTTGCAAGTTATGCAGTTGYTG
CCTCAAGCTTGCTGCCTCCCTCTGCCAAAACTAGTATGGACAAAATAACCCATGTCAATGCATTWTGGTAATGTG
ATGATTCAAAGATATTTTTTGATTAATGTTAAAATCCAGCAAGCTTCAAATTTGAAGTTGTCTCATGGTTAAACA
ATGAGTATTGcttatgacattttttttttaaactaacaaATTCATTGCAAGGAGGCCCATCATTTACTAGTATGT
TTAAGATCTTAGTGAGATTTTGCTTCAGTGGTCTTCACAGATTTGCTGATCTGGTCAGAGTCGAACCTTCTAATG
CTATTCTGGTTAGATAAAAATCG**GTGAGCCTTTGTACAAATACACTTTTTTCATACTAGCTGTACTTTCTATAAC
TTTATGAATAGCCACCTACGCTTATCTGTTGTTGCATGAGAACCCTTGTAACAAATGGTTGaagggtttattttt
tttataatgtag**gTTGAGAATGTCTGTAGACCCTCTCATGTCAGCTTGGAGGCAAAACAAATCTTTGGCCAAAAG
CATTCAGTAATGTTCTGCATAACACATTTGCTAGCATCATATTGTTGGKACGCAGTGTGGGTATTCCTCTTTCAA
CTATTTCTTGGAGAATGCCCACTACATCTCCTTCACATGAGACGGCTGATTTGCGGATGTCTTAAATCCTGCCTA
ATATGTTTGTRCCTAAACCTCTTAAACTTCTTTTGTTCTATGAAAAAATTTACTCGCATTTGGCATCTTTAATAG
ACTCTTGCTTTTTACAATTGTATCCTGTATGTCTACATAAACTCAAATATGGTAATTGTACTGTCAAAAWtttga
tttaaattttcagCAACTACTGAATAAAGTGATTGGTTTAAAATTCTTTACTAAAAATTTACCTTACTTTTGTGG
TATGATTAAAGGCTGTTTAGCTTCCTCCTTTTGCGGATGCTTCATCGAAATGTTTCATCAGAACATTTTGACAAA
GCacatgtaagaaaaaaaaagaaggaattcATGGTTTTTAAACTTTCTTTTGTTTCGTAGAKCAGGCTggttgtt
taaatttttttctgCTGGGATAGACTTTTCACAAGCTACCTGCAAGATCTTCTGTGTCACAATCTACATTGTTCAA
GGCGGCGATCTTACTTAACAACTCTACATCTCATAAGGTTGTGATTTGATTACTCTTTATTCTTCCATATGCAAT
TTTGTGAAATGAAGAATTGTTTCAATCCTTTTCTTACAACTAAACTGCCCAGAACCTTTTGCCCACAAAGTGGCA
TCTTTCATTTACATGTACTCAAGCTCATTGTACTTTGTGATTAGAGGGttttctttttcccctcttcttttgtgt
gtgtgtgtgtgtgtgtgtgtgtgtgtgtgtgtgtgttattaGTATTCATTGTTAAAGGAGCAGTGTTAGTCAATT
ACTTTTGATCCCATATGTTAGTGCTGTtggttatttttcttcttccataTGCTGATTTGtgaaagaaattattat
tgctCCCTCTTCTTGACCAAAATTGACGGAGACCATTTTGTCGCATATTGCCATCTGTTCTTTTTTCATGTAGCT
TGCTAACTATTCAGCCGTTAGCTTATATCCCCAATGGCTTTCATATTCCTCTTTGGGACTGgagcttttgttttg**

tttttgttggtcCTTTTGCCGTAGAATTCCATAAATTTCAGTATCATTGATTATGTCTTCCATTTGAGCTTACAC
ATTTCACAAGCTTTCTGTCACTATCAGATCCATTTAATATTAAAGATCACAGAATTCCCATGATCCGTCTAGGCT
TCCTTTTAGTCTAAGCAATTTTAGTATTGAAGTCTCTTGTTATGgctcaaggaaaaaaaaattcatcaggACTTA
ATGGATTTCATGTCTTTCCGCAATTCTGATCTCAGATTTTCATTACTGGACTACAGACCACTTAAAGATTCCAAT
CCATTGTGTGGCCTTCTGATAACCAAAAGAACAGTAGCATCTCATCATACCAGCCAACAATGACTACCACTGAAG
TATAGGGAAAAAAGCCATTTACATYTACTTATTTTATCCATCTGTTGAAAAGACGAATTTATGTCATTGTCTAAC
AATAAATTATTTCCTGATTTATGATAAACATGTTGGGAAGGTTTAGCCTTGGTGTACATTTAATCTATGGGCACT
GAAGTGAAAGAGTGACAATCATCCTTGTATTGAGACAGGGTAAATAAAGCTTATAGATCTAATAGGCAAAGGATG
ATAATGTTCACTTTGACAAGTCATTAGAGGGAAAGAGGACATATAAGCGGCGAATTGGCACATATACTTAATTAT
TTCCTGATTTATGATAAACATGTTGGGAAGGTTTAGCCTTGGTGTACATTTACCATGTGGACTCAATAACAGTAC
TGgggaagttaaaaaaaaaaaaaaaatctgttttGTAGTGTACATTATATAGTTAAGTGTAGATGTTTTATAATCT
TATTGACATCTTTTGCAATCCTTACACCAAGTTTCAGGCCTAGATGCACATTCTTAGATACTTGAGTAGATTGAT
CATGTTAGAATGGTGTCTACTTAATTAAGTTCACAAGCTATTGTAGATAATATGTGATCAGTGATTGCGTTAGAT
CAAAAGTTCACAAGCTAATGAAGCTATTGTACATTATGCAGCTAAAGTTTAGATGTTTTATGATCTTATTGACAT
CATTTGCAATCCTTACACCAAGTTTCGGGCCTAGGTGCACATTCTCAGATACTTGGGTAGATTGATCATGTTAGA
ATGGTGTCTATTGAATTCAGTTCACAAGCTATTGTAGATAATATATGATCAGTGATTATGTAGATCAAAAGTTCT
CACTGATTATGTAGCTAAGTGGTTTGAAATATCACCAAGATAGCAGGATCTATAGAATATGGTGTATTAGTCTCA
GAGAGTTCTCCTTCTACGCTCGAGTATTAAGGTATACCTCAACTTCAGAAAGCTTAACTAAGCTGTGACCAATTG
CATAGCCTAGCAACACTGATTCtgattataaattttgtgaACTTTAGTCTGTTTGATAAATGAAATCTATCAAGT
TCATGTCTTAATGCATTTCACAAGTTGTTTKGTTTTATGCGGATTTGATGCTTCTTACGAGCTTGCGTTTCAGAA
TTTCTTCTGATAGCAATTTAGACATTGATGGAATCATAAATTATCTACTCCTAATTCATTTAATAAGTTGATGGT
TTTCATGCTAGTTGATGTTTCTTGCAATTTCATATGTTCAGAATTTTTCTAAATAGCATGGCAATTTGATCCAAA
GGTAAATATYCTACTGTGTTATTTAACATCAATGGTCCTTTACAATTTGATCGGTGCTTTTTAAGGCATCCCCTA
TTATAACAAGGACTTTTAGATGCCTAGCAAAacacacaaaatatatatatatatcaatatatagacaaataaata
aataaataagtagaaaaatgttttaaatatgCATTTCCTTTAGTAGACAATCTAATAGATACCTTAACCTCTGAA
GCTGGATTAAATTGTTGATGGTATTCCTCTTTAGTCCTGTTGTAACGCTCGATCACTGATTTCATGctgaaaatc
aaattaaaagcaTTGTTTTTAGTGGACATTCTACTTTACATCAAGATGAAAAAGAGAATAAAGTCCCTGAGAAAA
AATGTGCAACAACCATAACAAAGAATCATTTCACAAGCAGAAGAATTAGCTACTTTAATACATCATGAAAAYATA
CTTGGCAGCTAAAGCCACTTGCATATTCGCATCTATGTATTATCTTAACTTTGAAGAATCAAAGTAGAACTTTTC
tggaaatttatttttcaagaatTATATTACTAATCTTCTCATATGTGTTGATTATAGTTGTACCTTTAACTATAG
CTAGTTGCAAGTCCTATCTGTTGTTATCTATCAATTTGTTTACAGAAGATTAGGAGAAGCAACATGAAAAGATGA
AGCTGCTTCTCAACCCAAAAGAACCAACCATACAATCAGAGAACCATCTCCAATAAGCAAAGGCATTCACCATCA
TAAGCTTTCATTCAAGTTTTCTGAAAGCTGAAGTAAGCTGTTTTCAGACACCTTTGAGACGAACTTGAAAGGTaa
tgtttaataaatataatgtcTATCTGTTTTGAGATCTTTCGCAAGTGAGCCAGATTCTTTGTTTAGGAATCAATA
TCACTTATATCAGGGAAAATTAGATTTTTGAAGTGAACCTAGATGACAATTGTTGTGTTCAATAGTGAGCCAGGT
ATTTCATGTCAATAGTGCTCCTAGTGAAAACCTAAACCAATTGATGGTCTGGTGTGTTCTGTTTCTTAGACTATT
TTGGTAGCTTAAGCTTTATTTACATCTAATTGGCAAACCAACTCCCAGATAAAACAGAGTGTTCAtccaccatat
atatatatatactaacacTTATTTTGGTCAAGGATTAAATTGTACTTTGGTTTTATCTAGAGCCTAAATGAGTAT
AGAGGTGTATAGTTAATGTTACTCAACAGGCTTCTAAATTATGCTTCTTGTTGAAGGTTTCAGGTGAAAAGCTGA
GGCTGGAGAAAGACCAAGAGTTCTtgtgatctatatatatatatatatatatatatgtttatatatataattcat
gtcTGCTTTCTTGTTGTAATGTATAGTGAAAGATCTGTTGTAATGTATTGTAAAGAAAATCAAGTGTAGATTGAC
TTTGAATATTCTTGAAAGCAATAAAATCATATCTTGGTTTTTACTAAGTTGTATGATATTGTTGTGTCTTGTGTA
TACAGTTGCATGATGAATGAGTGGTTGGAGTTGATGGATTTGTAAATAATGAATTTTGATTGCTAAATTGAAGGA
TTTATAACCAACATCAAATGgttatatttttttgttgtttgtttcctGTGAAAGCTGTCTGAAATGTAAAACCTNN
NNNNNNNNNNNNNNNNNNNNNNNNNNNNNNNNNNNNNNNNNNNNNNNNNNNNNNNNNNNNNNNNNNNNNNNNNNNNNNNN
NNNNNNNNNNNNNNNNNNNNNNNNNNNNNNNNNNNNNNNNNNNNNNNNNNNNNNNNNNNNNNNNNNNNNNNNNNNNNNNN
NNNNNNNNNNNNNNNNNNNNNNNNNNNNNNNNNNNNNNNNNNNNNNNNNNNNNNNNNNNNNNNNNNNNNNNNNNNNNNNN
NNNNNNNNNNNNNNNNNNNNNNNNNNNNNNNNNNNNNNNNNNNNNNNNNNNNNNNNNNaaaagaaaagaaaaacttggac
agctaattttttcaatctaataaaATTGTGTATGAGTAATATTACTTTATTTACGtattatctatataaaattt
taaattattaacaaccAATTCAAATAGCATGCAATTGAAAGACAAATAATTGCTTAGGTAGTGTGATGCAAATAT
TATTGATATTCTCAAGCTATACAATAGGAATAAAGAGCATAGAATTATATATTATGCATTGGAATGATTACTACT
GGAATAAAGCTAAGTATCTTATGAAGGGAGATTCTAGGAATTTATTAATTAAGGGAAGCTTATAGATCCATGATA
CATGTAACCTTAGATCAGCTCTAGGAGTGCTTTCCCATAAGATTCTATTTTGTAGGACCTACAAGAGCTTTTCAT
GGTAACTAATTCATTGTARAAGTCCATTGTTAAGTACTTTGGAAGATGCTTTAAAAATTTGGATCTTTATGAAAT
TCATAAACTAATCAATTGATGATGATATGTAATAGTTGTTGAGTACGTCGAGTCTAATGTTTGCCATGGTGATAC
CTTGTGAAAGGGGGGGAATGCCCCTTGTGAGGTTTTCCAGGGGTAACATTCCAGGGGTAACACCCctggaagaaa
aaaattatactagGTCTAGGCTTATGTCAGAGTTTTGGCTTATTTTTACTCTTTTATGTAAACATTTCTATATTG
TAAATCCTAgtttttttgtattaaaaaaaaaaaaagatactaaTGCGGTCGTGGACATAGGTCACATATTGATTGTA
CTACATAAAATTCTGTCTTTGTGTATGtcagtttttttttgtttttgtcatgGCTCAAGTTCACTCTTGATTGCTT
GATTCTTAACAATGATATATGGATAAATATATTTGACAAAAGGAGATTCTTAAGTAAGCtttctaaatattttga
attcaaCTTCCCATTAATCMAAGTAGtggaattaaatatttcaaattcaatttttttacattaatcaaactcgcca
attttttttttaaacaaaatgaataaatcataatttatttaataaaaaaaaaacaagagcatAGAACCtgataaca
taaattaaaaaactaaaaaaaaaacatgcaaaaacataaactcaaataaaataaacaaatacataaaatatattt

atgatattaagGAGATTAACATTGTAGtagagtaaaaaaataataaaagtagcCTTTAATTATTATCACTATATA
ATTTTTGCATCTATTTTAAAAGAGACCACCAAGTTTTTACCCAATAATatttaagagaaaaagaagatcaGCAAT
CCAAATCCCTAAGAAGATTTATGATTGGAATACCTTCTCTTTATAGACAATACAAACAATGTATTATGAAtcact
taattatatatatatatatatatatatatatatatatataccaatacTAATTAAGTATCAttacagttatatata
tatatatatatatatatatatataactgtaaTGATACTTAATTAGTATTGGTGATCACCAACAAAATATCAATGA
GAAGGTGAACGTGATTAATAACAACATAATGTATTTTTGTCTTTAATTAGATGAACATAACTTCAAATCCAAGAA
TAATTTTGTTCctataatttcttcttcttttcctccaTTTCCCTCAAAACTAAATCTTCAtttagagatatat
atatataaacttattcCTAAAGTTGTAAGTTCTCTTCACCCTTTTTCTCAAACTACTTCAATTTTCCCTTCTTTT
CCTACATTTTCTAAAAAACTCTAAACTTAGAAGTTGATGAAATAAATGAagacttgaagaaaaaaaaaataatac
taatttttttttttagtttgattgGATGAACAAAGTCCTAGAGTTCTCTCTTCTTCYTTCCCATYCATCAACTTCT
TCAAYTCCCTCTTCTTTTCCTCCATAATctcaaaaaaatcataatttagaTGTTGATATAATAAATGAagacttg
aagaaaaaaaaaataatactaattttttttttagtttattgaaTGAACAAAGTCCCAGagttctctctttttctt
tctcaccCATCAACTTCTTCAACTCTCTTCTATTCCTTCATTTTCTCAAAAACTCCAAACCTAGAAATTGATGAA
ATAAATGAAGACTAAGAACCAAGGAAAAaaagtttgttgttgttgtttttttttttttttttttttgacaa
ggcGGATGACCCACCATACGCTATTTGCGTTTTAATCATGGAGGGAGTTGAACCCTCAATCTCTCGCATGGGGT
TAGATTTTTCATTAGAAGCTAAATCCATTGCTTCTAATGAATGtagtttattcattttcttataaaataaaaata
agttaaattttATTGAGTTTAKTGAATGAAGTCTTAGAATTCTCTCTTCTTTTCCATCCATGGACTTCATTAACT
CCCTCTTCTTTTCCTCCATTTTCTCCAAAACTCTAAACAAATGAAGACTCggaaaaaactaataaataaataaat
tttatcaacTTTGATCAAATTAGCAAAGTCCTAGGgtcctctcttcttctttcccatCCATCAATATCTTCAATT
GCCTCTTTCTTCCCTccatttttctcaaatattcCAAACTTAGAAATTGatgaaataaatgaagattaagGGRaa
aaaaacctaaaatttatTGAGTTTATTGAATGAACAGTCATAGAGTTAGCTCTTCTTTTCCATCCATGAACTTCA
YCAACTCCCTCTTCTTTTCTTYCATTTTCTCAAAAAACTACAAACTAATGATAACttgaggaaaaaaaaactaaat
tttattgaattcgATTGAATTAATAAACTCCTAGccctctctcttctttttcccATCCATCAACTTGTTGAACT
TCCTCTTCTTTTACTTCATTTTCTCAAAAAACTCCAAACTTAGAAGTTACTGAAATAGacgatgacttagaaaaa
ctaattaaaaaataatttttattgtgtttattgAATKAACAAAATCCTAGAGTTCTCCCTTCTTCTCCATCCATC
AATTTTTTCAACTTCCTCCTCTTTTActccatttttttcaaaaattccaAACTCACAAGTTGATGAATTAAATGAG
gacttttaaagaaaaaaaaagctatatatttttttattttgattgtatgaacgAAGTATTAGaattctctcttcttc
ttttccgtCCTTCAACTCCCTCATCTTTTCCTCCATTTTCTCAAAAACTCCAAACTAATGAAGACTTGGGGAgga
aaaaactaaattttattgaGCTTGATTGAATTAATGAAGTCTTAGAGTTTTTCATTCTAATTTCTCATACTTCAA
CTCTCCCTCCTTTTCGTCCATTTTCTCAAAAACTCCAActtttagaattttatgtaataattgAAGgcttaaaaa
aacacttaatttaaataagtttgtttgaaTGAACAAAGTCCTAGAgttctcttttcttgttttccCCTCGATAAA
CTTCTTCAAGTCTCTTCTTTTCCTCCATTTCTTAaaactttcaaattttaaaagttgATGAAATAAATGAGAACT
GAAAAAacgaaacaaaaacaaaaacaaaaacaattttgtTTGAGTTTAATTGAACGAACATTTCCTARAGCAACT
CTTTTTTATATActtcaatttctcaaatactcCCAACTAAGAAGTtggtaaaataaaacttaagagtcattaaaa
taaatgaagacttaaaaaaaaaaaaaaattgttgagtTTGATTGAATGAACAAATTAATKAAGGCTAAAAGACATT
CATAGCCAATTGGCATTGAATCCCTACGTGTGGGCAAAAASAAAGTCAAATCCCaccgcaaggtgagggcacgta
AAAAATTTAGATAGCCCACACACACCCCTGACTGTCCTAGAGCTCTCTCTTCTCCTTACCCATCCATCAACTTYT
TAAACTCTCTCTTCTTTCCTTCCATTTTCTTAAAAACTCCCAACTTAGAAAttgatgaaataaatgaaaactaag
ggaaaaaagataaattttattgaGTGTATTGAATGAACAAAGTCCTAGAGTTCTCTCTTCTTTATCATccattaa
tttcattaattccGTCTTCTAAAAAAACTACAAACTTATGAAaacttggaaaaataaataaataaattttattgt
gtttgatTGAATTAGCAAAGTCCTAGGCTCTTGTCTTCTTCTAAAAaactcttaatttttaaatttctctctctt
tcccatccatcaacttcttcaacaCCCTCTTCTTTCCTCCATTTTCTCAAAAACTCAAAGAAgttgacaaaataa
ataaagacttaaaaaaaaaaacatatagatTTTATTGAGTTTGATTGAATGAACAAAGTCCTAAAGTTCTCTCCT
CTTCATTCCCATccatcaacatcttcaacacCCTTCACTTCCTCCATTTTCTCAAAAACTCCAAACTTATAAGTT
GATGaaaaaaaaagcaacaacaacaacaacaaaatatcaacTTCTTGGATTCCCTCTTCTTTTCCTCCATTTTCtc
aaattaaaaatctaaactTACAAGTTGATGAAATAAAACTTAgaagttgataaaataaatgaagactttaaaaga
aaaaataaataaataaatgttattgaGTTTGATTGAATGAACAAATTAAGTCCTAgatttctctcttcttccttC
CCATCCATCAACTTCTTCAACTCCCTCTTCTTTTCCTCCAtttctcaaattaaaaaaccaaacttATGAGTTGA
TGAAATAGAACTTAGAAGgtgataaaataaatgaagactttaaaataaataaataaataaatgttattgaGTTTG
ATTGAATAAACAAATTAAGTCCTAGAGTTCTCTCTTCTTCCTTCCTATCCATCAACTTCTTGGACTCCCTCTTCT
TTTCCTCCATTATCTCAAAAACTCCAAAAATCTGAAGTTGATGAAATAAACAAagacttagaaaaaaaatatata
tttttttttattgagtttaaCTGATTGAATCAGCAAGACACTAGagttctctcttcttctttcccattcatcaac
atcttcaaaaaCCATAAACATTAAAGAGAAGAccatttagaaaaaaatgaaagtaaacCTATTCCTAATTAAATT
TCTCTTCTCTATTCTCCTCCACAAACTTCTTCACCTCCCTCTTCCTTTTCACCTTCATTCCCCCAAACCTTCAAA
CCTAGAAGMCAAAACACACAATAACATAATGACACACTTCATACAATGCTTGAGTTTGATTGAATAAACAATACT
AAACTTATTCCTAAATATCTCATTACCCTTTATCCTCAACCACCTTCAACTTCMtcctcatttctcctcaaaACT
ACTCCAAACATTGAAAARATCTCACACTTCAYGCAATGAAAAAACTATGAAGCTCAACCCTTCAAAGCTTTCAAT
GAAAAGATCACCCTTTCTCCTAACTTTCTCTTATCAAACATATTCTCAAACAGTACCATAATGATACAGTTCATG
CACAACCCTTCAAAACTTTCAAAAAGACacagaaagagagaagaaagaaatatataGAAACCTGGAACTAGAGAA
GTCATAGAGACGACCAGTGCTTGAGAAGATGATAAGGCCAACTTCAGCATCACAAAGGATTGAAAGCTCCTTAGC
CTTCTTCAGAAGTCCATTCCTTCTCTTAGAGAAAGTCACCTGCCTACTTGTTGTGTTATCAATCCTTCTTATCAC
TATCTTTCCTCTCCCCATGATCTTTCTCTGCCAACAAAGTTCACCCATCCATCAGATCTATCATTCAGTACTACA
TCCTTTCTCaatcttataatattttaatacaaatatatatatatatatatatagaaagagagaaatatatat

atatttatatatatatatagaaagagagaaggAGAGTGAGAGAAAAAGAATCCaaagagaaaggaaagaaaGGTG
CAAACATGGAAGCATGAGATCTGAAATAAATggatgaaagaaaagaagagaaaagaaggaaaCCTGTTTGTCTTT
CAGGGAAGTTGAAATCTTTTTTCTGATGGATGGAAACAGCAAGTTGAGAGCAGCTATTTCTTTTTATGGTaacat
tgttgttgttgtttttttttttttaattaaacatggGTTGGTTGTTCTGCTATTGGATGGTTCTTTTCATTTAGG
TCCTTGTTTGATTATGATTTCAGTTTGGGTACTTCTTTTGTGTAAATTTGTGAAGGAGAAGGGATTGAGAAATGG
GGTGCTTTTAAATTTGGAGTTTTTTAAGGGGGTATGTGAGAAATRtttttataatattttaggtTTGATAGCCGC
CTGGCCTATAGATCACTGACGCGTGTTTGTTGGAGTTGAGTATGAGTTAATTATGGATTAAGTAAGTCATTATAA
TATGGAAAACGAGGTTCATATATTTATCACTAGAGAAGTAGtgacactaataataataataataataata

>DroAGL17_1_Dioscorea_rotundata
ATGGGGAGAGGAAAGATAGTGATAAGAAGGATTGATAACACAACAAGTAGGCAGGTGACTTTCTCTAAGAGAAGG
AATGGACTTCTGAAGAAGGCTAAGGAGCTTTCAATCCTTTGTGATGCTGAAGTTGGCCTTATCATCTTCTCAAGC
ACTGGTCGTCTCTATGACTTCTCTAGTTCCAGCATGAAATCAGTGATCGAGCGTTACAACAGGACTAAAGAGGAA
TACCATCAACAATTTAATCCAGCTTCAGAGGTTAAGTTTTGGCAGAGGGAGGCAGCAAGCTTGAGGCAGCAACTG
CATAACTTGCAAGAAAGTCATCGGCAATTACTGGGAGAAGAGCTTTCTGGCCTAAACATTAAGGATCTACAAAAT
CTCGAGAACCAACTCGATATGAGTTTACACAGTGTGAGAATGAAAAAGGAGCAACTTCTGATTGAAGAAATTGAA
GAATTAAAGCAGAAGGGAAATCTACTCCATCTAGAAAACGTGGAACTCTACAAGAAGATTTATGACGTAAGACGC
ACAGGAGTTGACAATGCCCCAGTTATTCCACATGGGTTCAGTTTTATGGAGAACAGAGATGTCCGAGTTCACCTT
GAACTAAGCCAACCACAACATCAAAAGGATCGCGTGCAACCAAGGTCTTCAAAATTGGGGTGA

>dro_miR444
TTTTTTTTTCCTTTTTTTTTTTACGTGGTAGATCCGGTCTCCATCGTCCACTTTTTTAGTTTTTTGTTTGTGGAT
TTGGATCTCTGGTGCTTGGTTCTCGCCTATAATTAGCGCCTTAGAGGAGGAGAAGGAGTTGTCAATCTCCATCAA
TGGAAAGCTTTGTGGCTAGGGTTTGTTGGTTAGGGTTTGGATTCCCGCTTTTTTTTTTGTTCCTTTGATGGGGTT
TAAAACCTTTGTTTGATGTTTTTTGATGGAAGGCTGATGAGCTGCTGTAGGGTTTCAGTTTACGCAGATGGAGGC
AGTGAGTTTGAGGCAGCAAATGCATAACTTCAAGAATTATAAATGGGCAAAGTGATGGTACCGATGACTTTCTTG
CAAGTTATGCAGTTGTTGCCTCAAGCTTGCTGCCTCCCTCTGCCAAAACTAGTATGGACAAAATAACCCATGTCA
ATGCATTATGGTAATGTGATGATTCAAAGATATTTTTTGATTAATGTTAAAATCCAGCAAGCTTCAAATTTGAAG
TTGTCTCATGGTTAAACAATGAGTATTGCTTATGACATTTTTTTTTTAAACTAACAAATTCATTGCAAGGAGGCC
CATCATTTACTAGTATGTTTAAGATCTTAGTGAGATTTTGCTTCAGTGGTCTTCACAGATTTGCTGATCTGGTCA
GAGTCGAACCTTCTAATGCTATTCTGGTTAGATAAAAAATCGGTTGAGAATGTCTGTAGACCCTCTCATGTCAGCT
TGGAGGCAAAACAAATCTTTGGCCAAAAGCATTCAGTAATGTTCTGCATAACACATTTGCTAGCATCATATTGTT
GGGACGCAGTGTGGGTATTCCTCTTTCAACTATTTCTTGGAGAATGCCCACTACATCTCCTTCACATGAGACGGC
TGATTTGCGGATGTCTTAAATCCTGCCTAATATGTTTGTACCTAAACCTCTTAAACTTCTTTTGTTCTATGAAAA
AATTTACTCGCATTTGGCATCTTTAATAGACTCTTGCTTTTTACAATTGTATCCTGTATGTCTACATAAACTCAA
ATATGGTAATTGTACTGTCAAAATTTTGATTTAAATTTTCAGCAACTACTGAATAAAGTGATTGGTTTAAAATTC
TTTACTAAAAATTTACCTTACTTTTGTGGTATGATTAAAGGCTGTTTAGCTTCCTCCTTTTGCGGATGCTTCATT
GAAATGCTTCATCAGAACATTTTGACAAAGCACATGTAAGAAAAAAAAAAAGAAGGAATTCATGGTTTTTAAACT
TTCTTTTGTTTCGTAGAGCAGGCTGGTTGTTTAAATTTTTTCTGCTGGGATAGACTTTTCACAAGCTACCTGCAA
GATCTTCTGTGTCACAATCTACATTGTTCAAGGCGGCGATCTTACTTAACAACTCTACATCTCATAAGGTTGTGA
TTTGATTACTCTTTATTCTTCCATATGCAATTTTGTGAAATGAAGAATTGTTTCAATCCTTTTCTTACAACTAAA
CTGCCCAGAACCTTTTGCCCACAAAGTGGCATCTTTCATTTACATGTACTCAAGCTCATTGTACTTTGTGATTAG
AGGGTTTTCTTTTTCCCCTCTTCTTTTGTGTGTGTGTGTGTGTGTGTGTGTGTGTGTGTGTGTGTGTTATTAGTA
TTCATTGTTAAAGGAGCAGTGTTAGTCAATTACTTTTGATCCCATATGTTAGTGCTGTTGGTTATTTTTCTTCTT
CCATATGCTGATTTGTGAAAGAAATTATATTGCTCCCTCTTCTTGACCAAAATTGACGGAGACCATTTTGTCGC
ATATTGCCATCTGTTCTTTTTTCATGTAGCTTGCTAACTATTCAGCCGTTAGCTTATATCCCCAATGGCTTTCAT
ATTCCTCTTTGGGACTGGAGCTTTTGTTTTGTTTTTGTTGGTCCTTTTGCCGTAGAATTCCATAAATTTCAGTAT
CATTGATTATGTCTTCCATTTGAGCTTACACATTTCACAAGCTTTCTGTCACTATCAGATCCATTTAATATTAAA
GATCACAGAATTCCCATGATCCGTCTAGGCTTCCTTTTAGTCTAAGCAATTTTAGTATTGAAGTCTCTTGTTATG
GCTCAAGGAAAAAAAAAATTCATCAGGACTTAATGGATTTCATGTCTTTCCGCAATTCTGATCTCAGATTTTCATT
ACTGGACTACAGACCACTTAAAGATTCCAATCCATTGTGTGGCCTTCTGATAACCAAAAGAACAGTAGCATCTCA
TCATACCAGCCAACAATGACTACCACTGAAGTATAGGGAAAAAAGCCATTTACATTTACTTATTTTATCCATCTG
TTGAAAAGACGAATTTATGTCATTGTCTAACAATAAATTATTTCCTGATTTATGATAAACATGTTGGGAAGGTTT
AGCCTTGGTGTACATTTAATCTATGGGCACTGAAGTGAAAGAGTGACAATCATCCTTGTATTGAGACAGGGTAAA
TAAAGCTTATAGATCTAATAGGCAAAGGATGATAATGTTCACTTTGACAAGTCATTAGAGGGAAAGAGGACATAT
AAGCGGCGAATTGGCACATATACTTAATTATTTCCTGATTTATGATAAACATGTTGGGAAGGTTTAGCCTTGGTG
TACATTTACCATGTGGACTCAATAACAGTACTGGGGAAGTTAAAAAAAAAAAAAAATCTGTTTTGTAGTGTACAT
TATATAGTTAAGTGTAGATGTTTTATAATCTTATTGACATCTTTTGCAATCCTTACACCAAGTTTCAGGCCTAGA
TGCACATTCTTAGATACTTGAGTAGATTGATCATGTTAGAATGGTGTCTACTTAATTAAGTTCACAAGCTATTGT
AGATAATATGTGATCAGTGATTGCGTTAGATCAAAAGTTCACAAGCTAATGAAGCTATTGTACATTATGCAGCTA
AAGTTTAGATGTTTTATGATCTTATTGACATCATTTGCAATCCTTACACCAAGTTTCGGGCCTAGGTGCACATTC
TCAGATACTTGGGTAGATTGATCATGTTAGAATGGTGTCTATTGAATTCAGTTCACAAGCTATTGTAGATAATAT

ATGATCAGTGATTATGTAGATCAAAAGTTCTCACTGATTATGTAGCTAAGTGGTTTGAAATATCACCAAGATAGC
AGGATCTATAGAATATGGTGTATTAGTCTCAGAGAGTTCTCCTTCTACGCTCGAGTATTAAGGTATACCTCAACT
TCAGAAAGCTTAACTAAGCTGTGACCAATTGCATAGCCTAGCAACACTGATTCTGATTATAAATTTTGTGAACTT
TAGTCTGTTTGATAAATGAAATCTATCAAGTTCATGTCTTAATGCATTTCACAAGTTGTTTGGTTTTATGCGGAT
TTGATGCTTCTTACGAGCTTGCGTTTCAGAATTTCTTCTGATAGCAATTTAGACATTGATGGAATCATAAATTAT
CTACTCCTAATTCATTTAATAAGTTGATGGTTTTCATGCTAGTTGATGTTTCTTGCAATTTCATATGTTCAGAAT
TTTTCTAAATAGCATGGCAATTTGATCCAAAGGTAAATATCCTACTGTGTTATTTAACATCAATGGTCCTTTACA
ATTTGATCGGTGCTTTTTAAGGCATCCCCTATTATAACAAGGACTTTTAGATGCCTAGCAAAACACACAAAATAT
ATATATATATCAATATATAGACAAATAAATAAATAAATAAGTAGAAAAATGTTTTAAATATGCATTTCCTTTAGT
AGACAATCTAATAGATACCTTAACCTCTGAAGCTGGATTAAATTGTTGATGGTATTCCTCTTTAGTCCTGTTGTA
ACGCTCGATCACTGATTTCATGCTGAAAATCAAATTAAAAGCATTGTTTTTAGTGGACATTCTACTTTACATCAA
GATGAAAAAGAGAATAAAGTCCCTGAGAAAAAATGTGCAACAACCATAACAAAGAATCATTTCACAAGCAGAAGA
ATTAGCTACTTTAATACATCATGAAAACATACTTGGCAGCTAAAGCCACTTGCATATTCGCATCTATGTATTATC
TTAACTTTGAAGAATCAAAGTAGAACTTTTCTGGAAATTTATTTTTCAAGAATTATATTACTAATCTTCTCATAT
GTGTTGATTATAGTTGTACCTTTAACTATAGCTAGTTGCAAGTCCTATCTGTTGTTATCTATCAATTTGTTTACA
GAAGATTAGGAGAAGCAACATGAAAAGATGAAGCTGCTTCTCAACCCAAAAGAACCAACCATACAATCAGAGAAC
CATCTCCAATAAGCAAAGGCATTCACCATCATAAGCTTTCATTCAAGTTTTCTGAAAGCTGAAGTAAGCTGTTTT
CAGACACCTTTGAGACGAACTTGAAAGGTAATGTTTAATAAATATAATGTCTATCTGTTTTGAGATCTTTCGCAA
GTGAGCCAGATTCTTTGTTTAGGAATCAATATCACTTATATCAGGGAAAATTAGATTTTTGAAGTGAACCTAGAT
GACAATTGTTGTGTTCAATAGTGAGCCAGGTATTTCATGTCAATAGTGCTCCTAGTGAAAACCTAAACCAATTGA
TGGTCTGGTGTGTTCTGTTTCTTAGACTATTTTGGTAGCTTAAGCTTTATTTACATCTAATTGGCAAACCAACTC
CCAGATAAAACAGAGTGTTCATCCACCATATATATATATATACTAACACTTATTTTGGTCAAGGATTAAATTGTA
CTTTGGTTTTATCTAGAGCCTAAATGAGTATAGAGGTGTATAGTTAATGTTACTCAACAGGCTTCTAAATTATGC
TTCTTGTTGAAGGTTTCAGGTGAAAAGCTGAGGCTGGAGAAAGACCAAGAGTTCTTGTGATCTATATGATATATA
TATATATATATGTTTATATATATATAATTCATGTCTGCTTTCTTGTTGTAATGTATAGTGAAAGATCTGTTGTAATG
TATTGTAAAGAAAATCAAGTGTAGATTGACTTTGAATATTCTTGAAAGCAATAAAATCATATCTTGGTTTTTACC
AAGTTGTATGAAAAAAAAA

Stem-loop structure

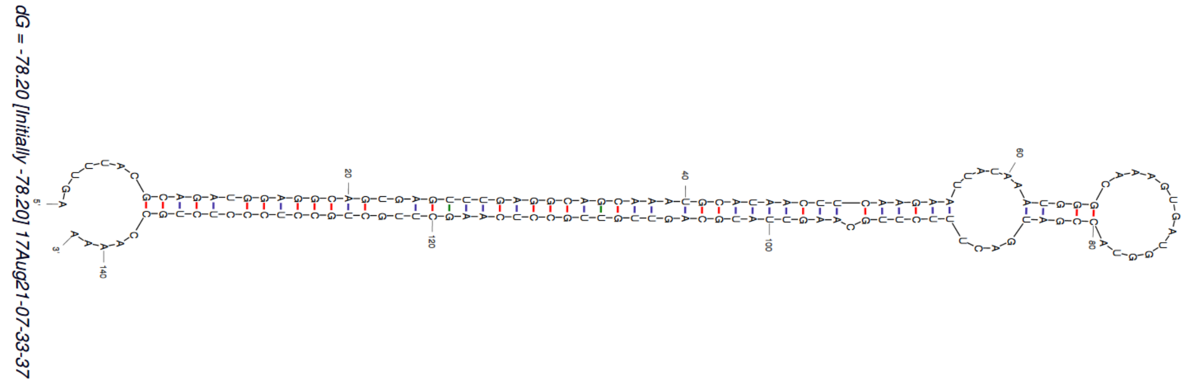


*DroAGL17_2*

>BDMI01000005.1

TGGTGCCTATACTTCCAATCCGGACTACATGGCGACCTGGTGTTTACTGATGGGAAAGCAAGACCATTTCA
TGGATCTGCTTGATAATTGTTATAAGGGCTCCATACTCGATTTTGATAGGTTTGGCAAGGTTAATGTTGTT
CATCTCCGGCAGCAACAGAAAGAGATTGCAGAGAAAGCTTTTGACCTGAGAATGAGGATGATTGCATACTG
GAGGAGTGTCGTACTAAGACTGGTTGATGGACCGGCATTGCACATTCTTTACAGTATTCAGAAATTGGTGG
AAGTTGAGATGGATGAAGAAATACTCAATGAGATTGCTGGGGATAGTGGGAAAGGTCTGGAGAAGATGCTG
GAGGAATCACCATCAGTTGCAAGGAAGCGTGATAGGCTTCGCAAGAGCGTTGAATTACTTGAGGAATCCAA
GCAGGTTGTTGCCACTATTATAGATCGCATTGCACTTCTTGATGATTAATTAGTtctatatatttcctttt
ttattgtatgtcattttagttgttttctcGCTTAAAATCTATCTTGtctggtaattaattaattactctgT
TTTGCTGTCTACCTTATTACTCTGTTATTGATTAATTTAGATGGGAAATCAAGTTACTTTCTATGTCAAGT
AccttaaccaaaaaaaaaacgagATTATGAAAACATTACTGTCATTTTTTAATATGCAACTAGTCAGCAAA
TACAGTCAACATAATCAAGCACTTAAAGGGATCAGCGATGATCTGAAACAACATGTCCTTTTACATCATGA
CTAAATATCCTACAAATGAAGTTGTGCAGATGTTTGAATATCTTCCTAAAGGAGTTGAAGCATGCTGCATC
AAGTCAATGAAATTTCTTTTAGTCAGAACTAAGAAAATACAGAAGCAATTCAGaagttgtttattttcaaa
ttaagtAAAATTACCCAAACTTTGGAGATATTGGTTGTGCCACAGCCacatcttgttgttgttgttgttgg

agatGGAGGGATATATTTTCATTCATGAGGCTATAAGAAGTATGCATTCCTCTGTTAACACCTTCATTTGA
TCCCCATGCATAAACCTGTAAACCATAAATCTATATTTAGTTTATCCTGTTAAAAGTACACAATGATTTTT
CAccttggttaatttttatatacctTGTTGTACATTTCCACGTTTTCTTGACTCATTACGTTTACCTTCT
TCTTAAGTTCCATGTTTTCTTGCTGAACAAGGTTCACCtgaaaattaaaagttatattCAAGTCAGTAATT
TCTCAAGTCTAGATCACCttatgcatttatatataaaaaaaagacagcATCAACCATATATGGTCCTCTTA
TTTATCAAAAAACCTATATATGGTCATATATTCATTTGTTTTCAATATTTctatacttaaaaattaaaaca
taaactaGTGTTTGTGCTctgtattttgaaaactacTTGCAACCATAGTTCATGTAATCTACACGTTTGCA
TATGCCTAGATCACATATGTTTTACATCATGACCAATATTAATTTGCATAGTTTAATTTTTGAACTATTTA
AAAATAGTCACTAAGGCGATTAAAAACTATGTTGTCAAAAACTAACGTTTACGATGTATATCACATTAata
gattttgattttaatttacaaaattgaAAAACCAATTAAGTAGCATTTAGTGATGAATCCTATATTGTACT
TTGTCGatccactatatatatatatatatatatatgtatgtatgtatcaaATATTTCTAATACAACCTTAT
GATGCAGTTCTTGAATTTCTGACCTCTGAGCTTGACCCTGCAAAGGTTAAAGACTATTAgaacagaaaata
aataataattatcaaaacaaagaaTGATCAACTAGTTTTATAATTGCAGAGCTACTAGCATCTTCACTCGA
TCATGACCTTCTCCTTGCGGACATTGTGTAAACTCCTTTCTAGTTGAACCTCTAGATCTTTTAGATCTTTA
ACACTTAACTGTGAAAGTCGTTCTCCCAACAATTTTCTGCAATAACATCAAAAATTAGCTGaaacttctaa
aaaaataatgtgaatttttaattgaactttttatgatgttattatatttcCTTGCTCATGAATGTTGAAATG
CTTGTACAAtcaaatagaagaaaattaatAGCAAACATATTCTTAAAAAATCATGCTTAGCAAGGCTTCAG
CATAATGCTAAGATTTAAGGCAATAGTGCAAGATAAAAGATGGTACCGATGAGCTTCATCCAGGCTATGAA
GTTGTTGCCTCAGGCTACCTGCCTCCTTTTGCCATATCTAATGAGATGCAATACAAAAGATTCATTCAACT
TATGAATTTCCAAATAAGGAGAAATATATGAAAGATTTAAGATCTAATTGTTTGGGTATATTGATCATGAA
GTATATAATTTGGCATATATAattggtgtttttatttttcccattaTGCATAGAAGAAAAGTCAGAAGAAA
GTTCACTAGCTAGTACAAGCAATTGATCACTAcaacacagacacacacacacatatatatatatatatatt
gtaagttGATTAAAAAATCACCACAATCTGTTCAAACATATAGCTggggaaaacaaaaaaacatgaattag
CAATTTTAAACATGCAACTTTAATTACCAGTTAGTTTGTAAATACTTGAAGTAAAtgtatgcatatttttt
ttttcctattatcatttttttttttttgacgaaATGTGGACAAACCACGTGTGAGGGTATGTGTACTAGC
CAGAAATTGACTGTCCAGTATGTACACTCTCAACCAGAAATGAGACACTGCAATTACAACCAAAGACCTAT
TACCACCACTGCTTTAATCAAGTTTTGAAATCTAAATTTTTGCATCTCACATTCCATCTTCACAATAAGAC
TAAATATTGCTGGGCTATAAACCCTTCGGTtactaataaaatttactccaattattttattaacatggCAT
GGTATACAATAAAACATCAACACTAATAACTTTGTGCCACAATAAGAAAATTCCAAGTAGCattggtttc
tttttattctttgaaTTGCCATCTATGTAAAACTGATTTCTTTCACTTCCTAAGATACTTCAAAATCCAAA
ATTTGTTAGCATTAAGCAATGCAATTATTTATTAATGGTACTCCTTTGACTAagataataacattttttt
ttacctttattTCTGAATGCATGTCCAACCCTTGTTTCTGCTCCACTTTTGTCTTGCCAAATCGATCAATA
ACTGATTTCATACTGAGCATAGAAACGTAAACTCTCATCACTTTCTATTCAAATGTAGATGACTAGGATAA
GACTCATAAATGCGTGCAGGTTTCCCTCTAGTTCACCATCttaatcattgaaaaaaaattacaaggtAAAT
CTACTTGAATGGCGTTGGTATAGAAACATgatacatgtatttttttaaaaataaaataaaatagataaact
acaaattttattaaaaaaaaaaaaacagatagaACCGAGTAGAAAAGAAATGATgcaagtattatatatata
gtttatgaGCACAgttaaagaatttaaaaataatttcaaataggttttcttttctatcTCCCGTAATAATG
TGCTATGCATAAACATATATTGAATTCAAGTTCACATATAAGAATAAACAATATTGAACTTGAATACTAGT
ATTGAATATTAAGTAGATAGACTTAATTATCCAATGCAAAATGCAAATACAGTCATACCCCCAATGCGAAA
AATGAAACATATATGTGTCTGCACACAATATTGTTTCAAAGAGaggaatatatataataactgtTCTAATG
TAGATGTAGTTTTGAATCTAAATGAATGTGTTCTAGGGACTTCAATTAGTTTCACAATcaaattatttaca
taaataagaTTATAAGCAATAACAAAATAGAATAACTAAATCACGGTCGCAATATTCCGGAAAGTTGATAT
AAAACTGTAACTCTTTAATCAAAATAATGCTTTCAAATTTTGTATActaaaagttattaatttacATTCAT
ATATAGTATCATCATGCTAGCTACAGTTTATTTACACCACATTATAAAAATTCACATAAATCACTTGacct
aaaaaaaaagtttgcatAGAACATTTAATTTATCAAGTCATAGCATAAGAGTTTGAGTACATAAATTATAT
CAACATATTTCATTAGGAATCACAATGAAcatcaataattatattttattaatctccgagccttcaatat
ctttatTTGTCATTCTCAACTCTCCATCTCCATTTAATTTACTCACACCGTCCATCGCAAATATTTTTGAA
agtgaaatttaattatgatatgaTTAATGCCTTTTTCCCTTTTGTACTTGGTACTCTctgtttctctcttt
tttctaataaatttgtggtttatccatttattcaaaaaaaaaaaaaaatgccttgacacataattttcaaa
ataatgtgAAAATCTTCCTCCTCTTCATTGTAGCAGCCTGACAGACGTCTGATGCATCATAATTGATGATG
AATTATATTACAATGTACATNNNNNNNNNNNNNNNNNNNNNNNNNNNNNNNNNNNNNNNNNNNNNNNNNNN
NNNNNNNNNNNNNNNNNNNNNNNNNNNNNNNNNNNNNNNNNNNNNNNNNNNNNNNNNNNNNNNNNNNNNNN
NNNNNNNNNNNNNNNNNNNNNNNNNNNNNNNNNNNNNNNNNNNNNNNNNNNNNNNNNNNNNNNNNNNNNNN
NNNNNNNNNNNNNNNNNNNNNNNNNNNNNNNNNNNNNNNNNNNNNNNNNNNttttaataatagacgacaagtgCC
CTTTTTacatgaatataaacagtacaaacagtactggaacccgaatgtacaatatatgtacagtatgagaa
tagtataagaatcccgggtgaacagtatatgaacagtacagtgaacagtactgtcgggggaggaatttgaa
cccatgacctcctccttatactttggtgtcataccattcaGGTTCttaggattatatatatatatatataa
attagaggtAATCGAAGCGAAACAGGCATCAAAGCATTGGCATACTTCGGAGCCAGGATGTGTAAATCACT
AGTGAGCATAAAATTGCAAAGTGCACCCAAAGCACAGTCATTCGGCTCAGAGCATGACTGTGATTAAAGAG
CACCAATGGCTTTAGCAATATTAATTTGCTAAAAGTCCAAAAACACGAAGATACAAGTCCTATAAGTGCCA

CCAGTCAATAGTACGATTGTACAGTATTTTTTTACCTATAATATAAATCCTGACAGTTAGGTTTTTGAAGA
GGACTTTTATTGAAAACTTCATCGGAGGCCAAGGTTTCAAATAGAGATCAAATGAAGCCAATCTAGGATGA
GTTGATGCTGCTATATCAGCCCTTTCGGTTCAAAATTGGCGTCAAGAACAACATCAATAAGTGAGTTCTAT
CATCCACATTAAGAGAAGCTAGCATTGAGGGAGCTTTTTGTATGTATTCATTAATGTCTCAAATCTTGGat
gttgttttataattattgtattatgaTAAGCTAAACCTTTAATGGTGCTTTGAATCGATGAATCTTGAGAT
GAACTTGTAGTTGATTTCCTTTTTCATTATATTGGCTATTTTGAGTTGTTCTTCTAAATCTTCTATGTTTG
TTTCATGCTTGTGTTTTGTTTGTGGGAAAGCCGGCTGTAAATACATATTTGATTTGTATGATTAAGATTGA
GAATCAATCATATATGATCACAACATAGAGAATAGAGGTTAAGTGCATGCCTAGAGATAGAGTATCGTGTC
GAGCTCTCTAAATTAAGGTTAAACCTACAAGAGGGATTCAGTACATAGTcttttaatgcaatcgtagagCA
ATTGAGTCGTCGTATATATGGATTAGAAAATAGCCACCTTCAAATAACGGTTAATCTACTGTAGAAGTCTC
CACGATTCAAGAATACTTAATATCAAACTAATATATTGCAAGGTAGTCTCAAGGGAATTCGTGTCCCAAGC
ACCATATTCTTATTCTGATTTTATCCTAAATTCACTATTTCAttggtttttagttttgaatttgagCAAA
CATCATTCAATGTGTCGGCTAAAAAATAGAGGAAAGATTAGTACTATGAGCGCTGATCCTCGTGAGATCAA
CTACCCTACATTAAGATACAGTACTTTACTACTTCTATGACTCATATACTTGCAGCTTCGCACTCATAGGA
AAGAGCCTATCAATTCCTTTCTGTCATTCAAagatgtaattttattattatgtctaTTATGACAAACTACA
ATATATACATTTCGTTTAGtgatttaatttcttgtttgCACCAGTCCATTAGCATTCactaaaattgatat
tttcttaTTAGGCTTCCAATAAAGCAACAGAAGATTACTTTCATTATGACTATATAGTGCGGAGCACACAT
ATATATCGGTAGGGGACTGTGACCGCCGTACGTTAAGCATTAATATCTTCATTAATTGAATATAATTAGAT
GCGTATATCCTTAAACACCATTGTATGACAGAGATAACTAAAGAGGCTTCAAAAGCATCAACATGAGCTAT
ATGTAGGTTGGTAGGAAAGTATCACTGCTTTCCATCACTGCCATTAatgagaatttatatttttgtttact
cCATGAATAAATTAGATGCTGCTGTCATTCATGATAGGTAGCTCATTTCACTTAGTTGAGTATaggaagaa
aattttgttatgGGGATTTTGCAGTTAAATCTTTTGGGTCtcttaattaaatatgttttaggTAAAGGATG
CATGTTCCTTCAATGAACTTAATTAATCTAAAATGATGGAGTGTGAGGATAAAATCAAagttacaaattaa
aatgaTGGTTCATAATAAGCAAGATCTTGAATCTTTACAAGGAGTGTTATATATTATTCGCAAATCTCATG
CTGCATCTTCCTAATTAATTAATGGCTTTGTTGGTGTAACTAAGATGATGTTCATGATTTTACTTTGCATC
TGACtcttagaaaatgaaaagaaaaaaaaaaaaatacttaattcaagAACTAGCTAGGTTAACAAGTTGAT
AACCtgagatatatatatctccaaAAACTAAGTGAACCTGTACTTGTTATGCAACTAATTAAGCTGAAGAT
CAGTGTAAACTTTTTGTtgagaatatataataataagtaCAACAAAGAGAAAGATATGAAACATTGAAAGA
AGCCCTGttcacatgaatatatatatgatagaaaGCTAGAGAATAAGATGAAGGGAGATACTTGGTGTTGG
CATAGTCATAGAGCCTGCCAGTGCTGGAGAAGACGATGAGACCGATTTCTGCATCACAAAGGATGGCAAGT
TCCTTTGCTTTCTTAAGCAGTCCATTCCTCCTCTTTGAGAATGTTACTTGCCTATTGCTAGAGTCTTCTAT
CCTTCTTATCTCAATCTTTCCCCTTCCCATCTTCTACTCTGTGTTTTTACACAAATCAATGGATCCAGTAG
CTAGATCTGTtaggaagaacaagaacaagaacaaatatatatgaatggTAACTATTGGTGATGAAATGACA
GGTGTTATTCGAGGAGAGGCTATGAATGAATGAAGATAAATAAGTGATGGGACTTGTAATAGAATCTGAAA
ACAACCCTTTATTTTTGGGTGGTTTTGTAGGAAAAGCTAAAGAGAGTTGTTGGTCTTCCTGTTGTAGTTTA
TGAATATTGTTGCAAGTTTCAAAAGCGCCACCAGTGCAATGGATCTCCGACGCGTATTacaatgaaaacat
atatatatatatatatatattatgtatatatatggctCGCTTGCCCAGGCGTGGTCAAGAAGAAACCTAGG
TTTGAAAGGATTTTGGGGTCCAACGGACCTATCAGTTTCTCAATTATCACCTCTTCAATAATGGATAATTT
ATTGGGGTTACTTTTGCCCTGGTTTGCCATGTAGTATATCTCACATTATTACACCAGGATCTAACAAATGG
GAGCATATTTAGTAAACACTGTTAGGATCTAACAAATGGGAGCATATTTAGTAAACACTGTTAGTGTTTGG
GAATAAACCTTggataaaagattttttttttcagggttATTTATGGATAAATTGTCAAATTAAACCACAAT
TTGAAAATGTTGATAGGTACAAGAATAGTTCTTCAAATCAACCTAAATATTTGTAGATAAgctgatataat
ttttttaaggcgATAAGTGTCCCtcataaaaaattgttaaaatgaTAGACAAATATGGAGGTACATTTGTT

>DroAGL17_2_Dioscorea_rotundata
ATGGGAAGGGGAAAGATTGAGATAAGAAGGATAGAAGACTCTAGCAATAGGCAAGTAACATTCTCAAAGAGGAGG
AATGGACTGCTTAAGAAAGCAAAGGAACTTGCCATCCTTTGTGATGCAGAAATCGGTCTCATCGTCTTCTCCAGC
ACTGGCAGGCTCTATGACTATGCCAACACCAATATGAAATCAGTTATTGATCGATTTGGCAAGACAAAAGTGGAG
CAGAAACAAGGGTTGGACATGCATTCAGAAATAAAGATATGGCAAAAGGAGGCAGGTAGCCTGAGGCAACAACTT
CATAGCCTGGATGAAGCTCATCGAAAATTGTTGGGAGAACGACTTTCACAGTTAAGTGTTAAAGATCTAAAAGAT
CTAGAGGTTCAACTAGAAAGGAGTTTACACAATGTCCGCAAGGAGAAGGGTCAAGCTCAGAGGTCAGAAATTCAA
GAACTGCATCATAAGGTGAACCTTGTTCAGCAAGAAAACATGGAACTTAAGAAGAAGGTAAACGTAATGAGTCAA
GAAAACGTGGAAATGTACAACAAGGTATATAAAAAATTAACCAAGGTTTATGCATGGGGATCAAATGAAGGTGTT
AACAGAGGAATGCATACTTCTTATAGCCTCATGAATGAAAATATATCCCTCCATCTCCAACAACAACAACAACAA
GATGTGGCTGTGGCACAACCAATATCTCCAAAGTTTGGGTAA

*DroAGL17_3*

>BDMI01000021.1

gaatattgATAAAGCATCTCACTTGAGTTCCATTTCATCTGAAGGGAGTTGAATTAATTTTCATCTTCTac
atttgtttctttgtttctttgtttctttgtttatttgtttattttgtttctttatgcaTAGATCTATTGGC
TTGCCAAAGAATGAGAAGAGAAGAGACACCAAGTTCATTGATTCTTAATTTTGATCCTTTTCttcatttct
atatatatatatatatatatatatatatatataatgaaacatCACACTAGAGCTCCATTTCATTTGAAGATAG
CTTAATTTTCATCTTCctttatttcattctttctttgttttgtgacAGTATTAATTCAAGCTGGttgatt
tcttcaattttttttatatagaggaattaattaattaacaaactaTTAATTCATGTGAAGGTATCTGGTT
TGCATCTTCCTTTATTAACTgtactttcttttcttcttttttttttgtgtcttgATGAATAGATCTGCTGC
aggttaataattaattttgtgaaAGGAAGAGCATGGGGAGAGGGAAGATTGTGATCCGTCGGATCGATAAT
TCGACAAGTAGGCAAGTGACTTTCTCTAAGAGGAGGAATGGTTTACTCAAAAAGGCGAAAGAGCTTGCCAT
TCTTTGCGATGCCGATATCGGTCTTATCGTCTTCTCTAGCACCGGCAGGCTCTATGATTACTCTAGCACCA
AGTATTACTtaactctttctttctttctactTCTCAATATTTCATGCTTTTCTTAGATGTACACCAagtta
gatttatatatatatatatatatgcaagaatTTATTGATTCTAGTATTACAAATATAatacatgaagaaga
agaagaagaagcttaaATGTTGAGTTCTAAGGTCTATTATGAGtccttaaaattattttcatgatGAATCT
TGACTTTATATTATTCCCAAATACATTCATATGTTAAAAAACCTAATATCTTTGTAGAGATGGTGCattaa
aaaagagtttttttttttaaaaaaaaaggtcacaTTCCATGCATTACAAAAATTGCATGAAATTAATTTC
ATCAATGAAGTGCTTCCTAATTTTTCTACTAATTTTTGCCCATATATTTAATGTagtatttatgatttata
tacttcatattaaaaaaaaaaaccgaagattaaaaatataaagaacttccaaacaaaacaacactttctttg
atttcttacATGAAAAATTCATATACCATAACGGACATATATTTCGCCAATGACCTCTTGGTCTATTGGAA
TAGAGTTTTTTGCATAGGGTATGAGACAGATTCAAACCTCATCATACGCAGGATGAGCGTGCAAGTACGTA
GAAGTATTAACTCCCATCTCATGTATGTATGTCCCTAACATGCTTGTCCACATTtgtcaaatttaaaaaaa
aaaaatttcccttgTTTCATGTGTTATATGTCAGAAGTATAAACTATAAACACTAAGTCATAGATCATTTT
CATGAACAGCATGAAGTCAATTATTGATCGATATAACAAGGCCAAAGAAGTGCATAATCAAGAGTTGAGCT
TGAGTTCAGAAATTAAGGTATGATAAATTTCATATTAGCATAATTAGaattttgaaatatctcatatatct
tttaacaaaattaacctATGAATTTGATctaaataatgaattaatttgattaatcaaTCTACTAAACATGC
TGATATGTTTTGGTCATTAGTTTTGGCAAAAGGAGGCAACGAGCTTGAGGCAACAACTTCATAACTTGCAA
GAAACTCATCGGTAAATCGCTAATTACTCTGTATTGCATAATGAATTCATCGAATATGAACATGAAATCAT
ACAgtaataatcaatttaatttcatAACTTAAATAGttgtaaataaataaccttTAATTTTTTGCATCAAA
TCTCACTCTTTATATGATCATGAGTTATACGAAGAACATTGATCACTGTTAATTCTTGCAGACAATTGCTG
GGCGAAGACCTTCTCGGTTTAACTGTCAAAGATCTACAAAGCATAGAGAATCAACTTGAAATAAGTCTTCG
AAATGTTCGAACAAAAAAGGTTTTCTGAAGAATCACATTATcattaaccaatttttttactGCATTGCACC
CTTTTTAATTAAACTCTGAGCACAAAATTTTGCAGGAACAAGTCTTGACTGAAGAAATCAACGAACTAAAC
CGAAAggttgaacatatatatatatatatatatatatgtatatgtatatatgtgatatcatgtttgatgtg
atAAATTTAAGATGAATGAGAAATTGAATTCTGTTGCAGGGGAAACTAATTCAACAAGAAAATATGGAACT
TTACAAGAAGGTACAGTTTGTGTGCCAGGAAAGCATGGGATTACAAAACAAGGTGTATTAATCTTAGAAAC
ATCatgatatatatgattataaaaCGTTATTAATGTTGAAGTTTGTGATCATTGAAGTGAATAAGAATTGA
AAAAAGTTTTTGATCATTATAGGTTTATGCAGTATGGGGATCAAATGTAGGAGTAATTACTAGAGGCTCAA
TGATTCCATGTGCATTCAGTATCACTGAAGATGCTCATGTGCCAGTTCATCTTGAACTGAGCCAACCTCAA
CCTCAACCTCAAAAACATGGACAAGAACAAACCGATATTTCACAAATGACGGCTCCGAAGCTTAGGTAAGT
TTTTGTGCTTGATTAGAacttaaaaatgatttttcttgtttggtttattaattgattgataAAAACACAGT
GTTTTTACTGATATATCAATAATGCAACAGACTTCAACTGCAATGTGATGGTGCTCTTCCACTATGAGAAG
GTGTTGTGCAAACttgcattaattaatttgtatttgatttgaatttgctGAAGGAACTATGTATTAATTTG
TTTGTGTGTTTAAGGCTTGTGTtgtaatgaaattgaaaattatgTGAAACTCTAGGAGGAAGAACTCAGTT
GCATGCATGGTTAAAAATTAACTTGTCTTATGAAGAAGTATGGTTCATTATTAGGCATTAATAATGATTAT
AAGAtgcatgtatttgtattttgtgCACTATAAAATTTCTTGTATCCatagtgtatatatacattaattat

>DroAGL17_3_Dioscorea_rotundata

ATGGGGAGAGGGAAGATTGTGATCCGTCGGATCGATAATTCGACAAGTAGGCAAGTGACTTTCTCTAAGAGGAGG
AATGGTTTACTCAAAAAGGCGAAAGAGCTTGCCATTCTTTGCGATGCCGATATCGGTCTTATCGTCTTCTCTAGC
ACCGGCAGGCTCTATGATTACTCTAGCACCAACATGAAGTCAATTATTGATCGATATAACAAGGCCAAAGAAGTG
CATAATCAAGAGTTGAGCTTGAGTTCAGAAATTAAGTTTTGGCAAAAGGAGGCAACGAGCTTGAGGCAACAACTT
CATAACTTGCAAGAAACTCATCGACAATTGCTGGGCGAAGACCTTCTCGGTTTAACTGTCAAAGATCTACAAAGC
ATAGAGAATCAACTTGAAATAAGTCTTCGAAATGTTCGAACAAAAAAGGAACAAGTCTTGACTGAAGAAATCAAC
GAACTAAACCGAAAGGGGAAACTAATTCAACAAGAAAATATGGAACTTTACAAGAAGGTACAGTTTGTGTGCCAG
GAAAGCATGGGATTACAAAACAAGGTTTATGCAGTATGGGGATCAAATGTAGGAGTAATTACTAGAGGCTCAATG
ATTCCATGTGCATTCAGTATCACTGAAGATGCTCATGTGCCAGTTCATCTTGAACTGAGCCAACCTCAACCTCAA
CCTCAAAAACATGGACAAGAACAAACCGATATTTCACAAATGACGGCTCCGAAGCTTAGGTAA

#### Dioscorea bulbifera

##### *DbuAGL17 and dbu-miR444*

>DbAGL17 [organism=Dioscorea bulbifera] AGL17-like MADS-box gene, partial genomic locus, start exon2

CAATTCTTTGCCTTTTTACATGCTGTTTGATTGGTTATTAATAATTCAAAATTTATATATAGATAACATGTAAAT
GAAGTGGTATTATTTAGACCACAATTTATTAGATTGGAAAAAATTAAAAATTTAAATTCTTTATAGGCATTAGCA
AAAGGATCACTCGAGAATCATCTATTTCCGAAATTAAAAAAATTCCAATATAATTTTAATAAGAAATAGTTAAGA
ATGTTGGTCTAATAGCGAAATAGTGCTTCAAGTTGCTTGATTCTAAGAACACCGATAGCAAATAGATATATATAT
TTCCTCCGTTTTATATTAACTGTCATATTTAAAAAAATTTTTAAAAATTTATTTATCATCTTTTAAAGATCTATA
AATATTAATCTCTTCTGTTTGTTTAGAAAAAGATATATATATTAATGAAAATAAATCTAACAAAAATAAAAATTA
CTTATCATAAAATTTTTTTAAAATTTTAAAAATTTTAACGACAGGTTAAAATTTAAAATTCAGATATGACAATTA
ACATGAAACAGAGAGAGTATATGTTTTGCCTTTTGAAACGTATTTCATAAGAAACAATCGACACAAACATAATCA
TTTGATGTTGGTTATGAATCATTCAATTTAGCCATCAAAATTCCTGATTTTCAAATTCTTCAACTCCAAATCACT
CATTCATCATGTAACTGTATATATATAGACACCCTACAATACCATGCAACTTGGTAAATACCAAGATATGACTTT
ATTACTTGCAAGAATATTCAAAGTCAATCTACACTTGATTTTCTTTACAATAGATTACAACAGAACTTTCACTAT
AAATTACAACAAGACAGCAGACAAGAATTATATATATACATATATTTCACATAGATCACAAGAACTCTTGCTCTT
TCTCCAGCCTCAGCTTTTCACCTGAAACCTTCAACAAGAAGCGTAAGTTAGAAGCCTCTTGAGTAATGAGATCAA
CATGGCACTTGTGGTTCTTCAACCATACACCTCTATATTCATTTTAGTCTCCATGTACAAAACAAAAGTACAATT
TGATCCTTGACCTTCAGCCATGAAACTTCAAGATAGTCCATAAACCATGAAAATGCAAGTAGTTTTGATCCCTAC
TTAAGAAATTTTTTCAGTCCTTAGACCAAAGCAATTGTATATAAATATATAGTGGGTGGCCACTCTGTTTTATCT
GGAAGTACATTTGCCAATTAGATGTCAAGAATATGTGCTAAGCACTAACACAGTTTTGATCGACAAAATAAATCT
TAAGCTGCCAAAAGAGTCGAACAAATAGAACACACCAGACCACAATTTGGTTTAGGTTTTCACTATGAGCATTCT
TGACATGAAATACCTGGCTCACTATTGAACACAGCAATAGTCATGGATCTTCTATGCTTTGATTTCAATTACCAC
TATCAGTCTAGTTTCACATCCAAAAATCTTTTGAATTTTCCCCAATACAACTGATATTGATTCCTAACAAAGAAT
CTGATTCACTTATGAAAGATCTCAGAACAGAAAGACATTATATTTATTATACATTACCTTTCAAGTTCATCTCAA
CGGTGTTTGTAAACAGCTTATCTTCAGCTTTCAGAACACTTGAATGAAAACTTATGATGGTAAACGCGTTTGCTT
ATTGGAGACGATTCTGTGATTGTATGGTTAGTTCTTTTGTGTGGGGAGCAGCTTCATCTTTAGATTTCATTTCAT
ATTGCTTCTCCTTGTCTCCTGAATACAAATTGGTAGATAATGACAGGTAGGACTTGCAACTAGCTATAGCTAAAG
GTACGACTATAATCAAAACATATGAGAAGATTAGTAGTATAATTTTTGAAACATAAATTTCCAGAGGGGTTCAAC
TTTGATTCTTCTAAGTTAAGATAATACACAGATGCGAATATGCAAGTGGCTTTAGCTGCCAAGTATGTTTTCATG
ATGAATTGAAGTAGCTAATTCTTCTGCTTATGAAATGGTTATTTCTATGTTATGGTTGTTGCACATCTTTTCTCA
GGGACTTAATTCTCTTTTTCATTTTGATGTAAAGTAGAATGTCCACTAAAAATAATGCTTTTAATTTGATTTTCA
GCATGAAATCAGTGATTGAGCGTTACAACAGGACTAAAGAGGAATACCATCAACAATTTAATCCAACTTCAGAGG
TTAAGGTATCTATCAGATTGTCTACCAAAAGAAATGCATATTTAAAACATTTATTTATTTATATACAGACATATA
TTTTTTGTGCGATTTGCTAGGTATCTAAAAGTCCTTGCTATAATAGGGGATGCCTTCAAAAGCATAGTGCAACAC
ATGCACGATCAAATTGTAAAGGACCATTGATGTTAAATAACACCTAGTAGTATATTTACCTTTGGATCAAATTGC
CATGCTATTTAGAAAAATTCTGAAAATATGAAATTGCGAGAAACATCAACTTGCATCAAAATCATCAACTCATTA
AATGAATTAGGGTAGATACTTTATGATTCCATCAATGTCTAAATTGCTGTCAGAAAAAATTCTGAAACGCAAGCT
CACAAGAAGCATCAAATCCACATGAAACCAAATAAGTTGTGAAATGCGTTAGGACATGAAATTGATAGGTTTTAT
TCATCAAAAAGACTAAAGTTCGCTAAATTTATAATCAGGATAAGCCATAAAGTGCTAGGCTATGCAATTGGTCAC
AGCTTAGTTGAGCTTTCTGAAGTTGAGGTATACCTTAATACTCAAGGGTGAAAGGAGAACTTTCTGAGACTAATA
TACCATATTCTATAGATCCTGCTAACTCAGTCATATTTCAAACCAGACAGCTACATTATCAGTGAGAACTTTTGA
TCTCCGTAATCACTGATCATATATTATCTACAATAGCTTGTGAACTTAATTCAGTGGGCACCATTCAAACATGAT
CAATCTACTCGAGCATCAACTTCCCCAGTATTGTTAATTGAGTCCGCACGGGAAGTGTCACCAAGGCTAAACCTT
CTCAACCTGTTTATCATAAATCAGGAAATAATAAAGCGACAATAAGATCATAAAACATCTATACTTAGCTACATA
ATGTACACTCCAAAACAGATTTTTTAAACTTCCCCAGTATAGTTATTGAGTCGCATGGGAAATCTACACCAAGAC
TAAACCTTCTCAACCTGTTTATCATAAATCAGGAAATAATAAAATGTATGTGCCAATTCGTCGCTTATATGTCCT
TTTCACCTCTAATGACTTGTTAAAGTGAACATTATCATCCTTTGCCTATTAGATCTATAAGTTTTGCTTACCCTG
TCTCACTAGAAGGATGATTGTCACTCTTTCACTTCAGTGCCCATAGATTATCTGTAGTAGGATTAAGTTAGGTTC
TTGTTAGACAATGAAATCAATTCGTGTTCTCAACAGATAGATGGAAGAAGTGAATAAGTAGATGTAAATGGCTTT
TTTCCCTTGCACTTCAGTAGTAGTCATTGTTGGCTGGTCTGATGAGATGCTATTGTTCCTTTGGTTATCAGAGGG
GCACATAATAGATAGGAATCTTTAAGCGGTCTGTAGCCAGTAATGAAAATCTGAGATCAGAATTGCGGAAAGACA
GGAAATCCACGAGGTCCTCATGAATTTTATTTTTCTTGAGCCATAACAAGAGATTTCAATGCTAAAATCGCTTAG
ACTAAAAGGAAGCCTAGACGGATCAGGGGAATTCTGTGATTTTAAATATTAAATGGATCAGAGAGTGACAAAAAG
CTCGTGAAATGTGTAGGCTCAAATGGAAGACATAATCAATGATACTGAAATTTTTGGAAATCCAAGGCGAAAAAA
CAAACAAAAGCAATATAAAAGCTCCAGCCCCAAAGAGGAAAATGAAAGCGATCGGGGATATAAGCTAGAGGCTGA
ATACTTGGCAAGCTGCACGAAAAAAAGAACAGATGGCAATATGCGACAAAATGGTCTCCATCAATTTTGTCCTAG
AAGAGGGAAAAACAATGTTTTGTTTCACAAAACAGCATATGGAAGAGAAAAATAACCAACAGCACTAACATATGG
GATCAAGCGTAGTTGACTAACACTACTCCTTTAACAATGAATACTAATAACCCCAAAAAAAAAAACACGGAGAGG
AAAAGAAAACCCTCTAATCACAAGTAAAATGAGCTTGAATACATGTAAATGAAAGATGCCATTTTGCGGCAAAA
GGTCCTGGGAAGTTTAGTTGAAAGAAAGGGATTGAAACAATTCTTCATTTCACAAAATTGCGTACGGAAGAATAA
AGAGTAATCAATTCACAACCTCATGAGATGTAGAGATGTTAAGTAAGACCGCTGCCTTGAACAATGTAGATTGTC
ACACAAGATCTTGCAGGAAGAAAGATACTAGCTATAGCCGTGTAAAAGTGCCAACGCAGCATGTGAAAAGTCTAT

CCCAGCAGAAAAAGAAAAATTAAACAACCAGCCTGACCTACGAAACAGAAGAAAGTTTACAAACCATGGGTTCAT
TTCTCTTACATATGCTTCGTCAAAATATTCTGATGAAGCATTTCGATGAAGCATCTTCAAAAGGAGGAAGCTAAA
CAGCTTTATTCATACCCCAAAAGTAAGGTAAAGATTTAGTAAAGAATTTCAAACCAATCAATTTACTCAGCAGTT
GCTGAAAATTTAAATCAAAGTTTTGACAGTACAATTACCCATTTAGATAATGTGCAAACTCTTCCATATAAAGAC
AGGAACATCATATTTGAGTTGATGTAGACAGATACAGGATATAAATGTAAAAAGCAAAAGAGTCTACCTAAGATC
CCAAATGCAAGTAAATTTTTTATAGAACTAAAGAAGTTAAGAGATTTAGGTACAAACATATTATGCAGGATT**TGA**
**GACATCTGCAAATCACCCATCTCATGTGAAGGAGATGTAATGGACAGCATTCACCTAGAAATAGTTGAAAGAGGA**
**GTACCCACACTGTGTCCCAAACAATATATAACGCTAGCAAATGTGTTATGCAGAACATTACTGAATGCTATAGGC**
**CAAAGATATGTTTTGCATCAAAGCTGACATGAGACGGTCTGCAGACATTCTCAAC**CTACATTATAAAAAAATATA
AACCCTTTAACCATTTGTTACAAGGGTTCTCATGTAAAAACAGATGTGCGTAGGTGGCTATTCATAAAGTTATAG
AAGATACAGCAAGTATGAAAAAAGTGTATTTGTCCAAAAGCTCACCGATTTTTATCTAACCAGAATAGCATTATA
AGGTTTGACTCCGACAAGATCAGCGAATCTGTGAAGATCACTGAAGCAAAATCTGACTAAGATCTCAAACATACT
AGCAAATGATGGGCCTCCTTGCAATAGATCATTTAGTTAAAAAAATGTCATAAGCAATACTCATTGTTTGACCGT
GAGGAAACTTCAAATTTTAAGCTTGTTGGATTTTAACATTAATCAAGAACAATCTTTGAATCATCACATTAC**CAT**
**AATGCATTGACATGGGTTATTTTGTCCACACTAGTTTTGGCAGAGGGAGGCAGCAAGCTTGAGGCAGCAACTGCA**
**TAACTTGCAAGAAAGTCATCGGTACCATCACTTTGCC**CTGGAATAAATTCAGCTTTATACATTAGAAATCTTTCC
TGATAGTAGAGCAATCACTTCACAACAAAGAGTGTTCAGCAGATCTATAACCAAAAGTAATAGAGCAAATGCCAT
ATCTCATTGTATAAAATTTCAAACTGAAAACTAGTTTTCCATCGAGAACATAGGATATCACACTTTGCTTTAAAA
CATAAAACAAACTTGTAAAAAAAATTCCCCATTTATCAGGCCAAGTTATTACCTTTGTTAAGCAGACATTGTCAG
TTAAAGCTACACTATAGGTCCTTCAAATCAAATGCAGTAACCTATTTGAGTTATATCTAATTAAAAAAAAAAATC
CTTCTCATAATTGAAATGATGTTTCATTATTACTTGCTAAACTTAGAGAAGTTGGAAGTGAGAAAGATAGGTTGG
CACTCAACTTCTTTATCAATCAAAACAGGTTTACTGTGAACCAATTTGATTTACCATGTTAGACGCACTTAAAAC
ACAGGTAATTAAAATTTGTCATTTAAAGTCACATCTAGAGGTAATACTATCGCAAAAATAAATCTAAATTATAGT
ATAAGTTTGAATATTAAAGAAATTGTGTAAAAGAGAAAAGATCAGTTTGACCATTTTGTTTAACTTCTCAATCTT
TAGTTTTATCATTACTATTATCGGTAAGCTTTTCAAAGTGATTTGACGTCAACAAAATTCCCAAACTAAAACTGT
GGGCTCAGACAAGCCTTAAAAACATATCGTAATGATATTGCATGCTTTCACTACATAAAATGATAATTGTCTAAG
ACTAATGGAATTGTTTTTAAGCAAGCTTGACTGCATTTATTGCAAATTTATAAAAACATATGAAGCAAAGACATT
CAAAATGCCCACTTAGTAATTTGCTTATTTTCTAGGAACATGAGAGCGATACAAAGGGTTAAACATCATCAAATG
TATGCATAGTACATTTTGTATCCTTCAATCAAAACCCTAATACTTATTGTCATACTATTGCTAATGGTAAGGTAT
ATTTAACTATGTTTTCATAAAATTCAGTGCTAACACCTTCCTCAAATCCCACAAATTATCACAATTAGGAATTCT
CTAGCACTAAAATTCCGCTCAGGCAAAGCAACTATGTATGAAGTCCGCATAATCATCCCCAAAAATCCTAAATGA
ATGCAATCTTATAAACTTCGATAAATATCAGCATATCACGCCAGCATATCTCTAACGCCTAACTGCAGGCAATTA
CTGGGAGAAGAGCTTTCTGGCCTAAACATTAAGGATCTACAAAATCTCGAGAGCCAACTCGATACGAGTTTACAC
AGTGTGAGAATGAAAAAGGTAAGAAAATAGCTCTGCTCTTAAGATTCAGCACTTAAAAAATAAAATTGCCACGAT
AATTGCTTCTTTTTCTTTTGATTTTTGTTTTTCTAGGAGCAACTTCAAGACAACCCCAAATAGTTGGGACTAACG
GCTTTTGTTGCTGTTGTTGTTGTAGGAGCAACTTCTGATTGAAGAAATTGAAGAATTAAAGCGAAAGGTCATTTA
AAGAAGTTTCATTGTGCGTCTTTAAACTTATGGTGATGAGATTAGCAGCCTAACTTTCTAGGATTTCCTCGCAGG
AAAATCTACTCCATCTCGAAAACGTGGAACTCTACAAGAAGGTAACCCTTTATTGAGAAGAAAATATAAAAAATA
TTATATTCATTAAATAAATGGGTATGAAGCTATAGCAAGACACTTAAAATTGCAAAGAACTACTGGGAACCTATA
TATCATAAAGGATAAATAAAGATTAAAAACATTAGGTTCTATTGTCTCACTTGCTATATGTGAATTTTGCATCAG
GTTTATGACGTAAGACGCACAGGAGTTGACAATGGTCCAGTTATTCCACATGGGTTCAGTTTTATGGAGAACACA
GATGTCCGAGTTCACCTTGAACTAAGCCAACCACAACATCAAAAGGATTGTGTGCAACCAGGGCCTTCAAAATTG
GGTTGATCTTAAATGGAATTCGAAACCTAGTACTTGAAGAAGCTTACAGCAGGATTGAACTATGCGATCAAGAAC
TAAAACCAAGGTTATGTTTTATTACTTCACAATAAACATTTACACTGTCTAAGTATGATTTGGTGTGTAAACAAG
AGAAGCAATGACAACTGAGTTCAATCACTTGTTTGCGGCAACAATATAGCACAACAACCAAGAAGTATGTATTAT
TCAACTGCAGCAATGATGTGAAAACTGATTATTTGCTTAATTATAATTGTATCATCAGATTACTAATACTGAAAC
ATGATTATTCTGATGCACATAAGGTGTCAAAGGGAGCCAGATCATACAAATGGCCATGACCATAGATGAAAACAC
ATCTTAACAAGTCTCATGCCACAGAATGACAAAAGATTACAAAATCATACGGGTGAAACAGGCATACCGAAGGAA
ATAGCAATGAGTTTCTTCGAATCTACCAACAATTTCTTACAAAATATTGCTATAAAGGAAAACACCACAAGATGC
ACAATTAGATAGTACTTGGCCATAGATTTCCCAAAGTACTTATTTACTTCCCAGTGAAATTACCATCTCATGAAC
TCCAAAATTCAAACACTGTCCTTGAAATTTCAAAAAGGACCACAAAAGGCCTTCAACAATAACTATCCAGAAATA
CTACATTTAATGTTTATACAAAACGCTGCTCTAGTTCTGTGATTCTAGAGCTATTCCAACTTTTCAAAAAACATA
GAAGCCACCCTGGCAATCATAAATCCATGTTTATCCCATTATCTCAAAGTAACTACAATAAATACCACCACAAGG
ATAACTATCACTCTTCTAGTATCACAAAATTGCCCCATTTTGTCTACCTAACTATAAATATACCATCCCCAATAA
TAATCACACTAAGTTGTCATGGCATTTCTAAAATTTATACAAGAACAAAATTCCATCAATAATTACCCACAGCAC
CTCTCAACAGCTGGGCTTGCCCCAGTTTTCAAATAAATAAATAAATAAAGAATAAATAGAAATCTAATGAAAGTC
ATCCCGCAATAATACAATATTTAAGCCTTAAACTCAACATACAGTATTTAGGTTCACAATCCTATGAAGGAAAAT
TTCAAGGTCAACAACCCAAGGTGTGATACAACAGCTTCATATGGAATTATTTCAAAAGCCTACCCAGGATATCTA
ACCTCTACTCCTTCAACCACCTCTCAGAGCACAGAAATGACATCCACAAAGATCTTCAACATGATGGATACGCAT
AAAACAAATAGCAAATGGTTGATTTATATTTTGGAGTTCACATGCCAAGATATGCTCTCCCAAATATTAAGGGTT
CTTAATGCTCATTGAATGCACTATTGGGGAAAACAACAAATATCTTGTTTCCTTCTGAACTTATAACTGATTGCA
ATCATTTAACAAATACAAAGAATTGTACATTAAAAGGCGTCTACACAACTAGGTATCTTCCACACGACCAATGAA
ATAAACCGTCCCAATCATTTAAACCTAACTTGATCAAAAGTTCTTTGTGAATCTGTAATTTGTAGGAATATTCTA

GAAGATGCCAAATGTCAGAATATCAATACCATTCTTTTTTAAGCAAAGTTAAACTATTTCTTACAAGTGTATGTT
AGTTGATTATGGGCTTCCATTGGACTCTTTGATTTAGCACTCACGCAATTTAACAACCAGCAGTAAGTTTTGCGA
CAAAAACAGGTAAACTGCAAATAGTGAAAGCTTCTGGATAAACATGATTGTGAGATCTATTACAATATTAACAGT
CACATGTGGAACTGAAGAGTTAGAAAACAGCAAAGCATGCTTCGAGCTGTGAAACCACCACTTTCGGACTTTAAT
GTTTCTTGAGAAAATGTAGATCATTTGATACTTAACCTATACATAGCGAAAATCAACAATTACCCTTCATTTATG
TATTAATTTTATAATTTCCTCCAACATTGTATGCCATCATCTTCTAGCCCAATAGAAGGCCATACATAAACTCCA
TAACAGTACCCACATCAACAATAAACTTAATTGATTAAAAGGACATAGTTCTGCGACTAAAAGGACATAGTCCTC
CAACATTGTACCCACATTAACAATAAACTTAATTGATTAACAGTACCCACCTCTACATCTCAGGGTATTAAAAAT
TATTCTATAACCAGATACAGATAACGAATTAATCATCAAAAAGGTTTGGAGACACTGAAGTGAAATGTGTTTGAA
GTGCAACTTTTAAGAGATAATACCAAAATATGTTGCCTTGGCATATCTTAAAGTGGTGTTCTAAGATATCATTTC
CACACAATATATCAAAAACATAAAGAACACACTCAAAAAATGCCAGTCAATCCCCATTTTTTGACTTAAGTATTT
CTGAGCACCCCAATTCAATATTTTTATTGACTACTTATGAACTGAATTCAATATTTAAAATGAACTCAAGCTGAA
AGAAATCATAC**CATTCATAATTCTACAAGTTATGCATTTGCTGCCTCAAACTCACTGCCTCCATCTGCGTAAA**CT
GGTTAATAAGAAAAATAAAGCTTAGTTCATGCAAAGCACTGAAAAATAATTCATTTGAGCATATTGACAAGACAA
TTATAACCAGGTATCTAATTTATTATGTGAATCATTAAGAAGGCAAATCATAGACAAAATAAGGGGTCATTGAAT
GAGTTAAAAGATGGGTGAATTAACAGAGCACAATATACACAACTAAGAGACATGGAGGACACCTACATACCTCTG
GATATAGCACAAAGATACAAAAAAAAGATTACCAGCTTCATACAAACAGCAATGTTACTATTTCACCAGAGCCAT
ATTCAAGATTGCTCAACTTAGATCGTACATTATAAGCCAGGGAACAAATTATTCCCATGACTTCCAGCCAAACAA
AATAATTTGAATAATTTGAAAAAGGAATAACATTTACTACAACCACAACAACAACAACAACAAGCCATTAGTCCC
AACTATTTGGGGCCGGCTACATGAACCCTCCTATTCTACAAAATTTATTAATTAGCTTAATATAATCATTAATGA
AGTCGAAGTTGATAATGACTTGGGATGAATTTTTCCTACTGATACAAATTTAAGAGTATCACTCTCCATTCATCG
TAGTCTCTTCTTTGCTTTTTTCCCTTTTTCTCTGCACCAACTTATATCACCCAATAATCTAACCTAGTGCATTCA
TAGGACTGCTAAATACGAAGCTTAACCATACTGCGCAGCATTTGCCAGTTTATCTTTCATATAAACCCACCATTT
CGCTACATAAATACAACCAGTATATTTGTATCCGATGGATGTACTTTTAAAGTTCGATTTGTAGTTATCTATATT
TCTTAACAGCAATAACGCCACAGGCCACTACTTCGTAATCTGTAGGTCTTCTTTCATAACTTATTTAAGATAAAT
ATAACGACATTTATAACAATGTCAAGCAAAAAGTTTATTTGGTTGTGTTTTTTTTTTTTCTTCTTTTGGAAGGAG
GGGGGGTAATATGAAGAGCAAAAAGTTGTATGAAAAAATAAATAAAAGACAAACATAAACGTGTTTGAATCCGAT
TTTAGACTGGGATATCCCAAAACACATGCCCAAATACCCACATGATGATCAGCTTAATACAATCTAATAGTAAAC
CAAACGCGATAAAATACGGATGACTTTATCCCCGCTCTACTTTAACCTATTATCCCAACCCCATTTGACCAGAAC
CCAAACAAGTCAAAAAGTTATTTGAGACACAACCTACATTTTTCCTATGCATTACGAACCAACAAATCTTTTTTT
TAAAAGATAAACATTTCAATAGCATTATAAGTAAATCTTAATATTCTACTTATTTCTCATCCTATTTATGTTCAA
CCACCACATACTCATTGGCCTCCTGATAGGTATACATCATTAACTCATTTGTCTTCTATTTATTCCTCTGACTAG
TGTTTGTTTTTATCATCCATACATTCGTTGAACCTGCATCCTTTACCAAAGCCTCTTGTCATCCTAGTTGACAAA
AAGCTATCCAGGAAGAACTTGATGATTTACATAAAACTCAAACATGGAAACTTGTGTCATTAGCCTTAGGTAAAA
AAATTATTAGTGCAAGTGGATATATAAGATTAAGACCAAATCCTATAAAGACTCGTCTTCTGGCTGAAGGTTTTT
CTCAAGAATATGGCATTAACTATGAGGAGACTTTGCTCTAGTAGCAAAATTGACTTGAATTTGTGTTTTGATAAC
CTTTACTTCTTCCAATCAATGGCTATTCTTTCAAATAGATGAAAAAAAATGCTTTCTTAAATGGAGAATTTACAT
GAAAAGATGGGCATGTGTCCTCATCCAGCTTTGCCCCACACTAAGGGACAAATATGTCTTTTTCATAAAGCACTA
TATAGTCTCAAACAATCACCTCGTGCTTGGTTTGACAAGTTCAGTTTAGCGTTACTCTAGCTTGACTTATTAACT
CACCTTATGGTTCAGATTTATTCACTCATGCCTCTTCTCGTGGCAAAGTTCTTTTGCTTTATTATGTGGATGATA
TGATCGTTACTAAAGATGACTCGGATGGCATTGTAAGATCACATTTCAAATTTATTTGAAATGAATGATTTAGGT
CCTCTCCATTATTCTCTTAGCATTGAAGTCGGCTCACCAGCACATATTTGTCTCAAACCGAATATGCACTTGATC
TACTTAATAGAACAAAGCAGGTCTTCCTGATGACAATACACTAACTACTCCTTTTGAGTACACTATTAAACTCCC
TCCAGATGATAGTGAACTTTTGAACGATCCCACTCGACATCGAGAAGTTATTGGGTCTTTAGTGTATCTCACAGT
CAGTCATCCTGATATACAGTTTGTTGTACAAGTTTGTCAATTTAATTCTGCTCCTTATACTACTCATTGCACAAA
TGTTCTTCAGATTCAAGACACATGTCATTATAGATTATTACTTCCTTCATCTATGATTTTGCATACCTTTTCTAT
TATAGATTAGGGTGGAGATATCACAAATTGTCATTCCACTACTGGTTGGTGAATTTATTTGGAAGACCTTTTGGC
TCTTAGAAAAGCAAGAAACAGTGTTGTCAGATCCTCCATTAAAGTAAAGTATCAATTTATGGCATGTACCATAGC
AGAGGCTTCATTGCCTTTTAGATGCCTTGGGACTCTCTAATCAATACATATGCCTCTCCATTGTGATAATAAAAG
CGCGATCTCAATTGCTCACAATTCGATTTTTAATAAACGAACAAAGCATATAGAAATCGATTGTCATTTTGTGCG
CCAACATCATCTACATGACACTATTACTCTAACATGGTCAATCTTCTGAACAGATTCCTGATACATTCATAAGGA
CTTATAGATTGGTCTGTTTTCAATTTTTCCTTTTAAGGCAAACTCCTGATGTTCACTTCTTCTCCATCAACAGTT
TCAAGGAGATGTTAGAGTAATAATAATAATAATAATATATGTCTAAAGTATAAGGGTATTAATGTCTTTTAGGTT
AAGTTCAGAAAACCCTAATCCTACTCTTATATAAAGAAAGCTGCACCCCAATTGTATTTGAGTGTAAAAAACTAT
TCTTTTAGTTCTACAATAGTTTCTTCGATTGTTTCCTTTGCATCTAACATGGCATAAGAGCTTAGAATATTTGAT
TTGTTAGCAACGTGGCTAGTTTAGAATAGACTAGCATGTATCAAAGCGGCATACTAGAAAAACTAATAATCCCAC
GTTAATTAAATTGGAAAATATGAACTTGAATATATAAGCCTATGTTTCTCGTCCCATCAGCTTAAATTTTTGGGT
TGAATTGAGCCCAGAACATATTTCCTTTACATCTCAAAAAAGAACTCTCTATATAAAACTAGTAGAAGGTACTAA
AAAAAGATTGACTAAGGAACATTTGTCTAAGAGAAATCTTGATGTTGGGATCTATCCCAAGATAAATATAAAATT
TCCCTATGACATCATATTTAGAGGTAAATGGTACAAATTGATGATTCTACAAAGTTCTTTTTATCACAATATTGA
CCCAAACCAACTTACCCAGAAAAGGCAAATGTGCCAAAGGTAAAAAGGGTACTAGGAAAATAAAGTATATCACAA
GACAGCCAGGGATAAAAGAATCCATCATAAAATACTCCAAGATCTTTCAAATTACACCATATTTTAAAGTCCATC
ATGAAACCATTGTACAATCACATAAACTACCATAAGGCAGAAGCCCAAATCATGTAATACCCTATCTTCATCTTC

ATTCCATGAAGCAAACTATGAGATGATCAATAAGATCATAAATCCATAACCATAATGAACAAAAAAAGTAACATA
TAGCTGAAAAATGATTTCTTACCAATAACATATGCAGTAATAACTCACTTTAAAGGTTCAAAGCATACAAGCATC
ATGGTTGAGGAAATTGCTATTCAATGCAAATTGCACAAGGTAGCAAATCACAGACGGCGAGGGGGCAAAAGATAA
AAGAGCATCATCTTTTACAAGCACAGACGATATCAAGACAAAACAAAGATGGAGCAGCAAGAGACAATTCACAGA
TGAAACCAGTGTAAACATAAACCTCATGTATATCAATCACCTCAAAGCTTAAAACAAAATAAAACCATCATCACA
CACAACAAAACTTTGACTTTTCACTCAAAAAAAAAATAAAAAATAAAGGGATTAGTTTCAAGCGTGCCAAAACAA
AGATCTGAAACAAATAATGAGTAAACAGTGCAAGAGCCCACTAAAAACCTGAAATTTCAAAAGCAAAACCACAAT
GCAGTAACAAACAGTTAAAAAAACCAAAAGCCCAAACCTGCATTGAAACAAAGATAAAGAAAGTAGATAGATTCT
CACTAAC**CTGAAAACCCTCAGCTGCTTATCAGT**CTTCCATCAAAAAAGAACCTCATTATCTCCAAGAAAAACAGA
AGAAAAAGGGTTCTAAACCCCATCAAAAGAACAAGAAAGCGGGAAGTCCAAAC**CCTAACCAAACCCTAGCCAAAA**
**GCTTTCCGTTGAT**GGAGATTGACGACGACGACTCCTCCTCTAAGGCGCTAATTATAGGCAAGAAGCAAGCCCCAG
AGATCCAAATCCACGAACAAAAAACTCAAAAAGAGGACGATGGAGACCGGATCCAACACGTAAAAAAAAGAAAAG
AAAAAAAAAAAGAACAAAAAAAAGAAAAAAATAAAAAAAACAGAAAGAATCGTACCCGAAAATTGTAAGGAACGA
ATTCACAAAGAGGGTTCTGGAAAATATCTTGCGAATTAAATAGTACCTGCGGATTAAAATAAAGAAAGGTTGACT
GGCTGAGAATGAACCACGTCGCGCGAGACTCTTTTCGCACGCAGCCCCTTGCTGGACCTTCGAGACAAAATCGGG
TACCTTGGGGCACACCACAACGCCCCCAAAACACCCAAGACGTCGTTTCCCGACACGATGACGTGTCCCTCTCTT
CGGCGGTTGGATGGGTCACGACTGTTGATGCCCGAATCTTGGGTGACGTGGCTGCTCCTCGGTGTGGGTGTTTGG
TGCACCGTGTAATGCAGCAGTGACGTCATGGGTGACGTCATGTGCTAGTGTCCACGTGGACGTCTAGTTCAGCTG
CCCGGCTGTGTCTCACCAATTCCATCATTATTTTTTATTAAAGAGTTTTATTTTAATCAATAAAGATCTTTCCAG
TTGCATTAATATATAGAGTATTATCTGTGTATTATTATTATTATTATGAGCAAAAGTGGAAAAATGAGAAAATAA
ACGGGATGCATTATTTATTCTAATAATATTATTATTATCATTATCTTTACGGTCCATGTCGCTTTGAAATTTGGA
TTGTGTCGCTAAGTGCATGAATCTATAATTTAGAGATTAAATATTTAATTTTTACTTTTACTTGAAAATAAATGA
TAGTTATAATATTTTTTATTTTGTGTTGGGGAACTGATCACCAAATTAATCAATTAATTAATTAATTAATTATGG
AAACATCAGGCAAACAGGCACTGAAAGAAACTCATGCAAAAGCATTGGACTCAGCGAACAGTGAGGATAAAATAT
CAAGGGGCTGTTAAAAACAAATATTTCAAAAATCAGAAGGCTCTTCAAATTAGGTTGATCAAGTGCAAACATCTA
AAAACTTTTATGTACTTTTAAGAGAAGAAACAGAGCAAGTGACAAACTAATATTCTACATAACAATGATGTAACA
TCCTCCTGTTCATATGGCTCACAGATAAATTTTTTATTATTATTTTTTATTTTCAAAATTATGGATAAGCCACAA
ATACAACTCCTATTGTAGTAGTGGGTAATGACTCACAAAAACTAAACTAGATATTTTTATATGAAATTGATATTA
CATTTATTTAGTTGAGAAAAATTTATATGGAGCTGACTTTATTAATAATAATAATAATAATTAAAAGGATCATTT
TTAATTTTAAAATAATATTAAATCTATTGATATAGATTTTAAATACTACCAAGGAAATGATTTAAGTGGTAAAGA
CTTAAATGGGTTAAACAAATGGTCTCAAGTCAAACACTTATATATGTAAAAGAAAACCCACAAAGAAAATTTCAC
CCATTTTGGAACTTATAGTATTTCATTCATATTGATTTAGAACCTGGGACTTTTAAAAAAAACAAGTCGAGCAAA
ATAAATAAATAAATAAATAAATACCCAACTCAAGGTACAGGTGTGATTGGTGAAGCTTAGCCTCCTGTTTGTTTT
GAGTTGAGTTTCAAAACTCAAAAGAAAAACTTTTAACAATAGAATCTCTCATTTTTGGCTTAGAAGTTAGCTGAT
AGGATTAGATATTTTATTTTGTGAAACTTCATTCTTATTTTCTAAAAATAGAATGAAAAAAATGGTTAGATCTAA
ACATAAGCCAAAATACAAATAATTTAGTTCAAAACACACATTTTAGCCAATATAATTTTTTTTTTAAAACAAGAT
GTAAAATAAAACAAATAATGATCTATTTTGATAAACACTTGAAATGTTGCAACCAAGCACTTAGTATCAACCGCA
AACAACACAAGGAAATTAGGGTTTCAATTATAAGTTCCACAGAAATGAATCAGGCCTTATGCTCTCACCAGATGA
ATATAACATAGGCAAACATAGAAGTGACAAATCAATGAAAAGCATCATTCTCGTTGAACCGTTGAATGGAACTCT
GACCTTTGATGCACATTCAAAATCATTGACAAAAGCAATTGCGTTTGTAAAGCCTCATACATGCTTAAAAAAATA
AGTTAATAGCAGCAAAACACTAATAACACAAACATCTAATGCATGTTAGCAGCATAATCATCAAACTTAAAAATG
AACGAAAATTTAAAAAAAAAAACAGCATAACAATGCATCTTTATTTTTTTAATCTTATGCAGGTACAAGCTTCA
TCATAATTCCAATTGTTTCTTCTGTGGACCAAGAAAACATTCACCTTTTCAACAATTTTCTCAAAAGCAGAACGA
GGTGAGGGGTAAAACCATCCCACTTCAAAGCGTAAATCTATATGTTAAACAGTAGCAGGTTTCTGGCAGTTCTCA
AATGGTGAATCTAGATACCTCTAGGGATACAATAAAGTTTTAGCAGGATTAATTGAAAGGCAAACTAACAGCGAC
AAATAGAATTTACAACCAAGTTCCTTAAAAAATTACACCTTCAAGTGCTTAGAGAAATTGATGAATTGCAGGGAA
TGAGTGACAACGAAAGCAGATGCAGGAAAAATTAACTGGAACAAACCAGGAATACAACAAGGCCGCAGAGGCAAA
AGAATTGTTCAAAACATATTCTTTCTTTTGGAACAAATGGAGCATATATGATGATCAAAACAATGATAGAAACAT
GAATATAAATAATTGGTAAAAGATAGGTGATTTTGCCAATTGATAAAAGTTTACAAAGGCGAATTTCCAGTGGAT
GCAACAGGGACAGCATACTTCTTTAATATATAAGGATTTTCTTTCCAACATTACCAGAAGGGATATTTGTCAATC
CACTGAAAGCATGTATGAAAAATGTGAAATATCACAAAATTGAAGGAAATTCTTCTGCTTATTCTTTCTTAACTC
CCGAAACTTCTTCGTAGAGGTTTTTCCCATGCCCCACATTTCATAGGGAACCTCAAAGCAGCTCTCTTTCAGTAT
ATTCCATGTATATACCAATTTCATTCAGTTAATATATCTTTGGTGTCAATGGCCGATAAATAGTGGCATCTCAAA
GAATCACACATGATCCCTGTTATTCTATAATCTAAAGGCGGCATGAATCCTCACTTCATAATTGCATAACAAAAC
ATATGACATTTGACAAAGATCTCCGCTGCTTATTAATTTGCAATGATCCTTCCCATTGTAAGATTTGAAGCAGGC
TACTTACTAAGATTACCAAGACAGGCTGATTTCTACAATAACAATAAGCCGTTAATCCCATCTATTTGGGGTCGG
CTACTAAGGTAGGTTAATTTCTATTGAATAAAAAGAGAAAGAAAAGATTTGGGGTATGTGTACATCATAAACCTA
TAAATGCAACTATACATAACACGGTATATGGATGGGATCAGAAGAAATGATGCAGTCACACTAGTATAAAGAACT
ATAAGACGAGCTGACATTTGATAAACATGAGTAGAGAAACAAAGTACCTGTGGTTGACTCTAAAGCTACTCACCT
CTTAAGGCTTTTCTATGAGACAAAATTTCTAAGAGGAGCTCCTCTGGGTAAAAAAACCATGATGGTTATCATACT
ACTTGTGGACAAATTTCAGGTTGGTGAACTGCATGATCATAAAATATTTTCTAAGATTAGCTCATCTTAGACATT
TTGGCCCCTTTATTACTATTTTGTTTGTTCAAAACAAAACTAAACTTGTCGAGCAAGTACAAACATAACAATAG
CATATGAATTCTCAGACACCTCAACATGATAGAAATATATGAACTTCTCTGAACACCAAATCAGAACTGATAGTT

CATATCAGGATGGGAAATGCAGTATAAAAGCAACAAACTTTACAAAATATCAATTCAGATAGATTCAACTACAAA
ACAAAACTACATTACCATGAACTTCAATCACAAGAACAAAAAGGGAAAGCAGTATTAAAGCAAGAATTTTGTATA
CTGATAATGATTAAGAGATATAAAAACAGCATTGCTACACTAACCAAGCATTCATATAATCAAACAACAAAAGAA
AAGAACAGAAACTGACT

>DbAGL17 [organism=Dioscorea bulbifera] AGL17-like MADS-box gene, partial CDS
CTCTATGACTTCTCTAGTTCCAGCATGAAATCAGTGATTGAGCGTTACAACAGGACTAAAGAGGAATGCCATCAA
CAATTTAATCCAACTTCAGAGGTTAAGTTTTGGCAGAGGGAGGCAGCAAGCTTGAGGCAGCAACTGCATAACTTG
CAAGAAAGTCATCGGCAATTACTGGGAGAAGAGCTTTCTGGCCTAAACATTAAGGATCTACAAAATCTCGAGAGC
CAACTCGATACGAGTTTACACAGTGTGAGAATGAAAAAGGAGCAACTTCTGATTGAAGAAATTGAAGAATTAAAG
CGAAAGGAAAATCTACTCCATCTCGAAAACGTGGAACTCTACAAGAAGGTTTATGACGTAAGACGCACAGGAGTT
GACAATGGTCCAGTTATTCCACATGGGTTCAGTTTTATGGAGAACACAGATGTCCGAGTTCACCTTGAACTAAGC
CAACCACAACATCAAAAGGATTGTGT

>dbu-MIR444 [organism=Dioscorea bulbifera] partial transcript of MIR444, transcript variant 1a_1
ATCAACGGAAAGCTTTTGGCTAGGGTTTGGTTAGGGTTTGGACTTCCCGCTTTCTTGTTCTTTTGATGGGGTTTA
GAACCCTTTTTCTTCTGTTTCTCTTGGAGATAATGAGGTTCTTTTTTGATGGAAGACTGATAAGCAGCTGAGGGT
TTTCAGTTTACGCAGATGGAGGCAGTGAGTTTGAGGCAGCAAATGCATAACTTGTAGAATTATGAATGGGAAAAG
TGATGGTACCGATGACTTTCTTGCAAGTTATGCAGTTGCTGCCTCAAGCTTGCTGCCTCCCTCTGCCAAAACTAG
TGTGGACAAAATAACCCATGTCAATGCATTATGGTTTCAGGTGAAAAGCTGAGGCTGGAGAAAGAGCAAGAGTTC
TTGTGATCTATGTGATATATATGTATATATATAGTTCTTGTCTGCTGTCTTGTTGTAATTTATAGTGAAAGTTCT
GTTGTAATCTATTGTAAAGAAAATCAAGTGTAGATTGACTTTGAATATTCTTGCAAG

Stem-loop structure

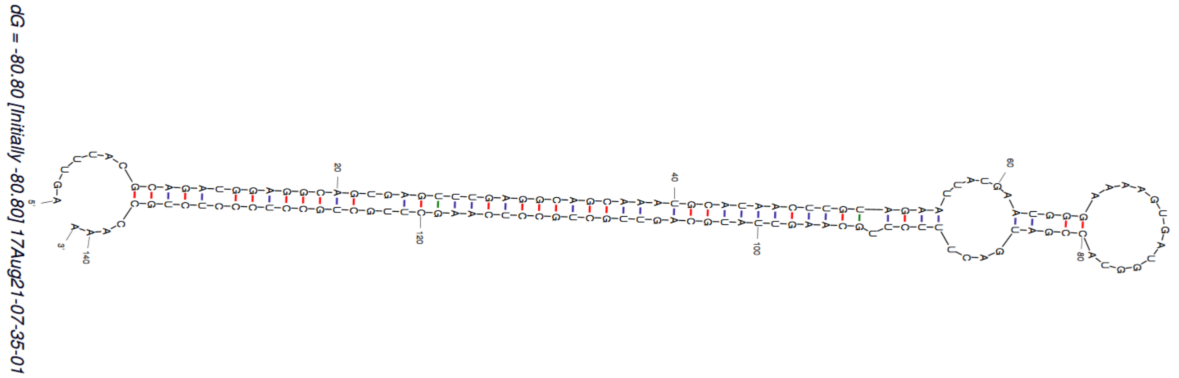


### Alismatales

#### Spirodela polyrhiza

*SpoAGL17*

>pseudo2
TCGCCGATTATTTATGCCCTGGCGCACGACGTCATCCGCTGGGTCCGGCATGTCCGCCATCGGACAGATCGAACA
TGGCCATTTGACGAACACACCCTTCCTCGTTGTCCGACCAGCCCTGGCGCGAACCCAGAAGGTTGCACGTGAGGT
CGATGCGAAGCTCCCTAGAAAGGCGACAAGTGGCTGCGATACCACTCCCTCAGCGCACAGTAATCTATCGGCTGC
GGACAGCGTGAGGGTGTTTTCGTCAAATATCAAAGGTGACCCGTGCGCCGTCAAGCCGTCGGAGGCGGACCCTGC
ACCAGACGGCGGGCGGCGGTGGGATCCACTCTCATTTTCGGATATATCTTCCCGAGTCGAAGGAAACCAGAAGCC
CCcTCGTCGTGGTACAAAACCTCGGGGTCGATTCGTCATGCTGTGAGGCGCGGGATGTGAGGAGAGGGCTTCATC
CAGTGGATCTCCGAGGCTCCGAAACTGAAAATGGAAAACCAGAGACCAGCCAGGGATTAAGGTTTCCATATATGT
GCATCACCAAACCGGGCAGTTCGAGTCTATTTTAAAGAGGAGGGAGGCCATCCTTTCCAGAaGGGACCTAAaTAA
AGCTAAGCAACAAAAACCGGTCCTCGCCTCCTCCTCCTCGCCCATCCCACTCTCTCTTCCCcTTAcACCtCTCTC
TNNNNNNNNNNNNNNNNNNNNNNNNNNNNNNNNNNNNNNNNNNNNNNNNNNNNNNNNNNNNNNNNNNNNNNNNNN
NNNNNNNNNNNNNNNNNNNNNNNNNNNNNNNNNNNNNNNNNNNNNNNNNNNNNNNNNNNNNNNNNNNNNNNNNNN
NNNNNNNNNNNNNNNNNNNNNNNNNNNNNNNNNNNNNNNNNNNNNNNNNNNNNNNNNNNNNNNNNNNNNNNNNNN
NNNNNNNNNNNNNNNNNNNNNNNNNNNNNNNNNNNNNNNCTCtCTCTCTCTCTCcTCTCTCTCTCTCTCTCTCTCT
CTCTCTCTCTCTCTCTCTCTCTCTCTCTCTCTCTCTCTCTcTTTTCCCTCCTTGTCCTCTGTGCCGGGAATTTCTTG
GTGAATCATCCCTTTCTGGTGGAGGTGAAGAGTAGTTCCGTCTCCGTGGCCGTCTCCTTCCTTGGTAAGGAAGAC
GTGCTCTCCCGAAATTCTTCCGTAGATCAGTTCCTTATTCGTTTCTGTGCTCGTCTGTCGTCTCGTTTCCTTCCT
CGGAGTCTTCCCAAACGGATCCGACAAGTGGCAAAGTTTTCATGGCTGATGATGATGATGATAGGGGTGGAAGAA
TGGGgAGGGGgAAGATTGTGATCCGCAGGATCGACAACTCGACCAGCCGGCAGGTGACCTTCTCCAAGCGCCGCA

ACGGGCTTCTGAAGAAGGCTAAGGAGCTTGCCATTCTCTGCGACGCTGAGGTTGGCGTCATGATCTTCTCCAGCA
CCGGACGCCTCTACGACTTCTCTAACACAAGGTACCAGCCCCTCCGCCCCCTGTCCCCCTTTTCTGCCCTTCTTG
TCCTTGTATCTTCTGTGTTTTTGTTCTAACTTAATCCTTTGCCCGTAATGCATCCAATCTTGGCTGTCGCAGGTT
TCTTTACCCTTCCTTAGTTTCCCTTACTCTTCATGTTTTCTTTGGAAACGAATCTATCCATAAGATCATGCTTGG
GATCAGCCATCGGCTGACTTACTGTTGCTGCCGAGCAGCTCCATGAGAAGAGAGGGCACTTGTTCATATTCAAAA
TACAGTTTCGTGAGGGAAGAAAGACATAAACTGAGGTTTTTTGGAGCCTGTTGGAAAGACCGAACACCCATCATT
CTTGATGACAGATTTATTGGATTTCACGCGGCTCCCTGCATTCCCACAATTAGTAACGTCCGTTCTTGGCACTCC
TCCCTTCCTTGTAGGCAGAACGCATTTTCCTATCATGGTTTCTGCCAGAGTCGAATAAATGTACTTGGCTCGCGA
CTTAACAGATTCATTGACGTCACAGGAATCATGACAAGTGGCTGAGGTAACAGACGTTTGAGCGTGAATCTACAT
ACTGAAGGAACAAATAATCTACAACTATTTGCACACATATTTATATCTATCCATCTGTCAATCTATCCAGCTGAC
CATCAGCCTATTTTCTCGCAATTTCTTGAGAGGATATAATTTTACGTGGTTAAATCAGTCCAGTGTCGTCGGTCG
CGACGCCTTTCCTGGATGAACCATTGAGTTGTTCTTTTGTGCTGTTCGGCCGAGAGGAGATGACAATGGCCTCAC
CTGCATTCATCACCAAGCTCATAGACATGAGGCTACAGACATGTAGAGAGGGGACGTCTTACAAGGATAATTTCG
TTACGGAAGCTTCATTGGACAACCGTATACAGCCGTTAGCTTCTTCTCAATGCGTAGGGCGGCTTGCGCGGGGTC
ATCCTTTCAAAGCTGCCGTCTATCTGCAATCTAATTTAGCTAAGGCTGTGGAGATCCATACGAAATAACCTAACA
TAATTGAGTGGATAGGACAGGAGATTATACACCTTTTAACTCAATCCAACTGGTAGCAGCTTCCACTGCACAGGA
AATATACACTGTCTTGATATGTTAGTCATATGAGTCTTCTGGAACTAGGACGGTAATGAATCTCATAGTTCACAT
CTAAAACCATCAAGCCGTCAGAAGGCCAGAGGATAGAGTGCATAGAGTAGAAGTGGACTGCCTCACGGTAGCTTC
ATCTGGGGAATACGTCGAGAAATCAAATAAGTTTCGCTACATATCTATTTTGCAGCGTTATTCCGGCATGCTTCG
TTGAACACCGCCTTGGTTACAATTGTTGTGTAAATGGATTGAAAGTGACTCACCCAGTGTTGATCATTGTAGTTA
AAGAGCCATCCTTTCTCTCAGGTAAAGTAGCAGGATAGTGCCATATCCTTGCGGGCAAAGGAATCATGACAAAAG
ATGGCTATCTTATCCGCAAATCATGACCTCGATGTTCATGTAGATGATCACTGGAAAGAGGGGTCATTATTCTTG
ATAGCGGACACTTGTCAACTTTCTCTTTTATATATAATTCTCCGCTTATTGTCGACTGAGTTTCTTCTTCTGGCC
ACTCACTTCCGATAGTGTTTTCTTACATTTTGTTGTGCATTATTAAAGAACTATTCTAAAAGAGGGTATCGAGTT
ACCAATTACCTATACATATTACAAGTATATAAGTTACGTTTGTGTCGATGCTGAGAGTAGGCAATAATTTGAAGG
AACATGTTGCCCATGGAAACTTTTTCTGGCCTGAATAGTTGGAAACTCATTCCTAAGTGCCCCTAAACGCTTCTT
TGGGATCTTCGTAAATCTTACGAGAAATCCATCAAAAGGCGACTGCCCTTTTCTTTCCTGGTTTTTGTGTCTCAG
CATTTACCCTAACCGGTGTTCTCGTATCCATAATCACTTTTTAAGGCTTGTTGTTCGTCCACGATACTGCTACTA
TCGATTCTCTAGATATCTGATACAATTCTTCCCTATTTTGGCTGTTTCCTTGGGTGTGGAATATTTTCAAAGAAC
TTTTGCAGCAAAGCGGCAAAAAAATCAAACAGTGGGATTCGTATGACGAGGATCCACGTTTCAGGAGCCAAGTAG
TACAGCTGGACAGTACAGCCGCATGTATTTTGCTGGAAATAGTTGCATTCATAATCTAGACTGGTAGATATTGAG
TTCAGATGCATGATAGCTTTCGCTACAGATGATCATGTTGCTTGATTTCGTAATTTCTAAAAGTTTCTTAAAAGT
TCCCCAGGGATGCACGTAGTAATATGTCTAACTCGTGGAGTATCATATAAAATTATGCCTAATTCATATTCTTCC
GACTAGCTGGAGTTTCCTGTGAAGAGAGAGTTGCACTCGACTAGGGACACTCTTAATACCAGCTCATCTATTTTT
ACGTTTGGATTTCCGAAACAGTTCCTTTTCTCGTTGCCCTTGCGATTCTATGCTTCATGCGTTCTGATCTAGTTC
GTATTTCAGTTCTCCATCACATCTATTTTTCGATATGTGGCCTCGTTCTTTCTCAAGTTCTAGTTCCTCTTTCAG
TTCTCCCTCGCATCCACTTGTCAATTTGTGCCCACACTCTCTCATTGATCTTTCTTTCTGCTCCATTCCCCTCTC
GTGCTGTTTCTCTCTTCCGCAGACTAATTTCCTTTGTCTCAGACTACTTATCTAGCTCCCTATACCCTTCCcTTC
ATCCCGCCACTCCTGATGACTCTCTCTCCCTCTcNNNNNNNNNNNNNNNNNNNNNNNNNNNNNNNNNNNNNNNNN
NNNNNNNNNNNNNNNNNNNNNNNNNNNNNNNNNNNNNNNNNNNNNNNNNNNNNNNNNNNNNNNNNNNNNNNNNNN
NNNNNNNNNNNNNNNNNNNNNNNNNNNNNNNNNNNNNNNNNNNNNNNNNNNNNNNNNNNNNNNNNNNNNNNNNNN
NNNNNNNNNNNNNNNNNNNNNNNNNNNNNNNNNNNNNNNNNNNNNNNNNNNNNNNNNNNNNNNNNNNNNNNNNNN
NNNNNNNNNNNNNNNNNNNNNNNNNNNNNNNNNNNNNNNNNNNNNNNNNNNNNNNNNNNNNNNNNNNNNNNNNNN
NNNNNNNNNNNNNNNNNNNNNNNNNNNNNNNNNNNNNNNNNNNNNNNNNNNNNNNNNNNNNNNNNNNNNNNNNNN
NNNNNNNNNNNNNNNNNNNNNNNNNNNNNNNNNNNNNNNNNNNNNNNNNNNNNNNNNNNNNNNNNNNNNNNNNNN
NNNNNNNNNNNNNNNNNNNNNNNNNNNNNNNNNNNNNNNNNNNNNNNNNNNNNNNNNNNNNNNNNNNNNNNNNNN
NNNNNNNNNNNNNNNNNNNNNNNNNNNNNNNNNNNNNNNNNNNNNNNNNNNNNNNNNNNNNNNNNNNNNNNNNNN
NNNNNNNNNNNNNNNNNNNNNNNNNNNNNNNNNNNNNNNNNNNNNNNNNNNNNNNNNNNNNNNNNNNNNNNNNNN
NNNNNNNNNNNNNNNNNNNNNNNNNNNNNNNNNNNNNNNNNNNNNNNNNNNNNNNNNNNNNNNNNNNNNNNNNNN
NNNNNNNNNNNNNNNNNNNNNNNNNNNNNNNNNNNNNNNNNNNNNNNNNNNNNNNNNNNNNNNNNNNNNNNNNNN
NNNNNNNNNNNNNNNNNNNNNNNNNNNNNNNNNNNNNNNNNNNNNNNNNNNNNNNNNNNNNNNNNNNNNNNNNNN
NNNNNNNNNNNNNNNNNNNNNNNNNNNNNNNNNNNNNNNNNNNNNNNNNNNNNNNNNNNNNNNNNNNNNNNNNNN
NNNNNNNNNNNNNNNNNNNNNNNNNNNNNNNNNNNNNNNNNNNNNNNNNNNNNNNNNNNNNNNNNNNNNNNNNNN
NNNNNNNNNNNNNNNNNNNNNNNNNNNNNNNNNNNNNNNNNNNNNNNNNNNNNNNNNNNNNNNNNNNNNNNNNNN
NNNNNNNNNNNNNNNNNNNNNNNNNNNNNNNNNNNNNNNNNNNNNNNNNNNNNNNNNNNNNNNNNNNNNNNNNNN
NNNNNNNNNNNNNNNNNNNNNNNNNNNNNNNNNNNNNNNNNNNNNNNNNNNNNNNNNNNNNNNNNNNNNNNNNNN
NNNNNNNNNNNNNNNNNNNNNNNNNNNNNNNNNNNNNNNNNNNNNNNNNNNNNNNNNNNNNNNNNNNNNNNNNNN
NNNNNNNNNNNNNNNNNNNNNNNNNNNNNNNNNNNNNNNNNNNNNNNNNNNNNNNNNNNNNNNNNNNNNNNNNNN
NNNNNNNNNNNNNNNNNNNNNNNNNNNNNNNNNNNNNNNNNNNNNNNNNNNNNNNNNNNNNNNNNNNNNNNNNNN
NNNNNNNNNNNNNNNNNNNNNNNNNNNNNNNNNNNNNNNNNNNNNNNNNNNNNNNNNNNNNNNNNNNNNNNNNNN
NNNNNNNNNNNNNNNNNNNNNNNNNNNNNNNNNNNNNNNNNNNNNNNNNNNNNNNNNNNNNNNNNNNNNNNNNNN
NNNNNNNNNNNNNNNNNNNNNNNNNNNNNNNNNNNNNNNNNNNNNNNNNNNNNNNNNNNNNNNNNNNNNNNNNNN
NNNNNNNNNNNNNNNNNNNNNNNNNNNNNNNNNNNNNNNNNNNNNNNNNNNNNNNNNNNNNNNNNNNNNNNNNNN

NNNNNNNNNNNNNNNNNNNNNNNNNNNNNNNNNNNNNNNNNNNNNNNNNNNNNNNNNNNNNNNNNNNNNNNNNNN
NNNNNNNNNNNNNNNNNNNNNNNNNNNNNNNNNNNNNNNNNNNNNNNNNNNNNNNNNNNNNNNNNNNNNNNNNNN
NNNNNNNNNNNNNNNNNNNNNNNNNNNNNNNNNNNNNNNNNNNNNNNNNNNNNNNNNNNNNNNNNNNNNNNNNNN
NNNNNNNNNNNNNNNNNNNNNNNNNNNNNNNNNNNNNNNNNNNNNNNNNNNNNNNNNNNNNNNNNNNNNNNNNNN
NNNNNNNNNNNNNNNNNNNNNNNNNNNNNNNNNNNNNNNNNNNNNNNNNNNNNNNNNNNNNNNNNNNNNNNNNNN
NNNNNNNNNNNNNNNNNNNNNNNNNNNNNNNNNNNNNNNNNNNNNNNNNNNNNNNNNNNNNNNNNNNNNNNNNNN
NNNNNNNNNNNNNNNNNNNNNNNNNNNNNNNNNNNNNNNNNNNNNNNNNNNNNNNNNNNNNNNNNNNNNNNNNNN
NNNNNNNNNNNNNNNNNNNNNNNNNNNNNNNNNNNNNNNNNNNNNNNNNNNNNNNNNNNNNNNNNNNNNNNNNNN
NNNNNNNNNNNNNNNNNNNNNNNNNNNNNNNNNNNNNNNNNNNNNNNNNNNNNNNNNNNNNNNNNNNNNNNNNNN
NNNNNNNNNNNNNNNNNNtCcTCtCTCtCtCTCTCtCTCTCtCTCTCtCTCTCTCTCTCtCTCTCTCTcTCTCtC
TCTCTCTCTCTCTCTCTCTCTCTCTCTCTCTCTCtCTCTCTCTCtctCTCcTCTCTCCTCTCTCTCcTCTCTCTC
tCacTCTCTCTCcTCTCTCcTCTCTCTCTCTCTCTCTCCcTCCcTCCCTCTCTCTcTTCCCCGCCTCCGAGGAAT
CCAATGGTCTTAAAGTTGCCGCGATATTATGCACGTCAGAGATATCTCCtGTTATCGACGGCCAAACCTAGGTCT
TTGTGAGGAGTTCATGAGGGTACGGCATCCACCATGTTTtCCCCTCTTTTAATATCTGAACGTTGCAAATGGAGA
GTTAGCACTAATGACGTTCACGCAGAAAaTTTTTAAAAAATATTTACATAGATGTCCCTTCTATCCTCTGCGATG
GGTGCTGTAATCTTTGACTTCTTCAGGTATCTGCGATGACCATCATCGCCCTGTTCCGCTCTGGAGTACATTTTA
AATTCTCGTCTTTAGCACTGATGTTACAATGAAATACAGAATGAATTAGATATTCGTATTATTTATATTTAAATG
AAGTGTATCATTGATAGAACTTTTAGCATGATCATTTCATTCTTCATCGGCGGAACTCATTCTCTTGACAAAATA
TGCCTCTAAAACTCAACATTTTTGGGAATTGCCATGGAGGGTTTGGTGAGTATTTAAGTATACCTACTTTTAGCT
TGCTTCATACATTATTTTATCGTGTCTCAAGCTAATTCCACAGCAAGGGTCACAGATTCACATCTACAGTAATGT
GAAAGCTTTTTCACCTCGCTTGTTTGAACTCGCGTGGAAAATCTTTAGTTGTAGGAATTTTTGCCCAGCACGCTT
TTCTAATTCCATTAAAGATGAAAATCTTTGTATTTTACCTTCCACCTTTATATTTTTTTtCATTGACGATTCCAC
CAAGCATATAACTGGCATCAATTTGAGAAGGTTGCTCGCTTAGAAATTTAAGATAAGCGCATTCTAAAACAGCAC
CCAATTAGAGCTATGGCCTCTCTCTTCTCAATAGTAGTATTTGATTAAAATTACACGATACATAAAATCTAGAGC
CAATGTGCACTCGGTCATACACAGAATTAGTCGCGTCAATAACCATTTGTTTTtCTTTCTCTAACACTTTTTtAG
GTCGTTTCAATCAGCCGGTTGCTGCTCTTCTACCGCTTTATCGCGCTACCAATTGATCCACACTTCTGCATAGGG
GCTGCATACTGTGGTTTAGCTTTCCCACCGATTGCACACTGACGAAGTTAGCCAAGCTTCAGGAGCATCAAGTTG
ACCATTACTCCTGAACAAAAAATTCTAGTTAACGCGGGATAAATGTGGCAAAGTATGATAATACAGCGATTTATA
GTCGAAATGCTGCTATTAGTACCATTAGTTAATGGTACCTCTGAATTATTGGTAGCATTAGTAAATTAGTCGAAA
TGATACTATTAGTATCATCAGTTTCTCTTATAATCGCGTGATTTGAGGTAGTCCTCCAAAACCGCGTGTTTCATT
AGGATTAGTTAATGTTGGCCGTTGTTCTGCAAGCTGAATCAATTCGGTCCCGACTCTGTAAGAACCCTAATGGCG
AATATCTTGGCCAATAAGTTGAAGAGTCGGATCAGTAGGCATATTAGTATTTGATGTAGGGGTTTTCTCGTCCCC
ACATGCTATGACAAGATACGCTATTCTCAGTTTAACTGTGTACTCAGTCACTAGTCGGATGGTCCAAAGAAAAAA
GGCGATGGCCTAGAGAAGGTGAAAAGAGAAAGAATAGCTTATAAATGTCTTATTGTTTGCAATAAGTCTTCCACC
GGGGGTGGTTCGTCGCGAAACGAGAATGGATGAGTTCCCCAACGCCTTTCTTTAAGCTTTACAGGTTCCTAGATG
TTGCATTTACAGGAGCACTGGATAAGTAATGACTGCCCATCACTGCTATCCAGCTTATCGGGGAAAAGAATAACA
TAATGGAGTTTTTCCTTGGATTTTTGACAGTATGAGGGCAGTGATGGAACGATACAACAAGGCCAAAGAAGATCA
TTGCCTGGACCTCAACGCAACTTCGGAGGCGAAGGTACTCTTCTTTGCACCTGTTTCTGCTAGCTTATGTGCACG
TCTTCAAGTTCATCTGTTTATTGATCGATGGTTTCCTTCGACTTTGGATGAGTAGTGCCATTGTAGCATCCGTGG
GCTCAGAAGCGGGAGAGTGCCGAGTCCCTGCATTCCCATAGCTTCTTAAAGAACCTATTGGAAGAACGATCATTC
CATCCACTTAAAAAAaCAGAAAATCAAGGTATTAGTCATAATATAGAAACTGAAACGTATAAAAGCATGAGCTCA
GGGATCTCCTAACAGAGGAAACGCTGAAAAAAAAAAAAaaCTGTTTTAAACTACAAGAAAGGGGGAAATTTTAGAA
GAGTTGCCGTCTCATCTAACTCTCATGAATGCGAAATGACGTCTGATTTTTCTGTCATTATTTTCTTTATCAGCG
TAAGCATTTGCATAGAAGATATATTTTCTAAATATAATTGATGTTGCCTCTTTTCTATTCTGATCTTCATGTAAT
CAAGGGCATATCTCAGTGGAATGCATGTAACTATTGAGAAAAGGTTAGAATCCTCAGCAATTATCTATTTCGCCA
TACCGGTACCTCTGAGCTGGTTCCTCAATAGCATGAGTCATCAATTAAAAAGAGTGAAAGCAAACCTTGCATGGC
TGACCTACGTCACAGAGGAAAAGAGAGACAAATCTATGACTCAGATTTGTAGATGCTACATCGGCACGAAAATAG
CTGATGCCGTTTCAACTTATCAAACTCTATCAATGCTTTTTAGGAGGATTAAGTACTGATATGCCTTTCTCCAGT
CAACGGAGGGGAAACCAATGGCAATTCCATCTGACTAACCCTCTTCACCGATCAAAATCAGATTTTCTGACTAGG
ATTAGCAAGCAGTATATTTGAGATATATCATCCATCGCAGTGTTACTTTGCAGAGGGTAAACGGCGATCCAAGCT
AACATATTATATCTTGTGACGCAAGAACGCAGGCTCTTGATTTTTtATATGCATTTGCCAAAGCATAGAATGTTC
ACTTGGGTAATATAAATATTAACGCTGTGTGCTTACAGTTCTGGCAAAAGGAGGCAGGAAGCTTAAGGCAGGAAC
TCCTAGCTCTGCAAGACAGCCATAGGTATCATCTTTCTCTCCGTTTATTACACTGCATTCGGAACCTCATGCCAA
GAGAATCTCGCAAGATGTGACGTGGCATGAAACAAGGTAGAATCACACGTGGAAAAGATCATCCATCCATCTCAT
CCTAAGCAGTATGGAAGTAAAATTAGGGCATCACCGGACGAACAATGTAATAAGGCCGATCAAATACCACAGATT
TTAAAGTATACATTTTCCAAGGCGCTGTAACAAGGTTCATCCGGTTGTCAAGTATCTCTTGAACTTCAGCAAGTA
GGTTAAAATAGACGTGCTACGGGTCGTGGTATCAGTAAGGCTGGTCCTTGCTTCGTGGGGGAACCCGTCTGTATG
GAAAAATCAAGCCCACTAGCACTGCTAGTTGCGATAGTTTGCCCTCTCTAACATATCAATCTATCTCGTTGATAA
TGTCAGGCAACTCAGGGGAGAAGAGCTTTCTGGCCTAAACATCAAAGATCTGCAAAACCTAGAGAACCAACTAGA
AATGAGCTTGCGAACAATTCGGACGAAGAAGGTGATAAGATGGGCAGGTCTATAGTGTTCTATGGCCCTCATGAC
AATGATTTCAAACTCTCTATTCACCTCTGGTTTTTTtACAGGAACAACTTTTGACCGATGAAATCCAAGAGCTCA
ATCGAAAGGTATTTTCCGAAAGAAACACTCGAGATCTACGTATTATTGCTTATGAAATGCTATTGATGGGTCATC
CGAAACTATCCTGGTCGTGTCTCAGGGAAATCTTATTCACCTGGAAAACACGGAACTGTACAAGCAGTTAAACCT

CATTCGTCAAGAAAACATGGAATTGTGTAAGAAGGTAAGTTTAGCACCCAATCTCAGATGCCTCCAGTGAAATTA
TAAAATCTATGATTTCCACTGTGATGTCCAATGAATTATTCATGAAGAAGTCTTCTCAGGTTTATGCATCAAGGG
AATCGAAGGGCGCCGAGGGATGCCCATCCTTTCAAAATAGTTTTAGCCTGACAACGGGACCGCATGTTCCCGTCG
ACCTTGAGCTAAGCCAGCCTCAGCACCAGACCTGTGATATGCGACAGGAACCTCCAAGATTGGGGTAATTTTCCC
TCACCAAGTGCTGAGAAGTATTTCGGGAGTATACAGCCTGCATAAATTTCTAATTCATAATCTATGATTCCTCAC
AGGCTAAAACTTAACTACGACGATGATCCGGCAAGTCATCAGTGGACCTCTCGGCGGTGATATCCCTTCTCTTTC
ATGAACCTGGCATCAACAATGGACAAGGTGTCATTGCATGACGTCAAATTATCAGAAAGCGGAGCCGGAGGAGAA
GAGCCACAAAGGTCGACGGTAATTTTTttTTTTtGGTTGTTACAAACCATGTAAAGCTAGTTATCTCATGGAATA
AGAAGGGTGCAGACATTGCCTCACATTTTCTTACTCATGTAATCCGATGCGTCATTGCCGATTAAAGTTTGGATC
CCCATTTTTTCAAACATATGTGAGGATACAAACGAAAGGAAGATAACAACCATCATCAGATATAAACCCCAGCAG
GCCTGACAATCGCTCAACAGGATCCTAAAGGTGCATGCATGTATCGTGTCATAGAATTCAGAGGAACGAGGTACA
TCATATTTCAACGAAAAATCTGAAATAAGAATCGAGCTTGTCTAAAAGGAGTCAGCTTTTACCGTCACACGGTGA
GGAAGTGCCACGCATAGAATTCACCCATTTAAAGGCGCACAAAAGACACGAAGTGCAGGGAACATTTTCTCTGAA
CCAGGAGAAACTCATGAACATTGGCGTTGCTGATCATTTTTCCTCTCAGAATTATATGACGGAGGGTCCTAAAGC
AAAGACAATGGCCGTCCAGAAGGATTCGTTAAAAAaTAAAAACCTTCCCATGAAGGGGCATCAAGGGCTACAATA
AAGTTAACTATCTATCCCAATATTGCCTTTCAAATCATGTATGGGATGACATCAAAGATTAATGAGGTCGCTAAG
TTTTTTTTCTCCCACCTGCAACAGCGTGATGTACCGATGCTAACTAAGCTTTCGGAATGAATCCGAAGATAATCG
CGATTAAGATCCAGATAATTTTAATCCATCACGCCTTCAAACTATTACCCTGGTGACATGAGCCGGATTTTAGCG
GTGCTAATTATCTCCAGGAAATCTAACCAGACATCAACCTGACCTTATCGTCTACATCCTTTGGATCCCGGAATG
ACGGCTGCCTTCATCTCGTCTTCGTCCCATGTGGGGTTATGTCGTTGTTTGAGTGTGTGGCCCACTCTATGGCAC
ATGCGAACCAAAAATTAGGGTAAAGGCATCTCGCATCCATCGTCGTGGCCGTTCATACCGAGTGCTGGAATTACG
GCAAGAGTCACGGTGGCATGTCAAAGACGAATGCCATCCCGTAATCGTTAAATAATAATCAGCGCCGCAGAACGA
TCCAAAGGAGGCCGGGGACGAGCTCCGATTATTATGGAGAGGTCCTTGAAGAGCTGGGGATGAGATCGTTGGCTT
CACTTATTTTAGGGT

>SpoAGL17
ATGGGGAGGGGGAAGATTGTGATCCGCAGGATCGACAACTCGACCAGCCGGCAGGTGACCTTCTCCAAGCGCCGC
AACGGGCTTCTGAAGAAGGCTAAGGAGCTTGCCATTCTCTGCGACGCTGAGGTTGGCGTCATGATCTTCTCCAGC
ACCGGACGCCTCTACGACTTCTCTAACACAAGTATGAGGGCAGTGATGGAACGATACAACAAGGCCAAAGAAGAT
CATTGCCTGGACCTCAACGCAACTTCGGAGGCGAAGTTCTGGCAAAAGGAGGCAGGAAGCTTAAGGCAGGAACTC
CTAGCTCTGCAAGACAGCCATAGGCAACTCAGGGGAGAAGAGCTTTCTGGCCTAAACATCAAAGATCTGCAAAAC
CTAGAGAACCAACTAGAAATGAGCTTGCGAACAATTCGGACGAAGAAGGAACAACTTTTGACCGATGAAATCCAA
GAGCTCAATCGAAAGGGAAATCTTATTCACCTGGAAAACACGGAACTGTACAAGCAGTTAAACCTCATTCGTCAA
GAAAACATGGAATTGTGTAAGAAGAAGTCTTCTCAGGTTTATGCATCAAGGGAATCGAAGGGCGCCGAGGGATGC
CCATCCTTTCAAAATAGTTTTAGCCTGACAACGGGACCGCATGTTCCCGTCGACCTTGAGCTAAGCCAGCCTCAG
CACCAGACCTGTGATATGCGACAGGAACCTCCAAGATTGGGGTAA

#### Zostera marina

##### *ZmaAGL17*

>scaffold_203
ATCAATAATATAAAATCAAAAAAAGGAATGAAAATGATGTGAAGAGACATCAATAGATCCTTAAAACTCCA
AATATAAACAAACATATTCATTCCCCTCTTCTACAAACCCATGTCATAAAATACCTTATTGGAGTCGTCTC
CCCCCTCCCCTCTTCTTCTGTTCTCTTCTCTTCGTTCCTTCCCCTTCCCCTCCATTCTTTTTCTTCTTCTT
CTTCGTGGTCTTTCTGCTACGCGTTCTTCTCATGATCTAATAAATCTGAATAAAGGAATAAGGAGAGAAAG
GAAAGATCTCCAGAAATTTTCTTCTCTGGTAGAACAAGTAAAGACTTAAGACACGATCTTACCTAGCCTTT
TTTGTTTGTTTTTCTTCTTGGTTTCTATTGTAGCTATGTGCAACAGATCGATGTTCTTGGTGGTTATAACG
TTGTTACTATATATATATTGCCAGTGGCTCTTATCCAAGATGCAGTTCCATCCGTTTTGTTATATCTTTTT
TCCCCTCCTGCTTATAAGTAGATCTGACTGATGTATCAACCTTTTGTTCTTTCTATCAGGGATGGGGAGAG
GTAAGATTATTATCCGAAGAATCGATAACTCAACAAGCAGACAGGTGACTTTTTCGAAGAGGAGGAATGGG
TTACTTAAGAAGGCCAAGGAACTTTCCATACTTTGCGATGCAGAAGTCGGTCTCATGGTTTTCTCAGGCAC
TGGAAGACTATATGACTTTGCAAGCACCAGGTATATACATACATATAATTTCTTCTCCTTCTTCTTTAGGG
TTTTACTTAATCTCGTCTTTTGAGAAGAAGAATAAGAAGAACAGATCGGTAGTTTTATATAATATTGTGCC
GAGATGATATTTTTCCGGCAACAGATGAGTAGTTTTATATAAATCGACAGACAAACTGAAACTACGTATAC
CATGCAACCAATCTCCATCTCCCTAGTTTCGGGTAACAACGATAGATTAATTGTAGAATGGTCGGAAATTC
TTTTTGCTAGCTAGCTAGCAACTTCTTTGTCCCCCAGTAATAGAGCTTATTATTCATGCATGTGATTTGAC
TGTAGTTATTTACTTCTCTTTTCAATATAAATGAAGATCTATCACTTATCGTCTATCTAGGAGCAAAATAC
AATAAAATATATATGTATTGGATTAACCACTCAAGTCATTTCTAATATCTGCAGCATGAAGTCTGTCATAG
ATCGCTACAACAAGGCAAAAGAGGATCATAATAATTTGACGAATGCAACTTCAGAAGATATGGTGATTAAT
ACCTATCCATGGTGACAAATGTTAATAATCACCAGTTTAATTTTAGTCAATATTTATTTAATTAATTGTTG
CAGTTCTGGCAAAGAGAAGCTGCTTGTATGAGGCAGAAATTGCAAATGCTGAGAGAAAATCAAAGGTGATT

TACATAATCATATAACTGTAATCTGTACATATAAGTTAACTAACAGTACATTTGCCGGCCGGCGGCCATGA
TGACAGACATTTGATGGGAGAAGAGCTATCTGGCTTGAGGCTGAAAGACTTACAGAATCTAGAAAATCAGC
TCGAGATGAGTTTACGTGGTATTCGTACCAAGAAGGTACTCTTTGAGTTTGATCAAGTTGTAAGTTGATGA
CAATTCACAAGTGCATGCATGAGTTGTTTAATTTCTTGCTTTCTTCCAGGATCAAATATTAACTAACGAGA
TCCAAGAACTCAGCCAAAAGGTACTTCCTGTTATAATTCACCTGTCATATATTACTGAATTCTTAATTTTA
CAGGGAAACCTTATTCACCAAGAGAATATAGAGCTGTACAAGGAGGTGCAGGTTTATAATCAGCATAAAGG
AGGAGGATCCAACGATCGGGCCGCAGATAAGTTTCCACAGGTTTGTCCTCACGGTTTATCTTCCAGAGATG
AAATGCGTGTTCCATTTCTTCATCTTGAACTATGTCAACCTCAACAACAGCAAAACTATGAAATGGCACCA
TCTGATCCGATATTAGGGTAATAATAACCAAAAGATTTTATAATTAATTTCTTCTTCATCCATCAATATAA
ATTACGAATTTCTTCTTAATTTGCAGATTACAACTAAATTGATCGATGGTCGGAATAAGAAGAAGAACGAT
CGACGCAATATCTATCGGAATATCTATCGGAATGATATATGGATTATATATTGGATCTGTGTAATTAATTA
ATTATATTGCTAAGAGGAATATCGGCCGTTCTTCAATTCCCTTTTTTTTCCTTAGTCAAAATTAAAGAATC
AATAAGAAGGGTGCGGGTGATCCAACACTGAGAGTTATTGTAACTACTGCTTGAGTAAATATATATGACTA
TCGATCTATCTATTTATCATTTATAACTACGTAACTACATATCTTTTAATTTATGTCCATTAATAACCATT
GACAATACCTGTACCATGGTAAAAAAATTCCATTACCTCTCTTATAAGAAATTAAATAGTCATTTTTTGGA
TCATATTCCATCAAAAATCCCTCATCGAACACCATGGTCCACGCTTCGTTTTGGTTCACCAGGATCCACCC
CGTTTTATGACCCAAACGTACGGGGTGGTGAAATTAATACTCTCCTCGAACACCTACTACTCATTACTAGC
TTTATGAGTTATAAATTAAACAATACAATCCATTTGACACAGAAAGATTAGCTTAATTGGTTAAAATAACT

>ZmaAGL17_Zosma203g00090.1
ATGGGGAGAGGTAAGATTATTATCCGAAGAATCGATAACTCAACAAGCAGACAGGTGACTTTTTCGAAGAG
GAGGAATGGGTTACTTAAGAAGGCCAAGGAACTTTCCATACTTTGCGATGCAGAAGTCGGTCTCATGGTTT
TCTCAGGCACTGGAAGACTATATGACTTTGCAAGCACCAGCATGAAGTCTGTCATAGATCGCTACAACAAG
GCAAAAGAGGATCATAATAATTTGACGAATGCAACTTCAGAAGATATGTTCTGGCAAAGAGAAGCTGCTTG
TATGAGGCAGAAATTGCAAATGCTGAGAGAAAATCAAAGACATTTGATGGGAGAAGAGCTATCTGGCTTGA
GGCTGAAAGACTTACAGAATCTAGAAAATCAGCTCGAGATGAGTTTACGTGGTATTCGTACCAAGAAGGAT
CAAATATTAACTAACGAGATCCAAGAACTCAGCCAAAAGGGAAACCTTATTCACCAAGAGAATATAGAGCT
GTACAAGGAGGTGCAGGTTTATAATCAGCATAAAGGAGGAGGATCCAACGATCGGGCCGCAGATAAGTTTC
CACAGGTTTGTCCTCACGGTTTATCTTCCAGAGATGAAATGCGTGTTCCATTTCTTCATCTTGAACTATGT
CAACCTCAACAACAGCAAAACTATGAAATGGCACCATCTGATCCGATATTAGGATTACAACTAAATTGA

#### Zostera muelleri

##### *ZmuAGL17*

>Zmu_v1_s8905_15862
TCCTCCTCCCCTCCCATCCCATCACCTCCTTCCTTCTTCCTCTTGCTCTTTCTGCTATGTTCTTCCCATGA
TCTAATAAATCTGAATAAAGGAATAAGGAGATCGAAAGGAAAGGTCTCCAGAAAATTAATTTCTTCTCTTG
TAGAAGTAAGGACTTAAGACACGACCCTAGCTACCCTTTTTTTCTCTTTTTCTTGGTTTGTATTGTAGCTA
TGCGCCACAGATCGATGTTCTTGATTCTTGCTGATGATAAACGTTGTTACTATTGCCTATTGCTGTTCTTC
AAGATGCAGATCGATCCCTTTTGTTATTATATNNNNNNNNNNNNNNNNNNNNNNNNNNNNNNNNNNNNNNN
NNNNNNNNNNNNNNNNNNNNNNNNNNNNNNNNNNNNNNNNNNNNNNNNNNNNNNNNNNNNNNNCCTTTTC
CTTTCTCCTCCTCCTCCTCCTCCTCCTCCTCCTTCTTCTTTTTTGCCTTCTTTTAGGGATGGGGAGAGGGA
AGATTGTAATCCGAAGAATCGATAACTCAACAAGCAGACAGGTGACTTTTTCTAAGAGGAGGAATGGGCTA
CTGAAGAAGGCCAGGGAGCTAGCTATACTTTGCGATGCAGAAGTTGGTCTCATGATTTTCTCAAGCACTGG
AAGACTCTATGATTTCGCAAGCACCAGGTATACATATTTTCTTCTTCTTCTTCTTCTTTAGGGTTTTAATT
AATCTCGTCTTTTGAGAAGAAGAAGAATAATAAGAAAACTACTCTCAGTCTCGATTATCATCTCCTGTAAG
ATGGTGCCAAGAAGTTATTTTTTGCTTTTTTTATATATACATACGAATCGACAAATATATTGAAACTACGT
ACCATGCAACCAATCTCTATCAGAATGATCGGAAATTCCTTTTACTAGCCTGCTACCTAGCTATTCCTTTG
TCCCCCAACTTATAGCTTTGTTTATACCCGAAATAGTTATCACTTTATTCACTTGTCGACAAACTTCTCTC
TCTCTATCTATATATCTACCTAAATATAAATATATATAAATACTTTTTTGGGAGATGAGATGGATCAGATA
TGAAGCAGGTGAATTCTGATGCTCCTATTGATGAATGATTTACTTTGTCCGCCGTGTGCAGCATGAAGTCT
GTTATAGATCGCTACAACAAGGCAAAAGAAGAGCAGAATAATGTGACGAATGCAACTTCAGAAGACAAGGT
GATTAATAATCATGCATGTTGAAAGTTAATATATTGGCAGGCAGGGTAGGTAGGTGACCAATAATGACTGG
TTTTGATTCAGTATTTATCTGTTGCAGTTCTGGCAAAGAGAAGCTGCTTGTTTGAGGCAGCAATTGCATAC
TCTGAGAGAAAATCACAGGTGATGTACATATGCATTCAGTTGAAAGTCTGTGAATTGATAACTGTAATCTA
AGCTGACAGACCTGTCCGGCCATGAGCCCATCATGATCTCGCAGACACTTGATGGGAGAAGAGCTATCTGG
CTTGAGGGTTAACGACTTACAGAATCTAGAAAATCAACTCGAGATGAGTCTACGTGGTGTTCGTACCAGGA
AGGTACTGTTACTGTTACTGTTACTGTCTATCACTGTCACTGTCACTGTCACTGTCATTCTTCCTAAACTA
CTACTATGTTCGATCTTAATTTCGTCTTTCTTCATCCAGGATGAAATATTAACTAATGAGATTCAAGAGCT
CAACCGGAAGGTACATTCCGATTCAACGTCTAAGTTAACATATCACTGCTTCTTCACAGCCCATTTACAGT

GCTACTGAATTGGATCTATTATTTCAGGGAAACATTATCCACCAGGAGAATCTAGAACTGTACAAGAAGGT
ACTTACTTAAATTAATCATACCAAAACCCTAAGGATTATATATCACCATGGATGAAATTTTTGATCAACTC
TTATTTGTTTAGGTTTATAACAATAGGGGAGGATCCAACGACCAGGCAGCAGCAGACAAGTTTCCACAGGT
CTCGCATGGTATTTCTACCAGAGATGACATGCGTGTTCCATTCATTCATCTTGAGCTAAGCCAACCCCAAC
ATCAACAAACCTATGAAATGTCACTATCTGATCCAAAGTTAGGGTATTACTGATCCAATATTATATAAATT
ACTATACGGGCGGGAACCTTTATCCTCATCTGTGCTGAACATCCCCATTTGATTTGCAGATTGCAACTAAA
TTGATCGATCGATGGTCGGAGTAAAAAGAAAAGCAACAACGACGACGAAGACGTAATATCATTCGGAATGA
TATATGGATTATGGAGTACGTACGTATGAATATGATGATATCTGTGTACTTATATTGCTAAGAAGAACACC
TGCCGTACTTCAATTCCTTTTTTTTTCCTTAGTCAAAATTAAAGAGTCAATAAGCGTGCCGGTGATCCAAG
ACTGCGAGTTATTGTAACGACTGCTTGAGTAAATATGAATATCTATCTGTATTCTATCTGTCAATCTATCA
TTAATACGTACGTAACCATACTTTTATGTCTATTAACGACCATTTACTTACCGGCGCCGGTGAAACCATTT
TTTGTTTTCTTGTCACATTTTGCATCACATTAAGGAAGAATCGATTATGATTTGAGAACTGTTGGGGAAGA
ACATCAAAAGTCATTAATCCGTCATTAAACACCTTGGGTGAAAAAACAAAACCATGTATCTCTCGTCAATC
AGAAAATTACAAATATTTGA

>Zmu_AGL17_1

ATGGGGAGAGGGAAGATTGTAATCCGAAGAATCGATAACTCAACAAGCAGACAGGTGACTTTTTCTAAGAGGAGG
AATGGGCTACTGAAGAAGGCCAGGGAGCTAGCTATACTTTGCGATGCAGAAGTTGGTCTCATGATTTTCTCAAGC
ACTGGAAGACTCTATGATTTCGCAAGCACCAGCATGAAGTCTGTTATAGATCGCTACAACAAGGCAAAAGAAGAG
CAGAATAATGTGACGAATGCAACTTCAGAAGACAAGTTCTGGCAAAGAGAAGCTGCTTGTTTGAGGCAGCAATTG
CATACTCTGAGAGAAAATCACAGACACTTGATGGGAGAAGAGCTATCTGGCTTGAGGGTTAACGACTTACAGAAT
CTAGAAAATCAACTCGAGATGAGTCTACGTGGTGTTCGTACCAGGAAGGATGAAATATTAACTAATGAGATTCAA
GAGCTCAACCGGAAGGGAAACATTATCCACCAGGAGAATCTAGAACTGTACAAGAAGGTTTATAACAATAGGGGA
GGATCCAACGACCAGGCAGCAGCAGACAAGTTTCCACAGGTCTCGCATGGTATTTCTACCAGAGATGACATGCGT
GTTCCATTCATTCATCTTGAGCTAAGCCAACCCCAACATCAACAAACCTATGAAATGTCACTATCTGATCCAAAG
TTAGGGTATTACTGA

#### Triglochin maritima

##### *TmaAGL17 and TmAGL17asRNA1-1*

>TmAGL17 [organism=Triglochin maritima] AGL17-like MADS-box gene, partial genomic DNA, genome walking (start exon3)

ATCTGATCGTCTTGTAAATTTGATGGAGTTCTTCGTGGAACTCAAAAACTCCGTGCACCTACAGACAATTAAATC
GTTAAACAAACACTAAACACAATTTACAAACATTGTAATTATTTAATTAAATGTTTAGATTTTATCATCTCACGT
TCTCTACCCGTTCGGGGTCTGTACACGTTACGTTTCGGAGCCAAAGTGTGGTGAACAATCCAACACACCGTGATG
TTACAGTATCATTCTATATATAATTGTGTGTTCTCTTCTAGAGTGACACCTAAGAGTGGTCGCTCAAACTGTAGA
TGATCTCTTCTATGAGACCAATGCTGAATGAATTTGTGATTTTCCACCGACCAATTAACGCATGCTCTCCCTAAG
CTGATTTCACTTCATGTCTCTCCATCGGCTCTGGAATGTATTGTTGTAGACTGAACTGCCTCATGACTCGGTCTG
GACAATGCATAACCATCTCTTCAAAATGTATAAGTTATGTACAACTCCTCCAAATCTCAGAACCTTCTGTACATA
GTGGTGGTAACTGACTCAGATCCAAACGCTCATACGGATCCCATGTGACCTGGAATACCTCCTACACATCGACCT
TATTCGGATAAAAATCAAGTGTTGATGTCTTACCAGTCCCTTGAATGTTCTTGGGCATCTCCAACGAGAACCTAA
TGGTGCTCTATCGTCCTCCGGGTACAGTAACAGTCTGCTCTTCAAAGGCCTCACAACGTGGAAATGTTCCCAGCC
CACAACTGTAGAAACATAAGAAATCCGCACAAATGTTTGGTCGTGATCAATGCAGCTTCGTCCAAACTGTGATAT
AAGTGAGCCAGAGCCGCACTACCCCAACTATACTCCGATGCAACAACCAAATCTTTGAGCAAAAGAATGTACTTC
AAGTGAACCATTGTGCCGCTCTTATCAGCCAAAATATTGCATCCCAGCTGGAACAGAATAAAAGCTCTAGTGTGT
CTGGTAATCGTAACCATGTTAGCATCGAAAGGAACCTCCTGAAACTGCATTTGCAACCAACGCAGAGTTATCGAA
CCCTTACCGCCCAAACCATCCGGACACTGGGCCAATAAATCCATGCAGACATCTCACAAATCGTGATTGTGTCGA
AGCCTTCCAGTTCCAGCAGAAACAACTTCCTCTCATCAGTTATCTGCCAAGCTTTGAGTTTGGATCCATGCTCAA
CACATTTTACTCCCTCGACATCTTTCTGAATAAAATAACATAAATTTTAGAATATTAATATGTACGTAAAAACAT
TGTATGAAAGTAGTAAAATACTTACGAAATCCTTGAACAAGTCAGATTTGTGGTCGACCTGCAAAACCAACACTG
ATCTGTCCACCGGACCTTGCGAAGCGTAGCTGCCTCTAATTGAATCAACACTAGAACCATCAAACTCCATCTGTA
CGTAACATGTTAAATATTAGAAAACGTTGAAGTGTTGCAAATTAAATGGTGCAAATCAAATATTAGAAAACAAAT
CAAGTGTTGCAAATCAAGTATTAGAAAATAAATCAAGTGTTGCAAACAAAATATTAGAAACCAAATTAAGTGTTG
CAAATCAATTAGGAGAGGATTTGGAACAATAACAAATCCGAGTCATAACAAAATATTTTACATTAATTCAAGTCT
TACACCACATCAATTCAACTCTTACATTTAATCAAGTCTTTATTTTCAAGTTTTACATTAATTCAAGTCTTACAT
TTATCCGAAATCCTACATAATAGTAACATAGACATTTTTAAACAATTTTACTATATAGATTCCTTGCATCATTTT
CTATTATGCCCCAAACCGCCACATCTCCCACACCGTACCCTTCTTCCAGAGTGCTCGTCCATCTCGTTTCGGATT
CGCGTTGATTTTGGCCGACTAGGAGGTCTTTTGGTTTGCGGATGCTTAATTTTGATGTTGCCGGAACTTCGGTTG
TAGGATTTGGACTTATATTCAAATATATTACGCAGTCCCAACCGGTAGACCTTACTGTACCATGCTCCGGCGAAT
GAGTGTTGGTTGTCAGCGTAGGGTCAGACACTTTATTTTCAGTTTCCACATACACTCCAACCACAGTCATCCGTT
TAACACAGACATCCGTTTAAACAAACACATCATTCCATAAATCGACATTAAATCTCACATACATTCGTTACTCTA

TTTTTTCGATTACAAAAATCAAACTTAAATTCCTACATACATTCAAAGTCACATACATTCAAACTCAAATGAATC
ATTTTTAAGTTTACATTCATTAAAACTAACATACATTCAAGCCAAAATAACAACTCTAATTTCAATCACAATAAT
CAAACTTAAATTTAATTATTTTTAAGATGACTACCGTTAACCTATAATCAACAAAAATGAAATCAATAATCAACT
AAGAAACAAGAAATAAATAATTTGACCTAAAAATCCAAAATCGAAATCGAGTTTGCACCAAAACGGGTTCTTAAA
CCAACAAAAAATACTACAATCATAGCTAAAACAACAATGTATATCATCATATACATAAAAAGACAAGAAAACGAT
GGAAACGAAAAAATGAAACCCTAACCTCCAATTCGCAAGAACTCAACGTGAGGGAGAAGGTGGCGTCATGCTGTT
TTTATAGCCCGACAAAACACGGTGTTAAGAACGCCGAGTTCACTTTTTTAAGTGAACTCGACGTTCTAGATGCCG
AATTTGCCTTTAGAACTCGACATCAAGGATGACAAGTTCTATTTAAGACATTGTTCTCCTGACGCACTAACCATG
TCATGCACGATCTGAGTTGTCAGGAGAAAAGTTTTAATAGAACTTGACACTTGTGATGTCAAGTTCTATTTGAAA
TGTTCTCCTGAGAACTCACTTTAAGAGAAATATGACAAGTGTTGTGCAATAAAACTAAAATGAACTTGACACTGG
TGATGTCAAGTTCTGCTTTAGAGAACTCAGCATCACAAGTGTAGAATCTCTCGAGCAGAATTCAATACTGATAAT
GTCGAATTCAGTATACACTAAAAAACAAACTCGCCGTCACGTGTCATCACATGGTGACAACAAGTTTAGTGTGAA
AAATATTTTTGAAAGAAATTTATTTTGACATTTTTTTATGGTGACAACAAGTTTAGTGTGAAAAAAAATTTTGAA
AAAAATTTATTTTTACAATTTTAAAAAAAAATATTTTGAAAAAAAATCCAAAAAGAAACTAGAGGCATATCCACT
TAGGATCTCGGAAACAAACGGTGGAATATTCCAGAATACGATGTTAAGATTTTTCTGATAATCAAATACGATGGC
AAGATTGGATGGGAAGAGGAATTAATAAGTAATGCATTCTCGCTACAAAACATCTGCAGCGGCAACAGACCGAGC
TGTTTCCACTTTCCACAATGCGCTACGAACTTCATTTACAAGAAAATCTTATTACAAGTTTAAGCACAAGGAACG
CCGGGACTTTGCCAGGTTTAAGCAAACAATGTAATATGAAAATTCATATATTGAGGTACTGAAATATGGCAGTGT
GGTATTAATTAAGATGGCACACATT**AGCGAACAACACCATTCATATTTGATATTATCAACTGGTTGACGCACATT**
**ATTTGTCAGAATGAAATTGAAACATAACTTTCATCGTCAACAACGAATGCCCCACAGGTGGACAAAAACAATGAC**
**GCAAAACTGCACAAAATTTACATATTTCGTTTGTTTTCTACCTTCTGAATCTGTAGCTGAAAATGAAATCAGAAC**
**TTATCTGCATTGTTTACCTACTCTCAGCTTTGGCAACATGAGGCTGCGAGCTTGAGACAACAGCTAAATGACTTG**
**CAAGAAAACCATCGGTATCTTTTCTAACCTCTTTCTGTGCTATCACCACAACAAT**CTGAGTACACAAGCATGATT
AGACATGTACATAAGTAGAAATTATGACAATATCAAATAATCTAATAGTCTCGACGAAAGTCAGATAATGCCAAA
CCGTAC**GTATTCCCTTCAAATTACTTCAGAGAAAAAGACTAGGCGAACAGGCTGTACTGCACCCATAACCATAGG**
**CGGCTCCAAGGTCAAAAACTGCAATAGTTGTTTAATACTTCAATTACAAAACTAACCAAGGTGCAAAAAAATATT**
**TCTGAAACAAGCAAATTGGTATAGATATAAGTTTATGGCCAGACTTGCATTTAATGACTAAGTTATATCA**TACAT
TTCAGAAAATGTATGCAGTCTATTATAGGATTTCATAACCAGCATGAAGCAAAACTTGGATGTTTTTTTAATTTT
AAAAAAAAGATTATTTTTACCAATATCATAAGTTTCAAATTATTAAGAGTAGACAACACATAAGAGGATCTCTAC
TTGCAACCAGGAAGCTGATGGGGGAAGAGCTCTCTGGCATGAATGTTAAAGACCTACAGAATCTAGAGAACAAAT
TGGAGATGAGCTTACGTGGTGTTCAAATGAAGAAGGTAATACTGCTTGGTCATGTAAAGGATATATACGATTGTA
AATTTTCAGTTATCTATCTATCTATCAAACACTTGAGGAAGTGATGTCTTACGTCCCTCATTTTTGCAGGACCAA
CTTTTGTCTGATGAAATCCAAGAACTTAATCGGAAGGTTTCTCACAAATGCCCCTATCTCCAACACTTAGCAATC
TGAGGGTACATATTCTTAATTTCACTAAATATCTGTTGCAGGGGAGCCTTGTTCACCAAGAAAACCTAGAGCTCT
GTAAGAAGGTAAACCTGATCCGTCAAGAAAACATGGAATTGTACAAGAAGGTATATTGATTGAAAAACAAAGATG
TGGGCCAGAACAAAATGCTCCATATTGACATGAGACCTGCTTATGCTAACAAGGATAAATTGTAAAATGTGAACC
TTAGGTTTATGGCACAAAGGAGCCGAATGGAGTCAACCGAAGTCCAGCTGTTTCTCAAGGCCTTAACATGACAGA
TAATATTCATACTTCTATACACCTCGGGTTAAGTCAGCCTCAGCAGCAAACCTTTGAGATGCTACCAAGCCCAGG
GTTAGGGTAATTTAACTTTTTCTCAACATTTTCTGAAAATGATACTTCAGTGGTCTCTTTTAAATTCTAAACTAT
TCATTGGGATTCAACAGATTACAACTACATTAAGAAGATTGTTCCTGAAGAAGAAACACCCTGGATACCTAACGC
GCAAGGGTGTTGTTTGCCTGTTATGGTACATGTTATCTTTAAAAATTCAGTGTATTAAAAAACATTACGGTTGCA
TATTTGGAAAGGAGTATACAATGTTCTGCTGTATCGATGGAAGTGATTATCCATGTAAAGCTGTGCAAGTAATAC
GAAGGCCTTGAGTAGGCTTGCTTTCCTTAATCCTCTAAATGTGTCTTGCGAAAAGTGTAAATTACGGGATTGCCC
GCTACCGTGCCTTTACACCTCCCACCTTTTAAAGTCTCTGTGTCTCTCTTCACCTTTCAATTAGTTTCGATTTTT
AACCCCTCCCTTAAAATTGTGTTGATTTTCTCTGTAAGAAAATGTGTTCAACATGGCAGACGTTGGAGGTTTCTA
GGGTCCTCTGCTTGCCTTCGGAGCTAGATGGACTTCTACAGTATATACACATTGAGGGTGAGGTCGTATATAGGT
TGAACATTGAGTGGATTAACCATGGATGAGTCAAAAAAGAGGATTCAAATAAAAAAGGTGGAAAGTTGTGAAAGA
TTATGGGATTTTGACAGTCCACCATAAACCAATGGACAAAATTAACAAACTTTGGTGAGAGGGGTGTAAACTAAA
ATAAAATGAAAAGTGATGAGGGCATTGAGATTGTTCAAAGGTGACCTGCAAAGTGAATGGAGGATTGGAGGCAAT
GGGGGGAGTACATTGTAATTTACTCTTGTACAATATGGAAGTATTATGAGTTTCCTTTGATCTACTAAAATTTCC
TAACTATGAATACAAACTACAAACGATCCCAATCATTTAAGAGTTGGACACTCAACTGTGTTTAGTTGAGTATTT
CTTTTGGGATATCTCAACATAAGTTGACTCCACTCATCAACAATTTCCTGCAAATGGTTAGGGTACAGCAAAGAC
TTATTAAAGAAAATAAGGACAGATATTACCTGCACGCTGCATCTCATACAGTGATTGAATGATGACTTTAAATCG
AAATGTGCCCATTATCAAATGAAGAGCCACCTGCATCACCAGATGAACTTCTGCGAAAATGAAATCCAATAACTA
ATTCATAAAGTAAGACATGTAAGAAAACTAAGGGACCAAAAGCATCTTCAGAGAAAATATGTCTCTTATTAATAC
TCTTGACATCGTTGCCAGAAGAAAAAAGAAAGAATATGCAAGGAACAAGTTGTCCGGACCTTATGTTTCAAACTT
TCAAACACAACAACTGAAAGTCTCAAAGTACATCTGAACCACAATTATGTAGCAAGTCCCAAAAGGCATCAAAGT
ACAACCAACAAGCACTTGGATTAGAAGGGGCAAGCTTCACCCTACTCAAACATAAAAAATCATATTCACCATCAG
GCAACAATTTTCGTTTAGAACCTCATCTGCAGGCACAACTTATCTCACATGTCAAACCCTCCCAGGACCAGAACG
TAGGTTCCAATGTGTGTATGTTTTACCACTTCACTAGAACCTAGACATATGTTTTACTGCTTCACTAAAACCTAG
ACATATGTTTTACTGCATCACTAAAACATAGACACATTTTATAACAGCCTACGAATAGCAGTAGCAAAGAAAGAG
GGAGGATGATCGTGATTTGCACTTTTATAATGTAATGTAATAAAATTTTCGTAATTGATTAATCCAAATAACTTC
AAAAGGGAAACTACATAGTCTAATATAATATTGTGATCATCAAAAAAATATCTCCTGACAAATAACAATATCATT

ACCAAGTACACATTCACAAGGCAGTCACTGGTTCTTCACTCTCTTCATGCAAAAAAGAAAAAAAGTTGTAATACA
TATATCTAAGAGAACTTATGCAGGCCTCTGGATGCAAGGACAGTTTATCAAGTGGCTGGGTTCAATCTCCCACAA
GGGAAGAGGGTAGTGGGAAACAAGAGCCTGTCAAACTACAGACCAACCGTTGGGAACAGAAAATATGAGTAATAC
GAACAAGAGTGTATTCTTTCTAATTCAAGGATCACATATCGAGTGAGATCGTCGTGTAGGGCAGTAGGCTATAGG
TTTGATTTTGTTAAATGACAAGGCTTAGGGTTTGGAGCACAGCTCATGGGTTGGATTATTGGAATTATGAGACTG
GTAATGAAATTGACTGGGTGATCTTAGGTTGACTTACGGGTTGTTGGTTTGAGCAATTTGTAGTCATTTACAAGC
AACTGAGGATGAAATCTAGCAACCTAGAAGTTGGAACTCCAGTAATTCAAGGGTATCAACATAAACGAATAGAGT
CAAAGATATTATATATTACACTAGGATCTCAAAAGTAGGGGAAATGTTGAAATCAAATTTCAAGATGTAGTTCTA
CCCTTCAACATGAGATGAACCTCTAGATTACCGATCATTATTATCTATGTAGATTTTCCATAGAAGTGGACCATG
CAATAGCATTTCACGAGGATGTATTCACAGGTGATCCTTCTATATCGGGTGTTCATGATACATTTACTCAAAACT
ATTACTAGTCACATTTGTATATCTGAGCAACATTTTTTTATCATCCTAATTAGAAGAAATTGATGCATAATCTAA
TGCTGGTCAAACAGTTTATGCTAGTAAATACTTTACGGATAAGACTCTAATTTACCCAGGAATGATATTGTTACT
TGTCTTAAACCATTAACTCTCATGACAAATTCTATAATACATGTCTAATATTTTATTTGGAGTACCCTAGCTGAG
CAATTATGATTTATTTTGGATACTTGTGGATATTTTATAGATACTATGAGATACTTCTAGAATTCTTTAATGTGC
CTCTACATAACAATTTAAGACAAAATAGAAATACATAATACATTGCCCATGACCGTTCAAATAATAATAATAATA
GCATTGTTCATGGAATTTGCATCTATAAAGGTCTATACTTGACTTGTGTAATTTTATTGTATTCGAGGGTTGAAA
GGCGTCTAGGCAGTCACTAGGCAGGGGAAGGGCACTTGGGTGCCTAGCTCACCTAGGTGACCTTCTCAATCCTTG
GGGCTTGGGCCAACTAGGCGGCCACCTTTTTAATCACTGATTGTATTACTAATTTTTTCATTTCAATCCATTTCA
ATTCACTTTGTAAATGTTTTCATGTATATGTCATACTAACTTTTTTTATTTTATGATAATATAACATGACAGTCA
TATACTCATATGTGTCCTACTTCTATAAAAAATTAGGAATCCATGTATTTATATCCTATTGCACTTGTATCTCAT
CTACATCCATCTACAACGTTTCACAAATCACTACAGTTAAGAAATGAAGACAATTTATATACCATTTTAACTTAG
TCCATTTCAATCCAAAATGTCAAAAATATAATTTGATTTTTGACTCATTGGAAGTACACATAAGCAATGAGGCTT
TCCAATGCCACATAGATATGTTTGTGTCTTTTGTCAATCATAGTGTACCTAGTGGATGTGAGTCAAGGGTCGAAT
CCCCAATGATTTGGTATTAAAGCCTCAGTTATGTCATCTCGTGTCCAAAATGAAGACTTGAAGAAAGTGTGGGGT
TGAAGAATAAGCGAAGGTATTAACTACATATAGATGTGACATTCGTTGGGCACGAAATTCATTAGAGTTCTATTG
GTTGGGTTGAGCAACAAGAAAGAACTTTCATCTCTTGTCTTGCCCCCATCTACTAAAGTGGACCACAAAAGGGTT
GTGAGGCAAGCGACAATAGGGATTGCATTGTGTATATTATGATGAGATAATTTTGACTATGAGAAAGTGCTCAAC
CAAGAGAATGGATGAAACCTTGAGTTAGTGATCTATTTTTTGAATATTATAAGCATTATTATCTTCTTTGAAGCA
TTTTCCCTCATGTCTCAATGCAATGGTTCTACTGAACATAAAATAAATAATAGAATCGAATGTACATTGTTATGA
GTTCTAGGGCCTTTAGATGTGCATAAAACCTACGGCAAGTAAGGTGTTTTACGTATTCATCCTTAAAAAGTCTCA
ATACTAACCTTTCAATTTACTCTCACCTTCCAATAATGGGAAGGCTTTGCAAGGTGAGAAAGAGCATTGGGTCTT
TTCAAAGGTGAAGCGTGCAAACTGAAATGGAAGCATAGGTGAGAAGGGGGCAAATTATAATCTACTTAAATGTTT
TCTTACTAATATGATCAGTGCGAAAAACAGACAAAATAGGAATGCAAGTTTTTTTTTGATCGGTAAAATAGGAAT
GCAAGTTATGACATGAACCCATTAATCTTTTATTACATTCCTAACTATTCAGTCATAATCTAATAATATATGATT
CCTTTTTAGCATGCCAGTGATGTACTATCTCATTGGTTTGTGATTTAGTGTCGTATTCTCACCACCTTGCGAAGT
TTGAATCCATCACTAAATCGCTAAAGGTGCATTCTTTTTGTCTTTCAATTTAAATTTTAGGAACATATAATTTTA
AAGCTTCATCATATCAACAGAAGAAATCTAACAGATTCCTCTTGCCAGTAACACTTTGTAAGGTAGCATTTTTTT
TAGTTCTCATTTTGTATTAATCCATATACCTAATAGAATATTTATACCCCTCTCCATGTCCAAACCACACTACAA
AGAGAAGATAGCTTAGACATGATACTGGAGAGAGATAATTTTCTCTAAGAATCCACGCTTGAATTGATTTCCTAA
CCCAAGACAATGCAAAAGAAAGGTGAACGGAATTCAAAACTATTACCACGAGTTTTTATGATGGCCCCATATTAG
GTTCATCTTGTAGGAGAATTTTTAGATTTTTGGGACATCACACTTAAGCTGAAACAATCACTCTCTAGATAGGGC
TTTCTATGCACAAAGGTAAATTATTTTGCGATGCATTAAGCATGTTGTTTAGTCCTACTTAAGTTTCCATGGCTA
GAAAACATATACCCATGCTCAAGTCATAATTACTTACAATTTAGTCAATCAGTTAGCGAATCTATTGAATTCTTG
CCATGACAAAAATGTGATCAATTTAAAGGATGATGAATCACTTCTCCTGGAAGAGAACTCAATCCAACTGATTCT
GAAAATTTTAAGTTCTTGAAATGAATAATAATATAGGTTATTGTGAACTTTACCTAACTCATGGTGTCATATATT
GTCGTCAAAAACACCTACGTCGCAATTGCTATATTGGAATAAATAACCTTAAAAAAGAATGAGTGATGGAATAGA
ATAGAGATAGAAACACACTTAAATCATGATTTTTTTTCTTGGGTTTTGTTGATATACACGAGGAAATGTTCTTAT
AAAGGAATAAGATTTTCATTTATTTGTTGGCTATCACCTTTTAACAAGGAAATGTGATCAATTTGTGGTGTGTTT
CCTAACATGCCCTTCAACTATAAGATATTTATGATTTAATTAATAGATAAACAGTGTGATATTCATCAAACGAAT
TATTTACCTTTTTATGAATTAGTCCTTTAAATTCTAATAAGTCAAACAGAAAGTCAAGAATACATTTCGTAAATG
ACTTTTACCTTGGTGCAGTGGCATAAGGTTACAAGTAAAAAAAATAAAGAATCACAATATTATATTGTTTTTTGT
ATTATCTTTTTGATGTTGCAATCCCATGACGTGAAAATCAATCTTGAATAGATAGATTATAAGAATTCGATGGTA
AAACAATCTTCCTTCTAAATAGCAAAATGGTTATGAAAACAGGTGAACTCAATTTTGAATAGGGAAAATTGATTA
AAAATTATTTGACCAATCTCTGCCTTAAAAACATATCTTCAATATGTCAAAATTACAAAAAGTAACAAAAATATT
CATAAACAATAATGTCAAATATGCATTTCAAAACATTGAAGGGTAAACATATATTTTTAAAATTGCATAGCAAGT
AGTTCAAAATATCGAAACAACCAATTACATAGAATGAGGGTAAATATGGAAATTCAAGAACTCCCAAAGCTCAAA
GTTGTAACTGCATGGAAAGAAATTGTGATTCCTAAAAGGAGGTGGGTATGGGATTTCCTCCGTATAAAGGAATTA
ATGACCCTTTCCATCCCTTAAAAGCACGCCAAATCTCAATTCATTCCTCTTATAAATTCTTATCACTTATGTACA
TCACATAACGCCAATGTTCAGGTAGAGCCTAGGTGATTGAGCTTGCCCAATCACTTAGGTGATTGAGCGGTTGCC
CAGGTGTGGGCAACCCTTTTTTTTAAATTAAAAAATAAGGGGAAAATACAAGAAAAATACAAATAAAATATTTAA
ATAAATCTTACATGTTTGGGCATAAAAAAGAAGGTCAAACCACTAACTCATTAGGGATTTATAGGAATGGAAGAG
TAGGGGAGGGGAGGGGAGGGGGTAAAACAAAAAAAAATGAAAATGAATGGCCAAGATTGCATCACAACAAAACAA
CTTACTATATAACCCTCTCTCTCCCTAACCCAGCCACCATCTCTCTTCCAAACCCTAACCAAGCCACCCTCTCTT

CCCCAAACCCTAACCCAGCCACCCTATTCTTCTGATCTTCTCTCCCACTATCTTCTCTCGTTGCTACAGTAGCAG
CCAGCAACGACCATCAACCAGCAGTGCACAACACATGGTCGCCTAGGCGACCATGGGCGGCTGGTCGCCTAGGTG
ACCATGGGCGGCCGCATAAACCGCACACCACTGTGTGCGGTGTGCAGATCGTCGCATATGAAACCACCCGACCGC
CCACCTGAACGTTGCATAACGTTGTTGCATTCCAATGTGTGACCGCACCAATGACACCCAATTGGGTGTGACTAC
GTTATGTGACCCCCAAAATTTGGCCGCACTCCAAAACATAAAAAATGAGGATGTTACAATGGATGTGGTCACATA
TGCGAATGGATCTAAAAAAATGAGGACATTCGAGAAAATGTTGGAGTAGTCCAGATCGAGAAGAAACTTATACAA
CATCATTTAAGATGGTTTGGACATATACAACCGAGACCTATGGATGCGTCAGTGATGATGGGTATGTTTGGCCAT
GAGGAATGTGCGAAAAGATGTACGGAAGACCAAGATTAAGTTGGGAGGAGACGGTCAAAAGAGATTTAAATGAGT
GAAACATCTCCAAAAATTTAACATTGGATAGGAGTGTATGGAAGAAGCCAAGAAGGCGATTCACATGCCTGAACT
GTGATTCTTTTTTAGTTTCATTTTTGGCCTACACCAACTAGTTTGGGATTAAAGGTTTTGTTATTGTTCTTGTTG
TTGTCACGCTAGCAGAATAAAATTGGCTCGTGATAGACTGGTCAAATGTTGATTTGAATTCTAACAATTTGAACT
TACACCTAAATTTCAGTTTAGTAAACCTAAAGAGAAAGCTTAAGCTTAAGTAAGCTCGATTATATTTGATACGTT
TACACCCTTAGGACTTCAAAAAGAAAACAGAAAATGTAATTGTTTTGTGCATTAAAAGTGTTTTTTGTCAAATTC
AATCTATCAAGAGACAAATGACATTTGATACATATAATTTTATATAACTCGCCTTATTGGATTAAAACTTGTTGA
GTTACAACTTCACAAAATTTTGATAGCATTTATTAGTCATTTTAGGACAAAGGAGGTGGGACTTGTGGGGGTTTA
AGCCCTATAGGGTAATAGGGCACAAGCCGCACAACGCATTTGCTAAATAAAAACTAAGTCTGTATAACCAAGACA
ATTGTGTATTATATGGTAATATAGTCTTCATAGTCCGTGCTAGGAACAAATCCAACATGATTAATACTACCGAAT
ATGGATTGGGTTGGATTCACTGGTAACTTTGTTGGCAAAATTCAAAAAGTTGTATATATTAGTTTCACAAAGAAG
TAACAATTTCATGTGAGCTTATAACATAGTGTTCTATCATGATTCCTACAACAAGATAACTGCATTCTTTTTTCC
CAAATTAACTTATTTTATGTGCAAGTATGGTTGTAGGTGGGCAAGTTTGATCCATGGCACCTACCACTAGGGGTG
GCAATGGAT

>TmAGL17 [organism=Triglochin maritima] AGL17-like MADS-box gene, partial CDS, splice variant 1
CGGAGGTGGGTCTTATAATCTTCTCCAGCACTAGTAGACTCTATGATTTCTCAAGTACTAGTATGAAGTCAGTAA
TCGATCGGTACAACAATGCAAAGGAAGAGCAACACCATTCGCTTAATACAGCATCAGAAATCAAGCTTTGGCAAC
ATGAGGCTACGAGCTTGAGACAACAGCTAAATGACTTGCAAGAAAACCATCGAAAGCTGATGGGGGAAGAGCTCT
CTGGCATGAGTGTCAAAGACCTACAGAATCTAGAGAACAAACTGGAGATGAGCTTACGTGGTGTTCAAATGAAGA
AGGACCAACTCTTGTCTAATGAAATCCAAGAACTTAATCGGAAGGGGAGCCTTGTTCACCTAGAAAACCTAGAGC
TCTGTAAGAAGGTAAACCTAATCCGTCAAGAAAACATGGAATTGTACAAGAAGGTTTACGGCACAAAGGAGCCGA
ATGGAGTCGACCGAAGTCCAGCTGTTTCTCAAGGCCTTCACATGATAGATGATATTCATACTTCCATA

>TmAGL17asRNA1-1 [organism=Triglochin maritima] antisense transcript 1 of an AGL17-like MADS-box gene, transcript variant 1
TGATATAACTTAGTCATTAAATGCAAGTCTGGCCATAAACTTATATCTATACCAATTTGCTTGTTTCAGAAATAT
TTTTTTGCACCTTGGTTAGTTTTGTAATTGAAGTATTAAACAACTATTGCAGTTTTTGACCTTGGAGCCGCCTAT
GGTTATGGGTGCAGTACAGCCTGTTCGCCTAGTCTTTTTCTCTGAAGTAATTTGAAGGGAATACATTGTTGTGGT
GATAGCACAGAAAGAGGTTAGAAAAGATACCGATGGTTTTCTTGCAAGTCATTTAGCTGTTGTCTCAAGCTCGCA
GCCTCATGTTGCCAAAGCTGAGAGTAGGTAAACAATGCAGATAAGTTCTGATTTCATTTTCAGCTACAGATTCAG
AAGGTAGAAAACAAACGAAATATGTAAATTTTGTGCAGTTTTGCGTCATTGTTTTTGTCCACCTGTGGGGCATTC
GTTGTTGACGATGAAAGTTATGTTTCAATTTCATTCTGACAAATAATGTGCGTCAAGCAGTTGATAATATCAAAT
ATGAATGGTGTTGTTCGCT

#### Acorales

##### Acorus gramineus

###### *AgAGL17*

>AgAGL17 [organism=Acorus gramineus] AGL17-like MADS-box gene, partial genomic DNA, gene-specific PCR on genomic DNA from leaves (start exon2)
TTCAGAGGTTAAGGTACCTAATGATTCTCCATCTATACCACATGTTATGCTGCATTTTTTTCATGGATATTCTAT
GTACAGAGATTGAATCGTCACTTCTATTATATACTTGGGATATTATAAGCTTTTCTAGAAGACAATTACTGATTA
ACATCTAGATTTGGTAGACCAATATTTAAGATGAACAGGAGCAATAATTAATCATACTGCATAACAACTATAACT
CAAAGAAGTGAGTGAGATTATGATAATTCCTTGTTACTAAGGGGCTGTTTGGCCAGAGGAATAAGGGTGGAATAG
GGGAGGAATAGAGGAGAATAAGGCAAAAAGGTGTTTGGGTAGAGGGAATAAGGGGGTATAGCTATGTGGAATAAA
GGGAGAATAAGCTGGTATTTGTATTCCAAATCCATCCATTGAATTGTGTTATTCCTCCACTGATATAAGTGGACC
CACCTATCAAATCACATATCATCACATCAAATCATAGTTTTGAATATCGATCGCGATCTCGATATATTGGTCGAT
ATATCGAAATCGCAGATGAAAATGCAAAAATCGAGGGATTTATTGAAGTTTTAGCTAAAATCGCTGACGTCAACG
ATATATCGACTATATCGACCTTATATCGTCGATATATCGACAAGCAATGTCCCAAACGCGACAGGCAAGGACCGG
CAGCCACCAAATCACATCCATCCAGCACAAAAAACGAGTAGACCCGGTGACCGGGTCAGAATCGGGAGACGGAGT
CGACGATGCCGGAGTAGTCGGATCCGGAGACGAAAGGCAGCGGCGGCTTCTCCTGGTACTTTCTGAGGATCTGAA
GGTAGTTGGCATCGTTGACGCTCGTCACCCTGACTTGAACTCGGACCAAGGTCGGCGATTGAAGGACCGGGACAG

GGTGGGGAGAGGGAGATCGTCAGGGGCGACGAGGGGTCGGCAATCGGGACCGTCGGATCGCCGATCGAACGGCAG
AGAAGGGCTTCCATGGATGGAGAGATGTAAAAGAGGCAGTGAAAGCGACTCCCCGTTTGTGTGTATATATATATG
TCATTAAATACTCACTTATATATATGTCATTAAATACTCACTTATATTGTTATTCATTTTTATTTTTTTTCACAA
TTTTATATCAGTTTTTCTGTTTTTTTCAGTTTTCTTTTTCAGTTTTTTTTTTAAAAAGAGCTAATCGATATTTCG
ATTGAACCCAAAAGCGATATATCGATATTTTCGATATTTAAAACACTGCATCAAATCACTTGTCATCACCTCTTA
ATCAACTTTTTACACCTTATTCACATCTTATTCTTCATCAACCAAACACTCCAACATTTATTATTACGTTTCCCA
ATAAGAATAACTGAAAACACAACCAAACACAAAATCTCTTATTTCATTGATAAAGTCAACAATTCCTGGAATAGG
CATAAATTCTAGAATAGTTTATGTCTCTATTCAAACGGGTCATAAGATATTCACAATAAAACTACAGGTCAAAAG
CTCCAAAGCAAAAAATAGAAAACTAGGATCAATTTCAGTGAAAGGATTTATATGAGTTATAGTACTCCTTAAAAA
TCCTTTATAGATAAAAATCTGAGGGAGATGACAATTAAGCCTCTTCGTACCCCATCCGATATCCCCATACGGCAT
ACAAATTGGATGGGAAGAACTTGGTGGGAGAGATCCTATGCGGGCCATACTAGCACATTTTCATGAGTCATCCCA
AGCTGTATATGCATGTTTGTTATTAAGAATGTTGGCCTAAGGATCTAGTAGATAGAAAATGATCGGTTGGCCAAA
AAAATCCAGAAATACCTGTGATGCAGAAAGTACCTTGTTAGTTGCTAGTCGGTTGACAGAAATATAGGCAGTGAC
GTGGACAGAGGATAATCTGATTGGCATTTTTATAAAAGATTCTATGTTCTGGTGATTTAGCTATATAAAGGAAAT
AAATAAAATGGATATAGAGTTTGTCAAATATTGGGATTTATGAGAAGAGGCTGTACTATCACTAATAATCATTTT
TGATAAATGGATGAGTGATGAGCATTAACTTGAGGAACATTGGGGCACGGACCTGTAGCTCAGTAGTTGATAAAA
GAATTTGCAATCAATAAGTCATGGGTTCGAGTCCTAGTATTTACCTACAAAAAAAAAAGGCACTGCAGGAACATT
ACATTGTACAAATAATAAAATCATTATTATTGGATTCATAGGGAAATAAAAAAAATTACCAAAACTAATGTTTTT
TCCTGAAATTAAATTCTGCTCAGTTTGTGCATAACGCATGCATTTTGTACCTTTGCATTTATATAATTTAAGACA
TCAACTCATACTGTTCAAATATTCGACAAATATTTTGTTATGTAAAGTTTACCGTGTAGTGAGGGCAGGGTAATT
AGGTTCCCATTTTATATTAAAGTATTACTAACAATCATTGAGCACGCTAGAGGAAAGATTTGAACCTTCAACCTT
GTGGTTAACAGCTATACGCTTTTAACCAACTGAGCTACTCCATCAAGATTCATTTGGGTAATTAATGTTCCTTTG
GGATGTTCGCTATTGAAAAAGGAGATGAGAGATTCATAAAAAAGTTCCAATACCAATACCATGCACTTCTGCAGA
GTAAGAAAGGGGTATGGAGCATAATGGATCAGTATGCCATTTTATACTTCCTGGATAGGAATTTGGGTTCTCTTA
CATCAATTTAAGCATACAAACACCTTGACTAGCTAGCCTTAGTGGGCCTAGGGTTCCTCAAAGACTTTAGTTCCT
TTAGTGTTTGTGGCTCAAAAATTTTGGTAGAACTTCAATAAATACAATTTCACAAAAAAATCCCAAGCAACAAAT
CTTCTTTTTGCAATTATATATATTTGATCCCCATCCTTTCAAGGAAATCTTCATAATTCACTTGTATGGGAACCT
TTTTTAAACTTATATTGATCAACAATTTTTTGTTCATGCAATTGCTAGTTCTGGCAAAGGGAGGCAGCAAGCTTG
CGGCAACAATTACATACCTTACAAGAAAGCCATAGGTATTATTTATCTCAATAATAATACATTTTGCCATGAATT
CATTGATCGCCTATTCATAATGTAAAAATATTGAAGTCTAGTTAGAGTATGCTTTATCAATAGGCAGTTACTGAC
AGCTAAGGTTATCTTATACTACCACCAAACATGCTTGATATGATATCTTGCTAATCTTTTGTTGATAAATTACAC
ACTTTGCAAGAAAACCTTAGGTATCATCTCAGTGCCCTACAATGCAAGTATTACGATACAAATTCATGGATCAGC
CCAATCATAAAGCTTATCAATAGACAATTTGTAGAACCAAAGGTTACTTATACTAACTTGAAACATGGTTTGTAA
CTCCGTATCATGTCATGTTCATTTTTTGGTTCATGCTGTGCAGCTTATATATTTGCTAACACACCCATATCTAAC
CCATATGTGTAGATTTGTATGATATGTGATTTTAATTGTTTTATTTAATCAAATTACTAGGATCTCAAACAACTG
GACTTACCCAGGAACCAAGCTTTTATCCCATTATTATAGCCAATCACCGACCCATCATTCTTTGGGGTAGATTAA
TATGAGGTTGATATCTAATCAGATTTAAATCAAGTACAAGCCCCGCCGTCCCTCTGTTTACTCATGTTCTGTCAC
TTATATTGAATAATCCTGGTGTTTCATATGCCCACCCAACCGGGGAGATTTTCTCAAATAAGATTAAAGCAGATT
GAATGAAATGGAGTTTTAGCCTAAAGGTTGATGACCGAACAAAGATAAACAAAGAAATTAAGGAGAAATACCTAT
CAGATAGAGGTTCAACAAGCTACATTATAAGGATATGGGTATTGGGCTATTAGAAAGCAACAGTTTGAAAAGTTG
CATATGGCTGAAAGGACATTAAATAGACAAATAGAATGACTAGGAAAGAGAAGGTTAGAAAGTGTGTTAAAAGAA
GCTCAGGAGTAGACCTACAAGAGATTGAATAACAGATCACACATGCACAATGACAACCAATTATTGCACTTGTGT
AACAAGGTGTGCCCAATAGAGCTAATAGTTTTTGGGAGCAAAACACAAATACTTCATGATGATGATGATGCGACT
TTATGGCACAGTAGTAGAATTTATGAAACTAACAGAAAACCATTGAGAAACTAACTACATAGAGTTGAGGGATCT
GATCCATAGAATAAAATATCCATAATCACTGGTTTACAAAAAGTTCTCACCATATAATAAAGTCAAAGATCAGAT
TTACGTTGTGATTTTAGTCCTATGTGATCCATGCATTGACTGTTCTCCACTGGCATATTACCTAAATTATATCCA
GAACTCTCTTATCTCTTAAAAAAAATGATAAGAATCTTACCACCAAGTTTTCATCATCTGTGGGATACGATATTG
GTAAAACAGGAGGAACAGAAAAGGAAAGGGAAAATGGTAGATATACCCTCCCCTGCCAATTTGCCTCTTCAGAAC
ACAGAATTCCACCAAGAATTTTATCTCCATCAAAATATTAATCTCGAATAGTAACCACCATGAATTGAGTGTAGA
ATCCCCACTATTGAACCAACAAACAATTCCACAAGAGAACCAACATCATCATTGAATTTATTTTGTTGAGCAATA
ACGACATACTCTACCATTTCAAGCATTTTCCTCAGTCTTCAATATCTTCTCCAAAGGGGCAGTCCAAATCAAACC
TTATGCTCCTCACCTCCACATCATTACATGAAGAGTACCCCAATCAAGCAAATCAACTGCCTAAGAAGTCCTACA
AAGCACCATGAACCTCTACCAAGAAGTTAGAAAATTCTGCATGCTTTAACTAGAGAATATCTATTATATAGGTCC
CATAAAGTACAGGCATCATCTCTTTTGGATACTTATAGTGGGATATTATAGAGACCAACAATAGTCTGCCGGGGA

GAAAAAATCAAACCTTCATGTTGATTTTTGAAGATTATCCATAGGTGAGTTGATAATTAGCAACAGCTATAAAATG
ATGGGATGAAGAGGAAAAACCCCCCCCCAAAAAACTGTCTAACCTCTGGTCCACAGAGTCAACCCATACCTTAGC
CCAGAAATGGCTCCATGACATAGTTGTCAAATTGCCAGCATGACAAAGTATCACACAAGTGGGAATTGAGGTCTA
CGCATGAGTATGCATTCTAGGTGGTGGCTAGTCTGCTAGGAAGCAAAATGCCTAGACCTTCATATTCCATCCACC
GCAGGATTGTCAAATCTTTTCTCTACATTATTTCATTTCATTACATCTAATCCCATTGACGGGTGAGATTGAATG
TATAAATATCAGCATGCTTGATTGATAGGTGCTACGCGTAACATGCAACAGGGAGAGCCTAGACATGCACAACAC
CAAAAATGATGGCTTAGTTTAATTAAGACCTAGGTGAACCTATAAGGTGCTCTAGGGGTCCTTGTGGACTAGGTA
TGACTGACCTAAGCATGGCCTTGACAAATATGCTCCATGAGATCAGAGAGTCAACAGGGAATTTAAAAAAAAAAT
GTGTTTGATAAATGGAGGAAAGACCGAAAAGGATTTGAATGGGATTAATGTAAATACAAAATCTGACACTATGAT
TGAAAGGATTAACTTTGAATGAAAACATTGGACAATATCCATTAAGGATCCCAAATTCCATGGTATCATAGGTAC
AAAGCCATCAGATAACAGGGAATTTTTTCTTAAAAAAAAGTTGTTCCCTCAAAACCTCTTCTTTTTCTGAAAAAA
AAAGTTTGAAATATAGATAATAAATGTTATCACGACTCAACCTCACACTCACCCCCACCCCAAATGAAAAAAAAA
ATTGCACGGGTGGTATATGAGGAGTAAATCCAATTTTCACATAAATGACTCCTACATCACCTAAAGATCTGATAG
TGTCTCAAGCATAATAAACTCTCCAAACAGATCAGTTATTGAGGCAAAACAATGTCCAAACCATGTCACATTCAG
TACTTCTTTCCTTCATTCATTGTAATACAACTCTTACCTTTCTCCTTTCTAAAAAAATACCTCTGTAATATATTC
ATCAGACGAAATCCACTATATAAATAAACAAAATTAAACCAAGGAGATATTGACTGTAGTAGAGTTTGCTACCAT
CTCTGCCCACGCTTATCACCCTCACCCCCACTCCTTCTATAATCAAAACTCTGTTTACAAATTCTACCCTCCATG
CCATTCACAATACATCTACCCCAATCTGAGATAAACTTTCAGTTGAATCCCAAATGTTGGGCAACAATGGTGTCA
TGCCTTGAAGGTAATATCTTTGAGGTACAAGGATGTCTAATATGGTAATGCCCTCAAGGGAACTCCTAAGGAAAT
TATCCAACTTTTTCACTAATACTTGCAGGCAACTCATGGGAGAGGAGCTCTCTGGTTTGAGTGTGAAAGATTTAC
AAAATCTAGAGAACCAACTGGAAATGAGCTTGCGTGGTGTAAGAATGAAAAAGGTCATCCTCAAGCTAATCAGTT
CACTTTGGTCTACAAGATCATATATGAGAAACATGCATATTTATCAAAACTTTTTAATTGCAGGACCAGATTTTG
ACTGATGAAATTCAAGAACTAAACCGGAAGGTCCTTCAGAATAGTGATTAAAATTTATTTATTGCAAAAGAATGA
TGAAATGCAAATTTCTAACATTTAATTTGTAATTTTTGTGCAGGGAAACCTCATTCATCAAGAAAATATGGAGCT
CTACAAGAAGGTAAACCTCATTCACAGAGAAAACATGGAATTGTATAAAAAGGTATTGAATTAGATAGTAAAAGG
AGCTAGAGAAATATCACAATATCTACCTTGTCTATAAAAAATTGAGTTGTGCGATAGCTTCTAAAATAATTTTTC
CTACAACAGATCAATGGAACAGGGGAGGTAAATGGAGGTATTAGCAAAAGTCCATTTCTTTCAAATGGTTTTAGC
ATCAGAGAGGACATGCACCCTCCCATCCATCTTGAACTAAGTCAACCTCAAACACAGAACTATGAAATGCCAGCA
AGAGCAACAAAGTTAGGGTAAATTTGCTTCATCAACACACGATAAAATTTCAAGAGCCTATTCTTTTCTGCTTTT
GATATGAATGTTATGATGAATATTGTGACAGATTACAGCTGCACTAGAATGAGGATTCATGAAGATGTCACGTTT
CAGCAATGCTTCCTATGGTTATATAATTATGGTACGTTGAACCATTCAAAAAAAATGATTATGCTATTTCAGAGT
GATGTATAAAACTTATATCAGGGTCAATCAACAGCACCAAATCTTCTAGATTTGATGAAAATGAACTACTTGTAA
ATGTAAATTAATGTGATAATAAAAGGCATAGACCACGCCTGGTGTTCGACATTTATTTGACATTGCCAAGTCCTA
GGTTCTAGCCTTCTAGATGCAGTGCAGAGATTTCTATGTACCTCAAATGTCCCTGCAAGAAAACTTATGATAATA
GAAACCAAACGAACTTACTAGTAAGTAAAATCATCATAGGCTAGCATCACATAAATATTGAAAACATAAGCACAT
AAAACTGTCATAAAGCATAGTGCAACCTTTGAGTTCACCACAGTCTGCATATTGATACATTGATAGGCAAAGAAA
AATGTGCAGTGAGACTGCAAATGATAAATGCACGTTTCAACATATAAAACTCAATCTGATAATAAAATAGAATAA
AGGTGATATAGAATTGTAATTCAACTGAATTTCTGTGTTTTGCATAAAATCTAGGTGGTTAAGTGGAACAAAATA
TTTTTAAGGACAGCCAATGAAGTGAAACAAAATATCCGAATAAGCGTATTACACCTTTTCTGGGATAGCTATTCA
CCACTTAACTATAACACTGAATATTATGGGGTATATTAGCGAACAAAACATGAATAACCAAAGAATTAATATTCT
ATTCCTAAAAACTGAGATACTTAAACCAAACATAATACAATAATTATAATAAGGAAAGAAATATGGCTGGTATAG
TAGTCACCTTACTAGAACTATGACATATGAAACCAGGACAGTTCCATGTTGGAGAAAATCCAGGTCCCCTCCAAA
ACCAACATAGAGATCTCTCCCTGCAACAGGCATCTCATCAGTGACCAGCACTGATGTATTCTTGTGTCATTGCTA
CACAGAGTGATGCAAATCAGAATGCATGAACATATTAAGAATAAAAAATTTAACTTATTTTTTCTTACATTAGAT
TTCCTTAAATAATGTTCTATGTCCATGATCACATCAGAAAGGTCATAACTATATGATATGAGATTAGCCATTTAA
ATGCAAGTCATGTTGATCCGATTTGGCTGGCAGATGAGATACATATTTTTGCCATATTCCCTCAGTAGCGTGATA
AGCTTTTGCATCGCTAGAATCCTATATCAAGAAAGTTCACAAGACCAATGGACCAAGACAGTTTTGCATCAAATA
CAAGAACTACAGTATAGTATTAGGCATAAGGTACTTTTGAGAACAATATAATAACGCTTTTGACAAGTTTATGTT
ACAAACATAGAAATTGCAGCCATTTAATACACACAGTAAATTTGTACTGTCTACACGAACTCCACTGCAAAGTTT
TTATAGTTCCAAAAAAAATTTCGTGAATTAATTAATTCTCAGAATACATGAAATTCTTTTGATGTGAAAGTACAGG
ACCATACAATTTATATTACAAAGTTTGAAGTTGTTTCCATTAATACATGCACTAAAATTTGCAGTGTCTGTAGAA
ACTCCAAAGCGAAGTTTTATTTACCAAAGCCCTTCCCTTGATATTTTAAGTCAGTCTATAGTATATAACAATTCT
CATTAAATTACAAACAACAAATCAATGAAATGATAAAGAATATATGATTATTCTGATATATCTATTTTGCTAGTT
TTCATTCAATGAACATTTACTATTGCATTTCTTTCATCTACTCCTATTTTTAAGTGAATAATCTCATGCGTATTG

AAACTCAATGATTAGCTGTCATGATTGTGTTTATACATAGGACTTGCAAGAAGACGTATTTCCGCAAAAATGTGT
ACAAGTGGGTGAACAAACTACTGATCAATCAGCATAAACATGCAAGCAAGCAAGCAGATTTGCTATGAACAATAA
ATTTATCATGGTTGACCACAAACTTTGTATGTACAGGAAGGAAGATAAACTGTAATAGTCAAATTTTGTCTCAGA
TGTTTCCTCAAAATATCCCATGGAAATGTATTTTCTAAGAAAATTGTCACCAATGCAACAATGTACTACTTCATG
CCAAATGATGCAAGCAAAAAACATTTTTTTTGGGTGAAATGTGAGAACACACCACTGTCACCACCAAAATTCTAT
TTATTAAAAAACGGAGTAGTTCCCTACAAGTGAAAAACATGAAATCTAGTATGACAAAATGTTTCCAACTAAAGA
ACAAACAAGAGAATCTCATGTATAAATTGGCAAGGTAACACCACAGGCTCGGAAAAAAAGAGAGACTGCAATGAA
AATACTAACTCTCTTGGCGAGGCAATTTAACATCACAAACAAAATTATCAACAAAGCCTATTCTCCATACTCCGG
GGTCACCATCATGCATCTTATATCATCATAAAGTGCAATCTAGCATCACTCATTACAGTTCATTCATATTTTTCA
GCTCCTACTCCCATCAAAGATATTTAGGTCTTCAAAGCTTCGCAGAATCTGACAGATCTATCTAAAACTGAGGAT
AGGCTTAACCCCCCTAACATCAGTCTTTGATTTCCTCATTTCAGCCATGGCACAACAATTCATGTACAGTTATCG
AACTGCCAACATTCACAACCCTATGTCCAGATTTAGTCTAAGCTGCTATAAAACAATCACATAATATTTCAAAAG
CACATTTTCATTTCATCTAACAGCTTTGCAGGAAGAGAACTTTGTCATTTTGCACAAAAGATGCCATAACAAATG
TGACAGCTGTGTCAGACCTATTCATGTGCAGTTACCGAGCTGACAACATTCAAAATCCTATGTCCAGATTGTGAG
GTCTAAGCTGCCATGCAACAATCACACAATACTTCAACAGCACATTTTCATTTCATCAAACCAACTTTGCAGGAG
GAACAAATGTGGCAGCAGACGTATCCTGTGTTTTCCAATACAGACTTGTAGAGACCAACAAAATCAAAATCCGCT
TGTGTGCTTTTTAAGTCAGAGTAGGCAACTGCCTCTCACCTGTTTTGATCAAAAGGTTGTTTTGGGAGTTGCAGA
GACTCCTCTGACTACAAAAAGAATCTAAACTACCTAAATCATAATCAACTATATCCTAACATCCAACATAAAAAC
AGAAAATCATTTCCGAATTATATCTCAATATTAACTAATCCTAATTATCCATGAAGGATATGATAAACCTAACAA
ATAAAATTAAAATACTAAAAATTGCAAATCATCCATAATCGTCCGTTTGGTGGTCCTCAATGCCGATCGAACGGC
CCATGGGTGTGCTACATCACTGTTCATATCTTGTAGCTTCTCGAAATGCCTAGGTTGAAACTAGGCAGGGAAGTG
AACTTGAATGGAGATTAGTGCTCTTTGTAACAGCTATTAAAAGAGGTAGGAGACTCTTTTTTTATTTTTTTTTGG
TTGGGATTGGGGGGGAGGAGCTTGGCAGCAAAGGTGTCTCAGCTGGATAACTCGTAGTTCTTTCAGGGAGTTGGG
CCATCTGGATAACTCGAAATGACTCCATATTCGAGGAAGCAGCCCTCTATTTTTGGAAAACATGTGGGAAAGTGT
GGCAAAGTTTTTGGAGACTGAGGAAGTAATGTATTGGAGCCTCTTCAATTAAGTTCTCCAGGGAGGGGATGCTG
ATTGTGATCCAATCATTAGTCTGGGATCTCTCTGAACCTGATTTTTGGGATAACTTTGTCTAGATGCTTTTACTT
GCTGTATATCCATTCCTGGAATTTATCCCTGACATTAATATATTCATGGAGGTAGACTCCAACCTTTTCTCAAGA
AAAAAAAAGGAACAAATTATCCAAATGTTAAAAATAAAAAAAAGGTAAACTCACCATTTCATAGTGATCATTGTA
TCATTTAGAAAGGGCAAATGGAAGGCGATGGCTGCACCACCACATTTCCTAAAGGAAATGGCACAGCCACTAGCA
AAAGAAATCATTCTTTAAATACATGCCTTAAATTTGGAAACATGGATCGGCATTCGGCAAAACTTATCAGTTATT
TCACAGATAGATATGAGAAGGAATCCAATGATAGGATTACTTGCATGTCCCTTCTTCTTCTTTTTTTTGAAAATT
GGTAGGAACTTTTATTAAGAGATAAAAACCCCGAACAAGAGGAGATGAGGGATCTCCATAACAAAAGAAAAATCT
GTTTAGGAGAAATCTCCCAATCAAGCAAAATTACAATACTAACTCCTCCCAAACCATGAAAGACCCATAATAGAT
GAAGGATTTCTTCTTGAAGAATCATTTTCACCCAATCTCTAAGCGTATATGAAATTAACACCTGAGTATTCTCCA
CATATACTCTCTTATTTTGGAAAAGAGTTTCATTCCGAGTAAACCATGGGCACCAACATATTGCTTGAGGAATCA
ACCATGGAAATTTAGCCCCATTACTACGGTCTCCTTTTTTTCTTCTATTCCGGAACCACTTCCACATATTGTCTA
TGTCTTGCACGTCCCTTCTTACAACTAAAAAAAACAACTCCGGATACCACACTGATCATTACAAAATAATCTCCC
TTGAAGCAAATTTTCCAGAAACTGAGAAAATAGTTTAAGGCAGAAGCTTGGCACTTAATCTCCTCTATAATTACT
GTCTTTTCTACCTGTACAAAGATAAATCAAGTTTCTAGACCCCCCGAGAAATAAAAACAACAGTTGCACTTATTC
CTTGGAATCCAATCCATAAACATCATCAGGAACCATGTTCCAAATCCACATTGACAAAAAAACTCATCAAATATA
TATTTTATTATAAAGAGTGGTATCCATGAATTCGACTAAACATCGACTTAAATCTAACAACTCTAAACCAATATA
AATAGGCATCTTTGTGTTAAAAAAAAATCCCATAGTACTTGTGGCAAACACGAAACACCGTGTTTGATTGGCATC
TGGTTCCCAACTTTTCTCAAGTTAACCAATATGGATTTGCTAGCAGGTGGTTATGATCAAAGAAAGGCACCCTTT
TGGGAACCTGAATCTGACAGGGTTTTAGTGATTTCAACTTTGTTTAGACATAATTTTTCCACTTGTCCTCGCTTC
AACAACTGTACACTTAACGCGAGGTTTTTCCCCCTAATTTCGCTTGCCTTTGCTTTCTATAGGGCTTGGGGCTGA
GACACTGACACCCATAGACAGATACAAGATCTCCAACACTCTACATAAAATGGTGCTGAGCCTTTGCCATCATAC
ACCACCCAGAATCACACTCTTCTTTTTTTTTCTTTTTTTCTTTCTTTTTTTGAGCAAATGTTAGTAATCATGTAA
TCATGTTATAAAGAATGCTTCCAAACAAAAGGAGATGAGAGATCTCCGAACAAAATTATGAGTGAAAAAAAAATA
AAAACTAAAGAAAACCCTCCCAAAGTTAGACAAACCTAACTATATCTCCTAACTATTTTTTTTTTGGTGGGTGTC
ATTTAACTTTTTGTTCTTTCTTAACCTTGCTCAAACATAATGGTGTTTGAAAACCATTCGTCAATGCAACACTGC
CACAAACCATATATGAATTACTCACTCACAAGTCACAACCATGGACAAACTTTACCTCTAAAACTTTTTGGCATG
CCCTCAAGTACCTGCACACCATTACCCAAATTTAAGTTCAATATCTGTCGACAACCAATTACCCATTACCCTCAA
CAGATGCAAATGCAATCATAAACTTATTGAAACAATAAAAATTAATAATAAAAAAACATATGTTTCATTTGCTTG
TTTTCCTCCCCTCCCTTTCCTTCTACCCTTTTTTTTATTTTAGTAAATCTTCTCTCTTTGCACATTGACCAAAGA

AATCCAACGAAGCAATGTCGAAAAAACAAAACACGAAATCCAACTCAACAGAACAAGTAACACCGTCACAGAAAC
TCACAATTGAGAAAGCGCGAAAGATTTTCACCCGTAAAAAAAAAAGATGTCCCCTCATTCGAAGATCGAATGTCC
TCTTCTTCGAGAAACTCGGGCAAAGAGAAATACTGTTAGGAGAGGAGCAGAAGCAGTAGAGAGAGAGAGAGAGAG
AGAGGGAGAGAGTTCATGATTAGCCTTGATCACCACGTAATCAATCAGCTTTTTAGGGTTTCACAAGTGGGCGAT
TATTTTAAAAAGAAAAAACTAATCGGCCGAACCCCGTGTTTCTTCGCGTCGGGGAACGGCTCGCTCTTGAGCCGA
TCGCGTGAGCCGATACAGAGTCTGTGATAGTCCAGCTGTCGAGCCTTGGCTCGCTTTAAAAGCAGTCCCGACCGT
TCGGTTGAACCATTACTCGGTATATATGATCGTACAAAAAAAAATCGATAAAATATTGGAAAAGATCGATATAGC
CCCTGAGAAGTTAATTAAATCGGTTTTAGTCCCCCATAAAATTAAAAAATCACCAAATGTCTCTAAAAAAAAAAA
AAATATGAATGGTTAATGGATTCAGTCCCTAAATAAGTTTATAAGGAATATTCCAGAAATATCTTATTTACGAAA
TAGAGCTCCAATAAATGACATTCAAATAAATCCTTACAAAATATGAAAAATCTGATTTAATCATTTTGATGACAA
AAATGAATCTTTCTTTTCCTGAAAATTCTCCTGATATTATATACTCACCACCAACGAAAAAAGAAAAAAAATCAA
TATAACTATGTGGAAACTTATGAAAATAAGAAAATGTCCTTGTAAAATCAGTAAATCCCCTTAAAATCAACCATA
GATCTGATATTTGTATTACATTAATTTATCAAGTAAATTTTAAAAAGGTATTACAATAATTTATCATGTATGGTT
AGAATCAACCACAAATCTACTTAATTTCAGTTATGATAGGAACTAAAATAAATATTATCTAATGAGATAGAAATC
TGACAGAATTTACCAATTTTGCATGAATATTTTCAAATTTTCATAAGTTTTTAGTGAGCTATATCGATTTTCCCA
TGAAATAAGAGGTGATGGCATTTTAGTAAGTTTTAATGTGTAATTAGGGTTAACAATGTACTTGATTAATTAGGG
TCACAGTAGGAACCAAAATAATATTTTTCTATTAACATATAGATTTGTGACAATTTTATAATTTTACAAGGATTA
AAATTAATTTAGCTAAATATGCATGGAGTTGGTGTTAATTTTTTCTTAAAACATCAATCAATACTGTATTTAACC
CAAAAAAACAGCTTGATTCTTTAAGTGTCGTTCGAAAGGACTGGAGAAGGCCACGTAGTTTGTTATTTCACCTTG
CATTTTAAGCTTATTATATTTGATTTGTGTCGCGCAAAAGAGACATATAAAAAGAAAAGAGGGATCGCTGAAAGG
AAAAGTACGGACGGTTTGGATTCTTTATGGTTGGGTCCACCCATCTCGGATAGCTCATCGAGATCTGACTGTCCG
TACGAATGTTTGACTCTGACCTTCCGTGGCCAAATATTGGGTTTGGGCGACGATGGCCTAACGCATTAATTATAC
TTCATTGAATTTACAAGGATGCATATAAAAATGTATGTTTTAGATTATAAAAAAAAAAATTGAGTGGAACGGCAC
TCTTTATCCAACCCATTCACGCCCAGCCCATTTTAAAAAATGAAAAAAAATACACATAGTTTCAATTACAACTTT
AATTACAGTTTATTTTATTTTTTTAATATATAAAACTATATATTTATCTTTTAAAGTGCATGCTATTATATGAAA
TAAAGTATATATTGAAATATATTAATGTATTTTTTTATGATAAAAAAAATAAAAAAAAAATAGACTGCAATATAT
ACTATGATAGTATAATATACACTACGTTATCATAACATACAGGCTGTAAAACAGACTGTAAAAATAGGCTGCGTG
TATCAGCACTCTAAAAAAAGAAAAAGAAAAGGAAAATGTTAAATTCAATCATTTCTACAAAACCAAAACTCGAAT
AATACGTGTTATCATTTTTTTAAAAAAGGTATATGTTTTAATAATAAATGTATATCACTTCATTTCAAGTTAAAT
GAACTTATTTTTAATAAGCGTAATCTTCATTCAAAAAAATTTTTTATTAAGTGTACTCTTTTTTTTATAAGATAT
ATACATCGTCTAACAACTAATGAGGTATATATTCCACTGGATTTTTAAAGAGATATATAAGACTGTATTTGAGAC
TGTAAAAAAAAACGAGTATATCAACACCCAAAGAAAAATGGGCGACAATAGGCTTTTCATCTCTTTCTTTCAAGC
ACTGTTATACTCAAGTCATTTTTACTAATCAAAGTTCTATATACTTCATTAGAAATTAAGTGAAACATAAACATC
GTTTAACAATGTATATATCTCATTAAAAAAAATTATATTTATTGGAAATAGGTTCATTCAACTCGAAATGGAACG
GTTCATTCAATTCGAAATGGAACGATATATGCTTTTATATTCGAAATGGTTCATTCAACTAGATGATATTTACTT
AAGAGTGGTAAACGAGCTGAGCTTTGACATCATGATATGTTCTAACTCGGCTCGTTTACCTTTAAACTTGGCTCG
GGCTCGACTCGATTTGGGACCGAGCTTAATTTTGTGTTCAAGCTCGGCTCAGTTCGGCTTGTTTCTAAAACGAGC
TCGAGTACAAGCTCAAAACAAGGCTCAAGCTCATGCTCATTTAGCTAAAAAAGAAACTTCTTTAACAATTAAAAC
ATGAAGAAAAAACACTATAACATGCTAAAAATAGAAAATCATATTCGAAAACCATCACTAAATAATCATATTAAA
AATTATACAACATTCCACACTTTTAAGTAAATGAGTTTGTATAGG

>AgAGL17 [organism=Acorus gramineus] AGL17-like MADS-box gene, partial CDS, splice variant 1, gene-specific PCR on cDNA from leaves

CAGCACCGGTCGCCTCTACGACTTCTCTAGCTCGAGCATGAAGACAGTTATTGATCGGTATAACAAATCAAAAGA
GGAGCACAACCATGTTGTCAACTCAACTTCAGAGGTTAAGTTCTGGCAAAGGGAGGCAGCAAGCTTGCGGCAACA
ATTACATACCTTACAAGAAAGCCATAGGCAACTCATGGGAGAGGAGCTCTCTGGTTTGAGTGTGAAAGATTTACA
AAATCTAGAGAACCAACTGGAAATGAGCTTGCGTGGTGTAAGAATGAAAAAGGACCAGATTTTGACTGATGAAAT
TCAAGAACTAAACCGGAAGGGAAACCTCATTCATCAAGAAAATATGGGGCTCTACAAGAAGGTAAACCTCATTCA
CAGAGAAAACATGGAATTGTATAAAAAAATCAATGGAACAGGGGAGGTAAATGGAGGTATTAGCAAAAGTCCATT
TCTTTCAAATGGTTTTAGC

### Marking of feature of AGL16-like genes and MIR824 genes

exons of AGL16-like genes
exons of miR824
mature miR824
target site

#### miR824 in Brassicales

##### Brassicaceae

###### Arabidopsis thaliana

*AGL16*

>Chr3
>AT3G57230
ATCCCAAATTGTCATATGTTCTTTCCTTATGTGGCTCTCTCTTCTCTTTCTCTTTCTCTTTCCTTTTTCTCTTTC
GATCAACAAAACCACATCTCTGAACTGAACCAATTTCTCTTCTCCCCCTTCCGGTTATCGGATTACCAGATCTCG
TTTCCCGCGATCTAGTTTATTCTTTGAAAAAGGTAAATGTTCAATTTTTTTATTTTATTCAAAATCTCCCCTGGA
TTCAGGAGATGATGATTTGAAATTGACGATGATGTTTGTGTGTAAAAGTGATAGAAGCAGAAATGGGAAGGGGCA
AGATCGCGATTAAGAGGATCAATAACTCTACGAGCCGTCAGGTTACGTTCTCGAAGCGAAGGAATGGATTGTTGA
AGAAAGCTAAGGAGCTTGCGATTCTCTGCGATGCTGAGGTTGGTGTCATCATCTTCTCCAGCACCGGTAGGCTCT
ACGATTTCTCCAGCTCCAGGTACCTTTTTTCTTTTCTCTTTGGTTGACTTTTTTTTTGGGTCTTGGTCCATCTAA
TTGGTTGATTGATTGATGTAGCAAATTCTAAATCTCGATTTTTTTTTTTCTTTCTTAAGTCGATTTCTTGTTAAT
TGTTTGCAAATTCAATTGCTTTAGAGAAGGAATCGTAGATCAACTTAGTCTGATAATTTGCAGAAGAAATCACTT
GGTTTGAATGATAATATGTTTTGTAAGGGGAATATGAGAGTGTGTTCTCTGTGTTGTTGGAGATGATTACGATAT
TTGTGTGGAACATAGATTGTGAGGGACTTGTGGTGTTTATTGAAACATACTAAATTTGGTAGAAACTTTATCAGA
AGAGGAAAGGAGGGGAGAAAGTGGTTAGTGGTTGAAGAAAACTCTAAACTCTAGAGAGGATTTAGAAAGAATCGT
CTCCTTTTCACTGTTCATTTACTTTATTGGCAACTAGTTGGTGGGTTTGATGTACCTAAAATGAGCTGCATTATC
TTCTCTTAATATATATTGTTAGTACACAAGTGGGGGAATCATTTCTGATTTCGTTGTCATCCATAATCAAGTTTT
TGTTTTTTTAATTCTTCAACTTGGTCATTTGTTAATGATATGTAGGAGTATAGTCTTTCTTTTTGGGAACATTTT
GCGATTTGTTAGAAAAATATTATATCTTCAATCCTTTCTGAATTTCTACTTTAGGAGTTCATTCATTTCTTTATT
TGATTGTATCTTTCGTGTTTACATAATTGTTTATTAAAAAAGGTTGAATAAAATCTGGCTCTTAGATCTGTTTCA
GAGGGAGAAAGCTTCCATTGTTCTGCAGAGCTTTTGTTGTTTCTCTGTTTTTTGAACTGAGACTCTGTTTTTTTC
TTTTTTTCTCTTCAATAAAGACAGCAAGATATCTATACTGTTTACTGATTAGAGTTGGCTCTATTTACATCAATT
TTGTCTGGCTTCTCTCATTCTCTGCTTAAACTTCATACATTATGTTAGATTTATTGCGAATCTGTAGTTGGGGTG
CTGAGAATCCTGTTCCATGTAGACAGAGATGGCTTAGGGTTAACTAAATCTATTTAAGTAAGAAAATTTGACACT
GAAAGATGTAAGGATAAATGTTGCAGTGGATGTTGCTTGGGTTACATTACCTATCAACCACACTCTACAACTATG
AAACTACTCTCTTTTTTGGCAGCATGAAATCGGTCATAGAGAGATACAGCGATGCCAAAGGAGAAACCAGTTCAG
AAAATGATCCCGCTTCAGAAATTCAGGTACTCCTTTTTTTTGTTACTCCTCCCTATGGATTTCAACGAGTAGACA
AAACAAGTAACATACTACAACAGAGAGACTGTTACTCGATTCTTGTACTAATATGTTAGCCTATTACAACTCATT
AGCATTTCTCAACTTTTAATTCCTCTCAAGCTGATCCTGATATCTAATCAGAGTTGCAGACCATAAATACCACCA
CACATCATATGTTGAAGTAGGGTTTGCAGTTTGCTATAAAACACAAGTCATGCCACCTGAACATCAGATTGCATA
CATAGTGTTGCATTTTATTTTGTCAAGAATCCGTTGTAGACTAGATAACATATTTATCCGAAAACAAAATACTTG
TTTACAGTTTCAAACACCTGAAAATCATTTCTACTATATTCTAGTTCTGGCAAAAGGAGGCTGCGATTCTAAAGC
GTCAGCTACATAACTTGCAAGAAAACCACCGGTATGTTTTGCATCCTCTGAGTTTGTCAAACACTCTCGATAGTA
GTAACAACTAGAGCAGGGAAATATGTACTAAAATGCAAAATGATAACATACGTGTAGTCCAAGGTTAGAAGATTT
TGCAGAACCCAAAACTATCCTACCATTTGCATTTGCTTAGTAAGGATAGTCGATATAAAGAATAGTAAGCCAAAT
GGATTAAGAGTTGTATACATAAACACATCTAAACCATCCATAAGAAGGAAATAACCAAACACTTTCACACTCCCC
ATCGTTTTCTTGTTTTGCTTTGGTGAATTAGGAAATGTATATAGTAACTCTGGAAAAGTATGCATATTCAGAAGA
GCTTGTACTTGACAGGCAAATGATGGGGGAGGAGCTCTCTGGACTAAGTGTAGAAGCTTTACAGAATTTGGAAAA
TCAGCTTGAATTGAGCCTTCGTGGCGTTCGAATGAAAAAGGTTTTGATGCTGTGACAAATAAGTCAAATTTTGGA
ACCATTTAGAAGAAGTAATGGATGTACTCTATCTACGCTCTTCTTTTGCCTTTTGCAGGATCAAATGTTAATCGA
AGAAATACAAGTACTTAACCGAGAGGTGGGACTATGGTTACTGTGATATAGGTTTTAGACGTTTTCCATAGTGTT
GGTAAGCACATTGCTAATTCTTTTTTTATTCGGTTGCAGGGGAATCTCGTTCACCAAGAGAATTTAGACCTCCAC
AAGAAAGTAAACCTAATGCACCAACAGAACATGGAACTACATGAAAAGGTACCTTTGGGTTTTGGGAAGTATCTG
GGAGCCAAGATACTCTTTAAAATTGCTAGAACATTTTTTTCTTTCTTATCTGATGATAATATGTGCTTATTCCAG
GTTTCAGAGGTCGAGGGTGTGAAAATCGCAAACAAGAAT<u>TCTCTTCTCACAAATGGTCTA</u>GACATGAGAGATACC
TCGAACGAACATGTCCATCTTCAGCTCAGCCAACCGCAGCATGATCATGAGACGCATTCAAAAGCTATCCAACTC
AACTATTTTTCCTTCATTGCATAATATAATTCGGTGTGCCAACACACTTATGTTGACCTCGTCGGAATCATATCA
CAATTCACTGTGTCAGCTTGCCTCTGCATAAGCGAAAATAAAAACATAAACATGATCAGTTTGCATTCCATATCT
ATCAAACACCAGCTTTGTAACTTTTAAAACTTTTTCTCCGTGCAAAGACCTTTGGTTTGGCGCTTAAGCATGTAG
TTTGATGATCAAAGGAATGGGTGTTTTAGCATAAAGTTGTCACCCTTCCGTTGCATTTT

*ath-miR824*

>Chr4

TGGGTAAGAAGCATGAGCGGCTTGAGCAGCGGCTTTATGTGAGAAATCGAAGCCCCACATGTCCCCAAATTCAAA
TTATAGATTAATCCAAATGAAATCTGATTAACGCACTAATCGATACTTAAGGTTTTAAATTACTCTCATTCTTCG
CTTACATTACTTATATAAAACCCTCTCTCGCATCCTTCCTCAGTT**AATCATCACTGCTCTCTTCTCCATCTCGCT**
**TTCACTCGCTCTCTCTCTTGCCCTAGCTTCCCTTTTCTCCTGAATCGGCCTCACATCACGGCTTTTTCCGACTGT**
**TTTCCGTCGAATTTCAATGGGATCTCGTGACGGCGATCCGCCATTTTCGAAATTCTTGAGCTGACTCGTCTCCAA**
**TAGTTCATCG**GTTAGTCTGATTCCGTGTATATCTATTCTTAACCTGCAATTTGATTTGATTGTACACATTTTTTG
TTGAGTCGTTTTTGTGATATTTGTTTTTATCCGTGAGATTAGTTTCGTAGTTATGGCGTTTATCATCGATAGATC
AGCTGCATGAGCTTTAAGATTATATAATTTCTGAGCTGCGGATCTAATTTTAGTAATGTATGGAATGATTTTGAG
TACTAATGTATCATAATTTGGTTTTGCATGTCCAG**ATTTGTGAGAGCTTTTAAAATCTTTAAGTGGACCTGTCTC**
**ATGTATTGAG**GTTTGTTTTCTCATCTATATTATTTGTGTTTGATGCATTTACATATATGTATATATCTTATGATC
TGTTTTATTCTACAAGTATTGTGAGTGTCATTTGTTGATTGTTACTTAGTTAGTCTATATATTCCTTTTACTTAG
TTAGTTATGAGCATAAGATAAAAACAAATAATAGACTCGACTCAATTGATCTGAAGAAACACAGAAAACAAATCT
ATACCCAGATATTTAGTTACTGTGGTTATTGATATCTGCACCTTCTTTAGTTAGCTACTGTGAATAAGAGACTCG
ACTCAGTTGATCTGAAGAAACATAGTTTGTTACTGTGGTTTTTTATATCTGCACCTTCTTTAGTTGTAATTCCTG
AGAAGATTTTGAAAGCATCATTAGATTCTTACTTGAGCCCACTTTAAGGTAAAAAGGAGCTCGTGGAAAGTCTAT
ACCCAAATTATATGAAATTAAGCAAAATATGCTTGTTCGTTCTTCTTCTTCGGCAAACTAACACTAATTCACCCA
CTAAAGAGTTTTCGAGTTTGTCTGAGTATGGTCTTTTTTTGTTTAACATGGTCTGTTAAGTAAAACCCTCAATAT
GGAACCTTCTGAGAATTCAAGCGTGATCATGATATCCTTGTCACTCAACTTTGATTCTTGGTCAAATAAGACTTT
TCAAAAAACTATATGATAGGCTCCTTTTAGAGACCATTTATTGGATAGCTTCATCTGTGTTTGCAGTCTCTAAAT
GTACAAAAATGCATTTGCACAATGTCTTCAAAGAAGAGAAAGTGAAGAAAACTTGCTTTACTATGAGGAGATTAA
GAAAAATTTTATTGATTGAATGCAAGACAATTTTGTCTCAAATCATTAAAATAAAAAATACTACTTTGCTGTTGC
ATAAACTAGCAATGCATAAATATCAGGTTTATATCTGCATATCTTTGTAATTGTATGCTGTTCGTTGTTTTTATA
ATATGAATTTTAAATCTGAAAATTCATCTTCCCTTGTTAATGCAG**TGGAAAACTGCTACAGTTGACTGAGATAAG**
**GTTCTGACAGTCCAGCTGAGATGCTGAGTATCACCATTTGTACTTGAGTTGTCTCTCATGTCTAGACCATTTGTG**
**AGAAGGGAGTTTTTGTTTACACCAATACCCCCCAGTCTCTAAATTTGTAAGAAGTATTATGCTCAATTAAGGAAC**
**ATAGCTAGTTCGACATACCATACTACCCTTTTTAAAACTATCCTATATGTTTGATGCTAGCATAGCGTAATTTTG**
**TGTTCTCATGGTGCAGCAGGGTTAGTTTATTGTGTACCTCTAGATATATATTCTCTTGCTGCGTAGGGTTCCCAA**
**GCTGCCAAAAACTTTTAAAAATTAGTGATCTGTTCCCCAAACCCCCATTCAATAAGAAAGGTCTACCTGTAGCTC**
**ACAGTCACAGCTATTAGAGCTGTCCTTGCTTCTTTGGGTTTACTGTTTTAGTTAATTAGTTACCCTATGAAAGTC**
**TGATCCTTCAGAAGTTAGGTATAGAAGTAAGTCGATTAAGTTCATTCATGACTTTTCCAGATGTTGCATACATAA**
**CTTTTTTTATTTGTATTCGAACTAGTCCAAGCCGATTCTCAAAATCATATAAAATCATTTACACCTTGGTCATGG**
**TAGCTAAGAATATTGTATCTAAAATTGGGGAGTGGGGAGATGTTTGGTTATATTCCCTTCTCATCGATGGTCTAG**
**ATGTGCGAGGTGACTCTCATGGAGGTAAAGAACAATGGTGATTTTGTGAAGAACCCAACGTAATGGTAATTCCTA**
**AAAAGGTTAGAAGTTTTTTCAGCTTGTTGTATTGCTAAAATGGGGTTGATGTACTCAACGACATCCAAGTGTACT**
**TGAGTGAGCTTTTTTGGGGTTGAGTACCTCGACCCATTATTCAAACTAATGTAAATGGTGAATGCAGCAGTGACT**
**TTGTTGCCTTTTGCAAGAACTAAAGAAGACAGAAACAGGTTGTAAAAGAGAGCCAAGTGTGTGTTTATGGTAGAA**
**AGAGCAAAGTGAACGAAAGGTGTACCTTTTTGACTTGTTGTCACTGGTTTTCTCCCACTTCATCCGTTTCATGCT**
**GCATCAGAAAACAACATAAGGAATGAATGACGTAACGCGAAGCATTAGGAGTTGCTTGTAAATTAATACATTGCC**
**ATTACTAACGTAATTCAGTAGATTCTAACTACAAATGAAGTCAATGTATCTATTTGTCTACTTTAGCCAATGTAT**
**GATAAGACCAAATAGTCTTCTCTTTTTTCAGAAACTCTCTAGGATTAAAAAGTTTGTGGGTGAAAGAAATATTAT**
**CGTGTGGATGATAAG**AATAATTGATCTTGTGTTAGTAAATTAGGAATAGATATACAAGTAGGTTTCTCTCTAAAT
AAAAAATAAAAGAGTTTAAATTGCATGCGTATAAAAGAAAAAAGTAAGAAGAAAATATGTTCCGGTTAATGGTTG
GGTGCATCCGAATCGAACCGGCGCAAACCAAAAAATCTAAAGGAGATTTGAGGTGATAAAAGGAAATCAGACATT

#### Arabidopsis lyrata

*AlMADS104*

>AL5G38720 | scaffold_5:18296101..18299840 forward

ATCCCAAATTGTCATATGTTCTTTCCTTATGTAGCTCTCTCTGCTCTTTCTCTTTCCTTTTTCTCTTTCGATCAA
CAAAACCACATCTCTGAACTGAACCATTTTTCTTCTCCTCTTCCGGTTATCGGATTACCAGATCTCGTTTCCCGC
GATCTGATTTATTCTTAGGTTTTTGACAAAAGGTAATGTTCGATTTTGAGTTTATTGAATTTTGGGAGATGATTT
TGAATTGACGATTATGTTTGTTGGGTGAAATTGATATAGCAGGAGCAGGAGAAGAAAATGGGAAGGGGAAAGATC
GCGATTAAGAGGATCAATAACTCTACGAGCCGTCAGGTTACGTTCTCCAAGCGAAGGAATGGATTGTTGAAGAAA
GCTAAGGAGCTTGCGATTCTCTGCGATGCTGAGGTTGGTGTTATCATCTTCTCTAGCACCGGTAGGCTCTACGAA
TTCTCCAGCTCCAGGTACTTTTTTTTTTCTTTCTCTTTGGTTGACTTTTTTTTTTTTTTGGTTTTGGTCCTTCTG
ATTGATTGATTCATGAAGCAAATTCTAAATCTCGAGTTTTATTTTTTCTTGAATCGATTTCTTGTTAATTTTTTT
TATCTGCAAATTCAATTGGTTTAGGGAAAGAATCGTAGATCAAATTAAAACAAAATCCAATATTCATAGATTATA
GGTTTTGTAAGTAGGGGGATATGATTACGATATTTGTGTGGAGCATAGATTGTGAGGGACTTGTGGTGTTTATTG
AAACATACTAAATTGGGTAGAAACTTTATCAGAAGAGGAAAGGAGGGGAGAAAGTGGTTAGTGGTTGAAGAAACT

CTAACTCTCTCCCTCATCAAGTCTTGAAATCACTTTTTTTTTTTTTTTTATAGTCTTCTTGGTTTGGGTAGAGAG
GATTTAGAAAGAATCGTCTCTTTTTCACTATTTCTTTTCTTTATTGGCAACTAGTTGGTGGGTTTGATGTACCTA
AAATGAGCTGCATTATCTTCTCTTACTATATTGTTAGTACACAAGTGGGGGAATCATTTCTGATTTCGTTGTCAT
CCATAATCAAGTTTTTGTTTTTTTTTTATTGTTCAACTTGGTCATTTGTTAATGATATGTACGAGTATAGTCTTT
CTTTTTGGAAACATTTTGCTATTCGTTAGAAAAATATTATATCATCAATCCCTTCTGAATTTCTACTTTAGGAAT
TCATTCATTTATTTATTTGAATGTATCATTTGTGTTTATTATTTATTAAAAAAAAAGGTTGAATAAAATCTGGCT
CTTAGATCTGTTTCAGAGGGAGAAAGCTTCCATTGTTGTGCAGAGCTTTTGTTGTTTCTCTGTTTTTAGAACTGA
GACTCTGTTTTTGTCTTACTTTTTCTCTTCAATAAAGACAGCAAAAAAAGAAAAAGAATTTGTAGCATCCATCAG
AGGTGTTACAGCAGAGAAGGAAATAATATTAAGTAGAAAAGATCGTCAGAAATAGATGTAATGGTTCCAAGAGAT
TGGGGCATTAGATGAGAAATATATTAGAGGCATGTTGTTAAAGGGTCTTTCACGTTTATGTCTCACTGATCCATA
TCTATACTCTTGTTTACTGATTAGAACTCTGCTTAAACTTCATACATCATATTAGATTTATTGCGAATCTGTATT
CCATGTATGCAAAGATGGCTTAGGGTAACAAAATCTATCTAAGTAAAAATTTTGACACTGAAAGATGTAAGGAGA
AATGTTGCAGTGGATGTTGCTTGGGTTACCTTACCTATCAACCACACTCTGCAACTATGCAACTACTCTTTTTTT
GGCAGCATGAAATCGGTCATAGAGAGATACAGCGATGCCAAAGGGGAAACCAGTTCAGAAAATGATCCCGCTTCA
GAAATTCAGGTACTCCTTTTTTGTTACTCCTCCCTATGGATTTCAACGAGGAATTCAGGGACCCAATTCTGTAAT
TGAGTAAAGAGTAGACAAAACAAGTAACATACTACAACAGAGAGACTGTTACTCGGTGCTTTTGCTAATGTTAGG
CTATTTCAACTTCATTAGCATTTCTCAACTTTTGATTCCTCTCAAGCTGATCCTGATTTCATATCGGACTTGCAG
AGCATAAATACCACACATCATATGTTGAAGTAGGGTTTGCTATAAAACACATGTCATGCAACGTGACCATCAGAT
TACATACATGATGTTGCATGTTCTTTTTCAAGAATCCGTTCTAGACTAGATAACATATTAATCCAAAAACAAAAT
ACTTTTTTTCTTAGTTTCAATCACCTGAAAACCTATTTCTACTATATTCTAGTTCTGGCAAAAGGAGGCTGCGAT
TCTAAAGCGACAGCTACATAACTTGCAAGAAAACCACCGGTATGTTTTGCATCCTCTGAGTTCTTCAAACACTCT
TGAACAGGGAATGATGTACTAAAATGCAAAATGATAACATATATGGAGTCCAAGGCTAGATTTTGCAGAACCCAA
AAGCATCCTACCACCTTCATAGGAATGGTTGATAGTAGTAATAACCAAAAAGCCAAATGGATGTTTTCCCCAAAG
AATTACCATTGTATTTCTACCACCTCCAGAGTTATATACATCCACACATCTAAAACATCCATAAGAAGGAAATAA
CCAAACACTTTCACACTCCCCATCTTTTTCTTGTTTTGTTTTGGTGAATTAGGAAATGTATATAGTAATTCTGGA
AAAGTATGCATATTCAGAAGAGCTTGTACTTGACAGGCAAATGATGGGGGAGGAGCTCTCTGGACTAAGTGTAGA
AGCTTTACAGAATTTGGAAAATCAGCTTGAATTGAGCCTTCGTGGGGTTCGAATGAAAAAGGTTTTGTTGCTGTG
ACAAATAAGTCACATTTTGGTACCATTTAGAAGAAGTAATGGATTTACTCTATCTACAATCTTCTTGCCTTTTGC
AGGATCAAATGTTAATCGAAGAAATACAAGTACTCAACCGAGAGGTGGGACTATGGTTACTGTAATATAGGTTTT
AGACATTTTCCACTGTGTTGGTAAGCACAATGCTGATTCTTATTTATTCGGTTGCAGGGGAATCTCGTGCACCAA
GAGAATTTAGACCTCCACAAGAAAGTAAACCTAATGCACCAACAGAACAAGGAACTACATGAAAAGGTACCTTTG
GGTTTTGGGAAGTATCTGGGAGCCAAGATTATCTTTAAATTGCTAGAAAATTTTGTTCTTATCTGAAGATAATAT
GTCGTTATTCCAGGTTTCAGAGGTTGAGGGTGTGAAAATCACAAACAAGAATTCTCTTCTCACAAATGGTCTAGA
CATGAGAGATAACTCAAGCGAACATGTCCATCTTCAGCTCAGCCAACCGCAGCATGATGAGACGCATGCAAAAGC
TATCCAACTCAACTATTTTTCCTTCATTGCATGATGATATATTCGGTGTGCCGACACACTTATGTTGACCTCATC
GGAATCATATCTCAATTAACTGTATCAGTTTGCAATCCCTTTCTATCAAACACCAGCTTTGAACCTTTTTAAACG
TTCTTCTCTCTGCAAAGAACCTTGGTTTGGCGTTTAAGCATGTACTTTGCAGGATAAAGGAATGTAGTGTTTTAA
CATAAAGTTTGTCTGTGTTATTCAAGAGGACTAATACTTTCCTTGATTTGTGACTTGTGATGATATGGTAAAGCA
AATTGTTTTTTTTTTTTTTTTTTTTTGACATTTTTTGGGGACATAGTAAAGCACATTGTATGAG

*aly-miR824*
>AL7G28800
**CTCCTCACTTGCTCTCTCCTCCATTTCGCTTTCACTCTCCCTTGCCCTAGCGTACATCTTCTTCTTAATCGGCCT**
**CACATGCCGGCTTTTTCCGACTGTTTTCTCTCGAATTTCAGTGGGATCTCGTGACGCCATTTTCGAAATTCTTGA**
**GCTGACGCGTTTCTATTAGCTCCTCG**GTTAGTCTGATTCCGTGTATCTCTTCTCTTAACCTGCAATTTGATTTGA
TTTTACAGATATTACATTTTTTTTTTTTTTTGTTTTTGTGATATCCTTTAGATTTCTCTTGCGGTTTGAGGATCC
AATTTTACTAATCTATGTATGGAATAGTAATGTATCAACATTTGGTTTTGCATGTGCAG**ATTGGTGAGAGCTGCT**
**TTCTCAATCTTTAAGTGGATCTCTCTCTCATGTTTTCAG**GTTTGGTTTCTCATCTTCTTTATTTGTGTTTGATGC
ATTTGCTTTTTAATTGTTTGGATCACACTTCTTTTTGTGTTGTTTTTGGGTATAGTTTTTTTCTTTTCTTAAGA
TGAAAATCACATAATAATGTATATTGTATACATATACATTATCTTATGATCCTTTTTATTCTAGAGGGATTGTGA
TTGTCATTTGTTGATTGTTACTTAGTTAGTCTATATTCCTTTTACTTAAGAATGTTGTAATAGTTATGAGCATAA
GATAAAACCAAATAATACCATTTTCTTGCACCTTCTTTAGTTATACTTCTAAAAAAGTTTGAAAGCATCATTAAA
TTGTTACTGGAACCTACTCTAAGGTAAAAAGGAACTCCTGGAACGTCTATACCCAATTTATATGGAATGTAGCAA
ATCATGTTTGTTCTTTCTTTTTCTTCGGCAAATTAACACTAAAGAGTTTTGAAGTTTGTCTGAGTATGGTCTTTT
TTTGTTTAACATGACCTGTTTTGTTAAACATTCACTATTGGCTCTCGATATGGAGATGATCTATTTAGAAACTTC
TGAGAATTTAAGCGTGATCTTTAGTTCCTTGTCATTCAACTTTGATTCTTTGTCAAACAAGACTTTTCAAAAAAC
TATATGATAGGAGAATATGCCCCTTTTAGAGAGCAGTTATATGATAGTTTCATTTGTGTTCACAGTCTCTAAATG
TACAAAAATGCATCTGCACAATGTTTTCAAAGAAGAGAAAGTGAAGAAAACTTGTTTTACTTTGGGGAAATTAAG
AATTTTTTTATTGATTGAATGCAAGTCAATTTTGTCTCAAATCAAAAAGAGTACTTTGCTGTTGCATAAACTAGC
AATGCATAAATTTCAGGTTTATATCTGCATATCTTTGTAGTTGTAGGCTGTTCGTTGTTTTTAATATGAATATGA
ATCTGAAAATTCATCTGCCTTTGTTAATGCAG**TGGAAAACTGCTACAGTTGACAGAGATAAGGTTCTGACAGTCC**
**AGCTGAGTATCACCGTTTGAAGACTGACAGGTGTACTTGAGTTGTCTCTCATGTCTAGACCATTTGTGAGAAGGG**

**AGTTTTTGTTTGCACCAATACCCACCAATCTCTAAGTTTGTAAGAAGTATTATGCTCAATTAAGGAAAATAGCTA
GTTCAACTTACCACACTACCCTTTTACAAACTATCCTATATGTTAGATGCTAGCATGGCGTAATTTTGTGTTCTC
ATTCTGCAGCAGGGTTAGTTTCTTGTGTACCTCTAGATATTTATTCTCTTGCTGCGTAGGGTTCCCAAGCTGCCA
AAAACTTTTCAAAATTAGAGATCTGTTCCCCACTAACCCCCATCCAATAAGGAAGGTCTACCTGTATGTAGCTCA
CAGCTATTAAAGCTGTGCTCGCGTCTTGGTTTTACTGTTTTAGTTACCCCATGAAATCTGATCCTTCAGAAGTTA
GGTATAGAAGTAAGTCGATTAAGTTCATTCATGACTTTTCCAGATGTTGCACAGATAACTTTTTTATTTGTATTA
TAACTAGTCCAAGCCGATTCTCGAAATCATATAAAATCATTTACACCTTGGTCATGGTAGCCAAGAATTTGTATC
TTACACTAGGTTTCCTAATAAATAAAAATGGGGAGTGGGGAGGTGTTTGGTTATATTCCCTTCTCATCGATGGTC
TAGATGTGCGAGGTGACTCTCATGGAGGTAAAGAAAAATGGTGATTTTGTGAAGAACCAACGTAATGGTAATTCC
TAAAAAGGTTAGAAGTTTTTCAGCTTGTTGTATTGCTAGATTTCAAGACCAGTTGATCTGCCAATGCAACTTAG
GCGATGCTTCCAAATGTACTTGTGTGAGCTTTTTTTGGGGTTGAGTACATCATCCCATTATTCAAACTAATGTAA
ATGGTGAATGCAGCAGTGATTTTGTTGCAAGGTGAAAGAATAGGTTTGTCTTTGCAAGAACTATAAGAAGACAG
AAGAAACAGGTTGTAAAAGAGAGCCAAGTGTGTGTTTATGGTAGAAAGAGCAAAGTGGACGAAAGGTGTACCTTT
TGTCTTTTGACTTGTTGTCACTCATTCTCTCCCACTCATCCATTTCATGCTGCATCAGAAAACAATATAAGGAAT
GAATGACGTAACGCCAAGCTTTAGGAGTTGCTTGTAAATTAATACATTCCATTACTAACGTAATTCACTAGATTC
TGACTACAAATGAAGTCAATGTATCTAACTATCTACTTTAACCAATGTATACTCTCTTTAAGTACTTTCATAAAT
ATGGATACAATTA**

#### Capsella rubella

*CrAGL16*

>Carubv10017951m.g | scaffold_5:11453570..11457320 forward

ATATGTTCTTTCCTTATGTGGCTCTGTCTGTCTCTCTCTCTTCTACTCCTCTCTCCTTCTCTTCTCTTTCGATCA
AACAACAACATAACCAAAACAAATCTCTGACCTGAACCATCTCTTATCTCACCCGGTTATCGGATTTTCCAGATC
TCTTTTCGATTTCATTTCCCGCGATCTTGGGTTTTTCAAAAAGGTAATGTTCGATTTTGTTTAATTGAATTCGGG
AGCTGATGGTTTGGATTTGAATTTGAAATTCACGATGATGATGATGTAGAAATGGGAAGGGGAAAGATCGCGATT
AAGAGGATCAATAACTCTACGAGCCGCCAGGTTACCTTCTCCAAGCGAAGGAATGGTTTGTTGAAGAAAGCAAAG
GAGCTTGCGATTCTCTGCGATGCTGAGGTTGGTGTTATCATCTTCTCTAGCACTGGTCGGCTCTACGATTTCTCC
AGCTCCAGGTACCTCTTTTTTCTCTTTCGTTGACTTTTTTTTTTTGGTATTGGTTTGGTCCATGTGATTCATTGA
TGAACCAATTTCGAAATCTTAAAATTCGATTTCTTGTTGGTTGTTCTATCTGCAAATTCATTTGGTTTAGGGGAA
GAATCGTAGATCAGCTCAGACTAAGTGAAGTGGTTGAAATAATCCCAAGTTCAATACGATTTGATAGAGATTAAA
AAAGGTTTTGAAATTAAGGGAACATGAGAGTGTGTTTTATTTCTCTGTTTTTGTATGTATGTTTCTATGTTGTGT
GGTGCGTAGATTGTGAGGGACTTGTGGTGTTTGCCAAAAACATACTAAATTTGGTAGAAACTTTATCAGAAGAGG
AAAGGGGGGGAGAGAAATAAAGTGGTTAGTGGTTGAAGAAACTCTTAAAACTCTCTCTCTCTCTCTCTCGCTCGC
TCATTAAAACCACTTTTTATTGTCTTCTTGGTTTGGTTAGAGAGGTTTTAGAAAGAATCGTCTCTTTTTGCGCTA
TTCCTTTTCTTTATTGGCAACTAGTTGGTGGGTTTGATGTACCTAAAATGAGCTGCAGCATTATCTTATGTTATT
AATACTTTAATATACACAAGTGGGGGAATCATTTCTGATTTCGTTGTCATCCATAATCAAGTTTTGCTCTTTTAT
TATTCAACTTGCTCATTTGTTTAATGATGATATGTAGAGGAGTTTAGTCTTTCTTTTTGGAAACATTTTGCCTAT
TGTTAGATAATAATCTCTCTGTTTGCATCATTTATCTATTAGAAAAGGTTGAATATAATCTGCCTCTTACATCTG
TCTCAGAGGGAGAAAGCTTCTGTTATTCTCTCTTTTTTTTTTTTGTTCCTTTCTTTCAGTTTTCTCTGTTTTTAG
AAGTGGGACTCTGCCCCTTTTCTTTTTCTTTTCAATAAAGACAGCAAAAACAAAAAGAAAAGAATTTGTACCATC
CATCAGAAGTGTTGCAGTATGAAAAGAAACAAAATATTAAGTACGGAAGATCTTCGGATATAATGTAATGGTTGC
AATTTCCAATGGTGTAAAATGGTCCTTCACATTTATGTCTCACTTACCAATATATCTATAACCTGTTTTTGTGAT
TAGAGTTGGCCACGTTCTCTGCTCAAGCTTCATACATCTTGTTTAGCGAATCTGTATCTGGGTTGCAGAGAACCA
TTCCATGTAGGCAATGATGGCTTAGGGTAGACTAAAATCCGTTTTTTTAAGTAAAACTTTTGACAATGAATTGTA
AGGAGAAATGTTGCAGTGTATGTTGCTTGGTTACCTTACCTACCAACCACACTCTGCAACTATGCAATTACCTTT
TTCGTTGGCAGCATGAAATCAGTCATAGAGCGATACACCGATGCCAAAGGGGAAACCAGTTCGGAAAATGACCCC
GCTTCAGAAATTCAGGTACTACCTTTTTTGTTGCTCCTCCTTATGGATTTTAACGAGTAATCAAGAGAACCAATT
ATGTGCTTGAGTTAAGAATATACCAAACAAGTAACATACAAAGACAGAGAGACTGTTATTGGTGCTTTTTACTGA
TGTTAGCCTATTACAACTTTATTGGCATTTCCCCACTTTTAATTCCTCTCCAGCTGATCCTGGTTTCAATATCAG
ACTTCTGAGTCTAAATACCACACATCATTTGTTGAAGTAAGTTTGCTATAAAACACAAGACATGCCCACCTGACC
ATCAGATTACATACATAATTTTGCATTTTCTTATTCAAGAATCCGTTCTAGACATAACATACAACATTATGTTTA
ATCCAAAGACAAAATACATGTTTTTCAATCACCTGAAAATCATTTCTACTATATTACAGTTCTGGCAAAAGGAGG
CTGCAATTCTAAAGCGACAGCTACATAACTTGCAAGAAAACCACCGGTATGTTTTGCATCCACTTAGATTGTCAA
ACACTCTCGATAGTGGTTACAACTAGAACAGGGAAGGATGTACTAAAATGCAAAATGAAAACTTATATGTAGTCT
AAGGCTGGATTTTTCAGAACCCAAAAGTATCCTACCATTTGCATAGGAATGGTCGATATTAAGAAGAAGAAAAAA
ACCAAACAGATTAAGAATGTTTTTCAACCTCTTAAGGTTTTACCAGGATACTGGATGTTTTCCCCAAAACCACCA
TTTTATCTCTACCACCTCCAGCATCTAAAGGCTAAACCATCCATAAGAAGGAAATAACCAAACACTTTCACACTC
ACACTTTCAAACACTTTGACAGGCAAATGATGGGGGAAGAGCTCTCTGGACTAAGTGTAGATGATTTACAGAAAT
TGGAAAATCAGCTTGAATTGAGCCTCCGTGGCGTTCGGATGAAAAAGGTTTTGATGCTGTGACGATAAAGTCAAT
TTTTGGAACCATTTAGAAGAAGTAATAGCTGTACTATATCTACTATGTTCTTGCCTTTTTCAGGATCAAATGTTA
ATCGAAGAAATACAAGAACTTAACCGAGAGGTAGGACAATGGTCACCATAATATAGATTTTAGACATTCTCCACT
CTGTTGGTAAGGAAGATGCTAATTTTTCTTATCTTTTCGGATGCAGGGGAATCTCGTGCACCAAGAGAATTTAGA

CCTCCACAAGAAAGTAAACTTAATGCACCAACAGAATCTGGAATTACATGAAAAGGTACCTTTGGGGTTTTTTTT
TGGAAGCATCTAAGAGCCAAGATTATTAGAATCTGTAACATGCTAGAACATTTTATATTTTCTTATCTAAGGATA
ATATGTGCATATTCCAGGTTTCAGAGGTTGAGGGTGTGAAAATCACAAACAAAAATTCTCTTCTCACAAATGGTC
TAGACATGAGAGATAACTCGAGCGAACATGTCCATCTTCAGCTCAGCCAACCGCAGCATGATGAGACACATGCAA
AAGCTATCCAACTCAACTACTTTTCCTTCATTGCATAATATATAAGTGGTGTGCATACGCACACACTTATGTTGA
CCTCTCGTTCGGAATCATATCTCAATTTAACTGTTGCAGCTTGTTACTGCATAAGCAAAGAAACATAAACATGAT
CAGTTTGCAATCCATTTCTATCAAACACCAGCTTTGTAACTCTTTCAAAAAACTTCTTTCTCCCTGCAAAATAAC
TTTGTAACTGTTGGTTTGGAGCTTAAGCCAATGAGTAGTTTGATGATCATAGGAAATGAGTGTTTTCACATAACC
TCGTAGACTTCCGTTGCATTCTAGTTTTCTCATCCAAAGCAATTTGTAAAATGTGAGTAAGTTAGCAGTAATAAA
A

*cru-miR824*

>scaffold_7

ATATAGAAGTAGTACATAAATATATACTTAATAGTAATAGAGAAAGACGCACGAGCCGCTTCAGTTGCGGCTTTA
TGCGAGACCTACATGTCGGCCTCAATTAAAAATTGAAATCTTCTCTTCTCTCTCTCTCTCTCTGATTAACGCA
CTAATCAATATAATTAAGATATAAAAATCTGTGGCATTACTTATATAAAACCCTCCCTCTCGTCTC**CTCACCTTA**
**ATCATCATCATCATCAGGACAGTGAGAGTGCTCGCTCTCTCTTCCCCTCTTTTAGCTCTCTCTCTATTTCCTCTC**
**TCTTGCCCTAGCTTCTTCATCTACCCCCTCAATTATTCAGCTACTATATAACTCATCGTCTTCACCTCTCACACG**
**AAACGCCCTTTTGTGTTGATTTTTCTGCCG**GTGAGATCTCGTCACGCGAGAATCGCCTTTTTTTCTTAAAAGTTG
TCAAATGTTCTTCATAGCGTCGATTACATCTCTGATTTAGTGTATCTCAACTTTGCGATTTGATTTTACAGATAT
TACATTTGGAATCATTTTTTGAAATTCCTTGAAATTCTCAAATAGAGTTTTTATGCAGTGATAGATCAGCTGCAT
GAGAGCTTTTTTTTAGATTTATAGCATCTCTGAGTTGCAGATCCAATTTTACTGTTCTGGACTTCTGGTAATCAT
CATCATTTTGGATAATTATGTATCTAAACTTTGGTTTGCATGTGCAG**ATTAATATGCCTATGGGTCGTGTTAGAG**
**AGCATGCCTATGGGTCGGTGTTAGAGAGCTTTCTCAATCAATCTTGAGTACTGGATCTGTCTCCTCTTGTTTCCA**
**G**GTTTGGATTCTCATCTTCACTTTCTTTTTTATTATTATTAGATGTATTTGCTAGTTTACAAATTTCGTACATTT
GAGATCATGTCTCAGTGTTTTTTTCTTCTTTTTCCGGAAGTTCAAAAACACATAATAATGTGTATATATATATGT
ATATCTTATGATCCTTGTCATTCTAATGGGATTGCTATTGTCATTTTGTTGATTGTTACTTAGTCTATTTGTATT
CCTCTTACTTAAGAACAGCTTAAATGCTTATGGAGCATAAAATAAAAGACTCTTTACCTATCTGTTAAACCAACT
AAATGAAGTCTTTACCCGAAATTATCTGAATCATACAAAACGTCTTTTGTTCTTATTCCTTTTGGATTTAACATA
GCTTATAATTAAGTTGGAACACTTACTATCTATTTGCAAGCTTTTGAGAATTGAGACTTGATCTTTGAGTTTCAT
GTCATCATTTTTTTTATCTTGTCTTTACTCAAACAATACCTTTCAAACACATATATGATAAAATTATAGGCTAAA
TCAACCCCATGAAGCTTATGAATGAGAGTTGAAAACCACCTTTGGAAGTAACACAGCCTCTTTTAGAGAGCAATA
TAGAATAGTCTTCATCTGTGTTCACAGCCTCTTTTAGAGAGCAATATAGAATAGTCTTCATCTGTGTTCACAGTC
TCTCTAGATGTACAATAATGCAATGCTGCTTATTTGAGAAAGCAAATGCCTTGTATAGAAACTCCAACACATCAA
AAAATGTTGGATGCTTTTTCGCACAATGTTTCGAACAAGAGAAAGTTGAGAAAATTAAGTATTTCATTGATTGGA
TATGAGCAAGTCATATTTTTCTTAGATCATGACAATCAAATAGGCTACTTGCGCGAACTTGCAATGCTGCAGGTT
TATATCTACATAAGTATATGCTGTTCTTGGTCATAATATTAATTTGAATTTGCCCTTGTTGATGCAG**TTGCAAAC**
**TGTTACAAATAATCGGGATGCTGGGGCTGATGTGGTTGATTAAACTGAAGACTGGCAGGTTTGCTCGAGTTATCT**
**CTCGTCTCTAGACCATTTGTGAGAAGGGATCTTTTATTTGCACCAATACCTCACCAGTCTCTAAATCTGCAAATA**
**AGTATTATGCTCACTTAAGGAAAATACCTTGTCAATATATCTCACCCTCTTTTAAAACTAATCTTAGAATTCAGA**
**TGCCTTTGGATCCTAAAATACATTGTTGTGCATATATGTAATCTACTGGCGCAGCAGGGTAAGTTTCTTGTGTAC**
**CTCTAGAAATTCTCTTACTGCTTAGGATTCCCGAGCTACCAAAACTTTCTTAAGGTCTACTCTGTAGGTCACAAC**
**TATTATTGCCGCACTCACCTTATTGTTAAGATGGTTATTTCTTCCAATCAATATCTGATCCTTCGGAAGTTAGGT**
**TTAAAAATAAGTAGATAATGTGCATTAGTGACTTTTCCTGATGCTGCACTCATAATTGTTTTGTTTGTGTTCCAA**
**CTCAATTCAAGATGATTCTCGAAATCATATAAATTCATTAACACCATGAACATGATAGCTAAGAATTTGTATGTC**
**TTACAACTAGTAGACACCCTAATTATAAACATAGGGACTGGTGAGGGGTTTGGTTATGTTCCCTTCTCATCGATG**
**GTCTAGATGTGCGAGGTGACTCTCATGGAGGTAAAGAAAAATGGTGAATTTGTGAAGCTCCCATCATGATGATAA**
**TTCCTAAAAAGGTTAGATTTCAGATTTTGAAATTGTTGTATTGCATGATTTCCAGATCAATAATGGATTTACTAA**
**TACACCACAGGTGATGGTTCCAAGTTATAGTTGTGTAATCCTTTTTTCTTTTTGTTGGGATTTAAGTATATCATC**
**CCACTTTTCAAACTAATGAAAATGATGGATGCAGTAGTGAAATTGTTGCAAGGTGAATATCTTTGTTGCCTTTTC**
**AAGAACCAACGAAGACAGAAACAAGTTGTAAAAGAATGCAAGTGTGAATACGGATGAAGGAGGAAAGTGGAGGGA**
**AGGTGTGTTACCTTTGACTTGTTGTCACTGATTATCTCACACTCATCCAATTCATGCTGCAAGAGGAAACAACTA**
**AAGGAATGAGTGACGTAACACCAAGCATCAGGAGTTTGCTTGTAGATTTGATAATACATTCCATTACTAACGTAA**
**TCTTTGTCATATTAGGTAGACTCTTGACTACAAATGAAGACTATGTAACTTAG**CTAATGTATATCATTAACTTTT
TAAAGTGTTTTCATGAATAACGTTAATACATGGAGTCAAACAAATGTGATGGTGTTTAGCGTGAATATATATAAT
ATCACTTCTAAGTTCTAATACAGTGCGACTATGGTTTCCAACAATATTTAACATAAAAATCATCATATCGATCAC

#### Camelina sativa

*CsaAGL16_1*

>contig_CM002734

aacaacaacaacagcttaacccaccaccaccaaatctctcagctgaaccgtttcttctctcacctccggttatcg
gattagccagatctcgtctcaatttcaaaaaaaaaaaaggtatataattaattcgattcttttgttgttgaattg

gggaatttgaatttgaaattgacgatgatgtttaatttgtggagttgaaagaaatgggaaggggaaagatcgcga
taaagaggatcaataactcaacgagccgtcaggttacgttctcgaagcgaaggaatggtttgttgaagaaagcta
aggagcttgcgattctctgcgatgctgaggttggtgttatcatcttctctagcaccggtcgcctctacgatttct
ccagctccaggtaccctttttttctttcttttggttggttgactttttttttttttttggggtattggtat
tggtttggttggttgattgaggaattaatttctaaatcttgatttcttgttaattgtttattgcaaattttcaa
ttggtttagggaaagactcgtagtagatcaattgaattggctgaaatttccaagtctaacttacggtttaattga
gagattcatagattctaaggtttttgtaattaagggaacatgataatgtgtgttgttgtgttttcttgttttgta
tgtatgtatgtttctatgtttgtttgagatgaggggagctttagattgtgaggtgtttctcgaaacatactaatt
ttggtagacacttttttcagaagaggaaaggtaggggtgggggtgggagagaaaaggaaagtggttagtggttgaa
gaaactcttcactctctctctctctctctctctgtcgcgcataaaccactttttttttagtatttgtcttgttggt
ttggttagagagggtttttttttgggaaagaatcgtctccttttccactcttccttttttcttttattggcaactag
ttggtgggtttgatgtacctaaaatgagctgcagcattatcttatgtttattaatatacacaagtgggatggggg
gaatcatttctgatttcgttgtcatccataatcaagttttgcttttaaattattccacttgctcatttgtttaat
gatatgtaggagtatagtctttcttttggaaacactttgcctattgttataaaaaaatcattatatcatcaatc
ccttccgatttctattttaattaggttgaatataatcttgctcttagatctgtttcagagggagaatgcttcta
tctttgttttgttatttctgtttttttgttcatttttttttagttccctggttttagaactgggttctgtcctt
ttcttttcttttttttgtttctctctttaatataaagatatcaaaaacaaaaaaacgaaaatgaatttgtaagttc
catcagaagtgttacagtcgaaacgaacaaaatattaagtagaaagatcgtcagaaataaatgtaatggtctaaa
atggtccttcacatttatgtctcactatgacactatctataccctgcttctttgttagcgaatctgtatttgggg
tgcagagaatcttcattccatgtaggcaaagatggctaagggtagactaaaatctgtttttaagtaaaaaaaatt
ctgacaatggaagttgtaaggagaaatgctgcagcggatgttgcttggttaccttacctaccaaccacactcttc
aactatgcaactaccttttttttggcagcatgaagtcggtcatagagagatacaccgatgccaaaggggaaaccag
ttcagaaaatgaccccgcttcagaaattcaggtactccctcctttgtggttactcctccctctatggatcccaac
gagtaattcaaagacctaattctgtaattgagtaacgaatataccaaacttagtaacatactaaaacagagagac
tgttattggtgcttttactgatgttagcctattacaacttcattagcatatctcaacattttaattcctcttcat
atcagacttgcagaccataaataccacacatcgttgctgacttaggatttgataaaacacaatacatgacaccag
accatcagattacatacataatgttgcattttcttattcaagaatcctattaatccaaaaacataatacttgttt
tctcagtttcaatcacctgaaaaatcatttctgttatattatagttctggcaaaaggaggctgcgattctaaagc
gacagctgcataacttgcaagaaaatcaccggtatgttttgcatcctctgagtttgtcaaacactctcgacagta
gtaacaactagaacagggaatgatgtactaacatgcaaaaactgataacgtatatgtagttcacggctagttttt
gcagaacccaaaagtatcctaccatttgtataggaagtcaataatattaagaacaaaaagccaaggattaagaat
ggttccccaaaatcaccattgtatctctaccacctccagagttatctatacctccacacatcttaaccatccata
agaaggaaataaccagaaactttgaaactccccatctatttgttttgttttgtttttttggtgaattaggaaatg
tatatagtagtagtaactctggaaaagtatgcatattcagaagagcttgtacttgacaggcaaatgatggggcaa
gaactctctggactaagtgtagacgatttacagaatttggaaaatcagcttgaactgagccttcgtggcgttcga
atgaaaaaggttttgatgctgtgacgatatgtcaatttttggaaccatttaagaagaagtaatagctgtactcta
tctactcttttcttgcgttttgcaggatcaaatgttaatccaagaaatacaagaactcaaccgagaggtgggact
ctatagttaccgtcatatatagattttagacattttcagagtgttagtaagcgcaatgctaatcctttatctttt
cggttgcagggaatctcgtgcaccaagagaatttagaccttcacaagaaagtaaacctaatgcaccaacagaat
atggaattgcatgaaaaggtacctttgaggttttgggaagcatctagcagccaaatgatacaacatttttttctcc
tatctgaggataatatgtgtgtgcttatccaggtttcagaggttgagggtgtgaaaatcacaaacaaaaattatc
ttctcacaaatggtctagacatgagagataactcaagcgaacatgtccatcttcagctcagccaaccgcagcatg
atgagacgcatgcaaaagctatccaactcaattacttttccttcattgcataatatttatggtgtgcgtatgtac
atacgcacacacttatgttgatgacctcgtcggaaacatatctcaattaactgtagcagcttgtcaatgcataag
caaagatacagaaacatgatcagtttgcaatccattttctatcaaacaccagctttgtaacttttttaaaaaaacat

*CsaAGL16_2*

>contig_CM002732

ggattagccagatctcgtcgtctcgatttcagaaaaaggtatttttttttattaattcgatttttttttaaaattga
attcgtgaatttgaatttgaaaattgacgatgatgatgatgatgtttaattttgtgggtgagaaattaaaaaaaca
ggagttgaaagaaatgggaaggggaaagatcgcgataaagaggatcaataactcaacgagccgtcaggttacgtt
ctcgaagcggaggaatggtttgttgaagaaagctaaggagcttgcgattctctgcgatgctgaggttggtgttat
catcttctctagcaccggtcgcctctacgatttctccagctcnnnnnnnnnnnnnnnnnnnnnnnnnnnnnnnnnn
nnnnnnnnnnnnnnnnnnnnnnnnnnnnnnnnnnnnnnnnnnnnnnnnnnnnnnnnnnnnnnnnnnnnnnnnnnnnnn
nnnnnnnnnnnnnnnnnnnnnnnnnnnnnnnnnnnnnnnnnnnnnnnnnnnnnnnnnnnnnnnnnnnnnnnnnnnnnnn
nnnnnnnnnnnnnnnnnnnnnnnnnnnnnnnnnnnnnnnnnnnnnnnnnnnnnnnnnnnnnnnnnnnnnnnnnnnnnnn
nnnnnnnnnnnnnnnnnnnnnnnnnnnnnnnnnnnnnnnnnnnnnnnnnnnnnnnnnnnnnnnnnnnnnnnnnnnnnnn
nnnnnnnnnnnnnnnnnnnnnnnnnnnnnnnnnnnnnnnnnnnnnnnnnnnnnnnnnnnnnnnnnnnnnnnnnnnnnnn
nnnnnnnnnnnnnnnnnnnnnnnnnnnnnnnnnnnnnnnnnnnnnnnnnnnnnnnnnnnnnnnnnnnnnnnnnnnnnnn
nnnnnnnnnnnnnnnnnnnnnnnnnnnnnnnnnnnnnnnnnnnnnnnnnnnnnnnnnnnnnnnnnnnnnnnnnnnnnnn
nnnnnnnnnnnnnnnnnnnnnnnnnnnnnnnnnnnnnnnnnnnnnnnnnnnnnnnnnnnnnnnnnnnnnnnnnnnnnnn
nnnnnnnnnnnnnnnnnnnnnnnnnnnnnnnnnnnnnnnnnnnnnnnnnnnnnnnnnnnnnnnnnnnnnnnnnnnnnnn

nnnnnnnnnnnnnnnnnnnnnnnnnnnnnnnnnnnnnnnnnnnnnnnnnnnnnnnnnnnnnnnnnnnnnnnnnnn
nnnnnnnnnnnnnnnnnnnnnnnnnnnnnnnnnnnnnnnnnnnnnnnnnnnnnnnnnnnnnnnnnnnnnnnnnnn
nnnnnnnnnnnnnnnnnnnnnnnnnnnnnnnnnnnnnnnnnnnnnnnnnnnnnnnnnnnnnnnnnnnnnnnnnnn
nnnnnnnnnnnnnnnnnnnnnnnnnnnnnnnnnnnnnnnnnnnnnnnnnnnnnnnnnnnnnnnnnnnnnnnnnnn
nnnnnnnnnnnnnnnnnnnnnnnnnnnnnnnnnnnnnnnnnnnnnnnnnnnnnnnnnnnnnnnnnnnnnnnnnnn
nnnnnnnnnnnnnnnnnnnnnnnnnnnnnnnnnnnnnnnnnnnnnnnnnnnnnnnnnnnnnnnnnnnnnnnnnnn
nnnnnnnnnnnnnnnnnnnnnnnnnnnnnnnnnnnnnnnnnnnnnnnnnnnnnnnnnnnnnnnnnnnnnnnnnnn
nnnnnnnnnnnnnnnnnnnnnnnnnnnnnnnnnnnnnnnnnnnnnnnnnnnnnnnnnnnnnnnnnnnnnnnnnnn
nnnnnnnnnnnnnnnnnnnnnnnnnnnnnnnnnnnnnnnnnnnnnnnnnnnnnnnnnnnnnnnnnnnnnnnnnnn
nnnnnnnnnnnnnnnnnnnnnnnnnnnnnnnnnnnnnnnnnnnnnnnnnnnnnnnnnnnnnnnnnnnnnnnnnnn
nnnnnnnnnnnnnnnnnnnnnnnnnnnnnnnnnnnnnnnnnnnnnnnnnnnnnnnnnnnnnnnnnnnnnnnnnnn
nnnnnnnnnnnnnnnnnnnnnnnnnnnnnnnnnnnnnnnnnnnnnnnnnnnnnnnnnnnnnnnnnnnnnnnnnnn
nnnnnnnnnnnnnnnnnnnnnnnnnnnnnnnnnnnnnnnnnnnnnnnnnnnnnnnnnnnnnnnnnnnnnnnnnnn
nnnnnnnnnnnnnnnnnnnnnnnnnnnnnnnnnnnnnnnnnnnnnnnnnnnnnnnnnnnnnnnnnnnnnnnnnnn
nnnnnnnnnnnnnnnnnnnnnnnnnnnnnnnnnnnnnnnnnnnnnnnnnnnnnnnnnnnnnnnnnnnnnnnnnnn
nnnnnnnnnnnnnnnnnnnnnnnnnnnnnnnnnnnnnnnnnnccccctttttttttttttttttggttggttgac
tttttttttttttttggtattggttggttggtccatgtgattttgattgaggaattaatttatctgcaaatttt
caattggtggtttagggaaagaatcgtagtagataaattgaattggctgaaaaatctaagtctaacttacggttt
aattgagagattcatagattctaaggtttttttgtaattaagggggaacatgagaatgtgtgttgttgtgttttc
ttgttttatttgtatgtacgtatgttttctatgttgtttgagaattgagatgagaggggatttagattgtgagg
tgtttctcgaaacatactaattttggtagacacttttttcagaagaggaaaggtaggggtgggggtgggggaga
gaaaaggaaagtggttagtggttgaagaaactcttaactctctctctctctctctctctctgtcgctcataaa
ccactttttagtatttgtcttgttggtttggttagagagggttttttttagaaagaatcgtctctttttccac
tattcctttttctttattggcaactagttggtgggtttgatgtacctaaaatgagctgcagcattatcttatgtt
tattnttgtgttttcttgttttgtatgtatgtatgtttctatgtttgtttgagatgaggggagctttagattgtg
aggtgtttctcgaaacatactaattttggtagacacttttttcagaagaggaaaggtaggggtgggggtgggagag
aaaaggaaagtggttagtggttgaagaaactcttcactctctctctctctctctctctgtcgcgcataaaccact
tttttttagtatttgtcttgttggtttggttagagagggttttttttagaaagaatcgtctcctttttccactct
tcctttttctttattggcaactagttggtgggtttgatgtacctaaaatgagctgcagcattatcttatgtttat
taatatacacaagtgggatgggggaatcatttctgatttcgttgtcatccataatcaagttttgcttttaaatt
attccacttgctcatttgtttaatgatatgtaggagtatcgtctttctttttggatacattttgcctattgttat
taaaaatacattatatcatcaatcccttctgatttcctattttagaaaaggttgaatataatctttgctcttag
atctgtttcagagggagaatgcttctatctttgtttttgttattctctgtttttttgttctcttttgttcagctct
ctgtttttagaactgggttctgtcctttttttttttctttttcttttcaatataaagatatcacaaacaaacacga
aaatgagtttgtaacttccatcagaagtgttacagtcgaaacgaacaatatattaagtagaaaagatcgtcagaa
ataaatgtaatggtctaaaatggtcctttacatttatgtctcactatgacactatcgataccctgcttctttgat
tagagttggccacattctctggcgaagcttcatacatcatgttagcgaatctgtatttggggtgcagagaatctt
cattccatgtaggcaaagatggcttagggtagactaaaatctgtattttaagtaaaaaaaatttgacagtggaag
ttgtaaggagaaatgttgcagcggatgttgcttggttaccttacctaccaacaacactcttcaactatgcaacta
cctttttttggcagcatgaagtcggtcatagagagatacagcgatgccaaaggggaaaccagttcagaaaatga
ccccgcttcggaaattcaggtactcactcctttcttgttgttcctccctatatggatctcaacgagtaattcagg
gaaccaatgctgtaattgagtaatgagtataccaaacttagtaacatactaaaacagagagactgttattggtgc
ttttactgatgttagcctattacaacttcattagcatatctcaacattttaagtcctcttcatatcagacttgca
gaccactaataccacatatcatatgctaaattagggtttgctataatacacaatacatgcaaccagaccatcaga
ttacatacataatgttgcattttcttattcaagaatccgttctagactagataacatattaatccaaaaacataa
tacacttgttttatcagtttcagtcacctgaaaatcatttctattatattctagttctggcaaaggaggctgcg
attctaaagcgacagctacataacttgcaagaaaaccaccggtatgttctgcatcctctgagtttgtcaaacact
ctcgatagtagtaacaactagaacgaggaaagatgtatgaacatgcaaaatgacaacgtatatgtagttcacggc
tagattctgcagaatcgaaatgtttcctaccatttgcataggaatggtcgatattaggaataaaaagcctaattg
attacgaaagttttccaacctcttgaacaccattgtatctctaccacctccggagttatatacctccatacatct
aaaccatccataagaaggaaataactaggaactttcacactccccatcatttttgtttgttttgtttttgatgaa
ttaggaaatgtatatagtagtagtaactctggaaaagtatgcatattcagaagagcttgtacttgacaggcaaat
gatggggcaagaactctctggactaagtgtagacgaattacagaatttggaaaatcagcttgagttgagccttcg
tggcgttcgaatgaaaaaggttttgatgctgtgacgatatgtcaattttgggaaccatttaagaagaagtaatag
ctgtactctaatctattcttttcttgccttttgcaggatcaaatgttaatccaagaaatacaagaactcaaccga
gaggtgggactatggttaccgtcatatagattttagacattttccacggtacagtgctaaatcctttttcttttttg
gttgcagggactcttgtgcaccaagagaatttagaccttcacaagaaagttaacctaatgcaccaacagaatat
ggaattgcatgaaaaggtacctttgaggttttgagaagcatctaggagtcacatgatagaacatttttttctcgta
tctgaggataatatgtgtgtgcttattgcaggtttcagaggttgagggtgtgaaaatcacaaacaaaaattctct
tctcacaaatggtctagacatgagagataactcaagcgaacatgtccatcttcagctcagccaaccgcagcatga
tgagacgcatgcaaaagctatccaactcaactactttttccttcattgcataatatatgtggtgtgcgtaaatacg
cacacacttatgttgatgacctcgtcggaatcatatctcaattaactgtagtagcttgtcactgcataagcaaag
atacataaacatgatcggtttgcaatccattttctatcaaacaccagctttgtaacttttaaaaaaaacttgttt

*CsaAGL16_3*

>NC_025690.1

CAACAACAGCTTAACCCACCACCACCAAATCTCTCAGCTGAACCGTTTCTTCTCTCACCTCCGGTTATCGGATTA
GCCAGATCTCGTCTCAATTTCAAAAAAAAAAAAGGTATATAATTAATTCGATTCTTTTGTTGTTGAATTGGGGAA
TTTGAATTTGAAATTGACGATGATGTTTAATTTGTGGAGTTGAAAGAAATGGGAAGGGGAAAGATCGCGATAAAG
AGGATCAATAACTCAACGAGCCGTCAGGTTACGTTCTCGAAGCGAAGGAATGGTTTGTTGAAGAAAGCTAAGGAG
CTTGCGATTCTCTGCGATGCTGAGGTTGGTGTTATCATCTTCTCTAGCACCGGTCGCCTCTACGATTTCTCCAGC
TCCAGGTACCCTTTTTTTCTTTTCTTTTTGGTTGGTTGACTTTTTTTTTTTTTTTTGGGGTATTGGTATTGGTT
TGGTTGGTTGATTGAGGAATTAATTTCTAAATCTTGATTTCTTGTTAATTGTTTATTTGCAAATTTTCAATTGGT
TTAGGGAAAGACTCGTAGTAGATCAATTGAATTGGCTGAAATTTCCAAGTCTAACTTACGGTTTAATTGAGAGAT
TCATAGATTCTAAGGTTTTTGTAATTAAGGGAACATGATAATGTGTGTTGTTGTGTTTTCTTGTTTTGTATGTAT
GTATGTTTCTATGTTTGTTTGAGATGAGGGGAGCTTTAGATTGTGAGGTGTTTCTCGAAACATACTAATTTTGGT
AGACACTTTTTCAGAAGAGGAAAGGTAGGGGTGGGGGTGGGAGAGAAAAGGAAAGTGGTTAGTGGTTGAAGAAAC
TCTTCACTCTCTCTCTCTCTCTCTCTCTGTCGCGCATAAACCACTTTTTTTTAGTATTTGTCTTGTTGGTTTGGT
TAGAGAGGGTTTTTTTTTGGGAAAGAATCGTCTCCTTTTCCACTCTTCCTTTTTCTTTATTGGCAACTAGTTGGT
GGGTTTGATGTACCTAAAATGAGCTGCAGCATTATCTTATGTTTATTAATATACACAAGTGGGATGGGGGGAATC
ATTTCTGATTTCGTTGTCATCCATAATCAAGTTTTGCTTTTAAATTATTCCACTTGCTCATTTGTTTAATGATAT
GTAGGAGTATAGTCTTTCTTTTTGGAAACACTTTGCCTATTGTTATAAAAAAATCATTATATCATCAATCCCTTC
CGATTTTCTATTTTAATTAGGTTGAATATAATCTTGCTCTTAGATCTGTTTCAGAGGGAGAATGCTTCTATCTTT
GTTTTTGTTATTTTCTGTTTTTTTGTTCATTTTTTTTTAGTTCCCTGGTTTTAGAACTGGGTTCTGTCCTTTTCTT
TTCTTTTTTTGTTTCTCTCTTTAATATAAAGATATCAAAAACAAAAAAACGAAAATGAATTTGTAAGTTCCATCA
GAAGTGTTACAGTCGAAACGAACAAAATATTAAGTAGAAAGATCGTCAGAAATAAATGTAATGGTCTAAAATGGT
CCTTCACATTTATGTCTCACTATGACACTATCTATACCCTGCTTCTTTGTTAGCGAATCTGTATTTGGGGTGCAG
AGAATCTTCATTCCATGTAGGCAAAGATGGCTAAGGGTAGACTAAAATCTGTTTTTAAGTAAAAAAAATTCTGAC
AATGGAAGTTGTAAGGAGAAATGCTGCAGCGGATGTTGCTTGGTTACCTTACCTACCAACCACACTCTTCAACTA
TGCAACTACCTTTTTTTGGCAGCATGAAGTCGGTCATAGAGAGATACACCGATGCCAAAGGGGAAACCAGTTCAG
AAAATGACCCCGCTTCAGAAATTCAGGTACTCCCTCCTTTGTGGTTACTCCTCCCTCTATGGATCCCAACGAGTA
ATTCAAAGACCTAATTCTGTAATTGAGTAACGAATATACCAAACTTAGTAACATACTAAAACAGAGAGACTGTTA
TTGGTGCTTTTACTGATGTTAGCCTATTACAACTTCATTAGCATATCTCAACATTTTAATTCCTCTTCATATCAG
ACTTGCAGACCATAAATACCACACATCGTTGCTGACTTAGGATTTGATAAAACACAATACATGACACCAGACCAT
CAGATTACATACATAATGTTGCATTTTCTTATTCAAGAATCCTATTAATCCAAAAACATAATACTTGTTTTCTCA
GTTTCAATCACCTGAAAAATCATTTCTGTTATATTATAGTTCTGGCAAAAGGAGGCTGCGATTCTAAAGCGACAG
CTGCATAACTTGCAAGAAAATCACCGGTATGTTTTGCATCCTCTGAGTTTGTCAAACACTCTCGACAGTAGTAAC
AACTAGAACAGGGAATGATGTACTAACATGCAAAAACTGATAACGTATATGTAGTTCACGGCTAGTTTTTGCAGA
ACCCAAAAGTATCCTACCATTTGTATAGGAAGTCAATAATATTAAGAACAAAAAGCCAAGGATTAAGAATGGTTC
CCCAAAATCACCATTGTATCTCTACCACCTCCAGAGTTATCTATACCTCCACACATCTTAACCATCCATAAGAAG
GAAATAACCAGAAACTTTGAAACTCCCCATCTATTTTGTTTTGTTTTGTTTTTTGGTGAATTAGGAAATGTATAT
AGTAGTAGTAACTCTGGAAAAGTATGCATATTCAGAAGAGCTTGTACTTGACAGGCAAATGATGGGGCAAGAACT
CTCTGGACTAAGTGTAGACGATTTACAGAATTTGGAAAATCAGCTTGAACTGAGCCTTCGTGGCGTTCGAATGAA
AAAGGTTTTGATGCTGTGACGATATGTCAATTTTTGGAACCATTTAAGAAGAAGTAATAGCTGTACTCTATCTAC
TCTTTTCTTGCGTTTTGCAGGATCAAATGTTAATCCAAGAAATACAAGAACTCAACCGAGAGGTGGGACTCTATA
GTTACCGTCATATATAGATTTTAGACATTTTCAGAGTGTTAGTAAGCGCAATGCTAATCCTTTATCTTTTCGGTT
GCAGGGGAATCTCGTGCACCAAGAGAATTTAGACCTTCACAAGAAAGTAAACCTAATGCACCAACAGAATATGGA
ATTGCATGAAAAGGTACCTTTGAGGTTTTGGGAAGCATCTAGCAGCCAAATGATACAACATTTTTTCTCCTATCT
GAGGATAATATGTGTGTGCTTATCCAGGTTTCAGAGGTTGAGGGTGTGAAAATCACAAACAAAAATTATCTTCTC
ACAAATGGTCTAGACATGAGAGATAACTCAAGCGAACATGTCCATCTTCAGCTCAGCCAACCGCAGCATGATGAG
ACGCATGCAAAAGCTATCCAACTCAATTACTTTTCCTTCATTGCATAATATTTATGGTGTGCGTATGTACATACG
CACACACTTATGTTGATGACCTCGTCGGAAACATATCTCAATTAACTGTAGCAGCTTGTCAATGCATAAGCAAAG
ATACAGAAACATGATCAGTTTGCAATCCATTTTCTATCAAACACCAGCTTTGTAACTTTTTAAAAAACATTTCCT

*csa-miR824a*

>contig_CM002738

tattatcagtttcgcacgcaagatgttacatttgtaaaacgtgttattgaaaagtagcagggaattgaacctagc
caatcatcaccacccaggttcattcaaacccaacatatcacatgtacaaaaaaaaagatgaaaccaatattttcg
aaattgggatattgtgaatatgatttctatgttaaaacttattcggaaaccaaggtcccactact**aaaaataata**
**ttttcatttacgctcactagtgacacattggctaagttaaatagtctcatttgtagtcagaaactacggaagatg**
**acaaagattacattagtaatggaatgatgatcaaatctacaagcaaactcctaatgcttggcgttacgtcactca**
**ttccttttgttgtttcctgttgcagcatgaattggatgagcgagagcatcagtgaccacaagtcaaaggtacaca**
**ttccgtccactctcctctttcatccatatacacgctttattttacaacctgtttttgtcttctttacttcttgca**
**aaggcaacaaacctattcaccttgcaacaaattcactactgcatccatcatttacattagtttgagttatgggat**
**gatttactcaatccccaaaaaagcttacacaactatagttggaaacatcatcacctgtattggaggatccattgg**

**cctggaaattatgcaatacaacaaattcacaagctgaaaatctaacccactttttaggaattaccattattatct**
**tgggaacttcacaaattcaccattttatctttacctccatgagagtcatctcgcacatctagaccatcgatgaga**
**agggaatataaccaaccacctcaccaatccccatttttatttattaggaaactagtgtaagacgatacaatttct**
**tagctaccatgtccgtgttgtaaatgattttatatgaattcgagaaccatcttaaattgagttggaacataaatg**
**gaaaaaaaaagttaatgagtgcagcatctggaaataccactaatgcacttttatctacttacttataattttaaa**
**acccaacttctgaaggatcagattttgatggaagaaataaatcaccttaactatgaggagtgcggctgtaatagc**
**tgtgacctacagagtagacctttcttatggatggggttcggctacagttgtttaatttcttaaagatttggtagc**
**ttgggaatcctacaaagtaagagaatatctagaggtacagaagaaactaaccctgacgtgccaagagattacata**
**ttcgcacaacaatgtattctaggctacggaagcatctaaaacataggatcggtttttagaaagggtagtgtgata**
**tgttgacaagctatattccttagctgagaaatagacttattgcagatatagagactggtggggtattggtgaaaa**
**caaaagatcccttctcacaaatggtctagacaagagagataactcgagcaaaccttgccaagtcttcggcttaat**
**caaccacagcggccccataccaaaatggaatctcagtcaatcttgtctcgactat**ctgtagcaaacaaaggaaaa
tgaatattcagattcatattcaaattcataatcataaaaacaagaacagcacacagcacacacttatgttgatat
aaacttggaattcatgcatagcaagttcatgcaattgcaaagcagcctttctaattttgtaagaaaaataggact
tgcttatgttcaatcaatgaaattcttagtttctcgacagtagaaacaagagttttctttactttctcttgtttg
aagacaataagcaaaacaagcatccaacattctcaatgtattggagttactattattctagccatatgtac**cttc**
**tcaaagaaactacattgcatatcagtggcatccggagactgtgaacacagatgaaaactatcctaaaagaggttg**
**tatcactcccaagggtgttttaatcaagattgagaaagctctcatcctctctctctctctctctctctctcagcc**
**atgggcatatcaaaat**ctgcacacatgcaaaccaaactttagatacatataattagtaaccaaataaaacataat
cctaaatcggatctgcagctctgaaaaatctgataagtaaaagctcatgcagctgatctatcatcatcattcatc
actaataattcctatttgaagcggaaacttatatagaaatcatcaaggggggggnnnnnnnnnnnnnnnnnnnnn
nnnnnnnnnnnnnnnnnnnnnnnnnnnnnnnnnnnnnnnnnnnnnnnnnnnnnnnnnnnnnnnnnnnn**ggaagtaatc**
**gacgcttcgaataacaacattaagagcttaggaggataaaatggcgattctccgtcacgagatcccaccggggca**
**aataaaatcaatagagaaaagctggtttgaagtgaggtgaaagatgatgttagctaagattgagaggagtagatg**
**aaaaaaaaaaaaaaaaagctagggcaagagagaaagaggggaaaaaaaagagagagagcagtgaggaggaggatt**
**aaggtgaggagacgaaatcgagagagggttttatataagtaatggcaaggaagaacgagctttaattcttttact**
**taattatagtaatagtactgattagtgcgtcgttaatctgagagagagagagag**atttctcaatagtttaattga
attgggaagggtggacatgatgtaggtctcgcatctgaagcggctcgtgcgtctttctctatttttatttttattt
ttattcttttactctagtaaagtacttccactactttatttatatcatcgctgacaattcggttttgtcgactct

*csa-miR824b*

>contig_CM002740
attcactactgcatccatcatttacattagtttgagtaatgggatgatttactcaatccccaaaaaagcttacac
aactataattggaaacatcacctgtgttggaggattcattggcatgaaaattatgcaatacaacaaattcacaag
ctgaaaatctaacccactttttaggaattaccattacta**tcttgggagcttcacaaattcatgagagtcacctcg**
**cacatctagaccatcgatgagaagggaatataaccaaccacctcaccaatccccatttttatttcttaggaaact**
**agtgtagctactagcgtaagacgatacaaattcttagaccttagttatcatgtccatggtgtaaacgattttata**
**tgaattcgagaatcatcttaaattgacttggaacacaaaggaaattaaggaaaagccactaatgcacttttatct**
**acttacttttaatatttaaatatagcttatcaaggatcagattttgatggaagaaataaacacgttaactatgag**
**gagtgcggctctaatagctgtgacctgcagagtagacctttcttataggatggggatcggctacagttctctaat**
**ttcttaaagttttggtagctcgggaatcctacacagtaagagaatatctagaggggtacagaagaaactaaccct**
**gatgtgccaagacattacatattcgcacaacaatgcattctaggctaggaaagcatctgaaacataggattggtt**
**ttaaaagtgtagtgtgagaataatacttagtgcagatttagagattggtgggggtattggtggaaacaaaagatc**
**ccttctcacaaatggtctagacaagagagataactcgagcaaacctgccaagtc**ttccgcttaatcaaccacagc
ggtcccatactaaaatggatttgcgccgcatctcagtcaatcttatctcgattatatgtacccgttttcaactgc
gttgacaaaggaaaatgaatattcagattcacattcatattgtgaccacgagcagcatacacacacttaagttga

*csa-miR824c*

>contig_CM002739
ccatcatttacattagtttgagtaatgggatgatgtactcaatccccaaaaagcttagacaactatacttggaaa
catcacctgtgttggaggatccattggcctggaaattatgcaatacaacaaattcacaagctgaaaatctaaccc
atttttaggaattaccattactatcttgggagcttcacaaat**tcatgagagtcacctcgcacatctagaccatc**
**gatgagaagggaatataaccaaccacctcaccaatccccatttttatttcttacgaaactagtgtagctactagc**
**gtaagacgatacaaattcttagttatcatgtccatggtttactaaacgattttatatgaattcgagaatcatctt**
**aaattgacttggaacacaaaggaaataaggaaaagccactaatgcacttttatctacttacttttaatatttaaa**
**catagcttctcaaggatcagattttgatggaagaaataaaacacgttaactatgaggagtgcggctctaatagct**
**gtgacctgcagagtagagctttcttatggatggggatcggctaaagttctctaatttcttaaagttttggtagct**
**cgggaatcctacacagtaagagaatatctagaggggtacagaagaaactaaccctgatgtgccaagagattacat**
**attcgcacaacaatgcattctaggctaggaaagcatctgaaacataggattggttttaaaagtgtagtgtgagaa**
**taatacttagtgcagatttagagattggtggggtattggtggaaacaaaagatcccttctcacaaatggtctaga**
**caagagagataactcgagcaaacctgccaagtc**ttccgcttaatcaactacagcggccccatacaaaaatggatt

tgcgccgcatctcagtcaatcttatctcgattatatgtacccgttttcaactgcgttgacaaaggaaaatgaata
ttcagattcagattcaaaattcatattgtgacatcgaacagcatacacttaagttgatataaacctgaaattcat

#### Leavenworthia alabamica

##### *LaAGL16_1*

>LA_scaffold2516

GTTTCCTTTCAATTCCCGCTAACTGTTTTTTGAATAAGGTAATGTTCAGATTTCGATTTTGACTTAATTTAATAT
CGTCTCCTGGATTCTGGAGAGTTGATTGATGTTTTTTTTTAATTGACGATGATTATGTTTGTTGTGTAAAAGAAC
TAAGAGTTGTAGATATGGGAAGGGGAAAGATCGCGATTAAGAGAATCAATAACTCAACGAGCCGTCAGGTTACGT
TTTCTAAGCGAAGAAATGGATTGTTGAAGAAAGCAAAGGAGCTTGCTATTTTGTGCGATGCAGAGGTTGGTGTAA
TCATCTTCTCAAGTACTGGCAGGCTCTACGATTTCTCCAGTTCCAGGTTCCTTTTTTTGGGTTGACTTTTGGTAT
TAGTTGTCATGTGATTGATGAGGAAACCAATTCTAGATCTCTAATTTCTTGTTAATTGTGCACATCATATCATCA
TCTACCCATTTATAGATACATTAACTTAGTCTGACAATCGCCGAAATCGAAATTTCTTGGTTTGGTTAACTATAT
GGTTTGATTGAGATTCATAGAATATATAAGTTTAGTTAAGGAATATGAGACTCTGCTTTCTTGGTTTATAATTAT
ATGTATGTTTGAGATAGAGGACATGGAAAAAACAAAGAATAAAAGGATATATAATTCGACAGTAGTAGAAAGATG
GTTTTTTTTTTGTTTGTGTGGAGCGTAGATTGTGAGTGAACTGTAAGTGTTTTTCAGAACATACTAAATTTGGTA
TAAAGTTTTTTCATAAGAGGAAAAAGGGTTAGTGGTTAAGAAACTCAAACTTTTGTCGCTCACCAAGTCATTAAC
CACTTATATTTTTTTATTTGTCTTCTTGCTTTGGTTTTGATAGAATCGTCTCTTTTCACTATTCCCTTTCTTTAT
TGGCTACTAGTTGGTGGGGTTGATGTACCTAGAATTAACTACATTATCTTATATATATAATAATTCACAAGTGGG
AGAATCATTTCTGATTTCTTTGTCATCCATAATCAAGTTATCTTTTTTTAAGTTATCAACATGTTCTTTGTTTAA
TGATATATAGGAGTATAGTCTTTCTTTTTAGAGATATTTTTAGAATTTTTTGAATATCATTATATCATCAATCCC
TTCTGAGTTTTTCTACTTTATGAGTTCTTTCAATTCTTTATTTGAATCTATCATTTTTGCTTACATAATAATTAT
CTGTTTTAAAAGGTTGAATAAGAATATAAGATCTGGCTCTTTAATCTGGCTTTTTTTTTTGTCAACAATCTGGCT
TTTCTTTCTCTTCAACAAAAAGGGGCAAAAGGAAGAAAAAGAGTTTCTAGAATATATCAGAAGTGTTATACGCAGA
AAAAAGTATAGATATTAAGTGGAAAAGATCATCAGAAATATATGTAATGGTTCCAAGAGATTTAGTCATTAGATA
AGGCAGATACTAGAAGCATTCTTTTTAAAGGGTCTTTCACGTTTATGTCTCACTAAGTCATATCTGCACTCTGGT
TTACTGATTAGAATTGGCTAGAATTACATCAACTTTGTCTGAGTTCTCTCATTATTTGCTTAAGTTTCATACTTC
ATGTTACATTTCTATTATAGTGAATCTGTGTTTTGGATGCAGAGAATCCCATTCCATGTAGGCAAAGATGGCTTA
GGGTAGACTAAATCTATTTAAGTAAAAAAAAATTTGACACTGAAAGATGTAAGGAGAAATATTGCAATTTATGTT
GCTTGGGTTACCTTACCTACCAACCACACTCTGCAACTACCTTTTTTGGCAGCATGCAATCGATCATAGAGAGAT
ACAACGATGCCAAAGGGGAAACCAGTTCAGAAAATGATCCTGCTTCAGAAATCAAGGTACTCCTTTATTTGTTAC
TCTTCCCTATGGATTTCAACGAGTAATTTAGGGATCTTTAATTAAGTAACGAATAGACTAAACAAGAGACATACT
GAAGCAGAGAGACTGTTACTTGGTGTTCTTTGTTAATGCTAGCCTATTGCAACTTCATCAACATTTCTCATCTTT
TAAATCCTCTTTAGTTGATCATGATTTCATATCAGACTTGCAGATCTAAATGCCACACATCATATGATGAAGTAG
GGTTTTTTATTCATGACATCTGACCTTCATATTACATATATAATATGTTGCATTTTATTTTTCAAGAATCCATTC
TAGACTAGATAACATAATATAATCCAAAAGCATCATACTTGTTTTCTCAGTTTCAATCACCTAAAAAAATAATTT
CTACTATATTTTAGTTCTGGCAAAAGGAGGCTGCGATTCTAAAGCAACAGCTACATAACTTGCAAGAAAACCACC
GGTATGTTTTGCATCCTCTTAGTAATAACAACTAGAACAGGGGATAACATGAATAGTCATATTACAAATAATCAA
ACACTTCCATACTCCACATCTCTGTTTAGGAAATGTATCTAATAGCTCTGGAGAAAAATGCATATTCAAATGAGC
TTGTACTTGAACAGGCAAATGATGGGGGAAGAGCTCTCTGGACTAAGTATAGAAGACTTACAGAATTTGGAAAAT
CAGCTTGAGTTGAGTTTCCGTGGCGTACGAATGAAAAAGGTAGGTTTTGATTCTCTGAAAACATATCTCAATTTT
TGGAAAAAAAGTAATGCGTTTACTCTATCTACTATTTTCTTTTCTTGACTTTTGTAGGATCAAATGCTAATTGAA
CAAATACAAGAACTCAACCGAGAGGTGCGACTCTCAGATTTATCCGCAGTGTTGGTAAGTACAATGCTAATTGTT
GCTTCTTCTTCTTTGTTCGGTTGCAGGGGAATCTCGTGCACCAAGAAAATTTAGACCTCCACAAGAAAGTAAACC
TAATGCACCAACAGAACATAAACTTACATGAAAAGGTACCTTTGGAATCTGTGGGGCAAGATGATGTTTAAAATG
GTATAACATTTTTGATCTTATCTGATGATCATCATCATATGTGCTTATTCCAGGTTTCAGAGGTCGAAGGAGTGA
ATATGGCAAACACAAATTTTCTTCTCACAAATGGTCTAGAAATGAGAGATAACTCATGCGAACATGTCAATCTTC
AGCTCAGCCAACCACAGCATGATGAGACACATGCAAAAGCTATCCAACTCAACTACTTCTCCTTCATTGCATGAT
AATATTTGGTGTGTCAACACACTTATGTTCACCTCAGCCAAATCATATCTCATTTGAATGTAGCAGCTTACCACT
GCATAAACATGATCCATATCAGACACCAGCTTTGTAACTTTTTCAGACATTTTTTTATCTCTGCTAAGAATCTTT

##### *LaAGL16_2*

>LA_scaffold2880

GCCATGTGTCAATATATAACAAGTGATGTATGTTTTATATAGTAAGAAACCCTTTTGCTTAATCTAAGATTCCCA
AATTTAGCATCATATGTTTTTTCCTTACATAGCTCTCTCCTCTATTTTGATTTCTATTTCTCTCTTAATTAAACA
CAAGAAGAATGGGAAGAGGAAAGATAGCAATAAAGAGGATTGATAACTCATCAAGTCGTCAAGTAACTTTCTCAA
AGCGAAGAAATGGATTGTTGAAGAAAGCAAAAGAGTTATCGATTCTATGTGATGCAGAGGTTGCTGTTATAATCT
TCTCCAGCACTTCTAAGCTTTACGATTTCTCCAGCACTAGGTATAATGTTACCTTTGACTCTTTTCTTTTTTTGT
AAATGTTCATTGATGATTCTAAATCTCCACTTTTTACAATTCATTTCTTGTTTTAATTCTAAACTGTAGATCACA
TATATAAACATTAATTTTAGTCTGTTACTGGATAAAAAAACACTACTTGTTTGGTTGGTGTTGACTTATATGAAA
GGTTCAGATTCATAGATTATAAGGTTTTGTAATAGGAATATGTATAAGGTTGTTGTTGAGATGGAGGACACAACA

AGGAAGATATAGAATTAGGACAATAATAGAATGACTTATTGTTTTTTTTTGTGGAATCATACTAAATTTGGCAGA
AGAGGAAAGTAGTTAGTAAGCATAAACCACTTTTTTTAATTTTATTTGTCTTCTTGGTTTGGTTTAAGAGGTTTT
AATTTTAGAAAGAATCGTCTCTTTTGAACTATTATTGACTTTCTTTATTGGCAACTAGTTGGTGGATGGATCGAT
GTACCTAAAATGAATGACCTTGCATTATCATTTCCTTTTAGTATTATTTTAAAAATACAAAAGTGGGGTGGCGAA
TCATTTCTGATTTAGTTGTCATCTACAATCAAGTATTTGCCATTATATTATTCATTTGGGAAACACTCGTTTTAA
GTGGTATATCATTTCTGCAAAATAATGATTTATATGATTATACTATAGTCTTTCATTTAATGCTTTATTAGAATA
GATTTATAATTAATTCAATAAGGTTGAATAGAACTTGTTGCAGTTCATAAGAAGTAACGCAGTGGTGGTAAAGGG
TCATTTACCTTTATATCCTATAGTCTATACTCCGTTTTGGTTATTAGACTTTCCCACATTTACATCAGTTTTCTC
ATTTTTTTGCTTAACCTTCATACATCAAGTTAGATTTTGATTGTAGCAAATCTATGTGTTTGGGTATAGAGAATC
ACATTCTATGTAGGCAAAGTTGGCTTTTAGGATATAAACTAAATGTATTTTAAGTAGAAATAGTTTGACACTGAA
AGATGTAATGATAAAGTGTTGCAGTTGATGTTGTTTGCAGATGTATAACTACTTTTTTTTGGCAGTATGAAATCT
GTAATAGAGAGATACATAGATTCCAAAGGAGAAACCAATTCAGAAAGCTACCCTGCTTCAGAAATCTTGGTACTC
CTTTTTTGTTCACTCTTCGTTACTCATCCATATCGATTTCAACGAGTAATTCATGGACTCAGTTCTCTAATTTTT
GAGTTAATGTATAGATAAAACAAGTAACGTACTANNNNNNNNNNNNNNNNNNNNNNNNNNNNNNNNNNNNNNNNN
NNNNNNNNNNNNNNNNNNNNNNNNNNNNNNNNNNNNNNNNNNNNNNNNNNNNNNNNNNNNNNNNNNNNNNNNNNN
NNNNNNNNNNNNNNNNNNNNNNNNNNNNNNNNNNNNNNNNNNNNNNNNAGGTTTTGTCATGATGTCTTCTCCAAAATCA
CAATTGTATTGTATCTCTACCACCTCCATAAGAAGGGAATACACCAAACAAAGGCCACAGTCTCCATTTTGTTAT
TCTTCTTGGTGAAAACTACGAAATGTATATAATAACTATGGCAATCTATGTATACTCAAAGAAGAGCTTGTACTT
GGACAGGCAAATGATGGGGGAAGAGCTCTCTGGACTGAGTGTAGAAGATTTGCAAAACTTAGAAAATCAGCTTGA
GTTAAGCCTTCATGGTGTACGAGTGAAAAAGGTTTTGATGCTGTGACATACAACAACTCTTATAATGAATCTACT
CTTCTTATCTACCTACTCTTTCTTTTTTCTTGGCTTTTTTAGGATCAAATATTACTCGAAGAAATACAAGAACTC
AACCGAGAGGTGAAGACTCTGCTTACTACCAAAGTGTTGGTACACACACAATGTATCTAATCCTGTCTTTTTTTT
TCCGGTTACAGGGGAATCTTGTGTACCAAGAGAACTTAGACCTTCACAAGAAAGTTAACCTAATGCACCAACAGA
ACATGGAATTACAAGAAAAGGTATCTTTGGTTTTAACAAGTAACTAGAACATGTTTTGTTTTTATTTGAGGATGA
ATATGTTACAATTCCAGGTTTCAGAGGGTTTGAGAAGTGCAAACAATGATTATCTTCTCACAAATGGTCTAGACA
GGAGAGATAACTCAACCGAACATGTCCATCTTCAACTTAACCAAACTCAGCATGATGAGACGGAGACAAAAGCTA
TTCAACTCACGTACTTCTCTTTCATTGCATAATGTATTCATACTTATGCTGACCATGGCATAATCATATTTTAAT
TAATTGTAGCAACTTCTGTGTAAACGAAACATAAACATGATCAGCCAATATTGATCTCTTTCCTAAAAAAAAAAT
AGAAAAAATAGTTTGTTTGGTAGCAGGCCCTGGCCATTGCCTATTGTTGTAAAAGGTCCAAAGACAAAAAAAAAA

*lal-miR824a*

>LA_scaffold423

AACTGAAGACTGACAGGTTTGCTCGAGTCGTCTCTCATGCCTAGACCATTTGTGAGAAGGGATTCATTGTT
TGTACCCATACCCCACTAATCTTGAAATCTGCAATAAGTATTATGCTAGCTTATTCAACATATTACATTAC
CCTTTTAAAGAACATCATATATTTCATATGCAATAGCCTGAAATACCTTGTTTGGTTATGTAGTTCTGTGT
TTACAGAACTGTTCCCCAAACATCTTCCAATAAAACAGATATTATTACTCTGTAGGTCAAAGTCTCATTAT
TTAGAACTGCTCTCACCTCTTAGTTAAGCTGTTTTCCTTCCATCAAAAATCTTATCCTTCTGAATTGAACT
TAGGTATAAAAGTAGATCAAGTGTATTCATGACTTTTACAGATGCTGCACACATAACTTTTTATAGAGTGT
CTGAGATTTTTCCATGATAGCTAAGAATCTATATCTTATACAATTAGCTACACTAGTTTTCATAATAAATA
AAAAATGTGGAGTGTGGAGATCTTGGGTTTATTCCCTTCTCATCGATGGTCTAGATGTGCGAGGTAACTCA
CTAACTCTCATAAAAGCTA

*lal-miR824b*

>LA_scaffold2880

GTTGAAGATGGACATGTTCGGTTGAGTTATCTCTCCTGTCTAGACCATTTGTGAGAAGATAATCATTGTTT
GCACTTCTCAAACCCTCTGAAACCTGGAATTGTAACATATTCATCCTCAAATAAAAACAAAACATGTTCTA
GTTACTTGTTAAAACCAAAGATACCTTTTCTTGTAATTCCATGTTCTGTTGGTGCATTAGGTTAACTTTCT
TGTGAAGGTCTAAGTTCTCTTGGTACACAAGATTCCCCTGTAACCGGAAAAAAAAAGACAGGATTAGATAC
ATTGTGTGTGTACCAACACTTTGGTAGTAAGCAGAGTCTTCACCTCTCGGTTGAGTTCTTGTATTTCTTCG
AGTAATATTTGATCCTAAAAAAGCCAAGAAAAAAGAAAGAGTAGGTAGATAAGAAGAGTAGATTCATTATA
AGAGTTGTTGTATGTCACAGCATCAAAACCTTTTTCACTCGTACACCATGAAGGCTTAACTCAAGCTGATT
TTCTAAGTTTTGCAAATCTTCTACACTCAGTCCAGAGAGCTCTTCCCCCATCATTTGCCTGTCCAAGTACA
AGCTCTTCTTTGAGTATACATAGATTGCCATAGTTATTATATACATTTCGTAGTTTTCACCAAGAAGAATA
ACAAAATGGAGACTGTGGCCTTTGTTTGGTGTATTCCCTTCTTATGGAGGTGGTAGAGATACAATACAATT
GTGATTTTGGAGAAGACATCATGACAAA

#### Brassica rapa

*BrAGL16_1*

>Brara.A00398

GTCCTGTCTCATTTAAGATCTGGTCTGCTTTCGTATTCTCTACGTGTGTTGTATGTCTGATCTTTGGGTGCTTTT
CGTCTCTGTCGGGATTCTGGGATTCCTTTCACTCTTTCTGGTTTGAGAAAATTAAAATTATTTGAAAATAAATAA
TTCTACTTTGGGTTTCTCACAATTATTGGGTTTATCTATTTATTTTATTCGTAGCGTATACATTTTTTCTTCTTT
TAAATAAAACAAAAAGGGTGCTACTTTATTAATGTTGTACGATACAACAATATTATAATGTCAAAATAAACCACT
TATAGAATGCCAAGTGTCAAGATATGACAAGAGATCTTATGTTTTATATAGTAAGAAGACTTTTGCCCTAAAATT
CCCAAACTAACCATAAACATATGTTCTTTCCTTTTGTGGGCGCTTCTCTGCTTTCTTCTCTGTCTCTTTCTTTCT
ATTTATATTTTCGATCATAAAACCCAAATTGATTTTCCTAGATCTCTTGACCTGTTTTCTCTTATTTGGAAAAAG
GTAATTGATTTCGATTTTTCTTAGTTTAATTGAAATATCTCGCCCTGGCGATGGTTTGACCATGTTTTTGAGTTT
TTTGAATTAGAGGTTGTAAGAAATGGGAAGGGGTAAGATCGCGATCAAGAGGATTGATAACTCGACGAGTCGTCA
GGTGACTTTCTCCAAGCGAAGGAACGGACTCTTGAAGAAAGCCAAGGAGCTTGCGATTCTCTGCGATGCTGAGGT
TGGTGTCATCATCTTCTCCAGCACCGGCAGGCTCTACGAGTTCTCCAGCTCCAGGTATACCCTTTTCTTCCTTTG
TTCTATGTTTGGTTGACTTTTTGTGGTATTGGTCCAGTGTGATTGATTAGAGAATCTAACCCATGATCCATCCGG
TTTTGTTGACCACAAACCAATTCCAAATTGTTTCCATTCTATTTCTAGTTATTTTTGTTGTTATCTGGAAAATTC
AATTGATTGTGTGCACATCCCTGAAGATGACGTGAGCAGCATTATTTTACCTTGATTCATATTATACGACAATGA
AACCTCAAAGCTGAAAATTGGTATTATGACTTGGTTGGTTTGACTATCTAGATGACTTGGCTGGAAAAAAATAAT
ATACAAAGCCAATACAGTTTGCTTGAGATTCATAGATTACAGGCTTTGTAACGGAATTATGAGATTGTGTTTGTG
GGTTTGTATGTTATGTGTGTATGCTATTGAGATGCAGGACATGACAAAAGTCAAATGGATTTAGGATTGGATATG
ACTAGAAATGATGTAGATTTGTGGTTGATGCGTAGATAGGAAGGGGGACTTGTAGTGTTTCTCAAAAAATACTAA
ATTTGGTAAAAACTGTTTTAAGAATGGAATAAAAAGTGGTTAGTGGTTAGTATGGTCAAGAAACTCTTAACTCTG
TCGCTAATCACCACATTTTTTTTTATGTCTTCTTGGTTAGAAAGAATAGTCTATTGAGCTATTTACTTTTTTTTT
ATTAGCAACTTGTTGTTGGTGGGTTGGATGTACCTATAATGATTTTCATGTTGAATAAGTGGGGCTGTGGGGGAA
TCATTTCTGCTTTCGTTCTCATCCATACTCAAGTTTTCCTATTTAATTCATCCCAATTGGGAAATAAAAGTATAA
AACCACCCCATTTTTAATACTCTACATTCAAGTTGGTCATTGTTTATTTGTTTATAAAGAGTAGATTCTTCTTCT
TTTGTTGGGGAAAATTAGTCCCTTCTGATTTTCTACATTAGGAGGTCTTCCTTAGAGTAAAACTGAAGTTAAACT
ATAAAGTAAAAATATCGTCAGAAATAAATGTAGCTTCAAAAGAGTTCATCCATGAGATTTAAGCCCATTAGATAT
CAGAATAAATGGTGTTAAACCATCCATGTCTCTCTAATTCATAACTTGCAAAGCTGATCACCATAGTCCATTAGA
TATCAGAATCAATAGTATCTGCAGATATACAACTTACCTTCTTTTTGTTTGGCAGCATGAAATCGATAATAGATA
GATACAGCGAGGCCAAATGTGAAACTAGTTCGGAAATCAACCCAGCTTCAGAAATCAAGGTACTTGGTGCTTTTA
GTCTATTACAGCTTCACTATCATGTTTTCAAATTTTGACACCTCTCAAGCTGATCCTGAGCTCATATTGGAAGTA
GGTTTTGCTATATCACACACACACATACAAGTTATGACACCTGACCATCATATTACATTCACTATACTGCATTTT
TCATTTTTTAAAGAATCCATTCTAGACTTGTTAGCACTTAGCATATCAATCTCAAAACCAAAAACAAAAGTTTAT
TTCCTCAGATTCAATCACCTGAAAATAATTCTGATGTGTTAATAGTTTTGGCAAAAGGAGGCCGCGATTCTAAAG
CGACAGCTACACAGCTTGCAAGAAAACCACCGGTATGTGTTTTGCATCCTCGTCAAACACTCTCGATATTAACAA
ACTAGAACACGGAACGATGTACTAAAATGTCTGCAGAAAAGCAAAACTATCCTAGCATTTGCATTTTTTTTAATA
ATGGGAACGTCGATGTTAAAAACAAAATTCCAAAATGATTTACCACCTCTTAAGGACTTTACCATGATGCTGGAT
ATCTTCCCCAAAATCACCACTGTATATATAGTTACTCTGGAAAAAGTTTGCATGTTCACAAGTGATGGAGGAAGA
GCTTGTACTTGGCAGGCAAATGATGGGGGAAGAGCTCTCTGGACTAAGTGTGGAAGATTTACAGAAACTGGAGAA
TCAGCTTGAACTGAGCCTTCATGATGTACGAATGAAAAAGGTATTGATATTGTGACATATATGTTGATATTTTTC
TTTTTCATCTAACGATTGGAAAAAAGAAGTAAAAACTGTACTCTACTCTCACACTTGTAGGATCAAATGTTACTA
GAAGAAATACAAGAGCTTAACCGCGAGGTGGGAATCTGATTACTCGAATATAGAAATTAGACATTTCCACAGTGC
TAATTCTTACATTTTTTTTGGTAGCAGGGGAATCTCTTACACAAAGAGAACTTAGACCTCCACAAGAAATTAAAC
CTAATGCGACAACAGAATATGGAGTTACATAAGAAGGTAATCTTTGTTTATTGGGAGGCAAAAATGCCTTTGAAA
AAAGCTACAACATGTGTTTTCTTATGAATTGTGAGGATAATGTGATATGGCTACCACTTGCAGGTTTCAGAGGTT
GAGGGTGTGAAAAGCTCAGAGAAAATTTCTCTTCTCACAAATGGTCTAGGCATGAGAGATAACTCGAGCGAACAC
GTCCATCTTCAGCTCAGCCAACCGCTGCAGCATGATGAGACGCACTCAAAAGCTATCCAACTAAGCTATTTCTCC
TTCATTTCATAATATATAATTCAATTTCAACTGTAGCAGCTTGCCACTGCATGAGCAAAAAAAAAAAACATGAGC
AGATTGTAATGTTTTCGATCAAAACACCAGCTTTGTATTTTTGTTTTGTTTTACTTATTTCTCTCAGCAAAGAAC
TCTGGTTTGACGTTTAGGTATGTACTGTGCGATGGTCAAAAAACGTAGTGTTCTTTCTTAGGTTCCTAGAAATGT
CAAAAATAATTTTAAGTATAATCAAACCTTAAGTTGTAGTCAAATTTCTCT

*BrAGL16_2*

>Brara.K00360

ATGGGAAGGGGAAAGATCGCGATTAAGAGGATCGATAACTCGACGAGTCGTCAGGTGACTTTCTCCAAGAGAAGG
AACGGCTTGTTGAAGAAAGCCAAGGAGCTTGCGATTCTTTGCGATGCTGAAGTTGGTGTTATCATCTTCTCTAGT
ACTGGCAGGCTCTACGATTTCTCCAGCTCCAGGTATCACTTGTTATCTCTTTGGTTGAATTTTTGATGCTATTGA
TTACTCCATCCATGATCCACCGACGGTTTATTGACCACAAACCAATCATAAATCTCCAGTTTTTTAATTCGATTT
TTCGCTATTTTTGTTCTCTGCAAATTCAATCCACTGAAGATAGACATAAACACCATCTTAGTCTTATGGCAATGG
CTGAAATTTGTACATCACTTGATTGGTTGACTATCTACATATATGAGAGCTGGAAAAAAAAACAAGGCCAGTACG
GTTGTGTTTAAGAGTCATAGATTATATGAGTTCTGTAACGTAATATGAGATTTTGATTTATGGTTTGTATATTAT
GTATGTTTCGATGCAGGACAGGACAAAAGTAAATTGGATAGGACTAGAATGATAGAGATTTGTGTGGAGTGTATA
GATCATAAGGTGGACTTGTAGTGTTCTCTCAAAACCATACTAAATTTAGTATAAAAGGGCAGAAGAAACTCTTCT
TTGGTTTGGTTTTGAGGTTTCTTAGAAAAAGAGTAGTCTCTCTTTCACTGTTTACTTTTTTCTTTATTGGCAACT

AGTTGGTGGGTTTGATGTACCTAAAATGAGCTGCATTATCTAATGGTTAATATATTAATACACTAGTGGGGAAT
CATTTTTGCTTTCGTGGTCATCCATAATCTATCTTTTTCCTATTTAATTTATCCAATTTGGAAAGAAACCATTCC
ATTTAAATGCTCAACTTGGTCATTGTTTAGTTATATATAAAGTATATTCTTTTTACATATCAATCATCAAATCC
CTTCTGATTTTCTACTTTAGAAGGAGTTCACTGAAGTAAAAAGAAAATAAACTATAGAGTAAAAACTAAAAAGAT
TGTCAGAAATAAATGTAGATCCAGAAGAGTTGATGGTTCCATGAGATTTAGATAGAATAGGCAATGGTTCTAAAG
GGACACATTTATGTCTCACTAGTTCATATTTGTCAGCTGACAACAATAATAACTTAACTCTGGTTTGGTGATTAG
AGTTGGCCACTTGTACATAAGTTTGGTCTTGTTTCTTTCATTCTCTACTTAAGCTTCATGCAGCATCATGTTATG
TTTAGAGTATGGAGAATCTGTATTTTTGGTGCAGAGAATTAATTTCCATGTAGGTAAAATTGGCTTAGGGTAGTT
TAAGTAAAAGTTTTGTCACCGAAACTATGTAAGTAGAAATGATGCAGTTGATGTTGCTCTAGTTACCTTACCTGC
CAAACTCACTCCTGCAGATATACAACTTAAATCTTCTTCTTTTTATGGCAGCATGAAATCGGTAATAGAGAGATA
CAGAGATGCCAAATGTGACACCAATTCGGAAATGAACCCAGCTTCAGAACTCAAGGTATTCCTCTTTTGTTTTTT
TTTTGATAAATTCCTCTTTTGGTTTAATGTTTATTACTCCTCTCTATGGATTTCAACGAATAATTGCTGCAGTTA
ATTTTCTAATGAGTTACGAATAGACCAAACAAGTAACATAGTAAAACAAAGAGATAGTTACTTGGTGCGTTTTAT
AATGTTAGCTTGTTACAGCTTCGTTAGCCTATTTCAACTTTTGACTCCTCTCAAGCTGCACCTGAACATAAGATT
ACATACATTACGTTGCATTATCTTTTTCAAGAATCCATTCTAGACTTGATAAGATGTTAATGCCGCAAAAAAAAA
AAATTGTTTTCTCAGTTTCAGTCACCTGAAAATAATTCTGCTATATTATAGTTTTGGCAAAACGAGGCTGCGATT
CTAAAGCGTCAGCTACATAACTTGCAAGAAAACCACCGGTATGTTCTTGCATCCTCTGAGATCGTCAAACACTCT
CGATATTAACATGACATGATGTACTAAAAATGTCAAAGGCTATAAGATTTTTCGGAAAAGTAAAACTATTCCAGC
ATTTGCATTTTTTCAATAAAAGAATGTGGATATCAAAAATAAAATGGAAAACTGTTTTAACACCTCTTAAGGACT
TTACCTGTTACCATGATGCTGGATATCTTCTCCAAAATCACCATTGTTATCTCAGCAAACTCGTGAGTTATATCT
CCACACATCTAGAGTTGTAGACCATCCACAAGAAGTGAATAACCAAACACTTCCACATTCCCTATCTCTCTTTTT
ATTTTTGTTTGGATGAATTAGGAAAGGTATATAATCACTCTAGAAAAGTTTGCATATTCAAAAGTGATGGGGAAA
GAGCTTGTAATTGATAGGCAAATGATGGGTGAAGAGCTCTCTGGACTAAGTGTGGAAGATCTTCAGAAACTGGAG
AATCAGCTTGAGATGAGCCTTCGTGACGTACGAATGAAAAAGGTATAGATAGGGCCTAGGTATAGATGTTGTGGC
AAATAAGTCAACTTTTTCATGTATGTTGTACTTACTCTTTTCTTTTATTGACGTTTGCAGGAACAAATGTTAGTC
GAAGAAATAAAAGAACTGAACCGAGAGGTAGGACTCTGATTAGGCATATCCACATTATTTAATACCTACAAATTG
CTAATTCTTATATTTTCGATTGCAGGGTAATCTCGTGCACCAAGAGAATTTAGAGCTCCATAAGAAAGTAAACCT
AATGCACCAGCAGCACATGGAGTTACATAAGAAGGTATTTTGAGTTTGGAAGGCAGATTAATCTTTCAAAAAACT
AACAATGTTTTTCTTCTTCTAAATTGAGGATAATGTAATCTGGCTACTACTTGCAGGTTTCAGAGGTTGAGAGTG
TGAAAAGCGCAGACAAAATTTCTCTTCTCACAAATGGTCTAGACATGGGAGGTAACTCGAGCGAACATGTCCATC
TTCAGCTCAGCCAACCCCAGCAGCATGACGAAACAAGTTCCAAAGCTATCCAACTAAGCTATTTTTCTTTCATTG
CGTGA

*bra-miR824a*
>Brara.A01447
**TTAGTATTCAATCTCCACACTCTCTTCTCTTCGTAGCTCTCTCTCTATCTTTTGCCCTAGCTAACTTGTTTTCTT**
**CTTCTTTTTCTCAATCCGCTTCACATCAATGCATCCTCACGATTCATGGGCTTTTCCGATTAAGTTTTCTCTGGG**
**ATCTCGAAGAATTGCCATTTTTTTAAATATGCTAAAGCAGACGAGTCGCTTCGATTACTTTTCATTTG**GTTAGTC
TGATTCTGTTTATATAGTTTCTTAACTTGCAATTTGTTTTATGAAATGTTTTGTGTTGTTTGTTAGATCTCTCCA
TATTTATATATTAGTTTGAGAAGTCAGATGTTTGCATGTACAG**ATTAATGCTCATGGATGAGAGGTTTGTCAACC**
**TTTAATGGATCTGTCTCCCCTTCTCAG**GTTTGGTTTCTCATCTCCATTTTATTTGTTAGATGTGTGTTATATAGA
ATGTCTCAGCCTTGATCGTACTTCTGTTGTGTTTTTTCTTTTGTTACTTCCGTTAAGATGCGTACCGTGTAAAAT
AAAAAGCACATAATAATGCAAGATCCCTTGTCTTTCTGCAAGTATTGTCTGATTGTCTAATTACTTAGTGTCTAT
GTTCCTTTCACTAATGAACAGCTTAATAGTGGTTAATTATGCGCATAAAATAAAAGCAAATTATTTTGTCTCTTG
ACCAGAGAAACTGATCAAAACTTAAATCACAGATATGAAACTGACTATCATGATGTGGTTTTCGATTCATTTTCT
CGCATCTTTTTTTTTTGTGTATACCTAAAAAAGAGTTTGAAAGATTCATTAGATTCTCACTTTAAGGTTAAAAAG
GTGTCTGTACCTAAATTACATCATATGTACGAAGTACTAACACATAATTTTTTTGTCTTCTTTTTGCAAATTAAC
ATTGCATCACCCACTAGAGAGTTCTGATGTTTTTTTTTTCTGATTGTAATGCTCTTTTTTCTGACTTCATGGCCT
GTACTAAGTTGAATTCTCACTATGCCTCTATTTGGAAGCTATTTTCAGAATTAAAGCTTGATCTTTGTGTTCCTT
GTCAATAAACGTTGATTTCTGAATAATATATTTTATACAACTATATCTATGATAAGTTAATAGTCCAAAGCAATC
CAGAAAAGCTCAAGGGTGATAGTTGAAAACACCTTAGGAAACATTCGTGTTCATAAGACTCTAGATGTTCAAAAA
TAAATTGTTACTTCTTTGAGAAAGCAGTGGCTTGAGTAGAAACTCCATTAAGTTTGTAATTAGGTTGGGTTCTTA
GATTTAAAAGGGAAGTGAATGGAGCTTGTTACACTGAAAAGAAACTAAGAATTCTGTTGACTAGAGACGAGCAAT
TCAATTTTGTGCATAATTCAGGTTTATGTCCACATATCTTTGTAACATGAATTTGAATCTAAAAATTCATCTGCG
TTTGTGTATGCAG**AGTGGTAAACTGCTGAGTTGAGATACTGATGATGGTTTTACAATGGGGCTGCCGTGGTTAAC**
**TAAACCAGAGACTGACAGGTTCGCTCGAGTTCTCTCTCATGTCTAGACCATTTGTGAGAAGGGATTTTCTGTTTG**
**CATCATTCACTCCCACCAATCTCTAAGTATGCAATAAGTATTATTCTCACTTAAGAAAAAAGCAATCCTATTATT**
**TGAGATGCTTCTACAGCTTGAAATATGTTGTTTAGTTCTTTATATGGTTACCAAAACGTTAAGAAACCATGTGAA**
**CATTCCCCCAAAAATTAAAGGTCTAGCCTGTAGGTCACAGTCATTAGAGCCACTCTCACTTCTCGTTTTTTTTTT**
**TGTCAAACCACTTCTCGTTTAAAATGTGTTACTTCTTCCATCAAAATCTGATCCTTGAGGAGTTAGGTCCATAAG**
**TAAGTACACAGAGTGAACACATAACTTGATAAAATCATTTACACTTTGGACAATTTCCCATGAGATGCCAAACTT**
**TTCAATGGAGTTTGAGTTTTTCAATGATAGCTAAAATTTGTATCTAGCACTACTTATAGCTACACTAGTATACTT**

**GATAATATTTTGAGAGGAGTGGGGGAGATGCTGTGTTATTATTCCCTTCTCATCGATGGTCTAGATGTGTGAGGT**
**GACTCTCATGGAAGTAAAGAGAAATGGTGATTTATAGGAAAGCATCCACCACAATGGTATTATCCTAAAAAGGGT**
**TAGAACATTGTCAGCTTATGGAATAATTGTTTTGTATTGCTAGATTTTGGACCAGTGGTCTGCTAATGTAACACA**
**TGTGATACTTTCGAGTATAGTTGTGTGAGCTTTATTTTTCAGGTTGAGTATAACAGCATGCCATTATTAATCAAC**
**GATGGATGCAGAAGCAGTTACTAAAGAATCAGAAAACAGAAAAAGTCGAAGTGTTTTAAACGATGAAAGAGCAAA**
**GTGAACGAAAGGTGTACCTTTTTTACTATATTCTCTCTCGTTCATCCATTTCATGCTGCAACCGAAAAGACCAAA**
**GGGAATGAATGGGTGTGTTTTAATTACATGACGTAACGCCAAGTATTTAGAGATTGCTTGTAAATTTATAAGTAC**
**TATATACAAATACTCGGTAGATTCTAACTCCCGAATGAAAGACTTTGTATCTATCTTCTTAATGGCACATGAATA**
**TAGTTATAGTTTATAAATGGATTTATAT**

*bra-miR824b*

>Scaffold70077
AGCTAGAGACTTTATGCCAGAATTAGAAGCCCTACATATCTCACtcaaatcaattaaatcatataattataatca
aaatctaaatAATGCACTAATCAATAATTAAGTTATAAATTAAGCTCAGTCTTCATTGCATTACTCGT**ATAAAAC**
**CCTCTCTAGCTTCCCTAGAATTCTCCGTGCTCTATTCCCTTCGTCGCTCTCCCTCAATAATATATCTCTTGCCCT**
**AGctgcttcttcttcttcttcttcACGATCCGCTTCACATCATGCATCCCCACGATTAACGTTTTTTTTCCCGAC**
**GGTTACCATTGATTTTTCAGTGGGATCTCGTGACGGAAAAGTCATCATTTCTCTAAAGTTCTTAAAGGAGAGGAG**
**TTCACTTCCTAAGCTTCGATTACTTCGTCCGTTAGTTTGATTCCGT**GTATAATTCTTGTTTATTAACTTTCAATT
TTACTTCACACACGACATTTTTTTTAGCTGAATCGTTTTCCGAGATCTCTCTAGTATTTTGAGAAGTCGCAATTA
TCATCTGATAGATCAGCTGCATGAGCATTACTTACCTGAATTTTATTAGCTGATCCAAATTTTACTGTTCCAGGA
AAGATGATGGATTTCTTCTTTTGGTTTGCATGTGCAG**ATCGATGCTCATGGGAAGAGAGATTATCTCTCAATCTA**
**ACAGATCTGTCTCCTCTTCTCAG**GTCCATTACTTTTGTTAGATGTACATGTTTCATAAGGTTTTAACAACCCTTG
ACTGTAGTACTTCTGTTGTGTTTTTTTTTTGGTATACAGTATCTGACTTTACAATAATGATTATCCTCGTCCTTC
TACAGCTATTGTGATTATAATGATTTTTTTTTCTAGAAGTATTGTGATTGTCATTTGGTGATTGTACTCTTTTTC
TTATTAACAGCTTAATAATAGTCATGAGCATATAATAAAAGCAACCATCGAAATTGATCCAAAGATAAAACACAG
ATAACAAAATGACTCATGTGTGTTTTTTTTAAATTGATTTTCTTGCATTTCTTTAGTGGTACCTGAAAAGAGTAT
TAAAGCATCATTATGAAAAAGGTTTACCATATACCCAAATTATATGAAATGTACAGATACATTTAGGTGGAGACA
TCTATCATCAGAGAATTAAAGCATGATCTTTGAGTTCCTTGTCAATAAACTTTGATTCTTCCTCGGTAACACTTT
CAAACAACTATATGATAAGAGAATATGCCAAATCGACCCAATAAAATTCAAGGGTTTGAAAACACTTTGGGAACG
ATAGAGCCTAGACATCAGTATTGGATAGTTTCATCTGTGTGTCCTGGCCTCCAGATGTCCAAAGGAACGAAATGT
ATCTTTTTGAGAAAGCAATAGCTTGGATGGAAACTTTATGAGATAAATACAGGTTAGGTGCTTGCTTGAACTAAC
AAGGAAATGAATGGAGCTAACAAGGAAATGAATGGAGCTTGTTTCATTGAATAGAAATGATGAGTTTCTATTGAT
TAATAGCAATTCATTTTTGCCTTCAATGATTAAATAGAAGTATATTTTGCTGTTGCATAAACTAGTTAATTTAAA
TCTAAAAATTCATCTGCCTTTGTTTATGCAG**TGCTACATTATAGATTTGGGGGCTGCCGTGGTTAACTAAACTGA**
**AGACTGGCGCAGGTTCCCTCGAGTTGTCTTTCATGTCTAGACCATTTGTGAGAAGGGATTTTTTGTTTGCACCGT**
**TCACCCCACTCTCAATCTGTAGTGAGTATTAAGGAAAATAGCTTGTTCAACATATCACACTACTCTTTTAGAAAC**
**AAAACTATATTTTAGATGCTTTTCCAGCTTGATATACGTTGTTTTGTTATGTATACCTGTATTCTCCCGGCTACC**
**AAAACGTTAAGAAATAAGGTGAACATCTCCCCAAAACCTGCATCCAATTAGAAAGGTCTAGGTCACAGCCATTAC**
**TTGCTATCTCTCGATTAAGCTATTTTATTTCTATCATCATCATCAGATCATTCAGGAGTTAAGTATAAAATAAAA**
**TACATTAAGTCCATTCATGACATTTCTGGATGCTGCATACATAATATTTTGTTTTTGGACTATGCCACACTTCGG**
**ACAGACATTTTTCCTACGAGATGTCAAACCTCTGTCTGGAGTGTTAGAGGTATTTCATGATAGCTAAGAACTTGA**
**ATCTTGCACTACTTAGCTACACTAGTTTACTTCATAGTAAATAACTTGGGGAGTGGGGAAGGTGCTGGGTTATTA**
**GTCCCTTCTCATCGATGGTCTAGATGTGCGAGGTGACTCTCATTGAGGTAAAAGAGATACGGTGACTATGGGAAG**
**CATTCACCATAATGGTATTTCTTAAAGGTTAGAAGATTTTCAACTAGTGAAATTACGGTATAATTTCTTGATTTT**
**GAATCAGTGATATGCTAATGCAACACTTGTGATACTTCCGAGTATAGCTGTGTGATATTTTCTTCAGGTTGAGTA**
**CACCACGCCATTATTCAAACTAAACCTAAACGATGGATACAGAAGCATGTCGGAATGACTTTATTGCAAGGTCCT**
**TTGCAAAAACTGAAGAAGTGCAAGAGCTAAAGAAGTCAGAAAAAAAGCAATGACATTAAGAAAATATTGCTTGTA**
**CATTGAATTGATACACATAGCAATACTTGGTAGATTCTAACTACAAATGAAAGACTGTGTATATATACTTAGCCA**
**ATGCTTCATATAAATAGAATTACTTCTTATACATGGACTATATCTATATCTCAGGACACGTATTAGTGTTGACTT**
**CTTAATGAGTTTACTTAAGCCTGCGAGTTGAATGTTGAATTGAACATACAAAAATACAGTGCGGTTTGTTATAGA**
**ATCAGTTAAAGTTTTGAAAATTTTACCAAATTAAACTAATTGTTCCGATTTGATTTTAATTTATTGAATCAAAAT**
**TTTCGGAGTGATTATGATTACAATTTTTTTTATGCTAA**TTCGGTTTGAAATTTGAATATGGACCACCATATGGTA
TCTTGGTGCGAAACGTAAATTTTGAAACCGTTATCTAATTAAAAAGGGGATTAAGGGGCTCTCCAAGGGTAAAAA

*bra-miR824c*

>A08
TATAATTCTAATTGAAATCTAATTAACGCAGTAATCGATAATTCAGTATTATAATGGGGGTCATGCTTCCTTTGC
ATTACTATATAAAACCCTCTCTCGCCTCCTTAGTAATCACCGTGCTCT**CTTCTCCCTTCTTAATTCGCTCTCACT**
**CTATGATATCTCGTTTATTAAG**GTGCTTTTTAAACTTTCAGTTTTATTTTCTGTTGTATTGTTTGAGACTCTCTC
TCACCAGTAGTTTGAGAAGGGGGAGGATGTGAGTTTATCGTTTAGTAGATCAGCTGCATGAGCTTTAATTATCAA
ACTTTCTGAATTGCAGATCCAAGTTTTATTGTTTTCCTTTCATTTTAGTACTAAAATGTATTAACATTAGTGAAA
TAAAAGAAGAGTATTAAATTTGGTTTGCATGTGTACGAATGAAGTTCATGAGGGGTGATAGATGGATCTGTCTCT

TTTTTTCAGGCTTGGTTTCTCATCCCCATTTACTTTTATTAAGATGCATATGTTTTATCCTTGATCGTATTTTTT
TTATGCTTTTTGGTATAAAATATCTGACTAAACAAATCTAATGCATGCATGATCGATCCGTGTTTCAAGTAGTGT
GATTGTCATTTTCCGTTGTAATTGTACTTAGTGTCCATGTTCCTTTTTACTTATGAACAGCTTAACAGTAGTTAT
GAGCATAAGATAAAAGCAAATTTATTTTTGTTCTTGGCTAACGAAACTGATAAAAAGAAAGAAACACATTTTTTT
TACTCTCTTGTGCTTTTTCATTTCGTTTTACTGTATCTTCTTTAGTCATACCTAAGAGAGTTTTGAAAGAATCAT
TAAAGTCACACTTTAAGGTACACATAAGTTTACTAAAAATTATATGCCCAAAATATATGAAATTAAACTTGTTTT
ACTGTTTGTTCAAAAAAAAAACTTGTTTTACTGAAGAAAAGTTAAGAATTCTATTGATTGGATACAAGCAATTCT
ATATTTTCTTAGATGACTAAAAATATAAAAGGTACTTTGCTGTTGCATAAACTTGCACCACATGAATTTTGATTC
ATATATATGTTTATATCCACATATCATGAATTTTGAATCTGAAAGTCATATGCCTTTATTAATTCAG**TGCTACCT**
**TTTAACATGGAGCTGCAATGGTTAAATAAACTGAAGAGAAGCAGGGTCGCTCGAGTTGTCTCTCATGTCTAGACC**
**ATTTGTGAGAAGGGATTTACTGTTTGCACCATCCACCTCACCAATCTCTAAATCTGCCATTAGTATTATGCTCAT**
**TTAAGAAAATTAGCATCTAGAAAATAGGTACATAAAGTGCATGAATGACTTTTGCAGATGCTGTACACATAATTT**
**TTTTTTATAGAATCAGTTCATGCTGGTTCTCCAAATCATATACAGTTTTGGACATATATTTTTCCTAAGAGATGT**
**CAAACTTCTATATGGATTGTTAGAAGGTTTTCATGATATCTTAGAATTTGTAGCTTGCACTACTTAGCTACATTA**
**GTTCACTTGTTAAAAGTTGGGAAGTGGGGGAGGTGCTGGGTTATTCCCTTCTCATCGATGGTCTAGATGTGCGAG**
**GTGACTCTCATAGTGTTAAAGAGAACCGGTAAAAGTGAGTGTGGTCACAACAAGCCTCAACAACAATAGTATTTC**
**CTGAAAAGGTTAGAAAATTTTCACCTTGTGAAATTATTATATTGCTAGATTTCTCGACCAGTTTTGGGAGTTGTT**
**TTTCAGGTTGAGTAGTACATCATGCCACTTTTCAAACTTAACGTAAATGATAGATGCAGAAGCAAGTAGCAATGA**
**TTTTATTGCAAGAACTAAAGAAGTCAGGCAAAAGAAGAAGAAACAAAGTGTGTTTATCGGTGACAGATCAATAAC**
**GTGACCGAAAGGTTTTACCATTTGACTTGTTGTCGTCGTTGATTCTCTCATTATATTCATCCATTTCATGCTGCA**
**AGAAAACCCAATGTATTAATACATTTCTTGTAAATTGATACGTACCATTACTCGGTAGATTATAACCACAAATGA**
**AAGACTATGTACATCTGTATACTTAGCCAATGAGAAGATGAATATAATTACTTCTTATCCATGAATGATAT**TAGC
CCAAAACAAATATAAGTGATGCTAGATAGCCTAAATAATAAATTTTATTTTTGTATAATTGTATGACATTCATTG
AAACTATTGTTTCCACAATTTAACATGTTTACCAAAAATATATATATATATTGTTTCCACTTTTTGTGAAGAAGT

#### Brassica oleraceae

##### *BoAGL16*

>C01

TTTCTTTCCATTTATATTTTCGATCATAAAACCCAAATTGATTTTCCTAGATCTCGCGCGACCTGTTTTCTCTTA
TTTGGAAAAAGGTAATTGATTTCGATTTTTCTTAGTTTAATTGAAATATCTCGCCCTGGCGATGGTTTGACCATG
TTTTTGAGTTTTTTGAATTAGAGGTTGTAAGAAATGGGAAGGGGTAAGATCGCGATCAAGAGGATTGATAACGCG
ACGAGTCGTCAGGTGACTTTCTCCAAGCGAAGGAACGGACTCTTGAAGAAAGCCAAGGAGCTTGCGATTCTCTGC
GACGCTGAGGTTGGTGTCATCATCTTCTCCAGCACCGGCAGGCTCTACGAGTTCTCCAGCTCCAGGTATACCCTT
TTCCATTTGCTTCCTTTGTTCTATGTTTGGTTGACTTTTTGTGGTATTGGTCCAGTGTGATTGATTAGAGAATCT
AACCCATGATCCATCCGGTTTTGTTGACCACAAACCAATTCCAAATTGTTTCCATTCTATTTCTAGTTATTTTTG
TTGTTATCTGGAAAACTCGTTTGATTGTGTGCACATCCCTGAAGATGACATAAGCAGCATTATTTTACCTTGATT
CATATTATATGACAGTGAAACCTCAAACCTGAAAATTGGTATTATGACTTGGTTGGTTTGACTATCTATATGAAT
TGGCTGGAAAAAAAAAATCACAAAGCCAATACAGTTTGCTTGAGATTCATAGATTACAGGCTTTGTAACGGAATT
ATGAGGTTTATGGGTTTGTATGTCATGTGTGTATGCTATTGAGATGCAGGACATGACAAAAGTCAAATGGATTTA
GGATTGGATATGACTAGGATGATGTAGATTTGTGGTTGATGCGTAGATAGGAAGGGACTTGTAGTGTTTCTTAAA
AATACTAAATTTGGTAAAAACTGTTTTAAGAATGGAATAAAAAGTGGTTAGTGGTTAGTATGGGTCAAGAAACTC
TTAACTCTGTCGCTAATCACCACATCTTTTTTTTTTATGTCTTCTTGGCTAGAAAGAATAGTCTATTGAGCTATT
TACTTTTTTTTATTAGCAACTTGTTGTTGGTGGGGTTGGATGTACCTATAATGATTTTAATGTTGAATAAGTGGG
GGAATCATTTCTGCTTTCGTTCTCATCCATACTCAACTTTTCCTATTTAATTCATCCCAATCGGGAAATAAAAGT
ATAAAACCACCCCATTTTTAATACTCTACATTCAAGTTGGTCATTGTTTATTTGTGTATAAAGAGTATATTCTTC
TTCTTTTGTTGGGAAAAATTAGTCCCTTCTGATTTTCTACATTAGGAGGTCTTCCTTAGAGTAAAACTGAAGTTA
AACTATAAAGTAAAAATATCGTCAGAAATAAATGTAGCTTCAAAAGAGTTCATCCATGAGATTTAGTCCATTAGA
TATCAGAATAAATGGTGTTTAACCATCTATGTCTCTCTAATTCATAACTTGCAAAGCTGATCACAATAGTCCATT
AGATATCAGAATCAATAGTATCTGCAGATATACAACTTACCTTCTTTTTGTTTGGCAGCATGAAATCGATAATAG
ATAGATACAGCGAGGCCAAATGTGAAACTAGTTCGGAAATCAACCCAGCTTTAGAAATCAAGGTACTTGGTGCTT
TTAGTCTATTACACAGCTTCACTATCATGTTTCAAATTTTGACACCTCTCAAGCTGATCCTGAGTTCATATTGGA
AGTAGGTTTTGCTATATCACACACACACATACAAGTTATGACACCTGACAATCATATTACATTCACTATACTGCA
TTTTTCATTTTTTAAAGAATCCATTCTAGACTTGTTAGCACTTAGCATATCAATCTCAAAACCAAAAACAAAAGT
TTATTTCCTCAGATTCAATCACCTCTGAAAATAGTTCTGATGTGTTATAGTTTTGGCAAAGGAGGCCGCGATTC
TAAAGCGACAGTTACACAACTTGCAAGAAAACCACCGGTATGTGTTTTGCATCCTCGTCAAACACTCTCGATATT
AACAAACTAGAACAGGGAACGATGTACTAAAATGTCTGCAGAAAAGCAAAACTATCCTAGCATTTGCATTTTTTT
TATAATGGGAACGTCGATGTAAAAACAAAATTCCAAAATGATTTACCACCTCTTAAGGACTTTACCATGATCCTG
GATATCTTCTCCAAAATCACCACTGTATATAGTTACTCTGGAAAAAGTTTGCATATTCACAAGTGATGGAGGAAG
AGCTTGTACTTGGCAGGCAAATGATGGGGGAAGAGCTCTCTGGACTAAGTGTGGAAGATTTACAGAAACTGGAGA
ATCAGCTTGAACTGAGCCTTCATGATGTACGAATGAAAAAGGTATTGATATTGTGTGACATATATGTCAGATAAA
TTTTTTTTCATCTAACTATTGGAAAAAAGACGTAAAAACTGTACTCTACTCTCACACTTGTAGGATCAAATGTTA
CTAGAAGAAATACAAGAGCTTAACCGCGAGGTGGGAATCTGATTACTCGAATATAGAAATTATTAGACATTTCCA

CAGTGCTAATTCTTATATTTTTGGTAGCAGGGAATCTCTTACACCAAGAGAACTTAGACCTCCACGAGAAATTA
AACCTAATGCGACAACAGAATATGGAGTTACATAAGAAGGTAATCTTTGTTTTTTAGGAGGCAAAATCGCCTTTG
AAAAAAAGCTACATCATGTGTTTTCTTATGAATTGTGAGGATAATGTGATATGGCCAAGACTTGCAGGTTTCAGA
GGTTGAGGGTGTGAAAAGCTCAGAGAAAATTTCTCTTCTCACAAATGGTCTAGGCATGAGAGATAACTCGAGCGA
ACATGTTCATCTTCAGCTCAGCCAACCGCTGCAGCATGATGAGACTCCTTCAAAAGCTATCCAACTAAGCTATTT
CTCCTTCATTTCATAATATATAATTCAATTACAATTGTAGCAGCTTGCCACTGCATAAGCAAAAAAAAACATGAG
CAGATTGTAATGTTTTCGATCAAAACACCAGCTTTGTATTTTTTTTGTTTTGTTTTACTTCTTTCTCCCAGCAAA
GAACTATGGTTTGACGTTTAGGTATGTACTGTGCGATGGTCAAAAAACGTAGTGTTCTTTCTTAGGTTCCGAGAA

*bol-miR824a*

>C01

ATTTACTTGCAACTTGAAGTTATGATGGTGATGTAACTTAACAATTTTTACTGATTTTGTAAAGATCAAAATTTT
ACTACATGCAAAAAAGTTAATAAGTAAAAGAAAACACAAAACTAAAATTATACAGTTTATTGAATAAATTAGCTA
AAACAATTATCCAAAATAGCTACGTTAATACAGTAGGCCTGGGCGTTCGGGTCGGGTTTTTCGGATTTCGGTTTT
TTTTTATAACACCTCCTAGGTCACATTCTAGTAGATTTACAAGTACGGGTCGGGTTCAGATATAACACATCGGAT
TCGGATCGGTTTTGTATCACATCATAGAACCCATAAAGTAACCATATATCATTCGGATTCGGATTATATCGGATC
GGTTCGGATATACATGAAATAAAATCTAAAATTTAAAAGCAAAACATAAGAAATATATACTTATGTATATATAAT
TAAGTATTTAAAGTAGTTATTTAAATTTTAAATACTTTTTGTTAGATACCATATCAAAATAAATATGAAATTGAA
TATCTGAAGTATATATTCATGTTTCATATAATTATATTGTATATTAGTTTGGACATTCGGAATGGTTTCTTCGGA
TATCTTTTCGGATTTTTTGGTTTTTTCGATTTTTCGGGTTATCCGTTCGGGTTCGGTTAATAACACTTCGAGTTC
AGATATGTTTTGTACCACTTTACAAGATCCATTCGGCTATTTTTTGCATTTCGGACCGGATACGGATCGGGTTTT
TCGATTCGGGTTCGGTTCGGATTATGGATTTTATGCCTATGCCTATTAATACATATCAATTTCAGGTTTCATCAT
ATGGTATTTTATGAATGTACATTTTAGTTATTCGATAAATATGTAATTGTAGTTTCAATTTGAAAAGATTATTAG
GAAAAAGAAATATTTTGTAGTTAGTAAAGTTATTTCCAACTAGTATTCCAAAGGATATCGGAGCCGAGCGTAGCC
ACTGGTCAAGTCCAAGTGTAAATAGCTAGAGCCGAAGAAAGTGGGCGATGATTTGTTTTATATATATATATATAT
GTATATATATATATATGAAGCCGCATAAGCTACAGCTTTATGCCAGAATTCGAAACCCTACATGTCTCACTCAAT
TCAATTGTAAAATTGAAATCTGATTAACGCACTAATCTATAATTAAGTTAAAAATTAATCTCATTCTTCCTTGCA
TTACTTATATAAAACCCTCTCTCGCCTCCTCCTTAGTATTCAA**TCTCCACACTCTCTTCTTTTCTTAGCTCTCGC**
**TCATATCTCTCTCTATGTTTTGCCCTAGTTAACTTGTTTTCTTCTTCTTTTTCTCAATCCGCTTCACATCAATGC**
**ATCCTCATGATTCATGTGCTTTTCCGATTAAGTTTTCTCTGGGATCTCGAAGAATTGCCAATTTCTTAAATATGT**
**TAAAGCAGACGAGTCGCTTCGATTACTTCATTTG**GTTAGTCTGATTCTGTTTATATAGTTTCTTAACTTGCAATT
TGTTTTATGAAATGTTTTGTGTTGTTTGTTAGATCTCTCCATATTTATATATTAGTTTGAGAAGTCAGATGTTTG
CATGTGCAG**ATTAATGCTCATGGATGAGAGGTTTGTCAACCTTTAATGGATCTGTCTCCCCTTCTCAG**GTTTGGG
TTCTCATCTCCATTTTTATTTGTTAGATGCATAGTGTGTGTTATATAGAATGTCTCAGCCTTGATCGTACTTCTG
TTGTGTTTTTTCTTTTGTTACTTCCGTTAAGATGCGTACTGTCTGATTGTCTAATTACTCAGTGTCTATGTTCCT
TTCACTAATGAACAGCTTAATAGTGGTTAATTAGGCGCATAAAATAAAAGCAAATTATTTTGTCTCTTGACCAGA
GAAACTGATCAAAACTTAGAAATGAAATTGACTCTCATGATGTGTTTTATTCATTTTCTCGCATCTTTTTTATGT
ATACCTAAAAAAGAGTTTGAAAGCTTAGATTCTCACTTTAAGGTTAAAAAGGTGTCTGTACGCAAATTACATCGT
ATGTACGAAGTACGAACACATCTTTTTCCTTCCTTTTGCAAATTAACGCTGCATCACCCACTAGAGAGTTCTGAT
GTTTATTTCTGATTGTAATGTTCTTTTTTCCGACTTCATGGACTGTACTAAGTTGAACTCTCACTATACCTCTAT
TTGGAAGCTATTTTCAGAATTAAAGCTTGATCTTTGTGTTCCTTGTCAATAAACGTTGATGTCTAAATATTATAT
TTTAGACAACTATATATATGATAAGTTAATAGTCCAAAGCAATCCAAAAAAAAGCTCGAGGGTGATAGTTGAAAA
CACCTTAGGAACCACTTTTAGACAGCAATATAGAATAGTTTCATTTGTGTTCATAAGCCTCTAGATGTTCAAAAA
TAAATTGTTATTTCTTTGAGAAAGCAGTGGCTTGAGTAGAAACTCCATTAAGTTTGTAATTAGGTTGGGTTCTTA
GATTAAAAAGGAAGTGAATGGAGCTTGTTACACTGGAAAGAAACTAAGAATTCTGTTGATTAGCGACGAGCAATT
CAATTTTGTGCATAATTCAGGTTTATGTCCACATATCTTTGTAACATGAATTTGAATCTAAAAATTCATCTGCCT
TTGTGTATGCAG**TGGTAAACTGCTGAGTTGAGATACGATTCCGATGATGGTTTTACTATGAGGCTGCCGTGGTTA**
**ACTAAACCAGAGACTGACAGGTTCGCTCGAGTTGTCTCTCATGTCTAGACCATTTGTGAGAAGGGATTTTCTGTT**
**TGCATCATTCACTCCCACCAATCTCTAAATATGCAATAAGTATTATTCTCACTTAAGAAAGAAACAATCCTATTA**
**TTTGAGATGCTTCTACAGCTTGGAATATGTTGTTTCGTTCTTTATATTGCTACCAAAACGTTAAGAAACCAGGTG**
**AACATTTCCCTTGAAAGGTCTAGCCTGTAGGTCATAGTCATTAGACCCACCTCTCACTTCTCGATTAAAATGTGT**
**TATTTCTTCCATCAAAATCTGATCCTTGAGGAGTTAGGTACATAAGTAAGTGCACAGAGTGAACACATAACTTGA**
**TAAAGTCATTTACACTTTGGACATTTTCCCAAGAGATGCCAAACTTCTCAATGGATTTTGAGTTTTTCAATGATA**
**GCTAAAATTTGTATCTAGCACTACTTATAGCTACACTAGTATACTTGATAATATTTGGGGGGAGTGGGGGAGATG**
**CCGTGTTATTATTCCCTTCTCATCGATGGTCTAGATGTGTGAGGTGACTCTCATGGAGGTAAAGAGAAATGGTGA**
**TTTATAGGAAAGCATCCACCACAATGGTATTATCCTAAAAGGGTTAGAAAATTTTCAGCTTTTGGAATTATTGTT**
**TTGTATTGCTAGATTTTGGACCAGTGGTCTGCTAATGCAACACATGTGATCCTTTCGAGTATAGTTGTGTAAGCT**
**TTATTTTTCAGGTTGAGTACAACAGCATGCCATTAGTCCATTACTAATCAACGATGGATGCACAAGCAATTACTA**
**AAGAAGTCAGAAAACAGAAAAAGTCGAGTGTTTTAAACGATGAAAGAGCAAAGTAAACGAAAGGTGTACCTTTTT**
**TACTATATTCTCTCTCGTTCATCCATTTCATGCTGCAACCGAAAAGACCAAAGGGAATGAATGGGTGTGTTTTAA**
**TACATGACGTAACGCCAAGTATTTAGAGATTGCTTGTAAATTCATACGTACTACAAATACTCGGTAGATTCTAAC**
**TCCCGAATGAAAGACTTTGTATCTATCTTCTTAATGGCACATGAATATAATTATACATGG**ATTTATACTAACGTT

AAACAAATATTAAGTGGACCATGGTTCCCACTAACATATCGTACATGAAAAATCATATTCTCAATAGACGTACTT
TGTATGTAATGAGGCAGGAGAACCCACCAACCCATGAAGTGGAATAATATGTAGATATGCAAAAGTATTTCCCAC

*bol-miR824b*

>C07

GAAGCCCTACATGTCACACTCAAATCAATTAAATCATATAATTATAGCCAAAATCTAATTAACGCACTAATCAAT
AATTAAGTTATAAATTAAGCTCAGTCTTCATTGCATTACTCATATAAAACCCTCTCTAGCTTCCCTAGTATTCTC
CGT**GCTCCTCTTCCCTTCTCCTCTCTCCCTCAATAATATCTCTCTTGCCCTAGCTTCTTCTTCTTCTTCTTC**
**ACGATCCGCTTCACATCATGCATCCACACGATTCACGGTTTTTCTCCTGACGGTTACCATTGATTTTTCAGTGGG**
**ATCTCGTGAAGGAAAATTCGCCATTTCTCTAAAGTTCTTAAAGGAGTGGAGTCACTTCCTAAGCTTCGATTACTT**
**CGTCGATTAGT**GTAATTCCGTATATAATTCTTGTTTATTAACTTTCAATTTTATTTTACACACATGACATTTTTT
TGCTGAAACTTTTTGTGAGATCTCTCTAGTATTTTGAGAAGTCGCAATTATCATCTGATAGATCAGCTGCATGAG
CATTACTTACCTGAATTTTATAGCTGATCCAAATTTTACTGTTCCAG**GAAAGATGATGGATTGCTTCTTTTGGTT**
**TGCATGTGCAGAACGATGCCCATGGGAAGAGAGATTATCTCT**CAATCTTTAACGGATCTGTATCCTCTTCTCAGG
TCCATTACTTTTGTTAGATGCACATGTTTAATAATGTTTTAACAACACTTGACTGTAGTACTTCTGTTTTGTTGT
TTTTTTGTTCTTTTTTTTTTTGGTGTATAGTATTTAACCTTACAATAATGATTATCCTCGTCCTTCTACAGCTAT
TGTGATTATAATGCATGATTCTTCCTTTTCTAGAAGTATTGTGATTGTCATTTGGTGATTGTATACTTAGTTAGT
GTCCTTGCTCTTTTTTTCTTATGAACAGCTTAATAGTAGTCATGAGCATATAATAAAAGCAATCATCGAAATTGA
TCGAAAGATAAAACACAGATAACAAAATGACTCTCCTGTTTTTTAAAAAAAAAAAAATTGTTTTTCTTGCATTTCT
TTAGTTGTACCTGAAAAGAGTATGAAAGCATCATTAATTCACACTTTAAGGTAAAAAAAAAGGTTTACCATATAC
CCAAATTATATGAAATGTACAGATACATTTAGGTGGAGATATCTATCACCAGAGAATTAAAGCTTGATCTTTGAG
TTCCTTGTCAATAAACTTTGATTCTTCCTCAGTAACACTTTCACACAACTATATGATAAGAGAATATGCCGAATA
GACCCAATAAAACTCAAGGGTTTGAAATGACTTTGGGAACGATAGAGCCTAGACATCAGTATTGGATAGTTTCAT
CTGTGTGTCTTGGCCTCTAGATGTCCAAAGGAACGAAATGTATCTTTTTTGAGAAAGCAATAGCTTGGATGGAAA
CTCTATTAGATTAATACAGGTTAGGTGCTTGCTTGAACTAACAAGAAAATGAATGGAACTTGTTTCATTGAATAG
AAATGATGAGTTTCTATTGATTAATAGCAATTCATTTTGCCTTCAATGATTAAATAGTAAGTGTATTTTGCTGTT
GCATAAACTAGTTAATTTAAATCTAAAAATTCATCTGCCTTTGTTTATGCAG**TGCTACATTATAGATTTGGGGGC**
**TGCCGTGGTTGACTAAACTGAAGACTGGCGCAGGTTCCCTCGAGTTGTCTTTCATGTCTAGACCATTTGTGAGAA**
**GGGATTTTTTGTTTGCACCGTTCACCCCACTTTCAATCTGTAGTGAGTATTAAGGAAAATAGCTTGTTCAACATA**
**TCACACTACTCTTTTAGAAACAATACTATATTTTAGATGCTTTTCCAGCTTGATATACGTTGTTTTGTTATGTAT**
**ATCTGTATTATCCCGGCTACCAAAACGTTAAGAAATAAGGTGAACATCCTAAAACCTGCATCCAATTAGAAAGGT**
**CTAGGTCACAGATATTACTTGTTACCTCTTGATTAAGCTATTTTATTTCTATCAGCATAATCAGATCATTCAGGA**
**GTTAAGTATAAACTAAAATACATTAAGTCCATTCATGACATTTCTGGGTGCTGCATACATAATATTTTCCTTGTT**
**TTTGGACTATGCCACACTTCGGACAGACATTTTGCCTAAGAGATGTCAAACCTCTGTCTGGAGTGTTAGAGGTAT**
**TTCATGATAGCTAAGAACTTGAATCTTGCACTACTTAGCTACACTAGATTACTTCATAGTAAATAGCTTGGGGAG**
**TGGGGGAAGGTGCTGGGTTATTATTTCCTTCTCATCGATGGTCTAGATGTGCGAGGTGACTCTCATTGAGGTAAA**
**AGAGATACGGTGACTATGGGAAGCATTCACCATAATGGTATTTCTTAAAAAGGTTAGAGATTTTCAACTTGTGAC**
**ATTATCGTAATATTGCTTGATTTTGATTCAGTGATCTGCTAATGCAACACTTGTG**ATACTTCCGAGTATAGCTAT
GTGAAGTTTTTTTCTTCAGGTTTGAGTACACCACACCATTATTCAAACTAAACGTAAACGATGAATACAGAAGCA
TGTAGCAATAACTCTGTTGCCAGGTCCTTTGCAAAAACTGAAGAAGTGCAAGAGCTAAGGAAGTCAGAAAAAAAA

*bol-miR824c*

>C08

AAGTGTACAGTGACTAGAGCAGCCTGAGTTACGGCTTTATGCCAGCATTAGAAGCCCTACATGTCTCACTCAATA
ATATAATTCTAATTGAAATCTAATCAACGCACTAATCGATAATTCAGTATTATAATTGAGGTCATGCTTCCTTTG
CATTACTATATAAAACCCTCTATCGCCTCCTTAGTAATCACCGTGCTCTCTT**CTCCCTTCCTAATTCGCTCTCGC**
**TCTATAATATCTCGTTTATTAAG**GTGCTTTTTAAACTTTCAGTTTTATTTTCTGTTGCATTGTTTGAGATCTCTC
TCACCAGTAGTTTGAGAGGGGGGGGGGGGGGGGGGTGATATNNNNNNNNNNNNNNNNNNNNNNNNNNNNNNNNNN
NNNNNGTTTGTGTAGGGAGGGGGGGGGGGGGGGGGGGGGGGGGGGGGGAATGTAAGTTTTATAGTTTAGTAGATCAG
CTGCAAGAGCTTTACTTATCAAACTTTCTGAATTGCAGATCCAATTTTATTGTTTTGCTTTTCTTTTTAGTACTA
AAATGTATTAACATTAGTGAAATAAAAGAAGAGTATATATTAAATTTGATTTGCATGTGTTAGGAATGAAGTTCA
TGAGGGGGTGATAGATGGATCTGTCTCTTTTTTTCAGGCTTGGTTTCTCATCCCCATTTACTTTTGTTAAGATGC
ATATGTTTTTATCCTTGATCGTATTTCTTATGCTTTTTGGTATAAAATATCTGACTAAATAAATCTAATGCATGC
ATGATCGATCCGTGTCTTTCAAGTAGTGTGATTGTCATTTTCCGTTGTAATTGTACTTAGTGTCCATGTTCCTTT
TTACATATGAACAGCTTAACAGTAGTTATGAGCATAAGATAAAAAGCAAATTTATTTTTGTTCTTGGCTAACGAA
ACTGATAAAAAGAAAGAAACATAATTTTTTTTTACTCTCATGTGCTTTTTCATTTCGTTTTACTGTATCTTCTTT
AGTTATACCTAAGAGAGTTTTGAAAGCATCATTAAAGTCACACTTTTAGGTACACAG**AAGTTTCGTACAAAAAAA**
**AAG**GTACACAGAAGTTTACTATAAATTTTGTGCCCAAAATATATGAAATTAAACTTGTTTTACTGATGAGAAGCT
AAGAATTCTATTGATTGGATACAATCAATTCGATATTTTCTTAGATGATTAAAATAAAAGGGTACTTTGCTGTTG
CATAAACTAGCAATACATGAATTTTGACATATGTTTATATTCACATATCTCTGCAACATGAATTTGAATCTGAAA
GTCATCTGCCTTTGTTAATGCAG**TGCTACCTTTTAGCATGGAGCTGCAATGGTTGAATAAACTGGAGACAAGCAG**
**GATCTCTCGAGTTGTCTCTCATGTCTAGACCATTTGTGAGAAGGGATTTACTGTTTGCACCATTCACCTCACCAA**

**TCTCTGAATCTGCAATTAGTATTATGCTCATTCAAGAAAATTAGCATCTAGAAAGTTGGTACATAAAGTGCATTT**
**ATGACTTTTGCTGATGCTGTACACATATTTTTTTTTTTTGTAGAATCAGTTCATGACAATTTTCCAAATCATATA**
**CAGTTTTGGACATACATTTTGCCTAAGAAATGTCAAACTTCTGTCTGGAGTGTTAGAAGGTTTTCATGATAGCTC**
**AGAATTTTATAGCTTGCCCTACTTAATTAGCTACATTAGTTCACTTGTTAAGAAGTTGGGAAGTGGGGGAGGTG**
**CTGGGTTTCCCTTCTCATCGATGGTCTAGATGTGCGAGGTGACTCTCATAGTGTTAAAGAGAAACG**GTATAAGTG
AGTGGTCACAACAAGCCTCCACATGCAACATTAGTATTTCCTGAAAAGGTTAGAAATTTTTCACATTGTGAAATT
ATTGTATTGCTAGATTTTCGACCAGTTTCGTGAGTTGTTTTTCAGGTTGATACATCATGCCACTTTTCAAACTTA

#### Sisymbrium irio

##### *SirAGL16*

##### >SI_scaffold669

TTTCTCTTTCTCTTTCTCTTTCTACTTGTAGTTTCGATCAACAAAACAAAACCACATTGATTTTCTTCTTACTTC
CCCGCGAGCAGATCTCGCTCGATCTGTTTTGTCTTAGATTCTGTCTGAACAAAAGGTAATTAATTTTTTTGTTTT
TATTTTGAGTTAATTGAATTATCTCTCTCTCTGGTTTGACTATGTTTTTGAACAAGAAGGGTGTGACGAAATGGG
AAGAGGGAAGATCGCGATCAAGAGGATCGATAACTCGACGAGCCGTCAGGTGACTTTCTCGAAGCGAAGGAATGG
ATTGTTGAAGAAAGCCAAGGAGCTGGCGATTCTCTGCGACGCAGAAGTCGGTGTCATCATCTTCTCCAGTACCGG
CAGGCTCTACGATTTCTCCAGCTCAAGGTATCCCTCCTCCTCCTCCTCCTCTTTTTCTCTTCGGTTGACTTTTGG
TCCTTTTGGGAATATATATATATGATCCATCATCATTCATCGCCTTTTGTTGACCACAAACCAATTCTAAATCTC
CATTTTATTTTttcaattcaattcaattcaattGTTGTGTGCACATTCTTGAAGACGATGACTTAAAAACCAATT
TTTATTAGTCTGATTCGTATTATATGACAGTGATTGAAATTTGTTACATCTCTTGGTTTGGTTTGACTTTCTATA
TGAACGGTCTGGAACAAAATAACCCGGTTTGCTTGAGTTGGTACGGAATTATGTTGTTGTGATATATATATGGGT
GGGTTTGTATGTTATGTGTGTTTCTGTGTTGTTGAGGTGCAGGACAGGACAAAAGTATAATGAATTTAGGATTTT
GATAGGACTAGAGAATGATAAAGATTTTTGTCAAGCGTTAGATCGAAACCATACTAAATTTGGTATAAACTATTT
CCAGAATGGAATAAAAAGGAGTCCAAAGTGGTTAGTGGATAGTGGTTAGTAGGGCAGGAGGAGGAAACTCTTTAA
CTCTCTTGCTTATCACCACTTTTTTTTTTTATAATGTCTTGTTGGTTTGGTTTGAGGTTCTTAGAAAGAATCGTCT
CTCTTCTGTTTTTTTTTTTCTTTTACATTATTAGCAACTAGTTGGTGGGGTTGATGTACCTAAAATGAGCTGCATT
ATTTTATGTTTAAATTTATTAATACACAAGTGGGGGGAATCATTTCTGCTTTCGTTGTCATCCATAATCATCTTT
TTTCCTATTTAATTTATCCATTTAGGAAATAGACCACCCCATTTTAATGCTCTACGTTCAACTTGGTCATTGTTT
ATACGAGTATATTCTTTCTTTTTTGGGAAAACACTATACGTATCAATCATCAATCTCTTCTCATTTTCTACTTTA
GGAGTAGTTCATTGAATTAAAACTGAAATTATACTATATAGTAAAGAAAAATCTTCAGAAAATATGTAAATTCAG
AAAAGTTATTCAAGAGATTGGGGGCTGATAACAATAATATCTATACTCGAGTTTGGTGATTAGTGTTGGCCAGTT
CTACATCAACTTTACTCCATTTCTCTCATTCTCTGCTAAAGCTTCATACGTCATGTTACATTTTGATTGTGGCGA
AACTGTATTTTGGCTGCAGAGAAACAAATTCCATGTAGGCAAAGATGGCTTAGGGTATAGCCTAAATCTATTTTA
AGTATTTTGACACTGAAAATATGTAAGGAGAAATGTTGCTTGAATGAACTTACCTGCCAATCACACTCTGCAGAT
ATTCAACTGACCTTCTTTTTGTCTGGTAGCATGAAATCGGTAATAGAGCGATACAGCGAGGCCAAAGGTGAAACC
AATTCGGAAATTAACCCAGCTTCAGAAATCAAGGTACTCTCTTTTTGGTTCTTTTTTTTACTCTTCTCTAAGGATT
TTAACTAGTAAGAGAATCAGTTTTTGTAACTGAGTAACGAGTAGGCCAAACAAGTAATATACTAAAACATAGAGT
TTGTTACTTGTAAATGTGTTAGCCTATTACAGCTTCATTAGTATATTTCAACTTTTAACTCCTCTGAAGCTGAAC
CTGATCTCATATTGGACTGGAGACCTTAAATACCACGCATTATATGTTGAATTAGGATTTGCTATAAAACACACA
CAAGTCATGACAACTGAGTGCCTGACCATCAGATTACATACATTATAGTGCAAATTCTTTTTTCAAGAAACCATT
CTAGACTAGATAACATATCAATCCCGAAAACTATAATAAAAAAAAAAACTTGTTTTCTCAGCTTCAGTCTCCTGAA
ATTAATTCTGCTATATTATAGTTTTGGCAAAAGGAGGCCGCAATTCTAAAGCGACAGCTACATAACTTGCAAGAA
AACCACCGGTATGTTTTGCATCTTCTGAGTTCGTCTCGATATTAACAAACTAGAACAGGGACGATGTACTAAAAT
GTCCAAGGCTAGAAGATTTTGCAGAAAGGCAAAACTATCCTAGCATTTGCATTTTTTTTTAATAGGAATGTCGATA
TTAAAATACAAAATTCCAAACTGATTTACAACCTCTTAAGGACTTTACCGTGATGCTGGATATCTTCCCCAAAAT
CACCATTGTATCTCTACAACCTTGAGAGTTAGCTCTCCACACATCTAGATCATCCACAAGAAGGGAATAACCAAA
CACTTCCACATTCCGGATCTCTCTTTCTTTTTATTTGGTGAATTAGGAAAGGTATATAATTACTCTGGAAAAGTC
ATATTCAAAAGTGATGGGGAAAGAGCTTGTACTTGGCAGGCAAATGATGGGGGAAGAGCTCTCTGGACTAAGTGT
GGAAGATTTACAGAAATTGGAAAGTCAGCTTGAGTTGAGCCTTCGTGACGTACGAATGAAAAAGGTATTGATGAT
GTTGTGACAAATAAGTCAGATACCTTTATATGTGACATTTGGAAAAAAAGTAATGACTGTACATTACTCTTCTCT
ACTATTGACGTTTGTAGGATCAAATGCTAGTCGAAGAAATACAAGAGCTTAACCGCGAGGTGGGAGTCTGATTAC
CGGAATATAGAAATTAGATAATGTTCACAGTGTTGGTACATACTAATGCTAATTCGTATATTTTTGGTTGCAGGG
GAATCTCGTGCACCAAGAGAATTTAGACCTCCACAAAAAAGTAAACCTAATGCAACAACAGAATATGGAGTTACA
TAAGAAGGTATAAATCTTTGAAAAAGCTACAGCATGTGTTTTCTTCTGAATTGAGGATAATGTGATCTGGTTACT
TGCAGGTTTCAGAGGTTGAGGGTGTGAAAAGCGCAGACAAAAATTCTCTTCTCACAAATGGTCTAGACATGAGAG
ATAACTCGAGCGAACATGTCCATCTTCAGCTCAGCCAACCGCAGCAGCATGATGAGACACATGCAAAAGCTATCC
AACTAAGCTATTTCTCCTTCATTGCATAATAATACAATTCAATTTTAACTGTAGCAGCTTGCCTCTGCATAAGCA
AAGAAATATAAACATGATCATTATTGTAATGTTTTTGATCAAACACCAGCTTTTATTTTTGTTTTGTTTTAACTC
TTTCTCTCAGCAAAGAACTTTGGTTTGGCGTTGAAACATAAAGAATGTAGTGGTCTTACTCAGCACTACAGTCAA
ACATTTCTTGATTTTTCTATCTATAAGTTTAGTCAACTTTTCCAAGCTATTTCTTTTGTGTAATTTAAATCCATT
TCTACAATAAAAGTTGATATTATCTTTAACACATGTGTAAGCCAATGTTGTGGGGGAGGAAATGGCCACAGGCC
TATAACAATTTGTGGTGTGAATCTGGAAAGGAGCTCACGTGGGCTTATCAGCCTCCGACGGTAAAAACAAAGCAT

*sir-miR824*

>Scaffold_189

GAGAAGCAATTCAAAAGTTTTTCCATCAATGATAAAGATAATATAGGCTACTTTGCTGTTGCATAATTCACGTTT
ATATCCACATCTTTGTAACATGAATTTGAATCTGATAATTCATCTGCCGTTGCCAATGCTGCAGTTAATCGAGAT
AAAATTCTGATGATCCAGCTGAGATACCGCCGTGGTTTTAGTACGGGACTGCCGTGCTTAACTAA**AGACTGACAG**
**GTCCGCTCGAGTTGTCTCTCATGTCTAGACCATTTGTGAGAAGGGATTTTTTGTTTGCACCGTTCACCCCACCTA**
**TCTCTAAATCTGAAATCAATAAGTATTATGCTCATTTAAGGAAAAACAATCCTAGATTTCAGATTCTTTCTACAG**
**CTTTTTAAATACGTTGTTTCGTTATGCATATCTGTATACTCCAGGCTACCAAAACGTTAAGAAACTGGGTGAACA**
**TTTCCCTAAACCCCATCCAATAAGGCAGGTCTAGCCTGTAGGTCACAGCCATTAGAGCCGCGTTCCCTTTCGATT**
**ATATAAGCTGTTTTATTTCTTCCATCAAAATCTGATCCTTGAGGAGTTAGTACATAAGTATGCATTCATGACTTT**
**TCCAGATCTGCATACACAACTTTTTCATTTGTATTTTGTCCGAATTAGATAAAATCATTTACATTTTGGATACAC**
**GTTTTTGCCTAAGAGATGACAAACTTCTGTCTGGAGTTTGAGAGTTTTTCATGATAGCCAAGAGTTTGTATCTTG**
**CACTAGTTAGCTACACTAGTTTACGTGATAATAATTTGGGGAGTGGGGGAGGTGCTGCGTCATTCCCTTCTCATC**
**GATGGTCTAGATGTGCGGGGTGACTCTCATGGAGGTAAAGAGAAATGGTGAT**TTTAGGAAGCATCCACGACTTCG
GTATTTCCTAAAAAGGTTTGAATATTTTCAGCTTGTGCAATTTATTACTAGATTTTGGAGCAGTGATCTGCTAAT
GCAGCACATGTGATACTTCCGAGTATAGTTGTGTGAACCTTTTGATTTCAGGTCTAGTACATCATCATTGCCATT

#### Schrenkiella parvula

*SpAGL16*

>EP_scaffold336473545

ATTTCGCTTTCTCTCACTTTCGATCAAAAATCACATTTTTTTCTCTCTTCTCGAGCAGATCTCGCGCGATCTGT
TTCCTCTTTGAGTTTTGGAAAAAAGGTAATGTTTTTCATTTTGGGTTTTGAGTTTATTGAATTATCTATCTCGTG
GCTGAATTTGCGGAGATGGTTTTAACGATGTTTTGTGATGTGAAAGGCTGAAGAAAATGGGAAGGGGAAAGATCG
CTATCAAGAGGATCGACAACTCCACGAGCCGTCAGGTTACTTTCTCCAAGCGAAGGAACGGGTTGTTGAAGAAAG
CTAAGGAACTTGCGATTCTCTGCGATGCAGAGGTCGGTGTCATCATCTTCTCCAGCACCGGCAGGCTGTACGATT
TCTCCAGCTCCAGGTACCCTTTTTTTGTCTCCTCTCTTTGGTTGACTTTTGGTATTGCTCCTTTGCGATTGATTA
GGGATTCACACATCCATGATCCACTGGGTTCCGTTGACCACAAATCATTTCTAAATCTCGATATTTTTCTAATCG
ATTTCTCGTTTTTTTTTAATCTGGAAATTCAGTTGTTGTGTCCACGTTCCTGAAGAAGAAATAAACACCAAATTT
TTTAAGTCTGATTTATACAACAAAGGCTGAAAATTGATACATCACTTGGTTGGTTGGATATCTATATGAAATGGG
GCTGAATAAACAAGGCCATTACAGTTTGCTTGAGATTCATAGATTACAAGTTTTGTAACGGAATAATGAGATTGT
GATTTATGGGTATGTATGACATGACAAAAGTAAAATGGATTTAGAATTGGATAGGACTAGAAAGATGGAGTCTTT
GTGTGAAGCGTAGATCGTGAGGGACTTGTAGTTGTTACTAAATTTGGTATGAACTATATTTCAGATGAAAAGGGG
TATAAAGTGGTTAGTGGTTAGTAGGGCAGAAGAAACTCTAACTCTCTCTCTCCCCCTTATCACCACTTTTTATGT
CTTCTTGGTTTGGTTTGAGGTTTTAGAAAGAATCGTCTCTGTTCACTATTCCTTTTTCTTTATTGGCAACTAGTT
GGTGGGTTGATATACCTAAAATGAGCTGCACTAATTAATGTTAAATATAATAATACACAAGTGGGGGAATCATTT
CTGCTTTCGTTATCATCCATAATCAACTTTTCCCATTTAATTATCCAATTAGGAAACAAACCCCCCAATTTTAAA
GCTCTACGTTCAACTTGGTCACTGTTTAGTGATATATAGGTGTATATTCTTTCTTTATTATACGTATCACAACCC
TTTCTGATTTTCTACTTTAGGAGGAGTTTTTCGTTGAAGTAAAAAACAAAATATACATTAGATTAGATGTAAGAA
GCAATGGTGTTAAAGAGTCCTTCACATTTATGTCTCACTAATTCGTATAGGAAAATCATTACTTGCCAGCTGATA
ACAATAATATATCTATACTCGGGTTTGGTGAAATGAGATGGCCACTTCTACATCAATTTCTCTCCTTTCTCTCAT
TCTCTACTTAAGCTTCATGCATCATGTTACATTGTGATTATGGCGAATCTGTATTTTGGGTGCAGAGAATCAAAT
CCATGTAGGCAAAGATGGCTTAAGGTAGCCTAAAATTATTCTTTAGTAAAAAATTTGACAATGAAAATATGTAAG
GAGAAATGTTTTAGTGGATGTCGCTTGGGTTACCTTACTTACCAACCACACTCTGCATATATACAACTACTTTCT
TCTTTTTTTGGCAGCATGAAATCGGTAATAGAGAGATACAGCGATGCAAAAGGGGACGCCAATTCCGAAAATAAC
CCAGCTTCAGAAATCAAGGTACTCCTTTTTTGGTTCACTTTTCTTACACTTTTCTATGGATTTCAACGAGTAGTT
CAGGGACTCTATTTTGTATATATGAGTAACGACCATCAAGTAACACTCTAAAACAGAGAGACAGTTACTTTGTGG
TTTTTATAGTGTTAGCCTATTACAACATCATTAGCATATTTCAACTTTTAGCTTCTCTCAACCTGATTCCGATCT
CATATTGGACTTGCAGACTTAAATACTACGCATTGTATGTGAAGTAGTATTTGCTATAAACACACACACACAAGT
CATGACACCTGACGTCCTATTACATACATTATGTTGCATTTCTTTTTTCAAGAATTCATTCAAGTATTGATAACA
TATTAATCCCAAAAAAAATATTGTTTCAATACCTGAAAATAATTTCTGCTATATTATAGTTTTGGCAAAAGGAAG
CCGCAATTCTAAAGAGACAGCTACATAACTTGCAAGAAAACCACCGGTATGTTTTGCATCCTCTGAGTTCGTCAG
ACACTCTCGATATTAACAACTAGAACAGGGAATGATGTACTAAATGCAAAATCATAACATATAGGCAAAGTGCAA
GGCTAGAAGATTTTGCAGAACCTAAAACTGTCCTATCATTTGCATCTCTTTTTAAAATTGGAATATCGATATTAA
AAATAAAATGCCAAACTGTTTTACCACCTCTTAAGGACTTTACCATGATGCTGGATATCTTCCCCAAAATCACCA
TTGTATCTCTACAACCTCGAGAGTTATATCTCTACACATCTAGACCATCTACAAGAGGGGAATAGCCAAACACTT
TCACACTTCCCATCTCTCTTTTTCGTTTTGTTTTGGTGAATTAGTTAACGTAAATAATTACTCTGGAAAAGTTTG
CATATTCAAAAGTTGATGCTGAAAGAGCTTGTACTTGACAGGCAAATGATGGGGGAAGAGCTCTCTGGACTAAGT
GTAGAAGATTTACAGAAATTGGAAAATCAGCTGGAATCGAGCCTTCGTGACGTAAGAATGAAAAAGGTATTGATG
CTGTGACAGATAAGTCGATATTTTTTTATATAACATTTGGAAGAAAATGGTAATGACTGTAATCTATATCTGTTC

TTTTCTTTTATTGACGTTTGTAGGATCAAATGTTAGTTGAGGAAATACGAGAACTCAACCGAGAGGTGGGACTCT
GGTTACCGGAATATACAAGCCACAGTGTTGGTACGTACACATTGCTAATTCTTTTTTGTTTTGGTTGCAGGGTAA
TCTTGTGCACCAAGAGAATTTAGACCTCCACAAGAAAGTAAACCTAATGCACCAACAGAATATGGAGTTACATAA
AAAGGTATCTTTGATTTTGGGAGAGGCAGGATTATCTTTAAGAAAGAAAGGTACAACATGTATTGTTTCTTCTAC
ATTGAGGATAATGTAATGTGGCTACCACTCCAGGTTTCAGAGGTTGAAGGTGTGAAAAGCGCAGAGAAAGATTCT
CTTCTCACAAATGGTCTAGGCATGAGAGATAACTCGAGCGAACATGTCCATCTTCAGCTCAGCCAACCGGAGCAG
CATGATGAGACACGAGCAAAAGCTATCCAGCTAAGCTATTTCTCTTTCATTGCATAATATAATTCAATTTAACCG
CAGCAGCTTGCTACTGCATAAGCGAAGCAAACACAAACATGATCAGATTGTAATCGATCTCGATCAGAAACCAGC
TTTGTATTTCTATTCTGTTTTACTCTGCAAAGAACTTTTGGTTTGGCGTTTAAGCATGTTCTGTGATGGACCTGA

*spa-miR824*

>CM001193.1

CTCATTCTTCCTTGCATTACTTATATAAAACCCTCTCTCGTCTCCTTAGTATTCTCCGTGCTCTCTCCTCCCTTC
GTCGCTCTCCCTCTCTATCCTCTCTCTTGCCCTAGctactacttcttcttcttcttcttctgaatccGCTTCTCA
TCAATGCATCTTCACGATTCaaagcttttattttatccCGATGGTTTCCCATTGGATTTTCAGTGGGATCTCGTG
ACGGAGGAAtcgccatttttttttcttaaagctgACCGGGTCTCTTCCTAAGCTTCGATTACTTCGTCAGTTAGTC
TGATTCCGTGTATCTCGTCTCTTAAGGTGCTTTTTAActtgcaattttattttacagatacgacattttgttttg
ttgaatcGTTTAAGATCTCTCTCTAGTAGTTTGAGAATTCGGAAAATGGgtttatcatttgattgatCAGCTGCA
TGCATGAGCTCTAGTCACCAAAATTTCTGAGCTGTGGATCTAAATTTTACTGTTTCAGAGAAAAAGATGgattgc
tttttttgtttttgtttttgtttgtgtgagtataaatgttttaaatttgggTTGCATGTGCAGATTGATGCTCGT
GGATGAGGAAATTCTCAATCTTTAATGGATCTATCTCCTTTTCTCAGGTTTGGTTTCTCatcttcatgtttttgt
tagatgcatatgttaaataaaatttcccATCCGTTGATGGTGctcttttgtgttgtttttttttgtccttggTA
TAAAAtatctgactttttttttttcttaagatgaaaaacacataataatgTATGATCCATGTCTTTCTAGAAGTAG
TGTGATTGTCAGTTGCTGATTGTAAACTTACTAGTATCTATAATCCTTTTACTTATGAAGAGCTTAATAGTAGTT
ATGAGCATGAAatgaaagcaaataaaagacATCTGTTTTCATAGCCTCTAAATgtccaaaaaatgaaatgtaacT
TAGAAAGCAATGGCTTGAATAGAAAGTCCATTAGAGAAAAAAGGTTGGGTGCTTAGATTAAGAAGGAAATGAATT
GAACTTGTTTCACTTgggagaaattaaaaattctgtTTATTGGATATAAGccattcaatttttcccTTAAATGA
CAAAGGGTGCTTCGCTGTTGCATAAACTAGTTTATTTGAATCTGAAACCCATCTGACTTTGTTAATGCAGTGCTA
CATTTTTGTATGGGGCTGCAGTGGTTGACTAAGTTGAAGACTGGCAGGTTCGCTCGAGTTGTCTCTTATGTCTAG
ACCATTTGTGAGAAGGGATACTTTGACTGCACCGTTCACCCCAATCTCTAAACCTGCAGAGTATTATGCTCAGTT
AAGGAAAATAGCTTGTTCAACATATCAAACTACCCTTTTAAAAGCAATCCTATATTCTAGATGCTTTTAGAGCTT
GAGATACGttgttttgttatgtataTCTGTACTCTCCCGGCTACCAAAACGTTAAGAACTAGTTGAATATTTCCC
CAAACCCCATCCAATAAGAAAGGTCTAGTCTGTAGGTCACACTCATTAGAGCCACGCTCACCCTCTCGATTAAGC
TGTTTAGTTTTTCCATCAAGATCTGATCCTTGAGGAGTTATGTATAAAAGTAAGAACATAAAGTGCATATATGCC
TTTTTCAGATGCTGcctttttcatttgtatttgGACTCGGTTCATGCTGATTctccaaattatataatatcattt
atacTTTGGACATACATTTAGCTTAAGAGATGTCAAACTTCTGTCTGGAGTGTTAAAGGTTTTCATGATACCTA
AGAATTTTGTATCTTGCACTAGTTAGCTACACTAgtttatttggtaaaaaaaattggggaGTGGGGGAGGTGCTG
GGTTATTTATTATTCCCTTCTCATCGATGGTCTAGATGTGCGAGGTGACTCTCATGGAGGTAAAGAGAAATGGTG
AGTTTGAGAAGCATCCACCATAATGGTATTTCCTAAGAAGGTTAGAAGATTTACAGCTTGTGGAATTAATTATTC
TACTGCTAGATTTAGGACCGGTGATCTGCTAATGCAACACATGATACTTCCAAGTATAGTCGtgtgagctttttt
ttttcaggttgaGTACATCATGCCATTATTCAAACTAACGTAGATGATGGATGCAGAAGCAAGTAGCAGTGACTT
GGTTGCAAGAACTAAAGAAgcaggaaaaaaaagcatagTGTGTTTATGAATGAAAGAGCGAAGTAAACGAAAGGG
TGTACCTTTTGACTTGTTGTCACCGATTCTCTCATTCATCCATGTCATGCTGCAGCAGAAAACACCAATGGGAAT
GGGTGTGTTTACATGGCGTAACGCCAAGCATTAAGCGATTGCTTGAAATCGATTCGTATGGTAGATTCTAACTGA
ATATATAAGACTATGTATCTATCTACTTGTCCAATGctaaatatgaatataattaCTTGTTTTATACATGGATTA
TATTAGCATTAAACAAGTATAAGTGATGCCAGATAgcgtaaataataaattatattttaatgtgaCATTCATTCA
GTGGGACCATGGTTTCCACTTAACATATTACATGAACATATTCACAATAGACGTGTATTTTGGGGTATACGGAGG
TAAGTGCTTGCGTGAGAACCCACCAACCCAcgaaatgaaaaggaaaaatatgtaGATATGCAATACttgaaaagt
aaattaaaaagaagctCGAAG

#### Eutrema salsugineum

*EsAGL16_1*

>scaffold_1

TCCTCTTTCTATTTCTATTTTTCTTTTTTTGGAACTTCTTCCAATTTCTCTCTCTCTTCTCTCTCTTTCGATCAA
AATCACATTTCTTTCTCCCCCGAGATCTCGCGCGACCTGTTTTCACTTAGATTTTTTGTTTTGTTTGTTGGATAA
GGTAATGTTTAAATTGTTGATTTTTGAGTTATTGAATATCTATCGCCTGGATTTGCGGGAAATGATTTGACGATG
TTAAAAAAAAAAAGGCTGCAGAAAATGGGAAGGGGAAAGATCGCGATCAAGAGGATCGATAACTCCACGAGCCGT
CAGGTCACTTTCTCGAAGCGAAGGAACGGATTGCTGAAGAAAGCAAAGGAGCTTGCGATTCTGTGCGATGCAGAG
GTCGGTGTCATCATCTTCTCCAGCACCGGCAGGCTCTACGATTTCTCCAGCTCCAGGTATACCCTTTTGTTTTCT
TCCTTCTTTTTTTGGGTTGACTTTTCCTTTTGGTCCACTCTGATTGATTTGGGATTCCATCGGGTTTTTTTGTTG
ACCGCAAGGCAAGTTCTGAATATCCAATTTTCTAATCGATTTTTCTCGTTTTTGATATCTGGACATTCATTCACT

TGCTGTGTGTACTGTACATCCCTTAAGGTGACATTAAACATCGAATTAGTGTGTTTCATAATTTTATATGGCAAT
GGGTGGAACTTAGTAGATCACTTGGTTGGTTGACTATATGATCAATGGGCTGAAAAAAACACTGTTTGTTTGAGA
TTCATGGATTATATAGCTTTTAGTAATGGAATATGAGATTTTGATTTGTGGGTATGTTGTATGTATGTTTCATAT
GTTATTGAGATGCAGGACAGGACAGGCCTAAATGATGATAGAATTGGATAGGACTAGAAAGATAGAGTTTTTTTG
GTGAAGCGTATAGATCTTCAGGGACTTGTTGTGTTTCTCGAAACATACTAAATTTGGTATTTTCAGAGGAAAAAA
GGGAGAGGAAGTGGTTAGTGGTTTGTAGGGCAGAAGAAACTCCAACTCTCTCTCTCTCTCTCTCTCTCTCTCTCT
CTCTCTCTCTCGCTTTATCTCTCTCGCTTTGTGTCTTCTTTCTTGGTTTGGTTTGAGGTTTTAGAAAAGAATCGT
CTTTGTTCGCTAATCTTTTTCTTTATTGGCAACTAGTTGGTGGGTTGATGTACCTAAAATGAGCTTGCATTATTT
TATGCTAATTTTATTTATTACCCAAGTGGGGGAATCATTTCTGCTTTCGTTGTCATCCATATTCAACTTTTTCCT
ATTTAATTATCCAATTAGGAACCCCTAATTTTAATGCTCTACTCTCAACTTGGTCATTATTTAGGAAGATAGTCT
TTCTTTTGGAAACATTATTTCATCCATTCCTTCTGATTTCCTACTACAACTTGGATAATTAAAAACATGGAGTTG
ATGGTTCTCAGAGATTGGGGGCATTAGATAAAAAAAAAAACAAAAAAAAACCATAGATAATAAGCAATGGTGTTA
ATTGGGTACTTCACACTATCGTCTCACTAGTTCATTTAGAAAAATCTTTACTTGCGAGCTGATAACTATAGTATT
TATAATCGGGTTTAGTGAATAAAGTTGGTCATCTACATCAAGATTGTCTAATTTCACTCATTCTCTACTTAAGCT
TTATACTTCATCTTACATTTATGATTATGGCGAATCTGTATTTTCGGTGCAGAGAATCCCGTTCCATGTAGGCAA
AAGATACTTAGGGTATTATAAATCTATTTAAGTCCAAGTTTTGACACTGAATATGTAAGGAGAAATGTTACTCTC
TCTGAACTCCGAAGATTTAAAACTATCTTCTTTTTTTTTTTTTTGGCAGCATGAAATCGGTGATAGAGAGATACAG
GGATGCCAAAGGGGATTCCAATTTGGAAAGTAGTCCAGCTTCAGAAATCAAGGTACTCCTTTTTATGACTCATCT
CTATGGATTTTTCAGAAACTCAATTTTGTAATGAGTAACAAAGAGACCGTTACTTGTTGTTAGCCTAATACAACT
CTATATAAGCATATTCTCAAGCTGATCCTGATTTCATATTGGAGTAGGATTTGCTATAAAACACAAGTCATCACA
CTTGACCATCAGATTACATTCATTATGTTGCATTTTCGTTTGCAAAAAAATCCATTCTAAACTTGATAACATATT
AATACCCAAAAGAACTTGATTTCTCAGTTTCAATCACCTGAAAATAACTTCTGCTATATTATAGTTTTGGCAAAA
GGAGTCCGCTATTCTAAAGCGACAGCTACATAACTTGCAAGAAAACCACCGGTATGTTTTGCATCCTCTGAGTTT
GTCAAACCCTCCCGATAGTAACACCTAGATCAGGGAACGATGTACTAATATTAAAATGGTAACATATATATGTAG
TCTAAGGCTTGAACGTTTAGCAGAACCCCAAAACTATCCTAGTATTTTGCATTTTTTTTTTCCAATAGAAATGTT
CATATATATAAAGCCAAACCAACAAAGAATGTTATAACCACCTCTTAAGGATTTTACCATGATGCTAGATGTCTT
CCCCAAAATCACAATTGTATCTCTACCGCCTCCAGAGTTATACCTCCACACATCTAAGACCATACATACGAAGGG
AATTACTAAACACTTCCACATTCCCCCATCATCTTCTTTGTTTTTTTCTTTTTATTAATTTAGACAAGGTATATA
ATTACTTTGGAAAATTTTGCATATTCAAAAGTGATGGGGAAACAAAGATCTTGTACTTGACAGGCAAATGATGGG
GGAAGAGCTCTCTGGACTAAGTGTAGAAGATTTACAGAAATTGGAGAATCAGCTTGAATTGAGCCTTCGTGGCGT
ACGAATGAAAAAGGTATTTGATGCTGTGACAAATAAGTTCTTTTTTTCCAAGGTGAAAAGTTTGGAAAAAAAAGT
AATGGCTGCACTCTATCTAAGCATTTATGTTTATCGACGATTGTAGGATCAAATGTTAGTCGAAGAAATCCAAGA
ACTGTACCGTGAGGTTGGACTCTGATTACAGGAATATAGGAATCAGACGTTTTCCACAGTGTTGGGTACGTAGAA
ATGCTAATTCTTATATTTTCGGTTGCAGGGGAATCTCGTGCACCAAGAGAATTTAGACCTCCACAAGAAAGTAAA
CTTAATGCACCAAAAAAATATAGAATTACAGAAAAAGGTATATATGATTTAGGGACTCTGGATTTTCTTTAAAAG
GGAAAGGTACCACATGTTTTTGCATTGAGGAGTAATATAATGTGGCTACTCACCACAGGTTTCAGAGGTTGAGGG
TGTGAAAAGCGCAGACAAAAGTTCTCTTCTCACAAATGGTCTAGACATGATCAGAGATAACTCGAGCGAACATGT
CCATCTTCAGCTCAGCCAACCGCAGCATGATGAGACACAGGCTAAAGCTATCCAACAAAGCTATTTTTCCTTCAT
TGCATAATATAGTTCAATCAATTAACTGTATGTAGCAGCTTGCCACTGCATAAGCAAAGAAACAGGAACATGTCT
GATAATTTCCATTTCGATCAAACACCAGCTTTGTTTCTTTTCTTAACTCTTCTTTCTCTTTGCATAGAACATTGG
TTTGGCGTTTAACTATGTACTGTGATGGTCAAAGAATGTAGTG

*esa-miR824*

>scaffold_1
CCTTAAACATCTTTCTTTTTTTTTTCATTTTTTTTTTTCACGTAGATCTATTTTTCTTCTAAAGAACTTAAAGCAC
ACAAAATGTCAAGTAGGCGAGGATAAGAAAATAATACTAATAGTTTCGGAGAGACCGAAGACAAAAAAAAAGAGA
GGGGGATGTGAAAAAAGGAACACAGAAAATGAGGGAGAAGTAGGAGAAAAAGAATGGATGAATTTGTCGGATAAT
TAGAAAAAAAAAACTAAAATTAAGCGTATACTATTTTTTTTTGTAACAAGCATACAGTATCTTCAACGATAGGAA
AGTAACAATGATTTATTATTGAGTCTTTATTATTTGACTAAAAAATAGACGAACTAGTCTTATTTGTCCTTAAAA
ATGAGTATATATACTTATGAGTTAAGAAAACGTATGAGTTCAGATTCATTCATCTATTTCAAGAAAATTGGTATA
TGTTCTTAAACATTCAATCAAAACATATGCAATATAATTGTTATTTGACTAAAAAAGTAGACGAACTACTCTTAT
TTTTCCTTAAAAATGAGTACATATACTTATGAGTTAAGAAAACTTATGAGTTTAGATTCATTCATCTATTTCAAG
AAAATTGGTATATGTTCTTAAACATTCAATCAAAACATATGCAAATGATTTATAATTCGACTAAAAAATAGACGA
ACTAGTCTTATTTTTCCTTAAAAATGAGTACATATACCTATAAGAAAACTTATGAGTTTAGATTCATTCATCTAT
TTCAAGAAAAATTGTATATGATTTTAAACATTCAATCAAAACGTATGCAAATAATCGCATACACATTGCTGTGTA
ATTTGAATAACAAACATAATAAACAATGATTTGAGTTTAAATATGTATGTATTCTCCTTTTCTTTCCGTGACCAA
TAATATGTATTATTATTATATAAAAGTATAGAACAATGCATTCTTATTTATTTGTTTGAAAAAATTGTGTGTCTG
ATAAAATTCAAATGATGATATGTGCAGATTTTTTTTTATATATTTCTTAGTTAGATACGTAAGTTTTTAAAAGAG
ATCATGAAAGGGATCACGTAAACAGAAAAGACACATGTTTTAATATATTCCAGGAAAATGATATCGAAGCCGAGT
GTAGCCACTGATAAAGTGTACAGTTACTACTCCGTAGAGAATGGAAGCCTGAGCTGCGGCTTTATGCGAGAATTT
TTGAAGCCCGACATGTCTCAATTAAATTAAGTATAATTTTATACTATTTGAAATCTGATTAACCCACTAATCAAT
GAATTATTAAGTTTTTAAACTAAGCTCATTCACATCACTTATATAATTACCCTCTCGCCTCCTTAGTATTCTCC**G**

TGCTCTCTCCTCCTTTCCCTCTCTCTCTCTCTCCCTTGCCCTAGCTTCTCCTTCTTCTTCTTCTTCTTCTCAATC
CGCTACATATCATGAATCCTTACGGTCCAGATTATGGCTTTTTATTTTTCCCGATGGTTTCACATCAGTGAGATC
TCGTGACTTGGGAATCGCCATTTTCGAAAGCTCATGCGTCTTCTTCACAAGCTTCGACTACTTCGTCGGTAAGTC
TGATTCCGCGTATCTCGTGTCTTTAGATACTCTTGTAGTTTAATTTAACATGCGGTTTATTTTTGTTAAATCGTT
TTATTGAATCCGTGAGAAATTAGTTTGAGAAGTCGAACTTAAATAGTTTTTGCTAAAATGGGGTTTATCATTGAT
AGACCAGCTGCATGAGCTTTAAATACCCGAATAATGAAATGCATATGTTTCAAGAGTTTTTGCTAGATTCATATG
TTTCAAGAAAATCTTCCCACCCTTGATCCTACTTCTTTTGTGTTGATTTTGGGTATAAAATATCGACATTTTTTG
TTTTCTGTATTGATTTTGGGTATAATCATGCATTGTGATTGTCATTTGCTGATTGTAATCACTAGTTAGTGTCTA
TGTTCCTTTTGCTTATGAACAGTTTAATTATTAGTTATGACTTATGAGCATAAAATCAAAGCAAAGTAAAGACTC
TTTATCTTCTCTTGACCAACGATAATAACACAGAAAACAGATTCACTCTCTTGTGTCTTTTTATGTTTTTTCATT
TTTCTTGCATCTTGTACCTAAAAAGAGTTTGAAAGCATTGTTAAATTCATAGAAAATGGTTCATAGAAAGTCTAT
GAAACGTACGAAACATCTTTGTTTCTTTTTTTTTTTTTTTATAAACAGTTTAACATGGCTATTAAGTTGAACACT
CACTATGGCTCTCGATGTGGCGATGATCTATTTGGAAGCTTCTGAGAATCAAAGCTTTGATCTTTCTGTTCCTTG
TCAATAAATGTTGGTTGTTCCTAAATAATACTTTTCAAACAACTATATGATAAGAGAATAGGCCAAATCAACCCA
ATAAAGCTCAAGGGTGAGAGTTGAAAACACCTACGGAGTGATACAGCCTCCTTTTGACAACAATATAGGATAGTT
TCATCTGTGTTCAAAAATGAAATGTAACTTCTTTAATTGAGAAAGCAATGTCTTGAATAGAAAAAGCATTTGATA
AAGAAAGTTGAGTGCTTAGATTAAGAAGGAAGTGAATGAAACTTGTTTTACTGAAGGTAAATCAAGAATTCTATT
GATTGGATAATAAGCAATTCAATACTTCATAAACGATTCAAATAAATAGGAGACTTTGCTGTTTCATAAACTAGC
AATACATGATCAGGTTCATATCACATATGTTTTTAACATAATTTATCTGCCTTTCCTTTGTTTATGCAGTGGCAA
ACCGTTCCACTTAATTGAGATTAGATTCTCACAGTCCAGCTGAAATACTGTGCATCAACCGTTTTAGCATGAGGC
TGCTCTGGTTGACTAAACCGAAGACTTGCAGGTTCGCTTGAGTTGTCTCTCATGTCTAGACCATTTGTGAGAAGG
GATTTTTATTTTATTTGCACCGTTCACACCACCAATTTCTAAATCTGCCATAAGTATTATATATGCCCAAGTTAG
GAAAATAGCTTGTTCAACATATCACACAACCCTTTTAATTAAAAACAATTCCATATATCAGATGCCTCTACGGCT
TGAAATATATTGTTTTCTTATGTAGTTATGTATACTCTTGGAGTAGTTTCTTGTGTATCTTTAGATATTCGCTAG
CCGCCTAGGATTTTTAAGCTACCAAAACGTTAAGAAATTAGTGAAATGTTCCCAAACCGCATTTTTTAAAAAGGT
CAGTTTATAGGTCACAGCCATTAGAGTCGCACTCACCTCCTTTTTAAGTTGTTTGATTTCTTCCATGGTCCATGA
AAATCTGATACTTTAGAAGTTAGGTATAAAGGTAAGTACTTAAGGTGCATTTCATGACTTTTCTTGATGCTGCGT
ACATAACTTTTTTCATTCGTATCAGCCGGGTTTAGTTCAAGCTGAGAACTCCAAATCATATAGAATTTGACACTT
TGGACATGCATTTTGCCTAGTAGATCTCAAAGTTCTGTCTGGAGTGTCCGAGGTTTTTCCATGATAGCTAAGAAT
TTTTTATCTTACACAATAAACTATACTAGTTTTCCTAAGAAAAAAAAATGGGGAATGGGGAGGTGTTGGGTAAAT
TCCCTTCTCATCGATGGTCTAGATTTGCGAGGTGACTCTCATGGAGGTAAAGAGAAATAGTGATTTTGAGAAGCA
TCCAACATAATGGTAATCTCTAAAAAAAGGTTAGAAGATTTTCAGCTTGTGAAATTGTTATTGCTAGATTTGTAG
ACCAGTGATCTGCTAATGCAAGACGTGAGATGCGTCCAAGTAAAGGTTGTGTGAGCTTTTTTTTCAGGTTGAGTA
ACAACAAGCCATTGTTGAAACTGACGTAGATGATGGATGTAGAGAAGCGAGTAGCAGTGATTCTGTTGCAAGGTC
CTTTGCAAGAACAAAAGAAGTAAGAATGCTGAACTTAAAGGAAAGGAAAGAAGGTGTGTTTATCGATGAAAGAGC
AAAGTGAACGAAAGGTTTACCTTCTGACTTGTTGCCACCGAATTCTCCCATTTATCTATTTCATGCTGCAACAGA
AAACACCAAAGGGGAATGGGTGTGTTTACATGACGTTAACGCCAAGAAATTAAGAGATTACTTGTAAAATTGATA
AGTTCCCTTACTCTGTTGATTCTGACTACTAATGAAAGATGTATCTATCTACCTAGCCAATGCTCAATATTTATA
CAATTATTTCTGTATTGATTATATTAGCACTAAATAATAAATTAATTTAATGTGGCATTCAGTGGGACCCATGGT
TCACAACAGACGTGTATAATTAAAAGAAGCTCGAAGCTTAAAGATACCACTTGTTTAGTCTTGAAGAAGAATCAA
AAACATCTTTACTCGTTGAAGATAATGAATCAAAGTTATTAACAGATTGTTTTTCTACAAAAGCTTCGACATTAT

#### Aethionema arabicum

##### *AaAGL16*

>Scaffold_817

GACTAATCTCTCCTCGAGTCTTCACCGTTTTCTTTTCTTCTTCCCTCAGTCATCGGAAGATCAGATCTCAGTTTC
GGTTTCGTCTTCACTTTCCCGCGGTCTAATACTCTAACTTGTTTTAATGTTGAAATTTCGATCTGGATTAGTGAA
TATTCCATGGATCTGAGAGATTTTTGATGATGTTTGTGGAATAAAACAGCAGAATGGGAAGGGGAAAGATCGCGA
TAAGGAGGATCGATAACTCGACGAGTCGTCAAGTGACTTTCTCGAAGAGAAGAAATGGATTGTTGAAGAAAGCTA
AGGAACTTTCTATCTTATGCGATGCAGAGGTTGGCGTTATCATCTTCTCAAGCACTGGCAGGCTTTACGATTTCT
CCAGCTCTAGGTAACTTCATTTCCCCTTGCTCTATCTGATTGGTAAATTACCTATTGCAAAATTGATATGTGGAA
ATTCAAATTGGTGTTTAAAAGAGGTGTAAATGGGTATGGGTTAAGGTTCTTAGGCCGGTTTATGTATGACGATGA
TGATGTTACATTGAAATTGAATGATTCGTTTTTGAGTTGGGAAAAAGGTCATATCATCTACGATCTACCCAATTA
GTATATTAGTATAAAAGATGAACAGAGTCATTTTAAAATCATATGACAATGGTGGTGACGTTGTGGATCACGTGG
GTTTGTTGACTTTCTGTTAATGGGCTGAGAAATTATATGAAGCCCGTTTACAAAAACACTATATTGTAATGGAAA
AGAATTAGATTGAAGAGCGGAGAGTCGCTTGTTACTTGTTAGGGACTTTGTTCTGTTTTTGGCAAACTTGCTAAA
TTTGATACAAAACTTTTATTTATTTGTTTATGTCTTAGATGGGTTCGGTAAGTTAGAGAGAAAGAATAGTCTTTA
TCTCTTGCCTTTCTTTTTATACTTAGATTTTATTATTTTAGCAACTTGTTGGTAGGTTGATGTACCTAAAGTAGC
TGCATTATTTACCTTTGATTATATTTAATTATCAAGTGGGAGAATCATTTCTCATCCCAAGTCTATTTAATTATG
CATTATAAATCCCATATTGGTCATTATTTATTAAACAATTTGGAGTTTCTAGAATCCATCAAAAGCTTTACGGCA
GAAGATGTAAGTTTTAAGTAGTGGACCAATTTCTAAAAATGGGAACACTCTTTCGAGTTTCATGTTGTTATATAT
ATCTCCAATTATATATTTATTTTTCATTTAAATCAAAGTTGAAAACAAATATATAGCATTATATCTCAATAATAT

TTATTATTTTAAAATAGAAAAAAACATTATCATTTTTTTAAACAAAAAAATAAATCATGTTTTTTAGATTTGAAC
AGTAGATGTTTTGAAAAAATCTATTAGAGTTAATATTTGTCAGAATCTCAATCCCACATTGGAAAGAGTCTAGCA
TTTAAATGAATATATAAAGGATATGGGCTACTCCGTTCATTGCCGGTTGGTTTTGAGTTGAAACCTAAAATGGTA
TCAGAGCCCAGGTATCCGTATGAATGTCATTGACTTTTTCGTCTGGGATGAATTGTACAAAAATGGTCTTCAAAG
ATGGACTATTTTGTGACATATTCTTGAAGAAAATGGAGAGTATAATTCATGTATAAATAAAATGAGAGTATGATT
AGAATCGTAAGTCGGTGTATATAATGAAAATTCCATTTCGTGCGGGCAAAAGATAGCTCGTAGTGTAACCTAAAT
CCTACACCATGAAAGCAAAAGATAGCTTACCAATACTACACTCAGCAAGTATACAACTCTTCAACTTTTTGGCAG
CATGAAATCAGTAATAGATAGATACAATGATGCAAAAGGGGAAACCAATTCATTGACCAACCCATCTTCAGAAAT
CAAGGTTACTCTCTAAATAACGCACTTTCTTCTTATTTATTCATAGATTAAACACATAATGTTGCATTTGGTTTT
CCCAGAATCTAGATCTAGGAATGTAGAAGAATCAAAGTAATTCAAGAGAAATAGATACTTGTTTTCTTAGTTTTA
ATCACCAAACCATCATTTCTGCTGTATTCTATAGTTTTGGCAAAAGGAGGCTGCTATTCTAAAGAGACAGCTACA
TACCTTGCAAGAGAACCACCGGTATGTTTTCTATCATCTTCTCCTTCCTCCTAGTTCATCAAACAGTCTCGACAG
TAATGCTACAACAGGGAATGATGTGATAAGAAGCAAAAATGATAAGATATGTAGACAGTAGTCAAAGGCTAAAAC
ATCTTCAGAACCTAAAACAATGCTACAATAAGGTAAATAAGAATGGACGATAGTAAGCATAAAAAAGCCAAACCG
ATAAAAAATGTTGCCACCTGTTTGGGATTACCATGATGCTGGAAGTCCTGCCCAAAATCACCATGGTATAAAGCA
TTATGAGAAGGAAAAAAACCAAAACACTTCCACGCTCTCCATTTCTTTTTTTTTCTTTTTGTGAATTAGGAAATGT
ATATTCCTAAGCTCCAATGTGAATTATTGTTGGAAACCTGAAGTAACACAGACAAAAGTTTGTCTTCTGTAACAA
GTTGGATATTCAAAAGTGATGGTGGAAGAGATTGTACTTGACAGGCAAATGATGGGGGAAGAGCTTTCTGGATTA
ACTGTAGAAGACTTGCAGAATTTGGAAAATCAGCTTGAAATGAGCCTTCGTGGTGTACGAATGAAGAAGGTTAGG
ATCCTGTAACAAATATAAGTCATTTTTGTAACACGTGGAAAAGTAATGGCTGTACTTTACGTACTCCTTTCTTCT
CTTCTATTTCTGTAGGATCAAATGTTGGTGGAAGAAATAAAAGAACTCAACAGAGAGGTGAAACTCAAACTTCCA
TTGCATAGACATTTACTAGAGTCGTGTGTCCTCACAATGCTAACCCCTAATATTTTGGACTGCAGGGAAATCTCG
CACACCAAGAGAATATAGAACTTCACAAGAAAGTATATCTAATGCAGCAAGAGAATATGGAACTATATAAAAAGG
TATTATGGGTTTTACGAAGTAGCAAGGAGCCAAGTAAATCTTTAGAAGGCTGGAATAATAAGTACAACTAATTAA
GGATAATGCTCATTCCAGGTTTCAGAGATTGAGGGTGTGAAAAGTGCAAACAAACATTCTCTTCTCACAAATGGT
CTAGACAAGAGAGATAGCTCGAGCGAACCTGTCAATCTTCAGCTCAGCCAGCCAGAGCAGTCCCACAATGAAGAG
AAGCAAACAAATGCCACCCAACTTGGCTACTTTTTCTTCATTGGATAGTATATTTTAAGCGAGGATACACTTTTA
AGTTGACTTCAGCAGATTCATATCTCCAATAACTGTAGCAGCTACCTCTGCATAAGCAAAGACACACAAGTCATT
GGATAGATGCGGTTTGGGACTTTTCTTTTTCGTAAAGACTTTGATTGGCGTTTAAGCATGTATTTGATTGATGAG

*aar-miR824*

>Scaffold_31

TTTTTTTTTATGAATATTGGAGCTGATCGAAACTTGGTTGCGATAATGAGCCGTTTGAGCTGCGCCTTTAAGGGG
AGCCCTACATGTTTTAATTTAATTAATTTAAAAATCTGATTAACGCACTAATCCATGATTAAAGTTTTAAATTAA
CCTCATTCTTCCTTTGCATTACTTATATAAAACCCTCTCCTCCCCTTCTTAATCATTCCTCATCT**CTCCTTCTTC**
**TTCTTCTCTCTCTCATTTGCCCTAGCTCTCCTTCTAATTCCGCTTCACACATACCTCCTCATTGTAATCGCCTTC**
**TCTTTCTCGTTGATTTTCAGTCGGATCTTGGTCGTGCTTGCCATGGAAGGTCGCCATTTCAGAAATTCTTGCGCA**
**AATGTTGCTTTCACCCTCTCGTCG**GTTAGTCTCGCTCCGTTTTCTCGATTTTTTTTATTTACGGTGCTTTAGAC
TTGCAATTTTAGCCTTGTAAGCGGTTATGGCATGTGGATTTTGTTAAATCTTTTGTGGTATTTGATTTGTCGATG
AAATCTCCTTTCGTTGATCGATCAACCGCATGGCATTTAGATATTATAAGAATTTTTTAGCTCCGAGCTTAATTT
TACAGTTCCAGAAATCGTGGTTAATGACAATTATGGTGTGCTATTTAGATTTTAGTCTGATGTCATTAAATTCGG
TTGCATGTGCTGCTGTGCAG**ATGATACCCGTCGGCGATGGGTGGGAGTTATCTTAATCTTGAATGGAAATGGAGC**
**TGTCTGCGTTCTCAG**GTTCGGTTTCTCACCTTCATTGATTGTCATTCTGAGATGGTTATGCTACATGAAAATTTA
AGGCTTATTTTGGTGTTTTTTTAAGATTAAGCAAACATATTAACACATGATTATAATGTGAGCCTTAGACGTATT
TGAAGCATTGTGACTAGCAGCATTTATTGATTGTTCTTTGGTTAGTCTATATTTGCTTTACATGAGAACTGCTTC
ATTACTGAGGAAAAAAAAAAAGAGAACTGCTTCATTGTTATCTGCATAGCACAAGATATCATAACAACATGGCCT
ATTAAGATAATCTATTTGGAACCTTGTAGATTATCCTTTTGATCTTTGGATTTGTTGTATTTTAACTGTGTTTGT
TCCTCAAAAGATATACTTTAAACAACTATATGATTAAAAAAATAAGCCAAACGAAACGATGAAGCTCAAGACTAG
AACCTGCCTTAAATAACATCCTGAGAGTTGAAAACACTTTGAAAGTGACGGTGTCCTCGGACCTTAAAAGAAATT
AGTTAAATCTCTCCCCTAGATAATGAATGTTTTACTACATTGACACTACTTTTATATTGCAATATAGGATATTTT
CATTTGTGTGCAGTCTCTGAATGTCCCAGATACACATGTCAACTAGTTTTAACTGGCTTGAATAGATCTAAAACG
TTAACAAGTTTGGGAATCCTATATCTGTATTAAGACTAAAACAAGTTTCCATGACCATGTGTGCTTTCTTAGAAT
GTCTTCAACTTAGAGAATGTGAAGGGAAGCTTGTTTTCCTGACAAAATAAGTTGAGTCACTAATTGGTTGGAACC
TTCCTTAAATCATTTGGAAAGTTACTTTTATGTTGCATAAACTATCAATGCACGAATGTCAGGTTTAAATCCACA
TTTAGTCAAGCTAGTATATCTATGCCTCTCTGTACATGAATCTTGTTTCATGTTGGTAATGTGGATTATTTGAAT
CTGATCATTCATCAGCCTTTGCTAATACGTTGCACTCTGATACAG**TTAATCGAGCTGTAATGTGGCTGCACATCA**
**CTCGTCTCATCAGTTTGGTGCTGCTGTGGTTGGCTGAGTTGAAGATTGACAGGTTAGCTCGAGTTATCTCTCATG**
**TCTAGACCATTTGTGAGAAGGGAATTTATATGCATCAACCCCACCAATCTCTTGAAAATAACTATCTTGCCTAAT**
**TAAAGAAACCCACCTACTCAACTTATTAAAGATAATCCTGGATTTCAAATACTCTTCTTAAGCATCAGGGTTACT**
**TTGTTGTGAGTTTCTAGATTCTTTTGTTGCAATTTATCTTCACACACTTCACCCAATGAGAAAAAGGTCAATTTA**
**TAGAGGTCACAGCCATTAGAGCGTCGGTTTCATCTGTTTTATATCTCAAATTGCATATCGATTCATGGTATCTCC**
**AAATTCTACATATAAAAGTTTTGTCATTTGTACATTTTAGACTCAGTTCTAGCCAATTTTTTGAAATTATATAAA**

**AAAAAAAGATTTCAAACTTGTCTTTGGAGTGTGTAAGATTTTTTCATAATATAATAAGAGAAGAGTGGCGTGGGG
GGAGGTGTGTGGTGTTAATCCCTTCTCATCGATGGTCTAAATGTGTGAGGTGACTCATGGTGTCATTTTGGGAAG
GTTTTATACACAGTAGATCGACATGAAAAAAGGTAAGAGAAAAATTTCAATTTCTGTGAATTTGATATATATGTA
TTGTGTGGATATGGTGATGAGTAATATATAAGTGGGCAGGTGAGGTGAGGTAGCTTTTTGGTATTGTGAAACTAA
TGTGTATGATGATGATAGATGCAGAAGCAAAGAAACGTTTTTATTGTTTGATTATTTGATATGATGTTAGAGGAA
AAAAAAGGTGTATGTGACGTAAAGCCAAAGAATAGGAGAGAGTTGGTTTTGAAGTTTATCGATTACGTAATGTCT
CTAAGTGGAAGATGATGAGTGGTGA**CGCTGTTTTTGTAACTGCTTTGATCTGGTCATGAATTGGATTTTTTGGTA
CAAGATTTGTGTATTTATGTTAATTTTGGGGTTTAATTGTTAATTGTATGTTTGGATTTATACAAATTGGGATGA
TGACAAATAAAATTTCACAATTTGTGTGTTAGTAACAAAAAAGAAAAATTTTACTGTTTTTTTTTGTGTGGCAAA

#### Cleomaceae

##### Tarenaya hassleriana

*ThAGL16a and ThAGL16b, tha-miR824a*

>gi|537644881|gb|AOUI01034514.1|

AAATGATACAAGTTGCAAAAAAAAAAAAAAAGCTTTTTGTTATAGGCCGAGCCTAATAAAATAATAAACGGGCCGA
TGATCTGTTAAATGGTACTATCCATCTGATGACGTGTACACGAAAACGCCCCACGCACGCTCCACGGTAGATATT
TAAAACCCGATCAAGCTCGCTTCTCCGCATAGGCGCATAGATATTTTCTTCCCTTTTTCTGATTCTCATAACTCT
CTCTTCGTGCATTTTCTCTGAGAATTTTCTCCAGATCCGTCGCCGAAGATATTTTTCTCTCTCGCCGGAGACAGA
TTCTTCGATTTTCTCAGGATTCCGGTTAGAGAATTCATGCCTTTTTTCGAATCTCAGATCGTTATTCTCGTGGAT
TTTCACTTTCTTAGGCGTAGGTTCCAATTATCTTTAGTTTTTGTTCGAGATTTTCTATCGTTTCTTTCCTGAATA
TCCTTGATTTTGTTTTCGTTTTCGTGGTTTTCACGTGTTCGTCTATATGTGTACTGTGTAGTTGCGACGCGACGT
GACTTTGTCTGTGTTTAGTGGTAGGGTTTAATCGTAGTGTCTAAGATCTGTTTGGTTGAGGAGAACGCGGGAGAA
AACGCGGTATTATTGCGTTTTAGGCTTTTAGCCCCTGATTCTGCATTCGTTGAACGTTTCAGGTTCTATGTTTTG
TTTACATTTCTCCTCTTAAACTTTGGCATTTCCTTTTATCCTTGAATGGATTTTTTCTTTCGGTTTCTGCTTTGT
TTCCATGTCAATCTGCTTTCGATTACTGCAAAGTATGATTGTTTTGGAATTTGGGTATTTGCATAATACATAATA
GCTTGAGTAGCTTGAGCTAGTCACAGCTTGTGTTATGGCTTCATTTTGTTAATGAGAAGATAAATCCTTACTTGT
TGCGCTCTGAAGTTAGGGTCATTTTGGTACATATGCAAGATAACTGGAGCTTAGATGAGTGGGAGATTACAATAT
CCACATTCTAGAATAATAGATTTGTGATAAAACAAAAATATGTAGTTGTTGCCCAAATTTGGGTAGTTCAAACGG
TGGAATGGAAGTGTAAAATGTTTGTCCCGCACCGAGTTTAAACCTCAAATTTGGAGAGGGAGTGTATAATCCAAA
TCACGTGTTTAGTTTCCGCCCTCAAAATTAGTTTGGACCTTGCTCCAGGATACCCAACTGTCCAAAAAAAATATG
TAGTTGTTACTTATTCGTGATGTTGAGCGATTAGTTCATTTTTAGTTTATTATTATTTTTACTGTGAGACAAGGG
GAATGTGTGCACAGGCCCATGGGAAAGATGAAATCTGAAGACGTTTTGTGTGAAACCAAACTATTTCTGAAGTGG
GTTACCATTGTGTAAAACAAAGGCTGCTCAAAATAGATTAATCAAATTACTTTGCCCAAAATAAATTTCTAGGAT
TCAAATCCTTGACCAACTTGCAGATCACTGTGATTCCAGATTTTGTTCGTTTTTTAACCAATTTTATAGAGCTGG
ATTGATGTTGCTGTGTTTACTGCTTTGCTTCAGTTAGAGAGGGAAAGCAAAGGCTTGTGATTTAGTTTATGTTGC
ACATAAAGCCTCGGACTGTTTTTTAAGTCAAATTAGACTTTTCCATCTTAGTTCTAGATATATACTGACAGTTAG
CTGGTAATATGGTTATCCATGCTTAAAATAAGATATGTAATGGCGCTTGAATCCGAGACACGCATACACTTCATT
TTTATGTATTACTGTAAGTAGATTTGTGTCTGTTATGCGTGCCCCCTGCAAGTATAAAATTATTATGGAGAGATC
TTAACTAAAGAGTTTGTGATCCATCTTAAAGAAGAGACAAAATTGTTATATTGACAGTTTTTAAGCTGTTTGCTT
GCATCTTTGTCGATTTCTCTTTTTGCTTGTTCATATTTTCTTTCCAAAAATTTCTGGAACTTCATTTTAATGAAT
TGACAAATCGAGTTTTCTCTTGTAGTCAGGCATTCTTACCACAAATAGAGATGGAGTCTTGGACCTCTTGGTATA
ATGCACATCGAGCTTTCTCTTGGCTTGTGATTTTAATGAATTGACAAATCATTGAAACGTTTTTAACAATGTCGC
TTGGAGATCAGCCGTCATGTAGTCAGGCATTCTTACCACAAATAGAGATGGAAATGAAAAAAATAGAATCTATCT
CTATTGCAGTGGAAACGACGCATTTTCGTTCTTCCTGAGAACATAGCAAGAGCGTTGAGTGTGTCACAGTTAACC
ATCTTTACGACCGTGGGTGTACATGTGCTTTACTGGTTCCTGCAACTTCTACTGATTGTGCTTTGTTTTGTAAAG
TTTATGTATATGTCTTCATATTCTACTCATCTGTGATAATTGTTGTTTCCATTTACTTCTACATCCTCTTTGAAA
TGACATTTGCTGCAGCAGCACAGTTTACTTTTTATTTATTTCACCTGATCTTATCTTTACATGAAACACTGCATT
CAGTTATCATCAAATACATGAATCTGAAAAAAAAAAACTTATATACCACATCTTAGATAGTAGAAGAAAATGTTTG
GAGTCTTGGTCTACTTTGTTAATGGAAGTAGCAATTTCCTGCTGCTGTTTTGGTCAATGTTGCAGCTAAAAGTTA
CTGCTCAGGAACAATGTCCGGGAAGAGGGAATAGTACCCTAACTTGGTGCCACTTGTCAATGGCTCGTAGATCTG
TTGCACAGGTACGCTTGATGCATCTCCTATATCTAGGCCATTTGTGAGAAGGGAATCCATGTTTGTACCATTTTT
GGCCCTTTCTTCGTGTACCTGGTGTTGAAGATTACTTGCTTTAGGGATCTCTTTGACATGTCATGCATATATTAG
TTCACATGATGGATACCTAGTAAACCATACTTGCTAATACAATTGATCAGAAAAACTCTTCTGCTGCAGCAAAAA
TATTAAAACTAGCTTCCTTTTTTTTTCTTGATGACTTCCGTTTACCTTTTTATAGAGTTCCACGTTCTCCTCGTGA
ATGAAAGTTCCCTGAAGAACAGAAATAAGACTAGCTCTTGCCAATGAAATTTTCAAGCAGATTTTGATGAGAACT
CCAAAAGCAATGATTACCTTTCTTTTCAATTCTTGTATCTCATCAATCATCCTCTGGTCCTGTGGAATTTGATAG
AATTGCACTATCAGCGTACTGAGTCATATGTTGCTGAGAATCATATTCTTTTCTTCATAGTGTCAGTCTAACCTT
TCTCATTCGAACCCCTCTAAGACTCATTTCAAGCTGGTTCTCCAAATTCTGTAATCCCTTTGCACCTAGTCCAGA
AAGCTCTTGTCCCATCATTTGCCTGTTTGCATTAAGAAAATAATTTCTAATACCAGGAAATTTCCAGTCTTTGCA
ACACCCATAGCCTTATTCCAAGTCTCTAGGATCCAGTGTTTTTATTTACAAGACGACATATCTTTGCTCTAGGAG
CTTTCTTAGAGGGCAGTCTCTCGTTTTGTGATCAAATGAAATGCCCTTTACCGGGAAACCATTTTCTAGTGCTTA

AAAAGCTCGGACCGTAAGGATTAGGGCTAATATGGCATTTACGTTTTGGACTTCTATTCCTTAGTTCTGATTCTT
AGTCCATTTGCTTTGAATGAAGCGAAACACTAATCTGTAGGCCACAGGAAAGAAGAAAAGTTGACTTACCTGTGA
TTTTCTTGCAAGCTGTCCAGTTGTTGCCTTAATGCCACCACCTCCCTTTGCCAGAACTAGAAAATAGGACGCAGA
TTTTCATGTGTACTGGAACTTAATAATGCTAAAATGCAAAAGAGACGATATTTTTCTAGTATATGCACCCTGCAA
TGGAATATGTGATAACATGATTACAAATATCCTGAAATCAGCAGAAAGATTGTAAAATTGTTAGATTTCACAAAT
TTAGAACAGATTTATTTATCATGGCCAGCTGATCATACTCAACAAACAACGTTTCAATCATATCCTTTATTTTGT
TATCTACACGTGCGCAGGAATCTAATATATAGTGCGGTAGATTGTTCTTTTTTTTTTAATATTCTGACCGATGTT
TAAATCTTACGGTGCAGCGTGCCTGGTTTTTGCAACCGGAACAGCAATGACTTCATGAGTCATGGTAACAACATC
TTGGCCTAGTAAGATCATCAGCTCAGTAGAATATATCATGCCTACATGGTTCTGCTATGTTATTTCTGCTGCACA
ATTACAAATGAATTATCCGGTGCCCCTAAGGTACCTGCACAACTAGTTAGCTAAAATTGTCTATCCTAGTTGTAG
GCGTACTACACATTTCCGTCCTTGCATGTTTTTCCTCCTTACCTAGACTAAGCCATTCTAACATTCAGGGAACCT
TCGTCATGACGGAAGTCTCCTGACATATCTAATAGATCGGAATTAAATACCCTGCCTTCAGCGGAATGGATCTGT
TTAAATTTCAGAGACCAGCACAGTTAGAGTGAAATTTCTGTTTATTTGAATAATATGAGAAACTTGTGAAAATAT
TCTGGTATTTAGACATCCATGCATTCACATGTTTCAATTGAATGCTAGAAATCCAAATACTGTGGTTGGAGTCAT
ATTTATTTTCGAGGTTAGTGATGAAAGTCAATAGCTTTTAAGATCCGAGTCATTCTATATCTTGTCACTGAAGAA
GAAAATAATTGCACAAAGCCAAGAGGTTTTTGCACCAGGACCAGCTTCATTAACACATGCAGTTATCTAAATTTT
CGGTAATAAATTAGCACGTTCTGTCATATATGCCATTGTCAGCGCTAGAATGATTCAGAAGTATGTTAGTATATA
CGCGAATGTTTCGGCAAGCTTAGAAGAACAGGAATGTGTCTGCACAGAATTCTCTTTTAATTGATGTTTTATTAG
TACCTTGACTTCAGATGCTGGGTCACTCATTTGATTATTTTCCTCCTTTGTCTTGTTGTACCGATCAATTGTGGA
CTTCATGCTGCCCAGACCAAGGGAAAACAAAGGAGAAAAAGGAACACTAACATCAGATCAAGCTGTTGATGGGTC
AGCAAGAAAGAAACTGAAGTCGTTCATATTTTTTTCATATGTAGATACCACAAGCAGTTATGGGTTTCATTCAAA
ATGACCGGGTGTATTCAAGGAGCATTAGATAATCACCATGGCTGAATGCTGTAAAGCTTATTATCTCATCTTCAG
CCTAACACAGGTAAGATTCTAAGCCTCAAGTCTTCTTTATATTGCGAATCAGAAAATGAGCTAGCTTAAGAGAAG
CCGAGGAAATCATTGGATGTTTAGTATAACTGGTAGATTAAGCTATTAGATGGAAAATAATCAGGGTCATGCTAG
CATTAGATAAGCACCAGAAGAAACCTAGCTTCTTTACAGCTTTTGTAGAAATCTGGATTGACATGGAAATCACCA
AAACAGTAGAAATCACCCAAAGCAGGTTTCAAGAAGATGAGCCGACAATGTGTTGGTGTGTAGAAGTATAGCAGA
TGAATAATACTGAGATTTTGTCCTTGTGATTACACCAGTGGCACGAGCTATACACTAAAGCATTCTAGGATTACC
TTGCTCTTTAATCCCAGGCAATGCAGAGAAAAACAATCGATAGAACCACTAGTTATATCAATGACCTACGTAGAA
TTGTAAGTCGCGAAGAACATGGTCTGACTATTACAATTGCTTCAGCGAGAGCTGTTGTTAGTGTTGGACTTCTCT
TTAGTTCTATTGGCTAAACAATTATTTGGTTATCCAAACTTGGATAATTCAACCGGTGAAATATATGCGCAAAAC
GTTTCTTCCGCCTCGAATTCGATCCTCAAGTTTGGAGAGGAGCGTATATTCCAAATCATGTAACTGACATCCGAT
CCCGAGATTAGTTTGGGTCTTGGTCCAGAACACTCGGTTACCAAAATAATTTTTTGATTGTCATTCTAAGCTTTG
GTCGATCAAACCGAACTCTAGTCGTTTTTTCCAATGAAAGGATGCAATATTTTTGTTCATCACTCACTCACTCGT
CTTTTCCAATGATTGAATTTTCTTTCTGTGATCGGTCTCTTCTAAATAAGGAATTATTCCCAAACTTGGTTTTAT
ATATCCAAGTCTTCGTATTATAGTCGGATTTAAATGGATGAGAAGCATGGTCGAACAGACCGAAAGAGCTCTTGT
TATACTTATCCTCGTGATAATTATTTATAACAGAAAATCCATATGATCAATGAGACTATGTAAGAAACATCGCAA
ATGCAGATGCATAAAAGAAATGGATGTTATTTCGCTAGGTTTTGTTATGCCTAGAAGTTTTTGTTAGCCAACCCA
TATACCACATGTATAGCAGACGAAAGACACCACGACGGTAAGCTACAAGAAGGATGGTGGGCGCATGCCAAGAAG
AGCAACACTCATAGTTAGGCCATTTGGTCAGAATTAGGCTAATTGAGCTTGGTATGGTTCATTTCAAAAGGCATT
ATTAATGGGCATTATATAAAAAATGTCATTCCTTTCCTATTTTGATTTCAACACATATCTCCAATTGTGACCTTA
TGACCCATGTGATCAGAAATATAGTCTGATCAAATCCTGGTCTTATTGACAATAATGTTAAATTTATGGTGATCA
TATTAATTCATTATGTTCATACATTGAAAATCTCATAAATCGTTTGATTAAAACATTATAAATTTAACGTTATGA
CCAATGAGATCAACCTGTGACCACACTATATTTCAGATCATTTGAGTAATAGGATCACAGTTAGAGAAAGTATGT
GTTGAAGTTAAAATAGAAAGGGAGAGTATTTTTTATAATGCCCTTTATTATGTCCATATATATAGAATTCTCTCG
TTAATAAGACATTTTTTTTTGGGATTTTGAAGAAATGGTAATAGTTTAGTTAGGTAAGCGCATTCGATGCTCTTA
AAGCTGAATATGGGACTCGGAAATTTCTATCAAAGTCGTTAGATTCAACTACATTTCTTCTATTTGGGTCAATAA
GAACCAGACTCGTTCGGAAGCCGCGACAGTCGAACAAAGTATGATATAATTGAAAGAGACATAGACAACATAGCC
AACAAATAAAGTTGTTGAGGTGACAGAATAGATTGAGAGTGACTCGGGGTCTTTTGCATTCGAATGCTAAAGTTC
ATTTTTGTTCCTTCGAGATGTCCTCATATACAGTAACGGGGAAGAAGAACAATGACACTTGCGCATCGTTTTGTT
CAAAGTGAGGGAACATCAATTGATGACTAACAGGAAGAAATGTTGTTTTAGTATGTGTAGAGGCGAATATCTTGG
CCACATTGTCACATGCAAAGGTATAGAGGTCGATCAAACCAAAATTAGCGCGGTGACAAAATGACCTATTGCGAA
AATTGCCCCATCGCATTCATAAGCTAAGCTTTTACAAGACCGACCAAGGTCAAATCTTAATATGAAAAAGAGTTC
ATGATAATCGTAGTTATCTAACAGTGGCAACATTATTTTTGTGGACACAAGTTTGTGGTCGCCCTATCCCGATGA
CCATTTACCAAGCGACACATTTAAACCAATTGTCAACTACTTCAGCTCAACAAATGGATCTAATAGAAGTAGATG
TGGATACGACATAATTGTTAACCCCATTTTATCCATTGTACTGTGAAATCTTCAGGACGAAAAGGTAGTCAGTCT
ATTTCACCCAATTAACCATTAGTAGATTGGCAAACGAATCTTTATTCTATGTTCATGTGATCTCCCGTTAAATTT
GTTAGGAGAGTTTCATAACCTTCTTATAGGTGGTCATTCAGGGATCAATAAAATTTACTGAAGGTTGAAGTGAGA
AATGTTCTAATGCGGTATGTAGAAGGATGTCATAATCCTTATGGATGTCTGTGATATATGCCAACGCTTCAAGGG
TTCAACACATTTATCAATAGGTTGATTGCCTTTACTAATTCCATAGAGAGTAAAAGTGATATCCCTATGGATTTT
ATGGAAGGTCTTCCTGTGTCTAAACGACTGGTCGATCGCCCAAAGCAAATGTGTTCATTTCTTGACACTGAAACA
TCTTTTATTCGGTATACGGATGGTAGAACTGTTCTTCTGGGATATCATGCGCCTCCACAAGATACTAGAATCTAT
TGTCTCGAATCGAGACATGGTTTCCTTAGTCAATTTTGAAAGACCCTTTTTGAACTCTACGAAATTAAATTGAA
GATGAGCACTATGTATCTTCCATAGATCGAATGAGTGAAATAGACAAAATATTGGTACAACACATCATTACATAG

ATCCATACCAATGATGCCTTTTCAGGCAGTTTATGAAAGAGAATCTCTGATTTTGTGACGACAAATCAGTCATGA
ATGTCTCCAAGACTTGCTTTCTAAAATGGTGTAAGAATAGGATCTTACATAGGTAACTCTAGATTAGCCTTAATC
GAGCACAAGTGATTACGAGGCATACCAATATGCAAAATGCCGATCCTAGGAATTTTGGAAAGTTGATTTAGCCTT
CTTGCGCATACGACCTTAACAGATGGGGTCATTAGCACGAAATTTTCTTGAAAGTTAGGGTCCAAGTATTTCGGT
CCCTGTGAGGTGGTCAAACGAATCAGTAAGTCAACTTATACATTGGCTTTGTCCCCCCACGACGACAATTCATCT
GATTTTCCATGTTTACCTTTTGAACCAATAGTGAGAATTACAAATTGTCGAACCGAGGAAATACCTCTGGGATTG
ACAGCAGATCTCGACGGCGAGTAGAACAGGAAAGCTTTCATGGGAAACGAGAATCTAACACTAATAGGGGTACAA
AGGTATTGATCCATTGGAGAGGATTTTCAGACTTTGAGGCAACTTGGGAACCTATTGAAGGTTTTTTAGAGCGAT
TCTATAAGTTTCACCTTGAGGATAATGTGAACATAGGAGGAGGGGTGTATTGTTAGGCGGCCCATGTACCACACA
TATAGGAGAAAGGGCAAGCAGTCTGAAAAACACAACGGGGTAGAAGACGGTAAGGCATAGAGAATGGTGGGCTCA
TGTAGAGAAGAGGAGTAGTTGGGGTCTCACTAGGCCATTTGGTCCTAATTAGGCCTTATGGTTTATTTTATTTTG
GATTTTATGTTTTATTGTTATTTGAAGCTTTAATTAATTATTAAACCATAACATCTCATCTAATGTATCTGTATA
AGAGTGTTTTCTCTCATTAATAAGACATTTTTGGGTTTTTACACGAGGTAGCAGTTGAGTTTGAGATTAAAATTG
AGGTTGAGTTAATAAGACAGTCTTTTGGTACCTTGTGTTACTTCATGGGTTATGGTTGATGATGATTTTCACGTT
TTAACAGTTTTTTCAAACGTTATCTTCGCTTCCTATGCAATAGGAAGTAACAATGACTAGTTGTATAGAAGTAAT
AACTACTTCCTCCATTTCAAAATAATTGTTGTTTTAGGTTATTTCACATTTTTTTAAAAAAAAGAGAAAAAGTCA
CATATACTCTTATTTAATGAAAATAATGTTATCATTAATAATTTATTGGAATTTGATAGGGATAAAATTGGAAAC
TATTGTTAGAAGTCGATGCATTACAAAGTTACTGAACCACGAGATTCTAAACTCTATGACGAGACTACATGGATT
ATATGTATATGAGCCAAAAAAAAAAAAAAAGTTAGCACCAAGTGGAGCCATCAAATCAACTCAAGCTTTATCGCTT
TCAGCGTATCTCAATTTGACATTTTAGAGAGAAAAAAGGAGTCTTAGAGACTTGGAAAGAGAAGCACAAAAACTA
GGCCAAGTGGCAATGCATGCGTTTAGCACAAACTTCCAAGGAAAGAGATTCCTAGGACTCAAATTGGTGTTTCGT
CCAATGAGCGTCACTTATCAGATTTTGTGGTTCATATCCTGCATGAGTGCATGCATGCATGCATGATACTTTCAC
GTGTTGCGAAAACGGAGATTCGTGTTATACTTCCACTTGTTTTCTCTTAACATCGTATGGTCACTTGAAAACTGT
TCGATAAATTGTTCATCCTAAAATTGCTTGACTAATAAAACACTAAATCTAGAATTATGAAGGAATGCCACCCGA
AATATGGAAAAACGCCTTGGTGATATATGAAGTGCAAACTATTTGTTTGAACTTCGTATAACCAATTGAATTTCC
AAAGCATTGTAACGTTACGAACTTACTTAAAAAGCGGTGTATCTTCTTGATTTTATATGGATAATAACATGAAAC
TCGGCTTGCAAAAACACTAAGCTCTTATATGGTCTTAAGGTTCATATATATGTGCAGTTCAAATGTACTATGCTA
CTATTCTTTTTTTATGATAGGACAATTGTTGGACTATGTGGAAGGCCATATCGCCAGTGGTTATGCCCAAACTCG
GGTTTGTGTTTAAACTTTCCACATTCCAACCAGTAGAATCAAGCCCGTGGGAATTTTCTATTATTTGTTCATCCT
ACTCTTTTGCATCTTTTTATTTTTATTTTTTTGCAATAGTTTTTATTCCTAGCGTTTATAAACCCACGCAAGAAT
ACACATATTGGAATTGATGGGGATTCTTCAATTTTCATTTTAGTTCTTATGATGGCCCCTAAACAAACGTTATTG
TTTTCATTTTCTCAAATATTTTTTATTTAGAATTTAAAAAATATAAATTTTTATTTTGTATTGGTTTGCAACCAA
TGCAAAACACTTCCCATGTAAAGCATGGAACTTATGCTGAATAACCAAAAAACTAAAAAATGAGACAAAGTTTCA
TAAAACCGATGCAGAAAAGCCCGAAATTTTGTTTATGATCCACCCATTATTTGTAAATACTTAGGAAAGATATAC
GTTTAGAAAATTCATATAACATATTTACTGGGCTTTGTTGATTTCCAAATATTACATTTGGAGAAGTTTGATTTA
GGAGAACAATTTGTTGTACAAAAGATATAAATTTTCATAGCCTAAGCAGAATTTGCATAGGCTATGAAGATCATG
GAGTTTAAGTATCCATTCTTAGCTGCAAATTCTTATACCGAGCATCCAAAGCATCGAGCAAAGTTGAAGGATGAA
ATGTGAAAACTACTTATAAGCTCATCAACTCTATGTATATATAGAGTTAAGGTTAACAACAAATGAGCCTAAAGC
TTTATACTAGGACTCAATGGTAAATGCCATTTAGGATAACATAGAGTGCATATGTCCCTTTAATATTTTTGCCTA
ATCATTCTATTTCCAAGATCAATGAAGTATGAGTCTATAGCCTAACTAATATAGATAGCCATTAGAAACAGTTCA
ATAGGATTCAAATATCAGCAAAAAAAAAACAATACATAAAATACGGCAATCTCATCGAGCTCATGAAAGCAGAAT
CACATAGTCCACAGACACTAAACCAATCAAATAACATATAGTAAAATCAAACGAACAGACAACCCTAGGATATGA
GCATGTATTACTACAAAACACGAACAAAACCTGACGAACACTGTATCTACGTATATATACAGTTACATATATGGG
CATATATATGTCTACTAAGCGCATTGATTTAGAGAAAAAGGAAACCGGAGGAAATACCTCGAGTTGGAGAAATCG
TAAAGCTTGCCTGTGCTGGAGAAGATGATGACACCGACCTCTGCGTCGCAGAGGATTGCGAGCTCCTTCGCCTTT
TTCAGCAATCCGTTCCTTCTCTTCGAGAACGTCACTTGACGACTCGTCGTGTCATCGATCCTCCGGATCACTATC
TTTCCCCTTCCCATTTCCGATCTGAGATTGCGATCTAGAATTCGAAAAAAAAAACACAAAAGTGCTCAGAATCCA
ATCGGAAGCAGACGGAGATTCGAAGATTACAGAGGACAGTATTCTATGAGATCGAAACAAAAATTTGAGTTCAGA
ATCCGGAAAAAAGTGCTCGAAGTCCAAATTGAAACAGACGGAGACTAAGATTACAGTTCAAGAATAATCGATAAG
ATCGCGAAAAAGCTGAAATGGAGAGAATCGAGGCTTACAGAGAAAAATGGTGAGGATTTCTGATTTTTGCAGCTC
ACATACGCGACGTATTTGGTCGCAGAAGATTCAAATCTTCGGAGAGGTGTGTGTGTGGCCATTTAAGGAAAAAAA
AGGAAGGAGCGTGATTCCGGAAACCGGAGCGGTTTCGAGAAAAACAGCTAACGGTATTCATTGACACGTGGTCGA
ATCATGAGACGGGAGAATTTTTTTAATGGTCAATGATGTTTGAAGCGAGATCGAGGAATCTTACCAAATGGGTAC
ATGACACGTGGTCATCTTAGTATTATTTCCTTTTTTGGAATCTATGGTTTTGTGCCCTTTTTTTCAAAATGCACA
GAGCATTATCCACAAACCATGAAGCATGTGTATTGTACCCGATAAAATTTTAAAATAAATATAATAATATAAGAT
TCAGTTTTCACTAGGTACACTTCTATGTGCAATTATATTTATTTATTTATTAAAAATATTTATACATGTAATGCA
AGAAAAGACATAATATTTATAGGCAGTTCTAACACAAAAATATAAATTCTTTGTAACACACACAAAAAAAAATATA
GGTACCTAAAATTATGCATTTGGAAAATATATCGCCGATTATGATAAGCTTTGTGTTTATCCTTTTTGTTCCGGT
AGTAGTGTCGTCTGTAAAAAAAAAAAAAGATTTTTCTTTTTGTCTTGCTAGAAGAAGTAACAGCAAACATTTCTTG
AAACAAAGGAAAATCGAAAATGAGCTGATTGCAGGGGAACGTTTTTAGCTGAACTAGAGGGGTTGCCAACTTGCC
ATAACACACCAAAACCAAGGTTTAACGAGCTCGGGGGTAATTTGGTAAAAGCGATATTGGCGGCTTTGCGTTTGA
TAAAACACAAAGCAGACGAAGGTCCGGGACTCTGCAATTCTTCTTCAAGTTTTGTAGGAGTTTTTCGGTCAAGAT
TTAGGCTCGATCGTGTTTTCTTTCTCGCGTTGTACACTTCGTGAATAAAAGAACTACCTTTCCCGAGACGTACGA

TGAACCCTAACGTTCGATTCAGAATTCGTAATTCCTTTCGTTTTCTTGCTCTATTTGAATAATGTTGTTTAATCA
TAGATCCTTAATTATGTATTTAAATTATGTACATGACGTATGGAAAAGAATTTTCATACTATGACTTTTGATATA
AAAAAAGGGATTATTTCTCTATTTGTACCTCGTAATGGATATTAAAATTGTTTACGTAGTTTCGAATTTATTGAG
TGCTCTAAATATATATATGCAATATACATTCTAAAAAATAATTTATTTCGGTTATTGTAAGATTGTCATAAAATT
ATTTACTAATTAAATAAATATAAAGTTATATATATTTAAAATTTTACTTAATCATAATTTTATATATTTTAAATA
TGTTTAATCACAGACATTTAAATGTATGTGAAAAACCGTATTTGTTGGGATGAAAATGTATGTTTCTTTAAATAA
TTATTTTTTATAAATTAATGGTTTATCAAAACATTTAAAGTTCACTATACTATATTCACAATTAATAGTAAATTG
AAATCGATTATACAATGCTTAGAATAATTAAGCGTTGATGTTTAAAAAAAAATTATCTTGAAAAAAAAAGTTACA
TAAACTCTTGTTAAGAGTAGGTTGTATCTTGACTAAGAACGATCTAATCAACAATTAATAATATTATTAGAATAG
AAAATTATCAATATTTCAAACAGCATGGTTCACATGATTCTTGCTTTGTAGCTGGAATAAGTTTCTTTTGAAATA
ACATTTTTTTGTAAATTGTTCCGGATTAAAATATAAAAATGAAATAAATACCAATATTTTATACTAAAATAAATA
ATATAAAAATTGATATAAATCCAACTTCGAACTCCCAGATTTCATTATAATATAATTAGATTCAATTGTAGATAT
ATTCATATAATTTTCCTTATATATATATATATGTGTGTGTGTGTGTGTGTGTGTGAATTAAAGGGTTTGCAGC
CAAGAAGACAAAGAAATAATAGGGATCGTTAACACTAGATACCCAACCATGTAGTCTATCAATTATTCCGCAATT
TTTTAAAATATTTCGTAATTTGTAATTTGTTGTTTTCCTGTTGGAAAGCTCTAAAAGGGAAAGTTTTATTCCACT
ACAAGCAATTATGGAAAGAATACTATATATATATATATATGAGATGGTTATGTCATGTGGCCTTGTATTTCCATC
TTCTTTAAATTAATTAATTGTCAATTTGCCTACCCACCTTGTATCCATTTTGCTTACTTACCCACCACCTTCTAC
GGAGGGAAAATATTTGACATATTCAAACAAAATAAAATTGGAAAATTAGGTTGCATATTATATCACTAATCAAAC
CAATTAGGTAATAAAGAGAATTTGTCTACCACTTTTAATGTTTAAACTTTAAATCTCTAAAAATTTATGAAACTA
TCTATATATCAAATCTTGTTATTCGATCGCATGGATCGAATCATCATGTCACGTTGTATTAGGAATGTATACAAA
TCTCTGATATGTTATCGGGATATATGATATAGATACATTTTAATTATTTAGTTGTTTAGTATACTTAAGATTTCG
TATTGTGTATATATATGTTGTTTGGTGAATAAAAACACGAGGTTTATTCCGTAAATATTGACAATATATCATAGT
TTGGTTATTATTGATTTCTAAAACAAATATTGTTCAACGTGTTCTTCGATTAGGGTTCTGAAACCCTAACTTTCC
ATTGCTGTTTTCGCACTCTATTTTCAGGTTTCGCCGAAAAAGATGACCGGAAAAACCAGAGAGAGAAAGATGAGA
GAGAAAATTGAGTAGAGAGAATGGAAAAATTTCTCTAGTTGTATTCCTCTGTTTCATGTACAGATGTTAAAATAC
ACTTCCGATTGTATAAACCGCAATATAACTAAGTAACAGATTTCTAAACAGATTGTAACTGTGTCATATACTAAT
ATACTCCCCTCAAACTTGAATCGACTGCACCTTCAGGGAGATACAAGCTTAAAATATGAAATTTCGACATTAGAA
CAGAAAACACCTGTAATATCAGCTAACTGATTTTCTGTCTTCACATGCATCATCCACAAGAAACCTGATTGAAGC
TTATCTCGAACTAGATGGCAATCAATGTCAATATGCTTCGTCCGTTCATGAAAAAATGGATTCGAAGCGATGTGT
ATGGCGGATTTGTTGTCACAAAAGAGATTGGCAGGTCCTTCCAAGGAAATCTGAAAGTCTGCCATCAAATTTGTT
AACCAAATTAACTCACAATGATCTGTATTCTGCCTCTGTGCTGCTCCTCGAAACCGTGGTCTGTTTCTTTGATTT
CCATGAAATCAAAGATTCTCTGAGAAAAATACAGAACCCAGACACTGAACGCCTCGTATTAGGACATGTTGTCCA
ATCTGCATCCGCAAAAGCATTGAAACGAAGAGCTGAAGATGCAGAATAAAACAGACCTTGCCCGGGGTTAGTTTT
CAGATACTGCACTACCCTCTGCGCCGCTTTCATATGTATATCTGTTGGAGCTGATAAAAACTGACTAAGCTTGTG
AGTAGCAAATGTAATGTCTGGTCTGGTTATGGTTAGGTATAATAACCTTCCCACAAGACGTCTGTAATTTGTTAT
GTCTTCGAGAGGAGTTCCAGCATCTTTACTCAAGTGCGTATTTGCATCACGAGGTGTAGAAACTGGTTTACATCC
CAATAAACCTGAATCTGATATTAAATCCAGACAATATTTCCTCTGATACAATGATATTCCCTTCTTCTGTCTTGC
TATCTCCAATCCAAGGAAGAATCTCAAATCTCCTAAGTCTTTAATTTTGAACTTCTGTCTCAGTAGATCCTTTAC
TTCTGAAACTGCATCAATGCTGTCACTAGCAATGACAATGTCATCCACATATACCAGTAGAACTACAAAACCAGA
TTTTGAACGTCTCACAACTTCGATTTCTTCATTTTCTTTTGTCCTATAAAATATAAAAAATTCATATGTATCTTT
GTATTCAAAGTTGGAATTTTAATACGTATGTAATGATTCTGATGATGCTAATGGTATCAAATAATCATTCATAAT
TATAACCTCATGCAAAACAATATGATATTATCTACTGATGGCTTATTAGTTAAAATTCTATTTTTTTAAGAGGAA
CCCACTAATATTCAAATTCCAGAAAATATTTATTGAGACAGAGAAAAATATTATATTACAACGATATTTTTGCAT
CGGGGAGAATGAGATGGTCTATGAGTAGTTTCCCATCCAAGTTCTTGGATCCAAGTCCCAAACTAAATTATTAGA
AAATTTAGAAAAAAGGTTTGAATATTGCACAAAAATTTTAAACTCATTGTATCTCAAGTGTTAAATCTTAATTAT
CCACTATGTACATAGACATAATGGATGTTTCGGGTCCGATTTTTTTTACATAATATACTGAAATTTTAGACCGAA
GTTTTTTTCGGTATAAGTTTTAGATCGAAGTTGGGTTCAACTACAACCAAGATTCATTATTCATATAAAAGACAT
ATAAATCCCGGTTAGCAACCAGTGAGAGAGGTCAGGAAGTTGTGTTTTGTATGGTTTATTAGCATTTGATTTGGA
TTTAATTTTTGTTAATAAAAACATCTATTTTCTAAAAAAACAATTTTTCTATTTTTGTATTCAGAATGATTTTAA
TTATTTTATTATAAAAAATTATATCCAATAAAAAATTTAGATGAAATCAGATTACTGTAGAAATTTACGAAATGA
TATTTTTATTGATAACCAAAATAACATATAAATAGATGAGAATCTTACATTGAATAAGTAAAGTAGACATAATGA
TAAATATTTCTAAATATATATTTTCCAGATTTGATAATATCTCTATTATACGATATATCTCGTAATATTCTTCAA
GATCCGGTATATCTCAACATCCCCTTCAAGTTGGAGCGTGGAAGATCACGAAACGCCCAACTTGCTGAAAAGATG
ATTAAACGCACGCCGCTTGAACCACATCACCTTGCGAACTTGTGTCAATAAAATACATAACTAAAAAAATCAAAC
AAACAACTGCGCTGCAACTTTGGCACCACGCACAAAAGAATCGTTCGATGATAACAAAAAAAGGACGACAATCGC
GATGTGGAAAGAAGTCTCGGGAATAACTCTGATTCATCCGAGATTGAAAGGCAACTACGCGAAAGCGTTTAATGA
CAACAAATTTTTTGGTAAAAATAAGATAATTGGCGGCTATGCGAATAATCTTTAGAAACATGTGGGATTATGAGA
GAAAAAAAAAAACAAAAAACAAAAACAAAATTACTCTCTTATTCTTAATAAAAAAAACTCGAAATGATATTACCA
CTGGATCGGTATATCATCATCTTCATCTTCACTCTCATTATCCATCATCATTTTTATGAAAAGAACAACAAATTA
GAAAAGAATCAGAAGCTCTGATACCATGTAGAAATTTACGGAATAATATTTTTATTGATAACCAAAATAACATAT
AAATAGATGACTTACATTGAATAAGTAAAGTAGACATAATGATAAATATTTCTAAATATATATTTTCCGGATTTG
ATAAGATATCCATTATACGATATATCTCAAAATATTCTTCAAGATCCGGTATATCTCTAACAATTACAACATAAT
TATATATATTATTATTTTGTTTGGTTCGATTTCATATTTGTAAGAAAATTAACCCAAAAAAAGAATATTATCACC

CAATCCCTGTAATGATATGATATTTTTATATAGACCGTATGGATCTTCGCCTATAAAAGATGGTGCAAATTAATT
CTTAAAATATTAGCCAATAAATGTAACTTCATAGTTAATGTTCTTTGGGGTATTAAATGCAACTGCTTGTTGTTC
ATGTAATCATCTTATATTAAAACTATATATATCAATGGTTAATCAATCCTCTAACCAAAAAAATGGTTATTTAAT
CGTTAGCTGCAACCCTATCGAACAATAGTTGTTACAAACAAAGCTATGCAACAATATTTTTCTAATAAATCATAC
TATAATCTTATACCGTTAGACCACATTGCATATGAATAAAAAGGTAAAATCTTATATAAAAAAAGAAGCATATAA
TAAATATACTGGCTCAGTGGCTTGGGTTTCCCTTTTCTCAATTTTTTTAAAGGAAAAAAAATGGAAAACAACTGC
CCCCAAGACAGAATTGGGCTCTAAATTTGGCCCATTAGTCTAGCCATTACATTTAAATAATCTATATAATATATT
GGGCTTTTAAGTGGCCCTTTAAATGGGCTTTCAATAGTACCTAAATGGGCTTAAACTAAAGCCTATAATATTTAA
CTGGTGTACTCGAATGACCATAATTTACGGAAGGTTATTAATTAGATGATATAAATTTCATAATGACATAGATTG
ATCACATGGACGAGGTCTAGTGTATTGTTTCGGTGGTGAACATTAATAATTAAATATAACACAAGACAATTTTTT
ACAATCTCCCATTTCTGAAATTCAAATTCAGATTCGGGTCATAAAGTTTGAGATGTGTTGTATTTTAATTTCATT
TATACCCAAGCTTGGTGGTATAGTGACAAGAGAATATGCAAAACATATTTGTATTCTAGATTCAAATCTAAACAT
GTAGAGAGACATTTATCTGGACTTTTCCGATCATTTGGAAGATGTAGTTGGGTCAGATTCGGTGAACTTCCGAAT
TATCAAAAAAAAAATTTGAGATGGATCGAAAAGAGTAGTTTGAGGACGATAGGACGAGAAAACCAGTCTAACCCTA
ATCCCGATTTTTTTAAACCTCGAGGTGATACGATATATCCATCTTATTAAGAAAAAAATCGAAAGCTCAAACCGA
GTTTCGAGTTTCGGACTAAAATGTACACCTCTAAAAACACACTTTATCTTTGATAATTCCTTTGATAAGAAGCAT
AACCCTCTCAAACATTTTTGAGATGATATGCATAATAAAAATATTTATTTTAATTATAAAAATATTAAATTTTAA
ATTCTTGAAATTTAATTTATCTATTAATAATATTAAAATATTAATATAAATAATTTTATTAGGGTGATTTTCCAA
ATAATATTTTGACTGGCCTTATTTGGATTCACATTAGATTTATAAACGAAATAATATAATTATTAAATATATTAA
TTATTAAATTTAACCATAAAAAACTTAACTAATTTAATAGATAATAAATGATTGAAAAAAGAAATAGAAAATAGA
CAAAAAATTGATAACTGTTTTCTTCTATTTAAGTAAATAAATTAATTATTTCAGTATCTGTAGTAGATCATGTAT
CTCTTGAATAGTTAGTTAATGTTGAAAAAGTTAATTAATTTTTAAAAATAAATCTTTTATTAATTATGGATATGA
TAAATCTTTGGGTTGTAACAGATATATTTGATTCTTTTTAAATAAACCTTTTAGTTTATTTAATAACGGATAAT
TATTTGACCCCACCAAAGCATCACGTGTAGGTAAAAACGGGTGGCATTGTCGGTACAATCGGAAAGAGGGGTAAA
AAATAAAAGTTCATAAAAAGTATAAATGAATTCTATTGGAAAGTTGTCTCCAAGTTTTATTTTATTTTATTTTAC
TTTTTCTAATCTTCCAACACATTCTCCAATCAGGACAGTAGTTTTCCAAACCCTCTGACCGAGGCAGAGACATAA
GATAAGTATACCAAAAATATCTTACCGGACATAGGAGCACGTGTCACTAGTCCTATAAATATCTTCTTATACAAT
CATTTCCTTTTGGTAATCGGAAATCAAGCATACAGCAAGACAGAACGCTAATCAAGCTGACACGTGTCATTTGCC
AACTCTCATTTTCACCAACTTTTGTTAGCGATTTAAAATAATCGTGGATTAGTCATATATTTTGTTTTCCATTTG
CATCAGGACATTATAATATGGTTGGTGTTTAGTATGACAAATTAACGTTTTATTTAAATTCAAATTTGGAAATCC
ATAGGATTCGTGCAATTAAAATTATAAAAAAAAAAAAATTGAAGTGTATTTGTCGTTGTTTGTGCTAAAGAGAGCA
GATACCCTAACATTGACAGATATGTACTGTCTTAAAAATAAAATAAAAATAAGATAATAGAGTATTACGAGGAAA
AAATTTAAAAATAGAAAGTGCGACGTGGCGAATTTATAATCGCTAAAAGAAAACTAATCAAAAGCATTTATTTTG
TAATCTCCAGCTACACTTTTCTCTCTCTTTAATATTTTTAAAATAAAATTTATTTTCTGAAATCGTCGGACTCGA
CTCGGCTCGACTCGGCTCAACTGACTATTTGGCTTCCAGATATTTTAGCATTAAAATACGAATTTTATATTCCTA
ATGATTATGTAATAATCCAATATATTTTGAGAAAAGAGTAGAATACTGAATGAAGCCCCTTTCTAAAGTATGGTG
TTTGCTTATTGTATTCAATCAACAAAACATGATTCGCTGTTATATATTCATAATTTCGTCGTATTTTTTAATTCT
TAATTCGAATACTTGTAATTATAAATTTCGTTGTATGATTTAATATTTGTGGATATTTATTTCTTAAATGATGAG
TCAAAAAGTGAAAATGTACATGAAAAAGTGTTAAATGGATTAAAAATAATATATATTATAATATTTTTGTAGATA
CCTTTTATAATATTTTGTAAACTCATAAGTATGGGCGGACCCAACATTATAGCAATGGCGTCACAAATTCTATTG
ATCTAATTTAAATATTTAATTTTATTTATAAATTTTATAAAATTCTAAGTAAATTTATTAAATATCCCTTATAAA
AATAATAAAATTAATCATGCCCCTATTACATATATTTCTAGATCCATCTCTACTCATAAAGTGTTAAAAAATAAA
TAATATACTGAATATATCTATATATCCGTTATTCGGCTACCTTCGTTATCTCACGATTTCTATGTTCTCCACTCA
CAATTTAATTTATGTTATATCTTTTTTATATTTATTCGTAAATTTATAAATAAATCTTTACATAAATTATTTGTA
CACACTATAAAATTATTTAAATACGATTTTTTAATATTTAAATATTTAAAATCTTTATATTTATTTTTGATGATA
TTGTGAAGAAAAACAAAGCTAAATCTCCAAGAAATAATATAATTGCAAATAAAAAATAAAAAATAAATCTACATT
TTAATTTAAGATTAATTTCTGTGACACAAAGGCCAAATAAAATAGAACTGACTAAGCAACAGCGCCGACCCAATC
AAATGTTTGCTACATGAAGCAAAGCCAATCAGCTGGGCCTTTGGGGGAAAATCCGAAACCCTAGATTATATTCTT
AGATATTTCATTTCATTAATTAAAAATCTGATTTAACGCACTAATAAGTAATTAAGATTTAAATTAAGTTCATTC
CCATCCCCCACACTCCTATATAAATATCCTCTGGCCTCGTAAATATCTTCGTTCACCTCTCTCCCATTCTATCTT
TTTAACTCCCTTGTCTCTAATTTGCCCTAGATGTTCTTCCGAATCTGCTCTAAAGCCTTCGCTGTTACGAATTCC
GCCTGATTTCCGGAGATTTATCCGCCGGATCTCGATTATGTTCGTGGTGAGAGATCGACATTTACAAATTCCTAC
GTGTACGATTCTCTTTTCGACGCCGCTTTGTTCGTCGGCGAATTTTGGTGAGTTTGCTTCTGATTTTTTTTTTGC
TTTTGTTTTCTCCGGTTCTTTCTATTTGCGGATTCGATTTACGACGATGATACGTGGTTTTCTGTTAAATCGTTT
TTGAGATCTGTTTGGGTGATGAGAATGCTCGAGAAAGTAGATCGCTAAAATGAAGGAAATGAGACTTGTTTTAGC
ATCGACGTATCAGCTGCATGACCTTATAATTTCGGACTTTCGCAGATAATGCTTCGATTATTTTTTTCTATTTCA
TTGTTGACGACAGCGATTGAAGTGCTTCCAATCCCCGTGTTATTAATTTGATTGCATGCGCAGATGATACCCGT
AGATGAGAATTATTTGGATCGATCTCAAGTGAACATGTCTCCATTTTCAGGTGAGATTCATCGTTTCGATTTTTC
GGGCGGATGCATTTCTTGAGCCTTTTCCCCTTTTTCATGGAGATACTGCGATCCATTTAAAAAAAATGTATATAT
CCAACTGTTTGTCATTTGCCTTATATCATATGATCCTTTGAGGCGTAGAAGCGTCTTCTTTGTCATTTGCTTGTG
TTGATCTGGCTCCTTTTTTTCTGGCTGTTTTAATTGGTAGACTATGTTTCCATTACAGGCACAACTTGATATATG
TGCTCATAAAATAAGGGCACATAACAAGATCTTGTGCATTGAAACTTTGAAAAGTGTCCATCATCTATGAGTCTA
TGACATGCAAAACTACTGTTTAGAGGAGTTTCCATCTATTTAATTAGAATCAACTACTGTTTTGAAAAAAAATAA

AAAATTCTCATTTATTTATTTATTTTATGCAAGCTCCATTTAAAAGTTCACAATCGGTGCTTCCTAAGATGTATA
TCTAACAATACATCTTGTTATTCACAATGTTCATGATCATTGATCATGATGCGCATTAATTGAGGACAGTCAGTA
GAACATGGTTGTTTTGTTAGGTTAGTTGTACTACAACTAATGCAAACATGCGATGACTACAATTTCTTTGCATCT
TCTCTTGACCGAAGTTTAATTAGCTTTTCGTTATGGTTTGAATCGTTTGATAGCAAATAACCATAGAAAGCATTG
TGGATTTTTTAATTACTAATGTTTAAGTTAAAAGGGATCTGTTGAAATACCACTGACTTAATTTGTGCCTTGTAC
TATCCACACCCAAACTAAAAGGAAATAATCGGTTTTCATAACATCCTTTACATTCTTAATTAATCACTCAATCTA
ATTAATTATGCTTGAGGTTGAGGTCCGTGAGAAAGTTTTATTGTCTAACGTTGCTCATTTAAAGTAAAGATTCTT
AATAATCTGGCTCTCATAATGGATCTTAGAGTGGAGCTTGTTCCACACATTCTTATCTCTCTCTGTTGCAATATT
TTCTCTAATAATTTGCTTAAAAAGAATAGATCATCATAGGTAACTTCTGTAAATTTTGCCCTATACTAATTAATA
CAATTTCCATGACCAAGAAATTGGTCTAGGTATGAAAATTTCAGTCCTTGACCTTGTTATAGGGTGGGAACTTGT
GTTGGTGCTATTATGGTGTGTTAAGCTCTTCCAGAAACTGCTGTTTAATGTTTTAACAGCAGTCACGACTCATGG
GAGACTAGGACTCTGAGAGTAACTTTGCATACTTGTTTTTGTCTTGAAGGTCCAATCATAACTTGACTGGAAGTA
TGGATAATGTAGAGTGGATATGATTGTCTATGATATGGATTTCTTGGGTCCATGTTTAAGACTTAAGTGTCATCT
TTACATTCCAACACTCCTTCCTACAAGTAATTTATCTATGGTTATAGTGTGGTTGGTCGTGTACTTTAGTTATAG
ATTTGTTGATACTGGGCCTTTCTTTTGGTTGTAGAGATATTCCTTAGATTGTATTCTAAATATGGCTTGATGGAT
ACCCCACAGTATGTGGTCTTGGCTTCATCTCATCTCCATTCTTTCACTCCCTGCTCTTTTTCTTACAAAGTAGGA
AGATATAATTAATCATCATATGGTAGAAGGACCAAAGGTTTCGACGTACGAGTGGTTATAAGTTAAAACTAATTT
TAGAAATTTAGTTTAGTATGAATTATTTGGTATTGATAATGGTGGAACTGCCATCAGAAAGTCATGTTCTGCTCC
ATTTTGAGTTTAAGAAGTAAATTCATTTTAATGAATCTTTAGGTATTAGTTTGGGTGAGATGATCATCAAAATAT
TACTGTCTGCTTCTTTTGTTTTCTAATTAGTTTATTCATCTGAACTGTTATCTTAGTGTTCATTGCATATGGCTT
TTAAACTGATAGACACTGATGTATTAACTTGCATTCCCTTGATTATGGCATACAAATTATTCAGTGAACAAGATA
GATATCGTGCTCTCATATATATTGAGCTCTTACAACATCATGGAGTGCTAAAAGGACTCAAATATACTTAACATA
ACCCTCTTTACGGTGTCTAGTAAAATTAAAAGGAAGCAAATCACTTACCCTTTTTCTTCCTTTAATTTGCTCGAG
CCTCAAGAGCCACCATCTAGTTGCTTATGTAAAAACATTTGCTGCTCTGAATAGATACCGTCAAAGGCTCTTTTA
CTTTGAGGGCCAGAGCCCTTCTCGTCTATCAAGGGCTGGTGCTACATTATCCTATTAGGAGTGCAAAATATTCCG
CTCAACCCTTGTACTTGAAAATTTCTTCATCCCATCTAAGCAAACTAAATACATTTATCCCGTAATTTGTGCATG
TTTGTCTTTGTTTGTTTTTCTTCCTCATTTCCATACAGTGACCATTTTAAATTTTGAGTATTTTACTTTAAGTAT
GGAAAACATTCCTCTATTACATTTGTAGTGTGATTTGGTTGTGTTGATTTCTGTGCGTTGTCGTTCGGGATTTTG
TGATTTCTATAGTTAACAGTGTTTGAATGTAGCACTGTTGTTCCCTCTCTGACCCAAACCGGAATGAGATTTAAT
GGTTTCCAATGTTTTTACAGTTTCTATTTTTTGTTCTCTGAGAAAAAAAAACCCTCCTCGACCATCAAATACACG
TTGAATGGCAAATTAAGGAAGTTCTTTTTTCAAATGAGTAATGTATTGGCCACAGAGGACATTGGAATCCACATT
AGTAATGTTGATACAAATGGATTTACTTTTGATCATGCTTTCTTTTATAATGCAGTGGCAATCTGCTGCAGTTA
AATCAAGGAGGATGAGATTCTGAAGAGTTGCAGGTGTCAATGCATACCACGTACACATTGAGTATGCTCTATGTG
ATGAAAGAAAAGTATCCGGGTTTGGTAGCCCCTGCGGCTGGTGCTTGCACGTGCCGTGGTAGGCTGAGGTGAAGA
TGGACAGGTTCTGTTGAGTCA**TCTCTTATGTCTAGACCATTTGTGAGAAGGGAATTCCTTTTTGCACTATCGGAA**
**CCCCCCAGCACTGAAACCTGGTATAGGTATTACCCTCGTTAAGAAAGCAACCCGCCTTTTAAATAGCAAATCCTA**
**GCTCCTAGATTAGTTCAATTTAGCAATACCTTTTTCTGTAGTTCCTTATTCTCTCGGCGCATTAGGTTTAGTTCC**
**TTGTGGAGTTCTAGATTTTCTTGGTGCATGAGATTCTCCTGCAGCAGGAAGAGTAAGAAATTAATTAGTTTCGGC**
**ATACGCACATAGACAAAAAATAGAAAAGTCTGTTCTAAGATAATTGCCACTAGAGGTCCACCTTTCGGTTCAGTT**
**CTTCTATTTCATCCATCAAAATCTGATCCTACAAAAGTCAAAAGAGAAGAAAGAGCAATAGATAGAGCACCAACC**
**CACTGCATTTCGAAAAAAGCCACAAACATGACTGATTAGTTATGGCGACCAAACCTTTTTCATTCGTACCCCTCG**
**AAGACTCAATTCAAGCTGGCTTTCCAAGCTCTGTAAATCTTTTATACTTAGCCCAGAAAGCTCTTCTCCCATCAT**
**TTTCCTGTCAGGTATATTTCATTAGGAGACTGTTTTTGAAGATGCATTTCCTTACGGAACACAAATTTCTGTCAC**
**CGTCTTGAGGATTTTGATAACTATATATGTTCCCAATACGGTTCTACATTTACCAATAAATTCTGGGGAGTGTT**
**TGGTTATTCCCTTCTCATGAATGGTCTAGATGTGTGAGGTGACTCTCA**AAAAGTGCAGAGATGCGACCGTGATTT
TGGAGAGGCTATAGGCATCTAGCATCTGATGTATTTTGCCCAGGTAATCCTAACCAGGTGAGTACATTTTCAGCC
GAAGAACTTTGCTATGCTGACGAAGTATTCTCGGGGATATTCCAATATATAAAGGTTTGTTATGGTAATGAGTTC
TGGTAATTGGACTCAACGGTCACATATACCCTATTATTAATAGTTTCCGAAGTCATATTATCCCCTAACTGGACT
TGCATTTATCTTTTTCATCTGCTGTTCTAACTGTTGTTTTCAGAATAATATGCATGCAATGTATTCGGAGGCAAG
ATATACAAAAACATACCGATGGTTTTCTTGCAAGTTATGTAGCTGTTGCCTCAGAATTGCTGCTTCCCTCTGCCA
GAACTAGAATATAGAACAAATTAAGTTCAGGTGGTTTACTAGAGCATAATATAGGAAACTGAGGATGATCCTTCC
ATATCTAAGTTGGTGAAAAAAAAAGATCGATGAGCTTTCTACATTTCCTGTTATTGTTGAAAACCCAAGTAGCAA
TTGCTGCGTTAGTAGAAATAATAGTATGCTTCAGTTGCTTGACCTCATCTCTGTTATGGCAGGTGTTTGCTTCAA
CAATATTGTGTGAGTAAATTAGGTCAGCAACAGGATCATTTCATTTGACCTTCCCCTCATTTTATTGCAAATAAT
TAGGTCAGTATTTCCTATCCATCTACAACATATACAGAAATGCATTATATCTGTACGGAATTCGGTTGTTTTGTT
TCGGTAAAATGCAAAAGCGGAAATCTAGTTTCCTATCTTGCTCGCATCATTTGGTACACACTAAAATCCTAAGCC
CTTTTAACAAGAACATGTTCCGGAGTTTTCTCCGCAAATGTGTATTGAATAGTAAAATGTGAGTATCGTACTGAT
CAGAACATTGTATGTTTCATCTTTTGCAAGTCAAATTAGATCATCTGAAGCAATTACCTTGATTTCGGATACTGT
GTCGCTTTCGCTATCCCCTTTTGTGTCACCGTATCGCTCGATTATCGATTTCATGCTGCAAGTGAAGATTTTAAC
ACATGAGAACGTTATCTTTCCCATTAATCAGCTCAAATCAGGTTTGTAAATTCATGTCCATGTCTGCTAATTCTA
TGTGTATATATATATATATATATATATCTGTTTGATTCTGCTGCATCTGGTCTGTGCATATGTGTCTTGTAGGATCT
CGTATGATTTAGTCAGTAAAGCTGGGTCAGAAACATAGAACAAGCGAGCCATCCTTCTTCTCCTTCTTTGGTTAT
ATCTTTTGGTTCTGTAATACCTTTTTTTTCTGTGTGTTGGCCTCAGTTGTTGCAGGGTCGGTAACCAAGCTACAA

GACGGGTCAAAACACAGGTATGTTATGTCTAACCTATAATCTGGAGCAGGTGGGTTTCACACCTTTGCATATATA
CTACAGAAGAAACTATGAAGAGATATCAAACACTTTGCCAAATTTGATTTTGGGATTTGTATTTTTTCCCTGGTG
GCACACCATTTCTTTTATGCGACGGAATCTACATAAAAACAATTAATTAGTGTTAGCCTGTGTTCAGAATTAATA
TATAGAGCTAATTTCGCTAATACCTGATTATTATAACCCCAGGTAGGTACAGCTACCCACCAACTAGTTGCCAAT
AAAAATATAAATAAGGGAAAAAATATTAAGAGGGAGTGAATGGTAAACACCGCACACAGACTATCCAGATTTTTT
GGTGTTACAATTTCAGTTTTTATCCCTTCAAATTTTTAAGTTGGAGTATCGAATCCAGATCTCGTGAGGGTCAAA
TATCAAACTCGGTTATCAGTTAGGCGTGGCGAAAGTTCGGGTGCTCATATCGCCCGATAGAAGGGCGAGTCGAGC
TGAACTGAACTCCGATAAGACTGAGGAAAGAGGGGGTGAAAGCACTGAGAGGACTTCGGGTTTACATATCCCCCA
CACCTACCTAAGAGCGCAAAGCAGAGGGAAAGCCTAAATTTGCAAGTTGATATCTATCCAGTTCTCTATAACCAC
TTCCTTGTCCTATCTTTTCTCTTTTCATCATTTCCTAAAAGCTCTTGATCCATTTTGGATCAGTGACTACACAGT
TTGTTTCTTTTTAGGATATGGATCGGTGTTAACAGGTTAGATTAGTGGTACCTCGCATAGTCACTGATTCAACAC
CCAAATCTCTAATTTTCTCCCAAGTTTGGGCTTTAGATGTACCATATGTTGACACATCTATCAATCTAGCAAGAA
ATTTGATGAATTATATCTAAAGTGAAAAAAGGTCCCCATTAAAGGAGTGAACTCACCCACCATTACATTTGTGTT
AGTGGTCAATCATCGAAGGATTTTATGATGGGTATAATTATATATATATATATATATATATATGTATATGCATAT
GTGTGTGTAATGATCCACTTGTTTCTTTTGATGGAATAATGATCCACTTGTGTATTGAAAAAGTAAATACACACT
CATTTCTCTATTTAGAAAGTTGTAGATACTACAGATATTTGAACTTTCTTTTTTTGTGGAAGGAGGACTCATTTA
ATGTATCATCATTGTCACATGTATGCACGCATCTATGCAGATGTCTATGAATACATACACACACATGCGCAGTTG
AAGTATACGTGTATCGGTGTATGCACATACACAGCACACACCATTTTATTCACCGTTCTCTACAGAGATATTTAC
TTCATAACAAAGCGATCTCATGAAATGCTATAACTCATCTCTGTCTTCCCCCAAACACTATCCTTGTGAAATTTT
AACGATAATTTGAATTGGAATACGAATTTATAATTTATCTACAAAATCTATTTTTTTTTTGGTGAAAACAAAATC
TAAATTGTTTATGCTCATGAAACTCTTCGATTATTAATTGATTTAAAAAAAACGATTCCCTCATCAATCATATGG
ACCAAACAAACAAGAGGACGAATTCACGCGCGATCGATCTAAAGTCAAACAAAAATATGAAAAAAGGAATGAAAA
AAAAGGCAAAAGTACCTGGAGCTGGAGAAATCGTAGAGCTTGCCGGTGCTGGAGAAGACGATGATTCCGACCTCC
GCGTCGCAGAGGATCGCAAGCTCCTTCGCCTTCTTTAACAAGCCACTCCTCCTCTTGGAGAACGTCACTTGTCGA
CTCGTTGAGTTATCGATCCTCCGAATCACTATCTTTCCCCTTCCCATTATTGCTCTCTGGATCATCATCGTCATC
ATCACCGATCACAAGAACAGAAATCAACCATATTCGAAATCGATATGAAAATCGAAACCAGATCGAATTCAACAT
TGCATCACAGTAAATCTAAAGTACCAATCGCGGATTCGCGAGAAAGTGACAGAGATTGGAGAGAGTGCGGAGACT
TTGCCCGTCCAGAAGTCGAGGAAAAAGAGAACATACAAAACGGCGAAGGATCAGAGGAACAGAGATGTCGTTTTA
ACTTTGTGGAGAGAGAGAGAGAAGAGAGAGAGAGAGATGCATTTGGGAATTTAGGGTAAAAACTTTTATTAAAAA

*tha-miR824b*

>scaffold00351

TCGGAAACAATGTTTGTGACGAATCACTGTTTCAGCTTTGATTCATATGGATTTGCTTCTGATCGTGGGTCTGCT
TTCTTTGCTAATTGCAGTGGCAATCTGCTGCAGTTAATCAAGGAGATGAGATTCTGAAGAGGTCAAGGTGATAAA
CGCGGCTAACACGGAAAATAGTGTGTTGAAGGAAAGGT**TCGTTCGAGTCATCTCTTATTTCTAGACCATTTGTGA**
**GAAGGGAATTCCTGTTTGCACTATTGACACCCCCTGGTGTAGGTATTTACCCTAACCTTTTAAAACACAATCCTA**
**GCTCCTAGATAGTTCAGTTTAACGATTCCCTTTACCCTTATTCCATATTCTCTTTAGACATTAGGTTTAGTTTCT**
**TGTGGAGTTCTAGATTTTCTTGGCGCATGAGATTAGTGAGTTTTGACACGCACACACAGACACAAAGGAAAGTCT**
**GTTCTAAGATAGTTGCATCACTAGAAGCCCTCCTTTTGGTTCAGTTCTTCTGTTTCTTCCAATTAAAATCTGATC**
**TTACAGAAGTGAAAAGAAAAGAAAGAGCACACCCACTGTTTTGGCAAAATTCACAAACATGACTGATTTGTCATG**
**GCATCCAAACATTTTTTCGTTTCCAAATTCTTTGAATCTATTACCACTTAGGCCAGAAAGCTCGTACTCGGTCCA**
**TTTGCCTGTCAGGTATTCAGTATGAGACTATTTGTTTCTTAAAATGCATTTGACAAATTTCTTCAGCTATATACG**
**TTCCTATAATACATAATAAAATCCGGGGAGGTGTTTGGTTATTCCCTTCTCATGAATGGTCTAGATATGCGCGGT**
**GACTCTGAAAAA**TGCTGAGGTTGAGACCGTGAGTTTTGGTCAGTGTATTATCAGCATCCAGCATCTCATGGTGTT
CGGCACAGGTAATAAATACTGAGACTAGGTAAGCACTTTTCCAGCTTTAGAACTTGCTAGGGCTGGACACCTGTT
TTCAGTGATAATCGATTTTTTTTGTCATTGCATGTTTGTAGGATCTTTTTAGGGAGTCCTGGTAAGATAGTTTTA

#### Caricaceae

##### Carica papaya

*CpAGL16*

>supercontig_177

ATTCTCGCGGTTAAAGTAAGTGAAAGTCTCTGCTATTTCTTTTAGATTCCTTTTTTCTCGCTCATAAAATCTCTT
TCTCTCTCTTTCTTATCAGATTTTGTCTGTTTTTTAATTTACTTTCGGAAATCTGGCACGAAAATAGTTTGATTT
CGACTTAGAATTTCGGTGTTCTTTTTGGAAGCTCGAAGAATATTCTGATTAGTATGGGGAGAGGGAAGATAGTGA
TAAGGAGGATTGATAATTCCACGAGCAGACAAGTAACTTTCTCCAAGAGAAGAAATGGATTGTTGAAGAAAGCGA
AGGAGCTCGCGATCCTCTGCGACGCGGAAGTCGGAGTCATGATCTTCTCCAGCACCGGGAAGCTCTACGATTTTG
CTAGCACCAGGTCATTTTTCCTCATCTAAGATCTCATGATTCCTTTCGTTCTTTTCATTTCTCTCTCTAGCATAT
ATACGATATCCATGGATTATACATATACACATTTATAACGAGCTCTTCTCATCTCTTTATGTACATAGTCAACTA
AGGTTTTTTGCTTATATACGGTATTAGGGACCACTGTCGTCGTTAGGGTTTAACCAAGCCGATCTAATTATTTCG

AGATAGTAGTTCGATTTCCGGAAGAAAAAAATAATTTAATGCTAAAAAACTGGAGCCTATTAATTGATATACTGT
TCTTTTCTCCTTTCGTGATTCTAAGAAACTAATAGGATATATGCATGTTACTGTTGATTTCTTTATGAGAATTAG
CATGATGTGCTTTTGAAGTTACCGAACTACTAAGCAGCGGCATTTTTGTTTTCTGTTGACATGAAATAGCTGAAA
GATTAGCTATATTTATTGTTAATTGTATATCAAACTATCTATATACAGTATTAACTTGCTATACATTGTGAAGTT
GCCAAAGATTTTTTTTTTAATTGAAAGAATTAGTAATGTTGCTTCATCTTATTAGTTCAAAATTTTAACTAAACT
AGAACGTAATAGACCTAGTTTGTCAATTTACCAAGTGAATCTATGAATACATGGTTATATGAGTGCTTGAGAAAC
TGTACTTAAACAAATTAGTTTAAGTCAAACAAATTAGTTTGTATAAGGATAATTCCAAGGCAAGGGATAATTCTT
ATGTTACATTAATATGGTACTATAATTTTATTATATAAAAATTGACATGGAATAAACTCACATCTAAAATATTTA
TAGATTTGTAATTTGATTCAATACATATACTAATATTCTAAGTTTCGTAGTTGAAGTGAGAGATAAGACTTGATA
TCAACTAGTCATCAACTCTTCGATAGTATTATGTTGATTGTCTTAAATACTTTCAGCTAAACATTTTATAATTCG
TTCTTTTATATATTGTTAAGTCTATCCAACACTTTCAAAAGAAAAAGAAAGTAAGTCTATCTAATGCGGCTCTCT
CCCTAAGCCTTTTCTTCCATATCCTAATTAAGAAGTACGTTACATAGCATCTCAATCTTCTCATTAATAAATTCC
TTTATGTCAGCTGTATAATCAAGGACAAGTAAAAGGAAATGGAGAAATATATATAGCTACATCTTACATATGGTA
AATACATTGCTAGACTCGGCTTTGGTTATTCTTAGCAACCCTGGTTATATTTTGGTATCCTATTCATCATGACTG
CTGGAACAAACAGTGCACGAAGTCTGAGGTGATTTATTGCTTAAGTTCTACACTGAAGCAAGTGATTGTTTTAGT
TTTTATAGAAGGTCAGGTTGCTGGTGATGAAAAAGTTTATTAGCATAGATGCAATAATTATAGAATTGGAGATAA
TAAAATAGGAAAAAGTTTTAATCATGTATCTATTTAATGTTTAAGTATGAAACTTGAAATGATAAAATATAATAG
AATTAGTTTTAAAACTTTAATACTAATCTTTGTATTTGGATTCCTTTAGGCAGTTTCTCATGATGACTGTGTGCA
TGGTAATAATTAGTCAAAATTTATCTGAGTCTTTTTTATGGTATACATAGGTGTAACACAATTATTATGAAAAA
TAACAATCATGAACAAACAAGTTTTCTCAGTTGATAGATAGACAAGTTCTACTCTTTATATGCTAAACTTTTCAA
AACAATATGAATTCCTGAATTTAAAGCAATATTAGTACTTTTAGGTTCAAAATCTCTTCTTCCTCGAAAAAGAAA
TCGACTATTCATTCCTTTTTTGAAAAGGAAGGTAACAAATCATTAACTATTCTAATATGATGGTAGAAGACTTTT
TAGAAACATGATACTAGAAGATGTATTGTATTTTAAATTCAAACACAAAGCGTTTCAAAGGGTTATGATCTTCTT
TCTAGTATTTTCTTTTTGAAGAGACAAAATTACATCCTCATAGAGTGAGAAGGGCATAATCAGTTTGCACCACCC
AAATACCAAAGAGGAATTAAAAATTCAATAACCCATATATTTTTCTTGAAAATACCCCAGTTCGTTGTTCTTCTT
TTTCTTACAATTTCTCCATCTTCTAAATCTTTAATCAAAGCAAACTTCCCTTCCTCTTTCTTTACATCACTTTTA
CATTAAGTATGACCTTTAAGACTTTTAGTAAGTTAACTAAGTTAATAAATAAATGTATGATCCAATACTAATATA
TATCTTATTCTGCATACTAGAAAATGACTTTTTGAAGTGAATGTTGTGAAAAACGTGAATAAAGGAATTTCCTAT
CCTGATATCTTTCCTAATCTGTTCAACCATATAATTATTAAATAGAATTAGCAAATTTTAAGATCAATTATACTG
TTATTTTTTTAAAAAATTATCGATTATACTGCTCGGAACCCCATATTTAGATATATCTTTTGCAAAAAAAGAAAA
GCCCCATGAATAAGATATGTTGTTATATACAAAAGAGGGAAACTACCTAATTAATTAATGAAATGTATTCTCAAG
ATAAATTTACGAGATATCATATATTATAGGTGTTTGATTTGGCACCGAGGAGCTTCCTTCGTAGAACATTCGTCA
CTTGACCTTAGTTTATATGTATTTTATTAGCCCATGGGTCCTCCATTAGTCTTTGACTAATTGCGTTTGATACAA
AGAGAAACAGTTAATCTTTTAACTAATTATTCTGGAAACAACAAGAAGGAACAATTAATGAATCTTCTGTCCAAC
ATTATAATCTTGTTTTGCCTCTCAAAGAAGGGAGAGTTCAGTTCTCTGTACTGCATAGGTACAGGGTCTCCTGGT
TGTACTCAGAAAGTAAGAAGTGCTTTCACGAAAAGAAAAAGTGGGTTGTGAAGGTGCCAACCGAAACAGGAGAGA
AAAAAAAGTTGAAGTTGAGGTAGGAATATAAAAAATTAAATGATGACTAAGGAATCATTATCCCAAGTCCAAACT
CATTGGATTTTGATAAGTTTCTATTAAGTTCTAAACATAAATTATTATCAAGAATTAGCATGCTAGTGTTAAAA
AACGATTTTTAATTTTAAGGATCTTTGAAACATGTATGATGACATTGCAATTTGATATCGTTTTCTGTAGGAGGG
AGCCTAAAAAATTCTGGCATTTTCCTGATTAAATTGTAATGTAAATTTTAAATATAACAAGTTGGAACTATATTG
TACATTGTAACTCTTATCATTTAATCATTTAATTATTTCTTGAATTATAAAGTTCAAAAATAAATTTTAGGTCTT
CAAATACACCTAATTTTATTGAACAAATGAGTCATTCACAAGACCTCTATAGATTTAGGTAAAACAATATGTCCA
CATTTCTATATAGAAAACTTTAGTATCATTATTAAAGGTTGAGAAAGCTAATCCATTGCAGAAGAATTGTCTTCC
CTATCTTTAACAACTTTGTTTCAATTCTTATAGGCAAAGACTTCAGGGTAAATCCCAAACATATAATGTCAATCT
AGAGAAATACCCTTTTTAAATTTGCTAGAAAAATTAGTGTTGGCAAAGTTTTGAAATATCAAGAAAAATGATAAT
TTACATAATTCTTTCGATTTCAAGCATAATTCTGATTATATATGTAGTTTTTCTTAAATAACTACATAGTATTTT
CTTTGATTTAAAAAATTACTTGTCCTGTCTCTCTCTGTGAAAACAAGTTTTACAGAATTTAAATCCAGTTGAAGC
AAATTTTAGTCAAATTCAATAAAAAAAAAACCTCTTTTTTATAATTTTAAAAATTTAACGAACTTACAAGTTTTAT
AAATTTTAAAACTAACATAACTTCATTATTATTTATATCTAATTCGATAAAATAAACTTATACTTAGCTGTCACA
TCCAGTTCATAGTCGTAATGAGTTGCTTTGTTTTGGAGAGTAATACACATACAATTCGCAAAGGATCGCCGAATA
ATTATGACAAGCAGTTTAAGAAAGCCACAAGTTTTTTAAGCATACTTTCCCGTATTATTACATAATAATACAAAA
AAATTAATTGTTGAAATATCTCTTTTCTCATCGTACATCTTGAGAGAATTCATGTCCGACTTAGAGGAAGAAACC
AGGGGTTCTATCCCGTGAGAGACTACTCCACGTTACTAAGAAAAGTTAAATAAAAGTTTTTAGATAGATTCAAAA
GTATCAATAAGTATGCTCATAATAATGTGTCCATGTAAATATAAATTAAAAAAGTTTTTAAAAAAATAGAATAATT
AAATCTTTACATCCCATTTTCTCTTGTGAAATCCGTCTTACATGTTCACAATTAATTTTCTCTCAGTTATCATCC
TTCGTTTTTGTATATACATACATACTAATTTTGGTGACGCATATTATTTGAACTTGGGATTTGGAACTTCCCAAG
TTCGGTAGCTATAACTGCATAACTTGATTTAGGGATGAATATGTAACTTGATGTCAATATTTGTTTTTAGGGTGA
ATTTAGCTGTAAGCAAGATATTAACAAAGTGCCGCCAATGTGCAGAATTCATGAGAAAATATTTTTATGAAGTTA
CACGAAACATGGTGATGGAAGAGAAAATGACTTTTTCTTTCTTTTAAAAAAATATTCAAGGTATGAAATTAATAAC
AATGGGGTCCGATAAAAAGAAACAAATGAACTCAAATACTCCTACTTATTTCTTTAAAATAACTAATAAAAGAGG
GGACAAAAACTTCTTTGAGTTAATGAAATTCTAAAACCATCCAATGATTATTTCTTAAAAAAAAAAAAAAAAAAA
AAACCATCCAATGCTCATCTTTACCAATGTTCAAGAAATAATTCAAGATATGTTGAGCAGAAACCAATTAAATCC
CACAACCTTTTTCACCTCTATACTGAAAGGATGAGATGATGAGGGAGATAGCTCTCTTGCAAACATATCTTTTAT

GTTTTAACCTTCTATTTATGGATATATGTTTTATGTTTGCATTTAGAGGGTGAAGAGCATAGACCGTACAATAAG
TTATAGTTTTGAAGGTTGTCTAGAGTGGTAGGTCAGAGTTGTAAATGAAGCTCAACATGCTTGGATGTCGCGTTT
ACTATGCAAATGAAGAAGTCATCTTGGAAGTCATCTTGGCATTTTAGTTAGACAAAAGGAGATACAAATGTAACC
AAGTTGTCAAAAAGCTTACAATCATGTGCCTCAGTTATTCTTTTTATATATTTTTAAAGACAGAATAATTTTTTT
AAAAAATGAAGAAAATGGGTTAAATGACATGTTCCTTTGCATTCCTTTCCTTAAGAATGAGATTTTGGATGTCTT
TTGATCCCACAAAGGGAATAAAGCTTTTAATCCAGATAGTTTTGCTATTGTAGTTATTGAGACATAATTGAGACA
ATTGTTTTGTTAGTCTAGAAAGAGATAGGAATCAATATTAGCTATTTAAAATATTTTTCTTTAAAAATAAATGTT
ACATCTCCTCAAATACACAATTTTTTGAGGCAAAAGGTCTTTAATAAATATCATTTTTGTGAAAAAAATATTGAA
ACCCATAATCATTTTTAATTGTGATGCACTTCTCTTCTTGTCTTTGGTATGCTACACTTGATTTGCTTGGCTTGA
GGTAGATGATAGTTGGTCAGATAGGTAATAATAAGTTTTGGATGTGGCAAGGTATCCAAAATGAACTAAGAGACC
AATAAAAGAAGGACGAGCCTTATCTCTATCATCATTTTCTTACTTTACTTGTTATTTAGAAGGAACTGAAGAGCT
TTATCAAGATATTATTTTTTGTCTTGTCTCTCTCGTTATGACACTTCCAATAAAATTAGAGACCTTAAGTTTTTA
AATATATTTTTTATTTAACAATCATCATTTTATGCATGACAAAAATCTCAGAAAACTTATAAAAAGTTTAACATA
ATTATAAATTCCTTTTTATTCCAGAAGGCACTTCATTAGTACCTCATTAATATTAACGAGTAAGAATAAGGCAAT
TGGGTAAGTAAGCTAACTATCAATGAGTAAAATTTTTTTTCTCAACAATATGTTTATGATGATTGAGCTAAAAAA
ATTAGTATAACACGAAAAATCATATTATATTTTTCTTAATTAATATTAATAAATAGAGTGATGGTTGTAGTCAGT
AGTGATTGAGACAATAATGATAGTAATAGAAATCATAACAATAGAGGTGAGTGGTAATAAGGTGCCAGTGAAATT
TTGGTGGAGCAAGAGAAAAAGCGGTGCCAATGGTAACAGCAGTAGCAACAACAATGATAGTAGCATAGTAGTGAT
CATGTAAATGATGGTGATAGTAATAATAACAATGATAATAATAATTACTAAAGCGATAAAGACGATAAAAATAAT
TTGCATTGTGGTGGTAGTTATTATAGTAGTAGCGGTGACAACATCGACATGCATGGTTGTTATAAAGGCAATAAG
GATGGAGGGCAACAACATTGGAGATAATAGTGTTGTCGTAATTGTGGTATTAGCGTTGATGCTAATGGTAGTAGC
AATGGTACCAACAATGATAGTAGCACTGGCAATGAGATAAGGTGATGACAATGGCGGCATAGTATTCTCGTGAAA
GGATAAAGTTTGGAAGAACAAAAAATTATCAAAGACATTTTTGTTCTTTTAGAAGACTAATCAAATAGGTAATGC
TTCTTTACTTCTTAAATTGAAGATTTTTAACAAGCACAATCACCTGCTTATATCTCCCTAAAAGATAATTTAAAT
TATCCTATTGATAATTTAAATTATCCTATTTTCCATAATTTCAAATATATCAAAAAATTTTAATTGCAATCTACC
ACAAACGCATTAAGTATTGATAGTTGAATAGTTAAGCCATTGAAGAATGTTGACTTTTGCTCTCTCACATGGTCA
AATTTATTTTCTTTGTTTCTCACTGTTCTCGGTTGAATACTCGTCTTCGATTATCTTATCACAATCTACCTTCTG
TTAAAATTGCTGATCCTTGGATATAAAGTTGTTGTGCATCATAATTTTCACTATCTCCAAATCCATTTAAGTACA
CAACCTAGATATCTAGCCATATTGCTTATCATCTTCCTTTAGTTGTACTTTTGGTCATCTTTATATATTGTTGGT
GAAGAATGGACATGAACGAGAATTTCTAAGTAGATTAAGAATTGTATTTCTAGATGCTCTAAACTTCAATGTAAC
AAAATAAAATAAAATAAATTTTTAGTAACCTTAGGATTCTATTAACTGTTCAAATTCTTTTAAGTTATTCTTTCC
ATCCTACTTGTATAGATATAGTTTCATTAGTGCTATATGTAGAACAAATCTATAAATAATATGAAGCTCATTTAC
AAATCTACACTTAGTTTAACTAGAAAAACGCATAAATTTTATTATCTTTATTTGTTTTAATTAGGAGAAAATTAA
ATTATATTAGGATGACAAAAAAATTCAGTATACATGAACTACTATAAGAACTTAATTTTATCATTTTCTCCTAGT
TAAAACAAGTATAAATAATAAAATATACACATTCTTAAATTTATCATTATGACAAAAACTCAATATACATGGACT
TCTATAAGAACTTAATTTTATCATTTTAAGTAGGATCCTTTCATCATTTTTTTAATTAAAATAAGTATATTTTCA
TAAATAAATGCAAAAAAATTTGAATTTATTTGAAAAATAGTGATGCAGAATAGGAAACTTCAAAAATAGACATAT
TTCAATAATAGAAGAATGACGAACCAAATAAGGGGAACAATTATTTTGAGAAATTTTACACCGAAAGGAATAAAA
AATGTTAATTTTTTTTTATAGCAAATTAAAATTTTGGAAAGTTTTATAATAAAAAGCTCTAATTCTTTTCATTTA
ACTTACTTTCCGATTTTAGCTTTTCTTTATTAGCAGTAAAAAGAAATGGGATGTTTCCTATTTACTAGCTTAGTC
ATAAGATAGGTTATTCGTAATTGTTATTCATATACTGAGATATTGTCTCAATTATTGAGATCTAAGATTTCTGAA
TTTTATTCTTAATAAAAATTCTCGAAATTTGTACTCAACCTCTCTTTGTTCTACATCAGATAGTATCAGAGCCTA
AAATGGAAATGCCATACCTTTAAATTTGCTCAAACTAAGTTTCTTGCCAAAAAGAAAAAAAATACTACTACAGCC
ACCAAGAGCAACAGGAACGAAAAGACAATTGTTGGCCAACGTGATTTCTAGCCACCACCAAGAGGTGGACGACAA
TGTTGCTTTTGTTCACTTCAGAGCATTCCTGTCAGCTTGTTCACCAAATATCGCTCATGGGGGCACCGAAAAGAT
TTCCTATATTAGGTTTTCTTTATTTACAATAAAGGAAATAGGAAGATTTTTCAGTTTTCTTCTATTTAATTTAT
TTTTCTATTTTATATGTATTCTATATTTTCTATTTTTATTGCATTTTAGTGCTTATAAATAAGGTACTTTAACTT
AGTTATTAGGTAGATTATTCATGATTATTATTCGTATATTAAGATATTATTTTGATTATTAAAACTCAAGATTTT
TAACTTTTGTTCTTAATAAAAATTCTCTCAATTTGTATTCAACCTCTCTTAGCCCTACGTTACTTAGCCTAGGAT
TTTTATGTTTATTATAAGTACCTATGTGATTTATCTTGCAAAATCAAGCCATGGGTTAATAAGATTATCATGAGA
GGTTTCTTAACAGCTTGGTCTTGTATGCAATCATGCGTATAGTGTGATGCATAAGAAACCTCTTCTTAGCATGAT
GCTCAATGGTTTCATAAATGTGGCTTCTTGTTAATTTTCTGTTCCTATACCAACTATTAGGGGTATCATATCTGG
CTTCTTGTTAGTGTTCTGTTTCTATACCAACTATTAGGCATCATACGTAAGAATTCCTTGAAACAACCCCAAATC
CCTAAATAAGCAAGTTAGCCCTTACTTTGTACTACATCAGTAGCTAACTACCATAATGGCCTATTCAGTTATGGT
CCTTTATACATCAATTAACCAACAGCTTGTTTTCTATACTTCTATTTGACCAAGTGCTAGAGTAACTATACCAGT
GCTGCAACAGCTAAAGCACCAATCCTGAGTTCTGATTCTAGCAAATTAGTGCTCAAGAGTTCTATCAGTATCCTC
CATAATTCTGGGTACTGTTGGGTAATGGTAGCAGTCCTAGGCAGTTCTGAAAGTAGTCAAAAGACTATGACCCTT
CTGGATTCCATTAGTTGCTTCGGCATCCCGTGGTTGTCTCAATGCCCCAGACATCTCTTGATCCACATGTTGTGC
CATAATCGAGATGGCTTTAGGGGTCCTAGCCAGCTTTCGGGTCTATGATTTTTTCCCAATGTTTGAGGTTAAGTC
CATGAATCGAGATGACTTTAGAGGAACAAATCTTTTGACCACTACCAAGTCCAAAAAGTTCTCTAATACTACATC
TTGAAATACTAAATAAAATGTTAGATGTTGATAACCTGGATTAGGAATATTTTGTAAAAGAACTCTATTAACGAT
TCGGAGGACCCTTACGGAAAGACTTCATAAAGGAGTTAAATTACACCAAACAAGTACCATTTTTGAGTACGTAAA
ACAACTACAACACACGATAAAATTCCTCAAGCCACAAATGCTTAATTAAACATTCGAACAGCTTTAACTGCATAA

ACAATCAGTAATAGAGTTGTCCATGAATCCTTATCTTCCCAACAATGATGCTCCATTATCAGAAAAATATTGATA
TAATGATAGACCCCTTAGGAGCATAGGAAATAAAGCAAAACCTAGAAGATCAGCAGTCTATTACAAATATGAAGA
TAGGTATTTTCCAGGGCACTGTTGTGCAAATAAAACAATAATGGCCATGGAAAAAATTGTTATGGAAAAAACTCT
CAAAGAAAAAATTTCTACAATTCAACCTTACGGACAAGGTTGATGCTCAAGAGGAGGGTATTGTGATGAACCCAT
TTAAGGTGTAATAAAAGGAAGGGTAAGTTATGAAATTCACATCAAAGAGCAAAATAATAAAACAAAACTAGGGTT
CGATTTTGTCAACCCTAATAATCTCTTCAATCTCACAGCCAATGCCTCTGCCATCTGCATTCCATCCCGCAGCCG
ACACCGCGTTCATTATCACCGCGCAATCCTGTCAAGCATGATTTTGCCAAGAAGAACGCTTTTGTTTACTGTATT
AATGCCTATAAAAGGCCATCTTCCATCACTTGTAAACACCAATCGAAATCAATAAAAATCTTATATCTATTTCTC
CCTTCATATCTTCTTCTTAGTATTCATTCTCTCATTCTTCTCTTTTTCTTCTCTCTTTTCTTTCTTCATATCAGA
GCCATGTCCAAGGCTTTTGATAAGGCTCTCTTGCTAATTGTGGAAGACTATTACTTCAATCACTGTTTTCTATAA
TAAGGCTAATCCCTGAGCCACTAAAGCAGCTATTTATATGGTGATTAGTTTGATGCAGAAACTTAACGTTTGTTA
CCAATCTTCTAGATGTTGGAGAAGGTTACTACAATCACTCTTTTTTAGATTGATAAATGGATAGGAGAGGGAGAG
ATACTTATAAGCAAGGATAAGTTTAGTTACTTGCCTTATCATGTAAGAATATAATTGTAAGCATGTATCTGTAAA
TTACTTCCAAAGAGATATTTTGTTTCTGTAGCAAAGCCATGAATGTTTATTTCAAGGCACAACTATATATTCTGG
TTGCATATTGACACCAACTGCAATACATGTGCTTTTCTGAAGATTCATCTTCTTCATGCCCTCCGGACAACTCTC
TTCTATTGGCTTATTTGTCAGGTGGACCTCAATTAATCTGAACTCTGATTTTCCCCGTCTATCTAACAACTGAGC
TACTGATGGATATTGTTGTCCATCAATATGTAATATTCTCTTTCCCATTTGGCTCCCAAAATGAAGAGAAAAGAA
GAATCTACAAAAGGTTAATGACCTGATCTAGATAAAATATGACTTCTGACAGGAAACTTCTATTAGAATATACTA
AAAGCATTATTTAAACTATCATCACTAACTCTAACTGGAACCTCCAGATTAAGACTTGTTGTGATTAGATCGCAT
CATTAGAAGTTAAAACCAAGAGAGCTTAGCTGTAAGTGATTTGTGATTAATATTCCTCATGAACTCTCAAATTTC
AGCTGCACAAAATCAAGCTAGTAACAAAGATAGAAAAAAGATTCACCATGGTGTCCCCTTATGGTGGCTCTGACA
GGAAACTGTCACAATGAAAACACGACTGCTCTTTATTATATATTTCAATTTGCATGGATAATATTTTATTGTTAG
CAGAATAGAAGAACAAGATGCTACCCAAACCTACTGAACTTCAACACTGTCCATAAAAAGCCAAGGAGAAGAAAA
CTTACTTCTAGGTTTGAGCTCCATAGGCACTGACAGATTTATATTTCCATGCTCCTGGAAATTTGTAACTATTAC
TTAATGGGATACCTTAACTTATATCTAGAAAGGAGAACAGAGAAGATAACGAACATCAAAAGGTCATAAGAACAT
ATCATACGTTGCATCTTCTGTCTTTGCTGTTAGTGACATTGTTTTCCTGATATAATTTGAATTTCTCTTACCAGC
ATGAAAACTGTAATTGAGCGGTACAATAGAACAAAAGAAGAAAACCAGCAGCTCGGTAACCAAACATCTGAAATA
AAGGTAATTTTCACATGTGATCCTTAAATTTCTTTGTCTTAACCTTAAAATATGATTTTGCTGCCACTGTGCTTG
ATGGGAAGATTGAAAGTAGGCACATCTATGCGGTTTCAATGTATATTGCATTCTTTAAAAACAATATTAAAAATA
TGCAAGAAGCAACAATCTCAAATTTGCTTCTGTTGTTCCACAAACTTATAACCTGTAGGAGAACAATGATGATCT
CTCCTAATGAACAGTGACTATGGCTGGAAAACATGTACCAAACAATATCGAACCAGAAACTGGAGCAATTTGATG
TATGTCTTAACTGAATCAACAGTCATTGCAACAGGAATGTTATAAAATAAGGAAGGCTAACTGTAGTGACAAGGA
AGATTCATGTTGTCTGTGCATGCCACTAACAACATGCCTACATATTTTAAACATGAATATTAAATTTCCTGGAAA
CAGCTAATAAATGTAGATGGGATGTAGGTAACTATCAATAACTTAATGCAGTCCAACATGGTAACGTCAAAATTA
TTAGACTGGGAAAATTGCAGGTGAATACCAAGATGGTAGCATTCCAACATTGTTTGCTGGACATATTATCAGTAT
TCGTAGTGTATGGTCGAACAAAGGATGTCCATATGGAAGCAGCAGAAGAATTGCTGTTTCTGAAACAAGGCCACC
GATTTCATGGAAGACTAAGTAAGCTAAACAAAGTAAGGCTCAAAAATACTGTAAAAGGTTGTGTGCACCAGTTTT
GAGCCAAAATGAGCCAAAAGAAAAGAAGGCAGTCATTCTGCCGCTAGTCATAAGGTTCACAAATGACAAACTCTA
GCAAACATCCCTTGAGTCCAACAAAAATGTCCATTTTGTAAAAGAAAAATAAAAAAAATTATAAATAGCTGTAAC
AATGAAAGTTTATAAAGGATTACTATTTTTTTTAAATACAATGCATAAAGTATCATTAAATACAAGCACCAAGAA
GGTACAAACCCATATATAAAAAATATTTATATATCAACTAAGATANNNNNNNNNNNNNNNNNNNNNNNNNNNNNN
NNNNNNNNNNCATCAACTTATCAATGTCAGAGTACATCACGACATGTTTCTAAGTGAAGAAAGCAATCTAGGAGG
TGCACTCACGTTCCAAATTTGCGTGAACTAGAAATTATCTCGTTGAGTTCTTCCTTGGTCAAATGTCTATAAAAA
GAACCAACTGTAATGTTCTTTGTTGAATCAAGGGACCAAATCGGTTGGTCATAATTCTGGTCAAGAGATACAGAA
GACAATGAGTTCAAGAGTGATTCGTATTTGTTAACTTCCCAATTATTGAGGTTCCCCAAGAGCTCTACATGCCAA
TCTCTTCCCTCTTTTCTATCGCTTAAGCCTTTTTAACCAATAGATCCTTATATCTATAATAGTGAAGATCTGAGA
GAACCTATCTCTCAACACTCCTCCTTTAGTCCAAACATCCATCCAAAAACCAATTCTCTCCTCATTCCCAATTCT
CTAATTGATACCTTCTTGAAATTTATCTCTTCCCATGTTTCTCTGTTTCCACAACCACTTCCCATGTGGCCTAGC
CAAAGCACCAGAGTATCTCCCCCTATCCATTGTACCAAAATTTGCCTCAATCGCACTCCTTCATAATATACCCCT
TTCTTTCATAAACATCCATATCCATTTTTCCTGAAGTGCCTGATTTATTTCTATTACAAATCTCAAACACAATCC
TCCCTCATTTAACGATCTTTTACTCACCTCCCATGCCACCAAGTGGTATTTCCTTCTTTTCTCTAAGTCACCCCA
TATAAGTCTGCAGTGAATTACTTTTAACTTCTTAGCAATAATTGTTGGAATTGTTAAAAATAATGTAGTAAACTG
GAAGATAGGATAACGTGCTCTTGATAAATGTTAGGCATTCACCTTTAGAGAGTAACTTCCTTCTCTCCTCTATAT
CCATCTATCCTTAGTTCTAGCCATTTCCCTTTCGCACCCTTCAAAAGTTTATTGTTCTTCTCTTTCCACACCACC
CAAAAAATAGTTAAAGGAACAAGATTTACAAATTTTTTCTTTGCACATATCTCATCTCATGCCCATAATTTATCT
CTAATTGTTCCTGTAATAAGCTAGTTAATCCCCAGCATACCATAAATCACTGTCTAAATGTTGTACATCATCGAA
CACTAGAGAAGAATGTGATTACAAGATTCAGTAGCTTTCTTTAACTAGAATTTTGCCCTTTTTATCAACTTATAA
ATGGTCAGAATGCATTCTCGTCTCGCTTCCCATGCAAAGAAAACAATTCTTAAGGGTGTTTCACCTTCCAAATTT
GATTAATTGGGAACTTGACCAGCCTCGCCTTATCCATTTTGACCAAATGATTATAAAAGACTAACAATGAACTCT
CCATCCTTATTCAGTTTCTGCATGATCTTGTCCTTGTTTTCACTCAAATGCATATTGAACAACAGCTAAAGCAAT
GTCTCATATTCTTCTACTTCCAATCGTGAGGTTCCTTGTCACATTCGTATTCCACTTCCTTCTACAATTCTCCCC
CTAAAGGCTTCCTCTACAACCATATTCTTCTCTTGATTAATAGCATAAATCCTCGGGAATTGATTTCTGAGACTA
CCCTTTCCTATACATTCATTATCCCAAAATTGGATTTTATTTTCCCTCTCAATCTCCATTGAATACACTCTAAAA

ATTGTTGTCTTCCATACCTATCTTCCTCCACAAACCCATCTTGCGAGGCCTCACTATCTCTCTAGTTAACCCTCC
CCCTTCGTCCACTCTATCAATAACTCTCTTCAACAACCTCCCCTCCTCTTTCAAATGCCTCCATATCAATTTTCC
TTGTAAGGGCCATACTCATCTCAAAAAATGATCTTAAGCCCAAACCCCCTTACTTCACCAGATGATATCTCCTAC
TCCTTTCATTATCCTCCCAAAAGAATTGACATTGAATATTTTTCAGGTTTTTTGCTACTTTACAGGCACTATCAA
GATAGATAAATAATAAATAGGAAGGTTTATGAGCTTGCTTTTAATGAGTTTTAACCTCCCACTCTTAGATAGGTA
ATTTCTTTTCCACCCAGTTAGTCCTCTCAAACATCTCAATCGCAGGGTCTCAAGTTCTCTCTCATCTTTATATTT
TGCCTCTAGTGGAAGTCTAAGATACTTAATAGGGACCTTTACAGATTTACAACTCAAAACCTCCACAACCCTAGC
CCCAAACCAAAAGATTTAATATCATCCATTCCTCAAGTTGACAGAAAAATCAGAGTATCATCAGCAAAAAATAAA
TGAGAGATCTCTATTTGATGGGACAACTCTTGCATTTTTACTTCAATACCTCCACCATAACTTTCTCCAACAATC
TGCTCAATGCTTCCATACTATAATAAAAAAGAAGTGTAGATAAAAAAATATCCTTGCCTCAATCTCTTGACACCC
TGAAAAGGAAGGCCCTCTATTTACCATCACCGCATAGAGGATGTGTGCCTACTCATGAACAAAGATTTTTAAAAT
CTTCCATGGAATATAAAGAAATACTCATAACCGACAAGCCCAATGTTTGATATCACCTAACTTTAACCTTATTAA
CATCCTCTTCCACTCCAAAAATCCTTCTATTCCTCTTTTCCCACACCACACAAAAGATAGTAAGAGGAATCAATC
CTAAATGTTTTTTTCCAATCACTGACACCCTTCCATGCTCATAACTCATTCCTAACATTATCCGCTATTACCTAA
CCGACGCTTAACAACCCATATGCCAGACACTATAGTTTATACGTTAAGGGGCACCATAGTAAAATATGATTGCAA
ATCTCTTCCACCTTAGAAAGGTAGCGCTCGTTCACCAAATATGATCGCATGTTTTGTTTTTAGGAATTCTCAATG
TCTGGAACATAGTTTTAAGCTTTTGAGCCCAGGCTTCTTCTAGCTTTGGTAACAATTAATATAGAATAAATTATG
ATTAATTTGGATCCTGATATATAATTTATTAGACATTATATTTAAAAAGTTTTAAAGCATCACACAACTAATTGA
AGAAAGTAAAACTTATTAAACAATAAAGTCTGTCTCAAAAGTAAAATTAGAAAGTTCTTCTTAGTTAACTAAAAC
TTATTGTACAACAAAGTCTGTCTCAAAAGTAAAATTAGAAAGTTCTTCATAATGCTTCTTAGTTAACCTTTCTCA
AATAAGATTAAACTAAGAAATACTGATGCTAATAAGATTAGGAAATAATACTACTAATATTAAATAAGAAAACTG
AATATTAATAAGAATAAAATCCTAAAGTTGCTAAAAGTATAACAATTTTTTTTCCGAATGATAATAAAACTATAA
CAATAGACTCCTATATAGTATAGTAAATTTATAGTTGATTGAATCATAATTTCTAATTTATATAAGTAACCAAGA
GCTAATCCATTCTGCATTATATAAAAGGACATAGGACCACCTGTTGCATACAGCAGATTTCCTCAACTAAAAACA
ATTGGTCCCTTTTCCCTAAAATCCTCACTTGTGAAGAAAGAGAGTTTTGAACATGGCTCAAACTGTGGTCAGTAT
AAAGATTTGTCCTAGAAAGAGTTTTAGTACTTCCTAGGAATATAGCACAACCCAAAATTTATTTCTACTTAAAAT
TCATATAGAATCCAAATTCAAGAAGTACTAGGAGTACCTAGAAACAAGGACATCCAAGTTTCAATTCTTTCTATC
ATGTCCCAAGTAAACTATAGTTTATCATCACGTATTTAAAATCAAATAGTTGACAGTCTAATTACTTCCTAGGAA
TATAGTATAACCCAAAATTTACTTCTACTTAAAATTCATATAGAATCCGAATTCAAAAAGTACTAGGAGTACCTA
GAAACAAGGACATCCAAGTTTCAATTCTTTCTATCATGTCCCAAGTAAACTATAGTTTATCATCACATATTTAAA
ATCAAATAGTTGACAGTCTAACTACTTCCTCGGTACACATCCTTCCTCTATATGGAACTTTTACCAATTAGAGAA
GACATGTTAAGTTTTTTTGAACTGTTTATGTTGATAAGCTCTTTTACTTAAAAATCATAAATAAACTGTAGAAAA
TAAGAAAAAAATAAAAAAAATAAAAACAGAAAACTGTGTGCCATATCCATTACTCATCTAAATCCTTCAATTTTT
TTTCCAGTTCTGGCAAAGGGAGGCAACAATGCTGAGGCAACAGCTGCAAAGCCTCCAAGAGAATCATCGGTATTA
TGTTTGGTTTTGCTTTTACTTTCAAGAATTTTTTCAATCCAATATCAATCAAAAAGCAGAAGAATTAGGTGCATA
CAGTGAAAAAAAATGCAGAACTAAATGTTATTTAGTGCATCAAGGACTTGTCACACATCGTATTGGACATAAACC
AAGGGACAATTTTTTTTGAATGTTATACACCAAAGCATACAGTCTATTCATCAACATCCAACCAGCAGATAAAAA
AAAAAAGCTTCTTTTCCTTTTGCCCGAAATCATGTTAAGAAAATGGAGACAACCTTTTCTCTAATTCATTATCAA
TAGTGAGGTTAAGAACATGTACACTGTCATTCTTATACTGATCAAACAACACGATAAAAATGCATCCATCAACAT
CTGACCTACAGACATAAAAAGAGGCTTTCTTCTCTTTGCCGGAAAGCATGCTAAGAACATGTAACTTCCTGATCT
CTGTACTTATTATTCTATTATTAAGTTAAGAACATATGCATTGTCAATATTAGTATCTCCTATATGTCAAAGGAT
AATTTTTTGACAGACTCTTAAAGTGAACATATCTGGACTGCTTAATGGGAAAATTATGGAATAGAATTTAGCCTG
TAAGCACTGATATTGGTGAACCAAATAGCCCATTAGTAATTTCTAAATCATATGCTTTTCTCAGTTTTCTGGTGA
ACGCAGAACAGGTCACATATTATTGTCTTCTAATTCCTCTTATGGTGTTATTCCATACATTTTAACATACGAGGC
AGTCTTCTAGGTAGTCTGGTCACTGCAAGACTGGGAAAATAATGTTATCAAGGAACTAAGGGGTGATACAGAGAT
CAGAGGTGATTTAGATTTCTGTTAAAATGTCTGTAAGGTTACGTAGGTCCTCATCATGCATAAGAGAATGATCCT
TAAAACAAAAACATAGGATTTGGTGTCAAAGGTCCCTCACAGTATGTAAGAGAAAAATACCATCTACAAAAACTC
TATGGTTCCTTTTTTTTTCCTAACAACCTGCAGTGTAGCCTAAGAGATGAGAGAGACAAGCTTCTTCCTTCTCTT
ATCTAGAAATGCCCTACCAAACTTATCAAATACCAAACTAGCCATGACCCAAACCCAATCCAAGGAACTGAAGTG
CAGCACACCAAAAAAGAAAGGAAAAAGTGCACCACGTTGAAAGGTAATTTCAAATGTCAGCTGCCCTCTTACAAA
GAAAGCACCTGTTAGCCAATCCCGAAAGATAATTATCAATAGTTAGGAATCTCAGCGGTTAGCCAAAAGCTGTTC
TCAAGGCCATATTTTACTTCCTAAGTGGGTTCTTGCTTTTGACAGTGCTGCATGAATCCCATTACAAGAAATACA
GGAGAAGCACATGATATTCAGGAGACTGACATATCCATCTTCTTCCAATCTTTCAAATTTCTAATCAGATCCACC
TCCCACACCACCTGACCAGAGGATATGCAATGGAGGTGACTCACTAATGCCTCCTTCCTCTTGCACAATAACAAC
ACTGAAGGAAAATGATGGTCTAAGGCCCTTCATCTTTCCCAAACTTCAACTTCCTACTAAACATGACACTCATCT
ATAAACAGTTGCAGAACTCATCCCTGCCTGCAATATACTCTTCTAAAAGTACAACCCATACTTGTAAGTCCCGGT
CCTCCAAACCCATCATCCAAACCAAATTTGGTTATAATCACACTCGATCAGAAGCTACCTTTATTGTGAATCTCC
AAATCCACTTCCCAATGAGTGTCTTGAAGAGTAATCATAACTGGATTCAAATCATGGCTGAATCCCATGAGACCT
AGTCCATCCTCTCTCTCCCTTCCTCCTGCGCTCTCCTCCAATTGCAAGAGTTGTCATTCACAAACAGCTAGGTGA
AGTGGCAATCGTTAGCCAAATAGAATGATTTTCCATTCATAGTGGTTGCTCGGTAGTGACTAGATTAGTGGCAAC
TTGCAAACAATCAGTAGTGGAGATAGGCAGGAATGGCTGAATAGCAGCTTCCCCAACTTTCCAAAAATTATAATA
TTAGCATTTTTACTTCACAGATACTCTTTAATTTAATTTTTTAATATTGAGAAAGTGTAATATACATTAACTCAC
GGCTCATTCCCCTGGCCACCTTCTGGGATTGGGAAGGAGAGGACTGGAGGGCTTTTGAAGGATTGAATCCTCTT

TTGACAGGTTTAGAGACTTGTGGTTCAATGTCCCAAGCTTTTTTATAAAGGACCAGCCATCGGGTTCCTGCGAGG
AGTCGAGGGAATTGATTGATGTTTTGCTTGAGATATAAAGAATTTTGTAGATAGGTTGGTGCTCATTTATATATT
TTCTTTTTCTTGCAGAAGAAATATGTATACACCACTAACTTAACAAAAGAGTTAATTTTCGTATGCTTTGATGAG
AGATCTTAGAAACTGTAGAAAAAATATTATGCTGAGGCAGGAATTCCCTTTTATCTTTATAAAAATTCTTCCTAT
CATTTTAGGATACTTAGCTTTTTTAGTTTTGGTCAAACAACAGTAACTTTGACTTGTTCACAACAGTGTAATTAT
TGTCAACATGAAAACTTCTTTTCCTACAAAAAAAAAATCCATAGTAACAACAACGGAGCCCCAGAAAGCAACATC
TTGCTCCCTACTACATAGACACAGCATGGCATATATCAAACTAAAAGGTACGGAGAAGATGTAACATGAAAAGTG
ACATCTCTTCAAATTTCCAATGCATTTCAGTAACATCTTTAAAAGATCAAAGTTTTGGAGTCAAATCACTTATTT
CCATACAACAATAACAGGCAGATGATGGGGGAAGAGCTTTCTGGCTTGAATGTTAAAGATTTACAGAACTTAGAG
AACCAACTGGAGATGAGCCTTCGAAGTGTTCGCATGAAAAAGGTCTGATAATGAAAATATAACCAAATAAAATTG
GAAACTGCTAGCAAACAAAGCGTATTCAGACTCACTTCAGTTATTGGTTTACTTTGTAGGACCAAATCTTGATGG
ATGAAATACAAGAACTAAACCGAAAGGTGAATTGAGAAAACAAAAGGTTTATTTTTTGTCTGACTTTTTTTAATC
CTGACTAACTAAATTTAGTATTGCAGGGGAACCTCATTCACGAAGAGAATGTTGAGCTCTATAAGAAGGTAAACC
TCATCCGTCAAGAAAACATGGAATTATATAAGAAGGTATCAGTTTTGGTAACTGATAAACACTGCAACCATGCAT
AAAACACTAGTCCAGTTCAGTCAGTAAACTTTGTTTCTATTTCAGTATGTTCCAGAGTCACTATAAGAACAGTGA
TTTGCATACCAGGTTTATGGAACAAGGGATGGGAATGGAGCAAACAGGAATTCCATTTTCAAAAATGGTCTAAGC
TTAGGAGATGACTTGCATGCACCTGTTCACCTCCAGCTAAGCCAACCAGAGCAGGCAAACTACGAGAGACCTTCA
AGGGCTACAAAACTGGAGTATTGTTCTTTCCTATTAAAACATTTTGGTTAT**GCCCAGATATTATTATGCCACAAA**
**ACTTACAGAGGCCTGTTTCCATTGTAATTGCAGCAAACTGCAACTGCATTAGCAAAGAAGATGCATTACAAAGGG**
**ATTAAATGAGCTCAATTTCTATCAAGAAGGATCCATGGATGAAGTTATGCTCCTAATACTTGGGATGTGGTATGT**
**GAACCACTAGCTATTCAAGAAAACCTTTTGGTTAAAAATGATTAAAGGAAGCAACTTTAGCAGGTGTGTGTGAAG**
**AGGGAAACCAGCTTCTTCTGTAAAGATAAGATTTAAGTGTAATAAAGGATGAAAAATAAATTTCACTGCATCTAC**
**CAGGCAATAATTTAGCAGTAATAAAATAAAGGGTGATCCTGAGAAAGATAATATGGTACCATCACAATATTACAA**
**GATCAGTAATTAAGCAAAACAAAAGTTTTCTGCAACAACATGATTCATGAATGCCCCATAAGCAACACTAACAAG**
**TAATAATCCGTGCCTCACTAAATAACTGTGTCAAATTAGTGTGTGACATACCTTAAACATTCAAAAGCGAATAGA**
**TTTTCCTTCAGATCAATAGACATCCCGGTAGATTAGAGCCCTG**CTGAAAGACAGATTCTGAATAATCAACTTATG
TAAAAACATATAAGCATAATGTAGCAATGCATTTAGGTTGGACTAGAAGTTAAATACCAACGAGAATCTGAAATA
CATAACCCTTCAAACAGAAGGAAAAATTTACAAAATTAATTCCACAAGGTATCCTCCCACTTTATACACATTCTT
TTTATCTCAAAAAAAGGTAACCTCCCATTTTATGTTCTTTTTATCTCTCGCTCTCTCGCACTTGCGCTCGCTCTC
TCTCTCTCTCTCTCCCTCTCTCTCTCTCCTCTCTCTCTCTCCTCTCTCTCTCTCTCCCTGTTTTAAGTT
TATTTTCTTATAATAAAAAGAGGAAACTACGAAAATCTGAAAGCCCTGATAAAGGTCAAGATTCATAACCAATTA
AAAATTGTACTTAGAAAAGAAGCCTTAC**ATCTGTATAGCCTGCAGATGTGAAAAATATCCTAACTTGACTTCTTA**
**CTACGTGGCCCGCGTGAAAGTAACTTTTAATCATAAACACTCCAACACTAAAGCCACGGAATTTACTCTCCAATG**
**TTGAACGCTCAAGTTAACTTTCT**CTGCCAGAAAAACAAGATCCATTTAAACTGAGGAAAGATCGCATAGCGATGA
CATTTAGATTACAAATAACCACAACTGTTAACTATTTGGCAAAGATATAACATAAGAGGATTGAGAAACCTAGGG
AATAAATAATTCTAAGCTATTCAATTTAGCTCAACCTCAAAAAGGATCTTCAATTTCTCTATTTCAAAAGAAAAA
AAGATCTTCAATTTCTCTATTTACCAATATGTTTCTCCTAGAAATATAAAGGGTCAAAGTTGACAGTTTTGGCTG
CCACTCATAATTTCCTAAGTTAAAATATATTTTGTTGCAATGCCACAAGAAACAGTGGTTTACATCAAAACTTCA
TGATGCAAATCTTTAGGTCTAACTGAACTCTTCCGACAAGCAAAGGCTGAAAAACATCAAGTCAGTTAAAGTAGC
AGGGAAGAAGGCAAGCCTCGCAATACACATGAGAGTTTCCATAAAAGTTTGATGTTTGGATCATTAGCGTATTAC
AAGATCTTTTTTTTTTTTTTATATAAAAAGAATATTAGAAGATTAAAACCAAATTTACATCCTCAAGAATCTTAC
TTTTATTCAATTAGACCAATGATTAAATGAGAAACAGAACTAAGGACTATCAAGAAAAGGAAAAGACAATGGATA
GAAAGAAACAACAGCCTGGAAACATTTAGTGTGTGAATCATCACCCACCACCTTCTAGGAAGAAAAGAGATCCTT
TCTTGAGCCTTCTACTTTTATCAGCTATTAGGAGACAAAGTAACAATTGAATTTAGCATGTCAATGCAAACCTAA
AAATAGGCACAGATATTCATGAACATGTTTGCATGAAGGCCCAATAATGGTAACAAATGGGTAAAAAATAAGGGG
GGAATTAAGTTCCTACTATTTGGAAAAATTTTGATACCATGTTAAATAATTAGCTGGGATCAGAGGCACGTGCTC
ATAGATCTAAGAAGCAAAAAGTGAATAGCATACTCTACTGCAGGGAATCATACAAAAGAGGGGCCTAAAGTATTT
CAAATAGACCAAAAAAAGCTTAAAGACAGATAGTAACTTAAATATATATGAAAAAGATAATCAAAATCTTGAAAC
TCAAAGAGTAAAGTTTTTCATTGATTAGAATCTGAGAAATCTAAACTTCAATATATTTAAATAGTTATCAATCGA
CCTCTATGCTATGCAGCAGAACTTCAAGTTCTAGCAGGTGTAATCAAAGCTTTTGTCATTGAAAACTTGCTGGTG
CTAGGAGAATTTACTAATAAATTCATTTTTGGCTGTGAGACAATGTACTAATAAATATTCAACTTTGGAATTGCA
GGCAGCCATCCTAACAGAGAATGCAACCTTGAAAGGAGCACTAGCTTGCTAGGAACGTGTTGCTAGAAAAGCTTG
ATCACTTTCAAAAACTCTAGCAATTTACAGTGACCACTAACATTTAAAATAGGGCTATATTACATTGTCCAGATT
TTATTGTGATATAATCATCTTCCTCTACAAGAGGGTACAACCCTCATCAAATGCACATTCTCCAACCTGTCGATT
TACTTCATGTCTCACAATACTGTGTACAGTTAAGAAGCTTGAGGCACTACAATATAGATTCCTTTGATTTGGGGG
ATCCAAAGTAAATACCACCTAGTGGCTTGGGAGGCTGTGGAACAACCTTTCTCTTTGGGGGTATCAATATATTTA
AATAGTTATCAATCGACCTCTATGCTATGCAGCAGAACTTCAAGTTCTAGCAGGTGTAATCAAAGCTTTTGTCAT
TGAAAACTTGCTGGTGCTAGGAGAATTTACTAATAAATTCATTTTTGGCTGTGAGACAATGTACTAATAAATATT
CAACTTTGGAATTGCAGGCAGCCATCCTAACAGAGAATGCAACCTTGAAAGGAGCACTAGCTTGCTAGGAACGTG
TTGCTAGAAAAGCTTGATCACTTTCAAAAACTCTAGCAATTTACAGTGACCACTAACATTTAAAATAGGGCTATA
TTACATTGTCCAGATTTTATTGTGATATAATCATCTTCCTCTACAAGAGGGTACAACCCTCATCAAATGCACATT
CTCCAACCTGTCGATTTACTTCATGTCTCACAATACTGTGTACAGTTAAGAAGCTTGAGGCACTACAATATAGAT
TCCTTTGATTTGGGGGATCCAAAGTAAATACCACCTAGTGGCTTGGGAGGCTGTGGAACAACCTTTCTCTTTGGG

GGTATCATGGGATTAGATCATTGATTACTTTCAATGATTACTATCATGGGATTAGATCATTGATTCCTTTGAAGA
GGTTGATGGAAGAAGAGATCCCAATTTCTTCCTTCATTGGCTTAATTCAATGGAAGAATACTTCAACTAGCATGA
GATTTCAAGGTGAGGTTGGTTAGACTTGTAAAAATATGGTAGTACATTATTGATGAAAACATTACAAGGATATAT
AGCTAACAATTGAAACATGGGATGAGATGAAAACTATTATTTGTGAAAAGTATTATTGAACCATGTTTCAATTGT
TAGCTATCTTCCAATTACTTTGACAAACTTCGTGACTAAGTTTATACTTTAGTTCAGGACAACATGTCAGTAACT
GAGTACACACTAAAGTTTGAATAATTAAAGATTCAAAGTCAACTAGTCGAAGATGATGGTCAAGCAAAATCTCAA
TTCAAGACTAGACTAAGAGTTAATATTAAAAAAGATTAATTTAATACCTTATGTCACTATAGCACAAATTTACCA
AGTGGCCCTTGATCTGGAAGAATATTCGAAATATCCAATCCCCATAAACATTGTGGAATTCATTATAAAGATACA
TTTGAATCAAAATGTAAGCTGAAGAATGGATCAATATCAAATTTCAAACTTCTTAAAGGTAAGGAAGTTGTGTGC
TTTAGATGTAGACAAAGTGAGAACATGAGTAATGAATGCCCAACAAAGAATAACTTGCATATTCGATTAAAAGAA
AAGGATGAGGTGCAAGAAAGAAATAATGGTTGTGTATCCTACGTGGATAATGACTTAGCTTGTAGTGAACAAGTA
CCTTTCTTTTGTCAAGCTGCATATCGTCAGACTCTAAAGTGAAAGAGTAAGAAGAATCACTATATCTTCCCATTG
CTCAAGATCTTTCCATATGAGTCAACTCTATAAATAGGCAAGAAAAACCAATATGGCTTCTTATTGCTCAATATC
TTTCATCATCCCATAGAGATTTAATACTCCTAAACAATTGTGTCTTTCTATTATTTTAGGTATTGAGGAGATCAA
CTAAACACAAAAGGTTGTTAAACTTTTATGGAAAAGCTCGAGTGTCAAGATTTTACATTATTTAGCACTTTAAAA
CTCGTGAACAAATTTTCTCCAATCAAGAAAAAATTATGTACAATGATTAACACTATATTTCTTAAATAAATTAGA
AATAGTAATTTTATTAAATGTAATACAACTATACTATTTATGAGTCTATTATTAAAAATATTATTATAGTTTTAG
TAATTTAGAATTTTATTCTTAATTATAGTTAAGAGTTTATTCTTATTAATATTATTTTCTATTGTATTAGGATTA
ATATTCTTAGTTCTACTAAATTATTATTTTCTACATTAGTTCTAGTTAGGAAAAGTTACTTACGGATTATTGTAA
ATAATTTTGCAATTTTATTTTTTATTCTTTTAAGTATTCAATAATATTTTGAGAGTTTTTCAATTTTGTAGTTGT
ATTATGTAATCAAACCTTTAAATATCATATTTAAAACTTCTTAAATGTGATGTTTAAGAAATCTTGGGTGTGACG
TCTAATTTTTTTACTCTCAAATCCTTAATCAGATAGATCAATTCTATTTGTCCTACATCATAAGTTATGCCTGGG
AGGATGCTGAAGCTTTGGAAGGATGAATGGTGTGGGGAAAGGACCTTACACCATCATTTTCCATTTGTGTGCAAG
AGGAAGGAGGCATTAGTGATTAACCTCTATTCTAAGTCGACTCTATACGTGGTGTAGGAAGTGGACCTATTTGGT
GCGAGTGGTGATCGTGTTGTAAGTTGGTAAAAGCATTCCGAAAATAGATTTTTGTAAATGTTGTTTTAAGATGAA
AAGCAGTATAATGTTTGGTAAAACTTACTGTAAAAAGTGCTTTTAATATTAAATATAATATTATATTTAGTTAAT
TATATTATAAAATAATTATAAATATTAAAGTAATAAAAATGATTACTATAATTTTTTAAATAATTTTTTAAAAAA
TACTCTAATCATTTAAATTATTTTAAAACATTTATTTTCATTTAATTATCTTTATTAATTTTCAATATAATATTA
ATTTTAAATAATTATAATTAGACTTTAAAACAAATATTATTTAAATAATTTATAATAAATGCAAAATAATGAATA
ACCAATTTTTATTTTAAAAATATAATTTATTAACTAATGAAACAAAAAGTTAATAAATATTCATTGAAAATATTC
TCTACCTCTTACGGCAGAAGTAATTTCTCTCAAAAACATCTATTTCCAGGTCTTGATTTTATCTTTATCAAAGAA
AAAATAAAAGATAACTTCTACATTTACTTAAAAACACTTTTTGTTTTTAGTGTCAAGCTCTTGAAGAAATTTCCT
ATTCCAAATTTGACAATAAGCAGAAGTTGACATCGTTAAATATCATTCCGCAAAAAAATTTCCACCTTAACAACT
TCAGAAAAAAAACTAAAATATACTTTTAAATTGCTGAAACAAAAATGCTGAAAAGAAACAAGATGCTTCTTGTTT
ATTACAATCAAGTTTATGGTTTTAGTAAATGATAAGTTAGTATATGTACTTATTATTGGCTGCAATATTCGGTTA
GGAAGAATTTGTTCCAAGTTAAAAAGATCCCCAAAAATATTTGGAATAGGAAAAAGAAAAACACACAAGCTTGAT
GTAATACCACTATTGTTTTTTATTACTCTGAAATGTTTAACTTTTAGCTGATTGTCACTTACATAAATCTTCTGA
AATGTTTCTTTACCAAGCACAGATGACCTTTACTAGGAGTGATGACCACTGGCTAAGATGCCAAGGTCCGGATGT
TTTGACTTGAGTTAATTTCATTAATATACATCTTTTGAAATTTTGTAAGTTAAACTAATATCATTTATATGAATT
TTTTCACAAAAAGAATATATCATTTATATACATTTTTTTTAGTGGAATACTAGTCATAGTTTTCGATAATTAATA
TCCTACTAAAAATATCATTTATAAATGCTAACAAATATTAGTAGTTTGTTAATCCTCTGATCTTAATAATATCAC
AGAATTATATTTTAAAATTATTGTTGTAATTGTAAAAAACAATTGTGAAATTTTCTTGCAGGAACTTGGATCCTA
TTGAATAGTTATCACATGTCCTGACGATAAAGATAACACCAATCAAGGAGGCATTTTGAGTACAAGGGAAGAATC
TTGAGGACCTTATTAATTTTCTAACTGAGCTGTAAATTCCTTAATTTGTTTCTAGGGTGGAACCTTCTTGGTGCA
TTGTTAAGATCTATTTTATCTTTTACTGTAAAAAACATTATGGGAAAAGGAGAGATGCAAAACTATAAACAGTTT
GCCTTATTGCAAAGCTTATTCATCAATTAGCAAATCATCTTTCTGTTGTACTCAATAATAACAATCACACACTGA
AAATACCTAAATGCCTTTATAAGACCTAAAGAAAAATGCATTTTTTCCACCTTAAGTTACTACACTTATGGTGTA
TGAACTCATGATTTGGCTGCACTACCGTAAGTTGAATGGGAGCAACAAAAGATATTTACTCTTCGGGATAGAGTG
GAAGAAAAGAGGAGATTCACAAAAGAGGAATATGATCATATATGTATATGTATGTATATATCTTAGTGTATTAAC
GGTATATGATGTAGATATAGCCCAAGGTAACTTGGTAACGATCAAATCATCATATCTGCATGTTTAACTATGATA
TTCACTAAAAATAGATCTCCAGAATTGCTATCACCGAATCTAACTGGTAATCAAATATGACATGTAATGACTTGC
GACAAAATCAACATCGATCTATAGAGTTTTAACGCTCAGTGGTGCAAAACAAGCTAAACCAACAATAAATTTAAA
CCTAAGAGTAATGTTTATCTTTTATCAGGTCATTGCAGCCTGAGTAAAACCACCTACCTTCACCCCTGCTTGTGA
TTATATATTTCACATAGATATCTCCAAGATGAAGATGTGATTTGAACATGAATGAGATTCAAGATGCTTGATCTG
TAATAACATGAACACAGCCAGTTTGCATCTCCGTTGTCAAGTATTAATACCCAAATATTCCTAAAGGTAAAGCTT
CTGGTCACCCCAAATTAGCCATAAGGCTTTTCTAAAGGATTTATTAAGCACGCCAAAAGGAACTAATATACAAGA
CCAAACTTTGGGCATTGGCAAGTTAGAATTATGCAATCTTCTTTTTCTTCAAAGGCAATTATTGGCAATTTAGAG
TTAAATGCCCACAGTTCAGAACCAATTCAGTGAACCTCCATTGAAATAACATGAGCTCAAAATTTTGTTCTAACT
TAATTCCACATAGATAACTCCTAAAAAAAAATAAATCTGAATACAAAATGACATTATATCCATGCATCTAGGAGC
TCAACCCCCTTTTTTCTTCATTTAACTTAAGATTTCAATGGTACTAATATATCATTTAGTAAAAAGCTGCTCTGT
TTAATTTGATAAACAGTGCATATCATGTGATTCTAGTTTCGGAGCTCATTTAATATGCCAGTATATGATCTATGA
TTAACACGAAAGTGGACCATGATGGGTTCACAGAATGAATCATGCTAAAAAAGACATTCTAAATATGTCCAACCT
GACTTGTCCCAGAATCTGAAGAACCATGATCCAGAAAACGTGTTCCCATGTGGTAGGGTTTACTAGGGAATTATG

ACAGAAAACTTAGAAGGTCATCATGAGATGATGTCAGTTAACATGACCTTGCGGAAAACTGTGGTTGATTAGTGA
GGAAAAGAGTAGAATAAGAGAAGGTATGTATATATACCATCGGAAGTGAATAAATGAAGGCATGGTTAGAGATTG
TTGCAGCCATAATGAAATCTCGCGAAATCATTGTGAGGGAAGGCCTTTAGACTAAGATGAAATTACTAGAGAAAG
GAATAACAAACAGGAACAAAAGGTTCAGGTGAACCCTTGGAAAAATATTAGGCTTGAAGGTTGGACATGAAGAAA
AAGTTCTATATTTTCAAAATGCACGGAGGCAATGAACAATCTCCTCAAGAAAAAAAAAAGGAAAGAAAAAGAAGA
AGCAACGAATAACTAATTATTTTCCAAGTCTTATGATAAATTGGTGAAGAATAATCAGCTCTTATGGCCAATATT
ACATATAGGTCAGACTTACAACAAACATATCACTTTATTTGTCTACCAAGCAAGTAACAGGGAACATGTTGGCAA
AATTTTGTAGAACCACAAACACAAGATGCAAATTTAGTGCTTCATCGATCACAAAGCAAATGAAAAGTAGAACAA
GAAACTTCTTTCCATGTAAATGCACAAAAATGAATGGGATGCATAGGAATGTGGATGTAATGTGTGGACATTTAT
GGGGTAATTCATACTACACACAATAAATTTTTCATTTTTCCCCCAAACAAAATAATAGATCATGGTTATGCGTAA
GAGAAACACAACTTGACCATTCTTCTACAATTTTATTTACTCATTATCTGTTAATTGAGATGCAGAATATCATGC
CCCTAACTGAAAGAAAAAAGAAGTATTTGCATCTTCAATGAAGGTAAAGTGGAAATGCTAAGTAAGCATTATGTA
ACATAC**CATTTTAATTTTCAGCTGAGACCTTGGACAGAGGAATAGCAGCAGCAATGATGGTTATCAAGCATTCAA**
**TTGGAATTGGTTGACTACCAGATGTACTTTCAGTTTCGGACAATTAAAC**CTGCAGTACCAACCAGAACTATAGGT
AAATGCCTTAATTAAAAACACATAAAAACACATGCACAACATCTAATGCCTATGTGAGTACTTCTTGGTTGTTTC
TACATTAAGCCATAGTGTAATTTGAATTTAAACTTGTCTATTTTGTTTTTTCTCCACTTTTTATTTCTTTCGAGG
ATTGTTATCCTCTGTTGACGTGCTTCATCTCTTTCTATAATTTTTTTCGGTAAGAAAAGAAAATTTTAATATATA
TCCTCATCTATTCATATATTTCCTATTATGGAAAAATAAAGTACTATTAGAAAGCAAGAGAAGAAAAGAACTTGA
GGGTAGGACTAGTGATGCAGAAGATGAATATCTTAAAATGCATTAATTTCTAATAAGATACATTTATATAATTGT
TAAATTTATGTATTATTCAGACATTTAGGTCATTTAGAAATTAGTATATATATATATGGAATAAGTTTATTATTT
CCTATTTAGTTGGGATCTTATTTAGTTATTCTTATATATAAGTTGTTTCAATTAATAAGCTCATTTTCTTACTCT
TCCAAATTTCTAGATAGACTTAAGAATTCCTATTTTGGTTGTATTTTTTACATATAGATAAGGTACCTAACTAGT
TATTAATTCATAATTGTTATTCAAATTTTAAGATATTGATTTGATTACTGAGATCTAAGATTTCAACTTTGATTC
TTAATAAATTTTCTTTAAGTTTTGTTCTTCAAGTTTTCTTTGTCTTCTATCAGTTAGTAATCCTTCAAAAATACA
AATACACAAAAGGAGAATAGGCCAAAACAACAAAATAAGCCGAATTATTAAAAAACGGCAAAGCAGTCCCCATGT
AAACTAGACAAGAAATTCGCCTTAAAGAGACTATTTGCCTTAGCCCAAAGAGACACCAAAAATGGCTATCTTGTA
CCAAAGAAAGAAAGGAGAGAAAGAAATAGTCCCTAAAATCCTAGCATTTTTCTCCAACCATAGCACCCATAACCC
AGCAAGAAACATCCCTCTTAACAGCAAATAGCCGTCTTTTTTCTTTGGAAAATCCATGAAAAGAAGAGAACAAAA
GCACTCATGATTCAAAGGAGAAACCCCATAAACACTTAAATATGCTTTCCGAGTTGATGCTCCAATGAAATATGT
AATGGTATCACCACTCCCTTTACATTAATAGAAGATAACAACCAATAAAAGACCAAATATAAAACTTAAAGTAGG
TATGTGTAGCCAATGTAAAATATGATACTTTCTCCACCTACATTGTCAAGTACAAAAATACTCCCATGTTACAGA
TCATAATGCTTTTCCGAAAAGATTTCTTAACCAAACCAATATGAATCAATACGTAAAGACCAGACATTCTGTATT
GGAAATTTAGAGATTACATTCCTATAATAGAGAACCAATTAACAAAAACAGCAAAAACAAAGACAAAATCATCCA
ACTGGGGATCAGCAAATGAATTCCGGAAAACTATTTAGACCAAGCTTATGCATCAAAAGCATCCACTAGCTATTA
GATGTAAAAGGAAAGAAAAGAAAGATATGTGGTCCCACACATACAGACACCAAGAGAGGTTATATCTATAACTAT
TATCCACAGATGACAACCAACATGCAACTACTTAAAAGTAACAAACAAGCAGTCGAAAAAAATATATATGTAGCT
TGAACATTTAGCTAACATCCATAAAGAAGGCGACGCAAATAACTCCAACTCAAAATATTAACATATTTGTATTAA
TTGACCTTAAGAAATTGAAGTTGCTTTGCTTTGATATACCATCTGATCGTTAATGTTACGACTGATATATTTAAC
AAAATGCCATAACAATACCACTACATATCTTTTGAATAAAGAACCCAGAGTGTTGGATCCTTCACATGCTATAAT
CAATTTAATCTCATTTTATAAAATCAACATAGAAGACATGTCAAAGATACAAAACCAATAAATAATAGAGACTAT
AGAATCTATAATGTTGCAATGTTCTACATATATATATATATATATATAAAAGAGGGTAATGGACTGCACATATTAAT
TCCACAAGCAACAAGTATATTTGCTCCTTCATATTCCAAAAAACAAATATCTCCTTGCAGATCTTTATATATGAA
TTGACAACTAATTATCCGCAGCTGCAAAAATTAACATGTCAATTCATATATTTTAGTTGTCTAAATAAATGTAGG
GATCTAATATTTTATGTGTTTGTCCATGCTGATTTTGATCAATAGAGGTATAGATTACCAGAAGCAAACAAATAA
ATGATAAGAATGATAATCCTGCAGCATTACATAAGCCTTAAAATATTGAAATGTATTGGAGAAAAAGTAAATTCG
CAAGATATAAAAGTCAGGAACTGTAGCCAAAAAGCGTCTATCATGAATGGTAAAAAATAAAATCCAAGAGCATTT
TAACAAGGCAATAAACAGCTCAAAACTAAAGTCATCTATGCAGTGATCTAACAAGTTAAAAAGGTGAACTCCTCA
ATACACTGTACAAAAGCAGCACCACACGGTACGCGTCTAAGAAACAACTGTATATACATCTAACAACGAGAAAAA
AAAAATCCTATATTAATTAAAAAGTAAATCGTACTACATACTCTAGAAAAAAGAAACAGCAGACATTTAAACGAT
ACGAACTGACAAATACGAAAAAGAATCATAATTCCGAC**CTGTAAGAAGAACGCGCACCCCTTCGAATAGAAACCA**
**ATATTAATTAGCAACTGTTAT**CTGCAAACATTTCCTCAACAAAACATGCCGTCAGAAATTTTCACGATGAAATAA
GCAAAAGACATAGTAAAAGCTTTTCCTTATTCACCAACAAAAAACCGAGAACTGCAATGTCTAGACAAAAGTAAG
GTTGGTTCTTAACCCGTTTCAATTTACTGTTGCTGCAGTTTCTCGGCAGCCAAACGTATTCCGATTCAAATTAAA
GTAGCAAAACCGTGTTCACGAACAAAAATATAAGAAGAAAAGAAACAAAAACACGAGAACGGCAAATCGCAACGA
AATTAATAAAATGCATAAGAGAAATTTCGACAAACTAAC**CAGTATGAAGCTTGATACCATGGATTCCAGCTTCAA**
**GTGTAAACAAGAAAAGATACAGAGGACTAAATTTAGAAAAGCAAATCAAACATCGCAACAAGGAATAAACATAAA**
**CCTTCGACC**AAGAATCGGAGAAAATTTGAGAAATGGATTGGTATGAGAGTGAGATCGTCTCGTACTGCAAAAAAC
AAGGAAAGTTGTAGAAAAAGAGAGAGAGGAAGAGAGAGAAAAAATATCTGGAGGAAAGGGGATTTAAATAAGGTA
ATGTGGGGGAGGGGGAGAGATTGGGTGGAACAAGATGAGCTATTTGCTTGNNNNNNNNNNNNNNNNNNNNNNNN
