## Supplementary Material for "Independent origin of *MIRNA* genes controlling homologous target genes by partial inverted duplication of antisense-transcribed sequences"

#### Supplementary Figures

#### Supplementary Figure Legends

**Supplementary Figure 1.** Fully resolved phylogeny of AGL17-like proteins as reconstructed using MrBayes (Ronquist and Huelsenbeck 2003). Numbers at the nodes represent posterior probabilities.

**Supplementary Figure 2.** Northern blot hybridization analysis of the presence of miR444 in different orders of monocotyledonous plants (A) as well as in different tissues of *Zea mays ssp. mays* (B). Abbreviations for species names, the orders to which the species belong and the tissues analysed in *Z. mays* are given on top of the corresponding gel. Analysis was done using total RNA isolated from leaves unless another tissue is indicated on top of the gel, and a probe which is complementary to miR444a.1 and miR444d.1 from *Oryza sativa*. RNA of *Oryza sativa* was used as a positive control and of *Arabidopsis thaliana* as a negative control. ath – *Arabidopsis thaliana*, osa – *Oryza sativa (Poales)*, zma – *Zea mays ssp. mays (Poales)*, mle – *Maranta leuconeura (Zingiberales)*, tbl – *Tradescantia blossfeldiana (Commelinales)*, hvi – *Hypoxis villosa (Asparagales)*, val – *Veratrum album (Liliales)*, mac – *Musa acuminata (Zingiberales)*, pda – *Phoenix dactylifera (Arecales)*, put – *Pandanus utilis (Pandanales)*, dbu – *Dioscorea bulbifera (Dioscoreales)*, eca – *Elodea canadensis (Alismatales)*, pna – *Pandanus natans (Pandanales)*, agr – *Acorus gramineus (Acorales).*

**Supplementary Figure 3.** Alignment of the stem-loop regions of *MIR444* transcripts identified in several orders of monocotyledonous plants. Names of transcripts are colored according to the orders to which the species belong in which the transcripts were found as follows: green, Poales; blue, Zingiberales and Commelinales; cyan, Asparagales and Liliales; purple, Arecales; red, Pandanales and Dioscoreales; orange, Alismatales and Acorales. The positions of the miRNAs and the miRNA*s are marked with boxes and labeled. The blue triangles indicate positions where parts of the alignment are not shown. osa – *Oryza sativa (Poales)*, zma – *Zea mays (Poales)*, epa - *Eichhornia paniculata*, mac – *Musa acuminata (Zingiberales)*, eol – *Elaeis oleifera (Arecales)*, egu - *Elaeis guineensis*, pda - *Phoenix dactylifera*, peq – *Phalaenopsis equestris (Asparagales)*, dbu – *Dioscorea bulbifera (Dioscoreales).*

**Supplementary Figure 4.** Alignment of the *MIR444* transcripts of variant 1 (A) and of variant 2 (B) identified in *Dioscorea bulbifera*. The *dbu-MIR444* transcripts were identified using RT-PCR on cDNA from *Dioscorea bulbifera* leaf material. The primers used for the RT-PCR are marked by arrows above the alignment. Exons are marked by cyan and purple bars above the alignment and numbered. Positions of the mature miRNAs and the miR* are boxed and labelled.

**Supplementary Figure 5.** GFP reporter gene assay to show the *dbu-MIR444* mediated down regulation of the *DbAGL17*-like target gene by transient expression in *Nicotiana benthamiana*. The *AGL17*-like target gene from *Dioscorea bulbifera* was fused with mGFP5ER in a binary vector. To show the *MIR444* mediated down regulation of the *AGL17*-like target gene, *Agrobacteria* carrying the *DbAGL17*-GFP fusion construct (*Agrobacteria* $OD_{600}$ = 0.5) were mixed with different amounts of *Agrobacteria* carrying the *dbu-MIR444* construct (Agrobacteria $OD_{600}$ = 0.01, 0.05, 0.1 and 0.5) and used for co-infiltration of *N. benthamiana* leaves. An untransformed leaf of *N. benthamiana* is shown as a negative control (A). The construct containing the *AGL17*-like gene fused to mGFP5ER was used to confirm the expression of the fusion protein (B). Increasing amounts of *Agrobacteria* containing the *dbu-MIR444* construct ($OD_{600}$ = 0.01, C; $OD_{600}$ = 0.05, D; $OD_{600}$ = 0.1, E; $OD_{600}$ = 0.5, F) were mixed with *Agrobacteria* containing the *DbAGL17*-GFP fusion construct ($OD_{600}$ = 0.5) and used for co-infiltration of *N. benthamiana* leaves. Tobacco leaves expressing the AGL17-GFP fusion protein (Agrobacteria $OD_{600}$ = 0.5) together with the *MIR160a* from *Arabidopsis thaliana* (Agrobacteria $OD_{600}$ = 0.5, G) or the empty vector used for miRNA cloning (Agrobacteria $OD_{600}$ = 0.5, H) show no reduced AGL17-GFP signal. Scale bars, 50 µm.

**Supplementary Figure 6.** Identification of cleavage products of the *DbAGL17a* transcript in *D. bulbifera* leaves. The *DbAGL17a* sequence is shown in blue from exon 2 to 5 with exons shaded alternately. The sequences of the mature miR444 species are also shown. Perfect base pairing between miR444 and *DbAGL17a* is shown by vertical lines. The directions of the sequences are indicated. The arrows indicate the cleavage sites identified by modified 5´RACE, where the second number indicates the total number of sequenced RACE products and the first number the number of clones with this cleavage site.

**Supplementary Figure 7.** Secondary structure prediction of the antisense transcripts to the *AGL17*-like gene of *Triglochin maritima*. Panel (A) shows the secondary structure of the transcript found in the 1KP project data where only the part aligning to *MIR444* sequences was used for prediction, panel (B) antisense transcript 1, variant 1, panel (C) antisense transcript 1, variant 2 and panel (D) antisense transcript 2. Prediction was done using mfold (Zuker 2003). The positions which are complementary to the *AGL17*-like gene are highlighted in orange. Delta free energy (dG) values of the structures are given.

**Supplementary Figure 8.** Alignment of the *AGL17*-like MADS-box transcripts identified in several orders of monocotyledonous plants. Names of transcripts are colored according to the orders to which the species belong in which the transcripts were found as follows: green, Poales; blue, Zingiberales and Commelinales; cyan, Asparagales and Liliales; purple, Arecales; red, Pandanales and Dioscoreales; orange, Alismatales and Acorales. Putative *AGL17*-like transcripts of *Smilax bona nox (Liliales), Xerophyllum asphodeloides (Liliales), Freycinetia multiflora (Pandanales)* and *Acorus americanus (Acorales)*; predictions of *AGL17*-like transcripts from sequenced plant species and experimentally determined *AGL17*-like transcripts (Db – *Dioscorea bulbifera, Dioscoreales*; Tm – *Triglochin maritima, Alismatales*; Ag – *Acorus gramineus, Acorales*) were aligned. The position of the miRNA target sites are marked with boxes and labelled. The blue triangles mark positions where part of the alignment is not shown. Os – *Oryza sativa (Poales)*, Zm – *Zea mays (Poales)*, Epa - *Eichhornia paniculata (Commelinales)*, Ma – *Musa acuminata (Zingiberales)*, Eo – *Elaeis oleifera (Arecales)*, Egu - *Elaeis guineensis (Arecales)*, Pd - *Phoenix dactylifera (Arecales)*, Deca – *Dendrobium catenatum (Asparagales)*, Pe – *Phalaenopsis equestris (Asparagales)*, Xvi – *Xerophyta viscosa (Pandanales)*, Zma – Zostera marina (Alismatales), Zmu – *Zostera muelleri (Alismatales)*, Spo – *Spirodela polyrhiza (Alismatales)*.

**Supplementary Figure 9.** Validation of miR824 mediated cleavage of *ThAGL16a* in *Tarenaya hassleriana*. Shown is the miRNA target site of *ThAGL16a* and the mature miR824. Perfect base pairing between miR824 and *ThAGL16a* is shown by vertical lines. The arrows indicate the cleavage sites identified by modified 5´RACE, where the second number indicates the total number of sequenced RACE products and the first number the number of clones with this cleavage site.

**Supplementary Figure 10.** Secondary structure prediction of the antisense transcript to the *AGL16*-like gene of *Carica papaya*. The positions which are complementary to the *AGL16*-like gene are highlighted in orange. Delta free energy (dG) values of the structures are given.

**Supplementary Figure 11.** Alignment of alleged miR444 from non-monocot species (Ge, et al. 2013; Lin and Lai 2013; Ruan, et al. 2009) with the most similar miR444 sequence from *Oryza sativa*. Identical nucleotides are indicated by a '*' at the corresponding alignment position.

**Supplementary Figure 12.** Multiple sequence alignment of ath-pre-miR824 and osa-pre-miR444d. Identical nucleotides are indicated by a '*' at the corresponding alignment positions. The sequence identity as calculated by dividing the number of alignment positions with identical nucleotides by the total number of alignment positions is given at the bottom.

**Supplementary Figure 13.** Scenarios for the origin of the *MIR444* (A) and *MIR824* (B) genes. The genomic loci of the (future) target genes and the *MIR* genes or antisense genes are depicted by double arrows representing double-stranded DNA; the arrow heads indicate the 3' ends. Exons are shown as boxes on the strand transcribed into RNA (e.g. coding strand), whereas introns are shown as lines. Exons of the (future) target genes are shown in black while exons of the *MIR* genes and the antisense genes are colored red. Regions which are reverse complementary to each other due to a partial inverted duplication are indicated by a color gradient. Shown are putative ancestral loci and the genomic locus of a representative target gene and its *MIR* gene as found today.

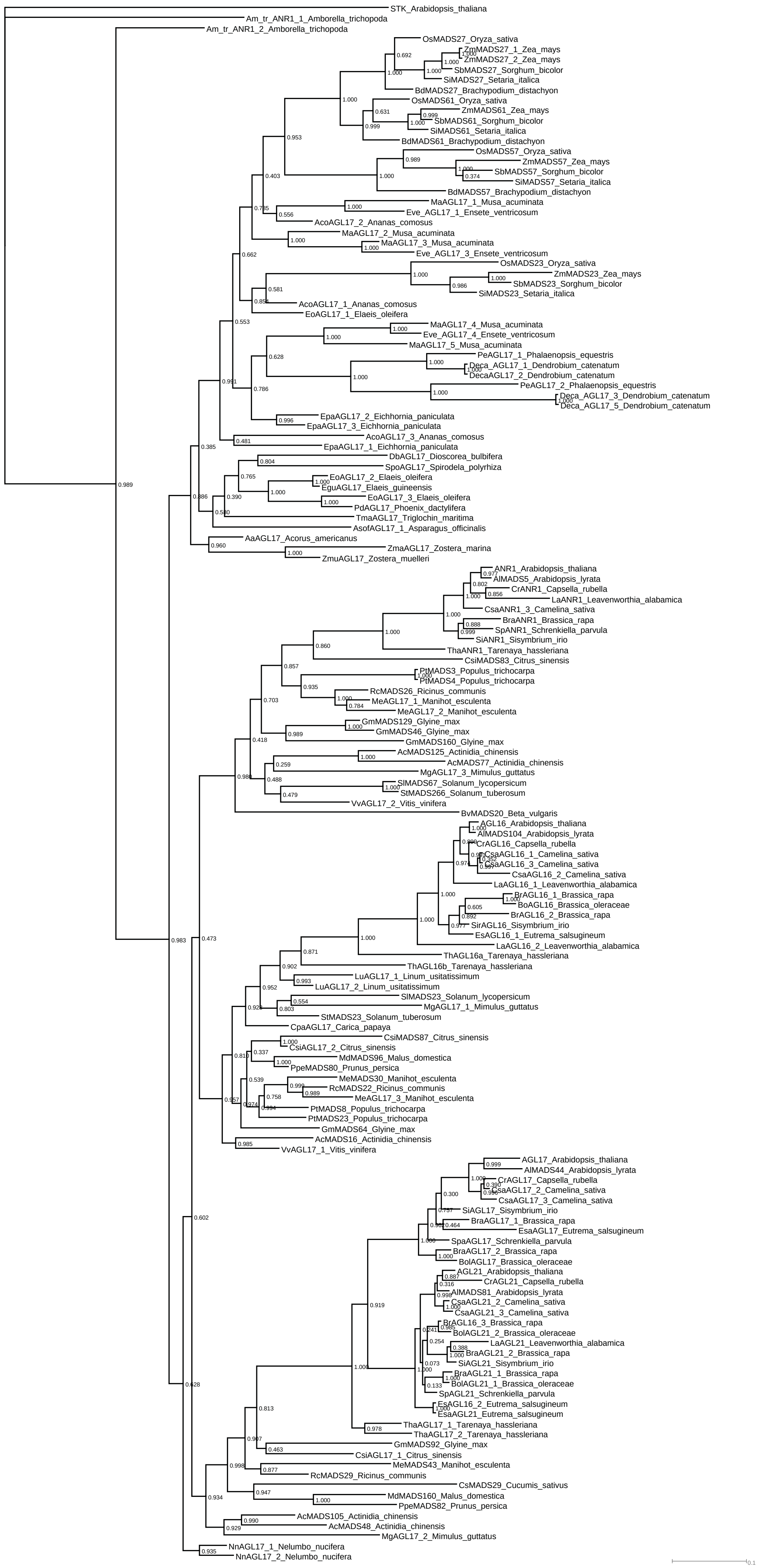

**Supplementary Figure 1.** Fully resolved phylogeny of AGL17-like proteins as reconstructed using MrBayes (Ronquist and Huelsenbeck 2003). Numbers at the nodes represent posterior probabilities.

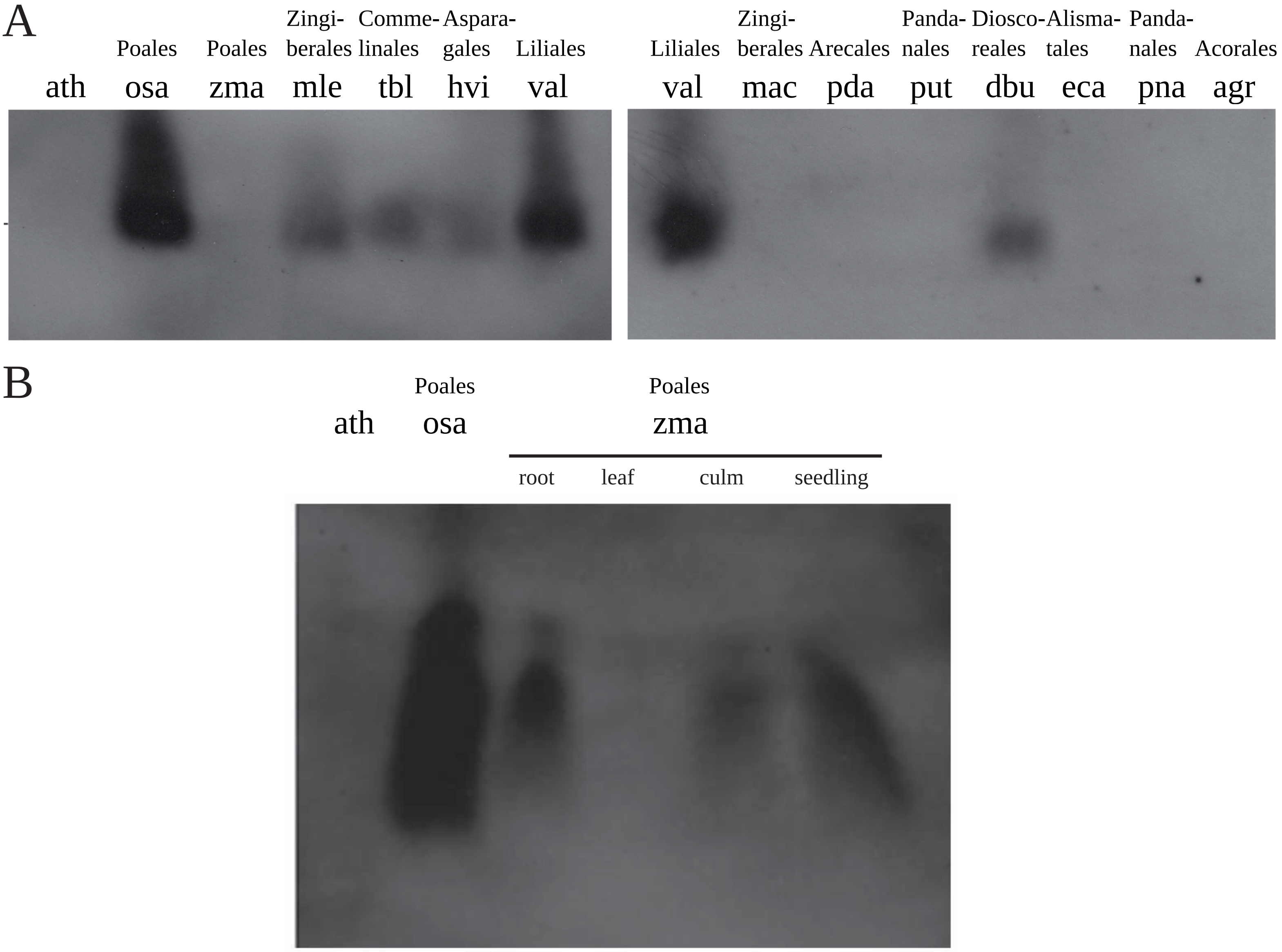


**Figure 2.** Northern blot hybridization analysis of the presence of miR444 in different orders of monocotyledonous plants (A) as well as in different tissues of *Zea mays ssp. mays* (B). Abbreviations for species names, the orders to which the species belong and the tissues analysed in *Z. mays* are given on top of the corresponding gel. Analysis was done using total RNA isolated from leaves unless another tissue is indicated on top of the gel, and a probe which is complementary to the miR444a.1 and miR444d.1 from *Oryza sativa*. RNA of *Oryza sativa* was used as a positive control and of *Arabidopsis thaliana* as a negative control. ath – *Arabidopsis thaliana*, osa – *Oryza sativa (Poales)*, zma – *Zea mays ssp. mays (Poales)*, mle – *Maranta leuconeura (Zingiberales)*, tbl – *Tradescantia blossfeldiana (Commelinales)*, hvi – *Hypoxis villosa (Asparagales)*, val – *Veratrum album (Liliales)*, mac – *Musa acuminata (Zingiberales)*, pda – *Phoenix dactylifera (Arecales)*, put – *Pandanus utilis (Pandanales)*, dbu – *Dioscorea bulbifera (Dioscoreales)*, eca – *Elodea canadensis (Alismatales)*, pna – *Pandanus natans (Pandanales)*, agr – *Acorus gramineus (Acorales)*

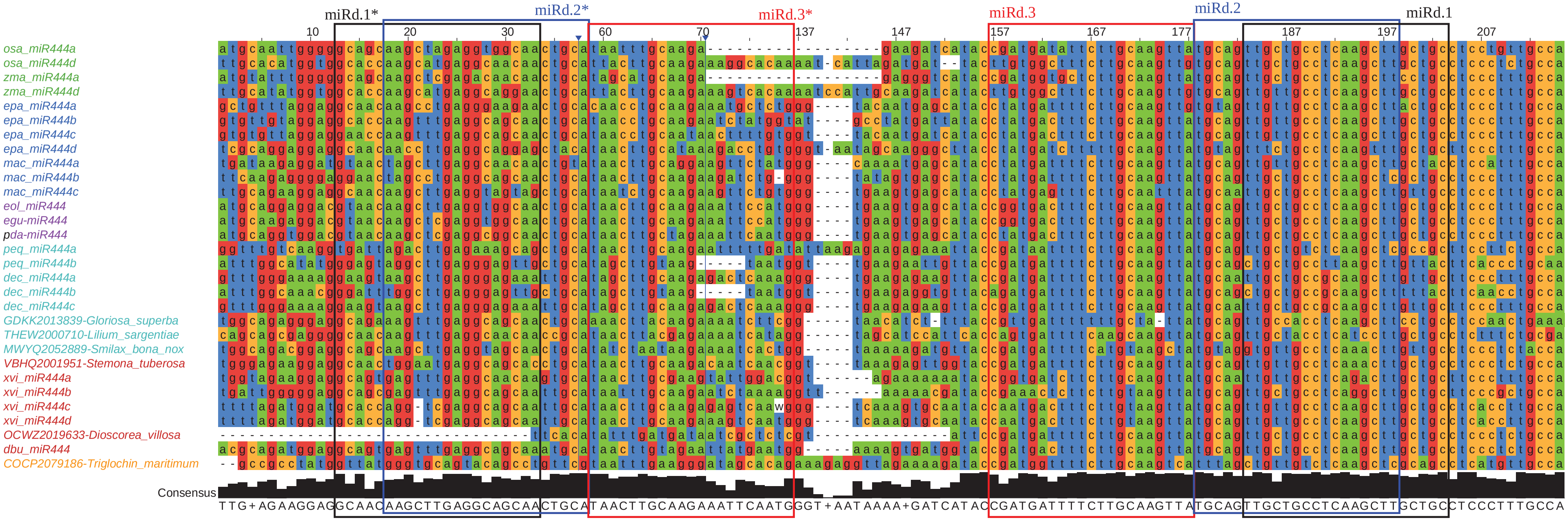

**Supplementary Figure 3.** Alignment of the stem-loop regions of *MIR444* transcripts identified in several orders of monocotyledonous plants. Names of transcripts are colored according to the orders to which the species belong in which the transcripts were found as follows: green, Poales; blue, Zingiberales and Commelinales; cyan, Asparagales and Liliales; purple, Arecales; red, Pandanales and Dioscoreales; orange, Alismatales and Acorales. The positions of the miRNAs and the miRNA*s are marked with boxes and labeled. The blue triangles indicate positions where parts of the alignment are not shown. osa – *Oryza sativa (Poales)*, zma – *Zea mays (Poales)*, epa - *Eichhornia paniculata*, mac – *Musa acuminata (Zingiberales)*, eol – *Elaeis oleifera (Arecales)*, egu - *Elaeis guineensis*, pda - *Phoenix dactylifera*, peq – *Phalaenopsis equestris (Asparagales)*, dbu – *Dioscorea bulbifera (Dioscoreales)*.

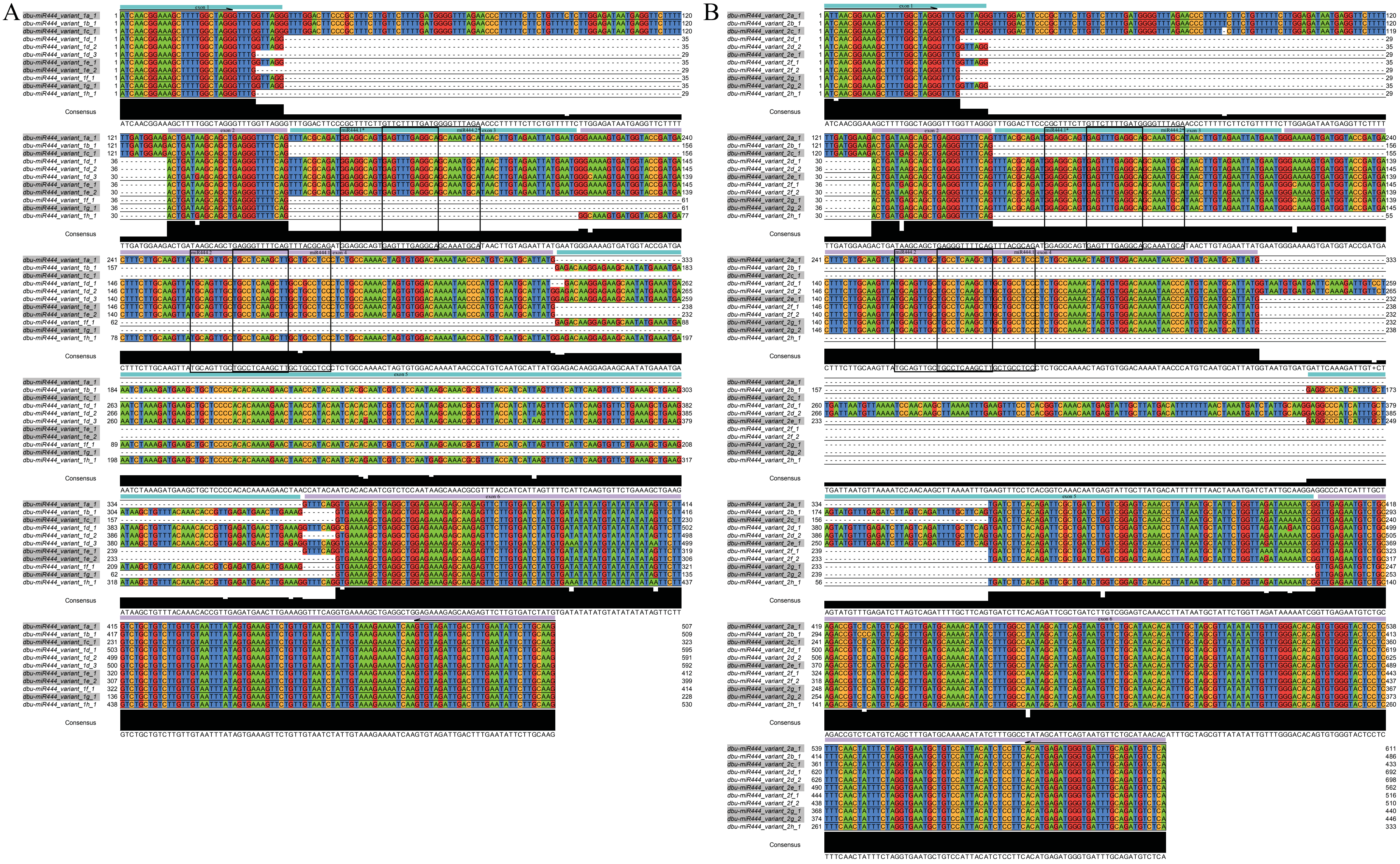

**Supplementary Figure 4.** Alignment of the *MIR444* transcripts of variant 1 (A) and of variant 2 (B) identified in *Dioscorea bulbifera*. The *dbu-MIR444* transcripts were identified using RT-PCR on cDNA from *Dioscorea bulbifera* leaf material. The primers used for the RT-PCR are marked by arrows above the alignment. Exons are marked by cyan and purple bars above the alignment and numbered. Positions of the mature miRNAs and the miR* are boxed and labelled.

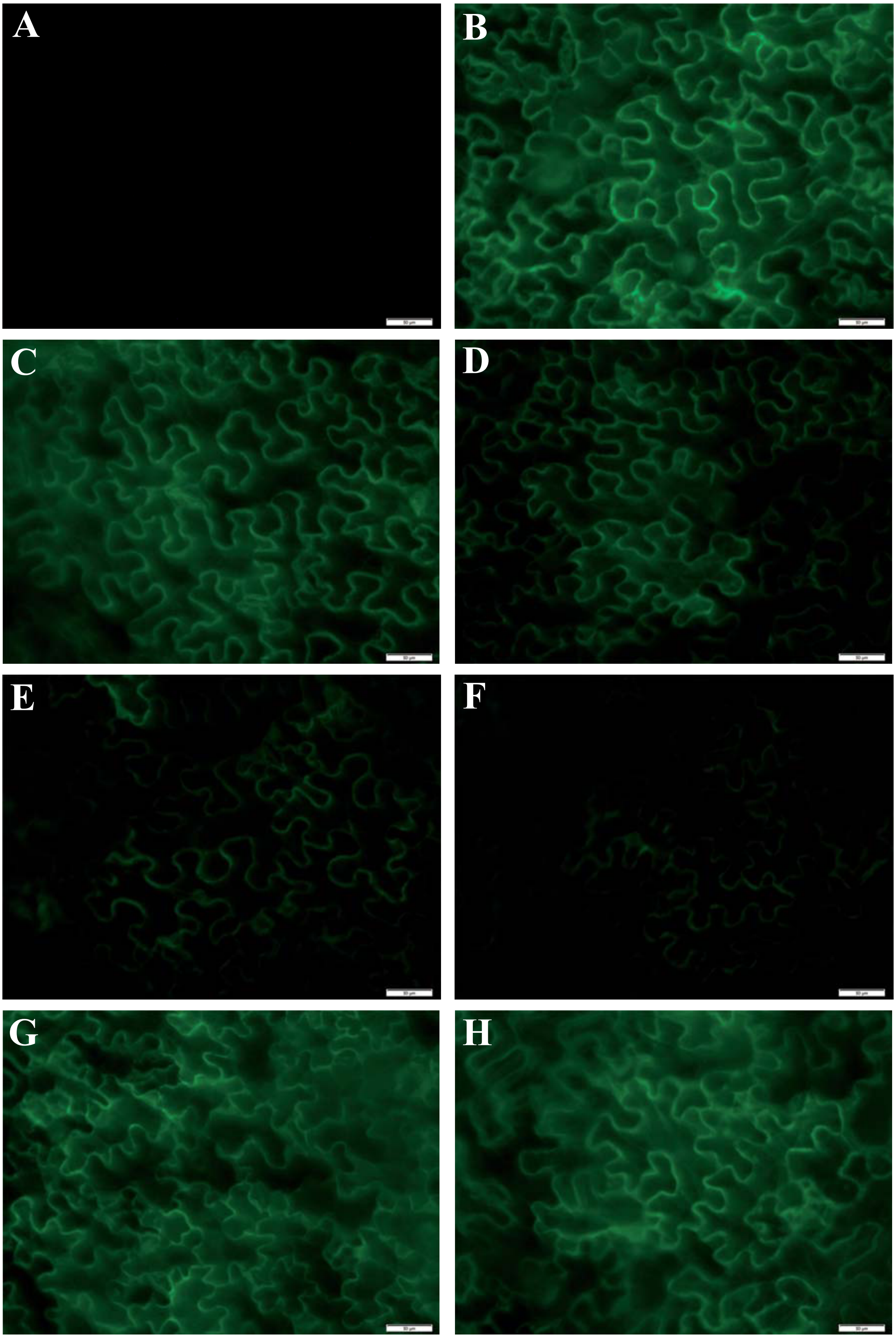

**Supplementary Figure 5 .** GFP reporter gene assay to show the *dbu-MIR444* mediated down regulation of the *DbAGL17*-like target gene by transient expression in *Nicotiana benthamiana*. The *AGL17*-like target gene from *Dioscorea bulbifera* was fused with mGFP5ER in a binary vector. To show the *MIR444* mediated down regulation of the *AGL17*-like target gene, *Agrobacteria* carrying the *DbAGL17*-GFP fusion construct (*Agrobacteria* $OD_{600}$ = 0.5) were mixed with different amounts of *Agrobacteria* carrying the *dbu-MIR444* construct (Agrobacteria $OD_{600}$ = 0.01, 0.05, 0.1 and 0.5) and used for co-infiltration of *N. benthamiana* leaves. An untransformed leaf of *N. benthamiana* is shown as a negative control (A). The construct containing the *AGL17*-like gene fused to mGFP5ER was used to confirm the expression of the fusion protein (B). Increasing amounts of *Agrobacteria* containing the *dbu-MIR444* construct ($OD_{600}$ = 0.01, C; $OD_{600}$ = 0.05, D; $OD_{600}$ = 0.1, E; $OD_{600}$ = 0.5, F) were mixed with *Agrobacteria* containing the *DbAGL17*-GFP fusion construct ($OD_{600}$ = 0.5) and used for co-infiltration of *N. benthamiana* leaves. Tobacco leaves expressing the AGL17-GFP fusion protein (Agrobacteria $OD_{600}$ = 0.5) together with the *MIR160a* from *Arabidopsis thaliana* (Agrobacteria $OD_{600}$ = 0.5, G) or the empty vector used for miRNA cloning (Agrobacteria $OD_{600}$ = 0.5, H) show no reduced AGL17-GFP signal. Scale bars, 50 µm.

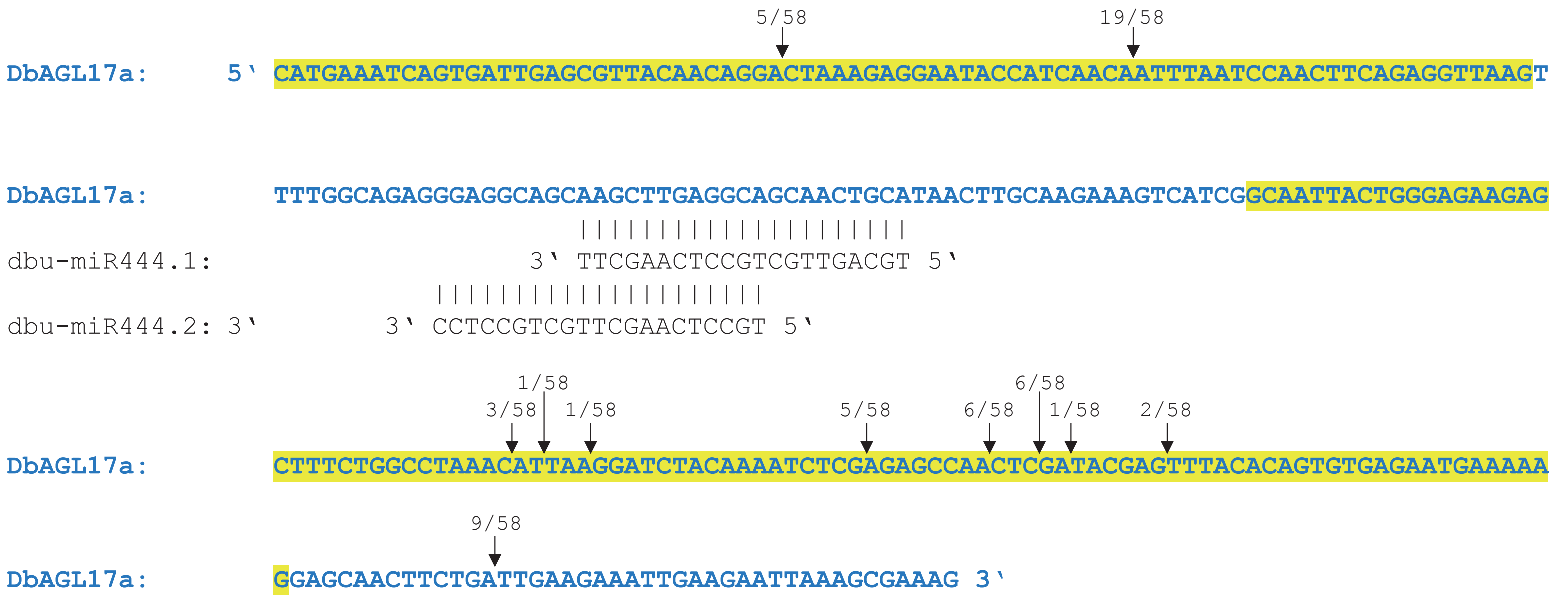


**Supplementary Figure 6.** Identification of cleavage products of the *DbAGL17a* transcript in *D. bulbifera* leaves. The *DbAGL17a* sequence is shown in blue from exon 2 to 5 with exons shaded alternately. The sequences of the mature miR444 species are also shown. Perfect base pairing between miR444 and *DbAGL17a* is shown by vertical lines. The directions of the sequences are indicated. The arrows indicate the cleavage sites identified by modified 5′RACE, where the second number indicates the total number of sequenced RACE products and the first number the number of clones with this cleavage site.

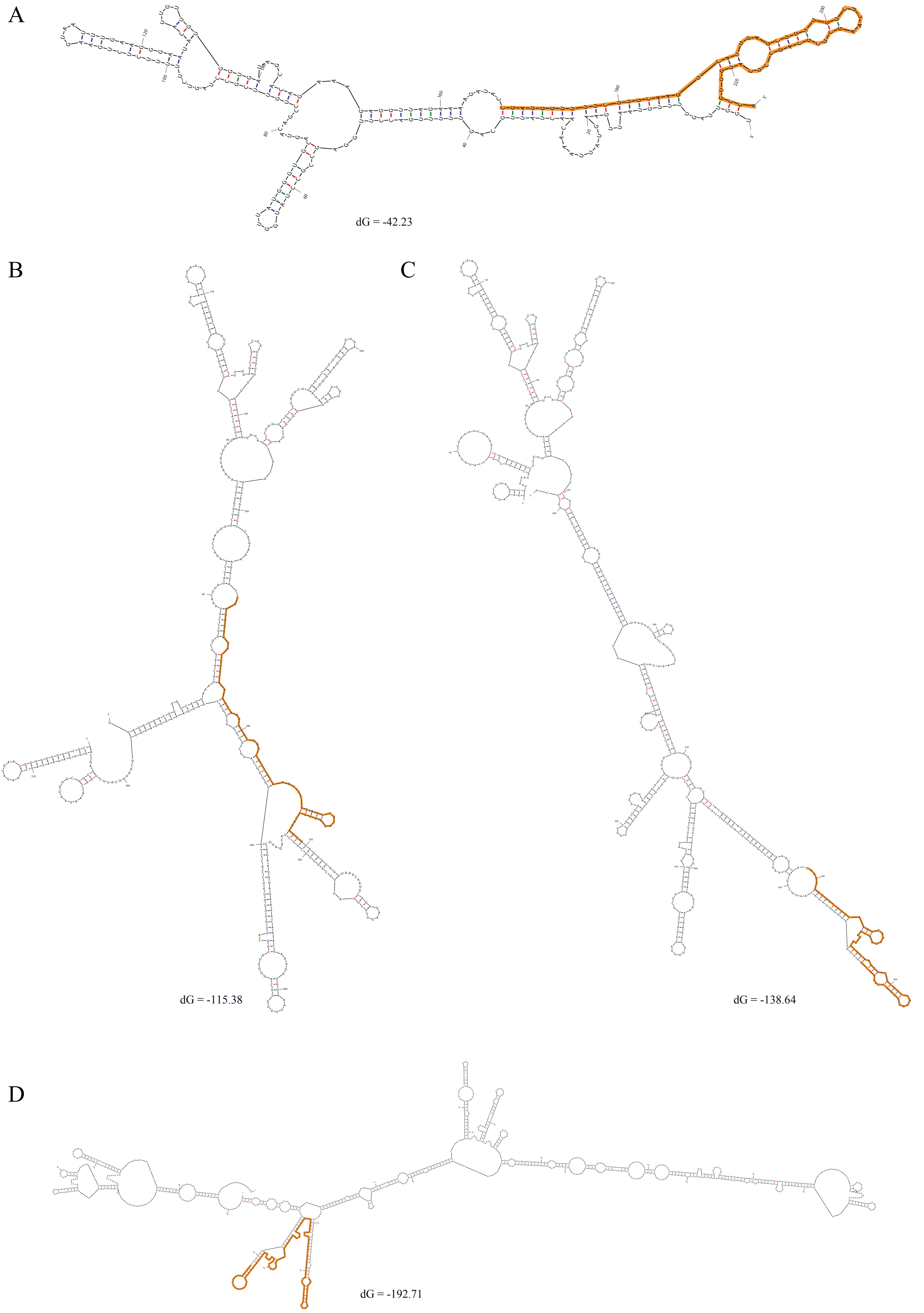


**Supplementary Figure 7.** Secondary structure prediction of the antisense transcripts to the *AGL17*-like gene of *Triglochin maritima*. Panel (A) shows the secondary structure of the transcript found in the 1KP project data where only the part aligning to *MIR444* sequences was used for prediction, panel (B) antisense transcript 1, variant 1, panel (C) antisense transcript 1, variant 2 and panel (D) antisense transcript 2. Prediction was done using mfold (Zuker 2003). The positions which are complementary to the *AGL17*-like gene are highlighted in orange. Delta free energy (dG) values of the structures are given.

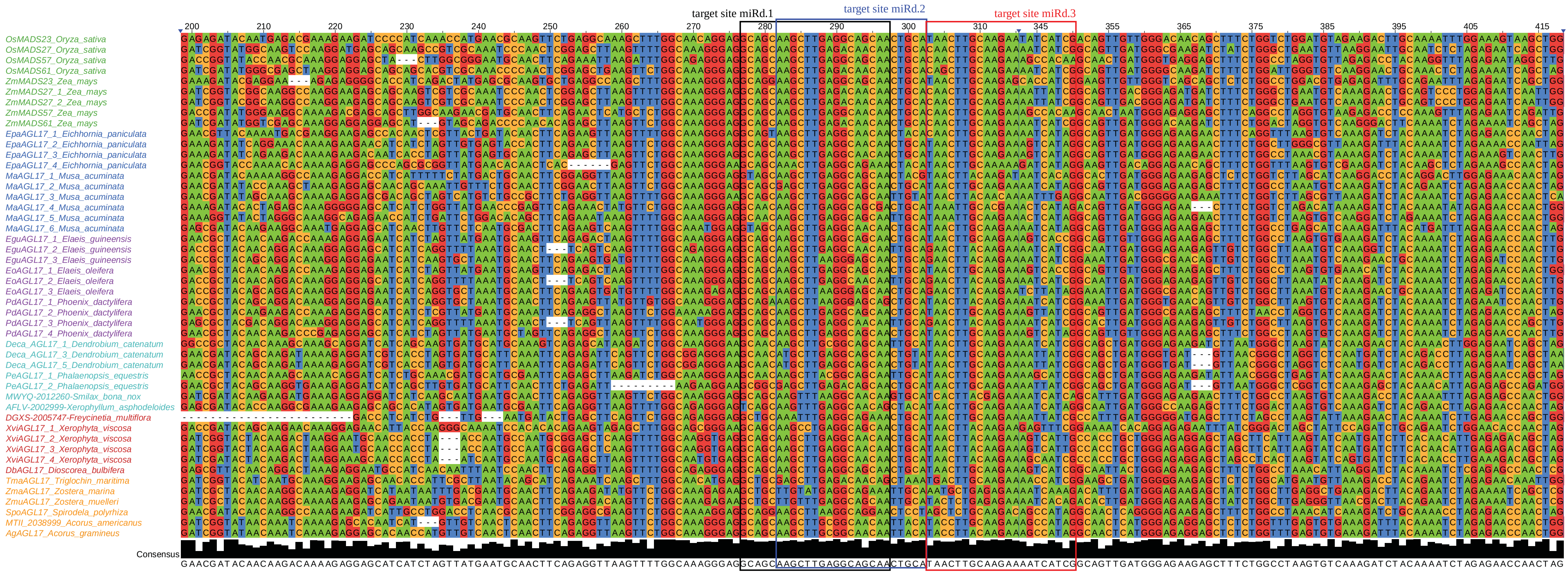

**Supplementary Figure 8.** Alignment of the *AGL17*-like MADS-box transcripts identified in several orders of monocotyledonous plants. Names of transcripts are colored according to the orders to which the species belong in which the transcripts were found as follows: green, Poales; blue, Zingiberales and Commelinales; cyan, Asparagales and Liliales; purple, Arecales; red, Pandanales and Dioscoreales; orange, Alismatales and Acorales. Putative *AGL17*-like transcripts of *Smilax bona nox (Liliales)*, *Xerophyllum asphodeloides (Liliales)*, *Freycinetia multiflora (Pandanales)* and *Acorus americanus (Acorales)*; predictions of *AGL17*-like transcripts from sequenced plant species and experimentally determined *AGL17*-like transcripts (Db – *Dioscorea bulbifera, Dioscoreales*; Tm – *Triglochin maritima, Alismatales*; Ag – *Acorus gramineus, Acorales*) were aligned. The position of the miRNA target sites are marked with boxes and labelled. The blue triangles mark positions where part of the alignment is not shown. Os – *Oryza sativa (Poales)*, Zm – *Zea mays (Poales)*, Epa - *Eichhornia paniculata (Commelinales)*, Ma – *Musa acuminata (Zingiberales)*, Eo – *Elaeis oleifera (Arecales)*, Egu - *Elaeis guineensis (Arecales)*, Pd - *Phoenix dactylifera (Arecales)*, Deca – *Dendrobium catenatum (Asparagales)*, Pe – *Phalaenopsis equestris (Asparagales)*, Xvi – *Xerophyta viscosa (Pandanales)*, Zma – Zostera marina (Alismatales), Zmu – *Zostera muelleri (Alismatales)*, Spo – *Spirodela polyrhiza (Alismatales)*.

```
                              12/16  1/16
                                 ↓    ↓
ThAGL16a:    5´ TCCCTTCTCACAAATGGTCTA 3´
                |||||||||||||||||||||
tha-miR824: 3´ AGGGAAGAGTGTTTACCAGAT 5´
```

**Supplementary Figure 9.** Validation of miR824 mediated cleavage of *ThAGL16a* in *Tarenaya hassleriana*. Shown is the miRNA target site of *ThAGL16a* and the mature miR824. Perfect base pairing between miR824 and *ThAGL16a* is shown by vertical lines. The arrows indicate the cleavage sites identified by modified 5´RACE, where the second number indicates the total number of sequenced RACE products and the first number the number of clones with this cleavage site.

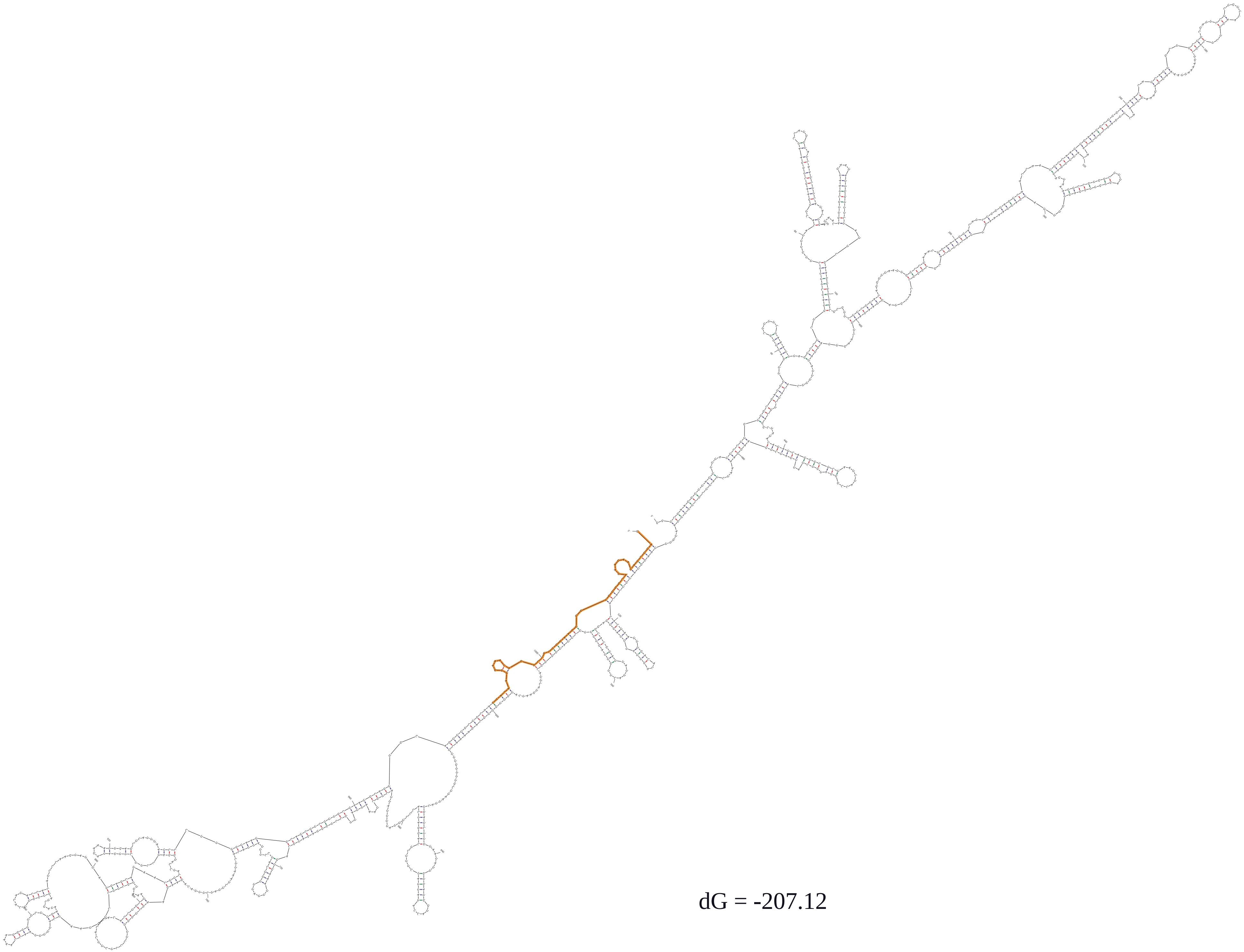


**Supplementary Figure 10.** Secondary structure prediction of the antisense transcript to the *AGL16*-like gene of *Carica papaya*. The positions which are complementary to the *AGL16*-like gene are highlighted in orange. Delta free energy (dG) values of the structures are given.

Alleged miR444 from *Fragaria×ananassa*

```
osa-miR444b.2   tgcagttgttg-tctcaagctt
fan-miR444      tgcagttgttggtcttcaact-
                *********** ***   * **
Ge, et al. 2013
```

Alleged miR444 from *Dimocarpus longan*

```
osa-miR444f     tgcagttgttgcctcaagctt
dlo-miR444b     tacaattctgtcgtcaagctt
                * ** ** *  * ********
Lin and Lai 2013
```

Alleged miR444s from *Gossypium hirsutum*

```
osa-miR444b.2   tgcagttgttgtctcaagctt
sRNA285490      tgcagttgttgtctcaagctt
                *********************

osa-miR444f     tgcagttgttgcctcaagctt
sRNA285485      tgcagttgttgcctcaagctt
                *********************

osa-miR444a.2   tgcagttgctgcctcaagctt
sRNA285470      tgcagttgctgcctcaagctt
                *********************
Ruan, et al. 2013
```

**Supplementary Figure 11.** Alignment of alleged miR444 from non-monocot species with the most similar miR444 sequence from *Oryza sativa*. Identical nucleotides are indicated by a ‘*’ at the corresponding alignment position.

CLUSTAL O(1.2.4) multiple sequence alignment

```
ath-MIR824      TATCACCATTTGTACTTGAGTTGTCTCATGTCTAGACCATTTGTGAGAAGGGAGTTTT   60
osa-MIR444d     ------------------AGTTATTGCACATG----------------------------   14
                                  **** *  * ****

ath-MIR824      TGTTTACACCAATACCCCCCAGTCTCTAAATTTGTAAGAAGTATTATGCTCAATTAAGGA   120
osa-MIR444d     ------------------------------------------------------------   14

ath-MIR824      ACATAGCTAGTTCGACATACCATACTACCCTTTTTAAAACTATCCTATATGTTTGATGCT   180
osa-MIR444d     ---------------------------------------------------GTGGCACCA   23
                                                                    * *   *

ath-MIR824      AGCATAGCGTAATTTTGTGTTCTCATGGTGCAGCAGGGTTAGTTTATTGTGTACCTCTAG   240
osa-MIR444d     AGCATGAGGCAAC-----------------------------------------------   36
                *****   * **

ath-MIR824      ATATATATTCTCTTGCTGCGTAGGGTTCCCAAGCTGCCAAAAACTTTTAAAAATTAGTGA   300
osa-MIR444d     ------------------------------------------------------AA----   38
                                                                       *

ath-MIR824      TCTGTTCCCCAAACCCCCATTCAATAAGAAAGGTCTACCTGTAGCTCACAGTCACAGCTA   360
osa-MIR444d     -----------------------------CTGCATTACTTGCAAGAAAGGCACAAAATCA   69
                                               *   *** ** *    *    ** *   *

ath-MIR824      TTAGAGCTGTCCTTGCTTCTTTGGGTTTACTGTTTTAGTTAATTAGTTACCCTATGAAAG   420
osa-MIR444d     TTAGATGATTACTTGTGGCTTTCTTGCAA--------GTTGTGCAGTTGCTGCCTCAAGC   121
                *****    * ****   ****      *        ***    **** *    * **

ath-MIR824      TCTGATCCTTCAGAAGTTAGGTATAGAAGTAAGTCGATTAAGTTCATTCATGACTTTTCC   480
osa-MIR444d     TTGCTGCCT------------------------------------------CCCTCTGCC   139
                *     ***                                            ** * **

ath-MIR824      AGATGTTGCATACATAACTTTTTTATTTGTATTCGAACTAGTCCAAGCCGATTCTCAAA   540
osa-MIR444d     AAAT--------------------------------------------------------   143
                * **

ath-MIR824      ATCATATAAAATCATTTACACCTTGGTCATGGTAGCTAAGAATATTGTATCTAAAATTGG   600
osa-MIR444d     ------------------------------------------------------------   143

ath-MIR824      GGAGTGGGGAGATGTTTGGTTATATTCCCTTCTCATCGATGGTCTAGATGTGCGAGGTGA   660
osa-MIR444d     ------------------------------------------------------------   143

ath-MIR824      CTCTCATGGAGGTAAAGAACAATGGTGAT                                  689
osa-MIR444d     -----------------------------                                  143
```

Sequence identity: 72/689 = 0.1045

**Supplementary Figure 12.** Multiple sequence alignment of ath-pre-miR824 and osa-pre-miR444d. Identical nucleotides are indicated by a ‘*’ at the corresponding alignment positions. The sequence identity as calculated by dividing the number of alignment positions with identical nucleotides by the total number of alignment positions is given at the bottom.

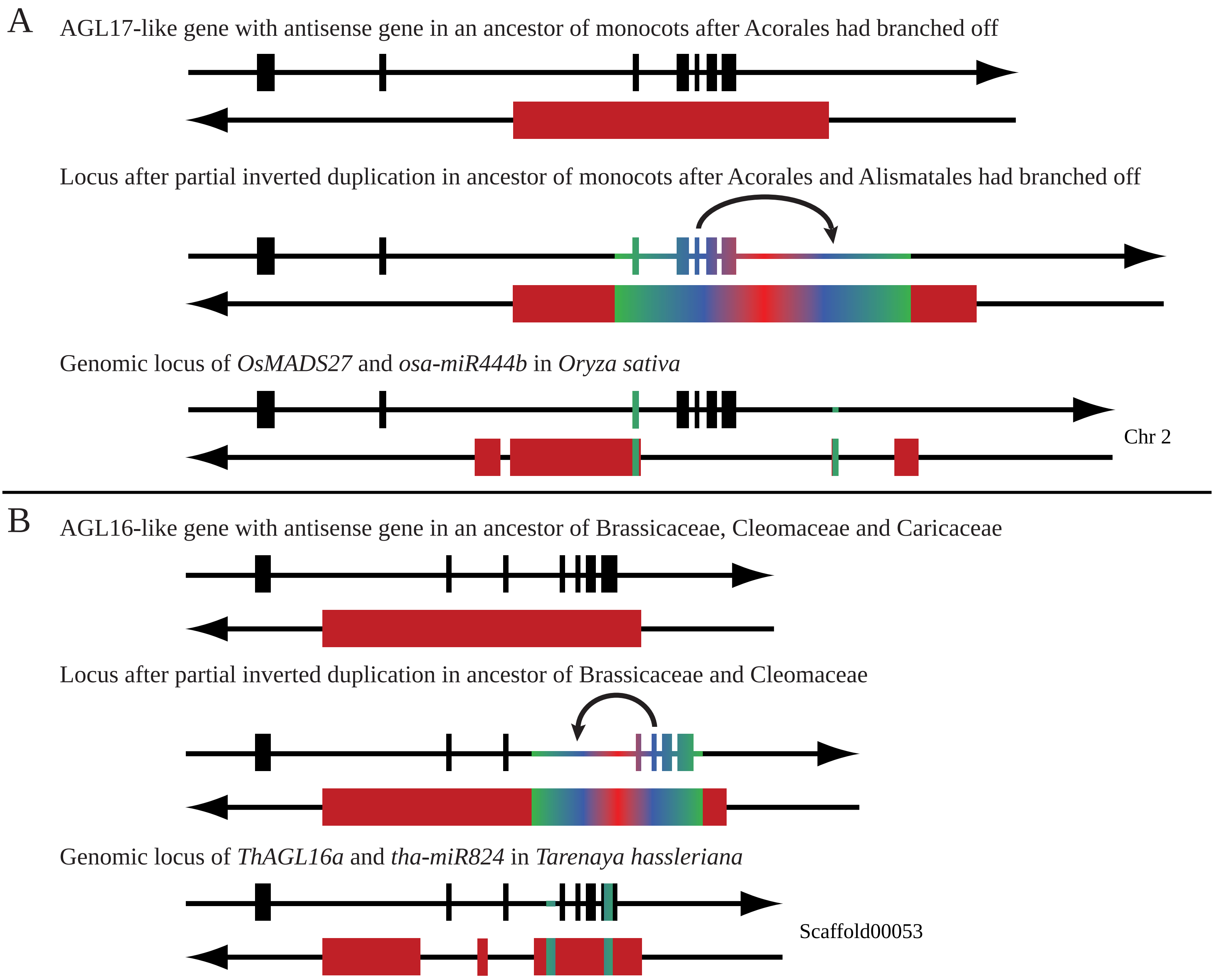


**Supplementary Figure 13.** Scenarios for the origin of the *MIR444* (A) and *MIR824* (B) genes. The genomic loci of the (future) target genes and the *MIR* genes or antisense genes are depicted by double arrows representing double-stranded DNA; the arrow heads indicate the 3' ends. Exons are shown as boxes on the strand transcribed into RNA (e.g. coding strand), whereas introns are shown as lines. Exons of the (future) target genes are shown in black while exons of the *MIR* genes and the antisense genes are colored red. Regions which are reverse complementary to each other due to a partial inverted duplication are indicated by a color gradient. Shown are putative ancestral loci and the genomic locus of a representative target gene and its *MIR* gene as found today.

#### Supplementary Tables

**Supplementary Table 1.** Primers used in this study.

| gene | name of primer | sequence (5´to 3´) |
|---|---|---|
| **General primers** | | |
| Oligo dT anchor primer for 5´RACE Tma | T125 | GCTCGCGAGCGCGTTTAAACGCGCACGCG TTTTTTTTTTTTTTTTTTVN |
| Anchor primer for 5´RACE Tma | T126 | GCTCGCGAGCGCGTTTAAAC |
| Oligo dT anchor primer for 3´RACE and 5´RACE Dbu | Oligo dT anchor primer | GACCACGCGTATCGATGTCGACTTTTTTTT TTTTTTTTV |
| Anchor primer for 3´RACE and 5´RACE Dbu | PCR anchor primer | GACCACGCGTATCGATGTCGAC |
| Degenerate MADS-box primer | LITU RQVT | GGCARGTSACMTWCTCSAARMGGAG |
| **Genome walking** | | |
| Dbu miR444 5´walking | DbumiRgenwalking5-1 | TGTTGCCTTGGCATATCTTAAAGTGGT |
| Dbu miR444 5´walking | DbumiRgenwalking5-2 | TGACTTAAGTATTTCTGAGCACCCCAATTC |
| Dbu miR444 5´walking | DbuAGL17GWalk3-1 | GCGAATCTGTGAAGATCACTGAAGCA |
| Dbu miR444 5´walking | DbuAGL17GWalk3-2 | AGCAATACTCATTGTTTGACCGTGAGG |
| Dbu miR444 5´walking | Dbu444Walk5-1 | CGCAGCATTTGCCAGTTTATCTTTCAT |
| Dbu miR444 5´walking | Dbu444Walk5-2 | AATAACGCCACAGGCCACTACTTCGTA |
| Dbu miR444 5´walking | Dbu444W5-1b | GGATTAGTTTCAAGCGTGCCAAAACA |
| Dbu miR444 5´walking | Dbu444W5-2b | AGTGCAAGAGCCCACTAAAAACCTGA |
| Tma AGL17 5´walking | tma444GenomeWalking3 -1 | TGTTGTGGTGATAGCACAGAAAGAGGTT |
| Tma AGL17 5´walking | tma444GenomeWalking3 -2 | TGCAAGTCATTTAGCTGTTGTCTCAAGC |
| Tma AGL17 3´walking | Tma444GenomeWalking 5-1 | CCCACAGGTGGACAAAAACAATGACG |
| Tma AGL17 3´walking | Tma444GenomeWalking 5-2 | CGTTTGTTTTCTACCTTCTGAATCTGTAGC |
| Tma AGL17 3´walking | TmaGenWalk5-1a | GAAAAGTGATGAGGGCATTGAGATTGT |
| Tma AGL17 3´walking | TmaGenWalk5-2a | GTGACCTGCAAAGTGAATGGAGGATT |
| Tma AGL17 3´walking | Tma444Walk5-1b | AGGGAGGATGATCGTGATTTGCACTTT |
| Tma AGL17 3´walking | Tma444Walk5-2b | TTCACAAGGCAGTCACTGGTTCTTCAC |
| Tma AGL17 3´walking | Tma444Walk5-1c | AAGCATAGGTGAGAAGGGGGCAAATTA |
| Tma AGL17 3´walking | Tma444Walk5-2c | TGATTCCTTTTTAGCATGCCAGTGATG |
| Agr AGL17 3´walking | AgrAGL17Walk3-1 | GGAACAGGGGAGGTAAATGGAGGTATT |
| Agr AGL17 3´walking | AgrAGL17Walk3-2 | GGTTTTAGCATCAGAGAGGACATGCAC |
| Agr AGL17 3´walking | AgrAGL17Walk3-1a | TGGGATCTCTCTGAACCTGATTTTTGG |
| Agr AGL17 3´walking | AgrAGL17Walk3-2a | CATGGAGGTAGACTCCAACCTTTTCTCA |
| **Genomic DNA** | | |
| Agr putative miR444 | Agr-miR444d1rc | GCAGCAAGCTTGCGGCAACAA |
| **5´RACE** | | |
| Dbu miR444 cDNA synthesis | Dbu444stemloop.rev | CCACACTAGTTTTGGCAGAGG |
| Dbu miR444 1. PCR | Dbu444RACE5-2 | GCATAACTTGCAAGAAAGTCATCGGTA |
| Dbu miR444 nested PCR | Dbu444RACE5-ExEx | CCATCACTTTTCCCATTCATAATTCT |
| Tma antisense transcript cDNA synthesis | Tma444-5RACE-1 | CAGGCTGTACTGCACCCATA |
| Tma antisense transcript 1. PCR | Tma444-5RACE-2 | TAGGCGGCTCCAAGGTCAAAAACT |
| Tma antisense transcript nested PCR | Tim-2_5-RACE-premiRNA-2 | CTTCAATTACAAAACTAACCAAGGTGC |
| **3´RACE** | | |

| | | |
|---|---|---|
| Dbu miR444 | miR444d1 | TTGCTGCCTCAAGCTTGCTGC |
| Agr miR444 | Acorus americanusd1.for | TTGTTGCCGCAAGCTTGCTGC |
| Tha miR824 (1. PCR) | ThmiR824nested.for | GCTGGCTTTCCAAGCTCTGT |
| Tha miR824 (nested PCR) | ThmiR824nested.new.for | GAGATGCGACCGTGATTTTG |
| **RT-PCR** | | |
| Tma AGL17 | tma-miR444d.1 | CTGTTGTCTCAAGCTCGCAGC |
| Dbu AGL17 | miR444rice | GCAGCAAGCTTGAGGCAGCAA |
| Dbu miR444 variant 1 | Dbu444 Exon1.for | ATCAACGGAAAGCTTTTGGCTAGG |
| Dbu miR444 variant 1 | DbuOwnSeqREV1 | CTTGCAAGAATATTCAAAGTCAATCTACAC |
| Dbu miR444 variant 2 | Dbu444 Exon1.for | ATCAACGGAAAGCTTTTGGCTAGG |
| Dbu miR444 variant 2 | Dbu444 clone8.rev | TGAGACATCTGCAAATCACCCATC |
| Agr AGL17 | A.gramineus.for | GTCGGCCTCATCATCTTCTC |
| Agr AGL17 | A.gramineus.rev | GGGTGCATGTCCTCTCTGAT |
| Tha AGL16-1 | ThaAGL16-1new.for | GACAAAGGAGGAAAATAATCAAATG |
| Tha AGL16-1 | ThaAGL16-1new.rev | GGAATAGTACCCTAACTTGGTG |
| Tha AGL16-2 (1. PCR) | ThaAGL16-2.for | AAGAAGGCGAAGGAGCTTGC |
| Tha AGL16-2 (1. PCR) | ThaAGL16-2.rev | GGCCACAGAGGACATTGGAA |
| Tha AGL16-2 (nested PCR) | ThaAGL16-2n.for | CACCGGCAAGCTCTACGATT |
| Tha AGL16-2 (nested PCR) | ThaAGL16-2n.rev | AATGCAGTGGCAATCTGCTG |
| **Modified 5´RACE** | | |
| Dbu AGL17 (cDNA synthesis) | Dbu cleavage cDNA | TCGGACATCTGTGTTCTCCA |
| Dbu AGL17 (1. PCR) | Dbu cleavage PCR1 | CATTGTCAACTCCTGTGCGTCTTACG |
| Dbu AGL17 (nested PCR) | Dbu cleavage nested | GAGTTCCACGTTTTCGAGATGGAG |
| Tha AGL16 (cDNA synthesis) | ThaAGL16-2.rev | GGCCACAGAGGACATTGGAA |
| Tha AGL16 (1. PCR) | ThaAGL16-2n.rev | AATGCAGTGGCAATCTGCTG |
| **MiR-RACE** | | |
| General primer | GeneRACER 5´primer | CGACTGGAGCACGAGGACACTGA |
| General primer | GeneRACER 5´nested primer | GGACACTGACATGGACTGAAGGAGTA |
| General primer | 3´Anchor miRACE primer | ATTCTAGAGGCCGAGGCGGCCGACATG |
| Dbu miR444.1 | Dbu-miR444.1 3miRACE | GGAGTAGAAATTGCTGCCTCAAGCTTG |
| Dbu miR444.1 | Dbu-miR444.1 5miRACE | TTTTTTTTTTGCAGCAAGCTTGAGGCA |
| Dbu miR444.1 | 444.1 5R short | TTTTTTTTTTGGAGGCAGCAAGCTTGA |
| Dbu miR444.1 | miR444.1 3R shifted | GGAGTAGAAATGCCTCAAGCTTGCTGC |
| Dbu miR444.2 | Dbu-miR444.2 3miRACE | GGAGTAGAAATGCAGTTGCTGCCTCAA |
| Dbu miR444.2 | Dbu-miR444.2 5miRACE | TTTTTTTTTTAAGCTTGAGGCAGCAAC |
| Dbu miR444.3 | Dbu-miR444.3 3miRACE | GGAGTAGAAACGATGACTTTCTTGCAA |
| Dbu miR444.3 | Dbu-miR444.3 5miRACE | TTTTTTTTTTTAACTTGCAAGAAAGTC |

**Supplementary Table 2.** Overview of the sequences submitted to GenBank including accession numbers and sequence descriptions.

| Accession number | Sequence description |
|---|---|
| KY094495 | AgAGL17, Acorus gramineus, AGL17-like MADS-box gene, partial CDS, splice variant 1 |
| KY172122 | AgAGL17, Acorus gramineus, AGL17-like MADS-box gene, partial CDS, splice variant 2 |
| KY319139 | AgAGL17, Acorus gramineus, AGL17-like MADS-box gene, partial genomic locus, starting from exon 2 |
| KY742743 | TmAGL17, Triglochin maritima, AGL17-like MADS-box gene, partial CDS, splice variant 1 |
| KY742744 | TmAGL17, Triglochin maritima, AGL17-like MADS-box gene, partial CDS, splice variant 2 |
| KY742745 | TmAGL17, organism=Triglochin maritima, AGL17-like MADS-box gene, partial genomic DNA |
| KY742746 | TmAGL17asRNA1-1, Triglochin maritima, antisense transcript 1 of an AGL17-like MADS-box gene, transcript variant 1 |
| KY742747 | TmAGL17asRNA1-2, Triglochin maritima, antisense transcript 1 of an AGL17-like MADS-box gene, transcript variant 2 |
| KY742748 | TmAGL17asRNA2, Triglochin maritima, antisense transcript 2 of an AGL17-like MADS-box gene |
| KY742749 | ThAGL16-1, Tarenaya hassleriana, AGL16 MADS-box gene, partial CDS, splice variant 1 |
| KY742750 | ThAGL16-1, Tarenaya hassleriana, AGL16 MADS-box gene, partial CDS, splice variant 2 |
| KY742751 | ThAGL16-2, Tarenaya hassleriana, AGL16 MADS-box gene, partial CDS |
| KY742752 | tha-MIR824, Tarenaya hassleriana, partial transcript of MIR824, splice variant 1 |
| KY742753 | tha-MIR824, Tarenaya hassleriana, partial transcript of MIR824, splice variant 2 |
| KY742754 | tha-MIR824, Tarenaya hassleriana, partial transcript of MIR824, splice variant 3 |
| KY742755 | tha-MIR824, Tarenaya hassleriana, partial transcript of MIR824, splice variant 4 |
| KY774629 | DbAGL17, Dioscorea bulbifera, AGL17-like MADS-box gene, partial CDS |
| KY774630 | DbAGL17, Dioscorea bulbifera, AGL17-like MADS-box gene, partial genomic locus |
| KY774631 | dbu-MIR444, Dioscorea bulbifera, partial transcript of MIR444, transcript variant 1a_1 |
| KY774632 | dbu-MIR444, Dioscorea bulbifera, partial transcript of MIR444, transcript variant 1b_1 |
| KY774633 | dbu-MIR444, Dioscorea bulbifera, partial transcript of MIR444, transcript variant 1c_1 |
| KY774634 | dbu-MIR444, Dioscorea bulbifera, partial transcript of MIR444, transcript variant 1d_1 |
| KY774635 | dbu-MIR444, Dioscorea bulbifera, partial transcript of MIR444, transcript variant 1d_2 |
| KY774636 | dbu-MIR444, Dioscorea bulbifera, partial transcript of MIR444, transcript variant 1d_3 |
| KY774637 | dbu-MIR444, Dioscorea bulbifera, partial transcript of MIR444, transcript variant 1e_1 |
| KY774638 | dbu-MIR444, Dioscorea bulbifera, partial transcript of MIR444, transcript variant 1e_2 |
| KY774639 | dbu-MIR444, Dioscorea bulbifera, partial transcript of MIR444, transcript variant 1f_1 |
| KY774640 | dbu-MIR444, Dioscorea bulbifera, partial transcript of MIR444, transcript variant 1g_1 |
| KY774641 | dbu-MIR444, Dioscorea bulbifera, partial transcript of MIR444, transcript variant 1h_1 |
| KY774642 | dbu-MIR444, Dioscorea bulbifera, partial transcript of MIR444, transcript variant 2a_1 |
| KY774643 | dbu-MIR444, Dioscorea bulbifera, partial transcript of MIR444, transcript variant 2b_1 |
| KY774644 | dbu-MIR444, Dioscorea bulbifera, partial transcript of MIR444, transcript variant 2c_1 |
| KY774645 | dbu-MIR444, Dioscorea bulbifera, partial transcript of MIR444, transcript variant 2d_1 |
| KY774646 | dbu-MIR444, Dioscorea bulbifera, partial transcript of MIR444, transcript variant 2d_2 |
| KY774647 | dbu-MIR444, Dioscorea bulbifera, partial transcript of MIR444, transcript variant 2e_1 |
| KY774648 | dbu-MIR444, Dioscorea bulbifera, partial transcript of MIR444, transcript variant 2f_1 |
| KY774649 | dbu-MIR444, Dioscorea bulbifera, partial transcript of MIR444, transcript variant 2f_2 |
| KY774650 | dbu-MIR444, Dioscorea bulbifera, partial transcript of MIR444, transcript variant 2g_1 |
| KY774651 | dbu-MIR444, Dioscorea bulbifera, partial transcript of MIR444, transcript variant 2g_2 |
| KY774652 | dbu-MIR444, Dioscorea bulbifera, partial transcript of MIR444, transcript variant 2h_1 |

#### Supplementary Data

see separate file

### Supplementary Methods

#### Access of genome data

*Monocot genomes*

The genome of *Z. mays* ssp. *mays* was assessed on Phytozome (www.phytozome.net) (Goodstein, et al. 2012). The genome sequences of *Elaeis guineensis* and *E. oleifera* (Singh, et al. 2013), *Ensete ventricosum*, *Ananas comosus* (Ming, et al. 2015), *Eichhornia paniculata*, *Dendrobium catenatum* (Zhang, et al. 2016), *Xerophyta viscosa* (Costa, et al. 2017), *Dioscorea rotundata* and *Asparagus officinalis* were downloaded from NCBI (Sayers, et al. 2012). The genome sequence of *P. dactylifera* (Al-Mssallem, et al. 2013) was downloaded from http://qatar-weill.cornell.edu/research/datepalmGenome/, the genome sequence of *Phalaenopsis equestris* (Cai, et al. 2015) was downloaded from ftp://ftp.genomics.org.cn/from_BGISZ/20130120/ and the genome sequence of *Spirodela polyrhiza* (Wang, et al. 2014) was downloaded from SpirodelaBase (https://www.waksman.rutgers.edu/spirodela/home). The genome of *Musa acuminata* (D'Hont, et al. 2012) was accessed on the website http://banana-genome.cirad.fr/. The genome of *Zostera muelleri* (Golicz, et al. 2015) was downloaded from http://www.appliedbioinformatics.com.au/index.php/Seagrass_Zmu_Genome. For the genome of *Z. marina* we assessed the sequence reads on the short read archive (SRA) at NCBI (Sayers, et al. 2012).

*Brassicales genomes*

The genomes of *A. lyrata*, *C. rubella*, *E. salsugineum*, *B. rapa* and *C. papaya* were accessed at Phytozome (Goodstein, et al. 2012). The genomes of *A. arabicum*, *C. sativa*, *B. oleracea* and *S. irio* were investigated on CoGe (https://genomevolution.org/coge/). Finally, the genomes of *L. alabamica* and *S. parvula* were downloaded from VEGI (Haudry, et al. 2013) and the genome of *T. hassleriana* was downloaded from NCBI (Sayers, et al. 2012).

#### Identification of *AGL17*-like genes

Known *AGL17*-like genes were taken from Gramzow and Theißen (2013), Gramzow and Theissen (2015) and Cai, et al. (2015). The *AGL17*-like genes from monocots were identified by BLAST searches of the genomes listed above using the coding sequence of *OsMADS23* as query. *AGL17*-like genes from *Tarenaya hassleriana*, *Capsella rubella*, *Leavenworthia alabamica*, *Camelina sativa*, *Brassica rapa*, *B. oleraceae*, *Schrenkiella parvula*, *Sisymbrium irio* and *Eutrema salsugineum* were identified by searches for syntenic genes on the Brassica Database (Cheng, et al. 2011) and by BLAST searches of the genomes on Phytozome (Goodstein, et al. 2012) and CoGe (Lyons, et al. 2008) with the *AGL17*-like genes from *A. thaliana* as queries.

**Alignments**

*MIR444* sequences of *O. sativa*, *Z. mays* ssp. *mays*, *M. acuminata*, *E. oleifera* and *P. equestris* and potential *MIR444* sequences identified from the 1KP data were aligned using MAFFT (Katoh, et al. 2002). Sequences of AGL17-like proteins as identified in genomes and transcriptomes of monocots were aligned using Probalign (Roshan and Livesay 2006). The resulting protein alignment was reverse translated into the corresponding nucleotide alignment using RevTrans (Wernersson and Pedersen 2003). Both alignments were visualized using Jalview (Waterhouse, et al. 2009).

**miRNA Northern blotting**

Plant material was obtained as follows: *Maranta leuconeura, Tradescantia blossfeldiana, Veratrum album, Pandanus utilis, Pandanus natans, Dioscorea bulbifera, Elodea canadensis, Acorus gramineus* and *Musa acuminata* plant material was provided by the Botanical Garden of the Friedrich Schiller University in Jena. The *O. sativa* cv. Dongjin seeds were obtained from the Rice T-DNA Insertion Sequence Database (RISD). *A. thaliana* Col-0, *O. sativa* cv. Dongjin*, Z. mays* ssp. *mays, Hypoxis villosa*, *Phoenix dactylifera* and *T. hassleriana* were grown in the greenhouse. *H. villosa* was grown in a greenhouse with no additional artificial light. *O. sativa* and *P. dactylifera* were grown under the following conditions: 8 h light with 24°C during the day and 20°C at night. The conditions for the *A. thaliana* were as follows: 16 h light at 19°C during the day and 17°C at night. *Z. mays* ssp. *mays* was grown with 16 h light, 19°C during the day and 10°C at night.

RNA was isolated as follows: Young leaf material and *Zea mays* root, leaf and culm material was harvested and ground under liquid nitrogen conditions either with a mortar and a pestle or using the Mixer Mill MM 400 (Retsch) with 35 ml grinding jars and 20 mm grinding balls. For *M. acuminata*, *P. dactylifera* and *Zea mays* 5 day old seedlings, a CTAB-based RNA isolation method (Chang, et al. 1993) was used with the following modification: For the first RNA precipitation one volume of isopropanol was used instead of 10 M LiCl. For all other samples, total RNA was isolated with QIAzol Lysis reagent (Qiagen) according to the manufacturer's instructions.

For Northern blotting 50 µg of total RNA was separated on an 8% denaturing polyacrylamide gel and transferred by semi-dry electro blotting (10-15V and 0.45 mA for 60-90 minutes using a Hoefer TE77 semi-dry transfer unit, Amersham Biosciences) to a positively charged nylon membrane (Biodyne Plus Membrane, Pall Gelman Laboratory). Membranes were cross-linked using freshly prepared cross-linking solution (0.16M l-ethyl-3-(3-dimethylaminopropyl) carbodiimide (EDC) (Sigma) in 0.13M 1-methylimidazole at pH 8 (pH adjusted with HCl)) for 1 hour at 60°C. Hybridizations were carried out at 60°C overnight in miRNA hybridization buffer (50% deionized formamide, 5x SSPE, 5x Denhardt's, 0.5% SDS, supplemented with 20 µg/ml boiled salmon sperm DNA) and 100 ng/ml probe. As a probe a 3´ DIG-labelled RNA oligo probe (Biomers) against the *O. sativa* mature miR444a.1/d.1 was used. Blots were washed twice for 5 min in 2x SSC, 0.1% SDS at 60°C. Probe detection was performed using an alkaline phosphate conjugated DIG antibody and the chemiluminescent substrate Disodium 3-(4-methoxyspiro {l,2-dioxetane-3,2'-(5'-chloro)tricyclo[3.3.1.13,7]decan}-4-yl) phenyl phosphate CSPD. Briefly, blots were incubated in washing buffer (0.3% Tween 20 in maleic acid buffer [0.1 M maleic acid, 0.15 M NaCl, pH 7.5]) for 5 minutes. After an incubation for 30 minutes in blocking solution (1% blocking reagent [Roche Life Science] in maleic acid buffer), the blots were incubated for 30 min in antibody solution (75 mU/ml anti-DIG antibody in blocking solution). The blots were washed twice for 15 min in washing buffer, and subsequently equilibrated with detection buffer (0.1 M Tris-HCl pH 9.5, 0.1 M NaCl) for 5 min. Finally, the blots were incubated with chemiluminescent substrate CSPD and exposed to CL-XPosure film (Perbio Science).

**5´ RACE**

*5´ RACE of MIR444 from Dioscorea bulbifera*

3 µg of DNase I digested total RNA was reverse transcribed using Transcriptor Reverse Transcriptase (Roche Life Science) according to the manufacturer's instructions. The cDNA was purified using the High Pure PCR purification kit (Roche Life Science). The cDNA was dA-tailed using Terminal Deoxylnucleotidyl Transferase (ThermoFisher Scientific). The first PCR amplification was carried out with an oligo(dT) anchor primer and a gene-specific primer using HotStarTaq *Plus* DNA Polymerase (Qiagen) and the following PCR program: 95°C for 5 min, 10 cycles of 94°C for 30 s, 55°C for 30 s, 72°C for 1 min, followed by 25 cycles of 94°C for 30s, 63.9°C for 30 s, 72°C for 1 min, and a final step of 72°C for 10 min. A nested PCR amplification was carried out with a PCR anchor primer and a nested gene-specific primer using HotStarTaq *Plus* DNA Polymerase (Qiagen) and the following PCR program: 95°C for 5 min, 35 cycles of 94°C for 30 s, 59.2° or 61.9°C for 30 s, 72°C for 1 min, and a final step of 72°C for 10 min. The PCR reactions were separated on a 1% agarose gel with ethidium bromide (EtBr) staining, all bands were excised from the gel and the DNA

was extracted using the QIAquick Gel Extraction kit (Qiagen) according to the manufacturer's instructions. The purified PCR products were ligated into the pGEM®-T vector (Promega) as recommended by the manufacturer and subsequently sequenced (Macrogen).

*5´ RACE of the antisense transcript to the AGL17-like gene from Triglochin maritima*

3 µg of DNase I digested total RNA was reverse transcribed using RevertAid H Minus Reverse Transcriptase (ThermoFisher Scientific) according to the manufacturer's instructions. The cDNA was purified using the High Pure PCR purification kit (Roche Life Science). The cDNA was dA-tailed using Terminal Deoxylnucleotidyl Transferase (ThermoFisher Scientific). The first PCR amplification was carried out with an oligo(dT) anchor primer and a gene-specific primer using DreamTaq DNA Polymerase (ThermoFisher Scientific) and the following PCR program: 95°C for 2 min, 10 cycles of 95°C for 15 s, 50°C for 45 s, 72°C for 2 min, followed by 25 cycles of 95°C for 30 s, 66.9°C for 45 s, 72°C for 2 min, and a final step of 72°C for 5 min. A nested PCR amplification was carried out with a PCR anchor primer and a nested gene-specific primer using DreamTaq DNA Polymerase (ThermoFisher Scientific) and the following PCR program: 95°C for 2 min, 35 cycles of 95°C for 15 s, 59.2° or 60.7°C for 1 min, 72°C for 2 min, and a final step of 72°C for 10 min. The PCR reactions were separated on a 1% agarose gel with ethidium bromide (EtBr) staining, all bands were excised from the gel and the DNA was extracted using the QIAquick Gel Extraction kit (Qiagen) according to the manufacturer's instructions. The purified PCR products were ligated with the pGEM®-T vector (Promega) as recommended by the manufacturer and subsequently sequenced (Macrogen). An overview of the primers used is provided in Supplementary Table 1.

**RT-PCR and 3´ RACE**

Approximately 2.5 µg of DNase I digested total RNA was reverse transcribed using an oligo(dT) anchor primer and Transcriptor Reverse Transcriptase (Roche Life Science) according to the manufacturer's instructions. For the PCRs two gene-specific primers (for RT-PCR) or a gene-specific primer and a PCR anchor primer (for 3´ RACE), and HotStarTaq *Plus* DNA Polymerase (Qiagen) were used according to the manufacturer's instructions. The PCR reactions were separated on a 1% agarose gel with ethidium bromide (EtBr) staining, all bands were excised from the gel and the DNA was extracted using the QIAquick Gel Extraction kit (Qiagen) according to the manufacturer's instructions. The purified PCR products were ligated into the pGEM®-

T vector (Promega) as recommended by the manufacturer and subsequently sequenced (Macrogen). An overview of the primers used is provided in Supplementary Table 1.
